# Proteomic profiling of extracellular vesicles distinguishes prostate cancer molecular subtypes

**DOI:** 10.1101/2025.07.31.667906

**Authors:** Megan L Ludwig, Ali T Arafa, Saasha Vinoo, Jason C Jones, Abderrahman Day, Hannah E Bergom, Zoi Sychev, Alec Horrmann, Nicholas M Levinson, Scott M Dehm, Emmanuel S Antonarakis, Justin Hwang, Justin M Drake

**Affiliations:** Department of Pharmacology, University of Minnesota, Minneapolis, MN, United States; Department of Medicine, Division of Hematology, Oncology, and Transplantation, University of Minnesota, Minneapolis, MN, United States; Department of Laboratory Medicine and Pathology, University of Minnesota, Minneapolis, MN, United States; Department of Urology, University of Minnesota, Minneapolis, MN, United States; Masonic Cancer Center, University of Minnesota, Minneapolis, MN, United States

## Abstract

Prostate cancer is the most common non-cutaneous cancer among men in the United States. Most prostate cancers are driven by androgen receptor (AR) signaling, but there are an increasing number of cases that lose AR and gain neuroendocrine (NE) features (AR−/NE+) or lack both (AR−/NE−). These latter subtypes are particularly aggressive and lethal. Extracellular vesicles (EVs) have shown great potential as biomarkers for noninvasive liquid biopsy assays, as EVs contain biomolecules from their cells of origin. Here, we used a shotgun proteomics approach with mass spectrometry to interrogate the global proteome of EVs isolated from prostate cancer cell lines reflecting diverse clinical subtypes, including AR−/NE+ and AR−/NE−models. We identified 3,952 EV proteins, which clustered largely by tumor subtype and provided enough proteomic coverage to derive classic gene signatures of AR or NE identity that are of high relevance for prostate cancer prognostication. EVs isolated from AR+ cells displayed high levels of proteins regulated by AR and mTOR signaling. EVs isolated from AR−/NE+ cells contained known NE markers such as SYP and CHGA, whereas EVs from AR−/NE−models were enriched in basal cell markers and proteins that regulate epithelial-to-mesenchymal transition (EMT). We integrated our cell line data with recently published EV proteomics data from 27 advanced prostate cancer patients and found 2,733 overlapping proteins including cell surface markers relevant to prostate cancer, AR activity indicators, and proteins enriched in specific subtypes (AR+, AR−/NE−, AR−/NE+). This approach is especially promising for rare cancer subtypes, such as prostate cancers that lose AR− related features and gain NE features, so as to optimize the use of these liquid biopsy samples for clinical decision making.

## Introduction

Prostate cancer is the second most common male cancer in the United States with an estimated 313,780 cases to be diagnosed in 2025^1^. The majority of prostate cancers are diagnosed as prostate adenocarcinomas and rely on androgen receptor (AR) signaling. Blocking AR signaling through androgen deprivation therapy has been an effective mainstay treatment of prostate cancer for decades^2^. However, resistance inevitably develops as castration resistant prostate cancer (CRPC). Most cases of CRPC retain adenocarcinoma histology and AR signaling^3, 4^, though an increasing number of CRPC tumors can develop into poorly differentiated carcinomas that are independent of AR signaling, termed treatment-emergent small cell carcinoma^5^. AR− independent tumors have been classified into two main molecular subtypes: neuroendocrine (NE) prostate cancer (NEPC, AR−/NE+) and double negative prostate cancer (DNPC, AR−/NE−). Both subtypes share loss of AR signaling, *RB* deletion, and *TP53* mutation^6^. NEPC additionally expresses NE genes such as synaptophysin (SYP) or chromogranin A (CHGA) which are not normally expressed in AR+ CRPC^7^. Both NEPC and DNPC exhibit minimal response to androgen deprivation therapy, lose PSMA expression making them unlikely responders to the recently developed PSMA-targeted radioligand therapy, and require rapid and aggressive therapy. Monitoring the emergence of ADT resistance during a patient’s treatment could catch transformation into aggressive disease and stratify patients to more aggressive or alternate forms of treatment.

Extracellular vesicles (EVs) have attracted significant interest as liquid biopsy biomarkers in disease settings. These particles represent nano-sized subpopulations of exosomes and microvesicles which originate from the endosomal system or the plasma membrane respectively and can range from 40-1,000nm in size^8^, but also includes larger vesicles of 1-10µm in size such as large oncosomes^9^. EVs can be found in all biofluids including blood, urine, and saliva^10^, and are more abundant than circulating tumor cells^11, 12^. The contents of EVs include proteins, miRNAs, mRNAs, and DNA, which are protected by a lipid bilayer membrane that stabilizes their contents^13^. As EVs carry markers from their cells of origin, there is much potential for using liquid biopsy assays to interrogate EVs for prognostic and predictive biomarkers.

Earlier work in breast cancer noted that EVs displayed proteome profiles that were indicative of their molecular subtypes^14^, though much of the work in EVs in prostate cancer has focused on RNA. mRNA expression levels of the TMPRSS2:ERG fusion in EVs isolated from urine is a component of the ExoDx Prostate (*IntelliScore*) (EPI) test that was developed to detect high-grade prostate cancer^15^. Proteome profiling of EVs from the 60 National Cancer Institute (NCI) cell lines only included two prostate cancer models, DU145 and PC3, which are both DNPC models^16^. Other proteome profiling work in prostate cancer has been restricted to a singular representative cell line^17, 18^, which has hindered the ability to make comparisons based on disease phenotypes.

In this study, we generated comprehensive proteomic profiles of EVs in multiple prostate cancer cell lines representing three clinically relevant molecular subtypes. Our results revealed a diversity of proteins that are carried by EVs, and that proteome profiles are largely distinguished by molecular subtypes. Individual proteins as well as signaling pathways are differentially enriched by prostate cancer subtype, including known therapeutic targets such as PSMA and TROP2. Remarkably, the proteomes of EVs carry proteins that indicate levels of AR signaling and NE identity from their cells of origin, further showcasing the potential of EVs to give information about signaling activity from tumors. While to date it has been a challenge to translate findings from EVs in cell lines to the clinic, we compare our results to our previous proteome profiling of EVs isolated from the plasma of advanced prostate cancer patients^19^. Our results suggest that with enough depth, profiling EVs isolated from patients can uncover significant protein markers relevant to cancer cells.

## Materials & Methods

### Cell Culture

Human prostate cancer cell lines LNCaP, C4–2, 22RV1, DU-145, PC3 and NCI-H660 cells were obtained from the American Type Culture Collections (ATCC). LASCPC-01 cells were a gift from Owen Witte at UCLA, EF1 cells were a gift from John K Lee at Fred Hutchinson, and LNCaP 95 were a gift from Scott Dehm at University of Minnesota. NCI-H660 and LASCPC-01 cells were grown in Advanced DMEM/F12 (Gibco), with 1X B27 Supplement (Gibco), 10 ng/mL EGF (PeproTech), 10ng/mL bFGF (PeproTech) 1% penicillin-streptomycin, and 1X Glutamax (Life Technologies). All other lines were grown in RPMI 1640 without phenol red (Gibco) supplemented with 10% FBS (Sigma Aldrich), 1% penicillin-streptomycin, and 1X Glutamax (Life Technologies). Cells were grown and maintained in a humified incubator at 37°C and 5% CO_2_.

### EV Isolation

FBS was spun at 100,000xg for 18 hours at 4°C to deplete bovine EVs. Cells were then plated at 5 million cells per 15cm dish and cultured in media with 5% EV-depleted FBS for 48 hours. As the media for LASCPC-01 and NCI-H660 contained no additional FBS, conditioned media was collected from LASCPC-01 in the same 48-hour window. Due to the slow growth of NCI-H660s, conditioned media was collected after 7 days post-plating. Conditioned media was then spun at 2,000xg for 30 minutes followed by 10,000xg for 30 minutes. Extracellular vesicles were then pelleted using the Beckman Coulter Optima XPN-100 ultracentrifuge and SW-32 rotor at 120,000xg for 70 minutes, washed with PBS, and ultra-centrifuged again at 120,000xg for 70 minutes. Biological replicates represent separate EV isolations. All spins took place at 4°C. Final extracellular vesicle pellets were resuspended in PBS and stored at −80°C.

### Mass Spectrometry

EV pellets were processed for mass spectrometry analysis as previously described^20^. EVs were lysed in buffer containing 8M urea, 2M thiourea, 400mM Tris pH 8.0, 20% acetonitrile, 10mM TCEP, 25mM chloroacetamide, and HALT protease (Roche) and phosphatase inhibitor cocktails (Thermo Scientific). Samples were then sonicated in a water bath for 10 minutes before incubating at 37°C for 30 minutes and then room temperature for 15 minutes to reduce and alkylate cysteines. Proteins were quantified by Bradford assay (BioRad) and 5µg of protein was digested with Lys-C (1:50) and trypsin (1:10) at 37°C overnight. Peptides were de-salted using MCX stage tips followed by C18 stage tips, and then quantified using Colormetric Peptide Assay kit (Thermo Fisher). 200 ng of peptides were analyzed by capillary LC-MS with a Dionex UltiMate 3000 RSLCnano system on-line with an Orbitrap Eclipse mass spectrometer (Thermo Scientific, Waltham MA) with FAIMS (high-field asymmetric waveform ion mobility) separation. Gradient separation was on a 40 cm, self-packed C18 capillary column with 100 um inner diameter (Dr. Maisch GmbH ReproSil-PUR 120 Å C18-AQ, 1.9 um particle size); the column was maintained at 55°C with a column heater from Sonation (Biberach, Germany). Peptides were separated with the following profile: 5% B solvent from 0 – 2 minutes, 8% B at 2.5 minutes, 21% B at 60 minutes, 35% B at 90 minutes and 90% B at 92 minutes with a flowrate of 350 nl/min from 0 – 2 minutes and 315 nl/minute from 2.5 – 92 minutes, where solvent A was 0.1% formic acid in water and solvent B was 0.1% formic acid in acetonitrile. The FAIMS nitrogen cooling gas setting was 5.0 L/min, the carrier gas was 4.6 L/Min and the inner and outer electrodes were set to 100 °C. The CV (compensation voltage) was scanned at −45, −60 and −70 for 1 second each with a data dependent acquisition method. The following MS parameters were employed: ESI voltage +2.1 kV, ion transfer tube 275 °C; no internal calibration; Orbitrap MS1 scan 120k resolution in profile mode from 400 – 1400 m/z with 50 msec injection time and 100% (4 × 10E5) automatic gain control (AGC); MS2 was triggered on precursors with 2 – 6 charges above 2.5 × 10E4 counts; MIPS (monoisotopic peak determination) was set to Peptide; MS2 settings (all CV’s) were: 1.6 Da quadrupole isolation window, 30% fixed collision energy, ion trap detection with 35 msec max injection time, 100% (1 × 10E4) AGC, 12 sec dynamic exclusion duration with +/-10 ppm mass tolerance and exclusion lists were not shared among CV’s.

Raw MS files were then searched using MaxQuant (v1.6.10.43) and Andromeda against the Uniprot human reference proteome database with canonical and isoform sequences (UP000005640, downloaded August 10^th^, 2021). Known contaminant sequences from the common Repository of Adventitious Proteins were removed as previously described^20^. The false discovery rate was set at 1%. Group-specific parameters included max missed cleavages of 2 and label free quantification (LFQ) with an LFQ minimum ratio count of 1. Global parameters included match between runs with a match time window and alignment time window of 5 and 20 minutes respectively, and match unidentified features selected. Quantitative, label-free proteomic data from MaxQuant was then imputed using random values generated from a normal distribution centered on the 1% quantile and the median standard deviation of all peptides. We then averaged the intensity of peptides that belonged to the same protein, requiring that the protein have at least two peptides. The data were then normalized by variance stabilization normalization (VSN)^21, 22^.

### Clustering & R Packages

Hierarchical clustering was performed using the Cluster 3.0 program with Pearson correlation and average linkage analysis. Java TreeView was used to visualize clustering results. Quantitative data for each peptide and protein can be found in Supplementary Tables 1&2. R scripts were run with RStudio (2023.06.1) and R 4.3.1. PCA plot was generated with the package “PCA tools”^23^ and ternary plot with package “Ternary”^24^.

### NanoTracking Analysis

The ZetaView Quatt PMX-430 (Particle Metrix GmbH, Innan am Ammersee, Germany) was used to analyze EV samples for size and particle concentration with ZetaView Software Suite (v1.3). Samples were diluted with PBS to manufacturer recommendations of optimal particles within reading frame. Measurements were collected across eleven positions using scatter with a 488nm laser wavelength at set temperature of 25°C, camera rate of 30 frames per second for 30 seconds, sensitivity of 85 and shutter speed of 200. Results were plotted using GraphPad Prism.

### Transmission Electron Microscopy

EVs isolated from DU145 and 22RV1 cell lines were applied to glow discharged 200 mesh copper grids for 2 minutes. Grids were washed with ddH_2_O and then floated on a droplet of freshly prepared 2% uranyl formate for 30 seconds before blotting away excess and air-drying. Samples were imaged using a FEI Tecnai G2 F30 Field Emission Gun Transmission Electron Microscope equipped with a Gatan 4k x 4k ultrascan CCD camera.

### Western Blotting

Whole cell lysates or EV samples were lysed with RIPA buffer (ThermoFisher) with added protease and phosphatase inhibitors. Lysates were then quantified using BCA assay (ThermoFisher) according to manufacturer recommendations. Samples were run and transferred onto PVDF 0.45um membrane (Millipore) and blocked with 5% BSA Fraction V. Primary antibodies included CD9 (Cell Signaling, 13174), ALIX (Cell Signaling, 2171), Calnexin (Cell Signaling, 2679), PSMA (Cell Signaling, 12815), TROP2 (abcam SP295). Secondary antibodies were Licor IR-conjugated, and membranes were imaged using the BioRad Chemi-Doc Imager.

### Gene Set Enrichment Analysis

Data was run in GSEA v4.3.2 (Broad Institute) against Hallmark and KEGG gene sets. 100 permutations were performed with the gene_set type given the limited number of samples and gene IDs compared to RNA-seq data. Additional parameters are weighted enrichment statistic and Signal2Noise metric for gene ranking, which are the GSEA default. GSEA results are listed in Supplementary Tables 3-5. Gene ontology and pathway enrichment was also calculated using DAVID^25^ using the default *Homo sapien* background or EnrichR^26^.

### Activity Scoring

AR activity scores were calculated by summative z-score of gene sets and then converted to a scale of 0 – 1, where 0 is the lowest score and 1 the highest as described^27^. The basal and luminal scores were similarly calculated. All gene sets are in Supplementary Table 6. The NE score was calculated by correlation with reference dataset as described^28^.

### Data Availability

Data files have been deposited at ProteomeXchange PXD058251 under MassIVE MSV000096519.

## Results

### Prostate cancer EVs cluster by subtype

We used unbiased mass spectrometry to characterize the proteome of EVs from clinically relevant subtypes that emerge in prostate cancer, such as those reliant on AR signaling (AR+), AR indifferent with NE features (AR−/NE+), and those without AR or NE features (AR−/NE−) (Figure 1a). We isolated EVs from prostate cancer cell lines of these differing subtypes - AR+ (LNCaP, LNCaP 95, C4-2, 22RV1), AR−/NE+ (NCI-H660, EF1, LASCPC-01), and AR−/NE− (PC3, DU145) – targeting small EVs with our differential ultracentrifugation protocol. We confirmed the presence of known EV markers CD9 and ALIX, the absence of a cell lysate marker Calnexin (Figure 1b), and an expected size range of small EVs (<200nm) for the majority of the population by NanoTracking Analysis (NTA) (Figure 1c, Supplementary Figure 1). However, the AR−/NE−isolations (PC3, DU145) contained an increased proportion of large EVs (>200nm) and so we will refer to these isolations as EVs rather than specifically small EVs. To further characterize the EV isolations, we performed transmission electron microscopy (TEM) (Figure 1d, Supplementary Figure 2) which showed intact vesicles with minimal impact on sphericity and few non-EV particles such as lipids or protein aggregates. We also observed larger EVs (>200nm) by TEM in DU145, consistent with the NTA, whereas this range of size was absent from 22RV1 EVs (Supplementary Figure 2).

**Figure 1.**
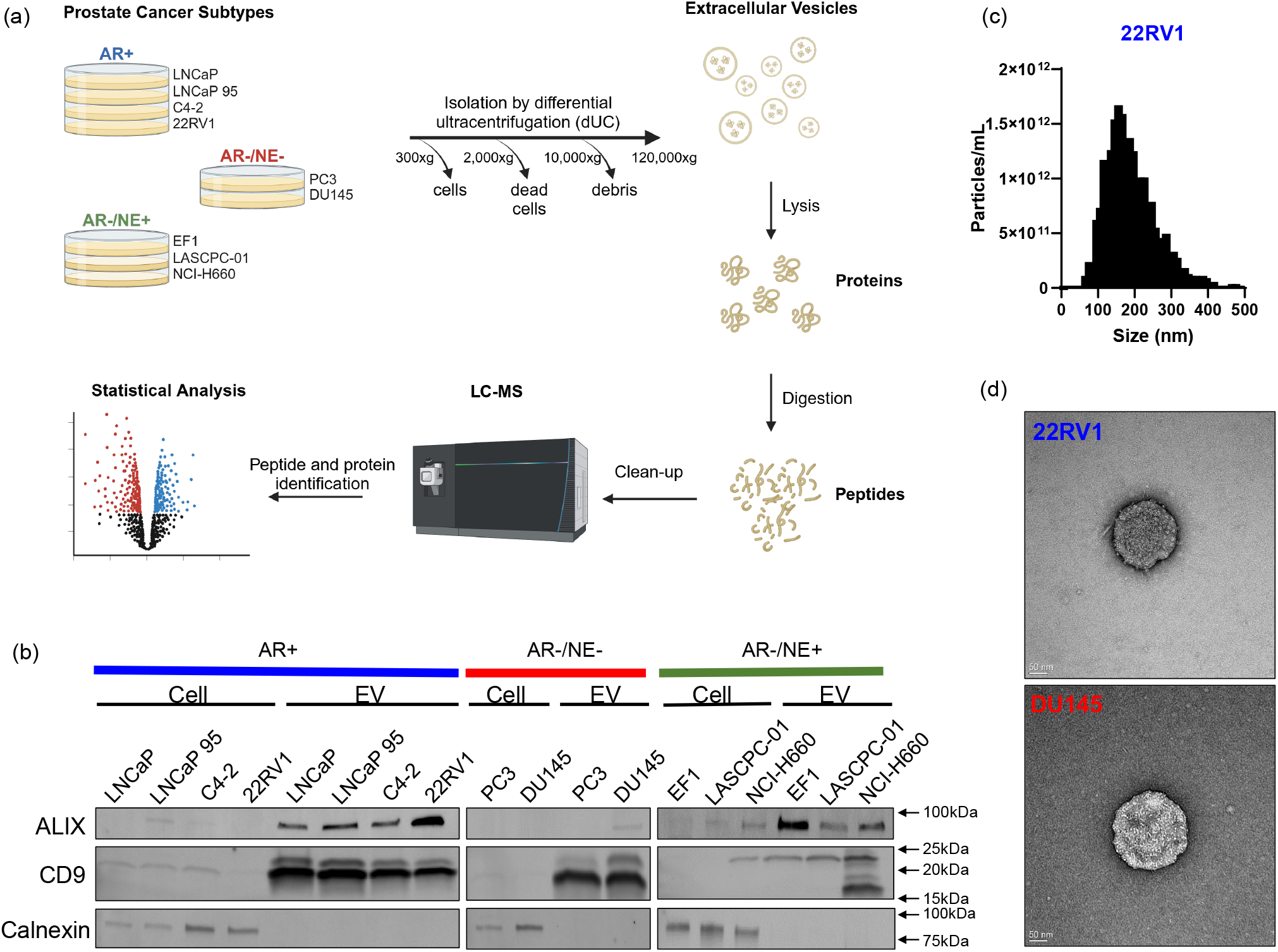
Extracellular vesicles (EVs) isolated from prostate cancer cell lines. a) Schematic of workflow from isolating EVs from prostate cancer cell lines of different subtypes by ultracentrifugation to preparing and running peptides on mass spectrometry. Created with Biorender.com. b) Western blot of cell or EV lysates indicating expression of EV markers ALIX and CD9 or cell lysate marker Calnexin. c) Nanotracking analysis (NTA) using ZetaView showing concentration of EVs from 22RV1 (AR+). EVs from the other cell lines are in Supplementary Figure 1. d) Transmission electron microscopy images of negatively-stained EVs from 22RV1 (top) and DU145 (bottom) with scale bar shown in the lower left

We then prepared the EV proteomes for quantitative mass spectrometry analysis, which identified over 2,000 proteins in each sample and 3,952 proteins across the entire cohort (Supplementary Table 7). Unsupervised hierarchical clustering of the complete EV proteomes showed that EVs clustered mainly by prostate cancer subtype (Figure 2a). Proteins were largely shared between EVs isolated from cell lines of the same subtype (Supplementary Figure 3), and 861 proteins (17.4%) were common to every EV proteome (Figure 2b). Unsurprisingly, these shared proteins were enriched for extracellular vesicle proteins (Supplementary Figure 4) and included canonical EV markers CD9 and CD81. Principal Component Analysis (PCA) demonstrated tight clustering of EVs isolated from AR+ cell lines, with the exception of LNCaP 95 which clustered more with EVs from AR-cell lines (Figure 2c). This observation was intriguing to us as LNCaP 95 cells were derived from LNCaP through androgen deprivation to exhibit resistance^29^, and while LNCaP 95 cells express variants of AR, its EV proteome is more similar to other AR-cell lines.

**Figure 2.**
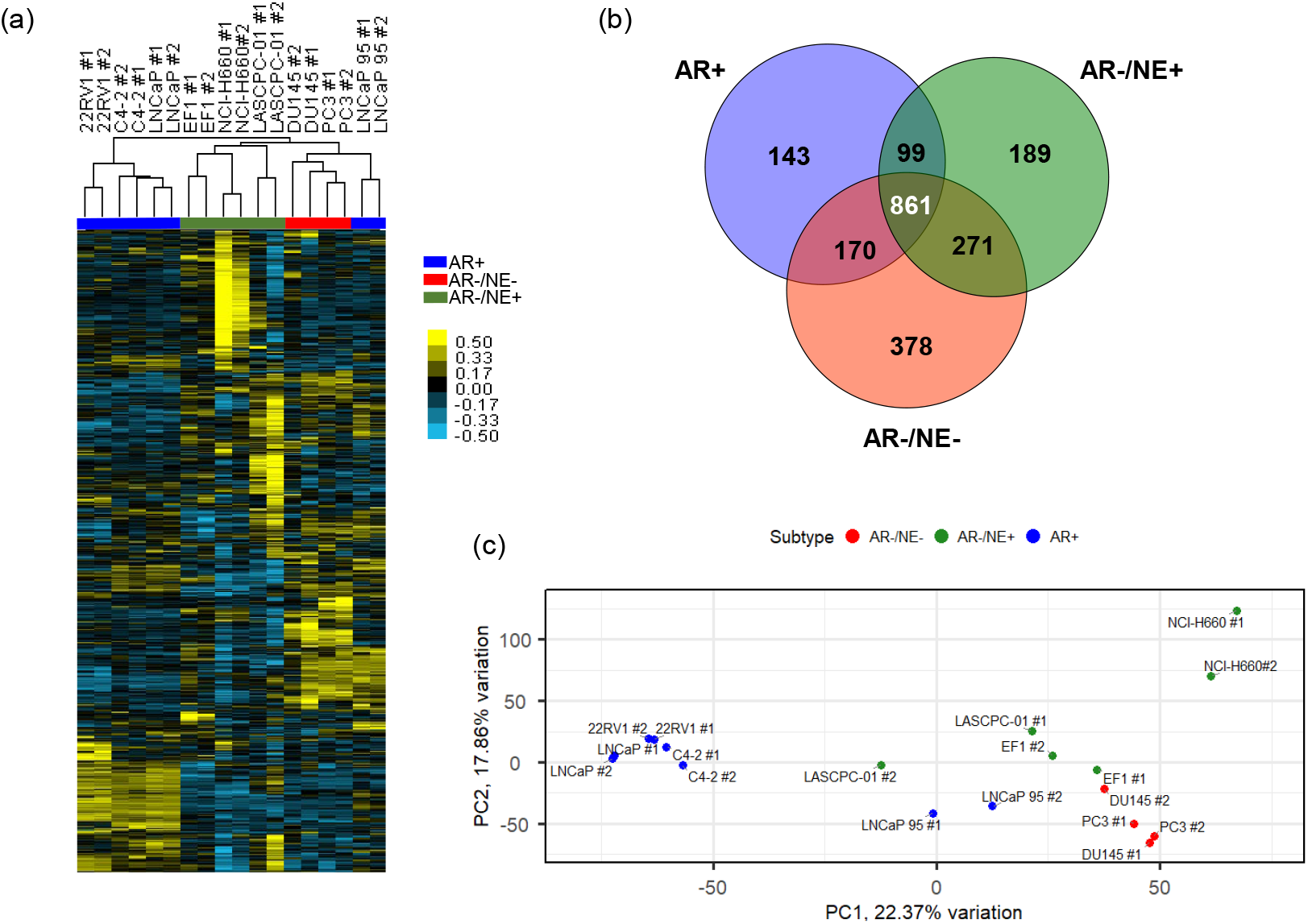
EV proteomes cluster by subtype of prostate cancer. a) Heatmap of unsupervised hierarchical clustering of 3,952 proteins identified by mass spectrometry in EVs. Biological replicates of EV isolations are indicated as #1 and #2. Samples are color coded by prostate cancer subtype: AR+ (blue), AR−/NE−(red), AR−/NE+ (green). b) Venn diagram of proteins found in all EV isolations of indicated subtype. c) Principal component analysis of EV proteomes for each replicate, with samples color coded by subtype.

### EVs contain prostate cancer markers from cell of origin

While the contents of EVs can reflect their cells of origin, EVs have differing enrichment of protein signatures and do not directly correlate with cell lysate expression^30^. We wanted to investigate markers of known relevance to prostate cancer, such as AR. While AR has been directly observed in EVs by immunoblotting^31^, we didn’t identify AR by mass spectrometry in our cohort and so turned to markers that indicate AR activity. We evaluated three signatures of AR activity^27, 32, 33^ (Supplementary Table 6, Supplementary Figure 5), as coverage of the proteins corresponding to these gene-based signatures was limited. Using the most complete gene signature^32^ (6/9, 67%), we observed that EVs isolated from AR+ models demonstrated the highest AR activity scores (Figure 3a). Notably, EVs from LNCaP 95 displayed the highest AR activity scores but with a distinct protein pattern compared to EVs from other AR+ cell lines (Figure 3b). Specifically, EVs from LNCaP 95 cells displayed elevated levels of RAB3B, FKBP5, and ACSL3, which were more commonly found in AR-models, rather than PSA (gene name: *KLK3*) and STEAP1/2. EVs from AR-models all displayed low AR activity as expected based on the independence from AR signaling in the cells of origin.

**Figure 3.**
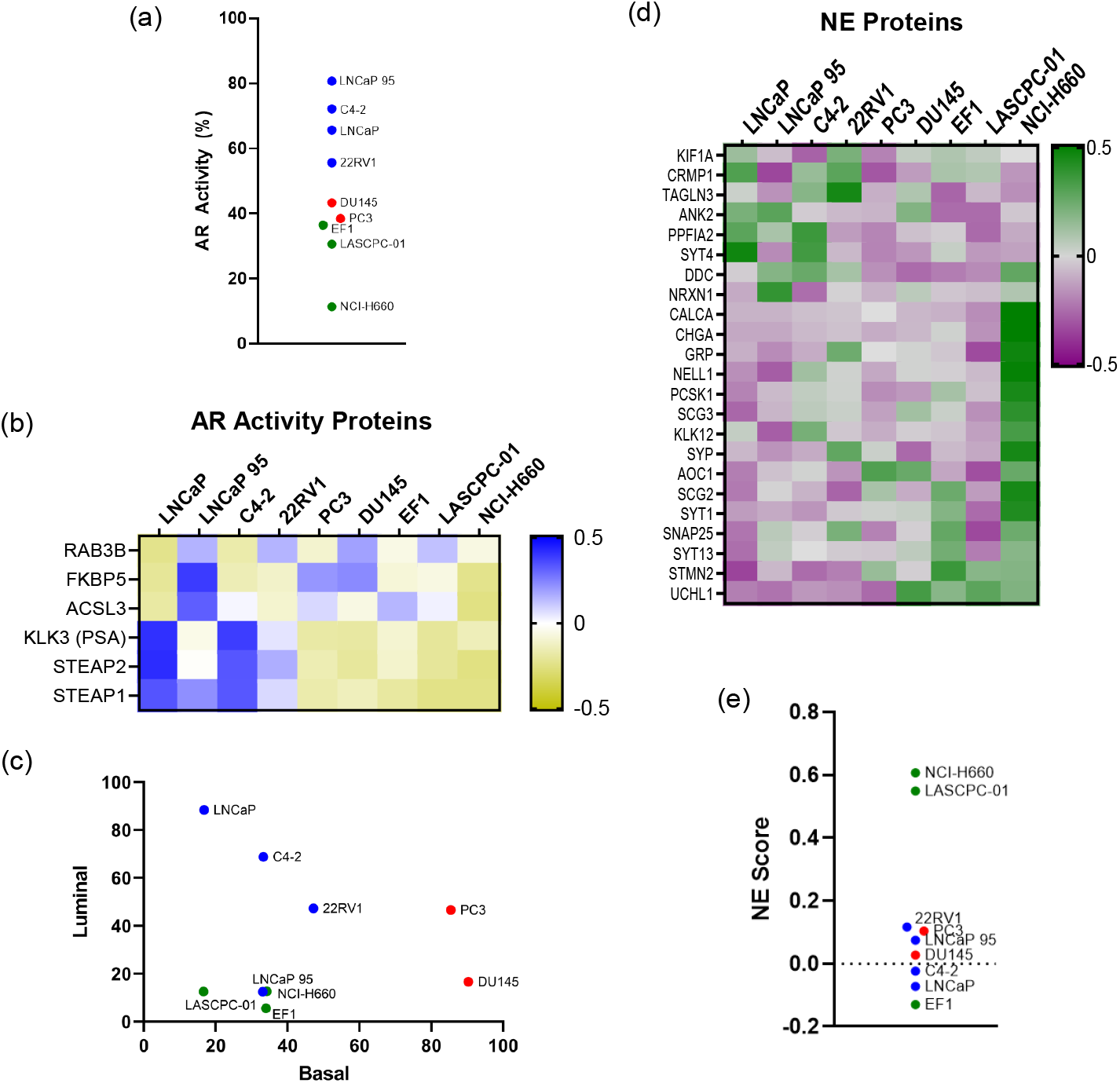
Prostate cancer EVs carry markers from cell of origin. a) AR activity scores as calculated for EV proteomes from most comprehensive signature, with average of replicates shown. b) Heatmap showing 6 proteins used for AR activity score in (a). c) Basal and luminal scores calculated for EV proteomes, with average of replicates shown. d) Heatmap of proteins with known association with neuroendocrine prostate cancer. e) Calculated neuroendocrine score by correlation with CRPC-NE samples ^28^ for EV proteomes, with average of replicates shown. Samples are color coded by subtype: AR+ (blue), AR−/NE+ (green), and AR−/NE−(red).

We then evaluated if EVs carried proteins associated with additional gene signatures relevant to prostate cancer, such as for luminal, basal, or NE cell identities. Prostate cancer cells that are reliant on AR signaling typically exhibit a luminal phenotype, while AR-independent cells display more basal (AR−/NE−) or neuroendocrine (AR−/NE+) features. Using luminal (33/71, 49.2%) and basal (9/30, 30%) gene signature scores^27^, we observed that EVs from AR+ cell lines predominantly contained luminal proteins as expected, while EVs from AR−/NE−were enriched for markers of basal identity (Figure 3c). We then examined the levels of NE-associated proteins^27^ and observed their presence predominantly in AR−/NE+ models (Figure 3d). Further, EVs from AR−/NE+ models NCI-H660 and LASCPC-01 scored highly using the Integrated Neuroendocrine Prostate Cancer score^28^ (20/70, 28.6%) (Figure 3e). However, EVs isolated from the EF1 AR−/NE+ cell line model scored low for this NE signature, despite clustering with the other AR−/NE+ in the total proteome (Figure 2a). This discrepancy may reflect further subtype heterogeneity in NEPC, as NCI-H660 and LASCPC-01 cells express ASCL1^34^, whereas EF1 expresses NEUROD1^35^. Neither transcription factor was detected directly in our mass spectrometry dataset, though similar to AR, the contents of EVs may retain specific subtype-defining proteins. These findings highlight the ability of EVs to carry distinct proteins indicative of luminal, basal, and NE cell identities, providing insights into the cellular origins and subtype-specific biology of prostate cancer.

We next examined cell surface markers associated with clinically relevant prostate cancer cell identities such as epithelial, adenocarcinoma, or NE. We identified 1,127 proteins that are classified as prostate cancer cell surface proteins^36^ (Supplementary Figure 6) whose levels clustered similarly to the total proteome (Figure 4a, Figure 2a). EpCAM was more highly enriched in EVs isolated from AR+ cell lines than those isolated from AR-cell lines (Figure 4b), highlighting potential limitations of using EpCAM to identify tumor-derived EVs in patient samples^11^. PSMA (gene name: *FOLH1*), a key target in prostate cancer therapy^37^, was enriched in EVs from AR+ cell lines, consistent with its relevance to the adenocarcinoma subtype. Similarly, STEAP1 and STEAP2 which have garnered interest as biomarkers for prostate cancer, are also highly enriched in EVs isolated from AR+ cell line models. Interestingly, TROP2 (gene name: *TACSTD2*), typically associated with epithelial and adenocarcinoma cells, was predominantly enriched in EVs from AR−/NE−models. Neuroendocrine cell surface markers NCAM1, DLL3, and CEACAM6 were present and enriched in EVs from AR−/NE+ cells. We then used a ternary plot to visualize how the cell surface markers clustered towards one specific subtype, plotting the average level for each protein by subtype where each axis of the triangle represents percentage of the total signal (Figure 4c) As was observed in Figure 4b, PSMA and STEAP1/2 expression were enriched in EVs from AR+ cell lines specifically, with 95-100% of PSMA and STEAP1/2 identified coming from the AR+ subtype (lower left point). TROP2 as well as ITGA3 and S100A16 were more specifically enriched in EVs from AR−/NE−models (top point), and rather than traditional neuroendocrine markers, CALR and HSPA5 were more specifically enriched in the AR−/NE+ models (lower right point). We further validated PSMA and TROP2 expression by subtype via Western blot (Supplementary Figure 7).

**Figure 4.**
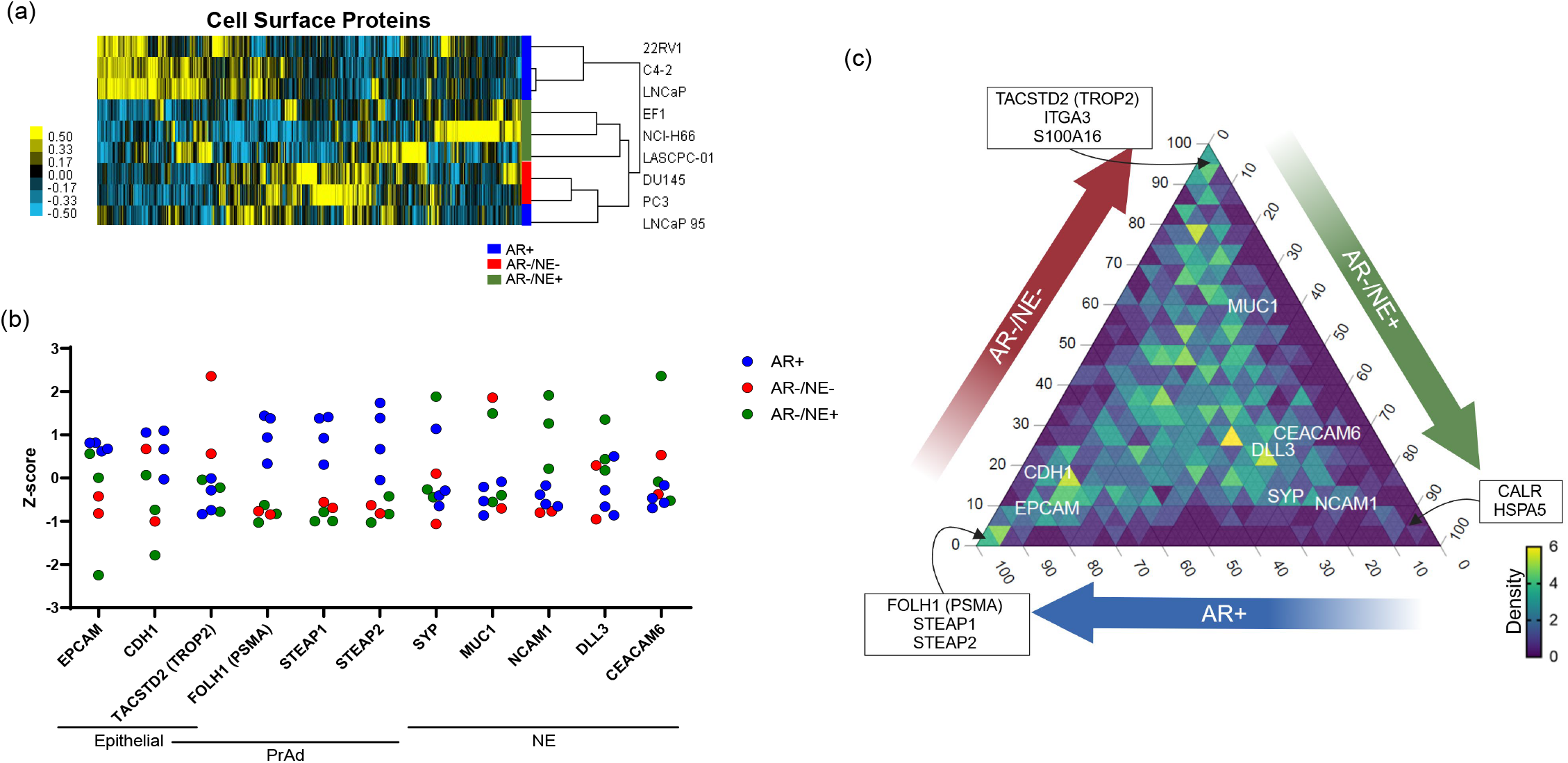
EVs exhibit cell surface markers of prostate cancer. a) Heatmap of unsupervised hierarchical clustering of 1,127 cell surface proteins found in the EV proteomes. b) Z-scores of cell surface markers with known relevance to prostate cancer cellular identities, such as epithelial, prostate adenocarcinoma (PrAd), and neuroendocrine (NE). Samples are color coded by subtype: AR+ (blue), AR−/NE+ (green), and AR−/NE−(red). c) Ternary plot of 1,127 cell surface proteins, plotting average protein level for each subtype. The percentage of total signal is represented on the edges of the triangle, 0-100. Density of proteins within an inner triangle is indicated by color, scaled from 0 (fewest proteins) to 6 (most proteins). Proteins more enriched to a specific subtype cluster to corners, whereas genes with similar expression in EVs across subtypes would cluster in the middle. Proteins of interest from (b) are labeled, with additional proteins of interest from each corner.

We then identified proteins and cell signaling networks that were enriched in each prostate cancer subtype. Comparing EVs isolated from AR+ vs. AR-cell lines, we observed an increase in proteins of known relevance to androgen-driven prostate cancer, including STEAP1 and PSMA (gene name: *FOLH1*), in EVs isolated from AR+ cell lines (Figure 5a). The most significantly enriched protein was RMC1, a regulator of CCZ1-MON1A/B function necessary for the localization of RAB7A which is involved in vesicle trafficking^38^. Gene set enrichment analysis (GSEA) revealed a global enrichment of proteins in the mTOR signaling pathway (Figure 5b), which is known to play a critical role in prostate cancer progression^39^ and may contribute to resistance to therapy. In EVs isolated from AR−/NE−models, we observed an enrichment of Vimentin and ICAM1 (Figure 5c) both of which are involved in epithelial-to-mesenchymal transition (EMT). This was further supported by GSEA, which identified the Hallmark EMT pathway as the most significantly enriched in EVs from AR−/NE−cell lines (Figure 5d), and that the EMT pathway has been previously noted to be enriched in AR−/NE−cells^40^. For EVs isolated from AR−/NE+ cell lines, we observed fewer significantly enriched proteins (Figure 5e), possibly reflecting greater heterogeneity within this subtype which can be visualized in the PCA plot (Figure 2c). Nevertheless, EVs still carried proteins of known importance to NEPC, such as UCHL1^41^. Overall, the most significantly enriched Hallmark pathway was oxidative phosphorylation (Figure 5f), which differs from expected as NEPC cells exhibit a glycolytic phenotype^42^. These findings highlight the potential of EVs to capture distinct protein signatures and signaling activities associated with different prostate cancer subtypes, providing insights into tumor biology and therapeutic resistance mechanisms.

**Figure 5.**
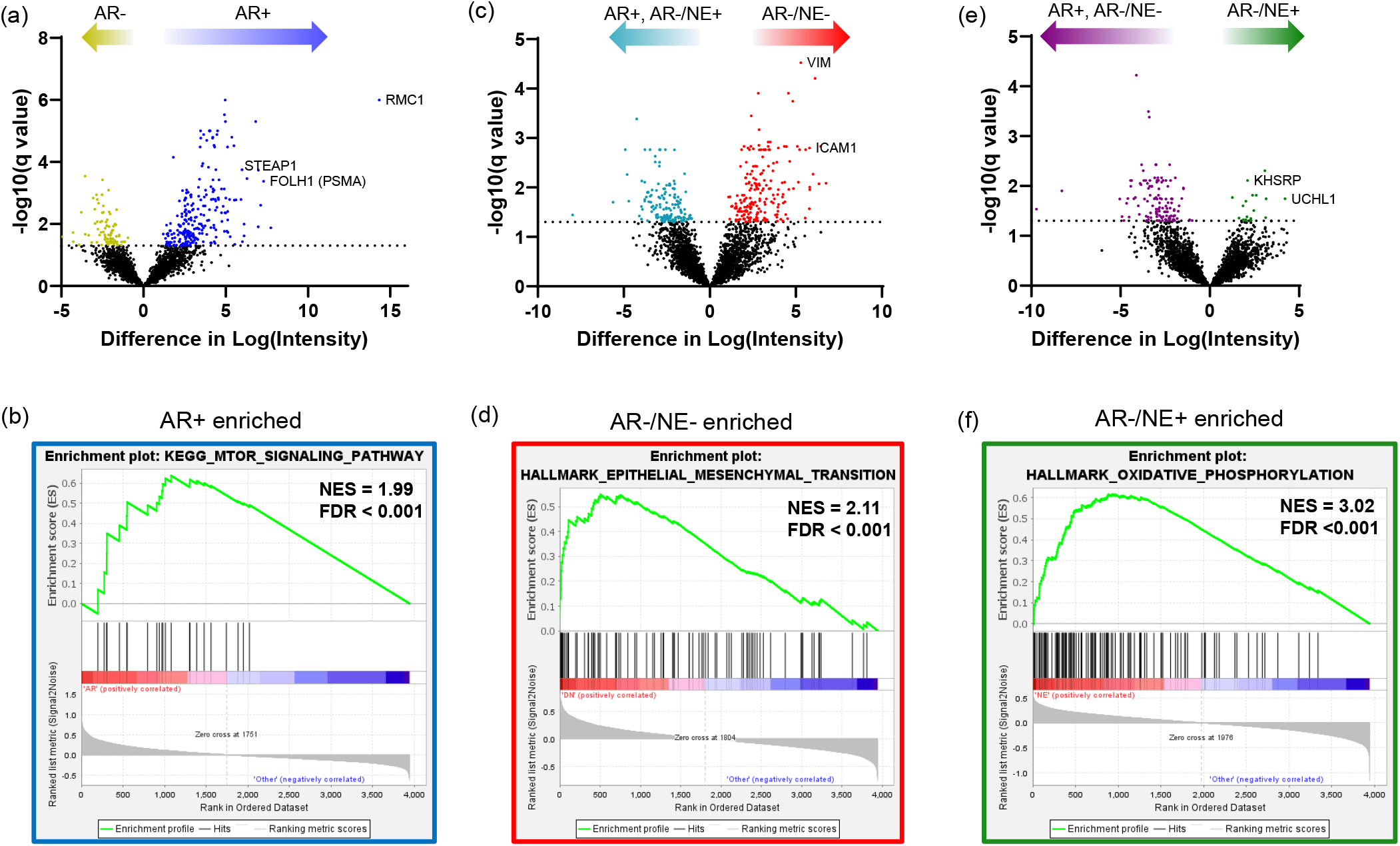
EVs display differential enrichment of proteins and pathways by subtype. a) Volcano plot comparing protein expression in EVs from AR+ cell lines to AR-, where significant proteins (q-value < 0.05) are colored for enrichment in AR+ (blue) or AR-(yellow). b) Gene set enrichment analysis (GSEA) of top enriched KEGG pathway in EVs from AR+ compared to AR−. c,d) As in (a,b), indicate enrichment of proteins in AR−/NE−EVs compared to the rest, with enrichment in AR−/NE−(red) and rest (teal) and most significant GSEA Hallmark pathway. e,f) As in (a-d), indicate enrichment of proteins in AR−/NE+ EVs compared to the rest, with enrichment in AR−/NE+ (green) and rest (purple) and most significant GSEA Hallmark pathway.

### Plasma-derived EVs from patients carry prostate cancer-relevant proteins

To determine whether traditional markers of advanced prostate cancer are detectable in EVs from patient samples, we analyzed recently published proteomic data from EVs isolated from the plasma of 27 metastatic castration resistant prostate cancer (mCRPC) patients^19^. This dataset identified over 5,000 proteins, including 4/9 (44%) genes from the AR activity signature. The levels of these proteins were similar across patients (Figure 6a), with higher levels of FKBP5 and ACSL3 compared to STEAP1/2. Calculated AR activity scores were low across much of the cohort (Supplementary Figure 8), which is likely due to the similar levels of these proteins. Given the rarity of AR-independent cases in patients, it is unlikely that so many of these patients exhibit AR−/NE+ or AR−/NE−cancer despite having low AR activity scores, similar to EVs from the AR-independent cell lines. Instead, it suggests that there was a lack of AR-independent models in the patient population for comparison, and clinical confirmation of histology is necessary to resolve this. Further, only 8/121 (6.6%) NE proteins were identified across the EV cohort with low correlation scores when compared to CRPC-NE models (Supplementary Figure 9). There were 25/71 (35%) luminal and 10/30 (30%) basal markers identified (Supplementary Figure 10), and the same patient’s EV proteome scored highest on both scales which may be indicative of the multiple cell types from which the EVs are being captured. Multiple surface markers relevant to prostate cancer were detected, including EpCAM, PSMA, and TROP2 (Figure 6b). While these EVs are not necessarily tumor-derived, the presence of these clinically relevant markers in the plasma of mCRPC patients is notable and highlights the potential for EV-based biomarker discovery in this patient population.

We then compared the total EV proteome from mCRPC patients to the EV proteome from prostate cancer cell lines. A total of 2,733 proteins overlapped between the datasets (Figure 7a), with levels varying across patients (Figure 7b). We used EnrichR on proteins that visually distinguished the three largest clusters of patients, which identified enrichments of Hallmark pathways such as mTOR, oxidative phosphorylation, and EMT pathways consistent with findings from our cell line analysis and supporting their relevance to prostate cancer progression. We next asked how these 2,733 shared proteins were represented in the cell line dataset and whether their levels were indicative of a specific prostate cancer subtype. The majority of these proteins were shared between all subtypes, as indicative of the density of the ternary plot being primarily in the middle (Figure 7c). In evaluating the 1,219 proteins that were identified in the EVs isolated from cell lines but not observed in this cohort of patients, we observed that many of the proteins not shared between datasets had been primarily enriched in the EVs from AR−/NE+ cell lines (Supplementary Figure 11). This concurs with our earlier analysis in which few markers of neuroendocrine prostate cancer were present in these patients.

Next, we further evaluated the proteins that were found to be significantly enriched in specific prostate cancer subtypes from the cell line EVs (Figures 5a, 5c, 5e). We asked if these proteins were also represented in the EVs from advanced prostate cancer patients and found that the majority of all three profiles were identified (AR+: 175/250, AR−/NE−: 161/201, AR−/NE+: 11/19). To further enhance possible prostate cancer-related signals, we removed proteins that have also been identified in EVs isolated from the plasma of healthy, non-cancer controls^43^. The average levels of these remaining proteins (AR+: 36, AR−/NE−: 27, AR−/NE+: 4) (Supplementary Tables 8-10, Supplementary Figure 12) are represented in Figure 7d. All patient EVs exhibited an elevated score of AR−/NE−subtype, which could be expected from mCRPC patients who had progressed on multiple lines of therapy including chemotherapies. We expected the AR+ score to be consistently elevated but instead found more variation from patient to patient, though comparing to a validated AR-sample is needed for improved context. Few AR−/NE+ markers were found, though it is notable that in some patients such as 5 and 6 that the levels of these proteins were higher than markers representing the other subtypes. This could indicate, along with expression of AR−/NE−enriched proteins, that these patients could be developing more AR-independent features or acquiring heterogeneity. Further exploration in a greater patient cohort with detailed review of tissue biopsies, specifically in patients with known clinical indications of AR-independent disease, is needed to clarify these findings.

**Figure 6.**
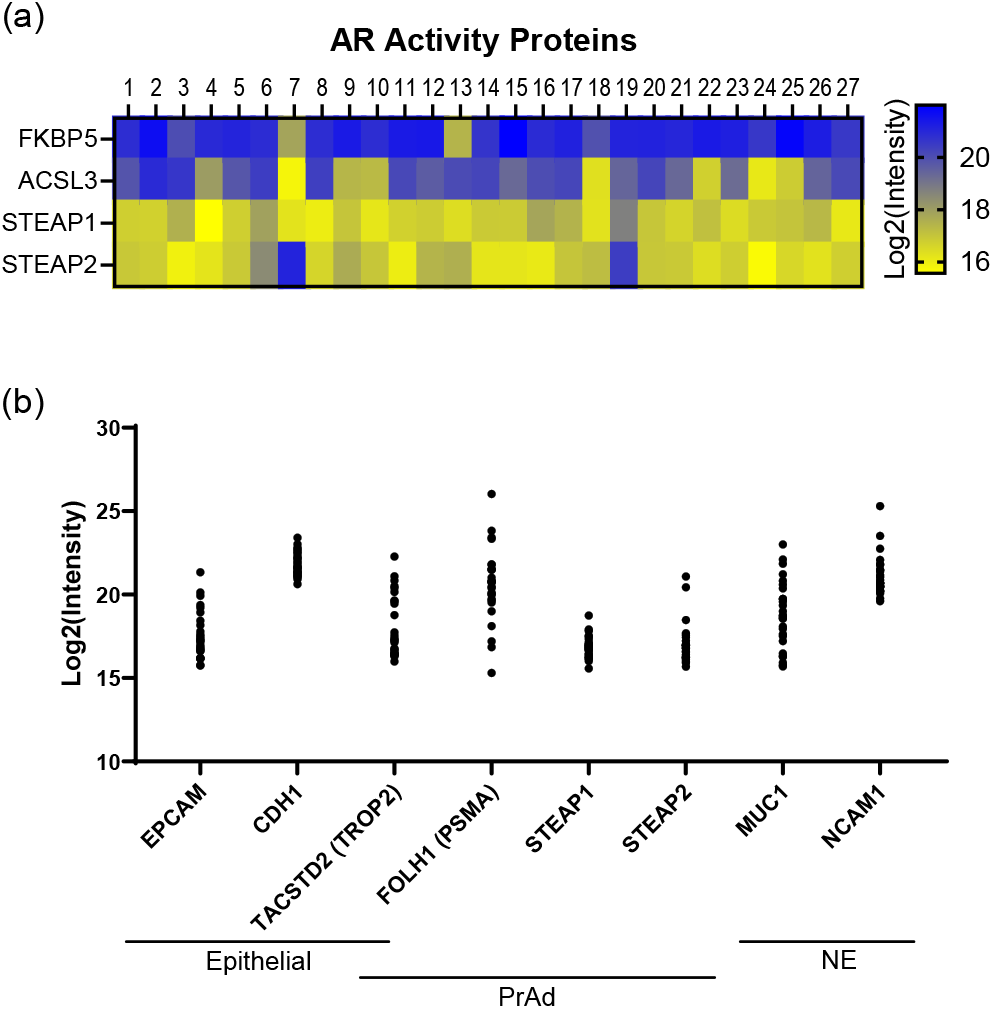
EVs from plasma of advanced prostate cancer patients carry signature proteins. a) Heatmap of 4 proteins found in AR activity score used in Figure 3a,3b for EVs isolated from the plasma of 27 advanced prostate cancer patients. b) Expression of cell surface proteins relevant to prostate cancer, similar to Figure 4b, of each patient.

**Figure 7.**
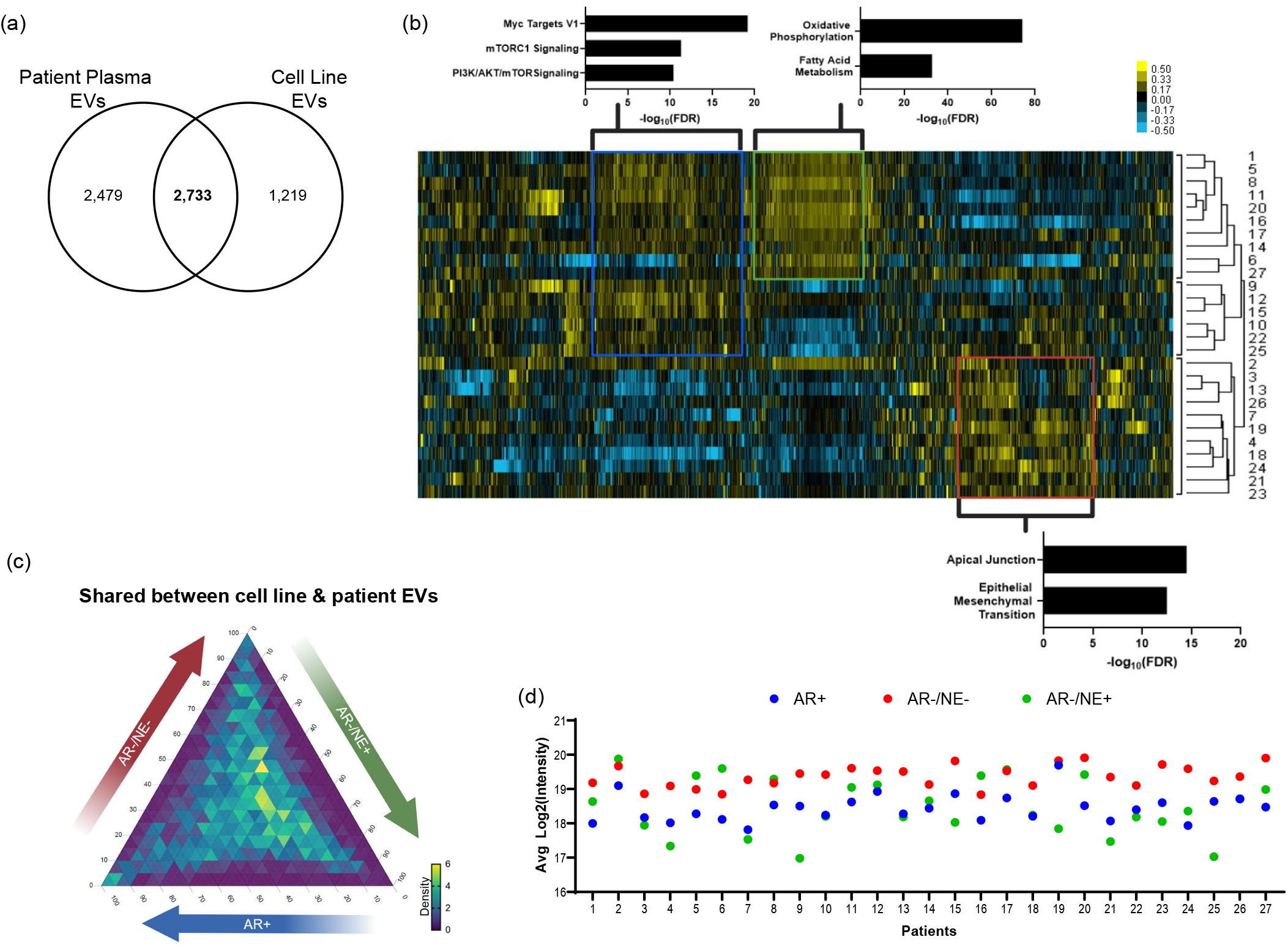
EVs from plasma of advanced prostate cancer patients overlap with cell line isolations. a) Venn diagram showing overlap of protein cohorts between plasma EVs from prostate cancer patients and the EVs isolated from prostate cancer cells. b) Heatmap of unsupervised hierarchical clustering of 2,733 proteins that overlapped between plasma and cell line EVs. Three largest clusters by dendrogram are indicated on the right. Genes in each indicated box were put into EnrichR, with the most significant Hallmark pathways indicated by graph. c) Ternary plot of 2,733 proteins that were shared between plasma and cell line EVs. The values plotted are average protein level for each subtype from cell line EVs. The percentage of total signal is represented on the edges of the triangle, 0-100. Density of proteins within an inner triangle is indicated by color, scaled from 0 (fewest proteins) to 6 (most proteins). d) Average expression of proteins associated with prostate cancer subtype for each patient, represented by 36 proteins for AR+ (blue), 27 proteins for AR−/NE−(red), and 4 genes for AR−/NE+ (green).

## Discussion

While it is known that EVs carry markers from their cells of origin, our results are the first to apply gene signatures relevant in prostate cancer to the proteome of EVs. That the EV proteomes largely reflected expectations on AR and NE activity, or basal and luminal identity, underscores the wealth of information carried in the EV proteome. Obtaining detailed information about the cell signaling activity occurring in the tumor cells of origin, beyond individual protein markers, could have immense clinical relevance. This is especially important in the case of AR, as we did not identify AR in our mass spectrometry dataset though other groups have observed AR in EVs by immunoblotting^31^. Given the nature of shotgun proteomics by mass spectrometry, this does not mean that AR protein was not present and could have been missed for technical reasons. More sensitive methods of identification, such as targeted mass spectrometry, may be needed to detect the AR protein. Targeted mass spectrometry requires the design, optimization, and validation of specific peptides that can be detected with higher sensitivity than shotgun proteomics approaches. This technique could be especially of use in detecting variants of AR, such as AR-v7 which is a known marker of resistance to ADT in prostate cancer. As nuclear proteins such as transcription factors are less commonly found in EVs, which carry largely membrane and cytoplasmic proteins^30^, more sensitive techniques need to be developed to detect these specific proteins. EVs isolated from cell lines displayed AR activity scores that correlated with expected based on the cells of origin, but it is yet unknown what the AR activity scores mean in EVs from patients. Detecting AR or AR-v7 levels could offer more direct measurements of AR activity and indicate burgeoning resistance to ADT. Identification of AR-v7 in CTCs has been informative for patient care^44^, and it is worthwhile exploring this concept in EVs for even earlier detection.

Our methods targeted small EVs (<200nm) and these were the majority of the EV population. However, EVs isolated from AR−/NE−cell lines contained more EVs of larger sizes, seen by both NTA and TEM. These larger vesicle populations have not previously been appreciated by NTA using the Nanosight for DU145 or PC3 EVs^45, 46^ despite using similar methods of ultracentrifugation as our protocol. The Nanosight has been shown to represent size of EVs more accurately than ZetaView^47^, which was used in this manuscript. However, this analysis focused on small EVs and TEM imaging confirmed the lack of larger vesicles (>200nm) suggesting that the Nanosight may not be optimal for identifying these larger EV sizes as compared to the ZetaView. Additional studies are warranted to determine if these larger EV sizes are of biological significance in AR-cells, indicating changes to mechanisms of EV sorting and packaging. Further separation of EVs by size, such as by size-exclusion chromatography or including gradients during differential ultracentrifugation, would help address this question as well as lead to further clarification on any changes in protein cargo carried by vesicles of different sizes. It is also important to note that EV purity is an important consideration. While our TEM imaging indicated minimal contamination by non-EV particles such as lipid or protein aggregates, our analysis by mass spectrometry could identify protein co-isolates alongside EV cargo. Comparative methods of isolation could help differentiate true protein cargo or with techniques such as immunogold labeling if the protein is projected to be on the surface of the EVs.

A limitation for the current clinical utilization of this protein information, both individual markers as well as activity scores, is the difficulty in isolating specific subsets of tumor-derived EVs. Our results suggest that using markers such as PSMA or EpCAM in prostate cancer patients to isolate or identify potential tumor-derived EVs are likely to miss the progression to AR-independence, given the lack of these markers in EVs isolated from AR-negative models. Specifically, EpCAM+ EVs have been used as an indicator of tumor-derived EVs^48^. However, our results indicate that using EpCAM as a singular marker in prostate cancer would miss EVs deriving from AR−/NE+ or AR−/NE−tumors. We anticipate that our results will help facilitate a more inclusive set of markers relevant for prostate cancer patients, such as including TROP2 or NCAM1, increasing the potential to find and identify these aggressive tumor subtypes. Importantly for developing EV-based tools, this may require tumor tissue in addition to blood from patients with these rare AR-independent disease types. Additionally, our approach did not give us information on how many EVs from the AR-positive models contained PSMA or EpCAM. It’s possible that there are a few EVs that carry a high load of PSMA or EpCAM, rather than more limited expression across most the EVs exported. These questions should be addressed for future exploration into isolating tumor-derived EVs using individual gene expression, and it may have relevance in developing companion assays to PSMA- or TROP2-targeting therapies in prostate cancer^37, 49^.

In analyzing total EVs without further enrichment that were isolated from advanced cases of prostate cancer patients, our results indicate the possibility of non-tumor EVs interfering with activity scoring. Unfortunately, the patient information to further investigate the EV proteome and its correlation with clinical outcomes such as survival, cancer progression, or tumor histology is not currently available. Patients received various ADT and chemotherapy treatments and exhibited multiple sites of metastasis, including bone, lymph node, and visceral, which may exhibit further differentiation of subtype by the EV proteome^19^. In addition, 25 of the 27 patients in this dataset identified as White with only 2 Black men. While our translational analysis of patient data is severely limited without histological confirmation or greater diversity, our results display the potential depth of information of cell signaling activity when the EV proteome is profiled. Gene signatures developed from RNA expression data on tumor lysates were informative in EVs from cell lines, however, these signatures appear to fall short with patient EVs without tumor-specific enrichment. Proteins enriched in the specific prostate cancer subtypes from cell lines could be more informative in establishing clinically relevant signatures. Moving forward, profiling patients with the rarer disease subsets of AR-independent prostate cancer needs to be prioritized as there is a lack of data available from these subsets. Our proteome profiles of the EVs from cell lines could be used as a starting point for investigating markers and activity relevant to these AR-independent cancers, such as NEPC activity or EMT signaling, though it is important to note that these conclusions originate from a handful of cell line models. As AR-independent cell lines are few in the prostate cancer field, it may be more effective to focus on collecting data from patients with histological confirmation of AR-independent disease. Progression samples in which patients are monitored over time in response to treatment could be critical in understanding the dynamics of EV cargo and identifying robust protein markers. It is also important to note that there is much left to be learned about the influence of other factors that affect EV biology, such as age and ethnicity, as well as length or timing of therapeutic treatment. These factors could affect the protein cargo of EVs and can only be addressed by profiling more and a wider variety of patients.

In looking at EVs isolated from NEPC patients, our results suggest that further differences within the NEPC subtype will need to be evaluated. EF1 cells are driven by NEUROD1 which exhibits a different transcriptional profile than the other two cell lines driven by ASCL1^50^. NEUROD1 and ASCL1 are key lineage transcription factors that can drive further subtyping of NEPC^51^, and the ASCL1-target DLL3 was more highly expressed in the EVs from ASCL1-driven NCI-H660 and LASCPC01 (Supplementary Table 2). However, AURKA is thought to be of more therapeutic benefit in the NEUROD1 subtype of NEPC^51^ though we observed higher AURKA expression in EVs from the ASCL1-driven cell lines. Further work is warranted for EVs from NEPC cell lines to investigate proteomic profile differences between these two NEPC subtypes. However, our work suggests that proteins involved in metabolism are enriched in these EVs from both NEPC subtypes. Proteins involved in oxidative phosphorylation were also specifically enriched in the NEPC subset, which differs from elevated glycolysis that is exhibited in NEPC tumors. Typically, it is primary prostate tumors that are seen to favor increased oxidative phosphorylation and lower levels of glycolysis^42^. More investigation is needed to determine if there are biological reasons for NEPC cells to export EVs enriched with proteins involved in oxidative phosphorylation, such as increasing glycolysis in the cells of origin or promoting primary prostate tumor formation in surrounding tissues, or if this is an artifact of cell culturing conditions. If biological, it could highlight the importance of monitoring the metabolic reprogramming occurring in NEPC cells^52^ and serve to further validate metabolism as a therapeutic vulnerability in the cells of origin.

Our data supports the utility of using the proteome profiles of EVs for subtyping clinically relevant prostate cancer subsets. We show that the proteome profiles carry a diverse wealth of information on the cell signaling activity from the cells of origin, including using classic gene set signatures on the EV proteome. Many of these markers can also be seen in EVs isolated from advanced prostate cancer patients. It has been difficult in translating protein markers and information gained from EVs isolated from cell lines to patients due to limited overlap in protein identification. Proteins such as albumin and immunoglobulins in plasma has made profiling the EV proteomes difficult for clinical samples, but recent work has overcome this challenge and identified over 5,000 proteins in patient EVs^19^. Our results indicate that with enough depth, key indicators of tumor status and cell signaling activity can be found. Future work should continue to focus on profiling patient samples, specifically from rare AR-independent subsets, to identify liquid biopsy markers of clinical relevance. Larger cohort sizes of patients followed over time will be critical to understanding how EV protein cargo changes and can be best utilized clinically for advancing therapeutic treatment. Comparing EV proteome profiles of patients who do or do not respond to the latest therapies would be informative in predicting patient response and may serve to match patients with future therapies in development. Even if individual markers such as PSMA, TROP2, or DLL3 are not the best for matching patients to the targeted therapies in development for prostate cancer, our results showcase that the depth of proteins that EVs carry can create informative activity scores that may lead to more robust predictions. Pairing informative, minimally invasive biomarker information from EVs alongside therapies as they’re developed can increase the speed and effectiveness in the clinic.

## Acknowledgements

We thank Dr. LeeAnn Higgens and the Center for Metabolomics and Proteomics at the University of Minnesota for providing services related to generation of quantitative proteomics data. The Orbitrap Eclipse instrumentation platform used in this work was purchased through High-end Instrumentation Grant S10OD028717 from the NIH. Portions of this work were conducted in the Minnesota Nano Center, which is supported by the National Science Foundation through the National Nanotechnology Coordinated Infrastructure (NNCI) under Award Number ECCS-2025124. Parts of this work were carried out in the Characterization Facility, University of Minnesota, which receives partial support from the NSF through the MRSEC (DMR-2011401) and the NNCI (ECCS-2025124). The schematic in Figure 1a was created with https://biorender.com.

## Disclosure Statement

HEB is a co-founder and CEO of Emergense. SMD has served as a paid consultant/advisor to Janssen, Bristol Myers Squibb, and Oncternal Therapeutics, and has served as principal investigator on grants awarded to the University of Minnesota by Janssen and Pfizer/Astellas. ESA reports grants and personal fees from Janssen, Sanofi, Bayer, Bristol Myers Squibb, Curium, MacroGenics, Merck, Pfizer, AstraZeneca, and Clovis; personal fees from Aadi Bioscience, Aikido Pharma, Astellas, Amgen, Blue Earth, Corcept Therapeutics, Exact Sciences, Hookipa Pharma, Invitae, Eli Lilly, Foundation Medicine, Menarini-Silicon Biosystems, Tango Therapeutics, Tempus and Z-alpha; grants from Novartis, Celgene, and Orion; and has a patent for an AR-V7 biomarker technology that has been licensed to Qiagen. JMD has no conflicts relevant to this work. However, he serves as a consultant and Chief Scientific Officer of Astrin Biosciences. The interest related to JMD has been reviewed and managed by the University of Minnesota in accordance with its Conflict-of-Interest policies. The other authors have no disclosures.

## Funding

ML and JJ were supported by the Targets of Cancer Training Program Fellowship (T32 CA009138). A.T.A. is supported by a Postdoctoral Fellowship, PF-23-1153194-01-CDP, Grant #: [https://doi.org/10.53354/ACS.PF-23-1153194-01-CDP.pc.gr.175399]. SMD is partially supported by NCI grants R01CA174777 and R01CA270539. ESA is partially supported by NCI grant P30 CA077598 and DOD grant W81XWH-22-2-0025. JMD is supported by the Masonic Cancer Center at the University of Minnesota and by the National Institutes of Health NCI R01CA269801. This work was also partially supported by a Masonic Cancer Center Cancer Research Translational Initiative (CRTI) pilot grant.

