## Supplemental Figures for "Proteomic profiling of extracellular vesicles distinguishes prostate cancer molecular subtypes"

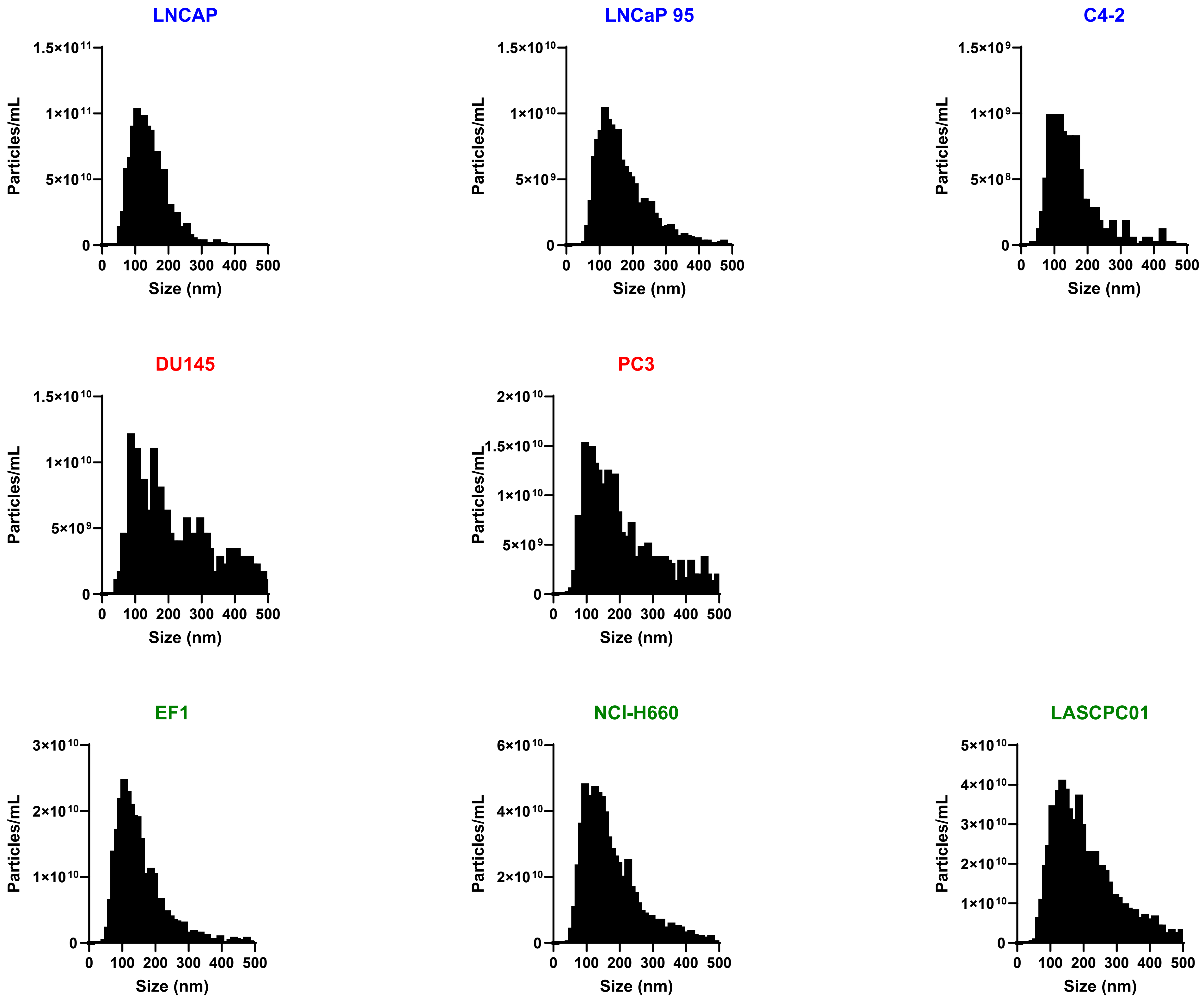


**Supplementary Figure 1.**
Nanotracking analysis (NTA) using ZetaView for EVs isolated from each indicated cell line.

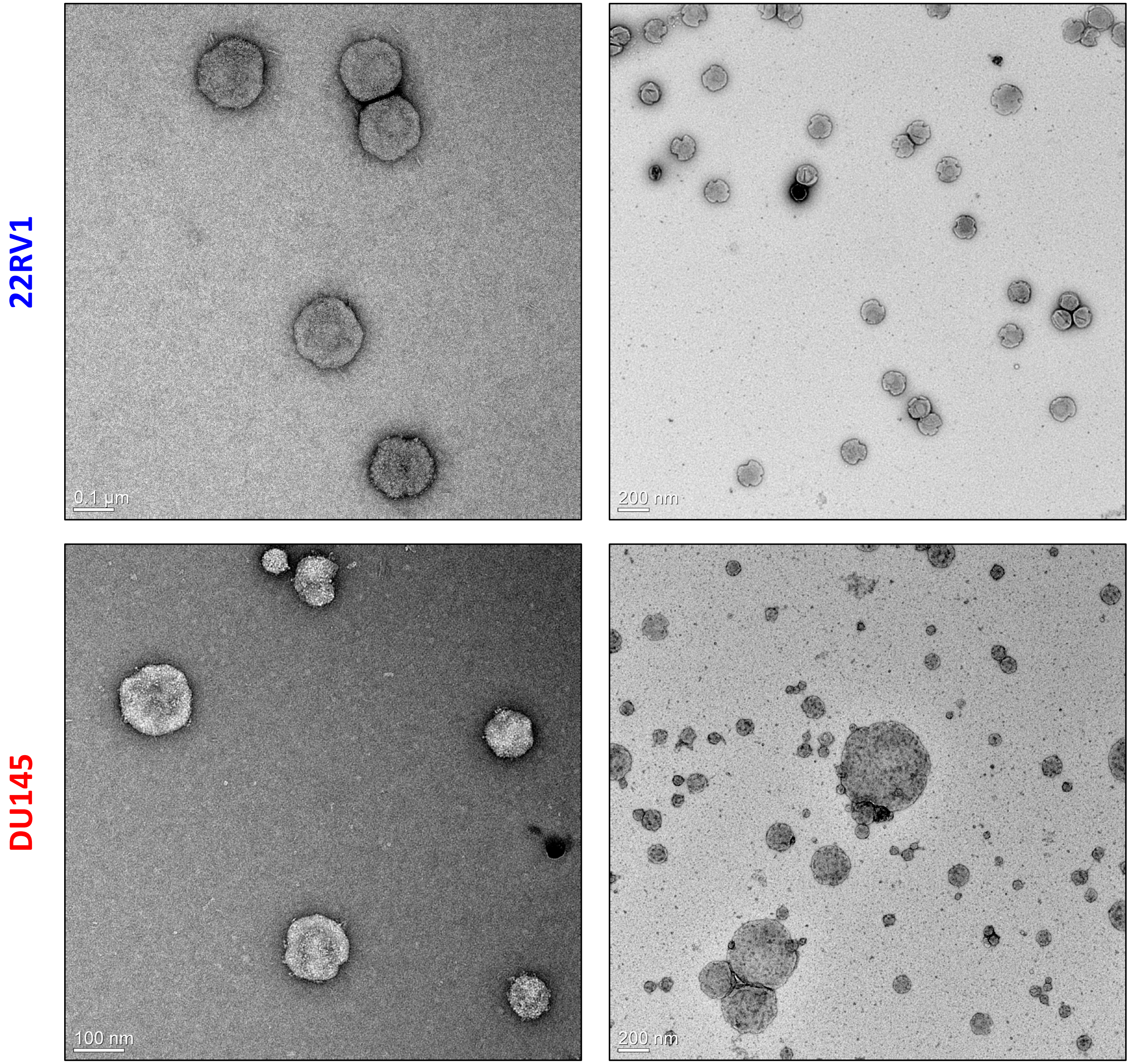


**Supplementary Figure 2.**
Transmission electron microscopy (TEM) images of EVs isolated from 22RV1 (AR+, top) or DU145 (AR-/NE-, bottom). Two images are shown for each cell line with differential zoomed out views, with scale bar noted in the bottom left of each image.

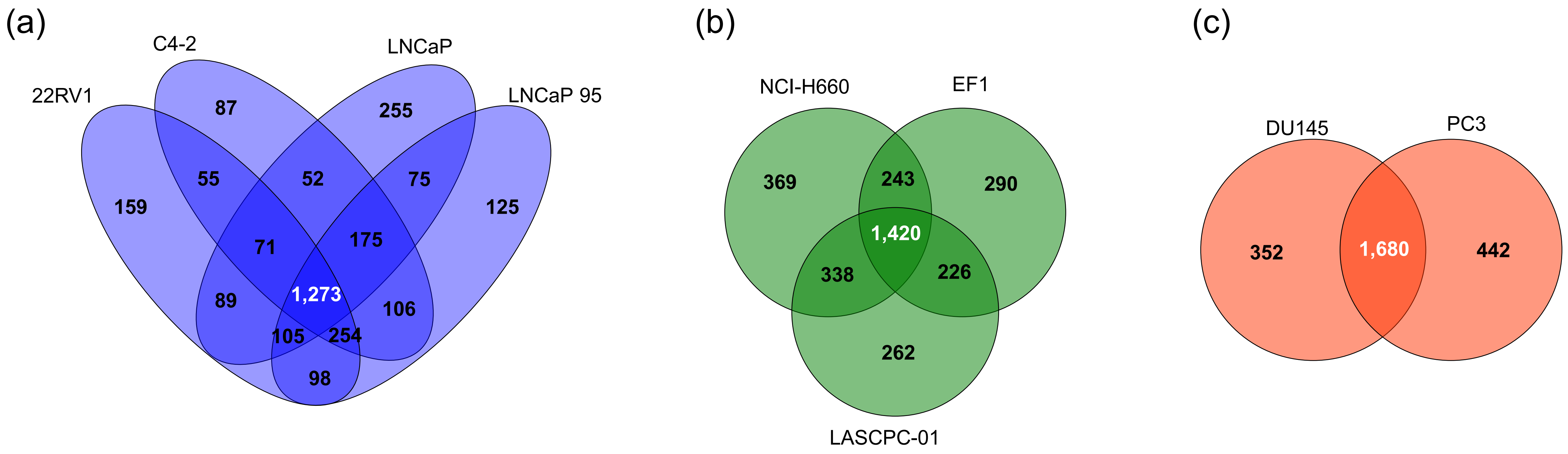


**Supplementary Figure 3.**
Venn diagram showing overlap of proteins identified from the EVs of cell lines with similar subtype, where a) AR+, b) AR-/NE+, c) AR-/NE-

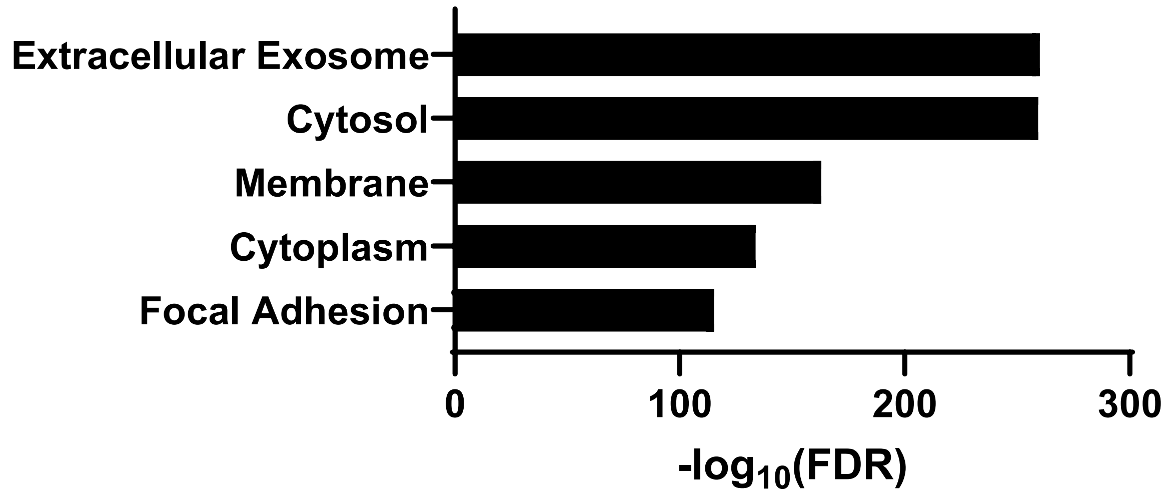


**Supplementary Figure 4.**
Enrichment analysis of gene ontology: cell compartment of 861 proteins common to all EV cell line proteomes (Figure 2b) using DAVID.

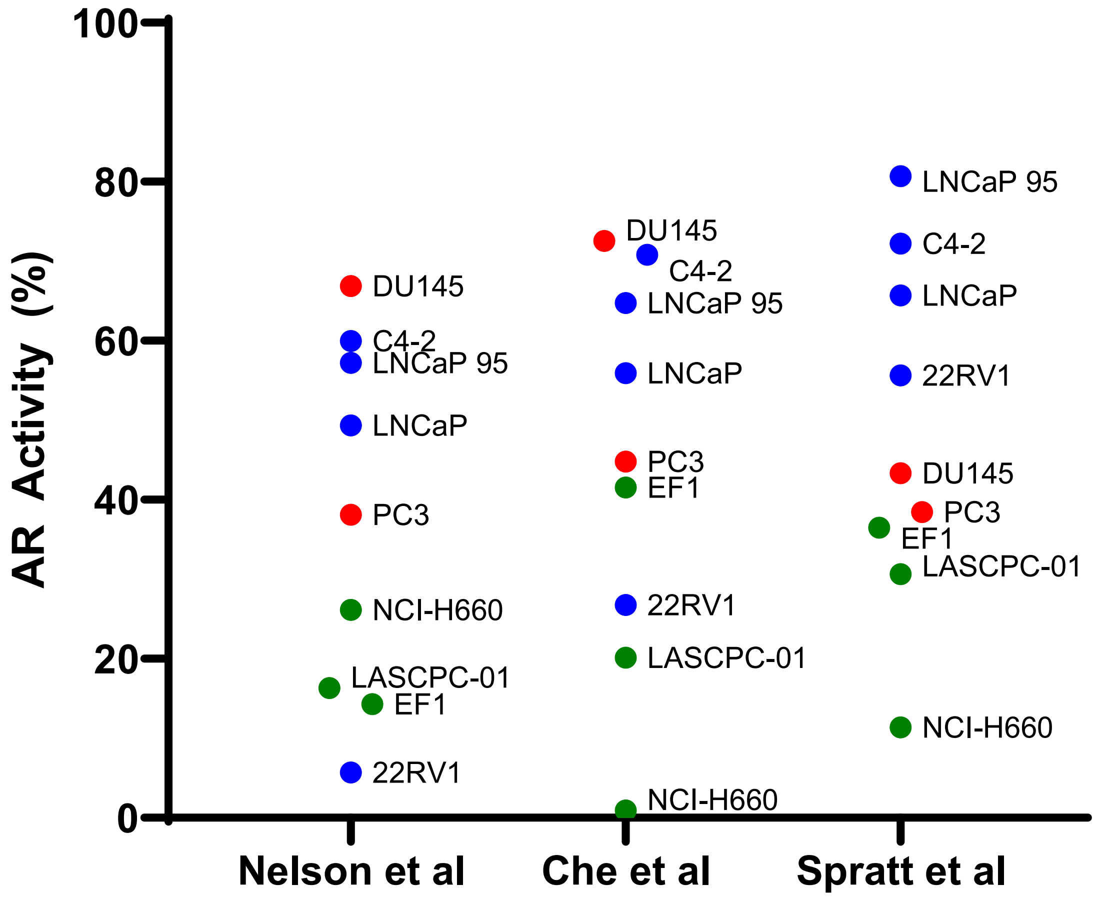


**Supplementary Figure 5.**

AR activity scores calculated from three gene signature lists. Samples are color coded by prostate cancer subtype: AR+ (blue), AR-/NE- (red), AR-/NE+ (green). Spratt et al[31] represented the most comprehensive gene list over the EV proteome, with 6/9 (66.7%) genes. The EV proteome contained 5/14 (35.7%) of the signature from Che et al[26] and 40/105 (38.1%) of the signature from Nelson et al[32].

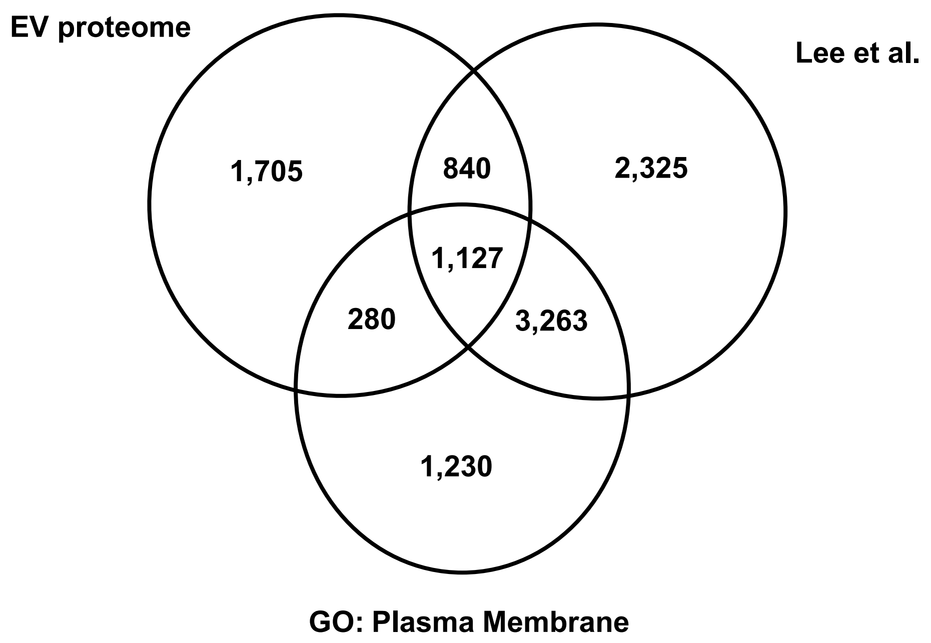


**Supplementary Figure 6.**
Venn diagram of proteins identified in the EVs from prostate cancer cells, the gene ontology term for plasma membrane (GO:0005886), and cell surface markers relevant to prostate cancer from Lee et al[35].

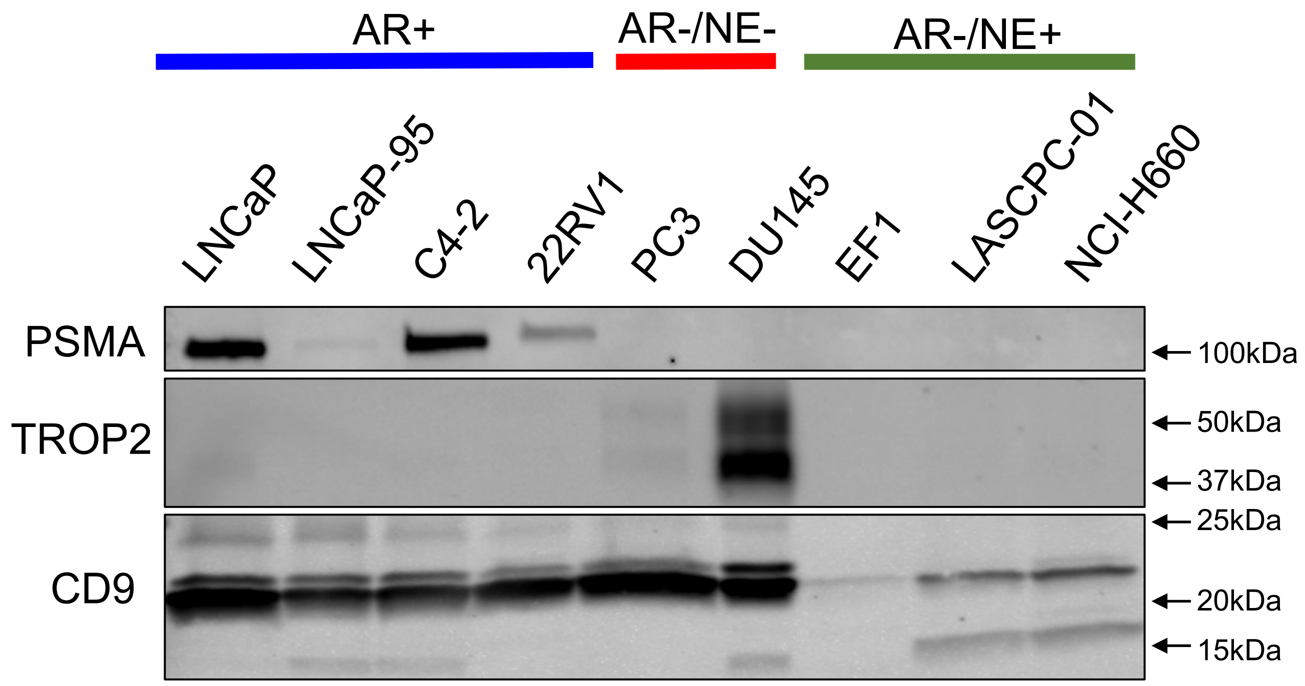


**Supplementary Figure 7.**

Western blot of 5µg of EV lysates from each cell line from indicated subtype, blotting for PSMA or TROP2 expression as well as CD9 as an EV marker. Molecular weights are indicated on the right side of image.

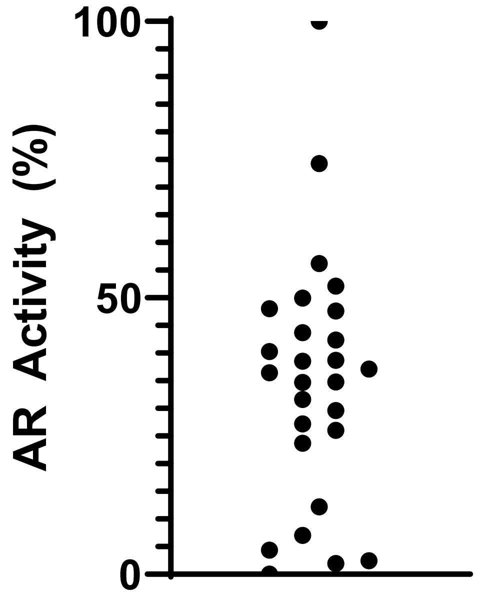


**Supplementary Figure 8.**
AR activity scores calculated using summative z-score with 4/9 proteins identified from gene signature from Spratt et al[31], similarly used in Figure 3a.

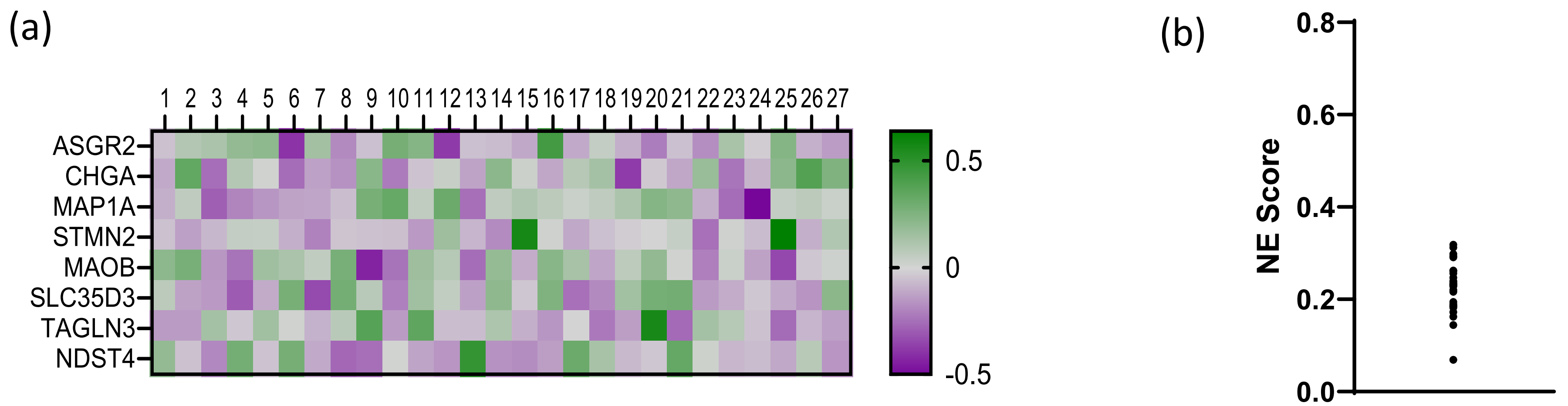


**Supplementary Figure 9.**
a) Heatmap of proteins associated with neuroendocrine prostate cancer[26] in EVs from plasma of advanced prostate cancer patients. b) Calculated neuroendocrine score by correlation with CRPC-NE samples[27] for EVs isolated from advanced prostate cancer patients.

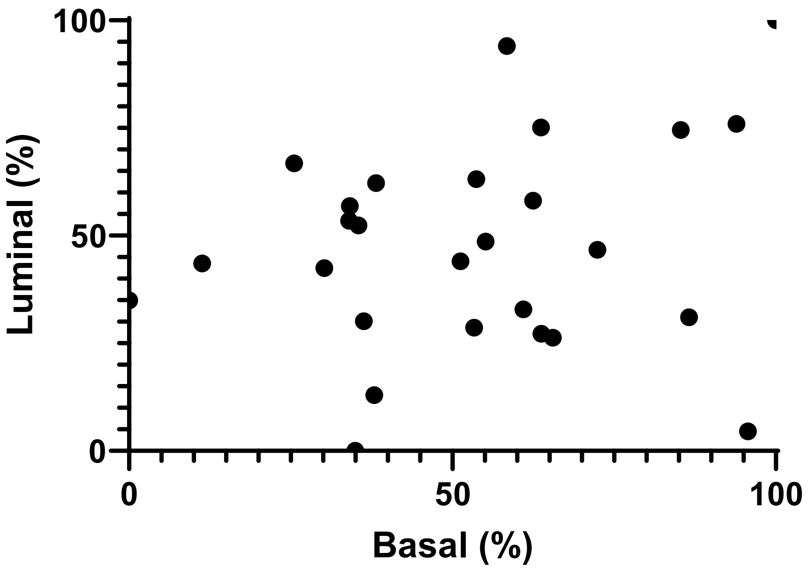


**Supplementary Figure 10.**
Basal and luminal scores[26] calculated for the EV proteomes of advanced prostate cancer patients.

**Not shared between cell line & patient EVs**

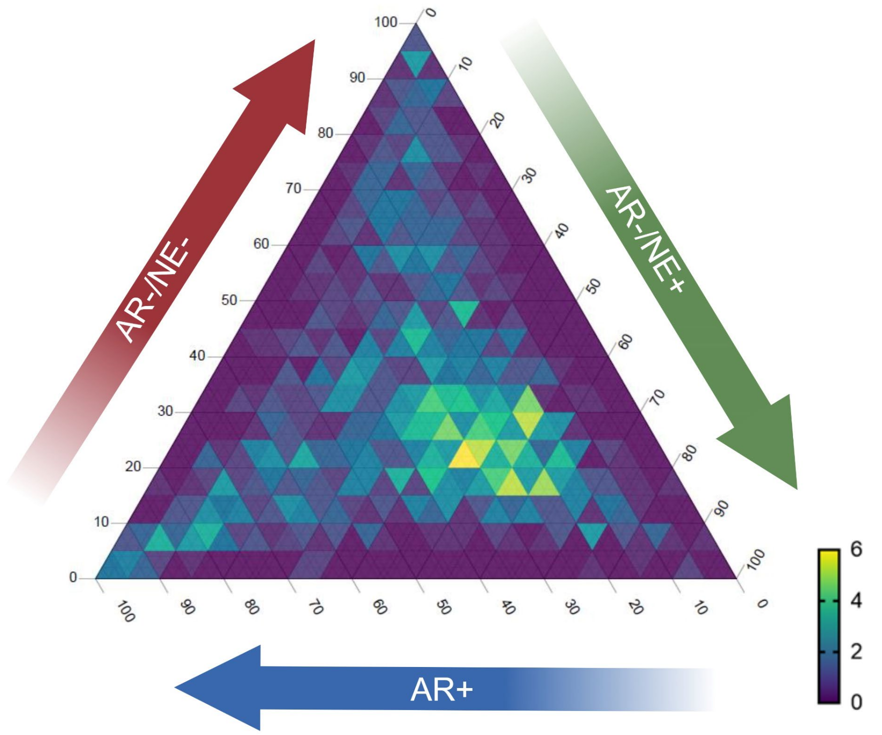


**Supplementary Figure 11.**
Ternary plot of 1,219 proteins that were found in EVs isolated from prostate cancer cell lines but not in plasma of patients. The values plotted are average protein level for each subtype from cell line EVs. The percentage of total signal is represented on the edges of the triangle, 0-100. Density of proteins within an inner triangle is indicated by color, scaled from 0 (fewest proteins) to 6 (most proteins).

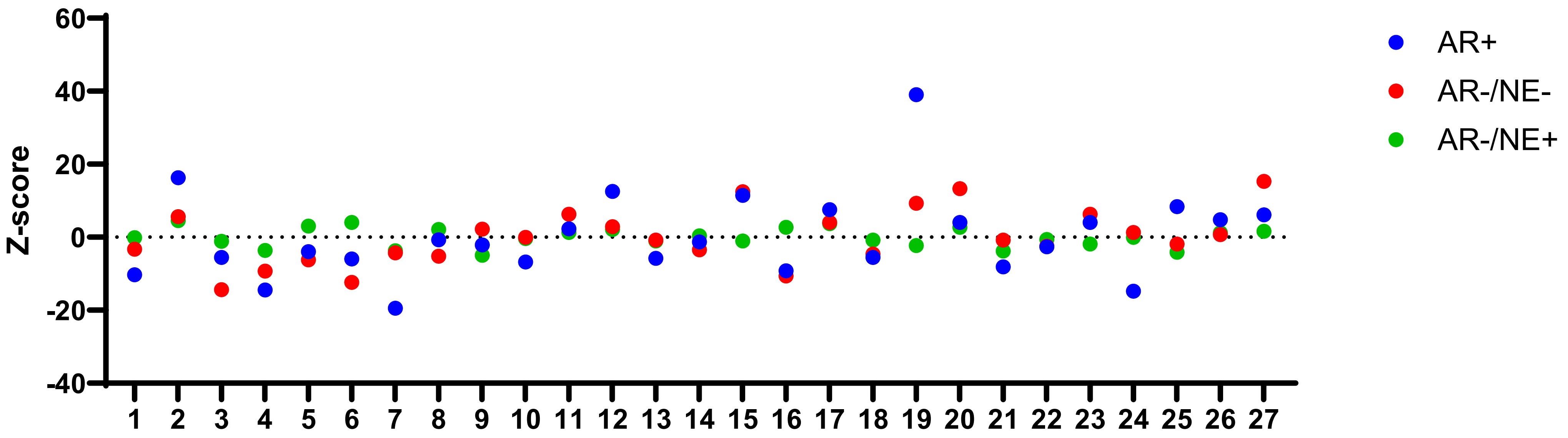


**Supplementary Figure 12.**

Summative z-score calculations for EV proteins associated with prostate cancer subtype for each patient, represented by 36 proteins for AR+ (blue), 27 proteins for AR-/NE- (red), and 4 genes for AR-/NE+ (green).
