## Supplemental Tables for "Proteomic profiling of extracellular vesicles distinguishes prostate cancer molecular subtypes"

| Supplementary Table # | Description |
|---|---|
| **STable 1** | Raw peptide intensities |
| **STable 2** | Proteome, imputed and normalized by VSN |
| **STable 3** | GSEA results for AR+ EVs |
| **STable 4** | GSEA results for AR-/NE- EVs |
| **STable 5** | GSEA results for AR-/NE+ EVs |
| **STable 6** | Gene signatures |
| **STable 7** | Count of peptides and proteins identified per EV sample |
| **STable 8** | AR+ enriched protein expression from plasma EVs of patients, log2(intensity) |
| **STable 9** | AR-/NE- enriched protein expression from plasma EVs of patients, log2(intensity) |
| **STable 10** | AR-/NE+ enriched protein expression from plasma EVs of patients, log2(intensity) |

| Peptide | Gene_Name | UniProt_ID | Description | Modifications | Intensity. 22RV1_1.11A | Intensity. 22RV1_2.12A | Intensity. C42_1.15A | Intensity. C42_2.16A | Intensity. DU145_1.3A | Intensity. DU145_2.4A | Intensity. EF1_1.7A | Intensity. EF1_2.8A | Intensity. H660_1.5A | Intensity. H660_2.6A | Intensity. LASCPC01_1.9A | Intensity. LASCPC01_2.10A | Intensity. LN95_1.17A | Intensity. LN95_2.18A | Intensity. LNCAP_1.13A | Intensity. LNCAP_2.14A | Intensity. PC3_1.1A | Intensity. PC3_2.2A |
|---|---|---|---|---|---|---|---|---|---|---|---|---|---|---|---|---|---|---|---|---|---|---|
| AAAAAAAA | SLC12A2; SLC1 | P55011; G3 | S12A2_HUMAN | Unmodified | 185760 | 0 | 0 | 0 | 0 | 0 | 0 | 0 | 0 | 0 | 0 | 0 | 0 | 0 | 1118100 | 528540 | 0 | 0 |
| AAAAAAAA | SLC12A2; SLC1 | P55011; G3 | S12A2_HUMAN | Unmodified | 0 | 0 | 0 | 0 | 0 | 0 | 0 | 0 | 0 | 0 | 0 | 0 | 0 | 0 | 0 | 0 | 826840 | 3911800 |
| AAAAAAAL | RPL4; RPL4; RP | P36578; H3E | RL4_HUMAN 6 | Unmodified | 10503000 | 7485900 | 4731100 | 3903000 | 1424000 | 735710 | 177760 | 226680 | 903540 | 254090 | 0 | 22108000 | 3051400 | 292960 | 6285700 | 2751500 | 123120 | 862980 |
| AAAAAAAL | RPL4; RPL4; RP | P36578; H3E | RL4_HUMAN 6 | Unmodified | 354130 | 358040 | 0 | 0 | 4219600 | 933670 | 620980 | 1760200 | 0 | 679520 | 0 | 0 | 6220700 | 1779100 | 0 | 0 | 334530 | 1715600 |
| AAAAAAGA | IPO9 | Q96P70 | IPO9_HUMAN I | Acetyl (Protein N | 0 | 0 | 0 | 0 | 1868700 | 0 | 1210300 | 79391 | 1954100 | 1934900 | 1632100 | 7288800 | 444060 | 0 | 0 | 0 | 215120 | 0 |
| AAAAAAGA | MAPK1 | P28482 | MK01_HUMAN | Acetyl (Protein N | 560810 | 0 | 331220 | 1012000 | 1019600 | 0 | 0 | 0 | 771610 | 340080 | 584850 | 598350 | 1514500 | 503730 | 433850 | 370280 | 0 | 0 |
| AAAAAAGE | QDPR; QDPR; C | P09417; D6F | DHPR_HUMAN | Acetyl (Protein N | 0 | 0 | 0 | 0 | 0 | 180410 | 0 | 0 | 0 | 66584 | 0 | 0 | 276290 | 154480 | 0 | 0 | 191900 | 146970 |
| AAAAARWN | WDR74; WDR7 | Q6RFH5; F5 | WDR74_HUMA | Acetyl (Protein N | 0 | 0 | 0 | 0 | 439870 | 0 | 189870 | 0 | 0 | 0 | 0 | 0 | 0 | 266200 | 0 | 0 | 247650 | 213900 |
| AAAAASAA | UNC119B | A6NIH7 | U119B_HUMAN | Unmodified | 0 | 992300 | 0 | 0 | 600220 | 86140 | 0 | 0 | 0 | 0 | 0 | 0 | 0 | 0 | 0 | 0 | 317510 | 1015600 |
| AAAAASRG | NDUFA2 | O43678 | NDUA2_HUMA | Acetyl (Protein N | 388370 | 482520 | 206100 | 397630 | 0 | 75367 | 238020 | 70557 | 0 | 388860 | 0 | 0 | 203420 | 0 | 1082300 | 261720 | 117820 | 141830 |
| AAAAELER | CHPF2; CHPF2 | Q9P2E5; G5 | CHPF2_HUMA | Unmodified | 21042 | 0 | 110680 | 0 | 13543 | 0 | 0 | 0 | 298530 | 45796 | 0 | 260290 | 0 | 0 | 19292 | 0 | 26067 | 461330 |
| AAAAELSLL | EEF1E1; EEF1E | O43324; D6 | MCA3_HUMAN | Acetyl (Protein N | 0 | 0 | 0 | 0 | 2129500 | 0 | 0 | 728000 | 795550 | 1002700 | 970780 | 9627900 | 3038000 | 1559400 | 0 | 327530 | 777070 | 1000100 |
| AAAAGRFP | ZNRF2 | Q8NHG8 | ZNRF2_HUMAN | Unmodified | 0 | 0 | 0 | 0 | 0 | 0 | 0 | 0 | 928050 | 0 | 0 | 0 | 0 | 0 | 0 | 0 | 0 | 0 |
| AAAAKPNN | SORD; SORD; S | Q00796; H0 | DHSO_HUMAN | Acetyl (Protein N | 0 | 0 | 0 | 4002500 | 0 | 0 | 0 | 0 | 0 | 0 | 8599200 | 20193000 | 0 | 0 | 4647000 | 7062800 | 0 | 0 |
| AAAAKPNN | SORD; SORD; S | Q00796; H0 | DHSO_HUMAN | Acetyl (Protein N | 0 | 0 | 980150 | 18846000 | 11323000 | 456950 | 762740 | 0 | 0 | 1797400 | 2362900 | 0 | 40787000 | 13141000 | 7796700 | 2301400 | 4217100 | 3932200 |
| AAAAQLSL | EPS15; EPS15 | P42566; B1A | EPS15_HUMAN | Acetyl (Protein N | 581750 | 2655100 | 0 | 0 | 0 | 0 | 0 | 0 | 0 | 0 | 0 | 0 | 0 | 0 | 0 | 151400 | 0 | 0 |
| AAAAVSES | LRRC47 | Q8N1G4 | LRC47_HUMAN | Acetyl (Protein N | 0 | 0 | 0 | 0 | 0 | 0 | 0 | 0 | 1427500 | 922120 | 0 | 1019200 | 0 | 0 | 0 | 0 | 0 | 0 |
| AAADLMAY | GNG2; GNG2; ( | P59768; G3 | GBG2_HUMAN | Oxidation (M) | 0 | 0 | 0 | 0 | 0 | 0 | 1018200 | 160470 | 0 | 306230 | 0 | 0 | 0 | 379970 | 0 | 0 | 4053200 | 11577000 |
| AAADSDPN | ZFPL1 | O95159 | ZFPL1_HUMAN | Oxidation (M) | 0 | 300220 | 0 | 0 | 0 | 0 | 0 | 0 | 1850200 | 379830 | 0 | 0 | 0 | 0 | 0 | 0 | 0 | 0 |
| AAADSDPN | ZFPL1 | O95159 | ZFPL1_HUMAN | Unmodified | 833270 | 0 | 0 | 0 | 0 | 0 | 0 | 0 | 3890600 | 0 | 0 | 0 | 0 | 0 | 0 | 0 | 0 | 0 |
| AAAEAGGD | PGP | A6NDG6 | PGP_HUMAN G | Acetyl (Protein N | 0 | 0 | 71060 | 0 | 0 | 26051 | 0 | 0 | 284800 | 0 | 0 | 1171200 | 0 | 0 | 0 | 36785 | 0 | 80089 |
| AAAEDVNV | PFDN4; PFDN4 | Q9NQP4; E9 | PFD4_HUMAN | Unmodified | 0 | 0 | 131220 | 156280 | 923480 | 315320 | 0 | 0 | 1694300 | 912600 | 1271500 | 6924400 | 1604100 | 515740 | 105940 | 125520 | 1625600 | 1007200 |
| AAAEDVNV | PFDN4; PFDN4 | Q9NQP4; E9 | PFD4_HUMAN | Unmodified | 0 | 0 | 0 | 0 | 1250900 | 0 | 0 | 0 | 0 | 0 | 0 | 0 | 571010 | 577860 | 0 | 0 | 1231100 | 591220 |
| AAAEDVNV | PFDN4; PFDN4 | Q9NQP4; E9 | PFD4_HUMAN | Unmodified | 0 | 0 | 0 | 0 | 0 | 0 | 0 | 0 | 0 | 0 | 0 | 0 | 96922 | 0 | 0 | 0 | 1292300 | 0 |
| AAAEELLAF | INPPL1 | O15357 | SHIP2_HUMAN | Unmodified | 232610 | 0 | 0 | 0 | 0 | 0 | 0 | 43876 | 1761500 | 538990 | 852810 | 3327500 | 0 | 0 | 0 | 0 | 0 | 0 |
| AAAEVAGQ | TFRC; TFRC; TF | P02786; H7 | TFR1_HUMAN | Unmodified | 1855300 | 1511800 | 2268700 | 1098300 | 0 | 0 | 0 | 26945 | 4012700 | 1461800 | 292470 | 1860600 | 0 | 0 | 2918000 | 2857000 | 1528900 | 1136900 |
| AAAEVNQD | FH | P07954 | FUMH_HUMAN | Unmodified | 0 | 0 | 0 | 0 | 0 | 0 | 0 | 0 | 676520 | 124190 | 391530 | 396350 | 0 | 0 | 0 | 0 | 0 | 0 |
| AAAFAGGD | TPBGL | P0DKB5 | TPBGL_HUMAN | Unmodified | 0 | 0 | 0 | 0 | 0 | 0 | 0 | 0 | 1160800 | 0 | 0 | 0 | 939350 | 0 | 0 | 0 | 0 | 0 |
| AAAFEEQEI | TLN1 | Q9Y490 | TLN1_HUMAN | Unmodified | 0 | 277300 | 1077300 | 1867300 | 6460200 | 2711800 | 2911800 | 1596300 | 0 | 817100 | 735220 | 6707100 | 2077400 | 3795400 | 1285700 | 499620 | 1870700 | 2376700 |
| AAAFEEQEI | TLN1 | Q9Y490 | TLN1_HUMAN | Unmodified | 0 | 0 | 0 | 0 | 2333600 | 557160 | 0 | 1742400 | 0 | 209150 | 0 | 0 | 781440 | 1462700 | 0 | 0 | 2214400 | 873440 |
| AAAFLAVLV | B3GALT6 | Q96L58 | B3GT6_HUMAN | Unmodified | 0 | 0 | 0 | 0 | 0 | 0 | 0 | 0 | 957720 | 0 | 0 | 0 | 0 | 0 | 0 | 0 | 0 | 0 |
| AAAGEFAD | CTNNA1; CTNN | P35221; E5R | CTNA1_HUMA | Unmodified | 465070 | 42951 | 786420 | 480110 | 0 | 0 | 0 | 0 | 0 | 0 | 333650 | 2765000 | 0 | 229160 | 1879000 | 1440200 | 0 | 0 |
| AAAGEFAD | CTNNA1; CTNN | P35221; E5R | CTNA1_HUMA | Unmodified | 414990 | 210660 | 483880 | 463320 | 0 | 0 | 0 | 0 | 0 | 0 | 0 | 2433300 | 60829 | 0 | 1791000 | 2411400 | 0 | 0 |
| AAAGELQE | EFHD2; EFHD2 | Q96C19; H0 | EFHD2_HUMA | Unmodified | 0 | 0 | 0 | 0 | 2533900 | 466240 | 0 | 0 | 653630 | 331610 | 0 | 840830 | 0 | 0 | 0 | 0 | 0 | 0 |
| AAAGELQE | EFHD2; EFHD2 | Q96C19; H0 | EFHD2_HUMA | Unmodified | 0 | 0 | 0 | 0 | 2001100 | 0 | 0 | 0 | 0 | 0 | 0 | 0 | 0 | 0 | 0 | 0 | 0 | 0 |
| AAAGGGG | RAB21 | Q9UL25 | RAB21_HUMAN | Acetyl (Protein N | 0 | 0 | 0 | 0 | 967130 | 340570 | 455060 | 270370 | 0 | 289460 | 0 | 0 | 454740 | 361970 | 0 | 0 | 315910 | 638940 |
| AAAGGQG | EGLN1 | Q9GZT9 | EGLN1_HUMA | Unmodified | 0 | 0 | 0 | 0 | 0 | 0 | 0 | 0 | 0 | 0 | 0 | 307310 | 0 | 0 | 0 | 0 | 0 | 0 |
| AAAGIQPP | CANT1 | Q8WVQ1 | CANT1_HUMA | Unmodified | 0 | 0 | 0 | 0 | 0 | 0 | 0 | 0 | 15232000 | 579080 | 0 | 0 | 0 | 0 | 0 | 0 | 0 | 0 |
| AAAGKAELI | PLEC | Q15149 | PLEC_HUMAN | Unmodified | 207240 | 0 | 0 | 1942200 | 222370 | 0 | 0 | 0 | 106220 | 32994 | 0 | 753020 | 0 | 0 | 0 | 0 | 300960 | 101920 |
| AAAGKVGD | TLN1 | Q9Y490 | TLN1_HUMAN | Unmodified | 0 | 0 | 72771 | 291640 | 5814800 | 1432400 | 10993000 | 3626500 | 106020 | 904840 | 114200 | 0 | 2417500 | 3311900 | 30972 | 0 | 5090900 | 1271200 |
| AAAGKVGD | TLN1 | Q9Y490 | TLN1_HUMAN | Unmodified | 0 | 0 | 0 | 0 | 0 | 0 | 1115500 | 312780 | 0 | 0 | 0 | 0 | 0 | 0 | 0 | 0 | 196500 | 0 |
| AAAGWRD | RRAS2; RRAS2; | P62070; E9P | RRAS2_HUMAN | Acetyl (Protein N | 378970 | 555630 | 0 | 0 | 3471500 | 0 | 45012 | 47949 | 0 | 409790 | 0 | 0 | 1341600 | 760150 | 901280 | 775030 | 347180 | 0 |
| AAAHLQGL | ACBD6; ACBD6 | Q9BR61; A0 | ACBD6_HUMA | Unmodified | 0 | 0 | 0 | 0 | 0 | 0 | 0 | 0 | 108000 | 0 | 72120 | 430090 | 0 | 0 | 0 | 0 | 0 | 0 |
| AAAKDTHE | RANBP1; RANB | P43487; C9J | RANG_HUMAN | Acetyl (Protein N | 621600 | 0 | 0 | 1415900 | 17183000 | 0 | 1201700 | 1103100 | 867780 | 0 | 3467700 | 11111000 | 997970 | 0 | 0 | 0 | 1464300 | 2853800 |
| AAALEAMK | STIP1; STIP1 | P31948; H0 | STIP1_HUMAN | Unmodified | 0 | 0 | 0 | 516060 | 4723900 | 671310 | 393240 | 312160 | 3659300 | 1985500 | 0 | 535840 | 5553600 | 3253000 | 335630 | 0 | 2875300 | 4263600 |
| AAALEAMK | STIP1; STIP1 | P31948; H0 | STIP1_HUMAN | Unmodified | 0 | 0 | 0 | 0 | 453430 | 0 | 247160 | 276040 | 0 | 0 | 0 | 206870 | 658580 | 258830 | 0 | 0 | 1440300 | 606780 |
| AAALEFLNI | STIP1; STIP1; S | P31948; F5H | STIP1_HUMAN | Unmodified | 1485200 | 568310 | 1720900 | 3066400 | 10204000 | 1136800 | 794080 | 0 | 27360000 | 6790700 | 7284300 | 73061000 | 20640000 | 4847900 | 3085000 | 3703500 | 5386200 | 9266200 |
| AAALEFLNI | STIP1; STIP1; S | P31948; F5H | STIP1_HUMAN | Unmodified | 0 | 0 | 0 | 578030 | 399340 | 382530 | 0 | 0 | 1936200 | 430180 | 1183600 | 474420 | 1264600 | 277610 | 0 | 0 | 308700 | 631750 |
| AAALLLQNI | ANK3; ANK3; A | Q12955; A0 | ANK3_HUMAN | Unmodified | 0 | 0 | 224450 | 210010 | 0 | 0 | 0 | 0 | 0 | 67465 | 0 | 0 | 2528600 | 2962400 | 2430100 | 2518700 | 0 | 0 |
| AAALLMLT | CCNDBP1 | O95273 | CCDB1_HUMA | Oxidation (M) | 930690 | 1285500 | 1013400 | 931730 | 0 | 0 | 0 | 0 | 0 | 0 | 0 | 380120 | 328550 | 150570 | 623130 | 2573300 | 0 | 0 |
| AAALRAEIT | DCTN1; DCTN1 | Q14203; A0 | DCTN1_HUMA | Unmodified | 0 | 0 | 0 | 0 | 0 | 0 | 0 | 0 | 0 | 21147 | 153720 | 355960 | 0 | 0 | 0 | 0 | 0 | 0 |
| AAALVLQTI | CTNND1; CTNN | O60716; A0 | CTND1_HUMA | Unmodified | 2966900 | 347210 | 1545800 | 10105000 | 21017000 | 3410600 | 0 | 0 | 0 | 0 | 4984100 | 11042000 | 16207000 | 13349000 | 11279000 | 7881000 | 9492900 | 20385000 |
| AAAMANNI | RBM39; RBM39 | Q14498; G3 | RBM39_HUMA | Unmodified | 529820 | 0 | 4181200 | 0 | 1772400 | 424490 | 0 | 0 | 1022600 | 0 | 2989700 | 246010 | 0 | 664210 | 0 | 0 | 0 | 0 |
| AAAMDVDT | RBX1 | P62877 | RBX1_HUMAN | Acetyl (Protein N | 0 | 0 | 0 | 0 | 0 | 102730 | 0 | 0 | 0 | 158130 | 0 | 0 | 0 | 0 | 0 | 0 | 286170 | 356540 |
| AAAMDVDT | RBX1 | P62877 | RBX1_HUMAN | Acetyl (Protein N | 0 | 0 | 0 | 0 | 128980 | 0 | 96854 | 85056 | 0 | 41518 | 0 | 0 | 0 | 118840 | 0 | 0 | 365120 | 191870 |
| AAAPAPEE | EEF1G | P26641 | EF1G_HUMAN | Unmodified | 0 | 0 | 0 | 0 | 1717800 | 0 | 0 | 171260 | 0 | 329380 | 499770 | 3896000 | 965800 | 745860 | 0 | 0 | 751210 | 1910400 |
| AAAPAPEE | EEF1G | P26641 | EF1G_HUMAN | Oxidation (M) | 0 | 0 | 0 | 0 | 0 | 0 | 0 | 0 | 0 | 0 | 152570 | 3119600 | 775220 | 0 | 0 | 0 | 0 | 0 |
| AAAPGVED | RRM2; RRM2; R | P31350; A0A | RIR2_HUMAN F | Unmodified | 0 | 0 | 0 | 0 | 0 | 0 | 0 | 0 | 0 | 0 | 0 | 761400 | 0 | 0 | 0 | 0 | 0 | 0 |
| AAAQLLQS | SEPTIN11; SEP | Q9NVA2; H0 | SEP11_HUMAN | Unmodified | 0 | 0 | 0 | 0 | 790610 | 0 | 0 | 0 | 0 | 157210 | 0 | 1368300 | 0 | 124160 | 177220 | 0 | 203280 | 0 |
| AAAQLLQS | SEPTIN11; SEP | Q9NVA2; H0 | SEP11_HUMAN | Unmodified | 163900 | 0 | 34849 | 1180500 | 4290200 | 1249100 | 441090 | 91036 | 755830 | 464050 | 154950 | 5039300 | 895920 | 527920 | 380310 | 484820 | 847590 | 1858500 |
| AAAQLLQS | SEPTIN11; SEP | Q9NVA2; H0 | SEP11_HUMAN | Unmodified | 0 | 0 | 0 | 0 | 530640 | 0 | 0 | 64139 | 0 | 0 | 0 | 0 | 0 | 0 | 0 | 0 | 431870 | 0 |
| AAAQVLNT | PKP4; PKP4 | Q99569; A0 | PKP4_HUMAN | Unmodified | 0 | 0 | 143650 | 215180 | 0 | 0 | 0 | 0 | 408670 | 22926 | 3943300 | 0 | 41061 | 0 | 180310 | 93426 | 0 | 0 |
| AAAQVLNT | PKP4; PKP4 | Q99569; A0 | PKP4_HUMAN | Unmodified | 0 | 0 | 0 | 152580 | 957910 | 171700 | 429440 | 74195 | 193090 | 781170 | 345040 | 0 | 1121400 | 508390 | 0 | 0 | 450660 | 584410 |
| AAAQYYLG | TTC4 | O95801 | TTC4_HUMAN | Unmodified | 0 | 0 | 0 | 0 | 0 | 0 | 0 | 0 | 0 | 0 | 0 | 0 | 362620 | 0 | 0 | 0 | 0 | 0 |
| AAARDVTLI | SARS1; SARS1 | P49591; Q5 | SYSC_HUMAN | Unmodified | 0 | 0 | 0 | 0 | 2280900 | 0 | 483270 | 175850 | 0 | 0 | 0 | 0 | 0 | 2482500 | 0 | 0 | 1145900 | 2600500 |
| AAARHSTLI | SMS | P52788 | SPSY_HUMAN | Acetyl (Protein N | 0 | 0 | 0 | 0 | 154750 | 0 | 150600 | 0 | 0 | 0 | 0 | 0 | 161280 | 103330 | 0 | 0 | 156200 | 255240 |
| AAARHSTLI | SMS | P52788 | SPSY_HUMAN | Acetyl (Protein N | 0 | 0 | 0 | 0 | 3568600 | 0 | 1541700 | 0 | 0 | 0 | 0 | 0 | 1323600 | 3986500 | 0 | 0 | 3221400 | 1405800 |
| AAARHSTLI | SMS | P52788 | SPSY_HUMAN | Acetyl (Protein N | 0 | 0 | 0 | 0 | 944880 | 0 | 1869400 | 0 | 0 | 174480 | 0 | 0 | 1706500 | 2049800 | 0 | 0 | 4290200 | 1145000 |
| AAASLHEL | ENTPD6; ENTP | O75354; H0 | ENTP6_HUMAN | Unmodified | 0 | 0 | 0 | 0 | 0 | 0 | 0 | 0 | 887070 | 0 | 126620 | 0 | 0 | 0 | 0 | 0 | 0 | 0 |
| AAASTDYYI | PDHA1 | P08559 | ODPA_HUMAN | Unmodified | 0 | 0 | 668040 | 0 | 0 | 0 | 0 | 0 | 1763800 | 0 | 37974 | 128010 | 790390 | 118690 | 315160 | 0 | 0 | 0 |
| AAASVTPP | TRAF2; TRAF2; | Q12933; B1 | TRAF2_HUMAN | Acetyl (Protein N | 0 | 0 | 0 | 0 | 27493000 | 7498400 | 0 | 462050 | 955330 | 817230 | 322000 | 1427300 | 555600 | 0 | 0 | 0 | 477600 | 791440 |
| AAATAEEPI | NAP1L4; NAP1L | Q99733; C9 | NP1L4_HUMAN | Unmodified | 0 | 0 | 0 | 0 | 0 | 0 | 0 | 0 | 465550 | 182370 | 0 | 2214700 | 0 | 0 | 259360 | 0 | 0 | 0 |
| AAATGPSF | PEPD; PEPD | P12955; A0A | PEPD_HUMAN | Acetyl (Protein N | 0 | 0 | 0 | 0 | 31530 | 0 | 1139900 | 0 | 0 | 0 | 0 | 429360 | 0 | 0 | 0 | 0 | 0 | 0 |
| AAATPESQ | MARCKSL1 | P49006 | MRP_HUMAN N | Unmodified | 10750000 | 5931200 | 29191000 | 47078000 | 13501000 | 689060 | 2282600 | 3869000 | 87062000 | 26615000 | 25336000 | 2.41E+08 | 69313000 | 38319000 | 29990000 | 41683000 | 29015000 | 2.79E+08 |
| AAATQPDA | SND1 | Q7KZF4 | SND1_HUMAN | Unmodified | 0 | 193330 | 370280 | 320630 | 70620 | 0 | 0 | 0 | 1055200 | 3416300 | 0 | 2411400 | 160460 | 0 | 632960 | 344770 | 0 | 0 |
| AAATQPDA | SND1 | Q7KZF4 | SND1_HUMAN | Unmodified | 0 | 0 | 0 | 1506000 | 0 | 0 | 0 | 0 | 0 | 0 | 0 | 0 | 725160 | 0 | 65506 | 0 | 0 | 0 |
| AAATTAQE | VIL1; VIL1 | P09327; B4D | VILI_HUMAN Vi | Unmodified | 464860 | 0 | 877220 | 0 | 0 | 69489 | 0 | 0 | 1067300 | 570280 | 837000 | 9955800 | 0 | 0 | 403100 | 160960 | 0 | 74349 |
| AAAVDFTA | ANK3; ANK3; A | Q12955; A0 | ANK3_HUMAN | Unmodified | 0 | 0 | 0 | 0 | 0 | 0 | 0 | 0 | 0 | 0 | 0 | 0 | 0 | 0 | 1115500 | 3042400 | 0 | 0 |
| AAAVDFTA | ANK3; ANK3; A | Q12955; A0 | ANK3_HUMAN | Unmodified | 0 | 69256 | 1970000 | 5032600 | 0 | 0 | 197710 | 0 | 0 | 0 | 0 | 0 | 8310900 | 4175800 | 4010700 | 2229300 | 0 | 0 |
| AAAVGTAN | VCL | P18206 | VINC_HUMAN | Unmodified | 0 | 0 | 0 | 0 | 1263400 | 230750 | 2134100 | 738150 | 0 | 695720 | 0 | 0 | 199160 | 154410 | 0 | 0 | 19227000 | 380660 |
| AAAVGTAN | VCL | P18206 | VINC_HUMAN | Unmodified | 0 | 0 | 0 | 0 | 0 | 0 | 0 | 0 | 0 | 0 | 0 | 0 | 0 | 0 | 0 | 0 | 796810 | 0 |
| AAAVPVEF | IDH3B; IDH3B; | O43837; A0 | IDH3B_HUMAN | Unmodified | 81239 | 0 | 0 | 0 | 0 | 0 | 0 | 0 | 666440 | 0 | 0 | 0 | 0 | 0 | 0 | 0 | 0 | 0 |
| AAAVSLEN | FMNL2; FMNL3 | Q96PY5; Q8 | FMNL2_HUMA | Unmodified | 0 | 0 | 0 | 0 | 911340 | 17342 | 0 | 0 | 32559 | 69009 | 46021 | 95628 | 0 | 0 | 0 | 0 | 24471 | 0 |
| AAAVSSGF | AGRN; AGRN; A | O00468; A0 | AGRIN_HUMAN | Unmodified | 1973700 | 779520 | 794320 | 0 | 135600 | 237650 | 357720 | 0 | 1011400 | 0 | 196860 | 68834 | 1358300 | 0 | 0 | 435780 | 0 | 0 |
| AAAVSSGF | AGRN; AGRN; A | O00468; A0 | AGRIN_HUMAN | Unmodified | 0 | 0 | 0 | 0 | 0 | 0 | 0 | 0 | 0 | 0 | 0 | 0 | 1012000 | 0 | 0 | 0 | 0 | 2345200 |
| AAAYLDPN | PTK2; PTK2; PT | Q05397; E5F | FAK1_HUMAN | Acetyl (Protein N | 0 | 0 | 0 | 0 | 282440 | 0 | 0 | 61239 | 0 | 0 | 0 | 0 | 0 | 0 | 0 | 0 | 442850 | 786980 |
| AACDLVRG | CSE1L | P55060 | XPO2_HUMAN | Unmodified | 0 | 0 | 0 | 0 | 0 | 0 | 192880 | 64432 | 0 | 0 | 0 | 0 | 57820 | 0 | 0 | 0 | 0 | 0 |
| AACHLKLE | UNC45A; UNC4 | Q9H3U1; A0 | UN45A_HUMA | Unmodified | 0 | 0 | 0 | 225570 | 0 | 81536 | 0 | 0 | 168010 | 0 | 0 | 0 | 443210 | 309500 | 0 | 0 | 132970 | 328190 |
| AACIAEQY | KIF20A | O95235 | KI20A_HUMAN | Unmodified | 1727000 | 0 | 0 | 716950 | 3286800 | 0 | 234200 | 38929 | 874890 | 496490 | 0 | 0 | 3009300 | 1754200 | 0 | 0 | 1699300 | 390070 |
| AACLESAQ | HDLBP; HDLBP | A0A024R4E | A0A024R4E5_H | Unmodified | 0 | 0 | 0 | 0 | 0 | 0 | 0 | 0 | 0 | 0 | 0 | 0 | 607600 | 805550 | 0 | 0 | 0 | 0 |
| AACNGPYD | GLOD4; GLOD4 | F6TLX2; I3L3 | F6TLX2_HUMA | Unmodified | 0 | 0 | 0 | 0 | 437390 | 179960 | 153770 | 0 | 96761 | 99512 | 1309900 | 0 | 816260 | 2610400 | 14424 | 0 | 0 | 423170 |
| AACNLLQR | PFKP; PFKP; PF | Q01813; A0 | PFKAP_HUMAN | Unmodified | 0 | 0 | 0 | 0 | 499080 | 0 | 0 | 0 | 342750 | 141870 | 363730 | 2309800 | 0 | 0 | 0 | 0 | 23508 | 376960 |
| AADAVEDL | PACSIN2; PACS | Q9UNF0; A0 | PACN2_HUMA | Unmodified | 49506000 | 37995000 | 22846000 | 12717000 | 0 | 0 | 0 | 0 | 1425000 | 0 | 358360 | 6324500 | 0 | 0 | 59807000 | 44431000 | 0 | 0 |
| AADAVEDL | PACSIN2; PACS | Q9UNF0; A0 | PACN2_HUMA | Unmodified | 22511000 | 12581000 | 8860000 | 20894000 | 0 | 0 | 0 | 0 | 1832100 | 135830 | 1170100 | 0 | 11790000 | 0 | 22428000 | 8177900 | 0 | 322290 |
| AADEEAFEI | CSE1L | P55060 | XPO2_HUMAN | Unmodified | 173780 | 176410 | 994670 | 447140 | 0 | 0 | 0 | 0 | 993860 | 599030 | 719080 | 15018000 | 0 | 0 | 506630 | 757850 | 0 | 0 |
| AADEEAFEI | CSE1L | P55060 | XPO2_HUMAN | Unmodified | 0 | 0 | 0 | 811430 | 0 | 0 | 0 | 0 | 1587100 | 0 | 299810 | 445610 | 55721 | 0 | 195160 | 0 | 159810 | 0 |
| AADEVAEG | FH | P07954 | FUMH_HUMAN | Oxidation (M) | 0 | 0 | 0 | 0 | 0 | 0 | 0 | 0 | 2042600 | 1170100 | 0 | 0 | 0 | 0 | 0 | 0 | 0 | 0 |
| AADFIDQAI | DUSP3; DUSP3 | P51452; K7E | DUS3_HUMAN | Unmodified | 0 | 0 | 0 | 0 | 0 | 0 | 0 | 0 | 0 | 0 | 0 | 0 | 0 | 0 | 0 | 0 | 0 | 0 |
| AADFQLHT | VDAC3 | Q9Y277 | VDAC3_HUMA | Unmodified | 0 | 0 | 0 | 0 | 0 | 0 | 401460 | 0 | 30393000 | 11064000 | 777210 | 0 | 0 | 0 | 0 | 0 | 928120 | 0 |
| AADIDQEVI | CAND1 | Q86VP6 | CAND1_HUMA | Unmodified | 172490 | 0 | 288260 | 169130 | 136610 | 513660 | 0 | 16645 | 666760 | 205070 | 374740 | 6052300 | 0 | 0 | 0 | 256850 | 0 | 451560 |
| AADIDQEVI | CAND1 | Q86VP6 | CAND1_HUMA | Unmodified | 0 | 0 | 0 | 268700 | 0 | 0 | 0 | 16263 | 329640 | 130570 | 28777 | 568730 | 0 | 87575 | 0 | 0 | 0 | 0 |
| AADKLIQNI | S100A16 | Q96FQ6 | S10AG_HUMAN | Unmodified | 0 | 0 | 0 | 0 | 5532600 | 0 | 0 | 0 | 0 | 231200 | 0 | 0 | 0 | 0 | 0 | 0 | 5007100 | 12624000 |
| AADKPADQ | CNOT10; CNOT | Q9H9A5; E9 | CNO10_HUMA | Acetyl (Protein N | 0 | 0 | 0 | 0 | 757760 | 171730 | 0 | 0 | 0 | 144840 | 0 | 0 | 0 | 0 | 0 | 0 | 187090 | 0 |
| AADLNGDL | RCN1; N/A | Q15293; E9F | RCN1_HUMAN | Unmodified | 0 | 0 | 0 | 0 | 0 | 0 | 0 | 0 | 283450 | 101980 | 1422600 | 0 | 0 | 0 | 0 | 0 | 0 | 0 |
| AADLNGDL | RCN1; N/A | Q15293; E9F | RCN1_HUMAN | Unmodified | 0 | 0 | 0 | 0 | 0 | 0 | 0 | 0 | 5898500 | 311120 | 0 | 0 | 0 | 0 | 0 | 0 | 0 | 0 |
| AADLNGDL | RCN1; N/A | Q15293; E9F | RCN1_HUMAN | Oxidation (M) | 0 | 0 | 0 | 0 | 629120 | 0 | 423950 | 0 | 15211000 | 0 | 762010 | 0 | 642170 | 0 | 795660 | 218010 | 0 | 0 |

| | | | | | | | | | | | | | | | | | | | | | | |
|---|---|---|---|---|---|---|---|---|---|---|---|---|---|---|---|---|---|---|---|---|---|---|
| AADLSKPID | WDR20; WDR2 | Q8TBZ3; H0 | WDR20_HUMA | Unmodified | 0 | 0 | 0 | 0 | 290570 | 0 | 0 | 38322 | 0 | 0 | 0 | 0 | 209650 | 194130 | 0 | 0 | 83902 | 113740 |
| AADMDLEV | PLXNA1 | Q9UIW2 | PLXA1_HUMAN | Unmodified | 0 | 0 | 0 | 51174 | 1079200 | 153330 | 51089 | 22769 | 208770 | 247380 | 93641 | 0 | 1034000 | 781890 | 0 | 0 | 1020700 | 1811300 |
| AADMGNP | SEL1L | Q9UBV2 | SE1L1_HUMAN | Oxidation (M) | 0 | 0 | 0 | 0 | 0 | 0 | 0 | 0 | 1930100 | 0 | 0 | 0 | 0 | 0 | 0 | 0 | 0 | 0 |
| AADMGNP | SEL1L | Q9UBV2 | SE1L1_HUMAN | Unmodified | 0 | 118340 | 0 | 0 | 0 | 0 | 0 | 0 | 1675500 | 0 | 0 | 0 | 0 | 0 | 0 | 0 | 0 | 0 |
| AADPPAEN | YBX1; YBX1 | P67809; H0 | YBOX1_HUMAN | Unmodified | 1981900 | 1272800 | 895380 | 1572000 | 1232000 | 233220 | 255640 | 473070 | 294680 | 0 | 569620 | 5198900 | 2748700 | 267610 | 318430 | 851940 | 607420 | 945790 |
| AADRLEEEI | TMCO3; TMCO | Q6UWJ1; A0 | TMCO3_HUMA | Unmodified | 0 | 0 | 0 | 0 | 0 | 0 | 0 | 0 | 3652300 | 760820 | 417400 | 0 | 0 | 0 | 0 | 0 | 0 | 0 |
| AADRLPNL | TOM1 | O60784 | TOM1_HUMAN | Unmodified | 8124500 | 7004800 | 3601800 | 5329700 | 3512200 | 576860 | 0 | 0 | 4847200 | 0 | 0 | 137860 | 7909000 | 3030300 | 7076500 | 6731700 | 0 | 2713900 |
| AADTDGD | CALML3 | P27482 | CALL3_HUMAN | Unmodified | 0 | 0 | 0 | 0 | 0 | 0 | 0 | 0 | 462490 | 147520 | 0 | 0 | 0 | 0 | 0 | 0 | 0 | 0 |
| AADTIEALQ | ATP11A; ATP11 | P98196; H0 | AT11A_HUMAN | Unmodified | 0 | 21541 | 0 | 0 | 1844600 | 390300 | 432640 | 355360 | 0 | 0 | 0 | 0 | 0 | 0 | 0 | 0 | 0 | 901710 |
| AAEAAAAP | BASP1 | P80723 | BASP1_HUMAN | Unmodified | 0 | 0 | 0 | 0 | 0 | 0 | 0 | 0 | 2956900 | 107760 | 0 | 0 | 0 | 0 | 0 | 0 | 0 | 0 |
| AAEAAAAP | BASP1 | P80723 | BASP1_HUMAN | Unmodified | 0 | 638710 | 0 | 0 | 20263000 | 5312400 | 2634400 | 1319600 | 60177000 | 5861800 | 0 | 0 | 394770 | 388940 | 0 | 0 | 1604700 | 13823000 |
| AAEAAAAP | BASP1 | P80723 | BASP1_HUMAN | Unmodified | 0 | 0 | 0 | 0 | 554500 | 0 | 0 | 0 | 463070 | 0 | 0 | 0 | 0 | 0 | 0 | 0 | 0 | 0 |
| AAEAAASA | WBP2; WBP2; V | Q969T9; A6 | WBP2_HUMAN | Oxidation (M) | 0 | 0 | 0 | 0 | 1703400 | 0 | 1090400 | 931670 | 0 | 0 | 0 | 0 | 0 | 1427400 | 0 | 0 | 1907900 | 0 |
| AAEAAASA | WBP2; WBP2; V | Q969T9; A6 | WBP2_HUMAN | Unmodified | 0 | 0 | 0 | 0 | 0 | 0 | 0 | 0 | 0 | 0 | 0 | 0 | 868050 | 987400 | 0 | 0 | 0 | 0 |
| AAEAAGGK | EXOC4 | Q96A65 | EXOC4_HUMA | Acetyl (Protein N | 0 | 0 | 0 | 0 | 933110 | 231980 | 180500 | 0 | 0 | 104980 | 0 | 0 | 127090 | 208800 | 0 | 0 | 920120 | 311740 |
| AAEAAINIL | PRKRA; PRKRA | O75569; A0 | PRKRA_HUMA | Unmodified | 0 | 0 | 0 | 0 | 0 | 0 | 0 | 410210 | 43812 | 0 | 0 | 281940 | 0 | 0 | 0 | 65804 | 0 | 0 |
| AAEADGPL | MPDU1; MPDU | O75352; J3 | MPU1_HUMAN | Acetyl (Protein N | 0 | 0 | 0 | 0 | 0 | 0 | 0 | 0 | 1877400 | 665820 | 321520 | 0 | 0 | 0 | 0 | 0 | 0 | 0 |
| AAEADGPL | MPDU1; MPDU | O75352; J3 | MPU1_HUMAN | Acetyl (Protein N | 635690 | 723300 | 0 | 0 | 0 | 0 | 0 | 0 | 7786200 | 525000 | 1006400 | 0 | 0 | 0 | 0 | 0 | 0 | 0 |
| AAEAEQEM | NF2; NF2 | P35240; A0 | MERL_HUMAN | Unmodified | 0 | 0 | 0 | 0 | 3578200 | 0 | 0 | 0 | 155680 | 0 | 0 | 0 | 1198600 | 600700 | 75529 | 0 | 881560 | 5418400 |
| AAEAFSELS | EPS8; EPS8; EP | Q12929; A0 | EPS8_HUMAN | Unmodified | 194500 | 0 | 181010 | 0 | 1092400 | 0 | 0 | 140260 | 0 | 54256 | 0 | 0 | 130030 | 523930 | 283580 | 0 | 0 | 0 |
| AAEALHGE | PDIA5 | Q14554 | PDIA5_HUMAN | Unmodified | 0 | 0 | 0 | 0 | 0 | 0 | 0 | 0 | 1109600 | 0 | 0 | 0 | 0 | 0 | 0 | 0 | 0 | 0 |
| AAECNIVV | DHX9 | Q08211 | DHX9_HUMAN | Unmodified | 0 | 0 | 0 | 0 | 0 | 0 | 55534 | 60806 | 0 | 0 | 0 | 1227600 | 0 | 0 | 0 | 0 | 0 | 0 |
| AAEDDEDE | PTMA; PTMA; P | P06454; H7 | PTMA_HUMAN | Unmodified | 0 | 0 | 0 | 0 | 0 | 0 | 0 | 0 | 0 | 0 | 0 | 0 | 0 | 0 | 0 | 0 | 0 | 0 |
| AAEDDEDE | PTMA; PTMA; P | P06454; H7 | PTMA_HUMAN | Unmodified | 31521 | 36385 | 0 | 210270 | 130270 | 76325 | 1337200 | 285640 | 2014400 | 0 | 17142 | 305080 | 46318 | 0 | 71179 | 0 | 46540 | 95931 |
| AAEDEEVP | DPYSL2 | A0A1C7CYX | A0A1C7CYX9_ | Unmodified | 0 | 0 | 0 | 0 | 583710 | 0 | 0 | 0 | 0 | 0 | 0 | 0 | 0 | 0 | 0 | 0 | 0 | 0 |
| AAEDLFVN | NCKAP1 | Q9Y2A7 | NCKP1_HUMA | Unmodified | 0 | 0 | 0 | 0 | 0 | 0 | 0 | 0 | 0 | 227560 | 83225 | 3092000 | 0 | 0 | 1761300 | 0 | 0 | 0 |
| AAEDRSKQ | TPM1; TPM1; T | P09493; A0 | TPM1_HUMAN | Unmodified | 0 | 0 | 0 | 0 | 752530 | 161550 | 0 | 0 | 0 | 0 | 0 | 0 | 0 | 0 | 0 | 0 | 0 | 0 |
| AAEEAKKF | PSAT1 | Q9Y617 | SERC_HUMAN | Unmodified | 0 | 0 | 0 | 0 | 0 | 0 | 0 | 129120 | 0 | 147090 | 0 | 0 | 0 | 3196900 | 0 | 0 | 1155700 | 0 |
| AAEEIGIK | MTHFD1; MTHF | P11586; V9 | C1TC_HUMAN | Unmodified | 0 | 0 | 701040 | 0 | 7615800 | 329260 | 2936200 | 1471600 | 621850 | 1107000 | 1309200 | 9372600 | 0 | 904350 | 225640 | 295300 | 5041200 | 7931500 |
| AAEEPSKV | MARCKS | P29966 | MARCS_HUMA | Unmodified | 70250 | 0 | 0 | 0 | 0 | 0 | 91035 | 125970 | 6682600 | 910850 | 31051 | 2190500 | 218680 | 0 | 0 | 0 | 395400 | 1569600 |
| AAEEVRHL | DBI; DBI; DBI | P07108; B8 | ACBP_HUMAN | Unmodified | 0 | 13962 | 0 | 0 | 0 | 0 | 0 | 0 | 22399 | 20471 | 53080 | 84869 | 48093 | 0 | 107120 | 0 | 0 | 0 |
| AAELEMEL | PFDN2 | Q9UHV9 | PFD2_HUMAN | Oxidation (M) | 0 | 208310 | 0 | 0 | 291300 | 0 | 0 | 0 | 786990 | 215600 | 874400 | 955140 | 777750 | 0 | 0 | 0 | 120950 | 1283200 |
| AAELEMEL | PFDN2 | Q9UHV9 | PFD2_HUMAN | Oxidation (M) | 0 | 0 | 0 | 0 | 1850400 | 120550 | 464400 | 0 | 0 | 259380 | 0 | 0 | 2749400 | 1361400 | 0 | 0 | 1475600 | 2002000 |
| AAELEMEL | PFDN2 | Q9UHV9 | PFD2_HUMAN | Unmodified | 0 | 0 | 0 | 0 | 493590 | 1257500 | 0 | 0 | 0 | 0 | 0 | 0 | 930090 | 584350 | 0 | 0 | 581320 | 3433100 |
| AAELFGLTA | PTRH1; PTRH1; | Q86Y79; A0 | PTH_HUMAN P | Unmodified | 0 | 79359 | 0 | 69395 | 27250 | 0 | 0 | 0 | 108760 | 80301 | 0 | 0 | 0 | 0 | 100930 | 0 | 0 | 0 |
| AAELIANSL | PHB; PHB; PHB | P35232; C9 | PHB_HUMAN P | Unmodified | 0 | 0 | 0 | 0 | 0 | 171700 | 0 | 0 | 1468800 | 613040 | 174960 | 0 | 0 | 0 | 0 | 0 | 0 | 0 |
| AAEMVPDL | TOM1L2; TOM1 | Q6ZVM7; F5 | TM1L2_HUMAN | 2 Oxidation (M) | 4428600 | 2434200 | 1947100 | 876200 | 0 | 0 | 0 | 0 | 0 | 0 | 0 | 0 | 1235300 | 129360 | 2598700 | 2443800 | 0 | 0 |
| AAEMVPDL | TOM1L2; TOM1 | Q6ZVM7; F5 | TM1L2_HUMAN | Oxidation (M) | 0 | 490910 | 0 | 0 | 0 | 0 | 0 | 0 | 0 | 0 | 0 | 0 | 1072300 | 0 | 469390 | 0 | 0 | 0 |
| AAEPVSRR | SEPTIN9; SEPT | Q9UHD8; K7 | SEPT9_HUMAN | Unmodified | 0 | 0 | 0 | 0 | 773380 | 0 | 0 | 0 | 0 | 0 | 0 | 0 | 712510 | 284870 | 0 | 0 | 1445300 | 2534700 |
| AAEQAGCN | TYSND1 | Q2T9J0 | TYSD1_HUMAN | Unmodified | 0 | 0 | 0 | 0 | 0 | 0 | 0 | 0 | 619420 | 42967 | 0 | 0 | 0 | 0 | 0 | 0 | 0 | 0 |
| AAEQYTPK | FASN; FASN | P49327; A0 | FAS_HUMAN F | Unmodified | 0 | 0 | 0 | 0 | 0 | 0 | 0 | 0 | 0 | 0 | 0 | 4245200 | 0 | 0 | 0 | 0 | 0 | 0 |
| AAESLADP | COPB2; COPB2 | P35606; H0 | COPB2_HUMA | Unmodified | 0 | 0 | 0 | 0 | 1067000 | 532050 | 846420 | 0 | 752130 | 597820 | 516600 | 2254400 | 1047200 | 851730 | 0 | 0 | 407810 | 1183300 |
| AAESPDQK | SLC16A1 | P53985 | MOT1_HUMAN | Unmodified | 0 | 0 | 0 | 8102800 | 0 | 0 | 7115400 | 3108000 | 226860 | 0 | 0 | 0 | 35420000 | 59164000 | 0 | 4515300 | 46533000 | 62368000 |
| AAESVSKP | RTF2; RTF2; RT | Q9BY42; A2 | RTF2_HUMAN | Unmodified | 0 | 0 | 0 | 0 | 150570 | 28329 | 0 | 0 | 89780 | 53224 | 81381 | 0 | 127610 | 0 | 0 | 0 | 97295 | 96021 |
| AAETYGVR | TAGLN3; TAGL | Q9UI15; H7 | TAGL3_HUMAN | Unmodified | 0 | 0 | 0 | 0 | 0 | 0 | 0 | 0 | 364100 | 84762 | 0 | 0 | 0 | 0 | 0 | 0 | 0 | 0 |
| AAEVWMD | GALNT4; GALN | Q8N4A0; Q1 | GALT4_HUMAN | Oxidation (M) | 0 | 0 | 0 | 0 | 0 | 0 | 0 | 0 | 3321800 | 362830 | 0 | 232380 | 0 | 0 | 0 | 0 | 0 | 0 |
| AAEVWMD | GALNT4; GALN | Q8N4A0; Q1 | GALT4_HUMAN | Unmodified | 0 | 0 | 0 | 0 | 0 | 0 | 198280 | 0 | 9875800 | 775590 | 0 | 0 | 0 | 0 | 0 | 0 | 0 | 0 |
| AAFAELHN | SPATS2L; SPAT | Q9NUQ6; B | SPS2L_HUMAN | Unmodified | 0 | 0 | 0 | 0 | 0 | 0 | 0 | 0 | 0 | 0 | 0 | 852610 | 0 | 0 | 15931 | 0 | 0 | 0 |
| AAFDDAIAE | YWHAE; YWHA | P62258; K7 | 1433E_HUMAN | Unmodified | 0 | 0 | 1459300 | 864810 | 0 | 260200 | 0 | 0 | 509830 | 0 | 315640 | 1865200 | 0 | 0 | 6992300 | 672250 | 0 | 0 |
| AAFERESD | MYDGF; MYDG | Q969H8; M0 | MYDGF_HUMA | Unmodified | 0 | 0 | 0 | 0 | 0 | 0 | 0 | 0 | 5728500 | 546130 | 110090 | 0 | 0 | 0 | 79382 | 0 | 0 | 0 |
| AAFERESD | MYDGF; MYDG | Q969H8; M0 | MYDGF_HUMA | Unmodified | 0 | 0 | 0 | 0 | 3465300 | 689410 | 3117600 | 2402400 | 62746000 | 56677000 | 1508500 | 0 | 781820 | 2664100 | 0 | 0 | 1568200 | 1882300 |
| AAFGKADD | TLN2; TLN2; TL | Q9Y4G6; H0 | TLN2_HUMAN | Unmodified | 0 | 0 | 0 | 0 | 0 | 0 | 0 | 0 | 993250 | 0 | 0 | 416670 | 55426 | 219030 | 0 | 0 | 449660 | 625170 |
| AAFGLSEA | SDHA; SDHA; S | P31040; D6 | SDHA_HUMAN | Unmodified | 0 | 0 | 0 | 0 | 0 | 0 | 442680 | 0 | 1950000 | 258330 | 53347 | 0 | 0 | 0 | 0 | 0 | 0 | 0 |
| AAFGQGS | LGALS3BP; LGA | Q08380; K7 | LG3BP_HUMAN | Oxidation (M) | 0 | 0 | 0 | 0 | 0 | 0 | 0 | 0 | 2413900 | 1515600 | 10272000 | 1471900 | 2056200 | 1243400 | 0 | 0 | 0 | 7396200 |
| AAFGQGS | LGALS3BP; LGA | Q08380; K7 | LG3BP_HUMAN | Unmodified | 0 | 0 | 0 | 0 | 0 | 0 | 0 | 0 | 5463000 | 0 | 0 | 0 | 0 | 0 | 0 | 0 | 0 | 2223700 |
| AAFIKAEK | PDCD10; PDCD | Q9BUL8; H7 | PDC10_HUMA | Unmodified | 0 | 0 | 0 | 0 | 710290 | 0 | 0 | 0 | 0 | 94594 | 59976 | 0 | 284210 | 169580 | 0 | 0 | 0 | 0 |
| AAFIKAEKE | PDCD10; PDCD | Q9BUL8; H7 | PDC10_HUMA | Unmodified | 0 | 0 | 0 | 0 | 1687300 | 0 | 0 | 0 | 0 | 0 | 0 | 0 | 533420 | 484050 | 140040 | 0 | 0 | 492640 |
| AAFIKAEKE | PDCD10; PDCD | Q9BUL8; H7 | PDC10_HUMA | Oxidation (M) | 0 | 0 | 0 | 0 | 0 | 0 | 0 | 0 | 0 | 0 | 0 | 0 | 1416300 | 0 | 0 | 0 | 0 | 0 |
| AAFKLDFLF | CELF1; CELF1; | G5EA30; F5 | G5EA30_HUMA | Acetyl (Protein N | 0 | 211350 | 0 | 0 | 0 | 0 | 50529 | 0 | 0 | 0 | 0 | 0 | 360220 | 113580 | 0 | 0 | 0 | 0 |
| AAFKTSYGI | ANXA7 | P20073 | ANXA7_HUMAN | Unmodified | 0 | 0 | 0 | 0 | 197130 | 0 | 188040 | 55880 | 0 | 784490 | 0 | 0 | 835920 | 0 | 0 | 0 | 547430 | 561260 |
| AAFKTSYGI | ANXA7 | P20073 | ANXA7_HUMAN | Unmodified | 0 | 0 | 0 | 0 | 1177300 | 0 | 179590 | 81177 | 258560 | 1148300 | 0 | 0 | 84541 | 0 | 0 | 0 | 700910 | 797790 |
| AAFPQYQP | CEMIP2; CEMIP | Q9UHN6; H | CEIP2_HUMAN | Oxidation (M) | 0 | 0 | 0 | 224650 | 1302200 | 98150 | 0 | 0 | 0 | 373150 | 0 | 0 | 0 | 0 | 722120 | 391820 | 51507 | 0 |
| AAFPQYQP | CEMIP2; CEMIP | Q9UHN6; H | CEIP2_HUMAN | Unmodified | 0 | 0 | 0 | 0 | 0 | 0 | 0 | 0 | 2790900 | 0 | 0 | 0 | 0 | 0 | 0 | 0 | 0 | 0 |
| AAFQLDCS | IGSF3 | O75054 | IGSF3_HUMAN | Unmodified | 589090 | 679780 | 0 | 441110 | 72250 | 0 | 0 | 0 | 397550 | 0 | 0 | 1026300 | 0 | 0 | 1341300 | 814520 | 0 | 0 |
| AAFRQVQ | VPS28; VPS28 | Q9UK41; E9 | VPS28_HUMAN | Unmodified | 26528000 | 46389000 | 0 | 655250 | 0 | 0 | 0 | 0 | 0 | 0 | 0 | 0 | 1389600 | 0 | 14744000 | 245870 | 0 | 0 |
| AAFRQVQ | VPS28; VPS28 | Q9UK41; E9 | VPS28_HUMAN | Unmodified | 0 | 0 | 0 | 1462400 | 8311900 | 1527300 | 3648300 | 1287100 | 0 | 3910000 | 0 | 0 | 26929000 | 16091000 | 0 | 0 | 4273500 | 8305900 |
| AAFSKDESI | NCAM1; NCAM | P13591; A0 | NCAM1_HUMA | Unmodified | 0 | 0 | 0 | 0 | 0 | 0 | 76544 | 0 | 5704900 | 3664500 | 153980 | 10616000 | 0 | 0 | 0 | 0 | 0 | 0 |
| AAFTAFEEA | CCDC124 | Q96CT7 | CC124_HUMA | Unmodified | 445740 | 0 | 448570 | 546820 | 0 | 0 | 0 | 0 | 212150 | 41952 | 0 | 383840 | 0 | 0 | 320960 | 714360 | 0 | 0 |
| AAFVLPEFA | SRM; SRM | P19623; K7 | SPEE_HUMAN | Unmodified | 0 | 326360 | 759280 | 462850 | 0 | 0 | 0 | 41607 | 5620500 | 204070 | 482980 | 6515100 | 346600 | 117670 | 1246200 | 1431600 | 0 | 24139 |
| AAFVLPEFA | SRM; SRM | P19623; K7 | SPEE_HUMAN | Unmodified | 0 | 0 | 0 | 157570 | 188810 | 0 | 0 | 0 | 885690 | 714380 | 162070 | 129140 | 1443400 | 234190 | 0 | 0 | 238790 | 434820 |
| AAFVSEQQ | ITGA3; ITGA3 | P26006; H0 | ITA3_HUMAN I | Unmodified | 639100 | 680120 | 409930 | 0 | 1.06E+08 | 57715000 | 0 | 0 | 1130600 | 123830 | 0 | 4631600 | 0 | 0 | 702550 | 519250 | 17139000 | 84658000 |
| AAFVSEQQ | ITGA3; ITGA3 | P26006; H0 | ITA3_HUMAN I | Unmodified | 0 | 0 | 0 | 0 | 2450600 | 495860 | 0 | 0 | 0 | 0 | 0 | 0 | 46819 | 0 | 0 | 0 | 2976600 | 3665400 |
| AAFVSEQQ | ITGA3; ITGA3 | P26006; H0 | ITA3_HUMAN I | Unmodified | 0 | 0 | 0 | 0 | 1387700 | 0 | 0 | 0 | 0 | 60334 | 0 | 0 | 0 | 0 | 0 | 0 | 1939100 | 1720800 |
| AAGAAGAA | C2orf72 | A6NCS6 | CB072_HUMAN | Unmodified | 170630 | 424280 | 250710 | 0 | 122000 | 58319 | 0 | 0 | 0 | 0 | 120280 | 459750 | 0 | 0 | 707540 | 547390 | 82810 | 172430 |
| AAGARPLT | BAG6; BAG6; B | P46379; A0 | BAG6_HUMAN | Unmodified | 0 | 0 | 0 | 0 | 0 | 0 | 0 | 0 | 276180 | 46414 | 19853 | 0 | 0 | 0 | 0 | 0 | 0 | 0 |
| AAGASDVV | FARSB | Q9NSD9 | SYFB_HUMAN | Unmodified | 0 | 246650 | 0 | 0 | 432910 | 5266100 | 851470 | 114220 | 290530 | 501720 | 608050 | 2634400 | 424350 | 0 | 0 | 286860 | 1093900 | 0 |
| AAGCDFTN | RIDA; RIDA; RID | P52758; H0 | RIDA_HUMAN | Unmodified | 0 | 0 | 0 | 0 | 0 | 0 | 0 | 141070 | 834140 | 518260 | 0 | 366900 | 0 | 0 | 409230 | 196550 | 275280 | 238930 |
| AAGDTTVIE | KTN1; KTN1; KT | Q86UP2; B7 | KTN1_HUMAN | Unmodified | 0 | 0 | 0 | 0 | 0 | 0 | 0 | 0 | 132690 | 0 | 158530 | 3231700 | 0 | 0 | 0 | 0 | 0 | 49179 |
| AAGDVDIG | TACSTD2 | P09758 | TACD2_HUMA | Unmodified | 0 | 0 | 0 | 223970 | 25776000 | 5622800 | 0 | 0 | 518450 | 142340 | 0 | 0 | 0 | 0 | 0 | 0 | 202970 | 442840 |
| AAGEPIKE | DNAJC2; DNAJ | Q99543; F2 | DNJC2_HUMA | Unmodified | 0 | 0 | 0 | 0 | 331360 | 0 | 190660 | 148000 | 0 | 64220 | 0 | 226480 | 559430 | 352550 | 0 | 0 | 189040 | 0 |
| AAGFKDPL | GART | P22102 | PUR2_HUMAN | Unmodified | 0 | 186480 | 0 | 173790 | 852970 | 0 | 703170 | 471480 | 462800 | 250700 | 591940 | 3794000 | 908860 | 350610 | 0 | 246840 | 500150 | 904090 |
| AAGFKDPL | GART | P22102 | PUR2_HUMAN | Unmodified | 0 | 0 | 0 | 0 | 0 | 0 | 0 | 577040 | 0 | 0 | 0 | 0 | 1142600 | 0 | 0 | 0 | 0 | 0 |
| AAGFNVLR | HSPA14 | Q0VDF9 | HSP7E_HUMA | Unmodified | 0 | 0 | 0 | 0 | 0 | 0 | 0 | 0 | 0 | 0 | 0 | 1481800 | 0 | 0 | 0 | 0 | 0 | 0 |
| AAGGAPSP | TBC1D10B | Q4KMP7 | TB10B_HUMAN | Unmodified | 39978 | 0 | 0 | 0 | 0 | 0 | 0 | 0 | 756740 | 109270 | 125930 | 0 | 0 | 0 | 55906 | 75401 | 0 | 0 |
| AAGGAVCE | MUC5B | Q9HC84 | MUC5B_HUMA | Unmodified | 0 | 0 | 0 | 0 | 0 | 0 | 0 | 0 | 0 | 0 | 0 | 0 | 119750 | 0 | 0 | 0 | 0 | 879290 |
| AAGGDHG | ADSL; ADSL; AD | P30566; B4 | PUR8_HUMAN | Acetyl (Protein N | 0 | 0 | 0 | 0 | 0 | 0 | 0 | 0 | 0 | 0 | 146910 | 811530 | 0 | 0 | 0 | 0 | 0 | 0 |
| AAGGDHG | ADSL; ADSL; AD | P30566; B4 | PUR8_HUMAN | Acetyl (Protein N | 80460 | 87166 | 179990 | 0 | 0 | 146590 | 152750 | 79520 | 0 | 85670 | 594040 | 7755400 | 651500 | 0 | 420570 | 418880 | 259340 | 299450 |
| AAGGLHLL | KIAA2013 | Q8IYS2 | K2013_HUMAN | Unmodified | 0 | 55267 | 0 | 0 | 0 | 715820 | 0 | 0 | 4248500 | 562760 | 0 | 1020700 | 0 | 0 | 0 | 0 | 0 | 0 |
| AAGGLHLL | KIAA2013 | Q8IYS2 | K2013_HUMAN | Unmodified | 0 | 0 | 0 | 0 | 0 | 0 | 0 | 0 | 3262800 | 0 | 0 | 0 | 0 | 0 | 0 | 0 | 0 | 0 |
| AAGHPGDI | TLN1 | Q9Y490 | TLN1_HUMAN | Unmodified | 122610 | 205950 | 44422 | 0 | 120050 | 0 | 27470 | 0 | 19106 | 17862 | 13056 | 517240 | 38303 | 0 | 128100 | 148200 | 27237 | 18665 |
| AAGHPGDI | TLN1 | Q9Y490 | TLN1_HUMAN | Unmodified | 0 | 0 | 0 | 74928 | 3650300 | 1178400 | 4244000 | 1189200 | 0 | 732240 | 0 | 0 | 756440 | 1460900 | 0 | 0 | 1650000 | 498730 |
| AAGIDEQEI | LAP3 | P28838 | AMPL_HUMAN | Unmodified | 0 | 0 | 123780 | 461580 | 886890 | 144730 | 198700 | 140580 | 1525900 | 117400 | 536100 | 1812200 | 587770 | 586080 | 0 | 320880 | 176750 | 657060 |
| AAGKGPLA | TMA7 | Q9Y2S6 | TMA7_HUMAN | Unmodified | 333790 | 702600 | 0 | 0 | 3706500 | 0 | 386470 | 283950 | 540990 | 2173500 | 0 | 0 | 4630800 | 3460200 | 0 | 0 | 1603700 | 6554000 |
| AAGKITDA | VPS33B; VPS33 | Q9H267; F5 | VP33B_HUMAN | Unmodified | 0 | 0 | 0 | 0 | 473760 | 0 | 150990 | 107370 | 0 | 57859 | 0 | 0 | 181830 | 157120 | 0 | 0 | 0 | 226200 |
| AAGMSSP | MYO9B; MYO9 | Q13459; M0 | MYO9B_HUMA | Unmodified | 0 | 0 | 0 | 0 | 0 | 0 | 0 | 0 | 0 | 0 | 0 | 876060 | 0 | 0 | 0 | 0 | 0 | 0 |
| AAGPELRV | C9orf64 | Q5T6V5 | QSPP_HUMAN | Unmodified | 0 | 0 | 0 | 0 | 5935500 | 0 | 0 | 529280 | 308090 | 486890 | 0 | 4830200 | 1406000 | 0 | 0 | 0 | 329780 | 72434 |
| AAGPISERI | SF3B4 | Q15427 | SF3B4_HUMAN | Acetyl (Protein N | 310170 | 580930 | 0 | 0 | 0 | 0 | 0 | 723430 | 0 | 0 | 0 | 0 | 0 | 0 | 0 | 0 | 0 | 0 |
| AAGSGELG | FLNB; FLNB | O75369; E7 | FLNB_HUMAN | Unmodified | 0 | 83394 | 0 | 0 | 4829500 | 1015100 | 0 | 0 | 1105300 | 1353700 | 221240 | 2464200 | 0 | 0 | 616080 | 0 | 501390 | 845930 |
| AAGSGELG | FLNB; FLNB | O75369; E7 | FLNB_HUMAN | Oxidation (M) | 0 | 0 | 369230 | 0 | 1247400 | 366330 | 0 | 0 | 169470 | 0 | 0 | 910380 | 254990 | 0 | 0 | 0 | 0 | 432890 |
| AAGSGELG | FLNB; FLNB | O75369; E7 | FLNB_HUMAN | Oxidation (M) | 0 | 0 | 0 | 0 | 1583700 | 191190 | 0 | 29363 | 0 | 0 | 0 | 337480 | 281880 | 375280 | 0 | 33264 | 336390 | 250640 |
| AAGSGELG | FLNB; FLNB | O75369; E7 | FLNB_HUMAN | Unmodified | 0 | 0 | 0 | 0 | 4446800 | 703800 | 0 | 0 | 0 | 0 | 0 | 447360 | 0 | 0 | 0 | 0 | 1159800 | 611510 |
| AAGSGELG | FLNB; FLNB | O75369; E7 | FLNB_HUMAN | Unmodified | 0 | 0 | 0 | 0 | 1746700 | 0 | 0 | 159140 | 0 | 0 | 0 | 0 | 0 | 0 | 0 | 0 | 1738100 | 0 |
| AAGTLYTY | EEF1G | P26641 | EF1G_HUMAN | Acetyl (Protein N | 0 | 0 | 0 | 0 | 0 | 58545 | 0 | 0 | 227980 | 0 | 127790 | 1250400 | 0 | 0 | 0 | 0 | 0 | 0 |
| AAGVNVEP | RPLP1 | P05386 | RLA1_HUMAN | Unmodified | 29065000 | 33153000 | 7042800 | 21639000 | 0 | 0 | 9296500 | 235120 | 5664400 | 4299100 | 0 | 0 | 33565000 | 15410000 | 46783000 | 33315000 | 5880800 | 6690200 |
| AAGVPSAT | HSPG2 | P98160 | PGBM_HUMAN | Unmodified | 0 | 0 | 2996200 | 0 | 1630900 | 1062100 | 0 | 6025.3 | 401100 | 709250 | 0 | 287570 | 0 | 1505700 | 0 | 0 | 0 | 1411400 |
| AAGVVLEM | RUVBL2; RUVB | Q9Y230; M0 | RUVB2_HUMA | Oxidation (M) | 0 | 0 | 13380 | 0 | 0 | 0 | 66954 | 33018 | 0 | 0 | 0 | 678430 | 0 | 0 | 0 | 0 | 0 | 0 |
| AAGVVLEM | RUVBL2; RUVB | Q9Y230; M0 | RUVB2_HUMA | Unmodified | 197580 | 97022 | 82316 | 102010 | 0 | 0 | 0 | 0 | 257530 | 23259 | 25304 | 1681600 | 0 | 0 | 124990 | 0 | 0 | 0 |
| AAGWSELS | PCBD2; PCBD2 | Q9H0N5; H | PHS2_HUMAN | Unmodified | 0 | 0 | 0 | 0 | 294690 | 87235 | 111570 | 0 | 715200 | 0 | 383060 | 335980 | 487120 | 0 | 0 | 0 | 0 | 0 |
| AAHIFFTDT | STXBP2; STXBP | Q15833; M0 | STXB2_HUMAN | Unmodified | 452740 | 446580 | 338380 | 2444000 | 2952600 | 863960 | 0 | 0 | 824210 | 1305000 | 729950 | 3086600 | 3736100 | 2404300 | 984010 | 1039500 | 2241000 | 2107400 |

| | | | | | | | | | | | | | | | | | | | | | | |
|---|---|---|---|---|---|---|---|---|---|---|---|---|---|---|---|---|---|---|---|---|---|---|
| AAHIFFTDT | STXBP2; STXBP | Q15833; M0 | STXB2_HUMAN | Unmodified | 229670 | 308020 | 45133 | 114150 | 0 | 0 | 0 | 0 | 0 | 0 | 0 | 0 | 66600 | 0 | 125920 | 105560 | 2015900 | 0 |
| AAHIFNEAL | MYO1B; MYO1B | O43795; E9F | MYO1B_HUMA | Unmodified | 172310 | 469970 | 0 | 705600 | 966270 | 285300 | 185180 | 0 | 0 | 253110 | 5075400 | 17398000 | 0 | 0 | 998070 | 150500 | 0 | 2906500 |
| AAHIFNEAL | MYO1B; MYO1B | O43795; E9F | MYO1B_HUMA | Unmodified | 0 | 0 | 0 | 0 | 3715000 | 0 | 363650 | 0 | 0 | 653520 | 0 | 0 | 0 | 0 | 0 | 0 | 2666100 | 0 |
| AAHLCAEA | PA2G4; PA2G4; | Q9UQ80; F8 | PA2G4_HUMAN | Unmodified | 0 | 0 | 0 | 0 | 4389900 | 485470 | 1361300 | 0 | 0 | 8145700 | 0 | 0 | 5104500 | 2056400 | 0 | 0 | 2977500 | 1190300 |
| AAHSEGNT | CCT2 | P78371 | TCPB_HUMAN | Unmodified | 0 | 0 | 0 | 158670 | 0 | 96761 | 0 | 0 | 718840 | 46235 | 221390 | 7320700 | 0 | 0 | 0 | 0 | 42343 | 0 |
| AAHSEGNT | CCT2 | P78371 | TCPB_HUMAN | Oxidation (M) | 0 | 0 | 483050 | 0 | 0 | 0 | 0 | 0 | 0 | 0 | 878060 | 1733900 | 0 | 0 | 0 | 1643000 | 0 | 0 |
| AAHVFFTD | STXBP1; STXBP | P61764; A0A | STXB1_HUMAN | Unmodified | 784070 | 80746 | 56837 | 677670 | 25936 | 0 | 2402300 | 0 | 1729300 | 445380 | 93490 | 1658800 | 393980 | 56875 | 479250 | 0 | 300640 | 404980 |
| AAIDWFDG | TAF15; FUS; TA | Q92804; P35 | RBP56_HUMAN | Unmodified | 595240 | 604910 | 0 | 0 | 2270100 | 0 | 0 | 0 | 2154500 | 1240900 | 778170 | 0 | 948400 | 1164800 | 0 | 0 | 385270 | 1112800 |
| AAIDWFDG | FUS; FUS | P35637; H3E | FUS_HUMAN R | Unmodified | 0 | 0 | 0 | 0 | 0 | 178780 | 0 | 166940 | 0 | 967730 | 0 | 0 | 0 | 0 | 0 | 0 | 0 | 0 |
| AAIEAFNK | SRP68 | Q9UHB9 | SRP68_HUMAN | Unmodified | 0 | 0 | 43562 | 166260 | 0 | 0 | 18694 | 0 | 0 | 112580 | 294750 | 3534800 | 0 | 0 | 0 | 81381 | 0 | 0 |
| AAIHYGIIDI | CRISPLD1; CRIS | Q9H336; B7 | CRLD1_HUMAN | Unmodified | 282630 | 0 | 0 | 0 | 0 | 0 | 0 | 0 | 1018300 | 166610 | 0 | 0 | 0 | 1249500 | 0 | 0 | 0 | 0 |
| AAIISAEGD | PHB; PHB; PHB | P35232; C9J | PHB_HUMAN P | Unmodified | 0 | 0 | 0 | 0 | 0 | 0 | 0 | 0 | 1402600 | 510660 | 0 | 0 | 0 | 0 | 0 | 0 | 0 | 0 |
| AAILPTSIFL | PRMT5; PRMT5 | O14744; H0 | ANM5_HUMAN | Unmodified | 0 | 0 | 0 | 0 | 0 | 0 | 0 | 0 | 0 | 0 | 0 | 925610 | 0 | 0 | 0 | 0 | 0 | 0 |
| AAILPTSIFL | PRMT5; PRMT5 | O14744; H0 | ANM5_HUMAN | Unmodified | 0 | 0 | 0 | 0 | 0 | 0 | 0 | 28635 | 0 | 0 | 0 | 12321000 | 0 | 0 | 0 | 0 | 0 | 0 |
| AAILQTEVD | ERC2; ERC1; EF | O15083; Q8 | ERC2_HUMAN | Unmodified | 0 | 0 | 0 | 0 | 171430 | 0 | 0 | 0 | 0 | 33922 | 0 | 0 | 0 | 0 | 0 | 0 | 0 | 74359 |
| AAIPSALDT | LGALS3BP | Q08380 | LG3BP_HUMAN | Unmodified | 0 | 233110 | 0 | 0 | 0 | 0 | 0 | 118680 | 0 | 372420 | 113830 | 0 | 142650 | 0 | 0 | 0 | 0 | 1791500 |
| AAIQSQLD | EXOC3; EXOC3 | O60645; D6 | EXOC3_HUMAN | Unmodified | 0 | 263650 | 0 | 0 | 252710 | 0 | 0 | 0 | 455760 | 0 | 0 | 1091200 | 0 | 0 | 0 | 132290 | 0 | 659400 |
| AAIQSQLD | EXOC3; EXOC3 | O60645; D6 | EXOC3_HUMAN | Unmodified | 33652 | 0 | 0 | 267840 | 4533700 | 531840 | 287200 | 0 | 723740 | 633580 | 0 | 0 | 1356800 | 814850 | 83230 | 0 | 451850 | 992710 |
| AAISDSAD | PARD3; PARD3; | Q8TEW0; Q5 | PARD3_HUMAN | Oxidation (M) | 0 | 0 | 2549400 | 0 | 0 | 0 | 0 | 0 | 0 | 0 | 0 | 0 | 0 | 0 | 0 | 0 | 2368100 | 0 |
| AAISNKITS | PAICS; PAICS | P22234; E9P | PUR6_HUMAN | Unmodified | 0 | 0 | 0 | 0 | 164550 | 0 | 153890 | 0 | 0 | 29641 | 0 | 0 | 689390 | 427690 | 0 | 0 | 427710 | 148080 |
| AAITETQRK | EHBP1 | Q8NDI1 | EHBP1_HUMAN | Unmodified | 0 | 0 | 0 | 0 | 1899500 | 139890 | 1162000 | 70761 | 565160 | 573030 | 38807 | 210400 | 0 | 324910 | 0 | 0 | 425290 | 1274000 |
| AAIVEKLSS | NUTF2; NUTF2 | P61970; H3E | NTF2_HUMAN | Unmodified | 0 | 0 | 0 | 0 | 0 | 0 | 634600 | 0 | 0 | 0 | 0 | 0 | 0 | 0 | 0 | 0 | 985250 | 96231 |
| AAIVFTDGF | MATN2; MATN2 | O00339; H0 | MATN2_HUMAN | Unmodified | 0 | 0 | 0 | 283250 | 167650 | 0 | 0 | 0 | 5208900 | 2036300 | 38727 | 357590 | 346730 | 0 | 104120 | 0 | 0 | 397850 |
| AAIVHNVD | RNF13; RNF13; | O43567; C9J | RNF13_HUMAN | Oxidation (M) | 0 | 0 | 0 | 0 | 0 | 0 | 0 | 0 | 1092200 | 0 | 0 | 0 | 0 | 0 | 0 | 0 | 0 | 0 |
| AAIYQTQAF | ARRDC3 | Q96B67 | ARRD3_HUMAN | Unmodified | 0 | 0 | 0 | 0 | 1145900 | 233540 | 786310 | 102870 | 301720 | 0 | 0 | 0 | 245080 | 0 | 0 | 0 | 0 | 0 |
| AAKDGANI | HSDL2 | Q6YN16 | HSDL2_HUMAN | Unmodified | 0 | 0 | 0 | 0 | 496110 | 0 | 0 | 0 | 688120 | 523380 | 0 | 0 | 0 | 0 | 0 | 0 | 0 | 0 |
| AAKGGELV | PLEC; PLEC | Q15149; H0 | PLEC_HUMAN | Unmodified | 0 | 0 | 0 | 0 | 0 | 0 | 0 | 0 | 0 | 0 | 0 | 0 | 0 | 0 | 0 | 0 | 1926400 | 0 |
| AAKLEILQC | NUMA1; NUMA | Q14980; F5H | NUMA1_HUMA | Unmodified | 129460 | 0 | 0 | 165470 | 0 | 0 | 0 | 0 | 0 | 0 | 180470 | 398320 | 0 | 0 | 249150 | 634610 | 0 | 0 |
| AAKQGDTII | NUP210 | Q8TEM1 | PO210_HUMAN | Unmodified | 0 | 0 | 0 | 0 | 0 | 0 | 0 | 0 | 0 | 674090 | 0 | 0 | 0 | 0 | 0 | 0 | 0 | 0 |
| AAKTQLAV | FKBP4; FKBP4 | Q02790; H0 | FKBP4_HUMAN | Unmodified | 0 | 116750 | 0 | 0 | 0 | 0 | 135400 | 130560 | 0 | 0 | 0 | 0 | 203260 | 14475 | 0 | 0 | 748770 | 67592 |
| AAKVLEQL | RPL11; RPL11 | P62913; Q5 | RL11_HUMAN | Unmodified | 0 | 0 | 0 | 0 | 0 | 0 | 564830 | 176240 | 0 | 0 | 0 | 0 | 143650 | 196020 | 0 | 0 | 76546 | 84629 |
| AALAGGTTI | DPYSL3 | Q14195 | DPYL3_HUMAN | Oxidation (M) | 0 | 0 | 0 | 0 | 4424000 | 0 | 3564200 | 382440 | 860760 | 0 | 41359000 | 86049000 | 0 | 0 | 0 | 0 | 927350 | 0 |
| AALAGGTTI | DPYSL3 | Q14195 | DPYL3_HUMAN | Unmodified | 0 | 0 | 0 | 0 | 0 | 0 | 0 | 0 | 0 | 0 | 1167900 | 2972000 | 0 | 0 | 0 | 0 | 0 | 0 |
| AALAGTAYI | N/A | N/A | N/A | Unmodified | 0 | 0 | 0 | 0 | 7028500 | 6838900 | 54815000 | 8260700 | 0 | 970850 | 0 | 0 | 1197500 | 36938000 | 0 | 0 | 6513400 | 4431900 |
| AALAPELAL | AHNAK2 | Q8IVF2 | AHNK2_HUMAN | Unmodified | 75752000 | 0 | 0 | 0 | 1347400 | 290850 | 0 | 0 | 0 | 0 | 0 | 0 | 1.03E+08 | 0 | 0 | 5470800 | 0 | 0 |
| AALAPLPPL | VTA1; VTA1 | Q9NP79; A0 | VTA1_HUMAN | Acetyl (Protein N | 2555300 | 1451900 | 1061000 | 1992300 | 6940700 | 1391500 | 1221900 | 560540 | 12421000 | 7223500 | 2895600 | 13972000 | 8212200 | 6559700 | 2224000 | 1165200 | 3242600 | 3071500 |
| AALCHFIVE | NAPB; NAPB | Q9H115; A0 | SNAB_HUMAN | Unmodified | 250050 | 314250 | 0 | 0 | 0 | 14226 | 0 | 0 | 918900 | 345210 | 0 | 570290 | 107990 | 0 | 449320 | 230780 | 0 | 0 |
| AALDEQFEI | HPGD; HPGD; | P15428; E9P | PGDH_HUMAN | Unmodified | 0 | 0 | 0 | 0 | 0 | 0 | 184160 | 150710 | 232710 | 0 | 0 | 0 | 684270 | 268980 | 0 | 0 | 331780 | 3798300 |
| AALEALGS | SHMT2; SHMT2 | P34897; G3 | GLYM_HUMAN | Unmodified | 0 | 0 | 0 | 0 | 0 | 0 | 0 | 0 | 4490700 | 0 | 0 | 0 | 0 | 0 | 0 | 0 | 0 | 0 |
| AALEALLG | FAM83F | Q8NEG4 | FA83F_HUMAN | Unmodified | 0 | 0 | 0 | 748970 | 0 | 0 | 322900 | 0 | 1380600 | 0 | 837910 | 0 | 0 | 0 | 1110900 | 863340 | 0 | 0 |
| AALEAQNA | CELF2; CELF2; | O95319; Q5 | CELF2_HUMAN | Unmodified | 0 | 0 | 0 | 0 | 0 | 0 | 1250000 | 1282500 | 0 | 0 | 0 | 254780 | 0 | 0 | 0 | 0 | 0 | 0 |
| AALEAQNA | CELF1; CELF1; | Q92879; G5 | CELF1_HUMAN | Unmodified | 2722400 | 1625500 | 1126900 | 1872900 | 0 | 0 | 1180600 | 0 | 0 | 237520 | 195940 | 2703900 | 4057800 | 1416400 | 1119200 | 898240 | 272010 | 384040 |
| AALEAQNA | CELF1; CELF1; | Q92879; G5 | CELF1_HUMAN | Oxidation (M) | 2388700 | 0 | 1884600 | 852580 | 0 | 0 | 382430 | 165350 | 0 | 54810 | 0 | 544400 | 1659600 | 758100 | 1656300 | 1194400 | 0 | 0 |
| AALEDSSG | EPS8; EPS8; EP | Q12929; A0 | EPS8_HUMAN | Unmodified | 0 | 0 | 0 | 0 | 692270 | 0 | 0 | 107350 | 0 | 0 | 0 | 0 | 0 | 0 | 0 | 0 | 0 | 0 |
| AALEELQR | TUBGCP3; TUB | Q96CW5; A0 | GCP3_HUMAN | Unmodified | 0 | 1011800 | 0 | 32753 | 0 | 0 | 0 | 0 | 0 | 2267600 | 0 | 639230 | 0 | 0 | 0 | 0 | 0 | 0 |
| AALEEVYPI | XRN2 | Q9H0D6 | XRN2_HUMAN | Unmodified | 0 | 0 | 0 | 0 | 0 | 634720 | 0 | 0 | 0 | 0 | 273280 | 2343400 | 0 | 0 | 0 | 0 | 86859 | 0 |
| AALEGFLA | APLP1; APLP1; | P51693; B7Z | APLP1_HUMAN | Unmodified | 159620 | 0 | 0 | 0 | 97964 | 0 | 0 | 0 | 12366000 | 369400 | 0 | 0 | 0 | 0 | 0 | 0 | 0 | 58137 |
| AALESQLR | CIT; CIT; CIT; CI | O14578; A0 | CTRO_HUMAN | Unmodified | 0 | 0 | 0 | 0 | 0 | 0 | 0 | 0 | 0 | 0 | 0 | 795430 | 0 | 0 | 0 | 0 | 0 | 0 |
| AALESQVRI | SPECC1; SPEC | Q5M775; J3 | CYTSB_HUMAN | Unmodified | 0 | 0 | 0 | 68031 | 508680 | 52473 | 0 | 0 | 0 | 121790 | 0 | 0 | 524290 | 504490 | 0 | 0 | 117810 | 255470 |
| AALEYLEDI | PRXL2A | Q9BRX8 | PXL2A_HUMAN | Unmodified | 0 | 0 | 0 | 0 | 0 | 0 | 397670 | 24462 | 5667200 | 777670 | 0 | 0 | 0 | 0 | 0 | 0 | 0 | 0 |
| AALEYLEDI | PRXL2A | Q9BRX8 | PXL2A_HUMAN | Unmodified | 0 | 0 | 0 | 0 | 0 | 0 | 0 | 0 | 0 | 1130700 | 0 | 0 | 0 | 0 | 0 | 0 | 0 | 0 |
| AALEYLEDI | PRXL2A | Q9BRX8 | PXL2A_HUMAN | Unmodified | 0 | 0 | 0 | 0 | 0 | 0 | 0 | 0 | 0 | 990520 | 0 | 0 | 0 | 0 | 0 | 0 | 0 | 0 |
| AALEYLGS | PTPRD; PTPRF; | P23468; P10 | PTPRD_HUMAN | Unmodified | 1741000 | 1085200 | 1487000 | 1613700 | 2203800 | 0 | 0 | 0 | 3251700 | 1008100 | 0 | 658510 | 831530 | 0 | 3286800 | 2055300 | 0 | 0 |
| AALFALQN | MTR; MTR; MTR | Q99707; A0 | METH_HUMAN | Unmodified | 0 | 0 | 0 | 12563 | 41541 | 0 | 33412 | 0 | 69838 | 86048 | 0 | 0 | 26137 | 0 | 16628 | 0 | 0 | 14335 |
| AALGEPVRI | LIN7C; LIN7C | Q9NUP9; G3 | LIN7C_HUMAN | Acetyl (Protein N | 0 | 759990 | 7888800 | 9663300 | 7850500 | 0 | 300930 | 157650 | 11583000 | 0 | 5030700 | 6889800 | 5123000 | 0 | 7245500 | 5091600 | 1728500 | 7225400 |
| AALGEPVRI | LIN7C; LIN7C | Q9NUP9; G3 | LIN7C_HUMAN | Acetyl (Protein N | 0 | 0 | 0 | 0 | 0 | 3111500 | 0 | 0 | 0 | 0 | 0 | 0 | 57852 | 66405 | 0 | 0 | 0 | 9088000 |
| AALGPLVT | UMPS; UMPS; U | P11172; F2Z | UMPS_HUMAN | Unmodified | 0 | 0 | 0 | 0 | 673030 | 0 | 0 | 0 | 414180 | 0 | 234960 | 1865600 | 0 | 0 | 0 | 0 | 0 | 0 |
| AALHLYSN | NAA10; NAA10; | P41227; F8W | NAA10_HUMAN | Unmodified | 0 | 3922000 | 2516700 | 0 | 1143300 | 0 | 4536600 | 0 | 0 | 0 | 1084200 | 1577900 | 0 | 0 | 4026600 | 3396100 | 0 | 0 |
| AALIYTCTV | ZNF706 | Q9Y5V0 | ZN706_HUMAN | Unmodified | 0 | 25614 | 0 | 0 | 0 | 0 | 0 | 0 | 819970 | 0 | 0 | 0 | 0 | 0 | 0 | 0 | 425590 | 1524600 |
| AALLDQAL | MACF1; MACF1 | H3BQK9; H3 | H3BQK9_HUM | Unmodified | 0 | 0 | 0 | 0 | 0 | 0 | 0 | 0 | 0 | 0 | 0 | 696240 | 0 | 0 | 116010 | 0 | 0 | 0 |
| AALLDQIR | WASL | O00401 | WASL_HUMAN | Unmodified | 1052800 | 1074700 | 0 | 1387100 | 0 | 0 | 0 | 0 | 6084200 | 411800 | 0 | 395230 | 0 | 0 | 1903400 | 1160400 | 0 | 0 |
| AALLDQIRE | WASL | O00401 | WASL_HUMAN | Unmodified | 1649800 | 1090200 | 694840 | 2168300 | 2542900 | 420040 | 517550 | 318030 | 0 | 0 | 0 | 0 | 3591800 | 3507400 | 2227900 | 827930 | 1072500 | 1999700 |
| AALLELWE | SPTAN1; SPTAN | Q13813; A0 | SPTN1_HUMAN | Unmodified | 0 | 0 | 0 | 0 | 0 | 0 | 0 | 0 | 0 | 0 | 0 | 3055500 | 0 | 0 | 0 | 0 | 0 | 0 |
| AALLNFDE | FHOD1 | Q9Y613 | FHOD1_HUMA | Unmodified | 0 | 0 | 0 | 0 | 1814100 | 0 | 0 | 71804 | 622550 | 0 | 0 | 0 | 0 | 0 | 0 | 0 | 419400 | 0 |
| AALLNQHY | EPRS1; EPRS1 | P07814; V9 | SYEP_HUMAN | Unmodified | 509940 | 323120 | 1275500 | 1095200 | 3394900 | 1750000 | 3537400 | 1370400 | 1976700 | 1661800 | 1791700 | 17037000 | 6689800 | 3457200 | 1049700 | 1057700 | 1925600 | 3781100 |
| AALMFANA | BCCIP | Q9P287 | BCCIP_HUMAN | Oxidation (M) | 0 | 0 | 0 | 28300 | 271880 | 58021 | 92781 | 0 | 0 | 0 | 210700 | 0 | 436710 | 0 | 0 | 0 | 46372 | 111330 |
| AALNAVRLI | SLC9A3R1; SLC | O14745; J3 | NHRF1_HUMAN | Unmodified | 0 | 0 | 0 | 0 | 6185700 | 840630 | 0 | 380790 | 0 | 1397800 | 0 | 0 | 14817000 | 14554000 | 0 | 0 | 2615600 | 3184700 |
| AALPAQELI | ACE; ACE; ACE; | P12821; J3K | ACE_HUMAN A | Unmodified | 1451000 | 895150 | 0 | 0 | 0 | 0 | 0 | 0 | 0 | 0 | 0 | 0 | 0 | 0 | 0 | 0 | 0 | 0 |
| AALPDLTK | DNAJC3; DNAJ | Q13217; X6 | DNJC3_HUMAN | Unmodified | 0 | 0 | 0 | 0 | 0 | 0 | 0 | 0 | 0 | 0 | 0 | 0 | 0 | 0 | 0 | 0 | 0 | 0 |
| AALPEGLPI | GDF15 | Q99988 | GDF15_HUMAN | Unmodified | 0 | 0 | 1160100 | 550500 | 579890 | 0 | 0 | 0 | 5165900 | 285390 | 0 | 0 | 0 | 0 | 2194400 | 1165100 | 766220 | 570190 |
| AALPEGLPI | GDF15 | Q99988 | GDF15_HUMAN | Unmodified | 0 | 0 | 0 | 0 | 0 | 0 | 0 | 0 | 0 | 847520 | 0 | 0 | 0 | 0 | 0 | 0 | 0 | 0 |
| AALPLTTSD | HUWE1 | Q7Z6Z7 | HUWE1_HUMA | Unmodified | 0 | 0 | 0 | 0 | 0 | 0 | 0 | 679510 | 0 | 0 | 0 | 3206400 | 0 | 0 | 0 | 79832 | 0 | 0 |
| AALPTPALS | PTPN23 | Q9H3S7 | PTN23_HUMAN | Unmodified | 1033200 | 0 | 0 | 0 | 0 | 0 | 0 | 0 | 466150 | 0 | 111690 | 484630 | 0 | 0 | 0 | 0 | 0 | 0 |
| AALQEELQ | FASN; FASN | P49327; A0A | FAS_HUMAN F | Unmodified | 570490 | 1034300 | 1139900 | 1492100 | 0 | 272430 | 0 | 122760 | 47866 | 0 | 3144600 | 34675000 | 0 | 0 | 0 | 73515 | 471600 | 0 |
| AALQELLSK | RPS25 | P62851 | RS25_HUMAN | Unmodified | 8862700 | 6946300 | 4414800 | 2941100 | 2713600 | 0 | 4210900 | 3082200 | 1019900 | 1048000 | 5000300 | 33981000 | 9540300 | 2978300 | 5224800 | 4047100 | 341380 | 2528000 |
| AALQELLSK | RPS25 | P62851 | RS25_HUMAN | Unmodified | 19753 | 0 | 0 | 0 | 0 | 0 | 238670 | 0 | 0 | 0 | 0 | 192310 | 888220 | 378450 | 0 | 0 | 0 | 0 |
| AALQLVVD | PAK4 | O96013 | PAK4_HUMAN | Unmodified | 494940 | 423240 | 270590 | 327240 | 0 | 0 | 0 | 0 | 0 | 0 | 126040 | 665460 | 0 | 0 | 492860 | 644130 | 0 | 0 |
| AALSALESF | PRKDC | P78527 | PRKDC_HUMAN | Unmodified | 0 | 0 | 0 | 0 | 0 | 0 | 0 | 0 | 0 | 0 | 0 | 1454100 | 0 | 0 | 0 | 0 | 0 | 0 |
| AALSASEGI | AIFM1; AIFM1; | O95831; A0 | AIFM1_HUMAN | Unmodified | 0 | 0 | 0 | 0 | 0 | 0 | 0 | 0 | 0 | 0 | 0 | 419700 | 0 | 0 | 0 | 0 | 0 | 0 |
| AALSEEELE | EIF4G1; EIF4G1 | Q04637; E7E | IF4G1_HUMAN | Unmodified | 0 | 0 | 256420 | 0 | 888210 | 0 | 0 | 0 | 0 | 64891 | 620870 | 2976100 | 0 | 0 | 0 | 0 | 436220 | 0 |
| AALSEEELE | EIF4G1; EIF4G1 | Q04637; E7E | IF4G1_HUMAN | Unmodified | 0 | 0 | 0 | 0 | 675820 | 38018 | 63630 | 0 | 246200 | 356770 | 1112100 | 2302200 | 221260 | 48819 | 0 | 0 | 0 | 438510 |
| AALTAEHF | COMMD9; COM | Q9P000; E9F | COMD9_HUMA | Acetyl (Protein N | 0 | 0 | 0 | 72495 | 691840 | 0 | 0 | 0 | 144480 | 0 | 120630 | 323640 | 491660 | 296520 | 135580 | 98726 | 594770 | 576640 |
| AALTERLAL | PSPH; PSPH | P78330; C9J | SERB_HUMAN | Unmodified | 0 | 0 | 0 | 0 | 1527300 | 0 | 0 | 0 | 0 | 0 | 0 | 0 | 3764600 | 0 | 0 | 0 | 314220 | 626820 |
| AALTGLLHI | HADHB; HADHI | P55084; C9J | ECHB_HUMAN | Unmodified | 0 | 0 | 0 | 0 | 0 | 0 | 0 | 0 | 557460 | 0 | 0 | 0 | 0 | 0 | 0 | 0 | 0 | 0 |
| AALTRDPQ | GPI; GPI; GPI; G | P06744; K7E | G6PI_HUMAN | Acetyl (Protein N | 2112900 | 2881500 | 4810300 | 0 | 63432000 | 9710100 | 12376000 | 7724000 | 27001000 | 26709000 | 34311000 | 1.46E+08 | 22523000 | 11665000 | 6664700 | 5859800 | 17216000 | 12956000 |
| AALTTLFK | CNDP2; CNDP2 | Q96KP4; J3 | CNDP2_HUMA | Acetyl (Protein N | 0 | 0 | 0 | 1112200 | 2051400 | 0 | 917400 | 720660 | 2415200 | 2655600 | 9556000 | 16883000 | 9681600 | 7477400 | 0 | 3816900 | 2051300 | 1160000 |
| AALTTLFKY | CNDP2; CNDP2 | Q96KP4; J3 | CNDP2_HUMA | Acetyl (Protein N | 0 | 0 | 0 | 0 | 0 | 0 | 652240 | 0 | 0 | 96686 | 0 | 0 | 2986500 | 2159900 | 0 | 0 | 1597400 | 0 |
| AALVAQNY | LXN | Q9BS40 | LXN_HUMAN L | Unmodified | 0 | 0 | 0 | 0 | 0 | 0 | 0 | 93797 | 0 | 0 | 0 | 0 | 0 | 0 | 863890 | 0 | 4324300 | 11829000 |
| AALVGGTTI | DPYSL5; DPYSL | Q9BPU6; E9 | DPYL5_HUMAN | Oxidation (M) | 0 | 0 | 0 | 0 | 0 | 0 | 116340 | 0 | 0 | 0 | 36757 | 1277400 | 203590 | 0 | 0 | 0 | 0 | 0 |
| AALVGGTTI | DPYSL5; DPYSL | Q9BPU6; E9 | DPYL5_HUMAN | Oxidation (M) | 0 | 0 | 0 | 291160 | 0 | 0 | 4813500 | 0 | 0 | 291920 | 0 | 0 | 0 | 0 | 0 | 0 | 0 | 0 |
| AALVVPAFE | LARGE2; LARGE | Q8N3Y3; E9 | LARG2_HUMAN | Unmodified | 0 | 0 | 727310 | 0 | 0 | 0 | 700010 | 168590 | 0 | 167070 | 138830 | 990740 | 0 | 0 | 0 | 0 | 0 | 397480 |
| AAMDNSEI | IMMT; IMMT; IM | Q16891; B9 | MIC60_HUMAN | Unmodified | 0 | 0 | 0 | 0 | 0 | 0 | 63131 | 0 | 300080 | 115880 | 0 | 0 | 0 | 0 | 0 | 0 | 0 | 0 |
| AAMEALVV | DHX9 | Q08211 | DHX9_HUMAN | Oxidation (M) | 0 | 0 | 0 | 0 | 395440 | 0 | 367160 | 0 | 0 | 0 | 0 | 385280 | 0 | 0 | 0 | 0 | 116800 | 0 |
| AAMEPIVIS | TLN1 | Q9Y490 | TLN1_HUMAN | Oxidation (M) | 124370 | 148460 | 336140 | 1025800 | 2396600 | 715760 | 820670 | 701060 | 124840 | 96601 | 168730 | 1424700 | 1880500 | 446710 | 199970 | 271240 | 0 | 1009500 |
| AAMEPIVIS | TLN1 | Q9Y490 | TLN1_HUMAN | Unmodified | 57562 | 119510 | 0 | 468920 | 3406600 | 525330 | 737030 | 282720 | 0 | 0 | 0 | 0 | 1339300 | 1840300 | 190570 | 104380 | 601120 | 936580 |
| AAMFPETLI | CTNNB1; CTNN | P35222; A0A | CTNB1_HUMAN | 2 Oxidation (M) | 1779200 | 403200 | 2241700 | 5410900 | 3959400 | 3033900 | 0 | 0 | 0 | 0 | 1358700 | 2286900 | 11881000 | 4211200 | 5760000 | 5787800 | 2306400 | 2962100 |
| AAMFPETLI | CTNNB1; CTNN | P35222; A0A | CTNB1_HUMAN | Oxidation (M) | 0 | 0 | 203440 | 3111200 | 6932500 | 1849100 | 0 | 0 | 0 | 0 | 0 | 610770 | 4555800 | 0 | 667230 | 1327700 | 919590 | 2722600 |
| AAMFPETLI | CTNNB1; CTNN | P35222; A0A | CTNB1_HUMAN | 2 Oxidation (M) | 0 | 0 | 0 | 0 | 6551500 | 0 | 0 | 0 | 0 | 0 | 0 | 0 | 0 | 7055600 | 0 | 0 | 2694700 | 0 |
| AAMFPETLI | CTNNB1; CTNN | P35222; A0A | CTNB1_HUMAN | 3 Oxidation (M) | 0 | 0 | 0 | 0 | 3472500 | 2991500 | 0 | 0 | 0 | 0 | 0 | 0 | 12547000 | 10637000 | 0 | 0 | 6160800 | 12074000 |
| AAMIVNQL | JUP; JUP; JUP; J | P14923; C9J | PLAK_HUMAN | Unmodified | 0 | 0 | 0 | 370240 | 0 | 0 | 0 | 0 | 0 | 154240 | 770700 | 213170 | 598640 | 252960 | 0 | 269580 | 0 | 0 |
| AAMIVNQL | JUP; JUP; JUP; J | P14923; C9J | PLAK_HUMAN | Oxidation (M) | 0 | 0 | 0 | 0 | 0 | 0 | 0 | 0 | 0 | 0 | 0 | 442390 | 0 | 0 | 380070 | 332220 | 0 | 0 |
| AAMIVNQL | JUP; JUP; JUP; J | P14923; C9J | PLAK_HUMAN | Oxidation (M) | 0 | 0 | 0 | 2979200 | 3315100 | 783320 | 0 | 0 | 1204700 | 290480 | 489960 | 9998600 | 0 | 1379100 | 3465700 | 3767600 | 1237200 | 1774800 |
| AAMIVNQL | JUP; JUP; JUP; J | P14923; C9J | PLAK_HUMAN | Unmodified | 287780 | 246540 | 0 | 313660 | 1214700 | 0 | 0 | 0 | 2305500 | 925430 | 1259100 | 3723100 | 0 | 0 | 1193600 | 0 | 198690 | 499240 |
| AAMIVNQL | JUP; JUP; JUP; J | P14923; C9J | PLAK_HUMAN | Unmodified | 0 | 0 | 0 | 0 | 1783200 | 90541 | 0 | 0 | 0 | 0 | 0 | 0 | 886040 | 0 | 0 | 0 | 482620 | 1020100 |
| AAMKGLGT | ANXA1; ANXA1 | P04083; Q5 | ANXA1_HUMAN | Oxidation (M) | 0 | 0 | 0 | 0 | 0 | 0 | 0 | 0 | 13904 | 0 | 0 | 2897700 | 0 | 0 | 0 | 0 | 0 | 0 |
| AAMRETWL | SPTBN1; SPTBN | Q01082; A0 | SPTB2_HUMAN | Oxidation (M) | 0 | 0 | 0 | 0 | 1539600 | 0 | 0 | 0 | 0 | 0 | 0 | 0 | 0 | 0 | 0 | 0 | 0 | 0 |
| AAMTDAM | RAI14 | Q9P0K7 | RAI14_HUMAN | Unmodified | 0 | 0 | 0 | 0 | 555850 | 0 | 60185 | 0 | 0 | 0 | 0 | 208390 | 0 | 0 | 0 | 0 | 226930 | 486600 |
| AANAITKN | TMOD3; TMOD3 | Q9NYL9; H0 | TMOD3_HUMA | Unmodified | 56648 | 52715 | 0 | 0 | 216050 | 0 | 0 | 0 | 0 | 0 | 0 | 0 | 0 | 61231 | 0 | 17253 | 53449 | 65335 |
| AANATTNP | LSM5; LSM5 | Q9Y4Y9; B8 | LSM5_HUMAN | Acetyl (Protein N | 0 | 0 | 0 | 0 | 5482400 | 789180 | 1951300 | 1133300 | 2244300 | 3543400 | 4065800 | 3578300 | 6127400 | 2320500 | 0 | 0 | 3254000 | 2096000 |

| | | | | | | | | | | | | | | | | | | | | | | |
|---|---|---|---|---|---|---|---|---|---|---|---|---|---|---|---|---|---|---|---|---|---|---|
| AANDAGYF | ACAA2; ACAA2; | P42765; A0A | THIM_HUMAN | Oxidation (M) | 0 | 0 | 0 | 0 | 0 | 0 | 0 | 0 | 1021000 | 647670 | 0 | 0 | 0 | 0 | 0 | 0 | 0 | 0 |
| AANEVSSA | LSAMP; LSAMP; | Q13449; F5H | LSAMP_HUMAN | Unmodified | 0 | 700910 | 0 | 0 | 0 | 0 | 0 | 0 | 2428200 | 300870 | 0 | 0 | 0 | 0 | 0 | 0 | 224100 | 0 |
| AANGVVLA | PSMA2; N/A; PS | P25787; A0A | PSA2_HUMAN | Unmodified | 0 | 97600 | 0 | 0 | 2050400 | 0 | 553250 | 235480 | 2018300 | 1445900 | 1789300 | 6127700 | 2003500 | 669980 | 320050 | 0 | 1270800 | 1380900 |
| AANILVGEN | YES1; SRC; YES | P07947; P12 | YES_HUMAN Ty | Unmodified | 698150 | 339010 | 441060 | 848010 | 478860 | 0 | 0 | 0 | 2967300 | 352680 | 327820 | 1683200 | 0 | 0 | 1678100 | 819860 | 1027300 | 538880 |
| AANKPKGC | RALA; RALA | P11233; C9J | RALA_HUMAN | Acetyl (Protein N | 0 | 0 | 0 | 0 | 3733900 | 551540 | 4020500 | 5068800 | 0 | 804210 | 0 | 0 | 7007800 | 14892000 | 0 | 0 | 5633700 | 1185700 |
| AANKSKGC | RALB; RALB; RA | P11234; C9J | RALB_HUMAN | Acetyl (Protein N | 0 | 0 | 0 | 0 | 275930 | 0 | 0 | 0 | 0 | 0 | 0 | 0 | 0 | 0 | 0 | 0 | 475390 | 360350 |
| AANLSRNG | DBNL; DBNL; D | Q9UJU6; F8V | DBNL_HUMAN | Acetyl (Protein N | 0 | 0 | 0 | 205470 | 0 | 0 | 0 | 0 | 0 | 0 | 0 | 0 | 0 | 1886800 | 0 | 0 | 989960 | 0 |
| AANNGALF | CYC1 | P08574 | CY1_HUMAN C | Unmodified | 0 | 0 | 0 | 0 | 0 | 0 | 0 | 0 | 7011000 | 0 | 0 | 0 | 0 | 0 | 0 | 0 | 0 | 0 |
| AANNIASLI | THUMPD3; THU | Q9BV44; H7 | THUM3_HUMA | Unmodified | 316330 | 247070 | 217490 | 227510 | 0 | 0 | 785760 | 141660 | 326220 | 0 | 85794 | 387470 | 195730 | 0 | 320020 | 392050 | 0 | 0 |
| AANPFQQK | TTYH3 | Q9C0H2 | TTYH3_HUMAN | Unmodified | 999120 | 397190 | 820390 | 971480 | 0 | 0 | 38943 | 301510 | 465480 | 1728700 | 0 | 3352600 | 23182 | 0 | 733540 | 2195400 | 0 | 0 |
| AANSLEAFI | HYOU1; HYOU1 | Q9Y4L1; A0A | HYOU1_HUMA | Unmodified | 147900 | 0 | 764470 | 55895 | 193590 | 105050 | 0 | 0 | 85802000 | 38950000 | 3163700 | 4387400 | 0 | 217000 | 0 | 0 | 11317 | 17980 |
| AANVLLSEI | STK24; STK24; | Q9Y6E0; H0 | STK24_HUMAN | Unmodified | 344040 | 685470 | 243750 | 597120 | 5485700 | 2798200 | 167760 | 793460 | 4851100 | 1377100 | 960410 | 4106700 | 2508100 | 2432400 | 595660 | 913870 | 1573700 | 3938200 |
| AANVLLSE | STK25; STK26; | O00506; Q9 | STK25_HUMAN | Unmodified | 32637 | 20025 | 615280 | 797870 | 5452100 | 1360300 | 262010 | 28348 | 705500 | 444060 | 1742900 | 4312900 | 5432500 | 3756700 | 953850 | 1353800 | 1861800 | 6276500 |
| AAPAQQTT | THUMPD1; THU | Q9NXG2; H3 | THUM1_HUMA | Acetyl (Protein N | 0 | 0 | 0 | 0 | 190430 | 0 | 0 | 0 | 110590 | 0 | 0 | 303540 | 0 | 0 | 0 | 0 | 0 | 0 |
| AAPDITGHI | NPTN; NPTN; N | Q9Y639; Q9 | NPTN_HUMAN | Unmodified | 106000 | 109140 | 817660 | 500060 | 0 | 889610 | 13067 | 307640 | 1947600 | 372490 | 0 | 7903400 | 3829200 | 3924700 | 1087000 | 746890 | 691340 | 4646300 |
| AAPDITGHI | NPTN; NPTN; N | Q9Y639; Q9 | NPTN_HUMAN | Unmodified | 0 | 0 | 0 | 0 | 655360 | 89731 | 63509 | 0 | 0 | 0 | 0 | 0 | 364480 | 160800 | 0 | 0 | 147860 | 582790 |
| AAPEASGTI | CORO1A | P31146 | COR1A_HUMA | Unmodified | 0 | 0 | 0 | 0 | 0 | 0 | 0 | 0 | 0 | 0 | 214140 | 705920 | 0 | 0 | 0 | 0 | 0 | 0 |
| AAPEASGTI | CORO1A | P31146 | COR1A_HUMA | Unmodified | 0 | 0 | 0 | 0 | 0 | 0 | 0 | 0 | 0 | 0 | 0 | 1781300 | 0 | 0 | 0 | 0 | 0 | 0 |
| AAPEASGTI | CORO1A | P31146 | COR1A_HUMA | Unmodified | 0 | 0 | 0 | 0 | 0 | 328740 | 0 | 0 | 1408700 | 0 | 5938500 | 11854000 | 0 | 0 | 0 | 0 | 0 | 0 |
| AAPEASSPI | NIBAN2 | Q96TA1 | NIBA2_HUMAN | Unmodified | 3147900 | 2831400 | 2329100 | 3417000 | 54578000 | 6982800 | 955680 | 537020 | 6136400 | 4685500 | 512320 | 2767000 | 9983900 | 8467100 | 3592000 | 2791700 | 13703000 | 18748000 |
| AAPEPARA | PPP3CB; PPP3C | P16298; Q5 | PP2BB_HUMAN | Acetyl (Protein N | 0 | 0 | 0 | 0 | 0 | 0 | 0 | 0 | 0 | 0 | 0 | 0 | 0 | 0 | 0 | 0 | 1038500 | 481350 |
| AAPFSLEYF | PPA1; PPA1 | Q15181; Q5 | IPYR_HUMAN I | Unmodified | 717460 | 559750 | 0 | 816780 | 338090 | 0 | 0 | 0 | 2497900 | 87309 | 5656000 | 34320000 | 403410 | 0 | 1203900 | 1115400 | 0 | 207660 |
| AAPFSLEYF | PPA1; PPA1 | Q15181; Q5 | IPYR_HUMAN I | Unmodified | 0 | 105660 | 0 | 0 | 732290 | 92464 | 112060 | 0 | 0 | 146470 | 71573 | 0 | 1187700 | 264650 | 0 | 0 | 305720 | 527940 |
| AAPGAPAE | PKP2 | Q99959 | PKP2_HUMAN | Acetyl (Protein N | 0 | 0 | 0 | 0 | 0 | 0 | 0 | 0 | 0 | 0 | 0 | 0 | 0 | 0 | 0 | 0 | 508990 | 0 |
| AAPGCTLW | BMX | P51813 | BMX_HUMAN C | Unmodified | 0 | 0 | 0 | 0 | 0 | 0 | 0 | 0 | 0 | 0 | 1793700 | 5336700 | 0 | 0 | 0 | 0 | 0 | 0 |
| AAPGPALC | PMM2; PMM2; F | O15305; H3 | PMM2_HUMAN | Acetyl (Protein N | 0 | 0 | 0 | 0 | 0 | 0 | 0 | 0 | 73192 | 49343 | 25438 | 63680 | 0 | 0 | 0 | 0 | 0 | 0 |
| AAPGPALC | PMM2; PMM2; F | O15305; H3 | PMM2_HUMAN | Acetyl (Protein N | 0 | 0 | 0 | 0 | 1156500 | 0 | 0 | 0 | 0 | 160840 | 0 | 0 | 1876900 | 2491300 | 0 | 0 | 970630 | 806130 |
| AAPGQEPP | QSOX1 | O00391 | QSOX1_HUMA | Unmodified | 0 | 0 | 75921 | 0 | 0 | 0 | 0 | 0 | 1292500 | 0 | 0 | 0 | 0 | 0 | 32983 | 0 | 0 | 0 |
| AAPLDSIHS | FASN; FASN | P49327; A0A | FAS_HUMAN Fa | Unmodified | 0 | 11512 | 0 | 0 | 0 | 0 | 0 | 0 | 0 | 0 | 0 | 2532500 | 0 | 0 | 1031900 | 0 | 0 | 0 |
| AAPLIPLSQ | EPS15L1; EPS1 | Q9UBC2; K7 | EP15R_HUMAN | Acetyl (Protein N | 593950 | 638040 | 0 | 0 | 0 | 0 | 0 | 0 | 0 | 0 | 0 | 317430 | 0 | 1039500 | 0 | 0 | 0 | 0 |
| AAPREEPLT | INPPL1; INPPL1 | O15357; A0A | SHIP2_HUMAN | Unmodified | 0 | 0 | 0 | 0 | 87746 | 0 | 40672 | 0 | 0 | 0 | 403550 | 1915800 | 0 | 0 | 0 | 0 | 73687 | 0 |
| AAPSGGVN | MIX23 | Q4VC31 | MIX23_HUMAN | Acetyl (Protein N | 0 | 0 | 0 | 0 | 0 | 0 | 0 | 0 | 0 | 69903 | 150710 | 0 | 0 | 0 | 0 | 0 | 0 | 0 |
| AAPSPQEA | CHMP4C | Q96CF2 | CHM4C_HUMA | Unmodified | 2535000 | 1629000 | 0 | 0 | 0 | 0 | 0 | 0 | 1290100 | 203810 | 0 | 60184 | 0 | 0 | 332590 | 575880 | 0 | 0 |
| AAPTASGK | GPRIN1 | Q7Z2K8 | GRIN1_HUMAN | Unmodified | 203400 | 0 | 0 | 0 | 1558900 | 0 | 1852000 | 1981800 | 4250700 | 3969900 | 0 | 0 | 1859300 | 1367200 | 0 | 0 | 5291800 | 7801000 |
| AAPVDLELI | PFDN1; PFDN1 | O60925; D6 | PFD1_HUMAN | Acetyl (Protein N | 0 | 0 | 0 | 0 | 1490200 | 0 | 0 | 51067 | 462270 | 305720 | 0 | 1920400 | 616340 | 429860 | 0 | 0 | 1252200 | 728640 |
| AAPVDLELI | PFDN1; PFDN1 | O60925; E5F | PFD1_HUMAN | Acetyl (Protein N | 0 | 2770400 | 0 | 0 | 0 | 0 | 0 | 0 | 0 | 0 | 0 | 0 | 0 | 0 | 0 | 0 | 3674400 | 0 |
| AAQALALLI | PSMD5; PSMD5 | Q16401; F2 | PSMD5_HUMA | Acetyl (Protein N | 0 | 0 | 50727 | 0 | 0 | 0 | 0 | 0 | 232100 | 0 | 99028 | 0 | 0 | 0 | 0 | 89083 | 0 | 0 |
| AAQALSVQ | SLC19A1 | P41440 | S19A1_HUMAN | Unmodified | 0 | 0 | 1481200 | 1684600 | 4339500 | 757970 | 311480 | 62112 | 707600 | 356280 | 505690 | 0 | 1777700 | 585850 | 1445400 | 1586000 | 463510 | 977470 |
| AAQAPSSF | MTHFD1; MTHF | P11586; V9 | C1TC_HUMAN | Unmodified | 0 | 0 | 0 | 0 | 0 | 0 | 0 | 0 | 268700 | 0 | 139190 | 2302000 | 0 | 0 | 0 | 0 | 0 | 0 |
| AAQAPSSF | MTHFD1; MTHF | P11586; V9 | C1TC_HUMAN | Unmodified | 0 | 585260 | 593170 | 2352100 | 11802000 | 3735000 | 3507900 | 0 | 6812700 | 3268500 | 4032700 | 21158000 | 4518500 | 3159200 | 1613100 | 2438100 | 3779200 | 8559000 |
| AAQAPSSF | MTHFD1; MTHF | P11586; V9 | C1TC_HUMAN | Unmodified | 0 | 0 | 0 | 0 | 6310700 | 0 | 0 | 0 | 0 | 3171000 | 0 | 0 | 0 | 0 | 0 | 0 | 5502400 | 0 |
| AAQASDLE | TALDO1 | P37837 | TALDO_HUMA | Unmodified | 0 | 0 | 0 | 0 | 0 | 0 | 0 | 0 | 0 | 0 | 0 | 796800 | 0 | 0 | 0 | 0 | 0 | 0 |
| AAQASDLE | TALDO1 | P37837 | TALDO_HUMA | Unmodified | 0 | 15481 | 0 | 46474 | 5909800 | 0 | 476370 | 0 | 0 | 1344800 | 0 | 0 | 1646900 | 1884000 | 36134 | 0 | 4895500 | 964130 |
| AAQCVTK | CPNE3; CPNE3 | O75131; A0A | CPNE3_HUMA | Acetyl (Protein N | 49774 | 785390 | 0 | 23631 | 268390 | 139750 | 357350 | 37539 | 1139400 | 100020 | 0 | 0 | 139640 | 111690 | 168960 | 0 | 642350 | 920550 |
| AAQELQEG | CBS; CBSL; CB | P35520; P0 | CBS_HUMAN C | Unmodified | 0 | 0 | 0 | 0 | 0 | 0 | 0 | 0 | 150620 | 0 | 439400 | 9108900 | 0 | 0 | 0 | 0 | 0 | 0 |
| AAQGEPQ | RAN; RAN; RAN | P62826; A0A | RAN_HUMAN G | Acetyl (Protein N | 1062100 | 140880 | 1947700 | 1565000 | 7944300 | 842200 | 233930 | 2188200 | 3405900 | 2173600 | 7357900 | 30220000 | 5503700 | 2134900 | 1538900 | 1864300 | 1984800 | 3609400 |
| AAQGEPQ | RAN; RAN | P62826; B5 | RAN_HUMAN G | Acetyl (Protein N | 0 | 0 | 0 | 0 | 466400 | 0 | 1413400 | 0 | 0 | 0 | 0 | 0 | 789820 | 386540 | 0 | 0 | 1087600 | 891680 |
| AAQGVGP | MZT2B; MZT2A; | Q6NZ67; Q6 | MZT2B_HUMAN | Acetyl (Protein N | 0 | 0 | 0 | 0 | 541600 | 0 | 1240200 | 317430 | 3121500 | 441000 | 0 | 0 | 831300 | 0 | 0 | 115830 | 563710 | 359390 |
| AAQIPESD | TIMM9; TIMM9; | Q9Y5J7; G3 | TIM9_HUMAN M | Acetyl (Protein N | 0 | 0 | 197620 | 0 | 0 | 0 | 0 | 0 | 629920 | 0 | 0 | 677130 | 0 | 0 | 0 | 0 | 0 | 0 |
| AAQKAAAF | TLN1 | Q9Y490 | TLN1_HUMAN | Unmodified | 0 | 0 | 0 | 0 | 1358400 | 0 | 288690 | 890010 | 0 | 267240 | 0 | 0 | 0 | 367370 | 0 | 0 | 242600 | 448920 |
| AAQKAAAF | TLN1 | Q9Y490 | TLN1_HUMAN | Unmodified | 0 | 0 | 0 | 0 | 0 | 0 | 1209200 | 744430 | 0 | 0 | 0 | 0 | 0 | 0 | 0 | 0 | 954610 | 377620 |
| AAQKINEG | NAPG; NAPG; N | Q99747; J3 | SNAG_HUMAN | Acetyl (Protein N | 0 | 0 | 0 | 0 | 0 | 114860 | 1554800 | 964620 | 0 | 231930 | 0 | 0 | 87861 | 922870 | 0 | 0 | 3515700 | 1379000 |
| AAQKINEG | NAPG; NAPG; N | Q99747; J3 | SNAG_HUMAN | Acetyl (Protein N | 0 | 0 | 0 | 0 | 420430 | 0 | 0 | 44280 | 0 | 0 | 0 | 0 | 0 | 0 | 0 | 0 | 4119600 | 1166400 |
| AAQLDGLE | LAMB2 | P55268 | LAMB2_HUMA | Unmodified | 0 | 0 | 0 | 0 | 0 | 0 | 0 | 0 | 1181500 | 0 | 0 | 0 | 0 | 0 | 0 | 0 | 0 | 0 |
| AAQLDGLE | LAMB2 | P55268 | LAMB2_HUMA | Unmodified | 0 | 0 | 0 | 0 | 0 | 0 | 0 | 7409.8 | 0 | 151230 | 0 | 0 | 82980 | 179140 | 0 | 0 | 0 | 203330 |
| AAQLQGLG | ALDH3B1; ALD | P43353; A0A | AL3B1_HUMAN | Unmodified | 0 | 0 | 0 | 0 | 809030 | 0 | 0 | 0 | 0 | 0 | 0 | 0 | 0 | 0 | 0 | 0 | 0 | 0 |
| AAQLQGLG | ALDH3B1; ALD | P43353; A0A | AL3B1_HUMAN | Unmodified | 0 | 0 | 0 | 0 | 3555600 | 1132900 | 0 | 0 | 7319000 | 0 | 0 | 0 | 0 | 0 | 0 | 0 | 0 | 0 |
| AAQLRPISL | EFCAB14; EFCA | O75071; A0 | EFC14_HUMAN | Unmodified | 0 | 0 | 0 | 0 | 0 | 0 | 0 | 0 | 0 | 1087300 | 0 | 0 | 0 | 0 | 0 | 0 | 0 | 0 |
| AAQLRPISL | EFCAB14; EFCA | O75071; A0 | EFC14_HUMAN | Unmodified | 0 | 0 | 0 | 0 | 0 | 0 | 0 | 0 | 32180000 | 4267300 | 0 | 0 | 0 | 0 | 0 | 0 | 0 | 0 |
| AAQLVDKD | NCLN; NCLN; N | Q969V3; A0 | NCLN_HUMAN | Unmodified | 0 | 0 | 0 | 0 | 0 | 0 | 0 | 0 | 53783000 | 3574200 | 0 | 0 | 0 | 0 | 0 | 0 | 0 | 0 |
| AAQNSGEA | LAMB1; LAMB1; | P07942; G3 | LAMB1_HUMA | Unmodified | 0 | 0 | 0 | 0 | 0 | 0 | 0 | 0 | 0 | 0 | 0 | 0 | 0 | 0 | 0 | 0 | 0 | 0 |
| AAQQAASS | CSNK1A1; CSN | P48729; U3 | KC1A_HUMAN | Unmodified | 0 | 0 | 0 | 0 | 0 | 0 | 0 | 0 | 0 | 0 | 0 | 0 | 0 | 0 | 0 | 0 | 0 | 0 |
| AAQRGMPS | SPAG16; SPAG | Q8N0X2; E7 | SPG16_HUMA | Oxidation (M) | 0 | 0 | 0 | 0 | 0 | 0 | 0 | 0 | 218600 | 0 | 0 | 0 | 1681300 | 0 | 0 | 0 | 868250 | |
| AAQSANIS | HMGCL | P35914 | HMGCL_HUMA | Unmodified | 0 | 0 | 0 | 46227 | 0 | 0 | 0 | 23179 | 5086800 | 573840 | 0 | 278590 | 0 | 0 | 0 | 0 | 92532 | 41901 |
| AAQSGPSE | ADAM15 | Q13444 | ADA15_HUMAN | Unmodified | 863450 | 1260500 | 872530 | 0 | 0 | 0 | 0 | 0 | 0 | 142170 | 242500 | 1358500 | 0 | 0 | 542170 | 674220 | 88313 | 0 |
| AAQSTAMN | NSF; NSF | P46459; I3L | NSF_HUMAN V | Unmodified | 0 | 0 | 0 | 0 | 385770 | 0 | 47948 | 0 | 0 | 139950 | 0 | 0 | 914920 | 907500 | 0 | 0 | 157600 | 903690 |
| AAQSTGAV | TRPM5; TRPM5; | Q9NZQ8; A0 | TRPM5_HUMA | Unmodified | 0 | 0 | 0 | 0 | 0 | 0 | 0 | 0 | 0 | 0 | 0 | 299190 | 0 | 0 | 0 | 0 | 0 | 0 |
| AAQSTGAV | TRPM4 | Q8TD43 | TRPM4_HUMA | Unmodified | 89256 | 0 | 0 | 0 | 0 | 0 | 0 | 0 | 0 | 0 | 0 | 0 | 0 | 0 | 1441000 | 918370 | 0 | 0 |
| AAQTAERLI | TRIP10; TRIP10; | Q15642; W4 | CIP4_HUMAN ( | Unmodified | 494040 | 0 | 0 | 0 | 7388400 | 0 | 0 | 0 | 0 | 0 | 0 | 0 | 719710 | 0 | 0 | 225930 | 649530 | 1537700 |
| AAQVAQDE | CCDC50 | Q8IVM0 | CCD50_HUMA | Unmodified | 14309000 | 23156000 | 16937000 | 13210000 | 0 | 0 | 0 | 19412 | 1548900 | 0 | 0 | 832510 | 0 | 0 | 30098000 | 20659000 | 0 | 0 |
| AAQVAQDE | CCDC50 | Q8IVM0 | CCD50_HUMA | Oxidation (M) | 0 | 0 | 0 | 0 | 0 | 0 | 0 | 0 | 0 | 0 | 0 | 0 | 1320700 | 36562 | 0 | 0 | 0 | 0 |
| AAQVAQDE | CCDC50 | Q8IVM0 | CCD50_HUMA | Oxidation (M) | 0 | 0 | 0 | 79629 | 0 | 0 | 0 | 0 | 0 | 0 | 0 | 0 | 653260 | 342270 | 0 | 0 | 0 | 0 |
| AAQVFYNN | ABCA2; ABCA2; | Q9BZC7; A0 | ABCA2_HUMA | Unmodified | 0 | 0 | 0 | 0 | 0 | 0 | 0 | 0 | 682140 | 84424 | 0 | 0 | 0 | 0 | 0 | 0 | 0 | 0 |
| AAQYACSL | PEX11B | O96011 | PX11B_HUMAN | Unmodified | 0 | 0 | 0 | 0 | 0 | 0 | 0 | 26546000 | 3068500 | 370630 | 0 | 0 | 0 | 188720 | 0 | 0 | 0 | 0 |
| AARDIFNK | VDAC2; VDAC2 | P45880; Q5 | VDAC2_HUMA | Unmodified | 0 | 0 | 0 | 0 | 0 | 0 | 155190 | 0 | 0 | 749510 | 0 | 0 | 0 | 0 | 0 | 0 | 0 | 0 |
| AARDNEVR | ARVCF; ARVCF; | O00192; E9 | ARVC_HUMAN | Unmodified | 0 | 594220 | 0 | 0 | 0 | 0 | 0 | 0 | 0 | 0 | 0 | 0 | 0 | 603980 | 0 | 0 | 0 | 60789 |
| AAREALLSS | CCT2; CCT2; C | P78371; F5 | TCPB_HUMAN | Unmodified | 0 | 0 | 0 | 0 | 0 | 0 | 894570 | 585000 | 0 | 1466100 | 0 | 0 | 7614200 | 2617000 | 0 | 0 | 188470 | 291730 |
| AAREECPV | TIGAR; TIGAR; | Q9NQ88; A0 | TIGAR_HUMAN | Unmodified | 41929 | 0 | 0 | 71648 | 411870 | 0 | 0 | 0 | 0 | 0 | 173700 | 1923900 | 680920 | 0 | 113080 | 0 | 343490 | 0 |
| AARESLTEI | RIPOR2; RIPOR | Q9Y4F9; B7 | RIPR2_HUMAN | Unmodified | 0 | 0 | 0 | 0 | 0 | 0 | 0 | 0 | 0 | 0 | 0 | 0 | 0 | 500190 | 0 | 0 | 0 | 0 |
| AAREVKLLL | GNAI2; GNAI1; | P04899; P63 | GNAI2_HUMAN | Unmodified | 0 | 0 | 0 | 0 | 514350 | 20801 | 1554500 | 176930 | 0 | 61121 | 0 | 0 | 155910 | 339880 | 0 | 0 | 2142700 | 2256800 |
| AARVIHVVT | CTNNA1; CTNN | P35221; G3 | CTNA1_HUMA | Unmodified | 0 | 0 | 0 | 299530 | 29386000 | 989130 | 0 | 0 | 0 | 460630 | 0 | 0 | 9204200 | 11159000 | 0 | 0 | 0 | 0 |
| AARVIHVVT | CTNNA1; CTNN | P35221; G3 | CTNA1_HUMA | Oxidation (M) | 0 | 0 | 0 | 0 | 0 | 0 | 0 | 0 | 0 | 0 | 0 | 0 | 0 | 2080200 | 0 | 0 | 0 | 0 |
| AASAAAAS | TRIM28 | Q13263 | TIF1B_HUMAN | Acetyl (Protein N | 398520 | 0 | 236900 | 302220 | 1452500 | 0 | 1317900 | 2424800 | 2621600 | 2026600 | 1423700 | 7629400 | 2341400 | 0 | 0 | 0 | 2361700 | 291590 |
| AASAAFQE | TBCD; TBCD; T | Q9BTW9; I3 | TBCD_HUMAN | Unmodified | 0 | 0 | 0 | 0 | 0 | 0 | 0 | 0 | 696620 | 43542 | 0 | 1955600 | 0 | 0 | 0 | 0 | 272800 | 0 |
| AASAAQRE | TLN1 | Q9Y490 | TLN1_HUMAN | Unmodified | 22438 | 0 | 50777 | 672870 | 8328300 | 2091800 | 4331100 | 3292400 | 85167 | 785900 | 222560 | 149970 | 2881900 | 2346400 | 22885 | 18537 | 1059300 | 2365500 |
| AASAATAA | AGPS; AGPS; A | O00116; A0 | ADAS_HUMAN | Unmodified | 0 | 0 | 0 | 0 | 0 | 0 | 0 | 0 | 2636800 | 486000 | 0 | 0 | 0 | 0 | 0 | 0 | 20671 | 0 |
| AASADSTT | MACROH2A1 | O75367 | H2AY_HUMAN | Unmodified | 10985000 | 7968500 | 294120 | 0 | 0 | 0 | 2942600 | 9339800 | 0 | 278300 | 0 | 845960 | 0 | 609840 | 0 | 0 | 28422 | 506890 |
| AASAFFTY | ACOT7 | O00154 | BACH_HUMAN | Unmodified | 12947 | 0 | 0 | 0 | 0 | 0 | 0 | 0 | 925740 | 896110 | 327380 | 1389400 | 0 | 0 | 0 | 0 | 915330 | 1720000 |
| AASAGVPY | SPTBN2; SPTBN | O15020; A4 | SPTN2_HUMAN | Unmodified | 0 | 0 | 0 | 0 | 0 | 0 | 0 | 0 | 0 | 0 | 0 | 1671300 | 683370 | 0 | 0 | 0 | 0 | 0 |
| AASAPYFE | TUBGCP2; TUB | Q9BSJ2; F2 | GCP2_HUMAN | Unmodified | 0 | 0 | 0 | 0 | 0 | 0 | 0 | 0 | 0 | 0 | 0 | 1374200 | 0 | 0 | 0 | 0 | 0 | 0 |
| AASATREQ | B4GALNT1; B4 | Q00973; F8 | B4GN1_HUMA | Unmodified | 527200 | 1011000 | 791360 | 902820 | 0 | 0 | 0 | 0 | 2228400 | 0 | 0 | 0 | 0 | 0 | 735170 | 1002400 | 0 | 0 |
| AASAYAVG | PRKAR2A; PRKA | P13861; H7 | KAP2_HUMAN | Unmodified | 0 | 0 | 569560 | 839460 | 1977300 | 1219500 | 169750 | 166840 | 1128000 | 382700 | 254320 | 1384400 | 5279700 | 5274500 | 905500 | 819780 | 873970 | 1834300 |
| AASCVLLH | AKR1A1; AKR1A | P14550; V9 | AK1A1_HUMAN | Acetyl (Protein N | 0 | 0 | 0 | 1207600 | 1971200 | 0 | 0 | 0 | 0 | 1551200 | 482180 | 3881400 | 2135400 | 1668300 | 1691500 | 1102300 | 1397800 | 276570 |
| AASCVLLH | AKR1A1; AKR1A | P14550; V9 | AK1A1_HUMAN | Acetyl (Protein N | 0 | 0 | 0 | 0 | 0 | 0 | 351750 | 0 | 0 | 0 | 0 | 0 | 343840 | 910280 | 0 | 0 | 2115800 | 271790 |
| AASDELSK | VCL | P18206 | VINC_HUMAN | Oxidation (M) | 0 | 0 | 0 | 0 | 0 | 0 | 0 | 0 | 0 | 0 | 0 | 496810 | 0 | 0 | 0 | 0 | 738880 | 88256 |
| AASDIAMT | YWHAE; YWHA | P62258; K7 | 1433E_HUMAN | Unmodified | 0 | 259330 | 698240 | 1353300 | 0 | 0 | 0 | 0 | 1094800 | 0 | 234780 | 10029000 | 46759 | 0 | 2178900 | 1726700 | 0 | 0 |
| AASDIAMT | YWHAE; YWHA | P62258; K7 | 1433E_HUMAN | Oxidation (M) | 681660 | 911820 | 2363000 | 935030 | 0 | 0 | 74846 | 0 | 182980 | 0 | 0 | 21071000 | 0 | 0 | 6279900 | 4724100 | 0 | 0 |
| AASEFESS | CLSTN1; CLSTN | O94985; Q5 | CSTN1_HUMA | Unmodified | 0 | 0 | 0 | 0 | 0 | 0 | 0 | 0 | 9770900 | 1566300 | 0 | 0 | 0 | 0 | 0 | 0 | 0 | 0 |
| AASELYTK | MYO5A; MYO5A | Q9Y4I1; G3 | MYO5A_HUMA | Acetyl (Protein N | 0 | 0 | 82775 | 205440 | 418620 | 0 | 0 | 0 | 0 | 0 | 0 | 64159 | 0 | 0 | 0 | 0 | 277010 | 0 |
| AASGALRN | CTNND1; CTNN | O60716; A0 | CTND1_HUMA | Unmodified | 0 | 0 | 92445 | 287690 | 9781400 | 2632100 | 0 | 0 | 0 | 0 | 0 | 0 | 8522800 | 12839000 | 0 | 0 | 4950500 | 16490000 |
| AASGALRN | CTNND1; CTNN | O60716; A0 | CTND1_HUMA | Unmodified | 0 | 0 | 0 | 0 | 6319800 | 1119900 | 31627 | 0 | 0 | 0 | 0 | 0 | 5326400 | 12522000 | 0 | 0 | 7636900 | 7423200 |
| AASGFNA | ANXA4; ANXA4; | P09525; B4 | ANXA4_HUMA | Unmodified | 0 | 0 | 0 | 0 | 259080 | 0 | 0 | 0 | 2002800 | 132500 | 365380 | 0 | 0 | 0 | 0 | 0 | 0 | 0 |
| AASGFNA | ANXA4; ANXA4; | P09525; B4 | ANXA4_HUMA | Oxidation (M) | 183050 | 351010 | 575400 | 170270 | 10650000 | 3214000 | 243210 | 2073300 | 11686000 | 8235600 | 14390000 | 6936100 | 2994800 | 2507700 | 635420 | 392660 | 2271300 | 3760300 |
| AASGFNA | ANXA4; ANXA4; | P09525; B4 | ANXA4_HUMA | Unmodified | 1821300 | 703330 | 1103500 | 1152500 | 81875000 | 7553600 | 7595100 | 3479000 | 1.49E+08 | 67332000 | 58527000 | 30947000 | 18389000 | 16746000 | 962960 | 787840 | 11126000 | 28059000 |
| AASGFNA | ANXA4; ANXA4; | P09525; B4 | ANXA4_HUMA | Unmodified | 0 | 0 | 0 | 119020 | 1115500 | 0 | 0 | 0 | 0 | 0 | 0 | 0 | 41333 | 0 | 150790 | 80413 | 378490 | 194820 |
| AASGPGPE | HSPG2 | P98160 | PGBM_HUMAN | Unmodified | 0 | 0 | 670080 | 0 | 0 | 0 | 0 | 0 | 0 | 0 | 0 | 0 | 0 | 0 | 0 | 0 | 0 | 0 |
| AASHRALD | REXO2; REXO2; | Q9Y3B8; F5 | ORN_HUMAN ( | Unmodified | 0 | 0 | 0 | 0 | 319200 | 0 | 0 | 0 | 0 | 181990 | 0 | 0 | 171650 | 92915 | 0 | 0 | 0 | 185420 |
| AASIFGGA | EIF4B; EIF4B; E | P23588; E7 | IF4B_HUMAN E | Unmodified | 542780 | 0 | 241960 | 391250 | 0 | 0 | 0 | 0 | 298790 | 177550 | 0 | 1517500 | 0 | 0 | 0 | 479290 | 0 | 0 |
| AASITSEVF | SUPT16H | Q9Y5B9 | SP16H_HUMAN | Unmodified | 0 | 0 | 0 | 0 | 572730 | 0 | 0 | 89204 | 0 | 446610 | 0 | 1229000 | 0 | 0 | 0 | 0 | 0 | 0 |

| | | | | | | | | | | | | | | | | | | | | | |
|---|---|---|---|---|---|---|---|---|---|---|---|---|---|---|---|---|---|---|---|---|---|
| AASLDGFY | SEZ6; SEZ6 | Q53EL9; H0Y | SEZ6_HUMAN S | Unmodified | 0 | 0 | 0 | 0 | 0 | 0 | 37107 | 0 | 2852700 | 239260 | 0 | 0 | 0 | 0 | 0 | 0 | 0 | 0 |
| AASLGSSQ | CAST; CAST; CA | A0A6Q8PG4 | A0A6Q8PG40_I | Unmodified | 0 | 0 | 0 | 0 | 237210 | 0 | 0 | 26505 | 0 | 794690 | 0 | 0 | 0 | 97188 | 0 | 0 | 246090 | 343560 |
| AASLGSSQ | CAST; CAST; CA | H0Y944; A0A | H0Y944_HUMA | Unmodified | 0 | 0 | 0 | 0 | 0 | 0 | 0 | 0 | 0 | 0 | 0 | 0 | 0 | 0 | 0 | 0 | 291670 | 0 |
| AASLVGKK | ITPA | Q9BY32 | ITPA_HUMAN I | Acetyl (Protein N | 0 | 0 | 0 | 0 | 803320 | 0 | 96275 | 427000 | 68448 | 217900 | 488860 | 2209800 | 462500 | 589730 | 0 | 0 | 974260 | 928760 |
| AASLVGKKI | ITPA | Q9BY32 | ITPA_HUMAN I | Acetyl (Protein N | 0 | 0 | 0 | 0 | 0 | 0 | 375940 | 200910 | 0 | 0 | 0 | 0 | 0 | 0 | 0 | 0 | 910200 | 0 |
| AASNIIFSN | DPP7; DPP7 | Q9UHL4; R4 | DPP2_HUMAN | Unmodified | 123740 | 72512 | 153430 | 0 | 0 | 0 | 0 | 0 | 526270 | 23807 | 0 | 168390 | 0 | 0 | 409320 | 119470 | 0 | 0 |
| AASNIIFSN | DPP7; DPP7 | Q9UHL4; R4 | DPP2_HUMAN | Unmodified | 0 | 0 | 0 | 0 | 0 | 0 | 0 | 0 | 2355400 | 0 | 0 | 0 | 0 | 0 | 0 | 0 | 0 | 0 |
| AASQGYTV | SEL1L | Q9UBV2 | SE1L1_HUMAN | Unmodified | 0 | 0 | 163140 | 0 | 0 | 0 | 357840 | 0 | 3816900 | 420570 | 0 | 0 | 0 | 0 | 150580 | 550100 | 0 | 0 |
| AASQILILK | ABCC4; ABCC4 | O15439; A0A | MRP4_HUMAN | Unmodified | 0 | 0 | 1433900 | 0 | 99450 | 0 | 145040 | 70192 | 0 | 0 | 0 | 0 | 40455 | 0 | 1500300 | 1733700 | 0 | 0 |
| AASQPGEL | CPE; CPE; CPE; | P16870; C9J | CBPE_HUMAN | Unmodified | 0 | 0 | 0 | 55651 | 0 | 0 | 0 | 0 | 20991000 | 2807800 | 0 | 371390 | 0 | 0 | 0 | 70802 | 0 | 236650 |
| AASQPGEL | CPE; CPE; CPE; | P16870; C9J | CBPE_HUMAN | Unmodified | 0 | 0 | 0 | 0 | 0 | 0 | 62015 | 5701100 | 0 | 3036600 | 0 | 0 | 0 | 0 | 0 | 0 | 0 | 0 |
| AASSAAQG | SCAMP2 | O15127 | SCAM2_HUMA | Unmodified | 0 | 0 | 0 | 1257800 | 545060 | 167960 | 0 | 0 | 0 | 0 | 0 | 211310 | 500290 | 0 | 613710 | 0 | 0 | 145230 |
| AASSIYINS | CAPN5; CAPN5 | O15484; E7E | CAN5_HUMAN | Unmodified | 0 | 310240 | 0 | 219070 | 0 | 111950 | 92800 | 50284 | 1129500 | 233450 | 0 | 731250 | 0 | 0 | 381620 | 381890 | 261260 | 587420 |
| AASSLLANI | PKP3; PKP3 | Q9Y446; H0Y | PKP3_HUMAN | Unmodified | 486130 | 378900 | 1051900 | 1388900 | 1326600 | 140690 | 0 | 0 | 1433200 | 398180 | 775300 | 6494300 | 2451700 | 329420 | 4579800 | 4221800 | 606880 | 341670 |
| AASSLNLSI | MAP4K4; MAP4 | O95819; C9J | M4K4_HUMAN | Unmodified | 0 | 0 | 0 | 356730 | 0 | 0 | 0 | 0 | 181800 | 0 | 0 | 1307600 | 0 | 0 | 523930 | 375450 | 0 | 0 |
| AASSQRAE | CHMP4C | Q96CF2 | CHM4C_HUMA | Unmodified | 1072800 | 955950 | 338820 | 0 | 1098300 | 0 | 0 | 0 | 443730 | 0 | 0 | 0 | 0 | 0 | 0 | 0 | 0 | 0 |
| AASSQRAE | CHMP4C | Q96CF2 | CHM4C_HUMA | Unmodified | 0 | 0 | 0 | 0 | 0 | 0 | 0 | 0 | 0 | 1634300 | 0 | 0 | 5055600 | 459130 | 0 | 0 | 0 | 0 |
| AASTDMAG | TPPP3 | Q9BW30 | TPPP3_HUMAN | Acetyl (Protein N | 0 | 0 | 0 | 0 | 0 | 0 | 507350 | 310460 | 0 | 0 | 0 | 0 | 0 | 0 | 0 | 0 | 0 | 0 |
| AASTDMAG | TPPP3 | Q9BW30 | TPPP3_HUMAN | Acetyl (Protein N | 0 | 0 | 0 | 0 | 0 | 0 | 0 | 134860 | 0 | 0 | 0 | 0 | 0 | 0 | 0 | 0 | 0 | 0 |
| AASTFNIRF | SEZ6L; SEZ6L; S | Q9BYH1; B0 | SE6L1_HUMAN | Unmodified | 0 | 0 | 0 | 0 | 0 | 0 | 0 | 0 | 4338500 | 640300 | 0 | 0 | 0 | 0 | 0 | 0 | 0 | 0 |
| AASWNQAI | SYT13; SYT13 | Q7L8C5; H0Y | SYT13_HUMAN | Unmodified | 0 | 0 | 0 | 0 | 0 | 0 | 0 | 0 | 714730 | 83911 | 0 | 0 | 0 | 0 | 0 | 0 | 0 | 0 |
| AASYTEEDI | DSG2 | Q14126 | DSG2_HUMAN | Unmodified | 0 | 0 | 0 | 0 | 38776000 | 1557900 | 0 | 0 | 562380 | 0 | 0 | 0 | 607460 | 0 | 627050 | 0 | 0 | 0 |
| AATAAADF | STRAP | Q9Y3F4 | STRAP_HUMAN | Unmodified | 295260 | 921960 | 875490 | 501740 | 4719100 | 704200 | 1686800 | 1524600 | 1377000 | 881330 | 1343600 | 14138000 | 6199800 | 1877100 | 730020 | 982620 | 2048800 | 1771500 |
| AATALKDV | PDIA6 | Q15084 | PDIA6_HUMAN | Unmodified | 0 | 0 | 0 | 0 | 248520 | 0 | 246170 | 0 | 1242300 | 4576000 | 85648 | 0 | 147330 | 0 | 0 | 0 | 61361 | 0 |
| AATALKDV | PDIA6 | Q15084 | PDIA6_HUMAN | Unmodified | 0 | 0 | 0 | 0 | 115940 | 0 | 105050 | 94721 | 0 | 1636000 | 0 | 520920 | 0 | 0 | 0 | 0 | 0 | 103680 |
| AATASAGA | SLC25A11; SLC | Q02978; I3L | M2OM_HUMAN | Acetyl (Protein N | 0 | 0 | 0 | 0 | 0 | 0 | 0 | 0 | 1002900 | 0 | 0 | 0 | 0 | 0 | 0 | 0 | 0 | 0 |
| AATASAGA | SLC25A11; SLC | Q02978; I3L | M2OM_HUMAN | Acetyl (Protein N | 0 | 0 | 0 | 0 | 0 | 0 | 875450 | 185520 | 0 | 679880 | 0 | 0 | 0 | 0 | 0 | 0 | 0 | 0 |
| AATDIARQ | SEPTIN6; SEPTI | Q14141; B1A | SEPT6_HUMAN | Acetyl (Protein N | 0 | 0 | 0 | 0 | 3662100 | 0 | 1771700 | 0 | 0 | 0 | 0 | 0 | 0 | 0 | 0 | 0 | 15943000 | 3853900 |
| AATDLERF | SEPTIN8; SEPTI | Q92599; A6N | SEPT8_HUMAN | Acetyl (Protein N | 0 | 0 | 0 | 0 | 2469900 | 189190 | 161560 | 0 | 0 | 157790 | 0 | 0 | 852680 | 0 | 0 | 0 | 549350 | 0 |
| AATFFGCIC | ADK; ADK; ADK; | P55263; A0A | ADK_HUMAN A | Unmodified | 0 | 0 | 92506 | 0 | 520910 | 0 | 134010 | 0 | 293630 | 0 | 0 | 2264900 | 0 | 0 | 0 | 263380 | 0 | 0 |
| AATFFGCIC | ADK; ADK; ADK; | P55263; A0A | ADK_HUMAN A | Unmodified | 0 | 0 | 0 | 92559 | 7452300 | 873110 | 1285300 | 0 | 0 | 0 | 0 | 0 | 1706400 | 2267700 | 0 | 0 | 5071900 | 5463300 |
| AATGANAE | UPP1; UPP1; UI | Q16831; C9I | UPP1_HUMAN | Acetyl (Protein N | 0 | 0 | 0 | 0 | 1151900 | 299460 | 0 | 0 | 0 | 254310 | 0 | 0 | 0 | 0 | 0 | 0 | 849850 | 490700 |
| AATIVATSE | PRKAR2A; PRKA | P13861; H7C | KAP2_HUMAN | Unmodified | 0 | 0 | 0 | 366340 | 0 | 0 | 0 | 0 | 0 | 598580 | 0 | 265930 | 4726200 | 992620 | 291890 | 0 | 0 | 0 |
| AATLAERSA | SERPINH1; SER | P50454; E9P | SERPH_HUMAI | Unmodified | 0 | 0 | 0 | 0 | 0 | 0 | 0 | 0 | 19408 | 752280 | 0 | 0 | 0 | 0 | 0 | 0 | 0 | 0 |
| AATLAQELE | SMC1A; SMC1A | Q14683; G8J | SMC1A_HUMA | Unmodified | 0 | 0 | 0 | 0 | 0 | 306120 | 0 | 0 | 0 | 0 | 0 | 0 | 0 | 0 | 0 | 0 | 0 | 0 |
| AATLILEPA | ACOT2; ACOT1; | P49753; Q86 | ACOT2_HUMAI | Unmodified | 0 | 0 | 0 | 0 | 0 | 699550 | 0 | 0 | 894580 | 0 | 0 | 0 | 0 | 0 | 0 | 0 | 0 | 0 |
| AATLILEPA | ACOT2; ACOT1; | P49753; Q86 | ACOT2_HUMAI | Acetyl (Protein N | 0 | 0 | 0 | 0 | 0 | 0 | 0 | 0 | 642370 | 0 | 0 | 0 | 0 | 0 | 0 | 0 | 0 | 0 |
| AATLMMLG | DNAJC7; DNAJC | Q99615; A0A | DNJC7_HUMAI | Oxidation (M) | 0 | 0 | 0 | 136230 | 0 | 0 | 0 | 0 | 0 | 0 | 0 | 589230 | 0 | 0 | 0 | 0 | 0 | 0 |
| AATQFWR | PDIA4; PDIA4 | P13667; A0A | PDIA4_HUMAN | Unmodified | 0 | 62253 | 0 | 0 | 0 | 0 | 0 | 0 | 2878700 | 513400 | 937850 | 1649500 | 0 | 0 | 0 | 0 | 0 | 0 |
| AATQLRGH | PDIA5 | Q14554 | PDIA5_HUMAN | Unmodified | 0 | 0 | 0 | 0 | 1.59E+08 | 8250100 | 4338500 | 0 | 0 | 178480 | 0 | 0 | 33773000 | 28470000 | 0 | 0 | 64840000 | 65944000 |
| AATQLRGH | PDIA5 | Q14554 | PDIA5_HUMAN | Unmodified | 0 | 0 | 0 | 0 | 861620 | 0 | 0 | 0 | 1530900 | 0 | 0 | 0 | 0 | 0 | 0 | 0 | 3632700 | 2627000 |
| AATSDLEH | NUCB2; NUCB2 | P80303; Q2L | NUCB2_HUMA | Unmodified | 117150 | 71075 | 973690 | 955370 | 0 | 0 | 0 | 0 | 31142000 | 4702600 | 939340 | 288520 | 794400 | 0 | 1113100 | 445190 | 0 | 0 |
| AATSDLEH | NUCB2; NUCB2 | P80303; Q2L | NUCB2_HUMA | Unmodified | 0 | 0 | 0 | 0 | 0 | 0 | 0 | 0 | 167040 | 1685000 | 0 | 0 | 190160 | 0 | 0 | 0 | 0 | 0 |
| AATSGVPSI | LSR; LSR | Q86X29; S4F | LSR_HUMAN Li | Unmodified | 10263000 | 6498300 | 18829000 | 16929000 | 3465400 | 0 | 0 | 0 | 52198000 | 13317000 | 2946600 | 33469000 | 4124300 | 1965400 | 17003000 | 17962000 | 236930 | 0 |
| AATSPALFN | DYNC1H1; DYN | Q14204; A0A | DYHC1_HUMA | Unmodified | 0 | 0 | 0 | 0 | 0 | 0 | 0 | 0 | 1862700 | 0 | 337790 | 3678600 | 0 | 0 | 0 | 0 | 0 | 0 |
| AATSRYTLN | VCAN; VCAN; V | P13611; Q86 | CSPG2_HUMAI | Unmodified | 249580 | 192030 | 0 | 501420 | 817850 | 0 | 186420 | 0 | 3693900 | 825660 | 437150 | 3437300 | 0 | 0 | 0 | 226670 | 222520 | 332680 |
| AATTGSGV | UBE2V1 | Q13404 | UB2V1_HUMAI | Acetyl (Protein N | 0 | 0 | 0 | 1303300 | 3323700 | 1072700 | 484220 | 1391600 | 906480 | 10512000 | 2176600 | 2221400 | 3856600 | 1738200 | 676510 | 1046700 | 4493200 | 5021800 |
| AATVGYGIL | ASRGL1; ASRGI | Q7L266; E9F | ASGL1_HUMAN | Unmodified | 0 | 0 | 0 | 0 | 0 | 0 | 0 | 0 | 1727800 | 0 | 0 | 1253100 | 0 | 0 | 0 | 0 | 0 | 0 |
| AATVTGKL | TFRC; TFRC | P02786; G3V | TFR1_HUMAN | Unmodified | 0 | 0 | 0 | 0 | 413780 | 0 | 1075800 | 147470 | 0 | 205800 | 0 | 0 | 0 | 387270 | 0 | 0 | 59742 | 0 |
| AATVTGKL | TFRC; TFRC | P02786; G3V | TFR1_HUMAN | Unmodified | 0 | 0 | 0 | 0 | 80462 | 0 | 298460 | 106460 | 0 | 1367700 | 0 | 0 | 0 | 0 | 0 | 0 | 207160 | 656540 |
| AAVAGKKP | RPL4; RPL4 | P36578; H3E | RL4_HUMAN 6 | Unmodified | 0 | 89837 | 29774 | 0 | 56922 | 0 | 0 | 0 | 0 | 0 | 0 | 278010 | 101100 | 0 | 16906 | 23671 | 0 | 0 |
| AAVATFLQ | UBA1; UBA1 | P22314; Q5J | UBA1_HUMAN | Unmodified | 1040100 | 542040 | 788960 | 911340 | 0 | 0 | 324660 | 72479 | 10609000 | 1312900 | 1776000 | 17370000 | 701400 | 295190 | 2112100 | 1223700 | 445190 | 613620 |
| AAVAWEAG | ADH5; ADH5; A | P11766; D6F | ADHX_HUMAN | Unmodified | 0 | 0 | 0 | 0 | 529500 | 0 | 0 | 0 | 0 | 561080 | 4637500 | 13929000 | 0 | 0 | 0 | 319550 | 0 | 0 |
| AAVAWEAG | ADH5; ADH5; A | P11766; D6F | ADHX_HUMAN | Unmodified | 0 | 0 | 0 | 0 | 1260600 | 436470 | 0 | 0 | 0 | 0 | 0 | 0 | 0 | 0 | 0 | 0 | 1072200 | 1240900 |
| AAVDAGFV | ETFA; ETFA; ETF | P13804; H0Y | ETFA_HUMAN | Unmodified | 0 | 0 | 0 | 0 | 0 | 0 | 0 | 7691.2 | 1161800 | 160990 | 0 | 0 | 0 | 0 | 151670 | 38655 | 0 | 0 |
| AAVDAMTR | DECR2; DECR2 | Q9NUI1; Q4 | DECR2_HUMAI | Unmodified | 0 | 74848 | 0 | 0 | 0 | 0 | 0 | 0 | 3357100 | 252780 | 134910 | 0 | 0 | 0 | 0 | 0 | 0 | 0 |
| AAVDLEKL | PFKL; PFKL | P17858; F8V | PFKAL_HUMAN | Acetyl (Protein N | 1232300 | 1964400 | 430830 | 4591300 | 850900 | 0 | 1860300 | 1737800 | 0 | 241100 | 0 | 0 | 1671500 | 669350 | 1118500 | 1430300 | 4612300 | 2010400 |
| AAVEALQS | SEPTIN8; SEPTI | Q92599; A6N | SEPT8_HUMAN | Unmodified | 0 | 0 | 140430 | 899100 | 2900800 | 0 | 135250 | 0 | 440860 | 264260 | 185170 | 1798700 | 576240 | 468360 | 309290 | 251660 | 447230 | 1507700 |
| AAVEDPRV | DLST | P36957 | ODO2_HUMAN | Unmodified | 0 | 0 | 0 | 0 | 0 | 0 | 0 | 0 | 0 | 183320 | 0 | 0 | 0 | 0 | 0 | 0 | 0 | 0 |
| AAVEEGIVL | HSPD1; HSPD1 | P10809; A0A | CH60_HUMAN | Unmodified | 22157 | 0 | 75440 | 0 | 0 | 0 | 0 | 172710 | 17589000 | 3267500 | 341800 | 318170 | 0 | 0 | 123990 | 0 | 0 | 0 |
| AAVENHVR | RPIA | P49247 | RPIA_HUMAN | Unmodified | 0 | 0 | 0 | 0 | 375740 | 0 | 0 | 0 | 0 | 277040 | 820260 | 0 | 454000 | 0 | 0 | 0 | 0 | 0 |
| AAVENLPTI | KPNB1; KPNB1 | Q14974; J3C | IMB1_HUMAN | Unmodified | 1412500 | 949600 | 260840 | 1239400 | 729470 | 144530 | 0 | 0 | 2393000 | 904200 | 1928100 | 5153200 | 1274000 | 189410 | 1859200 | 988220 | 339350 | 204040 |
| AAVESLGFI | PSMD1; PSMD1 | Q99460; A0A | PSMD1_HUMA | Unmodified | 0 | 0 | 0 | 0 | 0 | 0 | 0 | 0 | 0 | 867560 | 0 | 500210 | 0 | 0 | 0 | 0 | 0 | 0 |
| AAVEWFDC | EWSR1; EWSR1 | Q01844; H7I | EWS_HUMAN | Unmodified | 91359 | 0 | 0 | 0 | 144140 | 0 | 0 | 0 | 271460 | 0 | 1039600 | 6548600 | 0 | 0 | 0 | 39725 | 0 | 0 |
| AAVGRPLD | PDCD6IP; PDCI | Q8WUM4; F | PDC6I_HUMAN | Unmodified | 3023600 | 4087700 | 5553900 | 806370 | 259540 | 0 | 0 | 0 | 563410 | 0 | 0 | 1640800 | 0 | 0 | 3635300 | 5742200 | 52434 | 78878 |
| AAVGRPLD | PDCD6IP; PDCI | Q8WUM4; F | PDC6I_HUMAN | Unmodified | 57043000 | 84293000 | 7856300 | 17968000 | 3281800 | 775880 | 0 | 0 | 14356000 | 2922000 | 1525600 | 13061000 | 12022000 | 365050 | 3046400 | 25313000 | 336900 | 1093900 |
| AAVHLEGK | VCL | P18206 | VINC_HUMAN | Unmodified | 165160 | 0 | 612550 | 737150 | 230180 | 0 | 0 | 0 | 0 | 1743500 | 0 | 0 | 2416000 | 0 | 0 | 0 | 102990 | 0 |
| AAVIGDVIR | DDX21 | Q9NR30 | DDX21_HUMAI | Unmodified | 4690400 | 0 | 86043 | 0 | 0 | 0 | 34247 | 111000 | 0 | 0 | 231770 | 3322600 | 0 | 0 | 0 | 0 | 0 | 0 |
| AAVKGYLG | MDH2 | P40926 | MDHM_HUMAI | Unmodified | 0 | 0 | 0 | 0 | 0 | 0 | 204950 | 67144 | 0 | 0 | 0 | 0 | 0 | 0 | 0 | 0 | 0 | 0 |
| AAVLKPVLL | PDCD2L | Q9BRP1 | PDD2L_HUMAI | Acetyl (Protein N | 0 | 0 | 0 | 368250 | 5583800 | 605280 | 0 | 0 | 252490 | 688870 | 0 | 0 | 7543400 | 3335800 | 0 | 0 | 2514000 | 3132500 |
| AAVLQQVL | TPR; TPR | P12270; Q5S | TPR_HUMAN N | Acetyl (Protein N | 0 | 0 | 0 | 0 | 1145800 | 0 | 352050 | 0 | 172210 | 682500 | 135040 | 0 | 812950 | 0 | 0 | 0 | 868600 | 0 |
| AAVLQRVEI | LACTB2 | Q53H82 | LACB2_HUMAN | Acetyl (Protein N | 218700 | 201370 | 0 | 159810 | 2049200 | 0 | 0 | 0 | 20231000 | 5175800 | 0 | 746090 | 429440 | 0 | 0 | 0 | 1033200 | 1955900 |
| AAVMVHQI | CTNNB1; CTNN | P35222; A0A | CTNB1_HUMAI | Unmodified | 0 | 0 | 0 | 0 | 366190 | 0 | 0 | 0 | 0 | 0 | 0 | 686750 | 0 | 0 | 381530 | 253950 | 366580 | 644400 |
| AAVMVHQI | CTNNB1; CTNN | P35222; A0A | CTNB1_HUMAI | Unmodified | 1210800 | 206830 | 1322700 | 2820400 | 11401000 | 3341600 | 0 | 0 | 644630 | 132340 | 1250400 | 5065500 | 4020200 | 4438200 | 4045500 | 2917400 | 5981400 | 8068800 |
| AAVMVHQI | CTNNB1; CTNN | P35222; A0A | CTNB1_HUMAI | Oxidation (M) | 0 | 0 | 0 | 0 | 0 | 0 | 0 | 0 | 0 | 0 | 297560 | 425240 | 0 | 0 | 1056400 | 511390 | 1013900 | 1855400 |
| AAVMVHQI | CTNNB1; CTNN | P35222; A0A | CTNB1_HUMAI | Unmodified | 0 | 0 | 0 | 0 | 4339200 | 525300 | 0 | 0 | 0 | 41682 | 0 | 0 | 600830 | 1585600 | 0 | 0 | 1715700 | 3391300 |
| AAVMVYDE | ENAH; ENAH; E | Q8N8S7; A0 | ENAH_HUMAN | Unmodified | 0 | 0 | 0 | 206510 | 193530 | 156570 | 3077700 | 803110 | 360120 | 184950 | 233460 | 1837600 | 0 | 0 | 198700 | 119380 | 0 | 0 |
| AAVMVYDE | ENAH; ENAH; E | Q8N8S7; A0 | ENAH_HUMAN | Oxidation (M) | 0 | 0 | 0 | 0 | 183640 | 1130100 | 171510 | 165420 | 82103 | 48682 | 178590 | 1108400 | 327320 | 0 | 0 | 0 | 65557 | 0 |
| AAVMVYDE | ENAH; ENAH; E | Q8N8S7; A0 | ENAH_HUMAN | Unmodified | 0 | 0 | 0 | 0 | 0 | 0 | 0 | 0 | 0 | 0 | 0 | 2187700 | 0 | 0 | 0 | 0 | 0 | 0 |
| AAVMVYDE | ENAH; ENAH; E | Q8N8S7; A0 | ENAH_HUMAN | Oxidation (M) | 0 | 0 | 0 | 0 | 0 | 0 | 0 | 0 | 41926 | 0 | 117500 | 942670 | 0 | 0 | 0 | 0 | 0 | 0 |
| AAVPDAVG | ATP1A1; ATP1A | P05023; P50 | AT1A1_HUMAN | Unmodified | 0 | 0 | 16217000 | 7727700 | 853050 | 192940 | 204040 | 83635 | 4918300 | 966670 | 1705000 | 62431000 | 0 | 0 | 16519000 | 22713000 | 0 | 0 |
| AAVPDTVVI | VIL1 | P09327 | VILI_HUMAN Vi | Unmodified | 639090 | 0 | 0 | 0 | 0 | 0 | 0 | 0 | 4335700 | 2656900 | 3643700 | 12392000 | 0 | 0 | 0 | 0 | 320710 | 0 |
| AAVPSGAS | ENO1; ENO3; E | P06733; P13 | ENOA_HUMAN | Unmodified | 1821900 | 2163500 | 1182700 | 2206600 | 791290 | 2226600 | 1806700 | 1795200 | 7124500 | 4093700 | 7739300 | 16265000 | 2182500 | 1322300 | 2975900 | 2463400 | 3177900 | 2315000 |
| AAVPSGAS | ENO3; ENO2; E | P13929; P09 | ENOB_HUMAN | Unmodified | 1208100 | 0 | 0 | 0 | 2063000 | 0 | 0 | 0 | 6888000 | 1158200 | 286190 | 4924400 | 949630 | 0 | 0 | 0 | 1139500 | 2864700 |
| AAVPSGAS | ENO2; ENO2; E | P09104; F5H | ENOG_HUMAN | Unmodified | 0 | 0 | 0 | 0 | 0 | 0 | 0 | 0 | 0 | 0 | 0 | 1319900 | 0 | 0 | 0 | 0 | 1447900 | 1849100 |
| AAVPSGAS | ENO1; ENO1; E | P06733; K7E | ENOA_HUMAN | Unmodified | 15144000 | 29340000 | 18371000 | 40859000 | 2.73E+08 | 46119000 | 3963300 | 846520 | 1.63E+08 | 47271000 | 37569000 | 3.64E+08 | 1.44E+08 | 37766000 | 48413000 | 47709000 | 26420000 | 1.27E+08 |
| AAVPSGAS | ENO1; ENO1; E | P06733; K7E | ENOA_HUMAN | Unmodified | 0 | 0 | 0 | 207840 | 35982000 | 8245400 | 705150 | 0 | 6413100 | 12219000 | 0 | 1196100 | 20677000 | 4471400 | 0 | 0 | 11671000 | 15603000 |
| AAVPSIKFC | PGK1 | P00558 | PGK1_HUMAN | Unmodified | 0 | 0 | 0 | 0 | 5397100 | 0 | 8436800 | 6187500 | 0 | 3329100 | 0 | 0 | 8515500 | 11574000 | 0 | 0 | 1.2E+08 | 18623000 |
| AAVQELSS | PSEN1; PSEN1 | P49768; E7E | PSN1_HUMAN | Unmodified | 76796 | 0 | 484430 | 27222 | 361100 | 0 | 0 | 0 | 1680900 | 232360 | 0 | 3401900 | 0 | 0 | 1042200 | 1021300 | 0 | 0 |
| AAVQVLDD | MMUT | P22033 | MUTA_HUMAN | Unmodified | 0 | 0 | 0 | 0 | 0 | 0 | 0 | 0 | 0 | 247470 | 0 | 0 | 0 | 0 | 0 | 0 | 0 | 0 |
| AAVSHWQ | CTNNB1; CTNN | P35222; C9J | CTNB1_HUMAI | Unmodified | 1510400 | 888780 | 7121700 | 4185500 | 3675000 | 0 | 0 | 0 | 0 | 0 | 0 | 2208500 | 931440 | 0 | 8570500 | 11058000 | 0 | 4258500 |
| AAVVESHK | GSR; GSR; GSR | P00390; H0Y | GSHR_HUMAN | Unmodified | 0 | 0 | 0 | 0 | 766400 | 0 | 0 | 0 | 0 | 0 | 0 | 0 | 777520 | 634890 | 0 | 0 | 270420 | 0 |
| AAVVHHFE | TOR1AIP2 | Q8NFQ8 | TOIP2_HUMAN | Unmodified | 0 | 0 | 0 | 0 | 576740 | 0 | 775400 | 0 | 18498000 | 4973900 | 507380 | 834920 | 0 | 71669 | 0 | 0 | 0 | 237040 |
| AAVVPLGT | EBP; EBP; N/A; | Q15125; C9J | EBP_HUMAN 3 | Unmodified | 0 | 0 | 0 | 0 | 0 | 0 | 0 | 0 | 780800 | 150640 | 0 | 336150 | 0 | 0 | 0 | 0 | 0 | 0 |
| AAVYQPQP | FN1 | P02751 | FINC_HUMAN | Oxidation (M) | 0 | 0 | 4368500 | 3629900 | 0 | 0 | 0 | 0 | 0 | 0 | 0 | 0 | 1348600 | 1877900 | 0 | 1602400 | 0 | 0 |
| AAWAQGY | FURIN; FURIN; | P09958; A0A | FURIN_HUMAN | Unmodified | 0 | 0 | 0 | 0 | 0 | 896400 | 0 | 0 | 1327300 | 0 | 1214300 | 0 | 0 | 0 | 0 | 0 | 0 | 0 |
| AAWEAGKF | ACAT1; ACAT1; | P24752; A0A | THIL_HUMAN A | Unmodified | 0 | 0 | 0 | 0 | 0 | 0 | 0 | 0 | 411350 | 1185700 | 0 | 189230 | 991370 | 873690 | 0 | 0 | 0 | 452610 |
| AAWRVLSS | SFN | P31947 | 1433S_HUMAN | Unmodified | 0 | 0 | 0 | 26733000 | 3727500 | 0 | 0 | 0 | 4975100 | 0 | 0 | 0 | 0 | 0 | 0 | 3645500 | 7593500 |
| AAWRVLSS | SFN | P31947 | 1433S_HUMAN | Unmodified | 0 | 0 | 0 | 0 | 953110 | 347440 | 0 | 190630 | 0 | 133220 | 0 | 0 | 0 | 482010 | 0 | 0 | 333250 | 306820 |
| AAYDIEVN | FLOT1; FLOT1; | O75955; A2A | FLOT1_HUMAN | Unmodified | 0 | 0 | 1311600 | 1083300 | 3182400 | 1338700 | 320330 | 257580 | 2905700 | 825370 | 670650 | 2540200 | 947750 | 649370 | 1585400 | 864280 | 0 | 0 |
| AAYDIEVN | FLOT1; FLOT1; | O75955; A2A | FLOT1_HUMAN | Unmodified | 0 | 0 | 524520 | 626540 | 1968800 | 207050 | 133000 | 70510 | 1367800 | 822000 | 293340 | 2465900 | 0 | 0 | 128770 | 0 | 0 | 0 |
| AAYDIEVN | FLOT1; FLOT1; | O75955; A0A | FLOT1_HUMAN | Unmodified | 0 | 0 | 0 | 0 | 11761000 | 0 | 1714100 | 0 | 0 | 0 | 0 | 0 | 1245100 | 2134100 | 0 | 0 | 1861200 | 3919900 |
| AAYDREES | RALBP1 | Q15311 | RBP1_HUMAN | Unmodified | 0 | 0 | 276480 | 0 | 0 | 0 | 0 | 42436 | 643550 | 0 | 387670 | 753620 | 1007100 | 1023900 | 773320 | 946290 | 0 | 744680 |
| AAYEAELGI | LMNA; LMNA; L | P02545; A0A | LMNA_HUMAN | Unmodified | 0 | 0 | 0 | 0 | 8373100 | 637820 | 64290 | 0 | 156010 | 0 | 0 | 0 | 222740 | 110730 | 0 | 0 | 3102100 | 1615800 |
| AAYEAELGI | LMNA; LMNA; L | P02545; A0A | LMNA_HUMAN | Unmodified | 0 | 0 | 0 | 0 | 0 | 0 | 0 | 0 | 0 | 118620 | 0 | 0 | 0 | 0 | 0 | 0 | 363450 | 0 |
| AAYFGIYDT | SLC25A5 | P05141 | ADT2_HUMAN | Unmodified | 0 | 71349 | 0 | 0 | 194380 | 0 | 0 | 0 | 3184900 | 0 | 0 | 0 | 0 | 0 | 0 | 0 | 0 | 0 |
| AAYFGVYD | SLC25A4; SLC2 | P12235; P12 | ADT1_HUMAN | Unmodified | 0 | 0 | 36277 | 0 | 0 | 0 | 0 | 0 | 1063000 | 329680 | 0 | 0 | 0 | 0 | 0 | 0 | 0 | 0 |
| AAYIYFTDF | STXBP3 | O00186 | STXB3_HUMAN | Unmodified | 184480 | 88062 | 105170 | 543430 | 3769400 | 114420 | 700110 | 29742 | 0 | 410780 | 0 | 232300 | 4101200 | 6490000 | 413300 | 91180 | 2605700 | 5233800 |
| AAYKLVLIR | PGAM1; PGAM4 | P18669; Q8N | PGAM1_HUMA | Acetyl (Protein N | 0 | 63690 | 0 | 0 | 1497500 | 0 | 0 | 53518 | 0 | 0 | 0 | 0 | 0 | 0 | 0 | 25059 | 920200 | 964070 |

| | | | | | | | | | | | | | | | | | | | | | | |
|---|---|---|---|---|---|---|---|---|---|---|---|---|---|---|---|---|---|---|---|---|---|---|
| AAYLQETGI | ANXA1; ANXA1; | P04083; Q5T | ANXA1_HUMAN | Unmodified | 52303 | 4055700 | 26902 | 0 | 707390 | 0 | 371810 | 193300 | 0 | 472030 | 105360 | 4629800 | 0 | 0 | 69864 | 54598 | 630510 | 371720 |
| AAYLQETGI | ANXA1; ANXA1; | P04083; Q5T | ANXA1_HUMAN | Unmodified | 0 | 155510 | 0 | 0 | 1003400 | 394110 | 0 | 0 | 0 | 0 | 6806700 | 78675000 | 0 | 0 | 0 | 0 | 0 | 0 |
| AAYLQETGI | ANXA1; ANXA1; | P04083; Q5T | ANXA1_HUMAN | Unmodified | 1761100 | 0 | 0 | 0 | 41756000 | 10469000 | 1489400 | 0 | 652990 | 916930 | 4.41E+08 | 3.09E+09 | 0 | 0 | 0 | 0 | 5949800 | 7709500 |
| AAYPDLEN | RARS1; RARS1 | P54136; E5R | SYRC_HUMAN | Unmodified | 0 | 0 | 584280 | 0 | 4425800 | 804710 | 1353700 | 782120 | 1547300 | 962160 | 2468000 | 16775000 | 3833500 | 2862000 | 0 | 1024200 | 0 | 2052900 |
| AAYRAYQN | ECE1; ECE1 | P42892; A0A | ECE1_HUMAN | Unmodified | 0 | 0 | 0 | 0 | 6174200 | 1260800 | 0 | 0 | 0 | 3159200 | 0 | 0 | 0 | 0 | 0 | 0 | 0 | 714130 |
| AAYSTGGC | LRP11 | Q86VZ4 | LRP11_HUMAN | Unmodified | 0 | 0 | 0 | 42801 | 41445 | 0 | 0 | 0 | 619520 | 18304 | 0 | 0 | 0 | 0 | 0 | 0 | 0 | 0 |
| AAYSYRPG | PDCD6; PDCD6 | O75340; Q8 | PDCD6_HUMA | Acetyl (Protein N | 1657400 | 2263700 | 0 | 1444900 | 0 | 0 | 0 | 0 | 8023900 | 1182400 | 0 | 0 | 0 | 0 | 4352300 | 0 | 0 | 504880 |
| AAYSYRPG | PDCD6; PDCD6 | O75340; Q8 | PDCD6_HUMA | Acetyl (Protein N | 0 | 1281700 | 635140 | 1294400 | 2764000 | 0 | 543360 | 0 | 5274000 | 4774100 | 0 | 0 | 3063400 | 2055000 | 1787200 | 0 | 1120500 | 2292000 |
| AAYVLFYQI | USP11; USP4; U | P51784; Q13 | UBP11_HUMAN | Unmodified | 0 | 0 | 0 | 0 | 0 | 0 | 0 | 0 | 0 | 0 | 0 | 2487500 | 0 | 0 | 0 | 0 | 0 | 0 |
| AAYYPPSQ | PABPC1; PABP( | P11940; A0A | PABP1_HUMAN | Unmodified | 4502100 | 3822300 | 3139900 | 4408700 | 0 | 0 | 0 | 0 | 0 | 0 | 0 | 0 | 12853000 | 3790900 | 3983300 | 3567300 | 0 | 0 |
| ACAALGDV | SMYD4; SMYD4 | Q8IYR2; I3L4 | SMYD4_HUMA | Unmodified | 0 | 0 | 0 | 0 | 272530 | 0 | 0 | 107130 | 0 | 0 | 0 | 2007200 | 0 | 0 | 0 | 0 | 0 | 93252 |
| ACADATLS | GNB1; GNB1; G | P62873; F6U | GBB1_HUMAN | Unmodified | 4783400 | 2754600 | 2626500 | 6461100 | 4745600 | 2048000 | 1076300 | 755410 | 35632000 | 4913200 | 13381000 | 77016000 | 4554300 | 4047300 | 5152200 | 4646500 | 3654300 | 8830800 |
| ACADATLS | GNB1; GNB1; G | P62873; F6U | GBB1_HUMAN | Oxidation (M) | 0 | 0 | 593490 | 0 | 14895000 | 7919100 | 2992700 | 461220 | 0 | 17747000 | 1916600 | 0 | 12386000 | 20886000 | 0 | 0 | 30727000 | 59104000 |
| ACADATLS | GNB1; GNB1; G | P62873; F6U | GBB1_HUMAN | Unmodified | 0 | 0 | 0 | 0 | 5779400 | 398960 | 1855500 | 67623 | 696590 | 4767900 | 0 | 0 | 4966500 | 7503100 | 0 | 0 | 6632900 | 19147000 |
| ACALSQIN | CRYZL1; CRYZL | O95825; A6 | QORL1_HUMA | Unmodified | 0 | 0 | 0 | 0 | 525470 | 0 | 70266 | 0 | 207660 | 0 | 324410 | 272500 | 125440 | 0 | 0 | 0 | 350790 | 468080 |
| ACANPAAG | PGK1 | P00558 | PGK1_HUMAN | Unmodified | 0 | 0 | 0 | 0 | 5250400 | 0 | 0 | 0 | 0 | 1427900 | 0 | 0 | 10215000 | 0 | 0 | 0 | 11138000 | 22962000 |
| ACANPAAG | PGK1 | P00558 | PGK1_HUMAN | Unmodified | 0 | 0 | 0 | 0 | 18392000 | 1654400 | 6302000 | 0 | 0 | 8426400 | 0 | 0 | 20390000 | 6770000 | 0 | 0 | 59100000 | 1.07E+08 |
| ACARPLISV | RPL4; RPL4; RP | P36578; H3E | RL4_HUMAN 6 | Acetyl (Protein N | 6697500 | 1155300 | 1837200 | 1383200 | 1445200 | 0 | 1188100 | 0 | 0 | 403650 | 864410 | 12917000 | 1743500 | 1350100 | 1764600 | 2168100 | 366980 | 1092000 |
| ACDGVVH | C2orf72 | A6NCS6 | CB072_HUMAN | Unmodified | 192640 | 97283 | 93610 | 216120 | 500050 | 98220 | 0 | 0 | 0 | 68423 | 0 | 0 | 0 | 47104 | 467420 | 356980 | 35241 | 268990 |
| ACDSVTSN | MMS19; MMS19 | Q96T76; H7 | MMS19_HUMA | Unmodified | 0 | 0 | 0 | 0 | 0 | 0 | 0 | 0 | 0 | 0 | 0 | 1774600 | 0 | 0 | 0 | 0 | 0 | 0 |
| ACEFAGFQ | TLN1 | Q9Y490 | TLN1_HUMAN | Unmodified | 919320 | 0 | 0 | 205700 | 4657600 | 279740 | 4469000 | 1886800 | 450910 | 383900 | 85713 | 1381100 | 5481500 | 5744800 | 0 | 0 | 148530 | 3114200 |
| ACEFAGFQ | TLN1 | Q9Y490 | TLN1_HUMAN | Unmodified | 0 | 0 | 266490 | 0 | 0 | 0 | 983550 | 259650 | 0 | 0 | 0 | 0 | 0 | 399040 | 0 | 0 | 876120 | 0 |
| ACFHCETC | LASP1; LASP1; | Q14847; K7E | LASP1_HUMAN | Unmodified | 0 | 109630 | 77330 | 656400 | 0 | 1478200 | 60052 | 57145 | 0 | 0 | 0 | 0 | 0 | 0 | 58288 | 0 | 2864500 | 107280 |
| ACGADSYE | EGFR; EGFR; E | P00533; Q5 | EGFR_HUMAN | Unmodified | 493490 | 0 | 0 | 1320400 | 2309900 | 458660 | 0 | 0 | 248570 | 27834 | 0 | 0 | 571400 | 0 | 667210 | 682530 | 225770 | 479870 |
| ACGADSYE | EGFR; EGFR; E | P00533; Q5 | EGFR_HUMAN | Oxidation (M) | 0 | 0 | 755810 | 362580 | 826180 | 133900 | 0 | 0 | 0 | 0 | 0 | 0 | 0 | 0 | 177990 | 585580 | 0 | 162030 |
| ACGDSTLT | GNB2; GNB2; G | P62879; C9J | GBB2_HUMAN | Unmodified | 246430 | 4750400 | 5152500 | 7757600 | 2527800 | 1771800 | 269090 | 408000 | 8020100 | 766030 | 10232000 | 96973000 | 2260000 | 3170000 | 6271100 | 5483600 | 1677200 | 4497900 |
| ACGDSTLT | GNB2; GNB2; G | P62879; C9J | GBB2_HUMAN | Unmodified | 0 | 0 | 0 | 0 | 998040 | 366330 | 0 | 0 | 0 | 0 | 0 | 0 | 0 | 0 | 0 | 0 | 3362800 | 4041600 |
| ACGDSTLT | GNB2; GNB2; G | P62879; C9J | GBB2_HUMAN | Oxidation (M) | 0 | 0 | 0 | 0 | 4664000 | 0 | 1780900 | 498040 | 0 | 1312100 | 0 | 0 | 0 | 3416400 | 0 | 0 | 5784000 | 5008100 |
| ACGLNFAD | VAT1; VAT1 | Q99536; K7E | VAT1_HUMAN | Unmodified | 0 | 0 | 0 | 251580 | 564570 | 0 | 0 | 0 | 1512300 | 0 | 0 | 456650 | 0 | 0 | 0 | 0 | 0 | 0 |
| ACGLVASN | LGALS1; LGALS | P09382; F8V | LEG1_HUMAN | Acetyl (Protein N | 1689400 | 0 | 0 | 0 | 19011000 | 5228200 | 0 | 0 | 0 | 0 | 0 | 0 | 0 | 0 | 0 | 0 | 31011000 | 33028000 |
| ACGNFGIP | PAICS; PAICS; F | P22234; D6F | PUR6_HUMAN | Unmodified | 0 | 0 | 0 | 378780 | 0 | 369960 | 0 | 0 | 0 | 0 | 914350 | 10411000 | 0 | 0 | 0 | 0 | 165080 | 0 |
| ACGQIFCG | HGS; HGS; HGS | O14964; A0A | HGS_HUMAN H | Unmodified | 432830 | 253690 | 0 | 142290 | 0 | 226590 | 167870 | 0 | 478680 | 309970 | 0 | 260820 | 519370 | 0 | 86266 | 72330 | 309380 | 760260 |
| ACGVSRPV | SNU13; SNU13 | P55769; A0A | NH2L1_HUMAN | Unmodified | 122600 | 0 | 0 | 0 | 0 | 0 | 0 | 88771 | 0 | 231250 | 0 | 825660 | 444680 | 227630 | 0 | 0 | 82180 | 235330 |
| ACIDSNED | TPP2; TPP2; TP | P29144; A0A | TPP2_HUMAN | Unmodified | 0 | 0 | 0 | 0 | 0 | 0 | 0 | 0 | 0 | 0 | 0 | 3153900 | 0 | 0 | 0 | 0 | 0 | 0 |
| ACISPKPQI | LYN | P07948 | LYN_HUMAN T | Unmodified | 181990 | 170960 | 210500 | 336790 | 1117200 | 154660 | 512390 | 237360 | 647430 | 188770 | 0 | 0 | 0 | 283160 | 281930 | 373840 | 0 | 683990 |
| ACKVDSPT | KLC1; KLC1; KL | Q07866; G3 | KLC1_HUMAN | Unmodified | 29524 | 0 | 0 | 0 | 1144000 | 185770 | 553670 | 680350 | 343610 | 1055400 | 0 | 312630 | 589960 | 517520 | 22284 | 21168 | 1104500 | 1085900 |
| ACLDTAVEI | FASN; FASN | P49327; A0A | FAS_HUMAN F | Unmodified | 79530 | 0 | 0 | 923100 | 709560 | 0 | 412540 | 53473 | 202880 | 199850 | 839320 | 6806800 | 725350 | 380270 | 0 | 161840 | 591200 | 545090 |
| ACLDTAVEI | FASN; FASN | P49327; A0A | FAS_HUMAN F | Oxidation (M) | 0 | 0 | 0 | 206850 | 0 | 0 | 0 | 0 | 0 | 0 | 189570 | 4945600 | 436710 | 0 | 333060 | 261960 | 0 | 0 |
| ACLDYPVT | PTTG1IP; PTTG | P53801; A8 | PTTG_HUMAN | Unmodified | 0 | 0 | 0 | 0 | 0 | 0 | 0 | 0 | 5478900 | 1134000 | 0 | 752580 | 0 | 0 | 0 | 0 | 0 | 0 |
| ACLENLGL | SERPINB5 | P36952 | SPB5_HUMAN | Unmodified | 0 | 82025 | 177080 | 0 | 331640 | 0 | 0 | 0 | 0 | 0 | 0 | 0 | 0 | 0 | 113860 | 0 | 1770300 | 2458000 |
| ACLERIQEI | LMNB1; LMNB1 | P20700; E9P | LMNB1_HUMA | Unmodified | 0 | 0 | 128240 | 0 | 939060 | 233490 | 0 | 0 | 152090 | 1000100 | 830920 | 0 | 1846800 | 0 | 0 | 0 | 0 | 297070 |
| ACLGCMG | CRELD1; CRELD | Q96HD1; A0 | CREL1_HUMAN | Unmodified | 0 | 0 | 0 | 0 | 0 | 0 | 0 | 0 | 489630 | 146130 | 0 | 0 | 0 | 0 | 0 | 0 | 0 | 0 |
| ACLILRSIEI | SRP72; SRP72 | O76094; D6I | SRP72_HUMAN | Unmodified | 0 | 0 | 0 | 0 | 141660 | 0 | 2436500 | 0 | 0 | 0 | 0 | 0 | 371770 | 0 | 0 | 0 | 237940 | 385860 |
| ACLISLGYD | ACTN4; ACTN4 | O43707; F5 | ACTN4_HUMA | Unmodified | 0 | 130000 | 0 | 0 | 576980 | 359340 | 0 | 0 | 486350 | 0 | 0 | 9497800 | 0 | 0 | 0 | 0 | 0 | 0 |
| ACLYAGVK | GLUL; GLUL | P15104; A0A | GLNA_HUMAN | Unmodified | 0 | 0 | 0 | 0 | 757700 | 0 | 1174800 | 39989 | 250960 | 0 | 0 | 3637900 | 0 | 0 | 0 | 0 | 0 | 0 |
| ACNLMIFD | MANSC1 | Q9H8J5 | MANS1_HUMA | Unmodified | 0 | 0 | 0 | 0 | 0 | 0 | 0 | 0 | 1086800 | 343000 | 0 | 0 | 0 | 0 | 0 | 0 | 0 | 0 |
| ACNPLDAG | PTPRF; PTPRF; | P10586; H0 | PTPRF_HUMAN | Oxidation (M) | 0 | 0 | 0 | 0 | 0 | 0 | 0 | 0 | 0 | 0 | 0 | 0 | 0 | 0 | 1101600 | 357170 | 0 | 0 |
| ACNQLGQI | AP2A2; AP2A1; | O94973; O9 | AP2A2_HUMAN | Unmodified | 480160 | 347600 | 0 | 0 | 0 | 0 | 0 | 0 | 0 | 0 | 0 | 4173800 | 0 | 0 | 503020 | 0 | 0 | 0 |
| ACPDSLGS | ANO1; ANO1 | Q5XXA6; E9 | ANO1_HUMAN | Unmodified | 0 | 0 | 0 | 352680 | 0 | 0 | 50057 | 0 | 0 | 0 | 0 | 0 | 0 | 0 | 0 | 0 | 1847700 | 499750 |
| ACPPGTAA | IGF2R; IGF2R; I | P11717; A0A | MPRI_HUMAN | Unmodified | 0 | 0 | 0 | 0 | 0 | 0 | 0 | 0 | 6137500 | 0 | 0 | 0 | 0 | 0 | 0 | 78393 | 0 | 0 |
| ACQGQYE | TBC1D10A; N/A | Q9BXI6; H0 | TB10A_HUMAN | Unmodified | 193350 | 0 | 61616 | 0 | 624660 | 44963 | 0 | 0 | 64258 | 0 | 0 | 269150 | 0 | 0 | 114760 | 117040 | 0 | 67943 |
| ACQIFVRN | HNRNPM; HNR | P52272; M0 | HNRPM_HUMA | Unmodified | 0 | 0 | 0 | 10267000 | 1079400 | 0 | 5233000 | 0 | 0 | 283340 | 0 | 0 | 1451100 | 3508400 | 0 | 0 | 1244300 | 1427300 |
| ACQLTYPLI | TMEM59; TMEM | Q9BXS4; Q5 | TMM59_HUMA | Unmodified | 0 | 0 | 0 | 0 | 0 | 0 | 0 | 0 | 4398700 | 0 | 0 | 0 | 0 | 0 | 0 | 0 | 0 | 0 |
| ACQSIYPLH | RPS3A; RPS3A; | P61247; D6F | RS3A_HUMAN | Unmodified | 1264100 | 0 | 0 | 0 | 0 | 589940 | 866160 | 0 | 0 | 172600 | 0 | 5726000 | 0 | 0 | 0 | 0 | 0 | 0 |
| ACQSIYPLH | RPS3A; RPS3A; | P61247; D6F | RS3A_HUMAN | Unmodified | 8812700 | 3874400 | 4898900 | 4487200 | 6059100 | 895080 | 6072400 | 2796200 | 1538700 | 1361000 | 1988000 | 47774000 | 7773000 | 3384900 | 3936800 | 4451000 | 831640 | 4409500 |
| ACRDVDEC | GAS6 | Q14393 | GAS6_HUMAN | Unmodified | 0 | 0 | 0 | 0 | 0 | 0 | 0 | 0 | 772830 | 26887 | 22091 | 0 | 0 | 0 | 0 | 0 | 0 | 0 |
| ACSDELTV | SORL1; SORL1; | Q92673; E9 | SORL_HUMAN | Unmodified | 0 | 0 | 0 | 0 | 0 | 0 | 0 | 0 | 1156600 | 0 | 0 | 0 | 0 | 0 | 0 | 75244 | 0 | 0 |
| ACSFIREAA | NIT2; NIT2; NIT | Q9NQR4; F8 | NIT2_HUMAN ( | Unmodified | 0 | 0 | 0 | 0 | 269020 | 0 | 54274 | 0 | 0 | 274540 | 115380 | 0 | 725240 | 391540 | 0 | 0 | 425790 | 299920 |
| ACSLAASE | KIAA1522 | Q9P206 | K1522_HUMAN | Unmodified | 0 | 0 | 583360 | 0 | 2624300 | 0 | 0 | 0 | 0 | 0 | 0 | 497120 | 3624000 | 4961500 | 3706500 | 0 | 4210900 | 2969800 |
| ACTGDSW | SOGA1; SOGA1 | O94964; X6 | SOGA1_HUMA | Unmodified | 0 | 0 | 0 | 0 | 0 | 0 | 0 | 0 | 0 | 0 | 0 | 0 | 398900 | 147110 | 0 | 0 | 106090 | 0 |
| ACTILLR | CCT3; CCT3 | P49368; B4D | TCPG_HUMAN | Unmodified | 124800 | 0 | 0 | 0 | 0 | 0 | 0 | 0 | 0 | 0 | 4356100 | 22129000 | 0 | 0 | 0 | 0 | 0 | 0 |
| ACVMMQG | PNP; PNP; PNP | P00491; G3 | PNPH_HUMAN | Unmodified | 0 | 0 | 31175 | 0 | 0 | 0 | 0 | 18076 | 92439 | 0 | 0 | 3527000 | 0 | 0 | 0 | 0 | 0 | 0 |
| ACVQVLDP | CHID1; CHID1; | Q9BWS9; H | CHID1_HUMAN | Unmodified | 0 | 0 | 0 | 0 | 0 | 0 | 0 | 0 | 2028300 | 364480 | 0 | 0 | 0 | 0 | 0 | 0 | 0 | 0 |
| ACVVHGSI | ATP1A1; ATP1A | P05023; P50 | AT1A1_HUMAN | Unmodified | 316810 | 115920 | 2467800 | 6268000 | 0 | 0 | 372110 | 0 | 1205700 | 382860 | 1364900 | 10257000 | 5228600 | 1559900 | 2265000 | 2422800 | 436970 | 481650 |
| ACVVHGSI | ATP1A1 | P05023 | AT1A1_HUMAN | Unmodified | 0 | 0 | 0 | 0 | 9641900 | 0 | 0 | 0 | 0 | 1394300 | 0 | 0 | 5248900 | 4368800 | 0 | 0 | 0 | 1446800 |
| ACVVHGSI | ATP1A1 | P05023 | AT1A1_HUMAN | Oxidation (M) | 0 | 0 | 0 | 2102400 | 5805000 | 689460 | 0 | 0 | 0 | 1274700 | 0 | 606180 | 4967400 | 1273300 | 0 | 0 | 0 | 1295500 |
| ACVVHGSI | ATP1A1 | P05023 | AT1A1_HUMAN | Oxidation (M) | 0 | 0 | 0 | 0 | 0 | 0 | 0 | 0 | 0 | 763380 | 0 | 0 | 11818000 | 9296600 | 0 | 0 | 0 | 0 |
| ACYGVLR | RPS3; RPS3; RP | P23396; H0 | RS3_HUMAN 4 | Unmodified | 863160 | 583730 | 503510 | 450500 | 145750 | 0 | 238950 | 95867 | 291040 | 77296 | 295040 | 12452000 | 167500 | 120470 | 464680 | 450200 | 0 | 0 |
| ACYLSINP | ACTR1A; ACTR1 | P61163; R4 | ACTZ_HUMAN | Unmodified | 202900 | 0 | 0 | 203690 | 0 | 0 | 0 | 134960 | 0 | 0 | 0 | 4980100 | 0 | 0 | 184410 | 134550 | 0 | 0 |
| ADAELDDP | CACNA2D2; CA | Q9NY47; C9 | CA2D2_HUMA | Unmodified | 120360 | 0 | 0 | 0 | 0 | 0 | 0 | 0 | 1974100 | 0 | 0 | 339870 | 0 | 0 | 0 | 0 | 0 | 0 |
| ADAGFVYS | CASK; CASK; C | O14936; A0A | CSKP_HUMAN | Oxidation (M) | 0 | 0 | 0 | 0 | 0 | 0 | 0 | 0 | 0 | 0 | 0 | 0 | 0 | 0 | 2070300 | 1015600 | 0 | 0 |
| ADALQAGA | VAMP3; VAMP2 | Q15836; P6 | VAMP3_HUMA | Unmodified | 656880 | 582390 | 3795400 | 0 | 0 | 0 | 0 | 280600 | 4741900 | 1271500 | 0 | 0 | 0 | 0 | 1710400 | 1413500 | 0 | 360010 |
| ADASGFAP | MST1R; MST1R | Q04912; J3K | RON_HUMAN | Unmodified | 0 | 0 | 0 | 0 | 0 | 0 | 0 | 0 | 218770 | 0 | 44346 | 0 | 0 | 0 | 0 | 0 | 0 | 0 |
| ADATNVN | AHSA1; AHSA1 | O95433; G3 | AHSA1_HUMA | Unmodified | 0 | 120270 | 136610 | 318390 | 0 | 0 | 0 | 0 | 425210 | 0 | 399840 | 1747500 | 0 | 0 | 317270 | 299360 | 0 | 0 |
| ADDLDFET | EIF5A; EIF5AL1; | P63241; Q6I | IF5A1_HUMAN | Acetyl (Protein N | 5176100 | 3724100 | 689150 | 4984400 | 30597000 | 6892600 | 13447000 | 6985100 | 2605100 | 2954200 | 2629000 | 3238500 | 22107000 | 18351000 | 3956800 | 3149900 | 27662000 | 15330000 |
| ADDLDFET | EIF5A; EIF5AL1; | P63241; Q6I | IF5A1_HUMAN | Acetyl (Protein N | 0 | 0 | 0 | 0 | 0 | 0 | 0 | 0 | 701850 | 0 | 0 | 537980 | 0 | 0 | 0 | 0 | 2247600 | 2998600 |
| ADDLDFET | EIF5A; EIF5AL1; | P63241; Q6I | IF5A1_HUMAN | Acetyl (Protein N | 0 | 0 | 0 | 0 | 0 | 0 | 0 | 0 | 0 | 0 | 0 | 0 | 1125000 | 0 | 0 | 0 | 4971100 | 0 |
| ADDPSAAD | ETF1; ETF1 | P62495; D6F | ERF1_HUMAN | Acetyl (Protein N | 618420 | 397040 | 301370 | 1618000 | 4300900 | 812440 | 750940 | 317710 | 239270 | 619720 | 1989400 | 7412500 | 3787800 | 1565700 | 936430 | 577320 | 195280 | 1518800 |
| ADDVDQQ | LSM3 | P62310 | LSM3_HUMAN | Acetyl (Protein N | 0 | 0 | 0 | 0 | 0 | 0 | 0 | 0 | 807650 | 0 | 256010 | 0 | 0 | 0 | 0 | 0 | 0 | 0 |
| ADDYEQVK | ATP6V0D1; ATP | P61421; R4 | VA0D1_HUMAN | Unmodified | 0 | 0 | 0 | 0 | 168080 | 0 | 85477 | 46182 | 1533600 | 0 | 0 | 828450 | 417790 | 0 | 0 | 0 | 122770 | 75297 |
| ADEASELA | FASN; FASN | P49327; A0A | FAS_HUMAN F | Unmodified | 76310 | 0 | 0 | 0 | 0 | 0 | 0 | 0 | 0 | 0 | 0 | 1540300 | 0 | 0 | 0 | 0 | 0 | 0 |
| ADEASELA | FASN; FASN | P49327; A0A | FAS_HUMAN F | Unmodified | 0 | 0 | 0 | 0 | 0 | 0 | 0 | 0 | 0 | 0 | 0 | 0 | 0 | 0 | 0 | 0 | 0 | 0 |
| ADEATRRV | UFC1 | Q9Y3C8 | UFC1_HUMAN | Acetyl (Protein N | 0 | 0 | 0 | 0 | 10506000 | 394660 | 1931600 | 0 | 0 | 228370 | 0 | 0 | 4251700 | 3630800 | 0 | 0 | 7820300 | 4572600 |
| ADEDPIMG | NUTF2; NUTF2 | P61970; H3E | NTF2_HUMAN | 2 Oxidation (M) | 239130 | 569830 | 0 | 0 | 0 | 0 | 0 | 0 | 0 | 0 | 0 | 4190300 | 0 | 0 | 0 | 894200 | 0 | 0 |
| ADEEKLPP | PIN1; PIN1P1; F | Q13526; O1 | PIN1_HUMAN | Acetyl (Protein N | 2828700 | 2546700 | 2343100 | 1716500 | 0 | 0 | 270660 | 110340 | 1140700 | 534210 | 0 | 948650 | 725830 | 0 | 1469200 | 2141000 | 0 | 871160 |
| ADEEKLPP | PIN1; PIN1P1; F | Q13526; O1 | PIN1_HUMAN | Acetyl (Protein N | 6515500 | 5258200 | 0 | 3120200 | 242060 | 0 | 0 | 0 | 342450 | 36580 | 0 | 504740 | 286800 | 0 | 976270 | 1385000 | 239000 | 211020 |
| ADEHVIDQ | PRKAR2A; PRKA | P13861; H7 | KAP2_HUMAN | Unmodified | 0 | 0 | 724920 | 0 | 0 | 0 | 0 | 0 | 543960 | 0 | 0 | 3981300 | 0 | 0 | 2417200 | 933610 | 0 | 0 |
| ADEHVIDQ | PRKAR2A; PRKA | P13861; H7 | KAP2_HUMAN | Unmodified | 0 | 0 | 0 | 0 | 0 | 0 | 0 | 0 | 0 | 0 | 229970 | 0 | 0 | 0 | 973260 | 0 | 0 | 0 |
| ADEIAKAQ | HINT1; HINT1; | P49773; H0 | HINT1_HUMAN | Acetyl (Protein N | 0 | 0 | 0 | 0 | 13548000 | 1004400 | 37232000 | 30774000 | 0 | 7971000 | 0 | 0 | 13955000 | 21981000 | 55369 | 0 | 74206000 | 11434000 |
| ADELSEK | CAV1; CAV1; C | Q03135; C9J | CAV1_HUMAN | Acetyl (Protein N | 0 | 0 | 0 | 0 | 1758600 | 696520 | 233810 | 0 | 383750 | 218470 | 204570 | 0 | 0 | 0 | 0 | 0 | 148440 | 357930 |
| ADELSEKQ | CAV1; CAV1; C | Q03135; C9J | CAV1_HUMAN | Acetyl (Protein N | 0 | 0 | 0 | 0 | 8627900 | 944430 | 2262800 | 767830 | 0 | 0 | 148970 | 0 | 0 | 0 | 0 | 0 | 1493800 | 1112200 |
| ADESQEPC | MAN1C1 | Q9NR34 | MA1C1_HUMA | Unmodified | 0 | 0 | 0 | 0 | 0 | 0 | 0 | 0 | 1487200 | 0 | 0 | 0 | 0 | 0 | 0 | 0 | 0 | 0 |
| ADEVVWVF | DARS1; DARS1; | P14868; H7 | SYDC_HUMAN | Unmodified | 0 | 0 | 0 | 0 | 0 | 154790 | 0 | 0 | 0 | 99219 | 516420 | 3530200 | 0 | 0 | 0 | 0 | 0 | 0 |
| ADFDDRVS | CAPZA1 | P52907 | CAZA1_HUMAN | Acetyl (Protein N | 2147400 | 2843100 | 2576000 | 2149000 | 26068000 | 3961300 | 1162300 | 982920 | 3188400 | 2051400 | 3149200 | 14874000 | 9733300 | 4517600 | 3494500 | 1921700 | 8450000 | 7369100 |
| ADFDDRVS | CAPZA1 | P52907 | CAZA1_HUMAN | Acetyl (Protein N | 46504 | 0 | 0 | 86362 | 2442300 | 491590 | 459750 | 0 | 0 | 834950 | 0 | 0 | 2131800 | 835910 | 78435 | 57965 | 2209100 | 1322700 |
| ADFDNTVA | GSR | P00390 | GSHR_HUMAN | Unmodified | 967330 | 637580 | 624240 | 1368500 | 10970000 | 1100400 | 269610 | 244790 | 4817700 | 954050 | 2348600 | 10050000 | 3976100 | 721780 | 5036900 | 2708400 | 0 | 1040100 |
| ADFDTYDD | EIF4H; EIF4H; E | Q15056; A0A | IF4H_HUMAN E | Acetyl (Protein N | 5142000 | 1535700 | 418940 | 0 | 0 | 0 | 0 | 0 | 0 | 0 | 0 | 853710 | 0 | 0 | 2095000 | 937120 | 0 | 0 |
| ADFDTYDD | EIF4H; EIF4H | Q15056; A0A | IF4H_HUMAN E | Acetyl (Protein N | 943870 | 797180 | 1467100 | 3393900 | 3348500 | 0 | 169180 | 0 | 1013900 | 0 | 0 | 0 | 1673000 | 0 | 1165600 | 426860 | 0 | 0 |
| ADFIRESSE | PRRC2C; PRRC | Q9Y520; E7E | PRC2C_HUMA | Unmodified | 498300 | 369980 | 0 | 245680 | 0 | 0 | 0 | 0 | 0 | 0 | 0 | 0 | 237820 | 0 | 0 | 111340 | 0 | 0 |
| ADFLIFR | KIAA0319L; KIA | Q8IZA0; C9J | K319L_HUMAN | Unmodified | 243160 | 0 | 125660 | 149800 | 0 | 0 | 0 | 0 | 3190900 | 493810 | 291620 | 224930 | 0 | 0 | 255340 | 340890 | 0 | 0 |
| ADFQGISPI | DAD1; DAD1 | P61803; F5G | DAD1_HUMAN | Unmodified | 0 | 0 | 0 | 0 | 0 | 254250 | 0 | 0 | 1806600 | 0 | 0 | 0 | 0 | 0 | 0 | 0 | 0 | 0 |
| ADFSGMS | SERPINB6; SER | P35237; A0A | SPB6_HUMAN | Oxidation (M) | 0 | 0 | 0 | 0 | 0 | 0 | 0 | 0 | 0 | 0 | 0 | 3742300 | 0 | 0 | 0 | 0 | 0 | 0 |
| ADFSGMS | SERPINB6; SER | P35237; A0A | SPB6_HUMAN | Unmodified | 0 | 0 | 0 | 0 | 0 | 0 | 0 | 0 | 0 | 0 | 313050 | 1963500 | 719630 | 0 | 0 | 0 | 0 | 0 |
| ADGELNVD | PPP1CB; PPP1C | P62140; C9J | PP1B_HUMAN | Acetyl (Protein N | 0 | 0 | 0 | 0 | 1372000 | 0 | 0 | 0 | 0 | 0 | 0 | 2074100 | 0 | 0 | 0 | 0 | 0 | 0 |
| ADGETILK | SMS; SMS | P52788; H7 | SPSY_HUMAN | Unmodified | 192040 | 272560 | 684430 | 395940 | 239070 | 0 | 92745 | 52776 | 0 | 86336 | 726700 | 648500 | 0 | 334280 | 745610 | 0 | 216230 | 390440 |
| ADGEYWL | MFAP4; MFAP4 | P55083; K7E | MFAP4_HUMA | Oxidation (M) | 326010 | 0 | 656800 | 95012 | 405460 | 1365500 | 0 | 0 | 0 | 205630 | 0 | 212170 | 284580 | 0 | 3470700 | 1655800 | 0 | 0 |
| ADGGAEYA | CD46 | R4GN18 | R4GN18_HUM | Unmodified | 0 | 4290300 | 4006100 | 2565700 | 562000 | 0 | 0 | 0 | 3403300 | 725280 | 438080 | 14881000 | 0 | 0 | 5919800 | 3064500 | 147860 | 479180 |
| ADGGTQVI | DLD; DLD | P09622; E9P | DLDH_HUMAN | Unmodified | 1354200 | 1117300 | 571710 | 644000 | 0 | 0 | 62941 | 0 | 4625300 | 0 | 0 | 166680 | 0 | 0 | 37828 | 0 | 113910 | 0 |
| ADGKAGD | PCNP; PCNP; P | Q8WW12; A | PCNP_HUMAN | Acetyl (Protein N | 0 | 0 | 17138 | 0 | 224410 | 0 | 0 | 0 | 0 | 0 | 0 | 90958 | 0 | 0 | 0 | 0 | 201070 | 0 |

| | | | | | | | | | | | | | | | | | | | | | | |
|---|---|---|---|---|---|---|---|---|---|---|---|---|---|---|---|---|---|---|---|---|---|---|
| ADGNQLTL | CD109 | Q6YHK3 | CD109_HUMAI | Unmodified | 3239700 | 3945400 | 1211200 | 0 | 0 | 0 | 0 | 0 | 1690600 | 0 | 0 | 1014100 | 0 | 0 | 1285700 | 0 | 432140 | 0 |
| ADGPPHG( | PLXNA2 | O75051 | PLXA2_HUMAN | Unmodified | 0 | 0 | 0 | 0 | 0 | 0 | 524460 | 125200 | 814080 | 2038300 | 0 | 0 | 0 | 0 | 0 | 0 | 940410 | 424990 |
| ADGSGSVL | LRP1 | Q07954 | LRP1_HUMAN | Unmodified | 0 | 0 | 0 | 0 | 0 | 841460 | 580960 | 0 | 0 | 0 | 0 | 0 | 91385 | 0 | 0 | 0 | 722830 | 997490 |
| ADGSSLPE | PTK7 | Q13308 | PTK7_HUMAN | Unmodified | 0 | 0 | 0 | 0 | 0 | 0 | 100270 | 0 | 3336300 | 0 | 0 | 0 | 327530 | 0 | 883340 | 544640 | 0 | 0 |
| ADGSWRE( | BCAM; BCAM | P50895; A0A | BCAM_HUMAN | Unmodified | 1912300 | 482110 | 0 | 0 | 0 | 0 | 0 | 0 | 0 | 0 | 0 | 0 | 0 | 0 | 0 | 0 | 0 | 0 |
| ADGVHTVE | TMED1 | Q13445 | TMED1_HUMAI | Unmodified | 0 | 0 | 46668 | 0 | 0 | 0 | 0 | 0 | 2340500 | 117720 | 0 | 53979 | 0 | 0 | 0 | 0 | 0 | 0 |
| ADGYEPPV | RPS3A; RPS3A; | P61247; D6F | RS3A_HUMAN | Unmodified | 0 | 0 | 0 | 177020 | 0 | 0 | 0 | 0 | 0 | 0 | 960650 | 1022300 | 0 | 0 | 0 | 0 | 0 | 0 |
| ADGYVLEG | RPS8; RPS8 | P62241; Q5J | RS8_HUMAN 4 | Unmodified | 3868000 | 2471300 | 2267200 | 792660 | 0 | 0 | 0 | 64508 | 790540 | 0 | 156280 | 3616000 | 0 | 0 | 1702100 | 2111900 | 0 | 0 |
| ADHCIDSA | N/A | N/A | N/A | Unmodified | 0 | 6217000 | 0 | 0 | 0 | 687790 | 0 | 0 | 0 | 0 | 0 | 0 | 0 | 0 | 2254300 | 0 | 0 | 0 |
| ADHGDELF | CES2 | O00748 | EST2_HUMAN | Unmodified | 0 | 0 | 0 | 0 | 0 | 0 | 0 | 0 | 554250 | 0 | 0 | 0 | 0 | 0 | 0 | 0 | 0 | 0 |
| ADHGEPIG | HSP90AB1; HSI | P08238; Q58 | HS90B_HUMAI | Unmodified | 0 | 0 | 0 | 0 | 0 | 0 | 0 | 0 | 0 | 0 | 0 | 2403900 | 130010 | 0 | 0 | 0 | 0 | 0 |
| ADHGEPIG | HSP90AB1; HSI | P08238; Q58 | HS90B_HUMAI | Unmodified | 418790 | 0 | 465490 | 1869500 | 1793700 | 765810 | 318530 | 105510 | 2212600 | 228200 | 805330 | 5209100 | 1793400 | 317890 | 701210 | 563340 | 554690 | 784370 |
| ADHQPLTE | RPSA; RPSA; RF | P08865; A0A | RSSA_HUMAN | Unmodified | 4481500 | 0 | 0 | 0 | 0 | 0 | 0 | 0 | 1228200 | 0 | 1236700 | 17931000 | 0 | 0 | 0 | 658990 | 0 | 0 |
| ADHSAAVE | FARP2; FARP2 | O94887; H7( | FARP2_HUMAN | Unmodified | 0 | 0 | 356110 | 277130 | 0 | 0 | 0 | 0 | 0 | 0 | 0 | 252660 | 0 | 0 | 780850 | 386190 | 0 | 0 |
| ADHSFSDC | NAP1L4; NAP1L | Q99733; C9J | NP1L4_HUMAN | Acetyl (Protein N | 517660 | 0 | 0 | 917010 | 1348600 | 0 | 0 | 396530 | 786230 | 486470 | 257840 | 755280 | 2213100 | 984240 | 783040 | 457810 | 986920 | 1403000 |
| ADHSFSDC | NAP1L4; NAP1L | Q99733; C9J | NP1L4_HUMAN | Unmodified | 0 | 0 | 0 | 0 | 0 | 0 | 0 | 116240 | 0 | 0 | 0 | 0 | 258610 | 319860 | 0 | 0 | 465290 | 250200 |
| ADHTLSSLI | BROX; BROX | Q5VW32; F5 | BROX_HUMAN | Unmodified | 5382000 | 17285000 | 3948800 | 5478900 | 2626600 | 2043100 | 92957 | 0 | 1423200 | 2670500 | 0 | 4390500 | 1395500 | 1225100 | 16827000 | 2501900 | 653670 | 0 |
| ADHVQSLA | XPO7; XPO7 | Q9UIA9; H0\ | XPO7_HUMAN | Acetyl (Protein N | 0 | 148780 | 0 | 0 | 667520 | 0 | 0 | 0 | 0 | 0 | 0 | 711250 | 0 | 0 | 0 | 0 | 403080 | 0 |
| ADHWMLY | ACOT8; ACOT8 | O14734; A0A | ACOT8_HUMAI | Oxidation (M) | 0 | 0 | 0 | 0 | 0 | 0 | 0 | 0 | 345410 | 0 | 0 | 0 | 0 | 0 | 0 | 0 | 0 | 0 |
| ADHWMLY | ACOT8; ACOT8 | O14734; A0A | ACOT8_HUMAI | Unmodified | 0 | 0 | 0 | 0 | 0 | 0 | 0 | 0 | 188000 | 0 | 0 | 0 | 0 | 0 | 0 | 0 | 0 | 0 |
| ADIDNLPR | NAP1L1 | F8W543 | F8W543_HUM | Acetyl (Protein N | 229700 | 239730 | 522700 | 352000 | 0 | 0 | 0 | 0 | 122080 | 0 | 274880 | 1490800 | 268440 | 0 | 620590 | 0 | 0 | 0 |
| ADIDVSGP | AHNAK | Q09666 | AHNK_HUMAN | Unmodified | 0 | 0 | 0 | 0 | 1230300 | 0 | 0 | 0 | 0 | 0 | 0 | 0 | 0 | 0 | 0 | 0 | 133510 | 661890 |
| ADIEEIK | NUCB2; NUCB2 | P80303; Q2L | NUCB2_HUMA | Unmodified | 0 | 0 | 770070 | 325560 | 364940 | 0 | 0 | 0 | 1384500 | 276240 | 0 | 316020 | 102210 | 68868 | 989190 | 0 | 0 | 59137 |
| ADIEEIKSG | NUCB2; NUCB2 | P80303; Q2L | NUCB2_HUMA | Unmodified | 0 | 0 | 0 | 0 | 0 | 0 | 0 | 0 | 0 | 898610 | 0 | 0 | 0 | 0 | 0 | 0 | 0 | 0 |
| ADIFMFDEI | ABCE1; ABCE1 | P61221; D6F | ABCE1_HUMAN | Oxidation (M) | 144660 | 0 | 0 | 0 | 14228 | 0 | 0 | 0 | 0 | 0 | 0 | 0 | 0 | 0 | 420540 | 166910 | 138730 | 0 |
| ADIFVDPVI | GLG1 | Q92896 | GSLG1_HUMAI | Unmodified | 93118 | 0 | 0 | 0 | 0 | 0 | 0 | 0 | 8961000 | 245570 | 1825000 | 1590600 | 0 | 0 | 0 | 0 | 0 | 0 |
| ADIGVAMG | ATP1A1; ATP1A | P05023; P13 | AT1A1_HUMAN | Oxidation (M) | 0 | 176220 | 1994700 | 806480 | 0 | 0 | 0 | 0 | 0 | 168100 | 672820 | 1652400 | 247820 | 0 | 0 | 370170 | 0 | 0 |
| ADIIARLREI | AIP; AIP | O00170; E9F | AIP_HUMAN AI | Acetyl (Protein N | 0 | 0 | 0 | 0 | 1938300 | 1101900 | 46710 | 0 | 16903000 | 1173400 | 0 | 0 | 5182500 | 2419800 | 0 | 0 | 4959200 | 6249700 |
| ADIIYPGHC | LACTB2 | Q53H82 | LACB2_HUMAN | Unmodified | 0 | 0 | 0 | 0 | 0 | 0 | 0 | 0 | 4817900 | 231680 | 0 | 335340 | 0 | 0 | 245490 | 0 | 85308 | 0 |
| ADILEDKD( | HNRNPM; HNR | P52272; M0 | HNRPM_HUMA | Unmodified | 0 | 0 | 0 | 0 | 0 | 0 | 0 | 0 | 0 | 0 | 193190 | 1361300 | 0 | 0 | 0 | 0 | 0 | 0 |
| ADILQVGLI | PAIP1; PAIP1; P | Q9H074; D6 | PAIP1_HUMAN | Unmodified | 302210 | 312250 | 171750 | 0 | 0 | 0 | 0 | 0 | 0 | 0 | 0 | 0 | 0 | 0 | 200430 | 223340 | 0 | 0 |
| ADIQMPFT( | ALCAM; ALCAM | Q13740; F5( | CD166_HUMAI | Unmodified | 1811400 | 0 | 0 | 0 | 668760 | 0 | 0 | 159640 | 2390200 | 0 | 0 | 0 | 0 | 0 | 0 | 0 | 0 | 0 |
| ADIQMPFT( | ALCAM; ALCAM | Q13740; F5( | CD166_HUMAI | Oxidation (M) | 1995100 | 932750 | 310790 | 232960 | 0 | 0 | 0 | 0 | 0 | 0 | 0 | 0 | 0 | 0 | 928520 | 305040 | 0 | 0 |
| ADIQTER | RPS11; RPS11; | P62280; M0I | RS11_HUMAN | Acetyl (Protein N | 536750 | 0 | 0 | 0 | 0 | 0 | 0 | 0 | 0 | 59035 | 317090 | 3585800 | 0 | 0 | 0 | 0 | 0 | 0 |
| ADIQTERA | RPS11; RPS11; | P62280; M0I | RS11_HUMAN | Acetyl (Protein N | 0 | 0 | 0 | 0 | 7473300 | 1001200 | 453180 | 175220 | 0 | 255070 | 0 | 0 | 2983700 | 169740 | 0 | 0 | 0 | 1926700 |
| ADIQTERA | RPS11; RPS11; | P62280; M0I | RS11_HUMAN | Acetyl (Protein N | 0 | 0 | 0 | 0 | 3566500 | 575040 | 2389900 | 6696000 | 0 | 1184500 | 0 | 0 | 11421000 | 5998400 | 0 | 0 | 2036300 | 3520700 |
| ADISREQAI | TNS3 | Q68CZ2 | TENS3_HUMAN | Unmodified | 0 | 0 | 0 | 0 | 1302300 | 174340 | 0 | 0 | 0 | 25905 | 0 | 536960 | 0 | 0 | 0 | 0 | 938440 | 920140 |
| ADIVQQLL( | ANK3 | Q12955 | ANK3_HUMAN | Unmodified | 0 | 0 | 1464900 | 2494000 | 2805000 | 0 | 0 | 0 | 0 | 0 | 0 | 0 | 0 | 0 | 5009200 | 6030300 | 0 | 0 |
| ADIWSLGIT | STK24; STK24; | Q9Y6E0; H0\ | STK24_HUMAN | Oxidation (M) | 0 | 0 | 0 | 203900 | 1765100 | 509650 | 0 | 0 | 0 | 241440 | 854660 | 0 | 803300 | 411810 | 252050 | 0 | 360230 | 690780 |
| ADKDYHFK | NPM1; NPM1; N | P06748; A0A | NPM_HUMAN I | Unmodified | 0 | 131040 | 0 | 205320 | 1804300 | 809440 | 0 | 0 | 4517200 | 1584600 | 3422900 | 674470 | 2645700 | 0 | 0 | 0 | 83189 | 862300 |
| ADKEAAFD | RBBP4; RBBP4; | Q09028; E9F | RBBP4_HUMAN | Acetyl (Protein N | 0 | 0 | 0 | 0 | 2461100 | 0 | 987040 | 438850 | 479910 | 2828700 | 0 | 0 | 3521100 | 1843900 | 0 | 0 | 2744500 | 1384000 |
| ADKLTRIAI | ABCE1; ABCE1; | P61221; D6F | ABCE1_HUMAN | Acetyl (Protein N | 0 | 0 | 0 | 0 | 564880 | 196920 | 96889 | 345090 | 0 | 0 | 0 | 0 | 0 | 0 | 0 | 0 | 391480 | 0 |
| ADKNEVAA | MARS1; MARS1 | P56192; H0\ | SYMC_HUMAN | Unmodified | 380870 | 255050 | 555040 | 666290 | 183980 | 0 | 29312 | 51188 | 859790 | 102750 | 484730 | 2395400 | 0 | 110320 | 335410 | 472700 | 27650 | 62340 |
| ADKPDMGI | TMSB10 | P63313 | TYB10_HUMAN | Acetyl (Protein N | 13889000 | 8018100 | 772370 | 804750 | 32667000 | 8491700 | 2505400 | 5079300 | 4401200 | 6241100 | 833800 | 139870 | 1037100 | 1324800 | 0 | 106860 | 39187000 | 42930000 |
| ADKPDMGI | TMSB10 | P63313 | TYB10_HUMAN | Acetyl (Protein N | 16272000 | 10631000 | 0 | 989060 | 1.21E+08 | 20491000 | 6329100 | 5654300 | 39045000 | 22899000 | 2118700 | 1936100 | 3212800 | 5393100 | 701030 | 0 | 1.55E+08 | 1.41E+08 |
| ADKPDMGI | TMSB10 | P63313 | TYB10_HUMAN | Acetyl (Protein N | 0 | 0 | 0 | 0 | 0 | 0 | 0 | 860430 | 0 | 0 | 0 | 0 | 0 | 0 | 0 | 0 | 7157100 | 0 |
| ADKPDMGI | TMSB10 | P63313 | TYB10_HUMAN | Acetyl (Protein N | 0 | 0 | 0 | 0 | 794060 | 0 | 1091100 | 1206900 | 0 | 311960 | 344240 | 0 | 0 | 0 | 0 | 0 | 21220000 | 629820 |
| ADKRDPEF | NCLN; NCLN; N | Q969V3; A0A | NCLN_HUMAN | Unmodified | 0 | 0 | 0 | 0 | 0 | 0 | 0 | 0 | 5689900 | 987570 | 0 | 0 | 0 | 0 | 0 | 0 | 0 | 0 |
| ADLAAVEAI | DYNC1H1; DYN | Q14204; A0A | DYHC1_HUMA | Unmodified | 0 | 230600 | 0 | 0 | 594820 | 0 | 0 | 0 | 518290 | 53925 | 459100 | 5984200 | 123400 | 0 | 0 | 360590 | 142240 | 0 |
| ADLAECNII | KIF5B | P33176 | KINH_HUMAN | Acetyl (Protein N | 0 | 0 | 90855 | 0 | 886590 | 266570 | 938220 | 754850 | 388210 | 739240 | 0 | 1156000 | 0 | 486240 | 143880 | 74434 | 1230000 | 1304800 |
| ADLAEEYSI | UBE2L3 | P68036 | UB2L3_HUMAN | Unmodified | 284030 | 0 | 207060 | 0 | 0 | 0 | 0 | 0 | 0 | 0 | 70772 | 1018100 | 0 | 0 | 504250 | 663040 | 0 | 0 |
| ADLANEEK | RANBP3; RANB | Q9H6Z4; B7 | RANB3_HUMAI | Acetyl (Protein N | 0 | 0 | 0 | 0 | 586430 | 0 | 0 | 0 | 0 | 294850 | 518570 | 657970 | 624370 | 673370 | 0 | 0 | 0 | 0 |
| ADLATAPPI | LRRC47 | Q8N1G4 | LRC47_HUMAN | Unmodified | 0 | 0 | 0 | 0 | 0 | 0 | 0 | 94863 | 432370 | 52163 | 180810 | 3164900 | 0 | 0 | 377200 | 611790 | 148870 | 0 |
| ADLDHQR( | RRAS2; RRAS2 | P62070; E9P | RRAS2_HUMAN | Unmodified | 0 | 0 | 0 | 403780 | 282760 | 96807 | 0 | 0 | 0 | 0 | 0 | 0 | 0 | 85686 | 70857 | 0 | 90130 | 87784 |
| ADLDKLNII | PPP1CC; PPP1( | P36873; F8V | PP1G_HUMAN | Acetyl (Protein N | 0 | 281470 | 0 | 509270 | 1137400 | 381560 | 110420 | 0 | 0 | 0 | 236070 | 0 | 0 | 0 | 0 | 0 | 348400 | 0 |
| ADLDVSGP | AHNAK | Q09666 | AHNK_HUMAN | Unmodified | 0 | 0 | 0 | 0 | 2223300 | 0 | 0 | 0 | 0 | 0 | 0 | 0 | 0 | 0 | 165090 | 0 | 0 | 2778400 |
| ADLEAQRD | RAB14; RAB14 | P61106; X6R | RAB14_HUMAN | Unmodified | 1357800 | 874970 | 1957700 | 405530 | 137100 | 0 | 97472 | 29615 | 2034400 | 0 | 1946700 | 9529800 | 0 | 0 | 995890 | 1004700 | 0 | 0 |
| ADLEEQLSI | CAPZA2; CAPZ/ | P47755; F8V | CAZA2_HUMAN | Acetyl (Protein N | 1236400 | 0 | 0 | 0 | 2769600 | 0 | 456430 | 361430 | 1517800 | 515730 | 3500400 | 3729500 | 1466900 | 755290 | 889100 | 605970 | 828160 | 1109500 |
| ADLEEQLSI | CAPZA2; CAPZ/ | P47755; F8V | CAZA2_HUMAN | Acetyl (Protein N | 0 | 0 | 0 | 0 | 3079400 | 0 | 1970900 | 0 | 0 | 625860 | 0 | 0 | 2031100 | 2268500 | 0 | 0 | 3135100 | 1610500 |
| ADLELPWR | PSME4; PSME4 | Q14997; F6V | PSME4_HUMAI | Oxidation (M) | 0 | 0 | 0 | 0 | 0 | 0 | 0 | 41236 | 0 | 0 | 0 | 1537900 | 0 | 0 | 0 | 0 | 17602 | 0 |
| ADLEQRHF | UTRN | P46939 | UTRN_HUMAN | Unmodified | 0 | 0 | 190530 | 758860 | 1121700 | 0 | 0 | 0 | 2580800 | 203680 | 0 | 0 | 0 | 0 | 0 | 873650 | 0 | 0 |
| ADLGALEL\ | SORT1 | Q99523 | SORT_HUMAN | Unmodified | 0 | 0 | 0 | 0 | 0 | 0 | 0 | 0 | 5488300 | 0 | 861150 | 1827600 | 0 | 0 | 0 | 0 | 0 | 0 |
| ADLGALEL\ | SORT1 | Q99523 | SORT_HUMAN | Unmodified | 0 | 0 | 0 | 0 | 0 | 0 | 0 | 0 | 0 | 1282500 | 602950 | 0 | 0 | 0 | 0 | 0 | 0 | 0 |
| ADLHRQLC | SFT2D3 | Q587I9 | SFT2C_HUMAN | Acetyl (Protein N | 0 | 0 | 0 | 0 | 397240 | 0 | 538430 | 0 | 997920 | 119600 | 0 | 0 | 683720 | 770920 | 0 | 0 | 0 | 0 |
| ADLILPCAI | RNF141; RNF14 | Q8WVD5; ES | RN141_HUMAN | Unmodified | 188760 | 82961 | 0 | 0 | 0 | 0 | 0 | 0 | 0 | 0 | 0 | 0 | 117480 | 0 | 54164 | 39082 | 0 | 0 |
| ADLINNLG | HSP90AA2P; H: | Q14568; P0I | HS902_HUMAN | Unmodified | 4115100 | 6222500 | 17654000 | 17442000 | 36759000 | 7823100 | 3088500 | 1819200 | 39380000 | 13463000 | 50417000 | 3.66E+08 | 23608000 | 4851300 | 19537000 | 15940000 | 10373000 | 11590000 |
| ADLINNLG | HSP90AA2P; H: | Q14568; P0I | HS902_HUMAN | Unmodified | 0 | 0 | 0 | 0 | 395970 | 0 | 233350 | 340350 | 0 | 78827 | 0 | 422140 | 0 | 0 | 0 | 0 | 1632500 | 0 |
| ADLISEDV( | ARHGEF1; ARH | Q92888; M0 | ARHG1_HUMA | Unmodified | 14275 | 10450 | 23993 | 18702 | 239820 | 0 | 0 | 0 | 0 | 0 | 0 | 491860 | 0 | 0 | 25646 | 30463 | 0 | 9823.3 |
| ADLLLSTQI | SLC3A2; SLC3A | P08195; A0A | 4F2_HUMAN 4I | Unmodified | 38679000 | 36443000 | 23934000 | 18814000 | 2810800 | 1040600 | 0 | 161770 | 63766000 | 10046000 | 36805000 | 4.23E+08 | 0 | 0 | 69970000 | 49614000 | 693330 | 66245 |
| ADLLLSTQI | SLC3A2; SLC3A | P08195; A0A | 4F2_HUMAN 4I | Unmodified | 3443700 | 1048900 | 21971000 | 2014300 | 1991600 | 0 | 1087000 | 0 | 8276200 | 0 | 7659200 | 43230000 | 0 | 0 | 8347300 | 6908700 | 0 | 0 |
| ADLNQGIG | EFHD2 | Q96C19 | EFHD2_HUMAI | Unmodified | 0 | 0 | 0 | 0 | 0 | 307390 | 0 | 0 | 0 | 0 | 0 | 0 | 0 | 0 | 0 | 0 | 0 | 0 |
| ADLQNHLE | EEA1 | Q15075 | EEA1_HUMAN | Unmodified | 21559 | 0 | 0 | 0 | 0 | 0 | 0 | 0 | 0 | 0 | 0 | 571430 | 561310 | 0 | 0 | 0 | 0 | 0 |
| ADLRDNYS | EPRS1 | P07814 | SYEP_HUMAN | Unmodified | 0 | 0 | 0 | 22684 | 0 | 0 | 0 | 0 | 0 | 211100 | 0 | 0 | 480600 | 0 | 0 | 0 | 177190 | 0 |
| ADLSGMSC | SERPINB1 | P30740 | ILEU_HUMAN L | Unmodified | 0 | 0 | 0 | 0 | 100260 | 0 | 0 | 0 | 642140 | 35286 | 0 | 0 | 0 | 0 | 0 | 0 | 0 | 0 |
| ADLTALFLP | FLII | Q13045 | FLII_HUMAN Pr | Unmodified | 0 | 0 | 0 | 0 | 0 | 0 | 0 | 0 | 203210 | 0 | 85780 | 785150 | 0 | 0 | 0 | 204000 | 0 | 0 |
| ADLTEYLST | UQCRC1 | P31930 | QCR1_HUMAN | Unmodified | 0 | 0 | 0 | 0 | 0 | 0 | 0 | 0 | 8225700 | 611170 | 365210 | 790640 | 0 | 0 | 96576 | 0 | 0 | 0 |
| ADLVNRSI/ | PDCD6IP | Q8WUM4 | PDC6I_HUMAN | Unmodified | 0 | 0 | 0 | 2140600 | 0 | 0 | 0 | 0 | 0 | 175930 | 0 | 0 | 778600 | 330850 | 0 | 0 | 0 | 0 |
| ADLYIHVTY | BPNT2 | Q9NX62 | IMPA3_HUMAN | Unmodified | 0 | 0 | 0 | 0 | 0 | 0 | 197800 | 0 | 29030000 | 8818000 | 1079600 | 505090 | 337040 | 244380 | 0 | 0 | 0 | 0 |
| ADMEDLMS | MYH10 | P35580 | MYH10_HUMAI | Unmodified | 0 | 0 | 0 | 0 | 0 | 0 | 0 | 0 | 0 | 0 | 0 | 781050 | 0 | 0 | 54693 | 62365 | 0 | 0 |
| ADMETLQR | TARS1 | P26639 | SYTC_HUMAN | Unmodified | 0 | 0 | 0 | 0 | 0 | 389800 | 0 | 0 | 125980 | 0 | 361160 | 4080700 | 0 | 0 | 0 | 61297 | 0 | 0 |
| ADMGGAA | LAP3; LAP3 | P28838; H0\ | AMPL_HUMAN | Oxidation (M) | 0 | 0 | 38641 | 24647 | 0 | 0 | 0 | 25629 | 84812 | 260890 | 51879 | 613010 | 63440 | 0 | 0 | 10814 | 0 | 0 |
| ADMQNLVI | CAP1; CAP1; C/ | Q01518; Q5 | CAP1_HUMAN | Acetyl (Protein N | 0 | 0 | 0 | 0 | 1465400 | 131750 | 73404 | 10271 | 1689900 | 295220 | 381180 | 1641100 | 634470 | 2013500 | 0 | 0 | 472850 | 605430 |
| ADNEKLDN | KPNA4; KPNA4 | O00629; H7( | IMA3_HUMAN | Acetyl (Protein N | 0 | 0 | 0 | 0 | 3708900 | 520920 | 765550 | 0 | 0 | 0 | 0 | 0 | 3062500 | 2909900 | 92909 | 0 | 8502300 | 3752000 |
| ADNIEVFA( | TPP2; TPP2; TPI | P29144; A0A | TPP2_HUMAN | Unmodified | 0 | 0 | 0 | 0 | 0 | 0 | 0 | 0 | 0 | 0 | 0 | 4343400 | 0 | 0 | 145880 | 0 | 0 | 0 |
| ADNKQKGI | MYO1C; MYO1( | O00159; F5H | MYO1C_HUMA | Unmodified | 0 | 0 | 0 | 793300 | 994770 | 389430 | 0 | 0 | 0 | 0 | 0 | 0 | 892540 | 322850 | 0 | 0 | 298590 | 1069200 |
| ADPAAGPP | PRKCB | P05771 | KPCB_HUMAN | Acetyl (Protein N | 0 | 180170 | 0 | 449450 | 0 | 0 | 217630 | 387110 | 0 | 0 | 0 | 0 | 0 | 0 | 0 | 0 | 0 | 0 |
| ADPAAGPP | PRKCB | P05771 | KPCB_HUMAN | Acetyl (Protein N | 0 | 0 | 0 | 125090 | 0 | 0 | 1283500 | 939860 | 0 | 0 | 0 | 0 | 610050 | 867620 | 0 | 0 | 0 | 0 |
| ADPAAGPP | PRKCB | P05771 | KPCB_HUMAN | Acetyl (Protein N | 0 | 0 | 0 | 0 | 0 | 0 | 10266000 | 8073000 | 0 | 0 | 0 | 0 | 4769300 | 5635700 | 0 | 0 | 0 | 0 |
| ADPDNLQ/ | CAPN5; CAPN5 | O15484; E7E | CAN5_HUMAN | Unmodified | 0 | 0 | 382340 | 798620 | 0 | 0 | 0 | 0 | 498540 | 0 | 0 | 1406500 | 280390 | 115200 | 1126600 | 971790 | 0 | 0 |
| ADPEELFTI | STK24; STK24 | Q9Y6E0; B4I | STK24_HUMAN | Unmodified | 0 | 0 | 0 | 0 | 0 | 0 | 0 | 0 | 0 | 0 | 0 | 99570 | 0 | 0 | 0 | 383650 | 179330 | 296840 |
| ADPSLNPE | PACSIN2; PACS | Q9UNF0; A0 | PACN2_HUMAI | Unmodified | 12068000 | 8216000 | 12784000 | 8758700 | 0 | 0 | 0 | 0 | 0 | 0 | 0 | 1461000 | 4281900 | 1427000 | 16309000 | 12187000 | 0 | 0 |
| ADPSLNPE | PACSIN2; PACS | Q9UNF0; A0 | PACN2_HUMAI | Unmodified | 12582000 | 6312300 | 7880800 | 5283100 | 327550 | 115920 | 0 | 29107 | 262340 | 0 | 0 | 0 | 2073100 | 565630 | 13904000 | 13678000 | 196460 | 127790 |
| ADPVLYKD | EXT2 | Q93063 | EXT2_HUMAN | Unmodified | 0 | 0 | 0 | 0 | 0 | 0 | 149720 | 0 | 4644700 | 0 | 430960 | 1290200 | 0 | 0 | 0 | 0 | 46123 | 180040 |
| ADQAAGGI | COL2A1 | P02458 | CO2A1_HUMAI | Unmodified | 0 | 0 | 0 | 30547 | 0 | 0 | 0 | 0 | 255900 | 57094 | 6823200 | 113930 | 0 | 0 | 0 | 0 | 0 | 0 |
| ADQCYEDV | CORO1A; COR | P31146; H3E | COR1A_HUMA | Unmodified | 107680 | 106390 | 0 | 0 | 0 | 0 | 0 | 0 | 0 | 0 | 0 | 2163500 | 0 | 0 | 155770 | 0 | 0 | 0 |
| ADQEEQIH | LIMA1; LIMA1; L | Q9UHB6; F8 | LIMA1_HUMAN | Unmodified | 0 | 96930 | 0 | 447290 | 0 | 0 | 23814 | 150680 | 130800 | 0 | 0 | 0 | 0 | 0 | 828100 | 813990 | 14092 | 27628 |
| ADQELMTY | GNB1; GNB1 | P62873; B3K | GBB1_HUMAN | Unmodified | 0 | 0 | 0 | 0 | 0 | 0 | 0 | 0 | 2650700 | 0 | 0 | 0 | 0 | 0 | 0 | 0 | 0 | 0 |
| ADQELMTY | GNB1; GNB1 | P62873; B3K | GBB1_HUMAN | Oxidation (M) | 0 | 1122500 | 389100 | 0 | 0 | 0 | 0 | 0 | 2449000 | 660650 | 2358300 | 10618000 | 0 | 0 | 0 | 1281100 | 0 | 0 |
| ADQIETQQ | EHD1; EHD3; E | Q9H4M9; Q9 | EHD1_HUMAN | Unmodified | 69057 | 403010 | 325090 | 0 | 0 | 0 | 0 | 0 | 200290 | 0 | 0 | 3342200 | 0 | 0 | 704750 | 556090 | 0 | 0 |
| ADQLADES | SNAP25; SNAP2 | P60880; A0A | SNP25_HUMAN | Unmodified | 688550 | 357250 | 200720 | 0 | 884910 | 0 | 0 | 0 | 1529000 | 0 | 0 | 0 | 0 | 0 | 0 | 0 | 329520 | 0 |
| ADQLTEEQ | CALM2; CALM3 | P0DP24; P0I | CALM2_HUMAI | Unmodified | 4772800 | 5712800 | 3466800 | 1453000 | 4256100 | 880250 | 0 | 1102600 | 5889600 | 1096900 | 9454200 | 18161000 | 932500 | 882420 | 9928800 | 5593600 | 1003700 | 509030 |
| ADQLTEEQ | CALM2; CALM3 | P0DP24; P0I | CALM2_HUMAI | Acetyl (Protein N | 0 | 1688500 | 0 | 920770 | 16615000 | 1688800 | 1868100 | 63905 | 45265 | 2508900 | 0 | 0 | 8613700 | 5914400 | 0 | 0 | 15170000 | 10830000 |
| ADQLTEEQ | CALML3 | P27482 | CALL3_HUMAN | Acetyl (Protein N | 0 | 0 | 0 | 0 | 0 | 0 | 0 | 0 | 791850 | 146440 | 0 | 0 | 0 | 0 | 0 | 0 | 0 | 0 |
| ADQVDTQ( | EHD4 | Q9H223 | EHD4_HUMAN | Unmodified | 334280 | 215290 | 0 | 0 | 0 | 0 | 0 | 0 | 0 | 0 | 0 | 0 | 0 | 0 | 0 | 527010 | 0 | 0 |
| ADRAGVLA | GNB1; GNB1 | P62873; B3K | GBB1_HUMAN | Unmodified | 0 | 828910 | 373280 | 0 | 0 | 0 | 0 | 0 | 14748000 | 0 | 3405900 | 11546000 | 16249 | 0 | 0 | 0 | 0 | 0 |
| ADRAGVLA | GNB1; GNB1 | P62873; B3K | GBB1_HUMAN | Oxidation (M) | 248370 | 286800 | 0 | 1157000 | 34328000 | 1705900 | 13923000 | 0 | 4503900 | 8820700 | 1864000 | 0 | 10716000 | 288470 | 236820 | 0 | 23858000 | 35321000 |
| ADRAGVLA | GNB1; GNB1 | P62873; B3K | GBB1_HUMAN | Unmodified | 0 | 0 | 0 | 0 | 6905800 | 1485600 | 4288300 | 0 | 8725700 | 3296300 | 0 | 0 | 0 | 3876500 | 0 | 0 | 8006700 | 16962000 |
| ADRDESSP | RPS14; RPS14; | P62263; E5R | RS14_HUMAN | Unmodified | 608310 | 0 | 0 | 0 | 0 | 0 | 0 | 0 | 0 | 0 | 0 | 4657300 | 0 | 0 | 261250 | 396340 | 0 | 0 |
| ADRDESSP | RPS14; RPS14; | P62263; E5R | RS14_HUMAN | Unmodified | 0 | 0 | 0 | 0 | 0 | 0 | 640980 | 411690 | 0 | 0 | 0 | 0 | 0 | 0 | 0 | 0 | 0 | 0 |

| | | | | | | | | | | | | | | | | | | | | | | |
|---|---|---|---|---|---|---|---|---|---|---|---|---|---|---|---|---|---|---|---|---|---|---|
| ADRETVAVI | EIF5B; EIF5B | O60841; A0A | IF2P_HUMAN E | Unmodified | 279760 | 0 | 285310 | 0 | 230810 | 0 | 0 | 0 | 0 | 0 | 273690 | 1213700 | 0 | 0 | 374320 | 200250 | 153090 | 0 |
| ADRFLFEM | DAAM1 | Q9Y4D1 | DAAM1_HUMA | Unmodified | 65695 | 59700 | 0 | 0 | 330750 | 0 | 0 | 0 | 0 | 0 | 0 | 920610 | 0 | 0 | 0 | 0 | 0 | 0 |
| ADRQALNE | APLP1; APLP1; | P51693; B7Z | APLP1_HUMAN | Unmodified | 0 | 0 | 0 | 0 | 0 | 0 | 0 | 0 | 2564000 | 563970 | 0 | 0 | 0 | 0 | 0 | 0 | 0 | 0 |
| ADRQALNE | APLP1; APLP1; | P51693; B7Z | APLP1_HUMAN | Unmodified | 0 | 0 | 0 | 0 | 0 | 0 | 0 | 0 | 2739600 | 944950 | 0 | 0 | 0 | 0 | 0 | 0 | 0 | 0 |
| ADRTTAAH | ASNS | P08243 | ASNS_HUMAN | Unmodified | 0 | 74248 | 0 | 0 | 0 | 0 | 0 | 0 | 0 | 0 | 0 | 1420700 | 0 | 0 | 51724 | 136640 | 0 | 0 |
| ADSAVSQE | PACSIN3; PACS | Q9UKS6; E9I | PACN3_HUMAI | Unmodified | 4568800 | 4293700 | 1738200 | 2531300 | 970660 | 0 | 20214 | 12797 | 193820 | 0 | 517350 | 0 | 4652200 | 582100 | 2331100 | 1883600 | 171850 | 0 |
| ADSAVSQE | PACSIN3; PACS | Q9UKS6; E9I | PACN3_HUMAI | Unmodified | 28427000 | 19892000 | 12263000 | 9871000 | 0 | 0 | 0 | 0 | 1306300 | 0 | 397710 | 0 | 9082900 | 0 | 20147000 | 11456000 | 0 | 3816600 |
| ADSELQLVI | APRT; APRT; AP | P07741; H3E | APT_HUMAN A | Acetyl (Protein N | 0 | 0 | 0 | 0 | 1713500 | 0 | 0 | 0 | 0 | 0 | 543900 | 2385800 | 0 | 0 | 0 | 0 | 0 | 753410 |
| ADSELQLVI | APRT; APRT; AP | P07741; H3E | APT_HUMAN A | Acetyl (Protein N | 0 | 0 | 0 | 0 | 1152000 | 435770 | 165700 | 50556 | 0 | 0 | 0 | 0 | 0 | 670430 | 0 | 0 | 1035900 | 871690 |
| ADSGTAGC | PPP1R14B | Q96C90 | PP14B_HUMAN | Acetyl (Protein N | 319980 | 1273200 | 835930 | 1333600 | 2208800 | 0 | 0 | 0 | 559930 | 0 | 5460400 | 8385700 | 5214200 | 1486200 | 1375600 | 637390 | 2233600 | 2392700 |
| ADSPSIDYA | VAC14 | Q08AM6 | VAC14_HUMAN | Unmodified | 0 | 0 | 0 | 0 | 0 | 0 | 0 | 42954 | 0 | 0 | 0 | 428010 | 0 | 59003 | 0 | 0 | 0 | 0 |
| ADSVVVLL( | UPF1 | Q92900 | RENT1_HUMAN | Unmodified | 428390 | 149140 | 0 | 278140 | 0 | 0 | 0 | 0 | 0 | 0 | 0 | 0 | 0 | 0 | 1760400 | 0 | 0 | 0 |
| ADSYYEYLI | MAN1B1; MAN1 | Q9UKM7; H( | MA1B1_HUMAI | Unmodified | 0 | 0 | 0 | 0 | 0 | 0 | 0 | 0 | 5732100 | 510440 | 0 | 115740 | 0 | 0 | 0 | 0 | 0 | 0 |
| ADTDGGLII | MELTF | P08582 | TRFM_HUMAN | Unmodified | 0 | 0 | 0 | 0 | 0 | 0 | 64969 | 0 | 2200700 | 244220 | 702810 | 879670 | 0 | 0 | 0 | 0 | 0 | 0 |
| ADTDGGLII | MELTF | P08582 | TRFM_HUMAN | Unmodified | 0 | 0 | 0 | 0 | 0 | 651640 | 0 | 0 | 166500 | 238440 | 0 | 0 | 0 | 0 | 0 | 0 | 0 | 511000 |
| ADTIAKNEL | SEPTIN11; SEPT | Q9NVA2; D6 | SEP11_HUMAN | Unmodified | 0 | 0 | 0 | 0 | 0 | 0 | 0 | 0 | 0 | 0 | 0 | 0 | 0 | 0 | 0 | 0 | 0 | 0 |
| ADTLTLEER | SEPTIN9; SEPTI | Q9UHD8; K7 | SEPT9_HUMAN | Unmodified | 0 | 96545 | 0 | 0 | 0 | 0 | 0 | 1505100 | 581870 | 285540 | 476270 | 7120400 | 77301 | 0 | 647020 | 606540 | 0 | 0 |
| ADTLTLEER | SEPTIN9; SEPTI | Q9UHD8; K7 | SEPT9_HUMAN | Unmodified | 0 | 0 | 0 | 0 | 1664600 | 308430 | 180480 | 0 | 0 | 203150 | 0 | 0 | 446240 | 266530 | 0 | 0 | 1530200 | 1583700 |
| ADTLTPEEC | SEPTIN7; SEPTI | Q16181; E7E | SEPT7_HUMAN | Unmodified | 0 | 353000 | 1197000 | 1966400 | 3429600 | 5219500 | 1493700 | 686650 | 0 | 495160 | 3435300 | 3671800 | 383480 | 6016100 | 1535800 | 0 | 1438100 | 1295100 |
| ADTPSLGE | STXBP2; STXBP | Q15833; M0I | STXB2_HUMAN | Unmodified | 120250 | 152060 | 0 | 345040 | 0 | 0 | 0 | 0 | 296600 | 0 | 483290 | 2205800 | 220420 | 0 | 0 | 371110 | 0 | 0 |
| ADTQTYQP | ERH | P84090 | ERH_HUMAN E | Unmodified | 0 | 0 | 0 | 0 | 0 | 0 | 0 | 0 | 189900 | 0 | 0 | 2143000 | 0 | 0 | 0 | 0 | 0 | 0 |
| ADTQYILPN | DPP3; DPP3; DI | Q9NY33; G3 | DPP3_HUMAN | Acetyl (Protein N | 0 | 0 | 0 | 0 | 0 | 0 | 0 | 0 | 0 | 98738 | 0 | 287220 | 40299 | 0 | 0 | 0 | 0 | 0 |
| ADTVSKTEI | SEPTIN10; SEPT | Q9P0V9; B5I | SEP10_HUMAN | Unmodified | 0 | 0 | 0 | 0 | 7531200 | 1654200 | 242370 | 0 | 0 | 0 | 0 | 0 | 542100 | 239120 | 0 | 0 | 1537100 | 1398800 |
| ADVDVQPY | GTPBP4 | Q9BZE4 | GTPB4_HUMAN | Unmodified | 0 | 0 | 0 | 0 | 433980 | 0 | 0 | 561590 | 0 | 0 | 0 | 296290 | 0 | 0 | 0 | 0 | 0 | 0 |
| ADVDVSGF | AHNAK | Q09666 | AHNK_HUMAN | Unmodified | 0 | 0 | 0 | 0 | 1883000 | 205600 | 0 | 0 | 0 | 0 | 0 | 0 | 0 | 0 | 0 | 0 | 174060 | 1641700 |
| ADVDVSGF | AHNAK | Q09666 | AHNK_HUMAN | Oxidation (M) | 0 | 0 | 0 | 0 | 785480 | 0 | 0 | 0 | 0 | 0 | 0 | 0 | 0 | 0 | 0 | 0 | 0 | 0 |
| ADVEDGEE | RNF7; RNF7 | Q9UBF6; F8V | RBX2_HUMAN | Acetyl (Protein N | 0 | 0 | 0 | 109520 | 418600 | 0 | 0 | 0 | 0 | 0 | 116920 | 0 | 541250 | 0 | 0 | 153040 | 855830 | 1360400 |
| ADVFHAYL | CAND1; CAND1 | Q86VP6; A0A | CAND1_HUMA | Unmodified | 0 | 0 | 114970 | 0 | 0 | 0 | 0 | 0 | 605710 | 267280 | 248010 | 877590 | 0 | 0 | 522000 | 178830 | 0 | 0 |
| ADVFPGNE | PRKCA; PRKCA | P17252; J3KI | KPCA_HUMAN | Acetyl (Protein N | 0 | 0 | 0 | 0 | 7014900 | 0 | 4080900 | 0 | 0 | 198430 | 0 | 0 | 2160000 | 0 | 0 | 0 | 2049000 | 7196000 |
| ADVGDALE | CHPF2; CHPF2 | Q9P2E5; G5I | CHPF2_HUMAI | Unmodified | 0 | 0 | 73602 | 0 | 0 | 0 | 0 | 0 | 983620 | 0 | 0 | 0 | 0 | 0 | 0 | 0 | 0 | 0 |
| ADVGGEAA | BTBD17 | A6NE02 | BTBDH_HUMAI | Unmodified | 0 | 0 | 0 | 0 | 0 | 0 | 0 | 73096 | 1386200 | 0 | 0 | 0 | 0 | 0 | 0 | 0 | 0 | 0 |
| ADVLDLHE | RBM8A | Q9Y5S9 | RBM8A_HUMAI | Acetyl (Protein N | 1351100 | 0 | 0 | 0 | 0 | 0 | 0 | 228700 | 0 | 362430 | 0 | 0 | 1377300 | 0 | 0 | 726860 | 0 | 0 |
| ADVLDLHE | RBM8A | Q9Y5S9 | RBM8A_HUMAI | Acetyl (Protein N | 0 | 0 | 0 | 0 | 1015500 | 0 | 774130 | 0 | 0 | 563370 | 0 | 0 | 0 | 730680 | 0 | 0 | 0 | 0 |
| ADVQHLED | CEP55; CEP55 | Q53EZ4; H0 | CEP55_HUMAN | Unmodified | 232660 | 380140 | 379190 | 1080900 | 6125800 | 1623300 | 0 | 0 | 3581900 | 1332700 | 397980 | 284810 | 707010 | 314650 | 672540 | 286770 | 112990 | 600560 |
| ADVQSIIGL | TRIM28; TRIM28 | Q13263; M0I | TIF1B_HUMAN | Unmodified | 0 | 0 | 0 | 0 | 0 | 0 | 0 | 0 | 0 | 0 | 0 | 1442700 | 0 | 0 | 0 | 0 | 0 | 0 |
| ADVTEWR | MELTF | P08582 | TRFM_HUMAN | Unmodified | 0 | 0 | 0 | 0 | 0 | 0 | 0 | 0 | 1325500 | 80901 | 0 | 203720 | 0 | 0 | 0 | 0 | 0 | 0 |
| ADVVESWI | SPTAN1; SPTAN | Q13813; A0A | SPTN1_HUMAN | Unmodified | 0 | 0 | 0 | 0 | 0 | 0 | 0 | 0 | 0 | 0 | 0 | 1159600 | 0 | 0 | 0 | 0 | 0 | 0 |
| ADVYRDILA | TJP2; TJP2; TJP2 | Q9UDY2; A0 | ZO2_HUMAN T | Unmodified | 0 | 0 | 0 | 0 | 0 | 0 | 0 | 0 | 1757200 | 0 | 0 | 0 | 0 | 0 | 0 | 0 | 0 | 0 |
| ADYDTLSLI | PKP3; PKP3; PK | Q9Y446; E9F | PKP3_HUMAN | Unmodified | 180730 | 0 | 977460 | 1008700 | 0 | 0 | 277580 | 517330 | 464420 | 0 | 0 | 2523000 | 0 | 0 | 1863500 | 1083100 | 207820 | 0 |
| AEAAALTET | ABCC4 | O15439 | MRP4_HUMAN | Unmodified | 0 | 0 | 2949200 | 0 | 0 | 0 | 0 | 0 | 329480 | 53610 | 21556 | 362400 | 0 | 0 | 1804700 | 1426600 | 0 | 0 |
| AEAAASALA | CHMP2A; CHM | O43633; M0I | CHM2A_HUMA | Unmodified | 1558400 | 1833000 | 0 | 8119200 | 7896400 | 3519600 | 0 | 0 | 4109900 | 2091000 | 0 | 2298900 | 3044400 | 1767200 | 3936500 | 1731700 | 0 | 2385200 |
| AEAAGHRD | TNFRSF10B | O14763 | TR10B_HUMAN | Unmodified | 0 | 0 | 104920 | 0 | 2660200 | 294060 | 0 | 0 | 2149500 | 61740 | 121960 | 1545500 | 0 | 213610 | 243790 | 122430 | 51735 | 311640 |
| AEAALLLLP | COG3 | Q96JB2 | COG3_HUMAN | Acetyl (Protein N | 11894 | 0 | 0 | 0 | 0 | 0 | 0 | 0 | 0 | 0 | 0 | 234990 | 0 | 0 | 0 | 0 | 0 | 0 |
| AEAALLLLP | COG3 | Q96JB2 | COG3_HUMAN | Acetyl (Protein N | 0 | 0 | 37773 | 0 | 0 | 0 | 0 | 0 | 0 | 0 | 0 | 382140 | 1233900 | 1736000 | 0 | 0 | 0 | 0 |
| AEAARIEGE | MVP | Q14764 | MVP_HUMAN N | Unmodified | 1011600 | 886990 | 0 | 0 | 0 | 0 | 0 | 0 | 1916300 | 0 | 0 | 20255000 | 0 | 0 | 0 | 0 | 0 | 0 |
| AEAARQK | HNRNPA3 | A0A7I2V4G0 | A0A7I2V4G0_H | Unmodified | 0 | 82230 | 0 | 0 | 0 | 0 | 0 | 0 | 0 | 0 | 0 | 0 | 0 | 0 | 61246 | 0 | 0 | 0 |
| AEAAVESA | SHPK; N/A | Q9UHJ6; A0A | SHPK_HUMAN | Unmodified | 0 | 0 | 0 | 0 | 0 | 0 | 0 | 0 | 0 | 0 | 0 | 338020 | 0 | 0 | 0 | 0 | 0 | 0 |
| AEADMLVL | N/A | N/A | N/A | Oxidation (M) | 0 | 0 | 0 | 0 | 0 | 0 | 5253400 | 0 | 0 | 0 | 0 | 0 | 0 | 3945500 | 0 | 0 | 0 | 0 |
| AEADRIVFE | DCPS; DCPS | Q96C86; A0A | DCPS_HUMAN | Unmodified | 0 | 0 | 0 | 0 | 0 | 0 | 0 | 0 | 0 | 0 | 60600 | 209650 | 0 | 0 | 0 | 0 | 0 | 0 |
| AEAELLQQ | PLEC | Q15149 | PLEC_HUMAN | Unmodified | 0 | 0 | 0 | 0 | 230290 | 0 | 0 | 0 | 0 | 0 | 0 | 332840 | 0 | 0 | 0 | 0 | 318260 | 235070 |
| AEAESRAE | MVP | Q14764 | MVP_HUMAN N | Unmodified | 454400 | 0 | 0 | 0 | 516230 | 0 | 0 | 0 | 756650 | 0 | 0 | 2168900 | 0 | 0 | 0 | 0 | 0 | 0 |
| AEAETLVSF | CHDH | Q8NE62 | CHDH_HUMAN | Unmodified | 0 | 0 | 0 | 0 | 0 | 0 | 0 | 0 | 1808700 | 57958 | 27162 | 0 | 0 | 0 | 0 | 0 | 0 | 0 |
| AEAGDNLC | TUFM | P49411 | EFTU_HUMAN | Unmodified | 0 | 0 | 0 | 0 | 0 | 0 | 0 | 0 | 6607400 | 0 | 0 | 0 | 0 | 0 | 0 | 0 | 0 | 0 |
| AEAGPEGV | HYOU1; HYOU1 | Q9Y4L1; A0A | HYOU1_HUMA | Unmodified | 130240 | 0 | 483320 | 176510 | 0 | 0 | 19745 | 0 | 26337000 | 1934900 | 1809000 | 1427000 | 0 | 0 | 735420 | 212880 | 0 | 0 |
| AEAGPEGV | HYOU1; HYOU1 | Q9Y4L1; A0A | HYOU1_HUMA | Unmodified | 175260 | 0 | 0 | 511840 | 0 | 0 | 199820 | 52916 | 35617000 | 6080900 | 7055200 | 1773600 | 319880 | 611380 | 925330 | 507670 | 0 | 0 |
| AEAGRPYE | EFNB1 | P98172 | EFNB1_HUMAN | Unmodified | 306650 | 0 | 0 | 248290 | 42123 | 0 | 0 | 0 | 135810 | 287530 | 0 | 321580 | 0 | 0 | 261310 | 0 | 0 | 321240 |
| AEAGSGGF | DNAJB2; DNAJE | P25686; C9J | DNJB2_HUMAN | Unmodified | 729470 | 840720 | 0 | 0 | 0 | 0 | 0 | 0 | 0 | 0 | 0 | 0 | 0 | 0 | 734790 | 133830 | 0 | 0 |
| AEAGVPAE | FLNA; FLNA; FL | P21333; A0A | FLNA_HUMAN | Unmodified | 1729100 | 0 | 4331100 | 4426300 | 0 | 0 | 0 | 0 | 0 | 0 | 0 | 4882100 | 0 | 0 | 3479800 | 5173900 | 0 | 0 |
| AEALPLQR | ITGA6 | P23229 | ITA6_HUMAN I | Unmodified | 0 | 0 | 0 | 274390 | 0 | 344230 | 64726 | 0 | 160290 | 0 | 678450 | 4433900 | 0 | 0 | 1303200 | 1400400 | 97447 | 359700 |
| AEALTEPLN | CMTM6 | Q9NX76 | CKLF6_HUMAN | Unmodified | 52777 | 875050 | 0 | 0 | 377460 | 0 | 0 | 0 | 2759500 | 765300 | 3015600 | 6320800 | 0 | 0 | 0 | 64801 | 295280 | 0 |
| AEAQGSTT | NUP210 | Q8TEM1 | PO210_HUMAN | Unmodified | 0 | 0 | 0 | 0 | 0 | 0 | 49556 | 0 | 768620 | 0 | 0 | 0 | 0 | 0 | 0 | 0 | 0 | 0 |
| AEASIQAGA | AHNAK | Q09666 | AHNK_HUMAN | Unmodified | 0 | 0 | 0 | 0 | 5731300 | 1668300 | 0 | 0 | 0 | 0 | 0 | 0 | 130630 | 0 | 0 | 0 | 693600 | 2781700 |
| AEASIQAGA | AHNAK | Q09666 | AHNK_HUMAN | Unmodified | 0 | 0 | 0 | 0 | 1456900 | 0 | 0 | 0 | 0 | 0 | 0 | 0 | 0 | 0 | 0 | 0 | 890440 | 387030 |
| AEASIQAGA | AHNAK | Q09666 | AHNK_HUMAN | Unmodified | 0 | 0 | 0 | 396220 | 734730 | 0 | 0 | 0 | 0 | 0 | 0 | 0 | 0 | 0 | 0 | 208370 | 835190 | 833550 |
| AEASSGDH | FKBP4 | Q02790 | FKBP4_HUMAN | Oxidation (M) | 0 | 0 | 129080 | 0 | 0 | 0 | 0 | 0 | 29215 | 0 | 0 | 758930 | 0 | 0 | 145410 | 274120 | 0 | 0 |
| AEATNTGK | KIT | P10721 | KIT_HUMAN M | Unmodified | 0 | 0 | 0 | 0 | 0 | 0 | 0 | 0 | 0 | 45609 | 0 | 6922400 | 0 | 0 | 0 | 0 | 0 | 0 |
| AECLAMLG | DNAJC7; DNAJ( | Q99615; A0A | DNJC7_HUMAI | Unmodified | 0 | 0 | 0 | 0 | 0 | 0 | 0 | 0 | 0 | 0 | 0 | 0 | 118780 | 0 | 0 | 0 | 0 | 0 |
| AEDADMRI | SNAP25; SNAP2 | P60880; A0A | SNP25_HUMAN | Acetyl (Protein N | 0 | 0 | 0 | 0 | 0 | 0 | 0 | 0 | 0 | 0 | 0 | 0 | 0 | 0 | 0 | 0 | 0 | 0 |
| AEDAPLSS | SARG | Q9BW04 | SARG_HUMAN | Unmodified | 0 | 0 | 0 | 0 | 0 | 0 | 0 | 0 | 0 | 0 | 0 | 424450 | 0 | 65577 | 0 | 302940 | 0 | 0 |
| AEDFGNRL | MAN1B1; MAN1 | Q9UKM7; H( | MA1B1_HUMAI | Unmodified | 0 | 0 | 0 | 0 | 0 | 0 | 0 | 0 | 1004300 | 0 | 0 | 0 | 0 | 0 | 0 | 0 | 0 | 0 |
| AEDGATPS | MARCKS | P29966 | MARCS_HUMA | Unmodified | 1197600 | 1069600 | 2752600 | 2047800 | 212690 | 0 | 396330 | 323700 | 62183000 | 6748400 | 6885000 | 27386000 | 1010500 | 452390 | 3143000 | 3216000 | 733260 | 3468600 |
| AEDGATPS | MARCKS | P29966 | MARCS_HUMA | Unmodified | 1793000 | 777520 | 1754400 | 4692000 | 0 | 0 | 0 | 181320 | 38545000 | 0 | 4757400 | 23679000 | 0 | 0 | 13009000 | 2467600 | 1680300 | 2600700 |
| AEDGATPS | MARCKS | P29966 | MARCS_HUMA | Unmodified | 0 | 0 | 0 | 0 | 0 | 0 | 296710 | 32937 | 0 | 219510 | 0 | 0 | 0 | 44827 | 0 | 0 | 63489 | 438910 |
| AEDGSVID | ANXA2; ANXA2; | P07355; H0 | ANXA2_HUMAN | Unmodified | 357860 | 114250 | 40263 | 105450 | 801380 | 147040 | 719040 | 93150 | 4205000 | 0 | 0 | 83025 | 0 | 0 | 114870 | 44203 | 306350 | 514560 |
| AEDGSVID | ANXA2; ANXA2; | P07355; H0 | ANXA2_HUMAN | Unmodified | 413520 | 0 | 0 | 0 | 724640 | 404090 | 0 | 0 | 3604100 | 0 | 0 | 0 | 0 | 0 | 0 | 0 | 0 | 0 |
| AEDKQKLE | TM9SF2 | Q99805 | TM9S2_HUMAN | Unmodified | 0 | 0 | 0 | 0 | 0 | 0 | 0 | 0 | 0 | 837030 | 0 | 0 | 0 | 0 | 0 | 0 | 25879 | 0 |
| AEDMYSAC | GPRC5C; GPRC | Q9NQ84; A8 | GPC5C_HUMA | Unmodified | 0 | 0 | 493010 | 306430 | 0 | 0 | 15719 | 0 | 0 | 0 | 0 | 0 | 0 | 0 | 514180 | 561020 | 0 | 0 |
| AEDMYSAC | GPRC5C; GPRC | Q9NQ84; A8 | GPC5C_HUMA | Oxidation (M) | 0 | 0 | 323370 | 0 | 0 | 0 | 0 | 0 | 0 | 0 | 0 | 0 | 1011500 | 489880 | 350580 | 514150 | 0 | 0 |
| AEDNFEDII | DYNC1LI1; DYN | Q9Y6G9; E9I | DC1L1_HUMAN | Unmodified | 0 | 0 | 299220 | 127700 | 0 | 0 | 0 | 0 | 452630 | 0 | 331050 | 1975800 | 0 | 0 | 298410 | 249230 | 0 | 0 |
| AEDSGEYH | NPTN; NPTN | Q9Y639; Q9I | NPTN_HUMAN | Unmodified | 1772600 | 679550 | 3629600 | 3604800 | 1433400 | 349030 | 0 | 0 | 11269000 | 289350 | 6415700 | 39170000 | 1998100 | 0 | 5734200 | 4676700 | 190790 | 1239600 |
| AEDSSKVS | ADD3 | Q9UEY8 | ADDG_HUMAN | Unmodified | 0 | 0 | 0 | 0 | 0 | 0 | 0 | 92216 | 0 | 0 | 0 | 0 | 921160 | 1196000 | 0 | 0 | 720400 | 579080 |
| AEDYEENL | KIF23; KIF23; KI | Q02241; A0A | KIF23_HUMAN | Unmodified | 0 | 0 | 0 | 0 | 0 | 4409600 | 0 | 0 | 1637500 | 0 | 0 | 1066100 | 0 | 0 | 257170 | 0 | 0 | 0 |
| AEDYEENL | KIF23; KIF23; KI | Q02241; A0A | KIF23_HUMAN | Oxidation (M) | 0 | 0 | 0 | 0 | 0 | 0 | 0 | 0 | 722140 | 0 | 0 | 0 | 0 | 0 | 0 | 0 | 0 | 0 |
| AEDYEENL | KIF23; KIF23; KI | Q02241; A0A | KIF23_HUMAN | Oxidation (M) | 0 | 0 | 0 | 0 | 1404600 | 0 | 0 | 0 | 0 | 0 | 0 | 0 | 0 | 0 | 0 | 0 | 0 | 0 |
| AEDYEENL | KIF23; KIF23; KI | Q02241; A0A | KIF23_HUMAN | Unmodified | 0 | 0 | 0 | 0 | 809530 | 0 | 0 | 0 | 0 | 0 | 0 | 0 | 0 | 0 | 0 | 0 | 0 | 0 |
| AEDYGEHI | LLGL2; LLGL2 | Q6P1M3; J3( | L2GL2_HUMAN | Unmodified | 0 | 0 | 0 | 0 | 0 | 0 | 0 | 0 | 0 | 0 | 0 | 0 | 0 | 0 | 414790 | 0 | 0 | 0 |
| AEEASHWL | PLXNB2 | O15031 | PLXB2_HUMAN | Unmodified | 540000 | 296510 | 1000300 | 1058300 | 100530 | 0 | 0 | 0 | 953320 | 0 | 0 | 860200 | 0 | 0 | 1677900 | 1655500 | 0 | 0 |
| AEEDEDKE | ZDHHC5; ZDHI | Q9C0B5; H0 | ZDHC5_HUMA | Unmodified | 0 | 0 | 193310 | 0 | 0 | 0 | 0 | 0 | 0 | 184120 | 0 | 1750600 | 0 | 0 | 331760 | 247950 | 0 | 0 |
| AEEDEDKE | ZDHHC5; ZDHI | Q9C0B5; H0 | ZDHC5_HUMA | Unmodified | 1009700 | 507510 | 1709100 | 0 | 0 | 0 | 0 | 0 | 232080 | 73813 | 0 | 19530000 | 0 | 0 | 2606100 | 3308100 | 48555 | 0 |
| AEEDEILNF | CANX; CANX; C | P27824; A0A | CALX_HUMAN | Unmodified | 536480 | 0 | 131010 | 109170 | 0 | 0 | 0 | 0 | 0 | 0 | 630760 | 0 | 0 | 0 | 728420 | 713010 | 0 | 0 |
| AEEDKIPLL | RALB; RALB; RA | P11234; C9J | RALB_HUMAN | Unmodified | 244290 | 173200 | 906390 | 168590 | 620810 | 110970 | 0 | 0 | 0 | 0 | 1214000 | 17049000 | 0 | 211680 | 2613200 | 1082200 | 0 | 0 |
| AEEDKIPLL | RALB; RALB; RA | P11234; C9J | RALB_HUMAN | Unmodified | 361350 | 0 | 0 | 270810 | 0 | 0 | 0 | 0 | 0 | 0 | 945760 | 2697600 | 0 | 0 | 889350 | 1136800 | 0 | 0 |
| AEEDKLEQ | P4HA1 | P13674 | P4HA1_HUMAN | Unmodified | 0 | 0 | 0 | 0 | 0 | 0 | 22434 | 0 | 1076600 | 0 | 1194100 | 0 | 0 | 0 | 0 | 0 | 0 | 0 |
| AEEDVEPE | DBNL; DBNL; D | Q9UJU6; B4I | DBNL_HUMAN | Unmodified | 0 | 0 | 0 | 0 | 0 | 0 | 0 | 0 | 0 | 0 | 0 | 1261800 | 0 | 0 | 0 | 0 | 0 | 0 |
| AEEEVAKLE | ANKFY1; ANKF | Q9P2R3; I3L | ANFY1_HUMAN | Acetyl (Protein N | 75114 | 0 | 0 | 506490 | 3098900 | 907650 | 808390 | 452440 | 1742100 | 193480 | 193750 | 823090 | 618890 | 469000 | 0 | 246530 | 546330 | 775240 |
| AEEGIAAG | RPS12 | P25398 | RS12_HUMAN | Acetyl (Protein N | 0 | 323110 | 0 | 0 | 368120 | 0 | 174860 | 0 | 0 | 0 | 0 | 0 | 583750 | 117210 | 312590 | 0 | 0 | 0 |
| AEEIDAKAE | ATP6V1E2; ATP | Q96A05; P3 | VATE2_HUMAN | Unmodified | 71310 | 0 | 254300 | 64531 | 0 | 0 | 0 | 0 | 0 | 0 | 0 | 1553900 | 0 | 0 | 263340 | 216800 | 0 | 0 |
| AEEILEK | RPL11; EMP3; E | P62913; P54 | RL11_HUMAN ( | Unmodified | 0 | 2139700 | 3790700 | 0 | 0 | 0 | 0 | 0 | 385030 | 0 | 2007400 | 15925000 | 0 | 0 | 4707400 | 0 | 0 | 0 |
| AEELGLPIL | ACAA1; ACAA1; | P09110; C9J | THIK_HUMAN 3 | Unmodified | 0 | 0 | 61688 | 0 | 0 | 0 | 0 | 0 | 10409000 | 370680 | 765940 | 259950 | 0 | 0 | 0 | 0 | 0 | 0 |
| AEELHYPL( | RCN2; RCN2; R | Q14257; H0 | RCN2_HUMAN | Unmodified | 0 | 0 | 0 | 0 | 262580 | 0 | 61568 | 0 | 1753400 | 367590 | 0 | 0 | 0 | 0 | 0 | 0 | 145290 | 828420 |
| AEELVNTA | TMEM132A | Q24JP5 | T132A_HUMAN | Unmodified | 230350 | 366000 | 0 | 214870 | 0 | 0 | 0 | 0 | 0 | 0 | 0 | 455770 | 0 | 0 | 457910 | 201070 | 0 | 0 |
| AEEMDTED | EHBP1L1; EHBI | Q8N3D4; E9 | EH1L1_HUMAN | Unmodified | 26787 | 0 | 0 | 0 | 0 | 0 | 0 | 0 | 295400 | 0 | 0 | 0 | 0 | 0 | 0 | 0 | 0 | 0 |
| AEEQPQVE | CLIC1 | O00299 | CLIC1_HUMAN | Acetyl (Protein N | 0 | 0 | 0 | 4611900 | 62690000 | 1829600 | 7819600 | 6972600 | 11456000 | 9484900 | 3119800 | 6217400 | 31617000 | 28349000 | 5917700 | 6550100 | 54998000 | 38206000 |
| AEEQPQVE | CLIC1 | O00299 | CLIC1_HUMAN | Acetyl (Protein N | 0 | 0 | 0 | 0 | 2513100 | 0 | 0 | 0 | 0 | 0 | 0 | 0 | 0 | 0 | 0 | 0 | 12887000 | 889230 |
| AEEVATFFA | TOP1 | P11387 | TOP1_HUMAN | Unmodified | 0 | 0 | 0 | 0 | 0 | 0 | 0 | 0 | 0 | 0 | 0 | 0 | 0 | 0 | 0 | 0 | 0 | 0 |
| AEEVELYLE | TMEM87A | Q8NBN3 | TM87A_HUMAN | Unmodified | 0 | 0 | 0 | 0 | 0 | 0 | 0 | 0 | 982070 | 332030 | 581740 | 1844800 | 0 | 0 | 0 | 0 | 0 | 0 |
| AEEVVVR | ITGB4 | P16144 | ITB4_HUMAN I | Unmodified | 0 | 0 | 21137 | 0 | 286650 | 10903 | 0 | 1116600 | 310750 | 746150 | 18548 | 103800 | 0 | 0 | 45001 | 34850 | 227790 | 2025400 |
| AEEWGVQ | RALB; RALB | P11234; C9J | RALB_HUMAN | Unmodified | 218340 | 224610 | 653740 | 477460 | 1197800 | 0 | 0 | 46780 | 0 | 0 | 1355700 | 3171500 | 500620 | 0 | 1210500 | 863180 | 0 | 381030 |
| AEEYEFLTP | ARHGDIA; ARH | P52565; J3K | GDIR1_HUMAN | Unmodified | 610040 | 0 | 564820 | 579550 | 0 | 0 | 0 | 0 | 837220 | 164770 | 1305400 | 7815100 | 0 | 0 | 1310000 | 1245400 | 0 | 0 |

| | | | | | | | | | | | | | | | | | | | | | | |
|---|---|---|---|---|---|---|---|---|---|---|---|---|---|---|---|---|---|---|---|---|---|---|
| AEFFADVVI | ENOPH1; ENOF | Q9UHY7; D6 | ENOPH_HUMA | Unmodified | 0 | 0 | 0 | 0 | 0 | 0 | 0 | 40194 | 0 | 259450 | 603960 | 0 | 623090 | 425240 | 0 | 405620 | 0 | 0 |
| AEFGPPGP | RNPEP; RNPEP; | Q9H4A4; A6I | AMPB_HUMAN | Unmodified | 0 | 0 | 0 | 0 | 0 | 0 | 0 | 0 | 494880 | 119210 | 392980 | 2800600 | 0 | 65797 | 0 | 0 | 109720 | 315610 |
| AEFLDDQE | TRAFD1; TRAFD | O14545; F8V | TRAD1_HUMAN | Acetyl (Protein N | 13428 | 800940 | 0 | 0 | 0 | 198120 | 0 | 0 | 0 | 0 | 0 | 0 | 0 | 0 | 0 | 0 | 0 | 0 |
| AEFLDSIK | N/A | N/A | N/A | Unmodified | 48504000 | 47342000 | 51907000 | 36614000 | 30754000 | 34030000 | 14468000 | 0 | 34620000 | 24036000 | 22325000 | 0 | 34173000 | 29284000 | 47125000 | 46659 | 0 | 224130 |
| AEGAATEEI | BASP1; BASP1 | P80723; U3K | BASP1_HUMAN | Unmodified | 0 | 0 | 0 | 0 | 0 | 0 | 0 | 102480 | 4424700 | 0 | 0 | 0 | 0 | 0 | 0 | 0 | 0 | 0 |
| AEGAATEEI | BASP1 | P80723 | BASP1_HUMAN | Unmodified | 0 | 0 | 0 | 0 | 1670300 | 0 | 0 | 0 | 3988500 | 800340 | 0 | 0 | 0 | 0 | 0 | 0 | 125240 | 1311800 |
| AEGAATEEI | BASP1 | P80723 | BASP1_HUMAN | Unmodified | 0 | 0 | 0 | 0 | 1760500 | 0 | 116400 | 176870 | 286000 | 0 | 0 | 0 | 0 | 0 | 0 | 0 | 0 | 0 |
| AEGAERFA | ENDOD1 | O94919 | ENDD1_HUMA | Unmodified | 0 | 0 | 0 | 0 | 0 | 0 | 0 | 0 | 342710 | 62830 | 0 | 0 | 0 | 0 | 0 | 0 | 0 | 0 |
| AEGDVAAL | TPM4; TPM4; TF | P67936; A0A | TPM4_HUMAN | Unmodified | 0 | 0 | 0 | 0 | 943340 | 308980 | 0 | 0 | 0 | 0 | 0 | 314980 | 0 | 0 | 0 | 192920 | 0 | 0 |
| AEGERQPP | PTPA; PTPA; PTI | Q15257; A6F | PTPA_HUMAN S | Acetyl (Protein N | 0 | 0 | 0 | 524600 | 0 | 0 | 0 | 0 | 883950 | 0 | 0 | 2862500 | 0 | 0 | 0 | 0 | 0 | 0 |
| AEGILDVFC | PTPRA | P18433 | PTPRA_HUMAN | Unmodified | 0 | 0 | 333220 | 0 | 0 | 83128 | 0 | 0 | 0 | 0 | 704250 | 1912100 | 0 | 0 | 947140 | 695730 | 0 | 0 |
| AEGILLLVK | PARP4 | Q9UKK3 | PARP4_HUMAN | Unmodified | 0 | 0 | 0 | 0 | 647610 | 49066 | 0 | 0 | 626190 | 83177 | 0 | 568860 | 0 | 0 | 0 | 0 | 119010 | 0 |
| AEGLGDGY | HINT2 | Q9BX68 | HINT2_HUMAN | Unmodified | 0 | 0 | 0 | 0 | 0 | 0 | 0 | 0 | 2090100 | 0 | 0 | 0 | 0 | 0 | 0 | 0 | 0 | 0 |
| AEGLGDGY | HINT2 | Q9BX68 | HINT2_HUMAN | Unmodified | 0 | 0 | 0 | 0 | 0 | 0 | 0 | 0 | 4265300 | 357760 | 0 | 0 | 0 | 0 | 0 | 0 | 0 | 0 |
| AEGNNQAI | CHGA | P10645 | CMGA_HUMAN | Unmodified | 0 | 0 | 0 | 0 | 0 | 0 | 0 | 0 | 13616000 | 2172900 | 0 | 0 | 0 | 0 | 0 | 0 | 0 | 0 |
| AEGNPKPE | PXDN; PXDN | Q92626; H7( | PXDN_HUMAN | Unmodified | 0 | 0 | 0 | 0 | 0 | 0 | 0 | 0 | 841790 | 81094 | 0 | 0 | 0 | 0 | 0 | 0 | 0 | 0 |
| AEGNSRFT | COL2A1 | P02458 | CO2A1_HUMAI | Unmodified | 0 | 0 | 0 | 0 | 0 | 0 | 0 | 0 | 0 | 0 | 1693000 | 0 | 0 | 0 | 0 | 0 | 0 | 0 |
| AEGPDVAV | AHNAK | Q09666 | AHNK_HUMAN | Unmodified | 0 | 0 | 0 | 0 | 0 | 0 | 0 | 334360 | 0 | 0 | 0 | 0 | 0 | 0 | 0 | 0 | 0 | 0 |
| AEGPEVDV | AHNAK | Q09666 | AHNK_HUMAN | Unmodified | 0 | 0 | 359780 | 0 | 940440 | 0 | 0 | 0 | 0 | 0 | 0 | 0 | 0 | 0 | 233500 | 196260 | 145870 | 368900 |
| AEGPGLSR | FLNA; FLNA; FL | P21333; A0A | FLNA_HUMAN | Unmodified | 0 | 0 | 0 | 0 | 647900 | 0 | 0 | 0 | 0 | 0 | 0 | 0 | 0 | 0 | 0 | 0 | 825820 | 0 |
| AEGPGRGI | ATAD3A; ATAD3 | Q9NVI7; Q5S | ATD3A_HUMAN | Unmodified | 0 | 0 | 0 | 0 | 0 | 0 | 167410 | 0 | 2161500 | 556380 | 0 | 0 | 0 | 0 | 0 | 0 | 0 | 0 |
| AEGQRQAC | KRT18; KRT18 | P05783; F8V | K1C18_HUMAN | Unmodified | 0 | 0 | 0 | 0 | 0 | 90037 | 0 | 0 | 0 | 0 | 805020 | 0 | 0 | 0 | 0 | 0 | 0 | 0 |
| AEGSDVAN | PKM; PKM; PKM | P14618; B4D | KPYM_HUMAN | Oxidation (M) | 0 | 0 | 0 | 0 | 0 | 0 | 0 | 197590 | 1078000 | 0 | 0 | 0 | 0 | 0 | 0 | 599170 | 0 | 0 |
| AEGSEIRLA | P4HB; P4HB; P4 | P07237; A0A | PDIA1_HUMAN | Unmodified | 0 | 0 | 0 | 0 | 0 | 0 | 0 | 0 | 0 | 665710 | 0 | 0 | 0 | 0 | 0 | 0 | 0 | 0 |
| AEGTNFRT | PDCD6IP | Q8WUM4 | PDC6I_HUMAN | Unmodified | 173490 | 0 | 0 | 0 | 928070 | 116940 | 0 | 0 | 412990 | 126640 | 0 | 3251300 | 204470 | 102100 | 0 | 0 | 437270 | 475030 |
| AEHDQLLL | LMNB1; LMNB1 | P20700; A0A | LMNB1_HUMAI | Unmodified | 0 | 0 | 0 | 0 | 0 | 0 | 0 | 21497 | 0 | 0 | 0 | 969650 | 298300 | 201530 | 0 | 0 | 0 | 0 |
| AEHDSILAE | TARS1; TARS1; | P26639; D6F | SYTC_HUMAN | Unmodified | 0 | 0 | 0 | 0 | 0 | 0 | 0 | 0 | 98555 | 0 | 218950 | 1479400 | 0 | 0 | 0 | 112720 | 0 | 0 |
| AEHDSILAE | TARS1; TARS1; | P26639; D6F | SYTC_HUMAN | Unmodified | 0 | 0 | 0 | 183900 | 578150 | 99070 | 76787 | 37273 | 85230 | 84036 | 242440 | 281090 | 0 | 0 | 213100 | 0 | 96752 | 168220 |
| AEHLDHVN | PGM3; PGM3; F | O95394; A0A | AGM1_HUMAN | Unmodified | 65703 | 0 | 0 | 0 | 109950 | 0 | 0 | 26804 | 0 | 42886 | 0 | 0 | 349710 | 0 | 0 | 0 | 0 | 128780 |
| AEHLELLAE | PPP1R16A; PPP | Q96I34; A0A | PP16A_HUMAN | Acetyl (Protein N | 0 | 0 | 0 | 0 | 0 | 0 | 0 | 14617 | 0 | 0 | 0 | 0 | 0 | 0 | 0 | 0 | 195960 | 0 |
| AEHPTWGI | PTGS1; PTGS1; | P23219; A0A | PGH1_HUMAN | Unmodified | 0 | 0 | 0 | 0 | 0 | 0 | 0 | 0 | 0 | 0 | 0 | 1125700 | 0 | 0 | 0 | 0 | 0 | 0 |
| AEIDMLDIR | ANXA4; ANXA4 | P09525; Q6F | ANXA4_HUMAI | Oxidation (M) | 0 | 140520 | 193730 | 0 | 686560 | 0 | 0 | 0 | 1031500 | 0 | 331700 | 2980600 | 80077 | 0 | 0 | 399030 | 270400 | 0 |
| AEIDMLDIR | ANXA4; ANXA4 | P09525; Q6F | ANXA4_HUMAI | Unmodified | 0 | 283110 | 0 | 0 | 0 | 0 | 0 | 0 | 6471200 | 0 | 208500 | 553180 | 0 | 0 | 0 | 0 | 0 | 0 |
| AEIIHHLAD | ALDH18A1 | P54886 | P5CS_HUMAN | Unmodified | 0 | 0 | 0 | 0 | 0 | 0 | 0 | 0 | 800330 | 0 | 0 | 0 | 0 | 0 | 0 | 0 | 0 | 0 |
| AEIITVSDG | VPS25; VPS25 | Q9BRG1; K7 | VPS25_HUMAN | Unmodified | 2329100 | 2341800 | 5611000 | 0 | 0 | 0 | 0 | 0 | 0 | 0 | 0 | 7162400 | 0 | 0 | 3588500 | 2821300 | 0 | 0 |
| AEIKEMGEI | EPCAM; EPCAM | P16422; B5N | EPCAM_HUMA | Unmodified | 0 | 0 | 0 | 0 | 0 | 0 | 0 | 0 | 177750 | 666090 | 0 | 0 | 0 | 0 | 0 | 182390 | 0 | 0 |
| AEIKEMGEI | EPCAM; EPCAM | P16422; B5N | EPCAM_HUMA | Unmodified | 0 | 0 | 0 | 293710 | 3475500 | 202240 | 0 | 0 | 1092100 | 8807300 | 79588 | 0 | 11096000 | 7381200 | 0 | 0 | 301120 | 447540 |
| AEIKEMGEI | EPCAM; EPCAM | P16422; B5N | EPCAM_HUMA | Oxidation (M) | 0 | 0 | 0 | 103530 | 594850 | 149900 | 0 | 0 | 510150 | 1880100 | 0 | 116090 | 859930 | 396410 | 0 | 0 | 0 | 61246 |
| AEIKQGIRE | SDCBP; SDCBP | O00560; G5I | SDCB1_HUMAI | Unmodified | 0 | 0 | 0 | 0 | 0 | 0 | 0 | 0 | 149460 | 762060 | 0 | 0 | 0 | 0 | 0 | 0 | 0 | 0 |
| AEITELILQF | TMPPE | Q6ZT21 | TMPPE_HUMAI | Unmodified | 0 | 0 | 0 | 0 | 0 | 0 | 0 | 0 | 2015600 | 0 | 0 | 0 | 0 | 0 | 0 | 0 | 0 | 0 |
| AEKDEPGA | GANAB; GANAE | Q14697; F5H | GANAB_HUMA | Unmodified | 0 | 0 | 0 | 0 | 0 | 0 | 0 | 0 | 29874000 | 2904000 | 5879400 | 0 | 0 | 0 | 0 | 0 | 0 | 0 |
| AEKFDCHY | FHL1; FHL1; FH | Q13642; A0A | FHL1_HUMAN | Acetyl (Protein N | 0 | 0 | 0 | 0 | 0 | 0 | 418380 | 0 | 0 | 0 | 0 | 0 | 0 | 0 | 0 | 0 | 810580 | 0 |
| AELAELPLR | VPS35 | Q96QK1 | VPS35_HUMAN | Unmodified | 0 | 0 | 0 | 0 | 0 | 0 | 0 | 0 | 90850 | 73591 | 31140 | 2622800 | 0 | 0 | 35955 | 0 | 0 | 0 |
| AELANIERE | SLK; SLK | Q9H2G2; A0 | SLK_HUMAN S | Unmodified | 0 | 81254 | 201410 | 0 | 315590 | 0 | 0 | 0 | 916460 | 93861 | 553010 | 2119100 | 230630 | 194980 | 353650 | 247840 | 0 | 128930 |
| AELDDTPM | SFPQ | P23246 | SFPQ_HUMAN | Oxidation (M) | 167970 | 304950 | 4995200 | 0 | 403440 | 2268400 | 0 | 0 | 1305600 | 290420 | 87221 | 337560 | 0 | 775000 | 0 | 0 | 0 | 2027100 |
| AELDSEPEI | SH3BP4 | Q9P0V3 | SH3B4_HUMAI | Unmodified | 0 | 0 | 83349 | 68379 | 0 | 0 | 0 | 0 | 436430 | 0 | 0 | 253550 | 0 | 30101 | 131730 | 140980 | 0 | 0 |
| AELEIQK | APRT; EVI5; EVI | P07741; O6C | APT_HUMAN A | Unmodified | 0 | 0 | 73938 | 0 | 280970 | 0 | 0 | 0 | 1036400 | 95853 | 108550 | 361670 | 30153 | 0 | 949760 | 714210 | 0 | 0 |
| AELEQHLE | CCDC6 | Q16204 | CCDC6_HUMA | Unmodified | 83576 | 0 | 0 | 0 | 0 | 0 | 0 | 0 | 1118000 | 0 | 565230 | 3722200 | 0 | 0 | 0 | 0 | 439310 | 0 |
| AELGALPDI | NLN; NLN | Q9BYT8; E9F | NEUL_HUMAN | Unmodified | 0 | 0 | 0 | 19479 | 27119 | 11997 | 0 | 0 | 221520 | 13838 | 116330 | 301280 | 0 | 0 | 0 | 13760 | 0 | 0 |
| AELGEYIR | DYNC1H1; DYN | Q14204; A0A | DYHC1_HUMA | Unmodified | 100880 | 115940 | 0 | 105940 | 0 | 0 | 0 | 0 | 0 | 7722 | 152740 | 3060700 | 0 | 0 | 0 | 21568 | 761050 | 161070 |
| AELGIPLEE | IDI1 | Q13907 | IDI1_HUMAN Is | Unmodified | 0 | 0 | 0 | 0 | 0 | 0 | 15474 | 0 | 918400 | 0 | 279970 | 1136900 | 0 | 0 | 0 | 0 | 69679 | 0 |
| AELGLNEH | LZTFL1 | Q9NQ48 | LZTL1_HUMAN | Acetyl (Protein N | 0 | 0 | 0 | 0 | 564240 | 252830 | 97624 | 0 | 1637400 | 701980 | 0 | 591030 | 0 | 569660 | 0 | 0 | 206640 | 775320 |
| AELGTDLLS | SMC3 | Q9UQE7 | SMC3_HUMAN | Unmodified | 158270 | 0 | 0 | 0 | 0 | 0 | 0 | 25474 | 0 | 0 | 50915 | 90006 | 0 | 0 | 0 | 0 | 0 | 0 |
| AELHVLDVI | UTRN | P46939 | UTRN_HUMAN | Unmodified | 0 | 0 | 0 | 691200 | 1612700 | 0 | 0 | 0 | 0 | 0 | 0 | 0 | 0 | 0 | 1643100 | 1167900 | 0 | 0 |
| AELIVQPEL | DSP | P15924 | DESP_HUMAN | Unmodified | 0 | 0 | 0 | 88924 | 0 | 0 | 0 | 0 | 770770 | 0 | 0 | 383970 | 0 | 0 | 0 | 0 | 0 | 0 |
| AELLEQSR | N4BP1 | O75113 | N4BP1_HUMAI | Unmodified | 234530 | 600270 | 97425 | 212580 | 0 | 0 | 0 | 0 | 953120 | 33584 | 458130 | 281600 | 0 | 0 | 0 | 0 | 0 | 0 |
| AELLQSDEI | SLC29A2 | Q14542 | S29A2_HUMAN | Unmodified | 179580 | 0 | 300250 | 0 | 0 | 855890 | 0 | 0 | 0 | 0 | 314010 | 1297700 | 541350 | 0 | 388890 | 209240 | 0 | 0 |
| AELNEFLTF | RPS3; RPS3; RP | P23396; H0Y | RS3_HUMAN 4 | Unmodified | 1018700 | 2539000 | 2950500 | 0 | 0 | 0 | 0 | 0 | 4971900 | 0 | 959590 | 25474000 | 0 | 0 | 1852400 | 1432700 | 0 | 0 |
| AELNGKDG | GRB2; GRB2; G | P62993; J3Q | GRB2_HUMAN | Unmodified | 0 | 0 | 0 | 0 | 193920 | 0 | 129690 | 0 | 981610 | 53505 | 0 | 208620 | 0 | 0 | 905490 | 940510 | 0 | 0 |
| AELNSTTEE | KIF20A | O95235 | KI20A_HUMAN | Unmodified | 41559 | 0 | 713760 | 5414100 | 2674000 | 632100 | 23684 | 54206 | 1644400 | 257390 | 0 | 0 | 0 | 706890 | 465880 | 0 | 345910 | 636040 |
| AELQEVQIT | NDRG2; NDRG2 | Q9UN36; G3 | NDRG2_HUMA | Acetyl (Protein N | 0 | 0 | 0 | 1195000 | 0 | 0 | 0 | 0 | 0 | 0 | 2200600 | 0 | 1346700 | 0 | 1613400 | 1478400 | 0 | 0 |
| AELQMLLEI | ABI2; ABI2; ABI2 | Q9NYB9; E7I | ABI2_HUMAN A | Acetyl (Protein N | 78087 | 0 | 0 | 0 | 38739 | 0 | 0 | 0 | 0 | 152110 | 0 | 0 | 198840 | 0 | 0 | 0 | 426320 | 0 |
| AELQMLLEI | ABI1; ABI1; ABI1 | Q8IZP0; A0A | ABI1_HUMAN A | Acetyl (Protein N | 0 | 39493 | 0 | 16557 | 871840 | 40528 | 125610 | 0 | 17113 | 99521 | 0 | 0 | 264660 | 139940 | 148770 | 18456 | 0 | 15447 |
| AELQMLLEI | ABI1; ABI1; ABI1 | Q8IZP0; A0A | ABI1_HUMAN A | Acetyl (Protein N | 0 | 0 | 0 | 127610 | 823810 | 93883 | 106350 | 0 | 32285 | 68243 | 0 | 0 | 439500 | 391590 | 0 | 28433 | 463430 | 238580 |
| AELRDDPII | PSMD11 | O00231 | PSD11_HUMAI | Unmodified | 0 | 0 | 339210 | 0 | 1451300 | 827120 | 98256 | 37792 | 189010 | 279130 | 836380 | 13808000 | 0 | 0 | 409810 | 603580 | 344610 | 383560 |
| AELSEEALL | USP5; USP5 | P45974; F5H | UBP5_HUMAN | Acetyl (Protein N | 0 | 99597 | 0 | 30807 | 168210 | 0 | 69249 | 0 | 0 | 0 | 0 | 0 | 515120 | 0 | 37106 | 0 | 320720 | 143070 |
| AELSYRGP | CTTN; CTTN; C1 | Q14247; H0Y | SRC8_HUMAN | Oxidation (M) | 792560 | 0 | 767520 | 0 | 0 | 0 | 0 | 0 | 0 | 0 | 0 | 0 | 0 | 0 | 1348700 | 777160 | 0 | 0 |
| AELTSQES( | EPS15 | P42566 | EPS15_HUMAN | Unmodified | 621340 | 566160 | 239160 | 378620 | 80628 | 0 | 0 | 0 | 0 | 0 | 94252 | 93261 | 0 | 0 | 430520 | 308290 | 0 | 0 |
| AEMDEVER | DIAPH1; DIAPH | O60610; A0A | DIAP1_HUMAN | Unmodified | 0 | 0 | 0 | 0 | 0 | 0 | 0 | 0 | 0 | 0 | 0 | 777680 | 0 | 0 | 187350 | 0 | 0 | 0 |
| AEMDEVER | DIAPH1; DIAPH | O60610; A0A | DIAP1_HUMAN | Oxidation (M) | 25086 | 0 | 0 | 0 | 0 | 0 | 175250 | 13135 | 0 | 0 | 0 | 1073200 | 0 | 0 | 0 | 0 | 0 | 0 |
| AEMETLVR | DPYSL5; DPYSL | Q9BPU6; E9I | DPYL5_HUMAN | Unmodified | 0 | 0 | 839700 | 0 | 261530 | 0 | 0 | 0 | 264020 | 408330 | 0 | 668530 | 0 | 0 | 0 | 0 | 209790 | 0 |
| AEMGIQLIV | LAMP3; LAMP3 | Q9UQV4; E7 | LAMP3_HUMAN | Oxidation (M) | 0 | 0 | 0 | 0 | 758320 | 0 | 163590 | 0 | 1254800 | 178950 | 0 | 0 | 612760 | 0 | 0 | 0 | 0 | 0 |
| AEMGLGDI | SLC20A1 | Q8WUM9 | S20A1_HUMAN | Unmodified | 0 | 0 | 0 | 0 | 0 | 0 | 0 | 0 | 78264 | 0 | 0 | 766760 | 0 | 0 | 0 | 0 | 0 | 0 |
| AEMKSQPS | ITGA3 | P26006 | ITA3_HUMAN I | Oxidation (M) | 0 | 0 | 0 | 0 | 5070000 | 0 | 0 | 87714 | 0 | 0 | 0 | 0 | 0 | 0 | 0 | 0 | 3269000 | 4261700 |
| AEMKSQPS | ITGA3 | P26006 | ITA3_HUMAN I | Unmodified | 0 | 0 | 0 | 0 | 405510 | 1570200 | 0 | 0 | 0 | 0 | 51565 | 0 | 0 | 0 | 0 | 0 | 2576200 | 4741500 |
| AEMNGSKI | EPCAM; EPCAM | P16422; C9J | EPCAM_HUMA | Unmodified | 0 | 28404 | 0 | 0 | 0 | 0 | 0 | 0 | 318330 | 68939 | 0 | 159720 | 698890 | 357640 | 0 | 0 | 73801 | 497760 |
| AENDAQKF | ABCC4; ABCC4 | O15439; A0A | MRP4_HUMAN | Unmodified | 0 | 0 | 997150 | 0 | 0 | 0 | 0 | 0 | 0 | 0 | 0 | 0 | 0 | 0 | 0 | 0 | 0 | 0 |
| AENFFILR | ARPC4; ARPC4- | P59998; A0A | ARPC4_HUMAI | Unmodified | 783930 | 1158100 | 422860 | 0 | 0 | 0 | 0 | 61562 | 0 | 0 | 208910 | 4777900 | 0 | 0 | 777750 | 1252300 | 0 | 0 |
| AENGKLVIN | GAPDH | P04406 | G3P_HUMAN G | Unmodified | 212410 | 205790 | 0 | 0 | 0 | 0 | 1055200 | 0 | 0 | 0 | 0 | 1001800 | 0 | 0 | 0 | 97924 | 0 | 0 |
| AENGKLVIN | GAPDH | P04406 | G3P_HUMAN G | Unmodified | 2383700 | 0 | 0 | 4689200 | 34783000 | 5430900 | 3645600 | 776410 | 6193200 | 3801200 | 3062300 | 13344000 | 26434000 | 6982900 | 0 | 905600 | 7220700 | 16829000 |
| AENIPAVVI | SLC30A1 | Q9Y6M5 | ZNT1_HUMAN . | Unmodified | 0 | 0 | 0 | 1607700 | 0 | 0 | 365640 | 0 | 0 | 0 | 0 | 2183200 | 0 | 0 | 1508300 | 0 | 0 | 1083100 |
| AENLGEVLI | TM9SF4; TM9SF | Q92544; A0A | TM9S4_HUMAN | Unmodified | 0 | 0 | 209560 | 0 | 0 | 0 | 0 | 0 | 34927000 | 1692900 | 1016500 | 0 | 0 | 0 | 156570 | 100940 | 0 | 0 |
| AENLGEVLI | TM9SF4; TM9SF | Q92544; A0A | TM9S4_HUMAN | Oxidation (M) | 0 | 74806 | 0 | 0 | 0 | 0 | 0 | 0 | 7870800 | 2020300 | 0 | 0 | 0 | 57723 | 0 | 0 | 0 | 0 |
| AENLGEVLI | TM9SF4; TM9SF | Q92544; A0A | TM9S4_HUMAN | Unmodified | 0 | 0 | 0 | 71612 | 0 | 0 | 0 | 0 | 5780300 | 376270 | 0 | 0 | 102220 | 327650 | 63831 | 0 | 0 | 0 |
| AENMYAQI | MACF1; MACF1 | Q9UPN3; H3 | MACF1_HUMAI | Unmodified | 0 | 0 | 0 | 0 | 0 | 0 | 0 | 0 | 0 | 0 | 0 | 1787300 | 0 | 0 | 0 | 0 | 0 | 0 |
| AENPSLEN | KPNA3 | O00505 | IMA4_HUMAN I | Acetyl (Protein N | 0 | 106790 | 114970 | 115880 | 0 | 0 | 0 | 0 | 0 | 0 | 225640 | 338040 | 0 | 0 | 77222 | 134510 | 0 | 217310 |
| AENPSLEN | KPNA3 | O00505 | IMA4_HUMAN I | Acetyl (Protein N | 0 | 0 | 0 | 0 | 3294000 | 0 | 517060 | 0 | 0 | 0 | 0 | 0 | 1184200 | 0 | 0 | 0 | 1680100 | 1639700 |
| AENQPIRD | LRPPRC | P42704 | LPPRC_HUMAN | Unmodified | 0 | 0 | 0 | 0 | 0 | 0 | 0 | 0 | 3208700 | 860890 | 0 | 0 | 0 | 0 | 0 | 0 | 0 | 0 |
| AENQVLAM | BCAP31; BCAP3 | P51572; A0A | BAP31_HUMAN | Unmodified | 0 | 0 | 0 | 0 | 0 | 0 | 205800 | 0 | 0 | 0 | 0 | 0 | 0 | 0 | 0 | 0 | 0 | 122020 |
| AENSSLNLI | NSF; NSF | P46459; I3L( | NSF_HUMAN V | Unmodified | 0 | 0 | 677530 | 644670 | 504310 | 0 | 700530 | 188270 | 1370400 | 420790 | 0 | 1019600 | 2710800 | 1184000 | 514280 | 465570 | 704290 | 875380 |
| AENYDIPS/ | UBA1; UBA1 | P22314; Q5J | UBA1_HUMAN | Unmodified | 0 | 0 | 0 | 0 | 0 | 154020 | 0 | 0 | 79887 | 0 | 327730 | 1561100 | 0 | 0 | 0 | 0 | 0 | 685010 |
| AENYDIPS/ | UBA1; UBA1 | P22314; Q5J | UBA1_HUMAN | Unmodified | 0 | 0 | 0 | 0 | 0 | 0 | 0 | 0 | 944340 | 0 | 0 | 3979600 | 0 | 0 | 0 | 0 | 0 | 0 |
| AEPAKIEAF | PGK1 | P00558 | PGK1_HUMAN | Unmodified | 0 | 0 | 0 | 669210 | 5196800 | 1220600 | 873480 | 758970 | 894850 | 2540400 | 1461200 | 0 | 9485900 | 2218600 | 0 | 0 | 6506100 | 16579000 |
| AEPATGGK | GPRIN1 | Q7Z2K8 | GRIN1_HUMAN | Unmodified | 0 | 0 | 0 | 0 | 0 | 0 | 0 | 0 | 1681700 | 334610 | 0 | 0 | 0 | 0 | 0 | 0 | 0 | 0 |
| AEPEDHYF | ACTR3; ACTR3; | P61158; B4D | ARP3_HUMAN . | Unmodified | 0 | 0 | 310600 | 0 | 0 | 0 | 0 | 0 | 0 | 0 | 0 | 5586200 | 0 | 0 | 689950 | 621530 | 0 | 0 |
| AEPEKNGE | SLC16A3; SLC1 | O15427; J3Q | MOT4_HUMAN | Unmodified | 0 | 129860 | 0 | 197020 | 10018000 | 0 | 0 | 0 | 0 | 0 | 0 | 0 | 0 | 0 | 0 | 0 | 13209000 | 11366000 |
| AEPHFLSIL | DIAPH1; DIAPH | O60610; A0A | DIAP1_HUMAN | Unmodified | 0 | 0 | 0 | 0 | 0 | 0 | 0 | 34660 | 0 | 0 | 0 | 469180 | 0 | 0 | 478220 | 22526 | 0 | 0 |
| AEPLAVGKI | GPRIN1 | Q7Z2K8 | GRIN1_HUMAN | Unmodified | 0 | 0 | 0 | 0 | 0 | 0 | 0 | 0 | 948060 | 0 | 0 | 0 | 0 | 0 | 0 | 151620 | 34581 | 0 |
| AEPLQPDP | HEBP2; HEBP2 | Q9Y5Z4; Q5T | HEBP2_HUMAI | Acetyl (Protein N | 0 | 0 | 0 | 0 | 0 | 0 | 0 | 0 | 2869800 | 0 | 621950 | 0 | 0 | 0 | 0 | 0 | 0 | 0 |
| AEPPKAPE( | BASP1 | P80723 | BASP1_HUMAN | Unmodified | 522230 | 288760 | 0 | 0 | 73249000 | 16551000 | 2984400 | 2544000 | 59393000 | 11269000 | 0 | 0 | 507060 | 417330 | 0 | 0 | 8921100 | 21224000 |
| AEPPKTHV | HLA-C; HLA-C; | P10321; A0A | HLAC_HUMAN | Unmodified | 0 | 0 | 0 | 57189 | 0 | 0 | 0 | 0 | 8894300 | 0 | 0 | 0 | 0 | 0 | 0 | 0 | 0 | 0 |
| AEPQPPSG | GLO1 | Q04760 | LGUL_HUMAN | Acetyl (Protein N | 0 | 0 | 0 | 0 | 0 | 0 | 0 | 0 | 1096200 | 0 | 0 | 2668400 | 0 | 0 | 0 | 482720 | 0 | 0 |
| AEPSAATQ | RTN3; RTN3; RT | O95197; B7Z | RTN3_HUMAN | Acetyl (Protein N | 445410 | 0 | 2581000 | 1038500 | 0 | 0 | 854210 | 405070 | 1180800 | 0 | 782950 | 0 | 2295700 | 0 | 1071800 | 842040 | 0 | 5281400 |
| AEPSADLL\ | SIGIRR; SIGIRR | Q6IA17; C9J | SIGIR_HUMAN | Unmodified | 0 | 0 | 388520 | 0 | 0 | 0 | 0 | 0 | 0 | 0 | 0 | 229640 | 0 | 0 | 0 | 996490 | 0 | 0 |
| AEPVEVVAI | PARP1; PARP1 | P09874; A0A | PARP1_HUMAN | Unmodified | 409490 | 0 | 0 | 0 | 0 | 0 | 0 | 0 | 840610 | 139780 | 1744800 | 1268500 | 0 | 0 | 0 | 0 | 0 | 0 |
| AEPVSSTE/ | GPRIN1 | Q7Z2K8 | GRIN1_HUMAN | Unmodified | 0 | 0 | 0 | 0 | 0 | 0 | 0 | 0 | 898060 | 0 | 0 | 0 | 0 | 0 | 54117 | 33203 | 0 | 0 |
| AEQDVENI | DDX39A; DDX39 | O00148; K7E | DX39A_HUMAN | Acetyl (Protein N | 0 | 0 | 0 | 0 | 0 | 0 | 1364000 | 862360 | 0 | 0 | 0 | 0 | 800720 | 0 | 438190 | 0 | 0 | 0 |
| AEQEEEFIS | CCDC6 | Q16204 | CCDC6_HUMA | Unmodified | 0 | 0 | 0 | 0 | 0 | 0 | 0 | 0 | 313790 | 0 | 461070 | 1973800 | 0 | 0 | 0 | 0 | 0 | 0 |
| AEQEEEFIS | CCDC6 | Q16204 | CCDC6_HUMA | Unmodified | 0 | 0 | 0 | 0 | 0 | 0 | 0 | 0 | 1000200 | 0 | 234200 | 1889500 | 0 | 0 | 0 | 0 | 0 | 0 |

| | | | | | | | | | | | | | | | | | | | | | | |
|---|---|---|---|---|---|---|---|---|---|---|---|---|---|---|---|---|---|---|---|---|---|---|
| AEQEGGM | CD109 | Q6YHK3 | CD109_HUMAI | Unmodified | 0 | 0 | 0 | 0 | 0 | 0 | 0 | 0 | 1304100 | 0 | 0 | 0 | 0 | 0 | 0 | 0 | 0 | 0 |
| AEQEPTAE | ARHGDIA; ARH | P52565; J3K | GDIR1_HUMAN | Acetyl (Protein N | 17563000 | 0 | 9956300 | 17601000 | 68861000 | 7486900 | 30392000 | 36662000 | 9844400 | 0 | 18256000 | 70915000 | 60981000 | 29951000 | 22617000 | 11280000 | 1.1E+08 | 66962000 |
| AEQESLEF | MYO1D; MYO1I | O94832; J3Q | MYO1D_HUMA | Acetyl (Protein N | 1822300 | 872730 | 829940 | 0 | 0 | 0 | 0 | 0 | 170730 | 73217 | 0 | 555130 | 0 | 0 | 1125800 | 955070 | 0 | 0 |
| AEQEVISL | EVI5; EVI5 | O60447; A0A | EVI5_HUMAN E | Unmodified | 0 | 0 | 0 | 0 | 440360 | 0 | 218540 | 227850 | 0 | 0 | 0 | 0 | 0 | 163420 | 0 | 0 | 0 | 433670 |
| AEQINQAA | STOML2; STOM | Q9UJZ1; A0A | STML2_HUMAN | Unmodified | 0 | 0 | 0 | 0 | 0 | 0 | 0 | 0 | 8410800 | 1467200 | 0 | 0 | 0 | 0 | 0 | 0 | 0 | 0 |
| AEQNDFIPI | SCNN1A; SCNN | P37088; F5G | SCNNA_HUMA | Unmodified | 0 | 0 | 0 | 0 | 0 | 0 | 0 | 0 | 1283200 | 239690 | 0 | 0 | 0 | 0 | 0 | 0 | 0 | 0 |
| AEQPHQH | IDE; IDE; IDE; ID | P14735; A0A | IDE_HUMAN In | Oxidation (M) | 0 | 0 | 0 | 0 | 0 | 0 | 0 | 0 | 0 | 0 | 0 | 638570 | 0 | 0 | 0 | 0 | 0 | 0 |
| AEQPHQH | IDE; IDE; IDE; ID | P14735; A0A | IDE_HUMAN In | Unmodified | 0 | 0 | 0 | 0 | 0 | 0 | 0 | 0 | 0 | 0 | 109550 | 1654400 | 0 | 0 | 0 | 0 | 0 | 0 |
| AEQVALSR | G6PD; G6PD; G | P11413; E7E | G6PD_HUMAN | Acetyl (Protein N | 0 | 0 | 0 | 0 | 2033000 | 0 | 0 | 0 | 969760 | 0 | 312130 | 732260 | 290940 | 0 | 0 | 0 | 504760 | 0 |
| AEQVIFVR | NECTIN2; NEC | Q92692; K7E | NECT2_HUMAN | Unmodified | 0 | 0 | 0 | 0 | 0 | 0 | 0 | 0 | 138620 | 449330 | 831030 | 668150 | 0 | 0 | 0 | 0 | 0 | 0 |
| AERAEWIE | PLEK2; PLEK2 | Q9NYT0; H0 | PLEK2_HUMAN | Unmodified | 0 | 0 | 0 | 88373 | 0 | 60927 | 0 | 0 | 859420 | 181510 | 0 | 0 | 0 | 0 | 109390 | 35396 | 0 | 0 |
| AERELSEQI | EZR; EZR | P15311; E7E | EZRI_HUMAN E | Unmodified | 0 | 0 | 200230 | 0 | 432340 | 97409 | 0 | 0 | 2074600 | 0 | 9849.1 | 1872700 | 0 | 0 | 25745 | 0 | 190880 | 0 |
| AERGLEPSI | DIP2B | Q9P265 | DIP2B_HUMAN | Acetyl (Protein N | 0 | 0 | 0 | 1523300 | 0 | 0 | 0 | 19669 | 2214200 | 0 | 0 | 0 | 0 | 0 | 0 | 0 | 669190 | 3672700 |
| AERGYSFSI | PSMA2; N/A; PS | P25787; A0A | PSA2_HUMAN | Acetyl (Protein N | 0 | 0 | 355640 | 0 | 6144600 | 0 | 0 | 405130 | 0 | 93209 | 1177000 | 687720 | 5724100 | 0 | 0 | 0 | 0 | 66955 |
| AERPEDLN | POLE3 | Q9NRF9 | DPOE3_HUMAI | Acetyl (Protein N | 44874 | 0 | 0 | 0 | 1219000 | 116320 | 75152 | 0 | 1232400 | 339800 | 206080 | 835200 | 775420 | 0 | 0 | 0 | 803810 | 281180 |
| AERPEDLN | POLE3 | Q9NRF9 | DPOE3_HUMAI | Acetyl (Protein N | 0 | 0 | 0 | 0 | 0 | 0 | 934640 | 0 | 0 | 0 | 0 | 0 | 0 | 0 | 0 | 0 | 1646400 | 0 |
| AERPGLQN | LLGL2; LLGL2 | Q6P1M3; J3C | L2GL2_HUMAN | Unmodified | 425080 | 197330 | 0 | 315670 | 0 | 0 | 0 | 0 | 2910300 | 0 | 561010 | 1580400 | 0 | 0 | 395400 | 323660 | 0 | 1100000 |
| AERPGLQN | LLGL2; LLGL2 | Q6P1M3; J3C | L2GL2_HUMAN | Oxidation (M) | 153760 | 0 | 204940 | 0 | 0 | 0 | 0 | 0 | 457630 | 77588 | 266140 | 0 | 0 | 0 | 180600 | 0 | 44041 | 52496 |
| AERPKFEQI | EPHA7; EPHA7 | Q15375; A0A | EPHA7_HUMAN | Unmodified | 0 | 0 | 0 | 0 | 0 | 0 | 88966 | 0 | 514330 | 37834 | 0 | 0 | 0 | 0 | 0 | 0 | 0 | 0 |
| AERQTLIQH | APLP2 | Q06481 | APLP2_HUMAN | Unmodified | 0 | 0 | 0 | 0 | 348260 | 0 | 0 | 0 | 0 | 438240 | 0 | 0 | 0 | 0 | 0 | 0 | 0 | 0 |
| AERTSPSSI | SLC4A2; SLC4A | P04920; A0A | B3A2_HUMAN | Unmodified | 0 | 0 | 0 | 0 | 0 | 0 | 0 | 0 | 0 | 0 | 0 | 0 | 0 | 0 | 189430 | 0 | 0 | 0 |
| AERVHELN | LUC7L2; LUC7L | Q9Y383; A0A | LC7L2_HUMAN | Unmodified | 0 | 0 | 138470 | 0 | 0 | 69396 | 195860 | 146730 | 3800200 | 0 | 0 | 290060 | 0 | 208470 | 0 | 19511 | 0 | 0 |
| AERVLELVS | SHMT2; SHMT2 | P34897; H0Y | GLYM_HUMAN | Unmodified | 0 | 0 | 0 | 0 | 0 | 0 | 0 | 0 | 3405100 | 1075100 | 0 | 0 | 0 | 0 | 0 | 0 | 0 | 0 |
| AESDGSLLI | PPP4R3A; PPP4 | Q6IN85; Q5N | P4R3A_HUMAN | Unmodified | 0 | 189870 | 0 | 79998 | 115770 | 0 | 0 | 0 | 0 | 165380 | 0 | 865060 | 0 | 0 | 0 | 0 | 491530 | 0 |
| AESEEGPD | SEC23B; SEC23 | Q15437; A0A | SC23B_HUMAN | Unmodified | 0 | 0 | 0 | 0 | 0 | 0 | 0 | 0 | 0 | 0 | 0 | 2784800 | 0 | 0 | 96081 | 505270 | 0 | 0 |
| AESFFQTK | APEH; APEH; AI | P13798; C9J | ACPH_HUMAN | Unmodified | 0 | 0 | 586230 | 514780 | 1131200 | 365060 | 0 | 0 | 669870 | 0 | 1253400 | 5228600 | 1003600 | 0 | 0 | 0 | 889810 | 2725600 |
| AESFMFR | MTAP; MTAP; M | Q13126; J3Q | MTAP_HUMAN | Unmodified | 0 | 0 | 0 | 0 | 0 | 0 | 0 | 0 | 198220 | 0 | 90831 | 848440 | 0 | 0 | 0 | 0 | 0 | 0 |
| AESGPDLR | DSP | P15924 | DESP_HUMAN | Unmodified | 0 | 0 | 0 | 0 | 0 | 0 | 0 | 0 | 0 | 0 | 0 | 0 | 0 | 0 | 0 | 0 | 0 | 0 |
| AESLGIPQ | INPP5D | Q92835 | SHIP1_HUMAN | Unmodified | 0 | 0 | 0 | 0 | 0 | 0 | 0 | 13647 | 0 | 0 | 204300 | 1159500 | 0 | 0 | 0 | 0 | 0 | 0 |
| AESLQVTR | NSF; NSF | P46459; I3L | NSF_HUMAN V | Unmodified | 170200 | 0 | 222840 | 0 | 33477 | 10945000 | 2235800 | 0 | 620280 | 0 | 0 | 1237100 | 0 | 0 | 0 | 0 | 83641 | 160030 |
| AESQLNCL | FAM83F | Q8NEG4 | FA83F_HUMAN | Acetyl (Protein N | 0 | 0 | 505680 | 0 | 0 | 0 | 0 | 0 | 0 | 0 | 0 | 630150 | 339930 | 0 | 398910 | 395730 | 0 | 0 |
| AESSDKLYI | PARP1; PARP1; | P09874; A0A | PARP1_HUMAN | Acetyl (Protein N | 0 | 0 | 0 | 0 | 0 | 0 | 1385600 | 3684400 | 0 | 270370 | 0 | 0 | 549760 | 0 | 0 | 0 | 862720 | 429640 |
| AESSELVR | P4HTM; P4HTM | Q9NXG6; V9 | P4HTM_HUMAI | Unmodified | 0 | 0 | 0 | 0 | 0 | 0 | 90313 | 0 | 1498500 | 0 | 25081 | 24787 | 0 | 0 | 0 | 0 | 0 | 0 |
| AESVVLEHI | CHID1; CHID1; | Q9BWS9; E9 | CHID1_HUMAN | Unmodified | 0 | 0 | 0 | 0 | 0 | 0 | 0 | 0 | 1252900 | 0 | 0 | 0 | 0 | 0 | 0 | 0 | 0 | 0 |
| AETEALSK | HTATIP2; HTATI | Q9BUP3; E9I | HTAI2_HUMAN | Acetyl (Protein N | 0 | 0 | 0 | 0 | 0 | 0 | 0 | 0 | 0 | 201680 | 0 | 0 | 0 | 0 | 0 | 0 | 0 | 0 |
| AETEERSLI | CDV3 | Q9UKY7 | CDV3_HUMAN | Acetyl (Protein N | 0 | 0 | 0 | 416260 | 0 | 0 | 0 | 0 | 0 | 0 | 0 | 0 | 1732000 | 1312200 | 0 | 0 | 709340 | 0 |
| AETERSALI | RACGAP1; RAC | Q9H0H5; F8 | RGAP1_HUMAI | Unmodified | 0 | 0 | 0 | 0 | 1182000 | 83731 | 0 | 9316.3 | 0 | 167150 | 0 | 0 | 0 | 0 | 0 | 0 | 325600 | 71438 |
| AETLMTVTI | SEH1L; SEH1L | Q96EE3; K7E | SEH1_HUMAN | Oxidation (M) | 3375500 | 0 | 0 | 64690 | 0 | 0 | 0 | 0 | 280240 | 0 | 0 | 1307100 | 0 | 0 | 0 | 0 | 0 | 0 |
| AETNDSYH | TFCP2; TFCP2 | Q12800; F8V | TFCP2_HUMAN | Unmodified | 0 | 0 | 0 | 0 | 0 | 0 | 0 | 0 | 0 | 0 | 0 | 1129300 | 0 | 0 | 0 | 0 | 0 | 0 |
| AETTPSMIP | CLMP | Q9H6B4 | CLMP_HUMAN | Unmodified | 0 | 0 | 0 | 0 | 0 | 0 | 0 | 0 | 0 | 0 | 0 | 0 | 0 | 0 | 0 | 0 | 0 | 2513700 |
| AETTPSMIP | CLMP | Q9H6B4 | CLMP_HUMAN | Unmodified | 0 | 0 | 0 | 0 | 0 | 0 | 0 | 0 | 0 | 0 | 0 | 139710 | 0 | 0 | 0 | 0 | 148560 | 621350 |
| AETYDFLFK | RAB4B; RAB4B; | P61018; Q6F | RAB4B_HUMAN | Acetyl (Protein N | 0 | 0 | 0 | 0 | 0 | 0 | 0 | 0 | 78695 | 164300 | 0 | 157320 | 0 | 0 | 0 | 0 | 0 | 0 |
| AEVADEKLI | PRKCA; PRKCA | P17252; J3K | KPCA_HUMAN | Unmodified | 0 | 0 | 0 | 0 | 265240 | 0 | 537830 | 175610 | 0 | 0 | 0 | 0 | 0 | 0 | 0 | 0 | 121820 | 202790 |
| AEVAQLQE | PPFIBP2; PPFIB | Q8ND30; E9 | LIPB2_HUMAN | Unmodified | 0 | 98308 | 393930 | 93663 | 0 | 0 | 0 | 0 | 0 | 0 | 48574 | 114060 | 0 | 0 | 309040 | 481620 | 32799 | 0 |
| AEVARAQA | CCT6A | P40227 | TCPZ_HUMAN | Unmodified | 0 | 0 | 0 | 0 | 0 | 0 | 0 | 0 | 0 | 0 | 0 | 0 | 936270 | 493510 | 0 | 0 | 0 | 222460 |
| AEVEETLK | DYNLRB2; DYN | Q8TF09; Q9I | DLRB2_HUMAN | Acetyl (Protein N | 124990 | 0 | 0 | 0 | 1364300 | 0 | 484570 | 303480 | 0 | 759840 | 0 | 2012900 | 1181400 | 958640 | 222030 | 0 | 1765200 | 2215400 |
| AEVEETLKF | DYNLRB1; DYN | Q9NP97; B1 | DLRB1_HUMAN | Acetyl (Protein N | 531860 | 0 | 0 | 0 | 800790 | 0 | 1891300 | 360380 | 0 | 242650 | 22826 | 292530 | 458050 | 1171000 | 0 | 0 | 5459500 | 645560 |
| AEVEGKDL | HNRNPU; HNR | Q00839; A0A | HNRPU_HUMA | Unmodified | 0 | 0 | 0 | 0 | 0 | 0 | 278000 | 293900 | 0 | 171360 | 0 | 0 | 539140 | 0 | 0 | 0 | 0 | 0 |
| AEVGILTR | PKHD1L1; CEM | Q86WI1; Q9I | PKHL1_HUMAN | Unmodified | 163870 | 357600 | 871700 | 384390 | 0 | 0 | 0 | 38237 | 3577300 | 390480 | 0 | 907550 | 0 | 0 | 1478900 | 398880 | 0 | 0 |
| AEVMLGH | LRBA; LRBA; LR | P50851; A0A | LRBA_HUMAN | Oxidation (M) | 0 | 0 | 0 | 310390 | 0 | 0 | 0 | 0 | 0 | 0 | 0 | 334930 | 0 | 0 | 0 | 0 | 0 | 0 |
| AEVQVLVLI | RPL13A; RPL13 | P40429; A0A | RL13A_HUMAN | Acetyl (Protein N | 0 | 0 | 18742 | 347530 | 701450 | 0 | 0 | 0 | 0 | 0 | 0 | 0 | 66880 | 29091 | 0 | 0 | 91311 | 49710 |
| AEVQVLVLI | RPL13A; RPL13 | P40429; A0A | RL13A_HUMAN | Acetyl (Protein N | 3846100 | 0 | 0 | 21347 | 0 | 0 | 1935200 | 0 | 231030 | 448230 | 691760 | 0 | 665900 | 0 | 43544 | 17827 | 0 | 0 |
| AEVRGAEIE | MYDGF; MYDGI | Q969H8; M0 | MYDGF_HUMA | Unmodified | 0 | 0 | 0 | 0 | 1315400 | 0 | 338240 | 0 | 5049500 | 15431000 | 582450 | 0 | 803270 | 500730 | 0 | 0 | 0 | 220440 |
| AEVRGAEIE | MYDGF; MYDGI | Q969H8; M0 | MYDGF_HUMA | Oxidation (M) | 0 | 0 | 0 | 0 | 0 | 0 | 127180 | 0 | 491110 | 3132700 | 0 | 0 | 220080 | 129860 | 0 | 0 | 0 | 0 |
| AEVRLQFH | G6PD | P11413 | G6PD_HUMAN | Unmodified | 0 | 0 | 0 | 0 | 1568100 | 0 | 0 | 0 | 0 | 0 | 0 | 0 | 0 | 571380 | 0 | 0 | 224240 | 0 |
| AEVSIDQSI | CCDC50 | Q8IVM0 | CCD50_HUMA | Acetyl (Protein N | 0 | 0 | 6149200 | 3815500 | 0 | 0 | 0 | 0 | 606260 | 155040 | 0 | 735870 | 0 | 0 | 6427700 | 8950200 | 0 | 0 |
| AEVSIDQSI | CCDC50 | Q8IVM0 | CCD50_HUMA | Acetyl (Protein N | 12182000 | 5940400 | 0 | 9757900 | 0 | 210350 | 401520 | 205150 | 2046300 | 552630 | 10794000 | 0 | 11587000 | 4098100 | 7943800 | 7975600 | 1109300 | 0 |
| AEVTAPDVI | AHNAK2 | Q8IVF2 | AHNK2_HUMAI | Unmodified | 0 | 166320 | 0 | 0 | 0 | 0 | 0 | 326100 | 3599600 | 615580 | 0 | 0 | 2336500 | 0 | 0 | 0 | 0 | 0 |
| AEVTAPDVI | AHNAK2 | Q8IVF2 | AHNK2_HUMAI | Unmodified | 0 | 0 | 0 | 0 | 1340600 | 379710 | 0 | 0 | 0 | 0 | 0 | 48206 | 0 | 0 | 0 | 0 | 0 | 1344000 |
| AEVTEEVEI | SLK | Q9H2G2 | SLK_HUMAN S | Unmodified | 0 | 0 | 0 | 0 | 0 | 0 | 0 | 0 | 706830 | 0 | 1514300 | 2487600 | 0 | 0 | 673290 | 596150 | 0 | 0 |
| AEWLAVKD | CANT1 | Q8WVQ1 | CANT1_HUMAI | Unmodified | 0 | 326210 | 1892100 | 829890 | 86314 | 3582000 | 693550 | 326900 | 8188000 | 504410 | 0 | 0 | 373510 | 1930900 | 546910 | 126410 | 142180 | 1597900 |
| AEWLLAVR | AASDHPPT | Q9NRN7 | ADPPT_HUMAN | Unmodified | 0 | 0 | 0 | 0 | 493460 | 0 | 0 | 0 | 0 | 0 | 0 | 1779300 | 0 | 0 | 0 | 0 | 0 | 0 |
| AEYDLTTR | EIF3E; EIF3E; EI | P60228; A0A | EIF3E_HUMAN | Acetyl (Protein N | 0 | 0 | 0 | 50549 | 0 | 0 | 140840 | 174090 | 0 | 80547 | 134370 | 2707800 | 57537 | 0 | 0 | 0 | 0 | 0 |
| AEYEGDGII | PAICS; PAICS; P | P22234; D6F | PUR6_HUMAN | Unmodified | 0 | 0 | 259780 | 0 | 0 | 137740 | 0 | 0 | 615830 | 0 | 1590800 | 7710300 | 0 | 0 | 0 | 0 | 13294 | 0 |
| AFAAGADIK | ECHS1 | P30084 | ECHM_HUMAN | Unmodified | 148780 | 125690 | 167200 | 236720 | 508670 | 72125 | 1722600 | 306800 | 3380300 | 630730 | 0 | 0 | 1008700 | 407250 | 202370 | 182300 | 39700 | 510440 |
| AFAAQEDLI | RDX; RDX; RDX | P35241; A0A | RADI_HUMAN I | Unmodified | 2152300 | 1588200 | 2826700 | 5831800 | 28956000 | 7601200 | 926140 | 1285800 | 4544500 | 4078200 | 2608200 | 11346000 | 25801000 | 27631000 | 5586700 | 3725700 | 3829900 | 9390100 |
| AFAAQEDLI | RDX; RDX; RDX | P35241; A0A | RADI_HUMAN I | Unmodified | 0 | 0 | 0 | 0 | 40831000 | 4277400 | 192290 | 11320000 | 0 | 1457900 | 0 | 0 | 28173000 | 39404000 | 0 | 0 | 15388000 | 10250000 |
| AFAAVPTSI | SCRIB; SCRIB; S | Q14160; A0A | SCRIB_HUMAN | Unmodified | 0 | 0 | 0 | 0 | 0 | 0 | 0 | 0 | 4803800 | 0 | 0 | 0 | 0 | 0 | 0 | 0 | 0 | 0 |
| AFAAVPTSI | SCRIB; SCRIB; S | Q14160; A0A | SCRIB_HUMAN | Unmodified | 0 | 0 | 0 | 0 | 1094500 | 0 | 0 | 0 | 0 | 1014500 | 0 | 0 | 2601400 | 763230 | 0 | 0 | 3042300 | 7662600 |
| AFADAMEV | CCT4 | P50991 | TCPD_HUMAN | Unmodified | 0 | 0 | 0 | 0 | 514970 | 0 | 0 | 0 | 0 | 512050 | 0 | 0 | 1142000 | 649580 | 0 | 0 | 0 | 553350 |
| AFAEALLLK | HPGD; HPGD; H | P15428; E9P | PGDH_HUMAN | Unmodified | 0 | 0 | 0 | 0 | 0 | 0 | 0 | 0 | 0 | 0 | 0 | 0 | 475450 | 0 | 0 | 0 | 0 | 2302200 |
| AFAFQASQ | MKRN2; MKRN2 | Q9H000; C9 | MKRN2_HUMA | Unmodified | 676320 | 507830 | 342260 | 301250 | 0 | 0 | 0 | 68578 | 0 | 0 | 527550 | 568530 | 521760 | 197100 | 342390 | 0 | 0 | 0 |
| AFAHITGG | GART | P22102 | PUR2_HUMAN | Unmodified | 161740 | 365440 | 0 | 0 | 766620 | 0 | 1230700 | 144710 | 0 | 687250 | 1818100 | 9834800 | 1180000 | 726620 | 0 | 0 | 0 | 0 |
| AFAHITGG | GART | P22102 | PUR2_HUMAN | Unmodified | 0 | 0 | 0 | 0 | 2418300 | 2701700 | 1934500 | 0 | 0 | 1453400 | 0 | 0 | 4867800 | 1025600 | 0 | 0 | 4298800 | 4473100 |
| AFALEVIK | PPA2 | Q9H2U2 | IPYR2_HUMAN | Unmodified | 0 | 0 | 0 | 0 | 0 | 0 | 0 | 327000 | 5163700 | 931100 | 0 | 2582200 | 0 | 256330 | 0 | 0 | 0 | 0 |
| AFALNDLD | TTLL12 | Q14166 | TTL12_HUMAN | Unmodified | 0 | 0 | 0 | 0 | 0 | 0 | 0 | 0 | 0 | 0 | 0 | 0 | 0 | 0 | 0 | 0 | 0 | 0 |
| AFAMIIDKL | PCBP2; PCBP2; | Q15366; F8V | PCBP2_HUMAI | 2 Oxidation (M) | 5628200 | 20585000 | 2350000 | 12669000 | 1877100 | 0 | 0 | 0 | 0 | 541610 | 0 | 0 | 12581000 | 8312800 | 11140000 | 0 | 0 | 0 |
| AFAMIIDKL | PCBP2; PCBP2; | Q15366; F8V | PCBP2_HUMAI | Oxidation (M) | 1001700 | 2282100 | 0 | 2981500 | 0 | 0 | 0 | 0 | 0 | 0 | 0 | 0 | 3971800 | 2575500 | 2160000 | 0 | 0 | 0 |
| AFANLRK | PHGDH; PHGD | O43175; A0A | SERA_HUMAN | Acetyl (Protein N | 0 | 0 | 2855700 | 0 | 0 | 0 | 17855000 | 9889600 | 11129000 | 0 | 3072300 | 17385000 | 87646000 | 0 | 0 | 0 | 6358400 | 0 |
| AFASRVFPI | NSF; NSF | P46459; I3L | NSF_HUMAN V | Unmodified | 0 | 0 | 0 | 0 | 0 | 0 | 0 | 0 | 0 | 0 | 0 | 0 | 610940 | 0 | 0 | 0 | 0 | 0 |
| AFASRVFPI | NSF; NSF | P46459; I3L | NSF_HUMAN V | Oxidation (M) | 0 | 0 | 0 | 0 | 0 | 0 | 0 | 0 | 0 | 0 | 0 | 0 | 602360 | 470570 | 0 | 0 | 28080 | 0 |
| AFAVGVQC | IPO7 | O95373 | IPO7_HUMAN I | Unmodified | 0 | 0 | 187240 | 0 | 0 | 0 | 0 | 0 | 0 | 0 | 576710 | 5596800 | 0 | 0 | 0 | 199330 | 0 | 0 |
| AFCEPGNV | YARS1; YARS1 | P54577; A0A | SYYC_HUMAN | Unmodified | 0 | 0 | 62904 | 0 | 593190 | 89057 | 977270 | 298330 | 336880 | 275660 | 177090 | 860810 | 1916500 | 540160 | 222380 | 125730 | 1126600 | 426010 |
| AFCSFQIYA | CST3 | P01034 | CYTC_HUMAN | Oxidation (M) | 0 | 0 | 0 | 0 | 0 | 0 | 0 | 0 | 4971300 | 3202600 | 0 | 0 | 0 | 0 | 0 | 0 | 0 | 0 |
| AFCSFQIYA | CST3 | P01034 | CYTC_HUMAN | Unmodified | 0 | 0 | 0 | 0 | 0 | 0 | 0 | 0 | 1027100 | 298670 | 0 | 0 | 0 | 0 | 0 | 0 | 0 | 0 |
| AFDITYVR | LAMC1 | P11047 | LAMC1_HUMAI | Unmodified | 0 | 178370 | 0 | 257390 | 0 | 0 | 0 | 0 | 895190 | 0 | 252040 | 0 | 0 | 199040 | 115900 | 245770 | 0 | 802930 |
| AFDITYVRL | LAMC1 | P11047 | LAMC1_HUMAI | Unmodified | 0 | 0 | 0 | 0 | 612820 | 492620 | 0 | 0 | 0 | 0 | 0 | 0 | 556830 | 1254700 | 0 | 0 | 643720 | 3526200 |
| AFDLFNPN | MRPS23; MRPS | Q9Y3D9; J3C | RT23_HUMAN | Unmodified | 0 | 0 | 0 | 0 | 0 | 0 | 0 | 0 | 1517800 | 0 | 0 | 0 | 0 | 0 | 0 | 0 | 0 | 0 |
| AFDLIVDRI | NDUFB10; NDU | O96000; H3I | NDUBA_HUMA | Unmodified | 0 | 0 | 0 | 0 | 0 | 0 | 0 | 0 | 6541600 | 896550 | 0 | 989530 | 1182900 | 0 | 0 | 0 | 0 | 0 |
| AFDLVPPE | PGM2 | Q96G03 | PGM2_HUMAN | Unmodified | 0 | 0 | 0 | 0 | 0 | 0 | 0 | 0 | 0 | 0 | 0 | 2440000 | 704680 | 0 | 0 | 0 | 0 | 0 |
| AFDMISQR | GIPC1; GIPC1 | O14908; K7E | GIPC1_HUMAN | Oxidation (M) | 0 | 0 | 52397 | 0 | 566070 | 150690 | 0 | 0 | 0 | 74053 | 122450 | 191970 | 460660 | 147190 | 131490 | 53534 | 756540 | 644430 |
| AFDMISQR | GIPC1; GIPC1 | O14908; K7E | GIPC1_HUMAN | Unmodified | 0 | 0 | 0 | 0 | 2649300 | 0 | 0 | 0 | 0 | 0 | 0 | 0 | 0 | 0 | 0 | 0 | 1664100 | 3166700 |
| AFDMWAG | FAM3A; FAM3A; | P98173; D3D | FAM3A_HUMAI | Oxidation (M) | 0 | 0 | 0 | 0 | 0 | 0 | 0 | 0 | 1365300 | 246490 | 0 | 0 | 9585.5 | 0 | 0 | 0 | 0 | 0 |
| AFDPAELR | B4GALNT1; B4 | Q00973; F8V | B4GN1_HUMAI | Unmodified | 0 | 0 | 0 | 0 | 1564800 | 0 | 0 | 0 | 137150 | 826050 | 1611900 | 0 | 0 | 0 | 0 | 0 | 0 | 0 |
| AFDQGAD | UQCR10 | Q9UDW1 | QCR9_HUMAN | Unmodified | 0 | 0 | 0 | 0 | 0 | 0 | 380040 | 224910 | 1153600 | 63490 | 126290 | 207980 | 217880 | 0 | 0 | 0 | 291680 | 0 |
| AFDRHCNI | SNRPD2 | P62316 | SMD2_HUMAN | Unmodified | 353360 | 63194 | 0 | 0 | 4277600 | 401190 | 12121000 | 522630 | 338150 | 477980 | 523830 | 1381700 | 2509700 | 1041300 | 214180 | 86477 | 487270 | 2447000 |
| AFDRHCNI | SNRPD2 | P62316 | SMD2_HUMAN | Oxidation (M) | 122540 | 872820 | 148480 | 0 | 323000 | 59090 | 69085 | 100040 | 0 | 89712 | 0 | 471010 | 513780 | 197050 | 49541 | 659920 | 0 | 334620 |
| AFDSDGD | EFCAB14; EFCA | O75071; A0A | EFC14_HUMAN | Unmodified | 0 | 0 | 0 | 0 | 0 | 0 | 0 | 0 | 3774800 | 0 | 321740 | 143440 | 0 | 0 | 0 | 0 | 0 | 433130 |
| AFDSGIIPM | ACLY | P53396 | ACLY_HUMAN | Oxidation (M) | 0 | 0 | 0 | 0 | 0 | 0 | 0 | 392060 | 0 | 0 | 0 | 4702700 | 0 | 0 | 0 | 0 | 0 | 0 |
| AFDTAGNG | FASN; FASN | P49327; A0A | FAS_HUMAN F | Unmodified | 0 | 0 | 0 | 0 | 0 | 0 | 0 | 0 | 56741 | 0 | 0 | 7900400 | 0 | 0 | 403370 | 317870 | 0 | 0 |
| AFDTAGNG | FASN; FASN | P49327; A0A | FAS_HUMAN F | Unmodified | 0 | 0 | 0 | 0 | 1300900 | 0 | 0 | 0 | 0 | 225380 | 389060 | 0 | 1834900 | 399520 | 0 | 0 | 158060 | 2695500 |
| AFDYFNLA | SEL1L | Q9UBV2 | SE1L1_HUMAN | Oxidation (M) | 0 | 0 | 0 | 0 | 0 | 0 | 0 | 0 | 7761900 | 2323400 | 0 | 0 | 0 | 0 | 0 | 0 | 0 | 0 |
| AFDYFNLA | SEL1L | Q9UBV2 | SE1L1_HUMAN | Unmodified | 0 | 0 | 0 | 0 | 0 | 0 | 0 | 0 | 3656200 | 410390 | 0 | 0 | 0 | 0 | 0 | 0 | 0 | 0 |
| AFEADSTVI | UTRN | P46939 | UTRN_HUMAN | Unmodified | 0 | 0 | 0 | 465520 | 0 | 0 | 0 | 0 | 724230 | 190310 | 0 | 363100 | 0 | 0 | 950030 | 893900 | 0 | 0 |
| AFEDDDITI | OLA1; OLA1 | Q9NTK5; J3K | OLA1_HUMAN | Unmodified | 0 | 0 | 0 | 0 | 0 | 0 | 0 | 0 | 0 | 0 | 455300 | 948390 | 0 | 0 | 164720 | 0 | 0 | 0 |
| AFEEAEK | VCP; VCP; VCP; | P55072; C9J | TERA_HUMAN | Unmodified | 407850 | 52172 | 343840 | 475910 | 320640 | 0 | 0 | 65306 | 352750 | 0 | 0 | 1279900 | 3268400 | 617680 | 270020 | 260050 | 61630 | 968800 |
| AFEECMQK | ENTPD3 | O75355 | ENTP3_HUMAN | Unmodified | 0 | 0 | 0 | 0 | 0 | 0 | 0 | 0 | 217980 | 46493 | 0 | 0 | 0 | 0 | 0 | 0 | 0 | 0 |
| AFEEEYYR | STEAP2 | Q8NFT2 | STEA2_HUMAN | Unmodified | 0 | 0 | 989700 | 0 | 0 | 0 | 0 | 0 | 152590 | 123230 | 411930 | 0 | 0 | 0 | 3026300 | 2329100 | 0 | 0 |
| AFEEQER | ABCC1; ABCC1 | P33527; I3L | MRP1_HUMAN | Unmodified | 0 | 0 | 0 | 0 | 53175 | 0 | 0 | 0 | 0 | 23714 | 163230 | 1358100 | 0 | 0 | 558150 | 241560 | 0 | 0 |

| | | | | | | | | | | | | | | | | | | | | | | |
|---|---|---|---|---|---|---|---|---|---|---|---|---|---|---|---|---|---|---|---|---|---|---|
| AFEESLSTL | ECE1; ECE1; EC | P42892; B4D | ECE1_HUMAN | Unmodified | 272690 | 330250 | 496490 | 381710 | 1843400 | 938510 | 192560 | 0 | 9431000 | 2848800 | 0 | 382700 | 0 | 0 | 0 | 432960 | 0 | 0 |
| AFELMHSG | ADH5 | P11766 | ADHX_HUMAN | Unmodified | 1626600 | 0 | 0 | 0 | 1178400 | 61509 | 0 | 38473 | 0 | 81857 | 222060 | 1435000 | 0 | 0 | 0 | 0 | 454730 | 334460 |
| AFENLLGQ | LMO7; LMO7; LI | Q8WWI1; E9 | LMO7_HUMAN | Unmodified | 0 | 0 | 0 | 0 | 0 | 0 | 0 | 0 | 0 | 0 | 0 | 488750 | 0 | 0 | 0 | 0 | 1223900 | 0 |
| AFEPATGR | ALDH9A1 | P49189 | AL9A1_HUMAN | Unmodified | 507000 | 31046 | 438220 | 0 | 467260 | 0 | 185880 | 51896 | 1443800 | 358210 | 235430 | 1221400 | 834350 | 462450 | 380560 | 394020 | 153840 | 265790 |
| AFEPATGR | ALDH9A1 | P49189 | AL9A1_HUMAN | Unmodified | 0 | 0 | 0 | 0 | 1372800 | 331570 | 813440 | 221770 | 0 | 974920 | 0 | 0 | 2439400 | 2426700 | 0 | 0 | 1549700 | 1308500 |
| AFEPATGR | ALDH9A1 | P49189 | AL9A1_HUMAN | Unmodified | 0 | 0 | 0 | 0 | 0 | 0 | 0 | 0 | 0 | 0 | 0 | 0 | 0 | 0 | 0 | 0 | 2499700 | 0 |
| AFEPLRSFA | PLXNB2 | O15031 | PLXB2_HUMAN | Unmodified | 515890 | 303670 | 0 | 1740100 | 1639000 | 602760 | 154020 | 0 | 985390 | 529490 | 748160 | 152630 | 2119900 | 469650 | 860320 | 603200 | 870580 | 407440 |
| AFEPYLEILI | SGPL1 | O95470 | SGPL1_HUMAN | Unmodified | 1724900 | 0 | 0 | 114320 | 1817700 | 0 | 0 | 0 | 4538000 | 7006400 | 0 | 0 | 2089900 | 0 | 0 | 0 | 0 | 92725 |
| AFESDVFH | TRAFD1; TRAFD | O14545; F8V | TRAD1_HUMAN | Unmodified | 0 | 1104500 | 0 | 0 | 0 | 0 | 0 | 0 | 0 | 0 | 0 | 0 | 0 | 0 | 0 | 0 | 0 | 0 |
| AFEVPARAI | VIL1; VIL1; VIL1 | P09327; H7C | VILI_HUMAN Vi | Unmodified | 0 | 0 | 0 | 0 | 0 | 0 | 126430 | 0 | 471490 | 10663000 | 1899600 | 0 | 0 | 0 | 0 | 0 | 0 | 0 |
| AFEVSENG | FASN; FASN | P49327; A0A | FAS_HUMAN F | Unmodified | 209900 | 0 | 339430 | 282840 | 0 | 0 | 0 | 0 | 199860 | 0 | 1805700 | 27089000 | 0 | 0 | 460200 | 1052200 | 0 | 0 |
| AFFAGSQR | GANAB; GANAE | Q14697; F5H | GANAB_HUMA | Unmodified | 201360 | 147520 | 0 | 136300 | 0 | 0 | 0 | 0 | 84689000 | 5464400 | 0 | 1909600 | 0 | 0 | 221000 | 819830 | 0 | 0 |
| AFFAGSQR | GANAB; GANAE | Q14697; F5H | GANAB_HUMA | Unmodified | 272300 | 0 | 0 | 0 | 0 | 837360 | 0 | 0 | 2147200 | 20241000 | 0 | 0 | 0 | 0 | 0 | 0 | 0 | 0 |
| AFFAHDGV | TRPM5; TRPM5; | Q9NZQ8; A0 | TRPM5_HUMAN | Unmodified | 0 | 0 | 0 | 0 | 0 | 0 | 0 | 0 | 0 | 0 | 0 | 1513900 | 0 | 0 | 0 | 0 | 0 | 0 |
| AFFAQDGV | TRPM4 | Q8TD43 | TRPM4_HUMAN | Unmodified | 119850 | 0 | 154640 | 0 | 0 | 0 | 0 | 0 | 0 | 0 | 0 | 0 | 0 | 0 | 119240 | 164610 | 0 | 0 |
| AFFDLIDAL | PGAP1; PGAP1; | Q75T13; B4D | PGAP1_HUMAN | Unmodified | 0 | 0 | 0 | 0 | 0 | 0 | 136190 | 0 | 1827500 | 216680 | 0 | 0 | 0 | 0 | 0 | 0 | 0 | 0 |
| AFFESHPAI | NPEPPS; NPEPI | P55786; E9P | PSA_HUMAN P | Unmodified | 0 | 221690 | 0 | 580520 | 444500 | 0 | 0 | 49095 | 1838000 | 281650 | 177760 | 8637100 | 348260 | 367520 | 830360 | 769660 | 383040 | 613200 |
| AFFESHPAI | NPEPPS; NPEPI | P55786; E9P | PSA_HUMAN P | Unmodified | 0 | 0 | 0 | 0 | 133820 | 104010 | 0 | 0 | 0 | 0 | 0 | 0 | 56171 | 0 | 0 | 0 | 81931 | 151500 |
| AFFPCFDTI | RNPEP; RNPEP; | Q9H4A4; A6I | AMPB_HUMAN | Unmodified | 0 | 0 | 0 | 0 | 0 | 3855000 | 0 | 0 | 0 | 0 | 1719900 | 8031400 | 0 | 0 | 276170 | 0 | 0 | 0 |
| AFFSEVER | PA2G4; PA2G4 | Q9UQ80; F8 | PA2G4_HUMAN | Unmodified | 2107300 | 3351500 | 1196300 | 789760 | 0 | 0 | 260010 | 129920 | 0 | 446860 | 1153400 | 8175700 | 0 | 0 | 1067800 | 1510800 | 73049 | 0 |
| AFFSEVERF | PA2G4; PA2G4 | Q9UQ80; F8 | PA2G4_HUMAN | Unmodified | 0 | 0 | 0 | 0 | 0 | 0 | 0 | 0 | 0 | 0 | 0 | 1356600 | 726750 | 866270 | 0 | 0 | 0 | 0 |
| AFGDVFSV | PICK1; PICK1 | Q9NRD5; F6 | PICK1_HUMAN | Unmodified | 222870 | 31929 | 106840 | 891980 | 0 | 0 | 0 | 0 | 0 | 0 | 0 | 241370 | 0 | 0 | 76919 | 0 | 0 | 0 |
| AFGFMTR | PCBD1 | P61457 | PHS_HUMAN P | Unmodified | 0 | 0 | 0 | 0 | 0 | 0 | 0 | 0 | 2135400 | 123610 | 0 | 0 | 174400 | 0 | 0 | 203030 | 0 | 0 |
| AFGFSHLE | OTUB1; OTUB1 | Q96FW1; J3I | OTUB1_HUMAI | Unmodified | 493450 | 380530 | 241430 | 935010 | 991560 | 364510 | 42026 | 0 | 1457400 | 343540 | 1484500 | 6612700 | 1262800 | 281890 | 1335300 | 689350 | 501740 | 652260 |
| AFGPGLEG | FLNC | Q14315 | FLNC_HUMAN | Unmodified | 0 | 0 | 0 | 0 | 883360 | 0 | 0 | 0 | 0 | 0 | 0 | 0 | 0 | 0 | 0 | 0 | 0 | 0 |
| AFGPGLQG | FLNA; FLNA | P21333; Q60 | FLNA_HUMAN | Unmodified | 1525600 | 2081000 | 2002600 | 4026400 | 1740800 | 495310 | 679970 | 665040 | 135000 | 213490 | 206200 | 4128900 | 1904900 | 1983500 | 2496600 | 2328200 | 608200 | 560140 |
| AFGPGLQG | FLNA; FLNA | P21333; Q60 | FLNA_HUMAN | Unmodified | 0 | 0 | 1095300 | 4002500 | 29158000 | 8553200 | 30309000 | 16607000 | 0 | 1413000 | 0 | 0 | 24112000 | 25811000 | 1008200 | 589790 | 18060000 | 9176200 |
| AFGSVATRI | GALNT16 | Q8N428 | GLT16_HUMAN | Unmodified | 0 | 0 | 0 | 0 | 0 | 0 | 0 | 0 | 2654600 | 652310 | 0 | 0 | 0 | 0 | 0 | 0 | 0 | 0 |
| AFGYYGPL | SRSF3; SRSF3 | P84103; A0A | SRSF3_HUMAN | Unmodified | 5248800 | 5773500 | 2935400 | 2481800 | 0 | 0 | 0 | 52278 | 0 | 0 | 0 | 4303200 | 0 | 0 | 2868000 | 2154700 | 0 | 0 |
| AFHAALRN | ARPC5L | Q9BPX5 | ARP5L_HUMAN | Unmodified | 0 | 0 | 0 | 0 | 441480 | 0 | 38303 | 31008 | 0 | 64032 | 0 | 0 | 1052500 | 0 | 0 | 0 | 90763 | 0 |
| AFHDLKVA | ITGB4 | P16144 | ITB4_HUMAN I | Unmodified | 0 | 0 | 0 | 0 | 0 | 0 | 0 | 0 | 0 | 0 | 0 | 0 | 336020 | 0 | 0 | 0 | 1040700 | 0 |
| AFHNEAQV | TCP1; TCP1; TC | P17987; F5H | TCPA_HUMAN | Unmodified | 0 | 0 | 0 | 0 | 0 | 0 | 0 | 76324 | 271070 | 88724 | 373470 | 2112000 | 0 | 0 | 0 | 143460 | 0 | 0 |
| AFHNEAQV | TCP1; TCP1; TC | P17987; F5H | TCPA_HUMAN | Unmodified | 0 | 0 | 0 | 0 | 1539900 | 175380 | 0 | 0 | 1002200 | 1715300 | 2026600 | 12003000 | 0 | 739980 | 0 | 174640 | 0 | 0 |
| AFHPFIAGF | HDLBP; HDLBP | Q00341; A0A | VIGLN_HUMAN | Unmodified | 344450 | 451120 | 406670 | 350630 | 0 | 0 | 0 | 0 | 0 | 0 | 0 | 1591300 | 304870 | 0 | 415480 | 604350 | 0 | 0 |
| AFHSSLHE | ATP2B4 | P23634 | AT2B4_HUMAN | Unmodified | 0 | 0 | 0 | 0 | 2868000 | 612740 | 656590 | 286570 | 0 | 57073 | 0 | 0 | 0 | 0 | 0 | 0 | 1168100 | 2506000 |
| AFHYDMKT | VNN1 | O95497 | VNN1_HUMAN | Unmodified | 0 | 0 | 56415 | 241090 | 0 | 328930 | 0 | 0 | 0 | 0 | 0 | 0 | 18681 | 79892 | 100400 | 0 | 0 | 99497 |
| AFIDPLGLP | FOLH1B; FOLH | Q9HBA9; Q0 | FOH1B_HUMAI | Unmodified | 39964000 | 57116000 | 1.46E+08 | 2.17E+08 | 0 | 0 | 0 | 0 | 127970 | 47914 | 0 | 0 | 891260 | 47340 | 2.02E+08 | 1.56E+08 | 0 | 0 |
| AFIDVTAAA | ITGA6; ITGA6 | P23229; H7E | ITA6_HUMAN I | Unmodified | 0 | 598740 | 1024200 | 2066000 | 0 | 0 | 0 | 0 | 0 | 0 | 0 | 6509100 | 0 | 0 | 6226700 | 1886600 | 0 | 0 |
| AFIHWVSQ | QARS1; QARS1 | P47897; H7C | SYQ_HUMAN G | Oxidation (M) | 0 | 0 | 0 | 0 | 0 | 0 | 0 | 0 | 0 | 0 | 0 | 3772400 | 0 | 0 | 0 | 0 | 21279 | 0 |
| AFIHWVSQ | QARS1; QARS1 | P47897; H7C | SYQ_HUMAN G | Unmodified | 0 | 0 | 0 | 0 | 0 | 0 | 0 | 0 | 0 | 0 | 0 | 543800 | 0 | 0 | 0 | 0 | 0 | 0 |
| AFIITGQTY | ADSL; ADSL; AD | P30566; A0A | PUR8_HUMAN | Unmodified | 0 | 0 | 0 | 0 | 183550 | 1854900 | 325090 | 134240 | 0 | 0 | 0 | 3259300 | 0 | 0 | 1468800 | 159490 | 0 | 2242200 |
| AFIITLNSF | ATP6V0D2 | Q8N8Y2 | VA0D2_HUMAN | Unmodified | 0 | 0 | 0 | 0 | 0 | 0 | 0 | 0 | 0 | 218680 | 225150 | 730770 | 0 | 0 | 0 | 0 | 0 | 0 |
| AFIPQLLSR | IDE; IDE; IDE; ID | P14735; A0A | IDE_HUMAN In | Unmodified | 0 | 0 | 81942 | 0 | 0 | 0 | 0 | 0 | 0 | 657860 | 154780 | 6785000 | 0 | 0 | 0 | 123680 | 0 | 0 |
| AFITNIPFD | HNRNPM; HNR | P52272; M0I | HNRPM_HUMA | Unmodified | 0 | 0 | 670320 | 0 | 112190 | 0 | 0 | 311000 | 1196100 | 616820 | 0 | 6709300 | 0 | 0 | 0 | 0 | 0 | 0 |
| AFITNIPFD | HNRNPM; HNR | P52272; M0I | HNRPM_HUMA | Unmodified | 160040 | 138680 | 0 | 108850 | 3020300 | 545940 | 8582600 | 234830 | 429390 | 3040600 | 67774 | 42017 | 9857100 | 4937400 | 0 | 0 | 813800 | 3974000 |
| AFITNIPFD | HNRNPM; HNR | P52272; M0I | HNRPM_HUMA | Unmodified | 0 | 0 | 0 | 0 | 391100 | 0 | 1658700 | 0 | 0 | 147600 | 0 | 0 | 560980 | 430970 | 0 | 0 | 186530 | 0 |
| AFKDIDIED | EIF3A | Q14152 | EIF3A_HUMAN | Oxidation (M) | 0 | 109750 | 0 | 129090 | 0 | 0 | 0 | 0 | 0 | 0 | 0 | 0 | 669310 | 539870 | 107670 | 0 | 0 | 0 |
| AFKDTGKTI | RPS20; RPS20; | P60866; E5R | RS20_HUMAN | Acetyl (Protein N | 289370 | 492080 | 0 | 863330 | 2146600 | 0 | 1473900 | 1046300 | 215370 | 1758500 | 171070 | 1462200 | 7312800 | 2047700 | 0 | 0 | 466740 | 1566600 |
| AFKLFDDD | CETN2 | P41208 | CETN2_HUMAI | Unmodified | 0 | 0 | 0 | 0 | 595490 | 0 | 623040 | 0 | 0 | 107520 | 0 | 0 | 0 | 0 | 0 | 0 | 690730 | 640450 |
| AFKVAYSLI | MFGE8; MFGE8 | Q08431; H0Y | MFGM_HUMAN | Unmodified | 0 | 0 | 0 | 0 | 5120600 | 0 | 1019800 | 0 | 0 | 0 | 0 | 0 | 1113800 | 742340 | 0 | 0 | 3274100 | 1053200 |
| AFLDALQN | TRAP1 | Q12931 | TRAP1_HUMAN | Unmodified | 0 | 0 | 0 | 0 | 0 | 0 | 0 | 0 | 3602600 | 0 | 0 | 318540 | 0 | 0 | 0 | 0 | 0 | 0 |
| AFLDELK | FOLH1; FOLH1; | Q04609; E9F | FOLH1_HUMAI | Unmodified | 0 | 0 | 76893000 | 11675000 | 0 | 0 | 0 | 0 | 0 | 0 | 0 | 0 | 227020 | 0 | 31347000 | 25428000 | 45508 | 0 |
| AFLDELKAE | FOLH1; FOLH1; | Q04609; E9F | FOLH1_HUMAI | Unmodified | 3550900 | 6324700 | 1264200 | 20069000 | 0 | 0 | 0 | 0 | 0 | 0 | 0 | 0 | 3417400 | 1725800 | 5983000 | 7547900 | 0 | 0 |
| AFLDELKAE | FOLH1; FOLH1; | Q04609; E9F | FOLH1_HUMAI | Unmodified | 1325500 | 1137500 | 124250 | 8862600 | 0 | 0 | 0 | 0 | 0 | 0 | 0 | 0 | 1097500 | 794690 | 1288000 | 1600800 | 0 | 0 |
| AFLDEMK | LDOC1 | O95751 | LDOC1_HUMAI | Unmodified | 0 | 615200 | 1570900 | 0 | 0 | 0 | 0 | 0 | 0 | 0 | 234490 | 1027200 | 0 | 0 | 4262800 | 1960100 | 0 | 0 |
| AFLDFHALI | PTGES2; PTGES | Q9H7Z7; A0 | PGES2_HUMAN | Unmodified | 0 | 0 | 0 | 0 | 0 | 0 | 0 | 0 | 318310 | 0 | 0 | 1084200 | 30524 | 0 | 0 | 0 | 0 | 0 |
| AFLDFYHSI | ARHGEF1; ARH | Q92888; M0 | ARHG1_HUMA | Unmodified | 0 | 0 | 43555 | 0 | 5072600 | 0 | 0 | 0 | 0 | 0 | 185130 | 619780 | 0 | 0 | 0 | 0 | 1283700 | 10806000 |
| AFLDRPSQ | TXNIP | Q9H3M7 | TXNIP_HUMAN | Unmodified | 0 | 0 | 0 | 0 | 0 | 0 | 0 | 0 | 0 | 0 | 0 | 0 | 0 | 0 | 0 | 0 | 186460 | 448410 |
| AFLEENNLI | PPP5C; PPP5C; | P53041; A8M | PPP5_HUMAN | Unmodified | 50021 | 0 | 0 | 0 | 0 | 0 | 352500 | 47000 | 0 | 0 | 2009200 | 6563900 | 0 | 0 | 22280 | 0 | 60598 | 0 |
| AFLEENNLI | PPP5C; PPP5C; | P53041; A8M | PPP5_HUMAN | Unmodified | 0 | 0 | 0 | 0 | 0 | 0 | 0 | 0 | 0 | 0 | 0 | 0 | 399640 | 0 | 0 | 0 | 0 | 0 |
| AFLEENNLI | PPP5C; PPP5C; | P53041; A8M | PPP5_HUMAN | Unmodified | 0 | 0 | 0 | 0 | 638620 | 0 | 0 | 0 | 0 | 0 | 0 | 0 | 509150 | 0 | 0 | 0 | 244140 | 0 |
| AFLEETNAI | DOCK10; DOCI | Q96BY6; A0 | DOC10_HUMA | Unmodified | 0 | 0 | 0 | 0 | 1368500 | 206750 | 60733 | 28330 | 2915700 | 0 | 62538 | 0 | 0 | 0 | 0 | 0 | 33597 | 136200 |
| AFLFQDIEF | NELL1; NELL1; | Q92832; J3K | NELL1_HUMAN | Unmodified | 0 | 0 | 0 | 0 | 0 | 0 | 0 | 0 | 2709400 | 1580700 | 0 | 0 | 0 | 0 | 0 | 0 | 0 | 0 |
| AFLHVPAK | NID1 | P14543 | NID1_HUMAN | Unmodified | 0 | 0 | 0 | 0 | 0 | 0 | 0 | 0 | 522400 | 223760 | 0 | 0 | 0 | 0 | 0 | 0 | 0 | 0 |
| AFLIEEQK | RPL34 | P49207 | RL34_HUMAN | Unmodified | 2724100 | 3370500 | 1406500 | 1832000 | 0 | 0 | 209870 | 376490 | 531010 | 331380 | 0 | 7580100 | 881090 | 0 | 962930 | 1162900 | 103550 | 0 |
| AFLIEEQKI | RPL34 | P49207 | RL34_HUMAN | Unmodified | 0 | 0 | 0 | 186370 | 171020 | 0 | 460870 | 234050 | 0 | 0 | 0 | 0 | 2242900 | 1547000 | 0 | 0 | 76073 | 215580 |
| AFLLESLLK | CHAMP1 | Q96JM3 | CHAP1_HUMAI | Unmodified | 0 | 0 | 0 | 0 | 0 | 0 | 211790 | 0 | 187470 | 330980 | 0 | 519030 | 330340 | 103460 | 0 | 0 | 37859 | 0 |
| AFLLTPR | APOM; APOM | O95445; Q5 | APOM_HUMAN | Unmodified | 4317800 | 12763000 | 4954800 | 3732200 | 574260 | 964350 | 450670 | 114530 | 186800 | 0 | 0 | 0 | 2206400 | 2892700 | 2898600 | 2694900 | 109930 | 1318400 |
| AFLMQHAF | LONP1; LONP1 | P36776; K7E | LONM_HUMAN | Oxidation (M) | 0 | 0 | 0 | 0 | 0 | 0 | 0 | 0 | 1845400 | 317760 | 0 | 0 | 0 | 0 | 0 | 0 | 0 | 0 |
| AFLMQHAF | LONP1; LONP1 | P36776; K7E | LONM_HUMAN | Unmodified | 0 | 0 | 0 | 0 | 0 | 0 | 0 | 0 | 4790000 | 0 | 0 | 0 | 0 | 0 | 0 | 0 | 1089800 | 0 |
| AFLQENTH | APOBEC3B; AP | Q9UH17; P3 | ABC3B_HUMAI | Unmodified | 1199700 | 1434500 | 1284900 | 523870 | 0 | 0 | 0 | 10084 | 105690 | 0 | 131650 | 170860 | 0 | 0 | 1793200 | 1475300 | 0 | 0 |
| AFLTLAEDI | RAB10 | P61026 | RAB10_HUMAN | Unmodified | 7755000 | 6727400 | 2183400 | 7497100 | 1851600 | 52399 | 1117600 | 0 | 3850100 | 3947100 | 938570 | 3341100 | 2858500 | 103460 | 7451900 | 4083300 | 604600 | 2194300 |
| AFLTLAEDI | RAB10 | P61026 | RAB10_HUMAN | Unmodified | 1180400 | 1038700 | 1766200 | 904210 | 2013500 | 0 | 604330 | 0 | 1260800 | 458360 | 0 | 3904100 | 1629800 | 969980 | 5067900 | 2264600 | 116700 | 822580 |
| AFMIIQEQF | CDCP1 | Q9H5V8 | CDCP1_HUMA | Oxidation (M) | 97211 | 256270 | 537460 | 624640 | 0 | 0 | 0 | 0 | 0 | 0 | 337490 | 3072900 | 0 | 0 | 911430 | 890440 | 396890 | 0 |
| AFMIIQEQF | CDCP1 | Q9H5V8 | CDCP1_HUMA | Unmodified | 0 | 0 | 0 | 0 | 0 | 0 | 0 | 0 | 0 | 0 | 0 | 2730300 | 0 | 0 | 461020 | 0 | 231640 | 0 |
| AFMTADLP | CLTCL1; CLTC; | P53675; Q00 | CLH2_HUMAN | Oxidation (M) | 0 | 465490 | 0 | 0 | 0 | 0 | 0 | 0 | 0 | 20622 | 0 | 0 | 49508 | 26652 | 228450 | 0 | 0 | 0 |
| AFNDPFIQI | HSPH1; HSPH1 | Q92598; R4 | HS105_HUMAN | Unmodified | 0 | 0 | 653820 | 932250 | 362190 | 0 | 0 | 0 | 1221000 | 435570 | 1782200 | 3905800 | 813680 | 266990 | 1455400 | 0 | 0 | 615950 |
| AFNEFLER | TNFAIP2; TNFAI | Q03169; H0 | TNAP2_HUMAN | Unmodified | 410630 | 0 | 0 | 0 | 0 | 0 | 0 | 0 | 136310 | 0 | 0 | 859330 | 0 | 0 | 0 | 0 | 0 | 0 |
| AFNFLNEIK | VAMP7 | P51809 | VAMP7_HUMAI | Unmodified | 532820 | 0 | 792020 | 205820 | 0 | 0 | 0 | 0 | 0 | 8566.7 | 0 | 1251100 | 0 | 0 | 0 | 0 | 100590 | 0 |
| AFNKPHLA | MISP | Q8IVT2 | MISP_HUMAN | Unmodified | 0 | 0 | 0 | 0 | 1210000 | 0 | 0 | 0 | 0 | 0 | 0 | 0 | 0 | 0 | 0 | 0 | 0 | 0 |
| AFNLVKDS | U2AF2; U2AF2 | P26368; K7E | U2AF2_HUMAN | Unmodified | 0 | 106910 | 0 | 0 | 291740 | 750870 | 1237000 | 923740 | 0 | 0 | 0 | 286230 | 111610 | 840360 | 0 | 0 | 565550 | 773120 |
| AFNNKGD | CNTN1 | Q12860 | CNTN1_HUMA | Unmodified | 0 | 0 | 0 | 0 | 0 | 0 | 0 | 0 | 2416200 | 0 | 0 | 0 | 0 | 0 | 0 | 0 | 0 | 0 |
| AFNSSSFN | RANGAP1 | P46060 | RAGP1_HUMAI | Unmodified | 0 | 0 | 0 | 0 | 0 | 0 | 0 | 0 | 0 | 0 | 0 | 597700 | 0 | 0 | 0 | 29735 | 0 | 0 |
| AFNVISAW | STEAP3; STEAP | Q658P3; B8 | STEA3_HUMAN | Unmodified | 0 | 0 | 88229 | 0 | 0 | 0 | 0 | 0 | 448240 | 87990 | 252310 | 341380 | 0 | 0 | 0 | 248670 | 0 | 0 |
| AFPALHFL | TXNDC15; TXNI | Q96J42; H0Y | TXD15_HUMAN | Unmodified | 0 | 0 | 0 | 0 | 0 | 0 | 0 | 0 | 2391600 | 315220 | 0 | 0 | 0 | 0 | 0 | 0 | 0 | 0 |
| AFPALTSLD | CD14; CD14 | P08571; D6F | CD14_HUMAN | Unmodified | 0 | 0 | 0 | 0 | 0 | 0 | 0 | 0 | 732150 | 0 | 0 | 0 | 0 | 0 | 0 | 0 | 0 | 0 |
| AFPAQTAS | BAIAP2; BAIAP2 | Q9UQB8; I3I | BAIP2_HUMAN | Unmodified | 0 | 0 | 595740 | 850570 | 1805700 | 418090 | 0 | 90769 | 0 | 0 | 0 | 954260 | 1132200 | 485470 | 931510 | 1004300 | 0 | 504650 |
| AFPCWDEF | NPEPPS; NPEPI | P55786; A0A | PSA_HUMAN P | Unmodified | 0 | 0 | 457700 | 0 | 0 | 0 | 0 | 0 | 0 | 0 | 0 | 1284200 | 176370 | 0 | 0 | 0 | 0 | 0 |
| AFPEHLYQ | CRMP1; CRMP1 | Q14194; E9F | DPYL1_HUMAN | Unmodified | 0 | 0 | 0 | 0 | 0 | 0 | 1779000 | 430170 | 0 | 282520 | 0 | 0 | 0 | 0 | 0 | 0 | 0 | 0 |
| AFPHRPDA | VPS33B; VPS33 | Q9H267; F5I | VP33B_HUMAN | Acetyl (Protein N | 157250 | 196550 | 147700 | 323980 | 3726400 | 389780 | 1750800 | 300050 | 0 | 473670 | 414090 | 1979300 | 0 | 879370 | 228590 | 194950 | 572570 | 422140 |
| AFPHRPDA | VPS33B; VPS33 | Q9H267; F5I | VP33B_HUMAN | Acetyl (Protein N | 87263 | 0 | 65540 | 0 | 676590 | 0 | 69855 | 232130 | 258500 | 0 | 0 | 0 | 261310 | 150630 | 0 | 97529 | 151960 | 0 |
| AFPLAEWQ | METTL26; METT | Q96S19; H3I | MTL26_HUMAN | Unmodified | 0 | 464920 | 567620 | 0 | 0 | 0 | 0 | 0 | 148890 | 149900 | 79763 | 756410 | 0 | 0 | 431410 | 0 | 0 | 0 |
| AFPMPGFD | SHMT2 | P34897 | GLYM_HUMAN | Oxidation (M) | 0 | 0 | 0 | 0 | 0 | 202460 | 0 | 0 | 2798100 | 1086900 | 93627 | 154330 | 0 | 146640 | 0 | 0 | 0 | 0 |
| AFPMPGFD | SHMT2 | P34897 | GLYM_HUMAN | Unmodified | 0 | 0 | 0 | 0 | 0 | 0 | 0 | 0 | 9482900 | 1839600 | 0 | 0 | 0 | 0 | 0 | 0 | 0 | 0 |
| AFPNPYAD | LANCL1; LANCI | O43813; E9F | LANC1_HUMAI | Unmodified | 474340 | 699640 | 590380 | 574740 | 0 | 0 | 0 | 92668 | 1013700 | 471240 | 0 | 1881200 | 497010 | 0 | 583750 | 419650 | 154840 | 407940 |
| AFQNVFAP | GNS; GNS; GNS | P15586; H7C | GNS_HUMAN N | Unmodified | 0 | 261140 | 0 | 0 | 0 | 0 | 0 | 67250 | 5622200 | 873210 | 0 | 187290 | 506060 | 0 | 0 | 0 | 0 | 0 |
| AFQNVFAP | GNS; GNS; GNS | P15586; H7C | GNS_HUMAN N | Unmodified | 0 | 0 | 0 | 0 | 357170 | 0 | 329420 | 0 | 0 | 1240000 | 0 | 0 | 285570 | 606370 | 0 | 0 | 129830 | 877090 |
| AFQSVLEV | HTT; HTT; HTT | P42858; A0A | HD_HUMAN Hu | Unmodified | 0 | 0 | 0 | 0 | 0 | 81999 | 0 | 0 | 0 | 0 | 0 | 676240 | 0 | 0 | 0 | 0 | 0 | 0 |
| AFQVFEEAI | GAS6 | Q14393 | GAS6_HUMAN | Unmodified | 46143 | 0 | 0 | 156970 | 0 | 0 | 98994 | 137340 | 1818600 | 232420 | 0 | 0 | 0 | 0 | 0 | 0 | 0 | 0 |
| AFQYVETH | PRDX3 | P30048 | PRDX3_HUMAN | Unmodified | 0 | 0 | 0 | 0 | 0 | 0 | 1431300 | 435050 | 30351000 | 10322000 | 0 | 0 | 0 | 2356500 | 0 | 0 | 0 | 124050 |
| AFQYVHEH | C1GALT1 | Q9NS00 | C1GLT_HUMAN | Unmodified | 0 | 0 | 0 | 0 | 0 | 0 | 0 | 0 | 1502600 | 0 | 0 | 0 | 0 | 0 | 0 | 0 | 0 | 0 |
| AFRDLYSE | GPC1; GPC1; G | P35052; H7C | GPC1_HUMAN | Unmodified | 374360 | 155290 | 240980 | 0 | 0 | 0 | 0 | 0 | 0 | 0 | 0 | 0 | 0 | 0 | 419550 | 339310 | 118490 | 0 |
| AFRNIPGIT | RPL4; RPL4 | P36578; H3E | RL4_HUMAN 60 | Unmodified | 0 | 0 | 0 | 0 | 307290 | 0 | 824240 | 50668 | 0 | 157860 | 0 | 0 | 0 | 212890 | 0 | 0 | 0 | 0 |
| AFRSSTIQD | WASF2 | Q9Y6W5 | WASF2_HUMAI | Unmodified | 158530 | 0 | 101610 | 0 | 2872200 | 0 | 69319 | 881230 | 0 | 484170 | 0 | 21669 | 0 | 1231500 | 172000 | 138920 | 1876000 | 2669800 |
| AFRSSTIQD | WASF1; WASF1 | Q92558; Q5 | WASF1_HUMAI | Unmodified | 0 | 0 | 0 | 0 | 876940 | 0 | 285110 | 0 | 0 | 0 | 0 | 0 | 0 | 0 | 0 | 0 | 311630 | 230660 |
| AFRTATQL | CCT7; CCT7 | Q99832; F8V | TCPH_HUMAN | Unmodified | 0 | 0 | 0 | 0 | 976790 | 0 | 450180 | 192610 | 0 | 654610 | 82192 | 262270 | 1616300 | 1013400 | 0 | 0 | 338530 | 725170 |
| AFSDLTSR | ELOVL7; ELOVL | A1L3X0; D6F | ELOV7_HUMAN | Acetyl (Protein N | 0 | 0 | 0 | 0 | 0 | 0 | 0 | 0 | 2079900 | 359680 | 124040 | 0 | 0 | 0 | 0 | 0 | 0 | 0 |
| AFSDPFVE | HSPA4; HSPA4; | P34932; A0A | HSP74_HUMAN | Unmodified | 0 | 0 | 672360 | 630810 | 2000700 | 542770 | 552990 | 198330 | 1990800 | 924140 | 1901700 | 16399000 | 1873200 | 793400 | 824000 | 867520 | 415960 | 793140 |
| AFSGSGNF | UFD1; UFD1 | Q92890; C9J | UFD1_HUMAN | Unmodified | 0 | 0 | 0 | 227450 | 0 | 0 | 0 | 0 | 0 | 0 | 0 | 0 | 0 | 0 | 0 | 0 | 0 | 0 |

| | | | | | | | | | | | | | | | | | | | | | | |
|---|---|---|---|---|---|---|---|---|---|---|---|---|---|---|---|---|---|---|---|---|---|---|
| AFSGYLGTI | ESD; ESD; ESD; | P10768; X6R | ESTD_HUMAN | Unmodified | 0 | 0 | 0 | 1789900 | 3785200 | 1059400 | 1018100 | 523740 | 1952300 | 1451700 | 1149200 | 5350400 | 2650700 | 1840300 | 962700 | 1071700 | 1688400 | 1703500 |
| AFSGYLGTI | ESD; ESD; ESD; | P10768; X6R | ESTD_HUMAN | Unmodified | 0 | 0 | 0 | 0 | 470870 | 0 | 289340 | 399060 | 0 | 0 | 0 | 0 | 0 | 648410 | 0 | 0 | 1272600 | 0 |
| AFSIAQDAL | CLMN | Q96JQ2 | CLMN_HUMAN | Unmodified | 0 | 0 | 37893 | 39350 | 0 | 0 | 0 | 0 | 184800 | 0 | 177030 | 1264300 | 0 | 0 | 0 | 0 | 0 | 0 |
| AFSITQGLL | PFAS; PFAS | O15067; J3K | PUR4_HUMAN | Unmodified | 0 | 0 | 0 | 0 | 0 | 0 | 11357 | 0 | 0 | 0 | 601260 | 2981200 | 0 | 0 | 0 | 0 | 0 | 0 |
| AFSMDDR | TOLLIP; TOLLIP; | Q9H0E2; E9I | TOLIP_HUMAN | Unmodified | 1108400 | 1894300 | 477980 | 0 | 0 | 0 | 0 | 0 | 0 | 0 | 0 | 0 | 0 | 0 | 1729400 | 1866500 | 0 | 0 |
| AFSMDDRL | TOLLIP; TOLLIP; | Q9H0E2; E9I | TOLIP_HUMAN | Oxidation (M) | 0 | 3149000 | 524580 | 1500400 | 0 | 0 | 0 | 0 | 0 | 0 | 0 | 0 | 0 | 0 | 3726100 | 1591900 | 0 | 0 |
| AFSMDEHN | GPRC5B | Q9NZH0 | GPC5B_HUMA | Unmodified | 0 | 0 | 373920 | 727910 | 327630 | 135470 | 0 | 0 | 526750 | 99550 | 762840 | 636160 | 419370 | 0 | 525030 | 450090 | 0 | 0 |
| AFSMDEHN | GPRC5B | Q9NZH0 | GPC5B_HUMA | Oxidation (M) | 0 | 0 | 0 | 405650 | 0 | 0 | 0 | 0 | 321690 | 41075 | 0 | 0 | 8370.2 | 0 | 208400 | 234340 | 0 | 0 |
| AFSMDEPV | GPRC5C; GPRC | Q9NQ84; A8 | GPC5C_HUMA | Oxidation (M) | 0 | 0 | 1782100 | 1641900 | 350960 | 0 | 0 | 0 | 1344900 | 234000 | 0 | 0 | 738700 | 0 | 1166300 | 1159500 | 0 | 315940 |
| AFSMDEPV | GPRC5C; GPRC | Q9NQ84; A8 | GPC5C_HUMA | Unmodified | 0 | 0 | 2076600 | 3249200 | 690960 | 0 | 0 | 0 | 2167000 | 319940 | 0 | 0 | 1442100 | 0 | 1961500 | 1381800 | 0 | 604660 |
| AFSPTTVN1 | SEC61A1; SEC6 | P61619; B4D | S61A1_HUMAN | Unmodified | 0 | 0 | 0 | 0 | 0 | 0 | 0 | 0 | 1992900 | 0 | 0 | 319390 | 0 | 0 | 0 | 0 | 170800 | 0 |
| AFSVFLFN1 | IDI1; IDI1 | Q13907; C9J | IDI1_HUMAN Is | Unmodified | 1657400 | 0 | 15893 | 0 | 1882500 | 555500 | 710820 | 0 | 3037000 | 1585400 | 2182800 | 8004100 | 1772100 | 0 | 0 | 1130100 | 0 | 0 |
| AFSVNIFK | LAMP1 | P11279 | LAMP1_HUMAN | Unmodified | 0 | 0 | 0 | 0 | 555780 | 0 | 947330 | 569980 | 5581500 | 4905900 | 360130 | 0 | 176590 | 0 | 143170 | 0 | 0 | 0 |
| AFSYYGPLI | SRSF7; SRSF7; | Q16629; A0A | SRSF7_HUMAN | Unmodified | 1526900 | 1197000 | 489330 | 209760 | 0 | 0 | 0 | 0 | 0 | 0 | 0 | 1024300 | 0 | 0 | 405080 | 648250 | 0 | 0 |
| AFTDLNSIN | PROM1 | O43490 | PROM1_HUMA | Unmodified | 0 | 0 | 0 | 0 | 0 | 0 | 0 | 0 | 0 | 0 | 795800 | 5846100 | 46603 | 0 | 0 | 0 | 0 | 0 |
| AFTELQAK | PFDN1; PFDN1 | O60925; E5F | PFD1_HUMAN | Unmodified | 499810 | 635260 | 598570 | 603550 | 2188100 | 0 | 689480 | 600740 | 1415100 | 1379400 | 786950 | 5243300 | 1679100 | 535330 | 0 | 0 | 1744000 | 1007500 |
| AFTELQAK\ | PFDN1; PFDN1 | O60925; E5F | PFD1_HUMAN | Unmodified | 0 | 0 | 0 | 0 | 0 | 0 | 0 | 0 | 0 | 0 | 0 | 0 | 0 | 0 | 0 | 0 | 1889800 | 0 |
| AFTGREFDI | ATP2A2; ATP2A | P16615; H7C | AT2A2_HUMAN | Unmodified | 116610 | 164610 | 474250 | 272450 | 0 | 288120 | 93703 | 73454 | 765570 | 158360 | 0 | 0 | 0 | 0 | 320460 | 185500 | 0 | 322950 |
| AFTHTAQYI | ATIC; ATIC | P31939; C9J | PUR9_HUMAN | Unmodified | 0 | 0 | 0 | 0 | 10143000 | 0 | 3482600 | 0 | 3353100 | 1290900 | 1380500 | 9559900 | 9825800 | 8026600 | 664870 | 1707400 | 6148700 | 5294200 |
| AFTHTAQYI | ATIC; ATIC | P31939; C9J | PUR9_HUMAN | Unmodified | 0 | 0 | 0 | 0 | 0 | 0 | 0 | 0 | 0 | 0 | 0 | 0 | 0 | 0 | 0 | 0 | 2260200 | 0 |
| AFTKPEEAC | GART; GART; G | P22102; C9J | PUR2_HUMAN | Unmodified | 0 | 0 | 0 | 0 | 240490 | 0 | 0 | 0 | 0 | 0 | 0 | 0 | 483310 | 0 | 0 | 0 | 1061700 | 229850 |
| AFTPFSGPI | PEPD; PEPD; PE | P12955; A0A | PEPD_HUMAN | Unmodified | 0 | 0 | 0 | 0 | 1937700 | 0 | 857330 | 1359400 | 259500 | 646370 | 0 | 1283200 | 0 | 336320 | 74513 | 0 | 3087500 | 1503600 |
| AFTSTLEK | KIAA2013 | Q8IYS2 | K2013_HUMAN | Unmodified | 45872 | 23757 | 87059 | 44714 | 0 | 0 | 0 | 20831 | 15044000 | 2816500 | 94925 | 1786000 | 0 | 0 | 50266 | 44269 | 77156 | 259640 |
| AFTTTLNLL | NRDC; NRDC; I | O43847; G3\ | NRDC_HUMAN | Unmodified | 0 | 0 | 0 | 0 | 0 | 0 | 0 | 0 | 763160 | 0 | 0 | 1742800 | 0 | 0 | 0 | 0 | 192450 | 0 |
| AFTYINLDK | TFRC; TFRC | P02786; G3\ | TFR1_HUMAN | Unmodified | 2559300 | 4688400 | 0 | 1279500 | 0 | 0 | 1170600 | 0 | 3067300 | 1827500 | 0 | 2171700 | 463020 | 734070 | 0 | 0 | 318770 | 0 |
| AFVAIGDYI | RPS2; RPS2; RP | P15880; E9P | RS2_HUMAN 4 | Unmodified | 4386400 | 3123100 | 2602400 | 2620800 | 0 | 0 | 5858800 | 102790 | 1455100 | 628130 | 2964700 | 62417000 | 21671 | 0 | 3196300 | 2567900 | 224870 | 1561900 |
| AFVDFLSDI | C1QBP | Q07021 | C1QBP_HUMA | Unmodified | 0 | 0 | 0 | 0 | 0 | 0 | 0 | 0 | 4182100 | 0 | 2032000 | 0 | 0 | 0 | 0 | 0 | 64264 | 0 |
| AFVDFLSDI | C1QBP | Q07021 | C1QBP_HUMA | Unmodified | 0 | 0 | 0 | 196260 | 3757900 | 0 | 1309100 | 0 | 25408000 | 1915000 | 2856300 | 1938900 | 2973500 | 1864300 | 0 | 124460 | 0 | 455250 |
| AFVDNYKV | ABR; ABR; ABR; | Q12979; A0A | ABR_HUMAN A | Unmodified | 0 | 2480700 | 700590 | 0 | 188380 | 0 | 147520 | 76372 | 0 | 0 | 0 | 424300 | 80353 | 0 | 283490 | 251310 | 989970 | 465200 |
| AFVEAQNK | PREP | P48147 | PPCE_HUMAN | Unmodified | 0 | 0 | 0 | 0 | 0 | 0 | 0 | 0 | 0 | 0 | 0 | 1560700 | 0 | 0 | 0 | 0 | 0 | 0 |
| AFVELTK | PGM3; PGM3; F | O95394; J3K | AGM1_HUMAN | Unmodified | 136030 | 16664 | 65489 | 132600 | 841570 | 199130 | 62590 | 374740 | 237750 | 299350 | 143800 | 1349300 | 1857100 | 1447400 | 0 | 118680 | 959240 | 1362400 |
| AFVENYPQ | VPS45; VPS45; | Q9NRW7; A0 | VPS45_HUMAN | Unmodified | 152500 | 219380 | 113360 | 134750 | 694750 | 98964 | 304320 | 133200 | 253230 | 151390 | 0 | 757490 | 533470 | 340480 | 125170 | 111160 | 90420 | 187710 |
| AFVHWYVC | TUBA3C; TUBA | P0DPH7; Q7 | TBA3C_HUMAN | Oxidation (M) | 480720 | 447960 | 259710 | 66339 | 1623200 | 0 | 1310700 | 0 | 202050 | 467670 | 0 | 7899000 | 0 | 0 | 1602900 | 1616800 | 289330 | 2381500 |
| AFVPPLAEI | ARAP1; ARAP1; | Q96P48; F8V | ARAP1_HUMAN | Unmodified | 0 | 0 | 0 | 0 | 0 | 0 | 0 | 0 | 0 | 0 | 0 | 344400 | 0 | 0 | 0 | 0 | 0 | 0 |
| AFVRDPAP | TOR2A; TOR2A | Q8N2E6; Q5 | TOR2X_HUMAN | Unmodified | 0 | 0 | 0 | 0 | 0 | 0 | 0 | 0 | 1382800 | 1210100 | 0 | 0 | 0 | 0 | 0 | 0 | 0 | 0 |
| AFVRPSGTI | PGM3; PGM3; F | O95394; J3K | AGM1_HUMAN | Unmodified | 242380 | 0 | 115010 | 0 | 0 | 0 | 26164 | 0 | 635410 | 59686 | 96233 | 2562000 | 0 | 66463 | 578590 | 630980 | 0 | 56637 |
| AFVRRDWI | PALS2; PALS2; I | Q9NZW5; B8 | PALS2_HUMAN | Unmodified | 0 | 0 | 0 | 0 | 1778100 | 0 | 0 | 0 | 0 | 0 | 181460 | 0 | 0 | 0 | 0 | 0 | 366460 | 792510 |
| AFVYLSNLI | KIF1A; KIF1A; KI | Q12756; A0A | KIF1A_HUMAN | Unmodified | 0 | 0 | 0 | 0 | 0 | 0 | 0 | 0 | 209110 | 0 | 0 | 212270 | 0 | 0 | 88940 | 64349 | 0 | 0 |
| AFYAELYHI | PSME2; PSME2; | Q9UL46; A0A | PSME2_HUMA | Unmodified | 171570 | 198510 | 0 | 359620 | 828850 | 0 | 0 | 0 | 90436 | 428890 | 18732 | 730290 | 852560 | 0 | 0 | 0 | 263840 | 935140 |
| AFYAPVHAI | SLC12A2; SLC1 | P55011; G3X | S12A2_HUMAN | Unmodified | 0 | 0 | 0 | 329400 | 0 | 0 | 0 | 0 | 398200 | 0 | 0 | 0 | 0 | 0 | 0 | 0 | 0 | 0 |
| AFYNNVLG | OAT | P04181 | OAT_HUMAN C | Unmodified | 0 | 0 | 0 | 0 | 0 | 0 | 0 | 0 | 538950 | 301250 | 0 | 0 | 0 | 20574 | 0 | 0 | 0 | 0 |
| AFYPEEISS | HSPA1L; HSPA | P34931; P0D | HS71L_HUMAN | Oxidation (M) | 12618000 | 8179000 | 2707700 | 9439000 | 17020000 | 2021600 | 8960000 | 2661400 | 12627000 | 14163000 | 2716500 | 10156000 | 26101000 | 12243000 | 4268600 | 3301400 | 8263400 | 8435800 |
| AFYPEEISS | HSPA1L; HSPA | P34931; P0D | HS71L_HUMAN | Unmodified | 1996300 | 486120 | 104390 | 423750 | 20686000 | 0 | 4625000 | 67043 | 42220000 | 13321000 | 0 | 1663000 | 8725600 | 1292700 | 528480 | 130880 | 2601800 | 5731600 |
| AGAAERELI | ENO2 | P09104 | ENOG_HUMAN | Unmodified | 0 | 0 | 0 | 0 | 888410 | 0 | 386370 | 303760 | 3566500 | 1861300 | 0 | 1543500 | 0 | 0 | 0 | 0 | 1773500 | 2561300 |
| AGAAGALP | NXPH4 | O95158 | NXPH4_HUMA | Unmodified | 0 | 0 | 0 | 0 | 0 | 0 | 0 | 0 | 968990 | 120630 | 156900 | 0 | 0 | 0 | 0 | 0 | 0 | 0 |
| AGAAGGPE | PCNP | Q8WW12 | PCNP_HUMAN | Unmodified | 0 | 0 | 152880 | 43301 | 30626 | 64411 | 0 | 0 | 923970 | 155740 | 257640 | 0 | 141150 | 58257 | 0 | 17880 | 52836 | 48249 |
| AGAAGTAE | BCAM; BCAM; E | P50895; A0A | BCAM_HUMAN | Unmodified | 0 | 2589400 | 0 | 0 | 44957 | 0 | 0 | 0 | 716650 | 0 | 0 | 0 | 0 | 0 | 0 | 0 | 0 | 0 |
| AGAALVGV | C2orf72 | A6NCS6 | CB072_HUMAN | Unmodified | 0 | 0 | 0 | 0 | 0 | 0 | 0 | 0 | 0 | 0 | 0 | 0 | 0 | 0 | 486430 | 310370 | 0 | 0 |
| AGAAPVAP | ARHGEF2; ARH | Q92974; V9C | ARHG2_HUMA | Unmodified | 76500 | 65377 | 0 | 0 | 0 | 0 | 0 | 0 | 218130 | 14828 | 33353 | 1216100 | 0 | 0 | 16291 | 35368 | 0 | 0 |
| AGADEERA | CCT2; CCT2; C | P78371; F5G | TCPB_HUMAN | Unmodified | 212810 | 131060 | 351800 | 280550 | 0 | 1248500 | 71206 | 1988100 | 1559600 | 66884 | 95209 | 7835500 | 712140 | 147500 | 311140 | 223460 | 95685 | 410130 |
| AGADEERA | CCT2; CCT2; C | P78371; F5G | TCPB_HUMAN | Unmodified | 0 | 0 | 0 | 0 | 13476000 | 1729100 | 6829800 | 0 | 0 | 3594600 | 0 | 0 | 12529000 | 13477000 | 0 | 0 | 5657200 | 14613000 |
| AGAEGFVR | HTATIP2 | Q9BUP3 | HTAI2_HUMAN | Unmodified | 96577 | 55015 | 181930 | 229390 | 0 | 0 | 0 | 0 | 1161600 | 50222 | 0 | 716700 | 0 | 20045 | 0 | 271770 | 0 | 0 |
| AGAEGFVR | HTATIP2 | Q9BUP3 | HTAI2_HUMAN | Unmodified | 0 | 0 | 0 | 0 | 5433900 | 778630 | 1917700 | 398390 | 11795000 | 3831000 | 231100 | 316580 | 0 | 0 | 0 | 0 | 1010300 | 1292300 |
| AGAFEHLP | TPBG | Q13641 | TPBG_HUMAN | Unmodified | 0 | 0 | 0 | 0 | 0 | 0 | 0 | 0 | 0 | 0 | 0 | 2867700 | 0 | 0 | 0 | 0 | 0 | 0 |
| AGAGGGN | PPP2R2A; PPP2 | P63151; E5R | 2ABA_HUMAN | Acetyl (Protein N | 0 | 0 | 0 | 0 | 712260 | 0 | 66006 | 0 | 0 | 0 | 0 | 589370 | 562860 | 461200 | 0 | 0 | 1049000 | 813610 |
| AGAGLSSL | DDAH2 | O95865 | DDAH2_HUMA | Unmodified | 372720 | 0 | 0 | 0 | 4482600 | 429420 | 0 | 0 | 1753100 | 0 | 454160 | 710400 | 414160 | 373060 | 0 | 0 | 8199900 | 8826700 |
| AGAGSATL | MDH2; MDH2 | P40926; G3X | MDHM_HUMAN | Unmodified | 0 | 0 | 0 | 0 | 1398600 | 196040 | 1111800 | 223560 | 20507000 | 5889800 | 468500 | 226760 | 1543800 | 816080 | 0 | 0 | 355620 | 798450 |
| AGAGSATL | MDH2; MDH2 | P40926; G3X | MDHM_HUMAN | Oxidation (M) | 0 | 0 | 0 | 0 | 0 | 0 | 460340 | 33953 | 4046700 | 2123100 | 0 | 0 | 567330 | 255160 | 107330 | 0 | 0 | 0 |
| AGAGVPGE | AHNAK2 | Q8IVF2 | AHNK2_HUMA | Unmodified | 0 | 0 | 0 | 0 | 8013600 | 1341300 | 0 | 0 | 0 | 0 | 0 | 0 | 0 | 0 | 0 | 0 | 1892800 | 4786400 |
| AGAHLQG | GAPDH; GAPDH | P04406; E7E | G3P_HUMAN G | Unmodified | 0 | 0 | 0 | 0 | 0 | 0 | 0 | 0 | 0 | 0 | 0 | 0 | 0 | 0 | 0 | 0 | 0 | 0 |
| AGAHLQG | GAPDH; GAPDH | P04406; E7E | G3P_HUMAN G | 2 Oxidation (M) | 0 | 0 | 0 | 0 | 0 | 0 | 0 | 0 | 0 | 0 | 0 | 0 | 0 | 0 | 0 | 0 | 859870 | 0 |
| AGAHLQG | GAPDH; GAPDH | P04406; E7E | G3P_HUMAN G | Oxidation (M) | 0 | 0 | 0 | 0 | 0 | 0 | 0 | 0 | 0 | 0 | 0 | 0 | 0 | 0 | 0 | 0 | 1843200 | 887500 |
| AGAHLQG | GAPDH; GAPDH | P04406; E7E | G3P_HUMAN G | Unmodified | 0 | 0 | 0 | 289050 | 0 | 0 | 0 | 0 | 0 | 0 | 0 | 0 | 0 | 0 | 193420 | 0 | 777850 | 564840 |
| AGAIAPCE\ | RPLP0; RPLP0P | P05388; Q8 | RLA0_HUMAN | Unmodified | 0 | 0 | 0 | 19984000 | 1021200 | 0 | 0 | 413580 | 1730100 | 835540 | 3045500 | 52853000 | 2728200 | 2852300 | 0 | 0 | 0 | 2499600 |
| AGAISASGI | AHNAK | Q09666 | AHNK_HUMAN | Unmodified | 0 | 0 | 0 | 0 | 2237400 | 492310 | 0 | 0 | 0 | 0 | 0 | 119390 | 0 | 0 | 142420 | 0 | 377660 | 635780 |
| AGAISASGI | AHNAK | Q09666 | AHNK_HUMAN | Unmodified | 0 | 0 | 0 | 0 | 911910 | 0 | 0 | 0 | 0 | 0 | 0 | 0 | 0 | 0 | 0 | 0 | 343000 | 0 |
| AGAKPDQI | UPF1 | Q92900 | RENT1_HUMAN | Unmodified | 2340100 | 452300 | 0 | 0 | 0 | 0 | 0 | 0 | 0 | 0 | 0 | 0 | 0 | 0 | 213080 | 173390 | 0 | 0 |
| AGALQCSF | TLN1 | Q9Y490 | TLN1_HUMAN | Unmodified | 221480 | 162780 | 158300 | 1015600 | 5169200 | 792590 | 3953800 | 1795800 | 405550 | 660480 | 210360 | 1336500 | 2965900 | 4812200 | 277510 | 576720 | 2922500 | 2341200 |
| AGARKVIGI | PRMT1; PRMT1; | Q99873; E9F | ANM1_HUMAN | Unmodified | 0 | 0 | 0 | 0 | 0 | 0 | 263050 | 99297 | 0 | 0 | 0 | 0 | 0 | 0 | 0 | 0 | 481670 | 0 |
| AGASCPSG | SRGAP2; SRGA | O75044; Q5 | SRGP2_HUMA | Unmodified | 0 | 0 | 0 | 267120 | 1014400 | 217200 | 0 | 0 | 0 | 0 | 0 | 801120 | 720380 | 395670 | 0 | 186470 | 223690 | 283160 |
| AGASLWG | ITGA5 | P08648 | ITA5_HUMAN I | Unmodified | 687110 | 817980 | 714590 | 956310 | 2935700 | 1036000 | 0 | 0 | 0 | 0 | 0 | 0 | 421580 | 145800 | 1966900 | 1538800 | 1203400 | 873410 |
| AGATSEGV | DYNC1LI1; DYN | Q9Y6G9; E9I | DC1L1_HUMAN | Unmodified | 0 | 0 | 0 | 0 | 0 | 0 | 0 | 0 | 0 | 0 | 0 | 0 | 186910 | 140900 | 0 | 0 | 0 | 0 |
| AGATSEGV | DYNC1LI1; DYN | Q9Y6G9; E9I | DC1L1_HUMAN | Unmodified | 0 | 0 | 0 | 0 | 0 | 0 | 0 | 0 | 81155 | 143320 | 0 | 3929600 | 378170 | 275810 | 0 | 0 | 0 | 508420 |
| AGAVAERL | HEXA; HEXA; N | P06865; H3E | HEXA_HUMAN | Unmodified | 0 | 0 | 0 | 0 | 0 | 0 | 15430 | 0 | 579790 | 115490 | 275970 | 0 | 68730 | 0 | 0 | 0 | 0 | 0 |
| AGAVAERL | HEXA; HEXA; N | P06865; H3E | HEXA_HUMAN | Unmodified | 0 | 0 | 0 | 0 | 0 | 0 | 0 | 0 | 0 | 303070 | 0 | 0 | 0 | 0 | 0 | 0 | 0 | 0 |
| AGAVEKGV | ENO1; ENO1; E | P06733; K7E | ENOA_HUMAN | Unmodified | 0 | 0 | 0 | 0 | 356500 | 64470 | 0 | 81898 | 115600 | 80487 | 0 | 834340 | 0 | 0 | 0 | 0 | 908990 | 208020 |
| AGAVGAHL | SEC11C; SEC1 | Q9BY50; K7E | SC11C_HUMA | Unmodified | 0 | 0 | 0 | 0 | 0 | 0 | 0 | 0 | 2073800 | 0 | 610600 | 0 | 93224 | 85815 | 0 | 0 | 0 | 0 |
| AGAVGAHL | SEC11C; SEC1 | Q9BY50; K7E | SC11C_HUMA | Unmodified | 0 | 0 | 0 | 0 | 0 | 0 | 0 | 0 | 3209000 | 417190 | 556370 | 0 | 0 | 0 | 0 | 0 | 0 | 0 |
| AGAVLGRT | PLXNA4 | Q9HCM2 | PLXA4_HUMAN | Unmodified | 438730 | 903120 | 0 | 0 | 0 | 0 | 762560 | 0 | 0 | 0 | 0 | 0 | 0 | 0 | 0 | 0 | 0 | 0 |
| AGAVNPTV | DPP4; DPP4; DI | P27487; A0A | DPP4_HUMAN | Unmodified | 800610 | 612660 | 13126000 | 12327000 | 754200 | 99427 | 61637 | 80887 | 1371300 | 66883 | 0 | 657390 | 380270 | 0 | 11660000 | 6037500 | 4891800 | 3953700 |
| AGAWELR | DLL3 | Q9NYJ7 | DLL3_HUMAN | Unmodified | 0 | 0 | 0 | 0 | 0 | 0 | 0 | 0 | 1597000 | 0 | 0 | 0 | 0 | 0 | 0 | 0 | 82407 | 0 |
| AGAYDFPS | CAMK2D; CAMI | Q13557; Q1 | KCC2D_HUMA | Unmodified | 56863 | 46216 | 0 | 0 | 0 | 0 | 45935 | 55438 | 302270 | 79634 | 0 | 214700 | 0 | 0 | 0 | 0 | 0 | 0 |
| AGCECLNE | TXNL1; TXNL1 | O43396; K7E | TXNL1_HUMAN | Unmodified | 0 | 0 | 0 | 40161 | 0 | 0 | 0 | 0 | 0 | 0 | 0 | 925790 | 0 | 21542 | 0 | 0 | 238390 | 0 |
| AGCSNIAY | SLC5A3 | P53794 | SC5A3_HUMAN | Unmodified | 0 | 0 | 0 | 0 | 0 | 0 | 0 | 0 | 0 | 34592 | 0 | 383880 | 0 | 0 | 0 | 791530 | 0 | 0 |
| AGCSPEH | DLL3 | Q9NYJ7 | DLL3_HUMAN | Unmodified | 0 | 0 | 0 | 0 | 0 | 0 | 0 | 0 | 559010 | 0 | 0 | 0 | 0 | 0 | 0 | 0 | 0 | 0 |
| AGDAGIFV | SCRIB; SCRIB; | Q14160; A0A | SCRIB_HUMAN | Unmodified | 0 | 0 | 0 | 0 | 0 | 0 | 0 | 0 | 0 | 0 | 0 | 1368900 | 0 | 0 | 0 | 0 | 0 | 0 |
| AGDEFVEK | HADH; HADH; I | Q16836; A0A | HCDH_HUMAN | Unmodified | 867030 | 0 | 0 | 0 | 344430 | 2726700 | 921100 | 310950 | 3391200 | 104790 | 20406 | 232400 | 281440 | 91498 | 364190 | 458780 | 0 | 0 |
| AGDEIDEP | ZNF512; ZNF51 | Q96ME7; G3 | ZN512_HUMAN | Unmodified | 337440 | 262550 | 0 | 0 | 0 | 0 | 0 | 0 | 0 | 0 | 0 | 0 | 0 | 0 | 0 | 0 | 0 | 0 |
| AGDELTKL | PACSIN1; PACS | Q9BY11; F6 | PACN1_HUMA | Unmodified | 4454800 | 4223800 | 110370 | 245670 | 0 | 0 | 0 | 0 | 49612 | 0 | 0 | 0 | 52707 | 0 | 751660 | 492220 | 0 | 0 |
| AGDELTKM | PACSIN2 | Q9UNF0 | PACN2_HUMA | Unmodified | 10063000 | 6180600 | 3458800 | 5319400 | 0 | 0 | 0 | 0 | 1092800 | 0 | 0 | 677000 | 1636000 | 1075000 | 15715000 | 7225700 | 0 | 0 |
| AGDELTKM | PACSIN2 | Q9UNF0 | PACN2_HUMA | Oxidation (M) | 8748600 | 6219900 | 2683200 | 4847700 | 0 | 0 | 0 | 117910 | 0 | 0 | 81050 | 444930 | 1021100 | 234390 | 12357000 | 11002000 | 0 | 62102 |
| AGDISCNA | ITGA2 | P17301 | ITA2_HUMAN I | Unmodified | 2227900 | 681420 | 1308100 | 827150 | 1882000 | 383290 | 0 | 0 | 0 | 0 | 0 | 0 | 0 | 0 | 2099900 | 1422100 | 765740 | 1513400 |
| AGDKDDIT | JUP | P14923 | PLAK_HUMAN | Unmodified | 201290 | 0 | 1074500 | 1676400 | 0 | 0 | 0 | 0 | 1731900 | 0 | 484620 | 4363600 | 0 | 0 | 3023900 | 2352600 | 0 | 0 |
| AGDKPHM | SDHB; SDHB | P21912; A0A | SDHB_HUMAN | Unmodified | 0 | 0 | 0 | 0 | 0 | 0 | 15267 | 0 | 1606800 | 0 | 0 | 0 | 0 | 0 | 0 | 0 | 0 | 0 |
| AGDKPHM | SDHB; SDHB | P21912; A0A | SDHB_HUMAN | Oxidation (M) | 0 | 0 | 0 | 0 | 0 | 0 | 0 | 13081 | 277890 | 36785 | 26296 | 21602 | 0 | 0 | 0 | 0 | 0 | 0 |
| AGDLANEL | TNS3 | Q68CZ2 | TENS3_HUMAN | Unmodified | 0 | 0 | 0 | 0 | 780590 | 0 | 0 | 0 | 0 | 0 | 0 | 0 | 0 | 0 | 0 | 0 | 970360 | 490620 |
| AGDLLEDS | HDGF | P51858 | HDGF_HUMAN | Unmodified | 0 | 0 | 0 | 0 | 0 | 0 | 0 | 0 | 0 | 0 | 1512200 | 988440 | 0 | 0 | 0 | 0 | 0 | 0 |
| AGDLRDTG | XRCC6; XRCC6 | P12956; B1A | XRCC6_HUMA | Unmodified | 0 | 0 | 0 | 0 | 0 | 0 | 0 | 0 | 727190 | 0 | 498450 | 314890 | 0 | 0 | 0 | 0 | 0 | 0 |
| AGDMVYIV | SMPDL3B; SMP | Q92485; F8V | ASM3B_HUMA | Unmodified | 0 | 0 | 0 | 0 | 0 | 0 | 0 | 0 | 2041800 | 0 | 0 | 0 | 0 | 0 | 0 | 0 | 0 | 0 |
| AGDMVYIV | SMPDL3B; SMP | Q92485; F8V | ASM3B_HUMA | Oxidation (M) | 2095700 | 5762300 | 4790800 | 7995600 | 927190 | 0 | 0 | 0 | 1875100 | 562450 | 983660 | 3343400 | 2769700 | 1180600 | 9743100 | 6022200 | 163950 | 0 |
| AGDNILAVI | ERMP1; ERMP1 | Q7Z2K6; A0A | ERMP1_HUMA | Unmodified | 0 | 0 | 0 | 0 | 0 | 0 | 0 | 0 | 3871600 | 244020 | 0 | 386580 | 0 | 0 | 0 | 0 | 116940 | 0 |
| AGDPKLPV | ACOT8; ACOT8; | O14734; E9F | ACOT8_HUMA | Unmodified | 0 | 0 | 0 | 0 | 0 | 0 | 0 | 49299 | 16911000 | 460800 | 0 | 0 | 0 | 0 | 0 | 0 | 0 | 0 |
| AGDPVILY\ | TM9SF1; TM9SF | O15321; E9F | TM9S1_HUMAN | Unmodified | 0 | 0 | 0 | 0 | 371230 | 0 | 0 | 212400 | 53896000 | 8948200 | 0 | 1383400 | 0 | 0 | 0 | 0 | 0 | 0 |
| AGDREDITI | CTNNB1; CTNN | P35222; A0A | CTNB1_HUMA | Unmodified | 3173400 | 1206800 | 2495600 | 5047100 | 0 | 253310 | 0 | 0 | 883850 | 0 | 1258200 | 7815400 | 0 | 64801 | 7293000 | 10231000 | 747670 | 395820 |
| AGDREVQF | PSMC3; PSMC3 | P17980; E9P | PRS6A_HUMAN | Unmodified | 0 | 0 | 0 | 67434 | 5092000 | 418500 | 1654400 | 0 | 0 | 0 | 0 | 0 | 4886600 | 4675400 | 0 | 0 | 4100800 | 5613000 |
| AGDRLVEV | SLC9A3R2 | Q15599 | NHRF2_HUMA | Unmodified | 0 | 387630 | 0 | 1489500 | 0 | 0 | 0 | 0 | 806320 | 0 | 0 | 450300 | 0 | 0 | 505820 | 527960 | 0 | 0 |
| AGDSRNAI | KCNJ5 | P48544 | KCNJ5_HUMAN | Oxidation (M) | 0 | 0 | 0 | 0 | 0 | 0 | 49422 | 0 | 801630 | 0 | 0 | 0 | 36167 | 0 | 0 | 0 | 0 | 0 |
| AGDTGVKL | HIP1 | O00291 | HIP1_HUMAN I | Unmodified | 7698300 | 55801000 | 14850000 | 21098000 | 17858000 | 4785000 | 632540 | 0 | 10871000 | 7943100 | 840520 | 4393400 | 35538000 | 3493900 | 15647000 | 12659000 | 2316800 | 8136800 |
| AGDTLSGT | CARM1 | Q86X55 | CARM1_HUMA | Unmodified | 235640 | 0 | 0 | 0 | 252180 | 0 | 0 | 0 | 0 | 0 | 0 | 1655000 | 186420 | 0 | 0 | 0 | 0 | 0 |

| | | | | | | | | | | | | | | | | | | | | | | |
|---|---|---|---|---|---|---|---|---|---|---|---|---|---|---|---|---|---|---|---|---|---|---|
| AGDTTKFD | INF2; INF2; INF | Q27J81; A0A | INF2_HUMAN I | Unmodified | 0 | 0 | 0 | 0 | 0 | 0 | 152830 | 0 | 661940 | 308560 | 338960 | 3484400 | 0 | 1692700 | 0 | 167530 | 0 | 0 |
| AGDTVIPLY | ADH5; ADH5 | P11766; D6F | ADHX_HUMAN | Unmodified | 0 | 0 | 1218100 | 313040 | 3920500 | 176140 | 188550 | 435190 | 0 | 136820 | 1286700 | 6599600 | 409780 | 332690 | 57224 | 82042 | 629160 | 2667900 |
| AGEAGKLE | CORO1B | Q9BR76 | COR1B_HUMA | Unmodified | 1003000 | 655310 | 0 | 1349700 | 719540 | 0 | 0 | 0 | 2561500 | 150500 | 1280400 | 2696400 | 951250 | 0 | 2852900 | 867650 | 209510 | 0 |
| AGEARPGP | SQSTM1; SQST | Q13501; E9F | SQSTM_HUMA | Unmodified | 1992500 | 2540100 | 4271700 | 6079700 | 7629800 | 385380 | 1226500 | 370220 | 23897000 | 579930 | 0 | 1954300 | 3909900 | 443020 | 2081600 | 1769900 | 2189300 | 4985900 |
| AGEDAVW | QSOX2 | Q6ZRP7 | QSOX2_HUMA | Unmodified | 0 | 0 | 0 | 0 | 0 | 357230 | 0 | 0 | 23701000 | 1433200 | 0 | 675010 | 0 | 0 | 0 | 0 | 0 | 649870 |
| AGEDPYRC | GALNT16 | Q8N428 | GLT16_HUMAN | Unmodified | 0 | 0 | 0 | 0 | 0 | 0 | 0 | 0 | 2977400 | 0 | 0 | 0 | 0 | 0 | 0 | 0 | 0 | 0 |
| AGEELLKM | PACSIN3 | Q9UKS6 | PACN3_HUMA | Oxidation (M) | 927970 | 2125000 | 0 | 0 | 0 | 0 | 0 | 0 | 0 | 0 | 0 | 0 | 0 | 0 | 0 | 0 | 0 | 0 |
| AGEGGGVI | DDC; DDC; DD | P20711; A0A | DDC_HUMAN / | Unmodified | 0 | 0 | 0 | 0 | 0 | 0 | 0 | 0 | 1718100 | 2658500 | 0 | 0 | 0 | 0 | 0 | 0 | 0 | 0 |
| AGEHGVW | MARVELD3; MA | Q96A59; J3K | MALD3_HUMA | Unmodified | 440140 | 448200 | 0 | 615030 | 0 | 0 | 0 | 0 | 0 | 0 | 0 | 621890 | 0 | 0 | 0 | 273570 | 0 | 0 |
| AGELTEDE | RPS18 | P62269 | RS18_HUMAN | Unmodified | 198750 | 496280 | 186240 | 238110 | 130200 | 0 | 0 | 41771 | 641770 | 0 | 1898900 | 8338700 | 0 | 0 | 196370 | 0 | 0 | 0 |
| AGELTEDE | RPS18 | P62269 | RS18_HUMAN | Unmodified | 0 | 747300 | 0 | 0 | 0 | 0 | 0 | 0 | 355190 | 0 | 0 | 2154100 | 0 | 0 | 0 | 0 | 88084 | 0 |
| AGELTPEEE | CALML5 | Q9NZT1 | CALL5_HUMAN | Acetyl (Protein N | 0 | 85390 | 0 | 0 | 0 | 0 | 135800 | 0 | 0 | 0 | 0 | 0 | 0 | 0 | 0 | 0 | 414060 | 0 |
| AGENRWG | ANXA3; ANXA3; | P12429; D6F | ANXA3_HUMA | Unmodified | 0 | 0 | 0 | 0 | 1166300 | 2031300 | 0 | 0 | 0 | 0 | 0 | 66291 | 0 | 0 | 0 | 0 | 180110 | 797140 |
| AGEQKPSS | COPS7B; COPS | Q9H9Q2; J3I | CSN7B_HUMA | Acetyl (Protein N | 0 | 31986 | 0 | 0 | 108370 | 40189 | 0 | 0 | 31303 | 104380 | 80499 | 0 | 44565 | 0 | 56052 | 0 | 345020 | 0 |
| AGEQLAPF | ECPAS; ECPAS | Q5VYK3; J3K | ECM29_HUMA | Unmodified | 0 | 0 | 0 | 0 | 0 | 0 | 0 | 0 | 75133 | 0 | 59685 | 1499700 | 0 | 0 | 0 | 75396 | 0 | 0 |
| AGEVDVEC | CCNB2; CCNB2 | O95067; H1 | CCNB2_HUMA | Unmodified | 0 | 79789 | 0 | 0 | 235160 | 103700 | 86554 | 141270 | 0 | 98295 | 82225 | 226830 | 64209 | 39216 | 195060 | 97286 | 0 | 0 |
| AGEYIFER | ALDH6A1 | Q02252 | MMSA_HUMAN | Unmodified | 0 | 0 | 0 | 0 | 0 | 0 | 0 | 0 | 1013300 | 0 | 0 | 0 | 0 | 0 | 0 | 0 | 0 | 0 |
| AGFAFRQT | MYO1D; MYO1D | O94832; J3Q | MYO1D_HUMA | Unmodified | 257770 | 153940 | 216050 | 615780 | 241380 | 0 | 0 | 0 | 379140 | 561780 | 0 | 0 | 497290 | 254580 | 0 | 0 | 164520 | 710560 |
| AGFAGDD/ | POTEJ; POTEE; | P0CG39; Q6 | POTEJ_HUMAN | Unmodified | 1.26E+08 | 1.3E+08 | 1.66E+08 | 91290000 | 7.74E+08 | 2E+08 | 55748000 | 46583000 | 2.83E+08 | 51643000 | 1.08E+08 | 8.33E+08 | 1.96E+08 | 1.88E+08 | 1.91E+08 | 2.19E+08 | 1.17E+08 | 1.83E+08 |
| AGFAGDD/ | POTEE; POTEI; | Q6S8J3; P0C | POTEE_HUMAN | Unmodified | 0 | 0 | 0 | 0 | 0 | 1007900 | 959440 | 0 | 0 | 658920 | 0 | 0 | 1187800 | 586010 | 0 | 0 | 4891100 | 558460 |
| AGFAGDD/ | POTEE; POTEI; | Q6S8J3; P0C | POTEE_HUMAN | Unmodified | 0 | 0 | 1220800 | 2141900 | 1.02E+09 | 3.58E+08 | 85646000 | 23672000 | 1017000 | 88072000 | 4219900 | 0 | 4.95E+08 | 2.99E+08 | 0 | 0 | 2.12E+08 | 2.66E+08 |
| AGFAGDQI | ACTR1A; ACTR1 | P61163; P42 | ACTZ_HUMAN | Unmodified | 287460 | 0 | 0 | 0 | 4969200 | 1679600 | 6566200 | 4220300 | 134030 | 1831400 | 0 | 10788000 | 2614000 | 2302600 | 0 | 0 | 1536100 | 6882900 |
| AGFAGDQI | ACTR1A; ACTR1 | P61163; P42 | ACTZ_HUMAN | Unmodified | 0 | 0 | 0 | 0 | 48504 | 0 | 0 | 0 | 0 | 0 | 0 | 0 | 0 | 0 | 0 | 0 | 881500 | 0 |
| AGFFGTVV | DNAJC11 | Q9NVH1 | DJC11_HUMAN | Unmodified | 0 | 0 | 0 | 0 | 33189 | 0 | 0 | 0 | 2193400 | 715620 | 0 | 0 | 0 | 0 | 53974 | 63448 | 0 | 0 |
| AGFGGVLL | NAALADL2 | Q58DX5 | NADL2_HUMA | Unmodified | 0 | 0 | 0 | 0 | 0 | 0 | 0 | 0 | 0 | 0 | 0 | 0 | 456920 | 0 | 0 | 0 | 0 | 0 |
| AGFIESTFK | EIF2AK3 | Q9NZJ5 | E2AK3_HUMAN | Unmodified | 0 | 0 | 0 | 37036 | 0 | 0 | 0 | 0 | 967910 | 0 | 0 | 0 | 0 | 0 | 0 | 0 | 0 | 0 |
| AGFLAFTQI | DNAJC13; DNA | O75165; A0/ | DJC13_HUMAN | Unmodified | 0 | 0 | 0 | 0 | 0 | 0 | 0 | 0 | 0 | 0 | 15029 | 2582100 | 0 | 0 | 0 | 0 | 0 | 0 |
| AGFLDLKD | TLN1 | Q9Y490 | TLN1_HUMAN | Unmodified | 0 | 0 | 0 | 0 | 2121300 | 263380 | 5921700 | 179500 | 0 | 244630 | 0 | 0 | 830610 | 1383500 | 0 | 0 | 606820 | 729110 |
| AGFLDLKD | TLN1 | Q9Y490 | TLN1_HUMAN | Unmodified | 0 | 0 | 0 | 0 | 485250 | 0 | 1798700 | 0 | 0 | 0 | 0 | 0 | 292120 | 284660 | 0 | 0 | 232550 | 342450 |
| AGFPLAVH | PECR | Q9BY49 | PECR_HUMAN | Unmodified | 44847 | 625380 | 0 | 34031 | 200560 | 4124000 | 93399 | 27732 | 2931000 | 844090 | 0 | 0 | 0 | 0 | 0 | 0 | 223750 | 230240 |
| AGFRPVEA | CASC3 | O15234 | CASC3_HUMA | Unmodified | 637710 | 652360 | 0 | 0 | 0 | 0 | 27151 | 0 | 0 | 0 | 0 | 83556 | 77480 | 0 | 0 | 184190 | 0 | 0 |
| AGFTAAYSI | IGF2R | P11717 | MPRI_HUMAN | Unmodified | 846790 | 115210 | 0 | 0 | 0 | 0 | 0 | 0 | 32975000 | 318930 | 3272700 | 0 | 0 | 0 | 0 | 0 | 0 | 0 |
| AGFYYTGV | BIRC2; BIRC3; | Q13490; Q1 | BIRC2_HUMAN | Unmodified | 0 | 0 | 0 | 0 | 840320 | 164870 | 0 | 0 | 0 | 0 | 0 | 0 | 0 | 0 | 0 | 0 | 0 | 0 |
| AGFYYTGV | BIRC2; BIRC3; | Q13490; Q1 | BIRC2_HUMAN | Unmodified | 0 | 0 | 0 | 0 | 1958100 | 0 | 0 | 0 | 0 | 0 | 0 | 0 | 0 | 0 | 0 | 0 | 0 | 0 |
| AGGAAVVI | RCC2 | Q9P258 | RCC2_HUMAN | Unmodified | 0 | 0 | 0 | 0 | 811330 | 230180 | 543290 | 244970 | 3068000 | 2183400 | 4712700 | 14138000 | 391850 | 246040 | 0 | 0 | 0 | 0 |
| AGGADAV( | EHD4 | Q9H223 | EHD4_HUMAN | Unmodified | 6563100 | 7215000 | 4701000 | 5420400 | 3001000 | 2563700 | 0 | 0 | 1391900 | 192170 | 601300 | 5549300 | 2724500 | 0 | 9039000 | 7345600 | 1246600 | 3379500 |
| AGGADAV( | EHD4 | Q9H223 | EHD4_HUMAN | Unmodified | 0 | 0 | 0 | 0 | 928090 | 775010 | 0 | 0 | 0 | 0 | 0 | 0 | 1352300 | 0 | 0 | 0 | 1397000 | 1851300 |
| AGGADAV( | EHD4 | Q9H223 | EHD4_HUMAN | Unmodified | 0 | 0 | 0 | 0 | 5289500 | 0 | 0 | 0 | 0 | 823020 | 0 | 0 | 2030200 | 4237700 | 0 | 0 | 4820300 | 4917400 |
| AGGAGVP/ | OXCT1 | P55809 | SCOT1_HUMA | Unmodified | 0 | 0 | 0 | 0 | 1466300 | 0 | 654950 | 0 | 6273100 | 0 | 0 | 0 | 0 | 0 | 0 | 0 | 0 | 0 |
| AGGANYD/ | COTL1; COTL1 | Q14019; H3I | COTL1_HUMAN | Unmodified | 0 | 0 | 0 | 0 | 635380 | 0 | 0 | 0 | 456280 | 0 | 0 | 0 | 0 | 0 | 0 | 0 | 205630 | 453320 |
| AGGEAGVT | EIF2S3; EIF2S3I | P41091; Q2\ | IF2G_HUMAN E | Acetyl (Protein N | 105210 | 0 | 0 | 0 | 0 | 0 | 0 | 0 | 0 | 0 | 415460 | 2658800 | 0 | 0 | 0 | 0 | 0 | 0 |
| AGGEAGVT | EIF2S3; EIF2S3I | P41091; Q2\ | IF2G_HUMAN E | Acetyl (Protein N | 0 | 0 | 0 | 0 | 396360 | 49197 | 0 | 0 | 0 | 0 | 0 | 7937300 | 0 | 0 | 0 | 0 | 512200 | 0 |
| AGGEAGVT | EIF2S3; EIF2S3I | P41091; Q2\ | IF2G_HUMAN E | Unmodified | 0 | 0 | 277850 | 164890 | 0 | 0 | 0 | 0 | 0 | 0 | 0 | 853960 | 157140 | 0 | 0 | 514830 | 0 | 101570 |
| AGGEAGVT | EIF2S3; EIF2S3I | P41091; Q2\ | IF2G_HUMAN E | Acetyl (Protein N | 0 | 171600 | 0 | 6459700 | 10812000 | 983670 | 0 | 0 | 0 | 6799500 | 559900 | 531590 | 12923000 | 5005900 | 771870 | 168480 | 4855900 | 5234700 |
| AGGFLMKK | PGK1 | P00558 | PGK1_HUMAN | Unmodified | 0 | 0 | 0 | 0 | 2400000 | 609230 | 0 | 0 | 1397000 | 2163300 | 0 | 0 | 0 | 2114500 | 0 | 407010 | 7568500 | 12310000 |
| AGGFLMKK | PGK1 | P00558 | PGK1_HUMAN | Oxidation (M) | 0 | 0 | 0 | 0 | 0 | 0 | 207860 | 0 | 0 | 0 | 0 | 0 | 0 | 0 | 195640 | 0 | 1613500 | 0 |
| AGGFLMKK | PGK1 | P00558 | PGK1_HUMAN | Unmodified | 0 | 0 | 0 | 0 | 0 | 0 | 128880 | 0 | 0 | 0 | 0 | 0 | 0 | 0 | 0 | 0 | 1611100 | 0 |
| AGGGAGD | RAB32 | Q13637 | RAB32_HUMAN | Unmodified | 0 | 0 | 0 | 0 | 27795000 | 1585100 | 0 | 0 | 325260 | 305450 | 0 | 0 | 0 | 0 | 0 | 0 | 10984000 | 9095200 |
| AGGGPQEI | LRP11; LRP11 | Q86VZ4; Q5 | LRP11_HUMAN | Unmodified | 0 | 0 | 0 | 0 | 0 | 0 | 0 | 0 | 0 | 0 | 0 | 0 | 0 | 0 | 0 | 0 | 0 | 0 |
| AGGGPQEI | LRP11; LRP11 | Q86VZ4; Q5 | LRP11_HUMAN | Oxidation (M) | 0 | 0 | 0 | 0 | 0 | 0 | 0 | 0 | 755300 | 58970 | 0 | 0 | 0 | 0 | 0 | 0 | 0 | 0 |
| AGGIETIAN | HSPH1; HSPH1 | Q92598; R4 | HS105_HUMAN | Unmodified | 218100 | 0 | 0 | 0 | 0 | 0 | 0 | 0 | 895880 | 0 | 737700 | 6006400 | 71751 | 0 | 0 | 0 | 0 | 0 |
| AGGIETIAN | HSPH1; HSPH1 | Q92598; R4 | HS105_HUMAN | Unmodified | 0 | 988870 | 0 | 0 | 0 | 0 | 1207200 | 0 | 0 | 0 | 0 | 0 | 6867400 | 0 | 0 | 0 | 0 | 0 |
| AGGIETIAN | HSPA4; HSPA4; | P34932; A0A | HSP74_HUMAN | Unmodified | 163510 | 0 | 73007 | 92595 | 0 | 0 | 0 | 25596 | 721800 | 0 | 999960 | 8252700 | 0 | 0 | 622720 | 575760 | 0 | 0 |
| AGGIETIAN | HSPA4; HSPA4; | P34932; A0A | HSP74_HUMAN | Unmodified | 0 | 0 | 0 | 0 | 1183900 | 0 | 0 | 0 | 0 | 604620 | 0 | 140890 | 1916000 | 259220 | 0 | 0 | 981960 | 219310 |
| AGGLDWPI | PKP3; PKP3; PK | Q9Y446; E9F | PKP3_HUMAN | Unmodified | 93374 | 0 | 0 | 679010 | 0 | 0 | 0 | 0 | 0 | 0 | 1844700 | 2883000 | 372150 | 334770 | 0 | 589040 | 0 | 0 |
| AGGLGTN( | SNX33 | Q8WV41 | SNX33_HUMAN | Unmodified | 0 | 0 | 0 | 0 | 0 | 0 | 0 | 0 | 0 | 0 | 0 | 0 | 0 | 0 | 1960100 | 0 | 0 | 0 |
| AGGMHHI( | MCEE | Q96PE7 | MCEE_HUMAN | Oxidation (M) | 0 | 0 | 0 | 0 | 0 | 0 | 0 | 0 | 0 | 0 | 0 | 824990 | 0 | 0 | 0 | 0 | 0 | 0 |
| AGGNQAA | SLC38A10 | Q9HBR0 | S38AA_HUMAN | Unmodified | 0 | 0 | 0 | 0 | 0 | 0 | 0 | 0 | 13052000 | 0 | 0 | 0 | 0 | 0 | 0 | 0 | 0 | 0 |
| AGGPEGPF | TTLL12 | Q14166 | TTL12_HUMAN | Unmodified | 0 | 0 | 120240 | 0 | 0 | 0 | 0 | 0 | 0 | 0 | 254330 | 1896000 | 0 | 0 | 0 | 0 | 0 | 0 |
| AGGPGLEF | FLNA; FLNA; FL | P21333; A0A | FLNA_HUMAN | Unmodified | 0 | 0 | 803610 | 8133900 | 0 | 0 | 0 | 0 | 0 | 0 | 0 | 0 | 0 | 0 | 2417100 | 836550 | 0 | 0 |
| AGGPGLEF | FLNB; FLNB | O75369; E7E | FLNB_HUMAN | Unmodified | 0 | 0 | 912200 | 541230 | 910640 | 0 | 175740 | 0 | 284980 | 206580 | 127200 | 2453800 | 101610 | 0 | 905590 | 838830 | 0 | 0 |
| AGGPLGLS | SCRIB; SCRIB; | Q14160; A0/ | SCRIB_HUMAN | Unmodified | 0 | 0 | 1786500 | 4963100 | 12237000 | 2164600 | 0 | 0 | 61312000 | 21574000 | 2832500 | 11387000 | 13563000 | 4503200 | 0 | 0 | 25505000 | 20907000 |
| AGGPTTPL | LMNB1; LMNB1 | P20700; A0A | LMNB1_HUMA | Unmodified | 0 | 0 | 0 | 0 | 0 | 0 | 0 | 0 | 0 | 0 | 0 | 2959800 | 0 | 0 | 0 | 0 | 0 | 0 |
| AGGQAPR( | PROSER2; PRO | Q86WR7; D3 | PRSR2_HUMAN | Unmodified | 0 | 0 | 0 | 221620 | 3111300 | 0 | 152850 | 0 | 439280 | 617850 | 481050 | 0 | 0 | 0 | 61386 | 66645 | 0 | 323150 |
| AGGRIAVPI | SFTPA2; SFTPA | Q8IWL1; Q8I | SFPA2_HUMAN | Unmodified | 0 | 0 | 0 | 0 | 0 | 0 | 0 | 0 | 0 | 1363700 | 0 | 0 | 0 | 0 | 0 | 0 | 0 | 0 |
| AGGSAALS | H1-10 | Q92522 | H1X_HUMAN H | Unmodified | 0 | 0 | 0 | 0 | 0 | 0 | 0 | 418330 | 0 | 0 | 0 | 0 | 206190 | 0 | 0 | 0 | 0 | 0 |
| AGGSASAN | PSMB1 | P20618 | PSB1_HUMAN | Unmodified | 60411 | 132020 | 633460 | 0 | 1089100 | 42567 | 57427 | 20884 | 3182800 | 389770 | 0 | 1679200 | 0 | 0 | 458180 | 92027 | 190000 | 1075900 |
| AGGSASAN | PSMB1 | P20618 | PSB1_HUMAN | Oxidation (M) | 579870 | 0 | 0 | 933310 | 1250000 | 268370 | 0 | 140550 | 876130 | 644530 | 1290900 | 11008000 | 884110 | 692610 | 0 | 1219500 | 544800 | 670280 |
| AGGSFDLR | N/A; SYNGR2; | A8MWL6; O | SNG2L_HUMA | Unmodified | 46222 | 167800 | 0 | 1620500 | 292080 | 0 | 42888 | 32824 | 2489600 | 272640 | 252880 | 6994500 | 608050 | 54069 | 1393500 | 2853000 | 0 | 21138 |
| AGGTGGA | PCDHA4; PCDH | Q9UN74; A0 | PCDA4_HUMA | Unmodified | 0 | 0 | 0 | 0 | 1435500 | 260030 | 1313600 | 0 | 1772900 | 566680 | 0 | 0 | 0 | 0 | 0 | 0 | 591010 | 854680 |
| AGGTGPGI | APEH; APEH; AI | P13798; C9J | ACPH_HUMAN | Unmodified | 0 | 0 | 0 | 0 | 0 | 0 | 0 | 0 | 722070 | 0 | 0 | 815480 | 733660 | 736740 | 0 | 0 | 1331700 | 637910 |
| AGGTPRKA | H1-2 | P16403 | H12_HUMAN H | Unmodified | 0 | 0 | 0 | 0 | 0 | 0 | 678940 | 27029000 | 0 | 0 | 0 | 0 | 0 | 0 | 0 | 0 | 0 | 0 |
| AGGTSDSF | SYTL4 | Q96C24 | SYTL4_HUMAN | Unmodified | 0 | 0 | 0 | 0 | 898770 | 0 | 224660 | 0 | 0 | 212080 | 0 | 139160 | 0 | 0 | 0 | 0 | 184430 | 303340 |
| AGHFDKEI | ACAT2 | Q9BWD1 | THIC_HUMAN / | Unmodified | 0 | 0 | 0 | 0 | 5280200 | 0 | 604860 | 148130 | 0 | 521140 | 0 | 0 | 961350 | 0 | 0 | 0 | 0 | 0 |
| AGHMVPSI | SCPEP1; SCPEI | Q9HB40; I3L | RISC_HUMAN I | Oxidation (M) | 8151.8 | 0 | 0 | 0 | 58574 | 67078 | 61881 | 11507 | 236770 | 153290 | 0 | 287530 | 0 | 77066 | 0 | 95932 | 0 | 0 |
| AGHMVPSI | SCPEP1; SCPEI | Q9HB40; I3L | RISC_HUMAN I | Unmodified | 0 | 0 | 0 | 0 | 100430 | 0 | 21188 | 0 | 910980 | 328940 | 0 | 0 | 0 | 0 | 0 | 0 | 144810 | 94547 |
| AGHYLELG | MAN1A2; MAN1 | O60476; H0 | MA1A2_HUMA | Unmodified | 1583100 | 0 | 0 | 16585000 | 3156200 | 2852800 | 916950 | 307000 | 11486000 | 472400 | 0 | 3682100 | 248670 | 0 | 3719800 | 146480 | 1355900 | 3462400 |
| AGIAAKLAK | CAPN2 | P17655 | CAN2_HUMAN | Acetyl (Protein N | 0 | 0 | 0 | 0 | 0 | 0 | 0 | 109490 | 0 | 0 | 0 | 209700 | 0 | 0 | 0 | 0 | 2110100 | 0 |
| AGIIASARA | EEF2 | P13639 | EF2_HUMAN E | Unmodified | 194000 | 0 | 104970 | 0 | 2670400 | 490190 | 412700 | 21050 | 228590 | 412290 | 0 | 0 | 663170 | 0 | 196810 | 0 | 622520 | 869340 |
| AGIICQLN/ | MCM4; MCM4; | P33991; A0A | MCM4_HUMAN | Unmodified | 0 | 0 | 0 | 0 | 302800 | 0 | 265320 | 103580 | 154260 | 136040 | 0 | 903470 | 554500 | 0 | 0 | 0 | 0 | 0 |
| AGIISTVEVI | NPEPPS; NPEPI | P55786; A0A | PSA_HUMAN P | Unmodified | 0 | 0 | 0 | 0 | 0 | 0 | 0 | 0 | 0 | 0 | 0 | 4518700 | 0 | 0 | 0 | 0 | 0 | 0 |
| AGIPNQIPK | FREM1; FREM1 | Q5H8C1; F8 | FREM1_HUMA | Unmodified | 0 | 0 | 0 | 0 | 0 | 0 | 0 | 0 | 566270 | 349790 | 0 | 631790 | 0 | 0 | 0 | 0 | 0 | 0 |
| AGIPQHHP | NECTIN1 | Q15223 | NECT1_HUMA | Oxidation (M) | 0 | 0 | 0 | 0 | 0 | 0 | 0 | 0 | 0 | 0 | 0 | 800030 | 0 | 0 | 0 | 0 | 0 | 0 |
| AGIPQHHP | NECTIN1 | Q15223 | NECT1_HUMA | Unmodified | 0 | 0 | 0 | 189930 | 0 | 0 | 0 | 0 | 0 | 0 | 0 | 1684800 | 588240 | 0 | 0 | 0 | 0 | 0 |
| AGIQQVYTI | CUL1; CUL1; CI | Q13616; A0/ | CUL1_HUMAN | Unmodified | 0 | 0 | 0 | 0 | 4500500 | 2666700 | 0 | 31174 | 99537 | 168900 | 344240 | 1352200 | 0 | 0 | 0 | 0 | 1088700 | 996850 |
| AGITAAINL | GPR107 | Q5VW38 | GP107_HUMA | Unmodified | 0 | 0 | 0 | 176440 | 311230 | 1348300 | 228380 | 78727 | 0 | 0 | 0 | 162210 | 377700 | 0 | 0 | 149780 | 0 | 48753 |
| AGITTIEAV | TPM3; TPM3; TF | D6R904; A0/ | D6R904_HUM/ | Acetyl (Protein N | 0 | 0 | 1419400 | 2067200 | 61148000 | 15520000 | 3582400 | 8122400 | 5695200 | 3815300 | 1439200 | 16606000 | 19933000 | 12087000 | 132320 | 4361700 | 12500000 | 9176700 |
| AGITTIEAV | TPM3; TPM3; TF | D6R904; A0/ | D6R904_HUM/ | Acetyl (Protein N | 0 | 0 | 0 | 0 | 20868000 | 0 | 1956400 | 2819400 | 0 | 583560 | 0 | 0 | 5604000 | 0 | 0 | 0 | 11562000 | 1667400 |
| AGITTTLNS | MCM5; MCM5 | P33992; B1A | MCM5_HUMAN | Unmodified | 0 | 0 | 0 | 0 | 632430 | 66748 | 191920 | 0 | 178260 | 0 | 0 | 692250 | 0 | 0 | 0 | 0 | 56874 | 70129 |
| AGIVTSLQ/ | MCM2; MCM2 | P49736; H0 | MCM2_HUMAN | Unmodified | 0 | 436240 | 0 | 14415 | 172800 | 0 | 0 | 0 | 692790 | 370300 | 293090 | 1849600 | 383720 | 0 | 0 | 266090 | 62198 | 334770 |
| AGKEESGV | SLC29A1 | Q99808 | S29A1_HUMAN | Unmodified | 6224500 | 7983500 | 8943700 | 4296700 | 0 | 0 | 0 | 0 | 3006800 | 418840 | 2511000 | 48453000 | 0 | 0 | 14390000 | 18197000 | 66123 | 0 |
| AGKFPSLLT | RPL10A | P62906 | RL10A_HUMAN | Oxidation (M) | 0 | 0 | 0 | 0 | 176440 | 0 | 0 | 0 | 0 | 0 | 0 | 9676600 | 0 | 0 | 0 | 0 | 217110 | 0 |
| AGKFPSLLT | RPL10A | P62906 | RL10A_HUMAN | Unmodified | 1391100 | 721000 | 115340 | 0 | 0 | 0 | 0 | 0 | 0 | 0 | 0 | 12427000 | 0 | 0 | 395130 | 278540 | 0 | 0 |
| AGKFPSLLT | RPL10A | P62906 | RL10A_HUMAN | Oxidation (M) | 0 | 0 | 0 | 0 | 0 | 0 | 0 | 0 | 0 | 0 | 0 | 4457500 | 0 | 786570 | 1144200 | 0 | 0 | 0 |
| AGKFPSLLT | RPL10A | P62906 | RL10A_HUMAN | Unmodified | 598740 | 0 | 0 | 408690 | 0 | 0 | 258400 | 0 | 0 | 0 | 0 | 4573000 | 0 | 0 | 239700 | 40998 | 0 | 0 |
| AGKGGPTF | CHMP4B | Q9H444 | CHM4B_HUMA | Unmodified | 3276100 | 5213200 | 0 | 3774100 | 9759700 | 1706500 | 202870 | 19061 | 5045600 | 3463300 | 535550 | 0 | 7693400 | 2208600 | 758930 | 370470 | 665600 | 2558200 |
| AGKITFAEK | TFRC; TFRC | P02786; G3\ | TFR1_HUMAN | Unmodified | 0 | 0 | 0 | 0 | 0 | 0 | 0 | 0 | 1301100 | 270880 | 0 | 0 | 0 | 0 | 0 | 0 | 0 | 0 |
| AGKLDPHL | MYH9; MYH10 | P35579; P35 | MYH9_HUMAN | Unmodified | 406840 | 232070 | 1524900 | 1562400 | 0 | 0 | 0 | 0 | 1729500 | 730610 | 999930 | 10273000 | 72701 | 55687 | 1539900 | 1324500 | 89371 | 0 |
| AGKLDPHL | MYH9; MYH10 | P35579; P35 | MYH9_HUMAN | Unmodified | 0 | 0 | 0 | 145690 | 0 | 122560 | 47142 | 0 | 0 | 380560 | 0 | 0 | 0 | 83030 | 0 | 0 | 120450 | 155540 |
| AGKLIENVI | NAPG; NAPG | Q99747; J3K | SNAG_HUMAN | Unmodified | 0 | 0 | 0 | 0 | 361490 | 0 | 128030 | 0 | 0 | 0 | 0 | 0 | 256550 | 0 | 0 | 0 | 0 | 417080 |
| AGKPVICA | PKM; PKM; PKM | P14618; B4I | KPYM_HUMAN | 2 Oxidation (M) | 152620 | 157590 | 266880 | 779130 | 815860 | 217540 | 182930 | 0 | 0 | 129520 | 758320 | 4958300 | 269760 | 0 | 250400 | 338500 | 152250 | 403990 |
| AGKPVICA | PKM; PKM; PKM | P14618; B4I | KPYM_HUMAN | Oxidation (M) | 208260 | 179980 | 0 | 0 | 7271600 | 701860 | 283820 | 0 | 993220 | 292800 | 1050700 | 7784600 | 1511600 | 522850 | 417380 | 179850 | 1382900 | 2370500 |
| AGKPVICA | PKM; PKM; PKM | P14618; B4I | KPYM_HUMAN | Unmodified | 0 | 0 | 0 | 0 | 130730 | 15999 | 0 | 0 | 939500 | 0 | 0 | 0 | 356750 | 0 | 0 | 0 | 698270 | 758650 |
| AGKTLQIFN | CLTCL1; CLTC; | P53675; Q0 | CLH2_HUMAN | Unmodified | 0 | 0 | 0 | 0 | 241630 | 0 | 402300 | 34294 | 0 | 0 | 0 | 0 | 0 | 0 | 0 | 0 | 130730 | 141880 |
| AGKTLQIFN | CLTCL1; CLTC; | P53675; Q0 | CLH2_HUMAN | Oxidation (M) | 0 | 0 | 0 | 0 | 0 | 0 | 262860 | 162700 | 0 | 0 | 0 | 149440 | 145420 | 67731 | 0 | 0 | 82425 | 0 |
| AGKVDCQ/ | DNAJC10; DNA | Q8IXB1; A0A | DJC10_HUMAN | Unmodified | 0 | 0 | 0 | 0 | 0 | 0 | 0 | 0 | 5619900 | 3586400 | 292250 | 215310 | 0 | 0 | 0 | 0 | 0 | 0 |
| AGKVEAQH | PYCR3; PYCR3 | Q53H96; A0 | P5CR3_HUMA | Unmodified | 0 | 0 | 0 | 0 | 145860 | 42798 | 58912 | 0 | 0 | 0 | 0 | 382310 | 0 | 0 | 0 | 144020 | 0 | 0 |
| AGLAASLA( | SOD3 | P08294 | SODE_HUMAN | Unmodified | 75949 | 61212 | 0 | 0 | 0 | 0 | 0 | 0 | 9125700 | 2248900 | 0 | 0 | 0 | 0 | 0 | 0 | 0 | 0 |
| AGLAPLEVI | FLNB; FLNB | O75369; E7E | FLNB_HUMAN | Unmodified | 0 | 0 | 509540 | 263760 | 3412300 | 1101500 | 0 | 0 | 1084100 | 132490 | 0 | 3682300 | 305640 | 187760 | 685030 | 630610 | 840910 | 421580 |

| | | | | | | | | | | | | | | | | | | | | | | |
|---|---|---|---|---|---|---|---|---|---|---|---|---|---|---|---|---|---|---|---|---|---|---|
| AGLAPLEVI | FLNB; FLNB | O75369; E7E | FLNB_HUMAN | Unmodified | 0 | 0 | 227270 | 0 | 0 | 1979600 | 0 | 0 | 0 | 0 | 0 | 0 | 1321800 | 0 | 0 | 0 | 1376100 | 0 |
| AGLELLSD | EXOSC4; EXOS | Q9NPD3; E9 | EXOS4_HUMAN | Acetyl (Protein N | 65120 | 63263 | 61790 | 211160 | 669060 | 0 | 0 | 49969 | 146360 | 0 | 0 | 533470 | 1097100 | 327510 | 0 | 67525 | 618470 | 0 |
| AGLESGAE | DKC1 | O60832 | DKC1_HUMAN | Unmodified | 0 | 0 | 0 | 0 | 0 | 0 | 0 | 0 | 0 | 0 | 0 | 0 | 0 | 0 | 0 | 0 | 0 | 0 |
| AGLGHPAA | DDAH1 | O94760 | DDAH1_HUMA | Acetyl (Protein N | 592550 | 343770 | 646960 | 855850 | 450590 | 850940 | 0 | 0 | 2426500 | 1988500 | 0 | 2993800 | 3930100 | 354870 | 1131600 | 1217100 | 457290 | 2759900 |
| AGLGSGLS | ACADVL; ACAD | P49748; G3V | ACADV_HUMAN | Unmodified | 0 | 0 | 0 | 0 | 420670 | 0 | 0 | 0 | 14502000 | 0 | 0 | 0 | 0 | 0 | 0 | 0 | 375190 | 0 |
| AGLLAGDR | SLC9A3R1; SLC | O14745; J3Q | NHRF1_HUMAN | Unmodified | 0 | 0 | 996490 | 2032200 | 2535700 | 447930 | 0 | 191490 | 1143900 | 0 | 1192400 | 1838900 | 4158300 | 5635400 | 2139900 | 2357700 | 352630 | 315200 |
| AGLLAGDR | SLC9A3R1; SLC | O14745; J3Q | NHRF1_HUMAN | Unmodified | 0 | 0 | 0 | 0 | 12394000 | 4780500 | 693130 | 134940 | 0 | 1381000 | 183340 | 0 | 52065000 | 61032000 | 0 | 0 | 5516000 | 4387000 |
| AGLLEPEVI | CPT2; CPT2; CF | P23786; A0A | CPT2_HUMAN | Unmodified | 0 | 0 | 0 | 0 | 0 | 0 | 0 | 0 | 1407100 | 21862 | 0 | 140130 | 0 | 0 | 0 | 0 | 0 | 0 |
| AGLLFIFNF | GBE1; GBE1 | Q04446; E9F | GLGB_HUMAN | Unmodified | 0 | 0 | 0 | 0 | 0 | 0 | 0 | 0 | 192370 | 191010 | 20735 | 166720 | 0 | 167040 | 0 | 0 | 0 | 0 |
| AGLLQRAL | CLTC; CLTC | Q00610; A0A | CLH1_HUMAN | Unmodified | 0 | 0 | 0 | 0 | 961020 | 0 | 2547600 | 0 | 0 | 0 | 0 | 0 | 322360 | 253610 | 0 | 0 | 1564000 | 217330 |
| AGLLQRAL | CLTC; CLTC | Q00610; A0A | CLH1_HUMAN | Unmodified | 99622 | 83443 | 0 | 390400 | 129840 | 104430 | 243430 | 0 | 0 | 60775 | 0 | 0 | 103690 | 55046 | 0 | 0 | 120860 | 153680 |
| AGLLQVYI | EMG1; EMG1; E | Q92979; A0A | NEP1_HUMAN | Unmodified | 0 | 0 | 0 | 0 | 0 | 0 | 0 | 110400 | 0 | 0 | 53713 | 4521500 | 0 | 0 | 0 | 0 | 0 | 0 |
| AGLLTLED | MBTPS1 | Q14703 | MBTP1_HUMAN | Unmodified | 0 | 0 | 0 | 0 | 188050 | 0 | 0 | 0 | 863470 | 330180 | 65589 | 0 | 0 | 0 | 0 | 0 | 0 | 0 |
| AGLLVLAVL | IPO4; IPO4 | Q8TEX9; H0Y | IPO4_HUMAN I | Unmodified | 0 | 0 | 19512 | 0 | 0 | 0 | 0 | 0 | 0 | 0 | 0 | 418610 | 0 | 0 | 0 | 42488 | 0 | 0 |
| AGLNCSTE | EIF2S1; EIF2S1; | P05198; H0Y | IF2A_HUMAN E | Oxidation (M) | 0 | 0 | 0 | 0 | 0 | 0 | 0 | 0 | 0 | 0 | 0 | 435860 | 0 | 0 | 0 | 0 | 0 | 0 |
| AGLNCSTE | EIF2S1; EIF2S1; | P05198; H0Y | IF2A_HUMAN E | Unmodified | 0 | 0 | 0 | 0 | 0 | 0 | 0 | 0 | 0 | 0 | 143560 | 2322100 | 0 | 0 | 0 | 0 | 0 | 0 |
| AGLNSLEA | TPM4; TPM4; TF | P67936; A0A | TPM4_HUMAN | Acetyl (Protein N | 1376700 | 9211900 | 633340 | 0 | 21740000 | 2239300 | 1012400 | 414250 | 0 | 90714 | 521930 | 570720 | 1127300 | 1222600 | 990430 | 814700 | 0 | 0 |
| AGLNSLEA | TPM4; TPM4; TF | P67936; A0A | TPM4_HUMAN | Acetyl (Protein N | 0 | 0 | 0 | 0 | 3563100 | 0 | 245430 | 226540 | 0 | 0 | 477620 | 0 | 445270 | 558800 | 0 | 0 | 0 | 0 |
| AGLQFPVG | H2AX; H2AC6; I | P16104; Q93 | H2AX_HUMAN | Unmodified | 9.31E+08 | 5.4E+08 | 4043200 | 1854900 | 23431000 | 6849200 | 10252000 | 77017000 | 16806000 | 2093100 | 7730800 | 5.04E+09 | 4593900 | 1327900 | 1443000 | 2354600 | 384290 | 6118700 |
| AGLQFPVG | H2AX; H2AC6; I | P16104; Q93 | H2AX_HUMAN | Unmodified | 0 | 0 | 0 | 0 | 0 | 0 | 219670 | 412760 | 0 | 0 | 0 | 0 | 0 | 0 | 0 | 0 | 0 | 0 |
| AGLQVYNK | CD59; CD59; N | P13987; A0A | CD59_HUMAN | Unmodified | 33188000 | 840450 | 27570000 | 34187000 | 1.93E+08 | 25725000 | 13249000 | 8047500 | 25558000 | 17193000 | 2292700 | 31074000 | 1.19E+08 | 93413000 | 43692000 | 29446000 | 2.34E+08 | 7.77E+08 |
| AGLQVYNK | CD59; CD59; N | P13987; A0A | CD59_HUMAN | Unmodified | 0 | 0 | 0 | 0 | 0 | 0 | 0 | 0 | 0 | 0 | 0 | 0 | 0 | 0 | 0 | 0 | 1073100 | 0 |
| AGLRVSAP | AHNAK | Q09666 | AHNK_HUMAN | Unmodified | 0 | 0 | 0 | 0 | 1228600 | 161060 | 0 | 0 | 0 | 0 | 0 | 0 | 0 | 0 | 0 | 0 | 311080 | 688920 |
| AGLSQSLF | UPF1 | Q92900 | RENT1_HUMAN | Unmodified | 1988800 | 1961100 | 1847700 | 418600 | 0 | 0 | 0 | 0 | 0 | 0 | 130450 | 1511400 | 0 | 0 | 1276600 | 1096000 | 0 | 0 |
| AGLSSGFI | HSPG2; HSPG2 | P98160; A0A | PGBM_HUMAN | Unmodified | 0 | 0 | 1869900 | 582550 | 0 | 331940 | 0 | 0 | 0 | 0 | 0 | 0 | 0 | 0 | 0 | 409170 | 0 | 499670 |
| AGLTLVNLI | N/A | N/A | N/A | Unmodified | 831120 | 1322800 | 309540 | 127740 | 0 | 0 | 0 | 0 | 0 | 0 | 0 | 0 | 69572 | 0 | 332340 | 366140 | 0 | 0 |
| AGLVDDFE | ATP5PD; ATP5P | O75947; F5H | ATP5H_HUMAN | Unmodified | 0 | 190920 | 0 | 0 | 0 | 0 | 1115500 | 285830 | 5092200 | 0 | 0 | 0 | 0 | 411240 | 0 | 0 | 135700 | 219060 |
| AGLVELLLF | SNTB2; SNTB2 | Q13425; A0A | SNTB2_HUMAN | Unmodified | 150950 | 457600 | 1257700 | 862070 | 1501100 | 118230 | 0 | 0 | 2138300 | 675680 | 535140 | 1333600 | 1641700 | 791900 | 5148600 | 3638700 | 539000 | 856370 |
| AGLVELLLF | SNTB2; SNTB2 | Q13425; A0A | SNTB2_HUMAN | Unmodified | 0 | 0 | 0 | 1933900 | 5783300 | 1010600 | 0 | 0 | 0 | 1416900 | 0 | 0 | 3384600 | 1526100 | 0 | 0 | 2359800 | 4289600 |
| AGLVIGKG | FUBP1; KHSRP; | Q96AE4; Q92 | FUBP1_HUMAN | Unmodified | 68918 | 0 | 0 | 0 | 1139000 | 302050 | 362770 | 284230 | 443560 | 1923200 | 0 | 0 | 1256100 | 369600 | 0 | 0 | 188440 | 465410 |
| AGLYGLPR | FASN; FASN; FA | P49327; A0A | FAS_HUMAN Fa | Unmodified | 527240 | 767410 | 571050 | 2686300 | 1506800 | 313050 | 202710 | 0 | 926760 | 244250 | 1842100 | 22270000 | 1774100 | 0 | 3282000 | 2272600 | 158210 | 407190 |
| AGMERELE | OS9; OS9; OS9 | Q13438; B4E | OS9_HUMAN P | Oxidation (M) | 0 | 0 | 0 | 134030 | 0 | 0 | 0 | 0 | 2964000 | 312940 | 0 | 0 | 0 | 0 | 0 | 0 | 0 | 0 |
| AGMERELE | OS9; OS9; OS9 | Q13438; B4E | OS9_HUMAN P | Unmodified | 0 | 0 | 0 | 0 | 0 | 0 | 0 | 0 | 7081200 | 753480 | 0 | 0 | 0 | 0 | 0 | 0 | 0 | 52054 |
| AGMGAGA | N/A | N/A | N/A | Oxidation (M) | 0 | 0 | 105330 | 450610 | 0 | 0 | 0 | 0 | 0 | 0 | 0 | 0 | 388870 | 0 | 1952400 | 123270 | 0 | 0 |
| AGMIFYR | SHMT1 | P34896 | GLYC_HUMAN | Unmodified | 0 | 0 | 0 | 0 | 0 | 0 | 0 | 0 | 0 | 0 | 0 | 784840 | 0 | 0 | 0 | 0 | 0 | 0 |
| AGMIGFSN | N/A | N/A | N/A | 2 Oxidation (M) | 0 | 0 | 0 | 0 | 6788600 | 0 | 0 | 0 | 0 | 370970 | 0 | 0 | 0 | 0 | 0 | 0 | 6660100 | 3008600 |
| AGMSAEQ | GSTK1; GSTK1 | Q9Y2Q3; E9I | GSTK1_HUMAN | Unmodified | 0 | 0 | 0 | 0 | 0 | 0 | 0 | 0 | 1749500 | 292200 | 0 | 0 | 0 | 0 | 14125 | 0 | 0 | 0 |
| AGMTHIVR | RPL3; RPL3; RP | P39023; H7C | RL3_HUMAN 60 | Unmodified | 825410 | 730450 | 843370 | 293190 | 0 | 0 | 0 | 0 | 267130 | 0 | 278160 | 10286000 | 0 | 0 | 408150 | 331070 | 0 | 0 |
| AGMVNAW | MFGE8; MFGE8 | Q08431; H0Y | MFGM_HUMAN | Oxidation (M) | 0 | 1170800 | 333000 | 4888200 | 3985800 | 345560 | 0 | 0 | 1916000 | 900370 | 0 | 490260 | 393040 | 0 | 4464400 | 702360 | 256590 | 0 |
| AGMVNAW | MFGE8; MFGE8 | Q08431; H0Y | MFGM_HUMAN | Oxidation (M) | 0 | 0 | 1498500 | 0 | 11460000 | 0 | 0 | 0 | 5731900 | 1301100 | 0 | 2304200 | 0 | 0 | 22790000 | 5499100 | 0 | 0 |
| AGNASKDE | WARS1; WARS1 | P23381; G3V | SYWC_HUMAN | Unmodified | 0 | 0 | 373270 | 198320 | 0 | 107510 | 97226 | 0 | 1575100 | 301320 | 139770 | 296840 | 11886 | 0 | 526960 | 662520 | 146290 | 262450 |
| AGNEKEEG | TOP1; TOP1 | P11387; A0A | TOP1_HUMAN | Unmodified | 0 | 0 | 0 | 0 | 0 | 0 | 0 | 0 | 0 | 0 | 0 | 880000 | 0 | 0 | 0 | 0 | 0 | 0 |
| AGNFYVPA | RPL7; RPL7; RP | P18124; C9J | RL7_HUMAN 60 | Unmodified | 2271000 | 2157600 | 1220500 | 1082900 | 0 | 0 | 606830 | 767610 | 0 | 0 | 468270 | 21638000 | 388560 | 0 | 2054900 | 1700300 | 0 | 338910 |
| AGNGQWA | SF3B3 | Q15393 | SF3B3_HUMAN | Unmodified | 0 | 0 | 0 | 71665 | 0 | 0 | 0 | 0 | 0 | 0 | 0 | 4869200 | 0 | 0 | 0 | 0 | 0 | 0 |
| AGNILLNTE | STK3; STK4; STI | Q13188; Q13 | STK3_HUMAN S | Unmodified | 0 | 0 | 0 | 0 | 764610 | 0 | 0 | 0 | 0 | 149660 | 432110 | 1341800 | 0 | 0 | 0 | 0 | 144000 | 0 |
| AGNILLTEP | TAOK1; TAOK3; | Q7L7X3; Q9H | TAOK1_HUMAN | Unmodified | 579590 | 913180 | 862760 | 830810 | 1107200 | 753420 | 406040 | 415190 | 853690 | 463580 | 142890 | 1700000 | 438460 | 0 | 651610 | 720830 | 504010 | 621430 |
| AGNLGGG | RPL22; RPL22; I | P35268; K7E | RL22_HUMAN ( | Unmodified | 3763200 | 5009300 | 0 | 94455 | 392680 | 0 | 0 | 111810 | 1813600 | 182690 | 4010000 | 42681000 | 0 | 0 | 0 | 0 | 0 | 0 |
| AGNLGGG | RPL22; RPL22; I | P35268; K7E | RL22_HUMAN ( | Unmodified | 124670 | 86482 | 0 | 214900 | 3080100 | 563990 | 3097500 | 2188300 | 0 | 691880 | 624560 | 349080 | 0 | 0 | 49589 | 33995 | 338630 | 1717100 |
| AGNNMLL | FLNA; FLNA; FL | P21333; A0A | FLNA_HUMAN | Unmodified | 0 | 0 | 0 | 2962700 | 0 | 0 | 497760 | 351050 | 936780 | 105090 | 0 | 2001100 | 0 | 0 | 0 | 0 | 133960 | 0 |
| AGNNMLL | FLNA; FLNA; FL | P21333; A0A | FLNA_HUMAN | Oxidation (M) | 36620 | 2329100 | 1823300 | 2128100 | 340990 | 0 | 0 | 218820 | 230540 | 44967 | 0 | 1535700 | 1179700 | 291150 | 1867500 | 1327300 | 231110 | 202560 |
| AGNNMLL | FLNA; FLNA; FL | P21333; A0A | FLNA_HUMAN | Unmodified | 0 | 0 | 0 | 0 | 17130000 | 0 | 14351000 | 329300 | 254560 | 494460 | 0 | 0 | 7105400 | 11547000 | 0 | 0 | 11959000 | 10698000 |
| AGNNMLL | FLNA; FLNA; FL | P21333; A0A | FLNA_HUMAN | Oxidation (M) | 0 | 0 | 0 | 372520 | 1659300 | 0 | 3042100 | 426780 | 0 | 0 | 0 | 0 | 2420200 | 3037600 | 0 | 0 | 2302800 | 1037300 |
| AGNREHES | NAGPA | Q9UK23 | NAGPA_HUMA | Unmodified | 0 | 0 | 0 | 0 | 0 | 0 | 0 | 0 | 673740 | 308350 | 0 | 0 | 0 | 0 | 0 | 0 | 0 | 0 |
| AGPAAAFR | TMEM256; TME | Q8N2U0; K7 | TM256_HUMAN | Acetyl (Protein N | 0 | 0 | 0 | 0 | 0 | 0 | 0 | 0 | 534140 | 244990 | 0 | 0 | 0 | 0 | 62378 | 0 | 0 | 0 |
| AGPESDAC | ATP5MK | Q96IX5 | ATPMK_HUMAN | Unmodified | 0 | 0 | 59798 | 0 | 0 | 0 | 0 | 0 | 1972100 | 155190 | 0 | 0 | 0 | 0 | 0 | 0 | 0 | 0 |
| AGPESDAC | ATP5MK | Q96IX5 | ATPMK_HUMAN | Unmodified | 0 | 0 | 0 | 50342 | 201460 | 91046 | 461020 | 391620 | 6050400 | 1269600 | 165530 | 215120 | 232230 | 79826 | 0 | 0 | 0 | 229020 |
| AGPGPGP | RAB43; RAB43 | Q86YS6; C9J | RAB43_HUMAN | Acetyl (Protein N | 44395 | 84083 | 39859 | 210060 | 298510 | 0 | 157260 | 0 | 0 | 541100 | 0 | 0 | 203100 | 0 | 96733 | 0 | 93814 | 189620 |
| AGPPASIVF | PAK4; PAK4 | O96013; M0I | PAK4_HUMAN | Oxidation (M) | 322740 | 420920 | 0 | 696140 | 0 | 0 | 0 | 0 | 0 | 0 | 0 | 561010 | 386390 | 0 | 391110 | 591860 | 134680 | 0 |
| AGPSAQEP | SPAG9; SPAG9 | O60271; A0A | JIP4_HUMAN C | Unmodified | 0 | 0 | 0 | 0 | 156420 | 0 | 0 | 74552 | 0 | 0 | 0 | 806270 | 100840 | 0 | 0 | 43330 | 170400 | 710280 |
| AGPSRTVG | GPD2 | P43304 | GPDM_HUMAN | Unmodified | 0 | 0 | 0 | 0 | 18272000 | 2830600 | 1801200 | 614020 | 0 | 2390600 | 0 | 0 | 1362500 | 0 | 0 | 0 | 5666000 | 7009500 |
| AGPTSHVA | SIGIRR; SIGIRR | Q6IA17; C9J | SIGIR_HUMAN | Unmodified | 0 | 0 | 0 | 0 | 0 | 0 | 0 | 0 | 0 | 0 | 0 | 77044 | 0 | 0 | 20922 | 0 | 22370 | 0 |
| AGQAVDDI | PGD; PGD; PGD | P52209; K7E | 6PGD_HUMAN | Unmodified | 455460 | 0 | 1080200 | 1078500 | 17095000 | 2056900 | 2793800 | 79320 | 6720600 | 6240700 | 6492600 | 27585000 | 5570500 | 6104000 | 5641300 | 1240600 | 2680600 | 2904400 |
| AGQCVIGL | CNN2; CNN2; ( | Q99439; H3I | CNN2_HUMAN | Unmodified | 0 | 0 | 0 | 0 | 2581600 | 0 | 0 | 0 | 96492 | 0 | 0 | 0 | 0 | 0 | 0 | 0 | 174500 | 206990 |
| AGQDVVLI | LRP11; LRP11 | Q86VZ4; Q5Y | LRP11_HUMAN | Unmodified | 0 | 0 | 0 | 0 | 0 | 0 | 0 | 0 | 1899900 | 595220 | 0 | 0 | 0 | 0 | 0 | 0 | 0 | 0 |
| AGQEDPV | BRK1 | Q8WUW1 | BRK1_HUMAN | Acetyl (Protein N | 0 | 0 | 241450 | 455560 | 0 | 0 | 0 | 0 | 391510 | 136630 | 0 | 0 | 442260 | 360900 | 0 | 489600 | 0 | 0 |
| AGQEDPV | BRK1 | Q8WUW1 | BRK1_HUMAN | Acetyl (Protein N | 0 | 0 | 0 | 370980 | 19694000 | 1705100 | 3524200 | 0 | 0 | 0 | 0 | 0 | 6520300 | 3385200 | 0 | 0 | 23828000 | 8808700 |
| AGQEHLLR | UAP1L1; UAP1; | Q3KQV9; Q1 | UAP1L_HUMAN | Unmodified | 0 | 0 | 0 | 1392100 | 0 | 0 | 0 | 0 | 0 | 304820 | 1288900 | 331390 | 0 | 0 | 162820 | 79852 | 0 | 0 |
| AGQERPSV | ANAPC7 | Q9UJX3 | APC7_HUMAN | Unmodified | 0 | 0 | 0 | 0 | 0 | 0 | 0 | 0 | 21152 | 0 | 0 | 664970 | 0 | 0 | 0 | 0 | 0 | 0 |
| AGQMVTV | LMNB2 | Q03252 | LMNB2_HUMA | Oxidation (M) | 0 | 0 | 0 | 0 | 0 | 0 | 0 | 0 | 0 | 0 | 0 | 1224200 | 0 | 0 | 0 | 0 | 0 | 0 |
| AGQPEAGF | TYSND1 | Q2T9J0 | TYSD1_HUMAN | Unmodified | 0 | 0 | 0 | 0 | 0 | 0 | 0 | 0 | 495990 | 0 | 0 | 0 | 0 | 0 | 0 | 0 | 0 | 0 |
| AGQPLESD | TBC1D24; TBC1 | Q9ULP9; A0A | TBC24_HUMAN | Unmodified | 0 | 0 | 0 | 76908 | 117570 | 1197000 | 0 | 70442 | 0 | 0 | 35178 | 0 | 822570 | 268040 | 0 | 309560 | 0 | 478180 |
| AGQPLPLTI | ITSN2; ITSN2; IT | Q9NZM3; A0 | ITSN2_HUMAN | Unmodified | 0 | 0 | 0 | 0 | 362800 | 0 | 0 | 0 | 0 | 0 | 0 | 0 | 0 | 0 | 0 | 0 | 124540 | 0 |
| AGQPLQLL | MPST; MPST | P25325; B1A | THTM_HUMAN | Unmodified | 0 | 0 | 0 | 0 | 0 | 0 | 0 | 0 | 483200 | 42550 | 10746 | 1028900 | 856530 | 0 | 0 | 0 | 0 | 0 |
| AGQQGLS | MMUT | P22033 | MUTA_HUMAN | Unmodified | 0 | 0 | 0 | 0 | 0 | 0 | 0 | 0 | 1510800 | 0 | 0 | 0 | 0 | 0 | 284660 | 121190 | 0 | 0 |
| AGQQGLS | MMUT | P22033 | MUTA_HUMAN | Unmodified | 0 | 0 | 0 | 0 | 0 | 0 | 0 | 0 | 1748900 | 351980 | 0 | 0 | 0 | 0 | 0 | 0 | 0 | 0 |
| AGQSAAGA | FLNA; FLNA; FL | P21333; A0A | FLNA_HUMAN | Unmodified | 0 | 0 | 142780 | 131520 | 0 | 0 | 0 | 0 | 0 | 0 | 0 | 0 | 0 | 0 | 233400 | 141260 | 0 | 0 |
| AGQSVIGL | CNN3; CNN3 | Q15417; E9F | CNN3_HUMAN | Oxidation (M) | 0 | 0 | 0 | 0 | 654800 | 0 | 0 | 0 | 0 | 0 | 0 | 552970 | 0 | 0 | 0 | 0 | 0 | 0 |
| AGQTPQEF | FAR1; FAR1 | Q8WVX9; E9 | FACR1_HUMAN | Unmodified | 0 | 0 | 0 | 0 | 0 | 0 | 323150 | 0 | 0 | 0 | 0 | 0 | 0 | 0 | 0 | 0 | 0 | 0 |
| AGQTQPNI | AAK1; AAK1; AA | Q2M2I8; A0A | AAK1_HUMAN | Unmodified | 0 | 0 | 0 | 0 | 1214300 | 174350 | 0 | 0 | 0 | 0 | 0 | 0 | 1482500 | 2704700 | 0 | 0 | 395080 | 917250 |
| AGQVFLEE | CNP | P09543 | CN37_HUMAN | Unmodified | 1541700 | 0 | 5082800 | 0 | 38210000 | 5632800 | 6712000 | 3882700 | 5539500 | 5051700 | 33840000 | 55187000 | 22056000 | 14223000 | 5539100 | 6712100 | 10130000 | 27357000 |
| AGQVFLEE | CNP | P09543 | CN37_HUMAN | Unmodified | 0 | 0 | 0 | 0 | 0 | 0 | 0 | 0 | 0 | 0 | 0 | 0 | 448110 | 373070 | 0 | 0 | 776290 | 0 |
| AGQVVTIW | LMNA; LMNA; L | P02545; A0A | LMNA_HUMAN | Unmodified | 0 | 0 | 0 | 0 | 0 | 0 | 0 | 0 | 0 | 0 | 0 | 0 | 0 | 0 | 0 | 0 | 1552400 | 0 |
| AGRCADY | PSAT1 | Q9Y617 | SERC_HUMAN | Unmodified | 0 | 0 | 0 | 0 | 0 | 0 | 1450700 | 552110 | 0 | 462850 | 0 | 0 | 0 | 0 | 0 | 0 | 1470400 | 0 |
| AGRGGFKI | ACAA1; ACAA1; | P09110; F2Z | THIK_HUMAN 3 | Unmodified | 0 | 0 | 0 | 18364 | 0 | 0 | 0 | 0 | 73765 | 562190 | 0 | 0 | 0 | 0 | 0 | 0 | 0 | 0 |
| AGRLGSTV | HNRNPM; HNR | P52272; M0R | HNRPM_HUMA | Unmodified | 0 | 0 | 0 | 0 | 252950 | 0 | 509320 | 142640 | 0 | 0 | 70059 | 0 | 0 | 0 | 0 | 0 | 0 | 376880 |
| AGRLPACV | ACTR3; ACTR3; | P61158; F8V | ARP3_HUMAN | Acetyl (Protein N | 1255900 | 348530 | 519150 | 833880 | 4201200 | 1171400 | 2328500 | 1328500 | 436160 | 597080 | 675380 | 2730800 | 4596100 | 5911000 | 878030 | 541140 | 6924000 | 1829700 |
| AGRQGEITI | RELT | Q969Z4 | TR19L_HUMAN | Unmodified | 0 | 0 | 0 | 71508 | 226590 | 0 | 130680 | 0 | 0 | 0 | 0 | 0 | 203440 | 225330 | 0 | 0 | 106650 | 118800 |
| AGRSEAVV | SND1; SND1 | Q7KZF4; H7C | SND1_HUMAN | Unmodified | 333530 | 232840 | 493020 | 643160 | 0 | 0 | 0 | 0 | 998850 | 138540 | 772700 | 3201300 | 397340 | 159520 | 679540 | 425160 | 232380 | 69840 |
| AGRSMQA | NSF; NSF | P46459; I3L4 | NSF_HUMAN V | Acetyl (Protein N | 0 | 0 | 0 | 0 | 0 | 0 | 0 | 0 | 0 | 0 | 0 | 0 | 488580 | 807140 | 0 | 0 | 621780 | 0 |
| AGRTALHH | ANKRD28; ANK | O15084; B4D | ANR28_HUMAN | Unmodified | 160280 | 0 | 0 | 0 | 10271000 | 0 | 39212 | 0 | 2116400 | 1313200 | 0 | 322260 | 0 | 0 | 0 | 0 | 0 | 80223 |
| AGRVLVFVI | PFN2; PFN2 | P35080; C9J | PROF2_HUMAN | Unmodified | 0 | 0 | 0 | 0 | 0 | 0 | 0 | 0 | 0 | 124990 | 0 | 0 | 0 | 0 | 0 | 0 | 258830 | 0 |
| AGSDGAKI | CLIC1 | O00299 | CLIC1_HUMAN | Oxidation (M) | 0 | 0 | 0 | 0 | 721750 | 0 | 537140 | 0 | 0 | 118020 | 0 | 0 | 0 | 0 | 0 | 0 | 2535100 | 0 |
| AGSDGESI | CLIC4 | Q9Y696 | CLIC4_HUMAN | Unmodified | 0 | 140300 | 0 | 572480 | 2933100 | 954360 | 168200 | 0 | 867250 | 361530 | 578140 | 2054500 | 1377900 | 930930 | 815710 | 520990 | 986960 | 2281600 |
| AGSDGESI | CLIC4 | Q9Y696 | CLIC4_HUMAN | Oxidation (M) | 0 | 0 | 0 | 83371 | 196050 | 0 | 90387 | 0 | 978030 | 0 | 0 | 0 | 314700 | 671230 | 0 | 0 | 337910 | 794700 |
| AGSGLRQY | PSMC5; PSMC5 | P62195; J3Q | PRS8_HUMAN | Unmodified | 0 | 0 | 0 | 0 | 3287300 | 129560 | 210130 | 0 | 0 | 620510 | 0 | 0 | 1253600 | 1369200 | 0 | 0 | 408680 | 711690 |
| AGSGSQVF | SDC4 | P31431 | SDC4_HUMAN | Unmodified | 1265800 | 1267700 | 3312700 | 4048700 | 0 | 0 | 0 | 0 | 1730100 | 0 | 0 | 0 | 0 | 0 | 1463200 | 1805200 | 0 | 250530 |
| AGSIEDTS | PODXL2 | Q9NZ53 | PDXL2_HUMAN | Unmodified | 0 | 0 | 0 | 0 | 0 | 0 | 0 | 0 | 939250 | 0 | 0 | 0 | 0 | 0 | 0 | 0 | 0 | 0 |
| AGSLKDPE | TAOK1; TAOK1 | Q7L7X3; J3Q | TAOK1_HUMAN | Unmodified | 0 | 361820 | 284740 | 645590 | 0 | 0 | 0 | 0 | 857760 | 0 | 735790 | 778960 | 0 | 0 | 471020 | 554420 | 0 | 0 |
| AGSMKDPI | MYO6; MYO6; M | Q9UM54; A0 | MYO6_HUMAN | Unmodified | 0 | 0 | 0 | 0 | 0 | 0 | 0 | 0 | 416740 | 0 | 0 | 301880 | 0 | 0 | 429760 | 291720 | 0 | 0 |
| AGSMKDPI | MYO6; MYO6; M | Q9UM54; A0 | MYO6_HUMAN | Oxidation (M) | 0 | 0 | 2578000 | 2206500 | 0 | 0 | 0 | 0 | 0 | 0 | 0 | 1789300 | 198100 | 0 | 3661600 | 3913300 | 0 | 0 |
| AGSNMLLI | FLNB; FLNB; FL | O75369; A0A | FLNB_HUMAN | 2 Oxidation (M) | 0 | 0 | 0 | 457180 | 4752700 | 888330 | 0 | 115790 | 0 | 0 | 0 | 1761600 | 3123500 | 340530 | 0 | 396000 | 2134800 | 927290 |
| AGSNMLLI | FLNB; FLNB; FL | O75369; A0A | FLNB_HUMAN | Oxidation (M) | 0 | 0 | 0 | 0 | 3242100 | 616800 | 0 | 0 | 0 | 0 | 58265 | 0 | 1656700 | 686910 | 0 | 0 | 4353900 | 1052200 |
| AGSNMLLI | FLNB; FLNB; FL | O75369; A0A | FLNB_HUMAN | Unmodified | 0 | 0 | 0 | 0 | 8348900 | 0 | 0 | 0 | 0 | 0 | 0 | 0 | 0 | 0 | 0 | 0 | 0 | 0 |
| AGSNMLLI | FLNB; FLNB; FL | O75369; A0A | FLNB_HUMAN | 2 Oxidation (M) | 0 | 0 | 0 | 0 | 0 | 0 | 0 | 0 | 0 | 0 | 0 | 0 | 0 | 0 | 0 | 0 | 2118100 | 0 |
| AGSNMLLI | FLNB; FLNB; FL | O75369; A0A | FLNB_HUMAN | Oxidation (M) | 0 | 0 | 0 | 0 | 0 | 0 | 0 | 0 | 0 | 0 | 0 | 0 | 0 | 0 | 0 | 0 | 1927100 | 0 |
| AGSPTAPV | PTPRJ; PTPRJ; P | Q12913; A0A | PTPRJ_HUMAN | Unmodified | 322680 | 378520 | 0 | 0 | 0 | 0 | 0 | 0 | 3607700 | 0 | 440840 | 1444400 | 0 | 0 | 0 | 244910 | 0 | 0 |
| AGSPTLLN | STT3B | Q8TCJ2 | STT3B_HUMAN | Oxidation (M) | 0 | 0 | 0 | 0 | 0 | 0 | 0 | 0 | 1913100 | 0 | 0 | 0 | 0 | 0 | 0 | 0 | 0 | 0 |
| AGSPTLLN | STT3B | Q8TCJ2 | STT3B_HUMAN | Unmodified | 0 | 0 | 0 | 0 | 0 | 0 | 0 | 0 | 3432900 | 0 | 0 | 0 | 0 | 0 | 0 | 86438 | 0 | 0 |
| AGSQLGP | KIF23; KIF23; KI | Q02241; A0A | KIF23_HUMAN | Unmodified | 0 | 0 | 0 | 0 | 25068000 | 3659400 | 0 | 0 | 0 | 25583 | 0 | 0 | 0 | 0 | 0 | 0 | 0 | 23927 |

| | | | | | | | | | | | | | | | | | | | | | | |
|---|---|---|---|---|---|---|---|---|---|---|---|---|---|---|---|---|---|---|---|---|---|---|
| AGSRLETV( | MRPS23; MRPS | Q9Y3D9; J3C | RT23_HUMAN | Acetyl (Protein N | 0 | 0 | 0 | 0 | 0 | 0 | 0 | 0 | 438830 | 0 | 10431 | 0 | 0 | 0 | 0 | 0 | 0 | 0 |
| AGSRYEDF | UQCRC2; UQC | P22695; H3E | QCR2_HUMAN | Unmodified | 0 | 0 | 0 | 0 | 97794 | 0 | 0 | 0 | 20186000 | 6154700 | 0 | 448610 | 822960 | 1544500 | 0 | 35487 | 0 | 0 |
| AGSRYEDF | UQCRC2; UQC | P22695; H3E | QCR2_HUMAN | Unmodified | 0 | 0 | 0 | 0 | 0 | 0 | 0 | 0 | 0 | 2887900 | 0 | 0 | 90291 | 0 | 0 | 0 | 0 | 0 |
| AGSSESIAA | BAG6; BAG6; B/ | P46379; A0A | BAG6_HUMAN | Unmodified | 0 | 0 | 0 | 0 | 0 | 0 | 0 | 0 | 0 | 0 | 87054 | 1502300 | 0 | 0 | 0 | 0 | 0 | 0 |
| AGSSGNS( | PIP5K1A; PIP5K | Q99755; A6F | PI51A_HUMAN | Unmodified | 0 | 0 | 0 | 0 | 676360 | 0 | 0 | 0 | 234010 | 0 | 80110 | 585210 | 1346200 | 129700 | 0 | 0 | 0 | 0 |
| AGSSKYQV | LARS1; LARS1; | Q9P2J5; A0A | SYLC_HUMAN | Unmodified | 0 | 0 | 0 | 0 | 138720 | 0 | 163300 | 30656 | 0 | 86985 | 0 | 0 | 67010 | 0 | 0 | 0 | 51191 | 87635 |
| AGSSSLEA' | TPM1; TPM1; TF | H0YKJ4; H0Y | H0YKJ4_HUMA | Acetyl (Protein N | 0 | 0 | 0 | 0 | 364950 | 70289 | 0 | 0 | 0 | 0 | 0 | 770150 | 287350 | 0 | 0 | 0 | 12818 | 0 |
| AGSTALYW | OSTF1 | Q92882 | OSTF1_HUMAN | Unmodified | 0 | 0 | 0 | 180820 | 0 | 0 | 0 | 0 | 456960 | 0 | 401970 | 1839400 | 628260 | 150410 | 192490 | 467650 | 0 | 0 |
| AGSVSLDS | FMNL1; FMNL1 | O95466; A0A | FMNL1_HUMAI | Unmodified | 0 | 0 | 0 | 0 | 6399600 | 718250 | 0 | 0 | 0 | 127530 | 0 | 0 | 0 | 0 | 0 | 0 | 107040 | 0 |
| AGTEEAIVY | NIT2 | Q9NQR4 | NIT2_HUMAN ( | Unmodified | 0 | 0 | 0 | 0 | 0 | 0 | 0 | 0 | 0 | 0 | 0 | 0 | 1234900 | 0 | 0 | 0 | 0 | 0 |
| AGTEEAIVY | NIT2 | Q9NQR4 | NIT2_HUMAN ( | Unmodified | 0 | 0 | 143460 | 236170 | 0 | 0 | 0 | 0 | 689850 | 439010 | 233900 | 1379600 | 1877300 | 1662100 | 161300 | 113120 | 1325700 | 883260 |
| AGTFQAFE( | SPTAN1; SPTAN | Q13813; A0A | SPTN1_HUMAN | Unmodified | 0 | 0 | 351470 | 0 | 728470 | 347280 | 0 | 0 | 1413500 | 279670 | 1209300 | 3515000 | 0 | 0 | 889710 | 538830 | 0 | 0 |
| AGTGVDN\ | PHGDH; PHGD | O43175; A0A | SERA_HUMAN | Unmodified | 0 | 0 | 0 | 1070600 | 348950 | 0 | 0 | 53492 | 0 | 226660 | 13093000 | 1548900 | 1585900 | 553240 | 0 | 1415400 | 0 | 0 |
| AGTGVDN\ | PHGDH; PHGD | O43175; A0A | SERA_HUMAN | Unmodified | 3784500 | 3414800 | 11668000 | 16041000 | 3415400 | 1132900 | 2359200 | 1075800 | 16775000 | 8852200 | 4105300 | 30552000 | 41744000 | 13655000 | 13685000 | 15556000 | 985060 | 3634200 |
| AGTNMMM | FLNC | Q14315 | FLNC_HUMAN | 2 Oxidation (M) | 0 | 0 | 0 | 0 | 0 | 1657700 | 0 | 0 | 0 | 0 | 0 | 0 | 0 | 0 | 0 | 0 | 353300 | 0 |
| AGTNMMM | FLNC | Q14315 | FLNC_HUMAN | Oxidation (M) | 0 | 0 | 0 | 0 | 1801700 | 0 | 0 | 0 | 0 | 0 | 0 | 0 | 0 | 0 | 0 | 0 | 1464000 | 0 |
| AGTNMMM | FLNC | Q14315 | FLNC_HUMAN | Unmodified | 0 | 0 | 0 | 0 | 1190300 | 0 | 0 | 0 | 221500 | 0 | 0 | 0 | 324500 | 0 | 176780 | 0 | 1029300 | 244480 |
| AGTQIENID | ACTN4; ACTN4 | O43707; K7E | ACTN4_HUMAI | Unmodified | 2703100 | 2576600 | 3477500 | 4851600 | 86066000 | 21243000 | 5984100 | 1432700 | 10948000 | 11457000 | 7233100 | 60951000 | 19564000 | 16504000 | 6599400 | 5532100 | 18861000 | 10510000 |
| AGTQIENIE | ACTN1; ACTN1 | P12814; G3\ | ACTN1_HUMAI | Unmodified | 921990 | 800750 | 1207100 | 3872500 | 32278000 | 16503000 | 5516700 | 370720 | 3548500 | 1718500 | 6706800 | 22397000 | 8663600 | 9333100 | 4322600 | 2988200 | 20706000 | 18242000 |
| AGTQLLAG | ITGAV | P06756 | ITAV_HUMAN I | Unmodified | 1745700 | 591340 | 1355400 | 1414000 | 1249400 | 684850 | 0 | 0 | 3596000 | 218420 | 0 | 6318400 | 0 | 0 | 1869100 | 2015900 | 290320 | 0 |
| AGTRTDSR | PRKAR1A; PRK/ | P10644; K7E | KAP0_HUMAN | Unmodified | 0 | 0 | 0 | 0 | 5012700 | 0 | 1352700 | 574190 | 1554400 | 740400 | 0 | 0 | 1041800 | 926110 | 0 | 0 | 7955800 | 7303700 |
| AGTSPGTP | LRP1 | Q07954 | LRP1_HUMAN | Unmodified | 0 | 0 | 0 | 0 | 3438800 | 0 | 0 | 0 | 1647000 | 0 | 0 | 0 | 0 | 0 | 0 | 0 | 0 | 0 |
| AGTTATYE( | PKP2 | Q99959 | PKP2_HUMAN | Unmodified | 0 | 0 | 0 | 0 | 0 | 0 | 0 | 0 | 0 | 0 | 0 | 0 | 0 | 0 | 0 | 0 | 754160 | 1199400 |
| AGTVLQEA | CORO2A | Q92828 | COR2A_HUMAI | Unmodified | 0 | 0 | 0 | 0 | 0 | 0 | 0 | 0 | 0 | 0 | 0 | 0 | 0 | 0 | 799530 | 435020 | 0 | 0 |
| AGTWLAVH | SUSD2 | Q9UGT4 | SUSD2_HUMAI | Unmodified | 0 | 0 | 0 | 0 | 0 | 0 | 0 | 0 | 3833400 | 129290 | 0 | 0 | 0 | 0 | 0 | 0 | 0 | 0 |
| AGVAFFHL | TTC9C; TTC9C; | Q8N5M4; E9 | TTC9C_HUMAN | Unmodified | 0 | 0 | 0 | 0 | 0 | 0 | 0 | 21799 | 581290 | 0 | 188780 | 2224600 | 0 | 0 | 0 | 0 | 0 | 0 |
| AGVALPFEI | WWP2; WWP2; | O00308; H3I | WWP2_HUMAN | Unmodified | 5351100 | 7089200 | 3339200 | 1455600 | 0 | 117440 | 0 | 0 | 490250 | 0 | 0 | 0 | 0 | 342630 | 3660300 | 2972600 | 0 | 0 |
| AGVAPLQV | FLNA; FLNA | P21333; Q6( | FLNA_HUMAN | Unmodified | 111510 | 271920 | 141070 | 868750 | 15650000 | 4652300 | 324420 | 260680 | 1434600 | 1128100 | 361710 | 3025900 | 2973300 | 4004700 | 229490 | 258830 | 8745200 | 5386300 |
| AGVAPLQV | FLNA; FLNA | P21333; Q6( | FLNA_HUMAN | Unmodified | 0 | 0 | 0 | 0 | 2120600 | 0 | 0 | 453710 | 0 | 0 | 0 | 0 | 0 | 0 | 0 | 0 | 6743000 | 649770 |
| AGVENGKF | FLNB; FLNB | O75369; E7E | FLNB_HUMAN | Unmodified | 0 | 0 | 0 | 0 | 2062700 | 402410 | 35065 | 0 | 0 | 150710 | 0 | 0 | 0 | 0 | 0 | 10748 | 1195400 | 630510 |
| AGVFSDLS | AOC1; AOC1; A | P19801; C9J | AOC1_HUMAN | Unmodified | 0 | 0 | 0 | 0 | 0 | 0 | 0 | 0 | 4103100 | 1078200 | 89906 | 574510 | 0 | 0 | 0 | 0 | 0 | 62606 |
| AGVGEPPE | PSME4; PSME4 | Q14997; F8V | PSME4_HUMAI | Unmodified | 0 | 0 | 0 | 0 | 0 | 0 | 0 | 0 | 151540 | 0 | 0 | 1265300 | 0 | 0 | 0 | 0 | 0 | 0 |
| AGVHIKLPI | CKMT1A | P12532 | KCRU_HUMAN | Unmodified | 0 | 0 | 0 | 0 | 0 | 0 | 0 | 0 | 0 | 0 | 0 | 0 | 0 | 0 | 0 | 0 | 0 | 0 |
| AGVHIKLPI | CKB; CKB; CKB | P12277; H0Y | KCRB_HUMAN | Unmodified | 61706 | 73219 | 2303300 | 3271700 | 52367 | 0 | 514780 | 0 | 2386300 | 347960 | 170450 | 10222000 | 197200 | 0 | 2627500 | 511080 | 0 | 0 |
| AGVIFPVGI | MACROH2A2; N | Q9P0M6; O7 | H2AW_HUMAN | Unmodified | 17092000 | 10179000 | 0 | 0 | 0 | 0 | 0 | 1169700 | 0 | 0 | 0 | 94490000 | 902250 | 0 | 0 | 0 | 0 | 0 |
| AGVIGLTQ | HSD17B8 | Q92506 | DHB8_HUMAN | Unmodified | 0 | 0 | 0 | 0 | 0 | 0 | 0 | 0 | 706860 | 0 | 0 | 444480 | 0 | 0 | 0 | 0 | 0 | 0 |
| AGVLAGHE | GNB2; GNB4; G | P62879; Q9I | GBB2_HUMAN | Unmodified | 6010400 | 3729400 | 1315700 | 1151100 | 0 | 0 | 0 | 0 | 88340 | 0 | 0 | 45869000 | 0 | 0 | 0 | 2501600 | 0 | 0 |
| AGVLAGHE | GNB2; GNB1; G | P62879; P62 | GBB2_HUMAN | Oxidation (M) | 0 | 0 | 0 | 0 | 12282000 | 0 | 4733500 | 0 | 6044800 | 2233300 | 4779400 | 939460 | 0 | 0 | 1684800 | 0 | 8226400 | 16174000 |
| AGVLAGHE | GNB2; GNB1; G | P62879; P62 | GBB2_HUMAN | Unmodified | 0 | 0 | 0 | 0 | 584590 | 0 | 519810 | 0 | 9846200 | 384440 | 761230 | 0 | 381930 | 0 | 3206900 | 0 | 643190 | 838140 |
| AGVLAHLE | MYH9; MYH10 | P35579; P35 | MYH9_HUMAN | Unmodified | 182570 | 0 | 0 | 214110 | 0 | 0 | 0 | 0 | 110620 | 0 | 0 | 0 | 0 | 0 | 823530 | 808020 | 0 | 0 |
| AGVLAHLE | MYH9; MYH10 | P35579; P35 | MYH9_HUMAN | Unmodified | 227480 | 487640 | 1319200 | 851920 | 0 | 0 | 0 | 0 | 0 | 0 | 0 | 7121200 | 0 | 0 | 2163000 | 2074300 | 0 | 0 |
| AGVMALAN | RTRAF; RTRAF; | Q9Y224; G3' | RTRAF_HUMAN | 2 Oxidation (M) | 262690 | 0 | 73740 | 1617000 | 1195400 | 172840 | 0 | 0 | 0 | 216340 | 310040 | 160140 | 3709400 | 480360 | 476070 | 69340 | 0 | 2188300 |
| AGVNFSEF | PDCD6; PDCD6 | O75340; A0A | PDCD6_HUMA | Unmodified | 11632000 | 12037000 | 3603300 | 4507600 | 1756000 | 1908800 | 87758 | 0 | 27642000 | 6905100 | 701950 | 7970600 | 0 | 0 | 14060000 | 8542500 | 896220 | 0 |
| AGVNTVTTI | RPL7A; RPL7A; | P62424; Q5T | RL7A_HUMAN | Unmodified | 4326800 | 3668900 | 3039400 | 2414200 | 422880 | 0 | 924830 | 970770 | 2108500 | 264200 | 2448200 | 42756000 | 0 | 421980 | 3209700 | 1755700 | 0 | 539830 |
| AGVNTVTTI | RPL7A; RPL7A; | P62424; Q5T | RL7A_HUMAN | Unmodified | 5608500 | 4636300 | 3677400 | 3006000 | 1648200 | 0 | 646550 | 3472300 | 1461300 | 1126300 | 0 | 66523000 | 2253600 | 2216500 | 3320800 | 4185100 | 470540 | 1280000 |
| AGVPGVAA | PLK1; PLK1 | P53350; I3L3 | PLK1_HUMAN | Unmodified | 25722 | 0 | 0 | 23814 | 0 | 0 | 0 | 0 | 1092900 | 829420 | 228530 | 544460 | 0 | 0 | 322850 | 0 | 0 | 0 |
| AGVSAEPT | PRKG2 | Q13237 | KGP2_HUMAN | Unmodified | 0 | 0 | 0 | 0 | 0 | 0 | 10783000 | 5344900 | 0 | 0 | 0 | 0 | 0 | 0 | 0 | 0 | 0 | 0 |
| AGVSFSGH | DBN1; DBN1; D | Q16643; D6I | DREB_HUMAN | Acetyl (Protein N | 0 | 0 | 0 | 0 | 0 | 0 | 0 | 0 | 0 | 0 | 0 | 2976600 | 0 | 0 | 1291600 | 1488700 | 0 | 0 |
| AGVSISVVI | CPOX | P36551 | HEM6_HUMAN | Unmodified | 0 | 0 | 0 | 0 | 165720 | 0 | 0 | 91441 | 2862800 | 1887800 | 326120 | 1100800 | 0 | 0 | 0 | 0 | 0 | 280430 |
| AGVSVYGI | CMBL | Q96DG6 | CMBL_HUMAN | Unmodified | 0 | 0 | 0 | 0 | 0 | 0 | 0 | 0 | 0 | 0 | 0 | 0 | 0 | 0 | 421400 | 0 | 0 | 0 |
| AGVVGPEL | PLEC; PLEC | Q15149; H0Y | PLEC_HUMAN | Unmodified | 0 | 0 | 0 | 0 | 1471700 | 0 | 0 | 0 | 271190 | 427640 | 0 | 501610 | 209550 | 0 | 0 | 108740 | 8165700 | 3132800 |
| AGVYNLTK | PECR | Q9BY49 | PECR_HUMAN | Unmodified | 0 | 0 | 0 | 0 | 1944400 | 652010 | 0 | 0 | 2477500 | 331400 | 0 | 679740 | 0 | 35878 | 0 | 0 | 5560900 | 8468300 |
| AGWLDKNI | ARAP1; ARAP1 | Q96P48; E7E | ARAP1_HUMAN | Unmodified | 0 | 0 | 0 | 0 | 0 | 51971 | 0 | 0 | 0 | 193110 | 0 | 0 | 142790 | 124630 | 0 | 0 | 189970 | 0 |
| AGYAFEYLI | IREB2; IREB2; IF | P48200; H0Y | IREB2_HUMAN | Unmodified | 72722 | 0 | 0 | 0 | 0 | 0 | 0 | 0 | 0 | 0 | 0 | 0 | 209080 | 0 | 284510 | 0 | 0 | 0 |
| AGYAFRQA | MYO1B; MYO1E | O43795; E9F | MYO1B_HUMA | Unmodified | 0 | 0 | 0 | 0 | 1356100 | 626180 | 0 | 0 | 0 | 0 | 0 | 0 | 0 | 0 | 2284700 | 0 | 732750 | 0 |
| AGYIIPLQG | GAA | P10253 | LYAG_HUMAN | Unmodified | 18893 | 973030 | 0 | 0 | 0 | 0 | 0 | 0 | 11504000 | 0 | 0 | 0 | 0 | 0 | 39478 | 0 | 0 | 0 |
| AGYPQYVS | LRPPRC; LRPPF | P42704; B8Z | LPPRC_HUMAN | Unmodified | 0 | 0 | 0 | 0 | 0 | 0 | 0 | 0 | 1870900 | 527460 | 0 | 0 | 0 | 0 | 0 | 0 | 0 | 0 |
| AGYPTGTG | ELN; ELN; ELN; | P15502; B3K | ELN_HUMAN E | Unmodified | 0 | 0 | 0 | 0 | 0 | 0 | 424250 | 201330 | 0 | 0 | 0 | 0 | 0 | 0 | 0 | 0 | 0 | 0 |
| AHAAASIK | ANK2; ANK2; AI | A0A5F9ZGX3 | A0A5F9ZGX3_I | Acetyl (Protein N | 235250 | 146910 | 0 | 2151000 | 3096100 | 1267700 | 0 | 29604 | 753730 | 118980 | 0 | 1625000 | 2612000 | 4980000 | 560840 | 5285500 | 0 | 1217900 |
| AHAAIRENI | RPL5; RPL5; RP | P46777; A0A | RL5_HUMAN 6( | Unmodified | 2536200 | 2049700 | 1453000 | 1660300 | 3911200 | 967350 | 1912900 | 1673600 | 581970 | 835650 | 890470 | 9341400 | 2973700 | 1713200 | 1562700 | 1078700 | 527970 | 1676900 |
| AHAAQVTC | WDR77; WDR7 | Q9BQA1; H0 | MEP50_HUMAN | Unmodified | 0 | 0 | 0 | 0 | 0 | 0 | 0 | 0 | 0 | 0 | 0 | 5239600 | 0 | 0 | 0 | 0 | 0 | 0 |
| AHAAVWN | AIP; AIP | O00170; E9F | AIP_HUMAN AH | Unmodified | 214930 | 198330 | 0 | 0 | 2794900 | 0 | 1087900 | 333800 | 953610 | 532280 | 0 | 1835000 | 3439500 | 1043200 | 0 | 322360 | 2247400 | 3435900 |
| AHAHLDTG | GANAB; GANAE | Q14697; F5H | GANAB_HUMA | Unmodified | 0 | 0 | 0 | 0 | 0 | 0 | 0 | 0 | 6650100 | 289020 | 0 | 0 | 0 | 0 | 339420 | 0 | 0 | 0 |
| AHAQLVRE | GNAQ; GNAQ | P50148; B1A | GNAQ_HUMAN | Unmodified | 0 | 0 | 0 | 0 | 2020300 | 516380 | 579800 | 526970 | 0 | 454390 | 299540 | 118640 | 0 | 279690 | 0 | 0 | 117960 | 128450 |
| AHAQLVRE | GNAQ; GNAQ | P50148; B1A | GNAQ_HUMAN | Unmodified | 0 | 0 | 0 | 0 | 3183000 | 0 | 4183400 | 285340 | 0 | 844420 | 0 | 0 | 0 | 756130 | 0 | 0 | 0 | 0 |
| AHAVREKD | HS2ST1 | Q7LGA3 | HS2ST_HUMAN | Unmodified | 0 | 0 | 0 | 0 | 0 | 0 | 0 | 43676 | 0 | 1847200 | 0 | 0 | 556300 | 0 | 0 | 0 | 583640 | 590210 |
| AHAWPSP\ | GPRC5A | Q8NFJ5 | RAI3_HUMAN F | Unmodified | 0 | 0 | 0 | 0 | 7698600 | 2126200 | 0 | 0 | 0 | 0 | 0 | 0 | 0 | 0 | 53678 | 66344 | 0 | 40423 |
| AHAWPSP\ | GPRC5A | Q8NFJ5 | RAI3_HUMAN F | Unmodified | 0 | 0 | 0 | 0 | 4486200 | 1320800 | 0 | 0 | 210460 | 0 | 0 | 0 | 186780 | 0 | 0 | 0 | 0 | 0 |
| AHAWPSP\ | GPRC5A | Q8NFJ5 | RAI3_HUMAN F | Unmodified | 0 | 0 | 0 | 0 | 4809700 | 1557900 | 0 | 0 | 0 | 0 | 0 | 0 | 0 | 0 | 0 | 0 | 0 | 0 |
| AHCQTSG\ | MMUT | P22033 | MUTA_HUMAN | Unmodified | 0 | 0 | 0 | 0 | 969800 | 0 | 0 | 0 | 1766500 | 0 | 0 | 0 | 801020 | 618260 | 0 | 0 | 1012200 | 0 |
| AHDERSEA | NADK2; NADK2 | Q4G0N4; A0 | NAKD2_HUMAI | Unmodified | 0 | 0 | 0 | 0 | 0 | 0 | 0 | 0 | 518400 | 0 | 0 | 0 | 0 | 0 | 0 | 0 | 0 | 0 |
| AHDGGIYA | WDR1 | O75083 | WDR1_HUMAN | Unmodified | 6822300 | 1547700 | 5238700 | 5126500 | 33982000 | 10833000 | 3027400 | 36353 | 19757000 | 2744200 | 8559000 | 59585000 | 26196000 | 8924700 | 9006300 | 6333900 | 4657200 | 5391500 |
| AHDGGIYA | WDR1 | O75083 | WDR1_HUMAN | Unmodified | 427260 | 511890 | 0 | 479010 | 51553000 | 7636900 | 4167000 | 726780 | 1009300 | 4399900 | 67344 | 0 | 9027100 | 5535100 | 0 | 216460 | 9775000 | 9897900 |
| AHDQKVE( | TSPAN1 | O60635 | TSN1_HUMAN | Unmodified | 5628100 | 2644600 | 0 | 3749500 | 0 | 78993 | 0 | 0 | 10316000 | 1655200 | 0 | 0 | 0 | 0 | 3004700 | 0 | 0 | 0 |
| AHEASSAV | TRAF3; TRAF3; T | Q13114; A6N | TRAF3_HUMAN | Unmodified | 0 | 0 | 0 | 0 | 12078000 | 1572800 | 23560 | 0 | 0 | 0 | 0 | 156190 | 63319 | 0 | 0 | 0 | 59263 | 131260 |
| AHEEQDEE | KIF14 | Q15058 | KIF14_HUMAN | Unmodified | 0 | 0 | 109300 | 93650 | 0 | 0 | 0 | 0 | 0 | 0 | 3867400 | 582240 | 0 | 0 | 142150 | 0 | 0 | 0 |
| AHEFLRLS( | LSS; LSS | P48449; A0A | LSS_HUMAN La | Unmodified | 0 | 0 | 0 | 0 | 0 | 0 | 984950 | 139230 | 0 | 189800 | 0 | 0 | 0 | 0 | 0 | 0 | 175940 | 0 |
| AHEILPNLV | CCT8; CCT8 | P50990; H7( | TCPQ_HUMAN | Unmodified | 77929 | 0 | 0 | 62655 | 0 | 0 | 366950 | 406010 | 232390 | 538860 | 2092300 | 5186600 | 3750300 | 195790 | 0 | 569870 | 919360 | 1429500 |
| AHEPTYFT\ | FLNA; FLNA; FL | P21333; A0A | FLNA_HUMAN | Unmodified | 0 | 0 | 0 | 0 | 4001100 | 0 | 355370 | 0 | 0 | 0 | 0 | 1566200 | 785030 | 0 | 0 | 0 | 5369900 | 773710 |
| AHEWARAI | SCRN1 | Q12765 | SCRN1_HUMAI | Unmodified | 0 | 0 | 0 | 0 | 1443900 | 0 | 0 | 0 | 0 | 0 | 0 | 0 | 0 | 0 | 0 | 0 | 754190 | 1892200 |
| AHFFSDKD | TLL2 | Q9Y6L7 | TLL2_HUMAN T | Unmodified | 0 | 0 | 0 | 23355 | 0 | 0 | 37950 | 0 | 227670 | 95791 | 0 | 0 | 114550 | 0 | 0 | 16351 | 15698 | 0 |
| AHFLDADP | PAFAH1B3; PAF | Q15102; M0I | PA1B3_HUMAN | Oxidation (M) | 0 | 0 | 0 | 0 | 0 | 0 | 0 | 0 | 2604900 | 0 | 1288800 | 7143300 | 0 | 0 | 0 | 0 | 0 | 0 |
| AHFLDADP | PAFAH1B3; PAF | Q15102; M0I | PA1B3_HUMAN | Unmodified | 0 | 138300 | 0 | 0 | 0 | 0 | 0 | 0 | 4649500 | 249150 | 1170200 | 1493500 | 0 | 0 | 0 | 0 | 0 | 0 |
| AHFLVTQAI | CAPN7 | Q9Y6W3 | CAN7_HUMAN | Unmodified | 47575 | 0 | 0 | 0 | 879780 | 0 | 0 | 0 | 465320 | 0 | 0 | 961480 | 470280 | 206310 | 0 | 0 | 0 | 0 |
| AHFPLDVQ | GAA | P10253 | LYAG_HUMAN | Oxidation (M) | 81557 | 0 | 0 | 0 | 0 | 0 | 0 | 0 | 1036900 | 196630 | 28977 | 155640 | 0 | 0 | 0 | 0 | 0 | 0 |
| AHFPLDVQ | GAA | P10253 | LYAG_HUMAN | Unmodified | 0 | 0 | 0 | 0 | 0 | 0 | 0 | 0 | 1383400 | 0 | 0 | 0 | 0 | 0 | 0 | 0 | 0 | 0 |
| AHFPLDVQ | GAA | P10253 | LYAG_HUMAN | Unmodified | 0 | 0 | 0 | 0 | 0 | 0 | 0 | 0 | 1692500 | 0 | 0 | 0 | 0 | 0 | 0 | 0 | 0 | 0 |
| AHFQNFPN | TNPO1; TNPO1 | Q92973; A0A | TNPO1_HUMAI | Unmodified | 0 | 0 | 427600 | 0 | 0 | 0 | 587770 | 34752 | 0 | 0 | 254310 | 3877700 | 361260 | 0 | 589880 | 0 | 0 | 0 |
| AHFSGQEL | EHBP1 | Q8NDI1 | EHBP1_HUMAI | Unmodified | 0 | 0 | 0 | 0 | 0 | 0 | 0 | 0 | 0 | 0 | 0 | 0 | 0 | 0 | 0 | 0 | 0 | 0 |
| AHFSPSNII | QSOX1 | O00391 | QSOX1_HUMAI | Unmodified | 0 | 0 | 545310 | 243300 | 0 | 209860 | 0 | 0 | 3949400 | 638950 | 703100 | 0 | 0 | 0 | 381940 | 99679 | 0 | 466070 |
| AHFSPSNII | QSOX1 | O00391 | QSOX1_HUMAI | Unmodified | 0 | 0 | 476780 | 363400 | 0 | 0 | 0 | 0 | 3166600 | 271430 | 0 | 0 | 0 | 0 | 0 | 0 | 0 | 0 |
| AHGAPEPE | NFKBIB; NFKBII | Q15653; I3L4 | IKBB_HUMAN N | Unmodified | 0 | 0 | 0 | 0 | 0 | 0 | 0 | 0 | 0 | 0 | 0 | 76081 | 0 | 0 | 0 | 0 | 0 | 0 |
| AHGGISLSI | ATP13A3; ATP1 | Q9H7F0; H7 | AT133_HUMAN | Unmodified | 0 | 0 | 0 | 0 | 193130 | 135570 | 0 | 0 | 10851000 | 2346200 | 0 | 224970 | 0 | 0 | 0 | 0 | 76721 | 157360 |
| AHGGNSN | C1QL1 | O75973 | C1QRF_HUMAI | Unmodified | 0 | 0 | 0 | 0 | 0 | 0 | 0 | 0 | 0 | 2542700 | 0 | 0 | 0 | 0 | 0 | 0 | 0 | 0 |
| AHGGYSVF | ATP5F1B; ATP5I | P06576; H0Y | ATPB_HUMAN | Unmodified | 191090 | 159660 | 376600 | 0 | 76519 | 0 | 130450 | 63555 | 26879000 | 1929200 | 137190 | 425390 | 0 | 93048 | 80668 | 0 | 0 | 122650 |
| AHGGYSVF | ATP5F1B; ATP5I | P06576; H0Y | ATPB_HUMAN | Unmodified | 0 | 0 | 0 | 0 | 1347500 | 0 | 5873600 | 0 | 0 | 13629000 | 0 | 0 | 2174500 | 3455600 | 0 | 0 | 533540 | 2088800 |
| AHGHCLHI | AGPS; AGPS; A( | O00116; A0A | ADAS_HUMAN | Unmodified | 0 | 0 | 0 | 0 | 0 | 0 | 0 | 0 | 636520 | 0 | 0 | 0 | 0 | 0 | 0 | 0 | 0 | 0 |
| AHGPGALN | RHOQ; RHOQ | P17081; F8V | RHOQ_HUMAN | Acetyl (Protein N | 0 | 0 | 0 | 0 | 0 | 0 | 509810 | 0 | 0 | 0 | 0 | 0 | 0 | 141150 | 0 | 0 | 0 | 0 |
| AHGPGLE( | FLNB; FLNB | O75369; E7E | FLNB_HUMAN | Unmodified | 0 | 0 | 1933400 | 1745500 | 11805000 | 2492800 | 269460 | 125850 | 660900 | 381920 | 1012700 | 22298000 | 2093100 | 781830 | 1469400 | 1478500 | 2724800 | 1953900 |
| AHGVSSYD | LDLR; LDLR; LD | P01130; J3KI | LDLR_HUMAN | Unmodified | 0 | 0 | 3912200 | 206040 | 114140 | 3455400 | 0 | 0 | 978240 | 0 | 0 | 0 | 0 | 0 | 1039000 | 0 | 0 | 0 |
| AHHEVCPK | TRAF2 | Q12933 | TRAF2_HUMAN | Unmodified | 0 | 0 | 0 | 0 | 1651000 | 534990 | 0 | 0 | 0 | 0 | 0 | 0 | 723180 | 51131 | 0 | 0 | 0 | 46834 |
| AHHGEAGI | STC2 | O76061 | STC2_HUMAN | Unmodified | 0 | 0 | 0 | 0 | 0 | 0 | 0 | 0 | 540760 | 61323 | 0 | 0 | 0 | 0 | 21670 | 0 | 0 | 0 |
| AHIANPSG | FLNB; FLNB | O75369; E7E | FLNB_HUMAN | Unmodified | 0 | 0 | 0 | 0 | 20338000 | 2688400 | 0 | 166640 | 0 | 0 | 0 | 4603000 | 1899500 | 1568300 | 0 | 0 | 11288000 | 0 |
| AHIAQLCEI | CLTCL1; CLTC; | P53675; Q0( | CLH2_HUMAN | Unmodified | 2141400 | 1333200 | 0 | 0 | 0 | 0 | 0 | 0 | 0 | 47521 | 0 | 5210800 | 0 | 0 | 902430 | 1555800 | 0 | 0 |
| AHILACAPS | MOV10; MOV1( | Q9HCE1; Q5 | MOV10_HUMA | Unmodified | 6488400 | 1375100 | 372850 | 1084000 | 0 | 0 | 0 | 0 | 116850 | 0 | 0 | 0 | 0 | 0 | 932020 | 2389000 | 0 | 0 |
| AHIPNFTFN | NDST1; NDST1 | P52848; E7E | NDST1_HUMAI | Unmodified | 0 | 0 | 0 | 0 | 0 | 0 | 0 | 0 | 663600 | 44650 | 0 | 98812 | 0 | 0 | 45727 | 0 | 0 | 0 |
| AHIRQDLIH | CARMIL1; CARM | Q5VZK9; A0A | CARL1_HUMAN | Unmodified | 0 | 0 | 0 | 48248 | 1478200 | 31352 | 0 | 0 | 0 | 0 | 16737 | 766340 | 0 | 0 | 0 | 0 | 93786 | 845100 |
| AHIVFDFH( | ADSS2 | P30520 | PURA2_HUMAI | Unmodified | 0 | 0 | 0 | 0 | 0 | 0 | 0 | 0 | 0 | 0 | 0 | 1443300 | 0 | 0 | 0 | 0 | 0 | 0 |
| AHIVFDFH( | ADSS2 | P30520 | PURA2_HUMAI | Unmodified | 0 | 0 | 0 | 117060 | 0 | 0 | 0 | 0 | 218270 | 0 | 955930 | 34264000 | 115570 | 0 | 0 | 0 | 69667 | 49088 |

| | | | | | | | | | | | | | | | | | | | | | | |
|---|---|---|---|---|---|---|---|---|---|---|---|---|---|---|---|---|---|---|---|---|---|---|
| AHLGSCTL | ZNF512; ZNF51 | Q96ME7; G3 | ZN512_HUMAN | Unmodified | 202530 | 102610 | 0 | 0 | 0 | 0 | 0 | 0 | 0 | 0 | 0 | 0 | 0 | 0 | 0 | 0 | 0 | 0 |
| AHLGTALK | RPS23; RPS23; | P62266; D6F | RS23_HUMAN | Unmodified | 987560 | 865450 | 784350 | 671150 | 1071400 | 235770 | 550770 | 905480 | 363150 | 3202200 | 203930 | 4694500 | 1508300 | 559740 | 522870 | 591100 | 122930 | 480750 |
| AHLGTALK | RPS23; RPS23; | P62266; D6F | RS23_HUMAN | Unmodified | 0 | 0 | 0 | 0 | 0 | 0 | 684600 | 209070 | 0 | 0 | 0 | 0 | 1068900 | 354840 | 0 | 0 | 125820 | 201260 |
| AHLLADMA | SHMT2; SHMT2 | P34897; H0Y | GLYM_HUMAN | Unmodified | 0 | 0 | 0 | 0 | 0 | 0 | 494580 | 0 | 19166000 | 1952300 | 0 | 563890 | 0 | 0 | 0 | 0 | 0 | 0 |
| AHLMEIQVI | RPL3; RPL3; RP | P39023; B5N | RL3_HUMAN 6 | Oxidation (M) | 848740 | 650500 | 504940 | 0 | 242440 | 0 | 307880 | 147130 | 505410 | 0 | 324880 | 5441300 | 439530 | 0 | 527130 | 411420 | 19138 | 0 |
| AHLMEIQVI | RPL3; RPL3; RP | P39023; B5N | RL3_HUMAN 6 | Unmodified | 464900 | 1136700 | 0 | 0 | 0 | 0 | 0 | 0 | 381400 | 0 | 0 | 8249200 | 0 | 0 | 0 | 270950 | 0 | 0 |
| AHLRAAQD | EEA1 | Q15075 | EEA1_HUMAN | Unmodified | 0 | 826870 | 0 | 1128400 | 7775100 | 834800 | 1289500 | 0 | 0 | 0 | 0 | 0 | 2815900 | 0 | 0 | 0 | 0 | 2077500 |
| AHLRGFAN | STK25; STK25; | O00506; E7E | STK25_HUMAN | Acetyl (Protein N | 0 | 0 | 0 | 0 | 808630 | 0 | 0 | 0 | 0 | 185360 | 0 | 0 | 386180 | 0 | 0 | 0 | 305800 | 518390 |
| AHLTNQYN | PABPC4; PABP | Q13310; P11 | PABP4_HUMAN | Unmodified | 5436800 | 4844700 | 4431500 | 1719900 | 0 | 0 | 0 | 19915 | 0 | 0 | 0 | 1302100 | 793520 | 0 | 1630800 | 1594600 | 0 | 0 |
| AHLTNQYN | PABPC4; PABP | Q13310; P11 | PABP4_HUMAN | Oxidation (M) | 0 | 2186800 | 0 | 0 | 0 | 0 | 0 | 0 | 0 | 0 | 0 | 0 | 0 | 0 | 0 | 0 | 0 | 0 |
| AHLVAVFN | ANXA11; ANXA | P50995; H0Y | ANX11_HUMAN | Unmodified | 240080 | 3564500 | 1259500 | 237400 | 3814400 | 762060 | 0 | 140740 | 52459000 | 12640000 | 3438000 | 9085200 | 5617000 | 2710400 | 2328000 | 275600 | 0 | 8463400 |
| AHLVAVFN | ANXA11; ANXA | P50995; H0Y | ANX11_HUMAN | Unmodified | 0 | 0 | 0 | 0 | 7579500 | 0 | 1346700 | 0 | 287800 | 2553600 | 0 | 0 | 1618900 | 2139300 | 0 | 0 | 2195300 | 5854000 |
| AHLVAVFN | ANXA11; ANXA | P50995; H0Y | ANX11_HUMAN | Oxidation (M) | 0 | 0 | 0 | 0 | 0 | 0 | 0 | 0 | 0 | 1499900 | 0 | 0 | 753910 | 1047000 | 0 | 0 | 363600 | 1214100 |
| AHMEDSCI | PLXNB2 | O15031 | PLXB2_HUMAN | 2 Oxidation (M) | 407160 | 0 | 0 | 507020 | 0 | 0 | 0 | 0 | 0 | 0 | 0 | 391690 | 0 | 0 | 1249400 | 1066300 | 0 | 0 |
| AHMGMFTI | CLTCL1; CLTC; | P53675; Q00 | CLH2_HUMAN | 2 Oxidation (M) | 0 | 1426400 | 806160 | 0 | 0 | 0 | 0 | 11680 | 0 | 0 | 0 | 3660200 | 0 | 0 | 0 | 80718 | 0 | 0 |
| AHNNELEP | POLR2A; POLR2 | P24928; A0A | RPB1_HUMAN | Unmodified | 0 | 0 | 486680 | 0 | 0 | 0 | 0 | 0 | 0 | 137810 | 133710 | 698250 | 0 | 0 | 0 | 0 | 0 | 0 |
| AHNQDLGI | COL4A2; COL4 | P08572; A0A | CO4A2_HUMAN | Unmodified | 0 | 0 | 0 | 0 | 0 | 377650 | 0 | 0 | 284120 | 62941 | 0 | 0 | 0 | 0 | 0 | 0 | 0 | 0 |
| AHNTNDFV | SORL1; SORL1; | Q92673; E9F | SORL_HUMAN | Unmodified | 0 | 0 | 0 | 0 | 0 | 0 | 0 | 0 | 1029700 | 0 | 0 | 0 | 0 | 0 | 0 | 0 | 0 | 0 |
| AHPFFESV | PDPK2P; PDPK | Q6A1A2; O1 | PDPK2_HUMAN | Unmodified | 0 | 152320 | 1772200 | 580910 | 0 | 0 | 0 | 0 | 1258900 | 0 | 0 | 0 | 0 | 0 | 1642500 | 1601200 | 0 | 505150 |
| AHPQGLVL | AMDHD2; AMD | Q9Y303; H3I | NAGA_HUMAN | Unmodified | 17294 | 3565400 | 0 | 25009 | 0 | 0 | 0 | 0 | 0 | 0 | 0 | 263350 | 0 | 0 | 240830 | 192010 | 0 | 0 |
| AHQANQL | DYNC1H1; DYN | Q14204; A0 | DYHC1_HUMA | Unmodified | 0 | 0 | 0 | 0 | 0 | 0 | 0 | 0 | 0 | 0 | 0 | 317230 | 0 | 0 | 0 | 0 | 0 | 0 |
| AHQDALPF | CRAT; CRAT; CI | P43155; A0A | CACP_HUMAN | Unmodified | 0 | 0 | 0 | 0 | 0 | 0 | 0 | 0 | 3148900 | 1415400 | 0 | 0 | 0 | 0 | 0 | 0 | 0 | 0 |
| AHQDIHTQ | GOLIM4; GOLIN | O00461; F8V | GOLI4_HUMAN | Unmodified | 0 | 50959 | 0 | 0 | 0 | 269480 | 0 | 96813 | 11078000 | 72682 | 0 | 595260 | 0 | 209130 | 0 | 0 | 0 | 116100 |
| AHQDIHTQ | GOLIM4 | O00461 | GOLI4_HUMAN | Unmodified | 0 | 0 | 0 | 0 | 427700 | 0 | 0 | 944290 | 2102800 | 11321000 | 0 | 0 | 0 | 0 | 0 | 0 | 0 | 0 |
| AHQDIHTQ | GOLIM4 | F8W785 | F8W785_HUM | Unmodified | 0 | 0 | 0 | 0 | 0 | 0 | 0 | 0 | 0 | 3392600 | 0 | 0 | 0 | 0 | 0 | 0 | 0 | 0 |
| AHQFSIK | CDC42BPB; CD | Q9Y5S2; H0 | MRCKB_HUMA | Unmodified | 41226 | 0 | 0 | 0 | 381040 | 0 | 0 | 0 | 162250 | 0 | 0 | 0 | 0 | 82952 | 213770 | 0 | 388420 | 391760 |
| AHQITDESI | SNAP23; SNAP | O00161; H3I | SNP23_HUMAN | Unmodified | 735100 | 458900 | 0 | 874860 | 0 | 0 | 0 | 0 | 0 | 0 | 0 | 1684900 | 220820 | 0 | 0 | 622770 | 0 | 0 |
| AHQITDESI | SNAP23; SNAP | O00161; H3I | SNP23_HUMAN | Unmodified | 0 | 680830 | 2028600 | 1683900 | 712560 | 153260 | 86122 | 91401 | 938390 | 65761 | 2396900 | 9078000 | 607120 | 0 | 4453600 | 3584000 | 217920 | 0 |
| AHQITDESI | SNAP23; SNAP | O00161; H3I | SNP23_HUMAN | Unmodified | 0 | 0 | 0 | 0 | 3860900 | 0 | 0 | 0 | 0 | 0 | 0 | 0 | 2691800 | 1610500 | 0 | 0 | 524450 | 1844900 |
| AHQLEEDI | TAX1BP1; TAX1 | Q86VP1; B8 | TAXB1_HUMAN | Unmodified | 91682 | 0 | 0 | 152620 | 2268400 | 0 | 2313200 | 0 | 2515100 | 939900 | 96528 | 797520 | 1660900 | 0 | 0 | 0 | 796080 | 5503700 |
| AHQLEEDI | TAX1BP1; TAX1 | Q86VP1; B8 | TAXB1_HUMAN | Unmodified | 0 | 0 | 0 | 0 | 75697 | 0 | 444460 | 0 | 0 | 278840 | 0 | 0 | 41926 | 56180 | 0 | 0 | 150070 | 0 |
| AHQTGIHA | TWF2; TWF2; T\ | Q6IBS0; D6F | TWF2_HUMAN | Acetyl (Protein N | 0 | 390180 | 0 | 0 | 649670 | 227440 | 315940 | 0 | 415330 | 1581000 | 785880 | 6556700 | 0 | 168120 | 597920 | 417600 | 1997500 | 408830 |
| AHQTGIHA | TWF2; TWF2; T\ | Q6IBS0; D6F | TWF2_HUMAN | Acetyl (Protein N | 0 | 0 | 0 | 0 | 0 | 0 | 565500 | 0 | 0 | 286100 | 0 | 335810 | 0 | 0 | 0 | 0 | 3378900 | 0 |
| AHQVVED | PPP1CA; PPP1 | P62136; P62 | PP1A_HUMAN | Unmodified | 1323400 | 1164200 | 941180 | 1720500 | 2800300 | 1249300 | 618080 | 318710 | 8734500 | 3061200 | 3549700 | 26898000 | 422800 | 119270 | 3174100 | 2953600 | 582070 | 400160 |
| AHQVVED | PPP1CA; PPP1 | P62136; P62 | PP1A_HUMAN | Unmodified | 0 | 0 | 0 | 0 | 0 | 0 | 0 | 0 | 662240 | 0 | 0 | 3161400 | 0 | 0 | 0 | 0 | 0 | 0 |
| AHRLLGHV | ST13; ST13 | P50502; H7 | F10A1_HUMAN | Oxidation (M) | 0 | 0 | 0 | 0 | 0 | 0 | 693060 | 0 | 0 | 0 | 0 | 0 | 4770900 | 6299500 | 0 | 0 | 1567300 | 2349600 |
| AHRLLGHV | ST13; ST13 | P50502; H7 | F10A1_HUMAN | Unmodified | 0 | 0 | 0 | 0 | 0 | 0 | 561470 | 0 | 0 | 708250 | 0 | 0 | 0 | 1711500 | 0 | 0 | 0 | 988810 |
| AHRLSAEE | PCBD1 | P61457 | PHS_HUMAN P | Unmodified | 0 | 0 | 0 | 337110 | 304780 | 0 | 0 | 0 | 726080 | 382030 | 0 | 0 | 3568800 | 542410 | 0 | 0 | 280520 | 303800 |
| AHRWQDN | CACNA2D2; CA | Q9NY47; C9 | CA2D2_HUMA | Unmodified | 0 | 0 | 0 | 0 | 0 | 0 | 0 | 0 | 1831100 | 195420 | 0 | 0 | 0 | 0 | 0 | 0 | 181330 | 0 |
| AHSDGGD | CNTN1; CNTN1 | Q12860; H0 | CNTN1_HUMA | Unmodified | 0 | 0 | 0 | 0 | 189590 | 0 | 0 | 0 | 172680 | 0 | 0 | 0 | 0 | 0 | 0 | 0 | 0 | 316690 |
| AHSIQIMK | RPL18A; RPL18 | Q02543; M0 | RL18A_HUMAN | Unmodified | 0 | 0 | 0 | 0 | 0 | 0 | 0 | 0 | 0 | 0 | 0 | 1526600 | 0 | 0 | 0 | 0 | 43449 | 17254 |
| AHSIQIMK | RPL18A; RPL18 | Q02543; M0 | RL18A_HUMAN | Unmodified | 355970 | 434220 | 0 | 271980 | 0 | 0 | 817020 | 0 | 246070 | 114700 | 157850 | 5323400 | 305950 | 159830 | 142950 | 133370 | 214270 | 0 |
| AHSIQIMK | RPL18A; RPL18 | Q02543; M0 | RL18A_HUMAN | Oxidation (M) | 0 | 0 | 0 | 0 | 0 | 0 | 0 | 0 | 0 | 0 | 397340 | 2411800 | 236140 | 0 | 0 | 0 | 0 | 0 |
| AHSLLFEN | PHACTR4 | Q8IZ21 | PHAR4_HUMA | Unmodified | 0 | 0 | 0 | 175400 | 866420 | 0 | 0 | 0 | 2283000 | 0 | 117760 | 293910 | 0 | 90421 | 0 | 0 | 0 | 514840 |
| AHSPVAVQ | STK26; STK26; | Q9P289; B4 | STK26_HUMAN | Acetyl (Protein N | 0 | 0 | 367490 | 0 | 0 | 0 | 0 | 815500 | 0 | 2694600 | 569270 | 560130 | 167650 | 0 | 507220 | 392290 | 0 | 2481300 |
| AHSQTDRA | HLA-A; HLA-A; | A0A0G2JL56 | A0A0G2JL56_H | Unmodified | 0 | 0 | 0 | 0 | 0 | 0 | 0 | 0 | 1675200 | 0 | 0 | 527090 | 0 | 0 | 0 | 0 | 0 | 0 |
| AHSSMVG | PGK1 | P00558 | PGK1_HUMAN | Oxidation (M) | 0 | 674020 | 1059200 | 1381800 | 582510 | 254800 | 0 | 0 | 2464300 | 528150 | 2427900 | 20854000 | 0 | 0 | 1434000 | 2248500 | 759800 | 2050700 |
| AHSSMVG | PGK1 | P00558 | PGK1_HUMAN | Unmodified | 269360 | 385640 | 370310 | 4609000 | 2243600 | 275910 | 435140 | 113050 | 11695000 | 1375900 | 2629900 | 42222000 | 2710800 | 360590 | 4105200 | 2822400 | 1546400 | 4648400 |
| AHTCFNRL | NEDD4L; NEDD | Q96PU5; K7 | NED4L_HUMA | Unmodified | 1042600 | 0 | 0 | 0 | 0 | 0 | 0 | 0 | 0 | 0 | 0 | 0 | 0 | 0 | 0 | 0 | 0 | 0 |
| AHTDFFEA | COPS2; COPS2 | P61201; B4 | CSN2_HUMAN | Unmodified | 0 | 0 | 0 | 0 | 0 | 0 | 0 | 19569 | 172600 | 0 | 0 | 2192600 | 190380 | 127310 | 0 | 0 | 79022 | 0 |
| AHTDSLNT | PLXNB2 | O15031 | PLXB2_HUMAN | Unmodified | 1370300 | 0 | 3515800 | 4482300 | 0 | 0 | 0 | 0 | 5822900 | 446290 | 0 | 15096000 | 0 | 0 | 13080000 | 5825800 | 0 | 0 |
| AHTDVGP | PTPRF; PTPRF | P10586; H0 | PTPRF_HUMAN | Unmodified | 2205200 | 0 | 373000 | 280080 | 0 | 0 | 0 | 260610 | 0 | 0 | 0 | 0 | 0 | 0 | 442420 | 0 | 0 | 0 |
| AHTMTDD | CLTC; CLTC; CI | Q00610; J3K | CLH1_HUMAN | Unmodified | 2830500 | 2662600 | 686240 | 1052600 | 918260 | 655370 | 597450 | 0 | 466020 | 392820 | 0 | 3020900 | 798130 | 186900 | 1891700 | 1844800 | 403060 | 519310 |
| AHTMTDD | CLTC; CLTC; CI | Q00610; J3K | CLH1_HUMAN | Oxidation (M) | 3059500 | 2697700 | 349090 | 1586200 | 0 | 0 | 0 | 0 | 0 | 0 | 0 | 2519100 | 198900 | 0 | 2743700 | 2420200 | 0 | 0 |
| AHTSRFISA | PTPRD; PTPRF; | P23468; P10 | PTPRD_HUMAN | Unmodified | 0 | 0 | 0 | 0 | 323660 | 52002 | 46168 | 0 | 0 | 0 | 0 | 0 | 79841 | 136890 | 0 | 0 | 77495 | 43241 |
| AHTSSTQL | SEC23IP | Q9Y6Y8 | S23IP_HUMAN | Unmodified | 0 | 0 | 0 | 0 | 0 | 0 | 0 | 0 | 0 | 0 | 0 | 1042200 | 0 | 0 | 0 | 0 | 0 | 0 |
| AHTTPDHP | DIP2A; DIP2A | Q14689; A0 | DIP2A_HUMAN | Unmodified | 0 | 0 | 101200 | 138170 | 0 | 0 | 0 | 0 | 856460 | 236260 | 30963 | 433480 | 0 | 0 | 363770 | 203070 | 0 | 0 |
| AHVDEFKS | UGP2; UGP2; U | Q16851; E7E | UGPA_HUMAN | Unmodified | 0 | 0 | 0 | 0 | 125540 | 0 | 208910 | 159660 | 9650.2 | 57431 | 0 | 0 | 0 | 135030 | 0 | 0 | 206200 | 0 |
| AHVENTER | LAMC1 | P11047 | LAMC1_HUMA | Unmodified | 0 | 0 | 0 | 0 | 208740 | 0 | 0 | 0 | 223990 | 287440 | 65370 | 0 | 365940 | 185290 | 0 | 0 | 56597 | 1613900 |
| AHVIVMAA | VCP; VCP; VCP; | P55072; C9J | TERA_HUMAN | Oxidation (M) | 50335 | 44638 | 0 | 0 | 0 | 0 | 0 | 0 | 0 | 0 | 174880 | 709200 | 0 | 0 | 78369 | 112740 | 0 | 0 |
| AHVLAASV | CTNNA1; CTNN | P35221; E5R | CTNA1_HUMA | Unmodified | 0 | 0 | 3000300 | 3898200 | 2001000 | 0 | 0 | 0 | 0 | 0 | 2206000 | 6251100 | 1274800 | 1258800 | 6587700 | 5692200 | 163900 | 0 |
| AHVLAASV | CTNNA1; CTNN | P35221; E5R | CTNA1_HUMA | Unmodified | 933360 | 1106400 | 0 | 6086200 | 6702200 | 2230500 | 0 | 0 | 1143400 | 211410 | 2714600 | 6175800 | 5926400 | 3545400 | 2496700 | 4198000 | 0 | 0 |
| AHVLAASV | CTNNA1; CTNN | P35221; E5R | CTNA1_HUMA | Unmodified | 0 | 0 | 0 | 0 | 2609400 | 0 | 0 | 0 | 0 | 0 | 0 | 0 | 1576600 | 1282600 | 0 | 0 | 0 | 0 |
| AHVVPCFD | FLNA; FLNA | P21333; Q6 | FLNA_HUMAN | Unmodified | 0 | 0 | 0 | 0 | 2528500 | 325270 | 76379 | 0 | 476250 | 556610 | 0 | 0 | 546730 | 1345400 | 81386 | 0 | 3071700 | 0 |
| AHVVPCFD | FLNA; FLNA | P21333; Q6 | FLNA_HUMAN | Unmodified | 0 | 0 | 0 | 0 | 0 | 0 | 0 | 0 | 0 | 0 | 0 | 0 | 0 | 0 | 0 | 0 | 712410 | 23076 |
| AHYEPADR | MAP4K4; MAP4 | O95819; C9 | M4K4_HUMAN | Unmodified | 0 | 0 | 0 | 0 | 0 | 0 | 0 | 0 | 125040 | 9671.6 | 0 | 173540 | 0 | 0 | 0 | 0 | 38429 | 0 |
| AHYEPADR | MAP4K4; MAP4 | O95819; C9 | M4K4_HUMAN | Unmodified | 0 | 0 | 0 | 0 | 408920 | 0 | 0 | 0 | 0 | 0 | 0 | 0 | 0 | 0 | 99801 | 0 | 57413 | 236110 |
| AHYGGFTV | FGB; FGB | P02675; D6F | FIBB_HUMAN F | Unmodified | 0 | 1651400 | 0 | 0 | 0 | 0 | 0 | 0 | 0 | 0 | 0 | 0 | 0 | 211330 | 0 | 0 | 0 | 0 |
| AHYGGFTV | FGB; FGB | P02675; D6F | FIBB_HUMAN F | Unmodified | 0 | 5791200 | 0 | 0 | 0 | 2956900 | 0 | 0 | 0 | 0 | 0 | 0 | 0 | 3202000 | 0 | 0 | 0 | 0 |
| AIAALISEK | DNAJC10; DNA | Q8IXB1; A0A | DJC10_HUMAN | Unmodified | 0 | 329980 | 0 | 0 | 11219000 | 4132100 | 0 | 0 | 19227000 | 681520 | 665470 | 119950 | 0 | 0 | 446030 | 0 | 1599600 | 3763300 |
| AIAALISEKI | DNAJC10; DNA | Q8IXB1; A0A | DJC10_HUMAN | Unmodified | 0 | 0 | 0 | 0 | 0 | 0 | 0 | 0 | 10076000 | 1965100 | 718220 | 0 | 0 | 0 | 0 | 0 | 0 | 0 |
| AIAALISEKI | DNAJC10; DNA | Q8IXB1; A0A | DJC10_HUMAN | Unmodified | 0 | 0 | 0 | 0 | 0 | 0 | 0 | 0 | 7456400 | 5442500 | 0 | 0 | 0 | 0 | 0 | 0 | 284640 | 0 |
| AIADHLFW | LTV1; N/A | Q96GA3; A0 | LTV1_HUMAN | Unmodified | 915530 | 624660 | 804280 | 932640 | 0 | 0 | 0 | 0 | 0 | 225910 | 95034 | 0 | 649260 | 0 | 0 | 846030 | 0 | 0 |
| AIADTGAN | CCT8; CCT8 | P50990; H7 | TCPQ_HUMAN | Unmodified | 0 | 0 | 1288600 | 1251300 | 7229400 | 1093200 | 1717000 | 1149200 | 4241000 | 2418400 | 7810400 | 35003000 | 15415000 | 4687100 | 0 | 982200 | 2740400 | 4744100 |
| AIADTGAN | CCT8; CCT8 | P50990; H7 | TCPQ_HUMAN | Oxidation (M) | 0 | 0 | 0 | 0 | 1667000 | 0 | 1456800 | 0 | 0 | 394930 | 0 | 0 | 2311100 | 0 | 0 | 0 | 488860 | 1098700 |
| AIADTGAN | CCT8; CCT8 | P50990; H7 | TCPQ_HUMAN | Unmodified | 0 | 0 | 0 | 0 | 0 | 61366 | 0 | 0 | 0 | 0 | 324890 | 0 | 1106800 | 994420 | 0 | 0 | 0 | 0 |
| AIADWYNE | RO60; RO60 | P10155; H0 | RO60_HUMAN | Unmodified | 6009300 | 4832300 | 1989200 | 0 | 0 | 0 | 220460 | 0 | 0 | 0 | 94065 | 841590 | 268930 | 462260 | 3792300 | 1795900 | 0 | 0 |
| AIAELGIYP | ATP5F1B; ATP5 | P06576; H0 | ATPB_HUMAN | Unmodified | 0 | 0 | 255160 | 0 | 232070 | 0 | 0 | 158810 | 41915000 | 6128200 | 1312400 | 750600 | 0 | 0 | 0 | 0 | 0 | 0 |
| AIAFLQQPF | GART | P22102 | PUR2_HUMAN | Unmodified | 0 | 383550 | 0 | 970250 | 2845000 | 919390 | 694070 | 379190 | 1124800 | 3077300 | 0 | 5911100 | 2220400 | 1584000 | 706470 | 638610 | 559150 | 1540200 |
| AIAGVINQF | BPNT1; BPNT1; | O95861; F8V | BPNT1_HUMAN | Unmodified | 0 | 0 | 0 | 1255100 | 1180000 | 320890 | 0 | 36892 | 1068200 | 590080 | 2084000 | 3247700 | 1954700 | 0 | 616420 | 640380 | 625060 | 1350100 |
| AIAHYEQS | NAPA; NAPB; N | P54920; Q9 | SNAA_HUMAN | Unmodified | 334630 | 284660 | 373440 | 178570 | 0 | 0 | 0 | 0 | 490890 | 0 | 0 | 2400700 | 0 | 0 | 406100 | 399530 | 44979 | 78177 |
| AIAHYEQS | NAPA; NAPB; N | P54920; Q9 | SNAA_HUMAN | Unmodified | 2794700 | 2296200 | 101100 | 1890300 | 5069500 | 87993 | 2822500 | 1269600 | 3633800 | 1509000 | 829730 | 4040700 | 3805800 | 780800 | 982400 | 1331500 | 2598100 | 311760 |
| AIAKGVGII | ATP1A1; ATP1A | P05023; P50 | AT1A1_HUMAN | Unmodified | 0 | 0 | 0 | 0 | 562610 | 0 | 165540 | 129690 | 0 | 133080 | 0 | 757790 | 1547500 | 1118200 | 104470 | 75630 | 2105700 | 186120 |
| AIAKGVGII | ATP1A1; ATP1A | P05023; P13 | AT1A1_HUMAN | Unmodified | 0 | 0 | 0 | 0 | 0 | 0 | 0 | 0 | 0 | 0 | 0 | 0 | 0 | 0 | 0 | 0 | 2127100 | 0 |
| AIALQPENI | RMDN3; RMDN | Q96TC7; H0 | RMD3_HUMAN | Unmodified | 0 | 0 | 32143 | 45908 | 0 | 0 | 0 | 0 | 148900 | 0 | 55291 | 281200 | 0 | 0 | 0 | 0 | 0 | 0 |
| AIANECQA | VCP; VCP; VCP; | P55072; C9J | TERA_HUMAN | Unmodified | 0 | 0 | 0 | 0 | 0 | 0 | 0 | 0 | 1506600 | 79651 | 159560 | 12350000 | 0 | 0 | 0 | 202210 | 0 | 0 |
| AIAQLNSK | MME; MME; MM | P08473; A0A | NEP_HUMAN N | Unmodified | 0 | 0 | 21923000 | 32616000 | 41804 | 0 | 10486 | 166680 | 12611000 | 2096900 | 510000 | 680320 | 10588000 | 3371400 | 5399200 | 27139000 | 82333 | 258660 |
| AIAQLNSK | MME; MME; MM | P08473; A0A | NEP_HUMAN N | Unmodified | 0 | 0 | 0 | 601950 | 0 | 0 | 0 | 0 | 0 | 658810 | 0 | 0 | 6168500 | 3208600 | 0 | 0 | 0 | 0 |
| AIASIPTER | ITGA2; ITGA2; I | P17301; E9P | ITA2_HUMAN I | Unmodified | 2797900 | 1771300 | 4003600 | 2819500 | 0 | 347290 | 0 | 0 | 710090 | 78544 | 0 | 1321900 | 112880 | 0 | 4398300 | 4133300 | 414760 | 0 |
| AIATKCGIL | ATP2B1; ATP2B | P20020; E7E | AT2B1_HUMAN | Unmodified | 0 | 0 | 0 | 0 | 0 | 0 | 0 | 0 | 0 | 0 | 0 | 0 | 400380 | 382240 | 0 | 0 | 0 | 51040 |
| AIATLMSDE | SLC4A4 | Q9Y6R1 | S4A4_HUMAN | Oxidation (M) | 0 | 0 | 0 | 0 | 0 | 0 | 0 | 0 | 0 | 0 | 0 | 0 | 0 | 0 | 0 | 934040 | 0 | 0 |
| AIAVTVQEN | TLN1 | Q9Y490 | TLN1_HUMAN | Unmodified | 0 | 0 | 0 | 0 | 3017300 | 0 | 2057200 | 398970 | 301760 | 230490 | 0 | 185150 | 0 | 2374700 | 0 | 0 | 1280400 | 1169000 |
| AIAVTVQEN | TLN1 | Q9Y490 | TLN1_HUMAN | Oxidation (M) | 0 | 0 | 0 | 0 | 1110800 | 0 | 1025300 | 347090 | 0 | 174940 | 0 | 356030 | 450450 | 0 | 170230 | 0 | 471900 | 732880 |
| AICIPEGAK | DARS2; DARS2; | Q6PI48; A0A | SYDM_HUMAN | Unmodified | 0 | 0 | 0 | 0 | 119590 | 0 | 64447 | 18328 | 1163200 | 382400 | 0 | 0 | 0 | 0 | 0 | 0 | 0 | 0 |
| AIDEDLTAL | TNIK | Q9UKE5 | TNIK_HUMAN | Unmodified | 1093300 | 702490 | 0 | 231050 | 0 | 0 | 0 | 110830 | 905820 | 0 | 863710 | 2172100 | 271560 | 0 | 287280 | 166060 | 0 | 0 |
| AIDGINQR | CAMP; CAMP | P49913; J3K | CAMP_HUMAN | Unmodified | 0 | 0 | 234100 | 494360 | 0 | 123900 | 0 | 44930 | 6591300 | 266500 | 99886 | 281440 | 1512700 | 0 | 476850 | 395540 | 190350 | 0 |
| AIDGINQR | CAMP; CAMP | P49913; J3K | CAMP_HUMAN | Unmodified | 0 | 0 | 0 | 0 | 0 | 0 | 0 | 0 | 0 | 141270 | 0 | 0 | 0 | 0 | 0 | 0 | 0 | 0 |
| AIDGLDRN | AKR1C3; AKR1 | P42330; A0A | AK1C3_HUMAN | Unmodified | 0 | 0 | 0 | 0 | 6603300 | 0 | 0 | 0 | 0 | 0 | 0 | 0 | 0 | 0 | 0 | 0 | 0 | 0 |
| AIDGLNLLF | GALNS | P34059 | GALNS_HUMA | Unmodified | 0 | 0 | 0 | 0 | 0 | 0 | 0 | 0 | 601360 | 30055 | 0 | 0 | 0 | 0 | 0 | 0 | 0 | 0 |
| AIDLFTDAI | ST13P4; ST13; | Q8IZP2; P50 | ST134_HUMAN | Unmodified | 3409400 | 0 | 0 | 4196100 | 3899400 | 789730 | 2050100 | 908430 | 10729000 | 6635500 | 13736000 | 23803000 | 17165000 | 7589100 | 0 | 0 | 1264000 | 2625000 |
| AIDLFTDAI | ST13P4; ST13; | Q8IZP2; P50 | ST134_HUMAN | Unmodified | 0 | 0 | 0 | 378630 | 1227800 | 0 | 2367200 | 99979 | 1089300 | 4490100 | 0 | 0 | 11849000 | 7700900 | 0 | 0 | 1756600 | 1032200 |
| AIDLFTDAI | ST13P4; ST13; | Q8IZP2; P50 | ST134_HUMAN | Unmodified | 0 | 0 | 0 | 0 | 4033100 | 0 | 11753000 | 0 | 0 | 3485500 | 0 | 521900 | 5708100 | 17052000 | 0 | 0 | 4915400 | 1757100 |
| AIDPKLSTT | PPP3CA | Q08209 | PP2BA_HUMAN | Unmodified | 0 | 0 | 0 | 0 | 609320 | 0 | 387020 | 507390 | 0 | 0 | 0 | 0 | 831710 | 1286300 | 0 | 0 | 412310 | 474760 |
| AIDSSNLK | PCID2 | Q5JVF3 | PCID2_HUMAN | Unmodified | 0 | 0 | 0 | 0 | 0 | 0 | 0 | 0 | 0 | 0 | 26279 | 1250600 | 0 | 0 | 259600 | 98432 | 0 | 0 |
| AIDTIYQTT | ADAM10 | O14672 | ADA10_HUMAN | Unmodified | 6566400 | 6190600 | 11172000 | 14168000 | 1727100 | 3112600 | 826780 | 142610 | 2350500 | 0 | 68888 | 6606200 | 4057400 | 438490 | 20411000 | 17393000 | 0 | 1344900 |
| AIDTIYQTT | ADAM10 | O14672 | ADA10_HUMAN | Oxidation (M) | 0 | 0 | 0 | 0 | 3470000 | 0 | 597430 | 0 | 0 | 0 | 0 | 0 | 2915900 | 0 | 0 | 0 | 492720 | 1818700 |
| AIDTSGME | EFR3A; EFR3A | Q14156; A0 | EFR3A_HUMAN | Unmodified | 0 | 0 | 161280 | 0 | 1311600 | 171180 | 87482 | 0 | 374820 | 100610 | 51663 | 0 | 0 | 0 | 161600 | 0 | 275490 | 0 |
| AIDTSGME | EFR3A; EFR3A | Q14156; A0 | EFR3A_HUMAN | Unmodified | 0 | 0 | 0 | 0 | 856520 | 0 | 129520 | 62849 | 0 | 0 | 0 | 0 | 0 | 0 | 0 | 0 | 230740 | 618170 |
| AIDTTQLFS | PCK2; PCK2 | Q16822; H0 | PCKGM_HUMA | Unmodified | 0 | 0 | 0 | 0 | 0 | 0 | 0 | 0 | 802930 | 0 | 0 | 0 | 0 | 0 | 0 | 0 | 0 | 0 |

| | | | | | | | | | | | | | | | | | | | | | | |
|---|---|---|---|---|---|---|---|---|---|---|---|---|---|---|---|---|---|---|---|---|---|---|
| AIEASNAYS | LAMA5 | O15230 | LAMA5_HUMAN | Unmodified | 346850 | 37564 | 268260 | 47289 | 278550 | 37755 | 0 | 0 | 272680 | 0 | 0 | 0 | 0 | 0 | 255050 | 50640 | 549260 | 367810 |
| AIEAVSPGL | ATP7A | Q04656 | ATP7A_HUMAN | Unmodified | 0 | 0 | 332980 | 330070 | 0 | 0 | 0 | 0 | 501180 | 0 | 0 | 752680 | 0 | 0 | 326400 | 487510 | 0 | 0 |
| AIEDEGGN | SAFB; SAFB | Q15424; K7E | SAFB1_HUMAN | Unmodified | 0 | 0 | 0 | 0 | 0 | 817910 | 0 | 0 | 354860 | 165070 | 1452100 | 2097800 | 362440 | 0 | 0 | 0 | 223400 | 0 |
| AIEEELQEI | MATN2; MATN2 | O00339; H0 | MATN2_HUMAN | Unmodified | 0 | 0 | 0 | 0 | 0 | 0 | 0 | 0 | 1601000 | 493020 | 401800 | 0 | 0 | 0 | 0 | 0 | 0 | 0 |
| AIEEGVNHI | GNPDA2; GNPD | Q8TDQ7; P4 | GNPI2_HUMAN | Oxidation (M) | 0 | 0 | 0 | 0 | 0 | 0 | 0 | 0 | 0 | 0 | 819720 | 2099000 | 0 | 0 | 0 | 589100 | 0 | 0 |
| AIEGFNQS | UAP1 | Q16222 | UAP1_HUMAN | Unmodified | 0 | 0 | 1976700 | 795140 | 5203200 | 1048000 | 0 | 0 | 312130 | 131970 | 252230 | 1024600 | 1729900 | 957700 | 714220 | 397000 | 349930 | 912280 |
| AIEHADGG | GSS; GSS | P48637; A0A | GSHB_HUMAN | Unmodified | 0 | 0 | 0 | 0 | 5552500 | 809580 | 677610 | 391240 | 413830 | 1262700 | 1689900 | 2284900 | 0 | 0 | 24087 | 0 | 3599700 | 2407600 |
| AIEINPDSA | ST13P4; ST13; | Q8IZP2; P50 | ST134_HUMAN | Unmodified | 0 | 570940 | 4477100 | 1008800 | 0 | 0 | 0 | 0 | 3949900 | 238120 | 2042600 | 14454000 | 305380 | 0 | 1620700 | 1613600 | 0 | 0 |
| AIEIYTDMG | NAPA; NAPA; N | P54920; M0 | SNAA_HUMAN | Unmodified | 0 | 0 | 0 | 0 | 0 | 0 | 0 | 0 | 0 | 169390 | 70941 | 873850 | 0 | 142620 | 0 | 0 | 78090 | 0 |
| AIEIYTDMG | NAPA; NAPA; N | P54920; M0 | SNAA_HUMAN | Unmodified | 35666 | 82208 | 0 | 0 | 0 | 0 | 91406 | 0 | 0 | 128070 | 0 | 69409 | 346420 | 177860 | 0 | 0 | 97050 | 0 |
| AIEKETELD | TAX1BP1; TAX1 | Q86VP1; B8 | TAXB1_HUMAN | Unmodified | 0 | 0 | 0 | 0 | 537560 | 86713 | 655060 | 183170 | 0 | 338700 | 0 | 0 | 308080 | 102340 | 0 | 0 | 224130 | 653380 |
| AIEKLAVEA | CKB; CKB | P12277; G3 | KCRB_HUMAN | Unmodified | 0 | 44268 | 0 | 0 | 0 | 0 | 7925000 | 0 | 0 | 0 | 0 | 0 | 1276700 | 785580 | 0 | 0 | 0 | 0 |
| AIEKNLKED | GNAO1; GNAO | P09471; A0A | GNAO_HUMAN | Unmodified | 0 | 0 | 0 | 0 | 0 | 0 | 293000 | 0 | 0 | 108790 | 0 | 0 | 0 | 0 | 0 | 0 | 0 | 0 |
| AIELEAIKPT | NOSTRIN; NOS | Q8IVI9; E9PD | NOSTN_HUMA | Unmodified | 3844000 | 1305600 | 3599000 | 5046800 | 0 | 0 | 0 | 0 | 0 | 0 | 0 | 0 | 1504500 | 2352100 | 9130700 | 4172400 | 0 | 0 |
| AIELEAIKPT | NOSTRIN; NOS | Q8IVI9; E9PD | NOSTN_HUMA | Unmodified | 2153300 | 1699100 | 1829600 | 6970100 | 0 | 0 | 0 | 0 | 0 | 0 | 0 | 0 | 2269600 | 1087300 | 7195600 | 2101400 | 0 | 0 |
| AIELLEK | LIN7A; LIN7C; L | O14910; Q9 | LIN7A_HUMAN | Unmodified | 631490 | 473420 | 2893000 | 2529900 | 4043200 | 1306600 | 111280 | 149050 | 9714600 | 2849600 | 2389800 | 5883300 | 3608300 | 2820600 | 3227800 | 2202900 | 1029000 | 4675100 |
| AIELLQEFS | SRP72; SRP72 | O76094; D6 | SRP72_HUMAN | Unmodified | 0 | 0 | 0 | 0 | 459500 | 1299000 | 0 | 0 | 0 | 0 | 1040900 | 4003500 | 0 | 0 | 0 | 0 | 0 | 0 |
| AIELNPANA | SGTA; SGTA; SG | O43765; K7E | SGTA_HUMAN | Unmodified | 0 | 0 | 0 | 0 | 0 | 0 | 0 | 80883 | 0 | 0 | 531200 | 2423700 | 550700 | 258070 | 0 | 0 | 0 | 0 |
| AIELNPANA | SGTA; SGTA; SG | O43765; K7E | SGTA_HUMAN | Unmodified | 0 | 0 | 0 | 0 | 0 | 0 | 3402300 | 537040 | 0 | 0 | 0 | 0 | 5944100 | 1297300 | 0 | 0 | 2602400 | 0 |
| AIELSLQEQ | STAM2 | O75886 | STAM2_HUMAN | Unmodified | 0 | 0 | 0 | 0 | 0 | 0 | 0 | 0 | 0 | 0 | 0 | 0 | 0 | 0 | 0 | 0 | 0 | 857590 |
| AIELSLQEQ | STAM2 | O75886 | STAM2_HUMAN | Unmodified | 0 | 88310 | 0 | 0 | 917390 | 0 | 42918 | 155590 | 0 | 0 | 0 | 0 | 1010000 | 331310 | 0 | 0 | 724640 | 1350500 |
| AIELWYHD | XPO7; XPO7; XP | Q9UIA9; E5F | XPO7_HUMAN | Unmodified | 0 | 0 | 0 | 0 | 0 | 0 | 0 | 0 | 0 | 0 | 0 | 1811200 | 0 | 116910 | 0 | 50234 | 0 | 0 |
| AIEMLGGEI | FH | P07954 | FUMH_HUMAN | Oxidation (M) | 101000 | 0 | 0 | 0 | 0 | 0 | 242690 | 0 | 4433600 | 656780 | 0 | 1880000 | 185590 | 0 | 96601 | 0 | 124790 | 0 |
| AIENELLAR | GSS; GSS; GSS | P48637; A0A | GSHB_HUMAN | Unmodified | 0 | 0 | 0 | 0 | 0 | 0 | 0 | 0 | 2356000 | 0 | 1520100 | 5654500 | 0 | 0 | 879830 | 0 | 0 | 0 |
| AIEPNDYT | GGCT; N/A | O75223; M0 | GGCT_HUMAN | Unmodified | 0 | 0 | 0 | 0 | 91177 | 0 | 0 | 0 | 1725800 | 363060 | 964910 | 5191200 | 292760 | 191310 | 0 | 274950 | 37763 | 0 |
| AIEPPPLDA | DHX9 | Q08211 | DHX9_HUMAN | Unmodified | 0 | 0 | 0 | 0 | 0 | 0 | 0 | 0 | 0 | 0 | 0 | 11003000 | 0 | 0 | 0 | 0 | 0 | 0 |
| AIEPQKEEA | STX4; STX4 | Q12846; C9J | STX4_HUMAN | Unmodified | 0 | 118110 | 85583 | 162200 | 5123900 | 877590 | 306170 | 211320 | 2133900 | 1073000 | 1930800 | 5353200 | 329380 | 0 | 188890 | 296920 | 1807800 | 2921000 |
| AIEPQKEEA | STX4; STX4 | Q12846; C9J | STX4_HUMAN | Unmodified | 178430 | 325620 | 0 | 0 | 1118200 | 0 | 115520 | 53618 | 0 | 0 | 0 | 0 | 478370 | 0 | 0 | 316210 | 638560 | 819620 |
| AIEQADLLC | AP1S2 | P56377 | AP1S2_HUMAN | Unmodified | 0 | 0 | 0 | 0 | 0 | 0 | 0 | 0 | 0 | 0 | 0 | 0 | 0 | 0 | 0 | 0 | 0 | 0 |
| AIEQADLLC | AP1S1; AP1S1; | P61966; A0A | AP1S1_HUMAN | Unmodified | 0 | 0 | 369300 | 0 | 171560 | 0 | 0 | 0 | 343470 | 96421 | 484380 | 2524500 | 0 | 0 | 0 | 367260 | 0 | 0 |
| AIEQLKEQ | EIF2A; EIF2A; EI | Q9BY44; F8 | EIF2A_HUMAN | Unmodified | 0 | 0 | 0 | 0 | 228060 | 0 | 0 | 73470 | 0 | 0 | 0 | 0 | 0 | 178910 | 0 | 0 | 186820 | 219340 |
| AIEQSFDQI | STX2 | P32856 | STX2_HUMAN | Unmodified | 0 | 0 | 0 | 0 | 0 | 0 | 0 | 0 | 0 | 0 | 0 | 1673000 | 0 | 0 | 0 | 0 | 0 | 0 |
| AIEQSFDQI | STX2 | P32856 | STX2_HUMAN | Unmodified | 38159 | 62649 | 276010 | 0 | 0 | 0 | 0 | 0 | 0 | 0 | 0 | 10274000 | 0 | 0 | 361320 | 297990 | 0 | 0 |
| AIETFSGKV | IGF2BP1 | Q9NZI8 | IF2B1_HUMAN | Unmodified | 519650 | 557580 | 0 | 0 | 0 | 0 | 1715900 | 811590 | 229510 | 0 | 1583800 | 0 | 0 | 964560 | 0 | 0 | 0 | 211520 |
| AIETTDIISR | PGAM5; PGAM | Q96HS1; F5 | PGAM5_HUMA | Unmodified | 0 | 0 | 0 | 0 | 0 | 0 | 0 | 0 | 2289200 | 299390 | 272490 | 1551400 | 0 | 0 | 0 | 0 | 0 | 0 |
| AIETTDVTR | USP47 | Q96K76 | UBP47_HUMAN | Unmodified | 0 | 0 | 0 | 0 | 0 | 0 | 0 | 0 | 0 | 0 | 0 | 504840 | 0 | 0 | 0 | 0 | 0 | 0 |
| AIFIFLSNA | TOR1B | O14657 | TOR1B_HUMAN | Unmodified | 0 | 0 | 0 | 0 | 0 | 0 | 0 | 0 | 0 | 120690 | 0 | 0 | 0 | 0 | 0 | 0 | 0 | 0 |
| AIFLADGK | CORO1B; COR | Q9BR76; F5 | COR1B_HUMA | Unmodified | 1173700 | 645110 | 724110 | 2115600 | 3142900 | 1014400 | 257180 | 219340 | 913030 | 475130 | 809670 | 7938100 | 4365000 | 2272700 | 2923300 | 2443900 | 506240 | 1693800 |
| AIFLADGK | CORO1B; COR | Q9BR76; F5 | COR1B_HUMA | Unmodified | 0 | 0 | 0 | 12831 | 2208500 | 255910 | 253690 | 0 | 0 | 179830 | 0 | 65598 | 3726200 | 1976600 | 0 | 0 | 1124800 | 1230000 |
| AIFLADGN | CORO1C; COR | Q9ULV4; B4 | COR1C_HUMA | Unmodified | 410790 | 275450 | 301750 | 694060 | 31305000 | 5894800 | 228410 | 0 | 6732200 | 3108300 | 3388200 | 10742000 | 4609700 | 1051300 | 731800 | 381660 | 3140900 | 3744000 |
| AIFLADGN | CORO1C; COR | Q9ULV4; B4 | COR1C_HUMA | Oxidation (M) | 0 | 0 | 0 | 0 | 5107800 | 510090 | 436030 | 0 | 0 | 0 | 0 | 0 | 0 | 0 | 0 | 0 | 1075100 | 1053100 |
| AIFQEVHV | MMS19; MMS1 | Q96T76; Q5 | MMS19_HUMA | Unmodified | 0 | 0 | 0 | 0 | 0 | 0 | 0 | 0 | 0 | 0 | 0 | 841680 | 0 | 0 | 0 | 0 | 0 | 0 |
| AIFQTIQNR | IPO7 | O95373 | IPO7_HUMAN | Unmodified | 78063 | 289870 | 0 | 0 | 0 | 0 | 0 | 25539 | 219910 | 84745 | 327120 | 2810100 | 0 | 0 | 0 | 169350 | 0 | 0 |
| AIFQTIQNR | IPO7 | O95373 | IPO7_HUMAN | Unmodified | 0 | 0 | 252160 | 1213500 | 4114000 | 0 | 0 | 0 | 2033400 | 838380 | 3843400 | 7249000 | 2402100 | 1095600 | 0 | 0 | 2311500 | 1458400 |
| AIFSASIDD | VLDLR; VLDLR; | P98155; A0A | VLDLR_HUMAN | Unmodified | 0 | 0 | 0 | 0 | 455120 | 0 | 157530 | 0 | 0 | 427470 | 0 | 0 | 0 | 201890 | 0 | 0 | 0 | 0 |
| AIFTGHSA | RBBP7; RBBP7; | Q16576; Q5 | RBBP7_HUMAN | Unmodified | 0 | 0 | 0 | 0 | 0 | 0 | 0 | 0 | 0 | 0 | 0 | 0 | 1366800 | 0 | 0 | 0 | 0 | 0 |
| AIFTGYYGK | CNP | P09543 | CN37_HUMAN | Unmodified | 978820 | 720570 | 756210 | 1598900 | 2049300 | 0 | 0 | 423740 | 3749700 | 986890 | 11810000 | 43962000 | 0 | 110800 | 1696100 | 0 | 237480 | 447820 |
| AIFVTDR | MYO1D; MYO1 | O94832; J3 | MYO1D_HUMA | Unmodified | 79000 | 4067800 | 124430 | 0 | 0 | 0 | 0 | 0 | 424070 | 0 | 0 | 1165400 | 0 | 0 | 3241500 | 2961600 | 0 | 0 |
| AIFVTDRHL | MYO1D; MYO1 | O94832; J3 | MYO1D_HUMA | Unmodified | 1498400 | 1119800 | 645190 | 0 | 0 | 0 | 32478 | 0 | 448310 | 0 | 147970 | 0 | 0 | 276610 | 358560 | 0 | 0 | 0 |
| AIGGIILTAS | PGM1; PGM1 | P36871; A0A | PGM1_HUMAN | Unmodified | 0 | 0 | 0 | 0 | 156170 | 58826 | 181160 | 0 | 0 | 69157 | 0 | 325900 | 113490 | 145450 | 0 | 77904 | 94437 | 147680 |
| AIGHVIEEG | SEPTIN3; SEPT | Q9UH03; B1 | SEPT3_HUMAN | Unmodified | 0 | 0 | 55813 | 247130 | 277860 | 0 | 732070 | 452260 | 215420 | 40218 | 0 | 0 | 350610 | 0 | 138740 | 157760 | 0 | 45646 |
| AIGISNFNI | AKR1B1; AKR1B | P15121; E9P | ALDR_HUMAN | Unmodified | 0 | 0 | 0 | 0 | 3662700 | 0 | 0 | 0 | 0 | 0 | 0 | 0 | 0 | 0 | 0 | 0 | 0 | 0 |
| AIGISNFNI | AKR1B1; AKR1B | P15121; E9P | ALDR_HUMAN | Oxidation (M) | 0 | 0 | 0 | 0 | 12128000 | 924280 | 953080 | 0 | 0 | 0 | 3004000 | 9239100 | 606880 | 0 | 0 | 326440 | 2251600 | 1295800 |
| AIGKDNFT | DPYSL3; DPYSL | Q14195; H0 | DPYL3_HUMAN | Unmodified | 0 | 0 | 0 | 0 | 0 | 226690 | 0 | 30454 | 0 | 0 | 434200 | 469690 | 0 | 0 | 0 | 0 | 0 | 0 |
| AIGNTELEN | MTREX; MTREX | P42285; H0 | MTREX_HUMAN | Unmodified | 0 | 0 | 0 | 0 | 0 | 0 | 24766 | 132780 | 345130 | 0 | 619000 | 1328800 | 0 | 473510 | 0 | 0 | 62558 | 0 |
| AIGPHDVL | SF3B1; SF3B1 | O75533; H7 | SF3B1_HUMAN | Unmodified | 671360 | 647680 | 77436 | 70171 | 80943 | 0 | 161640 | 0 | 37931 | 45265 | 0 | 2178500 | 0 | 0 | 243690 | 209320 | 0 | 0 |
| AIGQAGSL | ALPL; ALPL | P05186; B1A | PPBT_HUMAN | Unmodified | 1147400 | 0 | 0 | 0 | 0 | 0 | 0 | 0 | 3483100 | 0 | 0 | 0 | 0 | 0 | 0 | 0 | 0 | 0 |
| AIGSASEG | PSMA5 | P28066 | PSA5_HUMAN | Unmodified | 791450 | 0 | 0 | 2814800 | 6227000 | 2412700 | 646880 | 389710 | 11966000 | 3432900 | 8615600 | 34046000 | 8632400 | 0 | 1815000 | 1998200 | 3368700 | 5629200 |
| AIGTFLFGA | PLPP1 | O14494 | PLPP1_HUMAN | Unmodified | 0 | 0 | 510710 | 211470 | 0 | 0 | 0 | 0 | 0 | 0 | 0 | 845400 | 0 | 0 | 4758600 | 806740 | 0 | 0 |
| AIGVKPPR | VCP; VCP; VCP; | P55072; C9J | TERA_HUMAN | Unmodified | 0 | 0 | 0 | 0 | 9696700 | 0 | 3075300 | 619860 | 0 | 3879600 | 166520 | 0 | 13694000 | 6361400 | 0 | 0 | 2061500 | 661270 |
| AIGVLTSG | PFKP; PFKL | Q01813; P17 | PFKAP_HUMAN | Unmodified | 0 | 0 | 0 | 143280 | 0 | 0 | 0 | 0 | 195380 | 257310 | 0 | 3696700 | 0 | 0 | 0 | 0 | 0 | 0 |
| AIGVLTSG | PFKP; PFKL | Q01813; P17 | PFKAP_HUMAN | Oxidation (M) | 0 | 0 | 0 | 0 | 492290 | 0 | 651840 | 761290 | 0 | 166660 | 0 | 0 | 0 | 0 | 0 | 466450 | 155980 | 0 |
| AIHKIDIPS | ABAT; ABAT; AB | P80404; H3 | GABT_HUMAN | Unmodified | 0 | 0 | 0 | 0 | 0 | 0 | 0 | 0 | 3210900 | 488200 | 0 | 0 | 0 | 0 | 0 | 0 | 0 | 0 |
| AIHNKVNI | SEPTIN2; SEPT | Q15019; H7 | SEPT2_HUMAN | Unmodified | 0 | 0 | 0 | 0 | 1632700 | 0 | 0 | 0 | 0 | 0 | 0 | 0 | 0 | 0 | 0 | 0 | 2855100 | 0 |
| AIHVHQFG | SOD3 | P08294 | SODE_HUMAN | Unmodified | 0 | 0 | 0 | 0 | 0 | 0 | 0 | 0 | 2295200 | 0 | 0 | 0 | 0 | 0 | 0 | 0 | 0 | 0 |
| AIHYALNC | CD9; CD9; CD9 | P21926; A0A | CD9_HUMAN | Unmodified | 1531500 | 2636900 | 327960 | 9621200 | 0 | 0 | 0 | 0 | 670830 | 2737300 | 0 | 0 | 1632400 | 1022700 | 33287000 | 558990 | 530450 | 0 |
| AIHYALNC | CD9; CD9; CD9 | P21926; A0A | CD9_HUMAN | Unmodified | 23050000 | 0 | 4200900 | 19771000 | 3379500 | 0 | 116910 | 0 | 14142000 | 2203900 | 857050 | 1080800 | 8576000 | 4351300 | 27953000 | 12049000 | 0 | 3856300 |
| AIHYALNC | CD9; CD9; CD9 | P21926; A0A | CD9_HUMAN | Unmodified | 0 | 2085800 | 0 | 8127700 | 9171000 | 0 | 0 | 0 | 0 | 4807100 | 0 | 0 | 36639000 | 47387000 | 0 | 0 | 1596300 | 9203500 |
| AIIEEGDKE | DNAJC13; DNA | O75165; A0 | DJC13_HUMAN | Unmodified | 0 | 0 | 0 | 0 | 420480 | 0 | 0 | 0 | 0 | 0 | 0 | 2317800 | 0 | 0 | 0 | 0 | 59801 | 0 |
| AIIEEYLHL | EIF4G1; EIF4G1 | Q04637; E7E | IF4G1_HUMAN | Oxidation (M) | 0 | 0 | 0 | 0 | 227880 | 0 | 0 | 0 | 0 | 0 | 581260 | 2082100 | 408340 | 0 | 0 | 0 | 255580 | 100710 |
| AIIENLIQK | CPD; CPD | O75976; J3 | CBPD_HUMAN | Unmodified | 0 | 169910 | 707760 | 568760 | 1469100 | 0 | 0 | 0 | 49981000 | 5656500 | 673720 | 809200 | 0 | 0 | 0 | 0 | 0 | 0 |
| AIIESDQEQ | SCRN1 | Q12765 | SCRN1_HUMA | Unmodified | 0 | 0 | 0 | 0 | 2457800 | 0 | 0 | 696210 | 2138500 | 0 | 0 | 0 | 0 | 122110 | 0 | 0 | 0 | 219540 |
| AIIIFVPVPQ | RPS7; RPS7 | P62081; B5 | RS7_HUMAN 4 | Unmodified | 1773900 | 1963200 | 0 | 890480 | 945600 | 131770 | 2700400 | 0 | 853070 | 670220 | 5003100 | 17606000 | 922000 | 0 | 1738100 | 669440 | 356270 | 639470 |
| AIIKPQYVD | ABCE1; ABCE1; | P61221; H0 | ABCE1_HUMAN | Unmodified | 79605 | 0 | 0 | 0 | 1663600 | 184170 | 0 | 94643 | 0 | 137850 | 595000 | 4114100 | 973870 | 266190 | 0 | 0 | 404100 | 400500 |
| AIILNTPHN | KYAT3; KYAT3 | Q6YP21; A0 | KAT3_HUMAN | Unmodified | 15797 | 0 | 0 | 0 | 201830 | 0 | 0 | 28544 | 99706 | 0 | 0 | 768140 | 0 | 0 | 0 | 80543 | 0 | 231120 |
| AIIRDRAALI | CHMP2B; CHM | Q9UQN3; C | CHM2B_HUMA | Unmodified | 0 | 0 | 0 | 0 | 3360000 | 322170 | 75702 | 18612 | 27896 | 700450 | 32222 | 0 | 1938400 | 1258800 | 0 | 0 | 115830 | 1273200 |
| AIIRGALCH | TTC4 | O95801 | TTC4_HUMAN | Unmodified | 0 | 0 | 0 | 0 | 0 | 0 | 105580 | 0 | 0 | 45339 | 0 | 0 | 203940 | 179270 | 0 | 0 | 435170 | 210380 |
| AIKELEEW | CLTA; CLTA; CL | P09496; C9 | CLCA_HUMAN | Unmodified | 0 | 100880 | 0 | 0 | 63451000 | 7609900 | 13073000 | 0 | 0 | 8520300 | 0 | 0 | 39611000 | 21188000 | 0 | 0 | 13406000 | 27814000 |
| AIKELGDH | FTH1; FTH1; FT | P02794; G3 | FRIH_HUMAN | Unmodified | 0 | 0 | 0 | 0 | 0 | 0 | 0 | 29107 | 0 | 758670 | 0 | 0 | 0 | 0 | 0 | 0 | 0 | 49240 |
| AIKELGDH | FTH1; FTH1; FT | P02794; G3 | FRIH_HUMAN | Unmodified | 0 | 0 | 0 | 0 | 53250 | 0 | 905610 | 357270 | 0 | 2703800 | 0 | 0 | 104400 | 440750 | 0 | 0 | 580070 | 212260 |
| AIKNDSVV | CCT7 | Q99832 | TCPH_HUMAN | Oxidation (M) | 0 | 0 | 0 | 0 | 0 | 0 | 0 | 0 | 0 | 227730 | 271730 | 1677900 | 232750 | 133020 | 0 | 71311 | 0 | 0 |
| AIKVEQATK | RBMX; RBMX; R | P38159; H3 | RBMX_HUMAN | Unmodified | 174830 | 126850 | 0 | 0 | 0 | 0 | 282630 | 170550 | 2509600 | 588490 | 3103200 | 9356700 | 336230 | 0 | 0 | 60602 | 0 | 0 |
| AILAARSSY | LZTR1 | Q8N653 | LZTR1_HUMAN | Oxidation (M) | 7387100 | 8539400 | 8743700 | 8091900 | 2838100 | 2336000 | 3158200 | 3228500 | 3307900 | 1863700 | 2164500 | 6322100 | 5090600 | 6831900 | 7398300 | 5290300 | 1763500 | 2886600 |
| AILDAEPGF | TBC1D10A; TB | Q9BXI6; B5 | TB10A_HUMAN | Unmodified | 589270 | 0 | 865820 | 2468300 | 10148000 | 0 | 698290 | 281830 | 2457200 | 1300800 | 0 | 2797100 | 14233000 | 9524700 | 0 | 0 | 6232000 | 9790800 |
| AILGATEVK | FUCA2 | Q9BTY2 | FUCO2_HUMA | Unmodified | 0 | 0 | 0 | 0 | 0 | 0 | 417700 | 27431 | 639950 | 942680 | 67036 | 3837700 | 0 | 0 | 0 | 0 | 0 | 130990 |
| AILGSVER | TARS1; TARS3; | P26639; A2 | SYTC_HUMAN | Unmodified | 0 | 0 | 0 | 0 | 0 | 0 | 0 | 0 | 0 | 0 | 0 | 6268500 | 0 | 0 | 839980 | 0 | 0 | 0 |
| AILILDNDG | COPZ1; COPZ1 | P61923; F8 | COPZ1_HUMA | Unmodified | 500810 | 1806400 | 784510 | 1025500 | 993630 | 382720 | 871100 | 24812 | 3116100 | 1942600 | 4470900 | 19779000 | 3697700 | 1001800 | 833310 | 1036000 | 783770 | 2275400 |
| AILNDTKLT | RABEP1 | Q15276 | RABE1_HUMAN | Unmodified | 0 | 18577 | 0 | 0 | 1848600 | 126580 | 49308 | 36164 | 147370 | 239750 | 102660 | 222810 | 357540 | 152270 | 398870 | 278980 | 0 | 106860 |
| AILNGIDSI | SMC3 | Q9UQE7 | SMC3_HUMAN | Unmodified | 0 | 0 | 0 | 0 | 250450 | 151280 | 0 | 98012 | 309020 | 136860 | 0 | 719460 | 95050 | 0 | 0 | 34365 | 58051 | 85967 |
| AILPCQDT | LTA4H | P09960 | LKHA4_HUMAN | Unmodified | 131450 | 0 | 0 | 0 | 0 | 0 | 0 | 0 | 0 | 0 | 0 | 7456700 | 0 | 0 | 0 | 0 | 0 | 0 |
| AILQATLRE | AIMP1; AIMP1; | Q12904; A0 | AIMP1_HUMAN | Unmodified | 0 | 0 | 0 | 0 | 1341200 | 0 | 447010 | 383240 | 0 | 444140 | 223610 | 3083500 | 1601700 | 546580 | 0 | 0 | 213440 | 457720 |
| AILQATLRE | AIMP1; AIMP1; | Q12904; A0 | AIMP1_HUMAN | Unmodified | 0 | 0 | 0 | 0 | 0 | 0 | 327850 | 278550 | 102730 | 0 | 0 | 1526300 | 762590 | 333790 | 0 | 0 | 0 | 239710 |
| AILQFYPK | ACE; ACE; ACE; | P12821; J3K | ACE_HUMAN A | Unmodified | 879780 | 805350 | 0 | 0 | 0 | 0 | 0 | 0 | 377290 | 0 | 0 | 0 | 0 | 0 | 0 | 0 | 0 | 0 |
| AILSMNGL | DIP2B | Q9P265 | DIP2B_HUMAN | Oxidation (M) | 0 | 0 | 0 | 0 | 0 | 0 | 0 | 0 | 0 | 0 | 0 | 2121300 | 0 | 0 | 0 | 0 | 0 | 0 |
| AILVDLEPG | TUBB2A; TUBB; | Q13885; P07 | TBB2A_HUMAN | Oxidation (M) | 1677400 | 749300 | 3918900 | 2714900 | 4034000 | 1465000 | 1147800 | 886420 | 31394000 | 6796900 | 22281000 | 1.52E+08 | 2280300 | 2180500 | 4554800 | 3803200 | 686070 | 2434700 |
| AILVDLEPG | TUBB2A; TUBB; | Q13885; P07 | TBB2A_HUMAN | Unmodified | 107660 | 97328 | 101610 | 2214200 | 1164500 | 305250 | 73758 | 15960 | 33426000 | 2606000 | 2476800 | 21527000 | 1706000 | 74068 | 570970 | 327270 | 85720 | 392710 |
| AILVDLEPG | TUBB2A; TUBB; | Q13885; P07 | TBB2A_HUMAN | Oxidation (M) | 0 | 0 | 0 | 0 | 0 | 0 | 3417600 | 0 | 0 | 2108200 | 0 | 0 | 14876000 | 3225400 | 0 | 0 | 3522300 | 7981700 |
| AIMAHMTV | FLOT1; FLOT1; | O75955; A0 | FLOT1_HUMAN | Unmodified | 0 | 0 | 0 | 0 | 0 | 0 | 0 | 0 | 1386300 | 0 | 0 | 0 | 0 | 0 | 0 | 0 | 0 | 0 |
| AIMAHMTV | FLOT1; FLOT1; | O75955; A0 | FLOT1_HUMAN | 2 Oxidation (M) | 0 | 0 | 160770 | 203450 | 0 | 0 | 0 | 0 | 1228900 | 0 | 1557500 | 297850 | 0 | 0 | 261380 | 145830 | 0 | 0 |
| AIMAHMTV | FLOT1; FLOT1; | O75955; A0 | FLOT1_HUMAN | Unmodified | 0 | 0 | 0 | 0 | 0 | 0 | 0 | 0 | 796050 | 213240 | 0 | 0 | 0 | 0 | 0 | 0 | 0 | 0 |
| AIMAHMTV | FLOT1; FLOT1; | O75955; A0 | FLOT1_HUMAN | Unmodified | 0 | 0 | 0 | 0 | 457570 | 326740 | 0 | 0 | 0 | 0 | 0 | 0 | 0 | 0 | 0 | 0 | 0 | 0 |
| AIMAQLPQ | CTNNA1; CTN | P35221; G3 | CTNA1_HUMA | Unmodified | 0 | 480080 | 1501200 | 6515900 | 12187000 | 4518800 | 0 | 0 | 1281600 | 547670 | 445760 | 7587900 | 10056000 | 10685000 | 3768900 | 2051200 | 0 | 0 |
| AIMAQLPQ | CTNNA1; CTN | P35221; G3 | CTNA1_HUMA | Oxidation (M) | 22238 | 0 | 2369400 | 5145500 | 3502600 | 1092600 | 0 | 0 | 0 | 0 | 536090 | 1867700 | 255450 | 5786000 | 4081800 | 5576800 | 0 | 0 |
| AIMAQLPQ | CTNNA1; CTN | P35221; G3 | CTNA1_HUMA | Unmodified | 0 | 171450 | 0 | 0 | 826360 | 0 | 0 | 0 | 0 | 0 | 0 | 0 | 1840600 | 5238900 | 0 | 0 | 0 | 0 |
| AIMGSGKE | SMS; SMS | P52788; H7 | SPSY_HUMAN | Unmodified | 255030 | 0 | 148740 | 112970 | 62363 | 12669 | 0 | 0 | 494440 | 0 | 470920 | 4195800 | 0 | 45009 | 235070 | 73752 | 0 | 0 |
| AIMKLDIEQ | GPD2; GPD2 | P43304; F5 | GPDM_HUMAN | Oxidation (M) | 0 | 0 | 0 | 0 | 285000 | 0 | 221220 | 0 | 0 | 476920 | 0 | 0 | 264300 | 180190 | 176610 | 0 | 0 | 62412 |
| AIMTYVSSF | ACTN4; ACTN1 | O43707; P12 | ACTN4_HUMA | Oxidation (M) | 179590 | 57379 | 837650 | 1813300 | 18002000 | 8128200 | 745270 | 0 | 4460900 | 1805400 | 4268000 | 21117000 | 8086400 | 5867400 | 5051700 | 4245800 | 5211400 | 5336700 |
| AIMTYVSSF | ACTN4; ACTN1 | O43707; P12 | ACTN4_HUMA | Unmodified | 305890 | 0 | 0 | 772300 | 24582000 | 8989800 | 879160 | 0 | 4746600 | 0 | 921760 | 6757500 | 3279000 | 2893000 | 2438800 | 0 | 155400 | 5007300 |
| AIMTYVSSF | ACTN4; ACTN1 | O43707; P12 | ACTN4_HUMA | Unmodified | 0 | 0 | 0 | 0 | 11967000 | 3523300 | 2282300 | 0 | 178220 | 1126800 | 0 | 0 | 8307600 | 10251000 | 0 | 0 | 19020000 | 6480900 |

| | | | | | | | | | | | | | | | | | | | | | | |
|---|---|---|---|---|---|---|---|---|---|---|---|---|---|---|---|---|---|---|---|---|---|---|
| AIMTYVSSF | ACTN4; ACTN1; | O43707; P12 | ACTN4_HUMAN | Oxidation (M) | 0 | 0 | 0 | 0 | 19095000 | 1684700 | 1092100 | 0 | 0 | 1561700 | 834990 | 0 | 8224000 | 11079000 | 0 | 0 | 12644000 | 2040800 |
| AIMVKGVD | ANXA1; ANXA1; | P04083; Q5T | ANXA1_HUMAN | Oxidation (M) | 0 | 0 | 0 | 0 | 9846300 | 1031000 | 1175000 | 0 | 0 | 196650 | 1310700 | 419980 | 0 | 0 | 0 | 0 | 2397900 | 508350 |
| AINCATSGY | FASN; FASN | P49327; A0A | FAS_HUMAN Fa | Unmodified | 0 | 0 | 0 | 0 | 0 | 0 | 0 | 0 | 0 | 0 | 0 | 7478700 | 0 | 0 | 0 | 0 | 0 | 0 |
| AINEAYKED | ANXA6; ANXA6; | P08133; E7E | ANXA6_HUMAN | Unmodified | 0 | 117730 | 2121200 | 1513200 | 1758200 | 49589 | 272220 | 21876 | 17724000 | 3874300 | 0 | 0 | 1459500 | 262550 | 3015500 | 613980 | 184240 | 1594500 |
| AINEFFR | MFAP3L; MFAP3 | O75121; P55 | MFA3L_HUMAN | Unmodified | 0 | 0 | 0 | 0 | 0 | 0 | 0 | 0 | 186470 | 0 | 0 | 2180000 | 0 | 0 | 0 | 0 | 0 | 0 |
| AINEFFRTE | MFAP3L; MFAP3 | O75121; P55 | MFA3L_HUMAN | Unmodified | 0 | 0 | 0 | 0 | 0 | 0 | 0 | 0 | 0 | 0 | 0 | 3230500 | 0 | 0 | 736000 | 661160 | 0 | 0 |
| AINFNFGY | QARS1; QARS1; | P47897; A0A | SYQ_HUMAN G | Unmodified | 0 | 0 | 0 | 0 | 385070 | 0 | 161020 | 257370 | 0 | 0 | 0 | 6500300 | 0 | 0 | 0 | 0 | 589690 | 0 |
| AINIEMYQK | RUFY2; RUFY2 | Q8WXA3; H0 | RUFY2_HUMAN | Unmodified | 0 | 0 | 0 | 256780 | 849410 | 0 | 2123600 | 0 | 0 | 0 | 0 | 0 | 729610 | 2560100 | 0 | 0 | 0 | 438060 |
| AINKKEGIC | LCP1; LCP1 | P13796; Q5T | PLSL_HUMAN I | Unmodified | 0 | 0 | 0 | 0 | 0 | 0 | 0 | 0 | 0 | 0 | 0 | 0 | 0 | 0 | 0 | 0 | 521420 | 0 |
| AINPINTFTI | ARHGAP1 | Q07960 | RHG01_HUMA | Unmodified | 0 | 0 | 0 | 0 | 1882200 | 324590 | 629360 | 495950 | 734180 | 468540 | 479030 | 728330 | 0 | 0 | 254820 | 0 | 0 | 492420 |
| AINQGGLT | PSMA6; PSMA6 | P60900; G3V | PSA6_HUMAN | Unmodified | 375160 | 293010 | 1056500 | 1064400 | 1635100 | 528850 | 0 | 0 | 4339400 | 0 | 3732600 | 31457000 | 1329800 | 540860 | 871740 | 1036100 | 391860 | 827020 |
| AINQGGLT | PSMA6; PSMA6 | P60900; G3V | PSA6_HUMAN | Unmodified | 0 | 0 | 0 | 156140 | 8727100 | 2873600 | 1517000 | 1339000 | 172740 | 3082700 | 604190 | 0 | 6230700 | 3021000 | 0 | 0 | 2618300 | 4039300 |
| AINQGGLT | PSMA6; PSMA6 | P60900; G3V | PSA6_HUMAN | Unmodified | 0 | 0 | 0 | 0 | 5368700 | 671130 | 2991200 | 1406400 | 0 | 4068600 | 0 | 0 | 5659700 | 7501000 | 0 | 0 | 11123000 | 6053800 |
| AINQIAAAL | NCKAP1 | Q9Y2A7 | NCKP1_HUMA | Unmodified | 0 | 0 | 0 | 0 | 0 | 0 | 0 | 0 | 0 | 0 | 227410 | 1711700 | 0 | 0 | 0 | 0 | 0 | 0 |
| AINQNLNS | KIDINS220; KID | Q9ULH0; E9 | KDIS_HUMAN I | Unmodified | 0 | 0 | 374970 | 0 | 0 | 0 | 0 | 0 | 0 | 0 | 0 | 1524900 | 0 | 0 | 348210 | 275690 | 0 | 0 |
| AINQQTGA | KHSRP; KHSRP | Q92945; M0 | FUBP2_HUMAN | Unmodified | 0 | 0 | 0 | 0 | 0 | 0 | 67132 | 0 | 2322100 | 402770 | 461590 | 5036600 | 0 | 0 | 0 | 0 | 0 | 0 |
| AINRKEGIC | PLS3; PLS3 | P13797; A0A | PLST_HUMAN I | Unmodified | 0 | 0 | 0 | 0 | 7131000 | 0 | 0 | 0 | 0 | 0 | 0 | 0 | 932660 | 3112700 | 0 | 0 | 8101500 | 1910200 |
| AINSSGLAI | ASAP2 | O43150 | ASAP2_HUMAN | Unmodified | 0 | 0 | 0 | 0 | 1106800 | 0 | 0 | 0 | 1109300 | 0 | 0 | 177120 | 502330 | 0 | 0 | 0 | 590950 | 1265200 |
| AINTLNGLF | ELAVL1; ELAVL1 | Q15717; Q1 | ELAV1_HUMAN | Unmodified | 241770 | 0 | 0 | 478560 | 0 | 0 | 89120 | 89860 | 524500 | 1259700 | 0 | 3413700 | 0 | 0 | 202960 | 504020 | 0 | 0 |
| AINTQEAP | HIP1R | O75146 | HIP1R_HUMAN | Unmodified | 0 | 0 | 0 | 0 | 134710 | 0 | 0 | 0 | 0 | 0 | 0 | 726460 | 0 | 0 | 95309 | 0 | 0 | 0 |
| AINTQEVA | HIP1; HIP1 | O00291; C9J | HIP1_HUMAN I | Unmodified | 0 | 0 | 0 | 0 | 847590 | 389300 | 0 | 173720 | 480530 | 0 | 0 | 218390 | 504350 | 392600 | 0 | 0 | 0 | 0 |
| AINTQEVA | HIP1; HIP1 | O00291; C9J | HIP1_HUMAN I | Unmodified | 0 | 0 | 0 | 0 | 0 | 0 | 0 | 0 | 0 | 0 | 0 | 0 | 0 | 0 | 0 | 0 | 0 | 0 |
| AINVQEEK | SPTAN1; SPTAN | Q13813; A0A | SPTN1_HUMAN | Unmodified | 76528 | 0 | 472030 | 0 | 383870 | 66246 | 0 | 18409 | 657980 | 277330 | 165350 | 2722800 | 193970 | 101560 | 463620 | 0 | 14831 | 529680 |
| AIPADSPTD | FLVCR1; FLVCR | Q9Y5Y0; H7 | FLVC1_HUMAN | Unmodified | 0 | 0 | 415350 | 0 | 461560 | 0 | 0 | 68130 | 737730 | 171240 | 671990 | 1798400 | 637180 | 0 | 0 | 607600 | 0 | 0 |
| AIPDTRHD | GALNT2 | Q10471 | GALT2_HUMAN | Unmodified | 0 | 0 | 0 | 0 | 0 | 0 | 45310 | 0 | 973940 | 80499 | 0 | 0 | 0 | 0 | 0 | 64836 | 0 | 0 |
| AIPDTRHD | GALNT2 | Q10471 | GALT2_HUMAN | Unmodified | 0 | 0 | 0 | 0 | 0 | 0 | 0 | 0 | 1029200 | 0 | 0 | 0 | 0 | 0 | 0 | 0 | 0 | 0 |
| AIPELTKLL | CTNNB1; CTNN | P35222; A0A | CTNB1_HUMA | Unmodified | 0 | 0 | 0 | 0 | 1175200 | 0 | 0 | 0 | 0 | 108040 | 0 | 0 | 0 | 0 | 0 | 0 | 2669000 | 1717900 |
| AIPNNQVL | EDF1 | O60869 | EDF1_HUMAN | Unmodified | 997040 | 1070100 | 730060 | 1053400 | 0 | 0 | 0 | 0 | 946770 | 0 | 336380 | 1411600 | 0 | 0 | 1468700 | 1187000 | 0 | 0 |
| AIPQLQGY | RPL6 | Q02878 | RL6_HUMAN 6 | Unmodified | 6269000 | 5207000 | 3766600 | 2681100 | 1270700 | 0 | 439870 | 0 | 1162200 | 979080 | 1934400 | 60100000 | 2226500 | 824530 | 3417700 | 3142900 | 446530 | 598950 |
| AIPSDGNF | DDC; DDC; DD | P20711; A0A | DDC_HUMAN | Oxidation (M) | 0 | 0 | 0 | 0 | 0 | 0 | 0 | 0 | 614670 | 0 | 0 | 0 | 0 | 0 | 0 | 0 | 0 | 0 |
| AIPVSPSAV | TJP1; TJP1; TJP1 | Q07157; A0A | ZO1_HUMAN T | Unmodified | 105770 | 0 | 1729800 | 3061700 | 14950000 | 2914400 | 454740 | 252450 | 1386300 | 154950 | 422020 | 2756000 | 10772000 | 3309200 | 2489600 | 2718500 | 1393000 | 1849300 |
| AIQASLAK | ABCB6 | Q9NP58 | ABCB6_HUMA | Unmodified | 126020 | 82234 | 167620 | 231570 | 136370 | 659220 | 0 | 0 | 1223700 | 62813 | 845590 | 376770 | 503260 | 0 | 241090 | 119550 | 76190 | 550880 |
| AIQDGTIVL | FAM3C | Q92520 | FAM3C_HUMA | Unmodified | 0 | 0 | 0 | 0 | 0 | 0 | 0 | 0 | 20368000 | 1256000 | 0 | 0 | 0 | 0 | 0 | 0 | 0 | 0 |
| AIQDGTIVL | FAM3C | Q92520 | FAM3C_HUMA | Oxidation (M) | 0 | 0 | 0 | 0 | 0 | 0 | 0 | 0 | 4805200 | 1063800 | 1716700 | 1202500 | 111500 | 0 | 0 | 0 | 0 | 0 |
| AIQDGTIVL | FAM3C | Q92520 | FAM3C_HUMA | Unmodified | 0 | 0 | 0 | 0 | 0 | 0 | 0 | 0 | 1303200 | 0 | 0 | 0 | 0 | 0 | 0 | 0 | 0 | 0 |
| AIQDGTIVL | FAM3C | Q92520 | FAM3C_HUMA | Oxidation (M) | 0 | 0 | 0 | 0 | 0 | 0 | 0 | 0 | 0 | 0 | 0 | 0 | 0 | 0 | 0 | 0 | 0 | 0 |
| AIQDGTIVL | FAM3C | Q92520 | FAM3C_HUMA | Unmodified | 0 | 0 | 0 | 0 | 714450 | 0 | 123750 | 0 | 3315300 | 0 | 0 | 0 | 0 | 0 | 0 | 0 | 0 | 0 |
| AIQDMFPN | TOLLIP; TOLLIP | Q9H0E2; F2 | TOLIP_HUMAN | 2 Oxidation (M) | 1717100 | 0 | 1110100 | 581670 | 0 | 0 | 0 | 0 | 0 | 65404 | 0 | 0 | 0 | 0 | 3017200 | 2492800 | 0 | 0 |
| AIQDMFPN | TOLLIP; TOLLIP | Q9H0E2; F2 | TOLIP_HUMAN | Oxidation (M) | 911440 | 771940 | 319300 | 580640 | 0 | 0 | 0 | 0 | 109660 | 0 | 0 | 0 | 0 | 0 | 1154000 | 680820 | 0 | 0 |
| AIQDPAFS | HUWE1; HUWE | Q7Z6Z7; A0A | HUWE1_HUMA | Unmodified | 0 | 0 | 0 | 0 | 0 | 0 | 0 | 0 | 194040 | 0 | 0 | 1821800 | 0 | 0 | 0 | 0 | 0 | 0 |
| AIQDYLST | INPP5D | Q92835 | SHIP1_HUMAN | Unmodified | 0 | 0 | 0 | 30274 | 688660 | 13408 | 0 | 0 | 95155 | 0 | 607500 | 1647100 | 0 | 0 | 79287 | 20748 | 31979 | 52610 |
| AIQERLFK | ANLN | Q9NQW6 | ANLN_HUMAN | Unmodified | 0 | 0 | 0 | 0 | 2734600 | 300760 | 0 | 0 | 0 | 0 | 0 | 0 | 0 | 0 | 0 | 0 | 1266600 | 699240 |
| AIQGGTSH | EPRS1 | P07814 | SYEP_HUMAN | Unmodified | 158300 | 108940 | 1554300 | 1172000 | 1767700 | 384390 | 310880 | 201100 | 1830200 | 1036800 | 2874800 | 17792000 | 3477900 | 1470000 | 1348800 | 1883500 | 499630 | 1214300 |
| AIQKGNME | CHMP1B | Q7LBR1 | CHM1B_HUMA | Unmodified | 0 | 0 | 0 | 0 | 48920000 | 0 | 0 | 0 | 0 | 532090 | 0 | 0 | 330710 | 0 | 0 | 0 | 0 | 141060 |
| AIQLEYSEA | PSMD3 | O43242 | PSMD3_HUMA | Unmodified | 0 | 83974 | 0 | 0 | 0 | 0 | 0 | 0 | 0 | 0 | 176380 | 2375300 | 0 | 0 | 256960 | 231550 | 0 | 0 |
| AIQLNSNS | ANAPC7 | Q9UJX3 | APC7_HUMAN | Unmodified | 0 | 11375 | 0 | 63954 | 0 | 0 | 48508 | 30019 | 0 | 5719.8 | 26966 | 628860 | 29140 | 25628 | 0 | 0 | 28687 | 0 |
| AIQLQPSA | TTC13; TTC13; | Q8NBP0; E5 | TTC13_HUMAN | Unmodified | 0 | 0 | 0 | 0 | 56717 | 0 | 0 | 0 | 2443500 | 0 | 0 | 0 | 0 | 0 | 0 | 0 | 0 | 0 |
| AIQLSGAE | COPS4; COPS4 | Q9BT78; D6 | CSN4_HUMAN | Unmodified | 0 | 0 | 0 | 0 | 0 | 0 | 0 | 0 | 250050 | 0 | 18170 | 741800 | 0 | 0 | 0 | 0 | 0 | 0 |
| AIQLTYNPD | FGG; FGG; FGG | P02679; C9J | FIBG_HUMAN | Oxidation (M) | 0 | 9517300 | 0 | 0 | 0 | 0 | 0 | 0 | 0 | 0 | 0 | 0 | 0 | 6714900 | 0 | 0 | 0 | 0 |
| AIRDGVIEA | PSMD3 | O43242 | PSMD3_HUMA | Unmodified | 0 | 200900 | 612620 | 501400 | 1939100 | 185180 | 1571400 | 404820 | 3013300 | 827510 | 855960 | 9821300 | 1889800 | 1822400 | 0 | 0 | 494220 | 2269000 |
| AIRDYQGP | ACLY; ACLY | P53396; K7E | ACLY_HUMAN | Unmodified | 0 | 0 | 0 | 0 | 0 | 0 | 533270 | 0 | 0 | 270460 | 0 | 0 | 0 | 0 | 0 | 0 | 0 | 0 |
| AIRGIGTDE | ANXA3; ANXA3; | P12429; D6F | ANXA3_HUMAN | Unmodified | 247340 | 407280 | 191320 | 0 | 1986700 | 707380 | 205880 | 210880 | 0 | 95908 | 0 | 0 | 0 | 0 | 448350 | 245540 | 536150 | 690790 |
| AIRGSNEY | SPRY1; SPRY1; | O43609; D6 | SPY1_HUMAN | Unmodified | 0 | 0 | 0 | 0 | 136470 | 250290 | 3649300 | 0 | 48978 | 0 | 83579 | 0 | 216390 | 667080 | 0 | 0 | 1123200 | 493910 |
| AIRILVQER | RTRAF; RTRAF; | Q9Y224; G3 | RTRAF_HUMAN | Unmodified | 0 | 0 | 0 | 0 | 708120 | 0 | 295800 | 0 | 0 | 0 | 0 | 0 | 385180 | 313600 | 0 | 0 | 537040 | 0 |
| AIRPQIDLK | PRKDC | P78527 | PRKDC_HUMA | Unmodified | 0 | 0 | 0 | 0 | 0 | 0 | 0 | 0 | 0 | 0 | 0 | 392780 | 0 | 0 | 0 | 0 | 0 | 0 |
| AISAHFDD | MACROH2A2; | Q9P0M6; A0 | H2AW_HUMAN | Unmodified | 291040 | 276800 | 0 | 0 | 175950 | 0 | 264240 | 1217900 | 0 | 0 | 0 | 2845700 | 119160 | 0 | 0 | 0 | 0 | 1339100 |
| AISCHFAS | PHYH; PHYH | O14832; B1 | PAHX_HUMAN | Unmodified | 0 | 0 | 0 | 0 | 0 | 0 | 0 | 0 | 813460 | 0 | 0 | 0 | 0 | 0 | 0 | 0 | 0 | 0 |
| AISCHFAS | PHYH; PHYH | O14832; B1 | PAHX_HUMAN | Unmodified | 0 | 0 | 0 | 0 | 0 | 0 | 0 | 0 | 614520 | 1665300 | 0 | 0 | 0 | 0 | 0 | 0 | 0 | 0 |
| AISEQTGKE | ACLY; ACLY | P53396; K7E | ACLY_HUMAN | Unmodified | 0 | 0 | 0 | 0 | 3298900 | 0 | 11600000 | 5218000 | 0 | 339170 | 0 | 0 | 1299400 | 538940 | 0 | 0 | 1333800 | 608590 |
| AISGLEQD | ZPR1; ZPR1; ZP | O75312; H7 | ZPR1_HUMAN | Unmodified | 0 | 0 | 0 | 0 | 0 | 0 | 0 | 0 | 0 | 0 | 0 | 611720 | 0 | 0 | 0 | 0 | 0 | 0 |
| AISGLEQD | ZPR1; ZPR1; ZP | O75312; H7 | ZPR1_HUMAN | Unmodified | 0 | 0 | 2044000 | 1009800 | 0 | 0 | 33101 | 0 | 0 | 0 | 260310 | 823230 | 0 | 0 | 0 | 715700 | 0 | 0 |
| AISGLEQD | ZPR1; ZPR1; ZP | O75312; H7 | ZPR1_HUMAN | Unmodified | 0 | 0 | 0 | 0 | 0 | 0 | 0 | 0 | 0 | 0 | 0 | 0 | 590200 | 79982 | 0 | 0 | 0 | 250200 |
| AISHEHSP | PPP2R1A; PPP2 | P30153; C9J | 2AAA_HUMAN | Unmodified | 1375900 | 0 | 2585400 | 4243100 | 7170500 | 0 | 1326700 | 0 | 4243600 | 1971400 | 10623000 | 28731000 | 8378600 | 2722200 | 3622300 | 3646000 | 3081900 | 7023700 |
| AISHEHSP | PPP2R1A; PPP2 | P30153; C9J | 2AAA_HUMAN | Unmodified | 0 | 0 | 133310 | 934070 | 0 | 0 | 0 | 0 | 1169500 | 0 | 0 | 7176700 | 386510 | 0 | 455760 | 713180 | 316520 | 0 |
| AISQILQES | LRSAM1; LRSAM | Q6UWE0; A0 | LRSM1_HUMA | Unmodified | 0 | 0 | 0 | 0 | 584750 | 0 | 0 | 0 | 163860 | 0 | 0 | 0 | 0 | 0 | 0 | 0 | 0 | 0 |
| AISQSGVAI | CEL | P19835 | CEL_HUMAN B | Unmodified | 0 | 0 | 0 | 0 | 0 | 0 | 0 | 0 | 5862600 | 1926600 | 4798200 | 0 | 0 | 0 | 0 | 0 | 0 | 0 |
| AISQSGVAI | CEL | P19835 | CEL_HUMAN B | Unmodified | 0 | 0 | 0 | 0 | 0 | 0 | 0 | 0 | 0 | 1127600 | 0 | 0 | 0 | 0 | 0 | 0 | 0 | 0 |
| AISRWSSL | CCT3; CCT3; C | P49368; Q5 | TCPG_HUMAN | Unmodified | 0 | 0 | 0 | 0 | 0 | 0 | 0 | 0 | 0 | 0 | 0 | 0 | 0 | 0 | 0 | 0 | 0 | 0 |
| AISSSAISK | CPSF7; CPSF7 | Q8N684; F5 | CPSF7_HUMAN | Unmodified | 954850 | 0 | 422930 | 0 | 1382500 | 30243 | 1875700 | 734640 | 0 | 0 | 0 | 183810 | 0 | 525780 | 0 | 0 | 0 | 0 |
| AISSSAISR | CPSF6; CPSF6 | Q16630; F8 | CPSF6_HUMAN | Unmodified | 1550400 | 1367700 | 703150 | 993200 | 0 | 0 | 0 | 0 | 0 | 0 | 0 | 0 | 0 | 0 | 305100 | 0 | 0 | 0 |
| AISSYFVST | MACROH2A1; | O75367; B4 | H2AY_HUMAN | Oxidation (M) | 0 | 0 | 0 | 0 | 0 | 0 | 0 | 391010 | 0 | 0 | 0 | 0 | 0 | 0 | 0 | 0 | 0 | 0 |
| AISTNSELS | COLEC12 | Q5KU26 | COL12_HUMAN | Unmodified | 113220 | 131170 | 201090 | 936800 | 0 | 0 | 0 | 0 | 0 | 0 | 0 | 0 | 0 | 0 | 283440 | 248680 | 0 | 0 |
| AITDGQAG | YTHDF3; YTHD | Q7Z739; A0A | YTHD3_HUMAN | Unmodified | 83080 | 34126 | 0 | 0 | 0 | 331580 | 0 | 0 | 328770 | 0 | 0 | 0 | 0 | 0 | 0 | 0 | 101190 | 0 |
| AITDNDMQ | CRISPLD1; CRI | Q9H336; E5 | CRLD1_HUMA | Unmodified | 0 | 0 | 0 | 0 | 1196300 | 0 | 0 | 0 | 811700 | 0 | 0 | 231600 | 0 | 0 | 0 | 0 | 0 | 378000 |
| AITEIYLTR | PLXNB2; PLXNE | O15031; A6 | PLXB2_HUMAN | Unmodified | 1084600 | 157230 | 1256400 | 695520 | 0 | 0 | 0 | 0 | 225330 | 161230 | 0 | 11141000 | 0 | 0 | 3743700 | 4158500 | 0 | 0 |
| AITEVTTTL | FARP2 | O94887 | FARP2_HUMAN | Unmodified | 18670 | 0 | 189680 | 57045 | 0 | 0 | 0 | 0 | 0 | 0 | 0 | 44085 | 0 | 0 | 914730 | 535470 | 0 | 0 |
| AITEVTTTL | FARP2 | O94887 | FARP2_HUMAN | Unmodified | 0 | 0 | 0 | 74775 | 0 | 0 | 0 | 0 | 0 | 0 | 0 | 0 | 0 | 0 | 428260 | 0 | 0 | 0 |
| AITGASLAD | RPL24; RPL24; | P83731; C9J | RL24_HUMAN | Oxidation (M) | 1772000 | 1649200 | 790820 | 0 | 0 | 0 | 743450 | 314920 | 0 | 0 | 614730 | 3217900 | 841190 | 0 | 791500 | 617440 | 0 | 0 |
| AITGASLAD | RPL24; RPL24; | P83731; C9J | RL24_HUMAN | Unmodified | 2042500 | 2852900 | 529140 | 0 | 806740 | 0 | 143640 | 280830 | 845370 | 0 | 1097700 | 6784600 | 4148200 | 0 | 1097000 | 546520 | 0 | 0 |
| AITGFDDP | DSP | P15924 | DESP_HUMAN | Unmodified | 0 | 0 | 0 | 0 | 219760 | 0 | 0 | 0 | 0 | 0 | 0 | 0 | 302980 | 0 | 0 | 0 | 839940 | 0 |
| AITGIQAFT | GCDH | Q92947 | GCDH_HUMAN | Unmodified | 289530 | 456110 | 1236500 | 1030800 | 1102600 | 0 | 605820 | 67628 | 4864000 | 450140 | 0 | 881230 | 0 | 340270 | 900210 | 492860 | 209620 | 0 |
| AITIAGIPQ | PCBP2; PCBP2; | Q15366; F8 | PCBP2_HUMAN | Unmodified | 23392000 | 6132300 | 2315400 | 11075000 | 1286100 | 141590 | 1698900 | 137180 | 3613300 | 799510 | 931570 | 2902600 | 3847100 | 1553800 | 5183700 | 5296800 | 412330 | 1058300 |
| AITIAGVPQ | PCBP1 | Q15365 | PCBP1_HUMAN | Unmodified | 5305600 | 5014500 | 1494300 | 3451100 | 0 | 0 | 540070 | 1080300 | 0 | 0 | 2482300 | 11961000 | 0 | 0 | 2912800 | 2768400 | 1516200 | 1003600 |
| AITIANQTN | DPYSL2; DPYSL | Q16555; A0A | DPYL2_HUMAN | Unmodified | 275160 | 0 | 0 | 0 | 431930 | 355850 | 892000 | 344240 | 1242400 | 617010 | 171310 | 3001000 | 150750 | 71078 | 104160 | 125190 | 124520 | 305030 |
| AITIASQTN | DPYSL3 | Q14195 | DPYL3_HUMAN | Unmodified | 0 | 0 | 0 | 0 | 0 | 0 | 160770 | 0 | 286510 | 73701 | 7874600 | 38784000 | 66991 | 0 | 0 | 0 | 0 | 0 |
| AITPLQVH | PAN2 | Q504Q3 | PAN2_HUMAN | Unmodified | 2204000 | 1689400 | 626150 | 2548800 | 0 | 0 | 0 | 0 | 0 | 0 | 0 | 0 | 0 | 0 | 520390 | 415130 | 0 | 0 |
| AITPQAQE | AKAP12 | Q02952 | AKA12_HUMAN | Unmodified | 0 | 0 | 0 | 0 | 623660 | 0 | 0 | 0 | 0 | 0 | 0 | 0 | 0 | 0 | 341560 | 0 | 141800 | 532950 |
| AITQEQCE | GAA; GAA; GAA | P10253; I3L | LYAG_HUMAN | Unmodified | 0 | 0 | 0 | 0 | 0 | 0 | 0 | 0 | 1820400 | 0 | 0 | 35845 | 0 | 0 | 0 | 0 | 0 | 0 |
| AITRNPLVA | STUB1 | Q9UNE7 | CHIP_HUMAN | Unmodified | 0 | 0 | 0 | 1281900 | 643340 | 0 | 0 | 0 | 1180200 | 0 | 0 | 802720 | 1753500 | 822170 | 0 | 0 | 338560 | 0 |
| AITSAYYR | RAB25; RAB11A | P57735; P62 | RAB25_HUMAN | Unmodified | 1504200 | 1255200 | 2515300 | 1743400 | 991280 | 392100 | 0 | 0 | 499760 | 103830 | 2020800 | 16718000 | 161950 | 0 | 3138200 | 3120300 | 0 | 0 |
| AITTGLQEY | TSNAX; TSNAX; | Q99598; Q5 | TSNAX_HUMAN | Unmodified | 0 | 0 | 0 | 0 | 0 | 0 | 0 | 0 | 1316900 | 0 | 667450 | 1889300 | 0 | 0 | 0 | 0 | 460210 | 0 |
| AITTQNVN | SMAD3 | P84022 | SMAD3_HUMA | Unmodified | 0 | 0 | 0 | 1088200 | 466960 | 0 | 0 | 415480 | 0 | 284390 | 171560 | 363060 | 0 | 0 | 239650 | 1193100 | 522670 | 1596600 |
| AITVFSPDG | PSMA7; PSMA8 | O14818; Q8 | PSA7_HUMAN | Unmodified | 0 | 0 | 0 | 0 | 0 | 0 | 0 | 0 | 5553300 | 0 | 2377100 | 7096900 | 0 | 0 | 0 | 1451900 | 0 | 0 |
| AITVFSPDG | PSMA7; PSMA8 | O14818; Q8 | PSA7_HUMAN | Unmodified | 1459500 | 0 | 1460400 | 2889600 | 3196400 | 0 | 1967700 | 0 | 10826000 | 1184100 | 6231500 | 25455000 | 3894500 | 1361800 | 3490000 | 2026800 | 0 | 0 |
| AITVHPEK | LRP1 | Q07954 | LRP1_HUMAN I | Unmodified | 0 | 23670 | 37935 | 28441 | 0 | 34272 | 0 | 26819 | 278160 | 37348 | 17332 | 0 | 0 | 390060 | 26576 | 19972 | 54845 | 111000 |
| AIVAESLNN | RFC1; RFC1 | P35251; H0 | RFC1_HUMAN | Unmodified | 0 | 0 | 0 | 0 | 0 | 0 | 0 | 254930 | 0 | 0 | 0 | 0 | 0 | 0 | 0 | 0 | 0 | 0 |
| AIVAGDQN | FHL1; FHL1; FH | Q13642; A0A | FHL1_HUMAN | Unmodified | 0 | 0 | 0 | 62814 | 2958100 | 261670 | 2789000 | 2274400 | 0 | 0 | 90095 | 0 | 0 | 0 | 298940 | 0 | 1410400 | 2631300 |
| AIVAIENPA | RPSA; RPSA; RF | P08865; A0A | RSSA_HUMAN | Unmodified | 6191800 | 5656300 | 2949500 | 2656400 | 677290 | 0 | 356550 | 204370 | 0 | 703350 | 6851200 | 36675000 | 2812500 | 878480 | 3075600 | 3650300 | 260790 | 0 |
| AIVAIENPA | RPSA; RPSA; RF | P08865; F8W | RSSA_HUMAN | Unmodified | 0 | 0 | 0 | 0 | 2829600 | 0 | 1017200 | 441490 | 0 | 1270200 | 0 | 0 | 5941200 | 3815000 | 0 | 0 | 649270 | 2100100 |
| AIVAIENPA | RPSA; RPSA; RF | P08865; F8W | RSSA_HUMAN | Unmodified | 0 | 0 | 0 | 0 | 2089900 | 0 | 3348400 | 1750900 | 0 | 874840 | 0 | 0 | 3794100 | 3619900 | 0 | 0 | 1812100 | 1638100 |
| AIVDALPPP | PPA1 | Q15181 | IPYR_HUMAN I | Unmodified | 576390 | 0 | 0 | 675080 | 0 | 0 | 605480 | 274910 | 0 | 0 | 540480 | 3492100 | 795240 | 760290 | 0 | 658600 | 0 | 627600 |
| AIVDGNLK | FLNC; FLNB; FL | Q14315; O7 | FLNC_HUMAN | Unmodified | 0 | 0 | 0 | 0 | 0 | 0 | 0 | 0 | 0 | 0 | 0 | 524370 | 0 | 0 | 0 | 0 | 0 | 0 |
| AIVDTVDNI | ADGRL2; ADGR | O95490; A0A | AGRL2_HUMAN | Unmodified | 0 | 0 | 0 | 0 | 0 | 0 | 0 | 0 | 696670 | 326750 | 0 | 125960 | 0 | 0 | 0 | 0 | 0 | 0 |
| AIVDVHFDI | AGRN; AGRN; A | O00468; A0A | AGRIN_HUMAN | Unmodified | 2478800 | 0 | 2995800 | 563510 | 0 | 442870 | 0 | 0 | 2189400 | 459020 | 0 | 0 | 0 | 279490 | 2286500 | 2131600 | 306240 | 94532 |
| AIVERAAQL | RPL32; RPL32; | P62910; D3Y | RL32_HUMAN | Unmodified | 0 | 0 | 0 | 0 | 0 | 0 | 403440 | 125000 | 0 | 148420 | 0 | 0 | 676600 | 340910 | 0 | 0 | 0 | 0 |
| AIVNSALK | SUN1; SUN1; S | O94901; H0 | SUN1_HUMAN | Unmodified | 151260 | 0 | 0 | 50228 | 315820 | 0 | 447060 | 83168 | 18754000 | 1000800 | 393300 | 152940 | 1664700 | 0 | 0 | 0 | 63657 | 0 |
| AIVQLVNEF | PAFAH1B3; PAF | Q15102; M0 | PA1B3_HUMAN | Unmodified | 763530 | 712900 | 808040 | 850630 | 643230 | 0 | 984420 | 511330 | 4310200 | 2903700 | 1893100 | 13697000 | 1160000 | 687200 | 678660 | 588720 | 0 | 0 |
| AIVQQWLE | EEF1E1; EEF1E1 | O43324; D6 | MCA3_HUMAN | Unmodified | 0 | 0 | 220050 | 0 | 0 | 0 | 0 | 28898 | 0 | 0 | 0 | 6162300 | 0 | 0 | 0 | 0 | 0 | 0 |
| AIVRHDFS | EXOC7; EXOC7 | Q9UPT5; B4 | EXOC7_HUMA | Unmodified | 0 | 0 | 0 | 0 | 1759900 | 0 | 0 | 0 | 549900 | 522840 | 0 | 0 | 0 | 0 | 0 | 0 | 76951 | 267030 |

| | | | | | | | | | | | | | | | | | | | | | | |
|---|---|---|---|---|---|---|---|---|---|---|---|---|---|---|---|---|---|---|---|---|---|---|
| AIVVDPVH( | LDLR; LDLR; LD | P01130; J3K | LDLR_HUMAN | Oxidation (M) | 0 | 0 | 0 | 201200 | 0 | 0 | 0 | 0 | 740150 | 0 | 0 | 0 | 0 | 0 | 0 | 0 | 0 | 0 |
| AIVVTDGEF | ME1 | P48163 | MAOX_HUMAN | Unmodified | 0 | 0 | 0 | 0 | 0 | 0 | 0 | 0 | 0 | 0 | 0 | 0 | 0 | 0 | 0 | 0 | 0 | 0 |
| AIVWGMQ1 | ACLY | P53396 | ACLY_HUMAN | Oxidation (M) | 0 | 0 | 0 | 0 | 0 | 0 | 0 | 0 | 0 | 0 | 0 | 3654800 | 0 | 0 | 0 | 0 | 0 | 0 |
| AIVWGMQ1 | ACLY | P53396 | ACLY_HUMAN | Unmodified | 48929 | 0 | 0 | 217350 | 1270000 | 0 | 0 | 0 | 431110 | 0 | 341120 | 6291100 | 326340 | 0 | 179220 | 0 | 0 | 0 |
| AIWAALQT( | UGGT1 | Q9NYU2 | UGGG1_HUMA | Unmodified | 0 | 0 | 0 | 0 | 0 | 0 | 0 | 0 | 11860000 | 2558500 | 0 | 0 | 0 | 0 | 0 | 0 | 0 | 0 |
| AIWHAFTAI | SWAP70; SWAF | Q9UH65; E9 | SWP70_HUMA | Unmodified | 0 | 0 | 0 | 0 | 691350 | 0 | 0 | 0 | 0 | 0 | 0 | 0 | 0 | 0 | 0 | 0 | 0 | 715600 |
| AIWNVINW | SOD2 | P04179 | SODM_HUMAN | Unmodified | 0 | 0 | 0 | 0 | 38348 | 0 | 0 | 0 | 10309000 | 557380 | 76862 | 42455 | 0 | 0 | 0 | 0 | 369530 | 529800 |
| AIWNVINW | SOD2 | P04179 | SODM_HUMAN | Oxidation (M) | 0 | 0 | 0 | 0 | 0 | 0 | 259400 | 0 | 0 | 1875800 | 0 | 0 | 0 | 0 | 0 | 0 | 0 | 0 |
| AIWVTGW( | ST14 | Q9Y5Y6 | ST14_HUMAN | Unmodified | 0 | 0 | 266910 | 0 | 0 | 0 | 0 | 0 | 0 | 0 | 0 | 0 | 0 | 0 | 974210 | 687410 | 0 | 0 |
| AIYFDGDF( | GFM1; GFM1; G | Q96RP9; C9 | EFGM_HUMAN | Unmodified | 0 | 0 | 0 | 0 | 0 | 0 | 0 | 0 | 803960 | 0 | 0 | 0 | 0 | 0 | 0 | 0 | 0 | 0 |
| AIYHDLEQ | PACSIN2; PACS | Q9UNF0; A0 | PACN2_HUMA | Unmodified | 38538000 | 32932000 | 25975000 | 35837000 | 0 | 0 | 104410 | 0 | 1350300 | 0 | 685550 | 5286000 | 8566500 | 1317000 | 62334000 | 28887000 | 0 | 0 |
| AIYHDLEQ | PACSIN2; PACS | Q9UNF0; A0 | PACN2_HUMA | Unmodified | 1446200 | 0 | 0 | 9733800 | 0 | 0 | 0 | 0 | 0 | 0 | 0 | 0 | 4953400 | 2636900 | 0 | 510160 | 0 | 0 |
| AIYHDLEQ | PACSIN2; PACS | Q9UNF0; A0 | PACN2_HUMA | Unmodified | 0 | 0 | 0 | 3000600 | 4380600 | 402890 | 0 | 0 | 0 | 1097100 | 0 | 0 | 74358000 | 9604700 | 0 | 0 | 436860 | 1323300 |
| AIYHTLNLC | ATP6V0A1; ATP | Q93050; B7 | VPP1_HUMAN | Unmodified | 0 | 0 | 162020 | 0 | 0 | 0 | 225050 | 85631 | 2131400 | 1757800 | 452180 | 0 | 0 | 0 | 0 | 112240 | 257000 | 0 |
| AIYKGPGSI | COMT; COMT | P21964; A0A | COMT_HUMAN | Unmodified | 660180 | 636140 | 0 | 1079600 | 1080800 | 173040 | 767540 | 262080 | 113010 | 0 | 189520 | 0 | 5029400 | 4799600 | 514990 | 718950 | 0 | 355140 |
| AIYPFDGHI | FNBP1L; FNBP1 | Q5T0N5; A0 | FBP1L_HUMAN | Oxidation (M) | 633910 | 578320 | 0 | 36810 | 196550 | 0 | 70824 | 138330 | 0 | 0 | 0 | 0 | 0 | 363340 | 2634200 | 0 | 533840 | 607650 |
| AIYPFDGHI | FNBP1L; FNBP1 | Q5T0N5; A0 | FBP1L_HUMAN | Unmodified | 390200 | 17289 | 0 | 0 | 0 | 0 | 6590500 | 0 | 566700 | 0 | 0 | 0 | 0 | 0 | 98454 | 0 | 57019 | 187450 |
| AIYQATYR | PSMB5 | P28074 | PSB5_HUMAN | Unmodified | 0 | 0 | 155930 | 137620 | 0 | 0 | 0 | 0 | 378410 | 0 | 0 | 2406900 | 0 | 0 | 0 | 0 | 0 | 0 |
| AIYQVYNAI | ABRAXAS2 | Q15018 | ABRX2_HUMAN | Unmodified | 0 | 0 | 0 | 0 | 92683 | 0 | 0 | 0 | 0 | 0 | 0 | 683650 | 108330 | 0 | 0 | 0 | 0 | 0 |
| AKADAECY | ERLIN2; ERLIN2 | O94905; E5F | ERLN2_HUMAN | Unmodified | 229000 | 0 | 0 | 0 | 158870 | 0 | 144440 | 61286 | 4542700 | 1039000 | 27674 | 34943 | 0 | 204680 | 0 | 0 | 0 | 0 |
| AKADAEYY | ERLIN1; ERLIN1 | O75477; B0 | ERLN1_HUMAN | Unmodified | 0 | 0 | 0 | 0 | 0 | 0 | 0 | 383780 | 4962600 | 0 | 282820 | 0 | 0 | 0 | 0 | 0 | 0 | 0 |
| AKAEASSG | FKBP4 | Q02790 | FKBP4_HUMAN | Unmodified | 0 | 0 | 21619 | 0 | 0 | 0 | 0 | 0 | 0 | 0 | 0 | 147680 | 0 | 0 | 0 | 17324 | 0 | 0 |
| AKAEASSG | FKBP4 | Q02790 | FKBP4_HUMAN | Unmodified | 287460 | 202610 | 438890 | 0 | 1051000 | 300070 | 43937 | 0 | 1266500 | 216010 | 627210 | 11967000 | 6184600 | 2715700 | 624330 | 964860 | 278580 | 1807800 |
| AKAEEMLS | GRB2; GRB2; G | P62993; J3Q | GRB2_HUMAN | Unmodified | 1519700 | 901520 | 1000700 | 830740 | 52806 | 419190 | 37160 | 0 | 0 | 146770 | 0 | 1051100 | 0 | 0 | 1468800 | 1173500 | 0 | 0 |
| AKAGAGSA | MDH2; MDH2 | P40926; G3 | MDHM_HUMAN | Unmodified | 0 | 0 | 0 | 0 | 974190 | 116760 | 749090 | 120010 | 2261700 | 2002600 | 0 | 0 | 704190 | 531290 | 0 | 0 | 196190 | 307360 |
| AKAGAGSA | MDH2; MDH2 | P40926; G3 | MDHM_HUMAN | Oxidation (M) | 0 | 0 | 0 | 0 | 0 | 0 | 147680 | 0 | 0 | 439620 | 0 | 0 | 0 | 0 | 0 | 0 | 0 | 0 |
| AKAQALVQ | LAMTOR2 | Q9Y2Q5 | LTOR2_HUMAN | Unmodified | 0 | 0 | 0 | 0 | 2260300 | 395560 | 2830300 | 0 | 381530 | 1606400 | 0 | 0 | 2916000 | 3402500 | 0 | 0 | 2458100 | 1836200 |
| AKAQDGA | ATP2B1; ATP2B | P20020; E7E | AT2B1_HUMAN | Unmodified | 0 | 0 | 0 | 0 | 0 | 0 | 0 | 0 | 0 | 0 | 0 | 0 | 1425300 | 0 | 0 | 0 | 0 | 0 |
| AKDDSELE | GLG1 | Q92896 | GSLG1_HUMA | Unmodified | 0 | 0 | 0 | 0 | 0 | 0 | 0 | 62866 | 4490500 | 2483500 | 936630 | 2271700 | 0 | 114190 | 0 | 0 | 0 | 0 |
| AKDGTHVL | YOD1 | Q5VVQ6 | OTU1_HUMAN | Unmodified | 0 | 0 | 0 | 0 | 523930 | 0 | 244660 | 0 | 0 | 0 | 0 | 0 | 1008000 | 0 | 0 | 0 | 0 | 0 |
| AKDLFIGD | B3GNT2 | Q9NY97 | B3GN2_HUMA | Unmodified | 0 | 0 | 0 | 0 | 0 | 0 | 44644 | 0 | 0 | 338060 | 0 | 0 | 0 | 0 | 0 | 0 | 0 | 0 |
| AKDLIVTPA | BCAT1; BCAT1 | P54687; F5H | BCAT1_HUMAN | Unmodified | 0 | 0 | 0 | 0 | 681380 | 0 | 0 | 0 | 0 | 0 | 611290 | 1762500 | 0 | 0 | 0 | 0 | 0 | 0 |
| AKDLPTFK | PIP4K2B | P78356 | PI42B_HUMAN | Unmodified | 0 | 0 | 0 | 0 | 209940 | 0 | 0 | 61334 | 0 | 0 | 0 | 0 | 267390 | 0 | 0 | 0 | 0 | 308620 |
| AKEDFLVY | GOLIM4; GOLIM | O00461; F8V | GOLI4_HUMAN | Unmodified | 0 | 0 | 0 | 0 | 0 | 0 | 0 | 0 | 382070 | 382110 | 0 | 0 | 0 | 0 | 0 | 0 | 0 | 0 |
| AKEDFLVY | GOLIM4; GOLIM | O00461; F8V | GOLI4_HUMAN | Unmodified | 0 | 0 | 0 | 0 | 0 | 0 | 873910 | 0 | 31980000 | 11205000 | 1293300 | 0 | 0 | 0 | 0 | 0 | 0 | 1954800 |
| AKEDIVTEQ | AKAP2; PALM2A | Q9Y2D5; C9 | AKAP2_HUMAN | Unmodified | 0 | 0 | 0 | 0 | 2285900 | 272550 | 0 | 0 | 0 | 0 | 0 | 0 | 0 | 0 | 0 | 0 | 485500 | 637540 |
| AKEEAERLE | RDX; RDX; RDX | P35241; A0A | RADI_HUMAN | Unmodified | 0 | 209610 | 27375 | 178410 | 229800 | 0 | 0 | 0 | 1122900 | 0 | 0 | 0 | 1079200 | 544320 | 203010 | 0 | 0 | 126340 |
| AKEELERQ | EZR; EZR | P15311; E7E | EZRI_HUMAN | Unmodified | 1038600 | 0 | 487220 | 585180 | 0 | 0 | 70495 | 0 | 15188000 | 1316400 | 1309900 | 3062100 | 818160 | 301590 | 845240 | 824420 | 162290 | 517170 |
| AKEEMMQ( | KIF14 | Q15058 | KIF14_HUMAN | Oxidation (M) | 0 | 0 | 0 | 0 | 306310 | 0 | 0 | 0 | 0 | 0 | 0 | 0 | 0 | 0 | 0 | 0 | 0 | 0 |
| AKEEMMQ( | KIF14 | Q15058 | KIF14_HUMAN | Unmodified | 0 | 0 | 0 | 0 | 2900700 | 0 | 0 | 0 | 246020 | 0 | 0 | 0 | 0 | 0 | 0 | 31319 | 905830 | 238410 |
| AKEEQELQ | GPM6B | B7Z613 | B7Z613_HUMA | Unmodified | 573170 | 546980 | 187980 | 0 | 0 | 0 | 347700 | 209890 | 638260 | 167630 | 95023 | 221900 | 0 | 0 | 158570 | 315220 | 0 | 0 |
| AKEEQELQ | GPM6B | B7Z613 | B7Z613_HUMA | Unmodified | 0 | 0 | 0 | 0 | 27707 | 0 | 1187100 | 0 | 81614 | 0 | 0 | 0 | 0 | 0 | 0 | 0 | 0 | 0 |
| AKEEQQDD | DCTN1; DCTN1 | Q14203; A0 | DCTN1_HUMA | Unmodified | 0 | 0 | 0 | 0 | 0 | 0 | 0 | 33189 | 0 | 0 | 0 | 740920 | 0 | 0 | 0 | 0 | 82863 | 60724 |
| AKEERLAE | INCENP | Q9NQS7 | INCE_HUMAN | Unmodified | 0 | 39868 | 0 | 0 | 458210 | 55600 | 0 | 0 | 985700 | 0 | 15432 | 540500 | 0 | 38943 | 19068 | 20264 | 78799 | 85849 |
| AKEERLEA | N/A | N/A | N/A | Unmodified | 0 | 0 | 0 | 0 | 3510700 | 1212600 | 0 | 0 | 616320 | 14419 | 256950 | 2702500 | 0 | 0 | 0 | 392010 | 520880 | 355220 |
| AKEFTNVYI | PABPC4; PABP | Q13310; P1 | PABP4_HUMAN | Unmodified | 27027000 | 11257000 | 0 | 19004000 | 0 | 0 | 0 | 0 | 0 | 0 | 0 | 0 | 5629600 | 0 | 0 | 0 | 143230 | 0 |
| AKEILQEEE | ATP6V1A | P38606 | VATA_HUMAN | Unmodified | 0 | 0 | 0 | 76298 | 301780 | 0 | 397000 | 0 | 0 | 211030 | 17008 | 30259 | 521540 | 656100 | 82482 | 23034 | 131490 | 0 |
| AKEILTKES | PPP2CB; PPP2C | P62714; P67 | PP2AB_HUMAN | Unmodified | 0 | 0 | 0 | 0 | 100170 | 0 | 0 | 0 | 0 | 0 | 0 | 0 | 109140 | 64121 | 0 | 0 | 0 | 91498 |
| AKELDPTN | STIP1 | P31948 | STIP1_HUMAN | Unmodified | 0 | 0 | 0 | 0 | 2307500 | 256970 | 367100 | 0 | 0 | 824780 | 0 | 0 | 3975200 | 1879600 | 0 | 0 | 2942200 | 3483100 |
| AKELDPTN | STIP1 | P31948 | STIP1_HUMAN | Oxidation (M) | 0 | 0 | 0 | 0 | 0 | 0 | 0 | 0 | 0 | 569590 | 0 | 0 | 2194300 | 2629200 | 0 | 0 | 2165800 | 2594100 |
| AKELDPTN | STIP1 | P31948 | STIP1_HUMAN | Oxidation (M) | 0 | 0 | 0 | 0 | 0 | 0 | 0 | 0 | 0 | 203460 | 0 | 0 | 0 | 0 | 0 | 0 | 2936000 | 666550 |
| AKELDPTN | STIP1 | P31948 | STIP1_HUMAN | Unmodified | 0 | 0 | 0 | 0 | 2466400 | 0 | 0 | 0 | 0 | 1269200 | 0 | 0 | 2834900 | 2141900 | 0 | 0 | 12475000 | 2570500 |
| AKELPTLKD | PIP4K2A; PIP4K | P48426; S4R | PI42A_HUMAN | Unmodified | 0 | 0 | 0 | 0 | 534340 | 0 | 295960 | 320350 | 0 | 0 | 0 | 0 | 584390 | 374390 | 0 | 0 | 236720 | 512970 |
| AKENSEES | DAAM1 | Q9Y4D1 | DAAM1_HUMA | Unmodified | 202630 | 0 | 0 | 265590 | 0 | 0 | 0 | 0 | 761570 | 181770 | 0 | 1491600 | 0 | 0 | 377930 | 361820 | 0 | 0 |
| AKENSEES | DAAM1 | Q9Y4D1 | DAAM1_HUMA | Unmodified | 0 | 0 | 0 | 0 | 2837600 | 0 | 2195500 | 0 | 0 | 844090 | 223060 | 0 | 2641500 | 1667700 | 0 | 0 | 0 | 0 |
| AKEQYAVV | ATRN | O75882 | ATRN_HUMAN | Unmodified | 0 | 0 | 0 | 0 | 0 | 0 | 0 | 0 | 203110 | 749830 | 112920 | 0 | 0 | 0 | 0 | 0 | 0 | 0 |
| AKESWEMI | FKBP5 | Q13451 | FKBP5_HUMAN | Unmodified | 0 | 0 | 0 | 113410 | 419270 | 0 | 63880 | 34215 | 94414 | 0 | 0 | 528810 | 598770 | 147750 | 0 | 55300 | 0 | 76295 |
| AKESWEMI | FKBP4 | Q02790 | FKBP4_HUMAN | Unmodified | 195540 | 0 | 110580 | 204440 | 207490 | 0 | 0 | 0 | 275090 | 117590 | 1266300 | 7370200 | 847660 | 345160 | 325610 | 237560 | 123170 | 65541 |
| AKESWEMI | FKBP4 | Q02790 | FKBP4_HUMAN | Oxidation (M) | 0 | 0 | 0 | 0 | 0 | 0 | 0 | 0 | 0 | 0 | 0 | 2478000 | 0 | 0 | 0 | 0 | 54981 | 0 |
| AKESWEMI | FKBP4 | Q02790 | FKBP4_HUMAN | Oxidation (M) | 0 | 0 | 0 | 161580 | 532010 | 0 | 695240 | 211440 | 0 | 587720 | 638200 | 0 | 2352400 | 1810200 | 0 | 0 | 673410 | 385480 |
| AKESWEMI | FKBP4 | Q02790 | FKBP4_HUMAN | Unmodified | 0 | 0 | 0 | 0 | 911380 | 0 | 1622500 | 164220 | 0 | 572040 | 0 | 0 | 4284600 | 825030 | 0 | 0 | 1225300 | 0 |
| AKFDGIFES | EPS15L1; EPS1 | Q9UBC2; M | EP15R_HUMAN | Unmodified | 0 | 654910 | 0 | 0 | 117240 | 15975 | 25556 | 0 | 0 | 114670 | 0 | 20419 | 214500 | 0 | 0 | 0 | 74553 | 21014 |
| AKFEELNM | HSPA5; HSPA5 | P11021; A0A | BIP_HUMAN Er | Oxidation (M) | 0 | 0 | 909480 | 1984900 | 539870 | 0 | 0 | 238870 | 72042000 | 12163000 | 15494000 | 4768800 | 0 | 0 | 3258900 | 1566400 | 0 | 0 |
| AKFEELNM | HSPA5; HSPA5 | P11021; A0A | BIP_HUMAN Er | Unmodified | 0 | 0 | 0 | 0 | 501800 | 0 | 369440 | 0 | 2.58E+08 | 7796800 | 4917600 | 1333500 | 315910 | 237340 | 0 | 0 | 0 | 0 |
| AKFENLCK | HSP90AB1; HSI | P08238; Q5 | HS90B_HUMA | Unmodified | 0 | 0 | 1125400 | 0 | 7504800 | 0 | 0 | 581160 | 2799900 | 2160800 | 1135900 | 12488000 | 12995000 | 4763900 | 1284000 | 0 | 2533800 | 3474100 |
| AKFFPEDV | RDX; RDX; RDX; | P35241; E9P | RADI_HUMAN | Unmodified | 302240 | 0 | 0 | 1371800 | 0 | 0 | 0 | 0 | 382440 | 0 | 0 | 1539600 | 0 | 0 | 1743600 | 321570 | 0 | 0 |
| AKFVGDPA | DAG1 | Q14118 | DAG1_HUMAN | Unmodified | 0 | 834600 | 0 | 0 | 768920 | 0 | 183990 | 0 | 0 | 0 | 0 | 0 | 372440 | 408180 | 0 | 0 | 154320 | 620420 |
| AKFYPEDV | EZR; EZR | P15311; E7E | EZRI_HUMAN | Unmodified | 28180 | 251820 | 0 | 405070 | 1367900 | 368890 | 0 | 0 | 718150 | 918690 | 0 | 333830 | 183780 | 585300 | 579790 | 0 | 716790 | 0 |
| AKFYPEDV | MSN; MSN | P26038; V9 | MOES_HUMAN | Unmodified | 0 | 390470 | 0 | 4319800 | 0 | 0 | 0 | 0 | 156560 | 0 | 0 | 0 | 0 | 0 | 243820 | 0 | 0 | 0 |
| AKGETAYLI | CXADR | P78310 | CXAR_HUMAN | Unmodified | 918620 | 0 | 364780 | 0 | 10965000 | 1768100 | 0 | 68366 | 0 | 406450 | 291100 | 4784800 | 344810 | 0 | 0 | 196860 | 0 | 0 |
| AKGEVDVD | AHNAK | Q09666 | AHNK_HUMAN | Unmodified | 0 | 0 | 0 | 0 | 4639200 | 0 | 0 | 0 | 0 | 0 | 0 | 0 | 0 | 0 | 0 | 229610 | 1811500 | 2676700 |
| AKGILFVG | PGD; PGD; PGD | P52209; K7E | 6PGD_HUMAN | Oxidation (M) | 0 | 0 | 0 | 0 | 2525500 | 0 | 0 | 0 | 0 | 0 | 0 | 0 | 1142700 | 0 | 0 | 0 | 0 | 0 |
| AKGILFVG | PGD; PGD; PGD | P52209; K7E | 6PGD_HUMAN | Unmodified | 0 | 0 | 0 | 0 | 4406500 | 0 | 0 | 0 | 0 | 0 | 0 | 0 | 0 | 3252400 | 0 | 0 | 440030 | 3297500 |
| AKGIRHEVI | GSTO1; GSTO1 | P78417; Q5 | GSTO1_HUMA | Unmodified | 0 | 0 | 0 | 0 | 4528000 | 0 | 861140 | 121930 | 0 | 195950 | 131750 | 0 | 0 | 0 | 0 | 0 | 1883000 | 1428300 |
| AKGPGGTS | RAB11FIP1 | Q6WKZ4 | RFIP1_HUMAN | Unmodified | 46330 | 153470 | 0 | 541010 | 132180 | 0 | 170710 | 108300 | 0 | 0 | 327970 | 562340 | 292130 | 244590 | 137850 | 166610 | 0 | 0 |
| AKGPLASQ | NT5E; NT5E | P21589; Q9 | 5NTD_HUMAN | Unmodified | 0 | 0 | 0 | 0 | 68202000 | 12996000 | 0 | 0 | 0 | 0 | 0 | 320900 | 0 | 0 | 0 | 0 | 14612000 | 32610000 |
| AKGPSENV | MEAK7; MEAK7 | Q6P9B6; H3 | MEAK7_HUMA | Unmodified | 0 | 163040 | 0 | 300420 | 26175000 | 0 | 0 | 0 | 3799200 | 2425000 | 0 | 1932000 | 0 | 1113800 | 527210 | 528310 | 4707600 | 5523300 |
| AKGPVHIVI | EMC1; EMC1 | Q8N766; H7 | EMC1_HUMAN | Unmodified | 0 | 0 | 0 | 0 | 0 | 0 | 0 | 0 | 1940200 | 0 | 0 | 0 | 0 | 0 | 0 | 0 | 0 | 0 |
| AKGVVIGT( | ATP2C1; ATP2C | P98194; B4E | AT2C1_HUMAN | Unmodified | 0 | 0 | 0 | 0 | 367590 | 0 | 155260 | 0 | 1462200 | 660290 | 0 | 0 | 139460 | 96467 | 0 | 0 | 707060 | 0 |
| AKHPMDTE | AP2A2; AP2A2 | O94973; A0 | AP2A2_HUMAN | Unmodified | 266410 | 0 | 0 | 0 | 1402000 | 313710 | 168450 | 78812 | 514220 | 282670 | 0 | 0 | 0 | 0 | 118610 | 126680 | 408060 | 301640 |
| AKIAEQVAS | CTNNA1; CTNN | P35221; G3 | CTNA1_HUMA | Unmodified | 3128900 | 2823800 | 2278900 | 1354200 | 23843000 | 4861700 | 251050 | 45437 | 0 | 714040 | 0 | 0 | 17458000 | 16645000 | 0 | 230920 | 0 | 0 |
| AKIDNYIPIF | ATP6V0D1; ATP | P61421; R4 | VA0D1_HUMAN | Unmodified | 0 | 0 | 0 | 0 | 0 | 0 | 0 | 0 | 639900 | 0 | 0 | 560850 | 0 | 0 | 0 | 0 | 0 | 0 |
| AKIELLGSY | ACP1 | P24666 | PPAC_HUMAN | Unmodified | 1772200 | 1205700 | 4324900 | 997640 | 7133700 | 494410 | 883370 | 877170 | 1476900 | 571030 | 1483600 | 9454200 | 5111800 | 2730300 | 1188300 | 1279500 | 4373000 | 4295600 |
| AKIIGFGSA | AP2A2; AP2A2 | O94973; A0 | AP2A2_HUMAN | Unmodified | 0 | 0 | 0 | 0 | 1695400 | 0 | 0 | 0 | 0 | 1032500 | 0 | 0 | 2559000 | 0 | 0 | 0 | 1133200 | 1507900 |
| AKINEAVE( | EPRS1; EPRS1 | P07814; V9 | SYEP_HUMAN | Unmodified | 203160 | 0 | 0 | 51830 | 1203700 | 0 | 0 | 0 | 86420 | 663180 | 0 | 837330 | 2345200 | 1561600 | 0 | 28035 | 444300 | 968650 |
| AKINQGWL | FERMT2; FERM | Q96AC1; A0 | FERM2_HUMA | Unmodified | 0 | 232250 | 0 | 0 | 1270600 | 526200 | 0 | 0 | 140440 | 0 | 0 | 0 | 0 | 0 | 0 | 77544 | 243870 | 583610 |
| AKISKPAPY | PRDX4; PRDX4 | Q13162; A6 | PRDX4_HUMA | Unmodified | 0 | 0 | 0 | 0 | 0 | 0 | 338090 | 0 | 17574000 | 38564000 | 7054700 | 0 | 0 | 0 | 0 | 0 | 0 | 0 |
| AKISSPTET | S100A11 | P31949 | S10AB_HUMAN | Acetyl (Protein N | 0 | 166810 | 0 | 0 | 1946700 | 93866 | 0 | 0 | 0 | 334170 | 0 | 0 | 0 | 650430 | 0 | 158570 | 9129200 | 701290 |
| AKKDDTD | CANX; CANX; C | P27824; A0A | CALX_HUMAN | Unmodified | 0 | 0 | 0 | 75655 | 0 | 0 | 0 | 0 | 2617200 | 88983 | 0 | 0 | 0 | 0 | 0 | 0 | 164920 | 0 |
| AKLAEQAEI | YWHAB; YWHA | P31946; Q4 | 1433B_HUMAN | Oxidation (M) | 197670 | 0 | 0 | 64488 | 13477000 | 2803000 | 4923000 | 1279100 | 140230 | 2590700 | 101380 | 92191 | 5259300 | 8341900 | 0 | 0 | 14624000 | 27088000 |
| AKLAEQAEI | YWHAB; YWHA | P31946; Q4 | 1433B_HUMAN | Unmodified | 0 | 68649 | 0 | 230190 | 34226000 | 1852200 | 5491300 | 368140 | 1737200 | 1122400 | 0 | 0 | 8104600 | 8461500 | 0 | 0 | 28577000 | 41763000 |
| AKLAEQAEI | YWHAB; YWHA | P31946; Q4 | 1433B_HUMAN | 2 Oxidation (M) | 0 | 0 | 0 | 0 | 494240 | 65196 | 123570 | 0 | 0 | 80631 | 0 | 0 | 0 | 194390 | 0 | 0 | 0 | 0 |
| AKLAEQAEI | YWHAZ; YWHA | P63104; E5R | 1433Z_HUMAN | Unmodified | 24720 | 0 | 0 | 466570 | 36090000 | 4951600 | 21458000 | 3509700 | 6287400 | 13134000 | 1271200 | 688810 | 15015000 | 15817000 | 0 | 0 | 90674000 | 91202000 |
| AKLAEQAEI | YWHAZ; YWHA | P63104; E5R | 1433Z_HUMAN | 2 Oxidation (M) | 0 | 0 | 0 | 0 | 222590 | 53657 | 1505700 | 676000 | 0 | 770490 | 0 | 0 | 225270 | 1803500 | 0 | 0 | 6820600 | 4044800 |
| AKLAEQAEI | YWHAZ; YWHA | P63104; E5R | 1433Z_HUMAN | Oxidation (M) | 0 | 0 | 0 | 84216 | 11946000 | 5183700 | 9779900 | 6145700 | 1176300 | 5331800 | 596340 | 330010 | 6516700 | 13035000 | 0 | 0 | 44673000 | 31923000 |
| AKLAEQAEI | YWHAQ; YWHA | P27348; E9P | 1433T_HUMAN | Unmodified | 0 | 0 | 0 | 168750 | 12552000 | 2537800 | 2131400 | 854820 | 1138000 | 2480500 | 0 | 280690 | 5023500 | 8278200 | 0 | 25186 | 25090000 | 17046000 |
| AKLAEQAEI | YWHAQ; YWHA | P27348; E9P | 1433T_HUMAN | Oxidation (M) | 0 | 0 | 0 | 0 | 5594900 | 0 | 1941100 | 1229900 | 75731 | 999110 | 236420 | 153010 | 5855300 | 3062400 | 0 | 0 | 7557900 | 5881400 |
| AKLAEQAEI | SFN | P31947 | 1433S_HUMAN | Oxidation (M) | 0 | 89728 | 0 | 0 | 15251000 | 2854500 | 0 | 0 | 425710 | 2900200 | 0 | 0 | 0 | 0 | 0 | 275590 | 7252500 | 4364800 |
| AKLAEQAEI | SFN | P31947 | 1433S_HUMAN | Unmodified | 0 | 0 | 0 | 0 | 32916000 | 3072700 | 0 | 0 | 1573500 | 2334600 | 705270 | 0 | 0 | 0 | 0 | 0 | 14703000 | 9875300 |
| AKLAEQAEI | SFN | P31947 | 1433S_HUMAN | 2 Oxidation (M) | 0 | 0 | 0 | 0 | 1157200 | 551700 | 0 | 0 | 0 | 231490 | 0 | 0 | 0 | 0 | 0 | 0 | 775110 | 308310 |
| AKLDENSA | SPTAN1; SPTAN | Q13813; A0 | SPTN1_HUMAN | Unmodified | 0 | 0 | 0 | 0 | 0 | 0 | 0 | 0 | 0 | 0 | 0 | 983160 | 0 | 0 | 0 | 0 | 0 | 0 |
| AKLDGAQL | AHNAK2 | Q8IVF2 | AHNK2_HUMA | Unmodified | 0 | 0 | 0 | 0 | 1527700 | 0 | 0 | 0 | 0 | 126510 | 0 | 0 | 0 | 0 | 0 | 0 | 0 | 445250 |
| AKLDGARL | AHNAK2 | Q8IVF2 | AHNK2_HUMA | Unmodified | 0 | 0 | 0 | 0 | 798810 | 0 | 0 | 25590 | 48805 | 0 | 0 | 0 | 447590 | 0 | 0 | 0 | 0 | 1076000 |
| AKLDGAWI | AHNAK2 | Q8IVF2 | AHNK2_HUMA | Unmodified | 0 | 0 | 0 | 0 | 1305000 | 0 | 0 | 0 | 0 | 0 | 0 | 0 | 0 | 0 | 0 | 0 | 433840 | 2577400 |
| AKLDSLQD | NUCB2; NUCB2 | P80303; Q2 | NUCB2_HUMA | Oxidation (M) | 0 | 0 | 1118000 | 1700900 | 0 | 0 | 1145300 | 664700 | 59643000 | 20607000 | 2438900 | 80464 | 491490 | 1815600 | 1004100 | 2558800 | 0 | 0 |
| AKLDSLQD | NUCB2; NUCB2 | P80303; Q2 | NUCB2_HUMA | Unmodified | 0 | 0 | 0 | 622930 | 0 | 0 | 551240 | 53088 | 1.69E+08 | 19774000 | 0 | 0 | 1156200 | 1807200 | 0 | 0 | 0 | 0 |
| AKLEETITQ | CACNA2D1; CA | P54289; H0 | CA2D1_HUMA | Unmodified | 477480 | 442710 | 0 | 0 | 0 | 0 | 687010 | 19837 | 5546400 | 1067800 | 0 | 0 | 0 | 0 | 0 | 0 | 69976 | 189120 |
| AKLEQLFQI | PLEC; PLEC | Q15149; H0 | PLEC_HUMAN | Unmodified | 0 | 0 | 0 | 0 | 228960 | 0 | 0 | 0 | 596710 | 0 | 0 | 0 | 0 | 0 | 0 | 0 | 1496100 | 0 |
| AKLHELNQ | SPTAN1; SPTAN | Q13813; A0 | SPTN1_HUMAN | Unmodified | 0 | 0 | 0 | 0 | 0 | 0 | 0 | 0 | 185460 | 206220 | 0 | 515710 | 0 | 0 | 0 | 0 | 289960 | 0 |

| | | | | | | | | | | | | | | | | | | | | | | |
|---|---|---|---|---|---|---|---|---|---|---|---|---|---|---|---|---|---|---|---|---|---|---|
| AKLITDLQD | KIF5B | P33176 | KINH_HUMAN | Unmodified | 56505 | 0 | 0 | 0 | 3099800 | 460540 | 1116800 | 756850 | 1237800 | 1056200 | 0 | 309620 | 2167600 | 1017600 | 0 | 100600 | 1072800 | 1430300 |
| AKLMGFFD | POGLUT3; POG | Q7Z4H8; H0 | PLGT3_HUMAN | Oxidation (M) | 0 | 0 | 0 | 0 | 0 | 0 | 0 | 0 | 0 | 533750 | 0 | 0 | 0 | 0 | 0 | 0 | 0 | 0 |
| AKLMLGWE | UXS1 | Q8NBZ7 | UXS1_HUMAN | Oxidation (M) | 0 | 0 | 0 | 0 | 0 | 0 | 0 | 30500 | 0 | 932430 | 0 | 0 | 0 | 0 | 4646600 | 0 | 0 | 0 |
| AKLNVEPD | SLITRK5 | O94991 | SLIK5_HUMAN | Unmodified | 0 | 72786 | 0 | 0 | 0 | 0 | 0 | 0 | 230200 | 264980 | 0 | 0 | 0 | 0 | 0 | 0 | 0 | 0 |
| AKLQELEG | MYH10 | P35580 | MYH10_HUMA | Unmodified | 0 | 0 | 0 | 0 | 5388100 | 542900 | 289770 | 105880 | 0 | 35510 | 0 | 865770 | 508350 | 537130 | 0 | 101490 | 301170 | 159820 |
| AKLQQCKA | KIF20A | O95235 | KI20A_HUMAN | Unmodified | 0 | 0 | 0 | 0 | 572980 | 0 | 0 | 0 | 0 | 115430 | 0 | 0 | 1152200 | 1430700 | 0 | 0 | 471500 | 154290 |
| AKLSSDVL | LONP1; LONP1 | P36776; K7E | LONM_HUMAN | Unmodified | 0 | 0 | 0 | 0 | 131090 | 12340 | 236720 | 0 | 2526700 | 902890 | 0 | 0 | 186550 | 283180 | 0 | 0 | 0 | 200820 |
| AKLTMLNT | SH3GL1; SH3G | Q99961; M0 | SH3G1_HUMA | Oxidation (M) | 957580 | 0 | 0 | 514740 | 0 | 0 | 0 | 70804 | 0 | 61699 | 0 | 0 | 141460 | 68099 | 575190 | 675840 | 0 | 0 |
| AKLTMLNT | SH3GL1; SH3G | Q99961; M0 | SH3G1_HUMA | Unmodified | 584280 | 204580 | 73698 | 824880 | 64697 | 0 | 89392 | 27132 | 0 | 71449 | 0 | 0 | 229160 | 205080 | 487120 | 479570 | 30459 | 0 |
| AKMDAEQD | NUCB1; NUCB | Q02818; C9J | NUCB1_HUMA | Unmodified | 0 | 0 | 0 | 0 | 0 | 0 | 0 | 0 | 4900000 | 554840 | 0 | 1301200 | 126940 | 0 | 0 | 0 | 24627 | 0 |
| AKMDAEQD | NUCB1; NUCB | Q02818; C9J | NUCB1_HUMA | Oxidation (M) | 0 | 0 | 0 | 0 | 0 | 132870 | 0 | 400380 | 2566700 | 955360 | 1347000 | 717930 | 0 | 0 | 215890 | 287330 | 0 | 0 |
| AKMEDTLK | GARS1; GARS1 | P41250; F8V | GARS_HUMAN | Unmodified | 0 | 261640 | 0 | 0 | 0 | 0 | 0 | 0 | 0 | 0 | 309930 | 678600 | 0 | 0 | 0 | 0 | 0 | 0 |
| AKNEILDEV | OPA1; OPA1; O | O60313; E5 | OPA1_HUMAN | Unmodified | 0 | 0 | 0 | 0 | 0 | 0 | 261470 | 0 | 535490 | 791080 | 2206100 | 15540000 | 0 | 0 | 0 | 0 | 156280 | 0 |
| AKNEPVDR | LRP1; LRP1; LR | Q07954; Q6 | LRP1_HUMAN | Unmodified | 0 | 0 | 136820 | 0 | 155680 | 0 | 0 | 0 | 4777600 | 55072 | 0 | 0 | 0 | 0 | 173860 | 0 | 0 | 0 |
| AKNEQEHH | GALNT7; GALN | Q86SF2; E9F | GALT7_HUMAN | Unmodified | 0 | 0 | 0 | 0 | 0 | 0 | 0 | 0 | 82814 | 0 | 0 | 0 | 83877 | 21833 | 0 | 0 | 0 | 21172 |
| AKNGHLAV | MYO1C; MYO1 | O00159; F5 | MYO1C_HUMA | Unmodified | 72738 | 18329 | 0 | 0 | 1679200 | 0 | 66755 | 21378 | 0 | 0 | 0 | 79078 | 701970 | 752250 | 0 | 0 | 538590 | 709230 |
| AKPAEAPA | DLST; DLST | P36957; Q8 | ODO2_HUMAN | Unmodified | 0 | 0 | 0 | 0 | 0 | 0 | 0 | 0 | 0 | 0 | 0 | 0 | 0 | 0 | 0 | 0 | 0 | 0 |
| AKPAMPQD | ACLY | P53396 | ACLY_HUMAN | Unmodified | 0 | 0 | 0 | 0 | 926100 | 0 | 413420 | 155630 | 473030 | 165220 | 261460 | 529870 | 288380 | 0 | 0 | 0 | 0 | 0 |
| AKPAMPQD | ACLY | P53396 | ACLY_HUMAN | Oxidation (M) | 0 | 0 | 0 | 0 | 112200 | 0 | 0 | 0 | 42262 | 0 | 0 | 850150 | 177360 | 0 | 117310 | 0 | 0 | 0 |
| AKPAMPQD | ACLY | P53396 | ACLY_HUMAN | Unmodified | 0 | 0 | 0 | 0 | 0 | 0 | 1265100 | 330750 | 0 | 0 | 0 | 0 | 0 | 0 | 0 | 0 | 339540 | 0 |
| AKPEGALQ | EPCAM; EPCAM | P16422; C9J | EPCAM_HUMA | Unmodified | 33777000 | 24742000 | 23163000 | 9539900 | 16447000 | 3874200 | 0 | 0 | 57827000 | 38241000 | 17980000 | 38248000 | 40825000 | 17184000 | 9602700 | 4329000 | 2557300 | 3304300 |
| AKPEGALQ | EPCAM; EPCAM | P16422; C9J | EPCAM_HUMA | Unmodified | 0 | 0 | 0 | 0 | 729550 | 0 | 0 | 0 | 0 | 1884400 | 0 | 0 | 1917000 | 3368600 | 0 | 0 | 734270 | 611560 |
| AKPLTDQE | PYGL; PYGL | P06737; E9P | PYGL_HUMAN | Acetyl (Protein N | 62744 | 271350 | 0 | 0 | 3125600 | 975200 | 574220 | 280070 | 387590 | 144980 | 0 | 53131 | 0 | 0 | 0 | 119530 | 1049400 | 5113300 |
| AKPLTDSE | PYGB | P11216 | PYGB_HUMAN | Acetyl (Protein N | 0 | 0 | 0 | 0 | 1597800 | 0 | 113680 | 0 | 0 | 745360 | 86570 | 230560 | 240820 | 0 | 0 | 0 | 2910100 | 19562000 |
| AKPPPGGG | PSMD3 | O43242 | PSMD3_HUMA | Oxidation (M) | 0 | 0 | 0 | 0 | 0 | 0 | 0 | 0 | 0 | 0 | 359460 | 1198100 | 0 | 0 | 241420 | 0 | 0 | 0 |
| AKPSPAPP | PEX19; PEX19; | P40855; Q5 | PEX19_HUMAN | Unmodified | 0 | 0 | 0 | 0 | 0 | 0 | 0 | 0 | 291110 | 49427 | 18820 | 0 | 227900 | 0 | 0 | 71201 | 92505 | 184400 |
| AKPTSDKP | ACSL4; ACSL4; | O60488; D6 | ACSL4_HUMAN | Unmodified | 27165 | 0 | 50024 | 88728 | 664190 | 0 | 0 | 31208 | 172020 | 0 | 1542700 | 5157400 | 169590 | 73358 | 72705 | 0 | 482310 | 872870 |
| AKPYEFNW | PLCB4; PLCB4; | Q15147; A0 | PLCB4_HUMAN | Acetyl (Protein N | 0 | 0 | 225220 | 677180 | 0 | 0 | 0 | 192590 | 325840 | 0 | 0 | 0 | 956910 | 510160 | 511960 | 596680 | 2940100 | 3124600 |
| AKPYEGSIL | GLUD1 | P00367 | DHE3_HUMAN | Unmodified | 0 | 0 | 0 | 0 | 393460 | 0 | 533130 | 0 | 1891400 | 992210 | 0 | 0 | 0 | 0 | 0 | 0 | 0 | 231540 |
| AKQHLEQE | KIF14 | Q15058 | KIF14_HUMAN | Unmodified | 0 | 0 | 0 | 0 | 484820 | 0 | 72654 | 0 | 0 | 20029 | 0 | 0 | 0 | 30456 | 0 | 0 | 228270 | 0 |
| AKQHLEQE | KIF14 | Q15058 | KIF14_HUMAN | Unmodified | 0 | 0 | 0 | 0 | 1693700 | 0 | 0 | 0 | 0 | 0 | 0 | 0 | 0 | 0 | 0 | 0 | 503740 | 0 |
| AKQIVWNG | PGK1 | P00558 | PGK1_HUMAN | Unmodified | 0 | 0 | 0 | 0 | 0 | 0 | 0 | 0 | 0 | 212050 | 0 | 0 | 0 | 0 | 0 | 0 | 619630 | 755170 |
| AKQIVWNG | PGK1 | P00558 | PGK1_HUMAN | Unmodified | 0 | 0 | 0 | 0 | 0 | 0 | 0 | 0 | 0 | 0 | 0 | 0 | 73226 | 2943000 | 0 | 0 | 7899800 | 0 |
| AKQTLENE | MYH9 | P35579 | MYH9_HUMAN | Unmodified | 683390 | 938310 | 447420 | 3921100 | 47814000 | 6683300 | 7625500 | 2697800 | 2305400 | 3624000 | 317090 | 4528700 | 14570000 | 15542000 | 2629700 | 2281400 | 23789000 | 17497000 |
| AKSESLELF | JOSD1; JOSD1; | Q15040; B0 | JOS1_HUMAN | Unmodified | 127250 | 0 | 0 | 0 | 1277400 | 0 | 410210 | 318870 | 0 | 0 | 0 | 482980 | 681080 | 0 | 0 | 0 | 609630 | 1433000 |
| AKSETVLTC | ITGA3 | P26006 | ITA3_HUMAN I | Unmodified | 135400 | 165630 | 0 | 158720 | 8385900 | 3446500 | 99894 | 139020 | 299680 | 0 | 38860 | 1702200 | 0 | 0 | 164000 | 89165 | 7162100 | 15439000 |
| AKSFEGNN | NRP1; NRP1; N | O14786; E9F | NRP1_HUMAN | Unmodified | 0 | 0 | 0 | 0 | 0 | 0 | 0 | 0 | 590010 | 423060 | 0 | 0 | 0 | 0 | 0 | 0 | 2390800 | 4147600 |
| AKSPQPPV | HNRNPU; HNR | Q00839; A0 | HNRPU_HUMA | Unmodified | 0 | 0 | 0 | 0 | 0 | 0 | 1041100 | 0 | 0 | 0 | 0 | 0 | 0 | 0 | 0 | 0 | 0 | 0 |
| AKSSMFRA | ARHGAP29; AR | Q52LW3; F8 | RHG29_HUMA | Oxidation (M) | 0 | 0 | 0 | 0 | 0 | 0 | 0 | 0 | 0 | 0 | 0 | 0 | 0 | 0 | 0 | 0 | 238280 | 107960 |
| AKSSMFRA | ARHGAP29; AR | Q52LW3; F8 | RHG29_HUMA | Unmodified | 0 | 0 | 0 | 0 | 126160 | 0 | 0 | 0 | 0 | 0 | 0 | 0 | 0 | 0 | 0 | 0 | 1032300 | 435910 |
| AKTASLFEC | MYOF | Q9NZM1 | MYOF_HUMAN | Unmodified | 0 | 0 | 0 | 0 | 1693200 | 0 | 0 | 0 | 0 | 0 | 0 | 0 | 0 | 0 | 0 | 0 | 0 | 0 |
| AKTDAGGE | ATG3 | Q9NT62 | ATG3_HUMAN | Unmodified | 263360 | 0 | 185140 | 223740 | 261810 | 0 | 0 | 0 | 661790 | 150280 | 414300 | 1820600 | 790090 | 237390 | 185700 | 233440 | 151580 | 297270 |
| AKTDAGGE | ATG3 | Q9NT62 | ATG3_HUMAN | Unmodified | 0 | 0 | 0 | 0 | 0 | 438510 | 0 | 0 | 0 | 397760 | 0 | 0 | 4096800 | 0 | 0 | 0 | 2416100 | 0 |
| AKTGGAYG | FASN; FASN | P49327; A0A | FAS_HUMAN F | Unmodified | 0 | 0 | 0 | 0 | 0 | 0 | 0 | 135840 | 0 | 0 | 0 | 1564700 | 0 | 0 | 0 | 0 | 0 | 0 |
| AKTQDGVA | ATP2B4 | P23634 | AT2B4_HUMAN | Unmodified | 0 | 0 | 0 | 0 | 3636500 | 0 | 0 | 0 | 614870 | 0 | 0 | 2659800 | 0 | 0 | 0 | 0 | 0 | 0 |
| AKVASLEES | CKAP4 | Q07065 | CKAP4_HUMAN | Unmodified | 0 | 0 | 0 | 0 | 175040 | 0 | 354930 | 18766 | 352240 | 567410 | 0 | 0 | 0 | 477190 | 0 | 0 | 79785 | 234840 |
| AKVDEFPL | RPL10; RPL10; | P27635; X1V | RL10_HUMAN | Oxidation (M) | 0 | 2948300 | 0 | 0 | 538200 | 0 | 0 | 0 | 0 | 0 | 0 | 6598300 | 0 | 0 | 3244500 | 503200 | 0 | 0 |
| AKVEQVLSI | VAPB; VAPB | O95292; E5F | VAPB_HUMAN | Acetyl (Protein N | 0 | 298130 | 0 | 0 | 7637500 | 1195100 | 2648800 | 1513200 | 2060700 | 4032600 | 517090 | 3723300 | 3596400 | 3267000 | 0 | 0 | 1196100 | 2764100 |
| AKVEVSQP | AHNAK2 | Q8IVF2 | AHNK2_HUMA | Unmodified | 0 | 0 | 0 | 0 | 1070600 | 0 | 0 | 0 | 0 | 0 | 0 | 0 | 0 | 0 | 0 | 0 | 0 | 0 |
| AKVIAAEGE | STOM | P27105 | STOM_HUMAN | Unmodified | 0 | 0 | 0 | 0 | 4009800 | 556920 | 377780 | 358880 | 0 | 0 | 0 | 68655 | 0 | 613930 | 0 | 0 | 0 | 0 |
| AKVIAAEGE | STOM | P27105 | STOM_HUMAN | Unmodified | 0 | 0 | 0 | 0 | 6403500 | 484090 | 0 | 0 | 0 | 0 | 0 | 0 | 0 | 0 | 0 | 0 | 0 | 0 |
| AKVLLSALE | MTHFD1; MTHF | P11586; V9G | C1TC_HUMAN | Unmodified | 0 | 0 | 0 | 0 | 683500 | 0 | 719190 | 0 | 0 | 0 | 0 | 0 | 0 | 171340 | 0 | 0 | 735910 | 221630 |
| AKVTEELAA | FKBP15 | Q5T1M5 | FKB15_HUMAN | Unmodified | 0 | 0 | 0 | 0 | 774590 | 0 | 0 | 0 | 0 | 187050 | 0 | 0 | 0 | 0 | 0 | 0 | 0 | 0 |
| AKVTGDEW | GYS1 | P13807 | GYS1_HUMAN | Unmodified | 0 | 0 | 0 | 0 | 1668200 | 0 | 0 | 0 | 0 | 0 | 0 | 2296100 | 0 | 0 | 0 | 0 | 824210 | 1297600 |
| AKVTGPRL | FLNC | Q14315 | FLNC_HUMAN | Unmodified | 0 | 0 | 0 | 0 | 2636700 | 0 | 0 | 0 | 0 | 0 | 0 | 0 | 0 | 0 | 0 | 0 | 2465300 | 0 |
| AKVTGPRL | FLNA; FLNA; FL | P21333; A0A | FLNA_HUMAN | Unmodified | 0 | 0 | 0 | 0 | 4609700 | 366060 | 0 | 0 | 0 | 110740 | 0 | 0 | 328570 | 338700 | 0 | 0 | 16131000 | 1496600 |
| AKVTLNGV | ICAM1; ICAM1 | P05362; E7E | ICAM1_HUMAN | Unmodified | 0 | 0 | 0 | 0 | 1297600 | 439050 | 0 | 0 | 0 | 0 | 0 | 0 | 0 | 0 | 0 | 0 | 446150 | 0 |
| AKWDAWN | ECI2; ECI2; ECI | O75521; F8V | ECI2_HUMAN E | Unmodified | 0 | 0 | 61560 | 0 | 0 | 0 | 494440 | 0 | 8614000 | 2947600 | 358860 | 0 | 211500 | 0 | 0 | 0 | 0 | 406510 |
| AKWDAWN | DBI; DBI; DBI | P07108; B8Z | ACBP_HUMAN | Unmodified | 953950 | 629510 | 2640300 | 686760 | 0 | 0 | 0 | 0 | 1496900 | 241120 | 1677100 | 2822000 | 94024 | 0 | 4331500 | 2671500 | 0 | 0 |
| AKWDTANI | ITGB3 | P05106 | ITB3_HUMAN I | Unmodified | 3940300 | 7096400 | 4930600 | 14463000 | 1820100 | 0 | 2106300 | 974170 | 0 | 0 | 0 | 0 | 3751200 | 1990900 | 6369200 | 6057500 | 174670 | 968820 |
| AKYDAIFDS | EPS15; EPS15 | P42566; B1A | EPS15_HUMAN | Unmodified | 326160 | 2343700 | 0 | 72931 | 175600 | 181790 | 55845 | 0 | 1263400 | 90224 | 0 | 62711 | 211210 | 153120 | 200270 | 160430 | 60878 | 103490 |
| AKYGEHEA | UTRN; UTRN | P46939; Q5 | UTRN_HUMAN | Acetyl (Protein N | 0 | 0 | 0 | 0 | 2421100 | 336030 | 0 | 0 | 0 | 460580 | 0 | 0 | 0 | 0 | 0 | 0 | 3956300 | 3174100 |
| AKYHGNVI | FASN; FASN | P49327; A0A | FAS_HUMAN F | Unmodified | 0 | 0 | 0 | 0 | 413810 | 0 | 0 | 0 | 0 | 0 | 0 | 0 | 0 | 0 | 0 | 0 | 0 | 0 |
| AKYPDYEV | PHPT1 | Q9NRX4 | PHP14_HUMAN | Unmodified | 0 | 0 | 0 | 145950 | 1477300 | 0 | 881990 | 224630 | 0 | 1153200 | 306200 | 1804300 | 1900500 | 305070 | 1431100 | 709950 | 1337200 | 0 |
| AKYPIVGYT | NENF | Q9UMX5 | NENF_HUMAN | Unmodified | 0 | 0 | 0 | 0 | 0 | 0 | 0 | 0 | 7212500 | 4738000 | 0 | 0 | 0 | 0 | 0 | 0 | 0 | 83556 |
| AKYPIVGYT | NENF | Q9UMX5 | NENF_HUMAN | Unmodified | 0 | 0 | 0 | 0 | 0 | 0 | 0 | 0 | 0 | 852210 | 0 | 0 | 0 | 0 | 0 | 0 | 0 | 0 |
| AKYPSLGQ | ENSA; ENSA; E | O43768; A0 | ENSA_HUMAN | Oxidation (M) | 0 | 0 | 297890 | 241600 | 1999100 | 0 | 110930 | 0 | 420640 | 492670 | 0 | 0 | 1144800 | 181400 | 0 | 0 | 1200400 | 2202400 |
| AKYPSLGQ | ENSA; ENSA; E | O43768; A0 | ENSA_HUMAN | Unmodified | 0 | 0 | 0 | 341090 | 1617300 | 0 | 0 | 0 | 0 | 0 | 0 | 0 | 814260 | 751780 | 0 | 0 | 698290 | 1810800 |
| ALAAAGYD | H1-3; H1-4; H1 | P16402; P10 | H13_HUMAN H | Unmodified | 37169000 | 19785000 | 417390 | 0 | 10208000 | 2555700 | 9677300 | 3.92E+08 | 1113200 | 684710 | 258310 | 1672600 | 0 | 0 | 215480 | 0 | 0 | 1851100 |
| ALAAAGYD | H1-3; H1-4; H1 | P16402; P10 | H13_HUMAN H | Unmodified | 0 | 0 | 0 | 0 | 138590 | 103270 | 933200 | 17735000 | 90176 | 41499 | 0 | 0 | 0 | 0 | 0 | 0 | 0 | 221620 |
| ALAAAGYD | H1-3; H1-4; H1 | P16402; P10 | H13_HUMAN H | Unmodified | 0 | 0 | 0 | 0 | 1642400 | 108060 | 6414000 | 2.99E+08 | 0 | 21115 | 0 | 0 | 400590 | 0 | 0 | 0 | 672900 | 1041900 |
| ALAAGGVG | ARPC5 | O15511 | ARPC5_HUMAN | Unmodified | 1592300 | 1359400 | 797370 | 1076000 | 2712800 | 528230 | 0 | 0 | 1446200 | 836040 | 1228200 | 22074000 | 3477600 | 1330800 | 903790 | 722850 | 567110 | 774090 |
| ALAAGGYD | H1-5 | P16401 | H15_HUMAN H | Unmodified | 40636000 | 25765000 | 49023 | 0 | 1528200 | 285960 | 1300600 | 35663000 | 276860 | 0 | 0 | 1456100 | 489220 | 0 | 0 | 0 | 92777 | 0 |
| ALAAGGYD | H1-5 | P16401 | H15_HUMAN H | Unmodified | 102600 | 0 | 0 | 164780 | 856810 | 788310 | 3852800 | 1867100 | 219760 | 1169800 | 1862000 | 0 | 358930 | 10492000 | 0 | 0 | 67202 | 0 |
| ALAAGGYD | H1-5 | P16401 | H15_HUMAN H | Unmodified | 0 | 0 | 0 | 0 | 239770 | 0 | 676990 | 37248000 | 0 | 49972 | 0 | 0 | 128330 | 0 | 0 | 0 | 22512 | 0 |
| ALAAKPGLI | NAGLU | P54802 | ANAG_HUMAN | Unmodified | 0 | 0 | 0 | 0 | 0 | 0 | 53287 | 0 | 1557600 | 0 | 0 | 0 | 0 | 0 | 0 | 0 | 0 | 0 |
| ALAAPAAEE | SLC7A5 | Q01650 | LAT1_HUMAN L | Unmodified | 4448500 | 1850800 | 7607500 | 1182200 | 0 | 0 | 98956 | 0 | 1768100 | 0 | 1803800 | 38371000 | 0 | 0 | 3597400 | 4566500 | 0 | 0 |
| ALAAPAAEE | SLC7A5 | Q01650 | LAT1_HUMAN L | Unmodified | 731200 | 0 | 0 | 0 | 162660 | 40524 | 0 | 0 | 1105300 | 0 | 111000 | 3948800 | 0 | 0 | 537110 | 0 | 484360 | 30740 |
| ALAAPAAEE | SLC7A5 | Q01650 | LAT1_HUMAN L | Unmodified | 16526000 | 14338000 | 3238900 | 7755700 | 15385000 | 5456100 | 3040300 | 1760200 | 8744700 | 4407200 | 16813000 | 67292000 | 17831000 | 3779600 | 7563000 | 7691700 | 3933000 | 2434800 |
| ALAAQNIVE | UAP1 | Q16222 | UAP1_HUMAN | Unmodified | 0 | 0 | 77347 | 346260 | 0 | 0 | 0 | 523210 | 0 | 0 | 70341 | 353080 | 0 | 0 | 0 | 0 | 0 | 0 |
| ALAAVPSA | SCRIB; SCRIB; | Q14160; A0 | SCRIB_HUMAN | Unmodified | 296110 | 129960 | 1392100 | 1584800 | 1694700 | 146410 | 0 | 0 | 7246100 | 2899600 | 244860 | 976040 | 1950900 | 498600 | 1048000 | 438110 | 2273400 | 8049200 |
| ALADENEF | GCN1 | Q92616 | GCN1_HUMAN | Unmodified | 115320 | 80945 | 0 | 0 | 0 | 0 | 0 | 0 | 0 | 0 | 0 | 1242700 | 0 | 0 | 508290 | 225830 | 0 | 0 |
| ALADLVHS | FECH; FECH | P22830; A0A | HEMH_HUMAN | Unmodified | 0 | 0 | 0 | 0 | 0 | 0 | 0 | 0 | 1544700 | 302090 | 0 | 0 | 0 | 220610 | 0 | 0 | 84577 | 0 |
| ALADVATVI | NCSTN; NCSTN | Q92542; H0 | NICA_HUMAN | Unmodified | 357580 | 346050 | 727010 | 487370 | 2946900 | 740680 | 0 | 0 | 16728000 | 7174100 | 1845800 | 13774000 | 0 | 0 | 1187200 | 832280 | 0 | 0 |
| ALADYIR | ERP44 | Q9BS26 | ERP44_HUMAN | Unmodified | 92480 | 0 | 84724 | 0 | 0 | 0 | 0 | 0 | 50583000 | 3817200 | 1956900 | 2928800 | 0 | 0 | 0 | 306290 | 0 | 0 |
| ALADYIRQ | ERP44 | Q9BS26 | ERP44_HUMAN | Unmodified | 0 | 0 | 0 | 0 | 333490 | 137830 | 115670 | 1091800 | 1092400 | 20466000 | 153670 | 0 | 522340 | 621900 | 0 | 0 | 0 | 716300 |
| ALAEENRN | MGAT5 | Q09328 | MGT5A_HUMA | Oxidation (M) | 0 | 0 | 0 | 0 | 0 | 0 | 0 | 0 | 1005700 | 362010 | 0 | 0 | 0 | 0 | 0 | 0 | 0 | 0 |
| ALAEEWGC | RAP2A | P10114 | RAP2A_HUMAN | Oxidation (M) | 139360 | 0 | 0 | 0 | 0 | 0 | 0 | 28880 | 404110 | 0 | 172460 | 567530 | 0 | 0 | 0 | 0 | 563740 | 0 |
| ALAEEWSC | RAP2B | P61225 | RAP2B_HUMAN | Oxidation (M) | 217530 | 214160 | 2159800 | 3554100 | 781670 | 509490 | 431350 | 438190 | 207840 | 361720 | 778460 | 875450 | 1079700 | 843290 | 1703000 | 614060 | 2514900 | 1728700 |
| ALAEEWSC | RAP2B | P61225 | RAP2B_HUMAN | Unmodified | 0 | 0 | 61011 | 1178000 | 625250 | 88141 | 105310 | 0 | 282790 | 0 | 0 | 62631 | 154160 | 477470 | 115870 | 0 | 1658400 | 4279600 |
| ALAEGGSIL | LAMB2 | P55268 | LAMB2_HUMAN | Unmodified | 0 | 276820 | 261030 | 102840 | 0 | 0 | 0 | 0 | 841570 | 0 | 0 | 0 | 0 | 0 | 0 | 0 | 0 | 0 |
| ALAEGVLLF | GSS; GSS; GSS | P48637; A0A | GSHB_HUMAN | Unmodified | 300440 | 261320 | 0 | 335770 | 0 | 0 | 29162 | 25671 | 104430 | 0 | 772050 | 6421600 | 0 | 0 | 727930 | 716000 | 0 | 0 |
| ALAEIAKAE | SFPQ | P23246 | SFPQ_HUMAN | Unmodified | 0 | 0 | 0 | 0 | 0 | 0 | 0 | 0 | 901480 | 0 | 202550 | 1730500 | 0 | 0 | 0 | 0 | 0 | 0 |
| ALAEIAKAE | SFPQ | P23246 | SFPQ_HUMAN | Oxidation (M) | 0 | 0 | 0 | 0 | 404260 | 0 | 0 | 0 | 0 | 0 | 165810 | 4399200 | 0 | 0 | 0 | 0 | 0 | 0 |
| ALAENSGV | CCT8; CCT8 | P50990; H7 | TCPQ_HUMAN | Unmodified | 76101 | 0 | 120300 | 181680 | 0 | 0 | 153140 | 76199 | 415010 | 169560 | 0 | 3600000 | 1008100 | 194940 | 123390 | 189450 | 0 | 0 |
| ALAENSGV | CCT8; CCT8 | P50990; H7 | TCPQ_HUMAN | Unmodified | 0 | 0 | 0 | 0 | 0 | 0 | 112670 | 0 | 0 | 112260 | 0 | 0 | 263060 | 303460 | 0 | 0 | 0 | 311630 |
| ALAERFHIS | PDIA5 | Q14554 | PDIA5_HUMAN | Unmodified | 0 | 0 | 0 | 0 | 0 | 0 | 0 | 0 | 11381000 | 13923000 | 119830 | 0 | 0 | 285120 | 0 | 0 | 0 | 186140 |
| ALAESWNA | RHEB; RHEB | Q15382; C9J | RHEB_HUMAN | Unmodified | 3350300 | 1428200 | 2424400 | 1544700 | 3397700 | 2525700 | 0 | 1114900 | 3427000 | 817140 | 741190 | 3247900 | 882060 | 459550 | 3564400 | 2030700 | 390680 | 4152500 |
| ALAETSYVK | MAGEA4; MAGE | P43358; P43 | MAGA4_HUMA | Unmodified | 535150 | 912560 | 0 | 0 | 0 | 0 | 0 | 0 | 0 | 0 | 0 | 0 | 0 | 0 | 0 | 340430 | 0 | 0 |
| ALAFHPIEP | STRN | O43815 | STRN_HUMAN | Unmodified | 0 | 0 | 0 | 0 | 16303000 | 3324400 | 1286300 | 0 | 524210 | 406850 | 0 | 1667800 | 6909300 | 8407600 | 0 | 0 | 7434400 | 8733900 |
| ALAFHPVEI | STRN3 | Q13033 | STRN3_HUMAN | Unmodified | 0 | 0 | 0 | 0 | 0 | 412340 | 0 | 0 | 934880 | 0 | 365780 | 3502700 | 0 | 0 | 0 | 0 | 0 | 0 |
| ALAGCDFL | TALDO1; TALDO | P37837; F2Z | TALDO_HUMAN | Unmodified | 0 | 0 | 0 | 0 | 0 | 276020 | 0 | 14920 | 0 | 0 | 231860 | 1607600 | 452580 | 172650 | 0 | 0 | 0 | 382720 |
| ALAGDVRD | QSOX2 | Q6ZRP7 | QSOX2_HUMA | Unmodified | 0 | 0 | 104650 | 102620 | 0 | 0 | 0 | 0 | 54458000 | 2312800 | 424890 | 0 | 0 | 0 | 0 | 0 | 0 | 0 |
| ALAGVTYA | BAIAP2; BAIAP2 | Q9UQB8; I3 | BAIP2_HUMAN | Unmodified | 243160 | 167050 | 946540 | 949140 | 4216100 | 1221200 | 477580 | 50759 | 241740 | 367070 | 180420 | 1888700 | 2963500 | 3329500 | 1417500 | 1554900 | 1087100 | 847330 |
| ALAHLLEAE | PCSK1N | Q9UHG2 | PCS1N_HUMA | Unmodified | 0 | 0 | 0 | 0 | 0 | 0 | 0 | 0 | 16227000 | 713770 | 0 | 0 | 0 | 0 | 0 | 0 | 0 | 0 |
| ALALECER | UACA; UACA; U | Q9BZF9; H0 | UACA_HUMAN | Unmodified | 0 | 0 | 0 | 90454 | 0 | 0 | 0 | 0 | 0 | 0 | 0 | 0 | 0 | 0 | 0 | 64600 | 373730 | 370540 |
| ALALSFHG | TOR3A; TOR3A | Q9H497; Q5 | TOR3A_HUMAN | Unmodified | 0 | 0 | 0 | 0 | 0 | 0 | 0 | 0 | 647060 | 231130 | 0 | 0 | 0 | 0 | 0 | 0 | 0 | 45858 |
| ALAMAESTI | HUWE1 | Q7Z6Z7 | HUWE1_HUMA | Unmodified | 0 | 0 | 0 | 0 | 0 | 0 | 74405 | 0 | 126230 | 426190 | 47668 | 940020 | 0 | 53350 | 0 | 0 | 0 | 0 |
| ALANMHES | EXOC4 | Q96A65 | EXOC4_HUMA | Unmodified | 0 | 0 | 0 | 0 | 0 | 0 | 0 | 0 | 312990 | 115570 | 0 | 0 | 0 | 0 | 0 | 0 | 0 | 0 |

| | | | | | | | | | | | | | | | | | | | | | | |
|---|---|---|---|---|---|---|---|---|---|---|---|---|---|---|---|---|---|---|---|---|---|---|
| ALANSLAC | ALDOA; ALDOA | P04075; J3KI | ALDOA_HUMAI | Unmodified | 0 | 0 | 0 | 0 | 0 | 0 | 0 | 0 | 0 | 0 | 0 | 0 | 0 | 0 | 0 | 0 | 2534500 | 0 |
| ALANVNIGS | RPLP1 | P05386 | RLA1_HUMAN ( | Unmodified | 6887900 | 5381600 | 2149400 | 0 | 1312300 | 0 | 0 | 0 | 0 | 0 | 0 | 6387400 | 3398300 | 0 | 5084100 | 3923900 | 1051400 | 0 |
| ALAPEYAK | P4HB; P4HB; P4 | P07237; A0A | PDIA1_HUMAN | Unmodified | 0 | 304840 | 608080 | 0 | 2847100 | 0 | 13061000 | 9235700 | 79520000 | 59091000 | 4225100 | 2244100 | 3906500 | 4912700 | 1024100 | 0 | 2668100 | 1995400 |
| ALAPLLLAF | RANGAP1 | P46060 | RAGP1_HUMAI | Unmodified | 0 | 28080 | 0 | 0 | 0 | 0 | 0 | 0 | 448060 | 0 | 64225 | 779580 | 1196700 | 133400 | 56088 | 38060 | 0 | 0 |
| ALAPMYYR | RAB22A | Q9UL26 | RB22A_HUMAN | Oxidation (M) | 291510 | 286390 | 0 | 0 | 0 | 0 | 0 | 0 | 0 | 0 | 0 | 263390 | 0 | 0 | 565350 | 329520 | 0 | 0 |
| ALAPTWEQ | TXNDC5 | Q8NBS9 | TXND5_HUMAI | Unmodified | 0 | 0 | 233940 | 0 | 5220200 | 1113900 | 3363100 | 0 | 2.17E+08 | 46449000 | 5249900 | 2137400 | 2903200 | 2115900 | 801120 | 614850 | 176150 | 2589000 |
| ALAQAPAP | SARG | Q9BW04 | SARG_HUMAN | Oxidation (M) | 0 | 0 | 0 | 589240 | 0 | 0 | 0 | 0 | 0 | 0 | 0 | 0 | 1190600 | 912620 | 0 | 0 | 0 | 0 |
| ALAQAPAP | SARG | Q9BW04 | SARG_HUMAN | Oxidation (M) | 0 | 0 | 0 | 0 | 0 | 0 | 0 | 0 | 0 | 0 | 0 | 0 | 2853700 | 2234700 | 0 | 0 | 0 | 0 |
| ALAQAPAP | SARG | Q9BW04 | SARG_HUMAN | Unmodified | 0 | 0 | 0 | 0 | 0 | 0 | 0 | 0 | 0 | 0 | 0 | 0 | 4443200 | 6624200 | 0 | 0 | 0 | 0 |
| ALAQEDQC | SPTBN2 | O15020 | SPTN2_HUMAN | Unmodified | 0 | 0 | 0 | 0 | 0 | 0 | 0 | 0 | 0 | 0 | 0 | 0 | 0 | 235580 | 216930 | 0 | 0 | 0 |
| ALAQEDQC | SPTBN2 | O15020 | SPTN2_HUMAN | Unmodified | 0 | 0 | 0 | 0 | 0 | 0 | 204560 | 338120 | 0 | 0 | 0 | 0 | 707980 | 833910 | 0 | 0 | 560500 | 281210 |
| ALAQEWG | RAP2C; RAP2C | Q9Y3L5; A0A | RAP2C_HUMAI | Unmodified | 0 | 0 | 0 | 0 | 0 | 0 | 0 | 0 | 393090 | 0 | 571050 | 131540 | 106670 | 0 | 44607 | 0 | 0 | 0 |
| ALAQEWG | RAP2C; RAP2C | Q9Y3L5; A0A | RAP2C_HUMAI | Oxidation (M) | 583500 | 390030 | 0 | 0 | 0 | 0 | 0 | 16344 | 350860 | 0 | 300880 | 2862000 | 199580 | 200110 | 424760 | 230190 | 153870 | 111370 |
| ALAQREFY | CEMIP2; CEMIP | Q9UHN6; F5 | CEIP2_HUMAN | Unmodified | 265600 | 0 | 0 | 558510 | 3472800 | 1544800 | 283560 | 0 | 4798700 | 2637800 | 0 | 0 | 0 | 0 | 0 | 0 | 0 | 0 |
| ALAREVDLI | FGA; FGA | P02671; A0A | FIBA_HUMAN F | Unmodified | 0 | 0 | 0 | 0 | 0 | 0 | 0 | 0 | 0 | 0 | 0 | 0 | 0 | 828310 | 0 | 0 | 0 | 0 |
| ALASIDSK | VCL | P18206 | VINC_HUMAN | Unmodified | 0 | 0 | 0 | 0 | 0 | 0 | 0 | 0 | 0 | 0 | 0 | 1082100 | 0 | 0 | 0 | 0 | 0 | 0 |
| ALASIDSKL | VCL | P18206 | VINC_HUMAN | Unmodified | 0 | 0 | 0 | 0 | 0 | 282470 | 0 | 443850 | 0 | 0 | 48631 | 1134700 | 174940 | 0 | 0 | 0 | 3222900 | 0 |
| ALASLGKPI | ELAC2; ELAC2; | Q9BQ52; E7I | RNZ2_HUMAN | Unmodified | 0 | 0 | 0 | 0 | 0 | 0 | 0 | 0 | 0 | 0 | 0 | 177090 | 0 | 0 | 0 | 0 | 0 | 0 |
| ALASQLQD | VCL | P18206 | VINC_HUMAN | Unmodified | 1031500 | 1650700 | 484680 | 0 | 0 | 0 | 0 | 0 | 4109700 | 0 | 0 | 3320500 | 0 | 0 | 1102200 | 1888400 | 662620 | 0 |
| ALASQLQD | VCL | P18206 | VINC_HUMAN | Unmodified | 0 | 0 | 0 | 0 | 305470 | 0 | 0 | 0 | 0 | 230380 | 0 | 0 | 0 | 0 | 0 | 0 | 1391300 | 0 |
| ALASTCKPI | NIBAN1; NIBAN | Q9BZQ8; H0 | NIBA1_HUMAN | Unmodified | 0 | 0 | 320590 | 483710 | 241360 | 0 | 208770 | 0 | 2208300 | 0 | 0 | 0 | 0 | 0 | 235630 | 290570 | 24444 | 25406 |
| ALAVGGLG | ARPC5L | Q9BPX5 | ARP5L_HUMAN | Unmodified | 611010 | 0 | 1176100 | 1355700 | 2079100 | 0 | 0 | 0 | 301530 | 979810 | 0 | 0 | 0 | 0 | 2233900 | 0 | 0 | 0 |
| ALAVNPRD | TLN1 | Q9Y490 | TLN1_HUMAN | Unmodified | 0 | 0 | 0 | 0 | 4819400 | 3095500 | 24138 | 0 | 556060 | 1421200 | 0 | 2462500 | 2495000 | 0 | 0 | 0 | 855810 | 4243900 |
| ALAVNPRD | TLN1 | Q9Y490 | TLN1_HUMAN | Unmodified | 0 | 0 | 0 | 0 | 8588100 | 574950 | 208220 | 0 | 0 | 230370 | 0 | 0 | 1166300 | 703060 | 0 | 0 | 2196500 | 2262700 |
| ALCADLSPI | XPNPEP1; XPNI | Q9NQW7; Q | XPP1_HUMAN | Unmodified | 0 | 0 | 0 | 0 | 287940 | 0 | 143060 | 103370 | 0 | 0 | 165780 | 0 | 443830 | 177100 | 0 | 0 | 267810 | 203150 |
| ALCATRQE | FASN; FASN | P49327; A0A | FAS_HUMAN F | Unmodified | 480910 | 0 | 1691800 | 1873700 | 1779700 | 615870 | 677520 | 0 | 0 | 0 | 1692700 | 25037000 | 3097200 | 1302900 | 1825500 | 1051500 | 1058000 | 1047700 |
| ALCDNVDT | CARS1; CARS1 | P49589; B4D | SYCC_HUMAN | Unmodified | 0 | 0 | 0 | 0 | 0 | 0 | 0 | 0 | 0 | 0 | 0 | 0 | 0 | 0 | 0 | 0 | 0 | 0 |
| ALCDPLEE | GCN1 | Q92616 | GCN1_HUMAN | Unmodified | 0 | 0 | 65922 | 0 | 0 | 0 | 0 | 75982 | 0 | 0 | 32967 | 549820 | 0 | 0 | 0 | 57292 | 87251 | 0 |
| ALCGLDES | LONP1; LONP1 | P36776; K7E | LONM_HUMAN | Unmodified | 0 | 0 | 14149 | 0 | 30978 | 0 | 0 | 0 | 1621900 | 265650 | 308930 | 0 | 0 | 0 | 0 | 0 | 0 | 0 |
| ALCLFEMP | AOC1 | P19801 | AOC1_HUMAN | Unmodified | 0 | 0 | 0 | 0 | 0 | 0 | 0 | 0 | 1367200 | 0 | 0 | 146350 | 0 | 0 | 0 | 0 | 0 | 0 |
| ALCPDHDS | SLC39A6 | Q13433 | S39A6_HUMAN | Unmodified | 0 | 0 | 0 | 0 | 715990 | 0 | 0 | 0 | 0 | 35980 | 0 | 0 | 0 | 0 | 0 | 0 | 769800 | 2161900 |
| ALCQELKE | DLG5 | Q8TDM6 | DLG5_HUMAN | Unmodified | 455080 | 0 | 0 | 0 | 0 | 0 | 0 | 0 | 0 | 0 | 391340 | 0 | 1328800 | 0 | 0 | 0 | 904610 | 1027600 |
| ALDAAFYK | RO60; RO60 | P10155; G5E | RO60_HUMAN | Unmodified | 26055000 | 20466000 | 7330200 | 7654700 | 88590 | 0 | 719610 | 395620 | 390580 | 110200 | 249760 | 3747700 | 1574500 | 889240 | 14757000 | 8905300 | 58903 | 42208 |
| ALDDISESI | REXO2; REXO2; | Q9Y3B8; F5C | ORN_HUMAN ( | Unmodified | 0 | 326750 | 0 | 464310 | 1120100 | 0 | 0 | 0 | 667780 | 288350 | 1334600 | 3374500 | 2700600 | 440200 | 755090 | 690240 | 402280 | 631260 |
| ALDDMISTI | CCT3; CCT3; C | P49368; Q5S | TCPG_HUMAN | Unmodified | 0 | 0 | 0 | 0 | 0 | 0 | 869090 | 243800 | 434470 | 250750 | 595310 | 4894200 | 1389700 | 419960 | 0 | 0 | 0 | 0 |
| ALDDMISTI | CCT3; CCT3; C | P49368; Q5S | TCPG_HUMAN | Oxidation (M) | 0 | 391850 | 0 | 0 | 631330 | 0 | 474430 | 129760 | 0 | 583500 | 0 | 4791600 | 411820 | 558530 | 0 | 0 | 774730 | 0 |
| ALDDMISTI | CCT3; CCT3; C | P49368; Q5S | TCPG_HUMAN | Unmodified | 0 | 0 | 0 | 0 | 877440 | 0 | 0 | 0 | 566840 | 736940 | 1839000 | 4806800 | 1021400 | 0 | 0 | 0 | 653370 | 460720 |
| ALDDMISTI | CCT3; CCT3; C | P49368; Q5S | TCPG_HUMAN | Oxidation (M) | 0 | 0 | 0 | 0 | 0 | 0 | 0 | 0 | 0 | 0 | 0 | 0 | 0 | 0 | 0 | 0 | 0 | 0 |
| ALDEGDIAI | PSMC2; PSMC2 | P35998; C9J | PRS7_HUMAN | Unmodified | 0 | 0 | 0 | 0 | 118860 | 0 | 0 | 144410 | 1132600 | 0 | 667130 | 3537400 | 0 | 0 | 553400 | 0 | 0 | 0 |
| ALDELFEAI | INF2; INF2; INF | Q27J81; A0A | INF2_HUMAN I | Unmodified | 0 | 0 | 0 | 0 | 0 | 0 | 0 | 0 | 449040 | 0 | 230910 | 2081200 | 0 | 0 | 0 | 0 | 0 | 0 |
| ALDENAKL | SHTN1 | A0MZ66 | SHOT1_HUMAI | Unmodified | 0 | 0 | 0 | 0 | 3237300 | 239730 | 882400 | 506310 | 0 | 1366100 | 0 | 0 | 0 | 0 | 0 | 0 | 982020 | 0 |
| ALDEYYDK | ATP6V1A | P38606 | VATA_HUMAN | Unmodified | 161790 | 227430 | 262180 | 0 | 0 | 0 | 0 | 0 | 611430 | 0 | 361820 | 3851900 | 53569 | 0 | 370010 | 355210 | 0 | 0 |
| ALDFAVGE | CST3 | P01034 | CYTC_HUMAN | Unmodified | 0 | 0 | 874510 | 325510 | 0 | 310500 | 0 | 0 | 1812200 | 3583200 | 0 | 0 | 0 | 0 | 278460 | 153220 | 256760 | 0 |
| ALDFIASK | ACTN4; ACTN1 | O43707; P12 | ACTN4_HUMAI | Unmodified | 1094600 | 0 | 1776200 | 1512400 | 15600000 | 6211200 | 1649700 | 1547800 | 4484700 | 1698300 | 10034000 | 33604000 | 4187000 | 4209100 | 2512700 | 3074600 | 5268300 | 3993600 |
| ALDGAFTEI | TLN1 | Q9Y490 | TLN1_HUMAN | Unmodified | 281800 | 318850 | 883070 | 2044800 | 3846900 | 924690 | 1509800 | 843210 | 585810 | 429960 | 233190 | 1866500 | 3859400 | 2586300 | 997290 | 829420 | 1256600 | 2598900 |
| ALDGDFSE | TLN2; TLN2; TLI | Q9Y4G6; H0 | TLN2_HUMAN | Unmodified | 0 | 0 | 0 | 0 | 0 | 0 | 0 | 44809 | 0 | 53933 | 0 | 404470 | 0 | 0 | 0 | 0 | 528490 | 556500 |
| ALDGDKAF | QSOX2 | Q6ZRP7 | QSOX2_HUMAI | Unmodified | 0 | 0 | 0 | 0 | 0 | 0 | 0 | 0 | 1193100 | 1769900 | 0 | 0 | 0 | 0 | 0 | 0 | 0 | 0 |
| ALDGLWNI | H6PD | O95479 | G6PE_HUMAN | Unmodified | 0 | 0 | 0 | 0 | 0 | 0 | 0 | 0 | 1081300 | 0 | 0 | 0 | 0 | 0 | 0 | 0 | 0 | 0 |
| ALDIAENEI | AHCY | P23526 | SAHH_HUMAN | Oxidation (M) | 124670 | 0 | 0 | 0 | 1382400 | 0 | 0 | 45979 | 0 | 313950 | 400930 | 3965600 | 0 | 0 | 0 | 0 | 1179600 | 1569100 |
| ALDIAENEI | AHCY | P23526 | SAHH_HUMAN | Unmodified | 0 | 0 | 0 | 0 | 1820500 | 0 | 0 | 0 | 0 | 0 | 0 | 0 | 0 | 0 | 0 | 0 | 741140 | 0 |
| ALDIAENEI | AHCY | P23526 | SAHH_HUMAN | 2 Oxidation (M) | 0 | 0 | 191420 | 0 | 57377 | 0 | 0 | 0 | 0 | 0 | 336180 | 5478000 | 141540 | 0 | 497580 | 247120 | 407520 | 0 |
| ALDIDLDKI | AP3D1; AP3D1 | O14617; A0A | AP3D1_HUMAN | Unmodified | 0 | 0 | 0 | 0 | 0 | 0 | 112640 | 0 | 2124800 | 0 | 0 | 4271100 | 150910 | 0 | 170780 | 54201 | 0 | 0 |
| ALDIDSDEI | ARHGEF26 | Q96DR7 | ARHGQ_HUMA | Unmodified | 0 | 0 | 0 | 0 | 455300 | 0 | 0 | 0 | 0 | 0 | 0 | 0 | 0 | 0 | 0 | 20795 | 187260 | 0 |
| ALDIPLVK | PRPF8; PRPF8 | Q6P2Q9; I3L | PRP8_HUMAN | Unmodified | 0 | 0 | 28059 | 0 | 0 | 0 | 0 | 0 | 0 | 0 | 0 | 719070 | 0 | 0 | 0 | 0 | 0 | 0 |
| ALDKKLEDI | STXBP2; STXBP | Q15833; R4C | STXB2_HUMAN | Unmodified | 0 | 0 | 0 | 0 | 251120 | 0 | 0 | 0 | 0 | 72630 | 0 | 0 | 0 | 237040 | 0 | 0 | 1071100 | 803560 |
| ALDLDSSC | STIP1; STIP1 | P31948; H0Y | STIP1_HUMAN | Unmodified | 0 | 0 | 240800 | 551600 | 2785800 | 713780 | 200530 | 0 | 1294700 | 843090 | 1322500 | 8489800 | 4037200 | 0 | 273830 | 380560 | 2734500 | 3376900 |
| ALDLDSSC | STIP1; STIP1 | P31948; H0Y | STIP1_HUMAN | Unmodified | 0 | 123730 | 0 | 0 | 569600 | 349480 | 35280 | 0 | 901740 | 684220 | 262470 | 0 | 2973400 | 1302300 | 0 | 0 | 2735800 | 3380800 |
| ALDLFSDN | PDIA6 | Q15084 | PDIA6_HUMAN | Unmodified | 0 | 0 | 0 | 38333 | 0 | 0 | 0 | 0 | 1173600 | 909040 | 30338 | 0 | 19628 | 0 | 276260 | 0 | 0 | 0 |
| ALDLVSDPI | MINDY3; MIND | Q9H8M7; X6 | MINY3_HUMAN | Oxidation (M) | 659000 | 1535900 | 175210 | 131470 | 0 | 35277 | 0 | 0 | 0 | 0 | 0 | 0 | 0 | 0 | 605080 | 241700 | 0 | 0 |
| ALDLVSDPI | MINDY3; MIND | Q9H8M7; X6 | MINY3_HUMAN | Unmodified | 0 | 71603 | 0 | 0 | 0 | 0 | 0 | 0 | 0 | 0 | 0 | 0 | 0 | 0 | 0 | 24774 | 0 | 0 |
| ALDQASEE | METAP2; METAI | P50579; F8V | MAP2_HUMAN | Unmodified | 337260 | 0 | 2835700 | 0 | 0 | 0 | 0 | 0 | 0 | 0 | 0 | 2556300 | 0 | 0 | 0 | 0 | 0 | 0 |
| ALDQFVNF | SMC3 | Q9UQE7 | SMC3_HUMAN | Unmodified | 135710 | 0 | 0 | 0 | 174730 | 0 | 293950 | 41944 | 95504 | 31692 | 36877 | 193960 | 115030 | 0 | 63229 | 0 | 0 | 0 |
| ALDRCSEG | NONO; NONO; | Q15233; A0A | NONO_HUMAN | Oxidation (M) | 0 | 0 | 0 | 0 | 0 | 0 | 0 | 0 | 0 | 3047700 | 0 | 524960 | 0 | 0 | 0 | 0 | 0 | 0 |
| ALDSEVDP | SIGIRR; SIGIRR | Q6IA17; H0Y | SIGIR_HUMAN | Unmodified | 744670 | 1117000 | 2033800 | 2896500 | 0 | 0 | 0 | 0 | 4788300 | 378910 | 0 | 1500900 | 0 | 0 | 1909500 | 1832900 | 0 | 0 |
| ALDSVVSD | DAGLB; DAGLB | Q8NCG7; E7 | DGLB_HUMAN | Unmodified | 0 | 742470 | 0 | 0 | 449270 | 0 | 10226 | 0 | 513140 | 0 | 498500 | 537000 | 171590 | 0 | 0 | 0 | 365670 | 220650 |
| ALDTMNFD | PABPC4; PABP( | Q13310; P11 | PABP4_HUMAN | Oxidation (M) | 57117000 | 38784000 | 20203000 | 15155000 | 0 | 0 | 0 | 0 | 0 | 0 | 292770 | 6669100 | 105300 | 0 | 11010000 | 22360000 | 0 | 0 |
| ALDTMNFD | PABPC4; PABP( | Q13310; P11 | PABP4_HUMAN | Unmodified | 11927000 | 12491000 | 2255600 | 2584400 | 0 | 0 | 0 | 0 | 0 | 0 | 0 | 534800 | 0 | 0 | 2788300 | 2596100 | 0 | 0 |
| ALDTVLFGI | STIM1; STIM1; S | Q13586; E9F | STIM1_HUMAN | Unmodified | 0 | 0 | 31393 | 0 | 0 | 0 | 0 | 30704 | 5520100 | 0 | 0 | 208770 | 0 | 0 | 0 | 25861 | 0 | 0 |
| ALDVGSGS | PCMT1; PCMT1 | P22061; C9J | PIMT_HUMAN I | Unmodified | 0 | 0 | 0 | 0 | 531630 | 0 | 0 | 89444 | 3762100 | 1614500 | 541420 | 2934500 | 0 | 0 | 0 | 290170 | 868400 | 766550 |
| ALDVMVST | S100A4 | P26447 | S10A4_HUMAN | Oxidation (M) | 439160 | 419170 | 480400 | 438160 | 1405100 | 376310 | 1005900 | 688420 | 0 | 412180 | 0 | 577500 | 418210 | 499200 | 760430 | 392580 | 609960 | 450730 |
| ALDVMVST | S100A4 | P26447 | S10A4_HUMAN | Unmodified | 0 | 0 | 0 | 109940 | 350890 | 63975 | 579160 | 111030 | 0 | 0 | 0 | 212000 | 195850 | 356770 | 0 | 0 | 0 | 360490 |
| ALDVSASD | APEH; APEH; AI | P13798; C9J | ACPH_HUMAN | Unmodified | 0 | 0 | 0 | 0 | 249120 | 0 | 95846 | 0 | 597130 | 118130 | 1591300 | 5473700 | 687910 | 230680 | 0 | 0 | 330160 | 0 |
| ALDVSASD | APEH; APEH; AI | P13798; C9J | ACPH_HUMAN | Unmodified | 0 | 0 | 377970 | 603720 | 801310 | 0 | 0 | 0 | 162050 | 129920 | 0 | 2652700 | 1173700 | 510760 | 880650 | 629060 | 537700 | 1510300 |
| ALEAFETFK | SORD | Q00796 | DHSO_HUMAN | Unmodified | 0 | 0 | 0 | 517270 | 0 | 0 | 0 | 0 | 0 | 0 | 0 | 1017500 | 515920 | 330210 | 540280 | 425160 | 0 | 0 |
| ALEAFETFK | SORD | Q00796 | DHSO_HUMAN | Unmodified | 0 | 0 | 0 | 439820 | 0 | 0 | 0 | 0 | 0 | 140030 | 228110 | 2813900 | 1551100 | 1065000 | 1125300 | 914940 | 237230 | 0 |
| ALEAHKRA | PPP2R5D; PPP2 | Q14738; H0Y | 2A5D_HUMAN | Unmodified | 0 | 0 | 0 | 0 | 1044800 | 0 | 0 | 0 | 0 | 261280 | 0 | 0 | 1233800 | 0 | 0 | 0 | 0 | 0 |
| ALEALVAK | AKR1A1 | P14550 | AK1A1_HUMAN | Unmodified | 0 | 46793 | 463250 | 1240700 | 1672000 | 318760 | 371070 | 181530 | 3156000 | 3937000 | 247000 | 5019200 | 135510 | 1514600 | 1779600 | 450590 | 690010 | 898420 |
| ALEAQNEI | MYO5C | Q9NQX4 | MYO5C_HUMA | Unmodified | 505020 | 0 | 0 | 9676.8 | 166430 | 0 | 9587.2 | 92402 | 73304 | 38401 | 0 | 632650 | 0 | 9356 | 0 | 344020 | 0 | 0 |
| ALEATTEHI | TLN1 | Q9Y490 | TLN1_HUMAN | Unmodified | 944550 | 372590 | 718470 | 1400500 | 2050200 | 0 | 417830 | 0 | 917410 | 521800 | 0 | 0 | 906620 | 605960 | 0 | 515300 | 333390 | 1004500 |
| ALEAVFGK | RBMX; RBMX; R | P38159; H3E | RBMX_HUMAN | Unmodified | 152150 | 543190 | 0 | 485300 | 301820 | 0 | 168490 | 21538 | 276450 | 1371100 | 2722600 | 5156600 | 284950 | 0 | 0 | 54545 | 0 | 0 |
| ALEDAFLAI | PPM1G | O15355 | PPM1G_HUMA | Unmodified | 0 | 0 | 0 | 0 | 0 | 0 | 0 | 0 | 0 | 0 | 0 | 0 | 0 | 0 | 0 | 0 | 0 | 0 |
| ALEDDIINH | DST; DST; DST; | Q03001; A0A | DYST_HUMAN | Unmodified | 0 | 0 | 0 | 0 | 674120 | 0 | 0 | 0 | 0 | 522820 | 0 | 1079800 | 0 | 0 | 170110 | 257420 | 314090 | 0 |
| ALEDFADN | EXOC7; EXOC7 | Q9UPT5; B4I | EXOC7_HUMAI | Unmodified | 0 | 0 | 0 | 0 | 463020 | 311370 | 711820 | 159340 | 1637600 | 197600 | 1944900 | 0 | 0 | 0 | 0 | 52056 | 109180 | 114600 |
| ALEDMFDA | CPT2; CPT2; CF | P23786; A0A | CPT2_HUMAN | Unmodified | 0 | 0 | 0 | 0 | 722390 | 0 | 0 | 0 | 1807100 | 0 | 0 | 0 | 0 | 0 | 0 | 0 | 0 | 0 |
| ALEDMNRE | ZPR1; ZPR1 | O75312; H7I | ZPR1_HUMAN | Unmodified | 0 | 0 | 0 | 0 | 0 | 0 | 0 | 0 | 0 | 0 | 66471 | 411810 | 0 | 0 | 0 | 0 | 0 | 0 |
| ALEDSSFLI | LMO7; LMO7; LI | Q8WWI1; E9 | LMO7_HUMAN | Unmodified | 63704 | 0 | 0 | 0 | 0 | 0 | 0 | 0 | 835260 | 853770 | 0 | 0 | 0 | 0 | 0 | 0 | 1301800 | 991500 |
| ALEEALEAK | MYH10 | P35580 | MYH10_HUMAI | Unmodified | 0 | 0 | 265870 | 167800 | 0 | 0 | 0 | 0 | 0 | 0 | 155380 | 7736000 | 0 | 0 | 0 | 335940 | 0 | 0 |
| ALEEALEAK | MYH10 | P35580 | MYH10_HUMAI | Unmodified | 0 | 0 | 0 | 0 | 1023500 | 0 | 0 | 0 | 0 | 0 | 0 | 0 | 0 | 0 | 0 | 0 | 0 | 0 |
| ALEEAMEQ | MYH9 | P35579 | MYH9_HUMAN | Unmodified | 0 | 0 | 0 | 0 | 0 | 0 | 0 | 0 | 0 | 0 | 0 | 2749800 | 0 | 0 | 0 | 0 | 0 | 0 |
| ALEEAMEQ | MYH9 | P35579 | MYH9_HUMAN | Unmodified | 0 | 499120 | 0 | 483420 | 0 | 0 | 0 | 0 | 602140 | 0 | 107760 | 3359100 | 0 | 0 | 1100900 | 920120 | 0 | 0 |
| ALEEAMEQ | MYH9 | P35579 | MYH9_HUMAN | Oxidation (M) | 0 | 0 | 0 | 0 | 0 | 0 | 0 | 0 | 143360 | 0 | 0 | 1564200 | 0 | 0 | 0 | 0 | 0 | 0 |
| ALEEAMEQ | MYH9 | P35579 | MYH9_HUMAN | Unmodified | 0 | 0 | 0 | 1740100 | 33835000 | 6319600 | 1253600 | 243850 | 2244000 | 4824800 | 735040 | 1915900 | 16876000 | 9281800 | 0 | 351480 | 7991000 | 14376000 |
| ALEEAMEQ | MYH9 | P35579 | MYH9_HUMAN | Oxidation (M) | 149170 | 63101 | 82463 | 412270 | 2212700 | 1634400 | 247560 | 179670 | 0 | 2633900 | 0 | 0 | 2173700 | 939190 | 265610 | 243230 | 660140 | 1525300 |
| ALEEDIENI | MACF1; MACF1 | H3BQK9; A0 | H3BQK9_HUM | Unmodified | 0 | 0 | 0 | 290510 | 7529300 | 1664900 | 0 | 0 | 1813400 | 1691000 | 0 | 1945700 | 0 | 538990 | 596710 | 0 | 1550100 | 2835300 |
| ALEEDLNQ | STK10 | O94804 | STK10_HUMAN | Unmodified | 0 | 0 | 0 | 0 | 1152000 | 269140 | 212080 | 978660 | 852840 | 835390 | 427180 | 232690 | 0 | 596110 | 0 | 0 | 281500 | 917990 |
| ALEEERRH | CTNND1; CTNI | O60716; A0A | CTND1_HUMAI | Unmodified | 0 | 0 | 0 | 0 | 0 | 0 | 0 | 19615 | 0 | 2561100 | 0 | 0 | 0 | 0 | 0 | 0 | 639110 | 4491100 |
| ALEEKPNN | IFIT3; IFIT3; IFIT | O14879; A0A | IFIT3_HUMAN I | Unmodified | 0 | 0 | 0 | 0 | 0 | 0 | 428770 | 0 | 0 | 0 | 126460 | 0 | 0 | 0 | 0 | 0 | 0 | 4575600 |
| ALEELATK | TPPP3 | Q9BW30 | TPPP3_HUMAN | Unmodified | 358480 | 418370 | 49072 | 0 | 0 | 0 | 4041300 | 2399400 | 0 | 48556 | 0 | 0 | 0 | 0 | 0 | 0 | 0 | 0 |
| ALEENNNF | STMN2; STMN2 | Q93045; E5F | STMN2_HUMAI | Unmodified | 544330 | 714300 | 510260 | 619150 | 7960500 | 690190 | 5532800 | 4698800 | 10362000 | 11545000 | 4570000 | 18419000 | 2388500 | 1406200 | 441650 | 439760 | 2672100 | 5950400 |
| ALEENNNF | STMN2; STMN2 | Q93045; E5F | STMN2_HUMAI | Unmodified | 0 | 0 | 0 | 0 | 0 | 0 | 1129900 | 1064300 | 0 | 328600 | 1369300 | 0 | 0 | 0 | 0 | 0 | 0 | 0 |
| ALEENNNF | STMN3; STMN3 | Q9NZ72; A0A | STMN3_HUMAI | Unmodified | 53539 | 0 | 0 | 0 | 0 | 0 | 171040 | 52315 | 284830 | 0 | 0 | 0 | 0 | 0 | 0 | 0 | 0 | 0 |
| ALEENNNF | STMN3; STMN3 | Q9NZ72; A0A | STMN3_HUMAI | Unmodified | 0 | 0 | 0 | 0 | 0 | 0 | 6240800 | 1340800 | 0 | 310330 | 0 | 0 | 0 | 0 | 0 | 0 | 0 | 0 |
| ALEEPAND | ESAM | Q96AP7 | ESAM_HUMAN | Unmodified | 0 | 0 | 0 | 0 | 265100 | 0 | 367020 | 479280 | 0 | 0 | 0 | 0 | 0 | 1586600 | 0 | 0 | 228770 | 851830 |
| ALEEPAVDI | MX1; MX1; MX1 | P20591; A0A | MX1_HUMAN I | Unmodified | 0 | 0 | 0 | 0 | 972800 | 0 | 0 | 0 | 14230000 | 1202000 | 0 | 1108600 | 0 | 0 | 0 | 0 | 0 | 0 |
| ALEEQHSLI | TXNDC11 | Q6PKC3 | TXD11_HUMAN | Unmodified | 0 | 0 | 0 | 0 | 0 | 0 | 0 | 0 | 2408900 | 103420 | 0 | 0 | 0 | 0 | 0 | 0 | 0 | 0 |
| ALEEVPIHI | CARMIL1; CARI | Q5VZK9; A0A | CARL1_HUMAN | Unmodified | 0 | 0 | 0 | 0 | 1730900 | 0 | 0 | 0 | 0 | 0 | 641790 | 0 | 317290 | 0 | 0 | 0 | 0 | 0 |
| ALEGCSPD | TMPPE | Q6ZT21 | TMPPE_HUMAN | Unmodified | 0 | 0 | 0 | 0 | 0 | 0 | 0 | 0 | 1319400 | 390120 | 0 | 0 | 0 | 0 | 0 | 0 | 0 | 0 |
| ALEGLQYPI | NID1 | P14543 | NID1_HUMAN I | Unmodified | 0 | 0 | 100370 | 0 | 168540 | 0 | 0 | 0 | 6147400 | 3730300 | 124430 | 0 | 0 | 0 | 0 | 0 | 2335600 | 1416200 |
| ALEHFPML | GPD2; GPD2 | P43304; F5G | GPDM_HUMAN | Unmodified | 0 | 0 | 0 | 0 | 0 | 0 | 0 | 0 | 1799400 | 0 | 0 | 0 | 932260 | 0 | 0 | 0 | 0 | 0 |

| | | | | | | | | | | | | | | | | | | | | | | |
|---|---|---|---|---|---|---|---|---|---|---|---|---|---|---|---|---|---|---|---|---|---|---|
| ALEHFTDL\ | CLTC; CLTC | Q00610; A0/ | CLH1_HUMAN | Unmodified | 4892000 | 7272400 | 3758700 | 1523800 | 1242200 | 782780 | 438520 | 114320 | 506370 | 299790 | 0 | 21733000 | 430600 | 194070 | 5921500 | 5784100 | 451300 | 0 |
| ALEHFTDL\ | CLTC; CLTC | Q00610; A0/ | CLH1_HUMAN | Unmodified | 41417000 | 19150000 | 18262000 | 10314000 | 0 | 0 | 0 | 0 | 3011000 | 232370 | 118780 | 1.77E+08 | 0 | 0 | 12788000 | 18760000 | 0 | 0 |
| ALEHSALAI | CTPS2; CTPS1; | Q9NRF8; P1 | PYRG2_HUMAN | Unmodified | 0 | 0 | 0 | 0 | 0 | 0 | 0 | 0 | 0 | 0 | 0 | 2794300 | 0 | 0 | 0 | 0 | 0 | 0 |
| ALEHSALAI | CTPS1; CTPS1; | P17812; B4E | PYRG1_HUMAN | Unmodified | 0 | 91107 | 0 | 0 | 338090 | 0 | 0 | 0 | 0 | 319960 | 617900 | 4928400 | 0 | 0 | 444700 | 212320 | 206580 | 0 |
| ALEHVDGT | ITGB4 | P16144 | ITB4_HUMAN I | Unmodified | 0 | 0 | 0 | 0 | 988690 | 81064 | 0 | 0 | 3214000 | 0 | 121590 | 147110 | 0 | 0 | 0 | 0 | 577950 | 425080 |
| ALEIADFSC | SCRIB; SCRIB; | Q14160; A0/ | SCRIB_HUMAN | Unmodified | 135040 | 0 | 236790 | 0 | 0 | 0 | 0 | 0 | 2066100 | 563170 | 223780 | 2897600 | 0 | 0 | 0 | 0 | 0 | 0 |
| ALEIADFSC | SCRIB; SCRIB; | Q14160; A0/ | SCRIB_HUMAN | Unmodified | 1330900 | 908440 | 519510 | 3203500 | 2642400 | 0 | 816690 | 0 | 19452000 | 6866100 | 879350 | 2129600 | 4229500 | 1913200 | 5855300 | 989010 | 8416100 | 4874400 |
| ALEIIPR | CCT7 | Q99832 | TCPH_HUMAN | Unmodified | 307210 | 245090 | 635090 | 721700 | 1384000 | 0 | 662860 | 489090 | 2264700 | 1351600 | 3239000 | 18405000 | 2542000 | 1330200 | 829150 | 533370 | 590020 | 0 |
| ALEIIPRQL | CCT7 | Q99832 | TCPH_HUMAN | Unmodified | 0 | 0 | 0 | 0 | 0 | 0 | 3690000 | 0 | 0 | 0 | 0 | 0 | 6515700 | 0 | 0 | 0 | 6065500 | 0 |
| ALEIYQTK | KLC1; KLC1; KL | Q07866; G3\ | KLC1_HUMAN | Unmodified | 0 | 0 | 0 | 0 | 757890 | 0 | 0 | 0 | 966000 | 205480 | 407920 | 2331500 | 0 | 0 | 0 | 187940 | 64638 | 230550 |
| ALELDGQS | STUB1; STUB1 | Q9UNE7; H3 | CHIP_HUMAN | Unmodified | 0 | 0 | 0 | 0 | 0 | 0 | 0 | 0 | 0 | 0 | 0 | 1837300 | 1755800 | 0 | 0 | 1259000 | 475250 | 0 |
| ALELDHK | DNAJC7; DNAJ( | Q99615; A0/ | DNJC7_HUMAN | Unmodified | 0 | 0 | 0 | 0 | 0 | 0 | 0 | 0 | 0 | 0 | 0 | 0 | 1404400 | 0 | 0 | 0 | 0 | 0 |
| ALELDHKN | DNAJC7; DNAJ( | Q99615; A0/ | DNJC7_HUMAN | Unmodified | 0 | 0 | 0 | 0 | 204590 | 0 | 0 | 0 | 0 | 0 | 0 | 0 | 173890 | 111180 | 0 | 0 | 0 | 0 |
| ALELDPDN | SGTA; SGTA; S( | O43765; A0/ | SGTA_HUMAN | Unmodified | 0 | 0 | 0 | 390630 | 0 | 0 | 161790 | 106230 | 748150 | 306940 | 724670 | 3309900 | 465740 | 338070 | 0 | 377830 | 0 | 0 |
| ALELDPDN | SGTA; SGTA; S( | O43765; A0/ | SGTA_HUMAN | Unmodified | 0 | 0 | 0 | 0 | 1125400 | 0 | 1048500 | 618980 | 0 | 725620 | 0 | 0 | 573990 | 541400 | 0 | 0 | 620170 | 724540 |
| ALELDPNL\ | MYH14; MYH10 | Q7Z406; P35 | MYH14_HUMA | Unmodified | 0 | 0 | 350980 | 223430 | 0 | 0 | 0 | 0 | 0 | 0 | 0 | 1948900 | 0 | 0 | 821400 | 550740 | 0 | 0 |
| ALELDQER | RDX; RDX; RDX | P35241; A0/ | RADI_HUMAN I | Unmodified | 0 | 1158900 | 0 | 136800 | 577660 | 126080 | 0 | 0 | 1626000 | 181270 | 1322000 | 3515300 | 168570 | 0 | 0 | 0 | 0 | 0 |
| ALELDQERI | RDX; RDX; RDX | P35241; A0/ | RADI_HUMAN I | Unmodified | 0 | 0 | 532490 | 631960 | 518790 | 97316 | 0 | 0 | 462390 | 0 | 144000 | 1582800 | 170950 | 163600 | 0 | 0 | 0 | 89812 |
| ALELDSNL\ | MYH9 | P35579 | MYH9_HUMAN | Unmodified | 762330 | 972830 | 1001200 | 1498200 | 274070 | 0 | 0 | 120450 | 0 | 1041400 | 975840 | 9735300 | 133110 | 0 | 2471400 | 1996600 | 0 | 1881000 |
| ALELDSNL\ | MYH9 | P35579 | MYH9_HUMAN | Unmodified | 0 | 0 | 249830 | 418270 | 8705300 | 3311100 | 8397600 | 3826500 | 0 | 5785100 | 146220 | 0 | 1877600 | 3036200 | 0 | 0 | 6922100 | 4510300 |
| ALELDSNN | FKBP4 | Q02790 | FKBP4_HUMAN | Unmodified | 522020 | 307520 | 904320 | 1403500 | 2159500 | 1013100 | 2429100 | 1886100 | 1242600 | 0 | 3636100 | 31428000 | 10547000 | 6418100 | 946430 | 948590 | 2538800 | 1984000 |
| ALELENDRI | LMNB2 | Q03252 | LMNB2_HUMAN | Unmodified | 101800 | 0 | 50971 | 135060 | 1085100 | 310130 | 0 | 0 | 1505500 | 709150 | 616230 | 2658300 | 337220 | 75002 | 55672 | 0 | 0 | 120860 |
| ALELEQER | MSN | P26038 | MOES_HUMAN | Unmodified | 557400 | 4686000 | 0 | 926060 | 592860 | 0 | 0 | 22701 | 0 | 0 | 0 | 49496 | 0 | 0 | 0 | 1262600 | 143490 | 0 |
| ALELEQERI | MSN | P26038 | MOES_HUMAN | Unmodified | 0 | 0 | 0 | 0 | 0 | 186960 | 0 | 0 | 0 | 0 | 0 | 0 | 0 | 0 | 0 | 0 | 0 | 0 |
| ALELGTVM | PLCG2; PLCG2 | P16885; A0/ | PLCG2_HUMAI | Oxidation (M) | 0 | 0 | 0 | 0 | 0 | 0 | 0 | 0 | 0 | 0 | 0 | 1364900 | 0 | 0 | 0 | 0 | 0 | 0 |
| ALELNMLSI | PSAT1 | Q9Y617 | SERC_HUMAN | Oxidation (M) | 0 | 0 | 0 | 1299300 | 1304400 | 0 | 397220 | 277020 | 370700 | 335600 | 1160300 | 9281800 | 1921000 | 1890800 | 0 | 0 | 464950 | 502130 |
| ALELNMLSI | PSAT1 | Q9Y617 | SERC_HUMAN | Unmodified | 0 | 0 | 0 | 0 | 2942400 | 474000 | 0 | 0 | 1234300 | 543210 | 747990 | 4461300 | 8347200 | 2246400 | 0 | 0 | 803440 | 828400 |
| ALELTGLK | NCL; NCL; NCL | P19338; C9J | NUCL_HUMAN | Unmodified | 0 | 0 | 97165 | 54943 | 1004000 | 30288 | 344780 | 0 | 2359300 | 1071100 | 5743400 | 1118600 | 545580 | 0 | 113020 | 0 | 196430 | 0 |
| ALENDPDC | PLS3; PLS3; PL! | P13797; U3K | PLST_HUMAN I | Oxidation (M) | 0 | 0 | 0 | 0 | 2262800 | 1091400 | 0 | 0 | 0 | 264530 | 0 | 0 | 485100 | 611560 | 0 | 0 | 3192300 | 527340 |
| ALENDPDC | PLS3; PLS3; PL! | P13797; U3K | PLST_HUMAN I | Oxidation (M) | 0 | 0 | 0 | 0 | 0 | 0 | 0 | 0 | 0 | 0 | 0 | 0 | 0 | 0 | 0 | 0 | 2844100 | 0 |
| ALENDPDC | LCP1; LCP1 | P13796; Q5T | PLSL_HUMAN I | Oxidation (M) | 0 | 0 | 0 | 0 | 0 | 0 | 0 | 0 | 0 | 0 | 0 | 0 | 0 | 0 | 0 | 0 | 1366300 | 0 |
| ALENPDTA | PTPN23 | Q9H3S7 | PTN23_HUMAN | Unmodified | 1001700 | 2159100 | 0 | 0 | 0 | 0 | 0 | 0 | 0 | 0 | 0 | 0 | 0 | 0 | 1641700 | 0 | 0 | 0 |
| ALENTAVK | TPP2; TPP2; TPI | P29144; A0/ | TPP2_HUMAN | Unmodified | 0 | 0 | 0 | 0 | 73605 | 0 | 0 | 7395.3 | 1075800 | 655090 | 0 | 1751600 | 0 | 0 | 90987 | 50168 | 84618 | 0 |
| ALENTAVK | TPP2; TPP2; TPI | P29144; A0/ | TPP2_HUMAN | Unmodified | 0 | 0 | 0 | 0 | 0 | 0 | 0 | 0 | 0 | 0 | 0 | 2081200 | 0 | 0 | 0 | 0 | 0 | 0 |
| ALEPQGLLI | AGRN; AGRN; / | O00468; A0/ | AGRIN_HUMAN | Unmodified | 740360 | 234180 | 2180500 | 292820 | 0 | 0 | 0 | 0 | 155970 | 0 | 0 | 0 | 0 | 0 | 841720 | 443320 | 0 | 0 |
| ALEPTSQKI | IMUP; N/A | Q9GZP8; K7I | IMUP_HUMAN | Unmodified | 0 | 0 | 0 | 0 | 0 | 0 | 0 | 0 | 0 | 358350 | 0 | 0 | 0 | 0 | 0 | 0 | 0 | 0 |
| ALEQALEK | ANPEP | P15144 | AMPN_HUMAN | Unmodified | 5197900 | 6520100 | 5053800 | 3415000 | 462730 | 15996 | 91289 | 31921 | 164060 | 95400 | 132650 | 1184700 | 1748700 | 0 | 7632300 | 5157900 | 3624600 | 9256200 |
| ALEQFATV\ | ACADVL; ACAD | P49748; J3Q | ACADV_HUMAI | Unmodified | 0 | 0 | 0 | 0 | 0 | 0 | 0 | 0 | 6964300 | 705130 | 0 | 0 | 0 | 0 | 0 | 0 | 0 | 0 |
| ALEQKPDD | SUGT1 | Q9Y2Z0 | SGT1_HUMAN | Unmodified | 0 | 0 | 0 | 0 | 527950 | 0 | 0 | 0 | 220770 | 96216 | 551120 | 2145100 | 144650 | 124000 | 0 | 122890 | 0 | 308070 |
| ALEQQVEE | MYH9 | P35579 | MYH9_HUMAN | Unmodified | 0 | 0 | 0 | 0 | 441090 | 0 | 0 | 0 | 0 | 0 | 242540 | 282530 | 0 | 0 | 0 | 0 | 0 | 0 |
| ALERFLQD' | PDIA3 | P30101 | PDIA3_HUMAN | Unmodified | 117210 | 0 | 0 | 0 | 10299000 | 1671800 | 11482000 | 84777 | 1211000 | 1.3E+08 | 3620100 | 0 | 10338000 | 10548000 | 115570 | 0 | 1668700 | 7878600 |
| ALERFLQD' | PDIA3 | P30101 | PDIA3_HUMAN | Unmodified | 0 | 0 | 0 | 0 | 3822400 | 0 | 408340 | 0 | 299230 | 4876000 | 570110 | 0 | 145890 | 0 | 0 | 0 | 980970 | 3517900 |
| ALERLEGVI | DYNC1H1; DYN | Q14204; A0/ | DYHC1_HUMA | Unmodified | 0 | 0 | 352080 | 228150 | 533410 | 327270 | 0 | 0 | 694990 | 1111100 | 1217400 | 10220000 | 0 | 0 | 930810 | 424020 | 0 | 372130 |
| ALESELQQI | PBXIP1 | Q96AQ6 | PBIP1_HUMAN | Unmodified | 279400 | 423990 | 633370 | 693140 | 0 | 0 | 0 | 33876 | 6407600 | 484710 | 1183100 | 0 | 957180 | 551520 | 1108400 | 2094000 | 1101000 | 0 |
| ALESLSCPI | H6PD | O95479 | G6PE_HUMAN | Unmodified | 0 | 0 | 0 | 0 | 0 | 0 | 0 | 0 | 1194000 | 397370 | 0 | 0 | 0 | 0 | 0 | 0 | 0 | 0 |
| ALESPERPI | PGK1 | P00558 | PGK1_HUMAN | Unmodified | 7584600 | 4461200 | 13597000 | 21568000 | 79459000 | 17720000 | 86202000 | 8964600 | 68859000 | 81212000 | 88371000 | 3.37E+08 | 1.41E+08 | 80045000 | 26263000 | 17477000 | 4.42E+08 | 3.43E+08 |
| ALESPERPI | PGK1 | P00558 | PGK1_HUMAN | Unmodified | 0 | 0 | 0 | 0 | 0 | 0 | 567690 | 0 | 0 | 0 | 0 | 0 | 432870 | 847510 | 0 | 0 | 3966500 | 0 |
| ALESSIAPI\ | RUVBL1; RUVBI | Q9Y265; E7E | RUVB1_HUMAI | Unmodified | 359750 | 0 | 132700 | 14765 | 0 | 0 | 0 | 0 | 254670 | 0 | 140620 | 9910100 | 0 | 0 | 786020 | 366120 | 0 | 0 |
| ALETCGGD | TSFM; TSFM; TS | P43897; F8V | EFTS_HUMAN I | Unmodified | 0 | 0 | 0 | 0 | 0 | 0 | 0 | 0 | 585630 | 128170 | 0 | 0 | 0 | 0 | 0 | 0 | 0 | 466220 |
| ALETLTGAL | DKC1; DKC1; D | O60832; H7I | DKC1_HUMAN | Unmodified | 2080000 | 1673800 | 1285700 | 2409200 | 0 | 0 | 0 | 0 | 0 | 0 | 0 | 0 | 0 | 0 | 0 | 2578800 | 0 | 0 |
| ALETLTGAL | DKC1; DKC1; D | O60832; H7I | DKC1_HUMAN | Unmodified | 0 | 0 | 70615 | 0 | 0 | 0 | 0 | 0 | 0 | 0 | 0 | 461780 | 0 | 0 | 0 | 114230 | 0 | 0 |
| ALEVAEYLT | ATG3; ATG3 | Q9NT62; F8\ | ATG3_HUMAN | Unmodified | 40113 | 0 | 29489 | 144150 | 474570 | 0 | 266380 | 0 | 315840 | 199390 | 113880 | 528440 | 906000 | 804540 | 0 | 71184 | 384700 | 508210 |
| ALEVIKPAH | EIF3A | Q14152 | EIF3A_HUMAN | Unmodified | 318930 | 227000 | 736650 | 274060 | 136140 | 0 | 0 | 0 | 205210 | 0 | 0 | 4520400 | 96483 | 0 | 338280 | 366380 | 0 | 0 |
| ALEVIKPAH | EIF3A | Q14152 | EIF3A_HUMAN | Unmodified | 382620 | 269920 | 0 | 1414700 | 330360 | 0 | 1356600 | 0 | 0 | 515050 | 154090 | 480330 | 1767600 | 603070 | 40910 | 0 | 90631 | 0 |
| ALEVSEDVI | MAGED2 | Q9UNF1 | MAGD2_HUMA | Unmodified | 0 | 0 | 0 | 400320 | 0 | 0 | 246130 | 0 | 0 | 382480 | 0 | 0 | 0 | 266490 | 0 | 0 | 237270 | 906990 |
| ALEVSEDVI | MAGED2 | Q9UNF1 | MAGD2_HUMA | Unmodified | 0 | 0 | 0 | 0 | 0 | 0 | 321950 | 66607 | 0 | 276760 | 0 | 47328 | 149240 | 294740 | 0 | 14115 | 573560 | 349880 |
| ALFDFNGN | CRK | P46108 | CRK_HUMAN A | Unmodified | 165650 | 75120 | 30573 | 0 | 0 | 0 | 0 | 0 | 160080 | 0 | 30212 | 239350 | 0 | 0 | 475500 | 307890 | 0 | 0 |
| ALFDFNGN | CRK | P46108 | CRK_HUMAN A | Unmodified | 511850 | 0 | 288510 | 737010 | 0 | 274640 | 0 | 0 | 0 | 383230 | 0 | 1494100 | 430610 | 0 | 0 | 561410 | 0 | 0 |
| ALFDYDK | DLG1; DLG4; D | Q12959; P78 | DLG1_HUMAN | Unmodified | 0 | 239390 | 0 | 0 | 10016000 | 2006000 | 0 | 55239 | 2437900 | 2216700 | 195710 | 0 | 2131600 | 2117200 | 0 | 0 | 0 | 6432900 |
| ALFDYNPN | MPP7; MPP7 | Q5T2T1; S4F | MPP7_HUMAN | Unmodified | 0 | 0 | 0 | 0 | 2412200 | 679670 | 0 | 0 | 482840 | 0 | 845600 | 696270 | 720780 | 865350 | 748920 | 0 | 205920 | 1208400 |
| ALFEEVPEL | ATIC; ATIC | P31939; H7( | PUR9_HUMAN | Unmodified | 0 | 0 | 0 | 0 | 0 | 134110 | 0 | 0 | 197280 | 0 | 142320 | 0 | 22621 | 272980 | 0 | 0 | 0 | 0 |
| ALFEEVPEL | ATIC; ATIC | P31939; H7( | PUR9_HUMAN | Unmodified | 363560 | 0 | 649630 | 2982500 | 5201700 | 1191500 | 521190 | 0 | 1605600 | 358880 | 6750800 | 23422000 | 7502600 | 1119600 | 7783700 | 4283000 | 553240 | 1779600 |
| ALFEEVPEL | ATIC; ATIC | P31939; H7( | PUR9_HUMAN | Unmodified | 0 | 0 | 0 | 703100 | 0 | 642710 | 0 | 0 | 168110 | 1446800 | 0 | 0 | 15956000 | 0 | 0 | 0 | 2617600 | 5114100 |
| ALFKPADVI | BTN3A1 | O00481 | BT3A1_HUMAN | Unmodified | 0 | 0 | 0 | 0 | 0 | 0 | 0 | 0 | 0 | 26888 | 67459 | 418450 | 29843 | 0 | 0 | 0 | 0 | 163790 |
| ALFLETEQL | ALCAM; ALCAM | Q13740; F5( | CD166_HUMAI | Unmodified | 16184000 | 16361000 | 5954000 | 5289100 | 20432000 | 2368500 | 4194800 | 1837500 | 16139000 | 6802100 | 0 | 3418800 | 5242600 | 3500600 | 0 | 0 | 7195700 | 17687000 |
| ALFLETEQL | ALCAM; ALCAM | Q13740; F5( | CD166_HUMAI | Unmodified | 2362600 | 2174800 | 1539200 | 1156800 | 2498400 | 376410 | 666500 | 313880 | 1768300 | 619370 | 60809 | 429540 | 1107000 | 252820 | 1005300 | 1013300 | 1067800 | 3822800 |
| ALFLIPR | TNIK; MINK1; M | Q9UKE5; Q8 | TNIK_HUMAN | Unmodified | 335350 | 0 | 0 | 0 | 0 | 0 | 0 | 0 | 2994700 | 394530 | 0 | 6679700 | 0 | 0 | 1402800 | 1572700 | 0 | 0 |
| ALFLYSR | ITGAV | P06756 | ITAV_HUMAN I | Unmodified | 0 | 0 | 0 | 0 | 0 | 0 | 0 | 27309 | 0 | 0 | 0 | 3171100 | 97620 | 141180 | 0 | 0 | 0 | 0 |
| ALFNAQFG | NT5DC2 | Q9H857 | NT5D2_HUMAI | Unmodified | 0 | 0 | 0 | 0 | 0 | 0 | 0 | 0 | 655300 | 214860 | 0 | 0 | 0 | 0 | 0 | 0 | 0 | 0 |
| ALFPTEIHS | MGAT4B; MGAT | Q9UQ53; E5 | MGT4B_HUMAI | Unmodified | 0 | 0 | 0 | 0 | 0 | 0 | 0 | 0 | 3065400 | 0 | 0 | 0 | 0 | 0 | 0 | 0 | 0 | 0 |
| ALFQDIKKF | FTL | P02792 | FRIL_HUMAN F | Unmodified | 0 | 0 | 0 | 0 | 0 | 0 | 0 | 0 | 4290800 | 114370 | 157840 | 0 | 0 | 0 | 0 | 0 | 0 | 0 |
| ALGAEIVR | CBS; CBSL | P35520; P0( | CBS_HUMAN ( | Unmodified | 0 | 6092800 | 0 | 0 | 0 | 0 | 525350 | 256870 | 128220 | 32160 | 0 | 15344000 | 28306 | 0 | 0 | 130360 | 0 | 0 |
| ALGAEIVRT | CBS; CBSL | P35520; P0( | CBS_HUMAN ( | Unmodified | 55125 | 17840 | 0 | 0 | 0 | 0 | 43391 | 0 | 1109300 | 169850 | 738000 | 2403300 | 0 | 0 | 114060 | 0 | 0 | 0 |
| ALGDLISAT | TLN1 | Q9Y490 | TLN1_HUMAN | Unmodified | 793210 | 803590 | 1774400 | 1508200 | 6969900 | 2096500 | 6896000 | 6222400 | 458530 | 1590500 | 0 | 5691100 | 2194100 | 2320100 | 1634600 | 1227500 | 5303300 | 332090 |
| ALGDLISAT | TLN1 | Q9Y490 | TLN1_HUMAN | Unmodified | 0 | 0 | 0 | 0 | 0 | 0 | 2724700 | 0 | 0 | 0 | 0 | 0 | 0 | 984570 | 0 | 0 | 1101100 | 0 |
| ALGFENAT( | LGALS3BP; LG/ | Q08380; K7I | LG3BP_HUMAI | Unmodified | 0 | 0 | 0 | 0 | 0 | 0 | 0 | 0 | 14341000 | 0 | 0 | 0 | 0 | 0 | 0 | 0 | 0 | 0 |
| ALGGHMTI | BPNT2 | Q9NX62 | IMPA3_HUMAN | Oxidation (M) | 0 | 0 | 0 | 0 | 0 | 0 | 0 | 0 | 0 | 2879700 | 0 | 0 | 0 | 0 | 0 | 0 | 0 | 0 |
| ALGHGVDI | PTGS1; PTGS1; | P23219; X6R | PGH1_HUMAN | Unmodified | 94314 | 90982 | 0 | 0 | 0 | 0 | 0 | 0 | 0 | 0 | 7953100 | 3870600 | 0 | 0 | 0 | 0 | 0 | 0 |
| ALGHIAVAI | PI4KA | P42356 | PI4KA_HUMAN | Unmodified | 27847 | 0 | 234330 | 0 | 0 | 0 | 0 | 0 | 0 | 0 | 0 | 475590 | 0 | 0 | 157360 | 188820 | 0 | 0 |
| ALGILGRSE | CIT; CIT | O14578; H7( | CTRO_HUMAN | Unmodified | 122660 | 436790 | 0 | 0 | 26714000 | 2055700 | 1708200 | 0 | 0 | 158990 | 0 | 0 | 816770 | 436800 | 0 | 0 | 2883500 | 1956400 |
| ALGILGRSE | CIT; CIT | O14578; H7( | CTRO_HUMAN | Oxidation (M) | 0 | 0 | 0 | 0 | 2311600 | 0 | 0 | 0 | 0 | 0 | 0 | 0 | 0 | 0 | 0 | 0 | 1243200 | 0 |
| ALGITEIFIK | SERPINI1; SERF | Q99574; H7( | NEUS_HUMAN | Unmodified | 0 | 0 | 0 | 0 | 0 | 0 | 0 | 0 | 2126000 | 263210 | 206480 | 0 | 0 | 0 | 0 | 0 | 0 | 0 |
| ALGKYGPA | EEF1B2; EEF1B | P24534; C9J | EF1B_HUMAN I | Unmodified | 0 | 0 | 0 | 0 | 3592400 | 0 | 0 | 0 | 0 | 0 | 0 | 0 | 0 | 0 | 0 | 0 | 0 | 0 |
| ALGLDSAN | FKBP5 | Q13451 | FKBP5_HUMAI | Unmodified | 0 | 0 | 0 | 0 | 3055700 | 453600 | 59884 | 31356 | 1435400 | 258300 | 0 | 1125900 | 1898400 | 1296900 | 0 | 0 | 395680 | 836010 |
| ALGLLCGK | NPC1 | O15118 | NPC1_HUMAN | Unmodified | 0 | 0 | 0 | 0 | 685150 | 279830 | 0 | 15685 | 1636900 | 0 | 0 | 398750 | 29350 | 0 | 0 | 0 | 0 | 265010 |
| ALGLNEND | MYO6; MYO6; M | Q9UM54; A0 | MYO6_HUMAN | Unmodified | 0 | 0 | 0 | 0 | 0 | 0 | 0 | 0 | 0 | 0 | 0 | 0 | 0 | 0 | 993080 | 838960 | 0 | 0 |
| ALGLNEND | MYO6; MYO6; M | Q9UM54; A0 | MYO6_HUMAN | Unmodified | 0 | 0 | 0 | 291110 | 0 | 0 | 112650 | 53185 | 0 | 0 | 0 | 0 | 119340 | 121860 | 320150 | 354280 | 110110 | 0 |
| ALGLVPQA | B3GNT8 | Q7Z7M8 | B3GN8_HUMAI | Unmodified | 0 | 0 | 0 | 0 | 0 | 0 | 0 | 0 | 1456700 | 211720 | 0 | 0 | 0 | 0 | 0 | 0 | 0 | 0 |
| ALGMKLPE | KARS1 | Q15046 | SYK_HUMAN L | Oxidation (M) | 0 | 0 | 0 | 0 | 0 | 0 | 0 | 0 | 0 | 0 | 0 | 557850 | 0 | 0 | 0 | 0 | 0 | 0 |
| ALGMKLPE | KARS1 | Q15046 | SYK_HUMAN L | Unmodified | 0 | 0 | 0 | 0 | 625750 | 0 | 0 | 0 | 803270 | 0 | 0 | 2285400 | 1435200 | 0 | 0 | 0 | 0 | 0 |
| ALGMKLPE | KARS1 | Q15046 | SYK_HUMAN L | Oxidation (M) | 0 | 0 | 0 | 159400 | 0 | 0 | 0 | 0 | 0 | 0 | 0 | 3148000 | 0 | 0 | 0 | 0 | 0 | 0 |
| ALGNICYD | RAP1GDS1; RAI | P52306; D6F | GDS1_HUMAN | Unmodified | 0 | 0 | 0 | 0 | 0 | 0 | 0 | 0 | 0 | 0 | 0 | 964060 | 0 | 0 | 0 | 0 | 0 | 0 |
| ALGNQQPS | GRP; GRP | P07492; H0 | GRP_HUMAN ( | Unmodified | 0 | 0 | 0 | 0 | 0 | 0 | 0 | 0 | 513740 | 0 | 0 | 0 | 0 | 0 | 0 | 0 | 0 | 0 |
| ALGNSEEA | UTRN; UTRN; U | P46939; Q5T | UTRN_HUMAN | Unmodified | 0 | 72758 | 295210 | 198140 | 0 | 0 | 0 | 0 | 615870 | 0 | 138400 | 0 | 0 | 0 | 564060 | 544470 | 0 | 0 |
| ALGPVESL( | LLGL1; LLGL1 | Q15334; A0/ | L2GL1_HUMAN | Unmodified | 0 | 0 | 0 | 0 | 944550 | 636340 | 0 | 0 | 0 | 401580 | 0 | 362320 | 0 | 233600 | 0 | 0 | 578220 | 0 |
| ALGQNPTN | MYL6B; MYL6; N | P14649; P60 | MYL6B_HUMAI | Unmodified | 2947800 | 4109700 | 5352700 | 7625700 | 52170000 | 17446000 | 1366600 | 1115700 | 16606000 | 7027900 | 8849100 | 53781000 | 15334000 | 8481900 | 6814800 | 5189100 | 5200700 | 8744500 |
| ALGSEVQD | SLK | Q9H2G2 | SLK_HUMAN S | Unmodified | 0 | 0 | 0 | 0 | 453600 | 0 | 0 | 0 | 4557800 | 205300 | 684180 | 1630000 | 888850 | 1223700 | 0 | 449010 | 0 | 350150 |
| ALGSEVQD | SLK | Q9H2G2 | SLK_HUMAN S | Unmodified | 0 | 0 | 0 | 0 | 1121000 | 0 | 0 | 0 | 0 | 185160 | 0 | 0 | 3598400 | 11155000 | 0 | 0 | 0 | 1062600 |
| ALGSGMA\ | TACSTD2 | P09758 | TACD2_HUMAI | Unmodified | 0 | 0 | 0 | 0 | 15214000 | 3722700 | 0 | 0 | 0 | 0 | 0 | 0 | 0 | 0 | 0 | 0 | 278240 | 977790 |
| ALGSGMA\ | TACSTD2 | P09758 | TACD2_HUMAI | Oxidation (M) | 0 | 0 | 0 | 0 | 6238000 | 936210 | 70660 | 0 | 0 | 0 | 16668 | 0 | 0 | 0 | 0 | 0 | 0 | 64897 |
| ALGSILAYL | PALD1 | Q9ULE6 | PALD_HUMAN | Unmodified | 0 | 0 | 0 | 0 | 0 | 0 | 0 | 0 | 0 | 0 | 29060 | 51109 | 0 | 0 | 0 | 0 | 0 | 0 |
| ALGSLHLP' | PRCP; PRCP; PI | P42785; A0/ | PCP_HUMAN L | Unmodified | 0 | 0 | 0 | 0 | 0 | 0 | 0 | 0 | 1237200 | 0 | 0 | 0 | 0 | 0 | 0 | 0 | 0 | 0 |
| ALGTEVIQL | AIFM1; AIFM1; / | O95831; A0/ | AIFM1_HUMAN | Unmodified | 0 | 0 | 0 | 0 | 0 | 0 | 0 | 0 | 959630 | 0 | 0 | 1491500 | 0 | 0 | 0 | 0 | 0 | 404410 |
| ALGTLVSH' | GCN1 | Q92616 | GCN1_HUMAN | Unmodified | 0 | 0 | 213710 | 0 | 0 | 0 | 0 | 0 | 0 | 0 | 0 | 1001400 | 0 | 0 | 0 | 0 | 0 | 0 |
| ALGVMDDI | MGAT1 | P26572 | MGAT1_HUMAI | Unmodified | 0 | 0 | 0 | 0 | 0 | 0 | 0 | 0 | 4461100 | 622040 | 125970 | 0 | 0 | 0 | 0 | 0 | 0 | 0 |
| ALGVSNFS | AKR1B10 | O60218 | AK1BA_HUMAN | Unmodified | 0 | 0 | 0 | 0 | 3310000 | 103230 | 10101000 | 4917900 | 0 | 0 | 0 | 0 | 0 | 0 | 0 | 0 | 0 | 0 |
| ALGYFAVV | OPA1; OPA1; OI | O60313; E5I | OPA1_HUMAN | Unmodified | 0 | 0 | 0 | 0 | 0 | 0 | 0 | 0 | 0 | 0 | 481990 | 6244000 | 0 | 0 | 0 | 0 | 0 | 0 |
| ALGYFHRA | RMDN1; RMDN | Q96DB5; H0 | RMD1_HUMAN | Unmodified | 0 | 0 | 0 | 0 | 394100 | 0 | 0 | 328190 | 0 | 333160 | 0 | 0 | 118600 | 0 | 0 | 0 | 0 | 0 |
| ALGYVLDR | INPP5A; INPP5/ | Q14642; H0\ | I5P1_HUMAN I | Unmodified | 0 | 0 | 0 | 0 | 0 | 0 | 0 | 0 | 0 | 0 | 0 | 0 | 0 | 0 | 558170 | 408570 | 0 | 0 |

| | | | | | | | | | | | | | | | | | | | | | | |
|---|---|---|---|---|---|---|---|---|---|---|---|---|---|---|---|---|---|---|---|---|---|---|
| ALGYVLDR | INPP5A; INPP5 | Q14642; H0 | I5P1_HUMAN I | Unmodified | 19036 | 0 | 0 | 179270 | 0 | 85872 | 0 | 0 | 0 | 130810 | 0 | 0 | 0 | 0 | 0 | 0 | 0 | 115150 |
| ALHARVFD | MYO1E; MYO1E | Q12965; H0 | MYO1E_HUMA | Unmodified | 0 | 557670 | 0 | 0 | 0 | 0 | 0 | 0 | 0 | 0 | 0 | 0 | 0 | 0 | 0 | 0 | 390080 | 0 |
| ALHDEEATI | B3GLCT | Q6Y288 | B3GLT_HUMAN | Unmodified | 0 | 0 | 0 | 0 | 704890 | 0 | 0 | 0 | 989040 | 0 | 0 | 0 | 0 | 0 | 0 | 0 | 0 | 0 |
| ALHFAISEY | CST4; CST1 | P01036; P01 | CYTS_HUMAN | Unmodified | 0 | 0 | 0 | 0 | 0 | 0 | 0 | 0 | 4354400 | 0 | 105290 | 0 | 0 | 0 | 0 | 0 | 0 | 0 |
| ALHHGIDLI | DARS1; DARS1 | P14868; H7E | SYDC_HUMAN | Unmodified | 0 | 0 | 0 | 0 | 221040 | 0 | 0 | 0 | 0 | 0 | 0 | 764610 | 0 | 0 | 0 | 0 | 0 | 0 |
| ALHLGNNV | LRRC8A | Q8IWT6 | LRC8A_HUMAN | Unmodified | 984180 | 0 | 0 | 0 | 0 | 0 | 0 | 0 | 0 | 0 | 0 | 4827700 | 0 | 0 | 0 | 0 | 0 | 0 |
| ALHLLEEYF | DLG1; DLG1; D | Q12959; A0A | DLG1_HUMAN | Unmodified | 49401 | 38223 | 84259 | 0 | 2408600 | 440750 | 0 | 0 | 0 | 111270 | 568160 | 0 | 1017300 | 762810 | 85684 | 53234 | 495270 | 1207600 |
| ALHSLLLR | PAFAH1B3; PAF | Q15102; M0 | PA1B3_HUMAN | Unmodified | 72733 | 85080 | 0 | 0 | 110870 | 0 | 0 | 0 | 580540 | 0 | 0 | 1747600 | 0 | 0 | 0 | 0 | 0 | 0 |
| ALHSLLLRL | PAFAH1B3; PAF | Q15102; M0 | PA1B3_HUMAN | Unmodified | 0 | 0 | 0 | 0 | 0 | 0 | 0 | 0 | 0 | 1080600 | 0 | 0 | 0 | 0 | 0 | 0 | 0 | 0 |
| ALHVPKAP | CCT8 | P50990 | TCPQ_HUMAN | Acetyl (Protein N | 0 | 0 | 0 | 0 | 235820 | 0 | 1558100 | 217300 | 0 | 247520 | 0 | 0 | 324590 | 743570 | 0 | 0 | 1728200 | 0 |
| ALHVPKAP | CCT8 | P50990 | TCPQ_HUMAN | Acetyl (Protein N | 0 | 0 | 0 | 0 | 175070 | 0 | 1130800 | 611970 | 0 | 195960 | 0 | 0 | 407010 | 439440 | 0 | 0 | 2776800 | 218600 |
| ALHYGREC | TLN1 | Q9Y490 | TLN1_HUMAN | Unmodified | 0 | 0 | 0 | 72208 | 4527900 | 0 | 136790 | 0 | 0 | 147990 | 0 | 0 | 1001000 | 636430 | 0 | 0 | 780890 | 1712900 |
| ALIAAQYSG | EEF1G | P26641 | EF1G_HUMAN | Unmodified | 295160 | 435270 | 5748400 | 2537200 | 8801000 | 3066400 | 418320 | 429300 | 3617000 | 922020 | 6599900 | 51049000 | 1051300 | 1180500 | 2390600 | 2157600 | 3566900 | 5976200 |
| ALIAEHQTF | MACF1; DST; D | Q9UPN3; Q0 | MACF1_HUMA | Unmodified | 0 | 0 | 0 | 0 | 267900 | 0 | 79398 | 60281 | 1503700 | 0 | 0 | 84244 | 0 | 0 | 60467 | 0 | 125330 | 0 |
| ALIAGGGAI | CCT4 | P50991 | TCPD_HUMAN | Unmodified | 0 | 0 | 0 | 0 | 0 | 0 | 0 | 0 | 0 | 0 | 0 | 0 | 0 | 0 | 0 | 0 | 0 | 0 |
| ALIAQLPSP | MACF1; MACF1 | Q9UPN3; H3 | MACF1_HUMA | Unmodified | 0 | 41229 | 0 | 0 | 0 | 0 | 0 | 0 | 542860 | 0 | 0 | 1321500 | 0 | 0 | 0 | 38362 | 0 | 0 |
| ALIASAGAD | STRN4; STRN4; | Q9NRL3; M0 | STRN4_HUMAN | Unmodified | 0 | 0 | 0 | 0 | 0 | 0 | 519070 | 950530 | 0 | 89579 | 0 | 556530 | 0 | 483330 | 0 | 0 | 413830 | 0 |
| ALIDQEVK | LYPLA1; LYPLA | O75608; B4 | LYPA1_HUMAN | Unmodified | 464330 | 787820 | 1703200 | 573130 | 1076800 | 1030900 | 226550 | 121010 | 1609600 | 345310 | 1557100 | 15267000 | 851660 | 804630 | 1488700 | 1101100 | 360850 | 363990 |
| ALIDTLLDG | C1orf43; C1orf | Q9BWL3; X6 | CA043_HUMAN | Unmodified | 0 | 0 | 0 | 0 | 0 | 0 | 0 | 0 | 27713 | 0 | 152650 | 1055500 | 0 | 0 | 0 | 0 | 0 | 0 |
| ALIDYENSN | SNX5; SNX5 | Q9Y5X3; A0A | SNX5_HUMAN | Unmodified | 0 | 0 | 0 | 457750 | 0 | 0 | 0 | 0 | 1959700 | 184820 | 0 | 1232900 | 776620 | 592320 | 0 | 0 | 0 | 0 |
| ALIEAEKVA | ERLIN2; ERLIN2 | O94905; E5F | ERLN2_HUMAN | Unmodified | 0 | 0 | 0 | 0 | 0 | 0 | 1630000 | 409840 | 0 | 1894600 | 0 | 0 | 0 | 0 | 0 | 0 | 0 | 0 |
| ALIEILATR | ANXA6; ANXA6; | P08133; E7E | ANXA6_HUMAN | Unmodified | 1298900 | 1423000 | 4728700 | 5671900 | 2719400 | 340980 | 491050 | 153500 | 39121000 | 7012300 | 307920 | 0 | 1958100 | 733130 | 7032900 | 3466300 | 419930 | 2291600 |
| ALIEILATRT | ANXA6; ANXA6; | P08133; E7E | ANXA6_HUMAN | Unmodified | 0 | 0 | 0 | 461690 | 7486600 | 1170900 | 1180500 | 0 | 2220400 | 6600700 | 0 | 0 | 5073700 | 2433600 | 0 | 0 | 1286300 | 8513700 |
| ALIEMEK | NONO; NONO; | Q15233; A0A | NONO_HUMAN | Unmodified | 0 | 0 | 0 | 0 | 0 | 0 | 1170000 | 813930 | 732350 | 608770 | 420320 | 1907400 | 890590 | 0 | 0 | 0 | 311280 | 1132700 |
| ALIEMEKQ | NONO; NONO; | Q15233; A0A | NONO_HUMAN | Unmodified | 0 | 0 | 0 | 0 | 0 | 0 | 285360 | 229570 | 0 | 112490 | 0 | 0 | 0 | 545340 | 0 | 0 | 0 | 0 |
| ALIEMEKQ | NONO; NONO; | Q15233; A0A | NONO_HUMAN | Oxidation (M) | 0 | 0 | 0 | 0 | 0 | 0 | 0 | 121850 | 0 | 0 | 0 | 0 | 0 | 0 | 0 | 0 | 0 | 205270 |
| ALIEMEKQ | NONO; NONO; | Q15233; A0A | NONO_HUMAN | Unmodified | 0 | 0 | 0 | 0 | 497990 | 0 | 635480 | 340690 | 0 | 0 | 0 | 0 | 591830 | 160720 | 0 | 0 | 259090 | 613490 |
| ALIESYQNL | ABI1; ABI1; ABI | Q8IZP0; A0A | ABI1_HUMAN A | Unmodified | 171450 | 0 | 111440 | 176680 | 0 | 0 | 0 | 73883 | 0 | 0 | 0 | 497370 | 472740 | 0 | 146280 | 254570 | 0 | 0 |
| ALIEVLQPL | WARS1 | P23381 | SYWC_HUMAN | Unmodified | 0 | 0 | 0 | 0 | 2827100 | 0 | 0 | 0 | 3318700 | 1370200 | 313670 | 4522600 | 0 | 775560 | 0 | 0 | 105510 | 0 |
| ALIEVLQPL | WARS1 | P23381 | SYWC_HUMAN | Unmodified | 0 | 0 | 0 | 0 | 1880400 | 0 | 381420 | 0 | 0 | 310890 | 0 | 0 | 0 | 592590 | 0 | 0 | 0 | 672490 |
| ALIEWIR | CPD | O75976 | CBPD_HUMAN | Unmodified | 0 | 0 | 0 | 0 | 0 | 0 | 0 | 0 | 10586000 | 126950 | 635000 | 2109900 | 0 | 0 | 0 | 0 | 0 | 0 |
| ALILVGGYG | GMPPB; GMPPE | Q9Y5P6; A0A | GMPPB_HUMA | Unmodified | 445460 | 629400 | 0 | 0 | 0 | 0 | 0 | 0 | 0 | 0 | 0 | 3475500 | 0 | 0 | 0 | 925990 | 0 | 0 |
| ALINSPEGA | DNASE2; DNAS | O00115; K7E | DNS2A_HUMA | Unmodified | 309550 | 0 | 0 | 361610 | 0 | 0 | 0 | 0 | 27772000 | 5956200 | 2582300 | 1626700 | 458850 | 228760 | 0 | 0 | 0 | 0 |
| ALIPLAKAE | TMEM132A | Q24JP5 | T132A_HUMAN | Unmodified | 0 | 0 | 0 | 0 | 0 | 0 | 0 | 0 | 2181500 | 0 | 0 | 0 | 0 | 0 | 0 | 0 | 0 | 0 |
| ALIPLALEG | UNC45A; UNC4 | Q9H3U1; A0 | UN45A_HUMA | Unmodified | 0 | 0 | 0 | 0 | 0 | 0 | 111740 | 0 | 237500 | 0 | 181840 | 1946800 | 0 | 0 | 0 | 0 | 0 | 0 |
| ALIQCAKDI | VCL | P18206 | VINC_HUMAN | Unmodified | 0 | 0 | 0 | 0 | 0 | 0 | 0 | 0 | 0 | 0 | 0 | 0 | 85110 | 177570 | 0 | 0 | 614180 | 0 |
| ALIQYFSPY | GPS1; GPS1; G | Q13098; A8 | CSN1_HUMAN | Oxidation (M) | 0 | 0 | 0 | 0 | 0 | 0 | 0 | 0 | 0 | 0 | 0 | 2344700 | 0 | 0 | 0 | 0 | 0 | 0 |
| ALISSNVQ | KIF20B | Q96Q89 | KI20B_HUMAN | Unmodified | 0 | 0 | 0 | 48453 | 1350600 | 0 | 0 | 0 | 0 | 0 | 0 | 0 | 0 | 0 | 0 | 0 | 731850 | 0 |
| ALIVEPSRE | DDX1; DDX1; D | Q92499; A0A | DDX1_HUMAN | Unmodified | 511690 | 0 | 611220 | 1266800 | 5128200 | 665410 | 2701100 | 199090 | 1110400 | 895990 | 1675900 | 10339000 | 8370800 | 2014300 | 1533900 | 734320 | 2154700 | 1714100 |
| ALKDEEKM | TPM3; TPM3; TF | P06753; J3K | TPM3_HUMAN | Unmodified | 34388 | 59611 | 0 | 0 | 49836000 | 10454000 | 480580 | 0 | 1496200 | 734780 | 211910 | 4476600 | 8375200 | 1113400 | 0 | 0 | 3575000 | 1927900 |
| ALKDEEKM | TPM3; TPM3; TF | P06753; J3K | TPM3_HUMAN | Oxidation (M) | 323420 | 171500 | 0 | 612360 | 20193000 | 5309900 | 111310 | 244870 | 1968800 | 1338500 | 1358900 | 3621200 | 8061800 | 1889600 | 0 | 367310 | 2417500 | 828600 |
| ALKDPNAF | EIF3M; EIF3M; E | Q7L2H7; H0 | EIF3M_HUMAN | Unmodified | 1328800 | 578570 | 680650 | 0 | 0 | 0 | 0 | 0 | 0 | 96805 | 663500 | 17140000 | 0 | 0 | 0 | 1989500 | 0 | 0 |
| ALKEDDPK | COPS2; COPS2 | P61201; H0 | CSN2_HUMAN | Unmodified | 0 | 0 | 0 | 0 | 0 | 0 | 0 | 135660 | 0 | 75795 | 0 | 0 | 356820 | 0 | 0 | 0 | 514600 | 378680 |
| ALKEGVLE | PHLDA1; PHLD | Q8WV24; R4 | PHLA1_HUMAN | Unmodified | 0 | 0 | 0 | 0 | 853150 | 0 | 0 | 0 | 365390 | 44039 | 0 | 0 | 0 | 0 | 0 | 0 | 3427800 | 1284000 |
| ALKPEVDKI | CTNNA1; CTNN | P35221; E5R | CTNA1_HUMA | Oxidation (M) | 892170 | 0 | 0 | 4458700 | 6133600 | 1831300 | 0 | 0 | 337060 | 0 | 1186500 | 5648200 | 4116600 | 7860500 | 3284700 | 4881900 | 0 | 0 |
| ALKPEVDKI | CTNNA1; CTNN | P35221; E5R | CTNA1_HUMA | Unmodified | 598790 | 924120 | 72718 | 2289800 | 10647000 | 2726500 | 0 | 0 | 614410 | 600380 | 1199700 | 0 | 4168500 | 3406000 | 859600 | 1370300 | 0 | 0 |
| ALKPGLSA | ENTPD6; ENTPI | O75354; Q5 | ENTP6_HUMAN | Unmodified | 0 | 0 | 0 | 103290 | 0 | 0 | 0 | 0 | 1885800 | 48142 | 0 | 816070 | 0 | 0 | 0 | 0 | 0 | 0 |
| ALKQISSN | ESD; ESD | P10768; U3 | ESTD_HUMAN | Acetyl (Protein N | 0 | 0 | 0 | 0 | 0 | 0 | 217880 | 266690 | 0 | 0 | 0 | 0 | 0 | 0 | 0 | 0 | 751710 | 0 |
| ALKVLDNY | CLIC1 | O00299 | CLIC1_HUMAN | Unmodified | 0 | 0 | 0 | 0 | 5390500 | 0 | 0 | 0 | 0 | 0 | 0 | 0 | 0 | 2625700 | 0 | 0 | 9134400 | 2579800 |
| ALKYSVDP | SLC4A7 | H7C3C4 | H7C3C4_HUM | Unmodified | 0 | 0 | 0 | 0 | 338210 | 0 | 0 | 0 | 0 | 0 | 0 | 0 | 0 | 0 | 0 | 0 | 664210 | 462910 |
| ALKYSVDP | SLC4A7 | H7C3C4 | H7C3C4_HUM | Oxidation (M) | 0 | 216480 | 0 | 0 | 0 | 0 | 0 | 0 | 0 | 0 | 0 | 841660 | 0 | 0 | 0 | 0 | 602700 | 266230 |
| ALLAEGVIL | MRPS23; MRPS | Q9Y3D9; J3 | RT23_HUMAN | Unmodified | 0 | 0 | 0 | 0 | 0 | 0 | 0 | 0 | 861250 | 0 | 0 | 0 | 0 | 0 | 0 | 0 | 0 | 0 |
| ALLAEHLLK | GLTP; GLTP; GL | Q9NZD2; F5 | GLTP_HUMAN | Acetyl (Protein N | 751440 | 553540 | 316280 | 488230 | 1006700 | 548470 | 149920 | 0 | 1858200 | 471740 | 926960 | 2140300 | 749960 | 248680 | 669630 | 591050 | 483990 | 1197600 |
| ALLAFQR | TTC7B; TTC7B | Q86TV6; Q6 | TTC7B_HUMAN | Unmodified | 0 | 0 | 172320 | 125200 | 0 | 0 | 0 | 0 | 0 | 0 | 0 | 0 | 0 | 0 | 265450 | 167650 | 0 | 0 |
| ALLAGGGF | EXT2 | Q93063 | EXT2_HUMAN | Unmodified | 0 | 0 | 0 | 0 | 0 | 0 | 0 | 0 | 5993700 | 0 | 0 | 0 | 0 | 0 | 0 | 0 | 0 | 1706200 |
| ALLALCGG | ANXA6; ANXA6; | P08133; E7E | ANXA6_HUMAN | Unmodified | 0 | 0 | 0 | 0 | 1905100 | 0 | 2453200 | 1080900 | 0 | 0 | 0 | 10988 | 1327100 | 1814400 | 0 | 0 | 2222600 | 6223800 |
| ALLANALTS | TMEM33; TMEM | P57088; D6 | TMM33_HUMA | Unmodified | 136260 | 0 | 0 | 0 | 0 | 0 | 0 | 73205 | 4724300 | 206310 | 73966 | 4043000 | 0 | 0 | 0 | 0 | 0 | 0 |
| ALLCYNRG | CHORDC1; CH | Q9UHD1; E9 | CHRD1_HUMA | Acetyl (Protein N | 0 | 0 | 0 | 0 | 990380 | 0 | 0 | 0 | 0 | 779590 | 0 | 0 | 1966200 | 0 | 0 | 0 | 7854200 | 2166800 |
| ALLDASETI | TRIM25; TRIM2 | Q14258; A0A | TRI25_HUMAN | Unmodified | 0 | 0 | 0 | 0 | 726520 | 0 | 0 | 0 | 0 | 0 | 0 | 0 | 0 | 0 | 0 | 0 | 132420 | 182920 |
| ALLDASETI | TRIM25; TRIM2 | Q14258; A0A | TRI25_HUMAN | Unmodified | 0 | 0 | 0 | 0 | 12524000 | 2611400 | 0 | 0 | 622550 | 298620 | 301540 | 2219300 | 1009800 | 699720 | 0 | 482830 | 1321200 | 2679500 |
| ALLDFEDK | MVP; MVP; MVP | Q14764; H3 | MVP_HUMAN | Unmodified | 0 | 0 | 0 | 0 | 0 | 0 | 0 | 533530 | 44660 | 0 | 0 | 1307000 | 0 | 0 | 0 | 0 | 0 | 0 |
| ALLDFEDKI | MVP; MVP; MVP | Q14764; H3 | MVP_HUMAN | Unmodified | 109800 | 109580 | 0 | 0 | 0 | 0 | 0 | 0 | 0 | 0 | 0 | 2629300 | 0 | 0 | 0 | 0 | 0 | 0 |
| ALLDPPLEL | SH3BP4 | Q9P0V3 | SH3B4_HUMAN | Unmodified | 0 | 0 | 0 | 0 | 1319800 | 0 | 0 | 0 | 903480 | 677360 | 143600 | 0 | 0 | 0 | 0 | 0 | 1296700 | 0 |
| ALLDRPLLI | TPBGL | P0DKB5 | TPBGL_HUMAN | Unmodified | 87138 | 0 | 0 | 0 | 0 | 0 | 0 | 0 | 2109600 | 170090 | 0 | 0 | 0 | 0 | 0 | 0 | 0 | 0 |
| ALLDSHSIF | CHAT | P28329 | CLAT_HUMAN | Unmodified | 0 | 0 | 0 | 0 | 0 | 0 | 0 | 0 | 0 | 0 | 105360 | 607230 | 0 | 0 | 0 | 0 | 0 | 0 |
| ALLDSLQL | GANAB | Q14697 | GANAB_HUMA | Unmodified | 0 | 0 | 0 | 0 | 0 | 0 | 0 | 0 | 2005100 | 7832800 | 119370 | 0 | 0 | 0 | 0 | 0 | 0 | 210100 |
| ALLEEYNK | DCTN3; DCTN3 | O75935; X6 | DCTN3_HUMA | Unmodified | 0 | 0 | 0 | 0 | 0 | 0 | 0 | 0 | 65142 | 269260 | 770260 | 1272400 | 0 | 0 | 0 | 0 | 0 | 855590 |
| ALLEQKIEE | TAOK1 | Q7L7X3 | TAOK1_HUMAN | Unmodified | 350640 | 0 | 0 | 0 | 624330 | 0 | 175460 | 0 | 1361000 | 0 | 0 | 857900 | 0 | 0 | 453690 | 0 | 458740 | 0 |
| ALLERTGY | HNRNPR; SYNC | O43390; O6 | HNRPR_HUMA | Unmodified | 0 | 0 | 0 | 0 | 3268400 | 0 | 1505200 | 307760 | 0 | 1128900 | 960220 | 0 | 3910800 | 2251300 | 0 | 0 | 548210 | 353200 |
| ALLEVVQS | PSMA7; PSMA8 | O14818; Q8 | PSA7_HUMAN | Unmodified | 470200 | 562890 | 1609900 | 1863300 | 8192700 | 1321100 | 2643600 | 465430 | 5330700 | 3247300 | 2768000 | 15635000 | 7248600 | 2303700 | 1256800 | 1469300 | 1670100 | 4459500 |
| ALLFIPR | HSP90AB1 | P08238 | HS90B_HUMAN | Unmodified | 3695900 | 0 | 8007300 | 7573100 | 8025700 | 2640400 | 0 | 143210 | 13305000 | 5487600 | 14267000 | 1.59E+08 | 7042400 | 1994400 | 11399000 | 8399000 | 1369200 | 4326400 |
| ALLFIPRRA | HSP90AB1 | P08238 | HS90B_HUMAN | Unmodified | 0 | 0 | 0 | 0 | 5528800 | 0 | 270470 | 0 | 0 | 1834100 | 0 | 0 | 3894700 | 3814300 | 0 | 0 | 1434800 | 2858000 |
| ALLFVPR | HSP90AA1 | P07900 | HS90A_HUMAN | Unmodified | 2654700 | 0 | 4044400 | 2664600 | 8053700 | 2213300 | 0 | 174800 | 12452000 | 7297200 | 10129000 | 84972000 | 2674700 | 707000 | 5457900 | 5988800 | 1406300 | 0 |
| ALLGYADN | GLOD4; GLOD4 | Q9HC38; F6 | GLOD4_HUMA | Unmodified | 0 | 0 | 0 | 0 | 0 | 0 | 0 | 0 | 113750 | 0 | 0 | 1389800 | 0 | 0 | 0 | 0 | 0 | 0 |
| ALLIIVGNP | MOV10; MOV10 | Q9HCE1; Q5 | MOV10_HUMA | Unmodified | 537770 | 1035500 | 0 | 0 | 0 | 0 | 0 | 0 | 0 | 0 | 0 | 0 | 0 | 0 | 988170 | 317430 | 0 | 0 |
| ALLILEGGQ | FAM20B; FAM2 | O75063; X6 | XYLK_HUMAN | Unmodified | 0 | 0 | 0 | 0 | 0 | 0 | 0 | 0 | 11532000 | 2495300 | 0 | 1324500 | 0 | 0 | 0 | 0 | 0 | 0 |
| ALLILEGGQ | FAM20B; FAM2 | O75063; X6 | XYLK_HUMAN | Unmodified | 0 | 0 | 0 | 0 | 0 | 0 | 0 | 0 | 0 | 361340 | 0 | 0 | 0 | 0 | 0 | 0 | 0 | 0 |
| ALLKLEKLD | GPS1; GPS1; G | Q13098; J3K | CSN1_HUMAN | Unmodified | 0 | 0 | 0 | 0 | 701310 | 0 | 0 | 0 | 0 | 0 | 0 | 0 | 439700 | 0 | 0 | 0 | 253520 | 609940 |
| ALLLGLAC | CARMIL1; CARI | Q5VZK9; A0A | CARL1_HUMAN | Unmodified | 0 | 0 | 0 | 0 | 370070 | 0 | 0 | 0 | 0 | 0 | 0 | 1444700 | 0 | 0 | 0 | 0 | 0 | 0 |
| ALLLSTYIK | AP2A2; AP2A1; | O94973; O9 | AP2A2_HUMAN | Unmodified | 0 | 0 | 253720 | 0 | 0 | 0 | 0 | 0 | 0 | 260350 | 0 | 2104000 | 0 | 0 | 247410 | 354150 | 0 | 0 |
| ALLLTGDF | MRPL46 | Q9H2W6 | RM46_HUMAN | Unmodified | 0 | 0 | 0 | 0 | 0 | 0 | 0 | 0 | 216950 | 77667 | 34462 | 0 | 0 | 0 | 0 | 0 | 233830 | 0 |
| ALLNHLDV | SLC39A14; SLC | Q15043; E5F | S39AE_HUMAN | Unmodified | 0 | 335360 | 292880 | 0 | 0 | 0 | 0 | 0 | 0 | 0 | 0 | 0 | 0 | 0 | 529420 | 488660 | 0 | 0 |
| ALLNNSHY | DLD; DLD; DLD | P09622; A0A | DLDH_HUMAN | Unmodified | 0 | 0 | 0 | 0 | 0 | 0 | 608760 | 53566 | 1998100 | 594650 | 0 | 0 | 0 | 0 | 0 | 0 | 0 | 240370 |
| ALLPELR | CHRNB1; DNAJ | P11230; Q8 | ACHB_HUMAN | Unmodified | 0 | 0 | 0 | 0 | 0 | 0 | 0 | 0 | 2820400 | 844250 | 0 | 0 | 0 | 0 | 0 | 0 | 0 | 0 |
| ALLQEFDN | KIF23; KIF23; KI | Q02241; A0A | KIF23_HUMAN | Unmodified | 327450 | 181010 | 465300 | 1264900 | 95591000 | 14509000 | 3071000 | 89296 | 3996300 | 11131000 | 9115300 | 9009300 | 847650 | 702240 | 1137200 | 243940 | 3878400 | 899740 |
| ALLQEFDN | KIF23; KIF23; KI | Q02241; A0A | KIF23_HUMAN | Unmodified | 0 | 0 | 0 | 0 | 1586000 | 0 | 0 | 0 | 0 | 0 | 0 | 0 | 0 | 0 | 0 | 0 | 1223300 | 0 |
| ALLQEFDN | KIF23; KIF23; KI | Q02241; A0A | KIF23_HUMAN | Oxidation (M) | 0 | 0 | 0 | 0 | 1925100 | 267980 | 184260 | 0 | 790420 | 515450 | 0 | 0 | 399500 | 188320 | 0 | 0 | 962720 | 0 |
| ALLQEFDN | KIF23; KIF23; KI | Q02241; A0A | KIF23_HUMAN | Oxidation (M) | 0 | 0 | 0 | 0 | 1880500 | 0 | 0 | 0 | 0 | 135750 | 0 | 0 | 0 | 0 | 0 | 0 | 3347200 | 0 |
| ALLQEFDN | KIF23; KIF23; KI | Q02241; A0A | KIF23_HUMAN | Unmodified | 0 | 24932 | 0 | 0 | 2828900 | 343980 | 178750 | 0 | 0 | 0 | 0 | 0 | 39448 | 0 | 0 | 0 | 2375500 | 102770 |
| ALLQEMPL | RO60 | P10155 | RO60_HUMAN | Unmodified | 2785100 | 4150300 | 141210 | 742510 | 0 | 0 | 0 | 0 | 695370 | 10099 | 0 | 283510 | 0 | 0 | 1593600 | 354560 | 0 | 0 |
| ALLQEMPL | RO60 | P10155 | RO60_HUMAN | Oxidation (M) | 19603000 | 16012000 | 2814700 | 5735000 | 0 | 0 | 0 | 196370 | 0 | 0 | 0 | 0 | 413450 | 0 | 7523100 | 7790300 | 0 | 0 |
| ALLQEMPL | RO60 | P10155 | RO60_HUMAN | Oxidation (M) | 345750 | 123530 | 0 | 495420 | 0 | 0 | 435980 | 0 | 0 | 316320 | 36412 | 0 | 1865700 | 1845400 | 0 | 95038 | 154220 | 0 |
| ALLQEMPL | RO60 | P10155 | RO60_HUMAN | Unmodified | 0 | 0 | 0 | 0 | 0 | 0 | 65312 | 0 | 0 | 43459 | 0 | 0 | 374410 | 215940 | 0 | 0 | 0 | 0 |
| ALLQMVEA | SLC4A2; SLC4A | P04920; A0A | B3A2_HUMAN | Oxidation (M) | 0 | 0 | 902870 | 2032600 | 0 | 0 | 1368400 | 0 | 0 | 0 | 0 | 0 | 0 | 0 | 2000400 | 140000 | 0 | 0 |
| ALLQMVQ | DNM2; DNM2 | P50570; K7E | DYN2_HUMAN | Oxidation (M) | 119960 | 1027600 | 16856 | 52122 | 495450 | 0 | 469870 | 0 | 0 | 98119 | 0 | 0 | 904140 | 650490 | 60254 | 29022 | 877710 | 92472 |
| ALLQQQPE | SRP68 | Q9UHB9 | SRP68_HUMAN | Unmodified | 0 | 0 | 0 | 0 | 0 | 0 | 0 | 0 | 82205 | 192570 | 0 | 1361400 | 0 | 0 | 0 | 0 | 0 | 0 |
| ALLQQQPE | SRP68 | Q9UHB9 | SRP68_HUMAN | Unmodified | 51715 | 38472 | 24554 | 0 | 528490 | 0 | 0 | 0 | 0 | 36740 | 0 | 2285200 | 0 | 116160 | 19158 | 19773 | 1309700 | 504360 |
| ALLQSSASI | PA2G4 | Q9UQ80 | PA2G4_HUMAN | Unmodified | 1362100 | 1832500 | 463450 | 1445900 | 554060 | 0 | 0 | 52054 | 782220 | 533580 | 792230 | 7053300 | 474330 | 262460 | 845740 | 648550 | 35996 | 0 |
| ALLQSSASI | PA2G4 | Q9UQ80 | PA2G4_HUMAN | Unmodified | 604740 | 356640 | 314870 | 201580 | 446990 | 0 | 0 | 0 | 240310 | 82140 | 89323 | 3657900 | 0 | 228130 | 639200 | 427540 | 0 | 0 |
| ALLRGVFV | ACAA2 | P42765 | THIM_HUMAN | Acetyl (Protein N | 0 | 0 | 0 | 0 | 0 | 0 | 125410 | 0 | 0 | 588440 | 0 | 0 | 0 | 0 | 0 | 0 | 0 | 0 |
| ALLRLQDT | P4HA1 | P13674 | P4HA1_HUMAN | Unmodified | 0 | 0 | 0 | 0 | 420820 | 0 | 0 | 50810 | 0 | 2503100 | 78481 | 0 | 0 | 346420 | 0 | 0 | 430070 | 2687500 |
| ALLRLQDT | P4HA1 | P13674 | P4HA1_HUMAN | Unmodified | 0 | 0 | 0 | 0 | 0 | 0 | 0 | 0 | 0 | 1435100 | 0 | 530380 | 0 | 0 | 0 | 0 | 279870 | 0 |
| ALLRLSQE | HSPA4; HSPA4; | P34932; A0A | HSP74_HUMAN | Unmodified | 0 | 0 | 0 | 0 | 0 | 0 | 318000 | 142570 | 120450 | 0 | 0 | 0 | 617590 | 597660 | 0 | 0 | 165990 | 0 |
| ALLRLYQE | HSPH1; HSPA4 | Q92598; O9 | HS105_HUMAN | Unmodified | 0 | 0 | 0 | 0 | 0 | 0 | 0 | 0 | 0 | 0 | 0 | 0 | 0 | 0 | 0 | 0 | 419740 | 0 |
| ALLRRPTVS | CCNB2; CCNB | O95067; H1 | CCNB2_HUMA | Acetyl (Protein N | 0 | 0 | 0 | 0 | 1968700 | 284800 | 0 | 196490 | 0 | 619770 | 0 | 0 | 169510 | 0 | 0 | 0 | 0 | 539180 |
| ALLSAPWY | HEXA; HEXA; HI | P06865; H3 | HEXA_HUMAN | Unmodified | 0 | 2321300 | 0 | 0 | 0 | 0 | 0 | 0 | 0 | 589270 | 679740 | 0 | 0 | 0 | 0 | 0 | 0 | 0 |
| ALLSAVTR | CTNNA1; CTNN | P35221; P26 | CTNA1_HUMA | Unmodified | 289900 | 144710 | 1594600 | 1089200 | 519810 | 0 | 0 | 0 | 191450 | 58474 | 588850 | 4441800 | 689040 | 160520 | 4368700 | 3105300 | 0 | 0 |
| ALLSTLHTV | GAK; GAK; GAK | O14976; A0A | GAK_HUMAN C | Unmodified | 0 | 89156 | 42949 | 19323 | 0 | 0 | 0 | 0 | 118100 | 283220 | 32091 | 1321200 | 130910 | 0 | 105100 | 65233 | 79275 | 0 |
| ALLSVHDT | MPP7; MPP7 | Q5T2T1; S4F | MPP7_HUMAN | Unmodified | 0 | 0 | 0 | 389920 | 4823000 | 886650 | 75906 | 93895 | 514780 | 941080 | 119990 | 562810 | 1632200 | 2397600 | 481860 | 507490 | 1119600 | 2185600 |

| | | | | | | | | | | | | | | | | | | | | | | |
|---|---|---|---|---|---|---|---|---|---|---|---|---|---|---|---|---|---|---|---|---|---|---|
| ALLSYDGLI | EPDR1 | Q9UM22 | EPDR1_HUMAN | Unmodified | 0 | 0 | 0 | 0 | 0 | 0 | 0 | 0 | 1917100 | 0 | 941360 | 650640 | 0 | 0 | 0 | 0 | 0 | 0 |
| ALLTPVAIA | GSR | P00390 | GSHR_HUMAN | Unmodified | 0 | 0 | 0 | 0 | 2562200 | 116880 | 0 | 134650 | 686460 | 133230 | 0 | 2078100 | 2513000 | 177300 | 0 | 518560 | 334530 | 417560 |
| ALLTPVAIA | GSR | P00390 | GSHR_HUMAN | Unmodified | 475180 | 279390 | 0 | 2833400 | 16589000 | 3353900 | 774840 | 689470 | 0 | 1269400 | 1446300 | 5060500 | 6677700 | 0 | 1670900 | 1863600 | 1262700 | 1284100 |
| ALLVTASQ( | CAP1; CAP1; C/ | Q01518; Q5 | CAP1_HUMAN | Unmodified | 125300 | 0 | 191980 | 51998 | 131640 | 0 | 0 | 0 | 1146400 | 0 | 577840 | 9558800 | 0 | 1678800 | 574410 | 490740 | 89919 | 0 |
| ALLVTASQ( | CAP1; CAP1; C/ | Q01518; Q5 | CAP1_HUMAN | Unmodified | 0 | 0 | 0 | 0 | 1136900 | 0 | 0 | 0 | 211160 | 2055500 | 0 | 0 | 0 | 0 | 0 | 0 | 0 | 0 |
| ALLYLCGG | ANXA2P2; ANX/ | A6NMY6; P0 | AXA2L_HUMAN | Unmodified | 0 | 0 | 0 | 0 | 12725000 | 2662500 | 0 | 1814300 | 2406800 | 2832700 | 0 | 0 | 0 | 0 | 0 | 0 | 5280400 | 4316400 |
| ALMAAEDK | TPM1; TPM1; TF | H0YL42; B7Z | H0YL42_HUMA | Unmodified | 0 | 0 | 60117 | 0 | 0 | 0 | 0 | 0 | 0 | 0 | 0 | 0 | 557590 | 0 | 0 | 319710 | 0 | 0 |
| ALMDEIEHI | CNPY1 | A0A1B0GVE( | A0A1B0GVE0_ | Oxidation (M) | 0 | 0 | 0 | 0 | 0 | 0 | 0 | 0 | 0 | 0 | 1098800 | 166920 | 0 | 0 | 0 | 0 | 0 | 0 |
| ALMDEIEHI | CNPY1 | A0A1B0GVE( | A0A1B0GVE0_ | Unmodified | 0 | 0 | 0 | 0 | 0 | 0 | 0 | 0 | 0 | 0 | 1669700 | 723200 | 0 | 0 | 0 | 0 | 0 | 0 |
| ALMDEVVK | PGK1 | P00558 | PGK1_HUMAN | Oxidation (M) | 573860 | 473590 | 10310000 | 1317300 | 4171500 | 631300 | 671550 | 567890 | 2359000 | 2335000 | 3115800 | 7687700 | 1874100 | 2422000 | 1032400 | 1684500 | 7397400 | 6434000 |
| ALMDEVVK | PGK1 | P00558 | PGK1_HUMAN | Unmodified | 783280 | 576530 | 2182100 | 2366800 | 0 | 3030000 | 3990100 | 2284900 | 10409000 | 6332500 | 9233200 | 33808000 | 13434000 | 10558000 | 0 | 2050900 | 25989000 | 61180000 |
| ALMDEVVK | PGK1 | P00558 | PGK1_HUMAN | Oxidation (M) | 0 | 0 | 0 | 0 | 208360 | 0 | 0 | 263090 | 0 | 897860 | 0 | 266160 | 1512200 | 631840 | 0 | 0 | 1811300 | 1713800 |
| ALMDEVVK | PGK1 | P00558 | PGK1_HUMAN | Unmodified | 0 | 0 | 0 | 0 | 2260600 | 0 | 497140 | 281500 | 0 | 1135300 | 0 | 472750 | 1832200 | 831660 | 0 | 0 | 4057100 | 5843100 |
| ALMDEVVK | PGK1 | P00558 | PGK1_HUMAN | Unmodified | 0 | 0 | 0 | 0 | 0 | 0 | 0 | 0 | 0 | 0 | 0 | 0 | 0 | 890500 | 0 | 0 | 4451100 | 1813800 |
| ALMELGGC | FAM171A1 | Q5VUB5 | F1711_HUMAN | Unmodified | 0 | 0 | 0 | 0 | 1049000 | 0 | 0 | 0 | 0 | 0 | 0 | 183200 | 0 | 0 | 0 | 0 | 0 | 0 |
| ALMGSPQL | JUP; JUP; JUP; J | P14923; C9J | PLAK_HUMAN | Oxidation (M) | 1444100 | 0 | 5262600 | 4788500 | 0 | 1005800 | 0 | 0 | 1797200 | 273370 | 1682800 | 11081000 | 4044200 | 0 | 11090000 | 10472000 | 0 | 0 |
| ALMGSPQL | JUP; JUP; JUP; J | P14923; C9J | PLAK_HUMAN | Unmodified | 0 | 0 | 330070 | 0 | 1126000 | 564860 | 0 | 0 | 3964300 | 0 | 0 | 1109200 | 0 | 0 | 0 | 0 | 0 | 0 |
| ALMLQGVD | HSPD1; HSPD1 | P10809; A0A | CH60_HUMAN | 2 Oxidation (M) | 0 | 0 | 0 | 0 | 0 | 95767 | 410290 | 0 | 744800 | 1027100 | 0 | 0 | 1210200 | 667350 | 0 | 0 | 0 | 278610 |
| ALMTGSLP | PIGG; PIGG; PI( | Q5H8A4; D6 | PIGG_HUMAN | Unmodified | 0 | 0 | 0 | 0 | 0 | 0 | 0 | 0 | 1843300 | 0 | 0 | 0 | 0 | 0 | 0 | 0 | 0 | 0 |
| ALMTGSLP | PIGG; PIGG; PI( | Q5H8A4; D6 | PIGG_HUMAN | Oxidation (M) | 0 | 0 | 0 | 0 | 0 | 0 | 0 | 0 | 1040600 | 0 | 0 | 0 | 0 | 0 | 0 | 0 | 0 | 0 |
| ALMTVGKE | NARS1; NARS1 | O43776; K7 | SYNC_HUMAN | Oxidation (M) | 68813 | 0 | 304430 | 0 | 563010 | 0 | 0 | 85457 | 0 | 0 | 390620 | 4094100 | 0 | 0 | 394600 | 525700 | 0 | 0 |
| ALMTVGKE | NARS1; NARS1 | O43776; K7 | SYNC_HUMAN | Unmodified | 0 | 0 | 0 | 0 | 0 | 0 | 0 | 0 | 0 | 532800 | 0 | 680200 | 0 | 0 | 0 | 0 | 0 | 0 |
| ALNEACES | GLG1; GLG1 | Q92896; H3 | GSLG1_HUMAI | Unmodified | 0 | 0 | 0 | 0 | 0 | 0 | 0 | 0 | 2446600 | 0 | 0 | 1896300 | 0 | 0 | 0 | 0 | 0 | 0 |
| ALNENLQC | PARP4 | Q9UKK3 | PARP4_HUMAN | Unmodified | 0 | 0 | 0 | 0 | 0 | 0 | 0 | 0 | 0 | 0 | 0 | 1244200 | 0 | 0 | 0 | 0 | 0 | 0 |
| ALNFGGIG | ECE1; ECE1; EC | P42892; B4E | ECE1_HUMAN | Unmodified | 138980 | 93219 | 18602000 | 564460 | 0 | 14927000 | 0 | 0 | 2362200 | 637460 | 0 | 31857 | 0 | 61881 | 436800 | 0 | 0 | 0 |
| ALNFGGIG | ECE1; ECE1; EC | P42892; B4E | ECE1_HUMAN | Unmodified | 1391900 | 646230 | 716530 | 0 | 4637600 | 826100 | 0 | 0 | 42996000 | 1996200 | 0 | 0 | 585770 | 0 | 1059900 | 1039200 | 78648 | 0 |
| ALNFNLDN | FSD1L; FSD1L; | Q9BXM9; Q8 | FSD1L_HUMAN | Unmodified | 0 | 0 | 0 | 0 | 0 | 0 | 0 | 0 | 0 | 0 | 0 | 915640 | 0 | 0 | 0 | 0 | 0 | 0 |
| ALNGAEPN | LAMTOR1; LAM | Q6IAA8; H0Y | LTOR1_HUMAN | Unmodified | 0 | 52357 | 151890 | 83729 | 0 | 0 | 0 | 0 | 0 | 0 | 0 | 1210100 | 0 | 0 | 153050 | 0 | 0 | 0 |
| ALNGDLVV | DIS3L2; DIS3L2 | Q8IYB7; C9J | DI3L2_HUMAN | Unmodified | 0 | 0 | 0 | 0 | 0 | 0 | 0 | 0 | 0 | 0 | 0 | 566680 | 0 | 0 | 0 | 0 | 0 | 0 |
| ALNGFTPLI | ANK3; ANK2; AI | Q12955; Q0 | ANK3_HUMAN | Unmodified | 0 | 0 | 0 | 0 | 0 | 0 | 0 | 41967 | 0 | 0 | 0 | 0 | 1029700 | 973920 | 1753300 | 403600 | 0 | 0 |
| ALNLAYSSI | CYFIP1 | Q7L576 | CYFP1_HUMAN | Unmodified | 228530 | 247830 | 0 | 0 | 0 | 0 | 0 | 11372 | 0 | 0 | 0 | 2153700 | 0 | 0 | 539990 | 306150 | 0 | 0 |
| ALNLGYALI | COL6A3; COL6/ | P12111; E7E | CO6A3_HUMAI | Unmodified | 0 | 27254 | 0 | 0 | 0 | 0 | 608310 | 0 | 0 | 185180 | 0 | 0 | 173700 | 577620 | 0 | 0 | 0 | 261660 |
| ALNRPPTYI | EIF5; EIF5; EIF5 | P55010; H0Y | IF5_HUMAN Eu | Unmodified | 2797800 | 0 | 684080 | 275220 | 770060 | 0 | 185040 | 112170 | 0 | 0 | 0 | 654190 | 0 | 0 | 0 | 0 | 340900 | 394010 |
| ALNSIIDVY | S100A8 | P05109 | S10A8_HUMAN | Unmodified | 0 | 0 | 0 | 2251300 | 0 | 0 | 0 | 0 | 0 | 0 | 0 | 0 | 0 | 0 | 0 | 45120 | 0 | 0 |
| ALNVEPDG | PRDX5 | P30044 | PRDX5_HUMAI | Unmodified | 0 | 291980 | 90106 | 793570 | 6050400 | 1254500 | 3736500 | 0 | 2268600 | 9766500 | 0 | 248530 | 10379000 | 3902300 | 2632700 | 0 | 21278000 | 6995000 |
| ALNVTPDY | TTC5; TTC5 | Q8N0Z6; H9 | TTC5_HUMAN | Unmodified | 0 | 0 | 0 | 0 | 906530 | 164480 | 610170 | 279460 | 0 | 0 | 0 | 0 | 406210 | 256040 | 0 | 0 | 363140 | 0 |
| ALPAVQQN | UBA1 | P22314 | UBA1_HUMAN | Unmodified | 445060 | 0 | 0 | 645350 | 0 | 0 | 0 | 44459 | 2728900 | 615060 | 1457300 | 4915600 | 0 | 0 | 706730 | 557850 | 0 | 0 |
| ALPAVQQN | UBA1 | P22314 | UBA1_HUMAN | Unmodified | 575750 | 321910 | 1773500 | 937070 | 1972200 | 0 | 0 | 775850 | 6451700 | 2654000 | 4172800 | 19656000 | 2187300 | 1087400 | 2337000 | 1815700 | 481820 | 1442800 |
| ALPCIVDVI | HSDL2 | Q6YN16 | HSDL2_HUMAI | Unmodified | 0 | 0 | 0 | 0 | 0 | 398990 | 557420 | 31621 | 13224000 | 5212700 | 0 | 0 | 0 | 0 | 0 | 0 | 0 | 0 |
| ALPEFFNG | SMARCC2; SMA | Q8TAQ2; Q9 | SMRC2_HUMA | Unmodified | 103600 | 0 | 0 | 0 | 1292600 | 0 | 26458 | 0 | 172640 | 55127 | 82118 | 1731700 | 32849 | 0 | 0 | 0 | 131850 | 0 |
| ALPELTKLL | JUP; JUP; JUP; J | P14923; C9J | PLAK_HUMAN | Unmodified | 0 | 0 | 0 | 0 | 2087300 | 0 | 0 | 0 | 0 | 0 | 0 | 0 | 2038000 | 1702500 | 0 | 0 | 998420 | 2354200 |
| ALPESLGQ | DDAH1 | O94760 | DDAH1_HUMA | Unmodified | 671020 | 833670 | 475280 | 453300 | 0 | 105190 | 0 | 0 | 1020700 | 356770 | 0 | 999100 | 0 | 140470 | 863460 | 1679400 | 0 | 0 |
| ALPFWNEE | PGAM1; PGAM4 | P18669; Q8I | PGAM1_HUMA | Unmodified | 1015800 | 817140 | 1204100 | 2651900 | 20011000 | 3908400 | 673220 | 0 | 3785200 | 1328300 | 16778000 | 22585000 | 8885000 | 2868300 | 493920 | 2119800 | 3095200 | 9956100 |
| ALPFWNEE | PGAM1; PGAM4 | P18669; Q8I | PGAM1_HUMA | Unmodified | 0 | 0 | 0 | 0 | 23829000 | 4378500 | 605710 | 18973 | 0 | 2392700 | 0 | 0 | 26589000 | 0 | 0 | 0 | 12278000 | 15695000 |
| ALPFWNEE | PGAM1; PGAM4 | P18669; Q8I | PGAM1_HUMA | Unmodified | 0 | 0 | 0 | 0 | 0 | 990040 | 0 | 0 | 0 | 406940 | 0 | 0 | 2381300 | 788840 | 0 | 0 | 2477600 | 1427000 |
| ALPILEELLI | TMTC3 | Q6ZXV5 | TMTC3_HUMAI | Unmodified | 0 | 0 | 0 | 0 | 0 | 0 | 0 | 0 | 0 | 555120 | 0 | 0 | 0 | 0 | 0 | 0 | 0 | 0 |
| ALPIYTTSA | ZDHHC20; ZDH | Q5W0Z9; B4 | ZDH20_HUMAI | Unmodified | 0 | 0 | 2504900 | 3884500 | 3937700 | 0 | 355730 | 0 | 951560 | 323680 | 869850 | 0 | 1031800 | 0 | 2098100 | 2938900 | 77556 | 508690 |
| ALPLALVLH | TYMP; TYMP; TY | P19971; C9J | TYPH_HUMAN | Unmodified | 0 | 0 | 60006 | 274940 | 171920 | 38729 | 0 | 0 | 0 | 0 | 0 | 0 | 172310 | 0 | 484860 | 195260 | 107440 | 58501 |
| ALPLALVLH | TYMP; TYMP; TY | P19971; C9J | TYPH_HUMAN | Unmodified | 0 | 0 | 0 | 0 | 139370 | 66367 | 0 | 0 | 0 | 0 | 0 | 0 | 161910 | 70070 | 0 | 0 | 32260 | 71218 |
| ALPLEAEVA | MAP4; MAP4 | P27816; E7E | MAP4_HUMAN | Unmodified | 0 | 0 | 0 | 0 | 0 | 0 | 0 | 0 | 0 | 0 | 690360 | 1358400 | 0 | 0 | 0 | 0 | 0 | 0 |
| ALPLPHFD | TPX2 | Q9ULW0 | TPX2_HUMAN | Unmodified | 0 | 0 | 0 | 0 | 325520 | 0 | 0 | 60097 | 0 | 91154 | 0 | 3430900 | 163990 | 0 | 0 | 0 | 80606 | 192640 |
| ALPLPTPEE | NHS; NHS; NHS | Q6T4R5; A0/ | NHS_HUMAN | Unmodified | 0 | 0 | 0 | 0 | 950510 | 105860 | 0 | 0 | 244360 | 123500 | 0 | 0 | 0 | 0 | 0 | 0 | 573190 | 648880 |
| ALPPAQAG | IRGQ | Q8WZA9 | IRGQ_HUMAN | Unmodified | 0 | 0 | 0 | 0 | 0 | 0 | 0 | 0 | 256770 | 0 | 305030 | 958270 | 0 | 0 | 0 | 0 | 0 | 0 |
| ALPSEELNI | EIF4G1; EIF4G1 | Q04637; E7E | IF4G1_HUMAN | Unmodified | 0 | 0 | 0 | 0 | 0 | 0 | 201060 | 331980 | 349320 | 0 | 144220 | 1931600 | 0 | 0 | 0 | 0 | 0 | 0 |
| ALPSEELNI | EIF4G1; EIF4G1 | Q04637; E7E | IF4G1_HUMAN | Unmodified | 0 | 0 | 0 | 0 | 0 | 176110 | 0 | 0 | 1040700 | 171240 | 1237200 | 2269500 | 0 | 0 | 0 | 0 | 61899 | 0 |
| ALPSQGLS | SGPL1; SGPL1; | O95470; H7 | SGPL1_HUMAN | Unmodified | 0 | 0 | 0 | 0 | 0 | 0 | 0 | 0 | 0 | 0 | 471470 | 0 | 0 | 0 | 0 | 0 | 45052 | 0 |
| ALPTSKPE( | ZC3H4; ZC3H4 | Q9UPT8; M0 | ZC3H4_HUMAI | Unmodified | 0 | 0 | 0 | 0 | 65715 | 0 | 0 | 56531 | 0 | 67127 | 255230 | 455310 | 0 | 65880 | 0 | 0 | 522890 | 395370 |
| ALQAAYGA | AKR7A2; AKR7A | O43488; H3I | ARK72_HUMAN | Unmodified | 148010 | 0 | 167600 | 0 | 0 | 82138 | 0 | 0 | 0 | 0 | 0 | 1823000 | 196300 | 75463 | 236240 | 235740 | 107760 | 224360 |
| ALQALQQE | VPS25; VPS25 | Q9BRG1; K7 | VPS25_HUMAN | Unmodified | 0 | 0 | 0 | 0 | 0 | 0 | 0 | 0 | 0 | 0 | 0 | 0 | 0 | 0 | 344340 | 0 | 58452 | 0 |
| ALQALQQE | VPS25; VPS25 | Q9BRG1; K7 | VPS25_HUMAN | Unmodified | 18513000 | 24888000 | 7189300 | 10856000 | 468090 | 0 | 0 | 0 | 1769900 | 0 | 0 | 6560200 | 0 | 0 | 31479000 | 18283000 | 89700 | 0 |
| ALQALQQE | VPS25; VPS25 | Q9BRG1; K7 | VPS25_HUMAN | Unmodified | 614810 | 0 | 0 | 657720 | 0 | 0 | 0 | 0 | 0 | 0 | 0 | 0 | 0 | 0 | 850680 | 0 | 0 | 0 |
| ALQAQEIE( | COLGALT1; CO | Q8NBJ5; M0 | GT251_HUMAN | Unmodified | 0 | 38947 | 0 | 47572 | 36721 | 231750 | 104690 | 0 | 6085900 | 276370 | 135450 | 244310 | 0 | 147870 | 0 | 0 | 0 | 123930 |
| ALQAQEIE( | COLGALT1; CO | Q8NBJ5; M0 | GT251_HUMAN | Unmodified | 0 | 0 | 0 | 0 | 0 | 0 | 0 | 0 | 1161100 | 4022300 | 0 | 688710 | 0 | 0 | 0 | 0 | 0 | 0 |
| ALQASALN | ALDOC; ALDOC | P09972; A8I | ALDOC_HUMA | Unmodified | 123010 | 0 | 0 | 0 | 0 | 0 | 0 | 0 | 463280 | 92597 | 0 | 5093000 | 0 | 0 | 588360 | 462580 | 0 | 0 |
| ALQATVGN | LAMP1 | P11279 | LAMP1_HUMAI | Unmodified | 1807300 | 1447900 | 3777100 | 2228300 | 901260 | 254170 | 101360 | 121790 | 1.51E+08 | 9809600 | 6293700 | 6460800 | 872740 | 1750900 | 3073500 | 2169700 | 0 | 720480 |
| ALQDEWD/ | PRPF19; PRPF1 | Q9UMS4; F5 | PRP19_HUMAN | Oxidation (M) | 0 | 0 | 0 | 0 | 0 | 0 | 0 | 0 | 0 | 0 | 0 | 0 | 0 | 0 | 0 | 0 | 0 | 0 |
| ALQDFKEG | SQOR; SQOR; I | Q9Y6N5; H3 | SQOR_HUMAN | Unmodified | 0 | 0 | 0 | 593490 | 0 | 0 | 0 | 0 | 5909400 | 4744700 | 0 | 0 | 0 | 0 | 0 | 0 | 0 | 0 |
| ALQDLENA | RPRD1A; RPRD | Q96P16; A0/ | RPR1A_HUMAN | Unmodified | 0 | 0 | 0 | 0 | 0 | 0 | 0 | 0 | 0 | 0 | 0 | 0 | 0 | 0 | 0 | 0 | 0 | 0 |
| ALQDLENA | RPRD1B; RPRD | Q9NQG5; A2 | RPR1B_HUMAI | Unmodified | 0 | 0 | 0 | 0 | 755270 | 0 | 0 | 0 | 0 | 269200 | 0 | 0 | 685410 | 116760 | 0 | 0 | 242180 | 328230 |
| ALQDLGLR | DDAH2; DDAH2 | O95865; H0 | DDAH2_HUMA | Unmodified | 518030 | 428180 | 620380 | 840760 | 1132200 | 0 | 114330 | 32544 | 4719600 | 601210 | 466370 | 1212200 | 0 | 0 | 502390 | 452230 | 440020 | 1051000 |
| ALQDLQLD | LZTFL1; LZTFL1 | Q9NQ48; H7 | LZTL1_HUMAN | Unmodified | 0 | 0 | 0 | 84735 | 179500 | 37627 | 0 | 0 | 0 | 120870 | 0 | 0 | 138370 | 0 | 0 | 0 | 0 | 408320 |
| ALQDMSSI | INPPL1; INPPL1 | O15357; A0/ | SHIP2_HUMAN | Oxidation (M) | 0 | 0 | 0 | 0 | 0 | 0 | 0 | 0 | 0 | 0 | 65898 | 371510 | 0 | 0 | 0 | 0 | 0 | 0 |
| ALQDMSSI | INPPL1; INPPL1 | O15357; A0/ | SHIP2_HUMAN | Unmodified | 0 | 0 | 47029 | 0 | 0 | 0 | 0 | 0 | 0 | 0 | 113180 | 472840 | 0 | 137370 | 57628 | 48415 | 0 | 0 |
| ALQDMSSI | INPPL1; INPPL1 | O15357; A0/ | SHIP2_HUMAN | Unmodified | 0 | 0 | 0 | 0 | 0 | 0 | 0 | 0 | 0 | 0 | 430920 | 2122000 | 0 | 0 | 0 | 0 | 75757 | 388640 |
| ALQDPALA | IQGAP3; IQGAP | Q86VI3; F2Z | IQGA3_HUMAN | Unmodified | 52311000 | 242000 | 437360 | 808390 | 785110 | 429620 | 0 | 0 | 306150 | 601540 | 438450 | 1403400 | 0 | 0 | 245240 | 94392 | 309120 | 0 |
| ALQEGDLA | EXOC1 | Q9NV70 | EXOC1_HUMAI | Unmodified | 0 | 0 | 0 | 127300 | 85040 | 170340 | 0 | 0 | 626030 | 0 | 144020 | 0 | 83739 | 32527 | 0 | 129090 | 58127 | 142110 |
| ALQEGEGE | CARS1; CARS1 | P49589; B4I | SYCC_HUMAN | Unmodified | 0 | 0 | 0 | 0 | 0 | 0 | 0 | 0 | 0 | 913840 | 0 | 15414000 | 3827200 | 0 | 0 | 521750 | 0 | 0 |
| ALQELLR | MPP2; MPP2; M | Q14168; A0/ | MPP2_HUMAN | Unmodified | 0 | 0 | 0 | 0 | 0 | 0 | 0 | 0 | 0 | 0 | 0 | 0 | 0 | 0 | 262530 | 0 | 0 | 0 |
| ALQFLEEV | CCT6A | P40227 | TCPZ_HUMAN | Unmodified | 0 | 0 | 751530 | 815990 | 3457100 | 877600 | 796940 | 457480 | 5683600 | 1765100 | 2288600 | 19431000 | 8384800 | 2070400 | 0 | 857010 | 1425400 | 2705200 |
| ALQGASQI | DNM1L | O00429 | DNM1L_HUMA | Unmodified | 0 | 0 | 0 | 0 | 0 | 0 | 0 | 0 | 0 | 0 | 0 | 4056800 | 0 | 0 | 294970 | 36520 | 0 | 0 |
| ALQGASQI | DNM1L | O00429 | DNM1L_HUMA | Unmodified | 0 | 0 | 0 | 0 | 595090 | 83450 | 2489500 | 0 | 43482 | 436940 | 84236 | 0 | 2380900 | 1196900 | 0 | 0 | 0 | 689820 |
| ALQGDYKD | TMCO3; TMCO3 | Q6UWJ1; A0 | TMCO3_HUMA | Unmodified | 0 | 0 | 0 | 0 | 0 | 341240 | 0 | 0 | 3294200 | 989830 | 0 | 0 | 0 | 0 | 0 | 0 | 0 | 0 |
| ALQGQISS | DST; DST; DST; | Q03001; A0 | DYST_HUMAN | Unmodified | 0 | 0 | 0 | 0 | 744580 | 0 | 0 | 0 | 0 | 0 | 0 | 0 | 0 | 0 | 0 | 0 | 0 | 0 |
| ALQHMTDF | AP2B1; AP2B1 | P63010; A0A | AP2B1_HUMAN | Oxidation (M) | 172660 | 260980 | 342940 | 419230 | 801440 | 479340 | 360630 | 152150 | 334040 | 504180 | 2253700 | 0 | 1233300 | 761790 | 305050 | 402250 | 915750 | 954670 |
| ALQHMTDF | AP2B1; AP2B1 | P63010; A0A | AP2B1_HUMAN | Unmodified | 0 | 139350 | 72061 | 127560 | 2864100 | 248560 | 635030 | 0 | 0 | 920510 | 0 | 913640 | 1320100 | 1031500 | 140820 | 83848 | 2197800 | 2111300 |
| ALQHSYLH | CDK4; CDK4 | P11802; F8V | CDK4_HUMAN | Unmodified | 151900 | 0 | 117910 | 0 | 0 | 0 | 0 | 0 | 939690 | 274810 | 0 | 141800 | 0 | 0 | 168020 | 122940 | 188350 | 162290 |
| ALQLAQRP | GBA; GBA; GBA | P04062; A0A | GLCM_HUMAN | Unmodified | 0 | 0 | 0 | 0 | 0 | 0 | 0 | 0 | 5698100 | 51421 | 0 | 0 | 0 | 0 | 0 | 0 | 0 | 0 |
| ALQLEEER | EZR; EZR | P15311; E7E | EZRI_HUMAN E | Unmodified | 0 | 236350 | 213110 | 0 | 0 | 0 | 0 | 0 | 4669700 | 0 | 1956700 | 15375000 | 0 | 0 | 636770 | 1262600 | 0 | 0 |
| ALQLEEERI | EZR; EZR | P15311; E7E | EZRI_HUMAN E | Unmodified | 0 | 0 | 0 | 214180 | 0 | 0 | 0 | 0 | 520770 | 26008 | 172900 | 951020 | 0 | 0 | 291010 | 0 | 0 | 0 |
| ALQLEVDR | PRC1 | O43663 | PRC1_HUMAN | Unmodified | 0 | 0 | 0 | 0 | 975790 | 660570 | 0 | 54853 | 1270900 | 857850 | 138860 | 0 | 794400 | 569940 | 990980 | 0 | 595990 | 0 |
| ALQLLHGG | CHAT; CHAT; C | P28329; A0A | CLAT_HUMAN | Unmodified | 0 | 0 | 0 | 0 | 0 | 0 | 0 | 0 | 0 | 166960 | 0 | 1993800 | 0 | 0 | 0 | 0 | 0 | 0 |
| ALQNAVTT | HGS; HGS; HG | O14964; A0/ | HGS_HUMAN I | Unmodified | 0 | 0 | 0 | 0 | 0 | 0 | 0 | 0 | 2236100 | 665080 | 0 | 1239500 | 0 | 0 | 0 | 0 | 0 | 0 |
| ALQNAVTT | HGS; HGS; HG | O14964; A0/ | HGS_HUMAN I | Unmodified | 0 | 0 | 0 | 0 | 17844000 | 0 | 2267400 | 0 | 0 | 688950 | 0 | 0 | 0 | 1710400 | 0 | 0 | 469960 | 0 |
| ALQPIVSEE | CRAT; CRAT; CI | P43155; A0A | CACP_HUMAN | Unmodified | 0 | 0 | 0 | 0 | 0 | 0 | 472300 | 124240 | 2366900 | 927800 | 0 | 0 | 0 | 0 | 0 | 0 | 17150 | 520000 |
| ALQPLEEG | MVP | Q14764 | MVP_HUMAN I | Unmodified | 4552800 | 447110 | 0 | 0 | 0 | 0 | 0 | 0 | 2837800 | 169420 | 0 | 28720000 | 0 | 0 | 0 | 0 | 0 | 0 |
| ALQPLNQR | CA14; CA14 | Q9ULX7; U3 | CAH14_HUMAI | Unmodified | 257770 | 120870 | 0 | 41096 | 0 | 0 | 0 | 0 | 0 | 0 | 0 | 5389600 | 0 | 0 | 48256 | 92675 | 0 | 0 |
| ALQQYTLEI | COPG1 | Q9Y678 | COPG1_HUMA | Unmodified | 152450 | 0 | 0 | 0 | 436260 | 119220 | 0 | 45155 | 0 | 0 | 363220 | 5486600 | 1254800 | 404220 | 285200 | 408820 | 0 | 772200 |
| ALQSGPPC | IGF2BP3 | O00425 | IF2B3_HUMAN | Unmodified | 546540 | 0 | 0 | 0 | 0 | 42616 | 0 | 0 | 778660 | 0 | 0 | 0 | 0 | 0 | 0 | 0 | 0 | 0 |
| ALQSGQC/ | PHGDH; PHGD | O43175; A0/ | SERA_HUMAN | Unmodified | 0 | 0 | 1537900 | 2108000 | 0 | 0 | 0 | 99404 | 0 | 0 | 0 | 8793300 | 0 | 0 | 2969800 | 1884700 | 0 | 0 |
| ALQSGQC/ | PHGDH; PHGD | O43175; A0/ | SERA_HUMAN | Unmodified | 0 | 0 | 0 | 0 | 0 | 0 | 0 | 0 | 0 | 0 | 0 | 0 | 2656900 | 3605100 | 0 | 0 | 0 | 0 |
| ALQSLACG | CUL3; CUL3 | Q13618; A0/ | CUL3_HUMAN | Unmodified | 0 | 0 | 0 | 0 | 0 | 56531 | 0 | 0 | 244070 | 41738 | 132940 | 1272300 | 0 | 0 | 0 | 215400 | 0 | 0 |
| ALQSLATVI | SNX9; SNX9; SI | Q9Y5X1; A0/ | SNX9_HUMAN | Unmodified | 2262200 | 7467400 | 0 | 1200100 | 0 | 0 | 0 | 0 | 0 | 0 | 0 | 0 | 199470 | 0 | 1839400 | 0 | 0 | 0 |
| ALQSPALG | IQGAP1; IQGAP | P46940; A0A | IQGA1_HUMAN | Unmodified | 0 | 0 | 0 | 0 | 2573200 | 1220400 | 0 | 0 | 0 | 0 | 0 | 0 | 0 | 0 | 0 | 0 | 324760 | 0 |
| ALQSQSWI | ECPAS; ECPAS; | Q5VYK3; J3K | ECM29_HUMAI | Unmodified | 357780 | 149010 | 0 | 0 | 0 | 0 | 0 | 0 | 0 | 0 | 23879 | 431800 | 0 | 32518 | 38768 | 116630 | 63867 | 564210 |
| ALQSVGQI | PSMC6; PSMC6 | P62333; H0 | PRS10_HUMAN | Unmodified | 76315 | 388480 | 163810 | 307490 | 2401300 | 405700 | 1614300 | 0 | 905180 | 750910 | 244860 | 1171800 | 195030 | 1292000 | 0 | 281810 | 1386300 | 0 |
| ALQTERND | TXLNA | P40222 | TXLNA_HUMAN | Unmodified | 0 | 0 | 0 | 0 | 0 | 0 | 0 | 0 | 0 | 0 | 0 | 0 | 0 | 0 | 0 | 0 | 0 | 1387500 |
| ALQVADFS | LRRC1; LRRC1 | Q9BTT6; Q5 | LRRC1_HUMAN | Unmodified | 0 | 0 | 0 | 0 | 0 | 0 | 0 | 0 | 0 | 0 | 0 | 1162100 | 0 | 0 | 0 | 0 | 0 | 0 |
| ALQVELVP | MYO1C | O00159 | MYO1C_HUMA | Acetyl (Protein N | 0 | 0 | 0 | 0 | 0 | 0 | 0 | 0 | 0 | 0 | 0 | 0 | 0 | 0 | 0 | 0 | 7012200 | 5166800 |
| ALQWHPDI | DNAJB2; DNAJE | P25686; C9J | DNJB2_HUMAN | Unmodified | 121080 | 124420 | 82972 | 0 | 0 | 0 | 0 | 0 | 0 | 0 | 0 | 0 | 0 | 0 | 264270 | 198190 | 0 | 0 |

| | | | | | | | | | | | | | | | | | | | | | | |
|---|---|---|---|---|---|---|---|---|---|---|---|---|---|---|---|---|---|---|---|---|---|---|
| ALQYAFFAI | EHHADH | Q08426 | ECHP_HUMAN | Unmodified | 0 | 0 | 0 | 0 | 0 | 0 | 0 | 0 | 1842400 | 0 | 0 | 0 | 0 | 0 | 0 | 0 | 0 | 0 |
| ALQYAFFAI | EHHADH | Q08426 | ECHP_HUMAN | Unmodified | 0 | 0 | 0 | 0 | 0 | 0 | 0 | 0 | 1696100 | 914360 | 0 | 0 | 0 | 0 | 0 | 0 | 0 | 0 |
| ALRASESGI | ADGRE5 | P48960 | AGRE5_HUMAI | Unmodified | 597400 | 0 | 0 | 384460 | 6584800 | 0 | 0 | 0 | 822830 | 0 | 35399 | 698320 | 940760 | 265190 | 716580 | 154230 | 4647400 | 0 |
| ALREMNWI | CDCA8 | Q53HL2 | BOREA_HUMAI | Oxidation (M) | 0 | 173560 | 0 | 0 | 4000700 | 820210 | 0 | 0 | 112920 | 0 | 0 | 0 | 0 | 1684100 | 0 | 0 | 294870 | 0 |
| ALREMNWI | CDCA8 | Q53HL2 | BOREA_HUMAI | Unmodified | 0 | 2092000 | 0 | 0 | 1053900 | 0 | 0 | 0 | 0 | 9872.2 | 0 | 0 | 137910 | 0 | 0 | 0 | 0 | 0 |
| ALRLDVGN | RPS8 | P62241 | RS8_HUMAN 4 | Unmodified | 1387800 | 285250 | 0 | 438220 | 0 | 0 | 0 | 0 | 0 | 0 | 0 | 1144800 | 343290 | 0 | 0 | 134880 | 0 | 0 |
| ALRLDVGN | RPS8 | P62241 | RS8_HUMAN 4 | Unmodified | 307180 | 292150 | 87000 | 408160 | 0 | 0 | 1304300 | 0 | 0 | 0 | 0 | 1705800 | 338260 | 0 | 0 | 0 | 0 | 0 |
| ALRLGDAIL | SNTB2; SNTB2; | Q13425; A0A | SNTB2_HUMAI | Unmodified | 0 | 0 | 0 | 360250 | 0 | 74149 | 0 | 0 | 0 | 191110 | 0 | 0 | 478160 | 0 | 0 | 0 | 0 | 504800 |
| ALRLGDAIL | SNTB2; SNTB2; | Q13425; A0A | SNTB2_HUMAI | Unmodified | 0 | 0 | 0 | 0 | 1874700 | 0 | 0 | 0 | 0 | 441680 | 0 | 0 | 2743700 | 0 | 0 | 0 | 0 | 2362100 |
| ALRNHLLE | TAOK1 | Q7L7X3 | TAOK1_HUMAI | Unmodified | 0 | 0 | 0 | 0 | 804800 | 0 | 509490 | 167660 | 0 | 0 | 0 | 0 | 0 | 209920 | 0 | 0 | 200580 | 0 |
| ALRTDYNA | HNRNPK; HNRI | P61978; Q5T | HNRPK_HUMA | Unmodified | 2389300 | 2519200 | 1337500 | 1602600 | 944740 | 0 | 466090 | 420610 | 3714200 | 856660 | 3253700 | 14850000 | 1733100 | 455700 | 1196500 | 797120 | 74741 | 1039500 |
| ALRTVWSS | SPINT2; SPINT2 | O43291; K7E | SPIT2_HUMAN | Unmodified | 0 | 0 | 0 | 0 | 4374500 | 0 | 0 | 0 | 0 | 16930000 | 0 | 0 | 919330 | 0 | 0 | 0 | 0 | 1096900 |
| ALRVLQAQ | TOMM34 | Q15785 | TOM34_HUMAI | Unmodified | 0 | 0 | 0 | 0 | 0 | 0 | 0 | 0 | 0 | 0 | 0 | 0 | 730650 | 0 | 0 | 0 | 851560 | 0 |
| ALRYPMAV | RPL36; RPL36; I | Q9Y3U8; J3K | RL36_HUMAN ( | Oxidation (M) | 1385500 | 4013800 | 1401700 | 1662400 | 623400 | 148110 | 143330 | 246240 | 0 | 0 | 475040 | 3454200 | 1430800 | 361370 | 1920100 | 1839900 | 58518 | 279850 |
| ALRYPMAV | RPL36; RPL36; I | Q9Y3U8; J3K | RL36_HUMAN ( | Unmodified | 2770800 | 3046800 | 867550 | 1312400 | 3386000 | 224550 | 541310 | 0 | 593010 | 86941 | 206680 | 7246300 | 3377400 | 891000 | 1518900 | 1248100 | 0 | 1354600 |
| ALSADQGS | PTGFRN | Q9P2B2 | FPRP_HUMAN | Unmodified | 5405400 | 4893600 | 25817000 | 24169000 | 1695500 | 2074500 | 141600 | 34542 | 226120 | 0 | 0 | 23132000 | 13101000 | 2580700 | 15904000 | 18222000 | 185590 | 480910 |
| ALSADQGS | PTGFRN | Q9P2B2 | FPRP_HUMAN | Unmodified | 0 | 0 | 0 | 0 | 0 | 0 | 0 | 0 | 0 | 0 | 0 | 0 | 11678000 | 8052700 | 0 | 0 | 0 | 0 |
| ALSAIADLL | CTNND1; CTNN | O60716; A0A | CTND1_HUMAI | Unmodified | 5422300 | 3775700 | 13116000 | 14096000 | 1896100 | 118800 | 419630 | 435190 | 0 | 50942 | 4408700 | 57011000 | 4126600 | 1729800 | 32001000 | 29803000 | 856170 | 2395000 |
| ALSAIADLL | CTNND1; CTNN | O60716; A0A | CTND1_HUMAI | Unmodified | 522330 | 436270 | 3584100 | 8015700 | 73122000 | 13415000 | 1389500 | 23284 | 0 | 152880 | 3918300 | 2478700 | 71461000 | 56063000 | 6729700 | 4571600 | 44339000 | 61146000 |
| ALSDADVQ | ATP6V1E1 | P36543 | VATE1_HUMAN | Acetyl (Protein N | 0 | 0 | 0 | 0 | 2163500 | 0 | 1268700 | 0 | 469500 | 272440 | 506300 | 4805100 | 2105100 | 1176800 | 0 | 0 | 583090 | 1179200 |
| ALSEALTEL | SHMT1 | P34896 | GLYC_HUMAN | Unmodified | 0 | 0 | 0 | 0 | 0 | 0 | 0 | 0 | 505220 | 353010 | 286950 | 4787600 | 0 | 0 | 0 | 0 | 0 | 0 |
| ALSEAQARI | PKN2; PKN2; PI | Q16513; B1A | PKN2_HUMAN | Unmodified | 0 | 0 | 0 | 0 | 2392900 | 0 | 661880 | 0 | 0 | 1759100 | 0 | 0 | 0 | 1462800 | 0 | 0 | 966040 | 1517800 |
| ALSEEKDVI | CEP55 | Q53EZ4 | CEP55_HUMAI | Unmodified | 0 | 0 | 108300 | 0 | 696010 | 0 | 0 | 0 | 0 | 192640 | 0 | 0 | 0 | 0 | 0 | 0 | 36641 | 0 |
| ALSEEKDVI | CEP55 | Q53EZ4 | CEP55_HUMAI | Unmodified | 0 | 0 | 0 | 0 | 3447200 | 0 | 260120 | 0 | 0 | 2035400 | 0 | 0 | 0 | 0 | 0 | 0 | 156470 | 0 |
| ALSEIAGMT | NDUFS1; NDUI | P28331; B4D | NDUS1_HUMA | Unmodified | 0 | 0 | 0 | 11209 | 0 | 57329 | 0 | 0 | 1305100 | 0 | 0 | 0 | 0 | 0 | 0 | 0 | 0 | 0 |
| ALSETVVEE | LRRC8A | Q8IWT6 | LRC8A_HUMAI | Unmodified | 0 | 0 | 0 | 0 | 0 | 356860 | 0 | 32110 | 0 | 0 | 0 | 1766200 | 0 | 0 | 0 | 0 | 0 | 0 |
| ALSEVSGEI | LRRC8C | Q8TDW0 | LRC8C_HUMAI | Unmodified | 0 | 0 | 0 | 0 | 0 | 54297 | 0 | 0 | 0 | 0 | 51011 | 375510 | 0 | 0 | 0 | 0 | 0 | 0 |
| ALSGNVIAV | NPTX1 | Q15818 | NPTX1_HUMAI | Unmodified | 0 | 0 | 0 | 0 | 0 | 0 | 0 | 0 | 1097500 | 1106000 | 0 | 0 | 0 | 0 | 0 | 0 | 0 | 0 |
| ALSGTSVAS | MBTPS1 | Q14703 | MBTP1_HUMAI | Unmodified | 0 | 0 | 0 | 0 | 0 | 0 | 0 | 0 | 269650 | 41809 | 0 | 0 | 0 | 0 | 0 | 0 | 0 | 0 |
| ALSGVLEQ | HBS1L; HBS1L; | Q9Y450; G5I | HBS1L_HUMAI | Unmodified | 328180 | 377530 | 193140 | 188350 | 301370 | 67383 | 0 | 0 | 368820 | 304960 | 243830 | 840860 | 0 | 0 | 556830 | 297140 | 129750 | 414990 |
| ALSIENSQL | LDOC1 | O95751 | LDOC1_HUMAI | Oxidation (M) | 2340800 | 2510200 | 8305500 | 3309100 | 0 | 0 | 0 | 0 | 0 | 0 | 0 | 0 | 0 | 0 | 8274400 | 5739400 | 0 | 0 |
| ALSIENSQL | LDOC1 | O95751 | LDOC1_HUMAI | Unmodified | 2706800 | 1327700 | 1615700 | 1969200 | 0 | 0 | 123880 | 0 | 0 | 0 | 0 | 0 | 26879 | 0 | 5633700 | 1002700 | 0 | 0 |
| ALSIFNELR | CDC23 | Q9UJX2 | CDC23_HUMA | Unmodified | 0 | 0 | 0 | 0 | 0 | 0 | 0 | 0 | 0 | 0 | 0 | 960870 | 0 | 0 | 0 | 0 | 0 | 0 |
| ALSIGFETC | CD44; CD44; C | P16070; E9P | CD44_HUMAN | Unmodified | 2343000 | 1122800 | 0 | 0 | 308230 | 0 | 0 | 0 | 0 | 59236 | 0 | 0 | 85667 | 0 | 0 | 0 | 0 | 669030 |
| ALSITLVQE | TNS3 | Q68CZ2 | TENS3_HUMAI | Unmodified | 0 | 0 | 0 | 0 | 3902600 | 427290 | 0 | 0 | 0 | 0 | 413790 | 227800 | 0 | 0 | 0 | 0 | 1764600 | 0 |
| ALSLGGQV | SV2A | Q7L0J3 | SV2A_HUMAN | Unmodified | 0 | 0 | 421150 | 4823700 | 0 | 134620 | 0 | 0 | 0 | 0 | 0 | 0 | 3288100 | 1313600 | 0 | 0 | 0 | 0 |
| ALSLGSLEF | ADGRG2 | Q8IZP9 | AGRG2_HUMA | Unmodified | 0 | 0 | 0 | 0 | 2534400 | 412530 | 0 | 0 | 3956200 | 1162200 | 0 | 0 | 0 | 1124300 | 0 | 0 | 0 | 0 |
| ALSLGSLEF | ADGRG2 | Q8IZP9 | AGRG2_HUMA | Oxidation (M) | 0 | 0 | 0 | 0 | 1771000 | 0 | 0 | 0 | 1933400 | 1067100 | 49264 | 0 | 0 | 0 | 0 | 0 | 0 | 0 |
| ALSMNTEN | SLC4A7 | H7C3C4 | H7C3C4_HUM. | Unmodified | 0 | 109580 | 0 | 0 | 4174400 | 0 | 55187 | 27878 | 0 | 0 | 350060 | 1571600 | 617410 | 0 | 0 | 0 | 1138800 | 3661600 |
| ALSMNTEN | SLC4A7 | H7C3C4 | H7C3C4_HUM. | Oxidation (M) | 0 | 0 | 0 | 0 | 196460 | 0 | 0 | 0 | 0 | 0 | 0 | 95995 | 0 | 0 | 0 | 0 | 0 | 250690 |
| ALSMNTEN | SLC4A7 | H7C3C4 | H7C3C4_HUM. | Unmodified | 0 | 0 | 0 | 0 | 0 | 0 | 0 | 0 | 0 | 0 | 0 | 0 | 0 | 0 | 0 | 0 | 0 | 0 |
| ALSMPLNG | CLIC4 | Q9Y696 | CLIC4_HUMAN | Acetyl (Protein N | 0 | 0 | 0 | 687500 | 9032600 | 887420 | 1076600 | 0 | 0 | 667040 | 197310 | 0 | 6589200 | 9343200 | 1094100 | 374890 | 9351300 | 10264000 |
| ALSMPLNG | CLIC4 | Q9Y696 | CLIC4_HUMAN | Acetyl (Protein N | 0 | 0 | 0 | 0 | 787510 | 0 | 0 | 0 | 0 | 0 | 0 | 0 | 0 | 357140 | 0 | 0 | 559690 | 0 |
| ALSNLESIP | UBQLN2; UBQL | Q9UHD9; Q9 | UBQL2_HUMAI | Unmodified | 0 | 0 | 0 | 0 | 0 | 0 | 0 | 0 | 0 | 0 | 0 | 1847600 | 0 | 0 | 0 | 0 | 0 | 0 |
| ALSNLVPK | SULF2 | Q8IWU5 | SULF2_HUMAN | Unmodified | 0 | 670550 | 0 | 0 | 0 | 0 | 0 | 0 | 0 | 0 | 0 | 0 | 66030 | 0 | 27557 | 0 | 0 | 3696300 |
| ALSPAELR | SCRIB; SCRIB; S | Q14160; A0A | SCRIB_HUMAN | Unmodified | 0 | 103400 | 549390 | 250350 | 0 | 0 | 0 | 0 | 2273400 | 0 | 271180 | 900270 | 0 | 181350 | 175490 | 0 | 0 | 0 |
| ALSPAELRA | SCRIB; SCRIB; S | Q14160; A0A | SCRIB_HUMAN | Unmodified | 0 | 0 | 0 | 0 | 50242 | 0 | 0 | 0 | 0 | 129850 | 0 | 0 | 0 | 0 | 0 | 0 | 290110 | 397020 |
| ALSPAFELS | NCSTN | Q92542 | NICA_HUMAN | Unmodified | 0 | 0 | 0 | 0 | 0 | 0 | 0 | 0 | 1272600 | 217540 | 0 | 0 | 0 | 0 | 0 | 0 | 0 | 0 |
| ALSPLEEW | VARS1; VARS1; | P26640; A2A | SYVC_HUMAN | Unmodified | 0 | 0 | 0 | 0 | 0 | 14264 | 0 | 0 | 409280 | 139850 | 0 | 10759000 | 0 | 0 | 0 | 0 | 0 | 0 |
| ALSQLAEVI | SNX1; SNX2; SN | Q13596; O6 | SNX1_HUMAN | Unmodified | 0 | 0 | 0 | 0 | 0 | 0 | 0 | 0 | 0 | 0 | 0 | 3121500 | 0 | 0 | 0 | 257350 | 0 | 0 |
| ALSRQEMC | HNRNPA2B1; H | P22626; A0A | ROA2_HUMAN | Unmodified | 60706 | 0 | 39536 | 0 | 125070 | 37076 | 74422 | 41576 | 0 | 175180 | 142520 | 279150 | 244000 | 0 | 0 | 0 | 0 | 71887 |
| ALSSAGPIF | MTSS2; MTSS2 | Q765P7; A0A | MTSS2_HUMAI | Unmodified | 0 | 0 | 0 | 0 | 155600 | 38077 | 0 | 8210.5 | 0 | 0 | 0 | 347240 | 0 | 0 | 0 | 0 | 0 | 0 |
| ALSSEGKP | SPTAN1; SPTAN | Q13813; A0A | SPTN1_HUMAI | Unmodified | 0 | 0 | 0 | 0 | 0 | 0 | 0 | 0 | 0 | 0 | 0 | 1354100 | 0 | 37183 | 0 | 0 | 0 | 0 |
| ALSSESFEF | PDK3 | Q15120 | PDK3_HUMAN | Unmodified | 0 | 0 | 0 | 0 | 109140 | 0 | 176960 | 22661 | 2573600 | 652890 | 0 | 0 | 0 | 0 | 0 | 0 | 17345 | 0 |
| ALSSEWKP | NCAM1; NCAM | P13591; A0A | NCAM1_HUMA | Unmodified | 0 | 0 | 231390 | 0 | 0 | 214570 | 0 | 11712 | 522480 | 161910 | 178060 | 2811200 | 0 | 150380 | 0 | 0 | 0 | 0 |
| ALSSLELSS | NCAPG | Q9BPX3 | CND3_HUMAN | Unmodified | 0 | 0 | 0 | 0 | 0 | 0 | 0 | 245170 | 0 | 0 | 0 | 0 | 0 | 0 | 0 | 0 | 0 | 0 |
| ALSSLLVH | RIPOR1; RIPOR | Q6ZS17; A0A | RIPR1_HUMAN | Unmodified | 0 | 0 | 0 | 0 | 0 | 0 | 0 | 0 | 0 | 0 | 0 | 0 | 0 | 0 | 0 | 0 | 0 | 0 |
| ALSSLQEEI | CCPG1 | Q9ULG6 | CCPG1_HUMA | Unmodified | 0 | 0 | 0 | 0 | 117230 | 0 | 0 | 0 | 1549000 | 92320 | 0 | 0 | 131250 | 0 | 0 | 0 | 0 | 0 |
| ALSSQHQA | HSPA5; HSPA5; | P11021; A0A | BIP_HUMAN Er | Unmodified | 0 | 0 | 0 | 0 | 0 | 224630 | 0 | 0 | 2756100 | 3352500 | 0 | 0 | 0 | 0 | 0 | 0 | 0 | 533910 |
| ALSSQHQA | HSPA5; HSPA5; | P11021; A0A | BIP_HUMAN Er | Unmodified | 0 | 0 | 0 | 0 | 0 | 0 | 0 | 0 | 0 | 2436500 | 0 | 0 | 0 | 0 | 0 | 0 | 0 | 0 |
| ALSSSADD | ANLN | Q9NQW6 | ANLN_HUMAN | Unmodified | 0 | 0 | 0 | 0 | 5304500 | 659670 | 631720 | 452380 | 0 | 0 | 0 | 0 | 0 | 0 | 0 | 0 | 2860000 | 1157300 |
| ALSTDPAAI | TLN1 | Q9Y490 | TLN1_HUMAN | Unmodified | 183380 | 0 | 0 | 0 | 3543800 | 1244600 | 0 | 0 | 0 | 363930 | 625770 | 6157000 | 0 | 0 | 245830 | 346320 | 1318500 | 0 |
| ALSTDSIER | PDK1 | Q15118 | PDK1_HUMAN | Unmodified | 0 | 0 | 0 | 0 | 0 | 0 | 0 | 48036 | 575700 | 131530 | 0 | 0 | 0 | 0 | 0 | 0 | 0 | 0 |
| ALSTMDLQ | CHMP1A; CHM | Q9HD42; F8 | CHM1A_HUMA | Unmodified | 1496100 | 0 | 0 | 0 | 8024100 | 1534600 | 851960 | 470970 | 3251900 | 1809400 | 495160 | 1125800 | 5053300 | 0 | 0 | 0 | 484650 | 1792200 |
| ALSTTASTA | EEFSEC; EEFSE | P57772; C9J | SELB_HUMAN | Unmodified | 119580 | 102650 | 0 | 206200 | 85128 | 0 | 393760 | 1819300 | 340480 | 64187 | 148210 | 428750 | 0 | 0 | 128400 | 0 | 0 | 110950 |
| ALSVETEK | RO60; RO60 | P10155; H0Y | RO60_HUMAN | Unmodified | 11684000 | 9685800 | 3812500 | 3660100 | 93097 | 18387 | 13491 | 0 | 186580 | 106890 | 0 | 1251100 | 0 | 320180 | 7841000 | 2972800 | 223490 | 353490 |
| ALSVGNIDI | STIP1; STIP1; ST | P31948; F5H | STIP1_HUMAN | Unmodified | 0 | 0 | 0 | 0 | 0 | 105320 | 0 | 0 | 464050 | 0 | 308340 | 1842800 | 0 | 0 | 0 | 0 | 0 | 0 |
| ALSVGNIDI | STIP1; STIP1; ST | P31948; F5H | STIP1_HUMAN | Unmodified | 0 | 96740 | 0 | 0 | 4942700 | 0 | 0 | 0 | 5774300 | 1979500 | 1146200 | 9114300 | 11406000 | 817560 | 0 | 0 | 3004300 | 521390 |
| ALSVGNIDI | STIP1; STIP1; ST | P31948; F5H | STIP1_HUMAN | Unmodified | 0 | 0 | 0 | 0 | 9942800 | 0 | 625680 | 0 | 0 | 2344600 | 0 | 0 | 16800000 | 8877800 | 0 | 0 | 31935000 | 9611200 |
| ALTADLPK | FARP2; FARP2 | O94887; H7 | FARP2_HUMAN | Unmodified | 0 | 0 | 0 | 0 | 0 | 0 | 207150 | 0 | 0 | 225520 | 0 | 0 | 0 | 309510 | 1185300 | 0 | 0 | 0 |
| ALTDAGCN | SCFD1; SCFD1; | Q8WVM8; A( | SCFD1_HUMAI | Unmodified | 0 | 0 | 0 | 19766 | 1767200 | 144070 | 160400 | 46094 | 106190 | 0 | 69297 | 453780 | 144230 | 185710 | 0 | 35209 | 0 | 0 |
| ALTDFFR | GCA; GCA; GCA | P28676; H7( | GRAN_HUMAN | Unmodified | 0 | 170350 | 0 | 0 | 0 | 0 | 0 | 0 | 1054800 | 0 | 0 | 626780 | 0 | 0 | 0 | 142420 | 0 | 39488 |
| ALTDMPQN | FGA | P02671 | FIBA_HUMAN F | 2 Oxidation (M) | 0 | 721810 | 0 | 0 | 0 | 0 | 0 | 0 | 0 | 0 | 0 | 0 | 0 | 0 | 0 | 0 | 0 | 0 |
| ALTDSFR | SRI; SRI; SRI | P30626; B4D | SORCN_HUMA | Unmodified | 1359000 | 1117000 | 1770300 | 1147000 | 314640 | 73841 | 30232 | 17195 | 11638000 | 809900 | 3554700 | 1618100 | 297540 | 107640 | 1262100 | 1078400 | 105420 | 326940 |
| ALTDYRAEI | PSMD11 | O00231 | PSD11_HUMAI | Unmodified | 0 | 0 | 0 | 498070 | 4469300 | 352920 | 2962500 | 0 | 853050 | 1504400 | 1315300 | 4026300 | 5567500 | 4054800 | 305370 | 0 | 1384300 | 1149500 |
| ALTGHLEE | ANXA1; ANXA1; | P04083; Q5T | ANXA1_HUMAI | Unmodified | 0 | 0 | 0 | 0 | 6751300 | 614590 | 0 | 0 | 0 | 220190 | 4756800 | 18101000 | 0 | 0 | 0 | 0 | 331540 | 691110 |
| ALTGHLEE | ANXA1; ANXA1 | P04083; Q5T | ANXA1_HUMAI | Oxidation (M) | 0 | 0 | 0 | 0 | 1843500 | 0 | 0 | 0 | 0 | 0 | 0 | 0 | 0 | 0 | 0 | 0 | 0 | 497050 |
| ALTHIDHSI | MARS1; MARS1 | P56192; F5H | SYMC_HUMAN | Unmodified | 0 | 0 | 0 | 0 | 0 | 0 | 0 | 0 | 0 | 0 | 0 | 387430 | 0 | 0 | 0 | 0 | 0 | 0 |
| ALTHSELD | HPN | P05981 | HEPS_HUMAN | Unmodified | 0 | 0 | 494430 | 293300 | 0 | 0 | 0 | 213190 | 0 | 0 | 0 | 0 | 0 | 0 | 748640 | 886670 | 0 | 0 |
| ALTLFDTDE | CHN1; CHN1; C | P15882; A0A | CHIN_HUMAN | Acetyl (Protein N | 0 | 16940 | 0 | 0 | 0 | 0 | 333680 | 0 | 0 | 0 | 0 | 0 | 0 | 0 | 0 | 0 | 232330 | 0 |
| ALTLIAGSP | CAND1; CAND1 | Q86VP6; A0A | CAND1_HUMA | Unmodified | 0 | 0 | 0 | 0 | 0 | 0 | 0 | 0 | 0 | 109290 | 0 | 6493400 | 0 | 0 | 0 | 456820 | 0 | 0 |
| ALTLLDYLL | EPN3; EPN3; EI | Q9H201; D6 | EPN3_HUMAN | Unmodified | 1494800 | 1393300 | 109370 | 92076 | 0 | 0 | 0 | 0 | 0 | 0 | 0 | 260230 | 54913 | 0 | 459440 | 1495500 | 0 | 0 |
| ALTNLPHTI | EIF3K; EIF3K; EI | Q9UBQ5; K7 | EIF3K_HUMAN | Unmodified | 0 | 0 | 0 | 0 | 0 | 0 | 0 | 245420 | 0 | 0 | 0 | 557460 | 601710 | 0 | 822250 | 0 | 0 | 0 |
| ALTPQCGS | ENDOD1 | O94919 | ENDD1_HUMA | Unmodified | 0 | 0 | 0 | 0 | 0 | 0 | 0 | 0 | 2759100 | 0 | 0 | 0 | 0 | 0 | 0 | 0 | 1046300 | 0 |
| ALTQPLGLI | MYADM; MYADI | Q96S97; A0A | MYADM_HUMA | Unmodified | 1348600 | 2395700 | 206510 | 367970 | 0 | 0 | 0 | 0 | 0 | 0 | 0 | 224910 | 0 | 0 | 3448100 | 3067000 | 0 | 0 |
| ALTQPSPVS | SLC4A2; SLC4A | P04920; C9J | B3A2_HUMAN | Unmodified | 0 | 0 | 682250 | 0 | 0 | 0 | 0 | 0 | 0 | 0 | 0 | 0 | 0 | 0 | 0 | 0 | 0 | 0 |
| ALTQPSPVS | SLC4A2; SLC4A | P04920; C9J | B3A2_HUMAN | Unmodified | 715310 | 0 | 745990 | 1542600 | 0 | 0 | 372370 | 0 | 531740 | 0 | 0 | 0 | 0 | 0 | 2071600 | 2195500 | 0 | 0 |
| ALTQTGGP | FLNA; FLNA | P21333; Q6( | FLNA_HUMAN | Unmodified | 0 | 0 | 719340 | 0 | 2552100 | 0 | 885490 | 828970 | 128780 | 0 | 0 | 695720 | 2976100 | 1336800 | 0 | 0 | 738740 | 0 |
| ALTSEIALL( | RPN1; RPN1 | P04843; B7Z | RPN1_HUMAN | Unmodified | 0 | 0 | 0 | 0 | 0 | 0 | 0 | 0 | 0 | 0 | 0 | 908940 | 0 | 0 | 0 | 98588 | 0 | 0 |
| ALTSELANA | MSN | P26038 | MOES_HUMAN | Unmodified | 1294300 | 2312800 | 1963400 | 4085200 | 2.95E+08 | 46547000 | 7358100 | 6186600 | 0 | 0 | 36683 | 156190 | 24448000 | 25668000 | 3061800 | 3180400 | 1.17E+08 | 1.05E+08 |
| ALTSELANA | MSN | P26038 | MOES_HUMAN | Unmodified | 271230 | 0 | 387790 | 1991200 | 1.42E+08 | 24115000 | 9761100 | 2742400 | 0 | 0 | 0 | 0 | 14596000 | 8760200 | 835500 | 15902 | 58937000 | 80327000 |
| ALTSFERDS | HADHA; HADHA | P40939; A0A | ECHA_HUMAN | Unmodified | 0 | 0 | 0 | 0 | 0 | 0 | 0 | 0 | 663330 | 536760 | 0 | 0 | 0 | 0 | 0 | 0 | 0 | 0 |
| ALTSFLPAP | SNW1; SNW1; S | Q13573; G3 | SNW1_HUMAN | Acetyl (Protein N | 100500 | 101070 | 23593 | 158330 | 1201400 | 199060 | 72064 | 57427 | 520220 | 910190 | 817940 | 388740 | 380030 | 137070 | 102160 | 51874 | 194980 | 534310 |
| ALTSPLQDI | CMIP; CMIP; CN | Q8IY22; A0A | CMIP_HUMAN | Acetyl (Protein N | 0 | 0 | 0 | 0 | 0 | 0 | 2431700 | 2669100 | 0 | 0 | 0 | 0 | 0 | 0 | 0 | 0 | 0 | 0 |
| ALTSVVNYI | CUL2; CUL2; CI | Q13617; Q5 | CUL2_HUMAN | Unmodified | 82428 | 0 | 0 | 0 | 648230 | 0 | 161480 | 106360 | 102740 | 193730 | 0 | 2171400 | 0 | 0 | 123420 | 0 | 757490 | 993390 |
| ALTTMGFR | SRI; SRI | P30626; C9J | SORCN_HUMA | Oxidation (M) | 0 | 1408800 | 627300 | 1005600 | 0 | 0 | 0 | 0 | 6080700 | 0 | 583890 | 2796400 | 0 | 0 | 1221200 | 786610 | 0 | 0 |
| ALTTMGFR | SRI; SRI | P30626; C9J | SORCN_HUMA | Unmodified | 0 | 670010 | 0 | 1056300 | 330930 | 0 | 0 | 0 | 10546000 | 564130 | 937450 | 5610400 | 255970 | 0 | 0 | 0 | 234970 | 752900 |
| ALTTMGFRI | SRI; SRI | P30626; C9J | SORCN_HUMA | Unmodified | 0 | 0 | 0 | 0 | 1885700 | 0 | 729490 | 35838 | 605990 | 1458700 | 82577 | 0 | 591380 | 770940 | 0 | 0 | 316820 | 721180 |
| ALTTMGFRI | SRI; SRI | P30626; C9J | SORCN_HUMA | Oxidation (M) | 0 | 0 | 0 | 0 | 433190 | 0 | 547550 | 0 | 0 | 1248500 | 0 | 0 | 480120 | 803600 | 0 | 0 | 0 | 0 |
| ALTVPELTQ | TUBB4A; TUBB3 | P04350; Q13 | TBB4A_HUMAN | Oxidation (M) | 0 | 555500 | 229040 | 969030 | 0 | 2232700 | 6283900 | 2840300 | 7239800 | 1364600 | 1238400 | 14582000 | 2338900 | 0 | 1132400 | 0 | 482380 | 0 |
| ALTVPELTQ | TUBB4A; TUBB3 | P04350; Q13 | TBB4A_HUMAN | Unmodified | 1146800 | 729060 | 307540 | 582900 | 7015500 | 1482000 | 19199000 | 2615800 | 7267000 | 3775700 | 2473500 | 16803000 | 2876100 | 1487200 | 2549200 | 651100 | 1106200 | 4603700 |
| ALTVPELTQ | TUBB6; TUBB6 | Q9BUF5; K7I | TBB6_HUMAN | Unmodified | 0 | 0 | 0 | 0 | 111440 | 0 | 0 | 0 | 0 | 0 | 0 | 0 | 0 | 0 | 0 | 0 | 36874 | 0 |
| ALTVPELTQ | TUBB2A; TUBB2 | Q13885; Q9I | TBB2A_HUMAN | Unmodified | 0 | 0 | 0 | 0 | 243290 | 141510 | 320250 | 210820 | 9907200 | 718160 | 1259300 | 14044000 | 0 | 0 | 48359 | 0 | 0 | 271990 |
| ALTVPELTQ | TUBB2A; TUBB2 | Q13885; Q9I | TBB2A_HUMAN | Oxidation (M) | 0 | 0 | 0 | 2656600 | 876210 | 63714 | 923360 | 36194 | 7706900 | 3689800 | 2660600 | 17809000 | 1783200 | 1658900 | 0 | 0 | 356700 | 1326700 |
| ALTVPELTQ | TUBB; TUBB; TU | P07437; Q5J | TBB5_HUMAN | Unmodified | 6210500 | 6324800 | 6528400 | 14917000 | 47206000 | 10773000 | 62540000 | 28668000 | 1.04E+08 | 28612000 | 32244000 | 3.69E+08 | 35706000 | 15807000 | 16795000 | 12352000 | 16194000 | 39165000 |
| ALVADSHP | SPTBN1; SPTBN | Q01082; A0A | SPTB2_HUMAN | Unmodified | 23227 | 0 | 0 | 0 | 0 | 0 | 0 | 82851 | 236420 | 111820 | 42711 | 1036200 | 0 | 0 | 0 | 0 | 0 | 0 |
| ALVADSHP | SPTBN1; SPTBN | Q01082; A0A | SPTB2_HUMAN | Unmodified | 0 | 0 | 0 | 0 | 0 | 0 | 0 | 0 | 88966 | 0 | 193640 | 752890 | 205570 | 0 | 0 | 0 | 0 | 664520 |
| ALVAIGTHD | FARSB | Q9NSD9 | SYFB_HUMAN | Unmodified | 0 | 0 | 0 | 0 | 0 | 0 | 0 | 58036 | 0 | 0 | 0 | 2894700 | 0 | 0 | 0 | 0 | 0 | 0 |
| ALVATPGKI | NCL; NCL; NCL | P19338; C9J | NUCL_HUMAN | Unmodified | 0 | 0 | 857960 | 522690 | 248330 | 0 | 480730 | 238660 | 0 | 0 | 0 | 1311300 | 1316500 | 658290 | 316000 | 324100 | 0 | 0 |

| | | | | | | | | | | | | | | | | | | | | | | |
|---|---|---|---|---|---|---|---|---|---|---|---|---|---|---|---|---|---|---|---|---|---|---|
| ALVATPGKI | NCL; NCL; NCL | P19338; C9J | NUCL_HUMAN | Unmodified | 0 | 0 | 0 | 0 | 910640 | 0 | 0 | 0 | 0 | 0 | 0 | 0 | 524490 | 0 | 0 | 0 | 387090 | 0 |
| ALVAYYQK | RPS16; RPS16; | P62249; M0I | RS16_HUMAN | Unmodified | 0 | 0 | 2804000 | 2126700 | 1774700 | 0 | 8745600 | 11764000 | 1384900 | 753890 | 2588200 | 38688000 | 0 | 0 | 3028600 | 1857900 | 0 | 2105700 |
| ALVDELEW | CNPY2; N/A; CI | Q9Y2B0; F8\ | CNPY2_HUMAI | Unmodified | 0 | 0 | 0 | 0 | 0 | 0 | 0 | 0 | 463830 | 619230 | 0 | 0 | 0 | 0 | 0 | 0 | 206550 | 0 |
| ALVDELEW | CNPY2; N/A; CI | Q9Y2B0; F8\ | CNPY2_HUMAI | Unmodified | 0 | 0 | 0 | 0 | 0 | 0 | 0 | 0 | 1.15E+08 | 34564000 | 4092900 | 534470 | 103390 | 0 | 58331 | 0 | 0 | 0 |
| ALVDGPCT | RPL14; RPL14 | P50914; E7E | RL14_HUMAN ( | Unmodified | 3115500 | 1747500 | 1001800 | 716020 | 0 | 0 | 0 | 0 | 0 | 0 | 0 | 18657000 | 0 | 0 | 1538200 | 1317600 | 0 | 0 |
| ALVDGPCT | RPL14; RPL14 | P50914; E7E | RL14_HUMAN ( | Unmodified | 2991800 | 2244000 | 650650 | 0 | 0 | 0 | 0 | 0 | 735470 | 0 | 1240300 | 12190000 | 0 | 0 | 0 | 567190 | 0 | 0 |
| ALVDHENV | PHGDH; PHGD | O43175; A0/ | SERA_HUMAN | Unmodified | 612120 | 0 | 2040900 | 3290700 | 0 | 0 | 966190 | 415890 | 3200600 | 660220 | 1919000 | 16966000 | 5582200 | 982060 | 3244100 | 2335400 | 986400 | 627570 |
| ALVDLADPI | PCSK1 | P29120 | NEC1_HUMAN | Unmodified | 0 | 0 | 0 | 0 | 0 | 0 | 134840 | 0 | 0 | 471600 | 0 | 0 | 0 | 0 | 0 | 0 | 0 | 0 |
| ALVDLQYIK | CNNM4 | Q6P4Q7 | CNNM4_HUMA | Unmodified | 0 | 0 | 0 | 497760 | 0 | 0 | 29729 | 0 | 0 | 18251 | 1698800 | 4017600 | 1553000 | 0 | 0 | 0 | 0 | 0 |
| ALVEDPRLI | CHST1 | O43916 | CHST1_HUMAI | Unmodified | 0 | 0 | 0 | 0 | 153100 | 0 | 0 | 255600 | 4684900 | 564580 | 0 | 354650 | 0 | 0 | 0 | 0 | 0 | 100890 |
| ALVEEALA( | MYG1; MYG1; N | Q9HB07; F8' | MYG1_HUMAN | Unmodified | 319110 | 746180 | 599870 | 0 | 0 | 0 | 0 | 0 | 244560 | 327660 | 0 | 3010400 | 0 | 0 | 343360 | 69810 | 0 | 0 |
| ALVEEEITR | GAK; GAK | O14976; D6I | GAK_HUMAN C | Unmodified | 0 | 0 | 0 | 0 | 0 | 0 | 0 | 0 | 0 | 0 | 0 | 1129400 | 0 | 0 | 0 | 0 | 0 | 0 |
| ALVEMQDV | TTYH3 | Q9C0H2 | TTYH3_HUMAN | Unmodified | 26799 | 0 | 14641 | 29855 | 0 | 0 | 0 | 0 | 695620 | 590150 | 13744 | 65885 | 0 | 0 | 302930 | 18444 | 0 | 0 |
| ALVERGGV | ACADVL; ACAD | P49748; G3\ | ACADV_HUMAI | Unmodified | 0 | 0 | 0 | 0 | 0 | 0 | 156870 | 26957 | 905430 | 2485500 | 0 | 0 | 0 | 0 | 0 | 0 | 0 | 343250 |
| ALVFQPVAI | PFKM; PFKM; PI | P08237; A0A | PFKAM_HUMAI | Unmodified | 0 | 0 | 0 | 0 | 0 | 0 | 0 | 0 | 0 | 0 | 0 | 1880100 | 0 | 0 | 0 | 0 | 0 | 0 |
| ALVFQPVAI | PFKM; PFKM; PI | P08237; A0A | PFKAM_HUMAI | Unmodified | 0 | 0 | 0 | 0 | 0 | 0 | 0 | 0 | 0 | 0 | 0 | 187360 | 0 | 0 | 0 | 0 | 0 | 0 |
| ALVGICTGI | RFTN1; RFTN1 | Q14699; G3; | RFTN1_HUMAN | Unmodified | 0 | 0 | 0 | 0 | 2459500 | 627730 | 0 | 0 | 0 | 0 | 0 | 0 | 0 | 0 | 0 | 0 | 546800 | 1241400 |
| ALVHDGDT | PIP5K1A; PIP5K | Q99755; A6I | PI51A_HUMAN | Unmodified | 435840 | 455380 | 509630 | 595550 | 0 | 0 | 0 | 0 | 374480 | 0 | 192380 | 6913900 | 0 | 0 | 697810 | 613510 | 0 | 0 |
| ALVHERDE | PSME2; PSME2; | Q9UL46; A0/ | PSME2_HUMAI | Unmodified | 0 | 0 | 0 | 0 | 0 | 0 | 0 | 0 | 1531500 | 0 | 862600 | 3486800 | 0 | 0 | 0 | 0 | 0 | 139100 |
| ALVITNSAR | ABCC1; ABCC1 | P33527; I3L4 | MRP1_HUMAN | Unmodified | 4841900 | 0 | 2854300 | 0 | 0 | 0 | 0 | 17377 | 0 | 0 | 0 | 2724600 | 0 | 0 | 228080 | 2820300 | 148190 | 0 |
| ALVIVMGK | PFN2; PFN2 | G5E9Q6; C9 | G5E9Q6_HUM/ | Oxidation (M) | 0 | 0 | 0 | 0 | 1317200 | 221490 | 0 | 0 | 0 | 190570 | 0 | 0 | 624800 | 383820 | 0 | 0 | 501270 | 0 |
| ALVIVMGKI | PFN2; PFN2 | G5E9Q6; C9 | G5E9Q6_HUM/ | Unmodified | 0 | 0 | 0 | 0 | 2923300 | 0 | 0 | 0 | 0 | 0 | 0 | 540240 | 299900 | 306650 | 0 | 0 | 1454700 | 0 |
| ALVLDCHY | C1QBP; C1QBP | Q07021; I3L: | C1QBP_HUMAI | Unmodified | 0 | 0 | 0 | 0 | 0 | 0 | 0 | 0 | 3056600 | 0 | 0 | 0 | 0 | 0 | 0 | 0 | 0 | 0 |
| ALVLELAK | INHBE | P58166 | INHBE_HUMAN | Unmodified | 23848 | 0 | 26443 | 0 | 0 | 0 | 0 | 0 | 1714100 | 0 | 57229 | 0 | 0 | 0 | 0 | 0 | 0 | 0 |
| ALVNQLHE | MCCC2; MCCC | Q9HCC0; A0 | MCCB_HUMAN | Unmodified | 61997 | 91182 | 0 | 0 | 0 | 0 | 0 | 0 | 1095000 | 188820 | 0 | 0 | 0 | 0 | 0 | 0 | 214390 | 0 |
| ALVPFEDLI | ECSIT; ECSIT | Q9BQ95; K7I | ECSIT_HUMAN | Unmodified | 0 | 0 | 0 | 0 | 0 | 0 | 0 | 0 | 923350 | 0 | 175250 | 44396 | 0 | 0 | 0 | 0 | 0 | 0 |
| ALVQNDTL | H1-10 | Q92522 | H1X_HUMAN H | Unmodified | 0 | 0 | 0 | 0 | 0 | 0 | 0 | 517670 | 0 | 0 | 0 | 0 | 0 | 0 | 0 | 0 | 0 | 0 |
| ALVQNDTL | H1-10 | Q92522 | H1X_HUMAN H | Unmodified | 0 | 0 | 0 | 0 | 0 | 0 | 0 | 840130 | 0 | 0 | 0 | 0 | 0 | 0 | 0 | 0 | 0 | 0 |
| ALVRLALEA | CACNA2D1; CA | P54289; E7E | CA2D1_HUMAI | Unmodified | 0 | 0 | 0 | 0 | 0 | 0 | 5039100 | 464570 | 0 | 489230 | 0 | 0 | 0 | 0 | 0 | 0 | 0 | 0 |
| ALVSAQWV | MPST; MPST | P25325; B1A | THTM_HUMAN | Unmodified | 0 | 0 | 0 | 0 | 0 | 0 | 0 | 0 | 174180 | 0 | 0 | 2873000 | 0 | 0 | 89815 | 35343 | 0 | 0 |
| ALVSAQWV | MPST; MPST | P25325; B1A | THTM_HUMAN | Unmodified | 100410 | 0 | 194460 | 413300 | 0 | 0 | 128590 | 0 | 5231700 | 601180 | 1357100 | 4732000 | 476700 | 287600 | 374730 | 267910 | 145750 | 236590 |
| ALVSEWKE | DDB1; DDB1; D | Q16531; A0/ | DDB1_HUMAN | Unmodified | 0 | 88078 | 0 | 0 | 766090 | 209980 | 120010 | 0 | 702590 | 537750 | 0 | 3204600 | 0 | 171710 | 0 | 111260 | 604200 | 529340 |
| ALVTGYFM | DHX15 | O43143 | DHX15_HUMAI | Oxidation (M) | 0 | 0 | 0 | 0 | 0 | 0 | 78931 | 0 | 0 | 0 | 0 | 4668000 | 0 | 0 | 0 | 0 | 0 | 0 |
| ALWADSGI | GNA13 | Q14344 | GNA13_HUMAI | Unmodified | 411660 | 0 | 0 | 0 | 0 | 0 | 0 | 0 | 772290 | 136340 | 0 | 5560900 | 0 | 0 | 0 | 0 | 0 | 0 |
| ALWADSGI | GNA13 | Q14344 | GNA13_HUMAI | Unmodified | 0 | 573390 | 0 | 0 | 0 | 0 | 0 | 0 | 0 | 0 | 0 | 1622600 | 0 | 0 | 0 | 0 | 0 | 0 |
| ALWCSEDI | STRAP | Q9Y3F4 | STRAP_HUMAN | Unmodified | 0 | 0 | 0 | 36095 | 0 | 0 | 0 | 0 | 0 | 0 | 0 | 1580200 | 581720 | 241570 | 265460 | 0 | 0 | 0 |
| ALWCSEDI | STRAP | Q9Y3F4 | STRAP_HUMAN | Unmodified | 0 | 0 | 0 | 0 | 0 | 0 | 0 | 0 | 0 | 0 | 0 | 0 | 613590 | 0 | 0 | 0 | 0 | 0 |
| ALWEDEG\ | GNAS; GNAS; G | Q5JWF2; P6: | GNAS1_HUMAI | Unmodified | 590790 | 511600 | 3331100 | 1313400 | 0 | 0 | 0 | 0 | 666500 | 0 | 855490 | 14210000 | 0 | 0 | 3394400 | 2872400 | 0 | 0 |
| ALWEDEG\ | GNAS; GNAS; G | Q5JWF2; P6: | GNAS1_HUMAI | Unmodified | 588040 | 0 | 951980 | 0 | 592360 | 307550 | 483190 | 92609 | 844000 | 484000 | 0 | 3325900 | 380250 | 0 | 2124900 | 49320 | 1730100 | 2073200 |
| ALWLAENC | SCRIB; SCRIB; S | Q14160; A0/ | SCRIB_HUMAN | Oxidation (M) | 0 | 0 | 0 | 0 | 203280 | 0 | 0 | 0 | 1965400 | 368520 | 364520 | 1221500 | 0 | 0 | 0 | 0 | 303120 | 0 |
| ALWLAENC | SCRIB; SCRIB; S | Q14160; A0/ | SCRIB_HUMAN | Unmodified | 0 | 0 | 0 | 0 | 0 | 0 | 0 | 0 | 1981300 | 171690 | 0 | 0 | 0 | 0 | 0 | 0 | 77935 | 0 |
| ALWLAENC | SCRIB; SCRIB; S | Q14160; A0/ | SCRIB_HUMAN | Oxidation (M) | 0 | 0 | 0 | 0 | 0 | 0 | 1772500 | 188240 | 0 | 2697500 | 0 | 0 | 2826800 | 1843400 | 0 | 0 | 8699300 | 6042800 |
| ALWLAENC | SCRIB; SCRIB; S | Q14160; A0/ | SCRIB_HUMAN | Unmodified | 0 | 0 | 0 | 0 | 2771800 | 0 | 668380 | 0 | 0 | 1275800 | 0 | 0 | 1093900 | 1400300 | 0 | 0 | 5912400 | 4373600 |
| ALWTGQYF | USP8 | P40818 | UBP8_HUMAN | Unmodified | 0 | 0 | 0 | 0 | 626560 | 91036 | 155220 | 0 | 0 | 38545 | 0 | 0 | 0 | 0 | 0 | 0 | 66599 | 0 |
| ALYDFHSEI | SNX33 | Q8WV41 | SNX33_HUMAI | Unmodified | 0 | 0 | 423420 | 0 | 0 | 0 | 0 | 0 | 0 | 0 | 0 | 0 | 0 | 0 | 236970 | 0 | 0 | 0 |
| ALYDFRSEI | SNX18 | Q96RF0 | SNX18_HUMAI | Unmodified | 224490 | 323270 | 478860 | 404550 | 0 | 0 | 0 | 0 | 0 | 0 | 0 | 0 | 109600 | 0 | 421330 | 356990 | 0 | 0 |
| ALYDTFSAI | PABPC4; PABP( | Q13310; P11 | PABP4_HUMAN | Unmodified | 52967000 | 18481000 | 2112100 | 12978000 | 884050 | 166310 | 929660 | 0 | 1229100 | 719160 | 0 | 657370 | 3277200 | 2136800 | 10041000 | 6071700 | 585430 | 397630 |
| ALYDYAGQ | PACSIN3; PACS | Q9UKS6; A0, | PACN3_HUMAI | Unmodified | 9792400 | 11813000 | 0 | 2304200 | 0 | 0 | 0 | 0 | 447390 | 0 | 0 | 0 | 552330 | 0 | 7234600 | 4021700 | 0 | 0 |
| ALYDYAGQ | PACSIN3; PACS | Q9UKS6; A0, | PACN3_HUMAI | Unmodified | 0 | 0 | 0 | 0 | 0 | 0 | 1828600 | 0 | 0 | 0 | 0 | 0 | 12274000 | 0 | 0 | 0 | 0 | 0 |
| ALYDYDGC | PACSIN1; PACS | Q9BY11; F6L | PACN1_HUMAI | Unmodified | 4400300 | 5174800 | 1559200 | 273360 | 0 | 0 | 0 | 0 | 0 | 0 | 0 | 0 | 0 | 0 | 2539600 | 1324500 | 0 | 0 |
| ALYDYDGC | PACSIN1; PACS | Q9BY11; F6L | PACN1_HUMAI | Unmodified | 2292700 | 2431300 | 0 | 0 | 0 | 0 | 0 | 0 | 0 | 0 | 0 | 0 | 0 | 0 | 416150 | 275240 | 0 | 0 |
| ALYDYEGQ | PACSIN2; PACS | Q9UNF0; H0 | PACN2_HUMAI | Unmodified | 48643000 | 35698000 | 50192000 | 21289000 | 0 | 0 | 0 | 0 | 1751500 | 98174 | 535400 | 4149200 | 4181400 | 1303100 | 96404000 | 53847000 | 0 | 247460 |
| ALYDYEGQ | PACSIN2 | Q9UNF0 | PACN2_HUMAI | Unmodified | 6862100 | 3226400 | 1179000 | 2359300 | 0 | 0 | 0 | 0 | 0 | 0 | 0 | 0 | 0 | 0 | 10544000 | 6057600 | 0 | 0 |
| ALYDYEGQ | PACSIN2 | Q9UNF0 | PACN2_HUMAI | Oxidation (M) | 3801200 | 2375400 | 0 | 10471000 | 907660 | 0 | 0 | 0 | 0 | 0 | 0 | 0 | 26322000 | 0 | 2545300 | 1805800 | 509810 | 0 |
| ALYDYEGQ | PACSIN2 | Q9UNF0 | PACN2_HUMAI | Unmodified | 3152200 | 3013100 | 0 | 5229400 | 0 | 0 | 0 | 0 | 0 | 0 | 0 | 0 | 16348000 | 3743800 | 0 | 0 | 0 | 1012200 |
| ALYEAGER | ANXA1 | P04083 | ANXA1_HUMAN | Unmodified | 1344300 | 1855900 | 1002600 | 2461100 | 402380 | 709310 | 40095 | 413680 | 88102 | 0 | 13485000 | 1.17E+09 | 919130 | 156340 | 3046300 | 2211200 | 46099 | 113200 |
| ALYEAGERI | ANXA1 | P04083 | ANXA1_HUMAN | Unmodified | 74035 | 123460 | 0 | 0 | 0 | 0 | 0 | 0 | 0 | 0 | 2039800 | 83301000 | 0 | 0 | 62524 | 104440 | 0 | 0 |
| ALYEGYAT\ | DPP3; DPP3; DI | Q9NY33; G3 | DPP3_HUMAN | Unmodified | 0 | 0 | 0 | 0 | 0 | 0 | 0 | 0 | 0 | 0 | 0 | 187240 | 0 | 0 | 0 | 0 | 0 | 0 |
| ALYEQSLHI | COPS2; COPS2 | P61201; B4D | CSN2_HUMAN | Unmodified | 0 | 0 | 0 | 0 | 556710 | 0 | 325270 | 0 | 222360 | 0 | 0 | 0 | 0 | 0 | 144920 | 203780 | 321940 | 562290 |
| ALYETELAD | LMNB1; LMNB1 | P20700; A0A | LMNB1_HUMAI | Unmodified | 0 | 0 | 0 | 0 | 359120 | 0 | 150860 | 240160 | 795600 | 395810 | 739420 | 3911600 | 94246 | 109260 | 0 | 0 | 39013 | 40517 |
| ALYFITPTSI | STXBP3 | O00186 | STXB3_HUMAN | Unmodified | 992980 | 506480 | 832890 | 2453700 | 12108000 | 2426700 | 2897100 | 4096500 | 2301600 | 2699000 | 2027000 | 9200300 | 0 | 3971300 | 2199700 | 2375400 | 9121600 | 12059000 |
| ALYGTTSET | MME; MME; MM | P08473; A0A | NEP_HUMAN N | Unmodified | 0 | 0 | 7623300 | 3021200 | 0 | 0 | 0 | 0 | 0 | 386670 | 0 | 0 | 761130 | 0 | 12627000 | 8492900 | 0 | 0 |
| ALYGTTSET | MME; MME; MM | P08473; A0A | NEP_HUMAN N | Unmodified | 0 | 0 | 21626000 | 21310000 | 0 | 0 | 619360 | 801400 | 6367600 | 1436700 | 0 | 0 | 4481400 | 2170400 | 30160000 | 23699000 | 0 | 348090 |
| ALYHTEER | PPA2; PPA2; PP | Q9H2U2; A0 | IPYR2_HUMAN | Acetyl (Protein N | 0 | 0 | 845820 | 517310 | 637890 | 0 | 91055 | 0 | 0 | 401600 | 0 | 729540 | 0 | 0 | 0 | 0 | 0 | 0 |
| ALYHTEER( | PPA2; PPA2; PP | Q9H2U2; A0 | IPYR2_HUMAN | Acetyl (Protein N | 0 | 0 | 0 | 0 | 378260 | 0 | 0 | 0 | 106070 | 74360 | 238960 | 744360 | 0 | 0 | 0 | 0 | 129410 | 270850 |
| ALYLIATNG | PAK2; PAK1; PA | Q13177; Q1: | PAK2_HUMAN | Unmodified | 0 | 0 | 139720 | 0 | 295000 | 0 | 0 | 0 | 649210 | 0 | 0 | 4539800 | 0 | 0 | 191100 | 214560 | 0 | 0 |
| ALYPSSVG( | MYO1B; MYO1E | O43795; H7( | MYO1B_HUMA | Unmodified | 0 | 0 | 0 | 0 | 2207700 | 0 | 0 | 0 | 1120000 | 0 | 2919400 | 11418000 | 0 | 0 | 0 | 0 | 0 | 0 |
| ALYPSSVG( | MYO1B; MYO1E | O43795; H7( | MYO1B_HUMA | Unmodified | 908050 | 0 | 0 | 0 | 20420000 | 4872500 | 2842000 | 1271400 | 1887900 | 1713300 | 5212900 | 13633000 | 0 | 1788600 | 0 | 0 | 4432100 | 5573300 |
| ALYPSSVG( | MYO1B; MYO1E | O43795; H7( | MYO1B_HUMA | Unmodified | 0 | 0 | 0 | 0 | 3339100 | 0 | 0 | 0 | 0 | 904010 | 0 | 0 | 1432800 | 0 | 0 | 0 | 4246500 | 0 |
| ALYSDPKSI | SCPEP1; SCPEI | Q9HB40; I3L | RISC_HUMAN I | Unmodified | 0 | 0 | 0 | 0 | 0 | 0 | 63545 | 95094 | 0 | 120910 | 0 | 1296600 | 0 | 0 | 0 | 0 | 0 | 0 |
| ALYSFQAR | NOSTRIN; NOS | Q8IVI9; A0A( | NOSTN_HUMA | Unmodified | 2769900 | 2113700 | 2469400 | 2568200 | 0 | 0 | 0 | 0 | 0 | 0 | 0 | 0 | 0 | 0 | 5650700 | 4409200 | 0 | 0 |
| ALYSFQAR( | NOSTRIN; NOS | Q8IVI9; A0A( | NOSTN_HUMA | Unmodified | 0 | 0 | 5612400 | 4557500 | 0 | 0 | 0 | 0 | 0 | 0 | 0 | 0 | 0 | 1404400 | 6629600 | 2348100 | 164100 | 0 |
| ALYSFQAR( | NOSTRIN; NOS | Q8IVI9; A0A( | NOSTN_HUMA | Unmodified | 0 | 0 | 0 | 4080200 | 0 | 0 | 0 | 0 | 0 | 0 | 0 | 0 | 0 | 0 | 0 | 0 | 0 | 0 |
| ALYSTAMES | PIP5K1A; PIPSL | Q99755; A2/ | PI51A_HUMAN | Unmodified | 0 | 0 | 0 | 539030 | 265990 | 0 | 0 | 0 | 282620 | 0 | 0 | 605680 | 350130 | 0 | 0 | 0 | 207450 | 617950 |
| ALYTATEK | CLIP1; CLIP1; C | P30622; F5H | CLIP1_HUMAN | Unmodified | 0 | 0 | 0 | 0 | 812530 | 85541 | 0 | 0 | 269220 | 0 | 38850 | 0 | 409740 | 0 | 0 | 0 | 127920 | 373510 |
| ALYTFEPR | OSTF1 | Q92882 | OSTF1_HUMAN | Unmodified | 229210 | 0 | 251180 | 245110 | 0 | 0 | 0 | 0 | 1557200 | 0 | 567520 | 4475400 | 0 | 0 | 0 | 1448000 | 0 | 0 |
| ALYYDLISS | SERPINF1; SER | P36955; I3L2 | PEDF_HUMAN | Unmodified | 0 | 0 | 0 | 0 | 0 | 0 | 0 | 0 | 2548600 | 279840 | 10326000 | 2086200 | 0 | 0 | 0 | 0 | 0 | 0 |
| ALYYLQIHF | DDB1; DDB1; D | Q16531; A0/ | DDB1_HUMAN | Unmodified | 0 | 0 | 0 | 237350 | 0 | 0 | 344600 | 0 | 645680 | 275020 | 175030 | 3967800 | 0 | 0 | 0 | 0 | 0 | 0 |
| AMAAEAEA | STOM | P27105 | STOM_HUMAN | Unmodified | 729910 | 734000 | 573700 | 1038100 | 577580 | 0 | 223600 | 802670 | 0 | 0 | 0 | 56636 | 152500 | 168900 | 679520 | 498870 | 0 | 26093 |
| AMAAEAEA | STOM | P27105 | STOM_HUMAN | Unmodified | 2168700 | 1975700 | 1091500 | 6145600 | 15429000 | 3402600 | 341970 | 2413600 | 0 | 28829 | 35220 | 24381 | 3151300 | 704700 | 1290800 | 1163800 | 536160 | 0 |
| AMAAEAEA | STOM | P27105 | STOM_HUMAN | Oxidation (M) | 352510 | 439960 | 261370 | 0 | 2041500 | 860640 | 197890 | 481470 | 0 | 0 | 0 | 0 | 1122800 | 0 | 418760 | 468880 | 0 | 194820 |
| AMAAEAEA | STOM | P27105 | STOM_HUMAN | Unmodified | 0 | 0 | 0 | 0 | 4013800 | 3585000 | 447900 | 110760 | 1027000 | 1509700 | 0 | 0 | 119750 | 0 | 0 | 0 | 0 | 0 |
| AMAAVPQD | NCLN; NCLN | Q969V3; K7E | NCLN_HUMAN | Unmodified | 0 | 0 | 0 | 318710 | 0 | 247900 | 1024500 | 500360 | 2235200 | 319300 | 0 | 0 | 554540 | 0 | 0 | 0 | 0 | 0 |
| AMADALLE | SHMT2; SHMT2 | P34897; H0Y | GLYM_HUMAN | Unmodified | 0 | 66910 | 0 | 0 | 0 | 0 | 221070 | 0 | 6541700 | 527920 | 0 | 388580 | 0 | 0 | 0 | 91656 | 0 | 0 |
| AMADALLE | SHMT2; SHMT2 | P34897; H0Y | GLYM_HUMAN | Oxidation (M) | 0 | 0 | 0 | 39276 | 0 | 0 | 0 | 0 | 1183100 | 0 | 0 | 0 | 0 | 0 | 0 | 0 | 0 | 0 |
| AMADELSE | CAV1; CAV1 | Q03135; E9F | CAV1_HUMAN | Unmodified | 0 | 0 | 0 | 0 | 9084200 | 1614300 | 0 | 0 | 0 | 0 | 0 | 0 | 0 | 0 | 30064 | 0 | 629130 | 1310800 |
| AMADELSE | CAV1; CAV1 | Q03135; E9F | CAV1_HUMAN | Oxidation (M) | 0 | 0 | 0 | 0 | 3149900 | 644990 | 0 | 0 | 0 | 0 | 90760 | 2454300 | 0 | 0 | 0 | 0 | 0 | 760830 |
| AMADELSE | CAV1; CAV1 | Q03135; E9F | CAV1_HUMAN | Unmodified | 0 | 0 | 0 | 0 | 1512400 | 0 | 0 | 0 | 0 | 0 | 0 | 0 | 0 | 96552 | 0 | 0 | 0 | 54111 |
| AMAQTAIK | CELF1; CELF1 | Q92879; G5I | CELF1_HUMAN | Unmodified | 1756100 | 1412500 | 869440 | 1715500 | 0 | 0 | 35138 | 0 | 0 | 0 | 0 | 613340 | 767320 | 0 | 407720 | 732350 | 0 | 0 |
| AMATKGGT | ANXA4; ANXA4 | B4E1S2; Q6I | B4E1S2_HUMA | Acetyl (Protein N | 0 | 0 | 0 | 0 | 9181300 | 0 | 0 | 833000 | 63580 | 1241800 | 0 | 0 | 0 | 2267700 | 8772.3 | 0 | 2616400 | 2973900 |
| AMATLLSK | SLC12A2; SLC1 | P55011; G3X | S12A2_HUMAN | Unmodified | 162890 | 0 | 0 | 354930 | 0 | 0 | 0 | 0 | 343250 | 0 | 0 | 561660 | 0 | 0 | 1541100 | 726480 | 0 | 446020 |
| AMAVLESLI | QARS1; QARS1; | P47897; H7( | SYQ_HUMAN G | Oxidation (M) | 0 | 0 | 0 | 0 | 0 | 0 | 0 | 0 | 0 | 0 | 0 | 1555600 | 0 | 0 | 0 | 0 | 0 | 0 |
| AMCSEIILR | PIGX; PIGX; PIG | Q8TBF5; H7( | PIGX_HUMAN I | Unmodified | 0 | 0 | 0 | 0 | 0 | 0 | 0 | 0 | 2558600 | 0 | 0 | 360370 | 0 | 0 | 0 | 0 | 0 | 0 |
| AMDDSSVT | CRAT; CRAT; CI | P43155; A0A | CACP_HUMAN | Unmodified | 0 | 0 | 0 | 0 | 0 | 0 | 0 | 0 | 984000 | 0 | 0 | 0 | 0 | 0 | 0 | 0 | 0 | 0 |
| AMDEEIVSI | SLC9A6; SLC9A | Q92581; A0/ | SL9A6_HUMAN | Unmodified | 0 | 0 | 0 | 0 | 0 | 0 | 0 | 0 | 2119600 | 394710 | 0 | 0 | 0 | 0 | 0 | 0 | 0 | 0 |
| AMDEIQPD | VPS28; VPS28; | Q9UK41; E9I | VPS28_HUMAN | Unmodified | 895980 | 1571900 | 384550 | 0 | 473390 | 0 | 0 | 0 | 0 | 0 | 0 | 1376700 | 323160 | 0 | 1720200 | 919470 | 80429 | 0 |
| AMDEIQPD | VPS28; VPS28; | Q9UK41; E9I | VPS28_HUMAN | Oxidation (M) | 1368200 | 2114100 | 1300800 | 0 | 0 | 0 | 0 | 0 | 0 | 0 | 0 | 1477100 | 0 | 0 | 3111000 | 2472300 | 0 | 0 |
| AMDEIQPD | VPS28; VPS28 | Q9UK41; E9I | VPS28_HUMAN | 2 Oxidation (M) | 2105500 | 1360100 | 587280 | 0 | 0 | 0 | 0 | 0 | 96075 | 0 | 0 | 465700 | 0 | 0 | 5842400 | 0 | 0 | 0 |
| AMDEIQPD | VPS28; VPS28 | Q9UK41; E9I | VPS28_HUMAN | Oxidation (M) | 318950 | 805870 | 267420 | 88810 | 0 | 0 | 0 | 0 | 0 | 0 | 0 | 115480 | 0 | 0 | 1284200 | 423870 | 0 | 0 |
| AMDFHWIE | ST3GAL4; ST3G | Q11206; E9F | SIA4C_HUMAN | Oxidation (M) | 0 | 0 | 0 | 0 | 0 | 0 | 0 | 0 | 293300 | 0 | 27239 | 0 | 0 | 0 | 0 | 0 | 0 | 0 |
| AMDFNGIL | HSPG2 | P98160 | PGBM_HUMAN | Oxidation (M) | 0 | 0 | 579770 | 0 | 0 | 31558 | 0 | 0 | 594690 | 0 | 0 | 582750 | 0 | 2410200 | 0 | 0 | 0 | 0 |
| AMDQEITVI | COPS3; COPS3 | Q9UNS2; H7 | CSN3_HUMAN | Oxidation (M) | 276860 | 0 | 0 | 0 | 349040 | 0 | 0 | 227380 | 0 | 0 | 0 | 0 | 618750 | 420250 | 0 | 349900 | 262180 | 0 |
| AMDQLAKT | ADIRF; ADIRF | Q15847; Q5 | ADIRF_HUMAN | Unmodified | 0 | 0 | 0 | 0 | 0 | 0 | 250450 | 0 | 0 | 103010 | 0 | 336710 | 0 | 408650 | 266190 | 0 | 1812000 | 1045800 |
| AMDTLGIE' | GNAO1; GNAO | P09471; A0A | GNAO_HUMAN | Oxidation (M) | 0 | 0 | 0 | 0 | 0 | 0 | 0 | 0 | 585320 | 0 | 0 | 0 | 0 | 0 | 0 | 0 | 0 | 0 |
| AMDTLGIE' | GNAO1; GNAO | P09471; A0A | GNAO_HUMAN | Unmodified | 0 | 0 | 0 | 0 | 365380 | 0 | 0 | 0 | 1333000 | 291740 | 0 | 0 | 0 | 0 | 0 | 0 | 0 | 0 |
| AMDVKLDF | GLG1; GLG1 | Q92896; H3I | GSLG1_HUMAI | Unmodified | 111950 | 0 | 0 | 162110 | 748440 | 340150 | 31571 | 15098 | 14588000 | 3320500 | 2262500 | 606020 | 183800 | 0 | 0 | 0 | 19247 | 212090 |
| AMDVYQK | STIP1; STIP1 | P31948; H0Y | STIP1_HUMAN | Unmodified | 0 | 25578 | 0 | 0 | 5345900 | 0 | 752980 | 541540 | 533150 | 1353500 | 695920 | 8588800 | 960800 | 1955500 | 9318.4 | 296940 | 1512100 | 5108600 |
| AMEAVAAQ | PGAM1; PGAM2 | P18669; P15 | PGAM1_HUMA | Unmodified | 1015300 | 912140 | 3064900 | 4778800 | 15242000 | 5106400 | 935430 | 1523200 | 2333100 | 1600200 | 5566600 | 40964000 | 22167000 | 8044800 | 3758700 | 3042500 | 11669000 | 26539000 |
| AMEAVAAQ | PGAM1; PGAM2 | P18669; P15 | PGAM1_HUMA | Oxidation (M) | 108430 | 0 | 5089800 | 2794100 | 1404500 | 22521000 | 2250200 | 214320 | 920400 | 78924 | 1957500 | 3672600 | 4144300 | 5331000 | 950620 | 683690 | 3437500 | 6989500 |

| | | | | | | | | | | | | | | | | | | | | | | |
|---|---|---|---|---|---|---|---|---|---|---|---|---|---|---|---|---|---|---|---|---|---|---|
| AMEAVAAQ | PGAM1 | P18669 | PGAM1_HUMA | Unmodified | 0 | 0 | 0 | 0 | 0 | 0 | 0 | 119360 | 0 | 0 | 0 | 255220 | 0 | 0 | 0 | 0 | 754440 | 0 |
| AMEGAGTD | ANXA6; ANXA6; | P08133; E7E | ANXA6_HUMAN | Unmodified | 0 | 0 | 0 | 0 | 0 | 0 | 108160 | 19815 | 101520 | 0 | 0 | 0 | 0 | 20040 | 0 | 0 | 0 | 123070 |
| AMEGAGTD | ANXA6; ANXA6; | P08133; E7E | ANXA6_HUMAN | Oxidation (M) | 0 | 0 | 0 | 0 | 2053900 | 2572900 | 0 | 60637 | 895270 | 1359000 | 0 | 0 | 0 | 0 | 0 | 0 | 1738700 | 4043400 |
| AMEGIFIKP | SDF2L1 | Q9HCN8 | SDF2L_HUMAN | Oxidation (M) | 0 | 0 | 542000 | 0 | 1288700 | 0 | 3154800 | 2073000 | 26515000 | 11327000 | 7392500 | 2822200 | 1402300 | 604230 | 481790 | 325060 | 332650 | 654180 |
| AMEGIFIKP | SDF2L1 | Q9HCN8 | SDF2L_HUMAN | Unmodified | 0 | 0 | 0 | 0 | 388060 | 0 | 1204100 | 127160 | 29531000 | 4707100 | 849270 | 683490 | 0 | 242600 | 0 | 0 | 0 | 703120 |
| AMEGIFMK | SDF2 | Q99470 | SDF2_HUMAN | Unmodified | 0 | 0 | 0 | 0 | 0 | 0 | 0 | 0 | 1895300 | 0 | 0 | 0 | 0 | 0 | 0 | 0 | 0 | 0 |
| AMEGIFMK | SDF2 | Q99470 | SDF2_HUMAN | Oxidation (M) | 0 | 0 | 0 | 0 | 0 | 0 | 0 | 0 | 588180 | 0 | 0 | 0 | 0 | 0 | 0 | 0 | 0 | 0 |
| AMEGIFMK | SDF2 | Q99470 | SDF2_HUMAN | Oxidation (M) | 0 | 0 | 0 | 0 | 0 | 0 | 0 | 0 | 5252700 | 461350 | 0 | 0 | 0 | 0 | 0 | 0 | 0 | 0 |
| AMEGIFMK | SDF2 | Q99470 | SDF2_HUMAN | 2 Oxidation (M) | 0 | 0 | 0 | 0 | 112160 | 0 | 0 | 0 | 1353300 | 970850 | 696970 | 0 | 0 | 0 | 0 | 0 | 0 | 984330 |
| AMEGIFMK | SDF2 | Q99470 | SDF2_HUMAN | 2 Oxidation (M) | 0 | 0 | 0 | 0 | 0 | 0 | 63216 | 0 | 0 | 2343100 | 0 | 0 | 0 | 0 | 0 | 0 | 0 | 0 |
| AMEQLTKS | PLCB3 | Q01970 | PLCB3_HUMAN | Oxidation (M) | 0 | 0 | 0 | 0 | 0 | 0 | 92503 | 54874 | 0 | 0 | 0 | 0 | 0 | 0 | 0 | 0 | 264900 | 1340200 |
| AMESHKDE | MEMO1 | Q9Y316 | MEMO1_HUMA | Oxidation (M) | 0 | 0 | 241580 | 0 | 0 | 0 | 59649 | 0 | 0 | 0 | 0 | 941700 | 0 | 0 | 0 | 0 | 0 | 64924 |
| AMESHKDE | MEMO1 | Q9Y316 | MEMO1_HUMA | Oxidation (M) | 426120 | 1055200 | 0 | 974780 | 0 | 559070 | 0 | 0 | 0 | 0 | 141420 | 405560 | 0 | 0 | 0 | 522220 | 979770 | 1585600 |
| AMEVDERP | PSMC3; PSMC3 | P17980; E9P | PRS6A_HUMAN | Oxidation (M) | 0 | 71016 | 1312500 | 1416300 | 2950500 | 916680 | 568220 | 455140 | 2315900 | 559490 | 1355600 | 6835500 | 4941200 | 1346300 | 1000600 | 217790 | 2119900 | 3041300 |
| AMEVDERP | PSMC3; PSMC3 | P17980; E9P | PRS6A_HUMAN | Unmodified | 162060 | 90630 | 0 | 615750 | 5116700 | 1444000 | 1100400 | 169450 | 4664100 | 1814300 | 0 | 1599200 | 1932600 | 2404000 | 0 | 0 | 2927100 | 8170200 |
| AMEWITAK | PFKP; PFKP; PF | Q01813; A0A | PFKAP_HUMAN | Oxidation (M) | 0 | 0 | 0 | 0 | 558840 | 184690 | 33979 | 0 | 165540 | 152190 | 0 | 2437700 | 126800 | 0 | 534290 | 75523 | 479720 | 326890 |
| AMEWITAK | PFKP; PFKP; PF | Q01813; A0A | PFKAP_HUMAN | Unmodified | 0 | 0 | 0 | 0 | 0 | 0 | 0 | 0 | 472300 | 0 | 0 | 1592600 | 0 | 0 | 0 | 0 | 0 | 0 |
| AMGAAQV | SORD; SORD | Q00796; A0A | DHSO_HUMAN | Oxidation (M) | 0 | 0 | 0 | 0 | 0 | 0 | 0 | 0 | 640810 | 428890 | 0 | 3930500 | 0 | 0 | 0 | 1966200 | 0 | 0 |
| AMGAAQV | SORD; SORD | Q00796; A0A | DHSO_HUMAN | Unmodified | 0 | 0 | 102560 | 215010 | 0 | 0 | 0 | 0 | 0 | 0 | 0 | 992700 | 0 | 0 | 266220 | 0 | 0 | 0 |
| AMGAAQV | SORD; SORD | Q00796; A0A | DHSO_HUMAN | Unmodified | 0 | 0 | 0 | 0 | 4471100 | 0 | 0 | 0 | 468970 | 0 | 0 | 0 | 4968100 | 5237300 | 0 | 0 | 1044100 | 0 |
| AMGDAEIK | COPB2; COPB2 | P35606; H0Y | COPB2_HUMA | Oxidation (M) | 0 | 0 | 0 | 172690 | 0 | 238660 | 192720 | 143950 | 0 | 0 | 0 | 931720 | 698730 | 576120 | 0 | 0 | 0 | 0 |
| AMGIETSDI | PLCB4; PLCB4 | Q15147; A0A | PLCB4_HUMAN | Oxidation (M) | 0 | 0 | 252820 | 117480 | 0 | 0 | 0 | 0 | 0 | 0 | 0 | 0 | 0 | 0 | 161640 | 401540 | 0 | 0 |
| AMGIMNSF | H2BC4; H2BC1 | P62807; P06 | H2B1C_HUMA | 2 Oxidation (M) | 1.08E+08 | 91698000 | 876540 | 812260 | 996980 | 695750 | 739600 | 767760 | 1398700 | 828080 | 1912200 | 2.51E+08 | 0 | 611640 | 1202500 | 641120 | 420310 | 602070 |
| AMGIMNSF | H2BC4; H2BC1 | P62807; P06 | H2B1C_HUMA | Oxidation (M) | 18340000 | 17295000 | 27416 | 52680 | 493370 | 73433 | 359250 | 12844 | 663490 | 102130 | 97610 | 40519000 | 0 | 0 | 121190 | 35370 | 0 | 95607 |
| AMGIMNSF | H2BC4; H2BC1 | P62807; P06 | H2B1C_HUMA | Unmodified | 646530 | 637720 | 0 | 0 | 0 | 0 | 0 | 0 | 137440 | 0 | 0 | 2430300 | 0 | 0 | 0 | 0 | 0 | 0 |
| AMGIMNSF | H2BC4; H2BC1 | P62807; P06 | H2B1C_HUMA | 2 Oxidation (M) | 3888300 | 2636400 | 0 | 0 | 1883800 | 0 | 1331400 | 0 | 0 | 0 | 0 | 789830 | 362040 | 0 | 0 | 0 | 0 | 0 |
| AMGIMNSF | H2BC4; H2BC1 | P62807; P06 | H2B1C_HUMA | 2 Oxidation (M) | 0 | 0 | 0 | 0 | 1942100 | 0 | 20351000 | 0 | 0 | 0 | 0 | 0 | 834470 | 580870 | 0 | 0 | 0 | 716120 |
| AMGLAALE | TTC13 | Q8NBP0 | TTC13_HUMAN | Unmodified | 0 | 182570 | 0 | 0 | 0 | 0 | 0 | 0 | 350280 | 80650 | 0 | 0 | 0 | 0 | 0 | 0 | 0 | 0 |
| AMGNATA | N/A | N/A | N/A | Unmodified | 0 | 0 | 0 | 0 | 0 | 0 | 2980300 | 2426800 | 0 | 0 | 0 | 0 | 0 | 0 | 0 | 0 | 0 | 0 |
| AMGNLQID | GNAI2 | P04899 | GNAI2_HUMAN | Oxidation (M) | 0 | 0 | 230260 | 0 | 0 | 0 | 0 | 38024 | 0 | 214660 | 982010 | 6382500 | 0 | 0 | 0 | 0 | 0 | 0 |
| AMGNLQID | GNAI2 | P04899 | GNAI2_HUMAN | Unmodified | 177110 | 0 | 0 | 0 | 0 | 0 | 0 | 0 | 0 | 0 | 0 | 914730 | 0 | 0 | 0 | 0 | 0 | 0 |
| AMGNLQID | GNAI2 | P04899 | GNAI2_HUMAN | Unmodified | 697160 | 1108800 | 0 | 1250400 | 0 | 0 | 0 | 0 | 4030900 | 0 | 1611500 | 8745600 | 0 | 0 | 542320 | 346390 | 813960 | 2323200 |
| AMGNLQID | GNAI2 | P04899 | GNAI2_HUMAN | Oxidation (M) | 3477600 | 8705300 | 0 | 4451800 | 1210500 | 0 | 0 | 335650 | 2497600 | 623450 | 9005800 | 48115000 | 0 | 0 | 3887700 | 3094000 | 804640 | 1018800 |
| AMGPKITLN | PPP2R1B | P30154 | 2AAB_HUMAN | Oxidation (M) | 0 | 0 | 0 | 0 | 238930 | 0 | 51508 | 0 | 0 | 0 | 0 | 0 | 589510 | 328880 | 0 | 0 | 22938 | 0 |
| AMGPLVLT | EIF5 | P55010 | IF5_HUMAN Eu | Oxidation (M) | 0 | 0 | 0 | 0 | 564760 | 0 | 0 | 0 | 0 | 129860 | 77175 | 735560 | 0 | 0 | 70961 | 54906 | 308150 | 235420 |
| AMHQAQTI | CELF1; CELF1 | Q92879; G5I | CELF1_HUMAN | 3 Oxidation (M) | 197920 | 89423 | 0 | 95448 | 0 | 0 | 0 | 0 | 0 | 0 | 0 | 0 | 0 | 0 | 83837 | 87656 | 0 | 0 |
| AMHQAQTI | CELF1; CELF1 | Q92879; G5I | CELF1_HUMAN | 2 Oxidation (M) | 614100 | 72613 | 155800 | 90411 | 0 | 0 | 0 | 0 | 0 | 0 | 0 | 27041 | 0 | 0 | 150600 | 97570 | 0 | 0 |
| AMHQAQTI | CELF1; CELF1 | Q92879; G5I | CELF1_HUMAN | Oxidation (M) | 707470 | 862950 | 35243 | 292670 | 0 | 0 | 47161 | 0 | 0 | 0 | 0 | 128800 | 347360 | 197100 | 0 | 0 | 0 | 0 |
| AMIAYWTN | CEL | P19835 | CEL_HUMAN B | Unmodified | 0 | 0 | 0 | 0 | 0 | 0 | 0 | 0 | 7485900 | 703800 | 0 | 0 | 0 | 0 | 0 | 0 | 0 | 0 |
| AMIAYWTN | CEL | P19835 | CEL_HUMAN B | Oxidation (M) | 0 | 0 | 0 | 0 | 0 | 0 | 0 | 0 | 2734300 | 1133900 | 6421900 | 0 | 0 | 0 | 0 | 0 | 0 | 0 |
| AMIENFNA | NAA11; NAA10; | Q9BSU3; P4 | NAA11_HUMAN | Unmodified | 0 | 0 | 0 | 0 | 488740 | 0 | 0 | 0 | 138880 | 60723 | 303020 | 5023900 | 183420 | 0 | 520130 | 0 | 0 | 62066 |
| AMIENFNA | NAA11; NAA10; | Q9BSU3; P4 | NAA11_HUMAN | Oxidation (M) | 0 | 0 | 0 | 0 | 158880 | 0 | 319290 | 40098 | 0 | 861740 | 0 | 1420500 | 0 | 0 | 0 | 0 | 90133 | 513750 |
| AMINAMDS | CIT; CIT; CIT; CI | O14578; A0A | CTRO_HUMAN | Unmodified | 0 | 0 | 0 | 0 | 1736900 | 280520 | 0 | 0 | 0 | 0 | 0 | 139160 | 0 | 0 | 0 | 0 | 918400 | 0 |
| AMISEQELI | N/A | N/A | N/A | Unmodified | 0 | 0 | 0 | 0 | 559080 | 0 | 0 | 103440 | 0 | 180390 | 0 | 0 | 129510 | 38438 | 0 | 0 | 379280 | 1893200 |
| AMISRWYF | APP; APP; APP | P05067; H7C | A4_HUMAN Am | Unmodified | 0 | 0 | 0 | 0 | 312880 | 132360 | 0 | 0 | 0 | 0 | 0 | 0 | 0 | 0 | 0 | 0 | 69822 | 405730 |
| AMKGFGTD | ANXA11 | P50995 | ANX11_HUMAN | Oxidation (M) | 0 | 0 | 0 | 0 | 0 | 0 | 181360 | 33398 | 0 | 73999 | 0 | 0 | 184980 | 202190 | 0 | 0 | 269100 | 725370 |
| AMKGFGTD | ANXA7 | P20073 | ANXA7_HUMAN | Oxidation (M) | 0 | 674880 | 0 | 0 | 0 | 0 | 1000500 | 416140 | 370320 | 604840 | 0 | 0 | 322810 | 104520 | 0 | 104540 | 1181100 | 2297200 |
| AMKGFGTD | ANXA7 | P20073 | ANXA7_HUMAN | Unmodified | 0 | 0 | 719060 | 0 | 0 | 0 | 0 | 0 | 0 | 85809 | 0 | 0 | 0 | 0 | 0 | 0 | 42877 | 362750 |
| AMKGLGTD | ANXA6; ANXA6 | P08133; E5R | ANXA6_HUMAN | Oxidation (M) | 0 | 0 | 0 | 0 | 0 | 0 | 250010 | 0 | 0 | 191500 | 0 | 0 | 0 | 0 | 0 | 0 | 302150 | 238810 |
| AMKQGFEF | SEPTIN9; SEPTI | Q9UHD8; K7 | SEPT9_HUMAN | 2 Oxidation (M) | 99951 | 0 | 0 | 0 | 0 | 0 | 130250 | 0 | 842270 | 276160 | 0 | 0 | 0 | 0 | 0 | 0 | 293470 | 0 |
| AMLALTSTL | OS9; OS9 | Q13438; B4E | OS9_HUMAN P | Unmodified | 0 | 0 | 0 | 0 | 0 | 0 | 0 | 0 | 0 | 0 | 0 | 0 | 0 | 0 | 0 | 0 | 0 | 0 |
| AMLENEK | MSN | P26038 | MOES_HUMAN | Unmodified | 88966 | 0 | 117120 | 0 | 9170500 | 43151 | 0 | 8198.5 | 220440 | 36034 | 20016 | 288830 | 1562500 | 470130 | 100950 | 91058 | 1530200 | 3405800 |
| AMLENEK | MSN | P26038 | MOES_HUMAN | Oxidation (M) | 0 | 76337 | 28046 | 74992 | 1289800 | 142490 | 0 | 0 | 0 | 91299 | 0 | 0 | 101180 | 72157 | 37023 | 36481 | 384900 | 1038100 |
| AMLHLQWI | ITGAV | P06756 | ITAV_HUMAN I | Oxidation (M) | 0 | 0 | 0 | 0 | 447840 | 0 | 0 | 0 | 1119800 | 0 | 0 | 2420400 | 0 | 171700 | 950350 | 461750 | 0 | 0 |
| AMLIEAGSI | TRRAP; TRRAP; | Q9Y4A5; F22 | TRRAP_HUMAN | Oxidation (M) | 0 | 0 | 0 | 0 | 0 | 0 | 0 | 0 | 0 | 0 | 0 | 768250 | 0 | 0 | 0 | 0 | 0 | 0 |
| AMLSGPG | ELOC; ELOC | Q15369; E5F | ELOC_HUMAN | Oxidation (M) | 1277500 | 0 | 0 | 0 | 0 | 0 | 0 | 0 | 1944700 | 0 | 0 | 7978300 | 0 | 0 | 790790 | 0 | 0 | 0 |
| AMLSGPG | ELOC; ELOC | Q15369; E5F | ELOC_HUMAN | Oxidation (M) | 0 | 0 | 0 | 0 | 21099000 | 0 | 6717400 | 0 | 0 | 2886600 | 1778000 | 0 | 1520900 | 16119000 | 0 | 0 | 16180000 | 21816000 |
| AMLSGPG | ELOC; ELOC | Q15369; E5F | ELOC_HUMAN | Unmodified | 0 | 0 | 0 | 0 | 6768800 | 0 | 0 | 0 | 0 | 0 | 0 | 0 | 3916100 | 0 | 0 | 0 | 3161300 | 4859200 |
| AMLSSTAV | DLG5 | Q8TDM6 | DLG5_HUMAN | Unmodified | 0 | 0 | 0 | 0 | 0 | 0 | 0 | 0 | 0 | 0 | 0 | 0 | 0 | 0 | 0 | 0 | 0 | 850620 |
| AMLWLSEK | PFKL | P17858 | PFKAL_HUMAN | Unmodified | 0 | 0 | 0 | 0 | 214970 | 0 | 295800 | 0 | 0 | 89260 | 0 | 0 | 0 | 0 | 0 | 0 | 0 | 502790 |
| AMMYSGEI | IMPDH2; IMPDH | P12268; A0A | IMDH2_HUMAN | Unmodified | 0 | 0 | 0 | 0 | 785800 | 0 | 0 | 0 | 549890 | 0 | 0 | 0 | 0 | 0 | 0 | 0 | 277190 | 0 |
| AMNGESLD | RBM3 | P98179 | RBM3_HUMAN | Oxidation (M) | 3268800 | 2001900 | 1393500 | 1938000 | 0 | 0 | 0 | 0 | 0 | 0 | 0 | 0 | 0 | 0 | 0 | 1936600 | 0 | 0 |
| AMNGESLD | RBM3 | P98179 | RBM3_HUMAN | Unmodified | 6319700 | 3199000 | 1575400 | 2629800 | 0 | 0 | 0 | 0 | 0 | 0 | 0 | 133830 | 0 | 0 | 2182100 | 2519600 | 0 | 0 |
| AMNNSWH | LIMS1; LIMS4; L | P48059; A0A | LIMS1_HUMAN | Oxidation (M) | 0 | 0 | 0 | 0 | 0 | 1213300 | 780150 | 0 | 0 | 0 | 0 | 0 | 241690 | 1317700 | 0 | 0 | 0 | 0 |
| AMNYNAKI | TMEM30A; TME | Q9NV96; A0 | CC50A_HUMA | Acetyl (Protein N | 0 | 0 | 0 | 0 | 1203200 | 0 | 500450 | 0 | 0 | 0 | 0 | 0 | 610700 | 0 | 0 | 0 | 0 | 0 |
| AMQADISQ | TMPO | P42166 | LAP2A_HUMAN | Oxidation (M) | 0 | 0 | 0 | 0 | 948640 | 0 | 4122800 | 105760 | 0 | 0 | 0 | 0 | 0 | 0 | 0 | 0 | 1170200 | 0 |
| AMQDAEVS | HSPA9; HSPA9; | P38646; H0Y | GRP75_HUMAN | Oxidation (M) | 0 | 0 | 0 | 0 | 0 | 0 | 0 | 0 | 118980 | 0 | 0 | 0 | 0 | 0 | 0 | 0 | 0 | 0 |
| AMQDAEVS | HSPA9; HSPA9; | P38646; H0Y | GRP75_HUMAN | Unmodified | 0 | 0 | 0 | 0 | 0 | 0 | 0 | 0 | 1671700 | 0 | 0 | 0 | 0 | 0 | 0 | 0 | 33747 | 0 |
| AMQDAEVS | HSPA9; HSPA9; | P38646; H0Y | GRP75_HUMAN | 2 Oxidation (M) | 0 | 0 | 0 | 0 | 0 | 0 | 0 | 0 | 2178000 | 614250 | 0 | 0 | 0 | 0 | 0 | 0 | 0 | 0 |
| AMQDAEVS | HSPA9; HSPA9; | P38646; H0Y | GRP75_HUMAN | Oxidation (M) | 0 | 0 | 0 | 0 | 0 | 0 | 0 | 0 | 4170300 | 312410 | 0 | 0 | 0 | 0 | 0 | 0 | 0 | 0 |
| AMQEQLEN | EHD4 | Q9H223 | EHD4_HUMAN | Oxidation (M) | 1339800 | 836350 | 908550 | 1556700 | 3022400 | 767910 | 0 | 0 | 0 | 50817 | 0 | 0 | 1971300 | 625240 | 2063200 | 1579000 | 1866400 | 3550200 |
| AMQEQLEN | EHD4 | Q9H223 | EHD4_HUMAN | Unmodified | 458730 | 376270 | 345530 | 730150 | 8295700 | 1226300 | 0 | 0 | 952820 | 0 | 0 | 605780 | 2143400 | 1128900 | 1126500 | 600080 | 2478700 | 6164100 |
| AMQEQLEN | EHD4 | Q9H223 | EHD4_HUMAN | Oxidation (M) | 0 | 0 | 0 | 0 | 0 | 0 | 0 | 0 | 0 | 0 | 0 | 0 | 0 | 0 | 0 | 0 | 954420 | 412280 |
| AMQEQLEN | EHD4 | Q9H223 | EHD4_HUMAN | Unmodified | 0 | 0 | 0 | 0 | 754290 | 0 | 0 | 0 | 0 | 0 | 0 | 0 | 0 | 0 | 0 | 0 | 818640 | 943060 |
| AMQGAGT | ANXA7 | P20073 | ANXA7_HUMAN | Unmodified | 0 | 0 | 0 | 0 | 0 | 0 | 0 | 0 | 1165200 | 0 | 0 | 0 | 0 | 0 | 17711 | 0 | 12130 | 0 |
| AMQGAGT | ANXA7 | P20073 | ANXA7_HUMAN | Oxidation (M) | 40267 | 107020 | 0 | 0 | 0 | 0 | 69147 | 0 | 5014900 | 9213300 | 0 | 0 | 433370 | 0 | 0 | 0 | 0 | 0 |
| AMQHAEV | PBDC1; PBDC1 | Q9BVG4; A6 | PBDC1_HUMA | Unmodified | 76447 | 56242 | 25976 | 0 | 31649 | 0 | 14709 | 0 | 0 | 0 | 51420 | 477310 | 0 | 0 | 67169 | 23246 | 35587 | 0 |
| AMQHISYL | CLSTN1; CLSTN | O94985; Q5 | CSTN1_HUMA | Unmodified | 0 | 0 | 0 | 0 | 0 | 0 | 0 | 0 | 9234800 | 720600 | 0 | 0 | 0 | 0 | 0 | 0 | 0 | 0 |
| AMQHISYL | CLSTN1; CLSTN | O94985; Q5 | CSTN1_HUMA | Oxidation (M) | 0 | 0 | 0 | 0 | 0 | 0 | 0 | 0 | 1613000 | 287020 | 0 | 1588200 | 0 | 0 | 0 | 0 | 10841 | 0 |
| AMQSLVKI | CHMP3 | Q9Y3E7 | CHMP3_HUMA | Unmodified | 0 | 0 | 0 | 0 | 0 | 0 | 0 | 0 | 322050 | 0 | 0 | 0 | 0 | 0 | 0 | 0 | 0 | 0 |
| AMQSLVKI | CHMP3 | Q9Y3E7 | CHMP3_HUMA | Oxidation (M) | 0 | 0 | 0 | 0 | 1108900 | 0 | 0 | 0 | 0 | 748240 | 0 | 0 | 327320 | 0 | 0 | 0 | 560320 | 0 |
| AMSDLQSL | DLG5; DLG5 | Q8TDM6; R4 | DLG5_HUMAN | Unmodified | 0 | 0 | 0 | 71381 | 553140 | 0 | 0 | 0 | 191070 | 0 | 0 | 0 | 0 | 52965 | 57370 | 0 | 478150 | 714360 |
| AMSEVTSLI | LSR; LSR | Q86X29; S4F | LSR_HUMAN Li | Oxidation (M) | 1298500 | 1602000 | 4055200 | 4089600 | 379410 | 191150 | 0 | 0 | 9842300 | 2020800 | 1037200 | 4461200 | 428120 | 0 | 3841100 | 4019600 | 36233 | 55374 |
| AMSEVTSLI | LSR; LSR | Q86X29; S4F | LSR_HUMAN Li | Unmodified | 2544000 | 0 | 2102700 | 321520 | 2098300 | 0 | 0 | 41831 | 39142000 | 4311800 | 2237900 | 0 | 136830 | 0 | 12353000 | 2673000 | 131350 | 0 |
| AMSLTIEPR | CDCP1 | Q9H5V8 | CDCP1_HUMA | Unmodified | 0 | 0 | 0 | 0 | 0 | 0 | 0 | 0 | 0 | 0 | 0 | 950730 | 0 | 0 | 0 | 0 | 0 | 0 |
| AMSLTIEPR | CDCP1 | Q9H5V8 | CDCP1_HUMA | Unmodified | 200870 | 0 | 530000 | 642890 | 409240 | 157690 | 0 | 0 | 846270 | 0 | 1022300 | 7615500 | 207980 | 0 | 995730 | 738480 | 482110 | 921850 |
| AMSLTIEPR | CDCP1 | Q9H5V8 | CDCP1_HUMA | Oxidation (M) | 302660 | 0 | 3477400 | 910300 | 0 | 0 | 0 | 0 | 541250 | 86017 | 935820 | 6245900 | 0 | 0 | 1629100 | 115660 | 0 | 266550 |
| AMSQDGA | UGP2; UGP2; U | Q16851; C9J | UGPA_HUMAN | Oxidation (M) | 0 | 0 | 0 | 0 | 3004100 | 212860 | 0 | 0 | 0 | 0 | 0 | 0 | 0 | 0 | 0 | 0 | 142400 | 1485600 |
| AMSRPILYS | DYNC1H1; DYN | Q14204; A0A | DYHC1_HUMA | Unmodified | 0 | 0 | 0 | 0 | 0 | 0 | 0 | 0 | 365090 | 0 | 0 | 459160 | 0 | 0 | 0 | 0 | 0 | 0 |
| AMSRPILYS | DYNC1H1; DYN | Q14204; A0A | DYHC1_HUMA | Oxidation (M) | 0 | 0 | 0 | 0 | 0 | 0 | 299560 | 0 | 456570 | 409840 | 145420 | 4773400 | 277640 | 151410 | 0 | 0 | 0 | 250390 |
| AMTEDGFL | ASNS; ASNS; AS | P08243; C9J | ASNS_HUMAN | Oxidation (M) | 0 | 0 | 0 | 0 | 0 | 0 | 0 | 0 | 0 | 0 | 657940 | 1136700 | 0 | 0 | 453470 | 219810 | 0 | 0 |
| AMTGVEQV | CCT3; CCT3 | P49368; B4D | TCPG_HUMAN | Unmodified | 0 | 0 | 0 | 0 | 0 | 0 | 0 | 0 | 607210 | 0 | 0 | 2877000 | 0 | 0 | 0 | 0 | 0 | 0 |
| AMTGVEQV | CCT3; CCT3 | P49368; B4D | TCPG_HUMAN | Oxidation (M) | 0 | 0 | 0 | 0 | 0 | 0 | 0 | 0 | 0 | 0 | 644110 | 4627000 | 342420 | 0 | 195640 | 0 | 0 | 0 |
| AMTLQEAE | PLXNB2; PLXNE | O15031; H0Y | PLXB2_HUMAN | Unmodified | 0 | 0 | 86441 | 976420 | 1660800 | 77212 | 0 | 0 | 1670500 | 131010 | 0 | 1133400 | 331120 | 0 | 1261900 | 163360 | 279710 | 1138300 |
| AMTLQEAE | PLXNB2; PLXNE | O15031; H0Y | PLXB2_HUMAN | Oxidation (M) | 663370 | 359520 | 930380 | 1258500 | 487390 | 0 | 0 | 0 | 226140 | 0 | 94205 | 5015800 | 416280 | 2085100 | 2790600 | 2703100 | 277660 | 319560 |
| AMTLQEAE | PLXNB2; PLXNE | O15031; H0Y | PLXB2_HUMAN | 2 Oxidation (M) | 0 | 0 | 0 | 0 | 0 | 2554000 | 0 | 211740 | 0 | 360710 | 0 | 0 | 1679800 | 0 | 0 | 0 | 1608700 | 2093800 |
| AMTLQEAE | PLXNB2; PLXNE | O15031; H0Y | PLXB2_HUMAN | Oxidation (M) | 0 | 0 | 0 | 0 | 5066800 | 0 | 0 | 0 | 249180 | 604970 | 0 | 0 | 1850000 | 0 | 0 | 0 | 5067300 | 9699400 |
| AMTPSRIAD | NCOA4; NCOA4 | Q13772; B4D | NCOA4_HUMA | Unmodified | 0 | 0 | 0 | 0 | 0 | 0 | 0 | 0 | 0 | 0 | 0 | 0 | 0 | 0 | 0 | 0 | 716510 | 0 |
| AMTPSRIAD | NCOA4; NCOA4 | Q13772; B4D | NCOA4_HUMA | Oxidation (M) | 0 | 0 | 0 | 0 | 0 | 0 | 671100 | 0 | 0 | 0 | 0 | 0 | 0 | 0 | 0 | 0 | 0 | 0 |
| AMVADNSI | RELL1; RELL1 | Q8IUW5; D6 | RELL1_HUMAN | Oxidation (M) | 0 | 0 | 0 | 217020 | 4695000 | 0 | 465010 | 826520 | 892350 | 1067900 | 0 | 0 | 0 | 0 | 0 | 0 | 0 | 0 |
| AMVAQPAE | NIBAN1 | Q9BZQ8 | NIBA1_HUMAN | Unmodified | 0 | 0 | 0 | 0 | 1540600 | 370410 | 179250 | 0 | 2303700 | 1402100 | 0 | 0 | 0 | 131760 | 0 | 0 | 56075 | 220930 |
| AMVASGSE | AK2; AK2; AK2; | P54819; G3V | KAD2_HUMAN | Unmodified | 148960 | 159850 | 0 | 65291 | 0 | 0 | 0 | 0 | 1656500 | 289020 | 359030 | 2770400 | 239880 | 0 | 175490 | 134670 | 18306 | 0 |
| AMVASGSE | AK2; AK2; AK2; | P54819; G3V | KAD2_HUMAN | Oxidation (M) | 208770 | 49753 | 30800 | 0 | 0 | 0 | 179820 | 12499 | 586880 | 75548 | 68977 | 807910 | 88266 | 0 | 100490 | 95343 | 0 | 0 |
| AMVDGSW | TNFSF11 | O14788 | TNF11_HUMAN | Unmodified | 0 | 0 | 0 | 0 | 0 | 0 | 0 | 0 | 0 | 0 | 0 | 239630 | 0 | 0 | 0 | 0 | 0 | 0 |
| AMVSEFLK | ANXA1; ANXA1; | P04083; Q5T | ANXA1_HUMAN | Acetyl (Protein N | 452000 | 933480 | 0 | 2898800 | 16106000 | 4811700 | 11226000 | 13794000 | 0 | 0 | 53264000 | 1.53E+08 | 2274800 | 3485100 | 0 | 956670 | 3111800 | 0 |
| AMVSEFLK | ANXA1; ANXA1; | P04083; Q5T | ANXA1_HUMAN | Acetyl (Protein N | 0 | 0 | 0 | 0 | 11941000 | 0 | 3378900 | 593530 | 0 | 0 | 5350700 | 13353000 | 915390 | 1972800 | 0 | 0 | 1304800 | 2628400 |
| AMVSNAQI | LRRFIP2; LRRFI | Q9Y608; Q3 | LRRF2_HUMAN | Unmodified | 0 | 132110 | 0 | 0 | 0 | 197900 | 0 | 0 | 0 | 0 | 0 | 0 | 651730 | 449220 | 0 | 0 | 276300 | 0 |
| ANAAAYLQ | ARVCF; ARVCF; | O00192; E9F | ARVC_HUMAN | Unmodified | 0 | 0 | 495940 | 1020900 | 510450 | 193330 | 0 | 0 | 248000 | 161770 | 89348 | 649710 | 739540 | 1664800 | 1244200 | 924050 | 248460 | 0 |
| ANALHATN | ANKFY1; ANKFY | Q9P2R3; I3L | ANFY1_HUMAN | Unmodified | 0 | 0 | 0 | 0 | 3687700 | 1690500 | 546620 | 0 | 325520 | 0 | 0 | 1725300 | 0 | 0 | 0 | 0 | 0 | 0 |

| | | | | | | | | | | | | | | | | | | | | | | |
|---|---|---|---|---|---|---|---|---|---|---|---|---|---|---|---|---|---|---|---|---|---|---|
| ANALLIREV | GNA11 | P29992 | GNA11_HUMAI | Unmodified | 87438 | 0 | 0 | 0 | 2280300 | 807830 | 440850 | 334760 | 291450 | 368020 | 633290 | 0 | 147460 | 0 | 0 | 0 | 830660 | 1909300 |
| ANALLIREV | GNA11 | P29992 | GNA11_HUMAI | Unmodified | 0 | 0 | 0 | 0 | 3713800 | 993860 | 1539800 | 0 | 0 | 1202200 | 0 | 0 | 0 | 0 | 0 | 0 | 3847800 | 4750400 |
| ANAPPTPLI | WIPF2 | Q8TF74 | WIPF2_HUMAN | Unmodified | 0 | 0 | 0 | 415200 | 0 | 0 | 0 | 0 | 130620 | 0 | 0 | 0 | 0 | 0 | 1704500 | 0 | 0 | 0 |
| ANAQAAAL | CAPG; CAPG; C | P40121; E7E | CAPG_HUMAN | Unmodified | 0 | 0 | 0 | 0 | 0 | 215100 | 0 | 0 | 0 | 0 | 0 | 0 | 0 | 0 | 0 | 0 | 205500 | 0 |
| ANASICFA\ | PRKRA; PRKRA; | O75569; A0/ | PRKRA_HUMAN | Oxidation (M) | 0 | 0 | 0 | 0 | 0 | 0 | 651770 | 310110 | 0 | 125930 | 0 | 0 | 0 | 0 | 0 | 0 | 0 | 0 |
| ANATGGG( | ALOX5 | P09917 | LOX5_HUMAN | Unmodified | 0 | 0 | 0 | 0 | 0 | 0 | 0 | 25826 | 945750 | 0 | 316810 | 1796400 | 0 | 0 | 0 | 0 | 0 | 0 |
| ANCERTFIA | NME1; NME1-N | P15531; J3KI | NDKA_HUMAN | Acetyl (Protein N | 0 | 0 | 0 | 136180 | 15925000 | 2403700 | 1713900 | 659570 | 0 | 2757700 | 2840100 | 0 | 28796000 | 6580600 | 0 | 0 | 11076000 | 10993000 |
| ANDDIIVN\ | PLS3; PLS3 | P13797; A0/ | PLST_HUMAN I | Unmodified | 0 | 0 | 0 | 0 | 201530 | 28534 | 0 | 0 | 134690 | 17346 | 0 | 0 | 0 | 0 | 0 | 0 | 33278 | 51115 |
| ANDDIIVN\ | PLS3; PLS3 | P13797; A0/ | PLST_HUMAN I | Unmodified | 0 | 0 | 0 | 0 | 7949400 | 1561100 | 224630 | 0 | 0 | 432620 | 0 | 0 | 1261900 | 1198200 | 0 | 0 | 2229000 | 2088000 |
| ANDDSTIPI | SLC20A2 | Q08357 | S20A2_HUMAN | Unmodified | 0 | 0 | 608840 | 0 | 1287300 | 0 | 0 | 0 | 0 | 0 | 1164200 | 1001700 | 0 | 0 | 447270 | 380750 | 0 | 0 |
| ANDDVAQI | POGLUT1 | Q8NBL1 | PGLT1_HUMAN | Unmodified | 0 | 0 | 0 | 0 | 0 | 0 | 0 | 0 | 928480 | 0 | 183440 | 0 | 0 | 0 | 0 | 0 | 0 | 0 |
| ANDGLKFS | PTPRF; PTPRF; I | P10586; H0Y | PTPRF_HUMAN | Unmodified | 0 | 0 | 0 | 0 | 4384000 | 0 | 0 | 0 | 0 | 0 | 0 | 0 | 0 | 0 | 0 | 0 | 0 | 0 |
| ANDIEKAV( | ARPC5; ARPC5 | O15511; B1/ | ARPC5_HUMAI | Unmodified | 0 | 0 | 0 | 0 | 470180 | 0 | 37553 | 198440 | 0 | 53398 | 0 | 0 | 338870 | 198430 | 0 | 0 | 255000 | 183420 |
| ANEALRLRI | PLEC | Q15149 | PLEC_HUMAN | Unmodified | 0 | 0 | 0 | 0 | 1653600 | 0 | 0 | 0 | 0 | 0 | 0 | 0 | 0 | 0 | 0 | 0 | 4328900 | 3488400 |
| ANEEIAQVI | TACC1; TACC1; | O75410; E7E | TACC1_HUMAI | Unmodified | 0 | 0 | 0 | 0 | 518580 | 0 | 0 | 0 | 0 | 0 | 0 | 0 | 0 | 0 | 0 | 0 | 0 | 358210 |
| ANEFHDVN | PRDX3 | P30048 | PRDX3_HUMAI | Unmodified | 0 | 0 | 0 | 0 | 0 | 0 | 0 | 0 | 932150 | 0 | 0 | 0 | 0 | 0 | 0 | 0 | 0 | 0 |
| ANEGTVGV | ALPL | P05186 | PPBT_HUMAN | Unmodified | 0 | 841270 | 0 | 0 | 0 | 0 | 0 | 0 | 923620 | 94251 | 0 | 0 | 0 | 0 | 0 | 0 | 0 | 0 |
| ANEHQGIG | UPF1 | Q92900 | RENT1_HUMAI | Unmodified | 1697500 | 1643600 | 4787900 | 2068500 | 0 | 0 | 0 | 0 | 411760 | 0 | 0 | 1775600 | 23686 | 0 | 1324300 | 1652100 | 0 | 0 |
| ANENASFL | CLIP1; CLIP1; C | P30622; F5H | CLIP1_HUMAN | Unmodified | 0 | 0 | 0 | 0 | 560850 | 0 | 0 | 0 | 0 | 0 | 0 | 0 | 578350 | 370990 | 0 | 0 | 251210 | 266820 |
| ANEPTHFT | FLNB; FLNB | O75369; E7E | FLNB_HUMAN | Unmodified | 0 | 0 | 0 | 260790 | 1313900 | 0 | 0 | 0 | 0 | 0 | 0 | 1917800 | 0 | 0 | 0 | 211080 | 288370 | 0 |
| ANEPVKTE | EPB41L1; EPB4 | H7C2K6; X6I | H7C2K6_HUM/ | Unmodified | 0 | 0 | 0 | 320360 | 1300800 | 0 | 0 | 0 | 0 | 0 | 0 | 0 | 2190900 | 0 | 0 | 0 | 453250 | 2145200 |
| ANESSPKP | AIF1L; AIF1L | Q9BQI0; A6F | AIF1L_HUMAN | Unmodified | 166710 | 156920 | 439940 | 0 | 0 | 0 | 0 | 0 | 413470 | 368920 | 0 | 0 | 4732800 | 0 | 714940 | 607480 | 0 | 0 |
| ANEVEQMI | DYNC1H1; DYN | Q14204; A0/ | DYHC1_HUMA | Unmodified | 0 | 0 | 0 | 0 | 0 | 0 | 0 | 0 | 144040 | 0 | 0 | 1180500 | 0 | 0 | 0 | 0 | 0 | 0 |
| ANFENLAK | CTTN; CTTN | Q14247; H0Y | SRC8_HUMAN | Unmodified | 830580 | 681470 | 0 | 899370 | 2359900 | 423830 | 0 | 48054 | 0 | 223900 | 145120 | 2274500 | 1116700 | 0 | 1223700 | 0 | 402050 | 580880 |
| ANFLNSNE | VIL1; VIL1; VIL1 | P09327; H7( | VILI_HUMAN Vi | Unmodified | 100180 | 0 | 9582200 | 7716900 | 1291000 | 9440500 | 707350 | 0 | 9619600 | 3890800 | 52587000 | 30111000 | 0 | 0 | 0 | 0 | 0 | 0 |
| ANFWYQPS | CARM1; CARM1 | Q86X55; K7E | CARM1_HUMA | Unmodified | 737240 | 934280 | 165380 | 415100 | 0 | 1004000 | 0 | 57275 | 76655 | 0 | 60567 | 3302800 | 0 | 0 | 350370 | 343150 | 0 | 0 |
| ANGHLLLN | LARS1; LARS1; | Q9P2J5; A0A | SYLC_HUMAN | Unmodified | 0 | 0 | 0 | 0 | 0 | 38691 | 0 | 32179 | 0 | 0 | 0 | 826670 | 0 | 0 | 0 | 0 | 0 | 0 |
| ANGIVVPS | SLC22A5; SLC2 | O76082; H7( | S22A5_HUMAN | Unmodified | 0 | 0 | 112940 | 0 | 0 | 0 | 0 | 0 | 348110 | 118480 | 0 | 227170 | 0 | 0 | 1158000 | 494260 | 0 | 0 |
| ANGTTVHV | RPL26; RPL26L | P61254; Q9U | RL26_HUMAN ( | Unmodified | 2523000 | 1693500 | 801660 | 662630 | 448130 | 1385200 | 215250 | 178180 | 117290 | 0 | 696300 | 14439000 | 334710 | 213810 | 1553800 | 1384000 | 808760 | 0 |
| ANHEEVLA | PNP | P00491 | PNPH_HUMAN | Unmodified | 0 | 0 | 290560 | 0 | 986770 | 211240 | 0 | 0 | 63628 | 152260 | 801830 | 4587100 | 0 | 0 | 0 | 399510 | 0 | 283100 |
| ANHFFTVTI | SFXN1; SFXN1; | Q9H9B4; D6 | SFXN1_HUMAN | Unmodified | 0 | 0 | 0 | 0 | 0 | 0 | 0 | 0 | 1430900 | 0 | 0 | 0 | 0 | 0 | 0 | 0 | 0 | 0 |
| ANHGPGM | PACSIN2 | A0A0U1RR2: | A0A0U1RR22_I | Oxidation (M) | 1385600 | 0 | 0 | 0 | 0 | 0 | 0 | 0 | 0 | 0 | 0 | 0 | 0 | 0 | 0 | 0 | 0 | 0 |
| ANHGPGM | PACSIN2 | A0A0U1RR2: | A0A0U1RR22_I | 2 Oxidation (M) | 4476500 | 2364500 | 0 | 0 | 0 | 0 | 0 | 0 | 0 | 0 | 0 | 0 | 0 | 0 | 0 | 0 | 0 | 0 |
| ANHGPGM | PACSIN2 | Q9UNF0 | PACN2_HUMAI | 2 Oxidation (M) | 0 | 3700700 | 11477000 | 24855000 | 0 | 0 | 0 | 0 | 0 | 0 | 409410 | 3281600 | 4363400 | 0 | 34329000 | 20505000 | 0 | 0 |
| ANHGPGM | PACSIN2 | Q9UNF0 | PACN2_HUMAI | Oxidation (M) | 0 | 0 | 0 | 2193800 | 0 | 0 | 0 | 0 | 0 | 0 | 0 | 0 | 0 | 0 | 2965800 | 514650 | 0 | 0 |
| ANHKIEELE | KIF14 | Q15058 | KIF14_HUMAN | Unmodified | 0 | 0 | 0 | 0 | 0 | 0 | 0 | 0 | 0 | 0 | 0 | 0 | 0 | 0 | 0 | 0 | 0 | 0 |
| ANHPMDAI | AP2A1 | O95782 | AP2A1_HUMAN | Unmodified | 580470 | 0 | 1485300 | 1816500 | 2869500 | 0 | 122060 | 203320 | 503540 | 2084200 | 0 | 2765800 | 1072800 | 4987900 | 267610 | 985470 | 3069700 | 5297800 |
| ANIAVQR | UBE2K; UBE2K; | P61086; D6F | UBE2K_HUMAI | Acetyl (Protein N | 298200 | 0 | 0 | 0 | 267340 | 0 | 0 | 35490 | 259150 | 242760 | 434510 | 1558800 | 0 | 0 | 278590 | 245760 | 99512 | 0 |
| ANIGDAQA | GPT2; GPT2 | Q8TD30; H3 | ALAT2_HUMAN | Oxidation (M) | 0 | 0 | 0 | 0 | 0 | 0 | 0 | 0 | 540310 | 0 | 0 | 0 | 0 | 0 | 0 | 0 | 0 | 0 |
| ANIIFNTSL( | SQOR; N/A | Q9Y6N5; H3 | SQOR_HUMAN | Unmodified | 0 | 0 | 0 | 0 | 0 | 0 | 0 | 0 | 431250 | 435380 | 0 | 0 | 0 | 0 | 0 | 0 | 0 | 0 |
| ANIKDLSQI | STXBP2; STXBP | Q15833; M0I | STXB2_HUMAN | Unmodified | 0 | 0 | 0 | 0 | 137700 | 0 | 0 | 0 | 873760 | 641920 | 0 | 425600 | 0 | 178030 | 0 | 0 | 463230 | 137630 |
| ANILVSGNI | POMGNT2 | Q8NAT1 | PMGT2_HUMAI | Unmodified | 0 | 0 | 0 | 0 | 0 | 0 | 0 | 0 | 412830 | 239770 | 0 | 0 | 0 | 0 | 0 | 0 | 0 | 0 |
| ANINIEK | RAB10 | P61026 | RAB10_HUMAN | Unmodified | 0 | 0 | 4361100 | 3373000 | 1791100 | 0 | 0 | 0 | 0 | 0 | 0 | 0 | 62548 | 4333700 | 4317500 | 3634000 | 0 | 0 |
| ANINVENA | RAB8A | P61006 | RAB8A_HUMAN | Unmodified | 5717300 | 4417500 | 4261400 | 7273400 | 8756700 | 1136200 | 28936 | 0 | 7558000 | 3563800 | 4126200 | 9009000 | 5964100 | 1102500 | 7671800 | 5813500 | 2273400 | 1541300 |
| ANINVENA | RAB8A | P61006 | RAB8A_HUMAN | Unmodified | 0 | 0 | 0 | 0 | 17848000 | 790010 | 1730300 | 0 | 0 | 375300 | 0 | 0 | 6547300 | 6647100 | 184190 | 0 | 2993700 | 9025100 |
| ANINVENA | RAB8A | P61006 | RAB8A_HUMAN | Unmodified | 0 | 0 | 0 | 0 | 736080 | 0 | 281350 | 0 | 0 | 0 | 0 | 0 | 0 | 0 | 0 | 0 | 220770 | 0 |
| ANIPIMDT( | ATP6V0A1; ATP | Q93050; K7E | VPP1_HUMAN | Unmodified | 0 | 0 | 0 | 0 | 0 | 0 | 0 | 80983 | 3209800 | 483410 | 0 | 335530 | 0 | 0 | 0 | 0 | 0 | 0 |
| ANIPIMDT( | ATP6V0A1; ATP | Q93050; K7E | VPP1_HUMAN | Oxidation (M) | 0 | 0 | 0 | 46477 | 0 | 0 | 0 | 289370 | 361050 | 668820 | 998150 | 910360 | 0 | 0 | 0 | 0 | 0 | 0 |
| ANIQAVSLI | CHMP2A; CHM | O43633; M0I | CHM2A_HUMA | Unmodified | 5479400 | 7802000 | 5193400 | 5096600 | 2110100 | 0 | 399630 | 284450 | 13116000 | 2977900 | 0 | 23124000 | 0 | 1141700 | 6379700 | 6690000 | 168540 | 210670 |
| ANIQAVSLI | CHMP2A; CHM | O43633; M0I | CHM2A_HUMA | Unmodified | 0 | 0 | 0 | 0 | 985970 | 0 | 0 | 346910 | 345370 | 904240 | 0 | 0 | 0 | 0 | 0 | 0 | 0 | 0 |
| ANIRNMSV | EEF2 | P13639 | EF2_HUMAN El | Oxidation (M) | 0 | 0 | 71222 | 0 | 0 | 324490 | 535680 | 155510 | 0 | 370170 | 0 | 0 | 514790 | 273640 | 0 | 0 | 465900 | 786480 |
| ANIRNMSV | EEF2 | P13639 | EF2_HUMAN El | Unmodified | 0 | 192970 | 392240 | 0 | 6183000 | 2943300 | 1083800 | 1079700 | 0 | 705430 | 0 | 0 | 1330100 | 2087300 | 0 | 282580 | 2439800 | 2709200 |
| ANIRNMSV | EEF2 | P13639 | EF2_HUMAN El | Oxidation (M) | 0 | 0 | 0 | 0 | 0 | 0 | 235300 | 0 | 52638 | 0 | 0 | 0 | 0 | 0 | 0 | 0 | 656950 | 0 |
| ANIRNMSV | EEF2 | P13639 | EF2_HUMAN El | Unmodified | 0 | 0 | 0 | 0 | 0 | 0 | 399830 | 0 | 0 | 0 | 0 | 0 | 0 | 0 | 0 | 0 | 1141000 | 0 |
| ANIVAYHD: | UBA2; UBA2 | Q9UBT2; K7I | SAE2_HUMAN : | Oxidation (M) | 0 | 0 | 0 | 0 | 0 | 0 | 0 | 0 | 0 | 0 | 0 | 1457600 | 0 | 0 | 0 | 0 | 0 | 0 |
| ANIVHLML: | CSE1L | P55060 | XPO2_HUMAN | Unmodified | 0 | 0 | 0 | 0 | 377320 | 0 | 240430 | 0 | 913170 | 196280 | 502960 | 7604000 | 50018 | 0 | 0 | 0 | 548350 | 181720 |
| ANIVHLML: | CSE1L | P55060 | XPO2_HUMAN | Oxidation (M) | 0 | 49832 | 0 | 488250 | 167990 | 0 | 0 | 156500 | 179970 | 381460 | 1326100 | 7955500 | 0 | 0 | 569080 | 219850 | 92448 | 0 |
| ANIWLVR | MBLAC2 | Q68D91 | MBLC2_HUMAI | Unmodified | 0 | 172470 | 795900 | 323100 | 0 | 0 | 0 | 0 | 98169 | 0 | 10509 | 4411500 | 0 | 0 | 544220 | 619140 | 0 | 0 |
| ANKEVPCY | IGF2R; IGF2R; I | P11717; A0A | MPRI_HUMAN ( | Unmodified | 0 | 0 | 0 | 127120 | 0 | 0 | 0 | 0 | 1365600 | 227100 | 0 | 0 | 0 | 0 | 0 | 0 | 0 | 0 |
| ANKITITND | HSPA6; HSPA1/ | P17066; P0D | HSP76_HUMAI | Unmodified | 235590 | 258570 | 0 | 335580 | 1302000 | 540620 | 1281700 | 648900 | 683430 | 1996100 | 0 | 529710 | 1579000 | 157580 | 88754 | 149340 | 851800 | 746500 |
| ANKLDHV\ | PRMT1; PRMT1; | Q99873; E9F | ANM1_HUMAN | Unmodified | 0 | 0 | 0 | 0 | 0 | 0 | 356450 | 70072 | 0 | 164380 | 398910 | 10765000 | 762800 | 0 | 0 | 0 | 389380 | 262160 |
| ANKQTAEV | BICD1; BICD2; | Q96G01; Q8 | BICD1_HUMAN | Unmodified | 0 | 0 | 0 | 0 | 707900 | 0 | 175830 | 0 | 0 | 0 | 0 | 0 | 0 | 0 | 0 | 0 | 270530 | 246200 |
| ANKVPDSV | BMX | P51813 | BMX_HUMAN ( | Unmodified | 0 | 0 | 0 | 0 | 0 | 0 | 0 | 0 | 0 | 0 | 163430 | 2918500 | 0 | 0 | 0 | 0 | 0 | 0 |
| ANLCTLAEI | FAS | P25445 | TNR6_HUMAN | Unmodified | 0 | 0 | 0 | 0 | 120890 | 26720 | 0 | 0 | 0 | 22583 | 0 | 0 | 216340 | 0 | 0 | 0 | 81146 | 47208 |
| ANLEAFTVI | KIF5B | P33176 | KINH_HUMAN | Unmodified | 240130 | 0 | 0 | 0 | 0 | 0 | 0 | 0 | 1485600 | 0 | 644010 | 4617700 | 0 | 0 | 210730 | 0 | 0 | 0 |
| ANLEGANL | KCTD9 | Q7L273 | KCTD9_HUMAI | Unmodified | 0 | 0 | 0 | 0 | 69731 | 0 | 0 | 15658 | 4352400 | 135560 | 73892 | 463560 | 0 | 0 | 0 | 0 | 242270 | 568740 |
| ANLEISYAK | NOSTRIN | Q8IVI9 | NOSTN_HUMA | Unmodified | 4320600 | 4302600 | 5517000 | 10675000 | 0 | 0 | 0 | 36806 | 1418400 | 477420 | 0 | 0 | 0 | 0 | 10935000 | 6948500 | 0 | 0 |
| ANLERTFIA | NME2; NME1-N | P22392; J3KI | NDKB_HUMAN | Acetyl (Protein N | 0 | 0 | 0 | 427880 | 92230000 | 21735000 | 9402500 | 2909300 | 191980 | 12733000 | 5217600 | 73012 | 91302000 | 37669000 | 0 | 0 | 81424000 | 69721000 |
| ANLHAEPD | SLITRK6 | Q9H5Y7 | SLIK6_HUMAN | Unmodified | 1902200 | 1791000 | 0 | 0 | 173160 | 0 | 0 | 0 | 4676700 | 3806500 | 0 | 0 | 0 | 0 | 0 | 232550 | 1383500 | 4233600 |
| ANLINNIFE | LNPEP | Q9UIQ6 | LCAP_HUMAN | Unmodified | 0 | 0 | 35357 | 0 | 0 | 0 | 0 | 0 | 0 | 47183 | 0 | 868230 | 0 | 0 | 427570 | 130020 | 0 | 0 |
| ANLISEDIE: | EPS8; EPS8; EP | Q12929; F5H | EPS8_HUMAN | Unmodified | 125840 | 142430 | 0 | 0 | 295490 | 0 | 0 | 0 | 178790 | 0 | 0 | 561700 | 37871 | 0 | 70464 | 105790 | 0 | 0 |
| ANLISEDIE: | EPS8; EPS8; EP | Q12929; F5H | EPS8_HUMAN | Unmodified | 0 | 0 | 0 | 0 | 2815300 | 20630 | 136580 | 0 | 0 | 0 | 0 | 0 | 0 | 0 | 0 | 0 | 680900 | 0 |
| ANLLEIAPH | GPD2; GPD2 | P43304; F5G | GPDM_HUMAN | Oxidation (M) | 15465 | 132900 | 0 | 0 | 0 | 0 | 0 | 0 | 482330 | 345360 | 0 | 0 | 0 | 0 | 0 | 21573 | 0 | 0 |
| ANLLFKGS' | TOMM40 | O96008 | TOM40_HUMAI | Unmodified | 0 | 0 | 0 | 0 | 0 | 0 | 0 | 0 | 0 | 148780 | 0 | 0 | 0 | 0 | 0 | 0 | 0 | 0 |
| ANLLNNEA | RECQL | P46063 | RECQ1_HUMAI | Unmodified | 0 | 0 | 0 | 0 | 438450 | 0 | 485850 | 32710 | 269880 | 158640 | 0 | 0 | 118600 | 0 | 0 | 0 | 71306 | 167630 |
| ANLLNNEA | RECQL | P46063 | RECQ1_HUMAI | Oxidation (M) | 67811 | 0 | 0 | 0 | 131770 | 0 | 0 | 155880 | 0 | 148920 | 0 | 0 | 55419 | 65943 | 0 | 0 | 72979 | 873640 |
| ANLPQSFQ | FLNA; FLNA | P21333; Q6( | FLNA_HUMAN | Unmodified | 379830 | 957880 | 112950 | 1950800 | 7647400 | 1286200 | 1061800 | 775290 | 0 | 0 | 0 | 0 | 2186800 | 1960800 | 826260 | 891410 | 3095200 | 0 |
| ANLPQSFQ | FLNA; FLNA | P21333; Q6( | FLNA_HUMAN | Unmodified | 0 | 0 | 0 | 0 | 3526800 | 0 | 0 | 120050 | 0 | 0 | 0 | 0 | 0 | 736040 | 0 | 0 | 5553400 | 1239400 |
| ANLQAALQ | VPS39 | Q96JC1 | VPS39_HUMAN | Unmodified | 0 | 0 | 0 | 0 | 0 | 0 | 0 | 0 | 0 | 0 | 0 | 365540 | 0 | 0 | 0 | 0 | 0 | 0 |
| ANLQIDQII | MYH9 | P35579 | MYH9_HUMAN | Unmodified | 305950 | 406760 | 592860 | 768450 | 0 | 181610 | 90595 | 64253 | 1893700 | 346620 | 631810 | 6670600 | 1603900 | 645920 | 955470 | 1131000 | 0 | 0 |
| ANLQIDQII | MYH9 | P35579 | MYH9_HUMAN | Unmodified | 110600 | 0 | 642010 | 1934000 | 22755000 | 6835100 | 859570 | 0 | 641680 | 2088900 | 1230300 | 0 | 6934300 | 3689700 | 581770 | 282170 | 6442700 | 7708300 |
| ANLQIDQII | MYH9 | P35579 | MYH9_HUMAN | Unmodified | 0 | 0 | 0 | 0 | 11304000 | 0 | 0 | 0 | 0 | 0 | 0 | 0 | 5976700 | 1156900 | 0 | 0 | 3338700 | 4168000 |
| ANLVFKEIE | SCP2; SCP2; S( | P22307; H0Y | SCP2_HUMAN | Unmodified | 0 | 0 | 0 | 0 | 0 | 0 | 0 | 0 | 2394500 | 3780000 | 0 | 0 | 0 | 0 | 0 | 0 | 0 | 0 |
| ANMHISES | C1orf21 | Q9H246 | CA021_HUMAN | Oxidation (M) | 74165 | 0 | 0 | 627700 | 150200 | 50505 | 0 | 0 | 0 | 0 | 0 | 0 | 117660 | 0 | 386700 | 358060 | 0 | 0 |
| ANMHISES | C1orf21 | Q9H246 | CA021_HUMAN | Unmodified | 74065 | 0 | 0 | 799490 | 0 | 0 | 0 | 0 | 0 | 0 | 0 | 0 | 309160 | 0 | 658790 | 282290 | 0 | 313340 |
| ANNGICFL | QARS1; QARS1; | P47897; A0A | SYQ_HUMAN G | Unmodified | 0 | 0 | 0 | 0 | 0 | 0 | 0 | 0 | 0 | 0 | 0 | 925400 | 0 | 0 | 0 | 0 | 0 | 0 |
| ANNIDYTVI | TPP2; TPP2; TPI | P29144; A0A | TPP2_HUMAN | Unmodified | 0 | 0 | 0 | 0 | 0 | 0 | 0 | 0 | 248900 | 146200 | 253290 | 2296200 | 38494 | 0 | 66607 | 57180 | 0 | 0 |
| ANNIDYTVI | TPP2; TPP2; TPI | P29144; A0A | TPP2_HUMAN | Unmodified | 0 | 0 | 0 | 143010 | 0 | 0 | 0 | 0 | 481560 | 0 | 830880 | 2762200 | 0 | 0 | 79381 | 79802 | 0 | 0 |
| ANNPEQNI | SND1 | Q7KZF4 | SND1_HUMAN | Unmodified | 376740 | 0 | 988960 | 1009900 | 0 | 0 | 259150 | 56906 | 0 | 0 | 458680 | 2866200 | 886920 | 291420 | 787500 | 803700 | 0 | 0 |
| ANNTFYGL | ALDH1A1 | P00352 | AL1A1_HUMAN | Unmodified | 0 | 0 | 0 | 0 | 0 | 0 | 0 | 0 | 0 | 0 | 0 | 21482000 | 0 | 0 | 0 | 0 | 0 | 0 |
| ANNYNEA\ | NCBP1 | Q09161 | NCBP1_HUMAI | Unmodified | 0 | 0 | 466330 | 0 | 0 | 0 | 0 | 0 | 0 | 0 | 0 | 1502000 | 0 | 0 | 0 | 0 | 0 | 0 |
| ANPALYVLI | PRPF8 | Q6P2Q9 | PRP8_HUMAN | Unmodified | 0 | 0 | 731640 | 0 | 0 | 0 | 0 | 0 | 0 | 0 | 0 | 579200 | 0 | 0 | 0 | 0 | 0 | 0 |
| ANPGAWIL | UGGT2; UGGT1 | Q9NYU1; Q9 | UGGG2_HUMA | Unmodified | 0 | 0 | 0 | 0 | 0 | 0 | 0 | 0 | 5091900 | 3243700 | 370630 | 605900 | 0 | 0 | 94848 | 0 | 0 | 0 |
| ANPHPFAC | SNX33; SNX33 | Q8WV41; H3 | SNX33_HUMAN | Unmodified | 1370000 | 773920 | 7171300 | 6974700 | 0 | 0 | 0 | 0 | 0 | 0 | 0 | 242030 | 384810 | 122710 | 14742000 | 8599700 | 0 | 93288 |
| ANPHPFAC | SNX33; SNX33 | Q8WV41; H3 | SNX33_HUMAN | Unmodified | 0 | 0 | 0 | 1007700 | 0 | 0 | 0 | 0 | 0 | 0 | 0 | 0 | 0 | 0 | 0 | 81509 | 0 | 0 |
| ANPNAASY | TMTC4 | Q5T4D3 | TMTC4_HUMAI | Unmodified | 0 | 0 | 0 | 0 | 0 | 354800 | 0 | 0 | 1838000 | 0 | 0 | 0 | 0 | 0 | 0 | 0 | 0 | 0 |
| ANPQVGV/ | XRCC5 | P13010 | XRCC5_HUMAI | Unmodified | 185250 | 0 | 79936 | 0 | 0 | 0 | 332250 | 428670 | 233100 | 177700 | 482810 | 11685000 | 0 | 0 | 94648 | 50036 | 0 | 0 |
| ANPRCPES | DGKH; DGKH | Q86XP1; A0/ | DGKH_HUMAN | Unmodified | 0 | 0 | 0 | 0 | 0 | 0 | 77800 | 0 | 0 | 0 | 0 | 0 | 0 | 323310 | 0 | 0 | 208510 | 0 |
| ANPTFHHV | SPRED2; SPREI | Q7Z698; H7( | SPRE2_HUMAN | Unmodified | 0 | 0 | 0 | 0 | 61357 | 0 | 0 | 0 | 0 | 0 | 0 | 0 | 0 | 0 | 0 | 0 | 239280 | 455310 |
| ANPTQLNT | TFCP2; TFCP2; | Q12800; F8\ | TFCP2_HUMAN | Unmodified | 0 | 0 | 0 | 0 | 0 | 0 | 0 | 0 | 235550 | 0 | 59008 | 547850 | 0 | 0 | 0 | 0 | 81608 | 0 |
| ANPVPDGI | ECE1; ECE1; EC | P42892; B4D | ECE1_HUMAN | Unmodified | 0 | 0 | 0 | 0 | 1669200 | 700550 | 0 | 0 | 1726900 | 1300400 | 0 | 0 | 0 | 0 | 0 | 0 | 0 | 0 |
| ANQVATGII | SCIN; SCIN; SC | Q9Y6U3; E5I | ADSV_HUMAN | Unmodified | 1237200 | 90647 | 0 | 0 | 197070 | 27110 | 23615 | 22182 | 1138500 | 310580 | 20396 | 690600 | 0 | 0 | 0 | 75508 | 138630 | 966170 |
| ANRFTVETI | FLNC | Q14315 | FLNC_HUMAN | Unmodified | 0 | 0 | 0 | 0 | 2942700 | 709050 | 0 | 0 | 0 | 0 | 0 | 0 | 0 | 0 | 0 | 0 | 2154800 | 0 |
| ANRGPAY( | TAGLN2; TAGLI | P37802; X6R | TAGL2_HUMAN | Acetyl (Protein N | 14655000 | 13140000 | 3798400 | 8655400 | 29314000 | 8982100 | 2284400 | 1735400 | 8046400 | 3437800 | 5068200 | 21077000 | 9795700 | 2822000 | 5520500 | 4608100 | 7748700 | 8921800 |
| ANRGPAY( | TAGLN2; TAGLI | P37802; X6R | TAGL2_HUMAN | Acetyl (Protein N | 0 | 0 | 0 | 0 | 9953700 | 3965000 | 1269000 | 820030 | 0 | 996120 | 0 | 0 | 2958500 | 0 | 0 | 0 | 2513900 | 6714400 |
| ANSALTPPI | SARG | Q9BW04 | SARG_HUMAN | Unmodified | 386380 | 0 | 246540 | 427580 | 0 | 0 | 0 | 0 | 0 | 233930 | 0 | 0 | 854180 | 0 | 246580 | 298870 | 0 | 0 |
| ANSALTPPI | SARG | Q9BW04 | SARG_HUMAN | Oxidation (M) | 0 | 0 | 0 | 0 | 0 | 0 | 0 | 0 | 0 | 0 | 0 | 0 | 1445400 | 0 | 0 | 0 | 0 | 0 |
| ANSALTPPI | SARG | Q9BW04 | SARG_HUMAN | Unmodified | 0 | 0 | 0 | 0 | 0 | 0 | 0 | 0 | 1037200 | 0 | 0 | 0 | 7634000 | 1055800 | 0 | 0 | 0 | 0 |
| ANSEHNGI | HSPA4L; HSPA | O95757; E9F | HS74L_HUMAN | Unmodified | 0 | 0 | 0 | 0 | 0 | 0 | 0 | 0 | 0 | 0 | 0 | 273190 | 0 | 0 | 0 | 0 | 0 | 0 |

| | | | | | | | | | | | | | | | | | | | | | | |
|---|---|---|---|---|---|---|---|---|---|---|---|---|---|---|---|---|---|---|---|---|---|---|
| ANSFTVSS | HDLBP; HDLBP | Q00341; A0 | VIGLN_HUMAN | Unmodified | 464580 | 507070 | 0 | 0 | 0 | 0 | 30754 | 0 | 1150700 | 328430 | 531300 | 5038200 | 1198800 | 0 | 521540 | 656630 | 511190 | 0 |
| ANSFTVSS | HDLBP; HDLBP | Q00341; A0 | VIGLN_HUMAN | Unmodified | 0 | 0 | 0 | 0 | 0 | 0 | 49233 | 0 | 0 | 37000 | 0 | 0 | 470030 | 246910 | 0 | 0 | 0 | 0 |
| ANSFVGTA | PDPK2P; PDPK | Q6A1A2; O1 | PDPK2_HUMAN | Unmodified | 0 | 0 | 1393000 | 1367100 | 0 | 0 | 0 | 0 | 0 | 0 | 0 | 0 | 1040100 | 0 | 0 | 881110 | 0 | 0 |
| ANSLPPGN | PROM1 | O43490 | PROM1_HUMA | Unmodified | 0 | 0 | 0 | 0 | 0 | 0 | 0 | 0 | 0 | 0 | 0 | 3936500 | 0 | 0 | 0 | 0 | 0 | 0 |
| ANSPSLFG | SH3KBP1; SH3 | Q96B97; Q5 | SH3K1_HUMAN | Unmodified | 0 | 6164800 | 1871100 | 829940 | 0 | 0 | 0 | 0 | 0 | 0 | 0 | 0 | 0 | 0 | 432750 | 0 | 0 | 0 |
| ANSVFEDL | ATP6AP2; ATP6 | O75787; A0 | RENR_HUMAN | Unmodified | 0 | 0 | 0 | 0 | 0 | 0 | 0 | 0 | 0 | 2313400 | 0 | 0 | 0 | 0 | 0 | 0 | 0 | 0 |
| ANSVKQEF | CAND1; CAND | Q86VP6; A0 | CAND1_HUMA | Unmodified | 0 | 77392 | 0 | 339220 | 2255100 | 469570 | 367730 | 321470 | 1538300 | 1020600 | 275830 | 3833900 | 851770 | 0 | 0 | 70793 | 583970 | 2225900 |
| ANSWQLV | DLGAP4; DLGA | Q9Y2H0; F8 | DLGP4_HUMA | Unmodified | 0 | 256620 | 0 | 2113300 | 0 | 0 | 0 | 0 | 0 | 0 | 114390 | 1244300 | 445550 | 269390 | 0 | 0 | 0 | 0 |
| ANTAQEQ | GJC1; GJC1; GJ | P36383; Q5 | CXG1_HUMAN | Unmodified | 0 | 0 | 0 | 0 | 7195900 | 1938800 | 0 | 0 | 0 | 0 | 0 | 0 | 0 | 0 | 0 | 542230 | 0 | 0 |
| ANTAYHLH | ERBIN | Q96RT1 | ERBIN_HUMAN | Unmodified | 0 | 0 | 0 | 0 | 1305800 | 107320 | 520570 | 0 | 118490 | 0 | 71473 | 208010 | 122740 | 0 | 26081 | 57195 | 0 | 180910 |
| ANTFVAEL | MDH2; MDH2 | P40926; G3 | MDHM_HUMAN | Unmodified | 111150 | 95756 | 236930 | 0 | 0 | 0 | 129810 | 78839 | 30125000 | 4746700 | 555990 | 6470400 | 373110 | 0 | 0 | 80814 | 0 | 0 |
| ANTLSQLS | IQGAP2; IQGAP | Q13576; E7 | IQGA2_HUMAN | Unmodified | 0 | 0 | 1151100 | 240470 | 0 | 0 | 0 | 0 | 0 | 0 | 0 | 0 | 0 | 0 | 1066500 | 872910 | 0 | 0 |
| ANTVLERV | TMEM9; TMEM | Q9P0T7; B1 | TMEM9_HUMA | Unmodified | 230370 | 0 | 977870 | 927780 | 0 | 0 | 0 | 0 | 5545200 | 441270 | 0 | 656820 | 0 | 0 | 685080 | 627330 | 67783 | 0 |
| ANTVTPAV | CDCA8 | Q53HL2 | BOREA_HUMA | Oxidation (M) | 0 | 0 | 0 | 2133600 | 615490 | 0 | 0 | 0 | 0 | 0 | 0 | 0 | 1125200 | 0 | 0 | 0 | 0 | 0 |
| ANVAVVSG | SDCBP | G5EA09 | G5EA09_HUMA | Unmodified | 95329 | 117140 | 377080 | 90012 | 0 | 0 | 0 | 0 | 571780 | 0 | 0 | 2650000 | 0 | 0 | 802380 | 867290 | 0 | 0 |
| ANVAVVSG | SDCBP | G5EA09 | G5EA09_HUMA | Oxidation (M) | 172560 | 7849900 | 0 | 584890 | 0 | 0 | 0 | 0 | 21139000 | 960820 | 856990 | 9824300 | 0 | 0 | 1046400 | 287940 | 0 | 0 |
| ANVAVVSG | SDCBP | G5EA09 | G5EA09_HUMA | Unmodified | 146400 | 0 | 84497 | 428310 | 0 | 0 | 0 | 0 | 19947000 | 0 | 0 | 629060 | 0 | 0 | 582650 | 225990 | 0 | 0 |
| ANVAVVSG | SDCBP | O00560 | SDCB1_HUMA | Unmodified | 683400 | 1293400 | 2033800 | 812970 | 0 | 0 | 0 | 0 | 4667700 | 895250 | 205090 | 8188400 | 0 | 0 | 3152900 | 2350400 | 0 | 0 |
| ANVAVVSG | SDCBP | O00560 | SDCB1_HUMA | Oxidation (M) | 13206000 | 15298000 | 0 | 20890000 | 0 | 0 | 0 | 0 | 52583000 | 0 | 2222700 | 3532900 | 0 | 0 | 28228000 | 0 | 0 | 0 |
| ANVAVVSG | SDCBP | O00560 | SDCB1_HUMA | Unmodified | 0 | 0 | 0 | 0 | 0 | 0 | 0 | 0 | 35856000 | 0 | 0 | 1227300 | 0 | 0 | 0 | 0 | 0 | 0 |
| ANVDISAP | AHNAK | Q09666 | AHNK_HUMAN | Unmodified | 135660 | 0 | 0 | 0 | 18856000 | 2452800 | 0 | 0 | 0 | 0 | 0 | 0 | 912270 | 672100 | 0 | 0 | 3615600 | 16758000 |
| ANVDISAP | AHNAK | Q09666 | AHNK_HUMAN | Unmodified | 0 | 0 | 0 | 0 | 7811400 | 0 | 0 | 0 | 0 | 0 | 0 | 0 | 0 | 0 | 0 | 0 | 1377000 | 0 |
| ANVIHATV | PCYOX1L; PCY | Q8NBM8; E7 | PCYXL_HUMAN | Unmodified | 0 | 0 | 0 | 0 | 273810 | 0 | 0 | 0 | 3472600 | 1042500 | 0 | 0 | 0 | 0 | 0 | 0 | 0 | 418620 |
| ANVINKQE | BIRC2; BIRC2; | Q13490; E9 | BIRC2_HUMAN | Unmodified | 0 | 0 | 0 | 0 | 362290 | 163360 | 0 | 0 | 160480 | 0 | 0 | 72442 | 0 | 0 | 0 | 0 | 266410 | 0 |
| ANVPNKV | CLTC; CLTC | Q00610; A0 | CLH1_HUMAN | Unmodified | 0 | 0 | 0 | 0 | 0 | 0 | 0 | 0 | 0 | 0 | 0 | 0 | 0 | 0 | 0 | 0 | 0 | 0 |
| ANVTPQSS | PLCB4; PLCB4 | Q15147; A0 | PLCB4_HUMAN | Unmodified | 0 | 0 | 740050 | 1550700 | 0 | 0 | 0 | 0 | 299890 | 0 | 0 | 0 | 1176900 | 1178900 | 544930 | 905430 | 2291700 | 5061900 |
| ANVTPQSS | PLCB4; PLCB4 | Q15147; A0 | PLCB4_HUMAN | Unmodified | 0 | 0 | 737440 | 981900 | 0 | 0 | 0 | 0 | 498340 | 0 | 0 | 0 | 684810 | 1238800 | 0 | 876590 | 1012700 | 2856500 |
| ANVTVLDT | MYOF; MYOF | Q9NZM1; H | MYOF_HUMAN | Unmodified | 4437800 | 5482600 | 1362800 | 3993300 | 2934400 | 0 | 0 | 0 | 0 | 0 | 0 | 0 | 0 | 0 | 1717400 | 7986200 | 0 | 452580 |
| ANVTVLDT | MYOF; MYOF | Q9NZM1; H | MYOF_HUMAN | Unmodified | 0 | 0 | 0 | 0 | 17547000 | 5410200 | 0 | 0 | 414090 | 80643 | 0 | 0 | 51074 | 0 | 0 | 0 | 334200 | 1580900 |
| ANVTVLDT | MYOF; MYOF | Q9NZM1; H | MYOF_HUMAN | Unmodified | 0 | 0 | 0 | 0 | 4646200 | 0 | 0 | 0 | 0 | 0 | 0 | 0 | 0 | 0 | 0 | 0 | 0 | 346670 |
| ANVVGNV | MOGS; MOGS; | Q13724; A0 | MOGS_HUMAN | Unmodified | 0 | 0 | 0 | 0 | 246730 | 0 | 0 | 0 | 8872800 | 0 | 0 | 0 | 0 | 0 | 0 | 0 | 188210 | 0 |
| ANWEAQQ | SEPTIN7; SEPT | Q16181; E7 | SEPT7_HUMAN | Unmodified | 66935 | 82720 | 0 | 0 | 2292200 | 506110 | 0 | 0 | 526670 | 223150 | 251150 | 2240800 | 510280 | 190720 | 0 | 0 | 0 | 615780 |
| ANWEAQQ | SEPTIN7; SEPT | Q16181; E7 | SEPT7_HUMAN | Unmodified | 0 | 0 | 0 | 0 | 7670200 | 0 | 282520 | 0 | 0 | 0 | 0 | 0 | 1955300 | 1033200 | 0 | 0 | 1455200 | 2055400 |
| ANWEAQQ | SEPTIN7; SEPT | Q16181; E7 | SEPT7_HUMAN | Unmodified | 0 | 0 | 0 | 0 | 23058000 | 2943100 | 1855400 | 735890 | 0 | 0 | 0 | 0 | 2490500 | 2879100 | 0 | 0 | 7993300 | 12090000 |
| ANWYFLLA | ACADM; ACAD | P11310; Q5 | ACADM_HUMA | Unmodified | 0 | 0 | 0 | 0 | 0 | 0 | 0 | 0 | 2521100 | 86670 | 23115 | 0 | 0 | 0 | 0 | 0 | 0 | 0 |
| ANYDVLES | PSMD9; PSMD | O00233; J3K | PSMD9_HUMA | Unmodified | 0 | 505050 | 0 | 156220 | 493990 | 0 | 88750 | 109380 | 362540 | 0 | 306670 | 1546300 | 0 | 0 | 199380 | 306330 | 130510 | 0 |
| ANYLIVED | DPY19L4 | Q7Z388 | D19L4_HUMAN | Oxidation (M) | 0 | 0 | 0 | 0 | 0 | 0 | 0 | 0 | 788870 | 0 | 0 | 0 | 0 | 0 | 0 | 0 | 256880 | 0 |
| ANYLIVED | DPY19L4 | Q7Z388 | D19L4_HUMAN | Unmodified | 0 | 0 | 0 | 0 | 0 | 0 | 0 | 0 | 468470 | 0 | 0 | 0 | 0 | 0 | 0 | 0 | 0 | 0 |
| ANYWWLR | ATIC; ATIC | P31939; H7 | PUR9_HUMAN | Unmodified | 0 | 0 | 277350 | 981490 | 0 | 0 | 0 | 0 | 848540 | 0 | 530230 | 5318300 | 512410 | 0 | 420870 | 525770 | 0 | 204080 |
| APAEILNG | MTHFD1; MTHF | P11586; V9 | C1TC_HUMAN | Unmodified | 48265 | 48301 | 419660 | 410220 | 2325300 | 383530 | 696470 | 613640 | 845340 | 1127100 | 623950 | 4858300 | 1324400 | 525840 | 329740 | 348910 | 1412500 | 1560300 |
| APAEILNG | MTHFD1; MTHF | P11586; V9 | C1TC_HUMAN | Unmodified | 0 | 0 | 0 | 0 | 169060 | 0 | 0 | 0 | 0 | 0 | 0 | 0 | 0 | 0 | 0 | 0 | 345830 | 159990 |
| APAEILNG | MTHFD1; MTHF | P11586; V9 | C1TC_HUMAN | Unmodified | 0 | 0 | 0 | 0 | 233950 | 0 | 729850 | 564170 | 0 | 0 | 0 | 0 | 0 | 0 | 0 | 0 | 3955200 | 2431700 |
| APAGQLPR | MCM3; MCM3; | P25205; A0 | MCM3_HUMAN | Unmodified | 447690 | 381180 | 0 | 167800 | 305250 | 0 | 52965 | 0 | 280020 | 0 | 159770 | 1107500 | 63307 | 0 | 193290 | 44101 | 0 | 65160 |
| APAGQLPR | MCM3; MCM3; | P25205; A0 | MCM3_HUMAN | Unmodified | 0 | 0 | 0 | 0 | 2072000 | 0 | 0 | 0 | 0 | 0 | 0 | 0 | 0 | 0 | 0 | 0 | 102050 | 0 |
| APAHHPAA | RRP12 | Q5JTH9 | RRP12_HUMAN | Unmodified | 0 | 157270 | 0 | 0 | 0 | 0 | 0 | 0 | 0 | 0 | 28500 | 166310 | 35136 | 89042 | 23067 | 16092 | 0 | 0 |
| APAHLSLP | GRN; GRN; GR | P28799; K7 | GRN_HUMAN | Unmodified | 0 | 0 | 0 | 0 | 304480 | 0 | 0 | 0 | 6710900 | 0 | 173500 | 0 | 0 | 0 | 0 | 0 | 68386 | 446070 |
| APAMFNIR | RPS3A; RPS3A; | P61247; H0 | RS3A_HUMAN | Unmodified | 1129000 | 1005500 | 535230 | 710900 | 0 | 0 | 0 | 0 | 750140 | 0 | 583090 | 0 | 617480 | 0 | 659700 | 0 | 0 | 330280 |
| APAMFNIR | RPS3A; RPS3A; | P61247; H0 | RS3A_HUMAN | Oxidation (M) | 122360 | 1063000 | 864460 | 848510 | 0 | 0 | 0 | 0 | 0 | 0 | 0 | 95699 | 0 | 0 | 657310 | 781370 | 0 | 24684 |
| APAMFNIR | RPS3A; RPS3A; | P61247; H0 | RS3A_HUMAN | Unmodified | 0 | 0 | 0 | 0 | 1453600 | 0 | 678420 | 137800 | 0 | 126680 | 0 | 0 | 483820 | 799750 | 0 | 0 | 112750 | 904120 |
| APAPKPEL | CD2AP | Q9Y5K6 | CD2AP_HUMA | Unmodified | 15619000 | 10182000 | 12715000 | 10238000 | 4001500 | 894150 | 299560 | 371740 | 1306700 | 289880 | 717300 | 3705700 | 8859000 | 2708200 | 22612000 | 13084000 | 0 | 0 |
| APAPKPEL | CD2AP | Q9Y5K6 | CD2AP_HUMA | Unmodified | 2930600 | 1873700 | 1871500 | 1988700 | 536930 | 50008 | 0 | 0 | 0 | 0 | 0 | 321700 | 1586800 | 460480 | 3046800 | 1806600 | 0 | 0 |
| APASVLPA | PRKAR2A; PRK | P13861; C9 | KAP2_HUMAN | Unmodified | 0 | 0 | 574900 | 624180 | 0 | 0 | 0 | 0 | 859550 | 96083 | 385660 | 2719500 | 778780 | 291070 | 1608900 | 1033000 | 82680 | 232810 |
| APDELHYT | RPN1; RPN1 | P04843; B7 | RPN1_HUMAN | Unmodified | 0 | 0 | 0 | 0 | 0 | 0 | 0 | 73472 | 3085800 | 6056800 | 0 | 590970 | 0 | 0 | 0 | 0 | 625270 | 0 |
| APDELHYT | RPN1; RPN1 | P04843; B7 | RPN1_HUMAN | Unmodified | 0 | 0 | 0 | 0 | 0 | 0 | 0 | 0 | 8199500 | 3045500 | 0 | 728110 | 0 | 0 | 0 | 0 | 0 | 0 |
| APDELWN | EXTL2; EXTL2; | Q9UBQ6; C | EXTL2_HUMAN | Unmodified | 0 | 0 | 0 | 0 | 0 | 0 | 0 | 0 | 5380800 | 0 | 762860 | 749630 | 0 | 0 | 0 | 0 | 0 | 0 |
| APDFLPLL | RP2 | O75695 | XRP2_HUMAN | Unmodified | 2989400 | 1991500 | 3672500 | 4155300 | 3780400 | 1750800 | 478470 | 532070 | 9471900 | 2774300 | 6933300 | 47914000 | 5694100 | 3593400 | 5870400 | 5807800 | 4320200 | 11322000 |
| APDFLSPS | SIGIRR; SIGIRR | Q6IA17; C9 | SIGIR_HUMAN | Unmodified | 0 | 0 | 0 | 0 | 93161 | 0 | 0 | 0 | 611630 | 308910 | 0 | 0 | 0 | 0 | 0 | 0 | 0 | 0 |
| APDFVFYA | RDX; EZR; MSN | P35241; P15 | RADI_HUMAN | Unmodified | 2027500 | 0 | 2119800 | 2754100 | 1793600 | 801130 | 820140 | 120210 | 6167200 | 383270 | 815130 | 25218000 | 413750 | 0 | 3177200 | 3730000 | 513170 | 0 |
| APDFVFYA | RDX; EZR; MSN | P35241; P15 | RADI_HUMAN | Unmodified | 0 | 0 | 0 | 0 | 980040 | 427880 | 0 | 33921 | 0 | 159450 | 0 | 0 | 0 | 239360 | 0 | 0 | 371450 | 0 |
| APDGLPAL | MANEAL | Q5VSG8 | MANEL_HUMA | Unmodified | 0 | 0 | 0 | 0 | 0 | 0 | 0 | 0 | 4256400 | 0 | 0 | 0 | 0 | 0 | 0 | 0 | 0 | 0 |
| APDPGFQE | DPP7; DPP7; D | Q9UHL4; R4 | DPP2_HUMAN | Unmodified | 0 | 0 | 0 | 0 | 0 | 60291 | 0 | 0 | 3288500 | 125730 | 352660 | 0 | 0 | 0 | 0 | 122530 | 0 | 0 |
| APDVTTLP | NCSTN; NCSTN | Q92542; H0 | NICA_HUMAN | Unmodified | 246680 | 337990 | 1746000 | 507050 | 891630 | 524390 | 171720 | 88213 | 27608000 | 2313500 | 1137500 | 8714000 | 240290 | 0 | 2885500 | 2200300 | 151180 | 196920 |
| APEATVTLP | ARRDC1; ARRD | Q8N5I2; Q5 | ARRD1_HUMA | Unmodified | 0 | 0 | 0 | 0 | 0 | 0 | 0 | 0 | 0 | 0 | 0 | 0 | 0 | 0 | 900910 | 360180 | 0 | 0 |
| APEEDAGG | PACSIN3; PACS | Q9UKS6; E9 | PACN3_HUMA | Unmodified | 0 | 2509500 | 0 | 0 | 128950 | 0 | 0 | 0 | 0 | 0 | 0 | 0 | 0 | 0 | 2022200 | 0 | 0 | 0 |
| APEEEVTF | UBR1; UBR1; U | Q8IWV7; A0 | UBR1_HUMAN | Unmodified | 0 | 0 | 0 | 0 | 0 | 0 | 0 | 24641 | 0 | 0 | 0 | 0 | 0 | 0 | 0 | 0 | 0 | 0 |
| APEFSMQG | EPCAM; EPCAM | P16422; B5 | EPCAM_HUMA | Oxidation (M) | 0 | 0 | 0 | 0 | 0 | 0 | 0 | 0 | 1321700 | 0 | 0 | 861110 | 0 | 0 | 577960 | 1118000 | 0 | 0 |
| APEFSMQG | EPCAM; EPCAM | P16422; B5 | EPCAM_HUMA | Unmodified | 68476 | 0 | 0 | 0 | 0 | 0 | 0 | 0 | 2707500 | 122950 | 145730 | 0 | 0 | 0 | 773790 | 82526 | 0 | 0 |
| APELALVR | PLCD3 | Q8N3E9 | PLCD3_HUMA | Unmodified | 259700 | 471780 | 73332 | 0 | 244500 | 0 | 0 | 0 | 0 | 81186 | 0 | 784000 | 0 | 0 | 121730 | 0 | 0 | 0 |
| APEPGPAG | SNX18 | Q96RF0 | SNX18_HUMA | Unmodified | 0 | 1404100 | 7366400 | 1120200 | 0 | 0 | 0 | 0 | 0 | 0 | 0 | 154600 | 0 | 0 | 6555100 | 5628000 | 0 | 0 |
| APEQADLT | RHBDF1 | Q96CC6 | RHDF1_HUMA | Unmodified | 311500 | 0 | 578860 | 553410 | 0 | 0 | 0 | 0 | 2271600 | 262490 | 0 | 0 | 221230 | 0 | 811300 | 675620 | 0 | 0 |
| APEQEQAA | BASP1 | P80723 | BASP1_HUMA | Unmodified | 0 | 0 | 0 | 0 | 751840 | 0 | 144790 | 119650 | 6647200 | 791610 | 0 | 0 | 0 | 0 | 43012 | 0 | 0 | 121700 |
| APESDTGD | EPB41L1; EPB4 | Q9H4G0; A0 | E41L1_HUMAN | Unmodified | 0 | 0 | 0 | 0 | 0 | 0 | 0 | 0 | 0 | 0 | 0 | 2160400 | 0 | 0 | 0 | 488320 | 0 | 0 |
| APEYLHR | G3BP2; G3BP2; | Q9UN86; A0 | G3BP2_HUMA | Unmodified | 0 | 0 | 0 | 0 | 0 | 0 | 0 | 0 | 0 | 0 | 0 | 313350 | 0 | 0 | 0 | 0 | 0 | 0 |
| APFDLFEN | HSP90AB1; HS | P08238; Q5 | HS90B_HUMA | Unmodified | 0 | 0 | 0 | 0 | 0 | 0 | 0 | 0 | 177320 | 0 | 3682800 | 2469300 | 0 | 0 | 0 | 174720 | 0 | 0 |
| APFDLFEN | HSP90AA1 | P07900 | HS90A_HUMA | Unmodified | 0 | 0 | 0 | 0 | 0 | 0 | 0 | 0 | 0 | 0 | 0 | 822840 | 0 | 0 | 0 | 0 | 0 | 0 |
| APFEEIAAH | EFR3B; EFR3B; | Q9Y2G0; E7 | EFR3B_HUMA | Unmodified | 0 | 0 | 0 | 49022 | 348860 | 197230 | 40230 | 17694 | 237840 | 149030 | 0 | 0 | 137000 | 78738 | 0 | 55319 | 141580 | 457440 |
| APFEEIAAQ | EFR3A; EFR3A | Q14156; A0 | EFR3A_HUMA | Unmodified | 0 | 0 | 142000 | 0 | 823060 | 0 | 104030 | 47756 | 180580 | 0 | 0 | 0 | 233610 | 212950 | 0 | 144600 | 196250 | 648110 |
| APGAEEYA | CNP | P09543 | CN37_HUMAN | Unmodified | 121270 | 0 | 0 | 0 | 1059300 | 0 | 0 | 1645700 | 1057600 | 0 | 5514400 | 6935600 | 0 | 0 | 0 | 0 | 35604 | 479660 |
| APGAEEYA | CNP | P09543 | CN37_HUMAN | Unmodified | 793540 | 671710 | 1466300 | 4730700 | 1569400 | 383700 | 353350 | 589020 | 3618600 | 480390 | 28898000 | 64454000 | 1657500 | 880730 | 4360400 | 2312400 | 369020 | 1435600 |
| APGAIGPY | RIDA | P52758 | RIDA_HUMAN | Unmodified | 0 | 0 | 0 | 0 | 0 | 0 | 282010 | 34335 | 1553500 | 604020 | 0 | 213190 | 0 | 0 | 0 | 0 | 0 | 0 |
| APGDEEAQ | FXYD6 | Q9H0Q3 | FXYD6_HUMA | Unmodified | 0 | 0 | 0 | 0 | 0 | 0 | 111480 | 200770 | 1876900 | 234120 | 809660 | 24158000 | 0 | 0 | 0 | 0 | 0 | 0 |
| APGDEEAQ | FXYD6 | Q9H0Q3 | FXYD6_HUMA | Unmodified | 0 | 0 | 0 | 0 | 0 | 0 | 0 | 212690 | 3386900 | 411200 | 2553800 | 35516000 | 0 | 0 | 0 | 0 | 0 | 0 |
| APGEQTVP | SUPT16H | Q9Y5B9 | SP16H_HUMA | Unmodified | 0 | 0 | 0 | 0 | 0 | 0 | 0 | 0 | 0 | 0 | 0 | 1184100 | 0 | 0 | 0 | 0 | 0 | 0 |
| APGFAQML | CCT8 | P50990 | TCPQ_HUMAN | Unmodified | 0 | 0 | 0 | 0 | 1753500 | 0 | 0 | 0 | 3346000 | 1200700 | 2460500 | 7344800 | 7399600 | 0 | 0 | 0 | 246130 | 1058000 |
| APGFAQML | CCT8 | P50990 | TCPQ_HUMAN | Oxidation (M) | 0 | 0 | 0 | 0 | 1432800 | 0 | 478630 | 297900 | 0 | 344600 | 0 | 0 | 0 | 0 | 0 | 0 | 113180 | 1296300 |
| APGFAQML | CCT8 | P50990 | TCPQ_HUMAN | Unmodified | 0 | 0 | 0 | 0 | 0 | 0 | 264660 | 139080 | 0 | 0 | 0 | 0 | 0 | 0 | 0 | 0 | 492860 | 0 |
| APGFARLP | GATD1; GATD1 | Q8NB37; H0 | GALD1_HUMA | Unmodified | 0 | 0 | 28143 | 133860 | 342670 | 110180 | 0 | 0 | 621500 | 210090 | 106490 | 0 | 123390 | 49675 | 0 | 62201 | 0 | 156810 |
| APGIILLDE | ABCB6 | Q9NP58 | ABCB6_HUMA | Unmodified | 510440 | 0 | 401770 | 783910 | 0 | 0 | 0 | 0 | 1402800 | 237330 | 0 | 931510 | 0 | 0 | 1456000 | 458800 | 0 | 0 |
| APGIILLDE | ABCB6 | Q9NP58 | ABCB6_HUMA | Unmodified | 0 | 0 | 0 | 0 | 799070 | 0 | 760360 | 0 | 0 | 0 | 0 | 0 | 814680 | 1225900 | 0 | 0 | 118570 | 0 |
| APGLLDSL | PROM2; PROM | Q8N271; F8 | PROM2_HUMA | Unmodified | 0 | 0 | 283410 | 0 | 0 | 0 | 0 | 0 | 0 | 0 | 0 | 0 | 0 | 0 | 2476000 | 206780 | 0 | 0 |
| APGPGLAQ | LRRC47 | Q8N1G4 | LRC47_HUMA | Unmodified | 77464 | 44227 | 65057 | 0 | 0 | 0 | 0 | 0 | 0 | 0 | 0 | 3136100 | 0 | 0 | 240920 | 199700 | 0 | 0 |
| APGQLALF | ATIC; ATIC | P31939; F8 | PUR9_HUMAN | Unmodified | 0 | 0 | 695280 | 856500 | 4064400 | 0 | 0 | 0 | 0 | 0 | 1667700 | 7969100 | 2895700 | 0 | 1564500 | 0 | 1160900 | 0 |
| APGQLALF | ATIC; ATIC | P31939; F8 | PUR9_HUMAN | Unmodified | 27172 | 0 | 0 | 270960 | 3149800 | 0 | 453000 | 28762 | 416300 | 1132800 | 0 | 0 | 8817800 | 3587000 | 0 | 0 | 2960300 | 1044300 |
| APGTNVA | RANBP2 | P49792 | RBP2_HUMAN | Unmodified | 149220 | 120010 | 88086 | 261780 | 0 | 0 | 0 | 0 | 0 | 0 | 0 | 569280 | 181470 | 0 | 78966 | 100720 | 0 | 0 |
| APGTPHSH | RPL18; RPL18; | Q07020; F8 | RL18_HUMAN | Unmodified | 2247000 | 1262500 | 444240 | 114360 | 0 | 0 | 0 | 0 | 0 | 0 | 214100 | 2310700 | 0 | 0 | 879770 | 614500 | 0 | 0 |
| APGVTDVL | IFIT2; IFIT2; IFIT | P09913; A0 | IFIT2_HUMAN | Unmodified | 0 | 0 | 0 | 0 | 1296900 | 0 | 0 | 0 | 0 | 0 | 0 | 0 | 0 | 0 | 0 | 0 | 0 | 0 |
| APHKPHEV | SLC4A2; SLC4A | P04920; A0 | B3A2_HUMAN | Unmodified | 0 | 0 | 1326900 | 1800000 | 264580 | 0 | 0 | 0 | 0 | 0 | 0 | 876220 | 0 | 0 | 3649100 | 1418800 | 0 | 0 |
| APHLAYLR | ACAT2 | Q9BWD1 | THIC_HUMAN | Unmodified | 0 | 0 | 0 | 0 | 2413500 | 0 | 220000 | 56403 | 0 | 586410 | 0 | 0 | 918340 | 445100 | 0 | 0 | 294710 | 677510 |
| APHLQLIE | EPB41L2; EPB4 | O43491; H0 | E41L2_HUMAN | Unmodified | 0 | 0 | 0 | 0 | 1986500 | 187690 | 0 | 0 | 0 | 0 | 0 | 0 | 0 | 0 | 0 | 0 | 362880 | 1264600 |
| APIIAVTR | PKM; PKM; PKM | P14618; B4 | KPYM_HUMAN | Unmodified | 3423900 | 2634000 | 10630000 | 4333000 | 361790 | 218850 | 113620 | 0 | 6819500 | 324280 | 21574000 | 2.29E+08 | 0 | 223170 | 14479000 | 9334200 | 205270 | 886310 |
| APIILVGTQ | RHOU | Q7L0Q8 | RHOU_HUMAN | Unmodified | 987990 | 1084300 | 730760 | 697120 | 770440 | 0 | 249230 | 80725 | 738040 | 131520 | 0 | 641050 | 890860 | 621620 | 693170 | 419350 | 0 | 0 |
| APILIATDV | DDX5; DDX17; | P17844; Q9 | DDX5_HUMAN | Unmodified | 345960 | 311830 | 309670 | 0 | 0 | 0 | 0 | 63959 | 1655900 | 0 | 899290 | 18380000 | 0 | 0 | 145020 | 0 | 0 | 0 |
| APIQWEER | IDH3A; IDH3A; | P50213; H0 | IDH3A_HUMAN | Unmodified | 0 | 0 | 0 | 0 | 0 | 0 | 0 | 0 | 533110 | 0 | 0 | 0 | 0 | 0 | 0 | 0 | 0 | 0 |
| APIQWEER | IDH3A; IDH3A; | P50213; H0 | IDH3A_HUMAN | Unmodified | 0 | 0 | 0 | 0 | 0 | 0 | 0 | 0 | 855280 | 145290 | 0 | 0 | 0 | 0 | 0 | 0 | 0 | 0 |
| APIRPDIVN | RPL4; RPL4; RP | P36578; H3 | RL4_HUMAN 6 | Unmodified | 667910 | 376370 | 0 | 259480 | 0 | 0 | 0 | 0 | 0 | 0 | 0 | 1777500 | 0 | 0 | 340920 | 284890 | 0 | 0 |
| APKIPMPD | AHNAK | Q09666 | AHNK_HUMAN | Unmodified | 0 | 0 | 0 | 0 | 1199500 | 0 | 0 | 0 | 0 | 0 | 0 | 0 | 0 | 0 | 0 | 0 | 243450 | 611180 |
| APKIPMPD | AHNAK | Q09666 | AHNK_HUMAN | Oxidation (M) | 0 | 0 | 0 | 0 | 1765800 | 0 | 0 | 0 | 0 | 0 | 0 | 0 | 0 | 0 | 0 | 0 | 552790 | 1591600 |
| APKISMPD | AHNAK | Q09666 | AHNK_HUMAN | Unmodified | 0 | 0 | 0 | 0 | 351680 | 0 | 0 | 0 | 0 | 0 | 0 | 0 | 0 | 0 | 0 | 0 | 706170 | 0 |
| APKMDVN | AHNAK | Q09666 | AHNK_HUMAN | Unmodified | 0 | 0 | 0 | 0 | 0 | 0 | 0 | 0 | 0 | 0 | 0 | 0 | 0 | 0 | 0 | 0 | 512640 | 0 |

| | | | | | | | | | | | | | | | | | | | | | | |
|---|---|---|---|---|---|---|---|---|---|---|---|---|---|---|---|---|---|---|---|---|---|---|
| APKPDGPG | FUS; FUS | P35637; H3E | FUS_HUMAN R | Unmodified | 109280 | 100590 | 117470 | 65446 | 35710 | 0 | 0 | 35997 | 853720 | 148230 | 2664200 | 5603300 | 0 | 0 | 114700 | 0 | 0 | 0 |
| APKPDGPG | FUS; FUS | P35637; H3E | FUS_HUMAN R | Oxidation (M) | 0 | 0 | 0 | 0 | 0 | 0 | 0 | 0 | 0 | 0 | 2953500 | 2055100 | 0 | 0 | 0 | 57913 | 0 | 0 |
| APLDHINLI | MGAM2; MGAM | Q2M2H8; O4 | MGAL_HUMAN | Unmodified | 0 | 0 | 0 | 0 | 0 | 0 | 0 | 0 | 0 | 0 | 474740 | 1194000 | 0 | 0 | 0 | 0 | 0 | 0 |
| APLDIPVPD | PSME1 | Q06323 | PSME1_HUMAI | Unmodified | 0 | 0 | 0 | 979290 | 0 | 0 | 0 | 0 | 5849200 | 1313800 | 6706100 | 10344000 | 63223 | 0 | 0 | 2418600 | 0 | 3140700 |
| APLDIPVPD | PSME1 | Q06323 | PSME1_HUMAI | Unmodified | 0 | 0 | 0 | 0 | 595090 | 0 | 234410 | 229900 | 0 | 724730 | 0 | 0 | 890090 | 0 | 0 | 0 | 380790 | 820420 |
| APLGGIHIS | DIP2A; DIP2A | Q14689; A0A | DIP2A_HUMAN | Unmodified | 0 | 0 | 0 | 0 | 0 | 0 | 0 | 0 | 1107000 | 363840 | 0 | 1085700 | 0 | 0 | 0 | 0 | 0 | 0 |
| APLKPYPVS | IARS1; IARS1; IA | P41252; A0A | SYIC_HUMAN I | Unmodified | 0 | 0 | 0 | 0 | 798910 | 0 | 0 | 0 | 1720000 | 442120 | 122870 | 11792000 | 0 | 0 | 0 | 0 | 0 | 1037800 |
| APLNVQFN | FLNB; FLNB | O75369; E7E | FLNB_HUMAN | Unmodified | 0 | 0 | 622600 | 0 | 715820 | 0 | 0 | 0 | 0 | 0 | 0 | 1994900 | 0 | 0 | 787710 | 437490 | 0 | 0 |
| APLRVQVQ | FLNA; FLNA; FL | P21333; A0A | FLNA_HUMAN | Unmodified | 0 | 0 | 380810 | 0 | 5905500 | 2152700 | 0 | 0 | 1023100 | 117130 | 191000 | 3694100 | 1829500 | 742310 | 0 | 536110 | 2168600 | 2682100 |
| APMAAQGI | GNA13 | Q14344 | GNA13_HUMAI | Unmodified | 0 | 0 | 0 | 0 | 0 | 0 | 0 | 0 | 823620 | 0 | 145800 | 4050300 | 0 | 0 | 0 | 0 | 0 | 0 |
| APMAAQGI | GNA13 | Q14344 | GNA13_HUMAI | 2 Oxidation (M) | 0 | 0 | 0 | 0 | 0 | 0 | 0 | 0 | 36665 | 0 | 0 | 422900 | 0 | 0 | 0 | 0 | 0 | 0 |
| APMAAQGI | GNA13 | Q14344 | GNA13_HUMAI | Oxidation (M) | 0 | 0 | 0 | 0 | 0 | 0 | 0 | 0 | 0 | 0 | 0 | 1544100 | 0 | 0 | 0 | 0 | 0 | 0 |
| APNDPQQ/ | TMEM260 | Q9NX78 | TM260_HUMAI | Unmodified | 0 | 0 | 0 | 0 | 0 | 0 | 0 | 0 | 2423500 | 0 | 0 | 0 | 0 | 0 | 0 | 0 | 0 | 0 |
| APNISMPD | AHNAK | Q09666 | AHNK_HUMAN | Unmodified | 0 | 0 | 0 | 0 | 0 | 0 | 0 | 0 | 0 | 0 | 0 | 0 | 0 | 0 | 0 | 0 | 0 | 0 |
| APNIYLVR | SYPL1 | Q16563 | SYPL1_HUMAN | Acetyl (Protein N | 0 | 0 | 0 | 0 | 0 | 0 | 0 | 0 | 0 | 0 | 0 | 0 | 0 | 0 | 0 | 447730 | 0 | 0 |
| APNTPDILE | ASS1 | P00966 | ASSY_HUMAN | Unmodified | 0 | 0 | 0 | 129660 | 0 | 17883 | 191160 | 131490 | 1573100 | 0 | 150610 | 0 | 531940 | 0 | 0 | 0 | 0 | 0 |
| APNYPTEG | NIF3L1; NIF3L1 | Q9GZT8; Q6 | NIF3L_HUMAN | Unmodified | 0 | 0 | 162380 | 185740 | 533700 | 0 | 50917 | 0 | 2576500 | 197160 | 0 | 2864200 | 272980 | 0 | 109070 | 178110 | 121870 | 0 |
| APPAAPTRI | FMNL1; FMNL1 | O95466; A0A | FMNL1_HUMAI | Unmodified | 0 | 0 | 0 | 0 | 2088200 | 0 | 0 | 0 | 0 | 0 | 0 | 0 | 0 | 0 | 0 | 0 | 0 | 0 |
| APPAPGPA | COPE; COPE; C | O14579; M0I | COPE_HUMAN | Unmodified | 0 | 0 | 215040 | 0 | 0 | 0 | 0 | 0 | 0 | 0 | 927210 | 3456900 | 0 | 0 | 0 | 0 | 0 | 0 |
| APPASKPTI | NCAM1 | A0A0D9SF3( | A0A0D9SF30_H | Unmodified | 0 | 0 | 0 | 0 | 0 | 0 | 0 | 0 | 733160 | 0 | 0 | 0 | 0 | 0 | 0 | 0 | 0 | 0 |
| APPILPPHL | PRKAB1; PRKAE | Q9Y478; F5H | AAKB1_HUMAN | Unmodified | 0 | 0 | 0 | 84548 | 148010 | 0 | 95730 | 0 | 256500 | 98580 | 84310 | 400180 | 97479 | 0 | 122400 | 107080 | 63456 | 298850 |
| APPPSLTD( | CHMP5 | Q9NZZ3 | CHMP5_HUMA | Unmodified | 1645000 | 584670 | 680950 | 1403600 | 0 | 0 | 0 | 0 | 2505400 | 247540 | 0 | 4933300 | 0 | 0 | 578150 | 282630 | 0 | 0 |
| APPPSLTD( | CHMP5 | Q9NZZ3 | CHMP5_HUMA | Unmodified | 0 | 0 | 301290 | 616250 | 9245900 | 2801100 | 259550 | 187680 | 0 | 4404300 | 3371600 | 2284600 | 6147900 | 3867200 | 0 | 0 | 326080 | 2405900 |
| APPPSLTD( | CHMP5 | Q9NZZ3 | CHMP5_HUMA | Unmodified | 708450 | 0 | 0 | 0 | 8061500 | 0 | 278750 | 0 | 0 | 2149600 | 0 | 3396700 | 2383200 | 2701300 | 0 | 0 | 294270 | 1514500 |
| APPSVFAE\ | GOT1 | P17174 | AATC_HUMAN | Unmodified | 1366000 | 1008200 | 71560 | 0 | 7537900 | 1906100 | 925740 | 296140 | 210090 | 0 | 16565000 | 46707000 | 6713900 | 3969500 | 0 | 170130 | 1979400 | 1881600 |
| APPVDDAE | BAZ1B | Q9UIG0 | BAZ1B_HUMAN | Unmodified | 0 | 0 | 0 | 0 | 0 | 0 | 0 | 183460 | 0 | 0 | 0 | 0 | 0 | 0 | 0 | 0 | 0 | 0 |
| APPVVGGF | GLCE; GLCE | O94923; H0Y | GLCE_HUMAN | Unmodified | 0 | 0 | 0 | 0 | 0 | 0 | 0 | 0 | 962310 | 0 | 937490 | 0 | 0 | 0 | 0 | 0 | 0 | 0 |
| APQEFMTS | ADPGK | Q9BRR6 | ADPGK_HUMAI | Unmodified | 0 | 0 | 0 | 0 | 0 | 0 | 0 | 0 | 2897500 | 177760 | 186450 | 206310 | 0 | 0 | 0 | 0 | 0 | 0 |
| APQETYAD | PSMC1; PSMC1 | P62191; G3V | PRS4_HUMAN | Unmodified | 0 | 0 | 0 | 0 | 0 | 0 | 0 | 0 | 762110 | 132330 | 0 | 1409300 | 0 | 0 | 649690 | 526510 | 0 | 0 |
| APQVDVDK | GARS1; GARS1 | P41250; F8V | GARS_HUMAN | Unmodified | 0 | 0 | 828720 | 0 | 0 | 58015 | 0 | 0 | 93800 | 0 | 81181 | 1583100 | 0 | 0 | 0 | 0 | 0 | 0 |
| APQVDVDK | GARS1; GARS1 | P41250; F8V | GARS_HUMAN | Unmodified | 0 | 0 | 0 | 0 | 0 | 0 | 0 | 40466 | 1216400 | 192670 | 622160 | 970460 | 0 | 0 | 0 | 0 | 0 | 0 |
| APQVVAEA | DKC1 | O60832 | DKC1_HUMAN | Unmodified | 194300 | 261190 | 185490 | 143770 | 31483 | 0 | 0 | 0 | 472680 | 321200 | 0 | 626520 | 88779 | 28057 | 0 | 105150 | 69622 | 19372 |
| APQYHEIGI | SLC4A7; SLC4A | Q9Y6M7; C9 | S4A7_HUMAN | Unmodified | 0 | 0 | 0 | 0 | 132750 | 0 | 0 | 0 | 0 | 0 | 0 | 0 | 0 | 0 | 0 | 0 | 150220 | 146010 |
| APREPLEP( | ADAM15 | Q13444 | ADA15_HUMAN | Unmodified | 22797 | 0 | 0 | 0 | 0 | 0 | 0 | 0 | 3452400 | 0 | 80738 | 0 | 0 | 0 | 25941 | 0 | 0 | 0 |
| APREPLEP( | ADAM15 | Q13444 | ADA15_HUMAN | Unmodified | 0 | 0 | 0 | 0 | 0 | 0 | 0 | 0 | 2003300 | 0 | 0 | 0 | 0 | 0 | 0 | 0 | 0 | 0 |
| APRGDVTA | MARCKSL1 | P49006 | MRP_HUMAN N | Unmodified | 157450 | 54137 | 2784900 | 3078900 | 575580 | 230520 | 51694 | 68095 | 3906900 | 384100 | 4740000 | 9857400 | 5059900 | 2977100 | 3300700 | 3803900 | 2117200 | 15381000 |
| APRPAPGG | ENDOD1 | O94919 | ENDD1_HUMA | Unmodified | 0 | 0 | 79720 | 0 | 0 | 0 | 0 | 12645 | 444520 | 37478 | 0 | 0 | 0 | 23725 | 0 | 79301 | 0 | 0 |
| APRPFGSV | PDLIM5; PDLIM | Q96HC4; A0 | PDLI5_HUMAN | Unmodified | 0 | 161630 | 0 | 0 | 0 | 0 | 0 | 43926 | 0 | 0 | 0 | 262420 | 70018 | 98319 | 0 | 98428 | 121150 | 931500 |
| APSAARSLI | CHMP2B; CHM | Q9UQN3; C9 | CHM2B_HUMA | Unmodified | 0 | 329060 | 0 | 990320 | 2132100 | 58013 | 429700 | 486200 | 0 | 203410 | 0 | 32681 | 1011900 | 1749100 | 509630 | 0 | 1156700 | 1247800 |
| APSDLYQII | CAPN1; CAPN1 | P07384; E9P | CAN1_HUMAN | Unmodified | 44755 | 0 | 89813 | 67332 | 90765 | 0 | 0 | 0 | 0 | 145520 | 310970 | 1034300 | 0 | 0 | 0 | 183840 | 0 | 0 |
| APSGAVLD | EPHB4; EPHB4 | P54760; Q96 | EPHB4_HUMAI | Unmodified | 0 | 0 | 0 | 0 | 0 | 0 | 0 | 0 | 0 | 0 | 0 | 0 | 0 | 0 | 0 | 0 | 0 | 0 |
| APSIIFIDEL | PSMC3; PSMC3 | P17980; E9P | PRS6A_HUMAN | Unmodified | 0 | 0 | 164250 | 231650 | 1327300 | 21439 | 0 | 0 | 1486200 | 770300 | 143890 | 0 | 1027700 | 869270 | 68220 | 43471 | 2076300 | 1178900 |
| APSIIFIDEL | PSMC3; PSMC3 | P17980; E9P | PRS6A_HUMAN | Unmodified | 0 | 0 | 0 | 174920 | 835830 | 0 | 1132500 | 0 | 298440 | 187830 | 0 | 0 | 0 | 0 | 443600 | 247170 | 3593100 | 557550 |
| APSLDIKGI | AHNAK | Q09666 | AHNK_HUMAN | Unmodified | 0 | 0 | 0 | 0 | 808410 | 343850 | 0 | 0 | 2229900 | 0 | 0 | 0 | 0 | 0 | 0 | 0 | 92994 | 218990 |
| APSLEDGS | SLC43A1; SLC4 | O75387; E9F | LAT3_HUMAN I | Oxidation (M) | 0 | 0 | 5095000 | 0 | 0 | 0 | 0 | 0 | 140190 | 52130 | 265380 | 1347800 | 0 | 0 | 5033200 | 6305100 | 0 | 0 |
| APSLEDGS | SLC43A1; SLC4 | O75387; E9F | LAT3_HUMAN I | Unmodified | 0 | 0 | 1223400 | 2494900 | 5863100 | 26969000 | 10010000 | 6156200 | 271290 | 0 | 0 | 0 | 1173900 | 10946000 | 2914600 | 756770 | 2662300 | 31354000 |
| APSPEKAEI | ASNS | P08243 | ASNS_HUMAN | Unmodified | 127250 | 82319 | 0 | 0 | 0 | 0 | 0 | 0 | 0 | 0 | 419730 | 3776900 | 0 | 0 | 508980 | 774730 | 0 | 0 |
| APSPLYSVI | GAA; GAA | P10253; I3L3 | LYAG_HUMAN | Unmodified | 0 | 0 | 0 | 0 | 0 | 0 | 0 | 0 | 2865700 | 700300 | 0 | 0 | 0 | 0 | 0 | 0 | 0 | 0 |
| APSRQDVY | CTNND1; CTNN | O60716; A0A | CTND1_HUMAI | Unmodified | 0 | 0 | 0 | 0 | 172200 | 0 | 0 | 0 | 0 | 0 | 628660 | 1659400 | 0 | 0 | 2911300 | 0 | 712880 | 325120 |
| APSVALPV/ | HSPB6; HSPB6 | O14558; A0A | HSPB6_HUMAI | Unmodified | 862990 | 361770 | 0 | 0 | 5055400 | 1409800 | 683670 | 0 | 1834600 | 763780 | 149500 | 0 | 0 | 0 | 0 | 0 | 0 | 0 |
| APSVANPV | PLXNA1 | Q9UIW2 | PLXA1_HUMAN | Unmodified | 0 | 0 | 17673 | 0 | 0 | 0 | 0 | 0 | 411800 | 206100 | 0 | 198800 | 0 | 0 | 0 | 59796 | 0 | 0 |
| APSVASSW | PLEKHA2; PLEK | Q9HB19; A0A | PKHA2_HUMAI | Unmodified | 0 | 233210 | 0 | 88739 | 0 | 247140 | 0 | 0 | 0 | 0 | 0 | 0 | 0 | 0 | 82881 | 0 | 229480 | 516230 |
| APSVSAKP( | HEXB | P07686 | HEXB_HUMAN | Unmodified | 0 | 0 | 0 | 0 | 0 | 418730 | 44986 | 0 | 258590 | 255700 | 0 | 0 | 0 | 0 | 0 | 0 | 0 | 0 |
| APTAAPPPI | WASL | O00401 | WASL_HUMAN | Unmodified | 0 | 0 | 0 | 0 | 0 | 0 | 0 | 0 | 0 | 0 | 0 | 0 | 0 | 0 | 569300 | 0 | 0 | 0 |
| APTLQQAY | SLC43A1; SLC4 | O75387; E9F | LAT3_HUMAN I | Unmodified | 73345 | 991980 | 1409600 | 0 | 0 | 0 | 0 | 27514 | 104870 | 0 | 366680 | 821730 | 0 | 0 | 2193300 | 1961000 | 0 | 0 |
| APTLQQAY | SLC43A1; SLC4 | O75387; E9F | LAT3_HUMAN I | Unmodified | 0 | 23772 | 829150 | 465960 | 0 | 0 | 9407.5 | 0 | 96819 | 0 | 0 | 0 | 0 | 0 | 951780 | 520610 | 0 | 0 |
| APTNIVYKII | RPA2; RPA2 | P15927; Q5T | RFA2_HUMAN | 2 Oxidation (M) | 67239 | 0 | 0 | 23441 | 0 | 0 | 0 | 0 | 0 | 0 | 0 | 619650 | 0 | 234450 | 0 | 0 | 0 | 0 |
| APVAGTCY | PEBP1 | P30086 | PEBP1_HUMAN | Unmodified | 1243400 | 0 | 0 | 4399900 | 0 | 0 | 0 | 0 | 1998800 | 0 | 571130 | 4509500 | 0 | 0 | 2597600 | 1441500 | 0 | 0 |
| APVDFGYV | SEPTIN9; SEPTI | Q9UHD8; K7 | SEPT9_HUMAN | Unmodified | 0 | 0 | 0 | 0 | 8164100 | 393140 | 0 | 0 | 0 | 953900 | 15964 | 0 | 0 | 0 | 0 | 0 | 3358900 | 6208400 |
| APVDITGQI | CARS1; CARS1 | P49589; B4D | SYCC_HUMAN | Unmodified | 0 | 521390 | 0 | 0 | 0 | 0 | 0 | 0 | 0 | 0 | 180580 | 899800 | 0 | 0 | 0 | 0 | 0 | 0 |
| APVEHVVA | NOB1; NOB1 | Q9ULX3; H3 | NOB1_HUMAN | Unmodified | 194620 | 0 | 0 | 0 | 597270 | 40565 | 0 | 0 | 0 | 0 | 0 | 0 | 212640 | 143440 | 0 | 0 | 0 | 0 |
| APVILGSPD | FBP1; FBP1 | P09467; A0A | F16P1_HUMAN | Unmodified | 0 | 0 | 0 | 0 | 0 | 0 | 0 | 0 | 928030 | 144970 | 0 | 0 | 0 | 0 | 0 | 0 | 12193 | 0 |
| APVLFFDRI | ABCC4 | O15439 | MRP4_HUMAN | Unmodified | 0 | 0 | 5366400 | 461040 | 0 | 0 | 0 | 0 | 0 | 0 | 0 | 1058600 | 0 | 0 | 5137300 | 3540100 | 0 | 0 |
| APVLLSSLD | PTGFRN | Q9P2B2 | FPRP_HUMAN | Unmodified | 652780 | 648000 | 4175600 | 634190 | 0 | 0 | 0 | 0 | 0 | 0 | 0 | 5649300 | 0 | 0 | 4476100 | 4070600 | 0 | 0 |
| APVLLSSLD | PTGFRN | Q9P2B2 | FPRP_HUMAN | Unmodified | 166610 | 397750 | 2123600 | 706930 | 0 | 0 | 0 | 0 | 0 | 495640 | 373130 | 3724800 | 0 | 0 | 1717400 | 1551500 | 0 | 0 |
| APVNTAELT | BCCIP | Q9P287 | BCCIP_HUMAN | Unmodified | 340780 | 451280 | 206790 | 515820 | 0 | 0 | 0 | 0 | 1026700 | 0 | 4850400 | 9284300 | 107920 | 0 | 298900 | 324740 | 0 | 0 |
| APVPGTPD | STOML2; STOM | Q9UJZ1; A0A | STML2_HUMAN | Unmodified | 0 | 0 | 0 | 0 | 0 | 0 | 0 | 0 | 1869200 | 260820 | 232350 | 189020 | 0 | 0 | 0 | 0 | 0 | 0 |
| APVPGTPD | STOML2; STOM | Q9UJZ1; A0A | STML2_HUMAN | Unmodified | 0 | 0 | 0 | 0 | 1012800 | 0 | 0 | 0 | 6740100 | 0 | 300690 | 0 | 0 | 0 | 0 | 0 | 0 | 0 |
| APVPTGEV | CANX; CANX; C | P27824; A0A | CALX_HUMAN | Unmodified | 0 | 0 | 0 | 0 | 0 | 0 | 0 | 208550 | 15200000 | 1756700 | 1177000 | 567560 | 0 | 0 | 0 | 0 | 0 | 0 |
| APVPTGEV | CANX; CANX; C | P27824; A0A | CALX_HUMAN | Unmodified | 0 | 0 | 0 | 0 | 0 | 0 | 0 | 0 | 0 | 2194000 | 0 | 0 | 0 | 0 | 0 | 0 | 0 | 0 |
| APVQPQQS | PSMD2; PSMD2 | Q13200; F8V | PSMD2_HUMA | Unmodified | 395710 | 0 | 814230 | 503180 | 627910 | 0 | 0 | 0 | 737100 | 0 | 567680 | 1945900 | 311860 | 0 | 409660 | 767600 | 0 | 364560 |
| APVVAIAVL | LLGL1; LLGL1 | Q15334; A0A | L2GL1_HUMAN | Unmodified | 1174200 | 1541900 | 998750 | 1683100 | 0 | 0 | 0 | 0 | 732230 | 0 | 0 | 1042000 | 0 | 0 | 1107600 | 2066200 | 0 | 0 |
| APVVGILVL | LLGL2; LLGL2 | Q6P1M3; J3( | L2GL2_HUMAN | Unmodified | 0 | 1037100 | 0 | 0 | 0 | 0 | 0 | 0 | 684930 | 0 | 0 | 0 | 0 | 0 | 0 | 335800 | 0 | 0 |
| APVYFYEF( | CES2; CES2 | O00748; J3Q | EST2_HUMAN ( | Unmodified | 0 | 0 | 23474 | 76108 | 0 | 16043 | 0 | 0 | 1535700 | 0 | 0 | 0 | 0 | 0 | 0 | 0 | 0 | 0 |
| APWELLELI | YARS1; YARS1; | P54577; A0A | SYYC_HUMAN | Unmodified | 0 | 693100 | 271840 | 285390 | 89518 | 0 | 0 | 0 | 511010 | 0 | 223970 | 1744000 | 0 | 0 | 630370 | 577350 | 0 | 0 |
| APWIEQEG | HLA-A; HLA-A; I | Q53Z42; A0A | Q53Z42_HUMA | Unmodified | 0 | 0 | 0 | 123250 | 0 | 0 | 0 | 0 | 15188000 | 742780 | 15395000 | 8235400 | 0 | 0 | 63257 | 106090 | 0 | 0 |
| APWIEQEG | HLA-A; HLA-A; I | Q53Z42; A0A | Q53Z42_HUMA | Unmodified | 0 | 0 | 611510 | 1424500 | 0 | 0 | 0 | 0 | 0 | 0 | 6053800 | 1.03E+08 | 0 | 0 | 0 | 381070 | 0 | 302420 |
| APWIEQEG | HLA-A; HLA-A; I | P04439; A0A | HLAA_HUMAN | Unmodified | 0 | 0 | 427560 | 542520 | 0 | 0 | 0 | 0 | 25641000 | 1928300 | 2402300 | 11576000 | 0 | 0 | 660430 | 742900 | 736650 | 0 |
| APWIEQEG | HLA-A; HLA-A; I | A0A0G2JL56 | A0A0G2JL56_H | Oxidation (M) | 0 | 0 | 0 | 0 | 0 | 0 | 0 | 0 | 0 | 1734800 | 0 | 0 | 954730 | 0 | 0 | 0 | 0 | 3376800 |
| APWIEQEG | HLA-A; HLA-A; I | A0A0G2JL56 | A0A0G2JL56_H | Unmodified | 0 | 0 | 0 | 0 | 0 | 0 | 0 | 0 | 0 | 708810 | 0 | 0 | 0 | 0 | 0 | 0 | 711550 | 3666400 |
| APWIEQEG | HLA-B; HLA-B; I | P01889; D5H | HLAB_HUMAN | Unmodified | 0 | 0 | 0 | 0 | 170440 | 88196 | 0 | 0 | 1798000 | 317940 | 450580 | 1385600 | 0 | 0 | 0 | 0 | 0 | 26588 |
| APWIEQEG | HLA-B; HLA-B; I | D5H3J5; A0A | D5H3J5_HUMA | Unmodified | 0 | 0 | 298500 | 0 | 48740000 | 13087000 | 283830 | 0 | 13346000 | 710800 | 1296000 | 1.36E+08 | 0 | 0 | 79428 | 0 | 0 | 1867700 |
| APWIEQEG | HLA-A; HLA-A; I | A0A1W2PR6 | A0A1W2PR61_ | Unmodified | 0 | 0 | 0 | 0 | 38073000 | 10269000 | 0 | 0 | 31711000 | 2098600 | 0 | 0 | 0 | 0 | 0 | 0 | 0 | 0 |
| APWIEQEG | HLA-A; HLA-A; I | A0A1W2PR6 | A0A1W2PR61_ | Unmodified | 0 | 0 | 0 | 0 | 29076000 | 4326300 | 0 | 0 | 0 | 1842400 | 0 | 0 | 0 | 0 | 0 | 0 | 0 | 0 |
| APWVEQE( | HLA-C; HLA-C; | P10321; Q5( | HLAC_HUMAN | Unmodified | 0 | 0 | 0 | 0 | 0 | 0 | 0 | 0 | 521730 | 173050 | 0 | 191960 | 0 | 0 | 0 | 0 | 0 | 0 |
| APWVEQE( | HLA-C; HLA-C; | P10321; Q5( | HLAC_HUMAN | Unmodified | 0 | 324350 | 0 | 143610 | 2942700 | 451930 | 0 | 0 | 55132000 | 3530500 | 4142000 | 23396000 | 0 | 0 | 257520 | 132950 | 222580 | 1502500 |
| AQAAAPAS | RPL29; RPL29 | P47914; A0A | RL29_HUMAN ( | Unmodified | 10526000 | 6173500 | 3089600 | 3642500 | 3529300 | 0 | 327230 | 1073600 | 1913600 | 287080 | 481680 | 24984000 | 5692800 | 0 | 5752800 | 4978600 | 224450 | 1053000 |
| AQAAAPAS | RPL29; RPL29 | P47914; A0A | RL29_HUMAN ( | Unmodified | 0 | 198770 | 0 | 0 | 784100 | 0 | 0 | 319400 | 0 | 59816 | 0 | 743060 | 1090500 | 0 | 0 | 0 | 0 | 0 |
| AQAAAPAS | RPL29; RPL29 | P47914; A0A | RL29_HUMAN ( | Unmodified | 0 | 0 | 0 | 0 | 1869800 | 0 | 1149500 | 1449000 | 0 | 367380 | 0 | 0 | 6027100 | 2431300 | 0 | 0 | 426360 | 1561600 |
| AQAALAVN | CCT6A | P40227 | TCPZ_HUMAN | Unmodified | 0 | 0 | 0 | 0 | 0 | 0 | 0 | 0 | 1573300 | 186570 | 0 | 5158500 | 618170 | 0 | 22612 | 0 | 0 | 0 |
| AQADLAYQ | FLOT1; FLOT1; I | O75955; A0A | FLOT1_HUMAN | Unmodified | 0 | 0 | 138150 | 0 | 1517200 | 0 | 0 | 0 | 1864400 | 854480 | 0 | 1457700 | 342490 | 0 | 1057000 | 259850 | 61228 | 73518 |
| AQAELNALI | TSG101; TSG10 | Q99816; F5H | TS101_HUMAN | Unmodified | 4202900 | 3389200 | 795880 | 2330200 | 506030 | 0 | 0 | 0 | 993090 | 43836 | 17388 | 1192400 | 0 | 0 | 7103100 | 5491800 | 0 | 0 |
| AQAELVGT. | ATP5F1D | P30049 | ATPD_HUMAN | Unmodified | 0 | 0 | 0 | 0 | 0 | 0 | 0 | 0 | 7553500 | 0 | 0 | 0 | 0 | 0 | 0 | 0 | 0 | 0 |
| AQAEQQHI | CNOT1; CNOT1 | A5YKK6; H3E | CNOT1_HUMA | Unmodified | 0 | 0 | 0 | 0 | 0 | 0 | 0 | 0 | 0 | 0 | 0 | 239040 | 0 | 0 | 0 | 20905 | 0 | 0 |
| AQAEVDRL | ERC2; ERC1; EF | O15083; Q8I | ERC2_HUMAN | Unmodified | 48060 | 0 | 0 | 116960 | 515930 | 93123 | 0 | 0 | 795510 | 102820 | 0 | 686310 | 185570 | 69321 | 207260 | 0 | 0 | 82630 |
| AQAEVEGL | PLEC; PLEC | Q15149; H0Y | PLEC_HUMAN | Unmodified | 0 | 0 | 0 | 0 | 0 | 0 | 0 | 60096 | 21410 | 0 | 0 | 3050500 | 0 | 570410 | 0 | 0 | 1031700 | 2236400 |
| AQAEVEGL | PLEC; PLEC | Q15149; H0Y | PLEC_HUMAN | Unmodified | 0 | 0 | 0 | 0 | 101320 | 0 | 0 | 0 | 0 | 0 | 0 | 0 | 0 | 0 | 0 | 0 | 1209500 | 538960 |
| AQAEVEGL | PLEC; PLEC | Q15149; H0Y | PLEC_HUMAN | Unmodified | 0 | 0 | 0 | 0 | 250620 | 0 | 0 | 0 | 0 | 0 | 0 | 0 | 0 | 0 | 0 | 0 | 1172400 | 0 |
| AQALAIETE | MVP | Q14764 | MVP_HUMAN N | Unmodified | 0 | 0 | 0 | 0 | 0 | 0 | 0 | 0 | 221110 | 0 | 0 | 6516200 | 0 | 0 | 0 | 0 | 0 | 831540 |
| AQALAIETE | MVP | Q14764 | MVP_HUMAN N | Unmodified | 3260600 | 1416100 | 0 | 0 | 7011200 | 1995000 | 0 | 0 | 0 | 2465600 | 0 | 9249100 | 547550 | 0 | 0 | 0 | 474980 | 2783700 |
| AQALRDNS | HSP90AA1 | P07900 | HS90A_HUMAI | 2 Oxidation (M) | 109970 | 0 | 229730 | 417860 | 8284700 | 1519900 | 1081200 | 30056 | 1014200 | 737090 | 320190 | 928250 | 1746100 | 1924800 | 295550 | 276380 | 2849700 | 3769900 |
| AQALRDNS | HSP90AA1 | P07900 | HS90A_HUMAI | Oxidation (M) | 265250 | 0 | 254800 | 478000 | 5773500 | 1011700 | 1216500 | 479770 | 1160600 | 1041000 | 410560 | 2043400 | 2507700 | 737030 | 226550 | 225820 | 3782700 | 4790700 |
| AQALRDNS | HSP90AA1 | P07900 | HS90A_HUMAI | Unmodified | 100780 | 0 | 0 | 443090 | 10411000 | 1214100 | 1386300 | 354720 | 2764600 | 1371100 | 0 | 1728800 | 1978300 | 960540 | 0 | 0 | 5047900 | 8467000 |
| AQALRDNS | HSP90AA1 | P07900 | HS90A_HUMAI | Unmodified | 0 | 0 | 0 | 2040000 | 40040000 | 6006800 | 8380900 | 2335100 | 6837900 | 8323500 | 1107600 | 5344900 | 12453000 | 4742700 | 0 | 0 | 18943000 | 37524000 |
| AQALRDNS | HSP90AA1 | P07900 | HS90A_HUMAI | 2 Oxidation (M) | 0 | 0 | 0 | 0 | 2387000 | 419800 | 0 | 0 | 0 | 618940 | 0 | 0 | 0 | 0 | 0 | 0 | 1598900 | 0 |
| AQALRDNS | HSP90AA1 | P07900 | HS90A_HUMAI | Oxidation (M) | 205030 | 154220 | 0 | 866730 | 16212000 | 2613800 | 349250 | 1547800 | 98498 | 3007100 | 764290 | 3627800 | 3729400 | 283200 | 0 | 203700 | 7764800 | 10374000 |
| AQALRDNS | HSP90AB1; HSI | P08238; Q58 | HS90B_HUMAI | 3 Oxidation (M) | 0 | 0 | 232760 | 276350 | 3322700 | 532320 | 0 | 25937 | 184400 | 0 | 0 | 1005600 | 1338800 | 0 | 0 | 238060 | 1041700 | 1098000 |
| AQALRDNS | HSP90AB1; HSI | P08238; Q58 | HS90B_HUMAI | 2 Oxidation (M) | 0 | 0 | 0 | 600560 | 7271600 | 1489400 | 715680 | 277930 | 973000 | 835170 | 426600 | 3240300 | 3945500 | 921020 | 1021400 | 166600 | 3643800 | 3840400 |
| AQALRDNS | HSP90AB1; HSI | P08238; Q58 | HS90B_HUMAI | Oxidation (M) | 279520 | 253190 | 182310 | 987830 | 12225000 | 2021100 | 924480 | 485640 | 2671400 | 2631600 | 3616300 | 6375500 | 6541600 | 3950500 | 297640 | 205560 | 5468300 | 7206100 |

| | | | | | | | | | | | | | | | | | | | | | | |
|---|---|---|---|---|---|---|---|---|---|---|---|---|---|---|---|---|---|---|---|---|---|---|
| AQALRDNS | HSP90AB1; HSI | P08238; Q58 | HS90B_HUMAI | Unmodified | 0 | 59792 | 0 | 211170 | 15015000 | 988170 | 630470 | 0 | 3649200 | 985270 | 735830 | 1891100 | 4527300 | 1473100 | 143650 | 0 | 5599300 | 6885600 |
| AQALRDNS | HSP90AB1; HSI | P08238; Q58 | HS90B_HUMAI | Unmodified | 0 | 0 | 0 | 0 | 25584000 | 0 | 2021600 | 507320 | 2506300 | 2195300 | 720810 | 1649900 | 12538000 | 5015900 | 0 | 0 | 8924200 | 14032000 |
| AQALRDNS | HSP90AB1; HSI | P08238; Q58 | HS90B_HUMAI | Oxidation (M) | 0 | 0 | 0 | 0 | 10668000 | 819440 | 241500 | 494570 | 508340 | 1065000 | 1023200 | 956970 | 8507100 | 2093300 | 0 | 0 | 5876900 | 7111300 |
| AQALVQYLI | LAMTOR2 | Q9Y2Q5 | LTOR2_HUMAN | Unmodified | 71509 | 157690 | 61670 | 186670 | 917600 | 161550 | 782310 | 70874 | 780450 | 561610 | 295140 | 0 | 1604300 | 0 | 0 | 41814 | 325650 | 695200 |
| AQAREEQV | DST; DST; DST; | Q03001; A0/ | DYST_HUMAN | Unmodified | 0 | 0 | 0 | 0 | 0 | 0 | 0 | 0 | 0 | 0 | 0 | 560540 | 0 | 0 | 0 | 0 | 0 | 0 |
| AQASSIPV( | TBCB; TBCB; TE | Q99426; K7E | TBCB_HUMAN | Unmodified | 0 | 0 | 0 | 0 | 62021 | 0 | 36651 | 0 | 5938600 | 28600 | 665680 | 5460600 | 0 | 0 | 0 | 0 | 0 | 0 |
| AQASSIPV( | TBCB; TBCB; TE | Q99426; K7E | TBCB_HUMAN | Unmodified | 0 | 0 | 0 | 0 | 1440600 | 351420 | 0 | 0 | 222960 | 792920 | 0 | 0 | 0 | 0 | 0 | 0 | 427970 | 649000 |
| AQATPDHV | DIP2B | Q9P265 | DIP2B_HUMAN | Oxidation (M) | 497260 | 642700 | 576110 | 1911500 | 243500 | 0 | 185790 | 0 | 652850 | 182080 | 799230 | 6648700 | 0 | 0 | 0 | 0 | 0 | 260870 |
| AQATPDHV | DIP2B | Q9P265 | DIP2B_HUMAN | Unmodified | 0 | 20208 | 13609 | 52146 | 0 | 0 | 0 | 0 | 830320 | 0 | 69039 | 97996 | 0 | 0 | 75671 | 22191 | 0 | 0 |
| AQAVASTS | ZFAND2B; ZFAI | Q8WV99; B4 | ZFN2B_HUMAI | Unmodified | 938790 | 641740 | 0 | 0 | 0 | 0 | 0 | 0 | 0 | 0 | 0 | 0 | 0 | 0 | 581980 | 0 | 0 | 0 |
| AQAVASTS | ZFAND2B; ZFAI | Q8WV99; B4 | ZFN2B_HUMAI | Oxidation (M) | 284620 | 135000 | 0 | 0 | 0 | 0 | 0 | 0 | 0 | 0 | 0 | 0 | 0 | 0 | 0 | 0 | 0 | 0 |
| AQAVLEED | LONP1; LONP1 | P36776; K7E | LONM_HUMAN | Oxidation (M) | 0 | 0 | 0 | 0 | 0 | 0 | 0 | 13558 | 843150 | 0 | 0 | 0 | 0 | 0 | 0 | 0 | 38464 | 0 |
| AQAVLEED | LONP1; LONP1 | P36776; K7E | LONM_HUMAN | Unmodified | 0 | 0 | 0 | 0 | 0 | 0 | 0 | 0 | 3015800 | 155240 | 0 | 0 | 0 | 0 | 0 | 0 | 0 | 0 |
| AQAVLEED | LONP1; LONP1 | P36776; K7E | LONM_HUMAN | Unmodified | 0 | 0 | 0 | 0 | 0 | 0 | 0 | 0 | 602380 | 0 | 0 | 0 | 0 | 0 | 0 | 0 | 0 | 0 |
| AQAYQTGK | HSP90B2P; HSI | Q58FF3; P14 | ENPLL_HUMAN | Unmodified | 0 | 0 | 0 | 0 | 88561 | 0 | 0 | 0 | 639530 | 0 | 15965 | 0 | 0 | 0 | 0 | 0 | 0 | 0 |
| AQAYQTGK | HSP90B1; HSP! | P14625; H0Y | ENPL_HUMAN | Unmodified | 0 | 0 | 0 | 0 | 0 | 0 | 2251600 | 1925700 | 236150 | 6452000 | 0 | 0 | 293640 | 600180 | 0 | 0 | 0 | 349820 |
| AQAYQTGK | HSP90B1; HSP! | P14625; H0Y | ENPL_HUMAN | Unmodified | 0 | 0 | 0 | 0 | 139050 | 0 | 2525900 | 1179800 | 0 | 8827500 | 0 | 0 | 0 | 0 | 0 | 0 | 40010 | 314780 |
| AQCEDDLA | ASPH; ASPH | Q12797; A0/ | ASPH_HUMAN | Unmodified | 0 | 0 | 0 | 0 | 213870 | 0 | 0 | 0 | 3294100 | 388200 | 0 | 0 | 170040 | 294030 | 0 | 0 | 19026 | 0 |
| AQDDAFVI | B3GNT8 | Q7Z7M8 | B3GN8_HUMAI | Unmodified | 0 | 0 | 0 | 0 | 0 | 0 | 0 | 0 | 945690 | 238480 | 0 | 0 | 0 | 0 | 0 | 0 | 0 | 0 |
| AQDEAFAL | MRPL11; MRPL | Q9Y3B7; H0 | RM11_HUMAN | Unmodified | 0 | 0 | 0 | 0 | 212880 | 0 | 0 | 0 | 3131800 | 686440 | 0 | 273800 | 0 | 0 | 0 | 0 | 118880 | 0 |
| AQDGAAMI | ATP2B1; ATP2B | P20020; E7E | AT2B1_HUMAN | Oxidation (M) | 0 | 0 | 262440 | 0 | 0 | 0 | 0 | 0 | 2225100 | 472050 | 0 | 813240 | 0 | 0 | 285530 | 0 | 0 | 0 |
| AQDGAAMI | ATP2B1; ATP2B | P20020; E7E | AT2B1_HUMAN | Unmodified | 0 | 0 | 0 | 0 | 0 | 0 | 0 | 0 | 0 | 0 | 0 | 1619400 | 0 | 0 | 0 | 0 | 0 | 0 |
| AQDGAAMI | ATP2B1; ATP2B | P20020; E7E | AT2B1_HUMAN | Unmodified | 237650 | 390190 | 0 | 722490 | 845090 | 0 | 0 | 0 | 916780 | 134380 | 912430 | 6134300 | 2044600 | 1090100 | 927160 | 489470 | 167660 | 604680 |
| AQDGAAMI | ATP2B1; ATP2B | P20020; E7E | AT2B1_HUMAN | Oxidation (M) | 0 | 0 | 0 | 0 | 433620 | 0 | 0 | 0 | 0 | 0 | 688470 | 3909500 | 643280 | 361810 | 700750 | 2331600 | 0 | 129330 |
| AQDGAAMI | ATP2B1; ATP2B | P20020; E7E | AT2B1_HUMAN | 2 Oxidation (M) | 0 | 0 | 0 | 0 | 0 | 0 | 0 | 0 | 0 | 0 | 0 | 1111200 | 173660 | 200800 | 477500 | 286620 | 0 | 0 |
| AQDGDFIF | PTGFRN | Q9P2B2 | FPRP_HUMAN | Unmodified | 4200100 | 3974600 | 12008000 | 12237000 | 641880 | 0 | 0 | 0 | 0 | 74132 | 0 | 6801200 | 3189600 | 485420 | 6620100 | 7892800 | 0 | 208050 |
| AQDGDFIF | PTGFRN | Q9P2B2 | FPRP_HUMAN | Unmodified | 0 | 0 | 0 | 400030 | 0 | 0 | 0 | 0 | 0 | 0 | 0 | 0 | 341730 | 60225 | 0 | 0 | 0 | 0 |
| AQDIQCGL | NUDCD2; NUD | Q8WVJ2; E5I | NUDC2_HUMA | Unmodified | 0 | 0 | 0 | 0 | 0 | 0 | 0 | 0 | 0 | 0 | 0 | 1065800 | 0 | 0 | 0 | 0 | 0 | 0 |
| AQDLFGDI | MELTF | P08582 | TRFM_HUMAN | Unmodified | 0 | 0 | 0 | 0 | 3268700 | 0 | 0 | 0 | 3577800 | 561640 | 419250 | 1278300 | 0 | 0 | 0 | 0 | 157040 | 261750 |
| AQDLFGDI | MELTF | P08582 | TRFM_HUMAN | Unmodified | 0 | 0 | 0 | 0 | 1196700 | 1237100 | 0 | 0 | 0 | 22288 | 0 | 0 | 0 | 0 | 0 | 0 | 196480 | 39806 |
| AQDLSNLE | LZTFL1; LZTFL1 | Q9NQ48; H7 | LZTL1_HUMAN | Unmodified | 0 | 0 | 0 | 0 | 0 | 0 | 0 | 0 | 0 | 0 | 0 | 374620 | 0 | 0 | 0 | 0 | 0 | 0 |
| AQDPDQLF | DPP3; DPP3; DI | Q9NY33; G3 | DPP3_HUMAN | Unmodified | 0 | 0 | 0 | 0 | 0 | 0 | 0 | 0 | 108500 | 101410 | 0 | 1432300 | 0 | 0 | 0 | 238760 | 0 | 0 |
| AQDQGEKI | RPL11 | P62913 | RL11_HUMAN ( | Acetyl (Protein N | 729980 | 497230 | 0 | 0 | 88547 | 0 | 60075 | 0 | 0 | 0 | 69122 | 838060 | 0 | 0 | 0 | 0 | 0 | 0 |
| AQDQGEKI | RPL11 | P62913 | RL11_HUMAN ( | Acetyl (Protein N | 0 | 0 | 0 | 83851 | 476310 | 43287 | 396300 | 274850 | 0 | 0 | 0 | 0 | 413530 | 291990 | 0 | 0 | 129930 | 306130 |
| AQDRIVEVI | SLC9A3R1 | O14745 | NHRF1_HUMAI | Oxidation (M) | 111060 | 0 | 41497 | 50835 | 0 | 0 | 0 | 0 | 0 | 0 | 0 | 0 | 0 | 0 | 112380 | 184800 | 0 | 0 |
| AQDRLIEVI | SLC9A3R2; SLC | Q15599; H3I | NHRF2_HUMAI | Unmodified | 316170 | 329760 | 490790 | 687340 | 0 | 0 | 91407 | 23536 | 0 | 0 | 0 | 1454200 | 0 | 0 | 868670 | 480890 | 0 | 81738 |
| AQDRLIEVI | SLC9A3R2; SLC | Q15599; H3I | NHRF2_HUMAI | Unmodified | 0 | 0 | 0 | 263570 | 783840 | 0 | 0 | 0 | 0 | 0 | 0 | 0 | 573910 | 267540 | 0 | 0 | 0 | 0 |
| AQEAEAGV | EHBP1L1; EHBI | Q8N3D4; H0 | EH1L1_HUMAN | Unmodified | 0 | 241230 | 0 | 0 | 442120 | 76328 | 0 | 0 | 0 | 0 | 0 | 0 | 0 | 0 | 0 | 0 | 75223 | 210290 |
| AQEAEQLLI | LAMB2 | P55268 | LAMB2_HUMAI | Unmodified | 0 | 0 | 0 | 0 | 0 | 0 | 0 | 0 | 2343800 | 0 | 0 | 0 | 185470 | 0 | 0 | 0 | 0 | 0 |
| AQEAPGQ/ | SLC9A3R1; SLC | O14745; J3Q | NHRF1_HUMAI | Unmodified | 0 | 0 | 0 | 0 | 0 | 0 | 0 | 0 | 1467500 | 0 | 1719400 | 4956000 | 0 | 0 | 1807500 | 2218700 | 0 | 0 |
| AQEAPGQ/ | SLC9A3R1; SLC | O14745; J3Q | NHRF1_HUMAI | Unmodified | 0 | 0 | 3283100 | 1981800 | 0 | 0 | 0 | 0 | 1505200 | 0 | 430020 | 3393600 | 1319200 | 551480 | 3422300 | 5014600 | 0 | 0 |
| AQEAPGQ/ | SLC9A3R1 | O14745 | NHRF1_HUMAI | Unmodified | 0 | 0 | 0 | 0 | 0 | 0 | 0 | 0 | 0 | 0 | 0 | 0 | 861030 | 426290 | 0 | 0 | 0 | 0 |
| AQEEAEAEI | VGF | O15240 | VGF_HUMAN N | Unmodified | 0 | 0 | 0 | 0 | 0 | 0 | 0 | 0 | 339860 | 0 | 0 | 0 | 0 | 0 | 0 | 0 | 0 | 0 |
| AQEEAERLI | EZR; EZR | P15311; E7E | EZRI_HUMAN E | Unmodified | 177330 | 212450 | 67780 | 0 | 0 | 0 | 27551 | 13102 | 1719700 | 272900 | 3926100 | 6710000 | 0 | 0 | 195670 | 384860 | 0 | 0 |
| AQEEAPQC | EPB41L1; EPB4 | Q9H4G0; Q4 | E41L1_HUMAN | Unmodified | 0 | 0 | 410560 | 0 | 0 | 0 | 0 | 0 | 0 | 0 | 1253300 | 513930 | 0 | 0 | 0 | 0 | 0 | 0 |
| AQEEMISEI | CIT; CIT; CIT; CI | O14578; A0/ | CTRO_HUMAN | Unmodified | 0 | 0 | 0 | 183450 | 4842700 | 764050 | 0 | 14659 | 352520 | 178420 | 0 | 0 | 0 | 0 | 104630 | 57555 | 0 | 902810 |
| AQEEMISEI | CIT; CIT; CIT; CI | O14578; A0/ | CTRO_HUMAN | Oxidation (M) | 0 | 0 | 0 | 0 | 678600 | 0 | 0 | 0 | 0 | 0 | 0 | 0 | 0 | 0 | 0 | 0 | 53288 | 131890 |
| AQEENTWI | CANT1; CANT1 | Q8WVQ1; K7 | CANT1_HUMAI | Unmodified | 0 | 0 | 0 | 0 | 0 | 0 | 0 | 0 | 1301100 | 68636 | 0 | 0 | 0 | 0 | 0 | 0 | 0 | 0 |
| AQEENTWI | CANT1; CANT1 | Q8WVQ1; K7 | CANT1_HUMAI | Unmodified | 0 | 0 | 0 | 0 | 0 | 0 | 0 | 0 | 1964100 | 332530 | 0 | 0 | 0 | 0 | 0 | 0 | 0 | 0 |
| AQEFVNCK | GLRX | P35754 | GLRX1_HUMAN | Acetyl (Protein N | 0 | 0 | 0 | 0 | 0 | 0 | 0 | 0 | 0 | 0 | 0 | 2503100 | 332000 | 399710 | 0 | 0 | 1654700 | 703300 |
| AQEFVNCK | GLRX | P35754 | GLRX1_HUMAN | Acetyl (Protein N | 0 | 0 | 0 | 0 | 0 | 0 | 0 | 0 | 0 | 0 | 0 | 0 | 69827 | 0 | 0 | 0 | 2565000 | 249720 |
| AQEGGGSI | FAM98A; FAM9 | Q8NCA5; E9 | FA98A_HUMAN | Unmodified | 0 | 0 | 0 | 0 | 0 | 0 | 0 | 0 | 0 | 0 | 203160 | 2009200 | 490390 | 0 | 0 | 177780 | 0 | 0 |
| AQEILSQLF | GLRX | P35754 | GLRX1_HUMAN | Unmodified | 0 | 0 | 0 | 0 | 0 | 34404 | 0 | 0 | 0 | 0 | 2102200 | 5003500 | 0 | 0 | 0 | 0 | 0 | 104490 |
| AQEIQDES/ | IQGAP1; IQGAP | P46940; A0A | IQGA1_HUMAN | Unmodified | 0 | 0 | 0 | 0 | 3826000 | 0 | 0 | 0 | 0 | 0 | 0 | 0 | 0 | 0 | 0 | 0 | 0 | 0 |
| AQELGHSC | NUMA1; NUMA | Q14980; H0 | NUMA1_HUMA | Unmodified | 30006 | 0 | 0 | 0 | 0 | 0 | 0 | 0 | 0 | 0 | 181340 | 427120 | 0 | 0 | 30852 | 0 | 0 | 0 |
| AQELPAKD | SYT7 | O43581 | SYT7_HUMAN | Unmodified | 0 | 0 | 0 | 0 | 0 | 0 | 400160 | 439540 | 2124400 | 180580 | 0 | 0 | 0 | 0 | 0 | 0 | 202340 | 0 |
| AQETEAAP | MAGED2; MAGI | Q9UNF1; Q5 | MAGD2_HUMA | Unmodified | 148110 | 0 | 144230 | 285610 | 1076300 | 246600 | 364250 | 305970 | 1143900 | 678060 | 0 | 0 | 1269600 | 0 | 0 | 151230 | 304850 | 1753900 |
| AQFAQPEII | ECHS1 | P30084 | ECHM_HUMAN | Unmodified | 0 | 0 | 0 | 0 | 0 | 0 | 8314100 | 0 | 0 | 0 | 0 | 0 | 0 | 0 | 0 | 0 | 0 | 0 |
| AQFDTLQK | YES1; YES1 | P07947; J3Q | YES_HUMAN Ty | Unmodified | 441730 | 2237700 | 1051300 | 875880 | 549410 | 156290 | 110260 | 30114 | 1534400 | 715310 | 304670 | 3057600 | 225060 | 0 | 3697200 | 29481 | 863460 | 1178200 |
| AQFEGIVTI | HSPA9; HSPA9; | P38646; H0Y | GRP75_HUMAI | Unmodified | 0 | 0 | 0 | 0 | 0 | 0 | 0 | 31842 | 2531700 | 535720 | 112280 | 91150 | 0 | 0 | 0 | 0 | 0 | 0 |
| AQFEGIVTI | HSPA9; HSPA9; | P38646; H0Y | GRP75_HUMAI | Unmodified | 0 | 0 | 8203800 | 2445500 | 1848600 | 0 | 0 | 0 | 48781000 | 6079600 | 0 | 1341700 | 0 | 0 | 0 | 0 | 0 | 0 |
| AQFEQLCA | HSPA4L; HSPA | O95757; E9F | HS74L_HUMAI | Unmodified | 0 | 0 | 0 | 0 | 7328900 | 1062900 | 1008400 | 38208 | 929710 | 838180 | 174160 | 824180 | 1141100 | 449670 | 0 | 0 | 1497900 | 1068400 |
| AQFERDLQ | MYH9 | P35579 | MYH9_HUMAN | Unmodified | 956980 | 1042900 | 3631400 | 4901500 | 70099000 | 10467000 | 3317600 | 1031500 | 632040 | 7287600 | 1988000 | 4703800 | 26030000 | 15179000 | 1996300 | 1964400 | 9888700 | 21721000 |
| AQFERDLQ | MYH10 | P35580 | MYH10_HUMAI | Unmodified | 0 | 0 | 364600 | 692280 | 7230500 | 1846900 | 915900 | 185280 | 0 | 185170 | 0 | 899400 | 2704400 | 2216400 | 343210 | 227210 | 120800 | 655720 |
| AQFETLQQ | FYN | P06241 | FYN_HUMAN T | Unmodified | 0 | 0 | 0 | 0 | 0 | 0 | 0 | 0 | 5666500 | 0 | 0 | 0 | 0 | 0 | 0 | 0 | 151820 | 0 |
| AQFEYDPA | CASK; CASK; C/ | O14936; A0/ | CSKP_HUMAN | Unmodified | 368600 | 0 | 761060 | 1216600 | 1341100 | 538790 | 110580 | 59104 | 5519200 | 1261100 | 0 | 1136600 | 1929300 | 944240 | 1411600 | 890970 | 366310 | 1022100 |
| AQFVRNLV | ATP5MG; ATP5I | O75964; A0/ | ATP5L_HUMAN | Acetyl (Protein N | 0 | 0 | 0 | 0 | 0 | 0 | 0 | 0 | 156940 | 1242700 | 0 | 0 | 0 | 0 | 0 | 0 | 0 | 0 |
| AQGEPVAC | SEC31A; SEC31 | O94979; H7I | SC31A_HUMAN | Unmodified | 0 | 0 | 0 | 392230 | 0 | 0 | 0 | 0 | 347920 | 159400 | 319590 | 1735800 | 808450 | 0 | 0 | 0 | 147760 | 431260 |
| AQGGSSD! | GOLGA3 | Q08378 | GOGA3_HUMA | Unmodified | 0 | 0 | 0 | 0 | 0 | 0 | 0 | 0 | 0 | 0 | 23110 | 391810 | 0 | 0 | 65668 | 0 | 0 | 0 |
| AQGLEEGV | PTK7; PTK7; PTI | Q13308; H0 | PTK7_HUMAN I | Unmodified | 0 | 0 | 0 | 0 | 10893000 | 734610 | 3572900 | 376710 | 5031800 | 5180100 | 1872400 | 5328600 | 9699300 | 3334500 | 1968800 | 1090600 | 5793200 | 9383000 |
| AQGPAASA | BASP1 | P80723 | BASP1_HUMAN | Unmodified | 0 | 0 | 0 | 0 | 0 | 0 | 817860 | 529800 | 5632100 | 900130 | 0 | 0 | 0 | 0 | 0 | 0 | 0 | 0 |
| AQGPAASA | BASP1 | P80723 | BASP1_HUMAN | Unmodified | 0 | 0 | 0 | 0 | 6.66E+08 | 95046000 | 50503000 | 1.15E+08 | 1.31E+08 | 1.02E+08 | 0 | 0 | 12242000 | 15876000 | 0 | 0 | 2.53E+08 | 3.3E+08 |
| AQGPAIDL! | FMNL2; FMNL2 | Q96PY5; C9I | FMNL2_HUMAI | Unmodified | 0 | 0 | 0 | 0 | 2854100 | 413730 | 330590 | 0 | 912290 | 355150 | 410550 | 2173200 | 290380 | 0 | 0 | 0 | 560720 | 2467300 |
| AQGPGPEF | P4HTM; P4HTM | Q9NXG6; A0 | P4HTM_HUMAI | Unmodified | 0 | 0 | 0 | 0 | 0 | 0 | 0 | 0 | 1252300 | 0 | 0 | 0 | 0 | 0 | 0 | 0 | 0 | 0 |
| AQGPGPEF | P4HTM; P4HTM | Q9NXG6; A0 | P4HTM_HUMAI | Unmodified | 0 | 0 | 0 | 0 | 0 | 0 | 0 | 0 | 11904000 | 329030 | 0 | 0 | 0 | 0 | 0 | 0 | 0 | 0 |
| AQGSTSST | SIRPA | P78324 | SHPS1_HUMAI | Unmodified | 0 | 0 | 7866.5 | 0 | 0 | 0 | 0 | 0 | 0 | 0 | 0 | 0 | 0 | 0 | 0 | 0 | 0 | 1920400 |
| AQGTFNSP | ST14 | Q9Y5Y6 | ST14_HUMAN | Unmodified | 968180 | 689140 | 753340 | 1243100 | 0 | 0 | 0 | 0 | 868020 | 0 | 0 | 0 | 1616000 | 0 | 1756100 | 1185400 | 0 | 0 |
| AQGVLAAC | LAMB2 | P55268 | LAMB2_HUMAI | Unmodified | 0 | 0 | 0 | 0 | 0 | 0 | 0 | 0 | 1176200 | 105400 | 80436 | 59631 | 0 | 0 | 37164 | 123000 | 0 | 0 |
| AQGWAPLI | CPNE1; CPNE1 | Q99829; B0( | CPNE1_HUMAI | Unmodified | 0 | 0 | 129210 | 0 | 0 | 0 | 0 | 0 | 816430 | 0 | 0 | 842530 | 0 | 0 | 0 | 0 | 0 | 0 |
| AQGWAPLI | CPNE1; CPNE1 | Q99829; B0( | CPNE1_HUMAI | Unmodified | 0 | 0 | 172430 | 335720 | 0 | 395330 | 0 | 0 | 2520300 | 343950 | 0 | 8650000 | 0 | 0 | 416520 | 329100 | 0 | 609500 |
| AQHLSPAP | SARG | Q9BW04 | SARG_HUMAN | Unmodified | 1476800 | 986720 | 2622200 | 1992800 | 123070 | 0 | 0 | 0 | 0 | 0 | 0 | 0 | 13331000 | 8181100 | 2421200 | 1912000 | 0 | 0 |
| AQHQDPG | DNAJB2 | P25686 | DNJB2_HUMAN | Unmodified | 0 | 870020 | 0 | 1032900 | 370310 | 0 | 0 | 0 | 149850 | 97474 | 95553 | 34053 | 271540 | 0 | 0 | 0 | 22360 | 91700 |
| AQHTLPQD | LSM14B; LSM1 | Q9BX40; A0/ | LS14B_HUMAN | Unmodified | 928460 | 0 | 0 | 1563600 | 0 | 0 | 0 | 0 | 0 | 0 | 0 | 0 | 0 | 0 | 2243100 | 2170200 | 0 | 0 |
| AQHVFQH/ | BLMH; BLMH; B | Q13867; J3K | BLMH_HUMAN | Unmodified | 340150 | 0 | 163950 | 0 | 764290 | 0 | 128750 | 363220 | 3413500 | 324550 | 973890 | 11556000 | 744670 | 366510 | 594400 | 0 | 0 | 196390 |
| AQIALKLGV | MDH1; MDH1; I | P40925; C9J | MDHC_HUMAN | Unmodified | 0 | 0 | 0 | 0 | 375930 | 0 | 1011100 | 0 | 0 | 171900 | 0 | 0 | 276800 | 525000 | 0 | 0 | 1626000 | 330780 |
| AQIEKRIQE | HSPD1; HSPD1 | P10809; A0A | CH60_HUMAN | Unmodified | 0 | 0 | 0 | 0 | 0 | 0 | 228390 | 0 | 0 | 82912 | 0 | 0 | 28370 | 0 | 0 | 0 | 42990 | 0 |
| AQIFANTVI | KRT18; KRT18 | P05783; F8V | K1C18_HUMAN | Unmodified | 0 | 123820 | 40610 | 0 | 0 | 0 | 0 | 0 | 3414600 | 0 | 2298400 | 31207000 | 0 | 1704100 | 277890 | 93451 | 0 | 0 |
| AQIHDLVL | HSPA1A; HSPA | P0DMV8; P0 | HS71A_HUMAN | Unmodified | 15857000 | 20820000 | 11175000 | 7393600 | 2123100 | 0 | 81436 | 105530 | 30857000 | 2842800 | 2052600 | 20773000 | 2723400 | 345630 | 13879000 | 11710000 | 362550 | 577300 |
| AQIHDLVL | HSPA1A; HSPA | P0DMV8; P0 | HS71A_HUMAN | Unmodified | 3262300 | 3474600 | 3160600 | 1575500 | 1.01E+08 | 8077700 | 20023000 | 4311600 | 22761000 | 41796000 | 3756600 | 3984900 | 97158000 | 38634000 | 1503500 | 1037600 | 13424000 | 44732000 |
| AQIHDLVL | HSPA1A; HSPA | P0DMV8; P0 | HS71A_HUMAN | Unmodified | 0 | 0 | 0 | 0 | 8048600 | 464420 | 12523000 | 3304200 | 0 | 5607000 | 0 | 0 | 10684000 | 16858000 | 0 | 0 | 8376300 | 7824500 |
| AQIHGGILI | HSPG2 | P98160 | PGBM_HUMAN | Unmodified | 0 | 0 | 3912000 | 3550700 | 0 | 0 | 0 | 0 | 0 | 0 | 284870 | 0 | 668640 | 0 | 0 | 0 | 0 | 2493000 |
| AQIHGGILI | HSPG2 | P98160 | PGBM_HUMAN | Unmodified | 0 | 0 | 0 | 0 | 612720 | 701470 | 0 | 0 | 0 | 0 | 0 | 0 | 1280900 | 5163200 | 0 | 0 | 552090 | 11331000 |
| AQIHQFREI | CUL4B; CUL4B; | Q13620; A0/ | CUL4B_HUMAI | Unmodified | 0 | 0 | 0 | 0 | 1378100 | 5285300 | 2155100 | 1712300 | 95330 | 0 | 0 | 411180 | 0 | 3386000 | 528870 | 0 | 0 | 4403000 |
| AQIIELLCIV | CD2AP | Q9Y5K6 | CD2AP_HUMAI | Unmodified | 1364800 | 2910100 | 0 | 1217400 | 21061 | 11805 | 0 | 0 | 235920 | 18813 | 42090 | 168870 | 167320 | 0 | 6240500 | 1039700 | 0 | 0 |
| AQIIQRQL( | MICAL3; MICAL | Q7RTP6; J3Q | MICA3_HUMAN | Unmodified | 0 | 0 | 0 | 0 | 946070 | 114970 | 0 | 0 | 0 | 0 | 0 | 0 | 0 | 0 | 0 | 0 | 0 | 0 |
| AQILGGAN | UBE2T | Q9NPD8 | UBE2T_HUMAN | Unmodified | 0 | 0 | 0 | 0 | 0 | 0 | 0 | 0 | 0 | 0 | 0 | 758380 | 0 | 0 | 0 | 0 | 0 | 0 |
| AQILPIR | CLTC; CLTC; CI | Q00610; J3K | CLH1_HUMAN | Acetyl (Protein N | 0 | 0 | 0 | 0 | 0 | 0 | 0 | 0 | 0 | 0 | 0 | 7989300 | 0 | 0 | 0 | 0 | 0 | 0 |
| AQIQQFHS | KTN1 | Q86UP2 | KTN1_HUMAN | Unmodified | 0 | 0 | 0 | 0 | 0 | 0 | 327520 | 0 | 0 | 0 | 0 | 7686300 | 0 | 0 | 0 | 0 | 0 | 0 |
| AQITEEEAK | NF2; NF2 | P35240; A0A | MERL_HUMAN | Unmodified | 0 | 0 | 580340 | 0 | 0 | 0 | 0 | 0 | 233930 | 2991200 | 625900 | 0 | 486390 | 506320 | 0 | 0 | 623720 | 4356400 |
| AQITGYR | FN1 | P02751 | FINC_HUMAN I | Unmodified | 174960 | 1832900 | 8907200 | 0 | 0 | 1224400 | 485670 | 124020 | 655850 | 25602 | 87150 | 1200400 | 0 | 438750 | 242090 | 2291900 | 92338 | 113720 |
| AQIWDTAG | RAB25 | P57735 | RAB25_HUMAN | Unmodified | 0 | 0 | 405740 | 0 | 0 | 0 | 0 | 0 | 0 | 0 | 0 | 1342400 | 0 | 0 | 0 | 672580 | 0 | 0 |
| AQIWDTAG | RAB11A; RAB11 | P62491; Q15 | RB11A_HUMAN | Unmodified | 2118100 | 3012400 | 105750 | 4913000 | 565690 | 1280500 | 0 | 0 | 2534600 | 416690 | 3424600 | 15889000 | 0 | 0 | 4622400 | 5387300 | 0 | 0 |
| AQIWDTAG | RAB11A; RAB11 | P62491; Q15 | RB11A_HUMAN | Unmodified | 1735200 | 822600 | 2654700 | 9104500 | 8413600 | 2443300 | 953020 | 607650 | 869700 | 1008900 | 2120800 | 4268700 | 2724100 | 1603600 | 1453200 | 1186700 | 405570 | 1033200 |
| AQKDEEKI | TPM1; TPM1; TF | P09493; H0Y | TPM1_HUMAN | Unmodified | 0 | 0 | 0 | 0 | 2316100 | 653980 | 0 | 0 | 129390 | 109730 | 0 | 151500 | 1706100 | 0 | 348400 | 0 | 223350 | 152620 |
| AQKELEEQ | RDX; RDX; RDX | P35241; A0A | RADI_HUMAN I | Unmodified | 0 | 0 | 0 | 0 | 0 | 0 | 0 | 0 | 0 | 0 | 0 | 0 | 0 | 0 | 0 | 0 | 0 | 0 |
| AQKLPDLSI | PRRC2C; PRRC | Q9Y520; H7( | PRC2C_HUMAI | Unmodified | 379840 | 444630 | 0 | 230720 | 99716 | 0 | 0 | 0 | 346000 | 109510 | 0 | 0 | 690250 | 188310 | 0 | 153440 | 32469 | 267710 |
| AQLADSFH | SPTAN1; SPTAI | Q13813; A0/ | SPTN1_HUMAN | Unmodified | 0 | 0 | 0 | 0 | 0 | 0 | 0 | 0 | 0 | 0 | 0 | 0 | 0 | 0 | 0 | 0 | 0 | 0 |
| AQLADSFH | SPTAN1; SPTAI | Q13813; A0/ | SPTN1_HUMAN | Unmodified | 0 | 0 | 0 | 0 | 0 | 0 | 0 | 0 | 0 | 373900 | 0 | 1295400 | 0 | 0 | 0 | 0 | 0 | 0 |
| AQLDAAVTI | ARSA; ARSA | P15289; A0A | ARSA_HUMAN | Unmodified | 0 | 0 | 0 | 0 | 0 | 0 | 0 | 0 | 623030 | 0 | 0 | 2539200 | 0 | 0 | 0 | 0 | 0 | 0 |

| | | | | | | | | | | | | | | | | | | | | | | |
|---|---|---|---|---|---|---|---|---|---|---|---|---|---|---|---|---|---|---|---|---|---|---|
| AQLELEVSI | MVP | Q14764 | MVP_HUMAN N | Unmodified | 1581000 | 1702300 | 0 | 55308 | 0 | 245170 | 0 | 0 | 3987800 | 274140 | 0 | 20491000 | 63874 | 0 | 40434 | 0 | 0 | 0 |
| AQLELYPEI | FAM126A; FAM: | Q9BYI3; A0A | HYCCI_HUMAN | Unmodified | 0 | 0 | 0 | 295530 | 0 | 0 | 0 | 0 | 204670 | 0 | 0 | 292300 | 0 | 0 | 1033100 | 0 | 0 | 0 |
| AQLEPVASI | PLEC; PLEC | Q15149; H0Y | PLEC_HUMAN | Unmodified | 0 | 0 | 0 | 0 | 843870 | 145510 | 0 | 0 | 619800 | 0 | 177680 | 840250 | 0 | 0 | 0 | 0 | 411960 | 1242600 |
| AQLEPVASI | PLEC; PLEC | Q15149; H0Y | PLEC_HUMAN | Unmodified | 0 | 0 | 0 | 0 | 1040200 | 0 | 0 | 0 | 0 | 0 | 0 | 0 | 0 | 0 | 0 | 0 | 1693900 | 2586300 |
| AQLEQGG\ | CC2D1A; CC2D | Q6P1N0; A0A | C2D1A_HUMAI | Unmodified | 3281800 | 1943200 | 761120 | 746510 | 0 | 0 | 0 | 0 | 1062900 | 72480 | 356770 | 810350 | 105520 | 0 | 1189500 | 344510 | 0 | 0 |
| AQLETALK | MOV10; RPGRII | Q9HCE1; Q6 | MOV10_HUMA | Unmodified | 957010 | 934060 | 332010 | 463650 | 565440 | 77597 | 49829 | 0 | 627110 | 182560 | 894280 | 1009100 | 1353500 | 1140900 | 923430 | 2364600 | 270800 | 75762 |
| AQLFALTG\ | USP14; USP14; | P54578; J3K | UBP14_HUMAI | Unmodified | 0 | 0 | 449380 | 0 | 0 | 0 | 0 | 0 | 270650 | 0 | 0 | 12291000 | 0 | 0 | 737650 | 0 | 0 | 0 |
| AQLFALTG\ | USP14; USP14; | P54578; J3K | UBP14_HUMAI | Unmodified | 0 | 0 | 0 | 0 | 0 | 0 | 0 | 172250 | 0 | 336470 | 0 | 0 | 0 | 0 | 0 | 0 | 736680 | 589890 |
| AQLFETSSI | RAB24; RAB24 | Q969Q5; F8\ | RAB24_HUMAI | Unmodified | 722490 | 270050 | 0 | 343780 | 466820 | 0 | 0 | 2268900 | 0 | 0 | 0 | 394670 | 625090 | 0 | 248470 | 237750 | 0 | 72117 |
| AQLGEDIR | TFG; TFG; TFG; | Q92734; A0A | TFG_HUMAN P | Unmodified | 0 | 0 | 520530 | 0 | 133400 | 189610 | 224340 | 0 | 3096400 | 394390 | 183450 | 6030300 | 0 | 0 | 25419 | 674830 | 95032 | 90015 |
| AQLGGPEA | HSPB1; HSPB1; | P04792; A0A | HSPB1_HUMAI | Unmodified | 462480 | 259010 | 4947700 | 1585900 | 3539400 | 746800 | 0 | 0 | 6386300 | 216440 | 42704 | 662960 | 1680500 | 939070 | 3266100 | 1899900 | 0 | 0 |
| AQLGGPEA | HSPB1; HSPB1; | P04792; A0A | HSPB1_HUMAI | Unmodified | 1181400 | 1450100 | 334640 | 7917500 | 2498500 | 522740 | 658160 | 424370 | 22787000 | 5388300 | 191030 | 0 | 12465000 | 4374700 | 2194600 | 2158300 | 2005800 | 3396800 |
| AQLHDTNI | PPP1R12A; PPP | O14974; A0A | MYPT1_HUMAI | Unmodified | 0 | 0 | 0 | 0 | 4136300 | 0 | 0 | 0 | 0 | 0 | 0 | 0 | 0 | 0 | 0 | 0 | 1677500 | 0 |
| AQLHGASE | HSPG2 | P98160 | PGBM_HUMAN | Unmodified | 0 | 0 | 0 | 0 | 0 | 0 | 0 | 0 | 0 | 0 | 0 | 0 | 0 | 0 | 0 | 1498900 | 0 | 0 |
| AQLKPPATS | ITGA5; ITGA5 | P08648; H0Y | ITA5_HUMAN Ir | Unmodified | 0 | 0 | 616660 | 655180 | 4793100 | 960830 | 149470 | 71833 | 0 | 0 | 0 | 176380 | 0 | 187050 | 1081500 | 1590400 | 678650 | 0 |
| AQLLQPTLI | TRAP1; TRAP1 | Q12931; I3L( | TRAP1_HUMAN | Unmodified | 0 | 0 | 0 | 0 | 0 | 0 | 0 | 0 | 1475100 | 0 | 0 | 0 | 0 | 0 | 0 | 0 | 0 | 0 |
| AQLLVVGS | L1CAM | P32004 | L1CAM_HUMAI | Unmodified | 0 | 0 | 0 | 0 | 0 | 0 | 0 | 0 | 1713100 | 0 | 0 | 0 | 0 | 0 | 0 | 0 | 0 | 0 |
| AQLNIGNV | RPL8; RPL8; RP | P62917; E9P | RL8_HUMAN 6( | Oxidation (M) | 0 | 0 | 0 | 0 | 0 | 0 | 0 | 0 | 0 | 0 | 0 | 3906600 | 0 | 0 | 0 | 0 | 0 | 0 |
| AQMKNPIL | STXBP1; STXBP | P61764; A0A | STXB1_HUMAN | Oxidation (M) | 0 | 0 | 0 | 0 | 285360 | 0 | 1955500 | 0 | 0 | 499340 | 0 | 0 | 357740 | 409870 | 0 | 0 | 355120 | 289140 |
| AQMVLSR | SCRIB; SCRIB; S | Q14160; A0A | SCRIB_HUMAN | Unmodified | 0 | 23633 | 83320 | 0 | 0 | 301810 | 0 | 0 | 1183100 | 103990 | 62762 | 0 | 0 | 0 | 0 | 64524 | 88714 | 359090 |
| AQMVQEDI | MSN | P26038 | MOES_HUMAN | Oxidation (M) | 0 | 0 | 131710 | 255160 | 7117400 | 2572600 | 256210 | 303290 | 0 | 0 | 0 | 0 | 3375300 | 1247700 | 182540 | 186780 | 2796400 | 3791400 |
| AQMVQEDI | MSN | P26038 | MOES_HUMAN | Unmodified | 183220 | 237300 | 628960 | 2484500 | 56311000 | 13085000 | 1329000 | 734240 | 0 | 0 | 0 | 0 | 8559100 | 7747600 | 922310 | 629140 | 20458000 | 28218000 |
| AQMVQEDI | MSN | P26038 | MOES_HUMAN | Unmodified | 0 | 0 | 0 | 0 | 22796000 | 1881100 | 1053300 | 411910 | 0 | 0 | 0 | 0 | 2905200 | 3077100 | 0 | 0 | 16662000 | 12917000 |
| AQMVQEDI | MSN | P26038 | MOES_HUMAN | Oxidation (M) | 0 | 0 | 0 | 0 | 15780000 | 3004100 | 1328500 | 435450 | 0 | 0 | 80808 | 0 | 1962200 | 4148500 | 0 | 0 | 14056000 | 11396000 |
| AQNCNILL: | DAAM1 | Q9Y4D1 | DAAM1_HUMA | Unmodified | 0 | 0 | 0 | 0 | 0 | 247910 | 208490 | 0 | 256070 | 0 | 0 | 534030 | 184510 | 0 | 0 | 0 | 0 | 198120 |
| AQNDLIWI | PARP1; PARP1; | P09874; A0A | PARP1_HUMAN | Unmodified | 849130 | 385360 | 53180 | 0 | 74233 | 0 | 0 | 0 | 2402700 | 2656400 | 130970 | 2878500 | 41571 | 0 | 0 | 0 | 0 | 467900 |
| AQNDLIWI | PARP1; PARP1; | P09874; A0A | PARP1_HUMAN | Unmodified | 782940 | 0 | 163180 | 104660 | 812160 | 0 | 1835300 | 336350 | 1536600 | 2323700 | 1481100 | 5113200 | 2527300 | 808320 | 0 | 86725 | 800000 | 1258500 |
| AQNISSNA | CKAP5 | Q14008 | CKAP5_HUMAI | Unmodified | 0 | 0 | 0 | 0 | 0 | 0 | 0 | 0 | 0 | 0 | 0 | 279820 | 0 | 357960 | 0 | 0 | 0 | 0 |
| AQNKETSV | SLK | Q9H2G2 | SLK_HUMAN S | Unmodified | 625220 | 270070 | 0 | 0 | 671580 | 0 | 179990 | 0 | 0 | 298010 | 258490 | 0 | 1354100 | 1412000 | 0 | 0 | 118670 | 0 |
| AQNLPQSF | CNNM3 | Q8NE01 | CNNM3_HUMA | Unmodified | 0 | 0 | 0 | 159880 | 0 | 0 | 0 | 0 | 1737900 | 499410 | 0 | 0 | 397090 | 0 | 0 | 156450 | 0 | 0 |
| AQPAQPAD | PSMA1; PSMA1 | P25786; F5G | PSA1_HUMAN | Unmodified | 0 | 0 | 369710 | 0 | 0 | 0 | 0 | 0 | 289770 | 0 | 288840 | 1228200 | 0 | 0 | 0 | 138810 | 0 | 0 |
| AQPAQPAD | PSMA1; PSMA1 | P25786; F5G | PSA1_HUMAN | Unmodified | 0 | 0 | 0 | 0 | 8637200 | 1721600 | 569500 | 272770 | 3042500 | 2172600 | 1779100 | 6132200 | 0 | 0 | 0 | 0 | 4075000 | 4058100 |
| AQPAQPAD | PSMA1; PSMA1 | P25786; F5G | PSA1_HUMAN | Oxidation (M) | 90861 | 77474 | 0 | 367310 | 2875400 | 448390 | 168870 | 130700 | 747770 | 493660 | 1015800 | 2658400 | 2695000 | 1189500 | 0 | 381110 | 0 | 1260800 |
| AQPGVSET | ZDHHC5 | Q9C0B5 | ZDHC5_HUMA | Unmodified | 0 | 0 | 0 | 4885800 | 20398000 | 1775800 | 4710400 | 0 | 4651600 | 4639800 | 3022900 | 4366800 | 18307000 | 31565000 | 0 | 0 | 11499000 | 51340000 |
| AQPTPSSS | WDR5; WDR5; \ | P61964; V9C | WDR5_HUMAN | Unmodified | 0 | 0 | 0 | 0 | 0 | 0 | 420410 | 378670 | 0 | 149880 | 350630 | 934270 | 159370 | 0 | 0 | 0 | 0 | 0 |
| AQPVSCFT | CACNA2D2; CA | Q9NY47; C9 | CA2D2_HUMAI | Unmodified | 818070 | 0 | 0 | 0 | 183340 | 0 | 323940 | 60629 | 11843000 | 2490500 | 642170 | 2928200 | 0 | 0 | 252190 | 56863 | 0 | 0 |
| AQPVSCFT | CACNA2D2; CA | Q9NY47; C9 | CA2D2_HUMAI | Unmodified | 0 | 0 | 0 | 0 | 0 | 0 | 490860 | 0 | 0 | 822500 | 0 | 0 | 81934 | 0 | 0 | 0 | 0 | 0 |
| AQPVSCFT | CACNA2D2; CA | Q9NY47; C9 | CA2D2_HUMAI | Unmodified | 0 | 0 | 0 | 0 | 0 | 0 | 103770 | 0 | 0 | 231880 | 0 | 0 | 0 | 0 | 0 | 0 | 0 | 0 |
| AQQAADKY | MYH9; MYH9; M | P35579; Q5E | MYH9_HUMAN | Acetyl (Protein N | 0 | 0 | 0 | 0 | 1560800 | 0 | 1571600 | 548420 | 2357700 | 367830 | 0 | 630190 | 606000 | 1381800 | 0 | 0 | 5950100 | 1240800 |
| AQQALSEL | UBR4; UBR4 | Q5T4S7; A0A | UBR4_HUMAN | Unmodified | 0 | 0 | 0 | 0 | 109970 | 0 | 0 | 0 | 656650 | 0 | 137970 | 3646700 | 0 | 0 | 0 | 0 | 25240 | 67994 |
| AQQATPGC | HINT2 | Q9BX68 | HINT2_HUMAN | Unmodified | 0 | 0 | 0 | 0 | 0 | 0 | 152970 | 214290 | 2850700 | 792480 | 19734 | 0 | 0 | 0 | 0 | 0 | 0 | 0 |
| AQQEFAAG | SCAMP3 | O14828 | SCAM3_HUMA | Unmodified | 1730000 | 860030 | 2240400 | 1689400 | 901830 | 161000 | 0 | 108730 | 679560 | 0 | 0 | 2310800 | 0 | 493420 | 1648900 | 951650 | 0 | 0 |
| AQQEFATG | SCAMP1; SCAM | O15126; A0A | SCAM1_HUMA | Unmodified | 0 | 155300 | 586930 | 345340 | 645980 | 0 | 134470 | 0 | 860940 | 203560 | 0 | 560360 | 343210 | 315570 | 253340 | 342580 | 0 | 107890 |
| AQQELEEQ | MSN | P26038 | MOES_HUMAN | Unmodified | 0 | 0 | 157390 | 0 | 387580 | 0 | 0 | 18509 | 0 | 0 | 0 | 0 | 0 | 0 | 0 | 142160 | 87148 | 185090 |
| AQQEQELA | PRKCSH; PRKC | P14314; K7E | GLU2B_HUMAI | Unmodified | 539430 | 675030 | 1656600 | 321510 | 577010 | 0 | 0 | 63805 | 60364000 | 4674600 | 2610900 | 2065200 | 128030 | 0 | 382680 | 607270 | 0 | 0 |
| AQQEVLQK | N/A | N/A | N/A | Unmodified | 0 | 0 | 0 | 0 | 0 | 0 | 0 | 0 | 0 | 0 | 0 | 2357000 | 0 | 0 | 0 | 0 | 0 | 0 |
| AQQLAEVE | MVP | Q14764 | MVP_HUMAN N | Unmodified | 1149600 | 2096700 | 0 | 0 | 3197500 | 186550 | 0 | 0 | 2776200 | 315430 | 814930 | 18851000 | 0 | 0 | 0 | 0 | 70261 | 517400 |
| AQQLAEVE | MVP | Q14764 | MVP_HUMAN N | Unmodified | 2388000 | 1678400 | 0 | 0 | 1162200 | 16260 | 0 | 82430 | 1275000 | 38008 | 0 | 10707000 | 0 | 0 | 0 | 0 | 111160 | 0 |
| AQQNNVEI | RPS7; RPS7 | P62081; A0A | RS7_HUMAN 4 | Unmodified | 9265400 | 9917300 | 2857500 | 9691800 | 8339000 | 667670 | 4546900 | 4166300 | 1975100 | 1615600 | 5661000 | 10122000 | 14784000 | 6083400 | 4690700 | 2875000 | 1426600 | 7166100 |
| AQQNNVEI | RPS7; RPS7 | P62081; A0A | RS7_HUMAN 4 | Unmodified | 2617000 | 1709200 | 0 | 1855100 | 745040 | 211100 | 622100 | 0 | 0 | 340580 | 580720 | 9826200 | 601530 | 0 | 812200 | 2328500 | 101220 | 962720 |
| AQQNPTLL | DTNA; DTNB; D | Q9Y4J8; O6C | DTNA_HUMAN | Unmodified | 0 | 0 | 0 | 0 | 378330 | 0 | 0 | 0 | 1163800 | 471290 | 0 | 0 | 0 | 0 | 0 | 0 | 0 | 0 |
| AQQNPTLL | DTNA; DTNB; D | Q9Y4J8; O6C | DTNA_HUMAN | Unmodified | 0 | 0 | 0 | 0 | 713340 | 0 | 0 | 0 | 0 | 647410 | 0 | 0 | 1604300 | 0 | 0 | 0 | 0 | 544200 |
| AQQPAAEL | TRIM26 | Q12899 | TRI26_HUMAN | Unmodified | 1648500 | 2065700 | 1441500 | 822880 | 0 | 0 | 0 | 0 | 0 | 0 | 0 | 0 | 0 | 0 | 2403200 | 1441600 | 0 | 0 |
| AQQPAAEL | TRIM26 | Q12899 | TRI26_HUMAN | Oxidation (M) | 154830 | 33525 | 0 | 0 | 0 | 0 | 0 | 0 | 0 | 0 | 0 | 0 | 0 | 0 | 1801300 | 1093600 | 0 | 0 |
| AQQPSEVT | ASAP1; ASAP1; | Q9ULH1; H0 | ASAP1_HUMAN | Unmodified | 0 | 0 | 0 | 0 | 466230 | 0 | 0 | 0 | 0 | 78413 | 0 | 0 | 0 | 0 | 0 | 0 | 417760 | 409230 |
| AQQQKQE\ | TXNDC9; TXND | O14530; F8V | TXND9_HUMAI | Unmodified | 0 | 0 | 0 | 0 | 406620 | 0 | 0 | 0 | 0 | 0 | 0 | 1190100 | 0 | 152630 | 0 | 0 | 227020 | 329890 |
| AQQSLELIC | KTN1; KTN1; KT | Q86UP2; B7 | KTN1_HUMAN | Unmodified | 0 | 0 | 0 | 0 | 0 | 0 | 0 | 45711 | 0 | 80725 | 0 | 1043000 | 0 | 0 | 0 | 0 | 0 | 0 |
| AQQSYERL | FNBP1L | Q5T0N5 | FBP1L_HUMAN | Unmodified | 3670300 | 4440600 | 0 | 0 | 1991800 | 0 | 364830 | 310470 | 867330 | 355670 | 0 | 299620 | 4614500 | 1416300 | 867640 | 0 | 635570 | 2025100 |
| AQQVAVQE | FLOT1; FLOT1; I | O75955; A0A | FLOT1_HUMAN | Unmodified | 0 | 0 | 417990 | 316710 | 0 | 42554 | 424540 | 127600 | 3003300 | 507480 | 360860 | 2244400 | 56534 | 0 | 743600 | 0 | 0 | 0 |
| AQQVAVQE | FLOT1; FLOT1; I | O75955; A0A | FLOT1_HUMAN | Unmodified | 210770 | 207690 | 2668500 | 1493700 | 0 | 0 | 20440 | 21701 | 7570000 | 327190 | 654000 | 7672100 | 39398 | 0 | 3815300 | 2579100 | 0 | 0 |
| AQQVAVQE | FLOT1; FLOT1; I | O75955; A0A | FLOT1_HUMAN | Unmodified | 0 | 0 | 1304200 | 1512500 | 4092900 | 898650 | 0 | 74578 | 3726400 | 1852400 | 0 | 80660 | 2003700 | 192790 | 745950 | 38798 | 441210 | 1524200 |
| AQQVLHLQ | LZTS1 | Q9Y250 | LZTS1_HUMAN | Unmodified | 0 | 0 | 0 | 0 | 0 | 0 | 0 | 0 | 0 | 0 | 0 | 711180 | 0 | 0 | 0 | 0 | 0 | 0 |
| AQQVSQGI | VCL | P18206 | VINC_HUMAN ' | Unmodified | 0 | 605500 | 620210 | 0 | 0 | 0 | 0 | 0 | 0 | 136520 | 0 | 7630900 | 0 | 0 | 1980700 | 1100100 | 0 | 0 |
| AQQVSQGI | VCL | P18206 | VINC_HUMAN ' | Unmodified | 0 | 0 | 0 | 0 | 691110 | 431130 | 398810 | 0 | 0 | 1265100 | 0 | 2160600 | 0 | 0 | 0 | 0 | 6137900 | 0 |
| AQRAFPNP | LANCL1; LANCI | O43813; E9F | LANC1_HUMAI | Acetyl (Protein N | 0 | 0 | 0 | 0 | 425170 | 0 | 0 | 238340 | 0 | 0 | 0 | 0 | 0 | 0 | 0 | 125300 | 372320 | 0 |
| AQREHGVL | DDAH2; DDAH2 | O95865; Q5! | DDAH2_HUMA | Unmodified | 0 | 0 | 0 | 62916 | 622270 | 0 | 126690 | 147830 | 214020 | 104580 | 24693 | 44050 | 253060 | 141800 | 32334 | 19896 | 457770 | 997590 |
| AQRIEYDCI | RCC2 | Q9P258 | RCC2_HUMAN | Unmodified | 43229 | 0 | 0 | 66606 | 0 | 0 | 0 | 48951 | 1113500 | 311990 | 0 | 9845000 | 0 | 0 | 0 | 0 | 0 | 0 |
| AQRNNEDI | SSR4; SSR4 | P51571; A6N | SSRD_HUMAN | Unmodified | 0 | 0 | 0 | 0 | 0 | 0 | 0 | 0 | 1836000 | 0 | 0 | 0 | 0 | 0 | 0 | 0 | 0 | 0 |
| AQSDGIW( | IDI1 | Q13907 | IDI1_HUMAN Is | Unmodified | 0 | 0 | 0 | 0 | 0 | 0 | 1379700 | 0 | 0 | 0 | 0 | 1862800 | 0 | 0 | 0 | 602080 | 282420 | 502780 |
| AQSINITELI | PFDN5; PFDN5 | Q99471; H3I | PFD5_HUMAN | Acetyl (Protein N | 0 | 0 | 0 | 0 | 1298800 | 140430 | 0 | 0 | 118300 | 437050 | 0 | 0 | 2698500 | 834620 | 0 | 0 | 1572600 | 774590 |
| AQSLLSTDI | PSMD11; PSMD | O00231; J3Q | PSD11_HUMAI | Unmodified | 0 | 0 | 17466 | 0 | 0 | 0 | 0 | 0 | 0 | 0 | 339980 | 2405800 | 0 | 0 | 0 | 0 | 0 | 0 |
| AQSLLSTDI | PSMD11; PSMD | O00231; J3Q | PSD11_HUMAI | Unmodified | 0 | 0 | 284470 | 879620 | 0 | 0 | 0 | 0 | 1929900 | 0 | 1444800 | 8200600 | 331610 | 0 | 527000 | 399080 | 0 | 0 |
| AQSLPSDQ | PAK6 | Q9NQU5 | PAK6_HUMAN | Unmodified | 0 | 0 | 0 | 0 | 965160 | 0 | 0 | 0 | 0 | 0 | 0 | 0 | 0 | 0 | 0 | 0 | 0 | 973180 |
| AQSLTTGQ | LCK; LCK; LCK; | P06239; E9P | LCK_HUMAN T | Unmodified | 0 | 0 | 0 | 59500 | 0 | 0 | 65234 | 0 | 28830 | 146920 | 0 | 1343700 | 128570 | 304950 | 0 | 0 | 0 | 0 |
| AQSNPYYN | LMO7; LMO7; LI | Q8WWI1; E9 | LMO7_HUMAN | Unmodified | 0 | 0 | 0 | 0 | 0 | 0 | 0 | 0 | 0 | 0 | 0 | 0 | 0 | 0 | 0 | 0 | 798670 | 542670 |
| AQSQTDRV | HLA-A; HLA-A; I | P04439; Q5S | HLAA_HUMAN | Unmodified | 0 | 0 | 0 | 0 | 0 | 0 | 0 | 0 | 35854000 | 1308300 | 0 | 0 | 0 | 0 | 0 | 0 | 0 | 0 |
| AQTAELFAI | DNAJC13 | O75165 | DJC13_HUMAN | Unmodified | 0 | 0 | 63299 | 152490 | 0 | 915170 | 297430 | 0 | 0 | 37665 | 63826 | 3498000 | 416760 | 294520 | 0 | 154110 | 172830 | 520340 |
| AQTEGINIS | RUVBL1; RUVBI | Q9Y265; H7( | RUVB1_HUMAI | Unmodified | 0 | 0 | 0 | 0 | 0 | 0 | 0 | 0 | 0 | 0 | 1676200 | 13719000 | 632910 | 0 | 0 | 0 | 0 | 0 |
| AQTEWNT( | SCAMP4 | Q969E2 | SCAM4_HUMA | Unmodified | 1310800 | 1694100 | 6312400 | 1399300 | 0 | 0 | 0 | 0 | 0 | 161800 | 0 | 915210 | 0 | 0 | 4224100 | 4667000 | 0 | 0 |
| AQTEWNT( | SCAMP4 | Q969E2 | SCAM4_HUMA | Unmodified | 6349900 | 7252300 | 14544000 | 16501000 | 332810 | 117850 | 0 | 0 | 2759000 | 0 | 297350 | 1572500 | 538550 | 182140 | 16421000 | 14971000 | 81241 | 68386 |
| AQTGEVAA | DIP2A; DIP2A | Q14689; A0A | DIP2A_HUMAN | Unmodified | 0 | 0 | 0 | 0 | 0 | 0 | 0 | 0 | 0 | 0 | 0 | 789920 | 0 | 0 | 0 | 0 | 0 | 0 |
| AQTKEQAD | MYH14; MYH14 | Q7Z406; M0( | MYH14_HUMAI | Unmodified | 0 | 0 | 0 | 0 | 905040 | 205450 | 575450 | 135590 | 0 | 307800 | 0 | 0 | 0 | 249130 | 0 | 0 | 793980 | 119720 |
| AQTLPTSV\ | SPTBN1; SPTBN | Q01082; A0A | SPTB2_HUMAN | Unmodified | 147440 | 0 | 0 | 0 | 599860 | 0 | 0 | 0 | 556150 | 0 | 0 | 3310400 | 0 | 449060 | 0 | 0 | 0 | 0 |
| AQTMSNS( | AIMP1 | Q12904 | AIMP1_HUMAN | Unmodified | 0 | 0 | 63423 | 0 | 0 | 171940 | 0 | 0 | 238550 | 0 | 0 | 1290200 | 299560 | 0 | 0 | 0 | 0 | 0 |
| AQTYELQE! | SEPTIN10; SEPT | Q9P0V9; B5I | SEP10_HUMAN | Unmodified | 0 | 0 | 0 | 0 | 3765800 | 393780 | 52886 | 106580 | 0 | 120220 | 0 | 287290 | 0 | 0 | 105730 | 79722 | 154780 | 0 |
| AQVADVVV | FASN; FASN | P49327; A0A | FAS_HUMAN F | Unmodified | 717490 | 755770 | 2060900 | 3979000 | 0 | 0 | 0 | 296780 | 1575500 | 872160 | 6879800 | 1.16E+08 | 0 | 224040 | 4935800 | 4905500 | 214370 | 631200 |
| AQVAMSTL | GTF2I; GTF2I; G | P78347; A0A | GTF2I_HUMAN | Acetyl (Protein N | 0 | 0 | 0 | 0 | 0 | 0 | 0 | 0 | 0 | 0 | 0 | 1089700 | 0 | 0 | 0 | 0 | 0 | 0 |
| AQVARPGC | HINT1; HINT1; I | P49773; H0Y | HINT1_HUMAN | Unmodified | 0 | 0 | 604230 | 0 | 5033100 | 568910 | 3208800 | 1344700 | 2375700 | 1633800 | 498780 | 73351000 | 8004200 | 5299000 | 959530 | 1286200 | 16164000 | 2274900 |
| AQVDPLVQ | PTPRB; PTPRB; | P23467; F8V | PTPRB_HUMAN | Unmodified | 855560 | 927990 | 0 | 0 | 0 | 0 | 0 | 0 | 0 | 0 | 0 | 0 | 0 | 0 | 0 | 0 | 0 | 0 |
| AQVEAVHK | EPB41L5 | Q9HCM4 | E41L5_HUMAN | Unmodified | 0 | 0 | 0 | 0 | 474620 | 0 | 0 | 0 | 0 | 0 | 0 | 0 | 0 | 217940 | 0 | 0 | 367540 | 972310 |
| AQVEEFLA( | PLOD1 | Q02809 | PLOD1_HUMAI | Unmodified | 0 | 0 | 316410 | 209980 | 142790 | 302540 | 50789 | 33941 | 24304000 | 4057800 | 1177700 | 412950 | 333670 | 167610 | 175700 | 305290 | 0 | 1042700 |
| AQVEEFLA( | PLOD1 | Q02809 | PLOD1_HUMAI | Unmodified | 0 | 0 | 0 | 0 | 0 | 0 | 0 | 0 | 2634400 | 967210 | 0 | 0 | 471020 | 0 | 0 | 0 | 0 | 0 |
| AQVEQELT | PLEC | Q15149 | PLEC_HUMAN | Unmodified | 0 | 0 | 0 | 199040 | 3495600 | 0 | 95924 | 0 | 0 | 0 | 0 | 0 | 0 | 0 | 0 | 0 | 5418200 | 5546000 |
| AQVIYTR | RPS25 | P62851 | RS25_HUMAN | Unmodified | 4115100 | 3786600 | 33604 | 645620 | 0 | 0 | 29924 | 0 | 422930 | 55456 | 119250 | 10676000 | 665740 | 0 | 809180 | 1094300 | 298410 | 0 |
| AQVLATIH( | PROSER2; PRO: | Q86WR7; D3 | PRSR2_HUMAN | Unmodified | 0 | 0 | 0 | 0 | 984850 | 0 | 0 | 0 | 0 | 0 | 0 | 0 | 0 | 0 | 0 | 0 | 0 | 0 |
| AQVLVEDIS | ARHGEF16; ARI | Q5VV41; B0( | ARHGG_HUMA | Unmodified | 0 | 0 | 0 | 0 | 0 | 0 | 0 | 0 | 3872000 | 509350 | 75784 | 0 | 0 | 0 | 55155 | 0 | 0 | 0 |
| AQVPFEQIL | CIB1 | Q99828 | CIB1_HUMAN ( | Unmodified | 0 | 132050 | 63522 | 21921 | 139220 | 73890 | 0 | 0 | 159970 | 0 | 34423 | 128930 | 0 | 0 | 0 | 27873 | 100490 | 0 |
| AQVQVESQ | TXNDC11 | Q6PKC3 | TXD11_HUMAN | Unmodified | 0 | 0 | 0 | 0 | 0 | 0 | 0 | 0 | 383680 | 0 | 115560 | 0 | 0 | 0 | 0 | 0 | 0 | 0 |
| AQVSLNVL | UTRN | P46939 | UTRN_HUMAN | Unmodified | 0 | 0 | 413930 | 223940 | 0 | 0 | 0 | 0 | 403400 | 0 | 0 | 0 | 0 | 0 | 1004300 | 1415100 | 0 | 0 |
| AQVSLNVL | UTRN | P46939 | UTRN_HUMAN | Unmodified | 0 | 0 | 0 | 0 | 2363000 | 0 | 0 | 0 | 0 | 2947900 | 0 | 0 | 0 | 2748700 | 0 | 0 | 1506600 | 1208100 |
| AQVTGDP( | IQGAP3; IQGAP | Q86VI3; F2Z | IQGA3_HUMAN | Unmodified | 0 | 0 | 0 | 0 | 2028300 | 0 | 0 | 0 | 1002700 | 447030 | 225160 | 1637500 | 0 | 0 | 0 | 0 | 0 | 0 |
| AQVTSLLGI | IKBKG; IKBKG; I | Q9Y6K9; C9J | NEMO_HUMAN | Unmodified | 0 | 0 | 0 | 0 | 9289100 | 0 | 0 | 0 | 0 | 609050 | 0 | 0 | 3904900 | 0 | 0 | 0 | 0 | 0 |
| AQVVSNLK | TLN1 | Q9Y490 | TLN1_HUMAN | Unmodified | 0 | 0 | 0 | 0 | 0 | 0 | 0 | 44674 | 401400 | 57461 | 33371 | 1963000 | 86554 | 0 | 711500 | 678100 | 84144 | 80202 |
| AQYDELAR | KRT18; KRT18 | P05783; F8V | K1C18_HUMAN | Unmodified | 0 | 302220 | 176250 | 186760 | 0 | 0 | 0 | 0 | 0 | 0 | 1076100 | 861070 | 0 | 0 | 0 | 161090 | 0 | 0 |
| AQYDELAR | KRT18; KRT18 | P05783; F8V | K1C18_HUMAN | Unmodified | 404310 | 0 | 582120 | 0 | 742600 | 0 | 0 | 52527 | 68124 | 0 | 73590 | 1036400 | 0 | 0 | 645650 | 0 | 663510 | 151810 |
| AQYEDIAQI | KRT1 | P04264 | K2C1_HUMAN | Unmodified | 5299600 | 4413700 | 2537700 | 2707300 | 4559300 | 0 | 0 | 924270 | 9409300 | 662520 | 1040900 | 5443500 | 1163600 | 815330 | 3989800 | 0 | 1177400 | 2139400 |
| AQYEQTLA | PACSIN3; PACS | Q9UKS6; E9I | PACN3_HUMAI | Unmodified | 41404000 | 38302000 | 14569000 | 7467200 | 801690 | 333050 | 0 | 0 | 1434100 | 0 | 0 | 0 | 1088000 | 0 | 37681000 | 26858000 | 0 | 0 |
| AQYEQTLA | PACSIN3; PACS | Q9UKS6; E9I | PACN3_HUMAI | Unmodified | 21988000 | 13217000 | 9617600 | 14203000 | 1280600 | 0 | 0 | 0 | 0 | 48667 | 0 | 0 | 13380000 | 1328800 | 12276000 | 4211000 | 574440 | 667470 |
| AQYVRLYP( | EDIL3 | O43854 | EDIL3_HUMAN | Unmodified | 0 | 0 | 0 | 0 | 33548000 | 17134000 | 2061100 | 662640 | 0 | 2856900 | 0 | 0 | 0 | 0 | 0 | 0 | 3286700 | 12622000 |

| | | | | | | | | | | | | | | | | | | | | | | |
|---|---|---|---|---|---|---|---|---|---|---|---|---|---|---|---|---|---|---|---|---|---|---|
| AQYYLPDG | ACTR1A; ACTR1 | P61163; R4G | ACTZ_HUMAN | Unmodified | 0 | 0 | 0 | 0 | 0 | 0 | 0 | 0 | 0 | 0 | 0 | 1299400 | 0 | 0 | 0 | 13883 | 0 | 0 |
| ARAEAEQM | FLOT1; FLOT1; | O75955; A0A | FLOT1_HUMAN | Unmodified | 34318 | 0 | 0 | 0 | 1860600 | 0 | 0 | 95486 | 50176 | 276600 | 22685 | 552180 | 0 | 0 | 598420 | 0 | 192220 | 467120 |
| ARAEAQEA | PCSK1N | Q9UHG2 | PCS1N_HUMAI | Unmodified | 701900 | 0 | 0 | 0 | 0 | 0 | 0 | 0 | 9898500 | 1324200 | 0 | 0 | 0 | 0 | 0 | 0 | 0 | 259820 |
| ARAHSIQIN | RPL18A; RPL18 | Q02543; M0 | RL18A_HUMAN | Unmodified | 0 | 177770 | 0 | 60649 | 0 | 0 | 194950 | 0 | 0 | 0 | 0 | 628740 | 65575 | 0 | 0 | 0 | 0 | 0 |
| ARASFPDQ | SIT1; SIT1 | Q9Y3P8; A0A | SIT1_HUMAN S | Unmodified | 0 | 0 | 0 | 0 | 0 | 0 | 0 | 0 | 367820 | 546330 | 0 | 0 | 0 | 0 | 0 | 0 | 0 | 0 |
| ARAYGPGII | FLNA; FLNA; FL | P21333; A0A | FLNA_HUMAN | Unmodified | 74438 | 193870 | 0 | 290700 | 2662600 | 690400 | 3235000 | 895330 | 0 | 234920 | 0 | 0 | 906080 | 1191200 | 0 | 0 | 1903600 | 422710 |
| ARAYGPGII | FLNA; FLNA; FL | P21333; A0A | FLNA_HUMAN | Oxidation (M) | 0 | 0 | 0 | 0 | 398260 | 0 | 616500 | 297340 | 0 | 12186 | 0 | 0 | 0 | 0 | 0 | 0 | 386250 | 0 |
| ARAYGPGII | FLNA; FLNA; FL | P21333; A0A | FLNA_HUMAN | Unmodified | 0 | 0 | 0 | 0 | 1405600 | 538950 | 1060300 | 0 | 0 | 0 | 0 | 0 | 0 | 0 | 0 | 0 | 661730 | 178730 |
| ARCAELEE | CIT; CIT; CIT; CI | O14578; A0A | CTRO_HUMAN | Unmodified | 0 | 0 | 0 | 0 | 3103700 | 504480 | 676180 | 227670 | 0 | 184520 | 0 | 0 | 472600 | 0 | 0 | 0 | 1810800 | 572860 |
| ARDCLIPM | ACAA1; ACAA1; | P09110; C9J | THIK_HUMAN 3 | Unmodified | 0 | 0 | 40398 | 0 | 0 | 0 | 0 | 0 | 1502700 | 0 | 0 | 0 | 0 | 0 | 0 | 0 | 0 | 0 |
| ARDCLIPM | ACAA1; ACAA1; | P09110; C9J | THIK_HUMAN 3 | Oxidation (M) | 0 | 0 | 0 | 0 | 0 | 0 | 64851 | 0 | 1348900 | 0 | 0 | 0 | 0 | 0 | 0 | 0 | 0 | 0 |
| ARDDLITDL | ATP6V1E1; ATP | P36543; C9J | VATE1_HUMAN | Unmodified | 0 | 0 | 0 | 0 | 0 | 0 | 679890 | 0 | 0 | 229230 | 1652500 | 2802000 | 768230 | 399980 | 0 | 0 | 0 | 0 |
| ARDGNDYS | EPB41L5 | Q9HCM4 | E41L5_HUMAN | Unmodified | 45901 | 0 | 0 | 0 | 0 | 0 | 0 | 0 | 0 | 0 | 0 | 0 | 0 | 0 | 939350 | 459600 | 0 | 0 |
| ARDHGLLI | MRPS18B | Q9Y676 | RT18B_HUMAN | Unmodified | 0 | 0 | 0 | 0 | 0 | 0 | 0 | 0 | 794490 | 0 | 0 | 0 | 0 | 0 | 0 | 0 | 0 | 0 |
| ARDMDLTE | CTNND1; CTNN | O60716; A0A | CTND1_HUMAI | Oxidation (M) | 0 | 0 | 0 | 250250 | 760950 | 0 | 0 | 0 | 0 | 0 | 0 | 0 | 566010 | 0 | 5256300 | 0 | 0 | 554190 |
| ARDPHSGI | CDK4; CDK4; C | P11802; F8V | CDK4_HUMAN | Unmodified | 210710 | 0 | 0 | 0 | 0 | 0 | 0 | 0 | 0 | 0 | 0 | 0 | 0 | 184660 | 0 | 0 | 135540 | 333030 |
| ARDRHFAC | CHID1; CHID1; | Q9BWS9; E9 | CHID1_HUMAN | Unmodified | 0 | 0 | 0 | 0 | 0 | 0 | 0 | 0 | 1696400 | 7630500 | 0 | 0 | 0 | 0 | 0 | 0 | 0 | 0 |
| ARDYDAM | RUVBL2; RUVBI | Q9Y230; M0 | RUVB2_HUMAI | Unmodified | 0 | 0 | 0 | 0 | 0 | 0 | 0 | 0 | 0 | 0 | 0 | 0 | 0 | 0 | 0 | 0 | 0 | 0 |
| ARDYDHLF | RAB35; RAB35; | Q15286; F5H | RAB35_HUMAN | Acetyl (Protein N | 0 | 3269500 | 3860800 | 3147700 | 4870600 | 1505200 | 3128000 | 0 | 0 | 719350 | 950170 | 3266600 | 0 | 1863600 | 5575400 | 4540300 | 1255800 | 806690 |
| ARDYDHLF | RAB35; RAB35 | Q15286; F5H | RAB35_HUMAN | Acetyl (Protein N | 0 | 0 | 0 | 0 | 308130 | 0 | 477570 | 9033.2 | 0 | 0 | 0 | 0 | 715390 | 290590 | 0 | 0 | 598940 | 222540 |
| AREAAIWEI | SLK | Q9H2G2 | SLK_HUMAN S | Unmodified | 0 | 0 | 139440 | 155610 | 0 | 0 | 0 | 0 | 0 | 0 | 0 | 2026600 | 74207 | 0 | 126440 | 323490 | 0 | 0 |
| AREAAIWEI | SLK | Q9H2G2 | SLK_HUMAN S | Unmodified | 0 | 0 | 0 | 0 | 1208800 | 173990 | 0 | 0 | 535520 | 0 | 0 | 0 | 4085000 | 0 | 0 | 0 | 140800 | 400820 |
| AREDEARLI | SLC9A3R2; SLC | Q15599; H3I | NHRF2_HUMAI | Unmodified | 0 | 0 | 0 | 0 | 14083000 | 0 | 0 | 0 | 0 | 742890 | 0 | 630870 | 9391600 | 1381300 | 554510 | 423240 | 10889000 | 9687200 |
| AREDIFME | AZGP1; AZGP1 | P25311; C9J | ZA2G_HUMAN | Unmodified | 0 | 0 | 120660 | 0 | 0 | 0 | 0 | 0 | 0 | 0 | 1927900 | 310080 | 0 | 0 | 408460 | 0 | 0 | 0 |
| AREDIFME | AZGP1; AZGP1 | P25311; C9J | ZA2G_HUMAN | Oxidation (M) | 0 | 557780 | 885940 | 0 | 0 | 0 | 0 | 0 | 0 | 0 | 2181700 | 0 | 0 | 0 | 346000 | 0 | 0 | 0 |
| AREEFRPTI | GNA13 | Q14344 | GNA13_HUMAI | Unmodified | 181260 | 203690 | 0 | 0 | 0 | 0 | 0 | 0 | 627930 | 0 | 0 | 4622200 | 0 | 0 | 0 | 0 | 0 | 0 |
| AREGDCGI | CPD | O75976 | CBPD_HUMAN | Unmodified | 0 | 0 | 24267 | 0 | 0 | 0 | 0 | 0 | 5690800 | 0 | 0 | 0 | 0 | 0 | 0 | 0 | 0 | 0 |
| ARELYHEEI | SCIN; SCIN; SC | Q9Y6U3; C9 | ADSV_HUMAN | Acetyl (Protein N | 0 | 0 | 0 | 0 | 0 | 0 | 0 | 0 | 0 | 377590 | 0 | 0 | 0 | 0 | 0 | 0 | 0 | 0 |
| ARELYHEEI | SCIN; SCIN; SC | Q9Y6U3; C9 | ADSV_HUMAN | Acetyl (Protein N | 0 | 0 | 0 | 0 | 0 | 0 | 0 | 0 | 0 | 1445400 | 0 | 0 | 0 | 0 | 0 | 0 | 0 | 0 |
| ARENPSEE | SF3A3 | Q12874 | SF3A3_HUMAN | Unmodified | 453640 | 0 | 0 | 0 | 0 | 0 | 0 | 0 | 0 | 0 | 0 | 976540 | 0 | 0 | 0 | 0 | 0 | 0 |
| AREQAEAE | TPM3; TPM3; TF | D6R904; A0A | D6R904_HUMA | Unmodified | 1971300 | 60207 | 0 | 2531700 | 8458200 | 2042600 | 0 | 0 | 4064000 | 0 | 1797600 | 7912100 | 6532900 | 961030 | 1873400 | 148660 | 0 | 0 |
| AREQAEAE | TPM3; TPM3; TF | D6R904; A0A | D6R904_HUMA | Unmodified | 502380 | 977020 | 945310 | 1686600 | 3913600 | 0 | 0 | 0 | 2216600 | 260010 | 1086300 | 20781000 | 3640100 | 440190 | 2111100 | 0 | 572470 | 281640 |
| ARESYVETE | CLTC; CLTC; CL | Q00610; J3K | CLH1_HUMAN | Unmodified | 2035700 | 6464900 | 0 | 3145000 | 3079800 | 1045700 | 0 | 0 | 577770 | 0 | 0 | 4044200 | 8513200 | 3419000 | 3271000 | 2429000 | 1685000 | 2458300 |
| ARETVENLI | SLC38A10; SLC | Q9HBR0; H0 | S38AA_HUMAN | Unmodified | 4268600 | 2517100 | 0 | 0 | 0 | 0 | 0 | 297670 | 2129000 | 0 | 146500 | 11392000 | 0 | 0 | 0 | 0 | 0 | 0 |
| ARFAKPVYI | HSD17B4; HSD | P51659; E7E | DHB4_HUMAN | Oxidation (M) | 0 | 0 | 0 | 0 | 0 | 0 | 0 | 0 | 2550300 | 1321000 | 0 | 0 | 112700 | 0 | 0 | 0 | 111440 | 359850 |
| ARFAKPVYI | HSD17B4; HSD | P51659; E7E | DHB4_HUMAN | Unmodified | 0 | 0 | 0 | 0 | 0 | 0 | 0 | 0 | 8081100 | 1211900 | 530340 | 0 | 0 | 0 | 0 | 0 | 0 | 0 |
| ARFASEYG | MANBA; MANBA | O00462; A0A | MANBA_HUMA | Unmodified | 77661 | 0 | 0 | 0 | 0 | 0 | 0 | 0 | 891620 | 1082200 | 0 | 0 | 0 | 0 | 0 | 0 | 0 | 0 |
| ARFEELCSI | HSPA6; HSPA1A | P17066; P0D | HSP76_HUMAN | Unmodified | 9462200 | 5552400 | 3007600 | 5067300 | 0 | 282460 | 208340 | 28096 | 16860000 | 2321900 | 1241500 | 7760800 | 1899900 | 773260 | 3998700 | 5680200 | 189690 | 991530 |
| ARFEELCSI | HSPA6; HSPA1A | P17066; P0D | HSP76_HUMAN | Unmodified | 0 | 0 | 0 | 703440 | 23245000 | 2356400 | 3063000 | 0 | 614910 | 19552000 | 0 | 0 | 34828000 | 17773000 | 0 | 0 | 3108000 | 7069700 |
| ARFEELNAI | HSPA8; HSPA2; | P11142; P54 | HSP7C_HUMAI | Unmodified | 63034000 | 83401000 | 83456000 | 64772000 | 0 | 0 | 594690 | 217360 | 1.43E+08 | 5392400 | 27046000 | 3.3E+08 | 7506900 | 1809500 | 65750000 | 57476000 | 307590 | 1702200 |
| ARFEELNAI | HSPA8; HSPA8; | P11142; E9P | HSP7C_HUMAI | Unmodified | 20986000 | 32372000 | 40950000 | 89144000 | 1.9E+08 | 42939000 | 22290000 | 236230 | 1.32E+08 | 1.62E+08 | 50320000 | 27247000 | 3.4E+08 | 88298000 | 31561000 | 17602000 | 21504000 | 55900000 |
| ARFEELNAI | HSPA2 | P54652 | HSP72_HUMAN | Unmodified | 47458 | 316290 | 0 | 0 | 2615900 | 498550 | 0 | 0 | 0 | 0 | 0 | 0 | 0 | 0 | 0 | 0 | 295860 | 1652500 |
| ARFELLCSI | HSPA14 | Q0VDF9 | HSP7E_HUMAN | Unmodified | 129370 | 0 | 0 | 0 | 139940 | 0 | 85106 | 0 | 0 | 74575 | 0 | 202160 | 215650 | 0 | 0 | 0 | 0 | 106720 |
| ARFVVTDG | CACNA2D1; CA | P54289; H0Y | CA2D1_HUMAI | Unmodified | 426590 | 0 | 0 | 0 | 427640 | 2932100 | 0 | 0 | 884190 | 130480 | 0 | 0 | 719350 | 0 | 0 | 0 | 241570 | 459110 |
| ARFVVTDG | CACNA2D1; CA | P54289; H0Y | CA2D1_HUMAI | Unmodified | 0 | 0 | 0 | 0 | 0 | 207560 | 8355500 | 3280700 | 1417200 | 2465400 | 0 | 0 | 0 | 237130 | 0 | 0 | 0 | 0 |
| ARGADRMS | ATIC | P31939 | PUR9_HUMAN | Oxidation (M) | 0 | 0 | 0 | 0 | 2493100 | 43475 | 0 | 0 | 0 | 0 | 24787 | 0 | 2990000 | 2623100 | 0 | 0 | 1353300 | 592320 |
| ARGCPEDA | IGF2R; IGF2R; I | P11717; A0A | MPRI_HUMAN | Unmodified | 0 | 0 | 121380 | 0 | 0 | 0 | 83389 | 0 | 9783700 | 1209800 | 592510 | 904030 | 561480 | 796150 | 116130 | 178770 | 21451 | 103230 |
| ARGCPEDA | IGF2R; IGF2R; I | P11717; A0A | MPRI_HUMAN | Unmodified | 0 | 0 | 0 | 0 | 0 | 0 | 60494 | 271250 | 0 | 551370 | 0 | 0 | 0 | 0 | 0 | 0 | 0 | 0 |
| ARGGDHVI | SLC38A10 | Q9HBR0 | S38AA_HUMAN | Unmodified | 160950 | 0 | 0 | 0 | 0 | 0 | 0 | 0 | 217720 | 0 | 0 | 0 | 0 | 0 | 0 | 0 | 0 | 0 |
| ARGGDHVI | SLC38A10 | Q9HBR0 | S38AA_HUMAN | Unmodified | 0 | 0 | 0 | 0 | 0 | 0 | 0 | 0 | 5448200 | 0 | 0 | 0 | 197380 | 0 | 0 | 0 | 0 | 0 |
| ARGGNIGD | VCP; VCP; VCP; | P55072; C9J | TERA_HUMAN | Unmodified | 743020 | 984460 | 0 | 0 | 0 | 0 | 0 | 0 | 125580 | 16953 | 0 | 1170000 | 0 | 0 | 0 | 1152400 | 0 | 0 |
| ARGGNIGD | VCP; VCP; VCP; | P55072; C9J | TERA_HUMAN | Oxidation (M) | 0 | 0 | 0 | 0 | 2724500 | 0 | 0 | 0 | 0 | 574680 | 0 | 0 | 0 | 0 | 0 | 0 | 0 | 0 |
| ARGITINAA | TUFM | P49411 | EFTU_HUMAN | Unmodified | 0 | 0 | 0 | 0 | 0 | 0 | 109830 | 693890 | 787150 | 0 | 0 | 0 | 0 | 0 | 0 | 0 | 0 | 0 |
| ARGLEVTD | AARS1; AARS1; | P49588; A0A | SYAC_HUMAN | Unmodified | 0 | 0 | 0 | 0 | 0 | 0 | 0 | 0 | 0 | 0 | 0 | 409980 | 147780 | 20853 | 0 | 0 | 0 | 0 |
| ARGLPDNI | UGP2; UGP2; U | Q16851; A0A | UGPA_HUMAN | Unmodified | 204440 | 0 | 0 | 0 | 6353900 | 0 | 903480 | 461150 | 3755000 | 3803900 | 969950 | 4008900 | 3103300 | 990070 | 210350 | 0 | 1771300 | 2580600 |
| ARGLPDNI | UGP2; UGP2; U | Q16851; A0A | UGPA_HUMAN | Unmodified | 0 | 0 | 0 | 0 | 267110 | 0 | 501170 | 0 | 0 | 0 | 0 | 0 | 0 | 0 | 0 | 0 | 505480 | 0 |
| ARGSAIDEI | TOR1B | O14657 | TOR1B_HUMAN | Unmodified | 0 | 70076 | 0 | 0 | 0 | 0 | 0 | 0 | 1503400 | 0 | 0 | 0 | 0 | 0 | 0 | 0 | 0 | 0 |
| ARGTFDNA | DYNC1H1; DYN | Q14204; A0A | DYHC1_HUMA | Unmodified | 0 | 0 | 74341 | 0 | 0 | 0 | 49866 | 0 | 98397 | 76615 | 37037 | 972120 | 0 | 0 | 0 | 0 | 28024 | 44462 |
| ARGTGELT | FBP1 | P09467 | F16P1_HUMAN | Unmodified | 0 | 0 | 0 | 0 | 155570 | 0 | 0 | 0 | 593710 | 1149300 | 0 | 0 | 0 | 0 | 0 | 0 | 705760 | 49924 |
| ARGTRLDG | ALPL | P05186 | PPBT_HUMAN | Unmodified | 354090 | 0 | 0 | 0 | 0 | 0 | 0 | 0 | 2608300 | 739840 | 0 | 0 | 0 | 0 | 0 | 0 | 0 | 0 |
| ARGVCSDV | MUC5B | Q9HC84 | MUC5B_HUMA | Unmodified | 0 | 0 | 0 | 0 | 0 | 0 | 0 | 0 | 0 | 0 | 0 | 0 | 0 | 0 | 0 | 0 | 0 | 346930 |
| ARHEVLLIS | HDLBP; HDLBP | Q00341; A0A | VIGLN_HUMAN | Unmodified | 88596 | 0 | 0 | 0 | 0 | 0 | 36969 | 0 | 0 | 0 | 0 | 584330 | 0 | 0 | 0 | 0 | 0 | 0 |
| ARHPQLLY | CSNK1A1; CSN | P48729; A0A | KC1A_HUMAN | Unmodified | 0 | 0 | 835440 | 622590 | 274930 | 0 | 60814 | 67206 | 631430 | 1107000 | 131010 | 1166400 | 0 | 38540 | 2182600 | 796430 | 108950 | 170000 |
| ARIGSDPL | SLC9A1 | P19634 | SL9A1_HUMAN | Unmodified | 0 | 0 | 0 | 0 | 0 | 0 | 0 | 0 | 0 | 0 | 0 | 0 | 0 | 1485200 | 0 | 0 | 0 | 0 |
| ARIHAETAL | USP8; USP8 | P40818; A0A | UBP8_HUMAN | Unmodified | 0 | 0 | 0 | 0 | 336580 | 0 | 45673 | 0 | 0 | 0 | 0 | 0 | 0 | 0 | 0 | 0 | 645880 | 0 |
| ARIHQQELI | UTRN | P46939 | UTRN_HUMAN | Unmodified | 0 | 0 | 0 | 0 | 2043800 | 335150 | 0 | 0 | 0 | 1966700 | 0 | 0 | 0 | 0 | 0 | 0 | 1767300 | 1041100 |
| ARIPVIAVD | EHHADH | Q08426 | ECHP_HUMAN | Unmodified | 0 | 0 | 0 | 0 | 0 | 0 | 0 | 0 | 3011300 | 1255500 | 0 | 0 | 0 | 0 | 0 | 0 | 0 | 0 |
| ARIVAEEFL | ARPC4; ARPC4- | P59998; H7C | ARPC4_HUMAI | Unmodified | 0 | 0 | 0 | 0 | 1350500 | 0 | 0 | 0 | 110760 | 0 | 0 | 434450 | 770250 | 0 | 0 | 0 | 0 | 428050 |
| ARIVEYEKE | ATP5PD | O75947 | ATP5H_HUMAN | Unmodified | 0 | 0 | 791590 | 0 | 820660 | 0 | 391600 | 543860 | 202180 | 109360 | 0 | 0 | 0 | 0 | 0 | 0 | 0 | 0 |
| ARLAALAD | SPTAN1; SPTAN | Q13813; A0A | SPTN1_HUMAN | Unmodified | 0 | 0 | 83192 | 129720 | 3146600 | 1058300 | 848320 | 0 | 1959100 | 1399200 | 707020 | 0 | 522480 | 313700 | 203460 | 97555 | 2824700 | 1022700 |
| ARLAEQAEI | YWHAG | P61981 | 1433G_HUMAN | Oxidation (M) | 557460 | 246370 | 1196700 | 1944700 | 9698800 | 1618200 | 1600200 | 640530 | 1451300 | 2926400 | 271340 | 4012600 | 7753700 | 9128000 | 1263200 | 1379000 | 8720000 | 7081000 |
| ARLAEQAEI | YWHAG | P61981 | 1433G_HUMAN | Unmodified | 0 | 0 | 144630 | 638210 | 9991700 | 164580 | 1944300 | 0 | 1210100 | 1569500 | 1070900 | 1751800 | 6321600 | 6671400 | 1147100 | 333700 | 8374600 | 7669700 |
| ARLAEQAEI | YWHAG | P61981 | 1433G_HUMAN | 2 Oxidation (M) | 0 | 0 | 52786 | 307900 | 2945100 | 0 | 535340 | 0 | 184740 | 280490 | 290300 | 410040 | 8158000 | 2268500 | 228010 | 0 | 1316700 | 1694800 |
| ARLAEQAEI | YWHAH; YWHA | Q04917; F8W | 1433F_HUMAN | Oxidation (M) | 0 | 0 | 0 | 295050 | 948030 | 431830 | 0 | 146780 | 0 | 179380 | 245090 | 423130 | 705870 | 467820 | 40601 | 240880 | 278450 | 701110 |
| ARLAEQAEI | YWHAH; YWHA | Q04917; F8W | 1433F_HUMAN | Unmodified | 0 | 0 | 0 | 511760 | 7324900 | 1085700 | 362470 | 55978 | 1631200 | 1829900 | 532120 | 2127400 | 6349100 | 0 | 146190 | 52332 | 1642100 | 5907800 |
| ARLEIEPEW | FKBP3 | Q00688 | FKBP3_HUMAN | Unmodified | 0 | 2729700 | 0 | 0 | 2268500 | 1049800 | 587240 | 0 | 1837500 | 1487900 | 0 | 5720600 | 3148500 | 1246600 | 796160 | 0 | 876690 | 2413300 |
| ARLEIEPEW | FKBP3 | Q00688 | FKBP3_HUMAN | Unmodified | 461160 | 422070 | 278360 | 440070 | 7967900 | 0 | 0 | 0 | 553930 | 575270 | 0 | 0 | 0 | 0 | 0 | 420120 | 271810 | 1277500 |
| ARLEPFGD | GALNT12 | Q8IXK2 | GLT12_HUMAN | Unmodified | 0 | 0 | 0 | 0 | 0 | 44739 | 0 | 0 | 437700 | 141890 | 0 | 0 | 0 | 0 | 0 | 0 | 0 | 0 |
| ARLIYEDYV | RGS19 | P49795 | RGS19_HUMAI | Unmodified | 0 | 0 | 177700 | 0 | 758240 | 0 | 127600 | 47079 | 0 | 0 | 0 | 0 | 0 | 0 | 0 | 0 | 847760 | 2282700 |
| ARLLELFTD | BROX; BROX; B | Q5VW32; F5 | BROX_HUMAN | 2 Oxidation (M) | 242650 | 166900 | 0 | 482810 | 0 | 0 | 0 | 0 | 0 | 470680 | 0 | 0 | 434120 | 360640 | 0 | 122810 | 0 | 0 |
| ARLPQPEG | PTK7; PTK7 | Q13308; H0Y | PTK7_HUMAN | Unmodified | 0 | 0 | 360170 | 0 | 2285900 | 955350 | 734980 | 452030 | 1926500 | 969030 | 411260 | 1770400 | 512430 | 1873800 | 957720 | 605650 | 1351000 | 2877000 |
| ARLPQPEG | PTK7; PTK7 | Q13308; H0Y | PTK7_HUMAN | Unmodified | 0 | 0 | 0 | 0 | 190730 | 0 | 243960 | 148020 | 0 | 0 | 0 | 0 | 0 | 328200 | 0 | 0 | 699960 | 0 |
| ARLSGYEE | CKAP5; CKAP5 | Q14008; E9F | CKAP5_HUMAI | Unmodified | 0 | 0 | 0 | 0 | 608100 | 0 | 230140 | 169700 | 0 | 346670 | 0 | 758170 | 650810 | 289330 | 75334 | 0 | 0 | 0 |
| ARLSNLAL | PDCD5; PDCD5 | O14737; K7E | PDCD5_HUMA | Unmodified | 0 | 0 | 0 | 0 | 229000 | 52076 | 0 | 0 | 0 | 0 | 0 | 0 | 813050 | 0 | 0 | 0 | 44939 | 66530 |
| ARLTQKYQ | VPS37B; VPS37 | Q9H9H4; F5 | VP37B_HUMAN | Unmodified | 0 | 0 | 0 | 0 | 2194300 | 0 | 0 | 0 | 0 | 406450 | 0 | 0 | 7959300 | 6841300 | 0 | 0 | 924210 | 1753600 |
| ARLVVAVEI | VANGL1 | Q8TAA9 | VANG1_HUMAI | Unmodified | 0 | 0 | 0 | 0 | 569690 | 55549 | 118670 | 0 | 0 | 75617 | 0 | 0 | 129500 | 98645 | 0 | 0 | 0 | 195210 |
| ARMQEAM | VCL | P18206 | VINC_HUMAN | 2 Oxidation (M) | 0 | 0 | 0 | 0 | 0 | 0 | 0 | 0 | 0 | 0 | 0 | 0 | 0 | 0 | 0 | 0 | 1042500 | 0 |
| ARMQEAM | VCL | P18206 | VINC_HUMAN | Oxidation (M) | 0 | 241030 | 0 | 0 | 0 | 0 | 0 | 0 | 0 | 0 | 0 | 0 | 0 | 0 | 0 | 157300 | 3327000 | 0 |
| ARMQEAM | VCL | P18206 | VINC_HUMAN | Unmodified | 0 | 0 | 0 | 0 | 207950 | 0 | 0 | 56784 | 0 | 0 | 0 | 0 | 0 | 0 | 0 | 0 | 1098100 | 956460 |
| ARNEYLLAI | SRGAP2; SRGA | O75044; P0D | SRGP2_HUMAI | Unmodified | 0 | 0 | 0 | 0 | 5032300 | 513440 | 1219300 | 0 | 512010 | 697500 | 0 | 2610200 | 4439300 | 1993800 | 0 | 0 | 1093600 | 1271700 |
| ARNGAGW | ITGB4 | P16144 | ITB4_HUMAN I | Unmodified | 0 | 0 | 0 | 0 | 0 | 0 | 0 | 0 | 0 | 0 | 0 | 0 | 0 | 0 | 0 | 0 | 1598300 | 0 |
| ARNLAYDT | PLOD1 | Q02809 | PLOD1_HUMAI | Unmodified | 0 | 0 | 0 | 100880 | 0 | 0 | 0 | 0 | 1060900 | 328440 | 0 | 0 | 0 | 0 | 0 | 0 | 0 | 0 |
| ARNQITQLI | CEP55 | Q53EZ4 | CEP55_HUMAN | Unmodified | 0 | 890350 | 2713100 | 3677800 | 37840000 | 6965600 | 2801300 | 656610 | 13680000 | 12755000 | 824700 | 4560800 | 2186600 | 3653900 | 5306900 | 312950 | 2277100 | 2305600 |
| ARPAEVGG | DUT; DUT; DUT | P33316; H0Y | DUT_HUMAN D | Unmodified | 0 | 0 | 0 | 0 | 0 | 0 | 0 | 0 | 0 | 95789 | 493400 | 1614100 | 0 | 0 | 0 | 0 | 0 | 385000 |
| ARPDDEEG | FLVCR1 | Q9Y5Y0 | FLVC1_HUMAN | Unmodified | 0 | 289200 | 896270 | 1172400 | 7603100 | 1011300 | 1537300 | 1056700 | 2183700 | 1463800 | 1377000 | 3670600 | 4566200 | 2520100 | 804730 | 855610 | 2531700 | 3556000 |
| ARPDDEEG | FLVCR1 | Q9Y5Y0 | FLVC1_HUMAN | Acetyl (Protein N | 0 | 0 | 0 | 1165800 | 2887400 | 465300 | 2160700 | 1468200 | 1003200 | 1938300 | 0 | 1816700 | 3031200 | 2786500 | 436280 | 1135800 | 1510800 | 1834600 |
| ARPEDVISE | MPST | P25325 | THTM_HUMAN | Unmodified | 0 | 0 | 0 | 0 | 0 | 57751 | 0 | 0 | 0 | 0 | 169170 | 1301800 | 0 | 0 | 0 | 0 | 0 | 43887 |
| ARPEDVVN | PARVB; PARVB | Q9HBI1; A0A | PARVB_HUMAN | Unmodified | 418740 | 496290 | 739090 | 1379500 | 113110 | 0 | 794930 | 515820 | 0 | 0 | 0 | 657680 | 695200 | 36401 | 1003200 | 1159800 | 131930 | 516080 |
| ARPEYMLP | PYGB | P11216 | PYGB_HUMAN | Oxidation (M) | 0 | 0 | 0 | 0 | 0 | 0 | 299060 | 70082 | 995740 | 0 | 0 | 491680 | 0 | 0 | 205600 | 0 | 0 | 45972 |
| ARPEYMLP | PYGB | P11216 | PYGB_HUMAN | Unmodified | 0 | 0 | 0 | 0 | 0 | 0 | 0 | 0 | 0 | 0 | 0 | 0 | 0 | 0 | 0 | 0 | 0 | 417030 |
| ARPFFNEF | ROBO1; ROBO1 | Q9Y6N7; A0A | ROBO1_HUMA | Unmodified | 0 | 0 | 0 | 0 | 0 | 0 | 144070 | 10254 | 526810 | 333340 | 0 | 0 | 0 | 0 | 0 | 0 | 159660 | 1648500 |
| ARPFPDGL | EEF2 | P13639 | EF2_HUMAN El | Unmodified | 0 | 0 | 487840 | 0 | 859640 | 81286 | 71687 | 0 | 3569200 | 264560 | 7319700 | 49159000 | 0 | 0 | 3386100 | 1829300 | 719650 | 0 |
| ARPGGNR | ADSS2 | P30520 | PURA2_HUMAI | Unmodified | 0 | 0 | 0 | 0 | 2177100 | 138780 | 147290 | 0 | 73777 | 123430 | 1544800 | 580570 | 1321600 | 0 | 0 | 0 | 68808 | 1527600 |
| ARPPGPHS | EFNB3 | Q15768 | EFNB3_HUMAN | Unmodified | 262130 | 287450 | 0 | 0 | 0 | 0 | 0 | 0 | 0 | 0 | 0 | 0 | 0 | 0 | 0 | 0 | 0 | 0 |
| ARPSQFPE | KIF2A; KIF2A | A0A6Q8PH5 | A0A6Q8PH57_ | Unmodified | 0 | 0 | 21688 | 49552 | 394680 | 39123 | 0 | 0 | 195860 | 0 | 63878 | 166140 | 307360 | 108460 | 24079 | 28714 | 0 | 58159 |
| ARQAAKTE | PSMA3; PSMA3 | P25788; G3V | PSA3_HUMAN | Unmodified | 0 | 0 | 0 | 725280 | 399790 | 1374800 | 0 | 13317 | 0 | 0 | 191330 | 45015 | 548130 | 2617500 | 16309 | 0 | 3976100 | 6777100 |
| ARQAAPCV | VCP; VCP; VCP; | P55072; C9J | TERA_HUMAN | Unmodified | 63380 | 0 | 0 | 2230900 | 21086000 | 4425700 | 7572500 | 89534 | 5727100 | 7455900 | 882620 | 2044200 | 26756000 | 35122000 | 0 | 0 | 6362200 | 4671700 |
| ARQEELYSI | PLEC; PLEC | Q15149; H0Y | PLEC_HUMAN | Unmodified | 0 | 0 | 0 | 0 | 175810 | 0 | 0 | 0 | 0 | 32408 | 0 | 0 | 0 | 0 | 0 | 0 | 1223500 | 1368600 |
| ARQEFEERI | SMC1A; SMC1A | Q14683; G8J | SMC1A_HUMA | Unmodified | 0 | 0 | 0 | 0 | 0 | 0 | 0 | 0 | 0 | 0 | 50574 | 244160 | 0 | 0 | 0 | 0 | 0 | 0 |

| | | | | | | | | | | | | | | | | | | | | | | |
|---|---|---|---|---|---|---|---|---|---|---|---|---|---|---|---|---|---|---|---|---|---|---|
| ARQENGMI | PAK4; PAK4 | O96013; M0 | PAK4_HUMAN | Unmodified | 333930 | 438700 | 456060 | 616890 | 76166 | 0 | 0 | 0 | 115830 | 0 | 126450 | 599780 | 0 | 0 | 531710 | 668890 | 0 | 191780 |
| ARQLLDQV | TMED4 | Q7Z7H5 | TMED4_HUMAI | Unmodified | 0 | 0 | 0 | 212110 | 0 | 0 | 0 | 0 | 80886000 | 2231200 | 2245200 | 725090 | 0 | 0 | 570740 | 259930 | 0 | 0 |
| ARQLLDQV | TMED4 | Q7Z7H5 | TMED4_HUMAI | Unmodified | 0 | 0 | 0 | 0 | 0 | 0 | 0 | 0 | 27983000 | 1216500 | 204380 | 1176700 | 0 | 0 | 0 | 0 | 0 | 0 |
| ARQPKPFF | PTGFRN | Q9P2B2 | FPRP_HUMAN | Unmodified | 1446400 | 313100 | 429180 | 6306200 | 4439900 | 1267100 | 1517400 | 0 | 0 | 0 | 0 | 4759200 | 15584000 | 10016000 | 1791800 | 1536100 | 265140 | 1416700 |
| ARQPKPFF | PTGFRN | Q9P2B2 | FPRP_HUMAN | Oxidation (M) | 3208800 | 1511600 | 1931700 | 9423000 | 2938000 | 0 | 1622800 | 81094 | 0 | 0 | 0 | 6077300 | 17206000 | 8055500 | 4379500 | 6625600 | 355410 | 362360 |
| ARQPLSEA | ANLN; ANLN | Q9NQW6; C | ANLN_HUMAN | Unmodified | 0 | 0 | 0 | 0 | 2873600 | 232650 | 218160 | 109220 | 0 | 88129 | 0 | 0 | 137560 | 0 | 0 | 0 | 1875400 | 310240 |
| ARQVLEHL | SNF8 | Q96H20 | SNF8_HUMAN | Unmodified | 70570 | 90647 | 0 | 45161 | 0 | 0 | 0 | 0 | 0 | 0 | 0 | 0 | 0 | 0 | 343350 | 180600 | 0 | 0 |
| ARQYPWG | SEPTIN6; SEPTI | Q14141; B1 | SEPT6_HUMAN | Unmodified | 0 | 0 | 0 | 165030 | 1007500 | 0 | 0 | 0 | 1054200 | 0 | 0 | 0 | 0 | 0 | 0 | 0 | 747070 | 0 |
| ARQYPWG | SEPTIN8; SEPTI | Q92599; Q9 | SEPT8_HUMAN | Unmodified | 0 | 0 | 0 | 0 | 20952000 | 9868400 | 873550 | 0 | 0 | 942980 | 0 | 1157800 | 1786900 | 0 | 0 | 0 | 2298200 | 8644000 |
| ARQYTSPEI | PDAP1; PDAP1 | Q13442; F8 | HAP28_HUMAI | Unmodified | 0 | 0 | 0 | 0 | 246510 | 0 | 0 | 634140 | 0 | 0 | 0 | 0 | 165300 | 0 | 0 | 0 | 1405000 | 316260 |
| ARRPESALI | VGF; VGF | O15240; Q0 | VGF_HUMAN | Unmodified | 256430 | 252400 | 0 | 0 | 0 | 0 | 0 | 0 | 4959000 | 219890 | 0 | 0 | 0 | 0 | 0 | 0 | 0 | 0 |
| ARRPQMDI | FMNL1; FMNL1 | O95466; K7 | FMNL1_HUMAI | Unmodified | 0 | 0 | 0 | 0 | 10476000 | 758100 | 0 | 65617 | 0 | 38967 | 0 | 0 | 128150 | 122540 | 0 | 0 | 576820 | 0 |
| ARSCGEVR | GPC6 | Q9Y625 | GPC6_HUMAN | Unmodified | 0 | 0 | 0 | 0 | 0 | 0 | 96417 | 62470 | 0 | 49091 | 0 | 0 | 3283300 | 1892700 | 0 | 0 | 105750 | 217420 |
| ARSEQFINI | CAPN1 | P07384 | CAN1_HUMAN | Unmodified | 0 | 0 | 0 | 0 | 0 | 0 | 0 | 0 | 514530 | 0 | 0 | 3275300 | 0 | 0 | 360370 | 231790 | 0 | 0 |
| ARSTVRDII | DYNLRB1; DYN | Q9NP97; B1 | DLRB1_HUMAI | Unmodified | 0 | 0 | 0 | 49024 | 1468300 | 0 | 231920 | 0 | 164940 | 1349200 | 137810 | 0 | 2530200 | 803010 | 0 | 0 | 567810 | 1652100 |
| ARSVDALD | LSR; LSR | Q86X29; S4 | LSR_HUMAN | Unmodified | 152770 | 0 | 155330 | 551280 | 369250 | 0 | 0 | 0 | 1577000 | 781160 | 407150 | 0 | 623170 | 250450 | 278200 | 164920 | 51898 | 92037 |
| ARSYELQE | SEPTIN11; SEP | Q9NVA2; D6 | SEP11_HUMAN | Unmodified | 52410 | 0 | 49638 | 310450 | 1004000 | 158440 | 41340 | 27849 | 326990 | 177150 | 81303 | 377210 | 0 | 106210 | 85803 | 67318 | 274140 | 185270 |
| ARSYELQE | SEPTIN11; SEP | Q9NVA2; D6 | SEP11_HUMAN | Unmodified | 0 | 0 | 0 | 0 | 9567000 | 939060 | 0 | 41714 | 0 | 1044200 | 0 | 0 | 1415200 | 866070 | 0 | 0 | 2392000 | 2472800 |
| ARTATMLC | PTPRF; PTPRF | P10586; A2 | PTPRF_HUMAN | Unmodified | 70383 | 0 | 238400 | 0 | 1017900 | 69374 | 0 | 0 | 875170 | 0 | 0 | 0 | 322390 | 0 | 0 | 0 | 0 | 0 |
| ARTATMLC | PTPRF; PTPRF | P10586; A2 | PTPRF_HUMAN | Oxidation (M) | 0 | 0 | 0 | 0 | 19935000 | 44007000 | 0 | 0 | 536700 | 228030 | 0 | 0 | 1424600 | 1405900 | 0 | 0 | 2490500 | 68613000 |
| ARTSSFAEI | GSK3A; GSK3A | P49840; A0 | GSK3A_HUMAI | Unmodified | 0 | 0 | 0 | 0 | 607290 | 0 | 94421 | 0 | 0 | 0 | 0 | 0 | 0 | 0 | 0 | 0 | 0 | 281320 |
| ARVECQSH | PSMA7; PSMA7 | O14818; H0 | PSA7_HUMAN | Unmodified | 0 | 0 | 0 | 0 | 0 | 0 | 0 | 0 | 271860 | 412710 | 373410 | 0 | 898350 | 0 | 0 | 0 | 0 | 0 |
| ARVITEEEK | RPL13; RPL13; | P26373; J3K | RL13_HUMAN | Unmodified | 0 | 0 | 0 | 0 | 0 | 0 | 1082300 | 851470 | 0 | 125620 | 0 | 0 | 867780 | 658920 | 0 | 0 | 186840 | 0 |
| ARVLKEEGI | TOMM34 | Q15785 | TOM34_HUMAI | Unmodified | 0 | 0 | 0 | 117580 | 573480 | 66740 | 67937 | 66595 | 0 | 79475 | 0 | 276040 | 633690 | 306400 | 110910 | 0 | 373460 | 997900 |
| ARVLQSFT | FLNB; FLNB | O75369; E7 | FLNB_HUMAN | Unmodified | 0 | 0 | 0 | 112920 | 2175400 | 700560 | 0 | 34521 | 0 | 112850 | 337600 | 322650 | 430730 | 393480 | 280790 | 228040 | 1208100 | 775710 |
| ARVPPEFL | MOGS; MOGS; | Q13724; A0 | MOGS_HUMAN | Unmodified | 0 | 82379 | 1216300 | 0 | 4416500 | 480570 | 11185000 | 2263500 | 3.06E+08 | 76953000 | 7038800 | 2132000 | 5345100 | 10729000 | 0 | 92258 | 722880 | 1637500 |
| ARVPQPDF | EHBP1L1; EHBI | Q8N3D4; E9 | EH1L1_HUMAN | Unmodified | 0 | 0 | 0 | 56246 | 2053600 | 60991 | 0 | 0 | 189130 | 38881 | 0 | 0 | 0 | 0 | 0 | 0 | 470380 | 0 |
| ARVYLDEN | INPP5A; INPP5 | Q14642; H0 | I5P1_HUMAN | Unmodified | 67574 | 0 | 0 | 0 | 0 | 0 | 0 | 0 | 0 | 0 | 0 | 160330 | 0 | 0 | 0 | 136870 | 0 | 0 |
| ARYEEVSVS | TXNDC17; TXNI | Q9BRA2; I3L | TXD17_HUMAN | Acetyl (Protein N | 1424300 | 0 | 1482400 | 2724500 | 5053500 | 0 | 0 | 0 | 8343600 | 915870 | 4825600 | 5227600 | 3254800 | 881860 | 3150600 | 2585900 | 761380 | 1165700 |
| ARYEEVSVS | TXNDC17; TXNI | Q9BRA2; I3L | TXD17_HUMAN | Acetyl (Protein N | 0 | 0 | 0 | 476290 | 34534000 | 2994800 | 2409900 | 0 | 1564600 | 4225200 | 913690 | 0 | 23502000 | 15247000 | 212540 | 0 | 17256000 | 23271000 |
| ARYFGDTD | DNAJB12; DNA. | Q9NXW2; V5 | DJB12_HUMAN | Unmodified | 0 | 0 | 0 | 0 | 0 | 0 | 0 | 0 | 711440 | 0 | 0 | 0 | 0 | 0 | 0 | 0 | 0 | 0 |
| ARYFGDTD | DNAJB12; DNA. | Q9NXW2; V5 | DJB12_HUMAN | Unmodified | 0 | 0 | 0 | 0 | 0 | 0 | 0 | 0 | 0 | 690200 | 36344 | 0 | 0 | 0 | 0 | 0 | 0 | 0 |
| ARYLAEKYE | EEF2 | P13639 | EF2_HUMAN | Unmodified | 0 | 0 | 0 | 0 | 0 | 0 | 0 | 0 | 0 | 0 | 0 | 0 | 0 | 0 | 0 | 0 | 281600 | 0 |
| ARYNAQCC | MDK; MDK; MDI | P21741; E9 | MK_HUMAN | Unmodified | 0 | 0 | 136530 | 36701 | 657310 | 0 | 0 | 0 | 13662000 | 146510 | 0 | 0 | 0 | 0 | 90576 | 15582 | 0 | 0 |
| ARYPAVDY | DLL1 | O00548 | DLL1_HUMAN | Unmodified | 0 | 0 | 0 | 0 | 0 | 0 | 0 | 0 | 0 | 0 | 0 | 1933000 | 0 | 0 | 0 | 0 | 0 | 0 |
| ASAAFSSV | TPD52; N/A | P55327; H0 | TPD52_HUMAN | Unmodified | 0 | 0 | 299010 | 223200 | 3737300 | 586900 | 1995800 | 1692600 | 3771500 | 8874600 | 0 | 4793700 | 3154700 | 3696000 | 0 | 0 | 407820 | 2248200 |
| ASAAPKPEI | PDLIM5; PDLIM | Q96HC4; H0 | PDLI5_HUMAN | Unmodified | 186830 | 0 | 0 | 0 | 0 | 0 | 0 | 51138 | 0 | 0 | 0 | 4244800 | 0 | 0 | 0 | 492250 | 0 | 0 |
| ASAAQLRI | UBE2M | P61081 | UBC12_HUMAI | Unmodified | 0 | 0 | 0 | 0 | 0 | 0 | 0 | 18202 | 0 | 0 | 0 | 0 | 0 | 0 | 0 | 0 | 127980 | 0 |
| ASADLMSY | GNG12 | Q9UBI6 | GBG12_HUMAI | Unmodified | 1023600 | 505140 | 224200 | 300190 | 187760 | 0 | 36815 | 0 | 1904900 | 44589 | 0 | 0 | 181520 | 563830 | 649610 | 809910 | 87772 | 436940 |
| ASADLMSY | GNG12 | Q9UBI6 | GBG12_HUMAI | Oxidation (M) | 170760 | 138880 | 0 | 0 | 0 | 0 | 0 | 0 | 254780 | 0 | 120660 | 0 | 0 | 0 | 236430 | 411480 | 0 | 0 |
| ASADLMSY | GNG12 | Q9UBI6 | GBG12_HUMAI | Oxidation (M) | 711470 | 0 | 0 | 0 | 15532000 | 3198200 | 538000 | 0 | 0 | 0 | 0 | 0 | 8352500 | 12360000 | 1848900 | 1187000 | 7003700 | 37982000 |
| ASADLMSY | GNG12 | Q9UBI6 | GBG12_HUMAI | Unmodified | 0 | 0 | 0 | 0 | 1758900 | 310290 | 0 | 0 | 0 | 0 | 0 | 0 | 0 | 0 | 0 | 0 | 288150 | 2589500 |
| ASADLMSY | GNG12 | Q9UBI6 | GBG12_HUMAI | Oxidation (M) | 0 | 0 | 0 | 0 | 5862100 | 0 | 0 | 0 | 675200 | 1978000 | 0 | 0 | 0 | 0 | 0 | 0 | 2061900 | 0 |
| ASAELALGI | GOT2 | P00505 | AATM_HUMAN | Unmodified | 0 | 0 | 0 | 0 | 0 | 0 | 0 | 0 | 8211100 | 3133200 | 0 | 0 | 0 | 0 | 0 | 0 | 0 | 0 |
| ASAETVDP | FSCN1 | Q16658 | FSCN1_HUMAN | Unmodified | 0 | 0 | 0 | 0 | 20276000 | 3079400 | 1803100 | 2160700 | 3419600 | 2607900 | 5154100 | 9735100 | 0 | 0 | 0 | 0 | 9377100 | 9242100 |
| ASAFALQE | DLST | P36957 | ODO2_HUMAN | Unmodified | 0 | 0 | 148550 | 0 | 0 | 0 | 1158900 | 151320 | 24521000 | 5298200 | 0 | 246320 | 0 | 0 | 0 | 543690 | 0 | 0 |
| ASAGDTQA | RAB3D | O95716 | RAB3D_HUMAI | Acetyl (Protein N | 0 | 210210 | 0 | 1862100 | 0 | 0 | 0 | 40528 | 0 | 0 | 0 | 48957 | 0 | 0 | 1569200 | 1422000 | 0 | 0 |
| ASAGHAVS | CTTN | Q14247 | SRC8_HUMAN | Unmodified | 1921200 | 2157900 | 0 | 2910200 | 18502000 | 2936200 | 0 | 0 | 1262100 | 509830 | 0 | 2062100 | 10869000 | 7003600 | 2352000 | 1779600 | 4986500 | 9130900 |
| ASAGHAVS | CTTN | Q14247 | SRC8_HUMAN | Unmodified | 0 | 0 | 0 | 0 | 3512200 | 0 | 0 | 0 | 0 | 0 | 164210 | 0 | 3938300 | 7338400 | 0 | 0 | 6784400 | 3244800 |
| ASAGPAPG | TBC1D10B | Q4KMP7 | TB10B_HUMAN | Unmodified | 125020 | 0 | 381850 | 0 | 6323500 | 0 | 0 | 66856 | 1787300 | 0 | 0 | 2395800 | 681150 | 0 | 1878700 | 1009000 | 0 | 0 |
| ASAGPQPL | TLN1 | Q9Y490 | TLN1_HUMAN | Unmodified | 206500 | 0 | 0 | 0 | 2465700 | 296750 | 2396500 | 573760 | 0 | 183410 | 0 | 0 | 1276400 | 2346200 | 0 | 107880 | 1345900 | 1458200 |
| ASAGSYISA | PSMB8; PSMB8 | P28062; X5I | PSB8_HUMAN | Unmodified | 0 | 0 | 0 | 0 | 0 | 0 | 0 | 0 | 0 | 0 | 0 | 0 | 0 | 0 | 0 | 0 | 0 | 0 |
| ASAGSYISA | PSMB8; PSMB8 | P28062; X5I | PSB8_HUMAN | Unmodified | 0 | 0 | 0 | 0 | 0 | 0 | 0 | 0 | 402420 | 215940 | 587450 | 2764900 | 0 | 0 | 0 | 0 | 0 | 0 |
| ASAILPGVL | NIBAN1 | Q9BZQ8 | NIBA1_HUMAN | Unmodified | 0 | 0 | 0 | 3210500 | 2179300 | 0 | 0 | 0 | 13207000 | 1556900 | 0 | 0 | 0 | 0 | 0 | 390270 | 0 | 0 |
| ASALIDRPA | EPB41L3; EPB4 | Q9Y2J2; A0 | E41L3_HUMAN | Unmodified | 0 | 0 | 0 | 39646 | 0 | 0 | 0 | 0 | 295040 | 0 | 0 | 0 | 127540 | 72706 | 63422 | 51361 | 176250 | 1205900 |
| ASALIDRPA | EPB41L3; EPB4 | Q9Y2J2; A0 | E41L3_HUMAN | Unmodified | 0 | 0 | 0 | 0 | 0 | 0 | 0 | 0 | 0 | 0 | 0 | 0 | 0 | 0 | 0 | 0 | 785140 | 4210500 |
| ASANMDLN | LAP3; LAP3 | P28838; H0 | AMPL_HUMAN | Unmodified | 0 | 0 | 0 | 0 | 0 | 0 | 0 | 0 | 317760 | 0 | 0 | 774590 | 79328 | 0 | 0 | 0 | 0 | 0 |
| ASAPAADP | PDLIM7; PDLIM | Q9NR12; H7 | PDLI7_HUMAN | Unmodified | 0 | 0 | 163190 | 209940 | 2319100 | 0 | 0 | 0 | 0 | 0 | 0 | 0 | 514840 | 239180 | 195610 | 139890 | 417010 | 1042900 |
| ASAPESGLI | ISOC1 | Q96CN7 | ISOC1_HUMAN | Unmodified | 0 | 124810 | 0 | 0 | 102340 | 0 | 704280 | 730660 | 0 | 608030 | 1076700 | 6448400 | 1030200 | 805810 | 0 | 279050 | 781380 | 0 |
| ASASAPAP | ITM2C; ITM2C | Q9NQX7; C8 | ITM2C_HUMAN | Unmodified | 0 | 0 | 0 | 0 | 3088300 | 1045900 | 0 | 0 | 27295000 | 0 | 0 | 0 | 0 | 0 | 0 | 0 | 0 | 513160 |
| ASASDGSS | GLIPR2; GLIPR2 | Q9H4G4; Q5 | GAPR1_HUMAI | Unmodified | 2468900 | 3225300 | 2696900 | 4631800 | 5177900 | 1027400 | 630960 | 1008500 | 1481000 | 603840 | 0 | 0 | 6623900 | 4292000 | 5928700 | 4174100 | 522280 | 2678900 |
| ASASDGSS | GLIPR2; GLIPR2 | Q9H4G4; Q5 | GAPR1_HUMAI | Unmodified | 0 | 0 | 0 | 0 | 16528000 | 0 | 265540 | 0 | 0 | 3371400 | 0 | 0 | 0 | 0 | 0 | 0 | 5488300 | 14317000 |
| ASASGAMA | VAPA; VAPA | Q9P0L0; J3Q | VAPA_HUMAN | Acetyl (Protein N | 0 | 0 | 0 | 0 | 20320000 | 0 | 2838000 | 1196800 | 0 | 1651600 | 0 | 0 | 5703800 | 0 | 0 | 0 | 0 | 0 |
| ASASGAMA | VAPA; VAPA | Q9P0L0; J3Q | VAPA_HUMAN | Acetyl (Protein N | 0 | 0 | 0 | 0 | 5868300 | 0 | 0 | 138770 | 0 | 0 | 0 | 0 | 1149000 | 0 | 0 | 0 | 0 | 1605300 |
| ASASGSGA | LMNA; LMNA | P02545; A0 | LMNA_HUMAN | Unmodified | 0 | 0 | 0 | 0 | 921980 | 179450 | 0 | 0 | 0 | 0 | 0 | 0 | 0 | 0 | 0 | 0 | 0 | 407770 |
| ASASYHISN | CAND1 | Q86VP6 | CAND1_HUMA | Acetyl (Protein N | 0 | 0 | 0 | 0 | 557910 | 0 | 0 | 0 | 0 | 0 | 0 | 0 | 0 | 0 | 0 | 0 | 526330 | 770950 |
| ASATAPAA | CCNDBP1; CCI | O95273; H3 | CCDB1_HUMA | Acetyl (Protein N | 3094000 | 2817000 | 1260900 | 1449600 | 0 | 0 | 0 | 0 | 0 | 0 | 0 | 0 | 0 | 0 | 5157700 | 6951800 | 40577 | 0 |
| ASATEDPVI | POC1B; POC1B | Q8TC44; F8 | POC1B_HUMA | Acetyl (Protein N | 229840 | 0 | 0 | 274310 | 0 | 0 | 0 | 0 | 750580 | 0 | 0 | 2545500 | 0 | 0 | 567850 | 57720 | 0 | 0 |
| ASAVESLRI | EVPL | Q92817 | EVPL_HUMAN | Unmodified | 0 | 0 | 0 | 0 | 3231600 | 8772400 | 1014500 | 314030 | 0 | 0 | 0 | 0 | 505980 | 3291100 | 0 | 0 | 626780 | 7227100 |
| ASAVGERL | HEXB; HEXB; H | P07686; H0 | HEXB_HUMAN | Unmodified | 0 | 0 | 0 | 173880 | 196580 | 0 | 0 | 0 | 862420 | 475070 | 0 | 0 | 1604000 | 592760 | 0 | 0 | 0 | 0 |
| ASAVSPAN | HCFC1; HCFC1 | P51610; A6 | HCFC1_HUMA | Acetyl (Protein N | 0 | 24826 | 0 | 0 | 77567 | 29290 | 54231 | 0 | 211560 | 137710 | 153470 | 644410 | 0 | 0 | 0 | 0 | 57251 | 77605 |
| ASAYVQAL | LYPLAL1 | Q5VWZ2 | LYPL1_HUMAN | Unmodified | 0 | 0 | 0 | 110130 | 1993000 | 0 | 0 | 0 | 274700 | 100800 | 0 | 0 | 448780 | 365670 | 318380 | 0 | 153580 | 115770 |
| ASAYVFASI | BPNT1; BPNT1 | O95861; A6 | BPNT1_HUMAN | Unmodified | 0 | 0 | 0 | 86169 | 346050 | 0 | 0 | 58943 | 0 | 0 | 1086000 | 1838300 | 425460 | 397210 | 0 | 0 | 244060 | 313040 |
| ASCILQLVK | AP3D1; AP3D1 | O14617; A0 | AP3D1_HUMAI | Unmodified | 981020 | 989650 | 191690 | 273690 | 0 | 0 | 0 | 0 | 0 | 0 | 70632 | 1290000 | 0 | 0 | 0 | 229120 | 0 | 0 |
| ASCSALMS | UTRN | P46939 | UTRN_HUMAN | Unmodified | 0 | 0 | 0 | 0 | 0 | 0 | 0 | 0 | 346660 | 0 | 0 | 0 | 0 | 0 | 0 | 0 | 0 | 0 |
| ASCSALMS | UTRN | P46939 | UTRN_HUMAN | Oxidation (M) | 0 | 0 | 513440 | 0 | 0 | 0 | 0 | 0 | 0 | 0 | 0 | 0 | 0 | 0 | 627650 | 438470 | 0 | 0 |
| ASDAFHFF | CLUH; CLUH; C | O75153; I3L | CLU_HUMAN | Unmodified | 0 | 0 | 0 | 0 | 178860 | 0 | 117200 | 0 | 0 | 0 | 311550 | 12103000 | 390670 | 112920 | 0 | 0 | 47488 | 56673 |
| ASDEVPLAI | SLC35B2 | Q8TB61 | S35B2_HUMAN | Unmodified | 0 | 0 | 0 | 0 | 0 | 0 | 0 | 1051100 | 0 | 0 | 0 | 3296200 | 0 | 0 | 103130 | 0 | 0 | 0 |
| ASDEVTRL | VCL | P18206 | VINC_HUMAN | Unmodified | 0 | 0 | 0 | 0 | 0 | 0 | 310080 | 195800 | 0 | 0 | 0 | 0 | 0 | 200360 | 0 | 0 | 679360 | 139470 |
| ASDHEGLL | RAB11FIP1 | Q6WKZ4 | RFIP1_HUMAN | Unmodified | 0 | 0 | 0 | 0 | 0 | 0 | 0 | 0 | 0 | 0 | 0 | 0 | 0 | 0 | 0 | 0 | 0 | 0 |
| ASDKETAQ | CCER2; CCER2 | I3L3R5; I3L3 | CCER2_HUMAI | Unmodified | 0 | 0 | 0 | 0 | 19821 | 0 | 4138200 | 16155000 | 40187 | 0 | 96885 | 0 | 0 | 0 | 0 | 0 | 0 | 0 |
| ASDKETAQ | CCER2; CCER2 | I3L3R5; I3L3 | CCER2_HUMAI | Unmodified | 0 | 0 | 0 | 0 | 329340 | 0 | 0 | 221640 | 927750 | 0 | 0 | 0 | 0 | 164700 | 0 | 0 | 0 | 0 |
| ASDPQSPP | TAOK1 | Q7L7X3 | TAOK1_HUMAI | Unmodified | 87421 | 123440 | 135590 | 199960 | 0 | 44595 | 0 | 0 | 302980 | 48980 | 148540 | 395600 | 0 | 0 | 407500 | 173790 | 81319 | 199240 |
| ASDTEAGK | IGSF3 | O75054 | IGSF3_HUMAN | Unmodified | 507750 | 219880 | 0 | 66302 | 0 | 0 | 0 | 0 | 383590 | 0 | 0 | 0 | 0 | 0 | 0 | 0 | 0 | 0 |
| ASDVHEVR | PKM; PKM; PKM | P14618; B4 | KPYM_HUMAN | Unmodified | 0 | 0 | 0 | 0 | 10822000 | 676500 | 9176300 | 3720600 | 0 | 586620 | 0 | 0 | 2441200 | 1186900 | 0 | 0 | 26561000 | 6925300 |
| ASDYGMKI | MLEC; MLEC | Q14165; F5 | MLEC_HUMAN | Oxidation (M) | 0 | 0 | 0 | 0 | 0 | 0 | 0 | 0 | 4612000 | 65081 | 285570 | 0 | 0 | 0 | 0 | 0 | 0 | 0 |
| ASDYGMKI | MLEC; MLEC | Q14165; F5 | MLEC_HUMAN | Unmodified | 0 | 0 | 0 | 0 | 0 | 0 | 0 | 0 | 14258000 | 933070 | 0 | 0 | 0 | 0 | 0 | 0 | 0 | 0 |
| ASEAASPLI | TNS3 | Q68CZ2 | TENS3_HUMAN | Unmodified | 0 | 0 | 0 | 0 | 443520 | 0 | 0 | 0 | 0 | 0 | 834110 | 0 | 0 | 0 | 0 | 0 | 141190 | 148340 |
| ASEAGEVP | COPS6; COPS6 | Q7L5N1; E7 | CSN6_HUMAN | Unmodified | 210660 | 431240 | 360450 | 338670 | 0 | 0 | 0 | 0 | 0 | 22910 | 1011200 | 4022700 | 0 | 0 | 397640 | 525120 | 0 | 0 |
| ASEAKEGE | CDC37; CDC37 | Q16543; K7 | CDC37_HUMA | Unmodified | 0 | 0 | 634810 | 0 | 0 | 0 | 0 | 0 | 0 | 0 | 0 | 916190 | 0 | 0 | 0 | 0 | 0 | 1231800 |
| ASEEEPEY | CHGB | P05060 | SCG1_HUMAN | Unmodified | 0 | 0 | 0 | 0 | 0 | 0 | 0 | 0 | 1125800 | 0 | 0 | 0 | 0 | 0 | 0 | 0 | 0 | 0 |
| ASEEHLKQ | NME2P1; NME2 | O60361; P2 | NDK8_HUMAN | Unmodified | 0 | 0 | 0 | 0 | 0 | 0 | 0 | 0 | 0 | 0 | 0 | 0 | 260420 | 146140 | 0 | 0 | 0 | 0 |
| ASEELQK | CACYBP; CACY | Q9HB71; B2 | CYBP_HUMAN | Acetyl (Protein N | 0 | 0 | 0 | 0 | 0 | 1002900 | 218860 | 90510 | 1568300 | 322960 | 0 | 2819200 | 555670 | 116540 | 0 | 0 | 55361 | 0 |
| ASEELQKD | CACYBP; CACY | Q9HB71; B2 | CYBP_HUMAN | Acetyl (Protein N | 0 | 0 | 0 | 0 | 0 | 0 | 0 | 502140 | 0 | 387620 | 0 | 966100 | 2076700 | 1351600 | 0 | 0 | 0 | 0 |
| ASEELQKD | CACYBP; CACY | Q9HB71; B2 | CYBP_HUMAN | Acetyl (Protein N | 0 | 0 | 0 | 0 | 3077600 | 0 | 1993100 | 0 | 0 | 1606100 | 0 | 0 | 11516000 | 3110500 | 0 | 0 | 4947300 | 670070 |
| ASEFYERN | GPSM1; GPSM1 | Q86YR5; A0 | GPSM1_HUMA | Unmodified | 0 | 0 | 0 | 0 | 1008900 | 0 | 0 | 395960 | 0 | 0 | 0 | 0 | 0 | 213460 | 0 | 0 | 454660 | 482220 |
| ASEGPAFFI | FARSB | Q9NSD9 | SYFB_HUMAN | Unmodified | 0 | 0 | 0 | 275980 | 0 | 0 | 0 | 0 | 0 | 0 | 805720 | 7367100 | 207200 | 0 | 0 | 0 | 0 | 0 |
| ASEIQPLQ | MFSD6; MFSD6 | Q6ZSS7; H7 | MFSD6_HUMAI | Unmodified | 0 | 0 | 0 | 0 | 0 | 0 | 0 | 0 | 133870 | 0 | 0 | 558350 | 0 | 0 | 0 | 0 | 0 | 0 |
| ASEKPLAA | MVD; MVD; MVI | P53602; H3 | MVD1_HUMAN | Acetyl (Protein N | 0 | 0 | 0 | 0 | 4309500 | 0 | 770750 | 404220 | 647450 | 958100 | 355200 | 2264000 | 283790 | 0 | 0 | 0 | 3543100 | 0 |
| ASELEQEQ | CALCA; CALCA | P01258; P0 | CALC_HUMAN | Unmodified | 0 | 0 | 0 | 0 | 0 | 0 | 0 | 0 | 839220 | 0 | 0 | 0 | 0 | 0 | 0 | 0 | 403940 | 0 |
| ASELEQEQ | CALCA | P06881 | CALCA_HUMAI | Unmodified | 0 | 0 | 0 | 0 | 0 | 0 | 0 | 0 | 791540 | 34667 | 21533 | 0 | 0 | 0 | 0 | 0 | 0 | 0 |
| ASELEQEQ | CALCA | P01258 | CALC_HUMAN | Unmodified | 0 | 0 | 0 | 202820 | 0 | 0 | 0 | 0 | 45028000 | 2478300 | 87470 | 506870 | 0 | 0 | 153940 | 0 | 0 | 0 |
| ASELEQEQ | CALCA | P01258 | CALC_HUMAN | Unmodified | 0 | 0 | 0 | 0 | 0 | 0 | 0 | 0 | 938820 | 4795100 | 0 | 0 | 0 | 0 | 0 | 0 | 0 | 0 |
| ASENAIVW | AP2M1; AP2M1 | Q96CW1; E9 | AP2M1_HUMAI | Unmodified | 0 | 0 | 0 | 0 | 0 | 0 | 0 | 0 | 0 | 0 | 0 | 0 | 0 | 0 | 0 | 0 | 0 | 0 |
| ASERWQPI | SF3A3 | Q12874 | SF3A3_HUMAN | Unmodified | 0 | 0 | 0 | 0 | 0 | 0 | 0 | 84660 | 0 | 206360 | 0 | 744390 | 2285600 | 0 | 0 | 0 | 0 | 0 |
| ASESISLK | LMO7; LMO7; L | Q8WWI1; E9 | LMO7_HUMAN | Unmodified | 0 | 0 | 0 | 0 | 1935000 | 136530 | 18202 | 0 | 22026 | 109990 | 154860 | 0 | 0 | 0 | 0 | 0 | 586600 | 769860 |
| ASESQILSI | MIA3; MIA3 | Q5JRA6; A0 | TGO1_HUMAN | Unmodified | 0 | 0 | 156450 | 0 | 0 | 0 | 0 | 0 | 4044600 | 350960 | 693970 | 0 | 0 | 0 | 0 | 0 | 0 | 0 |
| ASESSKPW | THRAP3; THRA | Q9Y2W1; A0 | TR150_HUMAN | Unmodified | 0 | 0 | 0 | 0 | 0 | 0 | 0 | 0 | 0 | 0 | 0 | 309240 | 0 | 0 | 0 | 0 | 0 | 0 |
| ASESTTPAF | MICALL1; MICA | Q8N3F8; B0 | MILK1_HUMAN | Unmodified | 452180 | 445890 | 392380 | 342330 | 0 | 113140 | 0 | 552790 | 186740 | 0 | 0 | 307880 | 650660 | 294010 | 302390 | 261990 | 0 | 227810 |

| | | | | | | | | | | | | | | | | | | | | | | |
|---|---|---|---|---|---|---|---|---|---|---|---|---|---|---|---|---|---|---|---|---|---|---|
| ASETPPPV/ | CC2D1A | Q6P1N0 | C2D1A_HUMAI | Unmodified | 3062700 | 0 | 843890 | 0 | 0 | 0 | 0 | 0 | 3158400 | 0 | 0 | 603610 | 0 | 0 | 1084400 | 1893200 | 0 | 0 |
| ASETQRSLI | CIT; CIT; CIT; CI | O14578; A0/ | CTRO_HUMAN | Unmodified | 0 | 0 | 0 | 0 | 0 | 0 | 0 | 0 | 0 | 0 | 0 | 0 | 0 | 0 | 0 | 0 | 0 | 0 |
| ASEVQEFT/ | SHCBP1 | Q8NEM2 | SHCBP_HUMA | Unmodified | 0 | 0 | 0 | 0 | 628600 | 453200 | 0 | 0 | 287370 | 0 | 0 | 0 | 0 | 0 | 0 | 0 | 0 | 0 |
| ASEVQLHL | ENTPD2 | Q9Y5L3 | ENTP2_HUMAN | Unmodified | 0 | 969430 | 0 | 135120 | 172100 | 0 | 0 | 0 | 0 | 0 | 0 | 549350 | 144140 | 149290 | 2235100 | 782580 | 0 | 0 |
| ASEWVQQ | AARS1; AARS1; | P49588; A0A | SYAC_HUMAN | Unmodified | 1254900 | 0 | 0 | 84035 | 305190 | 51696 | 30833 | 0 | 738070 | 123180 | 114000 | 815860 | 432900 | 161310 | 73498 | 24550 | 48414 | 121200 |
| ASEWVQQ | AARS1; AARS1; | P49588; A0A | SYAC_HUMAN | Oxidation (M) | 0 | 0 | 0 | 0 | 0 | 0 | 0 | 0 | 0 | 0 | 0 | 5413600 | 0 | 0 | 0 | 0 | 0 | 0 |
| ASEWVQQ | AARS1; AARS1; | P49588; A0A | SYAC_HUMAN | Unmodified | 0 | 0 | 0 | 0 | 4881600 | 0 | 0 | 0 | 641310 | 493120 | 141640 | 0 | 4545900 | 1620300 | 0 | 0 | 0 | 847340 |
| ASEWVQQ | AARS1; AARS1; | P49588; A0A | SYAC_HUMAN | Unmodified | 0 | 0 | 0 | 0 | 5851800 | 0 | 935270 | 128670 | 0 | 1689700 | 0 | 0 | 7826200 | 6427400 | 0 | 0 | 2080300 | 4162300 |
| ASFCLEDTI | LOXL2 | Q9Y4K0 | LOXL2_HUMAN | Unmodified | 0 | 0 | 0 | 0 | 0 | 0 | 0 | 0 | 341530 | 0 | 0 | 0 | 0 | 0 | 0 | 0 | 0 | 0 |
| ASFENNCE | EIF6; EIF6; EIF6 | P56537; A0A | IF6_HUMAN Eu | Unmodified | 795050 | 276440 | 0 | 0 | 314970 | 0 | 0 | 0 | 0 | 0 | 0 | 0 | 386320 | 0 | 0 | 0 | 0 | 217720 |
| ASFGSGPA | ADAMTS1; ADA | Q9UHI8; E5F | ATS1_HUMAN / | Unmodified | 0 | 0 | 511600 | 319140 | 0 | 0 | 0 | 0 | 0 | 0 | 0 | 0 | 0 | 0 | 0 | 0 | 0 | 0 |
| ASFHELYN | MYO6; MYO6; M | Q9UM54; A0 | MYO6_HUMAN | Unmodified | 0 | 0 | 0 | 0 | 0 | 0 | 0 | 0 | 0 | 0 | 0 | 387170 | 0 | 0 | 667370 | 292720 | 0 | 0 |
| ASFHELYN | MYO6; MYO6; M | Q9UM54; A0 | MYO6_HUMAN | Oxidation (M) | 23712 | 0 | 39841 | 0 | 0 | 0 | 0 | 0 | 0 | 0 | 65189 | 84889 | 0 | 0 | 483240 | 425460 | 0 | 0 |
| ASFLQLR | SPPL2B; SPPL2 | Q8TCT7; A0/ | SPP2B_HUMAN | Unmodified | 0 | 0 | 0 | 0 | 0 | 0 | 0 | 0 | 3567300 | 197720 | 0 | 838810 | 0 | 0 | 257420 | 132570 | 0 | 0 |
| ASFNHFDK | ACTN4; ACTN4; | O43707; F5C | ACTN4_HUMAI | Unmodified | 0 | 0 | 0 | 0 | 133240 | 0 | 0 | 0 | 380570 | 63719 | 69332 | 1276000 | 0 | 0 | 206820 | 0 | 0 | 0 |
| ASFPGPSE | GDF15; GDF15 | Q99988; A0/ | GDF15_HUMAI | Unmodified | 0 | 0 | 0 | 0 | 0 | 0 | 0 | 0 | 0 | 0 | 0 | 0 | 0 | 0 | 0 | 0 | 0 | 0 |
| ASFPQGPI | LYPLA1; LYPLA | O75608; A0/ | LYPA1_HUMAN | Unmodified | 166960 | 0 | 0 | 0 | 0 | 0 | 0 | 0 | 0 | 0 | 0 | 8271500 | 0 | 0 | 128380 | 123900 | 172040 | 0 |
| ASFPRWYF | SPINT2; SPINT2 | O43291; K7E | SPIT2_HUMAN | Unmodified | 345190 | 0 | 0 | 0 | 159450 | 0 | 0 | 0 | 527540 | 852220 | 0 | 0 | 66909 | 0 | 0 | 0 | 0 | 0 |
| ASFTPFVDI | TNIK | Q9UKE5 | TNIK_HUMAN T | Unmodified | 0 | 0 | 0 | 0 | 0 | 0 | 0 | 0 | 629770 | 427580 | 0 | 2015400 | 0 | 0 | 0 | 0 | 0 | 0 |
| ASFVDEHT | TXNRD2; TXNRI | Q9NNW7; A( | TRXR2_HUMAN | Unmodified | 0 | 0 | 0 | 0 | 0 | 0 | 0 | 9914.7 | 991730 | 247090 | 0 | 0 | 0 | 0 | 0 | 0 | 0 | 0 |
| ASFVTEVLA | ZW10; ZW10 | O43264; F5F | ZW10_HUMAN | Acetyl (Protein N | 0 | 0 | 169870 | 0 | 1484400 | 0 | 0 | 0 | 709970 | 523630 | 379930 | 1879500 | 0 | 0 | 0 | 0 | 815980 | 0 |
| ASFVTEVLA | ZW10; ZW10 | O43264; F5F | ZW10_HUMAN | Acetyl (Protein N | 0 | 0 | 0 | 35839 | 453590 | 78239 | 0 | 0 | 0 | 0 | 0 | 0 | 469020 | 282150 | 0 | 0 | 0 | 267040 |
| ASGADSKG | VCP; VCP; VCP; | P55072; A0A | TERA_HUMAN | Acetyl (Protein N | 3165300 | 2839600 | 543090 | 8698300 | 33125000 | 10894000 | 11217000 | 9998600 | 10265000 | 10748000 | 7285300 | 36000000 | 33371000 | 14316000 | 3346700 | 4318000 | 13719000 | 13084000 |
| ASGADSKG | VCP; VCP; VCP; | P55072; A0A | TERA_HUMAN | Acetyl (Protein N | 0 | 101560 | 0 | 0 | 0 | 0 | 1367500 | 1476400 | 0 | 0 | 0 | 0 | 196470 | 0 | 0 | 0 | 3192500 | 0 |
| ASGANYSF | DBNL; DBNL; D | Q9UJU6; B4I | DBNL_HUMAN | Unmodified | 0 | 0 | 0 | 0 | 0 | 0 | 44955 | 89275 | 0 | 52755 | 0 | 365570 | 0 | 377160 | 177450 | 116850 | 226170 | 218580 |
| ASGANYSF | DBNL; DBNL; D | Q9UJU6; B4I | DBNL_HUMAN | Unmodified | 0 | 0 | 0 | 0 | 0 | 0 | 661230 | 275530 | 0 | 0 | 0 | 0 | 0 | 112970 | 0 | 0 | 891470 | 0 |
| ASGAPAAA | FMN2 | Q9NZ56 | FMN2_HUMAN | Unmodified | 0 | 0 | 0 | 0 | 0 | 0 | 401370 | 107180 | 847340 | 729450 | 0 | 0 | 0 | 5400700 | 0 | 0 | 3042300 | 2807700 |
| ASGDFQTT | ITGAV | P06756 | ITAV_HUMAN I | Unmodified | 53538 | 0 | 0 | 322770 | 1205900 | 333810 | 31935 | 0 | 0 | 473800 | 0 | 0 | 794850 | 317660 | 0 | 0 | 601600 | 0 |
| ASGDSARP | SLC25A13 | Q9UJS0 | CMC2_HUMAN | Unmodified | 128820 | 83021 | 0 | 250020 | 0 | 0 | 0 | 0 | 8529700 | 0 | 307310 | 1339800 | 0 | 0 | 120740 | 68175 | 0 | 0 |
| ASGEHSPG | RNPEP; RNPEP | Q9H4A4; A6I | AMPB_HUMAN | Acetyl (Protein N | 0 | 0 | 0 | 0 | 0 | 0 | 0 | 0 | 0 | 0 | 0 | 3764700 | 0 | 0 | 0 | 71644 | 0 | 0 |
| ASGELMTV | GCLC; GCLC; G | P48506; A0A | GSH1_HUMAN | Unmodified | 0 | 0 | 0 | 0 | 0 | 0 | 0 | 0 | 0 | 0 | 0 | 878130 | 0 | 0 | 0 | 0 | 0 | 0 |
| ASGGSGG | SRP72; SRP72 | O76094; R4C | SRP72_HUMAN | Acetyl (Protein N | 0 | 0 | 0 | 0 | 0 | 0 | 0 | 0 | 0 | 0 | 0 | 449020 | 0 | 0 | 0 | 0 | 0 | 0 |
| ASGLAAGK | GART; GART; G/ | P22102; C9J | PUR2_HUMAN | Unmodified | 0 | 0 | 0 | 0 | 1362100 | 466510 | 1206500 | 0 | 0 | 169420 | 0 | 0 | 867540 | 845950 | 0 | 0 | 1790100 | 627780 |
| ASGLLLER | METTL26; METT | Q96S19; A0/ | MTL26_HUMAN | Unmodified | 428820 | 299220 | 0 | 294430 | 0 | 0 | 0 | 0 | 0 | 0 | 0 | 101490 | 0 | 0 | 749300 | 483870 | 0 | 0 |
| ASGNARIG | PRDX2; PRDX2 | P32119; A6N | PRDX2_HUMAI | Acetyl (Protein N | 0 | 0 | 340560 | 0 | 422230 | 0 | 1113300 | 752520 | 673330 | 431310 | 0 | 0 | 238550 | 400720 | 0 | 0 | 2266800 | 460520 |
| ASGNYATV | RPL8; RPL8; RP | P62917; E9P | RL8_HUMAN 6( | Unmodified | 3646100 | 3559000 | 2949000 | 0 | 2046900 | 785660 | 751590 | 618340 | 676810 | 284690 | 1449100 | 19923000 | 3991800 | 775170 | 0 | 2419100 | 291480 | 902150 |
| ASGNYATV | RPL8; RPL8; RP | P62917; E9P | RL8_HUMAN 6( | Unmodified | 3892300 | 3943300 | 2694600 | 532420 | 2862400 | 0 | 364030 | 637370 | 858510 | 457210 | 1587300 | 29340000 | 5606200 | 756700 | 2697200 | 2091000 | 243420 | 1665200 |
| ASGNYATV | RPL8; RPL8; RP | P62917; E9P | RL8_HUMAN 6( | Unmodified | 0 | 0 | 58319 | 0 | 646360 | 0 | 174080 | 142090 | 110150 | 41031 | 0 | 0 | 5277000 | 1399500 | 0 | 0 | 531010 | 487370 |
| ASGPGLNT | FLNA; FLNA | P21333; Q6( | FLNA_HUMAN | Unmodified | 2044000 | 2761000 | 0 | 4053300 | 7122800 | 1396800 | 3393100 | 0 | 964060 | 1836300 | 0 | 2894400 | 1787100 | 1413800 | 2242100 | 1893700 | 2556900 | 0 |
| ASGPPKGP | CRIP2; CRIP2 | P52943; H0Y | CRIP2_HUMAN | Unmodified | 0 | 0 | 0 | 0 | 486520 | 0 | 0 | 1286200 | 0 | 1020600 | 0 | 0 | 0 | 0 | 0 | 0 | 1872500 | 5522100 |
| ASGPPVSE | H1-3; H1-4; H1 | P16402; P10 | H13_HUMAN H | Unmodified | 46442000 | 21506000 | 398070 | 0 | 7180700 | 1914000 | 5784400 | 2.52E+08 | 609630 | 637980 | 1707600 | 1548500 | 764770 | 0 | 285280 | 219710 | 379580 | 3982500 |
| ASGPPVSE | H1-3; H1-4; H1 | P16402; P10 | H13_HUMAN H | Unmodified | 0 | 508460 | 0 | 0 | 155480 | 26642 | 1766600 | 33601000 | 95230 | 67451 | 0 | 0 | 0 | 0 | 0 | 0 | 0 | 200430 |
| ASGQAFEL | STMN1; STMN1 | P16949; A2A | STMN1_HUMAI | Unmodified | 0 | 0 | 0 | 0 | 0 | 0 | 0 | 48771 | 256870 | 374540 | 702540 | 440680 | 0 | 0 | 0 | 0 | 0 | 0 |
| ASGQAFEL | STMN1; STMN1 | P16949; A2A | STMN1_HUMAI | Unmodified | 0 | 0 | 0 | 0 | 0 | 0 | 0 | 307460 | 0 | 0 | 0 | 0 | 0 | 0 | 0 | 0 | 0 | 0 |
| ASGSPEPA | NCAM2; NCAM | O15394; H9I | NCAM2_HUMA | Unmodified | 0 | 1617700 | 4470500 | 5372600 | 0 | 0 | 0 | 0 | 0 | 912390 | 0 | 0 | 0 | 0 | 9605100 | 3767100 | 0 | 0 |
| ASGTLREYI | RPL18A; RPL18 | Q02543; M0I | RL18A_HUMAN | Unmodified | 0 | 0 | 0 | 0 | 0 | 0 | 56808 | 0 | 0 | 0 | 0 | 0 | 409690 | 443540 | 0 | 0 | 0 | 0 |
| ASGTTTTAV | MTAP; MTAP; N | Q13126; F8V | MTAP_HUMAN | Acetyl (Protein N | 67951 | 33043 | 0 | 0 | 534400 | 0 | 65862 | 56991 | 202090 | 0 | 0 | 64980 | 0 | 0 | 30977 | 63076 | 502400 | 0 |
| ASGVAVSD | CFL1; CFL1; CF | P23528; E9P | COF1_HUMAN | Acetyl (Protein N | 6935600 | 6800500 | 4313200 | 4651700 | 33283000 | 6005300 | 11656000 | 10941000 | 10315000 | 7738700 | 7118700 | 30137000 | 26082000 | 19332000 | 5972500 | 6268500 | 13918000 | 12223000 |
| ASGVAVSD | CFL1; CFL1; CF | P23528; E9P | COF1_HUMAN | Acetyl (Protein N | 0 | 0 | 0 | 0 | 26091000 | 4642300 | 20247000 | 9907800 | 0 | 2406500 | 0 | 0 | 11806000 | 11749000 | 0 | 0 | 61271000 | 9865200 |
| ASGVAVSD | CFL1; CFL1; CF | P23528; E9P | COF1_HUMAN | Acetyl (Protein N | 0 | 0 | 0 | 0 | 7056200 | 0 | 5749300 | 695470 | 0 | 831410 | 0 | 0 | 3168700 | 0 | 0 | 0 | 17883000 | 3061200 |
| ASGVAVSD | CFL1; CFL1; CF | P23528; E9P | COF1_HUMAN | Acetyl (Protein N | 0 | 0 | 0 | 0 | 0 | 0 | 0 | 0 | 0 | 930150 | 0 | 0 | 0 | 0 | 0 | 0 | 2460900 | 0 |
| ASGVAVSD | CFL1; CFL1; CF | P23528; E9P | COF1_HUMAN | Acetyl (Protein N | 0 | 6021800 | 0 | 0 | 1495200 | 0 | 0 | 0 | 0 | 594900 | 0 | 0 | 0 | 0 | 0 | 0 | 4316200 | 0 |
| ASGVEARQ | ERP29; ERP29 | P30040; F8V | ERP29_HUMAN | Unmodified | 0 | 0 | 0 | 0 | 92444 | 0 | 114950 | 34901 | 0 | 16577000 | 194130 | 0 | 43512 | 0 | 0 | 0 | 846030 | 625870 |
| ASGVPERY | TTLL12 | Q14166 | TTL12_HUMAN | Unmodified | 0 | 0 | 0 | 0 | 267530 | 0 | 0 | 0 | 377670 | 0 | 0 | 1272300 | 0 | 0 | 0 | 198850 | 0 | 0 |
| ASGVQVAD | DSTN; DSTN | P60981; F6R | DEST_HUMAN | Acetyl (Protein N | 0 | 0 | 0 | 0 | 10480000 | 0 | 2282500 | 0 | 34252 | 1119300 | 0 | 0 | 9645700 | 9837800 | 0 | 0 | 10510000 | 5201400 |
| ASGVQVAD | DSTN; DSTN | P60981; F6R | DEST_HUMAN | Acetyl (Protein N | 0 | 0 | 0 | 0 | 1010100 | 0 | 289970 | 0 | 0 | 120560 | 0 | 0 | 1488200 | 842540 | 0 | 0 | 1114200 | 632670 |
| ASGVTVND | CFL2; CFL2; CF | Q9Y281; F8V | COF2_HUMAN | Acetyl (Protein N | 1192500 | 854640 | 723260 | 747670 | 49401000 | 8364900 | 1018800 | 1065400 | 574050 | 354210 | 0 | 3082600 | 0 | 1325500 | 852310 | 947420 | 3178500 | 3891900 |
| ASGVTVND | CFL2; CFL2; CF | Q9Y281; F8V | COF2_HUMAN | Acetyl (Protein N | 0 | 0 | 0 | 0 | 19142000 | 4630500 | 0 | 799460 | 0 | 0 | 302400 | 0 | 0 | 1681900 | 0 | 0 | 6102100 | 0 |
| ASGVTVND | CFL2; CFL2; CF | Q9Y281; F8V | COF2_HUMAN | Acetyl (Protein N | 0 | 0 | 0 | 29025 | 12903000 | 528080 | 1076100 | 0 | 0 | 68538 | 0 | 0 | 0 | 0 | 22013 | 0 | 3109200 | 1158900 |
| ASHEEVEG | LGALS3BP | Q08380 | LG3BP_HUMAN | Unmodified | 0 | 0 | 196660 | 0 | 0 | 890280 | 0 | 124070 | 22087000 | 2477200 | 7210700 | 16312000 | 0 | 78659 | 137090 | 195830 | 0 | 263500 |
| ASHGYGGI | CTTN | Q14247 | SRC8_HUMAN | Oxidation (M) | 0 | 0 | 0 | 0 | 124100 | 0 | 0 | 0 | 0 | 0 | 0 | 0 | 1854900 | 54166 | 0 | 0 | 923210 | 1074200 |
| ASHGYGGI | CTTN | Q14247 | SRC8_HUMAN | Unmodified | 0 | 65894 | 0 | 0 | 4340700 | 773410 | 171220 | 62728 | 0 | 330960 | 0 | 0 | 3387700 | 2173500 | 0 | 0 | 1204500 | 1799300 |
| ASHNIGIAI | DBT; DBT; DBT | P11182; A0A | ODB2_HUMAN | Oxidation (M) | 106210 | 254820 | 0 | 0 | 0 | 0 | 0 | 0 | 425300 | 0 | 0 | 80273 | 0 | 0 | 150560 | 74607 | 0 | 0 |
| ASHPEDPA | DPP7; DPP7 | Q9UHL4; R4 | DPP2_HUMAN | Unmodified | 0 | 0 | 0 | 0 | 0 | 0 | 0 | 0 | 660700 | 0 | 0 | 0 | 0 | 0 | 0 | 0 | 0 | 0 |
| ASHTAPQV | ARPC2; ARPC2 | O15144; H7( | ARPC2_HUMAI | Unmodified | 2253400 | 0 | 0 | 250440 | 0 | 0 | 0 | 0 | 0 | 0 | 0 | 7778000 | 0 | 0 | 0 | 1450200 | 0 | 0 |
| ASHVTPQP | CCNDBP1 | O95273 | CCDB1_HUMA | Oxidation (M) | 0 | 445260 | 0 | 647550 | 0 | 0 | 0 | 0 | 0 | 0 | 0 | 0 | 0 | 0 | 1373700 | 228640 | 0 | 0 |
| ASIHEAWTI | ACTN4; ACTN1; | O43707; P12 | ACTN4_HUMAI | Unmodified | 1076100 | 1112900 | 1578700 | 846670 | 16069000 | 4788500 | 1263900 | 258230 | 3693900 | 2277400 | 1597100 | 27173000 | 1737600 | 0 | 2649400 | 3067700 | 4623800 | 739930 |
| ASIHEAWTI | ACTN4; ACTN4; | O43707; F5C | ACTN4_HUMAI | Unmodified | 0 | 0 | 0 | 162580 | 12857000 | 3584000 | 1230400 | 63144 | 387270 | 3611400 | 216290 | 2577500 | 1569600 | 923620 | 0 | 53346 | 6898000 | 235130 |
| ASIHEAWTI | ACTN4; ACTN4; | O43707; F5C | ACTN4_HUMAI | Oxidation (M) | 85210 | 0 | 0 | 216010 | 2797700 | 1980600 | 300120 | 218560 | 0 | 587130 | 248150 | 0 | 190560 | 121420 | 0 | 0 | 1601900 | 0 |
| ASIHEAWTI | ACTN1; ACTN1 | P12814; H9I | ACTN1_HUMAI | Unmodified | 45593 | 0 | 0 | 0 | 5403100 | 1541400 | 738710 | 0 | 0 | 297470 | 0 | 0 | 161270 | 147480 | 0 | 0 | 7534800 | 672920 |
| ASIHEAWTI | ACTN1; ACTN1 | P12814; H9I | ACTN1_HUMAI | Oxidation (M) | 0 | 0 | 0 | 0 | 295240 | 0 | 10429 | 0 | 0 | 0 | 0 | 0 | 0 | 0 | 0 | 0 | 415250 | 0 |
| ASIKKGEDI | MDH2; MDH2 | P40926; G3X | MDHM_HUMAI | Unmodified | 0 | 0 | 0 | 0 | 1537300 | 0 | 3275900 | 877190 | 269780 | 6835000 | 0 | 0 | 1976600 | 0 | 0 | 0 | 607500 | 791880 |
| ASILNTWIS | PAICS; PAICS | P22234; D6F | PUR6_HUMAN | Unmodified | 0 | 0 | 0 | 0 | 0 | 0 | 0 | 0 | 889710 | 212110 | 92588 | 8293800 | 0 | 0 | 427260 | 349020 | 580750 | 0 |
| ASILWLIGE | AP3B1 | O00203 | AP3B1_HUMAN | Unmodified | 0 | 0 | 0 | 0 | 109600 | 13021 | 191900 | 0 | 7660200 | 5356900 | 63081 | 1676700 | 149260 | 0 | 0 | 0 | 0 | 0 |
| ASINMLR | RPL7; RPL7; RP | P18124; C9J | RL7_HUMAN 6( | Oxidation (M) | 0 | 0 | 0 | 0 | 0 | 0 | 0 | 0 | 347150 | 0 | 0 | 3779500 | 0 | 0 | 0 | 0 | 16183 | 0 |
| ASINMLR | RPL7; RPL7; RP | P18124; C9J | RL7_HUMAN 6( | Unmodified | 869380 | 974700 | 460670 | 414530 | 83886 | 0 | 105550 | 82648 | 0 | 0 | 0 | 45160000 | 0 | 0 | 689360 | 496650 | 0 | 0 |
| ASINSLLSD | ATP7A | Q04656 | ATP7A_HUMAN | Unmodified | 0 | 0 | 253100 | 179290 | 0 | 0 | 35808 | 0 | 0 | 418680 | 0 | 0 | 0 | 0 | 0 | 346590 | 0 | 0 |
| ASINSLLSD | ATP7A | Q04656 | ATP7A_HUMAN | Unmodified | 0 | 0 | 137400 | 680700 | 0 | 0 | 0 | 0 | 0 | 0 | 0 | 0 | 0 | 55950 | 105080 | 190990 | 0 | 0 |
| ASIPFSVVG | SEPTIN2; SEPTI | Q15019; H7( | SEPT2_HUMAN | Unmodified | 521090 | 1421500 | 1011100 | 3339200 | 36000000 | 9627200 | 4909500 | 4368500 | 3252700 | 2393200 | 851280 | 10864000 | 9124000 | 3164500 | 2761000 | 1064300 | 19096000 | 16673000 |
| ASIPFSVVG | SEPTIN2; SEPTI | Q15019; H7( | SEPT2_HUMAN | Unmodified | 0 | 0 | 0 | 0 | 2982200 | 0 | 0 | 349350 | 0 | 0 | 0 | 0 | 0 | 0 | 0 | 0 | 2248400 | 0 |
| ASIPFSVVG | SEPTIN2; SEPTI | Q15019; H7( | SEPT2_HUMAN | Unmodified | 0 | 0 | 0 | 0 | 0 | 0 | 1436200 | 162320 | 0 | 0 | 0 | 0 | 0 | 0 | 0 | 0 | 1228400 | 289110 |
| ASISTENFR | EXT1; EXT1 | Q16394; H7( | EXT1_HUMAN I | Unmodified | 0 | 132550 | 0 | 279630 | 234750 | 0 | 2604500 | 46094 | 11098000 | 1292700 | 90316 | 294530 | 0 | 0 | 278730 | 240230 | 68171 | 378200 |
| ASITALEAK | MYH9 | P35579 | MYH9_HUMAN | Unmodified | 8.71E+08 | 168050 | 3E+08 | 4.84E+08 | 30273000 | 6927200 | 10229000 | 464070 | 2507600 | 1829700 | 1746000 | 13946000 | 1.57E+08 | 60732000 | 7.27E+08 | 5.54E+08 | 7416300 | 8749200 |
| ASITALEAKI | MYH9 | P35579 | MYH9_HUMAN | Unmodified | 0 | 0 | 0 | 0 | 380230 | 0 | 0 | 0 | 0 | 0 | 0 | 0 | 0 | 151710 | 0 | 0 | 525430 | 0 |
| ASITPGTILI | RPL6; RPL6; RP | Q02878; U3I | RL6_HUMAN 6( | Unmodified | 518650 | 564340 | 113240 | 77943 | 0 | 0 | 0 | 0 | 0 | 0 | 32606 | 2666400 | 0 | 0 | 298580 | 145730 | 0 | 0 |
| ASIWVGHR | ANXA3; ANXA3; | P12429; D6F | ANXA3_HUMAI | Acetyl (Protein N | 0 | 0 | 0 | 0 | 1.37E+08 | 15927000 | 0 | 0 | 0 | 182020 | 0 | 0 | 0 | 0 | 0 | 0 | 27480000 | 25888000 |
| ASKGLQDL | ADIRF; ADIRF | Q15847; Q5 | ADIRF_HUMAN | Acetyl (Protein N | 0 | 0 | 0 | 0 | 0 | 0 | 0 | 0 | 0 | 0 | 0 | 0 | 0 | 0 | 0 | 0 | 344850 | 0 |
| ASKLQNEL | MYH10 | P35580 | MYH10_HUMAI | Unmodified | 0 | 0 | 0 | 0 | 4366600 | 0 | 648400 | 0 | 0 | 0 | 0 | 0 | 1899200 | 1491200 | 0 | 0 | 324250 | 0 |
| ASKPLPPAI | SF3A1; SF3A1 | Q15459; H7( | SF3A1_HUMAN | Unmodified | 654870 | 526230 | 0 | 0 | 0 | 0 | 0 | 0 | 0 | 0 | 0 | 973550 | 0 | 0 | 0 | 413630 | 0 | 0 |
| ASLAETDKI | ATP6V1A | P38606 | VATA_HUMAN | Unmodified | 247990 | 0 | 53444 | 402110 | 473090 | 0 | 0 | 0 | 975470 | 0 | 0 | 0 | 1610000 | 191730 | 0 | 549020 | 206500 | 0 |
| ASLASLPPL | DCTN1; DCTN1 | Q14203; A0/ | DCTN1_HUMAI | Unmodified | 0 | 0 | 115470 | 0 | 1093600 | 0 | 326590 | 243850 | 171120 | 108860 | 0 | 3048400 | 767850 | 520470 | 0 | 77305 | 483440 | 1365800 |
| ASLAYFCQ | TXNDC16 | Q9P2K2 | TXD16_HUMAN | Unmodified | 312760 | 0 | 0 | 0 | 285240 | 0 | 0 | 0 | 834270 | 0 | 0 | 0 | 0 | 0 | 0 | 0 | 0 | 0 |
| ASLDRPFTI | RPN2; RPN2 | P04844; Q5J | RPN2_HUMAN | Unmodified | 0 | 0 | 0 | 0 | 0 | 0 | 0 | 0 | 443060 | 5441700 | 0 | 0 | 0 | 0 | 0 | 0 | 0 | 0 |
| ASLDSAGG | RTKN | Q9BST9 | RTKN_HUMAN | Unmodified | 0 | 0 | 0 | 0 | 0 | 0 | 0 | 0 | 314800 | 0 | 0 | 3784500 | 0 | 0 | 0 | 0 | 0 | 0 |
| ASLEDAPVI | ZC3HAV1; ZC3I | Q7Z2W4; C9 | ZCCHV_HUMA | Unmodified | 0 | 651030 | 615600 | 2682100 | 0 | 0 | 199850 | 0 | 1803200 | 0 | 0 | 0 | 0 | 148910 | 0 | 0 | 0 | 0 |
| ASLEDLGW | GDF15 | Q99988 | GDF15_HUMAI | Unmodified | 0 | 17952 | 42519 | 125300 | 0 | 0 | 0 | 0 | 797310 | 183030 | 0 | 0 | 0 | 0 | 365330 | 31361 | 0 | 0 |
| ASLENSLRI | KRT18; KRT18 | P05783; F8V | K1C18_HUMAI | Unmodified | 432440 | 309300 | 635640 | 0 | 0 | 562720 | 0 | 0 | 8612700 | 1828900 | 27069000 | 20484000 | 672040 | 290870 | 621540 | 742240 | 299750 | 0 |
| ASLGSLEG | AHNAK | Q09666 | AHNK_HUMAN | Unmodified | 85600 | 0 | 0 | 0 | 13270000 | 2175000 | 0 | 0 | 1044100 | 426260 | 0 | 0 | 0 | 207310 | 0 | 49900 | 4212100 | 4493900 |
| ASLGSLEG | AHNAK | Q09666 | AHNK_HUMAN | Unmodified | 0 | 0 | 0 | 0 | 0 | 0 | 0 | 0 | 98278 | 0 | 0 | 0 | 0 | 0 | 0 | 0 | 2715100 | 0 |
| ASLHALVG | ACOT2; ACOT1; | P49753; Q8( | ACOT2_HUMAI | Unmodified | 0 | 0 | 0 | 0 | 0 | 0 | 0 | 0 | 0 | 0 | 0 | 0 | 0 | 0 | 0 | 0 | 0 | 0 |
| ASLHGGEF | NID1 | P14543 | NID1_HUMAN | Unmodified | 0 | 65795 | 0 | 0 | 0 | 0 | 0 | 0 | 454400 | 0 | 0 | 254360 | 67075 | 0 | 0 | 0 | 0 | 41352 |
| ASLHGGEF | NID1 | P14543 | NID1_HUMAN | Unmodified | 0 | 0 | 0 | 0 | 0 | 0 | 0 | 0 | 17797000 | 660840 | 0 | 0 | 0 | 0 | 0 | 0 | 0 | 0 |
| ASLIEYIK | CPM; CPM | P14384; H0Y | CBPM_HUMAN | Unmodified | 0 | 0 | 0 | 0 | 2966100 | 0 | 0 | 336170 | 5542500 | 1773300 | 630640 | 418960 | 0 | 0 | 33914 | 0 | 157840 | 0 |
| ASLISAVSD | TSG101; TSG10 | Q99816; F5H | TS101_HUMAN | Unmodified | 34859000 | 31568000 | 48052000 | 35017000 | 329050 | 0 | 0 | 0 | 12883000 | 553410 | 2331600 | 14948000 | 1190300 | 0 | 70201000 | 44643000 | 0 | 0 |
| ASLISAVSD | TSG101; TSG10 | Q99816; F5H | TS101_HUMAN | Unmodified | 24184000 | 7589300 | 0 | 0 | 439480 | 0 | 0 | 47508 | 906370 | 534020 | 0 | 0 | 3557800 | 935550 | 0 | 0 | 57137 | 410810 |
| ASLLDPVPI | GCN1 | Q92616 | GCN1_HUMAN | Unmodified | 0 | 0 | 0 | 0 | 0 | 0 | 0 | 0 | 211920 | 0 | 201160 | 1254000 | 0 | 0 | 0 | 0 | 55716 | 0 |
| ASLLDPVPI | GCN1 | Q92616 | GCN1_HUMAN | Unmodified | 0 | 0 | 0 | 0 | 0 | 0 | 915730 | 400200 | 0 | 0 | 0 | 0 | 532180 | 0 | 0 | 0 | 0 | 0 |
| ASLLQNES | COPS4; COPS4 | Q9BT78; D6I | CSN4_HUMAN | Unmodified | 0 | 0 | 0 | 0 | 0 | 0 | 0 | 0 | 506770 | 184510 | 0 | 3960600 | 0 | 0 | 191200 | 446090 | 374760 | 0 |
| ASLLQSDR | DCTN4; DCTN4 | Q9UJW0; E5 | DCTN4_HUMAI | Acetyl (Protein N | 0 | 0 | 0 | 0 | 132440 | 0 | 0 | 0 | 287350 | 615100 | 90450 | 518600 | 1130200 | 1173400 | 0 | 26245 | 0 | 166330 |

| | | | | | | | | | | | | | | | | | | | | | | |
|---|---|---|---|---|---|---|---|---|---|---|---|---|---|---|---|---|---|---|---|---|---|---|
| ASLMNDDI | PACSIN2; PACS | Q9UNF0; B0 | PACN2_HUMAI | Unmodified | 11316000 | 12806000 | 7436200 | 9069800 | 3300200 | 563390 | 326330 | 248320 | 910480 | 707240 | 1099100 | 3328900 | 26560000 | 8760400 | 14199000 | 8778200 | 472860 | 1041200 |
| ASLMNDDI | PACSIN2; PACS | Q9UNF0; B0 | PACN2_HUMAI | Oxidation (M) | 16657000 | 14228000 | 7034200 | 5780500 | 1315000 | 335830 | 326690 | 0 | 197620 | 161690 | 194420 | 905260 | 17660000 | 7845300 | 17473000 | 7940400 | 222640 | 0 |
| ASLMNDDI | PACSIN2; PACS | Q9UNF0; B0 | PACN2_HUMAI | Unmodified | 62744 | 85132 | 0 | 31233 | 692640 | 0 | 368490 | 97409 | 0 | 0 | 0 | 0 | 4306600 | 4643400 | 150410 | 22241 | 264110 | 193400 |
| ASLMNDDI | PACSIN2; PACS | Q9UNF0; B0 | PACN2_HUMAI | Oxidation (M) | 0 | 386930 | 0 | 0 | 0 | 0 | 0 | 143060 | 0 | 0 | 0 | 0 | 1416500 | 759630 | 0 | 0 | 0 | 0 |
| ASLNGADI | HNRNPL; HNRI | P14866; B4D | HNRPL_HUMAI | Unmodified | 0 | 0 | 0 | 0 | 0 | 0 | 0 | 272910 | 0 | 0 | 181160 | 1922200 | 0 | 0 | 0 | 0 | 0 | 0 |
| ASLNPSDTI | AGPS; AGPS; A( | O00116; A0/ | ADAS_HUMAN | Unmodified | 0 | 0 | 0 | 0 | 0 | 0 | 0 | 0 | 36694000 | 3173800 | 0 | 0 | 0 | 0 | 0 | 0 | 0 | 0 |
| ASLNPSDTI | AGPS; AGPS; A( | O00116; A0/ | ADAS_HUMAN | Unmodified | 0 | 0 | 0 | 0 | 0 | 0 | 0 | 0 | 15275000 | 0 | 0 | 0 | 0 | 0 | 0 | 0 | 0 | 0 |
| ASLPTLGFT | ADSL; ADSL; AD | P30566; A0A | PUR8_HUMAN | Unmodified | 0 | 0 | 237900 | 51273 | 0 | 0 | 0 | 0 | 0 | 0 | 627450 | 5072800 | 0 | 0 | 592150 | 297130 | 0 | 0 |
| ASLPTLGFT | ADSL; ADSL; AD | P30566; A0A | PUR8_HUMAN | Unmodified | 0 | 0 | 281610 | 0 | 0 | 0 | 0 | 0 | 0 | 22066 | 0 | 7762600 | 0 | 0 | 524400 | 377740 | 0 | 0 |
| ASLPVIATV | RMC1; RMC1; F | Q96DM3; K7 | RMC1_HUMAN | Unmodified | 9.3E+08 | 1.05E+09 | 9.03E+08 | 1.05E+09 | 0 | 0 | 0 | 0 | 0 | 0 | 0 | 0 | 1.69E+09 | 6.6E+08 | 1.29E+08 | 8.03E+08 | 0 | 0 |
| ASLPVVRS | KIAA1217; KIAA | Q5T5P2; Q5 | SKT_HUMAN S | Unmodified | 0 | 0 | 0 | 0 | 0 | 0 | 0 | 0 | 0 | 55144 | 0 | 0 | 0 | 0 | 0 | 0 | 100790 | 103750 |
| ASLPYSQIS | PIGS | Q96S52 | PIGS_HUMAN ( | Unmodified | 0 | 0 | 0 | 0 | 0 | 0 | 0 | 0 | 4994800 | 785330 | 0 | 738370 | 0 | 0 | 0 | 0 | 0 | 0 |
| ASLRELALC | RNH1; RNH1 | P13489; H0Y | RINI_HUMAN R | Unmodified | 4899700 | 585820 | 0 | 2688400 | 0 | 33019 | 0 | 0 | 269170 | 78394 | 428100 | 0 | 1444000 | 0 | 486140 | 343320 | 455960 | 839380 |
| ASLRELALC | RNH1; RNH1 | P13489; H0Y | RINI_HUMAN R | Oxidation (M) | 0 | 0 | 0 | 163970 | 1714300 | 0 | 0 | 0 | 0 | 0 | 0 | 0 | 621010 | 0 | 0 | 0 | 973980 | 0 |
| ASLSKLGD | PGK1 | P00558 | PGK1_HUMAN | Oxidation (M) | 0 | 0 | 0 | 0 | 0 | 0 | 0 | 0 | 0 | 0 | 0 | 0 | 0 | 0 | 0 | 0 | 1297700 | 0 |
| ASLSLAPV | CCT2; CCT2; C( | P78371; F5G | TCPB_HUMAN | Acetyl (Protein N | 65096 | 0 | 0 | 0 | 0 | 0 | 0 | 0 | 88335 | 0 | 143930 | 442630 | 269980 | 0 | 0 | 64393 | 0 | 0 |
| ASLSLIEK | KPNA2; KPNA2; | P52292; A0A | IMA1_HUMAN I | Unmodified | 310540 | 589200 | 0 | 0 | 4715300 | 0 | 739890 | 9729.7 | 1457400 | 1096900 | 0 | 1906500 | 1656900 | 0 | 0 | 470590 | 2199100 | 59605 |
| ASLVALPEC | ACIN1; ACIN1; | Q9UKV3; E7I | ACINU_HUMAI | Unmodified | 0 | 0 | 0 | 0 | 0 | 0 | 0 | 0 | 0 | 655950 | 0 | 1360900 | 0 | 0 | 0 | 0 | 0 | 0 |
| ASLYNAVTI | PSAT1 | Q9Y617 | SERC_HUMAN | Unmodified | 0 | 0 | 463370 | 629580 | 0 | 0 | 0 | 0 | 0 | 0 | 819550 | 3585200 | 1358300 | 0 | 399130 | 755270 | 0 | 40975 |
| ASMGTLAF | CCT5; CCT5; C( | P48643; D6F | TCPE_HUMAN | Acetyl (Protein N | 0 | 144000 | 0 | 0 | 0 | 36691 | 0 | 0 | 0 | 0 | 0 | 224940 | 0 | 0 | 0 | 0 | 0 | 0 |
| ASMHPVTA | LTA4H | P09960 | LKHA4_HUMAI | Unmodified | 0 | 0 | 0 | 0 | 3842400 | 175630 | 0 | 0 | 392850 | 0 | 0 | 979240 | 0 | 0 | 0 | 0 | 361880 | 1277000 |
| ASMKGLGT | ANXA2P2; ANX/ | A6NMY6; P0 | AXA2L_HUMAI | Oxidation (M) | 0 | 0 | 0 | 0 | 1095100 | 114470 | 0 | 0 | 106850 | 36533 | 0 | 0 | 0 | 0 | 0 | 0 | 1270000 | 447450 |
| ASNEDVTK | PSIP1 | O75475 | PSIP1_HUMAN | Unmodified | 0 | 0 | 0 | 0 | 0 | 0 | 0 | 74011 | 0 | 78335 | 0 | 0 | 55801 | 0 | 0 | 0 | 41984 | 0 |
| ASNGDAW | HSPA9; HSPA9; | P38646; H0Y | GRP75_HUMAI | Unmodified | 0 | 0 | 0 | 0 | 0 | 0 | 0 | 0 | 357330 | 211080 | 178760 | 144640 | 0 | 0 | 0 | 0 | 0 | 0 |
| ASNIFGTPE | JPT1; JPT1; JPT1 | Q9UK76; J3K | JUPI1_HUMAN | Acetyl (Protein N | 0 | 0 | 0 | 0 | 256060 | 0 | 0 | 0 | 0 | 0 | 0 | 0 | 0 | 0 | 0 | 0 | 257980 | 200890 |
| ASNIITECK | CDC42BPA; CD | Q5VT25; A0/ | MRCKA_HUMA | Unmodified | 0 | 0 | 0 | 0 | 957280 | 272360 | 0 | 107930 | 0 | 0 | 0 | 743470 | 0 | 0 | 0 | 0 | 417270 | 1190000 |
| ASNKTTLQI | CBX3; CBX3; CI | Q13185; B82 | CBX3_HUMAN | Acetyl (Protein N | 0 | 0 | 0 | 0 | 0 | 0 | 0 | 801320 | 0 | 0 | 0 | 0 | 0 | 0 | 0 | 0 | 0 | 0 |
| ASNLENST | COPA; COPA; C | P53621; A0A | COPA_HUMAN | Unmodified | 0 | 0 | 0 | 0 | 0 | 0 | 0 | 0 | 0 | 0 | 1864900 | 7023900 | 0 | 0 | 0 | 0 | 0 | 0 |
| ASNLEPPA | N/A | N/A | N/A | Oxidation (M) | 0 | 0 | 0 | 0 | 0 | 0 | 0 | 0 | 3488000 | 0 | 0 | 0 | 0 | 0 | 0 | 0 | 0 | 0 |
| ASNLLLGFI | SORL1 | Q92673 | SORL_HUMAN | Unmodified | 0 | 0 | 0 | 0 | 0 | 0 | 0 | 0 | 1084400 | 0 | 0 | 0 | 0 | 0 | 0 | 0 | 0 | 0 |
| ASNLLRSS | CD2AP | Q9Y5K6 | CD2AP_HUMAI | Unmodified | 70202 | 0 | 109920 | 0 | 1257800 | 0 | 124790 | 0 | 0 | 132930 | 0 | 0 | 2601800 | 1717700 | 0 | 0 | 0 | 0 |
| ASNLLYGQ | DNAJC10; DNA | Q8IXB1; A0A | DJC10_HUMAI | Unmodified | 0 | 0 | 0 | 0 | 0 | 0 | 0 | 0 | 6587800 | 2247000 | 0 | 0 | 0 | 0 | 0 | 0 | 0 | 0 |
| ASNNTASI/ | GNG2; GNG2; ( | P59768; G3\ | GBG2_HUMAN | Acetyl (Protein N | 0 | 210130 | 0 | 0 | 0 | 0 | 634910 | 509550 | 0 | 0 | 0 | 0 | 0 | 444750 | 0 | 0 | 6957200 | 13039000 |
| ASNNTASI/ | GNG2; GNG2; ( | P59768; G3\ | GBG2_HUMAN | Acetyl (Protein N | 0 | 0 | 0 | 95405 | 535630 | 83826 | 364970 | 362480 | 324810 | 0 | 94763 | 580700 | 0 | 0 | 50287 | 0 | 1358300 | 5020500 |
| ASNNTASI/ | GNG2; GNG2; ( | P59768; G3\ | GBG2_HUMAN | Acetyl (Protein N | 0 | 0 | 0 | 0 | 439640 | 0 | 2443200 | 0 | 0 | 0 | 0 | 0 | 0 | 982390 | 0 | 0 | 8941100 | 8456400 |
| ASNPFLPG | EPN1 | Q9Y6I3 | EPN1_HUMAN | Unmodified | 827240 | 1314600 | 265810 | 707890 | 661530 | 0 | 0 | 0 | 512980 | 577470 | 151830 | 253290 | 5110400 | 2256000 | 0 | 1061700 | 1450900 | 7544900 |
| ASNPVIAA( | EPB41L4B | Q9H329 | E41LB_HUMAI | Unmodified | 0 | 0 | 901510 | 0 | 0 | 0 | 0 | 0 | 255050 | 178140 | 0 | 0 | 604920 | 531490 | 0 | 0 | 0 | 0 |
| ASNTADTLI | EXOC2 | Q96KP1 | EXOC2_HUMAI | Unmodified | 175110 | 99637 | 613850 | 984630 | 1268600 | 175160 | 0 | 0 | 948240 | 0 | 860730 | 4505700 | 90926 | 0 | 423940 | 949810 | 0 | 339290 |
| ASNTAEVFI | ACADVL | P49748 | ACADV_HUMAI | Unmodified | 0 | 0 | 0 | 0 | 0 | 0 | 0 | 0 | 8380400 | 4914400 | 0 | 0 | 0 | 0 | 0 | 0 | 0 | 0 |
| ASNVTNKT | HNRNPC; HNR | P07910; G3\ | HNRPC_HUMA | Acetyl (Protein N | 928980 | 337600 | 93606 | 252140 | 1349000 | 617300 | 0 | 852290 | 1285600 | 0 | 427530 | 1422000 | 2646800 | 547300 | 0 | 99908 | 176150 | 906170 |
| ASNVTNKT | HNRNPC; HNR | P07910; G3\ | HNRPC_HUMA | Acetyl (Protein N | 0 | 0 | 0 | 0 | 1154300 | 236480 | 1001900 | 866720 | 0 | 827810 | 0 | 0 | 1366900 | 720420 | 0 | 0 | 741160 | 446600 |
| ASNVTNKT | HNRNPC; HNR | P07910; G3\ | HNRPC_HUMA | Acetyl (Protein N | 0 | 0 | 0 | 0 | 0 | 0 | 311170 | 322500 | 0 | 0 | 0 | 0 | 0 | 0 | 0 | 94716 | 0 | 0 |
| ASPAAEDA | USP6NL; USP6I | Q92738; X6F | US6NL_HUMAI | Unmodified | 0 | 0 | 0 | 0 | 0 | 0 | 0 | 0 | 255760 | 0 | 0 | 0 | 0 | 0 | 0 | 0 | 0 | 822510 |
| ASPAAPAD | CAMSAP3 | Q9P1Y5 | CAMP3_HUMA | Unmodified | 0 | 0 | 0 | 0 | 794750 | 0 | 672350 | 503750 | 0 | 377990 | 0 | 0 | 0 | 0 | 0 | 0 | 156870 | 566160 |
| ASPAGGPL | FKBP10; FKBP1 | Q96AY3; K7E | FKB10_HUMAI | Unmodified | 0 | 0 | 0 | 0 | 0 | 0 | 0 | 0 | 0 | 0 | 734620 | 830090 | 0 | 0 | 0 | 0 | 0 | 0 |
| ASPATQPPI | KHDRBS1 | Q07666 | KHDR1_HUMAI | Unmodified | 0 | 0 | 0 | 0 | 0 | 0 | 0 | 0 | 0 | 1435800 | 2040000 | 4515000 | 0 | 0 | 0 | 0 | 0 | 0 |
| ASPFLLQYI | PRPSAP2; PRPS | O60256; E7E | KPRB_HUMAN | Unmodified | 0 | 0 | 0 | 17661 | 0 | 0 | 0 | 0 | 313130 | 0 | 0 | 313460 | 0 | 0 | 81575 | 0 | 0 | 0 |
| ASPGTPLSI | SLCO4A1; SLC( | Q96BD0; E1I | SO4A1_HUMAI | Unmodified | 0 | 0 | 0 | 751680 | 0 | 0 | 0 | 0 | 0 | 0 | 0 | 1924200 | 0 | 0 | 881540 | 840970 | 1828300 | 3957100 |
| ASPLLPANI | IARS2 | Q9NSE4 | SYIM_HUMAN I | Unmodified | 0 | 0 | 0 | 0 | 0 | 0 | 0 | 0 | 682380 | 180180 | 0 | 503800 | 0 | 0 | 0 | 0 | 0 | 0 |
| ASPLTHSPI | CNPY3 | Q9BT09 | CNPY3_HUMAI | Unmodified | 0 | 0 | 0 | 0 | 0 | 0 | 563880 | 361570 | 17110000 | 7981500 | 1063300 | 0 | 1011300 | 0 | 0 | 0 | 0 | 0 |
| ASPNTPPG | GCN1 | Q92616 | GCN1_HUMAN | Unmodified | 55135 | 115890 | 84533 | 38796 | 0 | 0 | 0 | 0 | 0 | 0 | 0 | 962100 | 0 | 0 | 180630 | 184010 | 0 | 0 |
| ASPQAADL | MAPK13 | O15264 | MK13_HUMAN | Unmodified | 0 | 0 | 0 | 0 | 977330 | 335870 | 225040 | 170260 | 884850 | 666030 | 299780 | 2420000 | 443990 | 0 | 0 | 0 | 317170 | 671970 |
| ASPQSEFM | ANLN; ANLN | Q9NQW6; H | ANLN_HUMAN | Unmodified | 1257000 | 410770 | 362460 | 0 | 8546100 | 0 | 0 | 0 | 299300 | 0 | 0 | 482650 | 0 | 0 | 0 | 0 | 807910 | 1020700 |
| ASPSTAGE | EIF2B4; EIF2B4 | Q9UI10; A0A | EI2BD_HUMAN | Unmodified | 339460 | 0 | 49625 | 0 | 85500 | 0 | 0 | 0 | 0 | 0 | 151200 | 454420 | 344090 | 82713 | 0 | 0 | 60516 | 289240 |
| ASPVNDIF( | ERLEC1 | Q96DZ1 | ERLEC_HUMAI | Unmodified | 0 | 0 | 0 | 0 | 0 | 0 | 0 | 0 | 7586400 | 557450 | 0 | 0 | 0 | 0 | 0 | 0 | 0 | 0 |
| ASPVPAPS | KIAA1522 | Q9P206 | K1522_HUMAI | Unmodified | 0 | 0 | 0 | 2834800 | 870160 | 152820 | 0 | 0 | 2253000 | 0 | 704340 | 2389600 | 1849700 | 724310 | 2979500 | 1640300 | 446450 | 491970 |
| ASPVPAPS | KIAA1522 | Q9P206 | K1522_HUMAI | Unmodified | 0 | 0 | 0 | 0 | 2343300 | 411770 | 0 | 0 | 0 | 0 | 0 | 0 | 3061400 | 2950300 | 0 | 0 | 1683900 | 1381200 |
| ASQDPFPA | PBK; PBK | Q96KB5; E5F | TOPK_HUMAN | Unmodified | 0 | 0 | 455520 | 0 | 0 | 0 | 0 | 286120 | 1523900 | 0 | 0 | 990040 | 0 | 0 | 0 | 756240 | 0 | 0 |
| ASQESEAR | SUN2 | Q9UH99 | SUN2_HUMAN | Unmodified | 0 | 44539 | 0 | 0 | 0 | 0 | 0 | 0 | 1586900 | 352540 | 95232 | 0 | 0 | 0 | 0 | 0 | 0 | 0 |
| ASQGGRGI | BBS4 | H3BN76 | H3BN76_HUM/ | Unmodified | 0 | 0 | 0 | 0 | 0 | 0 | 0 | 0 | 0 | 0 | 0 | 0 | 0 | 0 | 0 | 0 | 1788300 | 0 |
| ASQGPVYK | CARHSP1; CAR | Q9Y2V2; H3I | CHSP1_HUMAI | Unmodified | 0 | 0 | 0 | 0 | 599840 | 0 | 368510 | 739950 | 1056400 | 212300 | 0 | 483880 | 0 | 0 | 0 | 0 | 393820 | 536320 |
| ASQGPVYK | CARHSP1; CAR | Q9Y2V2; H3I | CHSP1_HUMAI | Unmodified | 0 | 0 | 0 | 0 | 0 | 0 | 859370 | 937330 | 120510 | 0 | 0 | 0 | 0 | 259430 | 0 | 0 | 725430 | 443890 |
| ASQKPQPN | USO1 | O60763 | USO1_HUMAN | Oxidation (M) | 0 | 0 | 0 | 0 | 1097900 | 100170 | 0 | 0 | 0 | 0 | 494890 | 7252500 | 0 | 1848900 | 0 | 0 | 0 | 0 |
| ASQLVSEK | SH3GL1 | Q99961 | SH3G1_HUMAI | Unmodified | 4376300 | 2130900 | 4719600 | 5088900 | 1043900 | 117760 | 663780 | 1694500 | 156290 | 60016 | 0 | 52189 | 3436100 | 1027900 | 6213400 | 2691500 | 285640 | 389960 |
| ASQLYSHN | MAP4K3; MAP4 | Q8IVH8; F5H | M4K3_HUMAN | Unmodified | 134020 | 7203900 | 0 | 0 | 0 | 0 | 0 | 0 | 0 | 0 | 0 | 1356100 | 0 | 0 | 82100 | 0 | 0 | 0 |
| ASQSQGIQ | ATP6V1G1; ATP | O75348; O9 | VATG1_HUMAI | Acetyl (Protein N | 0 | 0 | 0 | 0 | 2371500 | 202940 | 3614200 | 1207800 | 0 | 317870 | 176050 | 2761800 | 8243300 | 2698800 | 0 | 84152 | 271800 | 1399700 |
| ASQSQGIQ | ATP6V1G1; ATP | O75348; O9 | VATG1_HUMAI | Acetyl (Protein N | 0 | 0 | 0 | 0 | 0 | 0 | 0 | 409860 | 0 | 0 | 0 | 0 | 0 | 0 | 0 | 0 | 0 | 0 |
| ASQSQGIQ | ATP6V1G1; ATP | O75348; O9 | VATG1_HUMAI | Acetyl (Protein N | 0 | 0 | 1563500 | 0 | 0 | 0 | 2797900 | 727190 | 0 | 0 | 0 | 0 | 0 | 0 | 0 | 0 | 0 | 0 |
| ASQSRPNS | STK10 | O94804 | STK10_HUMAI | Unmodified | 0 | 0 | 0 | 0 | 2165600 | 621560 | 407540 | 0 | 239760 | 54746 | 0 | 937610 | 523240 | 692680 | 0 | 0 | 1663600 | 3953000 |
| ASQVKPGD | ADRA2A | P08913 | ADA2A_HUMAI | Unmodified | 0 | 41685 | 478500 | 428130 | 0 | 0 | 18048 | 95125 | 838240 | 300640 | 0 | 0 | 238750 | 362340 | 402750 | 1064100 | 0 | 0 |
| ASQVVFMS | KIF14 | Q15058 | KIF14_HUMAN | Unmodified | 0 | 0 | 0 | 0 | 2176400 | 0 | 132180 | 47160 | 0 | 0 | 0 | 0 | 0 | 0 | 0 | 0 | 839540 | 0 |
| ASRDEIFAC | MYH10 | P35580 | MYH10_HUMAI | Unmodified | 0 | 0 | 0 | 0 | 0 | 0 | 0 | 0 | 0 | 0 | 0 | 2534200 | 0 | 0 | 109110 | 187140 | 0 | 0 |
| ASREEILAQ | MYH9 | P35579 | MYH9_HUMAN | Unmodified | 0 | 1264100 | 0 | 0 | 306810 | 140660 | 0 | 0 | 0 | 0 | 0 | 0 | 201640 | 0 | 0 | 0 | 126960 | 101620 |
| ASRENPAM | SPAG9; SPAG9 | O60271; A0/ | JIP4_HUMAN C | Unmodified | 0 | 0 | 0 | 0 | 196680 | 0 | 58270 | 40770 | 0 | 0 | 0 | 0 | 57468 | 55822 | 0 | 0 | 0 | 246500 |
| ASRKEGTG | EIF3H; EIF3H; E | O15372; E5F | EIF3H_HUMAN | Acetyl (Protein N | 0 | 0 | 0 | 0 | 645520 | 76801 | 0 | 0 | 0 | 0 | 0 | 0 | 1429400 | 850540 | 0 | 0 | 0 | 480280 |
| ASRKLTELE | AFDN; AFDN; A | P55196; H0Y | AFAD_HUMAN | Unmodified | 0 | 0 | 0 | 0 | 1884700 | 71937 | 241190 | 0 | 0 | 84289 | 0 | 0 | 592210 | 1716300 | 0 | 0 | 716770 | 550540 |
| ASRVPSSD | TNKS1BP1 | Q9C0C2 | TB182_HUMAI | Unmodified | 0 | 28886 | 0 | 196350 | 118760 | 0 | 0 | 0 | 186470 | 126040 | 0 | 1050500 | 407130 | 73634 | 0 | 0 | 0 | 0 |
| ASSAQSGC | SND1 | Q7KZF4 | SND1_HUMAN | Acetyl (Protein N | 224920 | 288330 | 1330600 | 1870200 | 1329900 | 398600 | 0 | 312700 | 0 | 216690 | 4508100 | 9259000 | 3600400 | 1879000 | 1361500 | 2772200 | 1428900 | 0 |
| ASSAQSGC | SND1 | Q7KZF4 | SND1_HUMAN | Acetyl (Protein N | 0 | 0 | 0 | 0 | 3383100 | 0 | 5690500 | 0 | 0 | 690690 | 0 | 0 | 3697300 | 3634400 | 0 | 0 | 6902400 | 4700700 |
| ASSDIQVK | STMN1; STMN1 | P16949; A2A | STMN1_HUMAI | Acetyl (Protein N | 53864 | 0 | 0 | 375520 | 158700 | 0 | 2823300 | 4558200 | 17176000 | 9152900 | 3249100 | 5672200 | 1918900 | 873670 | 0 | 0 | 2400400 | 3324000 |
| ASSDIQVKI | STMN1; STMN1 | P16949; A2A | STMN1_HUMAI | Acetyl (Protein N | 0 | 0 | 0 | 0 | 0 | 0 | 5808900 | 6496700 | 0 | 2098600 | 0 | 0 | 520380 | 0 | 0 | 0 | 2525800 | 0 |
| ASSDIQVKI | STMN1; STMN1 | P16949; A2A | STMN1_HUMAI | Acetyl (Protein N | 0 | 0 | 0 | 0 | 0 | 0 | 0 | 892660 | 0 | 351080 | 0 | 0 | 0 | 0 | 0 | 0 | 0 | 0 |
| ASSEGGTA | ARPC1B; ARPC | O15143; C9J | ARC1B_HUMAI | Unmodified | 0 | 335600 | 0 | 75088 | 2024400 | 0 | 0 | 89785 | 6752500 | 102020 | 0 | 456700 | 0 | 0 | 130680 | 0 | 341330 | 153540 |
| ASSEGGTA | ARPC1B | O15143 | ARC1B_HUMAI | Unmodified | 0 | 0 | 0 | 0 | 0 | 0 | 0 | 68742 | 0 | 0 | 0 | 0 | 111270 | 64882 | 0 | 0 | 1539100 | 757520 |
| ASSELFSQI | HMGCS1 | Q01581 | HMCS1_HUMA | Unmodified | 0 | 0 | 777920 | 244590 | 2561400 | 301500 | 130970 | 976100 | 953860 | 672670 | 1608600 | 7144400 | 337220 | 0 | 748230 | 628950 | 0 | 0 |
| ASSEQAEQ | PEX14 | O75381 | PEX14_HUMAI | Acetyl (Protein N | 0 | 0 | 0 | 0 | 0 | 0 | 0 | 0 | 3853800 | 0 | 0 | 0 | 0 | 0 | 0 | 0 | 0 | 0 |
| ASSEQAEQ | PEX14 | O75381 | PEX14_HUMAI | Acetyl (Protein N | 0 | 0 | 0 | 0 | 0 | 0 | 0 | 0 | 18682000 | 4481700 | 0 | 0 | 0 | 0 | 0 | 0 | 0 | 0 |
| ASSGGQSI | NOSTRIN; NOS | Q8IVI9; A0A( | NOSTN_HUMA | Unmodified | 6089900 | 4569800 | 13622000 | 10727000 | 0 | 0 | 0 | 0 | 0 | 0 | 0 | 0 | 1153100 | 0 | 22650000 | 8846200 | 0 | 0 |
| ASSGGQSI | NOSTRIN; NOS | Q8IVI9; A0A( | NOSTN_HUMA | Unmodified | 0 | 0 | 0 | 0 | 0 | 0 | 0 | 0 | 0 | 0 | 0 | 0 | 365300 | 651540 | 0 | 0 | 0 | 0 |
| ASSHSVSI( | PTPRB; PTPRB; | P23467; F8V | PTPRB_HUMAI | Unmodified | 569500 | 301460 | 0 | 0 | 0 | 0 | 0 | 0 | 0 | 0 | 0 | 0 | 0 | 0 | 0 | 0 | 0 | 0 |
| ASSIENKQI | TRIO; TRIO; TRI( | O75962; E7E | TRIO_HUMAN T | Unmodified | 0 | 0 | 0 | 59214 | 0 | 61452 | 50954 | 0 | 388490 | 128940 | 69992 | 0 | 0 | 111340 | 0 | 0 | 167460 | 160890 |
| ASSIIDELF( | CLU; CLU; CLU | P10909; H0Y | CLUS_HUMAN | Unmodified | 252590 | 290410 | 0 | 0 | 0 | 0 | 0 | 0 | 133240 | 0 | 0 | 0 | 0 | 0 | 304340 | 46018 | 0 | 0 |
| ASSLIEEAK | TLN1 | Q9Y490 | TLN1_HUMAN | Unmodified | 26700000 | 29255000 | 466790 | 398920 | 1548200 | 491680 | 1236000 | 133700 | 0 | 99970 | 170900 | 787140 | 1516000 | 174460 | 293410 | 0 | 196810 | 0 |
| ASSLISLLK | EIF4G2; EIF4G2 | P78344; H0Y | IF4G2_HUMAN | Unmodified | 0 | 0 | 0 | 0 | 0 | 0 | 0 | 0 | 0 | 0 | 0 | 1806800 | 0 | 0 | 0 | 0 | 0 | 0 |
| ASSLSESSI | LIMA1; LIMA1; L | Q9UHB6; F8 | LIMA1_HUMAN | Unmodified | 0 | 0 | 0 | 0 | 0 | 0 | 0 | 0 | 0 | 0 | 0 | 0 | 0 | 0 | 0 | 0 | 0 | 0 |
| ASSPSGKM | TARS1; TARS1; | P26639; D6F | SYTC_HUMAN | Unmodified | 0 | 0 | 0 | 0 | 3551100 | 304190 | 0 | 170820 | 0 | 256840 | 0 | 0 | 1997200 | 1392200 | 0 | 0 | 818210 | 713670 |
| ASSPVPLPI | LGMN | Q99538 | LGMN_HUMAN | Oxidation (M) | 0 | 0 | 0 | 0 | 0 | 0 | 0 | 0 | 3006500 | 879780 | 0 | 0 | 0 | 0 | 0 | 0 | 0 | 0 |
| ASSQVNVE | RGS10 | O43665 | RGS10_HUMAI | Unmodified | 0 | 0 | 0 | 0 | 1002200 | 251830 | 396030 | 359140 | 0 | 205060 | 0 | 0 | 1559500 | 397690 | 64201 | 37942 | 349410 | 592340 |
| ASSRVSGS | STX7 | O15400 | STX7_HUMAN S | Unmodified | 0 | 0 | 0 | 0 | 833440 | 120390 | 0 | 0 | 0 | 195250 | 101210 | 0 | 0 | 0 | 0 | 0 | 0 | 0 |
| ASSSGNDI | DR1 | Q01658 | NC2B_HUMAN | Acetyl (Protein N | 0 | 234420 | 0 | 0 | 0 | 0 | 0 | 0 | 0 | 0 | 0 | 579590 | 1688600 | 0 | 83323 | 0 | 603080 | 0 |
| ASSSGSKA | CSNK1A1; CSN | P48729; A0A | KC1A_HUMAN | Acetyl (Protein N | 0 | 0 | 0 | 609670 | 318800 | 0 | 0 | 0 | 214150 | 288180 | 0 | 0 | 142340 | 0 | 0 | 0 | 0 | 0 |
| ASSSSMKII | SNX9; SNX9; SI | Q9Y5X1; A0/ | SNX9_HUMAN | Unmodified | 167220 | 376560 | 0 | 0 | 0 | 150840 | 0 | 0 | 0 | 0 | 0 | 0 | 96833 | 0 | 0 | 0 | 0 | 0 |
| ASSTSPVEI | ETFA; ETFA; ETI | P13804; H0Y | ETFA_HUMAN I | Unmodified | 0 | 0 | 0 | 0 | 0 | 0 | 0 | 20448 | 1881100 | 0 | 0 | 0 | 0 | 0 | 0 | 0 | 0 | 0 |
| ASSTSTPEF | ENAH; ENAH | Q8N8S7; A0 | ENAH_HUMAN | Unmodified | 118370 | 89422 | 275770 | 958530 | 814770 | 131740 | 165950 | 132500 | 1157400 | 261220 | 489110 | 5695500 | 640310 | 318600 | 775310 | 775880 | 913890 | 178650 |
| ASSVIFGLK | ILKAP; ILKAP | Q9H0C8; F8 | ILKAP_HUMAN | Unmodified | 0 | 268410 | 0 | 0 | 283140 | 0 | 1398000 | 717660 | 0 | 74956 | 0 | 838300 | 314760 | 0 | 0 | 11852000 | 114160 | 408450 |
| ASSVTTFTC | CRIP2; CRIP2 | P52943; H0Y | CRIP2_HUMAN | Unmodified | 0 | 0 | 0 | 0 | 0 | 0 | 0 | 0 | 3572800 | 135710 | 0 | 422070 | 0 | 0 | 0 | 0 | 0 | 0 |
| ASSVTTFTC | CRIP2; CRIP2 | P52943; H0Y | CRIP2_HUMAN | Unmodified | 0 | 0 | 0 | 0 | 0 | 0 | 345880 | 282940 | 0 | 169780 | 0 | 0 | 0 | 0 | 0 | 0 | 0 | 282300 |
| ASSVTTFTC | CRIP2; CRIP2 | P52943; H0Y | CRIP2_HUMAN | Unmodified | 0 | 0 | 0 | 0 | 0 | 0 | 188030 | 0 | 0 | 0 | 0 | 0 | 0 | 0 | 0 | 0 | 0 | 0 |
| ASTDTMGR | PLAU; PLAU | P00749; E7E | UROK_HUMAN | Oxidation (M) | 0 | 0 | 0 | 0 | 0 | 0 | 0 | 0 | 0 | 0 | 0 | 0 | 0 | 0 | 0 | 0 | 0 | 5495900 |

| | | | | | | | | | | | | | | | | | | | | | | |
|---|---|---|---|---|---|---|---|---|---|---|---|---|---|---|---|---|---|---|---|---|---|---|
| ASTEATELL | CIT; CIT; CIT; CI | O14578; A0A | CTRO_HUMAN | Unmodified | 18366 | 0 | 79808 | 225320 | 618530 | 199270 | 20353 | 0 | 561600 | 0 | 48356 | 354220 | 137460 | 0 | 64001 | 0 | 128260 | 110520 |
| ASTEATELL | CIT; CIT; CIT; CI | O14578; A0A | CTRO_HUMAN | Unmodified | 0 | 0 | 0 | 0 | 21972000 | 3026900 | 1113300 | 308120 | 0 | 462170 | 0 | 0 | 1524800 | 1168100 | 0 | 0 | 3910300 | 2428800 |
| ASTHCTYD | DNASE1L1 | P49184 | DNSL1_HUMAI | Unmodified | 0 | 0 | 0 | 0 | 0 | 0 | 0 | 0 | 1276500 | 0 | 74251 | 500340 | 191990 | 0 | 0 | 0 | 0 | 0 |
| ASTISAYK | STMN3 | Q9NZ72 | STMN3_HUMAI | Acetyl (Protein N | 1022500 | 1362200 | 646200 | 775530 | 1410200 | 62958 | 2033900 | 112230 | 518030 | 168700 | 237500 | 1254500 | 673510 | 240960 | 554090 | 676160 | 478440 | 0 |
| ASTISAYKE | STMN3 | Q9NZ72 | STMN3_HUMAI | Acetyl (Protein N | 0 | 0 | 0 | 0 | 0 | 0 | 920340 | 0 | 0 | 0 | 52589 | 0 | 0 | 0 | 0 | 0 | 0 | 0 |
| ASTLLRDEE | CHP1; CHP1; C | Q99653; H0Y | CHP1_HUMAN | Unmodified | 424230 | 294960 | 4574200 | 1305400 | 0 | 0 | 0 | 0 | 1213700 | 66600 | 0 | 5430000 | 0 | 0 | 2929800 | 3417800 | 0 | 0 |
| ASTLRLDFI | CACNA2D2; CA | Q9NY47; C9. | CA2D2_HUMAI | Unmodified | 398020 | 481520 | 75083 | 0 | 1972400 | 0 | 2436600 | 0 | 25074000 | 9559700 | 1606800 | 975250 | 0 | 0 | 0 | 0 | 269050 | 0 |
| ASTNIQDTE | PSMD10; PSMD | O75832; B1A | PSD10_HUMAN | Unmodified | 0 | 0 | 728710 | 1005000 | 1972800 | 536430 | 1270200 | 1006600 | 1761900 | 1498800 | 0 | 3205900 | 1959900 | 5797800 | 969810 | 954060 | 7309400 | 5033400 |
| ASTVEGGD | ACOX3 | O15254 | ACOX3_HUMAI | Acetyl (Protein N | 0 | 0 | 0 | 0 | 1586800 | 0 | 0 | 0 | 986780 | 0 | 34290 | 0 | 0 | 0 | 0 | 0 | 0 | 0 |
| ASVATLMK | MYO1B; MYO1E | O43795; E9F | MYO1B_HUMA | Unmodified | 0 | 152070 | 0 | 82704 | 16984000 | 5339300 | 1339000 | 1381500 | 2338600 | 986710 | 2778600 | 9170600 | 0 | 245260 | 204440 | 0 | 2428300 | 3596800 |
| ASVATLMK | MYO1B; MYO1E | O43795; E9F | MYO1B_HUMA | Oxidation (M) | 0 | 0 | 0 | 0 | 475960 | 0 | 127840 | 91892 | 0 | 63164 | 0 | 1299500 | 0 | 0 | 0 | 0 | 0 | 0 |
| ASVDELFAE | RAP2B | P61225 | RAP2B_HUMAN | Unmodified | 79643 | 0 | 3521700 | 1951100 | 41353 | 0 | 0 | 0 | 2307100 | 162170 | 536230 | 3194900 | 0 | 0 | 7320000 | 2579200 | 38908 | 37232 |
| ASVEERITQ | CHD4; CHD3; C | Q14839; Q1. | CHD4_HUMAN | Unmodified | 0 | 0 | 0 | 0 | 178460 | 0 | 148250 | 121760 | 0 | 99863 | 0 | 0 | 0 | 0 | 0 | 0 | 0 | 34394 |
| ASVEGGGF | ABCA3; ABCA3 | Q99758; H0Y | ABCA3_HUMAI | Unmodified | 0 | 0 | 44112 | 0 | 0 | 278020 | 0 | 0 | 625250 | 40930 | 0 | 0 | 0 | 0 | 0 | 0 | 0 | 0 |
| ASVGFGGS | UGDH | O60701 | UGDH_HUMAN | Unmodified | 0 | 0 | 0 | 0 | 4324500 | 688810 | 0 | 0 | 117880 | 0 | 0 | 0 | 701680 | 1428700 | 0 | 0 | 0 | 0 |
| ASVHCLET | THOC2; THOC2 | Q8NI27; H0Y | THOC2_HUMA | Unmodified | 0 | 0 | 0 | 0 | 0 | 0 | 177880 | 0 | 0 | 0 | 0 | 0 | 0 | 0 | 0 | 0 | 0 | 0 |
| ASVITQVFH | HDLBP; HDLBP | Q00341; A0A | VIGLN_HUMAN | Unmodified | 0 | 0 | 0 | 0 | 0 | 0 | 0 | 0 | 180580 | 0 | 0 | 1742200 | 0 | 0 | 0 | 0 | 117670 | 347680 |
| ASVKEQDY | YKT6; YKT6; YKT | O15498; A0A | YKT6_HUMAN S | Unmodified | 191940 | 154420 | 0 | 0 | 0 | 0 | 0 | 0 | 0 | 0 | 0 | 0 | 0 | 0 | 0 | 50947 | 0 | 0 |
| ASVLEPGRI | RTN4R; RTN4R; | Q9BZR6; H7 | RTN4R_HUMAN | Unmodified | 262970 | 249210 | 437790 | 234970 | 0 | 0 | 0 | 0 | 476980 | 58155 | 182720 | 0 | 0 | 34739 | 206890 | 122980 | 0 | 95002 |
| ASVPREPG | PFAS; PFAS | O15067; H0Y | PUR4_HUMAN | Unmodified | 79295 | 25128 | 70716 | 131170 | 1274100 | 0 | 0 | 0 | 306810 | 0 | 0 | 1972100 | 778100 | 189500 | 109870 | 103860 | 203580 | 567740 |
| ASVPTIQDC | TLN1 | Q9Y490 | TLN1_HUMAN T | Oxidation (M) | 0 | 32491 | 426910 | 0 | 5278200 | 943120 | 2634500 | 2746800 | 0 | 288140 | 189950 | 0 | 1529000 | 2182100 | 0 | 567790 | 1564800 | 0 |
| ASVPTIQDC | TLN1 | Q9Y490 | TLN1_HUMAN T | Unmodified | 0 | 263430 | 0 | 707560 | 7875600 | 867070 | 3777900 | 1770500 | 137570 | 279670 | 0 | 668670 | 2640500 | 4844900 | 379270 | 0 | 3376500 | 3329200 |
| ASVQFVDY | PLCB2 | Q00722 | PLCB2_HUMAN | Unmodified | 48914 | 0 | 80679 | 120110 | 0 | 0 | 0 | 0 | 0 | 0 | 158430 | 2111700 | 0 | 0 | 0 | 142280 | 0 | 7232.6 |
| ASVSQVEA | CKAP4 | Q07065 | CKAP4_HUMAN | Unmodified | 0 | 171150 | 0 | 0 | 979900 | 0 | 1119000 | 774410 | 5070200 | 2260500 | 125690 | 322630 | 836860 | 1610500 | 104870 | 0 | 92219 | 864060 |
| ASVSVTAEI | ICAM1; ICAM1 | P05362; E7E | ICAM1_HUMAN | Unmodified | 0 | 0 | 1176000 | 0 | 2631700 | 1648300 | 70987 | 0 | 264260 | 0 | 0 | 0 | 48319 | 424370 | 54142 | 0 | 215280 | 342400 |
| ASVTDGKT | RAB3B | P20337 | RAB3B_HUMAN | Acetyl (Protein N | 0 | 0 | 0 | 0 | 537560 | 0 | 0 | 0 | 74462 | 152080 | 205870 | 303440 | 709710 | 0 | 0 | 0 | 0 | 54280 |
| ASVTIQAR | CCDC6 | Q16204 | CCDC6_HUMA | Unmodified | 0 | 0 | 0 | 0 | 0 | 0 | 0 | 0 | 0 | 0 | 0 | 0 | 0 | 0 | 0 | 0 | 0 | 0 |
| ASVTLSEAE | EXOSC7 | Q15024 | EXOS7_HUMAN | Acetyl (Protein N | 0 | 0 | 0 | 0 | 304780 | 0 | 0 | 0 | 0 | 0 | 0 | 1352800 | 0 | 0 | 184670 | 292010 | 0 | 244410 |
| ASVTRAVF | GALM; GALM | Q96C23; B8Z | GALM_HUMAN | Acetyl (Protein N | 0 | 0 | 0 | 0 | 0 | 0 | 0 | 0 | 0 | 0 | 0 | 0 | 0 | 0 | 0 | 0 | 1369800 | 1428800 |
| ASVTVGGE | DTD1; DTD1; DT | Q8TEA8; A0A | DTD1_HUMAN | Unmodified | 0 | 0 | 0 | 0 | 0 | 0 | 0 | 0 | 432750 | 0 | 0 | 3335800 | 0 | 0 | 0 | 0 | 0 | 0 |
| ASVVTLPVY | DYNC1H1; DYN | Q14204; A0A | DYHC1_HUMA | Unmodified | 0 | 0 | 0 | 0 | 0 | 0 | 0 | 0 | 102990 | 27165 | 44363 | 0 | 0 | 0 | 0 | 0 | 0 | 0 |
| ASWGEFQ | VGF; VGF | O15240; Q0 | VGF_HUMAN N | Unmodified | 838910 | 0 | 0 | 0 | 0 | 0 | 0 | 0 | 4384700 | 220720 | 0 | 0 | 0 | 0 | 0 | 0 | 0 | 0 |
| ASWSSLSM | COX4I1; COX4I | P13073; H3B | COX41_HUMAI | Oxidation (M) | 0 | 0 | 0 | 0 | 0 | 0 | 0 | 0 | 2814300 | 0 | 0 | 471460 | 0 | 152200 | 0 | 0 | 0 | 0 |
| ASWTLLAR | CEMIP2; CEMIP | Q9UHN6; F5 | CEIP2_HUMAN | Unmodified | 0 | 310350 | 1162400 | 475600 | 2321600 | 791840 | 0 | 0 | 2915000 | 892000 | 0 | 172630 | 141320 | 120160 | 882900 | 575930 | 0 | 0 |
| ASWTRPEK | NCAM1; NCAM | P13591; A0A | NCAM1_HUMA | Unmodified | 0 | 0 | 0 | 0 | 0 | 0 | 0 | 0 | 579830 | 0 | 0 | 0 | 0 | 0 | 0 | 0 | 0 | 0 |
| ASWWPEF | ACOX3 | O15254 | ACOX3_HUMAI | Unmodified | 0 | 0 | 0 | 0 | 0 | 0 | 0 | 0 | 2112400 | 243830 | 45420 | 0 | 0 | 0 | 0 | 0 | 0 | 0 |
| ASYAVSETI | XPNPEP1; XPNI | Q9NQW7; Q | XPP1_HUMAN | Unmodified | 0 | 0 | 0 | 0 | 419010 | 107550 | 124870 | 120720 | 268380 | 0 | 598510 | 2500700 | 529970 | 221680 | 0 | 0 | 202290 | 292830 |
| ASYVASTKY | CTNNA1; CTNN | P35221; P26 | CTNA1_HUMAI | Unmodified | 0 | 400230 | 0 | 203030 | 7009700 | 0 | 0 | 0 | 0 | 53140 | 0 | 0 | 5833200 | 15639000 | 91351 | 296060 | 0 | 0 |
| ASYVAWGH | NIT2 | Q9NQR4 | NIT2_HUMAN ( | Unmodified | 323440 | 0 | 0 | 0 | 1246000 | 403000 | 0 | 0 | 2060900 | 910190 | 1378200 | 5541500 | 2477000 | 959410 | 1381400 | 596610 | 1580800 | 1596900 |
| ATAAELLK | PAK4; PAK4 | O96013; M0 | PAK4_HUMAN | Unmodified | 46967 | 276940 | 0 | 884280 | 1150700 | 0 | 76946 | 54910 | 145920 | 78852 | 0 | 540930 | 0 | 0 | 931190 | 0 | 480170 | 391340 |
| ATAAFILAN | IPO5; IPO5 | O00410; H0Y | IPO5_HUMAN I | Unmodified | 0 | 0 | 1373000 | 861110 | 0 | 0 | 0 | 0 | 1277300 | 738040 | 1054900 | 14573000 | 0 | 0 | 869140 | 536670 | 319120 | 0 |
| ATAAFSNV | TPD52L1; TPD5 | Q16890; E9F | TPD53_HUMAN | Unmodified | 0 | 0 | 0 | 0 | 0 | 266020 | 567050 | 0 | 0 | 182930 | 0 | 0 | 1003200 | 1896800 | 0 | 81295 | 386550 | 1405200 |
| ATAAGVKQ | PSMA6; PSMA6 | P60900; G3V | PSA6_HUMAN | Unmodified | 0 | 0 | 0 | 0 | 2201800 | 120870 | 430350 | 292740 | 0 | 440990 | 0 | 0 | 867130 | 489550 | 0 | 0 | 879130 | 884520 |
| ATAAPAGA | UQCRC2; UQC | P22695; H3B | QCR2_HUMAN | Unmodified | 0 | 0 | 0 | 0 | 0 | 0 | 0 | 0 | 13979000 | 4010400 | 0 | 0 | 0 | 0 | 0 | 0 | 0 | 0 |
| ATAATEEPF | TPP2; TPP2; TPI | P29144; A0A | TPP2_HUMAN T | Acetyl (Protein N | 0 | 0 | 0 | 0 | 0 | 0 | 0 | 0 | 0 | 0 | 389760 | 3597000 | 0 | 0 | 0 | 0 | 0 | 0 |
| ATAAVIFLH | LYPLA1; LYPLA1 | O75608; B4D | LYPA1_HUMAN | Unmodified | 0 | 178340 | 0 | 0 | 233770 | 100340 | 0 | 0 | 1211400 | 0 | 0 | 0 | 300470 | 0 | 177800 | 0 | 146270 | 0 |
| ATAAYEQLK | SACM1L; SACM | Q9NTJ5; A0A | SAC1_HUMAN | Acetyl (Protein N | 64557 | 0 | 0 | 0 | 276460 | 0 | 1469500 | 0 | 0 | 0 | 106270 | 450060 | 0 | 0 | 0 | 0 | 97789 | 309890 |
| ATADDELSI | GRB2; GRB2; G | P62993; J3Q | GRB2_HUMAN | Unmodified | 7866800 | 5136000 | 4729300 | 1125000 | 0 | 0 | 19568 | 0 | 841580 | 0 | 0 | 4322000 | 375490 | 0 | 472340 | 240770 | 0 | 64497 |
| ATADDELSI | GRB2; GRB2; G | P62993; J3Q | GRB2_HUMAN | Unmodified | 258790 | 222900 | 0 | 0 | 0 | 0 | 0 | 0 | 0 | 0 | 0 | 0 | 0 | 0 | 288410 | 306150 | 0 | 0 |
| ATADDELSI | GRB2; GRB2; G | P62993; J3Q | GRB2_HUMAN | Unmodified | 753910 | 289920 | 0 | 0 | 0 | 0 | 0 | 0 | 0 | 273690 | 0 | 0 | 322390 | 0 | 91879 | 0 | 0 | 0 |
| ATAEGSAG | RAB11FIP1 | Q6WKZ4 | RFIP1_HUMAN | Unmodified | 0 | 0 | 0 | 0 | 807410 | 0 | 0 | 0 | 0 | 0 | 0 | 0 | 0 | 0 | 0 | 0 | 0 | 0 |
| ATAEQISSC | DECR1; DECR1 | Q16698; E5F | DECR_HUMAN | Unmodified | 0 | 0 | 0 | 0 | 0 | 0 | 0 | 0 | 9498100 | 0 | 0 | 0 | 0 | 0 | 0 | 0 | 0 | 0 |
| ATAEVLNIG | PAICS; PAICS | P22234; E9P | PUR6_HUMAN | Acetyl (Protein N | 152860 | 0 | 1523900 | 795300 | 2005900 | 433280 | 1202100 | 46993 | 1273200 | 1043600 | 816200 | 3243400 | 4443400 | 0 | 597880 | 517280 | 6540400 | 4433400 |
| ATAEVLNIG | PAICS; PAICS | P22234; E9P | PUR6_HUMAN | Acetyl (Protein N | 0 | 0 | 0 | 146080 | 0 | 75099 | 1140000 | 79924 | 0 | 34288 | 0 | 0 | 0 | 0 | 0 | 0 | 6651800 | 0 |
| ATAFNEQV | RPA1 | P27694 | RFA1_HUMAN | Unmodified | 490100 | 316290 | 0 | 162330 | 0 | 0 | 0 | 0 | 0 | 69419 | 0 | 1669900 | 0 | 0 | 0 | 0 | 0 | 0 |
| ATAFNEQV | RPA1 | P27694 | RFA1_HUMAN | Unmodified | 98405 | 0 | 0 | 0 | 0 | 0 | 0 | 0 | 0 | 0 | 0 | 376500 | 0 | 0 | 0 | 0 | 0 | 0 |
| ATAFQREEI | RFFL; RFFL | Q8WZ73; C9 | RFFL_HUMAN E | Unmodified | 0 | 44741 | 261830 | 550170 | 687320 | 0 | 0 | 0 | 0 | 0 | 0 | 0 | 233010 | 0 | 0 | 176500 | 0 | 0 |
| ATAGAYIAS | PSMB5; PSMB5 | P28074; H0Y | PSB5_HUMAN | Unmodified | 285690 | 499170 | 0 | 0 | 0 | 0 | 0 | 0 | 153810 | 0 | 0 | 1569300 | 0 | 0 | 0 | 0 | 0 | 0 |
| ATAGAYIAS | PSMB5; PSMB5 | P28074; H0Y | PSB5_HUMAN | Unmodified | 0 | 0 | 0 | 0 | 0 | 0 | 0 | 0 | 664070 | 0 | 202920 | 2181600 | 0 | 0 | 0 | 0 | 0 | 0 |
| ATAGDTHL | HSPA1A; HSPA1 | P0DMV8; P0 | HS71A_HUMAN | Unmodified | 34912000 | 25964000 | 8252700 | 30669000 | 33225000 | 9548300 | 3109000 | 240470 | 88092000 | 38271000 | 10803000 | 18454000 | 49103000 | 27492000 | 17387000 | 8345400 | 3735900 | 20185000 |
| ATAGDTHL | HSPA1A; HSPA1 | P0DMV8; P0 | HS71A_HUMAN | Unmodified | 3146600 | 4119900 | 1028200 | 2685400 | 1505100 | 0 | 180010 | 0 | 11979000 | 946670 | 102620 | 1897200 | 1687300 | 0 | 1197000 | 851900 | 346420 | 282430 |
| ATAIRTEEEI | NF2; NF2 | P35240; A0A | MERL_HUMAN | Unmodified | 0 | 0 | 0 | 0 | 363650 | 0 | 0 | 0 | 39859 | 0 | 0 | 0 | 0 | 0 | 0 | 0 | 116150 | 304830 |
| ATALAIMGI | LRP1 | Q07954 | LRP1_HUMAN I | Unmodified | 0 | 0 | 0 | 0 | 0 | 0 | 0 | 0 | 1021200 | 0 | 0 | 0 | 0 | 0 | 0 | 0 | 0 | 0 |
| ATANSQVM | GPRC5C; GPRC | Q9NQ84; A8 | GPC5C_HUMA | Unmodified | 0 | 0 | 555120 | 1543900 | 0 | 0 | 0 | 0 | 692270 | 64308 | 0 | 508480 | 186260 | 30124 | 706240 | 2104100 | 0 | 0 |
| ATANSQVM | GPRC5C; GPRC | Q9NQ84; A8 | GPC5C_HUMA | Oxidation (M) | 22342 | 200420 | 140240 | 237290 | 0 | 0 | 64653 | 0 | 0 | 0 | 0 | 0 | 0 | 0 | 196090 | 209380 | 0 | 0 |
| ATANSQVM | GPRC5C; GPRC | Q9NQ84; A8 | GPC5C_HUMA | Unmodified | 0 | 0 | 0 | 250240 | 1206600 | 457800 | 0 | 0 | 1969400 | 316860 | 0 | 0 | 252890 | 1165300 | 0 | 0 | 0 | 1891200 |
| ATAPNKPP | LRP11 | Q86VZ4 | LRP11_HUMAN | Unmodified | 0 | 0 | 0 | 0 | 0 | 0 | 0 | 0 | 1843200 | 0 | 0 | 0 | 0 | 0 | 0 | 0 | 0 | 0 |
| ATAPQTQH | EEF1D; EEF1D; | P29692; E9P | EF1D_HUMAN | Unmodified | 0 | 0 | 418740 | 0 | 0 | 121180 | 0 | 0 | 5634400 | 88905 | 0 | 31967000 | 321340 | 0 | 430370 | 609590 | 0 | 0 |
| ATAPVPTV | RBM4; RBM4 | Q9BWF3; E9 | RBM4_HUMAN | Unmodified | 0 | 379060 | 599010 | 808320 | 0 | 0 | 0 | 0 | 0 | 0 | 0 | 0 | 0 | 0 | 536470 | 647100 | 0 | 0 |
| ATAPVSFN | BROX; BROX; BI | Q5VW32; F5 | BROX_HUMAN | Unmodified | 8356800 | 10361000 | 8748200 | 13836000 | 13472000 | 3585100 | 0 | 1123000 | 17157000 | 9859500 | 0 | 10286000 | 8404300 | 3875500 | 8558800 | 8402600 | 890470 | 2143500 |
| ATAPVSFN | BROX; BROX; BI | Q5VW32; F5 | BROX_HUMAN | Unmodified | 0 | 0 | 0 | 0 | 0 | 0 | 0 | 0 | 0 | 629400 | 0 | 0 | 0 | 1354900 | 0 | 0 | 0 | 0 |
| ATAPYNYS | RAB14; RAB14 | P61106; X6R | RAB14_HUMAN | Acetyl (Protein N | 0 | 0 | 0 | 637920 | 689940 | 0 | 660690 | 785030 | 939920 | 532970 | 924290 | 2519000 | 0 | 0 | 0 | 485320 | 0 | 0 |
| ATAQPPTE | GAP43 | P17677 | NEUM_HUMAN | Unmodified | 0 | 0 | 0 | 0 | 0 | 0 | 874230 | 1961200 | 0 | 0 | 0 | 0 | 0 | 0 | 0 | 0 | 0 | 0 |
| ATASPRSD | WWP1 | Q9H0M0 | WWP1_HUMAN | Acetyl (Protein N | 0 | 0 | 0 | 0 | 3200900 | 0 | 529480 | 0 | 0 | 0 | 0 | 0 | 1195200 | 0 | 0 | 0 | 1833600 | 1772700 |
| ATATQLLQI | STK3; STK3; STI | Q13188; A0A | STK3_HUMAN S | Unmodified | 394840 | 0 | 106170 | 106290 | 399640 | 0 | 0 | 83037 | 1594600 | 0 | 64333 | 427750 | 227750 | 185340 | 103730 | 123730 | 218330 | 137660 |
| ATATSCRP | HSPG2 | P98160 | PGBM_HUMAN | Unmodified | 0 | 0 | 311020 | 289660 | 0 | 0 | 0 | 0 | 217250 | 220190 | 0 | 0 | 230810 | 0 | 0 | 124310 | 0 | 271040 |
| ATATTSQP | ABR; ABR; ABR; | Q12979; A0A | ABR_HUMAN A | Unmodified | 0 | 0 | 0 | 0 | 548590 | 60958 | 119470 | 0 | 53652 | 0 | 0 | 0 | 163990 | 93495 | 256340 | 172270 | 126750 | 391970 |
| ATAVVDGA | PRDX2; PRDX2 | P32119; A6N | PRDX2_HUMAN | Unmodified | 1334500 | 3082500 | 6679800 | 5298100 | 833160 | 341800 | 401300 | 227210 | 19799000 | 2316100 | 17115000 | 2.75E+08 | 1820800 | 285090 | 7075300 | 7899000 | 857040 | 692130 |
| ATAVVDGA | PRDX2; PRDX2 | P32119; A6N | PRDX2_HUMAN | Unmodified | 1513000 | 0 | 0 | 1356300 | 0 | 0 | 0 | 0 | 3595500 | 1705100 | 1889300 | 12211000 | 1480700 | 0 | 0 | 517600 | 862250 | 0 |
| ATAVVDGA | PRDX2; PRDX2 | P32119; A6N | PRDX2_HUMAN | Unmodified | 0 | 0 | 0 | 0 | 0 | 0 | 1412400 | 2510200 | 0 | 794140 | 0 | 0 | 832830 | 0 | 0 | 0 | 2441600 | 177160 |
| ATCAPQHC | FLNA; FLNA; FL | P21333; A0A | FLNA_HUMAN | Unmodified | 0 | 0 | 0 | 0 | 961550 | 0 | 0 | 0 | 0 | 0 | 0 | 0 | 0 | 0 | 0 | 0 | 5993600 | 373380 |
| ATCIGNNS | PSMA4; PSMA4 | P25789; H0Y | PSA4_HUMAN | Unmodified | 0 | 0 | 0 | 145380 | 245980 | 0 | 0 | 0 | 646560 | 141970 | 357690 | 1948400 | 262610 | 0 | 0 | 0 | 0 | 0 |
| ATCLEEMTI | QTRT1 | Q9BXR0 | TGT_HUMAN Q | Unmodified | 0 | 48168 | 0 | 0 | 0 | 0 | 0 | 0 | 0 | 0 | 0 | 197690 | 0 | 0 | 0 | 0 | 0 | 0 |
| ATDAQLCL | NBEA; LRBA; LR | Q8NFP9; P5 | NBEA_HUMAN | Unmodified | 0 | 0 | 0 | 0 | 0 | 0 | 23905 | 12694 | 0 | 0 | 0 | 1803700 | 0 | 0 | 79328 | 0 | 0 | 0 |
| ATDEEPLGI | RTN4R; RTN4R; | Q9BZR6; H7 | RTN4R_HUMAN | Unmodified | 202290 | 200810 | 351770 | 0 | 309160 | 0 | 0 | 0 | 455440 | 0 | 0 | 0 | 0 | 0 | 114000 | 0 | 0 | 0 |
| ATDELATK | EFHD2 | Q96C19 | EFHD2_HUMAN | Acetyl (Protein N | 0 | 0 | 0 | 0 | 1206400 | 0 | 111150 | 42323 | 0 | 0 | 1235200 | 0 | 0 | 1906600 | 0 | 0 | 401080 | 0 |
| ATDFVADR | IDH2; IDH2 | P48735; H0Y | IDHP_HUMAN | Unmodified | 124290 | 0 | 19577 | 0 | 30075 | 0 | 0 | 0 | 4872500 | 183220 | 55004 | 579140 | 0 | 0 | 0 | 0 | 38272 | 0 |
| ATDFVADR | IDH2; IDH2 | P48735; H0Y | IDHP_HUMAN | Unmodified | 101240 | 0 | 0 | 67895 | 142170 | 51388 | 747570 | 259620 | 29373000 | 5793000 | 403480 | 665740 | 793970 | 252030 | 58396 | 0 | 25817 | 197050 |
| ATDFVVPG | IDH1; IDH1 | O75874; C9J | IDHC_HUMAN | Unmodified | 0 | 0 | 0 | 0 | 0 | 0 | 0 | 0 | 0 | 0 | 0 | 0 | 0 | 0 | 456530 | 232560 | 0 | 0 |
| ATDFVVPG | IDH1 | O75874 | IDHC_HUMAN | Unmodified | 0 | 0 | 937340 | 4200800 | 0 | 0 | 0 | 0 | 8691400 | 366750 | 7447600 | 33192000 | 0 | 0 | 5383900 | 11510000 | 0 | 0 |
| ATDGVTLT | PACSIN2; PACS | Q9UNF0; A0A | PACN2_HUMAI | Unmodified | 1926200 | 593620 | 6019000 | 611160 | 0 | 0 | 0 | 0 | 0 | 0 | 0 | 638290 | 0 | 0 | 6726300 | 3597300 | 0 | 0 |
| ATDHPHPS | CIT; CIT; CIT; CI | O14578; A0A | CTRO_HUMAN | Unmodified | 0 | 0 | 0 | 0 | 324330 | 0 | 0 | 0 | 105810 | 0 | 0 | 82282 | 0 | 0 | 73707 | 37116 | 0 | 0 |
| ATDLGGTS | ZC3HAV1; ZC3H | Q7Z2W4; C9 | ZCCHV_HUMA | Unmodified | 990070 | 878630 | 977600 | 1353200 | 667030 | 0 | 0 | 0 | 301480 | 0 | 0 | 107590 | 160970 | 0 | 876100 | 1063500 | 0 | 0 |
| ATDMPWK | CDC42BPA; CD | Q5VT25; A0A | MRCKA_HUMA | Unmodified | 0 | 0 | 0 | 0 | 0 | 0 | 0 | 0 | 0 | 0 | 0 | 0 | 0 | 0 | 0 | 0 | 0 | 0 |
| ATDSSDPLI | TRNT1 | Q96Q11 | TRNT1_HUMAN | Unmodified | 0 | 27419 | 0 | 0 | 0 | 0 | 0 | 0 | 0 | 0 | 0 | 222710 | 0 | 0 | 0 | 0 | 0 | 0 |
| ATDTLDRTI | GFM2 | Q969S9 | RRF2M_HUMAI | Unmodified | 0 | 0 | 0 | 0 | 36164 | 0 | 0 | 0 | 172940 | 0 | 0 | 0 | 0 | 0 | 0 | 0 | 0 | 0 |
| ATDVMIAGI | AHCY | P23526 | SAHH_HUMAN | Unmodified | 0 | 40390 | 0 | 0 | 292470 | 0 | 105470 | 188500 | 538630 | 131660 | 111300 | 4071700 | 0 | 0 | 0 | 0 | 277220 | 122190 |
| ATEAGSLE | ARHGEF2; ARH | Q92974; V9 | ARHG2_HUMA | Unmodified | 0 | 1056300 | 0 | 0 | 312830 | 2284400 | 118970 | 0 | 0 | 28557 | 145040 | 893290 | 0 | 0 | 451450 | 197250 | 0 | 1906300 |
| ATEARIISG | PSD3; PSD3 | Q9NYI0; E5R | PSD3_HUMAN | Unmodified | 0 | 0 | 0 | 0 | 743310 | 0 | 0 | 0 | 0 | 29462 | 0 | 0 | 0 | 206590 | 0 | 0 | 1324700 | 698630 |
| ATEDGTPYI | OSBP | P22059 | OSBP1_HUMAI | Unmodified | 0 | 0 | 0 | 0 | 603770 | 0 | 0 | 0 | 0 | 0 | 1660900 | 968900 | 0 | 0 | 0 | 0 | 0 | 0 |
| ATEDVLVK | CLSTN1; CLSTN | O94985; A0A | CSTN1_HUMAI | Unmodified | 0 | 0 | 0 | 0 | 0 | 0 | 0 | 0 | 0 | 0 | 0 | 0 | 0 | 0 | 0 | 0 | 0 | 0 |
| ATEEDSQV | GALNT1 | Q10472 | GALT1_HUMAN | Unmodified | 0 | 307970 | 0 | 288790 | 0 | 0 | 0 | 0 | 1435300 | 0 | 0 | 1360300 | 1205100 | 0 | 0 | 0 | 0 | 0 |
| ATEEDSQV | GALNT1 | Q10472 | GALT1_HUMAN | Unmodified | 0 | 0 | 100850 | 0 | 0 | 0 | 0 | 0 | 2111700 | 38607 | 559870 | 486630 | 0 | 0 | 0 | 30045 | 0 | 0 |
| ATEEFIIR | MVP; MVP; MVP | Q14764; H3I | MVP_HUMAN N | Acetyl (Protein N | 747790 | 862350 | 0 | 169760 | 0 | 0 | 0 | 52742 | 1079100 | 0 | 0 | 7657000 | 0 | 0 | 0 | 0 | 0 | 0 |
| ATEEPEDVI | PRRC2C; PRRC | Q9Y520; E7E | PRC2C_HUMAI | Unmodified | 1486000 | 1006600 | 246400 | 0 | 0 | 0 | 0 | 0 | 0 | 0 | 0 | 0 | 0 | 0 | 0 | 211830 | 0 | 0 |
| ATEEVSKNI | CAB39; CAB39 | Q9Y376; A0A | CAB39_HUMAN | Unmodified | 0 | 0 | 0 | 22991 | 0 | 0 | 0 | 0 | 0 | 0 | 0 | 0 | 0 | 419320 | 32052 | 0 | 0 | 449650 |
| ATEGKWEV | STXBP2; STXBP | Q15833; R4 | STXB2_HUMAN | Unmodified | 0 | 0 | 0 | 0 | 0 | 0 | 0 | 0 | 0 | 0 | 0 | 1312900 | 0 | 0 | 0 | 0 | 0 | 0 |
| ATEGMILTN | GPHN; GPHN; ( | Q9NQX3; G3 | GEPH_HUMAN | Acetyl (Protein N | 0 | 0 | 0 | 0 | 0 | 0 | 0 | 0 | 0 | 0 | 0 | 726100 | 0 | 0 | 0 | 0 | 0 | 0 |
| ATEGMVVA | PSMB7; PSMB7 | Q99436; Q5 | PSB7_HUMAN | Unmodified | 0 | 0 | 150190 | 0 | 0 | 0 | 0 | 0 | 0 | 0 | 817170 | 6397600 | 0 | 0 | 0 | 0 | 0 | 0 |

| | | | | | | | | | | | | | | | | | | | | | | |
|---|---|---|---|---|---|---|---|---|---|---|---|---|---|---|---|---|---|---|---|---|---|---|
| ATEGMVVA | PSMB7; PSMB7 | Q99436; Q5 | PSB7_HUMAN | Oxidation (M) | 0 | 0 | 0 | 0 | 0 | 0 | 0 | 0 | 0 | 0 | 155570 | 592720 | 0 | 0 | 0 | 111310 | 0 | 0 |
| ATEHPEPPI | SF3B2; SF3B2; | Q13435; E9I | SF3B2_HUMAN | Acetyl (Protein N | 0 | 0 | 0 | 0 | 0 | 0 | 0 | 0 | 0 | 0 | 0 | 3815500 | 603940 | 0 | 0 | 0 | 0 | 0 |
| ATEMLIFGK | ECI2; ECI2 | O75521; A0A | ECI2_HUMAN | Unmodified | 0 | 0 | 0 | 0 | 499330 | 0 | 0 | 0 | 4894300 | 1032900 | 0 | 0 | 0 | 0 | 0 | 0 | 0 | 0 |
| ATEMVEVG | CAVIN1 | Q6NZI2 | CAVN1_HUMAN | Oxidation (M) | 0 | 0 | 0 | 0 | 1109700 | 0 | 0 | 0 | 0 | 0 | 0 | 0 | 0 | 0 | 0 | 0 | 0 | 0 |
| ATENDIANI | HNRNPH3 | P31942 | HNRH3_HUMA | Unmodified | 472120 | 0 | 69916 | 59550 | 0 | 0 | 0 | 0 | 243360 | 47774 | 0 | 446440 | 0 | 0 | 204720 | 98444 | 0 | 0 |
| ATENDIYNI | HNRNPH1; HNI | P31943; P52 | HNRH1_HUMA | Unmodified | 739090 | 0 | 104420 | 221940 | 90651 | 132540 | 138160 | 45027 | 1023000 | 80596 | 239200 | 2677800 | 0 | 0 | 1013300 | 382700 | 0 | 0 |
| ATEPGLFG | PCDHB8; PCDI | Q9UN66; Q9 | PCDB8_HUMA | Unmodified | 0 | 294910 | 0 | 0 | 168080 | 36837 | 0 | 0 | 1310500 | 0 | 0 | 299820 | 0 | 69933 | 0 | 0 | 0 | 27128 |
| ATEPVIAFY | AK1; AK1 | P00568; Q5I | KAD1_HUMAN | Unmodified | 520240 | 0 | 0 | 1183600 | 8165400 | 0 | 3820100 | 3451400 | 4492500 | 3627100 | 0 | 5012200 | 4801000 | 3306100 | 1809600 | 1120300 | 3973700 | 2436700 |
| ATESGAQS | FKBP4; FKBP4 | Q02790; F5I | FKBP4_HUMAN | Unmodified | 0 | 0 | 0 | 0 | 0 | 0 | 0 | 0 | 0 | 0 | 423120 | 0 | 0 | 0 | 250650 | 0 | 0 | 0 |
| ATESGAQS | FKBP4; FKBP4 | Q02790; F5I | FKBP4_HUMAN | Oxidation (M) | 0 | 391750 | 178550 | 74900 | 0 | 0 | 0 | 0 | 0 | 0 | 524480 | 2326800 | 0 | 0 | 194820 | 337610 | 0 | 0 |
| ATESGAQS | FKBP4; FKBP4 | Q02790; F5I | FKBP4_HUMAN | Unmodified | 0 | 0 | 0 | 0 | 729830 | 0 | 0 | 90432 | 1277200 | 0 | 903950 | 7243700 | 2691200 | 2483700 | 0 | 0 | 470310 | 860190 |
| ATESGAQS | FKBP4; FKBP4 | Q02790; F5I | FKBP4_HUMAN | Oxidation (M) | 0 | 0 | 0 | 2381900 | 556120 | 0 | 0 | 0 | 0 | 0 | 3421800 | 15219000 | 5783500 | 1786000 | 0 | 1064800 | 496460 | 0 |
| ATESIEVDN | N/A | N/A | N/A | Unmodified | 0 | 0 | 0 | 0 | 0 | 0 | 0 | 0 | 74276000 | 0 | 0 | 0 | 0 | 0 | 0 | 41631000 | 0 | 0 |
| ATETSRPSL | INPP5D | Q92835 | SHIP1_HUMAN | Unmodified | 0 | 0 | 0 | 0 | 0 | 0 | 0 | 0 | 0 | 0 | 389160 | 4067400 | 112860 | 0 | 0 | 0 | 0 | 0 |
| ATETVELHK | SARNP; SARNP | P82979; F8V | SARNP_HUMAI | Acetyl (Protein N | 0 | 73368 | 0 | 0 | 1192500 | 0 | 0 | 224020 | 693180 | 728590 | 0 | 2562600 | 0 | 0 | 0 | 0 | 0 | 0 |
| ATEVAADAI | RPS6; RPS6; RP | P62753; A2A | RS6_HUMAN | Unmodified | 0 | 0 | 0 | 0 | 0 | 0 | 0 | 0 | 0 | 0 | 0 | 1443900 | 0 | 0 | 0 | 0 | 0 | 0 |
| ATEVHNLQ | GOLGA7B | Q2TAP0 | GOG7B_HUMA | Acetyl (Protein N | 0 | 0 | 0 | 0 | 0 | 0 | 0 | 0 | 765980 | 367080 | 654830 | 2084400 | 0 | 0 | 0 | 0 | 0 | 0 |
| ATEVHNLQ | GOLGA7B | Q2TAP0 | GOG7B_HUMA | Acetyl (Protein N | 0 | 208860 | 0 | 0 | 0 | 0 | 0 | 0 | 389120 | 0 | 972200 | 5119400 | 0 | 0 | 0 | 54485 | 0 | 0 |
| ATEVSKTPE | MAGED2; MAGI | Q9UNF1; Q5 | MAGD2_HUMA | Unmodified | 0 | 0 | 0 | 0 | 821230 | 0 | 526500 | 750190 | 0 | 421630 | 0 | 0 | 621440 | 1040700 | 0 | 0 | 192880 | 1247900 |
| ATEVTVAR | MACF1; MACF1 | Q9UPN3; H3 | MACF1_HUMA | Unmodified | 0 | 0 | 0 | 0 | 0 | 0 | 0 | 61729 | 217220 | 62376 | 77034 | 0 | 0 | 0 | 0 | 0 | 115680 | 21411 |
| ATFAEDSK | ECE1; ECE1; EC | P42892; B4E | ECE1_HUMAN | Unmodified | 0 | 0 | 0 | 0 | 789240 | 0 | 0 | 0 | 1213100 | 0 | 0 | 0 | 0 | 0 | 0 | 0 | 0 | 0 |
| ATFAEDSK | ECE1; ECE1; EC | P42892; B4E | ECE1_HUMAN | Unmodified | 0 | 0 | 0 | 0 | 0 | 0 | 0 | 0 | 0 | 345610 | 0 | 0 | 0 | 0 | 0 | 0 | 0 | 0 |
| ATFAFSPEE | CEMIP2; CEMIP | Q9UHN6; F5 | CEIP2_HUMAN | Unmodified | 1468800 | 432490 | 327680 | 248250 | 0 | 0 | 0 | 0 | 519170 | 200210 | 0 | 141550 | 0 | 0 | 512630 | 470120 | 0 | 0 |
| ATFAFSPEE | CEMIP2; CEMIP | Q9UHN6; F5 | CEIP2_HUMAN | Unmodified | 275910 | 0 | 358510 | 0 | 0 | 0 | 0 | 0 | 698800 | 0 | 0 | 0 | 0 | 0 | 828230 | 500440 | 0 | 0 |
| ATFDAISK | RPS2; RPS2; RP | P15880; E9P | RS2_HUMAN | Unmodified | 9846.2 | 2653200 | 175270 | 6951 | 2596200 | 0 | 6919100 | 4007600 | 0 | 0 | 0 | 66020000 | 4218500 | 131780 | 261900 | 221310 | 0 | 858500 |
| ATFDISLVV | NPEPPSL1; NPI | A6NEC2; P5 | PSAL_HUMAN | Unmodified | 0 | 0 | 0 | 0 | 724990 | 0 | 0 | 0 | 508080 | 458170 | 602850 | 0 | 0 | 0 | 0 | 0 | 554700 | 820520 |
| ATFGSGGH | CAPN15 | O75808 | CAN15_HUMAI | Unmodified | 132760 | 173810 | 0 | 0 | 0 | 0 | 0 | 0 | 0 | 0 | 0 | 0 | 0 | 0 | 0 | 82161 | 0 | 0 |
| ATFHTPFSI | ECI2; ECI2 | O75521; A0A | ECI2_HUMAN | Unmodified | 0 | 0 | 0 | 0 | 0 | 0 | 0 | 0 | 4275400 | 519550 | 0 | 0 | 0 | 0 | 1157700 | 408960 | 499970 | 0 |
| ATFIKVPQN | NCL; NCL; NCL | P19338; C9J | NUCL_HUMAN | Unmodified | 0 | 0 | 0 | 79267 | 1263200 | 0 | 1036400 | 589730 | 0 | 519940 | 0 | 4277000 | 2829200 | 1306000 | 0 | 0 | 444780 | 321650 |
| ATFISVQLK | PDCD6IP; PDCI | Q8WUM4; FI | PDC6I_HUMA | Acetyl (Protein N | 2903700 | 7911800 | 568530 | 1352500 | 2883100 | 0 | 357680 | 0 | 1499000 | 2475700 | 388000 | 650680 | 4832400 | 1923100 | 0 | 1452300 | 622720 | 1595700 |
| ATFISVQLK | PDCD6IP; PDCI | Q8WUM4; FI | PDC6I_HUMA | Acetyl (Protein N | 14082000 | 15413000 | 3045900 | 6655700 | 13671000 | 2247700 | 1936900 | 861650 | 6193100 | 8988000 | 2233800 | 4046900 | 19142000 | 7318300 | 7844300 | 6227100 | 2132600 | 5286600 |
| ATFNPAQD | PODXL | O00592 | PODXL_HUMAI | Unmodified | 0 | 403670 | 0 | 1488500 | 3323100 | 0 | 2097700 | 863660 | 24215000 | 4102700 | 4774100 | 1561900 | 6971400 | 5280900 | 981030 | 0 | 1091200 | 3621300 |
| ATFNPAQD | PODXL | O00592 | PODXL_HUMAI | Unmodified | 0 | 0 | 0 | 0 | 0 | 0 | 0 | 57151 | 0 | 188200 | 0 | 411450 | 145950 | 88141 | 0 | 0 | 0 | 0 |
| ATFSPIVTV | ANK3; ANK1; AI | Q12955; P16 | ANK3_HUMAN | Unmodified | 0 | 0 | 0 | 0 | 0 | 0 | 0 | 0 | 0 | 0 | 0 | 4001900 | 3514800 | 2781100 | 4459500 | 3405400 | 0 | 436530 |
| ATFVELSTK | AKR1B15; AKR1 | C9JRZ8; O6 | AK1BF_HUMAN | Acetyl (Protein N | 0 | 0 | 0 | 0 | 0 | 61304 | 3049600 | 3386600 | 0 | 0 | 0 | 0 | 0 | 0 | 0 | 0 | 0 | 46146 |
| ATFYGEQV | IQGAP1; IQGAP | P46940; H0 | IQGA1_HUMAN | Unmodified | 0 | 0 | 0 | 0 | 3611100 | 1017200 | 515240 | 168920 | 326050 | 0 | 0 | 615140 | 710820 | 382250 | 0 | 0 | 121200 | 367230 |
| ATGASRDM | DSG2 | Q14126 | DSG2_HUMAN | Oxidation (M) | 0 | 0 | 0 | 0 | 22386000 | 4139000 | 3517600 | 0 | 0 | 1216200 | 0 | 0 | 0 | 9538500 | 0 | 0 | 0 | 0 |
| ATGASRDM | DSG2 | Q14126 | DSG2_HUMAN | Unmodified | 0 | 0 | 0 | 0 | 7671700 | 0 | 0 | 0 | 0 | 0 | 0 | 0 | 0 | 0 | 0 | 0 | 0 | 0 |
| ATGDETGA | RPS3A; RPS3A; | P61247; D6F | RS3A_HUMAN | Unmodified | 82073 | 0 | 0 | 521380 | 0 | 0 | 0 | 0 | 2064200 | 0 | 167610 | 395720 | 0 | 0 | 68007 | 58233 | 32943 | 0 |
| ATGDETGA | RPS3A; RPS3A; | P61247; D6F | RS3A_HUMAN | Unmodified | 4249500 | 3287800 | 1359700 | 5176500 | 2112300 | 0 | 696900 | 785000 | 1911300 | 840720 | 118260 | 3900900 | 1859600 | 1029800 | 1456300 | 2106700 | 534570 | 1380900 |
| ATGDPTLLI | EDEM2 | Q9BV94 | EDEM2_HUMAI | Unmodified | 0 | 0 | 0 | 0 | 0 | 0 | 0 | 0 | 1253400 | 0 | 0 | 0 | 0 | 0 | 0 | 0 | 0 | 0 |
| ATGDPYYL | EDEM3 | Q9BZQ6 | EDEM3_HUMAI | Unmodified | 0 | 0 | 0 | 0 | 0 | 0 | 0 | 0 | 744240 | 0 | 0 | 0 | 0 | 0 | 0 | 0 | 0 | 0 |
| ATGEGFVE | RBM12 | Q9NTZ6 | RBM12_HUMAI | Unmodified | 0 | 0 | 0 | 0 | 0 | 0 | 0 | 0 | 0 | 0 | 318100 | 5119000 | 0 | 0 | 0 | 0 | 0 | 0 |
| ATGFDGGN | THOC2; THOC2 | Q8NI27; H0 | THOC2_HUMA | Unmodified | 0 | 0 | 0 | 0 | 0 | 0 | 0 | 0 | 0 | 0 | 0 | 430120 | 0 | 0 | 0 | 0 | 0 | 0 |
| ATGGYRTS | PAWR; PAWR; F | Q96IZ0; F8W | PAWR_HUMAN | Acetyl (Protein N | 0 | 0 | 0 | 0 | 2081600 | 0 | 0 | 0 | 0 | 0 | 0 | 0 | 0 | 692150 | 0 | 0 | 1054600 | 0 |
| ATGILLYGL | QARS1; QARS1 | P47897; F2Z | SYQ_HUMAN | Unmodified | 0 | 0 | 0 | 0 | 0 | 0 | 0 | 0 | 585850 | 0 | 721780 | 5909500 | 0 | 0 | 0 | 0 | 88298 | 0 |
| ATGIYSKTS | CPD | O75976 | CBPD_HUMAN | Unmodified | 0 | 0 | 0 | 0 | 695450 | 207670 | 0 | 0 | 0 | 1682500 | 0 | 0 | 84084 | 71046 | 0 | 0 | 0 | 0 |
| ATGIYSKTS | CPD | O75976 | CBPD_HUMAN | Unmodified | 0 | 13821000 | 0 | 126410 | 678530 | 0 | 0 | 0 | 9647700 | 3367300 | 163040 | 2188700 | 396200 | 370100 | 0 | 2814500 | 51529 | 0 |
| ATGLGEPV | NEDD4L; NEDD | Q96PU5; A0 | NED4L_HUMAI | Acetyl (Protein N | 0 | 0 | 0 | 0 | 6146700 | 1568100 | 0 | 251680 | 0 | 2845500 | 0 | 0 | 2951300 | 2622100 | 0 | 0 | 6592900 | 6514600 |
| ATGLHFRH | IQGAP1; IQGAP | P46940; A0A | IQGA1_HUMAN | Oxidation (M) | 835410 | 0 | 0 | 0 | 13415000 | 440100 | 329460 | 0 | 0 | 35768 | 0 | 0 | 458720 | 2364000 | 0 | 0 | 1159600 | 439670 |
| ATGLHFRH | IQGAP1; IQGAP | P46940; A0A | IQGA1_HUMAN | Unmodified | 0 | 0 | 0 | 0 | 13618000 | 0 | 518350 | 0 | 0 | 0 | 0 | 0 | 0 | 483720 | 0 | 0 | 172560 | 843440 |
| ATGPPVSEI | H1-5 | P16401 | H15_HUMAN | Unmodified | 39081000 | 13417000 | 0 | 0 | 1009800 | 0 | 525940 | 25208000 | 347200 | 304200 | 698870 | 235340 | 278730 | 0 | 0 | 0 | 117730 | 0 |
| ATGPPVSEI | H1-5 | P16401 | H15_HUMAN | Unmodified | 67843 | 57729 | 0 | 0 | 0 | 0 | 0 | 3739300 | 0 | 0 | 0 | 0 | 0 | 0 | 0 | 0 | 0 | 0 |
| ATGSPVSIF | SCYL1; SCYL1; | Q96KG9; E9I | SCYL1_HUMA | Unmodified | 0 | 0 | 0 | 0 | 0 | 0 | 0 | 0 | 238680 | 0 | 0 | 1656700 | 0 | 0 | 0 | 0 | 0 | 0 |
| ATGTSSGA | STIM1; STIM1; | Q13586; G0 | STIM1_HUMA | Unmodified | 0 | 0 | 0 | 0 | 0 | 0 | 0 | 0 | 1494100 | 304180 | 0 | 0 | 0 | 0 | 0 | 0 | 0 | 0 |
| ATGYPLAY | CAD; CAD | P27708; F8V | PYR1_HUMAN | Unmodified | 0 | 0 | 0 | 0 | 0 | 0 | 0 | 0 | 0 | 0 | 0 | 1188600 | 0 | 0 | 174730 | 208670 | 0 | 0 |
| ATHGGRID | APRT; APRT; AP | P07741; H3 | APT_HUMAN | Unmodified | 169070 | 620340 | 460620 | 546570 | 598650 | 0 | 214140 | 75052 | 0 | 0 | 1631200 | 3258000 | 710080 | 450270 | 637550 | 496050 | 0 | 0 |
| ATHGQTCA | VDAC2; VDAC2 | P45880; Q5J | VDAC2_HUMA | Acetyl (Protein N | 0 | 0 | 0 | 0 | 0 | 0 | 287020 | 148550 | 6201900 | 4203600 | 0 | 0 | 0 | 0 | 0 | 0 | 0 | 0 |
| ATHGQTCA | VDAC2; VDAC2 | P45880; Q5J | VDAC2_HUMA | Acetyl (Protein N | 0 | 0 | 0 | 0 | 0 | 0 | 0 | 0 | 10650000 | 3430300 | 0 | 0 | 0 | 0 | 0 | 0 | 0 | 0 |
| ATHIKLPRT | MTHFD1; MTHF | P11586; V9C | C1TC_HUMAN | Unmodified | 0 | 0 | 0 | 0 | 1005300 | 0 | 501140 | 0 | 0 | 805130 | 0 | 0 | 346200 | 410410 | 0 | 0 | 827580 | 1256900 |
| ATHRLLLLC | GNAS; GNAL; G | P63092; P38 | GNAS2_HUMA | Unmodified | 0 | 0 | 0 | 0 | 3615000 | 1211300 | 223330 | 76508 | 0 | 307570 | 0 | 0 | 2845600 | 1451700 | 0 | 0 | 733270 | 3834800 |
| ATHRLLLLC | GNAS; GNAL; G | P63092; P38 | GNAS2_HUMA | Unmodified | 0 | 0 | 0 | 0 | 2045800 | 489160 | 368270 | 0 | 0 | 0 | 0 | 0 | 1000100 | 880460 | 0 | 0 | 1359400 | 1430000 |
| ATIAGGGVI | H2AZ2; H2AZ1 | Q71UI9; P0C | H2AV_HUMAN | Unmodified | 20110000 | 8859300 | 0 | 0 | 2051300 | 364240 | 7347000 | 1.13E+08 | 175680 | 353390 | 0 | 1.3E+08 | 0 | 0 | 0 | 0 | 95083 | 1744300 |
| ATIDPELNE | MKLN1; MKLN1 | Q9UL63; C9 | MKLN1_HUMA | Unmodified | 0 | 0 | 0 | 0 | 446040 | 206120 | 0 | 0 | 0 | 0 | 248110 | 799980 | 0 | 0 | 0 | 0 | 0 | 92712 |
| ATIEGYYQ | PGM3; PGM3; F | O95394; J3K | AGM1_HUMAN | Unmodified | 0 | 174580 | 0 | 0 | 599400 | 71709 | 368110 | 78558 | 0 | 120950 | 0 | 0 | 905490 | 1298100 | 0 | 0 | 456140 | 916400 |
| ATIERDGY | ACO2; ACO2; A | Q99798; A2A | ACON_HUMAN | Unmodified | 0 | 0 | 0 | 0 | 0 | 0 | 0 | 0 | 3355700 | 0 | 0 | 0 | 0 | 0 | 0 | 0 | 0 | 0 |
| ATIFNLK | EIF2A; EIF2A; EI | Q9BY44; F8 | EIF2A_HUMAN | Unmodified | 139190 | 0 | 0 | 0 | 0 | 0 | 0 | 224700 | 0 | 0 | 0 | 5616300 | 685610 | 108420 | 0 | 0 | 0 | 0 |
| ATIGADFLT | RAB7A; RAB7A; | P51149; C9J | RAB7A_HUMAN | Unmodified | 10615000 | 13061000 | 11491000 | 14894000 | 29387000 | 4906900 | 7466800 | 5987500 | 27908000 | 14585000 | 9313000 | 61072000 | 18914000 | 18893000 | 15105000 | 13835000 | 4723500 | 11939000 |
| ATILCTSYV | RPA3; RPA3 | P35244; B5I | RFA3_HUMAN | Unmodified | 0 | 0 | 0 | 0 | 1376500 | 0 | 1733600 | 0 | 0 | 447930 | 0 | 0 | 4024600 | 0 | 0 | 0 | 1918100 | 1519300 |
| ATIPDWKL | PPP1R18; PPP1 | Q6NYC8; A0 | PPR18_HUMAN | Acetyl (Protein N | 0 | 0 | 0 | 0 | 876470 | 111350 | 39525 | 26415 | 0 | 18912 | 0 | 0 | 0 | 29168 | 0 | 0 | 367650 | 302530 |
| ATIQGVLR | AP2A1 | O95782 | AP2A1_HUMA | Unmodified | 140980 | 299470 | 0 | 223050 | 0 | 0 | 0 | 0 | 0 | 0 | 0 | 3532500 | 141300 | 0 | 0 | 0 | 0 | 0 |
| ATISALEAK | MYH10 | P35580 | MYH10_HUMA | Unmodified | 61582 | 239550 | 571520 | 607950 | 1824500 | 0 | 70502 | 142950 | 0 | 0 | 203710 | 1083600 | 1857800 | 898700 | 502330 | 764260 | 0 | 0 |
| ATISDEEIE | CHMP2B; CHM | Q9UQN3; C | CHM2B_HUMA | Unmodified | 2114900 | 1963400 | 541590 | 718310 | 0 | 0 | 0 | 0 | 3309300 | 317210 | 549370 | 3014900 | 0 | 0 | 1166100 | 0 | 0 | 1224500 |
| ATISDEEIE | CHMP2B; CHM | Q9UQN3; C | CHM2B_HUMA | Unmodified | 1346500 | 1145500 | 559060 | 1062500 | 13887000 | 1967000 | 0 | 0 | 355160 | 7038300 | 883430 | 550240 | 20107000 | 15190000 | 856840 | 438870 | 1907800 | 5712800 |
| ATISHVSPD | PTPRG | P23470 | PTPRG_HUMAI | Unmodified | 267690 | 328670 | 117800 | 418820 | 0 | 0 | 0 | 0 | 0 | 0 | 81920 | 0 | 33709 | 0 | 115910 | 272680 | 0 | 0 |
| ATISNDGA | CCT7; CCT7 | Q99832; F8V | TCPH_HUMAN | Unmodified | 665710 | 0 | 500790 | 1877400 | 8762200 | 801210 | 1285400 | 742860 | 3164600 | 3423400 | 3582900 | 40094000 | 16672000 | 10732000 | 1131900 | 1250400 | 1189500 | 2719300 |
| ATISNDGA | CCT7; CCT7 | Q99832; F8V | TCPH_HUMAN | Unmodified | 0 | 0 | 0 | 0 | 218040 | 0 | 1048100 | 0 | 0 | 147310 | 0 | 34031 | 751510 | 582510 | 0 | 0 | 1438900 | 292850 |
| ATISNPITG | P4HA1 | P13674 | P4HA1_HUMAI | Unmodified | 0 | 0 | 0 | 0 | 0 | 0 | 0 | 0 | 1215400 | 0 | 0 | 0 | 0 | 0 | 0 | 0 | 0 | 0 |
| ATISNPITG | P4HA1 | P13674 | P4HA1_HUMAI | Unmodified | 0 | 0 | 0 | 0 | 0 | 0 | 166670 | 0 | 0 | 351020 | 0 | 0 | 0 | 446680 | 0 | 0 | 302500 | 472720 |
| ATITLYGK | CEMIP2; CEMIP | Q9UHN6; F5 | CEIP2_HUMAN | Unmodified | 345750 | 0 | 772500 | 0 | 906410 | 151650 | 35977 | 12780 | 1520100 | 0 | 0 | 1542200 | 0 | 0 | 1196700 | 0 | 108680 | 408960 |
| ATITLYGKS | CEMIP2; CEMIP | Q9UHN6; F5 | CEIP2_HUMAN | Unmodified | 0 | 0 | 0 | 122020 | 2270600 | 349270 | 148820 | 44581 | 1291400 | 241780 | 0 | 0 | 89881 | 0 | 0 | 0 | 0 | 0 |
| ATKIDKEAC | COTL1 | Q14019 | COTL1_HUMAI | Acetyl (Protein N | 0 | 0 | 0 | 0 | 1603500 | 344130 | 141250 | 174800 | 689090 | 2071100 | 9337 | 57907 | 55981 | 165430 | 11033 | 0 | 2047100 | 1962200 |
| ATKSPEEG | CBR1 | P16152 | CBR1_HUMAN | Unmodified | 0 | 0 | 0 | 0 | 14664000 | 0 | 0 | 160940 | 0 | 784780 | 0 | 0 | 827990 | 0 | 0 | 0 | 6993100 | 20519000 |
| ATKVQDIK | GNAS; GNAS; G | Q5JWF2; P6 | GNAS1_HUMA | Unmodified | 0 | 0 | 0 | 0 | 799070 | 0 | 207190 | 197380 | 0 | 45216 | 0 | 0 | 418070 | 379940 | 0 | 0 | 1548300 | 400050 |
| ATLAENPN | PRR15L | Q9BU68 | PR15L_HUMAI | Unmodified | 0 | 0 | 0 | 0 | 0 | 0 | 0 | 0 | 0 | 0 | 0 | 0 | 1921000 | 2532500 | 0 | 899590 | 0 | 0 |
| ATLELTHN | GLO1 | Q04760 | LGUL_HUMAN | Unmodified | 0 | 0 | 0 | 0 | 0 | 0 | 0 | 0 | 4406500 | 0 | 0 | 0 | 0 | 0 | 0 | 0 | 0 | 0 |
| ATLENITNL | CZIB | Q9NWV4 | CZIB_HUMAN | Unmodified | 0 | 0 | 0 | 0 | 234030 | 0 | 0 | 0 | 1080800 | 0 | 0 | 2694200 | 0 | 0 | 653620 | 0 | 0 | 212480 |
| ATLENITNL | CZIB | Q9NWV4 | CZIB_HUMAN | Unmodified | 0 | 0 | 0 | 938150 | 7376600 | 2106700 | 5576200 | 0 | 0 | 1289900 | 0 | 0 | 5182000 | 2722200 | 0 | 0 | 6592200 | 8152900 |
| ATLKDQLIY | LDHA; LDHA; LI | P00338; F5C | LDHA_HUMAN | Acetyl (Protein N | 0 | 0 | 0 | 0 | 1819600 | 365950 | 0 | 0 | 0 | 464820 | 184750 | 576510 | 603730 | 0 | 0 | 0 | 1161100 | 2844700 |
| ATLKDQLIY | LDHA; LDHA; LI | P00338; F5C | LDHA_HUMAN | Acetyl (Protein N | 0 | 0 | 0 | 2112500 | 46491000 | 8921700 | 3499800 | 0 | 1675300 | 14859000 | 0 | 8419600 | 18434000 | 5439800 | 0 | 0 | 39931000 | 0 |
| ATLNAFLYF | ATAD3A; ATAD3 | Q9NVI7; Q5 | ATD3A_HUMA | Unmodified | 0 | 0 | 0 | 0 | 28551 | 0 | 0 | 0 | 1830200 | 0 | 207560 | 0 | 0 | 0 | 0 | 0 | 183750 | 0 |
| ATLPDADK | ACTN1; ACTN1 | P12814; A0 | ACTN1_HUMAI | Unmodified | 0 | 0 | 0 | 0 | 21722000 | 8310800 | 972880 | 0 | 3395700 | 536520 | 339380 | 2157700 | 906600 | 733510 | 0 | 108130 | 11702000 | 477640 |
| ATLPSPDK | UBA1; UBA1 | P22314; Q5J | UBA1_HUMAN | Unmodified | 493600 | 0 | 3067000 | 1292500 | 2826700 | 2999200 | 1728100 | 0 | 5080400 | 2201400 | 3457800 | 13305000 | 2727800 | 993370 | 2829400 | 1457600 | 1613700 | 4888100 |
| ATLQAALC | ARPC4; ARPC4 | P59998; A0 | ARPC4_HUMA | Unmodified | 0 | 448250 | 0 | 1467000 | 2643900 | 0 | 0 | 0 | 437220 | 0 | 0 | 0 | 1688700 | 446060 | 0 | 0 | 239320 | 1146200 |
| ATLRYFQH | HYOU1; HYOU | Q9Y4L1; J3C | HYOU1_HUMA | Unmodified | 0 | 0 | 0 | 0 | 117120 | 0 | 613070 | 0 | 849760 | 8295200 | 32190 | 0 | 294670 | 777560 | 0 | 0 | 93726 | 144180 |
| ATLWYVPL | ACOT7; ACOT7 | O00154; K7I | BACH_HUMAN | Unmodified | 347460 | 157880 | 238490 | 401450 | 5691300 | 625030 | 2798900 | 0 | 7355500 | 6059700 | 1154200 | 6915800 | 0 | 0 | 492620 | 350510 | 6508300 | 8397400 |
| ATLWYVPL | ACOT7; ACOT7 | O00154; K7I | BACH_HUMAN | Unmodified | 0 | 0 | 0 | 0 | 0 | 0 | 368660 | 0 | 0 | 0 | 0 | 0 | 0 | 0 | 0 | 0 | 880220 | 0 |
| ATMEAYSR | ANXA7 | P20073 | ANXA7_HUMAI | Unmodified | 692060 | 365440 | 882210 | 2364400 | 866210 | 544680 | 392350 | 179770 | 64274000 | 7740100 | 142220 | 1244700 | 1318600 | 985930 | 2840600 | 1476100 | 707920 | 904060 |
| ATMEAYSR | ANXA7 | P20073 | ANXA7_HUMAI | Oxidation (M) | 57893 | 75921 | 125350 | 137370 | 0 | 0 | 0 | 0 | 5008000 | 304990 | 52343 | 180100 | 52903 | 56807 | 162280 | 116440 | 33930 | 211190 |
| ATNDEIFSI | NCBP1 | Q09161 | NCBP1_HUMA | Unmodified | 0 | 0 | 0 | 0 | 0 | 0 | 0 | 0 | 0 | 0 | 0 | 2771100 | 0 | 0 | 0 | 0 | 0 | 0 |
| ATNFLAHE | EEF1D; EEF1D; | P29692; E9F | EF1D_HUMAN | Acetyl (Protein N | 0 | 0 | 540820 | 758340 | 16265000 | 3519900 | 554760 | 679490 | 1650700 | 1530500 | 2737500 | 38332000 | 7008400 | 4224700 | 633990 | 650900 | 15072000 | 9550000 |
| ATNFLAHE | EEF1D; EEF1D; | P29692; E9F | EF1D_HUMAN | Acetyl (Protein N | 0 | 0 | 0 | 0 | 0 | 0 | 0 | 0 | 0 | 0 | 0 | 44597 | 0 | 83666 | 0 | 0 | 1893800 | 0 |
| ATNIEVLSN | CSDE1; CSDE1 | O75534; E9F | CSDE1_HUMAI | Unmodified | 0 | 213330 | 0 | 0 | 0 | 0 | 0 | 0 | 0 | 0 | 267490 | 1999900 | 0 | 0 | 0 | 0 | 0 | 0 |
| ATNMTVSA | CPT2; CPT2; CF | P23786; A0A | CPT2_HUMAN | Unmodified | 0 | 0 | 0 | 0 | 0 | 0 | 0 | 0 | 1251900 | 0 | 0 | 0 | 0 | 0 | 0 | 0 | 0 | 0 |
| ATNSTEEEI | ARHGAP29; AR | Q52LW3; F8 | RHG29_HUMA | Unmodified | 0 | 0 | 0 | 0 | 0 | 165910 | 0 | 0 | 0 | 0 | 0 | 0 | 0 | 0 | 0 | 0 | 1215200 | 1807900 |
| ATNVCTRF | SMARCA5 | O60264 | SMCA5_HUMA | Unmodified | 287890 | 187720 | 0 | 0 | 115450 | 0 | 195060 | 183240 | 370550 | 0 | 0 | 0 | 0 | 49644 | 0 | 0 | 0 | 0 |
| ATNWGSLL | GSS; GSS; GSS | P48637; A0 | GSHB_HUMAN | Acetyl (Protein N | 0 | 0 | 268300 | 0 | 0 | 0 | 0 | 0 | 0 | 0 | 0 | 3322000 | 0 | 0 | 502910 | 303910 | 0 | 0 |
| ATNWGSLL | GSS; GSS; GSS | P48637; A0 | GSHB_HUMAN | Acetyl (Protein N | 0 | 0 | 0 | 0 | 70114 | 0 | 475330 | 0 | 345160 | 0 | 0 | 0 | 0 | 0 | 0 | 198850 | 0 | 0 |
| ATPEDNGF | ICAM1; ICAM1 | P05362; E7E | ICAM1_HUMAI | Unmodified | 303770 | 0 | 254550 | 0 | 9810900 | 2160500 | 0 | 0 | 0 | 0 | 0 | 0 | 0 | 0 | 0 | 0 | 1297200 | 2004500 |

| | | | | | | | | | | | | | | | | | | | | | | |
|---|---|---|---|---|---|---|---|---|---|---|---|---|---|---|---|---|---|---|---|---|---|---|
| ATPEQYQIL | FDPS; FDPS | P14324; A0A | FPPS_HUMAN | Unmodified | 519840 | 1921000 | 509190 | 610110 | 4100200 | 40387000 | 2930500 | 915650 | 5667800 | 2935600 | 3868200 | 18926000 | 2162100 | 268190 | 1055600 | 523420 | 5455800 | 725440 |
| ATPEQYQIL | FDPS; FDPS | P14324; A0A | FPPS_HUMAN | Unmodified | 0 | 0 | 0 | 0 | 2910800 | 0 | 0 | 0 | 0 | 28859 | 0 | 0 | 0 | 0 | 0 | 0 | 628700 | 0 |
| ATPEQYQIL | FDPS; FDPS | P14324; A0A | FPPS_HUMAN | Unmodified | 0 | 0 | 0 | 0 | 873030 | 0 | 0 | 0 | 0 | 501680 | 0 | 0 | 791880 | 144860 | 0 | 0 | 392620 | 81465 |
| ATPGNLGS | PIP4K2A | P48426 | PI42A_HUMAN | Acetyl (Protein N | 0 | 0 | 0 | 0 | 1064900 | 0 | 0 | 654800 | 0 | 269390 | 392820 | 1759700 | 0 | 1225500 | 0 | 0 | 527610 | 1285800 |
| ATPHLPLAT | N/A | N/A | N/A | Oxidation (M) | 3421400 | 2141000 | 539080 | 326870 | 0 | 0 | 0 | 0 | 0 | 0 | 0 | 0 | 0 | 0 | 3074100 | 1996700 | 0 | 0 |
| ATPQHTVS | SIRPA; SIRPB1 | P78324; Q5T | SHPS1_HUMAI | Unmodified | 0 | 0 | 0 | 0 | 0 | 0 | 0 | 0 | 0 | 0 | 509380 | 226090 | 113330 | 54362 | 0 | 0 | 0 | 0 |
| ATPYTFPG( | GALNT13 | Q8IUC8 | GLT13_HUMAN | Unmodified | 0 | 0 | 0 | 0 | 0 | 0 | 0 | 0 | 461040 | 130670 | 0 | 0 | 0 | 0 | 0 | 0 | 0 | 0 |
| ATPYTFPG( | GALNT1; GALN | Q10472; K7E | GALT1_HUMAN | Unmodified | 0 | 0 | 0 | 0 | 0 | 0 | 0 | 0 | 0 | 215100 | 223230 | 621220 | 0 | 0 | 0 | 0 | 0 | 0 |
| ATQADLME | CTNNB1; CTNN | P35222; A0A | CTNB1_HUMAI | Acetyl (Protein N | 182880 | 0 | 430400 | 719170 | 0 | 0 | 0 | 0 | 0 | 0 | 0 | 0 | 0 | 582150 | 939920 | 682960 | 784240 | 3894400 |
| ATQADLME | CTNNB1; CTNN | P35222; A0A | CTNB1_HUMAI | Acetyl (Protein N | 0 | 0 | 0 | 294620 | 2933200 | 0 | 0 | 0 | 0 | 0 | 0 | 207710 | 0 | 2520400 | 0 | 0 | 3119600 | 4805400 |
| ATQALVLAF | EIF4A1; EIF4A1; | P60842; J3K | IF4A1_HUMAN | Unmodified | 1006600 | 1130500 | 1708200 | 655180 | 0 | 0 | 293900 | 0 | 831750 | 0 | 906600 | 8345800 | 0 | 0 | 2209800 | 2262300 | 0 | 0 |
| ATQALVLAF | EIF4A1; EIF4A1; | P60842; J3K | IF4A1_HUMAN | Unmodified | 0 | 0 | 19812000 | 6958400 | 10622000 | 2844600 | 5300100 | 3210500 | 1886000 | 3749400 | 4150500 | 0 | 6230200 | 0 | 0 | 0 | 2412700 | 4300800 |
| ATQCVPKE | GRPEL1 | Q9HAV7 | GRPE1_HUMAI | Unmodified | 0 | 0 | 0 | 0 | 702850 | 32238 | 143250 | 0 | 990210 | 548800 | 0 | 0 | 129270 | 178210 | 0 | 0 | 88569 | 146290 |
| ATQEELK | DNAJA1 | P31689 | DNJA1_HUMAI | Unmodified | 0 | 0 | 0 | 0 | 0 | 21723000 | 0 | 0 | 0 | 0 | 0 | 0 | 0 | 0 | 0 | 0 | 0 | 0 |
| ATQEEVAVI | DOCK9; DOCK9 | Q9BZ29; A0A | DOCK9_HUMA | Unmodified | 845600 | 3938900 | 583830 | 0 | 0 | 0 | 0 | 0 | 0 | 0 | 0 | 0 | 1801900 | 665700 | 3504000 | 1420000 | 0 | 0 |
| ATQELIPIEI | MRPL40 | Q9NQ50 | RM40_HUMAN | Unmodified | 0 | 0 | 0 | 0 | 0 | 0 | 0 | 0 | 408780 | 254000 | 0 | 0 | 0 | 0 | 0 | 0 | 0 | 0 |
| ATQFTGAT( | DSG2 | Q14126 | DSG2_HUMAN | Unmodified | 0 | 0 | 0 | 415400 | 7592100 | 557590 | 0 | 0 | 2279500 | 258020 | 631940 | 0 | 0 | 0 | 316610 | 0 | 244020 | 1239800 |
| ATQFTGAT( | DSG2 | Q14126 | DSG2_HUMAN | Oxidation (M) | 0 | 0 | 1451600 | 734030 | 2314400 | 480910 | 63293 | 0 | 454190 | 0 | 868840 | 865960 | 0 | 0 | 626920 | 518380 | 0 | 281230 |
| ATQIPSYK | ACP3 | P15309 | PPAP_HUMAN | Unmodified | 0 | 0 | 0 | 0 | 104780 | 0 | 0 | 13644 | 396380 | 54521 | 260230 | 0 | 0 | 0 | 0 | 0 | 0 | 0 |
| ATQPETTEE | SHTN1 | A0MZ66 | SHOT1_HUMAI | Unmodified | 0 | 0 | 0 | 0 | 0 | 0 | 0 | 0 | 0 | 0 | 320910 | 0 | 0 | 0 | 0 | 0 | 0 | 0 |
| ATQQQHDI | PRKDC | P78527 | PRKDC_HUMAI | Unmodified | 0 | 0 | 592870 | 0 | 0 | 0 | 0 | 0 | 269460 | 29246 | 0 | 1855600 | 0 | 0 | 0 | 0 | 25702 | 0 |
| ATRFNLETE | OPA1; OPA1; OI | O60313; E5K | OPA1_HUMAN | Unmodified | 0 | 0 | 0 | 0 | 268510 | 0 | 0 | 0 | 0 | 0 | 398620 | 1499400 | 354490 | 0 | 0 | 0 | 0 | 0 |
| ATRPSDDP | PTPN23 | Q9H3S7 | PTN23_HUMAI | Unmodified | 422960 | 1163100 | 0 | 0 | 1500000 | 48711 | 0 | 0 | 550430 | 66824 | 0 | 0 | 0 | 0 | 0 | 0 | 0 | 0 |
| ATSFLLALE | RPN1; RPN1 | P04843; F8W | RPN1_HUMAN | Unmodified | 146500 | 0 | 22546 | 58483 | 0 | 0 | 47009 | 0 | 21218000 | 1872700 | 1422900 | 417110 | 0 | 0 | 495010 | 0 | 0 | 0 |
| ATSITVTGS | DAG1 | Q14118 | DAG1_HUMAN | Unmodified | 0 | 0 | 0 | 568620 | 0 | 0 | 0 | 0 | 0 | 0 | 0 | 0 | 0 | 0 | 458470 | 240180 | 0 | 0 |
| ATSKPMGC | CKAP5 | Q14008 | CKAP5_HUMAI | Unmodified | 0 | 0 | 0 | 104020 | 317380 | 1633600 | 435860 | 1833900 | 59740 | 14365 | 0 | 181480 | 59580 | 503430 | 0 | 0 | 19057 | 883170 |
| ATSNVFAM | MYL12A; MYL12 | P19105; O14 | ML12A_HUMAI | Unmodified | 0 | 0 | 0 | 86558 | 319040 | 0 | 0 | 0 | 487190 | 0 | 75324 | 537040 | 0 | 0 | 166250 | 145760 | 0 | 115090 |
| ATSNVFAM | MYL12A; MYL12 | P19105; O14 | ML12A_HUMAI | Oxidation (M) | 0 | 0 | 0 | 0 | 0 | 608350 | 0 | 32633 | 0 | 395820 | 1323400 | 8247200 | 2757100 | 0 | 2997000 | 0 | 0 | 620460 |
| ATSPQKSP: | PARVA | Q9NVD7 | PARVA_HUMAI | Acetyl (Protein N | 0 | 0 | 0 | 0 | 1764900 | 0 | 0 | 0 | 0 | 107000 | 0 | 0 | 0 | 0 | 0 | 0 | 984470 | 1819500 |
| ATSPVKSTT | ANLN | Q9NQW6 | ANLN_HUMAN | Unmodified | 0 | 0 | 0 | 0 | 277740 | 69187 | 118400 | 0 | 0 | 0 | 0 | 0 | 0 | 0 | 0 | 0 | 645780 | 153140 |
| ATSRGCITI | PGK1 | P00558 | PGK1_HUMAN | Unmodified | 0 | 0 | 0 | 0 | 328700 | 0 | 0 | 65891 | 0 | 129250 | 0 | 0 | 1302400 | 0 | 0 | 0 | 2182000 | 4097500 |
| ATSRGCITI | PGK1 | P00558 | PGK1_HUMAN | Unmodified | 0 | 0 | 0 | 0 | 0 | 0 | 0 | 0 | 0 | 0 | 0 | 0 | 464240 | 0 | 0 | 0 | 2445400 | 720310 |
| ATSSHPNS | TRIM25; TRIM25 | Q14258; A0A | TRI25_HUMAN | Unmodified | 0 | 0 | 34945 | 0 | 2382500 | 601990 | 0 | 0 | 28671 | 0 | 0 | 384270 | 262070 | 0 | 0 | 0 | 164380 | 334480 |
| ATSTATSGI | HSD17B4; HSD | P51659; E7E | DHB4_HUMAN | Unmodified | 0 | 0 | 0 | 0 | 0 | 0 | 0 | 0 | 860070 | 0 | 0 | 0 | 0 | 0 | 0 | 0 | 0 | 0 |
| ATSTRAQQ | CC2D1A; CC2D | Q6P1N0; A0A | C2D1A_HUMAI | Unmodified | 0 | 0 | 0 | 0 | 511430 | 143600 | 0 | 0 | 0 | 400820 | 0 | 0 | 395470 | 0 | 0 | 0 | 0 | 0 |
| ATSVALTW: | CNTN1 | Q12860 | CNTN1_HUMA | Unmodified | 0 | 416130 | 0 | 0 | 0 | 0 | 0 | 0 | 652540 | 0 | 0 | 0 | 0 | 0 | 0 | 0 | 0 | 0 |
| ATSVYLVQI | DIS3L2 | Q8IYB7 | DI3L2_HUMAN | Unmodified | 0 | 0 | 0 | 0 | 0 | 0 | 0 | 0 | 0 | 0 | 0 | 674990 | 0 | 0 | 0 | 239270 | 0 | 0 |
| ATTADGSSI | COPS4; COPS4 | Q9BT78; D6I | CSN4_HUMAN | Unmodified | 0 | 0 | 142180 | 0 | 0 | 0 | 0 | 0 | 170100 | 0 | 172680 | 2212700 | 0 | 0 | 0 | 257020 | 47664 | 0 |
| ATTADGSSI | COPS4; COPS4 | Q9BT78; D6I | CSN4_HUMAN | Unmodified | 0 | 0 | 57029 | 0 | 862560 | 239150 | 0 | 0 | 560400 | 541860 | 820360 | 183950 | 2864000 | 1008000 | 0 | 0 | 1005000 | 996740 |
| ATTATCTRF | CAMK2G; CAMI | Q13555; A0A | KCC2G_HUMA | Acetyl (Protein N | 0 | 0 | 68362 | 0 | 709530 | 0 | 0 | 0 | 0 | 0 | 0 | 0 | 0 | 0 | 0 | 0 | 3669000 | 4381000 |
| ATTGTPTAI | DPP9; DPP9; DI | Q86TI2; M0R | DPP9_HUMAN | Acetyl (Protein N | 0 | 0 | 0 | 0 | 1235800 | 0 | 0 | 0 | 0 | 0 | 0 | 0 | 0 | 0 | 0 | 0 | 1386300 | 1521600 |
| ATTLSNAVS | PICALM; PICALI | Q13492; H0Y | PICAL_HUMAN | Unmodified | 759280 | 1163900 | 0 | 504120 | 0 | 0 | 0 | 0 | 1832500 | 0 | 0 | 918830 | 0 | 0 | 0 | 570010 | 57828 | 0 |
| ATTPHPPPI | N/A | N/A | N/A | Unmodified | 0 | 0 | 0 | 0 | 0 | 0 | 0 | 0 | 0 | 0 | 0 | 0 | 28753000 | 297040 | 0 | 0 | 0 | 0 |
| ATTPVIMV( | POR; POR; POR | P16435; H0Y | NCPR_HUMAN | Oxidation (M) | 0 | 13754 | 0 | 0 | 0 | 0 | 0 | 0 | 1038900 | 160730 | 0 | 490650 | 0 | 0 | 0 | 0 | 0 | 0 |
| ATTVPDFQI | SH3GLB2; SH3( | Q9NR46; B7. | SHLB2_HUMAI | Unmodified | 0 | 0 | 0 | 0 | 208890 | 1873100 | 234530 | 0 | 735260 | 39281 | 0 | 366710 | 0 | 0 | 0 | 0 | 143730 | 97910 |
| ATTVPDGC | GPRC5A; GPRC | Q8NFJ5; F5C | RAI3_HUMAN F | Acetyl (Protein N | 0 | 0 | 0 | 0 | 7446200 | 2853900 | 0 | 0 | 0 | 0 | 0 | 0 | 0 | 0 | 0 | 0 | 0 | 0 |
| ATTVPDGC | GPRC5A; GPRC | Q8NFJ5; F5C | RAI3_HUMAN F | Unmodified | 0 | 0 | 0 | 0 | 0 | 0 | 0 | 0 | 0 | 0 | 0 | 0 | 0 | 0 | 10080000 | 0 | 0 | 0 |
| ATTVPDGC | GPRC5A; GPRC | Q8NFJ5; F5C | RAI3_HUMAN F | Acetyl (Protein N | 0 | 0 | 0 | 0 | 677720 | 0 | 0 | 0 | 0 | 21851 | 0 | 0 | 0 | 0 | 0 | 0 | 0 | 0 |
| ATTVSTQR | TOLLIP; TOLLIP; | Q9H0E2; E9I | TOLIP_HUMAN | Acetyl (Protein N | 1602100 | 2036300 | 0 | 673530 | 129980 | 0 | 0 | 0 | 822910 | 0 | 125010 | 816030 | 0 | 0 | 1987900 | 1365600 | 0 | 0 |
| ATTVSTQR( | TOLLIP; TOLLIP; | Q9H0E2; F2Z | TOLIP_HUMAN | Acetyl (Protein N | 13489000 | 0 | 2693800 | 15403000 | 9704600 | 3844900 | 4558200 | 0 | 5715400 | 7763900 | 2366200 | 607260 | 25852000 | 18770000 | 7771000 | 7186000 | 6890100 | 13900000 |
| ATTVSTQR( | TOLLIP; TOLLIP; | Q9H0E2; F2Z | TOLIP_HUMAN | Acetyl (Protein N | 0 | 0 | 0 | 0 | 0 | 0 | 0 | 0 | 0 | 2195700 | 0 | 0 | 36902000 | 17299000 | 0 | 0 | 4194900 | 0 |
| ATVAAFAAS | LIN7A; LIN7C; L | O14910; Q9I | LIN7A_HUMAN | Unmodified | 0 | 0 | 2748800 | 965750 | 628260 | 0 | 120370 | 70237 | 2595100 | 1125300 | 291280 | 4764400 | 81216 | 0 | 5038100 | 4357400 | 761060 | 0 |
| ATVAPEDV: | ACAT2 | Q9BWD1 | THIC_HUMAN / | Unmodified | 0 | 0 | 0 | 0 | 0 | 0 | 0 | 0 | 1049800 | 0 | 651550 | 1665800 | 0 | 0 | 604980 | 0 | 0 | 0 |
| ATVASSTQI | SEPTIN9; SEPTI | Q9UHD8; K7 | SEPT9_HUMAN | Unmodified | 0 | 0 | 0 | 0 | 0 | 0 | 0 | 0 | 0 | 0 | 0 | 0 | 0 | 0 | 0 | 0 | 806510 | 561690 |
| ATVDHNYY | DIPK2A; DIPK2A | Q8NDZ4; E7 | DIK2A_HUMAN | Unmodified | 341680 | 0 | 637730 | 0 | 0 | 0 | 0 | 0 | 41137000 | 7287000 | 0 | 1957400 | 0 | 0 | 0 | 47375 | 0 | 0 |
| ATVECVKD | CSDE1; CSDE1 | O75534; E9F | CSDE1_HUMAI | Unmodified | 0 | 0 | 0 | 0 | 0 | 0 | 0 | 0 | 0 | 0 | 0 | 0 | 1121900 | 0 | 0 | 0 | 0 | 0 |
| ATVEKPSNI | NUMBL; NUMB | Q9Y6R0; A0A | NUMBL_HUMA | Unmodified | 0 | 0 | 0 | 0 | 275970 | 65330 | 0 | 0 | 0 | 0 | 0 | 0 | 0 | 58807 | 0 | 0 | 0 | 0 |
| ATVEPETTP | EIF4E; EIF4E | P06730; D6F | IF4E_HUMAN E | Acetyl (Protein N | 0 | 0 | 0 | 0 | 0 | 0 | 0 | 0 | 0 | 0 | 1701100 | 3481500 | 0 | 0 | 74051 | 55693 | 0 | 0 |
| ATVEPETTP | EIF4E; EIF4E | P06730; D6F | IF4E_HUMAN E | Acetyl (Protein N | 969160 | 600190 | 0 | 1819200 | 0 | 0 | 0 | 1087300 | 2133700 | 0 | 3125500 | 12105000 | 4613000 | 0 | 0 | 733740 | 797520 | 0 |
| ATVFDDCK | STXBP3 | O00186 | STXB3_HUMAI | Unmodified | 0 | 0 | 393710 | 0 | 176450 | 0 | 0 | 0 | 57115 | 0 | 0 | 581280 | 0 | 71233 | 100620 | 0 | 23950 | 157620 |
| ATVFDDCK | STXBP3 | O00186 | STXB3_HUMAI | Unmodified | 0 | 0 | 0 | 0 | 1409200 | 564890 | 0 | 203760 | 130630 | 193810 | 66372 | 101640 | 1201800 | 412160 | 0 | 151920 | 373270 | 2094300 |
| ATVFLAMG | DNAJC3; DNAJ( | Q13217; X6F | DNJC3_HUMAI | Unmodified | 0 | 0 | 0 | 0 | 0 | 0 | 0 | 0 | 5462500 | 1307300 | 0 | 0 | 0 | 0 | 0 | 0 | 0 | 0 |
| ATVHIQVNI | CLSTN1; CLSTN | O94985; A0A | CSTN1_HUMAI | Unmodified | 530790 | 0 | 0 | 0 | 0 | 0 | 0 | 0 | 6320800 | 2224900 | 1592000 | 0 | 0 | 0 | 0 | 0 | 0 | 0 |
| ATVHIQVNI | CLSTN1; CLSTN | O94985; A0A | CSTN1_HUMAI | Unmodified | 0 | 0 | 0 | 0 | 0 | 0 | 82361 | 0 | 0 | 759980 | 0 | 0 | 0 | 0 | 0 | 0 | 0 | 0 |
| ATVLEDSYF | NEDD4; NEDD4 | P46934; H0Y | NEDD4_HUMA | Unmodified | 273140 | 539450 | 57589 | 0 | 348310 | 152620 | 0 | 0 | 415670 | 11211 | 0 | 0 | 0 | 0 | 481930 | 360480 | 0 | 0 |
| ATVLTTER | PRPF19; PRPF1 | Q9UMS4; F5 | PRP19_HUMAI | Unmodified | 0 | 130410 | 0 | 0 | 113990 | 0 | 0 | 0 | 0 | 0 | 0 | 1464000 | 0 | 0 | 0 | 0 | 179430 | 220090 |
| ATVMFECD | CD46 | P15529 | MCP_HUMAN I | Unmodified | 1215700 | 921140 | 411190 | 440710 | 1121500 | 0 | 0 | 0 | 1547500 | 166410 | 0 | 3567300 | 0 | 0 | 681350 | 448120 | 0 | 0 |
| ATVMFECD | CD46 | P15529 | MCP_HUMAN I | Oxidation (M) | 423090 | 119010 | 0 | 488780 | 0 | 0 | 0 | 0 | 0 | 0 | 0 | 0 | 0 | 0 | 0 | 603210 | 0 | 0 |
| ATVMLYDD | VASP; VASP; VA | P50552; K7E | VASP_HUMAN | Unmodified | 0 | 0 | 0 | 0 | 0 | 0 | 0 | 0 | 0 | 0 | 0 | 749050 | 0 | 0 | 0 | 0 | 0 | 0 |
| ATVMLYDD | VASP; VASP; VA | P50552; K7E | VASP_HUMAN | Unmodified | 0 | 0 | 0 | 94666 | 65012 | 0 | 0 | 0 | 177340 | 0 | 176800 | 3020500 | 39846 | 0 | 233990 | 108340 | 36210 | 0 |
| ATVMLYDD | VASP; VASP; VA | P50552; K7E | VASP_HUMAN | Oxidation (M) | 0 | 0 | 0 | 0 | 0 | 0 | 0 | 0 | 0 | 0 | 0 | 0 | 0 | 0 | 0 | 0 | 0 | 0 |
| ATVNTFGYI | SF3B1; SF3B1 | O75533; H7( | SF3B1_HUMAI | Unmodified | 395750 | 0 | 87771 | 0 | 368440 | 0 | 132130 | 0 | 0 | 368180 | 0 | 2196400 | 0 | 0 | 0 | 0 | 0 | 0 |
| ATVPKTEIR | RPS24; RPS24; | P62847; A0A | RS24_HUMAN | Unmodified | 0 | 0 | 0 | 0 | 1711900 | 0 | 1960000 | 974000 | 90820 | 0 | 0 | 11650 | 1369800 | 1044000 | 0 | 0 | 0 | 569890 |
| ATVQQLEG | FABP5 | Q01469 | FABP5_HUMAI | Acetyl (Protein N | 1185100 | 104890 | 0 | 0 | 4614300 | 929020 | 0 | 40797 | 425400 | 0 | 8082500 | 27620000 | 0 | 0 | 0 | 0 | 4156000 | 10479000 |
| ATVQQLEG | FABP5 | Q01469 | FABP5_HUMAI | Acetyl (Protein N | 0 | 0 | 0 | 0 | 2066500 | 0 | 0 | 0 | 0 | 0 | 0 | 0 | 0 | 0 | 0 | 0 | 1547200 | 2198200 |
| ATVQQLEG | FABP5 | Q01469 | FABP5_HUMAI | Acetyl (Protein N | 0 | 0 | 0 | 0 | 13940000 | 4367000 | 2433700 | 0 | 0 | 1114100 | 172320 | 0 | 0 | 0 | 0 | 0 | 35955000 | 49414000 |
| ATVSAPFGI | PLEC; PLEC | Q15149; H0Y | PLEC_HUMAN | Unmodified | 0 | 0 | 0 | 0 | 123860 | 0 | 0 | 0 | 0 | 39297 | 0 | 0 | 0 | 0 | 0 | 0 | 1482300 | 248450 |
| ATVTPSPVI | NUCKS1 | Q9H1E3 | NUCKS_HUMA | Unmodified | 0 | 0 | 0 | 53258 | 3066800 | 1027300 | 283450 | 305410 | 2245800 | 2190200 | 1464700 | 3012500 | 0 | 0 | 47177 | 128210 | 1309100 | 2968400 |
| ATVVVEATE | MAP1B; MAP1B | P46821; D6F | MAP1B_HUMAI | Acetyl (Protein N | 0 | 0 | 0 | 0 | 0 | 0 | 0 | 0 | 0 | 0 | 0 | 847900 | 0 | 0 | 0 | 0 | 0 | 0 |
| ATVWDEAE | PSMC3; PSMC3 | P17980; E9P | PRS6A_HUMAI | Acetyl (Protein N | 0 | 0 | 0 | 171030 | 369960 | 0 | 181990 | 0 | 232370 | 180520 | 162410 | 0 | 0 | 0 | 0 | 0 | 1259200 | 614670 |
| ATYIHNCLI | VTA1; VTA1; VTA | Q9NP79; Q5 | VTA1_HUMAN | Unmodified | 0 | 0 | 119680 | 166720 | 671710 | 0 | 44115 | 46584 | 397730 | 267170 | 167000 | 1166100 | 0 | 0 | 102880 | 91567 | 74466 | 99192 |
| ATYLEFIQQ | SEC23B; SEC23 | Q15437; A0A | SC23B_HUMAI | Acetyl (Protein N | 305660 | 0 | 0 | 0 | 0 | 0 | 0 | 0 | 0 | 0 | 0 | 2453200 | 0 | 0 | 0 | 0 | 0 | 0 |
| ATYLSVADI | TTYH2; TTYH2 | Q9BSA4; B4I | TTYH2_HUMAI | Unmodified | 0 | 0 | 0 | 0 | 97132 | 0 | 0 | 0 | 7213600 | 1359900 | 0 | 2256200 | 367850 | 0 | 0 | 0 | 0 | 0 |
| AVAAGNSC | TLN2; TLN1; TLI | Q9Y4G6; Q9 | TLN2_HUMAN | Unmodified | 0 | 0 | 0 | 0 | 1025200 | 234880 | 240810 | 421420 | 0 | 0 | 0 | 0 | 1615300 | 1315800 | 0 | 0 | 558100 | 846650 |
| AVAAGNSC | TLN1 | Q9Y490 | TLN1_HUMAN | Unmodified | 362710 | 0 | 0 | 0 | 1243800 | 0 | 2871200 | 2025600 | 0 | 0 | 0 | 1680900 | 4682800 | 6680500 | 0 | 796060 | 4034900 | 4260000 |
| AVAALLTIPI | CAND1; CAND1 | Q86VP6; A0A | CAND1_HUMA | Unmodified | 0 | 0 | 754800 | 669830 | 0 | 0 | 0 | 0 | 3257200 | 476860 | 1415900 | 8912400 | 676360 | 0 | 1510900 | 0 | 0 | 0 |
| AVAASKERI | H1-5 | P16401 | H15_HUMAN H | Unmodified | 0 | 0 | 0 | 0 | 0 | 0 | 178780 | 4341000 | 0 | 150110 | 0 | 0 | 0 | 0 | 0 | 0 | 0 | 0 |
| AVAASKER: | H1-3; H1-4; H1- | P16402; P10 | H13_HUMAN H | Unmodified | 0 | 0 | 0 | 0 | 0 | 0 | 1544200 | 83905000 | 0 | 0 | 0 | 0 | 0 | 0 | 0 | 0 | 0 | 374070 |
| AVADAIRTS | CCT4 | P50991 | TCPD_HUMAN | Unmodified | 0 | 0 | 0 | 0 | 2635900 | 559310 | 2368800 | 1636900 | 718510 | 1944300 | 0 | 0 | 8128800 | 5153500 | 0 | 0 | 2032500 | 2470700 |
| AVADAIRTS | CCT4 | P50991 | TCPD_HUMAN | Unmodified | 0 | 0 | 0 | 0 | 245350 | 0 | 1897900 | 556440 | 0 | 126090 | 0 | 0 | 156580 | 329880 | 0 | 0 | 885790 | 0 |
| AVAEFLNR( | TFAP2A; TFAP2A | P05549; A0A | AP2A_HUMAN | Unmodified | 0 | 0 | 0 | 0 | 0 | 0 | 0 | 390180 | 0 | 0 | 0 | 0 | 0 | 0 | 0 | 0 | 0 | 0 |
| AVAEQIPLL | TLN1 | Q9Y490 | TLN1_HUMAN | Unmodified | 1928500 | 3155800 | 5309600 | 5472900 | 2216600 | 114320 | 368510 | 785280 | 3417900 | 406750 | 1986100 | 11300000 | 1653700 | 329350 | 5314300 | 2715900 | 196010 | 1047100 |
| AVAEQIPLL | TLN1 | Q9Y490 | TLN1_HUMAN | Unmodified | 0 | 0 | 0 | 1084300 | 18474000 | 40472 | 14196000 | 0 | 0 | 0 | 0 | 0 | 13866000 | 26079000 | 0 | 0 | 7019800 | 9476300 |
| AVAFFLESI | PRKDC | P78527 | PRKDC_HUMAI | Oxidation (M) | 0 | 0 | 0 | 0 | 0 | 0 | 177180 | 0 | 0 | 0 | 0 | 0 | 0 | 0 | 0 | 0 | 0 | 80988 |
| AVAFQNPC | UQCRC2; UQCI | P22695; H3E | QCR2_HUMAN | Unmodified | 0 | 104910 | 169870 | 252210 | 0 | 0 | 0 | 0 | 40100000 | 2129800 | 888000 | 0 | 0 | 0 | 0 | 0 | 0 | 0 |
| AVAGDASE | ATP1A1 | P05023 | AT1A1_HUMAI | Unmodified | 432060 | 0 | 8235000 | 8105900 | 11790000 | 1493500 | 2205300 | 2143300 | 13241000 | 7350800 | 11067000 | 26536000 | 10915000 | 7275700 | 5986700 | 5169300 | 3508400 | 3532100 |
| AVAGEYSD | FSD1L; FSD1L; | Q9BXM9; Q8 | FSD1L_HUMAI | Unmodified | 0 | 0 | 0 | 0 | 0 | 0 | 836900 | 365940 | 0 | 0 | 0 | 0 | 0 | 0 | 0 | 0 | 0 | 0 |
| AVAGLLAD. | PSMA3; PSMA3 | P25788; G3V | PSA3_HUMAN | Acetyl (Protein N | 0 | 0 | 0 | 0 | 0 | 0 | 0 | 0 | 372980 | 291370 | 0 | 0 | 1420800 | 0 | 0 | 0 | 0 | 0 |
| AVAGNISDI | VCL | P18206 | VINC_HUMAN | Unmodified | 1595300 | 1835800 | 1225700 | 1057100 | 15417000 | 0 | 9145100 | 5531900 | 1551000 | 2709300 | 1178200 | 12256000 | 4267800 | 5821800 | 0 | 1769900 | 36897000 | 8113800 |
| AVAGNISDI | VCL | P18206 | VINC_HUMAN | Unmodified | 0 | 0 | 0 | 0 | 1579100 | 0 | 2087300 | 0 | 0 | 32172 | 0 | 0 | 0 | 0 | 0 | 0 | 11250000 | 2384900 |
| AVAGQQQ/ | TNIP1; TNIP1; T | Q15025; A0A | TNIP1_HUMAN | Unmodified | 0 | 0 | 0 | 0 | 0 | 0 | 0 | 0 | 0 | 0 | 0 | 0 | 0 | 0 | 137800 | 0 | 0 | 0 |
| AVAGWEE\ | DHX30; DHX30 | Q7L2E3; H7E | DHX30_HUMAI | Unmodified | 0 | 0 | 0 | 0 | 0 | 0 | 0 | 0 | 0 | 0 | 0 | 548420 | 0 | 0 | 0 | 0 | 0 | 0 |
| AVAHHTDC | PSMC5; PSMC5 | P62195; J3Q | PRS8_HUMAN | Unmodified | 0 | 0 | 0 | 0 | 0 | 0 | 0 | 0 | 0 | 0 | 0 | 0 | 0 | 0 | 0 | 0 | 0 | 0 |
| AVAHHTDC | PSMC5; PSMC5 | P62195; J3Q | PRS8_HUMAN | Unmodified | 0 | 0 | 0 | 0 | 0 | 0 | 315990 | 106990 | 0 | 249290 | 0 | 0 | 0 | 0 | 0 | 0 | 503190 | 398580 |
| AVAHHTTA | PSMC4 | P43686 | PRS6B_HUMAI | Unmodified | 0 | 0 | 0 | 156200 | 249650 | 301490 | 0 | 51823 | 335540 | 107490 | 155030 | 3019300 | 582640 | 177490 | 0 | 140300 | 128130 | 199120 |
| AVAHHTTA | PSMC4 | P43686 | PRS6B_HUMAI | Unmodified | 0 | 0 | 0 | 0 | 786750 | 0 | 5702000 | 0 | 0 | 0 | 0 | 0 | 3655000 | 6617500 | 0 | 0 | 6861500 | 5331000 |
| AVAIDLPGI | ABHD14B; ABH | Q96IU4; B4D | ABHEB_HUMAI | Unmodified | 659560 | 477130 | 738710 | 255950 | 197520 | 0 | 0 | 0 | 398130 | 147330 | 358450 | 1939000 | 0 | 0 | 555370 | 462130 | 314810 | 0 |
| AVALSSSVI | GRN; GRN; GRI | P28799; K7E | GRN_HUMAN F | Unmodified | 0 | 0 | 0 | 1049300 | 0 | 0 | 0 | 117270 | 895010 | 0 | 0 | 0 | 0 | 0 | 571950 | 0 | 0 | 0 |
| AVANETGA | VCP; VCP; VCP; | P55072; C9J | TERA_HUMAN | Oxidation (M) | 0 | 660860 | 0 | 414740 | 2691700 | 0 | 0 | 0 | 707400 | 2380100 | 0 | 420040 | 4862900 | 2066600 | 340750 | 100070 | 0 | 28912 |

| | | | | | | | | | | | | | | | | | | | | | | |
|---|---|---|---|---|---|---|---|---|---|---|---|---|---|---|---|---|---|---|---|---|---|---|
| AVANETGA | VCP; VCP; VCP; | P55072; C9J | TERA_HUMAN | Unmodified | 0 | 0 | 0 | 0 | 245420 | 0 | 0 | 0 | 41391 | 143230 | 0 | 0 | 0 | 0 | 0 | 0 | 0 | 0 |
| AVANQTSA | PSMC1 | P62191 | PRS4_HUMAN | Unmodified | 213470 | 0 | 493770 | 440010 | 603560 | 0 | 111650 | 0 | 1481600 | 0 | 0 | 280240 | 481380 | 183580 | 326040 | 334990 | 212940 | 269100 |
| AVANQTSA | PSMC1 | P62191 | PRS4_HUMAN | Unmodified | 0 | 0 | 0 | 0 | 18676000 | 3042100 | 9928100 | 172640 | 0 | 2388600 | 0 | 0 | 7449400 | 5738900 | 0 | 0 | 11378000 | 9284800 |
| AVANQTSA | PSMC1 | P62191 | PRS4_HUMAN | Unmodified | 0 | 0 | 0 | 0 | 116780 | 0 | 376990 | 0 | 0 | 0 | 0 | 0 | 167550 | 0 | 0 | 0 | 379020 | 70356 |
| AVANTMRT | CCT5; CCT5; C( | P48643; B7Z | TCPE_HUMAN | Unmodified | 0 | 0 | 0 | 0 | 1638700 | 0 | 1119000 | 322300 | 0 | 301290 | 0 | 0 | 1501700 | 1655800 | 0 | 0 | 1629600 | 1975200 |
| AVANYDSV | HSD17B4; HSD | P51659; E7E | DHB4_HUMAN | Unmodified | 0 | 0 | 0 | 0 | 0 | 0 | 345640 | 148950 | 17362000 | 4015700 | 1418400 | 308010 | 0 | 271140 | 0 | 0 | 0 | 0 |
| AVANYDSV | HSD17B4; HSD | P51659; E7E | DHB4_HUMAN | Unmodified | 0 | 0 | 0 | 0 | 0 | 0 | 0 | 0 | 596850 | 0 | 0 | 0 | 240780 | 111040 | 0 | 0 | 0 | 0 |
| AVAQALEVI | CCT3; CCT3 | P49368; B4D | TCPG_HUMAN | Unmodified | 0 | 0 | 279770 | 424550 | 0 | 0 | 0 | 85221 | 3505900 | 698400 | 4726900 | 37698000 | 1141000 | 334040 | 348960 | 333560 | 0 | 0 |
| AVAQQPEG | PROM2 | Q8N271 | PROM2_HUMA | Unmodified | 1555100 | 1315600 | 1458200 | 0 | 0 | 0 | 0 | 0 | 1680200 | 0 | 0 | 1184700 | 0 | 0 | 1854400 | 1193100 | 55270 | 0 |
| AVASAAAAI | TLN1 | Q9Y490 | TLN1_HUMAN | Unmodified | 0 | 341170 | 276830 | 518830 | 1286900 | 0 | 1876300 | 582040 | 0 | 0 | 0 | 1645600 | 805670 | 691050 | 612230 | 513260 | 781520 | 362790 |
| AVASEIFK | MYO1C; MYO1( | O00159; F5H | MYO1C_HUMA | Unmodified | 6674700 | 954160 | 8906200 | 8580500 | 15514000 | 7699800 | 559700 | 271030 | 552200 | 854660 | 0 | 2163300 | 4426100 | 5585800 | 15794000 | 0 | 2776400 | 5126900 |
| AVASQLDC | PSMC6; PSMC6 | P62333; A0A | PRS10_HUMAN | Unmodified | 0 | 0 | 0 | 0 | 0 | 0 | 513600 | 422480 | 0 | 0 | 0 | 1995700 | 1644600 | 660110 | 0 | 0 | 1086600 | 315580 |
| AVASQPDS | PANK4; PANK4; | Q9NVE7; H0 | PANK4_HUMAN | Unmodified | 3072200 | 0 | 0 | 3263300 | 8845500 | 3557300 | 3827100 | 7004200 | 4132500 | 3294400 | 3303200 | 2192500 | 0 | 0 | 1720900 | 1543500 | 5719300 | 5863100 |
| AVASSPAA | CAMSAP3 | Q9P1Y5 | CAMP3_HUMA | Unmodified | 0 | 0 | 0 | 0 | 0 | 0 | 0 | 0 | 0 | 0 | 0 | 0 | 440040 | 188640 | 0 | 0 | 0 | 0 |
| AVATEANN | VPS4B | O75351 | VPS4B_HUMAN | Unmodified | 0 | 0 | 0 | 0 | 6582200 | 0 | 0 | 0 | 3743600 | 2993600 | 1998000 | 2210400 | 5012100 | 3224700 | 0 | 0 | 1394200 | 1207700 |
| AVATEANN | VPS4A; N/A | Q9UN37; I3L | VPS4A_HUMAN | Oxidation (M) | 1005100 | 0 | 0 | 0 | 3170400 | 474500 | 915590 | 504000 | 0 | 1180000 | 856530 | 2027100 | 4127400 | 3098400 | 0 | 0 | 0 | 0 |
| AVATEANN | VPS4A; N/A | Q9UN37; I3L | VPS4A_HUMAN | Unmodified | 0 | 0 | 0 | 0 | 1494000 | 0 | 0 | 0 | 1236600 | 0 | 0 | 393950 | 642160 | 0 | 0 | 0 | 192140 | 0 |
| AVATGKMD | CRMP1; DPYSL | Q14194; Q1 | DPYL1_HUMAN | Oxidation (M) | 0 | 0 | 0 | 0 | 1503400 | 0 | 2139000 | 3843100 | 0 | 497710 | 3111300 | 8365000 | 0 | 0 | 0 | 0 | 458200 | 0 |
| AVATGKMD | CRMP1; DPYSL | Q14194; Q1 | DPYL1_HUMAN | Unmodified | 0 | 24911 | 0 | 0 | 1067200 | 0 | 3240000 | 831670 | 596390 | 729110 | 4267400 | 11033000 | 0 | 0 | 0 | 186210 | 159370 | 0 |
| AVAVGRPS | SEPTIN11; SEP | Q9NVA2; D6 | SEP11_HUMAN | Acetyl (Protein N | 651150 | 0 | 814460 | 874230 | 1388400 | 0 | 0 | 0 | 1270400 | 0 | 1400600 | 9095100 | 0 | 0 | 532840 | 738870 | 695060 | 0 |
| AVAVGRPS | SEPTIN11; SEP | Q9NVA2; D6 | SEP11_HUMAN | Acetyl (Protein N | 0 | 0 | 0 | 0 | 89960000 | 7863000 | 15587000 | 130160 | 0 | 9866900 | 608620 | 0 | 10608000 | 8735900 | 0 | 0 | 76130000 | 27026000 |
| AVAVVVDP | PSMD14 | O00487 | PSDE_HUMAN | Unmodified | 0 | 0 | 707870 | 0 | 0 | 0 | 0 | 0 | 788890 | 0 | 974400 | 10113000 | 0 | 0 | 0 | 699330 | 0 | 0 |
| AVCEADQS | TRAFD1; TRAFD | O14545; F8V | TRAD1_HUMAN | Unmodified | 1128900 | 571010 | 0 | 0 | 0 | 0 | 0 | 0 | 0 | 0 | 0 | 0 | 0 | 0 | 0 | 0 | 0 | 0 |
| AVCGLGSF | MMAB; MMAB; | Q96EY8; F5H | MMAB_HUMAN | Acetyl (Protein N | 537940 | 566660 | 170870 | 635960 | 137490 | 0 | 0 | 0 | 0 | 92798 | 347370 | 271380 | 589070 | 148210 | 0 | 284310 | 0 | 0 |
| AVCMLSNT | TUBA3C; TUBA | P0DPH7; Q7 | TBA3C_HUMAN | Unmodified | 0 | 0 | 0 | 943770 | 1315000 | 0 | 816710 | 0 | 13920000 | 1031400 | 1376100 | 7750200 | 0 | 0 | 0 | 0 | 0 | 1211400 |
| AVCMLSNT | TUBA3C; TUBA | P0DPH7; Q7 | TBA3C_HUMAN | Oxidation (M) | 2418300 | 0 | 29520 | 1289300 | 938080 | 0 | 0 | 0 | 4252000 | 1497700 | 3132000 | 24756000 | 1752500 | 0 | 1246300 | 2001000 | 345090 | 0 |
| AVCMLSNT | TUBA3C; TUBA | P0DPH7; Q7 | TBA3C_HUMAN | 2 Oxidation (M) | 0 | 0 | 0 | 0 | 29012000 | 1200200 | 51020000 | 75355 | 0 | 18831000 | 0 | 0 | 21852000 | 27957000 | 0 | 0 | 9334000 | 27690000 |
| AVCMLSNT | TUBA3C; TUBA | P0DPH7; Q7 | TBA3C_HUMAN | Oxidation (M) | 0 | 0 | 0 | 0 | 4796500 | 238100 | 4567400 | 0 | 0 | 3055200 | 0 | 0 | 7727400 | 2456900 | 0 | 0 | 1692300 | 4918500 |
| AVCMLSNT | TUBA1C; TUBA | Q9BQE3; A0 | TBA1C_HUMAN | Unmodified | 0 | 0 | 0 | 0 | 0 | 0 | 0 | 0 | 2238300 | 0 | 0 | 2394900 | 0 | 0 | 0 | 0 | 0 | 0 |
| AVCMLSNT | TUBA1C; TUBA | Q9BQE3; A0 | TBA1C_HUMAN | Oxidation (M) | 65968 | 0 | 0 | 0 | 0 | 0 | 55744 | 0 | 0 | 0 | 375320 | 0 | 271530 | 0 | 392830 | 59498 | 88818 | 0 |
| AVCMLSNT | TUBA1C; TUBA | Q9BQE3; F5I | TBA1C_HUMAN | 2 Oxidation (M) | 0 | 0 | 0 | 0 | 3397300 | 0 | 1477000 | 0 | 0 | 935700 | 0 | 0 | 0 | 0 | 0 | 0 | 0 | 0 |
| AVCSQEAM | APLP2 | Q06481 | APLP2_HUMAN | Unmodified | 0 | 0 | 0 | 0 | 0 | 0 | 0 | 0 | 926860 | 250930 | 0 | 0 | 0 | 0 | 0 | 0 | 0 | 0 |
| AVCVEAGM | PSMC3; PSMC3 | P17980; E9P | PRS6A_HUMAN | Unmodified | 0 | 0 | 0 | 0 | 0 | 0 | 0 | 0 | 659050 | 242840 | 0 | 1276500 | 154710 | 0 | 0 | 0 | 662420 | 0 |
| AVDATREA | LAMA5 | O15230 | LAMA5_HUMAN | Unmodified | 188970 | 27856 | 0 | 47183 | 79231 | 0 | 0 | 0 | 329760 | 84945 | 0 | 0 | 0 | 0 | 39719 | 60867 | 74311 | 28332 |
| AVDCLLDS | SEC62; SEC62; | Q99442; F8V | SEC62_HUMAN | Unmodified | 0 | 1334300 | 654160 | 1943400 | 0 | 392470 | 70903 | 0 | 986890 | 0 | 315180 | 1.66E+08 | 0 | 0 | 1726900 | 1421500 | 0 | 0 |
| AVDCLLDS | SEC62; SEC62; | Q99442; F8V | SEC62_HUMAN | Unmodified | 0 | 0 | 0 | 0 | 0 | 0 | 285940 | 104740 | 0 | 786050 | 0 | 0 | 0 | 0 | 0 | 0 | 0 | 0 |
| AVDDGVN | CCT8 | P50990 | TCPQ_HUMAN | Unmodified | 0 | 465390 | 0 | 0 | 1494100 | 0 | 0 | 0 | 156080 | 66766 | 797440 | 6618800 | 0 | 0 | 0 | 0 | 0 | 0 |
| AVDDLNLH | DYNC1H1; DYN | Q14204; A0A | DYHC1_HUMA | Unmodified | 0 | 0 | 272890 | 0 | 0 | 0 | 0 | 0 | 0 | 0 | 0 | 3254300 | 0 | 0 | 0 | 0 | 0 | 0 |
| AVDFAER | RPL8; RPL8; RP | P62917; E9P | RL8_HUMAN 60 | Unmodified | 2362900 | 1880000 | 1011300 | 724350 | 366460 | 0 | 0 | 0 | 1100200 | 72596 | 1207800 | 16457000 | 0 | 0 | 1413200 | 1135200 | 192900 | 1826400 |
| AVDGEWQ | IGSF3 | O75054 | IGSF3_HUMAN | Unmodified | 0 | 586360 | 0 | 0 | 0 | 0 | 0 | 0 | 0 | 0 | 0 | 778380 | 0 | 0 | 644000 | 107720 | 0 | 0 |
| AVDGEWQ | IGSF3 | O75054 | IGSF3_HUMAN | Unmodified | 957200 | 1076300 | 543290 | 441840 | 0 | 0 | 0 | 0 | 413610 | 0 | 0 | 2017900 | 0 | 0 | 1666800 | 1020000 | 157150 | 152610 |
| AVDHATNF | ACADVL; ACAD | P49748; J3K | ACADV_HUMAI | Unmodified | 0 | 0 | 0 | 0 | 0 | 0 | 0 | 0 | 0 | 0 | 0 | 0 | 0 | 0 | 0 | 0 | 0 | 0 |
| AVDIPHMD | RPL10A | P62906 | RL10A_HUMAN | Unmodified | 2499400 | 2564700 | 693680 | 0 | 1521600 | 0 | 643750 | 138560 | 852340 | 327120 | 3741900 | 11666000 | 679560 | 0 | 1381300 | 673340 | 0 | 734110 |
| AVDIPHMD | RPL10A | P62906 | RL10A_HUMAN | Oxidation (M) | 70913 | 855460 | 0 | 975630 | 232600 | 0 | 232590 | 252010 | 0 | 173770 | 763000 | 6157400 | 351400 | 145660 | 1159300 | 457300 | 0 | 0 |
| AVDIPHMD | RPL10A | P62906 | RL10A_HUMAN | Unmodified | 0 | 0 | 0 | 0 | 672080 | 0 | 303990 | 53490 | 0 | 0 | 1502600 | 4727400 | 584980 | 0 | 0 | 0 | 0 | 0 |
| AVDLGPPK | NIBAN2 | Q96TA1 | NIBA2_HUMAN | Unmodified | 3058600 | 4335000 | 3285000 | 6515500 | 87447000 | 13758000 | 1524800 | 0 | 10319000 | 8550300 | 0 | 2281900 | 19567000 | 14453000 | 5288400 | 3779600 | 27133000 | 26894000 |
| AVDNDPGI | AVIL; AVIL; AVIL | O75366; F8V | AVIL_HUMAN A | Unmodified | 0 | 0 | 0 | 0 | 0 | 0 | 0 | 0 | 0 | 0 | 0 | 28445000 | 305740 | 0 | 0 | 0 | 128810 | 0 |
| AVDNDPGI | AVIL; AVIL | O75366; F8V | AVIL_HUMAN A | Unmodified | 0 | 0 | 0 | 0 | 0 | 0 | 0 | 0 | 0 | 0 | 19514000 | 15635000 | 0 | 0 | 0 | 0 | 0 | 0 |
| AVDNQVYV | NIT2 | Q9NQR4 | NIT2_HUMAN ( | Unmodified | 0 | 0 | 0 | 0 | 512630 | 231880 | 0 | 0 | 872830 | 267550 | 824220 | 3386500 | 745910 | 405580 | 0 | 701580 | 0 | 359470 |
| AVDPDGD | SDF4; SDF4; SD | Q9BRK5; H0 | CAB45_HUMAN | Unmodified | 144980 | 58951 | 773310 | 576990 | 0 | 0 | 0 | 0 | 61491000 | 1541400 | 2441800 | 1578700 | 615490 | 164740 | 527640 | 651990 | 0 | 246790 |
| AVDPSSVA | NUP210 | Q8TEM1 | PO210_HUMAN | Unmodified | 0 | 0 | 0 | 0 | 0 | 0 | 0 | 0 | 5655700 | 1850000 | 0 | 261630 | 27257 | 0 | 0 | 0 | 0 | 0 |
| AVDPTSGQ | NUP210 | Q8TEM1 | PO210_HUMAN | Unmodified | 0 | 0 | 0 | 0 | 0 | 0 | 0 | 0 | 34699000 | 3587400 | 352360 | 0 | 0 | 0 | 0 | 0 | 0 | 0 |
| AVDSLVPIC | ATP5F1A; ATP5 | P25705; K7E | ATPA_HUMAN | Unmodified | 0 | 0 | 0 | 0 | 0 | 0 | 0 | 51413 | 20456000 | 1121500 | 775990 | 436850 | 0 | 0 | 0 | 0 | 0 | 0 |
| AVDSLVPIC | ATP5F1A; ATP5 | P25705; K7E | ATPA_HUMAN | Unmodified | 0 | 0 | 0 | 239130 | 352260 | 0 | 0 | 77935 | 5929300 | 1608000 | 0 | 0 | 206780 | 90928 | 0 | 0 | 0 | 0 |
| AVDSQILPK | RPL6 | Q02878 | RL6_HUMAN 60 | Unmodified | 3363100 | 4924100 | 3698100 | 1505400 | 1081800 | 144980 | 0 | 178540 | 821730 | 202630 | 3962000 | 32127000 | 923000 | 214980 | 3522800 | 2639400 | 416900 | 590210 |
| AVDTWSW | LGALS3BP | Q08380 | LG3BP_HUMAN | Unmodified | 0 | 0 | 0 | 0 | 227700 | 0 | 0 | 0 | 11151000 | 383610 | 2207200 | 9319000 | 0 | 0 | 0 | 0 | 0 | 0 |
| AVDTWSW | LGALS3BP | Q08380 | LG3BP_HUMAN | Unmodified | 0 | 0 | 0 | 0 | 0 | 2508200 | 2362400 | 0 | 32403000 | 14072000 | 31459000 | 0 | 6315000 | 7187800 | 406920 | 0 | 1504800 | 39716000 |
| AVDVFFPPI | CLTC; CLTC | Q00610; A0A | CLH1_HUMAN | Oxidation (M) | 0 | 0 | 0 | 0 | 0 | 0 | 0 | 0 | 0 | 0 | 0 | 671010 | 0 | 0 | 3682300 | 122960 | 0 | 0 |
| AVEALKEA( | DLG2; DLG1; D | Q15700; Q1 | DLG2_HUMAN | Unmodified | 0 | 0 | 0 | 0 | 0 | 0 | 0 | 0 | 0 | 0 | 0 | 0 | 0 | 0 | 0 | 0 | 845100 | 0 |
| AVEALKEA( | DLG1; DLG1; D | Q12959; H7 | DLG1_HUMAN | Unmodified | 0 | 0 | 0 | 0 | 348100 | 0 | 0 | 0 | 0 | 0 | 0 | 0 | 0 | 0 | 0 | 0 | 923050 | 0 |
| AVEATQVC | MBTPS2 | O43462 | MBTP2_HUMAN | Unmodified | 0 | 0 | 0 | 243480 | 0 | 0 | 0 | 0 | 561630 | 0 | 0 | 256170 | 0 | 0 | 0 | 0 | 0 | 0 |
| AVECVEST( | PHLDA3 | Q9Y5J5 | PHLA3_HUMAN | Unmodified | 249840 | 0 | 100610 | 0 | 0 | 0 | 0 | 0 | 0 | 0 | 0 | 0 | 0 | 93665 | 571360 | 659200 | 0 | 0 |
| AVEDEGLR | RASA3 | Q14644 | RASA3_HUMAN | Acetyl (Protein N | 0 | 0 | 0 | 0 | 176770 | 0 | 307070 | 156660 | 0 | 196220 | 0 | 0 | 0 | 0 | 0 | 0 | 916050 | 861670 |
| AVEDIHGLI | EIF3A | Q14152 | EIF3A_HUMAN | Unmodified | 0 | 0 | 0 | 0 | 0 | 0 | 0 | 0 | 0 | 0 | 0 | 0 | 0 | 0 | 0 | 0 | 0 | 0 |
| AVEEAERLS | USP8; USP8; US | P40818; H0Y | UBP8_HUMAN | Unmodified | 0 | 0 | 0 | 0 | 2204000 | 0 | 0 | 0 | 563370 | 317910 | 0 | 282670 | 108950 | 136530 | 132350 | 0 | 0 | 496910 |
| AVEEEDKM | PSMD14 | O00487 | PSDE_HUMAN | Unmodified | 0 | 0 | 0 | 0 | 3099100 | 0 | 686630 | 327300 | 0 | 202260 | 1278400 | 4786000 | 2680100 | 1791700 | 0 | 0 | 1044100 | 3381300 |
| AVEEEDKM | PSMD14 | O00487 | PSDE_HUMAN | Oxidation (M) | 191870 | 53696 | 175500 | 235550 | 1068800 | 343440 | 154460 | 366370 | 0 | 230290 | 0 | 2963700 | 1767200 | 1061600 | 123350 | 321190 | 317920 | 717760 |
| AVEEKIEWI | HSPA5; HSPA5; | P11021; A0A | BIP_HUMAN Er | Unmodified | 0 | 0 | 0 | 476790 | 1606900 | 0 | 6842900 | 168300 | 4.96E+08 | 1.09E+08 | 47176000 | 7556700 | 2266000 | 1629300 | 7297000 | 2086600 | 0 | 0 |
| AVEEKIEWI | HSPA5; HSPA5; | P11021; A0A | BIP_HUMAN Er | Unmodified | 0 | 0 | 0 | 0 | 0 | 0 | 5529800 | 0 | 1.84E+08 | 82861000 | 5469700 | 1290700 | 1430600 | 752460 | 0 | 0 | 159170 | 0 |
| AVEELQSIII | GLMN | Q92990 | GLMN_HUMAN | Acetyl (Protein N | 0 | 0 | 78088 | 0 | 539500 | 0 | 388380 | 0 | 190800 | 140360 | 199520 | 903380 | 609300 | 0 | 0 | 70626 | 313560 | 0 |
| AVEEMNGK | PABPC4; PABP( | Q13310; B1A | PABP4_HUMAN | Unmodified | 663880 | 566520 | 0 | 0 | 0 | 0 | 0 | 9433.4 | 161280 | 202550 | 0 | 0 | 0 | 0 | 0 | 114390 | 0 | 0 |
| AVEEMNGK | PABPC4; PABP( | Q13310; B1A | PABP4_HUMAN | Oxidation (M) | 0 | 0 | 0 | 0 | 0 | 0 | 0 | 0 | 0 | 0 | 0 | 0 | 0 | 0 | 0 | 0 | 0 | 0 |
| AVEEPLNAI | CD47; CD47 | Q08722; H7I | CD47_HUMAN | Unmodified | 2205700 | 0 | 0 | 1776400 | 0 | 0 | 542510 | 0 | 13440000 | 2011300 | 7312900 | 34482000 | 1201400 | 0 | 2573600 | 2126600 | 0 | 0 |
| AVEEPLNAI | CD47; CD47 | Q08722; H7I | CD47_HUMAN | Unmodified | 0 | 0 | 0 | 184020 | 0 | 0 | 0 | 185210 | 0 | 1844200 | 124620 | 0 | 1329900 | 511220 | 0 | 0 | 0 | 410010 |
| AVEGDFFR | FKRP | Q9H9S5 | FKRP_HUMAN | Unmodified | 0 | 0 | 0 | 0 | 0 | 0 | 0 | 341370 | 600100 | 0 | 0 | 0 | 0 | 0 | 0 | 0 | 0 | 0 |
| AVEHINKTI | ENO1; ENO1; E | P06733; K7E | ENOA_HUMAN | Unmodified | 0 | 0 | 0 | 0 | 2100800 | 0 | 2298700 | 2619700 | 0 | 0 | 0 | 0 | 368420 | 831690 | 0 | 0 | 20911000 | 1609400 |
| AVEHINKTI | ENO1; ENO1; E | P06733; K7E | ENOA_HUMAN | Unmodified | 0 | 0 | 0 | 0 | 347630 | 0 | 0 | 0 | 0 | 0 | 0 | 0 | 117320 | 0 | 0 | 0 | 2859400 | 0 |
| AVEILAQEN | SCP2; SCP2; S( | P22307; H0Y | SCP2_HUMAN | Unmodified | 0 | 0 | 0 | 0 | 0 | 0 | 0 | 0 | 984410 | 170190 | 0 | 0 | 0 | 0 | 0 | 0 | 0 | 0 |
| AVEILGNTE | TMEM265 | A0A087WTH | TM265_HUMAN | Unmodified | 379530 | 153030 | 0 | 0 | 0 | 0 | 0 | 0 | 0 | 0 | 0 | 0 | 0 | 0 | 0 | 0 | 0 | 0 |
| AVEIYIQGK | CACNA2D1 | P54289 | CA2D1_HUMAI | Unmodified | 1303000 | 1645600 | 0 | 113860 | 154460 | 0 | 5868300 | 1796400 | 4106100 | 1662100 | 0 | 212580 | 124440 | 0 | 0 | 0 | 0 | 0 |
| AVEKFGEEI | TXNRD1; TXNRI | Q16881; A0A | TRXR1_HUMAN | Unmodified | 0 | 0 | 0 | 0 | 12760000 | 0 | 0 | 0 | 0 | 0 | 0 | 0 | 0 | 0 | 0 | 0 | 1741600 | 0 |
| AVELAANTI | TKT; TKT | P29401; A0A | TKT_HUMAN Tr | Unmodified | 471730 | 1480300 | 0 | 0 | 23974000 | 10944000 | 4744400 | 2406600 | 634990 | 4135100 | 776230 | 18545000 | 249960 | 694850 | 1411100 | 347710 | 7257700 | 5954100 |
| AVELERQIE | RNF114; RNF11 | Q9Y508; A0A | RN114_HUMAN | Unmodified | 0 | 0 | 0 | 0 | 0 | 0 | 31548 | 0 | 0 | 0 | 0 | 0 | 2017900 | 1212500 | 0 | 0 | 171910 | 0 |
| AVELLGDIV | UQCRC1 | P31930 | QCR1_HUMAN | Unmodified | 0 | 0 | 0 | 0 | 0 | 0 | 0 | 0 | 35786000 | 4126600 | 69413 | 0 | 0 | 0 | 0 | 0 | 0 | 1585400 |
| AVELMDRT | CKAP5 | Q14008 | CKAP5_HUMAN | Unmodified | 152360 | 0 | 0 | 0 | 159440 | 0 | 0 | 0 | 335800 | 0 | 0 | 0 | 112160 | 61015 | 0 | 0 | 0 | 78995 |
| AVEMAAQF | PGP | A6NDG6 | PGP_HUMAN G | Unmodified | 116000 | 72587 | 175500 | 0 | 0 | 0 | 0 | 0 | 111190 | 0 | 0 | 1920400 | 0 | 0 | 143740 | 0 | 0 | 0 |
| AVENSSTAI | PSMA3 | P25788 | PSA3_HUMAN | Unmodified | 249190 | 195850 | 0 | 747230 | 2598700 | 771590 | 331820 | 0 | 8756200 | 2467000 | 2205600 | 15359000 | 1149800 | 709460 | 788490 | 939850 | 0 | 1095100 |
| AVENSSTAI | PSMA3 | P25788 | PSA3_HUMAN | Unmodified | 0 | 0 | 0 | 0 | 0 | 393000 | 0 | 0 | 0 | 623530 | 0 | 0 | 0 | 0 | 0 | 0 | 74687 | 0 |
| AVENSSTAI | PSMA3 | P25788 | PSA3_HUMAN | Unmodified | 0 | 0 | 0 | 0 | 144910 | 0 | 124520 | 84355 | 0 | 90295 | 0 | 0 | 0 | 119530 | 0 | 0 | 1036600 | 0 |
| AVEPGLVQ | CHPF2; CHPF2 | Q9P2E5; G5I | CHPF2_HUMAI | Unmodified | 0 | 0 | 0 | 0 | 0 | 0 | 135820 | 21959 | 617710 | 0 | 0 | 0 | 0 | 0 | 0 | 0 | 0 | 0 |
| AVEPLRTFS | NAGPA; NAGPA | Q9UK23; K7I | NAGPA_HUMA | Unmodified | 0 | 0 | 0 | 0 | 414250 | 0 | 0 | 0 | 1376400 | 769540 | 293000 | 0 | 0 | 0 | 0 | 0 | 0 | 0 |
| AVEQLKME | GNG4 | P50150 | GBG4_HUMAN | Oxidation (M) | 0 | 0 | 0 | 0 | 0 | 0 | 0 | 0 | 0 | 467910 | 0 | 0 | 0 | 0 | 0 | 0 | 0 | 0 |
| AVEQLKME | GNG4 | P50150 | GBG4_HUMAN | Unmodified | 0 | 0 | 0 | 0 | 1263800 | 0 | 2055000 | 420090 | 0 | 2210400 | 114230 | 0 | 0 | 0 | 0 | 0 | 0 | 0 |
| AVERGQLE | HP1BP3; HP1B | Q5SSJ5; B0( | HP1B3_HUMAN | Unmodified | 0 | 0 | 0 | 0 | 0 | 0 | 0 | 249730 | 0 | 0 | 0 | 0 | 0 | 0 | 0 | 0 | 0 | 0 |
| AVESDCYA | SEMA4C | Q9C0C4 | SEM4C_HUMAI | Unmodified | 0 | 0 | 0 | 0 | 0 | 0 | 0 | 0 | 2257400 | 0 | 0 | 155770 | 0 | 0 | 0 | 0 | 0 | 0 |
| AVESLKPSI | ASAP2; ASAP2 | O43150; A0A | ASAP2_HUMAN | Unmodified | 0 | 0 | 0 | 0 | 0 | 0 | 0 | 0 | 0 | 0 | 0 | 1208600 | 0 | 0 | 0 | 0 | 0 | 0 |
| AVESLRAIS | PPP2R1A; PPP2 | P30153; C9J | 2AAA_HUMAN | Unmodified | 0 | 0 | 0 | 0 | 21292000 | 2835800 | 9539700 | 0 | 0 | 3036800 | 0 | 0 | 15770000 | 21537000 | 0 | 0 | 12817000 | 17187000 |
| AVESLRQIS | PPP2R1B | P30154 | 2AAB_HUMAN | Unmodified | 0 | 0 | 0 | 0 | 1427800 | 0 | 841380 | 0 | 0 | 0 | 0 | 0 | 5151300 | 2780200 | 0 | 0 | 380970 | 1021900 |
| AVETELEY( | DAAM1 | Q9Y4D1 | DAAM1_HUMA | Unmodified | 64476 | 0 | 157930 | 199540 | 798470 | 0 | 833770 | 266860 | 728260 | 456770 | 438250 | 893940 | 0 | 491900 | 386740 | 196690 | 168830 | 0 |
| AVETLLDLII | RAB27B | O00194 | RB27B_HUMAN | Oxidation (M) | 21025 | 30843 | 161160 | 691450 | 37652 | 0 | 648150 | 0 | 0 | 0 | 0 | 0 | 195140 | 0 | 905710 | 580880 | 0 | 0 |
| AVETPPLSS | GAPVD1; GAPV | Q14C86; C9 | GAPD1_HUMAI | Unmodified | 0 | 0 | 0 | 0 | 0 | 0 | 0 | 0 | 0 | 0 | 0 | 501630 | 0 | 0 | 0 | 0 | 0 | 0 |
| AVETTAQSI | ATP6V1A | P38606 | VATA_HUMAN | Unmodified | 0 | 0 | 0 | 0 | 0 | 0 | 0 | 0 | 0 | 0 | 0 | 2250200 | 0 | 0 | 0 | 0 | 517730 | 760360 |
| AVEVATVVI | PTGFRN | Q9P2B2 | FPRP_HUMAN | Unmodified | 0 | 0 | 3853300 | 11519000 | 2814100 | 1265900 | 0 | 0 | 0 | 0 | 0 | 0 | 19756000 | 5554800 | 8871800 | 6438700 | 0 | 0 |
| AVEVATVVI | PTGFRN | Q9P2B2 | FPRP_HUMAN | Unmodified | 0 | 0 | 0 | 0 | 10634000 | 1750100 | 0 | 0 | 0 | 0 | 0 | 0 | 17160000 | 7835900 | 0 | 0 | 0 | 0 |
| AVEVAWET | OPA1; OPA1; OI | O60313; E5K | OPA1_HUMAN | Unmodified | 0 | 0 | 0 | 0 | 0 | 0 | 0 | 0 | 1616500 | 0 | 867430 | 4101100 | 51451 | 0 | 20568 | 0 | 0 | 0 |
| AVEVPAFAS | AVIL | O75366 | AVIL_HUMAN A | Unmodified | 0 | 0 | 0 | 0 | 212990 | 0 | 0 | 0 | 0 | 0 | 1246700 | 1687700 | 0 | 0 | 0 | 0 | 7202700 | 16308 |
| AVEVQGPS | AHNAK | Q09666 | AHNK_HUMAN | Unmodified | 0 | 0 | 0 | 288750 | 3127200 | 531480 | 0 | 0 | 0 | 0 | 0 | 244280 | 116060 | 61208 | 0 | 0 | 351550 | 579060 |
| AVEYFQDN | EXOC7; EXOC7 | Q9UPT5; B5I | EXOC7_HUMAI | Unmodified | 98408 | 60116 | 0 | 0 | 1206100 | 295330 | 0 | 0 | 1005300 | 451340 | 0 | 1449500 | 322650 | 0 | 350500 | 270710 | 327210 | 279530 |
| AVEYFQDN | EXOC7; EXOC7 | Q9UPT5; C9 | EXOC7_HUMAI | Unmodified | 0 | 0 | 0 | 0 | 0 | 0 | 0 | 0 | 0 | 0 | 0 | 0 | 0 | 0 | 0 | 0 | 649780 | 0 |

| | | | | | | | | | | | | | | | | | | | | | | |
|---|---|---|---|---|---|---|---|---|---|---|---|---|---|---|---|---|---|---|---|---|---|---|
| AVFDETYPI | AARS1; AARS1; | P49588; A0A | SYAC_HUMAN | Unmodified | 110940 | 0 | 0 | 0 | 0 | 0 | 0 | 0 | 0 | 0 | 0 | 4204700 | 0 | 0 | 273240 | 421000 | 0 | 0 |
| AVFDNLIQI | NRBP1; NRBP1 | Q9UHY1; F8 | NRBP_HUMAN | Unmodified | 0 | 74781 | 0 | 209200 | 230620 | 0 | 279020 | 0 | 238330 | 138540 | 0 | 1200200 | 74155 | 124470 | 243000 | 125810 | 0 | 0 |
| AVFLLTR | TRPM5; TRPM5; | Q9NZQ8; A0 | TRPM5_HUMAI | Unmodified | 4193200 | 0 | 2442900 | 1706000 | 0 | 0 | 0 | 0 | 0 | 0 | 0 | 658330 | 0 | 0 | 2383300 | 0 | 0 | 0 |
| AVFPLLYVF | EEF1AKNMT | Q8N6R0 | EFNMT_HUMAI | Unmodified | 0 | 0 | 0 | 0 | 0 | 0 | 0 | 0 | 0 | 0 | 0 | 913630 | 0 | 0 | 0 | 0 | 0 | 0 |
| AVFPSIVGF | POTEE; POTEI; I | Q6S8J3; P0C | POTEE_HUMAN | Unmodified | 0 | 0 | 1971100 | 1345200 | 2073700 | 290900 | 0 | 0 | 5150300 | 863020 | 885480 | 15125000 | 0 | 0 | 2475100 | 7264200 | 872590 | 0 |
| AVFPSIVGF | POTEE; POTEI; I | Q6S8J3; P0C | POTEE_HUMAN | Unmodified | 0 | 0 | 230230 | 410730 | 0 | 0 | 0 | 0 | 1077400 | 0 | 0 | 682060 | 223750 | 0 | 310060 | 97300 | 0 | 0 |
| AVFQANQE | ATP1A1 | P05023 | AT1A1_HUMAN | Unmodified | 4061300 | 2495600 | 41678000 | 31344000 | 25370000 | 4329900 | 844860 | 416610 | 26194000 | 9001900 | 49369000 | 2.16E+08 | 38471000 | 15115000 | 64231000 | 44895000 | 2658700 | 4062600 |
| AVFQANQE | ATP1A1 | P05023 | AT1A1_HUMAN | Unmodified | 837760 | 0 | 11440000 | 18339000 | 3096600 | 1069000 | 0 | 1096000 | 12876000 | 4798600 | 14323000 | 47121000 | 9308600 | 3688200 | 7073400 | 5822800 | 5123500 | 938970 |
| AVFQANQE | ATP1A1 | P05023 | AT1A1_HUMAN | Unmodified | 0 | 0 | 0 | 0 | 731920 | 0 | 0 | 0 | 0 | 0 | 0 | 0 | 3282700 | 2671400 | 0 | 0 | 463030 | 0 |
| AVFQDQFS | MARVELD2; MA | Q8N4S9; A1 | MALD2_HUMAI | Unmodified | 576500 | 437690 | 0 | 737360 | 0 | 0 | 0 | 0 | 349880 | 0 | 271800 | 892510 | 0 | 160800 | 873890 | 695420 | 0 | 0 |
| AVFVDLEPT | TUBA4A; TUBA4 | P68366; C9J | TBA4A_HUMAN | Unmodified | 199480 | 981380 | 455360 | 889620 | 0 | 79356 | 189930 | 65131 | 57157 | 248390 | 393060 | 3183400 | 487420 | 168810 | 1359400 | 639630 | 0 | 0 |
| AVFVDLEPT | TUBA4A; TUBA4 | P68366; C9J | TBA4A_HUMAN | Unmodified | 2630000 | 2713500 | 2207200 | 6083500 | 0 | 0 | 0 | 0 | 0 | 1207900 | 275570 | 1686500 | 668180 | 0 | 4125900 | 2280200 | 0 | 0 |
| AVFVDLEPT | TUBA1A; TUBA1 | Q71U36; Q9 | TBA1A_HUMAN | Unmodified | 9565500 | 10889000 | 3913000 | 11272000 | 10221000 | 2520400 | 3175900 | 2203200 | 67944000 | 9589900 | 28176000 | 2.12E+08 | 6521300 | 0 | 15645000 | 11329000 | 3195400 | 4019900 |
| AVFVDLEPT | TUBA1A; TUBA1 | Q71U36; Q9 | TBA1A_HUMAN | Unmodified | 42241 | 0 | 3068400 | 18171000 | 14017000 | 3825600 | 4364600 | 0 | 76415000 | 13587000 | 13584000 | 13237000 | 23139 | 2076200 | 0 | 3691700 | 3507200 | 5686300 |
| AVFVSEGK | CORO1A; COR( | P31146; H3E | COR1A_HUMAI | Unmodified | 0 | 53345 | 0 | 0 | 123600 | 0 | 14587 | 15296 | 844940 | 124630 | 2030200 | 12932000 | 0 | 0 | 0 | 0 | 10838 | 0 |
| AVGDGIVL( | LCP1; PLS3; PL | P13796; P13 | PLSL_HUMAN I | Unmodified | 434690 | 413490 | 0 | 0 | 8006300 | 2020200 | 479480 | 0 | 378210 | 306990 | 0 | 0 | 0 | 1672500 | 0 | 0 | 7901100 | 3332900 |
| AVGEEVWH | NCAM1; NCAM | P13591; A0A | NCAM1_HUMA | Unmodified | 0 | 0 | 0 | 0 | 0 | 0 | 0 | 0 | 291620 | 0 | 0 | 0 | 0 | 0 | 0 | 0 | 0 | 0 |
| AVGEIVQD' | CPNE8; CPNE8 | Q86YQ8; E7I | CPNE8_HUMAI | Unmodified | 0 | 0 | 0 | 0 | 1262900 | 338520 | 0 | 70429 | 391760 | 0 | 317890 | 0 | 0 | 221800 | 0 | 0 | 0 | 123210 |
| AVGESVQK | GALE; GALE; G/ | Q14376; Q5( | GALE_HUMAN | Unmodified | 0 | 0 | 393700 | 346030 | 0 | 0 | 0 | 0 | 0 | 0 | 0 | 0 | 0 | 0 | 593630 | 419970 | 0 | 0 |
| AVGFSSGT | P3H1 | Q32P28 | P3H1_HUMAN | Unmodified | 0 | 0 | 0 | 20825 | 0 | 0 | 44414 | 18004 | 10185000 | 709580 | 107230 | 162140 | 0 | 0 | 0 | 0 | 112340 | 0 |
| AVGGDHA\ | SNX11; SNX11 | Q9Y5W9; B4 | SNX11_HUMAN | Unmodified | 230100 | 0 | 0 | 0 | 824940 | 121610 | 110810 | 0 | 513470 | 204200 | 117780 | 151660 | 1102400 | 586180 | 266070 | 0 | 1832200 | 2643300 |
| AVGHPFVI( | XPO1; XPO1; XF | O14980; A0/ | XPO1_HUMAN | Unmodified | 0 | 0 | 0 | 0 | 0 | 0 | 0 | 0 | 0 | 0 | 0 | 0 | 0 | 0 | 0 | 0 | 0 | 0 |
| AVGIVGIER | VPS36; VPS36 | Q86VN1; A0. | VPS36_HUMAN | Unmodified | 1698200 | 4894100 | 3532200 | 4397100 | 226060 | 0 | 75070 | 61213 | 175220 | 0 | 86151 | 1595000 | 1706800 | 100400 | 5744900 | 4747500 | 231470 | 249530 |
| AVGIVGIER | VPS36; VPS36 | Q86VN1; A0. | VPS36_HUMAN | Unmodified | 0 | 0 | 0 | 0 | 0 | 0 | 0 | 0 | 0 | 0 | 0 | 0 | 9039500 | 1396500 | 0 | 0 | 456630 | 190320 |
| AVGIWHC( | RPL37A; RPL37. | P61513; M0I | RL37A_HUMAN | Unmodified | 622140 | 0 | 0 | 0 | 0 | 0 | 0 | 0 | 0 | 0 | 0 | 9991600 | 0 | 0 | 232360 | 106520 | 0 | 0 |
| AVGIWHC( | RPL37A; RPL37. | P61513; M0I | RL37A_HUMAN | Oxidation (M) | 354330 | 0 | 0 | 0 | 0 | 0 | 0 | 34481 | 0 | 0 | 0 | 1760300 | 0 | 0 | 0 | 0 | 0 | 0 |
| AVGKDNFT | DPYSL2; DPYSL | Q16555; A0/ | DPYL2_HUMAN | Unmodified | 0 | 0 | 0 | 0 | 3458600 | 0 | 0 | 0 | 0 | 0 | 0 | 0 | 0 | 0 | 0 | 0 | 0 | 669870 |
| AVGKDNFT | DPYSL2; DPYSL | Q16555; A0/ | DPYL2_HUMAN | Oxidation (M) | 0 | 0 | 0 | 0 | 0 | 0 | 1717400 | 0 | 0 | 467880 | 0 | 0 | 0 | 0 | 0 | 0 | 540710 | 827940 |
| AVGKEELG | HYOU1; HYOU1 | Q9Y4L1; E9F | HYOU1_HUMA | Oxidation (M) | 0 | 0 | 0 | 0 | 0 | 0 | 679300 | 0 | 0 | 5335000 | 0 | 0 | 0 | 0 | 0 | 0 | 0 | 0 |
| AVGKEELG | HYOU1; HYOU1 | Q9Y4L1; E9F | HYOU1_HUMA | Unmodified | 0 | 0 | 0 | 0 | 0 | 0 | 0 | 0 | 0 | 5724700 | 0 | 389570 | 0 | 0 | 0 | 0 | 0 | 0 |
| AVGKVIPEL | GAPDH; GAPDI | P04406; E7E | G3P_HUMAN G | Unmodified | 0 | 0 | 0 | 0 | 9041800 | 1019500 | 2184200 | 1970600 | 0 | 640060 | 0 | 0 | 5310700 | 4505600 | 0 | 0 | 8093100 | 7538800 |
| AVGPEITKT | PPP2R1A; PPP2 | P30153; B3K | 2AAA_HUMAN | Unmodified | 0 | 0 | 0 | 252350 | 0 | 386260 | 0 | 0 | 2535600 | 521100 | 0 | 0 | 3710000 | 1118800 | 0 | 0 | 780810 | 2957800 |
| AVGPEITKT | PPP2R1A; PPP2 | P30153; B3K | 2AAA_HUMAN | Oxidation (M) | 0 | 0 | 0 | 0 | 2127500 | 0 | 0 | 0 | 0 | 0 | 0 | 2262200 | 1424300 | 0 | 0 | 0 | 0 | 1708400 |
| AVGPHQFL | BPGM | P07738 | PMGE_HUMAN | Unmodified | 0 | 0 | 0 | 0 | 0 | 0 | 0 | 0 | 0 | 0 | 381660 | 0 | 0 | 0 | 0 | 0 | 0 | 0 |
| AVGRLEAV | CAP1; CAP1; C/ | Q01518; Q5 | CAP1_HUMAN | Unmodified | 0 | 0 | 0 | 0 | 115970 | 0 | 0 | 24052 | 1376500 | 0 | 269910 | 3837100 | 0 | 0 | 0 | 0 | 0 | 0 |
| AVGRLEAV | CAP1; CAP1; C/ | Q01518; Q5 | CAP1_HUMAN | Unmodified | 0 | 0 | 0 | 0 | 1520100 | 1533300 | 273490 | 51618 | 2595000 | 1683100 | 0 | 467620 | 1597200 | 1886200 | 0 | 0 | 766640 | 3041000 |
| AVGRLEAV | CAP1; CAP1; C/ | Q01518; Q5 | CAP1_HUMAN | Oxidation (M) | 22731 | 0 | 19625 | 73486 | 580210 | 0 | 50188 | 0 | 129050 | 0 | 72724 | 0 | 370260 | 137990 | 0 | 22921 | 148410 | 312960 |
| AVGSIAYLF | LDLR; LDLR; LD | P01130; J3K | LDLR_HUMAN | Unmodified | 0 | 0 | 0 | 0 | 0 | 0 | 0 | 0 | 190870 | 0 | 0 | 0 | 0 | 0 | 0 | 0 | 0 | 0 |
| AVGSIAYLF | LDLR; LDLR; LD | P01130; J3K | LDLR_HUMAN | Unmodified | 0 | 0 | 0 | 0 | 0 | 0 | 0 | 0 | 2149500 | 0 | 0 | 0 | 0 | 0 | 0 | 0 | 0 | 0 |
| AVGSISSTA | CLUH; CLUH; C | O75153; I3L | CLU_HUMAN C | Unmodified | 0 | 0 | 0 | 0 | 0 | 0 | 0 | 0 | 0 | 0 | 0 | 3399400 | 447050 | 0 | 0 | 0 | 0 | 141120 |
| AVGSVTVY | NUP210 | Q8TEM1 | PO210_HUMAN | Unmodified | 0 | 0 | 0 | 0 | 0 | 0 | 0 | 0 | 2002200 | 0 | 0 | 0 | 0 | 0 | 0 | 0 | 0 | 0 |
| AVGTQTLS( | SNX1; SNX1 | Q13596; H0 | SNX1_HUMAN | Unmodified | 0 | 160810 | 0 | 0 | 0 | 0 | 0 | 0 | 487970 | 0 | 0 | 1070500 | 0 | 0 | 0 | 0 | 0 | 0 |
| AVGVAVAT( | ATP2A3 | Q93084 | AT2A3_HUMAN | Unmodified | 0 | 87553 | 0 | 0 | 0 | 0 | 0 | 72297 | 0 | 0 | 0 | 2595300 | 0 | 0 | 0 | 0 | 543510 | 0 |
| AVGVPALG | ACY1; ABHD14 | Q03154; A0/ | ACY1_HUMAN | Oxidation (M) | 0 | 0 | 0 | 0 | 0 | 0 | 283230 | 0 | 3722400 | 1763500 | 205920 | 4929800 | 0 | 0 | 0 | 0 | 0 | 1147900 |
| AVGWGAT( | HIP1R | O75146 | HIP1R_HUMAN | Unmodified | 0 | 0 | 0 | 0 | 0 | 0 | 0 | 0 | 0 | 0 | 48529 | 267410 | 0 | 0 | 0 | 0 | 0 | 0 |
| AVGWNELE | PCBD1 | P61457 | PHS_HUMAN P | Unmodified | 0 | 3019300 | 3003100 | 1713000 | 0 | 0 | 0 | 0 | 3205600 | 0 | 0 | 4367200 | 0 | 0 | 2030500 | 2459600 | 0 | 0 |
| AVGWNELE | PCBD1 | P61457 | PHS_HUMAN P | Unmodified | 0 | 0 | 485210 | 2592900 | 3250100 | 748790 | 87201 | 0 | 14276000 | 4514400 | 0 | 0 | 122720 | 3570000 | 1238300 | 0 | 2514200 | 2634600 |
| AVHAHVNL | NIPSNAP3A | Q9UFN0 | NPS3A_HUMAI | Unmodified | 0 | 0 | 0 | 0 | 0 | 0 | 0 | 0 | 2112200 | 187140 | 0 | 0 | 0 | 0 | 0 | 90944 | 0 | 0 |
| AVHANPPT | MOGS; MOGS; | Q13724; A0/ | MOGS_HUMAN | Oxidation (M) | 0 | 0 | 0 | 0 | 0 | 0 | 719840 | 0 | 6661300 | 815790 | 0 | 0 | 0 | 0 | 1271800 | 0 | 0 | 0 |
| AVHANPPT | MOGS; MOGS; | Q13724; A0/ | MOGS_HUMAN | Unmodified | 0 | 0 | 0 | 0 | 0 | 0 | 0 | 0 | 8693100 | 723360 | 0 | 0 | 0 | 0 | 0 | 0 | 0 | 0 |
| AVHLQGHE | ELP2 | Q6IA86 | ELP2_HUMAN I | Unmodified | 0 | 0 | 0 | 34879 | 61570 | 0 | 0 | 0 | 173010 | 0 | 0 | 2649000 | 0 | 0 | 0 | 0 | 39924 | 0 |
| AVHNSVAA | ZCCHC3 | Q9NUD5 | ZCHC3_HUMA | Unmodified | 658360 | 580370 | 0 | 0 | 0 | 0 | 230680 | 175650 | 0 | 242830 | 0 | 935480 | 876370 | 373320 | 0 | 0 | 0 | 0 |
| AVHSFLWS | AOC1; AOC1; A | P19801; C9J | AOC1_HUMAN | Unmodified | 0 | 0 | 0 | 0 | 0 | 0 | 0 | 0 | 873460 | 577730 | 0 | 0 | 0 | 0 | 0 | 0 | 0 | 0 |
| AVHSPIRS( | MICAL3 | Q7RTP6 | MICA3_HUMAN | Unmodified | 0 | 0 | 0 | 0 | 888900 | 0 | 0 | 0 | 0 | 0 | 0 | 0 | 0 | 0 | 0 | 0 | 0 | 0 |
| AVHVESVQ | RANGRF | Q9HD47 | MOG1_HUMAN | Unmodified | 0 | 0 | 0 | 0 | 1353900 | 336020 | 0 | 0 | 0 | 0 | 0 | 599900 | 1767800 | 0 | 0 | 0 | 1646200 | 11041000 |
| AVIDLNNR | U2AF1; U2AF1L | Q01081; P0[ | U2AF1_HUMAN | Unmodified | 2198500 | 1453000 | 894000 | 847430 | 0 | 0 | 0 | 0 | 506910 | 0 | 364490 | 2420200 | 0 | 0 | 456410 | 444060 | 0 | 0 |
| AVIDQCNH | ITGA2; ITGA2; I | P17301; E9P | ITA2_HUMAN I | Unmodified | 0 | 0 | 0 | 0 | 0 | 0 | 0 | 0 | 0 | 0 | 0 | 0 | 0 | 0 | 0 | 0 | 2307700 | 761300 |
| AVIEHNLLS | COPS4; COPS4 | Q9BT78; D6I | CSN4_HUMAN | Unmodified | 0 | 513220 | 87312 | 91533 | 0 | 0 | 0 | 0 | 238100 | 0 | 0 | 3550200 | 0 | 0 | 0 | 168810 | 0 | 0 |
| AVIETADR | PPID | Q08752 | PPID_HUMAN I | Unmodified | 0 | 0 | 34447 | 0 | 0 | 83726 | 0 | 64280 | 103530 | 0 | 0 | 1139300 | 0 | 0 | 0 | 0 | 513250 | 18222 |
| AVIFCLSAD | DSTN; DSTN | P60981; F6R | DEST_HUMAN | Unmodified | 1113400 | 496390 | 329640 | 791900 | 2591500 | 422470 | 1055800 | 157680 | 316620 | 402830 | 1017300 | 4995000 | 3646400 | 4485900 | 336260 | 721150 | 2099200 | 1734800 |
| AVIFCLSAD | DSTN; DSTN | P60981; F6R | DEST_HUMAN | Unmodified | 2067700 | 1335500 | 0 | 0 | 10209000 | 749340 | 2773100 | 28689 | 70636 | 1753100 | 1228600 | 12815000 | 14089000 | 8401600 | 0 | 2827700 | 8158100 | 3615900 |
| AVIGSTYMI | MVP | Q14764 | MVP_HUMAN I | Oxidation (M) | 3105100 | 5164600 | 194800 | 1782400 | 0 | 0 | 0 | 0 | 1506600 | 0 | 0 | 5315100 | 0 | 0 | 0 | 3313700 | 0 | 0 |
| AVIHYRDD | ARPC2 | O15144 | ARPC2_HUMAI | Oxidation (M) | 0 | 0 | 0 | 0 | 54717 | 0 | 0 | 0 | 0 | 0 | 0 | 349800 | 0 | 0 | 172580 | 50717 | 0 | 0 |
| AVIHYRDD | ARPC2 | O15144 | ARPC2_HUMAI | Unmodified | 870530 | 1934600 | 1260200 | 0 | 0 | 0 | 0 | 0 | 2058400 | 0 | 759930 | 17284000 | 0 | 0 | 2180000 | 1693000 | 0 | 0 |
| AVIHYRDD | ARPC2 | O15144 | ARPC2_HUMAI | Oxidation (M) | 327190 | 513690 | 1808900 | 0 | 0 | 0 | 0 | 0 | 0 | 0 | 329280 | 7194900 | 0 | 0 | 158040 | 701320 | 0 | 0 |
| AVILIGGPQ | GMPPA; GMPP/ | Q96IJ6; C9J | GMPPA_HUMA | Unmodified | 0 | 0 | 158000 | 0 | 833370 | 2152800 | 5056300 | 1031000 | 0 | 125330 | 0 | 1139500 | 0 | 0 | 189620 | 297560 | 1355500 | 170550 |
| AVIMGAPG | AK3 | Q9UIJ7 | KAD3_HUMAN | Oxidation (M) | 0 | 0 | 0 | 0 | 0 | 0 | 0 | 0 | 382930 | 72170 | 0 | 0 | 0 | 0 | 0 | 0 | 0 | 0 |
| AVIMGAPG | AK3 | Q9UIJ7 | KAD3_HUMAN | Unmodified | 0 | 0 | 0 | 0 | 271620 | 0 | 0 | 0 | 2369400 | 321520 | 0 | 94877 | 0 | 0 | 0 | 0 | 0 | 0 |
| AVINSAGYI | NIBAN2 | Q96TA1 | NIBA2_HUMAN | Unmodified | 1411600 | 1093900 | 3070300 | 2345800 | 36998 | 257490 | 89266 | 0 | 9716200 | 937060 | 1749700 | 2078100 | 0 | 0 | 2468100 | 2146600 | 626310 | 452540 |
| AVIQVSQIV | DCXR; DCXR; D | Q7Z4W1; J3I | DCXR_HUMAN | Unmodified | 0 | 0 | 0 | 0 | 0 | 0 | 0 | 0 | 773030 | 0 | 0 | 1233200 | 0 | 0 | 0 | 0 | 0 | 0 |
| AVIVLNEGI | CPD | O75976 | CBPD_HUMAN | Unmodified | 0 | 113100 | 0 | 0 | 0 | 0 | 86139 | 84025 | 11721000 | 2424700 | 0 | 313430 | 0 | 0 | 0 | 0 | 0 | 0 |
| AVIVLNEGI | CPD | O75976 | CBPD_HUMAN | Unmodified | 0 | 0 | 0 | 0 | 0 | 0 | 0 | 0 | 1006300 | 688780 | 0 | 0 | 0 | 0 | 0 | 0 | 0 | 0 |
| AVIVLVENF | S100A16 | Q96FQ6 | S10AG_HUMAI | Unmodified | 0 | 0 | 0 | 0 | 2206100 | 275460 | 0 | 0 | 0 | 282890 | 0 | 0 | 0 | 0 | 0 | 0 | 1853200 | 2466300 |
| AVIVLVENF | S100A16 | Q96FQ6 | S10AG_HUMAI | Unmodified | 0 | 0 | 0 | 0 | 1647000 | 0 | 0 | 0 | 0 | 244510 | 0 | 0 | 0 | 0 | 0 | 0 | 12733000 | 4492400 |
| AVIVLVENF | S100A16 | Q96FQ6 | S10AG_HUMAI | Unmodified | 0 | 0 | 0 | 0 | 0 | 0 | 33986 | 0 | 0 | 0 | 0 | 0 | 0 | 0 | 0 | 0 | 2628400 | 0 |
| AVIYNPAT( | MYH10; MYH10 | P35580; E7E | MYH10_HUMAI | Unmodified | 0 | 0 | 0 | 112090 | 0 | 0 | 0 | 0 | 0 | 0 | 0 | 1042000 | 132660 | 0 | 0 | 0 | 0 | 0 |
| AVKASTEAT | CIT; CIT; CIT; CI | O14578; A0/ | CTRO_HUMAN | Unmodified | 0 | 0 | 0 | 0 | 0 | 0 | 0 | 0 | 0 | 0 | 0 | 0 | 0 | 0 | 0 | 0 | 1150600 | 0 |
| AVKEEGFPI | LRPPRC; LRPPF | P42704; B8Z | LPPRC_HUMAN | Unmodified | 0 | 0 | 0 | 0 | 0 | 0 | 0 | 0 | 3739800 | 963670 | 193010 | 0 | 0 | 966570 | 0 | 0 | 0 | 0 |
| AVKREDLFI | AKR1B10 | O60218 | AK1BA_HUMAN | Unmodified | 0 | 0 | 0 | 0 | 311990 | 0 | 567060 | 899660 | 0 | 99767 | 0 | 0 | 0 | 0 | 0 | 0 | 0 | 0 |
| AVKSSEHIN | BSG; BSG; BSG | P35613; A0A | BASI_HUMAN E | Oxidation (M) | 0 | 0 | 0 | 0 | 75156 | 0 | 141520 | 881610 | 0 | 0 | 0 | 0 | 0 | 35379 | 0 | 0 | 2122400 | 4562100 |
| AVKSSEHIN | BSG; BSG; BSG | P35613; A0A | BASI_HUMAN E | Unmodified | 0 | 0 | 0 | 0 | 0 | 0 | 1460400 | 568270 | 0 | 0 | 0 | 0 | 0 | 171760 | 0 | 174990 | 4205700 | 1260800 |
| AVKYVECS. | CDC42 | P60953 | CDC42_HUMA | Unmodified | 0 | 0 | 0 | 0 | 12513000 | 23213 | 14148000 | 8533600 | 0 | 1059400 | 0 | 0 | 0 | 14093000 | 0 | 0 | 33706000 | 16848000 |
| AVKYVECS. | CDC42 | P60953 | CDC42_HUMA | Unmodified | 0 | 0 | 0 | 0 | 0 | 0 | 289470 | 811520 | 0 | 0 | 0 | 0 | 0 | 0 | 0 | 0 | 2081800 | 611520 |
| AVLAELPQ( | CPNE2 | Q96FN4 | CPNE2_HUMAI | Unmodified | 0 | 0 | 77742 | 748140 | 7120700 | 1224700 | 6124900 | 0 | 2768900 | 2426700 | 0 | 59290 | 2072700 | 1772600 | 344010 | 168940 | 665720 | 4416100 |
| AVLAESYEF | ACO1 | P21399 | ACOC_HUMAN | Unmodified | 0 | 0 | 474070 | 0 | 0 | 0 | 0 | 559710 | 402990 | 0 | 241660 | 3792400 | 0 | 0 | 0 | 253370 | 0 | 0 |
| AVLASTPR | MMS19; MMS1! | Q96T76; Q5 | MMS19_HUMA | Unmodified | 79912 | 0 | 0 | 0 | 0 | 376770 | 0 | 0 | 35503 | 784750 | 0 | 1030300 | 0 | 0 | 0 | 0 | 0 | 0 |
| AVLCPPPVI | RAC3; RAC1 | P60763; P63 | RAC3_HUMAN | Unmodified | 6550800 | 0 | 0 | 7537000 | 1453700 | 203150 | 0 | 0 | 824490 | 188240 | 850450 | 4400800 | 1110400 | 746480 | 6003200 | 4071900 | 0 | 1606800 |
| AVLDALLE( | MAP1B; MAP1A | P46821; P78 | MAP1B_HUMAI | Unmodified | 0 | 163150 | 2356700 | 0 | 0 | 0 | 906320 | 1211000 | 1156400 | 923340 | 4600300 | 1135800 | 0 | 2084000 | 1091300 | 90151 | 0 | 3656100 |
| AVLDNPHH | TOR2A | Q5JU69 | TOR2A_HUMAN | Unmodified | 0 | 0 | 0 | 0 | 0 | 0 | 0 | 0 | 1843300 | 0 | 0 | 0 | 0 | 0 | 0 | 0 | 0 | 0 |
| AVLENNLG | MAT2B; MAT2B | Q9NZL9; H7 | MAT2B_HUMAN | Unmodified | 0 | 0 | 0 | 0 | 0 | 0 | 0 | 0 | 0 | 0 | 155310 | 3092400 | 0 | 0 | 0 | 90289 | 0 | 0 |
| AVLFCLSDI | CFL2; CFL2 | Q9Y281; F8V | COF2_HUMAN | Unmodified | 0 | 0 | 257390 | 0 | 2896600 | 1510400 | 0 | 0 | 0 | 0 | 0 | 774700 | 0 | 0 | 0 | 0 | 0 | 0 |
| AVLFCLSDI | CFL2; CFL2 | Q9Y281; F8V | COF2_HUMAN | Unmodified | 0 | 757510 | 0 | 13217 | 2647700 | 0 | 0 | 0 | 0 | 0 | 208840 | 0 | 347620 | 308360 | 338970 | 72817 | 233720 | 0 |
| AVLFCLSDI | CFL2; CFL2 | Q9Y281; F8V | COF2_HUMAN | Unmodified | 0 | 0 | 0 | 325920 | 43586000 | 7430600 | 601420 | 0 | 0 | 0 | 0 | 0 | 2762300 | 1642900 | 0 | 0 | 3642100 | 4775600 |
| AVLFCLSEI | CFL1; CFL1; CF | P23528; G3\ | COF1_HUMAN | Unmodified | 6097200 | 4548900 | 2857100 | 4302600 | 42679000 | 7749500 | 3184400 | 2567800 | 6541900 | 2773800 | 3013700 | 33109000 | 19534000 | 9531500 | 5761200 | 4608200 | 12196000 | 14437000 |
| AVLFCLSEI | CFL1; CFL1; CF | P23528; G3\ | COF1_HUMAN | Unmodified | 11832000 | 5545200 | 7138100 | 10626000 | 1.29E+08 | 13175000 | 10073000 | 6441800 | 13132000 | 6834000 | 6286900 | 70092000 | 58750000 | 28858000 | 8968700 | 10154000 | 36364000 | 35412000 |
| AVLFCLSEI | CFL1; CFL1; CF | P23528; G3\ | COF1_HUMAN | Unmodified | 0 | 0 | 0 | 0 | 5361300 | 930350 | 754390 | 0 | 692330 | 0 | 0 | 0 | 2398300 | 2135100 | 0 | 0 | 1647000 | 1665300 |
| AVLGEVKL( | EEF1G | P26641 | EF1G_HUMAN | Unmodified | 0 | 0 | 0 | 0 | 846970 | 355490 | 0 | 151920 | 0 | 253290 | 0 | 64408 | 653530 | 218620 | 0 | 0 | 1070900 | 687160 |
| AVLGEVVL\ | VARS1; VARS1; | P26640; A0A | SYVC_HUMAN | Unmodified | 0 | 0 | 0 | 0 | 0 | 0 | 0 | 0 | 0 | 0 | 0 | 13704000 | 0 | 0 | 0 | 0 | 0 | 0 |
| AVLGTSNFI | TFRC; TFRC | P02786; G3\ | TFR1_HUMAN ~ | Unmodified | 106340 | 0 | 0 | 1784800 | 1147300 | 584790 | 779280 | 256690 | 4000000 | 1881200 | 0 | 477130 | 0 | 0 | 3131800 | 0 | 0 | 0 |
| AVLIAGQP( | RUVBL2; RUVBI | Q9Y230; M0I | RUVB2_HUMAI | Unmodified | 745480 | 452220 | 527960 | 940210 | 3636300 | 740520 | 409860 | 397940 | 584580 | 633820 | 2143200 | 20102000 | 3716000 | 823720 | 513020 | 944470 | 393570 | 1506200 |
| AVLIASTRS | MICAL3; MICAL | Q7RTP6; A0/ | MICA3_HUMAN | Unmodified | 0 | 0 | 0 | 0 | 1123600 | 167280 | 0 | 0 | 0 | 0 | 0 | 0 | 0 | 0 | 0 | 0 | 0 | 0 |
| AVLIPKDD( | GALNT13; GALI | Q8IUC8; H7I | GLT13_HUMAN | Unmodified | 0 | 0 | 0 | 0 | 0 | 0 | 0 | 0 | 406410 | 0 | 0 | 0 | 0 | 0 | 0 | 0 | 0 | 0 |
| AVLLAGPP( | RUVBL1; RUVBI | Q9Y265; E7E | RUVB1_HUMAI | Unmodified | 474240 | 430380 | 408040 | 962830 | 4601600 | 1508200 | 567040 | 328740 | 594650 | 557120 | 3346300 | 15877000 | 4394700 | 2105600 | 425080 | 576800 | 529750 | 2958400 |
| AVLLDEDD | STXBP2; STXBP | Q15833; M0I | STXB2_HUMAN | Oxidation (M) | 0 | 0 | 0 | 0 | 2544700 | 0 | 1511800 | 0 | 0 | 0 | 0 | 0 | 1492600 | 1709200 | 0 | 0 | 603500 | 1365300 |
| AVLLDEDD | STXBP2; STXBP | Q15833; M0I | STXB2_HUMAN | Unmodified | 0 | 0 | 0 | 0 | 0 | 0 | 0 | 0 | 0 | 0 | 0 | 0 | 0 | 0 | 0 | 0 | 0 | 1320100 |
| AVLLDLEPF | TUBG1; TUBG2 | P23258; Q9I | TBG1_HUMAN | Unmodified | 0 | 0 | 0 | 566670 | 0 | 0 | 0 | 0 | 355010 | 0 | 391130 | 2775100 | 0 | 0 | 0 | 0 | 0 | 0 |
| AVLLGPPG. | AK2; AK2; AK2; | P54819; G3\ | KAD2_HUMAN | Unmodified | 0 | 0 | 0 | 132650 | 498310 | 0 | 0 | 0 | 1444400 | 913850 | 808300 | 1017000 | 1476600 | 0 | 0 | 0 | 78718 | 0 |
| AVLLLQEK | LSS; LSS | P48449; A0A | LSS_HUMAN La | Unmodified | 0 | 0 | 0 | 0 | 0 | 0 | 0 | 0 | 759600 | 0 | 0 | 0 | 0 | 0 | 0 | 0 | 0 | 0 |
| AVLLTQDTI | THUMPD3; THU | Q9BV44; H7 | THUM3_HUMA | Unmodified | 0 | 0 | 0 | 0 | 397460 | 0 | 0 | 204690 | 0 | 0 | 0 | 371880 | 0 | 0 | 274880 | 189600 | 0 | 0 |

| | | | | | | | | | | | | | | | | | | | | | | |
|---|---|---|---|---|---|---|---|---|---|---|---|---|---|---|---|---|---|---|---|---|---|---|
| AVLMIRTPE | CTNNA1; CTNN | P35221; G3X | CTNA1_HUMAI | Oxidation (M) | 0 | 0 | 0 | 0 | 863180 | 620800 | 0 | 0 | 0 | 0 | 0 | 0 | 3043200 | 0 | 0 | 0 | 0 | 0 |
| AVLNNVIF( | RAD50; N/A; RA | Q92878; A0/ | RAD50_HUMAI | Unmodified | 0 | 0 | 0 | 0 | 0 | 0 | 1457100 | 0 | 0 | 0 | 0 | 0 | 0 | 0 | 0 | 0 | 0 | 907690 |
| AVLNPLCQ | SEC23B; SEC2: | Q15437; Q1! | SC23B_HUMAI | Unmodified | 0 | 0 | 0 | 0 | 0 | 0 | 0 | 0 | 0 | 0 | 0 | 2706700 | 0 | 0 | 0 | 0 | 289690 | 0 |
| AVLNPLCQ | SEC23B; SEC2: | Q15437; Q1! | SC23B_HUMAI | Unmodified | 0 | 0 | 0 | 0 | 128210 | 0 | 0 | 0 | 0 | 0 | 0 | 0 | 0 | 724380 | 0 | 0 | 1909600 | 1532300 |
| AVLNPTPIK | RHOG | P84095 | RHOG_HUMAN | Unmodified | 536210 | 425810 | 890050 | 626740 | 540980 | 0 | 0 | 0 | 0 | 0 | 943010 | 4785800 | 233100 | 0 | 735790 | 791700 | 0 | 0 |
| AVLPLLDA( | TWF2; TWF2; T\ | Q6IBS0; D6F | TWF2_HUMAN | Unmodified | 40180 | 30272 | 0 | 0 | 0 | 0 | 0 | 0 | 112450 | 0 | 63544 | 281660 | 0 | 0 | 46683 | 0 | 0 | 0 |
| AVLPVAESI | EFL1 | Q7Z2Z2 | EFL1_HUMAN I | Unmodified | 0 | 0 | 0 | 0 | 203980 | 0 | 29444 | 0 | 0 | 42187 | 0 | 1171900 | 0 | 0 | 0 | 0 | 0 | 0 |
| AVLQAARM | HYOU1; HYOU1 | Q9Y4L1; J3Q | HYOU1_HUMA | Unmodified | 0 | 0 | 0 | 0 | 0 | 0 | 74558 | 0 | 0 | 525230 | 0 | 0 | 0 | 0 | 0 | 0 | 0 | 0 |
| AVLQEFGR | DHRS7; DHRS7 | Q9Y394; H0' | DHRS7_HUMAI | Unmodified | 494640 | 541770 | 295840 | 0 | 0 | 0 | 0 | 0 | 380690 | 17131 | 1324800 | 553830 | 0 | 0 | 452110 | 343370 | 0 | 0 |
| AVLQWTK | ERO1A; ERO1A; | Q96HE7; G3 | ERO1A_HUMAI | Unmodified | 0 | 0 | 0 | 0 | 270810 | 67249 | 111780 | 78315 | 9082600 | 886200 | 0 | 408460 | 146170 | 313370 | 0 | 0 | 88338 | 325640 |
| AVLRKAGG | APEH; APEH; AI | P13798; C9J | ACPH_HUMAN | Unmodified | 0 | 0 | 0 | 0 | 0 | 0 | 0 | 0 | 0 | 0 | 0 | 0 | 182710 | 346220 | 0 | 0 | 547520 | 328790 |
| AVLRYGLD: | SERPINF1; SER | P36955; I3L1 | PEDF_HUMAN | Unmodified | 0 | 0 | 0 | 0 | 0 | 0 | 83113 | 37249 | 0 | 341160 | 842490 | 0 | 0 | 0 | 0 | 0 | 0 | 0 |
| AVLSAEQLI | SERPINH1; SER | P50454; E9P | SERPH_HUMAI | Unmodified | 0 | 0 | 0 | 0 | 1574300 | 75452 | 0 | 0 | 2.2E+08 | 23708000 | 348310 | 0 | 184540 | 140320 | 0 | 0 | 0 | 0 |
| AVLSDGSE | OSMR | Q99650 | OSMR_HUMAN | Unmodified | 0 | 0 | 0 | 0 | 565670 | 120340 | 0 | 0 | 0 | 77441 | 0 | 0 | 0 | 0 | 0 | 0 | 0 | 0 |
| AVLTCSEQI | F11R; F11R | Q9Y624; A0/ | JAM1_HUMAN . | Unmodified | 5664100 | 2685400 | 3551000 | 4531400 | 1001500 | 0 | 0 | 0 | 5080800 | 0 | 760190 | 4447600 | 3204300 | 0 | 5963100 | 3369700 | 0 | 852340 |
| AVLTCSEQI | F11R; F11R | Q9Y624; A0/ | JAM1_HUMAN . | Oxidation (M) | 0 | 0 | 0 | 0 | 924190 | 0 | 0 | 0 | 0 | 0 | 0 | 0 | 1140500 | 0 | 0 | 0 | 708120 | 0 |
| AVLVDLEP( | TUBB4A; TUBB | P04350; Q3Z | TBB4A_HUMAN | Oxidation (M) | 2955900 | 929060 | 3332400 | 1806100 | 0 | 0 | 339060 | 347840 | 9107400 | 1009000 | 5187900 | 36678000 | 715210 | 224560 | 3864800 | 3584800 | 0 | 0 |
| AVLVDLEP( | TUBB4A; TUBB | P04350; Q3Z | TBB4A_HUMAN | Unmodified | 58012 | 92672 | 0 | 111800 | 210270 | 0 | 720900 | 538890 | 9640200 | 0 | 1800500 | 5043600 | 569990 | 59380 | 168330 | 85555 | 0 | 110640 |
| AVLWVSAD | NUMB; NUMB; | P49757; G3\ | NUMB_HUMAN | Unmodified | 0 | 0 | 0 | 280410 | 8653000 | 849240 | 0 | 0 | 0 | 641160 | 157390 | 0 | 1447200 | 0 | 0 | 0 | 498280 | 3067400 |
| AVLWVSAD | NUMB; NUMB; | P49757; G3\ | NUMB_HUMAN | Unmodified | 0 | 0 | 0 | 0 | 2854500 | 0 | 125250 | 18347 | 310920 | 0 | 0 | 0 | 557870 | 552650 | 0 | 0 | 606300 | 4474300 |
| AVMDHVSI | CTNNA1; CTNN | P35221; E5R | CTNA1_HUMAI | Oxidation (M) | 0 | 101530 | 0 | 21734 | 675100 | 0 | 0 | 0 | 0 | 0 | 0 | 0 | 0 | 17342 | 529660 | 0 | 0 | 0 |
| AVMDHVSI | CTNNA1; CTNN | P35221; E5R | CTNA1_HUMAI | Oxidation (M) | 0 | 0 | 0 | 0 | 3239300 | 510950 | 0 | 0 | 247090 | 220250 | 0 | 0 | 0 | 0 | 571880 | 0 | 0 | 0 |
| AVMEQANI | HSPA4L; HSPA | O95757; E9I | HS74L_HUMAN | Oxidation (M) | 0 | 0 | 0 | 0 | 0 | 0 | 0 | 0 | 0 | 0 | 738200 | 2182500 | 3189800 | 985200 | 0 | 0 | 0 | 0 |
| AVMGIFSL\ | NADSYN1; NAD | Q6IA69; E9P | NADE_HUMAN | Oxidation (M) | 0 | 0 | 0 | 0 | 0 | 0 | 0 | 0 | 0 | 0 | 0 | 716420 | 0 | 0 | 0 | 0 | 0 | 0 |
| AVMLPGAR | XRN2 | Q9H0D6 | XRN2_HUMAN | Unmodified | 0 | 0 | 0 | 0 | 0 | 0 | 0 | 0 | 0 | 0 | 0 | 508820 | 0 | 0 | 0 | 0 | 0 | 0 |
| AVMNFVVF | PLOD3; PLOD3 | O60568; H7( | PLOD3_HUMAI | Unmodified | 99512 | 0 | 0 | 0 | 0 | 0 | 0 | 0 | 6505000 | 0 | 0 | 0 | 0 | 0 | 0 | 0 | 0 | 0 |
| AVNAMILA( | BAIAP2L1 | Q9UHR4 | BI2L1_HUMAN | Unmodified | 0 | 0 | 0 | 333250 | 7105200 | 1402300 | 0 | 0 | 744890 | 327490 | 0 | 330380 | 2421800 | 2185700 | 393570 | 130950 | 472210 | 385450 |
| AVNAMILA( | BAIAP2L1 | Q9UHR4 | BI2L1_HUMAN | Oxidation (M) | 0 | 0 | 0 | 0 | 2621400 | 491200 | 0 | 45495 | 0 | 0 | 0 | 0 | 4266800 | 3637500 | 366330 | 223730 | 0 | 0 |
| AVNEKSCN | ENO1; ENO1 | P06733; A0A | ENOA_HUMAN | Unmodified | 0 | 0 | 0 | 0 | 0 | 0 | 0 | 0 | 0 | 0 | 0 | 0 | 0 | 0 | 0 | 0 | 1080100 | 0 |
| AVNPFLAD' | CKAP5 | Q14008 | CKAP5_HUMAI | Unmodified | 0 | 0 | 0 | 0 | 752410 | 0 | 691550 | 0 | 0 | 664650 | 0 | 0 | 1865200 | 631150 | 0 | 0 | 339150 | 822830 |
| AVNSKDDL | PAN3; PAN3 | Q58A45; A0/ | PAN3_HUMAN | Unmodified | 264860 | 313720 | 0 | 117210 | 0 | 0 | 0 | 0 | 0 | 0 | 0 | 0 | 41751 | 0 | 0 | 0 | 0 | 0 |
| AVNTAPVP | SYT7; SYT7 | O43581; F5I | SYT7_HUMAN ! | Unmodified | 0 | 0 | 0 | 260830 | 0 | 0 | 0 | 24614 | 904740 | 665480 | 0 | 116270 | 328710 | 0 | 296790 | 0 | 0 | 0 |
| AVNTAQGL | STX6 | O43752 | STX6_HUMAN ! | Unmodified | 279780 | 1668600 | 336600 | 0 | 103490 | 0 | 0 | 290180 | 0 | 127460 | 0 | 890110 | 238050 | 0 | 102670 | 118310 | 0 | 4024600 |
| AVNVYSTS' | MAPRE1 | Q15691 | MARE1_HUMAI | Acetyl (Protein N | 0 | 0 | 0 | 0 | 0 | 0 | 0 | 0 | 0 | 0 | 0 | 0 | 0 | 0 | 0 | 0 | 0 | 0 |
| AVNYFSK | CLTC; CLTC; CI | Q00610; J3K | CLH1_HUMAN | Unmodified | 8212300 | 0 | 4621900 | 5461600 | 1198300 | 0 | 372020 | 266400 | 2135800 | 571930 | 778450 | 73183000 | 1742200 | 499340 | 4682400 | 5244700 | 232730 | 1904700 |
| AVPEDENII | N/A | N/A | N/A | Oxidation (M) | 0 | 0 | 0 | 0 | 0 | 6100800 | 185470 | 0 | 0 | 0 | 0 | 0 | 0 | 0 | 0 | 0 | 0 | 335590 |
| AVPETRPNI | SNRPA; SNRPA | P09012; M0I | SNRPA_HUMAI | Acetyl (Protein N | 1022500 | 599810 | 469070 | 0 | 310920 | 0 | 136130 | 55171 | 583340 | 253880 | 1736800 | 14774000 | 105010 | 0 | 645790 | 453840 | 0 | 0 |
| AVPFPPTHI | PPP3CB; PPP3( | P16298; Q5I | PP2BB_HUMAN | Unmodified | 0 | 0 | 0 | 0 | 686430 | 0 | 0 | 0 | 0 | 0 | 0 | 0 | 0 | 0 | 0 | 0 | 0 | 1196300 |
| AVPGNPLG | PLA2G3 | Q9NZ20 | PA2G3_HUMAN | Unmodified | 0 | 0 | 0 | 0 | 0 | 0 | 0 | 0 | 307690 | 127840 | 0 | 0 | 0 | 0 | 0 | 0 | 0 | 0 |
| AVPLALALI | PSMD2; PSMD2 | Q13200; H7( | PSMD2_HUMA | Unmodified | 225050 | 0 | 111620 | 90145 | 0 | 0 | 0 | 0 | 99577 | 63773 | 50795 | 3361400 | 0 | 0 | 170780 | 133290 | 0 | 0 |
| AVPLAVPL( | THOC6 | Q86W42 | THOC6_HUMA | Unmodified | 0 | 0 | 0 | 0 | 0 | 0 | 0 | 0 | 201610 | 0 | 156760 | 1468300 | 0 | 0 | 0 | 0 | 0 | 0 |
| AVPLSVAL\ | GAS6 | Q14393 | GAS6_HUMAN | Unmodified | 0 | 0 | 294510 | 0 | 0 | 0 | 0 | 0 | 4304200 | 556470 | 0 | 83280 | 0 | 0 | 0 | 0 | 0 | 0 |
| AVPLSVAL\ | GAS6 | Q14393 | GAS6_HUMAN | Unmodified | 0 | 0 | 0 | 0 | 0 | 0 | 0 | 0 | 1135800 | 0 | 0 | 0 | 0 | 0 | 0 | 0 | 0 | 0 |
| AVPPTYADI | VDAC1; VDAC1 | P21796; C9J | VDAC1_HUMAI | Acetyl (Protein N | 0 | 0 | 0 | 147590 | 0 | 0 | 1054100 | 0 | 43122000 | 14905000 | 1244000 | 687940 | 283900 | 887420 | 0 | 0 | 311940 | 614730 |
| AVPREELF\ | AKR1A1; AKR1A | P14550; V9( | AK1A1_HUMAN | Unmodified | 0 | 0 | 264850 | 497100 | 1080000 | 285530 | 135110 | 68159 | 1369100 | 932390 | 0 | 1924600 | 1481100 | 677770 | 831550 | 406430 | 422410 | 1251300 |
| AVPSWFVR | IARS1; IARS1; I/ | P41252; A0A | SYIC_HUMAN I | Unmodified | 0 | 0 | 0 | 0 | 0 | 0 | 0 | 0 | 0 | 0 | 0 | 10435000 | 0 | 0 | 0 | 0 | 0 | 0 |
| AVPTDEAR, | RAB11A; RAB11 | P62491; Q1! | RB11A_HUMAN | Unmodified | 128270 | 0 | 139330 | 2077600 | 5345700 | 2802300 | 547980 | 159780 | 311760 | 1008200 | 1240500 | 95344 | 1760400 | 1275500 | 0 | 0 | 58609 | 166810 |
| AVPWVILSI | CANT1; CANT1 | Q8WVQ1; K7 | CANT1_HUMAI | Unmodified | 0 | 0 | 36498 | 0 | 0 | 0 | 0 | 0 | 14898000 | 1651400 | 629180 | 183900 | 120990 | 0 | 0 | 0 | 0 | 0 |
| AVQALCAV | UGDH | O60701 | UGDH_HUMAN | Unmodified | 168420 | 0 | 145780 | 491590 | 214770 | 298680 | 0 | 0 | 0 | 39136 | 0 | 261410 | 0 | 524890 | 0 | 111870 | 129870 | 0 |
| AVQDLCGV | RCC2 | Q9P258 | RCC2_HUMAN | Unmodified | 0 | 409890 | 0 | 228860 | 2301100 | 491000 | 274160 | 0 | 199880 | 375500 | 399880 | 5099300 | 0 | 0 | 81525 | 0 | 284180 | 6082700 |
| AVQDSKLD | EXTL3 | O43909 | EXTL3_HUMAN | Unmodified | 0 | 0 | 0 | 0 | 0 | 0 | 0 | 0 | 2875900 | 719770 | 464950 | 0 | 0 | 159710 | 45141 | 0 | 0 | 0 |
| AVQEIMQE | GART; GART; G/ | P22102; C9J | PUR2_HUMAN | Oxidation (M) | 0 | 0 | 0 | 0 | 453740 | 217620 | 206330 | 0 | 0 | 0 | 0 | 376700 | 622840 | 0 | 0 | 235920 | 750710 | 383460 |
| AVQEIMQE | GART; GART; G/ | P22102; C9J | PUR2_HUMAN | Unmodified | 112520 | 83377 | 292740 | 983730 | 2833900 | 522930 | 1168700 | 570480 | 703220 | 989670 | 0 | 3046500 | 1618000 | 1705000 | 299800 | 176180 | 1942800 | 2086500 |
| AVQFFVQA | DNAJC7; DNAJ( | Q99615; A0/ | DNJC7_HUMAI | Unmodified | 0 | 0 | 0 | 0 | 0 | 0 | 0 | 0 | 208900 | 189400 | 768190 | 1696800 | 1857500 | 0 | 0 | 0 | 0 | 0 |
| AVQFFVQA | DNAJC7; DNAJ( | Q99615; A0/ | DNJC7_HUMAI | Unmodified | 0 | 0 | 0 | 0 | 620650 | 0 | 733780 | 0 | 0 | 122280 | 0 | 0 | 835590 | 721070 | 0 | 0 | 233930 | 0 |
| AVQFFVQA | DNAJC7; DNAJ( | Q99615; A0/ | DNJC7_HUMAI | Oxidation (M) | 0 | 0 | 0 | 0 | 84779 | 0 | 1446500 | 89105 | 0 | 350760 | 0 | 0 | 654440 | 740400 | 0 | 0 | 411830 | 46801 |
| AVQFTEEK | SH3GLB2; SH3( | Q9NR46; Q9 | SHLB2_HUMAI | Unmodified | 0 | 0 | 0 | 0 | 793550 | 0 | 0 | 127320 | 914980 | 402130 | 181420 | 868020 | 0 | 950730 | 0 | 0 | 25298 | 375550 |
| AVQGAQQI | FLII | Q13045 | FLII_HUMAN Pr | Unmodified | 0 | 0 | 51628 | 0 | 0 | 425710 | 0 | 0 | 144910 | 0 | 138150 | 523880 | 0 | 0 | 0 | 122530 | 268590 | 0 |
| AVQGFFTSI | PSMD3 | O43242 | PSMD3_HUMA | Unmodified | 0 | 0 | 0 | 0 | 0 | 0 | 0 | 0 | 0 | 0 | 0 | 3816900 | 0 | 0 | 0 | 521080 | 0 | 0 |
| AVQIVSHPI | EDEM3; EDEM3 | Q9BZQ6; H0 | EDEM3_HUMAI | Unmodified | 0 | 0 | 45682 | 0 | 0 | 0 | 0 | 0 | 1743200 | 0 | 0 | 0 | 0 | 0 | 0 | 0 | 0 | 0 |
| AVQLYQQT | NAPG | Q99747 | SNAG_HUMAN | Unmodified | 0 | 0 | 0 | 0 | 0 | 0 | 149360 | 0 | 1116400 | 0 | 0 | 2094700 | 0 | 0 | 0 | 0 | 0 | 827810 |
| AVQLYQQT | NAPG | Q99747 | SNAG_HUMAN | Unmodified | 0 | 415860 | 0 | 2294200 | 2128900 | 623340 | 0 | 0 | 2720300 | 400410 | 1210800 | 3980500 | 1076300 | 0 | 0 | 0 | 0 | 1284200 |
| AVQLYQQT | NAPG | Q99747 | SNAG_HUMAN | Unmodified | 0 | 0 | 0 | 0 | 17194000 | 1799100 | 4426900 | 0 | 0 | 2388500 | 0 | 0 | 9686800 | 5584700 | 0 | 0 | 4966100 | 23316000 |
| AVQSLDKN | ARPC5; ARPC5 | O15511; B1/ | ARPC5_HUMAI | Oxidation (M) | 0 | 0 | 0 | 0 | 1086200 | 1018900 | 435060 | 178280 | 0 | 0 | 0 | 0 | 1108900 | 477550 | 0 | 0 | 434070 | 248880 |
| AVQTLQME | GFPT1; GFPT1; | Q06210; A0/ | GFPT1_HUMAN | Oxidation (M) | 0 | 0 | 16176 | 52791 | 0 | 0 | 0 | 0 | 537070 | 0 | 63930 | 74921 | 181380 | 0 | 109160 | 43177 | 0 | 0 |
| AVRASFEN | EIF6; EIF6; EIF6 | P56537; A0A | IF6_HUMAN Eu | Acetyl (Protein N | 0 | 0 | 0 | 0 | 1883500 | 517300 | 451170 | 259210 | 0 | 163930 | 0 | 0 | 1040900 | 1154800 | 0 | 0 | 3371800 | 2588300 |
| AVRPIEVTC | KLK14; KLK14 | Q9P0G3; A0. | KLK14_HUMAN | Unmodified | 0 | 0 | 0 | 0 | 0 | 0 | 0 | 0 | 4566700 | 383650 | 0 | 0 | 149350 | 90299 | 0 | 0 | 0 | 0 |
| AVRVFEFG( | CRYZ; CRYZ; CI | Q08257; A6I | QOR_HUMAN ( | Unmodified | 0 | 0 | 0 | 0 | 1255600 | 0 | 0 | 0 | 0 | 984530 | 0 | 0 | 666810 | 0 | 0 | 0 | 1178100 | 0 |
| AVSADPLPI | MARVELD2; MA | Q8N4S9; A1I | MALD2_HUMAI | Unmodified | 1816700 | 0 | 0 | 1143000 | 0 | 0 | 0 | 0 | 0 | 0 | 1063800 | 0 | 0 | 0 | 3782700 | 2959700 | 0 | 0 |
| AVSDSFGP | TMBIM1; TMBIN | Q969X1; F8\ | LFG3_HUMAN | Unmodified | 10048000 | 4940000 | 10412000 | 661070 | 0 | 0 | 0 | 0 | 0 | 0 | 0 | 0 | 0 | 0 | 13445000 | 9580800 | 0 | 0 |
| AVSDSFGP | TMBIM1; TMBIN | Q969X1; F8\ | LFG3_HUMAN | Unmodified | 3320200 | 5980400 | 2279600 | 1541400 | 0 | 0 | 0 | 0 | 0 | 0 | 0 | 0 | 0 | 0 | 16497000 | 7773900 | 0 | 0 |
| AVSDWIDE | ILF3; ILF3; ILF3; | Q12906; K7E | ILF3_HUMAN Ir | Unmodified | 1578600 | 2195200 | 1373600 | 1587500 | 0 | 0 | 0 | 0 | 918350 | 222370 | 1223800 | 3731700 | 1030400 | 0 | 831610 | 472310 | 0 | 0 |
| AVSDWLIA! | NCLN; NCLN; I | Q969V3; K7E | NCLN_HUMAN | Unmodified | 0 | 0 | 0 | 0 | 0 | 0 | 0 | 0 | 628240 | 123260 | 0 | 0 | 0 | 0 | 0 | 0 | 0 | 0 |
| AVSDYFGG | MELTF | P08582 | TRFM_HUMAN | Unmodified | 0 | 0 | 0 | 0 | 0 | 0 | 0 | 0 | 1859000 | 0 | 0 | 0 | 0 | 0 | 0 | 0 | 0 | 0 |
| AVSEAQEKI | PKN1 | Q16512 | PKN1_HUMAN | Unmodified | 0 | 0 | 0 | 0 | 194160 | 0 | 0 | 0 | 0 | 0 | 0 | 0 | 100960 | 0 | 0 | 0 | 110190 | 112250 |
| AVSEEQQP | ABCF1; ABCF1; | Q8NE71; A0, | ABCF1_HUMAN | Unmodified | 136200 | 0 | 0 | 0 | 514170 | 0 | 162060 | 0 | 74049 | 204180 | 149060 | 2167800 | 289450 | 0 | 218300 | 159650 | 0 | 481950 |
| AVSEGNLLI | PCID2 | Q5JVF3 | PCID2_HUMAN | Unmodified | 29563 | 0 | 0 | 0 | 0 | 0 | 0 | 0 | 0 | 0 | 0 | 2401100 | 0 | 0 | 0 | 32198 | 0 | 0 |
| AVSEKEVD: | TIMP2 | P16035 | TIMP2_HUMAN | Unmodified | 0 | 0 | 0 | 0 | 2493800 | 1830000 | 0 | 314360 | 0 | 1362500 | 0 | 0 | 0 | 0 | 0 | 0 | 1365800 | 4193600 |
| AVSESQLK | TSG101; TSG1( | Q99816; F5I | TS101_HUMAN | Acetyl (Protein N | 10679000 | 16740000 | 0 | 7442700 | 3369000 | 946140 | 487270 | 0 | 4637800 | 1476300 | 87809 | 1671700 | 5980000 | 3996200 | 14448000 | 10057000 | 537500 | 2248400 |
| AVSESQLKI | TSG101; TSG1( | Q99816; F5I | TS101_HUMAN | Acetyl (Protein N | 10544000 | 11184000 | 7529400 | 7229900 | 2943400 | 44624000 | 45228000 | 27733000 | 2564600 | 14252000 | 3423300 | 2010900 | 7541900 | 21126000 | 11784000 | 11107000 | 1443700 | 36028000 |
| AVSEVCRIF | RO60; RO60 | P10155; H0Y | RO60_HUMAN | Unmodified | 5631900 | 1404800 | 0 | 316740 | 0 | 0 | 0 | 0 | 0 | 0 | 0 | 0 | 20079 | 0 | 5525700 | 344310 | 0 | 0 |
| AVSEVCRIF | RO60; RO60 | P10155; H0Y | RO60_HUMAN | Unmodified | 0 | 1681400 | 0 | 2905100 | 0 | 0 | 0 | 0 | 0 | 0 | 0 | 0 | 1101300 | 0 | 0 | 0 | 0 | 0 |
| AVSHLVTW | SHPK; N/A | Q9UHJ6; A0/ | SHPK_HUMAN | Unmodified | 73240 | 0 | 0 | 0 | 0 | 0 | 0 | 0 | 0 | 0 | 0 | 1904100 | 0 | 0 | 421410 | 0 | 0 | 0 |
| AVSISPTN\ | PTPRJ; PTPRJ; P | Q12913; A0/ | PTPRJ_HUMAN | Unmodified | 0 | 0 | 0 | 0 | 149650 | 0 | 304590 | 10501 | 3349400 | 0 | 2058600 | 3308400 | 0 | 0 | 0 | 133970 | 0 | 0 |
| AVSNINTLL | UBR4 | Q5T4S7 | UBR4_HUMAN | Unmodified | 0 | 0 | 0 | 0 | 0 | 0 | 0 | 0 | 0 | 0 | 0 | 1365600 | 0 | 0 | 0 | 0 | 0 | 0 |
| AVSQVLDS | CASK; CASK; C/ | O14936; A0/ | CSKP_HUMAN | Unmodified | 0 | 0 | 0 | 0 | 0 | 0 | 0 | 0 | 1341200 | 140490 | 0 | 0 | 0 | 0 | 0 | 0 | 0 | 0 |
| AVSREDSV | HNRNPA3; HNI | P51991; A0A | ROA3_HUMAN | Unmodified | 0 | 0 | 0 | 0 | 0 | 0 | 381060 | 534360 | 0 | 44517 | 0 | 621500 | 0 | 0 | 0 | 0 | 0 | 0 |
| AVSSAIAQL | TLN1 | Q9Y490 | TLN1_HUMAN ' | Unmodified | 0 | 965390 | 0 | 167000 | 1839400 | 226960 | 216710 | 0 | 0 | 458940 | 0 | 30926 | 670060 | 267750 | 245090 | 0 | 344900 | 0 |
| AVSSPDRIL | BTN2A1 | Q7KYR7 | BT2A1_HUMAN | Unmodified | 0 | 0 | 0 | 0 | 331720 | 0 | 0 | 0 | 0 | 0 | 0 | 831220 | 0 | 0 | 0 | 0 | 99406 | 267860 |
| AVSTCAHLI | VPS35 | Q96QK1 | VPS35_HUMAN | Unmodified | 234710 | 0 | 0 | 830940 | 0 | 0 | 0 | 0 | 862230 | 0 | 2158300 | 2836300 | 163050 | 0 | 768040 | 320870 | 0 | 0 |
| AVSTTFGSI | DIP2B | Q9P265 | DIP2B_HUMAN | Unmodified | 24087 | 0 | 0 | 31433 | 0 | 0 | 0 | 0 | 0 | 68598 | 0 | 3052900 | 33524 | 0 | 0 | 23493 | 0 | 0 |
| AVSVEAAV' | TMEM132A | Q24JP5 | T132A_HUMAN | Unmodified | 0 | 0 | 0 | 0 | 0 | 0 | 0 | 0 | 1024600 | 0 | 0 | 0 | 0 | 0 | 0 | 0 | 0 | 0 |
| AVSVNEGY | MTMR12 | Q9C0I1 | MTMRC_HUMA | Unmodified | 0 | 0 | 0 | 105200 | 11374000 | 0 | 0 | 0 | 79087 | 33390 | 0 | 465700 | 90914 | 65186 | 0 | 63758 | 233750 | 236690 |
| AVSVTPIRD | MBNL1; MBNL1 | Q9NR56; A0, | MBNL1_HUMAI | Acetyl (Protein N | 5187400 | 4273800 | 0 | 4382300 | 1681100 | 0 | 0 | 0 | 0 | 0 | 0 | 1460000 | 4895400 | 0 | 1677800 | 1963200 | 0 | 0 |
| AVSWAQL( | USP43; USP43 | Q70EL4; V9( | UBP43_HUMAN | Unmodified | 0 | 0 | 0 | 0 | 0 | 0 | 0 | 45596 | 0 | 0 | 0 | 0 | 0 | 0 | 238780 | 92790 | 0 | 0 |
| AVTAADME | CAPN5; CAPN5 | O15484; E7E | CAN5_HUMAN | Unmodified | 0 | 0 | 0 | 0 | 0 | 244020 | 314320 | 51634 | 0 | 0 | 0 | 0 | 0 | 2678300 | 0 | 0 | 397130 | 514120 |
| AVTASNDE | COBL; COBL; C | O75128; J3K | COBL_HUMAN | Unmodified | 9300500 | 7454900 | 0 | 0 | 13217000 | 1166400 | 0 | 0 | 0 | 0 | 0 | 0 | 0 | 0 | 0 | 0 | 368750 | 1066900 |
| AVTDEEPFI | LRP1 | Q07954 | LRP1_HUMAN I | Unmodified | 0 | 205930 | 254430 | 0 | 0 | 0 | 0 | 0 | 1232500 | 0 | 0 | 0 | 0 | 0 | 85770 | 0 | 49637 | 0 |
| AVTDEEPFI | LRP1 | Q07954 | LRP1_HUMAN I | Unmodified | 0 | 0 | 0 | 0 | 2170500 | 833720 | 0 | 0 | 0 | 1028100 | 0 | 0 | 326220 | 0 | 0 | 0 | 179400 | 1337400 |
| AVTELNEPI | YWHAH | Q04917 | 1433F_HUMAN | Unmodified | 3451800 | 4131300 | 6052400 | 8132900 | 77588000 | 25567000 | 11654000 | 4208800 | 2444900 | 10159000 | 2827800 | 49713000 | 28025000 | 22733000 | 6569600 | 6926900 | 40376000 | 53779000 |
| AVTELNEPI | YWHAH | Q04917 | 1433F_HUMAN | Unmodified | 0 | 0 | 0 | 0 | 2385700 | 0 | 2164300 | 0 | 0 | 335170 | 0 | 0 | 0 | 1662700 | 0 | 0 | 3715300 | 2218900 |
| AVTEMNGF | PABPC4; PABP( | Q13310; P11 | PABP4_HUMAN | Unmodified | 7880700 | 5572800 | 0 | 4222400 | 503930 | 384400 | 0 | 0 | 380320 | 0 | 309550 | 3415500 | 959560 | 108180 | 4993800 | 2713100 | 197750 | 256640 |
| AVTEQGAE | YWHAQ; YWHA | P27348; E9P | 1433T_HUMAN | Unmodified | 294930 | 749240 | 1158900 | 725220 | 366290 | 15443 | 0 | 0 | 3070100 | 605780 | 7303100 | 25811000 | 248440 | 0 | 1781500 | 1958500 | 147720 | 342900 |
| AVTEQGAE | YWHAQ; YWHA | P27348; E9P | 1433T_HUMAN | Unmodified | 4152600 | 0 | 5913600 | 1168500 | 1.08E+08 | 33401000 | 13401000 | 1174300 | 23450000 | 14641000 | 35807000 | 56901000 | 9993600 | 35861000 | 9210600 | 8398200 | 88338000 | 89177000 |
| AVTEQGAE | YWHAQ; YWHA | P27348; E9P | 1433T_HUMAN | Unmodified | 0 | 0 | 0 | 0 | 2345500 | 0 | 0 | 0 | 0 | 0 | 0 | 0 | 3808400 | 1471800 | 0 | 0 | 7241200 | 1592700 |
| AVTEQGHE | YWHAB; YWHA | P31946; Q4\ | 1433B_HUMAN | Unmodified | 558470 | 1081000 | 0 | 0 | 1491700 | 0 | 426590 | 0 | 8744800 | 856570 | 16597000 | 21512000 | 787540 | 0 | 5352400 | 5334300 | 285220 | 2539900 |
| AVTEQGHE | YWHAB; YWHA | P31946; Q4\ | 1433B_HUMAN | Unmodified | 13986000 | 9560600 | 26680000 | 67175000 | 4.16E+08 | 60777000 | 64066000 | 3238300 | 57468000 | 47186000 | 30347000 | 1E+08 | 1.58E+08 | 1.34E+08 | 40633000 | 36125000 | 2.95E+08 | 4.28E+08 |
| AVTEQGHE | YWHAB; YWHA | P31946; Q4\ | 1433B_HUMAN | Unmodified | 0 | 0 | 0 | 0 | 5934600 | 0 | 0 | 0 | 0 | 0 | 0 | 0 | 6243900 | 4977700 | 0 | 0 | 12813000 | 10238000 |
| AVTEVLAR | SH3GL1; SH3G | Q99961; M0I | SH3G1_HUMAI | Unmodified | 49237000 | 36751000 | 25352000 | 24219000 | 2577200 | 0 | 1135400 | 1170500 | 2973400 | 1946800 | 956030 | 719670 | 36318000 | 12558000 | 35087000 | 30673000 | 501840 | 2139700 |

| | | | | | | | | | | | | | | | | | | | | | | |
|---|---|---|---|---|---|---|---|---|---|---|---|---|---|---|---|---|---|---|---|---|---|---|
| AVTEVLART | SH3GL1; SH3G | Q99961; M0 | SH3G1_HUMA | Unmodified | 0 | 0 | 0 | 0 | 1074700 | 0 | 1871800 | 0 | 0 | 339370 | 0 | 0 | 6317500 | 3880600 | 0 | 0 | 194980 | 408860 |
| AVTEVLART | SH3GL1; SH3G | Q99961; M0 | SH3G1_HUMA | Unmodified | 636610 | 0 | 0 | 0 | 2997400 | 320410 | 4869600 | 0 | 0 | 0 | 0 | 0 | 16390000 | 11868000 | 0 | 0 | 1125000 | 1354800 |
| AVTFIDLTTI | DERA; DERA; DI | Q9Y315; E9F | DEOC_HUMAN | Unmodified | 0 | 0 | 0 | 0 | 0 | 0 | 0 | 27016 | 0 | 0 | 0 | 3759100 | 0 | 0 | 0 | 140900 | 372210 | 0 |
| AVTFSPDL( | CAMSAP3 | Q9P1Y5 | CAMP3_HUMA | Unmodified | 0 | 0 | 0 | 389960 | 0 | 0 | 0 | 0 | 152830 | 0 | 0 | 0 | 0 | 0 | 0 | 0 | 0 | 0 |
| AVTFSPDL( | CAMSAP3 | Q9P1Y5 | CAMP3_HUMA | Unmodified | 0 | 0 | 0 | 0 | 480390 | 0 | 0 | 0 | 0 | 0 | 0 | 0 | 1426200 | 837520 | 0 | 0 | 0 | 0 |
| AVTGTHTL | EIF2B2; EIF2B2 | P49770; H0 | EI2BB_HUMAN | Unmodified | 0 | 0 | 0 | 0 | 1118200 | 1020700 | 2092800 | 1204500 | 0 | 57562 | 0 | 1080700 | 198890 | 0 | 0 | 0 | 233040 | 553490 |
| AVTGYNDP | DSP | P15924 | DESP_HUMAN | Unmodified | 0 | 0 | 0 | 0 | 0 | 0 | 0 | 0 | 357310 | 0 | 0 | 0 | 0 | 0 | 0 | 0 | 0 | 0 |
| AVTGYRDP | PLEC; PLEC | Q15149; H0 | PLEC_HUMAN | Oxidation (M) | 0 | 0 | 0 | 0 | 0 | 0 | 0 | 0 | 0 | 0 | 0 | 0 | 0 | 0 | 0 | 0 | 1193300 | 0 |
| AVTGYRDP | PLEC; PLEC | Q15149; H0 | PLEC_HUMAN | Unmodified | 0 | 0 | 0 | 0 | 0 | 0 | 197410 | 0 | 0 | 0 | 0 | 0 | 0 | 0 | 0 | 0 | 3865500 | 1411200 |
| AVTGYRDP | PLEC; PLEC | Q15149; H0 | PLEC_HUMAN | Unmodified | 0 | 0 | 0 | 0 | 1925200 | 0 | 0 | 0 | 0 | 439440 | 0 | 0 | 1114500 | 613000 | 0 | 0 | 12258000 | 10739000 |
| AVTHAIPAL | HTT; HTT; HTT; | P42858; A0A | HD_HUMAN Hu | Unmodified | 0 | 0 | 0 | 0 | 0 | 0 | 272820 | 0 | 113660 | 0 | 0 | 380850 | 0 | 0 | 0 | 19107 | 304210 | 0 |
| AVTHTSPEI | TBCD; TBCD; TI | Q9BTW9; I3L | TBCD_HUMAN | Unmodified | 240990 | 0 | 0 | 123630 | 0 | 0 | 0 | 0 | 113570 | 0 | 0 | 1405200 | 0 | 0 | 0 | 149210 | 0 | 0 |
| AVTHTSPEI | TBCD; TBCD; TI | Q9BTW9; I3L | TBCD_HUMAN | Unmodified | 0 | 0 | 0 | 0 | 0 | 0 | 0 | 0 | 0 | 0 | 82351 | 1507000 | 0 | 0 | 62413 | 91289 | 0 | 0 |
| AVTIFIR | CCT5; CCT5; C( | P48643; B7Z | TCPE_HUMAN | Unmodified | 0 | 0 | 0 | 0 | 0 | 0 | 0 | 0 | 86928 | 196170 | 2209700 | 11941000 | 0 | 0 | 37581 | 24283 | 0 | 0 |
| AVTIFIRGG | CCT5; CCT5; C( | P48643; B7Z | TCPE_HUMAN | Unmodified | 0 | 0 | 0 | 0 | 353150 | 0 | 0 | 0 | 0 | 0 | 0 | 0 | 402920 | 183060 | 0 | 0 | 0 | 0 |
| AVTITDLR | STAMBP | O95630 | STABP_HUMAN | Unmodified | 0 | 0 | 0 | 0 | 0 | 0 | 0 | 0 | 0 | 0 | 0 | 1537800 | 0 | 0 | 0 | 325700 | 0 | 0 |
| AVTKDEDE | CDV3 | Q9UKY7 | CDV3_HUMAN | Unmodified | 0 | 0 | 0 | 0 | 209310 | 0 | 0 | 0 | 190260 | 0 | 288970 | 616990 | 0 | 0 | 0 | 0 | 0 | 87940 |
| AVTKDEDE | CDV3 | Q9UKY7 | CDV3_HUMAN | Unmodified | 146480 | 118590 | 0 | 295520 | 3893900 | 0 | 141800 | 36132 | 929690 | 413220 | 1305800 | 3372900 | 2372400 | 860310 | 135950 | 284420 | 702390 | 1167100 |
| AVTKYTSAK | H2BC11; H2BC | P06899; O6 | H2B1J_HUMAN | Unmodified | 0 | 0 | 0 | 0 | 305110 | 0 | 1897900 | 8866900 | 0 | 16254 | 0 | 33664 | 54044 | 0 | 0 | 0 | 24367 | 880490 |
| AVTKYTSSK | H2BE1; H2BC4 | A0A2R8Y619 | H2BE1_HUMAN | Unmodified | 20238 | 14066 | 0 | 0 | 272480 | 38569 | 1467000 | 4084800 | 0 | 0 | 0 | 38744 | 0 | 0 | 0 | 0 | 0 | 468210 |
| AVTLECVS/ | HSPG2 | P98160 | PGBM_HUMAN | Unmodified | 0 | 0 | 1219500 | 887170 | 127300 | 785460 | 0 | 0 | 575630 | 229200 | 0 | 0 | 420060 | 524320 | 0 | 0 | 42482 | 1016800 |
| AVTLECVS/ | HSPG2 | P98160 | PGBM_HUMAN | Unmodified | 0 | 0 | 0 | 0 | 0 | 1262400 | 0 | 0 | 0 | 229330 | 348310 | 1514600 | 886100 | 1364900 | 0 | 0 | 200160 | 1970800 |
| AVTLHSAPI | MOGS; MOGS; | Q13724; A0/ | MOGS_HUMAN | Oxidation (M) | 0 | 0 | 0 | 0 | 0 | 0 | 0 | 0 | 0 | 1398000 | 0 | 0 | 0 | 0 | 0 | 0 | 0 | 0 |
| AVTPLGPPI | HLA-DQB1; HL/ | P01920; A0A | DQB1_HUMAN | Unmodified | 0 | 0 | 35156 | 0 | 284680 | 0 | 0 | 0 | 3034400 | 0 | 0 | 182650 | 0 | 0 | 0 | 0 | 0 | 0 |
| AVTPNGEIF | ARAP1; ARAP1; | Q96P48; F8V | ARAP1_HUMAN | Unmodified | 388390 | 0 | 223280 | 0 | 0 | 0 | 21292 | 0 | 0 | 83620 | 36103 | 601220 | 0 | 0 | 342820 | 261520 | 0 | 0 |
| AVTPPMPLI | PUF60; PUF60; | Q9UHX1; A0 | PUF60_HUMAN | Oxidation (M) | 0 | 0 | 0 | 0 | 0 | 0 | 0 | 0 | 1431400 | 1042500 | 658050 | 1275800 | 0 | 0 | 0 | 0 | 747430 | 0 |
| AVTPPMPLI | PUF60; PUF60; | Q9UHX1; A0 | PUF60_HUMAN | Unmodified | 0 | 0 | 0 | 0 | 0 | 0 | 0 | 0 | 307490 | 164310 | 0 | 0 | 0 | 0 | 0 | 0 | 0 | 0 |
| AVTPVPTKT | LARP1; LARP1; | Q6PKG0; H0 | LARP1_HUMAN | Unmodified | 0 | 375630 | 0 | 1018000 | 271520 | 0 | 46299 | 65210 | 319240 | 97677 | 354520 | 651150 | 782180 | 1088500 | 586060 | 566580 | 0 | 273170 |
| AVTQALNR | TLN1 | Q9Y490 | TLN1_HUMAN | Unmodified | 361860 | 1904500 | 709130 | 1139000 | 2102000 | 32243 | 362290 | 684410 | 341970 | 573240 | 136330 | 3140600 | 972540 | 249550 | 1577000 | 731910 | 148870 | 391310 |
| AVTSANIQE | ERBB2; ERBB2; | P04626; J3Q | ERBB2_HUMAN | Unmodified | 139240 | 89654 | 0 | 213930 | 61846 | 0 | 89916 | 0 | 238410 | 0 | 512790 | 0 | 339990 | 88306 | 706750 | 507750 | 30796 | 0 |
| AVTTPGKK( | NCL; NCL; NCL | P19338; C9J | NUCL_HUMAN | Unmodified | 0 | 0 | 0 | 0 | 341300 | 0 | 153930 | 250320 | 0 | 27314 | 0 | 0 | 706780 | 0 | 0 | 0 | 396630 | 268650 |
| AVTVYTNYI | AGL | P35573 | GDE_HUMAN ( | Unmodified | 0 | 0 | 0 | 136590 | 0 | 630600 | 0 | 0 | 0 | 0 | 0 | 810570 | 0 | 0 | 0 | 20988 | 0 | 0 |
| AVTVYTNYI | AGL | P35573 | GDE_HUMAN ( | Unmodified | 0 | 0 | 0 | 0 | 362710 | 0 | 0 | 0 | 0 | 0 | 1579200 | 2328800 | 0 | 0 | 0 | 0 | 0 | 0 |
| AVVDAGITT | PALS2; PALS2 | Q9NZW5; B8 | PALS2_HUMAN | Unmodified | 0 | 0 | 488150 | 562420 | 2687000 | 0 | 0 | 680180 | 0 | 568440 | 2239200 | 1969500 | 440650 | 583350 | 919340 | 536010 | 0 | 0 |
| AVVDAGITT | PALS2; PALS2 | Q9NZW5; B8 | PALS2_HUMAN | Unmodified | 0 | 0 | 0 | 0 | 690550 | 0 | 132910 | 0 | 0 | 0 | 0 | 0 | 38458 | 62554 | 0 | 0 | 729600 | 298630 |
| AVVGEKIMI | STXBP1; STXBP | P61764; A0A | STXB1_HUMAN | Unmodified | 0 | 0 | 0 | 0 | 0 | 0 | 177110 | 56615 | 0 | 0 | 0 | 0 | 0 | 0 | 0 | 0 | 46805 | 0 |
| AVVGEKIMI | STXBP1; STXBP | P61764; A0A | STXB1_HUMAN | Unmodified | 0 | 0 | 0 | 0 | 0 | 0 | 165720 | 45060 | 0 | 0 | 0 | 0 | 0 | 0 | 0 | 0 | 17273 | 0 |
| AVVGVVAG | RPL8; RPL8; RP | P62917; E9P | RL8_HUMAN 6( | Unmodified | 2907200 | 1585600 | 789020 | 416220 | 0 | 0 | 0 | 0 | 256350 | 0 | 1696600 | 15375000 | 0 | 0 | 1341400 | 1077400 | 0 | 0 |
| AVVGVVAG | RPL8; RPL8; RP | P62917; E9P | RL8_HUMAN 6( | Unmodified | 6283200 | 8389500 | 6443800 | 4515400 | 3003200 | 0 | 290070 | 207000 | 1343000 | 1094100 | 2673800 | 34200000 | 9153700 | 2043700 | 4078200 | 3384200 | 347850 | 1731400 |
| AVVHGILM | NPC2; NPC2; N | P61916; J3K | NPC2_HUMAN | Oxidation (M) | 62964 | 0 | 72224 | 58142 | 2148100 | 1241500 | 1116400 | 0 | 21333000 | 9556400 | 100580 | 0 | 683130 | 1032900 | 109830 | 0 | 0 | 182740 |
| AVVHGILM | NPC2; NPC2; N | P61916; J3K | NPC2_HUMAN | Unmodified | 0 | 0 | 0 | 0 | 0 | 0 | 0 | 0 | 4665000 | 544720 | 0 | 0 | 0 | 0 | 232870 | 0 | 0 | 0 |
| AVVISGAGI | ECH1; N/A; ECH | Q13011; M0 | ECH1_HUMAN | Unmodified | 0 | 1125900 | 0 | 0 | 0 | 0 | 0 | 6644.6 | 3428500 | 561130 | 79794 | 6135900 | 0 | 0 | 0 | 0 | 0 | 0 |
| AVVIVDDR( | SFPQ; NONO; N | P23246; Q15 | SFPQ_HUMAN | Unmodified | 0 | 0 | 0 | 0 | 0 | 0 | 0 | 0 | 0 | 0 | 0 | 456640 | 123170 | 0 | 0 | 0 | 0 | 0 |
| AVVKTFHET | CD81; CD81; C | P60033; A6N | CD81_HUMAN | Unmodified | 0 | 0 | 0 | 0 | 0 | 0 | 0 | 0 | 0 | 0 | 0 | 0 | 0 | 2960600 | 0 | 0 | 0 | 0 |
| AVVMISCN | M6PR; M6PR | P20645; H0 | MPRD_HUMAN | Oxidation (M) | 0 | 0 | 0 | 235630 | 0 | 0 | 0 | 0 | 3426300 | 288040 | 325300 | 375850 | 0 | 0 | 0 | 0 | 0 | 0 |
| AVVMISCN | M6PR; M6PR | P20645; H0 | MPRD_HUMAN | Unmodified | 0 | 0 | 0 | 0 | 0 | 0 | 0 | 0 | 5614200 | 291460 | 0 | 0 | 0 | 0 | 0 | 0 | 0 | 0 |
| AVVNDATIF | CUL5 | Q93034 | CUL5_HUMAN | Unmodified | 0 | 0 | 0 | 263660 | 1341900 | 0 | 0 | 227860 | 363490 | 348770 | 0 | 0 | 925330 | 809820 | 134890 | 0 | 946080 | 0 |
| AVVPEMEK | DHRS4L1; DHR | P0CG22; Q6 | DR4L1_HUMAN | Unmodified | 0 | 0 | 0 | 0 | 0 | 0 | 0 | 0 | 6803300 | 0 | 49255 | 0 | 0 | 0 | 0 | 0 | 0 | 0 |
| AVVPGPAE | EIF4E2; EIF4E2; | O60573; B9/ | IF4E2_HUMAN | Unmodified | 72607 | 0 | 0 | 0 | 175620 | 0 | 0 | 0 | 0 | 589110 | 0 | 1138800 | 0 | 0 | 0 | 0 | 84577 | 171730 |
| AVVPGVKP | EMC7; EMC7 | Q9NPA0; H0 | EMC7_HUMAN | Unmodified | 0 | 0 | 340930 | 0 | 103890 | 0 | 0 | 0 | 16509000 | 1167000 | 4659200 | 1277100 | 0 | 0 | 0 | 115460 | 0 | 0 |
| AVVPGVKP | EMC7; EMC7 | Q9NPA0; H0 | EMC7_HUMAN | Unmodified | 0 | 0 | 0 | 0 | 0 | 0 | 0 | 0 | 0 | 293940 | 0 | 0 | 0 | 0 | 0 | 0 | 0 | 72160 |
| AVVQVFEG | ATP6V1B2; ATP | P21281; H0 | VATB2_HUMAN | Unmodified | 0 | 0 | 0 | 0 | 967480 | 0 | 756600 | 331200 | 164550 | 76050 | 972030 | 2855600 | 390890 | 178080 | 76902 | 91940 | 0 | 634550 |
| AVVQVFEG | ATP6V1B2; ATP | P21281; H0 | VATB2_HUMAN | Unmodified | 0 | 0 | 0 | 509420 | 3095900 | 0 | 2255400 | 541720 | 1266700 | 314550 | 548320 | 0 | 2402700 | 1133200 | 0 | 612070 | 146600 | 1481400 |
| AVVRVSPE | COBL; COBL; C | O75128; A0/ | COBL_HUMAN | Unmodified | 0 | 0 | 0 | 0 | 502790 | 104890 | 242700 | 0 | 0 | 0 | 0 | 0 | 1139700 | 113620 | 0 | 0 | 114530 | 0 |
| AVVSGFDF | ARHGEF26 | Q96DR7 | ARHGQ_HUMA | Unmodified | 0 | 0 | 0 | 0 | 2962100 | 0 | 0 | 124060 | 0 | 298750 | 0 | 393600 | 467690 | 0 | 0 | 0 | 0 | 0 |
| AVVSQRLTI | HSP90B1; HSP9 | P14625; A0A | ENPL_HUMAN | Oxidation (M) | 0 | 0 | 2384400 | 809710 | 0 | 0 | 1077300 | 106880 | 0 | 40067000 | 0 | 0 | 0 | 0 | 190570 | 0 | 0 | 765190 |
| AVVSQRLTI | HSP90B1; HSP9 | P14625; A0A | ENPL_HUMAN | Unmodified | 0 | 0 | 0 | 0 | 0 | 0 | 0 | 0 | 0 | 7496800 | 0 | 0 | 0 | 0 | 0 | 0 | 0 | 0 |
| AVVSQRLTI | HSP90B1; HSP9 | P14625; A0A | ENPL_HUMAN | 2 Oxidation (M) | 0 | 0 | 0 | 0 | 0 | 0 | 0 | 0 | 0 | 18750000 | 0 | 0 | 0 | 0 | 0 | 0 | 0 | 0 |
| AVVSQRLTI | HSP90B1; HSP9 | P14625; A0A | ENPL_HUMAN | Oxidation (M) | 0 | 0 | 0 | 0 | 0 | 0 | 2262500 | 0 | 0 | 7572400 | 0 | 0 | 0 | 0 | 0 | 0 | 0 | 0 |
| AVVTGKMD | DPYSL2; DPYSL | Q16555; A0/ | DPYL2_HUMAN | Oxidation (M) | 0 | 0 | 0 | 0 | 3830500 | 594160 | 1779800 | 124280 | 176700 | 539150 | 0 | 391550 | 0 | 336630 | 0 | 0 | 437110 | 789050 |
| AVVTGKMD | DPYSL2; DPYSL | Q16555; A0/ | DPYL2_HUMAN | Unmodified | 0 | 0 | 0 | 0 | 2980300 | 0 | 0 | 511850 | 0 | 0 | 0 | 0 | 0 | 0 | 0 | 0 | 0 | 0 |
| AVVVCPKD | HNRNPU; HNR | Q00839; A0/ | HNRPU_HUMA | Unmodified | 583770 | 0 | 0 | 0 | 771340 | 277730 | 1890600 | 2587000 | 457540 | 454670 | 679950 | 2676700 | 1398300 | 487160 | 0 | 0 | 141780 | 835990 |
| AVVVCPKD | HNRNPU; HNR | Q00839; A0/ | HNRPU_HUMA | Unmodified | 0 | 0 | 0 | 0 | 0 | 0 | 195320 | 168560 | 0 | 76189 | 0 | 0 | 0 | 0 | 0 | 0 | 0 | 0 |
| AVVVDITEH | SLC31A1 | O15431 | COPT1_HUMAI | Unmodified | 0 | 0 | 0 | 0 | 0 | 0 | 0 | 0 | 0 | 0 | 0 | 0 | 0 | 0 | 423620 | 266070 | 0 | 0 |
| AVVVGFDP | PGP | A6NDG6 | PGP_HUMAN ( | Oxidation (M) | 120190 | 121660 | 0 | 449000 | 0 | 0 | 0 | 0 | 1097900 | 200930 | 857240 | 4203900 | 370150 | 0 | 505010 | 270800 | 0 | 0 |
| AVVVHAGE | SOD3 | P08294 | SODE_HUMAN | Unmodified | 629090 | 492640 | 0 | 0 | 885290 | 750610 | 1964400 | 272670 | 2418400 | 214540 | 0 | 124030 | 0 | 0 | 635850 | 0 | 66528 | 902660 |
| AVVVHAGE | SOD3 | P08294 | SODE_HUMAN | Unmodified | 0 | 0 | 0 | 0 | 0 | 0 | 0 | 0 | 14355000 | 2306500 | 0 | 0 | 0 | 0 | 0 | 0 | 0 | 173500 |
| AVVVHAGE | SOD3 | P08294 | SODE_HUMAN | Unmodified | 0 | 0 | 0 | 0 | 0 | 0 | 0 | 0 | 3266200 | 616870 | 0 | 0 | 0 | 0 | 0 | 0 | 0 | 0 |
| AVVVNAAQ | SLC25A11; SLC | Q02978; I3L | M2OM_HUMAN | Unmodified | 0 | 15267 | 0 | 0 | 56874 | 80141 | 0 | 0 | 1818700 | 557960 | 0 | 338660 | 0 | 108070 | 222670 | 375630 | 304010 | 123290 |
| AVVVPWGS | SPPL2A; SPPL2 | Q8TCT8; H0 | SPP2A_HUMAN | Unmodified | 0 | 0 | 0 | 0 | 7407100 | 0 | 2811900 | 649030 | 25144000 | 7393800 | 0 | 1486000 | 2292200 | 2129200 | 0 | 0 | 0 | 0 |
| AVVYSNTIC | GNAI1; GNAI1 | P63096; A0A | GNAI1_HUMAN | Unmodified | 0 | 79622 | 0 | 11362 | 0 | 0 | 0 | 0 | 68931 | 127730 | 151390 | 265260 | 145270 | 164650 | 131050 | 56913 | 0 | 0 |
| AVVYSNTIC | GNAI2 | P04899 | GNAI2_HUMAN | Oxidation (M) | 1584500 | 1485900 | 251360 | 2128800 | 2995100 | 494570 | 87922 | 159080 | 2103700 | 2874200 | 11858000 | 24411000 | 759490 | 343450 | 2064600 | 1316100 | 1664100 | 1544400 |
| AVVYSNTIC | GNAI2 | P04899 | GNAI2_HUMAN | Unmodified | 227000 | 3705500 | 0 | 699660 | 1602600 | 318960 | 0 | 0 | 4672700 | 855860 | 2278700 | 1255300 | 355820 | 0 | 867680 | 308540 | 488360 | 2280000 |
| AVYIGVHVI | SLC4A7; SLC4A | Q9Y6M7; C9 | S4A7_HUMAN | Unmodified | 0 | 0 | 0 | 1039000 | 475780 | 0 | 2640200 | 1193000 | 1107600 | 232560 | 672630 | 124490 | 0 | 0 | 444810 | 469110 | 1478000 | 346780 |
| AVYISVQDI | PBDC1 | Q9BVG4 | PBDC1_HUMAI | Unmodified | 0 | 0 | 0 | 0 | 357650 | 71540 | 0 | 0 | 0 | 0 | 0 | 2338300 | 329640 | 196350 | 0 | 0 | 1283500 | 0 |
| AVYLGELPI | UGGT1 | Q9NYU2 | UGGG1_HUMA | Oxidation (M) | 0 | 0 | 0 | 0 | 0 | 0 | 361480 | 0 | 4852400 | 2636300 | 0 | 0 | 0 | 0 | 0 | 0 | 0 | 0 |
| AVYLGELPI | UGGT1 | Q9NYU2 | UGGG1_HUMA | Unmodified | 0 | 0 | 0 | 0 | 0 | 0 | 0 | 0 | 9890500 | 704660 | 0 | 0 | 0 | 0 | 0 | 0 | 0 | 0 |
| AVYLGELPI | UGGT1 | Q9NYU2 | UGGG1_HUMA | Oxidation (M) | 0 | 0 | 0 | 0 | 0 | 0 | 0 | 0 | 0 | 1398900 | 0 | 0 | 0 | 0 | 0 | 0 | 0 | 0 |
| AVYLGELPI | UGGT1 | Q9NYU2 | UGGG1_HUMA | Unmodified | 0 | 0 | 0 | 0 | 0 | 0 | 0 | 0 | 0 | 1074800 | 0 | 0 | 0 | 0 | 0 | 0 | 0 | 0 |
| AVYLPNCD | IGFBP5 | P24593 | IBP5_HUMAN I | Unmodified | 0 | 0 | 0 | 0 | 0 | 0 | 0 | 135550 | 9889800 | 2028800 | 0 | 0 | 0 | 0 | 0 | 0 | 0 | 321980 |
| AVYPAFDK | ANKH | Q9HCJ1 | ANKH_HUMAN | Unmodified | 0 | 0 | 204360 | 0 | 0 | 0 | 0 | 0 | 2382400 | 0 | 0 | 123830 | 0 | 0 | 0 | 0 | 0 | 0 |
| AVYQARPQ | TRIM25; TRIM25 | Q14258; A0/ | TRI25_HUMAN | Unmodified | 0 | 0 | 0 | 0 | 532170 | 747620 | 68777 | 0 | 135620 | 0 | 0 | 0 | 0 | 684520 | 0 | 0 | 54273 | 800110 |
| AVYSDADIF | ABCC3 | O15438 | MRP3_HUMAN | Unmodified | 0 | 0 | 0 | 0 | 2411200 | 0 | 0 | 0 | 0 | 0 | 0 | 0 | 0 | 0 | 0 | 0 | 0 | 0 |
| AVYSRTFTV | MYO1C; MYO1( | O00159; F5F | MYO1C_HUMA | Unmodified | 0 | 0 | 0 | 0 | 1537700 | 0 | 289180 | 0 | 0 | 0 | 0 | 0 | 0 | 0 | 0 | 0 | 2932000 | 238520 |
| AVYTQDCP | AARS1; AARS1; | P49588; A0A | SYAC_HUMAN | Unmodified | 0 | 0 | 0 | 0 | 7416200 | 1728100 | 1009400 | 633630 | 2280700 | 1222600 | 1527000 | 4805400 | 8917500 | 6755900 | 1545500 | 2344900 | 2468100 | 2346000 |
| AWAEDVLA | PLCB2; PLCB2 | Q00722; Q9I | PLCB2_HUMAN | Unmodified | 0 | 0 | 0 | 0 | 0 | 0 | 0 | 0 | 0 | 0 | 472450 | 2791500 | 0 | 0 | 0 | 0 | 0 | 0 |
| AWAGIDLK | SPINT1; SPINT1 | O43278; H3I | SPIT1_HUMAN | Unmodified | 0 | 1915200 | 1763500 | 5150700 | 204840 | 0 | 0 | 0 | 3379100 | 628960 | 455080 | 0 | 1871300 | 926760 | 2890300 | 2614000 | 0 | 0 |
| AWDAHVT/ | RTN4IP1 | Q8WWV3 | RT4I1_HUMAN | Unmodified | 0 | 0 | 0 | 0 | 0 | 8255500 | 0 | 0 | 407320 | 109160 | 0 | 0 | 0 | 0 | 150540 | 0 | 0 | 0 |
| AWDIAIATN | FKBP4; FKBP4 | Q02790; F5H | FKBP4_HUMAN | Oxidation (M) | 44952 | 0 | 118430 | 0 | 0 | 0 | 204560 | 276950 | 0 | 0 | 1229500 | 3306800 | 2419600 | 532470 | 0 | 442960 | 0 | 0 |
| AWDIAIATN | FKBP4; FKBP4 | Q02790; F5H | FKBP4_HUMAN | Unmodified | 0 | 0 | 0 | 0 | 0 | 0 | 0 | 0 | 292120 | 0 | 418820 | 547720 | 1430300 | 0 | 0 | 0 | 89117 | 0 |
| AWDIGVAT | FKBP5 | Q13451 | FKBP5_HUMAN | Oxidation (M) | 0 | 0 | 0 | 0 | 0 | 0 | 0 | 0 | 0 | 0 | 0 | 0 | 1188200 | 0 | 0 | 0 | 0 | 0 |
| AWDVLLDI | MON2; MON2 | Q7Z3U7; A0/ | MON2_HUMAN | Unmodified | 148070 | 84105 | 0 | 38775 | 0 | 187560 | 0 | 0 | 0 | 737310 | 112230 | 1833300 | 0 | 0 | 37256 | 133430 | 189430 | 0 |
| AWEEYYK | FUBP1; KHSRP; | Q96AE4; Q9 | FUBP1_HUMAN | Unmodified | 0 | 0 | 0 | 0 | 221810 | 0 | 0 | 0 | 731600 | 449600 | 368610 | 3528600 | 0 | 0 | 0 | 0 | 99490 | 0 |
| AWEGNYL/ | MYO1D; MYO1I | O94832; J3C | MYO1D_HUMA | Unmodified | 0 | 0 | 5582600 | 29328000 | 0 | 0 | 0 | 0 | 3195600 | 0 | 1057300 | 1982000 | 0 | 0 | 0 | 0 | 0 | 2411900 |
| AWEGNYL/ | MYO1D; MYO1I | O94832; J3C | MYO1D_HUMA | Unmodified | 4324200 | 2677500 | 1284800 | 999140 | 0 | 0 | 0 | 0 | 0 | 0 | 388030 | 304640 | 0 | 0 | 1603100 | 1153300 | 0 | 624840 |
| AWENSPN | PRRC2C; PRRC | Q9Y520; H7 | PRC2C_HUMAI | Unmodified | 0 | 0 | 0 | 0 | 55792 | 104150 | 9028.6 | 69893 | 82328 | 0 | 0 | 18548 | 0 | 67240 | 0 | 0 | 77492 | 141060 |
| AWFREGFN | SMPDL3B; SMP | Q92485; F8V | ASM3B_HUMAI | Unmodified | 305940 | 0 | 0 | 0 | 1894200 | 619190 | 0 | 0 | 1517900 | 1037400 | 0 | 0 | 6469200 | 2847600 | 0 | 0 | 338400 | 482290 |
| AWGAIMTE | PACSIN1; PACS | Q9BY11; F6U | PACN1_HUMAI | Unmodified | 0 | 548250 | 0 | 0 | 0 | 0 | 0 | 579760 | 0 | 0 | 0 | 0 | 0 | 0 | 147020 | 0 | 143520 | 0 |
| AWGAIMTE | PACSIN1; PACS | Q9BY11; F6U | PACN1_HUMAI | Oxidation (M) | 13048000 | 12260000 | 0 | 2006700 | 0 | 0 | 43754 | 0 | 0 | 0 | 0 | 0 | 334360 | 0 | 1852800 | 990930 | 0 | 0 |
| AWGAIMTE | PACSIN1; PACS | Q9BY11; F6U | PACN1_HUMAI | Unmodified | 5528900 | 5610800 | 0 | 860560 | 0 | 0 | 0 | 0 | 0 | 0 | 0 | 1592500 | 0 | 0 | 512880 | 398630 | 0 | 0 |
| AWGAVVPI | CYRIB; CYRIB; ( | Q9NUQ9; E5 | CYRIB_HUMAN | Unmodified | 0 | 0 | 29866 | 0 | 0 | 0 | 0 | 24985 | 0 | 0 | 0 | 1781200 | 0 | 0 | 0 | 0 | 0 | 0 |
| AWGNNQD | SNAP25 | P60880 | SNP25_HUMAN | Unmodified | 149930 | 207710 | 0 | 0 | 0 | 0 | 39539 | 0 | 1095800 | 158900 | 0 | 0 | 0 | 0 | 0 | 0 | 0 | 0 |
| AWGPGLE( | FLNA; FLNA; FL | P21333; A0A | FLNA_HUMAN | Unmodified | 454510 | 0 | 712570 | 984170 | 1481100 | 403880 | 1797500 | 1638100 | 451380 | 196310 | 0 | 0 | 1308800 | 1495500 | 689830 | 885280 | 1891500 | 953360 |
| AWGPGLH | FLNB; FLNB | O75369; E7E | FLNB_HUMAN | Unmodified | 0 | 0 | 475100 | 252950 | 826940 | 238120 | 0 | 0 | 89194 | 0 | 111450 | 1315300 | 363120 | 0 | 233920 | 194000 | 242760 | 242190 |
| AWHAFFTA | PACSIN3; PACS | Q9UKS6; E9I | PACN3_HUMAI | Unmodified | 1547100 | 0 | 1003300 | 582780 | 0 | 0 | 0 | 0 | 0 | 0 | 0 | 235480 | 0 | 0 | 2016200 | 842210 | 0 | 0 |
| AWHAFFTA | PACSIN3; PACS | Q9UKS6; E9I | PACN3_HUMAI | Unmodified | 669310 | 1559400 | 75247 | 1207400 | 0 | 0 | 0 | 0 | 0 | 0 | 0 | 0 | 818700 | 217920 | 0 | 0 | 0 | 0 |
| AWHAFFTA | PACSIN3; PACS | Q9UKS6; E9I | PACN3_HUMAI | Unmodified | 0 | 0 | 0 | 832930 | 1430200 | 173730 | 0 | 0 | 0 | 137350 | 0 | 0 | 8914200 | 2323200 | 0 | 0 | 296440 | 399020 |
| AWITAPVAI | DSG2; DSG2 | Q14126; J3K | DSG2_HUMAN | Unmodified | 0 | 0 | 0 | 0 | 0 | 0 | 0 | 0 | 2281900 | 71912 | 0 | 351910 | 0 | 0 | 0 | 0 | 0 | 0 |
| AWITAPVAI | DSG2; DSG2 | Q14126; J3K | DSG2_HUMAN | Unmodified | 0 | 0 | 0 | 0 | 6632900 | 1452500 | 0 | 0 | 7447000 | 4219900 | 298060 | 0 | 0 | 1516000 | 0 | 0 | 305120 | 506140 |

| | | | | | | | | | | | | | | | | | | | | | | |
|---|---|---|---|---|---|---|---|---|---|---|---|---|---|---|---|---|---|---|---|---|---|---|
| AWKSGGAS | PCMT1; PCMT1 | P22061; F8W | PIMT_HUMAN I | Acetyl (Protein N | 0 | 0 | 0 | 0 | 492510 | 5929100 | 891440 | 0 | 0 | 540100 | 0 | 0 | 641380 | 735310 | 0 | 0 | 1753000 | 277200 |
| AWLMSDKT | MAN1A1 | P33908 | MA1A1_HUMAN | Unmodified | 0 | 0 | 0 | 0 | 0 | 0 | 0 | 0 | 555280 | 75906 | 195600 | 668310 | 0 | 0 | 0 | 0 | 0 | 0 |
| AWLTEKEE | UTRN | P46939 | UTRN_HUMAN | Unmodified | 0 | 0 | 0 | 868110 | 1960000 | 0 | 93106 | 0 | 0 | 757350 | 61049 | 0 | 276310 | 207540 | 0 | 0 | 442810 | 141210 |
| AWMAFMSI | PACSIN2; PACS | Q9UNF0; B0 | PACN2_HUMAN | Oxidation (M) | 1324100 | 349580 | 187820 | 251290 | 0 | 0 | 0 | 0 | 0 | 0 | 0 | 0 | 0 | 0 | 1649700 | 1989700 | 0 | 0 |
| AWMAFMSI | PACSIN2; PACS | Q9UNF0; B0 | PACN2_HUMAN | 2 Oxidation (M) | 475850 | 921190 | 0 | 0 | 0 | 0 | 0 | 0 | 0 | 0 | 236840 | 0 | 0 | 0 | 0 | 957090 | 0 | 0 |
| AWMAFMSI | PACSIN2; PACS | Q9UNF0; B0 | PACN2_HUMAN | 2 Oxidation (M) | 13128000 | 13786000 | 1860000 | 11499000 | 2997600 | 0 | 271080 | 0 | 0 | 790260 | 0 | 0 | 47099000 | 9136900 | 19624000 | 4938400 | 0 | 496790 |
| AWMAFMSI | PACSIN2; PACS | Q9UNF0; B0 | PACN2_HUMAN | Oxidation (M) | 3107900 | 0 | 0 | 0 | 364160 | 0 | 623900 | 0 | 0 | 480420 | 0 | 0 | 30593000 | 9312900 | 4614700 | 554110 | 208140 | 1029100 |
| AWMAFMSI | PACSIN2; PACS | Q9UNF0; B0 | PACN2_HUMAN | Unmodified | 212280 | 529760 | 0 | 289280 | 0 | 0 | 0 | 0 | 0 | 0 | 0 | 0 | 2304400 | 960930 | 485830 | 0 | 0 | 0 |
| AWNAYPY( | PITPNB; PITPN/ | P48739; Q0( | PIPNB_HUMAN | Unmodified | 0 | 0 | 0 | 0 | 0 | 0 | 0 | 0 | 0 | 213660 | 0 | 742250 | 668250 | 372560 | 572420 | 0 | 476210 | 0 |
| AWNDIKKV | RPL4; RPL4 | P36578; H3E | RL4_HUMAN 6( | Unmodified | 0 | 0 | 0 | 0 | 0 | 0 | 0 | 33029 | 0 | 0 | 0 | 313070 | 548560 | 155530 | 0 | 0 | 0 | 0 |
| AWNDTTKA | MAGED1 | Q9Y5V3 | MAGD1_HUMA | Unmodified | 0 | 0 | 0 | 0 | 0 | 0 | 0 | 0 | 0 | 0 | 0 | 0 | 1839200 | 0 | 0 | 0 | 286100 | 0 |
| AWNTMVLI | PYGL | P06737 | PYGL_HUMAN | Unmodified | 0 | 0 | 0 | 0 | 353800 | 0 | 0 | 0 | 0 | 0 | 0 | 0 | 0 | 0 | 0 | 0 | 0 | 239970 |
| AWNYELPA | PROM1; PROM1 | O43490; D6I | PROM1_HUMA | Unmodified | 0 | 0 | 0 | 0 | 0 | 0 | 0 | 0 | 0 | 0 | 0 | 9273800 | 0 | 0 | 0 | 0 | 0 | 0 |
| AWQYLSNS | TBC1D10B; TB( | Q4KMP7; H3 | TB10B_HUMAN | Unmodified | 0 | 0 | 0 | 0 | 875400 | 125840 | 113290 | 141470 | 0 | 173780 | 180650 | 629010 | 0 | 264490 | 0 | 0 | 0 | 289190 |
| AWRDPDEI | AKR1A1 | P14550 | AK1A1_HUMAN | Unmodified | 169190 | 0 | 153940 | 413110 | 80982 | 0 | 70735 | 0 | 1466000 | 173790 | 226440 | 1200100 | 0 | 0 | 1587200 | 550070 | 0 | 0 |
| AWSATFDE | SLC7A1 | P30825 | SL7A1_HUMAN | Unmodified | 299640 | 452550 | 1268500 | 1260600 | 0 | 0 | 0 | 0 | 1763100 | 2148000 | 0 | 6476500 | 0 | 0 | 14621000 | 2553100 | 0 | 0 |
| AWTVEQLR | FKBP3; FKBP3 | Q00688; G3 | FKBP3_HUMAN | Unmodified | 0 | 627720 | 0 | 0 | 0 | 0 | 0 | 0 | 475320 | 0 | 0 | 988240 | 0 | 0 | 922650 | 397240 | 0 | 0 |
| AWTVEQLR | FKBP3; FKBP3 | Q00688; G3 | FKBP3_HUMAN | Unmodified | 961430 | 944840 | 190830 | 165720 | 0 | 0 | 0 | 0 | 530040 | 0 | 279620 | 2692500 | 0 | 0 | 434840 | 520940 | 0 | 0 |
| AWTVEQLR | FKBP3; FKBP3 | Q00688; G3 | FKBP3_HUMAN | Unmodified | 1120000 | 0 | 0 | 0 | 0 | 0 | 0 | 0 | 3498600 | 0 | 1814000 | 4829500 | 0 | 0 | 0 | 0 | 0 | 0 |
| AWVESREK | CACYBP | Q9HB71 | CYBP_HUMAN | Unmodified | 51346 | 0 | 0 | 0 | 572760 | 0 | 213550 | 204540 | 0 | 192610 | 0 | 0 | 1332600 | 711920 | 24467 | 51248 | 364410 | 0 |
| AWVWNTH | RANBP1; RANB | P43487; F6V | RANG_HUMAN | Unmodified | 0 | 0 | 36062 | 0 | 0 | 0 | 0 | 0 | 0 | 0 | 0 | 1834800 | 0 | 0 | 860670 | 18903 | 0 | 0 |
| AYAALAALE | ILF3 | Q12906 | ILF3_HUMAN Ir | Unmodified | 2219400 | 2108100 | 1168700 | 650190 | 0 | 0 | 180860 | 0 | 1361800 | 277350 | 1305100 | 3894300 | 253300 | 0 | 559840 | 1103000 | 0 | 0 |
| AYAALTDEE | SEC63; SEC63 | Q9UGP8; A6 | SEC63_HUMAN | Unmodified | 0 | 0 | 0 | 0 | 0 | 629310 | 0 | 0 | 3776000 | 807290 | 0 | 0 | 0 | 0 | 0 | 0 | 0 | 0 |
| AYAALTDEE | SEC63; SEC63 | Q9UGP8; A6 | SEC63_HUMAN | Unmodified | 0 | 0 | 0 | 0 | 0 | 0 | 0 | 0 | 6947100 | 254430 | 0 | 291860 | 0 | 0 | 46306 | 0 | 0 | 0 |
| AYAANSHC | DPP3; DPP3; DI | Q9NY33; G3 | DPP3_HUMAN | Oxidation (M) | 40242 | 116800 | 0 | 0 | 602680 | 0 | 0 | 0 | 1257700 | 884260 | 1520100 | 4056100 | 1006700 | 153740 | 881150 | 629440 | 760810 | 0 |
| AYAANSHC | DPP3; DPP3; DI | Q9NY33; G3 | DPP3_HUMAN | Oxidation (M) | 0 | 0 | 0 | 0 | 0 | 0 | 0 | 0 | 0 | 0 | 630400 | 4646300 | 0 | 0 | 0 | 0 | 0 | 0 |
| AYADSYYYI | CCDC50 | Q8IVM0 | CCD50_HUMA | Oxidation (M) | 3347000 | 2596100 | 15372000 | 1636900 | 0 | 0 | 0 | 0 | 0 | 0 | 0 | 0 | 1031300 | 0 | 10695000 | 5521100 | 34980 | 0 |
| AYADSYYYI | CCDC50 | Q8IVM0 | CCD50_HUMA | Unmodified | 1914500 | 3000900 | 6566600 | 5455600 | 0 | 0 | 90953 | 76733 | 750050 | 120690 | 0 | 0 | 2550500 | 576300 | 9066700 | 2526600 | 0 | 252620 |
| AYADSYYYI | CCDC50 | Q8IVM0 | CCD50_HUMA | Oxidation (M) | 0 | 0 | 0 | 0 | 0 | 0 | 0 | 0 | 0 | 0 | 0 | 0 | 1315400 | 0 | 0 | 0 | 0 | 0 |
| AYADSYYYI | CCDC50 | Q8IVM0 | CCD50_HUMA | Unmodified | 0 | 0 | 0 | 0 | 0 | 0 | 0 | 0 | 0 | 0 | 0 | 0 | 744030 | 0 | 0 | 0 | 0 | 0 |
| AYALAFAEF | HSD17B4; HSD | P51659; A0A | DHB4_HUMAN | Unmodified | 0 | 0 | 56899 | 70569 | 111180 | 60938 | 205390 | 0 | 12905000 | 995730 | 1339100 | 340250 | 0 | 0 | 115740 | 94759 | 0 | 41839 |
| AYAPGGPA | COASY | Q13057 | COASY_HUMAN | Unmodified | 0 | 0 | 0 | 0 | 1379900 | 0 | 0 | 0 | 564590 | 0 | 0 | 878590 | 0 | 0 | 0 | 760240 | 0 | 2123200 |
| AYAQGISR | CS; CS; CS; CS; | O75390; F8V | CISY_HUMAN ( | Unmodified | 0 | 0 | 46473 | 0 | 0 | 0 | 0 | 136800 | 9929000 | 722870 | 153500 | 399000 | 0 | 0 | 73137 | 68163 | 5766.9 | 0 |
| AYAQQLAD | PACSIN3; PACS | Q9UKS6; E9I | PACN3_HUMAN | Unmodified | 10527000 | 11992000 | 5498900 | 3924400 | 0 | 0 | 0 | 0 | 0 | 118450 | 0 | 0 | 1871600 | 1674600 | 10120000 | 4717100 | 417890 | 0 |
| AYAQQLAD | PACSIN3; PACS | Q9UKS6; E9I | PACN3_HUMAN | Unmodified | 13732000 | 12815000 | 7477600 | 8128300 | 4006200 | 1125100 | 69905 | 0 | 646730 | 647450 | 0 | 0 | 18555000 | 8304800 | 7081700 | 5427800 | 1831700 | 2221500 |
| AYAQQLTE | PACSIN2; PACS | Q9UNF0; B0 | PACN2_HUMAN | Unmodified | 21492000 | 19233000 | 21984000 | 11959000 | 2135500 | 0 | 0 | 0 | 954570 | 529880 | 504900 | 4521600 | 17713000 | 5883600 | 42998000 | 24232000 | 0 | 926470 |
| AYAQQLTE | PACSIN2; PACS | Q9UNF0; B0 | PACN2_HUMAN | Unmodified | 47466000 | 40414000 | 25874000 | 40005000 | 13653000 | 0 | 0 | 0 | 1722800 | 885470 | 1927700 | 7504800 | 92059000 | 24404000 | 59939000 | 28060000 | 2050300 | 4884300 |
| AYARIGNS | STIP1 | P31948 | STIP1_HUMAN | Unmodified | 0 | 0 | 0 | 0 | 836740 | 0 | 419200 | 186900 | 0 | 0 | 0 | 0 | 460690 | 858170 | 0 | 0 | 2030200 | 0 |
| AYASNHPII | CPD | O75976 | CBPD_HUMAN | Unmodified | 0 | 0 | 0 | 0 | 826750 | 0 | 0 | 0 | 5386800 | 1927300 | 0 | 0 | 136220 | 0 | 0 | 0 | 0 | 0 |
| AYASPAKLE | GATD1; GATD1 | Q8NB37; E9I | GALD1_HUMAN | Unmodified | 0 | 0 | 0 | 0 | 299140 | 0 | 461610 | 348610 | 541200 | 0 | 0 | 688730 | 0 | 552950 | 412520 | 155380 | 0 | 325620 |
| AYAVILTTG | PTPRJ; PTPRJ; P | Q12913; A0A | PTPRJ_HUMAN | Unmodified | 1061600 | 548900 | 468880 | 653900 | 2880800 | 0 | 207710 | 0 | 8450600 | 900030 | 7133800 | 0 | 1106500 | 0 | 0 | 884120 | 0 | 841190 |
| AYAVILTTG | PTPRJ; PTPRJ; P | Q12913; A0A | PTPRJ_HUMAN | Unmodified | 0 | 0 | 0 | 0 | 7549100 | 1239000 | 61088 | 0 | 0 | 0 | 942960 | 0 | 1698000 | 1580400 | 0 | 0 | 2587600 | 2072800 |
| AYAYLFK | RAB2A; RAB2A | P61019; E9P | RAB2A_HUMAN | Acetyl (Protein N | 225780 | 0 | 190370 | 156430 | 376280 | 0 | 1198500 | 401180 | 2126300 | 2307400 | 380850 | 2567000 | 262320 | 447930 | 0 | 311400 | 0 | 0 |
| AYCERQGL | SUMO3; SUMO | P55854; P61 | SUMO3_HUMA | Unmodified | 0 | 0 | 0 | 0 | 3103400 | 0 | 145000 | 109240 | 0 | 0 | 217720 | 0 | 800880 | 0 | 0 | 0 | 2375200 | 2434000 |
| AYDATHLVI | ESD; ESD; ESD; | P10768; X6R | ESTD_HUMAN | Unmodified | 181380 | 0 | 0 | 0 | 0 | 0 | 229860 | 186140 | 276600 | 172510 | 216260 | 1229900 | 0 | 0 | 188860 | 115520 | 0 | 0 |
| AYDGRFKD | IDH2; IDH2 | P48735; H0Y | IDHP_HUMAN | Unmodified | 0 | 0 | 0 | 0 | 0 | 0 | 651210 | 0 | 456570 | 5541400 | 0 | 0 | 140810 | 207620 | 0 | 0 | 0 | 0 |
| AYDHLFK | RAB13 | P51153 | RAB13_HUMAN | Unmodified | 665290 | 1111200 | 1043700 | 0 | 654950 | 0 | 0 | 120990 | 0 | 84025 | 62351 | 402360 | 527730 | 799290 | 784070 | 757230 | 110870 | 145930 |
| AYDLLSK | PLCB2 | Q00722 | PLCB2_HUMAN | Unmodified | 0 | 0 | 0 | 0 | 0 | 0 | 267300 | 0 | 1371400 | 78637 | 863960 | 2903700 | 0 | 0 | 0 | 0 | 89468 | 0 |
| AYDPDYER | TJP2; TJP2; TJP2 | Q9UDY2; A0 | ZO2_HUMAN T | Unmodified | 0 | 0 | 0 | 52884 | 40884 | 0 | 0 | 0 | 1994100 | 80926 | 44953 | 333070 | 0 | 0 | 40830 | 39568 | 0 | 0 |
| AYDPDYER | TJP2; TJP2; TJP2 | Q9UDY2; A0 | ZO2_HUMAN T | Unmodified | 0 | 0 | 0 | 0 | 460440 | 0 | 0 | 0 | 0 | 202300 | 530180 | 0 | 0 | 0 | 0 | 0 | 19864 | 104140 |
| AYDYLVQN | TPP2; TPP2; TPI | P29144; A0A | TPP2_HUMAN | Unmodified | 0 | 0 | 0 | 40256 | 0 | 0 | 0 | 0 | 0 | 0 | 1949200 | 6617300 | 0 | 0 | 71893 | 51882 | 0 | 0 |
| AYEAAASAI | TMPO | P42166 | LAP2A_HUMAN | Unmodified | 1255000 | 0 | 0 | 0 | 0 | 0 | 341940 | 53454 | 0 | 0 | 0 | 903500 | 352320 | 0 | 0 | 0 | 0 | 350790 |
| AYEDGCKT | STIP1; STIP1; ST | P31948; F5H | STIP1_HUMAN | Unmodified | 0 | 0 | 0 | 0 | 231500 | 0 | 782220 | 227020 | 0 | 0 | 0 | 0 | 365900 | 539980 | 0 | 0 | 2957800 | 241450 |
| AYEDQTKP | AK3 | Q9UIJ7 | KAD3_HUMAN | Unmodified | 0 | 0 | 0 | 0 | 0 | 0 | 132470 | 78729 | 2449500 | 1140800 | 90259 | 0 | 0 | 0 | 0 | 0 | 0 | 0 |
| AYEDQTKP | AK3 | Q9UIJ7 | KAD3_HUMAN | Unmodified | 0 | 0 | 0 | 0 | 0 | 0 | 0 | 0 | 1324400 | 494760 | 0 | 0 | 0 | 0 | 0 | 0 | 0 | 0 |
| AYEFAER | CLTCL1; CLTC; | P53675; Q0( | CLH2_HUMAN | Unmodified | 8113700 | 6451000 | 7330700 | 1297600 | 2109300 | 0 | 1342900 | 0 | 44397 | 97750 | 216390 | 62538000 | 0 | 0 | 4287100 | 7017500 | 132120 | 3891300 |
| AYEFAERCI | CLTC; CLTC | Q00610; A0A | CLH1_HUMAN | Unmodified | 0 | 0 | 0 | 642150 | 0 | 0 | 0 | 0 | 0 | 414470 | 0 | 0 | 714840 | 0 | 0 | 0 | 0 | 0 |
| AYELALYLR | PFN2; PFN2 | G5E9Q6; C9 | G5E9Q6_HUM/ | Unmodified | 0 | 0 | 0 | 0 | 1262900 | 0 | 0 | 0 | 715290 | 0 | 0 | 0 | 0 | 294830 | 0 | 0 | 401470 | 0 |
| AYEQAGMI | NAPG; NAPG | Q99747; J3K | SNAG_HUMAN | Unmodified | 0 | 0 | 63901 | 0 | 3529700 | 84563 | 149540 | 0 | 0 | 0 | 0 | 0 | 0 | 29083 | 0 | 0 | 1323800 | 1457800 |
| AYEQAGMI | NAPG; NAPG | Q99747; J3K | SNAG_HUMAN | Oxidation (M) | 0 | 0 | 7128100 | 0 | 1651800 | 0 | 0 | 0 | 1678100 | 0 | 20603000 | 0 | 804500 | 658850 | 0 | 0 | 0 | 0 |
| AYESSRADI | NOXO1 | Q8NFA2 | NOXO1_HUMA | Unmodified | 0 | 0 | 0 | 0 | 0 | 775110 | 0 | 164890 | 0 | 161810 | 873100 | 1146000 | 0 | 0 | 0 | 0 | 0 | 0 |
| AYEVLKDEI | DNAJC10; DNA | Q8IXB1; A0A | DJC10_HUMAN | Unmodified | 40055 | 0 | 0 | 0 | 234030 | 0 | 148750 | 67061 | 2083000 | 81000 | 0 | 0 | 0 | 0 | 0 | 0 | 0 | 0 |
| AYEVLKDEI | DNAJC10; DNA | Q8IXB1; A0A | DJC10_HUMAN | Unmodified | 0 | 0 | 0 | 0 | 0 | 0 | 0 | 0 | 3480900 | 0 | 0 | 771630 | 0 | 0 | 0 | 0 | 0 | 0 |
| AYFETEK | GSPT1; GSPT1; | P15170; H3E | ERF3A_HUMAN | Unmodified | 0 | 0 | 0 | 0 | 75651 | 0 | 9641.1 | 0 | 345780 | 85542 | 0 | 1343500 | 0 | 15207 | 0 | 0 | 0 | 0 |
| AYFHLLNQ | PLS1; PLS3; PLS | Q14651; P13 | PLSI_HUMAN F | Unmodified | 1817500 | 710840 | 306290 | 378260 | 2583500 | 2178100 | 377960 | 0 | 2820400 | 0 | 192580 | 2896800 | 870080 | 204040 | 301860 | 472610 | 2007600 | 459980 |
| AYGDFAWS | SGPL1; SGPL1; | O95470; H7I | SGPL1_HUMAN | Unmodified | 0 | 0 | 0 | 0 | 0 | 0 | 0 | 0 | 1005500 | 52647 | 0 | 0 | 0 | 0 | 0 | 0 | 0 | 0 |
| AYGENIGY | ATP1B1 | P05026 | AT1B1_HUMAN | Unmodified | 1798900 | 1519900 | 15296000 | 13619000 | 68068000 | 0 | 21372000 | 8403600 | 29214000 | 9916500 | 19426000 | 38548000 | 40671000 | 27097000 | 11665000 | 11034000 | 15910000 | 25996000 |
| AYGFRGPG | CHGA; CHGA | P10645; G5E | CMGA_HUMAN | Unmodified | 0 | 0 | 0 | 0 | 0 | 0 | 0 | 0 | 1489000 | 1136200 | 0 | 0 | 0 | 0 | 0 | 0 | 0 | 0 |
| AYGFSCDG | C12orf57; C12o | Q99622; U3I | C10_HUMAN P | Unmodified | 148800 | 0 | 0 | 0 | 744380 | 0 | 462850 | 277880 | 0 | 215320 | 0 | 501420 | 900590 | 1738400 | 204100 | 113360 | 785420 | 617620 |
| AYGGSMC/ | RPL34 | P49207 | RL34_HUMAN ( | Unmodified | 2730800 | 1628000 | 2903500 | 2225400 | 0 | 0 | 0 | 249610 | 0 | 0 | 0 | 3831800 | 865090 | 0 | 2290800 | 1298800 | 0 | 0 |
| AYGPGIEPT | FLNA; FLNA; FL | P21333; A0A | FLNA_HUMAN | Oxidation (M) | 0 | 0 | 0 | 662570 | 0 | 0 | 913600 | 297700 | 0 | 0 | 0 | 0 | 611010 | 286070 | 0 | 442390 | 322040 | 0 |
| AYGPGIEPT | FLNA; FLNA; FL | P21333; A0A | FLNA_HUMAN | Unmodified | 446160 | 776010 | 0 | 524710 | 2432900 | 299370 | 1246300 | 363610 | 117160 | 0 | 0 | 0 | 530740 | 704940 | 0 | 0 | 947000 | 798350 |
| AYGPGIEPT | FLNA; FLNA; FL | P21333; A0A | FLNA_HUMAN | Unmodified | 308760 | 491840 | 0 | 1701300 | 0 | 0 | 0 | 343930 | 55035 | 0 | 0 | 596100 | 0 | 0 | 0 | 0 | 516660 | 0 |
| AYGPGLEK | FLNB; FLNB | O75369; E7E | FLNB_HUMAN | Unmodified | 0 | 0 | 0 | 0 | 211000 | 0 | 0 | 0 | 357050 | 0 | 0 | 978110 | 0 | 0 | 0 | 87071 | 16559 | 32617 |
| AYGQALAK | TKT; TKT | P29401; A0A | TKT_HUMAN Tr | Unmodified | 0 | 0 | 0 | 0 | 15837000 | 17853 | 165200 | 28940 | 91894 | 2313200 | 0 | 1677000 | 0 | 0 | 0 | 0 | 484630 | 1826400 |
| AYGQQLTD | PACSIN1; PACS | Q9BY11; F6I | PACN1_HUMAN | Unmodified | 7441700 | 6237700 | 1186800 | 1959000 | 0 | 0 | 0 | 0 | 0 | 0 | 0 | 485660 | 5017500 | 1268700 | 1876100 | 1342800 | 0 | 0 |
| AYGQQLTD | PACSIN1; PACS | Q9BY11; F6I | PACN1_HUMAN | Unmodified | 1854600 | 974430 | 0 | 0 | 36765 | 0 | 0 | 0 | 224740 | 319780 | 0 | 0 | 134200 | 0 | 319010 | 0 | 0 | 0 |
| AYHEQLSV | TUBA3C; TUBA | P0DPH7; Q7 | TBA3C_HUMAN | Oxidation (M) | 7053700 | 1449400 | 4759700 | 8397200 | 26682000 | 7581800 | 50400000 | 10232000 | 18013000 | 23271000 | 18824000 | 55209000 | 28353000 | 21247000 | 5986000 | 5971200 | 18768000 | 25201000 |
| AYHEQLSV | TUBA3C; TUBA | P0DPH7; Q7 | TBA3C_HUMAN | Unmodified | 0 | 0 | 0 | 0 | 14809000 | 1614400 | 23858000 | 331100 | 41695000 | 10976000 | 2792900 | 12049000 | 13417000 | 5738400 | 0 | 1041300 | 7196100 | 18580000 |
| AYHEQLSV | TUBA3C; TUBA | P0DPH7; Q7 | TBA3C_HUMAN | Oxidation (M) | 0 | 0 | 0 | 0 | 0 | 0 | 1174800 | 0 | 0 | 0 | 0 | 0 | 0 | 0 | 0 | 0 | 1185200 | 0 |
| AYHEQLSV | TUBA3C; TUBA | P0DPH7; Q7 | TBA3C_HUMAN | Unmodified | 0 | 0 | 0 | 0 | 0 | 0 | 2222100 | 0 | 0 | 0 | 0 | 0 | 0 | 0 | 0 | 0 | 0 | 0 |
| AYHEQLTV | TUBA1C; TUBA | Q9BQE3; A0 | TBA1C_HUMAN | Oxidation (M) | 881350 | 0 | 0 | 1015000 | 8268100 | 2471800 | 1653300 | 0 | 2683800 | 1506700 | 1203500 | 5002800 | 2095600 | 1967000 | 0 | 0 | 3440500 | 8601000 |
| AYHEQLTV | TUBA1C; TUBA | Q9BQE3; A0 | TBA1C_HUMAN | Unmodified | 0 | 0 | 0 | 0 | 5462100 | 0 | 0 | 0 | 0 | 685640 | 0 | 849300 | 1022600 | 0 | 0 | 0 | 1353200 | 3707900 |
| AYHLQDD( | ERLEC1 | Q96DZ1 | ERLEC_HUMAN | Unmodified | 0 | 0 | 0 | 0 | 0 | 0 | 0 | 0 | 8260900 | 1051500 | 315340 | 0 | 0 | 0 | 0 | 0 | 0 | 0 |
| AYHPHCFT | ZYX; ZYX | Q15942; H0 | ZYX_HUMAN Z | Unmodified | 0 | 0 | 0 | 0 | 0 | 0 | 0 | 0 | 0 | 0 | 0 | 0 | 0 | 0 | 0 | 0 | 1592800 | 0 |
| AYHSFLVEI | ARPC1B; ARPC | O15143; C9J | ARC1B_HUMAN | Acetyl (Protein N | 3176100 | 2048700 | 0 | 0 | 14268000 | 4839600 | 6809400 | 730660 | 0 | 1168400 | 0 | 12116000 | 4487900 | 5091500 | 0 | 0 | 9538500 | 7405900 |
| AYIATQGPL | PTPRF; PTPRF; | P10586; H0Y | PTPRF_HUMAN | Unmodified | 629010 | 805420 | 0 | 765090 | 0 | 0 | 0 | 0 | 1048400 | 0 | 223500 | 417150 | 0 | 0 | 2353800 | 1195000 | 0 | 268490 |
| AYIATQGPL | PTPRG | P23470 | PTPRG_HUMAN | Unmodified | 0 | 0 | 425420 | 154780 | 947530 | 243350 | 126930 | 0 | 0 | 0 | 235440 | 446950 | 0 | 0 | 0 | 538120 | 0 | 0 |
| AYIDSFR | DARS1; DARS1 | P14868; H7E | SYDC_HUMAN | Unmodified | 0 | 0 | 0 | 0 | 0 | 0 | 0 | 0 | 0 | 419090 | 124410 | 284170 | 4057300 | 72302 | 0 | 0 | 0 | 30671 | 159590 |
| AYIKDCVSI | VPS28; VPS28 | Q9UK41; E9I | VPS28_HUMAN | Unmodified | 0 | 0 | 0 | 450600 | 1204400 | 313380 | 313190 | 144450 | 0 | 665540 | 0 | 0 | 6560000 | 4601700 | 0 | 0 | 1005200 | 1124700 |
| AYIKDCVSI | VPS28; VPS28 | Q9UK41; E9I | VPS28_HUMAN | Unmodified | 0 | 0 | 0 | 0 | 0 | 0 | 0 | 0 | 0 | 0 | 0 | 0 | 2698600 | 1837700 | 0 | 0 | 1803900 | 799210 |
| AYILNLVK | CCT4 | P50991 | TCPD_HUMAN | Unmodified | 251980 | 0 | 0 | 0 | 0 | 0 | 0 | 378850 | 0 | 0 | 568320 | 23821000 | 1384900 | 0 | 0 | 372780 | 0 | 0 |
| AYINKVEEL | DBI; DBI; DBI | P07108; B8Z | ACBP_HUMAN | Unmodified | 284620 | 546630 | 544910 | 0 | 0 | 0 | 0 | 93045 | 1584800 | 0 | 1096500 | 4502300 | 0 | 0 | 673160 | 724450 | 0 | 0 |
| AYINKVEEL | DBI; DBI; DBI | P07108; B8Z | ACBP_HUMAN | Unmodified | 880500 | 891550 | 989580 | 0 | 0 | 118900 | 0 | 0 | 1348800 | 743250 | 913270 | 1827000 | 0 | 0 | 1482700 | 2396100 | 0 | 0 |
| AYIQENLEL | MTHFD1; MTHF | P11586; V9G | C1TC_HUMAN | Unmodified | 0 | 0 | 0 | 0 | 2065000 | 0 | 2440300 | 147140 | 813430 | 919610 | 472200 | 2356200 | 575360 | 0 | 0 | 0 | 3750400 | 661270 |
| AYIQENLEL | MTHFD1; MTHF | P11586; V9G | C1TC_HUMAN | Unmodified | 0 | 0 | 0 | 0 | 0 | 0 | 1281500 | 0 | 0 | 0 | 0 | 1552600 | 123500 | 0 | 0 | 0 | 1536300 | 103300 |
| AYIVQLQIE | SF1; SF1; SF1 | Q15637; A0A | SF01_HUMAN | Unmodified | 0 | 0 | 0 | 0 | 0 | 0 | 0 | 0 | 0 | 130390 | 0 | 670670 | 0 | 0 | 0 | 0 | 0 | 0 |
| AYKTTWGD | SNAP23; SNAP2 | O00161; H3I | SNP23_HUMAN | Unmodified | 0 | 0 | 0 | 0 | 220230 | 0 | 0 | 0 | 0 | 0 | 0 | 0 | 728650 | 0 | 0 | 0 | 762460 | 1540200 |
| AYKTTWGD | SNAP23; SNAP2 | O00161; H3I | SNP23_HUMAN | Unmodified | 0 | 0 | 0 | 0 | 0 | 0 | 0 | 0 | 0 | 0 | 0 | 0 | 0 | 0 | 0 | 0 | 1726400 | 0 |
| AYLEEECP/ | AZGP1; AZGP1 | P25311; C9J | ZA2G_HUMAN | Unmodified | 58170 | 0 | 603570 | 670870 | 0 | 29158 | 0 | 0 | 0 | 0 | 251910 | 807070 | 0 | 0 | 989560 | 238790 | 0 | 0 |
| AYLEGLCVI | HLA-C; HLA-C; | Q53YP1; A0/ | Q53YP1_HUM/ | Unmodified | 0 | 4906000 | 0 | 0 | 14274 | 0 | 0 | 0 | 453580 | 75163 | 140200 | 5565900 | 0 | 0 | 0 | 0 | 0 | 0 |
| AYLEGTCVI | HLA-C; HLA-C; | P10321; Q5( | HLAC_HUMAN | Unmodified | 0 | 0 | 0 | 0 | 660690 | 34118 | 0 | 0 | 8739200 | 816590 | 557500 | 7478600 | 0 | 0 | 0 | 0 | 0 | 2683100 |
| AYLEGTCVI | HLA-C; HLA-C; | P10321; Q5( | HLAC_HUMAN | Unmodified | 0 | 0 | 0 | 0 | 4049800 | 0 | 0 | 0 | 29186000 | 0 | 2106900 | 36925000 | 0 | 0 | 1295800 | 1480400 | 298230 | 782150 |
| AYLESFYK | ATP6V0D1; ATP | P61421; R4C | VA0D1_HUMAN | Unmodified | 123690 | 399250 | 181090 | 49330 | 0 | 0 | 274030 | 301910 | 1348000 | 849360 | 509350 | 0 | 0 | 0 | 1841500 | 558840 | 0 | 0 |
| AYLLGKED/ | SQSTM1; SQSTI | Q13501; C9J | SQSTM_HUMAN | Unmodified | 1032900 | 441810 | 441780 | 1379300 | 89438 | 0 | 150890 | 0 | 3461800 | 449050 | 0 | 259700 | 285010 | 867690 | 423430 | 1047400 | 227770 | 562300 |
| AYLLGKED/ | SQSTM1; SQSTI | Q13501; C9J | SQSTM_HUMAN | Unmodified | 0 | 0 | 0 | 198060 | 2620200 | 515250 | 688910 | 94609 | 2402700 | 644040 | 0 | 0 | 675400 | 680150 | 0 | 0 | 1312200 | 2733900 |
| AYLPTGKQ | ATG3 | Q9NT62 | ATG3_HUMAN | Unmodified | 0 | 0 | 0 | 0 | 0 | 0 | 208210 | 0 | 0 | 0 | 0 | 0 | 0 | 79488 | 0 | 0 | 368970 | 0 |
| AYLQQLR | ARPC3 | O15145 | ARPC3_HUMAN | Unmodified | 2064100 | 44418 | 395210 | 0 | 0 | 0 | 0 | 0 | 830330 | 0 | 188910 | 22097000 | 0 | 0 | 0 | 2222300 | 0 | 0 |
| AYLQQLRQ | ARPC3 | O15145 | ARPC3_HUMAN | Unmodified | 1146300 | 1271000 | 1202800 | 3200000 | 248460 | 0 | 0 | 0 | 2107700 | 0 | 982250 | 3627400 | 406060 | 0 | 976840 | 839990 | 0 | 0 |

| | | | | | | | | | | | | | | | | | | | | | | |
|---|---|---|---|---|---|---|---|---|---|---|---|---|---|---|---|---|---|---|---|---|---|---|
| AYLRDFMY | HADHB; HADHI | P55084; F5G | ECHB_HUMAN | Oxidation (M) | 0 | 0 | 0 | 0 | 0 | 0 | 0 | 0 | 1816600 | 1009000 | 0 | 0 | 0 | 0 | 0 | 0 | 0 | 0 |
| AYLSQGER | PLCB2; PLCB2 | Q00722; Q9I | PLCB2_HUMAN | Unmodified | 116440 | 0 | 202040 | 66927 | 101960 | 403960 | 104630 | 22446 | 2391300 | 428200 | 0 | 2891600 | 0 | 150650 | 0 | 0 | 0 | 0 |
| AYLSSGGG | ARVCF; ARVCF; | O00192; E9F | ARVC_HUMAN | Unmodified | 0 | 0 | 164660 | 820700 | 0 | 0 | 0 | 0 | 464540 | 0 | 0 | 161300 | 0 | 0 | 174450 | 535020 | 41281 | 0 |
| AYMAFGLV | N/A | N/A | N/A | Unmodified | 1681300 | 1509700 | 0 | 0 | 906280 | 0 | 0 | 0 | 0 | 0 | 0 | 0 | 0 | 0 | 3981000 | 580240 | 0 | 0 |
| AYNGENW | CRTAP; CRTAP | O75718; C9J | CRTAP_HUMAN | Unmodified | 0 | 0 | 0 | 0 | 0 | 0 | 0 | 0 | 895160 | 0 | 0 | 0 | 0 | 0 | 0 | 0 | 0 | 0 |
| AYNHYFDL | PALS2; PALS2 | Q9NZW5; B8 | PALS2_HUMAN | Unmodified | 0 | 0 | 471720 | 751270 | 2367500 | 899220 | 0 | 0 | 0 | 171010 | 1008300 | 117820 | 0 | 0 | 0 | 0 | 354370 | 2942800 |
| AYNTLIK | CRAT; CRAT; CI | P43155; A0A | CACP_HUMAN | Unmodified | 134930 | 445280 | 0 | 438050 | 63307 | 995470 | 47437 | 180950 | 951810 | 288740 | 0 | 0 | 465610 | 85982 | 135370 | 690690 | 101850 | 0 |
| AYNYVAQE | UGGT1 | Q9NYU2 | UGGG1_HUMA | Unmodified | 0 | 0 | 0 | 0 | 0 | 0 | 0 | 0 | 24449000 | 5383500 | 672360 | 532720 | 0 | 0 | 0 | 0 | 0 | 0 |
| AYPTSVCF | PPWD1; PPWD | Q96BP3; H0 | PPWD1_HUMA | Unmodified | 0 | 0 | 753900 | 594070 | 0 | 3278800 | 0 | 0 | 0 | 0 | 0 | 0 | 0 | 0 | 0 | 0 | 0 | 0 |
| AYQCVVLL | RARS1 | P54136 | SYRC_HUMAN | Unmodified | 0 | 0 | 0 | 0 | 190450 | 0 | 0 | 0 | 0 | 0 | 112370 | 925240 | 0 | 0 | 0 | 0 | 0 | 0 |
| AYQCVVLL | RARS1 | P54136 | SYRC_HUMAN | Unmodified | 0 | 0 | 0 | 0 | 0 | 0 | 0 | 0 | 206030 | 0 | 0 | 2755200 | 1013600 | 367260 | 0 | 0 | 0 | 0 |
| AYQDRLAY | IQGAP1; IQGAP | P46940; A0A | IQGA1_HUMAN | Unmodified | 156960 | 119610 | 0 | 386490 | 3206100 | 823620 | 498270 | 556670 | 945590 | 485950 | 0 | 0 | 1787400 | 0 | 124720 | 0 | 233880 | 432880 |
| AYQDRYQI | NDUFB10; NDU | O96000; H3I | NDUBA_HUMA | Unmodified | 0 | 0 | 0 | 0 | 305610 | 0 | 571240 | 0 | 491710 | 523310 | 0 | 0 | 0 | 513250 | 0 | 0 | 0 | 0 |
| AYQEELSRI | TRIOBP | Q9H2D6 | TARA_HUMAN | Unmodified | 0 | 0 | 92174 | 142800 | 705580 | 249200 | 105690 | 874000 | 0 | 141270 | 0 | 0 | 633050 | 653210 | 75981 | 97074 | 602370 | 1266900 |
| AYQGVAAP | VGF; VGF | O15240; Q0 | VGF_HUMAN N | Unmodified | 1234500 | 775850 | 0 | 0 | 0 | 0 | 1888900 | 2387600 | 11860000 | 4840500 | 0 | 0 | 0 | 0 | 0 | 0 | 0 | 0 |
| AYQHGGV | UPF1 | Q92900 | RENT1_HUMAN | Unmodified | 2015800 | 1334900 | 1078400 | 812130 | 0 | 0 | 0 | 0 | 0 | 0 | 0 | 0 | 273400 | 0 | 0 | 0 | 0 | 0 |
| AYQKQPTII | RPS11; RPS11; | P62280; M0I | RS11_HUMAN | Unmodified | 0 | 0 | 0 | 217820 | 2220700 | 187980 | 615730 | 399800 | 0 | 492300 | 364900 | 0 | 11644000 | 158650 | 0 | 0 | 755900 | 1475000 |
| AYQNWVK | ECE1; ECE1 | P42892; A0A | ECE1_HUMAN | Unmodified | 0 | 0 | 0 | 0 | 1547100 | 310050 | 0 | 0 | 5260400 | 610920 | 0 | 0 | 0 | 0 | 0 | 0 | 0 | 0 |
| AYQNYIK | MME; MME; MM | P08473; A0A | NEP_HUMAN N | Unmodified | 252850 | 193100 | 14497000 | 19983000 | 5950800 | 1780600 | 513690 | 239170 | 9795600 | 725420 | 3456300 | 23063000 | 6842300 | 2562800 | 18215000 | 14255000 | 2114300 | 4348300 |
| AYQNYIKK | MME; MME; MM | P08473; A0A | NEP_HUMAN N | Unmodified | 0 | 0 | 1099600 | 2521700 | 0 | 0 | 0 | 0 | 435120 | 170340 | 82222 | 1866400 | 0 | 0 | 2536200 | 1679600 | 0 | 0 |
| AYSDPSTG | PLEC; PLEC | Q15149; H0 | PLEC_HUMAN | Unmodified | 0 | 0 | 0 | 0 | 623260 | 0 | 0 | 0 | 0 | 0 | 0 | 0 | 0 | 0 | 0 | 0 | 2600800 | 2810600 |
| AYSDPSTG | PLEC; PLEC | Q15149; H0 | PLEC_HUMAN | Unmodified | 0 | 0 | 0 | 0 | 0 | 0 | 0 | 0 | 0 | 0 | 0 | 0 | 0 | 0 | 0 | 0 | 16866000 | 6387000 |
| AYSDQAIV | GCN1 | Q92616 | GCN1_HUMAN | Unmodified | 0 | 0 | 213570 | 0 | 0 | 0 | 0 | 0 | 0 | 0 | 0 | 1311300 | 0 | 0 | 0 | 0 | 0 | 0 |
| AYSEAHEIS | YWHAG | P61981 | 1433G_HUMAN | Unmodified | 0 | 85914 | 1573300 | 0 | 2230600 | 698700 | 201440 | 0 | 0 | 155640 | 2105600 | 7860800 | 0 | 2627600 | 117590 | 4674500 | 1974600 | 0 |
| AYSEAHEIS | YWHAG | P61981 | 1433G_HUMAN | Unmodified | 0 | 33338 | 0 | 293510 | 0 | 0 | 0 | 0 | 0 | 0 | 78884 | 790860 | 0 | 203310 | 0 | 0 | 0 | 0 |
| AYSMAKYL | PFN2; PFN2; PF | P35080; C9J | PROF2_HUMAN | Unmodified | 0 | 0 | 0 | 0 | 77760 | 0 | 0 | 0 | 0 | 0 | 0 | 0 | 0 | 24839 | 0 | 0 | 260950 | 0 |
| AYSNWPTY | GLRX3 | O76003 | GLRX3_HUMAN | Unmodified | 0 | 0 | 0 | 0 | 0 | 0 | 0 | 0 | 62254 | 0 | 0 | 2238400 | 0 | 0 | 0 | 0 | 0 | 0 |
| AYSQEEITC | GPRC5A | Q8NFJ5 | RAI3_HUMAN F | Unmodified | 0 | 0 | 0 | 0 | 5328000 | 0 | 0 | 0 | 0 | 0 | 0 | 0 | 0 | 0 | 0 | 0 | 0 | 0 |
| AYSSFNPA | CPE | P16870 | CBPE_HUMAN | Oxidation (M) | 0 | 0 | 0 | 0 | 173320 | 0 | 0 | 0 | 3997700 | 0 | 0 | 0 | 0 | 0 | 0 | 0 | 0 | 177030 |
| AYSSFNPA | CPE | P16870 | CBPE_HUMAN | Unmodified | 0 | 0 | 0 | 0 | 0 | 0 | 0 | 0 | 21719000 | 577990 | 0 | 0 | 0 | 0 | 0 | 0 | 0 | 0 |
| AYSSFNPA | CPE | P16870 | CBPE_HUMAN | Oxidation (M) | 0 | 0 | 0 | 0 | 0 | 0 | 240780 | 176680 | 556990 | 0 | 0 | 0 | 0 | 269400 | 0 | 0 | 0 | 0 |
| AYSSFNPA | CPE | P16870 | CBPE_HUMAN | Unmodified | 0 | 0 | 0 | 0 | 0 | 0 | 0 | 0 | 4643500 | 0 | 0 | 0 | 0 | 0 | 0 | 0 | 0 | 0 |
| AYTEVAAA( | EVPL; EVPL | Q92817; K7E | EVPL_HUMAN | Unmodified | 0 | 0 | 0 | 0 | 3443100 | 391280 | 0 | 0 | 375030 | 0 | 0 | 0 | 0 | 259040 | 0 | 0 | 0 | 0 |
| AYTGREFD | ATP2A3 | Q93084 | AT2A3_HUMAN | Unmodified | 0 | 0 | 0 | 0 | 0 | 0 | 0 | 0 | 0 | 0 | 0 | 1553000 | 0 | 0 | 0 | 0 | 0 | 0 |
| AYTNFDAE | ANXA2P2; ANX | A6NMY6; P0 | AXA2L_HUMAN | Unmodified | 0 | 0 | 0 | 0 | 293510 | 0 | 0 | 30152 | 496190 | 0 | 0 | 0 | 0 | 275270 | 0 | 0 | 0 | 19642 |
| AYTNFDAE | ANXA2P2; ANX | A6NMY6; P0 | AXA2L_HUMAN | Unmodified | 6176100 | 0 | 0 | 3854000 | 3.87E+08 | 80865000 | 11448000 | 91195 | 61399000 | 44685000 | 0 | 0 | 1719200 | 1726900 | 1283600 | 500110 | 1.15E+08 | 1.03E+08 |
| AYTNFDAE | ANXA2P2; ANX | A6NMY6; P0 | AXA2L_HUMAN | Unmodified | 0 | 0 | 0 | 122010 | 1.92E+08 | 33910000 | 7471500 | 105690 | 1095800 | 16181000 | 0 | 0 | 0 | 2223600 | 0 | 0 | 76271000 | 41705000 |
| AYVAFPDFI | MGAM2; MGAM | Q2M2H8; O4 | MGAL_HUMAN | Unmodified | 0 | 0 | 0 | 0 | 0 | 0 | 0 | 0 | 0 | 0 | 0 | 2035200 | 0 | 0 | 0 | 0 | 0 | 0 |
| AYVAVDGII | GLB1; GLB1 | P16278; K7E | BGAL_HUMAN | Unmodified | 0 | 0 | 0 | 0 | 0 | 0 | 0 | 127880 | 547150 | 0 | 0 | 0 | 0 | 0 | 0 | 0 | 0 | 0 |
| AYVESICSN | ZC3HAV1; ZC3I | Q7Z2W4; C9 | ZCCHV_HUMA | Unmodified | 559090 | 0 | 2133700 | 720170 | 533380 | 380910 | 0 | 0 | 0 | 0 | 0 | 687580 | 0 | 0 | 1308800 | 1044600 | 0 | 0 |
| AYVESICSN | ZC3HAV1; ZC3I | Q7Z2W4; C9 | ZCCHV_HUMA | Unmodified | 0 | 0 | 0 | 0 | 1133300 | 0 | 0 | 0 | 399230 | 0 | 0 | 0 | 589600 | 0 | 0 | 0 | 0 | 0 |
| AYVFERDQ | SCARB2; SCARI | Q14108; A0A | SCRB2_HUMAI | Unmodified | 1054700 | 565320 | 1061000 | 1844000 | 0 | 0 | 0 | 0 | 26424000 | 3767800 | 690850 | 1693100 | 0 | 0 | 2841400 | 1621800 | 0 | 0 |
| AYVRLAPD | RPL23A; RPL23 | P62750; K7E | RL23A_HUMAN | Unmodified | 0 | 0 | 0 | 0 | 1831900 | 0 | 447660 | 224060 | 0 | 270650 | 0 | 0 | 0 | 841260 | 0 | 0 | 340900 | 0 |
| AYVRLAPD | RPL23A; RPL23 | P62750; K7E | RL23A_HUMAN | Unmodified | 0 | 0 | 0 | 0 | 0 | 0 | 472530 | 0 | 0 | 0 | 0 | 0 | 775130 | 693770 | 0 | 0 | 0 | 0 |
| AYVSTLMG | EIF3F; EIF3F | O00303; H0 | EIF3F_HUMAN | Unmodified | 0 | 0 | 0 | 0 | 0 | 0 | 0 | 0 | 0 | 0 | 0 | 1955900 | 0 | 0 | 0 | 0 | 0 | 0 |
| AYVSTLMG | EIF3F; EIF3F | O00303; H0 | EIF3F_HUMAN | Oxidation (M) | 0 | 0 | 189690 | 0 | 0 | 0 | 0 | 0 | 0 | 0 | 0 | 2410500 | 0 | 0 | 0 | 219780 | 0 | 0 |
| AYVTKSPVI | CMBL | Q96DG6 | CMBL_HUMAN | Unmodified | 106790 | 0 | 0 | 0 | 205730 | 0 | 0 | 0 | 0 | 0 | 0 | 0 | 240460 | 0 | 0 | 0 | 980930 | 178330 |
| AYVVEPNL | PGAP4 | Q9BRR3 | PGAP4_HUMAI | Unmodified | 0 | 0 | 0 | 0 | 241730 | 0 | 0 | 38608 | 4302300 | 0 | 0 | 0 | 0 | 0 | 0 | 101290 | 121370 | 0 |
| AYVVLGQF | BANF1 | O75531 | BAF_HUMAN B | Unmodified | 72678 | 109890 | 0 | 0 | 160710 | 20266 | 0 | 0 | 138190 | 76702 | 0 | 701500 | 102540 | 0 | 204070 | 0 | 0 | 149920 |
| AYVVLGQF | BANF1 | O75531 | BAF_HUMAN B | Unmodified | 1728900 | 1353900 | 180880 | 234620 | 603270 | 148450 | 31994 | 0 | 1557000 | 0 | 705870 | 11010000 | 292320 | 116930 | 507790 | 525200 | 0 | 443290 |
| AYYPSIVAN | ABCC1; ABCC1 | P33527; I3L4 | MRP1_HUMAN | Unmodified | 0 | 0 | 0 | 0 | 0 | 0 | 0 | 0 | 0 | 0 | 0 | 1312100 | 0 | 0 | 0 | 0 | 0 | 0 |
| CAADAGRF | SUMF2; SUMF2 | Q8NBJ7; J3C | SUMF2_HUMAI | Unmodified | 0 | 0 | 0 | 0 | 0 | 0 | 0 | 0 | 10090000 | 1617100 | 0 | 0 | 0 | 0 | 0 | 0 | 0 | 0 |
| CAADLGLN | HINT1 | P49773 | HINT1_HUMAN | Unmodified | 886020 | 0 | 0 | 0 | 6099000 | 573500 | 4132400 | 3025600 | 3767400 | 3449900 | 0 | 7482600 | 8790000 | 5702400 | 0 | 1229700 | 8527200 | 7801100 |
| CAADVKAP | PLPBP | O94903 | PLPHP_HUMAN | Unmodified | 22068 | 0 | 0 | 0 | 0 | 0 | 0 | 0 | 0 | 0 | 0 | 0 | 0 | 0 | 0 | 0 | 498450 | 548900 |
| CADSSFTV | MUC5B | Q9HC84 | MUC5B_HUMA | Unmodified | 0 | 0 | 0 | 0 | 0 | 0 | 0 | 0 | 0 | 0 | 0 | 0 | 0 | 0 | 0 | 0 | 47930 | 3121700 |
| CADVTLEG | MESD | Q14696 | MESD_HUMAN | Unmodified | 0 | 0 | 0 | 0 | 0 | 0 | 0 | 0 | 1840300 | 0 | 0 | 0 | 0 | 0 | 0 | 0 | 0 | 0 |
| CADVTLEG | MESD | Q14696 | MESD_HUMAN | Unmodified | 0 | 0 | 0 | 0 | 0 | 0 | 0 | 0 | 1204600 | 87907 | 0 | 0 | 0 | 0 | 0 | 0 | 0 | 0 |
| CADYVVTG | PSAT1 | Q9Y617 | SERC_HUMAN | Unmodified | 0 | 0 | 0 | 286300 | 1173000 | 0 | 0 | 0 | 0 | 0 | 0 | 2797700 | 1142700 | 765980 | 0 | 363950 | 0 | 0 |
| CAEFELR | PLCG2; PLCG2 | P16885; A0A | PLCG2_HUMAI | Unmodified | 6217400 | 0 | 0 | 5869700 | 0 | 0 | 0 | 0 | 0 | 0 | 0 | 1208600 | 5796200 | 0 | 5986600 | 0 | 0 | 0 |
| CAEIFAR | FARSB | Q9NSD9 | SYFB_HUMAN | Unmodified | 0 | 0 | 0 | 0 | 0 | 0 | 0 | 0 | 103220 | 0 | 0 | 1726800 | 0 | 0 | 50099 | 53471 | 0 | 0 |
| CAFRTHTG | FSCN1; FSCN1 | Q16658; C9J | FSCN1_HUMAI | Unmodified | 0 | 0 | 0 | 0 | 282700 | 78201 | 0 | 0 | 0 | 0 | 0 | 0 | 0 | 0 | 0 | 0 | 0 | 290010 |
| CAFRTHTG | FSCN1; FSCN1 | Q16658; C9J | FSCN1_HUMAI | Unmodified | 0 | 0 | 0 | 0 | 0 | 0 | 0 | 0 | 0 | 0 | 0 | 0 | 0 | 0 | 0 | 0 | 1597500 | 0 |
| CAIGVTHF( | GLG1 | Q92896 | GSLG1_HUMAI | Unmodified | 0 | 0 | 0 | 0 | 0 | 0 | 0 | 0 | 8105000 | 1306100 | 0 | 0 | 0 | 0 | 0 | 0 | 0 | 0 |
| CALGVFR | PSMD2; PSMD2 | Q13200; C9J | PSMD2_HUMA | Unmodified | 47383 | 33830 | 0 | 0 | 0 | 0 | 0 | 0 | 0 | 0 | 0 | 726640 | 0 | 0 | 0 | 63385 | 43694 | 0 |
| CALGWDH | CTTN | Q14247 | SRC8_HUMAN | Unmodified | 0 | 0 | 192060 | 0 | 2052200 | 322080 | 0 | 44870 | 202940 | 107510 | 0 | 532490 | 1044700 | 837580 | 356670 | 570160 | 943590 | 1061400 |
| CALSSVDN | MEMO1 | Q9Y316 | MEMO1_HUMA | Unmodified | 0 | 0 | 0 | 0 | 0 | 0 | 105070 | 0 | 0 | 310530 | 0 | 0 | 505050 | 0 | 0 | 0 | 552710 | 0 |
| CANLFEAL | ABRACL | Q9P1F3 | ABRAL_HUMAN | Unmodified | 690250 | 0 | 0 | 0 | 100700 | 0 | 0 | 0 | 1361700 | 494270 | 35358 | 34908 | 0 | 0 | 0 | 0 | 120610 | 111940 |
| CANYVNG | MME; MME; MM | P08473; A0A | NEP_HUMAN N | Unmodified | 210520 | 89609 | 1017700 | 2112900 | 0 | 0 | 0 | 0 | 2293400 | 0 | 0 | 0 | 0 | 0 | 4611200 | 4521000 | 0 | 0 |
| CANYVNG | MME; MME; MM | P08473; A0A | NEP_HUMAN N | Oxidation (M) | 0 | 0 | 650780 | 1148400 | 0 | 0 | 0 | 0 | 0 | 0 | 0 | 0 | 0 | 0 | 1217300 | 1369500 | 0 | 0 |
| CAQGCICK | MT2A | P02795 | MT2_HUMAN M | Unmodified | 0 | 0 | 0 | 0 | 0 | 0 | 0 | 0 | 0 | 0 | 0 | 0 | 0 | 0 | 0 | 0 | 93188 | 0 |
| CAQGCICK | MT1X | P80297 | MT1X_HUMAN | Unmodified | 0 | 0 | 0 | 0 | 0 | 0 | 0 | 0 | 0 | 0 | 0 | 0 | 0 | 0 | 0 | 0 | 177600 | 0 |
| CAQIEAKF | NAP1L1; NAP1L | P55209; F8V | NP1L1_HUMAN | Unmodified | 0 | 0 | 0 | 0 | 415180 | 78779 | 528620 | 0 | 0 | 0 | 0 | 0 | 1250600 | 817210 | 0 | 0 | 1993000 | 1696800 |
| CASQAGM | CNN3; CNN3 | Q15417; E9F | CNN3_HUMAN | Unmodified | 0 | 0 | 0 | 0 | 325970 | 0 | 0 | 0 | 0 | 0 | 36679 | 404480 | 0 | 0 | 0 | 0 | 0 | 214030 |
| CASQSGM | CNN2; CNN2; ( | Q99439; H3I | CNN2_HUMAN | Unmodified | 0 | 0 | 0 | 0 | 943130 | 0 | 0 | 24291 | 0 | 0 | 0 | 0 | 157640 | 252360 | 0 | 0 | 483730 | 237350 |
| CATGNALC | ADAM9; ADAMS | Q13443; A0A | ADAM9_HUMAI | Unmodified | 2143900 | 517470 | 1120700 | 1049200 | 2065600 | 0 | 0 | 37768 | 418750 | 30614 | 0 | 155640 | 849520 | 330020 | 1497900 | 1748100 | 244560 | 119190 |
| CATITPDEA | IDH2 | P48735 | IDHP_HUMAN | Unmodified | 673660 | 708080 | 0 | 0 | 0 | 0 | 0 | 0 | 399490 | 24867 | 0 | 0 | 0 | 0 | 570370 | 0 | 0 | 0 |
| CATITPDEA | IDH2 | P48735 | IDHP_HUMAN | Unmodified | 0 | 0 | 0 | 0 | 0 | 0 | 297260 | 169020 | 9666400 | 2022800 | 0 | 0 | 0 | 0 | 0 | 0 | 0 | 0 |
| CATITPDEK | IDH1; IDH1 | O75874; C9J | IDHC_HUMAN | Unmodified | 0 | 0 | 0 | 691830 | 1128700 | 0 | 0 | 192200 | 863170 | 457660 | 0 | 2671800 | 7447000 | 4421800 | 972250 | 1048100 | 678090 | 596140 |
| CATITPDEK | IDH1; IDH1 | O75874; C9J | IDHC_HUMAN | Unmodified | 0 | 0 | 0 | 0 | 1918500 | 0 | 91275 | 177830 | 0 | 318790 | 0 | 0 | 3019700 | 3110800 | 0 | 0 | 1104000 | 1502300 |
| CATSKPAFI | ANXA1 | P04083 | ANXA1_HUMAI | Unmodified | 82579 | 0 | 0 | 0 | 11220000 | 2440100 | 217690 | 200350 | 0 | 0 | 13896000 | 4.28E+08 | 0 | 0 | 0 | 0 | 2030800 | 3103000 |
| CATVTENA | SPINT2; SPINT2 | O43291; K7E | SPIT2_HUMAN | Unmodified | 0 | 0 | 0 | 0 | 0 | 0 | 0 | 25087 | 7066300 | 106190 | 0 | 0 | 0 | 0 | 0 | 0 | 0 | 0 |
| CAVCLDTV | CIT; CIT; CIT; CI | O14578; A0A | CTRO_HUMAN | Unmodified | 0 | 0 | 0 | 0 | 0 | 0 | 0 | 0 | 0 | 0 | 0 | 0 | 0 | 0 | 0 | 0 | 907680 | 0 |
| CAVEHHAF | EPB41L5; EPB4 | Q9HCM4; Q | E41L5_HUMAN | Unmodified | 0 | 249580 | 0 | 3675000 | 0 | 0 | 0 | 0 | 0 | 50796 | 0 | 0 | 0 | 0 | 990390 | 849840 | 49633 | 0 |
| CAWCSKEI | ITGB5 | P18084 | ITB5_HUMAN I | Unmodified | 157690 | 0 | 0 | 0 | 364080 | 157410 | 0 | 0 | 0 | 145460 | 88811 | 0 | 865030 | 637110 | 0 | 0 | 761910 | 613350 |
| CCGGEDYI | TSPAN15; TSPA | O95858; H7( | TSN15_HUMAN | Unmodified | 424880 | 0 | 356090 | 530690 | 472570 | 0 | 0 | 0 | 470160 | 174340 | 0 | 498210 | 0 | 267400 | 1100600 | 875420 | 0 | 0 |
| CCGLVNG | TSPAN8 | P19075 | TSN8_HUMAN | Unmodified | 13104000 | 0 | 0 | 0 | 0 | 0 | 0 | 0 | 0 | 0 | 0 | 0 | 0 | 0 | 0 | 0 | 0 | 0 |
| CCILTEEGL | PHLDA1; PHLD. | Q8WV24; R4 | PHLA1_HUMAN | Unmodified | 0 | 0 | 0 | 0 | 469870 | 0 | 0 | 0 | 374450 | 0 | 0 | 0 | 0 | 0 | 0 | 0 | 1506200 | 404560 |
| CCNTCEDV | ERGIC3; ERGIC | Q9Y282; A0A | ERGI3_HUMAN | Unmodified | 0 | 0 | 0 | 0 | 0 | 0 | 0 | 0 | 320010 | 0 | 0 | 0 | 0 | 0 | 0 | 0 | 0 | 0 |
| CCSGAIIVL | PKM; PKM | P14618; B4E | KPYM_HUMAN | Unmodified | 0 | 0 | 0 | 0 | 261460 | 0 | 0 | 0 | 0 | 0 | 0 | 833770 | 0 | 0 | 0 | 0 | 291380 | 0 |
| CDAISVLEK | DYNC1LI1; DYN | Q9Y6G9; E9I | DC1L1_HUMAN | Unmodified | 0 | 0 | 0 | 47108 | 0 | 0 | 0 | 0 | 0 | 0 | 204680 | 1296000 | 0 | 0 | 0 | 83620 | 0 | 0 |
| CDAIVDLIH | NUP210 | Q8TEM1 | PO210_HUMAN | Unmodified | 0 | 50660 | 0 | 0 | 0 | 0 | 0 | 0 | 346630 | 125600 | 0 | 0 | 0 | 0 | 0 | 0 | 0 | 0 |
| CDAVSVLE | DYNC1LI2; DYN | O43237; B4E | DC1L2_HUMAN | Unmodified | 0 | 0 | 0 | 0 | 0 | 0 | 0 | 0 | 0 | 0 | 0 | 953590 | 0 | 0 | 0 | 0 | 0 | 0 |
| CDDLEALK | ITGB1; ITGB1; I | P05556; E7E | ITB1_HUMAN I | Unmodified | 625070 | 393830 | 746140 | 829790 | 1707700 | 0 | 0 | 0 | 2027200 | 57094 | 0 | 1948700 | 0 | 0 | 1081800 | 1047200 | 0 | 1818900 |
| CDDLEALK | ITGB1; ITGB1; I | P05556; E7E | ITB1_HUMAN I | Unmodified | 0 | 0 | 0 | 0 | 0 | 0 | 0 | 0 | 419950 | 0 | 0 | 0 | 0 | 0 | 0 | 0 | 159480 | 0 |
| CDEDEDIG | IGF2R | P11717 | MPRI_HUMAN | Unmodified | 0 | 0 | 86965 | 0 | 0 | 0 | 0 | 0 | 793840 | 58492 | 0 | 0 | 0 | 0 | 0 | 76158 | 0 | 0 |
| CDENILWL | PKM; PKM; PKM | P14618; B4E | KPYM_HUMAN | Unmodified | 0 | 0 | 702880 | 1805800 | 14714000 | 3258400 | 1306000 | 407180 | 2127000 | 913420 | 3794400 | 17505000 | 0 | 1386300 | 1246900 | 896730 | 4259700 | 8010200 |
| CDENILWL | PKM; PKM; PKM | P14618; B4E | KPYM_HUMAN | Unmodified | 0 | 0 | 0 | 0 | 856550 | 0 | 477980 | 0 | 0 | 0 | 0 | 0 | 0 | 0 | 0 | 0 | 3319900 | 801140 |
| CDENILWL | PKM; PKM; PKM | P14618; B4E | KPYM_HUMAN | Unmodified | 0 | 0 | 0 | 0 | 42204 | 0 | 95967 | 0 | 0 | 0 | 0 | 0 | 0 | 0 | 0 | 0 | 478280 | 0 |
| CDEPILSNI | PEBP1 | P30086 | PEBP1_HUMAN | Unmodified | 0 | 0 | 661510 | 0 | 0 | 0 | 0 | 0 | 382670 | 46339 | 437090 | 1996800 | 0 | 0 | 0 | 618100 | 0 | 0 |
| CDFPIMK | GLO1 | Q04760 | LGUL_HUMAN | Unmodified | 0 | 0 | 0 | 0 | 464070 | 0 | 0 | 0 | 327860 | 158160 | 249100 | 949440 | 942750 | 390590 | 152380 | 0 | 259530 | 533150 |
| CDFTEDQT | MYL6; MYL6; M | P60660; G3 | MYL6_HUMAN | Acetyl (Protein N | 0 | 0 | 0 | 0 | 4711100 | 1840700 | 796560 | 359960 | 1337100 | 859710 | 0 | 5311200 | 1801500 | 0 | 0 | 0 | 1289200 | 1340700 |
| CDFTEDQT | MYL6; MYL6; M | P60660; G8J | MYL6_HUMAN | Acetyl (Protein N | 0 | 0 | 0 | 0 | 10096000 | 430470 | 476300 | 0 | 0 | 0 | 0 | 0 | 3212500 | 1532100 | 0 | 0 | 3642400 | 2129600 |
| CDKEFMW | MTPN; MTPN | P58546; C9J | MTPN_HUMAN | Acetyl (Protein N | 0 | 0 | 0 | 0 | 1082500 | 0 | 0 | 0 | 0 | 0 | 0 | 0 | 0 | 0 | 0 | 0 | 1012900 | 0 |
| CDLEDERV | RAP1B; RAP1A; | P61224; P62 | RAP1B_HUMAN | Unmodified | 1294500 | 670930 | 6427600 | 0 | 0 | 0 | 303180 | 0 | 16564 | 0 | 861690 | 7561600 | 68670 | 0 | 4067900 | 3093900 | 0 | 53289 |
| CDLEDERV | RAP1B; RAP1A; | P61224; P62 | RAP1B_HUMAN | Unmodified | 0 | 0 | 0 | 1237800 | 0 | 0 | 0 | 0 | 158130 | 28025 | 0 | 859510 | 235260 | 362730 | 0 | 0 | 0 | 0 |
| CDLLQAMF | DIPK2A | Q8NDZ4 | DIK2A_HUMAN | Unmodified | 0 | 0 | 0 | 0 | 0 | 0 | 0 | 0 | 5654800 | 0 | 0 | 0 | 0 | 0 | 235460 | 0 | 0 | 0 |
| CDLVNYSY | TPP2; TPP2; TPI | P29144; A0A | TPP2_HUMAN | Unmodified | 0 | 0 | 648790 | 1233900 | 1415100 | 8971500 | 3494800 | 84587 | 0 | 0 | 0 | 1827400 | 0 | 3605300 | 343150 | 0 | 828380 | 5684200 |
| CDNCRRP( | FN1; FN1 | P02751; H0 | FINC_HUMAN | Unmodified | 0 | 0 | 484850 | 272990 | 0 | 0 | 0 | 0 | 0 | 0 | 0 | 0 | 0 | 0 | 0 | 143890 | 0 | 0 |
| CDPYAGKE | MYOF; MYOF | Q9NZM1; H0 | MYOF_HUMAN | Unmodified | 0 | 0 | 0 | 0 | 865310 | 434480 | 0 | 0 | 0 | 0 | 0 | 0 | 103860 | 0 | 0 | 0 | 0 | 416370 |

| | | | | | | | | | | | | | | | | | | | | | | |
|---|---|---|---|---|---|---|---|---|---|---|---|---|---|---|---|---|---|---|---|---|---|---|
| CDRVDQLT | VCL | P18206 | VINC_HUMAN ' | Unmodified | 0 | 0 | 0 | 0 | 0 | 0 | 0 | 25766 | 140420 | 269430 | 0 | 1025700 | 0 | 41700 | 1045000 | 0 | 0 | 0 |
| CDRVDQLT | VCL | P18206 | VINC_HUMAN ' | Unmodified | 0 | 0 | 0 | 0 | 0 | 0 | 0 | 0 | 1217700 | 0 | 307720 | 1918800 | 1176200 | 0 | 0 | 0 | 627320 | 131890 |
| CDVDIRK | ACTG1; ACTBL2 | P63261; Q56 | ACTG_HUMAN | Unmodified | 8245400 | 0 | 6962200 | 9741300 | 45948000 | 0 | 0 | 113730 | 7499800 | 1954500 | 722770 | 31839000 | 0 | 13126000 | 9131200 | 4986500 | 10437000 | 33541000 |
| CDVDIRKD | ACTG1; ACTB; / | P63261; P60 | ACTG_HUMAN | Unmodified | 0 | 0 | 0 | 0 | 1127900 | 0 | 0 | 0 | 0 | 0 | 0 | 0 | 0 | 0 | 0 | 0 | 568880 | 0 |
| CDVDIRKD | ACTG1; ACTB; / | P63261; P60 | ACTG_HUMAN | Oxidation (M) | 0 | 0 | 0 | 0 | 1693500 | 1656200 | 476990 | 0 | 0 | 1044700 | 0 | 0 | 583770 | 0 | 0 | 0 | 935850 | 1028400 |
| CDVIAQGIV | SUCLA2; SUCL | Q9P2R7; A0/ | SUCB1_HUMAI | Oxidation (M) | 0 | 0 | 0 | 0 | 0 | 0 | 0 | 0 | 0 | 0 | 0 | 336210 | 0 | 0 | 0 | 0 | 0 | 0 |
| CDVTDNQS | NUP93; NUP93 | Q8N1F7; H3 | NUP93_HUMAI | Unmodified | 0 | 0 | 228630 | 0 | 0 | 0 | 0 | 0 | 0 | 0 | 0 | 350080 | 0 | 0 | 0 | 0 | 0 | 0 |
| CDYENVPT | TXNRD1; TXNRI | Q16881; A0/ | TRXR1_HUMAN | Unmodified | 0 | 0 | 0 | 0 | 440990 | 0 | 0 | 0 | 690940 | 0 | 514760 | 0 | 0 | 0 | 0 | 0 | 0 | 0 |
| CEASAVPAI | LSAMP; LSAMP; | Q13449; F5H | LSAMP_HUMAI | Unmodified | 0 | 0 | 72150 | 0 | 0 | 0 | 0 | 0 | 1368500 | 0 | 0 | 0 | 0 | 0 | 0 | 0 | 0 | 0 |
| CEAVVADV | HSD17B4; HSD | P51659; E7E | DHB4_HUMAN | Unmodified | 0 | 0 | 0 | 0 | 0 | 0 | 0 | 0 | 3438500 | 0 | 0 | 0 | 0 | 0 | 0 | 0 | 0 | 0 |
| CECGSCV( | ITGB3; N/A | P05106; H3E | ITB3_HUMAN I | Unmodified | 248490 | 0 | 0 | 0 | 0 | 0 | 0 | 0 | 0 | 0 | 0 | 0 | 134470 | 330460 | 81132 | 155010 | 0 | 0 |
| CEDLETQT | PODXL | O00592 | PODXL_HUMAI | Unmodified | 0 | 0 | 0 | 167020 | 0 | 0 | 0 | 0 | 4501900 | 141230 | 695030 | 394820 | 173100 | 0 | 0 | 48361 | 0 | 201530 |
| CEESFVK | CD55; DENND1 | P08174; Q6F | DAF_HUMAN C | Unmodified | 461880 | 0 | 364250 | 0 | 112740 | 0 | 0 | 0 | 0 | 26325 | 0 | 2317500 | 0 | 0 | 509160 | 225390 | 0 | 0 |
| CEFAVGYQ | LARS1; LARS1; | Q9P2J5; A0A | SYLC_HUMAN | Unmodified | 0 | 0 | 0 | 0 | 0 | 0 | 0 | 0 | 0 | 0 | 202650 | 4695500 | 271520 | 0 | 0 | 0 | 0 | 0 |
| CEFDPLHT | COBLL1; COBLI | Q53SF7; A0/ | COBL1_HUMAI | Unmodified | 1008500 | 1348000 | 346520 | 583680 | 0 | 0 | 0 | 23856 | 0 | 0 | 102800 | 174460 | 377420 | 0 | 0 | 461590 | 0 | 0 |
| CEFFNAGC | CBS; CBSL; CB! | P35520; P0D | CBS_HUMAN C | Unmodified | 0 | 0 | 0 | 0 | 709870 | 0 | 61869 | 0 | 0 | 0 | 0 | 2594100 | 147440 | 0 | 0 | 0 | 205370 | 817230 |
| CEFQDAYV | HSPD1; HSPD1 | P10809; A0A | CH60_HUMAN | Unmodified | 0 | 0 | 0 | 0 | 0 | 0 | 0 | 0 | 9323700 | 208370 | 0 | 0 | 0 | 4296800 | 0 | 0 | 0 | 0 |
| CEFQDAYV | HSPD1; HSPD1 | P10809; A0A | CH60_HUMAN | Unmodified | 0 | 0 | 0 | 0 | 0 | 0 | 0 | 0 | 2794400 | 0 | 227340 | 102520 | 0 | 0 | 0 | 0 | 0 | 0 |
| CEGINISGI | RPL13A; RPL13 | P40429; A0A | RL13A_HUMAN | Unmodified | 1161500 | 1303200 | 313320 | 370930 | 0 | 0 | 0 | 0 | 0 | 0 | 234830 | 8439700 | 0 | 320480 | 904450 | 729290 | 0 | 0 |
| CEGIYRVS( | RALBP1 | Q15311 | RBP1_HUMAN | Unmodified | 0 | 0 | 0 | 462130 | 70338 | 0 | 26714 | 0 | 73085 | 52312 | 0 | 0 | 158850 | 0 | 0 | 0 | 96608 | 0 |
| CEHSINMC | TMEFF2 | Q9UIK5 | TEFF2_HUMAN | Unmodified | 122210 | 0 | 0 | 0 | 0 | 0 | 0 | 0 | 0 | 0 | 0 | 0 | 0 | 0 | 0 | 230680 | 0 | 0 |
| CELGPLHC | ITGA5 | P08648 | ITA5_HUMAN I | Unmodified | 0 | 0 | 0 | 0 | 0 | 0 | 0 | 0 | 0 | 0 | 0 | 0 | 0 | 0 | 0 | 408310 | 0 | 0 |
| CENVATLD | NRXN1; NRXN1 | Q9ULB1; A0/ | NRX1A_HUMAI | Unmodified | 0 | 0 | 0 | 0 | 0 | 0 | 0 | 0 | 145450 | 0 | 0 | 0 | 0 | 0 | 0 | 0 | 0 | 0 |
| CEQDRPSF | LAMP1 | P11279 | LAMP1_HUMAI | Unmodified | 0 | 0 | 0 | 0 | 0 | 0 | 0 | 0 | 1761900 | 0 | 0 | 0 | 245680 | 0 | 0 | 0 | 0 | 0 |
| CERHTSVQ | FN1 | P02751 | FINC_HUMAN I | Unmodified | 0 | 0 | 893650 | 0 | 0 | 0 | 0 | 0 | 0 | 0 | 0 | 0 | 0 | 0 | 0 | 0 | 0 | 0 |
| CERLPCHE | FBLN1 | B1AHL2 | B1AHL2_HUM/ | Unmodified | 0 | 0 | 0 | 0 | 0 | 0 | 0 | 0 | 121850 | 0 | 0 | 21906 | 0 | 0 | 0 | 0 | 0 | 0 |
| CESAFLSK | RPL24; RPL24; I | P83731; C9J | RL24_HUMAN ( | Unmodified | 1246100 | 106620 | 427210 | 517190 | 497570 | 158510 | 366050 | 367110 | 713700 | 52400 | 1319900 | 13750000 | 507270 | 322210 | 484910 | 488870 | 47927 | 367430 |
| CESAPGCC | CANX; CANX; C | P27824; A0A | CALX_HUMAN | Unmodified | 11208000 | 6625500 | 0 | 8150200 | 702480 | 1501200 | 0 | 0 | 2.61E+08 | 7907700 | 2197300 | 6097800 | 0 | 0 | 7221400 | 7616600 | 407430 | 0 |
| CESAPGCC | CANX; CANX; C | P27824; A0A | CALX_HUMAN | Unmodified | 0 | 0 | 0 | 0 | 0 | 0 | 0 | 0 | 1072200 | 1111000 | 0 | 0 | 0 | 0 | 0 | 0 | 0 | 0 |
| CESNLQMI | ITGA3 | P26006 | ITA3_HUMAN I | Unmodified | 0 | 0 | 0 | 16234 | 1060100 | 0 | 0 | 0 | 0 | 0 | 41407 | 699050 | 0 | 0 | 115950 | 0 | 220910 | 0 |
| CETQNPVS | CEACAM5; CEA | P06731; A0A | CEAM5_HUMAI | Unmodified | 0 | 0 | 0 | 0 | 223580 | 0 | 0 | 0 | 8034900 | 0 | 0 | 0 | 337330 | 0 | 0 | 0 | 0 | 98584 |
| CETSASELE | SLC19A1; SLC1 | P41440; H3E | S19A1_HUMAN | Unmodified | 161180 | 95340 | 632880 | 277530 | 0 | 0 | 0 | 0 | 0 | 0 | 0 | 486280 | 0 | 0 | 803780 | 437510 | 0 | 0 |
| CEVNGAG/ | GPX1; GPX1 | P07203; A0A | GPX1_HUMAN | Unmodified | 55632 | 0 | 136890 | 0 | 0 | 0 | 0 | 0 | 829900 | 56765 | 136360 | 951550 | 0 | 0 | 39762 | 50144 | 0 | 70408 |
| CEWLIQAP | NRP1; NRP1; N | O14786; Q5. | NRP1_HUMAN | Unmodified | 0 | 0 | 0 | 330870 | 1034500 | 381880 | 0 | 186670 | 0 | 0 | 0 | 276090 | 1206600 | 549900 | 0 | 0 | 316850 | 765330 |
| CFAFMEMI | AIDA | Q96BJ3 | AIDA_HUMAN / | 2 Oxidation (M) | 0 | 0 | 0 | 0 | 0 | 0 | 0 | 0 | 0 | 0 | 0 | 0 | 97600 | 0 | 239720 | 0 | 0 | 60094 |
| CFGGLQK | ESD; ESD | P10768; U3K | ESTD_HUMAN | Unmodified | 174350 | 0 | 0 | 417870 | 1597100 | 468960 | 1450200 | 1005400 | 603470 | 822440 | 755880 | 2014300 | 1740600 | 1512100 | 617340 | 294090 | 2517500 | 1343100 |
| CFIEEIPDE | TMED4; TMED9 | Q7Z7H5; Q9 | TMED4_HUMAI | Oxidation (M) | 0 | 20516 | 99106 | 0 | 0 | 0 | 0 | 0 | 7944900 | 937220 | 539180 | 0 | 0 | 0 | 0 | 0 | 0 | 0 |
| CFIEEIPDE | TMED4; TMED9 | Q7Z7H5; Q9 | TMED4_HUMAI | Unmodified | 0 | 0 | 0 | 0 | 0 | 0 | 0 | 0 | 5916300 | 0 | 0 | 0 | 0 | 0 | 0 | 0 | 0 | 0 |
| CFIEEIPDE | TMED9 | Q9BVK6 | TMED9_HUMAI | Oxidation (M) | 6041900 | 0 | 0 | 9977700 | 3028900 | 1431700 | 0 | 0 | 2672800 | 8320000 | 0 | 0 | 2421900 | 135630 | 0 | 0 | 1431400 | 1086300 |
| CFIVGADN | RPLP0; RPLP0P | P05388; Q8I | RLA0_HUMAN ( | Unmodified | 8338200 | 5012200 | 3012800 | 2889800 | 1329200 | 0 | 454410 | 423870 | 248060 | 161620 | 509730 | 14736000 | 660750 | 1183100 | 6890600 | 4694400 | 202740 | 398140 |
| CFLEADPYI | CD109 | Q6YHK3 | CD109_HUMAI | Unmodified | 0 | 0 | 0 | 0 | 0 | 0 | 0 | 0 | 0 | 0 | 0 | 0 | 0 | 0 | 0 | 0 | 0 | 0 |
| CFLSWLVK | UPF1 | Q92900 | RENT1_HUMAN | Unmodified | 0 | 0 | 317470 | 397920 | 0 | 0 | 0 | 0 | 204510 | 0 | 53559 | 0 | 0 | 0 | 571920 | 382190 | 0 | 0 |
| CFNTLTNS | ADAR; ADAR; AI | P55265; A0A | DSRAD_HUMAI | Unmodified | 114040 | 0 | 0 | 0 | 0 | 0 | 0 | 0 | 26433 | 71506 | 0 | 661280 | 0 | 0 | 0 | 0 | 0 | 0 |
| CFQPPIHQ | CYFIP1 | Q7L576 | CYFP1_HUMAN | Unmodified | 0 | 0 | 0 | 0 | 0 | 103290 | 0 | 34776 | 0 | 0 | 0 | 970110 | 231700 | 0 | 521090 | 0 | 378180 | 288390 |
| CFSIDNPG | WDR1 | O75083 | WDR1_HUMAN | Unmodified | 0 | 850680 | 0 | 0 | 0 | 785390 | 0 | 0 | 0 | 0 | 0 | 4262500 | 1508400 | 0 | 0 | 0 | 1016500 | 0 |
| CFSVLGFC | XRCC5 | P13010 | XRCC5_HUMAI | Unmodified | 92509 | 0 | 0 | 0 | 0 | 0 | 0 | 345900 | 0 | 120580 | 0 | 1384600 | 0 | 0 | 0 | 0 | 0 | 0 |
| CGDLEEEL | TPM4; TPM2; TF | P67936; P07 | TPM4_HUMAN | Unmodified | 0 | 0 | 0 | 0 | 890110 | 266290 | 0 | 0 | 0 | 0 | 241410 | 569890 | 266940 | 358560 | 0 | 0 | 0 | 0 |
| CGDLVFAK | HDGF | P51858 | HDGF_HUMAN | Unmodified | 0 | 0 | 0 | 0 | 490740 | 405310 | 125810 | 0 | 0 | 0 | 266070 | 1404700 | 442060 | 0 | 0 | 0 | 110520 | 90125 |
| CGDTIKIES | NRP1; NRP1; N | O14786; Q5. | NRP1_HUMAN | Unmodified | 0 | 0 | 0 | 0 | 603680 | 0 | 0 | 0 | 0 | 0 | 0 | 0 | 0 | 0 | 0 | 301940 | 217640 | 3337400 |
| CGEDYKLH | CLGN; CANX; C | O14967; P27 | CLGN_HUMAN | Unmodified | 0 | 0 | 0 | 0 | 0 | 0 | 0 | 0 | 917450 | 798070 | 0 | 0 | 0 | 0 | 0 | 0 | 0 | 0 |
| CGEMAQA/ | VARS1; VARS1 | P26640; A0A | SYVC_HUMAN | Unmodified | 0 | 0 | 0 | 0 | 0 | 0 | 0 | 0 | 0 | 0 | 0 | 601580 | 0 | 0 | 0 | 0 | 0 | 0 |
| CGFQDDV/ | MYO1D; MYO1I | O94832; J3Q | MYO1D_HUMA | Unmodified | 1751700 | 0 | 0 | 0 | 0 | 0 | 0 | 0 | 318770 | 0 | 0 | 0 | 0 | 0 | 0 | 0 | 0 | 0 |
| CGGIQYLQ | MYO6; MYO6; N | Q9UM54; A0 | MYO6_HUMAN | Unmodified | 0 | 0 | 333400 | 523080 | 0 | 0 | 0 | 0 | 233980 | 102250 | 0 | 265530 | 0 | 110800 | 808560 | 453950 | 0 | 0 |
| CGGVLIDH | KLK12; KLK12 | Q9UKR0; E9I | KLK12_HUMAN | Unmodified | 0 | 0 | 0 | 0 | 0 | 0 | 0 | 0 | 2146900 | 269080 | 0 | 86948 | 0 | 0 | 0 | 0 | 0 | 0 |
| CGIAVAPVS | DPP4; DPP4; DI | P27487; A0A | DPP4_HUMAN | Unmodified | 0 | 0 | 532350 | 865550 | 0 | 0 | 0 | 0 | 0 | 0 | 0 | 0 | 0 | 0 | 2437600 | 567800 | 0 | 0 |
| CGIAVAPVS | DPP4; DPP4; DI | P27487; A0A | DPP4_HUMAN | Unmodified | 0 | 0 | 0 | 0 | 0 | 0 | 0 | 0 | 0 | 0 | 0 | 0 | 0 | 0 | 702250 | 278960 | 0 | 0 |
| CGILHPGE | ATP2B1; ATP2B | P20020; E7E | AT2B1_HUMAN | Unmodified | 0 | 0 | 0 | 0 | 0 | 0 | 0 | 0 | 0 | 113610 | 0 | 0 | 2510500 | 3243800 | 0 | 0 | 416540 | 260730 |
| CGKGLEST | CSRP1; CSRP1; | P21291; E9P | CSRP1_HUMAI | Unmodified | 0 | 0 | 0 | 0 | 0 | 0 | 150890 | 0 | 0 | 0 | 0 | 0 | 0 | 0 | 0 | 0 | 606200 | 0 |
| CGLTDNEN | MUC5B | Q9HC84 | MUC5B_HUMA | Unmodified | 0 | 0 | 0 | 0 | 0 | 0 | 0 | 0 | 0 | 0 | 0 | 223010 | 0 | 0 | 0 | 0 | 0 | 535450 |
| CGLVIGRG | KHSRP; KHSRP | Q92945; M0I | FUBP2_HUMAN | Unmodified | 0 | 0 | 0 | 0 | 221990 | 0 | 161700 | 176720 | 0 | 274550 | 0 | 0 | 0 | 0 | 0 | 0 | 166240 | 281320 |
| CGNQAAIN | PPP2CB; PPP2( | P62714; P67 | PP2AB_HUMAN | Oxidation (M) | 0 | 0 | 0 | 0 | 0 | 0 | 0 | 0 | 1028700 | 222650 | 90812 | 726420 | 0 | 0 | 72570 | 0 | 0 | 0 |
| CGPGPLLP | MPPE1; MPPE1; | Q53F39; K7E | MPPE1_HUMAI | Unmodified | 0 | 0 | 0 | 0 | 0 | 0 | 0 | 0 | 1428800 | 334050 | 0 | 0 | 0 | 0 | 0 | 0 | 0 | 0 |
| CGPMVLD/ | SDHB; SDHB; S | P21912; A0A | SDHB_HUMAN | Oxidation (M) | 0 | 0 | 0 | 0 | 0 | 0 | 0 | 0 | 460170 | 577070 | 0 | 0 | 0 | 0 | 0 | 0 | 0 | 0 |
| CGQAVRLT | SDF2L1 | Q9HCN8 | SDF2L_HUMAN | Unmodified | 0 | 0 | 0 | 0 | 0 | 0 | 25101 | 0 | 0 | 393300 | 0 | 0 | 0 | 0 | 0 | 0 | 0 | 0 |
| CGQEEHD' | TES | Q9UGI8 | TES_HUMAN Te | Unmodified | 0 | 0 | 0 | 0 | 904950 | 0 | 0 | 0 | 0 | 0 | 0 | 233480 | 631330 | 474140 | 0 | 0 | 617350 | 1333700 |
| CGQGLRC' | IGFBP2 | P18065 | IBP2_HUMAN I | Oxidation (M) | 0 | 0 | 0 | 0 | 0 | 0 | 0 | 0 | 0 | 1682600 | 0 | 0 | 0 | 0 | 0 | 0 | 0 | 0 |
| CGQYIQK | MYOF; MYOF | Q9NZM1; H( | MYOF_HUMAN | Unmodified | 0 | 0 | 0 | 373950 | 2856500 | 106320 | 477960 | 0 | 0 | 0 | 0 | 86746 | 0 | 0 | 0 | 68934 | 0 | 0 |
| CGTGIVGV | PDLIM1 | O00151 | PDLI1_HUMAN | Unmodified | 0 | 0 | 0 | 0 | 0 | 0 | 0 | 0 | 0 | 0 | 0 | 503870 | 492430 | 0 | 0 | 0 | 3223600 | 2066900 |
| CGVISPRFI | RPS15A; RPS15 | P62244; H3E | RS15A_HUMAN | Unmodified | 0 | 0 | 0 | 0 | 0 | 0 | 705690 | 695190 | 0 | 136650 | 0 | 0 | 0 | 282210 | 0 | 0 | 0 | 375340 |
| CGVISPRFI | RPS15A; RPS15 | P62244; H3E | RS15A_HUMAN | Unmodified | 0 | 0 | 0 | 0 | 54793 | 0 | 407220 | 64691 | 0 | 0 | 0 | 0 | 80165 | 0 | 0 | 0 | 0 | 0 |
| CGVLGALR | THEM6 | Q8WUY1 | THEM6_HUMAI | Unmodified | 144980 | 0 | 0 | 0 | 0 | 0 | 0 | 0 | 730860 | 0 | 0 | 0 | 0 | 0 | 0 | 0 | 0 | 0 |
| CGVPDKFC | MMP14; MMP14 | P50281; A0A | MMP14_HUMA | Unmodified | 0 | 0 | 0 | 0 | 841710 | 963080 | 0 | 61202 | 0 | 0 | 0 | 0 | 0 | 0 | 0 | 0 | 341850 | 676570 |
| CGVPDQF( | MMP15 | P51511 | MMP15_HUMA | Unmodified | 0 | 0 | 0 | 0 | 0 | 0 | 0 | 0 | 0 | 0 | 0 | 0 | 1377100 | 0 | 0 | 0 | 0 | 0 |
| CGVPFSCC | TSPAN14; TSPA | Q8NG11; H7 | TSN14_HUMAN | Unmodified | 0 | 0 | 0 | 170770 | 0 | 0 | 0 | 0 | 0 | 0 | 0 | 678320 | 0 | 105090 | 183000 | 134930 | 0 | 0 |
| CGYAGSNI | ACTR2; ACTR2; | P61160; A0A | ARP2_HUMAN | Unmodified | 3009100 | 3433400 | 1269600 | 0 | 9155800 | 5638000 | 0 | 0 | 2812700 | 497820 | 1578800 | 21608000 | 8516100 | 4865000 | 4977800 | 5056000 | 4844900 | 4218900 |
| CGYHVVIG | STEAP2; STEAP | Q8NFT2; C9. | STEA2_HUMAN | Unmodified | 0 | 0 | 0 | 0 | 0 | 0 | 0 | 0 | 0 | 0 | 0 | 0 | 0 | 0 | 1187700 | 481110 | 0 | 0 |
| CGYHVVIG | STEAP2; STEAP | Q8NFT2; C9. | STEA2_HUMAN | Unmodified | 0 | 0 | 142240 | 472530 | 0 | 0 | 0 | 0 | 0 | 0 | 0 | 0 | 0 | 0 | 331130 | 252670 | 0 | 0 |
| CHAGHLN( | FGG; FGG; FGG | P02679; C9J | FIBG_HUMAN F | Unmodified | 0 | 0 | 0 | 0 | 0 | 0 | 0 | 0 | 0 | 145810 | 0 | 0 | 0 | 184680 | 0 | 0 | 0 | 104240 |
| CHAIIDEQI | LIMS1; LIMS4; L | P48059; A0A | LIMS1_HUMAN | Unmodified | 0 | 0 | 0 | 0 | 0 | 0 | 0 | 0 | 0 | 0 | 0 | 0 | 0 | 0 | 0 | 383970 | 0 | 0 |
| CHFDYNPY | PALS2; PALS2; I | Q9NZW5; B8 | PALS2_HUMAN | Unmodified | 0 | 0 | 0 | 0 | 670720 | 108760 | 0 | 0 | 0 | 0 | 0 | 0 | 0 | 189600 | 0 | 0 | 1136100 | 824740 |
| CHFDYNPY | PALS2; PALS2; I | Q9NZW5; B8 | PALS2_HUMAN | Unmodified | 0 | 0 | 0 | 0 | 0 | 0 | 0 | 0 | 3552700 | 0 | 0 | 0 | 0 | 0 | 0 | 0 | 606850 | 0 |
| CHGVPLVT | DPYSL5; DPYS | Q9BPU6; Q1 | DPYL5_HUMAN | Unmodified | 0 | 0 | 0 | 0 | 0 | 0 | 0 | 0 | 0 | 0 | 0 | 1692900 | 0 | 0 | 0 | 0 | 0 | 0 |
| CHLLVEHE | STK10; SLK; STI | O94804; Q9I | STK10_HUMAN | Unmodified | 0 | 0 | 0 | 424090 | 352230 | 0 | 0 | 300750 | 0 | 0 | 350430 | 1086800 | 0 | 0 | 0 | 0 | 243320 | 0 |
| CHSGECIT | LDLR; LDLR; LD | P01130; J3KI | LDLR_HUMAN | Unmodified | 0 | 0 | 0 | 0 | 283100 | 101890 | 63870 | 0 | 532110 | 108660 | 0 | 0 | 0 | 95181 | 0 | 13775 | 38684 | 155340 |
| CHTPPLYR | RPL18A; RPL18 | Q02543; M0I | RL18A_HUMAN | Unmodified | 248190 | 0 | 0 | 0 | 0 | 0 | 0 | 0 | 0 | 0 | 0 | 1231200 | 0 | 0 | 0 | 67485 | 0 | 0 |
| CIAYAESHI | GBE1; GBE1 | Q04446; E9F | GLGB_HUMAN | Unmodified | 0 | 0 | 0 | 0 | 0 | 0 | 0 | 0 | 0 | 127840 | 0 | 280440 | 0 | 0 | 0 | 0 | 256570 | 82642 |
| CICAEGYK | CRELD1; CRELI | Q96HD1; A0 | CREL1_HUMAN | Unmodified | 0 | 0 | 0 | 0 | 0 | 0 | 0 | 0 | 341120 | 29371 | 0 | 0 | 0 | 0 | 0 | 0 | 0 | 0 |
| CIDRSKPSI | DLG1; DLG1; D | Q12959; A0/ | DLG1_HUMAN | Unmodified | 0 | 0 | 0 | 0 | 7094800 | 0 | 0 | 0 | 0 | 1363800 | 0 | 0 | 0 | 0 | 0 | 0 | 3586500 | 5951100 |
| CIELCCGS' | ATP1A1 | P05023 | AT1A1_HUMAN | Unmodified | 306920 | 0 | 1499100 | 1194600 | 1951300 | 419210 | 691220 | 90213 | 781220 | 751010 | 954250 | 8545200 | 4174700 | 8890900 | 1575500 | 2066000 | 1378500 | 328500 |
| CIELSSGS\ | ATP1A3; N/A; A | P13637; A0A | AT1A3_HUMAN | Unmodified | 863660 | 0 | 0 | 0 | 951630 | 55046 | 586140 | 172760 | 270930 | 393350 | 0 | 0 | 273690 | 265680 | 0 | 70826 | 0 | 0 |
| CIESLIAVF( | S100A11 | P31949 | S10AB_HUMAN | Unmodified | 1375800 | 632350 | 479880 | 877800 | 0 | 0 | 0 | 0 | 3866400 | 0 | 0 | 0 | 0 | 0 | 493140 | 1100500 | 0 | 118660 |
| CIFGALHCI | CFC1; CFC1B | P0CG37; P0( | CFC1_HUMAN | Unmodified | 193160 | 113570 | 0 | 0 | 0 | 0 | 0 | 0 | 0 | 0 | 0 | 0 | 0 | 0 | 0 | 0 | 0 | 0 |
| CIGGVIFFH | ALDOC; ALDOC | P09972; J3Q | ALDOC_HUMA | Unmodified | 0 | 0 | 0 | 0 | 0 | 0 | 0 | 0 | 18636 | 254700 | 0 | 1399200 | 0 | 0 | 0 | 91743 | 517990 | 0 |
| CIGKPGGS | TIMM13; TIMM1 | Q9Y5L4; K7E | TIM13_HUMAN | Unmodified | 0 | 0 | 0 | 0 | 50703 | 0 | 0 | 0 | 93650 | 167990 | 0 | 2591000 | 1710400 | 0 | 0 | 0 | 61298 | 0 |
| CIIDILTEPK | FURIN; FURIN; | P09958; A0A | FURIN_HUMAN | Unmodified | 0 | 0 | 0 | 0 | 0 | 0 | 0 | 0 | 671950 | 598090 | 2201700 | 0 | 0 | 0 | 0 | 0 | 0 | 0 |
| CIIEVLSNA | CHGB; CHGB | P05060; A0A | SCG1_HUMAN | Unmodified | 0 | 0 | 0 | 0 | 0 | 0 | 0 | 0 | 969690 | 0 | 0 | 0 | 0 | 0 | 0 | 0 | 0 | 0 |
| CIIVEEGK | DSTN; DSTN | P60981; F6R | DEST_HUMAN | Unmodified | 0 | 0 | 0 | 0 | 0 | 0 | 0 | 0 | 195580 | 90841 | 374830 | 1257900 | 0 | 46972 | 0 | 0 | 0 | 0 |
| CILDLAHQ' | BCAT1 | P54687 | BCAT1_HUMAN | Unmodified | 18844 | 27113 | 76093 | 0 | 0 | 0 | 0 | 0 | 0 | 0 | 995410 | 5446000 | 0 | 0 | 0 | 0 | 0 | 0 |
| CILQDGRIF | SNRPB; SNRPN | P14678; P63 | RSMB_HUMAN | Unmodified | 35756 | 0 | 0 | 0 | 2102800 | 206780 | 892170 | 0 | 0 | 1658200 | 0 | 0 | 4352200 | 1503500 | 0 | 0 | 1102800 | 1160400 |
| CINRETGQ | CASK; CASK; C/ | O14936; A0/ | CSKP_HUMAN | Unmodified | 0 | 0 | 0 | 0 | 0 | 498940 | 0 | 210720 | 1198700 | 2032600 | 0 | 0 | 0 | 2593500 | 787570 | 493490 | 830870 | 1262400 |
| CIPALDSLT | HSPD1; HSPD1 | P10809; A0A | CH60_HUMAN | Unmodified | 0 | 0 | 0 | 0 | 0 | 0 | 0 | 0 | 2894900 | 0 | 0 | 0 | 0 | 0 | 0 | 0 | 0 | 0 |
| CIQVEITPT! | FAM126A; FAM | Q9BYI3; A0A | HYCCI_HUMAN | Unmodified | 0 | 0 | 0 | 0 | 0 | 0 | 0 | 0 | 0 | 0 | 0 | 0 | 0 | 0 | 0 | 0 | 0 | 228830 |
| CIRSTPEYF | ANXA6 | P08133 | ANXA6_HUMAI | Unmodified | 0 | 0 | 0 | 0 | 0 | 0 | 0 | 56353 | 0 | 1497300 | 0 | 0 | 0 | 1018300 | 0 | 0 | 1220200 | 0 |
| CISDLSLH\ | ITGA1 | P56199 | ITA1_HUMAN I | Unmodified | 0 | 0 | 0 | 0 | 0 | 0 | 688790 | 0 | 0 | 383090 | 0 | 0 | 0 | 0 | 0 | 0 | 2319500 | 2846800 |
| CISEVQANI | G6PD; G6PD | P11413; E7E | G6PD_HUMAN | Unmodified | 0 | 0 | 0 | 0 | 5000500 | 1625200 | 0 | 0 | 678670 | 0 | 0 | 2704500 | 1179100 | 0 | 0 | 0 | 952370 | 0 |
| CITPTGTHF | GRN; GRN | P28799; K7E | GRN_HUMAN F | Unmodified | 0 | 0 | 0 | 0 | 0 | 0 | 0 | 0 | 1791700 | 492900 | 0 | 0 | 0 | 0 | 0 | 0 | 0 | 623400 |
| CIVSEWIAE | PTGFRN | Q9P2B2 | FPRP_HUMAN | Unmodified | 0 | 0 | 35466 | 390510 | 0 | 0 | 119650 | 0 | 0 | 0 | 0 | 2206100 | 586560 | 112090 | 467610 | 292360 | 0 | 0 |
| CKAEHDQI | LMNB1; LMNB1 | P20700; A0A | LMNB1_HUMAI | Unmodified | 0 | 0 | 0 | 0 | 0 | 0 | 0 | 0 | 174090 | 0 | 293370 | 886730 | 195270 | 0 | 0 | 0 | 0 | 0 |
| CKDDEFTH | CALR; CALR; C/ | P27797; A0A | CALR_HUMAN | Unmodified | 0 | 0 | 0 | 0 | 0 | 0 | 0 | 0 | 1.73E+08 | 7926700 | 10015000 | 4855800 | 0 | 0 | 3940500 | 0 | 0 | 0 |

| | | | | | | | | | | | | | | | | | | | | | | |
|---|---|---|---|---|---|---|---|---|---|---|---|---|---|---|---|---|---|---|---|---|---|---|
| CKRQENEC | KRTCAP3; KRTC | Q53RY4; A0A | KCP3_HUMAN | Unmodified | 226670 | 262670 | 0 | 304720 | 0 | 0 | 20384 | 0 | 0 | 0 | 0 | 0 | 103000 | 70998 | 0 | 65121 | 0 | 0 |
| CKVAMANI | VCL | P18206 | VINC_HUMAN V | 2 Oxidation (M) | 0 | 0 | 0 | 0 | 0 | 0 | 0 | 0 | 0 | 0 | 0 | 0 | 0 | 0 | 0 | 0 | 1703100 | 0 |
| CLAAALIVL | PKM; PKM | H3BTN5; H3 | H3BTN5_HUM | Unmodified | 0 | 0 | 0 | 0 | 0 | 0 | 0 | 0 | 0 | 57680 | 153480 | 1096100 | 0 | 0 | 0 | 0 | 0 | 7425.5 |
| CLAALASLF | ILF3; ILF3; ILF3 | Q12906; K7E | ILF3_HUMAN In | Unmodified | 1282700 | 504110 | 269060 | 390400 | 0 | 0 | 16482 | 0 | 0 | 96480 | 253550 | 2264600 | 116470 | 38720 | 209110 | 381150 | 0 | 0 |
| CLADGNPF | MCAM | P43121 | MUC18_HUMA | Unmodified | 0 | 0 | 0 | 0 | 0 | 0 | 0 | 0 | 0 | 0 | 0 | 0 | 0 | 0 | 0 | 0 | 254240 | 526580 |
| CLAEGAGD | MELTF | P08582 | TRFM_HUMAN | Unmodified | 0 | 0 | 0 | 0 | 0 | 0 | 0 | 0 | 964920 | 481640 | 0 | 0 | 0 | 0 | 0 | 0 | 0 | 0 |
| CLATGPGI | FLNB; FLNB | O75369; E7E | FLNB_HUMAN | Unmodified | 0 | 0 | 0 | 0 | 2712400 | 913890 | 0 | 0 | 0 | 0 | 0 | 2140700 | 0 | 1109500 | 0 | 0 | 1918000 | 0 |
| CLATGPGI | FLNB; FLNB | O75369; E7E | FLNB_HUMAN | Unmodified | 0 | 0 | 0 | 0 | 0 | 0 | 0 | 0 | 0 | 0 | 0 | 0 | 0 | 0 | 0 | 0 | 589940 | 0 |
| CLAYDFYP | AZGP1 | P25311 | ZA2G_HUMAN | Unmodified | 0 | 0 | 0 | 0 | 0 | 0 | 0 | 0 | 0 | 0 | 976180 | 0 | 0 | 0 | 0 | 0 | 0 | 0 |
| CLDMIFNS | KIF23; KIF23; KI | Q02241; A0A | KIF23_HUMAN | Oxidation (M) | 0 | 0 | 0 | 0 | 1445300 | 329890 | 0 | 0 | 0 | 174400 | 0 | 0 | 0 | 0 | 0 | 0 | 592980 | 0 |
| CLDRSEVLI | GALNT7; GALNT | Q86SF2; H0Y | GALT7_HUMAN | Unmodified | 0 | 0 | 81495 | 0 | 0 | 0 | 0 | 0 | 1395700 | 0 | 0 | 0 | 0 | 131210 | 138100 | 0 | 0 | 0 |
| CLECQVMC | CIT; CIT; CIT; CI | O14578; A0A | CTRO_HUMAN | Unmodified | 0 | 0 | 0 | 0 | 0 | 0 | 95855 | 0 | 0 | 0 | 0 | 0 | 0 | 0 | 0 | 0 | 692690 | 0 |
| CLEEFELLC | EIF3CL; EIF3C; | B5ME19; Q9 | EIFCL_HUMAN | Unmodified | 63141 | 0 | 52110 | 143370 | 0 | 0 | 0 | 0 | 0 | 0 | 0 | 703710 | 107570 | 207450 | 0 | 0 | 0 | 0 |
| CLEGMILA | FNBP1L; FNBP1 | Q5T0N5; A0A | FBP1L_HUMAN | Unmodified | 0 | 0 | 0 | 0 | 0 | 0 | 0 | 0 | 0 | 0 | 0 | 0 | 197860 | 295600 | 0 | 0 | 0 | 0 |
| CLELFSELA | HSP90AB1 | P08238 | HS90B_HUMAI | Unmodified | 0 | 0 | 0 | 0 | 0 | 0 | 0 | 0 | 283680 | 0 | 397270 | 711740 | 0 | 0 | 0 | 0 | 0 | 0 |
| CLELFSELA | HSP90AB1 | P08238 | HS90B_HUMAI | Unmodified | 0 | 0 | 59653 | 0 | 890810 | 0 | 0 | 26743 | 698950 | 49066 | 388040 | 4083500 | 991770 | 465990 | 0 | 0 | 848070 | 0 |
| CLELFSELA | HSP90AB1 | P08238 | HS90B_HUMAI | Unmodified | 0 | 168630 | 0 | 849050 | 2587400 | 0 | 0 | 0 | 1065300 | 697210 | 0 | 5922500 | 1561700 | 1347300 | 224050 | 524660 | 1628100 | 0 |
| CLELFTELA | HSP90AA1 | P07900 | HS90A_HUMAI | Unmodified | 0 | 0 | 0 | 0 | 0 | 0 | 0 | 0 | 0 | 0 | 0 | 1009000 | 0 | 0 | 0 | 0 | 0 | 0 |
| CLELFTELA | HSP90AA1 | P07900 | HS90A_HUMAI | Unmodified | 0 | 0 | 0 | 0 | 425490 | 0 | 281850 | 0 | 0 | 0 | 0 | 3222500 | 0 | 0 | 210090 | 0 | 0 | 3652500 |
| CLFNHKFE | GLG1; GLG1 | Q92896; H3I | GSLG1_HUMAI | Unmodified | 0 | 0 | 0 | 0 | 0 | 0 | 0 | 0 | 3702900 | 943220 | 0 | 0 | 0 | 0 | 0 | 0 | 0 | 127530 |
| CLFNHKFE | GLG1; GLG1 | Q92896; H3I | GSLG1_HUMAI | Oxidation (M) | 0 | 0 | 0 | 0 | 0 | 0 | 0 | 0 | 443740 | 214270 | 0 | 0 | 0 | 0 | 0 | 0 | 0 | 0 |
| CLFQSPLF | CARM1 | Q86X55 | CARM1_HUMA | Unmodified | 0 | 0 | 0 | 0 | 0 | 0 | 0 | 0 | 0 | 0 | 0 | 2065600 | 0 | 0 | 0 | 0 | 0 | 0 |
| CLGNGNPI | ALCAM; ALCAM | Q13740; F5C | CD166_HUMAI | Unmodified | 5650600 | 1244600 | 184820 | 481760 | 143420 | 0 | 0 | 0 | 2337500 | 123480 | 0 | 0 | 0 | 0 | 0 | 814010 | 53489 | 0 |
| CLHPLANE | FHL1; FHL1; FH | Q13642; A0A | FHL1_HUMAN | Unmodified | 0 | 0 | 0 | 0 | 393000 | 0 | 927330 | 645180 | 0 | 903120 | 0 | 0 | 0 | 0 | 0 | 0 | 674250 | 0 |
| CLIEILASR | ANXA10; ANXA1 | Q9UJ72; P5C | ANX10_HUMAI | Unmodified | 514780 | 0 | 903200 | 1681400 | 0 | 0 | 113490 | 49733 | 0 | 1450200 | 0 | 0 | 0 | 0 | 1291300 | 738490 | 0 | 0 |
| CLIEILASRT | ANXA6; ANXA6; | P08133; E5R | ANXA6_HUMAI | Unmodified | 0 | 0 | 0 | 0 | 0 | 0 | 217520 | 0 | 564630 | 2679600 | 0 | 0 | 233230 | 191640 | 0 | 0 | 231640 | 534550 |
| CLIHDGAA | HSPG2 | P98160 | PGBM_HUMAN | Unmodified | 0 | 0 | 268990 | 256030 | 0 | 0 | 0 | 0 | 0 | 0 | 0 | 0 | 303590 | 1096200 | 0 | 304110 | 0 | 0 |
| CLIKNPAEF | MAP2K2; MAP2 | P36507; Q02 | MP2K2_HUMAI | Unmodified | 0 | 0 | 0 | 0 | 453810 | 0 | 1607400 | 132790 | 0 | 176670 | 0 | 0 | 734000 | 0 | 0 | 0 | 1140200 | 0 |
| CLKDEDPY | AP1B1; AP2B1; | Q10567; P63 | AP1B1_HUMAI | Unmodified | 0 | 0 | 0 | 547720 | 711730 | 192530 | 275660 | 147390 | 0 | 0 | 198990 | 1142200 | 924920 | 940090 | 437980 | 0 | 913590 | 941840 |
| CLKKDDES | KLC1; KLC1; KL | Q07866; G3V | KLC1_HUMAN | Unmodified | 13736000 | 0 | 0 | 0 | 494130 | 88582 | 0 | 0 | 0 | 0 | 0 | 172340 | 203790 | 0 | 0 | 0 | 255340 | 360940 |
| CLLDRVPEI | PRMT5; PRMT5; | O14744; H0Y | ANM5_HUMAN | Unmodified | 73714 | 0 | 0 | 0 | 0 | 0 | 0 | 0 | 0 | 0 | 0 | 1816600 | 0 | 0 | 0 | 0 | 0 | 0 |
| CLLEQLKP | SEZ6; SEZ6; SEZ | Q53EL9; C9J | SEZ6_HUMAN S | Unmodified | 0 | 0 | 0 | 0 | 0 | 0 | 0 | 0 | 1750000 | 0 | 0 | 0 | 0 | 0 | 0 | 0 | 0 | 0 |
| CLPARAPA | IGFBP6 | P24592 | IBP6_HUMAN I | Unmodified | 0 | 0 | 0 | 0 | 0 | 363550 | 0 | 0 | 0 | 55043 | 0 | 0 | 0 | 0 | 0 | 0 | 0 | 698900 |
| CLPRQDEE | IGFBP5; IGFBP5 | P24593; C9J | IBP5_HUMAN I | Unmodified | 0 | 0 | 0 | 0 | 0 | 0 | 0 | 0 | 6479700 | 1695000 | 0 | 0 | 0 | 0 | 0 | 0 | 0 | 0 |
| CLQLGSPG | KIAA2013 | Q8IYS2 | K2013_HUMAN | Unmodified | 0 | 0 | 0 | 0 | 0 | 0 | 0 | 0 | 12744000 | 562410 | 0 | 544670 | 0 | 0 | 0 | 0 | 0 | 0 |
| CLQNPSSL | GCN1 | Q92616 | GCN1_HUMAN | Unmodified | 0 | 0 | 0 | 0 | 0 | 0 | 756390 | 0 | 0 | 0 | 0 | 0 | 0 | 0 | 0 | 0 | 0 | 0 |
| CLQRQNRY | UBA1 | P22314 | UBA1_HUMAN | Unmodified | 0 | 0 | 0 | 0 | 2119600 | 399560 | 1061600 | 0 | 0 | 787570 | 0 | 0 | 3493900 | 1511000 | 0 | 0 | 3001200 | 4785800 |
| CLQSGTLF | CAPN1; CAPN1 | P07384; E9P | CAN1_HUMAN | Unmodified | 1004600 | 12837000 | 0 | 0 | 0 | 0 | 0 | 0 | 0 | 0 | 0 | 1249000 | 0 | 0 | 0 | 0 | 0 | 0 |
| CLQWTTVII | AKT3; AKT2; AK | Q9Y243; P31 | AKT3_HUMAN I | Unmodified | 0 | 0 | 0 | 0 | 0 | 0 | 0 | 0 | 0 | 0 | 0 | 0 | 0 | 0 | 0 | 674630 | 0 | 0 |
| CLREEIHKD | TMED10; TMED | P49755; G3V | TMEDA_HUMAI | Unmodified | 0 | 0 | 0 | 0 | 0 | 0 | 0 | 0 | 5546100 | 3328700 | 0 | 0 | 0 | 0 | 0 | 0 | 0 | 0 |
| CLSAAEEK | TPM3; TPM3; TF | J3KN67; A0A | J3KN67_HUMA | Unmodified | 0 | 0 | 0 | 0 | 20065 | 595450 | 0 | 176960 | 0 | 0 | 0 | 0 | 0 | 0 | 0 | 0 | 974330 | 58416 |
| CLSDPGPH | SLC9A1 | P19634 | SL9A1_HUMAN | Unmodified | 0 | 0 | 790850 | 121740 | 72857 | 0 | 0 | 0 | 0 | 0 | 0 | 0 | 1418800 | 309210 | 715130 | 485340 | 570680 | 74795 |
| CLSDPGPH | SLC9A1 | P19634 | SL9A1_HUMAN | Unmodified | 0 | 0 | 0 | 0 | 0 | 639330 | 0 | 0 | 0 | 0 | 436690 | 0 | 719850 | 779660 | 0 | 0 | 269770 | 0 |
| CLSEQIAD | TOR1AIP1; TOR | Q5JTV8; A0A | TOIP1_HUMAN | Unmodified | 0 | 0 | 0 | 0 | 0 | 0 | 0 | 0 | 3032600 | 0 | 0 | 0 | 0 | 0 | 0 | 0 | 0 | 0 |
| CLSHHVAD | TOR1AIP2 | Q8NFQ8 | TOIP2_HUMAN | Unmodified | 0 | 0 | 0 | 0 | 559040 | 0 | 130730 | 84637 | 2436900 | 653280 | 78117 | 0 | 0 | 0 | 0 | 0 | 0 | 0 |
| CLSSLKDEI | PGD; PGD | P52209; K7E | 6PGD_HUMAN | Unmodified | 0 | 0 | 0 | 136780 | 3345800 | 263170 | 314050 | 89951 | 0 | 642930 | 111110 | 0 | 1279100 | 1197600 | 0 | 0 | 596070 | 1144900 |
| CLSSSLR | LRP1 | Q07954 | LRP1_HUMAN I | Unmodified | 0 | 0 | 0 | 0 | 0 | 0 | 88554 | 0 | 288910 | 0 | 0 | 0 | 0 | 129460 | 0 | 0 | 0 | 0 |
| CLSTANQE | MYO1A; MYO1A | Q9UBC5; G3 | MYO1A_HUMA | Unmodified | 134850 | 121070 | 0 | 0 | 0 | 0 | 0 | 0 | 0 | 0 | 0 | 0 | 0 | 0 | 0 | 14380 | 0 | 0 |
| CLSVMEAK | AARS1; AARS1; | P49588; A0A | SYAC_HUMAN | Unmodified | 0 | 0 | 0 | 0 | 1024500 | 10074000 | 768220 | 244190 | 540560 | 0 | 0 | 0 | 1027500 | 2418100 | 0 | 0 | 270850 | 3314200 |
| CLTAIVK | ANXA1 | P04083 | ANXA1_HUMAI | Unmodified | 0 | 0 | 0 | 0 | 6449500 | 1447000 | 0 | 68913 | 0 | 286780 | 17580000 | 1.66E+08 | 0 | 0 | 0 | 0 | 761820 | 2193500 |
| CLTQSGIA | SRI; SRI; SRI | P30626; B4D | SORCN_HUMA | Unmodified | 1741300 | 1539300 | 291180 | 693740 | 0 | 0 | 0 | 0 | 14064000 | 0 | 576690 | 5941800 | 0 | 0 | 2489100 | 1358900 | 0 | 0 |
| CLTTDEYD | BLVRB; BLVRB; | P30043; M0F | BLVRB_HUMAI | Unmodified | 547190 | 290120 | 323600 | 1117000 | 2274500 | 0 | 0 | 0 | 0 | 0 | 0 | 0 | 2132500 | 1802900 | 1111800 | 1031000 | 2501800 | 1473000 |
| CLVDSAETI | LLGL2; LLGL2 | Q6P1M3; J3C | L2GL2_HUMAN | Unmodified | 54575 | 0 | 0 | 197190 | 0 | 0 | 0 | 7114.8 | 362430 | 86312 | 0 | 59500 | 0 | 0 | 190500 | 236070 | 0 | 0 |
| CLVGEFVS | APP; APP; APP; | P05067; H7C | A4_HUMAN Am | Unmodified | 0 | 0 | 0 | 0 | 0 | 170600 | 0 | 0 | 1066900 | 405670 | 78294 | 0 | 0 | 0 | 0 | 0 | 0 | 0 |
| CLVGEFVS | APLP2; APLP2 | Q06481; E9F | APLP2_HUMAN | Unmodified | 0 | 0 | 0 | 0 | 301340 | 144310 | 186930 | 0 | 632250 | 511740 | 0 | 0 | 184180 | 233500 | 0 | 0 | 0 | 0 |
| CLVLTGFG | VAT1; VAT1 | Q99536; K7E | VAT1_HUMAN S | Unmodified | 0 | 0 | 0 | 0 | 8500100 | 2255000 | 470200 | 296640 | 0 | 0 | 0 | 0 | 1626500 | 2006300 | 0 | 0 | 2192100 | 3997400 |
| CLVMKAEM | EPCAM; EPCAM | P16422; C9J | EPCAM_HUMA | Oxidation (M) | 0 | 0 | 0 | 0 | 0 | 0 | 0 | 0 | 0 | 524640 | 0 | 0 | 0 | 525700 | 0 | 0 | 0 | 0 |
| CLVMKAEM | EPCAM; EPCAM | P16422; C9J | EPCAM_HUMA | Unmodified | 0 | 0 | 0 | 0 | 141180 | 0 | 0 | 0 | 0 | 925040 | 0 | 0 | 615630 | 1026900 | 0 | 0 | 155590 | 124770 |
| CLYASVLTA | EEF2 | P13639 | EF2_HUMAN El | Unmodified | 0 | 0 | 0 | 679310 | 2362500 | 0 | 0 | 246880 | 1067900 | 535560 | 2096200 | 12522000 | 1866000 | 1494900 | 0 | 0 | 1668100 | 0 |
| CLYSLINEA | EHHADH | Q08426 | ECHP_HUMAN | Unmodified | 0 | 0 | 0 | 0 | 0 | 0 | 0 | 0 | 900860 | 0 | 0 | 0 | 0 | 0 | 0 | 0 | 0 | 0 |
| CMDVYTPK | CSNK2B; CSNK | P67870; Q5S | CSK2B_HUMAI | Unmodified | 782840 | 0 | 0 | 0 | 288450 | 86948 | 124690 | 52287 | 0 | 34009 | 0 | 312880 | 0 | 44574 | 0 | 0 | 251100 | 177840 |
| CMMAQYN | STIP1; STIP1 | P31948; H0Y | STIP1_HUMAN | Unmodified | 0 | 0 | 0 | 0 | 0 | 113900 | 0 | 0 | 0 | 0 | 31646 | 568340 | 0 | 0 | 0 | 0 | 0 | 0 |
| CMMAQYN | STIP1; STIP1 | P31948; H0Y | STIP1_HUMAN | Unmodified | 0 | 0 | 0 | 0 | 261140 | 0 | 0 | 0 | 1286300 | 23999 | 219410 | 4779100 | 0 | 0 | 0 | 0 | 64493 | 159210 |
| CMMAQYN | STIP1; STIP1 | P31948; H0Y | STIP1_HUMAN | Oxidation (M) | 0 | 0 | 20633 | 47160 | 0 | 0 | 0 | 0 | 76359 | 0 | 0 | 1005500 | 0 | 16413 | 27134 | 25854 | 0 | 0 |
| CMQLTDFII | RPL14; RPL14 | P50914; E7E | RL14_HUMAN ( | Oxidation (M) | 1415500 | 610800 | 0 | 0 | 503540 | 0 | 0 | 124770 | 0 | 0 | 0 | 5882300 | 0 | 0 | 0 | 767080 | 0 | 0 |
| CMVTAWSI | PTGFRN | Q9P2B2 | FPRP_HUMAN | Unmodified | 0 | 0 | 765500 | 663470 | 60581 | 0 | 0 | 0 | 0 | 0 | 0 | 1434100 | 0 | 0 | 1470900 | 1020600 | 1696600 | 0 |
| CMVTAWSI | PTGFRN | Q9P2B2 | FPRP_HUMAN | Oxidation (M) | 1410100 | 740230 | 1928300 | 714580 | 0 | 0 | 0 | 0 | 0 | 0 | 0 | 5849800 | 0 | 0 | 2333300 | 3118700 | 0 | 0 |
| CNAEEHVF | LAMP1 | P11279 | LAMP1_HUMAI | Unmodified | 0 | 0 | 0 | 0 | 0 | 0 | 0 | 0 | 3866500 | 741240 | 0 | 0 | 1149500 | 0 | 0 | 0 | 0 | 0 |
| CNDQDTR | FN1 | P02751 | FINC_HUMAN I | Unmodified | 0 | 0 | 0 | 0 | 246130 | 770180 | 0 | 0 | 0 | 206390 | 0 | 0 | 0 | 1012600 | 0 | 0 | 140040 | 1005100 |
| CNEIINWLI | HSPA8; HSPA8; | P11142; E9P | HSP7C_HUMAI | Unmodified | 17229000 | 0 | 8908600 | 16346000 | 18195000 | 5856900 | 3481100 | 777660 | 24422000 | 6156200 | 8673700 | 30109000 | 22403000 | 11348000 | 14529000 | 10187000 | 2867700 | 8943700 |
| CNEIINWLI | HSPA8; HSPA8; | P11142; E9P | HSP7C_HUMAI | Unmodified | 0 | 0 | 0 | 0 | 14263000 | 6860300 | 9901400 | 0 | 549130 | 8688600 | 0 | 0 | 36737000 | 24103000 | 0 | 0 | 7693400 | 9848100 |
| CNELQDIEI | SMARCA1; SMA | P28370; O6C | SMCA1_HUMA | Unmodified | 194140 | 0 | 0 | 0 | 0 | 0 | 0 | 0 | 0 | 367020 | 0 | 0 | 62723 | 0 | 58755 | 0 | 0 | 0 |
| CNEPAVWS | CLTC; CLTC | Q00610; A0A | CLH1_HUMAN | Unmodified | 8229600 | 2338500 | 2142100 | 1921500 | 0 | 0 | 0 | 0 | 31516 | 0 | 0 | 21143000 | 0 | 0 | 2912100 | 0 | 0 | 0 |
| CNIQYPATI | CEP55 | Q53EZ4 | CEP55_HUMAI | Unmodified | 0 | 191230 | 129490 | 0 | 117730 | 0 | 0 | 0 | 630550 | 0 | 0 | 441170 | 0 | 0 | 0 | 371160 | 143770 | 0 |
| CNIQYPATI | CEP55 | Q53EZ4 | CEP55_HUMAI | Unmodified | 67731 | 0 | 0 | 241540 | 6121400 | 1036300 | 118690 | 0 | 668790 | 1949100 | 0 | 174830 | 0 | 0 | 195400 | 59062 | 1327100 | 283610 |
| CNKEVYFA | CRIP1 | P50238 | CRIP1_HUMAN | Unmodified | 0 | 0 | 24254 | 0 | 0 | 0 | 535830 | 348710 | 0 | 0 | 0 | 0 | 0 | 0 | 0 | 0 | 0 | 0 |
| CNKEVYFA | CRIP1 | P50238 | CRIP1_HUMAN | Unmodified | 0 | 0 | 0 | 0 | 1035000 | 0 | 20855000 | 3044900 | 0 | 465450 | 0 | 0 | 0 | 0 | 0 | 0 | 264650 | 0 |
| CNLVPTDE | IARS1; IARS1; IA | P41252; A0A | SYIC_HUMAN I | Unmodified | 0 | 0 | 0 | 0 | 0 | 0 | 0 | 0 | 0 | 0 | 0 | 1870000 | 0 | 0 | 0 | 0 | 0 | 0 |
| CNNVLYIR | SNRPF | P62306 | RUXF_HUMAN | Unmodified | 856690 | 511800 | 0 | 0 | 0 | 0 | 0 | 0 | 0 | 0 | 0 | 77097 | 0 | 0 | 0 | 501070 | 0 | 0 |
| CNRAGKP\ | PKM; PKM; PKM | P14618; B4D | KPYM_HUMAN | Oxidation (M) | 241310 | 0 | 0 | 0 | 0 | 0 | 0 | 0 | 933060 | 0 | 0 | 0 | 0 | 0 | 197380 | 56473 | 1111600 | 1406800 |
| CNRAGKP\ | PKM; PKM; PKM | P14618; B4D | KPYM_HUMAN | Unmodified | 0 | 0 | 0 | 0 | 2048200 | 132750 | 0 | 0 | 0 | 40216 | 0 | 0 | 0 | 0 | 0 | 0 | 373050 | 783790 |
| CNSFIENS | RIPOR2; RIPOR | Q9Y4F9; B7Z | RIPR2_HUMAN | Unmodified | 0 | 0 | 0 | 0 | 0 | 0 | 0 | 0 | 0 | 0 | 0 | 0 | 227440 | 0 | 0 | 0 | 0 | 0 |
| CNSLSTLEI | LAMP2 | P13473 | LAMP2_HUMAI | Unmodified | 0 | 0 | 1148500 | 2136700 | 0 | 0 | 0 | 0 | 27043000 | 2402400 | 1239600 | 1316400 | 189490 | 0 | 1619900 | 1643800 | 0 | 0 |
| CNTCGEPI | ZYX; ZYX; ZYX | Q15942; H0Y | ZYX_HUMAN Zy | Unmodified | 0 | 0 | 0 | 0 | 121640 | 0 | 0 | 0 | 0 | 0 | 56105 | 0 | 0 | 0 | 0 | 0 | 243370 | 757700 |
| CNTCGEPI | ZYX; ZYX; ZYX | Q15942; H0Y | ZYX_HUMAN Zy | Unmodified | 0 | 86221 | 0 | 0 | 225340 | 0 | 0 | 0 | 0 | 0 | 0 | 0 | 73280 | 161030 | 505870 | 0 | 2772700 | 2662000 |
| CNTDDTIG | UBL5 | Q9BZL1 | UBL5_HUMAN | Unmodified | 275650 | 335170 | 351200 | 563630 | 831430 | 0 | 193970 | 0 | 248510 | 60019 | 0 | 131130 | 532610 | 207340 | 301800 | 269270 | 873590 | 1120100 |
| CNTDDTIG | UBL5 | Q9BZL1 | UBL5_HUMAN | Unmodified | 0 | 0 | 0 | 0 | 0 | 0 | 0 | 0 | 91892 | 116940 | 0 | 0 | 0 | 0 | 0 | 0 | 827960 | 0 |
| CNTPTYCD | VDAC3; VDAC3 | Q9Y277; E5F | VDAC3_HUMAI | Acetyl (Protein N | 0 | 0 | 0 | 0 | 0 | 0 | 0 | 0 | 918680 | 1940300 | 0 | 0 | 0 | 0 | 0 | 0 | 0 | 0 |
| CNVWILDG | DYNC1LI1; DYN | Q9Y6G9; O4 | DC1L1_HUMAI | Unmodified | 0 | 0 | 0 | 0 | 0 | 0 | 0 | 0 | 0 | 0 | 0 | 462710 | 1139600 | 0 | 73067 | 0 | 0 | 0 |
| CNYSQSEF | ZC3HAV1; ZC3H | Q7Z2W4; C9 | ZCCHV_HUMA | Unmodified | 0 | 0 | 0 | 0 | 2514700 | 856030 | 0 | 0 | 0 | 127260 | 0 | 0 | 930490 | 398240 | 0 | 0 | 438020 | 0 |
| CPEALFQP | ACTG1; ACTB; A | P63261; P60 | ACTG_HUMAN | Oxidation (M) | 0 | 0 | 0 | 0 | 0 | 0 | 0 | 0 | 183540 | 0 | 354460 | 0 | 0 | 0 | 0 | 0 | 0 | 0 |
| CPEALFQP | ACTG1; ACTB; A | P63261; P60 | ACTG_HUMAN | 2 Oxidation (M) | 1004000 | 606690 | 1143000 | 1017000 | 6279400 | 3654500 | 0 | 0 | 3744500 | 0 | 1643800 | 13296000 | 0 | 0 | 1175000 | 3922900 | 1856900 | 2996300 |
| CPEALFQP | ACTG1; ACTB; A | P63261; P60 | ACTG_HUMAN | Oxidation (M) | 672310 | 192190 | 0 | 498810 | 4736200 | 435870 | 0 | 0 | 4528800 | 0 | 0 | 738390 | 0 | 1106300 | 639680 | 458710 | 0 | 2324200 |
| CPEALFQP | ACTG1; ACTB; A | P63261; P60 | ACTG_HUMAN | Unmodified | 0 | 0 | 0 | 0 | 0 | 0 | 0 | 0 | 1298800 | 0 | 0 | 0 | 0 | 0 | 0 | 0 | 0 | 0 |
| CPENAFFLI | MYO1C; MYO1C | O00159; F5H | MYO1C_HUMA | Unmodified | 3315600 | 1662600 | 3754600 | 2516400 | 0 | 0 | 0 | 0 | 0 | 47626 | 0 | 807860 | 0 | 0 | 6739300 | 5797200 | 0 | 0 |
| CPFGALSIV | ABCE1; ABCE1; | P61221; D6F | ABCE1_HUMAI | Unmodified | 22982 | 0 | 51544 | 90363 | 853400 | 143500 | 0 | 44727 | 171970 | 0 | 256710 | 2360000 | 216700 | 0 | 0 | 235100 | 184120 | 274080 |
| CPFTGNVS | RPS11; RPS11 | P62280; M0C | RS11_HUMAN | Unmodified | 1251100 | 510730 | 0 | 0 | 0 | 0 | 0 | 0 | 0 | 0 | 241920 | 8964500 | 0 | 0 | 0 | 0 | 0 | 0 |
| CPKKDSDS | ITGB1; ITGB1; I | P05556; H7C | ITB1_HUMAN In | Unmodified | 59734 | 0 | 0 | 0 | 2905300 | 377720 | 0 | 0 | 99228 | 0 | 0 | 0 | 0 | 21993 | 0 | 0 | 1468100 | 2559000 |
| CPKVSVGA | AHNAK | Q09666 | AHNK_HUMAN | Unmodified | 0 | 172440 | 0 | 0 | 0 | 0 | 0 | 0 | 0 | 0 | 0 | 0 | 0 | 0 | 0 | 0 | 776110 | 0 |
| CPLSGACY | COPA; COPA; C | P53621; A0A | COPA_HUMAN | Unmodified | 0 | 0 | 0 | 0 | 0 | 0 | 0 | 0 | 0 | 0 | 0 | 1193500 | 0 | 0 | 0 | 0 | 0 | 0 |
| CPNLTHLN | ANP32A; ANP32 | P39687; H0Y | AN32A_HUMAI | Unmodified | 0 | 29007 | 0 | 0 | 1252500 | 450310 | 128710 | 0 | 1649100 | 346340 | 0 | 8384500 | 1513800 | 546040 | 0 | 0 | 273120 | 188280 |
| CPNLTYLN | ANP32E; ANP32 | Q9BTT0; Q5T | AN32E_HUMAI | Unmodified | 0 | 0 | 0 | 0 | 957590 | 0 | 0 | 0 | 102280 | 46413 | 0 | 1861700 | 59097 | 0 | 0 | 0 | 0 | 0 |
| CPNPPVQE | APOD; APOD; A | P05090; C9J | APOD_HUMAN | Unmodified | 0 | 0 | 0 | 0 | 0 | 0 | 0 | 0 | 59885000 | 4592500 | 0 | 0 | 0 | 0 | 0 | 0 | 0 | 0 |
| CPNPPVQE | APOD; APOD; A | P05090; C9J | APOD_HUMAN | Unmodified | 0 | 0 | 0 | 25784000 | 0 | 0 | 98350 | 0 | 13967000 | 495700 | 4247700 | 0 | 0 | 168360 | 0 | 0 | 0 | 0 |
| CPPGSPMI | ECE1; ECE1; EC | P42892; B4D | ECE1_HUMAN | Unmodified | 0 | 0 | 0 | 0 | 77034 | 0 | 0 | 0 | 382100 | 44785 | 0 | 66429 | 0 | 0 | 0 | 0 | 27252 | 0 |
| CPPGVVPA | FASN; FASN | P49327; A0A | FAS_HUMAN Fa | Unmodified | 0 | 74079 | 0 | 0 | 0 | 0 | 0 | 0 | 0 | 0 | 0 | 1544900 | 0 | 0 | 55627 | 78352 | 0 | 0 |
| CPQEAGAS | AKAP5 | P24588 | AKAP5_HUMAI | Unmodified | 0 | 0 | 0 | 132700 | 1874900 | 0 | 0 | 0 | 0 | 0 | 0 | 0 | 1777700 | 1491500 | 259200 | 154260 | 185810 | 468620 |
| CPRGFNGI | EDIL3 | O43854 | EDIL3_HUMAN | Unmodified | 0 | 0 | 0 | 0 | 1459900 | 0 | 0 | 0 | 0 | 0 | 0 | 0 | 0 | 0 | 0 | 0 | 362140 | 4699100 |
| CPSGDLPG | ADAM15 | Q13444 | ADA15_HUMAI | Unmodified | 4489100 | 1117100 | 182600 | 507150 | 0 | 0 | 0 | 0 | 534060 | 0 | 181700 | 862880 | 0 | 0 | 1925000 | 1072700 | 0 | 0 |

| | | | | | | | | | | | | | | | | | | | | | | |
|---|---|---|---|---|---|---|---|---|---|---|---|---|---|---|---|---|---|---|---|---|---|---|
| CPSPPMIN | PPT1; PPT1; PP | P50897; A0A | PPT1_HUMAN | Unmodified | 0 | 0 | 0 | 0 | 0 | 0 | 0 | 0 | 1086400 | 0 | 0 | 0 | 0 | 0 | 0 | 0 | 0 | 0 |
| CPTCPDAC | ITGB3; N/A | P05106; H3E | ITB3_HUMAN I | Unmodified | 852330 | 212650 | 212190 | 888210 | 668730 | 0 | 2576400 | 1063200 | 0 | 0 | 0 | 0 | 1727300 | 5082600 | 411410 | 364290 | 0 | 622290 |
| CPTQFPLIL | NIBAN2 | Q96TA1 | NIBA2_HUMAN | Unmodified | 0 | 0 | 0 | 0 | 3011900 | 1142300 | 0 | 0 | 1648300 | 870740 | 0 | 693050 | 0 | 0 | 0 | 0 | 0 | 0 |
| CPVDLSTA | ITGA2; ITGA2; I | P17301; D6F | ITA2_HUMAN I | Unmodified | 485340 | 206890 | 241690 | 905010 | 438050 | 0 | 0 | 0 | 0 | 0 | 0 | 0 | 0 | 0 | 1133100 | 740070 | 215590 | 545420 |
| CPVGYQLE | HEG1 | Q9ULI3 | HEG1_HUMAN | Unmodified | 0 | 0 | 141250 | 0 | 1206400 | 0 | 0 | 0 | 1365300 | 279500 | 0 | 0 | 0 | 0 | 293300 | 36379 | 0 | 398370 |
| CQADAQA | GALNT16 | Q8N428 | GLT16_HUMAN | Unmodified | 0 | 0 | 0 | 0 | 316890 | 0 | 0 | 0 | 1649800 | 533000 | 0 | 0 | 0 | 0 | 0 | 0 | 0 | 0 |
| CQCEEGF | LDLR; LDLR; LD | P01130; J3K | LDLR_HUMAN | Unmodified | 0 | 0 | 0 | 0 | 592580 | 0 | 225810 | 0 | 774380 | 0 | 1047100 | 0 | 0 | 0 | 0 | 0 | 0 | 0 |
| CQCHGHV | MEGF9 | Q9H1U4 | MEGF9_HUMA | Unmodified | 0 | 14960 | 0 | 0 | 0 | 0 | 0 | 0 | 159120 | 0 | 0 | 0 | 0 | 0 | 0 | 0 | 0 | 0 |
| CQEVISWL | HSPA1A; HSPA | P0DMV8; P0 | HS71A_HUMAN | Unmodified | 8532600 | 5156700 | 1104000 | 5185700 | 4613300 | 1482300 | 656140 | 0 | 12752000 | 2105700 | 0 | 3658800 | 10693000 | 3465200 | 3202600 | 0 | 1286700 | 4179200 |
| CQFAFYGC | SLC44A2 | Q8IWA5 | CTL2_HUMAN | Unmodified | 229400 | 390550 | 0 | 0 | 0 | 0 | 0 | 0 | 113580 | 0 | 0 | 0 | 0 | 0 | 321360 | 172930 | 0 | 0 |
| CQFQTKFP | GOLGA7 | Q7Z5G4 | GOGA7_HUMA | Unmodified | 0 | 0 | 0 | 259540 | 426440 | 0 | 0 | 0 | 0 | 0 | 0 | 651530 | 0 | 0 | 0 | 674360 | 0 | 0 |
| CQFTLKPIS | MCU; MCU | Q8NE86; S4 | MCU_HUMAN | Unmodified | 0 | 0 | 0 | 0 | 0 | 0 | 0 | 0 | 220290 | 54115 | 0 | 0 | 0 | 0 | 0 | 0 | 0 | 0 |
| CQGGDFT | PPIAL4C; PPIA; | A0A0B4J2A2 | PAL4C_HUMAN | Unmodified | 173230 | 0 | 0 | 0 | 0 | 0 | 0 | 0 | 281490 | 0 | 0 | 169530 | 0 | 0 | 161180 | 39225 | 0 | 0 |
| CQGGVNP | SORT1 | Q99523 | SORT_HUMAN | Unmodified | 0 | 0 | 0 | 0 | 0 | 0 | 633040 | 0 | 3118300 | 549320 | 0 | 0 | 208410 | 0 | 0 | 0 | 0 | 0 |
| CQGVVCA | CSDE1; CSDE1 | O75534; E9F | CSDE1_HUMA | Unmodified | 0 | 0 | 0 | 0 | 0 | 0 | 0 | 0 | 0 | 28265 | 0 | 593680 | 0 | 0 | 0 | 0 | 0 | 0 |
| CQKICDQV | ACTN1; ACTN1 | P12814; A0A | ACTN1_HUMA | Unmodified | 0 | 0 | 0 | 0 | 0 | 0 | 0 | 0 | 0 | 0 | 0 | 0 | 185860 | 0 | 0 | 0 | 270090 | 0 |
| CQKLELLL | ITGA3 | P26006 | ITA3_HUMAN I | Oxidation (M) | 0 | 0 | 0 | 0 | 3931600 | 2116800 | 0 | 0 | 0 | 0 | 0 | 0 | 137960 | 0 | 0 | 0 | 619210 | 540320 |
| CQKLELLL | ITGA3 | P26006 | ITA3_HUMAN I | Unmodified | 0 | 0 | 0 | 0 | 9325900 | 2738500 | 0 | 0 | 0 | 0 | 0 | 0 | 0 | 0 | 0 | 0 | 3614800 | 5972000 |
| CQLEINFN | ACTN2; ACTN4 | P35609; O4 | ACTN2_HUMA | Unmodified | 157320 | 90154 | 115500 | 340240 | 2386800 | 754090 | 360590 | 0 | 1137700 | 633400 | 324220 | 4444900 | 888260 | 601390 | 466120 | 494010 | 3936600 | 208830 |
| CQLLFALK | NUDT16L1; NU | Q9BRJ7; K7E | TIRR_HUMAN T | Unmodified | 0 | 0 | 0 | 43747 | 0 | 0 | 0 | 11540 | 0 | 0 | 0 | 172690 | 0 | 0 | 157290 | 112220 | 0 | 0 |
| CQNALQQ | GFPT1; GFPT1 | Q06210; A0 | GFPT1_HUMAN | Unmodified | 0 | 0 | 0 | 0 | 0 | 0 | 0 | 0 | 0 | 0 | 0 | 539960 | 1142100 | 434990 | 0 | 0 | 0 | 180420 |
| CQPIEFDA | ITGAV | P06756 | ITAV_HUMAN I | Unmodified | 1180900 | 0 | 0 | 333800 | 1239000 | 285780 | 0 | 0 | 0 | 0 | 0 | 831790 | 0 | 0 | 409620 | 226680 | 0 | 741450 |
| CQPSAITL | NPLOC4 | Q8TAT6 | NPL4_HUMAN | Unmodified | 0 | 0 | 0 | 0 | 293900 | 0 | 0 | 41632 | 0 | 0 | 0 | 0 | 0 | 0 | 0 | 0 | 490930 | 0 |
| CQQLAAYC | COPS4; COPS4 | Q9BT78; D6 | CSN4_HUMAN | Unmodified | 0 | 0 | 0 | 0 | 0 | 0 | 0 | 0 | 0 | 0 | 0 | 1165900 | 294680 | 0 | 0 | 0 | 0 | 0 |
| CQRVDLLG | ADAM15 | Q13444 | ADA15_HUMA | Unmodified | 1547700 | 370880 | 0 | 2473000 | 0 | 0 | 0 | 0 | 859630 | 0 | 0 | 1048000 | 194010 | 0 | 0 | 699010 | 0 | 80217 |
| CQVAFSYL | SH3KBP1; SH3 | Q96B97; Q5 | SH3K1_HUMA | Unmodified | 0 | 0 | 0 | 0 | 0 | 0 | 0 | 0 | 0 | 0 | 0 | 0 | 0 | 0 | 258540 | 189670 | 0 | 0 |
| CQVFEETQ | CCT7 | Q99832 | TCPH_HUMAN | Unmodified | 0 | 0 | 0 | 0 | 247630 | 0 | 0 | 0 | 0 | 0 | 0 | 2056300 | 0 | 0 | 0 | 0 | 0 | 0 |
| CQVFEETQ | CCT7 | Q99832 | TCPH_HUMAN | Unmodified | 0 | 0 | 0 | 0 | 0 | 188570 | 0 | 0 | 607320 | 294360 | 0 | 1770000 | 0 | 0 | 0 | 0 | 326150 | 0 |
| CQYYSVSF | DPP4; DPP4; D | P27487; A0A | DPP4_HUMAN | Unmodified | 139540 | 113110 | 669520 | 786220 | 0 | 0 | 0 | 0 | 0 | 0 | 0 | 0 | 0 | 0 | 2471300 | 2437400 | 0 | 0 |
| CRAPEVSQ | AP1B1 | Q10567 | AP1B1_HUMA | Unmodified | 0 | 0 | 0 | 0 | 107880 | 0 | 0 | 0 | 0 | 0 | 0 | 553230 | 95761 | 92224 | 0 | 0 | 159520 | 392660 |
| CRAPEVSQ | AP2B1; AP2B1 | P63010; A0A | AP2B1_HUMA | Unmodified | 188390 | 0 | 0 | 0 | 766860 | 0 | 801420 | 0 | 0 | 0 | 0 | 0 | 0 | 0 | 0 | 0 | 1036000 | 0 |
| CRDPTLGV | SMOC1 | Q9H4F8 | SMOC1_HUMA | Unmodified | 0 | 0 | 0 | 0 | 0 | 0 | 0 | 0 | 419910 | 0 | 0 | 0 | 0 | 0 | 553360 | 0 | 0 | 0 |
| CREMDEQ | TPM3; TPM3; T | J3KN67; A0A | J3KN67_HUMA | Unmodified | 143810 | 0 | 141990 | 0 | 0 | 0 | 32846 | 0 | 386920 | 0 | 336120 | 2253000 | 0 | 0 | 0 | 0 | 0 | 0 |
| CREMDEQ | TPM3; TPM3; T | J3KN67; A0A | J3KN67_HUMA | Unmodified | 0 | 0 | 0 | 0 | 1304300 | 1629600 | 416830 | 0 | 0 | 0 | 0 | 0 | 786740 | 0 | 0 | 0 | 0 | 0 |
| CRFPVVEN | CD46 | P15529 | MCP_HUMAN | Unmodified | 5027300 | 1949300 | 932340 | 733090 | 935170 | 0 | 0 | 0 | 1539900 | 604400 | 0 | 3019200 | 666220 | 0 | 2253400 | 952890 | 0 | 608410 |
| CRLDMIPD | MYOF | Q9NZM1 | MYOF_HUMAN | Oxidation (M) | 0 | 0 | 0 | 0 | 1096500 | 367900 | 0 | 0 | 0 | 0 | 0 | 0 | 0 | 0 | 0 | 0 | 0 | 143180 |
| CRLFVGNL | SFPQ | P23246 | SFPQ_HUMAN | Unmodified | 0 | 0 | 302410 | 800730 | 0 | 0 | 420880 | 0 | 0 | 88712 | 209940 | 196310 | 2922300 | 482030 | 0 | 0 | 0 | 0 |
| CRLVDADC | ADAM10 | O14672 | ADA10_HUMA | Unmodified | 379940 | 343510 | 0 | 1120200 | 110200 | 0 | 0 | 0 | 0 | 0 | 0 | 0 | 0 | 0 | 1018400 | 0 | 0 | 0 |
| CRPVNQE | HSPG2 | P98160 | PGBM_HUMAN | Unmodified | 0 | 0 | 727240 | 347910 | 501780 | 42960 | 0 | 29178 | 245460 | 0 | 523990 | 4394800 | 0 | 0 | 139240 | 291590 | 149240 | 208270 |
| CRSAGIKV | ATP1A1; ATP1A | P05023; P50 | AT1A1_HUMAN | Unmodified | 0 | 0 | 160310 | 0 | 3445400 | 0 | 896630 | 0 | 0 | 416360 | 0 | 0 | 6194300 | 2235200 | 0 | 0 | 2155200 | 547750 |
| CRSAGIKV | ATP1A1; ATP1A | P05023; P50 | AT1A1_HUMAN | Oxidation (M) | 0 | 0 | 0 | 0 | 735780 | 0 | 0 | 0 | 0 | 0 | 0 | 0 | 83031 | 1169500 | 0 | 0 | 289470 | 0 |
| CRSGFSLG | LRP1 | Q07954 | LRP1_HUMAN | Unmodified | 0 | 0 | 0 | 0 | 200610 | 0 | 0 | 0 | 199450 | 0 | 0 | 123540 | 0 | 0 | 0 | 0 | 0 | 290170 |
| CRTSGSVY | SRP14; SRP14 | P37108; H0 | SRP14_HUMAN | Unmodified | 559330 | 306680 | 158540 | 566990 | 0 | 0 | 182230 | 75578 | 0 | 0 | 0 | 0 | 0 | 0 | 595860 | 747380 | 0 | 0 |
| CRVDLPLA | DCTPP1; DCTP | Q9H773; H3 | DCTP1_HUMA | Unmodified | 0 | 0 | 0 | 0 | 0 | 0 | 0 | 0 | 0 | 0 | 26290 | 329460 | 0 | 0 | 0 | 0 | 0 | 0 |
| CRVIPSQE | SLC9A3R1 | O14745 | NHRF1_HUMA | Unmodified | 0 | 0 | 0 | 0 | 0 | 0 | 0 | 0 | 0 | 0 | 0 | 1632400 | 7530600 | 1962600 | 1840000 | 1928000 | 0 | 0 |
| CRVLTSGIF | GNAS; GNAS; G | Q5JWF2; P6 | GNAS1_HUMA | Unmodified | 55085 | 0 | 272110 | 149020 | 0 | 0 | 0 | 0 | 0 | 0 | 0 | 517170 | 0 | 0 | 455830 | 259700 | 0 | 177790 |
| CRVPCNW | MDK; MDK; MD | P21741; E9P | MK_HUMAN Mi | Unmodified | 135050 | 0 | 0 | 0 | 8182.9 | 0 | 0 | 0 | 396970 | 535490 | 0 | 0 | 0 | 0 | 166530 | 0 | 57735 | 158830 |
| CRVVGFAL | SNX33; SNX33 | Q8WV41; H3 | SNX33_HUMA | Oxidation (M) | 0 | 0 | 0 | 0 | 0 | 0 | 0 | 0 | 0 | 0 | 0 | 0 | 0 | 0 | 488000 | 402410 | 0 | 0 |
| CSACNVW | CSDE1; CSDE1 | O75534; E9F | CSDE1_HUMA | Unmodified | 0 | 0 | 0 | 0 | 85130 | 0 | 61303 | 112980 | 0 | 0 | 0 | 0 | 407940 | 314700 | 0 | 0 | 766440 | 192870 |
| CSASNRIIG | RPS21; RPS21; | P63220; Q9 | RS21_HUMAN | Unmodified | 0 | 0 | 0 | 0 | 0 | 0 | 117890 | 18600 | 0 | 0 | 0 | 0 | 0 | 559310 | 0 | 0 | 233410 | 236020 |
| CSATGSPA | HSPG2 | P98160 | PGBM_HUMAN | Unmodified | 0 | 0 | 1134400 | 519810 | 137210 | 0 | 0 | 0 | 0 | 63147 | 0 | 0 | 342000 | 589590 | 0 | 774720 | 65805 | 1566300 |
| CSDAAGYF | PCBP1 | Q15365 | PCBP1_HUMA | Unmodified | 13290000 | 3888900 | 2960600 | 1490900 | 0 | 0 | 0 | 0 | 0 | 0 | 0 | 3065200 | 0 | 0 | 1555300 | 4903800 | 0 | 0 |
| CSDIVFAR | PDCD6IP | Q8WUM4 | PDC6I_HUMAN | Unmodified | 13820000 | 9050900 | 3494800 | 9850300 | 4802700 | 2067800 | 189270 | 0 | 6153000 | 2641100 | 893660 | 4895000 | 9735100 | 6004000 | 6811500 | 6650700 | 780780 | 3389700 |
| CSDNTEVE | ANLN; ANLN | Q9NQW6; C | ANLN_HUMAN | Unmodified | 0 | 0 | 0 | 0 | 0 | 0 | 0 | 0 | 0 | 0 | 0 | 0 | 0 | 0 | 0 | 0 | 813210 | 0 |
| CSENKLPA | APEX1; APEX1; | P27695; A0A | APEX1_HUMAN | Unmodified | 0 | 0 | 0 | 0 | 0 | 0 | 0 | 0 | 0 | 0 | 0 | 2946300 | 0 | 0 | 0 | 0 | 0 | 0 |
| CSENNETI | SLC5A3 | P53794 | SC5A3_HUMAN | Unmodified | 0 | 0 | 0 | 0 | 0 | 0 | 0 | 0 | 0 | 0 | 0 | 0 | 0 | 0 | 366530 | 0 | 0 | 0 |
| CSFLHPEP | C6orf89 | Q6UWU4 | CF089_HUMAN | Unmodified | 0 | 0 | 0 | 0 | 0 | 0 | 0 | 0 | 869650 | 71874 | 0 | 0 | 0 | 0 | 0 | 0 | 59847 | 0 |
| CSGIASAA | QPRT | Q15274 | NADC_HUMAN | Unmodified | 0 | 0 | 0 | 0 | 0 | 0 | 0 | 0 | 624370 | 0 | 0 | 1296900 | 0 | 0 | 0 | 0 | 0 | 0 |
| CSGPGLEF | FLNA; FLNA | P21333; Q6 | FLNA_HUMAN | Unmodified | 0 | 0 | 0 | 1039400 | 457260 | 0 | 13389 | 0 | 11085 | 0 | 0 | 0 | 260540 | 207360 | 503460 | 511520 | 268180 | 142390 |
| CSGPGLG | FLNC | Q14315 | FLNC_HUMAN | Unmodified | 0 | 0 | 0 | 0 | 818840 | 243400 | 0 | 0 | 0 | 0 | 0 | 0 | 0 | 0 | 0 | 0 | 648250 | 178850 |
| CSGPGLPL | DPP4; DPP4; D | P27487; A0A | DPP4_HUMAN | Unmodified | 0 | 0 | 3331500 | 11612000 | 0 | 0 | 0 | 83988 | 0 | 0 | 0 | 0 | 0 | 0 | 11405000 | 6526200 | 1179000 | 0 |
| CSGPGLSF | FLNA; FLNA | P21333; Q6 | FLNA_HUMAN | Oxidation (M) | 117440 | 0 | 0 | 249750 | 450990 | 0 | 0 | 115870 | 0 | 0 | 0 | 0 | 444060 | 380880 | 0 | 282000 | 888540 | 0 |
| CSGPGLSF | FLNA; FLNA | P21333; Q6 | FLNA_HUMAN | Unmodified | 0 | 0 | 0 | 165410 | 0 | 0 | 1085800 | 700620 | 0 | 0 | 238870 | 0 | 1379500 | 0 | 0 | 0 | 1490200 | 213970 |
| CSGPGLSF | FLNA; FLNA | P21333; Q6 | FLNA_HUMAN | Oxidation (M) | 0 | 0 | 0 | 0 | 1081900 | 235500 | 4803200 | 2816400 | 0 | 0 | 0 | 0 | 1077600 | 3353800 | 0 | 0 | 6203700 | 610960 |
| CSGPGLSF | FLNA; FLNA | P21333; Q6 | FLNA_HUMAN | Unmodified | 92302 | 0 | 0 | 0 | 2737400 | 3656800 | 1853100 | 921290 | 0 | 0 | 0 | 0 | 817780 | 2775300 | 0 | 0 | 1945300 | 2406200 |
| CSGPGLSF | FLNA; FLNA | P21333; Q6 | FLNA_HUMAN | Oxidation (M) | 0 | 0 | 0 | 0 | 0 | 0 | 0 | 0 | 0 | 0 | 0 | 0 | 0 | 0 | 0 | 0 | 2977400 | 0 |
| CSGPLGIE | EDIL3 | O43854 | EDIL3_HUMAN | Unmodified | 1339300 | 0 | 0 | 0 | 45334000 | 39034000 | 1209300 | 305350 | 27434000 | 4236600 | 0 | 0 | 525490 | 0 | 0 | 0 | 4682700 | 27691000 |
| CSGPLGIE | EDIL3 | O43854 | EDIL3_HUMAN | Unmodified | 0 | 0 | 0 | 0 | 1.8E+08 | 72316000 | 23023000 | 0 | 1529700 | 7915700 | 0 | 0 | 0 | 0 | 0 | 0 | 57170000 | 1.07E+08 |
| CSGVACLV | WASL | O00401 | WASL_HUMAN | Unmodified | 385690 | 0 | 0 | 791860 | 0 | 0 | 0 | 75820 | 75767 | 0 | 0 | 0 | 0 | 0 | 290460 | 0 | 0 | 0 |
| CSHALIRG | DDAH2; DDAH2 | O95865; Q5 | DDAH2_HUMA | Unmodified | 0 | 0 | 0 | 0 | 0 | 0 | 0 | 0 | 0 | 260240 | 497020 | 0 | 0 | 0 | 0 | 421700 | 2654500 | 2960800 |
| CSHLLVK | PIN1; PIN1 | Q13526; K7E | PIN1_HUMAN | Unmodified | 1735600 | 1023500 | 458520 | 587040 | 0 | 0 | 16689 | 37333 | 125660 | 0 | 0 | 282110 | 442590 | 147240 | 800310 | 491520 | 316420 | 527110 |
| CSLCAFDA | KRTCAP3 | Q53RY4 | KCP3_HUMAN | Unmodified | 270990 | 0 | 0 | 0 | 0 | 0 | 0 | 0 | 0 | 0 | 0 | 105640 | 0 | 0 | 167860 | 146620 | 0 | 0 |
| CSLHSASP | KIAA1522 | Q9P206 | K1522_HUMAN | Unmodified | 0 | 0 | 282870 | 376880 | 435100 | 0 | 0 | 0 | 0 | 0 | 162160 | 592310 | 1369700 | 995960 | 819720 | 618170 | 396940 | 0 |
| CSLNPEW | PRKCB; PRKCB | P05771; I3L | KPCB_HUMAN | Unmodified | 0 | 0 | 0 | 0 | 137400 | 0 | 1487600 | 0 | 0 | 0 | 56694 | 0 | 0 | 1248200 | 0 | 0 | 0 | 749610 |
| CSNPLDTS | CTPS1; CTPS1; | P17812; B4E | PYRG1_HUMA | Unmodified | 0 | 0 | 227480 | 107210 | 0 | 0 | 0 | 0 | 0 | 0 | 244870 | 980540 | 0 | 0 | 133130 | 0 | 0 | 0 |
| CSPEIQGH | CHAT; CHAT | P28329; A0A | CLAT_HUMAN | Unmodified | 0 | 0 | 0 | 0 | 0 | 0 | 0 | 0 | 0 | 0 | 0 | 613950 | 0 | 0 | 0 | 0 | 0 | 0 |
| CSPTEAAV | HAPLN1 | P10915 | HPLN1_HUMA | Unmodified | 0 | 39229 | 0 | 132130 | 0 | 0 | 0 | 0 | 2574300 | 148250 | 0 | 0 | 0 | 0 | 365870 | 267300 | 0 | 0 |
| CSQAVYAA | CSRP1; CSRP1; | P21291; E9P | CSRP1_HUMA | Unmodified | 695600 | 512000 | 419830 | 799880 | 2485300 | 1275000 | 328710 | 148550 | 1472500 | 888260 | 0 | 1219500 | 1760500 | 1311100 | 702900 | 874790 | 851590 | 982210 |
| CSRFLNDT | MYO1B; MYO1E | O43795; E9F | MYO1B_HUMA | Unmodified | 0 | 0 | 0 | 0 | 413870 | 0 | 0 | 0 | 0 | 0 | 0 | 0 | 0 | 0 | 0 | 0 | 822120 | 0 |
| CSSILLHG | ATP1A1 | P05023 | AT1A1_HUMAN | Unmodified | 0 | 0 | 1031900 | 0 | 0 | 0 | 0 | 0 | 245960 | 112390 | 0 | 3049700 | 0 | 0 | 2039800 | 1822500 | 0 | 0 |
| CSSILLHG | ATP1A1 | P05023 | AT1A1_HUMAN | Unmodified | 0 | 0 | 0 | 1065600 | 0 | 0 | 0 | 0 | 1415700 | 0 | 1059100 | 7820100 | 0 | 0 | 1066100 | 1776300 | 0 | 0 |
| CSTPEEIK | DSTN; DSTN | P60981; F6R | DEST_HUMAN | Unmodified | 0 | 0 | 0 | 0 | 0 | 0 | 0 | 0 | 0 | 0 | 0 | 631580 | 433940 | 264300 | 0 | 0 | 417810 | 0 |
| CSTPSTDA | PTGFRN | Q9P2B2 | FPRP_HUMAN | Unmodified | 6870500 | 1842900 | 8191800 | 8279200 | 5198400 | 1747800 | 2811400 | 822420 | 0 | 0 | 0 | 0 | 36135000 | 25403000 | 11666000 | 9567100 | 714020 | 2610600 |
| CSVAVEMF | KIF23; KIF23; K | Q02241; A0 | KIF23_HUMAN | Unmodified | 0 | 0 | 102510 | 291430 | 0 | 287880 | 0 | 0 | 575120 | 0 | 135820 | 438000 | 0 | 0 | 0 | 0 | 0 | 0 |
| CSVCPDYD | SQSTM1; SQST | Q13501; E9F | SQSTM_HUMA | Unmodified | 0 | 0 | 0 | 0 | 0 | 0 | 0 | 0 | 0 | 0 | 0 | 0 | 0 | 0 | 0 | 0 | 698550 | 0 |
| CSVCSEPI | ZYX; ZYX | Q15942; H0 | ZYX_HUMAN Z | Oxidation (M) | 0 | 0 | 0 | 0 | 0 | 0 | 0 | 0 | 0 | 0 | 0 | 0 | 0 | 0 | 0 | 0 | 1521500 | 0 |
| CSVCSEPI | ZYX; ZYX | Q15942; H0 | ZYX_HUMAN Z | Unmodified | 0 | 0 | 0 | 0 | 0 | 0 | 0 | 0 | 0 | 0 | 0 | 0 | 0 | 0 | 0 | 0 | 1596600 | 4171100 |
| CSVIRDSL | PFN1; PFN1 | P07737; K7E | PROF1_HUMA | Oxidation (M) | 3171500 | 1659300 | 1201500 | 2486300 | 2501200 | 1864400 | 468460 | 0 | 2636500 | 2102100 | 2530200 | 10467000 | 4769100 | 1049200 | 2134300 | 1855000 | 2411900 | 2997300 |
| CSVIRDSL | PFN1; PFN1 | P07737; K7E | PROF1_HUMA | Unmodified | 930360 | 740370 | 0 | 1449600 | 5676200 | 642280 | 229180 | 0 | 2094200 | 766760 | 495850 | 4143400 | 4288800 | 827660 | 661000 | 138350 | 1506000 | 2046200 |
| CSVIRDSL | PFN1; PFN1 | P07737; K7E | PROF1_HUMA | Oxidation (M) | 100780 | 0 | 0 | 0 | 31128000 | 6152000 | 10634000 | 0 | 0 | 6889800 | 0 | 0 | 14862000 | 10889000 | 0 | 0 | 27832000 | 14603000 |
| CSVIRDSL | PFN1; PFN1 | P07737; K7E | PROF1_HUMA | Unmodified | 0 | 0 | 0 | 0 | 34490000 | 4656200 | 7621100 | 0 | 198680 | 2810700 | 0 | 0 | 13737000 | 10688000 | 0 | 0 | 30930000 | 16340000 |
| CSVLNSEE | GNPAT; GNPAT | O15228; A0 | GNPAT_HUMA | Unmodified | 0 | 0 | 0 | 0 | 0 | 565450 | 0 | 18204 | 1729800 | 350420 | 754560 | 0 | 366720 | 1395900 | 0 | 0 | 0 | 0 |
| CSVLPLSQ | GARS1; GARS1 | P41250; A0A | GARS_HUMAN | Oxidation (M) | 651160 | 0 | 175510 | 399140 | 1358700 | 314930 | 1085500 | 514900 | 1090500 | 917720 | 756860 | 5731800 | 1908000 | 2090500 | 416240 | 476630 | 1269500 | 922550 |
| CSVSLSNV | PLA2G4A | P47712 | PA24A_HUMAN | Unmodified | 0 | 0 | 0 | 0 | 0 | 0 | 0 | 0 | 0 | 0 | 0 | 553030 | 0 | 0 | 0 | 0 | 0 | 0 |
| CSWLVVQ | FDPS; FDPS | P14324; A0A | FPPS_HUMAN | Unmodified | 0 | 0 | 0 | 0 | 0 | 0 | 0 | 0 | 362260 | 160710 | 65350 | 2396400 | 130520 | 536350 | 0 | 0 | 0 | 0 |
| CSYQPTME | FLNA; FLNA; FL | P21333; A0A | FLNA_HUMAN | Oxidation (M) | 0 | 0 | 0 | 0 | 0 | 0 | 0 | 0 | 0 | 0 | 0 | 931000 | 0 | 0 | 0 | 0 | 0 | 0 |
| CSYQPTME | FLNA; FLNA; FL | P21333; A0A | FLNA_HUMAN | Unmodified | 1141300 | 0 | 0 | 3272300 | 0 | 0 | 0 | 0 | 0 | 0 | 0 | 0 | 0 | 0 | 0 | 965050 | 0 | 0 |
| CTAASASP | PTGFRN | Q9P2B2 | FPRP_HUMAN | Unmodified | 7545000 | 2090300 | 8478600 | 5057900 | 45918 | 1790300 | 2729400 | 0 | 658340 | 0 | 0 | 9155800 | 1431500 | 0 | 11760000 | 13449000 | 0 | 370000 |
| CTAASASP | PTGFRN | Q9P2B2 | FPRP_HUMAN | Unmodified | 5052200 | 2272600 | 3566600 | 1536100 | 0 | 0 | 0 | 0 | 0 | 0 | 0 | 3320900 | 1970600 | 401010 | 13575000 | 15797000 | 0 | 0 |
| CTACQESI | TRAF3; TRAF3; | Q13114; A6 | TRAF3_HUMAN | Unmodified | 0 | 0 | 0 | 0 | 749070 | 256190 | 0 | 0 | 0 | 0 | 0 | 0 | 0 | 0 | 0 | 0 | 0 | 0 |
| CTAQNPLG | IGSF21 | Q96ID5 | IGS21_HUMAN | Unmodified | 0 | 0 | 0 | 0 | 52639 | 0 | 0 | 0 | 329470 | 14910 | 0 | 0 | 0 | 0 | 0 | 0 | 0 | 0 |
| CTDPKFQG | ITGB1; ITGB1; | P05556; H7 | ITB1_HUMAN I | Unmodified | 0 | 0 | 0 | 0 | 3628300 | 1416300 | 0 | 0 | 0 | 0 | 0 | 0 | 0 | 575160 | 0 | 0 | 6486900 | 9224600 |
| CTDPKFQG | ITGB1; ITGB1; | P05556; H7 | ITB1_HUMAN I | Oxidation (M) | 0 | 0 | 0 | 0 | 875820 | 0 | 0 | 0 | 0 | 0 | 0 | 0 | 0 | 0 | 0 | 0 | 2002000 | 3684400 |
| CTDPKFQG | ITGB1; ITGB1; | P05556; H7 | ITB1_HUMAN I | Oxidation (M) | 0 | 0 | 0 | 0 | 0 | 0 | 0 | 0 | 0 | 0 | 0 | 0 | 0 | 0 | 0 | 0 | 1277500 | 0 |
| CTDPKFQG | ITGB1; ITGB1; | P05556; H7 | ITB1_HUMAN I | Unmodified | 0 | 0 | 0 | 0 | 443010 | 0 | 0 | 0 | 0 | 0 | 0 | 0 | 0 | 0 | 0 | 0 | 4733200 | 0 |
| CTESEEEE | USP14; USP14 | P54578; A6 | UBP14_HUMAN | Unmodified | 0 | 0 | 0 | 0 | 0 | 0 | 0 | 0 | 223710 | 0 | 0 | 1410400 | 0 | 0 | 0 | 310520 | 0 | 0 |
| CTEYLDRA | VPS4B; VPS4B; | O75351; K7E | VPS4B_HUMAN | Unmodified | 374080 | 194060 | 196680 | 0 | 0 | 38371 | 25154 | 13538 | 962430 | 143350 | 147720 | 0 | 0 | 0 | 389830 | 345580 | 0 | 0 |
| CTGGEVG | RPL12 | P30050 | RL12_HUMAN | Unmodified | 4004400 | 2158200 | 895450 | 1027200 | 660970 | 180400 | 198150 | 81519 | 261570 | 231710 | 581010 | 13904000 | 2500500 | 675640 | 1626500 | 1158800 | 517630 | 638540 |
| CTGTAANS | PRKCD; PRKCD | Q05655; A0 | KPCD_HUMAN | Unmodified | 0 | 0 | 0 | 0 | 243870 | 0 | 79416 | 32653 | 249840 | 163590 | 263640 | 254720 | 698550 | 226260 | 0 | 0 | 391090 | 1358400 |

| | | | | | | | | | | | | | | | | | | | | | | |
|---|---|---|---|---|---|---|---|---|---|---|---|---|---|---|---|---|---|---|---|---|---|---|
| CTHSSIEI | C1GALT1 | Q9NS00 | C1GLT_HUMAN | Unmodified | 0 | 0 | 0 | 0 | 0 | 0 | 0 | 136840 | 2003000 | 0 | 0 | 219770 | 0 | 0 | 0 | 0 | 0 | 0 |
| CTKEEHLC | TPM3; TPM3 | J3KN67; A0A | J3KN67_HUMA | Unmodified | 0 | 0 | 0 | 0 | 674530 | 224600 | 14412 | 0 | 67501 | 0 | 35072 | 0 | 350560 | 172080 | 0 | 36884 | 186250 | 139670 |
| CTLACQQF | RP2 | O75695 | XRP2_HUMAN | Unmodified | 189950 | 0 | 0 | 265670 | 0 | 0 | 0 | 0 | 426110 | 0 | 297730 | 1634500 | 303520 | 251440 | 337930 | 370030 | 0 | 0 |
| CTLEFQVIT | TMED7; TMED7 | Q9Y3B3; A0A | TMED7_HUMAI | Unmodified | 0 | 0 | 0 | 0 | 0 | 0 | 0 | 0 | 960600 | 0 | 0 | 0 | 0 | 0 | 0 | 0 | 0 | 0 |
| CTLHPNDS | LLGL1; LLGL1 | Q15334; A0A | L2GL1_HUMAN | Oxidation (M) | 0 | 33146 | 0 | 120370 | 0 | 38242 | 0 | 0 | 0 | 0 | 0 | 0 | 0 | 0 | 114780 | 122360 | 0 | 0 |
| CTLHPSDQ | LLGL2; LLGL2 | Q6P1M3; J3C | L2GL2_HUMAN | Unmodified | 0 | 0 | 157030 | 295550 | 0 | 0 | 0 | 0 | 1291500 | 200700 | 167920 | 0 | 188150 | 0 | 799580 | 680870 | 0 | 213150 |
| CTNCLEDE | CLDN4 | O14493 | CLD4_HUMAN | Unmodified | 229990 | 55286 | 0 | 0 | 0 | 0 | 0 | 0 | 0 | 0 | 0 | 0 | 0 | 0 | 103020 | 113970 | 0 | 0 |
| CTPGWTGI | MEGF10 | Q96KG7 | MEG10_HUMAI | Unmodified | 0 | 0 | 0 | 0 | 1199900 | 0 | 0 | 0 | 0 | 0 | 0 | 0 | 0 | 0 | 0 | 0 | 0 | 2794400 |
| CTSDTVFLE | KDF1 | Q8NAX2 | KDF1_HUMAN | Unmodified | 73862 | 35737 | 141090 | 110750 | 0 | 215480 | 0 | 0 | 286410 | 121960 | 691700 | 355180 | 129270 | 107810 | 202680 | 220940 | 0 | 0 |
| CTSLGIEVL | GGT7; GGT7 | Q9UJ14; A2A | GGT7_HUMAN | Unmodified | 0 | 0 | 0 | 0 | 0 | 0 | 0 | 0 | 900310 | 0 | 0 | 0 | 0 | 0 | 0 | 0 | 0 | 0 |
| CTSNFLSPI | SORT1 | Q99523 | SORT_HUMAN | Unmodified | 0 | 0 | 0 | 0 | 0 | 0 | 0 | 0 | 1778800 | 118660 | 0 | 0 | 0 | 0 | 0 | 0 | 122570 | 0 |
| CTTDENKV | GAR1 | Q9NY12 | GAR1_HUMAN | Unmodified | 0 | 0 | 0 | 0 | 0 | 0 | 0 | 0 | 0 | 0 | 0 | 532650 | 0 | 0 | 0 | 0 | 0 | 0 |
| CTTDHISA | ACO2; ACO2; A | Q99798; A2A | ACON_HUMAN | Unmodified | 0 | 0 | 0 | 0 | 0 | 0 | 0 | 0 | 942560 | 0 | 0 | 0 | 0 | 0 | 0 | 0 | 0 | 0 |
| CTTEAEQD | RELL1; RELL1 | Q8IUW5; D6 | RELL1_HUMAN | Unmodified | 66773 | 0 | 93505 | 0 | 0 | 0 | 14979 | 0 | 0 | 0 | 0 | 0 | 0 | 0 | 99852 | 179400 | 0 | 0 |
| CTVFHGAC | FASN; FASN | P49327; A0A | FAS_HUMAN F | Unmodified | 188570 | 0 | 0 | 0 | 0 | 0 | 0 | 0 | 0 | 0 | 0 | 19016000 | 0 | 0 | 845260 | 680150 | 0 | 0 |
| CTVSLTR | SPATS2L; SPAT | Q9NUQ6; B8 | SPS2L_HUMAN | Unmodified | 0 | 0 | 0 | 0 | 0 | 0 | 0 | 0 | 623130 | 0 | 25977 | 448750 | 0 | 0 | 0 | 0 | 0 | 0 |
| CTVSVETR | MYO1D; MYO1D | O94832; K7E | MYO1D_HUMA | Unmodified | 2813900 | 2327900 | 635200 | 0 | 0 | 0 | 0 | 0 | 0 | 0 | 0 | 0 | 0 | 0 | 0 | 1315700 | 0 | 638080 |
| CTVSVETRI | MYO1D; MYO1D | O94832; K7E | MYO1D_HUMA | Unmodified | 178720 | 140140 | 0 | 1640300 | 1646200 | 0 | 0 | 101350 | 0 | 0 | 0 | 8928800 | 3672000 | 2505200 | 825500 | 1106800 | 3394300 | 14024000 |
| CTWLIEGQ | ATRN | O75882 | ATRN_HUMAN | Unmodified | 0 | 0 | 0 | 96382 | 0 | 0 | 89313 | 0 | 1434100 | 0 | 0 | 0 | 0 | 0 | 0 | 0 | 0 | 382870 |
| CTWQLGRI | CBS; CBSL; CBS | P35520; P0D | CBS_HUMAN C | Unmodified | 77608 | 0 | 0 | 0 | 0 | 0 | 0 | 0 | 0 | 23232 | 119090 | 2852200 | 0 | 0 | 0 | 0 | 0 | 0 |
| CVANNQVI | LRPPRC | P42704 | LPPRC_HUMAN | Unmodified | 0 | 0 | 0 | 0 | 0 | 0 | 0 | 0 | 837810 | 0 | 0 | 0 | 0 | 0 | 0 | 0 | 0 | 0 |
| CVDLYNQA | GAS7 | O60861 | GAS7_HUMAN | Unmodified | 0 | 0 | 0 | 0 | 0 | 0 | 70976 | 0 | 0 | 0 | 0 | 1236600 | 0 | 0 | 0 | 0 | 0 | 0 |
| CVEENNG | SLC1A5; SLC1A | Q15758; M0 | AAAT_HUMAN | Unmodified | 138840 | 0 | 0 | 0 | 0 | 0 | 0 | 87777 | 190380 | 0 | 0 | 1137800 | 0 | 0 | 0 | 0 | 129790 | 0 |
| CVEPLGLEI | MFGE8; MFGE8 | Q08431; H0 | MFGM_HUMAN | Unmodified | 0 | 0 | 0 | 0 | 0 | 0 | 0 | 0 | 3088800 | 0 | 0 | 977960 | 0 | 0 | 0 | 0 | 0 | 0 |
| CVEQPELK | SORT1 | Q99523 | SORT_HUMAN | Unmodified | 0 | 0 | 637810 | 352330 | 0 | 0 | 48663 | 0 | 1688600 | 148400 | 189620 | 300080 | 0 | 0 | 331290 | 0 | 31237 | 0 |
| CVETCPPP | INSR | P06213 | INSR_HUMAN | Unmodified | 0 | 0 | 0 | 0 | 0 | 224310 | 0 | 0 | 380910 | 0 | 0 | 0 | 0 | 0 | 0 | 0 | 0 | 0 |
| CVFWDLGI | ADGRG2 | Q8IZP9 | AGRG2_HUMA | Unmodified | 0 | 0 | 0 | 0 | 0 | 0 | 0 | 0 | 1618700 | 130680 | 0 | 0 | 0 | 0 | 0 | 0 | 0 | 0 |
| CVGAELEY | C20orf27; C20 | Q9GZN8; H7 | CT027_HUMAN | Unmodified | 0 | 0 | 0 | 0 | 0 | 0 | 0 | 0 | 0 | 0 | 0 | 0 | 0 | 0 | 0 | 0 | 805980 | 608090 |
| CVHGTCLF | SLIT2; SLIT2; SL | O94813; X6F | SLIT2_HUMAN | Unmodified | 0 | 0 | 0 | 0 | 0 | 0 | 0 | 0 | 0 | 0 | 0 | 0 | 0 | 0 | 0 | 0 | 60484 | 411020 |
| CVIALQEK | CTNNA1; CTNN | P35221; G3 | CTNA1_HUMAI | Unmodified | 935240 | 738810 | 1671600 | 2841100 | 6650500 | 2731900 | 178980 | 76253 | 2008900 | 144780 | 1007300 | 0 | 3158100 | 5194400 | 5618600 | 5105300 | 151970 | 0 |
| CVILRNIDN | WDR1; WDR1 | O75083; D6 | WDR1_HUMAN | Unmodified | 0 | 0 | 0 | 0 | 18531000 | 3427100 | 4092300 | 0 | 0 | 1860200 | 0 | 0 | 9024300 | 13689000 | 0 | 0 | 14906000 | 6521900 |
| CVIMCQGS | SNRNP200 | O75643 | U520_HUMAN | Unmodified | 0 | 0 | 0 | 0 | 0 | 0 | 0 | 0 | 138060 | 142700 | 0 | 807410 | 0 | 0 | 0 | 0 | 18824 | 0 |
| CVINATGPI | GPD2; GPD2 | P43304; F5G | GPDM_HUMAN | Unmodified | 0 | 0 | 0 | 0 | 0 | 0 | 0 | 0 | 2243100 | 0 | 0 | 0 | 0 | 0 | 0 | 0 | 0 | 0 |
| CVLLSNLS | FASN; FASN | P49327; A0A | FAS_HUMAN F | Unmodified | 376550 | 0 | 0 | 368750 | 0 | 0 | 1094200 | 0 | 3138800 | 1725700 | 4528900 | 1.2E+08 | 656880 | 0 | 1729700 | 841300 | 1335500 | 0 |
| CVLNEGMI | DNAJA1 | P31689 | DNJA1_HUMAN | Unmodified | 0 | 0 | 0 | 0 | 7581500 | 998970 | 1567000 | 425580 | 0 | 4412500 | 0 | 0 | 11942000 | 7286400 | 0 | 0 | 1568500 | 2130300 |
| CVLNEGMI | DNAJA1 | P31689 | DNJA1_HUMAN | Oxidation (M) | 0 | 0 | 0 | 187520 | 314660 | 275970 | 199640 | 182530 | 0 | 1711800 | 0 | 0 | 1961100 | 2035300 | 0 | 0 | 597750 | 65753 |
| CVLPEEDS | STRAP | Q9Y3F4 | STRAP_HUMAN | Unmodified | 351790 | 225500 | 0 | 0 | 0 | 0 | 421470 | 323540 | 336710 | 0 | 615030 | 2399800 | 1514200 | 1323400 | 181970 | 282510 | 740850 | 1121800 |
| CVNLSIKDI | GSDMD; GSDM | P57764; G3 | GSDMD_HUMA | Unmodified | 0 | 0 | 0 | 0 | 0 | 2056500 | 0 | 0 | 0 | 0 | 0 | 0 | 0 | 0 | 0 | 0 | 0 | 5710700 |
| CVNSPGSF | FBLN1; FBLN1 | P23142; B1A | FBLN1_HUMAN | Unmodified | 0 | 0 | 0 | 0 | 0 | 0 | 0 | 0 | 3253400 | 134100 | 0 | 196200 | 561300 | 0 | 0 | 0 | 76651 | 0 |
| CVNSPGSF | FBLN1; FBLN1 | P23142; B1A | FBLN1_HUMAN | Unmodified | 0 | 0 | 0 | 0 | 0 | 0 | 0 | 0 | 0 | 468750 | 0 | 0 | 0 | 0 | 118570 | 0 | 0 | 0 |
| CVNTTLQIK | CAP1 | Q01518 | CAP1_HUMAN | Unmodified | 0 | 0 | 0 | 0 | 3976800 | 0 | 1459100 | 2424700 | 2642000 | 1333100 | 0 | 6839100 | 3422600 | 3607300 | 171640 | 129620 | 7464500 | 1272800 |
| CVNTTLQIK | CAP1 | Q01518 | CAP1_HUMAN | Unmodified | 0 | 0 | 0 | 0 | 0 | 0 | 0 | 0 | 0 | 0 | 0 | 0 | 0 | 0 | 0 | 0 | 208360 | 0 |
| CVQYLDRA | VPS4A; N/A | Q9UN37; I3L | VPS4A_HUMAN | Unmodified | 0 | 0 | 0 | 0 | 0 | 0 | 0 | 0 | 0 | 0 | 0 | 652890 | 0 | 0 | 247360 | 438940 | 0 | 0 |
| CVSSPHFC | PPP2R5D; PPP2 | Q14738; Q1 | 2A5D_HUMAN | Unmodified | 0 | 0 | 0 | 0 | 0 | 0 | 0 | 0 | 0 | 67536 | 0 | 1098700 | 0 | 0 | 0 | 0 | 0 | 0 |
| CVTVFSAPI | PPP5C; PPP5C; | P53041; A8N | PPP5_HUMAN | Oxidation (M) | 0 | 0 | 0 | 0 | 0 | 300200 | 0 | 21776 | 0 | 0 | 91350 | 439430 | 0 | 0 | 58815 | 0 | 0 | 25634 |
| CVTYTSLYK | TRIM26 | Q12899 | TRI26_HUMAN | Unmodified | 1580500 | 1203400 | 1387300 | 1705600 | 0 | 0 | 0 | 0 | 0 | 0 | 0 | 0 | 0 | 0 | 1962600 | 1394000 | 0 | 0 |
| CVVPGQAI | HSPG2 | P98160 | PGBM_HUMAN | Unmodified | 0 | 0 | 850060 | 815080 | 0 | 0 | 0 | 0 | 0 | 170000 | 0 | 0 | 778510 | 865300 | 1157900 | 237450 | 265890 | 2229400 |
| CVVTGEDC | NCAM1; NCAM | P13591; A0A | NCAM1_HUMA | Unmodified | 0 | 0 | 0 | 0 | 0 | 0 | 188160 | 155660 | 1126800 | 285460 | 0 | 0 | 0 | 0 | 0 | 0 | 0 | 0 |
| CVVVGDG | RHOQ; RAC2; R | P17081; P15 | RHOQ_HUMAN | Unmodified | 14311000 | 21257000 | 801410 | 15649000 | 22418000 | 9211200 | 11065000 | 2887000 | 2724000 | 1502500 | 6547200 | 30954000 | 13816000 | 23676000 | 25367000 | 21094000 | 21910000 | 21844000 |
| CVYKPMQF | FLNB; FLNB | O75369; E7E | FLNB_HUMAN | Unmodified | 0 | 0 | 198660 | 280100 | 3949500 | 1670000 | 0 | 0 | 0 | 0 | 0 | 1397000 | 454700 | 605560 | 145790 | 37768 | 1252300 | 1505000 |
| CVYYRKPLI | UBA1 | P22314 | UBA1_HUMAN | Unmodified | 0 | 0 | 0 | 128690 | 0 | 0 | 177360 | 0 | 0 | 236400 | 267790 | 0 | 662120 | 0 | 0 | 0 | 504530 | 962250 |
| CWAGSLW | WDR77 | Q9BQA1 | MEP50_HUMAI | Unmodified | 0 | 0 | 0 | 0 | 0 | 0 | 0 | 0 | 0 | 0 | 9001.6 | 416970 | 0 | 0 | 20230 | 10294 | 0 | 0 |
| CWALSPK | PTK7; PTK7 | Q13308; H0 | PTK7_HUMAN | Unmodified | 0 | 0 | 0 | 329900 | 2378800 | 685500 | 193730 | 181280 | 1352600 | 908590 | 324310 | 0 | 1573900 | 686150 | 708030 | 479080 | 649500 | 3011000 |
| CWGVDFQ | ADAM9; ADAM9 | Q13443; A0A | ADAM9_HUMA | Oxidation (M) | 0 | 0 | 0 | 0 | 1545000 | 0 | 0 | 0 | 0 | 33032 | 0 | 0 | 598560 | 559210 | 0 | 0 | 175740 | 0 |
| CWKFEHCI | CD59; CD59; N | P13987; A0A | CD59_HUMAN | Unmodified | 6193500 | 2934400 | 731060 | 9619900 | 4702700 | 2177900 | 949710 | 42643 | 3058600 | 915870 | 0 | 2864800 | 10145000 | 6273900 | 6402000 | 4885700 | 12206000 | 29778000 |
| CWVALGAI | MOGS; MOGS; | Q13724; A0A | MOGS_HUMAN | Unmodified | 0 | 0 | 0 | 0 | 0 | 0 | 0 | 0 | 422640 | 128960 | 0 | 0 | 0 | 0 | 0 | 0 | 0 | 0 |
| CYCGEGLS | DKK1 | O94907 | DKK1_HUMAN | Unmodified | 0 | 0 | 0 | 0 | 0 | 0 | 0 | 0 | 0 | 28973 | 0 | 60839 | 0 | 0 | 0 | 0 | 304400 | 724540 |
| CYEKFPLEI | LIMS2; LIMS1; L | Q7Z4I7; P48 | LIMS2_HUMAN | Unmodified | 0 | 0 | 0 | 0 | 237320 | 0 | 467020 | 531860 | 0 | 0 | 0 | 0 | 0 | 234070 | 0 | 0 | 0 | 0 |
| CYEKFPLEI | LIMS2; LIMS1; L | Q7Z4I7; P48 | LIMS2_HUMAN | Unmodified | 0 | 0 | 31590 | 0 | 0 | 0 | 530230 | 246950 | 0 | 0 | 0 | 0 | 0 | 0 | 0 | 0 | 0 | 0 |
| CYFQAQAF | MSMP | Q1L6U9 | MSMP_HUMAN | Unmodified | 0 | 0 | 0 | 0 | 0 | 0 | 0 | 0 | 0 | 0 | 0 | 0 | 0 | 0 | 0 | 0 | 234010 | 1326300 |
| CYGLHAFI | ACOX1 | Q15067 | ACOX1_HUMAI | Unmodified | 0 | 0 | 0 | 0 | 0 | 0 | 0 | 0 | 768150 | 0 | 0 | 0 | 845470 | 0 | 0 | 0 | 0 | 55203 |
| CYHGKTGF | RPL21; RPL21; | P46778; G3 | RL21_HUMAN | Unmodified | 0 | 0 | 0 | 0 | 0 | 0 | 369450 | 0 | 0 | 0 | 0 | 0 | 0 | 0 | 0 | 0 | 520240 | 0 |
| CYNDLLAS | CEP55; CEP55 | Q53EZ4; H0 | CEP55_HUMAN | Unmodified | 341880 | 198230 | 437910 | 1013900 | 5502300 | 996830 | 0 | 0 | 2011600 | 1123500 | 186680 | 785160 | 340410 | 364560 | 586620 | 313350 | 194520 | 756100 |
| CYPHPGSE | IGFBP2 | P18065 | IBP2_HUMAN | Oxidation (M) | 0 | 0 | 0 | 0 | 0 | 0 | 0 | 0 | 9849300 | 5023100 | 0 | 0 | 0 | 0 | 0 | 0 | 0 | 0 |
| CYQSEFGR | ANXA7 | P20073 | ANXA7_HUMAN | Unmodified | 0 | 0 | 0 | 0 | 298940 | 0 | 0 | 87573 | 3390000 | 0 | 0 | 85736 | 0 | 0 | 0 | 0 | 0 | 0 |
| CYQSEFGR | ANXA7 | P20073 | ANXA7_HUMAN | Unmodified | 1104900 | 562720 | 427500 | 1496700 | 296850 | 0 | 0 | 0 | 1.11E+08 | 8711400 | 855240 | 1509500 | 354120 | 0 | 1734000 | 1157600 | 539060 | 1443400 |
| CYRCGETC | CNBP | P62633 | CNBP_HUMAN | Unmodified | 0 | 0 | 0 | 0 | 494540 | 58831 | 271050 | 0 | 0 | 271180 | 0 | 0 | 1875900 | 1792900 | 0 | 0 | 1419200 | 259110 |
| CYSCGEFC | CNBP | P62633 | CNBP_HUMAN | Unmodified | 0 | 0 | 314750 | 456300 | 0 | 0 | 186260 | 0 | 0 | 161130 | 317250 | 757570 | 0 | 590470 | 0 | 341830 | 0 | 0 |
| CYVDGSEE | CXADR | P78310 | CXAR_HUMAN | Unmodified | 411770 | 0 | 0 | 0 | 1393300 | 324640 | 0 | 0 | 310450 | 0 | 0 | 7098100 | 0 | 0 | 0 | 0 | 0 | 0 |
| DAAAFLLS | CLUH; CLUH; C | O75153; K7E | CLU_HUMAN C | Unmodified | 0 | 0 | 0 | 0 | 0 | 0 | 0 | 0 | 0 | 12624 | 53384 | 402490 | 18764 | 0 | 0 | 29324 | 0 | 0 |
| DAAINSLLC | TOLLIP; TOLLIP | Q9H0E2; F2 | TOLIP_HUMAN | Oxidation (M) | 0 | 0 | 119320 | 46862 | 0 | 0 | 0 | 0 | 144280 | 0 | 0 | 0 | 61557 | 0 | 546860 | 191340 | 0 | 0 |
| DAALGPIRE | SPAST; SPAST; | Q9UBP0; A0 | SPAST_HUMAN | Unmodified | 0 | 0 | 282230 | 272370 | 1509300 | 454890 | 125630 | 69046 | 0 | 0 | 0 | 0 | 0 | 1386300 | 0 | 0 | 0 | 615790 |
| DAANLLND | KLC1; KLC1; KL | Q07866; G3 | KLC1_HUMAN | Unmodified | 0 | 0 | 0 | 0 | 0 | 0 | 0 | 0 | 113100 | 0 | 132610 | 975250 | 0 | 0 | 0 | 15651 | 0 | 0 |
| DAANLLND | KLC1; KLC1; KL | Q07866; G3 | KLC1_HUMAN | Unmodified | 109560 | 0 | 389160 | 471200 | 4468000 | 563790 | 911700 | 258060 | 2415700 | 961020 | 1135700 | 7948300 | 1166200 | 655170 | 1142600 | 287490 | 1069100 | 2263600 |
| DAARYHFF | TWF2; TWF2; T | Q6IBS0; D6F | TWF2_HUMAN | Unmodified | 0 | 0 | 0 | 0 | 381690 | 274230 | 0 | 0 | 0 | 0 | 1019800 | 0 | 0 | 0 | 0 | 0 | 314320 | 693470 |
| DAASLESQ | MYH10 | P35580 | MYH10_HUMAI | Unmodified | 0 | 0 | 0 | 755260 | 11679000 | 1918000 | 1906400 | 131420 | 451640 | 0 | 0 | 0 | 0 | 0 | 748420 | 665530 | 1833500 | 1362800 |
| DAASLESQ | MYH10 | P35580 | MYH10_HUMAI | Unmodified | 0 | 0 | 0 | 0 | 6529400 | 0 | 37288 | 0 | 0 | 0 | 0 | 0 | 3950500 | 0 | 0 | 0 | 3178900 | 0 |
| DAASVDKV | HNRNPDL; HNI | O14979; A0A | HNRDL_HUMA | Unmodified | 0 | 0 | 0 | 0 | 281120 | 0 | 337480 | 388950 | 0 | 300490 | 0 | 0 | 155090 | 173870 | 0 | 0 | 225010 | 130570 |
| DAATVTQM | LLGL1; LLGL1 | Q15334; A0A | L2GL1_HUMAN | Unmodified | 109870 | 0 | 350450 | 435740 | 181290 | 0 | 0 | 0 | 139530 | 0 | 0 | 3077000 | 0 | 236620 | 348160 | 221990 | 175220 | 1182400 |
| DACLREAV | NAPG; NAPG | Q99747; J3K | SNAG_HUMAN | Unmodified | 0 | 0 | 0 | 0 | 906430 | 127600 | 0 | 0 | 0 | 75816 | 0 | 0 | 355550 | 179790 | 0 | 0 | 0 | 576500 |
| DACLREAV | NAPG; NAPG | Q99747; J3K | SNAG_HUMAN | Unmodified | 0 | 0 | 0 | 0 | 1072800 | 0 | 0 | 0 | 0 | 118210 | 0 | 0 | 654360 | 711740 | 0 | 0 | 768800 | 639330 |
| DACVISSDI | TM9SF2 | Q99805 | TM9S2_HUMAN | Unmodified | 0 | 0 | 0 | 0 | 0 | 0 | 0 | 0 | 923970 | 0 | 0 | 1975600 | 0 | 0 | 0 | 0 | 0 | 0 |
| DADCRAVV | ECH1; N/A; ECH | Q13011; M0 | ECH1_HUMAN | Unmodified | 0 | 22900 | 0 | 93546 | 0 | 0 | 31489 | 0 | 98039 | 179320 | 0 | 0 | 0 | 0 | 0 | 0 | 0 | 0 |
| DADLTDTA | GFUS; GFUS; G | Q13630; E9F | FCL_HUMAN G | Unmodified | 0 | 0 | 0 | 0 | 0 | 0 | 0 | 0 | 704840 | 34378 | 0 | 842200 | 0 | 0 | 0 | 0 | 0 | 0 |
| DADLTDTA | GFUS; GFUS; G | Q13630; E9F | FCL_HUMAN G | Unmodified | 0 | 0 | 0 | 0 | 283040 | 0 | 0 | 0 | 0 | 224120 | 0 | 0 | 42044 | 0 | 0 | 0 | 227570 | 0 |
| DADPEVLL | TMEM260 | Q9NX78 | TM260_HUMAN | Unmodified | 0 | 0 | 0 | 0 | 0 | 0 | 0 | 0 | 1872100 | 0 | 0 | 0 | 0 | 0 | 0 | 0 | 0 | 0 |
| DADPILISLI | CORO1B | Q9BR76 | COR1B_HUMA | Unmodified | 725350 | 593010 | 765430 | 0 | 0 | 0 | 0 | 0 | 571870 | 0 | 1428500 | 6661800 | 0 | 0 | 1774500 | 1913200 | 0 | 0 |
| DADSITLFD | COPA; COPA; C | P53621; A0A | COPA_HUMAN | Unmodified | 0 | 0 | 0 | 0 | 0 | 0 | 0 | 0 | 0 | 0 | 0 | 4413000 | 0 | 0 | 311320 | 181430 | 0 | 0 |
| DADSQNPI | COPA; COPA; C | P53621; A0A | COPA_HUMAN | Unmodified | 0 | 0 | 0 | 0 | 0 | 0 | 0 | 0 | 63276 | 0 | 0 | 385980 | 0 | 0 | 0 | 0 | 0 | 0 |
| DADVQNF | ECI1; ECI1; ECI | P42126; Q9 | ECI1_HUMAN E | Unmodified | 0 | 0 | 0 | 0 | 0 | 0 | 228800 | 0 | 11259000 | 0 | 35523 | 0 | 0 | 404600 | 0 | 0 | 0 | 0 |
| DAEALSQR | MYH10 | P35580 | MYH10_HUMAI | Unmodified | 0 | 0 | 163330 | 965160 | 8766900 | 1938400 | 1114600 | 894990 | 116930 | 254050 | 498590 | 5313400 | 3760100 | 2209300 | 1064400 | 620430 | 1097400 | 2629100 |
| DAEDAIYGI | SRSF9; SRSF9; | Q13242; S4F | SRSF9_HUMAN | Unmodified | 179020 | 230300 | 98478 | 0 | 0 | 0 | 79798 | 0 | 176950 | 0 | 0 | 0 | 0 | 0 | 154410 | 0 | 0 | 0 |
| DAEDAVYG | SRSF1; SRSF1 | Q07955; J3K | SRSF1_HUMAN | Unmodified | 0 | 0 | 0 | 0 | 0 | 0 | 0 | 0 | 0 | 0 | 0 | 0 | 0 | 0 | 0 | 0 | 0 | 0 |
| DAEDAVYG | SRSF1; SRSF1 | Q07955; J3K | SRSF1_HUMAN | Unmodified | 1283700 | 1189500 | 255630 | 0 | 0 | 0 | 0 | 0 | 0 | 0 | 0 | 0 | 0 | 0 | 1063000 | 863400 | 0 | 0 |
| DAEGILEDI | CYRIB; CYRIB; | Q9NUQ9; E5 | CYRIB_HUMAN | Unmodified | 103440 | 90944 | 66104 | 66659 | 47866 | 0 | 0 | 0 | 1299100 | 0 | 426570 | 3471200 | 0 | 0 | 88115 | 87597 | 30882 | 39878 |
| DAEGILEDI | CYRIB; CYRIB; | Q9NUQ9; E5 | CYRIB_HUMAN | Unmodified | 235560 | 1867200 | 0 | 682360 | 115090 | 274700 | 0 | 0 | 9268400 | 0 | 986870 | 4428500 | 0 | 0 | 570160 | 302330 | 339300 | 192590 |
| DAEGLDEII | ATP2B4; ATP2B | P23634; H7E | AT2B4_HUMAN | Unmodified | 0 | 0 | 0 | 154640 | 117190 | 0 | 0 | 0 | 618650 | 0 | 0 | 0 | 0 | 0 | 378650 | 0 | 0 | 0 |
| DAEGLDEII | ATP2B4; ATP2B | P23634; H7E | AT2B4_HUMAN | Oxidation (M) | 78874 | 0 | 746510 | 465080 | 82785 | 0 | 183580 | 44231 | 309050 | 0 | 566810 | 779120 | 0 | 0 | 845660 | 1128900 | 0 | 0 |
| DAEIQERE | PRKG2 | Q13237 | KGP2_HUMAN | Unmodified | 0 | 0 | 0 | 0 | 0 | 0 | 715710 | 21719 | 0 | 0 | 0 | 0 | 0 | 0 | 0 | 0 | 0 | 0 |
| DAELQDQE | TNKS1BP1; TNK | Q9C0C2; A0 | TB182_HUMAN | Unmodified | 0 | 0 | 0 | 0 | 0 | 0 | 0 | 0 | 0 | 99228 | 0 | 610380 | 0 | 0 | 157160 | 216770 | 0 | 229690 |
| DAEMPATE | FLNA; FLNA; FL | P21333; A0A | FLNA_HUMAN | Unmodified | 326130 | 153510 | 0 | 336420 | 0 | 0 | 231880 | 217160 | 0 | 0 | 0 | 0 | 0 | 0 | 241520 | 136740 | 0 | 0 |
| DAERLLLNI | YES1; YES1 | P07947; J3Q | YES_HUMAN T | Unmodified | 291340 | 142040 | 455680 | 582490 | 2167200 | 1114800 | 161230 | 140780 | 1050000 | 1045700 | 902120 | 0 | 2877800 | 1010800 | 0 | 307830 | 2069200 | 4565000 |
| DAERQLLA | LYN | P07948 | LYN_HUMAN T | Unmodified | 0 | 0 | 0 | 692680 | 4911500 | 2778900 | 1431000 | 0 | 1593100 | 0 | 0 | 0 | 1379300 | 72184 | 0 | 0 | 0 | 0 |
| DAERQLLS | FYN; FYN; FYN; | P06241; E5R | FYN_HUMAN T | Unmodified | 0 | 0 | 0 | 0 | 0 | 0 | 0 | 133430 | 486670 | 134700 | 0 | 0 | 0 | 0 | 0 | 0 | 0 | 0 |
| DAESIHQY | NPEPPS; NPEPI | P55786; E9P | PSA_HUMAN P | Unmodified | 0 | 0 | 0 | 0 | 0 | 0 | 0 | 0 | 898060 | 36449 | 155200 | 3391400 | 0 | 0 | 246550 | 0 | 246020 | 816430 |
| DAEWTTVF | ARVCF; ARVCF; | O00192; E9F | ARVC_HUMAN | Unmodified | 0 | 0 | 1399000 | 892760 | 0 | 0 | 0 | 33137 | 0 | 96878 | 0 | 113250 | 0 | 0 | 1488800 | 1812500 | 0 | 0 |
| DAFADAVQ | KHSRP; KHSRP | Q92945; M0 | FUBP2_HUMAN | Unmodified | 0 | 0 | 0 | 0 | 588300 | 0 | 0 | 0 | 628900 | 0 | 270140 | 0 | 118430 | 0 | 92973 | 0 | 0 | 268080 |
| DAFDKGSL | PDCD6IP; PDCI | Q8WUM4; A | PDC6I_HUMAN | Unmodified | 1253200 | 1205400 | 0 | 264530 | 2324700 | 496890 | 731340 | 289020 | 830760 | 1731300 | 0 | 94487 | 3314800 | 3072400 | 153200 | 91370 | 721520 | 2002200 |
| DAFDRNPE | PGD | P52209 | 6PGD_HUMAN | Unmodified | 94485 | 166110 | 57761 | 203050 | 468490 | 136140 | 249280 | 0 | 652300 | 368820 | 134540 | 600710 | 138790 | 26133 | 1099200 | 167970 | 227820 | 0 |

| | | | | | | | | | | | | | | | | | | | | | | |
|---|---|---|---|---|---|---|---|---|---|---|---|---|---|---|---|---|---|---|---|---|---|---|
| DAFDTLFDI | PARVB; PARVA; | Q9HBI1; Q9I | PARVB_HUMAN | Unmodified | 94447 | 0 | 472610 | 142050 | 0 | 0 | 0 | 0 | 0 | 0 | 0 | 498630 | 0 | 0 | 554860 | 226780 | 0 | 0 |
| DAFQNAYL | ATP1A1 | P05023 | AT1A1_HUMAN | Unmodified | 571620 | 0 | 5473200 | 5656700 | 364840 | 93787 | 67297 | 24522 | 8805000 | 593580 | 11933000 | 49110000 | 0 | 161970 | 21266000 | 21119000 | 0 | 0 |
| DAGADLDK | NFKBIB; NFKBII | Q15653; I3L | IKBB_HUMAN I | Unmodified | 0 | 0 | 0 | 0 | 0 | 0 | 0 | 0 | 0 | 0 | 36635 | 414590 | 0 | 0 | 0 | 0 | 0 | 0 |
| DAGAGTEA | ERICH5 | Q6P6B1 | ERIC5_HUMAN | Unmodified | 0 | 0 | 0 | 0 | 0 | 0 | 0 | 0 | 0 | 0 | 0 | 0 | 0 | 0 | 0 | 0 | 501340 | 2305400 |
| DAGAGTEA | ERICH5 | Q6P6B1 | ERIC5_HUMAN | Unmodified | 0 | 0 | 0 | 0 | 0 | 0 | 107950 | 0 | 0 | 0 | 0 | 0 | 0 | 0 | 0 | 0 | 1714900 | 1524900 |
| DAGEGGLS | FLNA; FLNA | P21333; Q6( | FLNA_HUMAN | Unmodified | 0 | 0 | 0 | 0 | 541730 | 0 | 348090 | 147250 | 961410 | 0 | 0 | 425790 | 0 | 0 | 2594400 | 621900 | 0 | 0 |
| DAGEGLLA | FLNA; FLNA | P21333; Q6( | FLNA_HUMAN | Unmodified | 0 | 0 | 0 | 5742800 | 1159900 | 586830 | 1126600 | 0 | 0 | 0 | 0 | 0 | 0 | 0 | 0 | 0 | 428450 | 351300 |
| DAGEGLLA | FLNA; FLNA | P21333; Q6( | FLNA_HUMAN | Unmodified | 316440 | 553950 | 319680 | 707370 | 0 | 0 | 427500 | 0 | 396270 | 0 | 0 | 1506800 | 125840 | 0 | 539660 | 607600 | 99114 | 65029 |
| DAGEGLLA | FLNB; FLNB | O75369; E7E | FLNB_HUMAN | Unmodified | 0 | 0 | 0 | 0 | 2016400 | 251820 | 0 | 134860 | 0 | 0 | 338610 | 72027 | 0 | 0 | 1008700 | 0 | 0 | 0 |
| DAGFSSPD | WDR48 | Q8TAF3 | WDR48_HUMA | Unmodified | 0 | 0 | 0 | 0 | 0 | 0 | 0 | 0 | 0 | 0 | 0 | 888770 | 0 | 0 | 0 | 0 | 0 | 0 |
| DAGHPLYP | AKR1A1 | P14550 | AK1A1_HUMAN | Unmodified | 0 | 0 | 0 | 0 | 359600 | 0 | 0 | 0 | 1777500 | 475430 | 0 | 0 | 608550 | 0 | 0 | 0 | 0 | 303640 |
| DAGIEPGPI | LRPPRC; LRPPF | P42704; B8Z | LPPRC_HUMAN | Unmodified | 0 | 0 | 0 | 0 | 0 | 0 | 0 | 0 | 1074400 | 0 | 0 | 0 | 0 | 0 | 0 | 0 | 0 | 0 |
| DAGLWHG | DIP2B | Q9P265 | DIP2B_HUMAN | Oxidation (M) | 288940 | 0 | 43215 | 305720 | 206670 | 0 | 52069 | 0 | 412350 | 234290 | 51448 | 979980 | 0 | 0 | 331490 | 145250 | 297410 | 103260 |
| DAGLWHG | DIP2A; DIP2A | Q14689; A0A | DIP2A_HUMAN | Unmodified | 0 | 0 | 0 | 30751 | 0 | 0 | 11487 | 0 | 332610 | 227590 | 0 | 0 | 0 | 0 | 0 | 47256 | 0 | 0 |
| DAGMQLQ | GOT2 | P00505 | AATM_HUMAN | Unmodified | 0 | 0 | 0 | 0 | 0 | 0 | 0 | 0 | 1466800 | 67595 | 0 | 0 | 0 | 0 | 0 | 0 | 0 | 0 |
| DAGPLLISL | CORO1A | P31146 | COR1A_HUMAI | Unmodified | 0 | 0 | 158030 | 0 | 0 | 154490 | 38141 | 0 | 0 | 0 | 3234300 | 22964000 | 0 | 0 | 0 | 0 | 0 | 0 |
| DAGQCIDF | AIDA | Q96BJ3 | AIDA_HUMAN / | Unmodified | 0 | 0 | 0 | 2394900 | 722230 | 0 | 0 | 85843 | 0 | 0 | 0 | 0 | 1022400 | 1146800 | 360400 | 0 | 1182000 | 2680000 |
| DAGQIQSP | TLL2 | Q9Y6L7 | TLL2_HUMAN T | Unmodified | 0 | 0 | 0 | 0 | 0 | 0 | 0 | 54931 | 1309800 | 238270 | 254720 | 0 | 0 | 0 | 0 | 0 | 0 | 0 |
| DAGQISGL | HSPA9; HSPA9; | P38646; H0Y | GRP75_HUMAI | Unmodified | 0 | 0 | 0 | 332620 | 620980 | 0 | 0 | 74677 | 21780000 | 1727200 | 1011000 | 973500 | 505580 | 0 | 0 | 0 | 0 | 332750 |
| DAGQISGL | HSPA9; HSPA9; | P38646; H0Y | GRP75_HUMAI | Unmodified | 0 | 0 | 0 | 0 | 1022700 | 0 | 0 | 0 | 0 | 1456300 | 0 | 0 | 0 | 0 | 0 | 0 | 0 | 1053400 |
| DAGTIAGLI | HSPA8; HSPA8; | P11142; E9P | HSP7C_HUMAI | Unmodified | 62908000 | 72478000 | 60753000 | 81792000 | 1.83E+08 | 25856000 | 16027000 | 6521500 | 1.49E+08 | 74277000 | 53351000 | 2.16E+08 | 2.42E+08 | 75188000 | 48062000 | 49288000 | 15306000 | 40888000 |
| DAGTIAGLI | HSPA8; HSPA8; | P11142; E9P | HSP7C_HUMAI | Unmodified | 863000 | 3922500 | 0 | 0 | 2652900 | 0 | 1672000 | 0 | 0 | 52322 | 0 | 0 | 1704400 | 1928500 | 2553300 | 350570 | 867510 | 946750 |
| DAGTIAGLI | HSPA8; HSPA8; | P11142; E9P | HSP7C_HUMAI | Unmodified | 0 | 0 | 0 | 0 | 3229600 | 0 | 850700 | 0 | 0 | 595650 | 0 | 0 | 4075800 | 4225100 | 0 | 0 | 0 | 2191800 |
| DAGTIAGLI | HSPA5; HSPA5; | P11021; A0A | BIP_HUMAN Er | Unmodified | 0 | 0 | 612300 | 387200 | 5659300 | 0 | 2878000 | 674620 | 1.97E+08 | 41294000 | 6658000 | 3614600 | 1319500 | 1265500 | 1047500 | 0 | 0 | 5190400 |
| DAGTIAGLI | HSPA5; HSPA5; | P11021; A0A | BIP_HUMAN Er | Oxidation (M) | 0 | 1018600 | 3162500 | 898810 | 3110300 | 395710 | 2800700 | 2166900 | 64266000 | 30232000 | 10291000 | 3299000 | 2118500 | 1482800 | 2619000 | 1346400 | 0 | 1191700 |
| DAGTITGLI | HSPA2 | P54652 | HSP72_HUMAI | Unmodified | 2846000 | 2008400 | 0 | 500550 | 2775500 | 742920 | 0 | 0 | 0 | 0 | 0 | 2545300 | 0 | 0 | 0 | 219150 | 376110 | 504660 |
| DAGTITGLI | HSPA2 | P54652 | HSP72_HUMAI | Unmodified | 0 | 0 | 0 | 0 | 134320 | 0 | 0 | 0 | 0 | 0 | 0 | 0 | 0 | 0 | 0 | 0 | 0 | 0 |
| DAGVIAGLI | HSPA1L; HSPA1 | P34931; P0D | HS71L_HUMAI | Unmodified | 18857000 | 15748000 | 7842100 | 10202000 | 12526000 | 3841100 | 2660900 | 1069400 | 48783000 | 18987000 | 3877000 | 16839000 | 19531000 | 9531900 | 7962400 | 7065800 | 3007600 | 4881500 |
| DAGYEFDI( | PGAM1 | P18669 | PGAM1_HUMA | Unmodified | 0 | 0 | 0 | 0 | 0 | 0 | 0 | 0 | 430480 | 0 | 0 | 1849700 | 0 | 0 | 0 | 0 | 0 | 0 |
| DAHGQQM | NOTCH1; NOT( | P46531; A0A | NOTC1_HUMA | Oxidation (M) | 110320 | 136970 | 59711 | 254850 | 0 | 0 | 0 | 0 | 0 | 0 | 973920 | 2723500 | 0 | 0 | 336490 | 269870 | 0 | 0 |
| DAHNALLD | EIF3CL; EIF3C | B5ME19; Q9 | EIFCL_HUMAN | Unmodified | 752520 | 726650 | 1070400 | 962620 | 0 | 0 | 0 | 81929 | 491590 | 114080 | 2067200 | 12255000 | 283620 | 0 | 1319700 | 1121000 | 0 | 0 |
| DAHQGRP | MISP | Q8IVT2 | MISP_HUMAN I | Unmodified | 0 | 0 | 0 | 0 | 585050 | 294660 | 0 | 0 | 0 | 316450 | 0 | 327680 | 0 | 0 | 0 | 0 | 102310 | 0 |
| DAHSQGE\ | GLG1; GLG1; G | Q92896; H3I | GSLG1_HUMAI | Unmodified | 0 | 133520 | 0 | 0 | 1598200 | 371070 | 0 | 234970 | 21854000 | 4449900 | 4028300 | 3085700 | 482140 | 410340 | 300870 | 196290 | 0 | 665750 |
| DAIAILDGI | ANAPC7 | Q9UJX3 | APC7_HUMAN | Unmodified | 0 | 0 | 0 | 0 | 0 | 0 | 0 | 0 | 0 | 0 | 0 | 1295800 | 0 | 0 | 0 | 0 | 0 | 0 |
| DAICGQLQ | ADAM15 | Q13444 | ADA15_HUMAI | Unmodified | 2336300 | 508810 | 1390800 | 501270 | 0 | 0 | 0 | 0 | 1024600 | 0 | 0 | 2492500 | 58623 | 0 | 1363600 | 766310 | 0 | 0 |
| DAIEEKIIIAI | MYO1B; MYO1E | O43795; H7( | MYO1B_HUMA | Unmodified | 0 | 0 | 0 | 0 | 2167700 | 0 | 0 | 55893 | 0 | 0 | 0 | 0 | 0 | 0 | 0 | 0 | 1504000 | 0 |
| DAIHFYNK | STIP1 | P31948 | STIP1_HUMAN | Unmodified | 0 | 0 | 251960 | 0 | 1512700 | 0 | 0 | 0 | 1723500 | 497010 | 827540 | 10741000 | 975450 | 0 | 397170 | 278190 | 115680 | 0 |
| DAIILIFANK | ARF6 | P62330 | ARF6_HUMAN . | Oxidation (M) | 6180800 | 7753500 | 1543500 | 5646900 | 0 | 0 | 0 | 0 | 4963900 | 1259500 | 1560300 | 8104800 | 0 | 0 | 8059000 | 16519000 | 0 | 0 |
| DAIILIFANK | ARF6 | P62330 | ARF6_HUMAN . | Unmodified | 0 | 3038700 | 0 | 2444100 | 890140 | 788420 | 0 | 0 | 10980000 | 886640 | 0 | 1919000 | 0 | 0 | 0 | 0 | 198410 | 0 |
| DAIKENPEI | EXOC1 | Q9NV70 | EXOC1_HUMAI | Unmodified | 420990 | 0 | 0 | 0 | 795300 | 0 | 0 | 0 | 0 | 0 | 188300 | 889600 | 0 | 0 | 196500 | 83936 | 112110 | 331210 |
| DAIKPNLM | MTHFD1; MTHF | P11586; V9G | C1TC_HUMAN | Oxidation (M) | 0 | 0 | 0 | 0 | 0 | 0 | 0 | 0 | 0 | 0 | 0 | 3022600 | 0 | 0 | 0 | 0 | 0 | 0 |
| DAINQGMI | PDHB; PDHB; P | P11177; F8V | ODPB_HUMAN | Oxidation (M) | 0 | 0 | 0 | 0 | 0 | 0 | 0 | 0 | 992960 | 0 | 57606 | 183680 | 0 | 0 | 0 | 0 | 0 | 0 |
| DAINQGMI | PDHB; PDHB; P | P11177; F8V | ODPB_HUMAN | Unmodified | 0 | 0 | 0 | 0 | 0 | 0 | 0 | 0 | 4465100 | 220330 | 0 | 286460 | 0 | 0 | 0 | 0 | 0 | 0 |
| DAIQNPND | CYRIA; CYRIA; ( | Q9H0Q0; C9 | CYRIA_HUMAN | Unmodified | 0 | 0 | 0 | 0 | 0 | 0 | 0 | 0 | 0 | 0 | 0 | 5649500 | 0 | 0 | 0 | 0 | 0 | 0 |
| DAISGIGTD | ANXA6; ANXA6; | P08133; E5R | ANXA6_HUMAI | Unmodified | 1112200 | 915580 | 3207100 | 3831700 | 3615800 | 1085400 | 2315500 | 1233200 | 41824000 | 11047000 | 884650 | 0 | 2228600 | 1087500 | 3333600 | 1984600 | 828850 | 4661700 |
| DAKDYADS | RAB22A | Q9UL26 | RB22A_HUMAI | Unmodified | 358730 | 0 | 110370 | 179220 | 885630 | 0 | 0 | 0 | 693530 | 0 | 0 | 524560 | 83456 | 41278 | 0 | 0 | 339790 | 0 |
| DAKLYSNR | STIP1 | P31948 | STIP1_HUMAN | Unmodified | 0 | 0 | 0 | 0 | 0 | 0 | 0 | 40664 | 0 | 0 | 0 | 0 | 0 | 0 | 0 | 0 | 605770 | 0 |
| DALARVYG | RNASET2; RNA: | O00584; D6I | RNT2_HUMAN | Unmodified | 0 | 0 | 0 | 0 | 0 | 0 | 0 | 0 | 0 | 424560 | 0 | 0 | 0 | 0 | 0 | 0 | 0 | 0 |
| DALEKNQC | CEP55; CEP55 | Q53EZ4; H0Y | CEP55_HUMAI | Unmodified | 0 | 0 | 0 | 0 | 2727500 | 0 | 0 | 0 | 0 | 348340 | 0 | 0 | 0 | 0 | 0 | 0 | 65479 | 0 |
| DALEPGQR | APRT; APRT; AP | P07741; H3E | APT_HUMAN A | Unmodified | 137600 | 96597 | 1121700 | 520810 | 0 | 0 | 0 | 0 | 0 | 0 | 0 | 5644700 | 156230 | 0 | 1818100 | 947690 | 0 | 0 |
| DALHLLVF | ITGB5; ITGB5; I | P18084; H7( | ITB5_HUMAN I | Unmodified | 0 | 0 | 0 | 511630 | 321320 | 0 | 0 | 0 | 0 | 87102 | 0 | 0 | 0 | 97778 | 3124300 | 0 | 309010 | 0 |
| DALLDQIR | WASL | O00401 | WASL_HUMAN | Unmodified | 0 | 0 | 729630 | 0 | 0 | 2670400 | 0 | 0 | 0 | 186800 | 0 | 0 | 0 | 0 | 1952600 | 0 | 94520 | 0 |
| DALLDQIR( | WASL | O00401 | WASL_HUMAN | Unmodified | 0 | 0 | 0 | 0 | 0 | 0 | 0 | 0 | 0 | 0 | 0 | 0 | 319690 | 0 | 0 | 0 | 0 | 0 |
| DALLLIFAN | TRIM23 | P36406 | TRI23_HUMAN | Unmodified | 0 | 0 | 1306100 | 0 | 0 | 0 | 0 | 0 | 569980 | 0 | 217450 | 807370 | 0 | 0 | 2276200 | 1554800 | 0 | 0 |
| DALLSALSI | SPTBN1; SPTBN | Q01082; A0A | SPTB2_HUMAN | Unmodified | 0 | 0 | 0 | 0 | 0 | 0 | 0 | 0 | 511590 | 608540 | 0 | 1704700 | 0 | 0 | 0 | 0 | 0 | 0 |
| DALLVGVP | GNG10; DNAJC | P50151; A0A | GBG10_HUMAI | Unmodified | 1359200 | 1660300 | 478930 | 1001400 | 19028000 | 0 | 1987400 | 1889600 | 7478000 | 4792900 | 0 | 26973000 | 4046200 | 3095000 | 827110 | 921340 | 4084700 | 7270200 |
| DALMQPVF | VPS4B | O75351 | VPS4B_HUMAN | Unmodified | 0 | 0 | 0 | 0 | 0 | 0 | 0 | 0 | 228760 | 0 | 83102 | 288280 | 0 | 0 | 0 | 0 | 0 | 0 |
| DALNIETAI | ANXA2P2; ANX/ | A6NMY6; P0 | AXA2L_HUMAN | Unmodified | 0 | 0 | 0 | 80278 | 1868600 | 234210 | 304480 | 186470 | 4071500 | 760820 | 245820 | 960960 | 0 | 0 | 0 | 0 | 413510 | 0 |
| DALQAEGC | ABCA3; ABCA3 | Q99758; H0Y | ABCA3_HUMAI | Unmodified | 0 | 0 | 0 | 337500 | 0 | 0 | 0 | 0 | 966380 | 0 | 0 | 0 | 0 | 0 | 0 | 0 | 0 | 0 |
| DALRPPLQ | CKAP5 | Q14008 | CKAP5_HUMAI | Unmodified | 0 | 0 | 73779 | 0 | 0 | 0 | 0 | 0 | 0 | 0 | 298570 | 2560200 | 0 | 0 | 40370 | 96401 | 0 | 0 |
| DALSALAR | XYLT2; XYLT2; X | Q9H1B5; B4 | XYLT2_HUMAN | Unmodified | 164770 | 679530 | 0 | 159500 | 0 | 0 | 0 | 0 | 522210 | 0 | 0 | 0 | 0 | 0 | 572980 | 139500 | 41957 | 0 |
| DALSDLALI | CCT4 | P50991 | TCPD_HUMAN | Unmodified | 436200 | 367320 | 352630 | 314510 | 377050 | 140870 | 557620 | 60858 | 1128700 | 544730 | 3224900 | 20923000 | 536430 | 360400 | 965820 | 1085400 | 251130 | 227370 |
| DALSLFTVF | AMPD3; AMPD3 | Q01432; E9F | AMPD3_HUMA | Unmodified | 0 | 0 | 0 | 0 | 0 | 0 | 0 | 0 | 0 | 0 | 0 | 415740 | 0 | 0 | 0 | 0 | 0 | 0 |
| DALSSVQE | APOC3; APOC3 | P02656; B0Y | APOC3_HUMAI | Unmodified | 0 | 1039300 | 3486600 | 585280 | 0 | 0 | 561060 | 748940 | 0 | 0 | 2745400 | 0 | 0 | 0 | 1488300 | 587290 | 68024 | 216600 |
| DALTQINVI | ATP2B4 | P23634 | AT2B4_HUMAN | Unmodified | 0 | 105130 | 0 | 0 | 0 | 0 | 0 | 0 | 0 | 0 | 0 | 3480800 | 0 | 210110 | 0 | 0 | 0 | 0 |
| DALTQINVI | ATP2B4 | P23634 | AT2B4_HUMAN | Unmodified | 0 | 0 | 0 | 0 | 937170 | 0 | 323120 | 0 | 0 | 0 | 0 | 0 | 0 | 0 | 0 | 0 | 0 | 0 |
| DAMITVDD | EPS8; EPS8; EP | Q12929; F5H | EPS8_HUMAN | Unmodified | 0 | 0 | 0 | 0 | 855220 | 0 | 0 | 0 | 0 | 0 | 0 | 0 | 115530 | 0 | 0 | 0 | 0 | 0 |
| DAMMAMN | CIRBP; CIRBP; ( | Q14011; K7E | CIRBP_HUMAN | Oxidation (M) | 100290 | 0 | 142410 | 45519 | 0 | 0 | 0 | 0 | 0 | 0 | 0 | 94133 | 0 | 0 | 200830 | 88155 | 0 | 0 |
| DAMPSDAI | PPP3CA; PPP3( | Q08209; E7E | PP2BA_HUMAN | Unmodified | 0 | 0 | 0 | 0 | 0 | 0 | 0 | 0 | 0 | 0 | 0 | 850380 | 0 | 0 | 0 | 0 | 0 | 0 |
| DAMQYASE | CLTC; CLTC; CI | Q00610; J3K | CLH1_HUMAN | Unmodified | 1860900 | 2067300 | 1956600 | 0 | 0 | 0 | 0 | 0 | 0 | 0 | 0 | 13099000 | 0 | 0 | 0 | 2159900 | 0 | 0 |
| DANGNSF/ | RPS4Y2; RPS4X | Q8TD47; P6. | RS4Y2_HUMAN | Unmodified | 1670100 | 1113200 | 1060000 | 653670 | 203280 | 30571 | 32499 | 0 | 441900 | 49914 | 279480 | 17898000 | 0 | 0 | 1347100 | 888540 | 0 | 0 |
| DANGNSF/ | RPS4X | P62701 | RS4X_HUMAN . | Unmodified | 0 | 0 | 35595 | 31436 | 969540 | 156680 | 3518800 | 17134 | 0 | 606490 | 0 | 0 | 574890 | 586350 | 0 | 0 | 453980 | 740220 |
| DANLTLESI | CDC42BPB; CD | Q9Y5S2; H0Y | MRCKB_HUMA | Unmodified | 0 | 0 | 0 | 15879 | 0 | 0 | 0 | 15625 | 95580 | 0 | 0 | 0 | 0 | 0 | 34715 | 0 | 0 | 1732100 |
| DANLYISGI | ELAVL1; ELAVL | Q15717; M0I | ELAV1_HUMAN | Unmodified | 0 | 0 | 0 | 0 | 0 | 0 | 59716 | 0 | 0 | 0 | 315460 | 4238300 | 0 | 0 | 0 | 0 | 0 | 0 |
| DANNGNL( | SSB | P05455 | LA_HUMAN Lu | Unmodified | 0 | 0 | 0 | 712100 | 0 | 0 | 0 | 0 | 0 | 0 | 0 | 1408500 | 0 | 0 | 0 | 0 | 0 | 0 |
| DANSVTSS | BZW2; BZW2; B | Q9Y6E2; B5I | BZW2_HUMAN | Unmodified | 0 | 0 | 0 | 0 | 0 | 0 | 25568 | 0 | 0 | 0 | 0 | 1119000 | 0 | 0 | 0 | 0 | 85402 | 35052 |
| DANSVTSS | BZW2; BZW2; B | Q9Y6E2; B5I | BZW2_HUMAN | Unmodified | 0 | 0 | 0 | 0 | 0 | 0 | 0 | 0 | 0 | 0 | 0 | 324720 | 0 | 0 | 0 | 0 | 0 | 0 |
| DAPPPPPP | WIPF2; WIPF2 | Q8TF74; A8I | WIPF2_HUMAN | Unmodified | 236680 | 279200 | 638370 | 289120 | 0 | 0 | 0 | 0 | 0 | 0 | 0 | 0 | 0 | 0 | 913920 | 454530 | 0 | 0 |
| DAQEKLEL | TPM1; TPM3; TF | P09493; P06 | TPM1_HUMAN | Unmodified | 80419 | 68958 | 0 | 0 | 5669600 | 296040 | 210690 | 83277 | 48091 | 32857 | 0 | 68612 | 0 | 0 | 0 | 0 | 135700 | 0 |
| DAQELYAA | ANXA11 | P50995 | ANX11_HUMAN | Unmodified | 133740 | 0 | 0 | 0 | 0 | 427100 | 0 | 169710 | 27188000 | 515080 | 447340 | 6057300 | 0 | 0 | 559880 | 1301600 | 0 | 0 |
| DAQELYAA | ANXA11 | P50995 | ANX11_HUMAN | Unmodified | 1119800 | 784330 | 0 | 941010 | 1976600 | 0 | 0 | 0 | 90065000 | 1223900 | 3810900 | 3561500 | 308890 | 0 | 659530 | 763990 | 0 | 0 |
| DAQGLVLF | MVP; MVP; MVP | Q14764; H3I | MVP_HUMAN I | Unmodified | 0 | 0 | 0 | 0 | 0 | 0 | 0 | 0 | 0 | 0 | 0 | 615630 | 0 | 0 | 0 | 0 | 0 | 0 |
| DAQHYGG | GANAB; GANAE | Q14697; F5H | GANAB_HUMA | Unmodified | 0 | 0 | 0 | 0 | 0 | 88810 | 0 | 0 | 33857000 | 362860 | 0 | 1508600 | 0 | 0 | 0 | 0 | 0 | 0 |
| DAQHYGG | GANAB; GANAE | Q14697; F5H | GANAB_HUMA | Unmodified | 0 | 0 | 0 | 0 | 0 | 0 | 0 | 0 | 45850000 | 3389000 | 154280 | 0 | 0 | 0 | 0 | 0 | 0 | 0 |
| DAQHYGG | GANAB; GANAE | Q14697; F5H | GANAB_HUMA | Oxidation (M) | 0 | 0 | 0 | 199590 | 0 | 0 | 0 | 0 | 20388000 | 1834700 | 581160 | 0 | 0 | 0 | 233870 | 0 | 0 | 0 |
| DAQHYGG | GANAB; GANAE | Q14697; F5H | GANAB_HUMA | Unmodified | 0 | 0 | 0 | 0 | 0 | 0 | 0 | 0 | 8781100 | 13122000 | 0 | 0 | 0 | 0 | 184790 | 0 | 0 | 0 |
| DAQHYGG | GANAB; GANAE | Q14697; F5H | GANAB_HUMA | Oxidation (M) | 0 | 0 | 0 | 0 | 0 | 0 | 0 | 0 | 0 | 3689500 | 0 | 0 | 0 | 0 | 0 | 0 | 0 | 0 |
| DAQIGWIQ | RAB3GAP2 | Q9H2M9 | RBGPR_HUMAI | Unmodified | 4624000 | 0 | 0 | 668630 | 27962000 | 2979100 | 4728700 | 0 | 26928000 | 28298000 | 6630400 | 1720400 | 71291000 | 23539000 | 0 | 23844 | 4640600 | 10051000 |
| DAQPSFSA | MCM3; MCM3; | P25205; A0A | MCM3_HUMAN | Unmodified | 0 | 0 | 0 | 0 | 0 | 0 | 586450 | 273930 | 423840 | 0 | 0 | 853950 | 406480 | 0 | 0 | 0 | 0 | 0 |
| DAQQSSSF | UBE4A; UBE4A | Q14139; B7Z | UBE4A_HUMAI | Unmodified | 0 | 0 | 0 | 0 | 0 | 0 | 0 | 0 | 0 | 0 | 0 | 3576900 | 0 | 0 | 0 | 38219 | 0 | 0 |
| DAQQVEPE | PRRC2C; PRRC | Q9Y520; H7( | PRC2C_HUMAI | Unmodified | 735720 | 316880 | 104510 | 189440 | 49654 | 0 | 0 | 0 | 181320 | 0 | 311290 | 1102100 | 152160 | 0 | 92326 | 134450 | 0 | 68717 |
| DAQSNAAF | DHX9 | Q08211 | DHX9_HUMAN | Unmodified | 171940 | 55650 | 0 | 140020 | 2371000 | 129980 | 219290 | 0 | 387410 | 1146100 | 1039400 | 499820 | 2638100 | 374990 | 45613 | 48990 | 0 | 272900 |
| DAREPVVG | CHID1; CHID1; | Q9BWS9; H( | CHID1_HUMAN | Unmodified | 0 | 0 | 0 | 0 | 0 | 0 | 0 | 0 | 1678600 | 280670 | 191630 | 349440 | 0 | 0 | 0 | 0 | 0 | 0 |
| DARGYLQA | CDC42BPA; CD | Q5VT25; Q9 | MRCKA_HUMA | Unmodified | 173740 | 88392 | 0 | 77617 | 937740 | 0 | 0 | 0 | 0 | 402380 | 0 | 0 | 420910 | 228660 | 0 | 92699 | 212180 | 774510 |
| DARQVTSN | LRPAP1; LRPAP | P30533; A0A | AMRP_HUMAN | Unmodified | 0 | 0 | 0 | 0 | 0 | 0 | 0 | 0 | 0 | 4522700 | 0 | 0 | 0 | 0 | 0 | 0 | 0 | 0 |
| DASDDLDI | EIF2S2 | P20042 | IF2B_HUMAN E | Unmodified | 138680 | 0 | 128940 | 300410 | 572420 | 75986 | 0 | 52257 | 276620 | 47812 | 110010 | 1059100 | 230810 | 118790 | 310570 | 251460 | 116150 | 242960 |
| DASFHCA/ | BCAM; BCAM; E | P50895; A0A | BCAM_HUMAN | Unmodified | 0 | 0 | 0 | 0 | 0 | 0 | 0 | 0 | 0 | 0 | 0 | 0 | 0 | 0 | 0 | 0 | 0 | 0 |
| DASIVGFFI | PDIA3 | P30101 | PDIA3_HUMAN | Unmodified | 101820 | 131620 | 267900 | 955770 | 1634000 | 0 | 749570 | 0 | 34231000 | 13189000 | 6494200 | 805360 | 1015500 | 1542600 | 3029700 | 229950 | 116080 | 2109300 |
| DASLMVTN | CCT2; CCT2; C( | P78371; F5G | TCPB_HUMAN | Oxidation (M) | 352450 | 0 | 634980 | 595960 | 0 | 0 | 0 | 0 | 0 | 0 | 1535400 | 8253200 | 0 | 0 | 613740 | 573590 | 0 | 0 |
| DASLMVTN | CCT2; CCT2; C( | P78371; F5G | TCPB_HUMAN | Unmodified | 0 | 0 | 0 | 0 | 0 | 0 | 0 | 0 | 682640 | 0 | 0 | 875290 | 367450 | 0 | 0 | 0 | 0 | 0 |
| DASLVVTTS | AAMP; AAMP; A | Q13685; C9J | AAMP_HUMAN | Unmodified | 0 | 59437 | 0 | 0 | 385360 | 0 | 0 | 81318 | 0 | 0 | 0 | 0 | 92487 | 0 | 0 | 0 | 330590 | 311200 |
| DASNWSTI | AHSA1; AHSA1 | O95433; G3Y | AHSA1_HUMAI | Unmodified | 0 | 0 | 0 | 0 | 0 | 77793 | 0 | 0 | 42258 | 0 | 49369 | 582210 | 0 | 0 | 0 | 0 | 0 | 0 |
| DASQHNH | ATP8B1; ATP8B | O43520; A0A | AT8B1_HUMAN | Unmodified | 114900 | 0 | 0 | 0 | 0 | 0 | 0 | 0 | 0 | 0 | 0 | 1103500 | 0 | 0 | 0 | 0 | 0 | 0 |
| DASSFLAE\ | SLC3A2; SLC3A | P08195; A0A | 4F2_HUMAN 4I | Unmodified | 0 | 0 | 0 | 40360 | 0 | 0 | 0 | 0 | 0 | 0 | 19074 | 42623 | 0 | 0 | 388860 | 0 | 0 | 0 |
| DASSQGFL | CTH | P32929 | CGL_HUMAN C | Unmodified | 0 | 0 | 0 | 0 | 0 | 0 | 0 | 0 | 0 | 0 | 0 | 0 | 3517600 | 2615300 | 0 | 38949 | 678990 | 2853900 |
| DASVAEAW | SPTBN1; SPTBN | Q01082; A0A | SPTB2_HUMAN | Unmodified | 0 | 0 | 0 | 16482 | 0 | 0 | 0 | 0 | 114670 | 0 | 29632 | 540220 | 0 | 0 | 0 | 0 | 0 | 0 |
| DASVHTLLI | TNFRSF10B | O14763 | TR10B_HUMAN | Unmodified | 93615 | 116430 | 180840 | 284450 | 0 | 0 | 0 | 0 | 0 | 0 | 473580 | 177720 | 0 | 0 | 1502800 | 303000 | 0 | 0 |
| DASVPLID\ | EXOC4 | Q96A65 | EXOC4_HUMAI | Unmodified | 0 | 0 | 0 | 0 | 84290 | 0 | 0 | 102350 | 245550 | 1185500 | 744070 | 2425800 | 0 | 29331 | 0 | 49063 | 0 | 216510 |
| DASVPLID\ | EXOC4 | Q96A65 | EXOC4_HUMAI | Unmodified | 0 | 0 | 0 | 0 | 2459800 | 556640 | 0 | 183620 | 0 | 0 | 0 | 682880 | 0 | 0 | 0 | 0 | 0 | 575130 |

| | | | | | | | | | | | | | | | | | | | | | | |
|---|---|---|---|---|---|---|---|---|---|---|---|---|---|---|---|---|---|---|---|---|---|---|
| DATAERIDE | ZPR1; ZPR1; ZP | O75312; H7( | ZPR1_HUMAN | Unmodified | 0 | 0 | 94111 | 171460 | 0 | 0 | 0 | 0 | 0 | 0 | 481120 | 2419600 | 0 | 457890 | 0 | 0 | 0 | 0 |
| DATHDEAV | SNTB1; SNTB1 | Q13884; A0A | SNTB1_HUMAN | Unmodified | 0 | 0 | 0 | 0 | 55506 | 0 | 11919 | 0 | 2561300 | 0 | 0 | 0 | 0 | 0 | 0 | 0 | 39329 | 44264 |
| DATKEALH | UHRF1BP1; UH | Q6BDS2; H7 | URFB1_HUMAN | Oxidation (M) | 0 | 0 | 0 | 0 | 0 | 0 | 0 | 0 | 0 | 0 | 0 | 0 | 0 | 0 | 0 | 0 | 394240 | 0 |
| DATLTALDF | HADHA; HADHA | P40939; A0A | ECHA_HUMAN | Unmodified | 0 | 0 | 0 | 0 | 1230300 | 0 | 1065900 | 392750 | 6073300 | 1900500 | 0 | 0 | 672170 | 454560 | 0 | 0 | 122720 | 791460 |
| DATLTALDF | HADHA; HADHA | P40939; A0A | ECHA_HUMAN | Unmodified | 0 | 0 | 0 | 0 | 0 | 0 | 273940 | 36138 | 0 | 0 | 0 | 0 | 0 | 0 | 0 | 0 | 0 | 0 |
| DATNVGDE | ENO1; ENO1 | P06733; A0A | ENOA_HUMAN | Unmodified | 1511900 | 1044600 | 20192000 | 10376000 | 34047000 | 7822200 | 975500 | 1301500 | 53189000 | 6363800 | 64363000 | 2.19E+08 | 7727000 | 3887000 | 19806000 | 19858000 | 3784400 | 16824000 |
| DATNVGDE | ENO1; ENO1 | P06733; A0A | ENOA_HUMAN | Unmodified | 16717000 | 6949500 | 0 | 81603000 | 1.98E+09 | 2.72E+08 | 75633000 | 7486400 | 4.47E+08 | 4.76E+08 | 2.41E+08 | 2.03E+08 | 8.42E+08 | 3.51E+08 | 34059000 | 28346000 | 6.84E+08 | 8.67E+08 |
| DATNVGDE | ENO1; ENO1 | P06733; A0A | ENOA_HUMAN | Unmodified | 0 | 0 | 0 | 0 | 13607000 | 0 | 4951000 | 0 | 0 | 3537200 | 0 | 0 | 3938300 | 0 | 0 | 0 | 29434000 | 14385000 |
| DATNVGDE | ENO3; ENO3 | P13929; E5R | ENOB_HUMAN | Unmodified | 0 | 0 | 0 | 0 | 0 | 0 | 0 | 0 | 447140 | 2010900 | 70779 | 338950 | 0 | 0 | 0 | 0 | 0 | 623470 |
| DATNVGDE | ENO2 | P09104 | ENOG_HUMAN | Unmodified | 175760 | 0 | 0 | 0 | 3705600 | 925810 | 1812400 | 63976 | 3212200 | 2972300 | 119880 | 881640 | 638480 | 699410 | 0 | 0 | 4393300 | 10147000 |
| DATQITQGI | L1CAM | P32004 | L1CAM_HUMAI | Unmodified | 0 | 0 | 0 | 0 | 533610 | 0 | 0 | 82938 | 1107800 | 0 | 1561000 | 0 | 0 | 0 | 0 | 0 | 263820 | 160410 |
| DATQITQGI | L1CAM | P32004 | L1CAM_HUMAI | Unmodified | 0 | 0 | 0 | 0 | 0 | 0 | 0 | 0 | 0 | 0 | 0 | 0 | 0 | 0 | 0 | 0 | 0 | 0 |
| DATSRPTDI | SEC63 | Q9UGP8 | SEC63_HUMAN | Unmodified | 0 | 0 | 197350 | 0 | 0 | 0 | 0 | 0 | 660410 | 0 | 125090 | 811230 | 0 | 0 | 132350 | 143210 | 0 | 0 |
| DATVRAVP | MELTF | P08582 | TRFM_HUMAN | Unmodified | 0 | 0 | 0 | 0 | 5530900 | 192140 | 179190 | 127860 | 831830 | 122760 | 295740 | 1870200 | 0 | 245240 | 0 | 2113500 | 1013400 | 1143400 |
| DATWSPLN | WDR77; WDR7 | Q9BQA1; H0 | MEP50_HUMAN | Unmodified | 0 | 0 | 0 | 0 | 0 | 0 | 0 | 0 | 0 | 0 | 0 | 2832100 | 0 | 0 | 0 | 0 | 0 | 0 |
| DAVFIPAG | DYNC1LI1; DYN | Q9Y6G9; E9I | DC1L1_HUMAN | Unmodified | 0 | 0 | 0 | 0 | 0 | 0 | 0 | 0 | 730060 | 330110 | 476890 | 1733400 | 0 | 0 | 0 | 0 | 0 | 0 |
| DAVFIPAG | DYNC1LI1; DYN | Q9Y6G9; E9I | DC1L1_HUMAN | Unmodified | 0 | 0 | 0 | 0 | 608060 | 0 | 0 | 0 | 639560 | 0 | 0 | 1736500 | 0 | 0 | 0 | 0 | 70117 | 0 |
| DAVFIPAG | DYNC1LI2 | O43237 | DC1L2_HUMAN | Unmodified | 0 | 0 | 0 | 0 | 791530 | 303130 | 0 | 0 | 247650 | 358360 | 0 | 2213000 | 581250 | 995050 | 0 | 0 | 775470 | 0 |
| DAVFIPAG | DYNC1LI2 | O43237 | DC1L2_HUMAN | Unmodified | 0 | 0 | 0 | 0 | 0 | 0 | 0 | 0 | 0 | 0 | 0 | 0 | 0 | 0 | 0 | 0 | 0 | 0 |
| DAVGQPPF | SUN1; SUN1; S | O94901; H0 | SUN1_HUMAN | Oxidation (M) | 0 | 0 | 0 | 0 | 0 | 0 | 0 | 0 | 1058400 | 0 | 0 | 0 | 0 | 0 | 0 | 0 | 0 | 0 |
| DAVGQPPF | SUN1; SUN1; S | O94901; H0 | SUN1_HUMAN | Unmodified | 197200 | 0 | 0 | 0 | 0 | 0 | 0 | 0 | 5906200 | 0 | 0 | 0 | 0 | 0 | 0 | 0 | 0 | 0 |
| DAVITVPVF | HYOU1; HYOU | Q9Y4L1; J3Q | HYOU1_HUMA | Unmodified | 0 | 0 | 0 | 0 | 930970 | 183550 | 0 | 0 | 1600500 | 299670 | 250080 | 89711 | 0 | 0 | 44274 | 0 | 361390 | 486430 |
| DAVITVPVF | HYOU1; HYOU | Q9Y4L1; J3Q | HYOU1_HUMA | Unmodified | 85795 | 0 | 3278500 | 0 | 0 | 0 | 0 | 0 | 15420000 | 3362000 | 3672000 | 3091900 | 0 | 0 | 0 | 0 | 0 | 1046300 |
| DAVLLVFAI | ARF5; ARF3; AR | P84085; P61 | ARF5_HUMAN | Unmodified | 0 | 0 | 0 | 0 | 0 | 0 | 0 | 0 | 869400 | 345070 | 137520 | 0 | 0 | 0 | 0 | 0 | 0 | 0 |
| DAVQQDG | PPP3CB; PPP3C | P16298; Q5F | PP2BB_HUMAN | Unmodified | 0 | 0 | 0 | 0 | 362340 | 0 | 0 | 0 | 0 | 0 | 0 | 0 | 0 | 0 | 0 | 0 | 136720 | 1391400 |
| DAVRELDN | TAOK1; TAOK3; | Q7L7X3; Q9I | TAOK1_HUMAN | Unmodified | 36893 | 47435 | 0 | 196640 | 0 | 0 | 0 | 0 | 84153 | 0 | 58622 | 0 | 124670 | 0 | 141210 | 132020 | 21481 | 0 |
| DAVRELDN | TAOK1; TAOK3; | Q7L7X3; Q9I | TAOK1_HUMAN | Unmodified | 803350 | 598690 | 352970 | 996460 | 4524000 | 705740 | 1780300 | 1060900 | 2216700 | 1185400 | 1050200 | 5513900 | 2129700 | 1826600 | 1070300 | 2366100 | 8436900 | 27930000 |
| DAVRGND | MRE11; MRE11 | P49959; F5H | MRE11_HUMAI | Unmodified | 0 | 0 | 0 | 0 | 0 | 0 | 0 | 0 | 0 | 0 | 0 | 1144100 | 0 | 0 | 0 | 0 | 0 | 0 |
| DAVRIPTVD | KIAA1522 | Q9P206 | K1522_HUMAN | Unmodified | 42695 | 0 | 382680 | 591320 | 0 | 0 | 20890 | 0 | 747240 | 89296 | 218640 | 734110 | 0 | 132720 | 888860 | 708460 | 0 | 0 |
| DAVSNTTN | KTN1 | Q86UP2 | KTN1_HUMAN | Unmodified | 0 | 0 | 0 | 0 | 0 | 0 | 0 | 0 | 0 | 0 | 156130 | 1023800 | 0 | 0 | 0 | 0 | 0 | 0 |
| DAVTYTEH | H4C1 | P62805 | H4_HUMAN Hi | Unmodified | 44017000 | 43226000 | 607520 | 0 | 0 | 0 | 0 | 655390 | 553020 | 0 | 0 | 9.54E+08 | 0 | 0 | 0 | 259350 | 0 | 0 |
| DAVTYTEH | H4C1 | P62805 | H4_HUMAN Hi | Unmodified | 19427000 | 26765000 | 0 | 0 | 0 | 7621.6 | 0 | 0 | 0 | 0 | 0 | 4.62E+08 | 0 | 0 | 0 | 0 | 141960 | 1293100 |
| DAVVQHS | EXOC3; EXOC3 | O60645; D6I | EXOC3_HUMAI | Unmodified | 0 | 0 | 0 | 0 | 1357600 | 269160 | 0 | 0 | 0 | 273350 | 0 | 1459300 | 0 | 0 | 0 | 0 | 0 | 0 |
| DAVVTYTAI | SLC12A2; SLC1 | P55011; G3 | S12A2_HUMAN | Unmodified | 175740 | 73233 | 470230 | 98338 | 0 | 0 | 0 | 0 | 63069 | 0 | 0 | 573680 | 0 | 0 | 1227200 | 837350 | 0 | 0 |
| DAVVYPILV | HYOU1; HYOU | Q9Y4L1; E9F | HYOU1_HUMA | Unmodified | 0 | 0 | 110330 | 0 | 0 | 0 | 0 | 0 | 7593500 | 0 | 1663700 | 664380 | 0 | 0 | 657500 | 240630 | 0 | 0 |
| DAVVYPILV | HYOU1; HYOU | Q9Y4L1; E9F | HYOU1_HUMA | Unmodified | 0 | 0 | 51447 | 240210 | 0 | 0 | 0 | 0 | 3086000 | 1737100 | 0 | 0 | 0 | 129170 | 1508300 | 0 | 0 | 0 |
| DAWASPCI | SNTB2 | Q13425 | SNTB2_HUMAN | Unmodified | 110920 | 0 | 0 | 906910 | 0 | 0 | 0 | 0 | 0 | 84198 | 17005000 | 0 | 0 | 0 | 1048300 | 511380 | 0 | 0 |
| DAWEIPRE | HCK; HCK; HCK | P08631; J3K | HCK_HUMAN T | Unmodified | 0 | 0 | 0 | 0 | 0 | 0 | 0 | 0 | 409790 | 0 | 0 | 0 | 0 | 0 | 0 | 126930 | 0 | 0 |
| DAWEIPRE | HCK; HCK; HCK | P08631; J3K | HCK_HUMAN T | Unmodified | 0 | 0 | 0 | 0 | 0 | 0 | 0 | 0 | 0 | 0 | 2749300 | 1305100 | 0 | 0 | 0 | 0 | 0 | 0 |
| DAWEIPRE | YES1; SRC; YES | P07947; P12 | YES_HUMAN Ty | Unmodified | 1027200 | 1535500 | 2463300 | 2636700 | 2186600 | 845420 | 0 | 0 | 3137700 | 0 | 1268200 | 4193200 | 1150000 | 0 | 3123100 | 2173100 | 1423000 | 1986800 |
| DAWEIPRE | YES1; SRC; YES | P07947; P12 | YES_HUMAN Ty | Unmodified | 0 | 0 | 0 | 0 | 3225300 | 0 | 300470 | 0 | 0 | 647480 | 60337 | 0 | 851110 | 1467500 | 0 | 0 | 2450600 | 3857700 |
| DAWWAEIF | C2CD5; C2CD5 | Q86YS7; H0 | C2CD5_HUMA | Unmodified | 110300 | 122940 | 64214 | 118800 | 0 | 0 | 0 | 0 | 0 | 73042 | 108090 | 545940 | 0 | 0 | 258290 | 205710 | 0 | 0 |
| DAYERDLE | ANXA6; ANXA6; | P08133; E5R | ANXA6_HUMAN | Unmodified | 0 | 0 | 7484200 | 6554100 | 2646400 | 510900 | 1377000 | 0 | 99396000 | 16417000 | 0 | 0 | 2372600 | 559190 | 11264000 | 3552100 | 1734800 | 1698900 |
| DAYQVILD | API5; API5; API5 | Q9BZZ5; E9F | API5_HUMAN A | Unmodified | 0 | 0 | 0 | 0 | 578880 | 0 | 0 | 0 | 403240 | 150240 | 1644800 | 2774000 | 0 | 0 | 0 | 1029100 | 0 | 0 |
| DAYQVILD | API5; API5; API5 | Q9BZZ5; E9F | API5_HUMAN A | Unmodified | 0 | 0 | 0 | 120740 | 657290 | 0 | 929300 | 48204 | 0 | 0 | 0 | 0 | 889920 | 644180 | 0 | 0 | 0 | 722020 |
| DAYRPTTD | MACF1; MACF1 | Q9UPN3; H3 | MACF1_HUMAI | Unmodified | 0 | 26995 | 0 | 189650 | 0 | 0 | 0 | 0 | 653170 | 0 | 327000 | 1591500 | 0 | 0 | 119610 | 189060 | 0 | 0 |
| DAYSGGA | PSMB5 | P28074 | PSB5_HUMAN | Unmodified | 199170 | 0 | 110620 | 0 | 0 | 0 | 132710 | 0 | 0 | 0 | 0 | 6504400 | 0 | 0 | 0 | 274360 | 0 | 0 |
| DAYSGGA | PSMB5 | P28074 | PSB5_HUMAN | Unmodified | 258280 | 254170 | 0 | 129520 | 0 | 0 | 0 | 0 | 1043400 | 0 | 28773 | 6905700 | 0 | 5386400 | 0 | 567580 | 0 | 0 |
| DAYSLANN | CAPN7 | Q9Y6W3 | CAN7_HUMAN | Unmodified | 0 | 0 | 0 | 0 | 0 | 0 | 0 | 0 | 0 | 0 | 0 | 1249800 | 0 | 0 | 0 | 0 | 0 | 0 |
| DAYSSFGS | DDX3X; DDX3Y | O00571; O1 | DDX3X_HUMAI | Unmodified | 1213300 | 2466600 | 1095000 | 0 | 0 | 0 | 0 | 0 | 114920 | 0 | 0 | 320760 | 0 | 0 | 2585700 | 0 | 0 | 0 |
| DAYVSRLN | GSR; GSR; GSR | P00390; H0 | GSHR_HUMAN | Unmodified | 0 | 0 | 0 | 272980 | 3969300 | 1353800 | 143150 | 0 | 617900 | 543850 | 619700 | 0 | 3086400 | 1204600 | 0 | 0 | 394570 | 0 |
| DCAVIVTQI | PSMA6; PSMA6 | P60900; G3 | PSA6_HUMAN | Unmodified | 0 | 0 | 0 | 0 | 1264900 | 341920 | 167560 | 0 | 0 | 118150 | 225930 | 3909900 | 2053900 | 1425700 | 0 | 0 | 396730 | 1250400 |
| DCAVKPC | ERO1A; ERO1A | Q96HE7; G3 | ERO1A_HUMAI | Unmodified | 0 | 0 | 0 | 0 | 0 | 0 | 0 | 0 | 2838100 | 390620 | 177380 | 265940 | 0 | 0 | 0 | 0 | 0 | 0 |
| DCDRAIEIN | ST13P4; ST13; | Q8IZP2; P50 | ST134_HUMAN | Unmodified | 272440 | 0 | 373650 | 230370 | 0 | 0 | 0 | 0 | 572800 | 0 | 0 | 1887900 | 189890 | 0 | 316250 | 299580 | 0 | 0 |
| DCEDPNPI | AP1B1; AP2B1; | Q10567; P6 | AP1B1_HUMAN | Unmodified | 230610 | 187500 | 278370 | 373060 | 717700 | 347390 | 172960 | 0 | 875260 | 433610 | 775710 | 3748600 | 424890 | 335460 | 0 | 0 | 151400 | 0 |
| DCEECIQLI | STIP1; STIP1 | P31948; H0 | STIP1_HUMAN | Unmodified | 0 | 0 | 0 | 0 | 0 | 0 | 0 | 0 | 747060 | 0 | 700390 | 3651500 | 454770 | 0 | 0 | 0 | 0 | 0 |
| DCFGCLR | BANF1 | O75531 | BAF_HUMAN B | Unmodified | 0 | 0 | 0 | 0 | 0 | 0 | 0 | 0 | 471610 | 0 | 0 | 12969000 | 0 | 0 | 0 | 0 | 0 | 0 |
| DCFIEVNSI | ADAM9; ADAM9 | Q13443; A0A | ADAM9_HUMA | Unmodified | 862220 | 474960 | 1373600 | 489820 | 397930 | 0 | 0 | 0 | 213360 | 0 | 1066600 | 1572500 | 0 | 0 | 2200100 | 1718900 | 70657 | 0 |
| DCFIEVNSI | ADAM9; ADAM9 | Q13443; A0A | ADAM9_HUMA | Unmodified | 304860 | 0 | 77307 | 250420 | 0 | 0 | 0 | 0 | 165070 | 0 | 53169 | 524180 | 0 | 0 | 0 | 371130 | 0 | 0 |
| DCFIEVNSI | ADAM9; ADAM9 | Q13443; A0A | ADAM9_HUMA | Unmodified | 0 | 96508 | 0 | 0 | 1247600 | 3127200 | 0 | 0 | 0 | 0 | 828760 | 0 | 544740 | 0 | 0 | 0 | 0 | 605320 |
| DCFTCSN | FHL1; FHL1; FH | Q13642; A0A | FHL1_HUMAN | Unmodified | 40381 | 0 | 0 | 51835 | 105110 | 0 | 333490 | 192840 | 0 | 0 | 0 | 0 | 0 | 386600 | 0 | 22045 | 179730 | 132500 |
| DCGATWV | TPI1; TPI1; TPI1 | P60174; U3 | TPIS_HUMAN T | Unmodified | 2441700 | 0 | 1324600 | 1527700 | 3541300 | 0 | 0 | 0 | 1242500 | 393470 | 0 | 12457000 | 5192700 | 829360 | 2643900 | 983280 | 1243100 | 3512500 |
| DCGATWV | TPI1; TPI1; TPI1 | P60174; U3 | TPIS_HUMAN T | Unmodified | 3396300 | 2017500 | 2380900 | 4606800 | 7317700 | 3280200 | 168750 | 0 | 4516300 | 1100900 | 3016300 | 32644000 | 12270000 | 2909900 | 3757300 | 3748100 | 3649000 | 7312700 |
| DCGATWV | TPI1; TPI1; TPI1 | P60174; U3 | TPIS_HUMAN T | Unmodified | 0 | 0 | 0 | 0 | 96387000 | 11414000 | 8503400 | 0 | 0 | 7234000 | 0 | 0 | 66970000 | 48899000 | 0 | 0 | 68456000 | 1.13E+08 |
| DCGKLSKD | EPS15; EPS15 | P42566; B1A | EPS15_HUMAN | Unmodified | 0 | 0 | 0 | 0 | 137320 | 58172 | 35758 | 0 | 0 | 38745 | 0 | 0 | 870300 | 171350 | 0 | 0 | 195420 | 80168 |
| DCGSVDG | NPC2; NPC2; N | P61916; J3K | NPC2_HUMAN | Unmodified | 0 | 0 | 96449 | 105980 | 506420 | 313560 | 0 | 0 | 14805000 | 1394400 | 0 | 0 | 678600 | 152020 | 0 | 0 | 0 | 0 |
| DCGSVDG | NPC2; NPC2; N | P61916; J3K | NPC2_HUMAN | Unmodified | 0 | 0 | 0 | 0 | 0 | 0 | 0 | 0 | 0 | 1306100 | 0 | 0 | 1088500 | 0 | 0 | 0 | 0 | 0 |
| DCGVGFR | MDK; MDK; MD | P21741; E9P | MK_HUMAN Mi | Unmodified | 131910 | 0 | 242490 | 0 | 449940 | 8994000 | 0 | 0 | 11202000 | 188260 | 0 | 0 | 0 | 0 | 0 | 0 | 439100 | 0 |
| DCGVGFRI | MDK; MDK; MD | P21741; E9P | MK_HUMAN Mi | Unmodified | 0 | 0 | 0 | 0 | 0 | 0 | 0 | 0 | 7046100 | 3388600 | 0 | 0 | 0 | 0 | 0 | 0 | 0 | 0 |
| DCHEEVYA | ACBD3 | Q9H3P7 | GCP60_HUMAI | Unmodified | 0 | 0 | 0 | 0 | 0 | 0 | 0 | 0 | 0 | 0 | 0 | 608270 | 0 | 0 | 0 | 46572 | 0 | 0 |
| DCIGGCSD | GLRX | P35754 | GLRX1_HUMAN | Unmodified | 0 | 0 | 0 | 0 | 0 | 0 | 0 | 0 | 1716700 | 0 | 279300 | 0 | 0 | 0 | 0 | 0 | 3165700 | 0 |
| DCKVADGI | TPPP3 | Q9BW30 | TPPP3_HUMAN | Unmodified | 0 | 0 | 0 | 0 | 0 | 0 | 290650 | 274380 | 0 | 0 | 0 | 0 | 0 | 0 | 0 | 0 | 0 | 0 |
| DCLNVLNK | PFDN5; PFDN5 | Q99471; A0A | PFD5_HUMAN | Unmodified | 0 | 0 | 0 | 100440 | 1474200 | 150280 | 0 | 0 | 952380 | 204600 | 0 | 5925100 | 1316700 | 378400 | 0 | 0 | 1135200 | 565100 |
| DCPDNPHI | KHNYN; KHNYN | O15037; G3 | KHNYN_HUMA | Unmodified | 592820 | 0 | 117330 | 0 | 227100 | 84322 | 0 | 0 | 663100 | 0 | 0 | 0 | 130770 | 0 | 304290 | 168410 | 0 | 0 |
| DCPDSAEE | CYFIP1; CYFIP1 | Q7L576; A0A | CYFP1_HUMAN | Unmodified | 86100 | 0 | 0 | 0 | 0 | 0 | 0 | 0 | 0 | 0 | 30133 | 249590 | 0 | 0 | 0 | 0 | 0 | 0 |
| DCPLNAEA | AP1B1 | Q10567 | AP1B1_HUMAN | Unmodified | 0 | 0 | 0 | 0 | 0 | 0 | 0 | 0 | 0 | 0 | 0 | 637110 | 0 | 0 | 35202 | 0 | 0 | 0 |
| DCRDWSD | LDLR; LDLR; LD | P01130; J3K | LDLR_HUMAN | Unmodified | 72345 | 0 | 0 | 0 | 0 | 0 | 0 | 0 | 1545000 | 0 | 123600 | 477350 | 0 | 0 | 0 | 0 | 0 | 0 |
| DCRFLPG | ADAM9; ADAM9 | Q13443; A0A | ADAM9_HUMA | Unmodified | 0 | 0 | 0 | 176380 | 1557100 | 525460 | 0 | 0 | 0 | 0 | 0 | 0 | 1015800 | 525010 | 0 | 0 | 0 | 0 |
| DCRFLPG | ADAM9; ADAM9 | Q13443; A0A | ADAM9_HUMA | Unmodified | 0 | 0 | 0 | 0 | 3386600 | 382840 | 0 | 0 | 0 | 124880 | 0 | 0 | 1350200 | 1092200 | 0 | 0 | 231120 | 1272800 |
| DCRYALYD | DSTN; DSTN | P60981; F6R | DEST_HUMAN | Unmodified | 0 | 0 | 0 | 0 | 24214000 | 2691700 | 1908000 | 1602800 | 0 | 1816700 | 373110 | 0 | 19842000 | 11170000 | 0 | 0 | 7166300 | 9830600 |
| DCRYALYD | CFL1; CFL2; CF | P23528; Q9 | COF1_HUMAN | Unmodified | 141210 | 0 | 127380 | 687250 | 97816000 | 18428000 | 3274700 | 1589300 | 417420 | 3336300 | 1206500 | 0 | 26854000 | 9801200 | 0 | 0 | 13561000 | 27356000 |
| DCSLPYATI | FBLN1; FBLN1; | P23142; B1A | FBLN1_HUMAN | Unmodified | 0 | 0 | 0 | 0 | 0 | 0 | 0 | 474780 | 7714000 | 4082800 | 0 | 0 | 0 | 0 | 0 | 0 | 0 | 0 |
| DCTEALKLI | TOMM34 | Q15785 | TOM34_HUMAI | Unmodified | 0 | 0 | 0 | 0 | 411870 | 0 | 0 | 0 | 0 | 0 | 315470 | 492300 | 192930 | 86998 | 0 | 0 | 0 | 274220 |
| DCVGPEVE | PGK1 | P00558 | PGK1_HUMAN | Unmodified | 1199800 | 462390 | 1449900 | 1742600 | 8845400 | 0 | 3293900 | 147330 | 4878900 | 191470 | 0 | 26543000 | 12186000 | 0 | 0 | 2935400 | 31390000 | 48930000 |
| DCVGPEVE | PGK1 | P00558 | PGK1_HUMAN | Unmodified | 0 | 0 | 0 | 0 | 0 | 0 | 0 | 0 | 0 | 0 | 0 | 0 | 0 | 0 | 0 | 0 | 4798200 | 0 |
| DCVLLQED | KIAA2013 | Q8IYS2 | K2013_HUMAN | Unmodified | 0 | 0 | 0 | 0 | 0 | 0 | 0 | 0 | 13032000 | 1286600 | 0 | 0 | 0 | 0 | 0 | 0 | 0 | 0 |
| DCVSPSEY | VPS28; VPS28 | Q9UK41; E9I | VPS28_HUMAN | Unmodified | 15945000 | 5096700 | 8899700 | 7802000 | 1275200 | 0 | 0 | 0 | 1249500 | 833330 | 167800 | 1132500 | 5463300 | 3747400 | 13305000 | 9234500 | 308340 | 1424400 |
| DCVSPSEY | VPS28; VPS28 | Q9UK41; E9I | VPS28_HUMAN | Unmodified | 0 | 0 | 0 | 0 | 0 | 0 | 0 | 258540 | 0 | 277550 | 0 | 0 | 3060400 | 5035100 | 0 | 0 | 938880 | 0 |
| DDAMLLK | HSPD1; HSPD1 | P10809; A0A | CH60_HUMAN | Oxidation (M) | 0 | 0 | 0 | 0 | 2284100 | 617350 | 0 | 0 | 342940 | 0 | 0 | 0 | 0 | 0 | 0 | 0 | 237640 | 0 |
| DDANNDP | PAICS; PAICS; F | P22234; D6F | PUR6_HUMAN | Unmodified | 0 | 57853 | 0 | 0 | 0 | 0 | 0 | 0 | 113240 | 0 | 720140 | 3434400 | 0 | 0 | 0 | 0 | 0 | 0 |
| DDDGALH | SOD3 | P08294 | SODE_HUMAN | Unmodified | 0 | 0 | 0 | 0 | 0 | 0 | 0 | 0 | 1136800 | 0 | 0 | 0 | 0 | 0 | 0 | 0 | 0 | 0 |
| DDDHNGH | UBA6; UBA6 | A0AVT1; H0 | UBA6_HUMAN | Unmodified | 0 | 0 | 0 | 0 | 0 | 0 | 0 | 0 | 0 | 0 | 112990 | 644300 | 0 | 0 | 0 | 0 | 0 | 0 |
| DDDIAALV | ACTB; ACTB; AC | P60709; G5E | ACTB_HUMAN | Acetyl (Protein N | 7556400 | 7701600 | 2908300 | 14117000 | 1.03E+08 | 16915000 | 27130000 | 53670000 | 4802400 | 4562900 | 8850700 | 21759000 | 42129000 | 43825000 | 12305000 | 8216300 | 80539000 | 30505000 |
| DDDIAALV | ACTB; ACTB; AC | P60709; G5E | ACTB_HUMAN | Acetyl (Protein N | 96537 | 118560 | 28027 | 222120 | 5746400 | 976780 | 1393300 | 403080 | 1156200 | 517700 | 224320 | 619960 | 2932200 | 3260700 | 385580 | 63536 | 3890500 | 2085500 |
| DDDIAALV | ACTB; ACTB; AC | P60709; G5E | ACTB_HUMAN | Acetyl (Protein N | 0 | 0 | 0 | 0 | 0 | 0 | 2811300 | 2878900 | 0 | 0 | 0 | 0 | 0 | 0 | 0 | 0 | 10039000 | 1011900 |
| DDDIEEGD | MESD | Q14696 | MESD_HUMAN | Unmodified | 0 | 0 | 0 | 0 | 0 | 0 | 40420 | 0 | 4725800 | 0 | 0 | 0 | 0 | 0 | 0 | 0 | 0 | 0 |
| DDDKNND | MCFD2 | Q8NI22 | MCFD2_HUMA | Unmodified | 0 | 0 | 0 | 0 | 0 | 0 | 0 | 0 | 301710 | 0 | 0 | 0 | 0 | 0 | 67584 | 0 | 0 | 0 |
| DDEAEWQ | SEC63 | Q9UGP8 | SEC63_HUMAN | Unmodified | 0 | 0 | 0 | 0 | 0 | 0 | 0 | 0 | 0 | 202750 | 0 | 0 | 0 | 0 | 0 | 0 | 0 | 0 |
| DDEFTHLY | CALR; CALR; CA | P27797; A0A | CALR_HUMAN | Unmodified | 0 | 0 | 211520 | 0 | 0 | 0 | 0 | 0 | 40689000 | 2524500 | 1422900 | 1295000 | 0 | 0 | 2236000 | 313430 | 0 | 0 |
| DDEPVCGI | APMAP; APMAP | Q9HDC9; H0 | APMAP_HUMAI | Unmodified | 266580 | 0 | 0 | 0 | 119680 | 0 | 0 | 0 | 1492400 | 183310 | 0 | 0 | 0 | 0 | 0 | 0 | 0 | 0 |
| DDEVQVVF | RPL26; RPL26L | P61254; Q9L | RL26_HUMAN | Unmodified | 350770 | 0 | 273910 | 0 | 0 | 0 | 0 | 0 | 1177300 | 0 | 594460 | 4709600 | 0 | 0 | 199490 | 84146 | 0 | 0 |
| DDFKRELD | IMMT; IMMT; IM | Q16891; B9A | MIC60_HUMAN | Unmodified | 0 | 0 | 0 | 0 | 0 | 0 | 129390 | 0 | 1438500 | 83238 | 0 | 0 | 0 | 0 | 0 | 0 | 0 | 0 |
| DDGLFSGI | TAGLN2; TAGLN | P37802; X6R | TAGL2_HUMAN | Unmodified | 1139600 | 1075300 | 306410 | 323080 | 564010 | 149970 | 0 | 0 | 1246900 | 331990 | 1004800 | 4141700 | 144080 | 0 | 719330 | 840120 | 314220 | 134870 |
| DDGRGVSI | MAN2B1; N/A | O00754; M0 | MA2B1_HUMAI | Unmodified | 0 | 0 | 0 | 0 | 513410 | 0 | 0 | 0 | 908950 | 0 | 0 | 0 | 0 | 0 | 0 | 215170 | 0 | 0 |
| DDGVSIPG | PRMT5; PRMT5; | O14744; H0 | ANM5_HUMAN | Unmodified | 0 | 0 | 0 | 0 | 0 | 0 | 0 | 419930 | 0 | 0 | 1302700 | 845550 | 0 | 0 | 0 | 0 | 0 | 0 |
| DDIDLQNL | TSPAN14; TSPA | Q8NG11; A6 | TSN14_HUMAN | Unmodified | 18564 | 0 | 0 | 0 | 0 | 0 | 0 | 0 | 0 | 0 | 0 | 0 | 0 | 0 | 244290 | 34959 | 0 | 0 |
| DDIIENAPT | PRMT5; PRMT5; | O14744; H0 | ANM5_HUMAN | Unmodified | 0 | 0 | 0 | 0 | 0 | 0 | 0 | 0 | 0 | 0 | 0 | 5344200 | 0 | 0 | 353360 | 171880 | 0 | 0 |
| DDILNGSH | TLN1 | Q9Y490 | TLN1_HUMAN | Unmodified | 281030 | 719200 | 0 | 569650 | 0 | 0 | 0 | 0 | 0 | 0 | 0 | 1114800 | 92751 | 0 | 2300700 | 855050 | 0 | 0 |
| DDKHGSYI | TCP1; TCP1; TC | P17987; F5H | TCPA_HUMAN | Unmodified | 0 | 0 | 0 | 0 | 0 | 0 | 0 | 0 | 234000 | 48417 | 798960 | 2915500 | 0 | 0 | 0 | 0 | 106960 | 53885 |

| | | | | | | | | | | | | | | | | | | | | | | |
|---|---|---|---|---|---|---|---|---|---|---|---|---|---|---|---|---|---|---|---|---|---|---|
| DDLDEEED | PODXL | O00592 | PODXL_HUMAI | Unmodified | 0 | 0 | 259100 | 0 | 197320 | 0 | 0 | 0 | 306250 | 0 | 0 | 0 | 358990 | 0 | 0 | 0 | 607530 | 0 |
| DDLENVK | ITGB1; ITGB1; I | P05556; E7E | ITB1_HUMAN I | Unmodified | 100700 | 54052 | 565260 | 80974 | 974680 | 162500 | 74357 | 28012 | 374000 | 52749 | 0 | 316030 | 0 | 0 | 0 | 217260 | 505360 | 533390 |
| DDLGNTLE | MACROH2A1 | O75367 | H2AY_HUMAN | Unmodified | 0 | 0 | 0 | 0 | 0 | 0 | 0 | 0 | 0 | 0 | 305240 | 137140 | 0 | 0 | 0 | 0 | 0 | 0 |
| DDLVTVKTI | DLST; DLST; DL | P36957; Q86 | ODO2_HUMAN | Unmodified | 0 | 0 | 0 | 0 | 0 | 0 | 9405400 | 0 | 0 | 1500900 | 0 | 0 | 0 | 0 | 0 | 0 | 0 | 0 |
| DDNGVPF\ | ALDOC; ALDOC | P09972; J3Q | ALDOC_HUMA | Unmodified | 0 | 0 | 7770100 | 132470 | 0 | 0 | 0 | 0 | 0 | 0 | 253360 | 3203700 | 0 | 0 | 228020 | 221780 | 0 | 0 |
| DDPLTNLN | ACTN1; ACTN1; | P12814; G3\ | ACTN1_HUMAI | Unmodified | 0 | 0 | 0 | 0 | 0 | 0 | 0 | 0 | 0 | 0 | 113960 | 697540 | 8983.6 | 0 | 0 | 0 | 0 | 0 |
| DDPSKPVH | RPL3; RPL3; RP | P39023; H7C | RL3_HUMAN 6 | Unmodified | 5732600 | 4252700 | 6995000 | 1755300 | 408750 | 121230 | 1042900 | 185190 | 1154500 | 127350 | 3461600 | 86349000 | 137560 | 0 | 6009000 | 4223300 | 0 | 394500 |
| DDPVTNLN | ACTN4; ACTN4; | O43707; F5C | ACTN4_HUMAI | Unmodified | 0 | 0 | 524280 | 0 | 583350 | 202420 | 43671 | 43079 | 1363300 | 0 | 201460 | 3519800 | 361190 | 227140 | 1210100 | 193650 | 175910 | 0 |
| DDQALTRP | FGFRL1 | Q8N441 | FGRL1_HUMAN | Unmodified | 269770 | 146150 | 0 | 0 | 0 | 0 | 0 | 0 | 0 | 20944 | 0 | 0 | 0 | 0 | 0 | 764770 | 0 | 0 |
| DDREAQSI | AP1B1; AP2B1; | Q10567; P63 | AP1B1_HUMAN | Unmodified | 47827 | 43188 | 260980 | 92480 | 0 | 0 | 0 | 0 | 77201 | 0 | 0 | 504030 | 0 | 0 | 144310 | 178410 | 0 | 0 |
| DDSGQYY( | JAM3 | Q9BX67 | JAM3_HUMAN . | Unmodified | 0 | 0 | 0 | 0 | 0 | 0 | 0 | 0 | 235300 | 117800 | 471470 | 818940 | 0 | 0 | 0 | 0 | 0 | 0 |
| DDSNHTIG | RAB4A | P20338 | RAB4A_HUMAN | Unmodified | 58108 | 30763 | 219600 | 106420 | 0 | 0 | 0 | 0 | 0 | 0 | 0 | 614010 | 0 | 0 | 123350 | 100300 | 0 | 0 |
| DDSQFSEE | ELAPOR2; ELAP | A8MWY0; H7 | ELAP2_HUMAN | Unmodified | 0 | 0 | 0 | 0 | 0 | 0 | 0 | 0 | 715270 | 0 | 0 | 0 | 0 | 0 | 0 | 0 | 0 | 0 |
| DDSVEGLR | VPS36 | Q86VN1 | VPS36_HUMAN | Oxidation (M) | 466410 | 0 | 111370 | 250780 | 0 | 0 | 0 | 0 | 0 | 0 | 0 | 0 | 0 | 0 | 592750 | 242600 | 0 | 0 |
| DDTDDEIAI | CANX; CANX; C | P27824; A0A | CALX_HUMAN | Oxidation (M) | 0 | 0 | 0 | 0 | 0 | 0 | 0 | 0 | 6699800 | 0 | 440860 | 0 | 0 | 0 | 0 | 0 | 0 | 0 |
| DDTDDEIAI | CANX; CANX; C | P27824; A0A | CALX_HUMAN | Unmodified | 0 | 0 | 0 | 0 | 0 | 0 | 0 | 0 | 13082000 | 271810 | 0 | 0 | 0 | 0 | 0 | 0 | 0 | 0 |
| DDTHYYIQ | PLEK2; PLEK2 | Q9NYT0; H0 | PLEK2_HUMAN | Unmodified | 0 | 0 | 409040 | 0 | 0 | 0 | 0 | 0 | 383160 | 0 | 0 | 0 | 0 | 0 | 0 | 0 | 0 | 0 |
| DDTIYEDEI | UQCRB; UQCRI | P14927; B7Z | QCR7_HUMAN | Unmodified | 0 | 0 | 0 | 0 | 0 | 0 | 0 | 0 | 877330 | 0 | 148220 | 0 | 0 | 0 | 0 | 0 | 0 | 0 |
| DDTIYEDEI | UQCRB; UQCRI | P14927; B7Z | QCR7_HUMAN | Unmodified | 58362 | 228820 | 77088 | 0 | 0 | 0 | 0 | 0 | 908140 | 0 | 0 | 0 | 0 | 0 | 0 | 47598 | 0 | 0 |
| DDVAQTDL | SLC3A2; SLC3A | P08195; A0A | 4F2_HUMAN 4I | Unmodified | 450760 | 323020 | 3728700 | 0 | 0 | 0 | 0 | 0 | 1292800 | 295210 | 592920 | 7689500 | 0 | 0 | 3794300 | 576590 | 48871 | 0 |
| DDVAQTDL | SLC3A2; SLC3A | P08195; A0A | 4F2_HUMAN 4I | Unmodified | 0 | 0 | 0 | 0 | 0 | 0 | 0 | 0 | 0 | 0 | 0 | 0 | 11110000 | 0 | 1932800 | 0 | 0 | 0 |
| DDVFGYP( | USP9X | Q93008 | USP9X_HUMAN | Unmodified | 0 | 0 | 0 | 0 | 0 | 0 | 0 | 0 | 0 | 0 | 0 | 812900 | 0 | 0 | 0 | 0 | 0 | 0 |
| DEAAGGA | MARCKS | P29966 | MARCS_HUMA | Unmodified | 145820 | 0 | 0 | 0 | 0 | 0 | 0 | 0 | 8592200 | 1314800 | 0 | 0 | 0 | 0 | 173850 | 0 | 0 | 0 |
| DEAGHFLV | PCK2; PCK2 | Q16822; H0 | PCKGM_HUMA | Unmodified | 0 | 0 | 109810 | 0 | 0 | 386830 | 389680 | 183140 | 1535400 | 112820 | 0 | 0 | 0 | 0 | 46061 | 0 | 0 | 0 |
| DEAQNEGI | CCAR2; CCAR2 | Q8N163; G3 | CCAR2_HUMAI | Unmodified | 0 | 0 | 0 | 0 | 0 | 0 | 0 | 0 | 0 | 0 | 0 | 232360 | 996270 | 203810 | 0 | 0 | 209630 | 0 |
| DEAVGALH | GNE; GNE | Q9Y223; A0 | GLCNE_HUMA | Unmodified | 0 | 112530 | 54386 | 0 | 0 | 0 | 0 | 0 | 0 | 0 | 0 | 576140 | 0 | 0 | 20195 | 0 | 0 | 0 |
| DECCFDAN | ADAM10 | O14672 | ADA10_HUMAN | Unmodified | 0 | 0 | 3520500 | 0 | 0 | 0 | 0 | 0 | 213160 | 0 | 0 | 1590000 | 164210 | 92974 | 1326600 | 1731100 | 0 | 0 |
| DEDDEAYG | SLC44A4; SLC4 | Q53GD3; H0 | CTL4_HUMAN | Unmodified | 0 | 0 | 0 | 0 | 0 | 0 | 0 | 0 | 0 | 0 | 0 | 0 | 0 | 0 | 713470 | 595720 | 0 | 0 |
| DEDDEAYG | SLC44A4; SLC4 | Q53GD3; H0 | CTL4_HUMAN | Unmodified | 0 | 0 | 0 | 0 | 0 | 0 | 0 | 0 | 0 | 0 | 0 | 0 | 0 | 0 | 771420 | 0 | 0 | 0 |
| DEDEEDEE | CALR; CALR | P27797; K7E | CALR_HUMAN | Unmodified | 0 | 0 | 0 | 0 | 0 | 0 | 36007 | 84693 | 6880500 | 943410 | 0 | 0 | 0 | 0 | 0 | 0 | 0 | 0 |
| DEDKDKV( | ATP1B1 | P05026 | AT1B1_HUMAN | Unmodified | 0 | 0 | 0 | 0 | 1021200 | 0 | 0 | 0 | 2097100 | 1097300 | 187090 | 0 | 0 | 0 | 0 | 0 | 0 | 0 |
| DEDLFREW | BANF1 | O75531 | BAF_HUMAN B | Unmodified | 699720 | 2799800 | 697000 | 0 | 0 | 0 | 0 | 0 | 0 | 0 | 0 | 11625000 | 540650 | 0 | 1088500 | 680040 | 0 | 0 |
| DEEEEEVAI | CCER2 | I3L3R5 | CCER2_HUMAI | Unmodified | 0 | 0 | 0 | 0 | 10754 | 0 | 0 | 151380 | 0 | 0 | 0 | 0 | 0 | 0 | 0 | 0 | 0 | 0 |
| DEEKMELQ | TPM3; TPM3; TF | P06753; J3K | TPM3_HUMAN | Unmodified | 0 | 0 | 0 | 0 | 526860 | 0 | 0 | 0 | 0 | 0 | 0 | 0 | 0 | 0 | 0 | 47788 | 270950 | 110170 |
| DEGNYLDI | ANXA4; ANXA4 | P09525; Q6F | ANXA4_HUMAN | Unmodified | 0 | 0 | 0 | 0 | 0 | 0 | 0 | 0 | 1101400 | 0 | 558150 | 593250 | 0 | 0 | 0 | 0 | 0 | 0 |
| DEGSYSLEI | SDC1 | P18827 | SDC1_HUMAN | Unmodified | 0 | 0 | 0 | 0 | 1180300 | 138600 | 0 | 0 | 0 | 0 | 633590 | 0 | 0 | 0 | 0 | 0 | 674680 | 904460 |
| DEITFVSGA | ITGA6; ITGA6 | P23229; C9J | ITA6_HUMAN I | Unmodified | 0 | 0 | 399510 | 0 | 0 | 0 | 0 | 0 | 0 | 0 | 0 | 0 | 0 | 0 | 275110 | 0 | 0 | 0 |
| DEKPVEAIF | DNAJC3; DNAJ( | Q13217; X6F | DNJC3_HUMAN | Unmodified | 0 | 0 | 0 | 0 | 0 | 0 | 0 | 0 | 1701600 | 0 | 0 | 0 | 0 | 0 | 0 | 0 | 0 | 0 |
| DELEVIHLII | RO60 | P10155 | RO60_HUMAN | Unmodified | 1073000 | 1488400 | 632140 | 0 | 0 | 0 | 0 | 0 | 0 | 0 | 0 | 0 | 0 | 0 | 0 | 449330 | 0 | 0 |
| DELHIVEAE | NPM1; NPM1; N | P06748; A0A | NPM_HUMAN N | Oxidation (M) | 0 | 0 | 0 | 0 | 0 | 0 | 0 | 455000 | 1054000 | 0 | 79695 | 0 | 0 | 0 | 274930 | 0 | 370970 | 0 |
| DENLALYVI | TFRC; TFRC | P02786; G3\ | TFR1_HUMAN | Unmodified | 0 | 28104 | 263580 | 228370 | 0 | 0 | 0 | 0 | 991740 | 59019 | 0 | 27865 | 2866900 | 21291 | 689390 | 0 | 0 | 100420 |
| DENQSINH | ANXA7 | P20073 | ANXA7_HUMAN | Unmodified | 0 | 0 | 0 | 0 | 0 | 0 | 309950 | 102610 | 21272000 | 832140 | 406370 | 181990 | 0 | 0 | 0 | 0 | 0 | 0 |
| DENQSINH | ANXA7 | P20073 | ANXA7_HUMAN | Oxidation (M) | 0 | 0 | 0 | 0 | 17275 | 0 | 0 | 0 | 318180 | 0 | 0 | 0 | 17132 | 0 | 0 | 0 | 0 | 0 |
| DENSVELTI | GANAB; GANAE | Q14697; F5H | GANAB_HUMA | Unmodified | 0 | 0 | 0 | 0 | 0 | 0 | 0 | 0 | 2186200 | 0 | 0 | 0 | 0 | 0 | 0 | 0 | 0 | 0 |
| DENSVELTI | GANAB; GANAE | Q14697; F5H | GANAB_HUMA | Oxidation (M) | 0 | 0 | 0 | 0 | 0 | 0 | 0 | 0 | 679820 | 0 | 0 | 0 | 0 | 0 | 0 | 0 | 0 | 0 |
| DEPGAWEI | GANAB; GANAE | Q14697; F5H | GANAB_HUMA | Unmodified | 0 | 0 | 213580 | 0 | 0 | 0 | 0 | 0 | 8500800 | 328140 | 0 | 0 | 0 | 0 | 0 | 175210 | 0 | 0 |
| DEPGEQVE | HYOU1; HYOU1 | Q9Y4L1; E9F | HYOU1_HUMA | Unmodified | 0 | 0 | 0 | 0 | 0 | 0 | 0 | 0 | 0 | 0 | 685110 | 0 | 0 | 0 | 0 | 0 | 0 | 0 |
| DEPSVAAM | ACLY | P53396 | ACLY_HUMAN | Oxidation (M) | 0 | 0 | 0 | 0 | 0 | 0 | 0 | 0 | 0 | 0 | 0 | 1269400 | 0 | 0 | 0 | 0 | 0 | 0 |
| DESANQEE | VASP; VASP | P50552; K7E | VASP_HUMAN | Unmodified | 0 | 0 | 107200 | 0 | 0 | 0 | 0 | 0 | 0 | 0 | 0 | 674680 | 0 | 0 | 148570 | 0 | 0 | 0 |
| DESKEPIVE | NCAM1; NCAM | P13591; A0A | NCAM1_HUMA | Unmodified | 0 | 0 | 0 | 0 | 0 | 0 | 0 | 0 | 496110 | 411480 | 0 | 664980 | 446860 | 0 | 0 | 0 | 0 | 0 |
| DESTNVDN | ANXA11 | P50995 | ANX11_HUMAN | Unmodified | 0 | 0 | 0 | 0 | 0 | 0 | 0 | 0 | 1208000 | 111230 | 0 | 0 | 0 | 0 | 0 | 0 | 0 | 0 |
| DESVTAGT | GPC6; GPC6 | Q9Y625; A0 | GPC6_HUMAN | Unmodified | 558410 | 214810 | 448600 | 323930 | 0 | 0 | 0 | 0 | 155830 | 0 | 0 | 0 | 82620 | 0 | 0 | 155730 | 0 | 0 |
| DETGELSS | UPF1 | Q92900 | RENT1_HUMAN | Unmodified | 0 | 293470 | 0 | 123750 | 0 | 0 | 0 | 0 | 0 | 0 | 0 | 305610 | 0 | 0 | 0 | 329860 | 0 | 0 |
| DETNYGIP( | RACK1; RACK1; | P63244; D6F | RACK1_HUMAI | Unmodified | 0 | 0 | 0 | 0 | 0 | 0 | 0 | 0 | 0 | 0 | 0 | 4460100 | 0 | 0 | 0 | 0 | 0 | 0 |
| DETVSDCS | CAPZB; CAPZB; | P47756; B1A | CAPZB_HUMAN | Unmodified | 1059600 | 0 | 1825700 | 0 | 0 | 0 | 0 | 0 | 290630 | 272120 | 0 | 4836500 | 0 | 0 | 1745600 | 780310 | 0 | 0 |
| DEVAHTLTI | AKT1 | P31749 | AKT1_HUMAN | Unmodified | 0 | 0 | 414440 | 100970 | 0 | 0 | 0 | 0 | 0 | 0 | 0 | 739700 | 0 | 0 | 481010 | 0 | 0 | 0 |
| DEVAHTVTI | AKT2; AKT2 | P31751; M0F | AKT2_HUMAN | Unmodified | 0 | 0 | 0 | 0 | 0 | 0 | 0 | 0 | 0 | 0 | 0 | 0 | 0 | 0 | 0 | 183850 | 0 | 0 |
| DFAAEVVH | YARS1; YARS1 | P54577; A0A | SYYC_HUMAN | Unmodified | 294690 | 0 | 628020 | 901900 | 1625600 | 493990 | 398550 | 367610 | 2677100 | 1450600 | 871240 | 5190900 | 3530400 | 2564700 | 877580 | 789180 | 645930 | 703100 |
| DFAAEVVH | YARS1; YARS1 | P54577; A0A | SYYC_HUMAN | Unmodified | 0 | 0 | 0 | 0 | 695310 | 0 | 832060 | 0 | 0 | 522750 | 0 | 0 | 1054000 | 1322400 | 0 | 0 | 889810 | 267420 |
| DFADWYLE | IQGAP3; IQGAP | Q86VI3; F2Z | IQGA3_HUMAN | Unmodified | 568680 | 0 | 0 | 0 | 0 | 0 | 0 | 0 | 1085700 | 267550 | 0 | 387810 | 0 | 0 | 0 | 0 | 0 | 0 |
| DFAERGEA | SPON2; SPON2 | Q9BUD6; D6 | SPON2_HUMAI | Unmodified | 0 | 0 | 557290 | 228940 | 0 | 0 | 0 | 0 | 0 | 0 | 0 | 0 | 0 | 0 | 760160 | 265570 | 0 | 0 |
| DFAERGEA | SPON2; SPON2 | Q9BUD6; D6 | SPON2_HUMAI | Oxidation (M) | 72571 | 0 | 2195300 | 0 | 0 | 0 | 0 | 0 | 0 | 0 | 0 | 0 | 0 | 0 | 2516300 | 1719200 | 0 | 0 |
| DFAGIQVG | SARG | Q9BW04 | SARG_HUMAN | Unmodified | 0 | 0 | 0 | 0 | 0 | 0 | 0 | 0 | 0 | 0 | 370280 | 0 | 775970 | 1701500 | 0 | 0 | 424720 | 0 |
| DFAGIQVG | SARG | Q9BW04 | SARG_HUMAN | Unmodified | 0 | 0 | 0 | 0 | 0 | 0 | 0 | 0 | 0 | 0 | 0 | 0 | 2505000 | 2989500 | 0 | 0 | 0 | 0 |
| DFAGPYFV | TMEM222; TMEI | Q9H0R3; Q5 | TM222_HUMAN | Oxidation (M) | 0 | 160420 | 0 | 0 | 0 | 0 | 0 | 0 | 0 | 0 | 0 | 952780 | 0 | 0 | 0 | 2314300 | 0 | 0 |
| DFAIDIIK | PPA1 | Q15181 | IPYR_HUMAN I | Unmodified | 559610 | 546070 | 1564700 | 1087900 | 5556700 | 528620 | 230560 | 115960 | 885000 | 299100 | 8763100 | 30925000 | 3427800 | 1295400 | 2237700 | 1961200 | 1228500 | 2730600 |
| DFAPVIVEA | RPRD1A; RPRD | Q96P16; A0 | RPR1A_HUMAN | Unmodified | 0 | 825970 | 0 | 0 | 0 | 611180 | 4155000 | 0 | 72638 | 443430 | 214240 | 1290800 | 0 | 0 | 0 | 152050 | 1038000 | 0 |
| DFASEVSN | STAM | Q92783 | STAM1_HUMAN | Unmodified | 259160 | 254580 | 119020 | 52288 | 0 | 0 | 0 | 0 | 360340 | 0 | 0 | 0 | 0 | 0 | 62674 | 98695 | 0 | 0 |
| DFASFPSIN | POGLUT3 | Q7Z4H8 | PLGT3_HUMAN | Unmodified | 0 | 0 | 0 | 0 | 0 | 0 | 0 | 0 | 1488900 | 579720 | 0 | 0 | 20453 | 0 | 0 | 0 | 0 | 29121 |
| DFASFPSIN | POGLUT3 | Q7Z4H8 | PLGT3_HUMAN | Oxidation (M) | 0 | 0 | 0 | 0 | 0 | 0 | 0 | 0 | 90600 | 408600 | 0 | 28079 | 0 | 0 | 0 | 0 | 0 | 0 |
| DFASRGIEI | DLD; DLD | P09622; E9P | DLDH_HUMAN | Unmodified | 0 | 0 | 0 | 0 | 0 | 0 | 647980 | 0 | 0 | 364700 | 0 | 0 | 0 | 0 | 0 | 0 | 0 | 331090 |
| DFAVLEDH | CCDC50 | Q8IVM0 | CCD50_HUMA | Unmodified | 11096000 | 10607000 | 4272900 | 5151800 | 0 | 0 | 0 | 0 | 0 | 0 | 0 | 0 | 0 | 0 | 28180000 | 24930000 | 0 | 288160 |
| DFAYGETD | TTLL12 | Q14166 | TTL12_HUMAN | Unmodified | 0 | 0 | 314750 | 0 | 0 | 0 | 0 | 84300 | 0 | 0 | 0 | 5839000 | 0 | 0 | 428820 | 1081500 | 0 | 0 |
| DFDGLVQV | DARS2; DARS2; | Q6PI48; A0A | SYDM_HUMAN | Unmodified | 0 | 0 | 0 | 0 | 1707400 | 308410 | 0 | 0 | 1712800 | 0 | 327930 | 0 | 0 | 0 | 0 | 0 | 0 | 0 |
| DFDPAVTE | FBP1; FBP1; FB | P09467; Q5\ | F16P1_HUMAN | Unmodified | 0 | 0 | 0 | 0 | 412110 | 0 | 165780 | 21313 | 1579600 | 723570 | 188780 | 655320 | 260270 | 479340 | 0 | 0 | 356420 | 0 |
| DFDPVHGI | LYN | P07948 | LYN_HUMAN T | Unmodified | 612360 | 553050 | 567850 | 529890 | 0 | 0 | 68689 | 0 | 2092300 | 246890 | 288280 | 1112400 | 0 | 0 | 1277500 | 1283900 | 0 | 0 |
| DFDTALK | STIP1 | P31948 | STIP1_HUMAN | Unmodified | 0 | 0 | 29300 | 0 | 47778 | 0 | 0 | 0 | 300360 | 0 | 0 | 18068000 | 0 | 38643 | 0 | 0 | 7660400 | 5571800 |
| DFDTALKH | STIP1 | P31948 | STIP1_HUMAN | Unmodified | 0 | 0 | 0 | 0 | 0 | 0 | 0 | 136790 | 0 | 0 | 0 | 0 | 0 | 0 | 0 | 0 | 996870 | 0 |
| DFEDFRK | RACGAP1; RAC | Q9H0H5; F8 | RGAP1_HUMAI | Unmodified | 0 | 0 | 0 | 220370 | 3891600 | 0 | 119650 | 0 | 707640 | 594810 | 0 | 0 | 232840 | 0 | 0 | 0 | 422370 | 260540 |
| DFEDLAGK | THUMPD3 | Q9BV44 | THUM3_HUMA | Unmodified | 285260 | 422410 | 166650 | 0 | 0 | 0 | 0 | 0 | 0 | 0 | 99803 | 525810 | 0 | 0 | 746200 | 453330 | 0 | 0 |
| DFEEYPEH | XPO1; XPO1; XF | O14980; A0 | XPO1_HUMAN | Unmodified | 0 | 0 | 0 | 0 | 0 | 0 | 0 | 0 | 155690 | 76331 | 102170 | 2585500 | 0 | 0 | 0 | 28680 | 0 | 0 |
| DFEGCSVL | PTPN23; PTPN2 | Q9H3S7; C9 | PTN23_HUMAN | Unmodified | 501300 | 394620 | 0 | 0 | 0 | 0 | 0 | 20794 | 148790 | 43332 | 0 | 460440 | 0 | 0 | 0 | 0 | 0 | 0 |
| DFENSMN( | ECI2; ECI2; ECI | O75521; F8V | ECI2_HUMAN E | Unmodified | 0 | 0 | 0 | 0 | 0 | 0 | 0 | 0 | 1293200 | 330930 | 0 | 332070 | 0 | 0 | 0 | 0 | 0 | 339820 |
| DFEPILEQ( | ELP1; ELP1; ELF | O95163; A0 | ELP1_HUMAN I | Unmodified | 0 | 0 | 0 | 0 | 0 | 0 | 262590 | 69067 | 0 | 0 | 0 | 2604900 | 0 | 0 | 0 | 0 | 0 | 0 |
| DFEQPLAIS | PDLIM1 | O00151 | PDLI1_HUMAN | Unmodified | 0 | 0 | 0 | 0 | 0 | 1095600 | 619120 | 526900 | 0 | 0 | 0 | 0 | 0 | 0 | 0 | 0 | 635110 | 1202300 |
| DFESCLGA | ACO1 | P21399 | ACOC_HUMAN | Unmodified | 0 | 0 | 0 | 0 | 0 | 0 | 184620 | 263450 | 0 | 53938 | 0 | 846050 | 390450 | 335370 | 0 | 0 | 0 | 0 |
| DFESHITSY | EIF3CL; EIF3C; | B5ME19; Q9 | EIFCL_HUMAN | Unmodified | 602390 | 166660 | 0 | 0 | 0 | 0 | 0 | 0 | 0 | 0 | 0 | 1352200 | 0 | 0 | 213530 | 357550 | 0 | 0 |
| DFETIGQN | MYOF | Q9NZM1 | MYOF_HUMAN | Unmodified | 0 | 0 | 0 | 0 | 5619400 | 1612200 | 0 | 0 | 86220 | 224230 | 0 | 1284700 | 0 | 0 | 0 | 0 | 0 | 1054800 |
| DFETIGQN | MYOF | Q9NZM1 | MYOF_HUMAN | Unmodified | 0 | 0 | 0 | 0 | 2839100 | 0 | 0 | 0 | 0 | 0 | 0 | 0 | 0 | 0 | 0 | 0 | 0 | 0 |
| DFFLANASI | CAPN1 | P07384 | CAN1_HUMAN | Unmodified | 0 | 157350 | 0 | 195990 | 255580 | 0 | 0 | 0 | 702450 | 228480 | 674500 | 3961600 | 253380 | 0 | 442190 | 325520 | 0 | 118820 |
| DFFQSYGN | G3BP1; G3BP1; | Q13283; A0 | G3BP1_HUMAI | Unmodified | 0 | 0 | 0 | 0 | 0 | 0 | 0 | 0 | 283760 | 0 | 238890 | 2832400 | 0 | 0 | 0 | 0 | 0 | 0 |
| DFFRLPDP | SMURF1; SMUF | Q9HCE7; Q9 | SMUF1_HUMAI | Unmodified | 0 | 0 | 0 | 379790 | 1092100 | 172270 | 0 | 0 | 299600 | 173010 | 120010 | 538600 | 485630 | 0 | 0 | 0 | 276540 | 560130 |
| DFGDLSFT | ARG2 | P78540 | ARGI2_HUMAN | Unmodified | 0 | 0 | 0 | 0 | 0 | 0 | 0 | 0 | 1366800 | 0 | 0 | 0 | 0 | 0 | 0 | 0 | 0 | 0 |
| DFGHVQF\ | EDIL3 | O43854 | EDIL3_HUMAN | Unmodified | 2210800 | 1512900 | 0 | 219510 | 5.97E+08 | 1.45E+08 | 10583000 | 4850400 | 34738000 | 9231100 | 0 | 0 | 0 | 241900 | 0 | 0 | 14707000 | 1.95E+08 |
| DFGHVQF\ | EDIL3 | O43854 | EDIL3_HUMAN | Unmodified | 0 | 0 | 0 | 0 | 67152000 | 0 | 3696100 | 0 | 0 | 0 | 0 | 0 | 0 | 0 | 0 | 0 | 3486800 | 41922000 |
| DFGNYLFN | SYAP1 | Q96A49 | SYAP1_HUMAN | Unmodified | 0 | 0 | 0 | 65763 | 387380 | 27862 | 338080 | 0 | 0 | 14681 | 0 | 0 | 869360 | 646330 | 0 | 0 | 514160 | 1006200 |
| DFGNYLFN | SYAP1 | Q96A49 | SYAP1_HUMAN | Unmodified | 0 | 0 | 0 | 0 | 0 | 0 | 0 | 0 | 0 | 0 | 0 | 0 | 0 | 0 | 0 | 0 | 0 | 0 |
| DFGSFDK | SOD2; SOD2; S | P04179; G8J | SODM_HUMAN | Unmodified | 0 | 0 | 0 | 0 | 0 | 0 | 0 | 0 | 5277500 | 106990 | 131710 | 0 | 0 | 0 | 0 | 0 | 0 | 0 |
| DFHHHHFI | CPD | O75976 | CBPD_HUMAN | Unmodified | 0 | 0 | 0 | 0 | 0 | 0 | 0 | 0 | 4925500 | 991280 | 0 | 0 | 0 | 0 | 0 | 0 | 0 | 0 |
| DFHHHHFI | CPD | O75976 | CBPD_HUMAN | Unmodified | 0 | 0 | 0 | 0 | 0 | 0 | 0 | 0 | 1723800 | 912370 | 0 | 0 | 0 | 0 | 0 | 0 | 0 | 0 |
| DFHYIVFG | ITGA6; ITGA6 | P23229; C9J | ITA6_HUMAN I | Unmodified | 0 | 0 | 0 | 7158400 | 46115000 | 9247100 | 8506.6 | 0 | 0 | 0 | 1490100 | 6141200 | 3579500 | 14121000 | 8245700 | 9284600 | 47915000 | 1.45E+08 |
| DFHYIVFG | ITGA6; ITGA6 | P23229; C9J | ITA6_HUMAN I | Unmodified | 0 | 0 | 0 | 0 | 275970 | 0 | 0 | 0 | 0 | 0 | 0 | 0 | 0 | 0 | 0 | 0 | 1052300 | 656890 |
| DFIATQGPI | PTPRJ; PTPRJ; P | Q12913; A0 | PTPRJ_HUMAN | Unmodified | 0 | 0 | 0 | 0 | 337080 | 0 | 89221 | 9700 | 0 | 61617 | 738570 | 1063900 | 0 | 0 | 0 | 0 | 0 | 0 |
| DFIATQGPI | PTPRJ; PTPRJ; P | Q12913; A0 | PTPRJ_HUMAN | Unmodified | 0 | 0 | 0 | 0 | 231560 | 0 | 23040 | 0 | 281270 | 0 | 0 | 0 | 85711 | 59965 | 0 | 0 | 74473 | 85318 |
| DFIDLTLSK | GTPBP4 | Q9BZE4 | GTPB4_HUMAN | Unmodified | 8911 | 11515 | 51711 | 0 | 0 | 0 | 0 | 66428 | 0 | 0 | 0 | 0 | 0 | 0 | 315230 | 209450 | 0 | 0 |
| DFIDNLDRI | RABL6; RABL6; | Q3YEC7; F2 | RABL6_HUMAN | Unmodified | 0 | 0 | 0 | 0 | 0 | 0 | 0 | 16328 | 0 | 0 | 0 | 991790 | 0 | 0 | 0 | 0 | 0 | 0 |
| DFIDSLGLI | EIF3M; EIF3M | Q7L2H7; H0 | EIF3M_HUMAN | Oxidation (M) | 0 | 0 | 591130 | 548180 | 0 | 0 | 0 | 47211 | 0 | 0 | 550840 | 4546400 | 0 | 0 | 970280 | 168660 | 0 | 66258 |
| DFIDSVLVK | TOM1L2; TOM1 | Q6ZVM7; K7 | TM1L2_HUMAN | Unmodified | 412800 | 0 | 171670 | 0 | 0 | 0 | 0 | 0 | 0 | 0 | 0 | 0 | 0 | 0 | 741600 | 460710 | 0 | 0 |
| DFIIFEAAP( | PPP4C; PPP4C | P60510; H3E | PP4C_HUMAN | Unmodified | 0 | 0 | 0 | 0 | 0 | 0 | 0 | 0 | 0 | 0 | 209540 | 1812300 | 0 | 0 | 0 | 0 | 0 | 0 |

| | | | | | | | | | | | | | | | | | | | | | | |
|---|---|---|---|---|---|---|---|---|---|---|---|---|---|---|---|---|---|---|---|---|---|---|
| DFILPISDVI | MTHFD1L; MTH | Q6UB35; A0. | C1TM_HUMAN | Unmodified | 44937 | 0 | 119850 | 0 | 769240 | 73658 | 0 | 0 | 1022800 | 0 | 0 | 0 | 0 | 0 | 214560 | 127520 | 0 | 0 |
| DFINFISDK | TMX1 | Q9H3N1 | TMX1_HUMAN | Unmodified | 0 | 0 | 0 | 0 | 0 | 0 | 0 | 0 | 38949000 | 1275200 | 1310700 | 0 | 0 | 0 | 0 | 0 | 0 | 0 |
| DFINFISDK | TMX1 | Q9H3N1 | TMX1_HUMAN | Unmodified | 0 | 0 | 0 | 0 | 0 | 0 | 0 | 0 | 7087900 | 4486800 | 0 | 0 | 0 | 0 | 0 | 0 | 0 | 0 |
| DFISLGLQI | HSPG2 | P98160 | PGBM_HUMAN | Unmodified | 0 | 0 | 575570 | 386890 | 0 | 0 | 0 | 0 | 0 | 0 | 0 | 0 | 0 | 0 | 0 | 292970 | 0 | 0 |
| DFIWGHR( | POFUT2; POFU | Q9Y2G5; A0. | OFUT2_HUMAI | Unmodified | 0 | 0 | 0 | 0 | 0 | 0 | 0 | 0 | 1086900 | 0 | 0 | 0 | 0 | 353700 | 0 | 0 | 0 | 0 |
| DFKVQFVP | ABHD12; ABHD | Q8N2K0; A0. | ABD12_HUMAI | Unmodified | 0 | 0 | 0 | 46026 | 0 | 0 | 0 | 0 | 6389400 | 1491200 | 0 | 0 | 0 | 0 | 0 | 0 | 0 | 0 |
| DFLAGGIA/ | SLC25A6 | P12236 | ADT3_HUMAN | Unmodified | 0 | 0 | 0 | 0 | 0 | 0 | 624180 | 145480 | 17439000 | 3304700 | 840810 | 486120 | 0 | 0 | 0 | 0 | 0 | 0 |
| DFLAGGVA | SLC25A5 | P05141 | ADT2_HUMAN | Unmodified | 99907 | 0 | 0 | 0 | 0 | 0 | 7009400 | 3025000 | 32523000 | 11889000 | 2372300 | 881280 | 679410 | 1116600 | 0 | 0 | 0 | 328710 |
| DFLAGGVA | SLC25A4; SLC2 | P12235; V9C | ADT1_HUMAN | Unmodified | 230310 | 0 | 398930 | 0 | 3212900 | 153740 | 887320 | 128790 | 4873600 | 756340 | 768020 | 4115200 | 2504900 | 855660 | 729410 | 594070 | 454740 | 715490 |
| DFLDEHYD | SHCBP1 | Q8NEM2 | SHCBP_HUMA | Unmodified | 0 | 0 | 0 | 190480 | 1621500 | 600310 | 0 | 0 | 0 | 0 | 336600 | 584650 | 0 | 0 | 241930 | 0 | 0 | 0 |
| DFLDGVYA | FLNB; FLNB | O75369; E7E | FLNB_HUMAN | Unmodified | 0 | 0 | 0 | 0 | 0 | 0 | 0 | 0 | 0 | 0 | 0 | 134180 | 0 | 0 | 0 | 0 | 0 | 0 |
| DFLEAVNK | PSMC2; PSMC2 | P35998; C9J | PRS7_HUMAN | Unmodified | 245460 | 154450 | 323290 | 643520 | 0 | 129680 | 441120 | 260360 | 1431900 | 241450 | 269190 | 7211000 | 866620 | 546580 | 896070 | 640410 | 207730 | 0 |
| DFLGKSDP | CPNE1; CPNE1 | Q99829; Q5. | CPNE1_HUMAI | Unmodified | 0 | 0 | 0 | 22616 | 124000 | 68912 | 0 | 0 | 190450 | 59681 | 0 | 0 | 188180 | 0 | 0 | 0 | 0 | 339660 |
| DFLGKSDP | CPNE1; CPNE1 | Q99829; Q5. | CPNE1_HUMAI | Unmodified | 0 | 0 | 0 | 0 | 254170 | 74947 | 45598 | 0 | 0 | 160700 | 0 | 0 | 121850 | 77308 | 0 | 0 | 81637 | 482350 |
| DFLIPIGK | PDHB; PDHB; P | P11177; F8V | ODPB_HUMAN | Unmodified | 0 | 0 | 0 | 0 | 0 | 17889000 | 364920 | 0 | 3962700 | 982470 | 0 | 187820 | 423550 | 0 | 0 | 0 | 0 | 0 |
| DFLLKPELL | DDX39B; DDX3 | Q13838; O0 | DX39B_HUMAI | Unmodified | 0 | 0 | 0 | 0 | 0 | 0 | 0 | 0 | 0 | 0 | 262890 | 5394800 | 0 | 0 | 0 | 0 | 0 | 0 |
| DFLLNYPF | PRSS23 | O95084 | PRS23_HUMAI | Unmodified | 193770 | 213590 | 0 | 0 | 1723300 | 352590 | 0 | 0 | 0 | 0 | 71335 | 18926 | 0 | 0 | 0 | 0 | 25046 | 367620 |
| DFLLQQTM | GLO1 | Q04760 | LGUL_HUMAN | Unmodified | 0 | 0 | 665490 | 0 | 0 | 0 | 0 | 0 | 3736900 | 0 | 342400 | 47099 | 0 | 0 | 0 | 0 | 0 | 0 |
| DFLLQQTM | GLO1 | Q04760 | LGUL_HUMAN | Oxidation (M) | 91596 | 241720 | 149750 | 228760 | 54003 | 0 | 0 | 0 | 124230 | 0 | 1343000 | 5328900 | 202360 | 0 | 208090 | 1005300 | 0 | 20860 |
| DFLLSLK | KTN1 | Q86UP2 | KTN1_HUMAN | Unmodified | 0 | 0 | 0 | 0 | 0 | 0 | 0 | 36406 | 221900 | 82574 | 250050 | 1407300 | 0 | 0 | 0 | 0 | 0 | 0 |
| DFLLTAR | RPL38; RPL38; I | P63173; J3K | RL38_HUMAN ( | Unmodified | 0 | 0 | 966510 | 759280 | 568610 | 0 | 0 | 39945 | 380500 | 0 | 728360 | 10338000 | 853950 | 163220 | 1048600 | 1390000 | 0 | 0 |
| DFLLYLR | B3GNT2 | Q9NY97 | B3GN2_HUMAI | Unmodified | 0 | 0 | 0 | 0 | 0 | 0 | 0 | 0 | 1050600 | 0 | 0 | 0 | 0 | 0 | 0 | 0 | 0 | 0 |
| DFLMLAQT | OSBP | P22059 | OSBP1_HUMAI | Oxidation (M) | 0 | 0 | 0 | 0 | 0 | 0 | 0 | 0 | 0 | 0 | 0 | 1676200 | 0 | 0 | 0 | 23508 | 0 | 0 |
| DFLPLYFG\ | CNP; CNP | P09543; K7E | CN37_HUMAN | Unmodified | 0 | 0 | 0 | 45231 | 286840 | 0 | 0 | 0 | 0 | 0 | 0 | 0 | 0 | 194820 | 37683 | 0 | 78768 | 38297 |
| DFLPVDPA | PTPRF; PTPRF | P10586; A2A | PTPRF_HUMAI | Unmodified | 0 | 0 | 0 | 0 | 0 | 0 | 0 | 0 | 0 | 0 | 0 | 0 | 0 | 0 | 0 | 0 | 0 | 0 |
| DFLQEVCL | BAZ2A; BAZ2A; | Q9UIF9; H0Y | BAZ2A_HUMAI | Unmodified | 111330 | 0 | 0 | 0 | 0 | 0 | 0 | 0 | 0 | 0 | 0 | 0 | 0 | 0 | 0 | 0 | 0 | 0 |
| DFLTPPLLS | AP2A1 | O95782 | AP2A1_HUMAI | Unmodified | 0 | 0 | 81384 | 0 | 0 | 0 | 0 | 0 | 0 | 0 | 104610 | 1073400 | 0 | 0 | 0 | 0 | 0 | 0 |
| DFLVGQDF | MESD | Q14696 | MESD_HUMAN | Unmodified | 1926300 | 1213000 | 966080 | 824500 | 0 | 0 | 0 | 0 | 5400500 | 0 | 0 | 322880 | 0 | 0 | 432750 | 2084300 | 0 | 337280 |
| DFLVIPSPV | PI4KA | P42356 | PI4KA_HUMAN | Unmodified | 0 | 0 | 0 | 0 | 0 | 0 | 452600 | 0 | 0 | 0 | 0 | 416600 | 0 | 0 | 612910 | 270310 | 188710 | 0 |
| DFLVQIK | XPO1; XPO1; XF | O14980; A0/ | XPO1_HUMAN | Unmodified | 0 | 0 | 0 | 172910 | 389910 | 0 | 944430 | 0 | 1075300 | 277710 | 0 | 1626700 | 86255 | 700390 | 26086 | 0 | 158920 | 506460 |
| DFLVSISDF | SUCO; SUCO | Q9UBS9; B4 | SUCO_HUMAN | Unmodified | 0 | 0 | 0 | 0 | 0 | 0 | 0 | 0 | 503530 | 0 | 0 | 378720 | 0 | 0 | 0 | 0 | 0 | 0 |
| DFMIQGGI | PPIB; PPIB; PPII | P23284; A0A | PPIB_HUMAN F | Oxidation (M) | 0 | 0 | 0 | 0 | 0 | 0 | 0 | 0 | 1522100 | 0 | 386230 | 578020 | 0 | 0 | 0 | 0 | 0 | 0 |
| DFMIQGGI | PPIB; PPIB; PPII | P23284; A0A | PPIB_HUMAN F | Oxidation (M) | 1283400 | 231420 | 3201400 | 1341400 | 0 | 0 | 752220 | 649420 | 79217000 | 12546000 | 17343000 | 13742000 | 515910 | 1021300 | 3100000 | 2475800 | 0 | 121790 |
| DFMIQGGI | PPIB; PPIB; PPII | P23284; A0A | PPIB_HUMAN F | Unmodified | 169800 | 0 | 321600 | 490470 | 404140 | 0 | 0 | 89043 | 2.04E+08 | 8651500 | 8172500 | 4890500 | 0 | 0 | 1741000 | 586770 | 0 | 0 |
| DFMIQGGI | PPIL1 | Q9Y3C6 | PPIL1_HUMAN | Unmodified | 241360 | 0 | 0 | 0 | 0 | 0 | 0 | 0 | 0 | 0 | 0 | 565570 | 0 | 0 | 0 | 0 | 0 | 0 |
| DFMIQGGI | PPIL1 | Q9Y3C6 | PPIL1_HUMAN | Oxidation (M) | 114190 | 0 | 0 | 38790 | 0 | 0 | 0 | 0 | 0 | 0 | 0 | 0 | 0 | 0 | 108550 | 130860 | 0 | 0 |
| DFMIQGGI | PPIL1 | Q9Y3C6 | PPIL1_HUMAN | Oxidation (M) | 179360 | 34230 | 116050 | 208950 | 1150000 | 0 | 0 | 0 | 0 | 77070 | 211750 | 0 | 814790 | 0 | 115290 | 0 | 0 | 0 |
| DFMIQGGI | PPIL1 | Q9Y3C6 | PPIL1_HUMAN | Unmodified | 0 | 0 | 0 | 52803 | 505360 | 0 | 0 | 0 | 206490 | 81142 | 0 | 0 | 274240 | 0 | 0 | 0 | 0 | 0 |
| DFMLHSSS | IGSF3 | O75054 | IGSF3_HUMAN | Unmodified | 199170 | 303280 | 0 | 51315 | 0 | 0 | 0 | 0 | 168960 | 13689 | 0 | 2191000 | 0 | 0 | 327600 | 137530 | 0 | 0 |
| DFNDIRQD | MTA3; MTA2; M | Q9BTC8; O9 | MTA3_HUMAN | Unmodified | 160990 | 69003 | 0 | 258420 | 121320 | 30404 | 73688 | 67320 | 242150 | 219250 | 359760 | 1156700 | 61549 | 0 | 472410 | 0 | 33509 | 63151 |
| DFNHINVE | RPL9; RPL9; RP | P32969; A0A | RL9_HUMAN 6 | Unmodified | 294250 | 718800 | 183430 | 224250 | 0 | 0 | 0 | 0 | 0 | 0 | 0 | 2348500 | 0 | 0 | 426230 | 98119 | 0 | 0 |
| DFNIPGFP | QSOX1; QSOX1 | O00391; A8I | QSOX1_HUMAI | Unmodified | 0 | 0 | 24728 | 0 | 0 | 0 | 0 | 0 | 2757700 | 0 | 508890 | 0 | 0 | 0 | 0 | 0 | 0 | 0 |
| DFNLDGAF | UGGT1 | Q9NYU2 | UGGG1_HUMA | Unmodified | 0 | 0 | 0 | 0 | 0 | 0 | 0 | 0 | 727660 | 0 | 0 | 0 | 0 | 0 | 0 | 0 | 0 | 0 |
| DFNLDGAF | UGGT1 | Q9NYU2 | UGGG1_HUMA | Unmodified | 0 | 0 | 0 | 0 | 0 | 0 | 0 | 11527 | 1612900 | 0 | 0 | 0 | 0 | 0 | 0 | 0 | 0 | 0 |
| DFNLLRWL | GUSB | P08236 | BGLR_HUMAN | Unmodified | 0 | 0 | 0 | 0 | 0 | 0 | 0 | 0 | 34556 | 578760 | 0 | 0 | 0 | 194270 | 0 | 0 | 0 | 0 |
| DFNLPEYD | ERBIN | Q96RT1 | ERBIN_HUMAN | Unmodified | 131490 | 139870 | 64092 | 59678 | 0 | 0 | 0 | 0 | 1456300 | 0 | 166780 | 0 | 0 | 0 | 0 | 0 | 0 | 0 |
| DFNMPLTIS | PDLIM5 | Q96HC4 | PDLI5_HUMAN | Oxidation (M) | 0 | 0 | 0 | 0 | 972770 | 0 | 0 | 0 | 0 | 554480 | 0 | 824430 | 0 | 0 | 0 | 0 | 509710 | 0 |
| DFNQFIK | SMARCA1; SMA | P28370; O6 | SMCA1_HUMA | Unmodified | 619540 | 109840 | 78503 | 378720 | 844590 | 302330 | 0 | 80184 | 575080 | 447190 | 110600 | 3109000 | 829140 | 419660 | 156410 | 455660 | 617700 | 500450 |
| DFNRAFGF | PCBD1 | P61457 | PHS_HUMAN P | Unmodified | 0 | 0 | 0 | 0 | 1505100 | 265070 | 220980 | 0 | 951710 | 1555400 | 159810 | 0 | 2365100 | 489130 | 0 | 0 | 1455000 | 1693300 |
| DFPETNNII | GMPS | P49915 | GUAA_HUMAN | Unmodified | 0 | 0 | 0 | 0 | 0 | 0 | 0 | 226330 | 0 | 533370 | 0 | 1799200 | 911710 | 0 | 0 | 0 | 0 | 0 |
| DFPEYTFAI | PDIA4; PDIA4 | P13667; A0A | PDIA4_HUMAN | Unmodified | 0 | 0 | 0 | 0 | 334620 | 0 | 0 | 0 | 3113000 | 0 | 0 | 0 | 0 | 0 | 224290 | 0 | 0 | 0 |
| DFPLNDLL | DYNC1H1; DYN | Q14204; A0/ | DYHC1_HUMA | Unmodified | 0 | 0 | 18517 | 0 | 0 | 0 | 0 | 0 | 13117 | 23648 | 81222 | 374900 | 55585 | 0 | 50191 | 0 | 0 | 0 |
| DFPQYFAV | ANK3; ANK2; AI | Q12955; Q0 | ANK3_HUMAN | Unmodified | 0 | 0 | 0 | 2509800 | 1619300 | 0 | 0 | 0 | 0 | 0 | 0 | 1807800 | 6734900 | 0 | 3887300 | 2561000 | 226210 | 0 |
| DFQALNPI | GSTZ1; GSTZ1; | O43708; G3 | MAAI_HUMAN I | Unmodified | 0 | 0 | 0 | 0 | 0 | 0 | 0 | 0 | 1290800 | 129840 | 0 | 0 | 0 | 0 | 357330 | 0 | 0 | 0 |
| DFQDCPLC | GPR108; GPR1 | Q9NPR9; M0 | GP108_HUMAI | Unmodified | 0 | 70750 | 0 | 0 | 0 | 0 | 0 | 0 | 3193400 | 102830 | 75001 | 316460 | 0 | 0 | 202130 | 144960 | 0 | 0 |
| DFQEYVEP | DYNC1LI1; DYN | Q9Y6G9; E9I | DC1L1_HUMAI | Unmodified | 0 | 0 | 0 | 0 | 0 | 0 | 0 | 0 | 0 | 0 | 0 | 2762300 | 0 | 0 | 0 | 0 | 0 | 0 |
| DFQGTPTF | STXBP2; STXBP | Q15833; M0 | STXB2_HUMAI | Unmodified | 437000 | 0 | 346460 | 1018400 | 3479000 | 0 | 0 | 0 | 610260 | 316170 | 433510 | 2019200 | 1978600 | 1869300 | 633470 | 486610 | 1679400 | 1877500 |
| DFQPVLHL | COL18A1; COL | P39060; H7 | COIA1_HUMAI | Oxidation (M) | 0 | 0 | 0 | 0 | 0 | 0 | 293940 | 0 | 963630 | 0 | 0 | 0 | 0 | 0 | 0 | 230120 | 121900 | 117920 |
| DFQPVLHL | COL18A1; COL | P39060; H7 | COIA1_HUMAI | Unmodified | 0 | 0 | 0 | 0 | 0 | 0 | 0 | 0 | 1202000 | 19090 | 0 | 0 | 0 | 0 | 0 | 0 | 36643 | 0 |
| DFQPYFKG | PRXL2A | Q9BRX8 | PXL2A_HUMAI | Unmodified | 0 | 0 | 0 | 0 | 0 | 0 | 0 | 0 | 4653600 | 562300 | 0 | 119320 | 0 | 0 | 0 | 0 | 0 | 69961 |
| DFQPYFKG | PRXL2A | Q9BRX8 | PXL2A_HUMAI | Unmodified | 0 | 0 | 0 | 0 | 0 | 0 | 0 | 48413 | 5209400 | 227910 | 0 | 0 | 0 | 0 | 0 | 0 | 0 | 0 |
| DFQSGQH | NSF; NSF; NSF; | P46459; I3L | NSF_HUMAN V | Unmodified | 580350 | 208610 | 723000 | 876810 | 108870 | 0 | 55040 | 41977 | 1857700 | 725320 | 808320 | 1706700 | 2595200 | 1065000 | 1291000 | 1158300 | 307820 | 971810 |
| DFQYIDRD | EPN2; EPN2; EF | O95208; F6F | EPN2_HUMAN | Unmodified | 709530 | 536290 | 0 | 433390 | 0 | 0 | 0 | 0 | 386080 | 0 | 0 | 0 | 0 | 0 | 1302200 | 638660 | 71547 | 0 |
| DFQYIDRD | EPN3; EPN3; EF | Q9H201; D6 | EPN3_HUMAN | Unmodified | 317640 | 289360 | 0 | 130510 | 0 | 0 | 0 | 0 | 0 | 0 | 0 | 87335 | 0 | 0 | 208550 | 327270 | 0 | 0 |
| DFQYVDRI | EPN1 | Q9Y6I3 | EPN1_HUMAN | Unmodified | 4883000 | 4323300 | 963440 | 492750 | 0 | 0 | 0 | 79360 | 368730 | 0 | 138990 | 483500 | 0 | 0 | 0 | 2123800 | 539090 | 964220 |
| DFRFMATN | CAND1 | Q86VP6 | CAND1_HUMA | Oxidation (M) | 1026000 | 0 | 0 | 0 | 181700 | 0 | 0 | 0 | 0 | 0 | 0 | 0 | 0 | 0 | 0 | 0 | 0 | 135200 |
| DFSAFKDA | CNNM4 | Q6P4Q7 | CNNM4_HUMA | Unmodified | 0 | 0 | 0 | 68336 | 0 | 0 | 0 | 0 | 88768 | 34787 | 92388 | 1583200 | 86237 | 0 | 0 | 50641 | 0 | 0 |
| DFSALESQ | MYH9 | P35579 | MYH9_HUMAN | Unmodified | 554940 | 1133800 | 986340 | 1841200 | 51200000 | 4588900 | 0 | 131850 | 0 | 5174700 | 0 | 1028500 | 11724000 | 14059000 | 5603600 | 992810 | 16015000 | 9431800 |
| DFSALESQ | MYH9 | P35579 | MYH9_HUMAN | Unmodified | 0 | 0 | 0 | 0 | 847810 | 0 | 11554000 | 0 | 0 | 124330 | 0 | 0 | 3705400 | 4219200 | 0 | 0 | 43591000 | 4299600 |
| DFSALLQSI | KHNYN | O15037 | KHNYN_HUMA | Unmodified | 0 | 0 | 0 | 0 | 0 | 0 | 0 | 0 | 0 | 0 | 0 | 0 | 0 | 0 | 0 | 0 | 0 | 0 |
| DFSDKINR | PDCD6IP | Q8WUM4 | PDC6I_HUMAI | Unmodified | 1595100 | 1869400 | 1302000 | 774190 | 1862700 | 63514 | 0 | 0 | 0 | 293750 | 0 | 436880 | 460590 | 188980 | 2656000 | 2243600 | 0 | 0 |
| DFSESRNN | HNRNPL; HNRI | P14866; M0 | HNRPL_HUMAI | Unmodified | 48261 | 0 | 0 | 0 | 1629200 | 370200 | 1313500 | 0 | 255650 | 976220 | 1152600 | 0 | 4283000 | 0 | 0 | 0 | 245650 | 882990 |
| DFSGTSDP | SYT7 | O43581 | SYT7_HUMAN | Unmodified | 288730 | 226470 | 388870 | 312590 | 411400 | 0 | 508530 | 260390 | 3909600 | 671630 | 0 | 3617200 | 0 | 0 | 348470 | 307040 | 0 | 0 |
| DFSHPQMI | CCT5; CCT5; C | P48643; B7Z | TCPE_HUMAN | Unmodified | 0 | 0 | 0 | 0 | 0 | 0 | 0 | 0 | 0 | 112930 | 946890 | 2506000 | 485970 | 0 | 0 | 0 | 0 | 223930 |
| DFSPSGIF( | ATP5MF; ATP5I | P56134; G3 | ATPK_HUMAN | Unmodified | 59773 | 0 | 19642 | 0 | 0 | 0 | 0 | 0 | 3206800 | 816630 | 0 | 184840 | 0 | 0 | 0 | 0 | 0 | 0 |
| DFSPSSAK | VGF; VGF | O15240; Q0 | VGF_HUMAN N | Unmodified | 455370 | 394490 | 0 | 0 | 0 | 0 | 0 | 0 | 4869200 | 149930 | 0 | 177350 | 0 | 0 | 91755 | 0 | 0 | 0 |
| DFSRGLNV | CEMIP2; CEMIP | Q9UHN6; F5 | CEIP2_HUMAN | Unmodified | 0 | 519300 | 0 | 0 | 9229000 | 1283700 | 799570 | 0 | 0 | 1382800 | 286310 | 0 | 616260 | 455960 | 0 | 378970 | 0 | 0 |
| DFSRLEPL\ | IKBIP | Q70UQ0 | IKIP_HUMAN In | Unmodified | 0 | 0 | 0 | 0 | 0 | 0 | 0 | 0 | 4138800 | 777300 | 130640 | 0 | 0 | 0 | 0 | 0 | 0 | 0 |
| DFSRYNYL | MYO1B; MYO1E | O43795; E9I | MYO1B_HUMA | Unmodified | 338660 | 313580 | 97416 | 122290 | 1517300 | 0 | 0 | 0 | 879470 | 0 | 2230600 | 9920700 | 306550 | 0 | 521650 | 187540 | 241920 | 0 |
| DFSTYRPLI | MYORG | Q6NSJ0 | MYORG_HUMA | Unmodified | 38602 | 41427 | 0 | 0 | 0 | 0 | 0 | 0 | 454510 | 0 | 0 | 320520 | 0 | 0 | 0 | 0 | 0 | 0 |
| DFTATDLSI | ACAA2; ACAA2 | P42765; A0A | THIM_HUMAN | Unmodified | 0 | 3244800 | 0 | 0 | 0 | 0 | 80163 | 25020 | 10297000 | 504790 | 0 | 0 | 0 | 0 | 0 | 0 | 0 | 0 |
| DFTLNDFG | HMGCS1 | Q01581 | HMCS1_HUMA | Oxidation (M) | 0 | 0 | 0 | 0 | 0 | 0 | 0 | 0 | 64027 | 0 | 56392 | 448130 | 0 | 0 | 0 | 0 | 0 | 0 |
| DFTPVCTTI | PRDX6 | P30041 | PRDX6_HUMAI | Unmodified | 882080 | 0 | 1283500 | 1724500 | 317270 | 0 | 0 | 0 | 4575600 | 0 | 3227700 | 10704000 | 0 | 281940 | 3353100 | 0 | 632620 | 955310 |
| DFTPVCTTI | PRDX6 | P30041 | PRDX6_HUMAI | Unmodified | 21722 | 0 | 0 | 0 | 0 | 0 | 0 | 0 | 0 | 230350 | 0 | 0 | 1996200 | 0 | 0 | 0 | 1269800 | 1870500 |
| DFTQLFPR | MAPK13 | O15264 | MK13_HUMAN | Unmodified | 0 | 0 | 0 | 0 | 0 | 0 | 0 | 397470 | 654050 | 264240 | 0 | 534670 | 0 | 0 | 0 | 0 | 0 | 0 |
| DFTSLENT\ | CKAP4 | Q07065 | CKAP4_HUMAI | Unmodified | 0 | 0 | 92631 | 0 | 0 | 0 | 0 | 0 | 0 | 206370 | 0 | 0 | 0 | 0 | 193320 | 0 | 0 | 0 |
| DFTVSALH( | EIF4A2; EIF4A2 | Q14240; E7E | IF4A2_HUMAN | Unmodified | 30409 | 0 | 0 | 0 | 0 | 0 | 0 | 0 | 307330 | 0 | 244990 | 1096600 | 0 | 0 | 64985 | 46387 | 0 | 0 |
| DFTVSALH( | EIF4A2; EIF4A2 | Q14240; E7E | IF4A2_HUMAN | Oxidation (M) | 57850 | 0 | 103720 | 0 | 0 | 0 | 0 | 0 | 0 | 0 | 0 | 0 | 0 | 0 | 62163 | 87099 | 0 | 0 |
| DFTVSAMH | EIF4A1; EIF4A1; | P60842; J3Q | IF4A1_HUMAN | Oxidation (M) | 679220 | 514050 | 1151300 | 697830 | 0 | 0 | 0 | 28450 | 394990 | 0 | 694550 | 5417400 | 166020 | 0 | 1160900 | 1134700 | 0 | 77935 |
| DFTVSAMH | EIF4A1; EIF4A1; | P60842; J3Q | IF4A1_HUMAN | Unmodified | 274130 | 354520 | 0 | 488350 | 294570 | 0 | 0 | 0 | 1563400 | 57363 | 1618400 | 8062400 | 137500 | 94356 | 869900 | 0 | 51101 | 85575 |
| DFTVSAMH | EIF4A1; EIF4A1; | P60842; J3Q | IF4A1_HUMAN | 2 Oxidation (M) | 486550 | 0 | 0 | 0 | 0 | 0 | 0 | 0 | 0 | 0 | 126200 | 1438400 | 0 | 0 | 871520 | 0 | 0 | 0 |
| DFVAEPMG | BANF1 | O75531 | BAF_HUMAN B | Oxidation (M) | 0 | 0 | 0 | 0 | 0 | 0 | 0 | 0 | 0 | 0 | 1558500 | 6268700 | 0 | 0 | 0 | 0 | 0 | 0 |
| DFVNLHSF | VAV1; VAV1; VA | P15498; Q9 | VAV_HUMAN P | Unmodified | 0 | 0 | 0 | 0 | 0 | 0 | 0 | 0 | 0 | 0 | 0 | 672870 | 0 | 0 | 0 | 0 | 0 | 0 |
| DFVNYLVR | DHX9 | Q08211 | DHX9_HUMAN | Unmodified | 0 | 192270 | 0 | 0 | 0 | 0 | 0 | 0 | 0 | 0 | 0 | 10243000 | 0 | 0 | 0 | 110440 | 0 | 0 |
| DFVSALPD | ABCC1; ABCC1 | P33527; I3L | MRP1_HUMAN | Unmodified | 0 | 0 | 0 | 0 | 0 | 0 | 0 | 0 | 0 | 0 | 357340 | 3064800 | 0 | 0 | 0 | 351710 | 0 | 0 |
| DFVTTAEDI | PLAA; PLAA; PL | Q9Y263; H0 | PLAP_HUMAN I | Unmodified | 0 | 490110 | 411510 | 246050 | 0 | 0 | 202790 | 283210 | 185820 | 118500 | 0 | 1674800 | 0 | 269790 | 411990 | 28007 | 206220 | 290400 |
| DFVTVLAK | QSOX2; QSOX2 | Q6ZRP7; H0 | QSOX2_HUMAI | Unmodified | 30963 | 0 | 0 | 0 | 0 | 0 | 0 | 0 | 229670 | 535890 | 0 | 155290 | 0 | 0 | 0 | 0 | 0 | 0 |
| DFVWGLNI | PRKCD; PRKCD | Q05655; C9I | KPCD_HUMAN | Unmodified | 0 | 0 | 0 | 0 | 452240 | 0 | 0 | 0 | 320170 | 189310 | 0 | 1356100 | 0 | 0 | 0 | 0 | 374420 | 987170 |
| DFWNLHD | CASKIN2; CASK | Q8WXE0; J3 | CSKI2_HUMAN | Unmodified | 83943 | 0 | 0 | 409050 | 0 | 0 | 0 | 0 | 0 | 0 | 0 | 841560 | 0 | 0 | 655340 | 226910 | 0 | 0 |
| DFYELEPEI | PYGB | P11216 | PYGB_HUMAN | Unmodified | 0 | 0 | 0 | 0 | 0 | 0 | 0 | 0 | 690490 | 0 | 173820 | 0 | 245180 | 0 | 73671 | 0 | 558660 | 2260000 |
| DFYNPVVP | CLDN3 | O15551 | CLD3_HUMAN | Unmodified | 0 | 0 | 0 | 0 | 0 | 0 | 0 | 0 | 0 | 0 | 0 | 0 | 0 | 0 | 0 | 0 | 0 | 0 |
| DFYNPVVP | CLDN3 | O15551 | CLD3_HUMAN | Unmodified | 4173500 | 2023300 | 3749400 | 470610 | 0 | 0 | 0 | 0 | 0 | 434970 | 0 | 0 | 0 | 110240 | 7121500 | 5283100 | 0 | 0 |
| DFYQREFD | PI4KA; PI4KAP1 | P42356; Q8I | PI4KA_HUMAN | Unmodified | 0 | 0 | 581160 | 0 | 228620 | 0 | 0 | 0 | 0 | 0 | 0 | 0 | 0 | 0 | 126930 | 77167 | 0 | 0 |
| DGAASLSY | NUP210 | Q8TEM1 | PO210_HUMAI | Unmodified | 0 | 0 | 0 | 0 | 187820 | 106980 | 0 | 0 | 8599500 | 437070 | 0 | 347000 | 0 | 0 | 0 | 0 | 169290 | 0 |
| DGAASLSY | NUP210 | Q8TEM1 | PO210_HUMAI | Unmodified | 0 | 0 | 0 | 68162 | 0 | 0 | 0 | 0 | 364720 | 0 | 0 | 0 | 0 | 0 | 18933 | 0 | 0 | 0 |
| DGAASLSY | NUP210 | Q8TEM1 | PO210_HUMAI | Unmodified | 0 | 0 | 0 | 0 | 0 | 0 | 0 | 0 | 1299800 | 0 | 0 | 0 | 0 | 0 | 0 | 0 | 0 | 0 |
| DGAFAEFLI | ABCC1; ABCC1 | P33527; I3L | MRP1_HUMAN | Unmodified | 0 | 0 | 822280 | 0 | 0 | 0 | 0 | 0 | 0 | 0 | 962420 | 7024300 | 0 | 0 | 657860 | 1090300 | 0 | 0 |
| DGAGNSFI | IGF2R; IGF2R | P11717; A0A | MPRI_HUMAN ( | Unmodified | 0 | 0 | 0 | 0 | 0 | 0 | 0 | 0 | 43827000 | 1636700 | 0 | 5819600 | 0 | 0 | 1050200 | 654860 | 0 | 0 |
| DGALESQI | ABCC4; ABCC4 | O15439; A0/ | MRP4_HUMAN | Unmodified | 0 | 0 | 756790 | 0 | 0 | 0 | 0 | 0 | 0 | 0 | 0 | 0 | 0 | 0 | 0 | 509480 | 0 | 0 |

| | | | | | | | | | | | | | | | | | | | | | | |
|---|---|---|---|---|---|---|---|---|---|---|---|---|---|---|---|---|---|---|---|---|---|---|
| DGALESQD | ABCC4; ABCC4 | O15439; A0A | MRP4_HUMAN | Unmodified | 0 | 0 | 2009900 | 562160 | 0 | 0 | 0 | 0 | 0 | 0 | 0 | 507880 | 0 | 0 | 2296000 | 1606600 | 0 | 0 |
| DGALVGTA | CORO7; CORO | P57737; I3L1 | CORO7_HUMA | Unmodified | 1267700 | 4857400 | 4336600 | 3564400 | 0 | 0 | 2670000 | 1466400 | 374690 | 313830 | 872230 | 1111000 | 2431600 | 8009200 | 6587500 | 1305300 | 0 | 3640000 |
| DGANIVIAA | HSDL2 | Q6YN16 | HSDL2_HUMAI | Unmodified | 0 | 0 | 30783 | 0 | 498980 | 0 | 398500 | 148130 | 23361000 | 5304500 | 0 | 0 | 0 | 142150 | 24284 | 0 | 44683 | 0 |
| DGAPVATN | PIGS | Q96S52 | PIGS_HUMAN ( | Unmodified | 0 | 0 | 0 | 0 | 0 | 0 | 0 | 0 | 1999600 | 0 | 0 | 58083 | 0 | 0 | 0 | 0 | 0 | 0 |
| DGATLILSK | PLXNB2; PLXNE | O15031; A6( | PLXB2_HUMAN | Unmodified | 0 | 0 | 719140 | 0 | 0 | 0 | 0 | 0 | 2035300 | 369770 | 1190900 | 7889100 | 0 | 0 | 5934400 | 4088500 | 0 | 0 |
| DGAWGAF | FASN; FASN | P49327; A0A | FAS_HUMAN F | Unmodified | 0 | 0 | 925540 | 0 | 0 | 0 | 0 | 0 | 0 | 0 | 467510 | 34503000 | 0 | 0 | 938130 | 0 | 0 | 0 |
| DGDDAHE | ITGA6 | P23229 | ITA6_HUMAN I | Unmodified | 0 | 0 | 0 | 0 | 0 | 0 | 0 | 0 | 0 | 0 | 0 | 0 | 0 | 0 | 0 | 0 | 295530 | 4009700 |
| DGDFENP | ERP29; ERP29 | P30040; F8V | ERP29_HUMAN | Unmodified | 0 | 0 | 226880 | 0 | 0 | 32696 | 0 | 0 | 66058000 | 2445500 | 652250 | 2266400 | 0 | 0 | 268410 | 0 | 0 | 0 |
| DGDLVDW | MME; MME; MM | P08473; A0A | NEP_HUMAN N | Unmodified | 195920 | 272610 | 11812000 | 24739000 | 20453 | 0 | 0 | 0 | 23672000 | 644440 | 421110 | 559210 | 2436500 | 488860 | 77106000 | 29247000 | 30912 | 0 |
| DGDLYILA | HS2ST1 | Q7LGA3 | HS2ST_HUMAI | Unmodified | 0 | 0 | 0 | 0 | 0 | 0 | 0 | 0 | 214070 | 285310 | 0 | 0 | 0 | 0 | 0 | 0 | 0 | 0 |
| DGDRLWYI | PXDN | Q92626 | PXDN_HUMAN | Unmodified | 0 | 0 | 0 | 0 | 0 | 0 | 0 | 0 | 755360 | 464010 | 0 | 0 | 0 | 0 | 0 | 0 | 0 | 0 |
| DGDSYRSF | CAPZB; CAPZB; | P47756; A0A | CAPZB_HUMAI | Oxidation (M) | 0 | 521120 | 0 | 0 | 0 | 0 | 0 | 0 | 0 | 0 | 0 | 2549500 | 0 | 0 | 449610 | 647420 | 0 | 0 |
| DGDVIKTE | LACTB2 | Q53H82 | LACB2_HUMAI | Unmodified | 132410 | 94979 | 0 | 221750 | 0 | 0 | 0 | 0 | 25493000 | 437420 | 208710 | 735770 | 0 | 0 | 133320 | 202440 | 0 | 0 |
| DGDYWRLI | CPE | P16870 | CBPE_HUMAN | Unmodified | 0 | 0 | 0 | 0 | 0 | 0 | 0 | 0 | 216900 | 660510 | 0 | 0 | 0 | 0 | 0 | 0 | 0 | 47793 |
| DGEATLEVI | GAS6 | Q14393 | GAS6_HUMAN | Unmodified | 0 | 0 | 0 | 0 | 0 | 0 | 0 | 0 | 1279400 | 0 | 0 | 0 | 0 | 0 | 0 | 0 | 0 | 0 |
| DGEDQTQI | HLA-B; HLA-A; I | P01889; P04 | HLAB_HUMAN | Unmodified | 0 | 0 | 0 | 0 | 0 | 0 | 0 | 0 | 2767000 | 1970200 | 249100 | 459670 | 0 | 0 | 0 | 0 | 0 | 0 |
| DGEDQTQI | HLA-A; HLA-H; I | P04439; P01 | HLAA_HUMAN | Unmodified | 0 | 0 | 0 | 0 | 3665000 | 1247000 | 0 | 0 | 75449000 | 7154900 | 21417000 | 1.61E+08 | 0 | 0 | 329310 | 4412800 | 84624 | 0 |
| DGEDQTQI | HLA-B; HLA-B; I | P01889; D5I | HLAB_HUMAN | Unmodified | 571950 | 434840 | 0 | 0 | 0 | 0 | 0 | 0 | 4119700 | 1573200 | 2175500 | 78839000 | 0 | 0 | 0 | 0 | 0 | 0 |
| DGEDQTQI | HLA-B; HLA-B; I | P01889; D5I | HLAB_HUMAN | Unmodified | 820010 | 1668800 | 0 | 0 | 2919900 | 560900 | 0 | 0 | 30085000 | 343300 | 1832400 | 1.57E+08 | 0 | 0 | 288880 | 252360 | 82323 | 566510 |
| DGEEAGAY | PDIA3; PDIA3 | P30101; H7E | PDIA3_HUMAN | Unmodified | 0 | 0 | 0 | 0 | 0 | 0 | 0 | 0 | 4476200 | 146180 | 930850 | 751780 | 0 | 0 | 90166 | 111170 | 0 | 0 |
| DGEEYEWI | SLC4A7; SLC4A | Q9Y6M7; C9 | S4A7_HUMAN | Unmodified | 0 | 0 | 0 | 0 | 0 | 0 | 0 | 0 | 90755 | 0 | 0 | 515680 | 0 | 0 | 285150 | 372120 | 33348 | 0 |
| DGEIVGLS | TPP2; TPP2; TPI | P29144; A0A | TPP2_HUMAN | Unmodified | 0 | 0 | 0 | 0 | 0 | 0 | 0 | 0 | 395340 | 102780 | 0 | 6424800 | 0 | 0 | 0 | 76687 | 0 | 0 |
| DGEMDRN | ARVCF; ARVCF; | O00192; E9F | ARVC_HUMAN | Oxidation (M) | 165640 | 0 | 807710 | 555180 | 0 | 0 | 0 | 0 | 0 | 0 | 0 | 576240 | 0 | 0 | 952800 | 1066800 | 0 | 0 |
| DGENYVVL | NOMO1; NOMC | Q15155; P69 | NOMO1_HUMA | Unmodified | 0 | 0 | 0 | 0 | 0 | 0 | 0 | 0 | 1014400 | 0 | 0 | 80423 | 0 | 0 | 0 | 0 | 0 | 0 |
| DGEPEYNF | CAPN7; CAPN7 | Q9Y6W3; H7 | CAN7_HUMAN | Unmodified | 0 | 0 | 0 | 0 | 0 | 0 | 0 | 0 | 92228 | 0 | 0 | 280060 | 0 | 0 | 0 | 0 | 0 | 0 |
| DGEQIEQE | NCAM1; NCAM | P13591; A0A | NCAM1_HUMA | Unmodified | 0 | 0 | 0 | 0 | 0 | 0 | 0 | 0 | 551520 | 54339 | 0 | 0 | 0 | 0 | 0 | 0 | 0 | 0 |
| DGETPDPE | LTA4H | P09960 | LKHA4_HUMAI | Unmodified | 0 | 0 | 0 | 0 | 0 | 0 | 0 | 0 | 0 | 0 | 102440 | 289640 | 0 | 0 | 0 | 0 | 0 | 0 |
| DGETPLIK | KIDINS220; KID | Q9ULH0; E9 | KDIS_HUMAN I | Unmodified | 0 | 0 | 0 | 0 | 0 | 243080 | 15822 | 0 | 177940 | 0 | 0 | 229960 | 51217 | 0 | 49414 | 804630 | 0 | 0 |
| DGEVPPEE | FKBP10; FKBP1 | Q96AY3; H0 | FKB10_HUMAI | Unmodified | 0 | 0 | 0 | 0 | 4881400 | 0 | 0 | 0 | 0 | 797120 | 2914700 | 7244600 | 0 | 0 | 0 | 84501 | 0 | 0 |
| DGFDMFA | TOM1L2; TOM1 | Q6ZVM7; F5I | TM1L2_HUMAI | Oxidation (M) | 386770 | 395080 | 157990 | 167120 | 0 | 0 | 0 | 0 | 0 | 0 | 0 | 0 | 0 | 0 | 420430 | 294570 | 0 | 0 |
| DGFDMFA | TOM1L2; TOM1 | Q6ZVM7; F5I | TM1L2_HUMAI | Unmodified | 0 | 0 | 0 | 0 | 0 | 0 | 0 | 0 | 0 | 0 | 0 | 0 | 0 | 0 | 0 | 0 | 0 | 0 |
| DGFIDKED | MYL12A; MYL12 | P19105; O14 | ML12A_HUMAI | Unmodified | 0 | 0 | 0 | 0 | 0 | 0 | 0 | 0 | 0 | 0 | 37445 | 0 | 0 | 0 | 158060 | 115840 | 0 | 0 |
| DGFPSGTP | PAK2 | Q13177 | PAK2_HUMAN | Unmodified | 0 | 0 | 0 | 0 | 0 | 0 | 0 | 0 | 0 | 0 | 0 | 0 | 0 | 0 | 0 | 0 | 0 | 0 |
| DGFQPSR | TFRC; TFRC | P02786; G3V | TFR1_HUMAN | Unmodified | 776760 | 1086400 | 0 | 0 | 0 | 26166 | 16892 | 26045 | 2436300 | 367240 | 102990 | 1660100 | 0 | 0 | 1420300 | 1561600 | 0 | 0 |
| DGFRDFPA | GAA | P10253 | LYAG_HUMAN | Unmodified | 0 | 0 | 0 | 0 | 0 | 0 | 0 | 0 | 3693000 | 0 | 0 | 0 | 0 | 0 | 0 | 0 | 238320 | 0 |
| DGFRDFPA | GAA | P10253 | LYAG_HUMAN | Unmodified | 0 | 0 | 0 | 44255 | 1704800 | 0 | 0 | 0 | 8276100 | 407400 | 0 | 322490 | 0 | 0 | 0 | 0 | 0 | 72682 |
| DGFRDFPA | GAA | P10253 | LYAG_HUMAN | Oxidation (M) | 0 | 0 | 0 | 0 | 0 | 0 | 0 | 0 | 734100 | 27946 | 0 | 0 | 0 | 0 | 0 | 0 | 0 | 0 |
| DGFRFFTN | PNPO; PNPO; P | Q9NVS9; J3( | PNPO_HUMAN | Unmodified | 0 | 0 | 0 | 0 | 387590 | 0 | 0 | 0 | 0 | 0 | 137750 | 0 | 2443500 | 533620 | 0 | 0 | 257440 | 448290 |
| DGFVALSK | DDX1; DDX1; D | Q92499; A0A | DDX1_HUMAN | Unmodified | 446990 | 124970 | 0 | 646050 | 0 | 0 | 0 | 0 | 348320 | 81389 | 0 | 6467300 | 546540 | 169250 | 288890 | 226540 | 1377200 | 262980 |
| DGFVALSK | DDX1; DDX1; D | Q92499; A0A | DDX1_HUMAN | Unmodified | 0 | 0 | 0 | 0 | 492890 | 0 | 0 | 0 | 0 | 0 | 0 | 0 | 963370 | 0 | 0 | 0 | 3539400 | 397140 |
| DGFVTVDE | CALU | O43852 | CALU_HUMAN | Unmodified | 0 | 0 | 0 | 0 | 0 | 0 | 0 | 0 | 57411 | 61383 | 3040500 | 774690 | 0 | 0 | 428630 | 233110 | 0 | 0 |
| DGGAWGT | LGALS1 | P09382 | LEG1_HUMAN | Unmodified | 0 | 0 | 0 | 0 | 622990 | 237320 | 0 | 0 | 78092 | 0 | 158900 | 0 | 0 | 0 | 0 | 0 | 195690 | 627450 |
| DGGDKKEI | EPB41L2; EPB4 | O43491; E9F | E41L2_HUMAN | Unmodified | 0 | 0 | 0 | 0 | 3188900 | 0 | 0 | 0 | 0 | 0 | 0 | 0 | 0 | 0 | 0 | 0 | 0 | 398600 |
| DGGFCEV | PSAP; PSAP; PS | P07602; Q5E | SAP_HUMAN P | Unmodified | 67288 | 0 | 0 | 104420 | 500700 | 256910 | 279500 | 381340 | 6287900 | 5263500 | 38945 | 237380 | 916250 | 2146100 | 87505 | 53996 | 120580 | 525310 |
| DGGFCEV | PSAP; PSAP; PS | P07602; Q5E | SAP_HUMAN P | Unmodified | 0 | 0 | 0 | 0 | 0 | 0 | 0 | 0 | 1331700 | 557530 | 0 | 0 | 0 | 288780 | 0 | 0 | 0 | 159990 |
| DGGGTNSI | ABCC1; ABCC1 | P33527; I3L | MRP1_HUMAN | Unmodified | 0 | 0 | 815520 | 0 | 39216 | 0 | 0 | 0 | 130620 | 0 | 261330 | 5977000 | 0 | 0 | 621450 | 579530 | 0 | 0 |
| DGGGVRD | SLC12A2; SLC1 | P55011; G3 | S12A2_HUMAI | Unmodified | 0 | 2252300 | 1438400 | 743780 | 0 | 315520 | 0 | 0 | 238270 | 0 | 530630 | 1516000 | 0 | 0 | 2295100 | 2146700 | 311890 | 2387500 |
| DGGLICTS | CORO1A; COR( | P31146; H3E | COR1A_HUMA | Unmodified | 0 | 0 | 0 | 0 | 0 | 817380 | 0 | 129140 | 0 | 116320 | 0 | 1199900 | 0 | 0 | 0 | 103690 | 0 | 0 |
| DGGLICTS | CORO1A; COR( | P31146; H3E | COR1A_HUMA | Unmodified | 0 | 0 | 0 | 0 | 0 | 358530 | 0 | 0 | 0 | 0 | 88298 | 2158800 | 0 | 0 | 0 | 0 | 0 | 217800 |
| DGGNPFAI | SCAMP3 | O14828 | SCAM3_HUMA | Unmodified | 0 | 0 | 0 | 0 | 0 | 0 | 0 | 0 | 2610900 | 459560 | 345520 | 1013000 | 0 | 0 | 0 | 203310 | 0 | 0 |
| DGGNQEV | PRKAR2A; PRKA | P13861; H7( | KAP2_HUMAN | Unmodified | 338250 | 0 | 0 | 0 | 0 | 0 | 0 | 0 | 514950 | 153520 | 231110 | 1659800 | 0 | 0 | 1212200 | 739580 | 0 | 0 |
| DGGNQEV | PRKAR2A; PRKA | P13861; H7( | KAP2_HUMAN | Unmodified | 0 | 113510 | 0 | 0 | 0 | 0 | 0 | 0 | 0 | 0 | 0 | 0 | 267080 | 310110 | 0 | 0 | 0 | 0 |
| DGGPGPG | CELSR2 | Q9HCU4 | CELR2_HUMAI | Unmodified | 0 | 0 | 0 | 0 | 421060 | 0 | 0 | 0 | 0 | 0 | 0 | 0 | 335180 | 262370 | 0 | 0 | 0 | 0 |
| DGGPQPPI | PLEKHN1; PLEI | Q494U1; J3K | PKHN1_HUMA | Unmodified | 124040 | 258290 | 226830 | 178560 | 0 | 0 | 0 | 0 | 0 | 0 | 0 | 0 | 0 | 0 | 124420 | 151270 | 0 | 0 |
| DGGTDFPA | LRIG2; LRIG3; L | O94898; Q6I | LRIG2_HUMAN | Unmodified | 0 | 0 | 0 | 0 | 49129 | 0 | 0 | 0 | 633850 | 79087 | 32839 | 0 | 0 | 0 | 0 | 0 | 427620 | 0 |
| DGGVAIID | POFUT2 | Q9Y2G5 | OFUT2_HUMAI | Unmodified | 0 | 0 | 0 | 0 | 0 | 0 | 0 | 0 | 442650 | 0 | 0 | 0 | 0 | 0 | 0 | 0 | 0 | 0 |
| DGGVCLLS | IGF2R | P11717 | MPRI_HUMAN | Unmodified | 0 | 0 | 0 | 0 | 0 | 0 | 0 | 0 | 0 | 0 | 0 | 0 | 0 | 0 | 0 | 0 | 0 | 0 |
| DGGVQAC | GNAI3 | P08754 | GNAI3_HUMAI | Unmodified | 2027000 | 308810 | 1442400 | 1853800 | 222430 | 0 | 25373 | 32926 | 2940400 | 750920 | 1651900 | 10489000 | 139420 | 78924 | 2881900 | 2477700 | 384770 | 293590 |
| DGGVQWG | IGSF3 | O75054 | IGSF3_HUMAN | Unmodified | 176030 | 0 | 0 | 0 | 0 | 0 | 0 | 0 | 0 | 0 | 0 | 120420 | 0 | 0 | 437590 | 95029 | 89645 | 0 |
| DGHHHYE | GMPS | P49915 | GUAA_HUMAN | Unmodified | 0 | 0 | 0 | 0 | 820070 | 0 | 142610 | 0 | 0 | 0 | 0 | 0 | 0 | 0 | 0 | 0 | 230020 | 371360 |
| DGHSQLG | ARSD | P51689 | ARSD_HUMAN | Unmodified | 0 | 0 | 0 | 164580 | 0 | 0 | 0 | 0 | 469410 | 0 | 0 | 38443 | 0 | 0 | 0 | 0 | 0 | 0 |
| DGIDDESY | HDAC2 | Q92769 | HDAC2_HUMA | Unmodified | 0 | 0 | 0 | 0 | 0 | 0 | 0 | 0 | 0 | 0 | 0 | 2586500 | 0 | 0 | 0 | 0 | 0 | 0 |
| DGIEPGHI | MPST | P25325 | THTM_HUMAN | Unmodified | 0 | 0 | 104070 | 0 | 0 | 0 | 0 | 0 | 1942200 | 0 | 1028800 | 976790 | 0 | 0 | 809280 | 100750 | 0 | 0 |
| DGIEPMWI | EIF4E; EIF4E; EI | P06730; D6I | IF4E_HUMAN E | Oxidation (M) | 94463 | 0 | 0 | 0 | 0 | 0 | 0 | 0 | 0 | 0 | 182020 | 963830 | 0 | 0 | 0 | 0 | 0 | 0 |
| DGIILCELI | CNN3; CNN3 | Q15417; E9F | CNN3_HUMAN | Unmodified | 0 | 0 | 0 | 0 | 0 | 0 | 0 | 0 | 747220 | 23231 | 2962000 | 5094500 | 0 | 0 | 0 | 0 | 19779 | 0 |
| DGIILCELI | CNN3; CNN3 | Q15417; E9F | CNN3_HUMAN | Unmodified | 160280 | 0 | 0 | 0 | 460990 | 0 | 0 | 0 | 0 | 0 | 0 | 941730 | 0 | 0 | 0 | 0 | 73759 | 0 |
| DGISCEASI | BCAM; BCAM | P50895; A0A | BCAM_HUMAN | Unmodified | 1457400 | 0 | 0 | 0 | 276220 | 48450 | 0 | 0 | 0 | 0 | 0 | 0 | 0 | 0 | 0 | 0 | 55443 | 0 |
| DGISCEASI | BCAM; BCAM | P50895; A0A | BCAM_HUMAN | Unmodified | 1264900 | 804840 | 0 | 0 | 0 | 0 | 0 | 0 | 84043 | 0 | 0 | 26794 | 0 | 0 | 0 | 56664 | 0 | 0 |
| DGIVLGAD | PSMB7; PSMB7 | Q99436; Q5 | PSB7_HUMAN | Unmodified | 3.55E+08 | 2.87E+08 | 3.18E+08 | 7.32E+08 | 0 | 146510 | 4886300 | 0 | 2871000 | 0 | 987130 | 4486800 | 1.71E+08 | 56089000 | 4.76E+08 | 2.05E+08 | 355960 | 0 |
| DGIVMPTN | F11R; F11R | Q9Y624; A0A | JAM1_HUMAN | Unmodified | 1846800 | 1557800 | 2592600 | 2538000 | 5259100 | 961260 | 0 | 0 | 5126900 | 1063500 | 619380 | 3360300 | 5131500 | 2232800 | 2703500 | 1872600 | 0 | 0 |
| DGIVMPTN | F11R; F11R | Q9Y624; A0A | JAM1_HUMAN | Oxidation (M) | 0 | 1168800 | 1376400 | 1005200 | 2709100 | 408920 | 0 | 0 | 0 | 0 | 0 | 0 | 1740100 | 0 | 164330 | 1569600 | 0 | 487580 |
| DGIVPFLQ | SLC12A5; SLC1 | Q9H2X9; Q9 | S12A5_HUMAI | Unmodified | 374110 | 318380 | 341830 | 142250 | 0 | 0 | 0 | 0 | 666590 | 267210 | 0 | 3268000 | 0 | 0 | 2065500 | 1016700 | 0 | 0 |
| DGKALERF | PDIA3 | P30101 | PDIA3_HUMAN | Unmodified | 0 | 0 | 0 | 0 | 0 | 0 | 0 | 0 | 0 | 295100 | 0 | 0 | 23581 | 0 | 0 | 0 | 13872 | 33744 |
| DGKLDFEE | SLC25A24; SLC | Q6NUK1; J3I | SCMC1_HUMA | Unmodified | 0 | 0 | 0 | 0 | 0 | 0 | 0 | 0 | 531820 | 0 | 0 | 0 | 0 | 0 | 0 | 0 | 0 | 0 |
| DGKLHFGI | ASRGL1 | Q7L266 | ASGL1_HUMAI | Unmodified | 0 | 0 | 0 | 342170 | 0 | 12413000 | 1531300 | 115570 | 0 | 2451400 | 0 | 321500 | 0 | 0 | 0 | 133610 | 288870 | 0 |
| DGKLVVEC | FABP5; FABP5 | Q01469; I6L | FABP5_HUMAN | Unmodified | 0 | 0 | 0 | 0 | 844460 | 205720 | 0 | 0 | 0 | 0 | 79000 | 0 | 0 | 0 | 0 | 0 | 948180 | 2213200 |
| DGKVIIVDA | DIPK2A; DIPK2 | Q8NDZ4; E7 | DIK2A_HUMAN | Unmodified | 0 | 0 | 0 | 0 | 0 | 7304700 | 0 | 0 | 1122400 | 417120 | 0 | 0 | 0 | 0 | 0 | 0 | 16924 | 0 |
| DGKVIIVDA | DIPK2A; DIPK2 | Q8NDZ4; E7 | DIK2A_HUMAN | Unmodified | 0 | 0 | 0 | 0 | 0 | 0 | 0 | 0 | 4363100 | 684850 | 0 | 0 | 129540 | 0 | 0 | 0 | 0 | 0 |
| DGKYSQVL | RPS18; RPS18 | P62269; J3J | RS18_HUMAN | Unmodified | 0 | 430170 | 0 | 308840 | 0 | 0 | 0 | 0 | 0 | 0 | 0 | 2285500 | 127810 | 0 | 0 | 0 | 0 | 0 |
| DGLAFNAL | SPTBN4; ACTN | Q9H254; O4 | SPTN4_HUMAI | Unmodified | 0 | 430250 | 983270 | 2641600 | 2027600 | 1220600 | 0 | 0 | 4270100 | 0 | 1449500 | 19740000 | 0 | 0 | 1959000 | 1489900 | 216110 | 296090 |
| DGLEMEK | TPT1; TPT1; TPT | P13693; H0 | TCTP_HUMAN | Unmodified | 0 | 0 | 0 | 0 | 131620 | 62477 | 0 | 24112 | 0 | 0 | 521230 | 2370000 | 0 | 0 | 0 | 0 | 78396 | 110240 |
| DGLGFCAL | ACTN1; ACTN1 | P12814; G3 | ACTN1_HUMAI | Unmodified | 468910 | 0 | 0 | 1100700 | 1307900 | 0 | 0 | 0 | 0 | 0 | 1365600 | 5946700 | 0 | 0 | 0 | 652640 | 0 | 1103700 |
| DGLIIRKPV | RPL19; RPL19; I | P84098; J3K | RL19_HUMAN ( | Unmodified | 0 | 0 | 0 | 566340 | 0 | 0 | 0 | 0 | 152840 | 88667 | 0 | 2127000 | 1442100 | 0 | 0 | 0 | 0 | 213530 |
| DGLIVPIFQ | FUCA1 | P04066 | FUCO_HUMAN | Unmodified | 0 | 0 | 0 | 0 | 0 | 0 | 42120 | 0 | 0 | 260360 | 0 | 0 | 0 | 0 | 0 | 0 | 0 | 0 |
| DGLLENQT | FASN; FASN | P49327; A0A | FAS_HUMAN F | Unmodified | 0 | 0 | 0 | 0 | 0 | 0 | 45416 | 0 | 0 | 0 | 506220 | 22468000 | 0 | 0 | 0 | 706310 | 0 | 0 |
| DGLLPENT | G6PD; G6PD; G | P11413; E7E | G6PD_HUMAN | Unmodified | 0 | 0 | 0 | 0 | 0 | 0 | 0 | 0 | 287840 | 115530 | 0 | 492710 | 0 | 0 | 285080 | 32482 | 0 | 0 |
| DGLLYVYA | BMX | P51813 | BMX_HUMAN ( | Unmodified | 0 | 0 | 0 | 0 | 0 | 0 | 0 | 0 | 0 | 0 | 768880 | 0 | 215580 | 159890 | 726620 | 0 | 0 | 0 |
| DGLNDDD | YTHDF2; YTHD | Q9Y5A9; S4 | YTHD2_HUMAI | Unmodified | 724880 | 628370 | 355160 | 0 | 0 | 0 | 0 | 0 | 0 | 0 | 0 | 0 | 0 | 0 | 1781900 | 0 | 0 | 0 |
| DGLNQTTI | LARP4; LARP4 | Q71RC2; X6I | LARP4_HUMAN | Unmodified | 439940 | 324160 | 240360 | 425900 | 0 | 0 | 0 | 0 | 0 | 0 | 0 | 0 | 181100 | 0 | 123310 | 194270 | 0 | 0 |
| DGLPLGIG | SIGIRR; SIGIRR | Q6IA17; C9J | SIGIR_HUMAN | Unmodified | 0 | 0 | 146270 | 0 | 0 | 0 | 0 | 0 | 2617700 | 562590 | 0 | 0 | 0 | 0 | 921520 | 370880 | 0 | 0 |
| DGLTDVYN | ACAT1; ACAT1; | P24752; H0 | THIL_HUMAN A | Unmodified | 217880 | 0 | 270670 | 242780 | 321800 | 158070 | 0 | 160980 | 1804200 | 379110 | 0 | 0 | 611700 | 190630 | 294600 | 0 | 0 | 170360 |
| DGLVSLLT | GALK1; GALK1 | P51570; K7E | GALK1_HUMAI | Unmodified | 0 | 0 | 112250 | 119430 | 386220 | 174030 | 0 | 0 | 0 | 0 | 0 | 253860 | 412820 | 0 | 864260 | 201570 | 163710 | 257560 |
| DGMELVP | POGLUT3; POG | Q7Z4H8; H0 | PLGT3_HUMAI | Unmodified | 0 | 0 | 0 | 0 | 0 | 0 | 0 | 0 | 1470000 | 0 | 0 | 0 | 0 | 0 | 0 | 0 | 0 | 0 |
| DGMEYPFI | DNAJB11; N/A | Q9UBS4; H7 | DJB11_HUMAI | Oxidation (M) | 0 | 0 | 0 | 0 | 0 | 0 | 0 | 0 | 2566800 | 142390 | 1409600 | 653420 | 0 | 0 | 0 | 0 | 0 | 0 |
| DGMEYPFI | DNAJB11; N/A | Q9UBS4; H7 | DJB11_HUMAI | Unmodified | 0 | 0 | 0 | 0 | 0 | 0 | 0 | 0 | 2100200 | 0 | 1403500 | 0 | 0 | 0 | 0 | 0 | 0 | 0 |
| DGMSHLIQ | SLC12A7; SLC1 | Q9Y666; A0A | S12A7_HUMAI | Unmodified | 0 | 0 | 0 | 0 | 0 | 0 | 0 | 0 | 337780 | 0 | 0 | 1424700 | 0 | 0 | 322610 | 149200 | 0 | 0 |
| DGMSHLIQ | SLC12A7; SLC1 | Q9Y666; A0A | S12A7_HUMAI | Oxidation (M) | 0 | 0 | 109170 | 64463 | 0 | 0 | 0 | 0 | 0 | 0 | 30553 | 2192900 | 0 | 0 | 270840 | 646550 | 0 | 0 |
| DGNAIPLSI | PPM1L | Q5SGD2 | PPM1L_HUMAI | Unmodified | 0 | 0 | 0 | 0 | 601390 | 0 | 180170 | 0 | 7599100 | 0 | 0 | 0 | 0 | 0 | 0 | 0 | 0 | 0 |
| DGNASGT | EEF1A1P5; EEF | Q5VTE0; P68 | EF1A3_HUMAN | Unmodified | 33407000 | 8526100 | 2369100 | 9946800 | 0 | 1089700 | 869550 | 0 | 5367400 | 568430 | 2661400 | 12942000 | 2219600 | 998690 | 55315000 | 14959000 | 717920 | 2011300 |
| DGNASGT | EEF1A1P5; EEF | Q5VTE0; P68 | EF1A3_HUMAN | Unmodified | 12431000 | 10122000 | 0 | 1799100 | 1836200 | 694390 | 0 | 0 | 1839200 | 1026500 | 1459300 | 2208200 | 1584500 | 0 | 9194900 | 2774900 | 1696400 | 0 |
| DGNDLHM | DNAJA2 | O60884 | DNJA2_HUMAN | Unmodified | 0 | 0 | 0 | 0 | 0 | 0 | 0 | 0 | 369250 | 0 | 0 | 792820 | 0 | 0 | 0 | 0 | 0 | 344110 |
| DGNGYISA | CALM2; CALM3 | P0DP24; P0I | CALM2_HUMAI | Unmodified | 16052000 | 7960000 | 4561400 | 9912900 | 7554400 | 8961000 | 312360 | 325550 | 7553100 | 1548500 | 3988300 | 94447000 | 5374600 | 2267200 | 28629000 | 19072000 | 1753100 | 5550500 |
| DGNLASTL | TBL1XR1; TBL1 | Q9BZK7; O6 | TBL1R_HUMAI | Unmodified | 0 | 0 | 0 | 0 | 0 | 0 | 73699 | 0 | 220270 | 0 | 274090 | 835590 | 0 | 0 | 0 | 0 | 73670 | 0 |
| DGNLASTL | TBL1XR1; TBL1 | Q9BZK7; O6 | TBL1R_HUMAI | Unmodified | 0 | 0 | 0 | 0 | 754000 | 0 | 0 | 0 | 0 | 0 | 0 | 0 | 493630 | 0 | 0 | 0 | 754530 | 0 |
| DGNLHEGI | TJP2; TJP2; TJP2 | Q9UDY2; A0 | ZO2_HUMAN T | Unmodified | 0 | 0 | 0 | 0 | 2437500 | 906100 | 0 | 0 | 5961800 | 1435600 | 1768700 | 0 | 0 | 0 | 0 | 296130 | 0 | 0 |
| DGNLMETA | COBL; COBL; C | O75128; J3K | COBL_HUMAN | Unmodified | 0 | 163850 | 0 | 0 | 0 | 0 | 0 | 0 | 0 | 0 | 0 | 0 | 0 | 0 | 0 | 0 | 0 | 0 |
| DGNPDNE | DDX19B; DDX1 | Q9UMR2; Q9 | DD19B_HUMAI | Unmodified | 0 | 0 | 0 | 0 | 0 | 0 | 0 | 0 | 0 | 0 | 0 | 751870 | 0 | 0 | 0 | 0 | 0 | 0 |
| DGNRTIGT | TUFM | P49411 | EFTU_HUMAN | Unmodified | 0 | 0 | 0 | 0 | 793300 | 0 | 0 | 0 | 642850 | 508140 | 0 | 0 | 0 | 0 | 0 | 0 | 712740 | 0 |
| DGNVLLHE | CCT6A | P40227 | TCPZ_HUMAN | Unmodified | 254970 | 0 | 188760 | 2052700 | 13821000 | 0 | 5396300 | 76240 | 11533000 | 5142400 | 5639200 | 18511000 | 17417000 | 11775000 | 1519900 | 868660 | 6448300 | 14738000 |

| | | | | | | | | | | | | | | | | | | | | | | |
|---|---|---|---|---|---|---|---|---|---|---|---|---|---|---|---|---|---|---|---|---|---|---|
| DGNVLLHE | CCT6A | P40227 | TCPZ_HUMAN | Oxidation (M) | 138180 | 156820 | 244100 | 1055100 | 1008000 | 249410 | 403580 | 486350 | 0 | 1093700 | 3696500 | 11452000 | 4652600 | 3281300 | 604700 | 486670 | 1842900 | 3366200 |
| DGNVTCEF | FSCN1 | Q16658 | FSCN1_HUMAI | Unmodified | 0 | 0 | 0 | 0 | 0 | 0 | 0 | 0 | 107070 | 0 | 645720 | 1029600 | 0 | 0 | 0 | 0 | 0 | 0 |
| DGPGFYTT | HADHA; HADH/ | P40939; A0A | ECHA_HUMAN | Unmodified | 0 | 0 | 0 | 0 | 81668 | 1307400 | 0 | 0 | 1718400 | 262350 | 0 | 0 | 0 | 0 | 0 | 0 | 0 | 0 |
| DGPLGETV | UPF1 | Q92900 | RENT1_HUMAN | Unmodified | 381770 | 0 | 50836 | 106030 | 0 | 0 | 0 | 0 | 0 | 0 | 0 | 0 | 0 | 0 | 118260 | 86267 | 0 | 0 |
| DGPLNMIL | AHCY | P23526 | SAHH_HUMAN | Oxidation (M) | 0 | 0 | 194470 | 327550 | 0 | 0 | 0 | 0 | 106160 | 0 | 0 | 574880 | 0 | 0 | 1681400 | 152670 | 0 | 0 |
| DGQAMLW | RACK1; RACK1; | P63244; D6F | RACK1_HUMAI | Oxidation (M) | 0 | 0 | 1000300 | 0 | 0 | 0 | 0 | 0 | 0 | 0 | 544050 | 7505000 | 0 | 0 | 660740 | 564810 | 0 | 0 |
| DGQAMLW | RACK1; RACK1; | P63244; D6F | RACK1_HUMAI | Unmodified | 0 | 0 | 0 | 0 | 0 | 0 | 0 | 0 | 0 | 0 | 0 | 1171200 | 0 | 0 | 0 | 0 | 0 | 0 |
| DGQERDAI | FN1 | P02751 | FINC_HUMAN | Unmodified | 412180 | 0 | 1001700 | 0 | 180220 | 0 | 0 | 0 | 0 | 129680 | 208620 | 120980 | 0 | 0 | 0 | 268070 | 0 | 0 |
| DGQILPVPI | RCC2 | Q9P258 | RCC2_HUMAN | Unmodified | 0 | 0 | 0 | 0 | 0 | 0 | 0 | 265740 | 0 | 0 | 1269900 | 14090000 | 0 | 0 | 0 | 0 | 0 | 0 |
| DGQKITFH | DNAJA1 | P31689 | DNJA1_HUMAI | Unmodified | 0 | 0 | 0 | 0 | 6884000 | 423980 | 0 | 0 | 0 | 218060 | 0 | 0 | 0 | 0 | 0 | 0 | 1089700 | 0 |
| DGQLIYTPI | PSEN1; PSEN1; | P49768; G3\ | PSN1_HUMAN | Unmodified | 0 | 0 | 0 | 0 | 0 | 0 | 0 | 0 | 0 | 0 | 0 | 778940 | 0 | 0 | 0 | 0 | 0 | 0 |
| DGQLLIR | MOV10; MOV1C | Q9HCE1; Q5 | MOV10_HUMA | Unmodified | 3036100 | 5898500 | 405730 | 600370 | 0 | 0 | 0 | 0 | 0 | 0 | 0 | 630320 | 317540 | 120390 | 630640 | 2242400 | 0 | 0 |
| DGQVINET | VIM | P08670 | VIME_HUMAN | Unmodified | 0 | 0 | 0 | 0 | 427190 | 140370 | 101100 | 113380 | 0 | 33804 | 0 | 0 | 0 | 0 | 0 | 0 | 345340 | 1856900 |
| DGQYLLTG | LRBA; LRBA; LR | P50851; A0A | LRBA_HUMAN | Unmodified | 0 | 0 | 0 | 0 | 234120 | 0 | 0 | 0 | 0 | 0 | 0 | 1669400 | 0 | 0 | 0 | 0 | 0 | 0 |
| DGRGALQI | GAPDH; GAPDI | P04406; E7E | G3P_HUMAN G | Unmodified | 62470 | 0 | 968240 | 2457700 | 24874000 | 3259800 | 10152000 | 2756500 | 2704800 | 4047000 | 3190100 | 9300400 | 11079000 | 2783100 | 1332000 | 570100 | 31023000 | 17993000 |
| DGRGALQI | GAPDH; GAPDI | P04406; E7E | G3P_HUMAN G | Unmodified | 0 | 0 | 0 | 0 | 0 | 0 | 679330 | 198420 | 0 | 0 | 0 | 0 | 0 | 0 | 0 | 0 | 2232400 | 0 |
| DGRISVAG | GOT2 | P00505 | AATM_HUMAN | Unmodified | 2381400 | 0 | 0 | 0 | 0 | 0 | 0 | 0 | 0 | 5006300 | 0 | 866080 | 0 | 11496000 | 0 | 819240 | 91014 | 0 |
| DGRLEALLI | DIPK2A; DIPK2/ | Q8NDZ4; E7 | DIK2A_HUMAN | Unmodified | 0 | 39819 | 0 | 0 | 0 | 0 | 326100 | 114200 | 6583800 | 41846 | 0 | 0 | 0 | 79136 | 0 | 0 | 0 | 0 |
| DGRLEALLI | DIPK2A; DIPK2/ | Q8NDZ4; E7 | DIK2A_HUMAN | Unmodified | 0 | 0 | 0 | 0 | 0 | 0 | 0 | 0 | 3578000 | 318920 | 0 | 0 | 0 | 0 | 0 | 0 | 0 | 0 |
| DGRWTEL/ | LMAN2; LMAN2 | Q12907; D6I | LMAN2_HUMAI | Unmodified | 0 | 0 | 0 | 0 | 0 | 0 | 0 | 0 | 52117000 | 830540 | 970700 | 0 | 0 | 0 | 323440 | 0 | 0 | 0 |
| DGRYTDAT | DNAJC3; DNAJ( | Q13217; X6I | DNJC3_HUMAI | Unmodified | 0 | 0 | 0 | 0 | 0 | 0 | 0 | 0 | 4065000 | 0 | 0 | 0 | 0 | 0 | 0 | 0 | 0 | 0 |
| DGSCGVA | FLNA; FLNA; FL | P21333; A0A | FLNA_HUMAN | Unmodified | 0 | 0 | 449000 | 458110 | 0 | 0 | 0 | 0 | 0 | 0 | 0 | 343000 | 0 | 0 | 595940 | 0 | 0 | 0 |
| DGSDEDA/ | SORL1; SORL1; | Q92673; E9F | SORL_HUMAN | Unmodified | 0 | 0 | 0 | 0 | 0 | 0 | 0 | 0 | 952130 | 129840 | 0 | 0 | 0 | 0 | 0 | 0 | 0 | 0 |
| DGSDEPG1 | PRKCSH; PRKC | P14314; K7E | GLU2B_HUMAI | Unmodified | 0 | 0 | 0 | 0 | 0 | 0 | 0 | 0 | 9922700 | 0 | 0 | 522080 | 0 | 0 | 0 | 0 | 0 | 0 |
| DGSDVIYP/ | DNAJB1; DNAJE | P25685; M0( | DNJB1_HUMAI | Unmodified | 0 | 898110 | 57334 | 620080 | 0 | 0 | 0 | 0 | 1291900 | 0 | 314780 | 73903 | 0 | 0 | 0 | 155190 | 0 | 0 |
| DGSDYEG\ | GANAB; GANAE | Q14697; F5H | GANAB_HUMA | Oxidation (M) | 0 | 0 | 0 | 0 | 0 | 0 | 0 | 0 | 2246700 | 0 | 0 | 0 | 0 | 0 | 0 | 0 | 0 | 0 |
| DGSDYEG\ | GANAB; GANAE | Q14697; F5H | GANAB_HUMA | Unmodified | 0 | 0 | 0 | 0 | 0 | 0 | 0 | 0 | 407200 | 0 | 0 | 0 | 0 | 0 | 0 | 0 | 0 | 0 |
| DGSENLAC | WIPF2 | Q8TF74 | WIPF2_HUMAI | Unmodified | 2933800 | 2292700 | 2333300 | 2520500 | 595070 | 131670 | 0 | 0 | 0 | 0 | 0 | 375260 | 2138000 | 971800 | 5010200 | 2753000 | 0 | 437140 |
| DGSKEPIVI | NCAM2; NCAM: | O15394; H9I | NCAM2_HUMA | Oxidation (M) | 0 | 0 | 1001200 | 0 | 0 | 0 | 0 | 0 | 0 | 0 | 0 | 0 | 0 | 63017 | 1217500 | 905480 | 0 | 0 |
| DGSKEPIVI | NCAM2; NCAM: | O15394; H9I | NCAM2_HUMA | Unmodified | 0 | 0 | 620100 | 0 | 0 | 0 | 0 | 0 | 0 | 0 | 0 | 0 | 0 | 0 | 1753000 | 596780 | 0 | 0 |
| DGSNIVEA | RASEF | Q8IZ41 | RASEF_HUMAI | Unmodified | 54978 | 0 | 26690 | 54659 | 0 | 0 | 0 | 0 | 0 | 0 | 0 | 0 | 0 | 0 | 257420 | 68889 | 0 | 0 |
| DGSNKSG/ | TGOLN2; TGOL | O43493; A0/ | TGON2_HUMA | Unmodified | 122600 | 0 | 0 | 0 | 515010 | 0 | 144560 | 75390 | 1679200 | 1627600 | 65328 | 0 | 0 | 196260 | 0 | 0 | 21470 | 375500 |
| DGSPIADD | NLN; NLN; NLN | Q9BYT8; H0 | NEUL_HUMAN | Unmodified | 151840 | 0 | 0 | 0 | 0 | 0 | 0 | 0 | 588890 | 0 | 0 | 1353700 | 0 | 0 | 0 | 0 | 0 | 0 |
| DGSPILRDI | PLXNA2 | O75051 | PLXA2_HUMAI | Unmodified | 0 | 0 | 0 | 0 | 0 | 0 | 0 | 0 | 0 | 0 | 0 | 0 | 0 | 0 | 0 | 0 | 0 | 0 |
| DGSRADVI | MELTF | P08582 | TRFM_HUMAN | Unmodified | 0 | 0 | 0 | 0 | 0 | 155460 | 0 | 0 | 2920500 | 29147 | 156090 | 467460 | 0 | 0 | 0 | 0 | 0 | 0 |
| DGSSGGVI | PSMB9; PSMB9 | P28065; A0A | PSB9_HUMAN | Unmodified | 0 | 0 | 0 | 0 | 0 | 0 | 0 | 0 | 0 | 0 | 0 | 550630 | 0 | 0 | 0 | 0 | 0 | 0 |
| DGSTTAGN | ACAA1; ACAA1; | P09110; C9J | THIK_HUMAN | Unmodified | 0 | 0 | 0 | 0 | 953320 | 0 | 0 | 0 | 6386500 | 1565400 | 0 | 0 | 0 | 0 | 0 | 0 | 0 | 0 |
| DGSTTAGN | ACAA1; ACAA1; | P09110; C9J | THIK_HUMAN | Unmodified | 0 | 0 | 0 | 0 | 0 | 0 | 0 | 0 | 3703500 | 1425900 | 0 | 0 | 0 | 0 | 0 | 0 | 0 | 0 |
| DGSYAWEI | MESD | Q14696 | MESD_HUMAN | Unmodified | 20020 | 0 | 0 | 0 | 0 | 0 | 0 | 0 | 2344400 | 0 | 0 | 0 | 0 | 0 | 0 | 0 | 0 | 0 |
| DGTAGIPN | EIF2A; EIF2A; EI | Q9BY44; H7 | EIF2A_HUMAN | Unmodified | 89761 | 0 | 121250 | 52749 | 108540 | 0 | 26508 | 31782 | 116060 | 51606 | 177600 | 1216500 | 0 | 0 | 0 | 60480 | 271890 | 0 |
| DGTAGSHI | POFUT1 | Q9H488 | OFUT1_HUMAI | Oxidation (M) | 0 | 0 | 0 | 0 | 0 | 0 | 0 | 0 | 1794500 | 1015700 | 0 | 0 | 0 | 0 | 0 | 0 | 0 | 0 |
| DGTAGSHI | POFUT1 | Q9H488 | OFUT1_HUMAI | Unmodified | 0 | 0 | 0 | 0 | 0 | 0 | 0 | 0 | 7787400 | 638590 | 0 | 0 | 0 | 0 | 0 | 0 | 0 | 0 |
| DGTAPPPQ | PACSIN3; PACS | Q9UKS6; A0, | PACN3_HUMAI | Unmodified | 4099700 | 3506400 | 1918800 | 156850 | 0 | 0 | 0 | 0 | 0 | 0 | 0 | 0 | 0 | 0 | 3867500 | 1588800 | 0 | 0 |
| DGTFLVR | VAV1; PIK3R1; I | P15498; P27 | VAV_HUMAN P | Unmodified | 0 | 0 | 161020 | 0 | 264040 | 0 | 0 | 27293 | 86189 | 346680 | 547490 | 6054900 | 0 | 0 | 169110 | 244380 | 29434 | 0 |
| DGTFPLPIC | ICAM1; ICAM1 | P05362; E7E | ICAM1_HUMAI | Unmodified | 0 | 0 | 573230 | 1859100 | 3030800 | 1690700 | 0 | 0 | 99721 | 217980 | 0 | 0 | 0 | 0 | 1861400 | 2228900 | 75477 | 203680 |
| DGTFPLPIC | ICAM1; ICAM1 | P05362; E7E | ICAM1_HUMAI | Unmodified | 0 | 5641800 | 0 | 0 | 13109000 | 3519500 | 0 | 0 | 0 | 0 | 0 | 0 | 0 | 0 | 0 | 0 | 1744000 | 4761900 |
| DGTFPLPIC | ICAM1; ICAM1 | P05362; E7E | ICAM1_HUMAI | Unmodified | 0 | 0 | 0 | 0 | 0 | 4006900 | 0 | 0 | 0 | 0 | 0 | 0 | 29859 | 0 | 0 | 0 | 3224000 | 6158300 |
| DGTILCTLI | CNN2; CNN2; ( | Q99439; H3I | CNN2_HUMAN | Unmodified | 0 | 0 | 392890 | 0 | 1669300 | 0 | 0 | 0 | 0 | 0 | 0 | 0 | 0 | 0 | 0 | 0 | 0 | 0 |
| DGTISEDTI | TSG101; TSG1C | Q99816; F5H | TS101_HUMAI | Unmodified | 32151000 | 14825000 | 51490000 | 37066000 | 0 | 0 | 85378 | 0 | 2324500 | 0 | 0 | 4341000 | 0 | 0 | 30204000 | 77196000 | 0 | 0 |
| DGTISEDTI | TSG101; TSG1C | Q99816; F5H | TS101_HUMAI | Unmodified | 2155800 | 0 | 2472100 | 7504200 | 0 | 0 | 0 | 0 | 0 | 0 | 0 | 0 | 0 | 0 | 0 | 0 | 0 | 0 |
| DGTISEDTI | TSG101; TSG1C | Q99816; F5H | TS101_HUMAI | Unmodified | 754960 | 0 | 0 | 1091900 | 0 | 0 | 0 | 0 | 0 | 0 | 0 | 0 | 0 | 0 | 0 | 0 | 0 | 0 |
| DGTLFAW( | EIF2A; EIF2A; EI | Q9BY44; H7 | EIF2A_HUMAN | Unmodified | 0 | 0 | 0 | 0 | 0 | 0 | 0 | 0 | 0 | 0 | 0 | 2560600 | 0 | 0 | 0 | 0 | 0 | 0 |
| DGTMLINC | SLITRK6 | Q9H5Y7 | SLIK6_HUMAN | Oxidation (M) | 246870 | 195580 | 0 | 0 | 0 | 0 | 0 | 0 | 0 | 0 | 0 | 0 | 0 | 0 | 0 | 0 | 0 | 462950 |
| DGTSFGEY | KLC1; KLC1; KL | Q07866; G3\ | KLC1_HUMAN | Unmodified | 0 | 0 | 0 | 0 | 932810 | 0 | 332510 | 199750 | 0 | 217570 | 0 | 896790 | 0 | 0 | 0 | 0 | 412430 | 0 |
| DGTVLCEL | TAGLN2; TAGLI | P37802; X6R | TAGL2_HUMAI | Unmodified | 0 | 104320 | 0 | 52469 | 1242100 | 0 | 328570 | 0 | 0 | 322060 | 0 | 0 | 0 | 0 | 71145 | 0 | 814830 | 756470 |
| DGTVLCEL | TAGLN2; TAGLI | P37802; X6R | TAGL2_HUMAI | Unmodified | 2046900 | 1395300 | 52116 | 688780 | 20404000 | 3640900 | 2109100 | 0 | 1578100 | 4521400 | 1317000 | 1823700 | 5604400 | 1535000 | 774840 | 55840 | 14157000 | 13676000 |
| DGTVLCEL | TAGLN2; TAGLI | P37802; X6R | TAGL2_HUMAI | Oxidation (M) | 0 | 0 | 0 | 0 | 89558 | 0 | 138190 | 0 | 0 | 30079 | 0 | 0 | 124550 | 197560 | 0 | 0 | 3034100 | 275060 |
| DGTVTAGN | ACAA2; ACAA2; | P42765; A0A | THIM_HUMAN | Unmodified | 0 | 0 | 0 | 0 | 147520 | 0 | 0 | 0 | 3350600 | 1209800 | 0 | 0 | 0 | 0 | 0 | 0 | 0 | 0 |
| DGTYAVTYI | FLNB; FLNB | O75369; E7E | FLNB_HUMAN | Unmodified | 913420 | 2829200 | 0 | 296690 | 0 | 0 | 0 | 0 | 0 | 0 | 0 | 2068900 | 0 | 0 | 779460 | 583770 | 0 | 0 |
| DGVIEASIN | PSMD3 | O43242 | PSMD3_HUMA | Unmodified | 0 | 0 | 0 | 0 | 0 | 0 | 0 | 0 | 0 | 0 | 0 | 1329400 | 0 | 0 | 0 | 0 | 0 | 0 |
| DGVKFSAS | PCNA | P12004 | PCNA_HUMAN | Unmodified | 0 | 765670 | 0 | 0 | 0 | 0 | 5414700 | 3626100 | 19739000 | 36746000 | 1826400 | 561270 | 5444200 | 0 | 1373800 | 0 | 0 | 0 |
| DGVLEEQII | NCBP1; NCBP1 | Q09161; X6I | NCBP1_HUMAI | Unmodified | 0 | 0 | 0 | 0 | 0 | 0 | 121400 | 0 | 0 | 0 | 0 | 734230 | 0 | 0 | 1372600 | 0 | 0 | 0 |
| DGVLTLAN | MAP4; MAP4 | P27816; E7E | MAP4_HUMAN | Unmodified | 90021 | 0 | 0 | 0 | 0 | 0 | 0 | 0 | 0 | 0 | 0 | 1444900 | 0 | 0 | 0 | 0 | 0 | 0 |
| DGVMFQIC | EPDR1 | Q9UM22 | EPDR1_HUMAI | Unmodified | 0 | 0 | 0 | 0 | 0 | 0 | 0 | 0 | 732940 | 0 | 0 | 0 | 0 | 0 | 0 | 0 | 0 | 0 |
| DGVREVFE | RHOA; RHOA | P61586; A0A | RHOA_HUMAN | Unmodified | 0 | 903920 | 906580 | 1565500 | 3827100 | 1386800 | 477100 | 83089 | 7798100 | 3614000 | 2474400 | 6113700 | 2085700 | 1046100 | 1535700 | 0 | 896730 | 2738900 |
| DGVREVFE | RHOA; RHOA | P61586; A0A | RHOA_HUMAN | Oxidation (M) | 795620 | 329900 | 0 | 1368600 | 571570 | 4724600 | 61854 | 50949 | 1262300 | 1753000 | 4415800 | 6974200 | 275950 | 144910 | 1403300 | 1201600 | 180360 | 425460 |
| DGVREVFE | RHOA; RHOA | P61586; A0A | RHOA_HUMAN | Oxidation (M) | 0 | 0 | 0 | 0 | 1547800 | 0 | 211640 | 0 | 0 | 1288200 | 0 | 0 | 1023200 | 739990 | 0 | 0 | 0 | 472020 |
| DGVVEITGI | HSPB1; HSPB1; | P04792; A0A | HSPB1_HUMAI | Unmodified | 1188900 | 1444800 | 7427600 | 3297500 | 1931900 | 183020 | 300900 | 166140 | 15231000 | 3721100 | 0 | 0 | 3365600 | 330850 | 11692000 | 51683 | 991660 | 1171600 |
| DGVVFGVE | PSMA3 | P25788 | PSA3_HUMAN | Unmodified | 0 | 0 | 328480 | 345480 | 5429700 | 882980 | 136910 | 203850 | 4015000 | 563160 | 1100500 | 14834000 | 3519800 | 465350 | 264650 | 280670 | 1183200 | 6722800 |
| DGVWKPTI | ARPC1A; ARPC | Q92747; E9F | ARC1A_HUMAI | Unmodified | 1699900 | 1910400 | 1465000 | 969140 | 0 | 0 | 0 | 0 | 544140 | 0 | 681170 | 21579000 | 0 | 0 | 2996200 | 2122600 | 0 | 0 |
| DGVWNLQ | PRPF8 | Q6P2Q9 | PRP8_HUMAN | Unmodified | 0 | 0 | 0 | 0 | 0 | 0 | 0 | 0 | 0 | 0 | 0 | 1577800 | 0 | 0 | 0 | 0 | 0 | 0 |
| DGVYFLYE/ | ADSS2 | P30520 | PURA2_HUMAI | Unmodified | 0 | 0 | 0 | 0 | 0 | 0 | 0 | 0 | 1125400 | 0 | 0 | 6006600 | 0 | 0 | 98273 | 0 | 0 | 0 |
| DGVYFLYE/ | ADSS2 | P30520 | PURA2_HUMAI | Unmodified | 0 | 0 | 0 | 0 | 146420 | 0 | 0 | 0 | 0 | 0 | 1442000 | 12609000 | 0 | 0 | 0 | 0 | 0 | 0 |
| DGVYVLDL | ACLY | P53396 | ACLY_HUMAN | Unmodified | 0 | 0 | 0 | 0 | 0 | 1160800 | 0 | 81375 | 0 | 0 | 0 | 8733400 | 0 | 0 | 0 | 0 | 0 | 649510 |
| DGWHYGE | BAIAP2; BAIAP2 | Q9UQB8; I3I | BAIP2_HUMAN | Unmodified | 0 | 0 | 362130 | 0 | 0 | 0 | 0 | 0 | 697980 | 0 | 0 | 937490 | 0 | 0 | 1022000 | 483020 | 0 | 0 |
| DGWLYGEI | BAIAP2L1 | Q9UHR4 | BI2L1_HUMAN | Unmodified | 0 | 0 | 303390 | 0 | 114790 | 0 | 0 | 0 | 0 | 0 | 0 | 0 | 0 | 0 | 466610 | 189760 | 0 | 0 |
| DGWPAMC | DDX17; DDX17; | Q92841; A0/ | DDX17_HUMAI | Unmodified | 0 | 0 | 0 | 0 | 0 | 0 | 0 | 0 | 0 | 0 | 0 | 676130 | 0 | 0 | 0 | 0 | 0 | 0 |
| DGYADIVD | ACOT2; ACOT1; | P49753; Q8( | ACOT2_HUMAI | Unmodified | 0 | 1256800 | 0 | 0 | 0 | 0 | 0 | 0 | 1465900 | 371630 | 0 | 0 | 0 | 0 | 45913 | 1000400 | 0 | 0 |
| DGYIGPNC | MEGF9 | Q9H1U4 | MEGF9_HUMAI | Unmodified | 0 | 0 | 0 | 0 | 128150 | 0 | 0 | 0 | 862350 | 54122 | 100920 | 0 | 0 | 0 | 0 | 0 | 0 | 0 |
| DGYISNGE | PPP3R1; PPP3R | P63098; Q9( | CANB1_HUMAI | Unmodified | 0 | 0 | 0 | 0 | 0 | 96367 | 0 | 0 | 0 | 0 | 785790 | 3775500 | 0 | 0 | 173140 | 464040 | 0 | 0 |
| DGYNYTLS | S100A11 | P31949 | S10AB_HUMAI | Unmodified | 0 | 297800 | 0 | 0 | 6685700 | 2374200 | 0 | 0 | 12297000 | 140390 | 0 | 0 | 767000 | 0 | 1001300 | 1087800 | 258350 | 3544900 |
| DHADSNIV | RAB11A; RAB11 | P62491; Q15 | RB11A_HUMAI | Oxidation (M) | 0 | 0 | 539200 | 0 | 0 | 0 | 14022 | 0 | 0 | 0 | 392630 | 1987300 | 0 | 0 | 0 | 0 | 0 | 0 |
| DHADVSN( | ATP6V1B2 | P21281 | VATB2_HUMAI | Unmodified | 138540 | 0 | 124800 | 137820 | 200210 | 428080 | 0 | 0 | 0 | 0 | 271430 | 8896300 | 218810 | 0 | 263050 | 108690 | 0 | 0 |
| DHAFVTSS | PLCG2; PLCG2 | P16885; A0A | PLCG2_HUMAI | Unmodified | 0 | 0 | 0 | 0 | 0 | 0 | 0 | 0 | 0 | 0 | 0 | 2385200 | 0 | 0 | 0 | 0 | 0 | 0 |
| DHALLEEQ | ACTN4; ACTN4; | O43707; F5( | ACTN4_HUMAI | Unmodified | 0 | 0 | 0 | 0 | 0 | 0 | 0 | 0 | 63873 | 0 | 77484 | 369560 | 0 | 1816300 | 34333 | 43381 | 0 | 0 |
| DHASIQMI | RPS21; RPS21; | P63220; Q9E | RS21_HUMAN | Unmodified | 343620 | 872070 | 954900 | 426450 | 735370 | 430490 | 0 | 0 | 0 | 0 | 0 | 9111800 | 0 | 181770 | 1147400 | 928770 | 199030 | 2443200 |
| DHASIQMI | RPS21; RPS21; | P63220; Q9E | RS21_HUMAN | Oxidation (M) | 470800 | 237010 | 261630 | 0 | 0 | 27580 | 0 | 0 | 0 | 0 | 237570 | 3998900 | 0 | 0 | 585420 | 433090 | 0 | 0 |
| DHASIQMI | RPS21; RPS21; | P63220; Q9E | RS21_HUMAN | Unmodified | 1901600 | 2602600 | 0 | 1596600 | 924660 | 0 | 0 | 0 | 1464400 | 0 | 2855500 | 12830000 | 890310 | 94983 | 450280 | 934860 | 0 | 140820 |
| DHASIQMI | RPS21; RPS21; | P63220; Q9E | RS21_HUMAN | Oxidation (M) | 1263700 | 604040 | 0 | 0 | 0 | 0 | 0 | 0 | 0 | 154080 | 165800 | 1875200 | 0 | 0 | 17897 | 1189600 | 0 | 435110 |
| DHASIQMI | RPS21; RPS21; | P63220; Q9E | RS21_HUMAN | Oxidation (M) | 147080 | 0 | 0 | 168560 | 0 | 0 | 798780 | 0 | 0 | 0 | 0 | 0 | 3250500 | 2210100 | 0 | 0 | 1210400 | 545560 |
| DHASIQMI | RPS21; RPS21; | P63220; Q9E | RS21_HUMAN | Unmodified | 0 | 0 | 0 | 230060 | 1312800 | 0 | 1589300 | 0 | 1504100 | 266020 | 0 | 0 | 0 | 1989800 | 0 | 0 | 1036400 | 3475300 |
| DHASNIYK | CD151; CD151; | P48509; E9P | CD151_HUMAI | Unmodified | 2832800 | 0 | 0 | 58336 | 0 | 712370 | 228760 | 336310 | 200830 | 0 | 0 | 508300 | 0 | 0 | 0 | 155350 | 209720 | 0 |
| DHASNIYK | CD151; CD151 | P48509; K4D | CD151_HUMAI | Unmodified | 2676100 | 2172800 | 0 | 0 | 0 | 0 | 0 | 0 | 0 | 0 | 0 | 0 | 0 | 0 | 384340 | 625420 | 0 | 0 |
| DHASNIYK | CD151; CD151 | P48509; K4D | CD151_HUMAI | Unmodified | 393000 | 1320300 | 0 | 0 | 0 | 0 | 0 | 0 | 0 | 0 | 0 | 0 | 0 | 0 | 0 | 72187 | 0 | 0 |
| DHAVFTR | DNAJA1 | P31689 | DNJA1_HUMAI | Unmodified | 0 | 0 | 12002 | 0 | 100130 | 112820 | 0 | 11252 | 3172500 | 20413 | 0 | 0 | 930010 | 0 | 0 | 0 | 0 | 53311 |
| DHENIVIAK | P4HB; P4HB; P4 | P07237; A0A | PDIA1_HUMAN | Unmodified | 0 | 0 | 290210 | 0 | 0 | 0 | 0 | 0 | 12227000 | 556160 | 788620 | 763910 | 127780 | 0 | 269050 | 179750 | 0 | 0 |
| DHGGALGI | ACTN4; ACTN4 | O43707; F5( | ACTN4_HUMAI | Unmodified | 264890 | 173750 | 756340 | 0 | 281250 | 0 | 0 | 0 | 241660 | 0 | 0 | 1316000 | 0 | 0 | 0 | 218330 | 0 | 0 |
| DHGIRLLE/ | PLEC; PLEC | Q15149; H0 | PLEC_HUMAN | Unmodified | 0 | 0 | 0 | 0 | 0 | 0 | 0 | 0 | 0 | 0 | 0 | 0 | 0 | 1078100 | 0 | 0 | 2491200 | 1416100 |
| DHGVLESI | CANT1; CANT1 | Q8WVQ1; K7 | CANT1_HUMAI | Unmodified | 0 | 0 | 148120 | 141560 | 0 | 36219 | 0 | 0 | 16221000 | 694620 | 1334900 | 472120 | 0 | 0 | 0 | 0 | 0 | 122560 |
| DHGVNSFI | DPYSL2; DPYSL | Q16555; A0/ | DPYL2_HUMAI | Unmodified | 0 | 0 | 0 | 0 | 0 | 0 | 0 | 0 | 726960 | 44104 | 189110 | 127570 | 0 | 0 | 0 | 0 | 0 | 0 |
| DHGVNSFI | DPYSL2; DPYSL | Q16555; A0/ | DPYL2_HUMAI | Oxidation (M) | 0 | 0 | 361010 | 0 | 885560 | 0 | 120040 | 10696 | 337400 | 0 | 0 | 1100800 | 0 | 0 | 0 | 0 | 0 | 0 |
| DHGYSDVI | XXYLT1; XXYLT1 | Q8NBI6; A0/ | XXLT1_HUMAN | Unmodified | 0 | 0 | 0 | 0 | 0 | 0 | 0 | 0 | 283810 | 21516 | 0 | 0 | 0 | 0 | 0 | 0 | 0 | 0 |
| DHHLVIPS | MMS19; MMS1( | Q96T76; Q5 | MMS19_HUMA | Unmodified | 0 | 0 | 0 | 0 | 0 | 0 | 0 | 0 | 0 | 291390 | 0 | 559140 | 0 | 0 | 0 | 19490 | 223640 | 0 |
| DHINDIIK | IQGAP1; IQGAP | P46940; A0A | IQGA1_HUMAI | Unmodified | 78843 | 0 | 0 | 0 | 0 | 67509 | 0 | 0 | 97822 | 0 | 0 | 839900 | 0 | 0 | 0 | 0 | 13837 | 0 |
| DHINLPGF | PNP; PNP | P00491; G3\ | PNPH_HUMAN | Unmodified | 0 | 0 | 0 | 0 | 0 | 0 | 0 | 0 | 0 | 0 | 99383 | 675050 | 0 | 0 | 0 | 52534 | 0 | 0 |
| DHINLPGF | PNP; PNP | P00491; G3\ | PNPH_HUMAN | Unmodified | 0 | 0 | 0 | 0 | 0 | 0 | 0 | 0 | 0 | 0 | 0 | 1336200 | 0 | 0 | 0 | 0 | 0 | 0 |
| DHIWSQEE | DHTKD1 | Q96HY7 | DHTK1_HUMAI | Unmodified | 0 | 0 | 0 | 0 | 0 | 0 | 0 | 0 | 548880 | 0 | 0 | 0 | 0 | 0 | 0 | 0 | 0 | 0 |
| DHKLHIDH | SULF2 | Q8IWU5 | SULF2_HUMAI | Unmodified | 0 | 0 | 0 | 0 | 0 | 0 | 143890 | 340540 | 0 | 0 | 0 | 0 | 0 | 0 | 0 | 0 | 0 | 1177300 |
| DHLLSVSL | WDR1 | O75083 | WDR1_HUMAN | Unmodified | 0 | 0 | 0 | 1831200 | 56693000 | 2421600 | 4166700 | 0 | 0 | 4361100 | 0 | 197570 | 31081000 | 17372000 | 1139800 | 0 | 16439000 | 10779000 |

| | | | | | | | | | | | | | | | | | | | | | | |
|---|---|---|---|---|---|---|---|---|---|---|---|---|---|---|---|---|---|---|---|---|---|---|
| DHLVTAYN | FKBP3 | Q00688 | FKBP3_HUMAN | Unmodified | 1172000 | 1078700 | 1273300 | 1015700 | 318550 | 0 | 59284 | 31281 | 2476900 | 368350 | 1589700 | 6609400 | 299570 | 141550 | 2644800 | 1491100 | 0 | 314380 |
| DHLVTAYN | FKBP3 | Q00688 | FKBP3_HUMAN | Unmodified | 615480 | 527590 | 0 | 186660 | 0 | 0 | 0 | 0 | 252890 | 0 | 0 | 2761600 | 0 | 0 | 1155100 | 468900 | 0 | 0 |
| DHLYGTLD | DYNC1H1; DYN | Q14204; A0A | DYHC1_HUMA | Unmodified | 96545 | 0 | 0 | 0 | 0 | 0 | 0 | 0 | 0 | 56562 | 118110 | 5563700 | 0 | 39417 | 0 | 0 | 0 | 0 |
| DHNIPGEL | MYH10 | P35580 | MYH10_HUMA | Unmodified | 0 | 0 | 12376 | 0 | 3657800 | 0 | 0 | 0 | 0 | 0 | 0 | 0 | 1069500 | 74997 | 184130 | 0 | 0 | 0 |
| DHPESYHS | GNPDA1; GNPD | P46926; D6F | GNPI1_HUMAN | Oxidation (M) | 0 | 0 | 0 | 0 | 0 | 0 | 0 | 0 | 0 | 0 | 0 | 221920 | 0 | 0 | 0 | 0 | 0 | 0 |
| DHPLPEVA | COX4I1; COX4I | P13073; Q86 | COX41_HUMA | Unmodified | 36959 | 0 | 56757 | 0 | 0 | 0 | 0 | 0 | 8476000 | 1333100 | 519510 | 281880 | 0 | 0 | 0 | 0 | 0 | 0 |
| DHPPIPITD | PTPRF; PTPRF; | P10586; H0Y | PTPRF_HUMAN | Unmodified | 1151600 | 1380100 | 278410 | 170890 | 0 | 0 | 0 | 0 | 987430 | 837930 | 1322500 | 1175300 | 0 | 0 | 9396600 | 868510 | 0 | 0 |
| DHPPIPITD | PTPRF; PTPRF; | P10586; H0Y | PTPRF_HUMAN | Unmodified | 185530 | 176480 | 557970 | 0 | 685110 | 0 | 0 | 0 | 1328200 | 0 | 0 | 1414300 | 0 | 0 | 1320800 | 1489700 | 0 | 0 |
| DHQLLEPIF | TRIM29 | Q14134 | TRI29_HUMAN | Unmodified | 0 | 0 | 0 | 0 | 0 | 0 | 0 | 0 | 0 | 71460 | 89179 | 1503500 | 0 | 0 | 0 | 0 | 0 | 0 |
| DHQPCIIFN | PSMC6; PSMC6 | P62333; A0A | PRS10_HUMAN | Unmodified | 0 | 0 | 0 | 0 | 429100 | 0 | 0 | 0 | 84076 | 0 | 0 | 79901 | 0 | 0 | 0 | 0 | 0 | 0 |
| DHQYQFLE | TRIM28; TRIM28 | Q13263; M0 | TIF1B_HUMAN | Unmodified | 0 | 58587 | 170970 | 76057 | 0 | 0 | 0 | 0 | 2254600 | 0 | 491410 | 3862400 | 0 | 0 | 99357 | 101850 | 0 | 0 |
| DHQYQFLE | TRIM28; TRIM28 | Q13263; M0 | TIF1B_HUMAN | Unmodified | 0 | 0 | 0 | 0 | 0 | 0 | 0 | 0 | 574930 | 692590 | 0 | 0 | 117210 | 0 | 0 | 0 | 0 | 2204400 |
| DHRDNLEF | FASN; FASN | P49327; A0A | FAS_HUMAN F | Unmodified | 0 | 0 | 0 | 0 | 0 | 0 | 0 | 0 | 0 | 0 | 0 | 1947600 | 0 | 0 | 0 | 0 | 0 | 0 |
| DHRNDIML | KLK11; KLK11; | Q9UBX7; M0 | KLK11_HUMAN | Unmodified | 0 | 0 | 0 | 0 | 0 | 0 | 0 | 0 | 1030600 | 84695 | 0 | 0 | 0 | 0 | 0 | 0 | 0 | 0 |
| DHSFFIPDI | ZNF622 | Q969S3 | ZN622_HUMAN | Unmodified | 215390 | 686660 | 68375 | 227680 | 0 | 0 | 0 | 0 | 0 | 50297 | 1018300 | 13938 | 360860 | 0 | 531170 | 0 | 79598 | 388460 |
| DHSFFSNL | RIPOR2; RIPOR | B7Z6U4; F5C | B7Z6U4_HUMA | Unmodified | 0 | 0 | 0 | 0 | 0 | 0 | 0 | 0 | 138530 | 23359 | 410890 | 1508400 | 528190 | 300740 | 0 | 136220 | 0 | 0 |
| DHSFLGFE | SKAP2; SKAP2 | O75563; C9J | SKAP2_HUMAN | Unmodified | 0 | 0 | 0 | 0 | 0 | 0 | 181550 | 0 | 0 | 0 | 279770 | 995020 | 282770 | 373520 | 0 | 0 | 0 | 0 |
| DHSGGIGD | LACTB2 | Q53H82 | LACB2_HUMAN | Unmodified | 0 | 0 | 0 | 0 | 0 | 0 | 0 | 0 | 699240 | 127660 | 0 | 0 | 0 | 0 | 0 | 0 | 0 | 0 |
| DHSILHSRI | ADPGK; ADPGK | Q9BRR6; H3 | ADPGK_HUMA | Unmodified | 0 | 0 | 0 | 0 | 34177 | 0 | 0 | 0 | 2610700 | 0 | 0 | 0 | 0 | 0 | 0 | 0 | 0 | 0 |
| DHSKPISNI | TGOLN2; TGOL | O43493; A0A | TGON2_HUMA | Unmodified | 0 | 0 | 0 | 0 | 23148 | 0 | 0 | 0 | 4797600 | 97430 | 97549 | 334230 | 0 | 0 | 0 | 0 | 0 | 18486 |
| DHSPDLYS | ATP6AP2; ATP6 | O75787; A0A | RENR_HUMAN | Unmodified | 0 | 0 | 0 | 0 | 0 | 0 | 0 | 0 | 5585700 | 368930 | 0 | 0 | 0 | 0 | 646870 | 0 | 0 | 102970 |
| DHSPDLYS | ATP6AP2; ATP6 | O75787; A0A | RENR_HUMAN | Unmodified | 0 | 0 | 0 | 0 | 0 | 0 | 0 | 0 | 12491000 | 897860 | 357510 | 275260 | 0 | 0 | 0 | 0 | 0 | 0 |
| DHTLEDED | DRG1 | Q9Y295 | DRG1_HUMAN | Unmodified | 0 | 0 | 0 | 276980 | 0 | 1135700 | 0 | 0 | 133990 | 0 | 0 | 2271600 | 370150 | 751450 | 0 | 0 | 0 | 281700 |
| DHVFLCEG | JPT2; JPT2; JPT2 | Q9H910; H3 | JUPI2_HUMAN | Unmodified | 0 | 0 | 487660 | 289140 | 671480 | 0 | 0 | 0 | 414280 | 96289 | 212890 | 269550 | 305190 | 635150 | 211600 | 102090 | 79846 | 480000 |
| DHVTATQL | LRBA; LRBA; LR | P50851; A0A | LRBA_HUMAN | Unmodified | 0 | 0 | 0 | 0 | 398290 | 0 | 0 | 0 | 0 | 0 | 0 | 687640 | 0 | 0 | 0 | 0 | 0 | 0 |
| DHVVEGEF | FAM20B | O75063 | XYLK_HUMAN ( | Unmodified | 0 | 0 | 0 | 0 | 0 | 0 | 0 | 0 | 12808000 | 171650 | 0 | 988460 | 0 | 0 | 0 | 0 | 0 | 0 |
| DHVYGIHN | PROM1 | O43490 | PROM1_HUMA | Oxidation (M) | 0 | 0 | 0 | 0 | 0 | 0 | 0 | 0 | 0 | 0 | 0 | 607200 | 0 | 0 | 0 | 0 | 0 | 0 |
| DHVYGIHN | PROM1 | O43490 | PROM1_HUMA | Unmodified | 0 | 0 | 0 | 0 | 0 | 0 | 0 | 0 | 0 | 0 | 169490 | 5127400 | 0 | 0 | 0 | 0 | 0 | 0 |
| DHYSNGF( | CAPZA1 | P52907 | CAZA1_HUMAN | Unmodified | 363870 | 179410 | 0 | 0 | 1932800 | 0 | 0 | 424340 | 0 | 0 | 0 | 2575700 | 1230400 | 2347000 | 0 | 439470 | 998750 | 0 |
| DIAKASDE\ | VCL | P18206 | VINC_HUMAN ' | Unmodified | 0 | 20300 | 0 | 0 | 304140 | 88495 | 580780 | 444460 | 0 | 133300 | 0 | 0 | 149580 | 342150 | 0 | 0 | 1647900 | 284700 |
| DIALEITVTI | ITGA6 | P23229 | ITA6_HUMAN I | Unmodified | 35145 | 0 | 0 | 653420 | 1527800 | 655670 | 0 | 0 | 0 | 0 | 387570 | 515100 | 5241000 | 0 | 0 | 151950 | 2206500 | 22229000 |
| DIALVNLAI | TTC17 | Q96AE7 | TTC17_HUMAN | Unmodified | 0 | 0 | 0 | 0 | 0 | 0 | 0 | 0 | 1334800 | 0 | 0 | 0 | 0 | 0 | 0 | 0 | 0 | 0 |
| DIAWTEDS | WDR1; WDR1 | O75083; D6I | WDR1_HUMAN | Unmodified | 741200 | 0 | 833850 | 0 | 4981900 | 0 | 0 | 203650 | 949690 | 0 | 0 | 0 | 2150700 | 852910 | 768790 | 914410 | 259330 | 0 |
| DICNDVLSI | YWHAZ; YWHAZ | P63104; B0A | 1433Z_HUMAN | Unmodified | 539490 | 497500 | 105800 | 222610 | 0 | 0 | 318490 | 0 | 2194200 | 993400 | 55810 | 1985900 | 0 | 0 | 1495100 | 360100 | 250480 | 0 |
| DIDENAYAI | PSMD2 | Q13200 | PSMD2_HUMA | Unmodified | 0 | 0 | 0 | 0 | 405000 | 0 | 143990 | 34666 | 330780 | 82065 | 630500 | 3076500 | 117840 | 0 | 0 | 132760 | 207630 | 220330 |
| DIDEVSSLL | CCT8; CCT8 | P50990; H7C | TCPQ_HUMAN | Unmodified | 0 | 0 | 570790 | 0 | 0 | 0 | 0 | 0 | 1126200 | 0 | 0 | 0 | 0 | 0 | 0 | 377650 | 0 | 0 |
| DIDGHMVI | GPX4; GPX4; GI | P36969; A0A | GPX4_HUMAN | Unmodified | 1629200 | 1210000 | 0 | 0 | 157620 | 0 | 254240 | 103870 | 1265100 | 576520 | 293280 | 1635100 | 1892800 | 1057000 | 1292300 | 1263700 | 211580 | 0 |
| DIDGHMVI | GPX4; GPX4; GI | P36969; A0A | GPX4_HUMAN | Oxidation (M) | 0 | 19122 | 0 | 25499 | 0 | 0 | 0 | 0 | 0 | 0 | 0 | 0 | 0 | 0 | 443970 | 587390 | 0 | 0 |
| DIDISSPEF | AHNAK | Q09666 | AHNK_HUMAN | Unmodified | 0 | 0 | 0 | 0 | 9075100 | 0 | 0 | 0 | 0 | 0 | 0 | 0 | 0 | 0 | 0 | 0 | 0 | 8760600 |
| DIDNLVQR | LMAN1 | P49257 | LMAN1_HUMA | Unmodified | 0 | 0 | 161940 | 0 | 0 | 0 | 0 | 0 | 1.36E+08 | 6855800 | 5231400 | 0 | 0 | 0 | 0 | 0 | 0 | 0 |
| DIDPQNDL | DYNLRB2; DYN | Q8TF09; Q9I | DLRB2_HUMAN | Unmodified | 552480 | 0 | 0 | 0 | 0 | 0 | 0 | 0 | 0 | 0 | 1100500 | 3959300 | 0 | 0 | 0 | 0 | 0 | 0 |
| DIDSFVPSI | DLAT; DLAT; DL | P10515; A0A | ODP2_HUMAN | Unmodified | 380590 | 784440 | 79055 | 112040 | 250850 | 0 | 0 | 0 | 2302500 | 1054300 | 201160 | 263120 | 0 | 0 | 43267 | 26081 | 1197300 | 0 |
| DIDTLRDP( | IGF2R; IGF2R; I | P11717; A0A | MPRI_HUMAN ( | Unmodified | 0 | 0 | 3020900 | 1984200 | 365400 | 0 | 0 | 0 | 1.2E+08 | 1771800 | 8464800 | 16234000 | 0 | 0 | 1505300 | 2318800 | 0 | 0 |
| DIEAMDPSI | NSF; NSF; NSF | P46459; I3L( | NSF_HUMAN V | Oxidation (M) | 106200 | 0 | 0 | 0 | 0 | 0 | 0 | 0 | 0 | 0 | 0 | 564990 | 303280 | 0 | 0 | 508220 | 65330 | 0 |
| DIEAQLQH. | H6PD | O95479 | G6PE_HUMAN | Unmodified | 0 | 0 | 0 | 0 | 0 | 0 | 0 | 0 | 2866800 | 36172 | 0 | 0 | 0 | 0 | 0 | 0 | 0 | 0 |
| DIEDVFYK | SRSF1; SRSF1; | Q07955; J3K | SRSF1_HUMAN | Unmodified | 831660 | 1137800 | 1510100 | 680990 | 0 | 284670 | 0 | 0 | 0 | 0 | 213630 | 0 | 356820 | 0 | 792160 | 833160 | 0 | 0 |
| DIEDVFYKY | SRSF1; SRSF1; | Q07955; J3K | SRSF1_HUMAN | Unmodified | 0 | 0 | 0 | 0 | 0 | 0 | 0 | 0 | 0 | 242650 | 0 | 0 | 1389900 | 584430 | 0 | 0 | 0 | 0 |
| DIEEHLQR | STUB1; STUB1 | Q9UNE7; H3 | CHIP_HUMAN | Unmodified | 0 | 50531 | 385410 | 130710 | 0 | 0 | 0 | 0 | 290050 | 0 | 0 | 893960 | 121760 | 0 | 0 | 100140 | 0 | 0 |
| DIEGKLPQ1 | SMC4; SMC4 | Q9NTJ3; E9F | SMC4_HUMAN | Unmodified | 0 | 0 | 0 | 0 | 0 | 0 | 596160 | 69793 | 0 | 0 | 0 | 0 | 693860 | 686190 | 21285 | 0 | 0 | 201840 |
| DIEHALSVS | TOR1A; TOR1A | O14656; A0A | TOR1A_HUMAN | Unmodified | 0 | 0 | 0 | 0 | 87545 | 0 | 24266 | 0 | 6505700 | 1041500 | 88053 | 419930 | 0 | 0 | 0 | 0 | 0 | 0 |
| DIELQIR | DNM1L; DNM1I | O00429; F8V | DNM1L_HUMA | Unmodified | 0 | 0 | 0 | 0 | 0 | 0 | 0 | 0 | 1600700 | 244430 | 0 | 817140 | 0 | 0 | 0 | 0 | 0 | 0 |
| DIELVMSQ | NACA; NACA; N | E9PAV3; Q13 | NACAM_HUMA | Unmodified | 0 | 0 | 0 | 0 | 0 | 0 | 0 | 0 | 0 | 0 | 203440 | 986170 | 0 | 0 | 0 | 0 | 0 | 0 |
| DIEQSIKSE | ANXA4; ANXA4 | P09525; Q6F | ANXA4_HUMAN | Unmodified | 0 | 0 | 0 | 22772 | 163370 | 0 | 0 | 0 | 0 | 0 | 0 | 0 | 0 | 195060 | 0 | 0 | 0 | 43425 |
| DIEREDIEF | CCT4 | P50991 | TCPD_HUMAN | Unmodified | 144620 | 213260 | 0 | 62307 | 0 | 0 | 209580 | 0 | 0 | 0 | 117990 | 11186000 | 0 | 0 | 362180 | 455260 | 0 | 120880 |
| DIFAAVTQ\ | ENTPD8 | Q5MY95 | ENTP8_HUMAN | Unmodified | 0 | 0 | 0 | 0 | 0 | 0 | 0 | 0 | 520270 | 111150 | 0 | 0 | 0 | 0 | 0 | 0 | 182250 | 0 |
| DIFAMDDK | SUCLG2; SUCL | Q96I99; E9P | SUCB2_HUMA | Unmodified | 0 | 0 | 0 | 0 | 359410 | 0 | 0 | 0 | 2107800 | 0 | 0 | 0 | 0 | 0 | 0 | 0 | 0 | 1167400 |
| DIFGASDP | NEDD4L; NEDD | Q96PU5; A0 | NED4L_HUMA | Unmodified | 0 | 0 | 0 | 0 | 0 | 0 | 0 | 0 | 0 | 0 | 0 | 1225800 | 0 | 0 | 0 | 0 | 0 | 0 |
| DIFNKGFG | VDAC2; VDAC2 | P45880; Q5J | VDAC2_HUMA | Unmodified | 0 | 0 | 0 | 0 | 0 | 0 | 0 | 0 | 714290 | 1450800 | 0 | 0 | 0 | 0 | 0 | 0 | 0 | 0 |
| DIFQEIFDK | IDH2; IDH2 | P48735; H0Y | IDHP_HUMAN | Unmodified | 0 | 0 | 0 | 0 | 0 | 0 | 0 | 0 | 23157000 | 2393600 | 437660 | 478890 | 0 | 0 | 0 | 0 | 0 | 0 |
| DIFQEIYDK | IDH1 | O75874 | IDHC_HUMAN | Unmodified | 2027100 | 0 | 1203400 | 0 | 2497400 | 0 | 0 | 0 | 3149400 | 859420 | 3521600 | 7639500 | 6940100 | 2392300 | 4190600 | 2361900 | 0 | 1078300 |
| DIFSEVGP | CSTF2; CSTF2; | P33240; E7E | CSTF2_HUMAN | Unmodified | 0 | 0 | 0 | 0 | 0 | 0 | 0 | 0 | 0 | 0 | 143120 | 238350 | 0 | 0 | 0 | 0 | 0 | 0 |
| DIFYKPFH( | PLXNB2; PLXNE | O15031; E2F | PLXB2_HUMAN | Unmodified | 1715700 | 409150 | 420380 | 1585900 | 4061700 | 1824900 | 0 | 0 | 1689000 | 0 | 0 | 4683000 | 1876300 | 585900 | 1063300 | 1075700 | 1330300 | 1308600 |
| DIGAIAQV | DPYSL2; DPYSL | Q16555; A0A | DPYL2_HUMAN | Unmodified | 0 | 0 | 0 | 0 | 0 | 0 | 0 | 0 | 0 | 0 | 0 | 4423000 | 0 | 0 | 0 | 0 | 0 | 0 |
| DIGSKSDP | CPNE3; CPNE3 | O75131; A0A | CPNE3_HUMA | Unmodified | 0 | 0 | 0 | 0 | 0 | 473720 | 0 | 0 | 6611400 | 1650100 | 0 | 0 | 0 | 0 | 0 | 0 | 0 | 464160 |
| DIHDDQDY | FLOT1; FLOT1; | O75955; A0A | FLOT1_HUMAN | Unmodified | 239290 | 143190 | 5599000 | 1891900 | 7781800 | 4910400 | 2365300 | 394340 | 12899000 | 2559000 | 4412500 | 19661000 | 1245500 | 814020 | 10251000 | 2352700 | 1236500 | 0 |
| DIHFMPCS | GSPT1; GSPT1 | P15170; H3E | ERF3A_HUMAN | Unmodified | 0 | 0 | 0 | 0 | 2749100 | 659460 | 2129300 | 1960000 | 0 | 3092200 | 0 | 648200 | 1872100 | 1059300 | 0 | 0 | 1754400 | 5491900 |
| DIHTLAQLI | SRP72 | O76094 | SRP72_HUMAN | Unmodified | 166410 | 199850 | 74716 | 150630 | 469910 | 108010 | 0 | 0 | 171020 | 193460 | 0 | 578010 | 650380 | 0 | 270550 | 0 | 0 | 286630 |
| DIICQIAYAI | RPL5; RPL5; RP | P46777; Q5T | RL5_HUMAN 6( | Unmodified | 1817200 | 1211900 | 293260 | 175850 | 0 | 0 | 0 | 0 | 0 | 0 | 131190 | 19761000 | 0 | 0 | 564450 | 516670 | 0 | 0 |
| DIIDKQHTE | MYO1D; MYO1D | O94832; J3Q | MYO1D_HUMA | Unmodified | 188970 | 529600 | 0 | 181090 | 0 | 0 | 0 | 0 | 0 | 0 | 0 | 0 | 0 | 0 | 0 | 0 | 0 | 0 |
| DIIHDPGR( | RPL8; RPL8; RP | P62917; E9P | RL8_HUMAN 6( | Unmodified | 4390200 | 3691300 | 3409200 | 2150100 | 4004800 | 716480 | 1094200 | 718830 | 789970 | 486590 | 1291400 | 33654000 | 1893400 | 700580 | 3282700 | 3111500 | 259120 | 806620 |
| DIILATDLAI | PDE2A; PDE2A; | O00408; C9J | PDE2A_HUMAN | Unmodified | 0 | 0 | 0 | 0 | 5187500 | 1148600 | 316850 | 162760 | 0 | 403500 | 0 | 3338200 | 681790 | 443600 | 0 | 0 | 1101100 | 2121200 |
| DIILFSFK | VPS13C; VPS13 | Q709C8; A0 | VP13C_HUMAN | Unmodified | 0 | 0 | 0 | 0 | 0 | 0 | 0 | 0 | 0 | 0 | 0 | 478080 | 0 | 0 | 0 | 0 | 0 | 0 |
| DIILPFRVIP | AP2M1; AP2M1; | Q96CW1; C9 | AP2M1_HUMA | Unmodified | 0 | 0 | 476160 | 954020 | 388090 | 226990 | 174900 | 0 | 1046000 | 874420 | 700490 | 2172700 | 165110 | 0 | 0 | 0 | 123820 | 162400 |
| DIILQSNPL | MYO1E; MYO1F | Q12965; O0( | MYO1E_HUMA | Unmodified | 0 | 96424 | 164260 | 361080 | 185660 | 174550 | 36019 | 0 | 164040 | 7034.4 | 110660 | 412350 | 0 | 31068 | 0 | 19441 | 259990 | 120760 |
| DIINEEEVQ | AARS1; AARS1; | P49588; A0A | SYAC_HUMAN | Unmodified | 0 | 0 | 754360 | 363600 | 0 | 0 | 0 | 0 | 1539000 | 0 | 889830 | 6960000 | 399330 | 0 | 1320100 | 2096400 | 0 | 0 |
| DIIPHPSYL | PRSS8 | Q16651 | PRSS8_HUMAN | Unmodified | 992130 | 1274600 | 295840 | 21205000 | 0 | 0 | 0 | 0 | 25126000 | 7793800 | 135920 | 116110 | 6111100 | 1842400 | 9006200 | 3172700 | 0 | 0 |
| DIIRQPSEE | PEA15; PEA15 | Q15121; B1A | PEA15_HUMAN | Unmodified | 689390 | 417040 | 359880 | 557240 | 778860 | 277050 | 0 | 0 | 0 | 0 | 0 | 0 | 1332200 | 936820 | 387400 | 766140 | 0 | 840770 |
| DIISDTSGD | ANXA2P2; ANXA | A6NMY6; P0 | AXA2L_HUMAN | Unmodified | 0 | 0 | 0 | 0 | 2924200 | 0 | 0 | 164520 | 5151400 | 565610 | 0 | 345810 | 0 | 0 | 0 | 0 | 0 | 0 |
| DIISDTSGD | ANXA2P2; ANXA | A6NMY6; P0 | AXA2L_HUMAN | Unmodified | 998490 | 0 | 0 | 0 | 85625000 | 23642000 | 1590800 | 911140 | 26616000 | 6754600 | 4835300 | 0 | 0 | 0 | 67075 | 0 | 3854600 | 18335000 |
| DIISDTSGD | ANXA2P2; ANXA | A6NMY6; P0 | AXA2L_HUMAN | Unmodified | 0 | 0 | 0 | 0 | 2096400 | 409170 | 0 | 0 | 0 | 194250 | 0 | 0 | 0 | 0 | 0 | 0 | 1871600 | 2249200 |
| DIITRYQETI | ECH1; ECH1; E | Q13011; M0 | ECH1_HUMAN | Unmodified | 0 | 0 | 0 | 0 | 0 | 0 | 826300 | 0 | 1531400 | 0 | 0 | 0 | 0 | 0 | 0 | 0 | 0 | 0 |
| DIKAEELFL | SETD3; SETD3 | Q86TU7; Q6 | SETD3_HUMAN | Unmodified | 0 | 0 | 0 | 0 | 0 | 0 | 0 | 0 | 225350 | 0 | 0 | 199100 | 0 | 0 | 0 | 0 | 0 | 0 |
| DIKDTTVGI | PSMD7 | P51665 | PSMD7_HUMA | Unmodified | 0 | 0 | 0 | 0 | 0 | 0 | 0 | 0 | 297920 | 0 | 175600 | 1809800 | 0 | 0 | 186860 | 0 | 20803 | 0 |
| DIKEKLCY | POTEE; ACTG1; | Q6S8J3; P63 | POTEE_HUMAN | Unmodified | 0 | 0 | 0 | 0 | 356490 | 0 | 0 | 0 | 0 | 0 | 0 | 0 | 426410 | 78572 | 0 | 0 | 590410 | 295050 |
| DIKPAATHI | HINT3 | Q9NQE9 | HINT3_HUMAN | Unmodified | 0 | 0 | 230780 | 942610 | 3086900 | 0 | 366260 | 120360 | 2575800 | 603930 | 1003100 | 9921000 | 2085400 | 1776300 | 616780 | 899270 | 695900 | 4794900 |
| DIKPANVFI | NEK7; NEK6; NI | Q8TDX7; Q9 | NEK7_HUMAN | Unmodified | 422080 | 0 | 0 | 0 | 0 | 0 | 0 | 146200 | 0 | 0 | 0 | 3350000 | 0 | 0 | 483550 | 691060 | 0 | 0 |
| DIKPDNFLI | CSNK1A1; CSN | P48729; A0A | KC1A_HUMAN | Oxidation (M) | 435520 | 627080 | 0 | 1146400 | 0 | 0 | 0 | 0 | 0 | 0 | 0 | 1893300 | 0 | 0 | 2828600 | 1196600 | 0 | 0 |
| DIKPDNLLI | STK38 | Q15208 | STK38_HUMAN | Unmodified | 0 | 0 | 0 | 0 | 0 | 0 | 0 | 0 | 0 | 0 | 0 | 0 | 0 | 0 | 0 | 0 | 0 | 0 |
| DIKPENLLL | AURKB; AURKB | Q96GD4; J3I | AURKB_HUMA | Unmodified | 44301 | 38551 | 573220 | 108530 | 0 | 0 | 0 | 0 | 0 | 0 | 0 | 425370 | 0 | 0 | 729720 | 76893 | 0 | 0 |
| DIKPENLLV | DCLK1 | O15075 | DCLK1_HUMA | Unmodified | 0 | 0 | 0 | 0 | 0 | 0 | 0 | 0 | 0 | 0 | 0 | 0 | 0 | 0 | 0 | 0 | 0 | 0 |
| DIKPGNLLI | STK11; STK11; | Q15831; K7E | STK11_HUMAN | Unmodified | 0 | 0 | 0 | 0 | 0 | 0 | 0 | 0 | 88955 | 0 | 0 | 1035100 | 0 | 0 | 107110 | 0 | 0 | 0 |
| DIKPQLELII | MAP1B | P46821 | MAP1B_HUMA | Unmodified | 0 | 0 | 0 | 0 | 0 | 0 | 0 | 0 | 460650 | 0 | 247730 | 1277300 | 0 | 0 | 0 | 0 | 47731 | 0 |
| DIKSDNVLI | PAK2; PAK2 | Q13177; H7( | PAK2_HUMAN | Unmodified | 0 | 0 | 0 | 260030 | 0 | 0 | 0 | 0 | 0 | 0 | 0 | 422730 | 0 | 0 | 0 | 0 | 0 | 0 |
| DIKWDFTP | ATP6V1A; ATP6 | P38606; C9J | VATA_HUMAN ' | Unmodified | 0 | 0 | 0 | 0 | 0 | 0 | 0 | 0 | 0 | 0 | 0 | 1067700 | 691820 | 0 | 0 | 0 | 0 | 0 |
| DILACGRG | TJP2; TJP2; TJP2 | Q9UDY2; A0 | ZO2_HUMAN T | Unmodified | 0 | 0 | 17498 | 0 | 0 | 0 | 148880 | 99292 | 452710 | 0 | 38063 | 253190 | 0 | 0 | 0 | 0 | 0 | 0 |
| DILADLIPK | CS; N/A; CS; CS | O75390; F8V | CISY_HUMAN ( | Unmodified | 0 | 0 | 361530 | 114880 | 406940 | 0 | 391870 | 0 | 22442000 | 2672000 | 799950 | 580660 | 302970 | 0 | 117940 | 99743 | 0 | 799160 |
| DILADLIPKI | CS; N/A; CS; CS | O75390; F8V | CISY_HUMAN ( | Unmodified | 0 | 0 | 0 | 0 | 0 | 0 | 384580 | 0 | 0 | 476930 | 0 | 0 | 0 | 144530 | 0 | 0 | 0 | 162000 |
| DILAPNLQ\ | DNAAF5; DNAA | Q86Y56; H0Y | DAAF5_HUMAN | Unmodified | 0 | 0 | 20136 | 0 | 0 | 0 | 0 | 0 | 0 | 0 | 27994 | 337820 | 161670 | 0 | 0 | 0 | 0 | 0 |
| DILFPYIEEI | ENOPH1 | Q9UHY7 | ENOPH_HUMA | Unmodified | 0 | 0 | 0 | 0 | 0 | 0 | 0 | 0 | 1186800 | 355420 | 893740 | 6020500 | 546620 | 0 | 0 | 348850 | 0 | 0 |
| DILIQYDR | RPS16; RPS16 | P62249; M0F | RS16_HUMAN | Unmodified | 1332400 | 1454200 | 1482100 | 338400 | 808830 | 0 | 0 | 45736 | 157710 | 0 | 1775000 | 23652000 | 0 | 0 | 1470000 | 1504900 | 0 | 0 |
| DILIQYDRT | RPS16; RPS16 | P62249; M0F | RS16_HUMAN | Unmodified | 372310 | 0 | 181520 | 0 | 0 | 0 | 0 | 0 | 0 | 0 | 577140 | 1244200 | 0 | 0 | 294430 | 147400 | 0 | 0 |
| DILLPFQG( | CPM | P14384 | CBPM_HUMAN | Oxidation (M) | 0 | 0 | 0 | 0 | 0 | 0 | 0 | 0 | 533740 | 190360 | 0 | 0 | 0 | 0 | 0 | 0 | 0 | 0 |
| DILLRPELE | CYB5R3 | P00387 | NB5R3_HUMA | Unmodified | 0 | 0 | 132670 | 0 | 0 | 0 | 0 | 0 | 153230 | 0 | 0 | 1660300 | 0 | 0 | 0 | 0 | 0 | 0 |
| DILMHRWF | CNDP2 | Q96KP4 | CNDP2_HUMA | Oxidation (M) | 0 | 0 | 0 | 0 | 0 | 0 | 0 | 0 | 0 | 510730 | 0 | 0 | 7287600 | 7906700 | 0 | 0 | 0 | 0 |
| DILRAPVDI | CARS1; CARS1 | P49589; B4D | SYCC_HUMAN | Unmodified | 0 | 0 | 0 | 0 | 0 | 0 | 0 | 0 | 212640 | 0 | 271870 | 2442500 | 0 | 0 | 0 | 0 | 0 | 0 |
| DILVLPLDL | DHRS7; DHRS7 | Q9Y394; H0Y | DHRS7_HUMA | Unmodified | 0 | 0 | 2885900 | 0 | 61079 | 0 | 0 | 0 | 2398700 | 279820 | 1453400 | 3106300 | 0 | 0 | 0 | 751930 | 0 | 0 |
| DILVPSFDF | MGAT5 | Q09328 | MGT5A_HUMA | Unmodified | 0 | 0 | 0 | 0 | 0 | 0 | 0 | 0 | 8289800 | 1033900 | 1398100 | 1463800 | 0 | 0 | 0 | 0 | 0 | 0 |
| DIMEDAIDI | STXBP3 | O00186 | STXB3_HUMAN | Unmodified | 301700 | 179170 | 607900 | 289830 | 705560 | 0 | 0 | 38197 | 1308700 | 0 | 764350 | 3151000 | 212140 | 74890 | 814620 | 536920 | 0 | 569310 |
| DIMEDAIDI | STXBP3 | O00186 | STXB3_HUMAN | Oxidation (M) | 0 | 0 | 0 | 74278 | 203000 | 0 | 0 | 0 | 0 | 0 | 0 | 0 | 0 | 0 | 461360 | 974470 | 281240 | 0 |

| | | | | | | | | | | | | | | | | | | | | | | |
|---|---|---|---|---|---|---|---|---|---|---|---|---|---|---|---|---|---|---|---|---|---|---|
| DIMEDTIED | STXBP1; STXBP | P61764; A0A | STXB1_HUMAN | Oxidation (M) | 122510 | 22311 | 77431 | 0 | 53784 | 0 | 1265700 | 127550 | 407800 | 67681 | 0 | 1139900 | 0 | 0 | 172400 | 136340 | 0 | 112410 |
| DIMEDTIED | STXBP1; STXBP | P61764; A0A | STXB1_HUMAN | Unmodified | 0 | 0 | 0 | 0 | 0 | 0 | 701940 | 0 | 2993800 | 343590 | 0 | 1809700 | 0 | 0 | 480830 | 0 | 51524 | 324530 |
| DIMLATGK | FASN; FASN | P49327; A0A | FAS_HUMAN F | Unmodified | 174500 | 0 | 109620 | 40631 | 0 | 0 | 0 | 0 | 291100 | 74259 | 426450 | 11077000 | 0 | 0 | 430850 | 310510 | 0 | 0 |
| DIMLATGK | FASN; FASN | P49327; A0A | FAS_HUMAN F | Oxidation (M) | 0 | 0 | 0 | 0 | 83718 | 0 | 0 | 0 | 0 | 0 | 145110 | 2811600 | 0 | 0 | 0 | 272080 | 21547 | 0 |
| DINAYNCE | PRDX6 | P30041 | PRDX6_HUMAN | Unmodified | 0 | 0 | 676430 | 0 | 0 | 0 | 0 | 0 | 207660 | 108100 | 255220 | 2757800 | 0 | 0 | 432460 | 381350 | 185250 | 253850 |
| DINAYNCE | PRDX6 | P30041 | PRDX6_HUMAN | Unmodified | 0 | 0 | 0 | 0 | 0 | 0 | 0 | 0 | 0 | 0 | 941030 | 5211600 | 0 | 0 | 0 | 345970 | 0 | 0 |
| DINAYNCE | PRDX6 | P30041 | PRDX6_HUMAN | Unmodified | 0 | 0 | 0 | 0 | 0 | 0 | 0 | 0 | 0 | 0 | 0 | 6308200 | 0 | 0 | 0 | 0 | 0 | 0 |
| DINELNLPI | UBE2M | P61081 | UBC12_HUMAI | Unmodified | 1109800 | 930750 | 1020600 | 841610 | 1014800 | 0 | 941170 | 524240 | 3208300 | 1236400 | 1768000 | 5168200 | 1624300 | 1077100 | 1189900 | 1084400 | 1559700 | 1757400 |
| DINLQDED | PRPF8; PRPF8 | Q6P2Q9; I3L | PRP8_HUMAN | Unmodified | 196630 | 119850 | 49223 | 68918 | 0 | 0 | 0 | 0 | 0 | 0 | 165990 | 284610 | 0 | 0 | 0 | 78426 | 0 | 0 |
| DINQEVYN | ACO2; ACO2; A | Q99798; A2A | ACON_HUMAN | Unmodified | 0 | 0 | 0 | 0 | 315290 | 117220 | 309610 | 0 | 2982000 | 1308500 | 130490 | 18616 | 146210 | 237840 | 0 | 0 | 141000 | 158540 |
| DINRVYREI | ANXA1; ANXA1 | P04083; Q5T | ANXA1_HUMAN | Unmodified | 0 | 0 | 0 | 0 | 0 | 0 | 0 | 0 | 0 | 0 | 0 | 35822000 | 0 | 0 | 0 | 0 | 0 | 0 |
| DINSLYDVS | PLCG2; PLCG2 | P16885; A0A | PLCG2_HUMAI | Unmodified | 0 | 0 | 0 | 0 | 0 | 0 | 0 | 0 | 0 | 0 | 0 | 14106000 | 0 | 0 | 0 | 0 | 0 | 0 |
| DINWDSS( | UPF1 | Q92900 | RENT1_HUMAN | Unmodified | 1755800 | 2357800 | 441520 | 1301100 | 72230 | 0 | 0 | 0 | 969700 | 0 | 187130 | 794490 | 293100 | 71482 | 1291900 | 956590 | 33833 | 50601 |
| DIPFKENYI | MPPE1; MPPE1 | Q53F39; K7E | MPPE1_HUMAI | Unmodified | 0 | 0 | 0 | 0 | 0 | 0 | 0 | 0 | 462540 | 0 | 0 | 0 | 0 | 0 | 0 | 0 | 0 | 0 |
| DIPFSAIYFI | SLC25A13 | Q9UJS0 | CMC2_HUMAN | Unmodified | 0 | 1267300 | 0 | 0 | 0 | 0 | 0 | 0 | 1846800 | 0 | 0 | 44140 | 0 | 0 | 0 | 0 | 0 | 0 |
| DIPGLTDTT | RPS6; RPS6 | P62753; A2A | RS6_HUMAN 4 | Unmodified | 5340100 | 5290100 | 5586300 | 5091500 | 0 | 0 | 2190800 | 2217900 | 1523900 | 459080 | 0 | 40387000 | 0 | 0 | 2911200 | 0 | 0 | 0 |
| DIPGLTDTT | RPS6; RPS6 | P62753; A2A | RS6_HUMAN 4 | Unmodified | 0 | 0 | 0 | 0 | 0 | 0 | 0 | 0 | 0 | 0 | 0 | 2728800 | 0 | 0 | 0 | 0 | 0 | 0 |
| DIPKRPFSC | STK24; STK24; S | Q9Y6E0; H0Y | STK24_HUMAN | Unmodified | 0 | 0 | 0 | 0 | 2285900 | 306520 | 73244 | 0 | 0 | 574340 | 0 | 0 | 1256700 | 440140 | 0 | 0 | 736700 | 949050 |
| DIPNENEA( | AP1B1 | Q10567 | AP1B1_HUMAN | Unmodified | 0 | 0 | 0 | 0 | 0 | 0 | 0 | 0 | 0 | 0 | 0 | 0 | 0 | 0 | 0 | 0 | 0 | 0 |
| DIPNENEL( | AP2B1; AP2B1 | P63010; A0A | AP2B1_HUMAN | Unmodified | 149810 | 136520 | 216150 | 0 | 96025 | 0 | 0 | 20420 | 1016800 | 0 | 143590 | 781390 | 0 | 0 | 273560 | 225360 | 40102 | 0 |
| DIPNMFMD | ERLIN2; ERLIN2 | O94905; E5F | ERLN2_HUMAN | Oxidation (M) | 0 | 0 | 0 | 0 | 0 | 0 | 0 | 0 | 6128400 | 420170 | 0 | 0 | 0 | 0 | 0 | 0 | 0 | 0 |
| DIPNMFMD | ERLIN2; ERLIN2 | O94905; E5F | ERLN2_HUMAN | Unmodified | 0 | 0 | 0 | 0 | 0 | 0 | 0 | 0 | 7386500 | 0 | 30219 | 0 | 0 | 0 | 0 | 0 | 0 | 0 |
| DIPVVHQLI | NMT1; NMT1; N | P30419; A0A | NMT1_HUMAN | Unmodified | 0 | 0 | 0 | 0 | 0 | 0 | 0 | 0 | 192630 | 0 | 364400 | 1044000 | 0 | 0 | 0 | 0 | 0 | 0 |
| DIQAVMEE | NOSTRIN; NOS | Q8IVI9; A0A( | NOSTN_HUMA | Unmodified | 235030 | 0 | 247020 | 0 | 20071 | 46252 | 0 | 0 | 23694 | 0 | 28490 | 0 | 0 | 618610 | 787490 | 234700 | 0 | 0 |
| DIQDLKFG | PKM; PKM; PKM | P14618; B4D | KPYM_HUMAN | Oxidation (M) | 0 | 0 | 0 | 0 | 9761700 | 0 | 271960 | 0 | 0 | 209380 | 0 | 0 | 3247500 | 543570 | 0 | 0 | 2398200 | 4644900 |
| DIQDLKFG | PKM; PKM; PKM | P14618; B4D | KPYM_HUMAN | Unmodified | 0 | 130620 | 0 | 0 | 207060 | 0 | 0 | 0 | 0 | 0 | 0 | 0 | 39945 | 0 | 0 | 0 | 31223 | 0 |
| DIQDNKFIE | DLG3; DLG3 | Q92796; Q5J | DLG3_HUMAN | Unmodified | 0 | 0 | 0 | 1001300 | 668760 | 0 | 0 | 0 | 2553700 | 0 | 0 | 569050 | 0 | 0 | 0 | 0 | 0 | 0 |
| DIQDSLTVS | CAPZA2 | P47755 | CAZA2_HUMAN | Unmodified | 920760 | 0 | 0 | 0 | 0 | 0 | 0 | 0 | 0 | 0 | 236920 | 42390 | 0 | 0 | 0 | 728080 | 0 | 0 |
| DIQEDSGM | ZMPSTE24 | O75844 | FACE1_HUMAN | Unmodified | 0 | 0 | 0 | 0 | 0 | 183460 | 0 | 0 | 688280 | 0 | 0 | 0 | 0 | 0 | 0 | 0 | 0 | 0 |
| DIQEESTFS | LRPPRC; LRPPF | P42704; B8Z | LPPRC_HUMAN | Unmodified | 0 | 0 | 0 | 0 | 0 | 0 | 0 | 0 | 573840 | 0 | 0 | 0 | 0 | 0 | 0 | 0 | 0 | 0 |
| DIQEESTFS | LRPPRC; LRPPF | P42704; B8Z | LPPRC_HUMAN | Unmodified | 0 | 0 | 0 | 24468 | 0 | 0 | 0 | 0 | 752950 | 0 | 0 | 0 | 0 | 0 | 0 | 0 | 0 | 0 |
| DIQEHKFIE | DLG2; DLG2; D | Q15700; E9F | DLG2_HUMAN | Unmodified | 0 | 0 | 0 | 0 | 0 | 0 | 0 | 0 | 0 | 0 | 0 | 0 | 6586800 | 3844300 | 0 | 0 | 0 | 0 |
| DIQEHKFIE | DLG1; DLG1; D | Q12959; A0A | DLG1_HUMAN | Unmodified | 0 | 0 | 0 | 0 | 15441000 | 618910 | 0 | 0 | 548530 | 5008500 | 1620900 | 0 | 3818600 | 4682000 | 0 | 0 | 6713400 | 4343400 |
| DIQENDEE | PSMD11; PSMD | O00231; J3Q | PSD11_HUMAN | Unmodified | 0 | 0 | 0 | 0 | 0 | 0 | 0 | 0 | 0 | 0 | 604880 | 3595600 | 0 | 0 | 455180 | 0 | 0 | 0 |
| DIQLENYTI | RAET1L; ULBP2 | Q5VY80; Q9I | ULBP6_HUMAN | Unmodified | 121920 | 333350 | 0 | 838370 | 0 | 0 | 0 | 0 | 1311000 | 0 | 0 | 0 | 25657 | 0 | 524250 | 465590 | 0 | 0 |
| DIQQHKEE | GART; GART | P22102; A0A | PUR2_HUMAN | Unmodified | 0 | 0 | 0 | 117050 | 0 | 0 | 0 | 0 | 0 | 0 | 59298 | 2737000 | 0 | 0 | 396210 | 519200 | 0 | 0 |
| DIQQSLAD | EXOC3; EXOC3 | O60645; D6I | EXOC3_HUMAI | Unmodified | 0 | 0 | 1929100 | 0 | 0 | 0 | 0 | 0 | 0 | 0 | 0 | 999530 | 0 | 0 | 0 | 0 | 0 | 0 |
| DIQSLPQK | EHD4 | Q9H223 | EHD4_HUMAN | Unmodified | 1324200 | 942210 | 1079500 | 148800 | 0 | 0 | 352870 | 310030 | 0 | 0 | 0 | 2852400 | 162270 | 348420 | 2301200 | 1489500 | 0 | 0 |
| DIQTNVYIK | PSMD8; PSMD8 | P48556; K7E | PSMD8_HUMA | Unmodified | 0 | 0 | 0 | 0 | 0 | 47250 | 0 | 0 | 0 | 80812 | 0 | 3574200 | 72397 | 0 | 0 | 0 | 0 | 0 |
| DIQVPVCPI | ZFAND2B; ZFAN | Q8WV99; B4 | ZFN2B_HUMAN | Unmodified | 1747700 | 720390 | 72945 | 161120 | 0 | 0 | 0 | 0 | 79694 | 0 | 0 | 42968 | 0 | 0 | 146250 | 358210 | 89167 | 0 |
| DIQYPFLGI | NAA25 | Q14CX7 | NAA25_HUMAN | Unmodified | 0 | 0 | 0 | 0 | 0 | 0 | 0 | 0 | 0 | 0 | 0 | 0 | 0 | 0 | 0 | 0 | 0 | 0 |
| DIRAIYQVY | ABRAXAS2 | Q15018 | ABRX2_HUMAN | Unmodified | 0 | 0 | 0 | 0 | 490320 | 0 | 144280 | 0 | 0 | 62854 | 0 | 0 | 111140 | 0 | 0 | 0 | 370380 | 463810 |
| DIRDLLEED | PTPN23 | Q9H3S7 | PTN23_HUMAN | Unmodified | 0 | 0 | 0 | 0 | 0 | 0 | 0 | 0 | 0 | 0 | 0 | 0 | 0 | 0 | 0 | 0 | 0 | 0 |
| DIRDYNDA | MESD; MESD | Q14696; H0Y | MESD_HUMAN | Unmodified | 0 | 0 | 0 | 0 | 0 | 0 | 0 | 0 | 8527700 | 190160 | 248270 | 246280 | 0 | 0 | 0 | 0 | 0 | 0 |
| DIRDYNDA | MESD; MESD | Q14696; H0Y | MESD_HUMAN | Oxidation (M) | 0 | 0 | 0 | 0 | 0 | 0 | 0 | 0 | 700350 | 25267 | 0 | 28786 | 0 | 0 | 0 | 0 | 0 | 0 |
| DIREEAEKY | PSMA3; PSMA3 | P25788; H0Y | PSA3_HUMAN | Unmodified | 0 | 0 | 0 | 142990 | 571730 | 133240 | 85714 | 0 | 0 | 348140 | 0 | 0 | 753680 | 514280 | 0 | 0 | 287240 | 333570 |
| DIRESDYFT | STT3B | Q8TCJ2 | STT3B_HUMAN | Unmodified | 0 | 1073400 | 187040 | 86059 | 0 | 0 | 0 | 0 | 565280 | 0 | 0 | 0 | 0 | 581540 | 1879300 | 436420 | 0 | 0 |
| DIRESDYFT | STT3B | Q8TCJ2 | STT3B_HUMAN | Unmodified | 0 | 410560 | 362690 | 317480 | 0 | 0 | 0 | 0 | 6288400 | 1228900 | 0 | 0 | 112200 | 189820 | 220770 | 0 | 0 | 0 |
| DIRLQLEAC | PLEC; PLEC | Q15149; H0Y | PLEC_HUMAN | Unmodified | 0 | 0 | 0 | 0 | 0 | 0 | 0 | 0 | 0 | 0 | 0 | 0 | 0 | 0 | 0 | 0 | 327170 | 262300 |
| DIRNLEDIR | LAMC1 | P11047 | LAMC1_HUMAI | Unmodified | 0 | 301920 | 0 | 860580 | 135730 | 0 | 0 | 0 | 124380 | 22248 | 0 | 0 | 0 | 101900 | 0 | 0 | 132690 | 471680 |
| DIRPGAAFI | USP10; USP10 | Q14694; J3K | UBP10_HUMAN | Unmodified | 0 | 0 | 0 | 0 | 0 | 0 | 0 | 0 | 0 | 0 | 0 | 3442200 | 0 | 0 | 0 | 0 | 0 | 0 |
| DIRSDTSGI | ANXA7 | P20073 | ANXA7_HUMAN | Unmodified | 0 | 0 | 0 | 1226700 | 461750 | 530880 | 0 | 0 | 90605000 | 315460 | 1444900 | 0 | 0 | 0 | 652520 | 0 | 325470 | 1215800 |
| DISEASVFD | RPS26 | P62854 | RS26_HUMAN | Unmodified | 0 | 847430 | 86682 | 335950 | 0 | 0 | 244620 | 199850 | 101030 | 61910 | 168720 | 3866800 | 0 | 0 | 423140 | 419160 | 0 | 0 |
| DISEMFLQI | MUC1; MUC1; I | P15941; A0A | MUC1_HUMAN | Oxidation (M) | 0 | 0 | 0 | 0 | 41831 | 14772 | 0 | 0 | 7023500 | 10495 | 0 | 0 | 0 | 0 | 0 | 0 | 0 | 0 |
| DISEMFLQI | MUC1; MUC1; I | P15941; A0A | MUC1_HUMAN | Unmodified | 0 | 0 | 0 | 0 | 0 | 0 | 0 | 0 | 3164300 | 0 | 0 | 0 | 0 | 0 | 0 | 0 | 0 | 0 |
| DISGTSDPI | RASAL1 | O95294 | RASL1_HUMAN | Unmodified | 0 | 1416800 | 0 | 0 | 398730 | 0 | 0 | 0 | 2943700 | 0 | 0 | 1108700 | 0 | 0 | 0 | 0 | 65653 | 0 |
| DISLVPTRQ | PLEKHA6; PLEK | Q9Y2H5; A0A | PKHA6_HUMAN | Unmodified | 0 | 0 | 0 | 0 | 0 | 0 | 0 | 0 | 0 | 0 | 0 | 395040 | 0 | 0 | 254970 | 364440 | 0 | 0 |
| DISPFSYTA | ITGB5; ITGB5; I | P18084; V9C | ITB5_HUMAN I | Unmodified | 0 | 0 | 1159400 | 1047700 | 490880 | 0 | 0 | 0 | 235530 | 88944 | 0 | 3311500 | 0 | 0 | 1889600 | 889330 | 1041600 | 0 |
| DISPVLKDI | APRT; APRT; AP | P07741; H3E | APT_HUMAN A | Unmodified | 0 | 0 | 0 | 473870 | 0 | 0 | 0 | 0 | 0 | 0 | 0 | 12802000 | 0 | 13363000 | 0 | 5480600 | 0 | 0 |
| DISQAYYT | ANXA3; ANXA3; | P12429; D6F | ANXA3_HUMAN | Unmodified | 0 | 0 | 0 | 0 | 3443700 | 0 | 0 | 0 | 0 | 276160 | 0 | 0 | 0 | 0 | 0 | 0 | 0 | 485810 |
| DISQAYYT | ANXA3; ANXA3; | P12429; D6F | ANXA3_HUMAN | Unmodified | 0 | 0 | 0 | 0 | 35653000 | 4180600 | 0 | 0 | 0 | 0 | 0 | 1710100 | 0 | 0 | 0 | 0 | 2592000 | 3324800 |
| DISSALRVI | RNH1; RNH1; R | P13489; E9P | RINI_HUMAN R | Unmodified | 0 | 0 | 0 | 0 | 1072500 | 0 | 0 | 0 | 496840 | 0 | 0 | 0 | 994550 | 0 | 0 | 0 | 0 | 849410 |
| DISTNYYAS | HSP90B1; HSP | P14625; H0Y | ENPL_HUMAN | Unmodified | 0 | 0 | 960800 | 232250 | 635600 | 159400 | 0 | 494450 | 55094000 | 11714000 | 3808800 | 7693800 | 945550 | 574540 | 750790 | 74684 | 0 | 646030 |
| DISTNYYAS | HSP90B1; HSP | P14625; H0Y | ENPL_HUMAN | Unmodified | 0 | 162430 | 1137100 | 0 | 0 | 0 | 2916700 | 1665000 | 1.31E+08 | 54816000 | 14343000 | 9310400 | 2651900 | 1758100 | 804340 | 698040 | 0 | 3311900 |
| DISTTLNAD | HSPH1; HSPA4 | Q92598; O9 | HS105_HUMAN | Unmodified | 56021 | 24306 | 631880 | 719250 | 76223 | 0 | 0 | 34617 | 519740 | 74385 | 4793100 | 7893600 | 327100 | 0 | 895020 | 685050 | 0 | 0 |
| DITLTNDKI | KIF5B | P33176 | KINH_HUMAN | Unmodified | 0 | 0 | 0 | 0 | 0 | 0 | 0 | 0 | 0 | 0 | 0 | 4741600 | 0 | 0 | 0 | 0 | 0 | 0 |
| DITNQEEL | STEAP1 | Q9UHE8 | STEA1_HUMAN | Unmodified | 440750 | 458890 | 6075600 | 8932500 | 0 | 0 | 0 | 0 | 0 | 0 | 248580 | 0 | 4809200 | 1412300 | 10422000 | 6198800 | 0 | 0 |
| DITPLQVVL | MVP; MVP; MVP | Q14764; H3I | MVP_HUMAN I | Unmodified | 5764700 | 4505100 | 131610 | 238970 | 2185300 | 1365400 | 0 | 0 | 7780700 | 2577400 | 1250400 | 98481000 | 0 | 0 | 0 | 0 | 0 | 0 |
| DITSDSENS | FAS | P25445 | TNR6_HUMAN | Unmodified | 0 | 474860 | 79893 | 268490 | 2566200 | 314070 | 0 | 0 | 0 | 0 | 0 | 0 | 998100 | 364160 | 595920 | 0 | 0 | 0 |
| DITSDTSGI | ANXA1; ANXA1 | P04083; Q5T | ANXA1_HUMAN | Unmodified | 0 | 0 | 0 | 0 | 2629300 | 453440 | 0 | 41721 | 661430 | 100230 | 96350000 | 2.2E+09 | 0 | 0 | 0 | 613330 | 110730 | 336110 |
| DITSDTSGI | ANXA1; ANXA1 | P04083; Q5T | ANXA1_HUMAN | Unmodified | 132250 | 0 | 0 | 0 | 1.44E+08 | 28910000 | 7419700 | 0 | 1696800 | 496330 | 5.71E+08 | 3.83E+08 | 0 | 0 | 0 | 476910 | 5924300 | 15397000 |
| DITSDTSGI | ANXA1; ANXA1 | P04083; Q5T | ANXA1_HUMAN | Unmodified | 0 | 0 | 0 | 0 | 0 | 0 | 0 | 0 | 0 | 0 | 1155100 | 412710 | 0 | 0 | 0 | 0 | 0 | 0 |
| DITSSIHFE | SEPTIN9 | Q9UHD8 | SEPT9_HUMAN | Unmodified | 0 | 0 | 0 | 0 | 17511000 | 2022800 | 838550 | 0 | 371770 | 1466500 | 0 | 0 | 2272100 | 3969400 | 0 | 0 | 13951000 | 17283000 |
| DIVEMLFTC | OSTF1 | Q92882 | OSTF1_HUMAN | Unmodified | 42778000 | 62964000 | 20445000 | 73529000 | 59404000 | 46877000 | 10274000 | 211110 | 4.36E+08 | 33613000 | 10846000 | 33327000 | 38535000 | 36965000 | 46394000 | 22129000 | 9584200 | 53837000 |
| DIVEMLFTC | OSTF1 | Q92882 | OSTF1_HUMAN | Oxidation (M) | 203750 | 5.96E+08 | 62443000 | 75215000 | 1.6E+08 | 997790 | 5091600 | 2420600 | 36406000 | 0 | 1.26E+08 | 2.47E+08 | 491880 | 16546000 | 65494000 | 2732300 | 42346000 | 61965000 |
| DIVENYFMI | TRIM28; TRIM28 | Q13263; M0I | TIF1B_HUMAN | Unmodified | 0 | 0 | 0 | 0 | 550730 | 0 | 0 | 0 | 1593800 | 301950 | 0 | 0 | 202420 | 0 | 0 | 0 | 0 | 0 |
| DIVEYAELK | CYFIP1 | Q7L576 | CYFP1_HUMAN | Unmodified | 0 | 0 | 0 | 0 | 473090 | 0 | 0 | 0 | 0 | 0 | 0 | 2525600 | 0 | 0 | 0 | 823450 | 0 | 0 |
| DIVINANPA | AIMP2; AIMP2 | Q13155; A8N | AIMP2_HUMAN | Unmodified | 0 | 0 | 0 | 0 | 0 | 0 | 0 | 0 | 0 | 128000 | 767140 | 2816500 | 0 | 0 | 280890 | 0 | 0 | 0 |
| DIVNGLRS | CAPZB | B1AK88 | B1AK88_HUMA | Unmodified | 0 | 0 | 0 | 0 | 4550900 | 818730 | 1301700 | 706790 | 0 | 589270 | 0 | 0 | 2486400 | 1894400 | 0 | 0 | 1150900 | 2162900 |
| DIVPGDIVE | ATP2A2 | P16615 | AT2A2_HUMAN | Unmodified | 0 | 0 | 0 | 0 | 0 | 0 | 0 | 0 | 0 | 0 | 133020 | 586760 | 0 | 0 | 263130 | 24618 | 0 | 0 |
| DIVPGDIVE | ATP2A3 | Q93084 | AT2A3_HUMAN | Unmodified | 0 | 0 | 0 | 0 | 0 | 0 | 0 | 0 | 0 | 0 | 365570 | 778570 | 0 | 100530 | 0 | 0 | 102530 | 0 |
| DIVQFVPFI | CPNE9; CPNE8 | Q8IYJ1; Q86 | CPNE9_HUMAI | Unmodified | 2880100 | 3454200 | 5001800 | 1570600 | 0 | 0 | 199850 | 43849 | 11204000 | 914690 | 38810 | 7534100 | 0 | 0 | 6140900 | 4939600 | 0 | 0 |
| DIVQFVPFI | CPNE2 | Q96FN4 | CPNE2_HUMAI | Unmodified | 0 | 0 | 187450 | 0 | 0 | 0 | 0 | 0 | 2427700 | 0 | 0 | 47465 | 0 | 0 | 0 | 0 | 84240 | 0 |
| DIVQFVPFI | CPNE3 | O75131 | CPNE3_HUMAI | Unmodified | 0 | 0 | 0 | 0 | 0 | 0 | 0 | 0 | 1981500 | 1930500 | 0 | 0 | 0 | 0 | 0 | 0 | 0 | 0 |
| DIVQFVPYI | CPNE1; CPNE1 | Q99829; H0Y | CPNE1_HUMAI | Unmodified | 160260 | 150870 | 162310 | 0 | 87513 | 0 | 0 | 0 | 1361300 | 0 | 180700 | 1309000 | 0 | 0 | 356560 | 222260 | 0 | 103370 |
| DIVQFVPYI | CPNE1; CPNE1 | Q99829; H0Y | CPNE1_HUMAI | Unmodified | 0 | 0 | 0 | 0 | 683560 | 3747700 | 1022200 | 0 | 4278100 | 0 | 473940 | 1930600 | 0 | 0 | 0 | 0 | 319360 | 0 |
| DIVVLGVEI | PSMA7 | O14818 | PSA7_HUMAN | Unmodified | 0 | 0 | 0 | 116990 | 310600 | 0 | 0 | 0 | 631150 | 0 | 523700 | 2320500 | 288020 | 173910 | 0 | 0 | 0 | 242650 |
| DIVVQETMI | CALU; CALU | O43852; H0Y | CALU_HUMAN | Unmodified | 0 | 0 | 0 | 0 | 0 | 0 | 0 | 0 | 12543000 | 0 | 739830 | 0 | 0 | 0 | 0 | 0 | 0 | 0 |
| DIVVQETMI | CALU; CALU | O43852; H0Y | CALU_HUMAN | Oxidation (M) | 0 | 0 | 0 | 0 | 0 | 0 | 0 | 0 | 0 | 0 | 593000 | 0 | 0 | 0 | 0 | 0 | 0 | 0 |
| DIWPTRDE | ACO1 | P21399 | ACOC_HUMAN | Unmodified | 0 | 0 | 0 | 0 | 0 | 256050 | 86350 | 22169 | 504180 | 0 | 1166400 | 2856800 | 0 | 0 | 0 | 0 | 0 | 0 |
| DKADFCIIH | MYH9; MYH10 | P35579; P35 | MYH9_HUMAN | Unmodified | 172290 | 0 | 373100 | 0 | 0 | 0 | 211550 | 85116 | 0 | 0 | 0 | 3269800 | 0 | 0 | 358100 | 467090 | 0 | 0 |
| DKAPVQPQ | PSMD2; PSMD2 | Q13200; F8W | PSMD2_HUMA | Unmodified | 0 | 0 | 0 | 0 | 0 | 0 | 0 | 0 | 0 | 0 | 396320 | 0 | 0 | 0 | 0 | 0 | 0 | 0 |
| DKDDDEVI | EIF4G1; EIF4G1 | Q04637; C9J | IF4G1_HUMAN | Unmodified | 0 | 0 | 0 | 0 | 0 | 0 | 0 | 0 | 126980 | 0 | 0 | 586390 | 0 | 0 | 0 | 0 | 0 | 0 |
| DKDDDEVI | EIF4G1; EIF4G1 | Q04637; C9J | IF4G1_HUMAN | Unmodified | 0 | 0 | 0 | 0 | 69594 | 0 | 0 | 0 | 0 | 0 | 162010 | 501550 | 0 | 0 | 0 | 0 | 0 | 0 |
| DKDFAIDII | PPA1 | Q15181 | IPYR_HUMAN I | Unmodified | 0 | 0 | 0 | 0 | 1094700 | 0 | 0 | 637700 | 0 | 0 | 1171400 | 4815600 | 326780 | 0 | 0 | 0 | 0 | 0 |
| DKDFAIDII | PPA1 | Q15181 | IPYR_HUMAN I | Unmodified | 0 | 0 | 0 | 0 | 0 | 0 | 110830 | 0 | 0 | 0 | 0 | 0 | 839640 | 387230 | 0 | 0 | 0 | 1575700 |
| DKDISWFS | NCAM1; NCAM | P13591; A0A | NCAM1_HUMA | Unmodified | 0 | 0 | 0 | 0 | 0 | 0 | 0 | 0 | 431780 | 0 | 0 | 0 | 0 | 0 | 0 | 0 | 0 | 0 |
| DKDNFWM | ACTR2; ACTR2 | P61160; F5H | ARP2_HUMAN | Oxidation (M) | 231250 | 179190 | 0 | 0 | 873930 | 0 | 227940 | 0 | 0 | 0 | 0 | 297190 | 1329700 | 0 | 0 | 0 | 303000 | 0 |
| DKDNFWM | ACTR2; ACTR2 | P61160; F5H | ARP2_HUMAN | Unmodified | 141430 | 0 | 0 | 0 | 2377200 | 0 | 357490 | 0 | 489310 | 0 | 0 | 850290 | 1818500 | 804580 | 0 | 222350 | 232730 | 1287700 |
| DKDPPIPV | PDIA4; PDIA4 | P13667; A0A | PDIA4_HUMAN | Unmodified | 0 | 0 | 0 | 0 | 0 | 0 | 0 | 0 | 2246600 | 0 | 0 | 174780 | 0 | 0 | 0 | 0 | 0 | 0 |
| DKEAIQAYS | SCARB1 | B7ZKQ9 | B7ZKQ9_HUM | Oxidation (M) | 0 | 0 | 0 | 0 | 0 | 0 | 0 | 0 | 0 | 0 | 0 | 1057600 | 0 | 0 | 0 | 0 | 0 | 0 |
| DKEDQWLI | NCAM2; NCAM | O15394; H9I | NCAM2_HUMA | Unmodified | 0 | 0 | 0 | 745210 | 0 | 0 | 0 | 0 | 0 | 0 | 0 | 0 | 0 | 0 | 1260000 | 407840 | 0 | 0 |
| DKEDSDSN | AHNAK2 | Q8IVF2 | AHNK2_HUMAI | Unmodified | 157720 | 69662 | 0 | 104370 | 0 | 17782 | 0 | 0 | 0 | 0 | 0 | 674080 | 0 | 0 | 239680 | 304930 | 0 | 137070 |
| DKEIIAEYD | SRGAP2; SRGA | O75044; P0I | SRGP2_HUMAI | Unmodified | 109100 | 0 | 0 | 113750 | 4235100 | 655510 | 0 | 496570 | 0 | 0 | 0 | 1578100 | 2378700 | 2743500 | 48289 | 0 | 826670 | 2684600 |
| DKEIIAEYE | SRGAP1 | Q7Z6B7 | SRGP1_HUMAI | Unmodified | 0 | 0 | 0 | 0 | 0 | 0 | 712930 | 0 | 0 | 0 | 0 | 0 | 0 | 0 | 0 | 0 | 0 | 0 |
| DKESGGS( | SLC16A10; SLC | Q8TF71; H0Y | MOT10_HUMAI | Unmodified | 0 | 0 | 0 | 0 | 0 | 0 | 0 | 0 | 0 | 0 | 389250 | 1758600 | 0 | 0 | 0 | 0 | 0 | 0 |
| DKESGGS( | SLC16A10; SLC | Q8TF71; H0Y | MOT10_HUMAI | Unmodified | 282620 | 0 | 0 | 0 | 0 | 84862 | 0 | 0 | 265710 | 43447 | 0 | 272470 | 0 | 0 | 0 | 0 | 0 | 28893 |
| DKESGMSS | EMC1 | Q8N766 | EMC1_HUMAN | Oxidation (M) | 0 | 0 | 0 | 0 | 0 | 149400 | 0 | 0 | 2486500 | 447860 | 143720 | 277320 | 0 | 0 | 0 | 0 | 0 | 0 |
| DKESGMSS | EMC1 | Q8N766 | EMC1_HUMAN | Unmodified | 203300 | 0 | 0 | 0 | 0 | 0 | 0 | 0 | 1562500 | 0 | 0 | 0 | 0 | 0 | 0 | 0 | 0 | 0 |

| | | | | | | | | | | | | | | | | | | | | | | |
|---|---|---|---|---|---|---|---|---|---|---|---|---|---|---|---|---|---|---|---|---|---|---|
| DKESGMSS | EMC1 | Q8N766 | EMC1_HUMAN | Oxidation (M) | 0 | 0 | 0 | 0 | 0 | 0 | 0 | 0 | 992660 | 627450 | 0 | 0 | 0 | 0 | 0 | 0 | 0 | 0 |
| DKETTENEI | CHGB; CHGB | P05060; A0A | SCG1_HUMAN | Unmodified | 0 | 0 | 13318 | 0 | 0 | 0 | 0 | 17965000 | 2270900 | 516290 | 0 | 0 | 0 | 6.35E+08 | 0 | 0 | 0 | 0 |
| DKEVTALTS | CLIP2 | Q9UDT6 | CLIP2_HUMAN | Unmodified | 0 | 0 | 4007000 | 3842700 | 0 | 0 | 0 | 0 | 0 | 0 | 0 | 0 | 0 | 0 | 0 | 0 | 0 | 0 |
| DKFSFDLG | FKBP4; FKBP4 | Q02790; F5H | FKBP4_HUMAN | Unmodified | 0 | 0 | 132170 | 625860 | 0 | 0 | 0 | 0 | 0 | 166360 | 2486400 | 10822000 | 6197600 | 1252800 | 394170 | 284020 | 0 | 0 |
| DKFVEIGAI | PLCG2; PLCG2 | P16885; A0A | PLCG2_HUMAI | Unmodified | 0 | 0 | 0 | 0 | 894760 | 0 | 0 | 0 | 0 | 269960 | 0 | 4316900 | 0 | 0 | 0 | 0 | 0 | 0 |
| DKGDEEEE | CANX; CANX; C | P27824; A0A | CALX_HUMAN | Unmodified | 0 | 0 | 1771900 | 0 | 0 | 0 | 81730 | 57690 | 1155500 | 0 | 895000 | 1447400 | 0 | 0 | 1046600 | 0 | 0 | 0 |
| DKGEYTLV | FLNA; FLNA; FL | P21333; A0A | FLNA_HUMAN | Unmodified | 0 | 559940 | 902990 | 1522000 | 0 | 33235 | 257650 | 516440 | 292710 | 49658 | 0 | 1094300 | 864120 | 582410 | 907590 | 832600 | 0 | 0 |
| DKGGINLT | DRG1 | Q9Y295 | DRG1_HUMAN | Unmodified | 0 | 0 | 0 | 0 | 328180 | 0 | 0 | 88551 | 0 | 0 | 0 | 0 | 407070 | 316540 | 0 | 0 | 166940 | 142280 |
| DKGIFLHLS | PLOD3; PLOD3 | O60568; H7( | PLOD3_HUMAI | Unmodified | 0 | 0 | 0 | 0 | 0 | 0 | 0 | 0 | 1004200 | 0 | 0 | 0 | 0 | 0 | 0 | 0 | 0 | 140840 |
| DKGLQTSQ | CALR; CALR; C/ | P27797; A0A | CALR_HUMAN | Unmodified | 0 | 0 | 0 | 0 | 0 | 0 | 0 | 0 | 1012200 | 403770 | 92752 | 0 | 0 | 0 | 0 | 0 | 0 | 0 |
| DKGLQTSQ | CALR; CALR; C/ | P27797; A0A | CALR_HUMAN | Unmodified | 0 | 0 | 0 | 0 | 0 | 0 | 7971600 | 107110 | 0 | 50178000 | 269290 | 0 | 0 | 0 | 0 | 0 | 0 | 0 |
| DKGQDDFI | UNC13D; UNC: | Q70J99; K7E | UN13D_HUMA | Unmodified | 89662 | 113650 | 0 | 0 | 0 | 0 | 0 | 0 | 0 | 0 | 0 | 0 | 0 | 0 | 245680 | 169430 | 0 | 0 |
| DKGTGREC | HSPA9; HSPA9; | P38646; H0Y | GRP75_HUMAI | Unmodified | 0 | 0 | 0 | 0 | 0 | 0 | 0 | 0 | 606980 | 99169 | 0 | 0 | 0 | 0 | 0 | 0 | 0 | 0 |
| DKISSSSFII | TMTC3 | Q6ZXV5 | TMTC3_HUMAI | Unmodified | 0 | 0 | 0 | 0 | 0 | 535540 | 0 | 0 | 151590 | 0 | 0 | 35840 | 0 | 0 | 0 | 0 | 0 | 0 |
| DKITQYEK | EIF3J; EIF3J; EIF | O75822; H0Y | EIF3J_HUMAN I | Unmodified | 0 | 0 | 0 | 0 | 106740 | 0 | 0 | 0 | 319210 | 0 | 98079 | 1189500 | 0 | 0 | 0 | 43078 | 214040 | 352420 |
| DKKAEGAA | BASP1; BASP1 | P80723; U3K | BASP1_HUMAN | Unmodified | 12949 | 388250 | 0 | 0 | 13775000 | 3051600 | 393810 | 632190 | 4633500 | 1640600 | 0 | 0 | 204970 | 224590 | 0 | 0 | 748610 | 5489000 |
| DKKVELFH | PDIA5; PDIA5 | Q14554; C9J | PDIA5_HUMAN | Unmodified | 0 | 0 | 0 | 0 | 0 | 444840 | 205650 | 0 | 10220000 | 1917600 | 0 | 0 | 0 | 0 | 0 | 0 | 0 | 118260 |
| DKLDPDM/ | PLOD2; PLOD2 | O00469; E7E | PLOD2_HUMAI | Unmodified | 0 | 0 | 0 | 0 | 0 | 0 | 0 | 0 | 1541100 | 0 | 0 | 0 | 0 | 0 | 0 | 0 | 0 | 0 |
| DKLESEME | SFPQ; SFPQ | P23246; H0Y | SFPQ_HUMAN | Unmodified | 0 | 0 | 0 | 28021 | 0 | 0 | 0 | 0 | 970770 | 366710 | 2286700 | 3466300 | 0 | 0 | 99132 | 0 | 0 | 0 |
| DKLNTQST | NRP1; NRP1; N | O14786; E9F | NRP1_HUMAN | Unmodified | 0 | 0 | 0 | 0 | 13762000 | 2343900 | 105180 | 0 | 222170 | 0 | 0 | 0 | 525140 | 611500 | 0 | 0 | 3693600 | 25177000 |
| DKLPQPVQ | ITGB1; ITGB1; I | P05556; H7( | ITB1_HUMAN I | Unmodified | 3957700 | 905520 | 5000800 | 4169100 | 8187600 | 3093000 | 0 | 0 | 31004000 | 1250000 | 1686900 | 13800000 | 522090 | 0 | 5706700 | 7113900 | 1588200 | 7138500 |
| DKLPQSQL | RPS6KA1; RPS6 | Q15418; Q5 | KS6A1_HUMAN | Unmodified | 101380 | 0 | 0 | 142690 | 1686200 | 0 | 271980 | 152510 | 1266200 | 949400 | 0 | 1194200 | 576820 | 1039900 | 0 | 0 | 1095900 | 1770800 |
| DKLSELQLI | TMED9 | Q9BVK6 | TMED9_HUMAI | Unmodified | 0 | 0 | 0 | 0 | 0 | 0 | 0 | 0 | 21250000 | 1021400 | 0 | 1022700 | 0 | 0 | 206070 | 116580 | 0 | 0 |
| DKLTELQLF | TMED4 | Q7Z7H5 | TMED4_HUMAI | Unmodified | 78837 | 0 | 89624 | 0 | 0 | 0 | 134700 | 789640 | 22300000 | 1309700 | 802530 | 295080 | 0 | 0 | 127660 | 110940 | 0 | 82055 |
| DKMESFFL | MAN1B1; MAN1 | Q9UKM7; H( | MA1B1_HUMAI | Unmodified | 0 | 0 | 0 | 0 | 0 | 0 | 0 | 0 | 3585600 | 0 | 0 | 0 | 0 | 0 | 0 | 0 | 0 | 0 |
| DKMESFFL | MAN1B1; MAN1 | Q9UKM7; H( | MA1B1_HUMAI | Oxidation (M) | 0 | 0 | 0 | 0 | 0 | 0 | 471220 | 86412 | 5358200 | 242760 | 0 | 144730 | 0 | 790330 | 0 | 0 | 0 | 0 |
| DKNELVQK | YWHAZ; YWHA | P63104; E5R | 1433Z_HUMAN | Unmodified | 45206 | 0 | 0 | 0 | 2232400 | 0 | 1436500 | 0 | 893320 | 1372200 | 50006 | 0 | 0 | 506580 | 0 | 0 | 4646900 | 2541200 |
| DKNEQAFE | RPA1; RPA1 | P27694; I3L4 | RFA1_HUMAN | Unmodified | 0 | 0 | 0 | 0 | 0 | 0 | 0 | 77720 | 0 | 0 | 0 | 3757600 | 0 | 0 | 0 | 0 | 0 | 0 |
| DKNHPAV | GUSB | P08236 | BGLR_HUMAN | Oxidation (M) | 0 | 0 | 0 | 0 | 0 | 0 | 0 | 0 | 1810000 | 0 | 0 | 0 | 0 | 270490 | 0 | 0 | 0 | 0 |
| DKNHPAV | GUSB | P08236 | BGLR_HUMAN | Unmodified | 0 | 0 | 0 | 0 | 0 | 0 | 0 | 0 | 1298900 | 0 | 0 | 0 | 0 | 0 | 0 | 0 | 0 | 0 |
| DKNNKEPI | FERMT1 | Q9BQL6 | FERM1_HUMAI | Unmodified | 0 | 0 | 0 | 0 | 1409200 | 0 | 0 | 0 | 0 | 0 | 0 | 0 | 0 | 0 | 0 | 0 | 0 | 3939500 |
| DKNQDLC | PDIA5; PDIA5 | Q14554; H7( | PDIA5_HUMAN | Unmodified | 0 | 0 | 0 | 0 | 0 | 0 | 0 | 0 | 6157400 | 1874500 | 81999 | 0 | 0 | 0 | 20020 | 0 | 0 | 0 |
| DKNTPSPF | GCLC; GCLC; G | P48506; A0A | GSH1_HUMAN | Unmodified | 0 | 0 | 0 | 0 | 0 | 0 | 0 | 0 | 248370 | 0 | 0 | 1936000 | 0 | 0 | 0 | 0 | 0 | 0 |
| DKPELQFP | CNP; CNP; CNI | P09543; K7E | CN37_HUMAN | Unmodified | 145520 | 290040 | 143950 | 1057000 | 6468000 | 923860 | 0 | 0 | 488720 | 1015400 | 607640 | 1061200 | 4208500 | 1310200 | 1382600 | 142340 | 3274200 | 5592000 |
| DKPHVNV | TUFM; TUFM | P49411; H3E | EFTU_HUMAN | Unmodified | 0 | 0 | 0 | 0 | 0 | 0 | 0 | 184970 | 4613100 | 213610 | 126160 | 0 | 0 | 0 | 0 | 0 | 0 | 0 |
| DKPLKDVIL | PPIB; PPIB; PPII | P23284; A0A | PPIB_HUMAN F | Unmodified | 0 | 0 | 0 | 0 | 0 | 0 | 0 | 0 | 1842800 | 140080 | 256510 | 0 | 0 | 0 | 0 | 0 | 0 | 0 |
| DKPMYDEI | EHD3 | Q9NZN3 | EHD3_HUMAN | Oxidation (M) | 1078600 | 0 | 674540 | 1894200 | 0 | 0 | 0 | 0 | 0 | 0 | 0 | 0 | 0 | 120240 | 2746100 | 0 | 0 | 0 |
| DKPSVEPV | HBS1L; HBS1L; | Q9Y450; J3Q | HBS1L_HUMAI | Unmodified | 514350 | 0 | 525770 | 286800 | 0 | 643230 | 0 | 0 | 0 | 0 | 0 | 0 | 0 | 0 | 0 | 0 | 0 | 0 |
| DKPTYDEIF | EHD1; EHD1 | Q9H4M9; A0 | EHD1_HUMAN | Unmodified | 0 | 379940 | 724990 | 759760 | 1277100 | 0 | 1364400 | 0 | 0 | 0 | 0 | 11008000 | 0 | 0 | 3628100 | 1398200 | 0 | 1872600 |
| DKPVYDEL | EHD4 | Q9H223 | EHD4_HUMAN | Unmodified | 1707600 | 2612100 | 1264700 | 1989800 | 6403200 | 0 | 0 | 0 | 0 | 0 | 0 | 1311800 | 2665400 | 2362800 | 3868900 | 1757600 | 2126500 | 8466100 |
| DKQDEEQI | CALR; CALR | P27797; K7E | CALR_HUMAN | Unmodified | 0 | 0 | 0 | 0 | 0 | 0 | 0 | 0 | 764280 | 57818 | 0 | 0 | 0 | 0 | 0 | 0 | 0 | 0 |
| DKQGDITE | VPS36; VPS36 | Q86VN1; A0 | VPS36_HUMAN | Unmodified | 6278300 | 1278800 | 996350 | 1547000 | 0 | 0 | 0 | 0 | 470940 | 0 | 0 | 375730 | 2527100 | 748250 | 1458300 | 1070700 | 0 | 0 |
| DKQMELLE | HTATIP2; HTATI | Q9BUP3; E9I | HTAI2_HUMAN | Unmodified | 1895100 | 933070 | 0 | 0 | 0 | 0 | 0 | 25548 | 593550 | 203230 | 67594 | 15540000 | 0 | 0 | 0 | 0 | 0 | 0 |
| DKQPNAQI | CYFIP1 | Q7L576 | CYFP1_HUMAN | Unmodified | 491630 | 215510 | 562640 | 496530 | 0 | 0 | 0 | 0 | 0 | 0 | 0 | 3093900 | 0 | 0 | 796420 | 722990 | 0 | 0 |
| DKQTVAYR | SIAE | Q9HAT2 | SIAE_HUMAN S | Unmodified | 0 | 0 | 0 | 0 | 0 | 0 | 0 | 0 | 0 | 359920 | 0 | 0 | 0 | 0 | 0 | 0 | 0 | 0 |
| DKSKYDEIF | EHD2 | Q9NZN4 | EHD2_HUMAN | Unmodified | 0 | 0 | 0 | 0 | 721410 | 171180 | 0 | 0 | 0 | 338580 | 0 | 0 | 309040 | 0 | 0 | 0 | 0 | 0 |
| DKTIKMWE | PAFAH1B1; PAF | P43034; I3L3 | LIS1_HUMAN P | Oxidation (M) | 0 | 0 | 0 | 0 | 0 | 0 | 0 | 0 | 5242400 | 937140 | 0 | 319650 | 0 | 0 | 0 | 71812 | 0 | 0 |
| DKTYSYLNI | NPC2; NPC2; N | P61916; J3KI | NPC2_HUMAN | Unmodified | 0 | 0 | 0 | 0 | 0 | 0 | 112380 | 110740 | 0 | 864670 | 0 | 0 | 0 | 0 | 0 | 0 | 0 | 0 |
| DKVAELEN | CD58; CD58; C | P19256; H0Y | LFA3_HUMAN I | Unmodified | 0 | 0 | 0 | 0 | 1760100 | 0 | 0 | 0 | 512290 | 367330 | 264960 | 0 | 1777700 | 0 | 0 | 0 | 676470 | 0 |
| DKVFQGNI | EDIL3 | O43854 | EDIL3_HUMAN | Unmodified | 0 | 0 | 0 | 0 | 1.92E+08 | 376560 | 2403200 | 0 | 0 | 715870 | 0 | 0 | 0 | 0 | 0 | 0 | 1282700 | 50805000 |
| DKVGNVEY | ATP1B1 | P05026 | AT1B1_HUMAN | Unmodified | 129170 | 0 | 0 | 1798500 | 8036800 | 491000 | 1048800 | 0 | 7593700 | 1260800 | 827950 | 1171600 | 3269300 | 2215600 | 493540 | 218770 | 815750 | 2003300 |
| DKVMSEFN | CD63; CD63; C | P08962; F8V | CD63_HUMAN | Oxidation (M) | 705770 | 1872200 | 0 | 0 | 0 | 0 | 0 | 0 | 0 | 963460 | 56982 | 548390 | 0 | 0 | 155920 | 226000 | 0 | 0 |
| DKVMSEFN | CD63; CD63; C | P08962; F8V | CD63_HUMAN | Unmodified | 1760200 | 841550 | 0 | 0 | 0 | 0 | 0 | 0 | 342240 | 1048000 | 0 | 0 | 0 | 0 | 0 | 0 | 0 | 0 |
| DKVQAGD | RUVBL2; RUVBI | Q9Y230; M0I | RUVB2_HUMAI | Unmodified | 63605 | 75344 | 0 | 63183 | 1363100 | 0 | 262870 | 791180 | 0 | 114930 | 1734200 | 3103000 | 649880 | 733160 | 110850 | 65964 | 170040 | 1026200 |
| DKVQAGD | RUVBL2; RUVBI | Q9Y230; M0I | RUVB2_HUMAI | Unmodified | 0 | 0 | 0 | 0 | 740230 | 0 | 289010 | 140900 | 0 | 0 | 0 | 0 | 682430 | 256070 | 0 | 0 | 127610 | 0 |
| DKVSLIKDE | STXBP3 | O00186 | STXB3_HUMAN | Unmodified | 0 | 0 | 0 | 0 | 0 | 0 | 0 | 0 | 0 | 0 | 0 | 0 | 4487000 | 0 | 0 | 0 | 0 | 0 |
| DKVTYTLLF | SLC4A4 | Q9Y6R1 | S4A4_HUMAN | Unmodified | 0 | 0 | 0 | 0 | 5244200 | 456000 | 50053 | 40011 | 0 | 0 | 0 | 0 | 1594400 | 870310 | 0 | 0 | 0 | 692390 |
| DKWLAPD | PRMT1; PRMT1; | Q99873; E9F | ANM1_HUMAN | Unmodified | 0 | 0 | 0 | 0 | 0 | 0 | 0 | 0 | 0 | 0 | 1785800 | 509800 | 14172 | 0 | 0 | 0 | 0 | 0 |
| DKWSPALC | UBE2NL; UBE2I | Q5JXB2; P61 | UE2NL_HUMAI | Unmodified | 413720 | 1415800 | 0 | 0 | 0 | 0 | 0 | 0 | 341440 | 370640 | 0 | 9006900 | 0 | 0 | 0 | 1647800 | 85708 | 0 |
| DKYEPAAV | ATP1A1 | P05023 | AT1A1_HUMAN | Unmodified | 270690 | 0 | 208830 | 607120 | 0 | 0 | 0 | 0 | 486970 | 341480 | 693910 | 3543100 | 115020 | 0 | 2863100 | 1435700 | 20700 | 97705 |
| DKYEPAAV | ATP1A1 | P05023 | AT1A1_HUMAN | Unmodified | 0 | 0 | 1232900 | 390490 | 0 | 0 | 183290 | 0 | 578340 | 129840 | 3677100 | 8241900 | 0 | 0 | 735670 | 1154900 | 0 | 0 |
| DKYPNLQV | IMPDH2; IMPDI | P12268; A0A | IMDH2_HUMAI | Unmodified | 0 | 0 | 1862100 | 0 | 4177300 | 861740 | 7695400 | 1957600 | 2380900 | 2127500 | 1929700 | 5143500 | 1838300 | 0 | 0 | 0 | 3303000 | 0 |
| DLAAAIDRI | RARS2 | Q5T160 | SYRM_HUMAN | Oxidation (M) | 0 | 0 | 0 | 0 | 0 | 87129 | 0 | 0 | 0 | 0 | 0 | 0 | 0 | 0 | 0 | 0 | 0 | 0 |
| DLAALGDK | SPTAN1; SPTAN | Q13813; A0/ | SPTN1_HUMAI | Unmodified | 0 | 0 | 0 | 0 | 0 | 0 | 0 | 0 | 123960 | 0 | 68910 | 1745400 | 0 | 0 | 0 | 0 | 0 | 0 |
| DLAARNCL | PTK7; PTK7; PTI | Q13308; H0 | PTK7_HUMAN I | Unmodified | 0 | 0 | 0 | 0 | 5443300 | 24599000 | 1305300 | 189530 | 0 | 459380 | 158390 | 252630 | 924860 | 682490 | 0 | 0 | 541160 | 789150 |
| DLAARNCL | PTK7; PTK7; PTI | Q13308; H0 | PTK7_HUMAN I | Unmodified | 0 | 0 | 0 | 0 | 2379500 | 0 | 223560 | 1844900 | 0 | 857390 | 0 | 0 | 1582400 | 2949200 | 0 | 0 | 2820200 | 2963700 |
| DLAARNILV | EPHB3; EPHB1; | P54753; P54 | EPHB3_HUMAI | Unmodified | 0 | 0 | 0 | 0 | 0 | 0 | 0 | 0 | 0 | 0 | 0 | 0 | 0 | 0 | 0 | 0 | 276680 | 0 |
| DLADELAL | LDHA; LDHA; LI | P00338; F5G | LDHA_HUMAN | Unmodified | 82513 | 152450 | 105360 | 376150 | 1063900 | 368490 | 97111 | 91161 | 547620 | 770180 | 602120 | 902050 | 941780 | 142130 | 871560 | 145050 | 317110 | 931960 |
| DLADELAL | LDHA; LDHA; LI | P00338; F5G | LDHA_HUMAN | Unmodified | 0 | 0 | 0 | 22976 | 76281 | 0 | 0 | 0 | 0 | 40525 | 0 | 0 | 0 | 0 | 199180 | 0 | 0 | 0 |
| DLADKETLI | CIT; CIT; CIT; CI | O14578; A0/ | CTRO_HUMAN | Unmodified | 0 | 0 | 0 | 0 | 8957500 | 1058300 | 0 | 0 | 0 | 143090 | 0 | 0 | 0 | 0 | 0 | 0 | 0 | 0 |
| DLADYASK | UBE4A; UBE4A | Q14139; B7 | UBE4A_HUMAI | Unmodified | 534720 | 433890 | 0 | 0 | 0 | 0 | 0 | 0 | 0 | 0 | 0 | 460910 | 0 | 0 | 0 | 0 | 0 | 0 |
| DLAEDAPW | FLNC; FLNB; FL | Q14315; O7 | FLNC_HUMAN | Unmodified | 257840 | 720590 | 1423700 | 2397100 | 4651600 | 935400 | 527930 | 723100 | 352590 | 0 | 153350 | 1947000 | 2192500 | 1573600 | 1346500 | 1615900 | 815590 | 1118800 |
| DLAEDAPW | FLNC; FLNB; FL | Q14315; O7 | FLNC_HUMAN | Unmodified | 0 | 0 | 516510 | 0 | 2926900 | 0 | 739160 | 286950 | 261020 | 0 | 0 | 614150 | 0 | 0 | 0 | 0 | 0 | 0 |
| DLAEDLYD | PARVA; PARVA | Q9NVD7; E9 | PARVA_HUMAI | Unmodified | 0 | 0 | 0 | 0 | 274450 | 0 | 0 | 0 | 0 | 0 | 0 | 0 | 0 | 0 | 0 | 0 | 0 | 378380 |
| DLAEITTLD | ERP44 | Q9BS26 | ERP44_HUMAN | Unmodified | 0 | 0 | 0 | 0 | 0 | 0 | 2457100 | 475300 | 34687000 | 207030 | 1377800 | 925590 | 0 | 0 | 0 | 0 | 0 | 0 |
| DLAEITTLD | ERP44 | Q9BS26 | ERP44_HUMAN | Unmodified | 0 | 0 | 0 | 0 | 0 | 0 | 0 | 0 | 2313000 | 0 | 0 | 0 | 0 | 0 | 0 | 0 | 0 | 0 |
| DLAEVGEG | TNKS1BP1 | Q9C0C2 | TB182_HUMAN | Unmodified | 0 | 0 | 329460 | 196180 | 0 | 0 | 0 | 0 | 0 | 0 | 0 | 646580 | 97624 | 0 | 0 | 509140 | 0 | 0 |
| DLAFVDPD | CNNM4; CNNM | Q6P4Q7; Q9 | CNNM4_HUMA | Unmodified | 0 | 0 | 153320 | 0 | 0 | 0 | 0 | 0 | 0 | 0 | 0 | 2430200 | 0 | 0 | 0 | 0 | 0 | 0 |
| DLAFVDPE | CNNM3 | Q8NE01 | CNNM3_HUMA | Unmodified | 0 | 0 | 0 | 0 | 0 | 0 | 0 | 0 | 0 | 0 | 89023 | 333580 | 0 | 0 | 0 | 66455 | 0 | 0 |
| DLAGCIHG | IDH2 | P48735 | IDHP_HUMAN | Unmodified | 0 | 0 | 0 | 0 | 0 | 0 | 151290 | 0 | 19621000 | 2913100 | 0 | 0 | 0 | 0 | 0 | 0 | 122210 | 0 |
| DLAGSIIGK | HNRNPK; HNRI | P61978; S4F | HNRPK_HUMA | Unmodified | 0 | 0 | 0 | 1521900 | 1750300 | 686660 | 397900 | 359370 | 603050 | 820720 | 0 | 0 | 1662500 | 262650 | 0 | 0 | 123090 | 377470 |
| DLAGSIIGK | HNRNPK; HNRI | P61978; S4F | HNRPK_HUMA | Unmodified | 0 | 0 | 0 | 0 | 714440 | 78975 | 0 | 0 | 0 | 498670 | 0 | 0 | 0 | 0 | 0 | 0 | 702970 | 0 |
| DLAHFPAV | POGLUT2 | Q6UW63 | PLGT2_HUMAN | Unmodified | 0 | 0 | 0 | 0 | 0 | 0 | 0 | 0 | 917460 | 0 | 0 | 0 | 0 | 0 | 0 | 0 | 0 | 0 |
| DLAHTPSQ | CCNK; CCNK; ( | O75909; G3 | CCNK_HUMAN | Unmodified | 0 | 138710 | 296960 | 282630 | 0 | 0 | 0 | 0 | 0 | 0 | 0 | 866470 | 0 | 0 | 3034200 | 0 | 0 | 0 |
| DLAILQCH | LYPLA2; LYPLA | O95372; Q5 | LYPA2_HUMAN | Oxidation (M) | 253250 | 0 | 73236 | 0 | 0 | 0 | 86576 | 63435 | 565220 | 0 | 347200 | 4549000 | 0 | 0 | 321530 | 209430 | 0 | 0 |
| DLAILQCH | LYPLA2; LYPLA | O95372; Q5 | LYPA2_HUMAN | Oxidation (M) | 699700 | 384310 | 0 | 429700 | 1754300 | 0 | 2853400 | 0 | 0 | 0 | 1069900 | 1237700 | 0 | 980150 | 957670 | 211950 | 949580 | 3160800 |
| DLAKDITSI | ANXA1; ANXA1 | P04083; Q5T | ANXA1_HUMAI | Unmodified | 0 | 0 | 0 | 0 | 0 | 57655 | 0 | 13341 | 0 | 0 | 0 | 2050800 | 91984 | 0 | 0 | 0 | 66247 | 0 |
| DLALGLVP | MYO18A; MYO1 | Q92614; A0/ | MY18A_HUMAI | Unmodified | 0 | 0 | 0 | 60192 | 0 | 0 | 0 | 0 | 0 | 281140 | 0 | 1089300 | 349800 | 0 | 0 | 0 | 0 | 0 |
| DLAMADLE | TRAF2 | Q12933 | TRAF2_HUMAN | Oxidation (M) | 0 | 0 | 0 | 0 | 1642500 | 784510 | 0 | 0 | 0 | 0 | 67343 | 124610 | 0 | 0 | 0 | 0 | 0 | 0 |
| DLAMADLE | TRAF2 | Q12933 | TRAF2_HUMAN | Unmodified | 690400 | 946390 | 0 | 0 | 2728500 | 653660 | 0 | 0 | 0 | 0 | 0 | 4923600 | 0 | 0 | 0 | 0 | 0 | 0 |
| DLAPIGIR | HSD17B10 | Q99714 | HCD2_HUMAN | Unmodified | 103630 | 36256 | 0 | 0 | 0 | 0 | 0 | 0 | 4826700 | 441150 | 0 | 1010500 | 0 | 0 | 0 | 0 | 0 | 44452 |
| DLAPPSEP | TNC; TNC; TNC | P24821; F5H | TENA_HUMAN | Unmodified | 0 | 561760 | 0 | 0 | 0 | 0 | 0 | 0 | 840540 | 0 | 0 | 0 | 0 | 0 | 285390 | 0 | 1088700 | 0 |
| DLAPYLPS | GCN1 | Q92616 | GCN1_HUMAN | Unmodified | 0 | 0 | 0 | 0 | 0 | 0 | 0 | 0 | 0 | 0 | 0 | 1637200 | 0 | 0 | 0 | 0 | 0 | 0 |
| DLAQCVNE | VAV1; VAV1; VA | P15498; Q9 | VAV_HUMAN P | Unmodified | 0 | 0 | 0 | 0 | 0 | 0 | 0 | 0 | 0 | 0 | 0 | 417200 | 0 | 0 | 0 | 0 | 0 | 0 |
| DLAQINDL | MYO18A; MYO1 | Q92614; A0/ | MY18A_HUMAI | Unmodified | 0 | 0 | 0 | 0 | 0 | 0 | 0 | 0 | 0 | 0 | 0 | 0 | 0 | 0 | 0 | 0 | 0 | 0 |
| DLAQYDAA | NUCB1; NUCB | Q02818; H7I | NUCB1_HUMA | Unmodified | 0 | 0 | 0 | 0 | 0 | 0 | 0 | 0 | 1214100 | 0 | 309540 | 0 | 0 | 0 | 0 | 0 | 0 | 0 |
| DLASVNNL | SPTAN1; SPTAN | Q13813; A0/ | SPTN1_HUMAI | Unmodified | 0 | 1229600 | 0 | 0 | 297040 | 0 | 0 | 47523 | 0 | 155360 | 33628 | 2376200 | 0 | 0 | 0 | 0 | 0 | 0 |
| DLATVAFCI | CTBP2 | P56545 | CTBP2_HUMAN | Unmodified | 0 | 0 | 0 | 0 | 249690 | 0 | 0 | 391100 | 0 | 0 | 0 | 1317100 | 0 | 0 | 0 | 0 | 37693 | 648980 |
| DLAVAGPE | MVP | Q14764 | MVP_HUMAN N | Oxidation (M) | 0 | 0 | 0 | 0 | 0 | 0 | 0 | 0 | 0 | 0 | 0 | 2268500 | 382600 | 0 | 0 | 0 | 0 | 0 |
| DLAVAGPE | MVP | Q14764 | MVP_HUMAN N | Unmodified | 0 | 0 | 0 | 0 | 0 | 0 | 0 | 0 | 0 | 0 | 0 | 2511800 | 0 | 0 | 0 | 0 | 0 | 0 |
| DLCQRAEY | SULF2 | Q8IWU5 | SULF2_HUMAN | Unmodified | 0 | 0 | 0 | 0 | 0 | 0 | 0 | 0 | 0 | 0 | 0 | 0 | 0 | 0 | 0 | 0 | 0 | 1045700 |
| DLDADREV | RAB4A; RAB4A | P20338; A0A | RAB4A_HUMAI | Unmodified | 855510 | 528520 | 918230 | 1298000 | 0 | 0 | 0 | 0 | 605600 | 41124 | 226010 | 0 | 0 | 0 | 1011600 | 732230 | 0 | 0 |
| DLDDNLFG | TOR1A; TOR1A | O14656; A0/ | TOR1A_HUMAI | Unmodified | 0 | 0 | 0 | 0 | 0 | 0 | 0 | 0 | 2189000 | 0 | 0 | 0 | 0 | 0 | 0 | 0 | 0 | 0 |
| DLDDNLFG | TOR1A; TOR1A | O14656; A0/ | TOR1A_HUMAI | Unmodified | 0 | 0 | 0 | 0 | 0 | 0 | 0 | 0 | 708740 | 0 | 0 | 0 | 0 | 0 | 0 | 0 | 0 | 0 |
| DLDEVLQT | SBDS; SBDS; SI | Q9Y3A5; A0/ | SBDS_HUMAN | Unmodified | 1962900 | 2683600 | 550820 | 0 | 1990200 | 700430 | 502360 | 160550 | 660350 | 381270 | 751950 | 2729100 | 3139200 | 2226100 | 0 | 3613100 | 92337 | 1870600 |
| DLDFLHSV | CASK; CASK; C/ | O14936; A0/ | CSKP_HUMAN | Unmodified | 0 | 0 | 0 | 558030 | 1034700 | 132830 | 0 | 0 | 0 | 238250 | 0 | 0 | 2043900 | 1252100 | 0 | 0 | 897740 | 728120 |
| DLDFLHSV | CASK; CASK; C/ | O14936; A0/ | CSKP_HUMAN | Unmodified | 0 | 0 | 0 | 0 | 658460 | 0 | 158350 | 0 | 0 | 0 | 0 | 0 | 1071600 | 2389300 | 0 | 0 | 222470 | 0 |
| DLDGNGYI | ITGA5 | P08648 | ITA5_HUMAN I | Unmodified | 0 | 54239 | 0 | 0 | 0 | 0 | 0 | 0 | 0 | 0 | 58970 | 0 | 0 | 0 | 799710 | 811910 | 0 | 0 |
| DLDIFDR | MLEC; MLEC; M | Q14165; H0 | MLEC_HUMAN | Unmodified | 0 | 0 | 13087 | 280540 | 0 | 53081 | 0 | 0 | 1331700 | 0 | 0 | 0 | 0 | 0 | 0 | 190370 | 0 | 0 |

| | | | | | | | | | | | | | | | | | | | | | | |
|---|---|---|---|---|---|---|---|---|---|---|---|---|---|---|---|---|---|---|---|---|---|---|
| DLDIFDRV( | MLEC; MLEC; M | Q14165; H0' | MLEC_HUMAN | Unmodified | 0 | 0 | 0 | 0 | 0 | 0 | 0 | 0 | 1842300 | 0 | 0 | 0 | 0 | 0 | 0 | 0 | 0 | 0 |
| DLDIFDRV( | MLEC; MLEC; M | Q14165; H0' | MLEC_HUMAN | Unmodified | 0 | 0 | 0 | 0 | 392720 | 0 | 0 | 0 | 10274000 | 47278 | 0 | 0 | 0 | 0 | 0 | 0 | 0 | 0 |
| DLDIIDNYD | FLNB; FLNB | O75369; E7E | FLNB_HUMAN | Unmodified | 0 | 0 | 1200400 | 263580 | 1443600 | 995880 | 0 | 0 | 0 | 0 | 0 | 3561700 | 0 | 0 | 919050 | 559470 | 0 | 0 |
| DLDINRPG | NDRG4; NDRG | Q9ULP0; B7Z | NDRG4_HUMA | Unmodified | 0 | 0 | 0 | 0 | 1443700 | 0 | 246670 | 0 | 2600600 | 1112400 | 287140 | 1441700 | 0 | 0 | 0 | 0 | 334990 | 0 |
| DLDNAEEK | RABGAP1 | Q9Y3P9 | RBGP1_HUMAI | Unmodified | 0 | 0 | 0 | 0 | 2931500 | 265080 | 480530 | 0 | 0 | 659110 | 0 | 0 | 1713200 | 148920 | 0 | 0 | 204800 | 729800 |
| DLDPEREV | RAB4B; RAB4B | P61018; M0F | RAB4B_HUMAN | Unmodified | 244840 | 194250 | 0 | 0 | 0 | 0 | 0 | 0 | 339960 | 25193 | 0 | 1408000 | 0 | 0 | 297180 | 177000 | 0 | 0 |
| DLDPIGKAT | PDCD6IP | Q8WUM4 | PDC6I_HUMAN | Unmodified | 0 | 0 | 0 | 0 | 2531200 | 294700 | 308480 | 255460 | 0 | 692710 | 0 | 0 | 1548900 | 1328400 | 0 | 0 | 389420 | 2534400 |
| DLDQASLA | TLN1 | Q9Y490 | TLN1_HUMAN | Unmodified | 0 | 0 | 0 | 2099900 | 0 | 0 | 0 | 0 | 596200 | 0 | 353500 | 3156300 | 0 | 0 | 0 | 0 | 0 | 0 |
| DLDRIAGQ | RPS19; RPS19 | P39019; A0A | RS19_HUMAN | Unmodified | 2759600 | 2305800 | 1120200 | 999240 | 0 | 0 | 42524 | 0 | 766460 | 0 | 294820 | 22276000 | 785580 | 0 | 2022500 | 2086900 | 0 | 0 |
| DLDSLHRF | TXNDC5 | Q8NBS9 | TXND5_HUMAI | Unmodified | 0 | 0 | 0 | 0 | 0 | 0 | 0 | 0 | 14327000 | 100970 | 484740 | 0 | 0 | 0 | 0 | 0 | 0 | 0 |
| DLDTDFTN | CPD; CPD | O75976; J3Q | CBPD_HUMAN | Unmodified | 0 | 0 | 0 | 0 | 134220 | 0 | 0 | 0 | 1397800 | 227260 | 109540 | 0 | 0 | 0 | 0 | 0 | 0 | 0 |
| DLDVITIPSI | KIF2A; KIF2A; KI | O00139; A0A | KIF2A_HUMAN | Unmodified | 0 | 0 | 0 | 0 | 0 | 0 | 0 | 32293 | 0 | 0 | 66813 | 1582000 | 0 | 0 | 0 | 0 | 0 | 0 |
| DLDVITIPSI | KIF2A; KIF2A; KI | O00139; A0A | KIF2A_HUMAN | Oxidation (M) | 0 | 0 | 0 | 0 | 453320 | 0 | 1691700 | 560070 | 0 | 218240 | 0 | 0 | 237920 | 233090 | 0 | 0 | 0 | 0 |
| DLEADIIGD | ANXA6; ANXA6; | P08133; E5R | ANXA6_HUMAI | Unmodified | 0 | 0 | 1113700 | 0 | 0 | 0 | 955890 | 205760 | 8913500 | 1471500 | 0 | 0 | 0 | 0 | 1936400 | 0 | 0 | 0 |
| DLEAEHVE | BOLA2; BOLA2E | Q9H3K6; H3 | BOLA2_HUMAI | Unmodified | 3971000 | 4640300 | 2524700 | 1841300 | 0 | 0 | 317470 | 161320 | 2658100 | 0 | 8855500 | 42176000 | 353760 | 0 | 3418100 | 2956600 | 0 | 0 |
| DLEAHIDS/ | MYH9 | P35579 | MYH9_HUMAN | Unmodified | 0 | 152140 | 0 | 764090 | 5579500 | 562590 | 301350 | 309080 | 0 | 501960 | 1152800 | 3824500 | 0 | 1135700 | 891870 | 0 | 673220 | 1719400 |
| DLEALLNSI | LMNA; LMNA; L | P02545; A0A | LMNA_HUMAN | Unmodified | 0 | 0 | 0 | 0 | 6170000 | 991210 | 0 | 0 | 994970 | 175420 | 314710 | 1975600 | 1596200 | 0 | 0 | 0 | 602020 | 890450 |
| DLEALMFD | PGM1; PGM1 | P36871; A0A | PGM1_HUMAN | Oxidation (M) | 0 | 0 | 0 | 0 | 0 | 0 | 0 | 0 | 0 | 0 | 0 | 0 | 0 | 0 | 0 | 0 | 194910 | 426940 |
| DLEALMFD | PGM1; PGM1 | P36871; A0A | PGM1_HUMAN | Unmodified | 0 | 0 | 0 | 0 | 850440 | 89610 | 112320 | 0 | 903600 | 252440 | 112420 | 0 | 200300 | 225100 | 0 | 0 | 512250 | 834100 |
| DLEALQSS | PROM2 | Q8N271 | PROM2_HUMA | Unmodified | 1209400 | 608050 | 704980 | 854690 | 433400 | 0 | 0 | 0 | 0 | 174330 | 0 | 1120900 | 0 | 0 | 1612200 | 976020 | 0 | 0 |
| DLEAQIEA | MYH10 | P35580 | MYH10_HUMAI | Unmodified | 0 | 0 | 2100500 | 119560 | 0 | 0 | 0 | 0 | 115750 | 0 | 525520 | 2588700 | 0 | 0 | 4635100 | 349230 | 0 | 0 |
| DLEASIAR | DYNC1H1; DYN | Q14204; A0A | DYHC1_HUMA | Unmodified | 0 | 0 | 0 | 0 | 0 | 0 | 0 | 0 | 142440 | 48962 | 207630 | 1904700 | 0 | 0 | 39341 | 0 | 25270 | 27459 |
| DLEDIVGG | CD99L2; CD99I | Q8TCZ2; H0' | C99L2_HUMAN | Unmodified | 594300 | 0 | 739680 | 316470 | 0 | 0 | 0 | 0 | 0 | 0 | 0 | 0 | 0 | 0 | 0 | 0 | 0 | 0 |
| DLEDKEGE | DYNC1I2; DYN | Q13409; E7E | DC1I2_HUMAN | Unmodified | 0 | 0 | 0 | 0 | 1209400 | 0 | 48504 | 0 | 722400 | 128810 | 13804000 | 973660 | 0 | 0 | 0 | 60434 | 65413 | 0 |
| DLEDLFYK | SRSF9; SRSF9; | Q13242; S4F | SRSF9_HUMAN | Unmodified | 1895500 | 1962100 | 0 | 1386000 | 0 | 0 | 0 | 0 | 0 | 0 | 0 | 0 | 187890 | 0 | 1685900 | 1259800 | 0 | 0 |
| DLEEAEEY | TBCA; TBCA; TB | O75347; E5F | TBCA_HUMAN | Unmodified | 0 | 0 | 2200600 | 537320 | 0 | 0 | 0 | 0 | 1936500 | 0 | 2374900 | 8332800 | 469570 | 0 | 2523200 | 2914200 | 0 | 0 |
| DLEEAEEY | TBCA; TBCA; TB | O75347; E5F | TBCA_HUMAN | Unmodified | 0 | 0 | 0 | 481610 | 509080 | 0 | 0 | 0 | 962320 | 136390 | 391640 | 1564700 | 731850 | 247160 | 0 | 0 | 79850 | 0 |
| DLEEDHAC | EEF2 | P13639 | EF2_HUMAN EI | Unmodified | 510670 | 312670 | 380990 | 525820 | 292790 | 0 | 175500 | 187210 | 1481300 | 315640 | 1382400 | 11109000 | 505110 | 107810 | 500510 | 597080 | 0 | 0 |
| DLEEDHAC | EEF2 | P13639 | EF2_HUMAN EI | Unmodified | 0 | 0 | 0 | 0 | 2698500 | 0 | 464470 | 146600 | 1436600 | 306820 | 2875900 | 18104000 | 911490 | 441750 | 0 | 0 | 474950 | 818000 |
| DLEEEKTM | CD2AP | Q9Y5K6 | CD2AP_HUMAI | Unmodified | 0 | 0 | 0 | 0 | 0 | 0 | 0 | 0 | 0 | 0 | 0 | 0 | 326790 | 256740 | 0 | 0 | 0 | 0 |
| DLEEFFSTV | RBM39; RBM39 | Q14498; G3 | RBM39_HUMAI | Unmodified | 0 | 0 | 0 | 0 | 0 | 0 | 0 | 0 | 453200 | 0 | 261320 | 1905100 | 0 | 0 | 0 | 0 | 55073 | 0 |
| DLEEPINFF | IL1RAP; IL1RAP | Q9NPH3; C9 | IL1AP_HUMAN | Unmodified | 0 | 0 | 0 | 0 | 0 | 0 | 638020 | 28597 | 0 | 0 | 0 | 0 | 0 | 198220 | 0 | 234170 | 0 | 0 |
| DLEEVKVLI | CACYBP; CACY | Q9HB71; B2 | CYBP_HUMAN | Unmodified | 0 | 0 | 0 | 0 | 2000600 | 0 | 437260 | 0 | 0 | 450260 | 0 | 0 | 1473100 | 0 | 0 | 0 | 0 | 0 |
| DLEEWNQI | CLTB | P09497 | CLCB_HUMAN | Unmodified | 3487900 | 2881900 | 2149300 | 1536300 | 248510 | 0 | 0 | 0 | 467130 | 0 | 0 | 2686400 | 447300 | 0 | 1695600 | 1603200 | 0 | 0 |
| DLEEWNQI | CLTB | P09497 | CLCB_HUMAN | Unmodified | 1880700 | 956350 | 1433500 | 4296100 | 15014000 | 2299200 | 2530700 | 1767000 | 498820 | 660040 | 0 | 279480 | 34227000 | 11171000 | 882860 | 597930 | 2650400 | 6179800 |
| DLEFQLHE | DCP1A; DCP1A | Q9NPI6; A0A | DCP1A_HUMAI | Unmodified | 0 | 0 | 17290 | 0 | 0 | 0 | 0 | 0 | 0 | 24754 | 0 | 75475 | 0 | 0 | 48812 | 0 | 0 | 0 |
| DLEGENIEI | SYNCRIP; SYN( | O60506; B7Z | HNRPQ_HUMA | Unmodified | 0 | 0 | 0 | 0 | 0 | 0 | 334100 | 92382 | 0 | 639570 | 0 | 608650 | 0 | 0 | 0 | 0 | 0 | 0 |
| DLEGLSQR | MYH9 | P35579 | MYH9_HUMAN | Unmodified | 280460 | 0 | 2963500 | 2323400 | 26312000 | 3850800 | 13655 | 39917 | 1722500 | 1084400 | 58331 | 536490 | 7253100 | 3461400 | 2674700 | 1380100 | 13234000 | 6108100 |
| DLEGQLSD | HOMER1; HOM | Q86YM7; Q8 | HOME1_HUMA | Unmodified | 0 | 0 | 0 | 0 | 0 | 0 | 107760 | 0 | 0 | 154470 | 104090 | 222850 | 3666900 | 1510500 | 0 | 441780 | 0 | 0 |
| DLEGSDID | CSE1L | P55060 | XPO2_HUMAN | Unmodified | 0 | 0 | 86908 | 0 | 0 | 0 | 0 | 0 | 199700 | 193880 | 1564600 | 8480700 | 0 | 0 | 117300 | 143830 | 0 | 0 |
| DLEGTYLC | ICAM1; ICAM1 | P05362; E7E | ICAM1_HUMAN | Unmodified | 0 | 0 | 595190 | 322920 | 0 | 98044 | 0 | 0 | 0 | 0 | 0 | 0 | 0 | 0 | 429020 | 393900 | 178050 | 0 |
| DLEIERPIL( | NDRG3; NDRG | Q9UGV2; F8 | NDRG3_HUMA | Unmodified | 425010 | 305750 | 1041400 | 1705700 | 1650200 | 0 | 236370 | 229640 | 3468300 | 384370 | 13132000 | 73681000 | 0 | 0 | 2807500 | 0 | 369360 | 691670 |
| DLEKPFLLF | TUFM | P49411 | EFTU_HUMAN | Unmodified | 0 | 0 | 0 | 0 | 0 | 0 | 0 | 0 | 1876600 | 50101 | 36503 | 69437 | 0 | 0 | 0 | 0 | 0 | 0 |
| DLELAVPG | MTOR; MTOR | P42345; B1A | MTOR_HUMAN | Unmodified | 0 | 0 | 0 | 60871 | 0 | 294020 | 0 | 0 | 362120 | 41820 | 0 | 243940 | 0 | 0 | 146930 | 66205 | 168780 | 26498 |
| DLENLAGP | LRBA; LRBA; LR | P50851; A0A | LRBA_HUMAN | Unmodified | 0 | 0 | 0 | 0 | 0 | 0 | 0 | 0 | 0 | 0 | 0 | 417240 | 0 | 0 | 0 | 124550 | 0 | 0 |
| DLEPVLSV( | TOR1B | O14657 | TOR1B_HUMAI | Unmodified | 0 | 0 | 0 | 0 | 0 | 0 | 0 | 0 | 1165900 | 386050 | 0 | 0 | 25159 | 0 | 0 | 0 | 0 | 0 |
| DLEQQDC | CCDC50 | Q8IVM0 | CCD50_HUMA | Unmodified | 4392200 | 4310200 | 0 | 8665800 | 0 | 0 | 0 | 0 | 572220 | 199740 | 0 | 532110 | 4269300 | 1312200 | 10102000 | 8033600 | 0 | 853840 |
| DLERQSEQ | PFDN6 | O15212 | PFD6_HUMAN | Unmodified | 53095 | 96278 | 0 | 180120 | 0 | 0 | 0 | 0 | 0 | 0 | 641050 | 615290 | 0 | 0 | 103450 | 70004 | 0 | 0 |
| DLESIDPEF | WWP2 | O00308 | WWP2_HUMAI | Unmodified | 0 | 388770 | 0 | 140700 | 0 | 0 | 0 | 0 | 0 | 0 | 0 | 0 | 0 | 0 | 775240 | 66826 | 0 | 0 |
| DLESIDPEF | ITCH; ITCH; ITC | Q96J02; A0A | ITCH_HUMAN I | Unmodified | 202500 | 1018500 | 55293 | 208810 | 0 | 0 | 0 | 0 | 0 | 0 | 0 | 0 | 0 | 0 | 857950 | 124630 | 0 | 0 |
| DLESLSPRI | FAM131B | Q86XD5 | F131B_HUMAN | Unmodified | 0 | 0 | 0 | 3306500 | 0 | 0 | 0 | 0 | 0 | 0 | 0 | 0 | 0 | 0 | 0 | 0 | 0 | 0 |
| DLETQVIQI | MAPRE2 | Q15555 | MARE2_HUMAI | Unmodified | 0 | 0 | 0 | 2437200 | 0 | 0 | 0 | 0 | 1907400 | 0 | 0 | 0 | 0 | 0 | 0 | 0 | 0 | 0 |
| DLETSCSD | DCTN1; DCTN1 | Q14203; Q6 | DCTN1_HUMAI | Unmodified | 0 | 0 | 0 | 0 | 0 | 0 | 0 | 0 | 0 | 0 | 0 | 621270 | 0 | 0 | 0 | 0 | 0 | 0 |
| DLEVERQTI | CEP55; CEP55 | Q53EZ4; H0' | CEP55_HUMAN | Unmodified | 0 | 0 | 0 | 1845400 | 12991000 | 0 | 0 | 0 | 553900 | 1513900 | 0 | 0 | 0 | 0 | 0 | 0 | 3517600 | 2711300 |
| DLEYLDLK | PLEKHA7; PLEK | Q6IQ23; H0' | PKHA7_HUMAN | Unmodified | 0 | 0 | 0 | 0 | 0 | 0 | 0 | 0 | 0 | 0 | 0 | 0 | 865260 | 0 | 0 | 0 | 0 | 0 |
| DLFDPIIED | CKB; CKB; CKB | P12277; G3 | KCRB_HUMAN | Unmodified | 812100 | 0 | 10061000 | 7142300 | 0 | 0 | 0 | 171350 | 7724700 | 230410 | 4622900 | 81105000 | 0 | 0 | 34024000 | 17744000 | 0 | 0 |
| DLFEDELV | SYNCRIP; SYN( | O60506; B7Z | HNRPQ_HUMA | Unmodified | 0 | 0 | 0 | 0 | 0 | 0 | 0 | 0 | 295480 | 0 | 0 | 1172200 | 0 | 0 | 0 | 0 | 0 | 0 |
| DLFGDKEF | LIPA; LIPA; LIPA | P38571; Q5 | LICH_HUMAN I | Unmodified | 0 | 0 | 0 | 302940 | 215730 | 66333 | 166290 | 0 | 26541 | 271860 | 0 | 0 | 300510 | 457720 | 169410 | 0 | 0 | 60181 |
| DLFGKFGP | PABPC1; PABP( | P11940; Q9 | PABP1_HUMAN | Unmodified | 10017000 | 0 | 0 | 3846900 | 1181500 | 0 | 0 | 205980 | 0 | 1149600 | 0 | 0 | 4687200 | 0 | 0 | 0 | 779410 | 0 |
| DLFGKSDP | CPNE3; CPNE3 | O75131; A0 | CPNE3_HUMAI | Unmodified | 0 | 0 | 0 | 1735900 | 6959400 | 0 | 3361400 | 544110 | 5836000 | 8978200 | 92872 | 0 | 5998300 | 4824800 | 0 | 0 | 2470300 | 14801000 |
| DLFGLASN | APPL2; APPL2; | Q8NEU8; F8 | DP13B_HUMAN | Oxidation (M) | 0 | 0 | 0 | 306910 | 1185000 | 214440 | 0 | 79630 | 255440 | 0 | 0 | 118360 | 1537800 | 1322400 | 300440 | 182670 | 459510 | 0 |
| DLFGLASN | APPL2; APPL2; | Q8NEU8; F8 | DP13B_HUMAN | Unmodified | 0 | 0 | 0 | 182710 | 1248700 | 0 | 0 | 0 | 292680 | 157990 | 0 | 0 | 611950 | 1007500 | 0 | 2631700 | 0 | 0 |
| DLFNVDAF | NDC80 | O14777 | NDC80_HUMA | Unmodified | 23083 | 22196 | 0 | 81990 | 895730 | 67074 | 0 | 0 | 0 | 253920 | 0 | 93297 | 513980 | 462020 | 0 | 69108 | 311180 | 215590 |
| DLFPYEES | HDGF | P51858 | HDGF_HUMAN | Unmodified | 0 | 0 | 0 | 0 | 993230 | 381210 | 0 | 0 | 919310 | 196870 | 2589300 | 6523400 | 739590 | 190630 | 0 | 0 | 0 | 272250 |
| DLFQELTR | GRK6; GRK6; G | P43250; D6F | GRK6_HUMAN | Unmodified | 0 | 0 | 0 | 0 | 0 | 0 | 0 | 0 | 0 | 0 | 0 | 662510 | 0 | 0 | 173180 | 251020 | 0 | 0 |
| DLFQFNK | LLGL2; LLGL2; | Q6P1M3; J3 | L2GL2_HUMAN | Unmodified | 0 | 0 | 0 | 0 | 0 | 0 | 0 | 0 | 1481200 | 334850 | 395550 | 1916900 | 0 | 0 | 395650 | 0 | 274550 | 0 |
| DLFQVAFN | DYNC1H1; DYN | Q14204; A0A | DYHC1_HUMA | Unmodified | 0 | 0 | 0 | 0 | 0 | 0 | 0 | 0 | 253960 | 0 | 0 | 2553700 | 0 | 0 | 0 | 0 | 0 | 0 |
| DLFSMKPD | SLC12A7; SLC1 | Q9Y666; A0A | S12A7_HUMAN | Unmodified | 0 | 0 | 0 | 0 | 0 | 0 | 0 | 0 | 0 | 0 | 0 | 0 | 521950 | 428170 | 417790 | 79135 | 0 | 0 |
| DLFSMKPE | SLC12A7 | A0A0U1RR1 | A0A0U1RR18_I | Oxidation (M) | 0 | 0 | 0 | 0 | 0 | 0 | 222010 | 0 | 0 | 0 | 0 | 0 | 0 | 0 | 0 | 0 | 0 | 0 |
| DLGAQLDF | ACAP2; ACAP2; | Q15057; A0A | ACAP2_HUMAN | Unmodified | 77762 | 0 | 59696 | 128610 | 214950 | 61736 | 101520 | 0 | 220180 | 0 | 31753 | 252660 | 142650 | 137040 | 248110 | 178330 | 314530 | 0 |
| DLGDSHP | UBE3A; UBE3A; | Q05086; A0A | UBE3A_HUMAN | Unmodified | 0 | 29423 | 0 | 0 | 0 | 0 | 381080 | 131890 | 349130 | 221150 | 0 | 1367800 | 109280 | 330280 | 0 | 518180 | 452230 | 0 |
| DLGEELEAI | MYH9; MYH11 | P35579; P35 | MYH9_HUMAN | Unmodified | 0 | 0 | 571650 | 0 | 0 | 0 | 0 | 0 | 0 | 0 | 0 | 0 | 0 | 0 | 305570 | 376610 | 0 | 0 |
| DLGELLQA | CCDC22 | O60826 | CCD22_HUMA | Unmodified | 0 | 0 | 0 | 30002 | 73134 | 31236 | 0 | 0 | 60220 | 0 | 28906 | 55428 | 0 | 0 | 28956 | 36380 | 24349 | 36583 |
| DLGGFDEI | SDF4; SDF4; SD | Q9BRK5; H0 | CAB45_HUMAN | Unmodified | 0 | 0 | 0 | 0 | 0 | 0 | 0 | 0 | 11492000 | 1216300 | 1105100 | 0 | 307280 | 0 | 0 | 0 | 0 | 0 |
| DLGGFDEI | SDF4; SDF4; SD | Q9BRK5; H0 | CAB45_HUMAN | Unmodified | 0 | 0 | 1127000 | 599960 | 0 | 0 | 0 | 32592 | 22514000 | 3540600 | 1232000 | 623170 | 1695400 | 1502100 | 419390 | 349150 | 0 | 0 |
| DLGGFDEI | SDF4; SDF4; SD | Q9BRK5; H0 | CAB45_HUMAN | Unmodified | 0 | 0 | 0 | 0 | 0 | 0 | 0 | 0 | 0 | 3515100 | 0 | 0 | 0 | 0 | 0 | 0 | 0 | 0 |
| DLGGRDDI | EPHB3 | P54753 | EPHB3_HUMAN | Unmodified | 524150 | 337100 | 74834 | 0 | 0 | 0 | 0 | 0 | 0 | 0 | 0 | 0 | 0 | 0 | 0 | 0 | 0 | 0 |
| DLGHPVEE | USO1 | O60763 | USO1_HUMAN | Unmodified | 0 | 0 | 0 | 0 | 1494600 | 0 | 0 | 0 | 0 | 2855700 | 0 | 2457200 | 0 | 0 | 0 | 0 | 1078400 | 0 |
| DLGLAADL | SLC38A10 | Q9HBR0 | S38AA_HUMAN | Unmodified | 0 | 0 | 0 | 0 | 0 | 0 | 0 | 0 | 12177000 | 0 | 0 | 0 | 0 | 0 | 0 | 0 | 0 | 0 |
| DLGLAQDS | HIBADH | P31937 | 3HIDH_HUMAN | Unmodified | 0 | 0 | 0 | 0 | 0 | 0 | 196720 | 143080 | 2380400 | 0 | 0 | 0 | 0 | 290230 | 0 | 0 | 0 | 0 |
| DLGLAQDS | HIBADH | P31937 | 3HIDH_HUMAN | Unmodified | 0 | 0 | 0 | 0 | 0 | 0 | 0 | 0 | 0 | 1994200 | 0 | 0 | 0 | 0 | 0 | 0 | 0 | 0 |
| DLGLGVGE | PCDHB5 | Q9Y5E4 | PCDB5_HUMAI | Unmodified | 0 | 0 | 0 | 0 | 121120 | 0 | 0 | 77079 | 0 | 286410 | 0 | 2908500 | 290540 | 181620 | 0 | 0 | 0 | 0 |
| DLGLSESG | PDIA4; PDIA4 | P13667; A0A | PDIA4_HUMAN | Unmodified | 0 | 0 | 0 | 0 | 0 | 0 | 0 | 0 | 22305000 | 1591600 | 2814300 | 641690 | 0 | 0 | 0 | 172650 | 0 | 0 |
| DLGLSESG | PDIA4; PDIA4 | P13667; A0A | PDIA4_HUMAN | Unmodified | 0 | 0 | 2813300 | 897970 | 0 | 0 | 507360 | 378370 | 80689000 | 4140900 | 11725000 | 0 | 0 | 0 | 0 | 487940 | 0 | 0 |
| DLGNMEE | ALCAM; ALCAM | Q13740; F5 | CD166_HUMAI | Unmodified | 3306000 | 2986900 | 4693700 | 1878900 | 5782900 | 0 | 421750 | 188130 | 9593700 | 1159000 | 0 | 4621000 | 0 | 313610 | 3157300 | 2520800 | 1668600 | 11781000 |
| DLGQIEEN | DIP2B | Q9P265 | DIP2B_HUMAN | Unmodified | 0 | 0 | 0 | 0 | 0 | 0 | 0 | 0 | 0 | 0 | 0 | 2872200 | 0 | 0 | 0 | 0 | 0 | 0 |
| DLGQIEEN | DIP2B | Q9P265 | DIP2B_HUMAN | Unmodified | 128530 | 155520 | 338480 | 0 | 0 | 0 | 0 | 0 | 0 | 0 | 0 | 295190 | 0 | 0 | 505850 | 0 | 0 | 0 |
| DLGSDFQF | SOGA1; SOGA1 | O94964; X6F | SOGA1_HUMAI | Unmodified | 0 | 0 | 0 | 37631 | 0 | 0 | 0 | 0 | 0 | 0 | 0 | 0 | 990800 | 67847 | 0 | 0 | 0 | 0 |
| DLGTESQI | GDI2 | P50395 | GDIB_HUMAN | Unmodified | 2364700 | 1641500 | 2540300 | 6127100 | 4915700 | 1410800 | 481650 | 29691 | 16895000 | 3837900 | 8229400 | 21307000 | 8163100 | 2660400 | 5163300 | 5446800 | 1056100 | 1618600 |
| DLGTESQI | GDI2 | P50395 | GDIB_HUMAN | Unmodified | 0 | 0 | 0 | 0 | 11196000 | 0 | 0 | 0 | 0 | 1560000 | 0 | 0 | 2299700 | 4218300 | 0 | 0 | 0 | 6664100 |
| DLGTESQI | GDI2 | P50395 | GDIB_HUMAN | Unmodified | 0 | 0 | 0 | 0 | 15935000 | 802310 | 6492300 | 0 | 0 | 4295200 | 0 | 0 | 8132300 | 18472000 | 0 | 0 | 17624000 | 15034000 |
| DLGYIYFY( | USP5 | P45974 | UBP5_HUMAN | Unmodified | 0 | 2856500 | 0 | 0 | 0 | 0 | 0 | 0 | 670350 | 0 | 491610 | 2401500 | 0 | 734150 | 0 | 0 | 0 | 583180 |
| DLHDANT | SLC16A1; SLC1 | P53985; Q5 | MOT1_HUMAN | Unmodified | 185400 | 0 | 1683300 | 618080 | 0 | 0 | 132060 | 0 | 0 | 458190 | 3181100 | 6052900 | 0 | 0 | 1934200 | 2560100 | 0 | 0 |
| DLHDANT | SLC16A1; SLC1 | P53985; Q5 | MOT1_HUMAN | Unmodified | 34416000 | 1513400 | 18016000 | 12929000 | 1709400 | 189440 | 1173000 | 156540 | 4749900 | 1865700 | 8332300 | 43341000 | 925100 | 10486000 | 26368000 | 25515000 | 58915 | 0 |
| DLHDANT | SLC16A1; SLC1 | P53985; Q5 | MOT1_HUMAN | Unmodified | 794700 | 1612300 | 1867500 | 11118000 | 17931000 | 1672500 | 3246800 | 444200 | 6181000 | 733810 | 5862600 | 15046000 | 11828000 | 6227100 | 9059700 | 10200000 | 3938300 | 7136600 |
| DLHESSFS | DPYSL5 | Q9BPU6 | DPYL5_HUMAN | Unmodified | 0 | 0 | 0 | 0 | 0 | 0 | 0 | 0 | 0 | 0 | 0 | 2174500 | 0 | 0 | 0 | 0 | 0 | 0 |
| DLHFLMEC | SNX9; SNX9; SI | Q9Y5X1; A0A | SNX9_HUMAN | Oxidation (M) | 2557200 | 1774000 | 0 | 0 | 0 | 0 | 0 | 0 | 0 | 0 | 0 | 0 | 0 | 0 | 279770 | 877030 | 0 | 0 |
| DLHFLMEC | SNX9; SNX9; SI | Q9Y5X1; A0A | SNX9_HUMAN | Unmodified | 517150 | 0 | 0 | 0 | 0 | 0 | 53702 | 0 | 0 | 0 | 0 | 916160 | 0 | 0 | 0 | 0 | 0 | 0 |
| DLHMERVI | RHEB; RHEB | Q15382; C9 | RHEB_HUMAN | Unmodified | 0 | 0 | 0 | 0 | 5517200 | 0 | 1692400 | 176540 | 0 | 454980 | 0 | 0 | 0 | 0 | 0 | 0 | 659640 | 0 |
| DLHQGIEA | PACSIN3; PACS | Q9UKS6; E9 | PACN3_HUMAI | Unmodified | 8873300 | 5562500 | 1948400 | 467350 | 0 | 0 | 0 | 0 | 0 | 0 | 0 | 0 | 0 | 0 | 4328200 | 3459300 | 0 | 0 |
| DLHQGIEA | PACSIN3; PACS | Q9UKS6; E9 | PACN3_HUMAI | Unmodified | 33350000 | 34338000 | 9359900 | 6120500 | 0 | 0 | 0 | 0 | 0 | 0 | 0 | 0 | 0 | 0 | 27007000 | 13265000 | 0 | 0 |
| DLIADLKYE | ANXA6; ANXA6; | P08133; E5R | ANXA6_HUMAI | Unmodified | 0 | 114110 | 70066 | 104880 | 74860 | 0 | 118030 | 0 | 1354500 | 1219600 | 0 | 0 | 82711 | 31204 | 461510 | 75928 | 54989 | 42905 |
| DLIADLKYE | ANXA6; ANXA6; | P08133; E5R | ANXA6_HUMAI | Unmodified | 0 | 720830 | 0 | 3031600 | 1530300 | 628630 | 490330 | 0 | 14761000 | 9233400 | 227580 | 0 | 0 | 1083000 | 1594900 | 250820 | 931950 | 2117000 |
| DLIEDSSV( | LARP4; LARP4 | Q71RC2; X6 | LARP4_HUMAN | Unmodified | 268060 | 218340 | 0 | 311860 | 0 | 0 | 0 | 0 | 0 | 0 | 0 | 0 | 100660 | 0 | 0 | 141390 | 0 | 0 |
| DLIGIDNLV | FARP1; FARP1; | Q9Y4F1; C9 | FARP1_HUMAN | Unmodified | 0 | 0 | 18216 | 0 | 0 | 0 | 0 | 0 | 72672 | 0 | 730850 | 2946500 | 0 | 0 | 0 | 0 | 0 | 0 |
| DLIGIDNLV | FARP1 | Q9Y4F1 | FARP1_HUMAN | Unmodified | 541280 | 289030 | 0 | 276120 | 499820 | 396000 | 0 | 0 | 8354000 | 2926400 | 1319200 | 868610 | 0 | 0 | 489100 | 0 | 161830 | 373390 |
| DLIGIDNLV | FARP1 | Q9Y4F1 | FARP1_HUMAN | Unmodified | 0 | 0 | 0 | 0 | 878900 | 0 | 284890 | 0 | 29573 | 2017500 | 65173 | 0 | 104310 | 225390 | 0 | 0 | 1285700 | 576180 |
| DLIGSYAM | ARVCF; ARVCF; | O00192; E9F | ARVC_HUMAN | Oxidation (M) | 0 | 0 | 0 | 76678 | 0 | 0 | 0 | 0 | 0 | 0 | 0 | 0 | 0 | 0 | 158130 | 104700 | 0 | 0 |
| DLIGVQNL | SPTAN1; SPTAN | Q13813; A0A | SPTN1_HUMAN | Unmodified | 0 | 0 | 0 | 0 | 1279800 | 0 | 0 | 0 | 597080 | 308620 | 638230 | 4798600 | 0 | 0 | 203180 | 0 | 466150 | 490780 |

| | | | | | | | | | | | | | | | | | | | | | | |
|---|---|---|---|---|---|---|---|---|---|---|---|---|---|---|---|---|---|---|---|---|---|---|
| DLIGVQNLI | SPTAN1; SPTAN | Q13813; A0A | SPTN1_HUMAN | Unmodified | 0 | 0 | 0 | 0 | 292650 | 0 | 0 | 0 | 264100 | 120870 | 0 | 851240 | 79615 | 0 | 0 | 0 | 337580 | 0 |
| DLIHDQDE | NSFL1C; NSFL1 | Q9UNZ2; R4 | NSF1C_HUMAI | Unmodified | 205430 | 0 | 0 | 0 | 0 | 0 | 63684 | 0 | 0 | 0 | 368670 | 0 | 0 | 0 | 173160 | 173270 | 142340 | 0 |
| DLIKDLKSE | ANXA11 | P50995 | ANX11_HUMAN | Unmodified | 0 | 0 | 0 | 48681 | 3448000 | 283230 | 4521900 | 35536 | 612430 | 3518700 | 39695 | 0 | 1476500 | 3662700 | 0 | 0 | 6141500 | 8202500 |
| DLILNPR | SF3B3 | Q15393 | SF3B3_HUMAN | Unmodified | 456200 | 305270 | 0 | 158140 | 0 | 0 | 0 | 0 | 0 | 0 | 214240 | 2463600 | 0 | 0 | 0 | 0 | 200730 | 0 |
| DLILPTIQK | GCN1 | Q92616 | GCN1_HUMAN | Unmodified | 0 | 0 | 0 | 0 | 0 | 0 | 0 | 43884 | 0 | 0 | 69100 | 1060800 | 0 | 0 | 0 | 0 | 0 | 0 |
| DLILSVLQR | SYTL4 | Q96C24 | SYTL4_HUMAN | Unmodified | 0 | 0 | 0 | 0 | 0 | 0 | 0 | 0 | 208950 | 286550 | 0 | 0 | 0 | 0 | 0 | 0 | 0 | 0 |
| DLIQGKDLI | PDIA3 | P30101 | PDIA3_HUMAN | Unmodified | 0 | 145170 | 0 | 56999 | 0 | 0 | 0 | 0 | 0 | 2501700 | 0 | 0 | 22943 | 0 | 0 | 0 | 0 | 708490 |
| DLIQVVHE | MIA3 | Q5JRA6 | TGO1_HUMAN | Unmodified | 0 | 0 | 129520 | 0 | 0 | 0 | 0 | 0 | 1067600 | 0 | 437060 | 0 | 0 | 0 | 0 | 0 | 0 | 0 |
| DLISHDEM | TPT1; TPT1; TPT | P13693; A0A | TCTP_HUMAN | Oxidation (M) | 3575200 | 3566000 | 2128100 | 2336400 | 0 | 0 | 0 | 0 | 323910 | 135700 | 832310 | 11202000 | 167480 | 0 | 8932900 | 5758600 | 0 | 0 |
| DLISHDEM | TPT1; TPT1; TPT | P13693; A0A | TCTP_HUMAN | Unmodified | 587110 | 572790 | 0 | 540000 | 182240 | 0 | 0 | 0 | 1152400 | 0 | 0 | 2648700 | 0 | 0 | 1058800 | 491330 | 0 | 0 |
| DLIVCLFSK | MYO1D; MYO1D | O94832; J3Q | MYO1D_HUMA | Unmodified | 0 | 1477700 | 185320 | 0 | 0 | 0 | 0 | 0 | 137780 | 0 | 0 | 166260 | 0 | 0 | 1347500 | 904480 | 0 | 0 |
| DLIVCLFSK | MYO1D; MYO1D | O94832; J3Q | MYO1D_HUMA | Unmodified | 2020800 | 1305200 | 0 | 854500 | 310180 | 0 | 0 | 0 | 284980 | 119320 | 0 | 0 | 0 | 172390 | 0 | 512590 | 0 | 0 |
| DLIVSGDEI | GEMIN5 | Q8TEQ6 | GEMI5_HUMAN | Unmodified | 0 | 0 | 0 | 0 | 0 | 0 | 0 | 0 | 0 | 0 | 0 | 1417500 | 0 | 0 | 0 | 0 | 0 | 0 |
| DLIVTPATIL | BCAT1; BCAT1 | P54687; F5H | BCAT1_HUMAN | Unmodified | 0 | 0 | 41347 | 0 | 1361600 | 251480 | 0 | 0 | 0 | 0 | 8293400 | 40695000 | 0 | 0 | 0 | 0 | 0 | 0 |
| DLIYTLYYW | F3 | P13726 | TF_HUMAN Tis | Unmodified | 1187500 | 0 | 0 | 0 | 1553100 | 226650 | 0 | 0 | 0 | 0 | 0 | 0 | 0 | 0 | 0 | 0 | 0 | 0 |
| DLKAVKYVI | CDC42 | P60953 | CDC42_HUMA | Unmodified | 0 | 0 | 0 | 0 | 0 | 0 | 0 | 442070 | 0 | 0 | 0 | 0 | 0 | 1161000 | 0 | 0 | 2867200 | 908340 |
| DLKEEFTEA | TPP2; TPP2; TPI | P29144; A0A | TPP2_HUMAN | Unmodified | 0 | 0 | 0 | 0 | 0 | 0 | 0 | 0 | 0 | 0 | 723750 | 2196600 | 0 | 0 | 0 | 0 | 0 | 0 |
| DLKEEFTEA | TPP2; TPP2; TPI | P29144; A0A | TPP2_HUMAN | Unmodified | 58638 | 0 | 146270 | 131690 | 0 | 0 | 0 | 0 | 302280 | 514630 | 557120 | 6199500 | 306540 | 0 | 216250 | 121840 | 0 | 0 |
| DLKEIQHH | SH3GL1; SH3G | Q99961; M0I | SH3G1_HUMAI | Unmodified | 43512 | 0 | 0 | 0 | 158250 | 0 | 446700 | 136200 | 0 | 102050 | 0 | 0 | 1537400 | 759470 | 0 | 0 | 79920 | 0 |
| DLKLDNVM | PRKCB; PRKCA | P05771; P17 | KPCB_HUMAN | Oxidation (M) | 372590 | 100290 | 240350 | 652420 | 0 | 0 | 0 | 0 | 0 | 0 | 0 | 248380 | 0 | 0 | 479980 | 417390 | 0 | 0 |
| DLKNPDRV | UGDH | O60701 | UGDH_HUMAN | Unmodified | 0 | 1117900 | 0 | 147970 | 2381500 | 148800 | 0 | 0 | 0 | 0 | 0 | 0 | 405880 | 359020 | 0 | 0 | 0 | 0 |
| DLKPENILL | GRK6; GRK6; G | P43250; D6F | GRK6_HUMAN | Unmodified | 0 | 0 | 848860 | 0 | 0 | 0 | 0 | 0 | 1106800 | 0 | 0 | 0 | 0 | 0 | 0 | 852510 | 0 | 0 |
| DLKPENILL | GRK5 | P34947 | GRK5_HUMAN | Unmodified | 106980 | 0 | 26484 | 0 | 0 | 0 | 0 | 0 | 0 | 0 | 0 | 1354700 | 78227 | 0 | 0 | 0 | 34936 | 230620 |
| DLKPENILL | RPS6KA1; RPS6 | Q15418; P51 | KS6A1_HUMAN | Unmodified | 0 | 0 | 148890 | 0 | 0 | 0 | 0 | 0 | 0 | 0 | 0 | 746220 | 0 | 0 | 319260 | 300110 | 0 | 0 |
| DLKPGNLA | MAPK12; MAPK | P53778; O15 | MK12_HUMAN | Unmodified | 0 | 0 | 0 | 0 | 0 | 0 | 0 | 0 | 0 | 0 | 0 | 2298000 | 0 | 0 | 0 | 28232 | 0 | 0 |
| DLKPNGAN | HUWE1; HUWE | Q7Z6Z7; H0Y | HUWE1_HUMA | Unmodified | 0 | 0 | 0 | 0 | 0 | 0 | 0 | 0 | 0 | 0 | 0 | 616050 | 0 | 0 | 0 | 0 | 0 | 0 |
| DLKPQNLL | CDK1; CDK1; C | P06493; A0A | CDK1_HUMAN | Unmodified | 958810 | 1266200 | 1235400 | 2469700 | 0 | 0 | 0 | 0 | 3909000 | 45844 | 450370 | 4139600 | 87646 | 0 | 2390400 | 1814900 | 0 | 0 |
| DLKPQNLL | CDK5 | Q00535 | CDK5_HUMAN | Unmodified | 0 | 0 | 516800 | 0 | 0 | 0 | 0 | 0 | 0 | 0 | 0 | 1175500 | 0 | 0 | 0 | 571000 | 166070 | 0 |
| DLKPQNLL | CDK2; CDK2; C | P24941; G3V | CDK2_HUMAN | Unmodified | 674060 | 647890 | 0 | 135730 | 0 | 0 | 0 | 32151 | 0 | 0 | 0 | 961820 | 0 | 0 | 102090 | 227900 | 54136 | 0 |
| DLKPSNLLI | MAPK1 | P28482 | MK01_HUMAN | Unmodified | 63034 | 388570 | 196790 | 615850 | 0 | 0 | 0 | 51864 | 0 | 0 | 0 | 1416000 | 0 | 0 | 492960 | 0 | 0 | 0 |
| DLKPVLDS | TSG101; TSG10 | Q99816; F5H | TS101_HUMAN | Unmodified | 44886000 | 46761000 | 38874000 | 20699000 | 139800 | 131690 | 0 | 0 | 8184400 | 536220 | 0 | 12707000 | 499380 | 0 | 86259000 | 54966000 | 0 | 131750 |
| DLKPVLDS | TSG101; TSG10 | Q99816; F5H | TS101_HUMAN | Oxidation (M) | 0 | 0 | 152410 | 1846500 | 0 | 0 | 0 | 0 | 0 | 0 | 0 | 0 | 0 | 0 | 2913700 | 0 | 0 | 0 |
| DLKSELSGI | ANXA11 | P50995 | ANX11_HUMAN | Unmodified | 191080 | 0 | 287080 | 1245000 | 4256300 | 1198800 | 3985200 | 2323300 | 14609000 | 7647400 | 953100 | 259360 | 2679800 | 2956900 | 611370 | 314950 | 1706300 | 6346800 |
| DLKVENLLI | GAK | O14976 | GAK_HUMAN C | Unmodified | 0 | 0 | 0 | 0 | 0 | 0 | 0 | 0 | 0 | 0 | 0 | 1051900 | 0 | 0 | 0 | 0 | 0 | 0 |
| DLLDIANGI | DPY19L4 | Q7Z388 | D19L4_HUMAN | Unmodified | 0 | 0 | 0 | 0 | 0 | 0 | 0 | 0 | 367800 | 0 | 0 | 0 | 0 | 0 | 0 | 0 | 0 | 0 |
| DLLDTVLPI | CAND1; CAND1 | Q86VP6; A0A | CAND1_HUMA | Unmodified | 0 | 0 | 43064 | 0 | 0 | 0 | 0 | 0 | 507890 | 228510 | 122780 | 3036900 | 0 | 0 | 0 | 131250 | 0 | 0 |
| DLLEPESVI | DOCK10; DOCK | Q96BY6; A0A | DOC10_HUMA | Unmodified | 0 | 0 | 0 | 0 | 6569500 | 0 | 0 | 0 | 0 | 142680 | 0 | 0 | 0 | 0 | 0 | 0 | 0 | 1050000 |
| DLLEPGCS | PSMC1 | P62191 | PRS4_HUMAN | Unmodified | 0 | 0 | 41484 | 0 | 0 | 0 | 0 | 0 | 0 | 0 | 0 | 4729600 | 0 | 0 | 0 | 0 | 0 | 0 |
| DLLETVRVI | UTRN | P46939 | UTRN_HUMAN | Unmodified | 0 | 0 | 0 | 0 | 2707600 | 0 | 0 | 0 | 0 | 378140 | 0 | 0 | 1614700 | 457740 | 0 | 0 | 1776800 | 3966400 |
| DLLEVADVI | GRPEL1 | Q9HAV7 | GRPE1_HUMAI | Unmodified | 0 | 0 | 0 | 0 | 1453200 | 0 | 0 | 0 | 4341800 | 1232100 | 2667400 | 665570 | 421100 | 226990 | 0 | 0 | 0 | 0 |
| DLLFILTAK | DDB1; DDB1; D | Q16531; A0A | DDB1_HUMAN | Unmodified | 0 | 0 | 0 | 0 | 0 | 0 | 0 | 30992 | 0 | 87361 | 0 | 215850 | 0 | 0 | 0 | 0 | 0 | 0 |
| DLLFKDSTS | MELTF | P08582 | TRFM_HUMAN | Oxidation (M) | 0 | 0 | 0 | 0 | 1036400 | 0 | 0 | 0 | 0 | 276670 | 0 | 0 | 0 | 0 | 0 | 0 | 0 | 0 |
| DLLFQALGI | FARSB | Q9NSD9 | SYFB_HUMAN | Unmodified | 0 | 0 | 0 | 0 | 0 | 0 | 0 | 0 | 0 | 0 | 0 | 0 | 0 | 0 | 0 | 0 | 0 | 0 |
| DLLGLCEQ | XPO1; XPO1; XF | O14980; A0A | XPO1_HUMAN | Unmodified | 0 | 0 | 0 | 0 | 250100 | 0 | 110110 | 0 | 0 | 0 | 0 | 928330 | 0 | 0 | 0 | 0 | 0 | 0 |
| DLLHLPELI | PDCD11; PDCD | Q14690; A0A | RRP5_HUMAN | Oxidation (M) | 19039 | 0 | 0 | 18057 | 0 | 67078 | 178650 | 0 | 0 | 0 | 18586 | 108800 | 0 | 41721 | 0 | 0 | 0 | 47886 |
| DLLHNEDF | CPSF7 | Q8N684 | CPSF7_HUMAN | Unmodified | 1643100 | 1463900 | 1125500 | 417180 | 0 | 0 | 0 | 0 | 0 | 0 | 92805 | 167860 | 0 | 0 | 0 | 523430 | 0 | 0 |
| DLLHPSLEI | RPS27L; RPS27 | Q71UM5; C9 | RS27L_HUMAN | Unmodified | 834050 | 1033700 | 520740 | 603070 | 0 | 0 | 0 | 0 | 0 | 0 | 2015500 | 11661000 | 647640 | 195000 | 718990 | 659510 | 0 | 0 |
| DLLHPSLEI | RPS27L; RPS27 | Q71UM5; C9 | RS27L_HUMAN | Unmodified | 268130 | 0 | 0 | 0 | 95812 | 0 | 326940 | 0 | 0 | 154200 | 0 | 1127100 | 0 | 0 | 0 | 0 | 0 | 0 |
| DLLHPSPEI | RPS27 | P42677 | RS27_HUMAN | Unmodified | 616310 | 2270500 | 1810700 | 1540000 | 4149000 | 666010 | 761810 | 1037300 | 1319600 | 789450 | 5710200 | 23837000 | 4004900 | 2165000 | 1232300 | 1141000 | 265390 | 1385100 |
| DLLHPSPEI | RPS27 | P42677 | RS27_HUMAN | Unmodified | 67096 | 46773 | 0 | 64306 | 37662 | 0 | 0 | 0 | 59653 | 41962 | 535770 | 907290 | 26496 | 27147 | 0 | 37817 | 0 | 0 |
| DLLHPSPEI | RPS27 | P42677 | RS27_HUMAN | Unmodified | 0 | 44144 | 0 | 0 | 278680 | 53455 | 318510 | 261410 | 0 | 154970 | 0 | 0 | 680400 | 382050 | 0 | 0 | 203000 | 237730 |
| DLLIAYYDV | PDIA3 | P30101 | PDIA3_HUMAN | Unmodified | 0 | 0 | 0 | 0 | 0 | 0 | 0 | 0 | 2388000 | 1032900 | 241490 | 674590 | 0 | 0 | 0 | 0 | 0 | 0 |
| DLLIAYYDV | PDIA3 | P30101 | PDIA3_HUMAN | Unmodified | 0 | 0 | 0 | 0 | 0 | 0 | 231890 | 0 | 633580 | 613850 | 0 | 0 | 0 | 0 | 0 | 0 | 0 | 0 |
| DLLIYDSMI | SNTB1; SNTB1 | Q13884; A0A | SNTB1_HUMAN | Unmodified | 0 | 0 | 0 | 0 | 0 | 0 | 0 | 0 | 3497300 | 455380 | 0 | 0 | 0 | 0 | 0 | 0 | 0 | 0 |
| DLLLTGHEI | LLGL2; LLGL2 | Q6P1M3; J3 | L2GL2_HUMAN | Unmodified | 352720 | 285890 | 334690 | 186530 | 0 | 0 | 0 | 0 | 0 | 0 | 0 | 1868300 | 0 | 0 | 797620 | 185490 | 0 | 0 |
| DLLLTSSYL | SLC3A2; SLC3A | P08195; A0A | 4F2_HUMAN 4I | Unmodified | 16380000 | 13051000 | 21298000 | 12055000 | 23022000 | 7230300 | 2010600 | 220870 | 45026000 | 5986500 | 72480000 | 2.43E+08 | 0 | 0 | 55380000 | 36244000 | 1301400 | 1407400 |
| DLLLTVK | FBLN1 | B1AHL2 | B1AHL2_HUMA | Unmodified | 0 | 0 | 0 | 0 | 0 | 0 | 0 | 0 | 0 | 0 | 0 | 0 | 0 | 0 | 0 | 0 | 0 | 0 |
| DLLMKAETI | RACGAP1; RAC | Q9H0H5; F8 | RGAP1_HUMAI | Oxidation (M) | 0 | 0 | 0 | 0 | 7292900 | 648580 | 652570 | 57848 | 0 | 0 | 0 | 0 | 0 | 0 | 0 | 0 | 427020 | 2069500 |
| DLLMKAETI | RACGAP1; RAC | Q9H0H5; F8 | RGAP1_HUMAI | Unmodified | 0 | 0 | 0 | 0 | 9376500 | 1004400 | 0 | 0 | 0 | 513830 | 0 | 0 | 647460 | 111670 | 0 | 0 | 695870 | 132270 |
| DLLNTGGI | PKP2 | Q99959 | PKP2_HUMAN | Unmodified | 0 | 0 | 0 | 0 | 0 | 0 | 0 | 0 | 0 | 0 | 623870 | 0 | 0 | 0 | 0 | 0 | 0 | 0 |
| DLLPEDFV | ADAM9; ADAM9 | Q13443; A0A | ADAM9_HUMA | Unmodified | 0 | 0 | 0 | 122200 | 0 | 293640 | 79554 | 0 | 0 | 0 | 0 | 28243 | 0 | 96311 | 0 | 0 | 0 | 0 |
| DLLPGPVTI | YRDC | Q86U90 | YRDC_HUMAN | Unmodified | 0 | 0 | 0 | 0 | 0 | 0 | 0 | 0 | 645750 | 290940 | 54592 | 49998 | 0 | 323130 | 0 | 0 | 0 | 0 |
| DLLPLLPG1 | ITGB8 | P26012 | ITB8_HUMAN I | Unmodified | 0 | 0 | 0 | 0 | 0 | 0 | 0 | 0 | 0 | 0 | 0 | 0 | 0 | 0 | 0 | 0 | 0 | 239260 |
| DLLPYGFAI | SNRNP200; SN | O75643; A0A | U520_HUMAN | Oxidation (M) | 0 | 0 | 0 | 0 | 0 | 0 | 0 | 0 | 0 | 0 | 0 | 2864100 | 0 | 0 | 0 | 0 | 0 | 0 |
| DLLQDEPK | VPS37D; VPS37 | Q86XT2; C9J | VP37D_HUMAN | Unmodified | 1408700 | 880170 | 1033400 | 154580 | 0 | 0 | 0 | 0 | 0 | 0 | 0 | 0 | 0 | 0 | 2796600 | 1254500 | 0 | 0 |
| DLLQNLLK | CDK5 | Q00535 | CDK5_HUMAN | Unmodified | 0 | 0 | 0 | 0 | 0 | 0 | 0 | 0 | 0 | 0 | 0 | 1307500 | 0 | 0 | 740510 | 293630 | 0 | 0 |
| DLLQPHRL | POGLUT3; POG | Q7Z4H8; H0 | PLGT3_HUMAN | Unmodified | 0 | 0 | 0 | 0 | 172550 | 0 | 0 | 0 | 2354500 | 5437100 | 0 | 0 | 0 | 0 | 0 | 0 | 0 | 0 |
| DLLQSKIDI | NPTX1 | Q15818 | NPTX1_HUMAN | Unmodified | 0 | 0 | 0 | 0 | 0 | 0 | 0 | 0 | 15684000 | 673160 | 0 | 0 | 0 | 0 | 0 | 0 | 0 | 0 |
| DLLQSKIDI | NPTX1 | Q15818 | NPTX1_HUMAN | Unmodified | 0 | 0 | 0 | 0 | 0 | 0 | 0 | 0 | 35291000 | 6374500 | 0 | 0 | 0 | 0 | 0 | 0 | 18982 | 0 |
| DLLRNLLQ | PRKACA; PRKA | P17612; P22 | KAPCA_HUMAI | Unmodified | 0 | 0 | 0 | 0 | 286370 | 0 | 565100 | 0 | 0 | 481120 | 0 | 0 | 216850 | 169370 | 101670 | 0 | 156310 | 72021 |
| DLLRQLPA | ARHGAP29; ARI | Q52LW3; F8 | RHG29_HUMAI | Unmodified | 0 | 0 | 0 | 0 | 0 | 0 | 0 | 0 | 0 | 0 | 0 | 0 | 0 | 0 | 0 | 0 | 272840 | 0 |
| DLLSHENA | MCU; MCU | Q8NE86; S4I | MCU_HUMAN ( | Unmodified | 0 | 0 | 0 | 0 | 62253 | 0 | 0 | 0 | 835690 | 0 | 0 | 0 | 0 | 0 | 0 | 0 | 0 | 0 |
| DLLSSVSR | ANXA7 | P20073 | ANXA7_HUMAN | Unmodified | 385960 | 251740 | 651350 | 177940 | 0 | 0 | 430800 | 0 | 19558000 | 1683100 | 362400 | 2777100 | 0 | 0 | 994660 | 890730 | 0 | 0 |
| DLLSSVSRI | ANXA7 | P20073 | ANXA7_HUMAN | Unmodified | 0 | 0 | 0 | 0 | 0 | 0 | 0 | 0 | 456480 | 712640 | 0 | 0 | 0 | 0 | 0 | 0 | 0 | 0 |
| DLLVDQTIE | NUMBL; NUMB | Q9Y6R0; A0A | NUMBL_HUMA | Unmodified | 0 | 0 | 337350 | 387950 | 3740300 | 743610 | 134010 | 135070 | 838300 | 464940 | 0 | 0 | 1236400 | 583390 | 300050 | 405320 | 0 | 0 |
| DLLVEGLAI | DOP1B | Q9Y3R5 | DOP2_HUMAN | Unmodified | 65055 | 0 | 0 | 0 | 0 | 0 | 0 | 0 | 229870 | 0 | 43937 | 723410 | 0 | 0 | 156510 | 41793 | 0 | 0 |
| DLLVTGAYI | TMED10; TMED | P49755; G3V | TMEDA_HUMAI | Unmodified | 0 | 0 | 0 | 79591 | 5162300 | 684630 | 317170 | 254890 | 1.42E+08 | 2141000 | 7311900 | 2879100 | 0 | 0 | 634790 | 764540 | 3133400 | 3718500 |
| DLLVTGAYI | TMED10; TMED | P49755; G3V | TMEDA_HUMAI | Unmodified | 0 | 0 | 181520 | 0 | 3412700 | 761950 | 1171400 | 0 | 1.1E+08 | 48184000 | 13426000 | 0 | 7617700 | 7137100 | 158530 | 0 | 0 | 7193200 |
| DLLYFDGLI | PKP3 | Q9Y446 | PKP3_HUMAN | Unmodified | 134820 | 0 | 256580 | 83477 | 0 | 0 | 0 | 0 | 0 | 0 | 0 | 2562700 | 0 | 0 | 1401700 | 747020 | 0 | 0 |
| DLLYFDGLI | PKP3 | Q9Y446 | PKP3_HUMAN | Unmodified | 0 | 0 | 1354600 | 632890 | 0 | 0 | 0 | 0 | 0 | 0 | 265830 | 5004400 | 0 | 0 | 1433000 | 1221600 | 0 | 0 |
| DLMACAQ1 | DDX3X; DDX3Y | O00571; O15 | DDX3X_HUMAI | Oxidation (M) | 166760 | 120650 | 198640 | 158750 | 0 | 0 | 0 | 0 | 0 | 0 | 0 | 0 | 0 | 0 | 151150 | 134810 | 0 | 0 |
| DLMACAQ1 | DDX3X; DDX3Y | O00571; O15 | DDX3X_HUMAI | Unmodified | 351450 | 234570 | 0 | 632300 | 0 | 0 | 0 | 0 | 0 | 0 | 0 | 305420 | 95408 | 0 | 594780 | 293280 | 0 | 0 |
| DLMDEMVI | N/A | N/A | N/A | Oxidation (M) | 1.35E+09 | 3.27E+09 | 1.11E+09 | 0 | 0 | 0 | 0 | 47698000 | 0 | 8.53E+08 | 1.57E+09 | 6.94E+08 | 5.53E+08 | 2.93E+08 | 1.32E+09 | 1.36E+09 | 0 | 0 |
| DLMSKAEK | PGK1 | P00558 | PGK1_HUMAN | Unmodified | 0 | 0 | 0 | 0 | 2411900 | 523490 | 0 | 507360 | 101790 | 704090 | 93733 | 2765200 | 0 | 1792900 | 0 | 142670 | 0 | 5666400 |
| DLMSKAEK | PGK1 | P00558 | PGK1_HUMAN | Unmodified | 0 | 0 | 0 | 0 | 96168 | 34153 | 45936 | 19903 | 0 | 0 | 0 | 0 | 0 | 0 | 0 | 0 | 389370 | 133310 |
| DLMTDLK | ANXA6; ANXA6; | P08133; E7E | ANXA6_HUMAN | Unmodified | 0 | 0 | 394780 | 0 | 0 | 0 | 0 | 0 | 5627700 | 487130 | 0 | 0 | 0 | 0 | 897500 | 0 | 0 | 0 |
| DLMTDLKS | ANXA6; ANXA6; | P08133; E7E | ANXA6_HUMAN | Unmodified | 21373 | 26780 | 0 | 80785 | 57799 | 39022 | 0 | 0 | 1895800 | 341140 | 0 | 0 | 0 | 0 | 34258 | 0 | 34476 | 86948 |
| DLMTDLKS | ANXA6; ANXA6; | P08133; E7E | ANXA6_HUMAN | Oxidation (M) | 0 | 0 | 0 | 150560 | 0 | 0 | 0 | 0 | 346420 | 137400 | 0 | 0 | 0 | 0 | 0 | 0 | 0 | 0 |
| DLMVGDEA | ACTR2 | P61160 | ARP2_HUMAN | Oxidation (M) | 636900 | 746250 | 69784 | 651910 | 0 | 0 | 0 | 0 | 496610 | 36190 | 1122300 | 8003000 | 0 | 0 | 1392000 | 1202100 | 0 | 0 |
| DLNAVAFR | IDE; IDE; IDE; ID | P14735; A0A | IDE_HUMAN In | Unmodified | 0 | 0 | 119990 | 260650 | 0 | 0 | 0 | 0 | 222450 | 0 | 252470 | 4107300 | 54723 | 0 | 156660 | 235680 | 0 | 0 |
| DLNCVPEI | PRMT5; PRMT5; | O14744; G3 | ANM5_HUMAN | Unmodified | 0 | 0 | 0 | 0 | 0 | 0 | 0 | 0 | 0 | 63574 | 162340 | 1623900 | 0 | 0 | 0 | 0 | 0 | 0 |
| DLNFERGG | NDRG2; NDRG2 | Q9UN36; G3 | NDRG2_HUMA | Unmodified | 0 | 0 | 0 | 0 | 0 | 0 | 0 | 0 | 0 | 0 | 871820 | 0 | 0 | 0 | 555720 | 307110 | 0 | 0 |
| DLNGIDLTI | AIDA | Q96BJ3 | AIDA_HUMAN / | Unmodified | 884910 | 0 | 1432000 | 1784400 | 0 | 0 | 0 | 0 | 0 | 0 | 694660 | 1415500 | 0 | 0 | 7818200 | 3767800 | 0 | 0 |
| DLNGIDLTI | AIDA | Q96BJ3 | AIDA_HUMAN / | Unmodified | 0 | 0 | 619670 | 651590 | 0 | 0 | 0 | 0 | 0 | 0 | 0 | 301980 | 0 | 0 | 980590 | 922990 | 0 | 0 |
| DLNPDVN | ATP6V0A1; ATP | Q93050; B7Z | VPP1_HUMAN | Unmodified | 0 | 0 | 0 | 0 | 0 | 0 | 0 | 0 | 1190700 | 257290 | 0 | 0 | 0 | 0 | 0 | 0 | 0 | 0 |
| DLNPDVN | ATP6V0A1; ATP | Q93050; B7Z | VPP1_HUMAN | Unmodified | 27669 | 40373 | 230470 | 184920 | 82421 | 0 | 124250 | 81042 | 1943900 | 0 | 0 | 1418200 | 0 | 0 | 68673 | 43221 | 92551 | 210170 |
| DLNPFTPL | YRDC | Q86U90 | YRDC_HUMAN | Unmodified | 484330 | 0 | 122370 | 0 | 0 | 0 | 0 | 0 | 1205100 | 0 | 0 | 3002200 | 0 | 0 | 0 | 175960 | 0 | 0 |
| DLNQYRAD | TFRC; TFRC; TF | P02786; H7C | TFR1_HUMAN | Unmodified | 0 | 145020 | 341960 | 400680 | 0 | 0 | 0 | 0 | 1359300 | 183500 | 0 | 632390 | 0 | 0 | 462010 | 309340 | 0 | 0 |
| DLNRSFPD | CPD | O75976 | CBPD_HUMAN | Unmodified | 0 | 0 | 0 | 0 | 0 | 0 | 0 | 0 | 0 | 4035700 | 0 | 0 | 0 | 0 | 0 | 0 | 0 | 0 |
| DLPAHLQH | TRAF4; TRAF4; | Q9BUZ4; A0 | TRAF4_HUMAN | Unmodified | 0 | 0 | 0 | 554240 | 0 | 0 | 0 | 0 | 113690 | 0 | 100670 | 184920 | 403920 | 0 | 0 | 0 | 0 | 0 |
| DLPDNPER | BTN3A3; BTN3A | O00478; C9J | BT3A3_HUMAN | Unmodified | 0 | 0 | 0 | 0 | 0 | 0 | 0 | 208490 | 123610 | 0 | 192120 | 543440 | 0 | 0 | 0 | 0 | 0 | 0 |
| DLPDPLDQ | TRPM5; TRPM5; | Q9NZQ8; A0 | TRPM5_HUMAI | Unmodified | 0 | 0 | 0 | 0 | 0 | 0 | 0 | 0 | 0 | 0 | 0 | 1164600 | 0 | 0 | 0 | 0 | 0 | 0 |
| DLPDVQEL | SRP68 | Q9UHB9 | SRP68_HUMAN | Unmodified | 0 | 0 | 0 | 0 | 0 | 0 | 0 | 0 | 190700 | 0 | 212530 | 938540 | 0 | 0 | 0 | 0 | 0 | 0 |
| DLPELALD1 | PDCD4 | Q53EL6 | PDCD4_HUMA | Unmodified | 0 | 0 | 0 | 0 | 0 | 0 | 0 | 86303 | 0 | 0 | 0 | 1576200 | 0 | 0 | 0 | 0 | 0 | 0 |
| DLPELALD1 | PDCD4 | Q53EL6 | PDCD4_HUMA | Unmodified | 0 | 0 | 0 | 0 | 0 | 0 | 0 | 0 | 379360 | 254240 | 368010 | 873450 | 623800 | 0 | 0 | 0 | 0 | 0 |
| DLPGALDE | ARL8A | Q96BM9 | ARL8A_HUMAN | Unmodified | 0 | 0 | 0 | 0 | 0 | 0 | 442860 | 380500 | 431400 | 213340 | 0 | 0 | 195690 | 94212 | 0 | 0 | 41294 | 138520 |
| DLPGDARN | N/A | N/A | N/A | Oxidation (M) | 0 | 0 | 0 | 0 | 0 | 0 | 0 | 0 | 0 | 0 | 0 | 0 | 533840 | 257530 | 0 | 0 | 408940 | 1372400 |
| DLPLAQGII | ALDH7A1; ALDI | P49419; A0A | AL7A1_HUMAN | Unmodified | 0 | 0 | 0 | 0 | 0 | 0 | 0 | 0 | 0 | 0 | 244800 | 1533700 | 0 | 0 | 0 | 0 | 0 | 0 |
| DLPLAQGII | ALDH7A1; ALDI | P49419; A0A | AL7A1_HUMAN | Unmodified | 0 | 0 | 0 | 0 | 1595200 | 0 | 0 | 0 | 0 | 0 | 0 | 0 | 0 | 0 | 0 | 0 | 0 | 1478500 |

| | | | | | | | | | | | | | | | | | | | | | | |
|---|---|---|---|---|---|---|---|---|---|---|---|---|---|---|---|---|---|---|---|---|---|---|
| DLPLLLFR | PHGDH; PHGD | O43175; A0/ | SERA_HUMAN | Unmodified | 1019200 | 880350 | 2666700 | 1725600 | 86811 | 196400 | 132930 | 87228 | 7709600 | 1673600 | 1363900 | 76464000 | 356830 | 115200 | 9923200 | 10644000 | 101280 | 90649 |
| DLPNALDE | ARL8B | Q9NVJ2 | ARL8B_HUMAN | Unmodified | 82089 | 0 | 0 | 83963 | 0 | 0 | 0 | 283840 | 0 | 0 | 0 | 128650 | 712340 | 212630 | 0 | 0 | 0 | 0 |
| DLPPAALLY | FAM120A; FAM | Q9NZB2; A0. | F120A_HUMAN | Unmodified | 0 | 0 | 0 | 0 | 0 | 0 | 0 | 0 | 0 | 0 | 0 | 469720 | 0 | 0 | 0 | 90044 | 0 | 0 |
| DLPPILLVR | SLC12A2 | P55011 | S12A2_HUMAN | Unmodified | 49611000 | 52711000 | 73820 | 1405700 | 634750 | 87378 | 472250 | 148520 | 665070 | 178550 | 641800 | 4434100 | 6320300 | 0 | 36811000 | 36181000 | 0 | 1689100 |
| DLPQAMG | PAIP2B | Q9ULR5 | PAI2B_HUMAN | Unmodified | 304810 | 928580 | 0 | 0 | 3036000 | 669800 | 0 | 0 | 0 | 0 | 0 | 0 | 0 | 4755500 | 241180 | 0 | 0 | 0 |
| DLPRDIQFI | ALOX5 | P09917 | LOX5_HUMAN | Unmodified | 0 | 0 | 0 | 0 | 0 | 0 | 0 | 0 | 221390 | 0 | 188420 | 0 | 0 | 0 | 0 | 0 | 0 | 0 |
| DLPSLYPSI | GNPTAB; GNPT | Q3T906; H0 | GNPTA_HUMAI | Unmodified | 0 | 0 | 0 | 0 | 268520 | 0 | 0 | 0 | 480150 | 0 | 0 | 0 | 0 | 0 | 0 | 0 | 0 | 0 |
| DLPSLYPSI | GNPTAB; GNPT | Q3T906; H0 | GNPTA_HUMAI | Unmodified | 0 | 0 | 0 | 0 | 0 | 0 | 0 | 0 | 3503400 | 735090 | 199990 | 0 | 0 | 0 | 0 | 0 | 0 | 0 |
| DLPTFKDN | PIP4K2B | P78356 | PI42B_HUMAN | Unmodified | 0 | 0 | 0 | 776390 | 0 | 0 | 1306500 | 249490 | 1115500 | 0 | 0 | 0 | 1592000 | 1184700 | 0 | 0 | 0 | 0 |
| DLPVTEAVI | LRPPRC; LRPPI | P42704; B8Z | LPPRC_HUMAN | Unmodified | 0 | 0 | 0 | 0 | 0 | 0 | 629210 | 0 | 15318000 | 651950 | 0 | 302070 | 0 | 0 | 0 | 0 | 15084 | 0 |
| DLQANVEI | PFKM; PFKM | P08237; A0A | PFKAM_HUMAI | Unmodified | 304070 | 121220 | 461850 | 142470 | 0 | 0 | 0 | 0 | 0 | 0 | 0 | 4766100 | 0 | 0 | 975360 | 482450 | 0 | 0 |
| DLQEFIPLII | XPOT | O43592 | XPOT_HUMAN | Unmodified | 39597 | 0 | 20957 | 0 | 0 | 14194 | 0 | 0 | 89726 | 0 | 40410 | 431380 | 0 | 7353.8 | 76407 | 27461 | 0 | 18453 |
| DLQFLLR | MGAT5 | Q09328 | MGT5A_HUMAI | Unmodified | 0 | 0 | 0 | 0 | 0 | 0 | 0 | 0 | 3669100 | 0 | 267090 | 1368800 | 0 | 0 | 0 | 0 | 0 | 0 |
| DLQFVEVTI | FN1 | P02751 | FINC_HUMAN | Unmodified | 0 | 773240 | 5252100 | 889810 | 0 | 0 | 0 | 0 | 0 | 130160 | 0 | 1829700 | 191470 | 0 | 365240 | 1401000 | 0 | 0 |
| DLQGRDE | MYH9 | P35579 | MYH9_HUMAN | Unmodified | 0 | 15317 | 63471 | 0 | 0 | 0 | 0 | 0 | 0 | 0 | 10485 | 282650 | 0 | 0 | 81001 | 43248 | 28448 | 0 |
| DLQGRPSS | POFUT1 | Q9H488 | OFUT1_HUMAI | Unmodified | 0 | 0 | 280790 | 436300 | 247430 | 0 | 0 | 0 | 2461100 | 0 | 0 | 0 | 0 | 0 | 0 | 49161 | 0 | 0 |
| DLQHPNEI | COPB1; COPB1 | P53618; E9F | COPB_HUMAN | Unmodified | 0 | 0 | 0 | 0 | 0 | 0 | 30445 | 0 | 0 | 0 | 0 | 5663900 | 60049 | 0 | 197530 | 120140 | 0 | 0 |
| DLQIYSR | MYO1C; MYO1 | O00159; I3L | MYO1C_HUMA | Unmodified | 3979800 | 2431300 | 3404200 | 1268700 | 2048300 | 606530 | 29932 | 0 | 458480 | 1894500 | 49772 | 1027300 | 0 | 0 | 9489800 | 6255100 | 131200 | 158110 |
| DLQLLSQF | ADGRG1; ADGI | Q9Y653; H3I | AGRG1_HUMA | Unmodified | 0 | 0 | 0 | 130560 | 0 | 0 | 0 | 0 | 1731200 | 0 | 0 | 0 | 0 | 0 | 164090 | 138930 | 0 | 0 |
| DLQLQCEA | SLK | Q9H2G2 | SLK_HUMAN S | Unmodified | 0 | 0 | 0 | 0 | 0 | 0 | 0 | 0 | 0 | 0 | 0 | 1424700 | 0 | 88539 | 0 | 0 | 0 | 0 |
| DLQLQCEA | SLK | Q9H2G2 | SLK_HUMAN S | Unmodified | 0 | 0 | 64165 | 221230 | 0 | 0 | 0 | 0 | 0 | 0 | 559000 | 0 | 408520 | 0 | 60821 | 0 | 142610 | 0 |
| DLQLQILR | SUN1; SUN1; S | O94901; H0 | SUN1_HUMAN | Unmodified | 0 | 0 | 0 | 0 | 482910 | 197020 | 0 | 0 | 4995800 | 587430 | 666460 | 764240 | 464860 | 0 | 0 | 0 | 215450 | 0 |
| DLQMFLYH | SLC12A7; SLC1 | Q9Y666; A0/ | S12A7_HUMAN | Oxidation (M) | 0 | 0 | 486590 | 666380 | 0 | 0 | 0 | 0 | 0 | 0 | 35771 | 933130 | 0 | 0 | 1353000 | 1060800 | 0 | 0 |
| DLQNLMS | MME; MME | P08473; A0A | NEP_HUMAN | Oxidation (M) | 0 | 0 | 3155000 | 0 | 0 | 0 | 0 | 0 | 0 | 0 | 0 | 0 | 0 | 0 | 6400000 | 3685800 | 0 | 0 |
| DLQNLMS | MME; MME | P08473; A0A | NEP_HUMAN | Unmodified | 0 | 0 | 884640 | 249360 | 0 | 0 | 0 | 0 | 0 | 0 | 0 | 0 | 0 | 0 | 3712500 | 1526300 | 0 | 0 |
| DLQNVNIT | PHB; PHB; PHB | P35232; C9J | PHB_HUMAN | Unmodified | 0 | 0 | 0 | 184410 | 0 | 0 | 0 | 177040 | 1668000 | 517030 | 0 | 0 | 0 | 162050 | 0 | 0 | 0 | 0 |
| DLQQSIAR | PDCD6IP | Q8WUM4 | PDC6I_HUMAN | Unmodified | 0 | 0 | 0 | 497050 | 200400 | 0 | 43169 | 0 | 1221200 | 341910 | 82585 | 141890 | 251880 | 0 | 0 | 0 | 12695 | 235660 |
| DLQQSIARI | PDCD6IP | Q8WUM4 | PDC6I_HUMAN | Unmodified | 2.93E+08 | 2.07E+08 | 65993000 | 99150000 | 1.58E+08 | 28863000 | 3999800 | 3050900 | 1.11E+08 | 45526000 | 13711000 | 41019000 | 84549000 | 66118000 | 89627000 | 66056000 | 9885800 | 99381000 |
| DLQSNVEI | PFKP; PFKP; PF | Q01813; A0/ | PFKAP_HUMAI | Unmodified | 62817 | 210910 | 0 | 0 | 124230 | 0 | 173140 | 65987 | 0 | 0 | 0 | 7688000 | 0 | 0 | 822230 | 435440 | 139570 | 100120 |
| DLQSSDRL | G6PD; G6PD; G | P11413; E7E | G6PD_HUMAN | Unmodified | 0 | 0 | 0 | 0 | 0 | 0 | 0 | 0 | 0 | 0 | 0 | 1171700 | 0 | 0 | 0 | 0 | 0 | 0 |
| DLQTHGW | DHX29; DHX29 | Q7Z478; A0/ | DHX29_HUMAI | Unmodified | 0 | 0 | 0 | 0 | 0 | 0 | 0 | 0 | 0 | 0 | 0 | 608090 | 0 | 0 | 0 | 0 | 0 | 0 |
| DLQTRDE | MYH10 | P35580 | MYH10_HUMA | Unmodified | 0 | 0 | 189930 | 0 | 0 | 0 | 0 | 0 | 0 | 0 | 0 | 1541500 | 0 | 0 | 184890 | 313380 | 0 | 0 |
| DLQVQVR | GPR108; GPR1 | Q9NPR9; M | GP108_HUMAN | Unmodified | 0 | 0 | 0 | 0 | 0 | 0 | 0 | 0 | 270990 | 0 | 0 | 0 | 0 | 0 | 0 | 0 | 0 | 0 |
| DLQYVEK | OLFM1; OLFM1 | Q99784; F5I | NOE1_HUMAN | Unmodified | 0 | 0 | 0 | 0 | 300030 | 46331 | 0 | 18676 | 2005000 | 190750 | 0 | 0 | 0 | 300740 | 0 | 0 | 0 | 0 |
| DLREIQHH | SH3GL2 | Q99962 | SH3G2_HUMAI | Unmodified | 0 | 0 | 0 | 0 | 0 | 0 | 234720 | 13067 | 730690 | 322070 | 0 | 250320 | 0 | 0 | 0 | 0 | 0 | 0 |
| DLRHTENE | COASY; COASY | Q13057; K7E | COASY_HUMAI | Unmodified | 0 | 0 | 0 | 0 | 299090 | 36334 | 14130 | 31493 | 224430 | 0 | 215980 | 0 | 0 | 0 | 177720 | 0 | 0 | 0 |
| DLRNDEHT | RHOA; RHOA; F | P61586; C9J | RHOA_HUMAN | Unmodified | 0 | 0 | 0 | 0 | 0 | 0 | 0 | 0 | 990990 | 0 | 0 | 1089600 | 0 | 84053 | 0 | 0 | 264010 | 290310 |
| DLRPGDTV | CD276; CD276 | Q5ZPR3; A0/ | CD276_HUMAI | Unmodified | 2627900 | 1772100 | 6506800 | 7497800 | 0 | 0 | 0 | 0 | 1661200 | 0 | 0 | 5663200 | 0 | 0 | 6383100 | 7847200 | 0 | 0 |
| DLRPSALPI | TRRAP; TRRAP | Q9Y4A5; F2Z | TRRAP_HUMAN | Unmodified | 0 | 0 | 0 | 0 | 0 | 0 | 0 | 0 | 0 | 0 | 0 | 195670 | 0 | 0 | 0 | 0 | 0 | 0 |
| DLRREISYA | DNM2; DNM2 | P50570; K7E | DYN2_HUMAN | Unmodified | 267350 | 227440 | 0 | 306050 | 1379700 | 0 | 662810 | 0 | 635710 | 166390 | 0 | 0 | 1074400 | 1143100 | 162890 | 121360 | 770920 | 783000 |
| DLRSDEHV | RHOB | P62745 | RHOB_HUMAN | Unmodified | 0 | 250350 | 650260 | 3020300 | 1675000 | 0 | 521840 | 0 | 721110 | 0 | 0 | 1772700 | 2273600 | 750630 | 0 | 579240 | 151420 | 62571 |
| DLRTLLNET | PROM1 | O43490 | PROM1_HUMA | Unmodified | 0 | 0 | 0 | 0 | 0 | 0 | 0 | 16984 | 0 | 0 | 1081700 | 0 | 0 | 0 | 0 | 0 | 0 | 0 |
| DLSAENGL | CPD; CPD | O75976; J3C | CBPD_HUMAN | Unmodified | 0 | 0 | 0 | 0 | 401750 | 0 | 571380 | 0 | 12252000 | 552080 | 0 | 0 | 0 | 0 | 0 | 0 | 0 | 0 |
| DLSAGAVS | ASRGL1; ASRGI | Q7L266; A0/ | ASGL1_HUMAI | Unmodified | 0 | 0 | 0 | 0 | 0 | 0 | 0 | 0 | 1835300 | 718400 | 0 | 0 | 0 | 0 | 0 | 0 | 0 | 0 |
| DLSDGIHV | CKAP4 | Q07065 | CKAP4_HUMAI | Unmodified | 0 | 0 | 0 | 0 | 0 | 0 | 0 | 45748 | 927130 | 198850 | 0 | 0 | 0 | 0 | 0 | 0 | 0 | 0 |
| DLSDRSNI | SH3KBP1; SH3I | Q96B97; Q5. | SH3K1_HUMAN | Unmodified | 0 | 0 | 890560 | 1091200 | 2575500 | 0 | 0 | 0 | 0 | 0 | 0 | 0 | 1888100 | 0 | 1385000 | 1093300 | 0 | 0 |
| DLSEALPEI | MVB12B; MVB1 | Q9H7P6; F8 | MB12B_HUMAI | Oxidation (M) | 0 | 0 | 6990800 | 0 | 600360 | 0 | 0 | 0 | 3662500 | 1761100 | 522560 | 494670 | 649950 | 0 | 12325000 | 0 | 0 | 330740 |
| DLSEALPEI | MVB12B; MVB1 | Q9H7P6; F8 | MB12B_HUMAI | Unmodified | 0 | 0 | 0 | 0 | 146360 | 0 | 0 | 0 | 5506000 | 0 | 1142700 | 0 | 0 | 0 | 703890 | 0 | 0 | 0 |
| DLSEFLR | CPD; CPD | O75976; J3C | CBPD_HUMAN | Unmodified | 0 | 0 | 348190 | 0 | 0 | 0 | 0 | 0 | 11849000 | 781760 | 0 | 229440 | 0 | 0 | 0 | 230380 | 0 | 0 |
| DLSENND | GOLM1 | Q8NBJ4 | GOLM1_HUMA | Unmodified | 0 | 0 | 0 | 0 | 0 | 0 | 0 | 0 | 14766000 | 267420 | 0 | 0 | 0 | 0 | 0 | 0 | 0 | 0 |
| DLSFFGGL | P3H1; P3H1 | Q32P28; E2C | P3H1_HUMAN | Unmodified | 0 | 0 | 0 | 0 | 0 | 0 | 0 | 0 | 925490 | 0 | 0 | 0 | 0 | 0 | 0 | 0 | 0 | 0 |
| DLSFGHSR | PKP1 | Q13835 | PKP1_HUMAN | Unmodified | 0 | 55328 | 0 | 0 | 0 | 0 | 0 | 0 | 0 | 0 | 0 | 0 | 0 | 0 | 0 | 0 | 647200 | 726110 |
| DLSGLDAE | PTPN6; PTPN6 | P29350; U3K | PTN6_HUMAN | Unmodified | 0 | 0 | 0 | 0 | 0 | 0 | 0 | 0 | 237410 | 0 | 0 | 617290 | 0 | 0 | 0 | 87824 | 0 | 0 |
| DLSHIGDA | PCNA | P12004 | PCNA_HUMAN | Unmodified | 0 | 0 | 0 | 0 | 0 | 0 | 0 | 0 | 1920800 | 0 | 0 | 0 | 0 | 0 | 786960 | 0 | 0 | 0 |
| DLSKPIGAI | NBEA; LRBA; LF | Q8NFP9; P5 | NBEA_HUMAN | Unmodified | 0 | 0 | 0 | 0 | 0 | 0 | 0 | 0 | 0 | 0 | 0 | 1527700 | 0 | 0 | 0 | 0 | 0 | 0 |
| DLSLDDFK | PRDX3 | P30048 | PRDX3_HUMAI | Unmodified | 0 | 0 | 0 | 0 | 0 | 0 | 0 | 0 | 18806000 | 617450 | 0 | 1918300 | 0 | 0 | 0 | 0 | 0 | 0 |
| DLSLDDFK | PRDX3 | P30048 | PRDX3_HUMAI | Unmodified | 0 | 0 | 0 | 0 | 0 | 0 | 0 | 0 | 734120 | 432280 | 0 | 0 | 0 | 0 | 0 | 0 | 0 | 0 |
| DLSLEEIQK | STMN1; STMN2 | P16949; Q9 | STMN1_HUMAI | Unmodified | 697150 | 396790 | 479630 | 598500 | 7222400 | 1370800 | 3244300 | 5582800 | 35728000 | 15078000 | 14248000 | 17735000 | 5484200 | 1212700 | 458030 | 482700 | 1529900 | 3110700 |
| DLSLEEIQK | STMN1; STMN2 | P16949; Q9 | STMN1_HUMAI | Unmodified | 0 | 0 | 0 | 0 | 0 | 0 | 0 | 0 | 0 | 204410 | 1361400 | 0 | 0 | 0 | 0 | 0 | 0 | 0 |
| DLSLLQIQN | NAA16; NAA15 | Q6N069; Q9 | NAA16_HUMAN | Oxidation (M) | 0 | 0 | 0 | 0 | 0 | 0 | 0 | 0 | 0 | 0 | 0 | 2287900 | 0 | 0 | 0 | 0 | 0 | 0 |
| DLSLNKLG | GSR | P00390 | GSHR_HUMAN | Unmodified | 0 | 0 | 0 | 0 | 8647400 | 815750 | 0 | 0 | 0 | 0 | 0 | 0 | 4212600 | 2115500 | 0 | 0 | 1000600 | 0 |
| DLSLPPVD | NPEPPS; NPEPI | P55786; A0A | PSA_HUMAN P | Unmodified | 0 | 65317 | 0 | 0 | 0 | 0 | 0 | 0 | 0 | 22879 | 0 | 103020 | 0 | 0 | 931880 | 50681 | 13409 | 0 |
| DLSQKYTE | UACA; UACA; U | Q9BZF9; H0 | UACA_HUMAN | Unmodified | 0 | 0 | 0 | 0 | 0 | 0 | 0 | 0 | 120780 | 47993 | 0 | 0 | 0 | 0 | 0 | 0 | 537400 | 0 |
| DLSQNFPT | UGGT1 | Q9NYU2 | UGGG1_HUMA | Unmodified | 0 | 879070 | 918780 | 159750 | 0 | 0 | 88737 | 88578 | 9943200 | 3729500 | 165470 | 748000 | 286970 | 0 | 356840 | 358920 | 91159 | 374110 |
| DLSRFDAS | FASN; FASN; FA | P49327; A0A | FAS_HUMAN F | Unmodified | 435370 | 322500 | 4004600 | 1840200 | 7139400 | 3102600 | 4956800 | 193810 | 2148200 | 1346400 | 5641500 | 68072000 | 10724000 | 707000 | 5445800 | 2925500 | 3981100 | 3233900 |
| DLSSHQLN | LTA4H | P09960 | LKHA4_HUMAI | Unmodified | 0 | 0 | 0 | 0 | 0 | 0 | 0 | 0 | 1287800 | 0 | 2629700 | 15256000 | 0 | 0 | 0 | 0 | 0 | 30629 |
| DLSSHQLN | LTA4H | P09960 | LKHA4_HUMAI | Unmodified | 0 | 0 | 0 | 0 | 1587900 | 364780 | 224410 | 0 | 0 | 1525400 | 97363 | 0 | 953930 | 508630 | 0 | 0 | 0 | 1063200 |
| DLSTIEPLK | ANP32A; ANP3 | P39687; O4 | AN32A_HUMAN | Unmodified | 40776 | 82295 | 0 | 0 | 0 | 0 | 0 | 0 | 0 | 0 | 8461300 | 12841000 | 2544400 | 0 | 0 | 0 | 0 | 0 |
| DLSTVEAL | ANP32E; ANP3 | Q9BTT0; E9F | AN32E_HUMAN | Unmodified | 283150 | 240540 | 0 | 0 | 959880 | 0 | 0 | 316060 | 761380 | 818720 | 2240900 | 2861400 | 743050 | 0 | 0 | 102840 | 159260 | 369590 |
| DLSVKYPEI | TXNDC11 | Q6PKC3 | TXD11_HUMAN | Unmodified | 0 | 0 | 0 | 0 | 0 | 0 | 0 | 0 | 313320 | 0 | 0 | 0 | 0 | 0 | 0 | 0 | 0 | 0 |
| DLSWPSSY | TC2N | Q8N9U0 | TAC2N_HUMAI | Unmodified | 0 | 0 | 0 | 0 | 0 | 979810 | 0 | 181330 | 0 | 0 | 0 | 0 | 0 | 0 | 167670 | 114220 | 0 | 0 |
| DLTALSNM | ANPEP; ANPEP | P15144; H0 | AMPN_HUMAN | Oxidation (M) | 0 | 0 | 0 | 0 | 0 | 0 | 107880 | 0 | 0 | 0 | 0 | 0 | 0 | 0 | 0 | 293050 | 1422700 | 3852400 |
| DLTALSNM | ANPEP; ANPEP | P15144; H0 | AMPN_HUMAN | Unmodified | 0 | 0 | 0 | 0 | 904480 | 0 | 354190 | 0 | 0 | 608070 | 0 | 0 | 0 | 288740 | 0 | 0 | 969340 | 3872900 |
| DLTCNKSII | N/A | N/A | N/A | Oxidation (M) | 0 | 0 | 0 | 0 | 6754300 | 5728100 | 3751500 | 197230 | 0 | 1948400 | 0 | 0 | 2682900 | 0 | 0 | 0 | 0 | 21711000 |
| DLTDGHFE | SERPINB5 | P36952 | SPB5_HUMAN | Unmodified | 0 | 0 | 0 | 0 | 0 | 0 | 0 | 0 | 0 | 0 | 0 | 0 | 702910 | 482780 | 0 | 0 | 1463900 | 2161900 |
| DLTDYLMK | ACTG1; ACTC1 | P63261; P68 | ACTG_HUMAN | Oxidation (M) | 5258700 | 6129600 | 9546800 | 1296600 | 635280 | 0 | 531540 | 454350 | 1857200 | 725070 | 6505500 | 95140000 | 210910 | 147030 | 15691000 | 16982000 | 875360 | 0 |
| DLTDYLMK | ACTG1; ACTC1 | P63261; P68 | ACTG_HUMAN | Unmodified | 878450 | 1945500 | 1062800 | 421400 | 1146300 | 0 | 980270 | 80610 | 3860500 | 631860 | 2426500 | 44652000 | 0 | 0 | 2788700 | 1386500 | 1059100 | 0 |
| DLTEAVPRI | MTM1; MTM1 | Q13496; C9J | MTM1_HUMAN | Unmodified | 0 | 155520 | 63762 | 0 | 0 | 0 | 0 | 0 | 0 | 0 | 99981 | 225020 | 0 | 0 | 114430 | 114780 | 0 | 0 |
| DLTELVQR | FUT8; FUT8; FU | Q9BYC5; A0. | FUT8_HUMAN | Unmodified | 7390100 | 0 | 0 | 0 | 0 | 0 | 1608900 | 193350 | 3937000 | 506000 | 0 | 0 | 0 | 0 | 0 | 0 | 93165 | 0 |
| DLTELVQRI | FUT8; FUT8; FU | Q9BYC5; A0. | FUT8_HUMAN | Unmodified | 0 | 0 | 0 | 0 | 0 | 0 | 0 | 0 | 1683100 | 0 | 0 | 0 | 0 | 162310 | 0 | 0 | 94525 | 0 |
| DLTEVSTLK | APPL2; APPL2; | Q8NEU8; F8 | DP13B_HUMAI | Unmodified | 0 | 0 | 0 | 0 | 908270 | 0 | 0 | 0 | 555540 | 0 | 0 | 384790 | 787860 | 0 | 0 | 0 | 0 | 0 |
| DLTEYLSR | HNRNPDL; HNI | O14979; A0/ | HNRDL_HUMA | Unmodified | 0 | 0 | 0 | 0 | 0 | 0 | 0 | 0 | 0 | 0 | 536230 | 9333800 | 0 | 0 | 0 | 0 | 0 | 0 |
| DLTGDQSF | MYOF | Q9NZM1 | MYOF_HUMAN | Unmodified | 0 | 0 | 0 | 0 | 18444000 | 495850 | 74974 | 0 | 101910 | 0 | 384910 | 6704000 | 364920 | 287540 | 0 | 0 | 0 | 0 |
| DLTGDQSF | MYOF | Q9NZM1 | MYOF_HUMAN | Unmodified | 0 | 0 | 0 | 0 | 11864000 | 482810 | 0 | 0 | 0 | 0 | 0 | 0 | 0 | 0 | 0 | 0 | 802250 | 2728100 |
| DLTGQVPT | MAP1B; MAP1B | P46821; D6I | MAP1B_HUMAN | Unmodified | 0 | 0 | 0 | 0 | 0 | 0 | 0 | 0 | 553340 | 336750 | 371390 | 1219000 | 0 | 0 | 0 | 0 | 0 | 0 |
| DLTIGQMQ | PPME1 | Q9Y570 | PPME1_HUMAI | Oxidation (M) | 171020 | 228700 | 665420 | 0 | 0 | 0 | 0 | 0 | 0 | 0 | 0 | 1547900 | 0 | 0 | 91783 | 242530 | 0 | 0 |
| DLTIGQMQ | PPME1 | Q9Y570 | PPME1_HUMAI | Unmodified | 92352 | 141390 | 67037 | 0 | 137700 | 0 | 0 | 0 | 837630 | 69523 | 0 | 1435200 | 0 | 0 | 208390 | 99972 | 0 | 0 |
| DLTIQKADI | DARS1; DARS1 | P14868; H7C | SYDC_HUMAN | Unmodified | 0 | 0 | 0 | 0 | 0 | 0 | 0 | 0 | 0 | 0 | 0 | 1624400 | 445150 | 0 | 0 | 0 | 0 | 0 |
| DLTLLGAT | ATP11A; ATP11 | P98196; H0 | AT11A_HUMAN | Unmodified | 0 | 0 | 0 | 0 | 0 | 0 | 1870700 | 154940 | 0 | 277820 | 0 | 1018700 | 76474 | 0 | 942100 | 0 | 0 | 0 |
| DLTNVQNL | SPTAN1; SPTAN | Q13813; A0/ | SPTN1_HUMAN | Unmodified | 0 | 0 | 0 | 0 | 481120 | 0 | 0 | 0 | 0 | 332110 | 387100 | 1785400 | 0 | 0 | 159750 | 0 | 227570 | 0 |
| DLTNVQNL | SPTAN1; SPTAN | Q13813; A0/ | SPTN1_HUMAN | Unmodified | 0 | 0 | 0 | 0 | 441110 | 0 | 37703 | 0 | 0 | 0 | 159910 | 1507300 | 0 | 0 | 0 | 0 | 240320 | 0 |
| DLTQEMEV | UTRN | P46939 | UTRN_HUMAN | Unmodified | 0 | 0 | 0 | 350550 | 189140 | 0 | 0 | 26713 | 0 | 0 | 0 | 411060 | 0 | 0 | 702500 | 886870 | 0 | 0 |
| DLTQLFMF | DPP7; DPP7 | Q9UHL4; R4 | DPP2_HUMAN | Oxidation (M) | 0 | 0 | 0 | 0 | 0 | 0 | 0 | 0 | 1263600 | 103770 | 45339 | 299340 | 0 | 0 | 0 | 0 | 0 | 0 |
| DLTQLFMF | DPP7; DPP7 | Q9UHL4; R4 | DPP2_HUMAN | Unmodified | 0 | 0 | 0 | 0 | 0 | 119670 | 0 | 0 | 245050 | 0 | 0 | 0 | 0 | 0 | 0 | 0 | 0 | 0 |
| DLTQMADE | NAA10; NAA10 | P41227; C9J | NAA10_HUMAN | Unmodified | 0 | 0 | 0 | 0 | 0 | 0 | 0 | 0 | 0 | 0 | 0 | 760050 | 0 | 0 | 0 | 0 | 0 | 0 |
| DLTQTASS | MYOF; MYOF | Q9NZM1; H0 | MYOF_HUMAN | Unmodified | 0 | 0 | 0 | 0 | 2663000 | 1493400 | 547480 | 0 | 0 | 0 | 0 | 0 | 0 | 0 | 0 | 0 | 0 | 499820 |
| DLTSTMETI | N/A | N/A | N/A | 2 Oxidation (M) | 0 | 0 | 0 | 0 | 0 | 0 | 0 | 0 | 0 | 0 | 0 | 944570 | 0 | 14330000 | 206300 | 0 | 0 | 0 |
| DLTSVMR | SPTBN1; SPTBN | Q01082; A0/ | SPTB2_HUMAN | Unmodified | 0 | 0 | 0 | 0 | 143140 | 0 | 0 | 0 | 184000 | 124170 | 0 | 2656200 | 0 | 0 | 0 | 0 | 0 | 0 |
| DLTSVNILL | SPTBN2; SPTBN | O15020; Q0 | SPTN2_HUMAN | Unmodified | 0 | 0 | 0 | 0 | 373410 | 0 | 0 | 102490 | 0 | 0 | 0 | 485550 | 411030 | 139160 | 158270 | 82395 | 0 | 0 |
| DLTTAGAV | RPL18A; RPL18 | Q02543; M0 | RL18A_HUMAN | Unmodified | 4482600 | 2053100 | 1166900 | 342470 | 0 | 0 | 0 | 0 | 0 | 0 | 830940 | 18754000 | 0 | 0 | 1619000 | 2060500 | 0 | 0 |
| DLTTAGAV | RPL18A; RPL18 | Q02543; M0 | RL18A_HUMAN | Unmodified | 200350 | 0 | 0 | 0 | 0 | 0 | 98017 | 44101 | 0 | 0 | 0 | 222300 | 0 | 0 | 0 | 0 | 0 | 0 |
| DLTTFLESC | TAOK3; TAOK3 | Q9H2K8; G3 | TAOK3_HUMAN | Unmodified | 0 | 0 | 0 | 623140 | 1203300 | 0 | 802650 | 880200 | 132310 | 0 | 0 | 0 | 988730 | 1065100 | 508940 | 0 | 455280 | 415540 |
| DLTVANVG | PPM1L | Q5SGD2 | PPM1L_HUMAI | Unmodified | 0 | 0 | 0 | 0 | 0 | 0 | 323790 | 0 | 7488200 | 0 | 0 | 0 | 0 | 0 | 0 | 0 | 0 | 0 |
| DLVAIEAK | SPTBN1; SPTBN | Q01082; A0/ | SPTB2_HUMAN | Unmodified | 0 | 107480 | 0 | 0 | 256500 | 0 | 0 | 0 | 292520 | 0 | 444140 | 3280500 | 0 | 0 | 59056 | 0 | 0 | 0 |
| DLVAQTAQ | FAM91A1; FAM | Q658Y4; E7E | F91A1_HUMAN | Unmodified | 0 | 0 | 0 | 0 | 456350 | 0 | 0 | 0 | 217800 | 0 | 0 | 110730 | 0 | 0 | 48435 | 22500 | 0 | 419430 |
| DLVARESLI | AGRN | O00468 | AGRIN_HUMAN | Unmodified | 0 | 0 | 0 | 0 | 0 | 183950 | 0 | 0 | 0 | 73826 | 0 | 0 | 777850 | 573800 | 37435 | 0 | 0 | 628530 |
| DLVEISPNI | ADAM15 | Q13444 | ADA15_HUMAN | Unmodified | 0 | 490250 | 18204 | 0 | 0 | 0 | 0 | 0 | 0 | 0 | 0 | 0 | 0 | 0 | 1062500 | 0 | 0 | 0 |
| DLVEPWV | EPHB4; EPHB4 | P54760; Q9 | EPHB4_HUMAN | Unmodified | 637440 | 387560 | 0 | 0 | 0 | 0 | 0 | 0 | 0 | 0 | 0 | 0 | 0 | 0 | 142740 | 142570 | 0 | 0 |
| DLVGIENLI | FARP2 | O94887 | FARP2_HUMAN | Unmodified | 0 | 0 | 77888 | 0 | 0 | 0 | 0 | 0 | 0 | 0 | 0 | 0 | 0 | 0 | 0 | 0 | 0 | 0 |

| | | | | | | | | | | | | | | | | | | | | | | |
|---|---|---|---|---|---|---|---|---|---|---|---|---|---|---|---|---|---|---|---|---|---|---|
| DLVGIENLI | FARP2 | O94887 | FARP2_HUMAN | Unmodified | 0 | 0 | 82537 | 146670 | 0 | 0 | 0 | 0 | 0 | 0 | 0 | 0 | 0 | 0 | 676380 | 118430 | 0 | 0 |
| DLVHTEGC | HSPB11 | Q9Y547 | IFT25_HUMAN | Unmodified | 0 | 0 | 0 | 450270 | 0 | 0 | 0 | 0 | 1171900 | 665420 | 0 | 0 | 949320 | 0 | 0 | 1304200 | 0 | 0 |
| DLVKLQAG | ACSL4; ACSL3; | O60488; O9 | ACSL4_HUMAN | Unmodified | 0 | 0 | 0 | 0 | 0 | 0 | 0 | 0 | 0 | 0 | 0 | 0 | 0 | 0 | 0 | 0 | 1875400 | 0 |
| DLVNTANV | DSC2; DSC2 | Q02487; A0 | DSC2_HUMAN | Unmodified | 0 | 0 | 0 | 0 | 0 | 0 | 70860 | 0 | 2902900 | 0 | 0 | 0 | 0 | 0 | 0 | 0 | 125660 | 0 |
| DLVPDLSN | SDHB; SDHB; S | P21912; A0A | SDHB_HUMAN | Unmodified | 0 | 0 | 0 | 0 | 0 | 0 | 875150 | 0 | 5252500 | 1440400 | 0 | 0 | 0 | 0 | 0 | 0 | 0 | 0 |
| DLVQDPSL | GPHN; GPHN; | Q9NQX3; G3 | GEPH_HUMAN | Unmodified | 0 | 0 | 0 | 0 | 243290 | 0 | 0 | 0 | 0 | 0 | 235950 | 1323300 | 198470 | 0 | 0 | 0 | 0 | 0 |
| DLVSGLFSI | ENTPD2 | Q9Y5L3 | ENTP2_HUMAN | Unmodified | 0 | 34743 | 0 | 0 | 0 | 0 | 0 | 0 | 0 | 18748 | 0 | 0 | 0 | 0 | 99718 | 0 | 0 | 0 |
| DLVTQQLP | S100A14 | Q9HCY8 | S10AE_HUMAN | Unmodified | 0 | 0 | 0 | 0 | 0 | 0 | 0 | 0 | 237410 | 0 | 0 | 0 | 0 | 0 | 0 | 0 | 26737 | 529290 |
| DLVTVEQVI | FAM83F | Q8NEG4 | FA83F_HUMAN | Unmodified | 0 | 358730 | 521570 | 1510600 | 0 | 0 | 0 | 0 | 1512300 | 1045800 | 800910 | 0 | 707560 | 475430 | 2988800 | 661340 | 0 | 0 |
| DLVTVEQVI | FAM83F | Q8NEG4 | FA83F_HUMAN | Unmodified | 0 | 0 | 0 | 0 | 0 | 0 | 0 | 0 | 184820 | 888030 | 0 | 0 | 0 | 0 | 0 | 0 | 0 | 0 |
| DLVWLEEI | INCENP; INCEN | Q9NQS7; E9 | INCE_HUMAN I | Unmodified | 0 | 0 | 0 | 243860 | 89869 | 0 | 0 | 0 | 86090 | 0 | 0 | 0 | 28802 | 12468 | 166580 | 26719 | 0 | 0 |
| DLWASKNE | LIMA1; LIMA1; L | Q9UHB6; F8 | LIMA1_HUMAN | Unmodified | 1138400 | 1188800 | 0 | 3035700 | 1159600 | 0 | 0 | 0 | 0 | 0 | 0 | 2148800 | 0 | 0 | 4165200 | 869040 | 33196 | 0 |
| DLWLDGN | LRRC1 | Q9BTT6 | LRRC1_HUMAN | Unmodified | 0 | 0 | 0 | 0 | 0 | 0 | 0 | 0 | 0 | 0 | 0 | 0 | 647060 | 0 | 0 | 0 | 0 | 0 |
| DLYANTVLS | ACTG1; ACTB; | P63261; P60 | ACTG_HUMAN | Oxidation (M) | 96508000 | 1.01E+08 | 40928000 | 89908000 | 50362000 | 19633000 | 19281000 | 15952000 | 88520000 | 20551000 | 81424000 | 3.63E+08 | 32227000 | 12586000 | 1.65E+08 | 1.09E+08 | 10669000 | 25260000 |
| DLYANTVLS | ACTG1; ACTB; | P63261; P60 | ACTG_HUMAN | Unmodified | 3032500 | 10312000 | 1134900 | 7911300 | 19893000 | 0 | 0 | 0 | 56738000 | 3239300 | 3792500 | 14073000 | 3390700 | 0 | 10497000 | 2415400 | 0 | 9264100 |
| DLYANTVLS | ACTG1; ACTB; | P63261; P60 | ACTG_HUMAN | 2 Oxidation (M) | 9948300 | 6276100 | 0 | 35181000 | 3.69E+08 | 1.8E+08 | 74734000 | 31747000 | 3418500 | 30865000 | 18091000 | 6137300 | 1.99E+08 | 1.27E+08 | 20648000 | 5084500 | 93772000 | 76543000 |
| DLYANTVLS | ACTG1; ACTB; | P63261; P60 | ACTG_HUMAN | Oxidation (M) | 3085700 | 1650500 | 700120 | 16670000 | 9.46E+08 | 1.01E+08 | 93890000 | 5335500 | 7533600 | 26000000 | 7669600 | 528460 | 1.33E+08 | 1.64E+08 | 3650000 | 0 | 1.8E+08 | 2.03E+08 |
| DLYANTVLS | ACTG1; ACTB; | P63261; P60 | ACTG_HUMAN | Unmodified | 0 | 0 | 0 | 0 | 1.54E+08 | 12199000 | 11801000 | 0 | 6804800 | 5801700 | 0 | 0 | 29739000 | 25040000 | 0 | 0 | 17376000 | 43994000 |
| DLYANTVLS | ACTG1; ACTB; | P63261; P60 | ACTG_HUMAN | 2 Oxidation (M) | 0 | 0 | 0 | 0 | 10412000 | 0 | 8778300 | 0 | 0 | 0 | 0 | 0 | 0 | 10995000 | 0 | 0 | 17356000 | 0 |
| DLYEDELVI | HNRNPR; HNRI | O43390; B4I | HNRPR_HUMA | Unmodified | 0 | 0 | 0 | 0 | 0 | 0 | 0 | 0 | 0 | 0 | 0 | 308090 | 0 | 0 | 0 | 0 | 0 | 0 |
| DLYFLTNK | EFR3A; EFR3A | Q14156; A0 | EFR3A_HUMAN | Unmodified | 0 | 0 | 0 | 0 | 1486500 | 0 | 0 | 0 | 836570 | 0 | 150020 | 1333900 | 0 | 0 | 0 | 338420 | 0 | 0 |
| DLYHPICPI | DGKH; DGKH | Q86XP1; A0 | DGKH_HUMAN | Unmodified | 0 | 0 | 0 | 0 | 0 | 0 | 0 | 0 | 521100 | 0 | 0 | 0 | 210900 | 338840 | 0 | 44376 | 160070 | 125160 |
| DLYIDRPLF | WASHC2A; WA | Q641Q2; Q9 | WAC2A_HUMA | Unmodified | 174800 | 146390 | 256960 | 523420 | 2614000 | 400300 | 111690 | 0 | 1556400 | 153150 | 69939 | 520690 | 510220 | 256450 | 694310 | 245460 | 859580 | 798310 |
| DMAAAGLI | ANAPC7; ANAP | Q9UJX3; H0 | APC7_HUMAN | Unmodified | 0 | 0 | 0 | 0 | 0 | 0 | 0 | 0 | 0 | 0 | 0 | 418300 | 0 | 0 | 0 | 0 | 0 | 0 |
| DMAAFNEF | ME1 | P48163 | MAOX_HUMAN | Oxidation (M) | 0 | 0 | 0 | 408270 | 0 | 0 | 0 | 0 | 27009 | 0 | 464640 | 7317500 | 0 | 0 | 0 | 0 | 0 | 0 |
| DMADLPVF | REPS2 | Q8NFH8 | REPS2_HUMAN | Oxidation (M) | 0 | 0 | 0 | 0 | 0 | 0 | 0 | 0 | 0 | 0 | 0 | 0 | 209760 | 0 | 0 | 0 | 0 | 0 |
| DMAGAQA | DSG2 | Q14126 | DSG2_HUMAN | Unmodified | 0 | 0 | 0 | 0 | 0 | 173280 | 32893 | 0 | 1266700 | 209420 | 0 | 0 | 0 | 0 | 0 | 0 | 0 | 0 |
| DMAGAQA | DSG2 | Q14126 | DSG2_HUMAN | Oxidation (M) | 0 | 1405000 | 0 | 0 | 429080 | 0 | 0 | 612380 | 687120 | 0 | 2134200 | 1658500 | 297700 | 0 | 0 | 0 | 1148400 | 773120 |
| DMALNIGN | SNAP23; SNAP | O00161; H3I | SNP23_HUMAN | Oxidation (M) | 499420 | 822280 | 422810 | 2959600 | 5116600 | 1556400 | 0 | 158000 | 421740 | 0 | 1128200 | 0 | 2217000 | 2197600 | 1903900 | 1025900 | 2437700 | 5235000 |
| DMALNIGN | SNAP23; SNAP | O00161; H3I | SNP23_HUMAN | Unmodified | 0 | 0 | 0 | 0 | 3914200 | 0 | 0 | 31141 | 0 | 0 | 0 | 0 | 3152900 | 773940 | 0 | 0 | 1056300 | 4410900 |
| DMALTAFV | C3; C3 | P01024; M0 | CO3_HUMAN ( | Unmodified | 0 | 0 | 0 | 0 | 0 | 3803300 | 0 | 0 | 0 | 283710 | 0 | 0 | 0 | 1786500 | 0 | 0 | 0 | 1549900 |
| DMALYSLLI | PGAP4 | Q9BRR3 | PGAP4_HUMA | Unmodified | 0 | 0 | 0 | 0 | 76438 | 0 | 0 | 0 | 444840 | 0 | 0 | 0 | 0 | 0 | 0 | 0 | 0 | 0 |
| DMALYSLLI | PGAP4 | Q9BRR3 | PGAP4_HUMA | Oxidation (M) | 0 | 0 | 0 | 0 | 0 | 0 | 0 | 0 | 1243500 | 0 | 0 | 215600 | 0 | 0 | 0 | 0 | 0 | 0 |
| DMAQSIYR | SNW1; SNW1; | Q13573; G3 | SNW1_HUMAN | Unmodified | 0 | 0 | 0 | 0 | 498860 | 0 | 0 | 0 | 0 | 0 | 0 | 0 | 0 | 0 | 0 | 0 | 0 | 0 |
| DMDFLNK | PIP4K2C; PIP4K | Q8TBX8; H0 | PI42C_HUMAN | Unmodified | 0 | 0 | 0 | 0 | 812210 | 0 | 124060 | 55108 | 1019300 | 0 | 0 | 0 | 390550 | 381760 | 0 | 0 | 610610 | 1577800 |
| DMDLVEVN | ACAA2; ACAA2; | P42765; A0A | THIM_HUMAN | Oxidation (M) | 0 | 0 | 0 | 0 | 0 | 0 | 0 | 70995 | 363460 | 0 | 20150 | 78679 | 0 | 0 | 0 | 0 | 0 | 0 |
| DMEDPTPV | SMARCC1 | Q92922 | SMRC1_HUMA | Oxidation (M) | 0 | 0 | 0 | 0 | 0 | 0 | 0 | 0 | 160030 | 9371.4 | 782840 | 838310 | 0 | 0 | 0 | 0 | 0 | 0 |
| DMEESIR | PRKCSH; PRKC | P14314; K7E | GLU2B_HUMA | Unmodified | 0 | 0 | 0 | 0 | 0 | 0 | 0 | 0 | 49629000 | 483640 | 1498300 | 1148000 | 0 | 0 | 0 | 0 | 31943 | 45442 |
| DMELINNT | INMT-MINDY4 | F8WBC2 | F8WBC2_HUM | Oxidation (M) | 0 | 0 | 0 | 73203 | 0 | 0 | 911450 | 744990 | 985380 | 621000 | 0 | 0 | 0 | 67820 | 0 | 0 | 0 | 0 |
| DMELSFAV | MKRN1; MKRN | Q9UHC7; Q | MKRN1_HUMA | Oxidation (M) | 1561200 | 1340200 | 369000 | 738000 | 0 | 0 | 0 | 0 | 0 | 0 | 0 | 566910 | 0 | 0 | 0 | 407250 | 0 | 0 |
| DMELSFAV | MKRN1 | Q9UHC7 | MKRN1_HUMA | Unmodified | 0 | 0 | 0 | 0 | 0 | 0 | 0 | 0 | 0 | 0 | 0 | 0 | 558170 | 319630 | 0 | 0 | 0 | 0 |
| DMGTVVLG | GSPT1; GSPT2; | P15170; Q8 | ERF3A_HUMAN | Unmodified | 0 | 0 | 0 | 119450 | 141630 | 270510 | 0 | 0 | 0 | 130310 | 429960 | 3184000 | 305020 | 0 | 0 | 244770 | 0 | 0 |
| DMGTVVLG | GSPT1; GSPT1 | P15170; H3 | ERF3A_HUMAN | Unmodified | 0 | 0 | 0 | 0 | 0 | 0 | 199640 | 0 | 0 | 226820 | 0 | 0 | 1427400 | 706350 | 0 | 0 | 624100 | 0 |
| DMGTVVLG | GSPT1; GSPT1 | P15170; H3 | ERF3A_HUMAN | Oxidation (M) | 0 | 0 | 0 | 0 | 0 | 0 | 0 | 148390 | 0 | 0 | 0 | 0 | 1007600 | 0 | 0 | 0 | 0 | 0 |
| DMGYGNW | ADAR; ADAR; A | P55265; A0A | DSRAD_HUMA | Unmodified | 0 | 46707 | 0 | 0 | 0 | 0 | 132210 | 0 | 0 | 29866 | 0 | 785320 | 191740 | 0 | 0 | 0 | 0 | 196170 |
| DMILQFISR | DNM2 | P50570 | DYN2_HUMAN | Oxidation (M) | 799100 | 0 | 0 | 373750 | 0 | 0 | 0 | 0 | 0 | 0 | 0 | 1889600 | 0 | 0 | 502910 | 628900 | 0 | 0 |
| DMLDIFTR | SPPL2B; SPPL2 | Q8TCT7; A0 | SPP2B_HUMAN | Unmodified | 0 | 0 | 0 | 0 | 0 | 0 | 0 | 0 | 1028600 | 0 | 0 | 390460 | 0 | 0 | 0 | 0 | 0 | 0 |
| DMLEAGILI | CCT8; CCT8 | P50990; H7 | TCPQ_HUMAN | Unmodified | 0 | 0 | 0 | 0 | 0 | 0 | 0 | 0 | 400710 | 0 | 0 | 119310 | 0 | 0 | 0 | 0 | 0 | 272570 |
| DMLEAGILI | CCT8; CCT8 | P50990; H7 | TCPQ_HUMAN | Oxidation (M) | 0 | 0 | 0 | 0 | 1008400 | 0 | 0 | 0 | 1295500 | 74019 | 0 | 5359500 | 2762100 | 0 | 1116000 | 0 | 93472 | 0 |
| DMLLANKV | RPLP0; RPLP0P | P05388; Q8 | RLA0_HUMAN | Unmodified | 103960 | 0 | 0 | 0 | 0 | 0 | 0 | 0 | 0 | 0 | 0 | 1093200 | 0 | 0 | 0 | 0 | 0 | 0 |
| DMLLTLHQ | PACSIN3; PACS | Q9UKS6; E9 | PACN3_HUMA | Unmodified | 694490 | 1018000 | 0 | 635240 | 431950 | 0 | 0 | 0 | 170390 | 0 | 0 | 0 | 1289100 | 0 | 0 | 0 | 0 | 0 |
| DMLLTLHQ | PACSIN3; PACS | Q9UKS6; E9 | PACN3_HUMA | Oxidation (M) | 4375400 | 3160900 | 962170 | 1066700 | 0 | 0 | 0 | 0 | 0 | 0 | 0 | 0 | 1596500 | 393580 | 2068700 | 1568300 | 224620 | 0 |
| DMLYSAQD | XPR1 | Q9UBH6 | XPR1_HUMAN | Oxidation (M) | 0 | 0 | 0 | 0 | 0 | 0 | 0 | 0 | 0 | 0 | 0 | 1066400 | 0 | 0 | 0 | 87424 | 0 | 0 |
| DMNLTLEP | ACY1; ABHD14 | Q03154; A0 | ACY1_HUMAN | 2 Oxidation (M) | 0 | 0 | 91934 | 0 | 0 | 0 | 0 | 0 | 0 | 0 | 0 | 546980 | 0 | 0 | 0 | 133690 | 0 | 0 |
| DMNQVLD | WARS1; WARS1 | P23381; G3 | SYWC_HUMAN | Oxidation (M) | 0 | 0 | 0 | 0 | 0 | 0 | 0 | 0 | 0 | 332450 | 0 | 2780900 | 0 | 0 | 0 | 0 | 0 | 0 |
| DMPLIFIGG | TPST1; TPST1 | O60507; C9 | TPST1_HUMAN | Oxidation (M) | 0 | 0 | 0 | 0 | 0 | 0 | 0 | 0 | 1266300 | 0 | 0 | 0 | 0 | 0 | 0 | 0 | 0 | 0 |
| DMPLIFIGG | TPST1; TPST1 | O60507; C9 | TPST1_HUMAN | Unmodified | 0 | 0 | 0 | 0 | 0 | 0 | 0 | 0 | 424560 | 0 | 0 | 0 | 0 | 0 | 0 | 0 | 0 | 0 |
| DMPPAFIK | VCL | P18206 | VINC_HUMAN | Oxidation (M) | 227960 | 174290 | 341800 | 0 | 266280 | 0 | 0 | 0 | 0 | 0 | 0 | 709960 | 2813200 | 1068800 | 519610 | 924570 | 1035000 | 0 |
| DMQGLSLI | MVB12A; MVB1 | Q96EY5; M0 | MB12A_HUMA | Oxidation (M) | 1512700 | 1044700 | 16625000 | 6195400 | 0 | 283290 | 233090 | 0 | 0 | 0 | 233340 | 637010 | 719080 | 0 | 29480000 | 26776000 | 0 | 1097100 |
| DMQGLSLI | MVB12A; MVB1 | Q96EY5; M0 | MB12A_HUMA | Unmodified | 916720 | 1006400 | 1978400 | 1564100 | 0 | 0 | 0 | 0 | 377530 | 0 | 0 | 409130 | 0 | 0 | 2795600 | 1401000 | 0 | 0 |
| DMQGLSLI | MVB12A; MVB1 | Q96EY5; M0 | MB12A_HUMA | Oxidation (M) | 22655000 | 16924000 | 0 | 16791000 | 482460 | 0 | 0 | 0 | 1139800 | 0 | 0 | 0 | 3605000 | 463750 | 3951200 | 7012200 | 362740 | 254520 |
| DMQGLSLI | MVB12A; MVB1 | Q96EY5; M0 | MB12A_HUMA | Unmodified | 3070200 | 3251200 | 0 | 1743700 | 0 | 0 | 0 | 0 | 0 | 0 | 0 | 0 | 259450 | 0 | 0 | 0 | 0 | 0 |
| DMQNNRE | NOTCH1; NOT | P46531; A0A | NOTC1_HUMA | Unmodified | 0 | 0 | 0 | 0 | 0 | 0 | 0 | 0 | 1710900 | 0 | 0 | 1268100 | 34401 | 28914 | 0 | 0 | 0 | 0 |
| DMQNNRE | NOTCH1; NOT | P46531; A0A | NOTC1_HUMA | Oxidation (M) | 0 | 0 | 0 | 0 | 0 | 0 | 0 | 0 | 223420 | 0 | 241230 | 2663800 | 0 | 0 | 0 | 0 | 0 | 0 |
| DMRQTVA | EEF1A2; EEF1A | Q05639; Q5 | EF1A2_HUMAN | Unmodified | 0 | 67095 | 0 | 167530 | 14474000 | 2284700 | 2339100 | 614610 | 370140 | 3367800 | 426120 | 0 | 9884800 | 5188600 | 0 | 0 | 5093700 | 12993000 |
| DMRQTVA | EEF1A2; EEF1A | Q05639; Q5 | EF1A2_HUMAN | Oxidation (M) | 0 | 0 | 0 | 0 | 1113700 | 0 | 70870 | 0 | 0 | 0 | 82075 | 0 | 2203800 | 0 | 0 | 0 | 0 | 129300 |
| DMSMASW | HELZ2 | Q9BYK8 | HELZ2_HUMAN | Oxidation (M) | 0 | 0 | 0 | 0 | 0 | 0 | 0 | 0 | 0 | 0 | 1550000 | 0 | 0 | 0 | 0 | 0 | 0 | 0 |
| DMTGKEDI | COPG1 | Q9Y678 | COPG1_HUMA | Unmodified | 0 | 0 | 0 | 0 | 0 | 0 | 19878 | 0 | 0 | 0 | 0 | 523310 | 0 | 0 | 0 | 0 | 0 | 0 |
| DMTLIHGY | EXOC3; EXOC3 | O60645; H0 | EXOC3_HUMA | Oxidation (M) | 0 | 0 | 0 | 0 | 0 | 0 | 0 | 0 | 0 | 0 | 227700 | 719440 | 0 | 0 | 0 | 0 | 0 | 0 |
| DMTMFVTA | EIF3I; EIF3I; EIF | Q13347; A0 | EIF3I_HUMAN I | Oxidation (M) | 0 | 0 | 0 | 0 | 0 | 0 | 0 | 0 | 0 | 0 | 189080 | 1220800 | 0 | 0 | 0 | 0 | 0 | 0 |
| DMTSEQLD | ATP1A1 | P05023 | AT1A1_HUMAN | Oxidation (M) | 807040 | 656200 | 11628000 | 8858200 | 5904000 | 1081200 | 1119100 | 436350 | 4840100 | 1385100 | 16385000 | 51921000 | 6707300 | 3309500 | 13404000 | 13950000 | 850000 | 647980 |
| DMTSEQLD | ATP1A1 | P05023 | AT1A1_HUMAN | Unmodified | 0 | 0 | 699390 | 3694800 | 6062600 | 0 | 615960 | 0 | 10055000 | 608970 | 2178000 | 7739100 | 1987700 | 791520 | 2596300 | 2715600 | 0 | 0 |
| DMTSEQLD | ATP1A1 | P05023 | AT1A1_HUMAN | Oxidation (M) | 0 | 0 | 0 | 0 | 4523000 | 258320 | 539530 | 0 | 0 | 1646500 | 0 | 287750 | 10582000 | 3180000 | 0 | 0 | 920230 | 707400 |
| DMTSEQLD | ATP1A1 | P05023 | AT1A1_HUMAN | Oxidation (M) | 0 | 0 | 0 | 0 | 25855000 | 976190 | 0 | 0 | 0 | 938640 | 0 | 0 | 19013000 | 9723200 | 0 | 0 | 1522000 | 0 |
| DMVGIAQT | DDX17; DDX17 | Q92841; A0 | DDX17_HUMA | Unmodified | 0 | 0 | 0 | 0 | 4258600 | 0 | 0 | 59855 | 424520 | 0 | 732270 | 5279700 | 188690 | 0 | 0 | 0 | 0 | 0 |
| DMVGIAQT | DDX17; DDX17 | Q92841; A0 | DDX17_HUMA | Oxidation (M) | 0 | 0 | 0 | 0 | 0 | 185140 | 0 | 0 | 0 | 0 | 0 | 1091100 | 0 | 0 | 0 | 0 | 173030 | 237040 |
| DMVGLDAI | ACE; ACE; ACE | P12821; J3Q | ACE_HUMAN A | Unmodified | 0 | 0 | 0 | 0 | 0 | 0 | 0 | 0 | 328940 | 0 | 0 | 0 | 0 | 0 | 0 | 0 | 0 | 0 |
| DNAGAATE | ALDOC; ALDOC | P09972; A8 | ALDOC_HUMA | Unmodified | 0 | 0 | 0 | 206000 | 13085000 | 934080 | 251330 | 99306 | 3220300 | 1555100 | 943930 | 7704500 | 0 | 0 | 0 | 0 | 856500 | 2819000 |
| DNAGDRAI | TJP2; TJP2; TJP2 | Q9UDY2; A0 | ZO2_HUMAN T | Unmodified | 0 | 0 | 0 | 151730 | 0 | 0 | 0 | 0 | 1522300 | 104690 | 122370 | 892020 | 0 | 0 | 0 | 182740 | 0 | 0 |
| DNALLAQL | STXBP1; STXBP | P61764; A0A | STXB1_HUMAN | Unmodified | 0 | 0 | 51050 | 0 | 0 | 0 | 0 | 29767 | 92558 | 0 | 21305 | 202450 | 0 | 0 | 50529 | 25018 | 0 | 0 |
| DNALLAQL | STXBP1; STXBP | P61764; A0A | STXB1_HUMAN | Unmodified | 83899 | 64641 | 48739 | 48618 | 37943 | 0 | 0 | 0 | 386080 | 18203 | 124820 | 374320 | 0 | 0 | 123170 | 57692 | 0 | 0 |
| DNALLAQL | STXBP1; STXBP | P61764; A0A | STXB1_HUMAN | Unmodified | 0 | 0 | 0 | 0 | 0 | 0 | 0 | 0 | 1313800 | 0 | 911030 | 0 | 0 | 0 | 0 | 0 | 0 | 0 |
| DNALLAQL | STXBP1; STXBP | P61764; A0A | STXB1_HUMAN | Oxidation (M) | 0 | 0 | 0 | 0 | 753740 | 0 | 841750 | 0 | 6288400 | 1066000 | 1140900 | 3265000 | 0 | 0 | 0 | 0 | 0 | 0 |
| DNALLSAIE | NUP93; NUP93 | Q8N1F7; H3 | NUP93_HUMA | Unmodified | 0 | 0 | 0 | 0 | 0 | 0 | 0 | 0 | 0 | 0 | 0 | 0 | 0 | 0 | 0 | 0 | 0 | 0 |
| DNAMLEYL | EZR; EZR | P15311; E7E | EZRI_HUMAN E | Unmodified | 0 | 0 | 0 | 929440 | 23437000 | 3892000 | 387650 | 202340 | 17128000 | 4501800 | 1760900 | 12813000 | 1868400 | 2215000 | 0 | 0 | 2787200 | 236550 |
| DNAMLEYL | EZR; EZR | P15311; E7E | EZRI_HUMAN E | Oxidation (M) | 490710 | 270760 | 0 | 335810 | 3593500 | 993850 | 0 | 70234 | 1956100 | 1661300 | 1705800 | 7919100 | 1304600 | 1362800 | 476500 | 716300 | 1080000 | 0 |
| DNDFIYHD | PDCD6IP | Q8WUM4 | PDC6I_HUMAN | Unmodified | 1.66E+08 | 1.16E+08 | 87657000 | 64614000 | 10660000 | 1559900 | 185030 | 118280 | 45900000 | 6094100 | 7093100 | 27796000 | 13984000 | 4536900 | 90993000 | 64652000 | 469130 | 3188400 |
| DNDFIYHD | PDCD6IP | Q8WUM4 | PDC6I_HUMAN | Unmodified | 0 | 2801600 | 0 | 0 | 11029000 | 1346900 | 0 | 0 | 0 | 4640800 | 0 | 0 | 11966000 | 7177500 | 121830 | 93408 | 1082100 | 2664800 |
| DNDFLNEC | PIP4K2B | P78356 | PI42B_HUMAN | Unmodified | 0 | 0 | 0 | 0 | 0 | 0 | 0 | 57930 | 83140 | 0 | 346560 | 1253500 | 411790 | 0 | 250720 | 236710 | 213390 | 607870 |
| DNEESEQF | ABCC4; ABCC4 | O15439; A0 | MRP4_HUMAN | Unmodified | 0 | 0 | 845460 | 517250 | 0 | 0 | 0 | 0 | 0 | 0 | 0 | 139290 | 0 | 0 | 2319000 | 931900 | 0 | 0 |
| DNEETGFG | ANPEP | P15144 | AMPN_HUMAN | Unmodified | 0 | 0 | 0 | 0 | 0 | 0 | 0 | 0 | 0 | 0 | 0 | 0 | 0 | 0 | 0 | 0 | 631380 | 0 |
| DNEETGFG | ANPEP | P15144 | AMPN_HUMAN | Unmodified | 0 | 0 | 0 | 0 | 0 | 0 | 0 | 0 | 0 | 0 | 142180 | 0 | 0 | 0 | 0 | 0 | 11030000 | 15602000 |
| DNEETGFG | ANPEP | P15144 | AMPN_HUMAN | Unmodified | 0 | 0 | 0 | 0 | 0 | 0 | 0 | 0 | 0 | 0 | 0 | 0 | 0 | 0 | 0 | 0 | 3725600 | 2144000 |
| DNENVVNI | FGB | P02675 | FIBB_HUMAN F | Unmodified | 0 | 4737900 | 0 | 0 | 0 | 0 | 0 | 0 | 0 | 1237800 | 0 | 773360 | 0 | 0 | 0 | 0 | 53672 | 0 |
| DNERVVGF | TXNRD1; TXNRI | Q16881; A0 | TRXR1_HUMAN | Unmodified | 0 | 0 | 0 | 0 | 1.35E+08 | 4223500 | 5298800 | 0 | 11030000 | 12051000 | 13834000 | 2734800 | 46819000 | 10983000 | 0 | 0 | 25144000 | 49435000 |
| DNFFEVFTI | DNAJC2; DNAJ | Q99543; C9 | DNJC2_HUMA | Unmodified | 0 | 0 | 0 | 0 | 0 | 0 | 0 | 0 | 0 | 0 | 0 | 0 | 0 | 0 | 0 | 0 | 0 | 0 |
| DNFHGLAI | LMAN2; LMAN2 | Q12907; D6I | LMAN2_HUMA | Unmodified | 0 | 1148500 | 106330 | 127470 | 0 | 0 | 0 | 0 | 22912000 | 6867700 | 547110 | 436090 | 0 | 0 | 1048400 | 0 | 0 | 0 |
| DNFNVDEI | RAB12; RAB12 | Q6IQ22; A0A | RAB12_HUMAN | Unmodified | 0 | 0 | 0 | 0 | 0 | 0 | 0 | 0 | 0 | 0 | 181800 | 925870 | 0 | 0 | 0 | 934250 | 0 | 878010 |
| DNFQDTLC | ECE1; ECE1; EC | P42892; B4D | ECE1_HUMAN | Unmodified | 0 | 546730 | 637370 | 479090 | 2059200 | 420900 | 0 | 0 | 25474000 | 2618000 | 0 | 0 | 0 | 0 | 399000 | 534780 | 0 | 0 |
| DNFTAIPEC | DPYSL3; DPYSL | Q14195; H0 | DPYL3_HUMAN | Unmodified | 0 | 66289 | 0 | 0 | 0 | 0 | 0 | 452160 | 0 | 0 | 14082000 | 65249000 | 0 | 0 | 0 | 0 | 0 | 0 |
| DNFTLIPEG | DPYSL2; DPYSL | Q16555; A0 | DPYL2_HUMAN | Unmodified | 0 | 0 | 0 | 0 | 0 | 0 | 0 | 0 | 679370 | 0 | 0 | 3913800 | 0 | 0 | 127530 | 0 | 0 | 0 |
| DNFTLIPEG | DPYSL2; DPYSL | Q16555; A0 | DPYL2_HUMAN | Oxidation (M) | 0 | 0 | 0 | 0 | 2959400 | 285290 | 374880 | 70214 | 0 | 0 | 0 | 0 | 0 | 0 | 0 | 0 | 0 | 0 |
| DNFWMTR | ACTR2; ACTR2 | P61160; F5 | ARP2_HUMAN | Unmodified | 0 | 0 | 0 | 0 | 841230 | 235860 | 0 | 0 | 0 | 0 | 0 | 1166000 | 0 | 0 | 320500 | 0 | 163680 | 316930 |
| DNFWMTR | ACTR2; ACTR2 | P61160; F5 | ARP2_HUMAN | Oxidation (M) | 0 | 0 | 86027 | 0 | 0 | 6455900 | 0 | 0 | 0 | 0 | 0 | 584230 | 0 | 0 | 0 | 0 | 181990 | 0 |
| DNGLLAKP | OAT | P04181 | OAT_HUMAN C | Unmodified | 0 | 0 | 0 | 0 | 0 | 0 | 0 | 0 | 318980 | 0 | 0 | 0 | 0 | 0 | 0 | 0 | 0 | 0 |
| DNGPNYV | AOC1 | P19801 | AOC1_HUMAN | Unmodified | 0 | 0 | 0 | 0 | 94862 | 0 | 0 | 162380 | 1254100 | 228670 | 0 | 0 | 0 | 60103 | 0 | 0 | 124150 | 78025 |
| DNHLLGTF | HSPA5; HSPA5 | P11021; A0A | BIP_HUMAN Er | Unmodified | 0 | 0 | 0 | 1359000 | 3245600 | 1465900 | 4001600 | 0 | 67209000 | 57343000 | 9826200 | 1886300 | 179020 | 797940 | 0 | 0 | 685340 | 7808300 |
| DNHVVAA | QPRT | Q15274 | NADC_HUMAN | Unmodified | 0 | 0 | 0 | 0 | 0 | 0 | 45860 | 44317 | 0 | 0 | 204660 | 1740800 | 0 | 0 | 0 | 0 | 0 | 0 |
| DNIDITLQV | ARL8B; ARL8A | Q9NVJ2; Q9 | ARL8B_HUMAN | Unmodified | 0 | 0 | 0 | 0 | 310860 | 0 | 182630 | 0 | 0 | 153230 | 0 | 0 | 333670 | 316840 | 0 | 0 | 0 | 644990 |

| | | | | | | | | | | | | | | | | | | | | | | |
|---|---|---|---|---|---|---|---|---|---|---|---|---|---|---|---|---|---|---|---|---|---|---|
| DNIIPFLR | SLC12A6; SLC1 | Q9UHW9; Q | S12A6_HUMAN | Unmodified | 134970 | 98486 | 157630 | 0 | 0 | 0 | 0 | 0 | 0 | 47300 | 0 | 1576900 | 0 | 0 | 0 | 139360 | 0 | 0 |
| DNINIEEAA | RAB32 | Q13637 | RAB32_HUMAN | Unmodified | 0 | 0 | 0 | 0 | 0 | 0 | 0 | 0 | 0 | 0 | 0 | 0 | 0 | 0 | 0 | 0 | 0 | 0 |
| DNIQGITK | H4C7; H4C1 | Q99525; P62 | H4G_HUMAN H | Unmodified | 0 | 2258200 | 728050 | 946300 | 2428000 | 484510 | 733860 | 0 | 1270400 | 310880 | 2569300 | 66297000 | 2211200 | 423820 | 226790 | 0 | 608230 | 0 |
| DNIQGITKF | H4C1 | P62805 | H4_HUMAN Hi | Unmodified | 9.27E+08 | 5.92E+08 | 3409300 | 1155300 | 2996800 | 1762900 | 3557600 | 16495000 | 11394000 | 3849900 | 37018000 | 2.83E+09 | 1264300 | 0 | 1045800 | 1647100 | 0 | 1642300 |
| DNIQGITKF | H4C1 | P62805 | H4_HUMAN Hi | Unmodified | 1.44E+08 | 60061000 | 0 | 0 | 0 | 0 | 0 | 0 | 0 | 0 | 126400 | 8.39E+08 | 0 | 0 | 186770 | 235670 | 0 | 598000 |
| DNIRLRSSI | ALCAM; ALCAM | Q13740; F5C | CD166_HUMAI | Unmodified | 0 | 0 | 0 | 0 | 2734800 | 0 | 0 | 0 | 0 | 0 | 0 | 0 | 0 | 0 | 0 | 0 | 3632100 | 2314200 |
| DNKDLIVC | MYO1D; MYO1I | O94832; J3C | MYO1D_HUMA | Unmodified | 821420 | 998980 | 0 | 765620 | 0 | 0 | 0 | 0 | 0 | 0 | 0 | 0 | 0 | 0 | 0 | 0 | 0 | 0 |
| DNKPLVER | SRP68 | Q9UHB9 | SRP68_HUMAN | Unmodified | 179300 | 0 | 628440 | 0 | 40212 | 0 | 373600 | 0 | 0 | 0 | 646720 | 868710 | 657990 | 0 | 0 | 0 | 562780 | 0 |
| DNLAILEK | CAB39L; CAB39 | Q9H9S4; B7 | CB39L_HUMAN | Unmodified | 0 | 0 | 81562 | 399750 | 0 | 0 | 0 | 0 | 0 | 0 | 0 | 1684400 | 878200 | 0 | 126800 | 494420 | 0 | 0 |
| DNLAKQQ( | RAB35; RAB35 | Q15286; F5H | RAB35_HUMAN | Unmodified | 0 | 0 | 0 | 0 | 516450 | 0 | 0 | 0 | 0 | 0 | 0 | 0 | 422900 | 0 | 0 | 0 | 206590 | 366540 |
| DNLAQQSI | CHMP5 | Q9NZZ3 | CHMP5_HUMA | Unmodified | 0 | 0 | 0 | 0 | 0 | 0 | 0 | 0 | 4938800 | 0 | 0 | 0 | 0 | 0 | 299350 | 0 | 0 | 0 |
| DNLAQQSI | CHMP5 | Q9NZZ3 | CHMP5_HUMA | Oxidation (M) | 0 | 0 | 1120300 | 0 | 0 | 0 | 0 | 0 | 0 | 0 | 0 | 6106100 | 0 | 0 | 0 | 1031200 | 0 | 0 |
| DNLEFFLA( | FASN; FASN | P49327; A0A | FAS_HUMAN F | Unmodified | 117060 | 0 | 136130 | 71419 | 0 | 0 | 0 | 0 | 0 | 0 | 0 | 12078000 | 0 | 0 | 1102400 | 528130 | 0 | 0 |
| DNLENPDF | PLCG2; PLCG2 | P16885; A0A | PLCG2_HUMAI | Unmodified | 0 | 0 | 0 | 0 | 0 | 0 | 0 | 0 | 0 | 0 | 1163900 | 0 | 0 | 0 | 0 | 0 | 0 | 0 |
| DNLEWLAF | PSMD1; PSMD1 | Q99460; A0A | PSMD1_HUMA | Unmodified | 0 | 50321 | 0 | 0 | 0 | 0 | 0 | 0 | 0 | 0 | 0 | 801380 | 0 | 0 | 0 | 0 | 0 | 0 |
| DNLLDTYS. | QSOX2 | Q6ZRP7 | QSOX2_HUMAI | Unmodified | 0 | 0 | 0 | 0 | 0 | 0 | 0 | 0 | 1942200 | 0 | 0 | 225850 | 0 | 0 | 0 | 0 | 0 | 0 |
| DNLQILR | NAA16; NAA15; | Q6N069; Q9 | NAA16_HUMAN | Unmodified | 33561 | 291820 | 0 | 0 | 21601 | 0 | 0 | 0 | 0 | 0 | 0 | 3027900 | 0 | 0 | 30127 | 0 | 0 | 0 |
| DNLQLPLQ | NPC1 | O15118 | NPC1_HUMAN | Unmodified | 0 | 0 | 52061 | 93940 | 1442100 | 323200 | 0 | 0 | 5148700 | 731250 | 580380 | 193930 | 373240 | 97828 | 174990 | 74337 | 0 | 517020 |
| DNLSYIEHI | PEA15 | Q15121 | PEA15_HUMAN | Unmodified | 278430 | 259390 | 98799 | 124300 | 0 | 0 | 0 | 0 | 527930 | 39337 | 0 | 0 | 0 | 0 | 451190 | 344800 | 0 | 0 |
| DNLTLWTS | YWHAE | P62258 | 1433E_HUMAN | Unmodified | 0 | 0 | 0 | 0 | 0 | 0 | 0 | 0 | 1791900 | 0 | 0 | 1461300 | 0 | 0 | 2410100 | 850990 | 0 | 0 |
| DNLTLWTS | YWHAE | P62258 | 1433E_HUMAN | Oxidation (M) | 854360 | 637590 | 3089000 | 0 | 0 | 0 | 0 | 0 | 107310 | 0 | 0 | 5619300 | 0 | 0 | 6290800 | 8246900 | 0 | 0 |
| DNLTLWTS | YWHAG | P61981 | 1433G_HUMAN | Unmodified | 795560 | 766750 | 1175800 | 1222500 | 346890 | 0 | 67967 | 32150 | 444180 | 211250 | 693640 | 6649100 | 164540 | 0 | 3974500 | 2898400 | 192340 | 0 |
| DNLTLWTS | YWHAH | Q04917 | 1433F_HUMAN | Unmodified | 0 | 0 | 0 | 95111 | 0 | 0 | 0 | 0 | 412390 | 35380 | 156720 | 1873500 | 0 | 0 | 323590 | 239270 | 0 | 0 |
| DNLTLWTS | YWHAZ; YWHA | P63104; B7Z | 1433Z_HUMAN | Unmodified | 1562700 | 1806300 | 0 | 0 | 0 | 0 | 0 | 375540 | 8781500 | 1391300 | 0 | 13644000 | 0 | 0 | 3733900 | 1904600 | 517730 | 254330 |
| DNLTLWTS | YWHAB | P31946 | 1433B_HUMAN | Unmodified | 362230 | 806500 | 1248800 | 1762700 | 0 | 0 | 0 | 73084 | 2560300 | 0 | 374890 | 1558600 | 0 | 0 | 2879600 | 2707800 | 0 | 129440 |
| DNNLLGR | HSPA1L; HSPA( | P34931; P17 | HS71L_HUMAN | Unmodified | 172390 | 0 | 0 | 0 | 0 | 0 | 0 | 0 | 235320 | 52416 | 44634 | 458260 | 0 | 0 | 0 | 289720 | 0 | 0 |
| DNNMGLV | COPS3 | Q9UNS2 | CSN3_HUMAN | Unmodified | 0 | 116070 | 0 | 0 | 0 | 337050 | 0 | 274810 | 0 | 0 | 324470 | 1089200 | 501150 | 198520 | 0 | 0 | 0 | 0 |
| DNNQFAS/ | COPB2; COPB2 | P35606; H0Y | COPB2_HUMAI | Unmodified | 0 | 0 | 0 | 0 | 0 | 0 | 0 | 0 | 0 | 0 | 130910 | 2066600 | 0 | 0 | 0 | 0 | 0 | 0 |
| DNPGVVT( | UBA1; UBA1; UI | P22314; Q5J | UBA1_HUMAN | Unmodified | 0 | 0 | 0 | 313730 | 231070 | 0 | 0 | 0 | 637890 | 0 | 388640 | 5297000 | 0 | 0 | 331130 | 260370 | 0 | 0 |
| DNPSIFVH! | LRBA; LRBA; LR | P50851; A0A | LRBA_HUMAN | Oxidation (M) | 0 | 0 | 0 | 0 | 0 | 0 | 0 | 0 | 176940 | 0 | 0 | 1858600 | 0 | 0 | 0 | 0 | 0 | 0 |
| DNQHQDE | GOLIM4; GOLIN | O00461; F8V | GOLI4_HUMAN | Unmodified | 0 | 0 | 0 | 0 | 0 | 0 | 0 | 0 | 197720 | 38030 | 167800 | 39544 | 0 | 0 | 0 | 9774.9 | 0 | 0 |
| DNRGNLL( | FN1 | P02751 | FINC_HUMAN I | Unmodified | 0 | 306010 | 0 | 1837500 | 0 | 0 | 0 | 0 | 0 | 0 | 0 | 0 | 1371100 | 1135600 | 0 | 270500 | 223590 | 1474200 |
| DNRNLFEV | CACNA2D2; CA | Q9NY47; C9 | CA2D2_HUMAI | Unmodified | 73287 | 0 | 0 | 0 | 0 | 0 | 0 | 0 | 4692100 | 0 | 836390 | 1270600 | 0 | 0 | 0 | 0 | 0 | 0 |
| DNRNLFEV | CACNA2D2; CA | Q9NY47; C9 | CA2D2_HUMAI | Unmodified | 0 | 0 | 0 | 0 | 777510 | 0 | 0 | 0 | 5307700 | 853690 | 0 | 0 | 0 | 0 | 0 | 0 | 0 | 0 |
| DNRPGDKI | NPTX1 | Q15818 | NPTX1_HUMAN | Unmodified | 0 | 0 | 0 | 0 | 0 | 0 | 0 | 0 | 36231000 | 2292500 | 0 | 0 | 0 | 0 | 0 | 0 | 0 | 0 |
| DNRVVLFE | CPNE3; CPNE3 | O75131; H0Y | CPNE3_HUMAI | Oxidation (M) | 0 | 0 | 1207200 | 518010 | 913330 | 0 | 0 | 0 | 3744400 | 155330 | 0 | 0 | 0 | 0 | 0 | 849630 | 241250 | 119100 |
| DNRVVLFE | CPNE3; CPNE3 | O75131; H0Y | CPNE3_HUMAI | Unmodified | 0 | 0 | 0 | 0 | 0 | 0 | 0 | 0 | 1074800 | 0 | 0 | 0 | 0 | 0 | 0 | 0 | 0 | 0 |
| DNSDFDLL | EIF3D | O15371 | EIF3D_HUMAN | Unmodified | 0 | 0 | 0 | 0 | 0 | 0 | 0 | 0 | 0 | 0 | 551450 | 2969600 | 0 | 0 | 0 | 610790 | 0 | 0 |
| DNSHNEY\ | VANGL2 | Q9ULK5 | VANG2_HUMAI | Unmodified | 0 | 0 | 0 | 0 | 0 | 0 | 0 | 0 | 0 | 0 | 0 | 670120 | 0 | 0 | 0 | 0 | 0 | 0 |
| DNSIYVLAI | VNN2; VNN2; V | O95498; E9F | VNN2_HUMAN | Unmodified | 0 | 0 | 1985800 | 0 | 0 | 2266800 | 0 | 167890 | 0 | 0 | 0 | 0 | 0 | 0 | 214410 | 0 | 0 | 3713400 |
| DNSPVHLC | SLITRK6 | Q9H5Y7 | SLIK6_HUMAN | Unmodified | 229950 | 317640 | 0 | 0 | 0 | 0 | 0 | 0 | 192120 | 0 | 0 | 0 | 0 | 0 | 0 | 0 | 0 | 0 |
| DNSTMGYI | HSP90AA1 | P07900 | HS90A_HUMAI | Oxidation (M) | 0 | 0 | 0 | 0 | 0 | 0 | 0 | 0 | 365850 | 0 | 0 | 2667700 | 0 | 0 | 0 | 0 | 0 | 0 |
| DNSTMGYI | HSP90AA1 | P07900 | HS90A_HUMAI | Unmodified | 0 | 0 | 0 | 0 | 160300 | 0 | 0 | 0 | 1017100 | 0 | 0 | 6002700 | 0 | 0 | 260740 | 0 | 0 | 0 |
| DNSTMGYI | HSP90AA1 | P07900 | HS90A_HUMAI | Unmodified | 187410 | 0 | 0 | 0 | 0 | 0 | 39398 | 0 | 1219600 | 136240 | 429830 | 7311200 | 0 | 0 | 460360 | 0 | 134010 | 0 |
| DNSTMGYI | HSP90AA1 | P07900 | HS90A_HUMAI | Oxidation (M) | 54017 | 0 | 46976 | 0 | 0 | 0 | 0 | 0 | 0 | 0 | 0 | 2060900 | 0 | 0 | 232680 | 0 | 0 | 0 |
| DNSTMGYI | HSP90AB1; HSI | P08238; Q5 | HS90B_HUMAI | 2 Oxidation (M) | 0 | 0 | 34976 | 0 | 0 | 0 | 0 | 0 | 67332 | 0 | 0 | 2012100 | 0 | 0 | 0 | 214950 | 0 | 0 |
| DNSTMGYI | HSP90AB1; HSI | P08238; Q5 | HS90B_HUMAI | Oxidation (M) | 0 | 0 | 0 | 0 | 0 | 0 | 0 | 0 | 241660 | 0 | 0 | 2406000 | 0 | 0 | 0 | 0 | 0 | 0 |
| DNSTMGYI | HSP90AB1; HSI | P08238; Q5 | HS90B_HUMAI | Unmodified | 0 | 0 | 0 | 0 | 314540 | 0 | 0 | 0 | 669030 | 0 | 0 | 4856700 | 0 | 0 | 0 | 0 | 0 | 0 |
| DNSTMGYI | HSP90AB1; HSI | P08238; Q5 | HS90B_HUMAI | Oxidation (M) | 0 | 0 | 52652 | 0 | 1564800 | 2412400 | 70627 | 0 | 325100 | 0 | 0 | 640370 | 0 | 0 | 0 | 0 | 0 | 855480 |
| DNSTMGYI | HSP90AB1; HSI | P08238; Q5 | HS90B_HUMAI | Unmodified | 0 | 0 | 0 | 77176 | 106120 | 0 | 0 | 0 | 745450 | 0 | 107420 | 3513600 | 96524 | 0 | 166580 | 0 | 134080 | 0 |
| DNSVSQH/ | N/A | N/A | N/A | Unmodified | 0 | 0 | 0 | 0 | 0 | 0 | 0 | 0 | 0 | 0 | 0 | 2512100 | 0 | 0 | 0 | 0 | 0 | 0 |
| DNTAKLFD | EIF3I; EIF3I; EIF | Q13347; A0/ | EIF3I_HUMAN I | Unmodified | 8585000 | 0 | 0 | 0 | 0 | 0 | 0 | 0 | 0 | 0 | 0 | 0 | 0 | 0 | 170420 | 0 | 700490 | 0 |
| DNTIEHLLF | PPP2R1A; PPP2 | P30153; B3K | 2AAA_HUMAN | Unmodified | 0 | 0 | 0 | 0 | 0 | 0 | 0 | 0 | 0 | 0 | 0 | 0 | 0 | 0 | 76173 | 0 | 0 | 0 |
| DNTINLIHT | ARPC2; ARPC2 | O15144; H7( | ARPC2_HUMAI | Unmodified | 342250 | 0 | 467180 | 0 | 0 | 0 | 211900 | 0 | 0 | 1369400 | 0 | 5328600 | 0 | 0 | 612050 | 663150 | 0 | 0 |
| DNTINLIHT | ARPC2; ARPC2 | O15144; H7( | ARPC2_HUMAI | Unmodified | 3718200 | 2912900 | 575620 | 4304100 | 488630 | 0 | 373530 | 0 | 1752800 | 583060 | 1562900 | 7018500 | 2109800 | 0 | 4309600 | 4512000 | 0 | 348560 |
| DNTNEIYS( | ITGB1; ITGB1; I | P05556; H7( | ITB1_HUMAN I | Unmodified | 1083900 | 966470 | 2425000 | 2226600 | 12219000 | 0 | 86578 | 52246 | 7307600 | 475990 | 0 | 4139900 | 618780 | 399630 | 2223800 | 1921400 | 1361900 | 10127000 |
| DNTNEIYS( | ITGB1; ITGB1; I | P05556; H7( | ITB1_HUMAN I | Unmodified | 0 | 0 | 0 | 0 | 813580 | 0 | 0 | 0 | 0 | 0 | 0 | 0 | 0 | 0 | 0 | 0 | 2936100 | 0 |
| DNTQRFLQ | PLEC | Q15149 | PLEC_HUMAN | Unmodified | 0 | 0 | 0 | 0 | 720690 | 176430 | 0 | 0 | 0 | 58238 | 0 | 0 | 0 | 0 | 0 | 0 | 818840 | 713080 |
| DNTRPGAI | DDX1; DDX1; D | Q92499; A0/ | DDX1_HUMAN | Oxidation (M) | 0 | 131940 | 361550 | 423820 | 303850 | 0 | 150200 | 104490 | 0 | 0 | 597820 | 2209400 | 590540 | 272310 | 185770 | 0 | 0 | 187090 |
| DNTRPGAI | DDX1; DDX1; D | Q92499; A0/ | DDX1_HUMAN | Unmodified | 0 | 0 | 0 | 98720 | 0 | 0 | 0 | 0 | 0 | 0 | 0 | 1084300 | 270840 | 352600 | 0 | 0 | 0 | 0 |
| DNTVHDLF | GOLGA3 | Q08378 | GOGA3_HUMA | Unmodified | 0 | 0 | 0 | 0 | 324720 | 0 | 0 | 0 | 0 | 0 | 0 | 0 | 533630 | 438260 | 0 | 0 | 222880 | 741020 |
| DNTVIDHTI | KDM4A | O75164 | KDM4A_HUMA | Unmodified | 0 | 0 | 0 | 0 | 0 | 0 | 0 | 0 | 0 | 200620 | 0 | 1091400 | 0 | 0 | 0 | 0 | 0 | 0 |
| DNVAEQLF | PTGR2; PTGR2 | Q8N8N7; G3 | PTGR2_HUMAI | Unmodified | 0 | 0 | 0 | 207510 | 0 | 0 | 0 | 0 | 0 | 0 | 0 | 998540 | 0 | 0 | 0 | 225410 | 0 | 0 |
| DNVAGVTL | ATP6V1D; ATP6 | Q9Y5K8; G3 | VATD_HUMAN | Unmodified | 0 | 0 | 0 | 0 | 0 | 0 | 0 | 0 | 0 | 0 | 670430 | 1247400 | 0 | 0 | 0 | 1099200 | 0 | 0 |
| DNVCSPG( | PYCR2; PYCR2; | Q96C36; A0/ | P5CR2_HUMAI | Unmodified | 0 | 0 | 0 | 0 | 0 | 0 | 0 | 0 | 7631600 | 380300 | 0 | 0 | 0 | 0 | 0 | 0 | 0 | 0 |
| DNVDDPT( | LMAN2 | Q12907 | LMAN2_HUMAI | Unmodified | 0 | 0 | 0 | 0 | 0 | 0 | 0 | 0 | 8631100 | 1218300 | 245640 | 2004000 | 0 | 0 | 528880 | 0 | 0 | 0 |
| DNVFENNI | MICALL1; MICA | Q8N3F8; B0 | MILK1_HUMAN | Unmodified | 208610 | 0 | 0 | 112890 | 0 | 0 | 0 | 103240 | 0 | 0 | 0 | 0 | 912670 | 716110 | 0 | 0 | 0 | 0 |
| DNVSSPG( | PYCR1; PYCR1; | P32322; J3Q | P5CR1_HUMAI | Unmodified | 0 | 0 | 0 | 0 | 0 | 0 | 0 | 0 | 2511200 | 0 | 0 | 0 | 0 | 0 | 129230 | 0 | 0 | 0 |
| DNVYYYDE | CDH1 | P12830 | CADH1_HUMA | Unmodified | 0 | 0 | 0 | 0 | 0 | 0 | 0 | 0 | 0 | 0 | 0 | 0 | 0 | 0 | 14152000 | 8753700 | 0 | 0 |
| DNWEELYI | NPEPPS; NPEPI | P55786; A0A | PSA_HUMAN P | Unmodified | 0 | 0 | 0 | 0 | 65149 | 60238 | 0 | 0 | 0 | 0 | 59700 | 0 | 47918 | 0 | 493610 | 167080 | 0 | 109860 |
| DNYNADFI | SNX16; SNX16; | P57768; E5R | SNX16_HUMAN | Unmodified | 0 | 0 | 0 | 0 | 431300 | 0 | 0 | 0 | 0 | 42767 | 0 | 0 | 0 | 0 | 0 | 0 | 0 | 0 |
| DNYPPSFK | FUCA2 | Q9BTY2 | FUCO2_HUMA | Unmodified | 0 | 0 | 133480 | 31244 | 94147 | 0 | 0 | 0 | 1985100 | 233960 | 384780 | 60918 | 0 | 0 | 361710 | 0 | 85443 | 46810 |
| DNYPQSVI | MYO1C; MYO1( | O00159; F5H | MYO1C_HUMA | Unmodified | 2206500 | 2570700 | 5959900 | 5116600 | 7300600 | 1931600 | 60140 | 40201 | 0 | 0 | 295520 | 1206500 | 1951200 | 1110700 | 8955300 | 5302600 | 495190 | 1578800 |
| DNYPQSVI | MYO1C; MYO1( | O00159; F5H | MYO1C_HUMA | Unmodified | 0 | 0 | 0 | 0 | 875840 | 258800 | 191100 | 0 | 0 | 315890 | 0 | 0 | 0 | 464730 | 0 | 0 | 342460 | 478440 |
| DNYSPGW | EPRS1 | P07814 | SYEP_HUMAN | Unmodified | 0 | 0 | 0 | 0 | 0 | 0 | 0 | 0 | 0 | 0 | 0 | 2384900 | 0 | 0 | 0 | 0 | 0 | 0 |
| DNYVPEVS | RPS17; RPS17; | P08708; H0Y | RS17_HUMAN | Unmodified | 708140 | 1754300 | 26082 | 350740 | 0 | 0 | 0 | 0 | 0 | 0 | 0 | 145950 | 145290 | 0 | 1182300 | 114380 | 0 | 0 |
| DPADASEA | CHGB; CHGB | P05060; A0A | SCG1_HUMAN | Unmodified | 0 | 0 | 0 | 0 | 0 | 0 | 0 | 0 | 555490 | 0 | 0 | 0 | 0 | 138570 | 0 | 0 | 0 | 0 |
| DPAEGDG/ | GANAB; GANAE | Q14697; F5H | GANAB_HUMA | Unmodified | 0 | 0 | 0 | 0 | 0 | 0 | 0 | 0 | 3570600 | 683780 | 193480 | 0 | 0 | 0 | 0 | 0 | 0 | 0 |
| DPDASKPE | CALR; CALR; C/ | P27797; A0A | CALR_HUMAN | Unmodified | 0 | 0 | 0 | 0 | 0 | 0 | 0 | 0 | 1995100 | 431830 | 0 | 0 | 0 | 0 | 0 | 0 | 0 | 0 |
| DPDKQTQI | SETD2 | Q9BYW2 | SETD2_HUMAN | Unmodified | 0 | 0 | 0 | 0 | 23413000 | 2654000 | 1015700 | 722750 | 0 | 168580 | 0 | 393230 | 1015700 | 894950 | 0 | 0 | 10866000 | 47628000 |
| DPDTPILIIK | AVIL | O75366 | AVIL_HUMAN A | Unmodified | 0 | 0 | 0 | 0 | 0 | 0 | 0 | 0 | 0 | 0 | 1627400 | 8833800 | 0 | 0 | 0 | 0 | 0 | 0 |
| DPEGFVGI | P4HA1 | P13674 | P4HA1_HUMAI | Unmodified | 0 | 0 | 227400 | 0 | 0 | 0 | 0 | 0 | 1182300 | 136520 | 416030 | 225250 | 0 | 0 | 0 | 0 | 0 | 0 |
| DPELWGS\ | CLTC; CLTC | Q00610; A0/ | CLH1_HUMAN | Unmodified | 0 | 83125 | 0 | 0 | 0 | 0 | 0 | 0 | 0 | 0 | 0 | 103630 | 0 | 0 | 0 | 0 | 0 | 0 |
| DPENFPFV | RAB7A; RAB7A; | P51149; A0A | RAB7A_HUMAN | Unmodified | 0 | 0 | 1704700 | 0 | 0 | 0 | 0 | 246540 | 470930 | 345850 | 1196800 | 6377800 | 0 | 0 | 1280000 | 3131500 | 0 | 0 |
| DPENFPFV | RAB7A; RAB7A; | P51149; A0A | RAB7A_HUMAN | Unmodified | 1061200 | 972820 | 273150 | 468700 | 0 | 0 | 0 | 0 | 0 | 685680 | 317220 | 2156100 | 151270 | 0 | 1215900 | 619430 | 0 | 0 |
| DPFLNPDL | GNG13 | Q9P2W3 | GBG13_HUMAI | Oxidation (M) | 0 | 0 | 0 | 0 | 0 | 0 | 0 | 0 | 0 | 0 | 3391800 | 5886900 | 0 | 0 | 0 | 0 | 0 | 0 |
| DPFLNPDL | GNG13 | Q9P2W3 | GBG13_HUMAI | Unmodified | 0 | 0 | 0 | 0 | 0 | 0 | 0 | 0 | 0 | 0 | 312060 | 2421800 | 0 | 0 | 0 | 0 | 0 | 0 |
| DPGLVDQL | SARS1; SARS1 | P49591; Q5T | SYSC_HUMAN | Unmodified | 0 | 0 | 541090 | 0 | 803900 | 0 | 267740 | 46700 | 476810 | 0 | 0 | 2634500 | 0 | 0 | 0 | 0 | 0 | 0 |
| DPGTVANK | STAM; STAM | Q92783; A6N | STAM1_HUMAI | Unmodified | 0 | 0 | 0 | 0 | 0 | 0 | 0 | 21403 | 1752500 | 0 | 169820 | 0 | 0 | 0 | 0 | 43994 | 91187 | 35203 |
| DPGVITYDI | VARS1; VARS1; | P26640; A0A | SYVC_HUMAN | Unmodified | 0 | 0 | 79065 | 131090 | 0 | 0 | 0 | 0 | 0 | 0 | 0 | 1358200 | 112890 | 48887 | 50652 | 159220 | 0 | 0 |
| DPGVLSFLI | TEP1; TEP1; TEF | Q99973; G3 | TEP1_HUMAN | Unmodified | 0 | 0 | 0 | 0 | 0 | 0 | 0 | 0 | 0 | 0 | 0 | 465720 | 0 | 0 | 0 | 0 | 0 | 0 |
| DPHLACVA | CLTCL1; CLTC; | P53675; Q0( | CLH2_HUMAN | Unmodified | 923340 | 144040 | 228300 | 185240 | 0 | 0 | 0 | 10051 | 0 | 52974 | 77074 | 4855800 | 0 | 0 | 102470 | 207380 | 0 | 322640 |
| DPLLDDH( | MYO6; MYO6; N | Q9UM54; A0 | MYO6_HUMAN | Unmodified | 0 | 0 | 1431700 | 0 | 0 | 0 | 0 | 0 | 106420 | 0 | 0 | 416220 | 0 | 0 | 0 | 0 | 0 | 0 |
| DPLLLSGTI | ATP2B1 | P20020 | AT2B1_HUMAN | Unmodified | 0 | 0 | 0 | 0 | 0 | 0 | 0 | 0 | 0 | 0 | 165340 | 694920 | 0 | 0 | 102340 | 0 | 0 | 0 |
| DPLNPIKQI | PPA1; PPA1 | Q15181; Q5 | IPYR_HUMAN I | Unmodified | 0 | 0 | 0 | 0 | 0 | 0 | 0 | 0 | 0 | 0 | 409550 | 427280 | 0 | 0 | 0 | 0 | 0 | 0 |
| DPNAFLFD | EIF3M; EIF3M; E | Q7L2H7; H0 | EIF3M_HUMAN | Unmodified | 0 | 0 | 301750 | 0 | 0 | 0 | 0 | 0 | 0 | 0 | 0 | 1013500 | 0 | 0 | 0 | 0 | 0 | 0 |
| DPNIVIAK | PDIA3 | P30101 | PDIA3_HUMAN | Unmodified | 78696 | 0 | 707810 | 65592 | 161290 | 0 | 73534 | 0 | 13558000 | 1047400 | 2510300 | 2678400 | 206360 | 55189 | 459820 | 62470 | 0 | 0 |
| DPNNLAFN | MGAM; MGAM; | O43451; A0/ | MGA_HUMAN I | Unmodified | 0 | 0 | 0 | 0 | 0 | 0 | 0 | 0 | 5682200 | 0 | 0 | 0 | 0 | 0 | 0 | 0 | 0 | 0 |
| DPNNLFM\ | PSMD2; PSMD2 | Q13200; H7( | PSMD2_HUMA | Oxidation (M) | 0 | 0 | 0 | 0 | 0 | 1288100 | 0 | 0 | 629850 | 0 | 0 | 3564900 | 0 | 0 | 0 | 0 | 160330 | 1342500 |
| DPNNLLNI | PSMD12 | O00232 | PSD12_HUMAN | Unmodified | 0 | 0 | 0 | 0 | 0 | 0 | 0 | 0 | 0 | 0 | 0 | 3088400 | 0 | 0 | 65484 | 386070 | 0 | 0 |
| DPNVVGQI | SUCLG2; SUCL | Q96I99; E9P | SUCB2_HUMAI | Unmodified | 17933000 | 16839000 | 11305000 | 8074900 | 0 | 0 | 0 | 0 | 2059800 | 0 | 891490 | 739900 | 393750 | 0 | 15366000 | 12926000 | 0 | 0 |
| DPPIITPEL( | N/A | N/A | N/A | Unmodified | 0 | 0 | 0 | 0 | 0 | 0 | 0 | 26508 | 0 | 0 | 0 | 320580 | 0 | 0 | 0 | 0 | 0 | 0 |
| DPQALSEH | STIP1 | P31948 | STIP1_HUMAN | Unmodified | 0 | 0 | 0 | 0 | 0 | 0 | 0 | 0 | 0 | 54064 | 0 | 542200 | 55059 | 0 | 0 | 0 | 0 | 0 |
| DPQCNLH( | SMOC1 | Q9H4F8 | SMOC1_HUMA | Unmodified | 484870 | 287470 | 0 | 0 | 0 | 0 | 262340 | 0 | 0 | 23054 | 0 | 0 | 0 | 0 | 0 | 0 | 0 | 0 |
| DPQFSLIS/ | STAM2 | O75886 | STAM2_HUMAI | Unmodified | 0 | 0 | 305500 | 0 | 0 | 3843400 | 578210 | 18330 | 803580 | 0 | 95740 | 1414600 | 0 | 0 | 0 | 0 | 0 | 355380 |
| DPSKRPTA | STK39 | Q9UEW8 | STK39_HUMAN | Unmodified | 43722 | 0 | 0 | 535060 | 631420 | 0 | 323110 | 244620 | 639540 | 104540 | 298710 | 389190 | 3118300 | 1442200 | 332880 | 383130 | 68262 | 651560 |
| DPSQQELP | FASN; FASN | P49327; A0A | FAS_HUMAN F | Unmodified | 0 | 0 | 0 | 117050 | 13944 | 729770 | 126450 | 0 | 0 | 0 | 3094000 | 16547000 | 0 | 0 | 0 | 0 | 54715 | 0 |
| DPSVLAVR | PIK3C2B; PIK3( | O00750; F5( | P3C2B_HUMAI | Unmodified | 0 | 0 | 0 | 0 | 0 | 0 | 0 | 0 | 0 | 0 | 71163 | 322590 | 0 | 0 | 97549 | 166420 | 0 | 0 |
| DPSVTQVT | SCAMP1; SCAN | O15126; A0/ | SCAM1_HUMA | Unmodified | 0 | 0 | 4928600 | 13341000 | 0 | 0 | 0 | 0 | 4760200 | 756660 | 0 | 681960 | 0 | 0 | 7099100 | 8793700 | 0 | 0 |

| | | | | | | | | | | | | | | | | | | | | | | |
|---|---|---|---|---|---|---|---|---|---|---|---|---|---|---|---|---|---|---|---|---|---|---|
| DPTGMDPI | SPCS2; SPCS2; | Q15005; E9F | SPCS2_HUMAN | Oxidation (M) | 0 | 0 | 0 | 0 | 58239 | 0 | 0 | 0 | 757900 | 43586 | 86254 | 0 | 0 | 0 | 0 | 0 | 0 | 0 |
| DPTGMDPI | SPCS2; SPCS2; | Q15005; E9F | SPCS2_HUMAN | Unmodified | 0 | 0 | 0 | 0 | 116400 | 0 | 0 | 0 | 565970 | 0 | 0 | 0 | 0 | 0 | 0 | 0 | 0 | 0 |
| DPTGMDPI | SPCS2; SPCS2; | Q15005; E9F | SPCS2_HUMAN | Unmodified | 28544 | 0 | 0 | 0 | 0 | 0 | 0 | 0 | 442420 | 0 | 0 | 0 | 1645500 | 2920200 | 0 | 0 | 0 | 0 |
| DPTNPFRF | ADAM10 | O14672 | ADA10_HUMAN | Unmodified | 1092400 | 500950 | 10280000 | 1235500 | 616380 | 160390 | 222850 | 23575 | 465840 | 0 | 214390 | 2166500 | 644150 | 244970 | 9931000 | 3693300 | 0 | 0 |
| DPTPIEFSP | POMGNT1 | Q8WZA1 | PMGT1_HUMA | Unmodified | 0 | 719550 | 0 | 0 | 0 | 0 | 0 | 0 | 354340 | 0 | 0 | 0 | 0 | 0 | 0 | 0 | 0 | 0 |
| DPTWSETG | PAN3; PAN3 | Q58A45; A0A | PAN3_HUMAN | Unmodified | 376600 | 353560 | 0 | 162530 | 0 | 0 | 0 | 0 | 0 | 0 | 0 | 0 | 0 | 0 | 166390 | 0 | 0 | 0 |
| DPVPGYSV | NAMPT; NAMPT | P43490; A0A | NAMPT_HUMA | Unmodified | 30282 | 0 | 523720 | 77602 | 259770 | 0 | 0 | 0 | 196480 | 0 | 193560 | 3766300 | 0 | 0 | 455680 | 245100 | 42276 | 0 |
| DPVQEAW | PKM; PKM; PKM | P14618; B4D | KPYM_HUMAN | Unmodified | 70877 | 71853 | 658730 | 130640 | 0 | 0 | 0 | 36876 | 349220 | 83261 | 2785400 | 16203000 | 0 | 0 | 721700 | 875630 | 0 | 0 |
| DPVYTFSIS | NPLOC4; NPLO | Q8TAT6; I3L | NPL4_HUMAN | Unmodified | 0 | 55975 | 91619 | 23310 | 0 | 0 | 0 | 0 | 0 | 0 | 0 | 292240 | 33400 | 0 | 156690 | 77207 | 0 | 0 |
| DPYGGSEC | MARVELD2; MA | Q8N4S9; A1 | MALD2_HUMA | Unmodified | 228450 | 0 | 536210 | 912830 | 0 | 0 | 0 | 0 | 0 | 0 | 0 | 60939 | 0 | 0 | 932500 | 760010 | 0 | 0 |
| DPYQEEEW | ACP2; ACP2; AC | P11117; B7Z | PPAL_HUMAN | Unmodified | 0 | 0 | 38035 | 0 | 0 | 0 | 0 | 0 | 1686300 | 0 | 64554 | 52095 | 0 | 0 | 118810 | 0 | 0 | 0 |
| DQAAEGIN | MRPS22; MRPS | P82650; A0A | RT22_HUMAN | Unmodified | 0 | 0 | 0 | 0 | 0 | 0 | 0 | 0 | 1089800 | 0 | 0 | 0 | 0 | 0 | 0 | 0 | 0 | 0 |
| DQADKAST | MYH9 | P35579 | MYH9_HUMAN | Unmodified | 1410300 | 390640 | 0 | 248820 | 0 | 0 | 175590 | 62194 | 0 | 32477 | 0 | 0 | 0 | 221930 | 0 | 0 | 545530 | 447560 |
| DQAVMISG | MYO1C; MYO1C | O00159; I3L | MYO1C_HUMA | Oxidation (M) | 0 | 0 | 780510 | 0 | 0 | 0 | 0 | 0 | 0 | 0 | 0 | 0 | 0 | 0 | 783660 | 340450 | 0 | 0 |
| DQAVMISG | MYO1C; MYO1C | O00159; I3L | MYO1C_HUMA | Unmodified | 0 | 278070 | 736720 | 381510 | 0 | 0 | 0 | 0 | 0 | 0 | 0 | 0 | 0 | 0 | 897720 | 921920 | 0 | 0 |
| DQAVMISG | MYO1C; MYO1C | O00159; I3L | MYO1C_HUMA | Unmodified | 1608800 | 1992900 | 1771700 | 2492600 | 1304500 | 612380 | 0 | 0 | 444270 | 0 | 0 | 1720600 | 574750 | 1094100 | 5740600 | 4054700 | 402360 | 640660 |
| DQAVMISG | MYO1C; MYO1C | O00159; I3L | MYO1C_HUMA | Oxidation (M) | 614060 | 605980 | 1106800 | 925530 | 397480 | 195000 | 0 | 0 | 84832 | 29748 | 0 | 0 | 0 | 0 | 1996500 | 1767900 | 74472 | 0 |
| DQAVMISG | MYO1C; MYO1C | O00159; I3L | MYO1C_HUMA | Unmodified | 1687800 | 1917900 | 2060900 | 5533600 | 934560 | 319260 | 0 | 32135 | 331990 | 0 | 134300 | 0 | 0 | 0 | 2730900 | 1597600 | 260490 | 262950 |
| DQAVMISG | MYO1C; MYO1C | O00159; I3L | MYO1C_HUMA | Oxidation (M) | 776190 | 1186900 | 880370 | 2352200 | 0 | 0 | 0 | 0 | 0 | 0 | 0 | 110150 | 303300 | 0 | 2032200 | 2854900 | 184720 | 0 |
| DQCQVDS | WFDC2 | Q14508 | WFDC2_HUMA | Unmodified | 0 | 0 | 78490 | 0 | 0 | 0 | 0 | 0 | 1013200 | 235760 | 0 | 0 | 0 | 0 | 0 | 0 | 0 | 0 |
| DQDGEILLI | DKK3; DKK3 | Q9UBP4; F6 | DKK3_HUMAN | Unmodified | 0 | 422200 | 0 | 0 | 0 | 0 | 0 | 0 | 603380 | 0 | 2955100 | 0 | 0 | 0 | 514610 | 0 | 0 | 0 |
| DQDGKIGL | SDCBP; SDCBP | O00560; G5I | SDCB1_HUMA | Unmodified | 1820300 | 0 | 0 | 2545500 | 1986000 | 0 | 0 | 50752 | 12202000 | 3041900 | 257250 | 826480 | 2254000 | 1820000 | 689840 | 821460 | 0 | 0 |
| DQDGKIGL | SDCBP; SDCBP | O00560; G5I | SDCB1_HUMA | Unmodified | 0 | 0 | 0 | 0 | 26725000 | 9209900 | 3005000 | 63139 | 1869500 | 41661000 | 0 | 0 | 4432100 | 22006000 | 0 | 0 | 0 | 3638600 |
| DQDLEPGA | EPRS1 | P07814 | SYEP_HUMAN | Oxidation (M) | 0 | 0 | 0 | 0 | 151190 | 0 | 0 | 0 | 82968 | 0 | 325450 | 893250 | 0 | 0 | 0 | 0 | 0 | 0 |
| DQDLEPGA | EPRS1 | P07814 | SYEP_HUMAN | Unmodified | 0 | 0 | 0 | 0 | 0 | 0 | 0 | 0 | 163480 | 0 | 830820 | 1960100 | 0 | 0 | 0 | 0 | 0 | 0 |
| DQDPAAFC | MGAM2 | Q2M2H8 | MGAL_HUMAN | Unmodified | 0 | 0 | 0 | 0 | 0 | 0 | 0 | 0 | 0 | 0 | 0 | 1166900 | 0 | 0 | 0 | 0 | 0 | 0 |
| DQEEGVMI | SLC39A14; SLC | Q15043; E5F | S39AE_HUMAN | Unmodified | 383340 | 371660 | 1078400 | 350030 | 106180 | 0 | 0 | 0 | 376150 | 0 | 59380 | 1193700 | 0 | 0 | 873480 | 971450 | 384910 | 649140 |
| DQEEGVMI | SLC39A14; SLC | Q15043; E5F | S39AE_HUMAN | Oxidation (M) | 0 | 17479 | 68835 | 0 | 0 | 0 | 0 | 0 | 29172 | 0 | 0 | 59682 | 0 | 0 | 49315 | 73511 | 0 | 0 |
| DQELYFFH | TARS1 | P26639 | SYTC_HUMAN | Unmodified | 0 | 0 | 25055 | 121560 | 0 | 0 | 185920 | 0 | 97139 | 74274 | 0 | 53554 | 0 | 0 | 415180 | 0 | 0 | 0 |
| DQEWYDA | CCDC50 | Q8IVM0 | CCD50_HUMA | Unmodified | 887510 | 987520 | 867130 | 335750 | 0 | 0 | 0 | 0 | 0 | 71471 | 0 | 0 | 0 | 0 | 1742200 | 1454600 | 0 | 0 |
| DQEWYDA | CCDC50 | Q8IVM0 | CCD50_HUMA | Unmodified | 0 | 0 | 932830 | 119940 | 0 | 0 | 0 | 0 | 0 | 0 | 0 | 0 | 0 | 0 | 894420 | 408640 | 0 | 0 |
| DQFALAFH | EPS15; EPS15 | P42566; B1A | EPS15_HUMAN | Unmodified | 198880 | 448450 | 51881 | 136950 | 727430 | 94098 | 0 | 0 | 113360 | 526290 | 58576 | 314070 | 578200 | 284360 | 0 | 0 | 534980 | 379670 |
| DQFGFINY | CSDE1; CSDE1 | O75534; E9F | CSDE1_HUMA | Unmodified | 37091 | 23880 | 35779 | 84203 | 105020 | 21447 | 0 | 30416 | 56902 | 0 | 105410 | 2256000 | 0 | 0 | 133880 | 49424 | 0 | 64762 |
| DQFTDQQ | MYO1B; MYO1B | O43795; H7 | MYO1B_HUMA | Unmodified | 0 | 0 | 0 | 0 | 0 | 0 | 0 | 0 | 0 | 685890 | 0 | 572570 | 0 | 0 | 0 | 0 | 0 | 0 |
| DQGELERC | MYH9 | P35579 | MYH9_HUMAN | Unmodified | 22689 | 0 | 66100 | 0 | 4761300 | 567640 | 342510 | 0 | 195750 | 551980 | 51751 | 0 | 554450 | 1024300 | 131120 | 0 | 576490 | 731070 |
| DQGKPEV | CIT; CIT; CIT; CI | O14578; A0A | CTRO_HUMAN | Unmodified | 0 | 0 | 0 | 0 | 3299400 | 0 | 0 | 0 | 0 | 0 | 0 | 0 | 0 | 0 | 0 | 0 | 929000 | 570370 |
| DQGPPPSF | PNMA2 | Q9UL42 | PNMA2_HUMA | Unmodified | 1443600 | 0 | 0 | 0 | 0 | 0 | 179690 | 0 | 0 | 0 | 0 | 7050300 | 0 | 0 | 0 | 0 | 0 | 0 |
| DQGPTFDY | B4GALT3 | O60512 | B4GT3_HUMAN | Unmodified | 0 | 0 | 0 | 0 | 0 | 0 | 0 | 0 | 441440 | 188350 | 0 | 1159500 | 0 | 0 | 0 | 0 | 0 | 0 |
| DQGSFTEV | IGF2R; IGF2R | P11717; A0A | MPRI_HUMAN | Oxidation (M) | 0 | 0 | 0 | 0 | 0 | 0 | 38792 | 0 | 13186000 | 4522400 | 1619300 | 65199 | 1039600 | 0 | 0 | 0 | 0 | 0 |
| DQGTYEDY | MYL6; MYL6; MY | P60660; G3V | MYL6_HUMAN | Unmodified | 0 | 0 | 0 | 0 | 0 | 0 | 0 | 0 | 695500 | 0 | 0 | 8685500 | 0 | 0 | 0 | 1652700 | 0 | 0 |
| DQGTYEDY | MYL6; MYL6; MY | P60660; G3V | MYL6_HUMAN | Unmodified | 2109100 | 847610 | 1573500 | 3856700 | 11995000 | 4680600 | 629590 | 0 | 3883900 | 647830 | 2072800 | 6917300 | 3350100 | 1826700 | 4961800 | 2580800 | 1294500 | 2851900 |
| DQHMAIAV | CEL | P19835 | CEL_HUMAN B | Unmodified | 0 | 0 | 0 | 0 | 0 | 0 | 0 | 0 | 362000 | 141740 | 0 | 0 | 0 | 0 | 0 | 0 | 0 | 0 |
| DQHSFELD | ATL3; ATL3; ATL | Q6DD88; F5 | ATLA3_HUMAN | Unmodified | 159700 | 0 | 913070 | 0 | 465830 | 0 | 0 | 43198 | 138010 | 0 | 0 | 323580 | 0 | 0 | 0 | 323620 | 0 | 240540 |
| DQIAQLEA | LMNB1; LMNB1 | P20700; A0A | LMNB1_HUMA | Unmodified | 0 | 0 | 0 | 0 | 0 | 0 | 0 | 0 | 0 | 0 | 0 | 1198200 | 0 | 0 | 0 | 0 | 0 | 0 |
| DQIPSPGLI | ATP1B3; ATP1B | P54709; C9J | AT1B3_HUMAN | Oxidation (M) | 1264400 | 826600 | 2074400 | 2392400 | 0 | 0 | 0 | 0 | 1318400 | 969290 | 128200 | 14295000 | 0 | 0 | 27785000 | 93130 | 0 | 0 |
| DQIPSPGLI | ATP1B3; ATP1B | P54709; C9J | AT1B3_HUMAN | Unmodified | 0 | 0 | 0 | 0 | 0 | 0 | 0 | 0 | 602970 | 0 | 0 | 283180 | 0 | 0 | 458330 | 0 | 0 | 0 |
| DQITAGNA | PAICS; PAICS | P22234; E9P | PUR6_HUMAN | Unmodified | 0 | 0 | 0 | 0 | 0 | 0 | 0 | 0 | 497130 | 55028 | 195170 | 3820500 | 0 | 484370 | 43007 | 186770 | 0 | 0 |
| DQIYDIFQ | EIF4A1; EIF4A1; | P60842; J3K | IF4A1_HUMAN | Unmodified | 382190 | 486440 | 4587500 | 424980 | 0 | 0 | 282600 | 399250 | 485510 | 0 | 1008900 | 9789800 | 0 | 0 | 3292300 | 1942400 | 0 | 0 |
| DQKDIALEI | ITGA6 | P23229 | ITA6_HUMAN I | Unmodified | 0 | 0 | 0 | 0 | 966220 | 0 | 0 | 0 | 0 | 0 | 0 | 0 | 97204 | 0 | 0 | 0 | 348780 | 1495800 |
| DQLAAEAA | NOS1AP | O75052 | CAPON_HUMA | Unmodified | 0 | 0 | 0 | 0 | 0 | 0 | 0 | 0 | 206160 | 0 | 0 | 0 | 0 | 0 | 0 | 0 | 0 | 0 |
| DQLAAEAA | NOS1AP | O75052 | CAPON_HUMA | Unmodified | 0 | 0 | 200630 | 358910 | 0 | 0 | 0 | 0 | 1358900 | 517480 | 0 | 0 | 713970 | 743810 | 0 | 144130 | 0 | 0 |
| DQLAAEAA | NOS1AP | O75052 | CAPON_HUMA | Unmodified | 0 | 0 | 0 | 0 | 1484600 | 0 | 0 | 0 | 0 | 481310 | 0 | 0 | 2726000 | 1082900 | 0 | 0 | 725950 | 0 |
| DQLALAIGI | TSFM; TSFM | P43897; C9J | EFTS_HUMAN | Unmodified | 931160 | 1101500 | 0 | 273260 | 0 | 0 | 527850 | 187940 | 935850 | 846650 | 739400 | 1809500 | 948620 | 739600 | 1076500 | 552620 | 888880 | 805940 |
| DQLIYLLEC | VPS33B; VPS33 | Q9H267; G3 | VP33B_HUMAN | Unmodified | 0 | 0 | 0 | 0 | 0 | 0 | 0 | 723500 | 0 | 0 | 0 | 862530 | 0 | 0 | 0 | 0 | 0 | 0 |
| DQLIYNLLK | LDHA; LDHA; LI | P00338; F5G | LDHA_HUMAN | Unmodified | 1734300 | 748080 | 2528500 | 3847300 | 2554000 | 1314100 | 0 | 0 | 7205100 | 545440 | 9178600 | 2.44E+08 | 0 | 377630 | 7410500 | 7518700 | 541300 | 3837100 |
| DQLIYNLLK | LDHA; LDHA; LI | P00338; F5G | LDHA_HUMAN | Unmodified | 54823 | 31693 | 0 | 177020 | 4116300 | 2772100 | 918200 | 259200 | 1994800 | 721350 | 6054900 | 39308000 | 2448100 | 1236000 | 374520 | 1079600 | 1430500 | 9933800 |
| DQLKEVWI | NUCB2; NUCB2 | P80303; Q2L | NUCB2_HUMA | Unmodified | 0 | 0 | 159440 | 0 | 0 | 0 | 0 | 0 | 956900 | 636870 | 0 | 0 | 0 | 0 | 74186 | 27004 | 0 | 0 |
| DQLQTFSE | ACTR1A; ACTR1 | P61163; A0A | ACTZ_HUMAN | Unmodified | 0 | 0 | 679990 | 0 | 0 | 291000 | 0 | 87372 | 1248700 | 0 | 363220 | 25414000 | 0 | 0 | 0 | 1077300 | 1298200 | 807840 |
| DQLQTFSE | ACTR1A; ACTR1 | P61163; A0A | ACTZ_HUMAN | Unmodified | 0 | 0 | 0 | 906940 | 2691400 | 0 | 0 | 0 | 2122500 | 1111600 | 0 | 10177000 | 0 | 0 | 0 | 0 | 911480 | 1839800 |
| DQLQTFSE | ACTR1B | P42025 | ACTY_HUMAN | Unmodified | 0 | 0 | 0 | 0 | 1469900 | 7146100 | 2988600 | 0 | 909860 | 0 | 0 | 7453200 | 0 | 2921900 | 0 | 0 | 1443800 | 4629600 |
| DQLRPTQL | MYOF; MYOF | Q9NZM1; HC | MYOF_HUMAN | Unmodified | 0 | 1419100 | 0 | 0 | 373600 | 0 | 0 | 0 | 162420 | 70110 | 0 | 0 | 0 | 0 | 0 | 0 | 0 | 0 |
| DQLSDDVS | SCG2 | P13521 | SCG2_HUMAN | Unmodified | 0 | 0 | 0 | 0 | 0 | 0 | 0 | 0 | 863210 | 53916 | 0 | 0 | 0 | 0 | 0 | 0 | 0 | 0 |
| DQMSTSSV | STX6 | O43752 | STX6_HUMAN | Unmodified | 0 | 0 | 0 | 0 | 0 | 0 | 0 | 69999 | 656280 | 58168 | 0 | 0 | 0 | 30623 | 0 | 0 | 0 | 0 |
| DQNIFVQK | BLVRA | P53004 | BIEA_HUMAN E | Unmodified | 472350 | 0 | 0 | 483000 | 2928700 | 0 | 646500 | 0 | 430520 | 0 | 0 | 0 | 3116000 | 1939200 | 448350 | 593010 | 904410 | 0 |
| DQNVLSPV | SRGAP2; SRGA | O75044; P0 | SRGP2_HUMA | Unmodified | 0 | 33681 | 0 | 0 | 253590 | 55428 | 23098 | 0 | 0 | 0 | 0 | 0 | 0 | 0 | 0 | 0 | 0 | 0 |
| DQPPNSVE | CYRIB | Q9NUQ9 | CYRIB_HUMAN | Unmodified | 161050 | 207160 | 226630 | 656750 | 1240000 | 0 | 0 | 0 | 2968800 | 160130 | 794790 | 3101200 | 81573 | 0 | 661880 | 490780 | 92905 | 0 |
| DQPPNSVE | CYRIB | Q9NUQ9 | CYRIB_HUMAN | Unmodified | 0 | 0 | 0 | 0 | 6084000 | 23447 | 0 | 0 | 295710 | 792260 | 0 | 0 | 0 | 0 | 0 | 0 | 1409100 | 5253500 |
| DQPQFQN | RAD23A; RAD2 | P54725; A0A | RD23A_HUMA | Unmodified | 588870 | 157190 | 268660 | 0 | 0 | 0 | 0 | 0 | 0 | 0 | 0 | 0 | 0 | 0 | 0 | 0 | 0 | 0 |
| DQQLEALC | GOLGA3 | Q08378 | GOGA3_HUMA | Oxidation (M) | 0 | 0 | 0 | 0 | 0 | 0 | 0 | 0 | 55558 | 0 | 0 | 0 | 0 | 0 | 0 | 0 | 0 | 886690 |
| DQQNLPY | MARK2; MARK2 | Q7KZI7; E7E | MARK2_HUMA | Unmodified | 0 | 0 | 0 | 445980 | 0 | 0 | 0 | 0 | 0 | 0 | 0 | 793560 | 0 | 0 | 0 | 734160 | 0 | 0 |
| DQQQKPLY | GNA13 | Q14344 | GNA13_HUMA | Unmodified | 553850 | 462420 | 0 | 892190 | 445250 | 0 | 0 | 0 | 6954400 | 85395 | 1665500 | 25097000 | 0 | 0 | 214800 | 227000 | 447100 | 0 |
| DQSAVVVC | GTF2I; GTF2I | P78347; A0A | GTF2I_HUMAN | Unmodified | 0 | 0 | 0 | 0 | 0 | 0 | 0 | 0 | 0 | 0 | 0 | 3152000 | 0 | 0 | 0 | 0 | 0 | 0 |
| DQSVGDPI | SCARB2; SCARI | Q14108; A0A | SCRB2_HUMA | Unmodified | 0 | 0 | 0 | 0 | 0 | 57110 | 13516 | 45844 | 0 | 189710 | 0 | 1802900 | 0 | 287910 | 0 | 0 | 0 | 40182 |
| DQTASAPA | COBLL1; COBLI | Q53SF7; A0A | COBL1_HUMA | Unmodified | 287470 | 260220 | 161770 | 0 | 0 | 0 | 0 | 0 | 0 | 0 | 0 | 0 | 0 | 0 | 0 | 240560 | 0 | 0 |
| DQTMGLGI | AVIL | O75366 | AVIL_HUMAN A | Oxidation (M) | 0 | 0 | 0 | 0 | 0 | 0 | 0 | 0 | 52061 | 0 | 455380 | 3128600 | 0 | 0 | 0 | 0 | 0 | 0 |
| DQTPDENI | IGF2BP3 | O00425 | IF2B3_HUMAN | Unmodified | 585250 | 91143 | 0 | 0 | 0 | 0 | 0 | 0 | 0 | 0 | 0 | 1940700 | 0 | 0 | 91294 | 0 | 0 | 0 |
| DQTPLILSV | KIAA1549 | Q9HCM3 | K1549_HUMAN | Unmodified | 59719 | 34464 | 107170 | 66799 | 0 | 0 | 0 | 0 | 137170 | 18917 | 0 | 184040 | 0 | 0 | 192420 | 106690 | 0 | 0 |
| DQVACLTF | CDCP1 | Q9H5V8 | CDCP1_HUMA | Unmodified | 0 | 0 | 0 | 0 | 341510 | 0 | 0 | 119310 | 0 | 0 | 0 | 1466900 | 0 | 0 | 572130 | 241100 | 0 | 0 |
| DQVANSAF | HSP90AA1 | P07900 | HS90A_HUMAN | Unmodified | 1663600 | 1347900 | 4447100 | 2596100 | 943360 | 1011400 | 0 | 51355 | 16099000 | 1552900 | 20320000 | 1.45E+08 | 248540 | 0 | 5409500 | 4175700 | 0 | 968240 |
| DQVANSAF | HSP90AA1 | P07900 | HS90A_HUMAN | Unmodified | 0 | 0 | 9548900 | 367460 | 0 | 0 | 0 | 0 | 1025800 | 522080 | 0 | 1395900 | 0 | 0 | 33134 | 522070 | 0 | 0 |
| DQVANSAF | HSP90AA1 | P07900 | HS90A_HUMAN | Unmodified | 0 | 0 | 0 | 0 | 3100400 | 0 | 0 | 0 | 0 | 0 | 0 | 0 | 0 | 0 | 0 | 0 | 853710 | 2135600 |
| DQVEDYAL | MTR; MTR; MTR | Q99707; A0A | METH_HUMAN | Unmodified | 0 | 0 | 0 | 0 | 0 | 0 | 0 | 0 | 0 | 0 | 0 | 215670 | 0 | 0 | 0 | 0 | 0 | 0 |
| DQVELVEN | HINT3 | Q9NQE9 | HINT3_HUMAN | Unmodified | 0 | 0 | 0 | 0 | 0 | 0 | 0 | 0 | 545540 | 0 | 0 | 283980 | 0 | 0 | 0 | 0 | 0 | 0 |
| DQVGFVVS | EPB41L1 | A0A1B0GTW | A0A1B0GTW6_ | Unmodified | 2488700 | 2277000 | 0 | 0 | 0 | 0 | 0 | 0 | 0 | 0 | 0 | 0 | 0 | 0 | 3370100 | 0 | 0 | 0 |
| DQVLAMLE | CKAP5 | Q14008 | CKAP5_HUMA | Unmodified | 0 | 0 | 0 | 0 | 0 | 0 | 0 | 0 | 0 | 0 | 62917 | 202500 | 0 | 0 | 0 | 0 | 0 | 0 |
| DQVNTVGI | EPHA2 | P29317 | EPHA2_HUMA | Unmodified | 0 | 0 | 0 | 0 | 566140 | 0 | 0 | 0 | 0 | 0 | 268160 | 2045200 | 0 | 0 | 0 | 0 | 1129600 | 1220000 |
| DQVTAQEI | XRCC5 | P13010 | XRCC5_HUMA | Unmodified | 596630 | 104340 | 858360 | 0 | 0 | 0 | 0 | 1428800 | 0 | 0 | 737150 | 6923700 | 286960 | 0 | 173420 | 94381 | 0 | 0 |
| DQVYPQYV | ZC3HAV1; ZC3H | Q7Z2W4; C9 | ZCCHV_HUMA | Unmodified | 0 | 0 | 1354700 | 350170 | 655460 | 392940 | 0 | 0 | 752110 | 0 | 0 | 0 | 0 | 0 | 1501100 | 802720 | 0 | 0 |
| DQWEDRIC | EZR; EZR | P15311; E7E | EZRI_HUMAN E | Unmodified | 2166000 | 929250 | 645710 | 2562900 | 351440 | 0 | 0 | 0 | 15258000 | 366490 | 1426700 | 19136000 | 0 | 0 | 1810500 | 2068400 | 0 | 0 |
| DRAATSPA | DYNC1H1; DYN | Q14204; A0A | DYHC1_HUMA | Unmodified | 0 | 0 | 236290 | 0 | 0 | 471930 | 0 | 0 | 930040 | 0 | 0 | 0 | 328790 | 149770 | 0 | 125400 | 0 | 122610 |
| DRCPLPCII | RACGAP1 | Q9H0H5 | RGAP1_HUMA | Unmodified | 0 | 0 | 114980 | 225470 | 549740 | 201290 | 0 | 0 | 796050 | 0 | 206610 | 651530 | 0 | 0 | 0 | 39343 | 0 | 0 |
| DRDDFPVV | RRAS | P10301 | RRAS_HUMAN | Unmodified | 0 | 110070 | 1010000 | 391330 | 0 | 0 | 0 | 0 | 0 | 0 | 0 | 0 | 0 | 0 | 758600 | 355600 | 0 | 0 |
| DRDDFPVV | RRAS | P10301 | RRAS_HUMAN | Unmodified | 205920 | 376850 | 0 | 533740 | 0 | 0 | 0 | 39083 | 0 | 0 | 0 | 0 | 0 | 0 | 839610 | 515760 | 0 | 0 |
| DRDDIVKT | ATP2B1; ATP2B | P20020; E7E | AT2B1_HUMAN | Unmodified | 0 | 0 | 0 | 0 | 0 | 0 | 0 | 0 | 0 | 0 | 0 | 0 | 159700 | 104010 | 0 | 0 | 0 | 26482 |
| DRDEFPMI | RRAS2; RRAS2; | P62070; E9P | RRAS2_HUMAN | Oxidation (M) | 366320 | 351760 | 2950500 | 1248400 | 372710 | 50714 | 0 | 0 | 0 | 0 | 0 | 0 | 0 | 0 | 3257900 | 2762300 | 0 | 30935 |
| DRDEFPMI | RRAS2; RRAS2; | P62070; E9P | RRAS2_HUMAN | Oxidation (M) | 4329900 | 3108900 | 423870 | 931940 | 0 | 0 | 0 | 0 | 0 | 0 | 0 | 0 | 378620 | 36463 | 2236800 | 2285800 | 0 | 0 |
| DRDNDSDI | MYOF | Q9NZM1 | MYOF_HUMAN | Unmodified | 0 | 0 | 0 | 0 | 5097300 | 1573900 | 0 | 0 | 360240 | 0 | 0 | 0 | 0 | 0 | 0 | 0 | 0 | 0 |
| DRDPEAQF | PRMT5; PRMT5; | O14744; H0 | ANM5_HUMAN | Unmodified | 0 | 0 | 0 | 0 | 0 | 0 | 0 | 0 | 0 | 0 | 0 | 2780700 | 0 | 0 | 0 | 0 | 0 | 514490 |
| DRDSQRDI | FASN; FASN | P49327; A0A | FAS_HUMAN F | Unmodified | 37718 | 0 | 0 | 18011 | 0 | 0 | 0 | 0 | 0 | 0 | 27246 | 0 | 0 | 0 | 199240 | 0 | 0 | 0 |
| DRDVTFSP | MANF; MANF | P55145; H7 | MANF_HUMAN | Unmodified | 15971 | 0 | 520110 | 0 | 2819600 | 350290 | 1175700 | 231900 | 2.56E+08 | 43800000 | 21684000 | 5065500 | 1727700 | 1718900 | 0 | 633940 | 0 | 3482600 |
| DRDVTFSP | MANF; MANF | P55145; H7 | MANF_HUMAN | Unmodified | 0 | 0 | 0 | 0 | 423820 | 0 | 1074500 | 0 | 0 | 2594700 | 0 | 0 | 0 | 0 | 0 | 0 | 0 | 0 |
| DREEALHQ | DDX3X; DDX3Y | O00571; O1 | DDX3X_HUMA | Unmodified | 1638700 | 0 | 1221000 | 979690 | 0 | 0 | 0 | 0 | 0 | 0 | 0 | 2150600 | 0 | 0 | 1488400 | 1268300 | 0 | 0 |
| DREEILLQE | TOR2A | Q5JU69 | TOR2A_HUMA | Unmodified | 50958 | 73860 | 41028 | 0 | 0 | 0 | 0 | 0 | 22896000 | 1988600 | 249100 | 34193 | 0 | 0 | 127420 | 58506 | 0 | 0 |
| DREGFFTN | PFN2; PFN2; PF | P35080; G5E | PROF2_HUMA | Unmodified | 3296300 | 5129700 | 933570 | 1499300 | 3621000 | 872010 | 229730 | 132220 | 3555500 | 1356600 | 1954300 | 15577000 | 2454700 | 1698500 | 684290 | 1617500 | 2079900 | 1422800 |
| DREGFFTN | PFN2; PFN2; PF | P35080; G5E | PROF2_HUMA | Unmodified | 0 | 0 | 0 | 163230 | 3180500 | 0 | 0 | 0 | 402020 | 302370 | 179220 | 584780 | 550640 | 237920 | 297970 | 213380 | 1488200 | 335720 |
| DREGYTSF | GOLPH3 | Q9H4A6 | GOLP3_HUMA | Unmodified | 22241 | 0 | 0 | 0 | 0 | 0 | 0 | 0 | 250460 | 0 | 0 | 421480 | 68360 | 0 | 111080 | 45846 | 0 | 0 |
| DRESMGHI | HNRNPF; HNRI | P52597; A0A | HNRPF_HUMA | Unmodified | 0 | 0 | 0 | 0 | 1191600 | 387570 | 1579100 | 0 | 0 | 0 | 0 | 0 | 1376900 | 983690 | 0 | 0 | 438860 | 1880300 |
| DRETMGHI | HNRNPH1; HNI | P31943; P55 | HNRH1_HUMA | Oxidation (M) | 0 | 0 | 0 | 0 | 440770 | 0 | 909130 | 0 | 0 | 167300 | 0 | 0 | 663930 | 0 | 0 | 0 | 163730 | 505270 |
| DRETMGHI | HNRNPH1; HNI | P31943; P55 | HNRH1_HUMA | Unmodified | 0 | 0 | 0 | 0 | 264510 | 262780 | 0 | 0 | 0 | 325650 | 0 | 0 | 1213500 | 0 | 0 | 0 | 306490 | 1939200 |

| | | | | | | | | | | | | | | | | | | | | | | |
|---|---|---|---|---|---|---|---|---|---|---|---|---|---|---|---|---|---|---|---|---|---|---|
| DRETSKVIY | WASL | O00401 | WASL_HUMAN | Unmodified | 0 | 0 | 0 | 0 | 1010900 | 0 | 0 | 0 | 0 | 137310 | 0 | 0 | 0 | 0 | 0 | 0 | 48450 | 0 |
| DREVGIPPI | ACTR3; ACTR3 | P61158; B4D | ARP3_HUMAN | Unmodified | 626300 | 677500 | 924940 | 1002600 | 0 | 0 | 91699 | 24483 | 513710 | 0 | 738490 | 5919300 | 93760 | 60719 | 809520 | 1114500 | 0 | 0 |
| DREVNAVD | IDE; IDE; IDE; ID | P14735; A0A | IDE_HUMAN In | Unmodified | 0 | 0 | 0 | 0 | 0 | 82040 | 0 | 0 | 0 | 0 | 0 | 10172000 | 527360 | 119510 | 0 | 216520 | 637950 | 285270 |
| DRFGIDDQ | PIP4K2C; PIP4K | Q8TBX8; F8V | PI42C_HUMAN | Unmodified | 0 | 0 | 0 | 0 | 0 | 215720 | 0 | 0 | 462520 | 0 | 0 | 781580 | 215780 | 0 | 0 | 0 | 0 | 71900 |
| DRFNHFSL | GPI; GPI; GPI; G | P06744; K7E | G6PI_HUMAN ( | Unmodified | 0 | 0 | 0 | 0 | 2468100 | 0 | 1084900 | 0 | 0 | 477680 | 0 | 1965200 | 0 | 0 | 0 | 0 | 314170 | 0 |
| DRFQGRFD | ATP1B1 | P05026 | AT1B1_HUMAN | Unmodified | 0 | 0 | 3033000 | 1956000 | 4766600 | 373420 | 795690 | 291650 | 6117900 | 1249500 | 2748500 | 6376500 | 0 | 1297000 | 1959200 | 1711800 | 432890 | 147730 |
| DRFVNDYD | RCN2; RCN2 | Q14257; A8M | RCN2_HUMAN | Unmodified | 0 | 0 | 72767 | 0 | 0 | 0 | 0 | 0 | 4232700 | 0 | 0 | 0 | 0 | 0 | 0 | 0 | 0 | 0 |
| DRGFELTFI | DDOST; DDOST | P39656; U3K | OST48_HUMAN | Unmodified | 0 | 0 | 359930 | 159890 | 0 | 0 | 0 | 100470 | 16995000 | 684260 | 671630 | 1521900 | 0 | 0 | 1512300 | 759030 | 0 | 0 |
| DRGGAGKI | AHNAK2 | Q8IVF2 | AHNK2_HUMAI | Unmodified | 0 | 0 | 0 | 0 | 1007100 | 80701 | 0 | 0 | 0 | 0 | 0 | 0 | 0 | 0 | 0 | 0 | 0 | 608220 |
| DRGLTEFP | LRRC57; N/A; L | Q8N9N7; A0 | LRC57_HUMAN | Unmodified | 999800 | 590570 | 236390 | 558650 | 3810500 | 0 | 441070 | 620560 | 0 | 1100200 | 623930 | 3401800 | 0 | 0 | 486130 | 652240 | 1014600 | 2573500 |
| DRHFAGD | CHID1; CHID1; | Q9BWS9; E9 | CHID1_HUMAN | Unmodified | 0 | 0 | 0 | 0 | 0 | 0 | 0 | 0 | 9101800 | 244360 | 749480 | 0 | 0 | 0 | 0 | 0 | 0 | 0 |
| DRINDDLIS | DLG2; DLG1; D | Q15700; Q1 | DLG2_HUMAN | Unmodified | 0 | 0 | 776720 | 0 | 617630 | 390610 | 144660 | 0 | 0 | 0 | 812930 | 0 | 0 | 1016900 | 927650 | 0 | 0 | 422510 |
| DRIPVYSAI | ENDOD1 | O94919 | ENDD1_HUMA | Unmodified | 0 | 0 | 316560 | 350960 | 1853600 | 453490 | 0 | 0 | 1313200 | 416260 | 0 | 913060 | 1595900 | 613940 | 396010 | 463690 | 1532300 | 6210000 |
| DRISWAGD | MPZL1; MPZL1 | O95297; Q9 | MPZL1_HUMAN | Unmodified | 1975800 | 895950 | 2123200 | 0 | 2294300 | 663160 | 78057 | 0 | 1143600 | 0 | 0 | 16817000 | 0 | 404770 | 1328900 | 0 | 503850 | 1285800 |
| DRKESLDV | ASAH1; ASAH1; | Q13510; A0A | ASAH1_HUMAN | Unmodified | 0 | 0 | 0 | 0 | 0 | 0 | 0 | 0 | 7174500 | 417040 | 831040 | 0 | 137520 | 0 | 0 | 0 | 0 | 0 |
| DRLEFWCT | PSMD13; PSMD | Q9UNM6; J3 | PSD13_HUMAN | Unmodified | 28307 | 82209 | 33117 | 142380 | 0 | 0 | 352300 | 78616 | 0 | 0 | 310350 | 2837200 | 1720900 | 1260200 | 0 | 806120 | 0 | 1761200 |
| DRLHLQG | UBE2Z | Q9H832 | UBE2Z_HUMAN | Unmodified | 0 | 0 | 0 | 0 | 537300 | 0 | 0 | 0 | 0 | 0 | 0 | 0 | 0 | 0 | 0 | 0 | 429560 | 1280200 |
| DRLLELLQ | PROM2 | Q8N271 | PROM2_HUMA | Unmodified | 1659200 | 1065100 | 874910 | 901040 | 0 | 0 | 0 | 0 | 0 | 0 | 81406 | 1274000 | 0 | 0 | 1882700 | 1147900 | 0 | 0 |
| DRLLLATMI | ALDH1A1; ALDH | P00352; Q5 | AL1A1_HUMAN | Unmodified | 0 | 0 | 0 | 190980 | 0 | 0 | 0 | 0 | 0 | 0 | 0 | 5107000 | 0 | 0 | 0 | 349970 | 0 | 0 |
| DRLLLATMI | ALDH1A1; ALDH | P00352; Q5 | AL1A1_HUMAN | Oxidation (M) | 0 | 0 | 0 | 0 | 0 | 0 | 0 | 0 | 0 | 0 | 0 | 14091000 | 0 | 144080 | 0 | 0 | 0 | 0 |
| DRLLQSNP | MYO1C; MYO1 | O00159; I3L | MYO1C_HUMA | Unmodified | 1970000 | 2742200 | 3265400 | 7902000 | 3183900 | 3265000 | 0 | 0 | 2374000 | 0 | 343980 | 964420 | 1642800 | 884390 | 4639000 | 4214500 | 1218600 | 3038400 |
| DRLQYFRD | IQGAP1; IQGAP | P46940; A0A | IQGA1_HUMAN | Unmodified | 380210 | 423400 | 0 | 201620 | 2685800 | 780140 | 0 | 0 | 0 | 0 | 236760 | 985990 | 0 | 0 | 512560 | 378120 | 0 | 0 |
| DRLSLVLV | CDCP1 | Q9H5V8 | CDCP1_HUMA | Unmodified | 0 | 0 | 3134800 | 345970 | 12308000 | 1413300 | 0 | 0 | 0 | 418080 | 2776600 | 18865000 | 1764500 | 4243100 | 4566500 | 3056200 | 1657900 | 9488000 |
| DRLTCAYP | TPBG | Q13641 | TPBG_HUMAN | Unmodified | 112630 | 0 | 0 | 0 | 328080 | 0 | 0 | 0 | 0 | 0 | 0 | 3228900 | 0 | 0 | 0 | 0 | 0 | 55420 |
| DRLVAEFTI | STX7 | O15400 | STX7_HUMAN | Unmodified | 0 | 0 | 0 | 0 | 4539500 | 853300 | 0 | 0 | 1754300 | 0 | 0 | 0 | 429820 | 0 | 0 | 0 | 155300 | 364680 |
| DRLVLSYV | IGF2R; IGF2R; I | P11717; A0A | MPRI_HUMAN | Unmodified | 0 | 0 | 0 | 0 | 0 | 0 | 0 | 0 | 1031500 | 0 | 0 | 0 | 0 | 0 | 0 | 0 | 33525 | 0 |
| DRLVPGPV | LMAN2; LMAN2 | Q12907; D6 | LMAN2_HUMAI | Unmodified | 180030 | 0 | 358420 | 152830 | 0 | 0 | 0 | 0 | 76357000 | 5625800 | 1522700 | 1359300 | 0 | 0 | 496570 | 0 | 0 | 0 |
| DRLVTLEEI | NUCB2; NUCB | P80303; H0 | NUCB2_HUMA | Unmodified | 0 | 0 | 1321000 | 890620 | 0 | 0 | 0 | 0 | 67214000 | 1870700 | 635730 | 472670 | 353760 | 0 | 3542400 | 621190 | 0 | 0 |
| DRMVNSN | PDHA1; PDHA1 | P08559; Q5 | ODPA_HUMAN | Oxidation (M) | 0 | 0 | 0 | 0 | 0 | 0 | 424070 | 110850 | 1048200 | 0 | 0 | 0 | 0 | 0 | 0 | 0 | 0 | 0 |
| DRMVNSN | PDHA1; PDHA1 | P08559; Q5 | ODPA_HUMAN | Unmodified | 0 | 0 | 0 | 0 | 263340 | 0 | 0 | 0 | 1079900 | 0 | 0 | 0 | 0 | 0 | 0 | 0 | 0 | 0 |
| DRNLQEG | TMED1; TMED1 | Q13445; K7E | TMED1_HUMAI | Unmodified | 0 | 0 | 0 | 0 | 0 | 0 | 0 | 0 | 969900 | 94044 | 0 | 0 | 0 | 0 | 0 | 0 | 0 | 0 |
| DRPFKPYN | FARSA; FARSA; | Q9Y285; K7E | SYFA_HUMAN | Unmodified | 0 | 0 | 0 | 0 | 0 | 0 | 0 | 0 | 0 | 0 | 0 | 803060 | 0 | 0 | 0 | 0 | 0 | 0 |
| DRPFMGG | PTGES2; PTGES | Q9H7Z7; A0 | PGES2_HUMAI | Oxidation (M) | 0 | 0 | 0 | 0 | 0 | 0 | 0 | 0 | 0 | 0 | 0 | 299200 | 0 | 0 | 0 | 0 | 0 | 0 |
| DRPFQAQ | NOXO1 | Q8NFA2 | NOXO1_HUMA | Unmodified | 0 | 0 | 0 | 0 | 0 | 0 | 0 | 0 | 0 | 0 | 0 | 299510 | 0 | 0 | 0 | 0 | 0 | 0 |
| DRPLADW | PIK3C2B; PIK3C | O00750; F5 | P3C2B_HUMAI | Unmodified | 0 | 0 | 0 | 0 | 0 | 0 | 0 | 0 | 0 | 0 | 0 | 0 | 0 | 0 | 1409800 | 771730 | 0 | 0 |
| DRPQEAD | EIF3B; EIF3B | P55884; C9J | EIF3B_HUMAN | Unmodified | 2125100 | 1643500 | 0 | 2121100 | 1913200 | 0 | 0 | 0 | 1685700 | 252690 | 6372200 | 16343000 | 1132800 | 0 | 2629100 | 0 | 0 | 0 |
| DRPSFSEIA | PTK7; PTK7 | Q13308; H0 | PTK7_HUMAN | Unmodified | 0 | 281200 | 915070 | 1623300 | 18829000 | 108580 | 4701800 | 1444300 | 14448000 | 9644100 | 9061800 | 7206500 | 13467000 | 12192000 | 3378000 | 3351300 | 9810500 | 27669000 |
| DRPVSQPS | WNK1; WNK1; | Q9H4A3; F5 | WNK1_HUMAN | Unmodified | 0 | 0 | 0 | 0 | 0 | 0 | 0 | 0 | 0 | 0 | 0 | 608820 | 0 | 0 | 0 | 0 | 0 | 0 |
| DRRDHPLF | COX4I1; COX4I | P13073; Q8 | COX41_HUMAI | Unmodified | 113210 | 0 | 0 | 0 | 0 | 0 | 0 | 0 | 938930 | 458940 | 0 | 0 | 0 | 0 | 0 | 0 | 0 | 220300 |
| DRRYADLT | PGAM1 | P18669 | PGAM1_HUMA | Unmodified | 0 | 0 | 0 | 0 | 5857000 | 752360 | 103140 | 0 | 0 | 447450 | 0 | 0 | 971930 | 669930 | 0 | 0 | 2866200 | 4597600 |
| DRSAEETF | STXBP3 | O00186 | STXB3_HUMAN | Unmodified | 1591900 | 1347600 | 1416500 | 1024900 | 2323600 | 950900 | 108480 | 0 | 2240900 | 370590 | 782180 | 10034000 | 1846100 | 987810 | 2347700 | 2509600 | 561100 | 2357400 |
| DRSGVISD | PDCD6; PDCD6 | O75340; H0 | PDCD6_HUMA | Unmodified | 6463000 | 9674700 | 1207300 | 4389900 | 5022400 | 3988200 | 0 | 0 | 51365000 | 8887700 | 427490 | 826710 | 0 | 0 | 8186200 | 2379800 | 0 | 2288400 |
| DRSGYIDE | CALB2; CALB2 | P22676; H3E | CALB2_HUMAN | Unmodified | 0 | 0 | 0 | 0 | 0 | 0 | 404880 | 164740 | 0 | 0 | 0 | 0 | 0 | 0 | 0 | 0 | 0 | 0 |
| DRSGYIDE | CALB2 | P22676 | CALB2_HUMAN | Unmodified | 0 | 0 | 0 | 0 | 0 | 0 | 916350 | 0 | 0 | 0 | 0 | 0 | 0 | 0 | 0 | 0 | 0 | 0 |
| DRSSFYVN | PFN1; PFN1; PF | P07737; I3L | PROF1_HUMAN | Unmodified | 7109800 | 16195000 | 3615800 | 9360200 | 54999000 | 8700500 | 16977000 | 8063000 | 12200000 | 7064500 | 10090000 | 28007000 | 21733000 | 14365000 | 31600000 | 6460800 | 22369000 | 53658000 |
| DRTGFAVI | CDKN2C | P42773 | CDN2C_HUMA | Unmodified | 0 | 0 | 0 | 0 | 0 | 0 | 0 | 0 | 869620 | 0 | 0 | 0 | 0 | 0 | 0 | 0 | 0 | 0 |
| DRTLTPLG | PGAM5 | Q96HS1 | PGAM5_HUMA | Unmodified | 0 | 0 | 0 | 0 | 0 | 0 | 0 | 0 | 947400 | 523130 | 0 | 0 | 235880 | 464930 | 0 | 0 | 364740 | 0 |
| DRTQIAICP | ARPC1B; ARPC | O15143; C9 | ARC1B_HUMAI | Unmodified | 1687500 | 853760 | 0 | 954260 | 17811000 | 0 | 4649000 | 3265600 | 1533700 | 1291600 | 662620 | 14843000 | 4855100 | 4540400 | 1212300 | 1591200 | 8211800 | 5752000 |
| DRTQIALSI | ARPC1A; ARPC | Q92747; E9F | ARC1A_HUMAI | Unmodified | 248210 | 0 | 736190 | 0 | 0 | 0 | 0 | 0 | 181430 | 0 | 0 | 1804400 | 0 | 0 | 339100 | 638970 | 271070 | 0 |
| DRTQIALSI | ARPC1A; ARPC | Q92747; E9F | ARC1A_HUMAI | Unmodified | 962870 | 614170 | 405090 | 504860 | 0 | 0 | 0 | 0 | 159210 | 0 | 81595 | 5914900 | 0 | 0 | 894930 | 457360 | 0 | 0 |
| DRTQQSKI | FARP1 | A0A1B0GV6 | A0A1B0GV68_I | Unmodified | 62801 | 0 | 793300 | 0 | 0 | 0 | 21787 | 0 | 2018600 | 328010 | 55947 | 0 | 790370 | 0 | 953660 | 375290 | 0 | 102250 |
| DRTQQYD | ME2; ME2; ME2 | P23368; A0A | MAOM_HUMAN | Unmodified | 0 | 14348 | 0 | 0 | 0 | 0 | 32981 | 0 | 254540 | 0 | 0 | 0 | 0 | 23956 | 0 | 0 | 0 | 0 |
| DRTSGNVE | ADRM1 | Q16186 | ADRM1_HUMA | Unmodified | 150710 | 0 | 0 | 344150 | 656300 | 0 | 367660 | 74948 | 1518600 | 0 | 578750 | 1251600 | 1286200 | 731120 | 187290 | 76897 | 422780 | 678830 |
| DRVALSNM | NPEPPSL1; NPE | A6NEC2; P5 | PSAL_HUMAN | Oxidation (M) | 0 | 0 | 0 | 0 | 0 | 0 | 0 | 0 | 167030 | 0 | 0 | 709560 | 0 | 0 | 0 | 0 | 0 | 0 |
| DRVALSNM | NPEPPSL1; NPE | A6NEC2; P5 | PSAL_HUMAN | Unmodified | 0 | 0 | 0 | 0 | 0 | 0 | 0 | 0 | 129280 | 0 | 0 | 934430 | 190300 | 0 | 57473 | 0 | 0 | 0 |
| DRVDCGYI | TFF3 | Q07654 | TFF3_HUMAN | Unmodified | 0 | 0 | 0 | 0 | 0 | 0 | 0 | 0 | 11775000 | 2456800 | 0 | 0 | 0 | 0 | 0 | 0 | 0 | 0 |
| DRVEISPE | BRCC3; BRCC3 | P46736; A0A | BRCC3_HUMAI | Unmodified | 0 | 0 | 0 | 0 | 0 | 0 | 0 | 0 | 0 | 0 | 0 | 888010 | 0 | 0 | 0 | 0 | 0 | 0 |
| DRVETLRPI | MARK2; MARK2 | Q7KZI7; A0A | MARK2_HUMAI | Unmodified | 128330 | 91206 | 929480 | 198130 | 0 | 0 | 0 | 0 | 0 | 0 | 341830 | 1035200 | 0 | 0 | 880250 | 672280 | 0 | 0 |
| DRVHHEP | CALU | O43852 | CALU_HUMAN | Unmodified | 0 | 0 | 0 | 0 | 972750 | 0 | 0 | 0 | 30620000 | 757130 | 1025800 | 604360 | 0 | 0 | 0 | 0 | 0 | 0 |
| DRVHHEP | CALU | O43852 | CALU_HUMAN | Unmodified | 0 | 0 | 0 | 0 | 2673100 | 0 | 635400 | 0 | 6990600 | 6164900 | 0 | 0 | 876530 | 2457400 | 0 | 0 | 0 | 0 |
| DRVLAELP | RAB34; RAB34; | Q9BZG1; C9 | RAB34_HUMAN | Unmodified | 0 | 0 | 0 | 0 | 933680 | 190330 | 0 | 0 | 537530 | 0 | 0 | 0 | 0 | 0 | 0 | 0 | 0 | 0 |
| DRVLMYG | BSCL2; BSCL2; | Q96G97; E9 | BSCL2_HUMAN | Unmodified | 0 | 0 | 0 | 0 | 283500 | 0 | 0 | 0 | 391670 | 36738 | 0 | 0 | 0 | 0 | 0 | 0 | 52848 | 0 |
| DRVNDDLI | DLG3; DLG3 | Q92796; Q5 | DLG3_HUMAN | Unmodified | 29750 | 0 | 997900 | 915010 | 212510 | 0 | 34122 | 0 | 0 | 345430 | 521370 | 4597500 | 0 | 0 | 896010 | 0 | 205630 | 0 |
| DRVNDVC | SPTAN1; SPTAN | Q13813; A0 | SPTN1_HUMAN | Unmodified | 0 | 0 | 0 | 0 | 185000 | 0 | 0 | 0 | 0 | 0 | 121440 | 1502200 | 0 | 96087 | 163120 | 0 | 108980 | 0 |
| DRVSWDG | MPZL2 | O60487 | MPZL2_HUMAN | Unmodified | 619180 | 0 | 313830 | 0 | 923190 | 0 | 332830 | 0 | 0 | 0 | 90715 | 616650 | 0 | 0 | 586260 | 1611400 | 0 | 0 |
| DRVTDALN | HSPD1; HSPD1 | P10809; A0A | CH60_HUMAN | Unmodified | 20765 | 0 | 0 | 134670 | 0 | 0 | 54468 | 0 | 22536000 | 1439700 | 0 | 304940 | 721550 | 355570 | 112580 | 152640 | 0 | 0 |
| DRVTSAVE | USP5 | P45974 | UBP5_HUMAN | Unmodified | 0 | 0 | 0 | 0 | 0 | 149410 | 0 | 0 | 0 | 0 | 251860 | 921030 | 0 | 0 | 0 | 0 | 0 | 0 |
| DRWGQEV | DDR1; DDR1; D | Q08345; A0 | DDR1_HUMAN | Unmodified | 1217800 | 482490 | 270850 | 0 | 290600 | 0 | 0 | 0 | 3561200 | 462250 | 511930 | 643130 | 0 | 0 | 236690 | 0 | 455460 | 0 |
| DRYDSFIEL | FARSB | Q9NSD9 | SYFB_HUMAN | Unmodified | 0 | 0 | 241780 | 0 | 1478100 | 0 | 818270 | 469630 | 331020 | 0 | 0 | 2023300 | 731870 | 389520 | 0 | 0 | 0 | 0 |
| DRYGGQPI | HNRNPUL2; HN | Q1KMD3; H3 | HNRL2_HUMAI | Unmodified | 0 | 0 | 0 | 0 | 0 | 0 | 662760 | 0 | 0 | 0 | 0 | 0 | 1162900 | 0 | 0 | 0 | 0 | 0 |
| DRYLPDTLI | COMT; COMT | P21964; E7E | COMT_HUMAN | Unmodified | 2068400 | 0 | 270730 | 0 | 0 | 0 | 32866 | 0 | 1137400 | 22996 | 0 | 0 | 292610 | 0 | 1702400 | 2210600 | 0 | 0 |
| DSAAAVVV | RAB6C; RAB6D | Q9H0N0; Q5 | RAB6C_HUMAI | Unmodified | 0 | 0 | 0 | 0 | 0 | 0 | 0 | 0 | 0 | 0 | 0 | 0 | 0 | 0 | 1244200 | 624320 | 0 | 0 |
| DSAGEPLY | PLXNB2 | O15031 | PLXB2_HUMAN | Unmodified | 0 | 0 | 876400 | 33672 | 0 | 0 | 0 | 0 | 0 | 0 | 0 | 3204200 | 0 | 0 | 1907000 | 845940 | 0 | 0 |
| DSAIQQQV | DYNC1H1; DYN | Q14204; A0 | DYHC1_HUMA | Unmodified | 0 | 0 | 0 | 0 | 710180 | 0 | 0 | 0 | 315640 | 171760 | 0 | 1357400 | 183480 | 113200 | 42234 | 24526 | 82383 | 466990 |
| DSAQNSVI | TFRC; TFRC | P02786; G3 | TFR1_HUMAN | Unmodified | 4119000 | 2802100 | 6017100 | 5234600 | 1985900 | 893040 | 0 | 102360 | 29413000 | 2294500 | 3504000 | 4948200 | 839870 | 0 | 5085900 | 5273500 | 57623 | 815180 |
| DSAQNSVI | TFRC; TFRC | P02786; G3 | TFR1_HUMAN | Unmodified | 178170 | 175620 | 0 | 363250 | 247580 | 0 | 0 | 0 | 1134600 | 294080 | 0 | 0 | 137350 | 0 | 0 | 0 | 0 | 0 |
| DSAQNSVI | TFRC; TFRC | P02786; G3 | TFR1_HUMAN | Unmodified | 0 | 0 | 0 | 779400 | 17118000 | 4535500 | 3046800 | 40073 | 2273100 | 6988900 | 0 | 0 | 4610700 | 3962900 | 0 | 0 | 2507300 | 6672400 |
| DSAQRDDI | ATP1B1; ATP1B | P05026; V9 | AT1B1_HUMAN | Oxidation (M) | 0 | 0 | 1390800 | 316400 | 0 | 0 | 0 | 0 | 803210 | 0 | 0 | 5966200 | 0 | 0 | 1357600 | 1046700 | 0 | 210250 |
| DSAQRDDI | ATP1B1; ATP1B | P05026; V9 | AT1B1_HUMAN | Unmodified | 0 | 0 | 0 | 0 | 0 | 0 | 0 | 0 | 1259200 | 0 | 0 | 1740900 | 0 | 0 | 0 | 0 | 180670 | 0 |
| DSARYHFF | TWF1; TWF1 | Q12792; F8 | TWF1_HUMAN | Unmodified | 64192 | 0 | 0 | 0 | 249010 | 139690 | 0 | 0 | 0 | 82568 | 104720 | 288290 | 0 | 0 | 0 | 0 | 290650 | 0 |
| DSASFNPE | ACOX1; ACOX1 | Q15067; I3L | ACOX1_HUMAI | Unmodified | 0 | 0 | 0 | 0 | 35109 | 0 | 0 | 0 | 10053000 | 1589000 | 2489200 | 0 | 0 | 0 | 0 | 44941 | 0 | 0 |
| DSATAQAV | SNTB1; SNTB1 | Q13884; A0 | SNTB1_HUMAN | Unmodified | 97925 | 0 | 0 | 0 | 0 | 0 | 0 | 0 | 373300 | 0 | 0 | 0 | 0 | 0 | 0 | 0 | 313080 | 105770 |
| DSAVLDQS | RAB11FIP5; RAI | Q9BXF6; A0 | RFIP5_HUMAN | Unmodified | 0 | 0 | 0 | 0 | 639580 | 0 | 0 | 0 | 0 | 0 | 0 | 0 | 0 | 0 | 0 | 0 | 0 | 0 |
| DSAVNAIC | DKC1; DKC1 | O60832; H7 | DKC1_HUMAN | Unmodified | 156550 | 0 | 0 | 0 | 0 | 0 | 165190 | 143180 | 0 | 0 | 0 | 861130 | 0 | 0 | 0 | 0 | 0 | 0 |
| DSAYNTRT | ITGA1 | P56199 | ITA1_HUMAN I | Unmodified | 0 | 0 | 0 | 0 | 0 | 0 | 0 | 0 | 0 | 0 | 0 | 0 | 0 | 0 | 0 | 0 | 2489400 | 4152900 |
| DSCISLQK | CALCOCO2; CA | Q13137; D6 | CACO2_HUMA | Unmodified | 0 | 0 | 0 | 0 | 0 | 0 | 250700 | 0 | 603080 | 0 | 0 | 0 | 0 | 0 | 0 | 0 | 0 | 303280 |
| DSCPLDCK | SRSF3; SRSF3 | P84103; A0A | SRSF3_HUMAN | Unmodified | 929990 | 0 | 0 | 0 | 0 | 0 | 0 | 38042 | 0 | 0 | 0 | 321320 | 0 | 0 | 0 | 0 | 0 | 0 |
| DSCQGDS | KLK14; KLK14 | Q9P0G3; A0 | KLK14_HUMAN | Unmodified | 0 | 0 | 0 | 0 | 0 | 0 | 0 | 0 | 784850 | 0 | 0 | 0 | 0 | 0 | 0 | 0 | 0 | 0 |
| DSDDVPM | NRAS | P01111 | RASN_HUMAN | Unmodified | 0 | 0 | 0 | 0 | 84548 | 0 | 0 | 0 | 1652900 | 0 | 0 | 0 | 43896 | 0 | 0 | 0 | 0 | 0 |
| DSDDVPM | NRAS | P01111 | RASN_HUMAN | Oxidation (M) | 197420 | 0 | 0 | 0 | 0 | 0 | 0 | 0 | 926600 | 0 | 603110 | 906040 | 196330 | 0 | 98950 | 387560 | 0 | 0 |
| DSDDVPM | NRAS | P01111 | RASN_HUMAN | Oxidation (M) | 0 | 0 | 0 | 0 | 513780 | 0 | 0 | 0 | 0 | 395950 | 0 | 0 | 2214700 | 656480 | 0 | 0 | 1313500 | 211490 |
| DSDIIEDV | ENDOD1 | O94919 | ENDD1_HUMA | Unmodified | 0 | 0 | 0 | 0 | 0 | 0 | 0 | 0 | 1045000 | 0 | 0 | 0 | 0 | 0 | 0 | 0 | 0 | 0 |
| DSDLATQN | PDS5A; PDS5A | Q29RF7; H0 | PDS5A_HUMAN | Unmodified | 65046 | 0 | 86164 | 0 | 75237 | 0 | 0 | 0 | 0 | 0 | 0 | 445510 | 0 | 0 | 0 | 0 | 0 | 0 |
| DSDNYRVI | ERP44 | Q9BS26 | ERP44_HUMAN | Unmodified | 0 | 0 | 0 | 673850 | 0 | 0 | 0 | 79720 | 58602000 | 4174200 | 1470100 | 0 | 0 | 67089 | 0 | 0 | 0 | 0 |
| DSDSGLEQ | MLF2 | Q15773 | MLF2_HUMAN | Unmodified | 1568700 | 2536100 | 1298600 | 0 | 0 | 0 | 30880 | 0 | 0 | 0 | 0 | 0 | 0 | 0 | 563030 | 393060 | 0 | 0 |
| DSDSGLEQ | MLF2 | Q15773 | MLF2_HUMAN | Unmodified | 1950900 | 2689900 | 1697600 | 1999200 | 0 | 0 | 0 | 0 | 0 | 0 | 0 | 0 | 0 | 0 | 1189400 | 530660 | 0 | 0 |
| DSDSKLVE | CTNND1; CTNN | O60716; A0 | CTND1_HUMAI | Unmodified | 0 | 0 | 0 | 1596200 | 1341400 | 667910 | 0 | 32018 | 0 | 0 | 618630 | 0 | 2386600 | 1084200 | 243960 | 226580 | 1220700 | 1908200 |
| DSEADRFS | NUP210 | Q8TEM1 | PO210_HUMAN | Unmodified | 0 | 0 | 0 | 0 | 0 | 0 | 0 | 0 | 24430000 | 787000 | 4180100 | 0 | 0 | 0 | 0 | 0 | 0 | 0 |
| DSEDIYNL | CMBL | Q96DG6 | CMBL_HUMAN | Unmodified | 0 | 0 | 0 | 0 | 0 | 0 | 0 | 0 | 0 | 0 | 0 | 0 | 0 | 0 | 520910 | 0 | 0 | 0 |
| DSEDVPM | KRAS | P01116 | RASK_HUMAN | Unmodified | 0 | 0 | 0 | 0 | 0 | 0 | 0 | 0 | 1964500 | 353720 | 552970 | 0 | 0 | 0 | 0 | 200330 | 990810 | 0 |
| DSEDVPM | KRAS | P01116 | RASK_HUMAN | Oxidation (M) | 406790 | 709910 | 2297400 | 0 | 0 | 0 | 0 | 0 | 910150 | 506730 | 276520 | 2432300 | 0 | 0 | 2124900 | 988330 | 0 | 0 |
| DSEDVPM | KRAS | P01116 | RASK_HUMAN | Unmodified | 0 | 0 | 0 | 0 | 0 | 0 | 0 | 0 | 1086500 | 0 | 0 | 0 | 0 | 0 | 0 | 0 | 0 | 0 |
| DSEDVPM | KRAS | P01116 | RASK_HUMAN | Oxidation (M) | 823650 | 745610 | 0 | 635550 | 0 | 0 | 0 | 0 | 501590 | 0 | 902090 | 1592100 | 0 | 0 | 484100 | 821130 | 0 | 0 |
| DSEDVPM | KRAS | P01116 | RASK_HUMAN | Unmodified | 0 | 0 | 0 | 0 | 1489700 | 0 | 0 | 0 | 0 | 167410 | 0 | 0 | 1673000 | 0 | 0 | 0 | 107650 | 0 |
| DSEDVPM | KRAS | P01116 | RASK_HUMAN | Oxidation (M) | 0 | 0 | 0 | 134140 | 825360 | 0 | 0 | 40845 | 0 | 216450 | 0 | 0 | 3079700 | 381600 | 0 | 0 | 0 | 0 |
| DSEGYIYAI | ACY1; ACY1; AE | Q03154; F8V | ACY1_HUMAN | Unmodified | 0 | 0 | 0 | 0 | 0 | 0 | 0 | 0 | 0 | 0 | 0 | 1325800 | 0 | 0 | 0 | 0 | 0 | 0 |
| DSELGPGV | FMNL1; FMNL1 | O95466; A0 | FMNL1_HUMAI | Unmodified | 0 | 0 | 0 | 0 | 1969000 | 67951 | 313260 | 144240 | 0 | 430310 | 0 | 0 | 116980 | 0 | 65024 | 0 | 79323 | 159860 |
| DSELYQVLI | DPYSL5; DPYSL | Q9BPU6; E7 | DPYL5_HUMAN | Unmodified | 0 | 0 | 150900 | 0 | 0 | 0 | 0 | 0 | 0 | 0 | 0 | 3722100 | 0 | 0 | 0 | 0 | 0 | 0 |
| DSEQVAEL | USO1 | O60763 | USO1_HUMAN | Unmodified | 0 | 0 | 0 | 0 | 0 | 0 | 0 | 0 | 0 | 0 | 0 | 2838600 | 335370 | 0 | 0 | 0 | 204220 | 0 |

| | | | | | | | | | | | | | | | | | | | | | | |
|---|---|---|---|---|---|---|---|---|---|---|---|---|---|---|---|---|---|---|---|---|---|---|
| DSETGENII | HSPA9; HSPA9; | P38646; A0A | GRP75_HUMAI | Unmodified | 0 | 0 | 0 | 0 | 0 | 0 | 0 | 0 | 5067600 | 517370 | 0 | 0 | 0 | 0 | 0 | 0 | 0 | 0 |
| DSETSQRL | SHMT2 | P34897 | GLYM_HUMAN | Unmodified | 0 | 0 | 0 | 0 | 0 | 0 | 362120 | 0 | 332670 | 5314600 | 0 | 0 | 738140 | 0 | 0 | 0 | 0 | 0 |
| DSEVNFK | MARVELD2; MA | Q8N4S9; A1I | MALD2_HUMAI | Unmodified | 118060 | 146200 | 0 | 0 | 0 | 0 | 0 | 0 | 24173 | 10704 | 64647 | 464190 | 0 | 0 | 206340 | 192200 | 0 | 0 |
| DSEWIRSQ | CD151; CD151; | P48509; E9P | CD151_HUMAI | Unmodified | 2703500 | 1648100 | 2647700 | 1886500 | 0 | 198960 | 0 | 0 | 366900 | 0 | 0 | 233730 | 0 | 0 | 2096600 | 925420 | 111690 | 0 |
| DSFICSKQI | KIT | P10721 | KIT_HUMAN Ma | Unmodified | 0 | 0 | 0 | 0 | 0 | 0 | 0 | 0 | 894040 | 51454 | 0 | 0 | 0 | 0 | 0 | 0 | 0 | 0 |
| DSFSQLINI | ARL15; ARL15; | Q9NXU5; A0 | ARL15_HUMAN | Unmodified | 19280 | 0 | 38350 | 15894 | 0 | 0 | 0 | 0 | 46821 | 0 | 96387 | 178820 | 0 | 0 | 0 | 42244 | 0 | 0 |
| DSGAALGL | G6PC3; G6PC3 | Q9BUM1; K7 | G6PC3_HUMAI | Unmodified | 0 | 1323100 | 0 | 0 | 0 | 0 | 0 | 0 | 1533000 | 0 | 0 | 0 | 0 | 0 | 0 | 0 | 443620 | 0 |
| DSGAALGL | G6PC3; G6PC3 | Q9BUM1; K7 | G6PC3_HUMAI | Unmodified | 0 | 0 | 0 | 0 | 0 | 0 | 1980600 | 0 | 1560300 | 0 | 0 | 0 | 0 | 0 | 0 | 0 | 0 | 0 |
| DSGLPSQG | DLG1; DLG1; D | Q12959; A0A | DLG1_HUMAN | Unmodified | 0 | 0 | 1151100 | 0 | 4463800 | 1015400 | 0 | 0 | 4168900 | 1057100 | 1391100 | 2424000 | 0 | 0 | 0 | 864700 | 250210 | 1356900 |
| DSGNKPPG | GSTK1; GSTK1; | Q9Y2Q3; E9I | GSTK1_HUMAN | Unmodified | 0 | 0 | 0 | 0 | 0 | 0 | 72514 | 0 | 760790 | 231030 | 0 | 0 | 0 | 0 | 0 | 0 | 0 | 0 |
| DSGPAGQI | C6orf132 | Q5T0Z8 | CF132_HUMAN | Unmodified | 0 | 0 | 0 | 0 | 0 | 0 | 0 | 0 | 0 | 0 | 86445 | 331040 | 0 | 0 | 0 | 185960 | 0 | 0 |
| DSGQAFSL | RNF31; RNF31; | Q96EP0; H0Y | RNF31_HUMAN | Unmodified | 0 | 0 | 0 | 0 | 0 | 0 | 0 | 0 | 0 | 0 | 0 | 0 | 0 | 0 | 0 | 0 | 0 | 0 |
| DSGSDTAS | RAB11FIP1; RAI | Q6WKZ4; E7 | RFIP1_HUMAN | Unmodified | 151570 | 0 | 0 | 0 | 887140 | 0 | 0 | 0 | 0 | 0 | 0 | 727290 | 243550 | 0 | 0 | 0 | 0 | 0 |
| DSGSFLLR | DIPK2A | Q8NDZ4 | DIK2A_HUMAN | Unmodified | 842210 | 0 | 323270 | 0 | 471950 | 441750 | 0 | 25691 | 22332000 | 997480 | 0 | 728570 | 543240 | 0 | 0 | 178410 | 0 | 511420 |
| DSGSFLLRI | DIPK2A | Q8NDZ4 | DIK2A_HUMAN | Unmodified | 0 | 0 | 0 | 0 | 0 | 0 | 0 | 0 | 0 | 344700 | 0 | 0 | 0 | 0 | 0 | 0 | 0 | 0 |
| DSGSISLQI | SLK | Q9H2G2 | SLK_HUMAN S | Unmodified | 0 | 0 | 0 | 0 | 0 | 0 | 0 | 98555 | 266620 | 0 | 0 | 196210 | 0 | 239990 | 0 | 0 | 0 | 233930 |
| DSGSISLQI | SLK | Q9H2G2 | SLK_HUMAN S | Unmodified | 204390 | 202320 | 107520 | 149770 | 339650 | 89901 | 81957 | 0 | 327870 | 102790 | 0 | 1105100 | 548910 | 0 | 208330 | 292840 | 25271 | 167080 |
| DSGVQACI | GNAI1; GNAI1 | P63096; A0A | GNAI1_HUMAN | Unmodified | 0 | 0 | 0 | 0 | 0 | 0 | 0 | 0 | 517670 | 0 | 887900 | 9266400 | 302880 | 0 | 132290 | 328430 | 0 | 0 |
| DSGYPETL | COPE; COPE | O14579; M0 | COPE_HUMAN | Unmodified | 273530 | 0 | 31991 | 123060 | 0 | 89673 | 187200 | 0 | 1096900 | 61341 | 1130500 | 5120000 | 0 | 0 | 747160 | 309790 | 0 | 135760 |
| DSGYPLIIF | PAFAH2; PAFAH | Q99487; Q5 | PAFA2_HUMAN | Unmodified | 0 | 0 | 0 | 0 | 0 | 0 | 0 | 0 | 612430 | 92405 | 0 | 65145 | 0 | 0 | 0 | 0 | 0 | 0 |
| DSHDPLPS | PBXIP1 | Q96AQ6 | PBIP1_HUMAN | Unmodified | 0 | 0 | 0 | 0 | 0 | 0 | 0 | 0 | 5681400 | 336270 | 2093600 | 495750 | 0 | 0 | 0 | 0 | 0 | 222200 |
| DSHITNLK | CORO7; CORO | P57737; I3L1 | CORO7_HUMA | Unmodified | 0 | 0 | 0 | 0 | 0 | 0 | 0 | 0 | 0 | 0 | 0 | 0 | 0 | 0 | 0 | 0 | 0 | 0 |
| DSHRDVQ | TMED8 | Q6PL24 | TMED8_HUMAI | Unmodified | 0 | 0 | 0 | 0 | 0 | 0 | 0 | 0 | 0 | 0 | 715070 | 1252500 | 0 | 0 | 0 | 0 | 0 | 0 |
| DSHSLTTN | FGA; FGA | P02671; A0A | FIBA_HUMAN F | Oxidation (M) | 0 | 11746000 | 0 | 0 | 82904 | 0 | 34116 | 17682 | 0 | 0 | 0 | 0 | 0 | 30694 | 0 | 0 | 0 | 0 |
| DSIFEPAK | ADAR; ADAR; AI | P55265; A0A | DSRAD_HUMAI | Unmodified | 0 | 0 | 0 | 0 | 0 | 0 | 0 | 0 | 269640 | 0 | 0 | 1903200 | 0 | 0 | 0 | 0 | 0 | 0 |
| DSIFSNLTG | HADHA; HADHA | P40939; A0A | ECHA_HUMAN | Unmodified | 0 | 0 | 0 | 0 | 0 | 0 | 0 | 0 | 541460 | 87709 | 1160800 | 0 | 0 | 0 | 0 | 0 | 0 | 0 |
| DSIKLDDD | CAND1; CAND1 | Q86VP6; F5H | CAND1_HUMA | Unmodified | 0 | 0 | 0 | 43292 | 0 | 0 | 0 | 0 | 0 | 102990 | 293770 | 0 | 0 | 0 | 0 | 0 | 0 | 0 |
| DSIKLDDD | CAND1; CAND1 | Q86VP6; F5H | CAND1_HUMA | Unmodified | 147110 | 150410 | 300650 | 443360 | 666140 | 339180 | 174640 | 0 | 2024900 | 287020 | 190190 | 5480900 | 1641800 | 489050 | 315170 | 449800 | 159200 | 1039600 |
| DSILRSDE | FASN; FASN | P49327; A0A | FAS_HUMAN F | Unmodified | 0 | 0 | 351010 | 295580 | 0 | 0 | 0 | 0 | 0 | 0 | 0 | 3801200 | 0 | 0 | 198720 | 0 | 0 | 355890 |
| DSISIENK | FKBP9; FKBP9 | O95302; C9J | FKBP9_HUMAN | Unmodified | 0 | 0 | 0 | 0 | 0 | 0 | 0 | 0 | 11405000 | 276840 | 555140 | 674440 | 0 | 0 | 0 | 0 | 0 | 0 |
| DSIVAELDF | COPE; COPE | O14579; M0 | COPE_HUMAN | Unmodified | 0 | 0 | 0 | 0 | 0 | 0 | 0 | 0 | 0 | 0 | 0 | 1221100 | 0 | 0 | 0 | 0 | 0 | 0 |
| DSIVHQAG | ARPC2 | O15144 | ARPC2_HUMAI | Unmodified | 96624 | 0 | 0 | 0 | 660630 | 0 | 0 | 0 | 0 | 0 | 0 | 2192300 | 0 | 0 | 0 | 0 | 0 | 0 |
| DSIVHQAG | ARPC2 | O15144 | ARPC2_HUMAI | Oxidation (M) | 60690 | 0 | 0 | 0 | 0 | 0 | 0 | 0 | 0 | 0 | 65590 | 3574000 | 0 | 0 | 222480 | 522010 | 0 | 0 |
| DSIVHQAG | ARPC2 | O15144 | ARPC2_HUMAI | Unmodified | 0 | 0 | 0 | 0 | 0 | 0 | 0 | 0 | 175190 | 0 | 282210 | 3588000 | 0 | 0 | 221600 | 194360 | 0 | 0 |
| DSIVQGFQ | EFTUD2; EFTUD | Q15029; K7E | U5S1_HUMAN | Unmodified | 0 | 0 | 0 | 0 | 0 | 0 | 0 | 0 | 0 | 0 | 0 | 1063100 | 0 | 0 | 0 | 0 | 0 | 0 |
| DSKAEPHF | DIAPH1; DIAPH | O60610; A0A | DIAP1_HUMAN | Unmodified | 0 | 0 | 0 | 37854 | 0 | 0 | 0 | 0 | 147540 | 26519 | 62132 | 292430 | 0 | 0 | 70132 | 0 | 0 | 0 |
| DSKHSDYII | PTPRG | P23470 | PTPRG_HUMAN | Unmodified | 0 | 0 | 0 | 0 | 521720 | 0 | 0 | 0 | 0 | 0 | 0 | 0 | 0 | 0 | 0 | 0 | 0 | 0 |
| DSKPHFTT | DPY19L1; DPY1 | Q2PZI1; A0A | D19L1_HUMAN | Unmodified | 0 | 0 | 0 | 0 | 0 | 0 | 40104 | 0 | 4704900 | 683950 | 125820 | 0 | 0 | 0 | 0 | 0 | 108570 | 188570 |
| DSLAAASG | FLVCR1 | Q9Y5Y0 | FLVC1_HUMAN | Unmodified | 0 | 0 | 2872300 | 0 | 0 | 0 | 0 | 0 | 3121100 | 1364600 | 3891800 | 7387500 | 0 | 0 | 0 | 0 | 0 | 0 |
| DSLAAGAS | LRP11; LRP11 | Q86VZ4; Q5 | LRP11_HUMAN | Unmodified | 0 | 0 | 0 | 0 | 300200 | 0 | 0 | 0 | 1856800 | 0 | 0 | 0 | 0 | 0 | 139480 | 172610 | 60593 | 0 |
| DSLAAGAS | LRP11; LRP11 | Q86VZ4; Q5 | LRP11_HUMAN | Unmodified | 0 | 0 | 0 | 0 | 0 | 0 | 0 | 0 | 11860000 | 638170 | 0 | 0 | 0 | 0 | 0 | 0 | 0 | 0 |
| DSLAWLSL | DERA; DERA; DI | Q9Y315; G3 | DEOC_HUMAN | Unmodified | 0 | 0 | 44604 | 0 | 0 | 0 | 0 | 0 | 1328300 | 245920 | 0 | 840070 | 0 | 0 | 0 | 119060 | 0 | 0 |
| DSLDIEVK | SH3GL2; SH3G | Q99962; Q9 | SH3G2_HUMAI | Unmodified | 8918600 | 8503600 | 13067000 | 0 | 358840 | 0 | 0 | 0 | 0 | 0 | 0 | 0 | 1496000 | 0 | 15246000 | 15127000 | 0 | 0 |
| DSLDPSFTI | KHDRBS1 | Q07666 | KHDR1_HUMAI | Unmodified | 0 | 0 | 0 | 0 | 0 | 0 | 0 | 0 | 334340 | 66252 | 57793 | 75739 | 0 | 0 | 14452 | 0 | 0 | 0 |
| DSLDPSFTI | KHDRBS1 | Q07666 | KHDR1_HUMAI | Oxidation (M) | 0 | 0 | 0 | 0 | 70500 | 29702 | 52092 | 0 | 136650 | 223790 | 275640 | 960070 | 182750 | 0 | 59834 | 0 | 0 | 0 |
| DSLGAHSI | SEC14L1 | Q92503 | S14L1_HUMAN | Unmodified | 0 | 0 | 0 | 0 | 0 | 0 | 0 | 0 | 354000 | 50380 | 83724 | 1188900 | 0 | 0 | 0 | 0 | 0 | 272470 |
| DSLGTREIL | ACSL4; ACSL4; | O60488; D6I | ACSL4_HUMAN | Unmodified | 0 | 0 | 0 | 175150 | 1163600 | 348680 | 0 | 0 | 0 | 0 | 0 | 0 | 0 | 0 | 0 | 0 | 0 | 889040 |
| DSLHEKFP | IQGAP1; IQGAP | P46940; A0A | IQGA1_HUMAN | Unmodified | 163530 | 300690 | 205590 | 0 | 0 | 837790 | 0 | 0 | 209750 | 253570 | 0 | 6593400 | 0 | 62761 | 163740 | 646070 | 0 | 0 |
| DSLHSLMA | MYO10; MYO10 | Q9HD67; D6 | MYO10_HUMAI | Unmodified | 0 | 0 | 0 | 25737 | 0 | 0 | 0 | 0 | 81474 | 23186 | 0 | 0 | 0 | 0 | 61403 | 15605 | 0 | 0 |
| DSLIFLVDA | XRCC6 | P12956 | XRCC6_HUMAI | Unmodified | 0 | 0 | 0 | 0 | 0 | 0 | 0 | 0 | 158210 | 0 | 0 | 4499200 | 0 | 0 | 0 | 0 | 0 | 0 |
| DSLINLK | CISD2; CISD2 | Q8N5K1; D6 | CISD2_HUMAN | Unmodified | 10194000 | 15882000 | 19652000 | 7553600 | 13756000 | 1249500 | 0 | 4433200 | 6104800 | 589310 | 481940 | 0 | 0 | 2144900 | 1752500 | 2434700 | 222880 | 5313100 |
| DSLLAHSS | PTPRF; PTPRF; I | P10586; H0Y | PTPRF_HUMAN | Unmodified | 405130 | 327210 | 330100 | 288460 | 1385900 | 382850 | 0 | 0 | 1732800 | 0 | 330900 | 516070 | 340930 | 179040 | 774360 | 372730 | 0 | 0 |
| DSLLAHSS | PTPRF; PTPRF; I | P10586; H0Y | PTPRF_HUMAN | Oxidation (M) | 431320 | 462550 | 490900 | 329520 | 358110 | 191770 | 102000 | 121350 | 324450 | 69331 | 204010 | 357930 | 199770 | 76513 | 780720 | 788870 | 0 | 0 |
| DSLLQDGE | PFN1; PFN1 | P07737; K7E | PROF1_HUMAI | Oxidation (M) | 1282400 | 2687400 | 1390300 | 640710 | 0 | 0 | 0 | 156630 | 1318900 | 689750 | 0 | 7253400 | 277590 | 663620 | 3052400 | 2233400 | 0 | 0 |
| DSLLQDGE | PFN1; PFN1 | P07737; K7E | PROF1_HUMAI | Unmodified | 0 | 249410 | 16278 | 31108 | 41968 | 23848 | 0 | 19894 | 538420 | 0 | 39788 | 275980 | 0 | 0 | 0 | 25465 | 0 | 20443 |
| DSLPPSPLI | NHLRC2 | Q8NBF2 | NHLC2_HUMA | Unmodified | 0 | 0 | 0 | 0 | 0 | 0 | 0 | 0 | 0 | 0 | 0 | 982520 | 0 | 0 | 0 | 0 | 0 | 0 |
| DSLPSSDTI | MICAL3; MICAL | Q7RTP6; A0A | MICA3_HUMAN | Unmodified | 0 | 0 | 0 | 0 | 811560 | 0 | 0 | 0 | 0 | 0 | 0 | 0 | 0 | 0 | 0 | 37925 | 0 | 104710 |
| DSLSDDVV | HSDL2 | Q6YN16 | HSDL2_HUMAI | Unmodified | 0 | 0 | 0 | 0 | 0 | 0 | 0 | 0 | 5369300 | 308010 | 0 | 0 | 0 | 0 | 0 | 0 | 0 | 0 |
| DSLSPVLH | GPN1 | Q9HCN4 | GPN1_HUMAN | Unmodified | 0 | 1223800 | 0 | 0 | 52356 | 0 | 0 | 0 | 0 | 0 | 42621 | 0 | 0 | 0 | 0 | 249780 | 0 | 0 |
| DSLSVNEF | S100A13 | Q99584 | S10AD_HUMAN | Unmodified | 251320 | 284410 | 0 | 0 | 11925000 | 6032800 | 0 | 80951 | 1789900 | 388410 | 410470 | 1397100 | 0 | 0 | 0 | 0 | 2116500 | 16636000 |
| DSLSVNEF | S100A13 | Q99584 | S10AD_HUMAN | Unmodified | 0 | 0 | 0 | 0 | 4945400 | 478200 | 0 | 0 | 0 | 216860 | 0 | 0 | 0 | 0 | 0 | 0 | 5458700 | 6990000 |
| DSLVHSSP | PTGFRN | Q9P2B2 | FPRP_HUMAN | Unmodified | 578710 | 414290 | 1588300 | 815250 | 0 | 0 | 211520 | 0 | 0 | 0 | 0 | 6489200 | 0 | 131580 | 1725400 | 1410200 | 0 | 0 |
| DSLVYQYL | NHLRC2 | Q8NBF2 | NHLC2_HUMA | Unmodified | 0 | 0 | 0 | 0 | 0 | 689880 | 0 | 94455 | 0 | 0 | 0 | 2362400 | 0 | 0 | 0 | 0 | 0 | 0 |
| DSLYAQGK | RPL36A; RPL36 | P83881; H7E | RL36A_HUMAN | Unmodified | 56987 | 33725 | 59104 | 241880 | 432890 | 0 | 327750 | 668920 | 76705 | 56960 | 0 | 2121900 | 499480 | 600210 | 0 | 54260 | 0 | 688450 |
| DSMFGIAV | ITGA6 | P23229 | ITA6_HUMAN I | Oxidation (M) | 130970 | 238570 | 1448500 | 576280 | 4682200 | 332110 | 478030 | 403490 | 0 | 0 | 151740 | 1816900 | 3502800 | 1613600 | 1505300 | 1734500 | 2592500 | 8103600 |
| DSMFGIAV | ITGA6 | P23229 | ITA6_HUMAN I | Unmodified | 0 | 0 | 0 | 0 | 3916100 | 227060 | 0 | 0 | 0 | 0 | 0 | 0 | 943320 | 1019000 | 0 | 0 | 2097900 | 7581100 |
| DSMGHVG | SDCBP2 | Q9H190 | SDCB2_HUMAI | Oxidation (M) | 0 | 0 | 0 | 0 | 0 | 0 | 0 | 0 | 0 | 0 | 0 | 0 | 2739600 | 0 | 0 | 0 | 0 | 0 |
| DSMGHVG | SDCBP2 | Q9H190 | SDCB2_HUMAI | Unmodified | 0 | 0 | 0 | 0 | 0 | 0 | 0 | 0 | 152420 | 0 | 0 | 0 | 2745300 | 0 | 0 | 0 | 0 | 0 |
| DSMIWDC | FN1 | P02751 | FINC_HUMAN I | Oxidation (M) | 0 | 0 | 1258800 | 970150 | 0 | 0 | 0 | 0 | 0 | 0 | 0 | 0 | 108520 | 0 | 0 | 137600 | 0 | 0 |
| DSMSAAEV | EPB41L3; EPB4 | A0A0A0MRA | A0A0A0MRA8_I | Unmodified | 0 | 0 | 0 | 0 | 0 | 0 | 0 | 0 | 0 | 0 | 0 | 0 | 0 | 0 | 0 | 0 | 409610 | 2201800 |
| DSMSAAEV | EPB41L3; EPB4 | A0A0A0MRA | A0A0A0MRA8_I | Oxidation (M) | 0 | 0 | 427800 | 0 | 50553 | 0 | 0 | 0 | 29226 | 0 | 0 | 0 | 0 | 0 | 0 | 0 | 0 | 1776300 |
| DSNFAGDI | DDX42; DDX42 | Q86XP3; A0A | DDX42_HUMAI | Unmodified | 0 | 0 | 0 | 0 | 372190 | 95738 | 100920 | 0 | 0 | 217130 | 0 | 0 | 320510 | 88798 | 0 | 0 | 116090 | 98150 |
| DSNGAILV | RAB21 | Q9UL25 | RAB21_HUMAN | Unmodified | 156980 | 1172300 | 310240 | 291270 | 0 | 0 | 46217 | 0 | 442050 | 0 | 217810 | 910370 | 0 | 0 | 802160 | 898300 | 0 | 0 |
| DSNMPMD | F5; F5 | P12259; A0A | FA5_HUMAN C | 4 Oxidation (M) | 0 | 0 | 0 | 0 | 0 | 330070 | 0 | 0 | 0 | 110700 | 0 | 0 | 0 | 0 | 0 | 0 | 0 | 0 |
| DSNMVLSN | KIRREL1; KIRRE | Q96J84; Q5V | KIRR1_HUMAN | Unmodified | 0 | 0 | 0 | 0 | 354990 | 32650 | 0 | 0 | 111670 | 156440 | 0 | 367560 | 0 | 0 | 0 | 0 | 0 | 0 |
| DSNNLCLF | LGALS1 | P09382 | LEG1_HUMAN | Unmodified | 2238600 | 0 | 0 | 0 | 7762100 | 3662300 | 0 | 0 | 134820 | 0 | 300870 | 0 | 0 | 0 | 0 | 0 | 5455700 | 13739000 |
| DSNNLCLF | LGALS1 | P09382 | LEG1_HUMAN | Unmodified | 0 | 0 | 0 | 0 | 8894500 | 2518800 | 0 | 0 | 0 | 0 | 0 | 0 | 0 | 0 | 0 | 0 | 5146700 | 30384000 |
| DSPAFPDS | INCENP | Q9NQS7 | INCE_HUMAN I | Unmodified | 0 | 0 | 0 | 323970 | 403370 | 102850 | 107950 | 29210 | 0 | 0 | 0 | 100880 | 0 | 0 | 155870 | 124090 | 0 | 103690 |
| DSPLLLQQ | DCTN1; DCTN1 | Q14203; A0A | DCTN1_HUMAI | Oxidation (M) | 0 | 0 | 0 | 0 | 0 | 0 | 0 | 23636 | 0 | 0 | 0 | 350410 | 0 | 0 | 0 | 0 | 0 | 0 |
| DSPLLLQQ | DCTN1; DCTN1 | Q14203; A0A | DCTN1_HUMAI | Unmodified | 0 | 0 | 0 | 0 | 811550 | 0 | 247650 | 0 | 0 | 0 | 0 | 0 | 1316700 | 0 | 0 | 0 | 206860 | 650520 |
| DSPLQGSG | MAP4K4; MAP4 | O95819; C9J | M4K4_HUMAN | Unmodified | 170600 | 0 | 199990 | 391920 | 0 | 0 | 0 | 0 | 574760 | 0 | 955520 | 1687600 | 0 | 0 | 345140 | 239740 | 94319 | 113020 |
| DSPLQGSG | MAP4K4; MAP4 | C9J840; A0A | C9J840_HUMA | Unmodified | 0 | 0 | 0 | 0 | 0 | 0 | 0 | 0 | 0 | 0 | 0 | 0 | 271590 | 0 | 0 | 0 | 247250 | 922250 |
| DSPPTPTM | ZDHHC5; ZDHH | Q9C0B5; H0 | ZDHC5_HUMA | Unmodified | 0 | 0 | 110200 | 209630 | 0 | 0 | 0 | 0 | 0 | 134840 | 0 | 0 | 1813300 | 2366200 | 219840 | 354960 | 0 | 0 |
| DSPSVWA | PFN1; PFN1 | P07737; I3L3 | PROF1_HUMAI | Unmodified | 0 | 0 | 0 | 0 | 0 | 0 | 474860 | 0 | 0 | 0 | 0 | 0 | 399590 | 602880 | 0 | 0 | 2147300 | 0 |
| DSPTGANS | CAPN5; CAPN5 | O15484; E7E | CAN5_HUMAN | Unmodified | 0 | 0 | 0 | 0 | 0 | 0 | 169780 | 137790 | 0 | 0 | 0 | 145310 | 0 | 0 | 629210 | 630460 | 0 | 621810 |
| DSPTQERE | PPP2R5D; PPP2 | Q14738; H0Y | 2A5D_HUMAN | Unmodified | 0 | 0 | 0 | 379320 | 0 | 0 | 0 | 0 | 215890 | 0 | 0 | 9845300 | 533320 | 0 | 340290 | 365860 | 471450 | 0 |
| DSPYMAFII | FLNB; FLNB | O75369; E7E | FLNB_HUMAN | Oxidation (M) | 133300 | 0 | 847370 | 224900 | 154790 | 0 | 0 | 20142 | 0 | 0 | 0 | 1362000 | 0 | 0 | 372590 | 146930 | 0 | 0 |
| DSQAALPK | DBN1; DBN1; D | Q16643; D6I | DREB_HUMAN | Unmodified | 0 | 0 | 139750 | 95109 | 1200100 | 114390 | 438130 | 319540 | 187140 | 84596 | 861490 | 1295200 | 414330 | 417130 | 183070 | 112220 | 937210 | 2854800 |
| DSQHFENI | CCM2; CCM2; ( | Q9BSQ5; A0 | CCM2_HUMAN | Unmodified | 0 | 100140 | 70651 | 38362 | 49207 | 30519 | 0 | 0 | 0 | 0 | 40311 | 0 | 19945 | 22209 | 301540 | 209020 | 38242 | 0 |
| DSQICELK | CDNF; MANF | Q49AH0; P5 | CDNF_HUMAN | Unmodified | 0 | 0 | 0 | 0 | 0 | 0 | 0 | 0 | 4868400 | 172850 | 899440 | 396570 | 0 | 0 | 697250 | 208880 | 0 | 0 |
| DSQICELK | MANF | P55145 | MANF_HUMAN | Unmodified | 0 | 0 | 0 | 0 | 0 | 0 | 94954 | 117020 | 9873100 | 6660500 | 2799500 | 0 | 0 | 0 | 0 | 0 | 0 | 0 |
| DSQICELK | MANF | P55145 | MANF_HUMAN | Unmodified | 0 | 0 | 0 | 0 | 0 | 0 | 0 | 0 | 688580 | 605890 | 0 | 0 | 0 | 0 | 0 | 0 | 0 | 0 |
| DSQMVVD | FNBP1L; FNBP1 | Q5T0N5; A0A | FBP1L_HUMAN | Unmodified | 355400 | 577530 | 0 | 0 | 0 | 0 | 0 | 0 | 0 | 0 | 0 | 0 | 0 | 0 | 0 | 0 | 0 | 0 |
| DSQTQAIL | GPS1; GPS1; GI | Q13098; A8I | CSN1_HUMAN | Unmodified | 0 | 0 | 0 | 0 | 0 | 0 | 404170 | 0 | 0 | 0 | 37446 | 1134700 | 0 | 115140 | 0 | 0 | 0 | 0 |
| DSSAARNG | SDCBP; SDCBP | O00560; G5I | SDCB1_HUMAI | Unmodified | 0 | 0 | 0 | 0 | 4899200 | 2900600 | 0 | 0 | 3969700 | 12819000 | 1339200 | 0 | 2979700 | 4151600 | 0 | 0 | 380920 | 454770 |
| DSSDSGVF | KIDINS220 | Q9ULH0 | KDIS_HUMAN I | Unmodified | 0 | 0 | 0 | 0 | 454950 | 0 | 0 | 0 | 0 | 0 | 0 | 0 | 89802 | 60160 | 0 | 0 | 0 | 0 |
| DSSGNLH | ALDH16A1; ALD | Q8IZ83; F5H | A16A1_HUMAN | Unmodified | 0 | 0 | 0 | 0 | 0 | 0 | 0 | 0 | 0 | 0 | 0 | 202850 | 0 | 0 | 32241 | 0 | 0 | 0 |
| DSSHNELY | VANGL1 | Q8TAA9 | VANG1_HUMAI | Unmodified | 0 | 0 | 343650 | 304630 | 0 | 0 | 0 | 0 | 673700 | 56256 | 0 | 177740 | 0 | 0 | 0 | 370350 | 0 | 0 |
| DSSPHFQ | SUN2 | Q9UH99 | SUN2_HUMAN | Unmodified | 0 | 0 | 0 | 0 | 0 | 0 | 0 | 0 | 11127000 | 290640 | 10218 | 0 | 0 | 0 | 0 | 0 | 0 | 0 |
| DSSPPSLQ | FAM234B; FAM | A2RU67; Q6 | F234B_HUMAN | Unmodified | 0 | 0 | 0 | 0 | 0 | 0 | 0 | 0 | 1185700 | 186780 | 101170 | 269480 | 0 | 0 | 0 | 0 | 0 | 0 |
| DSSQDEQ | AVIL; AVIL | O75366; F8V | AVIL_HUMAN A | Unmodified | 0 | 0 | 0 | 0 | 0 | 0 | 0 | 0 | 0 | 0 | 0 | 9425300 | 0 | 0 | 0 | 0 | 0 | 0 |
| DSSQLGTD | EPB41L2; EPB4 | O43491; E9F | E41L2_HUMAN | Unmodified | 0 | 0 | 0 | 0 | 513720 | 0 | 0 | 180890 | 0 | 0 | 0 | 56467 | 0 | 0 | 0 | 61169 | 170780 | 26343 |
| DSSQTSRF | LARP1; LARP1; | Q6PKG0; H0 | LARP1_HUMAN | Unmodified | 66468 | 0 | 0 | 203660 | 0 | 0 | 321150 | 91554 | 0 | 70311 | 0 | 0 | 1292300 | 511490 | 0 | 0 | 208670 | 157890 |
| DSSRLPSE | BLVRB; BLVRB; | P30043; M0 | BLVRB_HUMAN | Unmodified | 5608900 | 1609400 | 2839700 | 7549400 | 254850 | 0 | 0 | 0 | 1887700 | 0 | 0 | 0 | 0 | 0 | 10092000 | 7744800 | 0 | 0 |
| DSSSSLTD | FARP2; FARP2 | O94887; H7 | FARP2_HUMAN | Unmodified | 18379 | 0 | 727840 | 991090 | 0 | 0 | 0 | 0 | 234340 | 0 | 0 | 381190 | 0 | 0 | 1045100 | 811470 | 0 | 202160 |
| DSSSTNLE | CAND1; CAND1 | Q86VP6; A0A | CAND1_HUMA | Unmodified | 0 | 0 | 0 | 0 | 0 | 0 | 0 | 0 | 758260 | 0 | 268330 | 757540 | 0 | 0 | 0 | 0 | 232000 | 717180 |
| DSSVFILGN | PLCG2; PLCG2 | P16885; A0A | PLCG2_HUMAI | Unmodified | 0 | 0 | 0 | 0 | 0 | 0 | 184530 | 0 | 699830 | 0 | 3816400 | 33677000 | 0 | 0 | 0 | 0 | 0 | 0 |
| DSTEAAICI | MVB12B; MVB1 | Q9H7P6; F8 | MB12B_HUMAI | Unmodified | 0 | 0 | 0 | 0 | 0 | 0 | 0 | 0 | 0 | 0 | 0 | 0 | 0 | 0 | 0 | 0 | 0 | 0 |
| DSTFLSTLE | NCLN; NCLN; N | Q969V3; A0A | NCLN_HUMAN | Unmodified | 0 | 0 | 0 | 0 | 0 | 0 | 0 | 0 | 4624100 | 0 | 0 | 489480 | 0 | 0 | 0 | 0 | 0 | 0 |

| | | | | | | | | | | | | | | | | | | | | | | |
|---|---|---|---|---|---|---|---|---|---|---|---|---|---|---|---|---|---|---|---|---|---|---|
| DSTGHVGF | SDCBP; SDCBP | O00560; G5I | SDCB1_HUMAI | Unmodified | 23309000 | 19637000 | 25794000 | 30714000 | 14087000 | 4957700 | 1016800 | 225570 | 95849000 | 30386000 | 3578800 | 26759000 | 8418500 | 4622300 | 44439000 | 23714000 | 445770 | 1133700 |
| DSTGHVGF | SDCBP; SDCBP | O00560; G5I | SDCB1_HUMAI | Unmodified | 202760 | 0 | 0 | 0 | 99200 | 0 | 161740 | 0 | 164000 | 750460 | 0 | 0 | 843000 | 0 | 0 | 0 | 11961 | 0 |
| DSTGHVGF | SDCBP; SDCBP | O00560; G5I | SDCB1_HUMAI | Unmodified | 488120 | 323750 | 0 | 235800 | 8013600 | 1747700 | 968210 | 0 | 3667000 | 16821000 | 0 | 0 | 7623900 | 3404300 | 224100 | 355750 | 0 | 0 |
| DSTGTYTC | NRCAM; NRCAI | Q92823; A0/ | NRCAM_HUMA | Unmodified | 0 | 0 | 0 | 0 | 0 | 0 | 0 | 0 | 371800 | 0 | 0 | 0 | 0 | 0 | 0 | 0 | 0 | 0 |
| DSTKDDNS | SCG3 | Q8WXD2 | SCG3_HUMAN | Unmodified | 0 | 0 | 0 | 0 | 0 | 0 | 216480 | 0 | 6713000 | 0 | 0 | 0 | 0 | 453830 | 0 | 0 | 0 | 0 |
| DSTLIMQLI | SFN; YWHAB; Y | P31947; P31 | 1433S_HUMAN | Oxidation (M) | 4836300 | 4885600 | 23764000 | 2708700 | 637120 | 554530 | 0 | 264740 | 6965700 | 899480 | 6084100 | 91060000 | 0 | 0 | 27738000 | 26542000 | 0 | 261360 |
| DSTLIMQLI | SFN; YWHAB; Y | P31947; P31 | 1433S_HUMAN | Unmodified | 0 | 0 | 898270 | 0 | 0 | 0 | 0 | 0 | 3718700 | 0 | 282890 | 4917300 | 0 | 0 | 2457700 | 0 | 0 | 0 |
| DSTQTPAV | CPT2; CPT2; CF | P23786; A0A | CPT2_HUMAN | Unmodified | 0 | 0 | 0 | 0 | 0 | 0 | 0 | 0 | 1971900 | 0 | 0 | 0 | 0 | 0 | 0 | 0 | 0 | 0 |
| DSTSELVPI | MELTF | P08582 | TRFM_HUMAN | Oxidation (M) | 0 | 0 | 0 | 0 | 3395300 | 696620 | 0 | 0 | 1962400 | 993850 | 0 | 0 | 0 | 22946 | 0 | 0 | 0 | 527050 |
| DSTSELVPI | MELTF | P08582 | TRFM_HUMAN | Unmodified | 0 | 0 | 0 | 0 | 384000 | 0 | 0 | 0 | 754150 | 0 | 0 | 0 | 0 | 0 | 0 | 0 | 0 | 0 |
| DSVFLSCS | WDR77; WDR7 | Q9BQA1; H0 | MEP50_HUMAI | Unmodified | 0 | 0 | 0 | 0 | 0 | 0 | 0 | 0 | 0 | 0 | 0 | 1617300 | 0 | 0 | 0 | 0 | 0 | 0 |
| DSVICLK | CD55; CD55; C | P08174; B1A | DAF_HUMAN C | Unmodified | 1177800 | 0 | 0 | 0 | 2852000 | 0 | 0 | 0 | 945940 | 0 | 1258500 | 5559500 | 590220 | 849610 | 0 | 0 | 3059500 | 10843000 |
| DSVVAGFC | EEF2 | P13639 | EF2_HUMAN EI | Unmodified | 696280 | 0 | 2843800 | 1781900 | 2839900 | 0 | 0 | 110530 | 270170 | 0 | 8038600 | 31907000 | 0 | 0 | 2618900 | 1942200 | 0 | 0 |
| DSVWGSG | PDCD4 | Q53EL6 | PDCD4_HUMA | Unmodified | 0 | 0 | 0 | 0 | 0 | 0 | 0 | 0 | 0 | 0 | 0 | 3621000 | 0 | 0 | 0 | 30687 | 0 | 0 |
| DSWVFGG | FOLH1 | Q04609 | FOLH1_HUMAI | Unmodified | 5632700 | 5153300 | 3281900 | 12323000 | 0 | 0 | 0 | 0 | 0 | 0 | 0 | 0 | 0 | 0 | 22544000 | 11258000 | 0 | 0 |
| DSWVFVG/ | FAM3A; FAM3A; | P98173; D3I | FAM3A_HUMAI | Unmodified | 0 | 0 | 0 | 0 | 0 | 0 | 0 | 0 | 2903000 | 0 | 0 | 0 | 0 | 0 | 0 | 0 | 0 | 0 |
| DSYDSYAT | CIRBP | Q14011 | CIRBP_HUMAN | Unmodified | 251990 | 241040 | 0 | 0 | 0 | 0 | 0 | 0 | 0 | 0 | 0 | 0 | 0 | 0 | 0 | 485010 | 0 | 0 |
| DSYLILETLI | PSMC3; PSMC3 | P17980; E9P | PRS6A_HUMAN | Unmodified | 54463 | 50886 | 50118 | 44485 | 42957 | 0 | 0 | 0 | 161440 | 0 | 274610 | 870140 | 21160 | 0 | 32901 | 118670 | 0 | 0 |
| DSYVGDEA | ACTG1; ACTC1; | P63261; P68 | ACTG_HUMAN | Unmodified | 64995000 | 79740000 | 1.04E+08 | 1.28E+08 | 7.58E+08 | 1.47E+08 | 19279000 | 14680000 | 1.47E+08 | 13822000 | 32300000 | 4.39E+08 | 1.79E+08 | 79748000 | 1.1E+08 | 91530000 | 53982000 | 1.49E+08 |
| DSYVGDEA | ACTG1; ACTC1; | P63261; P68 | ACTG_HUMAN | Unmodified | 2566500 | 0 | 8289900 | 3451900 | 71120000 | 25623000 | 1125500 | 5445100 | 13489000 | 9716600 | 41728000 | 35474000 | 84853 | 17512000 | 11852000 | 11931000 | 10083000 | 25411000 |
| DSYVGDEA | ACTG1; ACTC1; | P63261; P68 | ACTG_HUMAN | Unmodified | 0 | 0 | 0 | 0 | 24565000 | 2839000 | 5278500 | 2942900 | 0 | 1629900 | 0 | 0 | 10274000 | 5423400 | 0 | 0 | 6768200 | 5969900 |
| DTAALMDE | NOSTRIN; NOS | Q8IVI9; A0A( | NOSTN_HUMA | Unmodified | 466430 | 0 | 3146300 | 3499400 | 0 | 0 | 0 | 0 | 0 | 0 | 0 | 233760 | 379240 | 0 | 3217200 | 1227200 | 0 | 0 |
| DTAALMDE | NOSTRIN; NOS | Q8IVI9; A0A( | NOSTN_HUMA | Oxidation (M) | 0 | 0 | 1074500 | 0 | 0 | 0 | 0 | 0 | 0 | 0 | 0 | 105490 | 0 | 0 | 1982100 | 0 | 0 | 0 |
| DTAALMDE | NOSTRIN; NOS | Q8IVI9; A0A( | NOSTN_HUMA | Oxidation (M) | 0 | 0 | 0 | 1019900 | 0 | 0 | 0 | 0 | 0 | 0 | 0 | 0 | 0 | 0 | 3794100 | 2579500 | 0 | 0 |
| DTAAWTVG | KPNB1; KPNB1; | Q14974; J3Q | IMB1_HUMAN I | Unmodified | 448320 | 0 | 255620 | 0 | 0 | 0 | 0 | 0 | 0 | 0 | 0 | 3225200 | 0 | 0 | 0 | 542130 | 0 | 0 |
| DTANWLEI | CDH1 | P12830 | CADH1_HUMA | Unmodified | 0 | 0 | 0 | 0 | 0 | 0 | 0 | 0 | 747750 | 0 | 0 | 877420 | 0 | 0 | 432800 | 484730 | 0 | 0 |
| DTAVQGNI | PEF1 | Q9UBV8 | PEF1_HUMAN I | Unmodified | 800590 | 903260 | 1054100 | 1309600 | 1010400 | 489400 | 0 | 0 | 12776000 | 599650 | 408330 | 2558200 | 638040 | 467270 | 2690700 | 2294100 | 45041 | 254990 |
| DTCIVISGE | MYO1D; MYO1( | O94832; B0I | MYO1D_HUMA | Unmodified | 2892000 | 704140 | 355920 | 302420 | 0 | 0 | 0 | 0 | 0 | 0 | 0 | 347830 | 0 | 0 | 2494000 | 853920 | 73936 | 144450 |
| DTCLGPQA | ADAMTS13 | Q76LX8 | ATS13_HUMAN | Unmodified | 0 | 174810 | 5948100 | 938080 | 0 | 2175600 | 0 | 0 | 0 | 0 | 969170 | 0 | 0 | 0 | 1096900 | 253490 | 0 | 0 |
| DTCYSPKP | CD2AP | Q9Y5K6 | CD2AP_HUMAI | Unmodified | 121310 | 215430 | 0 | 539550 | 1524100 | 0 | 537930 | 158930 | 0 | 0 | 0 | 0 | 5792500 | 2850600 | 0 | 0 | 183830 | 0 |
| DTDALDLEI | ADRA2A | P08913 | ADA2A_HUMAI | Unmodified | 0 | 0 | 0 | 0 | 0 | 0 | 32399 | 0 | 426650 | 196870 | 0 | 0 | 0 | 0 | 0 | 940450 | 0 | 0 |
| DTDDVPMI | RAP1B; N/A; RA | P61224; A6N | RAP1B_HUMAN | Oxidation (M) | 729350 | 0 | 4155400 | 691080 | 0 | 0 | 0 | 0 | 0 | 0 | 378010 | 3554800 | 0 | 0 | 2298900 | 2233900 | 0 | 0 |
| DTDDVPMI | RAP1B; N/A; RA | P61224; A6N | RAP1B_HUMAN | Oxidation (M) | 5777500 | 2872300 | 1941400 | 4770500 | 0 | 0 | 0 | 0 | 0 | 0 | 2465400 | 12065000 | 812920 | 654700 | 6637100 | 6346400 | 0 | 0 |
| DTDGGPKE | SLC16A1 | P53985 | MOT1_HUMAN | Unmodified | 0 | 0 | 5247400 | 1937400 | 0 | 0 | 392240 | 0 | 0 | 0 | 0 | 0 | 1783400 | 1226200 | 6683400 | 5555300 | 1253800 | 3735900 |
| DTDIPLLNT | CDCP1 | Q9H5V8 | CDCP1_HUMA | Oxidation (M) | 0 | 0 | 806480 | 294950 | 157250 | 36507 | 0 | 0 | 0 | 39527 | 288440 | 1515500 | 0 | 0 | 762650 | 615480 | 379990 | 2483400 |
| DTDIVDEAI | ARPC3; ARPC3; | O15145; F8V | ARPC3_HUMAI | Unmodified | 0 | 229840 | 176590 | 168940 | 0 | 0 | 0 | 0 | 0 | 0 | 0 | 791160 | 0 | 0 | 41561 | 151880 | 0 | 0 |
| DTDNEEEIF | CALML3 | P27482 | CALL3_HUMAN | Unmodified | 0 | 0 | 0 | 0 | 0 | 0 | 0 | 0 | 181570 | 0 | 381160 | 23581 | 0 | 0 | 0 | 0 | 0 | 0 |
| DTDPDSKP | SEPTIN6; SEPTI | Q14141; Q9I | SEPT6_HUMAN | Unmodified | 0 | 52930 | 454670 | 194180 | 817360 | 740240 | 96318 | 0 | 0 | 0 | 0 | 1602000 | 0 | 0 | 320110 | 312350 | 244890 | 0 |
| DTDSEEEIF | CALM2; CALM3 | P0DP24; P0I | CALM2_HUMAI | Unmodified | 77195 | 43651 | 161950 | 81703 | 0 | 0 | 850330 | 400700 | 3695600 | 254490 | 30565 | 546960 | 0 | 119900 | 220110 | 117360 | 0 | 0 |
| DTDSEEEIF | CALM2; CALM3 | P0DP24; P0I | CALM2_HUMAI | Unmodified | 1504600 | 916000 | 1386600 | 617270 | 0 | 0 | 1390200 | 0 | 959980 | 233450 | 1263300 | 13351000 | 63759 | 0 | 3820100 | 2589900 | 0 | 0 |
| DTDSEEEIF | CALM2; CALM3 | P0DP24; P0I | CALM2_HUMAI | Unmodified | 1089000 | 1493700 | 1386400 | 0 | 458560 | 460650 | 623180 | 0 | 1924500 | 639310 | 21103 | 3019200 | 0 | 0 | 0 | 3063000 | 222270 | 168910 |
| DTDSINLYI | XPO1; XPO1; XF | O14980; A0/ | XPO1_HUMAN | Unmodified | 70663 | 0 | 0 | 21145 | 0 | 0 | 0 | 0 | 0 | 0 | 0 | 5777900 | 0 | 0 | 0 | 0 | 0 | 0 |
| DTDSSVAS | LSR; LSR | Q86X29; S4F | LSR_HUMAN Li | Unmodified | 0 | 0 | 0 | 4459200 | 262450 | 0 | 0 | 0 | 12466000 | 3590500 | 850640 | 6875400 | 0 | 0 | 2142100 | 6078100 | 65514 | 130720 |
| DTDSSVAS | LSR; LSR | Q86X29; S4F | LSR_HUMAN Li | Unmodified | 0 | 0 | 0 | 0 | 0 | 0 | 0 | 0 | 0 | 214980 | 179440 | 0 | 0 | 0 | 0 | 0 | 33885 | 0 |
| DTEDVPMII | RAP1A; RAP1A | P62834; A0A | RAP1A_HUMAN | Oxidation (M) | 0 | 0 | 787330 | 854670 | 0 | 0 | 0 | 0 | 0 | 0 | 568670 | 3304900 | 0 | 0 | 356840 | 0 | 0 | 0 |
| DTEEMLSK | CHMP4B | Q9H444 | CHM4B_HUMA | Unmodified | 273080 | 258530 | 105500 | 0 | 0 | 0 | 0 | 0 | 518950 | 0 | 0 | 144610 | 0 | 0 | 232810 | 154570 | 0 | 0 |
| DTEITCSER | EPCAM; EPCAM | P16422; B5N | EPCAM_HUMA | Unmodified | 620900 | 361900 | 3325700 | 279680 | 0 | 0 | 0 | 0 | 1569000 | 841010 | 34121 | 0 | 0 | 57925 | 2572500 | 1930200 | 0 | 0 |
| DTEITCSER | EPCAM; EPCAM | P16422; B5N | EPCAM_HUMA | Unmodified | 2492100 | 470000 | 9202400 | 456790 | 0 | 0 | 0 | 0 | 1475600 | 180740 | 811700 | 377650 | 0 | 184610 | 5270300 | 3031600 | 0 | 0 |
| DTEPGPAS | MTSS2; MTSS2 | Q765P7; A0/ | MTSS2_HUMAI | Unmodified | 0 | 0 | 0 | 0 | 0 | 65757 | 0 | 0 | 0 | 0 | 194250 | 890040 | 0 | 0 | 0 | 0 | 0 | 0 |
| DTEPLIQTA | CCT5; CCT5; C( | P48643; B7Z | TCPE_HUMAN | Unmodified | 0 | 0 | 0 | 0 | 0 | 0 | 0 | 56841 | 0 | 0 | 0 | 1143300 | 606920 | 0 | 0 | 0 | 0 | 0 |
| DTFADSLR | GNPTAB | Q3T906 | GNPTA_HUMAI | Unmodified | 0 | 0 | 0 | 0 | 205990 | 0 | 56069 | 28606 | 0 | 314880 | 1046500 | 0 | 0 | 0 | 0 | 0 | 0 | 0 |
| DTFFQNPC | CAPN5; CAPN5 | O15484; E7E | CAN5_HUMAN | Unmodified | 0 | 0 | 0 | 86237 | 0 | 0 | 0 | 0 | 156420 | 33482 | 29889 | 362790 | 0 | 0 | 267080 | 217840 | 0 | 0 |
| DTFNGNLF | MPI; MPI; MPI; I | P34949; H3E | MPI_HUMAN M | Unmodified | 0 | 0 | 0 | 0 | 0 | 0 | 0 | 0 | 0 | 0 | 0 | 0 | 0 | 0 | 0 | 0 | 0 | 0 |
| DTFNHLTT | RAB2A; RAB2A; | P61019; H7( | RAB2A_HUMAN | Unmodified | 1367500 | 1101900 | 569180 | 821450 | 0 | 0 | 0 | 0 | 12382000 | 1018200 | 1022700 | 7990300 | 0 | 0 | 1567200 | 1607800 | 0 | 0 |
| DTFYDNFE | APOBEC3B; AP( | Q9UH17; Q9 | ABC3B_HUMAI | Unmodified | 1053200 | 1113000 | 298400 | 0 | 0 | 0 | 0 | 0 | 0 | 0 | 0 | 0 | 0 | 0 | 3030200 | 877260 | 0 | 0 |
| DTGETSML | STEAP1; STEAP | Q9UHE8; C9 | STEA1_HUMAN | Oxidation (M) | 0 | 0 | 3416900 | 2344200 | 0 | 0 | 0 | 0 | 0 | 0 | 0 | 0 | 0 | 0 | 2689000 | 2462100 | 0 | 0 |
| DTGETSML | STEAP1; STEAP | Q9UHE8; C9 | STEA1_HUMAN | Unmodified | 701200 | 0 | 49510 | 5486700 | 152390 | 0 | 0 | 0 | 0 | 0 | 0 | 0 | 2627300 | 0 | 10021000 | 5091500 | 0 | 0 |
| DTGIFLDLI | XRCC6; XRCC6 | P12956; B1A | XRCC6_HUMAI | Oxidation (M) | 0 | 0 | 0 | 0 | 0 | 0 | 0 | 0 | 0 | 0 | 0 | 1710400 | 0 | 0 | 0 | 0 | 0 | 0 |
| DTGIFLDLI | XRCC6; XRCC6 | P12956; B1A | XRCC6_HUMAI | Unmodified | 0 | 0 | 0 | 0 | 0 | 0 | 0 | 0 | 349400 | 0 | 139730 | 778100 | 0 | 0 | 0 | 0 | 0 | 0 |
| DTGILDSIG | MBP; MBP; MBF | P02686; H7E | MBP_HUMAN I | Unmodified | 0 | 0 | 0 | 104190 | 514710 | 140550 | 0 | 0 | 0 | 0 | 0 | 0 | 935530 | 273160 | 22768 | 0 | 911110 | 242030 |
| DTGKTPVEI | RPS20; RPS20; | P60866; E5R | RS20_HUMAN | Unmodified | 5355100 | 3920700 | 2383000 | 3478200 | 2200600 | 0 | 0 | 126210 | 5170300 | 360960 | 7346400 | 32667000 | 4562300 | 491510 | 2600400 | 2998700 | 224020 | 1048700 |
| DTGKTPVEI | RPS20; RPS20; | P60866; E5R | RS20_HUMAN | Unmodified | 0 | 0 | 0 | 0 | 542770 | 0 | 0 | 207800 | 0 | 120280 | 0 | 368900 | 1539300 | 0 | 0 | 23571 | 0 | 724510 |
| DTGTAPFAI | COBLL1; COBLI | Q53SF7; A0/ | COBL1_HUMAI | Unmodified | 1773100 | 1330800 | 99303 | 855910 | 171360 | 0 | 0 | 0 | 104250 | 0 | 42060 | 176340 | 944260 | 131380 | 1132100 | 511830 | 664510 | 228550 |
| DTHEDHD | RANBP1; RANB | P43487; C9J | RANG_HUMAN | Unmodified | 0 | 0 | 11471000 | 0 | 0 | 457990 | 0 | 911180 | 4796600 | 0 | 13917000 | 0 | 0 | 0 | 5205300 | 13643000 | 0 | 0 |
| DTHLTITPT | SHPK; N/A | Q9UHJ6; A0/ | SHPK_HUMAN | Unmodified | 0 | 0 | 0 | 0 | 219700 | 0 | 0 | 0 | 0 | 0 | 0 | 2589900 | 0 | 0 | 0 | 0 | 0 | 83590 |
| DTIARALPF | PGAM1; PGAM4 | P18669; Q8I | PGAM1_HUMA | Unmodified | 0 | 0 | 37961 | 0 | 12678000 | 1623000 | 1543300 | 0 | 0 | 0 | 0 | 0 | 3685000 | 2062000 | 0 | 0 | 2904700 | 10067000 |
| DTIDFLNYT | EPS8; EPS8; EP | Q12929; A0/ | EPS8_HUMAN | Unmodified | 0 | 0 | 0 | 0 | 0 | 0 | 0 | 0 | 0 | 0 | 0 | 97668 | 0 | 0 | 0 | 0 | 0 | 0 |
| DTIEQYK | MYO1D; MYO1I | O94832; J3Q | MYO1D_HUMA | Unmodified | 511430 | 0 | 1108200 | 0 | 438720 | 2610900 | 0 | 0 | 0 | 0 | 121330 | 378020 | 0 | 0 | 1619200 | 1169400 | 0 | 0 |
| DTILHDNLI | GNA13 | Q14344 | GNA13_HUMAI | Unmodified | 0 | 0 | 215520 | 0 | 0 | 0 | 0 | 0 | 352030 | 0 | 0 | 1752000 | 0 | 0 | 0 | 0 | 0 | 0 |
| DTILQLNLK | GNAQ; GNA11; | P50148; P29 | GNAQ_HUMAN | Unmodified | 0 | 0 | 0 | 0 | 335800 | 0 | 0 | 49713 | 370650 | 0 | 555700 | 1547000 | 0 | 0 | 0 | 0 | 127430 | 0 |
| DTINLLDQI | GOLM1 | Q8NBJ4 | GOLM1_HUMA | Unmodified | 0 | 0 | 0 | 0 | 0 | 0 | 0 | 0 | 6334300 | 187600 | 0 | 305030 | 0 | 0 | 0 | 0 | 0 | 0 |
| DTINLLDQI | GOLM1 | Q8NBJ4 | GOLM1_HUMA | Unmodified | 0 | 0 | 0 | 0 | 0 | 0 | 282400 | 0 | 8980500 | 0 | 1158900 | 0 | 0 | 0 | 0 | 0 | 0 | 0 |
| DTIQHQQC | TJP2; TJP2; TJP2 | Q9UDY2; A0 | ZO2_HUMAN T | Unmodified | 142250 | 0 | 213310 | 0 | 74796 | 0 | 241630 | 151790 | 1559500 | 0 | 0 | 0 | 0 | 0 | 114730 | 118350 | 154660 | 0 |
| DTITPQQVI | PALS2; PALS2; I | Q9NZW5; B8 | PALS2_HUMAN | Unmodified | 0 | 0 | 0 | 0 | 0 | 0 | 0 | 0 | 0 | 0 | 0 | 1064700 | 0 | 0 | 0 | 0 | 0 | 0 |
| DTIVLLCKP | PDCD6IP | Q8WUM4 | PDC6I_HUMAN | Unmodified | 20424000 | 5580600 | 5075900 | 0 | 0 | 0 | 0 | 0 | 3126300 | 0 | 0 | 5493000 | 0 | 0 | 0 | 10637000 | 0 | 0 |
| DTKEIYTHF | GNAI3; GNAI2; | P08754; P04 | GNAI3_HUMAN | Unmodified | 0 | 0 | 0 | 0 | 1443500 | 0 | 779380 | 845000 | 0 | 0 | 0 | 0 | 1041600 | 1856500 | 0 | 0 | 1623700 | 3365000 |
| DTKTTVDAI | CHMP5 | Q9NZZ3 | CHMP5_HUMA | Unmodified | 0 | 0 | 0 | 0 | 283060 | 33073 | 0 | 0 | 0 | 0 | 0 | 0 | 3810100 | 0 | 0 | 0 | 22118 | 0 |
| DTKTTVDAI | CHMP5 | Q9NZZ3 | CHMP5_HUMA | Unmodified | 0 | 0 | 0 | 0 | 366310 | 88891 | 108820 | 66403 | 0 | 160480 | 0 | 0 | 91046 | 499650 | 0 | 0 | 82906 | 80830 |
| DTLFQDFK | MYO1D; MYO1I | O94832; J3Q | MYO1D_HUMA | Unmodified | 0 | 0 | 1344900 | 0 | 0 | 0 | 0 | 0 | 0 | 0 | 0 | 0 | 0 | 0 | 1383800 | 1115100 | 0 | 0 |
| DTLGLFLR | SNRNP200; SN | O75643; A0/ | U520_HUMAN | Unmodified | 0 | 0 | 0 | 0 | 0 | 0 | 0 | 0 | 175190 | 0 | 137630 | 4750700 | 0 | 0 | 0 | 0 | 0 | 0 |
| DTLGNTQII | FGFBP1 | Q14512 | FGFP1_HUMAN | Unmodified | 0 | 0 | 0 | 0 | 0 | 0 | 237290 | 653210 | 65140 | 32702 | 0 | 0 | 0 | 0 | 0 | 0 | 175460 | 500060 |
| DTLGPMQI | EIF3G; EIF3G; E | O75821; K7E | EIF3G_HUMAN | Unmodified | 74640 | 0 | 0 | 0 | 379700 | 0 | 61382 | 0 | 266440 | 0 | 440550 | 133470 | 0 | 0 | 0 | 0 | 124890 | 0 |
| DTLPKEVFI | PDCD6IP; PDCI | Q8WUM4; F8 | PDC6I_HUMAN | Unmodified | 0 | 0 | 0 | 216730 | 3763500 | 755850 | 0 | 199360 | 0 | 1881200 | 0 | 0 | 4250900 | 3070500 | 0 | 0 | 954590 | 1604800 |
| DTLPVGFIP | MVB12B; MVB1 | Q9H7P6; F8 | MB12B_HUMAI | Unmodified | 229600 | 664460 | 126790 | 187380 | 0 | 0 | 0 | 0 | 146630 | 0 | 0 | 0 | 0 | 0 | 0 | 147890 | 0 | 0 |
| DTLPVGFIP | MVB12B; MVB1 | Q9H7P6; F8 | MB12B_HUMAI | Unmodified | 0 | 0 | 523010 | 0 | 0 | 0 | 0 | 0 | 0 | 0 | 0 | 0 | 0 | 0 | 892950 | 0 | 0 | 0 |
| DTLQSELV( | DNM1L | O00429 | DNM1L_HUMA | Unmodified | 0 | 0 | 0 | 0 | 0 | 0 | 0 | 28674 | 41550 | 22204 | 127480 | 1815400 | 0 | 0 | 0 | 0 | 0 | 0 |
| DTLSIHYLI | IARS1; IARS1; I/ | P41252; A0A | SYIC_HUMAN I | Oxidation (M) | 26268 | 0 | 234100 | 0 | 30253 | 0 | 0 | 0 | 0 | 0 | 40566 | 5635700 | 0 | 0 | 139550 | 76685 | 0 | 0 |
| DTLSNPQS | NEDD4L; NEDD | Q96PU5; A0/ | NED4L_HUMAI | Unmodified | 2317000 | 2516000 | 1306900 | 2342800 | 0 | 0 | 0 | 0 | 0 | 0 | 0 | 637790 | 0 | 0 | 3395600 | 2079900 | 0 | 0 |
| DTLTSRPA( | FN1; FN1 | P02751; H0Y | FINC_HUMAN I | Unmodified | 0 | 0 | 0 | 0 | 0 | 0 | 0 | 0 | 0 | 0 | 0 | 0 | 0 | 0 | 0 | 0 | 0 | 0 |
| DTLTSRPA( | FN1; FN1 | P02751; H0Y | FINC_HUMAN I | Unmodified | 0 | 3122300 | 13542000 | 0 | 1476300 | 400200 | 0 | 0 | 0 | 0 | 0 | 2798700 | 833670 | 1141700 | 0 | 7869800 | 246320 | 0 |
| DTLVWDTP | ACSS1; ACSS1 | Q9NUB1; Q5 | ACS2L_HUMAN | Unmodified | 0 | 0 | 0 | 0 | 0 | 0 | 0 | 0 | 559510 | 0 | 0 | 0 | 0 | 0 | 0 | 0 | 0 | 0 |
| DTLYEAVRI | RPL10A | P62906 | RL10A_HUMAN | Unmodified | 283600 | 521140 | 81489 | 634930 | 0 | 0 | 0 | 0 | 185680 | 0 | 0 | 4625900 | 0 | 0 | 178080 | 235410 | 0 | 0 |
| DTMGDLPF | NPTX2 | P47972 | NPTX2_HUMAN | Unmodified | 0 | 0 | 0 | 0 | 0 | 0 | 0 | 0 | 2337200 | 0 | 0 | 0 | 0 | 0 | 0 | 0 | 0 | 0 |
| DTNGENIA | SND1 | Q7KZF4 | SND1_HUMAN | Unmodified | 307550 | 0 | 2449900 | 2833400 | 0 | 0 | 0 | 0 | 0 | 0 | 0 | 9040600 | 0 | 0 | 5998900 | 0 | 0 | 0 |
| DTNGSQFF | PPIB; PPIB; PPII | P23284; A0A | PPIB_HUMAN F | Unmodified | 141390 | 0 | 2784500 | 0 | 0 | 1594000 | 0 | 467920 | 1.13E+08 | 7329200 | 12837000 | 12271000 | 0 | 492350 | 1727000 | 2087100 | 1554600 | 0 |
| DTNKYEGV | FAM3C | Q92520 | FAM3C_HUMAI | Oxidation (M) | 0 | 0 | 0 | 0 | 0 | 0 | 0 | 0 | 4754000 | 0 | 0 | 0 | 0 | 0 | 0 | 0 | 0 | 0 |
| DTNKYEGV | FAM3C | Q92520 | FAM3C_HUMAI | Unmodified | 0 | 0 | 0 | 0 | 0 | 0 | 0 | 0 | 3354600 | 0 | 0 | 0 | 0 | 0 | 0 | 0 | 0 | 0 |
| DTPALFNF | GSK3B; GSK3B | P49841; A0A | GSK3B_HUMAI | Unmodified | 0 | 261490 | 0 | 0 | 0 | 0 | 0 | 0 | 0 | 0 | 0 | 0 | 0 | 0 | 337700 | 36018 | 0 | 0 |
| DTPQEMEC | CUL2; CUL2 | Q13617; A0/ | CUL2_HUMAN | Unmodified | 0 | 97472 | 0 | 0 | 0 | 0 | 0 | 0 | 0 | 0 | 0 | 459100 | 0 | 0 | 0 | 0 | 0 | 0 |
| DTPTQEDW | XPNPEP1; XPNI | Q9NQW7; Q | XPP1_HUMAN | Unmodified | 0 | 0 | 131660 | 0 | 0 | 0 | 0 | 0 | 0 | 0 | 254190 | 1322900 | 0 | 0 | 0 | 73167 | 0 | 0 |
| DTQALLSA | SLC26A6; SLC2 | Q9BXS9; C9. | S26A6_HUMAN | Oxidation (M) | 0 | 0 | 0 | 0 | 0 | 0 | 0 | 0 | 0 | 0 | 0 | 1506900 | 0 | 0 | 0 | 125240 | 0 | 0 |
| DTQALLSA | SLC26A6; SLC2 | Q9BXS9; C9. | S26A6_HUMAN | Unmodified | 0 | 0 | 0 | 0 | 0 | 0 | 0 | 0 | 53407 | 0 | 0 | 242290 | 0 | 0 | 0 | 0 | 0 | 0 |
| DTQDHLAV | LAMA5 | O15230 | LAMA5_HUMAI | Unmodified | 274130 | 148560 | 0 | 0 | 0 | 0 | 0 | 0 | 370500 | 4120900 | 0 | 0 | 0 | 0 | 98471 | 70418 | 0 | 0 |
| DTQEVPLEI | YTHDF1; YTHDI | Q9BYJ9; Q7Z | YTHD1_HUMAI | Unmodified | 1252800 | 57903 | 502350 | 0 | 0 | 0 | 3012600 | 0 | 0 | 0 | 275990 | 1125000 | 0 | 258990 | 495750 | 718580 | 0 | 0 |
| DTQFSYAV | IGSF8; IGSF8 | Q969P0; C9. | IGSF8_HUMAN | Unmodified | 23776000 | 20330000 | 8022400 | 8435800 | 3678600 | 1027800 | 398610 | 0 | 1621800 | 0 | 0 | 0 | 8567400 | 2006000 | 10907000 | 8199300 | 0 | 2863300 |
| DTQLQQIV | PPP3R1; PPP3R | P63098; F6U | CANB1_HUMAI | Unmodified | 391670 | 445630 | 461620 | 546660 | 503570 | 0 | 0 | 0 | 682560 | 96485 | 1491200 | 4571000 | 867250 | 0 | 776820 | 637410 | 0 | 0 |
| DTQSGSLL | SERPINH1; SER | P50454; E9P | SERPH_HUMAI | Unmodified | 0 | 0 | 0 | 0 | 0 | 0 | 0 | 0 | 6938600 | 322270 | 0 | 264130 | 0 | 0 | 0 | 0 | 0 | 0 |
| DTQVTRQA | STX4 | Q12846 | STX4_HUMAN | Unmodified | 0 | 0 | 0 | 0 | 0 | 0 | 0 | 0 | 0 | 0 | 0 | 0 | 0 | 0 | 0 | 0 | 872570 | 0 |
| DTQVTRQA | STX4 | Q12846 | STX4_HUMAN | Unmodified | 0 | 0 | 0 | 0 | 2076500 | 401900 | 92080 | 0 | 0 | 347470 | 0 | 0 | 822430 | 0 | 0 | 0 | 235840 | 1105300 |

| | | | | | | | | | | | | | | | | | | | | | | |
|---|---|---|---|---|---|---|---|---|---|---|---|---|---|---|---|---|---|---|---|---|---|---|
| DTSASAVA | HADHA; HADH | P40939; A0A | ECHA_HUMAN | Unmodified | 0 | 0 | 0 | 0 | 0 | 0 | 41480 | 0 | 3811900 | 0 | 0 | 0 | 0 | 0 | 0 | 0 | 0 | 0 |
| DTSCLAEQ | CNPY3 | Q9BT09 | CNPY3_HUMA | Unmodified | 0 | 0 | 0 | 0 | 0 | 0 | 0 | 0 | 1069900 | 0 | 0 | 0 | 0 | 0 | 0 | 0 | 0 | 0 |
| DTSEDIEEL | PSMD1; PSMD1 | Q99460; A0A | PSMD1_HUMA | Unmodified | 0 | 0 | 0 | 0 | 0 | 0 | 0 | 0 | 0 | 0 | 0 | 1551800 | 0 | 0 | 0 | 0 | 0 | 0 |
| DTSFEQHV | FASN; FASN | P49327; A0A | FAS_HUMAN F | Unmodified | 979580 | 0 | 2853900 | 745830 | 0 | 0 | 0 | 0 | 0 | 0 | 2331300 | 87119000 | 0 | 0 | 3344100 | 3178600 | 0 | 0 |
| DTSLEELQ | STMN3; STMN3 | Q9NZ72; A0 | STMN3_HUMA | Unmodified | 0 | 0 | 0 | 0 | 0 | 0 | 247800 | 0 | 281120 | 0 | 0 | 0 | 0 | 0 | 0 | 0 | 0 | 0 |
| DTSLFSDEI | ADAM10; ADAM | O14672; C9J | ADA10_HUMAN | Unmodified | 0 | 0 | 0 | 740670 | 1034200 | 117110 | 0 | 0 | 1590400 | 0 | 0 | 0 | 0 | 0 | 0 | 0 | 0 | 298680 |
| DTSLGRWF | HPN | P05981 | HEPS_HUMAN | Unmodified | 217790 | 211750 | 101910 | 65065 | 0 | 0 | 0 | 0 | 0 | 0 | 0 | 0 | 0 | 0 | 366050 | 805260 | 0 | 0 |
| DTSLYRPAI | PSMD2; PSMD2 | Q13200; H7 | PSMD2_HUMA | Unmodified | 0 | 0 | 0 | 0 | 0 | 0 | 0 | 0 | 0 | 0 | 0 | 0 | 0 | 0 | 0 | 0 | 0 | 0 |
| DTSLYRPAI | PSMD2; PSMD2 | Q13200; H7 | PSMD2_HUMA | Unmodified | 0 | 0 | 0 | 0 | 0 | 0 | 0 | 0 | 0 | 0 | 0 | 3299900 | 0 | 0 | 0 | 0 | 0 | 0 |
| DTSNDPG | OTULIN | Q96BN8 | OTUL_HUMAN | Unmodified | 0 | 0 | 0 | 0 | 0 | 0 | 0 | 0 | 0 | 677050 | 0 | 634590 | 0 | 0 | 0 | 0 | 0 | 0 |
| DTSNHFH | TIAL1; TIA1; TIA | Q01085; C9J | TIAR_HUMAN N | Unmodified | 0 | 0 | 0 | 0 | 0 | 0 | 0 | 0 | 0 | 0 | 0 | 898440 | 0 | 0 | 942660 | 0 | 0 | 0 |
| DTSPGGQ | EPHA1 | P21709 | EPHA1_HUMAN | Unmodified | 0 | 0 | 0 | 257820 | 0 | 0 | 0 | 0 | 0 | 0 | 0 | 0 | 0 | 0 | 482840 | 577540 | 0 | 0 |
| DTSSSTVV | TIA1; TIA1; TIA1 | P31483; E5R | TIA1_HUMAN N | Unmodified | 0 | 0 | 0 | 0 | 0 | 0 | 0 | 0 | 0 | 0 | 0 | 0 | 0 | 0 | 0 | 0 | 0 | 0 |
| DTSYLFITG | PCCB; PCCB; P | P05166; C9J | PCCB_HUMAN | Unmodified | 0 | 0 | 0 | 0 | 0 | 0 | 0 | 0 | 926520 | 0 | 0 | 0 | 0 | 0 | 0 | 0 | 0 | 0 |
| DTTAAHQA | SLC12A7; SLC1 | Q9Y666; A0A | S12A7_HUMAN | Unmodified | 34387 | 30182 | 83712 | 0 | 0 | 69799 | 83702 | 92737 | 0 | 0 | 0 | 707840 | 0 | 0 | 199000 | 113740 | 28365 | 0 |
| DTTALSFF | LRPPRC | P42704 | LPPRC_HUMAN | Unmodified | 0 | 0 | 0 | 0 | 0 | 0 | 364440 | 14484 | 187370 | 0 | 0 | 0 | 0 | 142750 | 0 | 0 | 0 | 0 |
| DTTHNLPE | PLXNB2 | O15031 | PLXB2_HUMAN | Unmodified | 1603000 | 711410 | 1198800 | 8641600 | 1906000 | 0 | 0 | 86865 | 1376000 | 0 | 0 | 2957300 | 0 | 0 | 3385800 | 3264100 | 1000900 | 0 |
| DTTINEIED | ATP6V1F | Q16864 | VATF_HUMAN | Unmodified | 0 | 0 | 0 | 0 | 0 | 0 | 0 | 0 | 0 | 0 | 0 | 785560 | 0 | 0 | 0 | 0 | 0 | 0 |
| DTTLNWK | DTYMK; DTYMK | P23919; H7 | KTHY_HUMAN | Unmodified | 23517 | 0 | 0 | 0 | 2309200 | 158290 | 625060 | 386050 | 230190 | 376110 | 91714 | 684990 | 841290 | 377790 | 0 | 0 | 0 | 592810 |
| DTTVGTLS | PSMD7 | P51665 | PSMD7_HUMA | Unmodified | 0 | 0 | 0 | 0 | 0 | 0 | 0 | 0 | 0 | 0 | 208970 | 2286300 | 0 | 0 | 0 | 0 | 0 | 0 |
| DTVATQLSI | PLIN3; PLIN3; P | O60664; K7E | PLIN3_HUMAN | Unmodified | 0 | 0 | 0 | 0 | 1604800 | 0 | 0 | 0 | 223240 | 0 | 0 | 0 | 0 | 0 | 0 | 0 | 0 | 1988800 |
| DTVTISGP | FASN; FASN | P49327; A0A | FAS_HUMAN F | Unmodified | 923410 | 568990 | 1111200 | 3179600 | 52903 | 0 | 697140 | 0 | 863950 | 204130 | 7411600 | 18895000 | 1199900 | 25902 | 8116400 | 2486600 | 418150 | 28896 |
| DTVVVQDL | CTSA; CTSA; CT | P10619; X6R | PPGB_HUMAN | Unmodified | 65256 | 105100 | 29778 | 34720 | 885070 | 793520 | 193910 | 0 | 748350 | 632130 | 124110 | 0 | 210440 | 108090 | 115970 | 27896 | 0 | 1422900 |
| DTVVVQDL | CTSA; CTSA; CT | P10619; X6R | PPGB_HUMAN | Unmodified | 0 | 0 | 0 | 0 | 0 | 151490 | 0 | 0 | 749630 | 56347 | 0 | 0 | 0 | 0 | 0 | 0 | 0 | 0 |
| DTYGVRPL | ASNS; ASNS; A | P08243; C9J | ASNS_HUMAN | Unmodified | 0 | 0 | 0 | 0 | 0 | 0 | 0 | 0 | 0 | 0 | 0 | 3227400 | 0 | 0 | 0 | 0 | 0 | 0 |
| DTYIENEK | RPN1; RPN1 | P04843; B7Z | RPN1_HUMAN | Unmodified | 0 | 0 | 263520 | 0 | 267260 | 0 | 0 | 18812 | 419620 | 0 | 37948 | 311790 | 148700 | 174480 | 0 | 216640 | 0 | 0 |
| DTYLSHFN | NNMT | P40261 | NNMT_HUMAN | Unmodified | 0 | 0 | 0 | 0 | 5300700 | 284360 | 0 | 0 | 0 | 0 | 0 | 0 | 0 | 0 | 0 | 0 | 0 | 0 |
| DTYSPDEH | GCNT2 | Q8N0V5 | GNT2A_HUMA | Unmodified | 0 | 0 | 0 | 0 | 0 | 0 | 0 | 0 | 0 | 0 | 0 | 244130 | 0 | 0 | 0 | 0 | 0 | 0 |
| DVAALNGL | EMC10; EMC10 | Q5UCC4; M | EMC10_HUMA | Unmodified | 1015500 | 0 | 0 | 0 | 0 | 0 | 0 | 0 | 1096100 | 0 | 0 | 0 | 0 | 0 | 0 | 0 | 0 | 0 |
| DVACGANI | RCC2 | Q9P258 | RCC2_HUMAN | Unmodified | 0 | 0 | 0 | 0 | 0 | 0 | 0 | 0 | 0 | 0 | 0 | 6801400 | 0 | 0 | 0 | 0 | 0 | 0 |
| DVACGANI | RCC2 | Q9P258 | RCC2_HUMAN | Unmodified | 0 | 0 | 0 | 0 | 0 | 0 | 0 | 55188 | 326940 | 0 | 0 | 8360000 | 0 | 0 | 0 | 0 | 0 | 0 |
| DVAEAKPE | IAH1; IAH1 | Q2TAA2; H7 | IAH1_HUMAN | Unmodified | 0 | 0 | 0 | 0 | 0 | 0 | 0 | 0 | 0 | 0 | 0 | 3802900 | 0 | 0 | 0 | 147270 | 0 | 0 |
| DVAEEIAN | SPTBN1; SPTBN | Q01082; A0 | SPTB2_HUMAN | Unmodified | 0 | 0 | 0 | 0 | 0 | 0 | 0 | 0 | 0 | 0 | 0 | 4514000 | 0 | 0 | 0 | 0 | 0 | 0 |
| DVAEYSEA | SLC26A6; SLC2 | Q9BXS9; A0 | S26A6_HUMAN | Unmodified | 0 | 0 | 0 | 0 | 0 | 0 | 0 | 0 | 0 | 0 | 0 | 2103800 | 0 | 0 | 0 | 0 | 0 | 0 |
| DVAFTVGE | FKBP5 | Q13451 | FKBP5_HUMAN | Unmodified | 268610 | 192030 | 0 | 0 | 211890 | 0 | 0 | 0 | 2641300 | 0 | 0 | 2782500 | 511260 | 0 | 203060 | 205160 | 0 | 0 |
| DVAGDASE | ATP1A3; N/A; A | P13637; A0A | AT1A3_HUMAN | Unmodified | 0 | 0 | 0 | 0 | 0 | 0 | 0 | 0 | 758780 | 0 | 0 | 0 | 0 | 0 | 0 | 0 | 0 | 0 |
| DVAGGLRS | TLN1 | Q9Y490 | TLN1_HUMAN | Unmodified | 0 | 114190 | 0 | 0 | 0 | 199630 | 210250 | 0 | 0 | 140930 | 334380 | 0 | 0 | 232240 | 0 | 0 | 108360 | 0 |
| DVAGLDTE | SPAG9; SPAG9 | O60271; A0 | JIP4_HUMAN C | Unmodified | 0 | 0 | 0 | 0 | 529220 | 0 | 0 | 78324 | 83552 | 51978 | 184550 | 262890 | 138670 | 80374 | 0 | 0 | 223580 | 1086800 |
| DVAIKEPLV | PRMT1; PRMT1; | Q99873; E9F | ANM1_HUMAN | Unmodified | 431990 | 0 | 0 | 0 | 0 | 0 | 0 | 0 | 0 | 0 | 0 | 69189000 | 0 | 0 | 0 | 560740 | 0 | 0 |
| DVAKALGD | TLN1 | Q9Y490 | TLN1_HUMAN | Unmodified | 0 | 0 | 0 | 0 | 0 | 0 | 368310 | 0 | 0 | 0 | 0 | 0 | 0 | 215130 | 0 | 0 | 207600 | 0 |
| DVAKGSEG | RAB11FIP1; RA | Q6WKZ4; E7 | RFIP1_HUMAN | Unmodified | 0 | 0 | 151330 | 0 | 4246100 | 175200 | 0 | 0 | 0 | 0 | 0 | 0 | 183030 | 194650 | 0 | 0 | 0 | 0 |
| DVAKINQD | PGM1 | P36871 | PGM1_HUMAN | Oxidation (M) | 0 | 0 | 0 | 0 | 0 | 0 | 147980 | 0 | 0 | 57066 | 0 | 0 | 0 | 0 | 0 | 0 | 414080 | 112420 |
| DVAPQAPV | HINT2 | Q9BX68 | HINT2_HUMAN | Unmodified | 0 | 0 | 76468 | 0 | 0 | 0 | 0 | 0 | 4047000 | 55335 | 0 | 0 | 0 | 0 | 0 | 0 | 0 | 0 |
| DVAPSTVKI | MAP4; MAP4 | P27816; E7E | MAP4_HUMAN | Unmodified | 0 | 0 | 0 | 0 | 0 | 0 | 0 | 0 | 3934300 | 340230 | 0 | 0 | 0 | 0 | 0 | 0 | 0 | 0 |
| DVAQIFNN | CAB39; CAB39 | Q9Y376; A0 | CAB39_HUMAN | Unmodified | 0 | 2080500 | 0 | 32291 | 0 | 139140 | 0 | 0 | 521000 | 118590 | 122190 | 648370 | 0 | 0 | 394340 | 195930 | 0 | 0 |
| DVAQIFNN | CAB39; CAB39 | Q9Y376; A0 | CAB39_HUMAN | Unmodified | 608440 | 745080 | 951860 | 1418600 | 2608700 | 1305000 | 66006 | 0 | 1794400 | 342320 | 1023300 | 7535400 | 138750 | 537870 | 1802400 | 1953900 | 347790 | 816780 |
| DVATVAFC | CTBP1; CTBP1; | Q13363; E9F | CTBP1_HUMAN | Unmodified | 0 | 54302 | 0 | 0 | 0 | 0 | 0 | 360930 | 0 | 0 | 0 | 2149700 | 1162400 | 531530 | 0 | 0 | 0 | 0 |
| DVATVCRQ | MAT2B; MAT2B | Q9NZL9; H7 | MAT2B_HUMAN | Unmodified | 0 | 0 | 0 | 0 | 0 | 0 | 0 | 261390 | 0 | 0 | 112130 | 0 | 0 | 0 | 0 | 0 | 0 | 633440 |
| DVAVVAGG | ATP6AP1; ATP6 | Q15904; A0 | VAS1_HUMAN | Unmodified | 784860 | 904780 | 0 | 0 | 0 | 0 | 111150 | 86293 | 5511800 | 647990 | 375690 | 144540 | 0 | 0 | 0 | 0 | 0 | 0 |
| DVAYQYVK | SSBP1; SSBP1; | Q04837; C9I | SSBP_HUMAN | Unmodified | 0 | 0 | 0 | 0 | 0 | 0 | 0 | 0 | 629390 | 0 | 0 | 228510 | 0 | 0 | 0 | 0 | 0 | 0 |
| DVCTELLPI | CBR1 | P16152 | CBR1_HUMAN | Unmodified | 359880 | 187820 | 855150 | 1044000 | 1020100 | 210660 | 1201100 | 7153.7 | 2561600 | 130130 | 1069300 | 10603000 | 167930 | 0 | 19740 | 1013800 | 245740 | 317410 |
| DVDFELIK | CCT5; CCT5; C | P48643; B7Z | TCPE_HUMAN | Unmodified | 0 | 0 | 0 | 0 | 0 | 0 | 0 | 0 | 2762900 | 0 | 3481900 | 15687000 | 2210100 | 315550 | 0 | 0 | 0 | 590970 |
| DVDGFHVI | MTHFD2; MTHF | P13995; A0A | MTDC_HUMAN | Unmodified | 0 | 0 | 0 | 0 | 0 | 0 | 0 | 0 | 779770 | 0 | 0 | 0 | 0 | 0 | 0 | 0 | 0 | 0 |
| DVDGLDRI | CTNNA1; CTNN | P35221; G3 | CTNA1_HUMA | Unmodified | 1273800 | 1701500 | 4852400 | 8355000 | 2695000 | 1199200 | 0 | 0 | 1367500 | 118030 | 2176700 | 12615000 | 1374100 | 914730 | 10935000 | 9920600 | 0 | 0 |
| DVDGVTDI | MTHFD1L; MTH | Q6UB35; Q5 | C1TM_HUMAN | Unmodified | 0 | 0 | 0 | 0 | 0 | 0 | 0 | 254640 | 500040 | 0 | 0 | 0 | 132250 | 0 | 0 | 0 | 0 | 420840 |
| DVDLEFLAI | VCP; VCP; VCP; | P55072; C9J | TERA_HUMAN | Unmodified | 2085700 | 740380 | 373710 | 2215400 | 10868000 | 344500 | 710850 | 892910 | 5440400 | 4703000 | 5513000 | 20557000 | 10024000 | 3698400 | 0 | 4187400 | 1640300 | 318910 |
| DVDNFKLP | MMP15; MMP1 | P51511; H3E | MMP15_HUMA | Unmodified | 1605400 | 1648900 | 2709900 | 1534100 | 0 | 0 | 0 | 0 | 0 | 0 | 0 | 0 | 0 | 0 | 6767100 | 4021500 | 0 | 0 |
| DVDPDVIQ | UTRN | P46939 | UTRN_HUMAN | Unmodified | 0 | 0 | 766380 | 366880 | 256900 | 0 | 0 | 0 | 354760 | 0 | 0 | 0 | 0 | 0 | 683170 | 792700 | 0 | 0 |
| DVDRIIGLD | MYH9 | P35579 | MYH9_HUMAN | Oxidation (M) | 131860 | 0 | 284740 | 661490 | 10113000 | 0 | 0 | 0 | 355970 | 0 | 437630 | 1524700 | 0 | 0 | 2746800 | 318780 | 3780700 | 0 |
| DVDVNLFE | MAN1B1; MAN1 | Q9UKM7; H | MA1B1_HUMA | Unmodified | 0 | 0 | 0 | 0 | 0 | 0 | 0 | 0 | 2306400 | 0 | 0 | 0 | 0 | 0 | 0 | 0 | 381830 | 0 |
| DVEDESTG | SERPINB5 | P36952 | SPB5_HUMAN | Unmodified | 0 | 0 | 0 | 0 | 0 | 0 | 0 | 18287 | 0 | 0 | 0 | 62841 | 0 | 677740 | 98266 | 0 | 247720 | 376100 |
| DVEDMFSF | ELAVL1; ELAVL | Q15717; M0 | ELAV1_HUMAN | Unmodified | 0 | 0 | 0 | 0 | 0 | 0 | 0 | 0 | 0 | 0 | 0 | 2135400 | 0 | 0 | 72122 | 0 | 0 | 0 |
| DVEEGDEK | SBDS; SBDS | Q9Y3A5; A0 | SBDS_HUMAN | Unmodified | 0 | 0 | 0 | 0 | 0 | 0 | 0 | 0 | 0 | 0 | 0 | 654380 | 0 | 0 | 0 | 0 | 0 | 0 |
| DVEFLAQLI | PIP4K2B; PIP4K | P78356; P48 | PI42B_HUMAN | Unmodified | 184430 | 1069400 | 0 | 0 | 1824800 | 0 | 703020 | 647950 | 0 | 0 | 0 | 1149200 | 1106000 | 1124300 | 0 | 0 | 0 | 0 |
| DVEGSTSP | CSDE1; CSDE1 | O75534; E9F | CSDE1_HUMA | Unmodified | 0 | 0 | 0 | 0 | 961800 | 0 | 62843 | 29391 | 0 | 254230 | 0 | 1557500 | 127680 | 0 | 0 | 0 | 0 | 0 |
| DVEHNVSF | P2RX4; P2RX4 | Q99571; F6S | P2RX4_HUMAN | Unmodified | 0 | 0 | 0 | 0 | 0 | 0 | 0 | 0 | 4090500 | 651980 | 0 | 0 | 0 | 0 | 0 | 0 | 0 | 0 |
| DVELLYPV | EPHX1 | P07099 | HYEP_HUMAN | Unmodified | 0 | 0 | 391890 | 0 | 0 | 0 | 0 | 0 | 194420 | 0 | 0 | 0 | 0 | 0 | 465240 | 194740 | 0 | 0 |
| DVEPEDPM | MLF2; MLF2; M | Q15773; F5 | MLF2_HUMAN | 2 Oxidation (M) | 5093500 | 6281200 | 929240 | 0 | 0 | 0 | 0 | 0 | 0 | 0 | 0 | 0 | 0 | 0 | 0 | 1432200 | 40547 | 0 |
| DVEPEDPM | MLF2; MLF2; M | Q15773; F5 | MLF2_HUMAN | Oxidation (M) | 92000 | 623530 | 0 | 0 | 0 | 0 | 0 | 0 | 0 | 0 | 0 | 0 | 0 | 0 | 0 | 0 | 0 | 0 |
| DVEREGLD | COLGALT1; CO | Q8NBJ5; M0 | GT251_HUMAN | Unmodified | 0 | 0 | 0 | 0 | 322570 | 0 | 0 | 0 | 6304000 | 0 | 0 | 108370 | 0 | 0 | 0 | 0 | 0 | 0 |
| DVERTDGA | SERPINH1; SER | P50454; E9P | SERPH_HUMA | Oxidation (M) | 0 | 0 | 0 | 0 | 0 | 0 | 0 | 0 | 7523500 | 1906100 | 0 | 0 | 0 | 0 | 0 | 0 | 0 | 0 |
| DVERTDGA | SERPINH1; SER | P50454; E9P | SERPH_HUMA | Unmodified | 0 | 0 | 0 | 0 | 0 | 748480 | 0 | 0 | 39635000 | 703980 | 0 | 0 | 0 | 0 | 0 | 0 | 0 | 0 |
| DVESPSWF | MYO1C; MYO1 | O00159; F5 | MYO1C_HUMA | Unmodified | 1336500 | 0 | 0 | 2862000 | 513860 | 805560 | 0 | 0 | 269310 | 40237 | 415880 | 0 | 126140 | 0 | 5338300 | 4316100 | 110240 | 80367 |
| DVEVTKEE | SLC25A13 | Q9UJS0 | CMC2_HUMAN | Unmodified | 0 | 0 | 0 | 0 | 0 | 0 | 78576 | 49604 | 4651900 | 0 | 0 | 0 | 0 | 0 | 0 | 0 | 1054000 | 0 |
| DVFFYQAD | TUBG1; TUBG2 | P23258; Q9 | TBG1_HUMAN | Unmodified | 0 | 0 | 0 | 0 | 0 | 0 | 0 | 0 | 0 | 0 | 321690 | 4255700 | 0 | 0 | 677160 | 0 | 0 | 0 |
| DVFHMVVE | PPA1; PPA1 | Q15181; Q5 | IPYR_HUMAN | Unmodified | 0 | 0 | 0 | 0 | 0 | 0 | 0 | 0 | 0 | 0 | 661590 | 1995700 | 0 | 0 | 0 | 0 | 0 | 0 |
| DVFHMVVE | PPA1; PPA1 | Q15181; Q5 | IPYR_HUMAN | Oxidation (M) | 0 | 0 | 0 | 471460 | 0 | 0 | 0 | 0 | 172870 | 0 | 2802500 | 16145000 | 0 | 0 | 1403200 | 206720 | 0 | 0 |
| DVFHMVVE | PPA1; PPA1 | Q15181; Q5 | IPYR_HUMAN | Oxidation (M) | 0 | 77848 | 0 | 482000 | 1713400 | 0 | 0 | 0 | 0 | 0 | 1774800 | 0 | 602070 | 0 | 0 | 137980 | 0 | 824190 |
| DVFHNHGI | SLC30A1 | Q9Y6M5 | ZNT1_HUMAN | Unmodified | 1979600 | 448480 | 5904100 | 8624000 | 5562100 | 0 | 1313600 | 216650 | 1693600 | 175450 | 1498100 | 25495000 | 912740 | 367480 | 8251800 | 6365700 | 0 | 526850 |
| DVFISAAEF | PSMB1 | P20618 | PSB1_HUMAN | Unmodified | 551910 | 0 | 0 | 1049600 | 1342100 | 0 | 0 | 391870 | 6826600 | 775330 | 2523300 | 9934500 | 1466000 | 0 | 932330 | 692810 | 0 | 0 |
| DVFNMDFF | PLCB2; PLCB2 | Q00722; Q9 | PLCB2_HUMAN | Oxidation (M) | 0 | 0 | 0 | 0 | 0 | 0 | 0 | 0 | 0 | 0 | 0 | 126730 | 0 | 0 | 37400 | 0 | 0 | 0 |
| DVFREATT | FLNB; FLNB | O75369; E7E | FLNB_HUMAN | Unmodified | 0 | 0 | 1667500 | 2052700 | 16658000 | 0 | 0 | 0 | 357890 | 0 | 1022100 | 1416200 | 633680 | 1875900 | 0 | 0 | 1248500 | 1836500 |
| DVFSKPVN | PTGFRN | Q9P2B2 | FPRP_HUMAN | Unmodified | 0 | 0 | 0 | 0 | 0 | 0 | 0 | 0 | 0 | 0 | 0 | 0 | 0 | 0 | 207050 | 0 | 0 | 0 |
| DVFSPIGEF | NPEPPS; NPEP | P55786; A0A | PSA_HUMAN P | Unmodified | 0 | 0 | 0 | 0 | 0 | 0 | 0 | 0 | 0 | 0 | 0 | 665050 | 0 | 0 | 219590 | 172090 | 0 | 0 |
| DVGAQILLI | CNDP2 | Q96KP4 | CNDP2_HUMA | Unmodified | 100520 | 0 | 549010 | 274980 | 442130 | 0 | 79201 | 0 | 344460 | 138630 | 995290 | 4766200 | 1144700 | 3144400 | 649060 | 507390 | 0 | 0 |
| DVGFRAIAI | GART | P22102 | PUR2_HUMAN | Unmodified | 0 | 0 | 0 | 0 | 0 | 0 | 0 | 0 | 0 | 522340 | 0 | 0 | 1081300 | 0 | 0 | 0 | 0 | 0 |
| DVGHRDQ | CTNNA1; CTNN | P35221; E5R | CTNA1_HUMA | Unmodified | 1085900 | 1084400 | 5022300 | 538450 | 11602000 | 1679900 | 0 | 0 | 0 | 0 | 1032300 | 6982800 | 5498500 | 3119300 | 5980200 | 8521300 | 0 | 0 |
| DVGHRDQ | CTNNA1; CTNN | P35221; E5R | CTNA1_HUMA | Unmodified | 0 | 0 | 0 | 0 | 964810 | 0 | 0 | 0 | 0 | 0 | 0 | 0 | 437180 | 490510 | 0 | 0 | 0 | 0 |
| DVGLSDEE | TMCO3 | Q6UWJ1 | TMCO3_HUMA | Unmodified | 0 | 0 | 0 | 0 | 0 | 0 | 0 | 0 | 4478700 | 0 | 0 | 0 | 0 | 0 | 0 | 0 | 0 | 0 |
| DVGRIVIGL | PPIC | P45877 | PPIC_HUMAN | Unmodified | 0 | 0 | 10961 | 0 | 14865 | 0 | 61519 | 0 | 53743 | 316120 | 0 | 0 | 0 | 19150 | 0 | 0 | 0 | 0 |
| DVGRPQP | TAGLN2; TAGL | P37802; X6R | TAGL2_HUMAN | Unmodified | 8051600 | 6372400 | 0 | 4736900 | 53941000 | 397680 | 10336000 | 0 | 9817500 | 8512800 | 9976500 | 23814000 | 19284000 | 0 | 5379300 | 2682500 | 865000 | 679560 |
| DVGSLDEK | S100A13 | Q99584 | S10AD_HUMAN | Unmodified | 0 | 0 | 130680 | 0 | 12022000 | 2213700 | 0 | 0 | 569700 | 0 | 0 | 457090 | 0 | 0 | 0 | 106020 | 6418700 | 14348000 |
| DVGVKTSE | ERBIN | Q96RT1 | ERBIN_HUMAN | Unmodified | 0 | 0 | 0 | 0 | 274850 | 23023 | 629920 | 0 | 0 | 0 | 30266 | 0 | 0 | 0 | 0 | 0 | 201760 | 0 |
| DVHEGQPI | CDH2 | P19022 | CADH2_HUMA | Unmodified | 0 | 0 | 0 | 0 | 0 | 0 | 0 | 0 | 870660 | 0 | 672860 | 0 | 0 | 0 | 0 | 0 | 113320 | 172000 |
| DVHNIYGL | GANAB; GANA | Q14697; F5 | GANAB_HUMA | Oxidation (M) | 271770 | 0 | 291670 | 0 | 0 | 0 | 535820 | 0 | 7456200 | 0 | 0 | 0 | 0 | 0 | 0 | 307710 | 0 | 0 |
| DVHNIYGL | GANAB; GANA | Q14697; F5 | GANAB_HUMA | Unmodified | 0 | 0 | 0 | 0 | 0 | 0 | 0 | 62713 | 54554000 | 320660 | 64087 | 0 | 0 | 0 | 0 | 0 | 0 | 0 |
| DVHNIYGL | GANAB; GANA | Q14697; F5 | GANAB_HUMA | Unmodified | 0 | 0 | 0 | 0 | 0 | 0 | 0 | 0 | 23376000 | 947170 | 104420 | 0 | 0 | 430740 | 0 | 0 | 0 | 0 |
| DVHSLSEY | LFNG | Q8NES3 | LFNG_HUMAN | Unmodified | 0 | 0 | 0 | 0 | 0 | 0 | 0 | 0 | 743170 | 36044 | 0 | 0 | 0 | 0 | 0 | 0 | 0 | 0 |
| DVIAQSQS | EIF4A3; EIF4A3 | P38919; I3L | IF4A3_HUMAN | Unmodified | 0 | 0 | 188630 | 0 | 0 | 0 | 0 | 96756 | 0 | 0 | 0 | 1800100 | 0 | 0 | 0 | 0 | 0 | 0 |
| DVIATDKE | MDH1; MDH1; | P40925; C9J | MDHC_HUMAN | Unmodified | 288580 | 0 | 865030 | 0 | 160160 | 0 | 90652 | 0 | 3020600 | 340750 | 4543300 | 27429000 | 0 | 0 | 871760 | 574770 | 0 | 599950 |
| DVIEEYFK | RPS12 | P25398 | RS12_HUMAN | Unmodified | 0 | 0 | 0 | 4389500 | 0 | 0 | 1005200 | 659460 | 1896100 | 601900 | 4573400 | 38187000 | 3492200 | 0 | 0 | 0 | 0 | 0 |
| DVIELTDDS | PDIA6 | Q15084 | PDIA6_HUMAN | Unmodified | 0 | 0 | 301310 | 0 | 0 | 0 | 0 | 0 | 46680000 | 1822700 | 4683100 | 3656000 | 0 | 0 | 769710 | 516210 | 0 | 0 |
| DVIILGDFG | EEPD1 | Q7L9B9 | EEPD1_HUMAN | Unmodified | 0 | 0 | 0 | 0 | 0 | 0 | 0 | 0 | 734010 | 128900 | 0 | 0 | 0 | 517570 | 1679300 | 716990 | 982080 | 2592900 |
| DVILLGDF | DNASE1L1; DN | P49184; A6 | DNSL1_HUMA | Unmodified | 90976 | 149650 | 0 | 50391 | 0 | 0 | 0 | 0 | 0 | 0 | 0 | 0 | 0 | 0 | 0 | 0 | 0 | 0 |
| DVIQAIKET | PLCB4; PLCB4 | Q15147; A0 | PLCB4_HUMAN | Unmodified | 0 | 0 | 0 | 0 | 0 | 0 | 0 | 0 | 0 | 0 | 0 | 0 | 0 | 0 | 0 | 0 | 704720 | 0 |
| DVIRRPTFV | ZC3HAV1; ZC3 | Q7Z2W4; C9 | ZCCHV_HUMA | Unmodified | 0 | 0 | 0 | 96935 | 2971900 | 0 | 0 | 0 | 0 | 0 | 0 | 0 | 127040 | 0 | 0 | 0 | 113910 | 0 |
| DVIYAVRD | PPAT | Q06203 | PUR1_HUMAN | Unmodified | 0 | 0 | 0 | 0 | 0 | 0 | 0 | 0 | 0 | 0 | 0 | 2760400 | 0 | 0 | 0 | 0 | 0 | 0 |
| DVKDTILH | GNA13 | Q14344 | GNA13_HUMA | Unmodified | 0 | 22771 | 0 | 0 | 0 | 0 | 0 | 0 | 571170 | 0 | 0 | 2506200 | 0 | 0 | 0 | 0 | 0 | 0 |
| DVKGESLF | EPCAM; EPCAM | P16422; B5 | EPCAM_HUMA | Unmodified | 27557000 | 18160000 | 12798000 | 27687000 | 19861000 | 3497700 | 0 | 0 | 64575000 | 50771000 | 13442000 | 29611000 | 76330000 | 26912000 | 32016000 | 34727000 | 1488300 | 2880500 |

| | | | | | | | | | | | | | | | | | | | | | |
|---|---|---|---|---|---|---|---|---|---|---|---|---|---|---|---|---|---|---|---|---|---|
| DVKPDNFL | CSNK1D; CSNK | P48730; P49 | KC1D_HUMAN | Oxidation (M) | 229520 | 312920 | 574260 | 464040 | 0 | 0 | 0 | 0 | 0 | 0 | 0 | 562450 | 0 | 0 | 369640 | 436520 | 0 | 0 |
| DVKPDNMI | ROCK2; ROCK1 | O75116; Q1 | ROCK2_HUMA | Unmodified | 39125 | 0 | 22224 | 49337 | 0 | 0 | 0 | 0 | 327160 | 0 | 0 | 633390 | 0 | 0 | 138840 | 67229 | 16583 | 0 |
| DVKPENFL | CSNK1G3 | Q9Y6M4 | KC1G3_HUMA | Unmodified | 280510 | 374310 | 494000 | 217460 | 0 | 0 | 0 | 0 | 0 | 0 | 0 | 3875500 | 0 | 0 | 0 | 473810 | 0 | 0 |
| DVKPENFL | CSNK1G2; CSN | P78368; K7E | KC1G2_HUMA | Unmodified | 188280 | 127530 | 338890 | 137150 | 0 | 0 | 0 | 0 | 383830 | 0 | 51648 | 0 | 0 | 0 | 624220 | 441380 | 0 | 0 |
| DVKPHCVL | CASK; CASK; C | O14936; A0 | CSKP_HUMAN | Unmodified | 144200 | 0 | 455440 | 781910 | 0 | 0 | 0 | 0 | 1599200 | 143810 | 0 | 556900 | 0 | 0 | 1049900 | 933180 | 0 | 89520 |
| DVKPHSLD | TM9SF1; TM9SF | O15321; E9F | TM9S1_HUMA | Unmodified | 0 | 0 | 0 | 0 | 0 | 0 | 0 | 0 | 43835000 | 1943100 | 0 | 2853100 | 0 | 0 | 0 | 75678 | 0 | 0 |
| DVKQFYDC | CD81; CD81; C | P60033; E9P | CD81_HUMAN | Unmodified | 0 | 221750 | 0 | 0 | 869210 | 0 | 0 | 0 | 0 | 907070 | 0 | 0 | 5000700 | 1089600 | 300350 | 0 | 586390 | 746600 |
| DVLAEIPEC | CPNE8; CPNE8 | Q86YQ8; E7I | CPNE8_HUMA | Oxidation (M) | 0 | 0 | 0 | 0 | 351630 | 75405 | 0 | 0 | 0 | 0 | 0 | 0 | 0 | 0 | 0 | 0 | 0 | 49437 |
| DVLAQLPS | TPBG | Q13641 | TPBG_HUMAN | Unmodified | 242630 | 204860 | 0 | 0 | 0 | 0 | 0 | 0 | 2217400 | 0 | 424130 | 20480000 | 0 | 0 | 0 | 0 | 0 | 0 |
| DVLDIEQFS | GRK6; GRK5; G | P43250; P34 | GRK6_HUMAN | Unmodified | 19015 | 0 | 29254 | 63069 | 256320 | 0 | 0 | 0 | 500750 | 0 | 1403900 | 2126700 | 0 | 0 | 51068 | 40255 | 100950 | 384210 |
| DVLEGYNC | KIF5B; KIF5C; K | P33176; O6 | KINH_HUMAN | Unmodified | 0 | 69737 | 0 | 0 | 1097200 | 0 | 0 | 0 | 1027600 | 0 | 0 | 5118000 | 0 | 135970 | 47071 | 0 | 0 | 1061400 |
| DVLETFTVK | CD9; CD9; CDS | P21926; A0A | CD9_HUMAN ( | Unmodified | 3.34E+08 | 2.32E+08 | 9.61E+08 | 4.94E+08 | 62817000 | 18621000 | 2778400 | 140130 | 3.84E+08 | 33488000 | 40121000 | 1.31E+08 | 1.2E+08 | 28879000 | 1.54E+09 | 9.89E+08 | 3078700 | 21832000 |
| DVLEVGEL | NAA50; NAA50; | Q9GZZ1; E7I | NAA50_HUMA | Unmodified | 0 | 0 | 0 | 0 | 141260 | 0 | 0 | 196980 | 0 | 0 | 387880 | 9270000 | 141240 | 0 | 153680 | 85603 | 0 | 0 |
| DVLFLKDC | PGK1 | P00558 | PGK1_HUMAN | Unmodified | 0 | 0 | 0 | 0 | 596150 | 46610 | 490950 | 323260 | 0 | 406870 | 0 | 0 | 545990 | 459480 | 0 | 0 | 4382300 | 1601500 |
| DVLILGGG | SMS; SMS | P52788; H7 | SPSY_HUMAN | Unmodified | 0 | 0 | 0 | 0 | 0 | 111500 | 0 | 0 | 0 | 0 | 20705 | 70456 | 66058 | 0 | 0 | 0 | 0 | 0 |
| DVLKQQLS | CEP55 | Q53EZ4 | CEP55_HUMA | Unmodified | 0 | 0 | 0 | 0 | 2070700 | 0 | 0 | 0 | 0 | 412850 | 0 | 38521 | 0 | 0 | 0 | 0 | 300720 | 0 |
| DVLLPWYN | IARS1; IARS1; I | P41252; A0A | SYIC_HUMAN | Unmodified | 0 | 0 | 0 | 0 | 0 | 0 | 0 | 0 | 0 | 0 | 0 | 5101800 | 0 | 0 | 0 | 0 | 0 | 0 |
| DVLLQVDD | MYH9 | P35579 | MYH9_HUMAN | Unmodified | 0 | 0 | 2233700 | 0 | 0 | 0 | 0 | 0 | 0 | 0 | 495630 | 8255800 | 0 | 0 | 1686800 | 1415900 | 0 | 0 |
| DVLLQVDD | MYH9 | P35579 | MYH9_HUMAN | Unmodified | 0 | 0 | 0 | 0 | 4671400 | 0 | 0 | 0 | 0 | 0 | 687710 | 0 | 1599600 | 0 | 0 | 0 | 489100 | 1190800 |
| DVLNDSEN | UTRN | P46939 | UTRN_HUMAN | Unmodified | 0 | 0 | 352690 | 201780 | 0 | 62130 | 0 | 0 | 0 | 117320 | 0 | 546880 | 86852 | 65937 | 579640 | 420080 | 0 | 0 |
| DVLQAETS | SNX6; SNX6 | Q9UNH7; A | SNX6_HUMAN | Unmodified | 0 | 0 | 31949 | 0 | 0 | 0 | 0 | 0 | 0 | 0 | 82567 | 898120 | 0 | 0 | 143610 | 58545 | 0 | 0 |
| DVLQEPKT | MME; MME; MM | P08473; A0A | NEP_HUMAN | Unmodified | 41949 | 0 | 75859000 | 1.28E+08 | 3262200 | 426930 | 0 | 0 | 57178000 | 6363600 | 1607400 | 871430 | 55795000 | 20764000 | 1.31E+08 | 98935000 | 0 | 0 |
| DVLQGELE | SPAG9; SPAG9; | O60271; H0 | JIP4_HUMAN C | Unmodified | 430700 | 0 | 0 | 0 | 0 | 0 | 0 | 0 | 226360 | 0 | 0 | 500660 | 0 | 0 | 0 | 0 | 0 | 0 |
| DVLSVAFS | RACK1; RACK1; | P63244; D6F | RACK1_HUMA | Unmodified | 0 | 0 | 0 | 0 | 0 | 0 | 0 | 0 | 0 | 0 | 0 | 2310000 | 0 | 0 | 0 | 0 | 0 | 0 |
| DVLSVAFS | RACK1; RACK1; | P63244; D6F | RACK1_HUMA | Unmodified | 269950 | 0 | 0 | 1777900 | 6036600 | 0 | 5482600 | 1913100 | 423000 | 1655100 | 2040000 | 161140 | 2177400 | 632710 | 0 | 0 | 1667400 | 2249000 |
| DVLSYHIPF | NCKAP1 | Q9Y2A7 | NCKP1_HUMA | Unmodified | 0 | 0 | 37715 | 0 | 0 | 0 | 0 | 0 | 0 | 0 | 0 | 0 | 0 | 0 | 145310 | 31631 | 0 | 0 |
| DVLTGQEF | GPD2; GPD2 | P43304; F5C | GPDM_HUMA | Unmodified | 103740 | 0 | 0 | 0 | 0 | 0 | 0 | 0 | 7482300 | 490040 | 0 | 0 | 0 | 0 | 0 | 0 | 0 | 0 |
| DVLTGQEF | GPD2; GPD2 | P43304; F5C | GPDM_HUMA | Unmodified | 0 | 0 | 0 | 0 | 0 | 501560 | 995520 | 0 | 1157100 | 3362300 | 0 | 1314200 | 0 | 0 | 0 | 0 | 0 | 0 |
| DVLTSTYG | EPB41L1; EPB4 | Q9H4G0; H | E41L1_HUMA | Unmodified | 0 | 0 | 0 | 2894000 | 3809900 | 732960 | 297350 | 0 | 0 | 0 | 2334200 | 2255600 | 1714800 | 4122700 | 1947300 | 1080800 | 1481500 | 0 |
| DVLYSQPP | REPS2 | Q8NFH8 | REPS2_HUMA | Unmodified | 0 | 0 | 0 | 0 | 0 | 0 | 0 | 0 | 147230 | 0 | 0 | 0 | 0 | 0 | 597810 | 275130 | 0 | 0 |
| DVMPGPR | ADGRE5 | P48960 | AGRE5_HUMA | Unmodified | 0 | 0 | 0 | 25984 | 873180 | 0 | 0 | 0 | 0 | 135710 | 0 | 0 | 0 | 0 | 0 | 0 | 0 | 1122100 |
| DVNAAIAA | TUBA4A | P68366 | TBA4A_HUMA | Unmodified | 2489600 | 2809800 | 6090300 | 7225200 | 10477000 | 2823400 | 11871000 | 6460300 | 266160 | 8210900 | 1082100 | 16854000 | 19813000 | 19498000 | 5342600 | 4927200 | 2837400 | 16065000 |
| DVNAAIATI | TUBA3C; TUBA | P0DPH7; Q7 | TBA3C_HUMA | Unmodified | 15805000 | 10139000 | 26796000 | 24088000 | 2.22E+08 | 28520000 | 1.95E+08 | 1.54E+08 | 1.36E+08 | 1.07E+08 | 77413000 | 5.94E+08 | 1.05E+08 | 53627000 | 10361000 | 20897000 | 51974000 | 1.4E+08 |
| DVNAAIATI | TUBA3C; TUBA | P0DPH7; Q7 | TBA3C_HUMA | Unmodified | 0 | 0 | 0 | 0 | 3718700 | 0 | 9852600 | 618830 | 0 | 621290 | 0 | 273100 | 852260 | 1074400 | 0 | 0 | 4104700 | 1195900 |
| DVNAAIATI | TUBA3C; TUBA | P0DPH7; Q7 | TBA3C_HUMA | Unmodified | 0 | 0 | 0 | 0 | 447800 | 0 | 1125500 | 808220 | 0 | 0 | 0 | 0 | 252790 | 350790 | 0 | 0 | 703050 | 0 |
| DVNGPEPL | CHP1; CHP1; C | Q99653; F5 | CHP1_HUMAN | Unmodified | 240090 | 143690 | 1122300 | 669930 | 0 | 0 | 0 | 0 | 0 | 0 | 0 | 2111100 | 490720 | 0 | 1437600 | 1125900 | 0 | 0 |
| DVNGPEPL | CHP1; CHP1; C | Q99653; F5 | CHP1_HUMAN | Unmodified | 0 | 0 | 0 | 0 | 1499700 | 480280 | 0 | 0 | 0 | 0 | 0 | 0 | 1168200 | 677680 | 0 | 0 | 157310 | 359260 |
| DVNLRPEC | ACAA1; ACAA1; | P09110; C9J | THIK_HUMAN | Oxidation (M) | 0 | 0 | 0 | 0 | 0 | 0 | 0 | 0 | 9442700 | 5516100 | 0 | 0 | 0 | 0 | 0 | 0 | 0 | 0 |
| DVNLRPEC | ACAA1; ACAA1; | P09110; C9J | THIK_HUMAN | Unmodified | 0 | 0 | 0 | 0 | 0 | 0 | 0 | 0 | 7872700 | 0 | 0 | 0 | 0 | 0 | 0 | 0 | 0 | 0 |
| DVNNWWI | POMT1; POMT1 | Q9Y6A1; A0 | POMT1_HUMA | Unmodified | 0 | 0 | 0 | 0 | 0 | 0 | 0 | 0 | 1040800 | 0 | 0 | 0 | 0 | 0 | 0 | 0 | 0 | 0 |
| DVNQQEF | RPS19; RPS19; | P39019; M0 | RS19_HUMAN | Unmodified | 3403100 | 2674900 | 2574700 | 2851900 | 0 | 0 | 0 | 0 | 1337500 | 474340 | 2747100 | 33684000 | 1603200 | 0 | 1034500 | 1856400 | 0 | 0 |
| DVNQQEF | RPS19; RPS19; | P39019; M0 | RS19_HUMAN | Unmodified | 41286 | 0 | 0 | 0 | 3594600 | 529330 | 3952700 | 0 | 0 | 1484500 | 0 | 0 | 2287900 | 2894800 | 0 | 0 | 740090 | 1985400 |
| DVNQQEF | RPS19; RPS19; | P39019; M0 | RS19_HUMAN | Unmodified | 23682 | 0 | 27831 | 35330 | 3170100 | 537750 | 2788200 | 0 | 0 | 1938800 | 0 | 70154 | 7450100 | 4833600 | 0 | 11208 | 1194300 | 2578500 |
| DVNVENVN | MTTP; MTTP; M | P55157; E9P | MTP_HUMAN | Unmodified | 0 | 0 | 0 | 0 | 76774 | 0 | 0 | 140300 | 189720 | 58377 | 20380 | 227260 | 0 | 0 | 0 | 0 | 48849 | 113200 |
| DVPAVEITH | MAOA | P21397 | AOFA_HUMAN | Unmodified | 0 | 0 | 544190 | 40220 | 0 | 0 | 0 | 0 | 0 | 0 | 0 | 0 | 0 | 0 | 399660 | 142320 | 0 | 0 |
| DVPAYSQD | RPN1; RPN1 | P04843; B7Z | RPN1_HUMAN | Unmodified | 0 | 0 | 0 | 0 | 0 | 0 | 0 | 0 | 393760 | 49510 | 0 | 1070800 | 0 | 0 | 0 | 0 | 0 | 0 |
| DVPFGFQT | SERPINB5; SER | P36952; C9J | SPB5_HUMAN | Unmodified | 278940 | 351170 | 0 | 0 | 0 | 0 | 0 | 0 | 0 | 0 | 0 | 0 | 0 | 737240 | 64151 | 70635 | 830120 | 302410 |
| DVPITSVRF | TMEM131 | Q92545 | TM131_HUMA | Unmodified | 0 | 0 | 0 | 0 | 0 | 0 | 0 | 0 | 2204100 | 0 | 0 | 0 | 0 | 0 | 0 | 0 | 0 | 0 |
| DVPIVHTEI | EPB41; EPB41; | P11171; A0A | EPB41_HUMA | Unmodified | 48409 | 69773 | 0 | 314970 | 268820 | 0 | 332370 | 354720 | 1042800 | 0 | 0 | 302250 | 333670 | 0 | 275620 | 180890 | 0 | 529640 |
| DVPLGTPL | DLAT; DLAT; DL | P10515; A0A | ODP2_HUMAN | Unmodified | 0 | 0 | 0 | 0 | 0 | 0 | 0 | 0 | 634500 | 0 | 0 | 0 | 0 | 0 | 0 | 0 | 0 | 0 |
| DVPLHALY | ASNS; ASNS | P08243; F8V | ASNS_HUMAN | Unmodified | 0 | 0 | 0 | 0 | 279430 | 0 | 0 | 0 | 824660 | 155480 | 582520 | 5484900 | 0 | 0 | 0 | 0 | 0 | 271810 |
| DVPLLPSD | MOV10; MOV1 | Q9HCE1; Q5 | MOV10_HUMA | Unmodified | 8702100 | 6452300 | 2739000 | 1722600 | 0 | 0 | 0 | 0 | 0 | 142930 | 0 | 0 | 0 | 0 | 3665400 | 4823700 | 0 | 0 |
| DVPVAEEV | AP3D1; AP3D1 | O14617; A0 | AP3D1_HUMA | Unmodified | 0 | 0 | 0 | 0 | 113170 | 0 | 25642 | 0 | 0 | 0 | 0 | 0 | 0 | 0 | 0 | 0 | 0 | 31879 |
| DVPVVAILC | PLA2G4A | P47712 | PA24A_HUMA | Unmodified | 0 | 0 | 0 | 0 | 0 | 0 | 0 | 0 | 0 | 0 | 293680 | 1502400 | 0 | 0 | 0 | 12886 | 0 | 0 |
| DVPYPPPL | PTGR2; N/A; PT | Q8N8N7; G3 | PTGR2_HUMA | Unmodified | 0 | 0 | 0 | 0 | 0 | 0 | 0 | 0 | 0 | 0 | 0 | 0 | 0 | 0 | 0 | 0 | 0 | 0 |
| DVQDSLTV | CAPZA1 | P52907 | CAZA1_HUMA | Unmodified | 1161600 | 1205000 | 958550 | 474480 | 1309200 | 0 | 0 | 0 | 737750 | 0 | 1003300 | 8518500 | 823270 | 0 | 974200 | 1724300 | 28444 | 990950 |
| DVQEIFR | DDX39B; DDX3 | Q13838; O0 | DX39B_HUMA | Unmodified | 0 | 291880 | 0 | 325540 | 0 | 0 | 0 | 47971 | 790700 | 76423 | 882400 | 9379900 | 0 | 0 | 0 | 0 | 0 | 0 |
| DVQEIFRLT | DDX39A; DDX3 | O00148; K7E | DX39A_HUMA | Unmodified | 0 | 0 | 0 | 0 | 0 | 0 | 645420 | 0 | 0 | 0 | 0 | 0 | 193230 | 0 | 0 | 0 | 0 | 0 |
| DVQELLTQ | SRP54; SRP54; | P61011; G3 | SRP54_HUMA | Unmodified | 0 | 0 | 0 | 0 | 0 | 0 | 0 | 0 | 0 | 0 | 0 | 1337600 | 292690 | 0 | 0 | 0 | 0 | 0 |
| DVQEQLSF | ROCK2; ROCK2 | O75116; Q1 | ROCK2_HUMA | Unmodified | 0 | 0 | 0 | 0 | 0 | 0 | 0 | 0 | 0 | 0 | 0 | 0 | 0 | 0 | 0 | 0 | 0 | 0 |
| DVQFEHG | TM9SF4; TM9SF | Q92544; A0 | TM9S4_HUMA | Unmodified | 0 | 0 | 0 | 0 | 0 | 0 | 0 | 0 | 3362700 | 107750 | 0 | 0 | 0 | 0 | 0 | 0 | 0 | 0 |
| DVQGTDAS | STOML2; STOM | Q9UJZ1; A0 | STML2_HUMA | Unmodified | 0 | 0 | 0 | 0 | 0 | 0 | 0 | 0 | 3185000 | 60594 | 255610 | 0 | 0 | 0 | 0 | 0 | 0 | 0 |
| DVQIGDIV | RPS11; RPS11 | P62280; M0 | RS11_HUMAN | Unmodified | 7695500 | 2496800 | 2382800 | 0 | 757850 | 0 | 760830 | 39098 | 0 | 514850 | 2364300 | 47969000 | 833190 | 0 | 1994500 | 1619800 | 463220 | 1151100 |
| DVQLIRSV | PLEC; PLEC | Q15149; H0 | PLEC_HUMAN | Unmodified | 0 | 0 | 0 | 0 | 153720 | 0 | 0 | 0 | 0 | 37919 | 0 | 0 | 0 | 0 | 0 | 0 | 374420 | 0 |
| DVQMLQD | CDC37; CDC37 | Q16543; K7E | CDC37_HUMA | Unmodified | 0 | 0 | 0 | 0 | 321640 | 0 | 0 | 0 | 354540 | 0 | 0 | 619910 | 48744 | 0 | 0 | 0 | 0 | 1111600 |
| DVQNFPA | DIAPH1; DIAPH | O60610; A0 | DIAP1_HUMA | Unmodified | 0 | 0 | 0 | 0 | 0 | 0 | 0 | 0 | 0 | 0 | 212490 | 394030 | 0 | 0 | 0 | 0 | 0 | 0 |
| DVQTALAL | HSD17B10; HS | Q99714; Q5 | HCD2_HUMAN | Unmodified | 0 | 0 | 0 | 0 | 0 | 0 | 1508100 | 224260 | 12436000 | 3475600 | 0 | 0 | 504220 | 0 | 0 | 0 | 0 | 350850 |
| DVRDIFER | PDCD11; PDCD | Q14690; A0 | RRP5_HUMAN | Unmodified | 0 | 0 | 0 | 0 | 0 | 0 | 205400 | 0 | 0 | 0 | 0 | 0 | 0 | 0 | 0 | 0 | 0 | 0 |
| DVRENPRA | HYOU1; HYOU | Q9Y4L1; E9F | HYOU1_HUMA | Unmodified | 0 | 22722 | 0 | 0 | 0 | 0 | 1306800 | 30798 | 0 | 10001000 | 414200 | 0 | 0 | 0 | 0 | 0 | 0 | 0 |
| DVRPYIVG | LRRC47 | Q8N1G4 | LRC47_HUMA | Unmodified | 0 | 0 | 276090 | 0 | 0 | 0 | 0 | 50939 | 0 | 0 | 0 | 957940 | 0 | 0 | 0 | 268140 | 0 | 0 |
| DVRPYIVG | LRRC47 | Q8N1G4 | LRC47_HUMA | Unmodified | 0 | 0 | 0 | 0 | 181760 | 0 | 0 | 0 | 0 | 0 | 0 | 0 | 336010 | 185810 | 0 | 0 | 0 | 134160 |
| DVSELTGFI | ATIC; ATIC | P31939; F8V | PUR9_HUMAN | Unmodified | 0 | 0 | 0 | 0 | 0 | 0 | 0 | 0 | 218550 | 0 | 0 | 173820 | 0 | 0 | 0 | 0 | 0 | 0 |
| DVSELTGFI | ATIC; ATIC | P31939; F8V | PUR9_HUMAN | Oxidation (M) | 76499 | 0 | 118040 | 0 | 0 | 0 | 0 | 0 | 99278 | 0 | 712120 | 2533400 | 0 | 0 | 219920 | 197280 | 0 | 0 |
| DVSGPMPI | VARS1; VARS1; | P26640; A0A | SYVC_HUMAN | Oxidation (M) | 0 | 0 | 47076 | 0 | 0 | 0 | 0 | 0 | 0 | 0 | 0 | 663740 | 0 | 0 | 0 | 30973 | 0 | 0 |
| DVSGPMPI | VARS1; VARS1; | P26640; A0A | SYVC_HUMAN | Unmodified | 0 | 0 | 0 | 0 | 0 | 0 | 0 | 0 | 0 | 0 | 0 | 2963200 | 0 | 0 | 102970 | 0 | 0 | 0 |
| DVSLGDWI | TNKS1BP1; TNI | Q9C0C2; A0 | TB182_HUMA | Unmodified | 45724 | 0 | 181950 | 140230 | 0 | 0 | 0 | 0 | 0 | 30750 | 61004 | 0 | 0 | 0 | 314330 | 0 | 103230 | 0 |
| DVSLLTQK | CMBL | Q96DG6 | CMBL_HUMAN | Unmodified | 91680 | 43948 | 0 | 0 | 37816 | 0 | 32813 | 0 | 370130 | 0 | 0 | 3561200 | 0 | 0 | 0 | 104780 | 491460 | 0 |
| DVSLSSLR | DIP2A; DIP2B; I | Q14689; Q9 | DIP2A_HUMA | Unmodified | 347250 | 448960 | 361650 | 557680 | 0 | 0 | 0 | 0 | 0 | 283810 | 193660 | 2955200 | 0 | 0 | 1102300 | 981140 | 0 | 0 |
| DVSSLFPD | AP2B1; AP2B1; | P63010; K7E | AP2B1_HUMA | Oxidation (M) | 0 | 0 | 0 | 0 | 0 | 0 | 0 | 0 | 0 | 0 | 0 | 0 | 0 | 0 | 0 | 0 | 297890 | 700820 |
| DVSSSTPP | SLC39A6 | Q13433 | S39A6_HUMA | Unmodified | 0 | 0 | 0 | 0 | 0 | 0 | 0 | 0 | 581210 | 0 | 0 | 0 | 0 | 0 | 0 | 0 | 0 | 0 |
| DVSSSVYS | KIT | P10721 | KIT_HUMAN Ma | Unmodified | 0 | 0 | 0 | 0 | 0 | 0 | 0 | 0 | 1688400 | 0 | 0 | 0 | 0 | 0 | 0 | 0 | 104730 | 227760 |
| DVSSVELLI | SPTBN1; SPTBN | Q01082; A0 | SPTB2_HUMA | Unmodified | 0 | 0 | 0 | 0 | 341080 | 0 | 0 | 0 | 0 | 0 | 0 | 1210300 | 0 | 0 | 0 | 0 | 426310 | 0 |
| DVSSVELLI | SPTBN1; SPTBN | Q01082; A0 | SPTB2_HUMA | Oxidation (M) | 0 | 0 | 0 | 0 | 758990 | 0 | 0 | 0 | 0 | 0 | 0 | 1052700 | 0 | 0 | 0 | 0 | 391680 | 0 |
| DVTADFEG | DPP7 | Q9UHL4 | DPP2_HUMAN | Unmodified | 88154 | 0 | 0 | 0 | 0 | 0 | 606610 | 0 | 739090 | 0 | 0 | 0 | 0 | 0 | 0 | 0 | 0 | 0 |
| DVTEESVTE | OCC1; C12orf7 | Q8TAD7; F8 | OCC1_HUMAN | Unmodified | 553660 | 555750 | 0 | 0 | 0 | 0 | 0 | 0 | 380360 | 0 | 0 | 272020 | 0 | 0 | 309070 | 160440 | 48216 | 322330 |
| DVTEESVTE | OCC1; C12orf7 | Q8TAD7; F8 | OCC1_HUMAN | Unmodified | 1656300 | 1868700 | 2031600 | 1714000 | 162300 | 33500 | 0 | 0 | 180370 | 21614 | 0 | 0 | 103290 | 0 | 0 | 1597000 | 69494 | 122250 |
| DVTEESVTE | OCC1; C12orf7 | Q8TAD7; F8 | OCC1_HUMAN | Unmodified | 3573200 | 4460800 | 0 | 0 | 0 | 0 | 0 | 0 | 1326500 | 0 | 0 | 0 | 189970 | 0 | 0 | 0 | 0 | 0 |
| DVTENLET | CREB3L4 | Q8TEY5 | CR3L4_HUMA | Unmodified | 0 | 0 | 0 | 0 | 0 | 4155800 | 0 | 0 | 453970 | 0 | 0 | 0 | 0 | 0 | 0 | 0 | 0 | 225060 |
| DVTGAEAL | SPTAN1; SPTAN | Q13813; A0 | SPTN1_HUMA | Unmodified | 0 | 0 | 0 | 0 | 0 | 0 | 0 | 0 | 0 | 0 | 131580 | 4591800 | 0 | 0 | 0 | 0 | 0 | 0 |
| DVTISKSEY | PHYH; PHYH; P | O14832; B1 | PAHX_HUMAN | Unmodified | 0 | 0 | 0 | 0 | 0 | 0 | 0 | 0 | 1791400 | 304640 | 0 | 0 | 0 | 0 | 0 | 0 | 0 | 0 |
| DVTNNVH | SEPTIN7; SEPTI | Q16181; E7E | SEPT7_HUMA | Unmodified | 0 | 0 | 909190 | 1916100 | 17212000 | 5516400 | 862070 | 0 | 1930200 | 77797 | 1267900 | 8076900 | 2929000 | 2920200 | 1001000 | 1420500 | 2946900 | 4005300 |
| DVTNNVH | SEPTIN7; SEPTI | Q16181; E7E | SEPT7_HUMA | Unmodified | 0 | 0 | 0 | 0 | 8630300 | 713700 | 600470 | 79719 | 0 | 198280 | 0 | 0 | 1546200 | 266230 | 0 | 0 | 484690 | 618420 |
| DVTPPPETI | MAP4; MAP4 | P27816; E7E | MAP4_HUMAN | Unmodified | 0 | 0 | 0 | 0 | 0 | 0 | 0 | 0 | 0 | 0 | 843200 | 1102500 | 0 | 0 | 0 | 0 | 0 | 0 |
| DVTQIFNN | CAB39L; CAB3 | Q9H9S4; B7 | CB39L_HUMA | Unmodified | 41259 | 56722 | 331770 | 424470 | 0 | 0 | 0 | 0 | 29292 | 0 | 227280 | 1234200 | 88142 | 39616 | 657730 | 668010 | 0 | 0 |
| DVTRGQA | CBR1; CBR1; C | P16152; E9P | CBR1_HUMAN | Unmodified | 0 | 0 | 0 | 842820 | 0 | 0 | 0 | 0 | 1481900 | 0 | 0 | 2711300 | 501330 | 0 | 1890700 | 829160 | 0 | 0 |
| DVTYEEAK | RAB14; RAB14 | P61106; X6R | RAB14_HUMA | Unmodified | 0 | 0 | 0 | 0 | 0 | 0 | 0 | 0 | 390520 | 0 | 0 | 2362000 | 0 | 0 | 0 | 0 | 0 | 0 |
| DVVAQQES | TAF2 | Q6P1X5 | TAF2_HUMAN | Unmodified | 0 | 0 | 0 | 0 | 870930 | 83061000 | 1.07E+08 | 66637000 | 0 | 0 | 0 | 0 | 10800000 | 1.04E+08 | 0 | 0 | 815680 | 2574000 |
| DVVEPLLRI | VARS1; VARS1 | P26640; A0A | SYVC_HUMAN | Unmodified | 0 | 0 | 563340 | 0 | 0 | 0 | 224160 | 0 | 0 | 81216 | 1467700 | 11157000 | 0 | 0 | 843670 | 399530 | 163290 | 0 |
| DVVFNYLH | UQCRC1 | P31930 | QCR1_HUMAN | Unmodified | 0 | 0 | 0 | 0 | 0 | 0 | 0 | 0 | 3874400 | 0 | 0 | 0 | 0 | 0 | 0 | 0 | 0 | 0 |
| DVVFNYLH | UQCRC1 | P31930 | QCR1_HUMAN | Unmodified | 0 | 0 | 0 | 0 | 0 | 0 | 0 | 0 | 23158000 | 0 | 0 | 0 | 0 | 0 | 0 | 0 | 913580 | 0 |
| DVVHLDSE | PDIA5; PDIA5 | Q14554; C9J | PDIA5_HUMA | Unmodified | 0 | 0 | 0 | 0 | 0 | 0 | 0 | 0 | 1582900 | 139830 | 0 | 0 | 0 | 0 | 0 | 0 | 0 | 0 |
| DVVICPDA | PARK7; PARK7; | Q99497; K7E | PARK7_HUMA | Unmodified | 1298400 | 230480 | 838860 | 921750 | 0 | 0 | 0 | 0 | 6065000 | 1614900 | 1829400 | 8792200 | 0 | 0 | 1627500 | 1720600 | 0 | 0 |
| DVVICPDA | PARK7; PARK7 | Q99497; K7E | PARK7_HUMA | Unmodified | 0 | 0 | 0 | 0 | 0 | 0 | 0 | 0 | 3088700 | 0 | 0 | 0 | 0 | 0 | 0 | 0 | 0 | 0 |
| DVVKVGAV | PDIA6 | Q15084 | PDIA6_HUMA | Unmodified | 0 | 0 | 0 | 0 | 347270 | 0 | 0 | 0 | 91743 | 319040 | 37528 | 0 | 0 | 0 | 0 | 0 | 0 | 187490 |
| DVVKVGAV | PDIA6 | Q15084 | PDIA6_HUMA | Unmodified | 0 | 0 | 0 | 0 | 0 | 0 | 1187400 | 0 | 1689900 | 16173000 | 0 | 0 | 0 | 0 | 0 | 0 | 408920 | 0 |
| DVVLPTYDI | POGLUT3 | Q7Z4H8 | PLGT3_HUMA | Oxidation (M) | 0 | 0 | 0 | 0 | 0 | 0 | 0 | 0 | 1731700 | 166370 | 0 | 0 | 0 | 3422900 | 0 | 0 | 0 | 0 |
| DVVLPTYDI | POGLUT3 | Q7Z4H8 | PLGT3_HUMA | Unmodified | 0 | 0 | 0 | 0 | 0 | 0 | 22998 | 0 | 3580700 | 0 | 0 | 0 | 0 | 0 | 0 | 0 | 0 | 21108 |
| DVVQAYPE | DHX9 | Q08211 | DHX9_HUMAN | Unmodified | 0 | 0 | 0 | 0 | 0 | 0 | 0 | 0 | 0 | 0 | 0 | 3148400 | 0 | 0 | 0 | 0 | 0 | 0 |
| DVVTAAGD | PSAP; PSAP; PS | P07602; Q5E | SAP_HUMAN P | Oxidation (M) | 0 | 0 | 202980 | 0 | 0 | 0 | 0 | 0 | 988920 | 92237 | 86369 | 0 | 0 | 0 | 0 | 104090 | 309040 | 0 |

| | | | | | | | | | | | | | | | | | | | | | | |
|---|---|---|---|---|---|---|---|---|---|---|---|---|---|---|---|---|---|---|---|---|---|---|
| DVVTAAGD | PSAP; PSAP; PS | P07602; Q5E | SAP_HUMAN P | Unmodified | 0 | 0 | 0 | 0 | 0 | 0 | 0 | 0 | 12525000 | 0 | 834790 | 0 | 0 | 0 | 0 | 0 | 0 | 0 |
| DVWEIPRE | FYN | P06241 | FYN_HUMAN T | Unmodified | 25897 | 0 | 0 | 0 | 7814.1 | 0 | 267440 | 0 | 1745300 | 829390 | 98947 | 88277 | 0 | 0 | 29579 | 0 | 70534 | 216610 |
| DVYDKVDY | FLOT2; FLOT2; I | Q14254; J3Q | FLOT2_HUMAN | Unmodified | 488840 | 587510 | 413140 | 1449300 | 1805600 | 443560 | 521020 | 15930 | 11443000 | 1308000 | 2083400 | 7064300 | 686460 | 322390 | 2486700 | 1703900 | 0 | 0 |
| DVYKEHFQ | NAMPT; NAMPT | P43490; A0A | NAMPT_HUMAI | Unmodified | 0 | 0 | 0 | 137770 | 3394800 | 1964800 | 146420 | 0 | 0 | 0 | 0 | 0 | 538080 | 594580 | 0 | 0 | 0 | 0 |
| DVYNSEYY | PTK7; PTK7; PTI | Q13308; H0Y | PTK7_HUMAN I | Unmodified | 0 | 0 | 824420 | 787010 | 393680 | 372260 | 0 | 0 | 4851100 | 718580 | 2462500 | 11857000 | 479220 | 0 | 2946200 | 1918100 | 229540 | 721490 |
| DVYSSTPTI | MAN1A2; MAN1 | O60476; Q5T | MA1A2_HUMAI | Unmodified | 0 | 0 | 0 | 0 | 0 | 0 | 0 | 0 | 5319200 | 0 | 0 | 0 | 0 | 0 | 0 | 0 | 0 | 0 |
| DVYVDSKD | NDUFAF4 | Q9P032 | NDUF4_HUMA | Unmodified | 0 | 0 | 0 | 0 | 0 | 0 | 100840 | 0 | 1688600 | 666090 | 0 | 0 | 0 | 0 | 0 | 0 | 705840 | 0 |
| DVYVGKPS | LFNG; LFNG | Q8NES3; A0, | LFNG_HUMAN | Unmodified | 0 | 0 | 969250 | 1153100 | 0 | 0 | 0 | 0 | 1456500 | 0 | 0 | 0 | 1765800 | 893770 | 0 | 552700 | 0 | 0 |
| DVYVPEASI | NUMB; NUMB; | P49757; G3\ | NUMB_HUMAN | Unmodified | 0 | 0 | 292900 | 0 | 26931000 | 2360500 | 0 | 0 | 2387600 | 646360 | 0 | 0 | 4813700 | 496540 | 1006800 | 699830 | 206130 | 7709100 |
| DWAESTLM | F10 | P00742 | FA10_HUMAN ( | Unmodified | 0 | 0 | 0 | 399030 | 359170 | 0 | 592950 | 61799 | 0 | 0 | 0 | 0 | 563530 | 512450 | 0 | 0 | 0 | 0 |
| DWAKPGP' | MTSS1; MTSS1; | O43312; H0Y | MTSS1_HUMAI | Unmodified | 0 | 0 | 0 | 0 | 0 | 0 | 197570 | 206860 | 753000 | 126090 | 0 | 186940 | 0 | 0 | 0 | 0 | 0 | 0 |
| DWEMHVH | LMAN2; LMAN2 | Q12907; D6I | LMAN2_HUMAI | Oxidation (M) | 0 | 0 | 0 | 0 | 0 | 0 | 0 | 0 | 9858200 | 2147700 | 0 | 0 | 0 | 0 | 0 | 83612 | 0 | 0 |
| DWEMHVH | LMAN2; LMAN2 | Q12907; D6I | LMAN2_HUMAI | Unmodified | 0 | 0 | 0 | 99587 | 0 | 0 | 457790 | 0 | 42333000 | 4693700 | 667450 | 338580 | 0 | 0 | 88928 | 72905 | 0 | 89447 |
| DWHPLAQI | DIP2A; DIP2A | Q14689; A0/ | DIP2A_HUMAN | Unmodified | 0 | 0 | 0 | 0 | 403260 | 0 | 0 | 0 | 0 | 109350 | 0 | 821870 | 0 | 0 | 575390 | 410180 | 0 | 0 |
| DWIGIFRV( | CALCOCO2; C/ | Q13137; D6I | CACO2_HUMA | Unmodified | 0 | 0 | 0 | 0 | 31597 | 0 | 227500 | 0 | 0 | 0 | 0 | 0 | 22828 | 20111 | 0 | 0 | 52699 | 0 |
| DWILPSDY | CALU; CALU | O43852; H0Y | CALU_HUMAN | Unmodified | 138040 | 0 | 282130 | 344220 | 198370 | 539160 | 170960 | 100910 | 40293000 | 3179700 | 7319000 | 1389200 | 310310 | 0 | 630410 | 273390 | 0 | 312050 |
| DWILPSDY | CALU; CALU | O43852; H0Y | CALU_HUMAN | Unmodified | 0 | 183920 | 0 | 0 | 538800 | 0 | 311710 | 0 | 230780 | 2973000 | 243150 | 0 | 270010 | 448110 | 0 | 0 | 0 | 833800 |
| DWKLDPVI | GLG1; GLG1 | Q92896; H3I | GSLG1_HUMAI | Unmodified | 0 | 0 | 0 | 0 | 0 | 0 | 0 | 0 | 1298800 | 0 | 0 | 481110 | 0 | 0 | 0 | 0 | 0 | 0 |
| DWKLDPVI | GLG1; GLG1 | Q92896; H3I | GSLG1_HUMAI | Unmodified | 0 | 0 | 0 | 0 | 0 | 0 | 0 | 41905 | 5905400 | 362080 | 1163800 | 2749300 | 0 | 0 | 0 | 114430 | 0 | 0 |
| DWKSDLSI | PTGFRN | Q9P2B2 | FPRP_HUMAN | Unmodified | 0 | 1270200 | 1319200 | 2337200 | 0 | 0 | 0 | 0 | 0 | 0 | 242020 | 4018500 | 0 | 0 | 3222200 | 2284700 | 0 | 0 |
| DWNTLIVG | PRMT5; PRMT5; | O14744; G3\ | ANM5_HUMAN | Unmodified | 0 | 0 | 0 | 0 | 0 | 0 | 0 | 10728 | 0 | 15714 | 0 | 2579600 | 0 | 0 | 0 | 0 | 0 | 0 |
| DWNVDLIP | GDI1; GDI2; GD | P31150; P50 | GDIA_HUMAN | Unmodified | 0 | 0 | 0 | 0 | 0 | 0 | 0 | 0 | 0 | 0 | 0 | 1347700 | 0 | 0 | 131980 | 0 | 0 | 0 |
| DWQEGGY | ACSL4; ACSL4 | O60488; H0Y | ACSL4_HUMAN | Unmodified | 0 | 0 | 0 | 0 | 0 | 0 | 0 | 124010 | 410050 | 0 | 2087300 | 12952000 | 88795 | 0 | 0 | 0 | 0 | 0 |
| DWQPHISF | DIP2B | Q9P265 | DIP2B_HUMAN | Unmodified | 1651200 | 649500 | 0 | 766010 | 0 | 368760 | 0 | 0 | 3643000 | 378180 | 1604100 | 12138000 | 265560 | 0 | 1945400 | 1647900 | 0 | 0 |
| DWQSETIE | COBLL1; COBLI | Q53SF7; A0/ | COBL1_HUMAI | Unmodified | 411390 | 291360 | 187660 | 517000 | 0 | 0 | 0 | 0 | 0 | 0 | 0 | 0 | 517410 | 335720 | 209210 | 176540 | 0 | 0 |
| DWQSETIE | COBLL1; COBLI | Q53SF7; A0/ | COBL1_HUMAI | Oxidation (M) | 0 | 1531300 | 0 | 0 | 0 | 0 | 0 | 0 | 0 | 0 | 0 | 0 | 0 | 349510 | 60307 | 1387800 | 0 | 0 |
| DWSHYFK | KRT18; KRT18 | P05783; F8V | K1C18_HUMAN | Unmodified | 43913 | 0 | 0 | 0 | 0 | 0 | 0 | 0 | 935170 | 0 | 753010 | 3763300 | 0 | 0 | 0 | 0 | 0 | 0 |
| DWSKVVLA | TPI1; TPI1; TPI1 | P60174; U3K | TPIS_HUMAN T | Unmodified | 0 | 0 | 0 | 0 | 0 | 0 | 0 | 0 | 0 | 0 | 0 | 0 | 0 | 0 | 0 | 0 | 1308600 | 0 |
| DWSNLAGI | PODXL2 | Q9NZ53 | PDXL2_HUMAN | Unmodified | 15303 | 0 | 589710 | 731330 | 290580 | 63264 | 0 | 0 | 4041100 | 652330 | 286870 | 0 | 0 | 435550 | 0 | 289070 | 0 | 135450 |
| DWSPTLYII | GPD2 | P43304 | GPDM_HUMAN | Unmodified | 0 | 0 | 0 | 0 | 0 | 0 | 0 | 0 | 9105400 | 269840 | 0 | 0 | 0 | 0 | 0 | 0 | 0 | 0 |
| DWTDTNY' | TSPAN6; TSPAN | O43657; A0/ | TSN6_HUMAN | Unmodified | 0 | 0 | 225470 | 0 | 0 | 0 | 0 | 0 | 381740 | 0 | 0 | 1593100 | 0 | 0 | 0 | 0 | 0 | 0 |
| DWTSDVN' | TNKS1BP1 | Q9C0C2 | TB182_HUMAN | Unmodified | 0 | 0 | 0 | 0 | 512390 | 0 | 0 | 0 | 0 | 0 | 0 | 0 | 1832500 | 1001400 | 0 | 0 | 0 | 0 |
| DWVCALG' | KIF21A; KIF21A | Q7Z4S6; A0/ | KI21A_HUMAN | Unmodified | 0 | 0 | 0 | 0 | 0 | 0 | 0 | 0 | 0 | 0 | 15883 | 126510 | 0 | 0 | 0 | 0 | 0 | 0 |
| DWVVAPIS' | CDH3 | P22223 | CADH3_HUMA | Unmodified | 0 | 4036700 | 0 | 0 | 23133000 | 2127700 | 0 | 0 | 471920 | 0 | 0 | 0 | 0 | 0 | 0 | 0 | 0 | 0 |
| DWYDVKAI | RPS3A; RPS3A; | P61247; H0Y | RS3A_HUMAN | Unmodified | 0 | 0 | 0 | 18614 | 322000 | 0 | 0 | 0 | 0 | 0 | 0 | 0 | 321200 | 65380 | 0 | 0 | 0 | 0 |
| DWYDVKAI | RPS3A; RPS3A; | P61247; H0Y | RS3A_HUMAN | Oxidation (M) | 0 | 0 | 0 | 0 | 0 | 0 | 1588000 | 58605 | 0 | 0 | 0 | 0 | 1212800 | 734180 | 0 | 0 | 0 | 490000 |
| DYAAYNVL | GLRX5 | Q86SX6 | GLRX5_HUMAN | Unmodified | 0 | 0 | 0 | 0 | 0 | 0 | 0 | 0 | 3436400 | 51363 | 0 | 0 | 0 | 0 | 0 | 0 | 0 | 0 |
| DYADSIHAI | RAB22A | Q9UL26 | RB22A_HUMAN | Unmodified | 2927300 | 2610500 | 2806600 | 2686500 | 0 | 0 | 0 | 0 | 4487400 | 289360 | 0 | 2563000 | 0 | 0 | 4616900 | 3927300 | 0 | 0 |
| DYAFVHFE | HNRNPR; HNRI | O43390; B4D | HNRPR_HUMA | Unmodified | 0 | 0 | 0 | 0 | 0 | 0 | 0 | 0 | 0 | 0 | 193120 | 2061800 | 0 | 0 | 0 | 0 | 0 | 0 |
| DYAFVHFE | HNRNPR; HNRI | O43390; B4D | HNRPR_HUMA | Unmodified | 0 | 0 | 0 | 0 | 0 | 0 | 683000 | 294410 | 468610 | 463650 | 0 | 2118500 | 534020 | 536990 | 0 | 0 | 0 | 0 |
| DYALHWL\ | RBBP7; RBBP7; | Q16576; C9J | RBBP7_HUMAN | Unmodified | 329460 | 0 | 172130 | 356490 | 0 | 0 | 0 | 0 | 2201700 | 0 | 1417000 | 9009600 | 834910 | 130390 | 567250 | 254840 | 82096 | 0 |
| DYAVSTVP' | DDAH1; DDAH1 | O94760; B4D | DDAH1_HUMA | Unmodified | 0 | 0 | 0 | 0 | 618700 | 209410 | 0 | 0 | 973230 | 279100 | 494350 | 1656300 | 1692800 | 461600 | 1138400 | 0 | 335670 | 863340 |
| DYAVVLR | FOLH1B; FOLH | Q9HBA9; Q0 | FOH1B_HUMAI | Unmodified | 1493200 | 550470 | 3341200 | 0 | 0 | 0 | 0 | 0 | 266180 | 16260 | 445940 | 395890 | 0 | 0 | 3315300 | 3330800 | 0 | 0 |
| DYCILYNP( | SPPL2B; SPPL2 | Q8TCT7; A0/ | SPP2B_HUMAN | Unmodified | 27057 | 0 | 0 | 0 | 0 | 0 | 0 | 0 | 6013100 | 0 | 0 | 0 | 0 | 0 | 0 | 0 | 0 | 0 |
| DYDAMGS( | RUVBL2; RUVBI | Q9Y230; M0I | RUVB2_HUMAI | Unmodified | 183350 | 98955 | 0 | 0 | 158970 | 0 | 0 | 0 | 73423 | 0 | 237450 | 514980 | 144680 | 0 | 0 | 0 | 0 | 0 |
| DYDDMSPI | HNRNPK; HNRI | P61978; Q5T | HNRPK_HUMA | Unmodified | 0 | 0 | 0 | 0 | 0 | 0 | 0 | 0 | 0 | 0 | 1544500 | 3042100 | 0 | 0 | 0 | 0 | 0 | 0 |
| DYDDMSPI | HNRNPK; HNRI | P61978; Q5T | HNRPK_HUMA | Oxidation (M) | 343720 | 0 | 0 | 0 | 0 | 0 | 0 | 0 | 0 | 0 | 252360 | 1010800 | 0 | 0 | 0 | 0 | 0 | 0 |
| DYDGWTPI | PPP1R12A; PPP | O14974; F8V | MYPT1_HUMAN | Unmodified | 0 | 0 | 0 | 0 | 0 | 0 | 0 | 0 | 0 | 0 | 79041 | 786320 | 85215 | 0 | 206170 | 185250 | 0 | 0 |
| DYDHLFK | RAB35; RAB35; | Q15286; F5H | RAB35_HUMAN | Unmodified | 0 | 1527500 | 1494900 | 0 | 0 | 0 | 0 | 0 | 0 | 0 | 0 | 1298600 | 0 | 0 | 2264700 | 1371500 | 0 | 0 |
| DYDSLAQP | COL6A2 | P12110 | CO6A2_HUMAI | Unmodified | 0 | 128980 | 100130 | 366900 | 604870 | 319780 | 2223800 | 0 | 0 | 1821600 | 0 | 0 | 1216600 | 2102000 | 144720 | 0 | 0 | 3069400 |
| DYDSTCVF | HINT3 | Q9NQE9 | HINT3_HUMAN | Unmodified | 0 | 0 | 0 | 0 | 0 | 0 | 0 | 0 | 0 | 0 | 0 | 1148000 | 141050 | 0 | 0 | 0 | 0 | 0 |
| DYDTLSK | BAIAP2L1 | Q9UHR4 | BI2L1_HUMAN | Unmodified | 0 | 0 | 0 | 295090 | 0 | 0 | 0 | 0 | 307260 | 0 | 0 | 0 | 777150 | 880510 | 0 | 0 | 0 | 0 |
| DYEAGQLY | LARP1 | Q6PKG0 | LARP1_HUMAN | Unmodified | 275840 | 0 | 390170 | 0 | 0 | 0 | 0 | 0 | 0 | 0 | 0 | 1493200 | 0 | 0 | 372640 | 242600 | 0 | 0 |
| DYEEIGPSI | ACTR3; ACTR3 | P61158; B4D | ARP3_HUMAN | Unmodified | 1049100 | 0 | 703430 | 770070 | 0 | 0 | 0 | 32088 | 736630 | 0 | 478680 | 6143400 | 0 | 0 | 917180 | 897220 | 0 | 0 |
| DYEEIGPSI | ACTR3; ACTR3 | P61158; B4D | ARP3_HUMAN | Oxidation (M) | 0 | 0 | 0 | 0 | 1966800 | 426040 | 869280 | 257780 | 0 | 329190 | 0 | 0 | 1485400 | 1499800 | 0 | 0 | 1124000 | 0 |
| DYEEVGAD | TUBA1C; TUBA1 | Q9BQE3; F5I | TBA1C_HUMAN | Unmodified | 0 | 0 | 0 | 0 | 4385400 | 1035000 | 191780 | 0 | 2775600 | 509390 | 0 | 0 | 0 | 0 | 0 | 0 | 665600 | 3135900 |
| DYEEVGVD | TUBA1A; TUBA1 | Q71U36; P68 | TBA1A_HUMAN | Unmodified | 513800 | 529510 | 239740 | 772620 | 10907000 | 1319900 | 7763200 | 11212000 | 15464000 | 5081400 | 4966100 | 5262100 | 3955100 | 2658800 | 845630 | 528410 | 2593300 | 6946700 |
| DYEFMWNI | CKB; CKB; CKB | P12277; H0Y | KCRB_HUMAN | Oxidation (M) | 0 | 0 | 0 | 0 | 0 | 0 | 0 | 0 | 0 | 0 | 0 | 108030 | 0 | 0 | 995120 | 0 | 0 | 0 |
| DYEILFK | GGH; GGH; GG | Q92820; A0/ | GGH_HUMAN ( | Unmodified | 85364 | 37558 | 194810 | 142820 | 0 | 0 | 0 | 0 | 60548000 | 6191400 | 3114600 | 2030200 | 60273 | 0 | 0 | 233970 | 0 | 0 |
| DYELQLVT' | PLEC; PLEC | Q15149; H0Y | PLEC_HUMAN | Unmodified | 0 | 0 | 0 | 0 | 0 | 827890 | 0 | 0 | 0 | 0 | 530610 | 85809 | 0 | 0 | 0 | 71980 | 0 | 597150 |
| DYENDED\ | TRAPPC3; TRAF | O43617; A0/ | TPPC3_HUMAN | Unmodified | 0 | 0 | 0 | 0 | 0 | 0 | 0 | 0 | 0 | 51638 | 0 | 462420 | 0 | 60234 | 0 | 0 | 78945 | 38265 |
| DYETATLSE | ACTN1; ACTN1; | P12814; A0A | ACTN1_HUMAI | Unmodified | 31462 | 0 | 183880 | 325830 | 6912700 | 1894400 | 1106400 | 518240 | 0 | 459770 | 356540 | 7136700 | 418360 | 0 | 700870 | 471400 | 3172000 | 0 |
| DYETATLSE | ACTN1; ACTN1; | P12814; A0A | ACTN1_HUMAI | Unmodified | 0 | 0 | 0 | 0 | 238370 | 0 | 330210 | 0 | 0 | 0 | 0 | 0 | 0 | 0 | 0 | 0 | 819500 | 0 |
| DYEVDATLI | COL1A2; COL1/ | P08123; A0A | CO1A2_HUMAI | Unmodified | 0 | 0 | 0 | 0 | 286920 | 919700 | 0 | 146140 | 406200 | 0 | 0 | 0 | 0 | 0 | 0 | 0 | 0 | 719610 |
| DYFEQYGK | HNRNPA1L2; H | Q32P51; P09 | RA1L2_HUMAN | Unmodified | 0 | 0 | 0 | 0 | 0 | 0 | 0 | 0 | 0 | 0 | 706890 | 1249400 | 0 | 0 | 0 | 0 | 0 | 0 |
| DYFGAHTY | PGD | P52209 | 6PGD_HUMAN | Unmodified | 0 | 0 | 0 | 668900 | 0 | 0 | 68518 | 0 | 1518400 | 229320 | 0 | 2717500 | 0 | 0 | 542270 | 385270 | 84042 | 106270 |
| DYFGLLFQI | EPB41L2; EPB4 | O43491; A0/ | E41L2_HUMAN | Unmodified | 0 | 0 | 80913 | 250230 | 4383100 | 744630 | 342040 | 76017 | 0 | 0 | 44872 | 345330 | 289680 | 234930 | 342800 | 239000 | 1448900 | 995670 |
| DYFGLLFQI | EPB41L2; EPB4 | O43491; A0/ | E41L2_HUMAN | Unmodified | 0 | 0 | 0 | 0 | 1139000 | 0 | 0 | 0 | 0 | 0 | 0 | 0 | 0 | 0 | 0 | 0 | 645020 | 161460 |
| DYFGLTYRI | EPB41L3; EPB4 | Q9Y2J2; A0A | E41L3_HUMAN | Unmodified | 0 | 0 | 0 | 0 | 0 | 0 | 98225 | 0 | 0 | 0 | 0 | 0 | 487770 | 460900 | 0 | 0 | 1330000 | 4606800 |
| DYFLFRDG | HSPE1; HSPE1 | P61604; B8Z | CH10_HUMAN | Unmodified | 0 | 0 | 0 | 0 | 0 | 0 | 0 | 0 | 1636500 | 0 | 0 | 0 | 570110 | 0 | 0 | 0 | 0 | 247150 |
| DYFNVPYP | NPEPPSL1; NPI | A6NEC2; P5! | PSAL_HUMAN I | Unmodified | 0 | 0 | 0 | 0 | 0 | 0 | 0 | 0 | 0 | 0 | 0 | 3264500 | 265180 | 572300 | 0 | 0 | 214230 | 0 |
| DYFQIHTP( | ELAPOR2; ELAP | A8MWY0; H7 | ELAP2_HUMAN | Unmodified | 0 | 0 | 0 | 0 | 0 | 0 | 39768 | 39043 | 2587100 | 417170 | 0 | 0 | 0 | 0 | 0 | 0 | 0 | 0 |
| DYFSKFGE' | HNRNPD; HNR | Q14103; H0Y | HNRPD_HUMA | Unmodified | 0 | 0 | 0 | 0 | 0 | 0 | 278280 | 0 | 0 | 289970 | 0 | 0 | 917560 | 0 | 0 | 0 | 0 | 79414 |
| DYFTKFGE\ | HNRNPAB; HNI | Q99729; D6I | ROAA_HUMAN | Unmodified | 0 | 32502 | 0 | 0 | 211280 | 0 | 0 | 0 | 0 | 214100 | 0 | 0 | 177870 | 0 | 0 | 0 | 0 | 57534 |
| DYGKESQA | RPS12 | P25398 | RS12_HUMAN | Unmodified | 0 | 0 | 0 | 0 | 1241100 | 108450 | 2613800 | 149630 | 0 | 1137400 | 0 | 0 | 2089300 | 491840 | 0 | 0 | 2456900 | 1137500 |
| DYGVLLEG | PRDX3 | P30048 | PRDX3_HUMAI | Unmodified | 273790 | 0 | 0 | 0 | 0 | 0 | 0 | 0 | 14454000 | 604250 | 158130 | 418940 | 0 | 0 | 0 | 163420 | 0 | 0 |
| DYGVQLVE | MRPL15; MRPL | Q9P015; E5F | RM15_HUMAN | Unmodified | 0 | 0 | 0 | 0 | 0 | 0 | 0 | 28890 | 2097600 | 57003 | 0 | 66920 | 0 | 0 | 0 | 0 | 0 | 0 |
| DYGVYLED | PRDX4; PRDX4 | Q13162; H7( | PRDX4_HUMAI | Unmodified | 346690 | 129050 | 550010 | 306770 | 0 | 0 | 0 | 0 | 1.62E+08 | 4260400 | 61002000 | 14340000 | 0 | 0 | 239760 | 477650 | 0 | 0 |
| DYHFKVDN | NPM1; NPM1; N | P06748; A0A | NPM_HUMAN N | Unmodified | 1820900 | 0 | 0 | 2043200 | 2499300 | 0 | 0 | 0 | 20956000 | 1365800 | 46960000 | 55928000 | 1710700 | 0 | 540190 | 0 | 0 | 0 |
| DYHMERPI | SLC26A6; SLC2 | Q9BXS9; C9. | S26A6_HUMAN | Unmodified | 0 | 0 | 0 | 0 | 0 | 0 | 0 | 0 | 0 | 0 | 0 | 993380 | 0 | 0 | 0 | 0 | 0 | 0 |
| DYHNLFSP | PXN; PXN; PXN | P49023; A0A | PAXI_HUMAN F | Unmodified | 0 | 0 | 0 | 0 | 0 | 0 | 0 | 0 | 0 | 0 | 0 | 0 | 0 | 0 | 0 | 0 | 1973600 | 0 |
| DYIALKEDL | HLA-A; HLA-A; I | Q53Z42; A0/ | Q53Z42_HUMA | Unmodified | 0 | 0 | 0 | 0 | 0 | 0 | 0 | 0 | 5155800 | 0 | 1561800 | 11765000 | 0 | 0 | 0 | 0 | 77951 | 820810 |
| DYIEFNK | AZGP1; AZGP1 | P25311; C9J | ZA2G_HUMAN | Unmodified | 0 | 0 | 872490 | 0 | 0 | 0 | 0 | 37868 | 0 | 0 | 6146900 | 952970 | 0 | 0 | 944540 | 160130 | 0 | 0 |
| DYIHINIGA | DDX5; DDX5; D | P17844; A0A | DDX5_HUMAN | Unmodified | 0 | 0 | 0 | 0 | 0 | 0 | 0 | 0 | 0 | 0 | 0 | 0 | 1616500 | 1030700 | 0 | 0 | 0 | 215560 |
| DYIKAEEDI | P4HA1 | P13674 | P4HA1_HUMAN | Unmodified | 0 | 0 | 0 | 0 | 0 | 0 | 0 | 0 | 3906300 | 2122300 | 339040 | 293060 | 0 | 134890 | 0 | 139570 | 0 | 0 |
| DYIPVDQEI | DYNC1H1; DYN | Q14204; A0/ | DYHC1_HUMA | Unmodified | 0 | 0 | 0 | 0 | 0 | 283300 | 0 | 0 | 0 | 0 | 0 | 1731900 | 0 | 0 | 0 | 0 | 0 | 0 |
| DYIPVDQEI | DYNC1H1; DYN | Q14204; A0/ | DYHC1_HUMA | Unmodified | 0 | 0 | 0 | 0 | 0 | 156330 | 0 | 140500 | 2612100 | 1016000 | 628850 | 6770100 | 0 | 0 | 682820 | 491190 | 600240 | 0 |
| DYIVVGSD! | SF3B3; SF3B3; | Q15393; J3Q | SF3B3_HUMAN | Unmodified | 15020000 | 11846000 | 6463200 | 6620400 | 0 | 183020 | 0 | 0 | 0 | 0 | 0 | 1441100 | 0 | 0 | 8564600 | 0 | 173510 | 192940 |
| DYKAEEDP | PITPNA; PITPN/ | Q00169; I3L | PIPNA_HUMAN | Unmodified | 0 | 0 | 0 | 0 | 578300 | 0 | 0 | 0 | 0 | 0 | 60212 | 0 | 0 | 0 | 0 | 0 | 369660 | 2080600 |
| DYKSSFAD | TOMM34 | Q15785 | TOM34_HUMAI | Unmodified | 0 | 0 | 0 | 0 | 0 | 0 | 0 | 0 | 0 | 0 | 0 | 0 | 0 | 0 | 0 | 0 | 234410 | 130750 |
| DYLALNED | HLA-G; HLA-G; | P17693; Q5F | HLAG_HUMAN | Unmodified | 364880 | 647480 | 460560 | 325220 | 0 | 254260 | 0 | 0 | 23853000 | 0 | 1093100 | 9434700 | 0 | 0 | 755800 | 667210 | 0 | 219330 |
| DYLEHITSI( | ERMP1; ERMP1 | Q7Z2K6; E7E | ERMP1_HUMAI | Unmodified | 65275 | 853220 | 465150 | 450730 | 0 | 0 | 0 | 209210 | 5200800 | 0 | 0 | 197180 | 0 | 0 | 719460 | 412500 | 0 | 0 |
| DYLHLPPEI | RPS10; RPS10-I | P46783; A0A | RS10_HUMAN | Unmodified | 0 | 0 | 0 | 0 | 0 | 0 | 0 | 0 | 0 | 0 | 0 | 2044000 | 0 | 0 | 0 | 120220 | 0 | 0 |
| DYLHYHIK | ARPC2; ARPC2 | O15144; H7( | ARPC2_HUMAI | Unmodified | 0 | 0 | 0 | 0 | 0 | 0 | 0 | 0 | 0 | 0 | 176310 | 815890 | 0 | 0 | 598500 | 0 | 0 | 0 |
| DYLHYIR | RPS11; RPS11; | P62280; M0I | RS11_HUMAN | Unmodified | 0 | 0 | 0 | 0 | 0 | 0 | 0 | 0 | 86361 | 0 | 195690 | 11250000 | 0 | 0 | 0 | 0 | 0 | 0 |
| DYLLCDYN | CAPZB; CAPZB; | P47756; A0A | CAPZB_HUMAN | Unmodified | 245270 | 0 | 0 | 0 | 0 | 0 | 0 | 0 | 0 | 0 | 0 | 2997800 | 0 | 0 | 227880 | 245630 | 0 | 0 |
| DYMFSGLK | RP2 | O75695 | XRP2_HUMAN | Unmodified | 0 | 0 | 0 | 0 | 0 | 0 | 0 | 0 | 287220 | 0 | 62096 | 919620 | 0 | 0 | 0 | 0 | 0 | 0 |
| DYMFSGLK | RP2 | O75695 | XRP2_HUMAN | Oxidation (M) | 0 | 98291 | 222780 | 0 | 0 | 0 | 0 | 0 | 0 | 0 | 1590400 | 1525400 | 0 | 0 | 348710 | 290090 | 0 | 0 |
| DYNEAYNY | DNAJC7; DNAJ( | Q99615; A0/ | DNJC7_HUMAN | Unmodified | 0 | 0 | 341390 | 361290 | 1037300 | 129940 | 601600 | 208970 | 658460 | 436380 | 161900 | 888950 | 1421400 | 862400 | 0 | 0 | 207290 | 200090 |
| DYNHLPTG | DMAC2L; DMA( | Q99766; A0/ | ATP5S_HUMAN | Unmodified | 0 | 0 | 0 | 0 | 0 | 0 | 0 | 39306 | 973030 | 239820 | 0 | 0 | 0 | 0 | 0 | 0 | 0 | 0 |
| DYNSSRED | CD82; CD82 | P27701; E9P | CD82_HUMAN | Unmodified | 0 | 0 | 0 | 0 | 0 | 0 | 0 | 0 | 0 | 0 | 0 | 0 | 0 | 0 | 182410 | 0 | 0 | 0 |
| DYNVTANS | LDHA; LDHA; LI | P00338; F5G | LDHA_HUMAN | Unmodified | 0 | 221650 | 351170 | 995920 | 1519200 | 425550 | 0 | 0 | 974570 | 148900 | 977710 | 20138000 | 251030 | 0 | 810870 | 563350 | 972440 | 1339500 |
| DYPVVSIED | ENO1; ENO1 | P06733; A0A | ENOA_HUMAN | Unmodified | 0 | 0 | 229220 | 1189600 | 29390000 | 2098900 | 555450 | 0 | 3979800 | 4130900 | 922180 | 3335100 | 8094500 | 6554800 | 1845000 | 222200 | 6379300 | 15958000 |
| DYQDNKAI | EIF1AY; EIF1AX; | O14602; P47 | IF1AY_HUMAN | Unmodified | 1912200 | 1932400 | 871320 | 0 | 0 | 0 | 16661 | 0 | 756080 | 153400 | 2184600 | 8494000 | 283750 | 0 | 1140300 | 1412500 | 0 | 0 |
| DYQDVRNI | ATP2B1; ATP2B | P20020; E7E | AT2B1_HUMAN | Unmodified | 174230 | 288250 | 2749800 | 6854600 | 279170 | 0 | 0 | 0 | 1651800 | 537100 | 1723500 | 10941000 | 1136100 | 0 | 10427000 | 4987800 | 0 | 0 |
| DYQRELEE. | MYH10 | P35580 | MYH10_HUMAI | Unmodified | 0 | 0 | 1028100 | 0 | 0 | 0 | 0 | 0 | 0 | 0 | 0 | 0 | 0 | 0 | 436500 | 377090 | 0 | 0 |
| DYQSQEPL | COBLL1; COBLI | Q53SF7; A0/ | COBL1_HUMAI | Unmodified | 1907000 | 1299200 | 728260 | 587880 | 270500 | 0 | 0 | 0 | 153160 | 0 | 0 | 0 | 2674200 | 1655600 | 564180 | 698430 | 0 | 0 |
| DYRQSSG/ | DDX3X; DDX3X; | O00571; A0/ | DDX3X_HUMAI | Unmodified | 0 | 0 | 145180 | 2115000 | 0 | 0 | 0 | 0 | 0 | 0 | 0 | 0 | 302790 | 68263 | 86683 | 72078 | 0 | 0 |
| DYSADTLAI | PLAU; PLAU | P00749; E7E | UROK_HUMAN | Unmodified | 0 | 0 | 0 | 0 | 1649400 | 1692300 | 0 | 0 | 473680 | 153520 | 0 | 0 | 0 | 0 | 0 | 0 | 1158100 | 2948700 |

| | | | | | | | | | | | | | | | | | | | | | | |
|---|---|---|---|---|---|---|---|---|---|---|---|---|---|---|---|---|---|---|---|---|---|---|
| DYSEADGL | EPB41L1; EPB4 | Q9H4G0; Q4 | E41L1_HUMAN | Unmodified | 0 | 0 | 0 | 0 | 0 | 3992400 | 0 | 0 | 0 | 0 | 543260 | 697900 | 0 | 0 | 0 | 610730 | 0 | 0 |
| DYSEADGL | EPB41L1; EPB4 | Q9H4G0; Q4 | E41L1_HUMAN | Unmodified | 117100 | 0 | 186140 | 438220 | 3657500 | 355630 | 47583 | 47376 | 0 | 72952 | 942120 | 0 | 911350 | 584180 | 330010 | 188070 | 509230 | 2175000 |
| DYSFTIQAY | CLSTN1; CLSTN | O94985; A0A | CSTN1_HUMAN | Unmodified | 0 | 0 | 0 | 0 | 873310 | 108540 | 54993 | 0 | 1035300 | 764150 | 0 | 0 | 380520 | 142030 | 0 | 0 | 163140 | 170480 |
| DYSFTIQAY | CLSTN1; CLSTN | O94985; A0A | CSTN1_HUMAN | Unmodified | 0 | 0 | 0 | 0 | 0 | 0 | 0 | 0 | 1563000 | 673450 | 0 | 0 | 0 | 0 | 0 | 0 | 0 | 0 |
| DYSGQGV | WDR1 | O75083 | WDR1_HUMAN | Unmodified | 89386 | 0 | 0 | 0 | 279830 | 0 | 24775 | 196740 | 776610 | 61447 | 438550 | 5668300 | 0 | 0 | 44698 | 65633 | 175060 | 32737 |
| DYSGQGV | WDR1 | O75083 | WDR1_HUMAN | Unmodified | 1535100 | 1339500 | 0 | 2131000 | 2734100 | 576180 | 781390 | 401410 | 2771800 | 132000 | 1899700 | 97593 | 1477100 | 0 | 734120 | 1122700 | 674640 | 0 |
| DYSNFDQE | PRKCD | Q05655 | KPCD_HUMAN | Unmodified | 0 | 0 | 18240 | 0 | 0 | 0 | 0 | 0 | 269320 | 0 | 67970 | 423460 | 0 | 0 | 0 | 0 | 341090 | 252790 |
| DYSSGFGC | CTTN | Q14247 | SRC8_HUMAN | Unmodified | 0 | 0 | 0 | 0 | 7659600 | 1918700 | 176620 | 202900 | 0 | 421730 | 0 | 0 | 5968200 | 4474500 | 0 | 0 | 3032600 | 2956800 |
| DYSVTANS | LDHB; LDHB; LI | P07195; A0A | LDHB_HUMAN | Unmodified | 384320 | 318650 | 371520 | 1069900 | 322860 | 539410 | 57476 | 0 | 0 | 169370 | 0 | 18132000 | 481860 | 139230 | 916890 | 484300 | 370460 | 524980 |
| DYTGEDVT | LGMN; LGMN; L | Q99538; G3 | LGMN_HUMAN | Unmodified | 0 | 0 | 0 | 0 | 362800 | 0 | 0 | 0 | 1073200 | 381810 | 236710 | 0 | 0 | 0 | 0 | 0 | 0 | 0 |
| DYTQMNEL | GPX1; GPX1; GI | P07203; A0A | GPX1_HUMAN | Unmodified | 0 | 0 | 0 | 0 | 0 | 0 | 0 | 0 | 0 | 0 | 0 | 1567100 | 0 | 0 | 0 | 0 | 77980 | 0 |
| DYTRPDLP | PTBP2; PTBP2; | Q9UKA9; A0 | PTBP2_HUMAN | Unmodified | 0 | 0 | 0 | 0 | 0 | 0 | 0 | 0 | 0 | 0 | 221740 | 1318600 | 0 | 0 | 0 | 0 | 0 | 0 |
| DYTSGAML | WARS1 | P23381 | SYWC_HUMAN | Unmodified | 0 | 0 | 0 | 0 | 0 | 0 | 0 | 0 | 615210 | 0 | 0 | 0 | 0 | 0 | 0 | 0 | 0 | 0 |
| DYTSGAML | WARS1 | P23381 | SYWC_HUMAN | Unmodified | 180650 | 0 | 0 | 167260 | 1232200 | 0 | 108880 | 0 | 1557800 | 397870 | 140670 | 553330 | 1041500 | 499930 | 0 | 0 | 75152 | 1114600 |
| DYTYEELLN | EIF2S2 | P20042 | IF2B_HUMAN E | Unmodified | 0 | 0 | 0 | 0 | 0 | 0 | 0 | 0 | 0 | 0 | 0 | 2371000 | 0 | 31358 | 0 | 0 | 0 | 0 |
| DYVAPTAN | SND1 | Q7KZF4 | SND1_HUMAN | Unmodified | 0 | 0 | 0 | 0 | 0 | 0 | 0 | 0 | 0 | 0 | 690540 | 4858300 | 0 | 0 | 0 | 1191000 | 0 | 0 |
| DYVAPTAN | SND1 | Q7KZF4 | SND1_HUMAN | Unmodified | 0 | 0 | 0 | 337770 | 0 | 0 | 0 | 0 | 202550 | 0 | 0 | 591160 | 0 | 0 | 0 | 0 | 225190 | 0 |
| DYVDLFR | PRKDC | P78527 | PRKDC_HUMA | Unmodified | 25537 | 0 | 42047 | 0 | 0 | 0 | 0 | 0 | 0 | 193730 | 22697 | 845690 | 0 | 0 | 0 | 16132 | 0 | 0 |
| DYVEINGEI | ST14 | Q9Y5Y6 | ST14_HUMAN S | Unmodified | 1196500 | 1610100 | 5959000 | 2836900 | 425120 | 42733 | 0 | 97800 | 331140 | 244810 | 0 | 2477700 | 1841200 | 516710 | 2712200 | 3254500 | 0 | 0 |
| DYVRQHFF | ACLY | P53396 | ACLY_HUMAN | Unmodified | 0 | 0 | 0 | 0 | 929560 | 0 | 2038100 | 0 | 177770 | 0 | 211650 | 0 | 0 | 0 | 0 | 0 | 0 | 0 |
| DYWETELL | CALCOCO2; CA | Q13137; D6I | CACO2_HUMA | Unmodified | 83732 | 0 | 0 | 0 | 268190 | 40350 | 0 | 0 | 237460 | 0 | 0 | 0 | 133680 | 0 | 0 | 0 | 72883 | 679570 |
| DYYECTGI | MME; MME; MM | P08473; A0A | NEP_HUMAN N | Unmodified | 0 | 0 | 1053700 | 565350 | 0 | 0 | 0 | 0 | 213370 | 0 | 0 | 0 | 0 | 0 | 1838800 | 1704900 | 0 | 0 |
| EAAAAGLP | CPD | O75976 | CBPD_HUMAN | Unmodified | 703070 | 0 | 0 | 0 | 0 | 0 | 0 | 0 | 10981000 | 535640 | 138330 | 2326800 | 0 | 0 | 0 | 0 | 0 | 0 |
| EAAAAGLP | CPD | O75976 | CBPD_HUMAN | Unmodified | 12522000 | 94032000 | 454930 | 0 | 0 | 0 | 0 | 0 | 0 | 0 | 0 | 0 | 0 | 0 | 0 | 0 | 0 | 0 |
| EAAAAVDIF | DIP2A; DIP2A | Q14689; A0A | DIP2A_HUMAN | Unmodified | 0 | 0 | 0 | 0 | 0 | 0 | 0 | 0 | 336300 | 173470 | 0 | 276390 | 0 | 0 | 0 | 65454 | 0 | 0 |
| EAAALGSR | ATP6V1G1 | O75348 | VATG1_HUMAN | Unmodified | 0 | 0 | 0 | 0 | 0 | 223950 | 1696400 | 0 | 0 | 0 | 0 | 0 | 0 | 2394900 | 0 | 0 | 655460 | 1036200 |
| EAAAQEAG | FMNL1; FMNL1 | O95466; K7E | FMNL1_HUMA | Unmodified | 0 | 0 | 0 | 0 | 1123300 | 0 | 0 | 0 | 0 | 0 | 0 | 0 | 0 | 0 | 0 | 0 | 0 | 0 |
| EAAAQEAG | FMNL1; FMNL1 | O95466; K7E | FMNL1_HUMA | Unmodified | 0 | 0 | 0 | 0 | 2596900 | 406640 | 0 | 0 | 0 | 0 | 0 | 0 | 0 | 0 | 0 | 0 | 77805 | 0 |
| EAAASGLPI | TXNDC12 | O95881 | TXD12_HUMAN | Oxidation (M) | 0 | 0 | 0 | 0 | 0 | 0 | 0 | 267710 | 1647300 | 2551800 | 0 | 0 | 0 | 0 | 0 | 0 | 0 | 0 |
| EAAASGLPI | TXNDC12 | O95881 | TXD12_HUMAN | Unmodified | 0 | 0 | 0 | 0 | 0 | 0 | 0 | 0 | 22907000 | 5621100 | 545810 | 0 | 0 | 0 | 0 | 0 | 0 | 0 |
| EAAASLSLT | SUN2 | Q9UH99 | SUN2_HUMAN | Unmodified | 0 | 0 | 0 | 0 | 0 | 0 | 0 | 0 | 4165700 | 0 | 0 | 0 | 0 | 0 | 0 | 0 | 0 | 0 |
| EAACLQQT | AGRN; AGRN; A | O00468; A0A | AGRIN_HUMAN | Unmodified | 434000 | 0 | 389980 | 466860 | 0 | 0 | 0 | 180450 | 267360 | 0 | 0 | 594480 | 0 | 0 | 300720 | 230410 | 0 | 0 |
| EAADAIDAE | JUP | P14923 | PLAK_HUMAN | Oxidation (M) | 0 | 0 | 3716300 | 0 | 0 | 0 | 0 | 0 | 0 | 0 | 0 | 6026000 | 0 | 0 | 4816800 | 3134000 | 0 | 0 |
| EAADTERL | CKAP4 | Q07065 | CKAP4_HUMA | Unmodified | 0 | 0 | 0 | 0 | 0 | 0 | 0 | 77066 | 1074900 | 0 | 0 | 148380 | 0 | 0 | 0 | 0 | 29189 | 1768900 |
| EAAEAIEAE | CTNNB1; CTNN | P35222; A0A | CTNB1_HUMA | Unmodified | 0 | 0 | 2359000 | 0 | 0 | 0 | 0 | 0 | 0 | 0 | 809130 | 8678100 | 0 | 0 | 18089000 | 14655000 | 0 | 0 |
| EAAENSLV | YWHAE; YWHA | P62258; K7E | 1433E_HUMAN | Unmodified | 313040 | 0 | 22369000 | 0 | 2501400 | 2228000 | 547380 | 0 | 9792000 | 3333500 | 8184600 | 1.18E+08 | 2184400 | 1645900 | 25868000 | 20588000 | 2734900 | 0 |
| EAAEVLQN | UBE2M; UBE2M | P61081; M0 | UBC12_HUMA | Unmodified | 0 | 205830 | 0 | 0 | 0 | 0 | 0 | 0 | 0 | 0 | 0 | 1297800 | 0 | 0 | 0 | 0 | 0 | 0 |
| EAAEVLQN | UBE2M; UBE2M | P61081; M0 | UBC12_HUMA | Unmodified | 497410 | 441080 | 748970 | 475250 | 228800 | 0 | 207990 | 58211 | 1685100 | 361250 | 563660 | 3452100 | 647860 | 0 | 859070 | 1144200 | 208450 | 92678 |
| EAAFDDAV | RBBP4; RBBP4; | Q09028; E9 | RBBP4_HUMA | Unmodified | 0 | 0 | 0 | 0 | 0 | 0 | 0 | 0 | 479180 | 0 | 1196200 | 3210600 | 0 | 0 | 0 | 0 | 0 | 0 |
| EAAFDDAV | RBBP4; RBBP4; | Q09028; E9 | RBBP4_HUMA | Unmodified | 37587 | 0 | 0 | 0 | 753070 | 0 | 0 | 0 | 1052800 | 680310 | 0 | 861890 | 0 | 0 | 0 | 0 | 0 | 0 |
| EAAFSLAE | ATP6V1D; ATP6 | Q9Y5K8; G3 | VATD_HUMAN | Unmodified | 0 | 0 | 0 | 0 | 0 | 0 | 0 | 0 | 0 | 0 | 0 | 2029300 | 0 | 0 | 0 | 0 | 0 | 0 |
| EAAGEGPA | APEX1; APEX1; | P27695; A0A | APEX1_HUMAN | Unmodified | 0 | 18906 | 0 | 0 | 0 | 0 | 0 | 42757 | 284020 | 0 | 911900 | 3905500 | 0 | 0 | 0 | 0 | 0 | 0 |
| EAAGEGPA | APEX1; APEX1; | P27695; A0A | APEX1_HUMAN | Unmodified | 0 | 0 | 0 | 0 | 1645700 | 0 | 472760 | 0 | 0 | 1180500 | 0 | 0 | 1464600 | 0 | 0 | 0 | 325490 | 0 |
| EAAHHAGF | SLC4A7; SLC4A | Q9Y6M7; C9 | S4A7_HUMAN | Unmodified | 0 | 0 | 0 | 0 | 0 | 0 | 0 | 0 | 0 | 0 | 0 | 0 | 0 | 0 | 0 | 0 | 0 | 0 |
| EAAHHAGF | SLC4A7; SLC4A | Q9Y6M7; C9 | S4A7_HUMAN | Unmodified | 0 | 0 | 819100 | 0 | 60194 | 0 | 0 | 0 | 26002 | 19507 | 148320 | 0 | 0 | 0 | 0 | 0 | 23776 | 0 |
| EAAIKEETE | TMCO3; TMCO3 | Q6UWJ1; A0 | TMCO3_HUMA | Unmodified | 0 | 0 | 0 | 0 | 0 | 0 | 0 | 0 | 1810500 | 227200 | 455560 | 1073200 | 0 | 0 | 0 | 0 | 0 | 0 |
| EAAIWELEE | SLK | Q9H2G2 | SLK_HUMAN S | Unmodified | 0 | 0 | 0 | 0 | 0 | 0 | 326710 | 175180 | 406950 | 0 | 310840 | 1522000 | 0 | 0 | 234200 | 213100 | 0 | 0 |
| EAALSTALS | LMNA; LMNA; L | P02545; A0A | LMNA_HUMAN | Unmodified | 0 | 0 | 0 | 0 | 5076500 | 970070 | 0 | 119930 | 756590 | 382020 | 322850 | 1457100 | 2722000 | 0 | 0 | 0 | 311700 | 1175700 |
| EAALYPHLI | SQSTM1 | Q13501 | SQSTM_HUMA | Unmodified | 564920 | 110050 | 1739100 | 0 | 0 | 0 | 0 | 0 | 0 | 0 | 0 | 54365 | 0 | 0 | 237620 | 188810 | 0 | 0 |
| EAANAMKL | ACSL4 | O60488 | ACSL4_HUMAN | Oxidation (M) | 0 | 0 | 0 | 0 | 0 | 0 | 0 | 0 | 0 | 0 | 970800 | 3102400 | 0 | 0 | 0 | 0 | 0 | 0 |
| EAANLLEG | GOLIM4; GOLIM | O00461; F8 | GOLI4_HUMAN | Unmodified | 0 | 0 | 0 | 0 | 0 | 0 | 0 | 0 | 1871200 | 0 | 0 | 0 | 0 | 0 | 0 | 0 | 0 | 0 |
| EAAQEAVK | HNRNPR; HNRI | O43390; B4 | HNRPR_HUMA | Unmodified | 0 | 0 | 0 | 0 | 613170 | 0 | 1140100 | 904930 | 0 | 0 | 0 | 0 | 401920 | 0 | 0 | 0 | 0 | 0 |
| EAAQKAVN | PABPC1; PABP | P11940; A0A | PABP1_HUMAN | Unmodified | 0 | 0 | 0 | 0 | 0 | 0 | 458500 | 192330 | 0 | 0 | 0 | 0 | 706930 | 408860 | 0 | 0 | 139360 | 0 |
| EAAQKVGA | PABPC4; PABP | Q13310; B1 | PABP4_HUMAN | Unmodified | 0 | 0 | 0 | 0 | 0 | 0 | 606470 | 247470 | 0 | 0 | 0 | 0 | 4689100 | 1443800 | 0 | 0 | 0 | 0 |
| EAARQNYV | ECI2; ECI2; ECI | O75521; F8 | ECI2_HUMAN E | Unmodified | 0 | 0 | 0 | 0 | 0 | 0 | 0 | 0 | 13414000 | 2095000 | 0 | 0 | 0 | 0 | 0 | 0 | 0 | 0 |
| EAARQQLL | PTPN23 | Q9H3S7 | PTN23_HUMAN | Unmodified | 341220 | 0 | 0 | 0 | 411930 | 0 | 0 | 0 | 0 | 0 | 0 | 0 | 0 | 0 | 0 | 0 | 0 | 110550 |
| EAASILQEL | PSMD12 | O00232 | PSD12_HUMAN | Oxidation (M) | 0 | 0 | 0 | 0 | 0 | 0 | 0 | 0 | 0 | 0 | 0 | 827630 | 0 | 0 | 0 | 0 | 0 | 0 |
| EAASLSEW | UTRN | P46939 | UTRN_HUMAN | Unmodified | 0 | 0 | 103180 | 416890 | 3652600 | 1656400 | 0 | 0 | 346900 | 839470 | 513430 | 0 | 1186200 | 796940 | 746140 | 96743 | 732470 | 788780 |
| EAASPELMI | AVIL | O75366 | AVIL_HUMAN A | Oxidation (M) | 0 | 0 | 0 | 0 | 0 | 0 | 0 | 0 | 0 | 0 | 207920 | 4812500 | 0 | 0 | 0 | 0 | 0 | 0 |
| EAASQGLLI | PPP1R16A; PPP | Q96I34; A0A | PP16A_HUMAN | Unmodified | 36287 | 0 | 0 | 0 | 135310 | 0 | 0 | 25362 | 2261700 | 93489 | 0 | 206130 | 141750 | 0 | 145840 | 100670 | 473740 | 2803500 |
| EAASRGILD | SVIP | Q8NHG7 | SVIP_HUMAN S | Unmodified | 0 | 0 | 0 | 0 | 2687100 | 0 | 2500600 | 1666000 | 0 | 5402000 | 0 | 0 | 12642000 | 10251000 | 0 | 0 | 3342300 | 10817000 |
| EAATLEVEF | HDGF | P51858 | HDGF_HUMAN | Oxidation (M) | 147590 | 0 | 0 | 0 | 680530 | 0 | 0 | 0 | 0 | 279960 | 1027900 | 4655000 | 1287700 | 0 | 0 | 0 | 0 | 0 |
| EAATLEVEF | HDGF | P51858 | HDGF_HUMAN | Unmodified | 40952 | 0 | 0 | 0 | 1350900 | 0 | 49018 | 0 | 1205500 | 257440 | 1012200 | 5984500 | 1395200 | 194950 | 0 | 0 | 150120 | 0 |
| EAATQAQQ | QARS1; QARS1 | P47897; F2Z | SYQ_HUMAN G | Unmodified | 0 | 0 | 52126 | 0 | 0 | 0 | 0 | 0 | 0 | 0 | 626330 | 3005900 | 2329900 | 0 | 0 | 0 | 0 | 0 |
| EAAVSHAG | NCKAP1 | Q9Y2A7 | NCKP1_HUMA | Unmodified | 0 | 87135 | 254030 | 154790 | 0 | 0 | 0 | 0 | 361970 | 70533 | 631120 | 3017400 | 0 | 0 | 0 | 1296900 | 0 | 0 |
| EAAWAISN | KPNA4; KPNA3; | O00629; O0 | IMA3_HUMAN I | Unmodified | 38775 | 0 | 0 | 0 | 0 | 0 | 0 | 0 | 1408100 | 0 | 0 | 589530 | 0 | 0 | 0 | 0 | 0 | 0 |
| EAAWAISN | KPNA4; KPNA3; | O00629; O0 | IMA3_HUMAN I | Unmodified | 746680 | 654370 | 197170 | 453410 | 618870 | 419320 | 40003 | 0 | 645080 | 0 | 254980 | 2005700 | 0 | 0 | 368630 | 483780 | 0 | 229940 |
| EAAYHPEV | TLN1 | Q9Y490 | TLN1_HUMAN | Unmodified | 0 | 170350 | 2940100 | 4510300 | 3344200 | 0 | 0 | 598600 | 1850200 | 491800 | 935600 | 5460500 | 1809500 | 1652200 | 1882100 | 2671300 | 323700 | 1076000 |
| EAAYHPEV | TLN1 | Q9Y490 | TLN1_HUMAN | Unmodified | 0 | 0 | 0 | 391010 | 10777000 | 2169300 | 4028100 | 2451500 | 1072100 | 300940 | 0 | 626980 | 7291200 | 7492600 | 0 | 0 | 2116000 | 4081100 |
| EACDQHGI | CPOX | P36551 | HEM6_HUMAN | Unmodified | 0 | 0 | 0 | 0 | 0 | 0 | 0 | 0 | 289160 | 91687 | 0 | 148530 | 0 | 28543 | 0 | 0 | 0 | 0 |
| EACGGHG | ACOX3 | O15254 | ACOX3_HUMA | Unmodified | 0 | 87224 | 0 | 0 | 0 | 30905 | 0 | 0 | 265950 | 0 | 0 | 0 | 0 | 0 | 0 | 0 | 0 | 0 |
| EACGVIVEI | RUVBL1 | Q9Y265 | RUVB1_HUMA | Unmodified | 0 | 0 | 0 | 0 | 0 | 0 | 0 | 0 | 179880 | 0 | 0 | 6684200 | 0 | 0 | 0 | 148150 | 0 | 0 |
| EACIPCGP | ELAPOR2; ELAP | A8MWY0; H | ELAP2_HUMAN | Unmodified | 0 | 0 | 0 | 0 | 0 | 0 | 0 | 0 | 468960 | 62711 | 17301 | 0 | 0 | 0 | 0 | 0 | 0 | 0 |
| EACLASNY | VPS37D | Q86XT2 | VP37D_HUMAN | Unmodified | 597810 | 0 | 0 | 0 | 0 | 0 | 0 | 0 | 0 | 0 | 0 | 0 | 0 | 0 | 1238500 | 495990 | 0 | 0 |
| EACNVPEP | IQGAP2; IQGAP | Q13576; E7E | IQGA2_HUMAN | Unmodified | 0 | 0 | 335930 | 228700 | 0 | 0 | 0 | 0 | 0 | 0 | 0 | 0 | 0 | 0 | 315520 | 0 | 0 | 0 |
| EADAIDDTI | DAAM1 | Q9Y4D1 | DAAM1_HUMA | Unmodified | 0 | 0 | 0 | 0 | 1890300 | 0 | 679430 | 181470 | 1383300 | 396970 | 216640 | 3502500 | 1083900 | 471360 | 719850 | 398680 | 0 | 355490 |
| EADAIDDTI | DAAM1 | Q9Y4D1 | DAAM1_HUMA | Unmodified | 0 | 0 | 0 | 0 | 1087400 | 0 | 493060 | 206660 | 0 | 179060 | 0 | 0 | 115210 | 299110 | 0 | 0 | 0 | 0 |
| EADAVFPR | MARVELD2; MA | Q8N4S9; A1 | MALD2_HUMA | Unmodified | 0 | 0 | 116750 | 92561 | 0 | 0 | 0 | 0 | 0 | 0 | 0 | 424790 | 0 | 0 | 135510 | 0 | 0 | 0 |
| EADDIVNW | P4HB; P4HB; P4 | P07237; A0A | PDIA1_HUMAN | Unmodified | 0 | 0 | 4370400 | 4341600 | 0 | 95555 | 0 | 0 | 20983000 | 1872200 | 3289700 | 2346800 | 1014200 | 0 | 467020 | 0 | 0 | 0 |
| EADDIVNW | P4HB; P4HB; P4 | P07237; A0A | PDIA1_HUMAN | Unmodified | 0 | 0 | 0 | 0 | 0 | 0 | 0 | 203100 | 6388400 | 135480 | 105460 | 0 | 0 | 0 | 995960 | 0 | 0 | 0 |
| EADESLNFI | TLN1 | Q9Y490 | TLN1_HUMAN | Unmodified | 300150 | 470160 | 0 | 766390 | 5287600 | 728500 | 1468500 | 242880 | 341030 | 532510 | 0 | 510610 | 2063700 | 2308600 | 452380 | 272440 | 844810 | 1690900 |
| EADGILKPL | AARS1; AARS1; | P49588; A0A | SYAC_HUMAN | Unmodified | 43161 | 0 | 0 | 0 | 0 | 0 | 0 | 0 | 305020 | 0 | 0 | 990760 | 0 | 0 | 0 | 98290 | 0 | 0 |
| EADGSETP | SND1 | Q7KZF4 | SND1_HUMAN | Unmodified | 0 | 0 | 0 | 524530 | 0 | 0 | 0 | 0 | 0 | 289340 | 706130 | 10228000 | 0 | 0 | 209400 | 203180 | 0 | 0 |
| EADIDGDG | CALM2; CALM3 | P0DP24; P0 | CALM2_HUMA | 2 Oxidation (M) | 405530 | 0 | 0 | 0 | 0 | 0 | 0 | 0 | 0 | 0 | 0 | 0 | 0 | 0 | 874270 | 450860 | 0 | 0 |
| EADLVFISV | UGDH; UGDH; | O60701; E7E | UGDH_HUMAN | Unmodified | 0 | 0 | 0 | 0 | 6546800 | 0 | 0 | 0 | 717520 | 0 | 318540 | 0 | 1141400 | 0 | 0 | 0 | 0 | 0 |
| EADQLKQD | NF2; NF2 | P35240; A0A | MERL_HUMAN | Unmodified | 0 | 0 | 0 | 206920 | 0 | 0 | 0 | 0 | 0 | 0 | 0 | 178690 | 93227 | 0 | 0 | 0 | 0 | 0 |
| EADYVAQA | HEBP1; HEBP1 | Q9NRV9; A0 | HEBP1_HUMAN | Unmodified | 0 | 0 | 0 | 249820 | 1070900 | 0 | 0 | 0 | 0 | 191110 | 262700 | 238230 | 2326100 | 1532200 | 285560 | 120710 | 357600 | 873250 |
| EADYVAQA | HEBP1; HEBP1 | Q9NRV9; A0 | HEBP1_HUMAN | Unmodified | 0 | 0 | 0 | 0 | 650880 | 0 | 0 | 0 | 0 | 216770 | 0 | 738480 | 86279 | 671280 | 157450 | 460400 | 0 | 859580 |
| EAEAAIYHL | PKM; PKM | P14618; B4D | KPYM_HUMAN | Unmodified | 0 | 1394400 | 402900 | 2100900 | 1577400 | 797500 | 0 | 0 | 2630700 | 203220 | 1237700 | 7744700 | 302670 | 0 | 3814900 | 0 | 187090 | 140730 |
| EAEAAIYHL | PKM; PKM | P14618; B4D | KPYM_HUMAN | Unmodified | 5077800 | 2818700 | 5395700 | 15579000 | 3965500 | 834850 | 522990 | 0 | 25691000 | 2554500 | 21575000 | 1.54E+08 | 1096400 | 117140 | 19549000 | 14562000 | 0 | 0 |
| EAEAAMFH | PKM; PKM | H3BTN5; H3 | H3BTN5_HUM | Unmodified | 0 | 0 | 0 | 0 | 0 | 0 | 0 | 0 | 135470 | 586090 | 0 | 353480 | 0 | 0 | 0 | 0 | 0 | 0 |
| EAEAGPAE | NBEAL2; NBEAL | Q6ZNJ1; H0 | NBEL2_HUMAN | Unmodified | 0 | 0 | 0 | 0 | 0 | 0 | 0 | 0 | 0 | 0 | 0 | 2012300 | 0 | 0 | 0 | 0 | 0 | 0 |
| EAEAMKAT | OSBP | P22059 | OSBP1_HUMA | Unmodified | 0 | 0 | 0 | 0 | 0 | 0 | 0 | 0 | 0 | 0 | 0 | 0 | 160540 | 0 | 0 | 0 | 0 | 0 |
| EAEDDDTG | TGOLN2; TGOL | O43493; A0A | TGON2_HUMA | Unmodified | 0 | 0 | 0 | 0 | 0 | 0 | 0 | 0 | 2686500 | 434830 | 2869100 | 0 | 0 | 0 | 0 | 0 | 0 | 0 |
| EAEDGFRK | PACSIN1; PACS | Q9BY11; Q9 | PACN1_HUMA | Unmodified | 8219300 | 0 | 0 | 0 | 0 | 0 | 0 | 0 | 0 | 0 | 0 | 0 | 0 | 0 | 5282800 | 0 | 0 | 0 |
| EAEDNIRE | PFDN1 | O60925 | PFD1_HUMAN | Oxidation (M) | 0 | 0 | 0 | 0 | 581790 | 0 | 0 | 0 | 0 | 0 | 196450 | 594900 | 238250 | 0 | 0 | 0 | 99606 | 215420 |
| EAEDNIRE | PFDN1 | O60925 | PFD1_HUMAN | Unmodified | 0 | 0 | 0 | 0 | 439940 | 0 | 0 | 0 | 926970 | 0 | 0 | 356240 | 161790 | 0 | 0 | 0 | 0 | 0 |
| EAEEREPKI | GLG1; GLG1 | Q92896; H3 | GSLG1_HUMA | Unmodified | 0 | 0 | 0 | 0 | 0 | 389600 | 0 | 0 | 702390 | 1053500 | 158770 | 0 | 0 | 0 | 0 | 0 | 0 | 0 |
| EAEESYMM | TMTC3 | Q6ZXV5 | TMTC3_HUMA | Unmodified | 0 | 0 | 0 | 0 | 0 | 0 | 0 | 0 | 1498600 | 215520 | 0 | 0 | 0 | 0 | 64895 | 0 | 0 | 0 |
| EAEEVYR | LUC7L2; LUC7L | Q9Y383; A0A | LC7L2_HUMAN | Unmodified | 0 | 0 | 0 | 19444 | 1531700 | 0 | 0 | 0 | 592950 | 55662 | 66530 | 273300 | 2188700 | 0 | 9384.8 | 0 | 1171100 | 586150 |
| EAEFLQK | PIP5K1A; PIP5K | Q99755; O6 | PI51A_HUMAN | Unmodified | 136020 | 120300 | 1071800 | 241470 | 240320 | 0 | 40056 | 44425 | 482080 | 202350 | 531540 | 1622700 | 114590 | 64877 | 365530 | 491160 | 198560 | 93868 |
| EAEGAPQV | CALD1; CALD1; | Q05682; C9 | CALD1_HUMA | Unmodified | 0 | 0 | 0 | 321740 | 665960 | 0 | 0 | 0 | 0 | 258290 | 227180 | 1037100 | 123830 | 201610 | 135660 | 353820 | 59412 | 105810 |
| EAEGSSAE | NASP; NASP; N | P49321; E9P | NASP_HUMAN | Unmodified | 0 | 0 | 0 | 0 | 0 | 0 | 0 | 0 | 0 | 354900 | 0 | 409740 | 150630 | 0 | 0 | 0 | 0 | 0 |
| EAEGVIVG | SORL1; SORL1; | Q92673; E9F | SORL_HUMAN | Unmodified | 0 | 0 | 0 | 0 | 0 | 0 | 0 | 0 | 3490000 | 107470 | 0 | 0 | 0 | 0 | 0 | 0 | 2634300 | 0 |
| EAEIHSKPV | PTGFRN | Q9P2B2 | FPRP_HUMAN | Unmodified | 1145500 | 937680 | 2535100 | 430480 | 0 | 0 | 0 | 0 | 0 | 0 | 0 | 6089600 | 0 | 0 | 4512500 | 3667700 | 0 | 0 |
| EAEKELHE | SEPTIN11; SEP | Q9NVA2; H0 | SEP11_HUMAN | Unmodified | 0 | 0 | 0 | 0 | 712300 | 0 | 0 | 0 | 0 | 0 | 0 | 0 | 0 | 0 | 0 | 0 | 174820 | 0 |
| EAEKLESE | SPTBN1; SPTBN | Q01082; A0A | SPTB2_HUMAN | Unmodified | 87426 | 0 | 0 | 0 | 0 | 0 | 0 | 0 | 720900 | 0 | 0 | 4353700 | 0 | 0 | 0 | 0 | 0 | 0 |
| EAELDVNE | PAFAH1B1; PAF | P43034; A0A | LIS1_HUMAN P | Unmodified | 0 | 0 | 0 | 0 | 736990 | 370330 | 0 | 65334 | 0 | 0 | 813830 | 2760100 | 492760 | 0 | 0 | 0 | 0 | 0 |
| EAELDVNE | PAFAH1B1; PAF | P43034; A0A | LIS1_HUMAN P | Unmodified | 0 | 0 | 0 | 0 | 2485900 | 0 | 0 | 0 | 0 | 310650 | 0 | 0 | 900440 | 1099100 | 0 | 0 | 523010 | 0 |

| | | | | | | | | | | | | | | | | | | | | | |
|---|---|---|---|---|---|---|---|---|---|---|---|---|---|---|---|---|---|---|---|---|---|
| EAELTRELE | MYO10; MYO10 | Q9HD67; A0 | MYO10_HUMA | Unmodified | 0 | 0 | 44209000 | 0 | 0 | 0 | 0 | 0 | 0 | 0 | 0 | 0 | 0 | 0 | 0 | 0 | 0 |
| EAEMEEHR | EIF4G1; EIF4G1 | Q04637; H7 | IF4G1_HUMAN | Unmodified | 0 | 26259 | 118430 | 0 | 737800 | 0 | 0 | 0 | 79875 | 0 | 132930 | 625610 | 98952 | 0 | 62426 | 72264 | 115910 | 706990 |
| EAENLMQE | IFIT1 | P09914 | IFIT1_HUMAN | Unmodified | 0 | 0 | 0 | 0 | 343570 | 0 | 0 | 0 | 0 | 0 | 0 | 0 | 38770 | 0 | 0 | 0 | 0 | 604350 |
| EAENPEGE | HDGF | P51858 | HDGF_HUMAN | Unmodified | 0 | 0 | 0 | 0 | 0 | 0 | 0 | 0 | 381020 | 0 | 0 | 0 | 1596600 | 0 | 0 | 0 | 0 | 0 |
| EAENPREV | SF3A1 | Q15459 | SF3A1_HUMAN | Unmodified | 1026100 | 636140 | 0 | 326750 | 0 | 0 | 256190 | 0 | 0 | 603090 | 214640 | 0 | 0 | 378900 | 0 | 0 | 497210 | 0 |
| EAEPDLLA | NCDN; NCDN | Q9UBB6; C9 | NCDN_HUMAN | Unmodified | 45427 | 23355 | 74493 | 0 | 0 | 0 | 0 | 0 | 0 | 0 | 0 | 0 | 0 | 0 | 0 | 95327 | 0 | 0 |
| EAEPHYMV | PTK7; PTK7; PTI | Q13308; H0 | PTK7_HUMAN | Oxidation (M) | 0 | 81834 | 15903 | 0 | 0 | 0 | 0 | 0 | 2501700 | 818690 | 648660 | 691730 | 0 | 0 | 3931100 | 0 | 0 | 0 |
| EAEPHYMV | PTK7; PTK7; PTI | Q13308; H0 | PTK7_HUMAN | Unmodified | 0 | 0 | 0 | 0 | 0 | 0 | 0 | 0 | 726400 | 0 | 0 | 0 | 1580100 | 0 | 0 | 0 | 0 | 0 |
| EAERDMLP | COBLL1; COBL | Q53SF7; A0 | COBL1_HUMA | Oxidation (M) | 889040 | 521610 | 0 | 0 | 0 | 0 | 0 | 0 | 0 | 0 | 0 | 0 | 627960 | 0 | 271670 | 280350 | 0 | 0 |
| EAERDMLP | COBLL1; COBL | Q53SF7; A0 | COBL1_HUMA | Unmodified | 198150 | 166010 | 81599 | 0 | 0 | 0 | 0 | 0 | 0 | 0 | 0 | 0 | 0 | 0 | 0 | 0 | 0 | 0 |
| EAERELQAI | SEPTIN10; SEP | Q9P0V9; B5 | SEP10_HUMAN | Unmodified | 0 | 0 | 0 | 0 | 6323500 | 0 | 0 | 0 | 0 | 0 | 0 | 0 | 0 | 0 | 0 | 0 | 631900 | 80293 |
| EAERKLEE | NUCB1; NUCB | Q02818; H7 | NUCB1_HUMA | Unmodified | 0 | 0 | 0 | 0 | 0 | 0 | 0 | 0 | 210240 | 0 | 0 | 0 | 0 | 0 | 0 | 0 | 0 | 0 |
| EAESCDCL | TUBB8; TUBB8; | Q3ZCM7; Q | TBB8_HUMAN | Unmodified | 0 | 0 | 0 | 0 | 5806900 | 0 | 5725000 | 0 | 14195000 | 3079400 | 435220 | 7142600 | 0 | 0 | 0 | 0 | 0 | 0 |
| EAESSPFV | HSP90B1; HSP | P14625; A0A | ENPL_HUMAN | Unmodified | 676560 | 296540 | 1917700 | 644410 | 0 | 97959 | 244900 | 82908 | 2.66E+08 | 19690000 | 28124000 | 28228000 | 0 | 0 | 1693000 | 1161600 | 0 | 0 |
| EAESSPFV | HSP90B1; HSP | P14625; A0A | ENPL_HUMAN | Unmodified | 0 | 0 | 0 | 0 | 347270 | 0 | 0 | 57165 | 0 | 2451900 | 0 | 0 | 0 | 61294 | 0 | 0 | 181930 | 0 |
| EAESSPFV | HSP90B1; HSP | P14625; A0A | ENPL_HUMAN | Unmodified | 0 | 81586 | 2842100 | 0 | 137750 | 0 | 104700 | 38832 | 2960200 | 18241000 | 373740 | 0 | 244070 | 0 | 297230 | 733510 | 944750 | 195330 |
| EAETKSPLV | ADD2 | P35612 | ADDB_HUMAN | Unmodified | 0 | 0 | 0 | 0 | 1430200 | 365880 | 149330 | 0 | 0 | 0 | 0 | 0 | 0 | 0 | 0 | 0 | 14568 | 0 |
| EAEVLPGL | SOGA1; SOGA1 | O94964; X6 | SOGA1_HUMA | Unmodified | 0 | 0 | 136750 | 0 | 0 | 0 | 0 | 0 | 0 | 0 | 0 | 0 | 443210 | 246960 | 0 | 0 | 0 | 0 |
| EAEVVLCG | ACAA2; ACAA2; | P42765; A0A | THIM_HUMAN | Unmodified | 0 | 0 | 0 | 0 | 0 | 0 | 0 | 72858 | 762040 | 0 | 0 | 0 | 0 | 0 | 0 | 0 | 0 | 0 |
| EAEYFELPE | KCTD12 | Q96CX2 | KCD12_HUMA | Unmodified | 0 | 0 | 0 | 0 | 0 | 0 | 0 | 0 | 1690600 | 244750 | 591580 | 0 | 0 | 0 | 0 | 0 | 0 | 0 |
| EAFEAYTD | PLXNB2; PLXNE | O15031; E2F | PLXB2_HUMAN | Unmodified | 580890 | 505970 | 899610 | 1083800 | 215880 | 0 | 0 | 0 | 1363500 | 0 | 0 | 303130 | 0 | 0 | 2147300 | 1932300 | 0 | 0 |
| EAFEEAGV | NUDT19 | A8MXV4 | NUD19_HUMA | Unmodified | 0 | 0 | 0 | 0 | 0 | 0 | 0 | 0 | 5725900 | 0 | 0 | 0 | 0 | 0 | 0 | 0 | 0 | 0 |
| EAFEISK | YWHAB; YWHA | P31946; P63 | 1433B_HUMAN | Unmodified | 0 | 0 | 891910 | 0 | 656170 | 0 | 6349.2 | 41606 | 1690100 | 0 | 3617700 | 3749100 | 0 | 0 | 988530 | 552740 | 11406 | 183900 |
| EAFELFDR | MYL6B; MYL6B | P14649; F8V | MYL6B_HUMAN | Unmodified | 0 | 0 | 0 | 0 | 345260 | 0 | 0 | 0 | 737590 | 0 | 165760 | 459290 | 0 | 0 | 0 | 47462 | 44167 | 66146 |
| EAFGDGYH | ABCA12 | Q86UK0 | ABCAC_HUMA | Unmodified | 0 | 0 | 0 | 0 | 0 | 0 | 0 | 0 | 394260 | 105550 | 0 | 0 | 51683 | 52530 | 0 | 0 | 0 | 0 |
| EAFGFIERG | CSDE1; CSDE1 | O75534; E9F | CSDE1_HUMA | Unmodified | 180160 | 105450 | 0 | 0 | 0 | 0 | 0 | 0 | 0 | 0 | 0 | 5988700 | 0 | 0 | 0 | 668230 | 0 | 0 |
| EAFIDWAR | ERO1B; ERO1B | Q86YB8; Q5 | ERO1B_HUMA | Unmodified | 0 | 0 | 0 | 0 | 0 | 0 | 0 | 0 | 1221900 | 0 | 0 | 0 | 0 | 0 | 0 | 0 | 0 | 0 |
| EAFLQEVW | VARS1; VARS1 | P26640; A0A | SYVC_HUMAN | Unmodified | 0 | 0 | 0 | 0 | 0 | 0 | 0 | 0 | 0 | 0 | 0 | 6119300 | 0 | 0 | 0 | 0 | 0 | 0 |
| EAFNIIGR | PIGS | Q96S52 | PIGS_HUMAN | Unmodified | 147960 | 0 | 0 | 0 | 0 | 235030 | 0 | 35104 | 5915200 | 727020 | 0 | 0 | 0 | 0 | 0 | 20769 | 0 | 0 |
| EAFNMIDQ | MYL12A; MYL12 | P19105; O1 | ML12A_HUMAN | Unmodified | 0 | 0 | 0 | 0 | 0 | 0 | 0 | 0 | 194970 | 0 | 0 | 1722900 | 0 | 0 | 0 | 230900 | 0 | 0 |
| EAFNMIDQ | MYL12A; MYL12 | P19105; O1 | ML12A_HUMAN | Oxidation (M) | 0 | 0 | 0 | 0 | 0 | 0 | 0 | 0 | 0 | 0 | 0 | 253490 | 0 | 0 | 158730 | 96258 | 0 | 0 |
| EAFPVGPE | ARVCF; ARVCF; | O00192; E9F | ARVC_HUMAN | Unmodified | 115250 | 251280 | 0 | 615900 | 70329 | 0 | 72984 | 67853 | 0 | 0 | 0 | 0 | 0 | 112500 | 0 | 440370 | 0 | 0 |
| EAFQALEK | TWF1; TWF1 | Q12792; F8V | TWF1_HUMAN | Unmodified | 525110 | 395660 | 346550 | 153010 | 0 | 0 | 0 | 0 | 0 | 0 | 337460 | 888690 | 48750 | 0 | 1103700 | 530130 | 0 | 0 |
| EAFQLFDR | MYL6; MYL6; M | P60660; G3 | MYL6_HUMAN | Unmodified | 498760 | 248610 | 400950 | 0 | 0 | 0 | 0 | 0 | 1142200 | 0 | 356130 | 6167600 | 0 | 0 | 532480 | 837440 | 0 | 0 |
| EAFQLFDR | MYL6; MYL6; M | P60660; G8J | MYL6_HUMAN | Unmodified | 2719200 | 2157700 | 4017100 | 5022600 | 67354000 | 9476200 | 1378600 | 523410 | 8084000 | 1854500 | 4194600 | 41505000 | 11729000 | 5821800 | 6327000 | 3432300 | 3425100 | 10881000 |
| EAFQLFDR | MYL6; MYL6; M | P60660; G8J | MYL6_HUMAN | Unmodified | 0 | 0 | 0 | 0 | 448390 | 0 | 0 | 0 | 0 | 0 | 0 | 0 | 0 | 0 | 0 | 0 | 0 | 114330 |
| EAFQPQEP | VCL | P18206 | VINC_HUMAN | Unmodified | 0 | 0 | 0 | 0 | 0 | 380780 | 0 | 0 | 566260 | 0 | 0 | 946150 | 0 | 0 | 0 | 87946 | 0 | 0 |
| EAFSDGITS | ASNS | P08243 | ASNS_HUMAN | Unmodified | 0 | 0 | 0 | 1.08E+09 | 8.74E+08 | 0 | 2.39E+08 | 0 | 0 | 0 | 3934800 | 0 | 8.05E+08 | 4.99E+08 | 0 | 0 | 0 | 0 |
| EAFSDGITS | ASNS | P08243 | ASNS_HUMAN | Unmodified | 0 | 0 | 0 | 0 | 0 | 0 | 0 | 62793 | 0 | 0 | 0 | 0 | 419830 | 0 | 0 | 0 | 430250 | 0 |
| EAFSLFDK | CALM2; CALM3 | P0DP24; P0 | CALM2_HUMA | Unmodified | 2544900 | 2246600 | 8192000 | 783290 | 623020 | 278640 | 0 | 53495 | 3012600 | 209500 | 2509100 | 29182000 | 120840 | 0 | 15227000 | 5596900 | 74969 | 175980 |
| EAFSLFDK | CALML3 | P27482 | CALL3_HUMAN | Unmodified | 0 | 0 | 0 | 0 | 46688 | 0 | 0 | 0 | 2416900 | 0 | 0 | 0 | 0 | 0 | 0 | 0 | 0 | 0 |
| EAFSLFDK | CALM2; CALM3 | P0DP24; P0 | CALM2_HUMA | Unmodified | 12295000 | 10966000 | 2304800 | 10114000 | 24343000 | 11834000 | 1449500 | 895630 | 16124000 | 5346200 | 22234000 | 62452000 | 4039000 | 2877800 | 20878000 | 12743000 | 7430100 | 5938300 |
| EAFTEAR | ITGA1 | P56199 | ITA1_HUMAN | Unmodified | 696220 | 1057000 | 0 | 1467000 | 0 | 0 | 0 | 0 | 0 | 0 | 0 | 0 | 355110 | 0 | 0 | 0 | 0 | 0 |
| EAFTNKPN | FLNB; FLNB | O75369; E7 | FLNB_HUMAN | Unmodified | 0 | 0 | 0 | 0 | 2628900 | 719380 | 0 | 319690 | 0 | 0 | 0 | 3281000 | 0 | 0 | 0 | 0 | 0 | 302330 |
| EAFVVEEW | COPB2; COPB2 | P35606; H0 | COPB2_HUMA | Unmodified | 0 | 0 | 104260 | 0 | 258150 | 0 | 0 | 29203 | 266720 | 201350 | 1294100 | 3843300 | 633440 | 0 | 270960 | 95793 | 0 | 0 |
| EAGAGGLA | FLNA; FLNA; FL | P21333; A0A | FLNA_HUMAN | Unmodified | 522760 | 740150 | 1945000 | 0 | 0 | 0 | 0 | 0 | 0 | 0 | 0 | 1183200 | 160060 | 0 | 1539000 | 1040900 | 0 | 0 |
| EAGAGGLA | FLNA; FLNA; FL | P21333; A0A | FLNA_HUMAN | Unmodified | 334350 | 1582200 | 0 | 2563800 | 0 | 0 | 599860 | 0 | 0 | 0 | 0 | 0 | 0 | 0 | 1298300 | 665690 | 0 | 0 |
| EAGDAESR | SFN | P31947 | 1433S_HUMAN | Unmodified | 0 | 0 | 0 | 0 | 6134600 | 2923200 | 0 | 0 | 1127600 | 5036200 | 0 | 0 | 186830 | 0 | 0 | 0 | 3948900 | 399930 |
| EAGDFHFC | PTPN23; PTPN2 | Q9H3S7; R4 | PTN23_HUMAN | Unmodified | 0 | 700610 | 0 | 0 | 0 | 0 | 0 | 0 | 0 | 0 | 0 | 0 | 0 | 0 | 0 | 0 | 0 | 205800 |
| EAGDFHFC | PTPN23; PTPN2 | Q9H3S7; R4 | PTN23_HUMAN | Unmodified | 362130 | 475480 | 0 | 0 | 279350 | 0 | 382680 | 0 | 0 | 0 | 0 | 0 | 0 | 0 | 0 | 0 | 0 | 0 |
| EAGEGGEA | MARCKS | P29966 | MARCS_HUMA | Unmodified | 0 | 0 | 734570 | 0 | 0 | 0 | 0 | 9044 | 22480000 | 2545800 | 648900 | 1102200 | 0 | 0 | 0 | 195600 | 0 | 0 |
| EAGELKPEI | COPB1 | P53618 | COPB_HUMAN | Unmodified | 89027 | 0 | 169940 | 0 | 0 | 0 | 0 | 0 | 1211400 | 0 | 1090200 | 7416300 | 331990 | 0 | 217650 | 295100 | 0 | 0 |
| EAGENWQ | CACNA2D1; CA | P54289; H0 | CA2D1_HUMA | Unmodified | 705190 | 644010 | 0 | 0 | 0 | 0 | 1299700 | 323740 | 2740300 | 315150 | 0 | 0 | 0 | 0 | 0 | 0 | 0 | 0 |
| EAGENWQ | CACNA2D1; CA | P54289; H0 | CA2D1_HUMA | Unmodified | 568830 | 248300 | 0 | 0 | 0 | 0 | 0 | 0 | 0 | 611490 | 0 | 0 | 0 | 0 | 0 | 0 | 0 | 0 |
| EAGEQGDI | G3BP1; G3BP1; | Q13283; A0 | G3BP1_HUMA | Unmodified | 1013100 | 578150 | 237120 | 0 | 31492 | 0 | 0 | 64138 | 127760 | 0 | 694030 | 1674300 | 0 | 0 | 0 | 0 | 0 | 0 |
| EAGFKRPV | TJP2; TJP2; TJP2 | Q9UDY2; A0 | ZO2_HUMAN T | Oxidation (M) | 0 | 0 | 0 | 220670 | 0 | 0 | 0 | 0 | 1803900 | 894810 | 0 | 168150 | 0 | 1261300 | 0 | 0 | 17635 | 0 |
| EAGFKRPV | TJP2; TJP2; TJP2 | Q9UDY2; A0 | ZO2_HUMAN T | Unmodified | 0 | 0 | 0 | 0 | 73360 | 0 | 0 | 0 | 2120200 | 0 | 0 | 0 | 0 | 0 | 0 | 0 | 0 | 0 |
| EAGFLRPV | TJP1; TJP1; TJP1 | Q07157; A0 | ZO1_HUMAN T | Unmodified | 65672 | 0 | 257920 | 597510 | 0 | 0 | 0 | 0 | 519370 | 0 | 90733 | 1106800 | 0 | 0 | 1161900 | 1138100 | 0 | 0 |
| EAGFPPGV | ALDH1A1 | P00352 | AL1A1_HUMAN | Oxidation (M) | 0 | 0 | 0 | 0 | 0 | 0 | 0 | 0 | 0 | 0 | 1613200 | 8278100 | 0 | 0 | 0 | 0 | 19783 | 0 |
| EAGHGTQ | ATP2B1; ATP2B | P20020; E7E | AT2B1_HUMAN | Unmodified | 0 | 0 | 0 | 0 | 0 | 0 | 0 | 0 | 0 | 0 | 572210 | 2506300 | 0 | 0 | 2745900 | 3116100 | 0 | 0 |
| EAGHGTQ | ATP2B1; ATP2B | P20020; E7E | AT2B1_HUMAN | Unmodified | 0 | 0 | 0 | 3652700 | 0 | 170870 | 0 | 0 | 1195400 | 339900 | 0 | 694490 | 2055800 | 744080 | 20054000 | 3215600 | 0 | 0 |
| EAGHGTTK | ATP2B4; ATP2B | P23634; H7E | AT2B4_HUMAN | Unmodified | 0 | 0 | 136120 | 110060 | 0 | 0 | 0 | 0 | 0 | 0 | 0 | 1037900 | 0 | 0 | 163200 | 0 | 0 | 0 |
| EAGIQPSLI | MELTF; MELTF | P08582; C9J | TRFM_HUMAN | Unmodified | 0 | 0 | 0 | 0 | 0 | 0 | 0 | 0 | 0 | 0 | 0 | 0 | 0 | 0 | 0 | 0 | 0 | 0 |
| EAGISKDRI | TALDO1; TALDC | P37837; F2Z | TALDO_HUMA | Unmodified | 0 | 0 | 0 | 0 | 412890 | 161540 | 0 | 23452 | 0 | 163650 | 0 | 0 | 330580 | 309500 | 0 | 0 | 150830 | 0 |
| EAGKEAAL | NCAPH; NCAPH | Q15003; E9F | CND2_HUMAN | Unmodified | 0 | 0 | 0 | 0 | 66836 | 0 | 1192600 | 69970 | 0 | 352970 | 0 | 0 | 0 | 0 | 0 | 0 | 0 | 265580 |
| EAGLAPVPI | UROD; UROD; U | P06132; A0A | DCUP_HUMAN | Oxidation (M) | 0 | 0 | 0 | 0 | 0 | 0 | 0 | 0 | 0 | 0 | 0 | 997400 | 0 | 0 | 0 | 0 | 0 | 0 |
| EAGMQPQI | HYOU1; HYOU1 | Q9Y4L1; E9F | HYOU1_HUMA | Oxidation (M) | 0 | 0 | 0 | 0 | 0 | 0 | 0 | 0 | 8721700 | 1296400 | 1303400 | 0 | 0 | 0 | 0 | 0 | 0 | 0 |
| EAGMQPQI | HYOU1; HYOU1 | Q9Y4L1; E9F | HYOU1_HUMA | Unmodified | 0 | 0 | 81966 | 0 | 0 | 0 | 1238300 | 0 | 19545000 | 2391400 | 1295000 | 0 | 0 | 0 | 0 | 0 | 0 | 0 |
| EAGNINQS | KIF23; KIF23; KI | Q02241; A0 | KIF23_HUMAN | Unmodified | 0 | 0 | 0 | 0 | 1864000 | 0 | 0 | 0 | 0 | 0 | 0 | 1359900 | 0 | 0 | 0 | 0 | 0 | 0 |
| EAGNINQS | KIF23; KIF23; KI | Q02241; A0 | KIF23_HUMAN | Oxidation (M) | 0 | 0 | 0 | 493420 | 643970 | 0 | 0 | 0 | 0 | 0 | 345680 | 1125900 | 0 | 0 | 0 | 0 | 0 | 0 |
| EAGNINTSI | KIF20A | O95235 | KI20A_HUMAN | Unmodified | 0 | 0 | 983080 | 2879000 | 4193400 | 0 | 0 | 0 | 1290700 | 604740 | 0 | 0 | 3214400 | 2840900 | 0 | 299120 | 1062300 | 0 |
| EAGNPGSL | ITGA3; ITGA3 | P26006; D6F | ITA3_HUMAN | Unmodified | 1271300 | 0 | 951140 | 767770 | 23944000 | 13816000 | 0 | 0 | 0 | 419710 | 1168300 | 8845900 | 0 | 0 | 1506800 | 0 | 7858700 | 18319000 |
| EAGNPGSL | ITGA3; ITGA3 | P26006; D6F | ITA3_HUMAN | Unmodified | 0 | 0 | 0 | 0 | 21290000 | 10643000 | 121580 | 0 | 0 | 302890 | 0 | 0 | 14478 | 121800 | 0 | 0 | 5446400 | 10933000 |
| EAGNPGSL | ITGA3; ITGA3 | P26006; D6F | ITA3_HUMAN | Unmodified | 0 | 0 | 0 | 0 | 59083000 | 18670000 | 0 | 0 | 0 | 0 | 0 | 0 | 0 | 0 | 0 | 0 | 20340000 | 35652000 |
| EAGQCFES | MOV10 | Q9HCE1 | MOV10_HUMA | Unmodified | 3213700 | 2682500 | 544650 | 1062600 | 0 | 0 | 0 | 0 | 0 | 0 | 0 | 0 | 0 | 0 | 1901300 | 0 | 0 | 0 |
| EAGSIVRLY | DLG1; DLG1; D | Q12959; H7 | DLG1_HUMAN | Unmodified | 0 | 0 | 0 | 0 | 548010 | 0 | 0 | 17230 | 0 | 1183300 | 0 | 0 | 0 | 0 | 0 | 0 | 799780 | 165130 |
| EAGSQKDE | TFRC; TFRC | P02786; G3 | TFR1_HUMAN | Unmodified | 1590700 | 1266000 | 1520900 | 1826900 | 0 | 0 | 0 | 0 | 18699000 | 218520 | 0 | 3980700 | 0 | 0 | 0 | 4401000 | 0 | 0 |
| EAGSQKDE | TFRC; TFRC | P02786; G3 | TFR1_HUMAN | Unmodified | 645340 | 740200 | 1035400 | 5462600 | 454330 | 0 | 0 | 0 | 28562000 | 0 | 1714000 | 844430 | 0 | 0 | 2362500 | 1347200 | 193680 | 0 |
| EAGVEMGI | DARS1; DARS1 | P14868; H7E | SYDC_HUMAN | Unmodified | 0 | 0 | 0 | 0 | 0 | 0 | 0 | 0 | 369940 | 0 | 0 | 3242900 | 0 | 0 | 59830 | 48195 | 0 | 0 |
| EAGVEMGI | DARS1; DARS1 | P14868; H7E | SYDC_HUMAN | Oxidation (M) | 0 | 0 | 74124 | 0 | 0 | 0 | 0 | 0 | 0 | 0 | 0 | 735060 | 0 | 0 | 0 | 47841 | 0 | 0 |
| EAGVFVPR | ACLY | P53396 | ACLY_HUMAN | Unmodified | 344990 | 268060 | 675830 | 635040 | 324050 | 151320 | 310840 | 46613 | 1688200 | 0 | 805590 | 13490000 | 110060 | 0 | 705030 | 755910 | 106420 | 0 |
| EAGVGNG | DCPS | Q96C86 | DCPS_HUMAN | Unmodified | 0 | 0 | 0 | 0 | 0 | 0 | 0 | 0 | 0 | 0 | 63358 | 795990 | 0 | 0 | 0 | 0 | 0 | 0 |
| EAGYEGPL | ACE; ACE | P12821; A0A | ACE_HUMAN A | Unmodified | 684430 | 0 | 0 | 0 | 0 | 0 | 0 | 0 | 0 | 0 | 0 | 0 | 0 | 0 | 0 | 0 | 0 | 0 |
| EAHQLFLEI | FEN1; FEN1 | P39748; I3L | FEN1_HUMAN | Unmodified | 684410 | 0 | 0 | 357180 | 547010 | 0 | 0 | 38084 | 0 | 0 | 2448000 | 7812900 | 552180 | 87068 | 0 | 0 | 0 | 0 |
| EAIDKAGY | ENO2; ENO2; E | P09104; F5H | ENOG_HUMAN | Unmodified | 0 | 0 | 0 | 0 | 297380 | 0 | 337850 | 28910 | 0 | 0 | 0 | 0 | 205860 | 0 | 0 | 0 | 728740 | 881180 |
| EAIDSPVSF | UPF1 | Q92900 | RENT1_HUMAN | Unmodified | 1329600 | 2078000 | 961940 | 1600800 | 0 | 0 | 0 | 0 | 215120 | 26622 | 123050 | 1372800 | 0 | 0 | 671080 | 953500 | 0 | 0 |
| EAIDSYIK | CLTC; CLTC | Q00610; A0 | CLH1_HUMAN | Unmodified | 7153900 | 5100900 | 7365500 | 2791200 | 1812300 | 454860 | 801720 | 263250 | 3001300 | 0 | 0 | 40099000 | 1193300 | 0 | 4426200 | 7219800 | 402860 | 0 |
| EAIEAIAES | PLCB2 | Q00722 | PLCB2_HUMAN | Unmodified | 0 | 0 | 0 | 0 | 0 | 0 | 0 | 0 | 0 | 0 | 572790 | 5079900 | 0 | 0 | 0 | 0 | 0 | 0 |
| EAIEGTYIDI | RPS11; RPS11 | P62280; M0 | RS11_HUMAN | Unmodified | 4083300 | 2857000 | 392810 | 3328900 | 2463300 | 432620 | 2033600 | 79440 | 70836 | 390430 | 1986900 | 34674000 | 0 | 621860 | 3649900 | 2783600 | 38515 | 1045100 |
| EAIEGTYIDI | RPS11; RPS11 | P62280; M0 | RS11_HUMAN | Unmodified | 305980 | 481330 | 0 | 0 | 323640 | 0 | 45211 | 309210 | 0 | 51671 | 212950 | 6658200 | 0 | 0 | 0 | 0 | 0 | 0 |
| EAIELREAV | RRBP1; RRBP1; | Q9P2E9; A0 | RRBP1_HUMAN | Unmodified | 0 | 0 | 0 | 0 | 158650 | 0 | 79694 | 13191 | 0 | 115580 | 0 | 0 | 1379300 | 489720 | 0 | 0 | 122980 | 330260 |
| EAIENATTN | CHMP4A; CHM | Q9BY43; E9F | CHM4A_HUMA | Unmodified | 2086200 | 2376000 | 334880 | 205170 | 0 | 0 | 0 | 0 | 2676400 | 0 | 0 | 1206100 | 0 | 0 | 634300 | 702070 | 0 | 0 |
| EAIESAIGG | DYNLT1 | P63172 | DYLT1_HUMAN | Unmodified | 0 | 0 | 0 | 0 | 0 | 0 | 0 | 0 | 0 | 368430 | 618720 | 8184800 | 0 | 0 | 0 | 119140 | 0 | 0 |
| EAIFQDTMI | SFTPB; SFTPB | P07988; H0 | PSPB_HUMAN | Unmodified | 0 | 0 | 0 | 0 | 407140 | 0 | 0 | 0 | 1077900 | 299880 | 0 | 0 | 0 | 278910 | 0 | 0 | 0 | 0 |
| EAIHSQLLE | PFDN1; PFDN1 | O60925; E5F | PFD1_HUMAN | Unmodified | 0 | 0 | 0 | 0 | 1195600 | 216810 | 0 | 136350 | 216830 | 557440 | 0 | 550220 | 371890 | 0 | 0 | 32338 | 583490 | 413230 |
| EAIHSQLLE | PFDN1; PFDN1 | O60925; E5F | PFD1_HUMAN | Unmodified | 0 | 0 | 0 | 0 | 0 | 0 | 0 | 0 | 0 | 0 | 0 | 0 | 0 | 0 | 0 | 0 | 286520 | 0 |
| EAIIPGSVY | UPF1 | Q92900 | RENT1_HUMAN | Unmodified | 1675400 | 2560600 | 1594600 | 906350 | 0 | 0 | 192460 | 0 | 0 | 0 | 0 | 1619300 | 0 | 0 | 0 | 1209400 | 0 | 0 |
| EAIKGVGTI | ANXA11 | P50995 | ANX11_HUMAN | Unmodified | 316740 | 0 | 0 | 79194 | 0 | 94538 | 0 | 0 | 16075000 | 4537900 | 387640 | 95854 | 372210 | 808480 | 0 | 0 | 93653 | 2378900 |
| EAIKSSLIIL | PSMA5 | P28066 | PSA5_HUMAN | Unmodified | 0 | 0 | 0 | 0 | 0 | 0 | 0 | 93379 | 0 | 0 | 0 | 0 | 0 | 0 | 0 | 0 | 508750 | 0 |
| EAILAIHK | ACTN4; ACTN4; | O43707; F5 | ACTN4_HUMA | Unmodified | 138900 | 162680 | 0 | 0 | 0 | 0 | 0 | 0 | 0 | 0 | 254240 | 2135200 | 0 | 0 | 205610 | 166950 | 0 | 0 |
| EAILDIITSR | ANXA6; ANXA6; | P08133; E5R | ANXA6_HUMA | Unmodified | 788690 | 1657800 | 8268700 | 9068500 | 2206500 | 1466600 | 820670 | 253640 | 1.16E+08 | 7156400 | 0 | 594470 | 1882800 | 823190 | 10386000 | 5208600 | 455510 | 3533600 |
| EAILDIITSR | ANXA6; ANXA6; | P08133; E5R | ANXA6_HUMA | Unmodified | 0 | 0 | 0 | 0 | 1490500 | 822100 | 453370 | 71964 | 0 | 3109300 | 0 | 0 | 1817100 | 647880 | 0 | 0 | 275790 | 2384500 |
| EAINLLEPM | PSMD1; PSMD1 | Q99460; A0 | PSMD1_HUMA | Oxidation (M) | 0 | 0 | 0 | 0 | 70551 | 0 | 0 | 0 | 0 | 20131 | 0 | 2668300 | 863100 | 0 | 0 | 0 | 0 | 0 |
| EAINVEQA | RAB7A; RAB7A; | P51149; C9J | RAB7A_HUMAN | Unmodified | 31404000 | 36760000 | 16812000 | 14327000 | 42534000 | 11666000 | 4750400 | 2280300 | 50832000 | 22042000 | 42159000 | 98830000 | 47231000 | 14205000 | 36658000 | 20174000 | 3733000 | 25073000 |
| EAINVEQA | RAB7A; RAB7A; | P51149; C9J | RAB7A_HUMAN | Unmodified | 0 | 0 | 0 | 0 | 0 | 359940 | 899510 | 0 | 0 | 301000 | 0 | 26388 | 938520 | 666030 | 0 | 0 | 645730 | 1351700 |
| EAIQAYSES | SCARB1 | B7ZKQ9 | B7ZKQ9_HUM | Oxidation (M) | 113940 | 103490 | 0 | 207210 | 0 | 0 | 0 | 0 | 0 | 0 | 0 | 1405000 | 0 | 0 | 328100 | 290090 | 0 | 0 |
| EAIQCVDN | MTREX | P42285 | MTREX_HUMA | Unmodified | 0 | 0 | 0 | 0 | 0 | 0 | 0 | 0 | 0 | 0 | 0 | 820820 | 0 | 0 | 0 | 0 | 0 | 0 |
| EAIQHPADI | CYRIB; CYRIB; | Q9NUQ9; E5 | CYRIB_HUMAN | Unmodified | 0 | 0 | 18531 | 0 | 36248 | 0 | 0 | 0 | 54158 | 0 | 0 | 169230 | 0 | 0 | 6785.4 | 8160 | 0 | 0 |
| EAIQHPADI | CYRIB; CYRIB; | Q9NUQ9; E5 | CYRIB_HUMAN | Unmodified | 73851 | 0 | 0 | 0 | 0 | 0 | 181760 | 0 | 415720 | 0 | 0 | 1443700 | 0 | 0 | 33559 | 0 | 0 | 75976 |

| | | | | | | | | | | | | | | | | | | | | | | |
|---|---|---|---|---|---|---|---|---|---|---|---|---|---|---|---|---|---|---|---|---|---|---|
| EAIQLIAR | PGM1; PGM1 | P36871; A0A | PGM1_HUMAN | Unmodified | 0 | 0 | 0 | 0 | 0 | 0 | 0 | 0 | 0 | 0 | 0 | 1930100 | 0 | 0 | 0 | 0 | 0 | 0 |
| EAIRELTGK | NPTX2 | P47972 | NPTX2_HUMAN | Unmodified | 0 | 0 | 0 | 0 | 0 | 0 | 66667 | 87871 | 3841400 | 164330 | 0 | 265050 | 0 | 0 | 0 | 0 | 0 | 35675 |
| EAISLVCEA | SHC1; SHC1; S | P29353; Q5T | SHC1_HUMAN | Unmodified | 0 | 0 | 0 | 0 | 0 | 0 | 0 | 0 | 328390 | 0 | 399920 | 884360 | 0 | 518440 | 0 | 0 | 732510 | 0 |
| EAIVNSCVF | DYNC1H1; DYN | Q14204; A0A | DYHC1_HUMA | Unmodified | 0 | 0 | 0 | 0 | 0 | 0 | 0 | 0 | 1783400 | 54986 | 312250 | 7757400 | 0 | 0 | 0 | 0 | 0 | 0 |
| EAKEPLWIF | CBS; CBSL; CBS | P35520; P0D | CBS_HUMAN C | Unmodified | 0 | 0 | 0 | 0 | 0 | 0 | 0 | 0 | 0 | 0 | 0 | 1216900 | 0 | 0 | 0 | 0 | 0 | 0 |
| EAKFPILSA | NT5E; NT5E | P21589; Q96 | 5NTD_HUMAN | Unmodified | 119030 | 103650 | 0 | 0 | 26474000 | 5257600 | 0 | 0 | 0 | 0 | 0 | 0 | 0 | 0 | 0 | 0 | 8153900 | 8434700 |
| EAKNSDVL | COLGALT1 | Q8NBJ5 | GT251_HUMAN | Unmodified | 0 | 0 | 0 | 0 | 0 | 0 | 70208 | 0 | 0 | 1780500 | 0 | 0 | 0 | 0 | 0 | 0 | 0 | 66237 |
| EAKPDELM | VAPA | Q9P0L0 | VAPA_HUMAN | Unmodified | 145740 | 0 | 137650 | 0 | 2574700 | 0 | 241710 | 65199 | 387440 | 57918 | 659000 | 933730 | 773310 | 0 | 89392 | 58811 | 94241 | 511250 |
| EAKPEDLM | VAPB | O95292 | VAPB_HUMAN | Unmodified | 0 | 0 | 0 | 0 | 0 | 0 | 0 | 0 | 1049000 | 0 | 0 | 2570700 | 0 | 0 | 0 | 0 | 0 | 0 |
| EALAEAALE | SLC9A3R1; SLC | O14745; J3Q | NHRF1_HUMA | Unmodified | 0 | 0 | 0 | 2951300 | 973780 | 0 | 0 | 0 | 1387200 | 0 | 0 | 8066600 | 0 | 0 | 4780500 | 3255100 | 205540 | 0 |
| EALDDVTLI | IDE; IDE; IDE; ID | P14735; A0A | IDE_HUMAN In | Unmodified | 0 | 0 | 0 | 0 | 0 | 0 | 0 | 0 | 0 | 0 | 0 | 3516500 | 43707 | 0 | 0 | 103480 | 0 | 0 |
| EALDLGITC | DPYSL5 | Q9BPU6 | DPYL5_HUMAN | Unmodified | 0 | 0 | 0 | 0 | 0 | 0 | 0 | 0 | 841640 | 0 | 2641400 | 8082100 | 0 | 0 | 0 | 0 | 0 | 0 |
| EALDVLGA | MRPS27; MRPS | Q92552; D6I | RT27_HUMAN : | Unmodified | 0 | 0 | 0 | 0 | 0 | 0 | 0 | 0 | 1208900 | 0 | 72053 | 379840 | 0 | 0 | 0 | 0 | 0 | 0 |
| EALEAVLIK | PHKA1; PHKA1; | P46020; A6N | KPB1_HUMAN | Unmodified | 0 | 0 | 25365 | 0 | 0 | 0 | 0 | 0 | 0 | 0 | 16387 | 354960 | 0 | 0 | 51993 | 0 | 0 | 0 |
| EALEEAEKA | LAMB1; LAMB1; | P07942; A0A | LAMB1_HUMA | Unmodified | 37264 | 85726 | 0 | 0 | 0 | 0 | 97224 | 0 | 0 | 0 | 0 | 0 | 0 | 0 | 0 | 0 | 369020 | 2601000 |
| EALEHKDR | TMOD3 | Q9NYL9 | TMOD3_HUMA | Unmodified | 0 | 0 | 0 | 0 | 4354000 | 629410 | 231870 | 0 | 0 | 161230 | 0 | 0 | 1808000 | 981110 | 0 | 0 | 0 | 0 |
| EALENANTI | CHMP4B | Q9H444 | CHM4B_HUMA | Unmodified | 4727000 | 7088300 | 4171600 | 2555200 | 629530 | 0 | 152280 | 56286 | 8023400 | 853840 | 1674000 | 6478800 | 542880 | 216890 | 4271000 | 4163100 | 0 | 0 |
| EALENSHTI | CHMP4C | Q96CF2 | CHM4C_HUMA | Unmodified | 5175700 | 4438100 | 665360 | 673390 | 0 | 0 | 58640 | 0 | 2663600 | 75163 | 0 | 449900 | 0 | 0 | 1951400 | 2980500 | 0 | 0 |
| EALEPSGEI | ERLIN1 | O75477 | ERLN1_HUMAN | Unmodified | 0 | 0 | 0 | 0 | 0 | 0 | 0 | 0 | 3473100 | 304840 | 598210 | 0 | 0 | 0 | 0 | 0 | 0 | 0 |
| EALESEFR | EXOC7; EXOC7 | Q9UPT5; C9. | EXOC7_HUMA | Unmodified | 0 | 422910 | 107700 | 0 | 0 | 0 | 0 | 0 | 0 | 0 | 1476900 | 709430 | 0 | 0 | 844450 | 0 | 0 | 0 |
| EALGDAQC | DNAJC7; DNAJ | Q99615; A0A | DNJC7_HUMA | Unmodified | 0 | 0 | 0 | 0 | 0 | 0 | 0 | 194360 | 793790 | 0 | 0 | 439780 | 0 | 0 | 0 | 0 | 0 | 0 |
| EALGHWL( | MOGS; MOGS; | Q13724; A0A | MOGS_HUMAN | Unmodified | 0 | 0 | 0 | 0 | 64738 | 13219 | 0 | 0 | 250020 | 255680 | 139860 | 37294 | 0 | 0 | 0 | 0 | 0 | 0 |
| EALGIPAAA | ATIC | P31939 | PUR9_HUMAN | Unmodified | 0 | 0 | 677070 | 851420 | 6369200 | 1897600 | 583710 | 489490 | 2337900 | 1142300 | 1724700 | 10108000 | 12887000 | 5272800 | 1252000 | 951390 | 2234500 | 2302400 |
| EALHERAN | GPD2; GPD2 | P43304; F5G | GPDM_HUMAN | Oxidation (M) | 0 | 0 | 110750 | 0 | 0 | 0 | 0 | 0 | 0 | 1584600 | 0 | 0 | 0 | 0 | 0 | 0 | 2448700 | 7392900 |
| EALKEAVILI | VPS4B; VPS4A; | O75351; Q9I | VPS4B_HUMAN | Unmodified | 0 | 0 | 0 | 0 | 1864800 | 0 | 0 | 0 | 0 | 638230 | 0 | 0 | 495940 | 607900 | 0 | 0 | 0 | 0 |
| EALLEEK | BAIAP2L1 | Q9UHR4 | BI2L1_HUMAN | Unmodified | 0 | 0 | 94734 | 0 | 2027500 | 1785300 | 0 | 11572 | 656590 | 271460 | 73453 | 486230 | 0 | 553260 | 229980 | 126130 | 0 | 43983 |
| EALLGVQEI | RCN2; RCN2 | Q14257; H0Y | RCN2_HUMAN | Unmodified | 206390 | 0 | 0 | 0 | 0 | 0 | 1637000 | 0 | 697150 | 182330 | 350070 | 44035 | 0 | 0 | 0 | 0 | 0 | 0 |
| EALLQASR | MSN | P26038 | MOES_HUMAN | Unmodified | 360280 | 1285400 | 1298800 | 238720 | 1118200 | 322820 | 528830 | 15316 | 1020900 | 0 | 82617 | 0 | 918010 | 26254 | 705300 | 349530 | 113390 | 424660 |
| EALLQASRI | MSN | P26038 | MOES_HUMAN | Unmodified | 0 | 0 | 736220 | 887590 | 18041000 | 7724100 | 0 | 53085 | 0 | 261250 | 133460 | 736620 | 3465300 | 2276500 | 787380 | 378450 | 2345200 | 14925000 |
| EALLQASRI | MSN | P26038 | MOES_HUMAN | Unmodified | 0 | 0 | 0 | 0 | 3819700 | 2330000 | 0 | 0 | 0 | 0 | 0 | 0 | 393080 | 0 | 0 | 0 | 698840 | 1443800 |
| EALLSIGK | FAM136A; FAM | Q96C01; E7I | F136A_HUMAN | Unmodified | 209770 | 0 | 0 | 0 | 61234 | 0 | 321210 | 86305 | 110750 | 86104 | 0 | 646160 | 201710 | 122510 | 0 | 0 | 189370 | 0 |
| EALLSSAVI | CCT2; CCT2; C( | P78371; F5G | TCPB_HUMAN | Unmodified | 0 | 0 | 0 | 0 | 1774800 | 0 | 237860 | 0 | 11339000 | 0 | 16080000 | 45561000 | 5561500 | 0 | 1748000 | 2284100 | 269090 | 3145400 |
| EALLVANG | ALDH16A1; ALD | Q8IZ83; F5H | A16A1_HUMAN | Unmodified | 5177400 | 0 | 0 | 0 | 0 | 0 | 0 | 0 | 0 | 92596 | 0 | 1109200 | 0 | 0 | 4326900 | 5182200 | 0 | 0 |
| EALNMLTW | CD109 | Q6YHK3 | CD109_HUMA | Unmodified | 0 | 0 | 0 | 0 | 0 | 0 | 0 | 0 | 1033600 | 0 | 0 | 0 | 0 | 0 | 0 | 0 | 0 | 0 |
| EALNPETIE | NCAM2; NCAM | O15394; H9I | NCAM2_HUMA | Unmodified | 0 | 813170 | 11417000 | 4911200 | 0 | 0 | 0 | 0 | 0 | 0 | 0 | 0 | 0 | 0 | 20230000 | 11696000 | 0 | 0 |
| EALNVFGN | GALE; GALE; GA | Q14376; Q5( | GALE_HUMAN | Unmodified | 0 | 0 | 0 | 0 | 2172100 | 945320 | 0 | 0 | 0 | 0 | 0 | 0 | 966410 | 0 | 0 | 0 | 1337900 | 0 |
| EALNVINTH | SRP72; SRP72; | O76094; D6I | SRP72_HUMAN | Unmodified | 0 | 0 | 0 | 0 | 726960 | 0 | 206700 | 110330 | 0 | 60592 | 0 | 1880800 | 1272000 | 0 | 0 | 0 | 162510 | 540280 |
| EALPGIIK | GALNT16 | Q8N428 | GLT16_HUMAN | Unmodified | 0 | 0 | 0 | 0 | 0 | 0 | 0 | 0 | 3438200 | 409580 | 0 | 0 | 0 | 0 | 0 | 0 | 19764 | 0 |
| EALPGSGC | ABHD14B; ABH | Q96IU4; F8V | ABHEB_HUMA | Unmodified | 0 | 0 | 127690 | 0 | 0 | 0 | 0 | 0 | 158100 | 29621 | 127530 | 978530 | 0 | 0 | 457050 | 196310 | 0 | 41917 |
| EALPSWLH | DAG1; DAG1 | Q14118; C9J | DAG1_HUMAN | Unmodified | 0 | 0 | 248480 | 383510 | 0 | 0 | 0 | 0 | 2440800 | 314960 | 192860 | 0 | 0 | 0 | 676660 | 0 | 0 | 0 |
| EALQDVED | YWHAE; YWHA | P62258; I3L( | 1433E_HUMAN | Unmodified | 0 | 0 | 9444000 | 10247000 | 10167000 | 2559300 | 936310 | 1200300 | 6586700 | 1172900 | 8166000 | 18765000 | 0 | 5796900 | 14393000 | 20109000 | 3363700 | 7113100 |
| EALQEADV | DLG5 | Q8TDM6 | DLG5_HUMAN | Unmodified | 0 | 0 | 338990 | 126850 | 1043900 | 60690 | 0 | 90614 | 201210 | 127340 | 0 | 0 | 0 | 0 | 160440 | 80488 | 691850 | 1350100 |
| EALQLMAT | PSMD1; PSMD1 | Q99460; A0A | PSMD1_HUMA | Oxidation (M) | 0 | 0 | 0 | 0 | 0 | 0 | 0 | 0 | 0 | 0 | 0 | 1059900 | 0 | 0 | 0 | 0 | 0 | 0 |
| EALQMLR | APOBEC3B; AP( | Q9UH17; P3 | ABC3B_HUMA | Oxidation (M) | 0 | 348120 | 378850 | 0 | 0 | 0 | 0 | 0 | 0 | 0 | 0 | 0 | 0 | 0 | 424930 | 290940 | 68760 | 0 |
| EALQNIND | DPYSL2 | A0A1C7CYX | A0A1C7CYX9_I | Unmodified | 0 | 0 | 0 | 0 | 4390700 | 711380 | 174430 | 0 | 712100 | 410670 | 2759800 | 0 | 0 | 0 | 0 | 0 | 0 | 254000 |
| EALQSDWI | UBXN6; UBXN6 | Q9BZV1; K7I | UBXN6_HUMA | Unmodified | 97463 | 209070 | 0 | 0 | 0 | 0 | 11052 | 0 | 146720 | 0 | 0 | 0 | 0 | 0 | 327960 | 105970 | 0 | 0 |
| EALQYFYFI | NUP93; NUP93 | Q8N1F7; H3 | NUP93_HUMA | Unmodified | 0 | 0 | 0 | 0 | 0 | 0 | 0 | 0 | 0 | 40039 | 0 | 1293300 | 0 | 0 | 0 | 0 | 0 | 0 |
| EALRAGLN | EIF2S1; EIF2S1; | P05198; H0Y | IF2A_HUMAN E | Unmodified | 195740 | 0 | 0 | 473060 | 884510 | 0 | 0 | 0 | 418720 | 0 | 0 | 0 | 1092200 | 428030 | 0 | 0 | 313060 | 0 |
| EALSAVLPF | PDE2A; PDE2A; | O00408; F5I | PDE2A_HUMA | Unmodified | 0 | 0 | 0 | 0 | 124240 | 0 | 0 | 0 | 0 | 0 | 475670 | 6751500 | 0 | 0 | 0 | 0 | 0 | 0 |
| EALSNKVD | MAGEA4; MAGE | P43358; C9J | MAGA4_HUMA | Unmodified | 544070 | 598690 | 0 | 0 | 0 | 0 | 0 | 0 | 302960 | 0 | 0 | 0 | 0 | 0 | 0 | 261450 | 0 | 0 |
| EALSTATV( | VPS13C; VPS13 | Q709C8; A0 | VP13C_HUMAN | Unmodified | 0 | 0 | 0 | 132600 | 420250 | 161110 | 0 | 117680 | 351840 | 134430 | 0 | 1830700 | 1331500 | 145760 | 69099 | 104790 | 748940 | 0 |
| EALTDADD | PLSCR3; PLSCF | Q9NRY6; I3L | PLS3_HUMAN I | Unmodified | 0 | 312050 | 0 | 20546 | 0 | 0 | 0 | 0 | 0 | 0 | 0 | 0 | 0 | 0 | 208090 | 0 | 0 | 0 |
| EALTYDGA | EIF4H; EIF4H; E | Q15056; A0A | IF4H_HUMAN E | Unmodified | 2192400 | 3218700 | 3659700 | 8723200 | 0 | 0 | 0 | 0 | 3067500 | 0 | 249440 | 3131300 | 0 | 0 | 6370500 | 4121300 | 0 | 0 |
| EALVDTLT( | IPO9 | Q96P70 | IPO9_HUMAN I | Unmodified | 0 | 0 | 0 | 0 | 34376 | 0 | 0 | 0 | 620500 | 721060 | 62388 | 308440 | 45873 | 0 | 651440 | 0 | 0 | 0 |
| EALVSVWL | CPE; CPE | P16870; H0Y | CBPE_HUMAN | Unmodified | 0 | 0 | 0 | 0 | 0 | 0 | 0 | 0 | 235530 | 152480 | 0 | 0 | 0 | 0 | 0 | 0 | 0 | 0 |
| EAMEDGEII | NCL; NCL; NCL | P19338; C9J | NUCL_HUMAN | Unmodified | 0 | 0 | 0 | 0 | 0 | 0 | 0 | 0 | 176660 | 0 | 500340 | 377830 | 0 | 0 | 0 | 0 | 0 | 0 |
| EAMGIYIST | AHSA1; AHSA1; | O95433; H0Y | AHSA1_HUMA | Oxidation (M) | 0 | 0 | 0 | 0 | 0 | 0 | 0 | 0 | 0 | 0 | 0 | 1200700 | 0 | 0 | 0 | 0 | 0 | 0 |
| EAMNDPLL | DHX15 | O43143 | DHX15_HUMA | Oxidation (M) | 0 | 0 | 0 | 0 | 0 | 0 | 0 | 0 | 0 | 0 | 0 | 1847100 | 0 | 0 | 0 | 0 | 0 | 0 |
| EAMNDPLL | DHX15 | O43143 | DHX15_HUMA | Unmodified | 203640 | 0 | 0 | 0 | 0 | 0 | 0 | 0 | 0 | 0 | 0 | 2388000 | 0 | 0 | 296620 | 251710 | 0 | 0 |
| EAMNHPGI | DHFR | P00374 | DYR_HUMAN D | Unmodified | 0 | 18925 | 0 | 0 | 82693 | 0 | 30647 | 23344 | 0 | 0 | 0 | 16708 | 61294 | 40050 | 23145 | 0 | 0 | 0 |
| EAMRIGAE | ENO1; ENO1; E | P06733; K7E | ENOA_HUMAN | Oxidation (M) | 0 | 0 | 0 | 0 | 10029000 | 2465300 | 387070 | 34115 | 0 | 2774800 | 0 | 0 | 5562300 | 1304800 | 0 | 0 | 2349500 | 7760000 |
| EAMRIGAE | ENO1; ENO1; E | P06733; K7E | ENOA_HUMAN | Unmodified | 0 | 0 | 0 | 0 | 32820000 | 7295600 | 487630 | 46383 | 1186900 | 8270600 | 0 | 1515800 | 10385000 | 7027500 | 0 | 0 | 10295000 | 13131000 |
| EAMSAYNS | AGL | P35573 | GDE_HUMAN ( | Oxidation (M) | 0 | 0 | 0 | 0 | 0 | 0 | 0 | 0 | 0 | 0 | 0 | 61874 | 0 | 0 | 0 | 0 | 0 | 0 |
| EAMSAYNS | AGL | P35573 | GDE_HUMAN ( | Unmodified | 0 | 0 | 0 | 0 | 0 | 0 | 0 | 0 | 0 | 0 | 0 | 609200 | 0 | 0 | 0 | 0 | 0 | 0 |
| EAMVESIEY | PLA2G4A | P47712 | PA24A_HUMAN | Unmodified | 0 | 0 | 0 | 0 | 0 | 0 | 0 | 0 | 0 | 0 | 0 | 1168900 | 0 | 0 | 0 | 0 | 0 | 0 |
| EANALAAG | SPTBN2 | O15020 | SPTN2_HUMAN | Unmodified | 0 | 0 | 773290 | 357830 | 0 | 0 | 0 | 0 | 0 | 0 | 0 | 1612000 | 0 | 0 | 875160 | 825260 | 0 | 0 |
| EANAPGPV | SUMF1; SUMF1 | Q8NBK3; F5( | SUMF1_HUMA | Unmodified | 57173 | 73218 | 0 | 0 | 0 | 0 | 139520 | 169240 | 632310 | 0 | 0 | 446190 | 0 | 0 | 0 | 0 | 0 | 98012 |
| EANEILQR | IMPDH2; IMPDI | P12268; P20 | IMDH2_HUMA | Unmodified | 326170 | 215420 | 363180 | 242980 | 445600 | 76916 | 285300 | 193540 | 1310100 | 308650 | 617620 | 3731100 | 749000 | 241630 | 122110 | 430300 | 730170 | 1359500 |
| EANELQQV | SPTAN1; SPTAN | Q13813; A0A | SPTN1_HUMA | Unmodified | 0 | 0 | 0 | 0 | 0 | 0 | 0 | 0 | 0 | 0 | 0 | 1556900 | 1228600 | 0 | 0 | 0 | 0 | 0 |
| EANFTVSSI | EIF4A3; EIF4A3 | P38919; I3L3 | IF4A3_HUMAN | Unmodified | 0 | 0 | 0 | 0 | 0 | 0 | 0 | 0 | 0 | 0 | 305630 | 1570200 | 0 | 0 | 0 | 0 | 0 | 0 |
| EANHDGDI | ATP2B1; ATP2B | P20020; F8V | AT2B1_HUMAN | Unmodified | 1505300 | 1348300 | 4267800 | 7240400 | 2344000 | 0 | 0 | 0 | 2804300 | 844430 | 6883500 | 19401000 | 11595000 | 294260 | 10122000 | 5787500 | 408640 | 0 |
| EANHNVSA | CYFIP1; CYFIP2 | Q7L576; Q9( | CYFP1_HUMAN | Unmodified | 206460 | 0 | 0 | 0 | 0 | 0 | 0 | 0 | 0 | 0 | 0 | 3337700 | 0 | 0 | 593580 | 0 | 0 | 0 |
| EANIQAVD | SLK | Q9H2G2 | SLK_HUMAN S | Unmodified | 0 | 0 | 0 | 0 | 0 | 0 | 0 | 0 | 265790 | 0 | 489420 | 1172800 | 0 | 0 | 0 | 201950 | 0 | 0 |
| EANIQAVD | SLK | Q9H2G2 | SLK_HUMAN S | Unmodified | 0 | 0 | 0 | 0 | 0 | 0 | 0 | 0 | 1352100 | 178960 | 1654400 | 2585600 | 1467000 | 1015500 | 0 | 314150 | 0 | 0 |
| EANIRVVSE | PARP1; PARP1 | P09874; A0A | PARP1_HUMAN | Unmodified | 0 | 0 | 0 | 0 | 0 | 50566 | 1072300 | 166350 | 0 | 1488000 | 0 | 0 | 1450300 | 740060 | 0 | 0 | 185000 | 1276600 |
| EANLAASF( | DLD; DLD | P09622; E9P | DLDH_HUMAN | Unmodified | 0 | 0 | 0 | 0 | 0 | 0 | 0 | 0 | 6280800 | 0 | 0 | 0 | 0 | 0 | 0 | 0 | 0 | 0 |
| EANLLNAV | NSFL1C | Q9UNZ2 | NSF1C_HUMA | Unmodified | 0 | 0 | 0 | 0 | 0 | 0 | 0 | 0 | 327230 | 0 | 0 | 1953000 | 0 | 0 | 0 | 0 | 0 | 0 |
| EANNFLWF | RPL7; RPL7 | P18124; A8M | RL7_HUMAN 6( | Unmodified | 5218000 | 4425700 | 3198200 | 1331400 | 0 | 0 | 0 | 172650 | 213060 | 317080 | 1551100 | 53396000 | 739470 | 0 | 4081500 | 2667500 | 0 | 0 |
| EANNYEED | SCG3 | Q8WXD2 | SCG3_HUMAN | Unmodified | 0 | 0 | 0 | 0 | 0 | 0 | 0 | 0 | 2880200 | 1032500 | 0 | 0 | 0 | 0 | 0 | 112730 | 0 | 0 |
| EANNYEED | SCG3 | Q8WXD2 | SCG3_HUMAN | Unmodified | 0 | 0 | 0 | 0 | 0 | 0 | 0 | 0 | 8201200 | 3583200 | 0 | 0 | 0 | 0 | 0 | 0 | 0 | 0 |
| EANSIIITPC | NNT; NNT; NNT | Q13423; E9F | NNTM_HUMAN | Unmodified | 0 | 0 | 0 | 0 | 0 | 0 | 0 | 0 | 1407200 | 376740 | 0 | 0 | 0 | 0 | 0 | 0 | 0 | 0 |
| EANSKADP | PACSIN2; PACS | Q9UNF0; A0 | PACN2_HUMA | Unmodified | 13918000 | 23510000 | 0 | 17010000 | 569780 | 0 | 459660 | 502560 | 74562 | 673270 | 675550 | 824340 | 20385000 | 4566600 | 4052300 | 5317900 | 307700 | 634440 |
| EANSKADP | PACSIN2; PACS | Q9UNF0; A0 | PACN2_HUMA | Unmodified | 5826400 | 4450800 | 0 | 3646500 | 0 | 0 | 0 | 0 | 638310 | 506070 | 1051100 | 0 | 1598600 | 325780 | 629010 | 1787600 | 0 | 0 |
| EAPAEGEA | MARCKS | P29966 | MARCS_HUMA | Unmodified | 1709600 | 1248600 | 0 | 4471900 | 276150 | 0 | 1176600 | 647740 | 88702000 | 14844000 | 19268000 | 39894000 | 957800 | 971280 | 4425900 | 4404100 | 726790 | 4161400 |
| EAPAEGEA | MARCKS | P29966 | MARCS_HUMA | Unmodified | 0 | 0 | 0 | 3112400 | 0 | 225910 | 2692300 | 2594100 | 51982000 | 24267000 | 7770000 | 7300900 | 7209300 | 4106400 | 0 | 0 | 6367400 | 18217000 |
| EAPAEGEA | MARCKS | P29966 | MARCS_HUMA | Unmodified | 0 | 0 | 0 | 0 | 0 | 0 | 1308000 | 1571500 | 19398000 | 11437000 | 1990000 | 3058800 | 3975600 | 0 | 0 | 0 | 1916300 | 6433900 |
| EAPCVLIYI | EPRS1; EPRS1 | P07814; V9C | SYEP_HUMAN | Unmodified | 0 | 0 | 0 | 0 | 1015400 | 0 | 1045500 | 0 | 0 | 712340 | 0 | 3542000 | 4654600 | 3809900 | 0 | 71366 | 1613100 | 1542800 |
| EAPEPGME | PFAS | O15067 | PUR4_HUMAN | Unmodified | 0 | 0 | 0 | 0 | 709530 | 0 | 0 | 0 | 0 | 0 | 0 | 3317100 | 165080 | 0 | 0 | 0 | 0 | 0 |
| EAPEPMELI | STAT1; STAT1; S | P42224; A0A | STAT1_HUMAN | Unmodified | 0 | 0 | 0 | 0 | 1038200 | 0 | 0 | 0 | 0 | 0 | 0 | 0 | 0 | 0 | 0 | 0 | 158740 | 2080900 |
| EAPNPIHLT | EIF3F; EIF3F; EI | O00303; A0A | EIF3F_HUMAN | Unmodified | 0 | 92112 | 0 | 0 | 0 | 0 | 0 | 0 | 0 | 0 | 0 | 14316000 | 0 | 0 | 109770 | 56849 | 0 | 0 |
| EAPPMEKP | RPS15; RPS15; | P62841; K7E | RS15_HUMAN | Oxidation (M) | 0 | 149870 | 0 | 0 | 116210 | 0 | 0 | 0 | 0 | 0 | 294720 | 3696400 | 0 | 0 | 0 | 0 | 0 | 238630 |
| EAPPMEKP | RPS15; RPS15; | P62841; K7E | RS15_HUMAN | Unmodified | 658420 | 383370 | 278530 | 0 | 0 | 0 | 0 | 0 | 82923 | 0 | 182550 | 8233400 | 94259 | 0 | 425180 | 471530 | 0 | 0 |
| EAPPPLSPI | PRKCSH; PRKC | P14314; K7E | GLU2B_HUMA | Oxidation (M) | 0 | 0 | 0 | 0 | 0 | 0 | 0 | 0 | 80350000 | 5361300 | 12379000 | 1388600 | 0 | 0 | 0 | 0 | 0 | 0 |
| EAPPPLSPI | PRKCSH; PRKC | P14314; K7E | GLU2B_HUMA | Unmodified | 0 | 0 | 0 | 0 | 0 | 0 | 0 | 0 | 1E+08 | 0 | 700590 | 0 | 0 | 0 | 0 | 0 | 0 | 0 |
| EAPSRSSA | ITGA5 | P08648 | ITA5_HUMAN I | Unmodified | 0 | 0 | 0 | 99342 | 3525900 | 448910 | 274290 | 113170 | 0 | 0 | 96943 | 341030 | 275670 | 0 | 0 | 44813 | 2412400 | 1369100 |
| EAPTAYSLL | PPAT | Q06203 | PUR1_HUMAN | Oxidation (M) | 0 | 0 | 0 | 0 | 0 | 0 | 27002 | 0 | 0 | 711300 | 323950 | 2943900 | 0 | 0 | 0 | 0 | 18490 | 0 |
| EAPTAYSLL | PPAT | Q06203 | PUR1_HUMAN | Unmodified | 0 | 0 | 0 | 0 | 0 | 0 | 0 | 0 | 0 | 0 | 0 | 1037000 | 0 | 0 | 0 | 0 | 0 | 0 |
| EAPVDVLT( | TMEM109 | Q9BVC6 | TM109_HUMAN | Unmodified | 0 | 0 | 514610 | 473060 | 0 | 0 | 0 | 0 | 41116000 | 4746100 | 2050300 | 2294600 | 0 | 0 | 396520 | 0 | 0 | 0 |
| EAPWQIQE | ABCC1; ABCC1 | P33527; I3L | MRP1_HUMAN | Unmodified | 0 | 0 | 0 | 0 | 0 | 0 | 26659 | 0 | 0 | 0 | 550160 | 3719900 | 0 | 0 | 0 | 0 | 0 | 0 |
| EAQAELED | RAI14 | Q9P0K7 | RAI14_HUMAN | Unmodified | 0 | 0 | 0 | 0 | 455050 | 196670 | 0 | 0 | 0 | 99220 | 0 | 0 | 0 | 0 | 0 | 0 | 269610 | 409380 |
| EAQAQHNI | AIDA | Q96BJ3 | AIDA_HUMAN / | Unmodified | 183350 | 0 | 2486000 | 0 | 930380 | 1352100 | 0 | 0 | 0 | 0 | 0 | 2350900 | 61143 | 0 | 770060 | 4808600 | 0 | 7948000 |
| EAQAQHNI | AIDA | Q96BJ3 | AIDA_HUMAN / | Unmodified | 0 | 0 | 0 | 0 | 0 | 0 | 213880 | 0 | 0 | 46462 | 0 | 0 | 0 | 1348100 | 0 | 0 | 2312500 | 1977700 |
| EAQAVPATI | PLEC; PLEC | Q15149; H0Y | PLEC_HUMAN | Unmodified | 0 | 0 | 0 | 0 | 537050 | 0 | 0 | 0 | 0 | 0 | 0 | 376990 | 0 | 327260 | 0 | 0 | 1795900 | 792770 |
| EAQAVPATI | PLEC; PLEC | Q15149; H0Y | PLEC_HUMAN | Unmodified | 0 | 0 | 0 | 0 | 0 | 0 | 0 | 0 | 0 | 0 | 0 | 0 | 0 | 0 | 0 | 0 | 1352200 | 0 |
| EAQDDLVK | EZR; EZR | P15311; E7E | EZRI_HUMAN E | Unmodified | 2879400 | 2449000 | 1620200 | 0 | 29461000 | 6579900 | 1486200 | 1343800 | 11739000 | 2936300 | 1348000 | 270840 | 15464000 | 8383600 | 3438600 | 3360400 | 2470500 | 8074100 |
| EAQDIKFTE | IGF2BP3 | O00425 | IF2B3_HUMAN | Unmodified | 819020 | 0 | 0 | 1183200 | 0 | 0 | 0 | 192460 | 0 | 0 | 366800 | 982690 | 1276400 | 433860 | 0 | 0 | 0 | 0 |
| EAQDTSDC | ATF6; ATF6; ATF | P18850; A0A | ATF6A_HUMAN | Unmodified | 0 | 0 | 0 | 0 | 0 | 0 | 0 | 0 | 0 | 0 | 0 | 0 | 0 | 0 | 0 | 0 | 0 | 0 |
| EAQELSQN | DDX21 | Q9NR30 | DDX21_HUMA | Unmodified | 0 | 0 | 0 | 0 | 0 | 0 | 0 | 647760 | 0 | 43253 | 0 | 0 | 0 | 0 | 0 | 0 | 0 | 0 |
| EAQERLTG | TXNDC12 | O95881 | TXD12_HUMAN | Unmodified | 0 | 0 | 0 | 0 | 99053 | 0 | 0 | 0 | 1008700 | 1147500 | 0 | 0 | 0 | 0 | 0 | 0 | 0 | 0 |

| | | | | | | | | | | | | | | | | | | | | | | |
|---|---|---|---|---|---|---|---|---|---|---|---|---|---|---|---|---|---|---|---|---|---|---|
| EAQLGNH( | TTC17; TTC17 | Q96AE7; A0/ | TTC17_HUMAN | Unmodified | 0 | 0 | 0 | 0 | 0 | 0 | 0 | 0 | 1395500 | 166890 | 0 | 0 | 0 | 0 | 0 | 0 | 0 | 0 |
| EAQLYAAQ | NASP; NASP; N. | P49321; E9P | NASP_HUMAN | Unmodified | 0 | 0 | 0 | 0 | 708980 | 0 | 913290 | 312030 | 777260 | 1580600 | 406210 | 1896700 | 9161000 | 0 | 0 | 290300 | 0 | 172720 |
| EAQMPAVII | CEL | P19835 | CEL_HUMAN B | Unmodified | 0 | 0 | 0 | 0 | 0 | 0 | 0 | 0 | 368400 | 28917 | 404460 | 0 | 0 | 0 | 0 | 0 | 0 | 0 |
| EAQMPAVII | CEL | P19835 | CEL_HUMAN B | Unmodified | 0 | 0 | 0 | 0 | 0 | 0 | 0 | 0 | 7445000 | 386590 | 1415200 | 0 | 0 | 0 | 0 | 0 | 0 | 0 |
| EAQPGQS( | LTBP1; LTBP1; I | Q14766; E7E | LTBP1_HUMAN | Unmodified | 0 | 43176 | 0 | 0 | 0 | 0 | 0 | 0 | 0 | 0 | 0 | 0 | 0 | 0 | 0 | 0 | 521720 | 3435400 |
| EAQQYSEA | GLIPR2; GLIPR2 | Q9H4G4; Q5 | GAPR1_HUMAI | Unmodified | 2132100 | 3058000 | 2804400 | 2744200 | 0 | 0 | 0 | 0 | 927770 | 0 | 0 | 0 | 0 | 0 | 5962500 | 3660600 | 0 | 0 |
| EAQRIAESI | ACTN4; ACTN4; | O43707; F5( | ACTN4_HUMAI | Unmodified | 37556 | 0 | 0 | 0 | 4973900 | 14843 | 27764 | 48308 | 0 | 3651800 | 0 | 156460 | 76705 | 0 | 405870 | 0 | 33968 | 103340 |
| EAQSQLIK | DNAJC3 | Q13217 | DNJC3_HUMAI | Unmodified | 0 | 0 | 0 | 0 | 0 | 0 | 0 | 0 | 7805200 | 247070 | 0 | 0 | 0 | 0 | 0 | 0 | 0 | 0 |
| EAQSQLIKS | DNAJC3 | Q13217 | DNJC3_HUMAI | Unmodified | 0 | 0 | 0 | 0 | 0 | 0 | 0 | 0 | 825890 | 232200 | 60023 | 57031 | 0 | 0 | 0 | 0 | 0 | 0 |
| EAQSRCGE | PHGDH; PHGD | O43175; A0/ | SERA_HUMAN | Oxidation (M) | 0 | 205170 | 557310 | 0 | 0 | 0 | 2884600 | 122980 | 0 | 312130 | 0 | 0 | 9854400 | 3676900 | 1443700 | 235520 | 777500 | 929340 |
| EAQSRCGE | PHGDH; PHGD | O43175; A0/ | SERA_HUMAN | Unmodified | 0 | 0 | 0 | 0 | 128200 | 0 | 0 | 0 | 0 | 0 | 0 | 0 | 3435400 | 2331100 | 0 | 24515 | 0 | 0 |
| EAQTSFLHI | DDX1; DDX1; D | Q92499; A0/ | DDX1_HUMAN | Unmodified | 1073800 | 981310 | 837530 | 4551300 | 1085900 | 322750 | 272960 | 0 | 874440 | 123900 | 1709400 | 5264100 | 3953000 | 879740 | 1851700 | 1310100 | 626020 | 1491000 |
| EAQTSFLHI | DDX1; DDX1; D | Q92499; A0/ | DDX1_HUMAN | Unmodified | 0 | 0 | 0 | 0 | 1796500 | 0 | 1304300 | 0 | 0 | 0 | 0 | 63873 | 2093000 | 3095000 | 0 | 0 | 1917000 | 1137300 |
| EAQVQYPL | ASNS; ASNS | P08243; F8V | ASNS_HUMAN | Oxidation (M) | 0 | 0 | 0 | 506490 | 1070500 | 0 | 0 | 0 | 792300 | 0 | 1149100 | 4257100 | 0 | 0 | 0 | 151060 | 227490 | 0 |
| EAQVYQAF | SUCLG2; SUCL | Q96I99; E9P | SUCB2_HUMAI | Unmodified | 0 | 0 | 0 | 0 | 0 | 0 | 0 | 0 | 2239200 | 0 | 0 | 0 | 0 | 0 | 0 | 0 | 78854 | 0 |
| EAQYNNFS | SCAMP4 | Q969E2 | SCAM4_HUMA | Unmodified | 0 | 0 | 789990 | 4849800 | 0 | 0 | 0 | 0 | 0 | 0 | 0 | 0 | 0 | 0 | 8958800 | 3638300 | 0 | 0 |
| EARAPASVI | PRKAR2A; PRK/ | P13861; C9J | KAP2_HUMAN | Unmodified | 0 | 0 | 0 | 519800 | 553020 | 0 | 0 | 0 | 300910 | 0 | 0 | 0 | 3070800 | 705230 | 0 | 0 | 215060 | 791310 |
| EARDIFYKP | PLXNB2; PLXNE | O15031; E2F | PLXB2_HUMAN | Unmodified | 0 | 0 | 0 | 0 | 864070 | 0 | 912660 | 0 | 0 | 0 | 0 | 0 | 493090 | 942690 | 0 | 0 | 0 | 2343600 |
| EARLSYLLN | GLG1; GLG1 | Q92896; H3I | GSLG1_HUMAI | Oxidation (M) | 0 | 0 | 0 | 0 | 0 | 0 | 0 | 0 | 83508 | 72564 | 0 | 0 | 0 | 0 | 0 | 0 | 0 | 0 |
| EARLVLDS\ | TBCA; TBCA; TE | O75347; E5F | TBCA_HUMAN | Unmodified | 0 | 0 | 0 | 0 | 1466400 | 0 | 0 | 18472 | 0 | 538130 | 0 | 0 | 3568900 | 1671200 | 0 | 0 | 514930 | 512420 |
| EARNVTME | SRP9 | P49458 | SRP09_HUMAN | Unmodified | 0 | 0 | 0 | 0 | 0 | 118300 | 2664600 | 1039200 | 0 | 117020 | 0 | 0 | 1167800 | 670030 | 0 | 0 | 1100500 | 805180 |
| EASAAEEAI | ENOPH1; ENOF | Q9UHY7; D6 | ENOPH_HUMA | Unmodified | 0 | 0 | 0 | 0 | 0 | 0 | 0 | 0 | 0 | 0 | 0 | 1343300 | 0 | 0 | 0 | 0 | 0 | 0 |
| EASADLSP' | AP3B1 | O00203 | AP3B1_HUMAN | Unmodified | 0 | 0 | 0 | 0 | 0 | 0 | 0 | 0 | 184880 | 0 | 279760 | 1254200 | 0 | 0 | 144030 | 87509 | 0 | 0 |
| EASDIILTDI | ATP2B3; ATP2B | Q16720; P2: | AT2B3_HUMAN | Unmodified | 2440300 | 0 | 0 | 0 | 714990 | 0 | 499250 | 0 | 0 | 349680 | 460290 | 11532000 | 0 | 0 | 0 | 0 | 205470 | 0 |
| EASDLLEQI | MPRIP; MPRIP; | Q6WCQ1; J3 | MPRIP_HUMAN | Unmodified | 0 | 0 | 118620 | 0 | 0 | 0 | 0 | 0 | 0 | 0 | 0 | 870350 | 0 | 0 | 0 | 0 | 0 | 0 |
| EASDPQPE | CAPZA1 | P52907 | CAZA1_HUMAN | Unmodified | 0 | 1413200 | 988520 | 0 | 10503000 | 801690 | 1444500 | 157530 | 2547500 | 1252100 | 2444800 | 8923500 | 9492100 | 5758200 | 1960400 | 1598200 | 970570 | 6983100 |
| EASDPQPE | CAPZA1 | P52907 | CAZA1_HUMAN | Unmodified | 0 | 0 | 0 | 0 | 0 | 0 | 0 | 0 | 0 | 0 | 0 | 0 | 0 | 0 | 0 | 0 | 724810 | 0 |
| EASELLR | MYO1C; UNC1: | O00159; Q7( | MYO1C_HUMA | Unmodified | 1935300 | 1996500 | 2190700 | 1219900 | 0 | 0 | 0 | 0 | 0 | 0 | 256470 | 141310 | 42551 | 0 | 4862900 | 4016000 | 0 | 0 |
| EASELLREL | MYO1C; MYO1( | O00159; F5F | MYO1C_HUMA | Unmodified | 0 | 0 | 0 | 498140 | 1190300 | 357390 | 235920 | 87631 | 55083 | 0 | 0 | 0 | 126110 | 133890 | 0 | 0 | 519970 | 234960 |
| EASFDGISI | PEPD; PEPD; PE | P12955; A0A | PEPD_HUMAN | Unmodified | 12944 | 0 | 586920 | 0 | 0 | 0 | 0 | 0 | 0 | 0 | 0 | 373300 | 0 | 0 | 0 | 0 | 0 | 713180 |
| EASFSPTDI | WDR33; WDR3: | Q9C0J8; C9J | WDR33_HUMA | Unmodified | 24039 | 0 | 0 | 177240 | 0 | 0 | 273540 | 0 | 0 | 221300 | 270690 | 429920 | 0 | 128790 | 30517 | 0 | 0 | 0 |
| EASHAGSV | MEMO1; MEMO | Q9Y316; B5I | MEMO1_HUMA | Unmodified | 202400 | 0 | 0 | 0 | 0 | 0 | 0 | 0 | 0 | 138890 | 0 | 0 | 0 | 0 | 914380 | 76256 | 0 | 0 |
| EASKQADV | PHGDH; PHGD | O43175; A0/ | SERA_HUMAN | Unmodified | 0 | 0 | 0 | 0 | 282970 | 0 | 6007800 | 2836500 | 0 | 809610 | 0 | 0 | 954950 | 8432600 | 0 | 0 | 2310600 | 560370 |
| EASLPLPG\ | PLS1; PLS1; PL! | Q14651; C9J | PLSI_HUMAN F | Unmodified | 0 | 0 | 0 | 0 | 0 | 0 | 0 | 0 | 0 | 0 | 250470 | 2514900 | 0 | 0 | 0 | 0 | 0 | 0 |
| EASMEGIV | NCAM1; NCAM | P13591; A0A | NCAM1_HUMA | Oxidation (M) | 0 | 0 | 0 | 0 | 0 | 119030 | 0 | 0 | 1814100 | 0 | 0 | 0 | 0 | 641090 | 0 | 0 | 0 | 0 |
| EASMEGIV | NCAM1; NCAM | P13591; A0A | NCAM1_HUMA | Unmodified | 0 | 0 | 0 | 571380 | 0 | 0 | 0 | 0 | 5457900 | 0 | 0 | 0 | 0 | 0 | 0 | 0 | 0 | 0 |
| EASMVITES | STOM | P27105 | STOM_HUMAN | Oxidation (M) | 2429500 | 2938600 | 0 | 2312600 | 10458000 | 1130200 | 0 | 659030 | 0 | 0 | 0 | 0 | 0 | 0 | 1127900 | 1282600 | 0 | 0 |
| EASMVITES | STOM | P27105 | STOM_HUMAN | Unmodified | 0 | 0 | 0 | 0 | 2040000 | 170440 | 375290 | 60644 | 0 | 0 | 0 | 0 | 846910 | 166800 | 0 | 0 | 0 | 0 |
| EASMVITES | STOM | P27105 | STOM_HUMAN | Oxidation (M) | 0 | 0 | 0 | 0 | 41691000 | 1025800 | 13320000 | 0 | 0 | 0 | 0 | 0 | 4218300 | 5388100 | 0 | 0 | 0 | 1946900 |
| EASMVITES | STOM | P27105 | STOM_HUMAN | Unmodified | 0 | 0 | 0 | 0 | 2232000 | 0 | 0 | 0 | 0 | 0 | 0 | 0 | 0 | 0 | 0 | 0 | 0 | 0 |
| EASNCFAII | ITM2B; ITM2B; I | Q9Y287; U3I | ITM2B_HUMAN | Unmodified | 0 | 0 | 0 | 0 | 213670 | 0 | 0 | 0 | 1224100 | 90584 | 0 | 0 | 0 | 0 | 0 | 0 | 0 | 0 |
| EASPNPED | PLXNB2 | O15031 | PLXB2_HUMAN | Unmodified | 561060 | 326850 | 917660 | 738610 | 0 | 0 | 0 | 0 | 1762000 | 91678 | 0 | 800160 | 0 | 0 | 2316300 | 1992400 | 0 | 0 |
| EASRSSPVI | CPNE3; CPNE3 | O75131; H0 | CPNE3_HUMAI | Unmodified | 0 | 507940 | 1179300 | 3081800 | 4177700 | 788830 | 5778600 | 5034900 | 14710000 | 10957000 | 267710 | 0 | 8911100 | 6344600 | 1222700 | 322080 | 5916400 | 9651600 |
| EASRSSPVI | CPNE3; CPNE3 | O75131; H0 | CPNE3_HUMAI | Unmodified | 0 | 0 | 0 | 0 | 0 | 647520 | 480130 | 137510 | 507510 | 690560 | 0 | 0 | 362780 | 53082 | 0 | 0 | 580970 | 0 |
| EASSTQDT | CSK | P41240 | CSK_HUMAN T | Unmodified | 105680 | 0 | 0 | 0 | 0 | 0 | 0 | 0 | 362700 | 640660 | 590230 | 2427400 | 0 | 0 | 0 | 0 | 0 | 0 |
| EASYSLIR | HP1BP3; HP1BI | Q5SSJ5; B0( | HP1B3_HUMAI | Unmodified | 1389400 | 974600 | 0 | 0 | 0 | 0 | 0 | 8689.9 | 0 | 86553 | 0 | 0 | 0 | 0 | 0 | 0 | 0 | 0 |
| EATADDLIK | DDB1; DDB1; D | Q16531; A0/ | DDB1_HUMAN | Unmodified | 0 | 0 | 0 | 0 | 0 | 0 | 0 | 0 | 0 | 0 | 0 | 0 | 0 | 0 | 0 | 0 | 0 | 0 |
| EATDAIGHI | ILF3 | Q12906 | ILF3_HUMAN Ir | Unmodified | 0 | 283540 | 231880 | 488440 | 0 | 0 | 0 | 0 | 0 | 0 | 0 | 1286400 | 0 | 0 | 0 | 0 | 0 | 0 |
| EATDEELEI | CUL4B; CUL4B; | Q13620; A0/ | CUL4B_HUMAI | Unmodified | 0 | 32848 | 27089 | 0 | 0 | 0 | 0 | 0 | 0 | 0 | 0 | 518150 | 0 | 0 | 0 | 17914 | 0 | 0 |
| EATDEELEF | CUL4A; CUL4A | Q13619; A0/ | CUL4A_HUMAI | Unmodified | 0 | 0 | 0 | 0 | 0 | 0 | 0 | 0 | 140760 | 34404 | 0 | 516310 | 0 | 0 | 0 | 0 | 0 | 0 |
| EATDEELEF | CUL4A; CUL4A | Q13619; A0/ | CUL4A_HUMAI | Unmodified | 0 | 0 | 0 | 0 | 47423 | 0 | 0 | 0 | 0 | 0 | 0 | 2143100 | 0 | 0 | 0 | 0 | 0 | 0 |
| EATDPRPC | CAPZA2; CAPZ/ | P47755; A0A | CAZA2_HUMAN | Unmodified | 2238300 | 1219600 | 856080 | 0 | 0 | 0 | 0 | 60114 | 1019300 | 0 | 521390 | 6700200 | 0 | 0 | 905980 | 573610 | 0 | 0 |
| EATDTPIVI( | DNAJC5 | Q9H3Z4 | DNJC5_HUMAI | Unmodified | 256590 | 0 | 2019400 | 0 | 0 | 0 | 0 | 0 | 0 | 0 | 0 | 1630600 | 0 | 0 | 0 | 0 | 0 | 0 |
| EATDVIIIHS | ANPEP | P15144 | AMPN_HUMAN | Unmodified | 0 | 0 | 0 | 0 | 0 | 0 | 0 | 0 | 102060 | 0 | 0 | 0 | 0 | 0 | 0 | 0 | 2600100 | 0 |
| EATELWGK | SCPEP1 | Q9HB40 | RISC_HUMAN I | Unmodified | 0 | 0 | 0 | 0 | 0 | 108110 | 0 | 0 | 529330 | 140880 | 176470 | 139380 | 0 | 0 | 0 | 0 | 0 | 0 |
| EATIPGHLI | FN1 | P02751 | FINC_HUMAN I | Unmodified | 0 | 335670 | 2059100 | 0 | 0 | 0 | 0 | 0 | 0 | 0 | 0 | 0 | 0 | 0 | 226870 | 401310 | 0 | 0 |
| EATMKGSS | DSG2 | Q14126 | DSG2_HUMAN | Unmodified | 0 | 0 | 0 | 0 | 3439800 | 0 | 0 | 27411 | 0 | 0 | 0 | 0 | 466420 | 294080 | 0 | 0 | 2606400 | 0 |
| EATNPPVI( | PDIA3 | P30101 | PDIA3_HUMAN | Unmodified | 29274 | 0 | 194010 | 0 | 0 | 0 | 0 | 0 | 4697600 | 0 | 364000 | 616540 | 0 | 0 | 167370 | 93919 | 0 | 0 |
| EATNPPVI( | PDIA3 | P30101 | PDIA3_HUMAN | Unmodified | 434550 | 0 | 587750 | 333430 | 442590 | 0 | 122830 | 116250 | 1.12E+08 | 8805500 | 543310 | 8862100 | 0 | 301790 | 1449600 | 1014100 | 0 | 433010 |
| EATNPPVI( | PDIA3 | P30101 | PDIA3_HUMAN | Unmodified | 0 | 0 | 0 | 0 | 0 | 0 | 0 | 0 | 23153000 | 1201900 | 1509000 | 2367600 | 126440 | 124160 | 174680 | 129950 | 0 | 0 |
| EATNTTSEI | TOM1L1; TOM1 | O75674; J3K | TM1L1_HUMAN | Unmodified | 5353400 | 7721700 | 4961300 | 5646700 | 0 | 0 | 0 | 122000 | 1896800 | 0 | 0 | 848820 | 4240100 | 2990600 | 6249500 | 5406100 | 880190 | 4222200 |
| EATPVVHE | PRRC2C; PRRC | Q9Y520; E7E | PRC2C_HUMAI | Unmodified | 0 | 0 | 0 | 459950 | 316040 | 0 | 0 | 0 | 0 | 0 | 0 | 0 | 0 | 0 | 0 | 0 | 0 | 0 |
| EATPYPALII | IDE; IDE; IDE; ID | P14735; A0A | IDE_HUMAN In | Unmodified | 0 | 0 | 0 | 0 | 0 | 0 | 0 | 0 | 0 | 0 | 0 | 4125200 | 0 | 0 | 0 | 203490 | 52986 | 0 |
| EATSILEEN | GCLC; GCLC; G | P48506; A0A | GSH1_HUMAN | Unmodified | 0 | 0 | 0 | 0 | 0 | 0 | 0 | 0 | 0 | 0 | 0 | 1371100 | 0 | 0 | 0 | 0 | 0 | 0 |
| EATSTFTNI | ITGB3 | P05106 | ITB3_HUMAN Ir | Unmodified | 290470 | 2023800 | 1218200 | 2474800 | 1511500 | 0 | 259690 | 217550 | 0 | 0 | 0 | 0 | 488490 | 0 | 2635200 | 407860 | 671440 | 90728 |
| EATSTFTNI | ITGB3 | P05106 | ITB3_HUMAN Ir | Unmodified | 243800 | 0 | 578000 | 2200000 | 4184100 | 0 | 12974000 | 5369200 | 342600 | 0 | 0 | 0 | 5545700 | 5291400 | 805880 | 575300 | 0 | 1557100 |
| EATTDFTVI | FLNB; FLNB | O75369; E7E | FLNB_HUMAN | Unmodified | 0 | 0 | 0 | 0 | 0 | 0 | 0 | 0 | 289160 | 0 | 0 | 7557200 | 0 | 0 | 308800 | 221160 | 251000 | 0 |
| EATTEFSVI | FLNA; FLNA | P21333; Q6( | FLNA_HUMAN | Unmodified | 950380 | 1391600 | 0 | 2581900 | 0 | 0 | 0 | 284040 | 1006100 | 0 | 0 | 220080 | 230810 | 147350 | 2776600 | 1326500 | 0 | 0 |
| EATVKPFAI | SUMF2; SUMF2 | Q8NBJ7; J3( | SUMF2_HUMAI | Unmodified | 0 | 0 | 0 | 0 | 0 | 0 | 28514 | 0 | 41085000 | 850990 | 501880 | 0 | 0 | 0 | 0 | 0 | 0 | 0 |
| EATVPCSA | PTK7 | Q13308 | PTK7_HUMAN I | Unmodified | 0 | 0 | 0 | 53616 | 0 | 0 | 49989 | 67871 | 1261200 | 39041 | 63702 | 105550 | 0 | 0 | 694230 | 426160 | 0 | 0 |
| EATWVVD\ | SCP2; SCP2 | P22307; H0 | SCP2_HUMAN | Unmodified | 35367 | 28643 | 0 | 0 | 0 | 0 | 181870 | 0 | 61312000 | 6715200 | 1178300 | 682660 | 372810 | 199930 | 44418 | 35101 | 0 | 80752 |
| EAVAHENI | NAPG; NAPG | Q99747; J3K | SNAG_HUMAN | Unmodified | 209950 | 164140 | 0 | 603580 | 145850 | 220200 | 149660 | 0 | 0 | 0 | 0 | 0 | 0 | 83351 | 1064200 | 725050 | 0 | 1323200 |
| EAVAMESY. | CCT2; CCT2 | P78371; F5G | TCPB_HUMAN | Unmodified | 0 | 0 | 0 | 0 | 2983700 | 42386 | 2282700 | 563480 | 0 | 763900 | 2740000 | 11703000 | 4694800 | 1666000 | 0 | 0 | 836850 | 2112600 |
| EAVAMESY. | CCT2; CCT2 | P78371; F5G | TCPB_HUMAN | Oxidation (M) | 62901 | 41404 | 121060 | 96243 | 277990 | 0 | 116670 | 63559 | 0 | 117420 | 0 | 2348300 | 640250 | 267050 | 129340 | 143790 | 0 | 234940 |
| EAVANKGP | CTSS; CTSS; C1 | P25774; U3I | CATS_HUMAN | Unmodified | 0 | 0 | 0 | 0 | 0 | 0 | 0 | 0 | 0 | 987770 | 0 | 0 | 0 | 0 | 0 | 0 | 0 | 0 |
| EAVAPVQE | DYNC1I2; DYN( | Q13409; E7E | DC1I2_HUMAN | Unmodified | 0 | 0 | 0 | 0 | 474020 | 0 | 0 | 0 | 1641600 | 281310 | 0 | 2022400 | 558870 | 0 | 0 | 0 | 0 | 0 |
| EAVAPVQE | DYNC1I2; DYN( | Q13409; E7E | DC1I2_HUMAN | Unmodified | 0 | 0 | 0 | 0 | 0 | 0 | 0 | 0 | 389540 | 0 | 927000 | 730580 | 0 | 0 | 0 | 0 | 34257 | 0 |
| EAVDGHTV | TRAF5 | O00463 | TRAF5_HUMAN | Unmodified | 0 | 0 | 0 | 0 | 0 | 0 | 0 | 0 | 0 | 0 | 0 | 316860 | 0 | 0 | 0 | 0 | 0 | 0 |
| EAVEAQIR | LRBA; LRBA; LR | P50851; A0A | LRBA_HUMAN | Unmodified | 0 | 0 | 0 | 0 | 0 | 0 | 0 | 0 | 0 | 0 | 0 | 0 | 0 | 0 | 0 | 0 | 0 | 0 |
| EAVEEPSSI | CHGA | P10645 | CMGA_HUMAN | Unmodified | 0 | 0 | 0 | 0 | 0 | 0 | 0 | 0 | 18381000 | 787720 | 0 | 0 | 0 | 0 | 0 | 0 | 0 | 0 |
| EAVEEPSSI | CHGA | P10645 | CMGA_HUMAN | Oxidation (M) | 0 | 0 | 0 | 0 | 116820 | 0 | 0 | 0 | 2179900 | 0 | 0 | 0 | 0 | 0 | 0 | 0 | 0 | 0 |
| EAVEFRQE | PACC1 | Q9H813 | PACC1_HUMAI | Unmodified | 0 | 0 | 0 | 0 | 0 | 0 | 0 | 0 | 382580 | 0 | 0 | 574560 | 0 | 0 | 73503 | 53603 | 0 | 0 |
| EAVEKEFEI | HSP90B1; HSP! | P14625; A0A | ENPL_HUMAN | Oxidation (M) | 0 | 0 | 0 | 0 | 0 | 0 | 429330 | 0 | 0 | 1711200 | 0 | 0 | 0 | 0 | 0 | 402970 | 508200 | 0 |
| EAVEKEFEI | HSP90B1; HSP! | P14625; A0A | ENPL_HUMAN | Unmodified | 0 | 0 | 0 | 0 | 0 | 0 | 165930 | 0 | 0 | 1800400 | 0 | 0 | 0 | 0 | 0 | 0 | 0 | 0 |
| EAVELPLTI | PSMC4 | P43686 | PRS6B_HUMAN | Unmodified | 0 | 0 | 1030200 | 0 | 543630 | 0 | 900270 | 56682 | 2070400 | 279790 | 934560 | 0 | 0 | 267880 | 0 | 409780 | 246480 | 665170 |
| EAVEVKGG | WDR44; WDR4 | Q5JSH3; H7I | WDR44_HUMA | Unmodified | 0 | 0 | 0 | 0 | 793010 | 0 | 0 | 0 | 0 | 0 | 0 | 0 | 841950 | 0 | 0 | 27883 | 421510 | 520230 |
| EAVGDLLD | EIF4G1; EIF4G1 | Q04637; C9J | IF4G1_HUMAN | Unmodified | 0 | 0 | 621700 | 685370 | 0 | 0 | 0 | 0 | 0 | 0 | 0 | 0 | 1771500 | 0 | 0 | 1119000 | 0 | 0 |
| EAVIAQISSI | ADAM10 | O14672 | ADA10_HUMAN | Unmodified | 0 | 0 | 5607700 | 3500500 | 1200700 | 527450 | 139640 | 40693 | 2353300 | 55740 | 0 | 2854500 | 1472600 | 531040 | 5334600 | 3768800 | 0 | 334540 |
| EAVILPIK | VPS4B; VPS4A; | O75351; Q9I | VPS4B_HUMAN | Unmodified | 228020 | 258670 | 1350700 | 468380 | 508510 | 369430 | 105670 | 0 | 1253800 | 155670 | 315200 | 2384700 | 213840 | 185810 | 550570 | 427240 | 296810 | 122060 |
| EAVILPIKFI | VPS4B; VPS4A; | O75351; Q9I | VPS4B_HUMAN | Unmodified | 1079600 | 1101000 | 283100 | 2981500 | 6733400 | 642400 | 198980 | 0 | 11246000 | 5133400 | 11048000 | 5375300 | 4841300 | 4714900 | 1057000 | 2332400 | 148240 | 0 |
| EAVILPIKFI | VPS4B; VPS4A; | O75351; Q9I | VPS4B_HUMAN | Unmodified | 976280 | 0 | 0 | 0 | 0 | 0 | 0 | 0 | 1856500 | 514050 | 0 | 2215000 | 0 | 0 | 698420 | 0 | 0 | 0 |
| EAVLLNLK | MRPL44 | Q9H9J2 | RM44_HUMAN | Unmodified | 173000 | 0 | 0 | 259740 | 261840 | 0 | 56597 | 82679 | 469640 | 160180 | 200670 | 467380 | 568000 | 0 | 272670 | 325450 | 0 | 0 |
| EAVLRAID( | CAMP; CAMP | P49913; J3K | CAMP_HUMAN | Unmodified | 0 | 0 | 0 | 0 | 0 | 0 | 0 | 0 | 0 | 380720 | 0 | 0 | 0 | 0 | 0 | 0 | 0 | 0 |
| EAVNLQQI | PGM3; PGM3; F | O95394; J3K | AGM1_HUMAN | Unmodified | 0 | 0 | 0 | 0 | 873910 | 142220 | 0 | 17123 | 0 | 642260 | 357820 | 0 | 2218000 | 1447300 | 0 | 0 | 589430 | 1488500 |
| EAVQGMV/ | CACNA2D2; CA | Q9NY47; C9 | CA2D2_HUMAI | Unmodified | 150290 | 208740 | 0 | 52683 | 0 | 61115 | 450770 | 221390 | 8273400 | 1631200 | 631160 | 782770 | 0 | 0 | 0 | 16318 | 0 | 0 |
| EAVRYINEI | TCP1; TCP1; TC | P17987; F5F | TCPA_HUMAN | Unmodified | 0 | 0 | 0 | 0 | 1830900 | 65312 | 0 | 0 | 0 | 0 | 0 | 0 | 3715800 | 0 | 0 | 0 | 2802500 | 1763600 |
| EAVSARGE | MMS19; MMS1! | Q96T76; H0 | MMS19_HUMA | Unmodified | 0 | 0 | 0 | 0 | 181400 | 24910 | 49016 | 0 | 0 | 0 | 0 | 0 | 0 | 0 | 42651 | 0 | 68268 | 181170 |
| EAVSGPMA | N/A; AAK1 | Q6ZSR9; A0/ | YJ005_HUMAN | Unmodified | 0 | 0 | 0 | 0 | 203490 | 0 | 0 | 0 | 0 | 0 | 0 | 0 | 950160 | 0 | 0 | 0 | 54129 | 388240 |
| EAVTEILGI | TSN; TSN; TSN | Q15631; H7( | TSN_HUMAN T | Unmodified | 0 | 0 | 0 | 0 | 0 | 0 | 0 | 44355 | 1056400 | 0 | 0 | 14316000 | 0 | 0 | 0 | 0 | 0 | 0 |
| EAVTFDQA | PLAA; PLAA; PL/ | Q9Y263; H0 | PLAP_HUMAN I | Unmodified | 0 | 0 | 0 | 0 | 0 | 0 | 0 | 92682 | 0 | 208930 | 0 | 969480 | 0 | 0 | 139320 | 51459 | 0 | 0 |
| EAVTGSRL( | LOXL2; LOXL2 | Q9Y4K0; H0 | LOXL2_HUMAN | Unmodified | 0 | 0 | 0 | 0 | 929490 | 0 | 741130 | 0 | 0 | 1903400 | 0 | 0 | 0 | 0 | 0 | 0 | 777770 | 1693600 |
| EAVTTIAAY | MYO1B; MYO1E | O43795; E9F | MYO1B_HUMA | Unmodified | 0 | 0 | 0 | 0 | 0 | 46659 | 0 | 0 | 226740 | 0 | 359450 | 4897300 | 0 | 0 | 0 | 0 | 0 | 0 |
| EAVTTIAAY | MYO1B; MYO1E | O43795; E9F | MYO1B_HUMA | Unmodified | 135360 | 0 | 0 | 0 | 559450 | 161470 | 0 | 0 | 0 | 0 | 608030 | 2861700 | 0 | 0 | 0 | 0 | 0 | 159950 |
| EAVVHGLN | ACSL4 | O60488 | ACSL4_HUMAN | Unmodified | 0 | 0 | 0 | 0 | 0 | 0 | 0 | 0 | 0 | 0 | 0 | 0 | 0 | 0 | 0 | 0 | 878240 | 0 |
| EAWFSPVI | SNTB1; SNTB1 | Q13884; A0/ | SNTB1_HUMAN | Unmodified | 0 | 0 | 0 | 0 | 0 | 0 | 0 | 0 | 8723800 | 0 | 0 | 0 | 0 | 0 | 0 | 0 | 0 | 0 |
| EAYDRDFL | MGAT1 | P26572 | MGAT1_HUMAI | Unmodified | 0 | 0 | 0 | 0 | 0 | 0 | 0 | 0 | 3292800 | 228800 | 0 | 0 | 0 | 0 | 0 | 0 | 0 | 0 |
| EAYDSTLPI | CDC42BPA; CD | Q5VT25; A0/ | MRCKA_HUMA | Unmodified | 0 | 0 | 0 | 430630 | 783960 | 0 | 0 | 0 | 627820 | 307200 | 384340 | 165050 | 0 | 0 | 691680 | 734750 | 672720 | 973320 |
| EAYEAGLIC | NDUFV1; NDUI | P49821; H0 | NDUV1_HUMA | Unmodified | 0 | 0 | 42979 | 0 | 572830 | 0 | 0 | 0 | 321060 | 0 | 0 | 1270500 | 0 | 0 | 67029 | 0 | 594230 | 0 |
| EAYEVKLG/ | PRSS8; PRSS8 | Q16651; H3I | PRSS8_HUMAN | Unmodified | 0 | 0 | 0 | 0 | 0 | 0 | 0 | 0 | 0 | 236970 | 0 | 0 | 0 | 133670 | 0 | 0 | 0 | 0 |

| | | | | | | | | | | | | | | | | | | | | | | |
|---|---|---|---|---|---|---|---|---|---|---|---|---|---|---|---|---|---|---|---|---|---|---|
| EAYFWLR | STT3B | Q8TCJ2 | STT3B_HUMAN | Unmodified | 0 | 0 | 0 | 0 | 0 | 0 | 0 | 0 | 958380 | 131380 | 0 | 5301300 | 0 | 0 | 0 | 0 | 0 | 0 |
| EAYGAVTQ | PMVK | Q15126 | PMVK_HUMAN | Unmodified | 0 | 865380 | 0 | 0 | 0 | 0 | 0 | 43460 | 461350 | 0 | 0 | 546900 | 0 | 0 | 0 | 0 | 49318 | 0 |
| EAYLVGLFE | H3-2; H3-3A; H | Q5TEC6; P84 | H3PS2_HUMAN | Unmodified | 714330 | 0 | 0 | 587070 | 0 | 0 | 4463400 | 0 | 0 | 0 | 0 | 1626700 | 0 | 0 | 0 | 624460 | 0 | 650710 |
| EAYMGNVL | ACAT1; ACAT1; | P24752; E9P | THIL_HUMAN A | Oxidation (M) | 89391 | 0 | 0 | 0 | 0 | 0 | 0 | 0 | 692280 | 0 | 0 | 0 | 0 | 0 | 0 | 0 | 0 | 0 |
| EAYMGNVL | ACAT1; ACAT1; | P24752; E9P | THIL_HUMAN A | Unmodified | 153160 | 0 | 0 | 0 | 0 | 0 | 676470 | 0 | 1843200 | 0 | 0 | 0 | 116740 | 0 | 79417 | 0 | 0 | 0 |
| EAYPEEAYI | ACLY | P53396 | ACLY_HUMAN | Unmodified | 0 | 0 | 0 | 0 | 0 | 0 | 0 | 377460 | 1645700 | 0 | 1960600 | 17751000 | 0 | 0 | 0 | 691050 | 0 | 0 |
| EAYPGDVF | ATP5F1A | P25705 | ATPA_HUMAN | Unmodified | 0 | 224970 | 0 | 0 | 0 | 0 | 0 | 0 | 32666000 | 1244800 | 0 | 0 | 0 | 0 | 381190 | 0 | 0 | 0 |
| EAYQRGAA | NIBAN1 | Q9BZQ8 | NIBA1_HUMAN | Unmodified | 688040 | 3549400 | 0 | 200950 | 353290 | 77913 | 0 | 92845 | 1518500 | 457740 | 0 | 0 | 29618 | 0 | 0 | 453130 | 37491 | 92430 |
| EAYYWLR | STT3A | P46977 | STT3A_HUMAN | Unmodified | 0 | 0 | 0 | 0 | 0 | 0 | 0 | 0 | 11731000 | 1253900 | 0 | 707270 | 0 | 0 | 0 | 0 | 0 | 0 |
| EAYYWLRH | STT3A | P46977 | STT3A_HUMAN | Unmodified | 0 | 0 | 0 | 0 | 0 | 0 | 0 | 0 | 674240 | 0 | 0 | 820500 | 0 | 0 | 0 | 0 | 0 | 0 |
| ECADEPVG | GLG1; GLG1; G | Q92896; H3I | GSLG1_HUMAI | Unmodified | 0 | 0 | 0 | 0 | 0 | 0 | 0 | 0 | 0 | 0 | 0 | 0 | 0 | 0 | 0 | 0 | 0 | 0 |
| ECAQLRQE | ITGB4 | P16144 | ITB4_HUMAN I | Unmodified | 0 | 354950 | 0 | 0 | 0 | 0 | 0 | 0 | 381760 | 0 | 0 | 0 | 0 | 0 | 0 | 0 | 855700 | 0 |
| ECDNALRE | TLN1 | Q9Y490 | TLN1_HUMAN | Unmodified | 230820 | 233230 | 190900 | 286160 | 243250 | 0 | 0 | 0 | 0 | 0 | 148550 | 1147800 | 0 | 139900 | 392560 | 364280 | 0 | 0 |
| ECEEEAINI | UTRN | P46939 | UTRN_HUMAN | Unmodified | 0 | 0 | 0 | 0 | 0 | 0 | 0 | 0 | 0 | 0 | 0 | 0 | 0 | 0 | 597440 | 618530 | 0 | 0 |
| ECFVCTAC | FHL2 | Q14192 | FHL2_HUMAN | Unmodified | 0 | 0 | 0 | 0 | 260110 | 0 | 0 | 0 | 0 | 0 | 0 | 145380 | 0 | 0 | 0 | 0 | 822750 | 347100 |
| ECFYQPMF | TMED5; TMED5 | Q9Y3A6; B1A | TMED5_HUMAI | Oxidation (M) | 0 | 0 | 0 | 0 | 0 | 0 | 0 | 0 | 9365100 | 2345300 | 0 | 0 | 0 | 0 | 0 | 0 | 0 | 0 |
| ECFYQPMF | TMED5; TMED5 | Q9Y3A6; B1A | TMED5_HUMAI | Unmodified | 0 | 0 | 0 | 0 | 0 | 0 | 0 | 0 | 13482000 | 1664800 | 108580 | 0 | 0 | 0 | 0 | 0 | 0 | 0 |
| ECGEHFEE | QSOX2; QSOX2 | Q6ZRP7; H0Y | QSOX2_HUMAI | Unmodified | 0 | 0 | 0 | 0 | 0 | 0 | 0 | 0 | 4148300 | 316570 | 238930 | 226520 | 0 | 0 | 0 | 0 | 0 | 0 |
| ECGPDNK | ITGA3 | P26006 | ITA3_HUMAN I | Oxidation (M) | 0 | 0 | 0 | 0 | 11314000 | 6754600 | 0 | 0 | 0 | 0 | 0 | 0 | 0 | 0 | 0 | 0 | 4114700 | 13382000 |
| ECGPDNK | ITGA3 | P26006 | ITA3_HUMAN I | Unmodified | 140960 | 0 | 0 | 0 | 21217000 | 7122200 | 0 | 0 | 0 | 0 | 96996 | 673370 | 211220 | 149580 | 0 | 78189 | 8826200 | 34048000 |
| ECGPDNK | ITGA3 | P26006 | ITA3_HUMAN I | Unmodified | 0 | 0 | 0 | 0 | 9427700 | 0 | 0 | 0 | 0 | 0 | 0 | 0 | 0 | 0 | 0 | 0 | 7953300 | 12691000 |
| ECGPDNK | ITGA3 | P26006 | ITA3_HUMAN I | Oxidation (M) | 0 | 0 | 0 | 0 | 1104500 | 0 | 0 | 0 | 0 | 0 | 0 | 0 | 0 | 0 | 0 | 0 | 847280 | 442210 |
| ECHLNADT | AP2B1; AP2B1 | P63010; A0A | AP2B1_HUMAN | Unmodified | 0 | 570590 | 0 | 0 | 1000700 | 929150 | 0 | 0 | 0 | 311140 | 0 | 0 | 727390 | 2161200 | 0 | 0 | 1116800 | 0 |
| ECIQEPSS | UTRN | P46939 | UTRN_HUMAN | Unmodified | 0 | 0 | 0 | 223520 | 298690 | 0 | 0 | 0 | 0 | 125680 | 0 | 0 | 247480 | 142310 | 272910 | 0 | 270200 | 359280 |
| ECISDNKS | MACF1; MACF1 | Q9UPN3; H3 | MACF1_HUMAI | Unmodified | 0 | 0 | 283810 | 220710 | 198290 | 1546200 | 265690 | 221140 | 0 | 0 | 0 | 0 | 94075 | 586490 | 77315 | 0 | 0 | 0 |
| ECISGYAR | CRELD2 | Q6UXH1 | CREL2_HUMAN | Unmodified | 0 | 0 | 82384 | 0 | 0 | 0 | 0 | 0 | 694130 | 48972 | 51787 | 0 | 0 | 0 | 0 | 77809 | 0 | 0 |
| ECLPLIIFLR | RPS4X | P62701 | RS4X_HUMAN | Unmodified | 561350 | 245580 | 90873 | 151220 | 0 | 0 | 0 | 0 | 91915 | 52025 | 84755 | 4814300 | 0 | 0 | 342910 | 0 | 0 | 0 |
| ECLPLIVFLI | RPS4Y2; RPS4Y | Q8TD47; P22 | RS4Y2_HUMAN | Unmodified | 0 | 0 | 0 | 0 | 0 | 0 | 0 | 0 | 0 | 0 | 0 | 1477800 | 0 | 0 | 0 | 0 | 0 | 0 |
| ECLQALEFI | PAK2; PAK2 | Q13177; H7C | PAK2_HUMAN | Unmodified | 272570 | 106110 | 188540 | 298260 | 0 | 0 | 0 | 0 | 843610 | 0 | 157100 | 3737400 | 0 | 0 | 506420 | 280910 | 0 | 0 |
| ECLQALEFI | PAK1; PAK1; PA | Q13153; E9F | PAK1_HUMAN | Unmodified | 0 | 0 | 0 | 0 | 0 | 0 | 0 | 0 | 0 | 0 | 0 | 655680 | 0 | 0 | 0 | 131820 | 0 | 0 |
| ECMAGTSC | KTN1; KTN1; KT | Q86UP2; G3 | KTN1_HUMAN | Unmodified | 0 | 0 | 0 | 0 | 0 | 0 | 0 | 0 | 0 | 0 | 59055 | 345440 | 0 | 0 | 0 | 0 | 0 | 0 |
| ECPIVGFRY | UTRN; UTRN; U | P46939; H0Y | UTRN_HUMAN | Unmodified | 0 | 0 | 0 | 0 | 461230 | 0 | 0 | 0 | 0 | 450620 | 0 | 0 | 0 | 565010 | 0 | 0 | 325600 | 0 |
| ECPPDTYLS | ELAPOR2; ELAP | A8MWY0; H7 | ELAP2_HUMAN | Unmodified | 0 | 0 | 0 | 0 | 0 | 0 | 0 | 0 | 1995300 | 856430 | 0 | 0 | 0 | 0 | 0 | 0 | 0 | 0 |
| ECPSVLEYI | PSAT1 | Q9Y617 | SERC_HUMAN | Unmodified | 0 | 35105 | 0 | 0 | 0 | 0 | 0 | 0 | 0 | 0 | 0 | 2388200 | 0 | 0 | 0 | 0 | 0 | 0 |
| ECREPELG | HSPBP1 | Q9NZL4 | HPBP1_HUMAI | Unmodified | 69007 | 0 | 0 | 0 | 0 | 0 | 0 | 0 | 290350 | 0 | 0 | 1557000 | 0 | 0 | 0 | 0 | 0 | 0 |
| ECRPCPAG | ELAPOR2; ELAP | A8MWY0; H7 | ELAP2_HUMAN | Unmodified | 0 | 0 | 0 | 0 | 0 | 0 | 0 | 0 | 6282400 | 290580 | 0 | 0 | 0 | 0 | 0 | 0 | 0 | 0 |
| ECRQSLSH | COPB1 | P53618 | COPB_HUMAN | Unmodified | 0 | 0 | 0 | 0 | 1155000 | 438070 | 86873 | 0 | 0 | 62675 | 0 | 0 | 230720 | 161920 | 0 | 0 | 343450 | 367310 |
| ECSIYLIGG | NIT2; NIT2 | Q9NQR4; H7 | NIT2_HUMAN ( | Unmodified | 0 | 0 | 0 | 0 | 0 | 0 | 0 | 0 | 0 | 0 | 0 | 909410 | 0 | 0 | 0 | 0 | 0 | 0 |
| ECTRGSAV | PSAP; PSAP; PS | P07602; Q5E | SAP_HUMAN P | Unmodified | 0 | 0 | 0 | 0 | 577790 | 802920 | 1950300 | 902070 | 1434200 | 6695500 | 89140 | 0 | 925730 | 2382200 | 0 | 0 | 449180 | 2286900 |
| ECVQCRAF | ITGB1; ITGB1; I | P05556; H7C | ITB1_HUMAN I | Unmodified | 0 | 0 | 0 | 0 | 418530 | 80006 | 91006 | 0 | 154500 | 0 | 0 | 22217 | 0 | 25318 | 58927 | 0 | 203120 | 1196200 |
| ECYQSHRD | PDCD6IP | Q8WUM4 | PDC6I_HUMAN | Unmodified | 23156000 | 5353400 | 0 | 1775500 | 9744600 | 0 | 0 | 0 | 4604800 | 3784700 | 0 | 1113000 | 18087000 | 5351400 | 7187400 | 5433700 | 4091800 | 10773000 |
| EDAALLPR | KLRG2 | A4D1S0 | KLRG2_HUMAN | Unmodified | 0 | 0 | 0 | 0 | 0 | 0 | 0 | 0 | 0 | 0 | 0 | 0 | 0 | 0 | 0 | 0 | 0 | 0 |
| EDAANNYA | TUBA3C; TUBA | P0DPH7; Q7 | TBA3C_HUMAN | Unmodified | 0 | 0 | 0 | 0 | 304880 | 111140 | 705740 | 780410 | 0 | 45935 | 0 | 0 | 0 | 0 | 0 | 0 | 96200 | 273150 |
| EDAENLLES | HSH2D; HSH2D | Q96JZ2; A0A | HSH2D_HUMA | Unmodified | 0 | 0 | 0 | 0 | 0 | 0 | 44057 | 0 | 29299 | 0 | 0 | 18336 | 0 | 0 | 952920 | 266700 | 0 | 0 |
| EDAGTYW | CEACAM1 | P13688 | CEAM1_HUMAI | Unmodified | 0 | 0 | 0 | 0 | 0 | 0 | 0 | 0 | 0 | 0 | 28174 | 410350 | 0 | 0 | 0 | 0 | 0 | 0 |
| EDALPGQK | BSG; BSG; BSG | P35613; A0A | BASI_HUMAN E | Unmodified | 0 | 0 | 451660 | 0 | 0 | 0 | 0 | 0 | 0 | 0 | 0 | 1443400 | 0 | 0 | 1469000 | 1417300 | 0 | 0 |
| EDAPIGPHI | EIF3A | Q14152 | EIF3A_HUMAN | Unmodified | 0 | 0 | 63679 | 0 | 0 | 0 | 140150 | 0 | 0 | 0 | 742710 | 3461700 | 0 | 0 | 0 | 93800 | 0 | 0 |
| EDAQEIAD | ANXA6 | E5RK69 | E5RK69_HUMA | Unmodified | 0 | 32258 | 191560 | 0 | 0 | 0 | 58495 | 28294 | 4207800 | 768820 | 0 | 0 | 0 | 44282 | 0 | 41407 | 0 | 0 |
| EDAQEIAD | ANXA6 | E5RK69 | E5RK69_HUMA | Unmodified | 0 | 0 | 0 | 0 | 0 | 0 | 0 | 0 | 781540 | 510320 | 0 | 0 | 0 | 0 | 0 | 0 | 0 | 0 |
| EDAQVAAE | ANXA6; ANXA6 | P08133; E7E | ANXA6_HUMAI | Unmodified | 0 | 0 | 0 | 0 | 0 | 0 | 0 | 142830 | 1449700 | 360700 | 0 | 0 | 0 | 0 | 0 | 0 | 0 | 0 |
| EDAVNWLS | PLCG2; PLCG2; | P16885; A0A | PLCG2_HUMAI | Unmodified | 0 | 0 | 0 | 0 | 0 | 0 | 0 | 0 | 0 | 0 | 0 | 2881000 | 0 | 0 | 0 | 0 | 0 | 0 |
| EDDEDKDE | CALR; CALR | P27797; K7E | CALR_HUMAN | Unmodified | 0 | 0 | 0 | 0 | 0 | 0 | 382840 | 234210 | 7474900 | 2210800 | 0 | 0 | 0 | 0 | 0 | 0 | 0 | 0 |
| EDDLNSFN | LTA4H | P09960 | LKHA4_HUMAN | Unmodified | 0 | 0 | 0 | 0 | 0 | 0 | 0 | 0 | 0 | 0 | 1050800 | 1463800 | 0 | 0 | 0 | 0 | 0 | 0 |
| EDDSIRPFK | EPHX1; EPHX1 | P07099; B1A | HYEP_HUMAN | Unmodified | 0 | 0 | 685370 | 0 | 0 | 0 | 0 | 0 | 7412400 | 212900 | 0 | 0 | 0 | 0 | 0 | 0 | 0 | 0 |
| EDDVGTGA | CCT3; CCT3 | P49368; B4D | TCPG_HUMAN | Unmodified | 0 | 0 | 0 | 0 | 0 | 0 | 0 | 0 | 0 | 0 | 13409000 | 0 | 0 | 0 | 0 | 0 | 0 | 0 |
| EDEISLEDL | ZC3H15 | Q8WU90 | ZC3HF_HUMAI | Unmodified | 0 | 0 | 0 | 0 | 0 | 0 | 0 | 0 | 0 | 0 | 0 | 1017700 | 0 | 0 | 0 | 0 | 0 | 0 |
| EDENVPFLI | RALA; RALA | P11233; H7C | RALA_HUMAN | Unmodified | 0 | 0 | 0 | 0 | 0 | 0 | 0 | 0 | 0 | 256640 | 446720 | 2972400 | 0 | 0 | 0 | 0 | 0 | 0 |
| EDENVPFLI | RALA; RALA | P11233; H7C | RALA_HUMAN | Unmodified | 47001 | 0 | 511440 | 150900 | 3137700 | 202910 | 0 | 15543 | 0 | 0 | 846270 | 6028000 | 0 | 0 | 2173500 | 966540 | 0 | 826840 |
| EDENVPFLI | RALA; RALA | P11233; H7C | RALA_HUMAN | Unmodified | 382960 | 244510 | 0 | 0 | 0 | 0 | 442560 | 70681 | 0 | 0 | 0 | 903300 | 0 | 0 | 212040 | 0 | 0 | 0 |
| EDENYYQK | NOSTRIN; NOS | Q8IVI9; A0A | NOSTN_HUMA | Unmodified | 357190 | 0 | 2156400 | 387310 | 0 | 0 | 0 | 0 | 0 | 0 | 0 | 0 | 0 | 0 | 1858500 | 621750 | 0 | 0 |
| EDEYFENI | MME; MME; MM | P08473; A0A | NEP_HUMAN N | Unmodified | 0 | 0 | 277920 | 0 | 0 | 0 | 0 | 0 | 0 | 0 | 0 | 0 | 0 | 0 | 656470 | 281600 | 0 | 0 |
| EDFDSLLQ | SLC3A2; SLC3A | P08195; A0A | 4F2_HUMAN 4I | Unmodified | 0 | 102950 | 2783600 | 0 | 0 | 0 | 0 | 0 | 0 | 0 | 1252600 | 7550300 | 0 | 0 | 2203100 | 1183900 | 0 | 0 |
| EDFDSLLQ | SLC3A2; SLC3A | P08195; A0A | 4F2_HUMAN 4I | Unmodified | 0 | 0 | 279170 | 274750 | 454820 | 0 | 1296700 | 0 | 0 | 0 | 0 | 1038500 | 459750 | 0 | 128410 | 117720 | 0 | 597400 |
| EDFGSPR | ITGB5 | P18084 | ITB5_HUMAN I | Unmodified | 0 | 0 | 65341 | 0 | 223990 | 100370 | 0 | 0 | 311640 | 0 | 413230 | 3092800 | 0 | 0 | 0 | 0 | 71378 | 190560 |
| EDFLVYKLE | GOLIM4; GOLIM | O00461; F8V | GOLI4_HUMAN | Unmodified | 0 | 0 | 0 | 0 | 1354800 | 340560 | 0 | 0 | 1321800 | 0 | 0 | 0 | 0 | 0 | 0 | 0 | 0 | 0 |
| EDFQRIPEL | CHP1; CHP1; C | Q99653; F5C | CHP1_HUMAN | Unmodified | 234680 | 4429300 | 225960 | 36498 | 0 | 454980 | 207380 | 0 | 0 | 48712 | 0 | 3545000 | 0 | 0 | 1575300 | 1168400 | 0 | 1590500 |
| EDGAISTIV | CCT8 | P50990 | TCPQ_HUMAN | Unmodified | 181120 | 83763 | 805780 | 314300 | 0 | 0 | 0 | 0 | 933280 | 0 | 1627100 | 27188000 | 357450 | 0 | 926340 | 896570 | 0 | 0 |
| EDGGFRHS | CKAP4 | Q07065 | CKAP4_HUMAN | Unmodified | 0 | 0 | 0 | 0 | 0 | 0 | 0 | 0 | 1484300 | 53721 | 0 | 0 | 0 | 0 | 0 | 0 | 0 | 0 |
| EDGLAQQ | FASN; FASN | P49327; A0A | FAS_HUMAN F | Unmodified | 0 | 0 | 3099200 | 216900 | 0 | 0 | 0 | 146220 | 0 | 0 | 8049600 | 55807000 | 0 | 0 | 2384700 | 2003100 | 0 | 0 |
| EDGNEEDK | YBX1; YBX1 | P67809; H0Y | YBOX1_HUMAN | Unmodified | 738630 | 517580 | 0 | 506910 | 0 | 32726 | 0 | 88850 | 82733 | 0 | 1494200 | 2112400 | 122800 | 0 | 201550 | 388500 | 16994 | 0 |
| EDGRVQAF | SPAG9; SPAG9 | O60271; A0A | JIP4_HUMAN C | Unmodified | 118790 | 0 | 0 | 64934 | 476770 | 156080 | 262250 | 0 | 0 | 0 | 40963 | 553780 | 183540 | 92353 | 53518 | 35862 | 900370 | 1130100 |
| EDHGRGYI | HNRNPU; HNR | Q00839; A0A | HNRPU_HUMA | Unmodified | 296300 | 116820 | 0 | 0 | 0 | 0 | 0 | 0 | 0 | 0 | 0 | 556700 | 177960 | 101890 | 0 | 36291 | 0 | 0 |
| EDIERMVQ | HSPA8; HSPA8; | P11142; E9P | HSP7C_HUMAI | Unmodified | 0 | 0 | 1174700 | 176320 | 899540 | 471610 | 387100 | 0 | 796790 | 695590 | 226980 | 161020 | 299430 | 0 | 0 | 0 | 430830 | 0 |
| EDITQSAQI | ILF3 | Q12906 | ILF3_HUMAN I | Unmodified | 1176900 | 756170 | 741440 | 706580 | 0 | 0 | 0 | 0 | 213520 | 0 | 820570 | 2437500 | 0 | 0 | 311690 | 447520 | 0 | 0 |
| EDIYAVEIV | HSPA4; HSPA4; | P34932; A0A | HSP74_HUMAN | Unmodified | 12116000 | 6060600 | 0 | 0 | 377870 | 0 | 138680 | 23136 | 725950 | 0 | 0 | 9558200 | 0 | 0 | 8087200 | 0 | 389620 | 0 |
| EDKAPQVD | GARS1; GARS1; | P41250; F8V | GARS_HUMAN | Unmodified | 66856 | 0 | 0 | 0 | 155940 | 0 | 0 | 0 | 173130 | 73435 | 269570 | 1300200 | 0 | 0 | 0 | 0 | 0 | 29762 |
| EDKGTGVV | LARS1; LARS1; | Q9P2J5; A0A | SYLC_HUMAN | Unmodified | 0 | 0 | 0 | 0 | 0 | 0 | 0 | 0 | 0 | 0 | 353410 | 3352200 | 0 | 0 | 0 | 0 | 0 | 0 |
| EDKLECSEI | CLTCL1; CLTC; | P53675; Q00 | CLH2_HUMAN | Unmodified | 1283100 | 226790 | 349040 | 476420 | 577150 | 0 | 0 | 571430 | 0 | 0 | 0 | 4333300 | 376570 | 496520 | 238760 | 666440 | 78523 | 0 |
| EDLDKVEP | DYNC1H1; DYN | Q14204; A0A | DYHC1_HUMA | Unmodified | 0 | 0 | 0 | 0 | 0 | 0 | 0 | 0 | 0 | 0 | 166230 | 1240700 | 0 | 0 | 0 | 0 | 0 | 0 |
| EDLGSLGK | GPRIN1 | Q7Z2K8 | GRIN1_HUMAN | Unmodified | 0 | 0 | 0 | 0 | 0 | 0 | 0 | 18201 | 356010 | 0 | 0 | 0 | 18051 | 0 | 0 | 0 | 45968 | 122940 |
| EDLLSPVQI | SLITRK5 | O94991 | SLIK5_HUMAN | Unmodified | 166630 | 138030 | 0 | 0 | 0 | 0 | 69787 | 0 | 772680 | 51328 | 0 | 0 | 0 | 0 | 0 | 0 | 0 | 0 |
| EDLNVNEV | RAB23 | Q9ULC3 | RAB23_HUMAN | Unmodified | 0 | 0 | 596810 | 504970 | 0 | 0 | 0 | 0 | 0 | 0 | 0 | 0 | 0 | 0 | 348080 | 0 | 0 | 0 |
| EDLRLPEGI | EIF5A; EIF5AL1; | P63241; Q6I | IF5A1_HUMAN | Unmodified | 114840 | 82599 | 0 | 0 | 0 | 0 | 0 | 0 | 0 | 30950 | 2137700 | 10464000 | 0 | 0 | 1568600 | 1413400 | 0 | 0 |
| EDLRSWTA | HLA-A; HLA-A; I | Q53Z42; A0A | Q53Z42_HUMA | Unmodified | 0 | 101450 | 0 | 0 | 0 | 0 | 0 | 0 | 3548300 | 496520 | 0 | 2406600 | 0 | 0 | 0 | 0 | 563390 | 0 |
| EDLRSWTA | HLA-A; HLA-A; I | Q53Z42; A0A | Q53Z42_HUMA | Oxidation (M) | 0 | 0 | 0 | 0 | 0 | 0 | 0 | 0 | 735070 | 164220 | 355690 | 2056000 | 0 | 0 | 22041 | 0 | 0 | 0 |
| EDLSWPGI | ATP6AP2; ATP6 | O75787; A0A | RENR_HUMAN | Unmodified | 0 | 0 | 0 | 0 | 0 | 0 | 0 | 0 | 3069400 | 0 | 0 | 273170 | 0 | 0 | 0 | 0 | 0 | 0 |
| EDLTEIRDN | RPLP0; RPLP0P | P05388; Q8N | RLA0_HUMAN ( | Oxidation (M) | 307450 | 497300 | 2219400 | 99127 | 0 | 0 | 0 | 0 | 0 | 0 | 0 | 1964700 | 0 | 0 | 4673800 | 0 | 0 | 0 |
| EDLTEIRDN | RPLP0; RPLP0P | P05388; Q8N | RLA0_HUMAN ( | Unmodified | 215880 | 171560 | 1454700 | 0 | 0 | 0 | 0 | 0 | 0 | 0 | 0 | 1519700 | 0 | 0 | 1769500 | 610750 | 114020 | 0 |
| EDLTEIRDN | RPLP0; RPLP0P | P05388; Q8N | RLA0_HUMAN ( | Oxidation (M) | 3378300 | 3390500 | 2449900 | 1860900 | 0 | 0 | 0 | 0 | 0 | 0 | 554960 | 12819000 | 0 | 0 | 11203000 | 5134100 | 0 | 0 |
| EDLYLKPIQ | EEF2 | P13639 | EF2_HUMAN EI | Unmodified | 876730 | 518140 | 1551200 | 324330 | 0 | 0 | 137780 | 0 | 662750 | 242810 | 979970 | 54894000 | 0 | 0 | 1719200 | 1714400 | 86613 | 0 |
| EDMAALEK | TUBA1A; TUBA4 | Q71U36; P68 | TBA1A_HUMAN | Unmodified | 553230 | 435820 | 1312900 | 178270 | 240500 | 0 | 207500 | 52852 | 1333100 | 556520 | 4384500 | 24365000 | 0 | 41473 | 1478300 | 1100700 | 26914 | 77265 |
| EDMAALEK | TUBA1A; TUBA4 | Q71U36; P68 | TBA1A_HUMAN | Oxidation (M) | 0 | 0 | 63347 | 0 | 0 | 0 | 0 | 40078 | 295100 | 0 | 688770 | 0 | 0 | 0 | 0 | 303220 | 0 | 21613 |
| EDQDKVAV | ACAT2 | Q9BWD1 | THIC_HUMAN | Unmodified | 4376900 | 1796200 | 686470 | 1094500 | 0 | 0 | 0 | 0 | 685700 | 0 | 0 | 1744800 | 0 | 0 | 0 | 0 | 0 | 0 |
| EDQTEYLEI | HSP90AB4P; HS | Q58FF6; P08 | H90B4_HUMAN | Unmodified | 0 | 0 | 441580 | 0 | 0 | 0 | 0 | 0 | 5289000 | 536570 | 5840700 | 39871000 | 0 | 0 | 361650 | 322620 | 0 | 0 |
| EDQTEYLEI | HSP90AB1; HSI | P08238; Q58 | HS90B_HUMAN | Unmodified | 106850 | 0 | 0 | 0 | 0 | 0 | 0 | 0 | 1079800 | 256240 | 0 | 7912000 | 0 | 0 | 343940 | 322770 | 0 | 0 |
| EDQVIQLM | RPN2; RPN2; RI | P04844; F2Z | RPN2_HUMAN | Unmodified | 0 | 0 | 0 | 0 | 0 | 0 | 0 | 0 | 381330 | 35181 | 0 | 0 | 0 | 0 | 0 | 0 | 0 | 0 |
| EDREGDDL | SELPLG; SELPL | Q14242; A0A | SELPL_HUMAN | Unmodified | 0 | 0 | 0 | 0 | 0 | 0 | 0 | 0 | 0 | 0 | 403020 | 1185300 | 0 | 0 | 0 | 0 | 0 | 0 |
| EDRPITIKD | VPS28; VPS28; | Q9UK41; E9I | VPS28_HUMAN | Unmodified | 1776700 | 1544200 | 8262700 | 876070 | 0 | 0 | 0 | 0 | 194620 | 0 | 0 | 0 | 0 | 0 | 6848300 | 4238800 | 0 | 0 |
| EDSLEAGLI | CHGA; CHGA | P10645; G5E | CMGA_HUMAN | Unmodified | 0 | 0 | 0 | 0 | 0 | 0 | 0 | 0 | 4583300 | 1132600 | 256000 | 0 | 0 | 0 | 0 | 0 | 0 | 0 |
| EDSQRPGA | HNRNPA1L2; H | Q32P51; P09 | RA1L2_HUMAN | Unmodified | 0 | 0 | 0 | 0 | 0 | 40173 | 18683 | 439950 | 1197400 | 0 | 4126300 | 4978300 | 167330 | 83318 | 256260 | 0 | 0 | 0 |
| EDSQRPGA | HNRNPA1L2; H | Q32P51; P09 | RA1L2_HUMAN | Unmodified | 0 | 0 | 0 | 0 | 0 | 0 | 0 | 0 | 0 | 0 | 2396500 | 0 | 0 | 0 | 0 | 0 | 93628 | 0 |
| EDSVKPGA | HNRNPA3; HNF | P51991; A0A | ROA3_HUMAN | Unmodified | 0 | 0 | 0 | 0 | 0 | 0 | 409120 | 16847 | 126390 | 72267 | 1038100 | 1306700 | 0 | 0 | 52582 | 0 | 0 | 68941 |
| EDTEEYNLI | HNRNPA3; HNF | P51991; A0A | ROA3_HUMAN | Unmodified | 0 | 0 | 0 | 0 | 211330 | 23094 | 110230 | 0 | 140480 | 0 | 233400 | 1541500 | 61702 | 0 | 0 | 31971 | 0 | 173170 |
| EDTESLEIF | ERP44 | Q9BS26 | ERP44_HUMAN | Unmodified | 0 | 0 | 0 | 0 | 0 | 0 | 0 | 0 | 21456000 | 359850 | 759960 | 409910 | 0 | 0 | 0 | 0 | 0 | 0 |
| EDTESLEIF | ERP44 | Q9BS26 | ERP44_HUMAN | Unmodified | 0 | 0 | 0 | 0 | 0 | 0 | 0 | 0 | 0 | 2044000 | 0 | 0 | 0 | 0 | 0 | 0 | 0 | 0 |
| EDTSEQVV | ABCC1; ABCC1 | P33527; I3L4 | MRP1_HUMAN | Unmodified | 0 | 0 | 0 | 0 | 0 | 0 | 0 | 0 | 0 | 0 | 0 | 3463700 | 0 | 0 | 0 | 0 | 0 | 0 |
| EDVFVHQT | YBX1; YBX3; YB | P67809; P16 | YBOX1_HUMAN | Unmodified | 0 | 0 | 789200 | 0 | 0 | 0 | 0 | 0 | 0 | 0 | 0 | 597510 | 0 | 0 | 0 | 0 | 0 | 0 |
| EEAEAPVEI | HYOU1; HYOU1 | Q9Y4L1; E9F | HYOU1_HUMA | Unmodified | 0 | 0 | 0 | 0 | 0 | 0 | 0 | 0 | 2020100 | 0 | 425740 | 0 | 0 | 0 | 0 | 0 | 0 | 0 |
| EEALELING | PIN1; PIN1; PIN | Q13526; K7E | PIN1_HUMAN F | Unmodified | 7712300 | 3961800 | 1332700 | 6257400 | 3056400 | 0 | 9435100 | 6471600 | 0 | 0 | 0 | 765540 | 5287200 | 4898600 | 10357000 | 7458100 | 322420 | 0 |

| | | | | | | | | | | | | | | | | | | | | | | |
|---|---|---|---|---|---|---|---|---|---|---|---|---|---|---|---|---|---|---|---|---|---|---|
| EEAPSLRP | FGB | P02675 | FIBB_HUMAN F | Unmodified | 0 | 299060 | 0 | 0 | 0 | 0 | 0 | 108170 | 0 | 0 | 0 | 0 | 0 | 0 | 0 | 0 | 0 | 0 |
| EEAQAEIEC | ATP6V1G1; ATP | O75348; A0A | VATG1_HUMAN | Unmodified | 0 | 0 | 0 | 0 | 0 | 0 | 0 | 0 | 0 | 0 | 121780 | 719620 | 0 | 0 | 0 | 0 | 0 | 0 |
| EEAQAEIEC | ATP6V1G1; ATP | O75348; A0A | VATG1_HUMAN | Unmodified | 0 | 0 | 0 | 0 | 0 | 0 | 0 | 0 | 0 | 0 | 0 | 456070 | 25074 | 0 | 0 | 266360 | 0 | 0 |
| EEASDYLEI | HSP90B1; HSP | P14625; A0A | ENPL_HUMAN | Unmodified | 0 | 0 | 2492500 | 0 | 0 | 0 | 0 | 0 | 83136000 | 4062000 | 15139000 | 16185000 | 0 | 0 | 0 | 0 | 0 | 0 |
| EEASDYLEI | HSP90B1; HSP | P14625; A0A | ENPL_HUMAN | Unmodified | 0 | 0 | 0 | 0 | 0 | 0 | 0 | 0 | 0 | 363630 | 0 | 0 | 0 | 0 | 0 | 0 | 34171 | 0 |
| EEASGSSV | XRCC5 | P13010 | XRCC5_HUMAI | Unmodified | 0 | 0 | 0 | 0 | 0 | 0 | 0 | 0 | 393010 | 0 | 0 | 2305100 | 0 | 0 | 0 | 0 | 0 | 0 |
| EEASNFR | PSMA3; PSMA3 | P25788; G3V | PSA3_HUMAN | Unmodified | 26435 | 100490 | 84836 | 41448 | 78609 | 0 | 0 | 0 | 367800 | 45119 | 0 | 1273200 | 0 | 0 | 111690 | 0 | 0 | 0 |
| EEDSANLK | CD2AP | Q9Y5K6 | CD2AP_HUMAI | Unmodified | 1049500 | 0 | 4176400 | 1210300 | 76133 | 0 | 0 | 0 | 214240 | 0 | 102040 | 531490 | 0 | 0 | 5423100 | 2886700 | 0 | 23915 |
| EEDSHTTLI | PLPP1 | O14494 | PLPP1_HUMAN | Unmodified | 0 | 0 | 2878400 | 602910 | 0 | 0 | 0 | 0 | 0 | 0 | 0 | 556030 | 0 | 0 | 884390 | 684000 | 0 | 0 |
| EEEAIQLDC | HSP90B1; HSP | P14625; A0A | ENPL_HUMAN | Unmodified | 0 | 0 | 0 | 0 | 0 | 0 | 0 | 0 | 0 | 0 | 3925700 | 0 | 0 | 0 | 0 | 0 | 0 | 0 |
| EEECHFYA | PRDX4; PRDX4 | Q13162; A6N | PRDX4_HUMAN | Unmodified | 0 | 0 | 0 | 0 | 0 | 0 | 0 | 0 | 3567700 | 0 | 580470 | 0 | 0 | 0 | 65655 | 37168 | 0 | 0 |
| EEEEFNTGI | SNRPD2; SNRP | P62316; K7E | SMD2_HUMAN | Unmodified | 0 | 0 | 0 | 0 | 0 | 0 | 0 | 0 | 258710 | 307520 | 467410 | 775500 | 0 | 0 | 19748 | 0 | 0 | 425740 |
| EEEEGISQE | HMGA1 | P17096 | HMGA1_HUMA | Unmodified | 0 | 0 | 0 | 0 | 0 | 0 | 0 | 0 | 0 | 0 | 353790 | 1143500 | 0 | 0 | 0 | 0 | 0 | 0 |
| EEEGSANR | CHGA; CHGA | P10645; G5E | CMGA_HUMAN | Unmodified | 0 | 0 | 0 | 0 | 0 | 0 | 0 | 0 | 3361900 | 0 | 0 | 0 | 0 | 0 | 0 | 0 | 0 | 0 |
| EEEGTGVV | IARS1; IARS1; IA | P41252; A0A | SYIC_HUMAN I | Unmodified | 0 | 0 | 0 | 0 | 0 | 0 | 0 | 0 | 0 | 0 | 0 | 4432200 | 0 | 0 | 0 | 0 | 0 | 0 |
| EEEIAALVIE | ACTG1; ACTG1 | P63261; A0A | ACTG_HUMAN | Acetyl (Protein N | 4651300 | 3742800 | 1484900 | 6825300 | 1.22E+08 | 19713000 | 20526000 | 23754000 | 3068200 | 6215200 | 2109000 | 7986900 | 57221000 | 48620000 | 6091400 | 4140200 | 94323000 | 30522000 |
| EEEIAALVIE | ACTG1; ACTG1 | P63261; A0A | ACTG_HUMAN | Acetyl (Protein N | 0 | 0 | 0 | 74463 | 3061500 | 0 | 400410 | 82766 | 656520 | 198360 | 35447 | 0 | 708420 | 837090 | 53216 | 16200 | 2112100 | 1102100 |
| EEEIAALVIE | ACTG1; ACTG1 | P63261; A0A | ACTG_HUMAN | Acetyl (Protein N | 0 | 0 | 0 | 0 | 0 | 0 | 0 | 0 | 0 | 0 | 0 | 0 | 0 | 0 | 0 | 0 | 5424700 | 0 |
| EEESQQQA | MYO6; MYO6; M | Q9UM54; A0 | MYO6_HUMAN | Unmodified | 0 | 0 | 1058500 | 120380 | 0 | 0 | 0 | 0 | 0 | 0 | 0 | 241500 | 0 | 0 | 681170 | 572760 | 0 | 0 |
| EEETSIDVA | SLC16A1 | P53985 | MOT1_HUMAN | Unmodified | 16645 | 0 | 0 | 0 | 0 | 0 | 192270 | 0 | 379880 | 153810 | 220690 | 133350 | 0 | 0 | 0 | 0 | 0 | 0 |
| EEETSIDVA | SLC16A1 | P53985 | MOT1_HUMAN | Unmodified | 0 | 0 | 2827500 | 419090 | 0 | 0 | 666090 | 0 | 634220 | 314640 | 1798600 | 1193800 | 97447 | 82455 | 1489100 | 1491200 | 0 | 0 |
| EEFASTCPI | CLIC1 | O00299 | CLIC1_HUMAN | Unmodified | 0 | 0 | 0 | 0 | 0 | 0 | 0 | 0 | 0 | 0 | 1420400 | 2771100 | 0 | 0 | 1347600 | 1251600 | 0 | 0 |
| EEFEHQQK | HSPA8; HSPA8; | P11142; E9P | HSP7C_HUMAI | Unmodified | 0 | 0 | 264590 | 0 | 0 | 0 | 0 | 0 | 83133 | 0 | 0 | 221820 | 27934 | 0 | 317940 | 0 | 0 | 0 |
| EEFTAFLHF | CALU; CALU | O43852; H0Y | CALU_HUMAN | Oxidation (M) | 0 | 0 | 179620 | 0 | 0 | 0 | 0 | 0 | 2909600 | 113920 | 668380 | 216140 | 0 | 0 | 217090 | 60503 | 17503 | 0 |
| EEFTAFLHF | CALU; CALU | O43852; H0Y | CALU_HUMAN | Unmodified | 0 | 0 | 0 | 0 | 0 | 0 | 0 | 0 | 1838100 | 0 | 0 | 0 | 0 | 0 | 0 | 0 | 0 | 0 |
| EEGGENLD | ANXA6; ANXA6; | P08133; E7E | ANXA6_HUMAN | Unmodified | 1.89E+09 | 0 | 0 | 0 | 0 | 0 | 555330 | 0 | 1423300 | 552920 | 0 | 0 | 0 | 0 | 0 | 0 | 0 | 0 |
| EEGSPLELE | SLC3A2; SLC3A | P08195; A0A | 4F2_HUMAN 4I | Unmodified | 290830 | 0 | 0 | 0 | 0 | 0 | 0 | 0 | 186780 | 0 | 0 | 0 | 0 | 0 | 388820 | 209590 | 0 | 0 |
| EEGSRDPA | PLEKHG3; PLEK | A1L390; G3V | PKHG3_HUMAI | Unmodified | 0 | 0 | 0 | 0 | 0 | 0 | 0 | 0 | 0 | 0 | 0 | 0 | 0 | 0 | 0 | 0 | 0 | 0 |
| EEGVFTVTF | CDCP1 | Q9H5V8 | CDCP1_HUMA | Unmodified | 0 | 0 | 2692700 | 0 | 929160 | 0 | 0 | 0 | 0 | 0 | 670460 | 5238800 | 0 | 0 | 1113300 | 1437200 | 0 | 287370 |
| EEGVLTLW | SLC25A11; SLC | Q02978; I3L | M2OM_HUMAN | Unmodified | 696430 | 680610 | 213180 | 0 | 0 | 0 | 0 | 0 | 1149700 | 0 | 0 | 1639300 | 600690 | 0 | 0 | 173200 | 0 | 0 |
| EEHEVAVL | IFITM3; IFITM3; | Q01628; A0A | IFM3_HUMAN I | Unmodified | 88970 | 0 | 15800000 | 3287900 | 0 | 0 | 0 | 0 | 0 | 0 | 0 | 0 | 0 | 0 | 6114700 | 5303500 | 0 | 0 |
| EEHLLEQL | TRIM26; TRIM26 | Q12899; A0A | TRI26_HUMAN | Unmodified | 424460 | 2708800 | 39948 | 559510 | 18673 | 0 | 13786 | 0 | 0 | 0 | 0 | 0 | 0 | 0 | 1777100 | 145630 | 0 | 0 |
| EEIFGPVQ | ALDH1A1 | P00352 | AL1A1_HUMAN | Oxidation (M) | 0 | 0 | 0 | 0 | 0 | 0 | 0 | 0 | 0 | 0 | 0 | 29744000 | 0 | 0 | 0 | 0 | 0 | 0 |
| EEIFGPVQ | ALDH1A1 | P00352 | AL1A1_HUMAN | Unmodified | 154450 | 0 | 0 | 0 | 0 | 0 | 0 | 0 | 0 | 0 | 0 | 41763000 | 0 | 0 | 0 | 0 | 0 | 0 |
| EEIPEEELAI | ATP2B1; ATP2B | P20020; E7E | AT2B1_HUMAN | Unmodified | 0 | 0 | 901790 | 0 | 0 | 0 | 0 | 0 | 0 | 0 | 0 | 0 | 0 | 0 | 0 | 0 | 0 | 0 |
| EEIPEEELAI | ATP2B1; ATP2B | P20020; E7E | AT2B1_HUMAN | Unmodified | 0 | 0 | 121760 | 0 | 0 | 0 | 0 | 0 | 364620 | 0 | 0 | 0 | 0 | 0 | 1061100 | 120280 | 0 | 0 |
| EEIQSAVTK | IQGAP3; IQGAP | Q86VI3; F2Z | IQGA3_HUMAN | Unmodified | 267940 | 0 | 91982 | 152670 | 56864 | 0 | 0 | 162860 | 159930 | 1047900 | 109670 | 300820 | 80875000 | 0 | 307690 | 41120 | 194940 | 0 |
| EEKEESDD | HSP90B1; HSP | P14625; A0A | ENPL_HUMAN | Unmodified | 0 | 0 | 102490 | 0 | 91090 | 0 | 0 | 21133 | 1641700 | 422790 | 496670 | 179110 | 0 | 0 | 69883 | 53549 | 0 | 0 |
| EELALLDGS | PFDN6; PFDN6 | O15212; A2A | PFD6_HUMAN | Unmodified | 0 | 0 | 0 | 0 | 0 | 0 | 0 | 0 | 0 | 0 | 0 | 1557800 | 0 | 0 | 0 | 0 | 0 | 0 |
| EELAQSQH | SLK | Q9H2G2 | SLK_HUMAN S | Unmodified | 0 | 0 | 104130 | 0 | 0 | 0 | 0 | 0 | 317160 | 0 | 0 | 891370 | 53398 | 0 | 65823 | 99966 | 0 | 0 |
| EELEEVIKD | PSMA5 | P28066 | PSA5_HUMAN | Unmodified | 0 | 0 | 368140 | 209630 | 0 | 0 | 0 | 0 | 620130 | 0 | 506700 | 2164900 | 0 | 0 | 0 | 0 | 0 | 0 |
| EELEQREAI | EIF3A | Q14152 | EIF3A_HUMAN | Unmodified | 0 | 0 | 0 | 121220 | 0 | 0 | 31867 | 0 | 84898 | 0 | 76654 | 1504300 | 0 | 0 | 924240 | 0 | 0 | 0 |
| EELESHRA | SLC4A7; SLC4A | Q9Y6M7; C9 | S4A7_HUMAN | Unmodified | 0 | 0 | 0 | 0 | 0 | 0 | 0 | 0 | 0 | 0 | 0 | 0 | 0 | 0 | 0 | 0 | 0 | 1622000 |
| EELGDAAA | AVIL | O75366 | AVIL_HUMAN A | Unmodified | 0 | 0 | 0 | 0 | 0 | 0 | 0 | 110180 | 0 | 0 | 946900 | 10438000 | 0 | 0 | 0 | 0 | 0 | 0 |
| EELGDAAA | AVIL | O75366 | AVIL_HUMAN A | Oxidation (M) | 0 | 0 | 0 | 0 | 0 | 0 | 0 | 0 | 0 | 0 | 268500 | 1454700 | 0 | 0 | 0 | 0 | 0 | 0 |
| EELGDEWL | DERA; DERA; DI | Q9Y315; G3V | DEOC_HUMAN | Unmodified | 0 | 0 | 0 | 0 | 0 | 0 | 0 | 0 | 0 | 0 | 0 | 1841000 | 0 | 0 | 0 | 0 | 0 | 0 |
| EELGLIEQA | PRPF8; PRPF8 | Q6P2Q9; I3L | PRP8_HUMAN | Unmodified | 22868 | 0 | 0 | 0 | 2877800 | 0 | 0 | 0 | 0 | 0 | 0 | 2893300 | 0 | 0 | 0 | 0 | 0 | 0 |
| EELIAELQD | PHGDH; PHGD | O43175; A0A | SERA_HUMAN | Unmodified | 0 | 0 | 0 | 64008 | 28204 | 0 | 0 | 0 | 1242100 | 49864 | 0 | 0 | 0 | 0 | 243830 | 0 | 0 | 0 |
| EELLFGGK | HK2; HK2 | P52789; E9P | HXK2_HUMAN | Unmodified | 0 | 0 | 0 | 0 | 0 | 0 | 0 | 0 | 0 | 0 | 0 | 1658000 | 0 | 432710 | 0 | 0 | 0 | 0 |
| EELMLRLQI | EZR; EZR | P15311; E7E | EZRI_HUMAN E | Unmodified | 0 | 0 | 0 | 154710 | 5357100 | 490400 | 67052 | 0 | 314960 | 465960 | 0 | 0 | 411470 | 368110 | 0 | 127440 | 0 | 611590 |
| EELMLRLQI | EZR; EZR | P15311; E7E | EZRI_HUMAN E | Oxidation (M) | 0 | 18476 | 0 | 0 | 2623700 | 2690600 | 0 | 0 | 605430 | 304950 | 0 | 0 | 705760 | 446560 | 0 | 0 | 178590 | 713120 |
| EELNAISGF | SF3A3 | Q12874 | SF3A3_HUMAN | Unmodified | 175320 | 113070 | 54126 | 72528 | 0 | 0 | 0 | 0 | 2398500 | 0 | 16921 | 1378500 | 0 | 0 | 0 | 311970 | 0 | 0 |
| EELQANGS | MARCKS | P29966 | MARCS_HUMA | Unmodified | 0 | 117640 | 2014400 | 775430 | 0 | 0 | 139460 | 0 | 3130100 | 314600 | 7414700 | 16016000 | 0 | 0 | 1687300 | 1316300 | 0 | 144060 |
| EELQANGS | MARCKS | P29966 | MARCS_HUMA | Unmodified | 0 | 0 | 0 | 0 | 0 | 0 | 0 | 0 | 0 | 0 | 1384200 | 447710 | 0 | 0 | 0 | 0 | 0 | 0 |
| EELQRQHD | NUCB2; NUCB2 | P80303; Q2L | NUCB2_HUMA | Unmodified | 0 | 0 | 0 | 0 | 0 | 0 | 0 | 0 | 1698500 | 0 | 0 | 0 | 0 | 0 | 0 | 0 | 0 | 0 |
| EELSNVLA | RPL36 | Q9Y3U8 | RL36_HUMAN ( | Unmodified | 0 | 0 | 0 | 0 | 0 | 0 | 0 | 0 | 0 | 0 | 0 | 1374100 | 0 | 0 | 0 | 0 | 379720 | 0 |
| EELSSGDS | AFDN; AFDN; A | P55196; Q5T | AFAD_HUMAN | Unmodified | 44589 | 0 | 0 | 0 | 0 | 0 | 0 | 0 | 0 | 0 | 282100 | 436030 | 0 | 0 | 0 | 76369 | 0 | 0 |
| EELSSGDS | AFDN; AFDN; A | P55196; Q5T | AFAD_HUMAN | Unmodified | 0 | 0 | 0 | 0 | 0 | 0 | 0 | 0 | 0 | 0 | 187010 | 0 | 441030 | 0 | 0 | 0 | 0 | 0 |
| EELVAAVEI | CTNNA1; CTNN | P35221; P26 | CTNA1_HUMAI | Unmodified | 215130 | 0 | 2713900 | 665220 | 0 | 0 | 0 | 0 | 455560 | 0 | 0 | 3076600 | 112330 | 0 | 3235100 | 1985300 | 0 | 0 |
| EELVLLVLG | RFFL; RFFL | Q8WZ73; C9 | RFFL_HUMAN I | Unmodified | 0 | 0 | 0 | 0 | 4691100 | 1792000 | 30319000 | 1800100 | 0 | 0 | 0 | 0 | 3618900 | 0 | 0 | 0 | 0 | 0 |
| EELYQNLTI | SPTAN1; SPTAN | Q13813; A0A | SPTN1_HUMAN | Unmodified | 0 | 0 | 0 | 0 | 0 | 0 | 0 | 0 | 246940 | 0 | 2332000 | 1371800 | 0 | 0 | 0 | 0 | 0 | 0 |
| EEMDRAQ | TSG101; TSG10 | Q99816; F5H | TS101_HUMAN | Unmodified | 11406000 | 7006700 | 8806200 | 7726800 | 0 | 0 | 0 | 0 | 4184300 | 0 | 0 | 3043800 | 0 | 0 | 25019000 | 10388000 | 0 | 0 |
| EEMDRAQ | TSG101; TSG10 | Q99816; F5H | TS101_HUMAN | Oxidation (M) | 8259000 | 7521600 | 17752000 | 1421300 | 0 | 188510 | 312730 | 0 | 474590 | 0 | 0 | 872990 | 0 | 313940 | 17674000 | 9676200 | 0 | 29958 |
| EEMDRAQ | TSG101; TSG10 | Q99816; F5H | TS101_HUMAN | Oxidation (M) | 98617 | 0 | 185280 | 163730 | 0 | 0 | 0 | 0 | 479160 | 0 | 0 | 252340 | 0 | 0 | 265700 | 165450 | 0 | 0 |
| EEMDRAQ | TSG101; TSG10 | Q99816; F5H | TS101_HUMAN | Unmodified | 191240 | 0 | 230810 | 159220 | 0 | 0 | 0 | 0 | 0 | 0 | 0 | 0 | 0 | 0 | 290250 | 155440 | 0 | 0 |
| EEMMQTLN | DAAM1 | Q9Y4D1 | DAAM1_HUMA | Unmodified | 0 | 0 | 0 | 0 | 113360 | 0 | 0 | 0 | 188610 | 0 | 0 | 697860 | 0 | 0 | 61613 | 0 | 0 | 0 |
| EEMQSNVE | NDRG1; NDRG | Q92597; A0A | NDRG1_HUMA | Unmodified | 0 | 0 | 222830 | 0 | 0 | 0 | 141280 | 25427 | 512130 | 0 | 0 | 1023200 | 0 | 0 | 201400 | 421670 | 0 | 0 |
| EENDIARGI | CEP55; CEP55 | Q53EZ4; H0Y | CEP55_HUMAN | Unmodified | 0 | 0 | 0 | 0 | 4481200 | 98907 | 0 | 0 | 1136800 | 343280 | 0 | 0 | 0 | 0 | 0 | 0 | 826270 | 1703000 |
| EENGVLVLI | PDIA4; PDIA4; F | P13667; C9J | PDIA4_HUMAN | Unmodified | 0 | 168330 | 0 | 281480 | 5042900 | 213160 | 0 | 62114 | 2714400 | 1006500 | 1204100 | 371140 | 1015300 | 790660 | 92701 | 69874 | 1398100 | 2772500 |
| EENLIPYSP | EMC1 | Q8N766 | EMC1_HUMAN | Unmodified | 0 | 0 | 0 | 0 | 0 | 0 | 0 | 0 | 931330 | 0 | 0 | 0 | 0 | 0 | 0 | 0 | 0 | 0 |
| EENPYAR | PTTG1IP; PTTG | P53801; A8M | PTTG_HUMAN | Unmodified | 0 | 0 | 3220100 | 0 | 0 | 0 | 0 | 0 | 0 | 0 | 0 | 0 | 0 | 0 | 1782100 | 1141600 | 0 | 0 |
| EENPYARFI | PTTG1IP; PTTG | P53801; A8M | PTTG_HUMAN | Unmodified | 349890 | 216450 | 6698200 | 2654400 | 7724500 | 2130400 | 584670 | 234990 | 3734200 | 760090 | 251000 | 0 | 606240 | 569360 | 2360000 | 1830500 | 0 | 1713400 |
| EENTIILQQ | NLN; NLN; NLN | Q9BYT8; E9F | NEUL_HUMAN | Unmodified | 140530 | 139720 | 0 | 116690 | 0 | 0 | 0 | 0 | 729660 | 0 | 65315 | 317850 | 42323 | 0 | 38341 | 17024 | 0 | 0 |
| EEPAAAGS | MARCKS | P29966 | MARCS_HUMA | Unmodified | 0 | 0 | 375300 | 0 | 0 | 0 | 0 | 0 | 4585300 | 286550 | 55071 | 420930 | 0 | 0 | 0 | 64628 | 113540 | 35553 |
| EEPAAAGS | MARCKS | P29966 | MARCS_HUMA | Unmodified | 0 | 0 | 0 | 0 | 0 | 0 | 0 | 0 | 536900 | 439330 | 0 | 0 | 0 | 0 | 0 | 0 | 808830 | 0 |
| EEPFFPPPE | HNRNPUL2; HN | Q1KMD3; H3 | HNRL2_HUMAI | Unmodified | 0 | 0 | 0 | 0 | 0 | 0 | 0 | 0 | 0 | 0 | 0 | 233770 | 0 | 0 | 0 | 0 | 0 | 0 |
| EEPGEDFP | TFRC; TFRC | P02786; G3V | TFR1_HUMAN | Unmodified | 0 | 0 | 278010 | 0 | 0 | 0 | 0 | 0 | 0 | 759780 | 0 | 0 | 0 | 0 | 0 | 0 | 0 | 0 |
| EEQELYQK | MYO6; MYO6; M | Q9UM54; A0 | MYO6_HUMAN | Unmodified | 0 | 0 | 535780 | 0 | 0 | 0 | 0 | 0 | 0 | 0 | 0 | 0 | 0 | 0 | 0 | 0 | 0 | 0 |
| EEQHQLAV | EIF3A | Q14152 | EIF3A_HUMAN | Unmodified | 361980 | 172490 | 0 | 279300 | 0 | 0 | 0 | 0 | 0 | 0 | 938060 | 4694600 | 0 | 0 | 795850 | 0 | 0 | 0 |
| EEQLFSSQ | TOR3A | Q9H497 | TOR3A_HUMAN | Unmodified | 0 | 0 | 0 | 0 | 0 | 0 | 0 | 0 | 1218200 | 0 | 0 | 0 | 0 | 0 | 0 | 0 | 0 | 0 |
| EEQNKMQI | CENPF | P49454 | CENPF_HUMAI | Oxidation (M) | 0 | 0 | 0 | 0 | 9855100 | 0 | 0 | 0 | 0 | 0 | 0 | 0 | 0 | 0 | 0 | 0 | 0 | 0 |
| EEQSQITS | AP2M1; AP2M1 | Q96CW1; E9 | AP2M1_HUMAI | Unmodified | 95685 | 0 | 0 | 0 | 0 | 0 | 0 | 0 | 0 | 0 | 0 | 1262300 | 0 | 0 | 0 | 0 | 490870 | 0 |
| EEQSQITS | AP2M1; AP2M1 | Q96CW1; E9 | AP2M1_HUMAI | Unmodified | 0 | 36324 | 0 | 0 | 0 | 0 | 0 | 0 | 402990 | 0 | 0 | 2110900 | 0 | 0 | 227200 | 347380 | 0 | 0 |
| EERQAHLTI | PABPC1; PABP( | P11940; A0A | PABP1_HUMAN | Unmodified | 0 | 0 | 0 | 0 | 0 | 0 | 0 | 0 | 45403 | 0 | 0 | 0 | 1004000 | 243300 | 0 | 0 | 28985 | 657660 |
| EESANNQV | JAG2 | Q9Y219 | JAG2_HUMAN I | Unmodified | 108350 | 0 | 0 | 0 | 0 | 0 | 0 | 0 | 763940 | 0 | 467560 | 4673800 | 0 | 0 | 0 | 155680 | 0 | 0 |
| EESDDEAA | HSP90B1; HSP | P14625; A0A | ENPL_HUMAN | Unmodified | 0 | 0 | 0 | 0 | 0 | 0 | 278290 | 619420 | 24351000 | 4369500 | 538560 | 745020 | 0 | 0 | 0 | 0 | 0 | 0 |
| EESDDEAA | HSP90B1; HSP | P14625; A0A | ENPL_HUMAN | Unmodified | 0 | 0 | 0 | 0 | 0 | 55909 | 0 | 188870 | 6222400 | 1580300 | 1721400 | 289590 | 0 | 0 | 0 | 0 | 0 | 0 |
| EESGKPGA | HNRNPA2B1; H | P22626; A0A | ROA2_HUMAN | Unmodified | 24599 | 0 | 0 | 0 | 0 | 990830 | 700800 | 487000 | 433670 | 176270 | 1576900 | 6077600 | 0 | 0 | 0 | 0 | 207590 | 30649 |
| EESGKPGA | HNRNPA2B1; H | P22626; A0A | ROA2_HUMAN | Unmodified | 0 | 0 | 0 | 0 | 1037300 | 12503 | 205430 | 314840 | 529260 | 0 | 559390 | 5498800 | 0 | 0 | 0 | 0 | 40479 | 96405 |
| EESGVSVS | SLC29A1 | Q99808 | S29A1_HUMAN | Unmodified | 1617900 | 484580 | 3501500 | 0 | 0 | 0 | 0 | 0 | 481520 | 0 | 837490 | 6707000 | 0 | 0 | 2386500 | 1797200 | 0 | 0 |
| EETFSTLK | RAB22A | Q9UL26 | RB22A_HUMAN | Unmodified | 0 | 0 | 309090 | 0 | 0 | 0 | 0 | 0 | 0 | 0 | 86929 | 0 | 0 | 0 | 3510000 | 0 | 0 | 0 |
| EETPGTEW | CLTB; CLTB | P09497; D6F | CLCB_HUMAN | Unmodified | 303710 | 136810 | 750100 | 175340 | 0 | 0 | 0 | 0 | 0 | 0 | 0 | 206890 | 0 | 0 | 333240 | 234670 | 0 | 0 |
| EETQPPVAI | SERBP1 | Q8NC51 | PAIRB_HUMAN | Unmodified | 0 | 0 | 0 | 0 | 0 | 0 | 0 | 0 | 0 | 0 | 505290 | 0 | 0 | 0 | 0 | 0 | 0 | 0 |
| EEVGEEAIV | MYH9; MYH9; M | P35579; Q5E | MYH9_HUMAN | Unmodified | 0 | 0 | 0 | 0 | 0 | 0 | 0 | 0 | 0 | 0 | 35839 | 1441600 | 0 | 0 | 0 | 0 | 0 | 0 |
| EEVGQYLK | PTRHD1 | Q6GMV3 | PTRD1_HUMAN | Unmodified | 0 | 0 | 489810 | 119280 | 0 | 319750 | 0 | 0 | 136240 | 334150 | 717630 | 2483400 | 0 | 0 | 385480 | 321930 | 171970 | 0 |
| EEVSGHLY | FAM234A; FAM2 | Q9H0X4; B4I | F234A_HUMAN | Unmodified | 0 | 0 | 0 | 0 | 0 | 0 | 0 | 0 | 0 | 0 | 0 | 0 | 0 | 0 | 367490 | 675370 | 0 | 0 |
| EEVVLLTH | GMPS | P49915 | GUAA_HUMAN | Unmodified | 29485 | 0 | 82663 | 0 | 177430 | 0 | 0 | 0 | 313110 | 0 | 289420 | 2534900 | 97537 | 0 | 72074 | 86183 | 100460 | 0 |
| EEVVLWMN | TM9SF3 | Q9HD45 | TM9S3_HUMAN | Unmodified | 0 | 0 | 0 | 0 | 0 | 0 | 0 | 0 | 3702200 | 0 | 0 | 0 | 0 | 0 | 0 | 0 | 0 | 0 |
| EFAAFFAK | EXOC4 | Q96A65 | EXOC4_HUMAI | Unmodified | 0 | 0 | 0 | 0 | 0 | 0 | 0 | 0 | 0 | 69208 | 0 | 606610 | 0 | 0 | 0 | 0 | 0 | 0 |
| EFAAGYNV | GPX4; GPX4; GI | P36969; A0A | GPX4_HUMAN | Unmodified | 710630 | 0 | 0 | 0 | 0 | 0 | 0 | 0 | 16302000 | 2150600 | 0 | 0 | 0 | 0 | 1286400 | 495810 | 0 | 0 |
| EFAAGYNV | GPX4; GPX4; GI | P36969; A0A | GPX4_HUMAN | Unmodified | 0 | 200830 | 0 | 0 | 0 | 0 | 241490 | 0 | 0 | 145410 | 0 | 0 | 234700 | 334690 | 0 | 0 | 236510 | 242400 |
| EFADSLGIF | RAB1C; RAB1A; | Q92928; P62 | RAB1C_HUMAI | Unmodified | 10095000 | 6295800 | 4269900 | 9740500 | 11334000 | 2066100 | 3661800 | 0 | 17866000 | 6576700 | 7289100 | 20828000 | 7152300 | 6487400 | 10027000 | 9322200 | 1970000 | 8320300 |
| EFAGEDTSI | XPO1; XPO1; XF | O14980; A0A | XPO1_HUMAN | Unmodified | 0 | 0 | 0 | 0 | 0 | 0 | 0 | 0 | 0 | 0 | 62028 | 5774200 | 0 | 0 | 0 | 0 | 0 | 0 |
| EFAGEDTSI | XPO1; XPO1; XF | O14980; A0A | XPO1_HUMAN | Unmodified | 0 | 0 | 0 | 0 | 45281 | 0 | 0 | 0 | 765740 | 0 | 0 | 1217900 | 0 | 0 | 0 | 0 | 0 | 0 |
| EFALADWK | PAN2 | Q504Q3 | PAN2_HUMAN | Unmodified | 322300 | 188940 | 163210 | 209050 | 0 | 0 | 0 | 0 | 0 | 0 | 0 | 154610 | 0 | 0 | 0 | 105090 | 0 | 0 |
| EFALTNPEI | SIL1; SIL1; SIL1 | Q9H173; D6 | SIL1_HUMAN N | Unmodified | 0 | 247300 | 0 | 0 | 0 | 0 | 247070 | 123850 | 2346300 | 1347800 | 0 | 0 | 316510 | 0 | 0 | 0 | 0 | 0 |
| EFCENLSAI | PPP2R1A; PPP2 | P30153; B3K | 2AAA_HUMAN | Unmodified | 0 | 0 | 0 | 0 | 0 | 0 | 0 | 0 | 0 | 0 | 0 | 348460 | 0 | 0 | 0 | 0 | 0 | 0 |
| EFDEDLED | SEPTIN3; SEPTI | Q9UH03; A0 | SEPT3_HUMAN | Unmodified | 0 | 0 | 0 | 0 | 0 | 0 | 0 | 0 | 0 | 0 | 0 | 323930 | 0 | 0 | 0 | 220930 | 0 | 0 |
| EFDEDSED | SEPTIN9; SEPTI | Q9UHD8; A0 | SEPT9_HUMAN | Unmodified | 261360 | 0 | 0 | 1081900 | 3711300 | 1428800 | 0 | 0 | 1718500 | 511570 | 1626200 | 9559000 | 775050 | 595350 | 638320 | 662880 | 1589300 | 3339100 |

| | | | | | | | | | | | | | | | | | | | | | | |
|---|---|---|---|---|---|---|---|---|---|---|---|---|---|---|---|---|---|---|---|---|---|---|
| EFDLTKEEI | OPA1; OPA1; OI | O60313; E5k | OPA1_HUMAN | Unmodified | 0 | 0 | 0 | 0 | 0 | 0 | 0 | 0 | 0 | 0 | 210420 | 213800 | 0 | 0 | 0 | 0 | 0 | 0 |
| EFDPTITDA | CALD1; CALD1; | Q05682; C9J | CALD1_HUMAI | Unmodified | 48432 | 0 | 526980 | 617900 | 435560 | 0 | 0 | 0 | 0 | 0 | 0 | 0 | 885000 | 592630 | 746250 | 588710 | 0 | 279510 |
| EFDSSTLNI | YIPF6 | Q96EC8 | YIPF6_HUMAN | Unmodified | 0 | 0 | 408210 | 0 | 0 | 0 | 0 | 0 | 0 | 0 | 0 | 920260 | 0 | 0 | 0 | 0 | 0 | 0 |
| EFEDRGFV | RARS1 | P54136 | SYRC_HUMAN | Unmodified | 0 | 0 | 119770 | 0 | 0 | 89555 | 0 | 0 | 193070 | 65598 | 560840 | 3472900 | 0 | 0 | 0 | 472500 | 0 | 209670 |
| EFEETAEK | BIRC5; BIRC5 | H3BLT4; A0A | H3BLT4_HUMA | Unmodified | 0 | 18940 | 0 | 0 | 0 | 0 | 0 | 0 | 436640 | 138430 | 0 | 0 | 0 | 0 | 0 | 0 | 494220 | 268770 |
| EFEGATLRY | APBB2; APBB2; | Q92870; H0Y | APBB2_HUMAN | Unmodified | 0 | 0 | 0 | 0 | 512950 | 680160 | 0 | 0 | 0 | 0 | 0 | 0 | 0 | 0 | 0 | 0 | 0 | 0 |
| EFEGLLAQI | LIMCH1; LIMCH | Q9UPQ0; D6 | LIMC1_HUMAN | Unmodified | 0 | 0 | 0 | 0 | 0 | 0 | 516250 | 0 | 0 | 0 | 0 | 0 | 343790 | 0 | 0 | 0 | 0 | 0 |
| EFEPLLNW | HSP90B1; HSP9 | P14625; A0A | ENPL_HUMAN | Unmodified | 0 | 0 | 234590 | 0 | 882310 | 0 | 1319200 | 0 | 1.86E+08 | 23032000 | 1769400 | 957490 | 0 | 680160 | 98334 | 0 | 0 | 0 |
| EFEPLLNW | HSP90B1; HSP9 | P14625; A0A | ENPL_HUMAN | Oxidation (M) | 554270 | 0 | 755640 | 0 | 0 | 0 | 4964500 | 652550 | 68501000 | 42851000 | 14801000 | 13213000 | 0 | 1090900 | 1346600 | 720390 | 0 | 0 |
| EFEPLLNW | HSP90B1; HSP9 | P14625; A0A | ENPL_HUMAN | Oxidation (M) | 23740 | 0 | 0 | 0 | 0 | 0 | 1283700 | 0 | 0 | 1169300 | 0 | 0 | 0 | 0 | 0 | 0 | 0 | 0 |
| EFEPLLR | INPPL1; INPPL1 | O15357; A0A | SHIP2_HUMAN | Unmodified | 0 | 0 | 0 | 0 | 0 | 0 | 0 | 0 | 0 | 0 | 0 | 702840 | 0 | 0 | 0 | 41337 | 0 | 0 |
| EFEPLLRVD | INPPL1; INPPL1 | O15357; A0A | SHIP2_HUMAN | Unmodified | 0 | 0 | 0 | 0 | 0 | 0 | 0 | 0 | 0 | 0 | 0 | 824070 | 0 | 0 | 0 | 0 | 0 | 0 |
| EFEQIEQLK | EMB; EMB | Q6PCB8; D6 | EMB_HUMAN E | Unmodified | 0 | 0 | 317540 | 0 | 0 | 0 | 0 | 0 | 0 | 0 | 0 | 0 | 0 | 0 | 747960 | 296290 | 114120 | 0 |
| EFESVLVDA | RPRD1B | Q9NQG5 | RPR1B_HUMAN | Unmodified | 0 | 0 | 0 | 0 | 0 | 0 | 0 | 0 | 0 | 0 | 0 | 0 | 0 | 0 | 0 | 0 | 0 | 0 |
| EFFESNIK | TSPAN14; TSPA | Q8NG11; A6 | TSN14_HUMAN | Unmodified | 554600 | 4238400 | 456590 | 108060 | 0 | 0 | 0 | 0 | 0 | 0 | 0 | 1468600 | 0 | 0 | 1619700 | 880700 | 0 | 0 |
| EFFNGKEP | HSPA5; HSPA5; | P11021; A0A | BIP_HUMAN Er | Unmodified | 66273 | 0 | 0 | 129460 | 0 | 0 | 0 | 0 | 9147600 | 184330 | 360600 | 662040 | 0 | 0 | 584280 | 197870 | 0 | 0 |
| EFFVDDSEI | DIP2B | Q9P265 | DIP2B_HUMAN | Unmodified | 0 | 0 | 0 | 0 | 1708800 | 0 | 0 | 0 | 4818300 | 2746800 | 5072000 | 8281400 | 6531100 | 2672300 | 0 | 0 | 2816800 | 9327100 |
| EFGADCKY | MDK; MDK; MDI | P21741; E9P | MK_HUMAN Mi | Unmodified | 439760 | 426560 | 1916700 | 800570 | 0 | 0 | 281360 | 160990 | 0 | 3428000 | 0 | 241980 | 227930 | 0 | 1038200 | 842350 | 335930 | 0 |
| EFGDKVNL | PRXL2A | Q9BRX8 | PXL2A_HUMAN | Unmodified | 0 | 0 | 0 | 0 | 0 | 0 | 1987500 | 0 | 6277200 | 4295300 | 0 | 0 | 0 | 0 | 0 | 0 | 0 | 0 |
| EFGFVIPER | OPLAH | O14841 | OPLA_HUMAN | Unmodified | 0 | 0 | 0 | 0 | 0 | 0 | 0 | 0 | 47999 | 0 | 46141 | 338690 | 0 | 0 | 0 | 0 | 0 | 44188 |
| EFGGGHIK | TWF2; TWF2; TV | Q6IBS0; D6F | TWF2_HUMAN | Unmodified | 0 | 0 | 0 | 298670 | 0 | 0 | 0 | 0 | 0 | 592030 | 0 | 459590 | 0 | 0 | 0 | 0 | 4404400 | 0 |
| EFGIADPDI | ACOX1; ACOX1 | Q15067; I3L | ACOX1_HUMAI | Oxidation (M) | 0 | 0 | 0 | 0 | 0 | 0 | 0 | 0 | 85683 | 0 | 13805 | 0 | 0 | 0 | 0 | 0 | 0 | 8024 |
| EFGNLTR | UQCRQ | O14949 | QCR8_HUMAN | Unmodified | 124520 | 0 | 0 | 0 | 131850 | 95608 | 0 | 0 | 908630 | 0 | 205540 | 0 | 0 | 0 | 0 | 0 | 0 | 0 |
| EFGPSHFS | GPX8; GPX8 | Q8TED1; E7E | GPX8_HUMAN | Unmodified | 0 | 1065600 | 0 | 0 | 0 | 0 | 0 | 0 | 1777400 | 0 | 1247800 | 0 | 0 | 0 | 0 | 0 | 0 | 0 |
| EFGPVVIDY | DYNC1H1; DYN | Q14204; A0A | DYHC1_HUMA | Unmodified | 0 | 0 | 532620 | 182700 | 770180 | 0 | 0 | 0 | 704770 | 0 | 406000 | 6450000 | 1019500 | 0 | 0 | 0 | 0 | 239300 |
| EFGSLPTTF | STX7 | O15400 | STX7_HUMAN S | Unmodified | 0 | 0 | 0 | 0 | 0 | 0 | 0 | 0 | 0 | 0 | 1576700 | 2065200 | 0 | 0 | 0 | 0 | 0 | 0 |
| EFGSLPTTF | STX7 | O15400 | STX7_HUMAN S | Unmodified | 146980 | 0 | 0 | 201670 | 1424000 | 278480 | 0 | 0 | 0 | 0 | 0 | 0 | 258010 | 237320 | 0 | 0 | 125520 | 250820 |
| EFGTNIK | HSP90B2P; HSI | Q58FF3; P14 | ENPLL_HUMAN | Unmodified | 199630 | 334680 | 414330 | 0 | 791210 | 101980 | 52767 | 27937 | 46747000 | 19823000 | 8666600 | 7858600 | 92548 | 0 | 551190 | 384820 | 63095 | 0 |
| EFGTNIKLC | HSP90B2P; HSI | Q58FF3; P14 | ENPLL_HUMAN | Unmodified | 0 | 0 | 0 | 0 | 0 | 0 | 0 | 0 | 1186000 | 1038100 | 0 | 0 | 0 | 0 | 0 | 0 | 0 | 0 |
| EFGTNIKLC | HSP90B1; HSP9 | P14625; A0A | ENPL_HUMAN | Unmodified | 0 | 0 | 0 | 0 | 713210 | 79488 | 440980 | 0 | 0 | 14296000 | 0 | 0 | 0 | 662720 | 0 | 0 | 252910 | 0 |
| EFGVTTGR | ADSS2 | P30520 | PURA2_HUMAI | Unmodified | 0 | 0 | 0 | 0 | 0 | 0 | 0 | 0 | 0 | 0 | 0 | 0 | 0 | 0 | 0 | 0 | 0 | 0 |
| EFHAPLIIDI | UAP1 | Q16222 | UAP1_HUMAN | Unmodified | 0 | 0 | 406840 | 507860 | 1285200 | 0 | 0 | 0 | 0 | 0 | 418250 | 0 | 1573800 | 0 | 0 | 254060 | 135260 | 367630 |
| EFHHGPDI | ERP44 | Q9BS26 | ERP44_HUMAN | Unmodified | 0 | 0 | 0 | 0 | 0 | 0 | 0 | 0 | 1757800 | 0 | 0 | 0 | 0 | 0 | 0 | 0 | 0 | 0 |
| EFHLNESG | SETSIP; SET; SE | P0DME0; Q0 | SETLP_HUMAN | Unmodified | 206270 | 120650 | 550290 | 740630 | 4047200 | 743540 | 394850 | 207790 | 8032500 | 6188600 | 12863000 | 24945000 | 5306100 | 941080 | 225870 | 241290 | 512060 | 523260 |
| EFHLNESG | SET; SET | Q01105; A0A | SET_HUMAN Pi | Unmodified | 0 | 0 | 0 | 0 | 1356800 | 0 | 0 | 133540 | 0 | 1109600 | 0 | 0 | 1515800 | 0 | 0 | 136480 | 92189 | 0 |
| EFHLNESG | SET; SET | Q01105; A0A | SET_HUMAN Pi | Unmodified | 0 | 0 | 0 | 0 | 488700 | 0 | 0 | 0 | 0 | 399230 | 0 | 0 | 552870 | 0 | 0 | 0 | 0 | 0 |
| EFIATTPSIT | EPB41L1; EPB4 | Q9H4G0; H7 | E41L1_HUMAN | Unmodified | 0 | 0 | 0 | 0 | 0 | 0 | 1501300 | 0 | 268040 | 0 | 218920 | 0 | 0 | 2923800 | 0 | 0 | 0 | 934400 |
| EFIDACLNI | STK26; STK26; S | Q9P289; B4E | STK26_HUMAN | Unmodified | 0 | 0 | 0 | 0 | 3088200 | 0 | 0 | 30365 | 0 | 0 | 0 | 0 | 3671500 | 5164000 | 0 | 0 | 2759900 | 2488900 |
| EFIEAEMK | NNT; NNT; NNT | Q13423; E9F | NNTM_HUMAN | Unmodified | 0 | 0 | 0 | 0 | 0 | 0 | 112120 | 0 | 6040700 | 831050 | 150240 | 0 | 0 | 0 | 0 | 0 | 0 | 0 |
| EFIEGVSQI | PPP3R1; PPP3R | P63098; F6U | CANB1_HUMAI | Unmodified | 0 | 0 | 0 | 0 | 0 | 0 | 0 | 0 | 0 | 0 | 0 | 1900300 | 0 | 0 | 0 | 0 | 0 | 0 |
| EFIEGVSQI | PPP3R1; PPP3R | P63098; F6U | CANB1_HUMAI | Unmodified | 216190 | 205230 | 0 | 321110 | 0 | 0 | 0 | 0 | 0 | 0 | 0 | 5185100 | 0 | 0 | 539720 | 451140 | 0 | 0 |
| EFIEKYDK | ANXA6; ANXA6; | P08133; E5R | ANXA6_HUMAI | Unmodified | 0 | 0 | 0 | 0 | 662340 | 0 | 122080 | 37569 | 1273400 | 531210 | 0 | 0 | 0 | 212180 | 0 | 0 | 101780 | 750480 |
| EFILNNEGI | FMNL2; FMNL2 | Q96PY5; C9I | FMNL2_HUMAI | Unmodified | 0 | 0 | 0 | 0 | 0 | 330520 | 0 | 0 | 322520 | 0 | 0 | 1831000 | 0 | 0 | 0 | 28335 | 199570 | 371320 |
| EFINFVK | NPC1 | O15118 | NPC1_HUMAN | Unmodified | 0 | 907710 | 0 | 0 | 301320 | 29974 | 1906500 | 587650 | 852860 | 132440 | 87732 | 0 | 0 | 36092 | 1524900 | 0 | 0 | 0 |
| EFIPDEPSK | USP54; USP54; | Q70EL1; A0A | UBP54_HUMAI | Unmodified | 124020 | 89684 | 228660 | 396070 | 0 | 0 | 0 | 0 | 0 | 0 | 0 | 0 | 0 | 0 | 188020 | 148830 | 0 | 0 |
| EFIQEPAK | PRMT5; PRMT5; | O14744; G3 | ANM5_HUMAN | Unmodified | 0 | 0 | 0 | 0 | 442820 | 0 | 0 | 0 | 285800 | 104760 | 717910 | 2900100 | 593490 | 0 | 0 | 0 | 0 | 249240 |
| EFITGDVEF | NAP1L4; NAP1L | Q99733; C9J | NP1L4_HUMAN | Unmodified | 0 | 0 | 0 | 0 | 0 | 0 | 0 | 0 | 317180 | 0 | 1364200 | 8235700 | 0 | 0 | 99000 | 72645 | 0 | 0 |
| EFKIPFEISK | MANBA; MANBA | O00462; A0A | MANBA_HUMA | Unmodified | 0 | 0 | 0 | 0 | 0 | 0 | 0 | 0 | 0 | 578720 | 0 | 0 | 0 | 0 | 0 | 0 | 0 | 0 |
| EFKRETGVI | HSPA9; HSPA9; | P38646; H0Y | GRP75_HUMAI | Unmodified | 0 | 0 | 0 | 0 | 558310 | 206900 | 0 | 105250 | 2891200 | 1500400 | 204820 | 0 | 269130 | 396420 | 0 | 0 | 189660 | 233040 |
| EFLALASSS | NCKAP1 | Q9Y2A7 | NCKP1_HUMAI | Unmodified | 0 | 0 | 215660 | 0 | 0 | 0 | 0 | 0 | 0 | 0 | 0 | 2656500 | 0 | 0 | 762120 | 0 | 0 | 0 |
| EFLHAQEE | PSMC4 | P43686 | PRS6B_HUMAN | Unmodified | 78896 | 0 | 0 | 222140 | 1502000 | 0 | 186190 | 264220 | 570390 | 101750 | 354870 | 3521400 | 2526300 | 0 | 152620 | 107420 | 803580 | 1791800 |
| EFLIALLEK | DNAJC13 | O75165 | DJC13_HUMAN | Unmodified | 0 | 0 | 0 | 0 | 0 | 0 | 0 | 0 | 16638 | 0 | 11580 | 422340 | 0 | 0 | 0 | 0 | 0 | 0 |
| EFLPEGQD | EIF4G1; EIF4G1 | Q04637; E7E | IF4G1_HUMAN | Unmodified | 540350 | 349920 | 21161000 | 0 | 50153000 | 2440300 | 0 | 1035200 | 6360500 | 6383300 | 0 | 53058000 | 0 | 0 | 0 | 31755000 | 5206500 | 14551000 |
| EFLQINSLC | GSTT2; GSTT2B | P0CG29; P0 | GST2_HUMAN | Unmodified | 0 | 0 | 0 | 0 | 0 | 0 | 0 | 0 | 0 | 0 | 0 | 1215600 | 0 | 0 | 0 | 0 | 0 | 0 |
| EFLSAKEET | SLC43A3; SLC4 | Q8NBI5; E9F | S43A3_HUMAN | Unmodified | 1120600 | 0 | 0 | 0 | 0 | 0 | 0 | 0 | 230560 | 98682 | 74901 | 0 | 0 | 0 | 205620 | 0 | 0 | 0 |
| EFLVAGGE | STRAP; STRAP | Q9Y3F4; H0Y | STRAP_HUMAN | Unmodified | 669620 | 550920 | 215420 | 713110 | 1321100 | 0 | 1590200 | 292450 | 1590400 | 1563100 | 783920 | 13469000 | 2370100 | 0 | 1970600 | 640380 | 726000 | 786340 |
| EFLVPSREF | DARS2; DARS2; | Q6PI48; A0A | SYDM_HUMAN | Unmodified | 0 | 0 | 0 | 0 | 0 | 0 | 0 | 0 | 1153900 | 92572 | 0 | 0 | 0 | 0 | 0 | 0 | 0 | 0 |
| EFLWRIEQI | MTCL1 | Q9Y4B5 | MTCL1_HUMAI | Unmodified | 0 | 0 | 0 | 0 | 308240 | 191560 | 0 | 0 | 0 | 0 | 0 | 0 | 19834 | 0 | 0 | 0 | 0 | 0 |
| EFMDRHGI | GART; GART; GA | P22102; C9J | PUR2_HUMAN | Unmodified | 0 | 0 | 0 | 0 | 1037100 | 0 | 0 | 0 | 0 | 0 | 0 | 0 | 0 | 0 | 0 | 0 | 573400 | 439880 |
| EFMDRHGI | GART; GART; GA | P22102; C9J | PUR2_HUMAN | Oxidation (M) | 89994 | 0 | 0 | 0 | 1145700 | 326500 | 383720 | 0 | 0 | 0 | 0 | 375210 | 837790 | 347010 | 0 | 53746 | 466700 | 0 |
| EFMPPGLQ | SARS1; SARS1 | P49591; Q5T | SYSC_HUMAN | Oxidation (M) | 0 | 0 | 294170 | 1024600 | 4760400 | 658450 | 1005000 | 0 | 2657800 | 2105000 | 1270100 | 5416800 | 6136900 | 5464500 | 0 | 0 | 0 | 3404000 |
| EFMPPGLQ | SARS1; SARS1 | P49591; Q5T | SYSC_HUMAN | Unmodified | 0 | 0 | 0 | 0 | 0 | 0 | 0 | 0 | 1917600 | 0 | 0 | 0 | 1404700 | 0 | 0 | 0 | 0 | 40281 |
| EFNEDGAL | SYNCRIP; SYNC | O60506; A0A | HNRPQ_HUMA | Unmodified | 0 | 0 | 89634 | 0 | 0 | 0 | 0 | 0 | 0 | 79887 | 970040 | 2059100 | 9974.1 | 0 | 0 | 127100 | 0 | 0 |
| EFNEDGAL | SYNCRIP; SYNC | O60506; A0A | HNRPQ_HUMA | Unmodified | 0 | 0 | 0 | 0 | 0 | 0 | 0 | 0 | 0 | 0 | 0 | 6647400 | 0 | 0 | 0 | 0 | 0 | 0 |
| EFNKYDTD | ACTR3; ACTR3 | P61158; B4D | ARP3_HUMAN | Unmodified | 0 | 1051900 | 0 | 1285600 | 2928700 | 1778600 | 0 | 194770 | 858500 | 214720 | 650080 | 6388000 | 0 | 0 | 1180100 | 2053100 | 460290 | 0 |
| EFNKYDTD | ACTR3; ACTR3 | P61158; B4D | ARP3_HUMAN | Unmodified | 0 | 0 | 0 | 0 | 3362700 | 253350 | 2299600 | 417730 | 0 | 199310 | 0 | 0 | 6100500 | 3081100 | 0 | 0 | 1310900 | 1436100 |
| EFNPTKPLI | ATXN2L; ATXN2 | Q8WWM7; H | ATX2L_HUMAN | Unmodified | 1095800 | 777720 | 0 | 236000 | 414750 | 0 | 291170 | 0 | 354520 | 0 | 288090 | 608940 | 777300 | 392940 | 0 | 214170 | 207370 | 347360 |
| EFPFDVQP | AIDA | Q96BJ3 | AIDA_HUMAN A | Unmodified | 0 | 0 | 0 | 0 | 0 | 0 | 0 | 0 | 0 | 0 | 319870 | 354160 | 0 | 0 | 0 | 826980 | 0 | 1943400 |
| EFPFDVQP | AIDA | Q96BJ3 | AIDA_HUMAN A | Unmodified | 452120 | 2246300 | 2280100 | 4293200 | 3106400 | 0 | 280330 | 122030 | 896780 | 477840 | 0 | 1405700 | 4654800 | 6055600 | 5776100 | 3338800 | 1733100 | 7977000 |
| EFPGFLEN | S100A10 | P60903 | S10AA_HUMAN | Unmodified | 0 | 0 | 0 | 0 | 1919100 | 0 | 0 | 0 | 0 | 0 | 0 | 0 | 0 | 0 | 0 | 0 | 0 | 0 |
| EFPGFLEN | S100A10 | P60903 | S10AA_HUMAN | Unmodified | 2169100 | 1640600 | 463370 | 639570 | 85801000 | 21279000 | 0 | 0 | 2938800 | 0 | 750130 | 0 | 0 | 565220 | 839300 | 406080 | 9222100 | 23089000 |
| EFPGFLEN | S100A10 | P60903 | S10AA_HUMAN | Oxidation (M) | 0 | 0 | 0 | 0 | 4398200 | 0 | 0 | 0 | 0 | 0 | 0 | 0 | 0 | 0 | 0 | 0 | 962510 | 896720 |
| EFPGLAGV | TXNDC5 | Q8NBS9 | TXND5_HUMAI | Unmodified | 0 | 0 | 0 | 0 | 0 | 0 | 914880 | 0 | 0 | 747550 | 0 | 0 | 0 | 0 | 0 | 0 | 0 | 0 |
| EFPTVPLVK | UGP2; UGP2; U | Q16851; E7E | UGPA_HUMAN | Unmodified | 0 | 0 | 0 | 103630 | 0 | 0 | 0 | 0 | 1107300 | 207880 | 0 | 6061400 | 0 | 0 | 0 | 0 | 0 | 0 |
| EFPVYLWQ | NIBAN1 | Q9BZQ8 | NIBA1_HUMAN | Unmodified | 0 | 0 | 0 | 124100 | 142340 | 20294 | 0 | 0 | 205390 | 437010 | 0 | 0 | 0 | 0 | 55982 | 0 | 0 | 0 |
| EFQASPLLI | CCDC22 | O60826 | CCD22_HUMA | Unmodified | 0 | 0 | 82325 | 226080 | 171800 | 0 | 0 | 0 | 493610 | 0 | 203180 | 1903800 | 0 | 0 | 0 | 228660 | 0 | 84313 |
| EFQAVVMA | EIF2B3; EIF2B3 | Q9NR50; A0 | EI2BG_HUMAN | Acetyl (Protein N | 6272000 | 0 | 0 | 0 | 0 | 0 | 156970 | 0 | 0 | 0 | 0 | 0 | 0 | 0 | 0 | 0 | 672600 | 0 |
| EFQDAGEC | HIBADH | P31937 | 3HIDH_HUMAI | Unmodified | 0 | 0 | 0 | 0 | 301570 | 0 | 0 | 0 | 3352800 | 0 | 0 | 0 | 0 | 0 | 0 | 0 | 0 | 0 |
| EFQDLNQE | RIPOR2; RIPOR | Q9Y4F9; B7Z | RIPR2_HUMAN | Oxidation (M) | 0 | 0 | 0 | 0 | 0 | 0 | 0 | 0 | 0 | 0 | 25838 | 33878 | 1644300 | 862790 | 0 | 0 | 0 | 0 |
| EFQIKEDIP | NMI | Q13287 | NMI_HUMAN N | Unmodified | 0 | 0 | 0 | 0 | 994450 | 0 | 0 | 53064 | 0 | 0 | 0 | 0 | 0 | 0 | 0 | 0 | 279760 | 917660 |
| EFQLGESV | GNA13; GNA12 | Q14344; Q0 | GNA13_HUMAI | Unmodified | 437970 | 376820 | 0 | 832220 | 419990 | 275270 | 0 | 0 | 3291200 | 765450 | 973840 | 15983000 | 0 | 0 | 484670 | 424370 | 0 | 0 |
| EFQLTEEN | MTPAP; MTPAP | Q9NVV4; Q5 | PAPD1_HUMAI | Unmodified | 0 | 0 | 0 | 0 | 0 | 0 | 0 | 0 | 569660 | 99023 | 0 | 0 | 0 | 0 | 0 | 0 | 0 | 0 |
| EFQSPDEE | SF3B1 | O75533 | SF3B1_HUMAN | Unmodified | 0 | 0 | 0 | 0 | 0 | 0 | 0 | 0 | 0 | 0 | 0 | 721450 | 0 | 0 | 0 | 0 | 0 | 0 |
| EFRFDLTQI | BMP8B; BMP8A | P34820; Q7Z | BMP8B_HUMAI | Unmodified | 0 | 0 | 0 | 0 | 0 | 0 | 79095 | 0 | 496100 | 0 | 0 | 0 | 0 | 0 | 0 | 0 | 0 | 0 |
| EFRFEPSS | NOMO1; NOMC | Q15155; P69 | NOMO1_HUMA | Unmodified | 0 | 0 | 0 | 215770 | 0 | 0 | 218510 | 170770 | 11257000 | 1220300 | 0 | 0 | 0 | 0 | 0 | 0 | 0 | 0 |
| EFRFEPSS | NOMO1; NOMC | Q15155; P69 | NOMO1_HUMA | Oxidation (M) | 0 | 0 | 0 | 0 | 0 | 0 | 0 | 90057 | 6777900 | 2410500 | 577110 | 0 | 0 | 0 | 0 | 0 | 0 | 0 |
| EFSEQFIR | COL21A1; COL | Q96P44; F5C | COLA1_HUMAI | Unmodified | 0 | 0 | 0 | 0 | 0 | 0 | 0 | 0 | 1016200 | 0 | 0 | 0 | 0 | 0 | 0 | 0 | 0 | 0 |
| EFSFHNFN | PCBD2; PCBD2 | Q9H0N5; HC | PHS2_HUMAN | Oxidation (M) | 0 | 0 | 78591 | 277560 | 0 | 0 | 0 | 0 | 0 | 324890 | 0 | 111840 | 320720 | 259460 | 330090 | 223570 | 0 | 0 |
| EFSGYVES | ANXA7 | P20073 | ANXA7_HUMAI | Unmodified | 904710 | 531260 | 1190300 | 781220 | 0 | 0 | 291360 | 28639 | 35783000 | 0 | 746580 | 4236200 | 192280 | 93066 | 3045000 | 1917600 | 249340 | 0 |
| EFSIDVGYE | ACTR3; ACTR3 | P61158; B4D | ARP3_HUMAN | Unmodified | 772180 | 1205600 | 0 | 0 | 0 | 0 | 0 | 0 | 670200 | 0 | 705370 | 6068100 | 0 | 0 | 0 | 1090700 | 0 | 0 |
| EFSIPRAHA | GPRC5A | Q8NFJ5 | RAI3_HUMAN F | Unmodified | 0 | 0 | 0 | 0 | 57975000 | 16956000 | 0 | 0 | 0 | 0 | 0 | 0 | 0 | 0 | 56325 | 0 | 222670 | 0 |
| EFSIPRAHA | GPRC5A | Q8NFJ5 | RAI3_HUMAN F | Unmodified | 0 | 0 | 0 | 0 | 17525000 | 3752100 | 0 | 0 | 0 | 0 | 0 | 0 | 0 | 0 | 0 | 0 | 0 | 0 |
| EFSITDVVP | HSPA4; HSPA4; | P34932; A0A | HSP74_HUMAI | Unmodified | 0 | 0 | 49154 | 0 | 0 | 0 | 0 | 0 | 327880 | 0 | 370170 | 722400 | 0 | 0 | 93872 | 56965 | 0 | 0 |
| EFSIYMTK | GOT2 | P00505 | AATM_HUMAN | Oxidation (M) | 0 | 0 | 0 | 0 | 0 | 0 | 0 | 0 | 658790 | 387570 | 0 | 0 | 0 | 0 | 0 | 80269 | 0 | 0 |
| EFSIYMTK | GOT2 | P00505 | AATM_HUMAN | Unmodified | 0 | 0 | 0 | 0 | 0 | 0 | 0 | 0 | 1485100 | 0 | 0 | 0 | 0 | 0 | 0 | 0 | 0 | 0 |
| EFSPFGSIT | PABPC4; PABP | Q13310; B1A | PABP4_HUMAN | Unmodified | 34005000 | 28494000 | 17961000 | 12929000 | 0 | 0 | 1233900 | 665250 | 0 | 0 | 0 | 1367300 | 0 | 1305700 | 8390200 | 8239700 | 0 | 0 |
| EFSPFGTIT | PABPC1; PABP | P11940; A0A | PABP1_HUMAN | Unmodified | 0 | 71284000 | 39491000 | 40292000 | 3486500 | 0 | 3017700 | 1470600 | 2396300 | 2056100 | 2039300 | 8649100 | 16954000 | 7428600 | 23941000 | 38666000 | 0 | 1074800 |
| EFSQNGEN | NOSTRIN; NOS | Q8IVI9; F8W | NOSTN_HUMA | Unmodified | 1303300 | 0 | 1151700 | 1955700 | 0 | 0 | 0 | 0 | 0 | 0 | 0 | 0 | 873590 | 706760 | 2359100 | 1233200 | 0 | 0 |
| EFSQSFDA | UCHL5; UCHL5 | Q9Y5K5; Q5 | UCHL5_HUMAI | Unmodified | 0 | 0 | 0 | 0 | 0 | 0 | 0 | 0 | 0 | 0 | 0 | 1222700 | 0 | 0 | 0 | 0 | 0 | 0 |
| EFSVNENQ | PLCG2 | P16885 | PLCG2_HUMAI | Unmodified | 0 | 0 | 0 | 0 | 0 | 0 | 0 | 0 | 0 | 0 | 3098000 | 11664000 | 0 | 2301400 | 0 | 0 | 0 | 0 |
| EFTAQNLG | EIF3H; EIF3H; E | O15372; B3k | EIF3H_HUMAN | Unmodified | 715560 | 286000 | 444150 | 823510 | 2405900 | 173750 | 738970 | 1440900 | 160380 | 599310 | 678380 | 9294700 | 4504700 | 767120 | 547540 | 613910 | 700790 | 1183000 |
| EFTEAVEAK | PHB; PHB; PHB | P35232; C9J | PHB_HUMAN F | Unmodified | 0 | 0 | 0 | 0 | 0 | 0 | 0 | 0 | 0 | 0 | 0 | 0 | 0 | 0 | 0 | 0 | 0 | 0 |
| EFTEREQG | UCHL1; UCHL1 | P09936; D6F | UCHL1_HUMAI | Unmodified | 45099 | 39381 | 0 | 24997 | 413780 | 205350 | 67126 | 0 | 13946000 | 236540 | 6495800 | 98539000 | 0 | 0 | 0 | 68328 | 0 | 0 |
| EFTEREQG | UCHL1; UCHL1 | P09936; D6F | UCHL1_HUMAI | Unmodified | 0 | 0 | 0 | 0 | 1606200 | 343170 | 110250 | 0 | 546640 | 349380 | 972400 | 0 | 0 | 0 | 0 | 0 | 0 | 0 |
| EFTKLEEVL | GSTO1; GSTO1 | P78417; Q5T | GSTO1_HUMAI | Unmodified | 0 | 0 | 0 | 0 | 0 | 0 | 0 | 0 | 0 | 0 | 0 | 2393700 | 0 | 0 | 0 | 0 | 0 | 0 |
| EFTKLEEVL | GSTO1; GSTO1 | P78417; Q5T | GSTO1_HUMAI | Unmodified | 0 | 0 | 0 | 0 | 664230 | 177260 | 0 | 0 | 0 | 0 | 713470 | 4684200 | 0 | 0 | 655180 | 0 | 0 | 0 |
| EFTMAEIEH | GARS1; GARS1; | P41250; A0A | GARS_HUMAN | Unmodified | 0 | 0 | 0 | 0 | 301900 | 658170 | 0 | 0 | 937980 | 129670 | 0 | 1339400 | 230400 | 0 | 16694 | 0 | 0 | 0 |
| EFTMAEIEH | GARS1; GARS1; | P41250; A0A | GARS_HUMAN | Oxidation (M) | 0 | 0 | 50038 | 0 | 0 | 0 | 0 | 0 | 0 | 0 | 147980 | 1975600 | 0 | 0 | 99866 | 104920 | 0 | 0 |
| EFTNVYIK | PABPC4L; PABP | P0CB38; Q1 | PAB4L_HUMAN | Unmodified | 94877000 | 69666000 | 34624000 | 30022000 | 907300 | 0 | 3447000 | 0 | 0 | 159130 | 0 | 0 | 10191000 | 3788000 | 22663000 | 38162000 | 2873100 | 1636700 |

| | | | | | | | | | | | | | | | | | | | | | | |
|---|---|---|---|---|---|---|---|---|---|---|---|---|---|---|---|---|---|---|---|---|---|---|
| EFTQGVKP | MAIP1 | Q8WWC4 | MAIP1_HUMAN | Unmodified | 0 | 0 | 0 | 0 | 0 | 0 | 136100 | 0 | 1527900 | 0 | 0 | 0 | 0 | 0 | 0 | 0 | 0 | 0 |
| EFTTGENFI | QSOX2 | Q6ZRP7 | QSOX2_HUMAI | Unmodified | 0 | 0 | 0 | 0 | 0 | 0 | 0 | 0 | 2187400 | 1182700 | 0 | 360320 | 0 | 0 | 0 | 0 | 42697 | 321080 |
| EFTTGENFI | QSOX2 | Q6ZRP7 | QSOX2_HUMAI | Unmodified | 81259 | 0 | 0 | 0 | 54191 | 0 | 0 | 0 | 12496000 | 1224500 | 373630 | 0 | 0 | 0 | 0 | 0 | 0 | 0 |
| EFVEACLNI | STK24; STK24; S | Q9Y6E0; H0\ | STK24_HUMAN | Unmodified | 160760 | 0 | 0 | 0 | 12203000 | 0 | 798480 | 560460 | 0 | 0 | 0 | 0 | 2997400 | 4750900 | 0 | 0 | 3129600 | 5275500 |
| EFVEAVLEL | MACROH2A1; M | O75367; B4I | H2AY_HUMAN | Unmodified | 665740 | 361760 | 0 | 0 | 0 | 0 | 0 | 0 | 0 | 0 | 0 | 0 | 0 | 0 | 0 | 0 | 0 | 0 |
| EFVEREVID | MRPL43; MRPL4 | Q8N983; H0 | RM43_HUMAN | Unmodified | 0 | 0 | 0 | 0 | 0 | 0 | 0 | 0 | 783170 | 0 | 0 | 0 | 0 | 0 | 0 | 0 | 0 | 54759 |
| EFVGNWN | MFGE8; MFGE8 | Q08431; H0\ | MFGM_HUMAN | Unmodified | 1496500 | 1734100 | 7695900 | 8423300 | 68353000 | 28154000 | 2464700 | 1206500 | 17320000 | 6780100 | 0 | 5907500 | 5563200 | 3490900 | 7153700 | 6846300 | 155330 | 18509000 |
| EFVGQLVA | VWA1; VWA1; V | Q6PCB0; J3( | VWA1_HUMAN | Unmodified | 91499 | 9899.2 | 165060 | 91397 | 0 | 0 | 0 | 0 | 461790 | 48967 | 20353 | 0 | 0 | 0 | 0 | 89197 | 0 | 0 |
| EFVISDRK | COTL1; COTL1 | Q14019; H3I | COTL1_HUMAN | Unmodified | 0 | 0 | 0 | 0 | 954530 | 699080 | 0 | 0 | 5328800 | 0 | 77418 | 920880 | 0 | 0 | 0 | 191730 | 0 | 0 |
| EFVISDRKE | COTL1; COTL1 | Q14019; H3I | COTL1_HUMAN | Unmodified | 0 | 0 | 0 | 0 | 3011600 | 562920 | 0 | 0 | 4734900 | 3233200 | 0 | 0 | 0 | 0 | 0 | 0 | 2435500 | 631100 |
| EFVSSDES | SSRP1 | Q08945 | SSRP1_HUMAN | Unmodified | 0 | 0 | 0 | 0 | 0 | 0 | 0 | 0 | 0 | 0 | 192900 | 423090 | 0 | 0 | 0 | 0 | 0 | 0 |
| EFVVIAPAA | RAB3GAP1 | Q15042 | RB3GP_HUMAN | Unmodified | 0 | 0 | 0 | 0 | 0 | 0 | 274720 | 0 | 0 | 1579800 | 0 | 2412200 | 0 | 0 | 126770 | 0 | 0 | 0 |
| EFWMDAN | ERBIN; ERBIN | Q96RT1; B4I | ERBIN_HUMAN | Oxidation (M) | 24943 | 0 | 0 | 156370 | 3899200 | 407720 | 0 | 0 | 0 | 0 | 0 | 0 | 0 | 681440 | 0 | 0 | 0 | 696970 |
| EFYGENIKE | ALDH3A2; ALDI | P51648; A0A | AL3A2_HUMAN | Unmodified | 223810 | 0 | 0 | 0 | 0 | 0 | 0 | 0 | 2088500 | 295120 | 0 | 0 | 0 | 0 | 0 | 0 | 0 | 0 |
| EFYGRGAP | NENF | Q9UMX5 | NENF_HUMAN | Unmodified | 0 | 0 | 0 | 0 | 0 | 0 | 0 | 0 | 1129000 | 704010 | 0 | 0 | 0 | 0 | 0 | 0 | 0 | 0 |
| EFYGRGAP | NENF | Q9UMX5 | NENF_HUMAN | Unmodified | 0 | 0 | 0 | 0 | 0 | 0 | 0 | 0 | 672510 | 855320 | 0 | 0 | 0 | 0 | 0 | 0 | 0 | 0 |
| EFYTVDGQ | CEMIP2; CEMIP | Q9UHN6; F5 | CEIP2_HUMAN | Unmodified | 0 | 0 | 620130 | 0 | 0 | 241730 | 0 | 0 | 1757200 | 0 | 0 | 150380 | 0 | 0 | 766450 | 637490 | 0 | 0 |
| EGAAHAFA | CAPZA2; CAPZ/ | P47755; A0A | CAZA2_HUMAN | Unmodified | 737280 | 0 | 1289500 | 0 | 0 | 0 | 465340 | 0 | 619190 | 0 | 91750 | 4134400 | 0 | 0 | 68939 | 1121700 | 0 | 0 |
| EGAAHAFA | CAPZA1 | P52907 | CAZA1_HUMAN | Unmodified | 423780 | 294320 | 868850 | 877590 | 0 | 0 | 0 | 0 | 802520 | 0 | 144430 | 2905400 | 0 | 0 | 0 | 427160 | 0 | 0 |
| EGAAHAFA | CAPZA1 | P52907 | CAZA1_HUMAN | Oxidation (M) | 338690 | 105490 | 857100 | 130500 | 0 | 0 | 0 | 0 | 0 | 0 | 0 | 1927300 | 0 | 0 | 0 | 781750 | 0 | 0 |
| EGAAPGAA | SHC1; SHC1; S | P29353; X6R | SHC1_HUMAN | Unmodified | 0 | 0 | 0 | 0 | 3117700 | 0 | 0 | 0 | 0 | 0 | 0 | 2913700 | 1509900 | 0 | 0 | 311510 | 0 | 1889500 |
| EGAAVIDV( | MTHFD2; MTHF | P13995; A0A | MTDC_HUMAN | Unmodified | 0 | 0 | 0 | 0 | 0 | 0 | 0 | 0 | 2064400 | 190960 | 0 | 0 | 0 | 0 | 0 | 0 | 0 | 0 |
| EGADEEEW | EHD4 | Q9H223 | EHD4_HUMAN | Unmodified | 0 | 263630 | 992290 | 0 | 418440 | 0 | 0 | 0 | 0 | 0 | 0 | 719690 | 0 | 0 | 1316700 | 963210 | 0 | 487240 |
| EGAEDKMT | BPNT2; BPNT2 | Q9NX62; H0 | IMPA3_HUMAN | Unmodified | 95889 | 0 | 0 | 0 | 0 | 0 | 27062 | 33923 | 2145200 | 1891000 | 0 | 0 | 0 | 0 | 0 | 0 | 0 | 0 |
| EGAFAYVH | SMCR8 | Q8TEV9 | SMCR8_HUMA | Unmodified | 0 | 1595900 | 122130 | 899350 | 0 | 0 | 0 | 0 | 0 | 0 | 0 | 117220 | 0 | 0 | 1551700 | 525630 | 0 | 0 |
| EGAFMVR | ITK; BMX | Q08881; P51 | ITK_HUMAN Ty | Unmodified | 0 | 0 | 0 | 0 | 0 | 0 | 0 | 0 | 1942400 | 247370 | 348690 | 1508300 | 0 | 0 | 0 | 0 | 0 | 0 |
| EGAFSNFP | DDX21; DDX50 | Q9NR30; Q9 | DDX21_HUMAI | Unmodified | 0 | 0 | 0 | 0 | 0 | 0 | 0 | 202460 | 0 | 0 | 668910 | 2122100 | 0 | 0 | 0 | 0 | 0 | 661540 |
| EGAKHFSG | CCT8; CCT8 | P50990; H7( | TCPQ_HUMAN | Unmodified | 0 | 0 | 0 | 0 | 0 | 0 | 422690 | 0 | 0 | 0 | 0 | 0 | 0 | 0 | 0 | 0 | 940360 | 0 |
| EGAKLECG | ALDH1A1 | P00352 | AL1A1_HUMAN | Unmodified | 0 | 0 | 0 | 0 | 0 | 0 | 0 | 0 | 0 | 0 | 3399800 | 55424000 | 0 | 0 | 0 | 0 | 0 | 0 |
| EGALCEEN | EEF2 | P13639 | EF2_HUMAN El | Unmodified | 1077300 | 0 | 772100 | 1221600 | 2301600 | 891690 | 118260 | 140060 | 1385700 | 258960 | 4101700 | 14369000 | 1855200 | 451300 | 0 | 847010 | 492160 | 1509800 |
| EGALCEEN | EEF2 | P13639 | EF2_HUMAN El | Oxidation (M) | 511050 | 0 | 339090 | 0 | 656140 | 434300 | 0 | 0 | 329390 | 9038.5 | 1088500 | 4063600 | 0 | 314140 | 781720 | 689020 | 198630 | 311650 |
| EGALDLSG | PCK2; PCK2 | Q16822; H0\ | PCKGM_HUMA | Unmodified | 0 | 0 | 0 | 0 | 0 | 0 | 0 | 0 | 3428000 | 0 | 0 | 0 | 0 | 0 | 0 | 0 | 0 | 0 |
| EGALSMAR | MYG1; MYG1; M | Q9HB07; F8\ | MYG1_HUMAN | Unmodified | 0 | 46046 | 33634 | 0 | 102090 | 0 | 24313 | 15184 | 314650 | 0 | 0 | 737930 | 56298 | 0 | 36732 | 0 | 0 | 0 |
| EGALTAFR | PSMD6; PSMD6 | Q15008; C9J | PSMD6_HUMAN | Unmodified | 0 | 0 | 0 | 0 | 0 | 0 | 0 | 0 | 0 | 0 | 0 | 0 | 0 | 0 | 0 | 0 | 0 | 0 |
| EGANREIL( | ARRB1; ARRB1 | P49407; H0\ | ARRB1_HUMAN | Unmodified | 0 | 0 | 0 | 159870 | 0 | 0 | 19499 | 0 | 0 | 42616 | 0 | 0 | 83925 | 0 | 270110 | 0 | 101800 | 63214 |
| EGAQFSSH | PLAA; PLAA | Q9Y263; H0\ | PLAP_HUMAN | Unmodified | 62460 | 75997 | 0 | 0 | 0 | 0 | 427330 | 18490 | 0 | 189230 | 0 | 5207500 | 849950 | 0 | 371670 | 120730 | 0 | 0 |
| EGAQYLMC | SLC12A2; SLC1 | P55011; G3\ | S12A2_HUMAN | Oxidation (M) | 76868 | 0 | 54605 | 0 | 0 | 0 | 0 | 0 | 0 | 0 | 0 | 0 | 0 | 0 | 120340 | 169210 | 0 | 0 |
| EGATPYNS | SLC6A6; SLC6A | P31641; A0A | SC6A6_HUMAN | Unmodified | 1281100 | 52589 | 0 | 0 | 0 | 0 | 0 | 0 | 0 | 0 | 0 | 6619500 | 0 | 0 | 149190 | 41447 | 0 | 0 |
| EGCGDDN | ITGA6 | P23229 | ITA6_HUMAN I | Unmodified | 0 | 0 | 198810 | 0 | 0 | 63690 | 0 | 0 | 0 | 0 | 60227 | 879610 | 152260 | 72337 | 502460 | 464990 | 271490 | 1409800 |
| EGCGDDN | ITGA6 | P23229 | ITA6_HUMAN I | Unmodified | 34397 | 137170 | 0 | 238300 | 6002800 | 2002100 | 0 | 0 | 0 | 0 | 160670 | 1962900 | 7137900 | 4450500 | 144300 | 544740 | 9068000 | 34948000 |
| EGCGDDN | ITGA6 | P23229 | ITA6_HUMAN I | Unmodified | 0 | 0 | 0 | 0 | 0 | 0 | 0 | 0 | 0 | 0 | 0 | 0 | 0 | 0 | 0 | 0 | 691480 | 333660 |
| EGCLAFGT | GEMIN5 | Q8TEQ6 | GEMI5_HUMAN | Unmodified | 0 | 0 | 0 | 0 | 0 | 0 | 0 | 0 | 0 | 0 | 0 | 612530 | 0 | 0 | 0 | 0 | 0 | 0 |
| EGCTEVSLI | HNRNPUL2; HN | Q1KMD3; H3 | HNRL2_HUMAN | Unmodified | 0 | 0 | 0 | 0 | 0 | 0 | 0 | 0 | 0 | 0 | 0 | 1481900 | 0 | 0 | 0 | 0 | 0 | 0 |
| EGDCGFGI | CPD | O75976 | CBPD_HUMAN | Unmodified | 0 | 0 | 0 | 0 | 0 | 0 | 0 | 0 | 2042000 | 91987 | 0 | 0 | 0 | 0 | 0 | 0 | 0 | 0 |
| EGDCGFGI | CPD | O75976 | CBPD_HUMAN | Unmodified | 0 | 0 | 51224 | 77746 | 0 | 0 | 0 | 0 | 3895300 | 0 | 0 | 0 | 0 | 0 | 15931 | 0 | 0 | 0 |
| EGDEQRTF | GARS1; GARS1 | P41250; A0A | GARS_HUMAN | Unmodified | 80757 | 85650 | 0 | 234430 | 0 | 0 | 124740 | 0 | 1606200 | 401040 | 771250 | 1571600 | 323310 | 0 | 343570 | 185470 | 0 | 0 |
| EGDFGCTV | ATP2B4 | P23634 | AT2B4_HUMAN | Unmodified | 0 | 0 | 0 | 0 | 145120 | 0 | 0 | 0 | 0 | 0 | 464650 | 2694400 | 0 | 0 | 138270 | 130660 | 0 | 0 |
| EGDLIAAQ/ | LMNA; LMNA; L | P02545; A0A | LMNA_HUMAN | Unmodified | 0 | 0 | 0 | 0 | 661280 | 559880 | 0 | 0 | 432500 | 0 | 102360 | 847570 | 295770 | 0 | 0 | 0 | 0 | 0 |
| EGDLIAAQ/ | LMNA; LMNA; L | P02545; A0A | LMNA_HUMAN | Unmodified | 0 | 0 | 0 | 0 | 776110 | 151340 | 177210 | 909370 | 0 | 0 | 0 | 0 | 0 | 119330 | 0 | 0 | 0 | 0 |
| EGDLPPLW | NDUFB9; NDUF | Q9Y6M9; E9 | NDUB9_HUMA | Unmodified | 0 | 0 | 0 | 0 | 0 | 0 | 0 | 0 | 1249600 | 0 | 0 | 0 | 0 | 0 | 0 | 0 | 0 | 0 |
| EGDNCRYS | MKRN1; MKRN | Q9UHC7; C9 | MKRN1_HUMA | Unmodified | 778080 | 0 | 0 | 0 | 0 | 0 | 0 | 0 | 0 | 0 | 0 | 0 | 569540 | 253480 | 0 | 0 | 0 | 0 |
| EGDPAIYAE | TES; TES | Q9UGI8; H7I | TES_HUMAN Te | Unmodified | 183210 | 94859 | 231060 | 298940 | 0 | 0 | 0 | 0 | 455980 | 0 | 0 | 2654200 | 0 | 0 | 680440 | 296400 | 50661 | 42512 |
| EGDPAIYAE | TES; TES | Q9UGI8; H7I | TES_HUMAN Te | Unmodified | 0 | 0 | 0 | 286190 | 1113600 | 0 | 0 | 0 | 319840 | 176930 | 0 | 574380 | 1088000 | 0 | 216640 | 161180 | 557260 | 1826600 |
| EGDQTSNI | DNAJB1; DNAJE | P25685; M0( | DNJB1_HUMAN | Unmodified | 935650 | 0 | 178190 | 659930 | 3861500 | 442330 | 440470 | 153970 | 1618000 | 662190 | 281150 | 670690 | 2048400 | 630100 | 386380 | 205870 | 1705700 | 1514000 |
| EGDTIIVPG | EIF5B; EIF5B | O60841; A0/ | IF2P_HUMAN E | Unmodified | 0 | 0 | 235720 | 1154200 | 1093600 | 0 | 0 | 0 | 299630 | 91113 | 619790 | 4352900 | 1776000 | 0 | 452380 | 355190 | 0 | 0 |
| EGDTMMG | PTGR1; PTGR1 | Q14914; F22 | PTGR1_HUMAN | Oxidation (M) | 0 | 0 | 0 | 0 | 419250 | 0 | 0 | 0 | 0 | 0 | 0 | 0 | 0 | 0 | 0 | 0 | 0 | 0 |
| EGDTMMG | PTGR1; PTGR1 | Q14914; F22 | PTGR1_HUMAN | Unmodified | 0 | 0 | 0 | 1163700 | 2623400 | 0 | 0 | 0 | 1161100 | 413190 | 0 | 0 | 181050 | 0 | 0 | 24562 | 0 | 179250 |
| EGDTVQLL | BCAM; BCAM; E | P50895; A0A | BCAM_HUMAN | Unmodified | 398160 | 618980 | 0 | 0 | 0 | 0 | 0 | 0 | 0 | 0 | 0 | 0 | 0 | 0 | 0 | 0 | 0 | 0 |
| EGDTVQLL | BCAM; BCAM; E | P50895; A0A | BCAM_HUMAN | Unmodified | 3953300 | 2372700 | 0 | 0 | 0 | 0 | 0 | 0 | 0 | 0 | 0 | 0 | 0 | 0 | 0 | 0 | 0 | 0 |
| EGDVLTLLE | RPS28 | P62857 | RS28_HUMAN | Unmodified | 0 | 0 | 219030 | 0 | 0 | 0 | 0 | 0 | 0 | 0 | 4301900 | 0 | 0 | 0 | 0 | 0 | 0 | 0 |
| EGDVLTLLE | RPS28 | P62857 | RS28_HUMAN | Unmodified | 0 | 932460 | 2006900 | 0 | 0 | 0 | 0 | 0 | 0 | 0 | 0 | 9027200 | 0 | 0 | 692260 | 862130 | 0 | 0 |
| EGDYVLFH | ACLY | P53396 | ACLY_HUMAN | Unmodified | 963970 | 535130 | 763950 | 1238000 | 475270 | 113390 | 469630 | 30480 | 2937400 | 246590 | 2536400 | 49779000 | 0 | 0 | 2491400 | 1637900 | 0 | 54352 |
| EGEDPNKA | ZNF622 | Q969S3 | ZN622_HUMAN | Unmodified | 0 | 436110 | 0 | 0 | 0 | 0 | 0 | 0 | 0 | 0 | 33003 | 76990 | 0 | 0 | 0 | 0 | 0 | 0 |
| EGEEAGPC | CDC37; CDC37 | Q16543; K7E | CDC37_HUMA | Unmodified | 79607 | 72155 | 181190 | 162750 | 409270 | 37426 | 0 | 0 | 0 | 0 | 0 | 3141300 | 124910 | 0 | 354660 | 279520 | 62737 | 185540 |
| EGEEFTVT( | KIT | P10721 | KIT_HUMAN M | Unmodified | 0 | 180010 | 20193000 | 0 | 0 | 872900 | 0 | 359500 | 494390 | 0 | 0 | 557240 | 0 | 0 | 0 | 0 | 0 | 0 |
| EGEEQLQT | POLR2B; POLR2 | P30876; C9J | RPB2_HUMAN | Unmodified | 0 | 0 | 0 | 0 | 0 | 0 | 0 | 0 | 0 | 0 | 0 | 245510 | 0 | 0 | 0 | 0 | 0 | 0 |
| EGEEVMFK | DYNC1H1; DYN | Q14204; A0/ | DYHC1_HUMA | Oxidation (M) | 0 | 0 | 0 | 0 | 0 | 0 | 0 | 0 | 0 | 0 | 0 | 439760 | 0 | 0 | 0 | 87106 | 0 | 0 |
| EGEEVMFK | DYNC1H1; DYN | Q14204; A0/ | DYHC1_HUMA | Unmodified | 0 | 0 | 0 | 61532 | 0 | 0 | 0 | 0 | 0 | 0 | 71342 | 1851600 | 0 | 0 | 91751 | 0 | 0 | 0 |
| EGEFVAQF | PA2G4 | Q9UQ80 | PA2G4_HUMAI | Unmodified | 3040200 | 3799900 | 1380300 | 811960 | 0 | 379380 | 425570 | 348990 | 0 | 1686600 | 1241500 | 26587000 | 2760400 | 806420 | 2905100 | 229960 | 1523400 | 2161900 |
| EGEGGLSL | MOV10; MOV10 | Q9HCE1; Q5 | MOV10_HUMAN | Unmodified | 875720 | 920000 | 0 | 0 | 0 | 0 | 0 | 0 | 0 | 0 | 0 | 0 | 282540 | 826440 | 0 | 0 | 0 | 0 |
| EGEGGQES | NIBAN1 | Q9BZQ8 | NIBA1_HUMAN | Unmodified | 0 | 0 | 151700 | 0 | 0 | 0 | 0 | 0 | 1775400 | 0 | 0 | 0 | 0 | 0 | 0 | 0 | 0 | 0 |
| EGEIIHLISK | CD2AP | Q9Y5K6 | CD2AP_HUMAI | Unmodified | 420010 | 337080 | 989020 | 271610 | 0 | 0 | 0 | 0 | 0 | 0 | 0 | 147800 | 0 | 0 | 902590 | 730420 | 0 | 0 |
| EGEPFELR | PTGFRN | Q9P2B2 | FPRP_HUMAN | Unmodified | 8364800 | 5801100 | 18372000 | 14166000 | 0 | 0 | 84869 | 0 | 0 | 0 | 0 | 22398000 | 749130 | 1958300 | 21927000 | 0 | 0 | 0 |
| EGEPGWVI | B3GAT3; B3GAT | O94766; E9F | B3GA3_HUMAI | Unmodified | 0 | 0 | 0 | 0 | 0 | 0 | 0 | 0 | 483790 | 0 | 102480 | 0 | 0 | 0 | 0 | 0 | 0 | 0 |
| EGEPRIDIN | PLS3; PLS3; PLS | P13797; H7( | PLST_HUMAN | Unmodified | 0 | 0 | 0 | 0 | 328180 | 118560 | 0 | 18354 | 0 | 0 | 0 | 0 | 0 | 0 | 0 | 0 | 77326 | 0 |
| EGESVQGS | SEC14L1 | Q92503 | S14L1_HUMAN | Unmodified | 0 | 0 | 0 | 0 | 28681 | 0 | 0 | 0 | 380170 | 0 | 0 | 120160 | 0 | 0 | 0 | 0 | 0 | 0 |
| EGETIIELK | CD47; CD47 | Q08722; A0/ | CD47_HUMAN | Unmodified | 0 | 0 | 939380 | 0 | 0 | 0 | 0 | 0 | 3528300 | 1548100 | 0 | 32513000 | 0 | 0 | 1799700 | 2565700 | 0 | 0 |
| EGEVLYIIEE | FNBP1L; FNBP1 | Q5T0N5; A0/ | FBP1L_HUMAN | Unmodified | 0 | 4789700 | 0 | 0 | 0 | 0 | 0 | 0 | 0 | 517470 | 0 | 0 | 0 | 584570 | 1737400 | 0 | 0 | 0 |
| EGFGHLSP | FGG; FGG; FGG | P02679; C9J | FIBG_HUMAN | Unmodified | 0 | 22632000 | 0 | 0 | 35749 | 0 | 0 | 0 | 0 | 0 | 0 | 0 | 0 | 3009000 | 0 | 0 | 109900 | 0 |
| EGFINYLTR | SPINT1; SPINT1 | O43278; H3I | SPIT1_HUMAN | Unmodified | 377810 | 574150 | 425670 | 0 | 0 | 0 | 0 | 0 | 1492100 | 0 | 317900 | 0 | 192820 | 0 | 1128100 | 1008200 | 0 | 0 |
| EGFIPSNY\ | LYN; LYN | P07948; E5R | LYN_HUMAN T | Unmodified | 1102500 | 866160 | 1030100 | 0 | 1379200 | 0 | 355810 | 175040 | 1863600 | 575010 | 119150 | 251520 | 313150 | 0 | 1238900 | 1885000 | 0 | 0 |
| EGFYQSPD | MEGF9 | Q9H1U4 | MEGF9_HUMAI | Unmodified | 0 | 0 | 0 | 0 | 0 | 0 | 0 | 0 | 2743700 | 161110 | 0 | 694020 | 0 | 0 | 0 | 0 | 0 | 0 |
| EGGAGAVE | CLASP2; CLASF | O75122; B3I | CLAP2_HUMAN | Unmodified | 0 | 0 | 0 | 0 | 0 | 0 | 0 | 0 | 0 | 0 | 0 | 1764100 | 0 | 0 | 0 | 0 | 0 | 71162 |
| EGGARLGA | STUB1; STUB1 | Q9UNE7; H3 | CHIP_HUMAN | Unmodified | 0 | 0 | 0 | 0 | 1860100 | 0 | 0 | 0 | 0 | 0 | 0 | 0 | 326910 | 719350 | 0 | 0 | 366830 | 1008300 |
| EGGFLLLH | FASN; FASN | P49327; A0A | FAS_HUMAN F | Unmodified | 69825 | 63946 | 212650 | 24982 | 0 | 0 | 0 | 0 | 52355 | 29010 | 68092 | 25751000 | 0 | 0 | 563370 | 310820 | 0 | 0 |
| EGGGDSS/ | MARCKSL1 | P49006 | MRP_HUMAN N | Unmodified | 3338900 | 1764200 | 10772000 | 21991000 | 2214000 | 0 | 634390 | 1000200 | 40624000 | 5177300 | 50074000 | 99987000 | 19784000 | 16540000 | 18422000 | 18119000 | 10733000 | 37453000 |
| EGGKELLS/ | PSD3; PSD3; PS | Q9NYI0; E5F | PSD3_HUMAN | Unmodified | 0 | 0 | 0 | 0 | 0 | 0 | 0 | 0 | 0 | 0 | 0 | 0 | 0 | 0 | 0 | 0 | 184050 | 0 |
| EGGLGPLN | PRDX2; PRDX2 | P32119; A6N | PRDX2_HUMAI | Unmodified | 1121600 | 817570 | 420170 | 1149100 | 541720 | 303970 | 277020 | 43364 | 9376600 | 1298900 | 2432200 | 15339000 | 1019500 | 0 | 1677200 | 1022700 | 549390 | 0 |
| EGGLGPLN | PRDX2; PRDX2 | P32119; A6N | PRDX2_HUMAI | Unmodified | 8380600 | 2261000 | 2403400 | 6950300 | 1987900 | 727660 | 578650 | 0 | 41839000 | 2822900 | 13187000 | 1.96E+08 | 4066800 | 1200000 | 7864800 | 6203700 | 1853300 | 2365100 |
| EGGLLTLA( | QDPR; QDPR; ( | P09417; H0\ | DHPR_HUMAN | Unmodified | 0 | 0 | 0 | 0 | 1180500 | 0 | 474350 | 457330 | 257860 | 348970 | 855990 | 3136800 | 1598100 | 975590 | 201720 | 198660 | 58176 | 490100 |
| EGGPNNH | CPE; CPE; CPE | P16870; C9J | CBPE_HUMAN | Unmodified | 0 | 0 | 0 | 0 | 0 | 0 | 0 | 58744 | 7379500 | 1720600 | 0 | 0 | 0 | 0 | 0 | 0 | 34102 | 0 |
| EGGPNPEF | TACO1 | Q9BSH4 | TACO1_HUMAI | Unmodified | 0 | 0 | 0 | 0 | 0 | 0 | 0 | 0 | 560340 | 0 | 0 | 0 | 0 | 0 | 0 | 0 | 0 | 0 |
| EGGSAGLII | PALS2; PALS2; | Q9NZW5; B8 | PALS2_HUMAN | Unmodified | 0 | 0 | 0 | 0 | 3804700 | 0 | 0 | 332530 | 968980 | 0 | 3350700 | 2347100 | 0 | 0 | 0 | 0 | 839080 | 0 |
| EGGSIPVTL | CNDP2 | Q96KP4 | CNDP2_HUMA | Unmodified | 0 | 0 | 0 | 0 | 0 | 0 | 0 | 29109 | 449780 | 0 | 2523000 | 9764000 | 0 | 0 | 0 | 0 | 0 | 0 |
| EGGTDSQF | P4HTM; P4HTM | Q9NXG6; V9 | P4HTM_HUMAI | Unmodified | 337700 | 0 | 0 | 0 | 0 | 0 | 0 | 321040 | 2073600 | 0 | 0 | 0 | 0 | 0 | 0 | 0 | 0 | 0 |
| EGGTVVYG | ALDH7A1; ALDI | P49419; A0A | AL7A1_HUMAN | Unmodified | 0 | 0 | 0 | 0 | 852990 | 0 | 0 | 0 | 0 | 0 | 0 | 0 | 0 | 0 | 0 | 0 | 0 | 0 |
| EGHGNKD | EPB41L5 | Q9HCM4 | E41L5_HUMAN | Unmodified | 0 | 0 | 0 | 0 | 181300 | 0 | 0 | 0 | 0 | 0 | 0 | 0 | 0 | 0 | 0 | 0 | 0 | 259490 |
| EGHLEKTE | HSPA13 | P48723 | HSP13_HUMAN | Unmodified | 0 | 0 | 0 | 0 | 0 | 0 | 0 | 0 | 5203400 | 1455900 | 0 | 0 | 0 | 0 | 0 | 0 | 0 | 0 |
| EGHLPDLL | SLC7A6 | Q92536 | YLAT2_HUMAN | Oxidation (M) | 0 | 0 | 0 | 0 | 0 | 0 | 0 | 0 | 0 | 0 | 0 | 1592500 | 0 | 0 | 0 | 0 | 0 | 0 |
| EGHLRVVE | CDKN2C | P42773 | CDN2C_HUMA | Unmodified | 0 | 0 | 0 | 0 | 0 | 0 | 136110 | 0 | 0 | 358350 | 0 | 0 | 0 | 0 | 0 | 0 | 218490 | 0 |
| EGHPVTSE | TST | Q16762 | THTR_HUMAN | Unmodified | 0 | 0 | 0 | 0 | 0 | 0 | 0 | 0 | 1274200 | 299740 | 0 | 106890 | 0 | 0 | 0 | 0 | 0 | 0 |
| EGHSYYYIS | EFNA1 | P20827 | EFNA1_HUMAN | Unmodified | 751440 | 936810 | 0 | 764030 | 0 | 105570 | 32504 | 345620 | 301820 | 0 | 0 | 0 | 50869 | 96062 | 76081 | 0 | 94224 | 295180 |
| EGHVEVVS | ANK3; ANK3; AI | Q12955; A0/ | ANK3_HUMAN | Unmodified | 0 | 0 | 913630 | 4456800 | 0 | 0 | 0 | 0 | 605200 | 0 | 0 | 280400 | 206120 | 0 | 9169000 | 5114700 | 367010 | 0 |
| EGHVEVVS | ANK3; ANK3; AI | Q12955; A0/ | ANK3_HUMAN | Unmodified | 0 | 0 | 0 | 2155800 | 0 | 0 | 0 | 0 | 0 | 0 | 0 | 0 | 5875900 | 3214500 | 1093400 | 0 | 0 | 0 |
| EGHVHIVEI | KIDINS220; KID | Q9ULH0; E9 | KDIS_HUMAN | Unmodified | 84589 | 0 | 249110 | 118680 | 0 | 0 | 0 | 0 | 0 | 0 | 0 | 874410 | 0 | 138940 | 127950 | 168860 | 0 | 0 |
| EGIAQTVFL | GSS; GSS; GSS | P48637; A0A | GSHB_HUMAN | Unmodified | 0 | 0 | 0 | 0 | 0 | 0 | 0 | 0 | 47353 | 0 | 25152 | 5807000 | 0 | 0 | 0 | 68480 | 0 | 0 |
| EGIAQTVFL | GSS; GSS | P48637; A0A | GSHB_HUMAN | Oxidation (M) | 0 | 0 | 0 | 0 | 1193100 | 0 | 0 | 0 | 0 | 447420 | 0 | 0 | 1100500 | 0 | 0 | 0 | 646920 | 0 |
| EGICAIGGI | LCP1; LCP1 | P13796; Q5I | PLSL_HUMAN | Unmodified | 0 | 0 | 0 | 0 | 0 | 0 | 0 | 0 | 341240 | 194620 | 0 | 0 | 0 | 0 | 0 | 0 | 280200 | 0 |
| EGICALGG | PLS3; PLS3 | P13797; A0A | PLST_HUMAN | Unmodified | 746080 | 0 | 0 | 314150 | 4292700 | 2779100 | 0 | 0 | 712280 | 0 | 0 | 0 | 690000 | 0 | 0 | 222170 | 1199700 | 2119400 |

| | | | | | | | | | | | | | | | | | | | | | | |
|---|---|---|---|---|---|---|---|---|---|---|---|---|---|---|---|---|---|---|---|---|---|---|
| EGICALGG | PLS3; PLS3 | P13797; A0A | PLST_HUMAN I | Unmodified | 0 | 0 | 0 | 0 | 6793800 | 0 | 0 | 0 | 0 | 0 | 0 | 0 | 1712900 | 0 | 0 | 0 | 5553600 | 0 |
| EGIDPTPYY | NARS1 | O43776 | SYNC_HUMAN | Unmodified | 0 | 0 | 0 | 0 | 0 | 3236500 | 0 | 0 | 236220 | 0 | 0 | 1424400 | 0 | 0 | 0 | 0 | 0 | 0 |
| EGIDPTPYY | NARS1 | O43776 | SYNC_HUMAN | Unmodified | 0 | 0 | 399910 | 743510 | 0 | 397630 | 0 | 362940 | 1928300 | 683260 | 1127200 | 9060200 | 0 | 0 | 786300 | 728560 | 0 | 0 |
| EGIESGDP | ASPH | Q12797 | ASPH_HUMAN | Oxidation (M) | 0 | 0 | 0 | 0 | 0 | 0 | 0 | 0 | 5048400 | 573940 | 0 | 0 | 184200 | 0 | 0 | 0 | 0 | 0 |
| EGIESGDP | ASPH | Q12797 | ASPH_HUMAN | Unmodified | 0 | 0 | 0 | 0 | 0 | 0 | 0 | 0 | 7986500 | 0 | 0 | 0 | 0 | 0 | 0 | 0 | 0 | 0 |
| EGIITIESQ | ITGB4; ITGB4 | P16144; J3K | ITB4_HUMAN I | Unmodified | 0 | 0 | 0 | 0 | 0 | 0 | 0 | 0 | 0 | 0 | 0 | 0 | 0 | 0 | 0 | 0 | 0 | 0 |
| EGILGQHQ | ITGA2; ITGA2; I | E9PB77; E7E | E9PB77_HUMA | Unmodified | 8979700 | 9511100 | 6954200 | 3281500 | 268620 | 109990 | 0 | 0 | 196990 | 0 | 0 | 3410300 | 0 | 0 | 17404000 | 9857500 | 101650 | 0 |
| EGILIKDFL | SHCBP1 | Q8NEM2 | SHCBP_HUMA | Unmodified | 0 | 775010 | 0 | 271760 | 14658000 | 1761200 | 0 | 0 | 0 | 0 | 0 | 0 | 0 | 0 | 0 | 368530 | 0 | 0 |
| EGILPERAE | ENSA; ENSA; E | O43768; A0A | ENSA_HUMAN | Unmodified | 0 | 0 | 0 | 0 | 373690 | 734690 | 53150 | 0 | 3616500 | 332860 | 0 | 0 | 0 | 0 | 0 | 0 | 1313000 | 1961800 |
| EGIPALDN | EEF2 | P13639 | EF2_HUMAN E | Unmodified | 72637 | 101890 | 0 | 98231 | 1263700 | 324690 | 248080 | 0 | 0 | 458110 | 248100 | 585230 | 532730 | 308450 | 182700 | 0 | 665350 | 602020 |
| EGIPGLYR | SLC25A24 | Q6NUK1 | SCMC1_HUMA | Unmodified | 0 | 0 | 0 | 233110 | 192850 | 1162200 | 0 | 151550 | 4008000 | 309550 | 0 | 350000 | 0 | 0 | 0 | 0 | 24371 | 200300 |
| EGIPLIILAG | IREB2 | P48200 | IREB2_HUMAN | Unmodified | 0 | 24733 | 80049 | 48953 | 335370 | 97385 | 64526 | 46086 | 0 | 0 | 117010 | 2770900 | 0 | 102700 | 0 | 73928 | 28566 | 182720 |
| EGITAIGGT | PLS1; PLS1 | Q14651; C9J | PLSI_HUMAN F | Unmodified | 245270 | 0 | 0 | 0 | 1376800 | 746420 | 0 | 0 | 2787300 | 0 | 2580900 | 4244200 | 831350 | 0 | 0 | 0 | 0 | 179410 |
| EGITTYFSG | DPP3; DPP3; D | Q9NY33; E9I | DPP3_HUMAN | Oxidation (M) | 0 | 0 | 0 | 0 | 0 | 0 | 35110 | 0 | 0 | 0 | 110750 | 879570 | 0 | 67074 | 0 | 0 | 0 | 0 |
| EGITTYFSG | DPP3; DPP3; D | Q9NY33; E9I | DPP3_HUMAN | Unmodified | 0 | 0 | 0 | 0 | 49286 | 0 | 0 | 0 | 0 | 0 | 0 | 257640 | 0 | 0 | 0 | 0 | 0 | 88624 |
| EGKLIMGI | ACLY | P53396 | ACLY_HUMAN | Unmodified | 0 | 0 | 0 | 0 | 0 | 0 | 231240 | 0 | 0 | 0 | 0 | 0 | 0 | 0 | 0 | 0 | 0 | 0 |
| EGKPCIIIK | ATP1B1; ATP1B | P05026; V9C | AT1B1_HUMAN | Unmodified | 256320 | 0 | 0 | 0 | 0 | 0 | 0 | 0 | 1943800 | 355620 | 1107400 | 3210300 | 0 | 0 | 0 | 0 | 0 | 0 |
| EGKPTIVEE | DNAJC8 | O75937 | DNJC8_HUMAI | Unmodified | 0 | 0 | 0 | 0 | 46228 | 0 | 0 | 0 | 870980 | 325200 | 0 | 3074900 | 0 | 0 | 0 | 0 | 0 | 0 |
| EGKRVLIAA | PGAM1 | P18669 | PGAM1_HUMA | Unmodified | 0 | 0 | 0 | 0 | 2465200 | 0 | 0 | 55977 | 0 | 203630 | 0 | 0 | 1492800 | 647570 | 0 | 0 | 768950 | 929680 |
| EGLDEVAG | CKAP5 | Q14008 | CKAP5_HUMAI | Unmodified | 0 | 0 | 0 | 0 | 0 | 0 | 0 | 0 | 0 | 0 | 383360 | 1276000 | 0 | 314280 | 0 | 0 | 0 | 0 |
| EGLDISHL | SLC12A2; SLC1 | P55011; G3 | S12A2_HUMAN | Unmodified | 0 | 0 | 0 | 0 | 0 | 0 | 721640 | 0 | 0 | 0 | 2227800 | 0 | 0 | 0 | 1790100 | 1719700 | 604250 | 2022600 |
| EGLDWDLI | COLGALT1; CO | Q8NBJ5; M0 | GT251_HUMAN | Unmodified | 0 | 83971 | 126510 | 252610 | 191800 | 395960 | 38521 | 0 | 2377100 | 0 | 45284 | 0 | 59159 | 0 | 86653 | 0 | 10723 | 254090 |
| EGLEFAVIR | PUS1; PUS1; P | Q9Y606; F5 | TRUA_HUMAN | Unmodified | 0 | 0 | 0 | 0 | 0 | 0 | 0 | 11142 | 262900 | 539490 | 0 | 304090 | 0 | 0 | 0 | 0 | 0 | 0 |
| EGLELLK | ENO1; ENO1 | P06733; A0A | ENOA_HUMAN | Unmodified | 409860 | 239200 | 1998400 | 915180 | 0 | 1787900 | 113590 | 0 | 0 | 1974500 | 9401200 | 30651000 | 1894500 | 835840 | 1923000 | 2050000 | 1240700 | 2325300 |
| EGLELPEDI | HSP90AB4P; H | Q58FF6; Q5 | H90B4_HUMAI | Unmodified | 4309400 | 2045000 | 0 | 9529000 | 8721700 | 2370900 | 1007400 | 335590 | 42736000 | 3986900 | 50999000 | 2.23E+08 | 3129800 | 2450000 | 10765000 | 9843400 | 897340 | 6277800 |
| EGLELPEDI | HSP90AB4P; H | Q58FF6; Q5 | H90B4_HUMAI | Unmodified | 622350 | 1305200 | 2906600 | 2477700 | 0 | 1034000 | 451990 | 0 | 8277700 | 879290 | 9769400 | 56886000 | 1556700 | 0 | 3175800 | 2370600 | 0 | 967800 |
| EGLENDLK | PDCD6IP | Q8WUM4 | PDC6I_HUMAN | Unmodified | 2601000 | 1907400 | 933010 | 1364700 | 68196 | 0 | 82910 | 0 | 1706900 | 392960 | 356040 | 777390 | 258410 | 0 | 727630 | 0 | 87989 | 112720 |
| EGLEYIPLR | DHCR24; DHC | Q15392; A0 | DHC24_HUMA | Unmodified | 0 | 0 | 0 | 0 | 244820 | 0 | 0 | 0 | 0 | 0 | 20491 | 1604700 | 0 | 439480 | 0 | 0 | 0 | 0 |
| EGLGVNEV | MYO6; MYO6; M | Q9UM54; A0 | MYO6_HUMAN | Unmodified | 379570 | 0 | 663200 | 2059900 | 0 | 0 | 0 | 0 | 980430 | 262190 | 0 | 859180 | 0 | 0 | 1913500 | 974270 | 0 | 0 |
| EGLHYTEV | TPP2; TPP2; TP | P29144; A0A | TPP2_HUMAN | Unmodified | 0 | 0 | 0 | 0 | 0 | 0 | 0 | 0 | 0 | 0 | 0 | 3400700 | 0 | 0 | 0 | 0 | 0 | 0 |
| EGLLDPNV | ERI3; ERI3; ERI | O43414; F6 | ERI3_HUMAN E | Unmodified | 749580 | 768200 | 0 | 0 | 0 | 0 | 0 | 0 | 113440 | 0 | 0 | 454560 | 0 | 0 | 364610 | 709820 | 0 | 0 |
| EGLLGLQN | CLASP1; CLAS | Q7Z460; O7 | CLAP1_HUMAN | Unmodified | 0 | 0 | 0 | 0 | 0 | 0 | 0 | 0 | 12545 | 0 | 169960 | 1192300 | 0 | 0 | 0 | 109360 | 0 | 0 |
| EGLLLWCC | ACTN2; ACTN4 | P35609; O4 | ACTN2_HUMAI | Unmodified | 0 | 0 | 0 | 226190 | 0 | 0 | 0 | 0 | 0 | 0 | 0 | 4191100 | 0 | 0 | 0 | 501920 | 0 | 0 |
| EGLLLWCC | ACTN2; ACTN4 | P35609; O4 | ACTN2_HUMAI | Unmodified | 37122 | 0 | 0 | 646030 | 1561400 | 473750 | 270760 | 0 | 480110 | 0 | 349590 | 2814900 | 684230 | 917760 | 0 | 0 | 680060 | 785940 |
| EGLLLYHT | TSPAN9; TSPAN | O75954; B5 | TSN9_HUMAN | Unmodified | 0 | 0 | 1000800 | 660830 | 0 | 0 | 0 | 0 | 0 | 0 | 0 | 0 | 0 | 0 | 431880 | 335550 | 0 | 0 |
| EGLMLDS | DYNC1H1; DYN | Q14204; A0 | DYHC1_HUMA | Oxidation (M) | 228970 | 0 | 0 | 0 | 0 | 0 | 0 | 0 | 379080 | 0 | 142340 | 834120 | 0 | 0 | 0 | 162310 | 0 | 0 |
| EGLMLDS | DYNC1H1; DYN | Q14204; A0 | DYHC1_HUMA | Unmodified | 0 | 0 | 0 | 0 | 87339 | 0 | 36085 | 0 | 281500 | 0 | 0 | 786220 | 0 | 0 | 0 | 0 | 0 | 0 |
| EGLNGGG | DNAJB6; DNAJ | O75190; A0 | DNJB6_HUMAI | Unmodified | 1391200 | 4788200 | 319070 | 0 | 0 | 0 | 0 | 0 | 0 | 0 | 0 | 0 | 0 | 0 | 2613000 | 1188400 | 0 | 0 |
| EGLPTEEIK | UTRN | P46939 | UTRN_HUMAN | Unmodified | 0 | 0 | 0 | 0 | 0 | 0 | 0 | 0 | 0 | 0 | 0 | 0 | 0 | 0 | 960980 | 0 | 0 | 0 |
| EGLPVALD | RTRAF | Q9Y224 | RTRAF_HUMAN | Unmodified | 332290 | 332370 | 1001000 | 805130 | 1586600 | 382780 | 490470 | 225170 | 0 | 252420 | 245970 | 6583900 | 2663900 | 915590 | 520290 | 445720 | 504320 | 1302300 |
| EGLQNME | STIP1; STIP1; S | P31948; F5 | STIP1_HUMAN | Unmodified | 166720 | 184950 | 0 | 516260 | 1422800 | 423130 | 49813 | 30649 | 4120800 | 911230 | 2580400 | 11628000 | 2270000 | 633580 | 410260 | 487920 | 879000 | 1681900 |
| EGLQNME | STIP1; STIP1; S | P31948; F5 | STIP1_HUMAN | Oxidation (M) | 65675 | 106760 | 0 | 0 | 0 | 0 | 0 | 0 | 0 | 0 | 653580 | 4142000 | 0 | 0 | 0 | 374760 | 132560 | 291540 |
| EGLQNME | STIP1 | F5GXD8 | F5GXD8_HUM | Unmodified | 0 | 0 | 0 | 0 | 0 | 0 | 0 | 0 | 0 | 0 | 0 | 0 | 0 | 5819500 | 0 | 0 | 0 | 0 |
| EGLTGTG | DNAJB2; DNAJ | P25686; C9J | DNJB2_HUMAI | Unmodified | 244510 | 184020 | 0 | 0 | 0 | 0 | 123710 | 58112 | 0 | 160710 | 0 | 47716 | 0 | 0 | 0 | 280080 | 306530 | 0 |
| EGLTSIEEV | PLEC; PLEC | Q15149; H0 | PLEC_HUMAN | Unmodified | 0 | 0 | 0 | 0 | 235050 | 0 | 0 | 0 | 0 | 87593 | 0 | 234540 | 0 | 0 | 0 | 0 | 1094300 | 291910 |
| EGLTSIEEV | PLEC; PLEC | Q15149; H0 | PLEC_HUMAN | Unmodified | 25405 | 15390 | 0 | 0 | 0 | 0 | 0 | 0 | 0 | 0 | 0 | 0 | 0 | 0 | 0 | 0 | 491730 | 0 |
| EGLVPVSE | USP10; USP10 | Q14694; J3K | UBP10_HUMAI | Unmodified | 0 | 0 | 0 | 0 | 0 | 0 | 0 | 0 | 0 | 0 | 343070 | 1958600 | 0 | 0 | 0 | 0 | 0 | 0 |
| EGMAALQS | ACADVL; ACAD | P49748; G3 | ACADV_HUMAI | Oxidation (M) | 0 | 0 | 0 | 0 | 0 | 0 | 0 | 0 | 496980 | 0 | 0 | 0 | 0 | 0 | 0 | 0 | 0 | 0 |
| EGMAALQS | ACADVL; ACAD | P49748; G3 | ACADV_HUMAI | Unmodified | 0 | 0 | 0 | 0 | 0 | 0 | 0 | 0 | 2488500 | 298660 | 0 | 0 | 0 | 0 | 0 | 0 | 0 | 0 |
| EGMEAAVE | XRN2 | Q9H0D6 | XRN2_HUMAN | Unmodified | 293020 | 124000 | 94212 | 114710 | 450980 | 2144900 | 116080 | 63221 | 424320 | 90515 | 0 | 545210 | 116370 | 0 | 260220 | 195310 | 567280 | 461700 |
| EGMELLAR | RAB3GAP2 | Q9H2M9 | RBGPR_HUMAI | Oxidation (M) | 0 | 0 | 0 | 0 | 91723 | 0 | 70494 | 19933 | 0 | 25192 | 0 | 0 | 0 | 0 | 0 | 0 | 156430 | 89892 |
| EGMFERIP | AGPS; AGPS; A | O00116; A0 | ADAS_HUMAN | Oxidation (M) | 0 | 0 | 0 | 0 | 0 | 0 | 0 | 0 | 2511500 | 0 | 0 | 0 | 0 | 0 | 0 | 0 | 0 | 0 |
| EGMLQHW | ACP2; ACP2; A | P11117; B7Z | PPAL_HUMAN | Unmodified | 0 | 0 | 0 | 0 | 0 | 0 | 0 | 0 | 8878600 | 0 | 0 | 0 | 0 | 0 | 0 | 0 | 0 | 0 |
| EGMQLISE | SPTBN1; SPTB | Q01082; A0 | SPTB2_HUMAN | Oxidation (M) | 0 | 0 | 0 | 0 | 0 | 0 | 0 | 0 | 0 | 0 | 0 | 1760100 | 853290 | 0 | 0 | 0 | 0 | 0 |
| EGMRANN | SND1 | Q7KZF4 | SND1_HUMAN | Unmodified | 0 | 0 | 0 | 0 | 0 | 0 | 0 | 0 | 0 | 0 | 0 | 0 | 793600 | 0 | 0 | 0 | 0 | 0 |
| EGMSNNS | GNG4 | P50150 | GBG4_HUMAN | Unmodified | 0 | 0 | 0 | 0 | 159910 | 0 | 296020 | 0 | 1577000 | 247120 | 0 | 0 | 0 | 0 | 0 | 0 | 0 | 0 |
| EGMTAFVE | ECHS1 | P30084 | ECHM_HUMAN | Unmodified | 0 | 0 | 0 | 0 | 0 | 0 | 101830 | 16214 | 1667900 | 42844 | 0 | 0 | 61519 | 43365 | 18552 | 0 | 0 | 0 |
| EGNAIFTF | SQOR; SQOR; | Q9Y6N5; H3 | SQOR_HUMAN | Unmodified | 0 | 0 | 0 | 0 | 0 | 81284 | 0 | 0 | 7054900 | 0 | 0 | 0 | 2332000 | 0 | 0 | 0 | 0 | 1035200 |
| EGNASGV | EEF1A2; EEF1A | Q05639; A0 | EF1A2_HUMAN | Unmodified | 10283000 | 7855700 | 27877000 | 9800000 | 667420 | 175000 | 0 | 0 | 1358500 | 96641 | 0 | 0 | 1336600 | 0 | 1.22E+08 | 33562000 | 183330 | 0 |
| EGNASGV | EEF1A2; EEF1A | Q05639; A0 | EF1A2_HUMAN | Unmodified | 1384900 | 1892400 | 0 | 1163900 | 0 | 154200 | 0 | 0 | 0 | 0 | 0 | 0 | 0 | 0 | 4736300 | 1354700 | 0 | 0 |
| EGNDLYHE | ATP5F1B; ATP5 | P06576; H0 | ATPB_HUMAN | Unmodified | 0 | 0 | 0 | 0 | 0 | 0 | 0 | 0 | 16082000 | 1372300 | 929640 | 137610 | 0 | 0 | 84508 | 0 | 0 | 685070 |
| EGNDLYHE | ATP5F1B; ATP5 | P06576; H0 | ATPB_HUMAN | Oxidation (M) | 68500 | 58109 | 0 | 107840 | 0 | 0 | 0 | 0 | 1418900 | 481200 | 128810 | 254130 | 0 | 0 | 74013 | 210170 | 0 | 0 |
| EGNEVFLP | GLCE; GLCE | O94923; H0 | GLCE_HUMAN | Unmodified | 0 | 0 | 0 | 0 | 0 | 0 | 0 | 0 | 663150 | 0 | 83224 | 196620 | 0 | 0 | 0 | 0 | 0 | 57041 |
| EGNGTVM | MYL6; MYL6; M | P60660; G3 | MYL6_HUMAN | Unmodified | 270240 | 0 | 0 | 0 | 10047000 | 2399500 | 0 | 0 | 103060 | 654980 | 91177 | 0 | 439280 | 387160 | 0 | 0 | 357050 | 1949400 |
| EGNGTVM | MYL6; MYL6; M | P60660; G3 | MYL6_HUMAN | Oxidation (M) | 0 | 0 | 0 | 0 | 1420600 | 0 | 0 | 0 | 0 | 0 | 0 | 0 | 3808600 | 196040 | 0 | 0 | 0 | 0 |
| EGNGTVM | MYL1; MYL3 | P05976; P08 | MYL1_HUMAN | Unmodified | 0 | 0 | 709730 | 945480 | 2000100 | 343890 | 0 | 0 | 2119200 | 0 | 0 | 12347000 | 574090 | 162100 | 1242800 | 1189500 | 0 | 317850 |
| EGNKPVLF | NRP1; NRP1; N | O14786; Q5 | NRP1_HUMAN | Unmodified | 629130 | 980790 | 0 | 1536300 | 2578200 | 846730 | 0 | 0 | 1067900 | 0 | 0 | 0 | 462470 | 0 | 1436300 | 2475000 | 528750 | 4943300 |
| EGNNPAE | HMGN2 | P05204 | HMGN2_HUM | Unmodified | 0 | 0 | 49873 | 0 | 0 | 0 | 0 | 0 | 99040 | 41505 | 27582 | 404240 | 0 | 0 | 22452 | 0 | 0 | 0 |
| EGNPEEDL | CAPG; CAPG | P40121; E7E | CAPG_HUMAN | Unmodified | 0 | 0 | 0 | 0 | 358270 | 0 | 0 | 0 | 0 | 0 | 0 | 0 | 0 | 0 | 0 | 0 | 77411 | 383380 |
| EGNQDKF | ITGA6 | P23229 | ITA6_HUMAN I | Unmodified | 0 | 0 | 0 | 0 | 6140600 | 0 | 0 | 0 | 0 | 0 | 80190 | 0 | 4707700 | 2607000 | 0 | 0 | 8367600 | 19308000 |
| EGNQEVPF | LRPPRC | P42704 | LPPRC_HUMAI | Unmodified | 0 | 0 | 0 | 0 | 0 | 0 | 0 | 0 | 2268300 | 69624 | 60107 | 25615 | 0 | 0 | 0 | 0 | 0 | 0 |
| EGNRIHFQ | HSD17B4; HSD | P51659; E7E | DHB4_HUMAN | Unmodified | 0 | 0 | 0 | 0 | 0 | 0 | 0 | 29543 | 834270 | 152850 | 0 | 62545 | 24173 | 0 | 0 | 0 | 0 | 176310 |
| EGNSPSFF | MOV10; MOV1 | Q9HCE1; Q5 | MOV10_HUMA | Unmodified | 3870800 | 2490600 | 341560 | 860720 | 0 | 0 | 0 | 0 | 117150 | 0 | 0 | 174050 | 607950 | 488550 | 1883400 | 2007400 | 0 | 0 |
| EGNVRVSR | GNB1; GNB1; G | P62873; B3K | GBB1_HUMAN | Unmodified | 0 | 0 | 0 | 0 | 0 | 122100 | 0 | 0 | 0 | 254750 | 0 | 0 | 0 | 0 | 0 | 0 | 434240 | 663070 |
| EGPAVVG | CAND1; CAND | Q86VP6; A0 | CAND1_HUMA | Unmodified | 0 | 0 | 438050 | 0 | 1127100 | 958860 | 0 | 0 | 2060700 | 0 | 2487800 | 6567200 | 0 | 0 | 0 | 1015400 | 298170 | 0 |
| EGPHYTPP | MRPL40 | Q9NQ50 | RM40_HUMAN | Unmodified | 0 | 0 | 0 | 0 | 0 | 0 | 0 | 0 | 505890 | 0 | 0 | 0 | 0 | 0 | 0 | 0 | 0 | 0 |
| EGPNHQLI | SEMA3C | Q99985 | SEM3C_HUMAI | Unmodified | 0 | 0 | 0 | 473430 | 0 | 0 | 0 | 0 | 424610 | 0 | 0 | 0 | 0 | 0 | 1200000 | 849760 | 0 | 0 |
| EGPPPASP | TKFC; TKFC; TK | Q3LXA3; I3L | TKFC_HUMAN | Unmodified | 0 | 0 | 274210 | 0 | 0 | 0 | 266270 | 0 | 0 | 0 | 297210 | 1780200 | 543570 | 0 | 278070 | 324200 | 0 | 0 |
| EGPPRGN | HSD17B8 | Q92506 | DHB8_HUMAN | Unmodified | 0 | 0 | 0 | 0 | 0 | 0 | 0 | 0 | 326450 | 0 | 0 | 0 | 0 | 0 | 0 | 0 | 76196 | 153790 |
| EGPVLILGF | PAM; PAM | P19021; H7E | AMD_HUMAN F | Unmodified | 0 | 0 | 0 | 0 | 0 | 0 | 0 | 162080 | 1218800 | 35465 | 1447500 | 955190 | 0 | 0 | 0 | 0 | 0 | 0 |
| EGPYDVVV | PARK7; PARK7 | Q99497; K7E | PARK7_HUMAN | Unmodified | 0 | 0 | 0 | 0 | 7474600 | 0 | 0 | 0 | 24197000 | 2515900 | 9734500 | 21621000 | 0 | 0 | 0 | 10719000 | 1735800 | 0 |
| EGPYDVVV | PARK7; PARK7 | Q99497; K7E | PARK7_HUMAN | Unmodified | 0 | 0 | 0 | 0 | 0 | 0 | 0 | 0 | 0 | 575560 | 0 | 0 | 3578900 | 0 | 0 | 0 | 0 | 0 |
| EGPYSISVL | FLNA; FLNA | P21333; Q6 | FLNA_HUMAN | Unmodified | 0 | 1957100 | 0 | 0 | 0 | 0 | 0 | 0 | 0 | 0 | 0 | 1185600 | 0 | 0 | 1739200 | 2310000 | 0 | 0 |
| EGQGNLP | NAE1 | Q13564 | ULA1_HUMAN | Unmodified | 0 | 0 | 0 | 0 | 634820 | 264160 | 58065 | 0 | 838570 | 60092 | 189190 | 562280 | 293630 | 107700 | 494080 | 0 | 46683 | 0 |
| EGQGNLP | NAE1 | Q13564 | ULA1_HUMAN | Oxidation (M) | 0 | 0 | 0 | 0 | 0 | 0 | 0 | 0 | 0 | 0 | 323880 | 600190 | 373920 | 0 | 0 | 0 | 0 | 0 |
| EGQLMAR | POGLUT3; POG | Q7Z4H8; H0 | PLGT3_HUMAN | Unmodified | 0 | 29240 | 0 | 0 | 545900 | 126570 | 147060 | 180070 | 3451800 | 403400 | 698120 | 3928100 | 1607400 | 594370 | 0 | 0 | 62580 | 0 |
| EGQLMARI | POGLUT3; POG | Q7Z4H8; H0 | PLGT3_HUMAN | Oxidation (M) | 0 | 0 | 0 | 0 | 0 | 0 | 0 | 0 | 0 | 924960 | 0 | 0 | 0 | 29438 | 0 | 0 | 0 | 0 |
| EGQNWLE | SEC11C; SEC1 | Q9BY50; K7E | SC11C_HUMAI | Unmodified | 0 | 0 | 0 | 0 | 43130 | 0 | 30483 | 0 | 2374700 | 230670 | 373950 | 0 | 0 | 0 | 0 | 0 | 0 | 0 |
| EGQTICVR | PRMT5; PRMT5 | O14744; H0 | ANM5_HUMAN | Unmodified | 0 | 0 | 0 | 0 | 0 | 0 | 0 | 0 | 194400 | 0 | 0 | 664370 | 0 | 0 | 0 | 0 | 0 | 0 |
| EGQTVSG | PNMA2 | Q9UL42 | PNMA2_HUMA | Unmodified | 3925700 | 3229400 | 0 | 82351 | 0 | 0 | 0 | 0 | 952230 | 1044800 | 0 | 0 | 0 | 0 | 0 | 0 | 1037200 | 1076600 |
| EGRLEESV | PPP3CA; PPP3 | Q08209; E9F | PP2BA_HUMAN | Unmodified | 157050 | 0 | 0 | 1108200 | 0 | 0 | 0 | 0 | 5132300 | 0 | 0 | 3140800 | 676090 | 0 | 917390 | 1016000 | 0 | 0 |
| EGRLQEVI | PSMD12 | O00232 | PSD12_HUMAI | Unmodified | 0 | 0 | 0 | 13712 | 178220 | 22653 | 225270 | 0 | 0 | 50912 | 0 | 0 | 46067 | 332220 | 18982 | 0 | 12337 | 0 |
| EGRWVIYN | USP5 | P45974 | UBP5_HUMAN | Unmodified | 0 | 0 | 0 | 0 | 616590 | 0 | 408640 | 212820 | 0 | 584980 | 0 | 0 | 298430 | 274160 | 0 | 0 | 229060 | 346740 |
| EGSALTEG | EPB41L3; EPB4 | Q9Y2J2; A0A | E41L3_HUMAN | Unmodified | 0 | 0 | 0 | 0 | 0 | 0 | 0 | 0 | 233900 | 39778 | 0 | 0 | 0 | 0 | 0 | 0 | 670950 | 6121500 |
| EGSASTEVI | SNRNP200; SN | O75643; A0 | U520_HUMAN | Unmodified | 127030 | 105390 | 0 | 0 | 0 | 0 | 0 | 0 | 0 | 0 | 0 | 5044300 | 0 | 0 | 0 | 45337 | 0 | 0 |
| EGSCPQV | WFDC2 | Q14508 | WFDC2_HUMA | Unmodified | 0 | 0 | 0 | 6197400 | 0 | 0 | 0 | 0 | 2950400 | 0 | 0 | 0 | 0 | 0 | 7163000 | 0 | 0 | 0 |
| EGSEAASK | MYO1B; MYO1 | O43795; H7 | MYO1B_HUMA | Oxidation (M) | 0 | 0 | 0 | 0 | 9840500 | 0 | 1078100 | 295090 | 1188900 | 700330 | 3211100 | 8284200 | 2206100 | 742300 | 1083600 | 0 | 2354700 | 2386800 |
| EGSEAASK | MYO1B; MYO1 | O43795; H7 | MYO1B_HUMA | Unmodified | 0 | 0 | 0 | 0 | 9845100 | 1209300 | 917570 | 23633 | 1966500 | 0 | 1075700 | 2355800 | 0 | 0 | 0 | 0 | 1859200 | 2839200 |
| EGSGDGG | TARS1; TARS1; | P26639; D6 | SYTC_HUMAN | Unmodified | 0 | 0 | 107190 | 144090 | 0 | 0 | 87314 | 0 | 416350 | 0 | 258890 | 6626500 | 0 | 0 | 0 | 475810 | 0 | 0 |
| EGSHSPLD | GPSM1; GPSM1 | Q86YR5; A0 | GPSM1_HUMA | Unmodified | 0 | 0 | 0 | 0 | 98517 | 16982 | 0 | 26902 | 457000 | 0 | 0 | 133920 | 0 | 0 | 0 | 0 | 169540 | 26623 |
| EGSIWNH | LMAN2; LMAN2 | Q12907; D6 | LMAN2_HUMAI | Unmodified | 0 | 0 | 0 | 262390 | 110260 | 0 | 265210 | 23315 | 1.03E+08 | 8467500 | 1918500 | 1911900 | 101730 | 177110 | 801040 | 809880 | 0 | 0 |
| EGSLGDEL | KCTD12 | Q96CX2 | KCD12_HUMAI | Unmodified | 0 | 0 | 0 | 0 | 0 | 0 | 0 | 0 | 1401300 | 0 | 810240 | 0 | 0 | 0 | 0 | 0 | 0 | 0 |
| EGSLPLQY | CXADR | P78310 | CXAR_HUMAN | Unmodified | 1734900 | 865670 | 786690 | 619480 | 13173000 | 3018200 | 1405500 | 832380 | 1550300 | 677250 | 1987900 | 23113000 | 0 | 0 | 1283800 | 1186800 | 0 | 0 |
| EGSRTDDE | HSP90B1; HSP | P14625; A0A | ENPL_HUMAN | Unmodified | 0 | 0 | 0 | 0 | 0 | 0 | 0 | 0 | 0 | 0 | 769440 | 102030 | 0 | 0 | 0 | 0 | 0 | 0 |
| EGSRTDDE | HSP90B1; HSP | P14625; A0A | ENPL_HUMAN | Unmodified | 0 | 0 | 0 | 0 | 77779 | 0 | 0 | 0 | 77631000 | 5581900 | 28185000 | 13420000 | 0 | 0 | 0 | 0 | 0 | 0 |
| EGSSAFLS | CHPF2; CHPF2 | Q9P2E5; H7 | CHPF2_HUMAI | Oxidation (M) | 0 | 0 | 0 | 0 | 0 | 0 | 0 | 0 | 3677000 | 0 | 0 | 211770 | 0 | 0 | 0 | 0 | 0 | 0 |
| EGSSAFLS | CHPF2; CHPF2 | Q9P2E5; H7 | CHPF2_HUMAI | Unmodified | 0 | 0 | 0 | 0 | 0 | 0 | 0 | 0 | 744540 | 0 | 172330 | 0 | 0 | 0 | 0 | 0 | 0 | 0 |
| EGSVMGV | DIP2B | Q9P265 | DIP2B_HUMAN | Unmodified | 0 | 136310 | 0 | 270580 | 0 | 0 | 0 | 0 | 0 | 205890 | 270610 | 1991400 | 0 | 0 | 914970 | 0 | 0 | 0 |
| EGSVTSVN | ACY1; ABHD14 | Q03154; A0 | ACY1_HUMAN | Unmodified | 73777 | 84028 | 301290 | 0 | 0 | 0 | 0 | 0 | 0 | 0 | 507570 | 4286800 | 0 | 0 | 421360 | 307810 | 623690 | 0 |

| | | | | | | | | | | | | | | | | | | | | | |
|---|---|---|---|---|---|---|---|---|---|---|---|---|---|---|---|---|---|---|---|---|---|
| EGSVVVDL | NNT; NNT; NNT | Q13423; E9F | NNTM_HUMAN | Unmodified | 0 | 0 | 0 | 0 | 0 | 0 | 0 | 0 | 5354700 | 0 | 0 | 0 | 0 | 0 | 0 | 0 | 0 | 0 |
| EGTDSSQG | CCT7; CCT7; C | Q99832; A0A | TCPH_HUMAN | Unmodified | 0 | 0 | 0 | 375890 | 0 | 0 | 0 | 0 | 901280 | 571940 | 0 | 665980 | 1011600 | 0 | 0 | 0 | 0 | 0 |
| EGTDSSQG | CCT7; CCT7; C | Q99832; A0A | TCPH_HUMAN | Oxidation (M) | 0 | 0 | 0 | 0 | 838740 | 0 | 542240 | 0 | 0 | 0 | 0 | 0 | 1672100 | 1971600 | 0 | 0 | 641140 | 1172200 |
| EGTEASASI | HM13; HM13 | Q8TCT9; A0A | HM13_HUMAN | Unmodified | 56382 | 0 | 17823 | 0 | 647310 | 219360 | 783140 | 532660 | 886830 | 199230 | 196620 | 489060 | 380300 | 155820 | 74828 | 146070 | 481880 | 1176300 |
| EGTEAWEA | DYNC1H1; DYN | Q14204; A0A | DYHC1_HUMA | Unmodified | 0 | 0 | 0 | 0 | 0 | 0 | 0 | 0 | 0 | 0 | 0 | 1449600 | 0 | 0 | 0 | 0 | 0 | 0 |
| EGTETFADI | TLN1 | Q9Y490 | TLN1_HUMAN | Unmodified | 0 | 0 | 0 | 0 | 0 | 242650 | 0 | 0 | 414430 | 284340 | 95075 | 1755800 | 0 | 0 | 0 | 0 | 122320 | 0 |
| EGTGSTATS | EIF3H; EIF3H; E | O15372; E5F | EIF3H_HUMAN | Unmodified | 26088 | 15673 | 0 | 97036 | 0 | 0 | 0 | 0 | 0 | 0 | 0 | 200300 | 44877 | 0 | 0 | 0 | 0 | 0 |
| EGTINVHD | MUC1; MUC1; M | P15941; B1A | MUC1_HUMAN | Unmodified | 0 | 0 | 0 | 0 | 0 | 0 | 0 | 0 | 9850100 | 748830 | 0 | 0 | 0 | 0 | 0 | 0 | 0 | 0 |
| EGTIQVQG | ABHD14B; ABH | Q96IU4; F8W | ABHEB_HUMAI | Unmodified | 77515 | 0 | 0 | 0 | 0 | 0 | 0 | 80613 | 0 | 0 | 0 | 348700 | 0 | 0 | 237050 | 71352 | 0 | 0 |
| EGTKFYNEI | PDCD6IP | Q8WUM4 | PDC6I_HUMAN | Unmodified | 953750 | 546680 | 0 | 498930 | 4009500 | 570280 | 293620 | 0 | 314410 | 3683500 | 0 | 0 | 4464100 | 2023900 | 140530 | 163260 | 645430 | 1975300 |
| EGTKFYNEI | PDCD6IP | Q8WUM4 | PDC6I_HUMAN | Unmodified | 0 | 0 | 0 | 676390 | 17203000 | 1578100 | 4820100 | 0 | 0 | 6286500 | 53195 | 0 | 15772000 | 10513000 | 0 | 0 | 8652200 | 6711000 |
| EGTLIVRQS | MAP4K4 | E7EN19 | E7EN19_HUMA | Unmodified | 0 | 0 | 0 | 0 | 92652 | 0 | 162750 | 94403 | 0 | 251450 | 0 | 0 | 0 | 422360 | 0 | 0 | 1182800 | 5803000 |
| EGTLQHAF | NCLN; NCLN; N | Q969V3; K7E | NCLN_HUMAN | Unmodified | 0 | 0 | 0 | 171590 | 615890 | 0 | 0 | 0 | 1830100 | 53392 | 0 | 0 | 0 | 0 | 0 | 0 | 0 | 0 |
| EGTQASEG | DBN1 | Q16643 | DREB_HUMAN | Unmodified | 0 | 0 | 0 | 0 | 0 | 0 | 0 | 0 | 0 | 0 | 0 | 4736500 | 0 | 0 | 0 | 0 | 0 | 0 |
| EGTRCIQVI | FAM126A; FAM | Q9BYI3; A0A | HYCCI_HUMAN | Unmodified | 0 | 0 | 0 | 0 | 34827 | 0 | 0 | 63287 | 219040 | 0 | 0 | 159630 | 0 | 0 | 0 | 0 | 0 | 1207000 |
| EGTSTLQS | SNX11; SNX11 | Q9Y5W9; B4 | SNX11_HUMAN | Unmodified | 0 | 0 | 0 | 0 | 0 | 0 | 125630 | 0 | 0 | 0 | 0 | 0 | 214240 | 165310 | 0 | 0 | 1727700 | 1201900 |
| EGVFPDNF | CD2AP | Q9Y5K6 | CD2AP_HUMAI | Unmodified | 5112800 | 7897000 | 2646700 | 3306100 | 221600 | 0 | 50915 | 170990 | 0 | 0 | 204080 | 396760 | 389150 | 0 | 11837000 | 3006700 | 0 | 0 |
| EGVFPDNF | CD2AP | Q9Y5K6 | CD2AP_HUMAI | Unmodified | 0 | 0 | 0 | 2072400 | 0 | 0 | 0 | 0 | 0 | 0 | 0 | 0 | 0 | 0 | 0 | 0 | 0 | 0 |
| EGVHGGLI | PFN1; PFN1 | P07737; K7E | PROF1_HUMAN | Unmodified | 2106200 | 1079400 | 1416100 | 2037700 | 15256000 | 4237200 | 1222400 | 1868700 | 5823400 | 2214200 | 3446600 | 12587000 | 7543200 | 2321100 | 1975700 | 1808900 | 5295900 | 6129500 |
| EGVHGGLI | PFN1; PFN1 | P07737; K7E | PROF1_HUMAN | Unmodified | 4565400 | 0 | 0 | 5941200 | 23519000 | 0 | 2241400 | 4679400 | 7687800 | 7577300 | 6595900 | 37947000 | 10442000 | 9102600 | 2550300 | 0 | 11969000 | 29008000 |
| EGVIEAGAI | ARFIP1 | P53367 | ARFP1_HUMAN | Oxidation (M) | 131930 | 0 | 191960 | 473000 | 0 | 0 | 0 | 0 | 0 | 0 | 0 | 43082 | 0 | 0 | 0 | 0 | 0 | 0 |
| EGVIGVTEE | SUN2 | Q9UH99 | SUN2_HUMAN | Unmodified | 0 | 0 | 0 | 0 | 0 | 0 | 0 | 0 | 52683000 | 2055800 | 0 | 0 | 0 | 0 | 0 | 0 | 0 | 0 |
| EGVIQNFK | AFDN; AFDN; A | P55196; Q5T | AFAD_HUMAN | Unmodified | 0 | 0 | 0 | 0 | 2046300 | 300650 | 0 | 0 | 926760 | 0 | 671080 | 0 | 1378000 | 0 | 0 | 0 | 0 | 0 |
| EGVKFDES | HSP90B1; HSP | P14625; A0A | ENPL_HUMAN | Unmodified | 0 | 0 | 183890 | 0 | 0 | 0 | 0 | 0 | 62329000 | 14734000 | 6881300 | 5791600 | 0 | 0 | 0 | 0 | 0 | 0 |
| EGVKFDES | HSP90B1; HSP | P14625; A0A | ENPL_HUMAN | Unmodified | 0 | 36435 | 0 | 0 | 330880 | 13303 | 493580 | 0 | 0 | 7849400 | 0 | 0 | 0 | 0 | 0 | 0 | 227560 | 0 |
| EGVKTENN | SUMO2 | P61956 | SUMO2_HUMA | Unmodified | 662050 | 660740 | 0 | 549320 | 146160 | 0 | 28042 | 0 | 361820 | 0 | 256290 | 3618000 | 0 | 154630 | 1386900 | 664080 | 0 | 0 |
| EGVKTENN | SUMO2 | P61956 | SUMO2_HUMA | Unmodified | 0 | 0 | 0 | 0 | 0 | 0 | 0 | 0 | 0 | 0 | 0 | 0 | 0 | 0 | 0 | 0 | 945260 | 0 |
| EGVLEKRSI | PHLDA1; PHLD | Q8WV24; R4 | PHLA1_HUMAN | Unmodified | 0 | 0 | 0 | 0 | 0 | 0 | 0 | 0 | 0 | 0 | 0 | 0 | 0 | 0 | 0 | 0 | 182540 | 0 |
| EGVLTGSPI | GCN1 | Q92616 | GCN1_HUMAN | Unmodified | 212980 | 113280 | 133880 | 0 | 0 | 0 | 0 | 0 | 187120 | 0 | 52734 | 1189200 | 0 | 0 | 166170 | 197430 | 0 | 62574 |
| EGVLVEHV | ENPP4 | Q9Y6X5 | ENPP4_HUMAN | Unmodified | 0 | 0 | 206010 | 0 | 0 | 0 | 90374 | 318010 | 4525000 | 803560 | 951390 | 680120 | 0 | 0 | 0 | 0 | 0 | 0 |
| EGVLVEHV | ENPP4 | Q9Y6X5 | ENPP4_HUMAN | Unmodified | 1936000 | 776500 | 0 | 1033700 | 0 | 0 | 0 | 0 | 415750 | 228950 | 0 | 0 | 433450 | 468150 | 0 | 0 | 0 | 0 |
| EGVREVFEI | RHOC; RHOC | P08134; Q5J | RHOC_HUMAN | Unmodified | 0 | 478500 | 0 | 1057100 | 748000 | 0 | 0 | 0 | 135630 | 0 | 0 | 0 | 3890600 | 507150 | 1805400 | 759160 | 628770 | 0 |
| EGVREVFET | RHOB | P62745 | RHOB_HUMAN | Unmodified | 0 | 0 | 1226300 | 901420 | 587740 | 0 | 0 | 0 | 892670 | 0 | 824520 | 5298000 | 598680 | 0 | 2403700 | 2098600 | 0 | 0 |
| EGVRYFVD | WWP1; WWP2 | Q9H0M0; O | WWP1_HUMAN | Unmodified | 0 | 229870 | 188960 | 198080 | 0 | 0 | 0 | 0 | 0 | 0 | 0 | 0 | 0 | 0 | 322830 | 319030 | 0 | 0 |
| EGVSAAFA | BZW1; BZW1; B | Q7L1Q6; C9 | BZW1_HUMAN | Unmodified | 240000 | 246100 | 77281 | 37607 | 1323200 | 0 | 321100 | 247840 | 88539 | 0 | 1269800 | 3276900 | 47082 | 371430 | 560200 | 564390 | 455540 | 1695200 |
| EGVVECSF | MDH2; MDH2 | P40926; G3X | MDHM_HUMAN | Unmodified | 29378000 | 46877000 | 170260 | 150700 | 0 | 0 | 0 | 0 | 22226000 | 1454500 | 920730 | 0 | 0 | 0 | 73291000 | 69417000 | 108600 | 0 |
| EGWKKSDI | CTNND1; CTNN | O60716; H0 | CTND1_HUMAI | Unmodified | 0 | 0 | 0 | 0 | 0 | 0 | 0 | 0 | 0 | 0 | 0 | 0 | 1099700 | 1979000 | 0 | 0 | 998070 | 2642900 |
| EGWPLDIR | ACO2; ACO2; A | Q99798; A2A | ACON_HUMAN | Unmodified | 73327 | 337850 | 18967 | 0 | 60402 | 532200 | 0 | 0 | 2397700 | 78887 | 501770 | 355290 | 0 | 0 | 738640 | 594290 | 116520 | 0 |
| EGWPSSAY | CBR1 | P16152 | CBR1_HUMAN | Unmodified | 131350 | 0 | 0 | 192690 | 6211200 | 630530 | 0 | 0 | 899300 | 365570 | 0 | 3730500 | 1218300 | 0 | 1297000 | 425200 | 2039700 | 4594200 |
| EGYAWAED | RTCB | Q9Y3I0 | RTCB_HUMAN | Unmodified | 0 | 0 | 276810 | 0 | 0 | 0 | 0 | 0 | 0 | 0 | 0 | 1687100 | 0 | 0 | 192790 | 113550 | 0 | 0 |
| EGYDFHSS | ACTR1A; ACTR1 | P61163; A0A | ACTZ_HUMAN | Unmodified | 0 | 359250 | 291040 | 259980 | 328890 | 0 | 273040 | 77530 | 0 | 332990 | 622220 | 14933000 | 157300 | 63518 | 903100 | 787150 | 207590 | 0 |
| EGYIPSNY | HCK; HCK; HCK | P08631; J3K | HCK_HUMAN T | Unmodified | 0 | 0 | 0 | 0 | 0 | 0 | 0 | 0 | 595100 | 0 | 90895 | 3180000 | 0 | 0 | 0 | 282850 | 0 | 0 |
| EGYSGVGL | APEX1; APEX1; | P27695; A0A | APEX1_HUMAN | Unmodified | 136600 | 0 | 295430 | 0 | 0 | 0 | 0 | 0 | 0 | 99791 | 0 | 6914900 | 0 | 0 | 353950 | 148760 | 0 | 0 |
| EGYVLTAVI | NOMO1; NOMO | Q15155; P69 | NOMO1_HUMA | Unmodified | 171540 | 0 | 0 | 0 | 0 | 135200 | 0 | 0 | 5712000 | 956380 | 426380 | 0 | 0 | 0 | 0 | 143690 | 0 | 0 |
| EHAFFRYID | PRKCB; PRKCB | P05771; I3L1 | KPCB_HUMAN | Unmodified | 0 | 0 | 0 | 0 | 567990 | 0 | 959480 | 0 | 0 | 0 | 0 | 0 | 517790 | 620460 | 0 | 0 | 0 | 0 |
| EHALAQAE | SCAMP1; SCAM | O15126; A0A | SCAM1_HUMA | Unmodified | 196380 | 763630 | 3078600 | 1676400 | 562920 | 0 | 0 | 37499 | 399190 | 9455.6 | 217390 | 1468100 | 0 | 0 | 1187100 | 1173000 | 168670 | 0 |
| EHALLAYTL | EEF1A2; EEF1A | Q05639; Q5 | EF1A2_HUMAN | Unmodified | 2.34E+08 | 2.81E+08 | 1.32E+08 | 1.58E+08 | 27011000 | 4593200 | 8159200 | 1943900 | 43367000 | 6268200 | 23280000 | 2.49E+08 | 24654000 | 7773100 | 2.3E+08 | 1.88E+08 | 7540100 | 12867000 |
| EHALLAYTL | EEF1A2; EEF1A | Q05639; Q5 | EF1A2_HUMAN | Unmodified | 506730 | 0 | 0 | 0 | 0 | 0 | 0 | 0 | 0 | 0 | 0 | 0 | 0 | 0 | 0 | 339890 | 0 | 0 |
| EHALTSGTI | ELOC; ELOC; E | Q15369; R4 | ELOC_HUMAN | Unmodified | 78760 | 119230 | 0 | 180790 | 227360 | 35263 | 0 | 0 | 635900 | 82448 | 205010 | 760610 | 157530 | 46723 | 45776 | 77082 | 42980 | 79832 |
| EHAMLAK | GSPT1; GSPT2; | P15170; Q8I | ERF3A_HUMAN | Oxidation (M) | 0 | 0 | 0 | 0 | 0 | 0 | 0 | 0 | 0 | 0 | 0 | 245180 | 0 | 0 | 0 | 0 | 0 | 0 |
| EHAPSIIFM | PSMC5; PSMC5 | P62195; J3Q | PRS8_HUMAN | Unmodified | 49267 | 0 | 0 | 0 | 0 | 0 | 0 | 0 | 808770 | 0 | 0 | 0 | 0 | 0 | 53825 | 0 | 0 | 0 |
| EHAPSIIFM | PSMC5; PSMC5 | P62195; J3Q | PRS8_HUMAN | Oxidation (M) | 0 | 0 | 98386 | 77214 | 0 | 0 | 0 | 0 | 0 | 0 | 0 | 954030 | 0 | 0 | 0 | 0 | 0 | 0 |
| EHASIDAQ | SIRT2 | Q8IXJ6 | SIR2_HUMAN N | Unmodified | 0 | 0 | 0 | 0 | 0 | 0 | 0 | 0 | 0 | 0 | 0 | 470080 | 0 | 0 | 0 | 0 | 0 | 0 |
| EHDPVGQI | CSE1L | P55060 | XPO2_HUMAN | Unmodified | 0 | 0 | 193140 | 266810 | 1432700 | 0 | 513230 | 9972 | 1761300 | 687950 | 1098500 | 7396600 | 0 | 324240 | 337380 | 106790 | 0 | 866220 |
| EHDQLIEK | XPO1; XPO1; XP | O14980; A0A | XPO1_HUMAN | Unmodified | 63435 | 639070 | 0 | 0 | 0 | 0 | 0 | 0 | 0 | 0 | 0 | 0 | 0 | 0 | 75687 | 64146 | 0 | 0 |
| EHDVKKED | NARS1 | O43776 | SYNC_HUMAN | Unmodified | 0 | 0 | 0 | 0 | 0 | 0 | 0 | 0 | 1010000 | 0 | 387790 | 12927000 | 0 | 84686 | 469890 | 280600 | 0 | 0 |
| EHDVQNTF | CBR4; CBR4 | Q8N4T8; H0 | CBR4_HUMAN | Unmodified | 0 | 0 | 0 | 0 | 0 | 0 | 0 | 0 | 1528500 | 0 | 0 | 0 | 0 | 0 | 0 | 0 | 0 | 0 |
| EHEDYCGA | TRAFD1; TRAFD | O14545; F8V | TRAD1_HUMAN | Unmodified | 175960 | 0 | 0 | 0 | 0 | 0 | 0 | 0 | 0 | 0 | 0 | 0 | 0 | 0 | 0 | 0 | 0 | 0 |
| EHEEPTTSE | IGFBP5; IGFBP5 | P24593; C9J | IBP5_HUMAN I | Unmodified | 0 | 0 | 0 | 0 | 0 | 0 | 0 | 0 | 1829800 | 372310 | 0 | 0 | 0 | 0 | 0 | 0 | 0 | 0 |
| EHELLEQQ | SF3B2; SF3B2; | Q13435; E9F | SF3B2_HUMAN | Unmodified | 190370 | 50870 | 0 | 0 | 0 | 0 | 0 | 0 | 0 | 0 | 0 | 486780 | 0 | 0 | 0 | 52083 | 0 | 0 |
| EHEVFQR | DNAJA2 | O60884 | DNJA2_HUMAN | Unmodified | 324660 | 516190 | 83314 | 185480 | 55714 | 0 | 0 | 0 | 738610 | 29684 | 64336 | 1399900 | 0 | 725040 | 1082000 | 45165 | 0 | 0 |
| EHEVTIFVR | ACLY; ACLY | P53396; K7E | ACLY_HUMAN | Unmodified | 35668 | 20499 | 97164 | 62001 | 0 | 0 | 0 | 23956 | 105340 | 0 | 67698 | 2788100 | 0 | 0 | 102400 | 84668 | 0 | 183920 |
| EHEVTIFVR | ACLY; ACLY | P53396; K7E | ACLY_HUMAN | Unmodified | 0 | 0 | 0 | 0 | 0 | 0 | 0 | 0 | 0 | 0 | 0 | 2144600 | 0 | 0 | 0 | 65951 | 0 | 0 |
| EHFAQFGH | SLIRP; SLIRP; S | Q9GZT3; H0 | SLIRP_HUMAN | Unmodified | 0 | 0 | 0 | 0 | 0 | 0 | 36118 | 0 | 860160 | 129750 | 0 | 0 | 0 | 0 | 0 | 0 | 0 | 0 |
| EHFAQFGH | SLIRP; SLIRP; S | Q9GZT3; H0 | SLIRP_HUMAN | Unmodified | 0 | 0 | 0 | 101430 | 0 | 0 | 300600 | 0 | 519760 | 434230 | 0 | 0 | 17191 | 28307 | 233710 | 0 | 0 | 0 |
| EHFGAREE | CLIP1; CLIP1; C | P30622; F5H | CLIP1_HUMAN | Unmodified | 460350 | 0 | 0 | 0 | 0 | 0 | 33895 | 0 | 0 | 0 | 39364 | 0 | 286440 | 0 | 22247 | 0 | 0 | 0 |
| EHFQDDVF | NAMPT; NAMPT | P43490; A0A | NAMPT_HUMAI | Unmodified | 0 | 0 | 0 | 542220 | 2151000 | 541880 | 0 | 25643 | 6960400 | 0 | 802200 | 3666900 | 365960 | 0 | 343620 | 284920 | 102650 | 133450 |
| EHFQSYDL | NAE1; NAE1; N | Q13564; J3Q | ULA1_HUMAN | Unmodified | 0 | 0 | 0 | 0 | 0 | 0 | 0 | 0 | 0 | 0 | 0 | 464130 | 0 | 0 | 0 | 0 | 0 | 0 |
| EHGAFDAV | MTHFD1; MTHF | P11586; V9G | C1TC_HUMAN | Unmodified | 0 | 0 | 0 | 0 | 0 | 0 | 0 | 0 | 0 | 0 | 0 | 0 | 0 | 0 | 0 | 0 | 0 | 0 |
| EHGFAGW | RAB32 | Q13637 | RAB32_HUMAN | Unmodified | 0 | 0 | 0 | 1184900 | 0 | 0 | 0 | 0 | 0 | 0 | 562430 | 0 | 319270 | 126120 | 0 | 0 | 0 | 232060 |
| EHGGSIVN | PECR | Q9BY49 | PECR_HUMAN | Unmodified | 0 | 0 | 0 | 0 | 0 | 0 | 133750 | 79183 | 2138200 | 0 | 0 | 0 | 0 | 0 | 0 | 0 | 0 | 0 |
| EHGIRFFET | RAB10; RAB13; | P61026; P51 | RAB10_HUMAN | Unmodified | 0 | 0 | 0 | 3009900 | 1908000 | 291980 | 0 | 0 | 581720 | 54115 | 0 | 0 | 1158500 | 802340 | 0 | 0 | 8826.4 | 1199700 |
| EHGLIFMET | RAB2A; RAB2B; | P61019; Q8 | RAB2A_HUMAN | Unmodified | 206670 | 211240 | 162570 | 230640 | 0 | 0 | 0 | 0 | 6836300 | 832170 | 513230 | 3654600 | 0 | 0 | 319630 | 228940 | 0 | 102670 |
| EHGLIFMET | RAB2A; RAB2B; | P61019; Q8 | RAB2A_HUMAN | Oxidation (M) | 0 | 0 | 0 | 0 | 0 | 0 | 0 | 0 | 2080200 | 464360 | 0 | 1481600 | 0 | 0 | 0 | 0 | 0 | 0 |
| EHGLKPVV | MELTF; MELTF | P08582; C9J | TRFM_HUMAN | Unmodified | 0 | 0 | 0 | 0 | 0 | 998220 | 0 | 0 | 4943400 | 0 | 445660 | 0 | 0 | 0 | 0 | 0 | 0 | 0 |
| EHGLNPD | ZC3HAV1; ZC3 | Q7Z2W4; C9 | ZCCHV_HUMA | Unmodified | 1260100 | 648420 | 1417500 | 1394900 | 560350 | 0 | 0 | 0 | 1089400 | 161560 | 0 | 3463300 | 305290 | 0 | 1597800 | 1688500 | 190810 | 0 |
| EHGQIEFY | VPS36; VPS36 | Q86VN1; A0 | VPS36_HUMAN | Unmodified | 514620 | 464940 | 725220 | 341360 | 0 | 462530 | 0 | 82696 | 0 | 0 | 141180 | 0 | 0 | 0 | 592870 | 499170 | 0 | 0 |
| EHHFEAIAL | AKR7A2; AKR7A | O43488; H3 | ARK72_HUMAN | Unmodified | 162650 | 0 | 179830 | 0 | 217830 | 0 | 111880 | 0 | 1064500 | 227520 | 164760 | 1868300 | 0 | 0 | 277500 | 277850 | 73926 | 0 |
| EHIMQMLC | IPO5; IPO5 | O00410; H0 | IPO5_HUMAN I | Oxidation (M) | 0 | 0 | 0 | 0 | 0 | 0 | 0 | 0 | 0 | 0 | 0 | 0 | 0 | 0 | 0 | 0 | 0 | 0 |
| EHINLGCD | VDAC1; VDAC1 | P21796; C9J | VDAC1_HUMAI | Oxidation (M) | 0 | 929040 | 0 | 0 | 0 | 0 | 0 | 29712 | 11680000 | 286280 | 320980 | 127020 | 0 | 0 | 0 | 0 | 0 | 0 |
| EHINLGCD | VDAC1; VDAC1 | P21796; C9J | VDAC1_HUMAI | Unmodified | 0 | 0 | 0 | 0 | 0 | 0 | 0 | 0 | 4998000 | 122040 | 0 | 0 | 0 | 0 | 0 | 0 | 0 | 0 |
| EHINRPAD | DNAJC16; DNA | Q9Y2G8; Q5 | DJC16_HUMAN | Unmodified | 0 | 0 | 0 | 99847 | 334030 | 94081 | 0 | 0 | 878340 | 75773 | 0 | 0 | 283090 | 137920 | 25043 | 16573 | 72284 | 244890 |
| EHINSVSAI | DYNC1H1; DYN | Q14204; A0A | DYHC1_HUMA | Unmodified | 0 | 0 | 0 | 0 | 0 | 0 | 0 | 0 | 55447 | 0 | 43981 | 622490 | 0 | 915520 | 0 | 0 | 0 | 0 |
| EHIPFFEDN | SERINC5; SERI | Q86VE9; A0A | SERC5_HUMAN | Oxidation (M) | 1490500 | 419560 | 4835900 | 0 | 0 | 0 | 0 | 0 | 0 | 0 | 0 | 0 | 0 | 0 | 11710000 | 5554600 | 0 | 0 |
| EHIPFFEDN | SERINC5; SERI | Q86VE9; A0A | SERC5_HUMAN | Unmodified | 0 | 0 | 580410 | 0 | 0 | 0 | 0 | 0 | 0 | 0 | 0 | 0 | 0 | 0 | 3942600 | 1995300 | 0 | 0 |
| EHIQLNESI | AGL | P35573 | GDE_HUMAN ( | Unmodified | 0 | 0 | 0 | 0 | 133130 | 0 | 0 | 0 | 0 | 0 | 0 | 336640 | 0 | 0 | 0 | 72005 | 20069 | 0 |
| EHIRTEVK | PRXL2A | Q9BRX8 | PXL2A_HUMAN | Unmodified | 0 | 0 | 0 | 0 | 0 | 0 | 0 | 23520 | 1622300 | 83004 | 0 | 0 | 0 | 0 | 0 | 0 | 0 | 0 |
| EHIYDEQPI | OCLN; OCLN; C | Q16625; A0A | OCLN_HUMAN | Unmodified | 1873000 | 1807800 | 4456900 | 3939400 | 2296400 | 0 | 0 | 0 | 5524500 | 823570 | 0 | 2964800 | 2970300 | 1495900 | 14389000 | 6720900 | 0 | 78355 |
| EHKAEILAL | CIT; CIT; CIT; C | O14578; A0A | CTRO_HUMAN | Unmodified | 0 | 0 | 0 | 216490 | 3243500 | 476080 | 110500 | 0 | 0 | 109370 | 110700 | 298670 | 320670 | 121730 | 0 | 34098 | 376310 | 351010 |
| EHKAEILAL | CIT; CIT; CIT; C | O14578; A0A | CTRO_HUMAN | Unmodified | 0 | 0 | 0 | 0 | 1922000 | 0 | 0 | 0 | 0 | 0 | 0 | 0 | 0 | 0 | 0 | 0 | 0 | 0 |
| EHLASKEVI | RAI14 | Q9P0K7 | RAI14_HUMAN | Unmodified | 0 | 0 | 0 | 0 | 603960 | 0 | 47772 | 62563 | 0 | 55874 | 0 | 0 | 0 | 0 | 0 | 0 | 286070 | 678020 |
| EHLDRLMN | NIBAN1 | Q9BZQ8 | NIBA1_HUMAN | Unmodified | 0 | 0 | 0 | 339880 | 291390 | 0 | 0 | 0 | 1870700 | 449290 | 0 | 0 | 0 | 0 | 0 | 0 | 0 | 0 |
| EHLELFWS | CLTCL1; CLTC; | P53675; Q0 | CLH2_HUMAN | Unmodified | 1447100 | 805580 | 466710 | 226680 | 0 | 0 | 0 | 0 | 0 | 0 | 0 | 6894800 | 0 | 0 | 804970 | 854180 | 0 | 0 |
| EHLELFWS | CLTCL1; CLTC; | P53675; Q0 | CLH2_HUMAN | Unmodified | 573800 | 202520 | 0 | 0 | 0 | 0 | 0 | 0 | 0 | 0 | 0 | 0 | 0 | 0 | 0 | 127200 | 0 | 0 |
| EHLHIVK | LMAN1 | P49257 | LMAN1_HUMAI | Unmodified | 22837 | 52569 | 173170 | 0 | 0 | 0 | 0 | 0 | 3004900 | 179070 | 127630 | 0 | 0 | 0 | 157250 | 44555 | 0 | 0 |
| EHLHIVKR | LMAN1 | P49257 | LMAN1_HUMAI | Unmodified | 0 | 0 | 0 | 0 | 0 | 0 | 0 | 341540 | 2751000 | 47530 | 95090 | 59968 | 0 | 0 | 37493 | 12009 | 0 | 0 |
| EHLLKYEY | WDR1; WDR1 | O75083; D6I | WDR1_HUMAN | Unmodified | 407270 | 0 | 0 | 803670 | 19326000 | 3573600 | 1921200 | 337960 | 339530 | 519280 | 0 | 123960 | 7238400 | 7247300 | 0 | 0 | 2009700 | 3186600 |
| EHLLLAR | TUFM | P49411 | EFTU_HUMAN | Unmodified | 0 | 0 | 0 | 277410 | 0 | 0 | 0 | 0 | 3682300 | 97009 | 150630 | 0 | 58012 | 0 | 0 | 0 | 0 | 0 |
| EHLLTNHLI | RO60 | P10155 | RO60_HUMAN | Unmodified | 1613200 | 0 | 0 | 0 | 0 | 0 | 0 | 0 | 0 | 0 | 0 | 0 | 0 | 0 | 980330 | 0 | 0 | 0 |
| EHLTSEAAS | UACA; UACA; U | Q9BZF9; H0 | UACA_HUMAN | Unmodified | 0 | 0 | 0 | 448280 | 574030 | 0 | 0 | 0 | 0 | 0 | 0 | 0 | 0 | 285790 | 0 | 0 | 1338600 | 1476800 |
| EHLYFETVI | NOMO1; NOMO | Q15155; P69 | NOMO1_HUMA | Unmodified | 0 | 0 | 0 | 0 | 0 | 0 | 0 | 0 | 13185000 | 2241900 | 0 | 390940 | 986950 | 0 | 0 | 0 | 0 | 0 |
| EHMGDILY | ATP6V1A | P38606 | VATA_HUMAN | Unmodified | 0 | 0 | 0 | 0 | 0 | 0 | 0 | 0 | 0 | 0 | 0 | 388080 | 0 | 0 | 0 | 0 | 0 | 0 |
| EHMQPTHI | YWHAG | P61981 | 1433G_HUMAN | Oxidation (M) | 251290 | 0 | 696920 | 0 | 0 | 0 | 0 | 95981 | 2304000 | 2793200 | 76331 | 2644000 | 0 | 0 | 783900 | 1042600 | 0 | 0 |
| EHNELQQI | DAAM1 | Q9Y4D1 | DAAM1_HUMA | Unmodified | 0 | 0 | 0 | 223430 | 494810 | 0 | 0 | 108810 | 0 | 6806.5 | 96828 | 953090 | 743270 | 0 | 0 | 0 | 94804 | 130510 |
| EHNGHITG | ARPC1A; ARPC | Q92747; E9F | ARC1A_HUMAN | Unmodified | 165380 | 473990 | 0 | 238500 | 0 | 0 | 0 | 0 | 528730 | 61806 | 0 | 1996900 | 0 | 0 | 0 | 0 | 0 | 0 |
| EHNGQVT | ARPC1B; ARPC | O15143; C9J | ARC1B_HUMAI | Unmodified | 1687400 | 1015400 | 0 | 0 | 0 | 0 | 0 | 0 | 0 | 0 | 0 | 4628900 | 0 | 0 | 0 | 16218 | 0 | 0 |
| EHPEAPVQ | TMEM237; TME | Q96Q45; F2 | TM237_HUMAN | Unmodified | 0 | 0 | 115310 | 134210 | 1338600 | 175450 | 0 | 0 | 68858 | 0 | 76227 | 320120 | 0 | 0 | 0 | 13963 | 4647600 | 245230 |
| EHPVLALP | POFUT1 | Q9H488 | OFUT1_HUMAN | Unmodified | 0 | 0 | 0 | 0 | 0 | 0 | 0 | 0 | 23692000 | 5521700 | 300240 | 0 | 0 | 0 | 0 | 724750 | 0 | 0 |
| EHPWEVMI | RPSA; RPSA; RP | P08865; A0A | RSSA_HUMAN | Oxidation (M) | 0 | 0 | 0 | 0 | 0 | 0 | 0 | 0 | 0 | 0 | 0 | 5873600 | 0 | 0 | 0 | 0 | 0 | 0 |

| | | | | | | | | | | | | | | | | | | | | | | |
|---|---|---|---|---|---|---|---|---|---|---|---|---|---|---|---|---|---|---|---|---|---|---|
| EHQISPGD | EHD1; EHD1; EI | Q9H4M9; C9 | EHD1_HUMAN | Unmodified | 0 | 0 | 0 | 380340 | 0 | 0 | 0 | 0 | 234500 | 0 | 3629300 | 1370000 | 0 | 0 | 0 | 0 | 0 | 0 |
| EHQISPGD | EHD1; EHD1; EI | Q9H4M9; C9 | EHD1_HUMAN | Unmodified | 0 | 0 | 0 | 0 | 179990 | 0 | 0 | 0 | 705060 | 0 | 226640 | 988550 | 0 | 0 | 0 | 0 | 0 | 0 |
| EHQPFSLQ | ITGA5; ITGA5 | P08648; H0 | ITA5_HUMAN I | Unmodified | 619220 | 379220 | 369680 | 400300 | 312940 | 0 | 0 | 0 | 0 | 0 | 0 | 0 | 0 | 0 | 1485300 | 1620800 | 74880 | 0 |
| EHQQALD\ | CDC16; CDC16 | Q13042; A0/ | CDC16_HUMA | Oxidation (M) | 0 | 0 | 0 | 0 | 0 | 0 | 0 | 0 | 1555300 | 0 | 0 | 307030 | 0 | 0 | 0 | 122020 | 0 | 213050 |
| EHRNLLQE | SMC4; SMC4 | Q9NTJ3; E9F | SMC4_HUMAN | Unmodified | 0 | 0 | 0 | 0 | 374620 | 0 | 168770 | 45178 | 0 | 91780 | 57398 | 0 | 246490 | 51172 | 0 | 0 | 73498 | 66524 |
| EHRPHTAV | TRIM26; TRIM2 | Q12899; Q5 | TRI26_HUMAN | Unmodified | 1087300 | 1536800 | 2167300 | 1819000 | 0 | 0 | 0 | 0 | 0 | 0 | 0 | 0 | 0 | 0 | 2258300 | 1685400 | 0 | 0 |
| EHSLIKPY( | LMAN2; LMAN2 | Q12907; D6I | LMAN2_HUMAI | Oxidation (M) | 0 | 0 | 0 | 0 | 1134500 | 0 | 0 | 0 | 64662000 | 2963400 | 0 | 0 | 0 | 0 | 0 | 0 | 0 | 0 |
| EHSLIKPY( | LMAN2; LMAN2 | Q12907; D6I | LMAN2_HUMAI | 2 Oxidation (M) | 0 | 0 | 0 | 0 | 0 | 0 | 0 | 0 | 63993000 | 12059000 | 1203500 | 0 | 0 | 0 | 0 | 0 | 0 | 0 |
| EHSLIKPY( | LMAN2; LMAN2 | Q12907; D6I | LMAN2_HUMAI | Unmodified | 0 | 0 | 0 | 0 | 0 | 0 | 0 | 0 | 11162000 | 150860 | 0 | 0 | 0 | 0 | 0 | 0 | 0 | 0 |
| EHSLQVSP | KIF20A | O95235 | KI20A_HUMAN | Unmodified | 0 | 12846 | 259580 | 682660 | 5642600 | 0 | 0 | 264460 | 757700 | 718200 | 0 | 0 | 1326400 | 1907000 | 288110 | 165540 | 2759000 | 602230 |
| EHSRQIND | EIF3I; EIF3I; EIF | Q13347; A0/ | EIF3I_HUMAN I | 2 Oxidation (M) | 0 | 0 | 0 | 0 | 0 | 0 | 0 | 0 | 0 | 0 | 0 | 0 | 564990 | 0 | 0 | 0 | 0 | 0 |
| EHSRQIND | EIF3I; EIF3I; EIF | Q13347; A0/ | EIF3I_HUMAN I | Oxidation (M) | 0 | 0 | 0 | 0 | 1111200 | 0 | 0 | 0 | 0 | 0 | 0 | 0 | 1263500 | 867780 | 0 | 0 | 0 | 822050 |
| EHSSTANII | SLC12A2 | P55011 | S12A2_HUMAN | Oxidation (M) | 0 | 0 | 995420 | 0 | 0 | 0 | 0 | 0 | 0 | 0 | 0 | 1239100 | 0 | 0 | 1565500 | 1317700 | 0 | 0 |
| EHSSTANII | SLC12A2 | P55011 | S12A2_HUMAN | Oxidation (M) | 0 | 0 | 0 | 0 | 338930 | 0 | 0 | 0 | 329840 | 0 | 191360 | 0 | 397130 | 0 | 774320 | 470610 | 382850 | 2370600 |
| EHSSTANII | SLC12A2 | P55011 | S12A2_HUMAN | Unmodified | 0 | 0 | 0 | 0 | 764130 | 0 | 0 | 0 | 0 | 0 | 0 | 0 | 0 | 0 | 114120 | 0 | 236900 | 2111100 |
| EHSSVLVM | PDIA5 | Q14554 | PDIA5_HUMAN | Oxidation (M) | 0 | 0 | 0 | 0 | 0 | 0 | 0 | 0 | 934910 | 125100 | 0 | 0 | 0 | 0 | 0 | 0 | 0 | 0 |
| EHTALLK | RPL35A; RPL35. | P18077; F8V | RL35A_HUMAN | Unmodified | 833540 | 961360 | 461640 | 388320 | 0 | 0 | 0 | 0 | 0 | 0 | 0 | 7472500 | 0 | 0 | 518580 | 447840 | 0 | 0 |
| EHTAYYIK | UQCRC1 | P31930 | QCR1_HUMAN | Unmodified | 0 | 0 | 0 | 0 | 0 | 0 | 0 | 0 | 3586100 | 127140 | 0 | 0 | 0 | 0 | 0 | 0 | 0 | 0 |
| EHTDTFNF | PTGFRN | Q9P2B2 | FPRP_HUMAN | Unmodified | 3836600 | 4226000 | 46194000 | 10285000 | 304190 | 152040 | 0 | 0 | 0 | 0 | 0 | 18156000 | 1125400 | 278050 | 8971300 | 10258000 | 0 | 0 |
| EHVEEISEL | RUVBL1; RUVBI | Q9Y265; H7 | RUVB1_HUMAI | Unmodified | 923270 | 462030 | 1372990 | 878670 | 4154100 | 978910 | 870580 | 175190 | 1075200 | 85145 | 2128700 | 20186000 | 5068100 | 2241900 | 655410 | 916630 | 561740 | 2083100 |
| EHVIEALRF | RPL10; RPL10L; | P27635; Q9( | RL10_HUMAN | Unmodified | 354460 | 314820 | 0 | 99263 | 0 | 0 | 0 | 0 | 0 | 0 | 228370 | 2916700 | 0 | 0 | 263720 | 150750 | 0 | 0 |
| EHVIHIIDF | CSNK1G1; CSN | Q9HCP0; U3 | KC1G1_HUMAI | Unmodified | 334680 | 407140 | 398510 | 374110 | 583570 | 137560 | 0 | 0 | 328550 | 0 | 340430 | 0 | 42609 | 0 | 769350 | 454010 | 145450 | 0 |
| EHVKPLFN | BLMH; BLMH; B | Q13867; K7E | BLMH_HUMAN | Unmodified | 0 | 0 | 0 | 0 | 0 | 0 | 32834 | 99185 | 1095400 | 0 | 0 | 1692000 | 0 | 0 | 0 | 0 | 0 | 0 |
| EHVMNEVI | NUCB2; NUCB2 | P80303; H0 | NUCB2_HUMA | Oxidation (M) | 196180 | 0 | 0 | 0 | 0 | 0 | 0 | 0 | 318910 | 0 | 0 | 0 | 0 | 0 | 0 | 0 | 0 | 0 |
| EHVMNEVI | NUCB2; NUCB2 | P80303; H0 | NUCB2_HUMA | Unmodified | 0 | 0 | 221340 | 0 | 0 | 0 | 0 | 0 | 6873600 | 88780 | 161040 | 0 | 0 | 0 | 462350 | 0 | 0 | 0 |
| EHVMNEVI | NUCB2; NUCB2 | P80303; H0 | NUCB2_HUMA | Oxidation (M) | 0 | 16831 | 182840 | 0 | 0 | 0 | 0 | 0 | 335220 | 0 | 49870 | 0 | 0 | 0 | 0 | 0 | 0 | 0 |
| EHVMNEVI | NUCB2; NUCB2 | P80303; H0 | NUCB2_HUMA | Oxidation (M) | 0 | 0 | 0 | 181580 | 0 | 0 | 0 | 0 | 5284800 | 1473900 | 0 | 0 | 455720 | 494620 | 0 | 0 | 0 | 0 |
| EHVMNEVI | NUCB2; NUCB2 | P80303; H0 | NUCB2_HUMA | Unmodified | 0 | 0 | 0 | 0 | 556830 | 0 | 0 | 0 | 5905800 | 276900 | 0 | 0 | 0 | 0 | 0 | 0 | 0 | 0 |
| EHVNPPAE | MGAT4B; MGAT | Q9UQ53; H0 | MGT4B_HUMAI | Unmodified | 0 | 0 | 0 | 0 | 1586800 | 0 | 96253 | 127500 | 2518300 | 1649900 | 0 | 0 | 0 | 496680 | 0 | 0 | 0 | 5349800 |
| EHVNVVFI( | GSPT1; GSPT2; | P15170; Q8I | ERF3A_HUMAN | Unmodified | 0 | 0 | 0 | 410120 | 2937900 | 221750 | 189020 | 0 | 1226200 | 387090 | 1266800 | 5258900 | 2834100 | 954140 | 206890 | 272540 | 396100 | 1556000 |
| EHWQPQE | RXYLT1; RXYLT | Q9Y2B1; G3 | RXLT1_HUMAN | Unmodified | 0 | 0 | 0 | 0 | 0 | 0 | 27544 | 0 | 1439800 | 53914 | 0 | 166360 | 0 | 0 | 0 | 0 | 0 | 0 |
| EHYPNGV( | CAPZA2; CAPZ/ | P47755; A0A | CAZA2_HUMAN | Unmodified | 648800 | 0 | 0 | 0 | 638430 | 249420 | 0 | 0 | 195930 | 0 | 0 | 1577900 | 504320 | 0 | 0 | 0 | 0 | 465540 |
| EHYTSLWD | STEAP2; STEAP | Q8NFT2; C9. | STEA2_HUMAN | Unmodified | 0 | 0 | 0 | 0 | 0 | 0 | 0 | 153720 | 811490 | 0 | 0 | 1366200 | 0 | 0 | 1733300 | 908240 | 562110 | 0 |
| EHYVDLK | NME1; NME1-N | P15531; J3K | NDKA_HUMAN | Unmodified | 203360 | 0 | 0 | 0 | 2056000 | 0 | 223010 | 109150 | 386000 | 0 | 0 | 761820 | 3246100 | 121320 | 0 | 789410 | 357170 | 1202200 |
| EHYVDLKD | NME1; NME1-N | P15531; J3K | NDKA_HUMAN | Unmodified | 0 | 0 | 0 | 210080 | 39945000 | 4713800 | 7609400 | 0 | 0 | 2516900 | 0 | 0 | 25048000 | 15451000 | 0 | 0 | 23499000 | 25145000 |
| EIADGLCLE | TPT1; TPT1; TPT | P13693; E9P | TCTP_HUMAN | Unmodified | 0 | 0 | 0 | 0 | 0 | 0 | 0 | 0 | 0 | 0 | 0 | 0 | 0 | 0 | 0 | 1961300 | 0 | 0 |
| EIADKTEKY | TAX1BP1; TAX1 | Q86VP1; B8 | TAXB1_HUMAN | Unmodified | 0 | 0 | 0 | 0 | 57781 | 0 | 592930 | 95605 | 0 | 29944 | 0 | 0 | 0 | 0 | 0 | 0 | 230550 | 98145 |
| EIADLGEAL | AARS1; AARS1; | P49588; A0A | SYAC_HUMAN | Unmodified | 0 | 0 | 0 | 0 | 0 | 0 | 0 | 0 | 248120 | 0 | 169220 | 602960 | 0 | 0 | 0 | 0 | 0 | 0 |
| EIADYLAAC | IST1; IST1; IST1 | P53990; H3E | IST1_HUMAN I | Unmodified | 806600 | 717560 | 1770600 | 2257300 | 0 | 0 | 0 | 0 | 7075100 | 0 | 2587300 | 32772000 | 197500 | 0 | 854980 | 1989400 | 0 | 0 |
| EIADYLAAC | IST1; IST1; IST1 | P53990; H3E | IST1_HUMAN I | Unmodified | 0 | 0 | 0 | 0 | 0 | 0 | 0 | 0 | 410590 | 457380 | 536380 | 0 | 0 | 0 | 0 | 0 | 0 | 0 |
| EIAEAYLGC | HSPA2 | P54652 | HSP72_HUMAI | Unmodified | 466090 | 440820 | 0 | 0 | 0 | 0 | 0 | 0 | 0 | 0 | 0 | 0 | 0 | 0 | 0 | 0 | 0 | 0 |
| EIAEAYLGK | HSPA8; HSPA8; | P11142; E9P | HSP7C_HUMAI | Unmodified | 29353000 | 30042000 | 52747000 | 34052000 | 72255000 | 10642000 | 29075000 | 11626000 | 76940000 | 26698000 | 13494000 | 81869000 | 88758000 | 55118000 | 34198000 | 35032000 | 17413000 | 22025000 |
| EIAEAYLGY | HSPA1A; HSPA | P0DMV8; P0 | HS71A_HUMAI | Unmodified | 0 | 0 | 0 | 0 | 4529900 | 0 | 1604800 | 0 | 0 | 3212600 | 0 | 0 | 0 | 0 | 0 | 0 | 1085800 | 0 |
| EIAENALGK | HNRNPH3 | P31942 | HNRH3_HUMA | Unmodified | 491540 | 389790 | 292260 | 0 | 332040 | 184190 | 650060 | 593320 | 0 | 217920 | 301330 | 2059900 | 538120 | 211040 | 310000 | 242390 | 102890 | 450090 |
| EIAENITK | RARS1 | P54136 | SYRC_HUMAN | Unmodified | 73707 | 0 | 141390 | 63655 | 181220 | 0 | 116970 | 53355 | 57184 | 84635 | 19304 | 1616300 | 345850 | 118220 | 74970 | 65002 | 40904 | 137630 |
| EIAETRFEV | TPR | P12270 | TPR_HUMAN N | Unmodified | 0 | 0 | 0 | 0 | 0 | 0 | 0 | 0 | 144740 | 0 | 0 | 598480 | 0 | 0 | 0 | 0 | 0 | 0 |
| EIAFLSEK | IKBIP | Q70UQ0 | IKIP_HUMAN Ir | Unmodified | 0 | 0 | 0 | 0 | 0 | 0 | 60338 | 0 | 720450 | 261770 | 0 | 0 | 0 | 0 | 0 | 0 | 0 | 0 |
| EIAFPKAEE | STAMBP; STAMI | O95630; C9. | STABP_HUMAN | Unmodified | 0 | 0 | 0 | 0 | 2561000 | 0 | 0 | 0 | 0 | 763020 | 0 | 0 | 2346100 | 1475600 | 0 | 0 | 0 | 0 |
| EIAKLDDFL | UBLCP1 | Q8WVY7 | UBCP1_HUMAI | Unmodified | 0 | 0 | 0 | 0 | 0 | 0 | 0 | 0 | 0 | 0 | 0 | 1999600 | 0 | 0 | 0 | 0 | 0 | 0 |
| EIAQDFK | H3-4; H3-3A; H | Q16695; P84 | H31T_HUMAN | Unmodified | 89520000 | 55612000 | 1616400 | 223490 | 239530 | 267080 | 402560 | 868510 | 8620500 | 1171500 | 4801100 | 9.54E+08 | 123090 | 0 | 271290 | 374860 | 158420 | 525470 |
| EIAQDFKTE | H3-4; H3-3A; H | Q16695; P84 | H31T_HUMAN | Unmodified | 1.03E+08 | 34853000 | 0 | 305170 | 0 | 0 | 0 | 107220 | 2863800 | 0 | 1619800 | 6.34E+08 | 42949 | 0 | 0 | 0 | 0 | 80460 |
| EIAQEFKTD | H3-2; H3-2 | Q5TEC6; A0/ | H3PS2_HUMAI | Unmodified | 80331000 | 5612500 | 512880 | 165020 | 0 | 0 | 0 | 0 | 0 | 0 | 732220 | 5.33E+08 | 70721 | 0 | 148490 | 279770 | 0 | 0 |
| EIARAVVEK | CUTA; CUTA; C | O60888; H0 | CUTA_HUMAN | Unmodified | 0 | 0 | 0 | 0 | 1247400 | 242820 | 0 | 0 | 0 | 462800 | 0 | 0 | 611460 | 447170 | 0 | 0 | 1801100 | 1246500 |
| EICDHQAE | SPECC1; SPEC | Q5M775; J3 | CYTSB_HUMAN | Unmodified | 0 | 0 | 0 | 0 | 180080 | 0 | 0 | 0 | 0 | 0 | 0 | 0 | 112460 | 270150 | 0 | 0 | 0 | 185900 |
| EICELTGID | MYO1B; MYO1E | O43795; E9F | MYO1B_HUMA | Unmodified | 361360 | 0 | 0 | 23016 | 904000 | 0 | 0 | 0 | 530330 | 0 | 2024300 | 10210000 | 0 | 0 | 0 | 316460 | 264960 | 0 |
| EIDDSIGK | GALNS | P34059 | GALNS_HUMAN | Unmodified | 0 | 0 | 83361 | 0 | 0 | 0 | 9246.2 | 0 | 239630 | 0 | 0 | 0 | 0 | 0 | 0 | 0 | 0 | 0 |
| EIDLVNRDI | CAV3; CAV1; C/ | P56539; Q0 | CAV3_HUMAN | Unmodified | 0 | 0 | 0 | 0 | 2414500 | 284530 | 88769 | 0 | 0 | 0 | 0 | 0 | 0 | 0 | 0 | 0 | 0 | 368200 |
| EIDSVKYLE | RAC2; RAC2; R/ | P15153; B1/ | RAC2_HUMAN | Unmodified | 0 | 0 | 0 | 0 | 0 | 0 | 1055500 | 0 | 0 | 0 | 0 | 0 | 0 | 0 | 0 | 0 | 583440 | 0 |
| EIDVSCVK | EPHB3 | P54753 | EPHB3_HUMAI | Unmodified | 0 | 0 | 0 | 0 | 0 | 0 | 0 | 0 | 0 | 0 | 0 | 656330 | 0 | 0 | 0 | 0 | 0 | 0 |
| EIEALNFK | MYO5C | Q9NQX4 | MYO5C_HUMAN | Unmodified | 0 | 0 | 0 | 0 | 0 | 0 | 0 | 0 | 0 | 0 | 0 | 0 | 0 | 0 | 0 | 0 | 0 | 0 |
| EIEDAEKYS | AHCYL1 | O43865 | SAHH2_HUMA | Oxidation (M) | 0 | 0 | 0 | 0 | 0 | 0 | 0 | 0 | 0 | 0 | 0 | 0 | 0 | 0 | 0 | 0 | 0 | 459040 |
| EIEDAEKYS | AHCYL1 | O43865 | SAHH2_HUMA | Unmodified | 0 | 0 | 0 | 0 | 3466300 | 0 | 0 | 0 | 0 | 0 | 0 | 0 | 0 | 305570 | 0 | 0 | 286720 | 648510 |
| EIEDPEDR | CANX; CANX; C | P27824; A0A | CALX_HUMAN | Unmodified | 0 | 0 | 0 | 0 | 0 | 0 | 61345 | 123780 | 7467900 | 1077200 | 466310 | 420170 | 0 | 0 | 0 | 0 | 0 | 0 |
| EIEELKQEL | AIMP1; AIMP1; | Q12904; A0/ | AIMP1_HUMAN | Unmodified | 0 | 0 | 0 | 0 | 670040 | 0 | 0 | 0 | 0 | 0 | 0 | 3477300 | 951310 | 0 | 0 | 0 | 0 | 0 |
| EIEELYANP | MGAM2 | Q2M2H8 | MGAL_HUMAN | Unmodified | 0 | 0 | 0 | 0 | 0 | 0 | 0 | 0 | 0 | 0 | 602240 | 0 | 0 | 0 | 0 | 0 | 0 | 0 |
| EIEEYELLH | ATP11C; ATP11 | Q8NB49; A0. | AT11C_HUMAN | Unmodified | 120490 | 130940 | 363690 | 1120100 | 0 | 0 | 0 | 0 | 1093000 | 329490 | 833010 | 1248500 | 89064 | 40845 | 1524600 | 536850 | 0 | 0 |
| EIEEYELLH | ATP11C; ATP11 | Q8NB49; A0. | AT11C_HUMAN | Unmodified | 0 | 0 | 218630 | 325230 | 0 | 0 | 0 | 0 | 734740 | 67648 | 159010 | 442280 | 0 | 29172 | 288630 | 0 | 0 | 0 |
| EIEIAEQDM | AHCYL1 | O43865 | SAHH2_HUMAI | Unmodified | 2840100 | 1228200 | 0 | 2130100 | 4723200 | 1361300 | 0 | 0 | 0 | 791260 | 0 | 0 | 1354100 | 1237900 | 1184600 | 514170 | 279310 | 0 |
| EIEIGVPNA | SPATA5; SPATA | Q8NB90; A0. | AFG2H_HUMAI | Unmodified | 44723 | 0 | 0 | 0 | 0 | 0 | 67030 | 0 | 106370 | 35636 | 109550 | 925140 | 0 | 0 | 0 | 0 | 0 | 0 |
| EIEILRNLYI | JAK1; JAK1; JAK | P23458; A0A | JAK1_HUMAN T | Unmodified | 0 | 0 | 0 | 0 | 1501600 | 0 | 0 | 0 | 0 | 0 | 0 | 0 | 0 | 0 | 0 | 0 | 0 | 0 |
| EIEKKLEEE( | SCP2; SCP2 | P22307; H0 | SCP2_HUMAN | Unmodified | 0 | 0 | 0 | 0 | 0 | 0 | 0 | 0 | 428770 | 2297000 | 0 | 0 | 0 | 0 | 0 | 0 | 0 | 0 |
| EIELENDLK | TBCE; TBCE; TB | Q15813; A0/ | TBCE_HUMAN | Unmodified | 0 | 0 | 0 | 0 | 0 | 0 | 0 | 0 | 0 | 0 | 0 | 978780 | 0 | 0 | 0 | 0 | 0 | 0 |
| EIEMHSINI | KDSR; KDSR; KI | Q06136; K7E | KDSR_HUMAN | Unmodified | 0 | 0 | 0 | 0 | 0 | 0 | 0 | 0 | 666260 | 0 | 0 | 0 | 0 | 0 | 0 | 0 | 0 | 0 |
| EIENEDMR | DIAPH1; DIAPH | O60610; A0/ | DIAP1_HUMAN | Unmodified | 0 | 88054 | 0 | 0 | 0 | 0 | 0 | 0 | 0 | 0 | 0 | 960390 | 0 | 0 | 219080 | 0 | 72537 | 0 |
| EIENLPLR | STEAP2; STEAP | Q8NFT2; B5I | STEA2_HUMAN | Unmodified | 669340 | 1005000 | 4036700 | 1474600 | 0 | 0 | 120600 | 108830 | 0 | 0 | 0 | 0 | 0 | 0 | 21711000 | 19724000 | 0 | 0 |
| EIENMILNE | ACOX1; ACOX1 | Q15067; I3L | ACOX1_HUMAI | Oxidation (M) | 0 | 0 | 0 | 0 | 573710 | 0 | 0 | 0 | 3478200 | 388860 | 0 | 0 | 0 | 0 | 0 | 0 | 293500 | 383530 |
| EIENMILNE | ACOX1; ACOX1 | Q15067; I3L | ACOX1_HUMAI | Unmodified | 0 | 0 | 0 | 0 | 0 | 0 | 0 | 0 | 1677100 | 0 | 0 | 0 | 0 | 0 | 0 | 0 | 0 | 0 |
| EIEPEPDFII | SMPDL3B; SMP | Q92485; Q5 | ASM3B_HUMAI | Unmodified | 4417500 | 0 | 8201000 | 6509800 | 224120 | 0 | 0 | 0 | 7735500 | 235910 | 3147700 | 5941400 | 836410 | 40845 | 31048000 | 11778000 | 0 | 25975 |
| EIEQEAAVE | UQCRFS1 | P47985 | UCRI_HUMAN | Unmodified | 0 | 0 | 0 | 0 | 0 | 0 | 0 | 0 | 7008100 | 211870 | 328320 | 1101000 | 0 | 0 | 0 | 0 | 0 | 0 |
| EIEQEAAVE | UQCRFS1 | P47985 | UCRI_HUMAN | Unmodified | 0 | 0 | 0 | 0 | 0 | 0 | 0 | 0 | 24603000 | 0 | 0 | 0 | 0 | 0 | 0 | 0 | 531230 | 0 |
| EIESSPQYF | ATXN2L; ATXN2 | Q8WWM7; F | ATX2L_HUMAN | Unmodified | 580350 | 177300 | 0 | 0 | 475020 | 0 | 0 | 0 | 423220 | 282920 | 0 | 0 | 0 | 0 | 0 | 0 | 0 | 0 |
| EIETLVEEK | P3H1; P3H1 | Q32P28; H7 | P3H1_HUMAN | Unmodified | 0 | 0 | 0 | 0 | 0 | 0 | 0 | 0 | 4747200 | 1396500 | 0 | 0 | 0 | 0 | 0 | 0 | 0 | 0 |
| EIETNLRSG | UTRN | P46939 | UTRN_HUMAN | Unmodified | 0 | 0 | 0 | 0 | 2151200 | 726820 | 0 | 0 | 136960 | 814220 | 0 | 0 | 0 | 914240 | 0 | 0 | 2055300 | 1498500 |
| EIEVDSSPS | RAP2C; RAP2A; | Q9Y3L5; P1 | RAP2C_HUMAI | Oxidation (M) | 109350 | 540510 | 73931 | 603240 | 0 | 0 | 0 | 0 | 0 | 0 | 0 | 0 | 0 | 906670 | 2120900 | 77789 | 0 | 0 |
| EIEVLERGD | PALM | O75781 | PALM_HUMAN | Unmodified | 0 | 0 | 0 | 0 | 2101800 | 421130 | 0 | 20111 | 248400 | 0 | 0 | 0 | 30554 | 0 | 0 | 0 | 0 | 411930 |
| EIEVLNK | OLA1; OLA1; OL | Q9NTK5; J3K | OLA1_HUMAN | Unmodified | 0 | 0 | 0 | 0 | 0 | 0 | 0 | 0 | 0 | 0 | 0 | 1842200 | 0 | 0 | 0 | 0 | 0 | 0 |
| EIEYVETVT | BAIAP2L1 | Q9UHR4 | BI2L1_HUMAN | Unmodified | 0 | 0 | 305020 | 349830 | 0 | 0 | 0 | 0 | 1607100 | 0 | 0 | 779590 | 337310 | 373640 | 491580 | 842260 | 0 | 0 |
| EIEYVETVT | BAIAP2L1 | Q9UHR4 | BI2L1_HUMAN | Unmodified | 0 | 0 | 0 | 0 | 5633700 | 2006800 | 0 | 0 | 0 | 0 | 0 | 0 | 1365400 | 1000900 | 0 | 0 | 0 | 0 |
| EIFAQEALA | CHDH | Q8NE62 | CHDH_HUMAN | Unmodified | 0 | 0 | 0 | 0 | 0 | 0 | 0 | 0 | 0 | 0 | 0 | 0 | 0 | 0 | 0 | 0 | 0 | 0 |
| EIFAQEALA | CHDH | Q8NE62 | CHDH_HUMAN | Unmodified | 0 | 0 | 0 | 0 | 0 | 0 | 0 | 0 | 1184200 | 0 | 0 | 0 | 0 | 0 | 0 | 0 | 0 | 0 |
| EIFDSRGNI | ENO1; ENO1; E | P06733; K7E | ENOA_HUMAN | Unmodified | 2515200 | 2034100 | 3387100 | 19177000 | 1.04E+08 | 11241000 | 1309900 | 63195 | 43159000 | 33092000 | 19496000 | 60440000 | 69978000 | 17507000 | 9620900 | 5490600 | 25058000 | 29278000 |
| EIFEQPESV | GFPT1; GFPT1; | Q06210; A0/ | GFPT1_HUMAN | Unmodified | 0 | 0 | 0 | 322800 | 0 | 0 | 0 | 0 | 0 | 268470 | 0 | 0 | 1053800 | 0 | 0 | 0 | 0 | 0 |
| EIFEQPESV | GFPT1; GFPT1; | Q06210; A0/ | GFPT1_HUMAN | Oxidation (M) | 0 | 0 | 0 | 0 | 0 | 0 | 0 | 0 | 147630 | 0 | 0 | 0 | 364480 | 0 | 106200 | 128740 | 0 | 0 |
| EIFFNILVP( | OS9; OS9 | Q13438; B4E | OS9_HUMAN P | Unmodified | 0 | 0 | 0 | 0 | 0 | 0 | 0 | 0 | 1978100 | 683830 | 0 | 0 | 0 | 0 | 0 | 0 | 28172 | 0 |
| EIFLRELISN | HSP90B1; HSP | P14625; P08 | ENPL_HUMAN | Unmodified | 177070 | 135610 | 384750 | 2942600 | 21180000 | 6815300 | 10120000 | 29399 | 33835000 | 1.3E+08 | 9922900 | 639030 | 28186000 | 17984000 | 685970 | 144750 | 4667300 | 9595400 |
| EIFLRELISN | HSP90B1; HSP | P14625; P08 | ENPL_HUMAN | Unmodified | 0 | 0 | 0 | 0 | 88159 | 0 | 627430 | 0 | 184120 | 10311000 | 0 | 0 | 237160 | 302140 | 0 | 0 | 41824 | 0 |
| EIFLRELISN | HSP90AA2P; H | Q14568; P0 | HS902_HUMAI | Unmodified | 0 | 136940 | 327030 | 2085600 | 16014000 | 5172800 | 4180400 | 175910 | 7534800 | 8630100 | 7587000 | 1563800 | 3550200 | 3752500 | 483760 | 244130 | 7013000 | 10681000 |
| EIFVDFAPF | NBEA; LRBA; LF | Q8NFP9; P5 | NBEA_HUMAN | Unmodified | 0 | 0 | 0 | 0 | 0 | 0 | 0 | 0 | 0 | 85104 | 0 | 686340 | 6628.7 | 0 | 0 | 0 | 0 | 0 |
| EIGALLQQI | TRNT1 | Q96Q11 | TRNT1_HUMAN | Unmodified | 0 | 0 | 0 | 0 | 0 | 0 | 0 | 0 | 0 | 0 | 0 | 678350 | 0 | 0 | 0 | 0 | 0 | 0 |
| EIGAVKYLE | RAC1 | P63000 | RAC1_HUMAN | Unmodified | 0 | 0 | 0 | 0 | 536370 | 0 | 207580 | 0 | 0 | 68601 | 0 | 0 | 1205100 | 1377000 | 0 | 0 | 1468600 | 846480 |
| EIGAVKYLE | RAC1 | P63000 | RAC1_HUMAN | Unmodified | 0 | 0 | 0 | 0 | 3390900 | 248330 | 2849400 | 153290 | 0 | 47999 | 0 | 124410 | 2460900 | 4272900 | 0 | 0 | 4558400 | 4519000 |
| EIGDLRAQ | DLG5 | Q8TDM6 | DLG5_HUMAN | Unmodified | 0 | 0 | 0 | 0 | 892860 | 0 | 0 | 0 | 0 | 0 | 0 | 0 | 0 | 0 | 0 | 0 | 518990 | 0 |
| EIGELTQLK | RSU1 | Q15404 | RSU1_HUMAN | Unmodified | 15947 | 0 | 0 | 204570 | 333510 | 564750 | 4422900 | 3214400 | 678770 | 206760 | 0 | 0 | 276390 | 5080900 | 120450 | 68818 | 686660 | 0 |
| EIGHGSFG | TAOK1; TAOK2; | Q7L7X3; Q9 | TAOK1_HUMAN | Unmodified | 412550 | 379840 | 451180 | 324490 | 0 | 0 | 0 | 0 | 0 | 0 | 0 | 2539500 | 0 | 0 | 721820 | 597020 | 0 | 0 |
| EIGILTTNK | ABI1; ABI2; ABI | Q8IZP0; Q9 | ABI1_HUMAN / | Unmodified | 395450 | 529830 | 858970 | 1155500 | 2409800 | 676770 | 135790 | 111550 | 1552000 | 243540 | 1299700 | 5956200 | 1170500 | 610690 | 1173500 | 894160 | 397830 | 1364900 |
| EIGILTTNKI | ABI1; ABI2; ABI | Q8IZP0; Q9 | ABI1_HUMAN / | Unmodified | 0 | 0 | 0 | 0 | 172560 | 45356 | 0 | 0 | 50565 | 0 | 0 | 0 | 0 | 0 | 0 | 0 | 413230 | 0 |
| EIGINEDQF | CFAP36; CFAP3 | Q96G28; B5 | CFA36_HUMAN | Unmodified | 351860 | 0 | 0 | 0 | 0 | 0 | 0 | 0 | 0 | 0 | 0 | 0 | 348710 | 0 | 0 | 0 | 0 | 0 |
| EIGNIISDAI | HSPD1; HSPD1 | P10809; A0A | CH60_HUMAN | Unmodified | 0 | 0 | 0 | 0 | 0 | 0 | 10904 | 0 | 6217800 | 0 | 777970 | 0 | 0 | 0 | 0 | 0 | 0 | 0 |
| EIGNIISDAI | HSPD1; HSPD1 | P10809; A0A | CH60_HUMAN | Unmodified | 0 | 0 | 0 | 0 | 0 | 0 | 0 | 0 | 4532500 | 229480 | 130480 | 0 | 0 | 0 | 0 | 0 | 0 | 0 |

| | | | | | | | | | | | | | | | | | | | | | | |
|---|---|---|---|---|---|---|---|---|---|---|---|---|---|---|---|---|---|---|---|---|---|---|
| EIGNLLHPS | SARS1; SARS1 | P49591; Q5T | SYSC_HUMAN | Unmodified | 0 | 0 | 0 | 321580 | 0 | 302110 | 0 | 0 | 1555100 | 0 | 763990 | 12110000 | 0 | 0 | 982530 | 2011700 | 245940 | 0 |
| EIGQSVDEY | SPTBN1; SPTBN | Q01082; A0A | SPTB2_HUMAN | Unmodified | 0 | 0 | 0 | 0 | 0 | 0 | 0 | 0 | 0 | 0 | 472540 | 2451300 | 0 | 0 | 0 | 0 | 0 | 0 |
| EIGSSTSSS | FRMD6 | Q96NE9 | FRMD6_HUMA | Unmodified | 0 | 0 | 0 | 0 | 541280 | 0 | 0 | 0 | 0 | 0 | 0 | 0 | 0 | 0 | 0 | 0 | 570150 | 2118900 |
| EIGTHKPLF | ACOX1 | Q15067 | ACOX1_HUMA | Unmodified | 0 | 0 | 0 | 0 | 0 | 0 | 0 | 0 | 385960 | 0 | 602120 | 0 | 0 | 0 | 0 | 0 | 53466 | 177810 |
| EIGWTDVG | PFKP; PFKP; PF | Q01813; A0A | PFKAP_HUMAN | Unmodified | 0 | 0 | 0 | 0 | 3753700 | 1114500 | 144420 | 0 | 790590 | 409050 | 1039900 | 0 | 0 | 0 | 0 | 0 | 1980000 | 2262100 |
| EIHDFVLEC | PI4KA | P42356 | PI4KA_HUMAN | Unmodified | 0 | 0 | 340750 | 0 | 0 | 0 | 0 | 0 | 0 | 0 | 0 | 869030 | 160810 | 0 | 401800 | 394260 | 0 | 0 |
| EIHKPGYLA | RDX; RDX; RDX; | P35241; E9P | RADI_HUMAN I | Unmodified | 489990 | 285940 | 1052700 | 716660 | 455810 | 32203 | 83536 | 0 | 681950 | 0 | 86704 | 5408400 | 0 | 0 | 2281300 | 1963100 | 0 | 0 |
| EIHKPGYLA | RDX; RDX; RDX; | P35241; E9P | RADI_HUMAN I | Unmodified | 538600 | 439340 | 601990 | 1346000 | 0 | 1181600 | 553570 | 0 | 1217600 | 0 | 293600 | 4801300 | 0 | 0 | 1226100 | 981560 | 0 | 0 |
| EIHPFIDDY | POLR1A | O95602 | RPA1_HUMAN | Unmodified | 0 | 0 | 0 | 0 | 1475200 | 0 | 0 | 0 | 16961000 | 14821000 | 0 | 0 | 383820 | 0 | 0 | 0 | 0 | 0 |
| EIHQDWAN | BRK1 | Q8WUW1 | BRK1_HUMAN | Unmodified | 0 | 0 | 0 | 570820 | 0 | 0 | 417240 | 0 | 608710 | 0 | 0 | 814800 | 688710 | 0 | 727960 | 698270 | 0 | 164770 |
| EIHQQRQE | SLC38A10; SLC | Q9HBR0; H0 | S38AA_HUMAN | Unmodified | 0 | 0 | 0 | 0 | 0 | 0 | 0 | 87321 | 4415500 | 0 | 0 | 0 | 0 | 0 | 0 | 0 | 0 | 0 |
| EIIDLVLDR | TUBA1A; TUBA1 | Q71U36; Q9 | TBA1A_HUMAN | Unmodified | 2568000 | 1941100 | 3658100 | 1280000 | 703380 | 915230 | 791110 | 317980 | 5535000 | 2498900 | 7221700 | 52419000 | 667180 | 279860 | 5753700 | 4699200 | 1410200 | 0 |
| EIIDLVLDRI | TUBA1A; TUBA1 | Q71U36; Q9 | TBA1A_HUMAN | Unmodified | 1491800 | 0 | 895460 | 0 | 0 | 0 | 829920 | 0 | 2290000 | 1787600 | 667460 | 19375000 | 0 | 0 | 1727700 | 1405300 | 258530 | 0 |
| EIIDPVLDR | TUBA4A; TUBA4 | P68366; C9J | TBA4A_HUMAN | Unmodified | 0 | 0 | 3320000 | 2267000 | 1053800 | 232800 | 0 | 0 | 0 | 380150 | 346850 | 8837000 | 135070 | 0 | 3301100 | 4682100 | 0 | 0 |
| EIIDPVLDRI | TUBA4A; TUBA4 | P68366; C9J | TBA4A_HUMAN | Unmodified | 626140 | 452200 | 786980 | 918660 | 296580 | 0 | 996790 | 0 | 0 | 585530 | 0 | 2867500 | 0 | 0 | 546890 | 511450 | 0 | 497020 |
| EIIHELEK | BAIAP2L1 | Q9UHR4 | BI2L1_HUMAN | Unmodified | 31779 | 0 | 0 | 0 | 1877300 | 0 | 0 | 0 | 513710 | 0 | 0 | 0 | 587850 | 931120 | 0 | 76270 | 101590 | 202880 |
| EIIIEERPGQ | MRI1 | Q9BV20 | MTNA_HUMAN | Unmodified | 0 | 0 | 0 | 93219 | 0 | 0 | 0 | 0 | 684650 | 0 | 377960 | 1238800 | 0 | 0 | 131550 | 0 | 0 | 0 |
| EIILDDDEC | AFDN; AFDN; A | P55196; Q5T | AFAD_HUMAN | Unmodified | 0 | 63489 | 0 | 259950 | 650700 | 0 | 0 | 0 | 146250 | 0 | 175660 | 0 | 1105900 | 1800800 | 0 | 0 | 0 | 0 |
| EIILVDDYSI | GALNT2 | Q10471 | GALT2_HUMAN | Unmodified | 0 | 268250 | 0 | 0 | 0 | 0 | 130670 | 24699 | 8557900 | 2471200 | 728260 | 1046500 | 391020 | 0 | 0 | 0 | 0 | 0 |
| EIILVDDYSI | GALNT2 | Q10471 | GALT2_HUMAN | Unmodified | 0 | 0 | 0 | 0 | 2581400 | 0 | 0 | 0 | 85615000 | 24881000 | 1640500 | 2244400 | 0 | 0 | 0 | 0 | 0 | 0 |
| EIILVDDYSI | GALNT2 | Q10471 | GALT2_HUMAN | Unmodified | 0 | 0 | 0 | 0 | 0 | 0 | 0 | 0 | 0 | 684670 | 0 | 0 | 0 | 465520 | 0 | 0 | 0 | 0 |
| EIISFGSGY | PCK1; PCK2; PC | P35558; Q16 | PCKGC_HUMA | Unmodified | 0 | 0 | 0 | 0 | 0 | 0 | 0 | 0 | 2421900 | 374520 | 0 | 0 | 0 | 0 | 0 | 0 | 0 | 0 |
| EIISNLAK | SF3B1 | O75533 | SF3B1_HUMAN | Unmodified | 0 | 0 | 186440 | 0 | 1790800 | 121790 | 274850 | 0 | 2662800 | 0 | 0 | 0 | 0 | 0 | 0 | 232770 | 266220 | 60950 |
| EIKDAISGI | ANXA6; ANXA6; | P08133; E5R | ANXA6_HUMAN | Unmodified | 77545 | 0 | 0 | 1022800 | 5879100 | 1421300 | 5338100 | 2075400 | 6207400 | 16550000 | 0 | 0 | 4892400 | 2645200 | 0 | 0 | 2773200 | 7451800 |
| EIKDILIQYD | RPS16; RPS16 | P62249; M0R | RS16_HUMAN | Unmodified | 0 | 0 | 62381 | 1144400 | 0 | 0 | 0 | 0 | 0 | 0 | 0 | 14271000 | 0 | 0 | 0 | 0 | 0 | 0 |
| EIKILNIFGV | TFRC; TFRC | P02786; G3V | TFR1_HUMAN | Unmodified | 0 | 0 | 0 | 0 | 0 | 0 | 137500 | 0 | 50718 | 90383 | 0 | 0 | 0 | 0 | 0 | 0 | 113830 | 217940 |
| EIKLENLEP | PTPRC; PTPRC; | P08575; E9P | PTPRC_HUMAN | Unmodified | 0 | 0 | 0 | 0 | 272640 | 207370 | 0 | 0 | 0 | 273120 | 0 | 0 | 377980 | 449540 | 0 | 0 | 0 | 0 |
| EIKNELLQA | BAG5 | Q9UL15 | BAG5_HUMAN | Unmodified | 0 | 0 | 0 | 0 | 0 | 0 | 0 | 0 | 1275700 | 0 | 0 | 0 | 0 | 0 | 784350 | 512900 | 0 | 0 |
| EIKTVESITD | LAMP1 | P11279 | LAMP1_HUMAN | Unmodified | 0 | 0 | 0 | 0 | 1431600 | 0 | 3174000 | 0 | 0 | 14703000 | 0 | 0 | 3085500 | 5613800 | 0 | 0 | 0 | 0 |
| EILDAFDK | PSAP; PSAP; PS | P07602; A0A | SAP_HUMAN P | Unmodified | 837230 | 425760 | 3498400 | 2371700 | 2348800 | 1000500 | 646100 | 0 | 2.25E+08 | 22257000 | 6573800 | 1770100 | 4693300 | 4142600 | 2338200 | 1097800 | 92227 | 1079900 |
| EILDESLRL | PDCD6IP | Q8WUM4 | PDC6I_HUMAN | Unmodified | 0 | 2849500 | 0 | 2588800 | 0 | 0 | 0 | 0 | 0 | 0 | 0 | 0 | 0 | 0 | 2222700 | 0 | 0 | 0 |
| EILDESLRL | PDCD6IP | Q8WUM4 | PDC6I_HUMAN | Unmodified | 15750000 | 20409000 | 0 | 13477000 | 5057100 | 0 | 0 | 0 | 7253100 | 2786400 | 0 | 0 | 11445000 | 6029900 | 5936900 | 0 | 0 | 0 |
| EILDKFTEE | ALDH9A1 | P49189 | AL9A1_HUMAN | Unmodified | 0 | 0 | 0 | 0 | 0 | 96817 | 0 | 0 | 1725700 | 214900 | 0 | 1247800 | 0 | 509550 | 0 | 0 | 203270 | 0 |
| EILDKFTEE | ALDH9A1 | P49189 | AL9A1_HUMAN | Unmodified | 0 | 0 | 0 | 0 | 805340 | 0 | 0 | 0 | 0 | 286480 | 0 | 0 | 1113100 | 465810 | 0 | 0 | 238050 | 0 |
| EILDKKVEK | HSP90AB1; HSI | P08238; Q58 | HS90B_HUMAN | Unmodified | 0 | 0 | 0 | 0 | 2877000 | 0 | 0 | 161510 | 0 | 678480 | 0 | 0 | 0 | 534730 | 0 | 0 | 1109400 | 1846400 |
| EILDSRGNI | ENO2; ENO2; E | P09104; F5H | ENOG_HUMAN | Unmodified | 0 | 0 | 0 | 0 | 0 | 0 | 901960 | 69779 | 1192800 | 1154000 | 0 | 0 | 94150 | 0 | 0 | 0 | 1607200 | 0 |
| EILEFPARD | DOCK7; DOCK7 | Q96N67; A0A | DOCK7_HUMA | Unmodified | 73224 | 42268 | 0 | 307290 | 0 | 0 | 0 | 23288 | 0 | 227400 | 0 | 0 | 0 | 0 | 166110 | 180650 | 0 | 0 |
| EILEQGLFS | HTATIP2; HTATI | Q9BUP3; E9I | HTAI2_HUMAN | Unmodified | 0 | 0 | 0 | 1260900 | 0 | 0 | 464330 | 0 | 10707000 | 0 | 706460 | 0 | 0 | 611040 | 0 | 0 | 0 | 986200 |
| EILEQGLFS | HTATIP2; HTATI | Q9BUP3; E9I | HTAI2_HUMAN | Unmodified | 0 | 0 | 0 | 0 | 495230 | 48234 | 416640 | 0 | 92100 | 1753400 | 0 | 0 | 168640 | 248110 | 0 | 0 | 475440 | 137040 |
| EILEQGLFS | HTATIP2; HTATI | Q9BUP3; E9I | HTAI2_HUMAN | Unmodified | 0 | 0 | 0 | 0 | 452700 | 0 | 144290 | 0 | 0 | 916320 | 0 | 0 | 214370 | 201370 | 0 | 0 | 207330 | 257390 |
| EILETLIK | GFPT1; GFPT1; | Q06210; A0A | GFPT1_HUMAN | Unmodified | 264880 | 541650 | 795840 | 532030 | 729180 | 0 | 217060 | 89830 | 1437500 | 705880 | 0 | 2344300 | 3826100 | 2363700 | 0 | 0 | 265630 | 713820 |
| EILGEIK | ITCH; ITCH; ITC | Q96J02; A0A | ITCH_HUMAN I | Unmodified | 0 | 492160 | 318990 | 64939 | 0 | 0 | 0 | 0 | 0 | 0 | 0 | 596420 | 0 | 0 | 0 | 807950 | 0 | 0 |
| EILGTAQSV | RPL12 | P30050 | RL12_HUMAN | Unmodified | 373710 | 0 | 20691 | 0 | 0 | 0 | 0 | 0 | 0 | 0 | 2577500 | 5188000 | 0 | 0 | 317520 | 232090 | 0 | 0 |
| EILGTAQSV | RPL12 | P30050 | RL12_HUMAN | Unmodified | 4535500 | 2045600 | 0 | 2198300 | 0 | 0 | 0 | 0 | 0 | 0 | 0 | 9791500 | 0 | 0 | 0 | 1252700 | 0 | 0 |
| EILGTCK | VCL | P18206 | VINC_HUMAN | Unmodified | 0 | 0 | 59899 | 0 | 0 | 0 | 0 | 0 | 1958100 | 80643 | 0 | 1547600 | 0 | 0 | 0 | 0 | 1714700 | 0 |
| EILIPVFK | ACLY | P53396 | ACLY_HUMAN | Unmodified | 0 | 0 | 0 | 0 | 0 | 0 | 0 | 0 | 0 | 0 | 196290 | 1386800 | 0 | 247480 | 0 | 0 | 47243 | 221040 |
| EILKEQENF | PARK7; PARK7; | Q99497; K7E | PARK7_HUMAN | Unmodified | 0 | 0 | 0 | 0 | 5079800 | 0 | 988330 | 0 | 0 | 1840900 | 0 | 60249 | 9510200 | 7720800 | 0 | 0 | 5304400 | 4374300 |
| EILLKGPDV | NDUFV1; NDUF | P49821; G3V | NDUV1_HUMA | Unmodified | 0 | 0 | 0 | 0 | 21761 | 0 | 103300 | 0 | 0 | 487240 | 279320 | 0 | 1584100 | 34947 | 0 | 0 | 0 | 0 |
| EILQEEEDL | ATP6V1A | P38606 | VATA_HUMAN | Unmodified | 0 | 0 | 0 | 53294 | 0 | 0 | 0 | 0 | 39206 | 0 | 0 | 0 | 364370 | 335080 | 149790 | 0 | 0 | 0 |
| EILSDDKHI | CLASP2; CLASP | O75122; B3K | CLAP2_HUMAN | Unmodified | 0 | 0 | 0 | 0 | 0 | 0 | 0 | 0 | 0 | 0 | 0 | 376770 | 0 | 0 | 0 | 0 | 0 | 0 |
| EILSEENEM | ACSL4; ACSL4; | O60488; D6I | ACSL4_HUMAN | Oxidation (M) | 0 | 0 | 0 | 0 | 0 | 0 | 0 | 0 | 0 | 0 | 0 | 1426200 | 0 | 0 | 0 | 0 | 0 | 0 |
| EILSEENEM | ACSL4; ACSL4; | O60488; D6I | ACSL4_HUMAN | Unmodified | 0 | 0 | 0 | 0 | 0 | 53307 | 0 | 0 | 0 | 0 | 461880 | 3301900 | 0 | 0 | 0 | 0 | 0 | 0 |
| EILSEVER | CCT3; CCT3 | P49368; B4D | TCPG_HUMAN | Unmodified | 0 | 0 | 0 | 0 | 0 | 0 | 0 | 0 | 0 | 0 | 0 | 18269000 | 0 | 0 | 0 | 0 | 0 | 0 |
| EILSLSASRI | IQGAP3; IQGAP | Q86VI3; F2Z | IQGA3_HUMAN | Unmodified | 0 | 0 | 0 | 0 | 1041200 | 0 | 0 | 0 | 551240 | 229540 | 0 | 341930 | 0 | 0 | 0 | 0 | 88596 | 0 |
| EILTLLQGV | TSN; TSN; TSN | Q15631; H7C | TSN_HUMAN T | Unmodified | 0 | 0 | 0 | 0 | 0 | 0 | 0 | 81631 | 0 | 0 | 540670 | 12672000 | 0 | 0 | 0 | 0 | 0 | 0 |
| EILTLLQGV | TSN; TSN; TSN | Q15631; H7C | TSN_HUMAN T | Unmodified | 0 | 0 | 0 | 0 | 0 | 0 | 0 | 0 | 0 | 0 | 144870 | 1572700 | 0 | 0 | 0 | 0 | 0 | 0 |
| EILVGDVG | CFL1; CFL1; CF | P23528; G3V | COF1_HUMAN | Unmodified | 25936000 | 20951000 | 17931000 | 19753000 | 1.04E+08 | 10665000 | 4581500 | 1080100 | 38009000 | 5038300 | 20375000 | 62633000 | 44405000 | 10343000 | 58660000 | 36905000 | 14941000 | 18809000 |
| EIMDDKSLI | IQGAP1; IQGAP | P46940; A0A | IQGA1_HUMAN | Unmodified | 0 | 0 | 0 | 0 | 3579000 | 0 | 0 | 0 | 0 | 0 | 0 | 0 | 0 | 0 | 0 | 0 | 75297 | 0 |
| EINGIHDES | LMO7; LMO7; LI | Q8WWI1; H0 | LMO7_HUMAN | Unmodified | 0 | 0 | 0 | 0 | 366480 | 56454 | 0 | 0 | 0 | 0 | 0 | 0 | 0 | 0 | 0 | 0 | 0 | 183820 |
| EINGISVAN | CASK; CASK; CA | O14936; A0A | CSKP_HUMAN | Unmodified | 0 | 0 | 538410 | 766200 | 0 | 0 | 0 | 0 | 0 | 0 | 0 | 813390 | 0 | 0 | 0 | 1155800 | 0 | 0 |
| EINLAPDSS | FN1; FN1 | P02751; H0Y | FINC_HUMAN I | Oxidation (M) | 0 | 0 | 3113900 | 0 | 0 | 0 | 0 | 0 | 0 | 782540 | 0 | 0 | 0 | 0 | 0 | 0 | 0 | 0 |
| EINNAHAIL | DNAJC5 | Q9H3Z4 | DNJC5_HUMAN | Unmodified | 176350 | 75027 | 4317600 | 0 | 0 | 0 | 0 | 0 | 0 | 0 | 0 | 5626200 | 0 | 35364 | 4018200 | 2290400 | 0 | 0 |
| EINNSHIQE | PLCB2; PLCB2 | Q00722; H0Y | PLCB2_HUMAN | Unmodified | 0 | 0 | 0 | 0 | 284090 | 0 | 0 | 0 | 0 | 0 | 748800 | 0 | 0 | 0 | 0 | 0 | 0 | 0 |
| EINTNQEAI | ATP6V0A1; ATP | Q93050; K7E | VPP1_HUMAN | Unmodified | 92613 | 75411 | 481300 | 177670 | 0 | 142940 | 261140 | 191790 | 5418500 | 834290 | 1461400 | 2202200 | 0 | 87059 | 228940 | 127250 | 106520 | 0 |
| EINTNQEAI | ATP6V0A1; ATP | Q93050; K7E | VPP1_HUMAN | Unmodified | 762340 | 541070 | 188810 | 981950 | 0 | 33441 | 127320 | 22712 | 1171900 | 284820 | 923960 | 397590 | 678850 | 164840 | 373710 | 230300 | 0 | 0 |
| EINTNQEAI | ATP6V0A1; ATP | Q93050; K7E | VPP1_HUMAN | Unmodified | 0 | 0 | 0 | 0 | 0 | 0 | 707840 | 54279 | 0 | 896420 | 0 | 0 | 0 | 0 | 0 | 0 | 0 | 0 |
| EIPAWVPFI | AZGP1; AZGP1 | P25311; C9J | ZA2G_HUMAN | Unmodified | 0 | 0 | 928430 | 857110 | 17119 | 0 | 0 | 0 | 376730 | 0 | 10337000 | 552420 | 0 | 0 | 1382000 | 409310 | 0 | 10014 |
| EIPETTPIEE | LARP4; LARP4; | Q71RC2; A0A | LARP4_HUMAN | Unmodified | 374960 | 0 | 0 | 0 | 0 | 312670 | 261290 | 0 | 0 | 0 | 0 | 320880 | 0 | 0 | 80295 | 0 | 0 | 0 |
| EIPFNEDPN | C2CD5; C2CD5 | Q86YS7; H0Y | C2CD5_HUMA | Unmodified | 114470 | 226130 | 0 | 409710 | 450640 | 0 | 123700 | 0 | 731530 | 195350 | 0 | 1827700 | 365200 | 0 | 0 | 474870 | 624280 | 1270700 |
| EIPQAERYC | CTNND1; CTNN | O60716; A0A | CTND1_HUMA | Unmodified | 826220 | 572510 | 711800 | 1416300 | 591160 | 0 | 0 | 0 | 0 | 0 | 901470 | 3650600 | 0 | 0 | 1547900 | 650790 | 554100 | 998070 |
| EIPQAERYC | CTNND1; CTNN | O60716; A0A | CTND1_HUMA | Unmodified | 263640 | 122760 | 0 | 0 | 0 | 0 | 0 | 0 | 0 | 0 | 0 | 0 | 0 | 0 | 572690 | 568850 | 0 | 595130 |
| EIQALKDTK | PPL; PPL; PPL | O60437; K7E | PEPL_HUMAN I | Unmodified | 0 | 0 | 0 | 0 | 341770 | 117940 | 0 | 0 | 0 | 0 | 0 | 0 | 97414 | 0 | 0 | 0 | 0 | 0 |
| EIQAQNFFS | NAGK; NAGK; N | Q9UJ70; C9J | NAGK_HUMAN | Oxidation (M) | 103400 | 0 | 0 | 0 | 0 | 0 | 0 | 0 | 0 | 0 | 106310 | 169470 | 0 | 0 | 0 | 14293 | 0 | 0 |
| EIQELQSKF | AFDN; AFDN; A | P55196; H0Y | AFAD_HUMAN | Unmodified | 0 | 0 | 0 | 0 | 0 | 0 | 0 | 0 | 0 | 87465 | 328240 | 1418200 | 0 | 36754 | 8177.1 | 0 | 0 | 0 |
| EIQEPDPTY | SMARCA5 | O60264 | SMCA5_HUMA | Unmodified | 0 | 0 | 0 | 0 | 0 | 0 | 0 | 0 | 0 | 0 | 174110 | 360280 | 0 | 0 | 0 | 0 | 0 | 0 |
| EIQGFFNIP | PRPSAP2; PRPS | O60256; E7E | KPRB_HUMAN | Unmodified | 0 | 81646 | 65188 | 79544 | 167470 | 48487 | 99517 | 28847 | 159860 | 169990 | 430800 | 1908000 | 116900 | 0 | 137900 | 201930 | 0 | 0 |
| EIQGFFSFF | PRPSAP1; PRPS | Q14558; C9J | KPRA_HUMAN | Unmodified | 0 | 0 | 0 | 0 | 36789 | 0 | 0 | 0 | 199700 | 0 | 0 | 204260 | 41511 | 0 | 0 | 0 | 0 | 0 |
| EIQHHLK | SH3GL2; SH3G | Q99962; Q9 | SH3G2_HUMA | Unmodified | 575950 | 0 | 0 | 547430 | 0 | 0 | 0 | 0 | 38883 | 0 | 0 | 0 | 125920 | 213470 | 448750 | 0 | 0 | 199190 |
| EIQKLETEL | NMI | Q13287 | NMI_HUMAN N | Unmodified | 0 | 0 | 0 | 0 | 1012600 | 0 | 0 | 0 | 0 | 54932 | 0 | 0 | 592300 | 0 | 0 | 0 | 264860 | 656240 |
| EIQKLPENS | ERO1B; ERO1B | Q86YB8; Q5T | ERO1B_HUMA | Unmodified | 0 | 0 | 0 | 0 | 0 | 120830 | 263220 | 0 | 0 | 566930 | 0 | 0 | 0 | 0 | 0 | 0 | 0 | 0 |
| EIQNAVNG | CLU; CLU; CLU | P10909; H0Y | CLUS_HUMAN | Unmodified | 0 | 0 | 0 | 0 | 2328600 | 2291500 | 0 | 0 | 0 | 979980 | 0 | 0 | 0 | 0 | 0 | 0 | 0 | 3033100 |
| EIQNLIK | TERF1; ISOC1; | P54274; Q96 | TERF1_HUMAN | Unmodified | 0 | 0 | 0 | 19045 | 0 | 0 | 0 | 0 | 310220 | 0 | 331280 | 3294700 | 789430 | 162470 | 0 | 0 | 0 | 0 |
| EIQSAFVSV | ECPAS; ECPAS | Q5VYK3; J3K | ECM29_HUMA | Unmodified | 0 | 0 | 0 | 0 | 1498600 | 0 | 724940 | 0 | 381330 | 128240 | 0 | 1159100 | 94096 | 96922 | 207680 | 258600 | 0 | 0 |
| EIQSTLEGL | LLGL2; LLGL2 | Q6P1M3; J3C | L2GL2_HUMAN | Unmodified | 593810 | 535140 | 238020 | 359830 | 297300 | 0 | 0 | 0 | 871760 | 123940 | 265140 | 561340 | 553970 | 0 | 580640 | 633930 | 81985 | 465660 |
| EIQTAVRLL | H2BC4; H2BC1 | P62807; P06 | H2B1C_HUMA | Unmodified | 4508400 | 1485200 | 0 | 0 | 14218000 | 15883000 | 8586000 | 10147000 | 0 | 1872300 | 265740 | 9161300 | 5414900 | 1405300 | 0 | 0 | 308280 | 12259000 |
| EIQTTTGN | RDH11 | Q8TC12 | RDH11_HUMA | Unmodified | 909340 | 579550 | 1259700 | 1980300 | 0 | 0 | 0 | 0 | 376880 | 88295 | 184930 | 1046500 | 0 | 0 | 154260 | 1036100 | 0 | 0 |
| EIQTTTGN | RDH11 | Q8TC12 | RDH11_HUMA | Unmodified | 0 | 0 | 0 | 0 | 260410 | 0 | 746950 | 180850 | 0 | 120180 | 0 | 0 | 0 | 3630100 | 0 | 0 | 0 | 233640 |
| EIQVGDVIF | ATP8B1; ATP8B | O43520; A0A | AT8B1_HUMAN | Unmodified | 104320 | 0 | 1240200 | 0 | 0 | 0 | 0 | 0 | 0 | 0 | 0 | 1097500 | 0 | 0 | 0 | 0 | 0 | 0 |
| EIQVIPLQR | IMPA1 | P29218 | IMPA1_HUMAN | Unmodified | 0 | 0 | 0 | 0 | 857290 | 555890 | 648970 | 89630 | 1919600 | 1161600 | 89449 | 1666000 | 1434600 | 681790 | 0 | 0 | 1322900 | 1461900 |
| EIQVQHPA | CCT3; CCT3; C | P49368; Q5S | TCPG_HUMAN | Unmodified | 0 | 0 | 117390 | 0 | 178810 | 0 | 0 | 0 | 434250 | 55195 | 441540 | 4132700 | 0 | 0 | 0 | 87937 | 0 | 0 |
| EIRAQLTEQ | SRGAP2; SRGA | O75044; P0D | SRGP2_HUMA | Unmodified | 0 | 0 | 0 | 0 | 1735500 | 0 | 249990 | 411560 | 0 | 284870 | 0 | 0 | 1290000 | 711140 | 0 | 0 | 271750 | 534310 |
| EIRAQLVEQ | SRGAP1 | Q7Z6B7 | SRGP1_HUMA | Unmodified | 0 | 0 | 0 | 0 | 868620 | 246460 | 382940 | 287490 | 0 | 91873 | 68450 | 0 | 229510 | 0 | 0 | 0 | 419420 | 615260 |
| EIRESTGA | PCBP1; PCBP3; | Q15365; P57 | PCBP1_HUMA | Unmodified | 2646000 | 0 | 0 | 2230800 | 1064900 | 0 | 0 | 241070 | 1708300 | 405960 | 0 | 0 | 2937800 | 1097000 | 0 | 0 | 0 | 0 |
| EIRESTGA | PCBP1; PCBP3; | Q15365; P57 | PCBP1_HUMA | Oxidation (M) | 2858900 | 1220800 | 1052300 | 4152000 | 0 | 0 | 293310 | 276250 | 0 | 0 | 782410 | 0 | 1187700 | 0 | 531920 | 0 | 0 | 0 |
| EIRGNPHLI | BMX | P51813 | BMX_HUMAN | Unmodified | 0 | 0 | 0 | 0 | 0 | 0 | 0 | 0 | 0 | 0 | 370740 | 0 | 0 | 0 | 0 | 0 | 0 | 0 |
| EIRPALELLI | GDI2 | P50395 | GDIB_HUMAN | Unmodified | 1241000 | 0 | 10424000 | 9544700 | 11456000 | 4090400 | 0 | 113450 | 41410000 | 4056200 | 25336000 | 63819000 | 11645000 | 3317800 | 2507500 | 15103000 | 0 | 5411700 |
| EIRTDDLCL | GALNT1 | Q10472 | GALT1_HUMAN | Unmodified | 0 | 0 | 0 | 0 | 163030 | 89058 | 0 | 0 | 3405700 | 0 | 0 | 695460 | 78646 | 0 | 0 | 0 | 0 | 0 |
| EISDGDVIIS | EGFR; EGFR; EC | P00533; Q5C | EGFR_HUMAN | Unmodified | 327920 | 269470 | 2002500 | 1454100 | 13761000 | 3315000 | 0 | 0 | 1154900 | 0 | 0 | 0 | 0 | 0 | 1150900 | 1369900 | 0 | 0 |
| EISESTPVE | LARP4B | Q92615 | LAR4B_HUMAN | Unmodified | 476920 | 0 | 0 | 0 | 0 | 0 | 0 | 0 | 0 | 0 | 0 | 0 | 0 | 0 | 0 | 0 | 0 | 0 |
| EISEVFPDC | HEXB; HEXB; HI | P07686; Q5U | HEXB_HUMAN | Unmodified | 483030 | 0 | 331280 | 1679800 | 393200 | 218960 | 0 | 0 | 7909700 | 1374500 | 263090 | 83894 | 1938400 | 1518900 | 1545000 | 214310 | 0 | 0 |
| EISFAYEVLS | DNAJA2 | O60884 | DNJA2_HUMAN | Unmodified | 1925800 | 2359600 | 1547500 | 317060 | 0 | 0 | 0 | 0 | 0 | 193180 | 252950 | 1570900 | 0 | 0 | 4194100 | 3649200 | 0 | 0 |
| EISGHTSGI | STRAP | Q9Y3F4 | STRAP_HUMAN | Unmodified | 0 | 0 | 0 | 0 | 0 | 0 | 109630 | 0 | 0 | 0 | 0 | 484420 | 0 | 0 | 0 | 0 | 37590 | 1734400 |
| EISLSAVRF | ROR2; ROR2 | Q01974; B1A | ROR2_HUMAN | Unmodified | 0 | 0 | 0 | 0 | 0 | 0 | 121230 | 0 | 0 | 968840 | 0 | 0 | 0 | 0 | 0 | 0 | 0 | 0 |
| EISLWFKPE | NME2; NME1-N | P22392; Q32 | NDKB_HUMAN | Unmodified | 215960 | 351050 | 230830 | 870800 | 21093000 | 1184600 | 1573500 | 0 | 579210 | 1423800 | 3611200 | 3381800 | 14645000 | 4028400 | 1185400 | 1172600 | 6671400 | 21996000 |
| EISNGKWL | ITGA6 | P23229 | ITA6_HUMAN I | Unmodified | 0 | 0 | 0 | 0 | 990710 | 109010 | 528410 | 18742 | 0 | 0 | 0 | 0 | 431320 | 504220 | 0 | 0 | 1731800 | 1625100 |
| EISNGKWL | ITGA6 | P23229 | ITA6_HUMAN I | Unmodified | 0 | 0 | 0 | 0 | 1510500 | 124100 | 562140 | 0 | 0 | 0 | 0 | 0 | 927010 | 922940 | 0 | 0 | 3102000 | 4243500 |
| EISNLLVAT | JAK1; JAK1; JAK | P23458; A0A | JAK1_HUMAN | Unmodified | 0 | 0 | 0 | 0 | 894250 | 202050 | 48312 | 25164 | 200580 | 67768 | 76450 | 532150 | 0 | 0 | 0 | 0 | 439560 | 181800 |
| EISPGSGP | EPB41L2; EPB4 | O43491; E9F | E41L2_HUMAN | Unmodified | 0 | 0 | 377830 | 345910 | 1607500 | 321800 | 0 | 84388 | 0 | 162280 | 0 | 748790 | 0 | 0 | 0 | 462650 | 328630 | 683910 |
| EISQDSLA | TJP1; TJP1; TJP1 | Q07157; A0A | ZO1_HUMAN T | Unmodified | 24435 | 17083 | 705520 | 280770 | 271210 | 52417 | 0 | 0 | 364290 | 0 | 0 | 1572100 | 264140 | 77404 | 311160 | 1271000 | 0 | 0 |

| | | | | | | | | | | | | | | | | | | | | | | |
|---|---|---|---|---|---|---|---|---|---|---|---|---|---|---|---|---|---|---|---|---|---|---|
| EISQDSLA | TJP1; TJP1; TJP1 | Q07157; A0 | ZO1_HUMAN T | Unmodified | 0 | 0 | 452410 | 2115400 | 24683000 | 6129400 | 942860 | 701900 | 580130 | 1977600 | 261940 | 216050 | 15043000 | 12163000 | 326870 | 571080 | 2887900 | 4859100 |
| EISQHNINI | SLC9A9; SLC9A | Q8IVB4; C9 | SL9A9_HUMAN | Unmodified | 0 | 0 | 0 | 0 | 38608 | 0 | 0 | 0 | 249290 | 19641 | 0 | 0 | 0 | 0 | 0 | 0 | 0 | 0 |
| EISSGLVEV | MGAT4A | Q9UM21 | MGT4A_HUMA | Unmodified | 0 | 0 | 0 | 0 | 0 | 0 | 0 | 0 | 687020 | 0 | 0 | 0 | 0 | 0 | 0 | 0 | 0 | 0 |
| EISYENTQI | UGGT1 | Q9NYU2 | UGGG1_HUMA | Unmodified | 0 | 0 | 0 | 0 | 194260 | 0 | 0 | 0 | 12546000 | 3542400 | 1116000 | 541360 | 0 | 0 | 0 | 102680 | 0 | 0 |
| EISYENTQI | UGGT1 | Q9NYU2 | UGGG1_HUMA | Unmodified | 0 | 0 | 0 | 0 | 0 | 0 | 0 | 0 | 0 | 899300 | 0 | 0 | 0 | 0 | 0 | 0 | 0 | 0 |
| EITALAPSTI | ACTG1; ACTC1; | P63261; P68 | ACTG_HUMAN | Oxidation (M) | 0 | 0 | 0 | 0 | 776230 | 0 | 0 | 168410 | 0 | 194870 | 0 | 0 | 0 | 0 | 0 | 0 | 252080 | 433390 |
| EITALAPSTI | ACTG1; ACTC1; | P63261; P68 | ACTG_HUMAN | Oxidation (M) | 39933000 | 36601000 | 49095000 | 50489000 | 6.8E+08 | 1.53E+08 | 1.38E+08 | 1.86E+08 | 47232000 | 37118000 | 79502000 | 2.94E+08 | 2.28E+08 | 1.72E+08 | 67920000 | 61994000 | 2.37E+08 | 1.81E+08 |
| EITALAPSTI | ACTG1; ACTC1; | P63261; P68 | ACTG_HUMAN | Unmodified | 43160000 | 41701000 | 42616000 | 88929000 | 1.99E+09 | 3.07E+08 | 3.46E+08 | 2.2E+08 | 1.98E+08 | 1.14E+08 | 72683000 | 5.43E+08 | 7.2E+08 | 4.56E+08 | 1.07E+08 | 53859000 | 5.79E+08 | 4.98E+08 |
| EITALAPSTI | ACTG1; ACTC1; | P63261; P68 | ACTG_HUMAN | Oxidation (M) | 0 | 178400 | 0 | 241430 | 7072500 | 0 | 5457600 | 10412000 | 218680 | 282190 | 0 | 788620 | 2955100 | 2895200 | 0 | 0 | 8680500 | 1886500 |
| EITALAPSTI | ACTG1; ACTC1; | P63261; P68 | ACTG_HUMAN | Unmodified | 0 | 0 | 0 | 0 | 15017000 | 1190500 | 18446000 | 10785000 | 0 | 664080 | 0 | 0 | 5882100 | 4470300 | 0 | 51537 | 34370000 | 4818900 |
| EITDGEEKT | CNPY4 | Q8N129 | CNPY4_HUMA | Unmodified | 0 | 0 | 0 | 0 | 0 | 0 | 0 | 0 | 1957700 | 0 | 145820 | 0 | 0 | 0 | 0 | 0 | 0 | 0 |
| EITEREMQI | CIT; CIT; CIT; CI | O14578; A0 | CTRO_HUMAN | Unmodified | 22587 | 0 | 0 | 0 | 904940 | 192620 | 0 | 0 | 0 | 33266 | 128610 | 376700 | 291530 | 53742 | 0 | 0 | 277370 | 256680 |
| EITGIGPST | VPS26A; VPS26 | O75436; S4 | VP26A_HUMAN | Unmodified | 0 | 0 | 0 | 0 | 0 | 0 | 0 | 0 | 237560 | 0 | 78118 | 908980 | 0 | 0 | 0 | 0 | 518490 | 0 |
| EITIHTKLP | PODXL | O00592 | PODXL_HUMA | Unmodified | 0 | 0 | 0 | 265300 | 0 | 0 | 0 | 0 | 13113000 | 168390 | 1976300 | 1045600 | 510940 | 0 | 0 | 79434 | 0 | 0 |
| EITIHTKLP | PODXL | O00592 | PODXL_HUMA | Unmodified | 0 | 0 | 0 | 0 | 0 | 0 | 0 | 0 | 2733400 | 350240 | 0 | 0 | 199010 | 0 | 0 | 0 | 0 | 0 |
| EITIINPAQF | GUF1 | Q8N442 | GUF1_HUMAN | Unmodified | 0 | 0 | 0 | 0 | 0 | 0 | 106820 | 26670 | 0 | 1101000 | 0 | 0 | 0 | 158830 | 0 | 0 | 0 | 136860 |
| EITLLMQTL | TXLNA | P40222 | TXLNA_HUMAN | Oxidation (M) | 0 | 0 | 0 | 0 | 328170 | 0 | 0 | 0 | 0 | 0 | 0 | 0 | 1510400 | 0 | 0 | 69469 | 774490 | 590430 |
| EITQDTNDI | SIRPA | P78324 | SHPS1_HUMA | Unmodified | 0 | 0 | 0 | 0 | 909580 | 0 | 44740 | 0 | 323200 | 2341500 | 0 | 1076900 | 0 | 0 | 48421 | 0 | 0 | 0 |
| EITQLSHLC | TYK2; TYK2; TY | P29597; E9P | TYK2_HUMAN I | Unmodified | 0 | 0 | 356070 | 843460 | 442200 | 0 | 0 | 0 | 304720 | 0 | 0 | 1401900 | 0 | 0 | 449440 | 353740 | 0 | 0 |
| EITTRYQLD | EPCAM; EPCAM | P16422; B5 | EPCAM_HUMA | Unmodified | 439060 | 146230 | 1453400 | 4478000 | 5566900 | 739380 | 0 | 0 | 4855100 | 17471000 | 2114000 | 0 | 22607000 | 6500300 | 0 | 175360 | 158720 | 867070 |
| EITYMHSE | GFPT1; GFPT2; | Q06210; O9 | GFPT1_HUMAN | Oxidation (M) | 75531 | 127170 | 0 | 628850 | 157390 | 0 | 0 | 67044 | 0 | 237150 | 0 | 844020 | 342980 | 0 | 538200 | 191390 | 0 | 0 |
| EIVAEAERL | EIF5 | P55010 | IF5_HUMAN Eu | Unmodified | 0 | 0 | 174310 | 99293 | 0 | 0 | 0 | 0 | 157500 | 0 | 0 | 1598400 | 0 | 89167 | 159940 | 183840 | 0 | 0 |
| EIVDPLYGI | ARHGAP17; AR | Q68EM7; C9 | RHG17_HUMA | Unmodified | 152090 | 297490 | 55618 | 465750 | 1804000 | 390510 | 36474 | 0 | 1333500 | 168350 | 588200 | 1384100 | 1271300 | 107110 | 149940 | 229270 | 377600 | 622150 |
| EIVEMNEIE | SLK | Q9H2G2 | SLK_HUMAN S | Unmodified | 0 | 0 | 0 | 90011 | 1233500 | 0 | 0 | 118370 | 349570 | 0 | 337960 | 868290 | 0 | 0 | 0 | 0 | 0 | 0 |
| EIVESGRG | HIP1R | O75146 | HIP1R_HUMAN | Unmodified | 0 | 0 | 0 | 171100 | 247740 | 0 | 0 | 0 | 0 | 202130 | 0 | 664560 | 1267000 | 756560 | 128810 | 0 | 181580 | 0 |
| EIVEVKEEN | HMGN5; HMGN | P82970; Q5 | HMGN5_HUMA | Unmodified | 108020 | 0 | 0 | 0 | 274120 | 0 | 0 | 13112 | 1769200 | 215920 | 1703200 | 1902400 | 765230 | 0 | 0 | 0 | 344720 | 0 |
| EIVHLQAG | TUBB4A; TUBB | P04350; P68 | TBB4A_HUMAN | Unmodified | 2189500 | 1013900 | 3536600 | 2087500 | 2788600 | 512960 | 4478100 | 1797400 | 14399000 | 4810200 | 12768000 | 1.43E+08 | 1235000 | 721650 | 4448300 | 3803800 | 898380 | 2193900 |
| EIVLADVID | PAICS; PAICS; F | P22234; D6F | PUR6_HUMAN | Unmodified | 323330 | 396720 | 166610 | 1109700 | 2734300 | 123940 | 481410 | 0 | 970160 | 153260 | 2871200 | 18475000 | 421720 | 0 | 595350 | 1279100 | 656720 | 0 |
| EIVLADVID | PAICS; PAICS; F | P22234; D6F | PUR6_HUMAN | Unmodified | 0 | 0 | 0 | 0 | 4544700 | 685580 | 1210500 | 0 | 448280 | 1487200 | 1847300 | 0 | 773530 | 1790000 | 0 | 0 | 1573200 | 1777900 |
| EIVPVLVSTI | ACAT2 | Q9BWD1 | THIC_HUMAN | Unmodified | 30954 | 0 | 0 | 0 | 0 | 0 | 0 | 0 | 1517000 | 315080 | 2511000 | 10956000 | 244030 | 0 | 0 | 0 | 0 | 330930 |
| EIVPVLVSTI | ACAT2 | Q9BWD1 | THIC_HUMAN | Unmodified | 0 | 0 | 0 | 102120 | 2322800 | 0 | 213850 | 64560 | 684510 | 364800 | 0 | 0 | 509140 | 146570 | 0 | 95199 | 0 | 0 |
| EIVRCYQSI | ANXA7 | P20073 | ANXA7_HUMA | Unmodified | 0 | 0 | 0 | 182900 | 4582800 | 123850 | 121800 | 0 | 919630 | 3146400 | 970600 | 376900 | 198510 | 338250 | 0 | 0 | 673460 | 2632100 |
| EIVRDIKEK | POTEJ; POTEE; | P0CG39; Q6 | POTEJ_HUMAN | Unmodified | 0 | 0 | 0 | 1604900 | 0 | 0 | 0 | 0 | 0 | 0 | 0 | 0 | 0 | 0 | 0 | 0 | 1534600 | 0 |
| EIVRDLDQI | SDF4; SDF4; SD | Q9BRK5; H0 | CAB45_HUMA | Unmodified | 0 | 78271 | 0 | 0 | 54033 | 0 | 0 | 44944 | 29021000 | 2452900 | 612010 | 437300 | 0 | 170550 | 0 | 0 | 0 | 0 |
| EIVRGYKW | PHPT1 | Q9NRX4 | PHP14_HUMA | Unmodified | 0 | 0 | 0 | 0 | 1449000 | 0 | 379470 | 53491 | 0 | 0 | 0 | 0 | 1706400 | 1253000 | 0 | 0 | 257260 | 458540 |
| EIVSSTRLD | EEA1 | Q15075 | EEA1_HUMAN | Unmodified | 0 | 0 | 0 | 0 | 1376400 | 0 | 144070 | 175190 | 0 | 89716 | 0 | 0 | 342670 | 0 | 0 | 0 | 60380 | 212910 |
| EIVVIHQDF | IARS1; IARS1; I | P41252; A0A | SYIC_HUMAN I | Unmodified | 0 | 0 | 416540 | 238870 | 0 | 585780 | 0 | 70035 | 937400 | 0 | 1469500 | 11860000 | 0 | 0 | 486280 | 273450 | 0 | 0 |
| EIVVLETLE | RCN1; N/A | Q15293; E9F | RCN1_HUMAN | Unmodified | 0 | 0 | 79662 | 0 | 0 | 0 | 0 | 0 | 1984600 | 272500 | 557850 | 62050 | 0 | 0 | 0 | 140760 | 0 | 0 |
| EIVVTDYSD | NNMT | P40261 | NNMT_HUMAN | Unmodified | 0 | 0 | 0 | 0 | 1197300 | 0 | 0 | 0 | 0 | 0 | 0 | 0 | 0 | 0 | 0 | 0 | 0 | 698110 |
| EIWDALK | UBTD1; UBTD2 | Q9HAC8; Q | UBTD1_HUMA | Unmodified | 185380 | 193560 | 253300 | 232770 | 574460 | 149560 | 0 | 0 | 950250 | 211920 | 439660 | 1997200 | 0 | 0 | 442610 | 404120 | 647230 | 1747400 |
| EIWGVEPS | PYGB | P11216 | PYGB_HUMAN | Unmodified | 0 | 0 | 335840 | 0 | 50626 | 72648 | 0 | 0 | 568230 | 0 | 76530 | 263560 | 0 | 0 | 0 | 0 | 759390 | 2328000 |
| EIWRTGQP | EVA1B | Q9NVM1 | EVA1B_HUMAN | Unmodified | 0 | 0 | 0 | 0 | 0 | 0 | 0 | 0 | 0 | 0 | 0 | 0 | 0 | 0 | 0 | 0 | 0 | 2028300 |
| EIYCPAPP | CD55; CD55; C | P08174; B1 | DAF_HUMAN C | Unmodified | 4691000 | 2293900 | 0 | 2396000 | 0 | 0 | 0 | 0 | 1461800 | 0 | 0 | 14722000 | 0 | 0 | 2640300 | 993830 | 678930 | 0 |
| EIYENMAP | PSMD2; PSMD2 | Q13200; H7 | PSMD2_HUMA | Unmodified | 0 | 0 | 0 | 0 | 434890 | 0 | 0 | 0 | 0 | 0 | 0 | 800690 | 0 | 0 | 0 | 0 | 0 | 0 |
| EIYENMAP | PSMD2; PSMD2 | Q13200; H7 | PSMD2_HUMA | Unmodified | 0 | 0 | 0 | 0 | 0 | 0 | 0 | 0 | 59193 | 0 | 0 | 740950 | 0 | 0 | 0 | 0 | 0 | 0 |
| EIYNQVNV | CYRIB; CYRIB; | Q9NUQ9; E | CYRIB_HUMAN | Unmodified | 0 | 0 | 0 | 0 | 0 | 502740 | 0 | 0 | 572480 | 0 | 316450 | 1490300 | 0 | 0 | 0 | 0 | 0 | 0 |
| EIYTHFTCA | GNAI3; GNAI2; | P08754; P04 | GNAI3_HUMAN | Unmodified | 6765600 | 4341700 | 285310 | 10471000 | 33090000 | 2307800 | 11633000 | 6083800 | 18975000 | 8380400 | 33540000 | 1.05E+08 | 27491000 | 38106000 | 11007000 | 6840000 | 27389000 | 62140000 |
| EIYYHGEPI | ARRB1; ARRB1; | P49407; H0 | ARRB1_HUMAN | Unmodified | 0 | 0 | 0 | 553690 | 0 | 0 | 0 | 0 | 289910 | 0 | 0 | 0 | 0 | 0 | 339670 | 181630 | 0 | 0 |
| EKADAIYNI | ECE1; ECE1; EC | P42892; B4 | ECE1_HUMAN | 2 Oxidation (M) | 0 | 0 | 0 | 0 | 0 | 0 | 0 | 0 | 1410200 | 0 | 0 | 0 | 0 | 0 | 0 | 0 | 0 | 0 |
| EKADAIYNI | ECE1; ECE1; EC | P42892; B4 | ECE1_HUMAN | Oxidation (M) | 0 | 0 | 0 | 0 | 1956500 | 0 | 0 | 0 | 2315000 | 0 | 0 | 0 | 0 | 676970 | 953870 | 624610 | 0 | 0 |
| EKADAIYNI | ECE1; ECE1; EC | P42892; B4 | ECE1_HUMAN | Unmodified | 0 | 1325600 | 0 | 147970 | 0 | 381090 | 0 | 0 | 1199100 | 0 | 0 | 0 | 0 | 0 | 0 | 0 | 0 | 0 |
| EKADAIYNI | ECE1; ECE1; EC | P42892; B4 | ECE1_HUMAN | Unmodified | 0 | 0 | 0 | 0 | 0 | 0 | 0 | 0 | 628610 | 0 | 0 | 0 | 0 | 0 | 0 | 0 | 0 | 0 |
| EKADAIYNI | ECE1; ECE1; EC | P42892; B4 | ECE1_HUMAN | 2 Oxidation (M) | 0 | 0 | 0 | 0 | 0 | 0 | 0 | 0 | 503880 | 591140 | 0 | 0 | 0 | 0 | 0 | 0 | 0 | 0 |
| EKAPSIIFID | PSMC3; PSMC3 | P17980; E9P | PRS6A_HUMAN | Unmodified | 0 | 0 | 0 | 0 | 1361600 | 156180 | 0 | 0 | 442220 | 420900 | 0 | 0 | 0 | 680420 | 0 | 0 | 1146800 | 901610 |
| EKDFIILSC | UPF1 | Q92900 | RENT1_HUMAN | Unmodified | 331730 | 0 | 0 | 0 | 0 | 98395 | 0 | 0 | 447930 | 0 | 0 | 94039 | 0 | 0 | 0 | 0 | 0 | 0 |
| EKDGDLYII | HS2ST1 | Q7LGA3 | HS2ST_HUMAN | Unmodified | 0 | 0 | 0 | 0 | 0 | 0 | 0 | 0 | 3096100 | 814540 | 0 | 0 | 0 | 0 | 0 | 0 | 0 | 0 |
| EKDGEFSV | EPHA2 | P29317 | EPHA2_HUMA | Oxidation (M) | 53328 | 67734 | 53651 | 0 | 1089600 | 0 | 1373100 | 51366 | 123830 | 121400 | 0 | 707710 | 515860 | 0 | 129600 | 0 | 266300 | 445680 |
| EKDILVLPL | DHRS7; DHRS7 | Q9Y394; H0 | DHRS7_HUMA | Unmodified | 548740 | 0 | 325710 | 0 | 0 | 0 | 0 | 0 | 3002300 | 0 | 989360 | 2738500 | 0 | 0 | 1075400 | 451960 | 0 | 0 |
| EKDIQEEST | LRPPRC; LRPPF | P42704; B8Z | LPPRC_HUMA | Unmodified | 0 | 0 | 0 | 0 | 0 | 0 | 0 | 0 | 2032300 | 80480 | 58489 | 0 | 0 | 0 | 0 | 0 | 211280 | 0 |
| EKDNIDITL | ARL8B; ARL8A | Q9NVJ2; Q9 | ARL8B_HUMAN | Unmodified | 0 | 0 | 0 | 0 | 1535500 | 236110 | 790950 | 0 | 882330 | 625680 | 317860 | 0 | 0 | 2225300 | 0 | 0 | 0 | 1813100 |
| EKDRFASA | PALS1 | Q8N3R9 | PALS1_HUMAN | Unmodified | 0 | 0 | 0 | 0 | 1215600 | 0 | 0 | 0 | 48727 | 77165 | 0 | 0 | 598290 | 195910 | 0 | 0 | 145420 | 620520 |
| EKDVQFEH | TM9SF4; TM9SF | Q92544; A0 | TM9S4_HUMAN | Unmodified | 0 | 0 | 0 | 0 | 0 | 0 | 0 | 0 | 4780200 | 593590 | 166060 | 459280 | 0 | 0 | 38142 | 19626 | 0 | 0 |
| EKDVQFEH | TM9SF4; TM9SF | Q92544; A0 | TM9S4_HUMAN | Unmodified | 0 | 0 | 0 | 0 | 0 | 0 | 0 | 0 | 2064100 | 0 | 0 | 22222000 | 0 | 0 | 0 | 0 | 0 | 0 |
| EKEAELKE | SEPTIN6; SEPTI | Q14141; Q9 | SEPT6_HUMAN | Unmodified | 0 | 0 | 0 | 0 | 1272900 | 0 | 0 | 0 | 112640 | 78265 | 0 | 0 | 0 | 0 | 0 | 0 | 49846 | 0 |
| EKEDAQEV | AKAP12 | Q02952 | AKA12_HUMAN | Unmodified | 0 | 0 | 0 | 0 | 635240 | 152570 | 38152 | 0 | 0 | 0 | 23122 | 69688 | 0 | 0 | 0 | 0 | 146590 | 993470 |
| EKEDAQEV | AKAP12 | Q02952 | AKA12_HUMAN | Unmodified | 0 | 0 | 0 | 0 | 1987900 | 0 | 0 | 0 | 0 | 0 | 0 | 0 | 0 | 0 | 0 | 0 | 640520 | 3208600 |
| EKEDDVPQ | LAP3; LAP3 | P28838; H0 | AMPL_HUMAN | Unmodified | 71307 | 84517 | 338740 | 654340 | 7632800 | 241990 | 1745400 | 1191400 | 2715100 | 1575600 | 1757500 | 6485300 | 6525500 | 3260800 | 707730 | 426840 | 2518800 | 4333400 |
| EKEDLLCG | IGF2R; IGF2R; I | P11717; A0A | MPRI_HUMAN | Unmodified | 111730 | 288500 | 0 | 458460 | 0 | 0 | 235760 | 66686 | 7471300 | 1030300 | 618720 | 0 | 1570400 | 271980 | 0 | 0 | 0 | 236070 |
| EKEDLLCG | IGF2R; IGF2R; I | P11717; A0A | MPRI_HUMAN | Unmodified | 0 | 0 | 0 | 0 | 0 | 0 | 0 | 41258 | 1688100 | 181400 | 119120 | 0 | 0 | 0 | 0 | 0 | 0 | 0 |
| EKEDPEPS | PCYOX1 | Q9UHG3 | PCYOX_HUMA | Unmodified | 0 | 0 | 26220 | 0 | 0 | 0 | 0 | 0 | 10029000 | 900620 | 209490 | 0 | 205460 | 173450 | 0 | 0 | 0 | 0 |
| EKEEFAVP | UBQLN2; UBQL | Q9UHD9; Q | UBQL2_HUMA | Unmodified | 764960 | 809500 | 564510 | 119640 | 0 | 0 | 0 | 0 | 0 | 241640 | 0 | 1469100 | 0 | 0 | 1097300 | 1524600 | 417590 | 0 |
| EKEELEQR | EIF3A | Q14152 | EIF3A_HUMAN | Unmodified | 0 | 0 | 0 | 0 | 0 | 0 | 0 | 0 | 0 | 0 | 0 | 1199600 | 0 | 0 | 0 | 108550 | 0 | 0 |
| EKEELMER | RDX; MSN; RDX | P35241; P26 | RADI_HUMAN | Unmodified | 0 | 0 | 0 | 0 | 0 | 0 | 0 | 0 | 0 | 0 | 0 | 707910 | 0 | 0 | 0 | 0 | 0 | 0 |
| EKEELMER | RDX; MSN; RDX | P35241; P26 | RADI_HUMAN | Unmodified | 0 | 0 | 0 | 0 | 637530 | 0 | 0 | 0 | 0 | 0 | 0 | 0 | 0 | 0 | 0 | 0 | 148810 | 0 |
| EKEFGDKV | PRXL2A | Q9BRX8 | PXL2A_HUMAN | Unmodified | 0 | 0 | 0 | 0 | 0 | 0 | 0 | 0 | 1794300 | 935020 | 0 | 0 | 0 | 0 | 0 | 0 | 0 | 0 |
| EKEKYQEE | LMAN1 | P49257 | LMAN1_HUMA | Unmodified | 0 | 0 | 0 | 0 | 0 | 0 | 0 | 0 | 2929100 | 3405500 | 0 | 0 | 0 | 0 | 0 | 0 | 0 | 427230 |
| EKENLSYD | HSPH1; HSPH1 | Q92598; R4 | HS105_HUMAN | Unmodified | 0 | 0 | 0 | 0 | 594230 | 0 | 0 | 0 | 0 | 336350 | 0 | 2849900 | 0 | 0 | 0 | 311020 | 436000 | 290940 |
| EKEPEEELY | FERMT3; FERM | Q86UX7; F5 | URP2_HUMAN | Unmodified | 652330 | 722800 | 1357600 | 2279400 | 1226100 | 0 | 2404000 | 978810 | 0 | 0 | 0 | 0 | 1958900 | 1834600 | 819550 | 904000 | 0 | 747030 |
| EKEPEYTLL | STAU1; STAU1 | O95793; Q5 | STAU1_HUMAN | Unmodified | 0 | 0 | 0 | 0 | 0 | 0 | 0 | 0 | 0 | 0 | 0 | 661990 | 0 | 0 | 0 | 0 | 0 | 0 |
| EKETGALY | UGGT1; UGGT1 | Q9NYU2; H7 | UGGG1_HUMA | Unmodified | 0 | 0 | 0 | 0 | 0 | 0 | 0 | 0 | 2294500 | 989960 | 0 | 0 | 0 | 0 | 0 | 0 | 0 | 0 |
| EKETLITIM | SCG3 | Q8WXD2 | SCG3_HUMAN | Unmodified | 0 | 0 | 0 | 0 | 0 | 0 | 0 | 0 | 12817000 | 1570700 | 0 | 0 | 0 | 0 | 0 | 0 | 0 | 0 |
| EKFAEASE | CLIP1; CLIP1; C | P30622; F5 | CLIP1_HUMAN | Unmodified | 0 | 0 | 0 | 0 | 0 | 0 | 0 | 0 | 472190 | 279590 | 0 | 0 | 325160 | 0 | 0 | 0 | 0 | 0 |
| EKFEEMIQ | CCT5; CCT5; C | P48643; B7Z | TCPE_HUMAN | Oxidation (M) | 0 | 0 | 0 | 0 | 0 | 0 | 0 | 0 | 60659 | 106550 | 490090 | 2805100 | 192860 | 0 | 0 | 0 | 0 | 0 |
| EKFEEMIQ | CCT5; CCT5; C | P48643; B7Z | TCPE_HUMAN | Unmodified | 29729 | 0 | 0 | 0 | 158710 | 0 | 178290 | 0 | 135090 | 0 | 716300 | 0 | 193350 | 0 | 0 | 0 | 0 | 163630 |
| EKFEGVEL | USP14; USP14 | P54578; A6 | UBP14_HUMA | Oxidation (M) | 0 | 0 | 0 | 0 | 0 | 0 | 0 | 0 | 0 | 0 | 0 | 3105000 | 0 | 0 | 0 | 0 | 0 | 0 |
| EKFEPTVFF | BZW2; BZW2; B | Q9Y6E2; Q7 | BZW2_HUMAN | Unmodified | 0 | 0 | 0 | 342060 | 0 | 0 | 0 | 0 | 0 | 0 | 0 | 0 | 621630 | 0 | 227810 | 0 | 0 | 0 |
| EKFHHLIPA | EEPD1 | Q7L9B9 | EEPD1_HUMAN | Unmodified | 0 | 0 | 0 | 0 | 0 | 0 | 202700 | 0 | 0 | 0 | 0 | 0 | 0 | 0 | 0 | 0 | 1549300 | 709760 |
| EKFPEADP | PNPT1; PNPT1 | Q8TCS8; H7 | PNPT1_HUMAN | Unmodified | 0 | 0 | 0 | 0 | 0 | 0 | 0 | 0 | 492150 | 153160 | 0 | 0 | 0 | 0 | 0 | 0 | 0 | 0 |
| EKGDYILIV | FLNC | Q14315 | FLNC_HUMAN | Unmodified | 0 | 0 | 0 | 56979 | 4234000 | 0 | 0 | 0 | 0 | 0 | 0 | 0 | 0 | 0 | 93278 | 0 | 5165800 | 516670 |
| EKGGELTL | DTX3L | Q8TDB6 | DTX3L_HUMAN | Unmodified | 0 | 0 | 0 | 0 | 0 | 0 | 189170 | 0 | 1054900 | 0 | 0 | 0 | 422410 | 322520 | 0 | 0 | 0 | 847630 |
| EKGGRSPD | PACSIN3; PACS | Q9UKS6; H0 | PACN3_HUMA | Unmodified | 291190 | 535020 | 0 | 296930 | 0 | 0 | 0 | 0 | 0 | 0 | 0 | 0 | 0 | 0 | 0 | 100940 | 0 | 0 |
| EKGNIQLS | IGF2R; IGF2R; I | P11717; A0A | MPRI_HUMAN | Unmodified | 0 | 0 | 0 | 0 | 0 | 0 | 0 | 0 | 264170 | 307690 | 53115 | 0 | 0 | 0 | 0 | 0 | 0 | 0 |
| EKGNIQLS | IGF2R; IGF2R; I | P11717; A0A | MPRI_HUMAN | Unmodified | 0 | 0 | 0 | 0 | 0 | 0 | 0 | 0 | 0 | 516620 | 104090 | 0 | 193780 | 199480 | 0 | 0 | 0 | 0 |
| EKGPTPEE | CHMP4A; CHM | Q9BY43; E9 | CHM4A_HUMA | Unmodified | 548790 | 2368800 | 364760 | 549460 | 1954900 | 60120 | 0 | 0 | 3100700 | 1834500 | 784280 | 786910 | 757140 | 577010 | 684980 | 937810 | 198650 | 184860 |
| EKHEMTAY | N/A | N/A | N/A | Unmodified | 18115000 | 49761000 | 5660500 | 3.79E+09 | 8.98E+08 | 2972800 | 0 | 348320 | 3.41E+09 | 13019000 | 2760500 | 18128000 | 0 | 0 | 10417000 | 0 | 97934000 | 4.46E+08 |
| EKHEMTAY | N/A | N/A | N/A | Oxidation (M) | 15341000 | 7841800 | 11258000 | 18152000 | 0 | 0 | 0 | 0 | 47161000 | 5928200 | 5465600 | 9603600 | 0 | 1811500 | 15598000 | 14202000 | 0 | 0 |
| EKIDLENTL | CCDC6 | Q16204 | CCDC6_HUMA | Unmodified | 0 | 64885 | 699260 | 0 | 0 | 739530 | 0 | 0 | 8851800 | 893090 | 979770 | 364810 | 2265200 | 0 | 834320 | 0 | 0 | 0 |
| EKIDQLQE | OPA1; OPA1; O | O60313; E5 | OPA1_HUMAN | Unmodified | 0 | 0 | 0 | 0 | 0 | 0 | 0 | 0 | 0 | 0 | 226910 | 1509100 | 0 | 0 | 0 | 0 | 192230 | 0 |
| EKIIQGQG | TIGAR | Q9NQ88 | TIGAR_HUMAN | Unmodified | 0 | 0 | 0 | 0 | 404720 | 84698 | 89669 | 219000 | 0 | 0 | 0 | 0 | 1061300 | 0 | 0 | 0 | 685640 | 249740 |
| EKIQEAKD | NAMPT; NAMPT | P43490; A0A | NAMPT_HUMA | Unmodified | 0 | 0 | 0 | 0 | 663840 | 76802 | 0 | 0 | 0 | 17048 | 0 | 0 | 0 | 0 | 0 | 0 | 0 | 0 |
| EKIQTGLK | IQGAP1; IQGAP | P46940; H0 | IQGA1_HUMAN | Unmodified | 0 | 0 | 86277 | 0 | 3095900 | 436820 | 60355 | 24766 | 0 | 32699 | 0 | 56813 | 359350 | 0 | 34583 | 0 | 50545 | 111050 |
| EKISVLQN | EEA1 | Q15075 | EEA1_HUMAN | Unmodified | 0 | 0 | 0 | 0 | 500270 | 0 | 0 | 0 | 0 | 0 | 0 | 0 | 0 | 0 | 0 | 0 | 0 | 0 |
| EKITAFVVE | ACADVL | P49748 | ACADV_HUMA | Unmodified | 0 | 0 | 0 | 0 | 0 | 0 | 0 | 0 | 192080 | 1834600 | 0 | 0 | 0 | 0 | 0 | 0 | 0 | 0 |
| EKKLTPITY | RAC1 | P63000 | RAC1_HUMAN | Oxidation (M) | 0 | 0 | 0 | 0 | 1513000 | 0 | 1572100 | 1296100 | 0 | 0 | 0 | 0 | 93746 | 928100 | 0 | 0 | 1384100 | 275720 |
| EKKLTPITY | RAC1 | P63000 | RAC1_HUMAN | Unmodified | 0 | 0 | 0 | 0 | 994580 | 0 | 440170 | 115290 | 0 | 164070 | 0 | 0 | 384270 | 365760 | 0 | 0 | 954020 | 0 |
| EKLAGLNQ | UTRN | P46939 | UTRN_HUMAN | Unmodified | 0 | 0 | 0 | 0 | 139400 | 0 | 0 | 0 | 0 | 77439 | 0 | 0 | 0 | 0 | 0 | 0 | 270990 | 113780 |
| EKLCYIALE | ACTRT1 | Q8TDG2 | ACTT1_HUMAN | Unmodified | 0 | 0 | 0 | 0 | 3253000 | 47295000 | 15611000 | 1331600 | 0 | 0 | 0 | 0 | 7058000 | 41973000 | 0 | 0 | 2657000 | 1.15E+08 |
| EKLCYVAL | POTEE; ACTG1; | Q6S8J3; P63 | POTEE_HUMAN | Oxidation (M) | 495960 | 0 | 65603 | 0 | 3141100 | 0 | 2621200 | 0 | 0 | 0 | 463580 | 1840300 | 4970500 | 2800000 | 232420 | 0 | 5264100 | 0 |
| EKLCYVAL | POTEE; ACTG1; | Q6S8J3; P63 | POTEE_HUMAN | Unmodified | 0 | 0 | 0 | 1442800 | 61660000 | 10434000 | 2551800 | 0 | 163800 | 6732000 | 0 | 0 | 19423000 | 2421800 | 0 | 0 | 26479000 | 18696000 |

| | | | | | | | | | | | | | | | | | | | | | | |
|---|---|---|---|---|---|---|---|---|---|---|---|---|---|---|---|---|---|---|---|---|---|---|
| EKLDCELQ | CLSTN1; CLSTN | O94985; A0A | CSTN1_HUMAI | Unmodified | 0 | 0 | 0 | 0 | 0 | 0 | 0 | 0 | 581540 | 0 | 0 | 0 | 0 | 0 | 0 | 0 | 0 | 0 |
| EKLEAYQH | IQGAP1; IQGAP | P46940; A0A | IQGA1_HUMAN | Unmodified | 1109100 | 1469800 | 0 | 1014900 | 1713900 | 304340 | 0 | 0 | 2116500 | 669040 | 407280 | 1885000 | 0 | 0 | 1463600 | 701020 | 0 | 0 |
| EKLEDPDP | AP3D1; AP3D1; | O14617; A0A | AP3D1_HUMAN | Unmodified | 0 | 0 | 0 | 0 | 0 | 0 | 0 | 0 | 0 | 0 | 0 | 996980 | 0 | 0 | 0 | 0 | 0 | 0 |
| EKLHIPWG | GNA13 | Q14344 | GNA13_HUMAI | Unmodified | 0 | 0 | 416940 | 0 | 0 | 0 | 0 | 0 | 0 | 0 | 1074200 | 0 | 0 | 0 | 467550 | 0 | 0 | 269960 |
| EKLIAPVAE | LDHB; LDHB; LI | P07195; A0A | LDHB_HUMAN | Unmodified | 0 | 0 | 0 | 0 | 4205600 | 0 | 0 | 0 | 0 | 0 | 0 | 0 | 1902700 | 510550 | 0 | 0 | 1113900 | 649990 |
| EKLMNVLS | FRYL; FRYL; FRY | O94915; H0' | FRYL_HUMAN I | Oxidation (M) | 0 | 0 | 0 | 0 | 0 | 0 | 0 | 0 | 0 | 0 | 0 | 641410 | 0 | 0 | 0 | 0 | 0 | 0 |
| EKLPANQY | TPP2; TPP2; TPI | P29144; A0A | TPP2_HUMAN ' | Unmodified | 0 | 0 | 0 | 0 | 0 | 0 | 84957 | 0 | 0 | 0 | 0 | 3152700 | 0 | 0 | 0 | 0 | 0 | 0 |
| EKLPGELEI | EIF3G; EIF3G; E | O75821; K7E | EIF3G_HUMAN | Unmodified | 962620 | 912950 | 1409200 | 2146800 | 1602200 | 262210 | 517780 | 469690 | 1217100 | 535060 | 2034400 | 5162900 | 3793800 | 1531200 | 1215100 | 1313400 | 980760 | 2405200 |
| EKLPLDINI | TTLL12 | Q14166 | TTL12_HUMAN | Unmodified | 0 | 0 | 712900 | 723430 | 0 | 0 | 193270 | 0 | 0 | 0 | 1073100 | 5221700 | 1362100 | 460130 | 929310 | 595010 | 157190 | 0 |
| EKLQIVEMI | SERPINH1; SER | P50454; E9P | SERPH_HUMAI | Unmodified | 0 | 0 | 0 | 0 | 321800 | 0 | 0 | 0 | 2262600 | 508180 | 0 | 0 | 0 | 0 | 0 | 0 | 0 | 0 |
| EKLTAASVC | SOD2; SOD2; S | P04179; F5C | SODM_HUMAN | Unmodified | 0 | 0 | 0 | 0 | 0 | 0 | 0 | 0 | 930000 | 1796800 | 0 | 0 | 0 | 0 | 0 | 0 | 0 | 0 |
| EKLTGTIQN | MMUT | P22033 | MUTA_HUMAN | Unmodified | 0 | 0 | 0 | 0 | 0 | 0 | 0 | 0 | 449210 | 138150 | 0 | 0 | 0 | 0 | 0 | 0 | 221290 | 0 |
| EKLVEENAI | UACA; UACA; U | Q9BZF9; H0' | UACA_HUMAN | Unmodified | 0 | 0 | 0 | 0 | 0 | 0 | 0 | 0 | 0 | 0 | 66319 | 0 | 855540 | 0 | 0 | 0 | 985980 | 0 |
| EKMNAKWI | ITGB1; ITGB1; I | P05556; H7( | ITB1_HUMAN I | Unmodified | 0 | 0 | 0 | 0 | 758950 | 0 | 0 | 0 | 0 | 0 | 0 | 0 | 0 | 0 | 0 | 0 | 1624800 | 4803700 |
| EKMNAKWI | ITGB1; ITGB1; I | P05556; H7( | ITB1_HUMAN I | Oxidation (M) | 0 | 0 | 0 | 0 | 3942300 | 651970 | 0 | 0 | 0 | 0 | 0 | 0 | 242780 | 144460 | 0 | 0 | 539080 | 2713500 |
| EKMNPLEQ | APLP1; APLP1; | P51693; B7Z | APLP1_HUMAN | Unmodified | 0 | 0 | 0 | 0 | 0 | 0 | 0 | 0 | 276500 | 0 | 0 | 0 | 0 | 0 | 146800 | 0 | 0 | 330500 |
| EKNAYQLT | CEP55 | Q53EZ4 | CEP55_HUMAN | Unmodified | 0 | 0 | 0 | 0 | 733660 | 91714 | 48932 | 356610 | 649300 | 1076000 | 15045 | 0 | 79889 | 0 | 0 | 0 | 271830 | 15709 |
| EKNDVEYY | CYLD; CYLD; C' | Q9NQC7; J3 | CYLD_HUMAN | Unmodified | 167850 | 204800 | 102070 | 0 | 382490 | 0 | 0 | 0 | 0 | 0 | 0 | 0 | 0 | 0 | 129350 | 103590 | 0 | 0 |
| EKNLLHVTI | HSP90B1; HSP | P14625; A0A | ENPL_HUMAN | Unmodified | 0 | 0 | 0 | 0 | 0 | 0 | 1256200 | 0 | 0 | 2301200 | 0 | 0 | 0 | 0 | 0 | 0 | 0 | 0 |
| EKNPLMYL | ITGA2 | P17301 | ITA2_HUMAN I | Unmodified | 0 | 183710 | 128300 | 107690 | 390370 | 0 | 0 | 0 | 0 | 0 | 0 | 0 | 0 | 0 | 0 | 0 | 0 | 195520 |
| EKNPLMYL | ITGA2 | P17301 | ITA2_HUMAN I | Oxidation (M) | 0 | 0 | 629440 | 0 | 0 | 0 | 0 | 0 | 0 | 0 | 0 | 0 | 0 | 0 | 102030 | 398940 | 0 | 0 |
| EKNQWQL | CNP; CNP | P09543; K7E | CN37_HUMAN | Unmodified | 0 | 52741 | 0 | 21863 | 5169300 | 1280100 | 935410 | 1049600 | 510850 | 63485 | 1040400 | 0 | 3248200 | 2959900 | 0 | 115430 | 2116200 | 5091800 |
| EKNQWQL | CNP; CNP | P09543; K7E | CN37_HUMAN | Unmodified | 0 | 0 | 0 | 0 | 3244000 | 0 | 752240 | 834800 | 560260 | 712090 | 1559600 | 748780 | 2992000 | 2324300 | 0 | 0 | 1414000 | 5459400 |
| EKPDDPLN | NDUFA11; NDU | Q86Y39; K7E | NDUAB_HUMA | Unmodified | 0 | 0 | 0 | 0 | 0 | 0 | 0 | 0 | 228590 | 65771 | 0 | 0 | 0 | 0 | 0 | 0 | 0 | 0 |
| EKPDQDAE | BASP1 | P80723 | BASP1_HUMAN | Unmodified | 0 | 14063 | 0 | 0 | 1666500 | 257830 | 0 | 0 | 10549000 | 0 | 0 | 0 | 0 | 0 | 0 | 0 | 57236 | 1652000 |
| EKPEEESLA | SEC16A; SEC1( | O15027; F1T | SC16A_HUMAN | Unmodified | 0 | 0 | 0 | 0 | 0 | 0 | 0 | 0 | 0 | 0 | 0 | 4264900 | 0 | 0 | 0 | 0 | 0 | 0 |
| EKPKEVAEI | EPB41L2; EPB4 | O43491; E9I | E41L2_HUMAN | Unmodified | 0 | 0 | 0 | 0 | 0 | 0 | 0 | 0 | 0 | 0 | 0 | 0 | 0 | 0 | 0 | 0 | 0 | 0 |
| EKPLALYVF | ALDH3A2; ALDI | P51648; A0A | AL3A2_HUMAN | Unmodified | 0 | 0 | 0 | 0 | 0 | 0 | 0 | 0 | 1088800 | 0 | 0 | 0 | 0 | 0 | 0 | 0 | 0 | 0 |
| EKPPGASV | USP43; USP43; | Q70EL4; V9C | UBP43_HUMAI | Unmodified | 0 | 0 | 0 | 296550 | 0 | 0 | 0 | 0 | 0 | 0 | 0 | 0 | 621510 | 142470 | 121930 | 91201 | 70703 | 0 |
| EKPPITVVG | PRPSAP1; PRPS | Q14558; O6I | KPRA_HUMAN | Unmodified | 0 | 0 | 647600 | 265520 | 269640 | 70752 | 0 | 0 | 1129900 | 0 | 2232700 | 8711900 | 242410 | 0 | 0 | 557640 | 0 | 0 |
| EKPQTLPS/ | PPP1R8 | Q12972 | PP1R8_HUMAI | Unmodified | 0 | 0 | 0 | 0 | 0 | 0 | 0 | 0 | 0 | 0 | 0 | 204550 | 0 | 0 | 0 | 0 | 0 | 0 |
| EKPSYDTE | CACYBP | Q9HB71 | CYBP_HUMAN | Unmodified | 0 | 0 | 60435 | 0 | 2374700 | 193730 | 319400 | 65913 | 6740000 | 422820 | 806220 | 4365500 | 2939100 | 1112500 | 0 | 0 | 479840 | 512140 |
| EKPSYDTE | CACYBP | Q9HB71 | CYBP_HUMAN | Oxidation (M) | 0 | 0 | 189960 | 402170 | 1957500 | 94662 | 325360 | 428750 | 1903800 | 476430 | 0 | 7977700 | 3045400 | 1072200 | 333920 | 310510 | 0 | 0 |
| EKPYFPIPE | HNRNPU; HNR | Q00839; A0A | HNRPU_HUMA | Unmodified | 4039000 | 0 | 0 | 0 | 0 | 0 | 4397600 | 0 | 3987300 | 871400 | 574380 | 5525900 | 547990 | 0 | 0 | 0 | 0 | 0 |
| EKPYFPIPE | HNRNPU; HNR | Q00839; A0A | HNRPU_HUMA | Unmodified | 1192400 | 775640 | 0 | 127270 | 0 | 0 | 0 | 0 | 2375600 | 0 | 0 | 0 | 0 | 0 | 0 | 0 | 0 | 0 |
| EKQDTFAL | ACAA1; ACAA1; | P09110; C9J | THIK_HUMAN 3 | Unmodified | 0 | 0 | 0 | 0 | 0 | 0 | 0 | 0 | 345880 | 346350 | 0 | 0 | 0 | 0 | 0 | 0 | 0 | 0 |
| EKQPVTGT | LMCD1; LMCD1 | Q9NZU5; H7 | LMCD1_HUMA | Unmodified | 0 | 0 | 0 | 0 | 0 | 0 | 29236 | 0 | 0 | 115250 | 329730 | 2401100 | 0 | 0 | 56934 | 0 | 120060 | 179080 |
| EKRDAFVY | TM9SF3; TM9SF | Q9HD45; Q5 | TM9S3_HUMA | Unmodified | 0 | 187120 | 0 | 0 | 0 | 0 | 126330 | 0 | 2057500 | 1485900 | 0 | 0 | 184180 | 139910 | 0 | 0 | 0 | 0 |
| EKRDLMAC | DDX3X; DDX3X | O00571; A0A | DDX3X_HUMAI | Unmodified | 0 | 85394 | 0 | 0 | 562950 | 106360 | 515100 | 206860 | 0 | 136510 | 0 | 0 | 0 | 481480 | 0 | 0 | 156310 | 0 |
| EKTELIQK | YWHAQ; YWHA | P27348; E9P | 1433T_HUMAN | Unmodified | 0 | 0 | 0 | 0 | 0 | 0 | 0 | 0 | 0 | 0 | 0 | 0 | 0 | 0 | 0 | 0 | 2548100 | 0 |
| EKTGQEYK | EPRS1; EPRS1 | P07814; V9C | SYEP_HUMAN | Unmodified | 0 | 0 | 0 | 0 | 335310 | 0 | 0 | 0 | 393220 | 104130 | 439040 | 0 | 599840 | 0 | 136050 | 0 | 89448 | 198880 |
| EKTHINIVV | EEF1A2; EEF1A | Q05639; Q5 | EF1A2_HUMAN | Unmodified | 0 | 0 | 0 | 0 | 2740500 | 279340 | 0 | 0 | 0 | 319630 | 0 | 0 | 4575900 | 1449000 | 0 | 0 | 1299600 | 1940800 |
| EKTHINIVV | EEF1A2; EEF1A | Q05639; Q5 | EF1A2_HUMAN | Unmodified | 0 | 0 | 0 | 0 | 1189200 | 0 | 1039800 | 0 | 0 | 0 | 0 | 0 | 1388100 | 1009200 | 0 | 0 | 3060300 | 1337400 |
| EKTPSPKEI | NUCKS1 | Q9H1E3 | NUCKS_HUMA | Unmodified | 0 | 0 | 0 | 0 | 0 | 0 | 0 | 0 | 0 | 265920 | 0 | 1220500 | 1090900 | 755380 | 0 | 0 | 0 | 0 |
| EKVDKDEV | MME | P08473 | NEP_HUMAN N | Unmodified | 0 | 0 | 390340 | 13858000 | 0 | 0 | 0 | 0 | 5312300 | 625110 | 0 | 292580 | 0 | 0 | 11633000 | 11082000 | 0 | 0 |
| EKVDLVLLC | ETFB | P38117 | ETFB_HUMAN I | Unmodified | 0 | 0 | 0 | 45434 | 133810 | 0 | 142290 | 0 | 1134500 | 172190 | 0 | 0 | 128990 | 0 | 0 | 16462 | 0 | 0 |
| EKVESELR | YWHAQ; YWHA | P27348; E9P | 1433T_HUMAN | Unmodified | 0 | 0 | 0 | 0 | 0 | 0 | 0 | 0 | 0 | 0 | 0 | 2846200 | 0 | 0 | 0 | 0 | 0 | 0 |
| EKVGEVTY | HNRNPM; HNR | P52272; M0I | HNRPM_HUMA | Oxidation (M) | 63395 | 0 | 0 | 0 | 0 | 0 | 1648100 | 23181 | 0 | 281020 | 0 | 0 | 255530 | 0 | 0 | 0 | 0 | 1302100 |
| EKVGEVTY | HNRNPM; HNR | P52272; M0I | HNRPM_HUMA | Unmodified | 12077000 | 0 | 0 | 17464 | 70556 | 0 | 158380 | 0 | 0 | 95748 | 182780 | 995920 | 0 | 0 | 60818 | 0 | 0 | 261640 |
| EKVNVDIIN | PSMD4; PIPSL; | P55036; A2A | PSMD4_HUMA | Unmodified | 0 | 0 | 0 | 0 | 1103000 | 0 | 0 | 0 | 0 | 0 | 0 | 677720 | 1244100 | 279410 | 0 | 0 | 0 | 890570 |
| EKVTNSTEI | EEA1 | Q15075 | EEA1_HUMAN I | Unmodified | 0 | 0 | 0 | 0 | 0 | 0 | 0 | 0 | 0 | 0 | 0 | 0 | 0 | 0 | 0 | 0 | 0 | 0 |
| EKWESLNA | UTRN; UTRN | P46939; Q5T | UTRN_HUMAN | Unmodified | 0 | 0 | 656220 | 1520000 | 3815000 | 0 | 0 | 0 | 0 | 0 | 0 | 0 | 654640 | 688740 | 1140900 | 1172000 | 0 | 1562800 |
| EKWVPEITI | CDC42; CDC42 | P60953; Q5J | CDC42_HUMA | Unmodified | 985760 | 1734200 | 0 | 715300 | 7372700 | 1650700 | 774440 | 0 | 1216600 | 1102700 | 766290 | 3064000 | 2955200 | 4197500 | 664880 | 571360 | 1440800 | 5629800 |
| EKYDNMAE | VPS28; VPS28; | Q9UK41; E9I | VPS28_HUMAN | Oxidation (M) | 2816200 | 1813200 | 864690 | 2640400 | 166240 | 234500 | 0 | 0 | 212840 | 0 | 0 | 279360 | 1057400 | 44054 | 2033100 | 1576500 | 0 | 0 |
| EKYDNMAE | VPS28; VPS28; | Q9UK41; E9I | VPS28_HUMAN | Unmodified | 3076100 | 3117700 | 636920 | 2547300 | 0 | 0 | 33445 | 0 | 0 | 436500 | 290220 | 123030 | 2045000 | 418580 | 2558800 | 35095 | 0 | 0 |
| EKYDTDFYI | DARS1 | P14868 | SYDC_HUMAN | Unmodified | 0 | 0 | 436900 | 93699 | 0 | 0 | 0 | 0 | 0 | 0 | 0 | 6165300 | 0 | 0 | 310470 | 0 | 0 | 0 |
| EKYEAIVEE | MYCBP; N/A | Q99417; A0A | MYCBP_HUMA | Unmodified | 80323 | 126850 | 0 | 0 | 388410 | 0 | 52550 | 0 | 175900 | 44608 | 168170 | 352210 | 209040 | 0 | 0 | 0 | 0 | 0 |
| EKYEITEQR | RPL6 | Q02878 | RL6_HUMAN 6( | Unmodified | 653830 | 973510 | 0 | 0 | 0 | 0 | 0 | 0 | 187500 | 0 | 0 | 0 | 0 | 0 | 0 | 0 | 0 | 0 |
| EKYEITEQR | RPL6 | Q02878 | RL6_HUMAN 6( | Unmodified | 0 | 0 | 0 | 0 | 0 | 0 | 0 | 0 | 0 | 0 | 0 | 0 | 0 | 0 | 0 | 436950 | 43543 | 0 |
| EKYGIVDYI | PDIA4; PDIA4 | P13667; A0A | PDIA4_HUMAN | Unmodified | 0 | 0 | 0 | 0 | 0 | 0 | 0 | 0 | 636260 | 0 | 0 | 0 | 0 | 0 | 0 | 0 | 0 | 0 |
| EKYGKDAT | ENO1; ENO1 | P06733; A0A | ENOA_HUMAN | Unmodified | 0 | 0 | 0 | 0 | 8279100 | 0 | 0 | 886310 | 0 | 0 | 0 | 0 | 1772800 | 0 | 0 | 0 | 3457400 | 0 |
| EKYIDQEEI | HSP90AA2P; HS | Q14568; P0I | HS902_HUMAI | Unmodified | 0 | 0 | 0 | 2950500 | 19309000 | 5329700 | 2329000 | 1114000 | 9671000 | 4681800 | 4042800 | 30530000 | 17166000 | 726820 | 0 | 1274800 | 5690700 | 14948000 |
| EKYPSHSFI | IMPA1; IMPA1; | P29218; E5R | IMPA1_HUMAN | Unmodified | 214280 | 0 | 0 | 76744 | 2200400 | 145820 | 482070 | 437460 | 1384600 | 1938400 | 0 | 1367400 | 1322300 | 349890 | 0 | 134540 | 1873200 | 3658400 |
| EKYQEEFEI | LMAN1 | P49257 | LMAN1_HUMAI | Unmodified | 0 | 0 | 545790 | 619640 | 0 | 0 | 124130 | 0 | 62870000 | 5222000 | 5654800 | 1027300 | 194050 | 218400 | 363860 | 268740 | 0 | 835370 |
| EKYQEEFEI | LMAN1 | P49257 | LMAN1_HUMAI | Unmodified | 0 | 0 | 0 | 0 | 0 | 0 | 0 | 0 | 1089900 | 426320 | 335810 | 0 | 0 | 0 | 0 | 0 | 0 | 0 |
| EKYSELVQI | HIP1 | O00291 | HIP1_HUMAN I | Unmodified | 0 | 0 | 0 | 0 | 0 | 0 | 0 | 0 | 143150 | 0 | 0 | 44712 | 1067600 | 793930 | 0 | 0 | 144310 | 107660 |
| ELAACQLR | KIF5B | P33176 | KINH_HUMAN | Unmodified | 0 | 0 | 0 | 0 | 666810 | 0 | 0 | 182230 | 0 | 0 | 0 | 0 | 343220 | 297330 | 0 | 0 | 283400 | 363170 |
| ELAAEVEV( | CYC1 | P08574 | CY1_HUMAN C | Oxidation (M) | 0 | 0 | 0 | 0 | 0 | 0 | 0 | 0 | 2319300 | 0 | 0 | 0 | 0 | 0 | 0 | 0 | 0 | 0 |
| ELAAEVEV( | CYC1 | P08574 | CY1_HUMAN C | Unmodified | 927050 | 0 | 363770 | 0 | 0 | 0 | 0 | 0 | 2604000 | 0 | 0 | 0 | 0 | 0 | 0 | 0 | 0 | 0 |
| ELAAEVEV( | CYC1 | P08574 | CY1_HUMAN C | 2 Oxidation (M) | 0 | 0 | 0 | 0 | 0 | 0 | 0 | 0 | 4626400 | 191190 | 0 | 0 | 0 | 0 | 0 | 0 | 0 | 0 |
| ELADESQTI | NSFL1C | Q9UNZ2 | NSF1C_HUMAI | Unmodified | 0 | 0 | 0 | 0 | 0 | 0 | 0 | 0 | 0 | 0 | 0 | 2382000 | 0 | 0 | 0 | 0 | 0 | 0 |
| ELADLMTR | JAK1; JAK1; JAK | P23458; A0A | JAK1_HUMAN 1 | Unmodified | 0 | 0 | 0 | 0 | 1601000 | 0 | 0 | 0 | 232630 | 130840 | 0 | 122580 | 0 | 0 | 136710 | 0 | 0 | 0 |
| ELAEAVAG( | UBA2; UBA2; UI | Q9UBT2; K7I | SAE2_HUMAN | Unmodified | 0 | 0 | 0 | 0 | 0 | 0 | 0 | 0 | 333200 | 0 | 612350 | 0 | 0 | 0 | 0 | 0 | 0 | 0 |
| ELAEDDSIL | ATP5ME | P56385 | ATP5I_HUMAN | Unmodified | 0 | 0 | 0 | 0 | 0 | 0 | 0 | 0 | 1014000 | 366070 | 481070 | 0 | 0 | 0 | 0 | 0 | 0 | 0 |
| ELAEDGYS | RPS3; RPS3; RF | P23396; H0' | RS3_HUMAN 4 | Unmodified | 1300300 | 1605900 | 745140 | 571660 | 0 | 222740 | 0 | 0 | 826760 | 0 | 3132700 | 46654000 | 0 | 0 | 1326500 | 1371800 | 0 | 0 |
| ELAEEEPHI | PEX19; PEX19; I | P40855; Q5( | PEX19_HUMAN | Unmodified | 85656 | 62964 | 282940 | 231670 | 698330 | 0 | 0 | 35583 | 630240 | 252020 | 179760 | 380220 | 1495400 | 1161300 | 189250 | 200100 | 383900 | 1113800 |
| ELAEQLGL! | EIF3G; EIF3G; E | O75821; K7E | EIF3G_HUMAN | Unmodified | 552660 | 409330 | 422630 | 646690 | 661590 | 0 | 0 | 142080 | 0 | 180420 | 0 | 3151900 | 1614700 | 1012700 | 399180 | 476610 | 724580 | 1357400 |
| ELAETLQQI | PTRHD1 | Q6GMV3 | PTRD1_HUMAI | Unmodified | 1359700 | 1014800 | 0 | 0 | 2443600 | 0 | 0 | 0 | 0 | 344930 | 1202800 | 6987200 | 0 | 0 | 121300 | 977410 | 783150 | 890500 |
| ELAFQISK | DDX39B; DDX3 | Q13838; O0( | DX39B_HUMAN | Unmodified | 0 | 0 | 0 | 0 | 296850 | 0 | 291180 | 196590 | 243480 | 232790 | 274660 | 3833100 | 0 | 0 | 0 | 0 | 27696 | 0 |
| ELAFRDSW | FAM3A; FAM3A; | P98173; D3I | FAM3A_HUMAI | Unmodified | 0 | 14632 | 0 | 97686 | 0 | 0 | 0 | 0 | 1206400 | 0 | 0 | 85879 | 0 | 0 | 44222 | 0 | 0 | 0 |
| ELAGHTGY | GNB1; GNB1; ( | P62873; B3K | GBB1_HUMAN | Unmodified | 4995200 | 1943800 | 2012900 | 2631400 | 378530 | 676930 | 537500 | 98710 | 26039000 | 1714500 | 8054200 | 1.18E+08 | 652970 | 597070 | 3269000 | 3874000 | 1931800 | 3684900 |
| ELAGQSRA | LAD1; LAD1 | O00515; E9I | LAD1_HUMAN | Unmodified | 0 | 0 | 0 | 0 | 85190 | 0 | 0 | 0 | 62983 | 0 | 0 | 0 | 154010 | 101400 | 0 | 0 | 0 | 500590 |
| ELAHILGFS | GRIK3 | Q13003 | GRIK3_HUMAN | Unmodified | 0 | 0 | 0 | 0 | 0 | 0 | 0 | 0 | 893090 | 34870 | 0 | 44379 | 0 | 0 | 0 | 162610 | 0 | 0 |
| ELAIERCFG | RABGAP1 | Q9Y3P9 | RBGP1_HUMAI | Unmodified | 0 | 0 | 0 | 0 | 163380 | 33071 | 78334 | 0 | 0 | 23905 | 0 | 0 | 37265 | 0 | 0 | 0 | 0 | 0 |
| ELAIQISK | ITGA1 | P56199 | ITA1_HUMAN I | Unmodified | 639410 | 579440 | 0 | 0 | 172040 | 0 | 145880 | 49421 | 145730 | 98580 | 407840 | 1303700 | 26991 | 0 | 0 | 0 | 51090 | 0 |
| ELAKVEEEI | TPD52; TPD52; | P55327; H0' | TPD52_HUMAN | Unmodified | 140640 | 0 | 0 | 213120 | 0 | 0 | 0 | 0 | 5103100 | 355550 | 409220 | 1755200 | 70757 | 0 | 371230 | 0 | 0 | 0 |
| ELALEEERL | BECN1; BECN1 | Q14457; K7E | BECN1_HUMAI | Unmodified | 22664 | 24516 | 0 | 0 | 350390 | 0 | 0 | 0 | 438210 | 0 | 0 | 61503 | 162940 | 0 | 66795 | 33292 | 112920 | 198410 |
| ELALEEERL | BECN1; BECN1 | Q14457; K7E | BECN1_HUMAI | Unmodified | 0 | 215790 | 0 | 62450 | 0 | 0 | 0 | 0 | 0 | 39523 | 0 | 0 | 113930 | 0 | 0 | 0 | 43947 | 183820 |
| ELALGQDR | DDT; DDTL; DD | P30046; A6N | DOPD_HUMAN | Unmodified | 821660 | 487640 | 923890 | 427410 | 61062 | 0 | 0 | 0 | 225790 | 0 | 521600 | 8657100 | 0 | 0 | 3822400 | 1874600 | 0 | 0 |
| ELALGQDR | DDT; DDT | P30046; J3K | DOPD_HUMAN | Unmodified | 0 | 0 | 0 | 0 | 247040 | 0 | 0 | 0 | 0 | 0 | 0 | 0 | 459540 | 0 | 0 | 0 | 279060 | 73769 |
| ELALNTAQI | SNX1; SNX1 | Q13596; H0' | SNX1_HUMAN | Unmodified | 0 | 0 | 0 | 115960 | 3409900 | 646590 | 0 | 174810 | 329360 | 143410 | 0 | 0 | 952090 | 388890 | 269730 | 0 | 200220 | 1003100 |
| ELALPGELT | VPS26A; VPS26 | O75436; S4I | VP26A_HUMAN | Unmodified | 0 | 0 | 0 | 0 | 3868300 | 0 | 0 | 0 | 1122500 | 232360 | 917720 | 4760900 | 0 | 0 | 1200800 | 519240 | 0 | 0 |
| ELALQPKD | GARS1; GARS1 | P41250; F8V | GARS_HUMAN | Unmodified | 117380 | 0 | 0 | 37581 | 89708 | 0 | 0 | 0 | 1588000 | 0 | 1483100 | 14607000 | 0 | 0 | 263310 | 829840 | 0 | 0 |
| ELALQPKD | GARS1; GARS1 | P41250; F8V | GARS_HUMAN | Unmodified | 0 | 0 | 0 | 0 | 0 | 0 | 156050 | 0 | 1199300 | 227760 | 0 | 5724700 | 300510 | 0 | 0 | 0 | 416420 | 0 |
| ELALVQTK | NEDD4L; NEDD | Q96PU5; A0A | NED4L_HUMAN | Unmodified | 0 | 0 | 0 | 0 | 0 | 0 | 0 | 0 | 0 | 0 | 0 | 267410 | 0 | 0 | 0 | 0 | 0 | 0 |
| ELAMPGED | TUFM | P49411 | EFTU_HUMAN | Oxidation (M) | 0 | 0 | 0 | 0 | 0 | 0 | 0 | 0 | 0 | 0 | 0 | 0 | 0 | 0 | 0 | 0 | 0 | 0 |
| ELAMPGED | TUFM | P49411 | EFTU_HUMAN | Unmodified | 0 | 0 | 0 | 0 | 263290 | 0 | 305120 | 133050 | 5053400 | 643390 | 0 | 0 | 0 | 0 | 0 | 0 | 0 | 0 |
| ELANFDPS | LSR; LSR | Q86X29; S4I | LSR_HUMAN L | Unmodified | 1314600 | 1115900 | 0 | 5065500 | 0 | 1829000 | 0 | 0 | 24654000 | 9593800 | 0 | 5225500 | 2401600 | 1278600 | 4749000 | 3303600 | 187350 | 395110 |
| ELAPAVSVI | COPG2 | Q9UBF2 | COPG2_HUMA | Unmodified | 0 | 0 | 0 | 0 | 0 | 0 | 0 | 0 | 106490 | 0 | 0 | 302020 | 0 | 0 | 0 | 61403 | 0 | 0 |
| ELAPLQELI | MOB1B; MOB1/ | Q7L9L4; Q9I | MOB1B_HUMA | Unmodified | 0 | 0 | 0 | 0 | 0 | 0 | 0 | 0 | 617220 | 0 | 1018000 | 5711800 | 0 | 0 | 0 | 1357200 | 0 | 0 |
| ELAPYDEN | RPS19; RPS19; | P39019; M0( | RS19_HUMAN | Unmodified | 3459200 | 8266300 | 0 | 1856400 | 0 | 0 | 0 | 0 | 1368900 | 0 | 935420 | 8970600 | 991290 | 0 | 1211400 | 1799800 | 0 | 0 |
| ELAPYDEN | RPS19 | P39019 | RS19_HUMAN | Unmodified | 0 | 0 | 0 | 0 | 2221800 | 0 | 0 | 0 | 520860 | 0 | 0 | 155090 | 0 | 0 | 0 | 0 | 0 | 0 |
| ELAQIAGRI | PPM1G | O15355 | PPM1G_HUMA | Unmodified | 0 | 0 | 0 | 0 | 0 | 0 | 0 | 0 | 0 | 0 | 0 | 236760 | 0 | 0 | 0 | 0 | 0 | 0 |
| ELAQLDQS | DIPK2A | Q8NDZ4 | DIK2A_HUMAN | Unmodified | 0 | 0 | 0 | 0 | 0 | 0 | 0 | 0 | 10070000 | 415460 | 0 | 328290 | 0 | 0 | 0 | 76710 | 263300 | 0 |
| ELAQQIQK | EIF4A1; EIF4A2 | P60842; Q14 | IF4A1_HUMAN | Unmodified | 1317200 | 1137400 | 2100000 | 782270 | 74897 | 0 | 0 | 245870 | 764600 | 232840 | 1011600 | 10059000 | 0 | 0 | 2348400 | 3250800 | 1168500 | 0 |
| ELAQQVQC | DDX17; DDX17 | Q92841; A0A | DDX17_HUMAN | Unmodified | 0 | 0 | 0 | 0 | 0 | 0 | 0 | 115080 | 462690 | 92426 | 634730 | 8153500 | 0 | 0 | 0 | 0 | 0 | 0 |
| ELAQTTRTA | CCT8; CCT8 | P50990; H7( | TCPQ_HUMAN | Oxidation (M) | 0 | 0 | 0 | 0 | 472590 | 73589 | 291660 | 316840 | 0 | 0 | 0 | 0 | 933830 | 401640 | 0 | 0 | 560340 | 631660 |
| ELAQTTRT/ | CCT8; CCT8 | P50990; H7( | TCPQ_HUMAN | Unmodified | 0 | 0 | 0 | 0 | 1038800 | 0 | 352830 | 237250 | 0 | 221580 | 0 | 0 | 884670 | 588310 | 0 | 0 | 706850 | 932940 |
| ELASALKS/ | ANXA2P2; ANX/ | A6NMY6; P0 | AXA2L_HUMAN | Unmodified | 0 | 0 | 0 | 0 | 2863800 | 0 | 0 | 0 | 0 | 0 | 0 | 0 | 0 | 0 | 0 | 0 | 1256200 | 1601900 |

| | | | | | | | | | | | | | | | | | | | | | | |
|---|---|---|---|---|---|---|---|---|---|---|---|---|---|---|---|---|---|---|---|---|---|---|
| ELASQPDV | TPI1 | P60174 | TPIS_HUMAN T | Unmodified | 0 | 0 | 0 | 0 | 1152100 | 0 | 0 | 0 | 416570 | 429680 | 5397500 | 557040 | 0 | 0 | 0 | 0 | 0 | 0 |
| ELASQPDV | TPI1 | P60174 | TPIS_HUMAN T | Unmodified | 4240500 | 10105000 | 318070 | 5514800 | 1.87E+08 | 13955000 | 41831000 | 0 | 6435800 | 31347000 | 3665200 | 8051700 | 51840000 | 51299000 | 3815900 | 2139200 | 1.25E+08 | 1.62E+08 |
| ELATLQR | IQGAP1; IQGAP | P46940; A0A | IQGA1_HUMAN | Unmodified | 349990 | 739420 | 32430 | 0 | 2480100 | 1427000 | 348700 | 207990 | 430600 | 663510 | 0 | 1900600 | 348940 | 0 | 0 | 0 | 183560 | 918240 |
| ELATVLSD( | NCKAP1 | Q9Y2A7 | NCKP1_HUMAI | Unmodified | 95270 | 0 | 411650 | 0 | 0 | 0 | 0 | 0 | 0 | 0 | 0 | 3778700 | 0 | 0 | 223380 | 184340 | 0 | 0 |
| ELAVEGMH | LARS1; LARS1; | Q9P2J5; A0A | SYLC_HUMAN | Unmodified | 79416 | 0 | 0 | 0 | 309940 | 0 | 0 | 26448 | 94878 | 0 | 0 | 559190 | 0 | 0 | 100650 | 0 | 54345 | 0 |
| ELAVFREK | IGFBP2; IGFBP2 | P18065; C9J | IBP2_HUMAN I | Unmodified | 0 | 0 | 0 | 0 | 278280 | 396190 | 95861 | 0 | 4098600 | 7092100 | 0 | 0 | 0 | 0 | 0 | 0 | 52293 | 584140 |
| ELAVQIQK | EIF4A3 | P38919 | IF4A3_HUMAN | Unmodified | 230030 | 0 | 0 | 164610 | 535810 | 125450 | 74277 | 15018 | 417290 | 108960 | 0 | 2593500 | 0 | 73102 | 0 | 0 | 400180 | 254960 |
| ELCFPETRI | RET; RET; RET; I | P07949; Q9E | RET_HUMAN P | Unmodified | 0 | 0 | 0 | 0 | 0 | 0 | 0 | 0 | 1661900 | 0 | 0 | 0 | 0 | 120760 | 0 | 0 | 0 | 0 |
| ELCGTEPH | CCER2 | I3L3R5 | CCER2_HUMAI | Unmodified | 0 | 0 | 0 | 0 | 0 | 0 | 0 | 1251500 | 0 | 0 | 0 | 0 | 0 | 0 | 0 | 0 | 0 | 0 |
| ELCHVCCI | ADAM10 | O14672 | ADA10_HUMAN | Unmodified | 533350 | 124340 | 580660 | 1104200 | 165470 | 0 | 82764 | 0 | 0 | 0 | 0 | 2496600 | 1037300 | 812620 | 786830 | 845490 | 0 | 80831 |
| ELCLHAAQ | GALNT3; GALNT | Q14435; E7E | GALT3_HUMAN | Unmodified | 0 | 0 | 0 | 156010 | 0 | 0 | 0 | 0 | 1668300 | 516490 | 364380 | 0 | 405740 | 490290 | 0 | 96096 | 0 | 149000 |
| ELCQGLG( | RNH1; RNH1 | P13489; H0Y | RINI_HUMAN R | Unmodified | 0 | 0 | 0 | 0 | 0 | 0 | 0 | 0 | 127260 | 0 | 0 | 3122300 | 0 | 0 | 0 | 0 | 0 | 0 |
| ELCSLASDI | ECPAS; ECPAS | Q5VYK3; J3K | ECM29_HUMAI | Unmodified | 0 | 0 | 0 | 140590 | 0 | 0 | 0 | 30005 | 0 | 0 | 0 | 574370 | 0 | 0 | 90868 | 0 | 0 | 0 |
| ELDATNISII | EPHA3; EPHA3 | P29320; C9J | EPHA3_HUMAN | Unmodified | 1790500 | 3828900 | 360130 | 262880 | 0 | 0 | 0 | 0 | 261110 | 0 | 0 | 0 | 0 | 0 | 0 | 178590 | 0 | 0 |
| ELDAYNPN | SORCS2; SORC | Q96PQ0; B5 | SORC2_HUMAI | Unmodified | 0 | 0 | 0 | 0 | 0 | 0 | 0 | 136040 | 4972900 | 730130 | 0 | 0 | 0 | 0 | 0 | 0 | 0 | 0 |
| ELDDATEAI | MYH10 | P35580 | MYH10_HUMAI | Unmodified | 0 | 0 | 0 | 0 | 0 | 0 | 0 | 0 | 0 | 0 | 278030 | 2192900 | 0 | 0 | 0 | 0 | 0 | 0 |
| ELDDISQEI | EPS15L1; EPS1 | Q9UBC2; M( | EP15R_HUMAN | Unmodified | 0 | 0 | 0 | 0 | 0 | 0 | 0 | 0 | 0 | 0 | 0 | 0 | 1858300 | 0 | 0 | 0 | 0 | 0 |
| ELDDYREE | OCLN; OCLN | Q16625; A0/ | OCLN_HUMAN | Unmodified | 0 | 0 | 0 | 800570 | 3012500 | 194210 | 0 | 0 | 2888600 | 0 | 0 | 0 | 1236100 | 1056400 | 2065300 | 309990 | 0 | 0 |
| ELDDYREE | OCLN; OCLN | Q16625; A0/ | OCLN_HUMAN | Oxidation (M) | 0 | 0 | 1106100 | 2200000 | 0 | 344550 | 0 | 0 | 0 | 0 | 0 | 0 | 1596600 | 2049100 | 7212000 | 529680 | 0 | 0 |
| ELDEEGNC | ROCK1; ROCK1 | Q13464; A0/ | ROCK1_HUMAI | Unmodified | 0 | 0 | 0 | 0 | 3900600 | 478920 | 0 | 0 | 228460 | 622050 | 0 | 0 | 1228800 | 0 | 0 | 0 | 1749900 | 2461800 |
| ELDEEHSQ | SLK | Q9H2G2 | SLK_HUMAN S | Unmodified | 69022 | 0 | 121100 | 130860 | 88031 | 194480 | 49869 | 22986 | 541040 | 217980 | 517300 | 4256400 | 250540 | 181560 | 163430 | 169390 | 23477 | 0 |
| ELDESLQV | CLU; CLU; CLU | P10909; H0Y | CLUS_HUMAN | Unmodified | 0 | 1200500 | 0 | 0 | 0 | 0 | 0 | 0 | 2628200 | 0 | 0 | 0 | 0 | 0 | 121550 | 0 | 0 | 0 |
| ELDFVSHH | NUCB1; NUCB1 | Q02818; H7I | NUCB1_HUMA | Unmodified | 0 | 0 | 0 | 0 | 93593 | 0 | 17017 | 0 | 1521700 | 0 | 0 | 168450 | 0 | 0 | 0 | 77397 | 0 | 0 |
| ELDGLWSF | GUSB; GUSB; G | P08236; F8W | BGLR_HUMAN | Unmodified | 0 | 0 | 0 | 0 | 0 | 0 | 0 | 0 | 26570000 | 1153100 | 0 | 188080 | 0 | 0 | 345530 | 0 | 0 | 0 |
| ELDGPVYE | SRGAP1; SRGA | Q7Z6B7; G5I | SRGP1_HUMAI | Unmodified | 0 | 679180 | 0 | 0 | 1159100 | 9318600 | 1446500 | 0 | 1404100 | 39010 | 19622000 | 4595500 | 0 | 0 | 0 | 0 | 875690 | 172420 |
| ELDIMEPK\ | PSMD2 | Q13200 | PSMD2_HUMA | Oxidation (M) | 0 | 0 | 0 | 239880 | 0 | 0 | 0 | 0 | 0 | 202810 | 85963 | 2434100 | 0 | 0 | 318120 | 267270 | 0 | 0 |
| ELDINNQV | GOLM2; GOLM2 | Q6P4E1; H0Y | GOLM2_HUMA | Unmodified | 0 | 0 | 0 | 0 | 0 | 0 | 0 | 0 | 8133700 | 898110 | 771330 | 861910 | 0 | 252680 | 0 | 0 | 0 | 0 |
| ELDLNSVLI | RNASET2; N/A; | O00584; H0Y | RNT2_HUMAN | Unmodified | 0 | 0 | 307970 | 0 | 0 | 0 | 0 | 0 | 2809500 | 0 | 979070 | 0 | 0 | 0 | 0 | 337840 | 0 | 0 |
| ELDLVSHH | NUCB2; NUCB2 | P80303; Q2L | NUCB2_HUMA | Unmodified | 0 | 0 | 203010 | 0 | 0 | 0 | 0 | 0 | 22363000 | 2934600 | 314380 | 0 | 0 | 0 | 425400 | 0 | 0 | 0 |
| ELDLVSHH | NUCB2; NUCB2 | P80303; Q2L | NUCB2_HUMA | Unmodified | 0 | 0 | 0 | 0 | 0 | 0 | 0 | 0 | 0 | 5057100 | 0 | 0 | 0 | 0 | 0 | 0 | 0 | 0 |
| ELDLYDNQ | PPP1R7; PPP1R | Q15435; C9J | PP1R7_HUMAN | Unmodified | 0 | 0 | 0 | 0 | 0 | 0 | 0 | 130660 | 0 | 594710 | 0 | 747010 | 0 | 0 | 0 | 0 | 0 | 0 |
| ELDPTNMT | STIP1 | P31948 | STIP1_HUMAN | Oxidation (M) | 0 | 0 | 0 | 0 | 2669400 | 0 | 0 | 0 | 0 | 661220 | 408420 | 1383900 | 3359100 | 0 | 0 | 0 | 1296500 | 2952700 |
| ELDPTNMT | STIP1 | P31948 | STIP1_HUMAN | Unmodified | 0 | 0 | 0 | 138130 | 0 | 0 | 0 | 0 | 1679500 | 0 | 0 | 0 | 1516800 | 0 | 0 | 26375 | 0 | 0 |
| ELDPTNMT | STIP1 | P31948 | STIP1_HUMAN | Oxidation (M) | 0 | 0 | 0 | 0 | 1301700 | 0 | 2860900 | 512520 | 0 | 0 | 1179000 | 0 | 0 | 10403000 | 0 | 0 | 1814200 | 0 |
| ELDPTNMT | STIP1 | P31948 | STIP1_HUMAN | Unmodified | 0 | 0 | 0 | 0 | 2832500 | 0 | 0 | 0 | 0 | 1785300 | 0 | 0 | 0 | 0 | 0 | 0 | 2521800 | 0 |
| ELDQGTPC | PACSIN2; PACS | Q9UNF0; A0 | PACN2_HUMAI | 2 Oxidation (M) | 2105600 | 1130700 | 0 | 1073700 | 0 | 0 | 0 | 0 | 0 | 0 | 0 | 0 | 2222900 | 2363000 | 2048900 | 455290 | 0 | 0 |
| ELDQGTPC | PACSIN2; PACS | Q9UNF0; A0 | PACN2_HUMAI | Oxidation (M) | 0 | 3345000 | 0 | 2136000 | 0 | 0 | 0 | 0 | 0 | 0 | 0 | 0 | 6417800 | 4860400 | 669450 | 0 | 0 | 0 |
| ELDQGTPC | PACSIN2; PACS | Q9UNF0; A0 | PACN2_HUMAI | Unmodified | 0 | 0 | 0 | 0 | 850840 | 0 | 0 | 0 | 0 | 0 | 0 | 0 | 6200500 | 2384200 | 1834600 | 0 | 0 | 0 |
| ELDQWIEQ | PPP2CA; N/A | P67775; E5R | PP2AA_HUMAN | Unmodified | 0 | 0 | 0 | 0 | 2240900 | 0 | 1456900 | 283180 | 3380600 | 0 | 0 | 2617600 | 0 | 2044100 | 0 | 0 | 2163700 | 0 |
| ELDQWIEQ | PPP2CA; N/A | P67775; E5R | PP2AA_HUMAN | Unmodified | 0 | 0 | 0 | 0 | 0 | 0 | 0 | 0 | 0 | 985030 | 0 | 0 | 789780 | 0 | 0 | 0 | 1965900 | 0 |
| ELDRYSLD | SHANK2; SHAN | Q9UPX8; A6I | SHAN2_HUMAI | Unmodified | 0 | 0 | 0 | 159720 | 0 | 0 | 0 | 0 | 0 | 0 | 0 | 0 | 222950 | 0 | 158630 | 154450 | 0 | 0 |
| ELDSGLAE | IST1; IST1; IST1 | P53990; H3E | IST1_HUMAN I | Unmodified | 0 | 524610 | 0 | 723910 | 1129700 | 191800 | 0 | 0 | 1471600 | 1162900 | 302410 | 577080 | 404710 | 213700 | 700960 | 70661 | 0 | 225380 |
| ELDSGLAE | IST1; IST1; IST1 | P53990; H3E | IST1_HUMAN I | Unmodified | 0 | 0 | 0 | 0 | 347410 | 0 | 0 | 0 | 0 | 0 | 0 | 0 | 0 | 418090 | 0 | 0 | 0 | 0 |
| ELDSNPFA | PNPO; PNPO | Q9NVS9; J3( | PNPO_HUMAN | Unmodified | 0 | 0 | 0 | 0 | 855600 | 0 | 0 | 0 | 0 | 118480 | 0 | 0 | 3647800 | 2351900 | 0 | 0 | 559990 | 1371100 |
| ELDTLNNE | EPS15; EPS15 | P42566; B1A | EPS15_HUMAN | Unmodified | 0 | 0 | 384090 | 93225 | 0 | 262950 | 398740 | 177200 | 0 | 0 | 0 | 906730 | 1416000 | 928120 | 0 | 0 | 0 | 1235700 |
| ELDTVTLED | PSMD3 | O43242 | PSMD3_HUMA | Unmodified | 0 | 0 | 0 | 0 | 457700 | 1283800 | 157950 | 0 | 0 | 0 | 612050 | 1646900 | 0 | 0 | 0 | 0 | 365130 | 0 |
| ELDVEEAH | DCPS | Q96C86 | DCPS_HUMAN | Unmodified | 0 | 0 | 0 | 0 | 0 | 0 | 0 | 0 | 271910 | 14829 | 1052800 | 2406100 | 0 | 0 | 0 | 0 | 0 | 0 |
| ELDVLQGR | NPTXR; NPTXR | O95502; A0/ | NPTXR_HUMAN | Unmodified | 0 | 0 | 0 | 0 | 0 | 0 | 0 | 0 | 6870200 | 1070400 | 375700 | 0 | 0 | 0 | 0 | 0 | 0 | 0 |
| ELDVSKNG | ERBIN; ERBIN | Q96RT1; B4I | ERBIN_HUMAN | Unmodified | 0 | 0 | 0 | 0 | 288570 | 0 | 0 | 0 | 3361100 | 916680 | 0 | 0 | 0 | 0 | 0 | 0 | 0 | 0 |
| ELDVVREIY | FMNL2 | Q96PY5 | FMNL2_HUMAI | Unmodified | 0 | 0 | 0 | 0 | 1554400 | 0 | 103650 | 131880 | 0 | 370540 | 0 | 0 | 0 | 0 | 0 | 0 | 0 | 1159700 |
| ELEAELEDE | MYH10 | P35580 | MYH10_HUMAI | Unmodified | 0 | 0 | 895880 | 0 | 0 | 443870 | 0 | 377740 | 185200 | 0 | 166310 | 4357900 | 0 | 0 | 0 | 0 | 0 | 0 |
| ELEAENYH | SPTBN1; SPTBN | Q01082; A0/ | SPTB2_HUMAN | Unmodified | 0 | 0 | 0 | 0 | 0 | 0 | 0 | 0 | 0 | 23892 | 0 | 2540200 | 0 | 0 | 0 | 0 | 0 | 0 |
| ELEAFREEL | LRPAP1 | P30533 | AMRP_HUMAN | Unmodified | 0 | 0 | 0 | 0 | 0 | 0 | 0 | 0 | 1498800 | 143930 | 0 | 0 | 0 | 0 | 0 | 0 | 0 | 0 |
| ELEALDEVI | NENF | Q9UMX5 | NENF_HUMAN | Unmodified | 0 | 0 | 0 | 0 | 0 | 0 | 0 | 174340 | 56105000 | 25789000 | 2474300 | 1355900 | 0 | 0 | 0 | 0 | 0 | 0 |
| ELEALDEVI | NENF | Q9UMX5 | NENF_HUMAN | Unmodified | 0 | 0 | 0 | 0 | 0 | 0 | 0 | 0 | 0 | 1091700 | 0 | 0 | 0 | 0 | 0 | 0 | 0 | 0 |
| ELEANVLA | SMC4; SMC4 | Q9NTJ3; E9F | SMC4_HUMAN | Unmodified | 0 | 0 | 0 | 0 | 0 | 0 | 0 | 0 | 355130 | 0 | 0 | 852330 | 0 | 0 | 0 | 0 | 0 | 0 |
| ELEAQLDD | CDC42BPB | Q9Y5S2 | MRCKB_HUMA | Unmodified | 0 | 0 | 0 | 0 | 1731200 | 0 | 0 | 0 | 0 | 0 | 0 | 0 | 0 | 0 | 0 | 0 | 0 | 2022800 |
| ELEASEELD | GLRX3 | O76003 | GLRX3_HUMAN | Unmodified | 0 | 0 | 0 | 0 | 0 | 0 | 0 | 0 | 0 | 0 | 388110 | 1369900 | 0 | 0 | 0 | 126840 | 0 | 0 |
| ELEDATETA | MYH9 | P35579 | MYH9_HUMAN | Unmodified | 0 | 0 | 0 | 0 | 0 | 0 | 0 | 0 | 197320 | 0 | 397200 | 1076700 | 0 | 0 | 0 | 0 | 0 | 0 |
| ELEDATETA | MYH9 | P35579 | MYH9_HUMAN | Oxidation (M) | 65559 | 0 | 0 | 0 | 0 | 0 | 0 | 0 | 0 | 0 | 0 | 1180700 | 0 | 0 | 0 | 49855 | 0 | 0 |
| ELEDATETA | MYH9 | P35579 | MYH9_HUMAN | Unmodified | 0 | 0 | 0 | 0 | 3207600 | 0 | 0 | 0 | 589710 | 894690 | 0 | 3572500 | 0 | 0 | 0 | 0 | 820590 | 0 |
| ELEDKVIEL | EPB41L3; EPB4 | Q9Y2J2; A0A | E41L3_HUMAN | Unmodified | 0 | 0 | 0 | 0 | 0 | 0 | 0 | 0 | 0 | 0 | 0 | 0 | 0 | 0 | 0 | 0 | 681110 | 0 |
| ELEDYIDNL | RAB11FIP2; RAI | Q7L804; Q6\ | RFIP2_HUMAN | Unmodified | 0 | 0 | 0 | 0 | 0 | 0 | 0 | 0 | 0 | 0 | 40538 | 64339 | 0 | 0 | 0 | 0 | 0 | 0 |
| ELEEDFIK | COTL1; COTL1 | Q14019; H3I | COTL1_HUMAN | Unmodified | 0 | 0 | 392810 | 0 | 706950 | 0 | 0 | 0 | 9961400 | 755090 | 0 | 4843500 | 0 | 0 | 650780 | 498990 | 100180 | 0 |
| ELEEDFIKS | COTL1; COTL1 | Q14019; H3I | COTL1_HUMAN | Unmodified | 0 | 0 | 0 | 0 | 48482 | 647750 | 0 | 0 | 5237100 | 2254000 | 0 | 893510 | 0 | 0 | 0 | 0 | 1902200 | 0 |
| ELEEDFIKS | COTL1; COTL1 | Q14019; H3I | COTL1_HUMAN | Unmodified | 0 | 0 | 0 | 0 | 630070 | 375810 | 0 | 0 | 2632500 | 1711200 | 0 | 0 | 0 | 0 | 0 | 0 | 287320 | 1773100 |
| ELEEEVNN | SEPTIN11; SEP1 | Q9NVA2; H0 | SEP11_HUMAN | Unmodified | 0 | 0 | 234500 | 336710 | 2845000 | 590360 | 199970 | 164880 | 259720 | 196510 | 388370 | 1437400 | 412290 | 308070 | 185010 | 258120 | 485920 | 665040 |
| ELEEEVNN | SEPTIN11; SEP1 | Q9NVA2; H0 | SEP11_HUMAN | Unmodified | 0 | 0 | 0 | 0 | 1526600 | 465800 | 0 | 0 | 2640500 | 0 | 0 | 0 | 0 | 0 | 0 | 0 | 352050 | 0 |
| ELEEEWVK | CKAP5 | Q14008 | CKAP5_HUMAN | Unmodified | 0 | 0 | 0 | 0 | 0 | 0 | 0 | 0 | 0 | 0 | 0 | 964970 | 0 | 0 | 0 | 0 | 0 | 0 |
| ELEEIVQPII | HSPA5; HSPA5; | P11021; A0A | BIP_HUMAN Er | Unmodified | 786140 | 0 | 6472200 | 1230800 | 2952000 | 424340 | 6926700 | 5486700 | 2.99E+08 | 79817000 | 32907000 | 7705100 | 2053900 | 2268900 | 6005900 | 1455300 | 338840 | 1331100 |
| ELEEIVQPII | HSPA5; HSPA5; | P11021; A0A | BIP_HUMAN Er | Unmodified | 0 | 0 | 0 | 0 | 0 | 0 | 1788900 | 0 | 0 | 3350200 | 0 | 0 | 0 | 0 | 0 | 0 | 0 | 0 |
| ELEEKVAEL | EPB41L2; EPB4 | O43491; A0/ | E41L2_HUMAN | Unmodified | 0 | 0 | 0 | 0 | 2784700 | 727640 | 0 | 296990 | 0 | 0 | 0 | 0 | 0 | 0 | 0 | 0 | 1636900 | 1498000 |
| ELEELQNN | SELENON; N/A; | Q9NZV5; H3 | SELN_HUMAN | Unmodified | 0 | 0 | 0 | 0 | 0 | 0 | 0 | 0 | 3517300 | 374210 | 0 | 0 | 0 | 0 | 0 | 0 | 0 | 0 |
| ELEESKYQ\ | AMPD3; AMPD3 | Q01432; E9F | AMPD3_HUMA | Unmodified | 0 | 0 | 0 | 0 | 0 | 0 | 0 | 0 | 0 | 0 | 0 | 1320600 | 0 | 0 | 0 | 0 | 0 | 0 |
| ELEESSFRI | INSR | P06213 | INSR_HUMAN I | Unmodified | 2.88E+08 | 2.4E+08 | 1.95E+08 | 99221000 | 0 | 0 | 0 | 0 | 1032600 | 61954 | 0 | 0 | 1476300 | 340090 | 3.5E+08 | 2.36E+08 | 38199 | 0 |
| ELEETLNK | CLASP2; CLASF | O75122; B3I | CLAP2_HUMAN | Unmodified | 0 | 0 | 0 | 0 | 0 | 0 | 0 | 0 | 0 | 0 | 0 | 1641500 | 0 | 0 | 0 | 0 | 0 | 0 |
| ELEEWYAR | CLTA; CLTA; CL | P09496; C9J | CLCA_HUMAN | Unmodified | 1653900 | 1144800 | 710160 | 0 | 114740 | 0 | 0 | 0 | 188880 | 0 | 0 | 16138000 | 0 | 0 | 653380 | 986420 | 0 | 0 |
| ELEEWYAR | CLTA; CLTA; CL | P09496; C9J | CLCA_HUMAN | Unmodified | 18778000 | 12298000 | 3374700 | 10039000 | 29239000 | 7338000 | 2966200 | 1086600 | 7814900 | 3234900 | 2015200 | 51518000 | 19292000 | 8078700 | 8131500 | 6988000 | 2912000 | 13374000 |
| ELEFLGWY | COPS6; COPS6 | Q7L5N1; E7I | CSN6_HUMAN | Unmodified | 0 | 0 | 0 | 0 | 0 | 0 | 0 | 0 | 0 | 0 | 0 | 1594300 | 0 | 0 | 0 | 0 | 0 | 0 |
| ELEFYLR | RPS8; RPS8 | P62241; Q5J | RS8_HUMAN 4 | Unmodified | 0 | 0 | 0 | 485990 | 0 | 0 | 162040 | 252960 | 326620 | 0 | 868350 | 2703500 | 0 | 0 | 0 | 0 | 0 | 0 |
| ELEFYLRK | RPS8; RPS8 | P62241; Q5J | RS8_HUMAN 4 | Unmodified | 706590 | 0 | 0 | 0 | 0 | 0 | 0 | 367750 | 0 | 0 | 0 | 6192300 | 0 | 203800 | 0 | 0 | 0 | 0 |
| ELEGILLPS | OS9; OS9 | Q13438; B4E | OS9_HUMAN P | Unmodified | 1378400 | 571510 | 0 | 368130 | 0 | 0 | 0 | 0 | 8184800 | 139140 | 737080 | 0 | 0 | 0 | 117740 | 276090 | 498360 | 0 |
| ELEGPVYS | SRGAP2; SRGA | O75044; Q5\ | SRGP2_HUMAI | Unmodified | 69829 | 36004 | 0 | 0 | 0 | 0 | 0 | 0 | 0 | 0 | 0 | 3132300 | 0 | 0 | 152780 | 242190 | 0 | 0 |
| ELEGWEPD | BIRC5; BIRC5 | O15392; A0/ | BIRC5_HUMAN | Unmodified | 0 | 0 | 0 | 631380 | 2326100 | 0 | 0 | 0 | 0 | 0 | 0 | 0 | 1447500 | 608190 | 0 | 0 | 167270 | 409080 |
| ELEGWEPD | BIRC5; BIRC5 | O15392; A0/ | BIRC5_HUMAN | Unmodified | 0 | 0 | 323750 | 941080 | 1148500 | 0 | 0 | 0 | 212030 | 0 | 0 | 0 | 1538300 | 281930 | 169170 | 94116 | 1303400 | 408740 |
| ELEIEER | EIF3A | Q14152 | EIF3A_HUMAN | Unmodified | 0 | 0 | 0 | 0 | 0 | 0 | 0 | 0 | 0 | 0 | 560220 | 1194600 | 0 | 0 | 0 | 1271500 | 0 | 0 |
| ELEISLR | FMNL2; FMNL3 | Q96PY5; Q8I | FMNL2_HUMAI | Unmodified | 0 | 0 | 0 | 0 | 0 | 0 | 0 | 0 | 0 | 0 | 0 | 532510 | 1387100 | 383190 | 0 | 0 | 0 | 0 |
| ELEKRASG | STMN1; STMN1 | P16949; A2A | STMN1_HUMAI | Unmodified | 0 | 0 | 0 | 0 | 0 | 0 | 327660 | 83055 | 0 | 74656 | 0 | 0 | 0 | 0 | 0 | 0 | 0 | 821510 |
| ELEKVCNP | HSPA8; HSPA8; | P11142; E9P | HSP7C_HUMAI | Unmodified | 0 | 0 | 0 | 0 | 1636400 | 0 | 7563800 | 2283000 | 0 | 0 | 0 | 0 | 0 | 5486700 | 0 | 0 | 9426800 | 0 |
| ELELLGGW | SH3BP4; SH3BI | Q9P0V3; C9. | SH3B4_HUMAN | Unmodified | 0 | 0 | 0 | 0 | 0 | 0 | 0 | 0 | 1215900 | 0 | 0 | 697120 | 0 | 0 | 0 | 0 | 695330 | 0 |
| ELENFRNLI | ERO1A | Q96HE7 | ERO1A_HUMAN | Unmodified | 0 | 0 | 0 | 0 | 248060 | 65283 | 297270 | 11981 | 3319900 | 7852600 | 899610 | 0 | 0 | 0 | 0 | 0 | 120250 | 149900 |
| ELENIIQETI | OS9; OS9; OS9 | Q13438; B4E | OS9_HUMAN P | Unmodified | 89346 | 0 | 0 | 0 | 0 | 0 | 0 | 0 | 2630900 | 0 | 481150 | 0 | 0 | 0 | 0 | 0 | 0 | 0 |
| ELENWAGS | CHMP4B | Q9H444 | CHM4B_HUMA | Unmodified | 0 | 0 | 0 | 0 | 470910 | 0 | 0 | 0 | 732690 | 0 | 0 | 0 | 0 | 0 | 0 | 0 | 0 | 0 |
| ELEPEAAEE | ACBD3 | Q9H3P7 | GCP60_HUMAI | Unmodified | 0 | 0 | 0 | 116140 | 0 | 0 | 0 | 0 | 0 | 0 | 600030 | 805580 | 0 | 0 | 0 | 0 | 0 | 0 |
| ELEQAIRG/ | PACSIN1; PACS | Q9BY11; F6U | PACN1_HUMAI | Unmodified | 1515200 | 969280 | 1080200 | 492290 | 0 | 0 | 0 | 0 | 0 | 0 | 0 | 0 | 0 | 0 | 452910 | 296320 | 341160 | 1690900 |
| ELEQGEPLI | FERMT1; FERM1 | Q9BQL6; G3 | FERM1_HUMAN | Unmodified | 64633 | 0 | 0 | 0 | 370060 | 97698 | 0 | 0 | 0 | 54128 | 0 | 143790 | 294310 | 0 | 0 | 0 | 241980 | 579200 |
| ELEQNANH | BAG5 | Q9UL15 | BAG5_HUMAN | Unmodified | 480470 | 313320 | 253660 | 2433100 | 0 | 0 | 1262100 | 0 | 8471400 | 0 | 0 | 1333100 | 0 | 0 | 925470 | 198910 | 0 | 0 |
| ELEQNANH | BAG5 | Q9UL15 | BAG5_HUMAN | Unmodified | 0 | 0 | 0 | 456010 | 4565800 | 0 | 1894500 | 0 | 4047500 | 5505100 | 0 | 0 | 4096200 | 2843000 | 0 | 0 | 0 | 1050400 |
| ELEQVCNP | HSPA1A; HSPA1 | P0DMV8; P0 | HS71A_HUMAN | Unmodified | 22269000 | 20823000 | 1919700 | 18335000 | 11942000 | 2447100 | 0 | 0 | 38078000 | 5605800 | 1477300 | 8866700 | 12666000 | 3268400 | 6386000 | 6051200 | 2658200 | 0 |
| ELEQVCNP | HSPA1A; HSPA1 | P0DMV8; P0 | HS71A_HUMAN | Unmodified | 0 | 0 | 0 | 0 | 24891000 | 0 | 0 | 0 | 0 | 13647000 | 0 | 0 | 0 | 16606000 | 0 | 0 | 22546000 | 14449000 |
| ELERHMED | ARHGEF7; ARH | Q14155; B1/ | ARHG7_HUMA | Unmodified | 0 | 0 | 0 | 0 | 515160 | 71654 | 244980 | 101090 | 264750 | 404730 | 96408 | 0 | 0 | 166610 | 0 | 0 | 290730 | 243010 |
| ELERLDLSI | FLRT1; FLRT1 | Q9NZU1; A0 | FLRT1_HUMAN | Unmodified | 0 | 0 | 0 | 0 | 0 | 0 | 0 | 0 | 0 | 3945800 | 0 | 0 | 0 | 0 | 0 | 0 | 0 | 0 |
| ELERVCNP | HSPA2 | P54652 | HSP72_HUMAN | Unmodified | 109830 | 0 | 0 | 0 | 1203900 | 707130 | 107720 | 90271 | 0 | 0 | 0 | 0 | 0 | 135970 | 0 | 0 | 472420 | 234100 |
| ELESQISEL | MYH9 | P35579 | MYH9_HUMAN | Unmodified | 555750 | 133790 | 0 | 124720 | 0 | 0 | 81451 | 0 | 0 | 79157 | 279910 | 688150 | 0 | 0 | 385720 | 513530 | 0 | 0 |
| ELESQISEL | MYH9 | P35579 | MYH9_HUMAN | Unmodified | 108140 | 0 | 0 | 491500 | 0 | 95898 | 0 | 0 | 1173000 | 0 | 0 | 531060 | 0 | 0 | 1170500 | 905430 | 0 | 0 |
| ELESQVSG | RRBP1; RRBP1; | Q9P2E9; A0/ | RRBP1_HUMAN | Unmodified | 0 | 0 | 0 | 0 | 0 | 0 | 0 | 0 | 0 | 0 | 0 | 0 | 853250 | 0 | 0 | 0 | 0 | 0 |
| ELESSEDRI | LUZP1; LUZP1 | Q86V48; E5I | LUZP1_HUMAN | Unmodified | 0 | 0 | 0 | 0 | 484070 | 0 | 0 | 0 | 0 | 0 | 0 | 0 | 118470 | 61208 | 0 | 0 | 130300 | 149910 |
| ELESVGIR | DRG2; DRG2; D | P55039; J3Q | DRG2_HUMAN | Unmodified | 941590 | 739560 | 580930 | 218550 | 0 | 143360 | 55687 | 119470 | 3401600 | 0 | 0 | 924440 | 0 | 0 | 110630 | 327320 | 0 | 0 |
| ELETILETQI | KIF20B | Q96Q89 | KI20B_HUMAN | Unmodified | 0 | 0 | 87315 | 275940 | 4744500 | 1883100 | 0 | 198720 | 0 | 0 | 469690 | 1003000 | 1636100 | 798840 | 0 | 0 | 514840 | 1053600 |

| | | | | | | | | | | | | | | | | | | | | | | |
|---|---|---|---|---|---|---|---|---|---|---|---|---|---|---|---|---|---|---|---|---|---|---|
| ELETVAAH( | NCLN; NCLN; N | Q969V3; K7E | NCLN_HUMAN | Unmodified | 0 | 0 | 0 | 0 | 0 | 0 | 0 | 0 | 15322000 | 0 | 0 | 747890 | 0 | 0 | 0 | 0 | 0 | 0 |
| ELETVCND | YWHAH; YWHA | Q04917; A2I | 1433F_HUMAN | Unmodified | 176420 | 127330 | 0 | 659310 | 4719500 | 733700 | 650380 | 30217 | 556700 | 1075200 | 358790 | 433100 | 2156200 | 1634100 | 755700 | 0 | 2068600 | 2273600 |
| ELETVDFK | SERPINB5; SER | P36952; C9J | SPB5_HUMAN | Unmodified | 0 | 0 | 28647 | 0 | 0 | 0 | 0 | 0 | 104830 | 79775 | 0 | 0 | 0 | 0 | 0 | 0 | 0 | 238950 |
| ELETVDFKI | SERPINB5; SER | P36952; C9J | SPB5_HUMAN | Unmodified | 65185 | 0 | 0 | 0 | 0 | 0 | 0 | 38059 | 572960 | 0 | 0 | 0 | 1021400 | 469300 | 0 | 0 | 446660 | 3292300 |
| ELEVAEGG | CDC37; CDC37 | Q16543; K7E | CDC37_HUMA | Unmodified | 0 | 0 | 0 | 0 | 23539 | 0 | 529620 | 151200 | 0 | 0 | 127460 | 397660 | 0 | 0 | 0 | 0 | 0 | 0 |
| ELEVAEGG | CDC37; CDC37 | Q16543; K7E | CDC37_HUMA | Unmodified | 129580 | 83319 | 0 | 0 | 2243900 | 0 | 0 | 0 | 455680 | 535110 | 1420700 | 5570500 | 0 | 0 | 612580 | 578440 | 0 | 0 |
| ELEVLLMCI | RPL32; RPL32; I | P62910; D3Y | RL32_HUMAN ( | Oxidation (M) | 0 | 0 | 0 | 0 | 711260 | 0 | 0 | 543820 | 0 | 949960 | 779250 | 6194500 | 293530 | 840620 | 0 | 0 | 416790 | 0 |
| ELEVLQAK | CLIP1; CLIP1 | P30622; A0A | CLIP1_HUMAN | Unmodified | 0 | 0 | 0 | 0 | 723820 | 0 | 0 | 0 | 0 | 0 | 0 | 0 | 319050 | 276810 | 0 | 0 | 112920 | 187830 |
| ELEVQGNG | PLXNA2 | O75051 | PLXA2_HUMAN | Unmodified | 127660 | 97286 | 40284 | 0 | 0 | 0 | 970090 | 0 | 0 | 0 | 72942 | 667010 | 0 | 0 | 98943 | 0 | 0 | 0 |
| ELEVQHPA | CCT8; CCT8 | P50990; H7C | TCPQ_HUMAN | Unmodified | 0 | 0 | 50237 | 0 | 0 | 0 | 0 | 0 | 0 | 0 | 409680 | 1598700 | 0 | 0 | 0 | 61525 | 0 | 0 |
| ELEYQANN | UXS1 | Q8NBZ7 | UXS1_HUMAN | Unmodified | 0 | 58812 | 0 | 0 | 0 | 0 | 0 | 0 | 1494100 | 0 | 108670 | 496140 | 0 | 0 | 557730 | 0 | 144070 | 0 |
| ELEYQANN | UXS1 | Q8NBZ7 | UXS1_HUMAN | Unmodified | 0 | 59687 | 0 | 0 | 0 | 0 | 0 | 0 | 1005400 | 754160 | 0 | 0 | 0 | 0 | 0 | 0 | 0 | 0 |
| ELFALHPSF | DOCK7; DOCK7 | Q96N67; A0A | DOCK7_HUMA | Unmodified | 0 | 108650 | 778680 | 0 | 0 | 0 | 0 | 0 | 0 | 148040 | 0 | 1342000 | 0 | 0 | 875840 | 906820 | 0 | 0 |
| ELFDELVK | ADK; ADK; ADK; | P55263; A0A | ADK_HUMAN A | Unmodified | 0 | 0 | 0 | 0 | 1519800 | 0 | 359880 | 98862 | 886580 | 294710 | 0 | 6397500 | 0 | 0 | 347990 | 0 | 595800 | 0 |
| ELFDVVANI | KIF23; KIF23; KI | Q02241; A0A | KIF23_HUMAN | Unmodified | 596880 | 972280 | 973250 | 3984900 | 48421000 | 5448100 | 1100400 | 0 | 10436000 | 8873600 | 3170500 | 2302500 | 2670700 | 1545600 | 4642900 | 844350 | 4832600 | 0 |
| ELFDVVANI | KIF23; KIF23; KI | Q02241; A0A | KIF23_HUMAN | Unmodified | 0 | 0 | 0 | 0 | 1119600 | 0 | 0 | 0 | 0 | 0 | 0 | 0 | 418060 | 0 | 0 | 0 | 0 | 0 |
| ELFEMAR | PSMC2; PSMC2 | P35998; C9J | PRS7_HUMAN | Unmodified | 0 | 0 | 0 | 0 | 0 | 0 | 0 | 0 | 0 | 0 | 0 | 1204500 | 0 | 0 | 0 | 0 | 0 | 0 |
| ELFESSTNI | MYO6; MYO6; N | Q9UM54; A0 | MYO6_HUMAN | Unmodified | 0 | 0 | 587970 | 479630 | 0 | 0 | 0 | 0 | 0 | 0 | 79230 | 586610 | 0 | 0 | 555620 | 557790 | 0 | 0 |
| ELFESSTNI | MYO6; MYO6; N | Q9UM54; A0 | MYO6_HUMAN | Unmodified | 77663 | 0 | 0 | 462820 | 54402 | 0 | 0 | 0 | 0 | 0 | 0 | 0 | 0 | 0 | 0 | 139050 | 52903 | 0 |
| ELFIQQER | LNPEP | Q9UIQ6 | LCAP_HUMAN | Unmodified | 172310 | 111580 | 1021500 | 375150 | 0 | 0 | 0 | 0 | 2810100 | 0 | 746750 | 1315800 | 0 | 0 | 334960 | 397140 | 0 | 0 |
| ELFLTHGNI | HAUS3; HAUS3 | Q68CZ6; A0A | HAUS3_HUMAI | Unmodified | 0 | 0 | 0 | 0 | 0 | 0 | 0 | 24861 | 264230 | 0 | 131850 | 432310 | 209230 | 93221 | 0 | 0 | 0 | 0 |
| ELFPIQME( | TOMM40; TOMI | O96008; K7E | TOM40_HUMAI | Unmodified | 46735 | 0 | 0 | 0 | 300430 | 0 | 0 | 0 | 5113500 | 0 | 0 | 0 | 138290 | 0 | 0 | 0 | 0 | 0 |
| ELFPIQME( | TOMM40; TOMI | O96008; K7E | TOM40_HUMAI | Oxidation (M) | 0 | 0 | 0 | 0 | 0 | 0 | 0 | 0 | 0 | 1110200 | 0 | 0 | 0 | 0 | 0 | 0 | 0 | 0 |
| ELFPVFTHL | C6orf89 | Q6UWU4 | CF089_HUMAN | Unmodified | 0 | 0 | 0 | 0 | 0 | 0 | 0 | 0 | 3196800 | 413120 | 0 | 0 | 0 | 0 | 0 | 0 | 0 | 0 |
| ELFREMESI | SCRN3 | Q0VDG4 | SCRN3_HUMAI | Unmodified | 0 | 0 | 0 | 100800 | 412830 | 58488 | 647560 | 0 | 687830 | 0 | 0 | 134910 | 788530 | 389230 | 77382 | 0 | 359810 | 461420 |
| ELFRVVPGI | ST3GAL1 | Q11201 | SIA4A_HUMAN | Oxidation (M) | 90169 | 0 | 0 | 0 | 735770 | 0 | 245400 | 0 | 0 | 0 | 0 | 0 | 0 | 0 | 0 | 0 | 0 | 1390000 |
| ELFSPLHAI | PAFAH1B2 | P68402 | PA1B2_HUMAN | Unmodified | 355590 | 361640 | 392790 | 363960 | 0 | 0 | 0 | 0 | 1197900 | 48235 | 1066800 | 37057000 | 29165 | 0 | 1495400 | 1303900 | 0 | 0 |
| ELFSQFGK | PABPC4; PABP( | Q13310; B1A | PABP4_HUMAN | Unmodified | 8421100 | 9188000 | 5523600 | 3525300 | 141410 | 0 | 242660 | 153600 | 916350 | 0 | 0 | 1207800 | 0 | 0 | 4726200 | 3947100 | 71345 | 0 |
| ELGDHVTN | FTH1; FTH1; FTI | P02794; G3V | FRIH_HUMAN F | Unmodified | 0 | 0 | 432000 | 447890 | 0 | 367230 | 3072900 | 107620 | 70631000 | 38988000 | 571830 | 0 | 2362400 | 3193500 | 309180 | 225000 | 1279000 | 2393000 |
| ELGEKLSKI | PDIA3 | P30101 | PDIA3_HUMAN | Unmodified | 0 | 0 | 0 | 0 | 0 | 0 | 366160 | 284140 | 0 | 1670200 | 0 | 0 | 59225 | 251070 | 0 | 0 | 0 | 0 |
| ELGENLDQ | IPO9 | Q96P70 | IPO9_HUMAN I | Unmodified | 0 | 0 | 0 | 0 | 0 | 0 | 0 | 0 | 0 | 0 | 0 | 931670 | 0 | 0 | 0 | 0 | 0 | 0 |
| ELGEYFLFI | DIAPH1; DIAPH | O60610; A0A | DIAP1_HUMAN | Unmodified | 0 | 0 | 137680 | 0 | 0 | 0 | 0 | 0 | 0 | 522380 | 899430 | 3854600 | 954590 | 0 | 0 | 0 | 0 | 0 |
| ELGEYGFH | ALDH1A1 | P00352 | AL1A1_HUMAN | Unmodified | 0 | 0 | 0 | 0 | 0 | 0 | 0 | 0 | 0 | 0 | 16331000 | 6.58E+08 | 0 | 0 | 0 | 0 | 0 | 0 |
| ELGFHSNS | LGR4 | Q9BXB1 | LGR4_HUMAN | Unmodified | 0 | 0 | 0 | 0 | 1651300 | 516500 | 0 | 0 | 0 | 0 | 150860 | 177400 | 413290 | 0 | 0 | 0 | 0 | 253890 |
| ELGGADNI | ARL15; ARL15; / | Q9NXU5; A0 | ARL15_HUMAN | Unmodified | 0 | 0 | 0 | 87307 | 0 | 0 | 0 | 0 | 0 | 0 | 0 | 1306700 | 0 | 0 | 0 | 0 | 0 | 0 |
| ELGGESNF | SH3GL1; SH3G | Q99961; M0I | SH3G1_HUMAI | Oxidation (M) | 15759000 | 21067000 | 12010000 | 14237000 | 2870000 | 0 | 739870 | 188480 | 719020 | 1369900 | 0 | 685430 | 14683000 | 6656500 | 18710000 | 13628000 | 148000 | 504600 |
| ELGGESNF | SH3GL1; SH3G | Q99961; M0I | SH3G1_HUMAI | Unmodified | 2717200 | 4532800 | 904160 | 2059800 | 1704100 | 0 | 0 | 0 | 1221300 | 355030 | 0 | 0 | 5357500 | 1845900 | 2565500 | 930370 | 0 | 486110 |
| ELGGESNF | SH3GL1; SH3G | Q99961; M0I | SH3G1_HUMAI | Oxidation (M) | 7545500 | 1434900 | 2916200 | 6946600 | 0 | 0 | 0 | 0 | 0 | 0 | 0 | 0 | 1693500 | 0 | 7417700 | 9082800 | 0 | 0 |
| ELGGESNF | SH3GL1; SH3G | Q99961; M0I | SH3G1_HUMAI | Unmodified | 7044300 | 0 | 0 | 2159500 | 0 | 0 | 0 | 0 | 70030 | 0 | 0 | 0 | 1009000 | 57307 | 0 | 1601800 | 0 | 562300 |
| ELGGLEGD | CNPY3 | Q9BT09 | CNPY3_HUMAI | Unmodified | 0 | 0 | 0 | 0 | 0 | 0 | 363920 | 86470 | 31687000 | 12291000 | 1321600 | 379430 | 0 | 0 | 0 | 0 | 0 | 0 |
| ELGGLEGD | CNPY3 | Q9BT09 | CNPY3_HUMAI | Unmodified | 0 | 0 | 0 | 0 | 0 | 0 | 1132800 | 648690 | 0 | 3563800 | 0 | 0 | 0 | 0 | 0 | 0 | 0 | 0 |
| ELGGLGMI | PRPF8; PRPF8 | Q6P2Q9; I3L | PRP8_HUMAN | 2 Oxidation (M) | 197690 | 0 | 72263 | 689680 | 0 | 0 | 0 | 0 | 697950 | 1003600 | 0 | 718400 | 0 | 0 | 94194 | 137240 | 0 | 0 |
| ELGIPASVI | STRN | O43815 | STRN_HUMAN | Oxidation (M) | 0 | 245310 | 0 | 0 | 0 | 0 | 729660 | 119970 | 0 | 0 | 0 | 1044300 | 0 | 0 | 0 | 0 | 0 | 0 |
| ELGISPIMT( | MICAL3; MICAL | Q7RTP6; A0A | MICA3_HUMAN | Unmodified | 0 | 0 | 0 | 180030 | 1332700 | 0 | 0 | 70442 | 0 | 0 | 0 | 0 | 0 | 0 | 0 | 0 | 0 | 0 |
| ELGITALHII | RPS14; RPS14; | P62263; E5R | RS14_HUMAN . | Unmodified | 209960 | 203500 | 0 | 0 | 232290 | 0 | 0 | 225360 | 0 | 0 | 955490 | 2297200 | 429750 | 262760 | 182340 | 171910 | 0 | 0 |
| ELGIWEPL/ | CCT3; CCT3 | P49368; B4D | TCPG_HUMAN | Unmodified | 552780 | 1332300 | 767970 | 616150 | 8129700 | 0 | 1596300 | 0 | 7042600 | 3445100 | 15841000 | 41176000 | 0 | 2266600 | 0 | 1571000 | 0 | 4098600 |
| ELGIWEPL/ | CCT3; CCT3 | P49368; B4D | TCPG_HUMAN | Unmodified | 0 | 0 | 0 | 0 | 1652300 | 281830 | 1336400 | 0 | 0 | 926400 | 0 | 0 | 1542200 | 2003100 | 0 | 0 | 1014000 | 1090200 |
| ELGLEKDD | LONP1; LONP1 | P36776; K7E | LONM_HUMAN | Unmodified | 0 | 0 | 0 | 0 | 0 | 0 | 0 | 0 | 2923600 | 694430 | 0 | 0 | 0 | 0 | 0 | 0 | 29552 | 0 |
| ELGLKESQI | LIMCH1; LIMCH | Q9UPQ0; D6 | LIMC1_HUMAN | Unmodified | 0 | 0 | 0 | 0 | 0 | 0 | 0 | 175510 | 0 | 0 | 0 | 0 | 2079400 | 0 | 0 | 0 | 0 | 0 |
| ELGLNPEK | ACAT2 | Q9BWD1 | THIC_HUMAN / | Unmodified | 0 | 94467 | 234150 | 0 | 0 | 0 | 323570 | 212400 | 0 | 579110 | 612930 | 10709000 | 0 | 0 | 0 | 273770 | 0 | 0 |
| ELGLPEELV | GMPS | P49915 | GUAA_HUMAN | Unmodified | 99365 | 0 | 82900 | 0 | 0 | 0 | 0 | 0 | 1609500 | 0 | 1106300 | 4463900 | 0 | 0 | 0 | 76220 | 0 | 79955 |
| ELGLPSGKI | PLCB4; PLCB4 | Q15147; A0A | PLCB4_HUMAN | Unmodified | 296960 | 0 | 1255000 | 3120200 | 361180 | 0 | 0 | 71448 | 0 | 172410 | 0 | 0 | 1441300 | 1251100 | 1854200 | 1771000 | 77083 | 2716300 |
| ELGLREENI | ALDH7A1; ALDI | P49419; A0A | AL7A1_HUMAN | Unmodified | 0 | 0 | 0 | 0 | 0 | 0 | 0 | 0 | 0 | 0 | 0 | 930550 | 0 | 0 | 0 | 0 | 0 | 0 |
| ELGNDAYK | STIP1 | P31948 | STIP1_HUMAN | Unmodified | 225340 | 114900 | 425000 | 604320 | 3824800 | 797040 | 150630 | 20911 | 362570 | 1152500 | 1443900 | 10285000 | 552800 | 1510300 | 571920 | 809910 | 951790 | 2374500 |
| ELGPLYTNI | B4GALT3 | O60512 | B4GT3_HUMAN | Unmodified | 0 | 0 | 0 | 0 | 0 | 0 | 0 | 0 | 0 | 0 | 0 | 0 | 2720900 | 0 | 0 | 0 | 0 | 0 |
| ELGPYSSK | CAPN2 | P17655 | CAN2_HUMAN | Unmodified | 1335800 | 0 | 282530 | 0 | 6872600 | 285400 | 10127000 | 120810 | 1299500 | 1184200 | 0 | 0 | 521440 | 1346500 | 0 | 0 | 3394900 | 4851900 |
| ELGQGSFG | INSR | P06213 | INSR_HUMAN I | Oxidation (M) | 1362300 | 1356900 | 0 | 0 | 0 | 0 | 0 | 0 | 0 | 0 | 0 | 0 | 0 | 0 | 0 | 0 | 0 | 0 |
| ELGQTLID( | UTRN | P46939 | UTRN_HUMAN | Unmodified | 0 | 0 | 25175 | 75934 | 0 | 0 | 0 | 0 | 0 | 0 | 0 | 0 | 0 | 0 | 520900 | 31572 | 0 | 0 |
| ELGRGLPTI | MMP14; MMP14 | P50281; A0A | MMP14_HUMA | Oxidation (M) | 0 | 0 | 0 | 0 | 2576500 | 0 | 0 | 0 | 0 | 0 | 0 | 0 | 0 | 0 | 0 | 0 | 534560 | 712750 |
| ELGRGLPTI | MMP14; MMP14 | P50281; A0A | MMP14_HUMA | Unmodified | 0 | 0 | 0 | 0 | 811500 | 0 | 0 | 0 | 0 | 255770 | 0 | 0 | 0 | 0 | 0 | 0 | 0 | 223830 |
| ELGSECGII | DDOST; DDOST | P39656; U3K | OST48_HUMAN | Unmodified | 0 | 0 | 0 | 0 | 0 | 0 | 0 | 0 | 1245600 | 0 | 0 | 0 | 0 | 0 | 0 | 0 | 0 | 0 |
| ELGSGQFG | BMX | P51813 | BMX_HUMAN ( | Unmodified | 0 | 0 | 0 | 0 | 0 | 0 | 0 | 0 | 0 | 0 | 3294200 | 12539000 | 0 | 0 | 0 | 0 | 0 | 0 |
| ELGSLPLPI | STX12; STX12 | Q86Y82; B1A | STX12_HUMAN | Unmodified | 0 | 0 | 913940 | 0 | 199710 | 0 | 0 | 26473 | 3144800 | 0 | 548610 | 769260 | 0 | 0 | 0 | 526820 | 0 | 0 |
| ELGSSTNA | SNRPB2 | P08579 | RU2B_HUMAN | Unmodified | 0 | 0 | 0 | 0 | 0 | 0 | 0 | 0 | 0 | 0 | 0 | 0 | 0 | 0 | 0 | 0 | 0 | 0 |
| ELGTVMR | CALM2; CALM3 | P0DP24; P0I | CALM2_HUMAI | Unmodified | 3800500 | 3610800 | 3032300 | 4143100 | 15113000 | 0 | 0 | 331460 | 7958300 | 4924200 | 5836200 | 22038000 | 4784400 | 1736600 | 6355500 | 3677400 | 4850300 | 8956500 |
| ELGVGIALF | FABP5 | Q01469 | FABP5_HUMAN | Unmodified | 66830 | 0 | 0 | 0 | 139770 | 0 | 0 | 25268 | 0 | 492400 | 6677600 | 2007500 | 0 | 0 | 0 | 0 | 1647600 | 747640 |
| ELGVGIALF | FABP5 | Q01469 | FABP5_HUMAN | Unmodified | 344990 | 0 | 0 | 0 | 7176800 | 1085800 | 0 | 124290 | 0 | 125440 | 2398000 | 4652700 | 0 | 0 | 0 | 0 | 16318000 | 15339000 |
| ELGVSTNA | UBLCP1 | Q8WVY7 | UBCP1_HUMAI | Unmodified | 166960 | 281050 | 0 | 0 | 0 | 0 | 0 | 0 | 0 | 0 | 0 | 1126600 | 0 | 0 | 0 | 0 | 0 | 0 |
| ELGYPREEI | PPP1R1B; PPP1 | Q9UD71; J3K | PPR1B_HUMAN | Unmodified | 0 | 0 | 0 | 0 | 0 | 0 | 0 | 0 | 0 | 0 | 0 | 4164100 | 0 | 0 | 0 | 0 | 0 | 0 |
| ELHEKFDLI | SEPTIN11; SEPT | Q9NVA2; H0 | SEP11_HUMAN | Unmodified | 0 | 0 | 0 | 0 | 5471100 | 0 | 292810 | 0 | 0 | 172590 | 0 | 315950 | 0 | 0 | 0 | 0 | 882270 | 1789700 |
| ELHEQLVAI | CPT2; CPT2; CF | P23786; A0A | CPT2_HUMAN | Unmodified | 0 | 0 | 0 | 0 | 0 | 0 | 0 | 0 | 1095400 | 0 | 0 | 0 | 0 | 0 | 0 | 0 | 0 | 0 |
| ELHEVCHL | RABEP1 | Q15276 | RABE1_HUMAN | Unmodified | 0 | 0 | 0 | 0 | 513480 | 0 | 0 | 0 | 0 | 65084 | 0 | 0 | 100360 | 144470 | 0 | 0 | 89066 | 126850 |
| ELHGQNP\ | CPSF6; CPSF6 | Q16630; F8V | CPSF6_HUMAN | Unmodified | 729890 | 1165400 | 0 | 0 | 56235 | 32943 | 0 | 0 | 0 | 0 | 0 | 0 | 0 | 0 | 0 | 0 | 0 | 0 |
| ELHINLIPN | HSP90AA2P; HS | Q14568; P07 | HS902_HUMAN | Unmodified | 0 | 0 | 951890 | 0 | 345300 | 0 | 0 | 0 | 1029200 | 0 | 628790 | 6197000 | 0 | 0 | 414150 | 235390 | 0 | 0 |
| ELHINLIPN | HSP90AA1; HSI | P07900; G3V | HS90A_HUMAN | Unmodified | 0 | 0 | 0 | 199230 | 0 | 95329 | 0 | 120200 | 2797600 | 325890 | 1394100 | 25497000 | 0 | 0 | 0 | 0 | 0 | 0 |
| ELHINLIPN | HSP90AA1; HSI | P07900; G3V | HS90A_HUMAN | Oxidation (M) | 0 | 0 | 0 | 0 | 6971500 | 0 | 0 | 0 | 0 | 7095100 | 0 | 0 | 3261100 | 0 | 0 | 0 | 1332200 | 5379100 |
| ELHINLIPN | HSP90AA1; HSI | P07900; G3V | HS90A_HUMAN | Unmodified | 0 | 0 | 0 | 0 | 2785100 | 0 | 136380 | 0 | 0 | 1181700 | 0 | 0 | 591710 | 220660 | 0 | 0 | 598250 | 1860000 |
| ELHIQGNR | RSU1 | Q15404 | RSU1_HUMAN | Unmodified | 268880 | 559170 | 150670 | 935700 | 39140 | 0 | 0 | 0 | 0 | 0 | 0 | 79945 | 0 | 0 | 206690 | 212370 | 0 | 0 |
| ELHIQGNR | RSU1 | Q15404 | RSU1_HUMAN | Unmodified | 0 | 0 | 0 | 0 | 7192600 | 0 | 13282000 | 108470 | 0 | 843680 | 0 | 0 | 5843200 | 5018000 | 227190 | 0 | 1551600 | 4252200 |
| ELHLDDHK | ATP12A | P54707 | AT12A_HUMAN | Unmodified | 0 | 0 | 0 | 0 | 0 | 0 | 0 | 0 | 764540 | 0 | 0 | 0 | 0 | 0 | 0 | 0 | 0 | 0 |
| ELHQLQNI | PIGO | Q8TEQ8 | PIGO_HUMAN | Unmodified | 0 | 323190 | 0 | 83977 | 0 | 1129900 | 0 | 0 | 2954300 | 436420 | 0 | 0 | 0 | 0 | 0 | 0 | 0 | 0 |
| ELHSAEEG. | DTNB; DTNB; D | O60941; E7E | DTNB_HUMAN | Unmodified | 0 | 0 | 273710 | 420830 | 98879 | 0 | 0 | 0 | 712390 | 102870 | 34771 | 0 | 0 | 0 | 554680 | 496570 | 0 | 0 |
| ELHSEFSE\ | HADHA; HADH/ | P40939; A0A | ECHA_HUMAN | Unmodified | 0 | 0 | 0 | 0 | 0 | 0 | 0 | 0 | 739440 | 0 | 0 | 0 | 0 | 0 | 0 | 0 | 0 | 0 |
| ELIATVR | FLOT2; FLOT2; I | Q14254; J3Q | FLOT2_HUMAN | Unmodified | 0 | 115010 | 0 | 0 | 47036 | 0 | 66411 | 564290 | 2750200 | 494030 | 98864 | 1635000 | 0 | 0 | 1269000 | 516710 | 0 | 0 |
| ELICHNQF | HSD17B4; HSD | P51659; E7E | DHB4_HUMAN | Unmodified | 0 | 0 | 0 | 0 | 0 | 0 | 625080 | 0 | 2820700 | 934810 | 0 | 0 | 131110 | 110710 | 0 | 0 | 0 | 0 |
| ELIECAR | TLN2; TLN1; TLI | Q9Y4G6; Q9 | TLN2_HUMAN | Unmodified | 0 | 0 | 0 | 294910 | 1211100 | 1185700 | 276700 | 174390 | 0 | 0 | 0 | 498000 | 573560 | 911510 | 397900 | 0 | 407980 | 666050 |
| ELIEELHPII | FRYL; FRYL; FR\ | O94915; F22 | FRYL_HUMAN I | Unmodified | 0 | 0 | 0 | 0 | 0 | 0 | 0 | 0 | 0 | 0 | 0 | 1093600 | 0 | 0 | 0 | 0 | 0 | 0 |
| ELIEIISGAA | ATP5F1C | P36542 | ATPG_HUMAN | Unmodified | 64271 | 0 | 0 | 0 | 0 | 0 | 24353 | 0 | 139380 | 312800 | 0 | 0 | 0 | 7700 | 0 | 0 | 0 | 0 |
| ELIEKMIAM | ERN1; ERN1 | O75460; A0A | ERN1_HUMAN | Unmodified | 0 | 0 | 0 | 628010 | 48686000 | 3.46E+08 | 57998000 | 15694000 | 0 | 38093 | 0 | 0 | 37847000 | 2.83E+08 | 0 | 0 | 0 | 2.63E+08 |
| ELIEKNSQL | TSC22D1; TSC2 | Q15714; A0A | T22D1_HUMAN | Unmodified | 0 | 0 | 0 | 0 | 0 | 0 | 0 | 0 | 0 | 0 | 0 | 0 | 51944 | 0 | 0 | 0 | 390090 | 0 |
| ELIELRNK | TFG; TFG; TFG; | Q92734; A0A | TFG_HUMAN P | Unmodified | 88020 | 10984000 | 244310 | 171330 | 2551000 | 135080 | 76289 | 67504 | 0 | 392570 | 0 | 94562 | 1370600 | 404730 | 912050 | 595000 | 768560 | 1225100 |
| ELIERIPELN | TFRC; TFRC | P02786; G3V | TFR1_HUMAN | Unmodified | 337900 | 2201900 | 4225500 | 2525200 | 774680 | 0 | 331470 | 0 | 9462200 | 2737100 | 0 | 3153400 | 330070 | 0 | 4125000 | 2371400 | 0 | 0 |
| ELIFEETAR | MAPK1 | P28482 | MK01_HUMAN | Unmodified | 0 | 324260 | 283390 | 0 | 539090 | 0 | 307170 | 326970 | 0 | 17845 | 0 | 2476400 | 0 | 0 | 841140 | 585070 | 0 | 0 |
| ELIFEETARI | MAPK1 | P28482 | MK01_HUMAN | Unmodified | 0 | 0 | 0 | 0 | 7161400 | 539270 | 4455400 | 0 | 161990 | 1536300 | 168920 | 143120 | 6755300 | 3690600 | 0 | 117360 | 3363100 | 5863100 |
| ELIGKHAIP | CTNND1; CTNN | O60716; A0A | CTND1_HUMAI | Unmodified | 0 | 0 | 0 | 0 | 422290 | 0 | 0 | 0 | 0 | 0 | 0 | 0 | 341690 | 925940 | 0 | 0 | 1376400 | 805300 |
| ELILSSNIG( | MTIF3 | Q9H2K0 | IF3M_HUMAN | Unmodified | 0 | 0 | 0 | 0 | 166870 | 0 | 0 | 0 | 1228000 | 238390 | 0 | 0 | 0 | 0 | 0 | 0 | 0 | 0 |
| ELIQDTSRF | SULT1A3; SULT | P0DMM9; P5 | ST1A3_HUMAN | Unmodified | 0 | 0 | 0 | 0 | 1436500 | 0 | 0 | 0 | 0 | 0 | 0 | 0 | 0 | 0 | 0 | 0 | 425190 | 0 |
| ELIQKELTIC | S100A6; S100A | P06703; R4C | S10A6_HUMAN | Unmodified | 0 | 0 | 0 | 0 | 1198200 | 199510 | 0 | 0 | 0 | 256110 | 0 | 0 | 0 | 0 | 0 | 0 | 936080 | 1410400 |
| ELISNASDA | HSP90AB4P; HS | Q58FF6; P14 | H90B4_HUMAN | Unmodified | 6258600 | 5458900 | 12499000 | 8262900 | 1273900 | 871240 | 1235400 | 635370 | 3.29E+08 | 17380000 | 1.02E+08 | 2.65E+08 | 1847600 | 0 | 16937000 | 12356000 | 395770 | 704580 |
| ELISNASDA | HSP90AB4P; HS | Q58FF6; P14 | H90B4_HUMAN | Unmodified | 0 | 0 | 0 | 0 | 9312000 | 0 | 141940 | 0 | 3699800 | 7366000 | 0 | 885070 | 287290 | 0 | 0 | 26662 | 1588600 | 0 |
| ELISNASDA | TRAP1 | Q12931 | TRAP1_HUMAN | Unmodified | 0 | 0 | 0 | 0 | 241430 | 663230 | 0 | 0 | 1839400 | 0 | 0 | 0 | 0 | 0 | 0 | 0 | 0 | 0 |
| ELISNSSDA | HSP90AA2P; HS | Q14568; P07 | HS902_HUMAN | Unmodified | 2495700 | 3235700 | 3249000 | 2830800 | 1630600 | 574500 | 0 | 0 | 18430000 | 1123400 | 11001000 | 1.03E+08 | 316580 | 0 | 4463500 | 3175800 | 323020 | 0 |
| ELISRLPEIY | EHD4 | Q9H223 | EHD4_HUMAN | Unmodified | 185780 | 502790 | 51326 | 503070 | 0 | 0 | 0 | 0 | 877330 | 237550 | 0 | 0 | 0 | 0 | 755060 | 217000 | 0 | 0 |
| ELITILEK | KPNB1; KPNB1 | Q14974; J3Q | IMB1_HUMAN I | Unmodified | 0 | 0 | 0 | 0 | 0 | 0 | 0 | 0 | 0 | 0 | 501500 | 0 | 0 | 0 | 0 | 0 | 0 | 0 |
| ELKEEIEEK | ROCK1; ROCK1 | Q13464; A0A | ROCK1_HUMAI | Unmodified | 0 | 0 | 0 | 0 | 424950 | 0 | 38036 | 60374 | 0 | 0 | 0 | 0 | 0 | 0 | 0 | 0 | 0 | 353470 |
| ELKELIQKE | S100A6; S100A | P06703; R4C | S10A6_HUMAN | Unmodified | 0 | 0 | 0 | 0 | 0 | 0 | 0 | 0 | 0 | 0 | 0 | 0 | 0 | 0 | 0 | 0 | 530760 | 580250 |
| ELKEQLGEI | DDX21 | Q9NR30 | DDX21_HUMAI | Unmodified | 0 | 0 | 0 | 0 | 0 | 0 | 119190 | 381890 | 0 | 0 | 0 | 0 | 0 | 0 | 241660 | 243310 | 0 | 0 |
| ELKIDIIPNF | HSP90AB1; HSI | P08238; Q58 | HS90B_HUMAN | Unmodified | 209610 | 0 | 0 | 536610 | 7049500 | 2080100 | 0 | 158750 | 13639000 | 3596000 | 2238200 | 7888700 | 21366000 | 3810200 | 238000 | 313200 | 3088900 | 4742700 |
| ELKIDIIPNF | HSP90AB1 | P08238 | HS90B_HUMAI | Oxidation (M) | 0 | 0 | 0 | 0 | 8571500 | 992220 | 768510 | 0 | 0 | 2801100 | 358620 | 0 | 0 | 0 | 0 | 0 | 3176500 | 3990000 |

| | | | | | | | | | | | | | | | | | | | | | | |
|---|---|---|---|---|---|---|---|---|---|---|---|---|---|---|---|---|---|---|---|---|---|---|
| ELKLTDYR( | PRDX4; PRDX4; | Q13162; A6N | PRDX4_HUMAN | Unmodified | 0 | 0 | 134890 | 0 | 0 | 0 | 0 | 0 | 1803400 | 5409500 | 40206 | 0 | 0 | 0 | 0 | 0 | 0 | 0 |
| ELKPMASG | SLC44A1 | Q8WWI5 | CTL1_HUMAN | Unmodified | 0 | 0 | 1192300 | 1960400 | 1135800 | 558670 | 89481 | 63477 | 0 | 1246500 | 1543100 | 2513400 | 1285400 | 347320 | 1279600 | 0 | 262080 | 473830 |
| ELKPMASG | SLC44A1 | Q8WWI5 | CTL1_HUMAN | Oxidation (M) | 299090 | 271430 | 1210600 | 3535300 | 412540 | 139740 | 11418 | 52969 | 672420 | 267480 | 324810 | 1621400 | 530140 | 0 | 2080800 | 1249400 | 149740 | 89900 |
| ELKREPAV( | ICAM1; ICAM1 | P05362; K7E | ICAM1_HUMAN | Unmodified | 0 | 0 | 0 | 1494800 | 22225000 | 4494700 | 0 | 0 | 0 | 0 | 0 | 0 | 0 | 0 | 531270 | 648730 | 604770 | 1420700 |
| ELKSLDQAI | SMC3 | Q9UQE7 | SMC3_HUMAN | Unmodified | 0 | 0 | 0 | 17111 | 285430 | 218890 | 0 | 0 | 0 | 178960 | 0 | 0 | 870510 | 317790 | 0 | 0 | 0 | 0 |
| ELKSPDEG | FOLH1B; FOLH | Q9HBA9; Q0 | FOH1B_HUMAN | Unmodified | 3233500 | 3863600 | 0 | 20242000 | 0 | 0 | 0 | 0 | 0 | 0 | 0 | 0 | 6869400 | 550640 | 4481700 | 5487000 | 0 | 0 |
| ELKSPDEG | FOLH1B; FOLH | Q9HBA9; Q0 | FOH1B_HUMAN | Unmodified | 0 | 0 | 0 | 0 | 0 | 0 | 0 | 0 | 0 | 0 | 0 | 0 | 1289100 | 1295000 | 0 | 0 | 0 | 0 |
| ELLAFQTSE | USP9X; USP9Y | Q93008; O0( | USP9X_HUMAN | Unmodified | 0 | 0 | 0 | 0 | 0 | 0 | 0 | 0 | 0 | 0 | 0 | 0 | 0 | 0 | 0 | 0 | 0 | 0 |
| ELLALDSVI | BAG3 | O95817 | BAG3_HUMAN | Unmodified | 0 | 0 | 0 | 140630 | 592130 | 0 | 0 | 0 | 0 | 0 | 0 | 0 | 0 | 0 | 0 | 0 | 715400 | 130560 |
| ELLAQRLQ( | CGN | Q9P2M7 | CING_HUMAN | Unmodified | 0 | 0 | 0 | 43284 | 555720 | 0 | 0 | 0 | 0 | 0 | 35507 | 0 | 211470 | 35787 | 0 | 0 | 0 | 0 |
| ELLAVACE( | FLOT2; FLOT2; | Q14254; J3Q | FLOT2_HUMAN | Unmodified | 0 | 0 | 400370 | 0 | 0 | 0 | 0 | 0 | 0 | 0 | 0 | 0 | 0 | 0 | 1013000 | 0 | 0 | 0 |
| ELLAVPDN | PSAT1 | Q9Y617 | SERC_HUMAN | Unmodified | 0 | 0 | 359580 | 777920 | 0 | 653170 | 0 | 0 | 1134800 | 2036200 | 1158000 | 4843200 | 0 | 0 | 774460 | 740300 | 0 | 0 |
| ELLDLLYK | ADA; ADA | P00813; F5G | ADA_HUMAN A | Unmodified | 0 | 0 | 804430 | 1413000 | 0 | 0 | 0 | 0 | 0 | 0 | 142090 | 988520 | 0 | 0 | 499240 | 306020 | 127490 | 170710 |
| ELLDTVTAF | SERPINF1; SER | P36955; I3L | PEDF_HUMAN | Unmodified | 0 | 0 | 0 | 0 | 0 | 0 | 190290 | 296290 | 1563300 | 789280 | 10869000 | 1659000 | 0 | 0 | 0 | 0 | 0 | 0 |
| ELLDYKGV( | PSAT1 | Q9Y617 | SERC_HUMAN | Oxidation (M) | 0 | 0 | 0 | 0 | 0 | 0 | 0 | 0 | 0 | 405080 | 0 | 0 | 0 | 1480500 | 0 | 0 | 0 | 0 |
| ELLDYKGV( | PSAT1 | Q9Y617 | SERC_HUMAN | Oxidation (M) | 0 | 0 | 0 | 0 | 1212100 | 0 | 1079300 | 0 | 0 | 586080 | 0 | 0 | 4440500 | 5805600 | 0 | 0 | 135310 | 0 |
| ELLEEMYEI | KIF20A; KIF20A | O95235; H0 | KI20A_HUMAN | Unmodified | 80991 | 87578 | 0 | 444850 | 707750 | 0 | 0 | 0 | 0 | 0 | 0 | 0 | 0 | 0 | 0 | 0 | 0 | 0 |
| ELLELASR | HDLBP; HDLBP | Q00341; A0A | VIGLN_HUMAN | Unmodified | 0 | 269760 | 183990 | 99051 | 0 | 0 | 0 | 0 | 125700 | 0 | 2021600 | 2087400 | 0 | 0 | 228900 | 230100 | 0 | 0 |
| ELLELEALS | MVP | Q14764 | MVP_HUMAN | Oxidation (M) | 161870 | 0 | 0 | 0 | 0 | 0 | 0 | 0 | 0 | 0 | 0 | 934480 | 386420 | 0 | 0 | 0 | 0 | 0 |
| ELLENAEK | NIN; GPC6; GP( | Q8N4C6; Q9 | NIN_HUMAN N | Unmodified | 772810 | 605620 | 0 | 0 | 0 | 0 | 0 | 0 | 0 | 0 | 0 | 0 | 0 | 257690 | 0 | 0 | 0 | 0 |
| ELLEPLFK | ERLEC1 | Q96DZ1 | ERLEC_HUMAN | Unmodified | 0 | 0 | 0 | 0 | 0 | 0 | 0 | 0 | 5727800 | 226050 | 295030 | 661940 | 148470 | 0 | 0 | 0 | 0 | 0 |
| ELLEQNPG | TBC1D10B; TB( | Q4KMP7; H3 | TB10B_HUMAN | Unmodified | 450380 | 318170 | 0 | 0 | 1347600 | 188440 | 216840 | 207460 | 4647200 | 323070 | 1226300 | 0 | 526120 | 194510 | 1929800 | 0 | 76861 | 181860 |
| ELLEREGLT | OS9 | Q13438 | OS9_HUMAN P | Unmodified | 0 | 0 | 0 | 0 | 0 | 0 | 0 | 0 | 997440 | 217570 | 0 | 0 | 0 | 0 | 0 | 0 | 0 | 0 |
| ELLEYDLQ( | UBR4 | Q5T4S7 | UBR4_HUMAN | Unmodified | 0 | 0 | 0 | 0 | 0 | 0 | 450590 | 0 | 165720 | 328900 | 420330 | 82342 | 403780 | 0 | 0 | 0 | 0 | 0 |
| ELLFHPALF | NRBP1; NRBP1 | Q9UHY1; F8 | NRBP_HUMAN | Unmodified | 105150 | 115350 | 0 | 252410 | 2077400 | 0 | 563590 | 0 | 611220 | 716510 | 0 | 2849200 | 1052100 | 535360 | 99314 | 238760 | 0 | 1338400 |
| ELLFLSNAI | DAP3; DAP3; D/ | P51398; V9G | RT29_HUMAN | Unmodified | 0 | 0 | 0 | 0 | 0 | 0 | 22950 | 0 | 1188800 | 0 | 265960 | 0 | 0 | 0 | 0 | 0 | 0 | 0 |
| ELLFYPPSN | NOMO1; NOMC | Q15155; P69 | NOMO1_HUMA | Unmodified | 0 | 0 | 0 | 0 | 0 | 0 | 0 | 0 | 2345500 | 0 | 0 | 0 | 0 | 0 | 0 | 0 | 0 | 0 |
| ELLGQGLLI | EIF3CL; EIF3C | B5ME19; Q9 | EIFCL_HUMAN | Unmodified | 0 | 0 | 0 | 0 | 0 | 0 | 0 | 0 | 0 | 0 | 1620400 | 0 | 0 | 0 | 0 | 0 | 0 | 0 |
| ELLHSQNT | TPR | P12270 | TPR_HUMAN N | Unmodified | 0 | 0 | 0 | 0 | 0 | 0 | 85791 | 0 | 0 | 0 | 419310 | 0 | 859860 | 136480 | 0 | 0 | 0 | 0 |
| ELLIFNK | PARP1; PARP1; | P09874; A0A | PARP1_HUMAN | Unmodified | 0 | 0 | 0 | 0 | 0 | 0 | 0 | 0 | 0 | 0 | 0 | 0 | 0 | 0 | 0 | 0 | 0 | 0 |
| ELLIIGGVA | DDX1; DDX1; D | Q92499; A0A | DDX1_HUMAN | Unmodified | 0 | 0 | 0 | 0 | 614880 | 650340 | 0 | 0 | 0 | 189970 | 751230 | 824830 | 0 | 0 | 0 | 0 | 0 | 370000 |
| ELLIQIFSTF | BRIX1 | Q8TDN6 | BRX1_HUMAN | Unmodified | 0 | 0 | 0 | 0 | 0 | 0 | 0 | 0 | 0 | 0 | 871340 | 247600 | 0 | 0 | 0 | 0 | 0 | 0 |
| ELLLPGNN | ICAM1; ICAM1 | P05362; K7E | ICAM1_HUMAN | Oxidation (M) | 0 | 0 | 0 | 0 | 645370 | 0 | 0 | 0 | 0 | 0 | 0 | 0 | 0 | 0 | 0 | 0 | 3767900 | 0 |
| ELLLPGNN | ICAM1; ICAM1 | P05362; K7E | ICAM1_HUMAN | Unmodified | 0 | 0 | 0 | 0 | 0 | 0 | 0 | 0 | 0 | 0 | 0 | 0 | 0 | 0 | 0 | 0 | 3858000 | 0 |
| ELLLQPVTI | ARPC4; ARPC4- | P59998; A0A | ARPC4_HUMAN | Unmodified | 1190300 | 1108900 | 0 | 0 | 0 | 0 | 0 | 0 | 388580 | 0 | 2174100 | 11156000 | 0 | 0 | 906220 | 0 | 0 | 0 |
| ELLLQPVTI | ARPC4; ARPC4- | P59998; A0A | ARPC4_HUMAN | Unmodified | 0 | 0 | 0 | 0 | 5354100 | 1772800 | 1161900 | 1258100 | 0 | 1144700 | 0 | 6683000 | 5744300 | 2777100 | 0 | 0 | 1347300 | 3364800 |
| ELLLQPVTI | ARPC4; ARPC4- | P59998; A0A | ARPC4_HUMAN | Unmodified | 0 | 0 | 0 | 0 | 0 | 0 | 328920 | 125840 | 0 | 0 | 0 | 0 | 0 | 0 | 0 | 0 | 231660 | 0 |
| ELLLTGPGI | LRRC47 | Q8N1G4 | LRC47_HUMAN | Unmodified | 0 | 0 | 0 | 0 | 0 | 0 | 0 | 0 | 650560 | 0 | 425350 | 0 | 0 | 0 | 0 | 0 | 0 | 0 |
| ELLPVLISAI | VCL | P18206 | VINC_HUMAN | Oxidation (M) | 0 | 0 | 487140 | 1328300 | 3933800 | 1572000 | 4652400 | 1267600 | 803080 | 1604200 | 1423200 | 8920500 | 2851800 | 3208900 | 1006500 | 1080500 | 13840000 | 3638700 |
| ELLPVLISAI | VCL | P18206 | VINC_HUMAN | Unmodified | 0 | 0 | 0 | 0 | 4157800 | 669940 | 2323000 | 83543 | 1553200 | 534290 | 254280 | 962170 | 1541300 | 2377800 | 0 | 0 | 8205100 | 2999500 |
| ELLQEQLE( | IKBKG; IKBKG; I | Q9Y6K9; A0A | NEMO_HUMAN | Unmodified | 0 | 0 | 0 | 0 | 218870 | 1080300 | 0 | 36896 | 0 | 0 | 0 | 0 | 967250 | 870830 | 0 | 0 | 0 | 0 |
| ELLQHPFLI | PAK2; PAK3 | Q13177; O7 | PAK2_HUMAN | Unmodified | 0 | 0 | 0 | 0 | 752720 | 215530 | 134070 | 184500 | 0 | 183740 | 0 | 602260 | 464390 | 404460 | 0 | 0 | 765410 | 509450 |
| ELLQTELS( | S100A1; S100A | P23297; Q5T | S10A1_HUMAN | Unmodified | 68283 | 147160 | 71636 | 279600 | 154840 | 0 | 149610 | 0 | 24602 | 17229 | 0 | 0 | 142850 | 0 | 353250 | 152230 | 0 | 0 |
| ELLQTPNFI | GPD1L; GPD1L | Q8N335; C9 | GPD1L_HUMAN | Unmodified | 0 | 0 | 109430 | 0 | 889690 | 237270 | 0 | 0 | 0 | 0 | 0 | 2567400 | 0 | 0 | 54545 | 0 | 0 | 0 |
| ELLRVAQIL | ATP5PO; ATP5P | P48047; A0A | ATPO_HUMAN | Unmodified | 0 | 0 | 0 | 0 | 0 | 0 | 323560 | 0 | 0 | 1150400 | 0 | 0 | 0 | 0 | 0 | 0 | 0 | 0 |
| ELLSNDESI | PSD3; PSD3; PS | Q9NYI0; E5R | PSD3_HUMAN | Unmodified | 0 | 0 | 0 | 0 | 712170 | 218640 | 0 | 0 | 219820 | 86186 | 0 | 277110 | 260810 | 45499 | 0 | 0 | 40596 | 457590 |
| ELLSQLEET | GOLGB1 | Q14789 | GOGB1_HUMA | Unmodified | 0 | 0 | 1761300 | 0 | 0 | 0 | 0 | 0 | 0 | 0 | 0 | 0 | 0 | 0 | 0 | 0 | 0 | 0 |
| ELLSSDAM | INPP5A; INPP5/ | Q14642; H0 | I5P1_HUMAN I | Unmodified | 0 | 114710 | 368390 | 326260 | 0 | 0 | 164610 | 0 | 158530 | 64522 | 0 | 1740900 | 350240 | 85717 | 0 | 710560 | 0 | 0 |
| ELLTLDEK | RPS9; RPS9; RP | P46781; F2Z | RS9_HUMAN 4 | Unmodified | 0 | 0 | 0 | 0 | 0 | 0 | 0 | 0 | 0 | 0 | 168220 | 2910900 | 0 | 0 | 0 | 0 | 0 | 0 |
| ELLTLDEKI | RPS9; RPS9; RP | P46781; F2Z | RS9_HUMAN 4 | Unmodified | 0 | 0 | 0 | 0 | 0 | 0 | 0 | 0 | 0 | 0 | 152020 | 7654300 | 0 | 0 | 0 | 0 | 0 | 0 |
| ELLTLDEKI | RPS9; RPS9; RP | P46781; F2Z | RS9_HUMAN 4 | Unmodified | 517300 | 0 | 314990 | 351170 | 102550 | 0 | 0 | 0 | 0 | 0 | 0 | 5783700 | 0 | 0 | 405960 | 339460 | 0 | 0 |
| ELLTLIR | ACOX1; ACOX1 | Q15067; K7E | ACOX1_HUMAN | Unmodified | 0 | 0 | 0 | 0 | 0 | 0 | 0 | 0 | 0 | 58850 | 262990 | 0 | 0 | 0 | 0 | 0 | 0 | 0 |
| ELLTRELPS | S100A4 | P26447 | S10A4_HUMAN | Unmodified | 0 | 0 | 0 | 0 | 2634100 | 0 | 828330 | 0 | 0 | 0 | 0 | 0 | 0 | 1010600 | 0 | 0 | 568280 | 1482200 |
| ELLTSFGPL | U2AF2; U2AF2 | P26368; K7E | U2AF2_HUMAN | Unmodified | 6038700 | 3012900 | 1422100 | 1748300 | 1462600 | 289590 | 1156600 | 651340 | 818530 | 1104300 | 2224500 | 8223200 | 2666100 | 1151800 | 622060 | 1706000 | 568580 | 1090300 |
| ELLTTMGDI | MYL12A; MYL12 | P19105; O14 | ML12A_HUMAN | Oxidation (M) | 0 | 0 | 0 | 0 | 0 | 0 | 0 | 0 | 0 | 0 | 0 | 1976200 | 0 | 0 | 772570 | 0 | 0 | 0 |
| ELLTTMGDI | MYL12A; MYL12 | P19105; O14 | ML12A_HUMAN | Oxidation (M) | 0 | 0 | 0 | 2326200 | 358100 | 0 | 0 | 0 | 0 | 0 | 0 | 1689600 | 0 | 0 | 2074500 | 0 | 0 | 1851500 |
| ELLTVGQEI | SNX9; SNX9; SN | Q9Y5X1; A0A | SNX9_HUMAN | Unmodified | 2477300 | 3504300 | 0 | 0 | 1119700 | 114690 | 0 | 0 | 0 | 0 | 216960 | 1691500 | 2025700 | 0 | 445360 | 647980 | 0 | 0 |
| ELLTVGQEI | SNX9; SNX9; SN | Q9Y5X1; A0A | SNX9_HUMAN | Unmodified | 511430 | 224860 | 151320 | 126370 | 0 | 0 | 0 | 29890 | 0 | 98956 | 0 | 0 | 0 | 0 | 120300 | 218220 | 0 | 0 |
| ELLVDCYKI | PPP1R14B; LIM | Q96C90; B5I | PP14B_HUMAN | Unmodified | 0 | 0 | 0 | 0 | 0 | 0 | 28584 | 0 | 0 | 134360 | 0 | 227440 | 36907 | 0 | 846380 | 0 | 516100 | 0 |
| ELLVIELSDI | CPE | P16870 | CBPE_HUMAN | Unmodified | 0 | 0 | 0 | 0 | 0 | 0 | 0 | 0 | 20390000 | 2415600 | 0 | 0 | 0 | 0 | 0 | 0 | 0 | 0 |
| ELLVPLTSS | PFDN5 | Q99471 | PFD5_HUMAN | Oxidation (M) | 0 | 0 | 0 | 0 | 0 | 0 | 0 | 0 | 0 | 0 | 0 | 3219700 | 24010 | 0 | 0 | 0 | 0 | 0 |
| ELMAQNQI | EXTL3 | O43909 | EXTL3_HUMAN | Unmodified | 0 | 0 | 0 | 0 | 59084 | 0 | 0 | 0 | 579020 | 59263 | 0 | 0 | 0 | 0 | 0 | 0 | 0 | 0 |
| ELMETMHF | VPS28; VPS28 | Q9UK41; E9I | VPS28_HUMAN | Unmodified | 0 | 0 | 0 | 0 | 0 | 0 | 0 | 0 | 0 | 0 | 0 | 0 | 0 | 0 | 2956800 | 0 | 0 | 0 |
| ELMETMHF | VPS28; VPS28 | Q9UK41; E9I | VPS28_HUMAN | Oxidation (M) | 433310 | 291920 | 782900 | 0 | 0 | 0 | 0 | 0 | 0 | 0 | 0 | 0 | 0 | 0 | 1262300 | 316290 | 0 | 0 |
| ELMETMHF | VPS28; VPS28 | Q9UK41; E9I | VPS28_HUMAN | 2 Oxidation (M) | 0 | 85719 | 45661 | 0 | 0 | 0 | 0 | 0 | 0 | 0 | 0 | 0 | 0 | 0 | 328260 | 151590 | 0 | 0 |
| ELMGDHVS | ACTL8 | Q9H568 | ACTL8_HUMAN | Unmodified | 0 | 0 | 0 | 0 | 0 | 0 | 0 | 0 | 0 | 0 | 261340 | 639540 | 0 | 0 | 0 | 0 | 0 | 0 |
| ELMNLTGT | TSG101; TSG10 | Q99816; F5H | TS101_HUMAN | Oxidation (M) | 3089600 | 1929300 | 2274400 | 214550 | 0 | 0 | 0 | 0 | 0 | 0 | 0 | 1519600 | 0 | 0 | 11715000 | 9823700 | 0 | 0 |
| ELMNLTGT | TSG101; TSG10 | Q99816; F5H | TS101_HUMAN | Unmodified | 2030700 | 877900 | 317150 | 73878 | 0 | 0 | 0 | 0 | 382340 | 0 | 0 | 631440 | 0 | 0 | 4571400 | 3223800 | 0 | 0 |
| ELMQVVLA | AP1S1; AP1S1; | P61966; A0A | AP1S1_HUMAN | Unmodified | 0 | 0 | 0 | 0 | 0 | 0 | 0 | 0 | 0 | 0 | 0 | 0 | 0 | 0 | 0 | 113740 | 0 | 0 |
| ELNACREQ | PPFIA2; PPFIA2 | O75334; G3 | LIPA2_HUMAN | Unmodified | 0 | 0 | 0 | 660990 | 0 | 0 | 0 | 0 | 0 | 0 | 0 | 0 | 412920 | 0 | 201140 | 191050 | 0 | 0 |
| ELNARPLE | SAR1B; SAR1B; | Q9Y6B6; D6I | SAR1B_HUMAN | Unmodified | 0 | 0 | 0 | 0 | 0 | 0 | 0 | 0 | 272470 | 599130 | 460530 | 0 | 0 | 0 | 98084 | 0 | 0 | 0 |
| ELNARPME | SAR1A; SAR1A | Q9NR31; H0 | SAR1A_HUMAN | Unmodified | 0 | 0 | 0 | 0 | 0 | 85539 | 328150 | 0 | 0 | 0 | 0 | 106370 | 0 | 0 | 0 | 0 | 0 | 1435800 |
| ELNARPME | SAR1A; SAR1A | Q9NR31; H0 | SAR1A_HUMAN | Oxidation (M) | 0 | 0 | 0 | 0 | 0 | 0 | 270090 | 0 | 0 | 19301 | 0 | 0 | 0 | 0 | 0 | 0 | 0 | 223810 |
| ELNEFREK | PFDN2 | Q9UHV9 | PFD2_HUMAN | Unmodified | 0 | 0 | 0 | 0 | 0 | 0 | 0 | 0 | 0 | 0 | 213130 | 1248800 | 590070 | 325570 | 0 | 0 | 0 | 0 |
| ELNELVSAI | PGM2 | Q96G03 | PGM2_HUMAN | Unmodified | 0 | 0 | 43017 | 170320 | 1213900 | 233250 | 0 | 0 | 200220 | 185930 | 97873 | 285020 | 4511900 | 565230 | 427020 | 77292 | 223130 | 1191300 |
| ELNELVSAI | PGM2 | Q96G03 | PGM2_HUMAN | Unmodified | 0 | 0 | 0 | 0 | 768820 | 0 | 287780 | 0 | 0 | 0 | 0 | 0 | 3383400 | 1838800 | 0 | 0 | 347420 | 491740 |
| ELNGTAPEI | PLAA; PLAA; PL/ | Q9Y263; H0 | PLAP_HUMAN | Unmodified | 0 | 0 | 0 | 0 | 0 | 0 | 0 | 0 | 0 | 0 | 0 | 629880 | 0 | 0 | 0 | 0 | 0 | 0 |
| ELNHMLSI | S100A16 | Q96FQ6 | S10AG_HUMAN | Unmodified | 0 | 0 | 0 | 0 | 0 | 0 | 0 | 0 | 0 | 0 | 0 | 0 | 0 | 0 | 0 | 0 | 0 | 0 |
| ELNHMLSI | S100A16 | Q96FQ6 | S10AG_HUMAN | Unmodified | 151540 | 0 | 0 | 0 | 15568000 | 256640 | 0 | 0 | 2375200 | 1401800 | 0 | 0 | 178150 | 0 | 0 | 0 | 8090500 | 19906000 |
| ELNHMLSI | S100A16 | Q96FQ6 | S10AG_HUMAN | Oxidation (M) | 0 | 0 | 0 | 0 | 6721900 | 0 | 0 | 0 | 112100 | 462370 | 0 | 0 | 0 | 0 | 0 | 33815 | 5009200 | 7030900 |
| ELNHMLSI | S100A16 | Q96FQ6 | S10AG_HUMAN | Unmodified | 0 | 63445 | 0 | 0 | 1375900 | 208440 | 0 | 0 | 212210 | 91790 | 0 | 0 | 0 | 0 | 0 | 0 | 1829200 | 1911600 |
| ELNITAAK | RPS7; RPS7; RP | P62081; A0A | RS7_HUMAN 4 | Unmodified | 0 | 2030100 | 1498100 | 1384700 | 0 | 0 | 0 | 0 | 0 | 0 | 0 | 0 | 0 | 0 | 1646300 | 1515700 | 0 | 0 |
| ELNKYSTHI | STXBP2; STXBP | Q15833; M0 | STXB2_HUMAN | Unmodified | 0 | 0 | 0 | 0 | 343830 | 0 | 86052 | 0 | 0 | 0 | 0 | 0 | 0 | 0 | 0 | 48949 | 280320 | 0 |
| ELNLQDFS | CARMIL1; CARI | Q5VZK9; A0A | CARL1_HUMAN | Unmodified | 0 | 0 | 0 | 0 | 0 | 0 | 0 | 0 | 0 | 0 | 0 | 1481500 | 0 | 0 | 0 | 0 | 0 | 0 |
| ELNLSFCEI | RANGAP1; RAN | P46060; H0 | RAGP1_HUMAN | Unmodified | 21680 | 0 | 0 | 0 | 122790 | 0 | 0 | 0 | 0 | 0 | 0 | 410890 | 0 | 0 | 33198 | 0 | 0 | 0 |
| ELNLVLDEI | MGAT4B; MGAT | Q9UQ53; H0 | MGT4B_HUMAN | Unmodified | 0 | 0 | 0 | 0 | 0 | 0 | 0 | 0 | 3421200 | 267560 | 0 | 650800 | 0 | 0 | 0 | 0 | 0 | 0 |
| ELNMIIMLP | SERPINB6; SER | P35237; A0A | SPB6_HUMAN | 2 Oxidation (M) | 0 | 0 | 0 | 0 | 0 | 0 | 0 | 0 | 0 | 0 | 0 | 674620 | 0 | 0 | 142310 | 0 | 0 | 0 |
| ELNNTCEP | HSPH1; HSPH1 | Q92598; Q5 | HS105_HUMAN | Unmodified | 0 | 0 | 166320 | 0 | 0 | 0 | 0 | 48865 | 436850 | 0 | 1115000 | 6085900 | 337550 | 0 | 259890 | 320120 | 0 | 189720 |
| ELNRFDGS | DNASE1L1; DN | P49184; A6C | DNSL1_HUMAN | Unmodified | 1178600 | 4173900 | 575650 | 690810 | 0 | 0 | 0 | 0 | 0 | 0 | 0 | 1325600 | 0 | 0 | 1456600 | 803490 | 0 | 0 |
| ELNSFLES( | TAOK1 | Q7L7X3 | TAOK1_HUMAN | Unmodified | 0 | 0 | 0 | 0 | 689660 | 0 | 339590 | 113210 | 0 | 417930 | 0 | 0 | 0 | 978650 | 0 | 0 | 0 | 141270 |
| ELNSPILFA | ADGRE5 | P48960 | AGRE5_HUMAN | Unmodified | 3050000 | 2002200 | 5303900 | 5972800 | 59041000 | 17698000 | 0 | 0 | 544020 | 0 | 1091500 | 2260700 | 4733200 | 0 | 4420500 | 4596700 | 5567800 | 57364000 |
| ELNVMFIET | RAB6A | P20340 | RAB6A_HUMAN | Oxidation (M) | 340570 | 0 | 0 | 0 | 776520 | 0 | 702560 | 269390 | 1245600 | 0 | 1132100 | 3475400 | 0 | 0 | 437390 | 356740 | 0 | 0 |
| ELNVMFIET | RAB6A | P20340 | RAB6A_HUMAN | Unmodified | 0 | 0 | 0 | 0 | 0 | 0 | 0 | 0 | 1398400 | 0 | 0 | 0 | 0 | 0 | 0 | 0 | 0 | 0 |
| ELNWDDYI | GPD2 | P43304 | GPDM_HUMAN | Unmodified | 0 | 0 | 0 | 0 | 0 | 0 | 0 | 0 | 1154100 | 0 | 0 | 0 | 0 | 0 | 0 | 0 | 0 | 0 |
| ELNYFAKAI | PGK1 | P00558 | PGK1_HUMAN | Unmodified | 0 | 0 | 199900 | 0 | 271830 | 0 | 646530 | 0 | 0 | 53081 | 0 | 0 | 153180 | 0 | 0 | 0 | 5537600 | 499590 |
| ELPAAVAP/ | PGLS; PGLS; P( | O95336; M0 | 6PGL_HUMAN | Unmodified | 0 | 0 | 0 | 0 | 0 | 0 | 0 | 0 | 0 | 466840 | 0 | 4051000 | 0 | 0 | 0 | 0 | 0 | 0 |
| ELPELLQR | PDCD6IP | Q8WUM4 | PDC6I_HUMAN | Unmodified | 1.05E+08 | 1.09E+08 | 52871000 | 44775000 | 13345000 | 6522800 | 488840 | 88641 | 36724000 | 12211000 | 7103500 | 17042000 | 19551000 | 9371000 | 60153000 | 56836000 | 332290 | 9260800 |
| ELPEQFFQI | LRRC1; LRRC1 | Q9BTT6; Q5 | LRRC1_HUMAN | Unmodified | 0 | 0 | 89127 | 0 | 0 | 0 | 0 | 0 | 1189700 | 0 | 0 | 2624400 | 0 | 0 | 0 | 0 | 0 | 0 |
| ELPGFLQS( | S100P | P25815 | S100P_HUMAN | Unmodified | 319700 | 189130 | 0 | 0 | 0 | 0 | 0 | 0 | 0 | 0 | 0 | 0 | 0 | 0 | 0 | 12266 | 0 | 0 |
| ELPGHTGY | GNB2; GNB4; G | P62879; Q9H | GBB2_HUMAN | Unmodified | 3139200 | 1118300 | 1158400 | 1816500 | 439940 | 497130 | 128040 | 52758 | 2759800 | 217330 | 5728500 | 40899000 | 346180 | 235570 | 2291500 | 2038400 | 643020 | 852120 |
| ELPGQTLH | SCRIB; SCRIB; S | Q14160; A0A | SCRIB_HUMAN | Unmodified | 0 | 0 | 0 | 0 | 0 | 0 | 0 | 0 | 2006700 | 338220 | 0 | 896740 | 0 | 0 | 0 | 442090 | 211430 | 636710 |
| ELPGQTLH | SCRIB; SCRIB; S | Q14160; A0A | SCRIB_HUMAN | Unmodified | 0 | 0 | 0 | 0 | 2188100 | 0 | 0 | 0 | 902650 | 883290 | 0 | 0 | 871820 | 536200 | 0 | 0 | 2783700 | 6730700 |
| ELPGQTLH | SCRIB; SCRIB; S | Q14160; A0A | SCRIB_HUMAN | Unmodified | 0 | 0 | 0 | 0 | 3023200 | 0 | 0 | 0 | 0 | 2664700 | 0 | 0 | 2415600 | 0 | 0 | 0 | 7577500 | 9889000 |
| ELPKADTN( | TGOLN2; TGOL | O43493; A0A | TGON2_HUMA | Unmodified | 0 | 101870 | 0 | 22032 | 269050 | 82256 | 69318 | 0 | 11117000 | 2839900 | 0 | 0 | 523170 | 115860 | 0 | 0 | 76915 | 197900 |
| ELPKADTN( | TGOLN2; TGOL | O43493; A0A | TGON2_HUMA | Unmodified | 0 | 0 | 0 | 0 | 0 | 0 | 0 | 0 | 0 | 167180 | 0 | 0 | 0 | 0 | 0 | 0 | 27255 | 0 |
| ELPKGPEQ | SLC38A10; SLC | Q9HBR0; H0 | S38AA_HUMAN | Unmodified | 0 | 0 | 0 | 0 | 0 | 0 | 701090 | 0 | 8307600 | 0 | 0 | 0 | 0 | 0 | 0 | 0 | 0 | 0 |
| ELPLAADAI | PKP3 | Q9Y446 | PKP3_HUMAN | Unmodified | 0 | 0 | 109130 | 7951600 | 0 | 0 | 0 | 0 | 2800100 | 0 | 0 | 5569900 | 24968 | 0 | 518000 | 4466100 | 0 | 0 |
| ELPLSPAFF | PBXIP1 | Q96AQ6 | PBIP1_HUMAN | Unmodified | 0 | 0 | 0 | 0 | 0 | 0 | 0 | 0 | 384950 | 11817 | 0 | 0 | 0 | 0 | 0 | 0 | 0 | 0 |
| ELPLSPAFF | PBXIP1 | Q96AQ6 | PBIP1_HUMAN | Unmodified | 0 | 0 | 0 | 0 | 0 | 0 | 0 | 0 | 5354800 | 495500 | 829670 | 0 | 0 | 0 | 0 | 0 | 0 | 0 |

| | | | | | | | | | | | | | | | | | | | | | | |
|---|---|---|---|---|---|---|---|---|---|---|---|---|---|---|---|---|---|---|---|---|---|---|
| ELPLSPAFF | PBXIP1 | Q96AQ6 | PBIP1_HUMAN | Unmodified | 0 | 0 | 0 | 0 | 0 | 0 | 0 | 0 | 4354000 | 2447000 | 1676300 | 0 | 0 | 0 | 0 | 0 | 0 | 918330 |
| ELPMRNLQ | EGFR; EGFR | P00533; C9J | EGFR_HUMAN | Unmodified | 0 | 0 | 0 | 0 | 1974400 | 0 | 0 | 0 | 0 | 0 | 0 | 0 | 0 | 0 | 0 | 0 | 0 | 0 |
| ELPNFWEC | CPD | O75976 | CBPD_HUMAN | Unmodified | 0 | 0 | 0 | 0 | 1326200 | 605460 | 0 | 0 | 10253000 | 4702700 | 0 | 207520 | 0 | 0 | 0 | 0 | 0 | 0 |
| ELPPDQAE | ACTN4; ACTN1; | O43707; P12 | ACTN4_HUMAI | Unmodified | 339000 | 0 | 256550 | 467150 | 1280100 | 648620 | 0 | 0 | 780400 | 0 | 746500 | 3515300 | 377100 | 218290 | 632710 | 497710 | 0 | 0 |
| ELPPGVEEI | MVP | Q14764 | MVP_HUMAN N | Unmodified | 0 | 3025700 | 0 | 0 | 960770 | 0 | 0 | 0 | 4401900 | 0 | 519420 | 30121000 | 0 | 0 | 0 | 0 | 0 | 0 |
| ELPPISPEH | DIS3L2 | Q8IYB7 | DI3L2_HUMAN | Unmodified | 139020 | 0 | 0 | 519580 | 0 | 0 | 1203200 | 0 | 0 | 1546600 | 0 | 797850 | 0 | 0 | 377500 | 0 | 0 | 0 |
| ELPPPAPP/ | SLC4A2; SLC4A | P04920; A0A | B3A2_HUMAN . | Unmodified | 110650 | 0 | 820170 | 608800 | 375420 | 0 | 0 | 0 | 0 | 0 | 172800 | 1214500 | 311860 | 0 | 1559100 | 691690 | 0 | 0 |
| ELPQFATGI | API5; API5 | Q9BZZ5; G3' | API5_HUMAN A | Unmodified | 0 | 0 | 0 | 0 | 0 | 0 | 0 | 0 | 309530 | 0 | 0 | 2914700 | 0 | 0 | 0 | 0 | 0 | 0 |
| ELPQLVAS\ | UBR4 | Q5T4S7 | UBR4_HUMAN | Unmodified | 0 | 0 | 0 | 0 | 0 | 0 | 0 | 0 | 71923 | 0 | 0 | 479240 | 0 | 0 | 0 | 0 | 0 | 0 |
| ELPQVSFVI | GLRX3 | O76003 | GLRX3_HUMAI | Unmodified | 0 | 0 | 0 | 0 | 0 | 0 | 0 | 0 | 0 | 0 | 461270 | 1323900 | 0 | 0 | 0 | 0 | 0 | 0 |
| ELPSFVGEI | S100A2; S100A | P29034; R4C | S10A2_HUMAN | Unmodified | 0 | 0 | 158450 | 196460 | 0 | 0 | 0 | 41351 | 0 | 0 | 0 | 0 | 0 | 0 | 366920 | 226700 | 516840 | 2553500 |
| ELPSFVGEI | S100A2; S100A | P29034; R4C | S10A2_HUMAN | Unmodified | 0 | 0 | 0 | 0 | 92089 | 0 | 0 | 0 | 0 | 0 | 0 | 0 | 0 | 0 | 0 | 0 | 193470 | 909430 |
| ELPSTFGQI | LRRC57; N/A; L | Q8N9N7; A0 | LRC57_HUMAN | Unmodified | 1543200 | 1599600 | 0 | 606570 | 0 | 0 | 0 | 0 | 0 | 0 | 690790 | 3682900 | 0 | 0 | 1263400 | 0 | 0 | 0 |
| ELPTLKDNI | PIP4K2A; PIP4K | P48426; S4F | PI42A_HUMAN | Unmodified | 84459 | 0 | 0 | 511440 | 2895600 | 699320 | 1430400 | 1154800 | 0 | 0 | 0 | 447610 | 1555800 | 1625800 | 0 | 0 | 929190 | 2965200 |
| ELPVLWHC | BYSL | Q13895 | BYST_HUMAN I | Unmodified | 0 | 0 | 0 | 0 | 0 | 0 | 0 | 0 | 0 | 0 | 0 | 68204 | 0 | 0 | 29511 | 16943 | 0 | 0 |
| ELPVNAQN | ADSS2 | P30520 | PURA2_HUMAI | Unmodified | 0 | 0 | 0 | 0 | 815180 | 147300 | 0 | 0 | 871330 | 0 | 2034200 | 0 | 0 | 0 | 0 | 0 | 0 | 393710 |
| ELQAAGKS | GPI; GPI | P06744; A0A | G6PI_HUMAN ( | Unmodified | 961960 | 1230900 | 784780 | 6056000 | 1620500 | 367670 | 0 | 66575 | 18157000 | 1001300 | 6685200 | 55974000 | 721200 | 278820 | 6007700 | 3220000 | 420260 | 0 |
| ELQAAGKS | GPI; GPI | P06744; A0A | G6PI_HUMAN ( | Unmodified | 0 | 0 | 0 | 2063400 | 39769000 | 1955600 | 2055100 | 278730 | 6424200 | 4602600 | 1697400 | 2436700 | 4323000 | 5792000 | 0 | 0 | 5814800 | 7622100 |
| ELQAANES | SWAP70; SWAF | Q9UH65; E7 | SWP70_HUMAI | Unmodified | 0 | 0 | 0 | 0 | 499430 | 0 | 0 | 0 | 0 | 41864 | 0 | 0 | 0 | 78424 | 0 | 0 | 0 | 994600 |
| ELQAHGAI | ARPC2; ARPC2; | O15144; G5I | ARPC2_HUMAI | Unmodified | 920230 | 922100 | 1675300 | 929030 | 2568700 | 685060 | 1552100 | 1099600 | 0 | 1306100 | 1565200 | 14303000 | 2847900 | 0 | 1496900 | 1218100 | 1434500 | 1132000 |
| ELQAHGAI | ARPC2; ARPC2; | O15144; G5I | ARPC2_HUMAI | Unmodified | 202690 | 176910 | 0 | 0 | 0 | 0 | 0 | 0 | 0 | 0 | 0 | 3522600 | 0 | 0 | 264290 | 324300 | 0 | 0 |
| ELQAQIAEL | MYH10 | P35580 | MYH10_HUMAI | Unmodified | 0 | 0 | 0 | 879640 | 728580 | 0 | 0 | 0 | 0 | 0 | 353040 | 1814700 | 0 | 0 | 1684800 | 487030 | 0 | 0 |
| ELQDAQEH | PTPN23 | Q9H3S7 | PTN23_HUMAN | Unmodified | 104530 | 100700 | 0 | 0 | 0 | 0 | 0 | 0 | 0 | 0 | 0 | 76268 | 0 | 0 | 0 | 0 | 0 | 0 |
| ELQDLALQ | CHGA; CHGA | P10645; G5E | CMGA_HUMAN | Unmodified | 0 | 295690 | 0 | 0 | 0 | 271460 | 371850 | 340750 | 1.4E+08 | 67400000 | 180850 | 0 | 0 | 0 | 0 | 0 | 0 | 0 |
| ELQDLALQ | CHGA; CHGA | P10645; G5E | CMGA_HUMAN | Unmodified | 0 | 0 | 0 | 50823 | 0 | 0 | 89808 | 0 | 0 | 1805600 | 0 | 0 | 0 | 0 | 0 | 0 | 0 | 0 |
| ELQDSTQM | CUL2; CUL2; CI | Q13617; Q5' | CUL2_HUMAN | Unmodified | 0 | 0 | 0 | 0 | 0 | 0 | 0 | 0 | 0 | 0 | 0 | 529190 | 0 | 0 | 0 | 0 | 0 | 0 |
| ELQEESGL' | NUDT1 | P36639 | 8ODP_HUMAN | Unmodified | 174850 | 320160 | 0 | 0 | 0 | 0 | 0 | 167150 | 691160 | 156120 | 0 | 1358300 | 0 | 0 | 0 | 244810 | 195080 | 178480 |
| ELQELNELI | ZC3H15; ZC3H' | Q8WU90; F8 | ZC3HF_HUMAI | Unmodified | 703940 | 0 | 1292800 | 0 | 423520 | 0 | 0 | 0 | 0 | 0 | 864790 | 0 | 0 | 0 | 0 | 0 | 0 | 797820 |
| ELQELSSSI | UFL1 | O94874 | UFL1_HUMAN | Unmodified | 0 | 0 | 0 | 0 | 0 | 0 | 99427 | 0 | 0 | 208370 | 0 | 0 | 143200 | 260320 | 0 | 0 | 0 | 0 |
| ELQELVQYI | VCP; VCP; VCP; | P55072; C9J | TERA_HUMAN ' | Unmodified | 2338200 | 3687900 | 2739500 | 2601000 | 1216100 | 329560 | 0 | 0 | 9222300 | 617360 | 6727000 | 46231000 | 1595700 | 421930 | 2611900 | 5169900 | 0 | 0 |
| ELQFYRNN | REXO2; REXO2; | Q9Y3B8; F5C | ORN_HUMAN ( | Unmodified | 0 | 0 | 0 | 48142 | 1147400 | 410630 | 99088 | 22202 | 64743 | 381440 | 85345 | 0 | 1722800 | 990360 | 0 | 0 | 530870 | 1033000 |
| ELQGQIAAI | YOD1 | Q5VVQ6 | OTU1_HUMAN | Unmodified | 964080 | 1201100 | 221220 | 280340 | 0 | 0 | 0 | 0 | 0 | 0 | 0 | 0 | 0 | 0 | 329380 | 185620 | 0 | 0 |
| ELQGQNVE | ANKRD13A | Q8IZ07 | AN13A_HUMAI | Unmodified | 1292900 | 1449800 | 0 | 1493500 | 277590 | 0 | 0 | 0 | 1072000 | 0 | 416590 | 284770 | 0 | 0 | 220450 | 184660 | 60470 | 20109 |
| ELQHAALG | SCAMP3 | O14828 | SCAM3_HUMA | Unmodified | 0 | 0 | 0 | 200100 | 0 | 0 | 0 | 0 | 266630 | 0 | 0 | 623490 | 0 | 0 | 0 | 0 | 27666 | 0 |
| ELQKPDSF | H6PD | O95479 | G6PE_HUMAN | Unmodified | 0 | 0 | 0 | 0 | 0 | 0 | 0 | 0 | 1940400 | 0 | 0 | 0 | 0 | 0 | 0 | 0 | 0 | 0 |
| ELQNAPDA | GPS1; GPS1; GI | Q13098; J3K | CSN1_HUMAN | Unmodified | 0 | 0 | 0 | 0 | 0 | 0 | 0 | 0 | 0 | 0 | 505030 | 754690 | 0 | 0 | 0 | 0 | 0 | 0 |
| ELQNAPDA | GPS1; GPS1; GI | Q13098; J3K | CSN1_HUMAN | Unmodified | 0 | 0 | 0 | 0 | 0 | 0 | 0 | 0 | 0 | 0 | 0 | 1610800 | 0 | 0 | 0 | 0 | 0 | 0 |
| ELQNEVEK | PABPN1; PABPI | Q86U42; B4I | PABP2_HUMAN | Unmodified | 0 | 0 | 0 | 0 | 549600 | 0 | 0 | 78205 | 710040 | 114910 | 249220 | 1799500 | 0 | 70185 | 0 | 0 | 265530 | 163860 |
| ELQNTVAN | SCAMP2; SCAM | O15127; H3I | SCAM2_HUMA | Unmodified | 1176800 | 588420 | 2166500 | 2466700 | 0 | 0 | 0 | 0 | 1082700 | 0 | 0 | 0 | 0 | 0 | 828730 | 4002900 | 0 | 0 |
| ELQPGSHI( | CHDH | Q8NE62 | CHDH_HUMAN | Unmodified | 0 | 0 | 0 | 0 | 97806 | 0 | 0 | 0 | 520970 | 0 | 0 | 0 | 36351 | 0 | 0 | 0 | 0 | 0 |
| ELQPGSHI( | CHDH | Q8NE62 | CHDH_HUMAN | Unmodified | 0 | 0 | 0 | 0 | 0 | 0 | 0 | 0 | 3896400 | 544380 | 0 | 0 | 0 | 61319 | 1946200 | 0 | 0 | 0 |
| ELQQAVLH | NUCB1 | Q02818 | NUCB1_HUMA | Unmodified | 0 | 0 | 0 | 0 | 0 | 0 | 0 | 0 | 1119200 | 0 | 329370 | 182570 | 0 | 0 | 0 | 0 | 0 | 0 |
| ELQQVREQ | GNPTAB | Q3T906 | GNPTA_HUMAI | Unmodified | 0 | 0 | 0 | 0 | 0 | 0 | 0 | 0 | 1754000 | 170570 | 84836 | 0 | 0 | 0 | 0 | 0 | 0 | 0 |
| ELQREPLTF | CIAPIN1; CIAPII | Q6FI81; H3B | CPIN1_HUMAN | Unmodified | 51669 | 45775 | 50770 | 31550 | 149230 | 0 | 0 | 0 | 124970 | 0 | 505820 | 2094400 | 0 | 0 | 167140 | 153450 | 63197 | 0 |
| ELQRQASP | RAB5C; N/A; RA | P51148; K7E | RAB5C_HUMAI | Unmodified | 0 | 0 | 0 | 810830 | 5226700 | 0 | 6281600 | 2047600 | 0 | 6968000 | 0 | 0 | 6352900 | 4815200 | 0 | 0 | 0 | 0 |
| ELQRQASP | RAB5A | P20339 | RAB5A_HUMAN | Unmodified | 0 | 0 | 0 | 0 | 2126700 | 0 | 1147600 | 498210 | 0 | 2032600 | 0 | 0 | 1773100 | 1525600 | 0 | 0 | 727780 | 1011400 |
| ELQRQASP | RAB5B | P61020 | RAB5B_HUMAN | Unmodified | 0 | 0 | 0 | 0 | 4529200 | 0 | 5148700 | 489760 | 0 | 3078500 | 0 | 0 | 6352900 | 4138500 | 0 | 0 | 1612000 | 1386000 |
| ELQSMADC | PYCR2; PYCR2; | Q96C36; A0/ | P5CR2_HUMAN | Unmodified | 0 | 0 | 0 | 0 | 78666 | 0 | 0 | 0 | 1136500 | 116740 | 0 | 0 | 0 | 0 | 0 | 695900 | 0 | 0 |
| ELQSMADC | PYCR1; PYCR1 | P32322; E2C | P5CR1_HUMAN | Unmodified | 0 | 0 | 0 | 0 | 0 | 0 | 0 | 0 | 737800 | 54763 | 0 | 0 | 0 | 0 | 0 | 0 | 0 | 0 |
| ELQTLHNLI | KIF5B; KIF5C; K | P33176; O6C | KINH_HUMAN | Unmodified | 97740 | 0 | 0 | 0 | 5917900 | 0 | 2016200 | 91123 | 1326000 | 694240 | 279220 | 2186200 | 0 | 0 | 827440 | 0 | 1605100 | 2513700 |
| ELQTVLSTY | ARHGEF7; ARH | Q14155; B1/ | ARHG7_HUMA | Unmodified | 17321000 | 251140 | 111890 | 193750 | 3026400 | 483420 | 2253800 | 110790 | 4561700 | 2683900 | 916700 | 3616700 | 1322900 | 879910 | 337000 | 233520 | 1774800 | 1040400 |
| ELQVLDGA | EXOC1 | Q9NV70 | EXOC1_HUMAI | Unmodified | 0 | 0 | 0 | 0 | 198380 | 0 | 0 | 23546 | 0 | 60324 | 0 | 320770 | 0 | 0 | 0 | 0 | 0 | 0 |
| ELQVPEEN | ARHGEF5; ARH | Q12774; A5Y | ARHG5_HUMA | Unmodified | 0 | 0 | 0 | 0 | 0 | 0 | 0 | 0 | 0 | 0 | 241890 | 212410 | 0 | 0 | 31447 | 0 | 0 | 0 |
| ELQYIDAISI | BAIAP2; BAIAP2 | Q9UQB8; I3I | BAIP2_HUMAN | Unmodified | 0 | 0 | 1394900 | 608410 | 686450 | 0 | 0 | 0 | 1329500 | 0 | 381570 | 2728500 | 348100 | 0 | 2049000 | 2443600 | 0 | 0 |
| ELQYIDAISI | BAIAP2; BAIAP2 | Q9UQB8; I3I | BAIP2_HUMAN | Unmodified | 0 | 0 | 0 | 0 | 11412000 | 4758100 | 0 | 0 | 2040300 | 1154600 | 16852 | 0 | 6054300 | 3949700 | 0 | 0 | 1378600 | 0 |
| ELRAEAERI | NUMA1; NUMA | Q14980; H0' | NUMA1_HUMA | Unmodified | 0 | 0 | 0 | 0 | 266840 | 0 | 0 | 0 | 267750 | 542800 | 0 | 0 | 1025200 | 0 | 0 | 0 | 401120 | 682630 |
| ELRAQHEE | GOLM2; GOLM2 | Q6P4E1; H0' | GOLM2_HUMA | Unmodified | 0 | 0 | 0 | 0 | 178760 | 0 | 69017 | 44801 | 445430 | 374620 | 43150 | 55877 | 244830 | 63513 | 0 | 0 | 0 | 145760 |
| ELRDEEQT/ | MCM6 | Q14566 | MCM6_HUMAN | Unmodified | 0 | 0 | 0 | 0 | 0 | 0 | 0 | 0 | 0 | 0 | 255920 | 770120 | 0 | 0 | 0 | 0 | 0 | 0 |
| ELRDFNLD | UGGT1 | Q9NYU2 | UGGG1_HUMA | Unmodified | 0 | 0 | 0 | 0 | 0 | 0 | 0 | 0 | 1738900 | 786710 | 0 | 298500 | 0 | 0 | 0 | 0 | 0 | 0 |
| ELRDHADS | RAB11A; RAB11 | P62491; Q15 | RB11A_HUMAN | Oxidation (M) | 1404700 | 1028500 | 3049500 | 2406500 | 0 | 0 | 0 | 0 | 0 | 0 | 1097200 | 10426000 | 0 | 0 | 2973900 | 4299500 | 0 | 0 |
| ELRDHADS | RAB11A; RAB11 | P62491; Q15 | RB11A_HUMAN | Oxidation (M) | 655060 | 763560 | 0 | 2573100 | 251940 | 0 | 185750 | 0 | 0 | 0 | 0 | 2141100 | 0 | 0 | 836480 | 834610 | 0 | 0 |
| ELRDQIVS\ | SHTN1 | A0MZ66 | SHOT1_HUMAI | Unmodified | 0 | 0 | 0 | 0 | 1978000 | 0 | 1003100 | 447270 | 1076100 | 1250600 | 0 | 1397400 | 1269400 | 1643100 | 0 | 0 | 502210 | 841460 |
| ELRDQIVS\ | SHTN1 | A0MZ66 | SHOT1_HUMAI | Unmodified | 0 | 0 | 0 | 0 | 0 | 0 | 0 | 0 | 0 | 0 | 167270 | 117230 | 353560 | 0 | 0 | 0 | 0 | 0 |
| ELREATSPK | EGFR; EGFR; E( | P00533; Q5( | EGFR_HUMAN | Unmodified | 0 | 0 | 0 | 0 | 4187900 | 0 | 0 | 0 | 0 | 0 | 0 | 0 | 0 | 0 | 0 | 0 | 0 | 0 |
| ELRENTQT | HNRNPK; HNRI | P61978; Q5T | HNRPK_HUMA | Unmodified | 1731500 | 1724500 | 473890 | 1340600 | 4991900 | 1124100 | 7029300 | 4424300 | 2665200 | 495310 | 3546700 | 27517000 | 9265900 | 7616700 | 602330 | 541190 | 1071800 | 2601300 |
| ELRETYNT( | ITSN2; ITSN2 | Q9NZM3; A0 | ITSN2_HUMAN | Unmodified | 0 | 0 | 0 | 0 | 731230 | 0 | 0 | 0 | 0 | 0 | 0 | 0 | 0 | 0 | 0 | 0 | 0 | 0 |
| ELREVIELPI | PSMC6; PSMC6 | P62333; A0A | PRS10_HUMAN | Unmodified | 16474 | 0 | 0 | 284070 | 0 | 0 | 0 | 0 | 97618 | 129890 | 296560 | 25028 | 0 | 0 | 30411 | 0 | 0 | 0 |
| ELREYLAQI | DIPK2A; DIPK2/ | Q8NDZ4; E7 | DIK2A_HUMAN | Unmodified | 0 | 0 | 0 | 1055900 | 0 | 0 | 914860 | 0 | 3360000 | 1749000 | 0 | 577710 | 0 | 392630 | 0 | 0 | 0 | 0 |
| ELRLQPSS1 | AOC1; AOC1; A | P19801; C9J | AOC1_HUMAN | Oxidation (M) | 0 | 0 | 0 | 0 | 0 | 0 | 0 | 0 | 0 | 401470 | 0 | 0 | 0 | 0 | 0 | 0 | 0 | 0 |
| ELRNQTPA | DIP2B | Q9P265 | DIP2B_HUMAN | Unmodified | 0 | 0 | 0 | 0 | 0 | 18526 | 0 | 0 | 0 | 0 | 0 | 0 | 305150 | 115030 | 0 | 0 | 102080 | 246960 |
| ELRNVLLNI | NSUN2 | Q08J23 | NSUN2_HUMA | Unmodified | 0 | 0 | 0 | 0 | 0 | 0 | 0 | 468000 | 0 | 0 | 0 | 0 | 0 | 0 | 0 | 0 | 0 | 0 |
| ELRQVFEG | LMAN1 | P49257 | LMAN1_HUMAI | Unmodified | 0 | 0 | 0 | 0 | 0 | 0 | 0 | 0 | 288590 | 115860 | 0 | 0 | 0 | 0 | 0 | 188250 | 0 | 0 |
| ELRQVNLQ | RASSF8; RASSF | Q8NHQ8; H( | RASF8_HUMAN | Unmodified | 0 | 0 | 0 | 0 | 0 | 0 | 0 | 0 | 0 | 0 | 0 | 0 | 0 | 0 | 0 | 0 | 553050 | 410090 |
| ELRVEQESI | ATXN10; ATXN1 | Q9UBB4; A0. | ATX10_HUMAN | Unmodified | 0 | 0 | 0 | 0 | 0 | 0 | 0 | 0 | 5034100 | 18146 | 27593 | 149440 | 727160 | 0 | 104820 | 81581 | 0 | 0 |
| ELSAERPLI | SCG3 | Q8WXD2 | SCG3_HUMAN | Unmodified | 0 | 0 | 0 | 0 | 0 | 0 | 0 | 214540 | 80146000 | 7121300 | 0 | 0 | 0 | 0 | 0 | 0 | 0 | 0 |
| ELSAERPLI | SCG3 | Q8WXD2 | SCG3_HUMAN | Unmodified | 0 | 0 | 0 | 0 | 0 | 0 | 0 | 0 | 1656100 | 3327500 | 0 | 0 | 0 | 0 | 0 | 0 | 0 | 0 |
| ELSANTAAI | SNX2 | O60749 | SNX2_HUMAN | Unmodified | 0 | 0 | 153050 | 241190 | 4856100 | 370800 | 1107800 | 1227900 | 1419300 | 180040 | 382960 | 5031700 | 4997000 | 1410800 | 0 | 771580 | 927720 | 1172500 |
| ELSAVTFPI | STAT1; STAT1; S | P42224; A0A | STAT1_HUMAN | Unmodified | 0 | 0 | 220760 | 0 | 0 | 0 | 0 | 0 | 980260 | 0 | 774080 | 0 | 0 | 0 | 0 | 0 | 0 | 0 |
| ELSAVTFPI | STAT1; STAT1; S | P42224; A0A | STAT1_HUMAN | Unmodified | 0 | 0 | 0 | 0 | 4244800 | 588610 | 174160 | 0 | 0 | 414720 | 0 | 0 | 0 | 607430 | 0 | 0 | 797720 | 9197300 |
| ELSCTELQI | TBC1D24; TBC1 | Q9ULP9; A0/ | TBC24_HUMAN | Unmodified | 0 | 0 | 0 | 0 | 0 | 0 | 0 | 0 | 372900 | 0 | 361350 | 893520 | 0 | 0 | 0 | 0 | 0 | 195250 |
| ELSDAVNV | TAX1BP1; TAX1I | Q86VP1; B8Z | TAXB1_HUMAN | Unmodified | 0 | 0 | 0 | 0 | 0 | 0 | 0 | 0 | 1166100 | 0 | 0 | 0 | 0 | 0 | 0 | 0 | 0 | 0 |
| ELSDDCFQ | PRCP; PRCP; PI | P42785; B7Z | PCP_HUMAN L | Unmodified | 0 | 0 | 0 | 0 | 0 | 0 | 0 | 0 | 1155700 | 0 | 0 | 0 | 0 | 0 | 0 | 0 | 0 | 0 |
| ELSDFISYL | PDIA3 | P30101 | PDIA3_HUMAN | Unmodified | 0 | 17061 | 114980 | 0 | 0 | 0 | 0 | 19063 | 29241000 | 113880 | 2238200 | 5420100 | 0 | 0 | 0 | 809280 | 0 | 0 |
| ELSDFISYL | PDIA3 | P30101 | PDIA3_HUMAN | Unmodified | 0 | 0 | 0 | 0 | 0 | 0 | 0 | 0 | 2257200 | 0 | 885930 | 662530 | 0 | 0 | 0 | 0 | 0 | 0 |
| ELSDIALR | ALDOC; ALDOC | P09972; J3Q | ALDOC_HUMA | Unmodified | 277950 | 0 | 0 | 0 | 80561 | 0 | 0 | 0 | 0 | 0 | 1287800 | 10287000 | 907050 | 308660 | 134720 | 0 | 525730 | 738490 |
| ELSDIALRI\ | ALDOC; ALDOC | P09972; J3Q | ALDOC_HUMA | Unmodified | 0 | 0 | 0 | 0 | 0 | 0 | 314940 | 113820 | 0 | 0 | 0 | 0 | 1509300 | 934000 | 0 | 0 | 2087700 | 2218800 |
| ELSDLESAI | CAD; CAD | P27708; F8V | PYR1_HUMAN | Unmodified | 0 | 139840 | 94690 | 72071 | 0 | 0 | 0 | 0 | 0 | 0 | 0 | 1448800 | 0 | 0 | 21859 | 0 | 0 | 0 |
| ELSEALGQI | LGALS3BP; LGA | Q08380; K7E | LG3BP_HUMAN | Unmodified | 0 | 0 | 0 | 0 | 0 | 0 | 0 | 0 | 1978300 | 0 | 0 | 6621500 | 0 | 0 | 0 | 0 | 0 | 0 |
| ELSEALTR | GARS1; GARS1; | P41250; A0A | GARS_HUMAN | Unmodified | 334520 | 134410 | 611790 | 718950 | 0 | 349200 | 208770 | 136050 | 4023100 | 994960 | 3904400 | 17559000 | 1246300 | 535360 | 779960 | 916750 | 312640 | 509450 |
| ELSEIMHE( | DOCK9; DOCK9 | A0A088AWN | A0A088AWN3_ | Unmodified | 0 | 0 | 0 | 0 | 614460 | 116710 | 59698 | 35655 | 200800 | 418420 | 0 | 0 | 0 | 208640 | 0 | 28985 | 113260 | 210810 |
| ELSEKLELA | OPTN | Q96CV9 | OPTN_HUMAN | Unmodified | 0 | 0 | 0 | 0 | 0 | 0 | 325250 | 0 | 179210 | 0 | 0 | 0 | 0 | 0 | 0 | 0 | 0 | 178720 |
| ELSELSLLS | ACP3 | P15309 | PPAP_HUMAN | Unmodified | 0 | 0 | 0 | 0 | 0 | 0 | 0 | 0 | 1836400 | 2092500 | 0 | 127120 | 0 | 0 | 0 | 0 | 0 | 0 |
| ELSEQIQR | EZR; EZR | P15311; E7E | EZRI_HUMAN E | Unmodified | 889290 | 1051700 | 1459400 | 787830 | 196510 | 0 | 0 | 0 | 12967000 | 444530 | 333050 | 24767000 | 0 | 0 | 1153100 | 2289800 | 65076 | 0 |
| ELSEVKDSI | SH3GL2 | Q99962 | SH3G2_HUMAI | Unmodified | 8157.8 | 13463 | 0 | 0 | 0 | 0 | 30504 | 0 | 0 | 0 | 0 | 0 | 0 | 0 | 0 | 0 | 0 | 0 |
| ELSEYVRN( | BZW1; BZW1 | Q7L1Q6; C9I | BZW1_HUMAN | Unmodified | 0 | 0 | 0 | 0 | 1662400 | 0 | 259130 | 0 | 0 | 0 | 0 | 0 | 1377900 | 463910 | 0 | 0 | 723840 | 1599400 |
| ELSFGRGS | PDIA6 | Q15084 | PDIA6_HUMAN | Unmodified | 0 | 0 | 0 | 0 | 0 | 0 | 0 | 0 | 4594600 | 0 | 0 | 0 | 0 | 0 | 0 | 0 | 0 | 0 |
| ELSGESDE | SH3KBP1; SH3I | Q96B97; Q5' | SH3K1_HUMAN | Unmodified | 0 | 209630 | 397390 | 598590 | 0 | 0 | 0 | 0 | 0 | 0 | 0 | 0 | 724930 | 0 | 465500 | 467720 | 0 | 0 |
| ELSGLGSA | ACADVL; ACAD' | P49748; G3\ | ACADV_HUMAI | Unmodified | 0 | 0 | 0 | 0 | 0 | 0 | 0 | 0 | 0 | 2261100 | 0 | 0 | 0 | 0 | 0 | 0 | 0 | 0 |
| ELSGSSPVI | GAA; GAA; GAA | P10253; I3L( | LYAG_HUMAN | Unmodified | 0 | 0 | 0 | 0 | 0 | 0 | 0 | 0 | 1994000 | 0 | 0 | 382810 | 0 | 0 | 0 | 0 | 0 | 0 |
| ELSLAGNEI | RNH1; RNH1 | P13489; H0Y | RINI_HUMAN R | Unmodified | 25792 | 0 | 0 | 0 | 98229 | 0 | 0 | 0 | 331350 | 0 | 434420 | 9911900 | 0 | 0 | 901470 | 585060 | 0 | 195290 |
| ELSLVMNE | N/A | N/A | N/A | Unmodified | 0 | 0 | 0 | 0 | 627620 | 0 | 19541000 | 3006900 | 0 | 979320 | 0 | 0 | 0 | 560100 | 0 | 0 | 60738000 | 2821800 |
| ELSNFYFSI | IARS2 | Q9NSE4 | SYIM_HUMAN I | Unmodified | 0 | 0 | 0 | 0 | 0 | 0 | 0 | 0 | 502900 | 0 | 0 | 0 | 0 | 0 | 0 | 0 | 357840 | 0 |
| ELSNSPLRI | SRRM2 | Q9UQ35 | SRRM2_HUMAI | Unmodified | 0 | 0 | 0 | 0 | 0 | 0 | 0 | 0 | 0 | 0 | 0 | 456490 | 309510 | 0 | 0 | 0 | 0 | 0 |
| ELSNTAAY( | SIAE | Q9HAT2 | SIAE_HUMAN S | Unmodified | 0 | 0 | 0 | 0 | 0 | 0 | 0 | 0 | 307930 | 66053 | 0 | 223430 | 0 | 0 | 109860 | 0 | 0 | 0 |
| ELSPDIAHL | LRRC47 | Q8N1G4 | LRC47_HUMAN | Unmodified | 144320 | 102330 | 239280 | 162410 | 0 | 0 | 0 | 27115 | 671510 | 0 | 136670 | 1565700 | 93924 | 0 | 367660 | 183200 | 0 | 0 |
| ELSSSTQGI | PLEKHG3; PLEK | A1L390; G3\ | PKHG3_HUMAI | Unmodified | 0 | 0 | 0 | 0 | 647320 | 0 | 0 | 0 | 0 | 0 | 0 | 0 | 0 | 0 | 0 | 0 | 613170 | 259980 |
| ELSSTEAQI | MYH14; MYH14 | Q7Z406; A0/ | MYH14_HUMAI | Unmodified | 0 | 0 | 0 | 0 | 446880 | 0 | 0 | 0 | 577260 | 417300 | 0 | 0 | 1836400 | 1108400 | 0 | 0 | 0 | 0 |
| ELSTTLNAI | HSPA4; HSPA4; | P34932; A0A | HSP74_HUMAN | Unmodified | 0 | 104550 | 376460 | 577280 | 0 | 0 | 350430 | 721210 | 2302000 | 130230 | 3019800 | 19737000 | 750320 | 0 | 553300 | 1000800 | 0 | 0 |
| ELSYMVVSI | CD109 | Q6YHK3 | CD109_HUMAI | Unmodified | 0 | 0 | 0 | 0 | 0 | 0 | 0 | 0 | 3455800 | 371890 | 0 | 0 | 0 | 0 | 0 | 0 | 0 | 0 |

| | | | | | | | | | | | | | | | | | | | | | | |
|---|---|---|---|---|---|---|---|---|---|---|---|---|---|---|---|---|---|---|---|---|---|---|
| ELTAPDENI | TOP1; TOP1 | P11387; A0A | TOP1_HUMAN | Unmodified | 830720 | 0 | 0 | 0 | 315720 | 0 | 0 | 579040 | 211080 | 93953 | 0 | 8010600 | 0 | 0 | 0 | 0 | 0 | 0 |
| ELTAVVQKI | RPS3; RPS3; RP | P23396; H0Y | RS3_HUMAN 4 | Unmodified | 0 | 1187900 | 0 | 0 | 0 | 0 | 0 | 0 | 0 | 0 | 0 | 1692700 | 0 | 166550 | 0 | 0 | 0 | 0 |
| ELTDEEAEI | NUDC; NUDC; | Q9Y266; A0A | NUDC_HUMAN | Unmodified | 0 | 0 | 765920 | 1084600 | 1470600 | 0 | 0 | 0 | 5658900 | 534590 | 3156200 | 12386000 | 3489100 | 605410 | 735350 | 972580 | 576420 | 1046900 |
| ELTDLDYRI | INPPL1; INPPL1 | O15357; A0A | SHIP2_HUMAN | Oxidation (M) | 0 | 0 | 0 | 152040 | 395610 | 0 | 0 | 0 | 0 | 392080 | 304150 | 476650 | 0 | 0 | 0 | 0 | 0 | 0 |
| ELTEEKESA | MDH1; MDH1; I | P40925; B8Z | MDHC_HUMAN | Unmodified | 0 | 0 | 0 | 0 | 7198200 | 2270200 | 2405100 | 452750 | 112880 | 1494000 | 151430 | 114290 | 3715300 | 2559200 | 0 | 0 | 4568600 | 4662600 |
| ELTIGSKLQ | S100A6; S100A | P06703; R4C | S10A6_HUMAN | Unmodified | 1097200 | 713350 | 0 | 0 | 1375400 | 439980 | 0 | 0 | 2951200 | 304690 | 64694 | 587950 | 0 | 0 | 0 | 0 | 780330 | 712850 |
| ELTLPVDST | KIAA0319L; KIA | Q8IZA0; C9J | K319L_HUMAN | Unmodified | 0 | 0 | 0 | 0 | 517170 | 0 | 116250 | 0 | 10121000 | 1004100 | 601340 | 658930 | 0 | 0 | 0 | 0 | 0 | 0 |
| ELTLQKGD | SORBS3; SORB | O60504; E5F | VINEX_HUMAN | Unmodified | 531700 | 0 | 0 | 0 | 242240 | 0 | 0 | 0 | 0 | 0 | 0 | 0 | 0 | 0 | 0 | 0 | 0 | 0 |
| ELTMASLPI | XRN2 | Q9H0D6 | XRN2_HUMAN | Oxidation (M) | 110920 | 105370 | 0 | 0 | 0 | 0 | 0 | 0 | 0 | 0 | 0 | 714900 | 0 | 0 | 423700 | 0 | 0 | 0 |
| ELTNLLGLI | UTRN | P46939 | UTRN_HUMAN | Unmodified | 0 | 0 | 0 | 0 | 0 | 0 | 0 | 0 | 0 | 0 | 0 | 0 | 0 | 0 | 0 | 465280 | 0 | 0 |
| ELTNQQEA | DCTN1; DCTN1 | Q14203; Q6 | DCTN1_HUMA | Unmodified | 0 | 0 | 0 | 0 | 0 | 0 | 0 | 0 | 0 | 0 | 0 | 3502700 | 0 | 0 | 0 | 0 | 0 | 0 |
| ELTPQVVS/ | VCL | P18206 | VINC_HUMAN | Unmodified | 935520 | 1045700 | 199230 | 191740 | 1412100 | 0 | 231870 | 0 | 0 | 992290 | 0 | 6164400 | 182350 | 91560 | 889330 | 782990 | 4996900 | 167460 |
| ELTQSELEL | CCNDBP1 | O95273 | CCDB1_HUMA | Unmodified | 333750 | 662480 | 273840 | 262990 | 0 | 0 | 0 | 0 | 0 | 0 | 0 | 0 | 475580 | 0 | 500550 | 547860 | 0 | 0 |
| ELTSQHPY | MGAT5 | Q09328 | MGT5A_HUMA | Unmodified | 0 | 0 | 0 | 0 | 0 | 0 | 0 | 0 | 2134800 | 490060 | 0 | 0 | 0 | 0 | 0 | 0 | 0 | 0 |
| ELTSTCSPI | HSPA4 | P34932 | HSP74_HUMAN | Unmodified | 0 | 0 | 0 | 0 | 0 | 0 | 0 | 0 | 0 | 0 | 890040 | 2336300 | 271870 | 0 | 0 | 0 | 0 | 0 |
| ELTTLLQK | SUN1; SUN1; S | O94901; H0Y | SUN1_HUMAN | Unmodified | 0 | 0 | 0 | 0 | 0 | 0 | 0 | 216270 | 2606300 | 196320 | 1170000 | 0 | 0 | 0 | 0 | 0 | 0 | 0 |
| ELTVSNND | RNH1; RNH1; R | P13489; E9P | RINI_HUMAN R | Unmodified | 0 | 0 | 0 | 0 | 188660 | 0 | 713330 | 1123300 | 424570 | 109650 | 932540 | 10585000 | 0 | 0 | 1282600 | 772250 | 270460 | 522090 |
| ELTVSNND | RNH1; RNH1 | P13489; E9P | RINI_HUMAN R | Unmodified | 0 | 0 | 0 | 0 | 0 | 0 | 0 | 0 | 0 | 0 | 0 | 0 | 0 | 654410 | 0 | 0 | 420720 | 2090100 |
| ELTYQNTD | LRRFIP1 | Q32MZ4 | LRRF1_HUMAN | Unmodified | 0 | 0 | 0 | 0 | 0 | 1365800 | 190060 | 0 | 588970 | 268780 | 0 | 646830 | 752050 | 0 | 0 | 0 | 0 | 592210 |
| ELVALMSAI | LTA4H | P09960 | LKHA4_HUMAN | Unmodified | 0 | 0 | 0 | 0 | 0 | 0 | 0 | 0 | 0 | 0 | 0 | 805690 | 0 | 0 | 0 | 0 | 0 | 0 |
| ELVALMSAI | LTA4H | P09960 | LKHA4_HUMAN | Unmodified | 0 | 0 | 0 | 0 | 0 | 0 | 0 | 0 | 0 | 0 | 0 | 2365900 | 0 | 0 | 0 | 0 | 0 | 524590 |
| ELVALMSAI | LTA4H | P09960 | LKHA4_HUMAN | Oxidation (M) | 0 | 0 | 0 | 0 | 0 | 0 | 0 | 0 | 0 | 0 | 0 | 397690 | 0 | 0 | 0 | 0 | 0 | 0 |
| ELVALMSAI | LTA4H | P09960 | LKHA4_HUMAN | Unmodified | 20567 | 0 | 0 | 0 | 2249000 | 0 | 58282 | 0 | 461720 | 257650 | 0 | 3778700 | 707420 | 0 | 0 | 0 | 196410 | 332320 |
| ELVALMSAI | LTA4H | P09960 | LKHA4_HUMAN | Oxidation (M) | 0 | 0 | 0 | 0 | 444930 | 0 | 0 | 0 | 0 | 0 | 395490 | 1110400 | 0 | 0 | 0 | 79783 | 0 | 0 |
| ELVDDSVN | PCMT1; PCMT1 | P22061; F6S | PIMT_HUMAN | Unmodified | 29196 | 0 | 0 | 0 | 0 | 0 | 0 | 0 | 255040 | 0 | 363070 | 1651900 | 0 | 0 | 0 | 0 | 63090 | 33109 |
| ELVDDSVN | PCMT1; PCMT1 | P22061; F6S | PIMT_HUMAN | Unmodified | 0 | 0 | 0 | 0 | 0 | 0 | 0 | 0 | 665690 | 0 | 0 | 912850 | 0 | 0 | 0 | 0 | 0 | 0 |
| ELVDQDIQ | NLGN1; NLGN1 | Q8N2Q7; J3 | NLGN1_HUMA | Unmodified | 179970 | 171540 | 0 | 0 | 0 | 0 | 0 | 0 | 0 | 0 | 0 | 0 | 0 | 0 | 0 | 0 | 0 | 0 |
| ELVDYFLN | MAN2B1 | O00754 | MA2B1_HUMA | Unmodified | 34557 | 28516 | 0 | 66860 | 0 | 0 | 0 | 0 | 1768900 | 377850 | 104000 | 43157 | 0 | 0 | 159900 | 22921 | 0 | 0 |
| ELVELNK | KIF4A | O95239 | KIF4A_HUMAN | Unmodified | 0 | 0 | 382070 | 0 | 778060 | 0 | 0 | 0 | 489420 | 65695 | 228030 | 151350 | 614550 | 502410 | 533820 | 152010 | 0 | 619590 |
| ELVELSQA | ARHGAP45; AR | Q92619; K7E | HMHA1_HUMA | Unmodified | 0 | 0 | 518370 | 0 | 788910 | 0 | 563440 | 70304 | 0 | 0 | 0 | 0 | 0 | 0 | 0 | 0 | 721780 | 321100 |
| ELVEPLTPS | EGFR; EGFR; E( | P00533; Q5 | EGFR_HUMAN | Unmodified | 0 | 0 | 3468800 | 0 | 0 | 0 | 0 | 0 | 0 | 0 | 0 | 0 | 0 | 0 | 1657300 | 1594500 | 0 | 0 |
| ELVEPVSR | ACADVL; ACAD | P49748; K7E | ACADV_HUMA | Unmodified | 0 | 489840 | 0 | 0 | 272530 | 1017900 | 162220 | 0 | 7367700 | 440190 | 171170 | 36197 | 0 | 22737 | 0 | 0 | 167960 | 0 |
| ELVFKEDG | EIF1AY; EIF1AX | O14602; P47 | IF1AY_HUMAN | Unmodified | 7826700 | 4458700 | 5641900 | 3911400 | 647380 | 0 | 490970 | 27079 | 1669000 | 238290 | 1524500 | 18799000 | 2090400 | 0 | 6795300 | 9871500 | 169450 | 707940 |
| ELVKEPLSN | VPS11; VPS11 | Q9H270; A0 | VPS11_HUMAN | Unmodified | 0 | 0 | 0 | 0 | 954680 | 109740 | 159860 | 218740 | 0 | 0 | 0 | 1928800 | 152820 | 151530 | 0 | 0 | 323650 | 292500 |
| ELVKLKETC | AKAP12 | Q02952 | AKA12_HUMAN | Unmodified | 0 | 0 | 0 | 0 | 0 | 0 | 0 | 0 | 0 | 0 | 0 | 0 | 0 | 0 | 0 | 0 | 230290 | 0 |
| ELVLDNCK | ANP32B | Q92688 | AN32B_HUMAN | Unmodified | 0 | 0 | 120400 | 0 | 0 | 0 | 0 | 0 | 0 | 71682 | 1220000 | 2692700 | 0 | 0 | 0 | 0 | 0 | 0 |
| ELVLDNSR | ANP32A; ANP3 | P39687; H7E | AN32A_HUMAN | Unmodified | 5510700 | 25276000 | 120260 | 0 | 0 | 765560 | 1094300 | 162440 | 899660 | 97833 | 4527400 | 16280000 | 914260 | 0 | 0 | 0 | 0 | 699570 |
| ELVLQREVI | SND1 | Q7KZF4 | SND1_HUMAN | Unmodified | 19135 | 0 | 191870 | 117100 | 857880 | 0 | 0 | 0 | 0 | 0 | 86683 | 155430 | 0 | 0 | 0 | 0 | 178540 | 341440 |
| ELVNEWSL | CHMP3 | Q9Y3E7 | CHMP3_HUMA | Unmodified | 2616600 | 3289500 | 1834200 | 2085900 | 1290600 | 479380 | 226640 | 0 | 5193900 | 1253400 | 472130 | 2315400 | 669120 | 899610 | 2774100 | 2878900 | 178460 | 352240 |
| ELVNNLAEI | EHD3 | Q9NZN3 | EHD3_HUMAN | Unmodified | 0 | 0 | 0 | 0 | 5284600 | 418620 | 0 | 0 | 0 | 323800 | 0 | 0 | 2945100 | 3143400 | 0 | 0 | 2005700 | 4699500 |
| ELVNNLGE | EHD1; EHD1; E | Q9H4M9; C9 | EHD1_HUMAN | Unmodified | 0 | 0 | 0 | 3231800 | 6388100 | 0 | 0 | 0 | 1378500 | 0 | 5367000 | 16048000 | 1559900 | 0 | 4203900 | 3261900 | 1131700 | 10542000 |
| ELVNNLGE | EHD1; EHD1; E | Q9H4M9; C9 | EHD1_HUMAN | Unmodified | 0 | 0 | 0 | 0 | 2289800 | 253760 | 0 | 0 | 0 | 48673 | 0 | 0 | 304090 | 216490 | 0 | 0 | 6097400 | 4531600 |
| ELVPGRPTI | ADAM15 | Q13444 | ADA15_HUMAN | Unmodified | 0 | 0 | 105240 | 0 | 0 | 0 | 0 | 92234 | 2324200 | 0 | 0 | 0 | 0 | 0 | 275860 | 0 | 0 | 0 |
| ELVPSSDPI | EMG1; EMG1; E | Q92979; V9 | NEP1_HUMAN | Unmodified | 0 | 0 | 0 | 0 | 0 | 0 | 0 | 0 | 0 | 0 | 0 | 726390 | 0 | 0 | 0 | 0 | 0 | 0 |
| ELVQDLLK | SEC24A | O95486 | SC24A_HUMAN | Unmodified | 0 | 0 | 0 | 0 | 0 | 0 | 0 | 55084 | 0 | 0 | 0 | 1619200 | 0 | 0 | 0 | 0 | 0 | 0 |
| ELVQPFSSI | DPP9; DPP9 | Q86TI2; M0F | DPP9_HUMAN | Unmodified | 0 | 482610 | 0 | 0 | 0 | 0 | 0 | 0 | 0 | 0 | 0 | 789980 | 0 | 0 | 0 | 0 | 0 | 0 |
| ELVQQYHC | BABAM2; BABA | Q9NXR7; F8 | BABA2_HUMAN | Unmodified | 0 | 0 | 58957 | 0 | 0 | 0 | 0 | 0 | 239310 | 82537 | 51303 | 983930 | 0 | 0 | 0 | 0 | 0 | 0 |
| ELVQSLK | P4HA2; MTRF1 | O15460; Q9 | P4HA2_HUMAN | Unmodified | 23728 | 0 | 0 | 67031 | 127890 | 235220 | 204710 | 545760 | 2664000 | 91823 | 0 | 0 | 0 | 159410 | 33489 | 0 | 231570 | 284050 |
| ELVQTVLAI | AP1S2; AP1S2; | P56377; H0 | AP1S2_HUMAN | Unmodified | 0 | 84885 | 0 | 94836 | 0 | 0 | 0 | 0 | 270280 | 0 | 36548 | 2317700 | 0 | 0 | 0 | 22909 | 0 | 0 |
| ELVRKYPQ | ASPH; ASPH; A | Q12797; E5F | ASPH_HUMAN | Unmodified | 0 | 0 | 0 | 0 | 535230 | 107300 | 38376 | 0 | 62137 | 649520 | 0 | 0 | 171410 | 471000 | 0 | 0 | 0 | 479270 |
| ELVRQLQLI | LRRC47 | Q8N1G4 | LRC47_HUMAN | Unmodified | 0 | 0 | 0 | 0 | 0 | 0 | 474520 | 268790 | 0 | 431530 | 0 | 0 | 823520 | 968430 | 0 | 0 | 0 | 0 |
| ELVSCSNC | SARS1; SARS1 | P49591; Q5 | SYSC_HUMAN | Unmodified | 0 | 0 | 0 | 54298 | 0 | 0 | 0 | 0 | 117820 | 0 | 0 | 539000 | 0 | 0 | 0 | 0 | 0 | 0 |
| ELVSDANQ | PPP2R1A; PPP2 | P30153; B3K | 2AAA_HUMAN | Unmodified | 159510 | 47597 | 456580 | 495610 | 1046300 | 497390 | 188900 | 271780 | 669800 | 472260 | 792120 | 3952400 | 952910 | 123630 | 684090 | 712780 | 1057100 | 813000 |
| ELVSLKQE | CKAP4 | Q07065 | CKAP4_HUMAN | Unmodified | 0 | 0 | 0 | 0 | 0 | 0 | 0 | 0 | 2063500 | 149480 | 0 | 721450 | 0 | 0 | 0 | 0 | 0 | 0 |
| ELVSSKVS | PLIN3; PLIN3; P | O60664; K7E | PLIN3_HUMAN | Unmodified | 0 | 0 | 0 | 0 | 749250 | 0 | 0 | 0 | 0 | 0 | 0 | 0 | 0 | 0 | 0 | 0 | 307430 | 1291600 |
| ELVSSSSS | SUB1; SUB1; SU | P53999; D6F | TCP4_HUMAN | Unmodified | 0 | 0 | 0 | 0 | 0 | 0 | 0 | 0 | 0 | 0 | 279450 | 236290 | 0 | 0 | 0 | 0 | 0 | 0 |
| ELVTGTLW | ARVCF; ARVCF; | O00192; E9F | ARVC_HUMAN | Unmodified | 0 | 0 | 470900 | 853200 | 0 | 0 | 0 | 0 | 601750 | 0 | 34935 | 196380 | 0 | 0 | 5981800 | 488320 | 0 | 0 |
| ELVTQQLPI | S100A13 | Q99584 | S10AD_HUMAN | Unmodified | 348890 | 495870 | 0 | 357000 | 60136000 | 9946100 | 79823 | 0 | 6113100 | 3898500 | 1064400 | 1397700 | 861720 | 819930 | 346610 | 190110 | 25475000 | 56344000 |
| ELVTQQLPI | S100A13 | Q99584 | S10AD_HUMAN | Unmodified | 0 | 0 | 0 | 0 | 4924300 | 522570 | 0 | 0 | 0 | 293140 | 0 | 0 | 0 | 0 | 0 | 0 | 10784000 | 12614000 |
| ELVVTQLG | PFKP; PFKP; PF | Q01813; A0 | PFKAP_HUMAN | Unmodified | 0 | 0 | 0 | 0 | 0 | 0 | 0 | 0 | 0 | 0 | 0 | 4538000 | 0 | 0 | 0 | 0 | 0 | 0 |
| ELVYPPDYI | XRCC6; XRCC6 | P12956; B1A | XRCC6_HUMA | Unmodified | 261190 | 0 | 0 | 0 | 0 | 0 | 0 | 0 | 0 | 0 | 1422400 | 5458700 | 0 | 0 | 50778 | 0 | 0 | 0 |
| ELVYPPDYI | XRCC6; XRCC6 | P12956; B1A | XRCC6_HUMA | Unmodified | 0 | 0 | 0 | 0 | 0 | 0 | 0 | 560050 | 0 | 0 | 0 | 0 | 791840 | 0 | 0 | 0 | 0 | 0 |
| ELWAVLNG | SRI; SRI | P30626; C9J | SORCN_HUMA | Unmodified | 605920 | 0 | 237890 | 0 | 0 | 0 | 0 | 0 | 7268400 | 173650 | 726510 | 2158900 | 0 | 0 | 676890 | 0 | 0 | 0 |
| ELWAVLNG | SRI; SRI | P30626; C9J | SORCN_HUMA | Unmodified | 0 | 0 | 0 | 0 | 0 | 0 | 0 | 0 | 0 | 246990 | 0 | 0 | 0 | 0 | 0 | 0 | 0 | 0 |
| ELWILNRYI | ANPEP | P15144 | AMPN_HUMAN | Unmodified | 0 | 0 | 0 | 0 | 334060 | 2175400 | 735980 | 0 | 0 | 246020 | 0 | 0 | 201240 | 3107700 | 0 | 0 | 1654400 | 3764200 |
| ELWLDRN | SCRIB; SCRIB; | Q14160; A0 | SCRIB_HUMAN | Unmodified | 0 | 0 | 0 | 0 | 0 | 0 | 0 | 0 | 2253800 | 112020 | 0 | 0 | 0 | 0 | 664140 | 206870 | 0 | 0 |
| ELWLDRN | SCRIB; SCRIB; | Q14160; A0 | SCRIB_HUMAN | Unmodified | 0 | 0 | 0 | 0 | 0 | 0 | 560140 | 0 | 2935100 | 202070 | 264850 | 1327300 | 0 | 0 | 0 | 0 | 0 | 381640 |
| ELYDKGGE | DNAJA1 | P31689 | DNJA1_HUMAN | Unmodified | 809470 | 0 | 0 | 0 | 0 | 0 | 0 | 0 | 0 | 0 | 0 | 0 | 0 | 0 | 0 | 0 | 0 | 0 |
| ELYDRYGE | DNAJA2 | O60884 | DNJA2_HUMAN | Unmodified | 2402400 | 1766300 | 1108500 | 0 | 0 | 0 | 66577 | 46828 | 188350 | 0 | 0 | 1064300 | 0 | 0 | 2269900 | 2124500 | 0 | 0 |
| ELYEALVK | UGT8 | Q16880 | CGT_HUMAN 2 | Unmodified | 0 | 79118 | 0 | 0 | 0 | 0 | 0 | 0 | 791980 | 0 | 339380 | 1377200 | 0 | 0 | 97116 | 0 | 0 | 0 |
| ELYERPPHI | MYO1D; MYO1 | O94832; J3 | MYO1D_HUMA | Unmodified | 16117000 | 12271000 | 2922000 | 5803600 | 0 | 728480 | 0 | 0 | 1515800 | 416600 | 0 | 0 | 0 | 0 | 5377700 | 4663600 | 17496 | 589610 |
| ELYGEVIR | INPP5D | Q92835 | SHIP1_HUMAN | Unmodified | 0 | 0 | 824250 | 0 | 0 | 0 | 0 | 0 | 0 | 0 | 0 | 1227100 | 0 | 0 | 0 | 0 | 0 | 0 |
| ELYIQHAKE | TFAM | Q00059 | TFAM_HUMAN | Unmodified | 0 | 0 | 0 | 0 | 0 | 0 | 0 | 0 | 564190 | 223270 | 0 | 0 | 0 | 0 | 0 | 0 | 0 | 0 |
| ELYLEDSPI | NUP210 | Q8TEM1 | PO210_HUMAN | Unmodified | 0 | 0 | 0 | 0 | 0 | 0 | 0 | 0 | 27902000 | 1452500 | 654520 | 0 | 0 | 0 | 0 | 0 | 0 | 0 |
| ELYLFDVLF | ASNS | P08243 | ASNS_HUMAN | Unmodified | 151890 | 0 | 194860 | 631920 | 0 | 0 | 0 | 0 | 499800 | 0 | 186620 | 4834100 | 0 | 0 | 952530 | 755560 | 0 | 0 |
| ELYLFDVLF | ASNS | P08243 | ASNS_HUMAN | Unmodified | 0 | 232690 | 131120 | 286520 | 0 | 0 | 0 | 0 | 593570 | 0 | 199860 | 3786200 | 0 | 0 | 1348900 | 542550 | 0 | 0 |
| ELYLSHNG | PPP1R7; PPP1R | Q15435; C9J | PP1R7_HUMAN | Unmodified | 0 | 0 | 0 | 0 | 235840 | 0 | 0 | 0 | 544710 | 0 | 0 | 1812100 | 0 | 0 | 136150 | 0 | 0 | 0 |
| ELYLSSSLK | USP8; USP8; U | P40818; H0 | UBP8_HUMAN | Unmodified | 0 | 0 | 0 | 0 | 1853500 | 0 | 216620 | 68326 | 0 | 0 | 0 | 0 | 981870 | 0 | 0 | 0 | 0 | 324590 |
| ELYSHAQF | TLL2 | Q9Y6L7 | TLL2_HUMAN T | Unmodified | 0 | 0 | 0 | 0 | 0 | 0 | 0 | 0 | 1442100 | 85160 | 0 | 0 | 0 | 0 | 0 | 0 | 0 | 0 |
| ELYVMEISI | CPD | O75976 | CBPD_HUMAN | Oxidation (M) | 0 | 0 | 218020 | 340310 | 0 | 0 | 0 | 0 | 24531000 | 3485600 | 0 | 0 | 0 | 0 | 0 | 191290 | 0 | 0 |
| ELYVMEISI | CPD | O75976 | CBPD_HUMAN | Unmodified | 0 | 0 | 0 | 0 | 0 | 0 | 0 | 0 | 52804000 | 2785700 | 0 | 0 | 0 | 0 | 0 | 0 | 0 | 0 |
| ELYYHGEPI | ARRB2; ARRB2; | P32121; K7E | ARRB2_HUMAN | Unmodified | 0 | 40958 | 130700 | 0 | 1263900 | 0 | 326280 | 128300 | 609630 | 0 | 0 | 981890 | 0 | 0 | 254850 | 293460 | 0 | 0 |
| EMAEERAE | DCTN1; DCTN1 | Q14203; Q6 | DCTN1_HUMA | Oxidation (M) | 0 | 0 | 0 | 285300 | 0 | 0 | 0 | 0 | 0 | 0 | 0 | 2748500 | 0 | 0 | 0 | 222860 | 0 | 0 |
| EMAKMVAI | VIL1; VIL1; VIL1 | P09327; H7 | VILI_HUMAN Vi | Unmodified | 0 | 0 | 0 | 0 | 590250 | 0 | 2699300 | 0 | 63535000 | 0 | 0 | 0 | 0 | 0 | 0 | 0 | 112510 | 189430 |
| EMALATNR | VPS37C; VPS37 | A5D8V6; F5 | VP37C_HUMAN | Unmodified | 5985500 | 7487300 | 4143100 | 2561100 | 54559 | 0 | 0 | 0 | 1918400 | 36448 | 0 | 124490 | 88010 | 0 | 6536200 | 7876600 | 97687 | 152280 |
| EMALATNR | VPS37C; VPS37 | A5D8V6; F5 | VP37C_HUMAN | Oxidation (M) | 1687300 | 1666900 | 1894400 | 758150 | 0 | 1256500 | 601410 | 0 | 412110 | 0 | 0 | 417440 | 0 | 0 | 2102500 | 2190200 | 0 | 0 |
| EMAQQELS | KIF14 | Q15058 | KIF14_HUMAN | Oxidation (M) | 0 | 0 | 0 | 0 | 885570 | 0 | 0 | 0 | 0 | 0 | 0 | 0 | 0 | 0 | 0 | 0 | 387650 | 244560 |
| EMAQQELS | KIF14 | Q15058 | KIF14_HUMAN | Unmodified | 0 | 0 | 0 | 0 | 1742200 | 244890 | 0 | 51894 | 131750 | 0 | 0 | 173140 | 0 | 165680 | 0 | 0 | 656000 | 223760 |
| EMDEAATA | EIF4G1; EIF4G1 | Q04637; C9J | IF4G1_HUMAN | Unmodified | 0 | 0 | 72270 | 0 | 0 | 0 | 0 | 0 | 142830 | 0 | 179980 | 1626700 | 0 | 0 | 103090 | 69369 | 0 | 0 |
| EMDEEDKA | TMA7 | Q9Y2S6 | TMA7_HUMAN | Unmodified | 0 | 0 | 0 | 0 | 215060 | 0 | 0 | 0 | 38895 | 0 | 0 | 0 | 215150 | 240090 | 0 | 0 | 153160 | 551580 |
| EMDEQIR | TPM3; TPM3; TF | J3KN67; A0A | J3KN67_HUMA | Unmodified | 0 | 0 | 67858 | 102540 | 24179 | 0 | 0 | 0 | 486020 | 23751 | 458380 | 2069400 | 0 | 0 | 150730 | 123340 | 0 | 0 |
| EMDEQIRLI | TPM3; TPM3; TF | J3KN67; A0A | J3KN67_HUMA | Oxidation (M) | 0 | 0 | 27715 | 39200 | 2265100 | 306950 | 0 | 47012 | 0 | 80527 | 0 | 0 | 513360 | 241000 | 51361 | 0 | 0 | 0 |
| EMDPVTQL | ALCAM; ALCAM | Q13740; F5 | CD166_HUMA | Oxidation (M) | 720340 | 1015400 | 27439 | 0 | 1708600 | 171070 | 0 | 0 | 1983600 | 0 | 0 | 0 | 0 | 0 | 233740 | 0 | 1053100 | 2670900 |
| EMDPVTQL | ALCAM; ALCAM | Q13740; F5 | CD166_HUMA | Unmodified | 38993 | 0 | 0 | 0 | 219840 | 0 | 0 | 0 | 1358600 | 0 | 0 | 0 | 0 | 0 | 0 | 0 | 0 | 245720 |
| EMDPVTQL | ALCAM; ALCAM | Q13740; F5 | CD166_HUMA | 2 Oxidation (M) | 2828500 | 3605800 | 164140 | 760840 | 1168200 | 64569 | 97019 | 0 | 195400 | 83508 | 0 | 98670 | 995350 | 0 | 2845800 | 276510 | 0 | 238800 |
| EMDQTMA | CMPK1; CMPK1 | P30085; Q5 | KCY_HUMAN U | Unmodified | 0 | 0 | 0 | 0 | 206820 | 0 | 0 | 0 | 203890 | 106320 | 0 | 334040 | 0 | 0 | 0 | 0 | 0 | 0 |
| EMDREASC | CES2; CES2 | O00748; J3C | EST2_HUMAN | Unmodified | 0 | 0 | 191620 | 0 | 0 | 0 | 0 | 0 | 525610 | 250560 | 0 | 0 | 0 | 26505 | 0 | 0 | 0 | 0 |
| EMDRETLII | CCT6A | P40227 | TCPZ_HUMAN | Unmodified | 0 | 0 | 143650 | 224170 | 0 | 0 | 0 | 0 | 2169100 | 421740 | 1887700 | 10546000 | 953270 | 194800 | 187670 | 236280 | 0 | 114920 |
| EMEAELED | MYH9 | P35579 | MYH9_HUMAN | Unmodified | 50161 | 0 | 0 | 0 | 1084900 | 276970 | 5359400 | 712600 | 1350400 | 177690 | 334390 | 1071400 | 0 | 0 | 245270 | 224060 | 87538 | 0 |
| EMEAELED | MYH9 | P35579 | MYH9_HUMAN | Oxidation (M) | 337290 | 297060 | 768580 | 341360 | 573050 | 0 | 0 | 0 | 591460 | 862480 | 289210 | 2960200 | 0 | 0 | 1457800 | 1622300 | 0 | 1801300 |
| EMEEPFEK | ADSL; ADSL; AD | P30566; A0A | PUR8_HUMAN | Unmodified | 0 | 0 | 0 | 0 | 155430 | 0 | 0 | 0 | 0 | 0 | 418920 | 1786000 | 0 | 0 | 0 | 0 | 0 | 112690 |
| EMEPLGWI | PRPF8 | Q6P2Q9 | PRP8_HUMAN | Oxidation (M) | 729500 | 0 | 0 | 0 | 0 | 0 | 0 | 0 | 0 | 0 | 0 | 838600 | 0 | 0 | 0 | 494070 | 0 | 0 |
| EMESRDEE | MYO18A; MYO1 | Q92614; A0 | MY18A_HUMA | Unmodified | 273770 | 346330 | 0 | 0 | 0 | 0 | 0 | 20368 | 0 | 0 | 0 | 288110 | 0 | 0 | 62094 | 24154 | 0 | 0 |
| EMFEDTVE | RBBP7; RBBP7 | Q16576; E9F | RBBP7_HUMAN | Unmodified | 0 | 0 | 0 | 0 | 0 | 0 | 0 | 0 | 0 | 0 | 0 | 1092700 | 0 | 0 | 0 | 0 | 0 | 0 |
| EMFEDTVE | RBBP7; RBBP7 | Q16576; E9F | RBBP7_HUMAN | Oxidation (M) | 0 | 0 | 0 | 0 | 0 | 142310 | 0 | 0 | 0 | 0 | 0 | 0 | 1415200 | 270800 | 0 | 0 | 0 | 0 |
| EMFEVTGL | CHID1 | Q9BWS9 | CHID1_HUMAN | Oxidation (M) | 0 | 0 | 0 | 0 | 0 | 0 | 0 | 0 | 3521800 | 0 | 0 | 0 | 0 | 0 | 0 | 0 | 0 | 0 |
| EMFEVTGL | CHID1 | Q9BWS9 | CHID1_HUMAN | 2 Oxidation (M) | 0 | 0 | 0 | 0 | 0 | 0 | 0 | 0 | 1940700 | 0 | 0 | 0 | 0 | 0 | 0 | 0 | 0 | 0 |
| EMFEVTGL | CHID1 | Q9BWS9 | CHID1_HUMAN | Unmodified | 0 | 0 | 0 | 0 | 0 | 0 | 0 | 0 | 834010 | 0 | 0 | 0 | 0 | 0 | 0 | 0 | 0 | 0 |
| EMFGSGTA | BCAT1 | P54687 | BCAT1_HUMAN | Oxidation (M) | 0 | 0 | 0 | 0 | 0 | 0 | 0 | 0 | 0 | 0 | 0 | 360270 | 0 | 0 | 0 | 0 | 0 | 0 |

| | | | | | | | | | | | | | | | | | | | | | | |
|---|---|---|---|---|---|---|---|---|---|---|---|---|---|---|---|---|---|---|---|---|---|---|
| EMGEAFAA | AP2A1 | O95782 | AP2A1_HUMAN | Oxidation (M) | 0 | 127290 | 142080 | 0 | 0 | 0 | 0 | 0 | 0 | 0 | 0 | 1074800 | 0 | 0 | 0 | 0 | 0 | 0 |
| EMGEMHRI | EPCAM; EPCAM | P16422; B5N | EPCAM_HUMA | Unmodified | 0 | 0 | 15027000 | 27111000 | 5222700 | 1140100 | 0 | 0 | 36942000 | 11407000 | 3331300 | 6915200 | 15239000 | 6820400 | 3928000 | 2929500 | 0 | 2019500 |
| EMGEMHRI | EPCAM; EPCAM | P16422; B5N | EPCAM_HUMA | Oxidation (M) | 480690 | 371690 | 2557900 | 526470 | 402180 | 79311 | 0 | 0 | 3992700 | 1261700 | 222070 | 1151100 | 1186200 | 0 | 925620 | 1884000 | 0 | 0 |
| EMGEMHRI | EPCAM; EPCAM | P16422; B5N | EPCAM_HUMA | 2 Oxidation (M) | 0 | 0 | 0 | 0 | 384460 | 0 | 0 | 0 | 0 | 149510 | 0 | 322490 | 0 | 0 | 0 | 2343800 | 0 | 0 |
| EMGIIPSW( | ECHDC1; ECHD | Q9NTX5; H0 | ECHD1_HUMA | Oxidation (M) | 0 | 0 | 0 | 0 | 3173700 | 0 | 500520 | 350830 | 0 | 0 | 352000 | 1896400 | 881960 | 0 | 0 | 0 | 902400 | 0 |
| EMGTPDVF | RPL31; RPL31; I | P62899; H7( | RL31_HUMAN ( | Unmodified | 160800 | 142480 | 139700 | 223300 | 0 | 0 | 0 | 0 | 0 | 0 | 0 | 0 | 347600 | 0 | 0 | 0 | 0 | 0 |
| EMGTPLAD | DPYSL5 | Q9BPU6 | DPYL5_HUMAN | Oxidation (M) | 0 | 0 | 0 | 0 | 0 | 0 | 0 | 67972 | 0 | 0 | 437740 | 2705900 | 0 | 0 | 0 | 0 | 0 | 0 |
| EMIDIYSTR | PSMD3 | O43242 | PSMD3_HUMA | Oxidation (M) | 177800 | 0 | 296910 | 770010 | 0 | 0 | 0 | 0 | 0 | 277320 | 0 | 3652200 | 0 | 0 | 415650 | 381050 | 64527 | 540500 |
| EMINNFR | CA12; CA12 | O43570; B3I | CAH12_HUMAI | Unmodified | 487740 | 0 | 0 | 0 | 0 | 0 | 0 | 0 | 0 | 0 | 0 | 299500 | 0 | 0 | 173920 | 168440 | 0 | 0 |
| EMKEIPFNI | C2CD5; C2CD5 | Q86YS7; H0 | C2CD5_HUMA | Oxidation (M) | 0 | 0 | 0 | 0 | 500160 | 0 | 149540 | 379210 | 0 | 309530 | 0 | 0 | 360970 | 214030 | 0 | 0 | 460220 | 624350 |
| EMKEIPFNI | C2CD5; C2CD5 | Q86YS7; H0 | C2CD5_HUMA | Unmodified | 0 | 0 | 0 | 0 | 502430 | 0 | 445870 | 57408 | 0 | 0 | 0 | 0 | 0 | 0 | 0 | 0 | 703440 | 186600 |
| EMKPGFYH | HS2ST1; HS2ST | Q7LGA3; K7I | HS2ST_HUMAN | Unmodified | 0 | 0 | 0 | 0 | 0 | 0 | 0 | 0 | 7116600 | 826720 | 0 | 0 | 0 | 0 | 0 | 0 | 0 | 0 |
| EMKPVIFLD | PRKDC | P78527 | PRKDC_HUMAI | Oxidation (M) | 0 | 0 | 0 | 0 | 0 | 0 | 0 | 0 | 0 | 0 | 0 | 102650 | 0 | 0 | 0 | 0 | 0 | 0 |
| EMLAVDAP | IDE; IDE; IDE; ID | P14735; A0A | IDE_HUMAN In | Unmodified | 0 | 0 | 0 | 0 | 393410 | 0 | 147770 | 0 | 0 | 0 | 0 | 1638700 | 0 | 0 | 0 | 0 | 0 | 230090 |
| EMLAVSVL | BPNT2 | Q9NX62 | IMPA3_HUMAN | Oxidation (M) | 0 | 0 | 0 | 0 | 0 | 0 | 0 | 0 | 315980 | 105070 | 0 | 0 | 0 | 0 | 0 | 0 | 0 | 0 |
| EMLLFPYDI | DOCK9; DOCK9 | Q9BZ29; A0A | DOCK9_HUMA | Oxidation (M) | 40978 | 35920 | 0 | 63061 | 0 | 0 | 0 | 0 | 83836 | 0 | 0 | 137090 | 0 | 0 | 101370 | 69927 | 0 | 0 |
| EMLQQSK | HSP90AB1; HSI | P08238; Q58 | HS90B_HUMAI | Unmodified | 287310 | 379740 | 1217900 | 909300 | 345470 | 0 | 0 | 0 | 2628500 | 498900 | 101290 | 12399000 | 0 | 0 | 2192800 | 637180 | 0 | 0 |
| EMLQQSK | HSP90AB1; HSI | P08238; Q58 | HS90B_HUMAI | Oxidation (M) | 0 | 0 | 132950 | 0 | 68752 | 0 | 36455 | 0 | 113310 | 131720 | 77054 | 4415900 | 22573 | 0 | 236160 | 93657 | 0 | 0 |
| EMMSPGLT | JAG1 | P78504 | JAG1_HUMAN I | 2 Oxidation (M) | 0 | 0 | 0 | 0 | 0 | 0 | 0 | 0 | 0 | 0 | 0 | 0 | 1228400 | 391190 | 0 | 78748 | 0 | 135270 |
| EMNDAAMI | CAP1; CAP1; CA | Q01518; Q5 | CAP1_HUMAN | 2 Oxidation (M) | 0 | 0 | 0 | 0 | 0 | 0 | 0 | 0 | 0 | 0 | 0 | 916170 | 0 | 0 | 0 | 0 | 0 | 0 |
| EMNDAAMI | CAP1; CAP1; CA | Q01518; Q5 | CAP1_HUMAN | Oxidation (M) | 0 | 0 | 0 | 0 | 0 | 0 | 0 | 0 | 0 | 0 | 0 | 3557100 | 0 | 0 | 0 | 0 | 0 | 0 |
| EMNDAAMI | CAP1; CAP1; CA | Q01518; Q5 | CAP1_HUMAN | 2 Oxidation (M) | 0 | 0 | 0 | 0 | 370010 | 0 | 218990 | 340960 | 248500 | 488760 | 0 | 657370 | 0 | 0 | 0 | 0 | 750760 | 0 |
| EMNIQQLT | CALB2; CALB2 | P22676; H3E | CALB2_HUMAN | Oxidation (M) | 0 | 0 | 138030 | 127490 | 0 | 0 | 500800 | 359480 | 62635 | 0 | 0 | 188570 | 0 | 0 | 123060 | 164650 | 0 | 0 |
| EMNPALGII | CCT5; CCT5; C( | P48643; B7Z | TCPE_HUMAN | Oxidation (M) | 0 | 0 | 218820 | 211030 | 542000 | 0 | 221220 | 112350 | 0 | 0 | 802810 | 4964100 | 1253900 | 824990 | 450720 | 0 | 464160 | 671410 |
| EMNPALGII | CCT5; CCT5; C( | P48643; B7Z | TCPE_HUMAN | Unmodified | 33758 | 0 | 0 | 412530 | 0 | 0 | 0 | 46672 | 0 | 171980 | 0 | 3595900 | 0 | 455090 | 114800 | 0 | 575880 | 726330 |
| EMPMQTLV | PSAP; PSAP; PS | P07602; A0A | SAP_HUMAN P | Unmodified | 0 | 0 | 0 | 0 | 0 | 0 | 492230 | 0 | 12408000 | 883020 | 0 | 0 | 0 | 0 | 0 | 0 | 0 | 0 |
| EMPNVFGI | EHD1; EHD1; E | Q9H4M9; C9 | EHD1_HUMAN | Oxidation (M) | 0 | 0 | 0 | 0 | 0 | 0 | 0 | 0 | 0 | 0 | 0 | 2326300 | 0 | 169150 | 0 | 0 | 0 | 0 |
| EMPNVFGI | EHD1; EHD1; E | Q9H4M9; C9 | EHD1_HUMAN | Unmodified | 0 | 0 | 119230 | 0 | 1095500 | 1395100 | 124330 | 0 | 0 | 0 | 0 | 0 | 485540 | 223040 | 0 | 0 | 567120 | 1137900 |
| EMPQIEPD | N/A | N/A | N/A | Oxidation (M) | 0 | 0 | 0 | 0 | 0 | 0 | 0 | 0 | 0 | 0 | 1369300 | 0 | 0 | 0 | 0 | 0 | 0 | 0 |
| EMPRGPTC | EPB41 | A0A2R8Y783 | A0A2R8Y783_H | Unmodified | 0 | 0 | 0 | 0 | 0 | 0 | 0 | 0 | 1125100 | 345880 | 0 | 0 | 1063800 | 647200 | 0 | 0 | 0 | 0 |
| EMPRGPTC | EPB41 | A0A2R8Y783 | A0A2R8Y783_H | Oxidation (M) | 0 | 0 | 70706 | 102730 | 0 | 0 | 0 | 0 | 0 | 360580 | 0 | 0 | 756310 | 360540 | 32851 | 60373 | 0 | 0 |
| EMQENDA | UQCRC1 | P31930 | QCR1_HUMAN | Unmodified | 0 | 0 | 0 | 0 | 0 | 0 | 0 | 0 | 1237700 | 0 | 0 | 0 | 0 | 0 | 0 | 0 | 0 | 0 |
| EMQKLPEA | NAPG; NAPG | Q99747; J3K | SNAG_HUMAN | Oxidation (M) | 0 | 0 | 0 | 0 | 2261500 | 0 | 0 | 1027000 | 0 | 0 | 0 | 2774200 | 1690400 | 0 | 0 | 0 | 2280900 | 0 |
| EMQKLPEA | NAPG; NAPG | Q99747; J3K | SNAG_HUMAN | Unmodified | 203120 | 0 | 0 | 0 | 2594300 | 0 | 0 | 49591 | 379720 | 292870 | 0 | 723010 | 609540 | 683800 | 0 | 0 | 944250 | 2257300 |
| EMQPTHPI | YWHAB; YWHA | P31946; P63 | 1433B_HUMAN | Unmodified | 813700 | 432650 | 567500 | 925190 | 0 | 0 | 0 | 0 | 3176300 | 23480 | 1000900 | 13096000 | 0 | 0 | 1305500 | 2210100 | 87524 | 0 |
| EMQPTHPI | YWHAB; YWHA | P31946; P63 | 1433B_HUMAN | Oxidation (M) | 0 | 0 | 0 | 0 | 0 | 0 | 0 | 0 | 0 | 106760 | 137240 | 1577100 | 0 | 0 | 0 | 27031 | 0 | 0 |
| EMQPVYR | XXYLT1; XXYLT1 | Q8NBI6; A0A | XXLT1_HUMAN | Unmodified | 0 | 0 | 80817 | 30707 | 0 | 103170 | 0 | 0 | 1665600 | 111550 | 0 | 0 | 0 | 0 | 0 | 0 | 0 | 0 |
| EMQQLSG( | SMC3 | Q9UQE7 | SMC3_HUMAN | Unmodified | 0 | 0 | 0 | 0 | 0 | 0 | 59121 | 48701 | 0 | 0 | 0 | 333850 | 0 | 0 | 0 | 0 | 0 | 0 |
| EMSEFLSR | MYO6; MYO6; M | Q9UM54; A0 | MYO6_HUMAN | Oxidation (M) | 0 | 0 | 0 | 0 | 0 | 0 | 0 | 0 | 0 | 214980 | 0 | 0 | 2824200 | 2724800 | 227240 | 223820 | 0 | 0 |
| EMSEFLSR | MYO6; MYO6; M | Q9UM54; A0 | MYO6_HUMAN | Unmodified | 0 | 0 | 0 | 0 | 0 | 0 | 0 | 0 | 291430 | 175000 | 0 | 0 | 1987400 | 2579400 | 59774 | 0 | 0 | 0 |
| EMSGDLEE | ANXA11; ANXA1 | P50995; H0Y | ANX11_HUMAN | 2 Oxidation (M) | 0 | 0 | 0 | 0 | 0 | 0 | 0 | 0 | 866210 | 229080 | 286090 | 261330 | 0 | 0 | 0 | 0 | 0 | 0 |
| EMSGDLEE | ANXA11; ANXA1 | P50995; H0Y | ANX11_HUMAN | Oxidation (M) | 0 | 0 | 0 | 0 | 0 | 1585100 | 63692 | 0 | 12472000 | 1443200 | 0 | 934430 | 0 | 0 | 0 | 0 | 163470 | 0 |
| EMSGDLEE | ANXA11; ANXA1 | P50995; H0Y | ANX11_HUMAN | Unmodified | 0 | 0 | 0 | 0 | 0 | 80105 | 0 | 0 | 18304000 | 0 | 932980 | 0 | 0 | 0 | 0 | 0 | 0 | 0 |
| EMSGDVRI | ANXA6; ANXA6; | P08133; E7E | ANXA6_HUMAN | Oxidation (M) | 422350 | 0 | 1116400 | 2830500 | 20708000 | 3900700 | 9836300 | 0 | 15405000 | 23827000 | 0 | 0 | 7226600 | 5981000 | 3098000 | 1039700 | 5698300 | 21711000 |
| EMSGDVRI | ANXA6; ANXA6; | P08133; E7E | ANXA6_HUMAN | Unmodified | 0 | 0 | 0 | 0 | 1742600 | 0 | 1713300 | 0 | 7034400 | 3198100 | 0 | 0 | 950260 | 0 | 0 | 0 | 1061500 | 3129500 |
| EMSNINHP | MON2; MON2; I | Q7Z3U7; A0 | MON2_HUMAN | Oxidation (M) | 0 | 0 | 0 | 0 | 551220 | 432870 | 0 | 0 | 0 | 982880 | 0 | 310980 | 0 | 0 | 0 | 0 | 694860 | 0 |
| EMTADVIEL | IARS2 | Q9NSE4 | SYIM_HUMAN I | Unmodified | 0 | 0 | 0 | 0 | 0 | 0 | 0 | 0 | 1542500 | 0 | 0 | 0 | 0 | 0 | 0 | 0 | 108360 | 0 |
| EMTFIFTDC | LFNG | Q8NES3 | LFNG_HUMAN | Unmodified | 0 | 0 | 0 | 0 | 0 | 0 | 0 | 0 | 128480 | 0 | 0 | 0 | 0 | 0 | 0 | 0 | 0 | 0 |
| EMTLASNR | VPS37B; VPS37 | Q9H9H4; F5 | VP37B_HUMAN | Oxidation (M) | 4634000 | 2426000 | 0 | 13654000 | 0 | 0 | 0 | 0 | 0 | 0 | 0 | 868310 | 0 | 393060 | 10899000 | 329790 | 0 | 0 |
| EMTLASNR | VPS37B; VPS37 | Q9H9H4; F5 | VP37B_HUMAN | Unmodified | 3548100 | 3213400 | 3350200 | 2457100 | 0 | 0 | 0 | 0 | 0 | 0 | 0 | 0 | 1768900 | 3198200 | 28451000 | 2387200 | 0 | 0 |
| EMTLASNR | VPS37B; VPS37 | Q9H9H4; F5 | VP37B_HUMAN | Oxidation (M) | 52720 | 0 | 277690 | 262560 | 11262000 | 2060800 | 3905800 | 74817 | 0 | 4487000 | 0 | 0 | 39418000 | 22847000 | 0 | 0 | 3263500 | 23408000 |
| EMTLASNR | VPS37B; VPS37 | Q9H9H4; F5 | VP37B_HUMAN | Unmodified | 0 | 0 | 0 | 0 | 0 | 0 | 0 | 0 | 0 | 0 | 0 | 0 | 11449000 | 9418500 | 0 | 0 | 0 | 5447500 |
| EMTLDEW | SERBP1; HABP4 | Q8NC51; Q5 | PAIRB_HUMAN | Oxidation (M) | 35513 | 0 | 107040 | 318200 | 0 | 0 | 0 | 0 | 0 | 0 | 668660 | 1114900 | 83769 | 0 | 0 | 0 | 0 | 0 |
| EMTRTGLA | TJP2; TJP2; TJP2 | Q9UDY2; A0 | ZO2_HUMAN T | Oxidation (M) | 0 | 0 | 0 | 0 | 806520 | 419310 | 0 | 0 | 0 | 594690 | 0 | 0 | 186980 | 194910 | 0 | 0 | 0 | 0 |
| EMTRTGLA | TJP2; TJP2; TJP2 | Q9UDY2; A0 | ZO2_HUMAN T | Unmodified | 0 | 0 | 0 | 0 | 2219300 | 0 | 0 | 0 | 0 | 815760 | 0 | 0 | 0 | 0 | 0 | 0 | 0 | 0 |
| EMVANWD | LARS1; LARS1; | Q9P2J5; A0A | SYLC_HUMAN | Oxidation (M) | 0 | 0 | 0 | 0 | 0 | 0 | 0 | 0 | 0 | 0 | 138340 | 1805300 | 0 | 0 | 0 | 0 | 0 | 0 |
| EMVECFNI | ECH1; ECH1; N | Q13011; M0 | ECH1_HUMAN | Oxidation (M) | 0 | 0 | 0 | 0 | 0 | 0 | 0 | 0 | 321580 | 30885 | 0 | 0 | 0 | 0 | 0 | 18607 | 0 | 0 |
| EMVELPLRI | VCP; VCP; VCP; | P55072; C9J | TERA_HUMAN | Oxidation (M) | 0 | 0 | 0 | 0 | 0 | 0 | 1310300 | 239170 | 0 | 1613300 | 0 | 5109200 | 7302000 | 3526100 | 0 | 0 | 1175700 | 0 |
| EMVELPLRI | VCP; VCP; VCP; | P55072; C9J | TERA_HUMAN | Unmodified | 138800 | 0 | 0 | 0 | 4464700 | 0 | 553470 | 25115 | 2496400 | 800430 | 0 | 2328500 | 2391000 | 1132900 | 0 | 0 | 644860 | 470380 |
| EMVKSKLP | EHD1; EHD1 | Q9H4M9; A0 | EHD1_HUMAN | Unmodified | 0 | 0 | 0 | 0 | 0 | 0 | 0 | 0 | 0 | 0 | 0 | 0 | 0 | 0 | 0 | 0 | 934900 | 0 |
| EMVLELIRD | FUBP1; FUBP1 | Q96AE4; E9I | FUBP1_HUMAN | Oxidation (M) | 0 | 0 | 87325 | 0 | 0 | 0 | 0 | 0 | 132360 | 0 | 759730 | 1336000 | 0 | 0 | 630970 | 0 | 0 | 0 |
| EMVLSEK | MAGT1; ZNF839 | Q9H0U3; A8 | MAGT1_HUMAI | Unmodified | 0 | 0 | 0 | 0 | 361280 | 0 | 386430 | 181200 | 8039700 | 117430 | 147890 | 1087300 | 0 | 0 | 131050 | 0 | 0 | 0 |
| EMVPEFVV | MRPL41 | Q8IXM3 | RM41_HUMAN | Oxidation (M) | 0 | 0 | 0 | 0 | 0 | 0 | 0 | 0 | 377260 | 125650 | 0 | 0 | 0 | 0 | 0 | 0 | 0 | 0 |
| EMVTSKLPI | EHD4 | Q9H223 | EHD4_HUMAN | Oxidation (M) | 1344900 | 1086700 | 936800 | 1928500 | 11605000 | 1977800 | 0 | 197140 | 324100 | 418050 | 0 | 1122300 | 5407200 | 3531400 | 1647800 | 1770000 | 8726100 | 12883000 |
| EMVTSKLPI | EHD4 | Q9H223 | EHD4_HUMAN | Unmodified | 664210 | 1066400 | 271640 | 1259300 | 24968000 | 2537600 | 213250 | 42467 | 838160 | 324240 | 0 | 1320600 | 5149600 | 2549900 | 2373700 | 936230 | 9135300 | 25020000 |
| EMVTSKLPI | EHD4 | Q9H223 | EHD4_HUMAN | Oxidation (M) | 0 | 0 | 0 | 0 | 0 | 0 | 0 | 0 | 0 | 0 | 0 | 0 | 0 | 0 | 0 | 0 | 651500 | 0 |
| EMYKTDLEI | ANXA2P2; ANXA | A6NMY6; P0 | AXA2L_HUMAN | Unmodified | 155880 | 0 | 0 | 54287 | 8168800 | 0 | 115080 | 0 | 2809700 | 0 | 0 | 0 | 0 | 0 | 0 | 0 | 1556200 | 2247100 |
| EMYKTDLEI | ANXA2P2; ANXA | A6NMY6; P0 | AXA2L_HUMAN | Oxidation (M) | 45064 | 0 | 0 | 0 | 4753500 | 1265200 | 67073 | 0 | 742630 | 177490 | 0 | 41217 | 0 | 0 | 0 | 0 | 0 | 579910 |
| EMYTTHED | CHSY3; CHSY1 | Q70JA7; Q86 | CHSS3_HUMAI | Unmodified | 0 | 0 | 0 | 0 | 0 | 0 | 0 | 0 | 339970 | 0 | 0 | 0 | 0 | 0 | 0 | 0 | 0 | 0 |
| ENAAAPSP | PALM | O75781 | PALM_HUMAN | Unmodified | 34851 | 0 | 812940 | 0 | 0 | 0 | 0 | 0 | 1137900 | 0 | 0 | 408200 | 0 | 0 | 258220 | 110940 | 156120 | 360260 |
| ENAAAPSP | PALM | O75781 | PALM_HUMAN | Unmodified | 0 | 0 | 0 | 0 | 139190 | 0 | 0 | 0 | 0 | 0 | 0 | 0 | 0 | 0 | 0 | 0 | 0 | 255680 |
| ENAAYFQF | DSP | P15924 | DESP_HUMAN | Unmodified | 0 | 0 | 0 | 0 | 0 | 0 | 0 | 0 | 0 | 186070 | 0 | 0 | 0 | 0 | 0 | 0 | 0 | 0 |
| ENAAYFQF | PLEC; PLEC | Q15149; H0 | PLEC_HUMAN | Unmodified | 0 | 0 | 0 | 0 | 870680 | 0 | 0 | 0 | 909920 | 0 | 0 | 0 | 4982700 | 0 | 0 | 0 | 1762900 | 901350 |
| ENAEQGEV | CD99 | P14209 | CD99_HUMAN | Unmodified | 2683800 | 2671100 | 11405000 | 5681600 | 1240500 | 615460 | 417140 | 85259 | 423890 | 58988 | 0 | 372580 | 309490 | 210490 | 22473000 | 13641000 | 566250 | 1159600 |
| ENAEQGEV | CD99 | P14209 | CD99_HUMAN | Oxidation (M) | 47699 | 0 | 1261300 | 4266300 | 0 | 0 | 0 | 0 | 0 | 0 | 0 | 0 | 0 | 0 | 15902000 | 21225000 | 0 | 0 |
| ENAEQGEV | CD99 | P14209 | CD99_HUMAN | Oxidation (M) | 0 | 67155 | 555020 | 887880 | 1894500 | 1615300 | 0 | 36358 | 0 | 0 | 0 | 0 | 722260 | 454670 | 345340 | 255820 | 308950 | 2789400 |
| ENAEQGEV | CD99 | P14209 | CD99_HUMAN | Unmodified | 908130 | 0 | 2868200 | 0 | 7175600 | 1723000 | 336380 | 173750 | 0 | 0 | 0 | 0 | 6401800 | 0 | 1461800 | 0 | 1274600 | 14366000 |
| ENAEQGEV | CD99 | P14209 | CD99_HUMAN | Oxidation (M) | 0 | 0 | 0 | 0 | 5107800 | 0 | 0 | 0 | 0 | 0 | 0 | 0 | 388270 | 0 | 0 | 0 | 1251800 | 4543700 |
| ENAEQGEV | CD99 | P14209 | CD99_HUMAN | Unmodified | 0 | 0 | 0 | 0 | 39558000 | 4609200 | 1536300 | 612560 | 0 | 0 | 0 | 0 | 10187000 | 6475800 | 0 | 0 | 17350000 | 83540000 |
| ENAEVGTT | DSC2; DSC2 | Q02487; A0A | DSC2_HUMAN | Unmodified | 0 | 0 | 0 | 0 | 236570 | 107960 | 0 | 0 | 1109600 | 170620 | 0 | 0 | 0 | 0 | 0 | 0 | 68260 | 0 |
| ENAGEDPC | DCD | P81605 | DCD_HUMAN I | Unmodified | 0 | 0 | 0 | 0 | 1867700 | 0 | 0 | 0 | 588290 | 0 | 0 | 0 | 0 | 0 | 0 | 0 | 0 | 0 |
| ENAIEDEEI | PDIA4; PDIA4; F | P13667; C9J | PDIA4_HUMAN | Unmodified | 0 | 0 | 0 | 0 | 0 | 0 | 0 | 613420 | 3136400 | 0 | 671090 | 0 | 0 | 0 | 211280 | 0 | 0 | 0 |
| ENAKNEEIL | GPX1; GPX1 | P07203; A0A | GPX1_HUMAN | Unmodified | 0 | 0 | 0 | 0 | 0 | 0 | 0 | 0 | 0 | 0 | 0 | 1252100 | 0 | 0 | 0 | 0 | 0 | 0 |
| ENALDRAE | TPM1; TPM1; TF | P09493; A0A | TPM1_HUMAN | Unmodified | 0 | 0 | 0 | 0 | 638530 | 156990 | 0 | 0 | 0 | 0 | 151910 | 882940 | 0 | 0 | 0 | 0 | 60914 | 110780 |
| ENALDRAE | TPM1; TPM1; TF | P09493; A0A | TPM1_HUMAN | Unmodified | 0 | 0 | 0 | 38822 | 2109600 | 305540 | 0 | 0 | 79812 | 0 | 1560900 | 6313200 | 0 | 36067 | 0 | 29240 | 321450 | 277070 |
| ENALMPNV | FAH | P16930 | FAAA_HUMAN | Oxidation (M) | 0 | 0 | 161410 | 0 | 0 | 0 | 0 | 0 | 0 | 0 | 329660 | 2248200 | 0 | 0 | 1374100 | 270520 | 0 | 0 |
| ENALSELD | VPS13C | Q709C8 | VP13C_HUMAN | Unmodified | 0 | 0 | 0 | 0 | 0 | 765370 | 344000 | 0 | 0 | 201720 | 183640 | 1709300 | 0 | 0 | 0 | 0 | 0 | 0 |
| ENALSQLD | VPS13A | Q96RL7 | VP13A_HUMAN | Unmodified | 0 | 0 | 0 | 0 | 0 | 0 | 0 | 0 | 0 | 0 | 0 | 4620600 | 2488700 | 0 | 6779700 | 0 | 0 | 0 |
| ENAPAIIFIC | PSMC4 | P43686 | PRS6B_HUMAN | Unmodified | 0 | 0 | 0 | 0 | 0 | 0 | 87160 | 0 | 0 | 42711 | 0 | 0 | 0 | 0 | 0 | 0 | 80170 | 0 |
| ENASAGVE | RAB13; RAB13 | P51153; A0A | RAB13_HUMAN | Unmodified | 0 | 0 | 0 | 0 | 977680 | 0 | 0 | 0 | 0 | 0 | 0 | 0 | 0 | 0 | 0 | 0 | 0 | 0 |
| ENASAGVE | RAB13; RAB13 | P51153; A0A | RAB13_HUMAN | Oxidation (M) | 0 | 0 | 0 | 196540 | 1480000 | 947270 | 0 | 0 | 0 | 0 | 0 | 0 | 880350 | 0 | 0 | 0 | 179800 | 449550 |
| ENAYDLEA | EIF3K; EIF3K; EI | Q9UBQ5; K7 | EIF3K_HUMAN | Unmodified | 0 | 0 | 0 | 0 | 0 | 0 | 0 | 0 | 0 | 0 | 307890 | 1593200 | 0 | 0 | 0 | 0 | 0 | 0 |
| ENDAHLVE | TMOD3; TMOD3 | Q9NYL9; H0 | TMOD3_HUMA | Unmodified | 247340 | 172070 | 474160 | 339640 | 3771500 | 733210 | 0 | 0 | 853020 | 0 | 0 | 1649800 | 870240 | 1171000 | 335700 | 376960 | 548050 | 0 |
| ENDAVTIQ | VPS35 | Q96QK1 | VPS35_HUMAN | Unmodified | 0 | 0 | 51923 | 0 | 0 | 0 | 0 | 0 | 0 | 163740 | 507410 | 535700 | 135850 | 0 | 0 | 0 | 0 | 0 |
| ENDAVWL | GIT1; GIT1; GIT | Q9Y2X7; J3Q | GIT1_HUMAN | Unmodified | 0 | 0 | 0 | 0 | 0 | 0 | 0 | 0 | 1148400 | 0 | 0 | 1924200 | 0 | 0 | 0 | 152750 | 0 | 0 |
| ENDFDRLV | COPE; COPE | O14579; M0 | COPE_HUMAN | Unmodified | 110160 | 0 | 0 | 0 | 1836300 | 271110 | 438980 | 47423 | 472680 | 407560 | 0 | 1187100 | 1456300 | 796510 | 200450 | 93677 | 584880 | 1351900 |
| ENDHLFLS | CALCOCO2; CA | Q13137; D6I | CACO2_HUMA | Unmodified | 0 | 0 | 0 | 180820 | 1036700 | 117280 | 189240 | 240710 | 2028900 | 214660 | 0 | 299830 | 332970 | 0 | 0 | 0 | 324020 | 935450 |
| ENDHLFLS | CALCOCO2; CA | Q13137; D6I | CACO2_HUMA | Unmodified | 0 | 0 | 0 | 0 | 479780 | 0 | 783050 | 0 | 0 | 0 | 0 | 0 | 0 | 0 | 0 | 0 | 359220 | 2677400 |
| ENDMSPSI | RAB11A; RAB11 | P62491; H3E | RB11A_HUMAN | Oxidation (M) | 0 | 0 | 1315600 | 0 | 0 | 0 | 0 | 0 | 0 | 0 | 0 | 10388000 | 0 | 0 | 492470 | 986720 | 0 | 0 |
| ENDPSSVL | RAB34; RAB34; | Q9BZG1; C9 | RAB34_HUMAN | Unmodified | 0 | 0 | 0 | 0 | 450020 | 0 | 0 | 0 | 882170 | 0 | 200060 | 1157300 | 0 | 0 | 0 | 0 | 0 | 0 |
| ENDRILVA( | GTPBP10; GTPE | A4D1E9; C9J | GTPBA_HUMAN | Unmodified | 0 | 0 | 0 | 0 | 0 | 110740 | 0 | 0 | 381210 | 0 | 0 | 0 | 0 | 0 | 0 | 0 | 0 | 238750 |
| ENDSRQKV | GALNT2 | Q10471 | GALT2_HUMAN | Unmodified | 0 | 0 | 0 | 0 | 103050 | 0 | 0 | 0 | 337930 | 76022 | 0 | 0 | 0 | 0 | 0 | 0 | 0 | 0 |
| ENDYYTPT( | STT3A | P46977 | STT3A_HUMAN | Unmodified | 0 | 0 | 0 | 0 | 0 | 138780 | 0 | 0 | 1634300 | 0 | 0 | 0 | 0 | 0 | 0 | 0 | 0 | 0 |
| ENDYYTPT( | STT3A | P46977 | STT3A_HUMAN | Unmodified | 0 | 305270 | 0 | 0 | 0 | 0 | 0 | 0 | 3329600 | 482320 | 146680 | 576550 | 0 | 0 | 83034 | 0 | 0 | 0 |
| ENDYYTPT( | STT3A | P46977 | STT3A_HUMAN | Unmodified | 0 | 0 | 0 | 0 | 0 | 0 | 0 | 0 | 1130400 | 0 | 0 | 0 | 0 | 0 | 0 | 0 | 0 | 0 |
| ENELRALEI | SMC2 | O95347 | SMC2_HUMAN | Unmodified | 0 | 0 | 11747 | 0 | 131200 | 31888 | 72665 | 0 | 81323 | 0 | 62644 | 0 | 65192 | 0 | 0 | 0 | 0 | 0 |
| ENELTYYC( | CD59; CD59; N | P13987; A0A | CD59_HUMAN | Unmodified | 0 | 292660 | 204100 | 0 | 718120 | 574280 | 0 | 0 | 701980 | 460020 | 0 | 556020 | 417370 | 520310 | 636970 | 459910 | 675390 | 1259600 |
| ENEQGEVD | CD99 | A0A096LP69 | A0A096LP69_H | Unmodified | 183030 | 205200 | 486080 | 356990 | 0 | 0 | 0 | 0 | 0 | 0 | 0 | 0 | 0 | 0 | 1800300 | 1117600 | 0 | 0 |
| ENEQGEVD | CD99 | A0A096LP69 | A0A096LP69_H | Oxidation (M) | 0 | 0 | 165070 | 0 | 0 | 0 | 150260 | 76515 | 0 | 0 | 0 | 0 | 0 | 0 | 0 | 588170 | 0 | 0 |

| | | | | | | | | | | | | | | | | | | | | | | |
|---|---|---|---|---|---|---|---|---|---|---|---|---|---|---|---|---|---|---|---|---|---|---|
| ENEQGEVD | CD99 | A0A096LP69 | A0A096LP69_H | Unmodified | 0 | 0 | 0 | 187760 | 494330 | 0 | 0 | 0 | 0 | 0 | 0 | 69554 | 0 | 0 | 0 | 0 | 0 | 1057800 |
| ENEQGEVD | CD99 | A0A096LP69 | A0A096LP69_H | Unmodified | 0 | 0 | 0 | 0 | 2033500 | 252030 | 62194 | 31864 | 0 | 0 | 0 | 0 | 221660 | 0 | 0 | 0 | 954620 | 3684400 |
| ENERFFGD | HYOU1; HYOU1 | Q9Y4L1; J3Q | HYOU1_HUMA | Oxidation (M) | 0 | 0 | 0 | 0 | 0 | 0 | 114230 | 0 | 1348700 | 225000 | 281540 | 0 | 0 | 0 | 105120 | 0 | 0 | 0 |
| ENERFFGD | HYOU1; HYOU1 | Q9Y4L1; J3Q | HYOU1_HUMA | Oxidation (M) | 0 | 0 | 0 | 146520 | 0 | 0 | 130560 | 0 | 2602100 | 4634800 | 426750 | 0 | 0 | 0 | 64077 | 0 | 0 | 0 |
| ENERFFGD | HYOU1; HYOU1 | Q9Y4L1; J3Q | HYOU1_HUMA | Unmodified | 0 | 0 | 0 | 0 | 219500 | 0 | 0 | 0 | 10393000 | 4641100 | 263170 | 0 | 468530 | 707760 | 0 | 0 | 0 | 0 |
| ENEVLEAW | SPTBN1; SPTBN | Q01082; A0A | SPTB2_HUMAN | Unmodified | 253700 | 0 | 0 | 0 | 159870 | 0 | 0 | 0 | 0 | 0 | 0 | 2728600 | 0 | 0 | 0 | 0 | 0 | 0 |
| ENEVQSLH | KTN1 | Q86UP2 | KTN1_HUMAN | Unmodified | 0 | 0 | 0 | 0 | 286750 | 0 | 106100 | 0 | 0 | 167870 | 0 | 0 | 779000 | 314650 | 181300 | 0 | 207170 | 382530 |
| ENFAGEAT | APOB | P04114 | APOB_HUMAN | Unmodified | 89186 | 530210 | 891520 | 0 | 0 | 745710 | 0 | 0 | 0 | 0 | 0 | 162050 | 0 | 0 | 686310 | 364970 | 0 | 0 |
| ENFCNIHV | CTPS1; CTPS1; | P17812; B4E | PYRG1_HUMAN | Unmodified | 0 | 0 | 0 | 0 | 0 | 0 | 0 | 0 | 0 | 0 | 0 | 3347500 | 0 | 0 | 0 | 0 | 0 | 0 |
| ENFFLGGV | PIK3C2A | O00443 | P3C2A_HUMAN | Unmodified | 0 | 0 | 0 | 0 | 0 | 0 | 0 | 0 | 0 | 0 | 0 | 516230 | 0 | 0 | 0 | 0 | 0 | 0 |
| ENFMTVDG | GCA; GCA; GCA | P28676; C9J | GRAN_HUMAN | Unmodified | 0 | 0 | 291680 | 0 | 0 | 0 | 0 | 0 | 1251700 | 0 | 0 | 791730 | 0 | 0 | 0 | 0 | 0 | 0 |
| ENFQNWLI | TAGLN2; TAGLN | P37802; X6R | TAGL2_HUMAN | Unmodified | 0 | 0 | 0 | 0 | 0 | 0 | 0 | 109060 | 0 | 238600 | 0 | 2911000 | 0 | 0 | 1261300 | 0 | 0 | 0 |
| ENFSCLTR | MDH1; MDH1; I | P40925; C9J | MDHC_HUMAN | Unmodified | 0 | 0 | 0 | 0 | 1273200 | 252120 | 161100 | 0 | 0 | 0 | 1393000 | 10631000 | 1264100 | 309700 | 598810 | 0 | 0 | 1015600 |
| ENFSCLTRI | MDH1; MDH1; I | P40925; C9J | MDHC_HUMAN | Unmodified | 672910 | 3663800 | 0 | 0 | 2144900 | 0 | 1022600 | 279990 | 207170 | 505900 | 0 | 0 | 1333600 | 1270800 | 0 | 0 | 1397800 | 1858900 |
| ENGELLPIL | GLRX3 | O76003 | GLRX3_HUMAN | Unmodified | 0 | 0 | 0 | 0 | 0 | 0 | 0 | 0 | 0 | 0 | 0 | 6794700 | 0 | 0 | 436270 | 687480 | 0 | 0 |
| ENGFIYFQI | BROX | Q5VW32 | BROX_HUMAN | Unmodified | 2723700 | 2038000 | 1082600 | 663570 | 0 | 0 | 0 | 0 | 1205200 | 0 | 0 | 3197600 | 0 | 0 | 1581300 | 1202400 | 0 | 0 |
| ENGFIYFQI | BROX | Q5VW32 | BROX_HUMAN | Unmodified | 0 | 340350 | 0 | 239070 | 0 | 0 | 0 | 0 | 0 | 254420 | 0 | 0 | 197240 | 0 | 0 | 0 | 0 | 31953 |
| ENGFTQHV | LARP1 | Q6PKG0 | LARP1_HUMAN | Unmodified | 117090 | 11745 | 0 | 184450 | 0 | 37263 | 0 | 0 | 0 | 0 | 140710 | 676840 | 70499 | 45046 | 0 | 106870 | 0 | 0 |
| ENGGYTGG | MOV10; MOV10 | Q9HCE1; Q5 | MOV10_HUMA | Unmodified | 323290 | 236270 | 173470 | 0 | 0 | 0 | 0 | 0 | 0 | 0 | 0 | 0 | 0 | 0 | 367930 | 385640 | 0 | 0 |
| ENGLEVLQ | DNAJC13 | O75165 | DJC13_HUMAN | Unmodified | 0 | 0 | 19561 | 0 | 0 | 0 | 0 | 0 | 0 | 0 | 0 | 737420 | 0 | 0 | 0 | 0 | 0 | 0 |
| ENGLRDILA | HSPA14 | Q0VDF9 | HSP7E_HUMAN | Unmodified | 0 | 0 | 0 | 0 | 10863000 | 36075000 | 8449600 | 0 | 236140 | 0 | 0 | 0 | 0 | 18834000 | 0 | 0 | 852620 | 28246000 |
| ENGLRDILA | HSPA14 | Q0VDF9 | HSP7E_HUMAN | Oxidation (M) | 0 | 0 | 0 | 0 | 0 | 29261000 | 0 | 0 | 67386 | 0 | 0 | 269160 | 0 | 0 | 51540 | 18530 | 0 | 0 |
| ENGNYIMY | ADAM10; ADAM | O14672; H0Y | ADA10_HUMAN | Unmodified | 0 | 0 | 0 | 0 | 865850 | 0 | 0 | 0 | 334920 | 0 | 0 | 975790 | 1109200 | 528730 | 0 | 0 | 0 | 299870 |
| ENGNYIMY | ADAM10; ADAM | O14672; H0Y | ADA10_HUMAN | Oxidation (M) | 0 | 0 | 3225100 | 1854000 | 521920 | 0 | 0 | 0 | 0 | 0 | 0 | 0 | 915360 | 543120 | 2127300 | 2390300 | 0 | 163450 |
| ENGNYIMY | ADAM10; ADAM | O14672; H0Y | ADA10_HUMAN | Unmodified | 0 | 0 | 0 | 0 | 1835000 | 117820 | 0 | 32329 | 0 | 0 | 0 | 0 | 0 | 0 | 0 | 0 | 0 | 0 |
| ENGRLPAL | MICAL3 | Q7RTP6 | MICA3_HUMAN | Unmodified | 0 | 0 | 0 | 0 | 914010 | 0 | 0 | 0 | 0 | 0 | 0 | 0 | 0 | 0 | 0 | 0 | 0 | 0 |
| ENGVTHPII | VAT1; VAT1; VAT | Q99536; K7E | VAT1_HUMAN S | Unmodified | 718110 | 385110 | 0 | 598700 | 16029000 | 5803300 | 0 | 0 | 6696900 | 0 | 0 | 8221400 | 1377300 | 1545000 | 673200 | 403430 | 1723000 | 5392200 |
| ENGVTHPII | VAT1; VAT1; VAT | Q99536; K7E | VAT1_HUMAN S | Unmodified | 156650 | 0 | 0 | 0 | 4329400 | 530970 | 0 | 0 | 0 | 0 | 487760 | 2805100 | 344540 | 0 | 340970 | 0 | 0 | 1268800 |
| ENGVTHPII | VAT1; VAT1; VAT | Q99536; K7E | VAT1_HUMAN S | Unmodified | 0 | 0 | 0 | 0 | 3420000 | 0 | 0 | 0 | 0 | 0 | 0 | 0 | 0 | 0 | 0 | 0 | 839540 | 0 |
| ENHLAASS | COBLL1; COBLI | Q53SF7; A0A | COBL1_HUMAI | Unmodified | 1195000 | 705200 | 451920 | 587310 | 0 | 0 | 0 | 0 | 0 | 19592 | 0 | 0 | 0 | 225290 | 482270 | 543150 | 0 | 565540 |
| ENHMQGK | KIF23; KIF23; KI | Q02241; A0A | KIF23_HUMAN | Unmodified | 909330 | 105660 | 2717400 | 3873600 | 5257800 | 1629800 | 37730 | 33027 | 30529 | 125530 | 0 | 21041 | 236060 | 471780 | 782680 | 0 | 919570 | 436530 |
| ENHMQGK | KIF23; KIF23; KI | Q02241; A0A | KIF23_HUMAN | Unmodified | 0 | 0 | 0 | 0 | 1017400 | 62054 | 0 | 0 | 0 | 0 | 0 | 0 | 0 | 0 | 0 | 0 | 1227700 | 57952 |
| ENHNRTQG | COPS7B; COPS | Q9H9Q2; J3I | CSN7B_HUMAI | Unmodified | 0 | 534010 | 0 | 0 | 635570 | 0 | 515170 | 208590 | 191390 | 177190 | 279460 | 668630 | 581700 | 182570 | 0 | 0 | 607930 | 687130 |
| ENIFAIQTLI | EPN2; EPN2; EF | O95208; F6F | EPN2_HUMAN | Unmodified | 925210 | 1024000 | 658450 | 341080 | 0 | 0 | 0 | 0 | 651470 | 0 | 0 | 0 | 0 | 0 | 1841600 | 1471200 | 0 | 0 |
| ENIIAVATTI | PPP2R2A; PPP2 | P63151; A0A | 2ABA_HUMAN | Unmodified | 0 | 0 | 0 | 0 | 426730 | 0 | 85979 | 0 | 0 | 514580 | 0 | 0 | 1201200 | 320880 | 0 | 0 | 226200 | 280090 |
| ENIKPNETS | ATXN2; ATXN2; | Q99700; A0A | ATX2_HUMAN A | Unmodified | 1000300 | 0 | 0 | 0 | 0 | 0 | 0 | 0 | 685180 | 118300 | 0 | 0 | 0 | 0 | 160750 | 0 | 0 | 0 |
| ENILEEFSK | HNRNPR; HNRI | O43390; B4D | HNRPR_HUMA | Unmodified | 0 | 0 | 0 | 0 | 0 | 0 | 0 | 0 | 0 | 0 | 0 | 0 | 0 | 0 | 0 | 0 | 0 | 0 |
| ENILFGCQI | ABCC1; ABCC1 | P33527; I3L4 | MRP1_HUMAN | Unmodified | 0 | 0 | 0 | 0 | 0 | 0 | 0 | 0 | 0 | 0 | 92549 | 1189900 | 0 | 0 | 90143 | 112700 | 0 | 0 |
| ENIQTLLK | RAI14 | Q9P0K7 | RAI14_HUMAN | Unmodified | 0 | 0 | 0 | 0 | 769330 | 112720 | 0 | 126930 | 125690 | 0 | 0 | 1073800 | 231900 | 208810 | 185480 | 123500 | 455930 | 588350 |
| ENIRPVWQ | TRIM26; TRIM26 | Q12899; A2A | TRI26_HUMAN | Unmodified | 416820 | 2447100 | 125730 | 956550 | 0 | 0 | 0 | 0 | 0 | 0 | 0 | 0 | 70403 | 30296 | 1794700 | 198770 | 0 | 0 |
| ENIRPVWQ | TRIM26; TRIM26 | Q12899; A2A | TRI26_HUMAN | Unmodified | 321870 | 1259200 | 0 | 1921100 | 0 | 0 | 0 | 0 | 0 | 0 | 0 | 0 | 1434000 | 0 | 1606900 | 173710 | 0 | 0 |
| ENITEDEAK | BROX; BROX; BI | Q5VW32; F5 | BROX_HUMAN | Unmodified | 986680 | 1638900 | 18478 | 455050 | 0 | 0 | 0 | 0 | 0 | 107570 | 0 | 1135300 | 123990 | 0 | 1682700 | 1013900 | 0 | 0 |
| ENITEDEAK | BROX; BROX; BI | Q5VW32; F5 | BROX_HUMAN | Unmodified | 553100 | 955880 | 0 | 535560 | 0 | 49287 | 0 | 0 | 0 | 31791 | 0 | 70728 | 0 | 0 | 0 | 64641 | 0 | 0 |
| ENITEDEAK | BROX; BROX; BI | Q5VW32; F5 | BROX_HUMAN | Unmodified | 0 | 0 | 0 | 0 | 308740 | 104610 | 0 | 0 | 0 | 0 | 0 | 0 | 0 | 188420 | 0 | 0 | 64005 | 49414 |
| ENIVEAIIHS | IPO7; IPO7 | O95373; E9F | IPO7_HUMAN I | Unmodified | 76507 | 304480 | 40320 | 5444200 | 48637 | 0 | 887910 | 0 | 200860 | 92271 | 259010 | 3152100 | 8222600 | 3135400 | 117910 | 89778 | 0 | 0 |
| ENIWDGVT | PRKCB | P05771 | KPCB_HUMAN | Unmodified | 440100 | 0 | 926470 | 1760000 | 0 | 0 | 1026100 | 912230 | 0 | 0 | 0 | 0 | 0 | 0 | 1025800 | 906070 | 0 | 0 |
| ENIWSASE | IQGAP2; IQGAP | Q13576; D6I | IQGA2_HUMAN | Unmodified | 0 | 0 | 0 | 13707 | 0 | 0 | 0 | 0 | 0 | 0 | 0 | 0 | 0 | 0 | 47367 | 37424 | 0 | 0 |
| ENKDDLNI | CD109 | Q6YHK3 | CD109_HUMAI | Unmodified | 0 | 0 | 0 | 0 | 0 | 0 | 0 | 0 | 1152500 | 0 | 0 | 0 | 0 | 0 | 0 | 0 | 0 | 0 |
| ENKEVVLQ | ANPEP | P15144 | AMPN_HUMAN | Unmodified | 0 | 885660 | 0 | 1098200 | 0 | 539230 | 0 | 0 | 0 | 0 | 1027000 | 0 | 0 | 0 | 178400 | 0 | 1.26E+08 | 1.36E+08 |
| ENKFIVRDF | ATP6V1C1; ATP | P21283; E7E | VATC1_HUMAN | Unmodified | 0 | 0 | 0 | 0 | 0 | 0 | 0 | 0 | 2019000 | 344730 | 84526 | 0 | 0 | 0 | 0 | 0 | 0 | 0 |
| ENKITITND | HSPA8; HSPA2; | P11142; P54 | HSP7C_HUMAI | Unmodified | 929210 | 2340200 | 793860 | 754960 | 7326100 | 2484400 | 974730 | 66684 | 1510400 | 2395800 | 232800 | 1249500 | 1147800 | 393890 | 1184700 | 899320 | 1065500 | 110740 |
| ENKPLPPVI | MAN1A2 | O60476 | MA1A2_HUMAI | Unmodified | 0 | 0 | 0 | 0 | 0 | 0 | 0 | 0 | 1632500 | 0 | 10678 | 117260 | 0 | 0 | 0 | 0 | 0 | 0 |
| ENKPSIIFID | VPS4B | O75351 | VPS4B_HUMAN | Unmodified | 732290 | 350890 | 171550 | 0 | 17912 | 0 | 0 | 46233 | 1787000 | 0 | 0 | 885590 | 0 | 0 | 0 | 0 | 0 | 0 |
| ENKPYALN | SCG2; SCG2 | P13521; C9J | SCG2_HUMAN | Unmodified | 0 | 0 | 0 | 0 | 246830 | 0 | 0 | 0 | 0 | 349740 | 0 | 0 | 0 | 32263 | 0 | 0 | 0 | 73617 |
| ENKQSLGL | CFAP20DC; CF | F5H9F7; F8V | F5H9F7_HUMA | Unmodified | 0 | 0 | 0 | 538980 | 10537000 | 3441300 | 472730 | 0 | 0 | 19042000 | 0 | 141280 | 10589000 | 12709000 | 0 | 0 | 239810 | 2135100 |
| ENLATVEGI | DCTN2; DCTN2 | Q13561; A8K | DCTN2_HUMAI | Unmodified | 0 | 0 | 0 | 0 | 115570 | 0 | 0 | 0 | 0 | 0 | 0 | 592010 | 71822 | 0 | 0 | 0 | 0 | 1792900 |
| ENLGQGW | EPHX1 | P07099 | HYEP_HUMAN | Oxidation (M) | 0 | 0 | 0 | 0 | 0 | 0 | 0 | 0 | 249480 | 0 | 0 | 387790 | 0 | 0 | 311460 | 0 | 0 | 0 |
| ENLGWQKI | DSP | P15924 | DESP_HUMAN | Unmodified | 0 | 0 | 0 | 72109 | 0 | 0 | 0 | 0 | 0 | 597780 | 0 | 0 | 0 | 0 | 0 | 139800 | 701680 | 0 |
| ENLKDTMT | CD151; CD151; | P48509; E9P | CD151_HUMAI | Unmodified | 585840 | 614710 | 0 | 0 | 1162700 | 0 | 0 | 0 | 130650 | 37537 | 0 | 192970 | 130100 | 0 | 128820 | 178170 | 184580 | 0 |
| ENLLAEKVI | TUSC3; TUSC3; | Q13454; D6I | TUSC3_HUMAI | Oxidation (M) | 0 | 0 | 0 | 0 | 0 | 0 | 0 | 0 | 107860 | 51606 | 0 | 0 | 0 | 0 | 0 | 0 | 0 | 0 |
| ENLLDFIK | P4HB; P4HB; P4 | P07237; A0A | PDIA1_HUMAN | Unmodified | 0 | 0 | 417290 | 274970 | 0 | 0 | 0 | 0 | 34115000 | 11460000 | 5838300 | 2173500 | 0 | 0 | 2561400 | 590740 | 0 | 0 |
| ENLLTYLPD | LRRC1 | Q9BTT6 | LRRC1_HUMAN | Unmodified | 11333 | 0 | 101250 | 168750 | 0 | 0 | 0 | 0 | 297640 | 0 | 24029 | 582210 | 0 | 0 | 67125 | 0 | 0 | 28899 |
| ENLQHTQA | TAOK3; TAOK3 | Q9H2K8; G3 | TAOK3_HUMAN | Unmodified | 199450 | 466770 | 993860 | 694730 | 213860 | 0 | 56635 | 26293 | 149570 | 0 | 214410 | 0 | 0 | 119410 | 731810 | 546740 | 0 | 99222 |
| ENLQHTQA | TAOK3; TAOK3 | Q9H2K8; G3 | TAOK3_HUMAN | Unmodified | 155860 | 179870 | 0 | 0 | 1267900 | 0 | 0 | 0 | 0 | 0 | 0 | 0 | 0 | 0 | 208360 | 115700 | 0 | 287760 |
| ENLQRTFLI | TAX1BP1; TAX1 | Q86VP1; B82 | TAXB1_HUMAN | Unmodified | 0 | 0 | 0 | 0 | 989740 | 0 | 0 | 0 | 0 | 0 | 0 | 0 | 0 | 0 | 0 | 0 | 904820 | 2647300 |
| ENLSDEDK | PLS1 | Q14651 | PLSI_HUMAN F | Unmodified | 112130 | 0 | 46137 | 0 | 91408 | 151480 | 0 | 0 | 390620 | 0 | 256790 | 1000900 | 0 | 0 | 120700 | 0 | 0 | 0 |
| ENLSYDLVI | HSPH1; HSPH1 | Q92598; R4( | HS105_HUMAN | Unmodified | 0 | 0 | 0 | 0 | 128900 | 0 | 204100 | 194800 | 638420 | 396250 | 1272200 | 4948600 | 0 | 0 | 743230 | 0 | 431400 | 0 |
| ENLYPYLGI | VPS45; VPS45; | Q9NRW7; A( | VPS45_HUMAN | Unmodified | 0 | 0 | 0 | 0 | 0 | 0 | 0 | 0 | 0 | 0 | 0 | 1156800 | 0 | 0 | 0 | 0 | 0 | 0 |
| ENLYTIQTL | EPN3; EPN3; EF | Q9H201; D6 | EPN3_HUMAN | Unmodified | 1255700 | 710090 | 360490 | 175390 | 0 | 0 | 0 | 0 | 0 | 0 | 181670 | 0 | 0 | 0 | 951070 | 1112500 | 0 | 0 |
| ENMPSHFK | PIP4K2A | P48426 | PI42A_HUMAN | Unmodified | 0 | 0 | 0 | 77502 | 0 | 0 | 221950 | 111360 | 0 | 0 | 190270 | 0 | 174170 | 0 | 0 | 0 | 0 | 296490 |
| ENMYAVQI | EPN1 | Q9Y6I3 | EPN1_HUMAN | Oxidation (M) | 1033400 | 923830 | 459890 | 225100 | 304510 | 0 | 107380 | 89681 | 69266 | 0 | 0 | 270120 | 0 | 0 | 454260 | 560420 | 300140 | 0 |
| ENMYAVQI | EPN1 | Q9Y6I3 | EPN1_HUMAN | Unmodified | 993460 | 0 | 597230 | 552960 | 3447100 | 0 | 0 | 0 | 0 | 0 | 0 | 0 | 0 | 0 | 1074100 | 0 | 1498700 | 0 |
| ENNAVYAF | SH3BGRL | O75368 | SH3L1_HUMAN | Unmodified | 199040 | 0 | 0 | 0 | 156910 | 0 | 0 | 0 | 0 | 32240 | 0 | 1430800 | 0 | 0 | 0 | 0 | 11944 | 0 |
| ENNFPPLP | SCAMP4; SCAM | Q969E2; C9J | SCAM4_HUMA | Unmodified | 1199500 | 1511000 | 2595600 | 3380300 | 674520 | 0 | 0 | 0 | 542920 | 140080 | 0 | 706470 | 1105700 | 328350 | 4112000 | 2742700 | 0 | 0 |
| ENNLKNVE | CLIP1; CLIP1; C | P30622; F5H | CLIP1_HUMAN | Unmodified | 0 | 0 | 0 | 0 | 793730 | 0 | 0 | 0 | 0 | 0 | 0 | 5790600 | 0 | 0 | 0 | 0 | 0 | 0 |
| ENNQVPAV | NID1 | P14543 | NID1_HUMAN I | Unmodified | 245440 | 0 | 171760 | 683160 | 0 | 0 | 5791900 | 726870 | 276630 | 338670 | 693330 | 0 | 2125800 | 6923800 | 0 | 190410 | 481540 | 3261800 |
| ENNTHPEV | STK26; STK26; | Q9P289; B4E | STK26_HUMAN | Unmodified | 0 | 0 | 0 | 0 | 172440 | 0 | 0 | 0 | 0 | 0 | 0 | 1231700 | 0 | 0 | 0 | 404700 | 0 | 0 |
| ENNYRGH( | THOC3 | Q96J01 | THOC3_HUMA | Unmodified | 0 | 0 | 0 | 0 | 0 | 0 | 0 | 0 | 0 | 0 | 0 | 371620 | 311800 | 201960 | 0 | 0 | 155830 | 196600 |
| ENPETEED' | ITGAV | P06756 | ITAV_HUMAN I | Unmodified | 0 | 0 | 1323700 | 0 | 0 | 0 | 0 | 0 | 600810 | 0 | 0 | 3484000 | 0 | 0 | 1646100 | 600740 | 0 | 0 |
| ENPFLRQE | CKAP5 | Q14008 | CKAP5_HUMAN | Unmodified | 0 | 0 | 95939 | 0 | 0 | 0 | 0 | 0 | 0 | 0 | 181830 | 410100 | 0 | 0 | 144610 | 0 | 0 | 0 |
| ENPGFDFS | NUDCD2; NUD | Q8WVJ2; E5I | NUDC2_HUMA | Unmodified | 0 | 0 | 0 | 0 | 2140000 | 7515100 | 0 | 386810 | 0 | 0 | 0 | 0 | 0 | 0 | 0 | 0 | 1173400 | 802740 |
| ENPGLTQD | PDCD10; PDCD | Q9BUL8; H7 | PDC10_HUMAI | Unmodified | 0 | 0 | 0 | 0 | 3664500 | 0 | 0 | 0 | 1503800 | 0 | 3999400 | 1671200 | 1888500 | 2191200 | 0 | 0 | 803830 | 2609400 |
| ENPGLTQD | PDCD10; PDCD | Q9BUL8; H7 | PDC10_HUMAI | Oxidation (M) | 0 | 0 | 0 | 494090 | 1681800 | 0 | 0 | 26177 | 0 | 402640 | 1166800 | 0 | 1781300 | 1004300 | 0 | 0 | 555950 | 1408600 |
| ENPGLTQD | PDCD10; PDCD | Q9BUL8; H7 | PDC10_HUMAI | Oxidation (M) | 0 | 0 | 0 | 0 | 347700 | 28261 | 114300 | 0 | 0 | 0 | 0 | 0 | 107210 | 164370 | 0 | 0 | 352940 | 303710 |
| ENPKVVNE | CISD2; CISD2 | Q8N5K1; D6 | CISD2_HUMAN | Unmodified | 0 | 0 | 0 | 0 | 624890 | 0 | 396950 | 55491 | 2498600 | 1861300 | 0 | 0 | 0 | 0 | 0 | 0 | 0 | 0 |
| ENPPVEDS | STT3B | Q8TCJ2 | STT3B_HUMAN | Unmodified | 0 | 0 | 0 | 0 | 0 | 0 | 0 | 0 | 3713300 | 140940 | 61553 | 0 | 0 | 0 | 0 | 0 | 0 | 0 |
| ENPRNFSD | TAGLN2; TAGLN | P37802; X6R | TAGL2_HUMAN | Unmodified | 1175500 | 0 | 381310 | 1071000 | 55252000 | 14453000 | 1290300 | 803180 | 1081600 | 2093100 | 644230 | 0 | 8332800 | 2136600 | 353800 | 196590 | 7135300 | 22441000 |
| ENPSATLEI | UPF1 | Q92900 | RENT1_HUMAN | Unmodified | 0 | 7474400 | 0 | 0 | 0 | 0 | 0 | 0 | 509440 | 0 | 0 | 3177900 | 0 | 768240 | 3643700 | 5406200 | 0 | 0 |
| ENPTGVVLI | CNK3/IPCEF1; | G9CGD6; Q6 | CNIPF_HUMAN | Unmodified | 0 | 0 | 0 | 0 | 0 | 0 | 0 | 0 | 176270 | 0 | 0 | 309700 | 0 | 0 | 23448 | 0 | 0 | 0 |
| ENPYYDSR | CLTC; CLTC | Q00610; A0A | CLH1_HUMAN | Unmodified | 2098100 | 0 | 1118600 | 641670 | 0 | 0 | 0 | 0 | 0 | 0 | 0 | 13634000 | 93727 | 0 | 1096000 | 1851400 | 0 | 0 |
| ENPYYDSR | CLTC; CLTC | Q00610; A0A | CLH1_HUMAN | Unmodified | 369210 | 365280 | 70039 | 761790 | 0 | 0 | 336690 | 131230 | 434900 | 108660 | 87299 | 0 | 0 | 0 | 154420 | 0 | 0 | 292030 |
| ENQELKDS | CALCOCO2; CA | Q13137; D6I | CACO2_HUMA | Unmodified | 0 | 0 | 0 | 0 | 440920 | 67495 | 1186400 | 158140 | 0 | 551320 | 0 | 75254 | 0 | 312030 | 318830 | 0 | 868090 | 1683200 |
| ENQGDET( | YBX1; YBX1 | P67809; H0Y | YBOX1_HUMAN | Unmodified | 0 | 0 | 138760 | 288540 | 189420 | 0 | 0 | 0 | 43791 | 0 | 124070 | 1868900 | 150640 | 0 | 48993 | 0 | 38223 | 47013 |
| ENQLQLSC | USP14; USP14 | P54578; A6N | UBP14_HUMAN | Unmodified | 0 | 0 | 0 | 0 | 679260 | 0 | 112880 | 0 | 0 | 0 | 613540 | 2268300 | 0 | 0 | 363560 | 0 | 0 | 0 |
| ENQQEEAF | ABAT; ABAT; AB | P80404; H3E | GABT_HUMAN | Unmodified | 0 | 0 | 0 | 0 | 0 | 0 | 0 | 0 | 1296900 | 418060 | 0 | 0 | 0 | 0 | 0 | 0 | 0 | 0 |
| ENQTAVDV | RHEB; RHEB | Q15382; C9J | RHEB_HUMAN | Unmodified | 939130 | 865800 | 1219700 | 1768700 | 2676700 | 1331500 | 0 | 0 | 2385200 | 490890 | 0 | 2167600 | 878790 | 453450 | 1288800 | 766860 | 0 | 778840 |
| ENQTAVDV | RHEB; RHEB | Q15382; C9J | RHEB_HUMAN | Unmodified | 1974800 | 1027300 | 928090 | 1439400 | 2296100 | 1219000 | 104600 | 0 | 1286200 | 0 | 319430 | 0 | 1524900 | 307340 | 1114400 | 945010 | 329820 | 1319000 |
| ENQVPGDI | CLGN; CLGN | O14967; D6I | CLGN_HUMAN | Unmodified | 0 | 0 | 0 | 0 | 0 | 0 | 72473 | 142390 | 11183000 | 1512200 | 650390 | 0 | 85088 | 98256 | 31039 | 0 | 0 | 0 |
| ENQWCEEI | SKP1; SKP1 | P63208; E5R | SKP1_HUMAN | Unmodified | 0 | 0 | 0 | 0 | 280820 | 44817 | 0 | 0 | 159950 | 22870 | 0 | 893690 | 162590 | 78416 | 0 | 0 | 69965 | 137790 |
| ENQWFDV( | HCFC1; HCFC1 | P51610; A6N | HCFC1_HUMA | Unmodified | 0 | 0 | 0 | 0 | 433790 | 0 | 0 | 0 | 185070 | 0 | 0 | 737340 | 0 | 0 | 0 | 0 | 487310 | 87252 |
| ENQWLGV | ITGA7; ITGA7 | Q13683; J3K | ITA7_HUMAN I | Unmodified | 0 | 0 | 0 | 0 | 0 | 0 | 378330 | 97286 | 0 | 0 | 0 | 0 | 0 | 0 | 0 | 0 | 0 | 0 |
| ENREAQMA | STMN1 | P16949 | STMN1_HUMAI | Unmodified | 0 | 0 | 0 | 0 | 0 | 0 | 0 | 0 | 2458900 | 844860 | 174260 | 0 | 0 | 0 | 0 | 0 | 0 | 0 |
| ENRLCYYI( | MANF | P55145 | MANF_HUMAN | Unmodified | 0 | 0 | 0 | 0 | 361370 | 0 | 343390 | 178700 | 10010000 | 8333800 | 2056800 | 0 | 613100 | 807990 | 0 | 0 | 88547 | 762560 |
| ENRLKDDI( | CIT; CIT; CIT; CI | A0A7I2V4H7 | A0A7I2V4H7_H | Unmodified | 0 | 0 | 0 | 0 | 678700 | 138580 | 234990 | 1719400 | 0 | 0 | 0 | 0 | 16776 | 0 | 0 | 0 | 228290 | 0 |
| ENRNHLLP | NTPCR | Q9BSD7 | NTPCR_HUMAI | Unmodified | 0 | 0 | 0 | 330970 | 0 | 0 | 2348500 | 0 | 0 | 0 | 0 | 0 | 6373400 | 1347400 | 0 | 0 | 0 | 0 |
| ENSDVIALV | CLIP1; CLIP1; C | P30622; F5H | CLIP1_HUMAN | Unmodified | 0 | 0 | 27639 | 0 | 0 | 0 | 0 | 0 | 162970 | 0 | 72247 | 307350 | 0 | 0 | 0 | 38660 | 98401 | 0 |
| ENSEESGE | DAAM1 | Q9Y4D1 | DAAM1_HUMA | Unmodified | 0 | 0 | 0 | 164700 | 0 | 0 | 482390 | 0 | 0 | 0 | 0 | 0 | 382360 | 0 | 0 | 0 | 0 | 0 |
| ENSESEGV | PPM1H | Q9ULR3 | PPM1H_HUMA | Unmodified | 0 | 0 | 0 | 0 | 0 | 903500 | 0 | 0 | 0 | 0 | 0 | 800850 | 0 | 0 | 0 | 0 | 0 | 0 |
| ENSIEILSS1 | CZIB | Q9NWV4 | CZIB_HUMAN ( | Unmodified | 0 | 0 | 0 | 0 | 0 | 460570 | 0 | 0 | 1160600 | 0 | 434350 | 1704400 | 0 | 0 | 0 | 0 | 0 | 0 |

| | | | | | | | | | | | | | | | | | | | | | | |
|---|---|---|---|---|---|---|---|---|---|---|---|---|---|---|---|---|---|---|---|---|---|---|
| ENSLITQFT | PARP4 | Q9UKK3 | PARP4_HUMAN | Unmodified | 14341 | 0 | 0 | 0 | 0 | 0 | 0 | 0 | 0 | 0 | 0 | 135260 | 18096 | 0 | 0 | 0 | 0 | 0 |
| ENSLITQFT | PARP4 | Q9UKK3 | PARP4_HUMAN | Unmodified | 0 | 0 | 0 | 0 | 0 | 0 | 0 | 27936 | 0 | 0 | 0 | 669010 | 0 | 0 | 0 | 0 | 0 | 0 |
| ENSLKTDD | SPTAN1; SPTAN | Q13813; A0/ | SPTN1_HUMAN | Unmodified | 0 | 477610 | 0 | 0 | 0 | 0 | 0 | 0 | 0 | 0 | 68822 | 833260 | 0 | 0 | 0 | 0 | 0 | 0 |
| ENSPFLNN | SLC12A7; SLC1 | Q9Y666; A0/ | S12A7_HUMAN | Unmodified | 0 | 0 | 148580 | 0 | 0 | 0 | 0 | 0 | 446170 | 0 | 61252 | 326290 | 0 | 0 | 0 | 773770 | 0 | 0 |
| ENSQMFQ( | CPD | O75976 | CBPD_HUMAN | Unmodified | 0 | 0 | 131390 | 275600 | 2009100 | 568410 | 189570 | 89986 | 27511000 | 6516100 | 0 | 198970 | 372650 | 272800 | 76794 | 0 | 35790 | 164830 |
| ENSRCPLN | UBR1; UBR1 | Q8IWV7; A0/ | UBR1_HUMAN | Unmodified | 0 | 0 | 0 | 0 | 0 | 0 | 0 | 0 | 0 | 0 | 0 | 481550 | 0 | 0 | 0 | 0 | 0 | 0 |
| ENSREALAI | SLC9A3R1; SLC | O14745; J3Q | NHRF1_HUMAI | Unmodified | 244180 | 206500 | 2521300 | 5463300 | 0 | 621920 | 0 | 0 | 263980 | 0 | 927780 | 3697300 | 5372400 | 2808400 | 1716800 | 765660 | 0 | 0 |
| ENSSSQDF | PVR; PVR | P15151; A0A | PVR_HUMAN P | Unmodified | 0 | 0 | 0 | 0 | 0 | 0 | 0 | 0 | 0 | 0 | 0 | 243330 | 0 | 0 | 0 | 0 | 0 | 0 |
| ENSSSVTV: | UMAD1 | C9J7I0 | UMAD1_HUMA | Oxidation (M) | 1446800 | 719580 | 1524900 | 0 | 0 | 0 | 0 | 0 | 0 | 0 | 86508 | 0 | 0 | 0 | 1993600 | 506440 | 0 | 0 |
| ENSSSVTV: | UMAD1 | C9J7I0 | UMAD1_HUMA | Unmodified | 2399900 | 1769800 | 2116300 | 1447900 | 0 | 0 | 0 | 0 | 101180 | 28843 | 0 | 0 | 0 | 0 | 3000400 | 2144600 | 0 | 0 |
| ENTEQRAE | PPFIA1; PPFIA1 | Q13136; A0/ | LIPA1_HUMAN | Unmodified | 103390 | 0 | 292130 | 503820 | 0 | 46061 | 0 | 0 | 681690 | 0 | 239500 | 187700 | 56558 | 0 | 390450 | 330720 | 39145 | 91846 |
| ENTHEATYI | PIK3C2B; PIK3C | O00750; F5C | P3C2B_HUMAN | Unmodified | 0 | 0 | 0 | 0 | 0 | 0 | 0 | 0 | 0 | 0 | 0 | 0 | 0 | 0 | 277130 | 0 | 0 | 0 |
| ENTLNQLV | ALDH6A1; ALDH | Q02252; G3\ | MMSA_HUMAN | Unmodified | 0 | 0 | 0 | 0 | 0 | 0 | 0 | 0 | 2633000 | 0 | 0 | 0 | 0 | 0 | 0 | 0 | 0 | 0 |
| ENTQPFVV | NIBAN1 | Q9BZQ8 | NIBA1_HUMAN | Unmodified | 0 | 0 | 254430 | 767030 | 817570 | 299890 | 808100 | 246880 | 8681100 | 2573500 | 152590 | 0 | 0 | 0 | 284070 | 217250 | 0 | 0 |
| ENTSDPSL' | LAMP1 | P11279 | LAMP1_HUMAI | Unmodified | 0 | 0 | 0 | 0 | 0 | 0 | 0 | 0 | 1358600 | 0 | 0 | 0 | 0 | 401820 | 0 | 0 | 0 | 0 |
| ENTTTIVFR | KNTC1; KNTC1 | P50748; E7E | KNTC1_HUMAI | Unmodified | 0 | 0 | 0 | 0 | 0 | 0 | 1693600 | 201430 | 1055400 | 3878900 | 0 | 20701000 | 0 | 2743900 | 263110 | 0 | 556270 | 1497700 |
| ENTVGLHR | PMPCA | Q10713 | MPPA_HUMAN | Unmodified | 0 | 0 | 0 | 0 | 0 | 0 | 0 | 0 | 0 | 27326000 | 0 | 0 | 0 | 0 | 0 | 0 | 0 | 0 |
| ENVFQEH( | CTTN | Q14247 | SRC8_HUMAN | Unmodified | 0 | 0 | 0 | 0 | 203570 | 0 | 0 | 0 | 0 | 10998 | 228130 | 223660 | 42419 | 25118 | 0 | 0 | 0 | 22850 |
| ENVIFFQQ( | UAP1 | Q16222 | UAP1_HUMAN | 2 Oxidation (M) | 0 | 0 | 0 | 272820 | 220820 | 0 | 0 | 19744 | 85115 | 0 | 0 | 227630 | 255940 | 0 | 200160 | 135120 | 55172 | 1354100 |
| ENVIFFQQ( | UAP1 | Q16222 | UAP1_HUMAN | Oxidation (M) | 0 | 0 | 0 | 0 | 129530 | 0 | 0 | 0 | 0 | 0 | 0 | 15578 | 40791 | 0 | 34717 | 0 | 0 | 0 |
| ENVLIGDG. | ACADM; ACADN | P11310; Q51 | ACADM_HUMA | Unmodified | 0 | 0 | 0 | 0 | 0 | 0 | 0 | 0 | 2743100 | 261350 | 0 | 0 | 0 | 0 | 0 | 0 | 251340 | 0 |
| ENVNATEN | IPO5; IPO5 | O00410; H0\ | IPO5_HUMAN I | Unmodified | 0 | 0 | 40984 | 186140 | 1708700 | 0 | 258550 | 0 | 360120 | 246840 | 872460 | 4560300 | 187850 | 0 | 0 | 177840 | 650610 | 658810 |
| ENVNLAQII | ARCN1; ARCN1 | P48444; Q6I | COPD_HUMAN | Unmodified | 0 | 0 | 0 | 0 | 0 | 197360 | 0 | 0 | 141550 | 0 | 0 | 1925100 | 0 | 0 | 111750 | 92077 | 0 | 0 |
| ENVSNCIQ | MCTS1 | Q9ULC4 | MCTS1_HUMAI | Unmodified | 200380 | 0 | 313420 | 458090 | 0 | 0 | 274880 | 0 | 596000 | 303770 | 0 | 1724000 | 880360 | 753160 | 371270 | 0 | 656460 | 1103300 |
| ENYAELLEI | COL6A1; COL6A | P12109; A0A | CO6A1_HUMAN | Unmodified | 0 | 0 | 0 | 0 | 0 | 0 | 0 | 0 | 714620 | 275340 | 0 | 0 | 0 | 0 | 20263 | 0 | 0 | 950470 |
| ENYLSEQD | AVIL | O75366 | AVIL_HUMAN A | Unmodified | 0 | 0 | 0 | 0 | 0 | 0 | 13735 | 0 | 0 | 0 | 80074 | 647620 | 0 | 0 | 0 | 0 | 0 | 0 |
| EPAAPVSIQ | LASP1; LASP1; I | Q14847; C9J | LASP1_HUMAN | Unmodified | 235620 | 210300 | 470060 | 0 | 0 | 0 | 0 | 0 | 167090 | 0 | 357870 | 2616900 | 56090 | 0 | 479230 | 338870 | 0 | 0 |
| EPAIHSEG( | GAA | P10253 | LYAG_HUMAN | Unmodified | 0 | 0 | 0 | 0 | 7485900 | 0 | 0 | 0 | 7350200 | 0 | 0 | 1112300 | 0 | 0 | 0 | 0 | 0 | 912980 |
| EPALNEAN | PSME1 | Q06323 | PSME1_HUMAI | Unmodified | 0 | 0 | 0 | 0 | 375300 | 0 | 0 | 0 | 3400900 | 204850 | 1873200 | 8196900 | 641310 | 0 | 0 | 0 | 0 | 762080 |
| EPAPDSGL | ISOC2 | Q96AB3 | ISOC2_HUMAN | Unmodified | 0 | 0 | 0 | 0 | 0 | 0 | 0 | 0 | 211770 | 0 | 0 | 0 | 0 | 0 | 0 | 0 | 0 | 0 |
| EPAVLELEC | CAP1 | Q01518 | CAP1_HUMAN | Unmodified | 0 | 0 | 346240 | 0 | 0 | 0 | 0 | 0 | 0 | 0 | 0 | 3072700 | 0 | 0 | 281840 | 329330 | 0 | 0 |
| EPCGGQN' | HAPLN1; HAPLI | P10915; D6F | HPLN1_HUMAN | Unmodified | 0 | 0 | 0 | 0 | 0 | 0 | 0 | 0 | 1326300 | 101950 | 0 | 0 | 0 | 0 | 0 | 0 | 0 | 0 |
| EPEEHQVE | GOLIM4; GOLIN | O00461; F8V | GOLI4_HUMAN | Unmodified | 0 | 0 | 0 | 0 | 0 | 0 | 0 | 0 | 295870 | 0 | 0 | 0 | 0 | 0 | 0 | 0 | 0 | 0 |
| EPELFQTV/ | EHD1; EHD1; EI | Q9H4M9; C9 | EHD1_HUMAN | Unmodified | 112730 | 40992 | 1392100 | 0 | 0 | 0 | 0 | 0 | 0 | 0 | 560510 | 8911800 | 0 | 0 | 1245300 | 270130 | 0 | 0 |
| EPELFQTV/ | EHD1; EHD1; EI | Q9H4M9; C9 | EHD1_HUMAN | Unmodified | 35719 | 0 | 20045 | 73870 | 498960 | 112440 | 0 | 0 | 0 | 52016 | 123840 | 0 | 174270 | 106080 | 0 | 0 | 392640 | 357470 |
| EPELGLEEI | HSPBP1 | Q9NZL4 | HPBP1_HUMAI | Unmodified | 0 | 0 | 0 | 0 | 0 | 0 | 490670 | 0 | 370760 | 0 | 1472600 | 441360 | 0 | 1038700 | 0 | 0 | 0 | 0 |
| EPEPPGVV | GALNT7; GALNT | Q86SF2; E9I | GALT7_HUMAN | Unmodified | 0 | 0 | 0 | 1063000 | 593420 | 956500 | 211110 | 52327 | 0 | 0 | 0 | 0 | 0 | 376770 | 0 | 0 | 630790 | 828170 |
| EPFDLGEPI | SH3GL1 | Q99961 | SH3G1_HUMAI | Unmodified | 2965700 | 0 | 0 | 445930 | 0 | 0 | 0 | 0 | 0 | 0 | 0 | 0 | 0 | 0 | 1887700 | 0 | 0 | 0 |
| EPFFHGHD | CSNK2A1; CSN | P68400; A0A | CSK21_HUMAN | Unmodified | 85202 | 0 | 62300 | 197890 | 0 | 0 | 0 | 0 | 0 | 0 | 0 | 759270 | 0 | 0 | 85697 | 0 | 0 | 0 |
| EPFHSILSV | NAXE; NAXE; N/ | Q8NCW5; A0 | NNRE_HUMAN | Unmodified | 0 | 0 | 0 | 0 | 0 | 0 | 0 | 0 | 0 | 0 | 0 | 634730 | 0 | 0 | 0 | 0 | 0 | 0 |
| EPFLQQLK | EIF3L; EIF3L; EII | Q9Y262; B0( | EIF3L_HUMAN | Unmodified | 0 | 99850 | 1124100 | 93618 | 0 | 0 | 0 | 0 | 452180 | 157810 | 2897600 | 11104000 | 107340 | 0 | 260250 | 459230 | 0 | 0 |
| EPGAGAQE | FAM171A1; FAN | Q5VUB5; Q5 | F1711_HUMAN | Unmodified | 0 | 0 | 0 | 0 | 0 | 0 | 50441 | 0 | 0 | 0 | 177850 | 824190 | 0 | 0 | 0 | 0 | 0 | 0 |
| EPGRPTPT' | SLC7A6; SLC7A | Q92536; H3I | YLAT2_HUMAN | Unmodified | 0 | 0 | 0 | 0 | 0 | 0 | 0 | 0 | 0 | 216540 | 0 | 14289000 | 0 | 0 | 0 | 1229300 | 0 | 0 |
| EPHFQSLLI | PALS2; PALS2; I | Q9NZW5; C9 | PALS2_HUMAN | Unmodified | 0 | 0 | 740510 | 46744 | 0 | 0 | 21288 | 0 | 0 | 0 | 565540 | 0 | 0 | 0 | 0 | 0 | 0 | 0 |
| EPHLNYSP | IGSF3 | O75054 | IGSF3_HUMAN | Unmodified | 922260 | 0 | 0 | 0 | 0 | 0 | 0 | 0 | 2584000 | 0 | 0 | 993170 | 306420 | 0 | 1894000 | 0 | 0 | 0 |
| EPIAASTNF | SPTAN1; SPTAN | Q13813; A0/ | SPTN1_HUMAN | Unmodified | 0 | 0 | 0 | 0 | 0 | 0 | 0 | 0 | 0 | 0 | 0 | 0 | 0 | 0 | 0 | 0 | 0 | 0 |
| EPIVGSTDY | SPTAN1; SPTAN | Q13813; A0/ | SPTN1_HUMAN | Unmodified | 0 | 0 | 0 | 0 | 0 | 0 | 0 | 0 | 526910 | 0 | 0 | 1919900 | 0 | 0 | 600100 | 0 | 0 | 0 |
| EPKLEAAVF | UGGT1; UGGT1 | Q9NYU2; H7 | UGGG1_HUMA | Unmodified | 0 | 0 | 0 | 0 | 4548200 | 0 | 0 | 0 | 0 | 477620 | 0 | 0 | 0 | 0 | 0 | 0 | 0 | 0 |
| EPLAAYPLK | GPX8; GPX8; GI | Q8TED1; E7I | GPX8_HUMAN | Acetyl (Protein N | 0 | 0 | 0 | 0 | 338400 | 0 | 0 | 0 | 0 | 0 | 0 | 0 | 0 | 0 | 0 | 0 | 326100 | 1041300 |
| EPLIYESDR | FMNL1 | K7EK60 | K7EK60_HUMA | Unmodified | 0 | 0 | 0 | 0 | 551700 | 0 | 0 | 0 | 0 | 0 | 0 | 0 | 1002700 | 0 | 0 | 0 | 0 | 0 |
| EPLLPPEDI | CDH1 | P12830 | CADH1_HUMA | Unmodified | 0 | 0 | 0 | 0 | 0 | 0 | 0 | 0 | 0 | 0 | 221830 | 0 | 0 | 0 | 683360 | 633980 | 0 | 0 |
| EPLLPPEDI | CDH1 | P12830 | CADH1_HUMA | Unmodified | 0 | 0 | 0 | 2267900 | 0 | 0 | 0 | 0 | 0 | 0 | 0 | 0 | 0 | 0 | 4945400 | 0 | 0 | 0 |
| EPLPPIQQI | XRCC5 | P13010 | XRCC5_HUMAI | Oxidation (M) | 0 | 0 | 0 | 0 | 0 | 0 | 0 | 0 | 0 | 0 | 0 | 8519300 | 0 | 0 | 0 | 0 | 0 | 0 |
| EPLYEGTAI | WWC2 | Q6AWC2 | WWC2_HUMAI | Unmodified | 65060 | 0 | 946340 | 823380 | 0 | 1638900 | 171270 | 122920 | 0 | 0 | 0 | 0 | 0 | 716160 | 0 | 0 | 0 | 1505700 |
| EPNSENVC | RAB10 | P61026 | RAB10_HUMAN | Unmodified | 375910 | 226560 | 0 | 0 | 0 | 0 | 0 | 0 | 124380 | 137320 | 0 | 745840 | 0 | 0 | 583450 | 0 | 0 | 0 |
| EPPAPAQQ | TMEM165 | Q9HC07 | TM165_HUMAN | Unmodified | 0 | 0 | 0 | 0 | 0 | 0 | 0 | 0 | 21836000 | 0 | 527530 | 5565100 | 0 | 0 | 0 | 0 | 0 | 0 |
| EPPLLLGVL | APMAP; APMAP | Q9HDC9; H( | APMAP_HUMAN | Unmodified | 0 | 0 | 0 | 0 | 0 | 0 | 0 | 0 | 1533600 | 0 | 0 | 551920 | 0 | 0 | 263740 | 0 | 0 | 0 |
| EPQGALER | DHTKD1; DHTK | Q96HY7; C9 | DHTK1_HUMAI | Unmodified | 2233600 | 2060700 | 7224100 | 6176800 | 0 | 0 | 123810 | 30496 | 491340 | 0 | 0 | 0 | 1813200 | 407570 | 7199200 | 6702700 | 0 | 0 |
| EPQPEVAA | SLC16A3; SLC1 | O15427; J3Q | MOT4_HUMAN | Unmodified | 152060 | 0 | 794070 | 0 | 0 | 0 | 0 | 0 | 893990 | 361310 | 47140 | 3286200 | 0 | 0 | 1024400 | 840160 | 0 | 0 |
| EPSAPSIPT | PDCD6IP | Q8WUM4 | PDC6I_HUMAN | Unmodified | 10781000 | 14644000 | 3993200 | 1099500 | 0 | 0 | 0 | 0 | 4274900 | 0 | 1609200 | 2842500 | 0 | 0 | 5099300 | 6186400 | 0 | 0 |
| EPSEVPTPI | HMGA1 | P17096 | HMGA1_HUMA | Unmodified | 0 | 0 | 0 | 0 | 0 | 0 | 0 | 254640 | 0 | 0 | 93293 | 0 | 0 | 0 | 0 | 0 | 0 | 0 |
| EPTGESSG | CEMIP2 | Q9UHN6 | CEIP2_HUMAN | Unmodified | 98400 | 0 | 0 | 0 | 1677900 | 0 | 0 | 0 | 2639900 | 169900 | 0 | 0 | 0 | 0 | 757000 | 0 | 0 | 0 |
| EPVAGDAV | VGF; VGF | O15240; Q0! | VGF_HUMAN N | Unmodified | 0 | 0 | 0 | 0 | 0 | 0 | 0 | 0 | 540540 | 0 | 0 | 295560 | 0 | 0 | 0 | 0 | 0 | 0 |
| EPVETAVDI | CIAPIN1; CIAPII | Q6FI81; H3B | CPIN1_HUMAN | Unmodified | 0 | 0 | 0 | 0 | 0 | 0 | 0 | 0 | 0 | 0 | 0 | 303020 | 0 | 0 | 0 | 0 | 0 | 0 |
| EPWLLPSQ | GANAB; GANAE | Q14697; F5H | GANAB_HUMA | Unmodified | 0 | 0 | 685970 | 0 | 0 | 0 | 0 | 0 | 3753200 | 608570 | 577220 | 0 | 0 | 0 | 0 | 828970 | 0 | 0 |
| EPWLLPSQ | GANAB; GANAE | Q14697; F5H | GANAB_HUMA | Unmodified | 567320 | 76448 | 199190 | 0 | 0 | 0 | 0 | 0 | 5815300 | 0 | 0 | 1179300 | 0 | 0 | 285510 | 168050 | 235890 | 0 |
| EQADFALE. | MYH14; MYH14 | Q7Z406; M0( | MYH14_HUMA | Unmodified | 366850 | 732410 | 3600500 | 617900 | 5732800 | 0 | 1193300 | 1056900 | 608920 | 2402500 | 1062800 | 21926000 | 861370 | 5592100 | 4594000 | 3237000 | 2458800 | 0 |
| EQADFAVE | MYH10; MYH11 | P35580; P35 | MYH10_HUMA | Unmodified | 0 | 0 | 405090 | 0 | 0 | 39179 | 0 | 58611 | 0 | 346960 | 263250 | 9249200 | 0 | 0 | 1970900 | 871320 | 56644 | 0 |
| EQAEAEVA: | TPM3; TPM3; TF | D6R904; A0/ | D6R904_HUMA | Unmodified | 0 | 0 | 0 | 0 | 623620 | 290090 | 0 | 0 | 1228300 | 0 | 1493900 | 5770800 | 1008100 | 119310 | 0 | 0 | 0 | 0 |
| EQAEAEVA: | TPM3; TPM3; TF | D6R904; A0/ | D6R904_HUMA | Unmodified | 475890 | 0 | 834290 | 1059800 | 2976400 | 1617100 | 0 | 0 | 2688800 | 959810 | 648020 | 9819000 | 1220800 | 303340 | 953730 | 0 | 383110 | 657630 |
| EQAELEAAI | PLEC | Q15149 | PLEC_HUMAN | Unmodified | 0 | 0 | 101300 | 0 | 0 | 0 | 20997 | 80376 | 95902 | 0 | 33845 | 435610 | 0 | 0 | 0 | 51294 | 0 | 0 |
| EQAIGEYEI | SHTN1 | A0MZ66 | SHOT1_HUMAN | Unmodified | 175810 | 68505 | 0 | 498380 | 1352300 | 298080 | 244800 | 138800 | 2226100 | 493640 | 0 | 600390 | 823560 | 781800 | 0 | 245580 | 461740 | 710540 |
| EQALQEAM | SWAP70; SWAF | Q9UH65; E7 | SWP70_HUMAI | Oxidation (M) | 0 | 0 | 0 | 0 | 0 | 0 | 0 | 0 | 0 | 525730 | 0 | 0 | 0 | 0 | 0 | 0 | 0 | 0 |
| EQAPDSVE | CYRIA | Q9H0Q0 | CYRIA_HUMAN | Unmodified | 0 | 0 | 0 | 0 | 0 | 0 | 0 | 0 | 0 | 0 | 138890 | 1338900 | 4278700 | 0 | 0 | 0 | 0 | 0 |
| EQCCYNC( | CNBP | P62633 | CNBP_HUMAN | Unmodified | 1266900 | 0 | 0 | 0 | 0 | 0 | 0 | 0 | 0 | 0 | 0 | 662970 | 0 | 0 | 0 | 0 | 0 | 0 |
| EQCEERIEI | GOLM1 | Q8NBJ4 | GOLM1_HUMA | Unmodified | 0 | 0 | 0 | 0 | 0 | 0 | 0 | 0 | 15284000 | 0 | 0 | 0 | 0 | 0 | 0 | 0 | 0 | 0 |
| EQDAVDQ\ | TRIM29; TRIM2( | Q14134; E9I | TRI29_HUMAN | Unmodified | 0 | 0 | 163850 | 0 | 0 | 0 | 0 | 0 | 0 | 0 | 0 | 1172600 | 0 | 0 | 54125 | 59813 | 0 | 0 |
| EQDQKWQ | LMO7; LMO7; LI | Q8WWI1; E9 | LMO7_HUMAN | Unmodified | 0 | 0 | 0 | 0 | 990800 | 195750 | 0 | 0 | 0 | 70519 | 0 | 0 | 0 | 0 | 0 | 0 | 327670 | 848120 |
| EQDYLCH\ | YKT6; YKT6; YK1 | O15498; A0/ | YKT6_HUMAN : | Unmodified | 417770 | 218720 | 101370 | 0 | 0 | 0 | 0 | 0 | 468400 | 0 | 0 | 1797900 | 0 | 0 | 405090 | 586050 | 0 | 0 |
| EQEAQFEK | CTNND1; CTNN | O60716; A0/ | CTND1_HUMAN | Unmodified | 0 | 0 | 0 | 0 | 0 | 0 | 0 | 0 | 855990 | 39326 | 145990 | 970980 | 15776 | 0 | 208330 | 21541 | 0 | 0 |
| EQEHMINV | ATP5PB; ATP5P | P24539; Q5( | AT5F1_HUMAN | Unmodified | 0 | 0 | 0 | 0 | 0 | 0 | 0 | 0 | 1585100 | 279340 | 0 | 0 | 0 | 0 | 0 | 108680 | 0 | 0 |
| EQEHMINV | ATP5PB; ATP5P | P24539; Q5( | AT5F1_HUMAN | Oxidation (M) | 0 | 206250 | 0 | 0 | 0 | 0 | 0 | 0 | 188340 | 61603 | 0 | 0 | 0 | 0 | 0 | 0 | 0 | 0 |
| EQEITAVQ/ | RRBP1; RRBP1; | Q9P2E9; A0/ | RRBP1_HUMAN | Unmodified | 0 | 0 | 0 | 0 | 0 | 0 | 0 | 39263 | 76232 | 42383 | 0 | 369930 | 0 | 0 | 0 | 0 | 0 | 0 |
| EQEPMPTV | JPT2; JPT2; JPT2 | Q9H910; A6I | JUPI2_HUMAN | Unmodified | 238380 | 208910 | 236150 | 546710 | 314600 | 189500 | 0 | 0 | 0 | 0 | 34891 | 997350 | 392520 | 0 | 539460 | 550680 | 0 | 0 |
| EQERLAYIN | STIP1 | P31948 | STIP1_HUMAN | Unmodified | 118140 | 0 | 235020 | 943630 | 5191400 | 1134200 | 540000 | 103000 | 2015600 | 2521100 | 3369700 | 0 | 7603300 | 3715900 | 0 | 145540 | 2019300 | 3206800 |
| EQEVNILK | MYH9 | P35579 | MYH9_HUMAN | Unmodified | 0 | 0 | 0 | 0 | 0 | 0 | 0 | 0 | 379370 | 0 | 384650 | 0 | 0 | 0 | 0 | 0 | 0 | 0 |
| EQEVPPVII | ACTL8 | Q9H568 | ACTL8_HUMAN | Unmodified | 0 | 0 | 0 | 0 | 0 | 0 | 0 | 0 | 0 | 0 | 0 | 877910 | 0 | 0 | 0 | 0 | 0 | 0 |
| EQFASTNI/ | RAP1GDS1; RAI | P52306; D6I | GDS1_HUMAN | Unmodified | 0 | 0 | 0 | 0 | 0 | 0 | 0 | 0 | 0 | 0 | 0 | 2146400 | 0 | 0 | 0 | 0 | 0 | 0 |
| EQFGQDGI | PTPRD; PTPRS; | P23468; Q13 | PTPRD_HUMAN | Unmodified | 54664 | 0 | 0 | 0 | 0 | 0 | 0 | 0 | 0 | 0 | 0 | 0 | 0 | 0 | 54427 | 91667 | 0 | 0 |
| EQFGQDGI | PTPRF; PTPRF; I | P10586; H0\ | PTPRF_HUMAN | Unmodified | 352760 | 369230 | 244860 | 250200 | 0 | 0 | 0 | 0 | 335190 | 0 | 491990 | 0 | 0 | 0 | 1291100 | 1095900 | 0 | 65921 |
| EQFLDGDC | CALR; CALR; C/ | P27797; A0A | CALR_HUMAN | Unmodified | 0 | 0 | 0 | 1022900 | 1687100 | 0 | 2148700 | 928280 | 7.3E+08 | 98002000 | 74274000 | 12344000 | 0 | 0 | 2116300 | 64889 | 191810 | 1958400 |
| EQFLDGDC | CALR; CALR; C/ | P27797; A0A | CALR_HUMAN | Unmodified | 1957300 | 322170 | 119540 | 0 | 5254800 | 0 | 24442000 | 980600 | 4344400 | 2.59E+08 | 506370 | 0 | 3718500 | 4221500 | 156150 | 0 | 0 | 8089200 |
| EQFLDGDC | CALR; CALR; C/ | P27797; A0A | CALR_HUMAN | Unmodified | 0 | 0 | 0 | 0 | 0 | 1449100 | 1768900 | 0 | 0 | 2717600 | 0 | 0 | 0 | 953800 | 0 | 0 | 0 | 0 |
| EQFNEFRD | RCN1 | Q15293 | RCN1_HUMAN | Unmodified | 0 | 0 | 0 | 0 | 0 | 0 | 0 | 0 | 1102500 | 61024 | 0 | 3641800 | 383920 | 0 | 0 | 35830 | 0 | 0 |
| EQFWEQAI | DBN1; DBN1 | Q16643; D6I | DREB_HUMAN | Unmodified | 0 | 0 | 207680 | 0 | 0 | 0 | 0 | 0 | 0 | 0 | 0 | 1716900 | 0 | 0 | 0 | 0 | 0 | 0 |
| EQGLKGTY | SLC25A1; SLC2 | P53007; B4I | TXTP_HUMAN T | Unmodified | 0 | 0 | 0 | 0 | 0 | 0 | 0 | 17949 | 0 | 318040 | 0 | 0 | 0 | 0 | 0 | 0 | 0 | 0 |
| EQGLRLLD | PLEC; PLEC | Q15149; H0\ | PLEC_HUMAN | Unmodified | 0 | 0 | 0 | 0 | 111210 | 0 | 0 | 0 | 0 | 0 | 0 | 0 | 0 | 0 | 0 | 0 | 224620 | 0 |
| EQGPREQE | PBXIP1 | Q96AQ6 | PBIP1_HUMAN | Unmodified | 0 | 0 | 0 | 0 | 0 | 0 | 0 | 0 | 7471700 | 325980 | 1169500 | 2318600 | 0 | 0 | 0 | 0 | 0 | 0 |
| EQGPYETY | RRM1; RRM1 | P23921; E9P | RIR1_HUMAN F | Unmodified | 0 | 0 | 166460 | 122480 | 238260 | 0 | 0 | 41477 | 247890 | 111060 | 1046000 | 2953200 | 182510 | 0 | 252410 | 174400 | 0 | 0 |
| EQGQAPITI | RHOG | P84095 | RHOG_HUMAN | Unmodified | 4652400 | 3100300 | 3388300 | 4584700 | 8474100 | 1623800 | 1680500 | 1194400 | 484500 | 564750 | 3681300 | 10598000 | 5202200 | 3358800 | 2740700 | 3073300 | 3731500 | 6575500 |
| EQGQNLAF | RAP1A | P62834 | RAP1A_HUMAN | Unmodified | 0 | 0 | 0 | 420670 | 0 | 0 | 0 | 0 | 0 | 0 | 0 | 0 | 336300 | 481320 | 0 | 0 | 0 | 261660 |
| EQGQNLAF | RAP1B; N/A; RA | P61224; A6N | RAP1B_HUMAN | Unmodified | 0 | 0 | 0 | 0 | 10131000 | 2542400 | 15627000 | 5096200 | 0 | 1227000 | 0 | 0 | 9176000 | 10664000 | 0 | 0 | 2732100 | 8615600 |
| EQGVLSFW | SLC25A6; SLC2 | P12236; P05 | ADT3_HUMAN | Unmodified | 0 | 0 | 0 | 0 | 0 | 0 | 0 | 41021 | 16496000 | 3570200 | 1613500 | 74676 | 0 | 0 | 0 | 0 | 0 | 0 |
| EQGVTFPA | STAM; STAM | Q92783; A6I | STAM1_HUMAN | Unmodified | 0 | 485270 | 175510 | 721760 | 2956800 | 0 | 758660 | 354360 | 106860 | 1895900 | 0 | 534940 | 1262800 | 0 | 293330 | 0 | 0 | 0 |
| EQGVTFPS | FASN; FASN | P49327; A0A | FAS_HUMAN F | Unmodified | 0 | 259610 | 157850 | 108110 | 0 | 0 | 0 | 0 | 0 | 0 | 337350 | 3533600 | 0 | 0 | 194300 | 176010 | 0 | 0 |
| EQGWDHII | AMMECR1 | Q9Y4X0 | AMMR1_HUMA | Unmodified | 0 | 0 | 149690 | 0 | 0 | 0 | 0 | 0 | 3204500 | 19947 | 0 | 0 | 545600 | 0 | 468430 | 169550 | 0 | 0 |
| EQHEQREA | GOLIM4; GOLIN | O00461; F8V | GOLI4_HUMAN | Unmodified | 0 | 0 | 0 | 0 | 0 | 0 | 0 | 0 | 2006700 | 0 | 0 | 0 | 0 | 0 | 0 | 0 | 0 | 0 |
| EQHLTFTR | ANK3; ANK3; AI | Q12955; B1A | ANK3_HUMAN | Unmodified | 0 | 0 | 0 | 0 | 0 | 0 | 0 | 0 | 0 | 0 | 0 | 0 | 0 | 0 | 153000 | 0 | 0 | 0 |
| EQHPDMSY | EPB41; EPB41L | P11171; Q9I | EPB41_HUMAN | Unmodified | 218930 | 0 | 134970 | 194710 | 174760 | 0 | 125360 | 131940 | 348720 | 84384 | 329960 | 321740 | 261240 | 0 | 114770 | 81094 | 127030 | 1510300 |

| | | | | | | | | | | | | | | | | | | | | | | |
|---|---|---|---|---|---|---|---|---|---|---|---|---|---|---|---|---|---|---|---|---|---|---|
| EQHQKLWI | PRKCSH; PRKC | P14314; K7E | GLU2B_HUMAI | Unmodified | 0 | 0 | 0 | 0 | 1048800 | 0 | 153940 | 0 | 0 | 3259300 | 0 | 0 | 0 | 0 | 0 | 0 | 45780 | 0 |
| EQIAGLSR | DAG1 | Q14118 | DAG1_HUMAN | Unmodified | 2766100 | 0 | 0 | 1359200 | 0 | 0 | 0 | 0 | 0 | 0 | 433330 | 83709 | 0 | 746260 | 829740 | 2292000 | 0 | 0 |
| EQIDIFEGII | SUCLG2; SUCL | Q96I99; E9P | SUCB2_HUMAI | Unmodified | 0 | 0 | 0 | 0 | 0 | 0 | 0 | 0 | 2236300 | 0 | 0 | 0 | 0 | 0 | 0 | 0 | 0 | 0 |
| EQIENYK | FUT8; FUT8 | Q9BYC5; A0. | FUT8_HUMAN . | Unmodified | 0 | 0 | 0 | 0 | 135440 | 0 | 105590 | 68061 | 19444000 | 2270900 | 0 | 197730 | 0 | 0 | 0 | 0 | 46658 | 0 |
| EQIGKVDEI | GAR1 | Q9NY12 | GAR1_HUMAN | Unmodified | 0 | 0 | 0 | 0 | 0 | 0 | 0 | 0 | 0 | 0 | 0 | 221670 | 0 | 0 | 0 | 0 | 0 | 0 |
| EQILDLSK | CCDC126; MG/ | Q96EE4; Q0! | CC126_HUMAI | Unmodified | 103000 | 0 | 0 | 269890 | 187050 | 0 | 194900 | 148660 | 2417900 | 0 | 98684 | 1022100 | 0 | 0 | 0 | 0 | 0 | 184420 |
| EQILEEFSK | SYNCRIP; SYN( | O60506; B7Z | HNRPQ_HUMA | Unmodified | 71100 | 0 | 524690 | 280990 | 0 | 0 | 0 | 0 | 943990 | 498980 | 2824600 | 19174000 | 184200 | 0 | 257250 | 285110 | 0 | 0 |
| EQILGDEAI | MOGS; MOGS; | Q13724; A0/ | MOGS_HUMAN | Unmodified | 0 | 0 | 0 | 0 | 0 | 0 | 0 | 0 | 2147600 | 493730 | 285330 | 72134 | 303680 | 0 | 0 | 0 | 0 | 0 |
| EQINQLTS\ | PXDN | Q92626 | PXDN_HUMAN | Unmodified | 0 | 0 | 0 | 0 | 0 | 0 | 0 | 0 | 1686600 | 0 | 0 | 0 | 0 | 0 | 0 | 0 | 0 | 0 |
| EQISDIDD/ | SUB1 | P53999 | TCP4_HUMAN . | Unmodified | 179890 | 0 | 0 | 142830 | 0 | 0 | 0 | 0 | 961390 | 174550 | 2262600 | 20781000 | 0 | 0 | 276640 | 185340 | 0 | 0 |
| EQISIGGLA | KIDINS220; KID | Q9ULH0; H7 | KDIS_HUMAN I | Unmodified | 0 | 0 | 525800 | 93022 | 0 | 0 | 0 | 0 | 0 | 0 | 142160 | 1689000 | 0 | 0 | 449490 | 644690 | 0 | 0 |
| EQITISQLSI | KIF23; KIF23; KI | Q02241; A0/ | KIF23_HUMAN | Unmodified | 218360 | 106420 | 0 | 742200 | 404020 | 0 | 0 | 0 | 387300 | 162740 | 141040 | 17959 | 0 | 0 | 0 | 0 | 21239 | 0 |
| EQITISQLSI | KIF23; KIF23; KI | Q02241; A0/ | KIF23_HUMAN | Unmodified | 0 | 0 | 0 | 0 | 10462000 | 2324600 | 0 | 0 | 3420100 | 1070200 | 1293500 | 0 | 0 | 0 | 0 | 0 | 0 | 0 |
| EQITISQLSI | KIF23; KIF23; KI | Q02241; A0/ | KIF23_HUMAN | Unmodified | 0 | 0 | 0 | 0 | 4611700 | 1809000 | 617510 | 0 | 0 | 0 | 0 | 148320 | 0 | 0 | 0 | 0 | 443890 | 0 |
| EQIVPKPEE | RPL17; RPL17; I | P18621; A0A | RL17_HUMAN ( | Unmodified | 14150000 | 8874500 | 5526200 | 5742900 | 3316100 | 839910 | 1216000 | 1247900 | 1850700 | 935120 | 14537000 | 70674000 | 4553000 | 1677200 | 6804500 | 5945200 | 509670 | 2246400 |
| EQIVPKPEE | RPL17; RPL17; I | P18621; A0A | RL17_HUMAN ( | Unmodified | 3357700 | 2541200 | 1683800 | 0 | 1486000 | 380320 | 0 | 261800 | 597640 | 1061100 | 1285500 | 15145000 | 2163900 | 536510 | 2138600 | 1588200 | 0 | 1490200 |
| EQIVVDLSI | UGDH | O60701 | UGDH_HUMAN | Unmodified | 573640 | 430070 | 462570 | 432290 | 626620 | 0 | 0 | 0 | 1300000 | 0 | 240830 | 1256000 | 0 | 0 | 327450 | 508690 | 132990 | 0 |
| EQIYDVYR | EIF4A3; EIF4A3 | P38919; I3L: | IF4A3_HUMAN | Unmodified | 996710 | 627190 | 0 | 0 | 0 | 0 | 0 | 39111 | 0 | 0 | 152240 | 4805100 | 0 | 0 | 0 | 0 | 205640 | 0 |
| EQKLGLEN | RO60 | P10155 | RO60_HUMAN | Unmodified | 716770 | 683400 | 0 | 830540 | 0 | 0 | 65365 | 17004 | 0 | 0 | 0 | 0 | 1377200 | 497110 | 0 | 0 | 0 | 0 |
| EQKVPEILC | CARS1; CARS1 | P49589; B4D | SYCC_HUMAN | Unmodified | 0 | 0 | 0 | 0 | 0 | 0 | 0 | 0 | 85335 | 0 | 388530 | 2301500 | 0 | 0 | 0 | 20687 | 0 | 0 |
| EQLAIAEFA | TCP1; TCP1; TC | P17987; F5H | TCPA_HUMAN | Unmodified | 431740 | 469240 | 499160 | 0 | 0 | 0 | 0 | 0 | 1213200 | 0 | 1143900 | 20338000 | 0 | 0 | 653620 | 971300 | 0 | 0 |
| EQLAQAMF | RTCB | Q9Y3I0 | RTCB_HUMAN | Unmodified | 358290 | 295070 | 434380 | 721550 | 1277300 | 144620 | 222120 | 0 | 639410 | 135020 | 1055600 | 8125400 | 1569600 | 572180 | 547170 | 337900 | 0 | 506110 |
| EQLAQAMF | RTCB | Q9Y3I0 | RTCB_HUMAN | Oxidation (M) | 0 | 0 | 0 | 0 | 0 | 0 | 0 | 0 | 0 | 0 | 870200 | 4179800 | 238970 | 0 | 0 | 82043 | 45980 | 0 |
| EQLAQAVA | HDLBP; HDLBP | Q00341; A0/ | VIGLN_HUMAN | Unmodified | 375120 | 434700 | 320350 | 267900 | 491360 | 131600 | 99788 | 38246 | 82368 | 163700 | 546310 | 1821800 | 964990 | 503720 | 264160 | 389170 | 25548 | 476200 |
| EQLDSSNS | USO1 | O60763 | USO1_HUMAN | Unmodified | 0 | 0 | 0 | 0 | 0 | 0 | 2116500 | 641480 | 0 | 0 | 348890 | 1737700 | 0 | 0 | 0 | 0 | 0 | 0 |
| EQLEAEAQ. | PBXIP1 | Q96AQ6 | PBIP1_HUMAN | Unmodified | 0 | 0 | 0 | 0 | 293120 | 0 | 80217 | 0 | 4985500 | 0 | 989190 | 279020 | 0 | 0 | 0 | 0 | 121470 | 109980 |
| EQLEEEEE/ | MYH9 | P35579 | MYH9_HUMAN | Unmodified | 0 | 0 | 492190 | 0 | 0 | 0 | 0 | 34269 | 263090 | 31957 | 288040 | 586300 | 0 | 16783 | 345150 | 458510 | 0 | 0 |
| EQLEKEEQ. | N/A | N/A | N/A | Unmodified | 0 | 0 | 0 | 0 | 0 | 0 | 0 | 0 | 0 | 1997300 | 0 | 0 | 0 | 0 | 0 | 0 | 0 | 0 |
| EQLFAEAS( | VWA1; VWA1 | Q6PCB0; J3( | VWA1_HUMAN | Unmodified | 0 | 0 | 0 | 0 | 0 | 0 | 0 | 0 | 0 | 0 | 0 | 0 | 0 | 0 | 0 | 0 | 0 | 0 |
| EQLGEEIDS | DDX21 | Q9NR30 | DDX21_HUMAI | Unmodified | 0 | 0 | 0 | 0 | 0 | 0 | 0 | 0 | 0 | 0 | 355200 | 787320 | 46191 | 0 | 0 | 0 | 0 | 0 |
| EQLGYIVFS | IDE; IDE; IDE; ID | P14735; A0A | IDE_HUMAN In | Unmodified | 0 | 0 | 0 | 0 | 0 | 0 | 0 | 236810 | 0 | 0 | 0 | 3859400 | 0 | 0 | 0 | 0 | 0 | 0 |
| EQLKEELNI | TAOK1 | Q7L7X3 | TAOK1_HUMAN | Unmodified | 135390 | 181310 | 59677 | 140300 | 555550 | 0 | 214180 | 113690 | 686490 | 197100 | 113280 | 622560 | 217380 | 0 | 110660 | 164150 | 0 | 0 |
| EQLLEKEEE | PPFIA2; PPFIA2 | O75334; G3\ | LIPA2_HUMAN | Unmodified | 0 | 0 | 0 | 0 | 0 | 0 | 0 | 0 | 0 | 0 | 0 | 0 | 0 | 0 | 314390 | 347680 | 0 | 0 |
| EQLLEREEE | PPFIA3; PPFIA1 | O75145; Q1: | LIPA3_HUMAN | Unmodified | 0 | 0 | 0 | 0 | 0 | 0 | 253750 | 0 | 449830 | 0 | 0 | 2552000 | 0 | 0 | 792670 | 446040 | 0 | 0 |
| EQLLQSNP | MYO1B; MYO1/ | O43795; Q9I | MYO1B_HUMA | Unmodified | 0 | 0 | 0 | 0 | 27562000 | 9588300 | 2419900 | 846810 | 4022100 | 0 | 10815000 | 47458000 | 2075500 | 0 | 1093600 | 636120 | 8968400 | 5334400 |
| EQLMREEA | CAPRIN1; CAPF | Q14444; G3\ | CAPR1_HUMAI | Unmodified | 0 | 0 | 0 | 0 | 85578 | 0 | 0 | 0 | 628310 | 0 | 361760 | 3032800 | 0 | 0 | 0 | 0 | 0 | 0 |
| EQLNEYKG | PLEC; PLEC | Q15149; H0\ | PLEC_HUMAN | Unmodified | 0 | 0 | 0 | 16836 | 0 | 0 | 0 | 0 | 0 | 0 | 0 | 0 | 253880 | 0 | 0 | 0 | 235200 | 316610 |
| EQLNRIEE( | SNAP23; SNAP: | O00161; H3I | SNP23_HUMAN | Unmodified | 3099200 | 2240000 | 1481300 | 2890100 | 6872000 | 1436000 | 0 | 0 | 1416500 | 228460 | 3000200 | 20719000 | 2891700 | 1891900 | 4223500 | 3154300 | 1307100 | 4128500 |
| EQLQAPVP | PBXIP1; PBXIP1 | Q96AQ6; Q5 | PBIP1_HUMAN | Unmodified | 0 | 0 | 0 | 0 | 0 | 0 | 0 | 0 | 2062800 | 0 | 168940 | 0 | 0 | 0 | 0 | 0 | 0 | 0 |
| EQLQPHEN | ITGAV | P06756 | ITAV_HUMAN I | Unmodified | 164080 | 98369 | 209930 | 0 | 0 | 0 | 0 | 0 | 0 | 0 | 79234 | 1850500 | 0 | 0 | 548250 | 231080 | 0 | 0 |
| EQLSAERQ | SMC3 | Q9UQE7 | SMC3_HUMAN | Unmodified | 313600 | 0 | 0 | 0 | 340130 | 114130 | 3804000 | 0 | 0 | 0 | 0 | 1222700 | 0 | 0 | 0 | 0 | 0 | 0 |
| EQLSDMMI | IQGAP1; IQGAP | P46940; A0A | IQGA1_HUMAN | Oxidation (M) | 0 | 0 | 0 | 0 | 655190 | 0 | 0 | 0 | 0 | 0 | 0 | 577980 | 0 | 0 | 0 | 0 | 0 | 0 |
| EQLTEEAR | DNAJB11; N/A | Q9UBS4; H7 | DJB11_HUMAN | Unmodified | 0 | 0 | 0 | 0 | 0 | 0 | 0 | 0 | 2261800 | 0 | 0 | 0 | 0 | 0 | 0 | 0 | 0 | 0 |
| EQLTEEARI | DNAJB11; N/A | Q9UBS4; H7 | DJB11_HUMAN | Unmodified | 40468 | 0 | 284530 | 122360 | 1085000 | 166070 | 532180 | 330420 | 61379000 | 8193200 | 7785900 | 2347100 | 123270 | 258850 | 258140 | 0 | 19965 | 251070 |
| EQLWLANI | IQGAP1; IQGAP | P46940; A0A | IQGA1_HUMAN | Unmodified | 239340 | 389580 | 0 | 274140 | 0 | 0 | 0 | 13677 | 0 | 0 | 0 | 2025900 | 0 | 0 | 645080 | 0 | 0 | 0 |
| EQMDNAV | NIBAN2 | Q96TA1 | NIBA2_HUMAN | Oxidation (M) | 0 | 0 | 209560 | 0 | 0 | 0 | 0 | 0 | 0 | 0 | 0 | 0 | 0 | 0 | 772780 | 0 | 0 | 0 |
| EQMDQISE | EXOC1 | Q9NV70 | EXOC1_HUMAI | Unmodified | 0 | 0 | 0 | 0 | 780050 | 0 | 0 | 0 | 649030 | 0 | 0 | 1006700 | 0 | 0 | 200110 | 0 | 189610 | 0 |
| EQMEAEIAI | KTN1 | Q86UP2 | KTN1_HUMAN | Oxidation (M) | 0 | 0 | 0 | 0 | 0 | 0 | 0 | 0 | 0 | 0 | 0 | 523160 | 0 | 0 | 0 | 0 | 0 | 0 |
| EQMEAEIAI | KTN1 | Q86UP2 | KTN1_HUMAN | Unmodified | 0 | 0 | 0 | 0 | 430400 | 0 | 216360 | 102910 | 0 | 94990 | 0 | 757380 | 0 | 0 | 0 | 0 | 532940 | 0 |
| EQMEEEQK | GNPTAB | Q3T906 | GNPTA_HUMAI | Oxidation (M) | 0 | 0 | 0 | 0 | 0 | 0 | 0 | 0 | 1612700 | 0 | 0 | 0 | 0 | 0 | 0 | 0 | 0 | 0 |
| EQMQPTHI | YWHAH; YWHA | Q04917; A2I | 1433F_HUMAN | Unmodified | 96652 | 94176 | 168330 | 485560 | 0 | 0 | 0 | 46912 | 728710 | 0 | 542280 | 4610800 | 0 | 0 | 486880 | 499240 | 0 | 0 |
| EQMVDASI | MGAT1 | P26572 | MGAT1_HUMAI | Unmodified | 0 | 0 | 0 | 0 | 0 | 0 | 0 | 0 | 545350 | 0 | 0 | 0 | 0 | 0 | 0 | 0 | 0 | 0 |
| EQNIVFNAI | LRPPRC | P42704 | LPPRC_HUMAN | Unmodified | 0 | 91038 | 0 | 0 | 0 | 0 | 0 | 34900 | 1310000 | 0 | 9116.4 | 0 | 0 | 0 | 0 | 0 | 2825600 | 0 |
| EQNQDSQ | OXR1; OXR1 | Q8N573; H0 | OXR1_HUMAN | Unmodified | 0 | 0 | 0 | 0 | 0 | 0 | 0 | 0 | 619670 | 0 | 0 | 1942500 | 0 | 0 | 0 | 0 | 0 | 0 |
| EQNSPIYIS | LIN7A; LIN7C; L | O14910; Q9I | LIN7A_HUMAN | Unmodified | 434610 | 0 | 1736400 | 0 | 0 | 0 | 0 | 0 | 0 | 0 | 0 | 0 | 0 | 0 | 3210500 | 2944200 | 0 | 0 |
| EQNYSDD\ | APP; APP; APP | P05067; E9P | A4_HUMAN Am | Unmodified | 0 | 0 | 0 | 0 | 0 | 0 | 0 | 0 | 1602300 | 0 | 0 | 0 | 0 | 0 | 0 | 0 | 0 | 0 |
| EQPEKEPEI | EIF3A | Q14152 | EIF3A_HUMAN | Unmodified | 0 | 0 | 0 | 0 | 0 | 0 | 0 | 0 | 0 | 0 | 0 | 14310000 | 0 | 0 | 0 | 0 | 0 | 0 |
| EQPFGEGV | UBA3; UBA3 | Q8TBC4; F8\ | UBA3_HUMAN | Unmodified | 0 | 0 | 0 | 0 | 0 | 0 | 0 | 0 | 407630 | 0 | 0 | 776190 | 0 | 0 | 0 | 52900 | 0 | 0 |
| EQPLDEELI | ATP1A1 | P05023 | AT1A1_HUMAN | Unmodified | 684520 | 384530 | 27340000 | 3615400 | 0 | 0 | 436950 | 0 | 8856500 | 1530600 | 13583000 | 61897000 | 445660 | 0 | 18711000 | 14103000 | 0 | 0 |
| EQPWVSV( | DYNC1H1; DYN | Q14204; A0/ | DYHC1_HUMA | Unmodified | 0 | 0 | 0 | 0 | 0 | 227810 | 0 | 78892 | 0 | 0 | 0 | 3561800 | 0 | 336990 | 116260 | 61503 | 0 | 172410 |
| EQQEAIEHI | SETSIP; SET; SE | P0DME0; Q0 | SETLP_HUMAN | Unmodified | 0 | 0 | 0 | 0 | 0 | 207380 | 0 | 0 | 0 | 0 | 7765400 | 0 | 463960 | 0 | 0 | 0 | 0 | 0 |
| EQQILTLFR | LACTB2 | Q53H82 | LACB2_HUMAN | Unmodified | 0 | 0 | 96725 | 0 | 0 | 0 | 0 | 0 | 5310000 | 0 | 0 | 873720 | 0 | 0 | 0 | 0 | 0 | 0 |
| EQQIVIQSS | HSPA9; HSPA9; | P38646; H0\ | GRP75_HUMAN | Oxidation (M) | 0 | 0 | 0 | 0 | 0 | 0 | 0 | 0 | 810320 | 0 | 0 | 0 | 0 | 0 | 0 | 0 | 0 | 0 |
| EQQIVIQSS | HSPA9; HSPA9; | P38646; H0\ | GRP75_HUMAN | Unmodified | 0 | 0 | 0 | 0 | 0 | 0 | 0 | 0 | 2180100 | 0 | 0 | 0 | 0 | 0 | 0 | 0 | 0 | 0 |
| EQQLAEIE/ | NOMO1; NOMC | Q15155; P6! | NOMO1_HUM/ | Unmodified | 0 | 0 | 0 | 119370 | 0 | 0 | 0 | 0 | 11603000 | 230720 | 972550 | 749110 | 0 | 0 | 0 | 0 | 237400 | 0 |
| EQQSLQRI | MIA3; MIA3 | Q5JRA6; A0/ | TGO1_HUMAN | Unmodified | 0 | 0 | 0 | 0 | 245950 | 0 | 149320 | 0 | 554290 | 704710 | 34425 | 0 | 0 | 0 | 0 | 0 | 170840 | 0 |
| EQQTVTKLI | UACA; UACA; U | Q9BZF9; H0\ | UACA_HUMAN | Unmodified | 0 | 0 | 0 | 0 | 0 | 0 | 724140 | 43874 | 0 | 0 | 0 | 0 | 0 | 0 | 0 | 0 | 114890 | 0 |
| EQRGTFNF | EMB; EMB | Q6PCB8; D6 | EMB_HUMAN E | Unmodified | 0 | 0 | 0 | 0 | 6545200 | 1599900 | 0 | 0 | 0 | 277650 | 0 | 0 | 0 | 960930 | 0 | 0 | 0 | 0 |
| EQRILNHVI | COMT; COMT; ( | P21964; F8V | COMT_HUMAN | Unmodified | 0 | 0 | 0 | 0 | 0 | 0 | 0 | 0 | 0 | 0 | 0 | 0 | 1578700 | 1968200 | 0 | 0 | 0 | 0 |
| EQRLNFGD | STUB1; STUB1; | Q9UNE7; H3 | CHIP_HUMAN | Unmodified | 324760 | 375230 | 220430 | 1079300 | 2015300 | 409210 | 516660 | 201070 | 937580 | 680660 | 670600 | 245750 | 4492200 | 2205500 | 314360 | 162730 | 1100300 | 2545100 |
| EQRLNFGD | STUB1; STUB1; | Q9UNE7; H3 | CHIP_HUMAN | Unmodified | 0 | 0 | 0 | 0 | 11225000 | 1431800 | 8955700 | 217230 | 0 | 642120 | 0 | 347410 | 20225000 | 17407000 | 0 | 0 | 8632700 | 13017000 |
| EQRNYQFD | SF3A1 | Q15459 | SF3A1_HUMAN | Unmodified | 0 | 0 | 0 | 0 | 0 | 0 | 342390 | 0 | 0 | 997400 | 0 | 0 | 973710 | 580020 | 0 | 0 | 0 | 145930 |
| EQSGTIYL( | NUP62; NUP62 | P37198; M0( | NUP62_HUMAI | Unmodified | 0 | 0 | 0 | 0 | 996440 | 125610 | 0 | 0 | 169880 | 0 | 0 | 733040 | 0 | 0 | 0 | 0 | 0 | 252000 |
| EQSIFGDH | DBN1; DBN1 | Q16643; D6I | DREB_HUMAN | Unmodified | 0 | 0 | 0 | 0 | 0 | 0 | 0 | 0 | 88803 | 0 | 69633 | 1255000 | 0 | 0 | 0 | 0 | 0 | 0 |
| EQSIFGDH | DBN1; DBN1 | Q16643; D6I | DREB_HUMAN | Unmodified | 0 | 0 | 0 | 0 | 0 | 0 | 0 | 0 | 65294 | 0 | 94826 | 1151100 | 0 | 0 | 0 | 0 | 0 | 0 |
| EQSILELGS | PSMD11; PSMD | O00231; J3Q | PSD11_HUMAN | Unmodified | 238020 | 21118 | 449840 | 200270 | 0 | 0 | 0 | 0 | 567710 | 0 | 776440 | 6132300 | 0 | 0 | 408040 | 345440 | 0 | 0 |
| EQTADGVA | NUDT5; NUDT5 | Q9UKK9; H0 | NUDT5_HUMAI | Unmodified | 0 | 0 | 246020 | 501880 | 2564300 | 0 | 0 | 0 | 4199400 | 987640 | 5897500 | 15133000 | 1885200 | 0 | 0 | 0 | 0 | 0 |
| EQTLQGAE | NECTIN2 | Q92692 | NECT2_HUMAN | Unmodified | 0 | 0 | 0 | 0 | 1340100 | 0 | 0 | 0 | 1037500 | 0 | 0 | 0 | 0 | 0 | 0 | 0 | 357450 | 931330 |
| EQTPWDSI | ERMP1; ERMP1 | Q7Z2K6; E7E | ERMP1_HUMAI | Unmodified | 0 | 0 | 0 | 0 | 0 | 0 | 0 | 0 | 1098200 | 0 | 0 | 117910 | 0 | 0 | 0 | 0 | 0 | 28990 |
| EQVANSAF | HSP90AB1; HSI | P08238; Q58 | HS90B_HUMAN | Unmodified | 4900900 | 7472400 | 13024000 | 8959100 | 5324000 | 2573900 | 173900 | 0 | 27037000 | 2206500 | 24193000 | 1.78E+08 | 4412700 | 1143900 | 13013000 | 14905000 | 747950 | 2069100 |
| EQVANSAF | HSP90AB1; HSI | P08238; Q58 | HS90B_HUMAN | Unmodified | 0 | 0 | 0 | 949750 | 5851200 | 2414700 | 0 | 0 | 1748600 | 2571900 | 2495800 | 1684700 | 5432700 | 0 | 368940 | 164720 | 0 | 95789 |
| EQVANSAF | HSP90AB1; HSI | P08238; Q58 | HS90B_HUMAN | Unmodified | 0 | 0 | 0 | 0 | 5532900 | 0 | 0 | 0 | 0 | 0 | 0 | 0 | 1458400 | 568770 | 0 | 0 | 874750 | 2134300 |
| EQVHQGG | SPC24; SPC24; | Q8NBT2; K7I | SPC24_HUMAN | Unmodified | 198310 | 0 | 0 | 940080 | 1003800 | 253730 | 633450 | 0 | 537540 | 377040 | 0 | 732050 | 2685400 | 0 | 458610 | 168170 | 0 | 519530 |
| EQVPGFTP | MTHFD1; MTHF | P11586; V9G | C1TC_HUMAN | Unmodified | 220520 | 0 | 1631700 | 548200 | 290520 | 0 | 0 | 0 | 3757800 | 814630 | 0 | 3639400 | 0 | 0 | 0 | 0 | 3283200 | 4605100 |
| EQVSEELVI | RABEP1 | Q15276 | RABE1_HUMAN | Unmodified | 0 | 0 | 0 | 0 | 0 | 0 | 0 | 0 | 0 | 0 | 0 | 4119300 | 0 | 0 | 0 | 0 | 0 | 0 |
| EQVTNVGG | SNCA; SNCA; S | P37840; A0A | SYUA_HUMAN | Unmodified | 0 | 0 | 0 | 375010 | 0 | 0 | 0 | 111820 | 0 | 0 | 556460 | 5680300 | 0 | 0 | 0 | 0 | 0 | 0 |
| EQVVEDRP | GOLM1 | Q8NBJ4 | GOLM1_HUMA | Unmodified | 0 | 0 | 0 | 0 | 0 | 0 | 0 | 0 | 1199100 | 198850 | 0 | 0 | 0 | 0 | 0 | 0 | 0 | 0 |
| EQVVKREE | AKR1B1; AKR1E | P15121; E9P | ALDR_HUMAN | Unmodified | 0 | 0 | 0 | 0 | 0 | 0 | 0 | 0 | 0 | 0 | 0 | 3431600 | 0 | 0 | 0 | 0 | 0 | 0 |
| EQWANLE( | SPTBN1; SPTBN | Q01082; A0/ | SPTB2_HUMAN | Unmodified | 0 | 0 | 0 | 0 | 0 | 0 | 0 | 0 | 0 | 59454 | 0 | 1494800 | 0 | 0 | 0 | 0 | 0 | 0 |
| EQWCSEHI | KIF20A; KIF20A | O95235; H0\ | KI20A_HUMAN | Unmodified | 0 | 0 | 0 | 0 | 20975 | 0 | 0 | 0 | 167880 | 20060 | 0 | 74977 | 0 | 0 | 136080 | 0 | 0 | 0 |
| EQWDTIEE | DECR1 | Q16698 | DECR_HUMAN | Unmodified | 0 | 0 | 0 | 0 | 0 | 0 | 0 | 0 | 12122000 | 414430 | 0 | 0 | 0 | 0 | 0 | 0 | 0 | 197740 |
| EQWTKYEE | UQCRB | P14927 | QCR7_HUMAN | Unmodified | 0 | 0 | 0 | 25949 | 0 | 0 | 0 | 0 | 8026500 | 3133900 | 0 | 0 | 0 | 0 | 0 | 0 | 0 | 0 |
| EQYAQEH( | PMVK | Q15126 | PMVK_HUMAN | Unmodified | 875520 | 0 | 0 | 225200 | 363250 | 0 | 16773 | 0 | 923810 | 121540 | 199100 | 891680 | 432540 | 0 | 0 | 351650 | 0 | 150450 |
| EQYAQEH( | PMVK | Q15126 | PMVK_HUMAN | Unmodified | 0 | 0 | 0 | 0 | 1233900 | 180890 | 1347500 | 0 | 0 | 425480 | 0 | 0 | 2111900 | 1442900 | 0 | 0 | 640410 | 4157100 |
| ERACYLSIN | ACTR1A; ACTR1 | P61163; R4G | ACTZ_HUMAN . | Unmodified | 0 | 0 | 0 | 0 | 0 | 0 | 0 | 0 | 0 | 0 | 0 | 4758900 | 0 | 0 | 279510 | 0 | 0 | 0 |
| ERAEQESAI | PLEC | Q15149 | PLEC_HUMAN | Unmodified | 0 | 0 | 0 | 0 | 270990 | 143280 | 0 | 0 | 0 | 74580 | 0 | 0 | 0 | 0 | 0 | 0 | 414280 | 343400 |
| ERAMAEGK | PPL; PPL; PPL | O60437; K7E | PEPL_HUMAN I | Unmodified | 0 | 0 | 0 | 0 | 842990 | 60680 | 81465 | 46675 | 0 | 0 | 0 | 0 | 0 | 0 | 0 | 0 | 709050 | 151950 |
| ERAMSTTSI | DBNL; DBNL; D | Q9UJU6; B4I | DBNL_HUMAN | Unmodified | 0 | 0 | 0 | 0 | 751330 | 127410 | 139090 | 86185 | 507920 | 67885 | 0 | 657180 | 36832 | 648900 | 0 | 0 | 0 | 805590 |
| ERAMSTTSI | DBNL; DBNL; D | Q9UJU6; B4I | DBNL_HUMAN | Oxidation (M) | 163890 | 136210 | 136800 | 207570 | 649920 | 0 | 0 | 0 | 0 | 49533 | 0 | 287440 | 637480 | 780230 | 0 | 269780 | 0 | 0 |
| ERARLQLEI | LMNA; LMNA; L | P02545; A0A | LMNA_HUMAN | Unmodified | 327180 | 576280 | 103700 | 997390 | 2240500 | 184000 | 174900 | 0 | 533260 | 2710000 | 0 | 0 | 61882 | 400220 | 0 | 125850 | 1552400 | 371300 |
| ERASLPTL( | ADSL; ADSL; AD | P30566; A0A | PUR8_HUMAN | Unmodified | 0 | 16695 | 0 | 28488 | 0 | 0 | 467480 | 40619 | 0 | 0 | 0 | 435810 | 157010 | 68700 | 81970 | 64018 | 0 | 0 |
| ERCRIEPHI | DSP | P15924 | DESP_HUMAN | Unmodified | 0 | 0 | 0 | 0 | 118900 | 0 | 0 | 0 | 0 | 738380 | 0 | 0 | 0 | 0 | 0 | 0 | 1000700 | 349740 |
| ERDFEDPY | SHB | Q15464 | SHB_HUMAN S | Unmodified | 0 | 0 | 0 | 0 | 598540 | 0 | 0 | 0 | 0 | 0 | 0 | 0 | 0 | 0 | 0 | 0 | 808530 | 1255700 |
| ERDFTSLEI | CKAP4 | Q07065 | CKAP4_HUMAN | Unmodified | 0 | 18607 | 0 | 0 | 0 | 0 | 0 | 0 | 3341200 | 347180 | 0 | 0 | 0 | 0 | 0 | 0 | 0 | 0 |
| ERDFTSLEI | CKAP4 | Q07065 | CKAP4_HUMAN | Unmodified | 0 | 0 | 0 | 0 | 66426 | 221960 | 0 | 0 | 2331900 | 1133600 | 0 | 0 | 0 | 0 | 0 | 0 | 0 | 0 |
| ERDGQSSF | CARMIL1; CARN | Q5VZK9; A0/ | CARL1_HUMAN | Unmodified | 0 | 46814 | 0 | 0 | 0 | 0 | 0 | 0 | 0 | 34180 | 173730 | 160820 | 0 | 0 | 0 | 0 | 0 | 0 |
| ERDSASFN | ACOX1; ACOX1 | Q15067; I3L( | ACOX1_HUMAI | Unmodified | 0 | 0 | 0 | 0 | 0 | 0 | 821640 | 0 | 17995000 | 2275200 | 5249500 | 0 | 0 | 0 | 0 | 0 | 0 | 0 |
| ERDVEELQ | CLIP1; CLIP1; C | P30622; F5H | CLIP1_HUMAN | Unmodified | 0 | 0 | 0 | 0 | 654760 | 0 | 155240 | 198110 | 0 | 0 | 0 | 246790 | 0 | 0 | 0 | 0 | 0 | 0 |
| ERDVILREN | UQCRC1 | P31930 | QCR1_HUMAN | Unmodified | 0 | 0 | 0 | 0 | 0 | 0 | 0 | 0 | 2309100 | 0 | 0 | 0 | 0 | 0 | 0 | 0 | 0 | 0 |

| | | | | | | | | | | | | | | | | | | | | | | |
|---|---|---|---|---|---|---|---|---|---|---|---|---|---|---|---|---|---|---|---|---|---|---|
| ERDVYEELI | IQGAP1; IQGAP | P46940; A0A | IQGA1_HUMAN | Unmodified | 3483200 | 0 | 0 | 0 | 0 | 0 | 0 | 0 | 653780 | 0 | 0 | 1245700 | 0 | 0 | 560550 | 603950 | 616470 | 0 |
| EREAELGAI | PABPC1; PABP( | P11940; Q9I | PABP1_HUMAN | Unmodified | 0 | 0 | 3411200 | 1660900 | 0 | 0 | 0 | 0 | 0 | 0 | 0 | 0 | 0 | 743460 | 1212800 | 658610 | 0 | 0 |
| EREAELGAI | PABPC1; PABP( | P11940; Q9I | PABP1_HUMAN | Unmodified | 0 | 0 | 364780 | 58647 | 1866400 | 0 | 999390 | 5374.9 | 21536 | 0 | 0 | 113310 | 3846000 | 817240 | 0 | 0 | 0 | 0 |
| EREEETPAY | OS9; OS9; OS9; | Q13438; B4E | OS9_HUMAN P | Unmodified | 0 | 0 | 0 | 0 | 0 | 0 | 0 | 0 | 8304400 | 0 | 0 | 249130 | 0 | 0 | 0 | 0 | 0 | 0 |
| EREEETPAY | OS9; OS9; OS9; | Q13438; B4E | OS9_HUMAN P | Oxidation (M) | 0 | 0 | 0 | 0 | 1311500 | 0 | 0 | 0 | 901690 | 0 | 0 | 0 | 0 | 0 | 0 | 0 | 0 | 0 |
| EREEETPAY | OS9; OS9; OS9; | Q13438; B4E | OS9_HUMAN P | Unmodified | 0 | 0 | 0 | 0 | 0 | 0 | 0 | 0 | 1809900 | 0 | 259090 | 0 | 0 | 0 | 0 | 0 | 0 | 0 |
| EREEHLLE( | TRIM26; TRIM2( | Q12899; A0A | TRI26_HUMAN | Unmodified | 0 | 0 | 0 | 0 | 0 | 0 | 0 | 0 | 0 | 0 | 0 | 0 | 0 | 0 | 565940 | 0 | 0 | 0 |
| EREELEQA( | SLC38A10 | Q9HBR0 | S38AA_HUMAN | Unmodified | 285320 | 276700 | 993520 | 41144 | 162160 | 0 | 0 | 0 | 8984600 | 343400 | 84937 | 274620 | 0 | 0 | 828430 | 723180 | 0 | 0 |
| EREENVKAI | CAND1; CAND1 | Q86VP6; A0A | CAND1_HUMA | Unmodified | 0 | 0 | 0 | 0 | 1644500 | 0 | 0 | 0 | 764100 | 1191400 | 276360 | 0 | 1907800 | 1624200 | 0 | 0 | 952200 | 1920300 |
| EREGLEND | PDCD6IP | Q8WUM4 | PDC6I_HUMAN | Unmodified | 27290000 | 32964000 | 5348600 | 3716800 | 1725500 | 934840 | 0 | 0 | 16101000 | 4066500 | 0 | 7072900 | 1962800 | 1894600 | 5493000 | 12986000 | 389080 | 1099600 |
| EREGLEND | PDCD6IP | Q8WUM4 | PDC6I_HUMAN | Unmodified | 0 | 0 | 0 | 60165 | 199090 | 0 | 233130 | 0 | 0 | 88477 | 0 | 0 | 280890 | 273600 | 0 | 0 | 132150 | 0 |
| EREKYDNM | VPS28; VPS28; | Q9UK41; E9I | VPS28_HUMAN | Unmodified | 7229700 | 9147800 | 0 | 6018400 | 3203700 | 0 | 85392 | 0 | 3775900 | 1388500 | 0 | 326020 | 10592000 | 786420 | 1320500 | 998390 | 135860 | 1090200 |
| EREKYDNM | VPS28; VPS28; | Q9UK41; E9I | VPS28_HUMAN | Oxidation (M) | 4758800 | 4987900 | 0 | 6678500 | 173780 | 0 | 0 | 0 | 0 | 105630 | 0 | 0 | 3179800 | 1153400 | 883180 | 1665000 | 0 | 492020 |
| ERELQHAA | SCAMP3 | O14828 | SCAM3_HUMA | Unmodified | 334540 | 385540 | 688080 | 495170 | 0 | 0 | 0 | 0 | 0 | 0 | 115580 | 1545000 | 0 | 0 | 249430 | 280770 | 0 | 0 |
| ERELQNTV. | SCAMP2; SCAM | O15127; H3I | SCAM2_HUMA | Unmodified | 0 | 595220 | 3058500 | 2085200 | 572220 | 0 | 0 | 0 | 901880 | 0 | 0 | 0 | 55441 | 0 | 1787200 | 2132800 | 0 | 0 |
| ERELQQLG | SLITRK6 | Q9H5Y7 | SLIK6_HUMAN | Unmodified | 4961000 | 4613600 | 0 | 0 | 0 | 0 | 0 | 0 | 9436900 | 6034000 | 0 | 0 | 0 | 0 | 0 | 197980 | 519350 | 1679800 |
| EREMQNLS | SCAMP1; SCAM | O15126; A0A | SCAM1_HUMA | Unmodified | 0 | 0 | 815150 | 321910 | 0 | 0 | 0 | 0 | 126790 | 0 | 0 | 381600 | 0 | 0 | 89648 | 360460 | 0 | 0 |
| EREPEPEP\ | MIA3 | Q5JRA6 | TGO1_HUMAN | Unmodified | 0 | 0 | 0 | 0 | 0 | 0 | 0 | 161040 | 7161400 | 767080 | 0 | 0 | 0 | 0 | 0 | 0 | 0 | 0 |
| EREQQIEEI | DBN1; DBN1 | Q16643; D6I | DREB_HUMAN | Unmodified | 0 | 0 | 133310 | 0 | 0 | 0 | 0 | 0 | 0 | 0 | 0 | 2673600 | 0 | 0 | 228830 | 0 | 0 | 0 |
| ERESLQQM | PRKCSH; PRKC | P14314; K7E | GLU2B_HUMAI | Unmodified | 98060 | 0 | 163570 | 71800 | 292180 | 209940 | 0 | 14318 | 2.79E+08 | 15931000 | 2963000 | 2743100 | 112020 | 103030 | 190230 | 177800 | 37310 | 103140 |
| ERESLQQM | PRKCSH; PRKC | P14314; K7E | GLU2B_HUMAI | Oxidation (M) | 33774 | 27909 | 0 | 0 | 107570 | 0 | 159150 | 38666 | 28290000 | 2424700 | 357780 | 667150 | 0 | 0 | 0 | 99693 | 0 | 0 |
| ERESLQQM | PRKCSH; PRKC | P14314; K7E | GLU2B_HUMAI | Unmodified | 0 | 0 | 0 | 0 | 0 | 0 | 228230 | 0 | 84524000 | 19996000 | 1019000 | 0 | 165650 | 0 | 0 | 0 | 0 | 299000 |
| ERESLQQM | PRKCSH; PRKC | P14314; K7E | GLU2B_HUMAI | Oxidation (M) | 0 | 0 | 0 | 0 | 46632 | 0 | 0 | 0 | 3836400 | 1598900 | 125610 | 0 | 0 | 0 | 0 | 0 | 0 | 0 |
| ERETAIRQL | STX12 | Q86Y82 | STX12_HUMAN | Unmodified | 0 | 0 | 0 | 0 | 0 | 0 | 179520 | 0 | 0 | 81746 | 0 | 0 | 168030 | 348860 | 0 | 0 | 0 | 0 |
| ERETALDIL | NF2; NF2 | P35240; A0A | MERL_HUMAN | Unmodified | 108370 | 0 | 646790 | 1015900 | 3824300 | 0 | 0 | 0 | 283650 | 0 | 0 | 496600 | 2149700 | 1342200 | 593980 | 790470 | 2660900 | 14230000 |
| EREVLQLQ | MRPS26 | Q9BYN8 | RT26_HUMAN : | Unmodified | 0 | 0 | 0 | 0 | 0 | 0 | 0 | 0 | 1635800 | 241180 | 0 | 0 | 163760 | 0 | 0 | 0 | 107590 | 0 |
| ERFGIDDQ | PIP4K2A | P48426 | PI42A_HUMAN | Unmodified | 0 | 274650 | 0 | 560090 | 0 | 0 | 0 | 0 | 0 | 0 | 383280 | 2859400 | 228520 | 0 | 198330 | 217690 | 116030 | 434310 |
| ERFGIDDQ | PIP4K2B; PIP4K | P78356; A0A | PI42B_HUMAN | Unmodified | 0 | 0 | 0 | 0 | 0 | 0 | 0 | 0 | 0 | 0 | 0 | 1160500 | 0 | 0 | 0 | 156460 | 0 | 0 |
| ERFNIDMPI | PRKCD; PRKCD | Q05655; A0A | KPCD_HUMAN | Unmodified | 0 | 0 | 0 | 0 | 0 | 0 | 0 | 0 | 283370 | 0 | 0 | 716630 | 0 | 0 | 0 | 0 | 0 | 0 |
| ERFQFPAQ | CDC42BPA; CD | Q5VT25; A0A | MRCKA_HUMA | Unmodified | 0 | 0 | 0 | 1435200 | 1869800 | 0 | 0 | 0 | 1195600 | 1002600 | 432230 | 0 | 1512300 | 2337200 | 842790 | 819480 | 2596600 | 2754500 |
| ERFSPLTTN | ATP5PO; ATP5P | P48047; A0A | ATPO_HUMAN | Unmodified | 0 | 0 | 0 | 0 | 25141 | 0 | 0 | 0 | 5227600 | 127160 | 95743 | 116180 | 0 | 0 | 0 | 0 | 0 | 0 |
| ERGALTVG | GFPT1; GFPT1; | Q06210; A0A | GFPT1_HUMAN | Unmodified | 178490 | 156250 | 191390 | 246630 | 0 | 0 | 0 | 0 | 0 | 0 | 0 | 1401700 | 0 | 0 | 238350 | 286670 | 0 | 0 |
| ERGALTVG | GFPT1; GFPT1; | Q06210; A0A | GFPT1_HUMAN | Unmodified | 0 | 0 | 0 | 0 | 0 | 0 | 0 | 0 | 0 | 480230 | 0 | 0 | 2870900 | 0 | 0 | 0 | 0 | 0 |
| ERGDFNHI | ATP1B1; ATP1B | P05026; V9C | AT1B1_HUMAN | Unmodified | 0 | 0 | 270030 | 0 | 774080 | 0 | 0 | 0 | 896170 | 0 | 0 | 3495200 | 0 | 0 | 0 | 622930 | 0 | 533760 |
| ERGDYVLA | FLNB; FLNB | O75369; E7E | FLNB_HUMAN | Unmodified | 0 | 0 | 0 | 0 | 19808000 | 3435700 | 2079100 | 0 | 0 | 401020 | 0 | 0 | 0 | 1010200 | 0 | 0 | 5245600 | 2575600 |
| ERGGNQTS | SMG9; SMG9 | Q9H0W8; M( | SMG9_HUMAN | Unmodified | 0 | 0 | 0 | 0 | 568380 | 0 | 0 | 0 | 0 | 0 | 0 | 176180 | 0 | 0 | 0 | 0 | 95759 | 0 |
| ERHGNVAS | CD2AP | Q9Y5K6 | CD2AP_HUMAI | Unmodified | 478920 | 587150 | 244450 | 655720 | 0 | 0 | 0 | 0 | 0 | 0 | 0 | 0 | 1098700 | 0 | 401350 | 283050 | 0 | 0 |
| ERHLEHEC | TRAF2; TRAF2; T | Q12933; B1A | TRAF2_HUMAN | Unmodified | 0 | 0 | 20709 | 0 | 754430 | 335180 | 0 | 0 | 0 | 0 | 0 | 0 | 0 | 0 | 0 | 0 | 0 | 0 |
| ERHPGSFD | RPS4X; RPS4Y1 | P62701; P22 | RS4X_HUMAN | Unmodified | 535740 | 711100 | 669000 | 216860 | 0 | 0 | 0 | 0 | 0 | 0 | 301820 | 6576300 | 0 | 0 | 776210 | 0 | 117560 | 0 |
| ERHTQLEQ | SBDS; SBDS | Q9Y3A5; A0A | SBDS_HUMAN | Unmodified | 63048 | 0 | 0 | 180790 | 404560 | 0 | 78640 | 0 | 66589 | 81925 | 0 | 0 | 268360 | 189150 | 0 | 0 | 0 | 314120 |
| ERHVSIQE/ | RAB21 | Q9UL25 | RAB21_HUMAN | Unmodified | 968230 | 3730100 | 2500800 | 5054800 | 15093000 | 2260700 | 2436000 | 0 | 7131300 | 3510700 | 3004200 | 11041000 | 3480300 | 3830800 | 2712600 | 2741100 | 478760 | 4674400 |
| ERIDFLPHY | SMCHD1 | A6NHR9 | SMHD1_HUMA | Unmodified | 100750 | 0 | 0 | 0 | 0 | 0 | 757880 | 110760 | 0 | 0 | 0 | 349020 | 0 | 0 | 0 | 0 | 0 | 0 |
| ERIDTRNEI | HSPA5; HSPA5; | P11021; A0A | BIP_HUMAN En | Unmodified | 382440 | 108700 | 3138400 | 0 | 2044900 | 0 | 6815900 | 48947 | 93615000 | 92844000 | 17572000 | 4083700 | 898370 | 407420 | 2994800 | 521220 | 93360 | 236730 |
| ERIEIEQNY | FNBP1L | Q5T0N5 | FBP1L_HUMAN | Unmodified | 2592000 | 2887600 | 0 | 0 | 1051900 | 0 | 0 | 0 | 0 | 480990 | 0 | 519410 | 1299100 | 0 | 76696 | 768950 | 1065100 | 1212800 |
| ERIGQFVF/ | AASDHPPT | Q9NRN7 | ADPPT_HUMAN | Unmodified | 0 | 0 | 0 | 0 | 0 | 0 | 0 | 0 | 0 | 0 | 385380 | 1689400 | 0 | 0 | 79787 | 0 | 0 | 0 |
| ERIGYPDDI | MME; MME; MM | P08473; A0A | NEP_HUMAN N | Unmodified | 0 | 0 | 9790400 | 7991100 | 632030 | 3464500 | 2286400 | 22516 | 2498800 | 395170 | 0 | 160260 | 1195800 | 0 | 13186000 | 10303000 | 200310 | 409520 |
| ERILQEHE( | EIF3A | Q14152 | EIF3A_HUMAN | Unmodified | 0 | 0 | 0 | 0 | 54787 | 0 | 0 | 0 | 0 | 25735 | 0 | 2675800 | 0 | 41697 | 0 | 0 | 43369 | 61258 |
| ERIPEAPAG | TLN1 | Q9Y490 | TLN1_HUMAN | Unmodified | 0 | 0 | 0 | 0 | 0 | 0 | 0 | 0 | 1205700 | 0 | 394500 | 3000000 | 0 | 1293600 | 0 | 0 | 0 | 0 |
| ERIQNAGG | PPM1A; PPM1B | P35813; O75 | PPM1A_HUMAI | Oxidation (M) | 0 | 0 | 0 | 214290 | 0 | 0 | 0 | 54689 | 75579 | 0 | 0 | 400510 | 0 | 0 | 1129000 | 792490 | 0 | 0 |
| ERIQNAGG | PPM1A; PPM1B | P35813; O75 | PPM1A_HUMAI | Unmodified | 0 | 0 | 0 | 588930 | 0 | 0 | 35863 | 0 | 440500 | 0 | 0 | 0 | 0 | 0 | 0 | 886340 | 105310 | 0 |
| ERISEQTY( | STXBP1; STXBP | P61764; A0A | STXB1_HUMAN | Unmodified | 0 | 62949 | 266650 | 337040 | 555620 | 242580 | 492560 | 253470 | 2267000 | 547260 | 241680 | 3819600 | 585070 | 136740 | 0 | 0 | 122130 | 397490 |
| ERIVAECN/ | CTNNA1; CTNN | P35221; P26 | CTNA1_HUMAI | Unmodified | 339860 | 0 | 0 | 1403700 | 441220 | 245150 | 0 | 0 | 63900 | 0 | 573180 | 1458500 | 554200 | 182350 | 1741300 | 2005800 | 36320 | 58289 |
| ERIVGWYH | PSMD7; PSMD7 | P51665; H3E | PSMD7_HUMA | Unmodified | 0 | 0 | 0 | 0 | 42630 | 0 | 0 | 0 | 68671 | 0 | 0 | 1767200 | 186810 | 0 | 0 | 41907 | 0 | 73305 |
| ERKEEDSH | PLPP1 | O14494 | PLPP1_HUMAN | Unmodified | 0 | 0 | 3620600 | 7109100 | 425140 | 0 | 0 | 0 | 0 | 0 | 0 | 3414000 | 0 | 0 | 13016000 | 7809200 | 0 | 0 |
| ERLAGLFNI | HYOU1; HYOU1 | Q9Y4L1; E9F | HYOU1_HUMA | Unmodified | 0 | 0 | 0 | 0 | 0 | 0 | 0 | 0 | 4656800 | 0 | 0 | 0 | 0 | 0 | 0 | 0 | 0 | 0 |
| ERLAGLFNI | HYOU1; HYOU1 | Q9Y4L1; E9F | HYOU1_HUMA | Unmodified | 0 | 0 | 0 | 0 | 187520 | 0 | 224050 | 0 | 3280100 | 1316900 | 0 | 296040 | 0 | 116480 | 122300 | 0 | 0 | 0 |
| ERLAILGIHI | ACTN1; ACTN1; | P12814; A0A | ACTN1_HUMAI | Unmodified | 0 | 0 | 0 | 0 | 447400 | 475300 | 0 | 0 | 0 | 0 | 113130 | 5720300 | 268660 | 0 | 202430 | 149890 | 462430 | 0 |
| ERLASLSSI | PPFIA1; PPFIA1 | Q13136; E9F | LIPA1_HUMAN | Unmodified | 0 | 0 | 0 | 491390 | 5201500 | 0 | 204030 | 0 | 1584200 | 0 | 0 | 469420 | 650960 | 0 | 0 | 0 | 335870 | 892990 |
| ERLDEPHF | PPM1L | Q5SGD2 | PPM1L_HUMAI | Unmodified | 0 | 0 | 0 | 0 | 0 | 0 | 78482 | 0 | 8397300 | 1616200 | 119540 | 0 | 0 | 0 | 0 | 0 | 0 | 0 |
| ERLDQPMT | UBA1; UBA1 | P22314; Q5J | UBA1_HUMAN | Unmodified | 184640 | 136130 | 376830 | 370450 | 435770 | 77585 | 356560 | 0 | 2108400 | 470290 | 786990 | 5249100 | 426960 | 0 | 655960 | 489670 | 279180 | 948740 |
| ERLDQPMT | UBA1; UBA1 | P22314; Q5J | UBA1_HUMAN | Oxidation (M) | 0 | 0 | 0 | 0 | 0 | 0 | 0 | 0 | 0 | 0 | 0 | 780180 | 0 | 0 | 0 | 0 | 0 | 0 |
| ERLDQPMT | UBA1; UBA1 | P22314; Q5J | UBA1_HUMAN | Unmodified | 0 | 0 | 0 | 0 | 2017000 | 0 | 147860 | 275890 | 0 | 358140 | 0 | 0 | 1557000 | 0 | 0 | 0 | 935310 | 4024900 |
| ERLEHPVLI | SUMF2; SUMF2 | Q8NBJ7; C9J | SUMF2_HUMAI | Unmodified | 0 | 0 | 0 | 0 | 0 | 0 | 0 | 0 | 31743000 | 1821500 | 0 | 0 | 0 | 0 | 0 | 46493 | 0 | 0 |
| ERLEQEQLI | ENAH; ENAH; E | Q8N8S7; A0A | ENAH_HUMAN | Unmodified | 76841 | 0 | 0 | 140490 | 163900 | 0 | 0 | 84751 | 480780 | 0 | 1467500 | 2314100 | 0 | 0 | 0 | 865980 | 75291 | 85622 |
| ERLEQQVP | RPL3; RPL3; RP | P39023; B5N | RL3_HUMAN 6( | Oxidation (M) | 9412300 | 6161000 | 689030 | 8046100 | 0 | 0 | 0 | 36993 | 453910 | 0 | 5911900 | 11689000 | 2572300 | 0 | 3062500 | 2665800 | 0 | 0 |
| ERLESLNIQ | EIF3A | Q14152 | EIF3A_HUMAN | Unmodified | 0 | 0 | 256020 | 0 | 0 | 0 | 0 | 0 | 0 | 0 | 0 | 3499900 | 0 | 0 | 160490 | 0 | 0 | 0 |
| ERLESLNIQ | EIF3A | Q14152 | EIF3A_HUMAN | Unmodified | 55697 | 0 | 337890 | 85073 | 0 | 0 | 123200 | 0 | 252070 | 137880 | 0 | 626840 | 0 | 0 | 188440 | 121630 | 0 | 0 |
| ERLFGSLQI | USP43; USP43 | Q70EL4; V9C | UBP43_HUMAI | Unmodified | 0 | 0 | 0 | 0 | 0 | 0 | 0 | 0 | 0 | 0 | 0 | 0 | 0 | 0 | 0 | 238070 | 0 | 0 |
| ERLLLAEK | VPS36 | Q86VN1 | VPS36_HUMAN | Unmodified | 1077500 | 1519900 | 2853400 | 0 | 223070 | 0 | 0 | 0 | 122900 | 124250 | 0 | 248820 | 3732000 | 1059700 | 1768600 | 1625000 | 0 | 0 |
| ERLSVYQAI | PLEC; PLEC | Q15149; H0Y | PLEC_HUMAN | Unmodified | 0 | 0 | 0 | 0 | 0 | 0 | 0 | 0 | 0 | 132730 | 0 | 2659900 | 0 | 0 | 0 | 0 | 572370 | 0 |
| ERMEPTYQ | STXBP2; STXBP | Q15833; M0I | STXB2_HUMAN | Unmodified | 73522 | 151860 | 147330 | 309900 | 144090 | 0 | 0 | 0 | 425220 | 105560 | 151830 | 952560 | 171070 | 39761 | 239290 | 155370 | 94725 | 267820 |
| ERMEVCEN | APLP2; APLP2 | Q06481; E9F | APLP2_HUMAN | Unmodified | 0 | 0 | 0 | 0 | 1201600 | 283510 | 217390 | 0 | 1124100 | 706070 | 196350 | 0 | 860680 | 725840 | 0 | 0 | 36192 | 429490 |
| ERMEVCEN | APLP2; APLP2 | Q06481; E9F | APLP2_HUMAN | Oxidation (M) | 39979 | 0 | 0 | 0 | 0 | 0 | 0 | 0 | 426170 | 350920 | 0 | 0 | 0 | 369030 | 0 | 0 | 0 | 0 |
| ERNDLIVFL | ITFG1 | Q8TB96 | TIP_HUMAN T-( | Unmodified | 0 | 0 | 588020 | 0 | 0 | 0 | 0 | 0 | 18240000 | 1729300 | 0 | 0 | 0 | 0 | 0 | 0 | 0 | 0 |
| ERNGLSLA | H1-5 | P16401 | H15_HUMAN H | Unmodified | 6430900 | 3326400 | 0 | 0 | 592620 | 0 | 517590 | 10682000 | 0 | 243990 | 0 | 0 | 0 | 0 | 0 | 0 | 0 | 0 |
| ERNPLTPA! | STS; STS | P08842; A0A | STS_HUMAN S | Unmodified | 0 | 0 | 0 | 0 | 0 | 0 | 0 | 0 | 565570 | 0 | 0 | 0 | 0 | 0 | 0 | 136570 | 0 | 0 |
| ERNQNEAL | TTC4 | O95801 | TTC4_HUMAN | Unmodified | 45734 | 156550 | 0 | 0 | 0 | 0 | 0 | 0 | 0 | 0 | 0 | 0 | 355150 | 0 | 0 | 323000 | 162910 | 152640 |
| ERNTDQAS | MYH9; MYH11; | P35579; P35 | MYH9_HUMAN | Unmodified | 0 | 0 | 1159300 | 1051000 | 6131300 | 2076500 | 1040900 | 0 | 932820 | 881340 | 0 | 10329000 | 949970 | 945280 | 1878500 | 1035200 | 1963500 | 0 |
| ERNTDQAS | MYH10 | P35580 | MYH10_HUMAI | Unmodified | 0 | 0 | 254900 | 211700 | 720170 | 219830 | 207490 | 0 | 0 | 33213 | 0 | 1182200 | 0 | 188890 | 406610 | 272440 | 103670 | 145940 |
| ERNVEDLS | ABCE1; ABCE1 | P61221; D6F | ABCE1_HUMAI | Unmodified | 4558300 | 4003600 | 3480300 | 2955100 | 640080 | 144010 | 610680 | 179790 | 398450 | 349450 | 345620 | 6517900 | 1588800 | 640910 | 1436400 | 1796900 | 152170 | 120630 |
| ERPAYAVH | COPA; COPA; C | P53621; A0A | COPA_HUMAN | Unmodified | 0 | 84465 | 102930 | 0 | 0 | 0 | 0 | 0 | 0 | 0 | 0 | 2261900 | 0 | 0 | 0 | 0 | 0 | 0 |
| ERPAYAVH | COPA; COPA; C | P53621; A0A | COPA_HUMAN | Oxidation (M) | 419320 | 0 | 0 | 0 | 0 | 0 | 0 | 0 | 0 | 0 | 0 | 474090 | 0 | 0 | 0 | 0 | 0 | 0 |
| ERPDLPPI( | SEC23A; SEC23 | Q15436; F5I | SC23A_HUMAN | Unmodified | 0 | 0 | 0 | 0 | 0 | 0 | 0 | 46523 | 0 | 0 | 0 | 840350 | 0 | 0 | 0 | 0 | 0 | 0 |
| ERPEAPDQ | AKT2; AKT2; AK | P31751; A8M | AKT2_HUMAN | Oxidation (M) | 0 | 0 | 795050 | 0 | 0 | 0 | 0 | 0 | 0 | 0 | 0 | 294910 | 735120 | 0 | 0 | 154610 | 0 | 0 |
| ERPETVLID | TAOK1 | Q7L7X3 | TAOK1_HUMAN | Unmodified | 508430 | 0 | 555760 | 1259600 | 0 | 0 | 0 | 0 | 1343000 | 0 | 745380 | 3795100 | 147730 | 0 | 0 | 577800 | 0 | 0 |
| ERPGLGSE | DDX42; DDX42 | Q86XP3; A0A | DDX42_HUMAI | Unmodified | 0 | 0 | 0 | 0 | 0 | 0 | 0 | 0 | 0 | 107220 | 109870 | 124660 | 0 | 0 | 53877 | 0 | 0 | 0 |
| ERPISLGIFI | SPAG9; SPAG9; | O60271; D6I | JIP4_HUMAN C | Unmodified | 0 | 0 | 0 | 0 | 115470 | 0 | 0 | 0 | 28493 | 0 | 23481 | 0 | 0 | 0 | 0 | 0 | 92945 | 788220 |
| ERPLCPLQI | CLUH; CLUH; C | O75153; I3L | CLU_HUMAN C | Unmodified | 0 | 0 | 0 | 0 | 0 | 0 | 0 | 0 | 0 | 0 | 120770 | 2456000 | 0 | 0 | 0 | 0 | 0 | 0 |
| ERPSSAIYP | ARHGEF2; ARH | Q92974; V9C | ARHG2_HUMA | Unmodified | 0 | 0 | 0 | 0 | 0 | 0 | 0 | 0 | 0 | 0 | 305450 | 1293500 | 0 | 0 | 0 | 53406 | 0 | 0 |
| ERPTPSLNI | EPRS1; EPRS1 | P07814; V9C | SYEP_HUMAN | Unmodified | 0 | 0 | 0 | 0 | 0 | 0 | 0 | 0 | 412060 | 0 | 0 | 10815000 | 0 | 0 | 0 | 0 | 0 | 0 |
| ERQAALQV | CAPG; CAPG | P40121; H7C | CAPG_HUMAN | Unmodified | 0 | 0 | 0 | 0 | 1613600 | 426970 | 0 | 0 | 0 | 0 | 345210 | 474350 | 0 | 0 | 0 | 0 | 1525000 | 2591900 |
| ERQDADG( | ROCK2; ROCK2 | O75116; Q1 | ROCK2_HUMAI | Oxidation (M) | 0 | 0 | 0 | 0 | 0 | 669150 | 0 | 0 | 0 | 0 | 0 | 0 | 0 | 0 | 0 | 0 | 808630 | 423570 |
| ERQEAEEAI | MSN | P26038 | MOES_HUMAN | Unmodified | 0 | 0 | 0 | 0 | 787190 | 188650 | 0 | 0 | 0 | 0 | 0 | 0 | 0 | 0 | 0 | 0 | 466680 | 189750 |
| ERQTNPSA | VCP; VCP; VCP; | P55072; C9J | TERA_HUMAN | Oxidation (M) | 173580 | 112950 | 177190 | 0 | 0 | 0 | 0 | 0 | 314200 | 0 | 0 | 3009900 | 0 | 0 | 149960 | 313350 | 0 | 0 |
| ERQTNPSA | VCP; VCP; VCP; | P55072; C9J | TERA_HUMAN | Unmodified | 101250 | 0 | 0 | 0 | 0 | 0 | 0 | 0 | 2039700 | 0 | 0 | 1995700 | 0 | 0 | 0 | 0 | 0 | 536360 |
| ERQVATAV( | ITGA5; ITGA5 | P08648; H0Y | ITA5_HUMAN I | Unmodified | 8859300 | 2638000 | 7738400 | 8845200 | 49303000 | 11264000 | 0 | 16534000 | 0 | 0 | 0 | 0 | 0 | 42095000 | 642300 | 416390 | 0 | 30401000 |
| ERQWEAW | CIT; CIT; CIT; CI | O14578; A0A | CTRO_HUMAN | Unmodified | 0 | 0 | 0 | 0 | 2219700 | 440470 | 0 | 0 | 0 | 0 | 0 | 0 | 0 | 0 | 0 | 0 | 264480 | 0 |
| ERSDALNS | CTNNA1; CTNN | P35221; G3X | CTNA1_HUMAI | Unmodified | 0 | 0 | 1550600 | 2537900 | 4546100 | 1550900 | 0 | 0 | 288670 | 149650 | 0 | 3864600 | 4237700 | 1991300 | 95842 | 3267500 | 0 | 0 |
| ERSDALNS | CTNNA1; CTNN | P35221; G3X | CTNA1_HUMAI | Unmodified | 0 | 0 | 0 | 0 | 2965400 | 374970 | 0 | 0 | 0 | 0 | 0 | 0 | 1595800 | 1468900 | 0 | 0 | 0 | 0 |
| ERSGPYGF | DNAJC13; DNA | O75165; H0Y | DJC13_HUMAN | Unmodified | 0 | 0 | 0 | 0 | 0 | 0 | 0 | 0 | 514390 | 0 | 0 | 258770 | 0 | 0 | 127770 | 0 | 0 | 0 |
| ERSGVSLA | H1-3; H1-4; H1 | P16402; P10 | H13_HUMAN H | Unmodified | 0 | 995880 | 0 | 0 | 1308500 | 0 | 1542100 | 68366000 | 0 | 41564 | 0 | 174610 | 0 | 0 | 0 | 0 | 0 | 0 |
| ERSGVSLA | H1-3; H1-4; H1 | P16402; P10 | H13_HUMAN H | Unmodified | 4672800 | 1440700 | 0 | 0 | 1475200 | 331840 | 3558100 | 1.3E+08 | 66414 | 130450 | 0 | 199900 | 0 | 0 | 0 | 0 | 0 | 0 |
| ERSGVVC( | CDCP1 | Q9H5V8 | CDCP1_HUMA | Unmodified | 145110 | 0 | 0 | 186880 | 609710 | 0 | 0 | 0 | 0 | 0 | 83559 | 1011800 | 120940 | 0 | 293580 | 262480 | 820890 | 1028900 |
| ERSLLHGD | SLC3A2; SLC3A | P08195; A0A | 4F2_HUMAN 4I | Unmodified | 10579000 | 18082000 | 5544700 | 13966000 | 18578000 | 3211300 | 1799000 | 0 | 34325000 | 3654400 | 73562000 | 59401000 | 564000 | 543590 | 12058000 | 5194500 | 1247300 | 296520 |
| ERSLLHGD | SLC3A2; SLC3A | P08195; A0A | 4F2_HUMAN 4I | Unmodified | 0 | 0 | 0 | 632500 | 12274000 | 2606200 | 0 | 0 | 0 | 756990 | 0 | 0 | 1499000 | 224870 | 0 | 0 | 0 | 2271200 |
| ERSPEVLS( | THOC6 | Q86W42 | THOC6_HUMA | Unmodified | 0 | 321160 | 0 | 0 | 0 | 0 | 0 | 0 | 0 | 0 | 0 | 318380 | 0 | 0 | 0 | 196900 | 0 | 0 |
| ERSRPFTV/ | LIMA1; LIMA1; L | Q9UHB6; F8 | LIMA1_HUMAN | Unmodified | 0 | 0 | 0 | 0 | 388680 | 0 | 0 | 0 | 0 | 0 | 0 | 0 | 167460 | 0 | 0 | 0 | 0 | 0 |
| ERSTSEPA\ | ASPH; ASPH; AS | Q12797; A0A | ASPH_HUMAN | Unmodified | 0 | 0 | 0 | 0 | 2578100 | 0 | 0 | 0 | 51380000 | 2047400 | 0 | 0 | 1345800 | 995140 | 0 | 0 | 0 | 733370 |
| ERTATGLTI | VPS13C | Q709C8 | VP13C_HUMAN | Unmodified | 0 | 0 | 0 | 0 | 0 | 0 | 0 | 0 | 55381 | 37389 | 54387 | 726850 | 0 | 0 | 0 | 0 | 0 | 0 |

| | | | | | | | | | | | | | | | | | | | | | | |
|---|---|---|---|---|---|---|---|---|---|---|---|---|---|---|---|---|---|---|---|---|---|---|
| ERTDDEQF | CAB39L; CAB39 | Q9H9S4; B7. | CB39L_HUMAN | Unmodified | 28031 | 0 | 200080 | 248440 | 0 | 0 | 0 | 0 | 112880 | 0 | 113090 | 541660 | 249810 | 412410 | 218400 | 262380 | 0 | 0 |
| ERTEIELSY | FNBP1; FNBP1 | Q96RU3; B7. | FNBP1_HUMAN | Unmodified | 0 | 0 | 0 | 0 | 1396400 | 2604100 | 87573 | 1005200 | 0 | 0 | 0 | 3380600 | 0 | 0 | 0 | 0 | 327900 | 3447800 |
| ERTESPSAS | COBLL1; COBL | Q53SF7; A0 | COBL1_HUMAN | Unmodified | 1111200 | 1020700 | 911880 | 0 | 0 | 0 | 0 | 0 | 0 | 0 | 0 | 0 | 1245500 | 815360 | 1203600 | 853490 | 0 | 0 |
| ERTESPSAS | COBLL1; COBL | Q53SF7; A0 | COBL1_HUMAN | Unmodified | 0 | 0 | 0 | 0 | 0 | 0 | 0 | 0 | 0 | 0 | 0 | 0 | 0 | 405840 | 0 | 0 | 0 | 0 |
| ERTEVEQA | TRIP10; TRIP10 | Q15642; W4 | CIP4_HUMAN | Unmodified | 0 | 0 | 0 | 0 | 0 | 0 | 0 | 0 | 0 | 0 | 0 | 0 | 0 | 0 | 0 | 0 | 0 | 0 |
| ERVCNLID | ALDH6A1; ALDH | Q02252; G3 | MMSA_HUMAN | Unmodified | 0 | 0 | 0 | 0 | 0 | 0 | 0 | 80874 | 524990 | 165980 | 0 | 0 | 0 | 0 | 0 | 0 | 0 | 0 |
| ERVEAGDV | RUVBL1; RUVB | Q9Y265; E7 | RUVB1_HUMAN | Unmodified | 136280 | 0 | 0 | 0 | 745420 | 0 | 0 | 0 | 423230 | 0 | 1782100 | 5115000 | 2185200 | 216220 | 0 | 0 | 0 | 0 |
| ERVEQMKE | INCENP | Q9NQS7 | INCE_HUMAN | Unmodified | 0 | 0 | 28089 | 271700 | 589390 | 0 | 0 | 0 | 1139800 | 0 | 0 | 0 | 291100 | 13094 | 72953 | 51386 | 162850 | 0 |
| ERVQKEEE | BTN2A1; BTN2A | Q7KYR7; H7 | BT2A1_HUMAN | Unmodified | 96655 | 0 | 0 | 557720 | 1058000 | 270930 | 123350 | 0 | 423910 | 142530 | 335050 | 1562500 | 1448000 | 0 | 292620 | 236090 | 441100 | 655210 |
| ERVTPPEG | PDLIM1 | O00151 | PDLI1_HUMAN | Unmodified | 0 | 0 | 0 | 0 | 2593900 | 0 | 0 | 0 | 331780 | 262640 | 259510 | 0 | 0 | 0 | 0 | 0 | 4266400 | 5969200 |
| ERWEQGQ | GYG1 | P46976 | GLYG_HUMAN | Oxidation (M) | 0 | 0 | 115880 | 0 | 0 | 411460 | 0 | 0 | 0 | 0 | 0 | 3613500 | 119840 | 0 | 0 | 0 | 0 | 0 |
| ERYGISSM | DARS1; DARS1 | P14868; H7 | SYDC_HUMAN | Unmodified | 0 | 0 | 0 | 0 | 0 | 0 | 0 | 0 | 534530 | 0 | 0 | 1675400 | 0 | 0 | 0 | 0 | 0 | 0 |
| ERYNFPNP | EIF3D | O15371 | EIF3D_HUMAN | Oxidation (M) | 0 | 0 | 8522500 | 0 | 0 | 0 | 0 | 0 | 348990 | 0 | 0 | 619680 | 0 | 0 | 0 | 301110 | 0 | 493420 |
| ERYSYVCP | ACTR3; ACTR3 | P61158; B4 | ARP3_HUMAN | Unmodified | 1216200 | 402080 | 0 | 2168800 | 5362300 | 905570 | 1259300 | 583070 | 931280 | 453860 | 0 | 11663000 | 2852200 | 3125400 | 355770 | 1313500 | 1427900 | 3266500 |
| ERYVVKN | NIBAN1 | Q9BZQ8 | NIBA1_HUMAN | Unmodified | 80732 | 0 | 0 | 1394800 | 4470600 | 245510 | 228880 | 0 | 6729200 | 2609300 | 0 | 0 | 0 | 0 | 0 | 394540 | 0 | 292800 |
| ESAALSEV | ARHGAP4; ARH | P98171; F8 | RHG04_HUMAN | Unmodified | 0 | 0 | 0 | 0 | 0 | 0 | 0 | 0 | 0 | 0 | 0 | 700090 | 0 | 0 | 0 | 0 | 0 | 0 |
| ESAELFQS | NAALADL2 | Q58DX5 | NADL2_HUMAN | Unmodified | 0 | 0 | 0 | 120880 | 0 | 0 | 0 | 0 | 0 | 0 | 0 | 0 | 0 | 0 | 236240 | 0 | 0 | 0 |
| ESAFEFLS | MDH1; MDH1; | P40925; B8 | MDHC_HUMAN | Unmodified | 403060 | 332870 | 277660 | 407780 | 3689000 | 599980 | 845080 | 0 | 3233900 | 1928900 | 3418300 | 13063000 | 1578400 | 767010 | 677290 | 408050 | 1165900 | 2401000 |
| ESAGNSQ | GGA3; GGA3 | Q9NZ52; J3 | GGA3_HUMAN | Unmodified | 0 | 0 | 93430 | 75294 | 0 | 0 | 0 | 0 | 0 | 0 | 0 | 0 | 0 | 0 | 0 | 66654 | 0 | 0 |
| ESALATAQ | AVIL | O75366 | AVIL_HUMAN | Unmodified | 0 | 0 | 0 | 46396 | 0 | 0 | 0 | 0 | 0 | 0 | 0 | 7000600 | 0 | 0 | 0 | 0 | 0 | 0 |
| ESALATAQ | AVIL | O75366 | AVIL_HUMAN | Unmodified | 0 | 0 | 0 | 0 | 0 | 0 | 0 | 0 | 0 | 0 | 4608700 | 85767000 | 0 | 0 | 0 | 0 | 0 | 0 |
| ESAPGALR | GGT7 | Q9UJ14 | GGT7_HUMAN | Unmodified | 0 | 0 | 0 | 0 | 0 | 0 | 0 | 0 | 12264000 | 0 | 0 | 0 | 0 | 0 | 0 | 0 | 0 | 89984 |
| ESAPGLIIA | SORL1 | Q92673 | SORL_HUMAN | Unmodified | 0 | 0 | 0 | 0 | 0 | 0 | 0 | 0 | 2174700 | 0 | 0 | 0 | 0 | 0 | 0 | 0 | 0 | 0 |
| ESAREEEE | VGF | O15240 | VGF_HUMAN | Unmodified | 0 | 0 | 0 | 0 | 0 | 441660 | 0 | 0 | 2976200 | 61141 | 64401 | 0 | 0 | 0 | 0 | 0 | 49373 | 0 |
| ESARQDEG | GAP43 | P17677 | NEUM_HUMAN | Unmodified | 0 | 0 | 0 | 0 | 0 | 0 | 8798200 | 12083000 | 0 | 0 | 0 | 0 | 0 | 0 | 0 | 0 | 0 | 0 |
| ESATEEKLT | DCTN2; DCTN2 | Q13561; A8 | DCTN2_HUMAN | Unmodified | 0 | 0 | 0 | 0 | 1626900 | 0 | 576410 | 521300 | 0 | 766080 | 0 | 0 | 1332700 | 777950 | 0 | 0 | 711530 | 1028400 |
| ESCDSALR | CAPZA1 | P52907 | CAZA1_HUMAN | Unmodified | 0 | 0 | 0 | 0 | 0 | 0 | 0 | 0 | 0 | 0 | 0 | 0 | 0 | 0 | 0 | 0 | 0 | 0 |
| ESCLEAYT | KPNB1; KPNB1 | Q14974; J3 | IMB1_HUMAN | Oxidation (M) | 0 | 0 | 0 | 0 | 0 | 0 | 0 | 0 | 0 | 0 | 0 | 1342800 | 0 | 0 | 0 | 612960 | 0 | 0 |
| ESDDPMA | HSP90B1; HSP9 | P14625; A0 | ENPL_HUMAN | Oxidation (M) | 0 | 0 | 0 | 0 | 0 | 0 | 0 | 0 | 0 | 946730 | 0 | 0 | 0 | 0 | 0 | 0 | 0 | 0 |
| ESDDPMA | HSP90B1; HSP9 | P14625; A0 | ENPL_HUMAN | Unmodified | 0 | 0 | 0 | 0 | 0 | 0 | 0 | 0 | 176920 | 356490 | 0 | 0 | 0 | 0 | 0 | 0 | 0 | 0 |
| ESDLNGAC | LMNB1; LMNB1 | P20700; A0 | LMNB1_HUMAN | Unmodified | 0 | 39363 | 0 | 0 | 691000 | 235500 | 421260 | 618020 | 313270 | 83405 | 127560 | 2305500 | 0 | 0 | 0 | 0 | 0 | 0 |
| ESDSVFVL | IGSF3 | O75054 | IGSF3_HUMAN | Unmodified | 2119600 | 1850300 | 642980 | 865060 | 1396400 | 782040 | 519540 | 214370 | 634550 | 166990 | 0 | 4478600 | 1405100 | 1286900 | 1949400 | 1274100 | 0 | 0 |
| ESDTGLAP | PSMC2; PSMC2 | P35998; C9 | PRS7_HUMAN | Unmodified | 0 | 0 | 222860 | 0 | 0 | 0 | 0 | 0 | 0 | 0 | 0 | 2748900 | 0 | 0 | 77452 | 223030 | 96091 | 0 |
| ESDVGFIP | HBS1L; HBS1L | Q9Y450; H0 | HBS1L_HUMAN | Unmodified | 235930 | 0 | 0 | 0 | 0 | 0 | 0 | 0 | 0 | 115970 | 199700 | 963530 | 0 | 0 | 297570 | 218850 | 0 | 0 |
| ESDVPLKT | MYDGF; MYDG | Q969H8; M0 | MYDGF_HUMA | Unmodified | 0 | 0 | 227900 | 0 | 0 | 0 | 0 | 30058 | 72744000 | 5359700 | 11179000 | 2634900 | 0 | 0 | 0 | 0 | 0 | 0 |
| ESDYFTPQ | STT3B | Q8TCJ2 | STT3B_HUMAN | Unmodified | 0 | 0 | 0 | 0 | 0 | 0 | 0 | 0 | 2376100 | 600670 | 0 | 0 | 0 | 0 | 0 | 0 | 0 | 0 |
| ESDYFTPQ | STT3B | Q8TCJ2 | STT3B_HUMAN | Unmodified | 0 | 0 | 0 | 0 | 0 | 0 | 0 | 0 | 15830000 | 1257600 | 0 | 1466700 | 0 | 0 | 0 | 0 | 0 | 0 |
| ESEAVEW | MSN | P26038 | MOES_HUMAN | Unmodified | 402340 | 0 | 0 | 0 | 34664000 | 0 | 484770 | 431760 | 0 | 0 | 0 | 0 | 0 | 5483600 | 0 | 916290 | 8505900 | 16887000 |
| ESEAVEW | MSN | P26038 | MOES_HUMAN | Oxidation (M) | 0 | 0 | 0 | 287330 | 0 | 0 | 0 | 0 | 0 | 0 | 0 | 0 | 507270 | 0 | 0 | 0 | 1200400 | 1209500 |
| ESEDKPEI | HSP90AA1 | P07900 | HS90A_HUMAN | Unmodified | 0 | 0 | 0 | 0 | 0 | 0 | 0 | 0 | 1993200 | 739530 | 2744400 | 9735600 | 0 | 0 | 0 | 0 | 0 | 0 |
| ESEDKPEI | HSP90AA1 | P07900 | HS90A_HUMAN | Unmodified | 0 | 0 | 65365 | 0 | 0 | 0 | 0 | 0 | 381560 | 0 | 3381300 | 10314000 | 0 | 0 | 210450 | 158440 | 0 | 0 |
| ESEITDED | SMARCA5 | O60264 | SMCA5_HUMA | Unmodified | 80745 | 77024 | 0 | 0 | 0 | 0 | 0 | 0 | 0 | 0 | 0 | 0 | 0 | 0 | 0 | 0 | 0 | 0 |
| ESEKELAL | M6PR; M6PR; M | P20645; F5 | MPRD_HUMAN | Unmodified | 0 | 207680 | 0 | 0 | 856500 | 0 | 0 | 0 | 0 | 326300 | 0 | 0 | 567060 | 0 | 0 | 122420 | 0 | 33690 |
| ESELIYVFK | MICALL1; MICA | Q8N3F8; B0 | MILK1_HUMAN | Unmodified | 258840 | 567350 | 170810 | 1087900 | 211590 | 62187 | 11529 | 0 | 129370 | 55186 | 156370 | 757450 | 372890 | 116860 | 289710 | 0 | 0 | 34477 |
| ESENVVCH | TXNDC9; TXND | O14530; F8 | TXND9_HUMAN | Unmodified | 0 | 0 | 0 | 0 | 0 | 0 | 0 | 0 | 0 | 0 | 97662 | 467990 | 0 | 0 | 0 | 48419 | 0 | 0 |
| ESEPQAAA | BASP1 | P80723 | BASP1_HUMAN | Unmodified | 137270 | 628620 | 0 | 0 | 4982700 | 904700 | 226100 | 201630 | 13730000 | 721010 | 0 | 0 | 0 | 61646 | 0 | 0 | 0 | 1649400 |
| ESEPQAAA | BASP1 | P80723 | BASP1_HUMAN | Unmodified | 0 | 49240 | 0 | 0 | 0 | 78457 | 0 | 0 | 1044400 | 372440 | 0 | 0 | 0 | 116100 | 0 | 0 | 308770 | 1032900 |
| ESEQRPPF | DDR1; DDR1; D | Q08345; A0 | DDR1_HUMAN | Unmodified | 504960 | 580960 | 207600 | 186600 | 0 | 0 | 0 | 0 | 638060 | 0 | 85664 | 482240 | 0 | 0 | 230180 | 285120 | 0 | 0 |
| ESERDRAM | OS9; OS9 | Q13438; B4 | OS9_HUMAN | Unmodified | 0 | 0 | 0 | 0 | 702230 | 0 | 0 | 0 | 2399200 | 685370 | 0 | 574550 | 0 | 0 | 0 | 0 | 0 | 0 |
| ESESAPGD | GRB2; GRB2 | P62993; J3 | GRB2_HUMAN | Unmodified | 1300900 | 916240 | 566430 | 0 | 0 | 0 | 0 | 0 | 0 | 0 | 0 | 915300 | 0 | 0 | 2085500 | 1660200 | 265100 | 399960 |
| ESESCDCL | TUBB2A; TUBB | Q13885; Q9 | TBB2A_HUMAN | Oxidation (M) | 0 | 0 | 0 | 0 | 0 | 0 | 0 | 223860 | 0 | 0 | 0 | 8439000 | 0 | 0 | 0 | 0 | 0 | 0 |
| ESESVDKV | HNRNPD; HNR | Q14103; H0 | HNRPD_HUMA | Unmodified | 0 | 0 | 0 | 0 | 0 | 0 | 0 | 0 | 103180 | 0 | 64160 | 280900 | 0 | 0 | 0 | 0 | 0 | 0 |
| ESETFPND | PLCG2; PLCG2 | P16885; A0 | PLCG2_HUMAN | Unmodified | 0 | 0 | 0 | 0 | 0 | 0 | 0 | 0 | 0 | 0 | 81730 | 525140 | 0 | 0 | 0 | 0 | 0 | 0 |
| ESEVLIGN | EXOC4 | Q96A65 | EXOC4_HUMAN | Unmodified | 0 | 0 | 0 | 0 | 0 | 0 | 0 | 0 | 0 | 0 | 0 | 762720 | 0 | 0 | 175120 | 0 | 0 | 0 |
| ESEVLIYAR | PIGX; PIGX; PIG | Q8TBF5; C9 | PIGX_HUMAN | Unmodified | 0 | 0 | 0 | 0 | 0 | 0 | 0 | 152050 | 6793800 | 704410 | 0 | 0 | 0 | 0 | 0 | 0 | 0 | 0 |
| ESEVLIYAR | PIGX; PIGX; PIG | Q8TBF5; C9 | PIGX_HUMAN | Unmodified | 0 | 0 | 0 | 19230 | 134380 | 0 | 0 | 0 | 2018100 | 0 | 0 | 0 | 97756 | 35061 | 55514 | 76144 | 0 | 0 |
| ESEYHYEY | ELAPOR1; ELAP | Q6UXG2; E9 | ELAP1_HUMAN | Unmodified | 174240 | 53104 | 0 | 0 | 0 | 0 | 0 | 0 | 552650 | 39206 | 0 | 0 | 0 | 0 | 0 | 0 | 0 | 0 |
| ESFAEMNR | COX4I1; COX4I | P13073; H3 | COX41_HUMAN | Unmodified | 561240 | 85333 | 0 | 0 | 0 | 2070600 | 0 | 0 | 689320 | 0 | 0 | 0 | 0 | 0 | 0 | 229590 | 0 | 0 |
| ESFDGSVR | NCL; NCL; NCL | P19338; C9 | NUCL_HUMAN | Unmodified | 0 | 0 | 0 | 0 | 0 | 0 | 0 | 0 | 25782 | 0 | 1608700 | 455340 | 0 | 0 | 0 | 0 | 0 | 0 |
| ESFESQFL | CUL1; CUL1; C | Q13616; A0 | CUL1_HUMAN | Unmodified | 0 | 0 | 0 | 0 | 0 | 0 | 0 | 0 | 0 | 0 | 0 | 583470 | 0 | 0 | 0 | 0 | 0 | 0 |
| ESGCSWH | ALDH16A1; ALD | Q8IZ83; F5 | A16A1_HUMAN | Unmodified | 0 | 0 | 0 | 0 | 0 | 0 | 0 | 0 | 0 | 0 | 0 | 1637500 | 0 | 0 | 0 | 0 | 0 | 0 |
| ESGEEFRM | LRPAP1; LRPAP | P30533; D6 | AMRP_HUMAN | Unmodified | 0 | 0 | 0 | 0 | 834420 | 0 | 0 | 0 | 224310 | 272690 | 0 | 0 | 0 | 0 | 0 | 0 | 0 | 0 |
| ESGGSGS | SLC16A10; SLC | Q8TF71; H0 | MOT10_HUMAN | Unmodified | 0 | 0 | 576390 | 0 | 0 | 0 | 0 | 0 | 0 | 0 | 0 | 2244100 | 0 | 0 | 219390 | 226180 | 0 | 0 |
| ESGHSNW | ATL3; ATL3; ATL | Q6DD88; F5 | ATLA3_HUMAN | Unmodified | 0 | 0 | 181570 | 91536 | 145530 | 0 | 0 | 0 | 0 | 0 | 0 | 0 | 0 | 0 | 719270 | 171480 | 0 | 116220 |
| ESGLQAAH | TTLL12 | Q14166 | TTL12_HUMAN | Unmodified | 0 | 0 | 0 | 0 | 0 | 0 | 0 | 0 | 0 | 0 | 363200 | 2956500 | 0 | 0 | 0 | 0 | 0 | 0 |
| ESGMSSLY | EMC1 | Q8N766 | EMC1_HUMAN | Oxidation (M) | 0 | 0 | 22693 | 0 | 0 | 0 | 0 | 0 | 264210 | 0 | 38943 | 0 | 0 | 0 | 0 | 0 | 0 | 0 |
| ESGPGIAP | ERBB3; ERBB3; | P21860; F8 | ERBB3_HUMAN | Unmodified | 375650 | 0 | 94172 | 235010 | 0 | 0 | 130020 | 0 | 409320 | 105170 | 230940 | 437810 | 109470 | 0 | 165620 | 180960 | 0 | 0 |
| ESGQVAII | STK3; STK3; STI | Q13188; Q8 | STK3_HUMAN | Unmodified | 0 | 0 | 0 | 0 | 892120 | 0 | 220570 | 0 | 353590 | 158850 | 0 | 0 | 0 | 0 | 0 | 0 | 209480 | 0 |
| ESGRFQD | DBNL; DBNL; D | Q9UJU6; B4 | DBNL_HUMAN | Unmodified | 1468400 | 912270 | 1391200 | 2187300 | 9082400 | 1319700 | 1990400 | 2507500 | 1040200 | 732500 | 1601600 | 6520900 | 10159000 | 8867200 | 2256100 | 1793600 | 5012800 | 6487100 |
| ESGVFEGI | SCARB1; SCARI | Q8WTV0; A0 | SCRB1_HUMAN | Unmodified | 3794700 | 1041300 | 1233600 | 1014400 | 2216700 | 230150 | 0 | 0 | 643450 | 84585 | 5718000 | 18164000 | 0 | 0 | 7070400 | 1306500 | 0 | 0 |
| ESGVFEGI | SCARB1; SCARI | Q8WTV0; A0 | SCRB1_HUMAN | Unmodified | 0 | 0 | 0 | 0 | 11918000 | 2911500 | 0 | 55696 | 158640 | 657790 | 1523600 | 0 | 0 | 0 | 0 | 0 | 931540 | 2258900 |
| ESGYCTGI | AGPS; AGPS; A | O00116; A0 | ADAS_HUMAN | Unmodified | 0 | 0 | 0 | 0 | 0 | 0 | 0 | 0 | 17501000 | 1609600 | 0 | 0 | 0 | 0 | 0 | 0 | 0 | 0 |
| ESGYNMIH | AGL | P35573 | GDE_HUMAN | Unmodified | 0 | 0 | 0 | 0 | 0 | 0 | 0 | 0 | 318080 | 0 | 0 | 0 | 0 | 0 | 168810 | 0 | 0 | 0 |
| ESGYNMIH | AGL | P35573 | GDE_HUMAN | Oxidation (M) | 63853 | 57128 | 308240 | 239840 | 0 | 0 | 40598 | 0 | 194070 | 434180 | 266040 | 2121300 | 0 | 0 | 90367 | 108980 | 99531 | 0 |
| ESHAKADS | PACSIN3; PACS | Q9UKS6; E9 | PACN3_HUMAN | Unmodified | 3361400 | 711160 | 0 | 2852500 | 76496 | 0 | 0 | 0 | 0 | 0 | 0 | 0 | 118230 | 0 | 0 | 46687 | 253710 | 0 |
| ESHATLVF | STAT3; STAT3; | P40763; A0 | STAT3_HUMAN | Unmodified | 86768 | 83536 | 0 | 0 | 0 | 0 | 0 | 0 | 0 | 0 | 0 | 401940 | 0 | 0 | 536830 | 297090 | 0 | 0 |
| ESHPGLFP | NID1 | P14543 | NID1_HUMAN | Unmodified | 0 | 0 | 0 | 0 | 0 | 0 | 0 | 0 | 0 | 0 | 0 | 0 | 0 | 0 | 0 | 0 | 0 | 125360 |
| ESHRLPVD | DSP | P15924 | DESP_HUMAN | Unmodified | 0 | 0 | 0 | 0 | 0 | 0 | 0 | 0 | 0 | 0 | 0 | 0 | 0 | 0 | 0 | 0 | 0 | 0 |
| ESIEEPSAK | SNRNP200; SN | O75643; A0 | U520_HUMAN | Unmodified | 0 | 0 | 50098 | 449310 | 0 | 0 | 0 | 0 | 416480 | 41829 | 0 | 1276000 | 115860 | 0 | 235780 | 0 | 0 | 0 |
| ESIEIINK | OAT | P04181 | OAT_HUMAN | Unmodified | 0 | 0 | 0 | 0 | 0 | 0 | 0 | 0 | 569450 | 0 | 0 | 0 | 0 | 0 | 0 | 0 | 0 | 0 |
| ESILQRPLS | OVCA2 | Q8WZ82 | OVCA2_HUMAN | Unmodified | 1433900 | 281480 | 393180 | 121760 | 0 | 0 | 0 | 0 | 0 | 0 | 0 | 2034100 | 0 | 0 | 720770 | 529540 | 0 | 0 |
| ESISDVGF | AGPS; AGPS; A | O00116; B7 | ADAS_HUMAN | Oxidation (M) | 0 | 0 | 0 | 0 | 0 | 0 | 1145600 | 674320 | 4828300 | 1798900 | 0 | 0 | 0 | 0 | 0 | 0 | 0 | 0 |
| ESISDVGF | AGPS; AGPS; A | O00116; B7 | ADAS_HUMAN | Unmodified | 0 | 0 | 0 | 0 | 0 | 0 | 1002500 | 0 | 9914400 | 1078600 | 0 | 0 | 0 | 0 | 0 | 0 | 0 | 0 |
| ESIVDVEG | DARS1; DARS1; | P14868; H7 | SYDC_HUMAN | Unmodified | 0 | 0 | 0 | 175400 | 0 | 0 | 0 | 0 | 0 | 0 | 294170 | 2176700 | 0 | 0 | 0 | 0 | 0 | 0 |
| ESIVDVEG | DARS1; DARS1; | P14868; H7 | SYDC_HUMAN | Unmodified | 43849 | 0 | 0 | 0 | 628690 | 0 | 127780 | 38967 | 1130800 | 317110 | 701620 | 10482000 | 0 | 0 | 0 | 0 | 0 | 0 |
| ESKADIFV | GLG1 | Q92896 | GSLG1_HUMAN | Unmodified | 0 | 0 | 0 | 0 | 0 | 0 | 0 | 0 | 29589000 | 3909600 | 2273800 | 0 | 0 | 0 | 0 | 0 | 36718 | 0 |
| ESKDPADE | STMN1 | P16949 | STMN1_HUMAN | Unmodified | 0 | 0 | 614510 | 0 | 1086700 | 0 | 0 | 0 | 10113000 | 252860 | 428150 | 4309500 | 0 | 0 | 0 | 0 | 761650 | 0 |
| ESKDQLSD | SCG2 | P13521 | SCG2_HUMAN | Unmodified | 0 | 0 | 0 | 0 | 0 | 0 | 0 | 62663 | 692020 | 270220 | 0 | 0 | 43874 | 0 | 0 | 0 | 0 | 0 |
| ESKEEETSI | SLC16A1 | P53985 | MOT1_HUMAN | Unmodified | 409610 | 372220 | 9390000 | 5936800 | 3366800 | 86606 | 1532100 | 40137 | 3180300 | 439860 | 3662100 | 14711000 | 925740 | 416060 | 10099000 | 11613000 | 0 | 1975900 |
| ESKEETPG | CLTB; CLTB | P09497; D6 | CLCB_HUMAN | Unmodified | 9372400 | 6657300 | 4842400 | 5435900 | 6018400 | 1033100 | 1340200 | 702030 | 0 | 0 | 278730 | 7781400 | 18237000 | 11646000 | 4409900 | 5906100 | 1438600 | 4116000 |
| ESKENQW | ITGA7; ITGA7 | Q13683; J3 | ITA7_HUMAN | Unmodified | 0 | 0 | 0 | 0 | 0 | 0 | 971990 | 346960 | 0 | 0 | 0 | 0 | 0 | 0 | 0 | 0 | 0 | 0 |
| ESKGPIVP | SLC12A2; SLC1 | P55011; G3 | S12A2_HUMAN | Unmodified | 0 | 0 | 0 | 0 | 1184000 | 250050 | 239720 | 98612 | 0 | 84656 | 0 | 0 | 1267400 | 1898200 | 0 | 0 | 872920 | 9022100 |
| ESKKEDLV | CFL1; CFL2; CF | P23528; Q9 | COF1_HUMAN | Unmodified | 47942000 | 69097000 | 9670100 | 38323000 | 2.3E+09 | 3.6E+08 | 1.44E+08 | 5203800 | 66551000 | 1.07E+08 | 40272000 | 81607000 | 6.76E+08 | 2.36E+08 | 54708000 | 47211000 | 5.96E+08 | 4.17E+08 |
| ESKLPGDK | CANX; CANX; C | P27824; A0 | CALX_HUMAN | Oxidation (M) | 0 | 0 | 0 | 0 | 0 | 0 | 96985 | 0 | 0 | 1451300 | 0 | 0 | 0 | 0 | 0 | 0 | 62560 | 0 |
| ESKLPGDK | CANX; CANX; C | P27824; A0 | CALX_HUMAN | Unmodified | 0 | 0 | 0 | 0 | 0 | 0 | 0 | 0 | 0 | 2536800 | 0 | 0 | 0 | 0 | 0 | 0 | 88510 | 0 |
| ESKTPAILE | CTNND1; CTNN | O60716; A0 | CTND1_HUMAN | Unmodified | 0 | 0 | 0 | 828620 | 8507000 | 749540 | 0 | 0 | 0 | 0 | 0 | 0 | 9164000 | 0 | 0 | 563800 | 3352900 | 5247900 |
| ESLAESAA | RANBP3; RANB | Q9H6Z4; B7 | RANB3_HUMAN | Unmodified | 0 | 0 | 0 | 0 | 0 | 0 | 0 | 0 | 0 | 0 | 0 | 0 | 0 | 0 | 0 | 0 | 0 | 0 |
| ESLCQAAL | MCCC1; MCCC | Q96RQ3; E9 | MCCA_HUMAN | Unmodified | 0 | 0 | 0 | 0 | 0 | 0 | 0 | 0 | 278370 | 0 | 0 | 0 | 0 | 0 | 0 | 0 | 0 | 0 |
| ESLDDLTN | IDE; IDE; IDE; ID | P14735; A0 | IDE_HUMAN | Unmodified | 0 | 0 | 0 | 0 | 0 | 0 | 0 | 0 | 0 | 0 | 0 | 0 | 0 | 0 | 0 | 0 | 0 | 0 |
| ESLDVYEL | ASAH1; ASAH1; | Q13510; A0 | ASAH1_HUMAN | Unmodified | 0 | 0 | 531560 | 436380 | 0 | 0 | 0 | 0 | 8362700 | 555950 | 947870 | 0 | 0 | 0 | 366320 | 371400 | 0 | 0 |
| ESLGPDSS | FGFRL1; FGFRL | Q8N441; D6 | FGRL1_HUMAN | Unmodified | 380330 | 0 | 549500 | 0 | 0 | 0 | 0 | 0 | 0 | 0 | 0 | 0 | 0 | 0 | 760200 | 970710 | 0 | 0 |
| ESLIDGIK | AHCY | P23526 | SAHH_HUMAN | Unmodified | 501060 | 67491 | 0 | 0 | 0 | 0 | 22172 | 0 | 1826200 | 0 | 1947600 | 20818000 | 0 | 0 | 1674600 | 1106300 | 30122 | 0 |
| ESLITLIEK | CPD | O75976 | CBPD_HUMAN | Unmodified | 9714 | 0 | 0 | 0 | 0 | 0 | 0 | 0 | 6758400 | 726120 | 114980 | 87477 | 0 | 0 | 0 | 0 | 0 | 0 |
| ESLKEEDE | PSMA3; PSMA3 | P25788; H0 | PSA3_HUMAN | Unmodified | 0 | 0 | 0 | 0 | 1993700 | 0 | 79311 | 38079 | 756570 | 457450 | 648200 | 1929400 | 924690 | 0 | 0 | 0 | 328160 | 0 |
| ESLKEEDE | PSMA3; PSMA3 | P25788; H0 | PSA3_HUMAN | Oxidation (M) | 0 | 0 | 0 | 0 | 0 | 0 | 0 | 0 | 0 | 0 | 0 | 0 | 0 | 0 | 0 | 0 | 0 | 0 |
| ESLLDGGN | AGRN | O00468 | AGRIN_HUMAN | Unmodified | 0 | 0 | 1136600 | 460650 | 94648 | 0 | 0 | 0 | 913970 | 99940 | 123690 | 143670 | 0 | 0 | 0 | 0 | 0 | 0 |
| ESLLLSEQ | CD320 | Q9NPF0 | CD320_HUMAN | Unmodified | 0 | 0 | 1064600 | 432830 | 1343400 | 0 | 0 | 0 | 816520 | 645640 | 0 | 612050 | 0 | 408210 | 803890 | 0 | 0 | 1437000 |
| ESLPVIAAP | ACBD3 | Q9H3P7 | GCP60_HUMAN | Oxidation (M) | 0 | 0 | 0 | 0 | 0 | 0 | 0 | 0 | 0 | 0 | 0 | 1938000 | 1785900 | 0 | 0 | 0 | 0 | 0 |
| ESLQQMAE | PRKCSH; PRKC | P14314; K7 | GLU2B_HUMAN | Unmodified | 643320 | 343460 | 0 | 0 | 0 | 0 | 0 | 75049 | 1328500 | 208720 | 997110 | 0 | 0 | 0 | 0 | 0 | 0 | 0 |

| | | | | | | | | | | | | | | | | | | | | | | |
|---|---|---|---|---|---|---|---|---|---|---|---|---|---|---|---|---|---|---|---|---|---|---|
| ESLQQMAE | PRKCSH; PRKC | P14314; K7E | GLU2B_HUMAI | Oxidation (M) | 0 | 0 | 0 | 0 | 0 | 0 | 0 | 0 | 87245 | 311060 | 617720 | 104200 | 0 | 0 | 0 | 0 | 0 | 0 |
| ESLQQMAE | PRKCSH; PRKC | P14314; K7E | GLU2B_HUMAI | Unmodified | 0 | 0 | 0 | 0 | 0 | 0 | 0 | 0 | 0 | 1409100 | 0 | 0 | 0 | 0 | 0 | 0 | 0 | 0 |
| ESLREVQLI | GALK1; GALK1 | P51570; K7E | GALK1_HUMAN | Unmodified | 0 | 0 | 0 | 0 | 1280900 | 224460 | 241270 | 15221 | 0 | 44800 | 123140 | 720180 | 1460600 | 996080 | 0 | 0 | 394430 | 622380 |
| ESLSAATA( | VPS13C | Q709C8 | VP13C_HUMAN | Unmodified | 0 | 0 | 0 | 0 | 0 | 0 | 0 | 0 | 0 | 0 | 0 | 915830 | 0 | 0 | 0 | 0 | 0 | 0 |
| ESLSEEEA( | SRRT; SRRT | Q9BXP5; H7 | SRRT_HUMAN | Unmodified | 0 | 0 | 0 | 71920 | 0 | 0 | 0 | 0 | 0 | 0 | 70734 | 993810 | 0 | 0 | 94737 | 214440 | 81781 | 59962 |
| ESLSGQAV | UQCRFS1P1; U | P0C7P4; P47 | UCRIL_HUMAN | Unmodified | 84241 | 0 | 0 | 0 | 0 | 0 | 0 | 0 | 4989200 | 0 | 97374 | 173650 | 0 | 0 | 0 | 54142 | 0 | 0 |
| ESLSGVCE | DEFA5 | Q01523 | DEF5_HUMAN | Unmodified | 0 | 0 | 0 | 0 | 0 | 0 | 0 | 0 | 9926600 | 0 | 0 | 0 | 0 | 0 | 0 | 0 | 0 | 0 |
| ESLTEINR | RIPOR2; RIPOR | Q9Y4F9; B7Z | RIPR2_HUMAN | Unmodified | 0 | 291890 | 0 | 0 | 0 | 0 | 0 | 0 | 0 | 0 | 0 | 2835600 | 0 | 0 | 83734 | 0 | 0 | 0 |
| ESLVISGLR | INSR | P06213 | INSR_HUMAN I | Unmodified | 241550 | 131020 | 501830 | 279330 | 147440 | 0 | 0 | 0 | 7356700 | 459180 | 0 | 0 | 0 | 0 | 574380 | 619710 | 0 | 0 |
| ESLVSFAM | DNAJC10; DNA | Q8IXB1; A0A | DJC10_HUMAN | Unmodified | 0 | 0 | 0 | 0 | 0 | 0 | 0 | 0 | 984350 | 184650 | 0 | 0 | 0 | 0 | 0 | 0 | 0 | 0 |
| ESLVVNYEI | MCM2; MCM2 | P49736; H0Y | MCM2_HUMAN | Unmodified | 0 | 0 | 0 | 0 | 0 | 0 | 0 | 0 | 0 | 0 | 332720 | 1997500 | 0 | 0 | 0 | 209480 | 0 | 0 |
| ESLWPFLID | MAT2B | Q9NZL9 | MAT2B_HUMAI | Unmodified | 0 | 0 | 0 | 0 | 872080 | 174660 | 68164 | 0 | 238490 | 234950 | 258200 | 985200 | 576860 | 360280 | 0 | 0 | 277160 | 315870 |
| ESLWPFLID | MAT2B | Q9NZL9 | MAT2B_HUMAI | Unmodified | 30871 | 0 | 0 | 0 | 0 | 0 | 0 | 0 | 0 | 0 | 142280 | 605370 | 0 | 0 | 0 | 0 | 0 | 0 |
| ESMAVLEK | CAB39; CAB39 | Q9Y376; A0A | CAB39_HUMAI | Oxidation (M) | 295780 | 0 | 304200 | 382620 | 228930 | 43676 | 0 | 0 | 143180 | 0 | 0 | 0 | 0 | 168140 | 495110 | 588880 | 70428 | 246020 |
| ESMAVLEK | CAB39; CAB39 | Q9Y376; A0A | CAB39_HUMAI | Unmodified | 141420 | 0 | 0 | 0 | 1608600 | 497570 | 64767 | 60476 | 0 | 0 | 0 | 0 | 291220 | 201640 | 0 | 0 | 22537 | 1085800 |
| ESMDSVKT | QSOX2; QSOX2 | Q6ZRP7; H0 | QSOX2_HUMAI | Oxidation (M) | 0 | 0 | 0 | 0 | 28158 | 0 | 20930 | 0 | 2761400 | 579570 | 0 | 0 | 0 | 0 | 0 | 0 | 0 | 0 |
| ESMDSVKT | QSOX2; QSOX2 | Q6ZRP7; H0 | QSOX2_HUMAI | Unmodified | 0 | 0 | 0 | 0 | 371920 | 0 | 0 | 0 | 516910 | 0 | 0 | 0 | 0 | 0 | 0 | 0 | 0 | 0 |
| ESMDSVKT | QSOX2; QSOX2 | Q6ZRP7; H0 | QSOX2_HUMAI | Unmodified | 0 | 0 | 0 | 0 | 0 | 0 | 0 | 0 | 1958700 | 0 | 0 | 0 | 0 | 0 | 0 | 0 | 0 | 0 |
| ESMQMQV | STOML2; STOM | Q9UJZ1; A0A | STML2_HUMAN | Unmodified | 0 | 0 | 0 | 0 | 0 | 0 | 0 | 0 | 602330 | 0 | 0 | 0 | 0 | 0 | 0 | 0 | 0 | 0 |
| ESNAEYLA | STEAP3; STEAP | Q658P3; B8 | STEA3_HUMAN | Unmodified | 0 | 0 | 0 | 0 | 0 | 0 | 0 | 0 | 0 | 0 | 728590 | 0 | 0 | 0 | 0 | 0 | 0 | 0 |
| ESNCYDPE | CLTC; CLTC | Q00610; A0A | CLH1_HUMAN | Unmodified | 789470 | 490390 | 233230 | 311110 | 0 | 0 | 0 | 0 | 0 | 0 | 0 | 3140500 | 0 | 0 | 301450 | 407440 | 0 | 0 |
| ESNQKLQE | SETD2; SETD2; | Q9BYW2; A0 | SETD2_HUMAN | Unmodified | 0 | 0 | 0 | 0 | 195660 | 63780 | 0 | 0 | 0 | 20608 | 0 | 0 | 0 | 0 | 0 | 0 | 171880 | 315260 |
| ESPEGSYTI | FNBP1L; FNBP1 | Q5T0N5; A0 | FBP1L_HUMAN | Unmodified | 350160 | 428360 | 0 | 0 | 0 | 0 | 0 | 0 | 0 | 0 | 0 | 0 | 0 | 0 | 62272 | 78134 | 0 | 0 |
| ESPESEGPI | VPS35 | Q96QK1 | VPS35_HUMAN | Unmodified | 103750 | 0 | 0 | 0 | 0 | 0 | 0 | 0 | 272640 | 0 | 253660 | 563460 | 0 | 0 | 0 | 0 | 0 | 0 |
| ESPFRHVF | TFRC; TFRC | P02786; G3 | TFR1_HUMAN | Unmodified | 0 | 0 | 23242 | 722470 | 4506100 | 2136700 | 4982100 | 0 | 4415200 | 14652000 | 689650 | 0 | 2712000 | 0 | 0 | 0 | 0 | 5607600 |
| ESPGLWG | AHNAK2 | Q8IVF2 | AHNK2_HUMAI | Unmodified | 20175 | 0 | 0 | 0 | 3726300 | 0 | 0 | 0 | 0 | 0 | 0 | 0 | 0 | 0 | 0 | 0 | 566990 | 2833600 |
| ESPQDSAI | RABGAP1 | Q9Y3P9 | RBGP1_HUMAI | Unmodified | 43412 | 33214 | 0 | 0 | 0 | 136190 | 0 | 0 | 114580 | 0 | 62483 | 432710 | 0 | 0 | 0 | 0 | 0 | 0 |
| ESPQEIAR | SORD; SORD | Q00796; A0A | DHSO_HUMAN | Unmodified | 0 | 0 | 2222100 | 1422400 | 0 | 20638 | 0 | 0 | 447360 | 50522 | 578380 | 4845300 | 0 | 0 | 2204700 | 1643500 | 0 | 0 |
| ESPYGLSFI | TJP1; TJP1; TJP1 | Q07157; A0A | ZO1_HUMAN T | Unmodified | 0 | 0 | 0 | 0 | 5423400 | 0 | 0 | 0 | 0 | 0 | 0 | 0 | 0 | 0 | 0 | 0 | 60585 | 33558 |
| ESQAKDVII | RPS12 | P25398 | RS12_HUMAN | Unmodified | 0 | 357390 | 0 | 231020 | 3022800 | 484520 | 1796100 | 561380 | 955480 | 887290 | 0 | 0 | 8914800 | 2664300 | 0 | 0 | 1695100 | 2059400 |
| ESQAYYQR | PRNP; PRNP | P04156; A2A | PRIO_HUMAN I | Unmodified | 915800 | 640450 | 1016200 | 0 | 0 | 0 | 0 | 0 | 0 | 0 | 0 | 0 | 0 | 0 | 529010 | 533890 | 0 | 0 |
| ESQFLKEEI | CTNNA1; CTNN | P35221; E5R | CTNA1_HUMAI | Unmodified | 972030 | 4303100 | 1364900 | 4461400 | 3597400 | 494250 | 0 | 0 | 539720 | 0 | 2549400 | 9736800 | 216290 | 0 | 9876900 | 7511600 | 0 | 0 |
| ESQIDVLQ | NECTIN3 | Q9NQS3 | NECT3_HUMAI | Unmodified | 0 | 0 | 0 | 0 | 0 | 0 | 0 | 0 | 1835600 | 0 | 0 | 0 | 512360 | 1073700 | 0 | 0 | 0 | 0 |
| ESQLFDPS | LIMCH1; LIMCH | Q9UPQ0; D6 | LIMC1_HUMAN | Unmodified | 0 | 0 | 0 | 0 | 0 | 0 | 0 | 819950 | 0 | 0 | 0 | 0 | 0 | 2737400 | 0 | 732410 | 0 | 0 |
| ESQSEREE | CDC42BPA; CD | Q5VT25; A0A | MRCKA_HUMA | Unmodified | 0 | 0 | 0 | 0 | 0 | 0 | 0 | 0 | 0 | 0 | 0 | 1435900 | 0 | 0 | 0 | 598670 | 0 | 0 |
| ESQSHPGI | PTPN11; PTPN1 | Q06124; A0A | PTN11_HUMAN | Unmodified | 291780 | 411090 | 19638000 | 171590 | 0 | 0 | 0 | 0 | 0 | 0 | 0 | 5164300 | 0 | 0 | 341690 | 333820 | 0 | 0 |
| ESQSPDTTI | TNPO1; TNPO1 | Q92973; A0A | TNPO1_HUMAI | Unmodified | 204910 | 0 | 527030 | 392310 | 353270 | 0 | 0 | 0 | 0 | 0 | 353110 | 2194200 | 494070 | 0 | 530030 | 295510 | 0 | 214300 |
| ESQTQDNI | UPF1 | Q92900 | RENT1_HUMAN | Unmodified | 0 | 1387800 | 956160 | 559990 | 0 | 0 | 0 | 0 | 145190 | 0 | 0 | 0 | 0 | 0 | 0 | 591690 | 0 | 0 |
| ESREAVEKI | HSP90B1; HSP | P14625; A0A | ENPL_HUMAN | Oxidation (M) | 0 | 0 | 0 | 0 | 108840 | 0 | 2176400 | 0 | 0 | 3867000 | 0 | 0 | 0 | 0 | 0 | 0 | 0 | 0 |
| ESREAVEKI | HSP90B1; HSP | P14625; A0A | ENPL_HUMAN | Unmodified | 0 | 0 | 0 | 0 | 604520 | 0 | 3419900 | 0 | 0 | 3728500 | 0 | 40033 | 0 | 0 | 0 | 0 | 0 | 0 |
| ESREVTVP | MCAM | P43121 | MUC18_HUMA | Unmodified | 0 | 0 | 0 | 1338100 | 1417600 | 0 | 387420 | 215570 | 0 | 252890 | 0 | 0 | 0 | 0 | 0 | 0 | 1110300 | 5750200 |
| ESRFDFIYD | MYO10; MYO10 | Q9HD67; D6 | MYO10_HUMAI | Unmodified | 0 | 0 | 0 | 0 | 0 | 0 | 0 | 0 | 0 | 0 | 0 | 0 | 0 | 0 | 307160 | 19819 | 0 | 0 |
| ESRKEELMI | DSTN; DSTN | P60981; F6R | DEST_HUMAN | Oxidation (M) | 0 | 303180 | 0 | 368110 | 7887400 | 334960 | 1217600 | 0 | 0 | 364930 | 15178 | 0 | 11125000 | 10289000 | 0 | 0 | 6565000 | 4401600 |
| ESRKEELMI | DSTN; DSTN | P60981; F6R | DEST_HUMAN | Unmodified | 0 | 0 | 0 | 0 | 0 | 0 | 0 | 0 | 0 | 0 | 0 | 0 | 1303500 | 379140 | 0 | 0 | 0 | 0 |
| ESRPENEEI | RP2 | O75695 | XRP2_HUMAN | Unmodified | 0 | 0 | 0 | 0 | 0 | 0 | 0 | 0 | 0 | 0 | 0 | 12038000 | 0 | 0 | 758710 | 0 | 0 | 0 |
| ESSDSFVPI | GALNT12 | Q8IXK2 | GLT12_HUMAN | Unmodified | 0 | 0 | 0 | 0 | 0 | 0 | 0 | 0 | 1570900 | 21332 | 0 | 0 | 0 | 0 | 0 | 0 | 0 | 0 |
| ESSETPDQ | CD44; CD44; C | P16070; H0 | CD44_HUMAN | Unmodified | 4032000 | 2268200 | 0 | 0 | 640800 | 0 | 0 | 0 | 0 | 0 | 0 | 0 | 0 | 0 | 0 | 0 | 426380 | 475150 |
| ESSETPDQ | CD44; CD44; C | P16070; H0 | CD44_HUMAN | Oxidation (M) | 1130200 | 327330 | 115560 | 0 | 161010 | 0 | 0 | 0 | 0 | 0 | 0 | 0 | 0 | 0 | 0 | 0 | 75266 | 0 |
| ESSETPDQ | CD44; CD44; C | P16070; H0 | CD44_HUMAN | Oxidation (M) | 0 | 0 | 0 | 0 | 4760100 | 0 | 0 | 0 | 0 | 0 | 0 | 0 | 0 | 0 | 0 | 0 | 0 | 4335200 |
| ESSETPDQ | CD44; CD44; C | P16070; H0 | CD44_HUMAN | Unmodified | 0 | 0 | 0 | 0 | 11457000 | 0 | 0 | 0 | 0 | 0 | 0 | 0 | 0 | 0 | 0 | 0 | 0 | 8528700 |
| ESSGKYTF | TMED2; TMED2 | Q15363; E7E | TMED2_HUMAI | Unmodified | 33698 | 0 | 0 | 0 | 0 | 0 | 0 | 0 | 6457200 | 364990 | 0 | 0 | 0 | 0 | 0 | 0 | 0 | 0 |
| ESSKVDAN | ABCC1; ABCC1 | P33527; I3L | MRP1_HUMAN | Unmodified | 0 | 0 | 875520 | 0 | 0 | 0 | 0 | 0 | 0 | 0 | 0 | 2834500 | 0 | 0 | 0 | 2193100 | 0 | 0 |
| ESSLLVSEL | KIF14 | Q15058 | KIF14_HUMAN | Unmodified | 0 | 0 | 0 | 0 | 1168800 | 0 | 0 | 0 | 73251 | 0 | 0 | 0 | 0 | 0 | 0 | 0 | 232620 | 297900 |
| ESSLQPAVI | MGAT4B; MGAT | Q9UQ53; E5 | MGT4B_HUMAI | Unmodified | 0 | 0 | 0 | 0 | 0 | 0 | 0 | 0 | 4807800 | 392600 | 0 | 0 | 84669 | 118330 | 1167500 | 1595500 | 0 | 0 |
| ESSLTEPVI | PPP2R5D; PPP2 | Q14738; E9F | 2A5D_HUMAN | Unmodified | 0 | 0 | 0 | 0 | 0 | 0 | 0 | 0 | 0 | 0 | 0 | 8276700 | 0 | 0 | 0 | 0 | 0 | 0 |
| ESSRSHSV | KIF23; KIF23; KI | Q02241; A0A | KIF23_HUMAN | Unmodified | 0 | 0 | 0 | 0 | 1194100 | 447400 | 18842 | 0 | 117980 | 206360 | 94619 | 80649 | 0 | 0 | 0 | 74181 | 0 | 0 |
| ESSSHHPG | FGA | P02671 | FIBA_HUMAN F | Unmodified | 0 | 306620 | 0 | 0 | 0 | 0 | 0 | 0 | 0 | 0 | 0 | 0 | 0 | 0 | 0 | 0 | 0 | 0 |
| ESSTPEEFS | CIT; CIT; CIT; CI | O14578; A0A | CTRO_HUMAN | Unmodified | 0 | 0 | 0 | 0 | 3888300 | 717270 | 0 | 0 | 64363 | 0 | 0 | 463490 | 0 | 0 | 97388 | 0 | 0 | 0 |
| ESSTVFELK | ELOB; ELOB; EL | Q15370; A0A | ELOB_HUMAN | Unmodified | 0 | 0 | 0 | 0 | 3659400 | 634130 | 0 | 0 | 0 | 0 | 0 | 3269700 | 2267400 | 16892 | 0 | 0 | 2064300 | 2233700 |
| ESSYGPEID | ST3GAL1; ST3G | Q11201; E5F | SIA4A_HUMAN | Unmodified | 0 | 0 | 319150 | 526600 | 60788 | 0 | 0 | 0 | 131160 | 0 | 23590 | 0 | 0 | 0 | 0 | 364120 | 0 | 0 |
| ESTAEKDEI | HSP90B2P; HSI | Q58FF3; P14 | ENPLL_HUMAN | Unmodified | 140110 | 0 | 1006400 | 0 | 0 | 0 | 495760 | 210770 | 91312000 | 4699700 | 7461900 | 14873000 | 0 | 122410 | 1300600 | 833570 | 0 | 0 |
| ESTELRAEE | PPFIA2; PPFIA2 | O75334; F8 | LIPA2_HUMAN | Unmodified | 0 | 0 | 0 | 329450 | 0 | 0 | 0 | 81782 | 0 | 0 | 0 | 0 | 0 | 0 | 0 | 0 | 0 | 0 |
| ESTESSNTI | CAMK2D; CAMK | Q13557; E9F | KCC2D_HUMA | Unmodified | 1588800 | 1011100 | 0 | 0 | 107730 | 19049 | 0 | 0 | 865260 | 34171 | 0 | 9168000 | 0 | 0 | 554530 | 362260 | 0 | 139620 |
| ESTGAQVQ | PCBP1; PCBP3; | Q15365; P57 | PCBP1_HUMAI | Unmodified | 15151000 | 14506000 | 4210700 | 9516700 | 0 | 0 | 0 | 0 | 2659900 | 0 | 2526900 | 15584000 | 3507500 | 0 | 5143700 | 3984200 | 0 | 0 |
| ESTGAQVQ | PCBP1; PCBP3; | Q15365; P57 | PCBP1_HUMAI | Oxidation (M) | 29511000 | 29603000 | 9096700 | 10350000 | 973350 | 264870 | 0 | 23423 | 217010 | 88025 | 7634000 | 20690000 | 887870 | 0 | 7047500 | 29559000 | 0 | 1551500 |
| ESTLGPDH | KLC4 | Q9NSK0 | KLC4_HUMAN | Unmodified | 0 | 0 | 0 | 0 | 0 | 0 | 0 | 0 | 257680 | 0 | 0 | 491580 | 0 | 0 | 0 | 0 | 0 | 0 |
| ESTSIHDMI | PPFIA2; PPFIA2 | O75334; G3 | LIPA2_HUMAN | Unmodified | 0 | 0 | 242430 | 236150 | 0 | 0 | 0 | 0 | 0 | 0 | 0 | 0 | 0 | 0 | 372440 | 0 | 0 | 0 |
| ESTTSEQS | ATP5F1C | P36542 | ATPG_HUMAN | Unmodified | 0 | 0 | 0 | 0 | 0 | 0 | 151760 | 40730 | 0 | 632120 | 0 | 0 | 0 | 32099 | 0 | 0 | 0 | 0 |
| ESTTTPGQ | SHC1; SHC1; S | P29353; X6R | SHC1_HUMAN | Unmodified | 0 | 0 | 0 | 0 | 0 | 0 | 605670 | 0 | 0 | 0 | 0 | 3423900 | 285630 | 0 | 0 | 0 | 0 | 0 |
| ESTTYISWK | VPS13C; VPS13 | Q709C8; A0 | VP13C_HUMAN | Unmodified | 0 | 0 | 0 | 24145 | 0 | 0 | 0 | 0 | 61351 | 0 | 51725 | 371410 | 0 | 0 | 31540 | 17658 | 0 | 0 |
| ESVDHLTIF | IARS1; IARS1; I | P41252; A0A | SYIC_HUMAN I | Unmodified | 0 | 0 | 226360 | 0 | 0 | 0 | 0 | 0 | 135550 | 0 | 0 | 5820400 | 0 | 0 | 96627 | 106380 | 0 | 0 |
| ESVELQNG | FLVCR1 | Q9Y5Y0 | FLVC1_HUMAN | Unmodified | 109370 | 196660 | 601410 | 538600 | 831870 | 0 | 0 | 65180 | 383270 | 0 | 330130 | 1093000 | 608230 | 121290 | 580270 | 734150 | 0 | 0 |
| ESVEQQAD | OPA1; OPA1; O | O60313; E5 | OPA1_HUMAN | Unmodified | 0 | 0 | 0 | 0 | 0 | 0 | 0 | 0 | 334420 | 0 | 0 | 4596600 | 0 | 0 | 0 | 0 | 0 | 0 |
| ESVFSPR | LAMP3; LAMP3 | Q9UQV4; E7 | LAMP3_HUMAI | Unmodified | 0 | 0 | 0 | 0 | 0 | 0 | 0 | 0 | 7422500 | 631160 | 0 | 0 | 0 | 0 | 0 | 0 | 0 | 0 |
| ESVLQFYPI | UBA2; UBA2 | Q9UBT2; K7 | SAE2_HUMAN | Unmodified | 0 | 0 | 0 | 0 | 427770 | 0 | 0 | 252060 | 0 | 0 | 0 | 1685600 | 0 | 0 | 0 | 0 | 433370 | 0 |
| ESVNAAFEI | ECHS1 | P30084 | ECHM_HUMAN | Oxidation (M) | 0 | 0 | 0 | 0 | 0 | 0 | 1423400 | 0 | 0 | 635380 | 0 | 0 | 0 | 0 | 0 | 0 | 158540 | 0 |
| ESVNYLVS | NIPSNAP3A | Q9UFN0 | NPS3A_HUMAI | Oxidation (M) | 0 | 0 | 0 | 0 | 0 | 0 | 0 | 0 | 4837800 | 735230 | 0 | 0 | 0 | 0 | 0 | 0 | 0 | 0 |
| ESVNYLVS | NIPSNAP3A | Q9UFN0 | NPS3A_HUMAI | Unmodified | 0 | 0 | 0 | 0 | 0 | 0 | 0 | 0 | 623160 | 0 | 0 | 0 | 0 | 0 | 0 | 0 | 0 | 0 |
| ESVPEFPLS | STMN1; STMN1 | P16949; A2A | STMN1_HUMAI | Unmodified | 0 | 0 | 720340 | 1351700 | 11896000 | 2249600 | 8762200 | 460030 | 37239000 | 22262000 | 22740000 | 1417000 | 7194700 | 0 | 1009200 | 855410 | 6826800 | 10771000 |
| ESVPLDDQ | PIGS | Q96S52 | PIGS_HUMAN ( | Unmodified | 0 | 0 | 0 | 0 | 0 | 0 | 0 | 0 | 378380 | 0 | 79072 | 0 | 0 | 0 | 0 | 0 | 40492 | 0 |
| ESVPLDDQ | PIGS | Q96S52 | PIGS_HUMAN ( | Unmodified | 0 | 0 | 0 | 0 | 0 | 0 | 203410 | 0 | 6718600 | 957380 | 0 | 0 | 0 | 0 | 0 | 0 | 0 | 0 |
| ESVPLDDQ | PIGS | Q96S52 | PIGS_HUMAN ( | Unmodified | 0 | 0 | 0 | 0 | 0 | 0 | 0 | 0 | 8717900 | 0 | 0 | 0 | 0 | 0 | 0 | 0 | 0 | 0 |
| ESVSPESLI | SYT4 | Q9H2B2 | SYT4_HUMAN | Unmodified | 296360 | 0 | 3183100 | 2704000 | 0 | 0 | 0 | 0 | 375290 | 0 | 77421 | 549610 | 302160 | 0 | 5452400 | 3977100 | 0 | 0 |
| ESVVDYCN | SEC62; SEC62 | Q99442; F8 | SEC62_HUMAN | Unmodified | 0 | 0 | 0 | 0 | 0 | 0 | 0 | 0 | 1176400 | 0 | 0 | 77678 | 0 | 0 | 0 | 0 | 0 | 0 |
| ESVVSINK( | JAK1; JAK1; JAK | P23458; A0A | JAK1_HUMAN | Unmodified | 0 | 0 | 0 | 0 | 324430 | 137360 | 100150 | 0 | 241580 | 452710 | 0 | 42949 | 0 | 0 | 0 | 32368 | 51722 | 103450 |
| ESWEMNSI | FKBP4 | Q02790 | FKBP4_HUMAN | Unmodified | 593580 | 318420 | 398240 | 992300 | 685970 | 0 | 0 | 0 | 390050 | 205550 | 558970 | 3063900 | 2740900 | 0 | 442410 | 548880 | 0 | 183230 |
| ESWEMNSI | FKBP4 | Q02790 | FKBP4_HUMAN | Unmodified | 0 | 0 | 0 | 0 | 0 | 0 | 0 | 0 | 0 | 0 | 0 | 0 | 881420 | 0 | 0 | 0 | 0 | 0 |
| ESYPVFYLF | ERP29; ERP29 | P30040; F8 | ERP29_HUMAN | Unmodified | 0 | 0 | 0 | 0 | 0 | 0 | 0 | 0 | 1.05E+08 | 2399400 | 998110 | 1125300 | 0 | 0 | 0 | 256550 | 0 | 0 |
| ESYPVFYLF | ERP29; ERP29 | P30040; F8 | ERP29_HUMAN | Unmodified | 0 | 0 | 0 | 0 | 1020100 | 0 | 649550 | 0 | 2.2E+08 | 1.07E+08 | 1211500 | 293700 | 0 | 0 | 0 | 0 | 0 | 0 |
| ESYPVFYLF | ERP29; ERP29 | P30040; F8 | ERP29_HUMAN | Unmodified | 0 | 0 | 0 | 0 | 0 | 0 | 1226500 | 0 | 0 | 5888500 | 0 | 0 | 0 | 0 | 0 | 0 | 0 | 0 |
| ESYSIYVYK | H2BC19P; H2B | Q6DRA6; P0 | H2B2D_HUMAI | Unmodified | 1.11E+08 | 0 | 0 | 0 | 0 | 0 | 0 | 15189000 | 0 | 0 | 0 | 4.89E+08 | 0 | 0 | 0 | 0 | 0 | 0 |
| ESYSVYVY | H2BC4; H2BC1 | P62807; O6 | H2B1C_HUMAI | Unmodified | 1.91E+08 | 86629000 | 2117200 | 893330 | 7113600 | 2527200 | 9397500 | 39191000 | 4787800 | 0 | 3874700 | 1.71E+09 | 835580 | 0 | 844190 | 1460800 | 373410 | 1894500 |
| ESYSVYVY | H2BC4; H2BC1 | P62807; O6 | H2B1C_HUMAI | Unmodified | 0 | 0 | 0 | 0 | 0 | 0 | 0 | 174520 | 0 | 0 | 0 | 1308200 | 0 | 0 | 0 | 0 | 346750 | 0 |
| ETAEAYLG | HSPA5; HSPA5; | P11021; A0A | BIP_HUMAN Er | Unmodified | 321610 | 0 | 0 | 0 | 78995 | 1871600 | 2966100 | 1074400 | 59945000 | 1515600 | 9902400 | 2193100 | 651980 | 0 | 0 | 0 | 219570 | 811910 |
| ETAEAYLG | HSPA5; HSPA5; | P11021; A0A | BIP_HUMAN Er | Unmodified | 0 | 0 | 0 | 105730 | 0 | 132120 | 109940 | 16802 | 991880 | 434620 | 0 | 50022 | 0 | 0 | 0 | 0 | 0 | 0 |
| ETAENYLGI | HSPA9; HSPA9; | P38646; H0 | GRP75_HUMAI | Unmodified | 1597100 | 4253300 | 69547000 | 0 | 3210100 | 24735000 | 2110100 | 1332300 | 919270 | 138710 | 0 | 0 | 0 | 0 | 14993000 | 2159600 | 564600 | 9655400 |
| ETAFEEDV | GPRC5B | Q9NZH0 | GPC5B_HUMAI | Unmodified | 0 | 0 | 688700 | 0 | 0 | 0 | 0 | 0 | 0 | 0 | 814290 | 1276600 | 0 | 0 | 279720 | 311150 | 0 | 0 |
| ETAHWKPF | NDUFS2; NDUF | O75306; A0A | NDUS2_HUMA | Unmodified | 0 | 0 | 0 | 119630 | 0 | 0 | 0 | 75796 | 1341700 | 0 | 0 | 0 | 0 | 198440 | 319020 | 0 | 104630 | 0 |
| ETAIELGYL | FH | P07954 | FUMH_HUMAN | Unmodified | 0 | 0 | 0 | 0 | 715730 | 0 | 0 | 0 | 1386700 | 850300 | 0 | 241640 | 2447500 | 1021600 | 96557 | 0 | 0 | 1231100 |
| ETAIQTEDS | COBLL1; COBL | Q53SF7; A0A | COBL1_HUMAI | Unmodified | 1591000 | 1915500 | 622060 | 444020 | 0 | 0 | 0 | 0 | 0 | 0 | 0 | 0 | 0 | 0 | 1005800 | 1001200 | 0 | 0 |
| ETALLIDPK | NPEPPSL1; NPI | A6NEC2; P5 | PSAL_HUMAN I | Unmodified | 0 | 40120 | 0 | 0 | 52601 | 0 | 0 | 0 | 0 | 171740 | 0 | 2091000 | 0 | 0 | 0 | 0 | 0 | 0 |
| ETCAWFTD | GFUS | Q13630 | FCL_HUMAN G | Unmodified | 0 | 0 | 0 | 0 | 0 | 0 | 0 | 0 | 0 | 283250 | 0 | 577200 | 401670 | 0 | 0 | 0 | 1397300 | 2566700 |
| ETCSLWPG | SRM; SRM | P19623; K7E | SPEE_HUMAN | Unmodified | 0 | 0 | 0 | 0 | 15478 | 0 | 0 | 0 | 687620 | 87228 | 0 | 166990 | 149440 | 0 | 0 | 0 | 0 | 50047 |
| ETCSLWPG | SRM; SRM | P19623; K7E | SPEE_HUMAN | Unmodified | 0 | 0 | 0 | 0 | 714150 | 0 | 0 | 0 | 1088900 | 137400 | 0 | 0 | 0 | 89082 | 0 | 116830 | 53182 | 78066 |
| ETCVSGED | AKAP12 | Q02952 | AKA12_HUMAN | Unmodified | 0 | 0 | 0 | 0 | 635740 | 0 | 0 | 0 | 0 | 0 | 0 | 0 | 0 | 0 | 0 | 0 | 277580 | 886970 |
| ETDLLLDD | PRDX5 | P30044 | PRDX5_HUMAI | Unmodified | 0 | 0 | 0 | 0 | 0 | 0 | 0 | 0 | 1161500 | 453530 | 302410 | 437790 | 0 | 0 | 0 | 0 | 0 | 0 |
| ETDLQELFI | EIF3G; EIF3G; E | O75821; K7E | EIF3G_HUMAN | Unmodified | 951730 | 891140 | 552810 | 777320 | 0 | 32576 | 0 | 0 | 410960 | 38662 | 354540 | 4153500 | 47110 | 29487 | 1101500 | 1046000 | 17963 | 0 |
| ETDRGMSA | SAMM50 | Q9Y512 | SAM50_HUMAI | Oxidation (M) | 0 | 0 | 0 | 0 | 0 | 0 | 0 | 0 | 1618600 | 0 | 0 | 0 | 0 | 0 | 35041 | 34777 | 0 | 0 |
| ETDRGMSA | SAMM50 | Q9Y512 | SAM50_HUMAI | Unmodified | 0 | 0 | 0 | 0 | 0 | 0 | 0 | 0 | 1895900 | 0 | 0 | 0 | 0 | 0 | 0 | 0 | 0 | 0 |

| | | | | | | | | | | | | | | | | | | | | | | |
|---|---|---|---|---|---|---|---|---|---|---|---|---|---|---|---|---|---|---|---|---|---|---|
| ETDVVNFD | CD2AP | Q9Y5K6 | CD2AP_HUMAI | Unmodified | 16266000 | 11966000 | 11487000 | 3896200 | 348520 | 0 | 565400 | 0 | 0 | 0 | 1242600 | 2597600 | 6159000 | 0 | 22437000 | 16760000 | 41344000 | 13050000 |
| ETEEITSLW | MESD | Q14696 | MESD_HUMAN | Unmodified | 0 | 0 | 0 | 1005200 | 0 | 0 | 0 | 0 | 716370 | 0 | 0 | 0 | 0 | 0 | 0 | 0 | 0 | 0 |
| ETEFKDDSI | CD2AP | Q9Y5K6 | CD2AP_HUMAI | Unmodified | 1518400 | 1224400 | 2080400 | 0 | 0 | 0 | 0 | 0 | 0 | 0 | 0 | 604040 | 0 | 0 | 2736300 | 2076600 | 0 | 0 |
| ETEGDVTS' | CCT8 | P50990 | TCPQ_HUMAN | Unmodified | 82534 | 0 | 576480 | 399560 | 0 | 0 | 42949 | 0 | 2195900 | 389080 | 437030 | 11023000 | 0 | 0 | 838490 | 763980 | 0 | 460700 |
| ETELSLQK | TRIM29; TRIM2 | Q14134; E9F | TRI29_HUMAN | Unmodified | 0 | 0 | 0 | 0 | 0 | 0 | 0 | 0 | 0 | 0 | 0 | 1713600 | 0 | 0 | 0 | 0 | 0 | 0 |
| ETEMALAK | MAP4; MAP4; M | P27816; E7E | MAP4_HUMAN | Unmodified | 0 | 0 | 0 | 0 | 0 | 0 | 0 | 0 | 0 | 0 | 0 | 1553900 | 0 | 0 | 0 | 0 | 0 | 0 |
| ETENDDLTI | IPO7 | O95373 | IPO7_HUMAN I | Unmodified | 0 | 0 | 0 | 0 | 563550 | 422240 | 0 | 66323 | 0 | 0 | 0 | 1789300 | 0 | 0 | 232510 | 205090 | 0 | 0 |
| ETEVIDPQD | SDC4 | P31431 | SDC4_HUMAN | Unmodified | 300570 | 192270 | 846000 | 0 | 0 | 0 | 0 | 0 | 0 | 0 | 0 | 0 | 0 | 0 | 290610 | 416880 | 0 | 0 |
| ETEVVQGK | TPBG | Q13641 | TPBG_HUMAN | Unmodified | 0 | 0 | 0 | 0 | 633300 | 0 | 0 | 0 | 0 | 203150 | 0 | 0 | 0 | 0 | 0 | 0 | 137950 | 0 |
| ETFASTASC | RNPEP; RNPEP | Q9H4A4; A6I | AMPB_HUMAN | Unmodified | 0 | 0 | 0 | 0 | 0 | 0 | 0 | 0 | 539530 | 87519 | 607210 | 1824300 | 0 | 0 | 0 | 104360 | 2079900 | 238510 |
| ETFASTASC | RNPEP; RNPEP | Q9H4A4; A6I | AMPB_HUMAN | Unmodified | 0 | 0 | 0 | 0 | 10522000 | 0 | 0 | 0 | 0 | 1230800 | 0 | 0 | 4182200 | 0 | 0 | 0 | 1236400 | 0 |
| ETFLTSPEE | AHSA1; AHSA1; | O95433; H0' | AHSA1_HUMAI | Unmodified | 0 | 0 | 0 | 0 | 0 | 0 | 0 | 0 | 751370 | 0 | 1873400 | 6160100 | 0 | 0 | 837330 | 0 | 0 | 0 |
| ETFLTSPEE | AHSA1; AHSA1; | O95433; H0' | AHSA1_HUMAI | Unmodified | 0 | 0 | 0 | 0 | 167510 | 0 | 0 | 0 | 0 | 0 | 0 | 0 | 141340 | 0 | 0 | 0 | 0 | 0 |
| ETFNHLTS\ | RAB2B; RAB2B | Q8WUD1; A( | RAB2B_HUMAI | Unmodified | 85791 | 82904 | 0 | 0 | 0 | 0 | 0 | 0 | 896770 | 0 | 251570 | 1707100 | 0 | 0 | 0 | 180750 | 0 | 0 |
| ETFNLYYMI | EPHA3; EPHA3 | P29320; C9J | EPHA3_HUMAI | Oxidation (M) | 815430 | 1023600 | 0 | 0 | 0 | 0 | 0 | 0 | 0 | 0 | 0 | 0 | 0 | 0 | 0 | 0 | 0 | 0 |
| ETFSAPSR/ | KIF14 | Q15058 | KIF14_HUMAN | Unmodified | 0 | 0 | 0 | 0 | 781470 | 0 | 0 | 0 | 0 | 0 | 0 | 0 | 0 | 0 | 0 | 0 | 430140 | 0 |
| ETGAASFL( | TPP2; TPP2; TPI | P29144; A0A | TPP2_HUMAN | Unmodified | 1457400 | 912930 | 0 | 1741400 | 1578300 | 714050 | 0 | 147550 | 0 | 0 | 0 | 10516000 | 0 | 0 | 2244600 | 0 | 0 | 111590 |
| ETGDPGG( | MOV10; MOV1( | Q9HCE1; Q5 | MOV10_HUMA | Unmodified | 4172100 | 6493800 | 2235300 | 1356800 | 0 | 0 | 112200 | 0 | 0 | 0 | 0 | 539980 | 0 | 0 | 4354000 | 3618300 | 0 | 0 |
| ETGEAGW\ | CD2AP | Q9Y5K6 | CD2AP_HUMAI | Unmodified | 0 | 0 | 200170 | 0 | 0 | 0 | 0 | 0 | 0 | 0 | 0 | 382820 | 0 | 0 | 584050 | 358530 | 0 | 0 |
| ETGEAGW\ | CD2AP | Q9Y5K6 | CD2AP_HUMAI | Unmodified | 5956400 | 5432900 | 6309600 | 2957400 | 0 | 0 | 0 | 36068 | 0 | 0 | 137520 | 0 | 0 | 0 | 12455000 | 6211200 | 0 | 0 |
| ETGFSHSQ | CHP1; CHP1; C | Q99653; F5( | CHP1_HUMAN | Unmodified | 200610 | 0 | 863780 | 258180 | 0 | 0 | 0 | 415040 | 53168 | 0 | 0 | 1063400 | 0 | 0 | 1370400 | 827550 | 0 | 0 |
| ETGHTNFG | YOD1 | Q5VVQ6 | OTU1_HUMAN | Unmodified | 1224300 | 935270 | 315000 | 448790 | 0 | 0 | 0 | 0 | 0 | 0 | 0 | 390300 | 491270 | 0 | 480700 | 410300 | 0 | 0 |
| ETGKPKGD | EWSR1; EWSR1 | Q01844; H7I | EWS_HUMAN F | Unmodified | 0 | 0 | 0 | 0 | 0 | 0 | 0 | 0 | 0 | 0 | 102910 | 2218000 | 0 | 0 | 0 | 0 | 0 | 0 |
| ETGQQFAV | CASK; CASK; C/ | O14936; A0/ | CSKP_HUMAN | Unmodified | 0 | 0 | 415070 | 0 | 231660 | 101750 | 2639400 | 0 | 2067600 | 146020 | 0 | 381820 | 91803 | 0 | 840900 | 611360 | 692600 | 313830 |
| ETGTEHSP | AAMDC; AAMD( | Q9H7C9; K4 | AAMDC_HUMA | Unmodified | 0 | 0 | 0 | 0 | 0 | 0 | 0 | 0 | 57920 | 17522 | 77917 | 221770 | 0 | 0 | 182310 | 0 | 0 | 0 |
| ETGVDLTK | HSPA9; HSPA9; | P38646; H0' | GRP75_HUMAI | Unmodified | 0 | 0 | 0 | 0 | 0 | 0 | 0 | 0 | 4630400 | 0 | 0 | 0 | 0 | 0 | 0 | 0 | 0 | 0 |
| ETGVDLTKI | HSPA9; HSPA9; | P38646; H0' | GRP75_HUMAI | Unmodified | 0 | 0 | 0 | 0 | 363410 | 623340 | 0 | 0 | 198980 | 0 | 2493200 | 0 | 0 | 0 | 0 | 0 | 416730 | 0 |
| ETGVPIAGF | MTHFD1; MTHF | P11586; V9( | C1TC_HUMAN | Unmodified | 0 | 109650 | 0 | 203360 | 712220 | 0 | 0 | 33602 | 899300 | 97251 | 225220 | 1465000 | 207940 | 0 | 171490 | 0 | 171660 | 175920 |
| ETGVPIAGF | MTHFD1; MTHF | P11586; V9( | C1TC_HUMAN | Unmodified | 0 | 0 | 0 | 0 | 2329400 | 342050 | 303770 | 121840 | 0 | 0 | 0 | 0 | 834310 | 658160 | 0 | 0 | 1605400 | 3090300 |
| ETGYTELVk | ASPH | Q12797 | ASPH_HUMAN | Unmodified | 0 | 0 | 0 | 0 | 0 | 0 | 0 | 0 | 0 | 1392200 | 0 | 0 | 0 | 0 | 0 | 0 | 0 | 0 |
| ETHKPLGD | ALDH4A1 | P30038 | AL4A1_HUMAN | Unmodified | 0 | 0 | 0 | 0 | 0 | 0 | 0 | 0 | 0 | 0 | 0 | 0 | 1267500 | 0 | 0 | 0 | 0 | 0 |
| ETHSDGPE | CACNA2D2; CA | Q9NY47; C9 | CA2D2_HUMAI | Unmodified | 490330 | 0 | 0 | 0 | 0 | 0 | 265070 | 0 | 3102400 | 758880 | 0 | 858190 | 0 | 0 | 0 | 0 | 0 | 0 |
| ETHVMDYF | PSME2; PSME2; | Q9UL46; A0/ | PSME2_HUMAI | Unmodified | 0 | 0 | 0 | 0 | 0 | 0 | 0 | 0 | 0 | 0 | 50748 | 526310 | 0 | 0 | 0 | 20660 | 0 | 31110 |
| ETIEDLPGK | ABCC4 | O15439 | MRP4_HUMAN | Unmodified | 0 | 0 | 2056500 | 0 | 0 | 0 | 0 | 0 | 0 | 0 | 0 | 0 | 0 | 12068 | 0 | 383260 | 0 | 0 |
| ETIEDLPGK | ABCC4 | O15439 | MRP4_HUMAN | Oxidation (M) | 0 | 0 | 0 | 209260 | 0 | 0 | 0 | 0 | 0 | 0 | 0 | 0 | 0 | 0 | 0 | 517150 | 0 | 0 |
| ETIEDLPGK | ABCC4 | O15439 | MRP4_HUMAN | Unmodified | 0 | 0 | 0 | 0 | 0 | 0 | 0 | 0 | 0 | 0 | 0 | 0 | 0 | 0 | 0 | 0 | 0 | 0 |
| ETIELVHTE | TWF2; TWF2; T\ | Q6IBS0; D6F | TWF2_HUMAN | Unmodified | 0 | 0 | 0 | 0 | 0 | 0 | 0 | 0 | 0 | 0 | 0 | 3031600 | 0 | 0 | 0 | 435160 | 0 | 0 |
| ETIEQEKQ/ | TMSB4X | P62328 | TYB4_HUMAN | Unmodified | 0 | 0 | 0 | 0 | 0 | 122480 | 0 | 495920 | 0 | 0 | 0 | 0 | 0 | 654830 | 0 | 0 | 2994300 | 5138200 |
| ETIEQEKRS | TMSB10 | P63313 | TYB10_HUMAN | Unmodified | 0 | 0 | 0 | 0 | 0 | 0 | 94409 | 148630 | 0 | 0 | 0 | 0 | 0 | 0 | 0 | 0 | 1560300 | 2344900 |
| ETIFSISK | OPA1; OPA1; OI | O60313; E5I | OPA1_HUMAN | Unmodified | 68546 | 57949 | 0 | 0 | 0 | 0 | 0 | 0 | 280690 | 0 | 379610 | 4381800 | 94038 | 0 | 0 | 0 | 0 | 0 |
| ETILCSSDK | CLSTN1; CLSTN | O94985; Q5 | CSTN1_HUMAI | Unmodified | 0 | 0 | 0 | 0 | 0 | 0 | 0 | 0 | 422890 | 0 | 0 | 0 | 0 | 0 | 0 | 0 | 0 | 0 |
| ETILSQK | NPTX1 | Q15818 | NPTX1_HUMAN | Unmodified | 1138800 | 696940 | 905720 | 1134400 | 0 | 814090 | 231820 | 213680 | 47593000 | 3435300 | 648240 | 0 | 45666000 | 4686700 | 0 | 3327000 | 0 | 0 |
| ETILSQKETI | NPTX1 | Q15818 | NPTX1_HUMAN | Unmodified | 0 | 0 | 0 | 0 | 0 | 0 | 0 | 270980 | 4126200 | 34571000 | 0 | 0 | 0 | 0 | 0 | 0 | 0 | 0 |
| ETIMNQEK | BTF3; BTF3; BTI | P20290; A0A | BTF3_HUMAN | Unmodified | 160390 | 0 | 120480 | 0 | 239260 | 4267500 | 54438 | 0 | 38450 | 66703 | 362730 | 1584200 | 0 | 0 | 0 | 29551 | 25813 | 23767 |
| ETIPDIDPN | COPA; COPA; C | P53621; A0A | COPA_HUMAN | Unmodified | 0 | 0 | 0 | 0 | 854600 | 0 | 0 | 0 | 260970 | 0 | 0 | 40226000 | 1079000 | 373150 | 0 | 0 | 0 | 0 |
| ETIPLQETS | PPT1; PPT1; PP | P50897; A0A | PPT1_HUMAN I | Unmodified | 0 | 0 | 0 | 0 | 0 | 0 | 560640 | 274290 | 2026000 | 0 | 0 | 0 | 614070 | 0 | 0 | 0 | 292210 | 949170 |
| ETIQDQLV( | TNFRSF10D | Q9UBN6 | TR10D_HUMAN | Unmodified | 0 | 0 | 0 | 0 | 0 | 178700 | 0 | 0 | 0 | 0 | 239740 | 969010 | 0 | 0 | 0 | 0 | 0 | 0 |
| ETIRELTAK | NPTX1 | Q15818 | NPTX1_HUMAN | Unmodified | 0 | 0 | 0 | 0 | 0 | 0 | 0 | 0 | 27790000 | 4183000 | 0 | 0 | 0 | 0 | 0 | 0 | 0 | 0 |
| ETKPEPMEI | STIP1; STIP1 | P31948; F5H | STIP1_HUMAN | Unmodified | 0 | 0 | 0 | 0 | 209710 | 0 | 21707 | 0 | 861920 | 0 | 2509300 | 2455500 | 121080 | 0 | 0 | 57194 | 0 | 90274 |
| ETKPEPMEI | STIP1; STIP1 | P31948; F5H | STIP1_HUMAN | Oxidation (M) | 0 | 0 | 65812 | 153200 | 0 | 0 | 0 | 0 | 245330 | 65895 | 308970 | 1103800 | 0 | 0 | 56299 | 54274 | 0 | 0 |
| ETKPEPMEI | STIP1; STIP1 | P31948; F5H | STIP1_HUMAN | Unmodified | 0 | 0 | 0 | 0 | 541980 | 0 | 85667 | 0 | 4952300 | 174950 | 4929500 | 25396000 | 0 | 0 | 0 | 0 | 101230 | 1148300 |
| ETKPEPMEI | STIP1; STIP1 | P31948; F5H | STIP1_HUMAN | Oxidation (M) | 0 | 0 | 453010 | 0 | 0 | 0 | 0 | 0 | 765410 | 164180 | 630880 | 4959100 | 724520 | 0 | 435980 | 687890 | 130260 | 339540 |
| ETKSEETLD | FKBP3 | Q00688 | FKBP3_HUMAN | Unmodified | 0 | 235810 | 0 | 0 | 0 | 0 | 0 | 38944 | 540420 | 0 | 135820 | 792970 | 0 | 0 | 581200 | 541140 | 0 | 0 |
| ETLAQLQQ | PFDN6 | O15212 | PFD6_HUMAN | Unmodified | 0 | 0 | 47793 | 0 | 0 | 0 | 0 | 0 | 309480 | 0 | 348890 | 4219600 | 0 | 0 | 125290 | 89576 | 0 | 0 |
| ETLATQLDI | ROCK1; ROCK1 | Q13464; A0/ | ROCK1_HUMAI | Unmodified | 0 | 0 | 0 | 0 | 0 | 0 | 0 | 0 | 0 | 0 | 0 | 1578100 | 0 | 0 | 0 | 0 | 0 | 0 |
| ETLGAQRE. | NPTX2 | P47972 | NPTX2_HUMAN | Unmodified | 0 | 0 | 0 | 0 | 0 | 0 | 0 | 0 | 1285000 | 3167600 | 0 | 0 | 0 | 0 | 0 | 0 | 0 | 0 |
| ETLIAEIEEL | TRIP11; TRIP11 | Q15643; H0' | TRIPB_HUMAN | Unmodified | 0 | 0 | 0 | 0 | 0 | 0 | 0 | 21833 | 0 | 0 | 0 | 357520 | 0 | 0 | 2463300 | 0 | 0 | 0 |
| ETLIQQLKE | KIF20B | Q96Q89 | KI20B_HUMAN | Unmodified | 67064 | 67238 | 0 | 0 | 1432700 | 0 | 0 | 0 | 0 | 0 | 0 | 0 | 0 | 0 | 0 | 0 | 418370 | 392810 |
| ETLITIMK | SCG3 | Q8WXD2 | SCG3_HUMAN | Unmodified | 0 | 0 | 0 | 0 | 0 | 0 | 0 | 0 | 7024200 | 955420 | 77791 | 620690 | 0 | 0 | 0 | 0 | 0 | 0 |
| ETLLAMFK | TARS1 | P26639 | SYTC_HUMAN | Unmodified | 0 | 0 | 66241 | 0 | 350980 | 59141 | 0 | 0 | 475200 | 62868 | 548810 | 1102700 | 0 | 0 | 119050 | 0 | 0 | 0 |
| ETLLNGVG | SNRNP200 | O75643 | U520_HUMAN | Oxidation (M) | 0 | 0 | 0 | 0 | 0 | 0 | 0 | 0 | 0 | 32403 | 0 | 2503700 | 0 | 0 | 0 | 0 | 0 | 0 |
| ETLPAEQDI | PSMA1; PSMA1 | P25786; F5G | PSA1_HUMAN | Unmodified | 0 | 0 | 148660 | 435350 | 910010 | 0 | 0 | 0 | 1230600 | 268700 | 783310 | 7550700 | 148150 | 0 | 351310 | 455880 | 102610 | 0 |
| ETLPLLLQH | MMS19; MMS1 | Q96T76; H0' | MMS19_HUMA | Unmodified | 0 | 0 | 0 | 0 | 0 | 0 | 0 | 0 | 111570 | 0 | 0 | 1254700 | 0 | 0 | 0 | 94658 | 0 | 0 |
| ETLQKLPEE | MAN1A1 | P33908 | MA1A1_HUMAI | Unmodified | 0 | 0 | 0 | 0 | 0 | 24225 | 57219 | 0 | 969950 | 369200 | 0 | 0 | 0 | 0 | 0 | 0 | 0 | 0 |
| ETLQKLPEE | MAN1A1 | P33908 | MA1A1_HUMAI | Unmodified | 0 | 0 | 0 | 0 | 0 | 0 | 0 | 0 | 5367600 | 479360 | 785140 | 810740 | 0 | 0 | 0 | 0 | 0 | 0 |
| ETLQQHKL | HSP90B2P; HSI | Q58FF3; P14 | ENPLL_HUMAN | Unmodified | 0 | 0 | 0 | 0 | 260870 | 0 | 0 | 0 | 246010 | 0 | 0 | 0 | 0 | 0 | 0 | 0 | 0 | 0 |
| ETLVYLTHL | XPO1; XPO1; XI | O14980; A0/ | XPO1_HUMAN | Unmodified | 735960 | 684440 | 375450 | 156890 | 0 | 29246 | 0 | 40595 | 594360 | 573470 | 311310 | 10542000 | 0 | 0 | 318740 | 0 | 0 | 0 |
| ETMQSLNE | KRT18; KRT18 | P05783; F8V | K1C18_HUMAN | Unmodified | 0 | 0 | 0 | 0 | 0 | 0 | 0 | 34548 | 475800 | 60145 | 0 | 122910 | 0 | 0 | 0 | 0 | 358170 | 44789 |
| ETMVTSTTE | PRKCSH; PRKC | P14314; K7E | GLU2B_HUMAI | Unmodified | 0 | 0 | 0 | 0 | 0 | 0 | 0 | 0 | 8202800 | 302310 | 0 | 0 | 0 | 0 | 0 | 0 | 0 | 0 |
| ETNISYSQE | AGPS; AGPS; A( | O00116; A0/ | ADAS_HUMAN | Unmodified | 0 | 0 | 0 | 0 | 0 | 0 | 0 | 0 | 17557000 | 0 | 290520 | 0 | 0 | 0 | 0 | 0 | 0 | 730960 |
| ETNLDSLPI | VIM | P08670 | VIME_HUMAN | Unmodified | 0 | 0 | 0 | 0 | 0 | 0 | 254860 | 0 | 0 | 80620 | 0 | 0 | 0 | 0 | 0 | 0 | 0 | 785970 |
| ETNNAAIIM | GMFB; GMFB | P60983; G3\ | GMFB_HUMAN | Unmodified | 0 | 73598 | 0 | 199760 | 0 | 0 | 68755 | 63294 | 296610 | 98219 | 176370 | 0 | 0 | 0 | 345670 | 41490 | 0 | 0 |
| ETNSGYSG | EXTL2 | Q9UBQ6 | EXTL2_HUMAN | Unmodified | 0 | 0 | 0 | 0 | 0 | 0 | 0 | 0 | 399870 | 0 | 0 | 153040 | 0 | 0 | 0 | 0 | 0 | 0 |
| ETNTVLAR | REPS1; REPS1; | Q96D71; H0 | REPS1_HUMAN | Unmodified | 0 | 0 | 0 | 0 | 477930 | 0 | 0 | 136190 | 0 | 122740 | 28179 | 0 | 0 | 0 | 530930 | 507370 | 0 | 0 |
| ETPAATEAF | BASP1 | P80723 | BASP1_HUMAN | Unmodified | 0 | 0 | 0 | 0 | 17878000 | 7201400 | 775660 | 1016000 | 27043000 | 2619900 | 0 | 0 | 486350 | 481060 | 249300 | 0 | 153980 | 12617000 |
| ETPALTINR | ACAA2; ACAA2; | P42765; A0A | THIM_HUMAN | Unmodified | 0 | 0 | 0 | 0 | 0 | 0 | 0 | 0 | 7778700 | 88226 | 0 | 0 | 0 | 0 | 0 | 0 | 0 | 0 |
| ETPATSAVT | FAM234B; FAM: | A2RU67; Q6 | F234B_HUMAN | Unmodified | 0 | 0 | 0 | 0 | 0 | 0 | 0 | 0 | 959020 | 111890 | 0 | 0 | 0 | 0 | 0 | 0 | 0 | 0 |
| ETPENLSN | PSD3; PSD3; PS | Q9NYI0; E5F | PSD3_HUMAN | Unmodified | 0 | 1326500 | 2814600 | 2146400 | 0 | 0 | 0 | 0 | 0 | 0 | 0 | 0 | 2322400 | 1965100 | 2373200 | 1509000 | 0 | 0 |
| ETPFLSNP( | NT5E; NT5E | P21589; Q9( | 5NTD_HUMAN | Unmodified | 0 | 0 | 0 | 0 | 845500 | 345580 | 0 | 0 | 0 | 0 | 0 | 0 | 0 | 0 | 0 | 0 | 0 | 503390 |
| ETPFLSNP( | NT5E; NT5E | P21589; Q9( | 5NTD_HUMAN | Unmodified | 0 | 0 | 0 | 0 | 45988000 | 0 | 182770 | 0 | 0 | 0 | 0 | 0 | 0 | 0 | 0 | 0 | 1875600 | 2338400 |
| ETPHSPGV | LIMA1; LIMA1; L | Q9UHB6; F8 | LIMA1_HUMAN | Unmodified | 757490 | 773390 | 1388800 | 910200 | 1243800 | 427670 | 0 | 0 | 0 | 0 | 0 | 1539400 | 794170 | 0 | 1026700 | 538070 | 0 | 182550 |
| ETPPDALIL | ABHD12; ABHD | Q8N2K0; A0. | ABD12_HUMAN | Unmodified | 14451 | 0 | 0 | 0 | 182430 | 25855 | 158630 | 0 | 598840 | 108990 | 47130 | 141900 | 0 | 0 | 0 | 0 | 110950 | 109370 |
| ETPPDALIL | ABHD12; ABHD | Q8N2K0; A0. | ABD12_HUMAN | Unmodified | 0 | 0 | 0 | 0 | 0 | 0 | 0 | 0 | 50293000 | 5042500 | 320490 | 1417100 | 0 | 0 | 0 | 0 | 0 | 0 |
| ETPQSLAFT | TJP2; TJP2; TJP2 | Q9UDY2; A0 | ZO2_HUMAN T | Unmodified | 0 | 0 | 0 | 0 | 0 | 0 | 0 | 0 | 7851500 | 0 | 0 | 4012500 | 0 | 0 | 0 | 0 | 0 | 0 |
| ETQALILAP | EIF4A3 | P38919 | IF4A3_HUMAN | Unmodified | 784610 | 552590 | 548540 | 0 | 0 | 0 | 0 | 0 | 106510 | 0 | 0 | 2720300 | 0 | 0 | 563150 | 763500 | 0 | 0 |
| ETQAPGEQ | RHBDF2; RHBD | K7EKI1; K7EI | K7EKI1_HUMAI | Unmodified | 0 | 0 | 0 | 0 | 797110 | 0 | 0 | 0 | 0 | 0 | 0 | 0 | 0 | 0 | 0 | 0 | 0 | 1385700 |
| ETQKSIYYI | HSP90AB1 | P08238 | HS90B_HUMAI | Unmodified | 0 | 0 | 0 | 0 | 402410 | 0 | 60910 | 0 | 0 | 51339 | 0 | 0 | 0 | 0 | 0 | 0 | 340770 | 61920 |
| ETQRLLEEE | CCDC124; CCI | Q96CT7; M0 | CC124_HUMAI | Unmodified | 584570 | 0 | 342090 | 2286100 | 152260 | 22585 | 0 | 0 | 691020 | 56443 | 0 | 0 | 0 | 67368 | 329820 | 215160 | 0 | 131900 |
| ETQTPVMA | DYNC1I2; DYN( | Q13409; E7E | DC1I2_HUMAN | Unmodified | 0 | 0 | 0 | 0 | 877720 | 0 | 0 | 0 | 1399500 | 0 | 0 | 1023000 | 794670 | 582000 | 0 | 0 | 0 | 1583900 |
| ETQTPVMA | DYNC1I2; DYN( | Q13409; E7E | DC1I2_HUMAN | Oxidation (M) | 0 | 0 | 0 | 0 | 0 | 0 | 0 | 218270 | 0 | 240070 | 0 | 3890600 | 695370 | 775530 | 0 | 196110 | 0 | 1255800 |
| ETRAQHLS | SARG | Q9BW04 | SARG_HUMAN | Unmodified | 0 | 0 | 0 | 0 | 1672200 | 434550 | 0 | 0 | 0 | 0 | 0 | 0 | 11100000 | 17465000 | 0 | 0 | 146250 | 0 |
| ETRLVLEVA | ATP5F1B | P06576 | ATPB_HUMAN | Unmodified | 0 | 0 | 0 | 0 | 0 | 0 | 0 | 0 | 162180 | 1123300 | 0 | 0 | 0 | 0 | 0 | 0 | 0 | 0 |
| ETRYGEVFI | AXL; AXL | P30530; M0I | UFO_HUMAN T | Unmodified | 0 | 0 | 0 | 0 | 1624400 | 0 | 0 | 0 | 0 | 0 | 0 | 0 | 0 | 0 | 0 | 0 | 0 | 0 |
| ETRYSYNTI | TMEM43 | Q9BTV4 | TMM43_HUMAI | Unmodified | 0 | 0 | 0 | 0 | 0 | 0 | 46359 | 0 | 1304200 | 695980 | 212350 | 0 | 0 | 0 | 0 | 0 | 0 | 0 |
| ETSDCSYLI | IGF2R; IGF2R | P11717; A0A | MPRI_HUMAN | Unmodified | 0 | 0 | 0 | 0 | 0 | 0 | 0 | 0 | 13033000 | 236990 | 0 | 1082500 | 0 | 0 | 0 | 0 | 0 | 0 |
| ETSLHSPSI | SYPL1 | Q16563 | SYPL1_HUMAN | Unmodified | 483320 | 0 | 807140 | 0 | 0 | 0 | 0 | 0 | 0 | 0 | 0 | 6560700 | 0 | 0 | 3764600 | 3343800 | 0 | 0 |
| ETSSPGTDI | SLC9A1 | P19634 | SL9A1_HUMAN | Unmodified | 0 | 0 | 565060 | 976490 | 302240 | 0 | 0 | 0 | 326440 | 0 | 0 | 0 | 784900 | 0 | 2751200 | 1628700 | 0 | 0 |
| ETSSPGTDI | SLC9A1 | P19634 | SL9A1_HUMAN | Unmodified | 0 | 0 | 0 | 0 | 848690 | 0 | 0 | 0 | 0 | 0 | 0 | 0 | 3205400 | 1198700 | 0 | 0 | 0 | 0 |
| ETTDTDTAI | ACTN4; ACTN4; | O43707; F5( | ACTN4_HUMAI | Unmodified | 158200 | 223750 | 0 | 132920 | 0 | 0 | 0 | 0 | 405300 | 85428 | 0 | 3325600 | 0 | 0 | 550130 | 674950 | 0 | 0 |
| ETTGTGPN | VPS26B | Q4G0F5 | VP26B_HUMAN | Unmodified | 0 | 0 | 123190 | 0 | 0 | 0 | 0 | 0 | 0 | 0 | 0 | 1718600 | 0 | 0 | 184070 | 0 | 0 | 0 |
| ETTVGAVT\ | TNPO3; TNPO3 | Q9Y5L0; C9J | TNPO3_HUMAI | Unmodified | 0 | 0 | 0 | 0 | 152090 | 0 | 0 | 0 | 0 | 132530 | 0 | 925590 | 0 | 0 | 26087 | 0 | 0 | 0 |
| ETTYYDVL( | DNAJA1 | P31689 | DNJA1_HUMAN | Unmodified | 636990 | 788040 | 3386300 | 0 | 0 | 0 | 0 | 0 | 0 | 0 | 0 | 0 | 0 | 0 | 2489400 | 164500 | 0 | 0 |
| ETTYYDVL( | DNAJA1 | P31689 | DNJA1_HUMAN | Unmodified | 461490 | 814890 | 1589200 | 0 | 0 | 0 | 0 | 0 | 0 | 0 | 0 | 466310 | 0 | 0 | 1960700 | 456620 | 0 | 0 |
| ETVDSFLDI | SLC1A4 | P43007 | SATT_HUMAN I | Unmodified | 762900 | 664180 | 321870 | 0 | 0 | 0 | 0 | 0 | 13493000 | 6411300 | 155350 | 18140000 | 0 | 0 | 829170 | 1426200 | 0 | 0 |
| ETVEEQAS | CUL4B; CUL4B; | Q13620; A0/ | CUL4B_HUMAI | Unmodified | 0 | 0 | 0 | 0 | 0 | 0 | 0 | 0 | 63546 | 0 | 134200 | 1067200 | 0 | 0 | 0 | 0 | 0 | 0 |
| ETVEEQVS | CUL4A; CUL4A | Q13619; A0/ | CUL4A_HUMAI | Unmodified | 0 | 0 | 0 | 0 | 0 | 0 | 0 | 0 | 0 | 0 | 0 | 977580 | 0 | 0 | 0 | 0 | 0 | 0 |
| ETVITMSGI | LRP1; LRP1 | Q07954; H0' | LRP1_HUMAN I | Unmodified | 0 | 0 | 0 | 0 | 0 | 0 | 0 | 0 | 247830 | 0 | 0 | 0 | 0 | 0 | 0 | 0 | 0 | 140110 |
| ETVLQQK | NPTX1 | Q15818 | NPTX1_HUMAN | Unmodified | 0 | 72788 | 0 | 0 | 0 | 26922 | 64561 | 46292 | 28654000 | 169060 | 23605 | 234620 | 0 | 0 | 237040 | 157210 | 156570 | 0 |
| ETVLQQKE | NPTX1 | Q15818 | NPTX1_HUMAN | Unmodified | 0 | 0 | 0 | 0 | 0 | 0 | 0 | 0 | 271920 | 951160 | 0 | 0 | 0 | 0 | 0 | 0 | 0 | 0 |

| | | | | | | | | | | | | | | | | | | | | | | |
|---|---|---|---|---|---|---|---|---|---|---|---|---|---|---|---|---|---|---|---|---|---|---|
| ETVLSALSF | PDCD6IP; PDCI | Q8WUM4; C | PDC6I_HUMAN | Unmodified | 7365600 | 8568000 | 3395500 | 798790 | 1365400 | 0 | 0 | 0 | 1415700 | 215850 | 0 | 2339200 | 0 | 0 | 5607900 | 6367000 | 557700 | 0 |
| ETVNVITLY | TSG101; TSG10 | Q99816; F5H | TS101_HUMAN | Unmodified | 14373000 | 47691000 | 41702000 | 25609000 | 0 | 0 | 0 | 0 | 10500000 | 705600 | 383990 | 11836000 | 1403100 | 607810 | 1.22E+08 | 78127000 | 0 | 0 |
| ETVQTTED( | VCL | P18206 | VINC_HUMAN | Unmodified | 0 | 0 | 0 | 0 | 0 | 0 | 0 | 0 | 0 | 0 | 532540 | 3735600 | 0 | 0 | 0 | 0 | 0 | 0 |
| ETVQTTED( | VCL | P18206 | VINC_HUMAN | Unmodified | 0 | 180330 | 462870 | 399850 | 0 | 0 | 0 | 0 | 678080 | 0 | 571030 | 0 | 0 | 0 | 651210 | 680620 | 59959 | 564890 |
| ETVREAEA | VCL | P18206 | VINC_HUMAN | Unmodified | 0 | 0 | 0 | 0 | 657710 | 0 | 56322 | 0 | 112750 | 60238 | 0 | 500760 | 0 | 0 | 119450 | 0 | 714850 | 0 |
| ETVSEESN | EEF2 | P13639 | EF2_HUMAN El | Unmodified | 0 | 0 | 0 | 699270 | 48559 | 0 | 208580 | 0 | 953150 | 0 | 1476200 | 31393000 | 0 | 0 | 199700 | 841330 | 0 | 399860 |
| ETVSQRPG | DCTN1; DCTN1 | Q14203; A0 | DCTN1_HUMA | Unmodified | 0 | 0 | 233470 | 376160 | 0 | 0 | 0 | 0 | 3369400 | 0 | 751750 | 7109700 | 445100 | 0 | 0 | 359050 | 0 | 0 |
| ETVTLEDGI | DPY19L4; DPY1 | Q7Z388; H0 | D19L4_HUMAN | Unmodified | 0 | 0 | 0 | 0 | 141090 | 0 | 0 | 0 | 1611600 | 93112 | 0 | 0 | 0 | 0 | 0 | 0 | 0 | 0 |
| ETVVEVPQ | VCP; VCP; VCP; | P55072; C9J | TERA_HUMAN | Unmodified | 0 | 6744500 | 0 | 313840 | 0 | 0 | 0 | 0 | 9314300 | 0 | 12111000 | 13728000 | 0 | 0 | 675230 | 3638800 | 0 | 0 |
| ETVYNLREI | GOLIM4; GOLIN | O00461; F8V | GOLI4_HUMAN | Unmodified | 0 | 0 | 0 | 0 | 0 | 0 | 0 | 0 | 28163000 | 0 | 0 | 0 | 0 | 0 | 0 | 0 | 0 | 0 |
| ETYDFDIAV | F10 | P00742 | FA10_HUMAN ( | Unmodified | 408520 | 2869800 | 954180 | 2241100 | 794700 | 0 | 0 | 68379 | 0 | 0 | 0 | 104800 | 254470 | 1254700 | 1269100 | 567930 | 366240 | 0 |
| ETYDFDIAV | F10 | P00742 | FA10_HUMAN ( | Unmodified | 0 | 0 | 0 | 0 | 1383700 | 918930 | 0 | 0 | 0 | 0 | 0 | 0 | 0 | 1361600 | 0 | 0 | 0 | 963290 |
| ETYEALLAR | SMC2 | O95347 | SMC2_HUMAN | Unmodified | 0 | 0 | 0 | 0 | 0 | 0 | 0 | 0 | 2934100 | 735800 | 0 | 318250 | 0 | 0 | 0 | 0 | 0 | 0 |
| ETYGAFLQI | EXOC7; EXOC7 | Q9UPT5; B4I | EXOC7_HUMAI | Unmodified | 0 | 0 | 0 | 0 | 3900800 | 284900 | 0 | 0 | 283580 | 0 | 74726 | 2059400 | 766830 | 649510 | 106380 | 104420 | 174650 | 0 |
| ETYGLVETF | ATPAF1; ATPAF | Q5TC12; H0 | ATPF1_HUMAN | Unmodified | 0 | 0 | 0 | 0 | 0 | 0 | 0 | 0 | 1723600 | 0 | 0 | 0 | 0 | 0 | 0 | 0 | 0 | 0 |
| ETYGSGTQ | VPS36; VPS36 | Q86VN1; A0 | VPS36_HUMAN | Unmodified | 2106200 | 1661500 | 2387500 | 1502400 | 0 | 0 | 0 | 0 | 551880 | 41863 | 232580 | 1226300 | 317310 | 49682 | 3380700 | 1738500 | 0 | 0 |
| ETYNALTN | RAB4A | P20338 | RAB4A_HUMAN | Unmodified | 162310 | 0 | 0 | 281580 | 0 | 198980 | 0 | 0 | 261030 | 0 | 0 | 500820 | 0 | 0 | 603060 | 314530 | 0 | 0 |
| ETYNSLAA | RAB4B; RAB4B | P61018; M0F | RAB4B_HUMAN | Unmodified | 136000 | 0 | 0 | 0 | 0 | 0 | 0 | 198470 | 341650 | 290980 | 0 | 199740 | 0 | 0 | 497480 | 161630 | 0 | 0 |
| ETYPNYLPL | TPP2; TPP2; TPI | P29144; A0A | TPP2_HUMAN | Unmodified | 0 | 0 | 0 | 0 | 0 | 0 | 0 | 0 | 590270 | 0 | 0 | 11087000 | 0 | 0 | 0 | 243560 | 118990 | 0 |
| EVAALCR | XRCC6; XRCC6 | P12956; B1A | XRCC6_HUMAI | Unmodified | 314760 | 67421 | 71924 | 28182 | 117960 | 56759 | 97863 | 109820 | 169750 | 122510 | 274380 | 3422200 | 0 | 0 | 55107 | 49378 | 0 | 55130 |
| EVAAQVKA | PARK7; PARK7; | Q99497; K7E | PARK7_HUMAN | Unmodified | 0 | 0 | 0 | 0 | 2810800 | 0 | 603910 | 408380 | 0 | 1037800 | 0 | 0 | 2830000 | 740480 | 0 | 0 | 2061100 | 2534700 |
| EVAATGTSI | SLC38A10; SLC | Q9HBR0; H0 | S38AA_HUMAN | Unmodified | 0 | 0 | 0 | 0 | 0 | 0 | 0 | 0 | 708130 | 0 | 0 | 117980 | 0 | 0 | 0 | 0 | 94965 | 0 |
| EVADAIRLN | SDF4; SDF4; SD | Q9BRK5; H0 | CAB45_HUMAN | Unmodified | 0 | 0 | 0 | 0 | 803500 | 156880 | 0 | 46406 | 1048400 | 0 | 0 | 0 | 0 | 0 | 97326 | 0 | 0 | 0 |
| EVADRWYS | GLIPR2 | Q9H4G4 | GAPR1_HUMAI | Unmodified | 1001100 | 996350 | 2938300 | 2113200 | 1274200 | 372910 | 460090 | 0 | 1409700 | 216720 | 0 | 737110 | 1709100 | 2594700 | 2319400 | 1947900 | 182340 | 721380 |
| EVAEIRQEI | SLC2A5 | P22732 | GTR5_HUMAN | Unmodified | 0 | 0 | 0 | 0 | 0 | 0 | 0 | 0 | 22009 | 0 | 0 | 0 | 0 | 0 | 0 | 169660 | 57536 | 0 |
| EVAENQQN | EPB41L2; EPB4 | O43491; E9F | E41L2_HUMAN | Unmodified | 0 | 0 | 0 | 0 | 720480 | 0 | 0 | 0 | 0 | 0 | 0 | 0 | 0 | 0 | 0 | 0 | 0 | 0 |
| EVAENQQN | EPB41L2; EPB4 | O43491; E9F | E41L2_HUMAN | Unmodified | 0 | 0 | 0 | 0 | 3032800 | 496580 | 0 | 0 | 0 | 0 | 0 | 0 | 293970 | 0 | 0 | 0 | 262070 | 1543300 |
| EVAGLWIK | GM2A; GM2A; ( | P17900; H0 | SAP3_HUMAN | Unmodified | 0 | 0 | 290650 | 136890 | 0 | 0 | 0 | 0 | 1837700 | 248370 | 456630 | 240480 | 746860 | 324070 | 253910 | 121750 | 0 | 322640 |
| EVAGPTEM | UTRN; UTRN; U | P46939; H0 | UTRN_HUMAN | Unmodified | 0 | 0 | 0 | 126440 | 0 | 0 | 0 | 0 | 218500 | 0 | 0 | 117330 | 0 | 0 | 303440 | 241370 | 0 | 0 |
| EVAIIVDQR | DGLUCY; DGLU | Q7Z3D6; H0 | GLUCM_HUMA | Unmodified | 0 | 0 | 0 | 0 | 0 | 0 | 0 | 0 | 0 | 207670 | 0 | 0 | 0 | 0 | 0 | 0 | 0 | 0 |
| EVAKPSPG | TP53I3; TP53I3 | Q53FA7; H7I | QORX_HUMAN | Unmodified | 0 | 0 | 0 | 0 | 635470 | 0 | 0 | 0 | 388100 | 1331100 | 0 | 743510 | 0 | 0 | 0 | 0 | 1635800 | 1667600 |
| EVALDLSQ | QSOX1 | O00391 | QSOX1_HUMAI | Unmodified | 0 | 0 | 0 | 0 | 0 | 0 | 0 | 0 | 1077900 | 0 | 107110 | 0 | 0 | 0 | 0 | 0 | 0 | 0 |
| EVANSTAN | TLN2; TLN1; TLI | Q9Y4G6; Q9 | TLN2_HUMAN | Unmodified | 635370 | 675930 | 1432600 | 3344400 | 9246800 | 2452900 | 6798100 | 3787000 | 995730 | 577310 | 634300 | 3342200 | 6998700 | 8323200 | 800220 | 1034100 | 4154300 | 6298000 |
| EVAQSLLN | SPC24; SPC24; | Q8NBT2; K7I | SPC24_HUMAN | Unmodified | 79957 | 0 | 0 | 170700 | 0 | 0 | 0 | 0 | 0 | 0 | 0 | 1239300 | 715060 | 475620 | 172960 | 122940 | 0 | 0 |
| EVARLEAPL | PSMD5; PSMD5 | Q16401; F2Z | PSMD5_HUMA | Unmodified | 0 | 0 | 0 | 0 | 0 | 0 | 0 | 0 | 0 | 0 | 0 | 0 | 0 | 0 | 0 | 0 | 0 | 0 |
| EVASEIWK | NCAM2; NCAM | O15394; H9I | NCAM2_HUMA | Unmodified | 120640 | 183780 | 1488700 | 0 | 0 | 138740 | 0 | 0 | 0 | 0 | 0 | 0 | 0 | 0 | 3849900 | 3278700 | 0 | 0 |
| EVATAIRGA | RPS2; RPS2; RP | P15880; E9P | RS2_HUMAN 4 | Unmodified | 205290 | 0 | 114600 | 553650 | 1392400 | 171680 | 9468300 | 2386900 | 0 | 0 | 1060900 | 1081300 | 393700 | 580770 | 0 | 0 | 237950 | 994240 |
| EVCEEAGV | NUDT3; RPS10- | O95989; A0 | NUDT3_HUMAI | Unmodified | 0 | 0 | 97732 | 141410 | 0 | 0 | 0 | 0 | 0 | 73618 | 146900 | 368190 | 0 | 0 | 38416 | 0 | 0 | 0 |
| EVCFACVD | CLTC; CLTC; CI | Q00610; J3K | CLH1_HUMAN | Unmodified | 1645700 | 1100000 | 0 | 393560 | 0 | 0 | 0 | 0 | 0 | 0 | 0 | 2141200 | 0 | 0 | 679380 | 1243400 | 0 | 0 |
| EVCFTIENK | SND1 | Q7KZF4 | SND1_HUMAN | Unmodified | 261390 | 185100 | 0 | 210230 | 0 | 0 | 0 | 0 | 778170 | 850010 | 0 | 747960 | 248710 | 151770 | 182550 | 0 | 831550 | 0 |
| EVCGFAPY | RPL36; RPL36 | Q9Y3U8; J3C | RL36_HUMAN ( | Unmodified | 1519700 | 79288 | 136140 | 637900 | 151510 | 235910 | 162670 | 0 | 0 | 0 | 154570 | 8279400 | 147820 | 55556 | 0 | 129830 | 0 | 0 |
| EVCRDFAV | CCDC50 | Q8IVM0 | CCD50_HUMA | Unmodified | 16808000 | 18183000 | 14571000 | 27399000 | 0 | 0 | 0 | 0 | 1213800 | 0 | 0 | 0 | 18278000 | 1866500 | 33613000 | 16819000 | 0 | 834270 |
| EVCRDFAV | CCDC50 | Q8IVM0 | CCD50_HUMA | Unmodified | 0 | 0 | 0 | 0 | 0 | 0 | 0 | 0 | 0 | 0 | 0 | 0 | 8738100 | 0 | 0 | 0 | 0 | 0 |
| EVDDFFEQ | SNX5; SNX5; SN | Q9Y5X3; Q5 | SNX5_HUMAN | Unmodified | 0 | 0 | 0 | 0 | 0 | 0 | 0 | 0 | 274520 | 0 | 0 | 900960 | 0 | 0 | 0 | 79004 | 0 | 0 |
| EVDDLGPE | DHX15 | O43143 | DHX15_HUMAI | Unmodified | 173120 | 85734 | 0 | 0 | 0 | 0 | 0 | 0 | 0 | 0 | 1267200 | 3744000 | 0 | 0 | 0 | 0 | 0 | 0 |
| EVDEDFDS | EFHD2; EFHD2 | Q96C19; H0 | EFHD2_HUMAI | Unmodified | 0 | 0 | 0 | 0 | 1064800 | 144220 | 0 | 0 | 1786700 | 0 | 345140 | 1605800 | 0 | 0 | 0 | 0 | 18577 | 117210 |
| EVDEDFDS | EFHD2; EFHD2 | Q96C19; H0 | EFHD2_HUMAI | Unmodified | 0 | 0 | 0 | 0 | 0 | 0 | 0 | 0 | 0 | 0 | 0 | 0 | 0 | 0 | 0 | 0 | 0 | 0 |
| EVDEKPAS | ARPC1A; ARPC | Q92747; E9F | ARC1A_HUMAI | Unmodified | 63709 | 47323 | 0 | 0 | 0 | 0 | 0 | 0 | 0 | 0 | 0 | 1576100 | 0 | 0 | 0 | 0 | 0 | 0 |
| EVDEQMLA | TUBB6; TUBB3; | Q9BUF5; Q1 | TBB6_HUMAN | Unmodified | 0 | 0 | 0 | 0 | 1298600 | 300940 | 586650 | 173230 | 2662600 | 353940 | 332840 | 8827700 | 0 | 0 | 0 | 0 | 145820 | 0 |
| EVDEQMLA | TUBB6; TUBB3; | Q9BUF5; Q1 | TBB6_HUMAN | Oxidation (M) | 0 | 0 | 0 | 0 | 367610 | 0 | 102360 | 71395 | 0 | 0 | 0 | 0 | 0 | 0 | 0 | 0 | 0 | 0 |
| EVDEQMLN | TUBB2A; TUBB; | Q13885; P07 | TBB2A_HUMAN | Oxidation (M) | 849080 | 556010 | 2403600 | 1288300 | 3631000 | 1306600 | 1681500 | 1159200 | 5306600 | 830230 | 2275900 | 15826000 | 1771900 | 865110 | 2437800 | 1958300 | 550870 | 1691700 |
| EVDEQMLN | TUBB2A; TUBB; | Q13885; P07 | TBB2A_HUMAN | Unmodified | 7007400 | 3825500 | 5461500 | 6396600 | 12653000 | 716780 | 6351600 | 3315200 | 45808000 | 5552400 | 22762000 | 2.59E+08 | 8007100 | 3664700 | 7458700 | 5043600 | 2225200 | 7501600 |
| EVDGEGKF | DPP3; DPP3; DI | Q9NY33; G3 | DPP3_HUMAN | Unmodified | 0 | 0 | 0 | 910490 | 0 | 209200 | 0 | 0 | 912180 | 0 | 1721200 | 7019600 | 0 | 0 | 447430 | 373740 | 0 | 0 |
| EVDGLDVS | DYNC1H1; DYN | Q14204; A0 | DYHC1_HUMA | Unmodified | 0 | 0 | 456280 | 0 | 0 | 0 | 0 | 18945 | 770560 | 0 | 625160 | 3252400 | 0 | 0 | 304320 | 234570 | 31094 | 0 |
| EVDGLLR | ARPC5L; DNAJ( | Q9BPX5; Q8 | ARP5L_HUMAN | Unmodified | 0 | 0 | 0 | 0 | 0 | 0 | 0 | 0 | 6731700 | 674640 | 167860 | 0 | 0 | 0 | 0 | 0 | 0 | 0 |
| EVDIGIPDA | VCP; VCP; VCP; | P55072; C9J | TERA_HUMAN | Unmodified | 0 | 0 | 0 | 0 | 0 | 0 | 0 | 0 | 658910 | 0 | 1713900 | 0 | 0 | 0 | 0 | 0 | 0 | 0 |
| EVDIGIPDA | VCP; VCP; VCP; | P55072; C9J | TERA_HUMAN | Unmodified | 824960 | 0 | 0 | 0 | 0 | 0 | 0 | 0 | 1555100 | 271050 | 2479400 | 4859100 | 0 | 0 | 0 | 1505000 | 278270 | 0 |
| EVDLKDYEI | FGA; FGA | P02671; A0A | FIBA_HUMAN F | Unmodified | 0 | 4063700 | 0 | 0 | 107740 | 0 | 0 | 0 | 0 | 0 | 0 | 0 | 0 | 0 | 0 | 0 | 0 | 0 |
| EVDLKDYEI | FGA; FGA | P02671; A0A | FIBA_HUMAN F | Unmodified | 0 | 439080 | 0 | 0 | 0 | 0 | 0 | 0 | 0 | 0 | 0 | 0 | 0 | 636460 | 0 | 0 | 0 | 0 |
| EVDNLYNII | CDCA8 | Q53HL2 | BOREA_HUMAI | Unmodified | 0 | 0 | 242320 | 1211000 | 4808300 | 232030 | 0 | 0 | 741990 | 1009700 | 0 | 0 | 2019800 | 0 | 313930 | 186840 | 529200 | 0 |
| EVDPSTGE | SQSTM1 | Q13501 | SQSTM_HUMAI | Oxidation (M) | 0 | 0 | 0 | 660240 | 2392500 | 0 | 0 | 0 | 2006200 | 0 | 0 | 0 | 0 | 0 | 0 | 0 | 0 | 0 |
| EVDPSTGE | SQSTM1 | Q13501 | SQSTM_HUMAI | Unmodified | 0 | 0 | 0 | 0 | 0 | 0 | 0 | 0 | 3952000 | 0 | 0 | 0 | 0 | 0 | 0 | 0 | 0 | 0 |
| EVDRYDAP | PACC1 | Q9H813 | PACC1_HUMAI | Unmodified | 0 | 0 | 0 | 0 | 0 | 0 | 0 | 0 | 6336400 | 0 | 0 | 2912500 | 0 | 0 | 0 | 0 | 0 | 0 |
| EVDSGNDI | TIMP2 | P16035 | TIMP2_HUMAN | Unmodified | 0 | 0 | 0 | 0 | 1545500 | 1449200 | 0 | 0 | 2166200 | 669550 | 556420 | 697220 | 0 | 0 | 263900 | 0 | 168750 | 1499000 |
| EVDSKSQV | PACS1; PACS1; | Q6VY07; B4I | PACS1_HUMAI | Unmodified | 0 | 0 | 0 | 0 | 850480 | 0 | 0 | 0 | 0 | 0 | 157600 | 0 | 185810 | 0 | 0 | 0 | 0 | 44886 |
| EVDVFSPLI | CNNM3 | Q8NE01 | CNNM3_HUMA | Unmodified | 0 | 0 | 342110 | 0 | 0 | 0 | 0 | 0 | 869320 | 135060 | 0 | 958780 | 0 | 0 | 369500 | 402380 | 0 | 0 |
| EVDVGLAA | ECH1; ECH1 | Q13011; M0I | ECH1_HUMAN | Unmodified | 0 | 0 | 0 | 0 | 0 | 0 | 0 | 0 | 4913700 | 1822500 | 0 | 0 | 236330 | 0 | 0 | 0 | 0 | 0 |
| EVDVRYTE | CMIP; CMIP; CN | Q8IY22; A0A | CMIP_HUMAN | Unmodified | 0 | 0 | 0 | 0 | 2368900 | 0 | 322920 | 209920 | 0 | 0 | 0 | 0 | 0 | 0 | 0 | 0 | 522560 | 1152100 |
| EVDYEEMK | DSG2 | Q14126 | DSG2_HUMAN | Unmodified | 0 | 32766 | 127800 | 0 | 623620 | 0 | 0 | 0 | 3049900 | 331860 | 217380 | 286710 | 0 | 0 | 0 | 0 | 0 | 0 |
| EVDYSGLR | CDV3; CDV3 | Q9UKY7; H0 | CDV3_HUMAN | Unmodified | 0 | 127770 | 0 | 223300 | 498220 | 0 | 0 | 0 | 0 | 0 | 0 | 1563700 | 294980 | 0 | 302510 | 286770 | 0 | 116500 |
| EVEAEFLR | IRAG2; IRAG2; I | Q12912; F5H | IRAG2_HUMAN | Unmodified | 0 | 0 | 0 | 0 | 0 | 0 | 0 | 0 | 0 | 0 | 783730 | 1382700 | 0 | 0 | 0 | 0 | 0 | 0 |
| EVEAVIPDH | HADHA; HADH | P40939; A0A | ECHA_HUMAN | Unmodified | 0 | 0 | 0 | 0 | 0 | 0 | 0 | 0 | 2393800 | 0 | 0 | 0 | 0 | 0 | 0 | 0 | 0 | 0 |
| EVEAVIPDH | HADHA; HADH | P40939; A0A | ECHA_HUMAN | Unmodified | 0 | 0 | 0 | 0 | 0 | 0 | 539600 | 0 | 6035500 | 636910 | 0 | 0 | 0 | 0 | 0 | 0 | 0 | 0 |
| EVEDKESE( | ZRANB2 | O95218 | ZRAB2_HUMAN | Unmodified | 0 | 0 | 0 | 0 | 97498 | 0 | 0 | 0 | 92009 | 20253 | 147210 | 380480 | 0 | 0 | 0 | 0 | 0 | 0 |
| EVEEDEYK | HSP90B1; HSP9 | P14625; A0A | ENPL_HUMAN | Unmodified | 0 | 0 | 515920 | 0 | 0 | 0 | 56797 | 0 | 31033000 | 856800 | 1165900 | 6488400 | 0 | 0 | 541130 | 247050 | 0 | 0 |
| EVEEDEYK | HSP90B1; HSP9 | P14625; A0A | ENPL_HUMAN | Unmodified | 0 | 0 | 0 | 0 | 0 | 0 | 0 | 100070 | 4024400 | 3610600 | 0 | 0 | 0 | 0 | 0 | 0 | 0 | 0 |
| EVEEEPGII | HYOU1; HYOU1 | Q9Y4L1; E9F | HYOU1_HUMA | Unmodified | 0 | 0 | 709400 | 0 | 394800 | 0 | 82773 | 46659 | 9377100 | 411430 | 1643100 | 961540 | 0 | 0 | 754900 | 423550 | 84919 | 0 |
| EVEEQHEV | CGN | Q9P2M7 | CING_HUMAN | Unmodified | 0 | 0 | 0 | 0 | 58018 | 0 | 0 | 0 | 58725 | 0 | 0 | 504360 | 0 | 0 | 0 | 0 | 0 | 0 |
| EVEERPAP | ARPC1B; ARPC | O15143; C9J | ARC1B_HUMAI | Unmodified | 0 | 0 | 0 | 0 | 0 | 0 | 0 | 0 | 0 | 0 | 0 | 599550 | 0 | 0 | 0 | 0 | 0 | 0 |
| EVEETDEM | DNAJA1 | P31689 | DNJA1_HUMAN | Oxidation (M) | 0 | 0 | 0 | 0 | 0 | 0 | 0 | 0 | 1620300 | 0 | 0 | 1583500 | 0 | 0 | 0 | 0 | 0 | 0 |
| EVEETDEM | DNAJA1 | P31689 | DNJA1_HUMAN | Unmodified | 0 | 0 | 0 | 0 | 0 | 0 | 0 | 0 | 3550100 | 0 | 0 | 0 | 0 | 0 | 0 | 0 | 0 | 0 |
| EVEHFLK | AREL1; SLC16A | O15033; O1 | AREL1_HUMAN | Unmodified | 0 | 0 | 0 | 0 | 251600 | 0 | 0 | 0 | 113510 | 33439 | 0 | 1119100 | 0 | 0 | 0 | 0 | 290980 | 271850 |
| EVEIAYSDV | CLIC4 | Q9Y696 | CLIC4_HUMAN | Unmodified | 162600 | 0 | 0 | 0 | 1084100 | 261600 | 0 | 0 | 0 | 206660 | 495820 | 0 | 0 | 0 | 0 | 178590 | 324750 | 316640 |
| EVELNELEF | SLC3A2; SLC3A | P08195; A0A | 4F2_HUMAN 4I | Unmodified | 257340 | 0 | 0 | 0 | 884640 | 0 | 217850 | 53848 | 985610 | 0 | 1748900 | 2268200 | 0 | 125880 | 0 | 0 | 297220 | 305500 |
| EVELNELEF | SLC3A2; SLC3A | P08195; A0A | 4F2_HUMAN 4I | Oxidation (M) | 0 | 0 | 0 | 0 | 917600 | 0 | 0 | 0 | 0 | 0 | 668360 | 377470 | 0 | 0 | 0 | 0 | 0 | 0 |
| EVELNELEF | SLC3A2; SLC3A | P08195; A0A | 4F2_HUMAN 4I | 2 Oxidation (M) | 0 | 193370 | 0 | 0 | 0 | 0 | 0 | 0 | 0 | 0 | 0 | 1785900 | 0 | 0 | 804930 | 287180 | 0 | 0 |
| EVENFEK | L1RE1 | Q9UN81 | LORF1_HUMAN | Unmodified | 0 | 662680 | 457880 | 0 | 0 | 0 | 0 | 0 | 0 | 0 | 0 | 0 | 0 | 0 | 0 | 0 | 0 | 0 |
| EVENSEDP | VCL | P18206 | VINC_HUMAN | Unmodified | 135980 | 249220 | 0 | 0 | 0 | 0 | 0 | 0 | 304910 | 134890 | 0 | 2307900 | 0 | 0 | 0 | 323420 | 0 | 0 |
| EVENSEDP | VCL | P18206 | VINC_HUMAN | Unmodified | 0 | 0 | 0 | 0 | 0 | 0 | 0 | 0 | 0 | 0 | 98368 | 0 | 0 | 0 | 0 | 0 | 313040 | 0 |
| EVEPALELL | GDI1 | P31150 | GDIA_HUMAN | Unmodified | 0 | 0 | 501880 | 271170 | 0 | 0 | 14490 | 0 | 2840200 | 199520 | 608240 | 2091900 | 0 | 0 | 513450 | 1124600 | 0 | 0 |
| EVEPEPTEI | EIF2S2 | P20042 | IF2B_HUMAN E | Unmodified | 0 | 0 | 307210 | 0 | 0 | 0 | 0 | 0 | 145440 | 0 | 126380 | 416680 | 0 | 0 | 188900 | 197640 | 0 | 0 |
| EVEPEPTEI | EIF2S2 | P20042 | IF2B_HUMAN E | Unmodified | 0 | 0 | 0 | 0 | 0 | 0 | 0 | 0 | 0 | 0 | 0 | 1466800 | 0 | 0 | 0 | 379400 | 0 | 0 |
| EVEQFTQV | NDUFB10; NDU | O96000; H3I | NDUBA_HUMA | Unmodified | 0 | 0 | 0 | 0 | 0 | 0 | 0 | 0 | 1402500 | 84964 | 0 | 0 | 596580 | 0 | 0 | 0 | 0 | 0 |
| EVEQLASK( | ACTRT1 | Q8TDG2 | ACTT1_HUMAN | Unmodified | 0 | 0 | 0 | 0 | 482890 | 5051600 | 632320 | 172750 | 0 | 0 | 0 | 0 | 0 | 0 | 0 | 0 | 246860 | 0 |
| EVERVVVD | CKMT1A; CKMT | P12532; C9J | KCRU_HUMAN | Unmodified | 0 | 0 | 678650 | 0 | 53523 | 0 | 0 | 34718 | 180860 | 35121 | 337070 | 0 | 0 | 0 | 106470 | 0 | 0 | 0 |
| EVESMGAH | UQCRC1 | P31930 | QCR1_HUMAN | Unmodified | 0 | 0 | 44309 | 0 | 0 | 0 | 0 | 0 | 18840000 | 607450 | 177460 | 508160 | 0 | 0 | 0 | 0 | 0 | 0 |
| EVESMGAH | UQCRC1 | P31930 | QCR1_HUMAN | Oxidation (M) | 0 | 0 | 0 | 0 | 0 | 0 | 0 | 0 | 3059700 | 0 | 0 | 0 | 0 | 0 | 0 | 0 | 0 | 0 |
| EVESMGAH | UQCRC1 | P31930 | QCR1_HUMAN | Oxidation (M) | 0 | 0 | 0 | 0 | 0 | 0 | 256420 | 0 | 912620 | 1634500 | 0 | 0 | 0 | 0 | 0 | 0 | 0 | 72805 |
| EVESRLSP( | PPP1R18; PPP1 | Q6NYC8; A0 | PPR18_HUMAN | Unmodified | 0 | 0 | 0 | 0 | 687920 | 62079 | 0 | 0 | 0 | 0 | 0 | 0 | 0 | 0 | 0 | 0 | 155780 | 342960 |
| EVESVDLPI | PON2; PON2; P | Q15165; G3 | PON2_HUMAN | Unmodified | 0 | 0 | 0 | 0 | 0 | 0 | 0 | 0 | 3614500 | 139070 | 0 | 0 | 0 | 0 | 0 | 0 | 0 | 294800 |
| EVETHANN | TAOK3; TAOK3 | Q9H2K8; G3 | TAOK3_HUMAN | Unmodified | 483240 | 365390 | 777230 | 0 | 2981400 | 445680 | 927920 | 0 | 192260 | 74723 | 262570 | 394870 | 1955300 | 1769800 | 743740 | 858690 | 439200 | 890760 |
| EVEVEVESI | SND1 | Q7KZF4 | SND1_HUMAN | Unmodified | 0 | 136520 | 0 | 0 | 0 | 0 | 0 | 0 | 0 | 0 | 0 | 2588900 | 0 | 0 | 423360 | 287650 | 0 | 0 |
| EVEVIGGAI | TK1 | P04183 | KITH_HUMAN T | Unmodified | 248620 | 209130 | 256870 | 271520 | 0 | 0 | 0 | 0 | 286820 | 69397 | 0 | 457580 | 0 | 0 | 765620 | 192900 | 0 | 0 |
| EVFAEAVR | RHOG | P84095 | RHOG_HUMAN | Unmodified | 0 | 0 | 0 | 0 | 1008900 | 0 | 142270 | 0 | 0 | 123830 | 193600 | 0 | 0 | 0 | 0 | 0 | 0 | 437980 |
| EVFDNDGI | PPP3CA | Q08209 | PP2BA_HUMAN | Unmodified | 323980 | 0 | 0 | 589220 | 592710 | 0 | 272710 | 164460 | 0 | 224030 | 0 | 0 | 1258200 | 906070 | 561740 | 340230 | 217850 | 378740 |
| EVFEDAAEI | NCL; NCL; NCL | P19338; C9J | NUCL_HUMAN | Unmodified | 127920 | 295830 | 126760 | 0 | 0 | 0 | 839920 | 533850 | 1414300 | 0 | 6404000 | 4461700 | 84492 | 0 | 0 | 0 | 0 | 0 |
| EVFEMATR | RHOA; RHOC; F | P61586; P08 | RHOA_HUMAN | Unmodified | 3052900 | 4152300 | 3094100 | 5262700 | 964500 | 0 | 76560 | 41782 | 18700000 | 0 | 5895500 | 41585000 | 1583000 | 502820 | 6299300 | 4228000 | 635560 | 1025500 |
| EVFEMATR | RHOA; RHOC; F | P61586; P08 | RHOA_HUMAN | Oxidation (M) | 1836400 | 2043700 | 1503200 | 1312900 | 370480 | 101250 | 0 | 0 | 3945300 | 272420 | 1266300 | 5598500 | 436870 | 0 | 2049500 | 1965600 | 148720 | 0 |
| EVFESSKMI | UGGT1 | Q9NYU2 | UGGG1_HUMA | Unmodified | 0 | 0 | 0 | 0 | 0 | 0 | 0 | 0 | 7183400 | 841280 | 0 | 0 | 0 | 0 | 0 | 0 | 0 | 0 |

| | | | | | | | | | | | | | | | | | | | | | | |
|---|---|---|---|---|---|---|---|---|---|---|---|---|---|---|---|---|---|---|---|---|---|---|
| EVFETATR | RHOB | P62745 | RHOB_HUMAN | Unmodified | 431810 | 353150 | 338500 | 900480 | 0 | 0 | 0 | 5888.6 | 368030 | 0 | 400250 | 3547700 | 96053 | 35248 | 0 | 695650 | 0 | 0 |
| EVFGDLLL | PLCG2; PLCG2 | P16885; A0A | PLCG2_HUMAN | Unmodified | 0 | 0 | 0 | 0 | 0 | 137790 | 0 | 0 | 0 | 0 | 2610100 | 11325000 | 0 | 35319 | 0 | 0 | 0 | 0 |
| EVFGDLLL | PLCG2; PLCG2 | P16885; A0A | PLCG2_HUMAN | Unmodified | 0 | 0 | 0 | 0 | 376020 | 0 | 0 | 0 | 0 | 0 | 0 | 21437000 | 0 | 0 | 0 | 0 | 0 | 1399200 |
| EVFIQTLDD | MME; MME; MM | P08473; A0A | NEP_HUMAN N | Oxidation (M) | 0 | 0 | 0 | 3468600 | 0 | 0 | 0 | 0 | 16707 | 51228 | 0 | 0 | 100430 | 0 | 8541100 | 322940 | 0 | 0 |
| EVFIQTLDD | MME; MME; MM | P08473; A0A | NEP_HUMAN N | Unmodified | 0 | 0 | 0 | 115370 | 0 | 0 | 0 | 0 | 27890 | 37998 | 0 | 0 | 0 | 0 | 434800 | 0 | 0 | 0 |
| EVFIQTLDD | MME; MME; MM | P08473; A0A | NEP_HUMAN N | Oxidation (M) | 0 | 0 | 605320 | 4288700 | 0 | 0 | 0 | 0 | 0 | 0 | 0 | 0 | 0 | 0 | 6028200 | 2889800 | 0 | 0 |
| EVFIRELISN | TRAP1 | Q12931 | TRAP1_HUMAN | Unmodified | 0 | 0 | 0 | 0 | 504090 | 0 | 309080 | 0 | 1088200 | 1524100 | 243160 | 0 | 151190 | 0 | 0 | 0 | 170360 | 0 |
| EVFLPSTPC | TMED4; TMED4 | Q7Z7H5; F8V | TMED4_HUMAN | Oxidation (M) | 0 | 0 | 0 | 0 | 0 | 109580 | 935450 | 0 | 18316000 | 1390200 | 0 | 0 | 0 | 0 | 0 | 0 | 0 | 708970 |
| EVFLPSTPC | TMED4; TMED4 | Q7Z7H5; F8V | TMED4_HUMAN | Unmodified | 0 | 0 | 0 | 0 | 0 | 0 | 0 | 0 | 37088000 | 742080 | 0 | 0 | 0 | 0 | 0 | 0 | 0 | 0 |
| EVFLPSTPC | TMED4; TMED4 | Q7Z7H5; F8V | TMED4_HUMAN | Oxidation (M) | 0 | 0 | 0 | 0 | 0 | 0 | 0 | 0 | 8700800 | 1196600 | 211010 | 0 | 0 | 0 | 0 | 0 | 0 | 0 |
| EVFLPSTPC | TMED4; TMED4 | Q7Z7H5; F8V | TMED4_HUMAN | Unmodified | 0 | 0 | 0 | 0 | 0 | 0 | 0 | 0 | 10761000 | 276800 | 0 | 0 | 113770 | 0 | 0 | 0 | 0 | 0 |
| EVFNILQAA | MET | P08581 | MET_HUMAN H | Unmodified | 0 | 0 | 0 | 0 | 0 | 0 | 0 | 0 | 3683700 | 0 | 0 | 0 | 0 | 0 | 0 | 0 | 0 | 0 |
| EVFPVLAAK | PDCD6IP; PDCI | Q8WUM4; F8 | PDC6I_HUMAN | Unmodified | 0 | 0 | 203850 | 0 | 0 | 3611100 | 0 | 435570 | 13460000 | 6237100 | 3067000 | 12081000 | 0 | 0 | 18387000 | 0 | 0 | 0 |
| EVFQIASNI | APPL1; APPL1 | Q9UKG1; C9 | DP13A_HUMAN | Unmodified | 548680 | 0 | 263740 | 0 | 1280900 | 1108300 | 0 | 100660 | 1490800 | 829890 | 246530 | 2028800 | 0 | 0 | 420350 | 269290 | 86491 | 0 |
| EVFQIASNI | APPL1 | Q9UKG1 | DP13A_HUMAN | Unmodified | 0 | 0 | 0 | 0 | 4286000 | 567110 | 727980 | 0 | 0 | 1490800 | 0 | 0 | 1826000 | 1324300 | 0 | 0 | 890160 | 2491700 |
| EVFQTASEI | MMS19; MMS19 | Q96T76; H7C | MMS19_HUMA | Unmodified | 0 | 0 | 0 | 0 | 0 | 0 | 0 | 0 | 0 | 0 | 0 | 470180 | 1682600 | 0 | 0 | 363840 | 0 | 0 |
| EVFRLPTDL | NUDCD1; NUD | Q96RS6; A0A | NUDC1_HUMA | Unmodified | 0 | 0 | 0 | 0 | 0 | 0 | 0 | 0 | 0 | 0 | 0 | 357920 | 732270 | 262000 | 0 | 0 | 0 | 0 |
| EVFRPLKPA | MAP4K4; MAP4 | O95819; H7C | M4K4_HUMAN | Unmodified | 0 | 0 | 411190 | 249290 | 316020 | 0 | 0 | 0 | 404010 | 188590 | 345260 | 652370 | 478520 | 0 | 349000 | 255090 | 286470 | 1736400 |
| EVFRPLKPA | MAP4K4 | C9J840 | C9J840_HUMA | Unmodified | 0 | 0 | 0 | 0 | 0 | 1328500 | 191980 | 0 | 2300400 | 230670 | 2258000 | 5229400 | 0 | 0 | 0 | 0 | 1373700 | 2933700 |
| EVFTRFCTC | CKB | P12277 | KCRB_HUMAN | Unmodified | 0 | 0 | 0 | 0 | 2330700 | 120870 | 7763200 | 0 | 0 | 976500 | 0 | 0 | 6521400 | 0 | 0 | 0 | 0 | 146920 |
| EVGEAFTIL | DNAJC7; DNAJC | Q99615; A0A | DNJC7_HUMAN | Unmodified | 0 | 0 | 0 | 0 | 0 | 0 | 0 | 0 | 704100 | 0 | 0 | 1245800 | 0 | 0 | 697130 | 0 | 0 | 0 |
| EVGEHVVS | FLNC | Q14315 | FLNC_HUMAN | Unmodified | 0 | 0 | 0 | 0 | 823220 | 585790 | 0 | 0 | 0 | 0 | 0 | 0 | 0 | 842900 | 0 | 0 | 1189900 | 0 |
| EVGQQLQI | SRGAP2; SRGA | O75044; P0D | SRGP2_HUMAN | Unmodified | 0 | 156210 | 0 | 0 | 1372100 | 0 | 0 | 0 | 0 | 0 | 0 | 1116200 | 0 | 0 | 0 | 45120 | 0 | 0 |
| EVGSHFDD | SOAT1; SOAT1 | P35610; B1A | SOAT1_HUMAN | Unmodified | 0 | 0 | 0 | 0 | 749450 | 0 | 0 | 0 | 0 | 0 | 0 | 0 | 544670 | 419170 | 0 | 0 | 0 | 1069500 |
| EVGSIIGKK | PCBP1; PCBP2; | Q15365; Q15 | PCBP1_HUMAN | Unmodified | 0 | 0 | 0 | 117470 | 1337500 | 256540 | 1222500 | 494570 | 0 | 287560 | 0 | 0 | 2501700 | 2528300 | 0 | 0 | 627610 | 747950 |
| EVGTPHGII | HPX | P02790 | HEMO_HUMAN | Unmodified | 0 | 0 | 0 | 0 | 0 | 0 | 0 | 0 | 0 | 0 | 0 | 14207000 | 0 | 0 | 0 | 0 | 0 | 0 |
| EVGVYEALI | MDH1; MDH1; I | P40925; B8Z | MDHC_HUMAN | Unmodified | 0 | 0 | 0 | 0 | 326230 | 0 | 501840 | 0 | 0 | 0 | 1261100 | 6007700 | 0 | 0 | 0 | 138920 | 0 | 0 |
| EVGVYEALI | MDH1; MDH1; I | P40925; B8Z | MDHC_HUMAN | Unmodified | 1329800 | 1092100 | 443290 | 1492800 | 10259000 | 1592100 | 189530 | 0 | 11735000 | 1661300 | 6411900 | 37626000 | 4569100 | 1666800 | 1384400 | 1045600 | 1035300 | 7225100 |
| EVHELYK | RO60; RO60; RO | P10155; H0Y | RO60_HUMAN | Unmodified | 418450 | 483920 | 216330 | 0 | 0 | 0 | 0 | 0 | 0 | 0 | 0 | 61397 | 0 | 0 | 653400 | 385990 | 0 | 0 |
| EVHGMAQI | SDF2 | Q99470 | SDF2_HUMAN | Unmodified | 0 | 0 | 0 | 0 | 764870 | 0 | 930490 | 275440 | 19521000 | 6379800 | 1382500 | 290060 | 0 | 0 | 0 | 0 | 0 | 2238200 |
| EVHGMAQI | SDF2 | Q99470 | SDF2_HUMAN | Oxidation (M) | 0 | 0 | 0 | 0 | 0 | 0 | 262630 | 203440 | 5310800 | 1259700 | 469640 | 154270 | 0 | 75979 | 0 | 0 | 0 | 0 |
| EVHHDIREI | CXADR | P78310 | CXAR_HUMAN | Unmodified | 1007800 | 695950 | 0 | 369260 | 5062400 | 657290 | 410520 | 111880 | 686930 | 243700 | 757280 | 4140900 | 0 | 0 | 1080500 | 1060800 | 243230 | 148620 |
| EVHKQVVE | LDHA | P00338 | LDHA_HUMAN | Unmodified | 0 | 0 | 0 | 0 | 2992300 | 0 | 0 | 0 | 0 | 255200 | 0 | 0 | 832810 | 0 | 0 | 0 | 1197800 | 2501900 |
| EVHKQVVE | LDHA | P00338 | LDHA_HUMAN | Unmodified | 0 | 0 | 0 | 0 | 3205400 | 212120 | 734140 | 0 | 0 | 695180 | 0 | 0 | 0 | 0 | 0 | 0 | 5266900 | 3709900 |
| EVHLGACG | CTNND1; CTNN | O60716; E9F | CTND1_HUMAN | Unmodified | 0 | 0 | 0 | 0 | 2584000 | 390800 | 0 | 0 | 1101100 | 0 | 127260 | 1377800 | 178750 | 1180100 | 0 | 1371800 | 25010 | 0 |
| EVHNLNQI | CEP55; CEP55 | Q53EZ4; H0Y | CEP55_HUMAN | Unmodified | 861580 | 1071600 | 1220000 | 3633100 | 9514500 | 2128800 | 0 | 249000 | 6673200 | 2771800 | 399710 | 1285400 | 434580 | 0 | 1400400 | 712060 | 440230 | 438990 |
| EVHNLNQI | CEP55; CEP55 | Q53EZ4; H0Y | CEP55_HUMAN | Unmodified | 787010 | 508240 | 1466900 | 1850700 | 6712000 | 677710 | 647980 | 456340 | 2727300 | 1447600 | 640460 | 851230 | 328310 | 0 | 997420 | 470920 | 106560 | 329990 |
| EVHVLCLG | ARL6; ARL6 | Q9H0F7; C9 | ARL6_HUMAN / | Unmodified | 0 | 0 | 0 | 0 | 0 | 0 | 0 | 0 | 0 | 0 | 0 | 273170 | 0 | 0 | 223540 | 74920 | 0 | 0 |
| EVIAGPLLR | NXN; NXN | Q6DKJ4; I3L4 | NXN_HUMAN N | Unmodified | 0 | 0 | 0 | 0 | 0 | 0 | 0 | 0 | 0 | 0 | 0 | 3923500 | 0 | 0 | 0 | 0 | 0 | 0 |
| EVIAVSCGI | ETFB | P38117 | ETFB_HUMAN I | Unmodified | 0 | 0 | 0 | 0 | 0 | 0 | 0 | 0 | 0 | 0 | 0 | 0 | 0 | 0 | 0 | 0 | 0 | 0 |
| EVIELPLTNI | PSMC6; PSMC6 | P62333; A0A | PRS10_HUMAN | Unmodified | 311470 | 111150 | 225690 | 395990 | 112840 | 0 | 0 | 0 | 1394200 | 83737 | 386820 | 3484200 | 0 | 0 | 260810 | 433740 | 0 | 0 |
| EVIELPVKH | PSMC5; PSMC5 | P62195; J3Q | PRS8_HUMAN | Unmodified | 0 | 0 | 0 | 0 | 0 | 0 | 0 | 0 | 1969400 | 0 | 0 | 2469800 | 2076700 | 1745200 | 0 | 0 | 0 | 0 |
| EVIHPLATS | STEAP1B; STEA | Q6NZ63; Q9 | STEAL_HUMAN | Unmodified | 0 | 0 | 0 | 0 | 0 | 0 | 0 | 0 | 0 | 0 | 0 | 439230 | 0 | 0 | 822070 | 225060 | 0 | 0 |
| EVIIAKDEV | AKT2; AKT2 | P31751; M0F | AKT2_HUMAN I | Unmodified | 0 | 0 | 148480 | 1099800 | 0 | 0 | 0 | 0 | 0 | 0 | 0 | 0 | 81872 | 0 | 1259300 | 1531400 | 0 | 0 |
| EVIKPYDW | TIPRL | O75663 | TIPRL_HUMAN | Unmodified | 0 | 0 | 0 | 0 | 0 | 0 | 100700 | 0 | 1195100 | 0 | 0 | 0 | 0 | 460510 | 0 | 61817 | 0 | 84804 |
| EVILCKDQI | SDCBP; SDCBP | O00560; G5I | SDCB1_HUMAN | Unmodified | 4978600 | 2923800 | 0 | 2519800 | 1116200 | 824860 | 224990 | 0 | 5496500 | 1833000 | 13237 | 1390700 | 887990 | 510120 | 494670 | 1980400 | 0 | 0 |
| EVILCKDQI | SDCBP; SDCBP | O00560; G5I | SDCB1_HUMAN | Unmodified | 707230 | 499290 | 0 | 1390700 | 1203500 | 0 | 0 | 0 | 1830200 | 1036000 | 0 | 0 | 1772100 | 0 | 0 | 0 | 0 | 99117 |
| EVINEVEK | SLK | Q9H2G2 | SLK_HUMAN S | Unmodified | 0 | 0 | 0 | 0 | 329280 | 0 | 31429 | 0 | 1127500 | 156090 | 0 | 977060 | 632600 | 0 | 0 | 0 | 37199 | 101870 |
| EVINEVEKA | SLK | Q9H2G2 | SLK_HUMAN S | Unmodified | 0 | 0 | 0 | 0 | 0 | 0 | 0 | 0 | 133390 | 236660 | 0 | 0 | 685670 | 237350 | 0 | 0 | 0 | 0 |
| EVINGNIK | EIF3G; EIF3G; E | O75821; K7E | EIF3G_HUMAN | Unmodified | 0 | 0 | 0 | 0 | 899220 | 153630 | 41134 | 0 | 171000 | 0 | 679390 | 2214400 | 813610 | 0 | 0 | 0 | 314510 | 465010 |
| EVIPVNVPE | EPRS1; EPRS1 | P07814; V9C | SYEP_HUMAN | Oxidation (M) | 0 | 0 | 0 | 0 | 0 | 0 | 0 | 0 | 220240 | 0 | 627980 | 2002700 | 0 | 0 | 0 | 0 | 0 | 0 |
| EVIQEWNL | TLN1 | Q9Y490 | TLN1_HUMAN | Unmodified | 0 | 119450 | 264970 | 140220 | 212430 | 0 | 1291600 | 425120 | 346870 | 0 | 0 | 593210 | 71065 | 0 | 207010 | 113900 | 0 | 111350 |
| EVIQEWNL | TLN1 | Q9Y490 | TLN1_HUMAN | Unmodified | 0 | 0 | 0 | 1188200 | 0 | 0 | 0 | 0 | 265970 | 0 | 443070 | 1989900 | 0 | 0 | 0 | 0 | 0 | 0 |
| EVIRNDGV | USO1 | O60763 | USO1_HUMAN | Unmodified | 268850 | 0 | 0 | 0 | 0 | 0 | 0 | 0 | 0 | 34112 | 112790 | 4685400 | 0 | 0 | 213460 | 0 | 0 | 0 |
| EVISRFDTP | MAN2B1 | O00754 | MA2B1_HUMAN | Unmodified | 105390 | 0 | 227450 | 166880 | 133920 | 0 | 0 | 61893 | 3287400 | 1249800 | 698510 | 295300 | 342850 | 390210 | 0 | 0 | 0 | 0 |
| EVITAQDTV | SMC2 | O95347 | SMC2_HUMAN | Unmodified | 0 | 0 | 0 | 0 | 586540 | 0 | 431940 | 181090 | 250810 | 132280 | 279410 | 2069800 | 0 | 0 | 0 | 0 | 0 | 0 |
| EVIVAKDEV | AKT1 | P31749 | AKT1_HUMAN I | Unmodified | 147840 | 129060 | 1251700 | 1655700 | 0 | 0 | 0 | 0 | 0 | 0 | 1507300 | 11198000 | 0 | 0 | 10538000 | 3122300 | 0 | 0 |
| EVKEQCEE | GOLM1 | Q8NBJ4 | GOLM1_HUMA | Unmodified | 0 | 0 | 0 | 0 | 0 | 0 | 0 | 0 | 1925800 | 542950 | 0 | 0 | 0 | 0 | 0 | 0 | 0 | 0 |
| EVKILEVHV | DNAJA2 | O60884 | DNJA2_HUMAN | Unmodified | 0 | 0 | 0 | 0 | 0 | 0 | 269760 | 2883300 | 0 | 230960 | 0 | 0 | 599570 | 639590 | 0 | 0 | 0 | 0 |
| EVKLLLLGA | GNAI3; GNAI2; | P08754; P04 | GNAI3_HUMAN | Unmodified | 0 | 0 | 0 | 0 | 805610 | 0 | 416070 | 360980 | 0 | 96380 | 0 | 65877 | 247020 | 333190 | 0 | 0 | 1072900 | 751920 |
| EVKLLLLGA | GNAI3; GNAI2; | P08754; P04 | GNAI3_HUMAN | Unmodified | 0 | 0 | 0 | 0 | 0 | 0 | 859900 | 317460 | 0 | 158370 | 0 | 0 | 672130 | 1222900 | 0 | 0 | 3008700 | 1807000 |
| EVKLQRQC | GNG11 | P61952 | GBG11_HUMAN | Unmodified | 0 | 0 | 0 | 0 | 294060 | 0 | 3192600 | 116960 | 0 | 0 | 0 | 0 | 0 | 0 | 0 | 0 | 124860 | 131550 |
| EVKPLGLT | PHYH; PHYH; P | O14832; B1A | PAHX_HUMAN | Oxidation (M) | 0 | 0 | 0 | 0 | 0 | 0 | 0 | 0 | 1460700 | 0 | 0 | 0 | 0 | 0 | 0 | 0 | 0 | 0 |
| EVKYEAPQ | TOM1 | O60784 | TOM1_HUMAN | Unmodified | 2050500 | 1980300 | 756590 | 1227600 | 0 | 0 | 0 | 0 | 482010 | 0 | 0 | 0 | 0 | 0 | 1279000 | 1134100 | 0 | 0 |
| EVKYEPFSF | USP14; USP14 | P54578; A6N | UBP14_HUMAN | Unmodified | 0 | 0 | 0 | 0 | 891980 | 0 | 0 | 0 | 290730 | 0 | 0 | 0 | 2505400 | 1114200 | 0 | 0 | 482230 | 0 |
| EVLAELEAL | PCK2; PCK2 | Q16822; H0Y | PCKGM_HUMA | Unmodified | 0 | 0 | 0 | 0 | 0 | 0 | 80651 | 0 | 2445700 | 0 | 0 | 0 | 0 | 0 | 0 | 0 | 0 | 0 |
| EVLAELEAL | PCK2; PCK2 | Q16822; H0Y | PCKGM_HUMA | Unmodified | 0 | 4745000 | 0 | 764900 | 0 | 0 | 0 | 0 | 1880200 | 0 | 0 | 0 | 0 | 0 | 0 | 0 | 0 | 0 |
| EVLAGRPLI | HNRNPU; HNR | Q00839; A0A | HNRPU_HUMA | Unmodified | 2494800 | 0 | 0 | 0 | 798700 | 0 | 9888900 | 0 | 586110 | 1268900 | 0 | 2983800 | 3891200 | 2801900 | 0 | 0 | 480930 | 0 |
| EVLAQTR | P4HTM; P4HTM | Q9NXG6; V9 | P4HTM_HUMAN | Unmodified | 243440 | 253980 | 266630 | 0 | 23464 | 0 | 18718 | 9776.2 | 1429300 | 0 | 0 | 0 | 0 | 25139 | 349290 | 285980 | 16642 | 34253 |
| EVLASDLV | NNT; NNT; NNT | Q13423; A0A | NNTM_HUMAN | Unmodified | 0 | 0 | 0 | 0 | 0 | 0 | 0 | 0 | 9130100 | 2534800 | 0 | 0 | 0 | 0 | 0 | 0 | 0 | 0 |
| EVLDEDTD | RABGAP1 | Q9Y3P9 | RBGP1_HUMAN | Unmodified | 0 | 0 | 0 | 0 | 344420 | 0 | 0 | 0 | 0 | 0 | 317760 | 1100700 | 0 | 0 | 0 | 0 | 0 | 0 |
| EVLDSFLDI | SLC1A5 | Q15758 | AAAT_HUMAN I | Unmodified | 2578900 | 1004200 | 2432000 | 0 | 0 | 0 | 0 | 0 | 677770 | 324960 | 611400 | 1.03E+08 | 0 | 0 | 5705300 | 4692500 | 0 | 0 |
| EVLEDFAEI | OPA1; OPA1; OI | O60313; E5K | OPA1_HUMAN | Unmodified | 0 | 0 | 0 | 0 | 0 | 0 | 0 | 0 | 0 | 0 | 633650 | 3916800 | 0 | 0 | 0 | 0 | 0 | 0 |
| EVLEDFAEI | OPA1; OPA1; OI | O60313; E5K | OPA1_HUMAN | Unmodified | 0 | 0 | 0 | 0 | 0 | 0 | 0 | 0 | 0 | 183680 | 0 | 1478300 | 0 | 0 | 0 | 0 | 0 | 0 |
| EVLEEISCY | CASK; CASK; C/ | O14936; A0A | CSKP_HUMAN | Unmodified | 116800 | 0 | 149640 | 811470 | 1751500 | 512350 | 0 | 270770 | 2180000 | 280430 | 0 | 0 | 1721500 | 1282500 | 1479600 | 849650 | 559960 | 1060000 |
| EVLEKQPVI | PSMB4 | P28070 | PSB4_HUMAN | Unmodified | 0 | 0 | 0 | 85383 | 0 | 0 | 0 | 0 | 0 | 0 | 0 | 334120 | 0 | 0 | 0 | 80488 | 0 | 0 |
| EVLELDSIR | ARL2 | P36404 | ARL2_HUMAN / | Unmodified | 0 | 0 | 343880 | 120400 | 0 | 0 | 53490 | 30850 | 0 | 0 | 418850 | 1647300 | 0 | 239520 | 0 | 0 | 0 | 0 |
| EVLEQVER | YES1; FYN; YES | P07947; P06 | YES_HUMAN Ty | Unmodified | 893170 | 0 | 1475200 | 70886 | 0 | 235790 | 0 | 104660 | 3725900 | 575690 | 613450 | 3281400 | 72982 | 0 | 2522600 | 2119400 | 375060 | 786850 |
| EVLEYNAIG | FDPS; FDPS; FD | P14324; A0A | FPPS_HUMAN | Unmodified | 187630 | 182240 | 0 | 0 | 0 | 0 | 0 | 0 | 0 | 0 | 0 | 7511300 | 0 | 0 | 0 | 0 | 0 | 0 |
| EVLGEHIVF | EIF1AD; EIF1AD | Q8N9N8; E9 | EIF1A_HUMAN | Unmodified | 142530 | 119350 | 86871 | 112860 | 177230 | 0 | 0 | 116150 | 209520 | 87464 | 436250 | 1172000 | 0 | 0 | 109230 | 110110 | 0 | 0 |
| EVLGSATR | UAP1 | Q16222 | UAP1_HUMAN | Unmodified | 0 | 0 | 0 | 0 | 0 | 0 | 0 | 0 | 2153700 | 0 | 0 | 826860 | 0 | 0 | 260690 | 0 | 0 | 0 |
| EVLLDIK | PACSIN1; PACS | Q9BY11; F6U | PACN1_HUMAN | Unmodified | 4969700 | 4024900 | 0 | 0 | 0 | 0 | 0 | 0 | 0 | 0 | 0 | 170790 | 0 | 150500 | 641300 | 502710 | 0 | 0 |
| EVLLEVK | SNTB1; SNTB2; | Q13884; Q13 | SNTB1_HUMAN | Unmodified | 417510 | 519230 | 2061900 | 0 | 0 | 0 | 0 | 0 | 0 | 0 | 0 | 0 | 0 | 0 | 997630 | 2764100 | 0 | 0 |
| EVLLEVQK | PACSIN2; PACS | Q9UNF0; A0 | PACN2_HUMAN | Unmodified | 21347000 | 17994000 | 12214000 | 9831000 | 803840 | 196320 | 0 | 0 | 1116100 | 166220 | 742550 | 6294100 | 13373000 | 1761100 | 29790000 | 30936000 | 176640 | 221980 |
| EVLLEVQKI | PACSIN2; PACS | Q9UNF0; A0 | PACN2_HUMAN | Unmodified | 0 | 0 | 0 | 0 | 0 | 0 | 0 | 0 | 0 | 0 | 0 | 0 | 417640 | 134060 | 0 | 0 | 0 | 0 |
| EVLLKTNLS | PPP4R3A | Q6IN85 | P4R3A_HUMAN | Unmodified | 0 | 0 | 0 | 174900 | 3063100 | 605940 | 0 | 447110 | 533670 | 480590 | 152910 | 883650 | 459220 | 220450 | 0 | 0 | 953200 | 1408700 |
| EVLLLAHNI | CEACAM6; N/A | P40199; M0C | CEAM6_HUMAN | Unmodified | 0 | 56247 | 0 | 0 | 0 | 0 | 0 | 0 | 5469400 | 613480 | 0 | 0 | 0 | 0 | 0 | 0 | 0 | 0 |
| EVLLLHNN | UBAP1; UBAP1 | Q9NZ09; A0A | UBAP1_HUMAN | Oxidation (M) | 1382700 | 1940200 | 0 | 0 | 0 | 0 | 411730 | 0 | 0 | 0 | 0 | 0 | 0 | 0 | 0 | 433280 | 0 | 0 |
| EVLLPEVGA | MOGS; MOGS; | Q13724; A0A | MOGS_HUMAN | Unmodified | 0 | 0 | 0 | 0 | 0 | 0 | 0 | 0 | 2150500 | 0 | 0 | 0 | 0 | 0 | 0 | 0 | 0 | 0 |
| EVLNMLK | GLG1 | Q92896 | GSLG1_HUMAN | Unmodified | 1273800 | 1701600 | 1234600 | 1204900 | 3080200 | 1026300 | 768560 | 544810 | 16469000 | 4643700 | 703850 | 2943900 | 1794400 | 1330600 | 1217000 | 806130 | 0 | 0 |
| EVLPALENI | PFDN2 | Q9UHV9 | PFD2_HUMAN | Unmodified | 433690 | 0 | 0 | 550940 | 394960 | 0 | 0 | 0 | 1128600 | 96513 | 3403000 | 12981000 | 1308700 | 377120 | 447270 | 438050 | 108570 | 585140 |
| EVLPALENI | PFDN2 | Q9UHV9 | PFD2_HUMAN | Unmodified | 0 | 0 | 0 | 0 | 2223900 | 478000 | 839760 | 0 | 270940 | 869040 | 0 | 0 | 4053300 | 1552500 | 0 | 0 | 4352300 | 2457600 |
| EVLPALENI | PFDN2 | Q9UHV9 | PFD2_HUMAN | Unmodified | 0 | 0 | 0 | 0 | 193810 | 0 | 394840 | 0 | 0 | 187640 | 0 | 0 | 505900 | 443850 | 0 | 0 | 0 | 470780 |
| EVLPGEIVE | PPAT | Q06203 | PUR1_HUMAN | Unmodified | 0 | 0 | 0 | 0 | 0 | 0 | 0 | 0 | 0 | 210770 | 293520 | 2405000 | 0 | 0 | 0 | 0 | 0 | 0 |
| EVLPGEIVE | PPAT | Q06203 | PUR1_HUMAN | Unmodified | 19223 | 0 | 0 | 0 | 0 | 0 | 114610 | 0 | 0 | 0 | 110640 | 763560 | 0 | 0 | 135670 | 60522 | 0 | 0 |
| EVLPHVPL | AUP1 | Q9Y679 | AUP1_HUMAN | Unmodified | 0 | 0 | 0 | 0 | 0 | 0 | 0 | 0 | 0 | 0 | 210730 | 636560 | 0 | 0 | 0 | 0 | 0 | 0 |
| EVLPLAFLG | ECPAS; ECPAS | Q5VYK3; J3K | ECM29_HUMAN | Oxidation (M) | 0 | 12850 | 0 | 0 | 90431 | 0 | 0 | 0 | 125030 | 0 | 0 | 1291900 | 0 | 0 | 0 | 0 | 935510 | 345510 |
| EVLQKALEI | STMN2; STMN2 | Q93045; E5F | STMN2_HUMAN | Unmodified | 0 | 0 | 0 | 0 | 0 | 0 | 1933300 | 1922600 | 0 | 167740 | 0 | 0 | 0 | 129130 | 0 | 0 | 1375300 | 0 |
| EVLQQLET | RABGGTA; RAB | Q92696; H0Y | PGTA_HUMAN | Unmodified | 0 | 0 | 0 | 0 | 0 | 0 | 0 | 0 | 0 | 0 | 0 | 886090 | 0 | 0 | 0 | 0 | 0 | 0 |
| EVLQQLET | RABGGTA; RAB | Q92696; H0Y | PGTA_HUMAN | Unmodified | 0 | 0 | 0 | 0 | 0 | 0 | 0 | 0 | 510220 | 0 | 0 | 1998300 | 0 | 0 | 0 | 0 | 0 | 0 |
| EVLRAEND | ABCC4; ABCC4 | O15439; A0A | MRP4_HUMAN | Unmodified | 0 | 0 | 268120 | 585510 | 0 | 0 | 0 | 0 | 0 | 137240 | 0 | 52303 | 0 | 0 | 150820 | 1099900 | 0 | 0 |
| EVLRENDLI | ERC1; ERC1; EF | Q8IUD2; K7E | RB6I2_HUMAN | Unmodified | 0 | 0 | 0 | 0 | 372320 | 0 | 99632 | 0 | 0 | 0 | 0 | 0 | 0 | 0 | 0 | 0 | 128560 | 77988 |
| EVLTEDK | UBA1 | P22314 | UBA1_HUMAN | Unmodified | 168490 | 0 | 395110 | 115260 | 415180 | 0 | 0 | 0 | 161700 | 0 | 0 | 1330400 | 0 | 2484000 | 219440 | 41616 | 0 | 185760 |
| EVLTGNDE | ATP6AP1; ATP6 | Q15904; A0A | VAS1_HUMAN | Unmodified | 0 | 0 | 135000 | 0 | 0 | 0 | 0 | 0 | 6086100 | 466380 | 0 | 1002700 | 0 | 0 | 0 | 141330 | 0 | 0 |
| EVMLILIR | SF3B1 | O75533 | SF3B1_HUMAN | Oxidation (M) | 120520 | 0 | 0 | 0 | 0 | 0 | 0 | 0 | 0 | 0 | 0 | 1006700 | 0 | 0 | 0 | 128420 | 0 | 0 |
| EVMLVGIGI | ATP5F1C | P36542 | ATPG_HUMAN | Unmodified | 0 | 0 | 0 | 0 | 0 | 0 | 0 | 0 | 5634100 | 151960 | 0 | 0 | 0 | 0 | 0 | 0 | 0 | 0 |
| EVMLVGIGI | ATP5F1C | P36542 | ATPG_HUMAN | Oxidation (M) | 197010 | 0 | 0 | 0 | 0 | 0 | 0 | 0 | 2958700 | 392020 | 211010 | 0 | 0 | 0 | 0 | 0 | 0 | 0 |
| EVMPLLLA | IPO4; IPO4 | Q8TEX9; H0Y | IPO4_HUMAN I | Oxidation (M) | 0 | 0 | 0 | 0 | 0 | 0 | 0 | 0 | 0 | 0 | 0 | 477200 | 0 | 0 | 79175 | 0 | 0 | 0 |
| EVMVDDR | RAB7A; RAB7A; | P51149; A0A | RAB7A_HUMAN | Unmodified | 591860 | 0 | 952830 | 0 | 0 | 0 | 0 | 0 | 215440 | 25558 | 572460 | 0 | 0 | 0 | 989430 | 0 | 0 | 0 |

| | | | | | | | | | | | | | | | | | | | | | | |
|---|---|---|---|---|---|---|---|---|---|---|---|---|---|---|---|---|---|---|---|---|---|---|
| EVMVDDRL | RAB7A; RAB7A; | P51149; A0A | RAB7A_HUMAN | 2 Oxidation (M) | 125120 | 1856100 | 0 | 1473200 | 0 | 40817 | 264440 | 0 | 679820 | 480190 | 119190 | 2256200 | 0 | 1477900 | 1921400 | 1212200 | 0 | 0 |
| EVNDLLQE | CENPF | P49454 | CENPF_HUMAN | Unmodified | 0 | 0 | 0 | 0 | 0 | 0 | 0 | 0 | 0 | 0 | 0 | 0 | 0 | 12636000 | 0 | 0 | 0 | 0 |
| EVNEVSQN | NIBAN1 | Q9BZQ8 | NIBA1_HUMAN | Unmodified | 0 | 0 | 0 | 0 | 1723800 | 852370 | 933610 | 209060 | 6426400 | 2804400 | 0 | 0 | 0 | 0 | 0 | 0 | 148140 | 327550 |
| EVNEVSQN | NIBAN1 | Q9BZQ8 | NIBA1_HUMAN | Unmodified | 0 | 0 | 0 | 0 | 833240 | 181250 | 1117500 | 499680 | 521240 | 1599900 | 0 | 0 | 0 | 0 | 0 | 0 | 0 | 0 |
| EVNKFQMA | SRP14 | P37108 | SRP14_HUMAN | Unmodified | 0 | 396560 | 0 | 797580 | 194460 | 0 | 0 | 0 | 0 | 51967 | 0 | 0 | 0 | 0 | 0 | 0 | 0 | 0 |
| EVNLAVQN | ALDH9A1 | P49189 | AL9A1_HUMAN | Unmodified | 219480 | 178060 | 352260 | 416470 | 972580 | 159310 | 364470 | 283190 | 1582000 | 1093500 | 405000 | 2468700 | 1492400 | 2720000 | 389060 | 313260 | 1338500 | 341820 |
| EVNLGLLAI | PARP4 | Q9UKK3 | PARP4_HUMAN | Unmodified | 90816 | 0 | 0 | 0 | 0 | 0 | 0 | 0 | 0 | 45635 | 120770 | 2380200 | 0 | 0 | 0 | 23516 | 0 | 0 |
| EVPAESVT | BPNT2; BPNT2 | Q9NX62; H0 | IMPA3_HUMAN | Unmodified | 0 | 0 | 0 | 0 | 0 | 0 | 0 | 0 | 569230 | 614790 | 0 | 0 | 0 | 0 | 0 | 0 | 0 | 0 |
| EVPAGKYP | NT5E; NT5E | P21589; H0Y | 5NTD_HUMAN | Unmodified | 314350 | 252220 | 0 | 0 | 2837800 | 0 | 0 | 0 | 0 | 0 | 0 | 684110 | 0 | 0 | 0 | 0 | 0 | 0 |
| EVPAGKYP | NT5E; NT5E | P21589; H0Y | 5NTD_HUMAN | Unmodified | 0 | 0 | 0 | 0 | 833260 | 746050 | 0 | 0 | 0 | 0 | 0 | 0 | 0 | 0 | 0 | 0 | 145810 | 354800 |
| EVPAVPETL | RPL7 | P18124 | RL7_HUMAN 6 | Unmodified | 1759300 | 1842100 | 1915000 | 499140 | 189250 | 0 | 0 | 0 | 0 | 0 | 1344200 | 6920700 | 316290 | 0 | 1382200 | 862230 | 0 | 0 |
| EVPAVPETL | RPL7 | P18124 | RL7_HUMAN 6 | Unmodified | 357630 | 269690 | 403210 | 266240 | 0 | 0 | 0 | 0 | 0 | 0 | 385710 | 1919000 | 0 | 0 | 417880 | 165590 | 0 | 0 |
| EVPCYVFD | IGF2R; IGF2R; I | P11717; A0A | MPRI_HUMAN | Unmodified | 0 | 0 | 0 | 0 | 0 | 0 | 0 | 0 | 1222400 | 217310 | 0 | 234910 | 0 | 0 | 0 | 0 | 0 | 0 |
| EVPCYVFD | IGF2R; IGF2R; I | P11717; A0A | MPRI_HUMAN | Unmodified | 0 | 0 | 0 | 0 | 0 | 303670 | 0 | 0 | 3041000 | 0 | 0 | 693180 | 0 | 0 | 79305 | 0 | 0 | 0 |
| EVPCYVFD | IGF2R; IGF2R; I | P11717; A0A | MPRI_HUMAN | Unmodified | 0 | 0 | 0 | 0 | 0 | 0 | 0 | 0 | 8523400 | 1214900 | 0 | 0 | 0 | 0 | 0 | 0 | 0 | 0 |
| EVPEEYSEF | GRK6; GRK6; G | P43250; D6F | GRK6_HUMAN | Unmodified | 0 | 0 | 83794 | 582190 | 0 | 0 | 0 | 0 | 258370 | 0 | 217690 | 0 | 0 | 0 | 0 | 0 | 0 | 0 |
| EVPEYLNH | CUL4A; CUL4A | Q13619; A0A | CUL4A_HUMAN | Unmodified | 0 | 0 | 0 | 0 | 0 | 0 | 0 | 0 | 0 | 214910 | 0 | 908630 | 0 | 0 | 0 | 0 | 0 | 0 |
| EVPGADRY | ARVCF; ARVCF; | O00192; E9F | ARVC_HUMAN | Unmodified | 152780 | 0 | 0 | 472760 | 0 | 0 | 0 | 0 | 257060 | 0 | 0 | 0 | 0 | 238500 | 0 | 0 | 0 | 0 |
| EVPIHRVAI | SND1; SND1 | Q7KZF4; H7 | SND1_HUMAN | Unmodified | 0 | 0 | 0 | 0 | 924280 | 520260 | 115320 | 0 | 0 | 97443 | 101310 | 0 | 837530 | 425920 | 0 | 0 | 233870 | 674560 |
| EVPIKEIVE | SLK | Q9H2G2 | SLK_HUMAN S | Oxidation (M) | 0 | 0 | 0 | 0 | 0 | 0 | 0 | 0 | 117780 | 560270 | 398360 | 1180500 | 241570 | 86261 | 0 | 0 | 0 | 0 |
| EVPIVQTET | EPB41L2; EPB4 | O43491; Q6 | E41L2_HUMAN | Unmodified | 0 | 0 | 0 | 0 | 11206000 | 0 | 0 | 0 | 0 | 0 | 0 | 0 | 180830 | 0 | 0 | 359770 | 2181900 | 6789600 |
| EVPNYKLIT | RPS25 | P62851 | RS25_HUMAN | Unmodified | 0 | 0 | 0 | 0 | 0 | 0 | 1230600 | 245590 | 0 | 0 | 0 | 0 | 688780 | 528530 | 0 | 0 | 0 | 0 |
| EVPVASGS | HTRA1; HTRA1 | Q92743; A0A | HTRA1_HUMAN | Unmodified | 0 | 0 | 0 | 0 | 0 | 0 | 0 | 0 | 2526000 | 0 | 0 | 0 | 0 | 0 | 0 | 0 | 0 | 1645300 |
| EVPVIFIHPI | RALGAPB; RAL | Q86X10; A2A | RLGPB_HUMAN | Unmodified | 53175 | 0 | 25129 | 32790 | 0 | 0 | 0 | 0 | 96765 | 0 | 0 | 919270 | 0 | 0 | 112400 | 94821 | 0 | 0 |
| EVQDGIAPI | USP5 | P45974 | UBP5_HUMAN | Unmodified | 1055100 | 1216600 | 534670 | 0 | 0 | 0 | 0 | 0 | 0 | 0 | 0 | 1425400 | 1032800 | 344600 | 0 | 458470 | 0 | 0 |
| EVQDLAFLI | RAP1GDS1 | P52306 | GDS1_HUMAN | Unmodified | 0 | 0 | 0 | 0 | 0 | 0 | 0 | 0 | 0 | 0 | 0 | 2472300 | 0 | 0 | 0 | 334510 | 1079400 | 0 |
| EVQDLFEA | UNC45A; UNC4 | Q9H3U1; A0 | UN45A_HUMAN | Unmodified | 465410 | 0 | 0 | 0 | 0 | 0 | 0 | 0 | 0 | 0 | 233370 | 4916400 | 1066600 | 0 | 0 | 0 | 0 | 0 |
| EVQEFYK | CD9; CD9; CD9 | P21926; A0A | CD9_HUMAN | Unmodified | 58383000 | 49270000 | 88368000 | 13155000 | 0 | 0 | 0 | 0 | 0 | 750790 | 667920 | 21867000 | 0 | 0 | 2.18E+08 | 1.43E+08 | 65187 | 0 |
| EVQEFYKD | CD9; CD9; CD9 | P21926; A0A | CD9_HUMAN | Unmodified | 99670000 | 85113000 | 2867100 | 34673000 | 3178100 | 0 | 0 | 0 | 10202000 | 5868200 | 1063900 | 17736000 | 2929000 | 529550 | 67220000 | 1.04E+08 | 326320 | 0 |
| EVQEVSRN | GOLIM4; GOLIM | O00461; F8V | GOLI4_HUMAN | Unmodified | 0 | 0 | 0 | 0 | 0 | 0 | 0 | 0 | 282890 | 0 | 0 | 0 | 0 | 0 | 0 | 0 | 0 | 0 |
| EVQGNESE | VIL1; VIL1 | P09327; C9J | VILI_HUMAN Vi | Unmodified | 139070 | 60775 | 0 | 0 | 0 | 0 | 0 | 0 | 815490 | 19937 | 407240 | 6099400 | 0 | 0 | 0 | 0 | 0 | 0 |
| EVQGNESE | VIL1; VIL1 | P09327; C9J | VILI_HUMAN Vi | Unmodified | 246600 | 114770 | 0 | 0 | 0 | 0 | 0 | 0 | 1159700 | 0 | 1222800 | 2400400 | 0 | 0 | 0 | 72316 | 0 | 0 |
| EVQHNCQ | GULP1 | Q9UBP9 | GULP1_HUMAN | Unmodified | 0 | 0 | 0 | 0 | 0 | 0 | 0 | 0 | 0 | 0 | 0 | 0 | 142520 | 112400 | 19642 | 91028 | 14524 | 0 |
| EVQILQQN | KIF14 | Q15058 | KIF14_HUMAN | Unmodified | 0 | 0 | 0 | 0 | 1413800 | 282400 | 106520 | 41770 | 0 | 30784 | 0 | 0 | 0 | 155600 | 0 | 0 | 320180 | 252730 |
| EVQLALPR | SIGIRR; SIGIRR | Q6IA17; C9J | SIGIR_HUMAN | Unmodified | 142610 | 0 | 704100 | 249490 | 1282500 | 201810 | 0 | 62298 | 1000700 | 0 | 0 | 1221400 | 136750 | 0 | 520240 | 507290 | 1247100 | 209490 |
| EVQLAQIFE | PDS5A; PDS5A | Q29RF7; H0Y | PDS5A_HUMAN | Unmodified | 0 | 0 | 0 | 0 | 0 | 0 | 0 | 0 | 0 | 0 | 79416 | 298720 | 0 | 0 | 0 | 0 | 0 | 0 |
| EVQLPDRG | IDE; IDE; IDE; ID | P14735; A0A | IDE_HUMAN In | Unmodified | 907450 | 24786 | 127490 | 301480 | 0 | 0 | 0 | 0 | 0 | 0 | 0 | 3565800 | 0 | 0 | 234320 | 234790 | 0 | 0 |
| EVQMGDF | FKBP10; FKBP1 | Q96AY3; K7E | FKB10_HUMAN | Oxidation (M) | 0 | 0 | 0 | 0 | 0 | 0 | 0 | 0 | 375380 | 45336 | 748570 | 85862 | 0 | 0 | 0 | 0 | 0 | 0 |
| EVQQGEEF | AK1; AK1; AK1 | P00568; H0Y | KAD1_HUMAN | Unmodified | 0 | 0 | 0 | 262960 | 0 | 0 | 0 | 19481 | 633310 | 59052 | 255260 | 0 | 1185400 | 0 | 0 | 0 | 0 | 0 |
| EVQQGEEF | AK1; AK1; AK1 | P00568; H0Y | KAD1_HUMAN | Unmodified | 0 | 0 | 12463000 | 0 | 0 | 134410 | 0 | 0 | 1504500 | 0 | 252960 | 28016 | 0 | 0 | 509250 | 0 | 0 | 0 |
| EVQQIQSN | KIF20B | Q96Q89 | KI20B_HUMAN | Unmodified | 0 | 0 | 0 | 0 | 1425400 | 0 | 0 | 0 | 0 | 0 | 0 | 354620 | 0 | 0 | 0 | 0 | 741210 | 0 |
| EVQTAVRLI | H2BE1; H2BU1 | A0A2R8Y61 | H2BE1_HUMAN | Unmodified | 1118000 | 0 | 0 | 0 | 0 | 0 | 451910 | 964850 | 0 | 0 | 0 | 0 | 0 | 0 | 0 | 0 | 0 | 0 |
| EVQTNDLK | RPS3A; RPS3A; | P61247; D6F | RS3A_HUMAN | Unmodified | 573730 | 389650 | 63185 | 303420 | 0 | 0 | 0 | 112720 | 3130600 | 6141000 | 151180 | 3626500 | 87097 | 0 | 450790 | 240110 | 239100 | 389590 |
| EVQTNDLK | RPS3A; RPS3A; | P61247; D6F | RS3A_HUMAN | Unmodified | 181670 | 259640 | 0 | 420380 | 0 | 54169 | 155820 | 159730 | 236030 | 128940 | 0 | 0 | 0 | 0 | 0 | 0 | 0 | 0 |
| EVQTNELK | EPB41L2; EPB4 | O43491; A0A | E41L2_HUMAN | Unmodified | 698870 | 123630 | 169750 | 275420 | 4936600 | 1741900 | 0 | 220610 | 51653 | 17493 | 0 | 0 | 1050200 | 361460 | 261840 | 120870 | 1123800 | 2559000 |
| EVQYDFSL | UBAP1 | Q9NZ09 | UBAP1_HUMAN | Unmodified | 1371000 | 2499300 | 0 | 0 | 0 | 0 | 0 | 0 | 0 | 0 | 0 | 286400 | 0 | 0 | 0 | 0 | 0 | 0 |
| EVQYHESD | AVIL; AVIL | O75366; F8V | AVIL_HUMAN A | Unmodified | 0 | 0 | 0 | 0 | 0 | 0 | 0 | 0 | 0 | 0 | 0 | 12928000 | 0 | 0 | 0 | 0 | 0 | 0 |
| EVQYHESD | AVIL; AVIL | O75366; F8V | AVIL_HUMAN A | Unmodified | 0 | 0 | 0 | 0 | 0 | 0 | 0 | 0 | 0 | 0 | 818610 | 346060 | 0 | 0 | 0 | 0 | 0 | 0 |
| EVQYLLNK | HYOU1; HYOU1 | Q9Y4L1; A0A | HYOU1_HUMA | Unmodified | 1529300 | 0 | 2699900 | 1488200 | 1303300 | 4725500 | 3131500 | 2831900 | 1335900 | 903100 | 0 | 0 | 0 | 6111500 | 7268800 | 6334900 | 353070 | 4271200 |
| EVRAAALV | CTNND1; CTNN | O60716; A0A | CTND1_HUMAN | Unmodified | 0 | 0 | 31934 | 129640 | 44797000 | 4072700 | 791020 | 0 | 0 | 50904 | 0 | 75627 | 12775000 | 29139000 | 80348 | 63727 | 49487000 | 53337000 |
| EVRVCEPC | HGS; HGS; HGS | O14964; A0A | HGS_HUMAN H | Unmodified | 165640 | 0 | 0 | 0 | 319990 | 0 | 0 | 0 | 0 | 282630 | 0 | 0 | 232460 | 124250 | 0 | 0 | 0 | 237580 |
| EVSDDEAE | HSP90AA1 | P07900 | HS90A_HUMAN | Unmodified | 0 | 0 | 100020 | 0 | 0 | 0 | 0 | 0 | 239100 | 0 | 115890 | 3939400 | 0 | 0 | 102300 | 24526 | 0 | 0 |
| EVSDGIIAP | ATIC | P31939 | PUR9_HUMAN | Unmodified | 0 | 0 | 247490 | 1183900 | 581600 | 0 | 138660 | 0 | 956330 | 147210 | 664730 | 1848000 | 0 | 0 | 1736000 | 789640 | 155850 | 0 |
| EVSFQSTG | C1QBP; C1QBP | Q07021; I3L | C1QBP_HUMAN | Unmodified | 0 | 0 | 0 | 0 | 0 | 0 | 0 | 0 | 882720 | 0 | 0 | 257540 | 0 | 0 | 0 | 0 | 0 | 0 |
| EVSMDDHI | ATP1A1; ATP1A | P05023; Q5T | AT1A1_HUMAN | Unmodified | 195850 | 0 | 0 | 0 | 968990 | 0 | 0 | 0 | 1963700 | 1022700 | 1362800 | 2598200 | 0 | 278620 | 787970 | 447480 | 224930 | 0 |
| EVSMDDHI | ATP1A1; ATP1A | P05023; Q5T | AT1A1_HUMAN | Unmodified | 430540 | 370780 | 0 | 4332600 | 7906300 | 473210 | 218070 | 0 | 5065500 | 1232100 | 8173500 | 19130000 | 3980200 | 1341700 | 1612800 | 1750400 | 433490 | 392650 |
| EVSMDDHI | ATP1A1; ATP1A | P05023; Q5T | AT1A1_HUMAN | Oxidation (M) | 0 | 0 | 0 | 1914700 | 1282400 | 0 | 359480 | 0 | 587940 | 328710 | 2336300 | 0 | 1420900 | 0 | 940000 | 1268100 | 0 | 0 |
| EVSMKAED | RP2 | O75695 | XRP2_HUMAN | Oxidation (M) | 0 | 0 | 0 | 0 | 0 | 0 | 0 | 0 | 0 | 0 | 149300 | 0 | 35772 | 0 | 0 | 0 | 0 | 0 |
| EVSMKAED | RP2 | O75695 | XRP2_HUMAN | Unmodified | 0 | 140650 | 16142 | 0 | 0 | 0 | 66202 | 32772 | 406400 | 0 | 0 | 1501000 | 0 | 66001 | 0 | 0 | 393240 | 472930 |
| EVSQPDW | PDIA4; PDIA4 | P13667; A0A | PDIA4_HUMAN | Unmodified | 58672 | 0 | 0 | 0 | 0 | 0 | 0 | 0 | 5467500 | 743050 | 810480 | 0 | 0 | 0 | 676070 | 241550 | 0 | 0 |
| EVSSATNA | SNRPA; SNRPA | P09012; M0 | SNRPA_HUMAN | Unmodified | 0 | 0 | 0 | 0 | 58461 | 32904 | 0 | 131710 | 329770 | 43875 | 89471 | 3468200 | 0 | 0 | 0 | 0 | 0 | 0 |
| EVSSEDIAD | PIP4K2C; PIP4K | Q8TBX8; F8V | PI42C_HUMAN | Unmodified | 0 | 0 | 0 | 0 | 0 | 845830 | 0 | 0 | 2967700 | 365680 | 0 | 0 | 740000 | 721110 | 259090 | 194250 | 641310 | 1190300 |
| EVSSEDIAD | PIP4K2C; PIP4K | Q8TBX8; F8V | PI42C_HUMAN | Oxidation (M) | 0 | 254980 | 0 | 387500 | 613260 | 0 | 0 | 0 | 494700 | 206960 | 0 | 1292000 | 306730 | 0 | 467670 | 329570 | 240230 | 0 |
| EVSSLEGSI | AKAP12 | Q02952 | AKA12_HUMAN | Unmodified | 0 | 0 | 0 | 0 | 0 | 0 | 0 | 0 | 0 | 0 | 0 | 0 | 0 | 0 | 0 | 0 | 184480 | 1906500 |
| EVSSSFDH | MTCH2; MTCH2 | Q9Y6C9; E9I | MTCH2_HUMA | Unmodified | 0 | 0 | 0 | 0 | 0 | 0 | 0 | 0 | 1563500 | 163810 | 688010 | 0 | 0 | 0 | 0 | 0 | 0 | 0 |
| EVSTTERTY | FARP1; FARP1 | Q9Y4F1; C9J | FARP1_HUMAN | Unmodified | 0 | 0 | 0 | 0 | 647000 | 0 | 0 | 0 | 0 | 0 | 0 | 0 | 0 | 0 | 0 | 0 | 753320 | 365630 |
| EVSTYIKK | EEF1A1P5; EEF | Q5VTE0; P68 | EF1A3_HUMAN | Unmodified | 12402000 | 12295000 | 5443200 | 11443000 | 6448000 | 1741600 | 1600500 | 791360 | 4073000 | 1096700 | 866920 | 18242000 | 11113000 | 3681600 | 10306000 | 9819700 | 3185600 | 5989400 |
| EVSVSAFTT | TSKU | Q8WUA8 | TSK_HUMAN T | Unmodified | 157190 | 0 | 0 | 0 | 28219 | 16819 | 0 | 0 | 1626800 | 139660 | 0 | 0 | 0 | 0 | 0 | 0 | 0 | 0 |
| EVTDEIVK | WARS1 | P23381 | SYWC_HUMAN | Unmodified | 0 | 0 | 0 | 0 | 0 | 0 | 0 | 0 | 1100600 | 0 | 0 | 0 | 0 | 0 | 0 | 0 | 0 | 0 |
| EVTDKEQE | EPB41L3; EPB4 | Q9Y2J2; A0A | E41L3_HUMAN | Unmodified | 0 | 0 | 0 | 0 | 0 | 0 | 108440 | 42704 | 0 | 0 | 32894 | 0 | 0 | 0 | 0 | 0 | 131900 | 680210 |
| EVTDKEQE | EPB41L3; EPB4 | Q9Y2J2; A0A | E41L3_HUMAN | Unmodified | 0 | 0 | 0 | 0 | 0 | 0 | 0 | 29629 | 0 | 0 | 0 | 0 | 0 | 0 | 0 | 0 | 0 | 990490 |
| EVTDMNLN | NIBAN2 | Q96TA1 | NIBA2_HUMAN | Oxidation (M) | 0 | 0 | 0 | 0 | 0 | 0 | 0 | 0 | 0 | 0 | 0 | 0 | 0 | 0 | 425150 | 0 | 0 | 0 |
| EVTDMNLN | NIBAN2 | Q96TA1 | NIBA2_HUMAN | Oxidation (M) | 0 | 0 | 0 | 0 | 2266900 | 0 | 0 | 107870 | 0 | 0 | 0 | 0 | 0 | 0 | 0 | 0 | 1622100 | 0 |
| EVTEFYVQ | ACAD11; ACAD | Q709F0; D6I | ACD11_HUMAN | Unmodified | 0 | 0 | 0 | 0 | 0 | 0 | 0 | 0 | 1270100 | 0 | 0 | 0 | 0 | 0 | 0 | 0 | 0 | 0 |
| EVTGIITQG | MFGE8 | Q08431 | MFGM_HUMAN | Unmodified | 4248800 | 3817900 | 12020000 | 16198000 | 11957000 | 15668000 | 229380 | 187110 | 12550000 | 900570 | 813940 | 17871000 | 27605000 | 0 | 14106000 | 10463000 | 0 | 3794400 |
| EVTGIITQG | MFGE8 | Q08431 | MFGM_HUMAN | Unmodified | 0 | 0 | 0 | 0 | 39989000 | 7035600 | 0 | 0 | 0 | 0 | 0 | 0 | 1465500 | 0 | 0 | 0 | 2928400 | 6179700 |
| EVTHNIHY | SEPTIN3; SEPTI | Q9UH03; A0 | SEPT3_HUMAN | Unmodified | 0 | 0 | 0 | 0 | 172700 | 0 | 434870 | 293380 | 0 | 0 | 0 | 0 | 580970 | 213160 | 0 | 0 | 0 | 0 |
| EVTLHLLP | MTMR12 | Q9C0I1 | MTMRC_HUMA | Unmodified | 0 | 0 | 0 | 0 | 0 | 0 | 29649 | 0 | 0 | 0 | 0 | 0 | 413390 | 307480 | 128130 | 34599 | 119130 | 590990 |
| EVTMELTK | TTYH2; TTYH2 | Q9BSA4; B4I | TTYH2_HUMAN | Unmodified | 0 | 0 | 0 | 0 | 0 | 0 | 0 | 0 | 0 | 0 | 0 | 0 | 0 | 0 | 0 | 0 | 0 | 0 |
| EVTPEGLQ | PAICS; PAICS; F | P22234; D6F | PUR6_HUMAN | Oxidation (M) | 0 | 39259 | 234360 | 138070 | 531360 | 0 | 0 | 0 | 0 | 0 | 1237400 | 6177800 | 605450 | 0 | 193060 | 199990 | 238500 | 458220 |
| EVTPEGLQ | PAICS; PAICS; F | P22234; D6F | PUR6_HUMAN | Unmodified | 0 | 0 | 0 | 0 | 0 | 0 | 404710 | 0 | 0 | 0 | 3499300 | 9152100 | 1058000 | 0 | 0 | 0 | 0 | 0 |
| EVTPEGLQ | PAICS; PAICS; F | P22234; D6F | PUR6_HUMAN | Unmodified | 0 | 0 | 0 | 0 | 0 | 353530 | 389680 | 67665 | 1235200 | 348600 | 200430 | 2284600 | 635620 | 0 | 0 | 0 | 937460 | 1953000 |
| EVTPEGLQ | PAICS; PAICS; F | P22234; D6F | PUR6_HUMAN | Oxidation (M) | 0 | 0 | 133420 | 0 | 254600 | 87290 | 117410 | 77766 | 0 | 111610 | 582000 | 4239700 | 924530 | 0 | 0 | 267060 | 605800 | 545380 |
| EVTVECKID | ATP1B3; ATP1B | P54709; C9J | AT1B3_HUMAN | Unmodified | 0 | 0 | 0 | 1499800 | 2644100 | 594550 | 113300 | 0 | 1396900 | 372920 | 3019600 | 3892200 | 2740300 | 2178500 | 819850 | 919700 | 356550 | 901030 |
| EVTYELDHI | CNOT1 | A5YKK6 | CNOT1_HUMA | Unmodified | 182280 | 0 | 0 | 0 | 67940 | 0 | 0 | 0 | 102980 | 0 | 0 | 2985400 | 234770 | 0 | 0 | 0 | 0 | 0 |
| EVVAGSHE | SPTBN1 | Q01082 | SPTB2_HUMAN | Oxidation (M) | 0 | 0 | 0 | 0 | 0 | 320380 | 0 | 0 | 0 | 0 | 0 | 1243900 | 0 | 500090 | 0 | 0 | 0 | 592670 |
| EVVCHASA | ACE; ACE; N/A | P12821; A0A | ACE_HUMAN A | Unmodified | 357450 | 102550 | 0 | 0 | 0 | 0 | 0 | 0 | 114290 | 0 | 0 | 0 | 0 | 0 | 119470 | 0 | 0 | 0 |
| EVVCTENL | PIGT; PIGT; PIG | Q969N2; A0 | PIGT_HUMAN | Unmodified | 0 | 0 | 0 | 0 | 0 | 0 | 0 | 124240 | 1592600 | 0 | 0 | 0 | 67783 | 0 | 0 | 0 | 0 | 0 |
| EVVDILTEQ | ULBP2 | Q9BZM5 | ULBP2_HUMAN | Unmodified | 0 | 0 | 474840 | 638400 | 0 | 0 | 0 | 0 | 602420 | 0 | 0 | 0 | 0 | 0 | 643440 | 469470 | 256050 | 0 |
| EVVDRGLQ | COLGALT1; CO | Q8NBJ5; M0 | GT251_HUMAN | Unmodified | 0 | 0 | 51431 | 70715 | 238490 | 38453 | 0 | 0 | 1715700 | 0 | 0 | 0 | 44257 | 44502 | 140490 | 64820 | 0 | 78087 |
| EVVEAHVD | GATD3B; GATD | A0A0B4J2D5 | GAL3B_HUMAN | Unmodified | 0 | 0 | 0 | 0 | 0 | 0 | 143760 | 0 | 926930 | 232420 | 0 | 39361 | 101750 | 0 | 0 | 0 | 0 | 0 |
| EVVEDINK | BAG5 | Q9UL15 | BAG5_HUMAN | Unmodified | 277850 | 0 | 294410 | 194240 | 0 | 32017 | 0 | 99108 | 1438600 | 161880 | 0 | 791090 | 0 | 0 | 284910 | 267200 | 0 | 0 |
| EVVEEAEN | PTMA; PTMA; P1 | P06454; H7 | PTMA_HUMAN | Unmodified | 183700 | 54620 | 0 | 0 | 0 | 0 | 142610 | 0 | 494210 | 0 | 2137400 | 16306000 | 0 | 0 | 701240 | 958500 | 0 | 0 |
| EVVETPLL | PSMC2; PSMC2 | P35998; C9J | PRS7_HUMAN | Unmodified | 0 | 0 | 0 | 0 | 0 | 0 | 0 | 0 | 0 | 0 | 0 | 1771700 | 0 | 0 | 0 | 87852 | 0 | 0 |
| EVVEVGQK | SLK | Q9H2G2 | SLK_HUMAN S | Unmodified | 0 | 54672 | 0 | 0 | 67273 | 0 | 0 | 0 | 163520 | 0 | 48999 | 553100 | 0 | 0 | 65072 | 0 | 25868 | 0 |
| EVVHTVSL | RUVBL2; RUVBI | Q9Y230; M0 | RUVB2_HUMAN | Unmodified | 0 | 71482 | 0 | 0 | 0 | 0 | 0 | 0 | 0 | 0 | 0 | 3813400 | 177010 | 0 | 0 | 0 | 0 | 0 |
| EVVIAQND | SEC31A; SEC31 | O94979; D6I | SC31A_HUMAN | Unmodified | 17134 | 0 | 0 | 0 | 234460 | 0 | 130040 | 29708 | 167640 | 0 | 130490 | 3426700 | 0 | 0 | 0 | 14252 | 422880 | 0 |
| EVVIFGAAS | HMGCL | P35914 | HMGCL_HUMA | Unmodified | 0 | 0 | 0 | 0 | 0 | 0 | 0 | 0 | 1677200 | 1262700 | 0 | 0 | 0 | 0 | 0 | 0 | 0 | 0 |
| EVVIVSATR | ACAT1; ACAT1; | P24752; E9P | THIL_HUMAN A | Unmodified | 0 | 0 | 0 | 57991 | 0 | 0 | 0 | 0 | 1797500 | 110480 | 0 | 224860 | 0 | 0 | 39431 | 0 | 213650 | 0 |
| EVVKKHSQ | HSP90AB1; HSI | P08238; Q5 | HS90B_HUMAN | Unmodified | 0 | 0 | 0 | 0 | 1067800 | 0 | 0 | 0 | 0 | 0 | 0 | 0 | 0 | 658360 | 0 | 0 | 2963100 | 622870 |
| EVVKTDSA | ZPR1; ZPR1; ZP | O75312; H7 | ZPR1_HUMAN | Unmodified | 0 | 0 | 0 | 0 | 0 | 0 | 0 | 0 | 0 | 0 | 0 | 859870 | 0 | 0 | 0 | 0 | 0 | 158060 |
| EVVLENQK | PALD1 | Q9ULE6 | PALD_HUMAN | Unmodified | 0 | 0 | 0 | 136400 | 47023 | 220610 | 98708 | 0 | 149720 | 0 | 148830 | 1044700 | 77390 | 0 | 0 | 0 | 0 | 21147 |
| EVVLQWFT | ANPEP | P15144 | AMPN_HUMAN | Unmodified | 0 | 0 | 0 | 0 | 0 | 0 | 0 | 0 | 0 | 0 | 60688 | 0 | 0 | 0 | 0 | 0 | 17787000 | 46341000 |
| EVVNKLIPD | RPS3A; RPS3A; | P61247; D6F | RS3A_HUMAN | Unmodified | 0 | 0 | 0 | 0 | 916610 | 0 | 1779200 | 359700 | 0 | 339210 | 0 | 0 | 1006500 | 288210 | 0 | 0 | 122150 | 494500 |
| EVVQNFAK | COTL1; COTL1 | Q14019; H3I | COTL1_HUMAN | Unmodified | 544920 | 0 | 1005200 | 1372000 | 23083000 | 7611200 | 3406300 | 2118400 | 14691000 | 11089000 | 234030 | 4293400 | 2702900 | 3403900 | 843080 | 0 | 17316000 | 11982000 |
| EVVSAQPA | GRN; GRN; GRI | P28799; K7E | GRN_HUMAN | Unmodified | 0 | 0 | 0 | 0 | 0 | 0 | 0 | 0 | 3085700 | 627260 | 0 | 133830 | 0 | 0 | 0 | 0 | 0 | 954710 |
| EVVSPQEFI | NCAM2; NCAM | O15394; H9I | NCAM2_HUMA | Unmodified | 0 | 357340 | 1643500 | 653570 | 107880 | 0 | 0 | 0 | 0 | 175320 | 318070 | 0 | 0 | 0 | 2397300 | 1997800 | 297130 | 301150 |
| EVVSPQEFI | NCAM2; NCAM | O15394; H9I | NCAM2_HUMA | Unmodified | 303520 | 374300 | 601030 | 2179800 | 0 | 0 | 0 | 0 | 0 | 0 | 0 | 0 | 0 | 0 | 3335600 | 2556400 | 0 | 0 |
| EVVSSTGVI | ROR1; ROR1 | Q01973; A0A | ROR1_HUMAN | Unmodified | 0 | 0 | 404320 | 0 | 1884900 | 323750 | 0 | 0 | 94306 | 0 | 0 | 414310 | 280410 | 0 | 1040800 | 894250 | 0 | 176260 |

| | | | | | | | | | | | | | | | | | | | | | | |
|---|---|---|---|---|---|---|---|---|---|---|---|---|---|---|---|---|---|---|---|---|---|---|
| EVVVSVPQ | NUP210 | Q8TEM1 | PO210_HUMAI | Unmodified | 0 | 1460100 | 0 | 96881 | 0 | 0 | 0 | 0 | 6560900 | 356830 | 0 | 355890 | 0 | 0 | 0 | 1530800 | 0 | 0 |
| EVWALVQA | FASN; FASN | P49327; A0A | FAS_HUMAN F | Unmodified | 0 | 0 | 0 | 0 | 0 | 0 | 0 | 0 | 0 | 0 | 0 | 2102400 | 0 | 0 | 42105 | 29633 | 0 | 0 |
| EVWDYVFF | LARS1; LARS1; | Q9P2J5; A0A | SYLC_HUMAN | Unmodified | 43540 | 0 | 91427 | 389370 | 528010 | 0 | 482900 | 0 | 770520 | 0 | 18377 | 5980400 | 302930 | 0 | 445690 | 157320 | 214070 | 408600 |
| EVWEELDG | NUCB1; NUCB: | Q02818; H7I | NUCB1_HUMA | Unmodified | 0 | 0 | 0 | 0 | 749730 | 963880 | 237310 | 176890 | 16594000 | 2452900 | 6666600 | 1283000 | 840650 | 0 | 41530 | 0 | 0 | 2156400 |
| EVWEETDG | NUCB2; NUCB: | P80303; Q2L | NUCB2_HUMA | Unmodified | 50464 | 0 | 0 | 596970 | 0 | 0 | 0 | 55179 | 50230000 | 3217300 | 424080 | 107880 | 540560 | 306180 | 522540 | 182640 | 0 | 0 |
| EVWEETDG | NUCB2; NUCB: | P80303; Q2L | NUCB2_HUMA | Unmodified | 0 | 0 | 0 | 0 | 0 | 0 | 1091100 | 0 | 0 | 2127400 | 0 | 0 | 850840 | 1290800 | 0 | 0 | 0 | 0 |
| EVWFFGLQ | RDX; RDX; RDX; | P35241; E9P | RADI_HUMAN I | Unmodified | 0 | 0 | 24446 | 33188 | 77951 | 0 | 0 | 0 | 71059 | 0 | 0 | 511940 | 0 | 0 | 644280 | 142300 | 0 | 0 |
| EVWYFGLH | EZR; EZR | P15311; E7E | EZRI_HUMAN E | Unmodified | 1013400 | 1177800 | 551920 | 528600 | 527530 | 0 | 164980 | 0 | 16888000 | 678390 | 2269100 | 18105000 | 0 | 100480 | 2822500 | 2098600 | 37319 | 0 |
| EVYDIAFSR | DCAF7; DCAF7 | P61962; A0A | DCAF7_HUMAI | Unmodified | 0 | 0 | 0 | 0 | 0 | 0 | 0 | 0 | 0 | 0 | 0 | 0 | 0 | 0 | 0 | 0 | 0 | 0 |
| EVYEEAGVI | NUDT4B; NUDT | A0A024RBG | NUD4B_HUMA | Unmodified | 0 | 0 | 0 | 0 | 0 | 0 | 0 | 0 | 0 | 0 | 54623 | 354670 | 0 | 0 | 0 | 0 | 0 | 0 |
| EVYEHVYE | LIN7C | Q9NUP9 | LIN7C_HUMAN | Unmodified | 0 | 0 | 0 | 0 | 0 | 0 | 0 | 0 | 1362400 | 0 | 0 | 1000100 | 0 | 0 | 0 | 0 | 0 | 0 |
| EVYELLDSF | PAICS; PAICS | P22234; E9P | PUR6_HUMAN | Unmodified | 0 | 231850 | 946170 | 904750 | 0 | 0 | 0 | 714340 | 1166700 | 521170 | 1973900 | 4659700 | 3512500 | 2280600 | 0 | 706490 | 1914400 | 2187300 |
| EVYFAERVI | CRIP1 | P50238 | CRIP1_HUMAN | Unmodified | 592930 | 369190 | 0 | 0 | 48385000 | 9284400 | 59678000 | 14987000 | 6637400 | 10378000 | 0 | 0 | 627250 | 698520 | 0 | 0 | 1453400 | 13156000 |
| EVYLDLVK | TOM1L1; TOM1 | O75674; J3K | TM1L1_HUMAN | Unmodified | 4843200 | 3998300 | 3613600 | 1917800 | 288220 | 0 | 0 | 0 | 885660 | 433590 | 199470 | 1518900 | 853620 | 789720 | 4385400 | 3283600 | 94307 | 0 |
| EVYLSGSFI | PRKAB1 | Q9Y478 | AAKB1_HUMAN | Unmodified | 1311300 | 0 | 0 | 0 | 0 | 0 | 0 | 0 | 0 | 275000 | 0 | 0 | 0 | 0 | 0 | 354380 | 0 | 0 |
| EVYPEETPE | FAM20B; FAM2( | O75063; X6F | XYLK_HUMAN ( | Unmodified | 0 | 0 | 0 | 0 | 0 | 0 | 0 | 0 | 5285000 | 273060 | 233930 | 0 | 0 | 0 | 263380 | 199010 | 0 | 0 |
| EVYPEETPE | FAM20B; FAM2( | O75063; X6F | XYLK_HUMAN ( | Unmodified | 0 | 0 | 0 | 0 | 0 | 0 | 0 | 0 | 2613600 | 163640 | 0 | 0 | 0 | 0 | 0 | 0 | 0 | 0 |
| EVYQQQQY | HNRNPAB; HNI | Q99729; D6I | ROAA_HUMAN | Unmodified | 137330 | 0 | 0 | 0 | 0 | 0 | 0 | 0 | 0 | 0 | 0 | 2329100 | 0 | 0 | 84735 | 70874 | 0 | 0 |
| EWAHATIIP | PPP2R1A; PPP2 | P30153; B3K | 2AAA_HUMAN | Unmodified | 0 | 0 | 0 | 0 | 0 | 0 | 0 | 0 | 0 | 0 | 0 | 1702800 | 0 | 0 | 0 | 95436 | 0 | 0 |
| EWAMADN | APLP1; APLP1; / | P51693; B7Z | APLP1_HUMAN | Unmodified | 0 | 0 | 0 | 0 | 0 | 0 | 0 | 0 | 666840 | 0 | 0 | 0 | 0 | 0 | 0 | 0 | 0 | 0 |
| EWEDSKRV | CHGA; CHGA | P10645; G5E | CMGA_HUMAN | Unmodified | 0 | 0 | 0 | 0 | 0 | 0 | 0 | 0 | 0 | 935130 | 0 | 0 | 0 | 0 | 0 | 0 | 0 | 0 |
| EWEEAELQ | APLP2 | Q06481 | APLP2_HUMAN | Unmodified | 132410 | 0 | 0 | 0 | 0 | 0 | 0 | 0 | 3969700 | 428190 | 486300 | 0 | 165370 | 0 | 0 | 0 | 0 | 0 |
| EWFLQAAK | GPI; GPI; GPI; G | P06744; K7E | G6PI_HUMAN ( | Unmodified | 0 | 0 | 0 | 0 | 0 | 0 | 0 | 203160 | 1494000 | 217470 | 1535700 | 13660000 | 0 | 0 | 1642800 | 1677200 | 0 | 0 |
| EWFLQAAK | GPI; GPI; GPI; G | P06744; K7E | G6PI_HUMAN ( | Unmodified | 0 | 1981500 | 0 | 6151500 | 12343000 | 6178400 | 2195400 | 378950 | 11793000 | 5008700 | 14804000 | 48028000 | 5154400 | 4085900 | 3440500 | 2633200 | 3045600 | 0 |
| EWIQNTLG | AGPS; AGPS; A( | O00116; A0/ | ADAS_HUMAN | Unmodified | 0 | 0 | 0 | 62220 | 0 | 0 | 320710 | 0 | 15665000 | 3033900 | 423370 | 0 | 0 | 0 | 0 | 9823.5 | 0 | 0 |
| EWIQNTLG | AGPS; AGPS; A( | O00116; A0/ | ADAS_HUMAN | Unmodified | 0 | 0 | 0 | 0 | 0 | 0 | 418240 | 0 | 0 | 1068700 | 0 | 0 | 0 | 0 | 0 | 0 | 171640 | 0 |
| EWLEDKG( | SSB; SSB; SSB | P05455; E9P | LA_HUMAN Lu | Oxidation (M) | 22649 | 0 | 0 | 607420 | 0 | 0 | 0 | 0 | 0 | 0 | 0 | 523550 | 0 | 0 | 379800 | 0 | 0 | 0 |
| EWMPVTK | RPS2; RPS2; RP | P15880; E9P | RS2_HUMAN 4 | Unmodified | 829020 | 489940 | 292330 | 452480 | 0 | 0 | 134050 | 86220 | 394170 | 0 | 1742800 | 8894600 | 0 | 52682 | 287620 | 297220 | 0 | 128890 |
| EWQAAWL | DERA; DERA; DI | Q9Y315; E9F | DEOC_HUMAN | Unmodified | 198040 | 0 | 0 | 0 | 462710 | 0 | 0 | 219950 | 0 | 0 | 0 | 642120 | 0 | 246390 | 0 | 218830 | 0 | 0 |
| EWQELDD/ | PSMD2 | Q13200 | PSMD2_HUMA | Unmodified | 0 | 0 | 271010 | 284720 | 733250 | 0 | 0 | 0 | 593740 | 144820 | 501610 | 2959800 | 559410 | 254090 | 204220 | 204530 | 0 | 0 |
| EWQELDD/ | PSMD2 | Q13200 | PSMD2_HUMA | Unmodified | 0 | 0 | 0 | 0 | 0 | 206710 | 0 | 31072 | 0 | 0 | 0 | 221140 | 2491900 | 891370 | 0 | 0 | 98911 | 1837400 |
| EWQSLAGS | EHHADH | Q08426 | ECHP_HUMAN | Unmodified | 0 | 0 | 0 | 0 | 0 | 0 | 0 | 0 | 3960300 | 846440 | 0 | 0 | 0 | 0 | 0 | 0 | 0 | 0 |
| EWSGLLEE | PFKP; PFKP; PF | Q01813; A0/ | PFKAP_HUMAN | Unmodified | 0 | 0 | 0 | 0 | 0 | 0 | 0 | 0 | 375300 | 0 | 513450 | 3331300 | 0 | 0 | 0 | 0 | 0 | 0 |
| EWSNSLEK | TRAF3; TRAF3; 1 | Q13114; A6I | TRAF3_HUMAN | Unmodified | 0 | 0 | 0 | 0 | 4097000 | 951280 | 0 | 0 | 0 | 0 | 46634 | 0 | 0 | 0 | 0 | 0 | 0 | 0 |
| EWTTTTGD | CANT1 | Q8WVQ1 | CANT1_HUMAI | Unmodified | 0 | 0 | 1712700 | 0 | 0 | 132240 | 0 | 0 | 15823000 | 2194500 | 745510 | 0 | 0 | 123510 | 0 | 0 | 0 | 336440 |
| EWVESMG( | ST13; ST13P5; ! | P50502; Q8I | F10A1_HUMAN | Unmodified | 0 | 0 | 0 | 0 | 0 | 0 | 0 | 0 | 0 | 0 | 159830 | 420020 | 0 | 0 | 0 | 0 | 0 | 0 |
| EWVLGRVE | PLXNB2 | O15031 | PLXB2_HUMAN | Unmodified | 0 | 0 | 856760 | 21659 | 0 | 0 | 0 | 0 | 0 | 35504 | 0 | 4005300 | 0 | 0 | 2252500 | 1359500 | 0 | 0 |
| EWYRLTEEI | IGSF3 | O75054 | IGSF3_HUMAN | Unmodified | 408250 | 201980 | 156300 | 219860 | 0 | 0 | 224110 | 0 | 234190 | 0 | 0 | 0 | 0 | 0 | 437440 | 214790 | 0 | 0 |
| EYAEDDNI | OTUB1; OTUB1 | Q96FW1; J3I | OTUB1_HUMAI | Unmodified | 0 | 0 | 243270 | 198040 | 2170100 | 722280 | 213570 | 0 | 553760 | 195590 | 1442400 | 6869900 | 1690600 | 305090 | 300260 | 421120 | 1296300 | 1714200 |
| EYAQNIWN | PYGL | P06737 | PYGL_HUMAN | Unmodified | 0 | 0 | 0 | 0 | 2563000 | 0 | 0 | 0 | 0 | 0 | 0 | 0 | 0 | 0 | 0 | 0 | 555810 | 3737100 |
| EYAQRIPS( | ITGA1 | P56199 | ITA1_HUMAN I | Unmodified | 1466500 | 1334800 | 0 | 0 | 87694 | 356100 | 0 | 57265 | 499530 | 0 | 0 | 0 | 0 | 51187 | 0 | 0 | 8570600 | 26140000 |
| EYAQRIPS( | ITGA1 | P56199 | ITA1_HUMAN I | Unmodified | 0 | 605040 | 0 | 1370500 | 163650 | 0 | 0 | 33495 | 0 | 103530 | 0 | 0 | 225730 | 168540 | 107600 | 0 | 1771700 | 3184400 |
| EYAQVFR | CTNNA1; CTNN | P35221; P26 | CTNA1_HUMAI | Unmodified | 156100 | 208440 | 0 | 0 | 0 | 0 | 0 | 0 | 0 | 0 | 0 | 1153200 | 0 | 0 | 898000 | 66462 | 0 | 0 |
| EYAQVFREI | CTNNA1; CTNN | P35221; P26 | CTNA1_HUMAI | Unmodified | 0 | 445490 | 0 | 0 | 1102700 | 1398800 | 0 | 7303.3 | 0 | 102290 | 0 | 1355000 | 241610 | 133900 | 3589900 | 1869400 | 0 | 0 |
| EYATEVDVI | AP1B1; AP2B1; | Q10567; P63 | AP1B1_HUMAN | Unmodified | 80998 | 0 | 0 | 0 | 90076 | 197830 | 0 | 10130 | 76448 | 0 | 0 | 1432300 | 0 | 0 | 351520 | 298000 | 110900 | 0 |
| EYDELAET( | KDM1A; KDM1A | O60341; R4( | KDM1A_HUMA | Unmodified | 0 | 0 | 0 | 0 | 0 | 0 | 0 | 108680 | 0 | 0 | 0 | 1152500 | 0 | 0 | 0 | 0 | 0 | 0 |
| EYDEPYVLI | ABCC4 | O15439 | MRP4_HUMAN | Unmodified | 0 | 0 | 416290 | 0 | 248760 | 0 | 0 | 0 | 0 | 0 | 0 | 0 | 0 | 0 | 1062000 | 969820 | 0 | 0 |
| EYDQALAD | PPID; PPID | Q08752; H0Y | PPID_HUMAN I | Unmodified | 925530 | 439090 | 0 | 0 | 0 | 0 | 0 | 76877 | 376090 | 261610 | 0 | 1158000 | 0 | 0 | 361440 | 0 | 0 | 271100 |
| EYDRLLEEI | BCAP31; BCAP: | P51572; A0A | BAP31_HUMAN | Unmodified | 0 | 0 | 443160 | 272780 | 0 | 26272 | 1083500 | 0 | 0 | 0 | 267810 | 960620 | 0 | 904690 | 289280 | 0 | 0 | 825560 |
| EYELESLFE | PEPD; PEPD; PE | P12955; A0A | PEPD_HUMAN | Unmodified | 93497 | 0 | 0 | 0 | 0 | 0 | 0 | 0 | 345100 | 145250 | 233840 | 796690 | 0 | 26275 | 0 | 128490 | 43794 | 0 |
| EYELLSDEL | KIF5B | P33176 | KINH_HUMAN | Unmodified | 0 | 0 | 0 | 0 | 0 | 0 | 0 | 0 | 0 | 0 | 0 | 1140200 | 0 | 0 | 0 | 0 | 0 | 0 |
| EYENIIALQI | NUCB2; NUCB: | P80303; Q2L | NUCB2_HUMA | Unmodified | 0 | 0 | 241270 | 395780 | 0 | 0 | 0 | 0 | 10733000 | 0 | 0 | 0 | 0 | 0 | 568540 | 0 | 0 | 0 |
| EYENIIALQI | NUCB2; NUCB: | P80303; Q2L | NUCB2_HUMA | Unmodified | 0 | 0 | 0 | 0 | 0 | 0 | 0 | 0 | 8912200 | 0 | 399350 | 230020 | 0 | 0 | 0 | 0 | 0 | 0 |
| EYEPEMGK | ATP2A1; ATP2A | O14983; P16 | AT2A1_HUMAN | Unmodified | 0 | 0 | 0 | 0 | 0 | 0 | 0 | 0 | 332160 | 70151 | 0 | 1904300 | 0 | 0 | 0 | 0 | 0 | 0 |
| EYERVTSGI | EREG | O14944 | EREG_HUMAN | Unmodified | 68261 | 168140 | 31254 | 191200 | 3359200 | 93258 | 0 | 32385 | 0 | 322160 | 0 | 0 | 0 | 52577 | 0 | 0 | 0 | 0 |
| EYEYSFR | MVB12B | Q9H7P6 | MB12B_HUMAI | Unmodified | 453870 | 651980 | 598290 | 387630 | 0 | 0 | 0 | 0 | 746890 | 0 | 0 | 226100 | 0 | 0 | 1067100 | 1449400 | 0 | 0 |
| EYEYSFRTE | MVB12B | Q9H7P6 | MB12B_HUMAI | Unmodified | 342220 | 460310 | 367000 | 699930 | 0 | 836580 | 0 | 66192 | 720420 | 0 | 0 | 0 | 82022 | 0 | 1086300 | 647580 | 38187 | 208260 |
| EYFGAFGEI | HNRNPDL; HNI | O14979; A0/ | HNRDL_HUMA | Oxidation (M) | 65309 | 103370 | 20636 | 142450 | 571850 | 0 | 415570 | 58114 | 224560 | 466760 | 0 | 818150 | 631900 | 139800 | 101300 | 32650 | 0 | 522140 |
| EYFGGFGE | HNRNPD; HNR | Q14103; H0Y | HNRPD_HUMA | Oxidation (M) | 0 | 0 | 0 | 0 | 0 | 0 | 0 | 0 | 191570 | 0 | 589910 | 603760 | 0 | 0 | 0 | 0 | 0 | 0 |
| EYFSWEGA | EEF1G | P26641 | EF1G_HUMAN | Unmodified | 0 | 0 | 396210 | 0 | 0 | 0 | 0 | 0 | 1003700 | 0 | 0 | 20519000 | 0 | 0 | 577120 | 525180 | 0 | 0 |
| EYFYYVDH | C8orf82; C8orf | Q6P1X6; J3K | CH082_HUMAI | Unmodified | 0 | 0 | 0 | 0 | 0 | 0 | 0 | 0 | 2739500 | 466500 | 0 | 0 | 0 | 0 | 0 | 0 | 0 | 0 |
| EYGEQIDP! | CNPY2; N/A; CI | Q9Y2B0; F8V | CNPY2_HUMAI | Unmodified | 0 | 0 | 0 | 0 | 0 | 0 | 0 | 0 | 0 | 0 | 0 | 0 | 0 | 0 | 0 | 0 | 0 | 0 |
| EYGEQIDP! | CNPY2; N/A; CI | Q9Y2B0; F8V | CNPY2_HUMAI | Unmodified | 0 | 0 | 0 | 0 | 315650 | 0 | 46656 | 90914 | 46729000 | 15219000 | 2785100 | 813880 | 0 | 373460 | 127430 | 0 | 645970 | 207200 |
| EYGETKGP | BROX; BROX | Q5VW32; F5 | BROX_HUMAN | Unmodified | 1214100 | 904840 | 0 | 1467600 | 1477700 | 376350 | 232180 | 0 | 0 | 1384700 | 0 | 0 | 1026200 | 502060 | 0 | 0 | 0 | 402660 |
| EYGGLDVL | CBR1; CBR1; C | P16152; E9P | CBR1_HUMAN | Unmodified | 85290 | 0 | 145580 | 263650 | 1691300 | 325630 | 634110 | 48448 | 0 | 1427600 | 65855 | 457340 | 394630 | 1261300 | 813120 | 172980 | 1179600 | 871140 |
| EYGMIYLGI | SND1 | Q7KZF4 | SND1_HUMAN | Unmodified | 0 | 50179 | 73951 | 90110 | 0 | 0 | 0 | 0 | 65115 | 0 | 100300 | 1125600 | 0 | 0 | 113700 | 59345 | 0 | 0 |
| EYHFISTEE | MPP1 | Q00013 | EM55_HUMAN | Unmodified | 0 | 0 | 0 | 0 | 0 | 0 | 0 | 0 | 619570 | 0 | 0 | 0 | 0 | 0 | 0 | 0 | 0 | 0 |
| EYIDPETK | CSNK1G1; CSN | Q9HCP0; P7 | KC1G1_HUMAI | Unmodified | 203770 | 196000 | 478930 | 23933 | 0 | 0 | 0 | 56377 | 434980 | 143360 | 263510 | 1581100 | 0 | 174320 | 343320 | 416790 | 0 | 0 |
| EYIEIITSSIK | BRK1 | Q8WUW1 | BRK1_HUMAN | Unmodified | 0 | 0 | 0 | 0 | 0 | 0 | 0 | 0 | 0 | 0 | 0 | 1078000 | 0 | 0 | 0 | 0 | 0 | 0 |
| EYIEIITSSIK | BRK1 | Q8WUW1 | BRK1_HUMAN | Unmodified | 0 | 0 | 0 | 0 | 0 | 0 | 0 | 0 | 498970 | 0 | 0 | 1264000 | 0 | 0 | 481950 | 382340 | 0 | 0 |
| EYIIPSPGYI | ABCC5; ABCC5 | O15440; C9J | MRP5_HUMAN | Unmodified | 0 | 0 | 0 | 0 | 0 | 0 | 0 | 0 | 0 | 0 | 0 | 1008600 | 0 | 0 | 0 | 0 | 0 | 0 |
| EYIIQVAAK | CNTFR | P26992 | CNTFR_HUMAI | Unmodified | 0 | 1209800 | 0 | 0 | 0 | 0 | 0 | 0 | 1453300 | 0 | 0 | 0 | 0 | 0 | 0 | 0 | 0 | 0 |
| EYILVEEAK | P4HA2; P4HA2; | O15460; E7E | P4HA2_HUMAI | Unmodified | 0 | 0 | 54289 | 0 | 167890 | 0 | 0 | 31769 | 3231500 | 1181400 | 0 | 0 | 0 | 0 | 0 | 0 | 0 | 0 |
| EYIPGQPPI | EPRS1; EPRS1 | P07814; V9G | SYEP_HUMAN | Unmodified | 55367 | 0 | 123970 | 0 | 0 | 0 | 0 | 0 | 423650 | 0 | 416060 | 1789900 | 0 | 0 | 117590 | 133550 | 0 | 0 |
| EYIPGQPPI | EPRS1; EPRS1 | P07814; V9G | SYEP_HUMAN | Unmodified | 0 | 0 | 0 | 0 | 0 | 0 | 0 | 0 | 0 | 0 | 0 | 0 | 2171100 | 0 | 0 | 0 | 0 | 0 |
| EYIPTVFDN | RHOG | P84095 | RHOG_HUMAN | Unmodified | 2903300 | 5235400 | 1157200 | 2960900 | 2962600 | 519670 | 0 | 211620 | 0 | 0 | 1637400 | 7526800 | 1141400 | 955880 | 3835000 | 2488000 | 796560 | 0 |
| EYIQLVEQL | VLDLR; VLDLR; | P98155; A0A | VLDLR_HUMAN | Unmodified | 0 | 0 | 0 | 0 | 0 | 0 | 0 | 0 | 1188100 | 0 | 0 | 0 | 442020 | 0 | 0 | 0 | 0 | 0 |
| EYIVEYSR | SORL1; SORL1; | Q92673; E9F | SORL_HUMAN | Unmodified | 0 | 0 | 0 | 0 | 0 | 0 | 0 | 0 | 1200700 | 0 | 0 | 334240 | 144940 | 0 | 0 | 0 | 0 | 0 |
| EYIVTQGPL | PTPRB; PTPRB | P23467; F8V | PTPRB_HUMAN | Unmodified | 693330 | 477580 | 0 | 0 | 0 | 0 | 0 | 38260 | 0 | 0 | 0 | 0 | 0 | 0 | 0 | 0 | 0 | 0 |
| EYKLVVLGS | RAP1B; RAP1A; | P61224; P62 | RAP1B_HUMAN | Unmodified | 0 | 0 | 0 | 173050 | 753930 | 233560 | 0 | 0 | 0 | 0 | 0 | 0 | 0 | 776810 | 0 | 0 | 297820 | 403270 |
| EYLAQLSNI | DIPK2A; DIPK2/ | Q8NDZ4; E7 | DIK2A_HUMAN | Unmodified | 0 | 0 | 0 | 0 | 0 | 0 | 0 | 0 | 1930500 | 83636 | 151460 | 0 | 0 | 0 | 0 | 0 | 438950 | 0 |
| EYLCLDNS | CEMIP2 | Q9UHN6 | CEIP2_HUMAN | Unmodified | 0 | 0 | 0 | 0 | 0 | 0 | 0 | 0 | 1623200 | 0 | 0 | 0 | 0 | 0 | 0 | 0 | 0 | 0 |
| EYLEGRNLI | SRGAP2; SRGA | O75044; Q5' | SRGP2_HUMAI | Unmodified | 0 | 0 | 0 | 0 | 2014200 | 307120 | 0 | 0 | 0 | 132100 | 0 | 0 | 965990 | 188610 | 0 | 0 | 373230 | 558440 |
| EYLERVDG | CUL1; CUL1; CI | Q13616; A0/ | CUL1_HUMAN | Unmodified | 0 | 0 | 0 | 0 | 1979600 | 0 | 0 | 0 | 0 | 270070 | 0 | 0 | 797230 | 0 | 0 | 0 | 0 | 255320 |
| EYLGAICS( | FN1; FN1 | P02751; H0Y | FINC_HUMAN I | Unmodified | 350630 | 321400 | 970210 | 1306500 | 0 | 421130 | 0 | 0 | 417090 | 240770 | 0 | 0 | 294170 | 732990 | 143890 | 503020 | 76755 | 431390 |
| EYLGAICS( | FN1; FN1 | P02751; H0Y | FINC_HUMAN I | Unmodified | 0 | 0 | 0 | 0 | 0 | 0 | 0 | 0 | 0 | 782840 | 0 | 0 | 2858500 | 6414700 | 0 | 0 | 671840 | 3730900 |
| EYLIAGKAE | TIMP2; TIMP2; T | P16035; K7E | TIMP2_HUMAN | Unmodified | 0 | 29935 | 0 | 72559 | 2062300 | 2284600 | 0 | 28083 | 1476400 | 375670 | 0 | 0 | 0 | 0 | 0 | 0 | 862690 | 1084600 |
| EYLLGSTAE | EEF1E1; EEF1E: | O43324; D6I | MCA3_HUMAN | Unmodified | 0 | 0 | 0 | 0 | 0 | 0 | 97225 | 0 | 402160 | 0 | 342220 | 4266500 | 0 | 0 | 0 | 0 | 0 | 0 |
| EYLLQYND | NDUFB4; NDUI | O95168; F2Z | NDUB4_HUMA | Unmodified | 38485 | 0 | 0 | 0 | 0 | 0 | 0 | 0 | 1934500 | 0 | 0 | 0 | 0 | 0 | 0 | 0 | 0 | 0 |
| EYLLSGDIS | PDCD4 | Q53EL6 | PDCD4_HUMA | Unmodified | 0 | 0 | 0 | 0 | 0 | 0 | 0 | 0 | 0 | 0 | 0 | 0 | 0 | 0 | 0 | 0 | 0 | 0 |
| EYLPIGGLA | GOT2 | P00505 | AATM_HUMAN | Unmodified | 0 | 0 | 0 | 0 | 0 | 0 | 0 | 0 | 4239000 | 1838000 | 222520 | 0 | 0 | 0 | 0 | 0 | 0 | 0 |
| EYLQTHWE | ENOPH1 | Q9UHY7 | ENOPH_HUMA | Unmodified | 0 | 0 | 0 | 0 | 369870 | 0 | 0 | 0 | 0 | 0 | 0 | 3471200 | 0 | 0 | 0 | 251650 | 181670 | 0 |
| EYLTKVDKI | CLTC; CLTC; CI | Q00610; J3K | CLH1_HUMAN | Unmodified | 4384900 | 1610800 | 58696 | 240950 | 0 | 0 | 0 | 0 | 0 | 0 | 0 | 14256000 | 0 | 0 | 203740 | 886180 | 0 | 0 |
| EYNEFAEVI | IPO7 | O95373 | IPO7_HUMAN I | Unmodified | 0 | 0 | 0 | 0 | 0 | 0 | 0 | 54671 | 0 | 72360 | 0 | 4222000 | 0 | 0 | 0 | 0 | 0 | 0 |
| EYNIGVLR | USP9X; USP9Y | Q93008; O0( | USP9X_HUMAN | Unmodified | 0 | 0 | 628680 | 324970 | 0 | 0 | 0 | 0 | 397810 | 0 | 0 | 788840 | 0 | 0 | 28623 | 437130 | 0 | 0 |
| EYPNSGPQ | SHPRH; SHPRH | Q149N8; A0/ | SHPRH_HUMA | Oxidation (M) | 2535600 | 8115200 | 2064000 | 4247000 | 4732500 | 1126400 | 0 | 0 | 31683000 | 2433900 | 348540 | 2150100 | 4996800 | 1430000 | 3518900 | 0 | 1624200 | 6285800 |
| EYPPITSDQ | OCLN; OCLN | Q16625; A0/ | OCLN_HUMAN | Unmodified | 1649900 | 363340 | 1158700 | 159640 | 0 | 0 | 0 | 0 | 839070 | 0 | 0 | 0 | 0 | 0 | 2206100 | 1498400 | 0 | 0 |
| EYQEDLALI | RBCK1; RBCK1 | Q9BYM8; A6 | HOIL1_HUMAN | Unmodified | 0 | 0 | 0 | 0 | 0 | 0 | 0 | 0 | 0 | 0 | 0 | 0 | 0 | 304540 | 0 | 0 | 0 | 0 |
| EYQEIENLI | ASDURF | L0R819 | ASURF_HUMAI | Unmodified | 0 | 0 | 0 | 0 | 122080 | 0 | 0 | 0 | 107780 | 0 | 0 | 799360 | 197980 | 0 | 0 | 0 | 0 | 363430 |
| EYQISAGDI | EHD4 | Q9H223 | EHD4_HUMAN | Unmodified | 741040 | 1219300 | 918160 | 1026400 | 0 | 0 | 0 | 0 | 0 | 0 | 0 | 734350 | 0 | 0 | 944310 | 810260 | 33185 | 0 |
| EYQLNDSA | GNAT3; GNAI1; | A8MTJ3; P63 | GNAT3_HUMAI | Unmodified | 0 | 0 | 0 | 0 | 0 | 295050 | 0 | 45389 | 339880 | 0 | 0 | 6978000 | 0 | 0 | 0 | 0 | 0 | 0 |
| EYQLNDSA | GNAI2 | P04899 | GNAI2_HUMAN | Unmodified | 0 | 0 | 0 | 1065300 | 0 | 0 | 0 | 0 | 4447300 | 1024500 | 1171300 | 15724000 | 0 | 0 | 832570 | 704680 | 0 | 0 |
| EYQLNDSA | GNAO1; GNAO | P09471; H3E | GNAO_HUMAN | Unmodified | 0 | 0 | 0 | 0 | 0 | 0 | 0 | 19414 | 1006700 | 109530 | 108790 | 0 | 0 | 0 | 0 | 30548 | 0 | 0 |
| EYQLNDSA | GNAI3 | P08754 | GNAI3_HUMAN | Unmodified | 0 | 0 | 0 | 0 | 0 | 0 | 0 | 0 | 0 | 0 | 0 | 1048100 | 0 | 0 | 0 | 0 | 0 | 0 |
| EYQLSDSA | GNA11; GNA14 | P29992; O95 | GNA11_HUMAI | Unmodified | 0 | 424890 | 0 | 0 | 814940 | 0 | 0 | 0 | 1759100 | 245360 | 425280 | 5958500 | 22076 | 0 | 0 | 0 | 218510 | 570140 |
| EYQLSDSTI | GNAQ; GNAQ | P50148; B1A | GNAQ_HUMAN | Unmodified | 306250 | 389140 | 0 | 1804800 | 436630 | 0 | 0 | 0 | 1393300 | 148250 | 523340 | 5104600 | 115020 | 0 | 128130 | 201660 | 0 | 0 |
| EYQTQLIQI | DYNC1H1; DYN | Q14204; A0/ | DYHC1_HUMA | Unmodified | 0 | 0 | 0 | 0 | 0 | 0 | 0 | 249920 | 497870 | 0 | 0 | 1041900 | 0 | 0 | 0 | 0 | 0 | 0 |
| EYRDLTTA( | RPL18A; RPL18. | Q02543; M0I | RL18A_HUMAN | Unmodified | 2146800 | 0 | 187990 | 471630 | 77104 | 0 | 0 | 0 | 264420 | 0 | 0 | 7339100 | 0 | 0 | 0 | 0 | 0 | 0 |
| EYRGQTPA | EPB41L5 | Q9HCM4 | E41L5_HUMAN | Unmodified | 0 | 0 | 1341400 | 1538600 | 922160 | 321440 | 0 | 0 | 2320300 | 1242600 | 446310 | 1028800 | 753310 | 402590 | 3424000 | 2181200 | 474810 | 320700 |

| | | | | | | | | | | | | | | | | | | | | | | |
|---|---|---|---|---|---|---|---|---|---|---|---|---|---|---|---|---|---|---|---|---|---|---|
| EYRQMVET | YWHAE; YWHAI | P62258; K7E | 1433E_HUMAN | Unmodified | 102350 | 42191 | 322040 | 1228400 | 1095200 | 1060900 | 248930 | 104850 | 978510 | 1193800 | 2589500 | 1337500 | 1044500 | 1077300 | 605280 | 328220 | 839920 | 233300 |
| EYRQMVET | YWHAE; YWHAI | P62258; K7E | 1433E_HUMAN | Oxidation (M) | 0 | 0 | 0 | 0 | 174000 | 0 | 31521 | 0 | 147680 | 116460 | 364950 | 543280 | 280040 | 172790 | 291500 | 223450 | 143640 | 0 |
| EYSIGTGST | EIF2AK2; EIF2AI | P19525; A0A | E2AK2_HUMAN | Unmodified | 68278 | 72006 | 0 | 0 | 257600 | 0 | 155740 | 134910 | 0 | 208110 | 0 | 415510 | 217850 | 240580 | 0 | 0 | 0 | 275350 |
| EYSNENAV | RIPK1 | Q13546 | RIPK1_HUMAN | Unmodified | 0 | 0 | 0 | 0 | 684940 | 0 | 0 | 52028 | 152740 | 0 | 0 | 0 | 0 | 601280 | 0 | 0 | 286860 | 0 |
| EYSSELNAI | RBM25; RBM25 | P49756; E9P | RBM25_HUMAN | Unmodified | 0 | 0 | 0 | 0 | 0 | 0 | 0 | 28489 | 0 | 0 | 0 | 833040 | 0 | 0 | 0 | 0 | 0 | 0 |
| EYSVLNAVI | UTRN; UTRN | P46939; Q5T | UTRN_HUMAN | Unmodified | 163700 | 0 | 552050 | 906860 | 402660 | 0 | 0 | 0 | 913010 | 414430 | 0 | 259250 | 0 | 0 | 952770 | 1004600 | 513560 | 0 |
| EYTAGREAI | P4HB; P4HB; P4 | P07237; A0A | PDIA1_HUMAN | Unmodified | 348530 | 0 | 463620 | 752230 | 7168700 | 2479700 | 21315000 | 252150 | 1.84E+08 | 1.36E+08 | 6174100 | 1591400 | 7281800 | 10880000 | 1647200 | 672800 | 1882200 | 4429200 |
| EYTAGREAI | P4HB; P4HB; P4 | P07237; A0A | PDIA1_HUMAN | Unmodified | 0 | 0 | 0 | 0 | 0 | 284740 | 3707200 | 0 | 37806000 | 16999000 | 0 | 0 | 998630 | 1117100 | 0 | 0 | 507930 | 1072600 |
| EYTEYNEEI | STAMBP; STAMI | O95630; C9J | STABP_HUMAN | Unmodified | 0 | 346240 | 0 | 0 | 804800 | 270710 | 0 | 0 | 534520 | 142550 | 0 | 354560 | 428660 | 0 | 0 | 151200 | 0 | 0 |
| EYTGFPDP | MYOF | Q9NZM1 | MYOF_HUMAN | Unmodified | 0 | 0 | 0 | 0 | 2756900 | 0 | 0 | 0 | 0 | 0 | 0 | 0 | 0 | 0 | 0 | 0 | 0 | 0 |
| EYTINIHKR | RPL31; RPL31; I | P62899; H7C | RL31_HUMAN ( | Unmodified | 0 | 0 | 45466 | 0 | 110950 | 0 | 43215 | 17541 | 0 | 0 | 106040 | 710060 | 0 | 0 | 0 | 0 | 109840 | 35960 |
| EYTLDVYR | YARS1; YARS1; | P54577; A0A | SYYC_HUMAN | Unmodified | 0 | 0 | 373880 | 0 | 0 | 0 | 0 | 0 | 1094300 | 0 | 385380 | 5517700 | 0 | 0 | 389740 | 534060 | 0 | 0 |
| EYTLDVYRI | YARS1; YARS1; | P54577; A0A | SYYC_HUMAN | Unmodified | 0 | 0 | 0 | 0 | 3466100 | 466160 | 671650 | 63530 | 0 | 705270 | 108440 | 0 | 2620500 | 2678600 | 0 | 0 | 1522800 | 3219300 |
| EYTLDVYRI | YARS1; YARS1; | P54577; A0A | SYYC_HUMAN | Unmodified | 0 | 0 | 0 | 0 | 0 | 0 | 0 | 0 | 0 | 0 | 0 | 0 | 315770 | 398740 | 0 | 0 | 0 | 0 |
| EYTVTEPEF | NID1 | P14543 | NID1_HUMAN | Unmodified | 0 | 0 | 0 | 0 | 136740 | 15987 | 0 | 0 | 1201100 | 0 | 0 | 0 | 0 | 0 | 0 | 0 | 52275 | 0 |
| EYVAYSHT( | ERGIC1 | Q969X5 | ERGI1_HUMAN | Unmodified | 0 | 0 | 0 | 0 | 0 | 0 | 0 | 0 | 2561000 | 109920 | 0 | 0 | 0 | 0 | 151000 | 822470 | 0 | 0 |
| EYVDLERE/ | CBFB; CBFB | Q13951; A0/ | PEBB_HUMAN | Unmodified | 0 | 0 | 0 | 0 | 0 | 0 | 0 | 0 | 0 | 0 | 0 | 0 | 0 | 0 | 0 | 0 | 0 | 0 |
| EYVDPNNII | AGPS; AGPS; A( | O00116; B7Z | ADAS_HUMAN | Unmodified | 0 | 0 | 0 | 25838 | 0 | 0 | 0 | 0 | 6579800 | 239570 | 0 | 0 | 90665 | 0 | 113000 | 0 | 0 | 0 |
| EYVDPNNII | AGPS; AGPS; A( | O00116; B7Z | ADAS_HUMAN | Unmodified | 0 | 0 | 0 | 0 | 0 | 0 | 2215200 | 209810 | 3503500 | 1875400 | 0 | 0 | 0 | 0 | 0 | 0 | 0 | 123430 |
| EYVESQLQ | TXNDC5 | Q8NBS9 | TXND5_HUMAN | Unmodified | 1.71E+08 | 0 | 0 | 0 | 0 | 0 | 0 | 2.89E+08 | 2819800 | 0 | 0 | 177030 | 0 | 0 | 0 | 0 | 52151 | 0 |
| EYVNASEV | ZNF185; ZNF18 | O15231; H0 | ZN185_HUMAN | Unmodified | 0 | 0 | 0 | 0 | 1236600 | 0 | 0 | 0 | 0 | 0 | 0 | 142460 | 0 | 92487 | 0 | 0 | 2019100 | 456860 |
| EYVNASEV | ZNF185; ZNF18 | O15231; H0 | ZN185_HUMAN | Oxidation (M) | 0 | 0 | 0 | 0 | 0 | 0 | 0 | 0 | 0 | 0 | 0 | 0 | 0 | 0 | 0 | 0 | 2250100 | 861940 |
| EYVNASEV | ZNF185; ZNF18 | O15231; H0 | ZN185_HUMAN | Unmodified | 0 | 0 | 0 | 0 | 0 | 0 | 0 | 0 | 0 | 0 | 0 | 0 | 0 | 0 | 0 | 0 | 1728800 | 0 |
| EYVRQLVY | ARHGAP39 | Q9C0H5 | RHG39_HUMAN | Unmodified | 0 | 0 | 0 | 0 | 0 | 0 | 0 | 0 | 0 | 210590 | 0 | 0 | 468440 | 0 | 0 | 0 | 582750 | 0 |
| EYWMDPE( | SUB1 | P53999 | TCP4_HUMAN | Unmodified | 0 | 0 | 0 | 0 | 0 | 0 | 0 | 0 | 0 | 0 | 0 | 774140 | 0 | 0 | 0 | 0 | 0 | 0 |
| EYYRKPVEI | MRPS7; MRPS7 | Q9Y2R9; J3C | RT07_HUMAN | Unmodified | 0 | 0 | 0 | 0 | 0 | 0 | 0 | 0 | 493410 | 0 | 0 | 0 | 197320 | 0 | 0 | 32208 | 0 | 0 |
| FAAATGATI | RPSA; RPSA; RF | P08865; A0A | RSSA_HUMAN | Unmodified | 6015900 | 6501900 | 5496100 | 5291300 | 1507800 | 0 | 0 | 0 | 2159300 | 384770 | 4923500 | 55170000 | 2221000 | 362420 | 4804200 | 3399000 | 0 | 634580 |
| FAADAVKLI | TALDO1 | P37837 | TALDO_HUMAN | Unmodified | 0 | 0 | 0 | 0 | 1204100 | 0 | 0 | 0 | 0 | 0 | 0 | 0 | 0 | 289220 | 0 | 0 | 0 | 0 |
| FAAEEFK | UFM1; UFM1 | P61960; H0Y | UFM1_HUMAN | Unmodified | 0 | 0 | 608330 | 0 | 0 | 0 | 0 | 0 | 0 | 0 | 0 | 0 | 0 | 0 | 93462 | 57459 | 0 | 0 |
| FAAEEFKVF | UFM1 | P61960 | UFM1_HUMAN | Unmodified | 0 | 0 | 0 | 0 | 0 | 0 | 0 | 0 | 0 | 0 | 0 | 0 | 0 | 0 | 1298800 | 271730 | 0 | 0 |
| FAAEHTIFA | HADH; HADH; I | Q16836; A0/ | HCDH_HUMAN | Unmodified | 0 | 0 | 0 | 0 | 0 | 0 | 0 | 0 | 10820000 | 0 | 0 | 853550 | 0 | 0 | 0 | 0 | 0 | 0 |
| FAAEHTIFA | HADH; HADH; I | Q16836; A0/ | HCDH_HUMAN | Unmodified | 0 | 0 | 0 | 0 | 0 | 0 | 0 | 0 | 1454600 | 0 | 0 | 0 | 0 | 0 | 0 | 0 | 0 | 0 |
| FAAGHDAE | C20orf27; C20 | Q9GZN8; H7 | CT027_HUMAN | Unmodified | 0 | 0 | 287770 | 294050 | 1297900 | 0 | 0 | 0 | 0 | 0 | 0 | 1164000 | 0 | 0 | 558150 | 139850 | 0 | 0 |
| FAAHWLGI | MGAM2 | Q2M2H8 | MGAL_HUMAN | Unmodified | 0 | 0 | 0 | 0 | 0 | 0 | 0 | 0 | 0 | 0 | 0 | 818360 | 0 | 29348 | 0 | 0 | 0 | 0 |
| FAAKGEGQ | EEF2 | P13639 | EF2_HUMAN EI | Unmodified | 73115 | 84564 | 0 | 0 | 85514 | 0 | 52651 | 0 | 0 | 58148 | 147560 | 890120 | 0 | 0 | 0 | 0 | 0 | 0 |
| FAAKGEGQ | EEF2 | P13639 | EF2_HUMAN EI | Unmodified | 0 | 0 | 0 | 30725 | 5505900 | 2104100 | 275230 | 29799 | 0 | 1627500 | 297010 | 306670 | 29504 | 202590 | 0 | 0 | 1628700 | 930640 |
| FAALDNEEI | ABCF1; ABCF1; | Q8NE71; A0/ | ABCF1_HUMAN | Unmodified | 0 | 0 | 0 | 0 | 573080 | 0 | 0 | 0 | 624840 | 0 | 1009700 | 5824300 | 0 | 0 | 0 | 0 | 455710 | 0 |
| FAANINKES | DARS1; DARS1; | P14868; H7C | SYDC_HUMAN | Unmodified | 0 | 0 | 1528400 | 0 | 0 | 0 | 0 | 0 | 0 | 0 | 0 | 0 | 3047000 | 0 | 0 | 0 | 1580200 | 0 |
| FAANPNQN | ELAVL1 | Q15717 | ELAV1_HUMAN | Unmodified | 0 | 0 | 0 | 0 | 0 | 67490 | 0 | 0 | 337120 | 0 | 0 | 1385800 | 0 | 0 | 0 | 194690 | 131120 | 0 |
| FAAQNLHC | ILKAP; ILKAP | Q9H0C8; E9 | ILKAP_HUMAN | Unmodified | 0 | 0 | 0 | 0 | 298680 | 0 | 129720 | 97311 | 0 | 0 | 0 | 2539400 | 358140 | 0 | 0 | 0 | 0 | 0 |
| FAATTVEEII | TNKS1BP1; TNI | Q9C0C2; E9 | TB182_HUMAN | Unmodified | 0 | 0 | 340150 | 549340 | 0 | 0 | 0 | 0 | 0 | 0 | 384620 | 0 | 0 | 0 | 579670 | 617230 | 0 | 0 |
| FACFERTVI | SLC25A3; SLC2 | Q00325; F8V | MPCP_HUMAN | Unmodified | 0 | 0 | 0 | 0 | 0 | 0 | 0 | 0 | 0 | 2352800 | 0 | 0 | 0 | 0 | 0 | 0 | 0 | 0 |
| FACHSASL | NONO; NONO; | Q15233; C9I | NONO_HUMAN | Unmodified | 0 | 0 | 0 | 0 | 367750 | 232930 | 225450 | 116370 | 473840 | 352270 | 515780 | 5462700 | 690160 | 76946 | 0 | 0 | 306240 | 554380 |
| FACNGTVII | EIF1B; EIF1; EIF | O60739; P41 | EIF1B_HUMAN | Unmodified | 0 | 813940 | 0 | 1164800 | 770550 | 0 | 0 | 170960 | 3850300 | 0 | 0 | 1088200 | 6376400 | 2314100 | 1680200 | 1412700 | 2318700 | 0 |
| FADDMYSL | ATP6AP2; ATP6 | O75787; A0/ | RENR_HUMAN | Oxidation (M) | 0 | 0 | 0 | 0 | 0 | 98702 | 0 | 0 | 952070 | 694580 | 0 | 0 | 0 | 0 | 0 | 0 | 0 | 0 |
| FADDTYTES | RAB1A; RAB1B; | P62820; Q9I | RAB1A_HUMAN | Unmodified | 0 | 0 | 0 | 0 | 0 | 0 | 0 | 0 | 0 | 0 | 460670 | 0 | 0 | 0 | 3067200 | 850700 | 0 | 0 |
| FADEFNK | BPHL | Q86WA6 | BPHL_HUMAN | Unmodified | 0 | 0 | 0 | 0 | 0 | 0 | 0 | 0 | 1689400 | 31661 | 0 | 0 | 0 | 0 | 0 | 0 | 0 | 0 |
| FADEHVPG | FLNB; FLNB; FL | O75369; A0/ | FLNB_HUMAN | Unmodified | 486670 | 373400 | 2186600 | 1931400 | 10186000 | 3609600 | 201510 | 108220 | 1211700 | 395160 | 1036900 | 8898500 | 638020 | 700840 | 2690500 | 1747200 | 2243800 | 821000 |
| FADFMVK | GARS1; GARS1 | P41250; F8V | GARS_HUMAN | Oxidation (M) | 0 | 125990 | 579910 | 0 | 699020 | 0 | 213500 | 0 | 1474300 | 0 | 0 | 5990700 | 932060 | 835670 | 738060 | 336530 | 0 | 0 |
| FADIVSILDI | EPHA2 | P29317 | EPHA2_HUMAN | Unmodified | 1902700 | 977400 | 1110200 | 528280 | 7308900 | 5119300 | 0 | 0 | 0 | 0 | 978590 | 18040000 | 0 | 0 | 1178700 | 647100 | 34449000 | 15492000 |
| FADIVSILDI | EPHA2 | P29317 | EPHA2_HUMAN | Unmodified | 0 | 0 | 0 | 0 | 207950 | 25725 | 0 | 0 | 0 | 0 | 0 | 0 | 0 | 0 | 0 | 0 | 2056900 | 138320 |
| FADLSEAAI | VIM; VIM; VIM | P08670; B0Y | VIME_HUMAN \ | Unmodified | 161990 | 0 | 0 | 279830 | 1420000 | 1035800 | 234970 | 250290 | 171230 | 0 | 0 | 0 | 0 | 0 | 0 | 0 | 2538100 | 3697600 |
| FADQEVR | PIK3C2A | O00443 | P3C2A_HUMAN | Unmodified | 0 | 0 | 137110 | 42812 | 110350 | 0 | 0 | 0 | 130390 | 0 | 0 | 510280 | 0 | 0 | 184190 | 130880 | 0 | 0 |
| FADQHVP( | FLNA; FLNA | P21333; Q6( | FLNA_HUMAN | Unmodified | 649640 | 2108600 | 187020 | 821770 | 7247400 | 400820 | 1020300 | 909390 | 0 | 0 | 927460 | 274390 | 3030400 | 619320 | 1007200 | 807490 | 2429500 | 0 |
| FADQHVP( | FLNA; FLNA | P21333; Q6( | FLNA_HUMAN | Unmodified | 0 | 0 | 0 | 0 | 0 | 0 | 620750 | 0 | 0 | 0 | 0 | 0 | 0 | 0 | 0 | 0 | 925250 | 0 |
| FAEAFEAIP | CCT8 | P50990 | TCPQ_HUMAN | Unmodified | 163960 | 219350 | 439220 | 0 | 1482500 | 375560 | 0 | 91023 | 366890 | 1161900 | 5247100 | 12285000 | 3221000 | 1017800 | 451910 | 485020 | 0 | 707960 |
| FAEAFEAIP | CCT8 | P50990 | TCPQ_HUMAN | Unmodified | 0 | 0 | 0 | 0 | 0 | 0 | 0 | 95432 | 0 | 1016100 | 0 | 0 | 192250 | 1646700 | 0 | 0 | 865100 | 3182900 |
| FAEAFEAIP | CCT8 | P50990 | TCPQ_HUMAN | Unmodified | 0 | 0 | 0 | 0 | 4519300 | 0 | 5535100 | 219930 | 0 | 0 | 0 | 0 | 3322000 | 4469500 | 0 | 0 | 8288000 | 3711700 |
| FAEALITFV | SLC44A1 | Q8WWI5 | CTL1_HUMAN ( | Unmodified | 1166400 | 968890 | 226940 | 277860 | 61372 | 0 | 0 | 0 | 462370 | 281870 | 0 | 442330 | 0 | 0 | 1761900 | 639140 | 0 | 0 |
| FAEDNFNN | RAB13 | P51153 | RAB13_HUMAN | Unmodified | 9521.2 | 2876800 | 0 | 0 | 0 | 0 | 0 | 0 | 241610 | 0 | 0 | 0 | 0 | 0 | 1193300 | 0 | 0 | 0 |
| FAEFDQIMI | MYO6; MYO6; I | Q9UM54; A0 | MYO6_HUMAN | Oxidation (M) | 0 | 0 | 671370 | 0 | 0 | 0 | 0 | 0 | 0 | 0 | 0 | 414660 | 0 | 0 | 670210 | 647590 | 0 | 0 |
| FAEFQYLQI | CCAR2; CCAR2 | Q8N163; G3 | CCAR2_HUMAI | Unmodified | 0 | 0 | 0 | 0 | 0 | 0 | 0 | 0 | 0 | 0 | 0 | 0 | 0 | 0 | 0 | 0 | 0 | 0 |
| FAEHGETL/ | NAA16; NAA15; | Q6N069; Q9 | NAA16_HUMAN | Unmodified | 0 | 0 | 0 | 0 | 742520 | 0 | 226850 | 204790 | 152100 | 98029 | 328730 | 3381100 | 284720 | 0 | 103150 | 0 | 0 | 22243 |
| FAEHLSAH | XPR1 | Q9UBH6 | XPR1_HUMAN | Unmodified | 0 | 0 | 116360 | 0 | 0 | 0 | 0 | 0 | 0 | 0 | 0 | 1905000 | 0 | 0 | 0 | 0 | 0 | 0 |
| FAEKYDSD | PDIA5 | Q14554 | PDIA5_HUMAN | Unmodified | 0 | 0 | 0 | 0 | 0 | 301130 | 0 | 0 | 7214900 | 10083000 | 0 | 0 | 0 | 0 | 0 | 0 | 0 | 180750 |
| FAELAQIYA | NIT2; NIT2 | Q9NQR4; H7 | NIT2_HUMAN ( | Unmodified | 0 | 0 | 2716400 | 0 | 0 | 0 | 0 | 0 | 986540 | 0 | 369440 | 878440 | 0 | 0 | 0 | 0 | 0 | 0 |
| FAELRSPNI | CNDP2 | Q96KP4 | CNDP2_HUMA | Unmodified | 0 | 0 | 0 | 938640 | 1432400 | 204680 | 0 | 0 | 0 | 0 | 582760 | 2856100 | 8272800 | 4809600 | 830880 | 421740 | 670000 | 503850 |
| FAELVYTGI | ASS1 | P00966 | ASSY_HUMAN | Unmodified | 104120 | 485400 | 0 | 356330 | 0 | 0 | 845430 | 0 | 1976900 | 470920 | 0 | 0 | 345510 | 765130 | 2018900 | 1363700 | 0 | 0 |
| FAELVYTGI | ASS1 | P00966 | ASSY_HUMAN | Unmodified | 0 | 16983 | 0 | 0 | 0 | 0 | 4776700 | 0 | 0 | 0 | 0 | 0 | 682980 | 3831200 | 0 | 0 | 0 | 547330 |
| FAEQDAKE | FKBP5 | Q13451 | FKBP5_HUMAN | Unmodified | 0 | 0 | 0 | 0 | 0 | 0 | 0 | 0 | 261320 | 0 | 0 | 0 | 2156400 | 0 | 0 | 0 | 90768 | 149400 |
| FAEQLIRVY | SAMHD1; SAMI | Q9Y3Z3; A0/ | SAMH1_HUMA | Unmodified | 0 | 0 | 0 | 0 | 0 | 0 | 647560 | 256620 | 0 | 0 | 0 | 514230 | 0 | 0 | 194270 | 0 | 105690 | 795800 |
| FAEQTLR | PLEC | Q15149 | PLEC_HUMAN | Unmodified | 0 | 0 | 0 | 0 | 0 | 0 | 0 | 0 | 0 | 0 | 0 | 0 | 0 | 0 | 0 | 0 | 244240 | 0 |
| FAEQTLRQI | PLEC | Q15149 | PLEC_HUMAN | Unmodified | 0 | 0 | 0 | 0 | 0 | 0 | 0 | 0 | 0 | 0 | 0 | 0 | 0 | 0 | 0 | 0 | 558560 | 0 |
| FAEVTQEVI | KIF23; KIF23; KI | Q02241; A0/ | KIF23_HUMAN | Unmodified | 1355900 | 1547400 | 4060400 | 13056000 | 28160000 | 8266500 | 807020 | 280080 | 18826000 | 7110500 | 10737000 | 15035000 | 3105200 | 1174800 | 5341700 | 2200000 | 5821900 | 2319000 |
| FAEVYFAQ | MLEC; MLEC; M | Q14165; F5C | MLEC_HUMAN | Unmodified | 511020 | 123460 | 0 | 0 | 0 | 0 | 0 | 0 | 11874000 | 5079400 | 0 | 0 | 0 | 0 | 0 | 141390 | 0 | 0 |
| FAFDYATK | IDH3B; IDH3B; | O43837; A0/ | IDH3B_HUMAN | Unmodified | 0 | 0 | 0 | 0 | 0 | 0 | 66804 | 41983 | 549980 | 155360 | 0 | 0 | 0 | 0 | 0 | 0 | 0 | 0 |
| FAFQAEVN | HSP90B1; HSP | P14625; A0A | ENPL_HUMAN | Unmodified | 743820 | 0 | 1876500 | 1762800 | 0 | 0 | 1141000 | 535920 | 2.93E+08 | 92970000 | 34359000 | 39174000 | 3877100 | 2507900 | 2624000 | 1915400 | 0 | 0 |
| FAFQAEVN | HSP90B1; HSP | P14625; A0A | ENPL_HUMAN | Unmodified | 0 | 0 | 0 | 0 | 1120000 | 0 | 2558300 | 73710 | 1048900 | 21090000 | 0 | 0 | 0 | 970940 | 0 | 0 | 0 | 2215300 |
| FAFQAEVN | HSP90B1; HSP | P14625; A0A | ENPL_HUMAN | 2 Oxidation (M) | 0 | 189490 | 0 | 0 | 0 | 0 | 817100 | 1208800 | 0 | 5223100 | 0 | 0 | 0 | 0 | 0 | 0 | 0 | 529550 |
| FAFQAEVN | HSP90B1; HSP | P14625; A0A | ENPL_HUMAN | Oxidation (M) | 0 | 0 | 0 | 0 | 0 | 0 | 3087300 | 433010 | 0 | 25060000 | 446730 | 0 | 136430 | 267790 | 0 | 0 | 0 | 0 |
| FAFSPLSEE | TMEM87A | Q8NBN3 | TM87A_HUMAN | Oxidation (M) | 0 | 0 | 0 | 0 | 0 | 0 | 0 | 0 | 0 | 0 | 0 | 0 | 0 | 0 | 0 | 0 | 0 | 0 |
| FAFSTEKEF | AHNAK2 | Q8IVF2 | AHNK2_HUMAI | Unmodified | 0 | 0 | 0 | 0 | 1155800 | 236090 | 0 | 0 | 0 | 0 | 0 | 0 | 0 | 117700 | 0 | 0 | 0 | 1482000 |
| FAGDKGYL | S100A10 | P60903 | S10AA_HUMAN | Unmodified | 0 | 0 | 0 | 0 | 699340 | 0 | 0 | 0 | 0 | 0 | 0 | 0 | 0 | 0 | 0 | 0 | 197950 | 0 |
| FAGDKGYL | S100A10 | P60903 | S10AA_HUMAN | Unmodified | 86336 | 199180 | 0 | 0 | 2146500 | 888430 | 0 | 0 | 68231 | 65891 | 0 | 0 | 0 | 0 | 0 | 0 | 1487100 | 1381100 |
| FAGGDYTT | EPRS1 | P07814 | SYEP_HUMAN | Unmodified | 0 | 0 | 0 | 46809 | 404120 | 0 | 99457 | 0 | 0 | 252480 | 110640 | 308040 | 1022100 | 109680 | 197940 | 0 | 0 | 283440 |
| FAGGDYTT | EPRS1 | P07814 | SYEP_HUMAN | Unmodified | 0 | 0 | 0 | 0 | 1815300 | 358220 | 2068900 | 0 | 0 | 1074600 | 0 | 0 | 2437400 | 0 | 0 | 0 | 1473700 | 1712900 |
| FAGGLHFS | AHNAK | Q09666 | AHNK_HUMAN | Unmodified | 0 | 0 | 0 | 0 | 17266000 | 1517800 | 0 | 0 | 0 | 0 | 0 | 0 | 808290 | 66371 | 0 | 0 | 3721200 | 8603900 |
| FAGHNFRN | ENO2; ENO2 | P09104; F5H | ENOG_HUMAN | Unmodified | 0 | 0 | 0 | 0 | 0 | 0 | 161740 | 0 | 2831700 | 0 | 0 | 2021900 | 0 | 0 | 0 | 0 | 291150 | 0 |
| FAGHSEAG | PAK4 | O96013 | PAK4_HUMAN | Unmodified | 221830 | 0 | 267580 | 0 | 0 | 11619 | 0 | 0 | 73624 | 0 | 95305 | 324560 | 0 | 0 | 331080 | 467200 | 0 | 0 |
| FAGIPLTPQ | CHST3 | Q7LGC8 | CHST3_HUMAI | Unmodified | 0 | 0 | 0 | 0 | 0 | 0 | 0 | 0 | 410590 | 0 | 0 | 79097 | 0 | 0 | 0 | 0 | 0 | 0 |
| FAGLHFFN | HADH; HADH; I | Q16836; A0/ | HCDH_HUMAN | Unmodified | 0 | 0 | 0 | 0 | 0 | 0 | 0 | 0 | 2944300 | 0 | 0 | 0 | 0 | 0 | 0 | 0 | 0 | 0 |
| FAGLSLVQ | VPS37B; VPS37 | Q9H9H4; F5 | VP37B_HUMAN | Oxidation (M) | 0 | 97442 | 0 | 0 | 0 | 0 | 0 | 0 | 0 | 0 | 0 | 0 | 0 | 0 | 244690 | 0 | 0 | 0 |
| FAGQMGIC | RAB35; RAB35 | Q15286; F5I | RAB35_HUMAN | Unmodified | 95207 | 146580 | 0 | 65215 | 0 | 0 | 0 | 0 | 356350 | 0 | 0 | 152600 | 0 | 0 | 281310 | 57403 | 0 | 0 |
| FAGQMGIC | RAB35; RAB35 | Q15286; F5I | RAB35_HUMAN | Oxidation (M) | 1793800 | 1685300 | 0 | 1826900 | 1254000 | 0 | 806770 | 0 | 1142200 | 0 | 2399300 | 2147700 | 643510 | 0 | 12015000 | 7191700 | 0 | 0 |
| FAGSESAVI | ITGA3 | P26006 | ITA3_HUMAN I | 2 Oxidation (M) | 28272 | 15516 | 0 | 0 | 22425000 | 16644000 | 0 | 0 | 593690 | 0 | 339280 | 1349000 | 0 | 0 | 225320 | 31027 | 6439200 | 9484000 |
| FAGSESAVI | ITGA3 | P26006 | ITA3_HUMAN I | Oxidation (M) | 72329 | 88326 | 0 | 134980 | 31847000 | 9078700 | 0 | 0 | 457760 | 118340 | 63793 | 232790 | 148090 | 0 | 49465 | 21257 | 8262800 | 20991000 |
| FAGSESAVI | ITGA3 | P26006 | ITA3_HUMAN I | Unmodified | 0 | 0 | 0 | 0 | 6013500 | 801700 | 0 | 0 | 0 | 0 | 0 | 0 | 0 | 0 | 0 | 0 | 970170 | 4858500 |
| FAGVFHVE | CD44; CD44; C | P16070; Q8( | CD44_HUMAN | Unmodified | 1412500 | 970650 | 0 | 0 | 1972100 | 271130 | 0 | 0 | 780910 | 226480 | 0 | 227500 | 0 | 0 | 78583 | 61226 | 233100 | 1319600 |
| FAHCTVLTI | ABCC4 | O15439 | MRP4_HUMAN | Unmodified | 0 | 0 | 814850 | 543990 | 0 | 0 | 0 | 0 | 0 | 0 | 0 | 0 | 0 | 0 | 669990 | 525490 | 0 | 0 |
| FAHEHEEN | ME1 | P48163 | MAOX_HUMAN | Unmodified | 0 | 0 | 0 | 0 | 55357 | 0 | 0 | 0 | 0 | 0 | 19125 | 212250 | 0 | 32469 | 0 | 0 | 0 | 0 |
| FAHIDGDH | DHX15 | O43143 | DHX15_HUMAI | Unmodified | 0 | 0 | 0 | 0 | 0 | 0 | 0 | 0 | 476310 | 122830 | 518760 | 9729300 | 0 | 0 | 0 | 0 | 0 | 0 |
| FAHTNVES | PDIA3 | P30101 | PDIA3_HUMAN | Unmodified | 0 | 0 | 0 | 0 | 0 | 0 | 0 | 0 | 11641000 | 257010 | 2093300 | 0 | 0 | 0 | 0 | 0 | 0 | 0 |
| FAHTNVES | PDIA3 | P30101 | PDIA3_HUMAN | Unmodified | 0 | 0 | 897780 | 0 | 0 | 0 | 0 | 0 | 4.28E+08 | 25342000 | 88478000 | 6946700 | 2021600 | 0 | 4383200 | 1383200 | 0 | 0 |
| FAIGSQVSI | MCM5; MCM5 | P33992; B1A | MCM5_HUMAN | Unmodified | 74996 | 116560 | 146650 | 100040 | 0 | 0 | 0 | 0 | 563130 | 147540 | 180830 | 1400000 | 0 | 0 | 97605 | 70654 | 35803 | 0 |
| FAIHGDPK/ | TPPP3 | Q9BW30 | TPPP3_HUMAN | Unmodified | 0 | 0 | 0 | 0 | 0 | 0 | 218640 | 111940 | 0 | 0 | 0 | 0 | 0 | 0 | 0 | 0 | 0 | 0 |
| FAIQDISVE | ACTN2; ACTN4 | P35609; O43 | ACTN2_HUMAI | Unmodified | 1060900 | 139630 | 0 | 395290 | 0 | 29109 | 0 | 0 | 364800 | 418640 | 871970 | 19984000 | 0 | 0 | 2886000 | 1975400 | 292600 | 0 |
| FAIQTDPNS | CDH2; CDH2; C | P19022; C9J | CADH2_HUMA | Unmodified | 0 | 0 | 0 | 0 | 0 | 0 | 0 | 0 | 4857100 | 0 | 0 | 2049100 | 0 | 0 | 0 | 0 | 0 | 0 |
| FAKPVYPG | HSD17B4; HSD | P51659; E7E | DHB4_HUMAN | Oxidation (M) | 0 | 0 | 0 | 0 | 0 | 0 | 0 | 0 | 2031700 | 0 | 0 | 0 | 0 | 0 | 0 | 0 | 718830 | 1176400 |
| FAKPVYPG | HSD17B4; HSD | P51659; E7E | DHB4_HUMAN | Unmodified | 0 | 0 | 0 | 0 | 0 | 0 | 0 | 0 | 10230000 | 142480 | 94446 | 0 | 0 | 0 | 0 | 0 | 184800 | 0 |

| | | | | | | | | | | | | | | | | | | | | | | |
|---|---|---|---|---|---|---|---|---|---|---|---|---|---|---|---|---|---|---|---|---|---|---|
| FALFDQDK | CPNE2 | Q96FN4 | CPNE2_HUMAI | Unmodified | 166060 | 0 | 0 | 0 | 2800100 | 488120 | 812750 | 321640 | 2847600 | 854950 | 0 | 0 | 0 | 253850 | 0 | 254960 | 0 | 0 |
| FALGIFAINI | IQGAP1; IQGAP | P46940; A0A | IQGA1_HUMAN | Unmodified | 0 | 0 | 21622 | 0 | 4725100 | 1341800 | 208640 | 0 | 147710 | 932650 | 0 | 0 | 301910 | 170270 | 183580 | 0 | 706270 | 0 |
| FALHSGYS | LRP11; LRP11 | Q86VZ4; Q5 | LRP11_HUMAN | Unmodified | 0 | 0 | 0 | 0 | 0 | 0 | 0 | 0 | 4474700 | 305080 | 0 | 0 | 0 | 0 | 0 | 0 | 0 | 0 |
| FALHSGYS | LRP11; LRP11 | Q86VZ4; Q5 | LRP11_HUMAN | Unmodified | 0 | 0 | 0 | 0 | 0 | 0 | 0 | 0 | 1695200 | 1363900 | 0 | 0 | 0 | 0 | 0 | 0 | 112070 | 0 |
| FALITWIGE | COTL1; COTL1 | Q14019; H3I | COTL1_HUMAN | Unmodified | 0 | 113460 | 0 | 436040 | 5938800 | 896610 | 585980 | 0 | 529010 | 7664200 | 0 | 22038 | 3239800 | 1212600 | 625660 | 0 | 3672400 | 1902000 |
| FALITWIGE | COTL1; COTL1 | Q14019; H3I | COTL1_HUMAN | Unmodified | 0 | 15766 | 0 | 0 | 17075000 | 1497900 | 5459900 | 0 | 0 | 8080000 | 0 | 0 | 2655300 | 7039400 | 0 | 0 | 19810000 | 7579900 |
| FALITWIGE | COTL1; COTL1 | Q14019; H3I | COTL1_HUMAN | Unmodified | 0 | 0 | 0 | 0 | 1206500 | 185750 | 2707500 | 0 | 0 | 472210 | 0 | 0 | 0 | 0 | 493710 | 175060 | 8131700 | 857550 |
| FALPYIRDV | UROD; UROD; l | P06132; A0A | DCUP_HUMAN | Unmodified | 0 | 0 | 0 | 0 | 0 | 0 | 82693 | 0 | 0 | 0 | 0 | 710970 | 0 | 0 | 0 | 0 | 386990 | 0 |
| FALTVVR | TIGAR | Q9NQ88 | TIGAR_HUMAN | Unmodified | 0 | 0 | 0 | 75689 | 100680 | 85493 | 0 | 9180.2 | 0 | 0 | 240260 | 1138200 | 125620 | 0 | 110360 | 70803 | 0 | 0 |
| FAMEPEEFI | PDIA4; PDIA4 | P13667; A0A | PDIA4_HUMAN | Unmodified | 0 | 0 | 0 | 0 | 0 | 0 | 0 | 0 | 2019800 | 0 | 0 | 0 | 0 | 0 | 0 | 0 | 0 | 0 |
| FAMEPEEFI | PDIA4; PDIA4 | P13667; A0A | PDIA4_HUMAN | Oxidation (M) | 0 | 0 | 0 | 0 | 0 | 0 | 0 | 0 | 2020600 | 477370 | 550450 | 0 | 0 | 0 | 0 | 0 | 0 | 0 |
| FAMEPEEFI | PDIA4; PDIA4 | P13667; A0A | PDIA4_HUMAN | Oxidation (M) | 0 | 0 | 0 | 0 | 193740 | 0 | 515070 | 0 | 15557000 | 6903800 | 6878100 | 0 | 0 | 440090 | 435340 | 0 | 0 | 133070 |
| FAMEPEEFI | PDIA4; PDIA4 | P13667; A0A | PDIA4_HUMAN | Unmodified | 0 | 0 | 0 | 0 | 0 | 0 | 0 | 0 | 10608000 | 1538000 | 388760 | 0 | 0 | 0 | 0 | 0 | 0 | 0 |
| FANDENVK | ABHD12; ABHD | Q8N2K0; A0, | ABD12_HUMAI | Unmodified | 0 | 0 | 0 | 0 | 926640 | 251090 | 0 | 173730 | 12663000 | 0 | 0 | 221650 | 0 | 0 | 0 | 0 | 904840 | 0 |
| FANIEEDTF | EFR3B; EFR3A; | Q9Y2G0; Q1 | EFR3B_HUMAN | Unmodified | 0 | 0 | 0 | 0 | 0 | 0 | 0 | 0 | 1239500 | 347080 | 228450 | 0 | 0 | 0 | 900410 | 693280 | 0 | 0 |
| FANPFPAA | PCCB; PCCB; P | P05166; C9J | PCCB_HUMAN | Unmodified | 0 | 0 | 0 | 0 | 0 | 0 | 0 | 0 | 1241100 | 0 | 0 | 0 | 659280 | 0 | 0 | 0 | 0 | 0 |
| FANSLVGV | SPTBN1; SPTBN | Q01082; A0/ | SPTB2_HUMAN | Unmodified | 0 | 0 | 0 | 0 | 0 | 0 | 0 | 0 | 0 | 0 | 0 | 1092100 | 0 | 0 | 0 | 559070 | 0 | 0 |
| FAPPLVIKE | OAT | P04181 | OAT_HUMAN C | Unmodified | 0 | 0 | 0 | 0 | 0 | 0 | 0 | 0 | 2308400 | 0 | 0 | 0 | 0 | 0 | 0 | 0 | 0 | 0 |
| FAPPLVIKE | OAT | P04181 | OAT_HUMAN C | Unmodified | 0 | 0 | 0 | 0 | 0 | 0 | 0 | 0 | 11883000 | 3234300 | 253580 | 0 | 0 | 0 | 0 | 0 | 0 | 228690 |
| FAPREGFIN | SPINT1; SPINT1 | O43278; H3I | SPIT1_HUMAN | Unmodified | 0 | 0 | 0 | 0 | 312250 | 0 | 0 | 0 | 293800 | 669450 | 0 | 0 | 955050 | 756730 | 0 | 0 | 0 | 0 |
| FAPSPTGFI | EARS2; EARS2 | Q5JPH6; H3I | SYEM_HUMAN | Unmodified | 114950 | 171510 | 0 | 0 | 0 | 0 | 0 | 0 | 647460 | 0 | 0 | 0 | 0 | 0 | 151970 | 95994 | 0 | 0 |
| FAQALGLTI | GTF2I; GTF2I | P78347; A0A | GTF2I_HUMAN | Unmodified | 0 | 0 | 0 | 0 | 0 | 346560 | 0 | 0 | 0 | 0 | 0 | 302750 | 0 | 0 | 0 | 0 | 0 | 0 |
| FAQDVSQD | SNF8 | Q96H20 | SNF8_HUMAN | Unmodified | 2905900 | 3327900 | 3008900 | 0 | 0 | 0 | 0 | 0 | 0 | 0 | 0 | 0 | 489740 | 0 | 4971600 | 4088400 | 0 | 0 |
| FAQDVSQD | SNF8 | Q96H20 | SNF8_HUMAN | Unmodified | 0 | 0 | 0 | 39206 | 399900 | 0 | 0 | 0 | 0 | 94399 | 0 | 0 | 596980 | 239950 | 0 | 0 | 0 | 154290 |
| FAQHGTFE | SFPQ; SFPQ | P23246; H0Y | SFPQ_HUMAN | Unmodified | 298300 | 0 | 0 | 0 | 0 | 0 | 0 | 0 | 0 | 0 | 0 | 5485500 | 0 | 0 | 0 | 391480 | 204660 | 0 |
| FAQIDEKTE | INCENP | Q9NQS7 | INCE_HUMAN I | Unmodified | 0 | 0 | 0 | 1568400 | 1653100 | 722160 | 33028 | 0 | 0 | 0 | 0 | 96710 | 1190500 | 668830 | 117900 | 107620 | 413390 | 401550 |
| FAQLNLAAI | CNTN2; CNTN2 | Q02246; A0/ | CNTN2_HUMA | Unmodified | 62609 | 0 | 0 | 5813200 | 0 | 0 | 4111100 | 0 | 0 | 0 | 0 | 0 | 0 | 0 | 0 | 0 | 0 | 0 |
| FAQMLEK | IDH2 | P48735 | IDHP_HUMAN | Oxidation (M) | 0 | 0 | 0 | 0 | 0 | 0 | 108920 | 26977 | 2654300 | 180240 | 0 | 0 | 0 | 0 | 94666 | 0 | 0 | 0 |
| FAQMLEK | IDH2 | P48735 | IDHP_HUMAN | Unmodified | 0 | 0 | 0 | 0 | 0 | 17980 | 87785 | 0 | 19503000 | 1752900 | 0 | 0 | 0 | 0 | 0 | 0 | 0 | 0 |
| FAQPGSFE | NONO; NONO; | Q15233; A0/ | NONO_HUMAN | Unmodified | 0 | 0 | 0 | 0 | 0 | 0 | 0 | 0 | 244970 | 0 | 73672 | 466100 | 0 | 0 | 0 | 0 | 0 | 0 |
| FAQPGSFE | NONO; NONO; | Q15233; A0/ | NONO_HUMAN | Oxidation (M) | 22783 | 97559 | 95473 | 0 | 0 | 0 | 0 | 48802 | 0 | 0 | 55546 | 925960 | 0 | 0 | 0 | 0 | 0 | 0 |
| FAQPGTFEI | PSPC1 | Q8WXF1 | PSPC1_HUMAN | Unmodified | 0 | 0 | 0 | 0 | 0 | 0 | 0 | 0 | 0 | 0 | 0 | 354270 | 0 | 0 | 0 | 0 | 0 | 0 |
| FAQQITGM | RAB12; RAB12 | Q6IQ22; A0A | RAB12_HUMAN | Unmodified | 0 | 0 | 130390 | 0 | 304370 | 0 | 0 | 0 | 245170 | 0 | 0 | 1246700 | 0 | 0 | 0 | 0 | 0 | 0 |
| FAQVSSSN | C1orf198; C1or | Q9H425; E5I | CA198_HUMAN | Unmodified | 0 | 0 | 0 | 0 | 2258300 | 108090 | 1260700 | 916400 | 1353900 | 1164600 | 3911300 | 5638000 | 979480 | 0 | 0 | 0 | 135640 | 645960 |
| FARHVEPG | HAGH; HAGH; I | Q16775; H3I | GLO2_HUMAN | Unmodified | 0 | 0 | 0 | 0 | 1051200 | 0 | 0 | 600010 | 0 | 524540 | 0 | 0 | 0 | 1170600 | 0 | 0 | 2374600 | 1944600 |
| FARLDIDPE | MTHFD1; MTHF | P11586; V9G | C1TC_HUMAN | Unmodified | 87367 | 63077 | 114100 | 0 | 78694 | 0 | 0 | 0 | 1528200 | 0 | 217580 | 2926500 | 0 | 0 | 0 | 185600 | 128870 | 0 |
| FASEFFPH | STEAP2; STEAP | Q8NFT2; C9. | STEA2_HUMAN | Unmodified | 2598700 | 3866900 | 29522000 | 8947300 | 0 | 0 | 0 | 0 | 0 | 0 | 0 | 155580 | 0 | 1375700 | 62768000 | 33972000 | 0 | 0 |
| FASEIAGVD | NDUFS1; NDUF | P28331; B4D | NDUS1_HUMA | Unmodified | 0 | 0 | 0 | 0 | 0 | 0 | 0 | 0 | 1145100 | 0 | 0 | 401600 | 0 | 0 | 0 | 0 | 0 | 0 |
| FASENDLPI | RANBP1; RANB | P43487; C9J | RANG_HUMAN | Unmodified | 918620 | 539680 | 1108800 | 1420700 | 1757600 | 673660 | 0 | 0 | 2496000 | 0 | 4484600 | 8229300 | 524210 | 0 | 2629500 | 1847200 | 0 | 0 |
| FASENDLPI | RANBP1; RANB | P43487; C9J | RANG_HUMAN | Unmodified | 84877 | 179600 | 0 | 264740 | 461880 | 0 | 0 | 0 | 390400 | 133530 | 601630 | 1838400 | 255570 | 0 | 0 | 0 | 0 | 0 |
| FASENDLPI | RANBP1; RANB | P43487; C9J | RANG_HUMAN | Unmodified | 0 | 0 | 99771 | 0 | 5519100 | 1376500 | 0 | 288800 | 0 | 421100 | 0 | 0 | 3009600 | 672350 | 0 | 0 | 1382600 | 908100 |
| FASLEFSPC | EMC1; EMC1 | Q8N766; Q5 | EMC1_HUMAN | Unmodified | 93991 | 0 | 0 | 0 | 0 | 0 | 191880 | 32823 | 12857000 | 2416600 | 2045400 | 563320 | 0 | 0 | 0 | 0 | 0 | 0 |
| FASLEFSPC | EMC1; EMC1 | Q8N766; Q5 | EMC1_HUMAN | Unmodified | 0 | 0 | 0 | 0 | 0 | 0 | 0 | 0 | 4062400 | 632640 | 223810 | 0 | 0 | 0 | 0 | 0 | 0 | 0 |
| FASLQPVSI | CYLD; CYLD | Q9NQC7; J3 | CYLD_HUMAN | Unmodified | 156480 | 207450 | 0 | 280010 | 0 | 0 | 313680 | 111530 | 0 | 0 | 0 | 0 | 432880 | 0 | 0 | 51690 | 0 | 0 |
| FASRIIDSLI | UGDH | O60701 | UGDH_HUMAN | Unmodified | 0 | 0 | 0 | 0 | 5526800 | 1006000 | 498850 | 0 | 0 | 566150 | 0 | 139440 | 73508 | 118090 | 0 | 0 | 2250400 | 1604100 |
| FASTVFPSD | ECPAS; ECPAS; | Q5VYK3; R4 | ECM29_HUMAI | Unmodified | 89865 | 141580 | 114100 | 61898 | 0 | 0 | 0 | 0 | 0 | 0 | 77285 | 2781400 | 0 | 0 | 0 | 170340 | 0 | 0 |
| FASVFGTM | EXT2 | Q93063 | EXT2_HUMAN I | Oxidation (M) | 0 | 0 | 0 | 0 | 273970 | 0 | 0 | 0 | 1019600 | 259780 | 0 | 0 | 109520 | 0 | 0 | 0 | 0 | 0 |
| FASVFGTM | EXT2 | Q93063 | EXT2_HUMAN I | Unmodified | 0 | 0 | 0 | 0 | 0 | 0 | 0 | 0 | 2405700 | 437240 | 0 | 0 | 0 | 0 | 0 | 0 | 0 | 0 |
| FASWALES | TRIM28 | Q13263 | TIF1B_HUMAN | Unmodified | 0 | 0 | 32764 | 49345 | 0 | 0 | 1138400 | 23567 | 293260 | 1110300 | 129200 | 468970 | 0 | 27796 | 0 | 0 | 145340 | 277890 |
| FASWALES | TRIM28 | Q13263 | TIF1B_HUMAN | Unmodified | 0 | 0 | 0 | 0 | 0 | 0 | 0 | 0 | 0 | 0 | 0 | 741120 | 0 | 0 | 0 | 0 | 0 | 0 |
| FASYLTFSP | ALDH18A1 | P54886 | P5CS_HUMAN | Unmodified | 0 | 0 | 0 | 0 | 0 | 0 | 0 | 0 | 1423500 | 0 | 0 | 0 | 0 | 0 | 0 | 0 | 0 | 25982 |
| FATCNPSTI | ALDH1A3; ALDI | P47895; H0 | AL1A3_HUMAN | Unmodified | 0 | 0 | 0 | 0 | 0 | 0 | 0 | 0 | 0 | 0 | 0 | 0 | 0 | 0 | 0 | 0 | 782460 | 419250 |
| FATCSDDG | WDR33; WDR3 | Q9C0J8; C9J | WDR33_HUMA | Unmodified | 0 | 0 | 0 | 0 | 0 | 0 | 0 | 0 | 91296 | 0 | 0 | 422250 | 0 | 0 | 0 | 0 | 0 | 0 |
| FATEAAITIL | TCP1; TCP1; TC | P17987; F5H | TCPA_HUMAN | Unmodified | 0 | 0 | 0 | 0 | 0 | 0 | 0 | 0 | 826330 | 0 | 1201900 | 3252900 | 0 | 0 | 0 | 0 | 0 | 0 |
| FATEAAITIL | TCP1; TCP1; TC | P17987; F5H | TCPA_HUMAN | Unmodified | 0 | 0 | 0 | 0 | 738470 | 0 | 608810 | 0 | 0 | 558370 | 0 | 0 | 2270700 | 1076900 | 0 | 0 | 0 | 2566400 |
| FATFVFNVI | NCS1 | P62166 | NCS1_HUMAN | Unmodified | 0 | 89124 | 0 | 30984 | 0 | 0 | 0 | 0 | 0 | 0 | 0 | 0 | 0 | 0 | 249590 | 250270 | 0 | 0 |
| FATGDIIQII | MPP1; MPP1; M | Q00013; C9J | EM55_HUMAN | Unmodified | 0 | 0 | 0 | 0 | 6612300 | 1174700 | 0 | 0 | 0 | 0 | 0 | 0 | 0 | 0 | 0 | 0 | 0 | 8315900 |
| FATHAAAL | SFPQ | P23246 | SFPQ_HUMAN | Unmodified | 0 | 0 | 0 | 0 | 0 | 0 | 0 | 0 | 205810 | 0 | 246930 | 2326700 | 0 | 0 | 0 | 0 | 0 | 0 |
| FATLAELVC | PTPN11; PTPN1 | Q06124; A0/ | PTN11_HUMAN | Unmodified | 0 | 0 | 38327 | 0 | 0 | 0 | 0 | 0 | 439050 | 57749 | 0 | 0 | 0 | 0 | 0 | 48346 | 0 | 0 |
| FATLAELVC | PTPN11; PTPN1 | Q06124; A0/ | PTN11_HUMAN | Oxidation (M) | 0 | 340310 | 0 | 0 | 0 | 0 | 0 | 0 | 0 | 151700 | 0 | 694720 | 0 | 0 | 1226500 | 0 | 0 | 0 |
| FATLQNEL | XPR1 | Q9UBH6 | XPR1_HUMAN | Unmodified | 0 | 0 | 252260 | 254340 | 0 | 0 | 0 | 0 | 0 | 0 | 522340 | 0 | 0 | 0 | 0 | 0 | 0 | 0 |
| FATLYSTR | ENDOD1 | O94919 | ENDD1_HUMA | Unmodified | 0 | 0 | 572390 | 408000 | 0 | 41321 | 0 | 0 | 5998300 | 576430 | 399840 | 1197300 | 0 | 0 | 431420 | 0 | 0 | 0 |
| FATNEAIQF | SEC16A; SEC1 | O15027; F1T | SC16A_HUMAN | Unmodified | 0 | 0 | 4020000 | 1078600 | 7660900 | 4123600 | 0 | 547480 | 0 | 57644 | 64518 | 1691500 | 1612400 | 17214000 | 5192000 | 424330 | 910940 | 2428700 |
| FATNEGFR | ABCA12 | Q86UK0 | ABCAC_HUMAI | Unmodified | 0 | 0 | 0 | 0 | 0 | 0 | 0 | 0 | 0 | 228290 | 0 | 0 | 0 | 0 | 0 | 0 | 0 | 0 |
| FATVEVTDI | HEBP1; HEBP1; | Q9NRV9; F5 | HEBP1_HUMAI | Oxidation (M) | 919960 | 0 | 0 | 0 | 564320 | 0 | 252820 | 42286 | 524390 | 0 | 180400 | 650380 | 0 | 0 | 0 | 0 | 0 | 0 |
| FAVALDSE | SUPT5H | O00267 | SPT5H_HUMAN | Unmodified | 42810 | 0 | 0 | 0 | 0 | 0 | 0 | 0 | 0 | 0 | 662310 | 868770 | 1116400 | 0 | 0 | 20681 | 0 | 0 |
| FAVAWYSL | IGSF3 | O75054 | IGSF3_HUMAN | Unmodified | 0 | 0 | 0 | 0 | 0 | 0 | 0 | 0 | 0 | 0 | 0 | 0 | 0 | 0 | 0 | 0 | 0 | 0 |
| FAVEEIIQK | LXN | Q9BS40 | LXN_HUMAN L | Unmodified | 0 | 0 | 0 | 0 | 0 | 0 | 0 | 0 | 0 | 0 | 0 | 0 | 0 | 0 | 0 | 0 | 595580 | 0 |
| FAVFLSEVS | LRRC8A; LRRC | Q8IWT6; Q8 | LRC8A_HUMAN | Unmodified | 0 | 0 | 0 | 0 | 0 | 0 | 0 | 0 | 0 | 0 | 0 | 681070 | 0 | 0 | 0 | 0 | 0 | 0 |
| FAVGIVIGR | FUBP1; FUBP3; | Q96AE4; Q9 | FUBP1_HUMAN | Unmodified | 0 | 0 | 0 | 0 | 0 | 0 | 0 | 0 | 0 | 0 | 0 | 621090 | 142950 | 0 | 0 | 0 | 0 | 0 |
| FAVGTAGL | B3GALT6 | Q96L58 | B3GT6_HUMAN | Unmodified | 0 | 0 | 0 | 0 | 0 | 0 | 0 | 0 | 3730200 | 0 | 0 | 0 | 0 | 0 | 0 | 0 | 0 | 0 |
| FAVGTAGL | B3GALT6 | Q96L58 | B3GT6_HUMAN | Unmodified | 0 | 0 | 0 | 0 | 0 | 0 | 0 | 10575 | 2404500 | 85133 | 0 | 0 | 0 | 0 | 0 | 0 | 0 | 0 |
| FAVLDFCYI | CYFIP1; CYFIP1 | Q7L576; H0 | CYFP1_HUMAN | Unmodified | 0 | 0 | 163720 | 17456 | 0 | 0 | 136470 | 0 | 0 | 0 | 0 | 0 | 0 | 0 | 85384 | 50062 | 0 | 16534 |
| FAVPTYAA | ADAM9; ADAM | Q13443; A0/ | ADAM9_HUMA | Unmodified | 4331700 | 3971200 | 5368700 | 7665400 | 984360 | 419550 | 0 | 0 | 609770 | 0 | 62719 | 332070 | 499020 | 65515 | 7776300 | 7876800 | 0 | 86288 |
| FAVPTYAA | ADAM9; ADAM | Q13443; A0/ | ADAM9_HUMA | Unmodified | 0 | 0 | 0 | 1893500 | 4312000 | 1333800 | 0 | 0 | 0 | 0 | 0 | 0 | 2596400 | 0 | 0 | 0 | 0 | 0 |
| FAVRDMRC | EEF1A2; EEF1A | Q05639; Q5 | EF1A2_HUMAN | Unmodified | 0 | 0 | 0 | 0 | 48713000 | 5476300 | 8274000 | 725390 | 0 | 8802300 | 0 | 0 | 30812000 | 24146000 | 0 | 0 | 22626000 | 36335000 |
| FAVRDMRC | EEF1A2; EEF1A | Q05639; Q5 | EF1A2_HUMAN | Oxidation (M) | 0 | 0 | 0 | 0 | 0 | 2146700 | 3398100 | 572000 | 0 | 2136600 | 0 | 0 | 9074200 | 7604600 | 0 | 0 | 0 | 5835800 |
| FAVYLPPK | ESD; ESD; ESD | P10768; X6R | ESTD_HUMAN | Unmodified | 0 | 0 | 641110 | 0 | 0 | 0 | 0 | 0 | 0 | 0 | 0 | 1833200 | 0 | 0 | 672580 | 0 | 0 | 0 |
| FAWGHDE | MAN1B1; MAN1 | Q9UKM7; H | MA1B1_HUMAI | Unmodified | 0 | 0 | 0 | 0 | 0 | 0 | 0 | 0 | 1054400 | 0 | 0 | 0 | 0 | 0 | 0 | 0 | 0 | 0 |
| FAYGYIEDL | SERPINB1 | P30740 | ILEU_HUMAN L | Unmodified | 0 | 88478 | 0 | 0 | 648700 | 0 | 0 | 0 | 1261800 | 455270 | 0 | 0 | 0 | 0 | 0 | 0 | 0 | 0 |
| FAYLGRLA | RPL13A; RPL13 | P40429; A0A | RL13A_HUMAN | Unmodified | 0 | 0 | 0 | 0 | 835270 | 192260 | 727250 | 0 | 0 | 0 | 0 | 0 | 464130 | 0 | 0 | 0 | 0 | 0 |
| FAYLGRLA | RPL13A; RPL13 | P40429; A0A | RL13A_HUMAN | Unmodified | 0 | 0 | 0 | 0 | 755450 | 200230 | 2367200 | 0 | 0 | 0 | 0 | 0 | 559990 | 866350 | 0 | 0 | 0 | 786690 |
| FAYPMPFC | CAPN7 | Q9Y6W3 | CAN7_HUMAN | Oxidation (M) | 0 | 0 | 0 | 0 | 0 | 0 | 0 | 0 | 108640 | 0 | 0 | 389930 | 0 | 0 | 0 | 0 | 0 | 0 |
| FCADHPFL | SERPINB6; SER | P35237; A0A | SPB6_HUMAN | Unmodified | 0 | 132920 | 0 | 0 | 0 | 0 | 0 | 0 | 0 | 0 | 0 | 2521300 | 0 | 0 | 304980 | 311360 | 0 | 0 |
| FCAFVQEA | ARHGEF1; ARH | Q92888; M0 | ARHG1_HUMA | Unmodified | 55844 | 0 | 0 | 0 | 1121400 | 364740 | 0 | 0 | 0 | 0 | 0 | 823670 | 0 | 0 | 0 | 0 | 503160 | 512270 |
| FCDNSSAI | PFAS | O15067 | PUR4_HUMAN | Unmodified | 0 | 0 | 52329 | 0 | 555380 | 186350 | 88050 | 31096 | 0 | 0 | 0 | 1392900 | 0 | 0 | 61710 | 29572 | 223100 | 259120 |
| FCDYGKAP | CNP | P09543 | CN37_HUMAN | Unmodified | 0 | 0 | 0 | 293500 | 1872500 | 917470 | 337030 | 245390 | 0 | 434690 | 1654600 | 2651700 | 1968000 | 1592300 | 107490 | 82006 | 2112400 | 4009300 |
| FCDYGKAP | CNP | P09543 | CN37_HUMAN | Unmodified | 0 | 892760 | 0 | 2154500 | 6762000 | 1831000 | 1062200 | 946770 | 1379500 | 2306100 | 3617100 | 4039600 | 3458800 | 4480700 | 0 | 0 | 5080700 | 6047100 |
| FCECDNFN | ITGB1; ITGB1; I | P05556; H7 | ITB1_HUMAN I | Unmodified | 282820 | 85196 | 0 | 0 | 0 | 0 | 0 | 0 | 731980 | 0 | 0 | 598260 | 0 | 0 | 0 | 0 | 0 | 0 |
| FCECDNFN | ITGB1; ITGB1; I | P05556; H7 | ITB1_HUMAN I | Unmodified | 578310 | 0 | 0 | 207980 | 2046900 | 0 | 0 | 0 | 0 | 0 | 0 | 0 | 456140 | 0 | 0 | 0 | 4232900 | 8246500 |
| FCEMLIK | MITD1; MITD1 | Q8WV92; H7 | MITD1_HUMAN | Oxidation (M) | 112070 | 0 | 0 | 0 | 0 | 0 | 0 | 0 | 0 | 0 | 0 | 1769700 | 0 | 0 | 0 | 75334 | 0 | 0 |
| FCEMLIK | MITD1; MITD1 | Q8WV92; H7 | MITD1_HUMAN | Unmodified | 0 | 0 | 0 | 0 | 0 | 0 | 0 | 0 | 0 | 0 | 0 | 974910 | 0 | 0 | 0 | 0 | 0 | 0 |
| FCENTQAG | GLG1; GLG1 | Q92896; H3I | GSLG1_HUMAI | Unmodified | 0 | 0 | 0 | 0 | 0 | 0 | 210910 | 0 | 6112100 | 0 | 0 | 1631300 | 0 | 0 | 0 | 0 | 0 | 0 |
| FCENTQAG | GLG1; GLG1 | Q92896; H3I | GSLG1_HUMAI | Unmodified | 0 | 0 | 0 | 0 | 0 | 0 | 17222 | 0 | 3388000 | 382490 | 721850 | 0 | 0 | 0 | 0 | 0 | 0 | 0 |
| FCETTIGSK | GPD1L; GPD1L | Q8N335; C9 | GPD1L_HUMAI | Unmodified | 31789 | 0 | 92692 | 0 | 0 | 0 | 197830 | 0 | 212050 | 0 | 0 | 412180 | 319770 | 1103500 | 35345 | 25511 | 61672 | 359810 |
| FCFLVEK | BAIAP2; BAIAP2 | Q9UQB8; I3I | BAIP2_HUMAN | Unmodified | 340860 | 0 | 0 | 0 | 3872600 | 999080 | 211260 | 107570 | 732900 | 337260 | 0 | 1239800 | 0 | 1999700 | 0 | 0 | 1043800 | 861650 |
| FCFSNEFS | TMED7; TMED7 | Q9Y3B3; A0/ | TMED7_HUMAI | Unmodified | 0 | 0 | 0 | 1423900 | 0 | 0 | 459820 | 39854 | 15785000 | 5704000 | 1688500 | 0 | 0 | 0 | 1581300 | 593250 | 0 | 0 |
| FCFSNRMS | TMED2; TMED2 | Q15363; E7E | TMED2_HUMAI | Oxidation (M) | 0 | 0 | 0 | 0 | 0 | 0 | 0 | 0 | 0 | 245560 | 0 | 0 | 0 | 0 | 0 | 0 | 0 | 0 |
| FCFTPHTEI | FASN; FASN | P49327; A0A | FAS_HUMAN F | Unmodified | 241100 | 0 | 348160 | 0 | 0 | 0 | 0 | 0 | 243120 | 0 | 0 | 18024000 | 328680 | 0 | 818760 | 456810 | 0 | 0 |
| FCGDKIPEI | TLL2 | Q9Y6L7 | TLL2_HUMAN T | Unmodified | 0 | 0 | 0 | 0 | 0 | 0 | 0 | 0 | 731680 | 0 | 0 | 0 | 0 | 0 | 0 | 0 | 0 | 0 |
| FCGKIAPPI | NRP1; NRP1; N | O14786; Q5. | NRP1_HUMAN | Unmodified | 289850 | 0 | 0 | 0 | 22283 | 0 | 0 | 0 | 0 | 0 | 0 | 0 | 0 | 0 | 0 | 0 | 230220 | 75502 |
| FCHGILTK | GLG1 | Q92896 | GSLG1_HUMAI | Unmodified | 0 | 0 | 0 | 0 | 0 | 0 | 0 | 118610 | 1927700 | 646670 | 0 | 0 | 0 | 0 | 0 | 0 | 0 | 0 |
| FCIWTESAI | RPL4; RPL4 | P36578; H3E | RL4_HUMAN 6 | Unmodified | 3517300 | 3018700 | 0 | 376710 | 0 | 0 | 0 | 0 | 0 | 0 | 708010 | 18876000 | 0 | 0 | 947020 | 792360 | 0 | 0 |
| FCIWTESAI | RPL4; RPL4 | P36578; H3E | RL4_HUMAN 6 | Unmodified | 123780 | 0 | 106190 | 0 | 0 | 0 | 0 | 0 | 0 | 0 | 0 | 1331600 | 0 | 0 | 0 | 0 | 0 | 0 |
| FCLDNGAK | PGK1 | P00558 | PGK1_HUMAN | Unmodified | 235800 | 91522 | 146390 | 68374 | 5139400 | 990370 | 924630 | 546900 | 2208000 | 1162300 | 1750200 | 11456000 | 536710 | 3239200 | 1071000 | 1850900 | 15968000 | 22037000 |
| FCNIMGSS | TMOD3; TMOD | Q9NYL9; H0 | TMOD3_HUMA | Unmodified | 0 | 0 | 0 | 0 | 863520 | 180940 | 0 | 0 | 0 | 0 | 0 | 0 | 641210 | 0 | 0 | 0 | 0 | 0 |
| FCNPGFPI | TM9SF2 | Q99805 | TM9S2_HUMAN | Unmodified | 47681 | 0 | 0 | 0 | 0 | 0 | 0 | 0 | 4466900 | 0 | 0 | 1426800 | 0 | 0 | 0 | 0 | 0 | 0 |
| FCNPGFPI | TM9SF2 | Q99805 | TM9S2_HUMAN | Unmodified | 0 | 0 | 0 | 0 | 177620 | 0 | 0 | 0 | 2020400 | 0 | 0 | 0 | 0 | 0 | 0 | 0 | 0 | 0 |
| FCSEAQFE | LRP1 | Q07954 | LRP1_HUMAN I | Unmodified | 0 | 0 | 0 | 0 | 0 | 0 | 0 | 0 | 391090 | 0 | 0 | 0 | 0 | 0 | 0 | 0 | 0 | 0 |

| | | | | | | | | | | | | | | | | | | | | | | |
|---|---|---|---|---|---|---|---|---|---|---|---|---|---|---|---|---|---|---|---|---|---|---|
| FCTELNQP | EEPD1 | Q7L9B9 | EEPD1_HUMAN | Unmodified | 0 | 0 | 0 | 31555 | 0 | 0 | 0 | 124420 | 0 | 0 | 0 | 0 | 154020 | 1005900 | 144390 | 73572 | 1160300 | 5248900 |
| FCTGLTQIE | CKB | P12277 | KCRB_HUMAN | Unmodified | 519020 | 147660 | 808850 | 1927800 | 434210 | 152030 | 685020 | 0 | 2834900 | 447500 | 701760 | 6908800 | 1158600 | 222350 | 2960700 | 1759400 | 45562 | 0 |
| FCTREGNC | ITGA6 | P23229 | ITA6_HUMAN I | Unmodified | 0 | 0 | 3091700 | 0 | 712400 | 186320 | 27315 | 0 | 0 | 28181 | 0 | 0 | 286130 | 150250 | 0 | 0 | 617010 | 2474100 |
| FCTREGNC | ITGA6 | P23229 | ITA6_HUMAN I | Unmodified | 0 | 0 | 0 | 0 | 17807000 | 746880 | 0 | 0 | 0 | 0 | 0 | 0 | 9440900 | 8178300 | 0 | 0 | 37448000 | 90813000 |
| FCVSDTEN | ECE1; ECE1; EC | P42892; B4D | ECE1_HUMAN | Oxidation (M) | 0 | 0 | 0 | 105910 | 3307800 | 0 | 0 | 0 | 2251700 | 1054700 | 0 | 0 | 0 | 0 | 110420 | 0 | 0 | 0 |
| FCVSDTEN | ECE1; ECE1; EC | P42892; B4D | ECE1_HUMAN | Unmodified | 0 | 0 | 0 | 0 | 0 | 0 | 0 | 0 | 281810 | 108440 | 0 | 0 | 0 | 0 | 0 | 0 | 0 | 0 |
| FCYEAVVR | PTPN23 | Q9H3S7 | PTN23_HUMAN | Unmodified | 475140 | 410330 | 0 | 36033 | 0 | 0 | 0 | 0 | 0 | 0 | 0 | 0 | 0 | 0 | 38419 | 0 | 0 | 0 |
| FDAGELITC | PHB; PHB; PHB | P35232; C9J | PHB_HUMAN F | Unmodified | 0 | 0 | 0 | 0 | 0 | 0 | 0 | 0 | 1591600 | 0 | 0 | 0 | 0 | 0 | 0 | 0 | 0 | 0 |
| FDAGELITC | PHB; PHB; PHB | P35232; C9J | PHB_HUMAN F | Unmodified | 0 | 0 | 0 | 0 | 1134200 | 0 | 0 | 1444700 | 1605600 | 101470 | 91472 | 195120 | 0 | 0 | 0 | 0 | 0 | 0 |
| FDAMPFTLI | PA2G4 | Q9UQ80 | PA2G4_HUMAN | Oxidation (M) | 194940 | 85867 | 9365.2 | 51629 | 0 | 0 | 0 | 0 | 132660 | 0 | 0 | 3949300 | 0 | 0 | 0 | 54631 | 0 | 0 |
| FDAMPFTLI | PA2G4 | Q9UQ80 | PA2G4_HUMAN | Unmodified | 0 | 0 | 0 | 0 | 0 | 0 | 0 | 0 | 0 | 0 | 0 | 2389800 | 0 | 0 | 0 | 0 | 0 | 0 |
| FDASFFGV | FASN; FASN; FA | P49327; A0A | FAS_HUMAN F | Unmodified | 577330 | 0 | 0 | 977830 | 0 | 0 | 0 | 0 | 0 | 0 | 0 | 27972000 | 0 | 0 | 0 | 1546600 | 0 | 0 |
| FDATSMHV | VIL1; VIL1 | P09327; B4D | VILI_HUMAN Vi | Unmodified | 0 | 0 | 0 | 0 | 0 | 0 | 0 | 0 | 705360 | 304250 | 0 | 4744000 | 0 | 0 | 0 | 0 | 0 | 0 |
| FDDAVVQS | HSPA8; HSPA8; | P11142; E9P | HSP7C_HUMAI | Unmodified | 1177700 | 797760 | 547490 | 956500 | 2045600 | 561270 | 197230 | 249200 | 3556100 | 1369400 | 2469100 | 6913400 | 1718700 | 708590 | 702310 | 434170 | 289160 | 844500 |
| FDDAVVQS | HSPA8; HSPA8; | P11142; E9P | HSP7C_HUMAI | Oxidation (M) | 585570 | 564750 | 523700 | 724610 | 1259800 | 327060 | 103920 | 15943 | 765510 | 229210 | 943840 | 1460400 | 887230 | 0 | 588080 | 571200 | 231450 | 172940 |
| FDDDKVSI | USP14; USP14 | P54578; A6N | UBP14_HUMAN | Unmodified | 114750 | 76094 | 47810 | 154050 | 304600 | 39593 | 0 | 0 | 177340 | 0 | 1820200 | 4307600 | 0 | 0 | 184830 | 263170 | 0 | 0 |
| FDDDVVSR | USP7; USP7; U | Q93009; A0A | UBP7_HUMAN | Unmodified | 40843 | 23470 | 36292 | 31826 | 0 | 0 | 0 | 26472 | 0 | 19713 | 157540 | 1267100 | 0 | 0 | 26727 | 0 | 0 | 0 |
| FDDGAGGI | PSMC2; PSMC2 | P35998; C9J | PRS7_HUMAN | Unmodified | 87964 | 0 | 0 | 0 | 0 | 0 | 0 | 0 | 255100 | 0 | 197710 | 2908300 | 0 | 0 | 210600 | 239680 | 0 | 0 |
| FDDHEVSD | USP8 | P40818 | UBP8_HUMAN | Unmodified | 51858 | 0 | 0 | 0 | 0 | 0 | 0 | 0 | 0 | 0 | 0 | 531660 | 0 | 0 | 0 | 0 | 0 | 0 |
| FDDPGLML | XRCC6; XRCC6 | P12956; B1A | XRCC6_HUMAI | 2 Oxidation (M) | 0 | 0 | 0 | 0 | 0 | 0 | 0 | 0 | 0 | 0 | 0 | 852580 | 0 | 0 | 0 | 0 | 0 | 0 |
| FDDPLLGP | LARS1; LARS1; | Q9P2J5; A0A | SYLC_HUMAN | Unmodified | 0 | 0 | 0 | 0 | 0 | 0 | 0 | 0 | 0 | 0 | 687460 | 5094400 | 0 | 0 | 172740 | 154210 | 0 | 0 |
| FDDTNPEK | QARS1; EPRS1; | P47897; P07 | SYQ_HUMAN G | Unmodified | 0 | 0 | 0 | 0 | 0 | 0 | 0 | 0 | 0 | 0 | 0 | 0 | 0 | 0 | 0 | 0 | 0 | 0 |
| FDDVVTNL | C1QL1 | O75973 | C1QRF_HUMAI | Unmodified | 0 | 0 | 0 | 0 | 101480 | 0 | 0 | 0 | 897290 | 156020 | 0 | 0 | 0 | 0 | 239340 | 0 | 0 | 0 |
| FDEFFRELL | GPC6 | Q9Y625 | GPC6_HUMAN | Unmodified | 153090 | 71493 | 12774 | 0 | 0 | 0 | 0 | 0 | 0 | 0 | 0 | 0 | 0 | 7882.3 | 0 | 0 | 0 | 0 |
| FDEGRNNI | P4HB; P4HB; P4 | P07237; A0A | PDIA1_HUMAN | Unmodified | 0 | 0 | 0 | 654110 | 404140 | 0 | 0 | 0 | 1088500 | 522080 | 190920 | 56193 | 0 | 623080 | 0 | 0 | 0 | 818030 |
| FDEGRNNI | P4HB; P4HB; P4 | P07237; A0A | PDIA1_HUMAN | Unmodified | 0 | 0 | 0 | 0 | 0 | 0 | 0 | 0 | 0 | 5181500 | 0 | 0 | 0 | 0 | 0 | 0 | 0 | 0 |
| FDEISFVNF | SSRP1 | Q08945 | SSRP1_HUMAN | Unmodified | 0 | 0 | 0 | 0 | 0 | 0 | 0 | 0 | 0 | 0 | 0 | 1055800 | 0 | 0 | 0 | 0 | 0 | 0 |
| FDENGLPT | CARS1 | P49589 | SYCC_HUMAN | Unmodified | 0 | 0 | 0 | 0 | 0 | 0 | 0 | 0 | 0 | 0 | 225170 | 844580 | 0 | 0 | 0 | 0 | 0 | 0 |
| FDENGLPT | CARS1 | P49589 | SYCC_HUMAN | Oxidation (M) | 0 | 0 | 0 | 0 | 0 | 0 | 0 | 0 | 0 | 0 | 0 | 176250 | 0 | 0 | 0 | 0 | 0 | 0 |
| FDENYAFLI | APOBEC3B; AP | Q9UH17; B0 | ABC3B_HUMAI | Unmodified | 782180 | 0 | 1138900 | 0 | 390320 | 0 | 0 | 0 | 0 | 0 | 0 | 562430 | 0 | 0 | 1639100 | 1549300 | 0 | 0 |
| FDEVLIR | MYO1C; MYO1C | O00159; F5H | MYO1C_HUMA | Unmodified | 207150 | 260880 | 412920 | 220870 | 0 | 0 | 0 | 0 | 72469 | 88057 | 0 | 0 | 0 | 0 | 754240 | 658940 | 0 | 0 |
| FDEVLVNH | HSPA4; HSPA4; | P34932; A0A | HSP74_HUMAN | Unmodified | 0 | 0 | 0 | 0 | 0 | 0 | 0 | 0 | 0 | 0 | 0 | 302240 | 0 | 0 | 0 | 47656 | 0 | 0 |
| FDFNAFR | MAN1C1 | Q9NR34 | MA1C1_HUMAI | Unmodified | 0 | 0 | 0 | 0 | 0 | 0 | 0 | 0 | 589430 | 91128 | 0 | 0 | 0 | 0 | 0 | 0 | 0 | 0 |
| FDFQAQSP | SORBS3; SORB | O60504; E5F | VINEX_HUMAN | Unmodified | 970890 | 0 | 0 | 0 | 0 | 0 | 0 | 0 | 0 | 141550 | 0 | 0 | 0 | 0 | 105820 | 0 | 0 | 0 |
| FDGAVEAV | MAN1A2; MAN1 | O60476; H0Y | MA1A2_HUMAI | Unmodified | 0 | 0 | 0 | 0 | 109530 | 0 | 0 | 0 | 2312800 | 402790 | 0 | 0 | 0 | 0 | 0 | 0 | 0 | 0 |
| FDGGEEVL | DECR1 | Q16698 | DECR_HUMAN | Unmodified | 0 | 0 | 0 | 0 | 0 | 0 | 0 | 0 | 1919000 | 0 | 0 | 0 | 0 | 0 | 0 | 0 | 0 | 0 |
| FDGGVEAI | MAN1A1 | P33908 | MA1A1_HUMAI | Unmodified | 0 | 0 | 0 | 0 | 0 | 0 | 0 | 0 | 1248200 | 254750 | 1939700 | 4635000 | 0 | 0 | 0 | 0 | 0 | 0 |
| FDGILTEGE | ERO1A | Q96HE7 | ERO1A_HUMAI | Unmodified | 0 | 0 | 0 | 0 | 0 | 0 | 0 | 0 | 549580 | 0 | 299390 | 0 | 0 | 0 | 0 | 0 | 43349 | 0 |
| FDGLLGFP | NUDT16L1; NU | Q9BRJ7; K7E | TIRR_HUMAN T | Unmodified | 0 | 0 | 0 | 0 | 0 | 0 | 0 | 0 | 0 | 0 | 0 | 1218700 | 87898 | 0 | 0 | 0 | 0 | 0 |
| FDINDATG | TRRAP; TRRAP; | Q9Y4A5; A0A | TRRAP_HUMAN | Unmodified | 0 | 207600 | 0 | 0 | 0 | 0 | 0 | 0 | 0 | 0 | 2873600 | 1323600 | 0 | 0 | 0 | 0 | 0 | 0 |
| FDLGLEDV | MAN1A2 | O60476 | MA1A2_HUMAI | Unmodified | 0 | 0 | 0 | 0 | 0 | 0 | 0 | 0 | 7754400 | 1138400 | 126990 | 279550 | 0 | 0 | 0 | 0 | 0 | 0 |
| FDLGQDVI | GDI1; GDI2; GD | P31150; P50 | GDIA_HUMAN | Unmodified | 0 | 0 | 0 | 26374 | 0 | 0 | 0 | 0 | 0 | 0 | 0 | 0 | 0 | 0 | 225570 | 0 | 0 | 0 |
| FDLHSSSH | PPP6R3; PPP6R | Q5H9R7; E9I | PP6R3_HUMAN | Unmodified | 0 | 0 | 0 | 0 | 0 | 0 | 0 | 0 | 0 | 0 | 88116 | 456160 | 0 | 0 | 0 | 50786 | 0 | 0 |
| FDLLASNFI | LARP4; LARP4 | Q71RC2; X6I | LARP4_HUMAN | Unmodified | 0 | 94438 | 24853 | 62042 | 0 | 0 | 0 | 0 | 0 | 0 | 0 | 0 | 0 | 0 | 0 | 20163 | 0 | 0 |
| FDLMYAK | TUBA3C; TUBA | P0DPH7; Q7 | TBA3C_HUMAN | Unmodified | 0 | 31756 | 132330 | 0 | 402490 | 0 | 427220 | 165020 | 2089400 | 357480 | 439070 | 21834000 | 0 | 0 | 226190 | 53189 | 309450 | 262180 |
| FDLMYAK | TUBA3C; TUBA | P0DPH7; Q7 | TBA3C_HUMAN | Oxidation (M) | 173040 | 0 | 1450200 | 0 | 411300 | 0 | 406150 | 296530 | 1009600 | 214710 | 267420 | 11060000 | 78088 | 0 | 463850 | 608540 | 106590 | 281340 |
| FDLNEPLH | UBA6; UBA6 | A0AVT1; H0Y | UBA6_HUMAN | Unmodified | 0 | 0 | 0 | 0 | 0 | 0 | 0 | 0 | 0 | 0 | 0 | 962780 | 0 | 0 | 0 | 48261 | 0 | 0 |
| FDMIVPILE | ILK; ILK | Q13418; A0A | ILK_HUMAN Int | Unmodified | 55904 | 164860 | 61527 | 111620 | 0 | 0 | 161730 | 0 | 295500 | 0 | 0 | 63245 | 65653 | 0 | 65748 | 50435 | 0 | 328120 |
| FDMIVPILE | ILK; ILK | Q13418; A0A | ILK_HUMAN Int | Oxidation (M) | 61030 | 186970 | 201810 | 412480 | 135420 | 21803 | 0 | 10818 | 37439 | 0 | 0 | 184090 | 0 | 263730 | 231500 | 140780 | 0 | 117290 |
| FDNLYGCF | AHCY | P23526 | SAHH_HUMAN | Unmodified | 0 | 0 | 0 | 0 | 0 | 0 | 0 | 0 | 0 | 0 | 0 | 570140 | 0 | 0 | 0 | 0 | 0 | 0 |
| FDNPAAVS | TMTC3; TMTC3 | Q6ZXV5; F8V | TMTC3_HUMAI | Unmodified | 0 | 0 | 0 | 0 | 0 | 0 | 0 | 0 | 740320 | 0 | 0 | 0 | 0 | 0 | 0 | 0 | 0 | 0 |
| FDNSYSLLI | TMED8 | Q6PL24 | TMED8_HUMAI | Unmodified | 33650 | 0 | 0 | 0 | 0 | 0 | 0 | 0 | 0 | 0 | 344260 | 1553000 | 0 | 0 | 0 | 0 | 0 | 0 |
| FDNSYSLW | ACBD3 | Q9H3P7 | GCP60_HUMAI | Unmodified | 0 | 0 | 0 | 0 | 0 | 0 | 0 | 72465 | 285060 | 0 | 434490 | 925710 | 230870 | 0 | 0 | 0 | 0 | 0 |
| FDQGDTTR | F11R | Q9Y624 | JAM1_HUMAN . | Unmodified | 243320 | 60560 | 260670 | 432800 | 0 | 0 | 130190 | 0 | 0 | 0 | 177680 | 0 | 0 | 0 | 227150 | 140100 | 0 | 0 |
| FDQLFDDE | SERBP1 | Q8NC51 | PAIRB_HUMAN | Unmodified | 0 | 0 | 0 | 0 | 0 | 0 | 0 | 0 | 0 | 0 | 122060 | 69018 | 0 | 0 | 0 | 0 | 0 | 0 |
| FDQLNIQR | OCRL; OCRL; O | Q01968; A0A | OCRL_HUMAN | Unmodified | 0 | 0 | 0 | 0 | 0 | 0 | 0 | 0 | 0 | 0 | 0 | 388960 | 0 | 0 | 0 | 0 | 0 | 0 |
| FDQPDDFK | HSPG2 | P98160 | PGBM_HUMAN | Unmodified | 0 | 0 | 779940 | 0 | 0 | 0 | 0 | 0 | 0 | 0 | 0 | 0 | 0 | 0 | 0 | 0 | 0 | 0 |
| FDQTGLMK | MAN2A1 | Q16706 | MA2A1_HUMAI | Unmodified | 0 | 0 | 0 | 0 | 0 | 0 | 0 | 32962 | 1441000 | 94266 | 0 | 708090 | 165320 | 21180 | 0 | 0 | 0 | 0 |
| FDRGYISPY | HSPD1; HSPD1 | P10809; A0A | CH60_HUMAN | Unmodified | 487490 | 0 | 484620 | 0 | 0 | 333500 | 645530 | 0 | 23103000 | 1570600 | 562350 | 638330 | 1299600 | 836200 | 645590 | 471060 | 88863 | 440670 |
| FDSASSSY | PIGS | Q96S52 | PIGS_HUMAN ( | Oxidation (M) | 0 | 0 | 0 | 0 | 0 | 0 | 0 | 0 | 1864500 | 110800 | 0 | 0 | 0 | 0 | 0 | 0 | 0 | 0 |
| FDSASSSY | PIGS | Q96S52 | PIGS_HUMAN ( | Unmodified | 0 | 0 | 0 | 0 | 0 | 0 | 0 | 0 | 7359500 | 612660 | 0 | 0 | 0 | 0 | 0 | 0 | 341240 | 0 |
| FDSDAASP | HLA-B; HLA-C; I | P01889; P10 | HLAB_HUMAN | Unmodified | 0 | 45734 | 34503 | 0 | 0 | 135280 | 0 | 0 | 4152700 | 566680 | 2534500 | 15363000 | 0 | 56692 | 51860 | 48232 | 0 | 18701 |
| FDSDAASP | HLA-C; HLA-C; | P10321; Q5( | HLAC_HUMAN | Unmodified | 0 | 0 | 0 | 0 | 0 | 0 | 0 | 0 | 2563900 | 336130 | 35700 | 0 | 0 | 0 | 0 | 0 | 0 | 0 |
| FDSDAASQ | HLA-A; HLA-A; I | P04439; Q9( | HLAA_HUMAN | Unmodified | 0 | 0 | 0 | 0 | 0 | 0 | 0 | 0 | 0 | 208560 | 388060 | 1915000 | 0 | 0 | 7908.7 | 0 | 0 | 0 |
| FDSDVGVY | HLA-DQB1; HLA | Q5SU54; A0 | Q5SU54_HUM | Unmodified | 0 | 0 | 0 | 0 | 0 | 0 | 0 | 0 | 1180000 | 0 | 0 | 0 | 0 | 0 | 0 | 0 | 0 | 0 |
| FDSPESHV | CBS; CBSL | P35520; P0D | CBS_HUMAN C | Unmodified | 0 | 0 | 0 | 0 | 0 | 0 | 0 | 0 | 0 | 0 | 0 | 1527500 | 0 | 0 | 0 | 0 | 0 | 0 |
| FDSSAPFL | CELSR3 | Q9NYQ7 | CELR3_HUMAN | Unmodified | 0 | 0 | 0 | 541250 | 98869 | 0 | 0 | 0 | 297450 | 0 | 0 | 0 | 0 | 0 | 0 | 0 | 392420 | 65410 |
| FDSSYDRD | FKBP9 | O95302 | FKBP9_HUMAN | Unmodified | 0 | 0 | 0 | 0 | 0 | 0 | 0 | 0 | 2113700 | 0 | 263920 | 151570 | 0 | 0 | 0 | 0 | 0 | 0 |
| FDTGNLCN | RPS4X | P62701 | RS4X_HUMAN | Oxidation (M) | 0 | 0 | 0 | 0 | 0 | 0 | 0 | 0 | 0 | 0 | 0 | 7046800 | 0 | 0 | 511350 | 448440 | 0 | 0 |
| FDTGNLCN | RPS4X | P62701 | RS4X_HUMAN | Unmodified | 0 | 0 | 0 | 0 | 0 | 0 | 0 | 0 | 0 | 0 | 0 | 868730 | 0 | 0 | 0 | 0 | 0 | 293120 |
| FDTGSGPA | AGRN; AGRN; A | O00468; A0A | AGRIN_HUMAN | Unmodified | 0 | 0 | 574140 | 0 | 0 | 241640 | 0 | 0 | 0 | 0 | 0 | 0 | 0 | 0 | 700480 | 803340 | 0 | 0 |
| FDTGSGPA | AGRN; AGRN; A | O00468; A0A | AGRIN_HUMAN | Unmodified | 988250 | 0 | 0 | 0 | 0 | 0 | 0 | 0 | 0 | 0 | 0 | 0 | 0 | 0 | 0 | 0 | 0 | 0 |
| FDTIYQILLK | TRIM25; TRIM25 | Q14258; A0A | TRI25_HUMAN | Unmodified | 0 | 147920 | 131440 | 0 | 284590 | 69290 | 0 | 0 | 0 | 0 | 84293 | 740530 | 0 | 0 | 104910 | 0 | 0 | 0 |
| FDTQLFHTI | RAB9A | P51151 | RAB9A_HUMAN | Unmodified | 0 | 0 | 132890 | 71554 | 0 | 0 | 0 | 0 | 93115 | 0 | 29937 | 80968 | 29128 | 0 | 228250 | 117620 | 0 | 34308 |
| FDTQYPYG | ERP29; ERP29 | P30040; F8V | ERP29_HUMAN | Unmodified | 0 | 0 | 0 | 0 | 0 | 0 | 0 | 0 | 19186000 | 391310 | 1008900 | 368260 | 0 | 0 | 0 | 0 | 0 | 0 |
| FDTQYPYG | ERP29; ERP29 | P30040; F8V | ERP29_HUMAN | Unmodified | 191720 | 0 | 0 | 0 | 0 | 0 | 0 | 0 | 43898000 | 7451600 | 1540200 | 0 | 0 | 0 | 96049 | 0 | 0 | 0 |
| FDTQYPYG | ERP29; ERP29 | P30040; F8V | ERP29_HUMAN | Unmodified | 323610 | 0 | 0 | 0 | 0 | 0 | 0 | 0 | 54447000 | 700820 | 140810 | 0 | 0 | 0 | 0 | 0 | 0 | 0 |
| FDVNTSAV | CLTC; CLTC | Q00610; A0A | CLH1_HUMAN | Unmodified | 0 | 859890 | 36102 | 0 | 0 | 0 | 0 | 0 | 0 | 0 | 0 | 0 | 0 | 0 | 593290 | 0 | 0 | 0 |
| FDVNTSAV | CLTC; CLTC | Q00610; A0A | CLH1_HUMAN | Unmodified | 0 | 2335200 | 0 | 147730 | 0 | 0 | 0 | 0 | 0 | 0 | 0 | 0 | 0 | 0 | 890120 | 0 | 0 | 0 |
| FDVQFPEN | DNAJA2 | O60884 | DNJA2_HUMAN | Unmodified | 0 | 105750 | 424270 | 29553 | 53036 | 0 | 0 | 0 | 0 | 0 | 0 | 289030 | 18600 | 0 | 520010 | 213090 | 0 | 15779 |
| FDVQSSLK | TPP2; TPP2; TPI | P29144; A0A | TPP2_HUMAN | Unmodified | 0 | 0 | 0 | 0 | 0 | 0 | 0 | 0 | 0 | 0 | 0 | 2380600 | 0 | 0 | 0 | 0 | 0 | 0 |
| FDVSGYPT | PDIA4; PDIA4 | P13667; A0A | PDIA4_HUMAN | Unmodified | 0 | 0 | 182610 | 0 | 0 | 0 | 0 | 0 | 6698000 | 419170 | 555160 | 0 | 0 | 0 | 0 | 167110 | 0 | 0 |
| FDVSSTSVI | NIBAN2 | Q96TA1 | NIBA2_HUMAN | Unmodified | 911260 | 0 | 0 | 531690 | 0 | 0 | 102190 | 0 | 1413600 | 0 | 0 | 2088800 | 0 | 0 | 1117400 | 1322800 | 0 | 0 |
| FDVTLLHTK | AVIL | O75366 | AVIL_HUMAN A | Unmodified | 0 | 0 | 0 | 0 | 0 | 0 | 385490 | 0 | 0 | 0 | 847060 | 20468000 | 0 | 371760 | 0 | 0 | 0 | 128320 |
| FDWRQQY | EMC1; EMC1 | Q8N766; Q5 | EMC1_HUMAN | Unmodified | 0 | 0 | 0 | 0 | 0 | 1112800 | 0 | 0 | 931820 | 100900 | 165610 | 1150900 | 0 | 0 | 0 | 0 | 0 | 0 |
| FEAHEVLN | XXYLT1; XXYLT1 | Q8NBI6; H7( | XXLT1_HUMAN | Unmodified | 0 | 0 | 0 | 0 | 0 | 0 | 0 | 0 | 753160 | 0 | 50407 | 0 | 0 | 0 | 0 | 0 | 0 | 0 |
| FEDPRDAE | SRSF9; SRSF9; | Q13242; S4F | SRSF9_HUMAN | Unmodified | 964480 | 1415900 | 379580 | 212440 | 0 | 0 | 0 | 0 | 0 | 0 | 0 | 298120 | 0 | 0 | 874810 | 617910 | 0 | 0 |
| FEEDFIEK | SNX33; SNX33 | Q8WV41; H3 | SNX33_HUMAN | Unmodified | 0 | 0 | 2454200 | 0 | 0 | 0 | 0 | 0 | 0 | 0 | 0 | 0 | 0 | 0 | 2523400 | 0 | 217550 | 0 |
| FEEDFIEKR | SNX33; SNX33 | Q8WV41; H3 | SNX33_HUMAN | Unmodified | 266410 | 0 | 625950 | 486610 | 0 | 0 | 0 | 0 | 0 | 0 | 0 | 0 | 0 | 0 | 2953300 | 1486000 | 0 | 0 |
| FEEDFISK | SNX18 | Q96RF0 | SNX18_HUMAN | Unmodified | 463390 | 189580 | 855870 | 506560 | 0 | 0 | 0 | 0 | 1981900 | 0 | 0 | 0 | 0 | 0 | 1265400 | 517350 | 0 | 0 |
| FEEDVEDG | SLC4A7; SLC4A | Q9Y6M7; C9 | S4A7_HUMAN | Unmodified | 0 | 0 | 394670 | 0 | 0 | 0 | 0 | 0 | 0 | 0 | 0 | 1205800 | 0 | 0 | 269360 | 225240 | 0 | 0 |
| FEEEFIK | SNX9; SNX9; SN | Q9Y5X1; A0A | SNX9_HUMAN | Unmodified | 0 | 0 | 0 | 0 | 0 | 0 | 0 | 0 | 0 | 0 | 0 | 0 | 0 | 0 | 0 | 0 | 0 | 0 |
| FEEESKEP | CLNS1A; CLNS | P54105; E9P | ICLN_HUMAN I | Unmodified | 0 | 0 | 0 | 0 | 0 | 0 | 0 | 0 | 0 | 0 | 0 | 7910900 | 0 | 0 | 0 | 0 | 0 | 1128400 |
| FEEFQNTL | TXLNA | P40222 | TXLNA_HUMAN | Unmodified | 0 | 0 | 0 | 0 | 0 | 0 | 0 | 0 | 0 | 0 | 0 | 524760 | 0 | 0 | 0 | 0 | 0 | 0 |
| FEELCSDLI | HSPA6; HSPA1 | P17066; P0D | HSP76_HUMAN | Unmodified | 1116700 | 390880 | 0 | 0 | 2008300 | 1380800 | 0 | 0 | 908070 | 155200 | 0 | 980220 | 0 | 0 | 0 | 0 | 1434200 | 1101700 |
| FEELCSDLI | HSPA6; HSPA1 | P17066; P0D | HSP76_HUMAN | Unmodified | 0 | 0 | 0 | 0 | 1784500 | 0 | 875130 | 16481 | 87941 | 2194700 | 0 | 0 | 2435500 | 1749000 | 0 | 0 | 267180 | 292510 |
| FEELNADLI | HSPA8; HSPA2; | P11142; P54 | HSP7C_HUMAI | Unmodified | 4881100 | 7315100 | 3956000 | 5397300 | 403810 | 0 | 241120 | 253280 | 7692300 | 769920 | 4848400 | 18594000 | 576780 | 279370 | 7474900 | 3815100 | 0 | 435040 |
| FEELNADLI | HSPA8; HSPA8; | P11142; E9P | HSP7C_HUMAI | Unmodified | 9640400 | 13916000 | 2924900 | 7222200 | 22790000 | 0 | 6268300 | 32496 | 28437000 | 27013000 | 0 | 8316700 | 30029000 | 22930000 | 6649300 | 5799200 | 6693200 | 5692000 |
| FEELNMDL | HSPA5; HSPA5 | P11021; A0A | BIP_HUMAN Er | Unmodified | 0 | 0 | 0 | 0 | 0 | 0 | 0 | 0 | 30552000 | 1141900 | 1457600 | 97357 | 0 | 0 | 0 | 0 | 0 | 0 |
| FEELNMDL | HSPA5; HSPA5 | P11021; A0A | BIP_HUMAN Er | Oxidation (M) | 0 | 0 | 0 | 0 | 0 | 0 | 0 | 0 | 8864600 | 1004500 | 2196300 | 1211900 | 0 | 0 | 329960 | 429990 | 0 | 0 |
| FEELNMDL | HSPA5; HSPA5 | P11021; A0A | BIP_HUMAN Er | Oxidation (M) | 0 | 0 | 0 | 0 | 0 | 0 | 0 | 0 | 0 | 1751300 | 0 | 0 | 0 | 0 | 0 | 0 | 0 | 0 |
| FEELNMDL | HSPA5; HSPA5 | P11021; A0A | BIP_HUMAN Er | Unmodified | 0 | 0 | 0 | 0 | 0 | 0 | 0 | 0 | 761330 | 981760 | 0 | 0 | 0 | 0 | 0 | 0 | 0 | 0 |
| FEELTNLIR | EIF3CL; EIF3C; | B5ME19; Q9 | EIFCL_HUMAN | Unmodified | 0 | 0 | 151390 | 0 | 0 | 0 | 0 | 0 | 222790 | 0 | 0 | 745890 | 0 | 0 | 0 | 186980 | 0 | 0 |
| FEESMSEK | GLG1; GLG1 | Q92896; H3I | GSLG1_HUMAI | Unmodified | 0 | 0 | 52342 | 0 | 0 | 0 | 0 | 0 | 2033700 | 0 | 0 | 537130 | 0 | 0 | 0 | 0 | 0 | 0 |
| FEETTADGI | FABP5; FABP5 | Q01469; I6L | FABP5_HUMAN | Unmodified | 0 | 0 | 0 | 0 | 0 | 0 | 0 | 0 | 0 | 0 | 335590 | 0 | 0 | 0 | 0 | 0 | 0 | 0 |
| FEGEPATH | SEPTIN6; SEPTI | Q14141; B1A | SEPT6_HUMAN | Unmodified | 0 | 0 | 0 | 0 | 0 | 0 | 0 | 0 | 0 | 0 | 0 | 2271000 | 551460 | 655780 | 0 | 0 | 0 | 1276800 |
| FEGGDRDI | PDIA4; PDIA4 | P13667; A0A | PDIA4_HUMAN | Unmodified | 0 | 0 | 0 | 0 | 148230 | 0 | 0 | 93151 | 37312000 | 3602900 | 5219100 | 194080 | 0 | 0 | 0 | 0 | 0 | 0 |
| FEGPQVQD | B3GAT3; B3GAT | O94766; G3 | B3GA3_HUMAI | Unmodified | 0 | 0 | 0 | 0 | 0 | 0 | 0 | 0 | 864280 | 0 | 0 | 0 | 0 | 0 | 0 | 0 | 0 | 0 |
| FEGQFSINI | ERGIC1; ERGIC | Q969X5; F22 | ERGI1_HUMAN | Unmodified | 0 | 0 | 484780 | 0 | 0 | 0 | 40926 | 0 | 1718300 | 0 | 0 | 0 | 0 | 0 | 281760 | 0 | 0 | 0 |

| | | | | | | | | | | | | | | | | | | | | | | |
|---|---|---|---|---|---|---|---|---|---|---|---|---|---|---|---|---|---|---|---|---|---|---|
| FEGQFSINI | ERGIC1 | Q969X5 | ERGI1_HUMAN | Unmodified | 0 | 0 | 0 | 0 | 0 | 0 | 0 | 0 | 17827000 | 550610 | 0 | 0 | 0 | 0 | 0 | 0 | 0 | 0 |
| FEHCNFNI | CD59; CD59; N | P13987; A0A | CD59_HUMAN | Unmodified | 9603700 | 3599800 | 15567000 | 4567700 | 752190 | 388600 | 0 | 0 | 5886200 | 14455 | 0 | 6399400 | 915870 | 390640 | 12845000 | 11598000 | 620700 | 3497800 |
| FEIWDTAG | RAB5B; RAB5C; | P61020; P51 | RAB5B_HUMAN | Unmodified | 1014500 | 0 | 0 | 0 | 0 | 0 | 0 | 0 | 0 | 0 | 0 | 4618000 | 0 | 0 | 1917400 | 0 | 0 | 0 |
| FELDTSER | RPN2 | P04844 | RPN2_HUMAN | Unmodified | 0 | 0 | 0 | 0 | 0 | 0 | 0 | 0 | 2620200 | 729370 | 0 | 0 | 0 | 0 | 138110 | 0 | 0 | 0 |
| FELLGCELI | MFGE8 | Q08431 | MFGM_HUMAN | Unmodified | 347460 | 0 | 114560 | 530990 | 0 | 0 | 0 | 0 | 0 | 0 | 0 | 0 | 0 | 0 | 822160 | 425610 | 0 | 0 |
| FELSCYSLA | ASS1; ASS1 | P00966; Q5T | ASSY_HUMAN | Unmodified | 0 | 0 | 0 | 0 | 0 | 0 | 0 | 0 | 0 | 0 | 0 | 0 | 0 | 0 | 2329900 | 1238000 | 0 | 0 |
| FENAAAGN | ARPC2; ARPC2; | O15144; G5I | ARPC2_HUMAN | Unmodified | 0 | 1282700 | 0 | 0 | 0 | 0 | 0 | 0 | 0 | 0 | 884360 | 3348100 | 0 | 0 | 5055000 | 1092800 | 0 | 365290 |
| FENAFLSH | ATP5F1A | P25705 | ATPA_HUMAN | Unmodified | 0 | 0 | 0 | 0 | 0 | 0 | 0 | 0 | 17844000 | 0 | 0 | 0 | 0 | 0 | 0 | 0 | 0 | 0 |
| FENAFLSH | ATP5F1A | P25705 | ATPA_HUMAN | Unmodified | 0 | 0 | 0 | 0 | 0 | 0 | 761980 | 0 | 8593800 | 355360 | 129310 | 0 | 0 | 0 | 0 | 0 | 0 | 0 |
| FENLGIQPI | PSMC3; PSMC3 | P17980; E9P | PRS6A_HUMAN | Oxidation (M) | 0 | 0 | 0 | 0 | 0 | 0 | 569180 | 360310 | 179510 | 112090 | 0 | 0 | 655790 | 314200 | 0 | 0 | 0 | 0 |
| FENWGACI | MDK; MDK; MD | P21741; E9P | MK_HUMAN Mi | Unmodified | 0 | 0 | 0 | 0 | 0 | 0 | 0 | 0 | 2251400 | 273610 | 0 | 0 | 0 | 0 | 0 | 0 | 0 | 0 |
| FEPIHGNFI | PRPF4 | O43172 | PRP4_HUMAN | Unmodified | 22173 | 0 | 0 | 0 | 133890 | 0 | 117630 | 0 | 0 | 0 | 0 | 946620 | 538860 | 0 | 0 | 0 | 406210 | 1005300 |
| FEQIVSILDI | EPHA3; EPHA3 | P29320; C9J | EPHA3_HUMAN | Unmodified | 2179000 | 3200500 | 0 | 0 | 0 | 0 | 0 | 23343 | 0 | 0 | 0 | 1484400 | 46110 | 0 | 0 | 0 | 0 | 0 |
| FEQYVEPYI | N/A | N/A | N/A | Unmodified | 0 | 0 | 1697400 | 1977100 | 0 | 0 | 0 | 0 | 0 | 0 | 0 | 0 | 1449200 | 0 | 0 | 3887000 | 0 | 0 |
| FESDPATH | SEPTIN11; SEPT | Q9NVA2; D6 | SEP11_HUMAN | Unmodified | 1192900 | 0 | 318780 | 142150 | 106900 | 0 | 0 | 0 | 360410 | 95221 | 650960 | 2272100 | 0 | 0 | 162480 | 0 | 0 | 0 |
| FEVIPQSIRI | TM9SF4; TM9SF | Q92544; A0A | TM9S4_HUMAN | Unmodified | 0 | 0 | 0 | 0 | 0 | 0 | 0 | 0 | 2236000 | 657940 | 0 | 4687400 | 0 | 0 | 0 | 0 | 0 | 0 |
| FEVIPQSIRI | TM9SF4; TM9SF | Q92544; A0A | TM9S4_HUMAN | Unmodified | 0 | 0 | 0 | 0 | 0 | 0 | 0 | 0 | 17276000 | 879170 | 88492 | 1528700 | 0 | 0 | 0 | 0 | 288280 | 0 |
| FFADLLDYI | HPRT1 | P00492 | HPRT_HUMAN | Unmodified | 0 | 0 | 0 | 0 | 457460 | 93412 | 652500 | 0 | 377590 | 574160 | 45824 | 416540 | 0 | 0 | 0 | 0 | 519580 | 356690 |
| FFAGIVWQ | AKT1; AKT1; AKT | P31749; A0A | AKT1_HUMAN | Unmodified | 0 | 0 | 797320 | 1270600 | 0 | 0 | 0 | 0 | 0 | 74165 | 0 | 161300 | 0 | 0 | 2706500 | 1959500 | 0 | 0 |
| FFAPAILSPI | RASA3 | Q14644 | RASA3_HUMAN | Unmodified | 691750 | 2089400 | 792280 | 0 | 0 | 0 | 209280 | 0 | 686330 | 288660 | 0 | 346940 | 0 | 0 | 1863600 | 2506500 | 0 | 0 |
| FFDANYDG | MAPRE3; MAPR | Q9UPY8; Q1 | MARE3_HUMAN | Unmodified | 0 | 0 | 0 | 0 | 0 | 0 | 0 | 0 | 0 | 0 | 0 | 2438600 | 0 | 128010 | 0 | 0 | 0 | 0 |
| FFDEESYSI | SSR4; SSR4 | P51571; A6N | SSRD_HUMAN | Unmodified | 0 | 0 | 0 | 0 | 0 | 0 | 0 | 0 | 40768000 | 412070 | 1792700 | 658790 | 0 | 0 | 0 | 0 | 0 | 0 |
| FFDEESYSI | SSR4; SSR4 | P51571; A6N | SSRD_HUMAN | Unmodified | 0 | 0 | 443830 | 182900 | 0 | 0 | 0 | 0 | 83799000 | 7943500 | 3375200 | 7049300 | 0 | 0 | 0 | 0 | 0 | 0 |
| FFDENESP | GOLM2; GOLM2 | Q6P4E1; A0A | GOLM2_HUMA | Unmodified | 0 | 0 | 0 | 0 | 0 | 0 | 0 | 0 | 2956900 | 177810 | 0 | 916910 | 0 | 0 | 0 | 0 | 0 | 0 |
| FFDGELTK | GNPDA1; GNPI | P46926; D6F | GNPI1_HUMAN | Unmodified | 0 | 0 | 0 | 0 | 0 | 0 | 0 | 0 | 0 | 0 | 623760 | 748990 | 0 | 0 | 308690 | 94028 | 0 | 0 |
| FFDHSGTL | SPCS2; SPCS2; | Q15005; E9F | SPCS2_HUMAN | Oxidation (M) | 0 | 0 | 0 | 0 | 0 | 269670 | 0 | 0 | 3454200 | 2639600 | 0 | 1015400 | 0 | 0 | 0 | 0 | 0 | 0 |
| FFDHSGTL | SPCS2; SPCS2; | Q15005; E9F | SPCS2_HUMAN | Unmodified | 0 | 0 | 0 | 0 | 0 | 0 | 0 | 0 | 7253700 | 654630 | 138500 | 0 | 0 | 0 | 0 | 0 | 0 | 0 |
| FFDHSGTL | SPCS2; SPCS2; | Q15005; E9F | SPCS2_HUMAN | Oxidation (M) | 0 | 0 | 0 | 0 | 0 | 0 | 793220 | 0 | 0 | 9427700 | 0 | 0 | 0 | 322000 | 0 | 0 | 0 | 0 |
| FFDHSGTL | SPCS2; SPCS2; | Q15005; E9F | SPCS2_HUMAN | Unmodified | 0 | 0 | 0 | 0 | 0 | 0 | 0 | 0 | 10863000 | 1722000 | 0 | 0 | 0 | 0 | 0 | 0 | 0 | 0 |
| FFDIDSGR | ATP2B3; ATP2B | Q16720; P23 | AT2B3_HUMAN | Unmodified | 0 | 0 | 469600 | 0 | 0 | 0 | 0 | 0 | 299490 | 0 | 0 | 1406800 | 0 | 0 | 475260 | 299440 | 0 | 0 |
| FFDKVQDL | ADGRE5 | P48960 | AGRE5_HUMAN | Unmodified | 0 | 0 | 0 | 0 | 1615200 | 0 | 0 | 35902 | 0 | 0 | 0 | 0 | 0 | 148900 | 0 | 0 | 0 | 0 |
| FFDTELENI | STK10 | O94804 | STK10_HUMAN | Unmodified | 0 | 0 | 0 | 0 | 0 | 0 | 0 | 291670 | 0 | 0 | 470390 | 447600 | 0 | 0 | 0 | 0 | 0 | 0 |
| FFEHFIEGC | OTUB1; OTUB1 | Q96FW1; J3I | OTUB1_HUMAN | Unmodified | 0 | 0 | 0 | 174980 | 0 | 0 | 0 | 0 | 0 | 141070 | 0 | 1469100 | 0 | 0 | 0 | 0 | 0 | 68061 |
| FFETSAK | RAB10; RAB3C; | P61026; Q96 | RAB10_HUMAN | Unmodified | 11397000 | 20027000 | 20438000 | 15744000 | 5761700 | 0 | 336050 | 279890 | 12367000 | 1930100 | 5062700 | 24462000 | 3436700 | 1913000 | 24398000 | 21843000 | 1241100 | 4403500 |
| FFEVILIDPF | RPL15; RPL15; | P61313; E7E | RL15_HUMAN | Unmodified | 8230300 | 7118200 | 1896500 | 2346000 | 3057100 | 0 | 4851200 | 0 | 613910 | 1328400 | 0 | 11147000 | 0 | 0 | 6528100 | 2876600 | 0 | 0 |
| FFGAENSV | PHYH; PHYH | O14832; B1A | PAHX_HUMAN | Unmodified | 0 | 0 | 0 | 0 | 0 | 0 | 0 | 0 | 18706000 | 2568700 | 0 | 1881000 | 0 | 0 | 0 | 109760 | 0 | 0 |
| FFGDSAAS | HYOU1; HYOU1 | Q9Y4L1; J3Q | HYOU1_HUMA | Unmodified | 0 | 0 | 577340 | 0 | 0 | 0 | 0 | 0 | 3251000 | 0 | 0 | 0 | 0 | 0 | 0 | 0 | 0 | 0 |
| FFGDSAAS | HYOU1; HYOU1 | Q9Y4L1; J3Q | HYOU1_HUMA | Unmodified | 0 | 0 | 0 | 0 | 0 | 0 | 0 | 0 | 5281600 | 389900 | 218510 | 462020 | 0 | 0 | 0 | 0 | 0 | 0 |
| FFGEIDPSL | CACNA2D1 | P54289 | CA2D1_HUMAN | Oxidation (M) | 1016000 | 1101700 | 0 | 0 | 0 | 0 | 0 | 0 | 994870 | 0 | 0 | 0 | 0 | 0 | 0 | 0 | 0 | 0 |
| FFGEIDPSL | CACNA2D1 | P54289 | CA2D1_HUMAN | Unmodified | 159410 | 0 | 0 | 0 | 0 | 0 | 0 | 0 | 1502300 | 0 | 0 | 0 | 0 | 0 | 0 | 0 | 0 | 0 |
| FFGGYVPE | MARS1; MARS1 | P56192; H0Y | SYMC_HUMAN | Oxidation (M) | 0 | 0 | 55886 | 0 | 0 | 0 | 0 | 0 | 0 | 0 | 99840 | 1490900 | 0 | 0 | 0 | 98200 | 0 | 0 |
| FFGGYVPE | MARS1; MARS1 | P56192; H0Y | SYMC_HUMAN | Unmodified | 0 | 0 | 0 | 0 | 0 | 0 | 0 | 0 | 0 | 0 | 0 | 150260 | 0 | 0 | 0 | 0 | 0 | 0 |
| FFGKDISTT | HSPH1; HSPH1 | Q92598; A0A | HS105_HUMAN | Unmodified | 0 | 0 | 0 | 136660 | 0 | 0 | 0 | 0 | 0 | 0 | 0 | 0 | 234400 | 0 | 0 | 0 | 403820 | 617990 |
| FFGNSWAI | AKR7A2; AKR7A | O43488; H3I | ARK72_HUMAN | Unmodified | 0 | 0 | 154250 | 417050 | 0 | 0 | 0 | 0 | 1214200 | 0 | 0 | 305010 | 0 | 0 | 0 | 0 | 0 | 0 |
| FFGSLPDS | HTATIP2; HTATI | Q9BUP3; E9I | HTAI2_HUMAN | Unmodified | 0 | 0 | 0 | 0 | 0 | 0 | 0 | 0 | 6630300 | 3671400 | 0 | 0 | 0 | 0 | 0 | 0 | 0 | 0 |
| FFHLAFEEE | EIF3I; EIF3I; EIF | Q13347; A0A | EIF3I_HUMAN | Unmodified | 0 | 0 | 0 | 0 | 2832500 | 0 | 331280 | 0 | 301440 | 716200 | 701910 | 898660 | 6162500 | 1718000 | 0 | 0 | 1045500 | 1877200 |
| FFHPEEWA | FUCA1 | P04066 | FUCO_HUMAN | Unmodified | 47260 | 0 | 0 | 0 | 0 | 0 | 0 | 0 | 2365900 | 282070 | 0 | 106780 | 0 | 0 | 0 | 32669 | 0 | 0 |
| FFHTGTNA | NDST1; NDST1 | P52848; E7E | NDST1_HUMAN | Unmodified | 0 | 0 | 0 | 0 | 53551 | 0 | 0 | 0 | 2550900 | 0 | 0 | 0 | 0 | 0 | 0 | 0 | 0 | 0 |
| FFKQPTFC | PRKCB; PRKCA | P05771; P17 | KPCB_HUMAN | Unmodified | 0 | 0 | 0 | 0 | 137320 | 0 | 1029500 | 0 | 0 | 0 | 0 | 0 | 0 | 436060 | 0 | 0 | 0 | 0 |
| FFLCQVAG | NCAM1; NCAM | P13591; A0A | NCAM1_HUMA | Unmodified | 0 | 0 | 0 | 0 | 0 | 0 | 0 | 0 | 5133900 | 94289 | 0 | 0 | 0 | 0 | 0 | 0 | 0 | 0 |
| FFLCQVAG | NCAM1; NCAM | P13591; A0A | NCAM1_HUMA | Unmodified | 0 | 0 | 0 | 0 | 0 | 375270 | 469910 | 0 | 194280 | 768330 | 274990 | 0 | 0 | 421970 | 0 | 0 | 0 | 0 |
| FFLGDLCS | NSDHL; NSDHL | Q15738; C9J | NSDHL_HUMA | Unmodified | 393270 | 0 | 0 | 82598 | 0 | 0 | 0 | 0 | 0 | 0 | 544700 | 0 | 0 | 0 | 52718 | 0 | 0 | 0 |
| FFLHGVTL | LAMA5 | O15230 | LAMA5_HUMAN | Unmodified | 304670 | 0 | 120140 | 219120 | 0 | 0 | 0 | 0 | 222960 | 0 | 0 | 0 | 0 | 0 | 161190 | 48683 | 0 | 0 |
| FFLQGIQLI | LAMP1 | P11279 | LAMP1_HUMAN | Unmodified | 0 | 0 | 38062 | 0 | 0 | 0 | 0 | 0 | 776860 | 545100 | 0 | 107070 | 0 | 0 | 400650 | 61268 | 0 | 0 |
| FFLQGIQLI | LAMP1 | P11279 | LAMP1_HUMAN | Unmodified | 649240 | 526540 | 465980 | 2093300 | 934630 | 0 | 171460 | 0 | 72379000 | 6999800 | 1373700 | 1287000 | 759240 | 1120200 | 2169700 | 399820 | 0 | 751590 |
| FFLSSGLID | TMEM43 | Q9BTV4 | TMM43_HUMA | Unmodified | 120220 | 182820 | 0 | 0 | 0 | 0 | 63387 | 0 | 1.15E+08 | 7095600 | 1423300 | 1633100 | 0 | 0 | 155660 | 91444 | 0 | 0 |
| FFMGNQV | XRCC5 | P13010 | XRCC5_HUMAN | Oxidation (M) | 0 | 0 | 0 | 0 | 0 | 0 | 0 | 314810 | 0 | 0 | 0 | 2143200 | 0 | 0 | 0 | 0 | 0 | 0 |
| FFMGTVVK | PLA2G4A | P47712 | PA24A_HUMAN | Unmodified | 0 | 0 | 0 | 0 | 0 | 0 | 0 | 0 | 0 | 0 | 364420 | 1637400 | 0 | 0 | 0 | 0 | 0 | 0 |
| FFNANQW | FUCA2 | Q9BTY2 | FUCO2_HUMA | Unmodified | 0 | 0 | 0 | 0 | 0 | 0 | 0 | 0 | 0 | 0 | 0 | 0 | 0 | 0 | 0 | 0 | 0 | 0 |
| FFNIPFLQL | RABL6; RABL6; | Q3YEC7; F2 | RABL6_HUMAN | Unmodified | 0 | 0 | 0 | 19219 | 11997 | 0 | 0 | 0 | 686040 | 120830 | 350470 | 1650800 | 0 | 0 | 0 | 14576 | 0 | 0 |
| FFNLNKLEI | ACO1 | P21399 | ACOC_HUMAN | Unmodified | 510130 | 354910 | 78557 | 128250 | 289550 | 105930 | 97247 | 14274 | 0 | 0 | 326960 | 3578400 | 256630 | 205200 | 145600 | 108830 | 220470 | 0 |
| FFPASADR | P4HB; P4HB; P | P07237; A0A | PDIA1_HUMAN | Unmodified | 0 | 0 | 0 | 0 | 0 | 0 | 0 | 26090 | 15439000 | 1533500 | 1029500 | 527870 | 0 | 0 | 0 | 362010 | 0 | 0 |
| FFPASADR | P4HB; P4HB; P | P07237; A0A | PDIA1_HUMAN | Unmodified | 0 | 398460 | 271990 | 0 | 480730 | 0 | 194520 | 128780 | 40784000 | 3574000 | 4117600 | 2672500 | 531840 | 0 | 1545700 | 198440 | 803720 | 222110 |
| FFPASADR | P4HB; P4HB; P | P07237; A0A | PDIA1_HUMAN | Unmodified | 0 | 0 | 0 | 0 | 0 | 0 | 2334100 | 443710 | 11767000 | 6840000 | 2349800 | 0 | 2181300 | 629170 | 0 | 0 | 0 | 0 |
| FFPEDVSEI | RDX; RDX; RDX | P35241; E9P | RADI_HUMAN | Unmodified | 0 | 0 | 0 | 0 | 0 | 0 | 0 | 0 | 0 | 0 | 0 | 19482 | 0 | 0 | 187170 | 0 | 0 | 0 |
| FFPFGLVQI | SIAE | Q9HAT2 | SIAE_HUMAN | Unmodified | 0 | 0 | 0 | 0 | 0 | 0 | 0 | 0 | 262250 | 51374 | 0 | 0 | 0 | 0 | 0 | 0 | 0 | 0 |
| FFPFGLVQI | SIAE | Q9HAT2 | SIAE_HUMAN | Unmodified | 0 | 0 | 0 | 0 | 0 | 0 | 0 | 25279 | 1546600 | 28305 | 794700 | 0 | 0 | 0 | 0 | 0 | 0 | 0 |
| FFPLESWQ | DDT; DDT; DDT | P30046; H7C | DOPD_HUMAN | Unmodified | 1946200 | 979410 | 1277200 | 606560 | 0 | 0 | 0 | 0 | 0 | 0 | 38764 | 14773000 | 0 | 0 | 5315700 | 3733100 | 0 | 0 |
| FFPLIEVNK | RPA1; RPA1 | P27694; I3L | RFA1_HUMAN | Unmodified | 0 | 0 | 0 | 0 | 0 | 0 | 0 | 0 | 0 | 0 | 348190 | 4105700 | 0 | 0 | 0 | 0 | 0 | 0 |
| FFPPDHTQ | FARP1; FARP1; | Q9Y4F1; A0A | FARP1_HUMAN | Unmodified | 0 | 0 | 0 | 0 | 0 | 0 | 0 | 0 | 0 | 0 | 0 | 1541500 | 0 | 0 | 0 | 1996000 | 0 | 0 |
| FFPPDPGQ | FARP2 | O94887 | FARP2_HUMAN | Unmodified | 0 | 0 | 0 | 0 | 0 | 0 | 0 | 0 | 0 | 0 | 0 | 0 | 0 | 0 | 296700 | 159170 | 0 | 0 |
| FFQEENTEI | PSMD12 | O00232 | PSD12_HUMAN | Unmodified | 0 | 0 | 0 | 0 | 642910 | 164200 | 299710 | 286580 | 0 | 0 | 269560 | 1823200 | 0 | 100170 | 0 | 0 | 0 | 0 |
| FFQEFKADI | UBXN6 | Q9BZV1 | UBXN6_HUMAN | Unmodified | 0 | 0 | 0 | 0 | 0 | 0 | 0 | 0 | 0 | 0 | 0 | 0 | 0 | 0 | 0 | 0 | 0 | 0 |
| FFQELPGSI | PDXDC1; PDXD | Q6P996; Q6I | PDXD1_HUMAN | Unmodified | 0 | 0 | 0 | 0 | 0 | 0 | 0 | 0 | 0 | 0 | 0 | 966340 | 0 | 0 | 0 | 0 | 0 | 0 |
| FFQLAAQV | PACS1; PACS1; | Q6VY07; B4I | PACS1_HUMAN | Unmodified | 0 | 0 | 0 | 0 | 857770 | 0 | 212480 | 0 | 0 | 0 | 0 | 0 | 0 | 0 | 0 | 100750 | 0 | 0 |
| FFQPTEMA | AP2A1 | O95782 | AP2A1_HUMAN | Oxidation (M) | 614940 | 651540 | 294110 | 784700 | 67575 | 0 | 0 | 0 | 395030 | 85392 | 310380 | 3409800 | 0 | 0 | 0 | 492450 | 11203 | 0 |
| FFQPTEMA | AP2A1 | O95782 | AP2A1_HUMAN | Unmodified | 0 | 0 | 0 | 0 | 0 | 0 | 0 | 0 | 26640 | 828930 | 0 | 0 | 0 | 0 | 0 | 0 | 1749600 | 1159900 |
| FFQPTEMA | AP2A2; AP2A2 | O94973; A0A | AP2A2_HUMAN | Oxidation (M) | 85873 | 74997 | 81534 | 0 | 0 | 0 | 0 | 71646 | 43960 | 0 | 157870 | 1603800 | 0 | 0 | 159700 | 266490 | 0 | 0 |
| FFQTACDV | IQGAP1; IQGAP | P46940; H0Y | IQGA1_HUMAN | Unmodified | 189690 | 135580 | 0 | 0 | 107700 | 156900 | 0 | 0 | 218570 | 0 | 0 | 734370 | 0 | 0 | 244790 | 160730 | 0 | 0 |
| FFQTCEK | XPR1 | Q9UBH6 | XPR1_HUMAN | Unmodified | 0 | 0 | 0 | 0 | 731170 | 0 | 0 | 0 | 0 | 0 | 239620 | 0 | 0 | 0 | 0 | 0 | 0 | 0 |
| FFREVLLEV | PACSIN2; PACS | Q9UNF0; A0 | PACN2_HUMAN | Unmodified | 31842000 | 18410000 | 15478000 | 37161000 | 11265000 | 1625500 | 1346700 | 36990 | 1078600 | 2323900 | 3028900 | 1473100 | 1.58E+08 | 52168000 | 34082000 | 21774000 | 2035200 | 6868700 |
| FFREVLLEV | PACSIN2; PACS | Q9UNF0; A0 | PACN2_HUMAN | Unmodified | 0 | 0 | 0 | 40373 | 580530 | 0 | 282070 | 0 | 0 | 0 | 0 | 0 | 6253800 | 3301900 | 41719 | 0 | 545420 | 323810 |
| FFRGATEL | HS2ST1 | Q7LGA3 | HS2ST_HUMAN | Unmodified | 0 | 0 | 0 | 0 | 0 | 0 | 58381 | 0 | 3665200 | 351160 | 0 | 0 | 0 | 0 | 0 | 0 | 0 | 0 |
| FFRNGDTA | P4HB; P4HB; P | P07237; A0A | PDIA1_HUMAN | Unmodified | 195410 | 0 | 0 | 195570 | 2990800 | 459730 | 3198100 | 99716 | 16291000 | 26951000 | 1723300 | 0 | 3131900 | 4377500 | 966660 | 536700 | 585070 | 2099400 |
| FFSEREASF | RPS6KA1; RPS6 | Q15418; Q5 | KS6A1_HUMAN | Unmodified | 0 | 0 | 76038 | 205760 | 927430 | 476860 | 658270 | 0 | 424420 | 591470 | 485660 | 404730 | 691010 | 700570 | 349580 | 132210 | 768170 | 1336900 |
| FFSFIEGCL | MAP4K4; MAP4 | O95819; A0A | M4K4_HUMAN | Unmodified | 0 | 0 | 0 | 0 | 0 | 0 | 0 | 0 | 0 | 326790 | 195360 | 507330 | 0 | 0 | 0 | 0 | 0 | 0 |
| FFSPLQK | DECR1; DECR1 | Q16698; E5F | DECR_HUMAN | Unmodified | 428250 | 0 | 0 | 0 | 184180 | 0 | 806470 | 209160 | 4298200 | 1417000 | 0 | 3425900 | 0 | 0 | 0 | 0 | 0 | 834290 |
| FFTDNVPA | MMS19; MMS19 | Q96T76; H0Y | MMS19_HUMA | Unmodified | 0 | 0 | 0 | 0 | 0 | 0 | 0 | 0 | 493510 | 0 | 0 | 2574000 | 0 | 0 | 0 | 0 | 0 | 0 |
| FFTDYGCD | SUSD2 | Q9UGT4 | SUSD2_HUMAN | Unmodified | 0 | 0 | 0 | 0 | 0 | 0 | 0 | 0 | 1417200 | 0 | 0 | 0 | 0 | 0 | 0 | 0 | 0 | 0 |
| FFTEEVDSI | ACAD9; ACAD9 | Q9H845; A0A | ACAD9_HUMAN | Unmodified | 0 | 0 | 0 | 0 | 682650 | 0 | 0 | 0 | 2427900 | 109090 | 0 | 96862 | 0 | 0 | 0 | 0 | 0 | 0 |
| FFTGQITAA | NNT; NNT; NNT | Q13423; E9I | NNTM_HUMAN | Unmodified | 0 | 0 | 0 | 0 | 0 | 0 | 0 | 0 | 2158400 | 253730 | 0 | 0 | 0 | 0 | 0 | 0 | 0 | 0 |
| FFTVKLPVA | RPN1 | P04843 | RPN1_HUMAN | Unmodified | 0 | 0 | 0 | 683570 | 0 | 0 | 250860 | 0 | 4.63E+08 | 14801000 | 3816000 | 0 | 0 | 0 | 0 | 0 | 0 | 0 |
| FFVADTAN | SUCLG2; SUCL | Q96I99; E9P | SUCB2_HUMAN | Unmodified | 0 | 0 | 0 | 0 | 0 | 0 | 0 | 0 | 4804100 | 1269000 | 0 | 0 | 0 | 0 | 0 | 0 | 0 | 0 |
| FFVAVPGQ | SPAG9; SPAG9 | O60271; A0A | JIP4_HUMAN C | Unmodified | 0 | 0 | 0 | 0 | 0 | 0 | 447980 | 0 | 685020 | 138360 | 309750 | 1625700 | 0 | 0 | 62975 | 218620 | 0 | 1708700 |
| FFVGGNW | TPI1 | P60174 | TPIS_HUMAN T | Unmodified | 4822000 | 3333000 | 3736900 | 8223100 | 95034000 | 10046000 | 5876500 | 1839300 | 4626300 | 8091400 | 4267800 | 44448000 | 48423000 | 29481000 | 6579000 | 7093700 | 37616000 | 69169000 |
| FFVSSSQG | BSG; BSG; BSG | P35613; A0A | BASI_HUMAN E | Unmodified | 3706000 | 1827900 | 18074000 | 189130 | 375580 | 0 | 0 | 497430 | 13468000 | 0 | 624540 | 39477000 | 0 | 0 | 21338000 | 25497000 | 279210 | 1062900 |
| FFVSSSQG | BSG; BSG; BSG | P35613; A0A | BASI_HUMAN E | Oxidation (M) | 275990 | 379340 | 0 | 3076200 | 0 | 0 | 0 | 0 | 4276000 | 0 | 0 | 0 | 0 | 0 | 3574700 | 1547500 | 0 | 0 |
| FFVTLLEAG | ARHGAP17 | Q68EM7 | RHG17_HUMAN | Unmodified | 316510 | 369150 | 260110 | 493510 | 905610 | 0 | 0 | 0 | 0 | 0 | 280350 | 1537800 | 1062700 | 1030600 | 372040 | 325330 | 163000 | 0 |
| FFVTTLPAF | TMX4 | Q9H1E5 | TMX4_HUMAN | Unmodified | 0 | 0 | 0 | 0 | 0 | 0 | 0 | 0 | 13704000 | 729050 | 0 | 274710 | 0 | 0 | 0 | 0 | 16412 | 0 |
| FFYAGTPP | ENDOD1 | O94919 | ENDD1_HUMA | Unmodified | 0 | 0 | 0 | 1631800 | 1028000 | 0 | 0 | 0 | 14926000 | 0 | 0 | 0 | 695650 | 362960 | 0 | 0 | 0 | 449130 |
| FFYLLEPGV | ST14 | Q9Y5Y6 | ST14_HUMAN | Unmodified | 0 | 0 | 2378000 | 1977000 | 0 | 0 | 0 | 0 | 1781300 | 248150 | 322490 | 0 | 1872100 | 1838000 | 2336000 | 2617800 | 0 | 0 |
| FFYSDQNV | TLN1 | Q9Y490 | TLN1_HUMAN | Unmodified | 0 | 0 | 0 | 0 | 9304200 | 0 | 0 | 0 | 0 | 0 | 0 | 0 | 2760400 | 2510600 | 0 | 0 | 0 | 7449000 |
| FGAAVLGN | FKBP15 | Q5T1M5 | FKB15_HUMAN | Unmodified | 0 | 0 | 0 | 0 | 0 | 0 | 0 | 0 | 0 | 0 | 283950 | 403920 | 0 | 0 | 0 | 0 | 0 | 0 |
| FGADDKNI | SYT4 | Q9H2B2 | SYT4_HUMAN | Unmodified | 0 | 0 | 0 | 0 | 0 | 0 | 0 | 0 | 0 | 0 | 0 | 0 | 0 | 0 | 581080 | 0 | 0 | 0 |
| FGALCEAE | AGRN; AGRN; A | O00468; A0A | AGRIN_HUMAN | Unmodified | 615290 | 159870 | 1194100 | 506610 | 0 | 0 | 0 | 0 | 838450 | 0 | 0 | 835430 | 238490 | 0 | 516380 | 410560 | 0 | 0 |
| FGALTAEK | LYPLA2; LYPLA | O95372; Q5I | LYPA2_HUMAN | Unmodified | 0 | 0 | 146980 | 0 | 0 | 0 | 0 | 0 | 123600 | 0 | 165710 | 7363900 | 0 | 0 | 238470 | 269360 | 67404 | 0 |
| FGAPPHAG | DARS1; DARS1 | P14868; H7E | SYDC_HUMAN | Unmodified | 0 | 0 | 0 | 0 | 0 | 0 | 0 | 0 | 306360 | 0 | 360740 | 6725300 | 0 | 0 | 0 | 289310 | 0 | 0 |
| FGAQLQCV | PLXNB2 | O15031 | PLXB2_HUMAN | Unmodified | 0 | 0 | 339850 | 0 | 0 | 0 | 0 | 0 | 0 | 0 | 0 | 0 | 0 | 0 | 0 | 741880 | 0 | 0 |
| FGATLEIVT | ETF1; ETF1 | P62495; B7Z | ERF1_HUMAN | Unmodified | 81925 | 107480 | 0 | 67475 | 464100 | 0 | 276230 | 0 | 0 | 160800 | 241900 | 0 | 382150 | 221670 | 0 | 0 | 859580 | 0 |

| | | | | | | | | | | | | | | | | | | | | | | |
|---|---|---|---|---|---|---|---|---|---|---|---|---|---|---|---|---|---|---|---|---|---|---|
| FGAVEGAV | PTGES2; PTGES | Q9H7Z7; A0 | PGES2_HUMAN | Unmodified | 0 | 0 | 49638 | 0 | 0 | 0 | 0 | 0 | 0 | 0 | 347450 | 0 | 0 | 0 | 286120 | 118810 | 0 | 0 |
| FGAVFLWD | WDR1; WDR1 | O75083; D6 | WDR1_HUMAN | Unmodified | 0 | 0 | 1393400 | 0 | 77172000 | 11786000 | 6700600 | 230140 | 8185000 | 5207600 | 5786400 | 15670000 | 49228000 | 26263000 | 9722300 | 10089000 | 14505000 | 24453000 |
| FGAVFLWD | WDR1; WDR1 | O75083; D6 | WDR1_HUMAN | Unmodified | 0 | 0 | 0 | 0 | 10892000 | 0 | 1557400 | 0 | 0 | 494930 | 0 | 0 | 3696300 | 1074500 | 0 | 0 | 0 | 0 |
| FGAVWTG | GANAB; GANAE | Q14697; F5 | GANAB_HUMA | Unmodified | 0 | 0 | 864810 | 0 | 0 | 0 | 0 | 0 | 60400000 | 7278800 | 2062900 | 2612000 | 0 | 0 | 0 | 0 | 0 | 0 |
| FGCPWGV | MAN2A1 | Q16706 | MA2A1_HUMA | Unmodified | 0 | 0 | 0 | 0 | 774600 | 0 | 0 | 0 | 4260000 | 759430 | 527240 | 2139600 | 0 | 0 | 0 | 0 | 0 | 2879700 |
| FGDEDLTW | C1orf198; C1or | Q9H425; E5 | CA198_HUMA | Unmodified | 0 | 1172500 | 0 | 0 | 0 | 0 | 0 | 120170 | 754780 | 0 | 0 | 2449500 | 0 | 0 | 1513600 | 112290 | 0 | 0 |
| FGDEFLGE | RASA3 | Q14644 | RASA3_HUMA | Unmodified | 0 | 0 | 0 | 0 | 0 | 90422 | 286430 | 0 | 0 | 657050 | 0 | 253000 | 0 | 0 | 63135 | 35349 | 1830400 | 2534000 |
| FGDFVLK | UMPS; UMPS; | P11172; F2Z | UMPS_HUMAN | Unmodified | 0 | 0 | 0 | 0 | 308970 | 58219 | 122510 | 56800 | 822090 | 224210 | 365330 | 1078400 | 0 | 123920 | 0 | 0 | 0 | 150840 |
| FGDILHVIN | DLG1; DLG1; D | Q12959; A0 | DLG1_HUMAN | Unmodified | 836210 | 0 | 0 | 0 | 1270900 | 366630 | 0 | 0 | 5762100 | 442490 | 672040 | 661580 | 404270 | 0 | 1495300 | 720770 | 0 | 0 |
| FGDLDEQE | DOCK8; DOCK | Q8NF50; Q9 | DOCK8_HUMA | Unmodified | 3064800 | 0 | 0 | 0 | 0 | 0 | 0 | 0 | 0 | 0 | 0 | 2345200 | 0 | 0 | 0 | 696720 | 0 | 0 |
| FGDLILK | PRPF8 | Q6P2Q9 | PRP8_HUMAN | Unmodified | 0 | 0 | 0 | 0 | 0 | 0 | 52801 | 0 | 0 | 0 | 133080 | 2715900 | 0 | 0 | 0 | 0 | 52295 | 0 |
| FGDPECQ | GART | P22102 | PUR2_HUMAN | Unmodified | 0 | 0 | 80722 | 250070 | 0 | 0 | 0 | 0 | 262170 | 0 | 294760 | 3397000 | 0 | 0 | 0 | 116940 | 0 | 0 |
| FGDPVVQS | HSPA1A; HSPA | P0DMV8; P0 | HS71A_HUMA | Oxidation (M) | 0 | 1778300 | 2251500 | 1319800 | 1772600 | 0 | 470590 | 0 | 3289400 | 752920 | 115440 | 1870900 | 1968700 | 0 | 1589400 | 1747200 | 0 | 1319000 |
| FGDPVVQS | HSPA1A; HSPA | P0DMV8; P0 | HS71A_HUMA | Unmodified | 2498500 | 3311300 | 1615000 | 1812100 | 5331000 | 1862900 | 355720 | 169320 | 11422000 | 1965500 | 1710100 | 6509300 | 4324000 | 2223400 | 1787400 | 1902200 | 598270 | 3043300 |
| FGDQDVW | MRPL46; MRPL | Q9H2W6; A | RM46_HUMAN | Unmodified | 0 | 0 | 0 | 0 | 0 | 0 | 0 | 0 | 945480 | 0 | 0 | 0 | 0 | 0 | 0 | 0 | 0 | 0 |
| FGDQLLQI | SDCBP2 | Q9H190 | SDCB2_HUMA | Unmodified | 243720 | 286990 | 0 | 0 | 0 | 0 | 162130 | 0 | 0 | 92096 | 0 | 0 | 0 | 0 | 596330 | 464000 | 0 | 0 |
| FGDQVLQI | SDCBP; SDCBP | O00560; G5 | SDCB1_HUMA | Unmodified | 46049000 | 37873000 | 28490000 | 58177000 | 10998000 | 5320000 | 584430 | 62979 | 1.95E+08 | 13385000 | 8726700 | 43855000 | 10194000 | 9072600 | 52608000 | 38342000 | 266950 | 2253600 |
| FGDYNKEV | EZR; EZR | P15311; E7E | EZRI_HUMAN E | Unmodified | 0 | 0 | 0 | 86264 | 2587100 | 419270 | 323120 | 197930 | 0 | 0 | 0 | 0 | 0 | 683150 | 0 | 0 | 3194400 | 577870 |
| FGEDVVEV | DOCK7; DOCK | Q96N67; A0 | DOCK7_HUMA | Unmodified | 0 | 0 | 0 | 31994 | 0 | 0 | 0 | 0 | 0 | 83563 | 0 | 2138900 | 0 | 0 | 0 | 238020 | 0 | 0 |
| FGEEIAR | PDCD6IP; PDCI | Q8WUM4; F | PDC6I_HUMA | Unmodified | 15282000 | 15918000 | 7261400 | 7168700 | 1179400 | 90666 | 223180 | 0 | 6338900 | 823610 | 868820 | 5199100 | 826060 | 322260 | 9597400 | 9382300 | 170740 | 111450 |
| FGEEIARL | PDCD6IP | Q8WUM4 | PDC6I_HUMA | Unmodified | 4124100 | 0 | 0 | 10360000 | 8197100 | 2550100 | 0 | 0 | 0 | 8535900 | 0 | 0 | 11444000 | 8178900 | 0 | 0 | 0 | 4409100 |
| FGEENIEV | TXNRD1; TXNR | Q16881; A0 | TRXR1_HUMA | Unmodified | 0 | 0 | 0 | 0 | 5183200 | 0 | 0 | 0 | 549780 | 851710 | 0 | 451010 | 2802500 | 0 | 0 | 0 | 0 | 0 |
| FGEGVSSP | VGF; VGF | O15240; Q0 | VGF_HUMAN N | Unmodified | 0 | 0 | 0 | 0 | 905280 | 66765 | 49243 | 0 | 14383000 | 1499400 | 0 | 0 | 0 | 0 | 0 | 0 | 0 | 538860 |
| FGELVMTK | HADHA; HADH | P40939; A0A | ECHA_HUMAN | Unmodified | 0 | 0 | 0 | 0 | 0 | 0 | 81764 | 0 | 3649500 | 185580 | 0 | 0 | 0 | 0 | 0 | 0 | 0 | 0 |
| FGEMQLDF | STT3B | Q8TCJ2 | STT3B_HUMA | Oxidation (M) | 0 | 0 | 0 | 0 | 0 | 0 | 0 | 0 | 2777500 | 577690 | 618530 | 0 | 0 | 0 | 0 | 0 | 0 | 0 |
| FGEMQLDF | STT3B | Q8TCJ2 | STT3B_HUMA | Unmodified | 0 | 0 | 0 | 0 | 0 | 0 | 0 | 0 | 6147700 | 609120 | 0 | 0 | 0 | 0 | 0 | 0 | 0 | 0 |
| FGEMQLDF | STT3B | Q8TCJ2 | STT3B_HUMA | Oxidation (M) | 0 | 0 | 0 | 0 | 0 | 0 | 0 | 0 | 4158800 | 0 | 0 | 0 | 0 | 0 | 0 | 0 | 0 | 0 |
| FGEMQLDF | STT3B | Q8TCJ2 | STT3B_HUMA | Unmodified | 0 | 0 | 0 | 0 | 0 | 0 | 0 | 0 | 3148100 | 0 | 0 | 0 | 0 | 0 | 0 | 0 | 0 | 0 |
| FGEPVLAG | PFAS | O15067 | PUR4_HUMAN | Unmodified | 0 | 0 | 0 | 0 | 0 | 0 | 0 | 0 | 0 | 0 | 0 | 2584800 | 0 | 0 | 0 | 0 | 0 | 0 |
| FGESIEDL | PIGX; PIGX; PIG | Q8TBF5; C9 | PIGX_HUMAN | Unmodified | 0 | 0 | 0 | 0 | 0 | 0 | 0 | 0 | 2266200 | 170200 | 0 | 0 | 0 | 0 | 0 | 0 | 0 | 0 |
| FGEVVDCT | HNRNPAB; HN | Q99729; D6 | ROAA_HUMAN | Unmodified | 118930 | 90270 | 153020 | 0 | 83903 | 72613 | 71587 | 0 | 542600 | 390900 | 209530 | 0 | 463060 | 182740 | 380820 | 0 | 86122 | 373520 |
| FGEVVDCT | HNRNPDL; HN | O14979; A0 | HNRDL_HUMA | Unmodified | 0 | 0 | 0 | 0 | 0 | 0 | 325720 | 0 | 565470 | 188660 | 809010 | 3186300 | 159010 | 0 | 0 | 0 | 0 | 0 |
| FGEVVDCT | HNRNPD; HNR | Q14103; H0 | HNRPD_HUMA | Unmodified | 0 | 0 | 0 | 0 | 0 | 61405 | 0 | 0 | 1950600 | 525240 | 874120 | 7220400 | 0 | 0 | 0 | 0 | 0 | 0 |
| FGEVVDCT | HNRNPD; HNR | Q14103; H0 | HNRPD_HUMA | Unmodified | 1153500 | 0 | 0 | 847290 | 1465500 | 346270 | 1466300 | 33664 | 3930000 | 3169600 | 10123000 | 25246000 | 3510600 | 0 | 0 | 476940 | 130680 | 373500 |
| FGFCPMA | FN1 | P02751 | FINC_HUMAN | 2 Oxidation (M) | 0 | 98454 | 3170100 | 340530 | 0 | 0 | 0 | 0 | 205910 | 0 | 0 | 0 | 0 | 311610 | 0 | 886890 | 0 | 0 |
| FGFCPMA | FN1 | P02751 | FINC_HUMAN | Oxidation (M) | 0 | 0 | 597400 | 651250 | 0 | 0 | 0 | 0 | 0 | 0 | 0 | 0 | 0 | 0 | 0 | 217060 | 0 | 0 |
| FGFCPMA | FN1 | P02751 | FINC_HUMAN | 2 Oxidation (M) | 0 | 0 | 0 | 0 | 0 | 560070 | 0 | 0 | 0 | 835710 | 0 | 0 | 0 | 4408200 | 0 | 0 | 0 | 2177700 |
| FGFEIIETK | NAA50; NAA50; | Q9GZZ1; A0 | NAA50_HUMA | Unmodified | 0 | 0 | 0 | 0 | 0 | 0 | 337580 | 80506 | 0 | 234960 | 0 | 7982200 | 149860 | 0 | 0 | 0 | 194130 | 0 |
| FGFELPQG | TXNIP | Q9H3M7 | TXNIP_HUMAN | Unmodified | 0 | 928670 | 135950 | 268100 | 7553.4 | 0 | 0 | 0 | 435970 | 0 | 256530 | 1893300 | 167700 | 0 | 246250 | 739900 | 70980 | 475570 |
| FGFGLLNA | PCSK1 | P29120 | NEC1_HUMAN | Unmodified | 0 | 0 | 0 | 0 | 0 | 0 | 0 | 0 | 2005100 | 584370 | 595920 | 0 | 0 | 0 | 0 | 0 | 0 | 0 |
| FGFPEGSV | RPS3; RPS3; RP | P23396; H0 | RS3_HUMAN 4 | Unmodified | 2145300 | 1424300 | 1171800 | 1935300 | 0 | 0 | 0 | 429380 | 385700 | 258780 | 902190 | 13913000 | 918710 | 0 | 1139700 | 1204400 | 0 | 0 |
| FGFSLPYV | B4GALT1 | P15291 | B4GT1_HUMA | Unmodified | 0 | 0 | 0 | 0 | 0 | 86412 | 0 | 0 | 0 | 185020 | 0 | 0 | 0 | 0 | 0 | 0 | 0 | 0 |
| FGFYEVFK | SLC25A3; SLC2 | Q00325; F8 | MPCP_HUMAN | Unmodified | 0 | 0 | 0 | 0 | 0 | 0 | 0 | 97890 | 6423100 | 8696000 | 253360 | 0 | 0 | 0 | 0 | 0 | 0 | 0 |
| FGGALDAA | ACLY | P53396 | ACLY_HUMAN | Unmodified | 0 | 2016700 | 201630 | 0 | 0 | 0 | 69142 | 0 | 291960 | 0 | 116710 | 6977900 | 0 | 0 | 316840 | 0 | 30573 | 0 |
| FGGEHVPN | FLNA; FLNA; FL | P21333; H0 | FLNA_HUMAN | Unmodified | 0 | 0 | 0 | 452160 | 9169100 | 3374600 | 0 | 0 | 0 | 0 | 524780 | 6222100 | 0 | 0 | 1018700 | 691080 | 7480500 | 696800 |
| FGGGNPEI | HADHA; HADH | P40939; A0A | ECHA_HUMAN | Unmodified | 0 | 0 | 0 | 0 | 0 | 0 | 0 | 0 | 2189200 | 0 | 0 | 0 | 0 | 0 | 0 | 0 | 0 | 0 |
| FGGNPGG | TARDBP; TARD | Q13148; A0 | TADBP_HUMA | Unmodified | 721260 | 0 | 0 | 0 | 210670 | 0 | 0 | 0 | 287440 | 0 | 0 | 77773 | 0 | 0 | 0 | 0 | 78904 | 0 |
| FGGPVHH | ELAVL1 | Q15717 | ELAV1_HUMA | Unmodified | 0 | 0 | 142430 | 204830 | 0 | 0 | 0 | 0 | 65635 | 0 | 74987 | 349350 | 209350 | 62176 | 97241 | 183980 | 0 | 0 |
| FGGSGSQ | PSMD2 | Q13200 | PSMD2_HUMA | Unmodified | 0 | 0 | 202720 | 145030 | 0 | 0 | 0 | 0 | 143080 | 0 | 275450 | 2207800 | 0 | 0 | 168630 | 155700 | 0 | 0 |
| FGHEFLEF | MAGOH; MAGC | P61326; Q9 | MGN_HUMAN | Unmodified | 309870 | 762460 | 0 | 0 | 0 | 0 | 0 | 0 | 0 | 0 | 0 | 855820 | 0 | 0 | 473770 | 0 | 0 | 0 |
| FGIAAKYQ | VDAC2; VDAC2 | P45880; A0A | VDAC2_HUMA | Unmodified | 0 | 0 | 0 | 0 | 0 | 0 | 0 | 0 | 568570 | 1477200 | 0 | 0 | 0 | 0 | 0 | 0 | 0 | 0 |
| FGIAVLGYL | ITGA2; ITGA2; I | P17301; E9P | ITA2_HUMAN I | Unmodified | 3262700 | 5234500 | 0 | 0 | 0 | 0 | 0 | 0 | 0 | 0 | 0 | 254340 | 0 | 0 | 5752400 | 2975000 | 0 | 0 |
| FGIAVLGYL | ITGA2; ITGA2; I | P17301; E9P | ITA2_HUMAN I | Unmodified | 0 | 0 | 310100 | 2296100 | 6896400 | 2536800 | 469430 | 0 | 0 | 0 | 0 | 0 | 1832600 | 575320 | 0 | 0 | 3295600 | 5137000 |
| FGIDDQDF | PIP4K2A | P48426 | PI42A_HUMAN | Unmodified | 0 | 0 | 0 | 0 | 145850 | 55107 | 0 | 0 | 222900 | 74103 | 225810 | 1199200 | 325390 | 81419 | 0 | 0 | 0 | 418890 |
| FGIDDQDY | PIP4K2B; PIP4K | P78356; A0A | PI42B_HUMAN | Unmodified | 0 | 0 | 0 | 0 | 0 | 0 | 0 | 0 | 634890 | 0 | 100160 | 726730 | 0 | 0 | 0 | 132000 | 0 | 0 |
| FGILSNCN | MKRN3 | Q13064 | MKRN3_HUMA | Unmodified | 488270 | 0 | 0 | 0 | 0 | 0 | 0 | 0 | 0 | 0 | 0 | 0 | 0 | 0 | 0 | 0 | 0 | 0 |
| FGILSNCN | MKRN1 | Q9UHC7 | MKRN1_HUMA | Unmodified | 5676000 | 2258300 | 688050 | 990900 | 0 | 0 | 0 | 0 | 0 | 0 | 0 | 0 | 819430 | 0 | 775310 | 893840 | 0 | 405540 |
| FGILSNCN | MKRN2; MKRN | Q9H000; C9 | MKRN2_HUMA | Unmodified | 336500 | 0 | 0 | 149610 | 0 | 0 | 0 | 0 | 0 | 0 | 0 | 0 | 36245 | 0 | 0 | 71859 | 0 | 0 |
| FGILSTGQ | VPS37D; VPS37 | Q86XT2; C9 | VP37D_HUMA | Unmodified | 886690 | 503710 | 0 | 0 | 0 | 0 | 0 | 25647 | 0 | 0 | 0 | 0 | 0 | 0 | 2279600 | 1368200 | 0 | 0 |
| FGILSTGQ | VPS37D; VPS37 | Q86XT2; C9 | VP37D_HUMA | Unmodified | 1764800 | 0 | 0 | 0 | 0 | 0 | 0 | 0 | 0 | 0 | 0 | 0 | 0 | 0 | 4026700 | 0 | 0 | 0 |
| FGIQAQMV | PSMB3; PSMB3 | P49720; A0A | PSB3_HUMAN | Oxidation (M) | 0 | 0 | 0 | 42513 | 0 | 0 | 0 | 0 | 0 | 67786 | 265700 | 1172200 | 0 | 50339 | 0 | 0 | 0 | 259060 |
| FGIYDIDN | CPNE1; CPNE1 | Q99829; Q5 | CPNE1_HUMA | Unmodified | 44709 | 0 | 0 | 13510 | 0 | 0 | 0 | 0 | 60910 | 0 | 0 | 516020 | 0 | 0 | 136710 | 117360 | 0 | 0 |
| FGKITNDF | EIF3B; EIF3B | P55884; C9 | EIF3B_HUMAN | Unmodified | 0 | 0 | 0 | 0 | 0 | 0 | 0 | 0 | 0 | 391580 | 0 | 0 | 0 | 492070 | 0 | 0 | 418400 | 302090 |
| FGKPDPIV | MYOF | Q9NZM1 | MYOF_HUMAN | Unmodified | 0 | 0 | 0 | 0 | 16066000 | 2970900 | 0 | 0 | 0 | 0 | 0 | 0 | 0 | 0 | 0 | 0 | 0 | 1966400 |
| FGLESHAH | ABCF1; ABCF1; | Q8NE71; H0 | ABCF1_HUMA | Unmodified | 0 | 0 | 117910 | 0 | 0 | 0 | 0 | 0 | 0 | 0 | 142500 | 6659400 | 112970 | 0 | 201080 | 187950 | 0 | 0 |
| FGLFSAEIK | PI4KA; PI4KAP2 | P42356; A4 | PI4KA_HUMAN | Unmodified | 0 | 0 | 162330 | 0 | 0 | 0 | 0 | 0 | 0 | 85120 | 0 | 430780 | 0 | 0 | 909360 | 407510 | 0 | 289650 |
| FGLGPAPF | NUDT19 | A8MXV4 | NUD19_HUMA | Unmodified | 0 | 0 | 0 | 0 | 0 | 0 | 0 | 0 | 1679200 | 321200 | 0 | 0 | 0 | 0 | 0 | 0 | 0 | 0 |
| FGLGSVAG | SLC25A13 | Q9UJS0 | CMC2_HUMAN | Unmodified | 0 | 0 | 0 | 0 | 0 | 0 | 0 | 0 | 818830 | 362970 | 0 | 0 | 0 | 0 | 0 | 0 | 0 | 0 |
| FGLIQSMK | IST1; IST1; IST1 | P53990; H3 | IST1_HUMAN I | Unmodified | 118360 | 0 | 0 | 0 | 537370 | 0 | 0 | 0 | 1881300 | 581140 | 0 | 4063600 | 0 | 0 | 0 | 95956 | 0 | 0 |
| FGLIQSMK | IST1; IST1; IST1 | P53990; H3 | IST1_HUMAN I | Oxidation (M) | 0 | 0 | 0 | 0 | 0 | 0 | 0 | 0 | 0 | 237940 | 0 | 5996800 | 0 | 0 | 0 | 0 | 0 | 0 |
| FGLLNIVM | GNPAT; GNPAT | O15228; Q5 | GNPAT_HUMA | Unmodified | 0 | 0 | 0 | 0 | 0 | 0 | 0 | 0 | 716000 | 39969 | 0 | 0 | 0 | 0 | 0 | 0 | 0 | 0 |
| FGLTFQSP | SPRED2; SPREI | Q7Z698; Q7 | SPRE2_HUMA | Unmodified | 116590 | 0 | 149070 | 281610 | 0 | 0 | 0 | 0 | 0 | 0 | 0 | 0 | 0 | 0 | 499660 | 615370 | 336320 | 451370 |
| FGLYLPLFK | SLC25A13 | Q9UJS0 | CMC2_HUMAN | Unmodified | 0 | 0 | 0 | 78376 | 0 | 0 | 0 | 0 | 1989000 | 544950 | 18777 | 0 | 32005 | 0 | 0 | 0 | 0 | 0 |
| FGMFEFLS | SLC25A1 | P53007 | TXTP_HUMAN | Oxidation (M) | 0 | 0 | 0 | 0 | 0 | 0 | 0 | 0 | 544920 | 0 | 0 | 0 | 0 | 0 | 0 | 0 | 0 | 0 |
| FGMKPYTS | PTGS1; PTGS1; | P23219; A0A | PGH1_HUMAN | Oxidation (M) | 0 | 0 | 0 | 0 | 0 | 0 | 0 | 0 | 0 | 0 | 509040 | 847390 | 0 | 0 | 0 | 0 | 0 | 0 |
| FGMTPSKG | VCP; VCP; VCP; | P55072; C9 | TERA_HUMAN | Oxidation (M) | 0 | 182580 | 87210 | 0 | 0 | 647220 | 350360 | 436160 | 0 | 0 | 0 | 0 | 324610 | 360630 | 0 | 0 | 459270 | 114340 |
| FGMTPSKG | VCP; VCP; VCP; | P55072; C9 | TERA_HUMAN | Unmodified | 0 | 0 | 0 | 0 | 0 | 0 | 0 | 0 | 0 | 0 | 0 | 77860 | 0 | 0 | 0 | 0 | 385230 | 1508100 |
| FGNAFLNR | EHD4; EHD1; E | Q9H223; Q9 | EHD4_HUMAN | Unmodified | 2423200 | 1822100 | 1748200 | 1268400 | 0 | 0 | 0 | 0 | 733910 | 0 | 976270 | 1876000 | 0 | 0 | 3097600 | 3064200 | 0 | 0 |
| FGNDGLHI | PUS1; PUS1; PU | Q9Y606; F5 | TRUA_HUMAN | Unmodified | 0 | 0 | 0 | 0 | 0 | 0 | 0 | 0 | 448760 | 0 | 0 | 936080 | 0 | 0 | 0 | 0 | 0 | 0 |
| FGNDVQHI | GRB2; GRB2 | P62993; J3K | GRB2_HUMAN | Unmodified | 0 | 0 | 1212900 | 0 | 243640 | 0 | 0 | 0 | 0 | 0 | 119770 | 473360 | 0 | 0 | 3001200 | 547130 | 0 | 0 |
| FGNELSAD | PLCB4; PLCB4 | Q15147; A0 | PLCB4_HUMA | Unmodified | 0 | 0 | 0 | 0 | 0 | 0 | 0 | 0 | 0 | 0 | 0 | 0 | 0 | 0 | 1171700 | 1462900 | 0 | 0 |
| FGNEVIPV | ACAT1; ACAT1; | P24752; A0A | THIL_HUMAN | Unmodified | 0 | 0 | 0 | 0 | 0 | 0 | 0 | 0 | 4831900 | 177270 | 0 | 0 | 0 | 0 | 0 | 0 | 0 | 0 |
| FGNPLLVQ | DYNC1H1; DYN | Q14204; A0 | DYHC1_HUMA | Unmodified | 135260 | 0 | 0 | 331940 | 0 | 0 | 0 | 0 | 1232200 | 651330 | 384870 | 1365400 | 0 | 0 | 1079600 | 132970 | 0 | 0 |
| FGNQADH | POFUT1 | Q9H488 | OFUT1_HUMA | Unmodified | 0 | 0 | 0 | 0 | 0 | 0 | 0 | 13999 | 17327000 | 2811300 | 1032300 | 572050 | 0 | 0 | 0 | 0 | 0 | 0 |
| FGNTTFKT | C2CD5 | Q86YS7 | C2CD5_HUMA | Unmodified | 0 | 73186 | 0 | 0 | 0 | 0 | 0 | 74436 | 377580 | 0 | 74672 | 224700 | 108890 | 0 | 0 | 0 | 181050 | 228870 |
| FGPALSVK | PABPC1; PABP | P11940; Q9 | PABP1_HUMA | Unmodified | 1.6E+08 | 1.04E+08 | 52081000 | 63157000 | 0 | 0 | 586280 | 453700 | 0 | 0 | 3325700 | 13680000 | 12478000 | 4012800 | 26045000 | 46245000 | 0 | 0 |
| FGPGVIEN | FKRP | Q9H9S5 | FKRP_HUMAN | Unmodified | 0 | 0 | 0 | 0 | 0 | 0 | 0 | 0 | 4448600 | 916590 | 0 | 0 | 195980 | 0 | 0 | 0 | 0 | 0 |
| FGPYYTEP | PSMB3; PSMB3 | P49720; A0A | PSB3_HUMAN | Unmodified | 0 | 0 | 0 | 0 | 0 | 0 | 0 | 0 | 323120 | 97712 | 523520 | 2781300 | 0 | 0 | 0 | 0 | 0 | 0 |
| FGQAEKTE | SH3GLB2; SH3 | Q9NR46; B7 | SHLB2_HUMA | Unmodified | 0 | 0 | 0 | 0 | 0 | 0 | 0 | 0 | 686970 | 267970 | 0 | 495650 | 0 | 0 | 0 | 0 | 0 | 0 |
| FGQFCTVG | PMPCA | Q10713 | MPPA_HUMAN | Unmodified | 0 | 0 | 0 | 0 | 0 | 0 | 0 | 0 | 1996800 | 0 | 245450 | 182540 | 0 | 0 | 0 | 0 | 0 | 0 |
| FGQGDLP | RANBP2 | P49792 | RBP2_HUMAN | Unmodified | 0 | 0 | 0 | 0 | 0 | 1201800 | 0 | 0 | 178030 | 0 | 0 | 1563900 | 101100 | 0 | 0 | 0 | 0 | 102580 |
| FGQGGAG | SFPQ | P23246 | SFPQ_HUMAN | Unmodified | 62770 | 17196 | 0 | 0 | 0 | 0 | 8452.3 | 0 | 508130 | 169970 | 1028300 | 2457700 | 0 | 0 | 0 | 0 | 61214 | 213530 |
| FGQIVNTL | EPHB2; EPHB2 | P29323; B1 | EPHB2_HUMA | Unmodified | 0 | 0 | 0 | 114510 | 214920 | 0 | 1001900 | 390950 | 946220 | 632260 | 0 | 0 | 306550 | 0 | 0 | 0 | 1760100 | 1636100 |
| FGQMLGS | DYNC1H1; DYN | Q14204; A0 | DYHC1_HUMA | Oxidation (M) | 0 | 0 | 0 | 0 | 395460 | 0 | 0 | 0 | 555960 | 330500 | 0 | 1507500 | 339220 | 278720 | 0 | 0 | 114150 | 286550 |
| FGQQNPR | ZC3H15; ZC3H | Q8WU90; F8 | ZC3HF_HUMA | Unmodified | 0 | 0 | 0 | 0 | 410260 | 0 | 91533 | 95295 | 0 | 31860 | 0 | 1424700 | 137850 | 129940 | 0 | 0 | 162140 | 255840 |
| FGQVYTEA | STT3A | P46977 | STT3A_HUMA | Unmodified | 0 | 0 | 0 | 0 | 72712 | 0 | 0 | 0 | 9327900 | 1085700 | 798860 | 405930 | 0 | 0 | 0 | 0 | 0 | 0 |
| FGQYAGH | CKAP5 | Q14008 | CKAP5_HUMA | Unmodified | 0 | 0 | 0 | 0 | 260260 | 520900 | 0 | 0 | 0 | 0 | 0 | 3146100 | 32048 | 0 | 0 | 0 | 50617 | 0 |
| FGQYNKE | SEC23B; SEC2 | Q15437; A0 | SC23B_HUMA | Unmodified | 215050 | 113240 | 0 | 0 | 0 | 0 | 0 | 0 | 0 | 0 | 0 | 3171300 | 0 | 0 | 93380 | 121880 | 40382 | 0 |
| FGRDEDPV | TXNL4B; TXNL4 | Q9NX01; H3 | TXN4B_HUMA | Unmodified | 309540 | 0 | 0 | 107150 | 0 | 0 | 255890 | 0 | 2266500 | 0 | 342520 | 0 | 149360 | 0 | 0 | 112190 | 36841 | 0 |
| FGRQTGG | ZNF330 | Q9Y3S2 | ZN330_HUMA | Unmodified | 0 | 0 | 0 | 0 | 0 | 0 | 0 | 0 | 0 | 0 | 0 | 0 | 237630 | 296910 | 0 | 0 | 208280 | 370240 |
| FGSAIAPL | ITGAV | P06756 | ITAV_HUMAN I | Unmodified | 0 | 0 | 234430 | 2186000 | 600340 | 313460 | 0 | 0 | 2636900 | 0 | 0 | 0 | 310290 | 0 | 1486700 | 0 | 0 | 0 |
| FGSAIPIPS | SNX9; SNX9; S | Q9Y5X1; A0 | SNX9_HUMAN | Unmodified | 33046000 | 15792000 | 0 | 0 | 1113200 | 0 | 0 | 0 | 1378300 | 428420 | 1618600 | 8735100 | 19893000 | 1435700 | 0 | 7622400 | 471650 | 0 |
| FGSALVPT | QTRT1; QTRT1 | Q9BXR0; K7 | TGT_HUMAN Q | Unmodified | 0 | 0 | 0 | 0 | 0 | 0 | 0 | 0 | 0 | 0 | 0 | 1679200 | 0 | 0 | 0 | 0 | 0 | 0 |
| FGSCQQG | ITGA6; ITGA6 | P23229; C9 | ITA6_HUMAN I | Unmodified | 2223200 | 480570 | 1388000 | 2981100 | 18023000 | 5627100 | 5246500 | 4453400 | 343430 | 22726 | 487490 | 7958700 | 14404000 | 13011000 | 3257000 | 2851900 | 42150000 | 69309000 |
| FGSCQQG | ITGA6; ITGA6 | P23229; C9 | ITA6_HUMAN I | Unmodified | 5278500 | 0 | 0 | 0 | 328640 | 0 | 0 | 0 | 0 | 0 | 0 | 0 | 506160 | 426240 | 0 | 0 | 4964100 | 0 |
| FGSLLPIHF | PTPA; PTPA; PT | Q15257; A6 | PTPA_HUMAN | Unmodified | 0 | 0 | 0 | 0 | 0 | 0 | 0 | 0 | 0 | 0 | 0 | 5201200 | 0 | 0 | 0 | 0 | 886770 | 843580 |
| FGSNINLE | PFDN4 | Q9NQP4 | PFD4_HUMAN | Unmodified | 0 | 0 | 0 | 0 | 0 | 0 | 0 | 156200 | 0 | 0 | 0 | 0 | 0 | 0 | 0 | 0 | 1333400 | 0 |
| FGSRHSAE | NNMT | P40261 | NNMT_HUMAN | Unmodified | 0 | 0 | 0 | 0 | 2006100 | 692140 | 0 | 0 | 0 | 0 | 0 | 381660 | 0 | 0 | 0 | 0 | 0 | 0 |
| FGSTADAL | MIA3 | Q5JRA6 | TGO1_HUMAN | Unmodified | 0 | 0 | 0 | 0 | 0 | 0 | 0 | 0 | 884300 | 0 | 726540 | 0 | 0 | 0 | 0 | 0 | 0 | 0 |

| | | | | | | | | | | | | | | | | | | | | | | |
|---|---|---|---|---|---|---|---|---|---|---|---|---|---|---|---|---|---|---|---|---|---|---|
| FGSVPFTK | EXOC7; EXOC7 | Q9UPT5; B4I | EXOC7_HUMAI | Unmodified | 0 | 0 | 124500 | 431100 | 1275200 | 297090 | 0 | 110570 | 638900 | 372590 | 151870 | 932590 | 533870 | 182360 | 171870 | 161440 | 0 | 0 |
| FGSVPFTKI | EXOC7; EXOC7 | Q9UPT5; B4I | EXOC7_HUMAI | Unmodified | 0 | 0 | 85744 | 0 | 3207200 | 0 | 0 | 409660 | 0 | 0 | 97946 | 168310 | 0 | 795490 | 0 | 0 | 628110 | 1123300 |
| FGSYAAIQE | PLD1 | Q13393 | PLD1_HUMAN | Unmodified | 0 | 0 | 0 | 0 | 0 | 0 | 0 | 0 | 452740 | 0 | 0 | 700280 | 0 | 0 | 0 | 0 | 0 | 0 |
| FGTAIAAVK | ITGA1 | P56199 | ITA1_HUMAN I | Unmodified | 1664200 | 1904000 | 45671 | 144630 | 840390 | 269160 | 52477 | 0 | 1336900 | 162550 | 114340 | 174470 | 68864 | 0 | 80729 | 61077 | 1514000 | 10289000 |
| FGTFGGLG | AHNAK | Q09666 | AHNK_HUMAN | Unmodified | 39254 | 0 | 0 | 0 | 3932200 | 804640 | 0 | 117000 | 0 | 0 | 298210 | 58509 | 184110 | 316380 | 0 | 0 | 1327500 | 3096600 |
| FGTLLSGS | PLAA; PLAA; PL | Q9Y263; H0Y | PLAP_HUMAN I | Unmodified | 0 | 0 | 0 | 0 | 0 | 0 | 0 | 0 | 0 | 0 | 0 | 2099200 | 0 | 0 | 0 | 0 | 0 | 0 |
| FGTVAVPN | TXNDC15; TXNI | Q96J42; H0Y | TXD15_HUMAN | Oxidation (M) | 0 | 0 | 0 | 0 | 0 | 0 | 0 | 0 | 2409900 | 0 | 0 | 0 | 0 | 0 | 0 | 0 | 0 | 0 |
| FGTVAVPN | TXNDC15; TXNI | Q96J42; H0Y | TXD15_HUMAN | Unmodified | 0 | 0 | 0 | 0 | 0 | 0 | 0 | 0 | 1343200 | 0 | 0 | 0 | 0 | 0 | 0 | 0 | 0 | 0 |
| FGTVLTEH | DNPH1; DNPH1 | O43598; H0Y | DNPH1_HUMA | Unmodified | 517680 | 354170 | 761060 | 361620 | 0 | 0 | 34349 | 37202 | 210490 | 0 | 1478500 | 5308900 | 0 | 0 | 1257000 | 811010 | 27976 | 101880 |
| FGVEQDRN | CTTN | Q14247 | SRC8_HUMAN | Oxidation (M) | 631770 | 0 | 392410 | 382170 | 2116400 | 406380 | 0 | 70941 | 496590 | 36901 | 0 | 660130 | 791640 | 329750 | 555580 | 518660 | 316800 | 383260 |
| FGVEQDRN | CTTN | Q14247 | SRC8_HUMAN | Unmodified | 1112700 | 1042200 | 554060 | 602410 | 4581800 | 724570 | 0 | 22712 | 979190 | 143540 | 107870 | 2091800 | 1747300 | 811770 | 0 | 0 | 452370 | 1644800 |
| FGVEQDVD | PKM; PKM; PKM | P14618; B4D | KPYM_HUMAN | Oxidation (M) | 447030 | 986980 | 1092000 | 2349500 | 38139000 | 4182200 | 2934800 | 0 | 1937200 | 2354900 | 2763900 | 6152300 | 5262600 | 2426600 | 11015000 | 945840 | 9994000 | 16210000 |
| FGVEQDVD | PKM; PKM; PKM | P14618; B4D | KPYM_HUMAN | Unmodified | 0 | 0 | 0 | 0 | 4019900 | 0 | 0 | 0 | 528640 | 159150 | 0 | 0 | 413600 | 0 | 0 | 0 | 0 | 1886600 |
| FGVIFAGA | PSAT1 | Q9Y617 | SERC_HUMAN | Unmodified | 328860 | 343500 | 827870 | 2727700 | 0 | 0 | 0 | 163520 | 0 | 0 | 3248700 | 11265000 | 1953700 | 1073700 | 2296500 | 3046100 | 0 | 0 |
| FGVPVIAD | IMPDH2; IMPDH | P12268; A0A | IMDH2_HUMAI | Unmodified | 0 | 0 | 704710 | 0 | 2871100 | 0 | 2507200 | 649630 | 2285900 | 1344100 | 1796600 | 4049900 | 1750900 | 720080 | 0 | 0 | 2143400 | 2317700 |
| FGVQMDR | CTTN | Q14247 | SRC8_HUMAN | Oxidation (M) | 357440 | 793090 | 244630 | 0 | 0 | 0 | 0 | 0 | 0 | 0 | 0 | 639810 | 0 | 0 | 2229300 | 993150 | 0 | 0 |
| FGVQMDR | CTTN | Q14247 | SRC8_HUMAN | Oxidation (M) | 1348000 | 1307000 | 0 | 0 | 1722400 | 164140 | 0 | 0 | 0 | 0 | 0 | 0 | 1586000 | 0 | 0 | 0 | 402520 | 0 |
| FGVQMDR | CTTN | Q14247 | SRC8_HUMAN | Unmodified | 640240 | 744670 | 0 | 523670 | 4649300 | 0 | 0 | 0 | 348580 | 0 | 0 | 860170 | 1806600 | 2160100 | 324020 | 0 | 1376800 | 0 |
| FGVQSERC | CTTN | Q14247 | SRC8_HUMAN | Unmodified | 1727100 | 1107800 | 458890 | 656600 | 3802700 | 0 | 0 | 0 | 897560 | 91916 | 0 | 1976500 | 1396400 | 757060 | 1467800 | 1374500 | 527430 | 1958000 |
| FGVQSERC | CTTN | Q14247 | SRC8_HUMAN | Unmodified | 0 | 0 | 0 | 263670 | 2973400 | 0 | 0 | 0 | 0 | 0 | 0 | 2115700 | 3184000 | 0 | 168950 | 0 | 0 | 2901700 |
| FGVQTDRC | CTTN | Q14247 | SRC8_HUMAN | Unmodified | 0 | 40659 | 0 | 0 | 682680 | 0 | 0 | 0 | 0 | 0 | 0 | 0 | 0 | 0 | 0 | 0 | 108850 | 323870 |
| FGVSTGRE | AHNAK | Q09666 | AHNK_HUMAN | Unmodified | 0 | 0 | 0 | 0 | 18398000 | 2428300 | 85504 | 0 | 2348100 | 36782 | 0 | 0 | 276850 | 259510 | 0 | 0 | 5111800 | 6943500 |
| FGVTDFPS | QSOX1 | O00391 | QSOX1_HUMAI | Unmodified | 0 | 0 | 0 | 0 | 0 | 0 | 0 | 0 | 1030000 | 0 | 0 | 0 | 0 | 0 | 0 | 0 | 0 | 0 |
| FGVVLDEIK | COPE; COPE; C | O14579; M0I | COPE_HUMAN | Unmodified | 609070 | 317040 | 0 | 768440 | 314910 | 0 | 234290 | 0 | 2399100 | 946750 | 2901500 | 16420000 | 0 | 0 | 665770 | 467720 | 309610 | 297830 |
| FGVVVVGV | BLVRA; BLVRA | P53004; C9J | BIEA_HUMAN E | Unmodified | 599120 | 1938700 | 789220 | 2069500 | 183420 | 0 | 0 | 243130 | 1426700 | 0 | 266960 | 2051700 | 631850 | 95802 | 1226900 | 972100 | 36043 | 0 |
| FGVYDIDN | CPNE3; CPNE3 | O75131; A0A | CPNE3_HUMAI | Unmodified | 267910 | 224570 | 2430300 | 1840600 | 1148000 | 0 | 0 | 193750 | 15206000 | 1762300 | 407560 | 1334300 | 348040 | 0 | 2631400 | 1899700 | 0 | 2924700 |
| FGWDCHG | IARS1; IARS1; I | P41252; A0A | SYIC_HUMAN I | Unmodified | 0 | 0 | 0 | 0 | 0 | 0 | 0 | 0 | 1062300 | 278300 | 6722300 | 1454600 | 0 | 0 | 0 | 0 | 0 | 0 |
| FGYDEIDN | ACOX1 | Q15067 | ACOX1_HUMAI | Unmodified | 0 | 0 | 0 | 0 | 0 | 0 | 0 | 0 | 1972500 | 0 | 0 | 0 | 0 | 0 | 0 | 0 | 0 | 0 |
| FGYQFTQC | TMOD3; TMOD3 | Q9NYL9; H0 | TMOD3_HUMA | Unmodified | 0 | 0 | 0 | 0 | 0 | 0 | 0 | 0 | 0 | 0 | 0 | 0 | 0 | 0 | 0 | 0 | 0 | 0 |
| FGYVDFES | NCL; NCL; NCL | P19338; C9J | NUCL_HUMAN | Unmodified | 0 | 0 | 0 | 0 | 0 | 0 | 0 | 246780 | 1616400 | 693570 | 5121000 | 4425800 | 0 | 0 | 0 | 0 | 0 | 0 |
| FHALGPIYY | RAB21 | Q9UL25 | RAB21_HUMAN | Unmodified | 374730 | 2832100 | 2166300 | 0 | 0 | 0 | 0 | 0 | 990600 | 0 | 282050 | 11318000 | 0 | 0 | 294550 | 280720 | 0 | 0 |
| FHCATGYC | SEZ6; SEZ6; SE | Q53EL9; H0Y | SEZ6_HUMAN | Unmodified | 120650 | 0 | 0 | 0 | 0 | 0 | 0 | 0 | 3881600 | 523760 | 0 | 0 | 0 | 0 | 0 | 0 | 0 | 0 |
| FHDFLGDS | PRDX6 | P30041 | PRDX6_HUMAN | Unmodified | 252680 | 419610 | 342450 | 1190200 | 331430 | 0 | 0 | 0 | 2197500 | 93852 | 1317600 | 4081200 | 4164300 | 328690 | 5484700 | 886440 | 174610 | 1553200 |
| FHDFLGDS | PRDX6 | P30041 | PRDX6_HUMAN | Unmodified | 0 | 0 | 0 | 0 | 0 | 0 | 0 | 0 | 0 | 0 | 0 | 0 | 1847500 | 0 | 0 | 0 | 913610 | 1921400 |
| FHDFLGDS | PRDX6 | P30041 | PRDX6_HUMAN | Unmodified | 0 | 0 | 0 | 0 | 1754500 | 0 | 0 | 0 | 0 | 0 | 0 | 0 | 3636700 | 2508200 | 0 | 0 | 7913900 | 8591500 |
| FHDIISDAE | P4HA1 | P13674 | P4HA1_HUMAN | Unmodified | 0 | 0 | 0 | 0 | 0 | 0 | 0 | 200700 | 3766100 | 0 | 680120 | 0 | 0 | 0 | 0 | 0 | 674330 | 0 |
| FHEAWQK | CTPS2 | Q9NRF8 | PYRG2_HUMAN | Unmodified | 0 | 0 | 0 | 0 | 0 | 0 | 0 | 0 | 0 | 0 | 0 | 251130 | 0 | 0 | 0 | 0 | 0 | 0 |
| FHEFHSPA | EHD1; EHD3; E | Q9H4M9; Q | EHD1_HUMAN | Oxidation (M) | 3592400 | 5813800 | 0 | 0 | 866430 | 0 | 2953500 | 0 | 0 | 0 | 796940 | 8477100 | 0 | 0 | 8519500 | 5308500 | 259010 | 0 |
| FHEFHSPA | EHD4 | Q9H223 | EHD4_HUMAN | Oxidation (M) | 1578200 | 0 | 0 | 976320 | 0 | 0 | 0 | 0 | 0 | 0 | 0 | 0 | 0 | 0 | 4822200 | 3148500 | 0 | 0 |
| FHEICSNL | ARL6IP1; N/A; A | Q15041; H3I | AR6P1_HUMAN | Unmodified | 0 | 0 | 0 | 0 | 0 | 0 | 0 | 0 | 788900 | 0 | 105850 | 1674600 | 0 | 0 | 0 | 0 | 0 | 0 |
| FHELHRDL | PACSIN3; PACS | Q9UKS6; E9I | PACN3_HUMAI | Unmodified | 1777100 | 890610 | 177000 | 145330 | 0 | 0 | 0 | 0 | 0 | 0 | 0 | 0 | 0 | 0 | 201850 | 0 | 0 | 0 |
| FHELHRDL | PACSIN3; PACS | Q9UKS6; E9I | PACN3_HUMAI | Unmodified | 1381900 | 904780 | 0 | 6411400 | 976190 | 0 | 142680 | 0 | 0 | 0 | 0 | 0 | 4858800 | 309750 | 0 | 0 | 0 | 0 |
| FHGTVKAE | GAPDH | P04406 | G3P_HUMAN G | Unmodified | 0 | 0 | 0 | 0 | 2688900 | 252680 | 0 | 0 | 160020 | 0 | 0 | 0 | 842090 | 502370 | 0 | 0 | 396120 | 1291000 |
| FHHLIPAH | EEPD1 | Q7L9B9 | EEPD1_HUMAN | Unmodified | 0 | 0 | 0 | 0 | 1574400 | 0 | 0 | 0 | 0 | 234360 | 0 | 0 | 0 | 1013500 | 0 | 0 | 5521800 | 9826700 |
| FHHTFSTEI | PDIA4; PDIA4 | P13667; A0A | PDIA4_HUMAN | Unmodified | 0 | 0 | 0 | 0 | 296640 | 0 | 0 | 283530 | 44781000 | 10670000 | 9095100 | 5958700 | 0 | 0 | 0 | 0 | 45052 | 57577 |
| FHHTIGGS | RPL15; RPL15 | P61313; E7E | RL15_HUMAN | Unmodified | 0 | 0 | 0 | 0 | 0 | 0 | 0 | 0 | 0 | 0 | 0 | 1763200 | 0 | 0 | 0 | 0 | 0 | 0 |
| FHILVANR | TOM1L2; TOM1 | Q6ZVM7; K7 | TM1L2_HUMAN | Unmodified | 1671700 | 2325800 | 303960 | 1305700 | 0 | 0 | 0 | 0 | 0 | 0 | 0 | 0 | 0 | 0 | 2810500 | 856260 | 0 | 0 |
| FHISEFPTL | PDIA5 | Q14554 | PDIA5_HUMAN | Unmodified | 0 | 0 | 0 | 0 | 0 | 0 | 0 | 0 | 3452800 | 283580 | 0 | 0 | 0 | 0 | 0 | 0 | 0 | 0 |
| FHIYNMGN | GRK5 | P34947 | GRK5_HUMAN | Oxidation (M) | 0 | 0 | 0 | 0 | 0 | 0 | 0 | 0 | 0 | 0 | 0 | 1230600 | 0 | 0 | 0 | 0 | 0 | 0 |
| FHIYNMGN | GRK5 | P34947 | GRK5_HUMAN | Unmodified | 0 | 0 | 0 | 0 | 0 | 0 | 0 | 0 | 0 | 0 | 0 | 813660 | 0 | 0 | 0 | 0 | 0 | 0 |
| FHKEIIHELI | BAIAP2L1 | Q9UHR4 | BI2L1_HUMAN | Unmodified | 0 | 0 | 0 | 0 | 237690 | 0 | 0 | 0 | 0 | 0 | 0 | 0 | 64521 | 139050 | 0 | 0 | 0 | 0 |
| FHLALENAI | FUT11 | Q495W5 | FUT11_HUMAN | Oxidation (M) | 0 | 0 | 0 | 0 | 0 | 0 | 0 | 0 | 295680 | 0 | 0 | 0 | 0 | 0 | 0 | 0 | 0 | 0 |
| FHLALENAI | FUT11 | Q495W5 | FUT11_HUMAN | Unmodified | 0 | 0 | 0 | 0 | 0 | 0 | 0 | 0 | 573420 | 0 | 0 | 0 | 0 | 0 | 0 | 0 | 0 | 0 |
| FHMVDGN | AHSA1; AHSA1; | O95433; H0Y | AHSA1_HUMAI | Oxidation (M) | 63415 | 0 | 34381 | 0 | 0 | 0 | 185150 | 18842 | 0 | 67702 | 2735700 | 1911200 | 0 | 0 | 168050 | 396550 | 0 | 0 |
| FHMVDGN | AHSA1; AHSA1; | O95433; H0Y | AHSA1_HUMAI | Unmodified | 0 | 0 | 0 | 0 | 676630 | 0 | 894160 | 26158 | 198930 | 0 | 190120 | 890760 | 0 | 0 | 0 | 0 | 0 | 0 |
| FHMYEGYF | PNP; PNP; PNP | P00491; G3V | PNPH_HUMAN | Oxidation (M) | 0 | 0 | 0 | 0 | 0 | 0 | 0 | 0 | 68473 | 0 | 0 | 3718200 | 0 | 0 | 0 | 0 | 0 | 0 |
| FHNELNLR | IPO8 | O15397 | IPO8_HUMAN I | Unmodified | 127750 | 0 | 642700 | 0 | 0 | 0 | 0 | 0 | 720300 | 0 | 587070 | 0 | 0 | 0 | 0 | 0 | 0 | 0 |
| FHNPQIHF | CDH1; CDH1; C | P12830; J3Q | CADH1_HUMA | Unmodified | 51780 | 230040 | 0 | 0 | 0 | 0 | 0 | 0 | 0 | 0 | 0 | 0 | 0 | 0 | 0 | 0 | 0 | 0 |
| FHNWFDD | STT3A; STT3A; S | P46977; E9P | STT3A_HUMAN | Unmodified | 387870 | 0 | 0 | 0 | 0 | 0 | 0 | 0 | 3598200 | 311770 | 0 | 0 | 0 | 0 | 0 | 0 | 0 | 0 |
| FHPDTDDV | NUCB1 | Q02818 | NUCB1_HUMA | Unmodified | 264960 | 0 | 335590 | 182290 | 317400 | 385000 | 326850 | 0 | 9995700 | 1755200 | 4097700 | 1687700 | 0 | 0 | 0 | 250980 | 0 | 484770 |
| FHPDTDDV | NUCB1 | Q02818 | NUCB1_HUMA | Unmodified | 0 | 0 | 0 | 0 | 956800 | 0 | 2451700 | 2124800 | 1091500 | 2360500 | 873730 | 0 | 425380 | 0 | 0 | 0 | 0 | 1226900 |
| FHPEHFVC | PXN; PXN; PXN | P49023; A0A | PAXI_HUMAN F | Unmodified | 0 | 0 | 0 | 0 | 0 | 0 | 0 | 0 | 0 | 0 | 0 | 0 | 0 | 0 | 0 | 0 | 202770 | 156450 |
| FHPEPYGL | CTNND1; CTNN | O60716; A0A | CTND1_HUMAI | Unmodified | 1172800 | 892580 | 3636900 | 2029700 | 0 | 0 | 0 | 0 | 0 | 0 | 1216000 | 6746700 | 0 | 0 | 7863400 | 11397000 | 0 | 0 |
| FHPGLGYE | PTGFRN | Q9P2B2 | FPRP_HUMAN | Unmodified | 10350000 | 8699600 | 27585000 | 15714000 | 127460 | 0 | 0 | 0 | 0 | 0 | 0 | 17849000 | 872090 | 0 | 21419000 | 23187000 | 0 | 131000 |
| FHPGLGYE | PTGFRN | Q9P2B2 | FPRP_HUMAN | Unmodified | 0 | 0 | 0 | 2175200 | 0 | 0 | 0 | 0 | 0 | 0 | 0 | 0 | 0 | 0 | 0 | 0 | 0 | 0 |
| FHPIQGHR | HNRNPA0 | Q13151 | ROA0_HUMAN | Unmodified | 0 | 0 | 18121 | 19619 | 0 | 0 | 0 | 0 | 203820 | 179590 | 15103 | 119470 | 300480 | 185560 | 0 | 12520 | 31594 | 133640 |
| FHPNGSTL | SNRNP40 | Q96DI7 | SNR40_HUMAI | Unmodified | 0 | 0 | 0 | 0 | 0 | 0 | 0 | 0 | 0 | 0 | 0 | 1859300 | 0 | 0 | 0 | 44875 | 113110 | 0 |
| FHPSIQSF | MUL1 | Q969V5 | MUL1_HUMAN | Unmodified | 0 | 0 | 0 | 709600 | 0 | 0 | 0 | 0 | 2965000 | 266000 | 0 | 0 | 0 | 0 | 0 | 0 | 0 | 0 |
| FHQLDIDD | CBR3; CBR1; C | O75828; P16 | CBR3_HUMAN | Unmodified | 0 | 481790 | 997920 | 1051900 | 441230 | 0 | 0 | 0 | 2412300 | 0 | 426250 | 5761700 | 330360 | 0 | 2658400 | 1023300 | 368540 | 501100 |
| FHQNQLLS | ZFP36L1; ZFP36 | Q07352; G3 | TISB_HUMAN n | Unmodified | 0 | 461510 | 0 | 0 | 0 | 0 | 0 | 0 | 0 | 0 | 0 | 0 | 0 | 0 | 0 | 0 | 0 | 0 |
| FHSAYYSSI | IDE; IDE; IDE; ID | P14735; A0A | IDE_HUMAN In | Oxidation (M) | 0 | 0 | 0 | 0 | 0 | 0 | 0 | 0 | 388680 | 1662300 | 0 | 982950 | 0 | 0 | 0 | 0 | 0 | 0 |
| FHTEALTAL | ETF1; ETF1; ETF | P62495; B7Z | ERF1_HUMAN | Unmodified | 629610 | 361680 | 881940 | 628730 | 0 | 35836 | 46666 | 17249 | 1390500 | 0 | 2802000 | 11553000 | 91452 | 29407 | 1147800 | 315050 | 0 | 0 |
| FHTFPQTAI | CALCA | P01258 | CALC_HUMAN | Unmodified | 0 | 0 | 0 | 0 | 0 | 0 | 0 | 0 | 3189400 | 1445100 | 0 | 0 | 0 | 0 | 0 | 0 | 259380 | 0 |
| FHTITTSYY | RAB10 | P61026 | RAB10_HUMAN | Unmodified | 590340 | 535050 | 795330 | 156650 | 0 | 0 | 0 | 0 | 194510 | 119580 | 143110 | 2594800 | 0 | 0 | 739580 | 782640 | 0 | 0 |
| FHVEEEGK | PGK1; PGK2 | P00558; P07 | PGK1_HUMAN | Unmodified | 0 | 423190 | 1507400 | 1557300 | 739820 | 430620 | 0 | 0 | 5822000 | 377320 | 4430500 | 23322000 | 899070 | 375910 | 2513500 | 0 | 890220 | 1819300 |
| FHVEEEGK | PGK1 | P00558 | PGK1_HUMAN | Unmodified | 11426 | 0 | 0 | 63991 | 4169900 | 746390 | 0 | 189560 | 49678 | 1617800 | 36190 | 4875600 | 3804100 | 2693400 | 0 | 203140 | 6443300 | 13193000 |
| FHVLVASQ | TOM1; TOM1; T | O60784; V9 | TOM1_HUMAN | Unmodified | 71369 | 502880 | 0 | 193370 | 29718 | 0 | 0 | 0 | 326020 | 0 | 0 | 135630 | 85396 | 0 | 378580 | 0 | 0 | 0 |
| FHVQLSNP | ITGB4 | P16144 | ITB4_HUMAN I | Unmodified | 0 | 0 | 0 | 0 | 745060 | 0 | 0 | 0 | 0 | 0 | 0 | 0 | 0 | 0 | 0 | 0 | 479690 | 1030500 |
| FHVQNLSC | ADRB2 | P07550 | ADRB2_HUMAI | Unmodified | 0 | 0 | 1155200 | 808860 | 0 | 0 | 0 | 0 | 160020 | 0 | 0 | 0 | 0 | 0 | 1596200 | 902400 | 222920 | 0 |
| FIAAQGPK | PTPRA; PTPRE; | P18433; P23 | PTPRA_HUMAN | Unmodified | 0 | 0 | 149670 | 0 | 0 | 0 | 0 | 0 | 0 | 9496 | 0 | 854290 | 0 | 0 | 0 | 144380 | 0 | 0 |
| FIAAQGPK | PTPRA | P18433 | PTPRA_HUMAN | Unmodified | 438640 | 0 | 0 | 0 | 0 | 154670 | 0 | 0 | 747800 | 76674 | 977120 | 2459000 | 363140 | 0 | 1138700 | 1300000 | 0 | 324430 |
| FIAAQGPK | PTPRE; PTPRE | P23469; S4F | PTPRE_HUMAN | Unmodified | 0 | 0 | 0 | 0 | 936990 | 564630 | 0 | 0 | 0 | 0 | 0 | 0 | 0 | 0 | 0 | 0 | 0 | 0 |
| FIADGCK | BAIAP2L1 | Q9UHR4 | BI2L1_HUMAN | Unmodified | 0 | 0 | 0 | 0 | 0 | 2403800 | 0 | 0 | 307590 | 8923.8 | 0 | 0 | 1441800 | 368600 | 0 | 190970 | 1458600 | 0 |
| FIADGCKE | BAIAP2L1 | Q9UHR4 | BI2L1_HUMAN | Unmodified | 0 | 0 | 0 | 0 | 1262400 | 297160 | 53455 | 0 | 0 | 110460 | 0 | 0 | 688320 | 1354900 | 0 | 0 | 231210 | 129430 |
| FIADQLDH | MRPL57 | Q9BQC6 | RT63_HUMAN | Unmodified | 0 | 0 | 0 | 0 | 0 | 0 | 100870 | 0 | 822390 | 0 | 0 | 0 | 0 | 141490 | 0 | 0 | 80824 | 0 |
| FIAGTGCL | PGP | A6NDG6 | PGP_HUMAN C | Unmodified | 60102 | 0 | 0 | 0 | 0 | 0 | 0 | 0 | 0 | 0 | 561440 | 1615500 | 0 | 0 | 0 | 276120 | 0 | 0 |
| FIASTGMDI | PRPF19; PRPF1 | Q9UMS4; H | PRP19_HUMAN | Unmodified | 0 | 0 | 0 | 0 | 0 | 0 | 0 | 65878 | 113580 | 0 | 0 | 0 | 0 | 0 | 0 | 0 | 291220 | 0 |
| FIATLQYIV | PREP; PREP | P48147; A0A | PPCE_HUMAN | Unmodified | 0 | 0 | 0 | 0 | 0 | 0 | 4243500 | 0 | 0 | 0 | 203510 | 667840 | 0 | 0 | 0 | 0 | 0 | 181180 |
| FIAVGYVD | HLA-A; HLA-A; | P04439; A0A | HLAA_HUMAN | Unmodified | 0 | 0 | 0 | 0 | 0 | 0 | 0 | 13081 | 8670700 | 2129400 | 2598500 | 23692000 | 0 | 0 | 135420 | 130820 | 0 | 182280 |
| FIAVGYVD | HLA-C; HLA-C; | Q53YP1; A0 | Q53YP1_HUMA | Unmodified | 0 | 0 | 0 | 0 | 4550300 | 131950 | 0 | 0 | 248130 | 769230 | 3271700 | 4775900 | 0 | 0 | 0 | 0 | 0 | 0 |
| FIAVGYVD | HLA-A; HLA-A; | A0A0G2JL56 | A0A0G2JL56_H | Unmodified | 0 | 0 | 0 | 0 | 0 | 0 | 0 | 0 | 1865500 | 283210 | 0 | 772520 | 0 | 2351500 | 0 | 0 | 0 | 0 |
| FIAVGYVD | HLA-A; HLA-A; | A0A0G2JL56 | A0A0G2JL56_H | Oxidation (M) | 0 | 0 | 0 | 0 | 0 | 0 | 0 | 0 | 7188700 | 2727300 | 605150 | 959500 | 544130 | 0 | 0 | 0 | 0 | 9697200 |
| FIAVGYVD | HLA-A; HLA-A; | A0A0G2JL56 | A0A0G2JL56_H | Unmodified | 0 | 0 | 198550 | 0 | 0 | 0 | 0 | 0 | 8103300 | 0 | 0 | 0 | 0 | 0 | 0 | 0 | 0 | 0 |
| FIAVGYVD | HLA-A; HLA-A; | P04439; Q9 | HLAA_HUMAN | Unmodified | 0 | 0 | 0 | 0 | 812110 | 0 | 0 | 0 | 511880 | 178980 | 0 | 1566500 | 0 | 0 | 0 | 0 | 780760 | 2093400 |
| FICASNQN | THNSL1 | Q8IYQ7 | THNS1_HUMAI | Unmodified | 0 | 0 | 168190 | 0 | 0 | 0 | 0 | 0 | 577810 | 0 | 0 | 0 | 0 | 0 | 0 | 0 | 0 | 0 |
| FICEQDHQ | HSPH1; HSPH1 | Q92598; A0 | HS105_HUMAN | Unmodified | 0 | 0 | 677630 | 1327800 | 185330 | 53293 | 220620 | 0 | 1419600 | 0 | 1739700 | 11354000 | 600740 | 192310 | 729020 | 0 | 0 | 0 |
| FICTTSAIQ | ACLY; ACLY | P53396; K7E | ACLY_HUMAN | Unmodified | 0 | 0 | 0 | 0 | 1042100 | 0 | 4327300 | 1107300 | 0 | 378210 | 0 | 0 | 299450 | 0 | 0 | 0 | 269170 | 0 |
| FIDADVEEF | DOP1B | Q9Y3R5 | DOP2_HUMAN | Unmodified | 0 | 0 | 0 | 0 | 0 | 0 | 0 | 0 | 0 | 0 | 0 | 829440 | 0 | 0 | 77482 | 0 | 0 | 0 |
| FIDDVVSA | PCYOX1L; PCY | Q8NBM8; E7 | PCYXL_HUMAN | Unmodified | 0 | 0 | 0 | 0 | 0 | 0 | 0 | 185800 | 1088200 | 120310 | 103190 | 0 | 0 | 0 | 0 | 21979 | 0 | 0 |
| FIDTGFSLP | WWP1; ITCH; W | Q9H0M0; Q | WWP1_HUMAN | Unmodified | 2248200 | 1313700 | 0 | 0 | 0 | 0 | 0 | 0 | 0 | 0 | 0 | 0 | 0 | 0 | 2584400 | 1558000 | 0 | 0 |
| FIDTGFSLP | WWP1; ITCH; W | Q9H0M0; Q | WWP1_HUMAN | Unmodified | 0 | 0 | 102790 | 0 | 72596 | 10611 | 0 | 0 | 34003 | 0 | 122500 | 129150 | 0 | 0 | 12012000 | 3300800 | 0 | 0 |
| FIDTGFTLP | WWP2 | O00308 | WWP2_HUMAN | Unmodified | 1147400 | 684280 | 0 | 0 | 0 | 0 | 0 | 0 | 0 | 0 | 0 | 0 | 0 | 0 | 1077700 | 663420 | 0 | 0 |
| FIDTTSK | RPL3L; RPL3; R | Q92901; P3 | RL3L_HUMAN | Unmodified | 4431100 | 3687500 | 2636700 | 2042800 | 2347900 | 361550 | 835580 | 670650 | 914050 | 341340 | 1521000 | 34531000 | 1226900 | 709160 | 2353100 | 2698700 | 199840 | 1376100 |
| FIDTTSKFG | RPL3; RPL3 | P39023; G5E | RL3_HUMAN 6 | Unmodified | 0 | 0 | 0 | 0 | 262110 | 0 | 403490 | 45364 | 0 | 0 | 0 | 0 | 296570 | 283500 | 0 | 0 | 0 | 223290 |
| FIEAAQR | COPS4; COPS4 | Q9BT78; D6 | CSN4_HUMAN | Unmodified | 15118 | 0 | 0 | 0 | 200340 | 0 | 193760 | 0 | 121740 | 89060 | 110790 | 3098900 | 18675 | 0 | 0 | 183160 | 191220 | 0 |
| FIEAGQFN | DLG3; DLG3 | Q92796; Q5 | DLG3_HUMAN | Unmodified | 0 | 0 | 0 | 679540 | 0 | 0 | 0 | 0 | 1567500 | 0 | 0 | 1789900 | 0 | 0 | 0 | 307140 | 20979 | 0 |
| FIEAGQYN | DLG1; DLG1; D | Q12959; A0 | DLG1_HUMAN | Unmodified | 0 | 0 | 0 | 0 | 0 | 0 | 0 | 0 | 3550300 | 0 | 0 | 3346500 | 0 | 0 | 170810 | 90992 | 0 | 0 |
| FIEAGQYN | DLG1; DLG1; D | Q12959; A0 | DLG1_HUMAN | Unmodified | 0 | 0 | 0 | 753480 | 10524000 | 7437000 | 0 | 0 | 1611300 | 3372300 | 0 | 0 | 2768900 | 1651000 | 0 | 0 | 2896200 | 5381900 |
| FIEDELQIP | ADSS2 | P30520 | PURA2_HUMAI | Unmodified | 0 | 0 | 0 | 0 | 684170 | 0 | 0 | 95171 | 0 | 0 | 3791600 | 13704000 | 0 | 0 | 0 | 0 | 0 | 0 |

| | | | | | | | | | | | | | | | | | | | | | | |
|---|---|---|---|---|---|---|---|---|---|---|---|---|---|---|---|---|---|---|---|---|---|---|
| FIEDLDENF | IL1RAPL1; IL1R/ | Q9NZN1; A0 | IRPL1_HUMAN | Unmodified | 230400 | 0 | 510550 | 582630 | 0 | 0 | 0 | 0 | 0 | 0 | 0 | 0 | 0 | 0 | 1237500 | 1329200 | 0 | 0 |
| FIEDRITEES | NAA25 | Q14CX7 | NAA25_HUMAI | Unmodified | 0 | 0 | 0 | 0 | 0 | 0 | 0 | 0 | 0 | 0 | 0 | 0 | 0 | 0 | 0 | 0 | 0 | 653150 |
| FIEEHATK | PDIA4; PDIA4 | P13667; A0A | PDIA4_HUMAN | Unmodified | 97488 | 0 | 263660 | 0 | 0 | 0 | 0 | 115330 | 11039000 | 3885300 | 1967300 | 0 | 1552500 | 0 | 172330 | 0 | 0 | 0 |
| FIEEHATKL | PDIA4; PDIA4 | P13667; A0A | PDIA4_HUMAN | Unmodified | 0 | 0 | 0 | 0 | 0 | 0 | 0 | 0 | 0 | 1006100 | 0 | 0 | 0 | 0 | 0 | 0 | 0 | 0 |
| FIEFEDSQE | SPAG9; SPAG9 | O60271; A0/ | JIP4_HUMAN C | Unmodified | 0 | 0 | 0 | 0 | 462550 | 0 | 0 | 0 | 0 | 0 | 0 | 486210 | 243590 | 0 | 0 | 0 | 0 | 1345100 |
| FIEIAAR | HNRNPU; HNR | Q00839; A0/ | HNRPU_HUMA | Unmodified | 967890 | 519620 | 0 | 0 | 721370 | 0 | 695620 | 794170 | 410380 | 312940 | 141840 | 2595700 | 1179000 | 0 | 446910 | 52531 | 0 | 0 |
| FIEVPCSED | OGFOD3; OGF( | Q6PK18; J3C | OGFD3_HUMA | Unmodified | 0 | 0 | 0 | 0 | 0 | 0 | 0 | 0 | 729830 | 0 | 0 | 0 | 0 | 0 | 0 | 0 | 0 | 0 |
| FIFDCVSQI | PGP | A6NDG6 | PGP_HUMAN ( | Unmodified | 0 | 0 | 147390 | 0 | 0 | 0 | 0 | 0 | 0 | 0 | 86074 | 851950 | 0 | 0 | 0 | 0 | 0 | 0 |
| FIFVGTLK | OTUD7B | Q6GQQ9 | OTU7B_HUMAI | Unmodified | 84039 | 0 | 502530 | 112890 | 1744300 | 365410 | 0 | 0 | 395770 | 110480 | 85147 | 249670 | 1936100 | 658120 | 133350 | 157110 | 340440 | 956830 |
| FIGELGKLD | EIF4G2; EIF4G2 | P78344; H0\ | IF4G2_HUMAN | Unmodified | 0 | 0 | 273170 | 904190 | 944750 | 0 | 889230 | 0 | 355080 | 0 | 0 | 3555700 | 745340 | 1598700 | 286510 | 852290 | 0 | 605890 |
| FIGSPPGY\ | CLPB; CLPB; CI | Q9H078; A0/ | CLPB_HUMAN | Unmodified | 0 | 0 | 0 | 0 | 0 | 0 | 0 | 0 | 1324300 | 0 | 0 | 1847100 | 0 | 0 | 0 | 0 | 0 | 0 |
| FIGVEAGG | CEMIP2; CEMIP | Q9UHN6; F5 | CEIP2_HUMAN | Unmodified | 92172 | 36299 | 0 | 0 | 0 | 0 | 204320 | 89341 | 596350 | 331770 | 0 | 372570 | 0 | 0 | 0 | 69070 | 0 | 0 |
| FIGVEAGG | CEMIP2; CEMIP | Q9UHN6; F5 | CEIP2_HUMAN | Unmodified | 0 | 0 | 446910 | 0 | 1498700 | 0 | 76096 | 0 | 0 | 0 | 0 | 0 | 0 | 0 | 0 | 0 | 0 | 0 |
| FIHDQTSPI | SLC25A1; SLC2 | P53007; B4D | TXTP_HUMAN 1 | Unmodified | 0 | 0 | 0 | 0 | 0 | 0 | 0 | 0 | 4088100 | 926420 | 104090 | 0 | 0 | 0 | 0 | 0 | 0 | 0 |
| FIHGSKEDI | ZC3H10 | Q96K80 | ZC3HA_HUMAI | Unmodified | 0 | 0 | 0 | 0 | 0 | 0 | 0 | 0 | 0 | 0 | 0 | 0 | 0 | 0 | 0 | 0 | 0 | 0 |
| FIHQQPQS | PXN; PXN; PXN; | P49023; F5H | PAXI_HUMAN F | Unmodified | 0 | 28468 | 71763 | 64431 | 365710 | 0 | 0 | 0 | 0 | 0 | 0 | 0 | 112360 | 88527 | 0 | 0 | 401380 | 1003100 |
| FIHQSDLK\ | ABCC1; ABCC1 | P33527; I3L4 | MRP1_HUMAN | Unmodified | 0 | 0 | 0 | 0 | 0 | 0 | 0 | 361170 | 0 | 0 | 0 | 1693400 | 0 | 0 | 148130 | 95268 | 147490 | 0 |
| FIHVSHLN/ | NDUFA9; N/A | Q16795; A0/ | NDUA9_HUMA | Unmodified | 0 | 0 | 0 | 0 | 0 | 0 | 104050 | 0 | 504540 | 188670 | 0 | 0 | 0 | 0 | 0 | 0 | 155300 | 0 |
| FIIDEELFG( | NEDD4; NEDD4 | P46934; H0\ | NEDD4_HUMA | Unmodified | 0 | 1936500 | 582220 | 0 | 0 | 0 | 0 | 0 | 0 | 0 | 0 | 0 | 0 | 0 | 0 | 611070 | 48854 | 0 |
| FIIHAPPGE | CAPZA2; CAPZ/ | P47755; A0A | CAZA2_HUMAI | Unmodified | 13776000 | 12573000 | 0 | 4899300 | 1035400 | 0 | 719620 | 0 | 7524400 | 461430 | 3385200 | 25626000 | 821030 | 788870 | 15806000 | 3647500 | 0 | 659530 |
| FIIPNVVK | LDHA; LDHA; LI | P00338; F5G | LDHA_HUMAN | Unmodified | 98756 | 1582800 | 4111500 | 3962600 | 11739000 | 2815000 | 880430 | 376040 | 3149900 | 4016700 | 1616700 | 2.2E+08 | 3608300 | 1913800 | 8725700 | 8742700 | 8016000 | 11661000 |
| FIIPQIVK | LDHB; LDHB; LI | P07195; A0A | LDHB_HUMAN | Unmodified | 1526100 | 2022800 | 1314100 | 0 | 209780 | 0 | 0 | 133720 | 0 | 0 | 9216600 | 70240000 | 909490 | 967090 | 1782900 | 1803600 | 0 | 0 |
| FIIPSPK | APOB | P04114 | APOB_HUMAN | Unmodified | 159700 | 0 | 1497800 | 0 | 0 | 0 | 0 | 0 | 0 | 0 | 0 | 0 | 0 | 0 | 0 | 335320 | 0 | 0 |
| FIIQSEKPPI | IDE; IDE; IDE; ID | P14735; A0A | IDE_HUMAN In | Unmodified | 0 | 0 | 0 | 0 | 0 | 0 | 0 | 0 | 0 | 0 | 0 | 5203400 | 0 | 0 | 802670 | 0 | 75248 | 0 |
| FIITALPTIY | TMX1 | Q9H3N1 | TMX1_HUMAN | Unmodified | 0 | 0 | 0 | 0 | 0 | 0 | 0 | 0 | 1.11E+08 | 3894100 | 3265600 | 3328800 | 0 | 0 | 0 | 0 | 0 | 0 |
| FIITALPTIY | TMX1 | Q9H3N1 | TMX1_HUMAN | Unmodified | 0 | 0 | 0 | 0 | 0 | 0 | 0 | 0 | 5271200 | 213300 | 0 | 0 | 0 | 0 | 0 | 0 | 0 | 0 |
| FIKDNWEE | NPEPPS; NPEPI | P55786; A0A | PSA_HUMAN P | Unmodified | 0 | 0 | 100120 | 0 | 0 | 0 | 0 | 0 | 0 | 0 | 0 | 618060 | 0 | 0 | 0 | 0 | 0 | 49418 |
| FIKGTDYQI | YARS1; YARS1; | P54577; A0A | SYYC_HUMAN | Unmodified | 0 | 0 | 0 | 0 | 224470 | 0 | 190640 | 105410 | 0 | 80008 | 0 | 0 | 293960 | 262980 | 0 | 0 | 116340 | 0 |
| FIKPLSNAY | COPS3 | Q9UNS2 | CSN3_HUMAN | Unmodified | 0 | 0 | 0 | 476730 | 0 | 0 | 0 | 0 | 774930 | 0 | 0 | 2170300 | 0 | 0 | 0 | 0 | 0 | 0 |
| FIKWDDET | PLCB2; PLCB2 | Q00722; Q9I | PLCB2_HUMAI | Unmodified | 0 | 0 | 0 | 0 | 0 | 0 | 0 | 0 | 0 | 0 | 0 | 2045700 | 0 | 0 | 0 | 0 | 378710 | 0 |
| FILAPAKQE | P4HA1 | P13674 | P4HA1_HUMAI | Unmodified | 0 | 0 | 0 | 0 | 0 | 0 | 0 | 0 | 14553000 | 3724600 | 3132000 | 643560 | 0 | 0 | 0 | 0 | 287190 | 751890 |
| FILHHSDP/ | TXNDC11 | Q6PKC3 | TXD11_HUMAI | Unmodified | 0 | 0 | 0 | 0 | 0 | 0 | 0 | 0 | 2014900 | 0 | 0 | 0 | 0 | 0 | 0 | 0 | 0 | 0 |
| FILIQNR | AP2S1; AP2S1; | P53680; M0( | AP2S1_HUMAI | Unmodified | 1002600 | 963120 | 894520 | 727980 | 0 | 0 | 0 | 0 | 1123200 | 131610 | 669760 | 6329400 | 0 | 0 | 1301800 | 988160 | 0 | 0 |
| FILNLPTFS | PSMB2; PSMB2 | P49721; A0A | PSB2_HUMAN | Unmodified | 226050 | 0 | 0 | 0 | 0 | 0 | 0 | 0 | 0 | 0 | 65416 | 3257400 | 0 | 0 | 0 | 385000 | 0 | 0 |
| FILPVNEGK | SBDS; SBDS | Q9Y3A5; A0/ | SBDS_HUMAN | Unmodified | 0 | 511330 | 0 | 0 | 0 | 0 | 0 | 539740 | 267650 | 0 | 0 | 1641500 | 0 | 0 | 0 | 529230 | 272520 | 0 |
| FILQEDGSI | GALNT12 | Q8IXK2 | GLT12_HUMAI | Unmodified | 0 | 0 | 0 | 0 | 0 | 0 | 0 | 0 | 1390900 | 0 | 0 | 0 | 0 | 0 | 0 | 0 | 0 | 0 |
| FILSCADDI | PAFAH1B1; PAF | P43034; I3L3 | LIS1_HUMAN P | Unmodified | 143160 | 0 | 271000 | 372050 | 254430 | 242630 | 201540 | 92831 | 123330 | 0 | 0 | 2396600 | 121410 | 0 | 179710 | 113490 | 260260 | 417700 |
| FIMDLVSSI | MME; MME | P08473; A0A | NEP_HUMAN N | Oxidation (M) | 0 | 0 | 4908700 | 2173600 | 0 | 0 | 0 | 0 | 0 | 0 | 0 | 0 | 0 | 0 | 19633000 | 12132000 | 0 | 0 |
| FIMDLVSSI | MME; MME | P08473; A0A | NEP_HUMAN N | Unmodified | 0 | 0 | 290390 | 488120 | 0 | 0 | 0 | 0 | 0 | 0 | 0 | 0 | 0 | 0 | 2643800 | 954180 | 0 | 0 |
| FIMESGAK | RPS3; RPS3; RP | P23396; H0\ | RS3_HUMAN 4 | Unmodified | 1587000 | 1487500 | 623320 | 773190 | 1144600 | 53704 | 1117900 | 870400 | 810610 | 0 | 1468700 | 30799000 | 267830 | 270950 | 1486000 | 1259200 | 128650 | 391340 |
| FIMESGAK | RPS3; RPS3; RP | P23396; H0\ | RS3_HUMAN 4 | Oxidation (M) | 542560 | 384980 | 188850 | 210120 | 235510 | 24848 | 232520 | 80619 | 0 | 44849 | 157160 | 0 | 0 | 0 | 0 | 316770 | 11168 | 163760 |
| FINDMVK | ELAPOR2; ELAP | A8MWY0; H7 | ELAP2_HUMAI | Unmodified | 0 | 0 | 244600 | 0 | 117230 | 0 | 24069 | 0 | 3843200 | 613090 | 719980 | 261280 | 0 | 0 | 0 | 0 | 43978 | 99447 |
| FINDQYEK | SEPTIN9; SEPTI | Q9UHD8; K7 | SEPT9_HUMAI | Unmodified | 57064 | 61877 | 204620 | 344340 | 7478600 | 1036700 | 702220 | 464590 | 164180 | 1015500 | 1332200 | 4958200 | 1130500 | 688550 | 230930 | 380340 | 4295300 | 4318200 |
| FINDQYEK' | SEPTIN9; SEPTI | Q9UHD8; K7 | SEPT9_HUMAI | Unmodified | 0 | 0 | 0 | 0 | 0 | 0 | 0 | 0 | 0 | 0 | 0 | 0 | 0 | 0 | 0 | 0 | 4283100 | 989040 |
| FINEDRLPI | RFC5; RFC5; RF | P40937; C9J | RFC5_HUMAN | Unmodified | 0 | 0 | 0 | 0 | 204580 | 0 | 0 | 0 | 642080 | 156390 | 0 | 143930 | 139370 | 170370 | 0 | 0 | 0 | 134100 |
| FINLNGLQI | ARHGAP4; ARH | P98171; E9P | RHG04_HUMAI | Unmodified | 0 | 0 | 0 | 0 | 0 | 0 | 0 | 0 | 0 | 0 | 85353 | 1010700 | 920120 | 0 | 0 | 0 | 0 | 0 |
| FINLYGLQI | SRGAP1; SRGA | Q7Z6B7; G5I | SRGP1_HUMAI | Unmodified | 0 | 215100 | 0 | 0 | 0 | 0 | 0 | 0 | 0 | 0 | 79891 | 1103500 | 404160 | 0 | 0 | 0 | 0 | 0 |
| FINNNAVTI | CUL1; CUL1; CI | Q13616; A0/ | CUL1_HUMAN | Unmodified | 0 | 0 | 0 | 0 | 470940 | 0 | 0 | 0 | 0 | 44321 | 584400 | 1364300 | 711640 | 180590 | 0 | 74502 | 608920 | 882140 |
| FINVPPQIA | BCCIP | Q9P287 | BCCIP_HUMAI | Oxidation (M) | 0 | 0 | 0 | 0 | 0 | 0 | 0 | 0 | 0 | 0 | 0 | 1971800 | 0 | 0 | 0 | 1041500 | 0 | 0 |
| FIPCSPFSD | DPYSL3 | Q14195 | DPYL3_HUMAI | Unmodified | 0 | 0 | 0 | 0 | 29089 | 0 | 0 | 0 | 17520 | 0 | 4030400 | 29111000 | 0 | 0 | 0 | 0 | 0 | 34630 |
| FIPCSPFSD | DPYSL3 | Q14195 | DPYL3_HUMAI | Unmodified | 0 | 0 | 0 | 0 | 0 | 0 | 0 | 0 | 0 | 0 | 1069800 | 2619500 | 0 | 0 | 0 | 0 | 0 | 0 |
| FIPDPNAEI | CLGN | O14967 | CLGN_HUMAN | Unmodified | 0 | 0 | 0 | 0 | 0 | 0 | 0 | 0 | 2934700 | 0 | 0 | 0 | 0 | 0 | 0 | 0 | 0 | 0 |
| FIPINGYPIF | ROR1; ROR1 | Q01973; A0/ | ROR1_HUMAN | Unmodified | 0 | 0 | 0 | 0 | 0 | 0 | 0 | 0 | 0 | 0 | 0 | 1235100 | 0 | 0 | 0 | 0 | 0 | 0 |
| FIPLIPIPER | CNTN1 | Q12860 | CNTN1_HUMA | Unmodified | 0 | 0 | 0 | 0 | 0 | 0 | 0 | 0 | 5835600 | 512730 | 0 | 0 | 0 | 0 | 0 | 0 | 0 | 0 |
| FIPLSEPAP | COPS8; COPS8 | Q99627; H7( | CSN8_HUMAN | Unmodified | 0 | 174370 | 1788100 | 0 | 727330 | 0 | 0 | 48171 | 844790 | 260650 | 2285700 | 8768400 | 2334900 | 1423000 | 0 | 950690 | 266980 | 796400 |
| FIPNQTEAI | EPB41L4B | Q9H329 | E41LB_HUMAI | Oxidation (M) | 872400 | 1606400 | 2355700 | 2063700 | 0 | 0 | 0 | 0 | 0 | 0 | 0 | 0 | 0 | 0 | 10321000 | 2269900 | 0 | 0 |
| FIPNTDDQI | CANT1 | Q8WVQ1 | CANT1_HUMAI | Unmodified | 0 | 0 | 0 | 0 | 0 | 0 | 0 | 0 | 2287900 | 0 | 0 | 0 | 0 | 0 | 0 | 0 | 0 | 0 |
| FIPRDSTEA | MVB12B; MVB1 | Q9H7P6; F8\ | MB12B_HUMAI | Unmodified | 509030 | 184700 | 524860 | 1200900 | 444720 | 0 | 0 | 0 | 1241700 | 569180 | 0 | 471120 | 803790 | 0 | 410920 | 274550 | 0 | 137660 |
| FIPRENGV\ | FLNA; FLNA; FL | P21333; A0A | FLNA_HUMAN | Unmodified | 0 | 0 | 91804 | 36727 | 573670 | 658730 | 0 | 0 | 0 | 71947 | 57375 | 575930 | 946920 | 0 | 142060 | 104180 | 201920 | 0 |
| FIPVKPCFY | SCAMP4; SCAM | Q969E2; C9J | SCAM4_HUMA | Unmodified | 0 | 0 | 0 | 1228000 | 0 | 0 | 0 | 0 | 0 | 0 | 0 | 0 | 0 | 0 | 2140300 | 578820 | 0 | 0 |
| FIPYTEEFSI | SEC23IP | Q9Y6Y8 | S23IP_HUMAN | Unmodified | 0 | 0 | 0 | 0 | 0 | 0 | 0 | 0 | 0 | 0 | 0 | 897400 | 0 | 0 | 0 | 0 | 0 | 0 |
| FIQENIFGI( | PDIA3 | P30101 | PDIA3_HUMAN | Oxidation (M) | 0 | 0 | 0 | 508040 | 0 | 0 | 413850 | 413610 | 66231000 | 6787600 | 19945000 | 18419000 | 310690 | 527660 | 5280300 | 0 | 0 | 0 |
| FIQENIFGI( | PDIA3 | P30101 | PDIA3_HUMAN | Unmodified | 0 | 0 | 0 | 0 | 856570 | 0 | 169280 | 0 | 71982000 | 2986000 | 2575300 | 910500 | 0 | 98505 | 0 | 0 | 771550 | 893950 |
| FIQENIFGI( | PDIA3 | P30101 | PDIA3_HUMAN | Unmodified | 0 | 0 | 0 | 0 | 0 | 0 | 0 | 0 | 17247000 | 6417600 | 794550 | 0 | 0 | 0 | 0 | 0 | 345370 | 1222100 |
| FIQENIFGI( | PDIA3 | P30101 | PDIA3_HUMAN | Oxidation (M) | 0 | 0 | 0 | 0 | 0 | 0 | 953740 | 0 | 10382000 | 11847000 | 2377000 | 0 | 0 | 0 | 0 | 0 | 0 | 0 |
| FIQGNQK | MYO1B; MYO1E | O43795; E9F | MYO1B_HUMA | Unmodified | 0 | 0 | 0 | 0 | 9244900 | 0 | 0 | 879160 | 0 | 0 | 0 | 2935900 | 0 | 0 | 0 | 0 | 1696600 | 1249800 |
| FIQQTYPS( | PDCD6IP; PDCI | Q8WUM4; F8 | PDC6I_HUMAI | Unmodified | 30038000 | 25303000 | 7768400 | 11346000 | 2104300 | 937060 | 0 | 0 | 10524000 | 2487100 | 1653200 | 229090 | 4901200 | 1251100 | 14517000 | 12561000 | 0 | 1427200 |
| FIQQTYPS( | PDCD6IP; PDCI | Q8WUM4; F8 | PDC6I_HUMAI | Unmodified | 704670 | 0 | 487540 | 4627100 | 49389000 | 14122000 | 5508500 | 3124800 | 812380 | 17184000 | 580400 | 679610 | 67147000 | 35789000 | 0 | 44936 | 9312900 | 34550000 |
| FIQQTYPS( | PDCD6IP; PDCI | Q8WUM4; F8 | PDC6I_HUMAI | Unmodified | 0 | 0 | 0 | 0 | 2126700 | 0 | 0 | 0 | 0 | 752570 | 0 | 0 | 3226300 | 1520000 | 0 | 0 | 798630 | 1348500 |
| FIREVTPYII | SNTB2; SNTB2; | Q13425; H7I | SNTB2_HUMAI | Unmodified | 0 | 0 | 513820 | 800160 | 1744600 | 305370 | 0 | 80557 | 383070 | 269740 | 0 | 0 | 1090200 | 411700 | 589970 | 445120 | 411830 | 898670 |
| FIRGVVDSI | TRAP1; TRAP1; | Q12931; I3L( | TRAP1_HUMAI | Unmodified | 0 | 0 | 0 | 0 | 0 | 0 | 0 | 0 | 357340 | 0 | 0 | 0 | 0 | 0 | 0 | 0 | 0 | 0 |
| FIRPLHEGT | FAM3A; FAM3A; | P98173; D3I | FAM3A_HUMAI | Unmodified | 0 | 0 | 0 | 0 | 0 | 0 | 0 | 0 | 6549600 | 1130100 | 666910 | 0 | 0 | 0 | 0 | 0 | 0 | 0 |
| FISADVHGI | UBA6 | A0AVT1 | UBA6_HUMAN | Unmodified | 0 | 2006200 | 0 | 167710 | 398740 | 0 | 0 | 0 | 0 | 0 | 658300 | 2908800 | 3174500 | 1610100 | 1276600 | 0 | 188870 | 0 |
| FISANLPCI | PTPRD; PTPRF; | P23468; P10 | PTPRD_HUMAI | Unmodified | 1229300 | 666770 | 855390 | 782430 | 1733800 | 964970 | 99981 | 0 | 1634700 | 578820 | 1309800 | 0 | 935470 | 434520 | 2844900 | 1779700 | 0 | 270570 |
| FISAPTKTPI | COLGALT1; CO | Q8NBJ5; M0 | GT251_HUMAI | Unmodified | 0 | 0 | 0 | 0 | 0 | 0 | 0 | 17385 | 3386800 | 143060 | 0 | 0 | 0 | 0 | 0 | 0 | 0 | 0 |
| FISDKDASI | PDIA3 | P30101 | PDIA3_HUMAN | Unmodified | 0 | 0 | 0 | 492940 | 9783200 | 506300 | 7017100 | 0 | 21263000 | 68555000 | 2106400 | 0 | 5424500 | 9514500 | 122760 | 0 | 1626000 | 11632000 |
| FISELAR | CYFIP1; CYFIP2 | Q7L576; Q9( | CYFP1_HUMAI | Unmodified | 424710 | 603560 | 803100 | 0 | 70750 | 3426200 | 2062500 | 61601 | 1318400 | 424960 | 946230 | 11123000 | 87728 | 4080700 | 1041300 | 1770800 | 552900 | 3791200 |
| FISGHTSEI | MOGS; MOGS; | Q13724; A0/ | MOGS_HUMAI | Unmodified | 0 | 103800 | 0 | 0 | 0 | 0 | 0 | 15111 | 3150700 | 199440 | 0 | 0 | 0 | 0 | 0 | 0 | 0 | 0 |
| FISGHTSEI | MOGS; MOGS; | Q13724; A0/ | MOGS_HUMAI | Unmodified | 0 | 0 | 0 | 0 | 0 | 0 | 0 | 0 | 3839400 | 0 | 1025800 | 0 | 0 | 0 | 0 | 0 | 0 | 0 |
| FISNGPFKI | CDH11; CDH11 | P55287; H3E | CAD11_HUMAI | Unmodified | 0 | 0 | 0 | 0 | 0 | 45484 | 0 | 0 | 0 | 0 | 0 | 0 | 0 | 0 | 0 | 0 | 73137 | 1004500 |
| FISVGYVDI | HLA-E; HLA-B; I | P13747; P01 | HLAE_HUMAN | Unmodified | 59915 | 946800 | 0 | 0 | 439000 | 0 | 161440 | 0 | 3609300 | 0 | 39895 | 578790 | 0 | 0 | 76549 | 81008 | 0 | 0 |
| FISVGYVDI | HLA-B; HLA-C; I | P01889; P10 | HLAB_HUMAN | Unmodified | 0 | 0 | 0 | 668290 | 0 | 0 | 0 | 0 | 8162500 | 0 | 0 | 1218400 | 0 | 0 | 0 | 0 | 0 | 0 |
| FITDNTVEE | SMARCA5 | O60264 | SMCA5_HUMA | Unmodified | 782490 | 595800 | 0 | 0 | 0 | 0 | 0 | 0 | 0 | 0 | 0 | 1251600 | 0 | 0 | 0 | 0 | 0 | 0 |
| FITEIWHPN | UBE2G1; UBE2( | P62253; I3L2 | UB2G1_HUMAI | Unmodified | 152920 | 0 | 213790 | 204060 | 316850 | 0 | 0 | 0 | 166290 | 14479 | 1388300 | 3376700 | 145770 | 0 | 343460 | 350630 | 0 | 0 |
| FITHAPPGE | CAPZA1 | P52907 | CAZA1_HUMAI | Unmodified | 5551100 | 5165300 | 2256300 | 3774800 | 2477600 | 650560 | 223320 | 0 | 7804500 | 462490 | 3191800 | 38588000 | 1821200 | 543200 | 6171800 | 4272000 | 590690 | 874810 |
| FITPEDLSK | HSPA4L; HSPA4 | O95757; E9F | HS74L_HUMAI | Unmodified | 0 | 1177900 | 0 | 0 | 244420 | 285550 | 976130 | 0 | 321210 | 0 | 510830 | 5788600 | 2397900 | 850060 | 1549100 | 0 | 492340 | 112780 |
| FITTVGIDFI | RAB27A; RAB27 | P51159; O0( | RB27A_HUMAI | Unmodified | 0 | 0 | 0 | 1353500 | 0 | 0 | 1933000 | 259410 | 0 | 0 | 0 | 216880 | 879320 | 1255200 | 0 | 0 | 0 | 0 |
| FITYPFDQN | PAF1 | Q8N7H5 | PAF1_HUMAN I | Unmodified | 0 | 0 | 0 | 0 | 163570 | 0 | 407470 | 0 | 756980 | 210560 | 81833 | 0 | 720850 | 0 | 0 | 0 | 0 | 0 |
| FIVDGWHE | HADH; HADH; I | Q16836; A0/ | HCDH_HUMAI | Oxidation (M) | 0 | 0 | 0 | 0 | 0 | 0 | 0 | 0 | 2414700 | 303350 | 0 | 0 | 0 | 0 | 0 | 0 | 0 | 0 |
| FIVDGWHE | HADH; HADH; I | Q16836; A0/ | HCDH_HUMAI | Unmodified | 0 | 0 | 0 | 0 | 0 | 0 | 0 | 0 | 1708800 | 0 | 0 | 0 | 0 | 0 | 0 | 0 | 0 | 0 |
| FIVGSDRAI | MESD | Q14696 | MESD_HUMAN | Unmodified | 0 | 0 | 0 | 0 | 0 | 0 | 0 | 0 | 0 | 0 | 0 | 0 | 0 | 0 | 0 | 0 | 0 | 0 |
| FIVLSNNYL | NCAM1; NCAM | P13591; A0A | NCAM1_HUMA | Unmodified | 0 | 0 | 0 | 0 | 0 | 0 | 100190 | 0 | 378230 | 288940 | 0 | 0 | 0 | 0 | 0 | 0 | 0 | 0 |
| FIVYSYK | GMFB; GMFB | P60983; G3\ | GMFB_HUMAN | Unmodified | 409030 | 279530 | 603570 | 749370 | 1997300 | 381450 | 197250 | 0 | 902760 | 451730 | 0 | 7188500 | 1576000 | 901620 | 931950 | 835790 | 947760 | 1218600 |
| FIYGNQDLI | ITCH; ITCH; ITC | Q96J02; A0A | ITCH_HUMAN I | Unmodified | 3838500 | 5280800 | 1257800 | 2264500 | 0 | 0 | 0 | 15518 | 628070 | 0 | 0 | 217110 | 0 | 0 | 2809100 | 2469600 | 0 | 0 |
| FIYMAVQH | PTPN11 | Q06124 | PTN11_HUMAI | Oxidation (M) | 0 | 0 | 0 | 0 | 0 | 13792 | 0 | 0 | 1160000 | 0 | 0 | 0 | 0 | 0 | 0 | 0 | 178720 | 212200 |
| FIYYPNPTF | PLXNA2 | O75051 | PLXA2_HUMAI | Unmodified | 0 | 0 | 0 | 0 | 0 | 0 | 0 | 0 | 1532400 | 0 | 0 | 0 | 0 | 0 | 0 | 0 | 0 | 0 |
| FKAEAPLPS | AHNAK | Q09666 | AHNK_HUMAN | Unmodified | 0 | 0 | 0 | 0 | 0 | 0 | 77381 | 0 | 0 | 0 | 0 | 0 | 0 | 211940 | 0 | 0 | 0 | 1729700 |
| FKDDVMP/ | TM9SF3; TM9SF | Q9HD45; Q5 | TM9S3_HUMAI | Oxidation (M) | 188710 | 75394 | 0 | 81641 | 0 | 0 | 51572 | 0 | 12010000 | 500240 | 573280 | 1902200 | 0 | 0 | 262280 | 173040 | 0 | 0 |
| FKDDVMP/ | TM9SF3; TM9SF | Q9HD45; Q5 | TM9S3_HUMAI | Unmodified | 0 | 0 | 0 | 0 | 0 | 0 | 0 | 0 | 21769000 | 434910 | 204600 | 493780 | 0 | 0 | 0 | 0 | 0 | 0 |
| FKDIFQEIFI | IDH2; IDH2 | P48735; H0\ | IDHP_HUMAN | Unmodified | 0 | 0 | 0 | 39108 | 0 | 0 | 0 | 0 | 6474400 | 1124000 | 147490 | 134920 | 0 | 0 | 0 | 0 | 0 | 0 |
| FKDIFQEIYI | IDH1 | O75874 | IDHC_HUMAN | Unmodified | 359410 | 200660 | 35490 | 487860 | 366990 | 91563 | 188920 | 16768 | 697170 | 584730 | 574550 | 1413200 | 2110900 | 784530 | 586110 | 492110 | 140410 | 128590 |
| FKDPLKDG | ATP6V1A | P38606 | VATA_HUMAN ' | Unmodified | 0 | 0 | 0 | 0 | 387160 | 0 | 172630 | 97570 | 0 | 85695 | 165280 | 0 | 1039200 | 0 | 0 | 0 | 142920 | 322830 |
| FKDPNCVG | DAZAP1; DAZAI | Q96EP5; K7E | DAZP1_HUMAI | Unmodified | 0 | 0 | 0 | 0 | 0 | 0 | 0 | 0 | 0 | 0 | 0 | 6261800 | 0 | 0 | 0 | 0 | 0 | 0 |
| FKDTYFVT( | F10 | P00742 | FA10_HUMAN ( | Unmodified | 17876 | 22553 | 0 | 1678600 | 0 | 0 | 0 | 0 | 0 | 0 | 0 | 0 | 0 | 0 | 642840 | 140250 | 0 | 0 |
| FKEANNFL' | RPL7; RPL7 | P18124; A8N | RL7_HUMAN 6( | Unmodified | 1877200 | 1357100 | 0 | 559780 | 62963 | 0 | 138200 | 0 | 240870 | 44764 | 0 | 2575900 | 0 | 0 | 127030 | 243130 | 0 | 46805 |
| FKEEGVRD | IARS1; IARS1; I/ | P41252; A0A | SYIC_HUMAN I | Unmodified | 149990 | 281710 | 0 | 0 | 149360 | 0 | 0 | 0 | 0 | 0 | 102400 | 1186700 | 0 | 0 | 0 | 159770 | 53068 | 0 |
| FKEGAEME | SIL1; SIL1; SIL1 | Q9H173; A0I | SIL1_HUMAN N | Oxidation (M) | 0 | 0 | 0 | 0 | 0 | 0 | 0 | 0 | 198450 | 0 | 0 | 0 | 0 | 0 | 0 | 0 | 0 | 0 |
| FKESFAEMI | COX4I1; COX4I | P13073; H3E | COX41_HUMAI | Unmodified | 0 | 0 | 0 | 0 | 0 | 0 | 0 | 99054 | 875840 | 0 | 0 | 0 | 0 | 0 | 0 | 0 | 0 | 0 |
| FKEVAEAYE | DNAJB2; DNAJE | P25686; C9J | DNJB2_HUMAI | Unmodified | 297630 | 286880 | 0 | 0 | 0 | 0 | 0 | 0 | 0 | 0 | 0 | 0 | 0 | 102360 | 260250 | 224920 | 0 | 0 |

| | | | | | | | | | | | | | | | | | | | | | | |
|---|---|---|---|---|---|---|---|---|---|---|---|---|---|---|---|---|---|---|---|---|---|---|
| FKFTIGDHS | WDR1; WDR1 | O75083; D6I | WDR1_HUMAN | Unmodified | 0 | 52637 | 0 | 416830 | 0 | 0 | 0 | 0 | 622090 | 409840 | 684060 | 2332500 | 750890 | 0 | 0 | 0 | 0 | 0 |
| FKGFDPNC | AGPS; AGPS; A( | O00116; A0A | ADAS_HUMAN | Unmodified | 0 | 0 | 0 | 0 | 0 | 0 | 1174700 | 0 | 707750 | 2722700 | 0 | 0 | 0 | 0 | 0 | 0 | 0 | 0 |
| FKGFPQPIL | MYOF; MYOF | Q9NZM1; H( | MYOF_HUMAN | Unmodified | 0 | 186190 | 0 | 0 | 503180 | 115300 | 0 | 0 | 0 | 0 | 0 | 0 | 0 | 0 | 0 | 0 | 117450 | 105460 |
| FKGPFTDV | VAPA | Q9P0L0 | VAPA_HUMAN | Unmodified | 153060 | 112050 | 0 | 0 | 12298000 | 2655900 | 4671700 | 1806200 | 1046700 | 2753400 | 2299400 | 5214500 | 8410200 | 5293400 | 147780 | 145550 | 1498200 | 4856500 |
| FKGPGDTS | PRKACA; PRKA( | P17612; A0A | KAPCA_HUMAN | Unmodified | 0 | 0 | 0 | 0 | 633550 | 0 | 1382000 | 0 | 0 | 955950 | 0 | 0 | 0 | 0 | 0 | 0 | 0 | 0 |
| FKGQILMPI | RPL32; RPL32; | P62910; D3Y | RL32_HUMAN ( | Unmodified | 386250 | 1202000 | 0 | 625600 | 0 | 0 | 0 | 0 | 0 | 0 | 0 | 3686900 | 751010 | 0 | 234140 | 0 | 0 | 0 |
| FKIGDQEFI | CRKL | P46109 | CRKL_HUMAN | Unmodified | 1069400 | 1166800 | 0 | 66634 | 0 | 0 | 0 | 0 | 0 | 0 | 0 | 482290 | 0 | 0 | 2190000 | 94281 | 0 | 0 |
| FKIPGSPPE | GDI2; GDI2; GD | P50395; V9G | GDIB_HUMAN | Unmodified | 151780 | 89555 | 92135 | 0 | 94296 | 0 | 0 | 0 | 1688300 | 68315 | 174280 | 2263800 | 0 | 0 | 0 | 192500 | 36783 | 0 |
| FKITIIPQDF | ABCC1; ABCC1 | P33527; I3L4 | MRP1_HUMAN | Unmodified | 97851 | 42375 | 0 | 350320 | 53756 | 19281 | 0 | 0 | 40222 | 96643 | 66789 | 894240 | 0 | 0 | 381360 | 463480 | 37236 | 0 |
| FKLDLDFPI | GSTM3; GSTM3 | P21266; A0A | GSTM3_HUMAN | Unmodified | 0 | 0 | 0 | 0 | 0 | 0 | 0 | 0 | 0 | 0 | 0 | 0 | 448620 | 0 | 0 | 0 | 0 | 77869 |
| FKLEESYDI | SERPINB6; SER | P35237; A0A | SPB6_HUMAN | Oxidation (M) | 0 | 0 | 0 | 0 | 0 | 0 | 0 | 0 | 0 | 0 | 393380 | 2194000 | 0 | 0 | 0 | 0 | 0 | 0 |
| FKLEESYTL | SERPINB1 | P30740 | ILEU_HUMAN L | Unmodified | 0 | 0 | 0 | 0 | 0 | 0 | 0 | 67967 | 474450 | 0 | 0 | 0 | 0 | 0 | 0 | 0 | 0 | 0 |
| FKLEWLGN | ATP1B1; ATP1B | P05026; V9G | AT1B1_HUMAN | Unmodified | 0 | 0 | 0 | 832950 | 138830 | 0 | 0 | 0 | 525450 | 405900 | 422360 | 1051300 | 0 | 0 | 324600 | 212980 | 32933 | 129660 |
| FKLITEDVQ | RPS3A; RPS3A; | P61247; D6F | RS3A_HUMAN | Unmodified | 2121900 | 1448300 | 0 | 865570 | 1579400 | 0 | 2561300 | 1416300 | 933650 | 975090 | 2158000 | 6495900 | 1144300 | 869170 | 655340 | 587000 | 179790 | 1486500 |
| FKLPPGEYI | CAPN2 | P17655 | CAN2_HUMAN | Unmodified | 0 | 0 | 0 | 0 | 0 | 0 | 0 | 0 | 951950 | 0 | 0 | 0 | 0 | 0 | 0 | 0 | 517050 | 0 |
| FKLPTKNEF | IDE; IDE; IDE; ID | P14735; A0A | IDE_HUMAN In | Unmodified | 0 | 0 | 0 | 0 | 3952000 | 0 | 0 | 0 | 1493400 | 0 | 1260800 | 2004600 | 16711 | 0 | 0 | 0 | 0 | 24559 |
| FKLVFLGE( | RAB6A; RAB6B; | P20340; Q9N | RAB6A_HUMAN | Unmodified | 0 | 0 | 0 | 0 | 3071000 | 0 | 2153100 | 16442 | 0 | 2307600 | 543900 | 952640 | 760260 | 0 | 0 | 133460 | 248550 | 101170 |
| FKMPEINIK | AHNAK | Q09666 | AHNK_HUMAN | Unmodified | 0 | 0 | 0 | 0 | 1033000 | 0 | 0 | 0 | 0 | 0 | 0 | 0 | 0 | 0 | 0 | 0 | 626700 | 546130 |
| FKMPEMHF | AHNAK | Q09666 | AHNK_HUMAN | Unmodified | 0 | 0 | 0 | 0 | 1410200 | 0 | 0 | 0 | 88351 | 0 | 0 | 0 | 0 | 0 | 0 | 0 | 0 | 0 |
| FKMPEMHF | AHNAK | Q09666 | AHNK_HUMAN | Unmodified | 0 | 0 | 0 | 0 | 1811900 | 0 | 48532 | 31491 | 0 | 201580 | 0 | 0 | 0 | 0 | 0 | 0 | 902250 | 1144100 |
| FKMPEMNI | AHNAK | Q09666 | AHNK_HUMAN | Oxidation (M) | 0 | 0 | 0 | 0 | 4806100 | 0 | 0 | 0 | 0 | 0 | 0 | 0 | 0 | 0 | 0 | 0 | 0 | 0 |
| FKMPEMNI | AHNAK | Q09666 | AHNK_HUMAN | Unmodified | 0 | 0 | 0 | 0 | 6396700 | 435350 | 0 | 0 | 0 | 0 | 0 | 0 | 0 | 0 | 0 | 0 | 3347300 | 3277100 |
| FKMPEMNI | AHNAK | Q09666 | AHNK_HUMAN | Oxidation (M) | 0 | 0 | 0 | 200640 | 6025700 | 0 | 0 | 0 | 0 | 0 | 466160 | 0 | 0 | 0 | 0 | 0 | 1141800 | 1368700 |
| FKMPEMNI | AHNAK | Q09666 | AHNK_HUMAN | Unmodified | 0 | 0 | 0 | 0 | 2111900 | 0 | 646010 | 0 | 0 | 0 | 0 | 0 | 0 | 0 | 0 | 0 | 1385600 | 0 |
| FKMPFLSIS | AHNAK | Q09666 | AHNK_HUMAN | Unmodified | 0 | 0 | 0 | 0 | 6148200 | 764940 | 0 | 0 | 0 | 0 | 0 | 0 | 0 | 0 | 0 | 0 | 1752900 | 3408200 |
| FKMPSFGV | AHNAK2 | Q8IVF2 | AHNK2_HUMAN | Unmodified | 0 | 0 | 0 | 0 | 9632700 | 1286300 | 0 | 0 | 0 | 0 | 0 | 0 | 0 | 0 | 0 | 0 | 3127300 | 12428000 |
| FKMPSFGV | AHNAK2 | Q8IVF2 | AHNK2_HUMAN | Oxidation (M) | 0 | 0 | 0 | 0 | 2043800 | 253970 | 0 | 92482 | 0 | 0 | 0 | 0 | 0 | 0 | 0 | 0 | 434650 | 1188000 |
| FKMPSFGV | AHNAK2 | Q8IVF2 | AHNK2_HUMAN | Unmodified | 0 | 0 | 0 | 0 | 669150 | 0 | 0 | 57044 | 0 | 0 | 0 | 0 | 0 | 0 | 0 | 0 | 122920 | 635630 |
| FKMPSMNI | AHNAK | Q09666 | AHNK_HUMAN | Unmodified | 11595 | 33758 | 0 | 0 | 1323500 | 478600 | 86040 | 233650 | 0 | 0 | 0 | 0 | 0 | 0 | 0 | 0 | 529400 | 1153700 |
| FKNEEEVF | ALDH7A1; ALDI | P49419; A0A | AL7A1_HUMAN | Unmodified | 0 | 0 | 0 | 0 | 0 | 0 | 937200 | 0 | 0 | 0 | 0 | 0 | 0 | 0 | 0 | 0 | 1280300 | 2247600 |
| FKNENFEII | SRP54; SRP54; | P61011; G3V | SRP54_HUMAN | Unmodified | 0 | 0 | 0 | 0 | 0 | 0 | 52898 | 0 | 0 | 0 | 0 | 725430 | 0 | 40195 | 0 | 0 | 0 | 0 |
| FKNQQAIQ | PHLDA3 | Q9Y5J5 | PHLA3_HUMAN | Unmodified | 0 | 0 | 0 | 0 | 882600 | 0 | 0 | 259590 | 0 | 0 | 0 | 0 | 1048500 | 0 | 0 | 0 | 0 | 0 |
| FKPGKWMI | MTHFD1; MTHF | P11586; V9G | C1TC_HUMAN | Oxidation (M) | 0 | 230220 | 0 | 0 | 1011400 | 0 | 1063600 | 0 | 701180 | 0 | 0 | 0 | 549590 | 0 | 0 | 0 | 2346500 | 387780 |
| FKPGKWMI | MTHFD1; MTHF | P11586; V9G | C1TC_HUMAN | Unmodified | 0 | 0 | 0 | 0 | 783840 | 0 | 444880 | 0 | 0 | 174880 | 0 | 0 | 0 | 0 | 0 | 0 | 520090 | 333730 |
| FKPGYLEA | PPA2; PPA2; PP | Q9H2U2; D6 | IPYR2_HUMAN | Unmodified | 0 | 0 | 0 | 0 | 0 | 0 | 0 | 0 | 1043300 | 0 | 147390 | 362130 | 0 | 0 | 0 | 0 | 0 | 0 |
| FKPNTPIGI | COBLL1; COBL | Q53SF7; A0A | COBL1_HUMAN | Oxidation (M) | 1785600 | 1237000 | 753970 | 0 | 0 | 0 | 0 | 0 | 0 | 0 | 0 | 0 | 0 | 603460 | 0 | 1101600 | 0 | 0 |
| FKPPPPNS | GCLC; GCLC; G | P48506; A0A | GSH1_HUMAN | Unmodified | 0 | 0 | 0 | 0 | 0 | 0 | 0 | 0 | 0 | 0 | 0 | 1701700 | 0 | 0 | 0 | 0 | 0 | 0 |
| FKPSLFQH | MGAT4B; MGAT | Q9UQ53; H( | MGT4B_HUMAN | Unmodified | 0 | 0 | 0 | 0 | 0 | 0 | 0 | 0 | 638810 | 0 | 0 | 0 | 0 | 0 | 0 | 0 | 0 | 0 |
| FKQISQAYE | DNAJA1 | P31689 | DNJA1_HUMAN | Unmodified | 180360 | 131080 | 0 | 231560 | 156220 | 0 | 0 | 0 | 241270 | 1650100 | 0 | 0 | 768930 | 413630 | 0 | 0 | 58707 | 0 |
| FKSDQNLC | IPO7 | O95373 | IPO7_HUMAN I | Unmodified | 0 | 0 | 0 | 0 | 0 | 0 | 0 | 0 | 0 | 0 | 0 | 1009000 | 0 | 0 | 0 | 0 | 0 | 0 |
| FKVDLTPYI | GSTZ1; GSTZ1; | O43708; G3 | MAAI_HUMAN I | Unmodified | 133930 | 112150 | 94210 | 62699 | 0 | 0 | 146300 | 64771 | 5536900 | 882580 | 0 | 800360 | 0 | 0 | 102500 | 112400 | 0 | 0 |
| FKVEYHAG | ADK; ADK; ADK; | P55263; A0A | ADK_HUMAN A | Unmodified | 0 | 0 | 0 | 0 | 525260 | 665820 | 0 | 0 | 0 | 0 | 0 | 659260 | 144080 | 190410 | 0 | 0 | 428970 | 0 |
| FKYGARNP | CIT | O14578 | CTRO_HUMAN | Unmodified | 0 | 0 | 0 | 0 | 4849100 | 0 | 0 | 0 | 0 | 0 | 0 | 0 | 0 | 0 | 0 | 0 | 968110 | 0 |
| FLAAVAHSI | MOV10; MOV1( | Q9HCE1; Q5 | MOV10_HUMA | Unmodified | 443440 | 159810 | 457320 | 42007 | 0 | 0 | 0 | 0 | 0 | 0 | 0 | 98714 | 0 | 0 | 712040 | 887860 | 0 | 0 |
| FLAEEGFYI | STT3A; STT3A; S | P46977; E9P | STT3A_HUMAN | Unmodified | 0 | 0 | 0 | 0 | 289350 | 0 | 0 | 0 | 12457000 | 775610 | 0 | 0 | 0 | 0 | 0 | 0 | 0 | 0 |
| FLAEEQAG | TRIM41; TRIM4 | Q8WV44; H( | TRI41_HUMAN | Unmodified | 117870 | 238730 | 153220 | 0 | 40939 | 0 | 23974 | 0 | 0 | 0 | 0 | 0 | 0 | 0 | 379850 | 77977 | 514030 | 0 |
| FLAITTDCL | JUP | P14923 | PLAK_HUMAN . | Unmodified | 84853 | 122110 | 0 | 0 | 0 | 0 | 0 | 0 | 0 | 966770 | 0 | 0 | 0 | 0 | 1022400 | 0 | 0 | 0 |
| FLALGDSG | RAB27A; RAB27 | P51159; H3E | RB27A_HUMAN | Unmodified | 85512 | 558060 | 2566100 | 1470700 | 632340 | 120790 | 742280 | 96411 | 362280 | 0 | 0 | 0 | 0 | 0 | 2385700 | 1437300 | 0 | 0 |
| FLALLREEG | NSF; NSF | P46459; I3L( | NSF_HUMAN V | Unmodified | 0 | 0 | 0 | 0 | 0 | 0 | 0 | 0 | 0 | 0 | 0 | 0 | 145780 | 213460 | 0 | 0 | 0 | 0 |
| FLANVNDR | SLC7A1 | P30825 | SL7A1_HUMAN | Unmodified | 148090 | 231920 | 2059400 | 0 | 0 | 0 | 0 | 0 | 0 | 0 | 0 | 24249000 | 0 | 0 | 3323000 | 0 | 0 | 0 |
| FLAPFAAHI | TMEM132A | Q24JP5 | T132A_HUMAN | Unmodified | 0 | 0 | 0 | 0 | 0 | 0 | 0 | 0 | 1101700 | 0 | 0 | 0 | 0 | 0 | 0 | 0 | 0 | 0 |
| FLAPYTTFS | PAAF1; PAAF1; | Q9BRP4; F5I | PAAF1_HUMAN | Unmodified | 0 | 0 | 0 | 0 | 0 | 0 | 413770 | 0 | 0 | 0 | 0 | 0 | 0 | 0 | 0 | 0 | 501660 | 0 |
| FLASVLHN | MRPL43; MRPL | Q8N983; H0 | RM43_HUMAN | Unmodified | 0 | 24898 | 0 | 0 | 0 | 0 | 0 | 0 | 314870 | 0 | 0 | 0 | 0 | 0 | 0 | 11563 | 0 | 0 |
| FLATEVSQI | CNNM4 | Q6P4Q7 | CNNM4_HUMA | Unmodified | 0 | 127010 | 0 | 0 | 154370 | 0 | 0 | 0 | 827900 | 0 | 1891000 | 14944000 | 0 | 0 | 0 | 991340 | 0 | 0 |
| FLATPPNGI | DNAJC13; DNA | O75165; A0A | DJC13_HUMAN | Unmodified | 8121.2 | 0 | 0 | 0 | 0 | 0 | 0 | 0 | 0 | 0 | 0 | 2436400 | 0 | 0 | 0 | 0 | 0 | 0 |
| FLATTPNSL | FN1; FN1; FN1 | P02751; H0Y | FINC_HUMAN I | Unmodified | 862280 | 2885700 | 3381600 | 0 | 0 | 0 | 0 | 0 | 887000 | 0 | 0 | 1581700 | 0 | 0 | 1286400 | 2345400 | 0 | 0 |
| FLAVGLVDI | SF3B3 | Q15393 | SF3B3_HUMAN | Unmodified | 747060 | 0 | 173040 | 0 | 0 | 0 | 0 | 0 | 184530 | 0 | 543370 | 5909300 | 0 | 0 | 0 | 521110 | 0 | 0 |
| FLCCQQSC | BMX | P51813 | BMX_HUMAN ( | Unmodified | 0 | 0 | 0 | 0 | 0 | 0 | 0 | 0 | 0 | 0 | 116710 | 662210 | 0 | 0 | 0 | 0 | 0 | 0 |
| FLCFGSEG | RO60 | P10155 | RO60_HUMAN | Unmodified | 10769000 | 16749000 | 0 | 2958100 | 0 | 0 | 0 | 0 | 0 | 0 | 0 | 70314 | 276210 | 0 | 5417700 | 3395100 | 0 | 0 |
| FLCHSNAL | WWP2 | O00308 | WWP2_HUMAN | Unmodified | 0 | 0 | 185820 | 0 | 0 | 0 | 0 | 0 | 0 | 0 | 0 | 0 | 0 | 0 | 763140 | 713170 | 0 | 0 |
| FLDAGHKL | PDIA3 | P30101 | PDIA3_HUMAN | Unmodified | 0 | 0 | 0 | 0 | 0 | 0 | 318430 | 0 | 3014700 | 4905800 | 0 | 0 | 231050 | 276470 | 0 | 0 | 0 | 0 |
| FLDAGHKL | PDIA3 | P30101 | PDIA3_HUMAN | Unmodified | 0 | 0 | 0 | 0 | 239100 | 0 | 225080 | 441410 | 0 | 11056000 | 0 | 0 | 566360 | 0 | 0 | 0 | 0 | 0 |
| FLDASGAK | BZW1; BZW1; B | Q7L1Q6; C9I | BZW1_HUMAN | Unmodified | 0 | 0 | 0 | 0 | 0 | 0 | 0 | 0 | 0 | 0 | 0 | 3069200 | 0 | 0 | 0 | 0 | 0 | 0 |
| FLDDLGLK | DSG2 | Q14126 | DSG2_HUMAN | Unmodified | 281280 | 0 | 2783400 | 0 | 0 | 0 | 0 | 0 | 0 | 0 | 0 | 1223800 | 0 | 0 | 2689500 | 3617400 | 0 | 0 |
| FLDDLGNA | BAG2 | O95816 | BAG2_HUMAN | Unmodified | 0 | 0 | 0 | 186690 | 0 | 1571100 | 0 | 0 | 712390 | 569470 | 0 | 1764300 | 1153600 | 658470 | 218040 | 118830 | 691390 | 0 |
| FLDGDELTI | CLIC5; CLIC6; ( | Q9NZA1; Q9 | CLIC5_HUMAN | Unmodified | 0 | 0 | 0 | 0 | 335300 | 0 | 336740 | 0 | 150980 | 104690 | 0 | 440080 | 257040 | 0 | 0 | 0 | 131110 | 204260 |
| FLDGDRLTI | CLIC3 | O95833 | CLIC3_HUMAN | Unmodified | 78119 | 0 | 0 | 0 | 1509800 | 506850 | 0 | 0 | 0 | 0 | 0 | 0 | 0 | 0 | 0 | 0 | 0 | 0 |
| FLDGELPLI | VPS37B; VPS37 | Q9H9H4; F5 | VP37B_HUMAN | Unmodified | 452760 | 4967500 | 778820 | 3949600 | 1346000 | 0 | 252640 | 0 | 0 | 288620 | 15132 | 51310 | 6354100 | 2635600 | 16018000 | 648570 | 288000 | 1074800 |
| FLDGELPLI | VPS37B; VPS37 | Q9H9H4; F5 | VP37B_HUMAN | Unmodified | 5148800 | 1639100 | 662610 | 2782900 | 0 | 0 | 0 | 0 | 0 | 0 | 0 | 0 | 770720 | 0 | 11507000 | 2197800 | 0 | 0 |
| FLDGIYVSE | RPL9; RPL9; RP | P32969; H0Y | RL9_HUMAN 6( | Unmodified | 3308900 | 6227300 | 2056700 | 2186700 | 3774800 | 679240 | 1358600 | 1046900 | 784690 | 575700 | 2383000 | 28217000 | 2393500 | 1266600 | 4263500 | 2501500 | 392310 | 3855000 |
| FLDGNELTI | CLIC1 | O00299 | CLIC1_HUMAN | Unmodified | 1754100 | 1377300 | 190480 | 1739500 | 36339000 | 8798300 | 2015600 | 0 | 11703000 | 5495100 | 1630400 | 5877100 | 13434000 | 18529000 | 2475900 | 2611900 | 13191000 | 15711000 |
| FLDGNELTI | CLIC1 | O00299 | CLIC1_HUMAN | Unmodified | 0 | 0 | 0 | 0 | 802540 | 0 | 0 | 0 | 0 | 0 | 0 | 0 | 0 | 1949100 | 0 | 0 | 3650700 | 0 |
| FLDGNELTI | CLIC1 | O00299 | CLIC1_HUMAN | Unmodified | 0 | 0 | 0 | 0 | 129310 | 0 | 269700 | 0 | 0 | 0 | 0 | 0 | 96235 | 305610 | 0 | 0 | 1570300 | 141190 |
| FLDGNEMI | CLIC4 | Q9Y696 | CLIC4_HUMAN | Oxidation (M) | 0 | 0 | 47350 | 0 | 4880200 | 2225200 | 0 | 0 | 0 | 3616100 | 0 | 2995500 | 2621400 | 5623800 | 0 | 0 | 3836800 | 2476900 |
| FLDGNEMI | CLIC4 | Q9Y696 | CLIC4_HUMAN | Unmodified | 0 | 0 | 0 | 0 | 1406500 | 21449 | 0 | 0 | 0 | 0 | 0 | 0 | 0 | 0 | 0 | 0 | 1021400 | 76131 |
| FLDINLYK | CTPS2 | Q9NRF8 | PYRG2_HUMAN | Unmodified | 0 | 0 | 0 | 0 | 0 | 0 | 0 | 0 | 0 | 0 | 0 | 1537000 | 0 | 0 | 0 | 0 | 0 | 0 |
| FLDKPEDVI | EPB41L1; EPB4 | Q9H4G0; H7 | E41L1_HUMAN | Unmodified | 308280 | 193200 | 615150 | 1473500 | 3605000 | 614750 | 0 | 96910 | 210960 | 195600 | 1510300 | 2316000 | 1925900 | 1918500 | 869280 | 951040 | 1396200 | 2494000 |
| FLDKPEDVI | EPB41L1; EPB4 | Q9H4G0; H7 | E41L1_HUMAN | Unmodified | 0 | 0 | 0 | 0 | 0 | 0 | 0 | 0 | 0 | 0 | 0 | 0 | 0 | 0 | 0 | 0 | 981110 | 0 |
| FLDLVPSL( | APOBEC3B; AP( | Q9UH17; P3 | ABC3B_HUMAN | Unmodified | 223640 | 314720 | 0 | 0 | 0 | 0 | 0 | 0 | 0 | 0 | 0 | 0 | 0 | 0 | 0 | 94667 | 19150 | 0 |
| FLDQNLQK | DSP | P15924 | DESP_HUMAN | Unmodified | 0 | 0 | 0 | 0 | 0 | 0 | 0 | 0 | 978110 | 202440 | 0 | 0 | 0 | 0 | 0 | 0 | 0 | 0 |
| FLDSTGSR | BZW2; BZW2; B | Q9Y6E2; Q7 | BZW2_HUMAN | Unmodified | 89847 | 115430 | 0 | 0 | 0 | 0 | 0 | 0 | 0 | 0 | 201770 | 1387900 | 0 | 0 | 13266 | 0 | 0 | 0 |
| FLDTSHYS | EXOC4 | Q96A65 | EXOC4_HUMAN | Unmodified | 0 | 0 | 0 | 378840 | 0 | 0 | 0 | 0 | 520040 | 0 | 60974 | 1307400 | 0 | 0 | 0 | 140780 | 0 | 0 |
| FLDVLELAE | MACF1; MACF1 | Q9UPN3; H3 | MACF1_HUMAN | Unmodified | 0 | 0 | 0 | 0 | 0 | 0 | 0 | 0 | 0 | 0 | 0 | 0 | 0 | 0 | 0 | 0 | 0 | 0 |
| FLEAGIYEV | CDH2; CDH2; ( | P19022; C9J | CADH2_HUMA | Unmodified | 0 | 24937 | 0 | 0 | 0 | 0 | 0 | 0 | 887900 | 276850 | 433610 | 233180 | 0 | 0 | 0 | 0 | 0 | 0 |
| FLEDALGQ | RIPOR1; RIPOR | Q6ZS17; A0A | RIPR1_HUMAN | Unmodified | 0 | 56767 | 52561 | 258160 | 0 | 0 | 0 | 82999 | 412420 | 0 | 0 | 759100 | 140050 | 0 | 144720 | 306200 | 0 | 0 |
| FLEDDPSD | EPHB3 | P54753 | EPHB3_HUMAN | Unmodified | 0 | 0 | 0 | 0 | 0 | 0 | 0 | 0 | 0 | 0 | 0 | 1207800 | 0 | 0 | 0 | 0 | 47308 | 0 |
| FLEDDPSD | EPHB3 | P54753 | EPHB3_HUMAN | Unmodified | 1179000 | 1431900 | 0 | 0 | 0 | 0 | 0 | 0 | 0 | 0 | 0 | 5743000 | 0 | 0 | 0 | 0 | 0 | 0 |
| FLEDTFGNI | MAN2B1; N/A | O00754; M0I | MA2B1_HUMAN | Unmodified | 125950 | 0 | 247810 | 130420 | 0 | 0 | 0 | 0 | 3265100 | 255900 | 1413500 | 0 | 51587 | 0 | 263970 | 119350 | 0 | 0 |
| FLEEEVYPL | ADSL; ADSL; AD | P30566; A0A | PUR8_HUMAN | Oxidation (M) | 0 | 0 | 0 | 552920 | 466520 | 0 | 0 | 0 | 304170 | 0 | 1402900 | 2988300 | 0 | 0 | 703250 | 0 | 0 | 0 |
| FLEEQEITD | PFKFB2 | O60825 | F262_HUMAN ( | Unmodified | 0 | 330190 | 0 | 0 | 0 | 0 | 0 | 0 | 232990 | 0 | 0 | 1048200 | 311390 | 0 | 0 | 0 | 0 | 0 |
| FLEEYLSST | DAD1; DAD1; D | P61803; F5G | DAD1_HUMAN | Unmodified | 0 | 0 | 0 | 0 | 0 | 0 | 0 | 0 | 2944200 | 0 | 0 | 0 | 0 | 0 | 0 | 0 | 0 | 0 |
| FLEEYLSST | DAD1; DAD1; D | P61803; F5G | DAD1_HUMAN | Unmodified | 0 | 0 | 0 | 0 | 0 | 0 | 0 | 0 | 908270 | 0 | 0 | 233430 | 0 | 0 | 0 | 0 | 0 | 0 |
| FLEFQYLTC | DSP | P15924 | DESP_HUMAN | Unmodified | 0 | 0 | 0 | 0 | 0 | 0 | 0 | 0 | 3977100 | 0 | 0 | 0 | 181880 | 110600 | 0 | 0 | 0 | 0 |
| FLEFTTLSA | RFTN1; RFTN1; | Q14699; C9J | RFTN1_HUMAN | Unmodified | 0 | 0 | 0 | 0 | 0 | 0 | 0 | 0 | 0 | 0 | 74399 | 0 | 0 | 0 | 0 | 0 | 17533 | 0 |
| FLEGEVPLE | VPS37C; VPS37 | A5D8V6; F5H | VP37C_HUMAN | Oxidation (M) | 0 | 943560 | 53136 | 395350 | 162710 | 22128 | 0 | 0 | 37525 | 146140 | 0 | 0 | 1103500 | 359750 | 563780 | 0 | 33725 | 148490 |
| FLEGTSCIA | PLEC; PLEC | Q15149; H0Y | PLEC_HUMAN | Unmodified | 0 | 0 | 163280 | 0 | 0 | 0 | 0 | 0 | 0 | 0 | 0 | 0 | 0 | 87760 | 0 | 0 | 2963100 | 2003400 |
| FLEHLSGA( | PFKP | Q01813 | PFKAP_HUMAN | Unmodified | 0 | 0 | 0 | 0 | 839600 | 0 | 47616 | 0 | 0 | 0 | 0 | 1273100 | 87218 | 0 | 0 | 0 | 180010 | 0 |
| FLEIGKFDL | FASN; FASN | P49327; A0A | FAS_HUMAN F | Oxidation (M) | 0 | 0 | 0 | 0 | 0 | 0 | 0 | 0 | 0 | 0 | 0 | 2381400 | 0 | 0 | 0 | 0 | 0 | 0 |
| FLEMCNDL | HSPA4; HSPA4; | P34932; A0A | HSP74_HUMAN | Oxidation (M) | 0 | 0 | 0 | 0 | 0 | 0 | 0 | 0 | 0 | 0 | 0 | 2009100 | 0 | 0 | 0 | 0 | 0 | 0 |
| FLEMHQL | PSAT1 | Q9Y617 | SERC_HUMAN | Oxidation (M) | 0 | 0 | 173140 | 0 | 700100 | 267570 | 0 | 0 | 0 | 0 | 0 | 1592000 | 799850 | 603440 | 0 | 242090 | 514250 | 282500 |
| FLEMHQL | PSAT1 | Q9Y617 | SERC_HUMAN | Unmodified | 0 | 0 | 0 | 0 | 1686200 | 0 | 0 | 88053 | 959470 | 404770 | 514310 | 1985300 | 1750300 | 1085900 | 0 | 0 | 340290 | 830990 |
| FLENALAVS | CLUH; CLUH; C | O75153; I3L | CLU_HUMAN C | Unmodified | 0 | 0 | 0 | 0 | 0 | 0 | 0 | 0 | 0 | 0 | 0 | 1198600 | 0 | 0 | 0 | 0 | 0 | 0 |
| FLEPTLR | DARS1; DARS1 | P14868; H7E | SYDC_HUMAN | Unmodified | 0 | 0 | 0 | 0 | 0 | 0 | 0 | 0 | 0 | 0 | 0 | 5520300 | 0 | 0 | 0 | 116610 | 0 | 0 |
| FLEQECNV | SFTPB; SFTPB | P07988; H0Y | PSPB_HUMAN | Unmodified | 0 | 0 | 0 | 0 | 0 | 0 | 0 | 0 | 1700000 | 775840 | 0 | 0 | 0 | 0 | 0 | 0 | 0 | 0 |
| FLERAAVEI | KPNB1; KPNB1 | Q14974; J3Q | IMB1_HUMAN I | Unmodified | 0 | 0 | 0 | 0 | 1380500 | 262610 | 437240 | 0 | 0 | 0 | 0 | 0 | 1005100 | 0 | 0 | 0 | 1140800 | 259800 |
| FLESGGKD | GULP1 | Q9UBP9 | GULP1_HUMAN | Unmodified | 0 | 0 | 0 | 247760 | 101810 | 22289 | 2323700 | 0 | 0 | 0 | 0 | 0 | 228660 | 120870 | 28983 | 88693 | 0 | 76032 |
| FLESGGKD | GULP1 | Q9UBP9 | GULP1_HUMAN | Unmodified | 449040 | 0 | 0 | 0 | 171490 | 282880 | 0 | 0 | 355610 | 174030 | 0 | 0 | 1249500 | 1650100 | 0 | 0 | 98417 | 0 |
| FLESGGQD | P4HB; P4HB; P4 | P07237; A0A | PDIA1_HUMAN | Oxidation (M) | 0 | 0 | 0 | 0 | 609360 | 0 | 0 | 396920 | 40831000 | 5492700 | 3587900 | 0 | 0 | 0 | 0 | 0 | 0 | 0 |
| FLESGGQD | P4HB; P4HB; P4 | P07237; A0A | PDIA1_HUMAN | Unmodified | 0 | 0 | 0 | 0 | 0 | 0 | 0 | 0 | 22989000 | 1793000 | 0 | 0 | 0 | 0 | 0 | 0 | 0 | 0 |

| | | | | | | | | | | | | | | | | | | | | | | |
|---|---|---|---|---|---|---|---|---|---|---|---|---|---|---|---|---|---|---|---|---|---|---|
| FLESGGQD | P4HB; P4HB; P4 | P07237; A0A | PDIA1_HUMAN | Oxidation (M) | 0 | 0 | 0 | 0 | 2030500 | 0 | 6228200 | 0 | 5373900 | 10506000 | 0 | 0 | 0 | 2008000 | 0 | 0 | 0 | 0 |
| FLESGGQD | P4HB; P4HB; P4 | P07237; A0A | PDIA1_HUMAN | Unmodified | 0 | 0 | 0 | 0 | 0 | 0 | 0 | 0 | 4164200 | 2774500 | 0 | 0 | 0 | 0 | 0 | 0 | 0 | 0 |
| FLESKGYDI | GFPT1; GFPT1; | Q06210; A0A | GFPT1_HUMAN | Unmodified | 0 | 0 | 0 | 0 | 0 | 0 | 0 | 0 | 0 | 313160 | 0 | 252050 | 0 | 0 | 0 | 0 | 0 | 0 |
| FLESPDFQI | PPP2R5D; PPP2 | Q14738; Q1 | 2A5D_HUMAN | Unmodified | 0 | 0 | 0 | 0 | 0 | 0 | 0 | 0 | 0 | 0 | 0 | 4164100 | 0 | 0 | 475530 | 458530 | 0 | 0 |
| FLESQEFQI | PPP2R5E | Q16537 | 2A5E_HUMAN | Unmodified | 0 | 0 | 0 | 0 | 0 | 0 | 0 | 0 | 0 | 0 | 0 | 2084800 | 0 | 0 | 0 | 0 | 0 | 0 |
| FLETELVEP | ERBB4; ERBB4; | Q15303; A0A | ERBB4_HUMAN | Unmodified | 0 | 0 | 0 | 0 | 0 | 0 | 0 | 0 | 0 | 0 | 0 | 1092400 | 0 | 0 | 0 | 0 | 0 | 0 |
| FLEVIKPFC | SEC61A1; SEC6 | P61619; B4D | S61A1_HUMAN | Unmodified | 0 | 0 | 0 | 0 | 0 | 0 | 0 | 0 | 1482600 | 1001800 | 194180 | 0 | 0 | 0 | 0 | 0 | 0 | 0 |
| FLFASVMAI | SORT1 | Q99523 | SORT_HUMAN | Unmodified | 0 | 0 | 0 | 0 | 0 | 0 | 0 | 0 | 3173400 | 384700 | 412900 | 0 | 0 | 0 | 0 | 0 | 0 | 0 |
| FLFDSVSS | IPO5; IPO5; IPO | O00410; E7E | IPO5_HUMAN I | Unmodified | 0 | 0 | 0 | 0 | 1089900 | 0 | 0 | 0 | 0 | 0 | 23053 | 9232600 | 0 | 0 | 0 | 0 | 78295 | 0 |
| FLFENQTP | U2SURP; U2SU | O15042; E7E | SR140_HUMAN | Unmodified | 0 | 0 | 0 | 0 | 0 | 0 | 0 | 0 | 0 | 0 | 0 | 2341700 | 0 | 0 | 0 | 0 | 0 | 0 |
| FLFILLGPK | SLC4A4 | Q9Y6R1 | S4A4_HUMAN | Unmodified | 0 | 0 | 0 | 0 | 0 | 0 | 0 | 0 | 0 | 0 | 0 | 0 | 0 | 0 | 203760 | 265510 | 0 | 0 |
| FLFLLLGPA | SLC4A7; SLC4A | Q9Y6M7; C9 | S4A7_HUMAN | Unmodified | 0 | 0 | 55413 | 0 | 0 | 0 | 0 | 0 | 69739 | 0 | 60068 | 1528900 | 0 | 0 | 0 | 131700 | 0 | 0 |
| FLFLLLGPS | SLC4A2; SLC4A | P04920; A0A | B3A2_HUMAN | Oxidation (M) | 98439 | 164870 | 257570 | 262100 | 0 | 0 | 0 | 0 | 0 | 0 | 20899 | 319540 | 0 | 0 | 6247200 | 465230 | 0 | 0 |
| FLFNLPLNI | EXOC2 | Q96KP1 | EXOC2_HUMAI | Unmodified | 0 | 0 | 86696 | 357830 | 0 | 0 | 0 | 0 | 1418500 | 311550 | 219310 | 637460 | 0 | 0 | 0 | 276950 | 0 | 0 |
| FLFNLPLNI | EXOC2 | Q96KP1 | EXOC2_HUMAI | Unmodified | 0 | 0 | 0 | 0 | 271960 | 80641 | 31890 | 0 | 0 | 84545 | 22639 | 0 | 67050 | 29655 | 0 | 0 | 55776 | 73419 |
| FLFSLFGQI | AIMP2; AIMP2 | Q13155; A8I | AIMP2_HUMAN | Unmodified | 0 | 0 | 192750 | 0 | 0 | 0 | 0 | 0 | 322360 | 177650 | 215260 | 5126400 | 0 | 0 | 0 | 182730 | 0 | 0 |
| FLFVDADQ | UGGT1 | Q9NYU2 | UGGG1_HUMA | Unmodified | 0 | 0 | 0 | 0 | 0 | 0 | 0 | 0 | 2730600 | 0 | 0 | 0 | 0 | 0 | 0 | 0 | 0 | 0 |
| FLFVDADQ | UGGT1 | Q9NYU2 | UGGG1_HUMA | Unmodified | 0 | 0 | 0 | 0 | 0 | 0 | 0 | 0 | 0 | 2729500 | 0 | 0 | 0 | 0 | 0 | 0 | 0 | 0 |
| FLGCVDIK | PSMD13; PSMD | Q9UNM6; A0 | PSD13_HUMAN | Unmodified | 0 | 0 | 0 | 0 | 0 | 0 | 0 | 0 | 0 | 0 | 0 | 1717100 | 0 | 0 | 125190 | 164700 | 0 | 0 |
| FLGDIEVW | TARS1 | P26639 | SYTC_HUMAN | Unmodified | 0 | 0 | 0 | 0 | 0 | 0 | 0 | 0 | 576260 | 0 | 661810 | 4313900 | 0 | 0 | 1065500 | 826570 | 0 | 0 |
| FLGDKVTS | HSPG2 | P98160 | PGBM_HUMAN | Unmodified | 0 | 0 | 0 | 196250 | 0 | 0 | 0 | 0 | 0 | 0 | 0 | 0 | 0 | 0 | 128460 | 415010 | 0 | 0 |
| FLGDYVEN | VPS13C | Q709C8 | VP13C_HUMAN | Unmodified | 2709000 | 0 | 0 | 0 | 0 | 0 | 0 | 0 | 0 | 0 | 0 | 1292700 | 0 | 0 | 3256700 | 2444600 | 0 | 0 |
| FLGGRLAM | DNAH11; DNAH | Q96DT5; A0A | DYH11_HUMAI | 2 Oxidation (M) | 0 | 0 | 0 | 0 | 0 | 0 | 0 | 0 | 0 | 867340 | 0 | 0 | 0 | 0 | 0 | 0 | 0 | 0 |
| FLGLTDTAS | RAB20 | Q9NX57 | RAB20_HUMAN | Unmodified | 0 | 0 | 208610 | 0 | 0 | 0 | 0 | 0 | 0 | 0 | 0 | 1996700 | 0 | 0 | 108180 | 194870 | 0 | 0 |
| FLGLTERD | EPHX1 | P07099 | HYEP_HUMAN | Unmodified | 15760 | 0 | 0 | 517090 | 959530 | 0 | 0 | 0 | 805140 | 125700 | 0 | 0 | 0 | 0 | 741730 | 229220 | 0 | 754430 |
| FLGPAEHL | PROM2; PROM2 | Q8N271; F8 | PROM2_HUMA | Unmodified | 8078700 | 5104200 | 4099600 | 8972600 | 0 | 0 | 0 | 0 | 211080 | 40162 | 879570 | 4110500 | 966310 | 551020 | 5214100 | 6944900 | 130370 | 0 |
| FLGQATVAI | RAB11FIP5; RAI | Q9BXF6; A0A | RFIP5_HUMAN | Unmodified | 0 | 0 | 0 | 111720 | 11356000 | 424760 | 353330 | 0 | 0 | 214330 | 0 | 0 | 2053600 | 1390400 | 0 | 0 | 436400 | 2722100 |
| FLGQNYYK | AMPD3 | Q01432 | AMPD3_HUMA | Unmodified | 0 | 0 | 1846800 | 181480 | 0 | 2284200 | 224470 | 134880 | 2203500 | 0 | 0 | 1503600 | 118400 | 0 | 438670 | 0 | 0 | 958490 |
| FLGSDEED | NUP88; NUP88 | Q99567; J3K | NUP88_HUMAI | Unmodified | 0 | 0 | 0 | 0 | 329260 | 0 | 0 | 121520 | 0 | 153650 | 0 | 1442600 | 0 | 0 | 0 | 0 | 0 | 0 |
| FLGSGGFI | PLOD1; PLOD1 | Q02809; Q5 | PLOD1_HUMAI | Unmodified | 0 | 0 | 0 | 0 | 0 | 0 | 0 | 0 | 1062000 | 0 | 0 | 0 | 0 | 0 | 0 | 0 | 0 | 0 |
| FLGSTEVE | GULP1; GULP1 | Q9UBP9; B8 | GULP1_HUMAI | Unmodified | 0 | 0 | 0 | 0 | 0 | 0 | 0 | 0 | 0 | 0 | 0 | 0 | 431490 | 444360 | 0 | 0 | 0 | 0 |
| FLGTEPEP | TST | Q16762 | THTR_HUMAN | Unmodified | 0 | 0 | 0 | 0 | 0 | 74387 | 0 | 0 | 5662400 | 408860 | 313420 | 0 | 0 | 0 | 0 | 0 | 0 | 0 |
| FLGTTCYDI | PRPF4 | O43172 | PRP4_HUMAN | Unmodified | 0 | 0 | 0 | 0 | 0 | 0 | 0 | 0 | 0 | 0 | 0 | 789550 | 0 | 0 | 0 | 0 | 0 | 0 |
| FLGVTNTIC | NDST1; NDST1 | P52848; E7E | NDST1_HUMAI | Unmodified | 0 | 0 | 0 | 0 | 0 | 0 | 0 | 0 | 426540 | 142050 | 0 | 0 | 0 | 0 | 0 | 0 | 0 | 0 |
| FLHKHDLD | PPP1CA; PPP1C | P62136; P36 | PP1A_HUMAN | Unmodified | 0 | 0 | 0 | 335610 | 462130 | 103720 | 0 | 0 | 653120 | 228880 | 0 | 1124800 | 0 | 0 | 213690 | 243710 | 0 | 0 |
| FLHKHDLD | PPP1CA; PPP1C | P62136; P36 | PP1A_HUMAN | Unmodified | 0 | 0 | 0 | 0 | 0 | 2830700 | 1015200 | 0 | 0 | 120970 | 0 | 0 | 696330 | 0 | 0 | 0 | 191090 | 0 |
| FLHLNTVQ | PIGG; PIGG; PI | Q5H8A4; E7 | PIGG_HUMAN | Unmodified | 0 | 0 | 0 | 0 | 0 | 0 | 0 | 0 | 4268900 | 0 | 0 | 0 | 0 | 0 | 0 | 0 | 0 | 0 |
| FLHNPDAA | SELENBP1; SEL | Q13228; A6 | SBP1_HUMAN | Unmodified | 0 | 0 | 589370 | 1537900 | 0 | 0 | 0 | 0 | 543330 | 0 | 0 | 0 | 0 | 0 | 1219100 | 1638000 | 0 | 0 |
| FLHQDIDS | IGF2R; IGF2R | P11717; A0A | MPRI_HUMAN | Unmodified | 291830 | 137140 | 1621900 | 536970 | 234620 | 0 | 0 | 0 | 1.02E+08 | 2466900 | 2719800 | 9252700 | 168220 | 50866 | 824590 | 877880 | 0 | 128330 |
| FLHQERMI | APLP1; APLP1; | P51693; B7Z | APLP1_HUMAN | Unmodified | 0 | 0 | 0 | 0 | 0 | 0 | 0 | 0 | 1007000 | 0 | 0 | 0 | 0 | 0 | 0 | 0 | 0 | 0 |
| FLHSEALLA | WDR46; WDR4 | O15213; H0 | WDR46_HUMA | Unmodified | 0 | 0 | 79659 | 0 | 0 | 0 | 0 | 0 | 0 | 0 | 0 | 1603800 | 0 | 0 | 0 | 0 | 0 | 0 |
| FLHTYSAAT | PIGO | Q8TEQ8 | PIGO_HUMAN | Unmodified | 0 | 0 | 0 | 0 | 0 | 0 | 0 | 0 | 7226500 | 751200 | 0 | 235950 | 0 | 0 | 0 | 0 | 448570 | 0 |
| FLICTDVAA | DDX1; DDX1; D | Q92499; A0A | DDX1_HUMAN | Unmodified | 262370 | 89924 | 129310 | 259660 | 97205 | 0 | 81645 | 0 | 0 | 0 | 515880 | 2238100 | 0 | 82194 | 0 | 211810 | 68440 | 57856 |
| FLIDGFPR | CMPK1; CMPK1 | P30085; Q5 | KCY_HUMAN U | Unmodified | 0 | 0 | 47967 | 0 | 205230 | 0 | 0 | 0 | 1661100 | 181570 | 293290 | 2316000 | 0 | 0 | 0 | 57059 | 0 | 0 |
| FLIDGFPR | CMPK1; CMPK1 | P30085; Q5 | KCY_HUMAN U | Unmodified | 0 | 0 | 0 | 0 | 9425400 | 2238600 | 2925300 | 266250 | 0 | 3118200 | 137630 | 0 | 19576000 | 8799200 | 0 | 0 | 10939000 | 14636000 |
| FLIDKNGC | GPX4; GPX4; GI | P36969; A0A | GPX4_HUMAN | Unmodified | 0 | 0 | 0 | 0 | 0 | 0 | 320600 | 257520 | 0 | 456010 | 0 | 0 | 674240 | 1415900 | 0 | 0 | 781800 | 376090 |
| FLIHEQQE | PLCG2; PLCG2 | P16885; A0A | PLCG2_HUMAI | Unmodified | 0 | 0 | 0 | 0 | 0 | 0 | 0 | 0 | 0 | 0 | 5662600 | 66497000 | 0 | 0 | 0 | 0 | 120580 | 0 |
| FLIKNCNDI | YWHAH; YWHA | Q04917; A2 | 1433F_HUMAN | Unmodified | 0 | 0 | 0 | 0 | 0 | 0 | 317540 | 222080 | 0 | 0 | 0 | 0 | 152940 | 0 | 0 | 0 | 1143400 | 0 |
| FLILLGSPK | HPSE; HPSE; H | Q9Y251; D6 | HPSE_HUMAN | Unmodified | 0 | 0 | 0 | 0 | 0 | 0 | 0 | 0 | 1284000 | 0 | 0 | 0 | 0 | 0 | 0 | 0 | 0 | 0 |
| FLILNPSK | MARK2; MARK2 | Q7KZI7; E7E | MARK2_HUMAI | Unmodified | 0 | 0 | 288490 | 168180 | 467490 | 0 | 302600 | 89252 | 0 | 145230 | 0 | 972300 | 362040 | 0 | 560470 | 518210 | 0 | 0 |
| FLINLEGGI | IARS2 | Q9NSE4 | SYIM_HUMAN I | Unmodified | 0 | 0 | 0 | 0 | 0 | 0 | 240950 | 0 | 8582700 | 346540 | 335310 | 1008400 | 0 | 0 | 0 | 70635 | 0 | 0 |
| FLIPNASQ | YWHAZ; YWHA | P63104; B0A | 1433Z_HUMAN | Unmodified | 10794000 | 12514000 | 14564000 | 29924000 | 46264000 | 15957000 | 28746000 | 20812000 | 1.46E+08 | 76295000 | 37763000 | 1.53E+08 | 34877000 | 1678000 | 24125000 | 18692000 | 1.19E+08 | 46481000 |
| FLIPNASQ | YWHAZ; YWHA | P63104; B0A | 1433Z_HUMAN | Unmodified | 0 | 0 | 0 | 0 | 2752900 | 3405600 | 25020000 | 2412700 | 0 | 16117000 | 0 | 0 | 10042000 | 31710000 | 0 | 0 | 77885000 | 22649000 |
| FLIQNVLR | IARS1; IARS1; I | P41252; A0A | SYIC_HUMAN I | Unmodified | 0 | 0 | 200860 | 0 | 0 | 0 | 0 | 0 | 0 | 0 | 355720 | 10772000 | 0 | 0 | 18631 | 0 | 0 | 0 |
| FLISPEFFT | HECW2; HECW | Q9P2P5; A0 | HECW2_HUMA | Unmodified | 0 | 0 | 0 | 0 | 0 | 0 | 0 | 0 | 1134100 | 180210 | 0 | 0 | 0 | 0 | 0 | 0 | 0 | 0 |
| FLITHNPTN | PTP4A1; PTP4A | Q93096; Q1 | TP4A1_HUMAN | Unmodified | 3029700 | 3373900 | 4010700 | 2348600 | 1168700 | 601210 | 387720 | 202380 | 5407500 | 1011600 | 5916600 | 29723000 | 1733500 | 624720 | 5343000 | 7127400 | 386550 | 982310 |
| FLITHNPTN | PTP4A2; PTP4A | Q12974; E9 | TP4A2_HUMAN | Unmodified | 0 | 0 | 0 | 1888700 | 772110 | 0 | 465950 | 0 | 0 | 1184100 | 0 | 0 | 2127900 | 875720 | 0 | 0 | 1499300 | 974400 |
| FLITHNPTN | PTP4A3; PTP4A | O75365; E5 | TP4A3_HUMAN | Unmodified | 0 | 0 | 0 | 0 | 0 | 0 | 0 | 0 | 2992200 | 0 | 0 | 0 | 0 | 0 | 0 | 0 | 0 | 0 |
| FLIVAHDD | FSCN1 | Q16658 | FSCN1_HUMAI | Unmodified | 0 | 0 | 0 | 183510 | 824830 | 85404 | 0 | 0 | 788070 | 98333 | 3115500 | 5704400 | 1153600 | 832990 | 0 | 0 | 107440 | 0 |
| FLIVGPQK | NDST2; N/A | P52849; S4 | NDST2_HUMAI | Unmodified | 0 | 0 | 0 | 0 | 0 | 0 | 0 | 0 | 1491500 | 329990 | 0 | 0 | 0 | 0 | 0 | 0 | 0 | 0 |
| FLIWDTAG | RAB22A; RAB31 | Q9UL26; Q1 | RB22A_HUMAN | Unmodified | 128360 | 5383100 | 257590 | 0 | 2929000 | 0 | 39285 | 0 | 443150 | 225540 | 0 | 3069200 | 948850 | 0 | 5280100 | 3031800 | 0 | 0 |
| FLIWDTAG | RAB22A | Q9UL26 | RB22A_HUMAN | Unmodified | 0 | 0 | 0 | 663630 | 646130 | 248190 | 0 | 0 | 0 | 1054000 | 0 | 0 | 67694 | 265940 | 0 | 0 | 492530 | 362400 |
| FLIYAQDKE | CDH2; CDH2; ( | P19022; C9 | CADH2_HUMA | Unmodified | 0 | 0 | 0 | 0 | 0 | 0 | 0 | 0 | 1147100 | 446010 | 409440 | 0 | 0 | 233640 | 0 | 0 | 0 | 0 |
| FLKPIIPNLI | PCDHB5 | Q9Y5E4 | PCDB5_HUMAI | Unmodified | 0 | 0 | 0 | 0 | 829150 | 0 | 0 | 0 | 3944000 | 724360 | 0 | 2134800 | 564330 | 201070 | 0 | 1163500 | 0 | 0 |
| FLKPVELRI | TJP1; TJP1; TJP1 | A0A087X0K | A0A087X0K9_H | Oxidation (M) | 0 | 0 | 0 | 0 | 1534300 | 237790 | 150300 | 0 | 0 | 59517 | 0 | 0 | 0 | 0 | 0 | 0 | 420830 | 349900 |
| FLKPYTLEE | ATP1B3; ATP1B | P54709; C9 | AT1B3_HUMAN | Unmodified | 0 | 182420 | 10880000 | 2165300 | 45035 | 0 | 0 | 0 | 4832100 | 0 | 3209800 | 40167000 | 0 | 1232300 | 19118000 | 13818000 | 0 | 0 |
| FLLADNLY | VBP1 | P61758 | PFD3_HUMAN | Unmodified | 0 | 0 | 0 | 0 | 979500 | 0 | 0 | 0 | 2055800 | 63738 | 0 | 4565400 | 1985900 | 0 | 0 | 0 | 655730 | 0 |
| FLLAVSR | ELP2 | Q6IA86 | ELP2_HUMAN I | Unmodified | 0 | 0 | 0 | 0 | 0 | 0 | 0 | 0 | 0 | 17260 | 0 | 1687300 | 0 | 0 | 0 | 0 | 0 | 0 |
| FLLDAVFA | USP47 | Q96K76 | UBP47_HUMAI | Unmodified | 60463 | 0 | 0 | 48288 | 105930 | 37535 | 89517 | 17390 | 332760 | 234080 | 287100 | 1269100 | 0 | 76230 | 106890 | 177730 | 128000 | 91877 |
| FLLDHQGE | ARHGAP1 | Q07960 | RHG01_HUMAI | Unmodified | 0 | 227190 | 0 | 251350 | 2358700 | 539070 | 1344100 | 0 | 1084600 | 608640 | 391460 | 383740 | 1678000 | 1601000 | 188700 | 232480 | 474700 | 673530 |
| FLLENSVLK | RABGGTA; RAB | Q92696; H0 | PGTA_HUMAN | Unmodified | 0 | 0 | 0 | 0 | 0 | 0 | 0 | 0 | 0 | 0 | 0 | 684030 | 0 | 0 | 0 | 174410 | 0 | 0 |
| FLLESFNNI | CUL3 | Q13618 | CUL3_HUMAN | Unmodified | 78655 | 0 | 97689 | 49991 | 246810 | 0 | 0 | 0 | 178550 | 1296300 | 407280 | 3131900 | 0 | 0 | 148460 | 106430 | 0 | 409850 |
| FLLEYIAPM | PLOD1 | Q02809 | PLOD1_HUMAI | Unmodified | 0 | 0 | 0 | 0 | 0 | 0 | 0 | 0 | 473800 | 91828 | 0 | 0 | 0 | 0 | 0 | 0 | 0 | 0 |
| FLLFSSLVT | NOMO1; NOMC | Q15155; P6 | NOMO1_HUMA | Unmodified | 0 | 0 | 0 | 0 | 0 | 0 | 0 | 0 | 2306800 | 416490 | 491720 | 410850 | 0 | 0 | 0 | 69685 | 0 | 0 |
| FLLGYFPW | TBC1D15; TBC | Q8TC07; C9 | TBC15_HUMAN | Unmodified | 0 | 0 | 0 | 0 | 0 | 0 | 0 | 0 | 25091 | 0 | 0 | 286340 | 0 | 0 | 0 | 76169 | 0 | 0 |
| FLLINRNEE | SLC2A1; SLC2A | P11166; A0A | GTR1_HUMAN | Unmodified | 2160900 | 2448800 | 7423600 | 3070000 | 0 | 0 | 0 | 90265 | 4725100 | 220750 | 0 | 42947000 | 0 | 0 | 9446500 | 7353400 | 0 | 0 |
| FLLINRNEE | SLC2A1; SLC2A | P11166; A0A | GTR1_HUMAN | Unmodified | 0 | 1166400 | 0 | 1335500 | 1180300 | 752930 | 0 | 0 | 1027800 | 102320 | 167930 | 0 | 226340 | 583130 | 585040 | 722770 | 360450 | 223550 |
| FLLLGLRPF | CCM2; CCM2; ( | Q9BSQ5; A0 | CCM2_HUMAN | Unmodified | 0 | 126170 | 86066 | 0 | 0 | 0 | 0 | 0 | 227790 | 221600 | 0 | 170950 | 0 | 0 | 0 | 318750 | 263610 | 0 |
| FLLLQQK | WDR26; WDR2 | Q9H7D7; C9 | WDR26_HUMA | Unmodified | 0 | 0 | 0 | 0 | 0 | 0 | 0 | 0 | 0 | 0 | 0 | 763860 | 0 | 0 | 0 | 0 | 33803 | 93649 |
| FLLPETK | SLC22A10; CAI | Q63ZE4; Q8 | S22AA_HUMAN | Unmodified | 0 | 0 | 0 | 0 | 0 | 0 | 0 | 0 | 20621000 | 744920 | 475800 | 452370 | 0 | 0 | 59011 | 52693 | 0 | 0 |
| FLLTFQRED | ADCY6 | O43306 | ADCY6_HUMAI | Unmodified | 167250 | 294580 | 123400 | 67714 | 358930 | 0 | 0 | 0 | 165640 | 15825 | 32950 | 234720 | 0 | 0 | 176690 | 209860 | 0 | 0 |
| FLLTKMEQ | HELLS; HELLS; | Q9NRZ9; A0 | HELLS_HUMAN | Oxidation (M) | 0 | 0 | 0 | 0 | 0 | 0 | 3786700 | 2055200 | 310370 | 0 | 0 | 0 | 1494400 | 2877900 | 0 | 74235 | 0 | 0 |
| FLLYNRSPI | APOM; APOM | O95445; Q5 | APOM_HUMAN | Unmodified | 517080 | 781390 | 0 | 1390100 | 541840 | 0 | 0 | 0 | 0 | 0 | 0 | 0 | 669540 | 598440 | 591960 | 78828 | 74180 | 0 |
| FLMANGQI | GDI1; GDI2; GD | P31150; P50 | GDIA_HUMAN | Oxidation (M) | 376220 | 0 | 0 | 0 | 0 | 0 | 0 | 388560 | 0 | 676260 | 0 | 4444000 | 465770 | 0 | 1413900 | 835270 | 699630 | 333910 |
| FLMANGQI | GDI1; GDI2; GD | P31150; P50 | GDIA_HUMAN | Unmodified | 0 | 0 | 0 | 408140 | 1176400 | 0 | 1027400 | 0 | 2307700 | 1093200 | 864560 | 5825100 | 497810 | 0 | 0 | 0 | 0 | 0 |
| FLMGTNSP | SUMF2; SUMF2 | Q8NBJ7; J3C | SUMF2_HUMAI | Oxidation (M) | 0 | 0 | 0 | 0 | 0 | 0 | 0 | 0 | 1440200 | 0 | 0 | 0 | 0 | 0 | 0 | 0 | 0 | 0 |
| FLMGTNSP | SUMF2; SUMF2 | Q8NBJ7; J3C | SUMF2_HUMAI | Unmodified | 0 | 0 | 0 | 0 | 0 | 0 | 0 | 0 | 6619900 | 0 | 0 | 0 | 0 | 0 | 0 | 0 | 0 | 0 |
| FLMSLVNQ | EIF3F; EIF3F | O00303; H0 | EIF3F_HUMAN | Oxidation (M) | 0 | 0 | 0 | 0 | 0 | 0 | 0 | 0 | 0 | 0 | 0 | 6575400 | 0 | 0 | 0 | 0 | 0 | 0 |
| FLMSLVNQ | EIF3F; EIF3F | O00303; H0 | EIF3F_HUMAN | Unmodified | 0 | 0 | 0 | 0 | 0 | 0 | 0 | 0 | 0 | 0 | 0 | 3494000 | 0 | 0 | 0 | 0 | 0 | 0 |
| FLNAENAQ | RANBP1; RANB | P43487; F6 | RANG_HUMAN | Unmodified | 10530000 | 613380 | 1315400 | 3590900 | 1537400 | 53959 | 0 | 111600 | 4307500 | 338200 | 5457800 | 15988000 | 0 | 0 | 1639000 | 2885500 | 0 | 0 |
| FLNDTSLPI | MYO1B; MYO1E | O43795; E9 | MYO1B_HUMA | Unmodified | 0 | 273850 | 0 | 0 | 789770 | 301470 | 0 | 57207 | 841830 | 0 | 1380100 | 12925000 | 0 | 0 | 402080 | 422500 | 251250 | 0 |
| FLNDTSLPI | MYO1B; MYO1E | O43795; E9 | MYO1B_HUMA | Unmodified | 0 | 0 | 0 | 0 | 695180 | 0 | 0 | 0 | 0 | 0 | 0 | 0 | 0 | 0 | 0 | 0 | 0 | 0 |
| FLNDTTKP | BCCIP | Q9P287 | BCCIP_HUMAN | Unmodified | 0 | 49711 | 100560 | 167110 | 0 | 0 | 25755 | 15375 | 0 | 0 | 555680 | 3298200 | 264750 | 0 | 128030 | 134790 | 0 | 0 |
| FLNEFFLN | CARS1; CARS1 | P49589; B4D | SYCC_HUMAN | Unmodified | 0 | 0 | 0 | 0 | 0 | 0 | 0 | 0 | 0 | 0 | 0 | 1804200 | 0 | 0 | 0 | 0 | 0 | 0 |
| FLNFFKDN | GFRA1 | P56159 | GFRA1_HUMAI | Unmodified | 0 | 0 | 0 | 0 | 0 | 0 | 148460 | 0 | 0 | 0 | 0 | 0 | 0 | 0 | 0 | 0 | 0 | 0 |
| FLNRHDLD | PPP1CB | P62140 | PP1B_HUMAN | Unmodified | 436650 | 605690 | 349880 | 350430 | 0 | 0 | 51955 | 267410 | 872310 | 199160 | 358740 | 2568700 | 396800 | 108790 | 952150 | 1176900 | 202800 | 0 |
| FLNRHDLD | PPP1CB | P62140 | PP1B_HUMAN | Unmodified | 0 | 0 | 0 | 0 | 766280 | 554870 | 648020 | 0 | 0 | 521730 | 0 | 0 | 12052000 | 317500 | 0 | 0 | 1491900 | 0 |
| FLNSGGFI | PLOD3 | O60568 | PLOD3_HUMAI | Unmodified | 0 | 0 | 0 | 0 | 0 | 137830 | 0 | 0 | 1695900 | 180790 | 0 | 0 | 0 | 0 | 0 | 0 | 0 | 0 |
| FLPEEEKLT | KIFBP; KIFBP; K | Q96EK5; A0A | KBP_HUMAN K | Unmodified | 0 | 60447 | 0 | 0 | 0 | 405940 | 0 | 0 | 0 | 0 | 0 | 1274500 | 0 | 666660 | 0 | 0 | 0 | 0 |
| FLPFKVVE | HSPA5; HSPA5; | P11021; A0A | BIP_HUMAN Er | Unmodified | 0 | 0 | 0 | 0 | 0 | 0 | 5269300 | 2426900 | 204140 | 8707900 | 0 | 0 | 470480 | 424150 | 0 | 0 | 0 | 0 |
| FLPFSFEHI | ATP6V0A1; ATP | Q93050; B7 | VPP1_HUMAN | Unmodified | 255360 | 0 | 694250 | 92648 | 0 | 0 | 0 | 0 | 971900 | 235230 | 0 | 3084600 | 0 | 0 | 429570 | 322620 | 0 | 0 |
| FLPGGTLC | ADAM9; ADAM9 | Q13443; A0A | ADAM9_HUMA | Unmodified | 2607700 | 1484000 | 2490800 | 4637200 | 931060 | 0 | 0 | 0 | 1616800 | 168660 | 235790 | 966810 | 560270 | 0 | 5227500 | 4197300 | 283500 | 0 |
| FLPGMAIG | SRM; SRM | P19623; K7E | SPEE_HUMAN | Unmodified | 3372000 | 0 | 0 | 0 | 0 | 247620 | 0 | 78391000 | 967110 | 0 | 0 | 1275800 | 0 | 6.03E+08 | 0 | 0 | 2177300 | 0 |
| FLPGQGLV | EFNB2 | P52799 | EFNB2_HUMAI | Unmodified | 0 | 0 | 0 | 0 | 0 | 0 | 0 | 0 | 0 | 0 | 0 | 0 | 0 | 0 | 0 | 977470 | 0 | 0 |
| FLPLFDR | HSPE1; HSPE1- | P61604; S4 | CH10_HUMAN | Unmodified | 0 | 0 | 381550 | 0 | 0 | 0 | 0 | 60314 | 63987000 | 1304900 | 1298100 | 3257600 | 0 | 0 | 0 | 425150 | 0 | 91948 |
| FLPLFDRV | HSPE1; HSPE1- | P61604; S4 | CH10_HUMAN | Unmodified | 1586200 | 861530 | 646240 | 1685800 | 4329400 | 1052600 | 2636300 | 48753 | 2.64E+08 | 55371000 | 10347000 | 1967300 | 13497000 | 5134600 | 1157000 | 763420 | 883550 | 1732800 |
| FLPLFDRV | HSPE1; HSPE1- | P61604; S4 | CH10_HUMAN | Unmodified | 0 | 0 | 0 | 0 | 0 | 0 | 7938200 | 0 | 0 | 13153000 | 408060 | 0 | 0 | 18153000 | 0 | 0 | 0 | 0 |
| FLPSELRDI | TLN1 | Q9Y490 | TLN1_HUMAN | Unmodified | 318950 | 0 | 737860 | 1179600 | 6581900 | 1536200 | 6494600 | 3285100 | 1559500 | 569420 | 509350 | 782790 | 2292100 | 6225200 | 715330 | 629990 | 2404000 | 4121500 |
| FLPSYQAVI | AGA; AGA | P20933; H0 | ASPG_HUMAN | Unmodified | 0 | 0 | 0 | 0 | 0 | 0 | 0 | 0 | 7962500 | 107850 | 0 | 0 | 0 | 0 | 0 | 0 | 0 | 0 |

| | | | | | | | | | | | | | | | | | | | | | | |
|---|---|---|---|---|---|---|---|---|---|---|---|---|---|---|---|---|---|---|---|---|---|---|
| FLPSYQAVI | AGA; AGA | P20933; H0Y | ASPG_HUMAN | Oxidation (M) | 0 | 0 | 0 | 0 | 0 | 0 | 0 | 0 | 688620 | 200680 | 40602 | 0 | 0 | 0 | 0 | 0 | 0 | 0 |
| FLPTENEVK | FMNL2; FMNL2 | Q96PY5; C9I | FMNL2_HUMAI | Unmodified | 0 | 0 | 0 | 0 | 103300 | 0 | 0 | 0 | 583070 | 362520 | 421390 | 1229200 | 0 | 0 | 0 | 0 | 113690 | 0 |
| FLQDYFDG | PDIA3 | P30101 | PDIA3_HUMAN | Unmodified | 1089100 | 0 | 1014600 | 1022900 | 1116600 | 0 | 543110 | 0 | 2.13E+08 | 33362000 | 45704000 | 14957000 | 3123000 | 0 | 1642400 | 1690700 | 0 | 0 |
| FLQDYFDG | PDIA3 | P30101 | PDIA3_HUMAN | Unmodified | 0 | 0 | 0 | 117970 | 0 | 0 | 0 | 0 | 28738000 | 1391500 | 6853500 | 987220 | 316820 | 0 | 0 | 0 | 0 | 0 |
| FLQEFYQD | MCM7 | P33993 | MCM7_HUMAN | Unmodified | 1597400 | 0 | 0 | 0 | 729190 | 370110 | 818290 | 0 | 745160 | 363430 | 431840 | 1291700 | 0 | 0 | 73869 | 1068800 | 192680 | 326830 |
| FLQEHGSD | FKBP3 | Q00688 | FKBP3_HUMAN | Unmodified | 3816800 | 3413500 | 2212800 | 2737100 | 2586300 | 714550 | 515430 | 527710 | 5056600 | 1562300 | 3729000 | 11580000 | 1777200 | 1238700 | 3372100 | 3157200 | 0 | 2209500 |
| FLQESNVL' | STAT3; STAT3; S | P40763; A0A | STAT3_HUMAN | Unmodified | 0 | 0 | 100400 | 0 | 0 | 0 | 50370 | 0 | 0 | 0 | 0 | 0 | 0 | 0 | 349090 | 303270 | 0 | 0 |
| FLQESNVL' | STAT3; STAT3; S | P40763; A0A | STAT3_HUMAN | Unmodified | 59300 | 41400 | 0 | 0 | 117070 | 0 | 0 | 0 | 0 | 0 | 0 | 785090 | 166980 | 0 | 142430 | 184050 | 0 | 0 |
| FLQFAYFK | CRTAP; CRTAP | O75718; C9J | CRTAP_HUMAN | Unmodified | 0 | 0 | 0 | 0 | 0 | 0 | 0 | 60408 | 6990400 | 2084500 | 0 | 155430 | 0 | 0 | 0 | 0 | 0 | 273580 |
| FLQKEEQR | SLK | Q9H2G2 | SLK_HUMAN S' | Unmodified | 0 | 0 | 0 | 0 | 692570 | 0 | 0 | 0 | 299880 | 270040 | 0 | 0 | 1893500 | 975100 | 0 | 0 | 125990 | 204590 |
| FLQLTPER | CCS; CCS; CCS | O14618; E9F | CCS_HUMAN ( | Unmodified | 41638 | 0 | 161210 | 148000 | 0 | 0 | 0 | 0 | 357300 | 0 | 307730 | 1481300 | 0 | 0 | 172710 | 101920 | 0 | 0 |
| FLQVGGNT | HCK; HCK; HCK | P08631; J3K | HCK_HUMAN T | Unmodified | 0 | 0 | 0 | 25596 | 351770 | 0 | 0 | 0 | 0 | 0 | 1686900 | 6814600 | 305620 | 276820 | 0 | 0 | 425800 | 0 |
| FLRASEEHI | NME2; NME1-N | P22392; J3K | NDKB_HUMAN | Unmodified | 0 | 0 | 0 | 1157400 | 80846000 | 9214300 | 142250 | 4776600 | 373850 | 4309800 | 4575800 | 437810 | 48443000 | 16253000 | 0 | 0 | 38100000 | 50198000 |
| FLRASEEHI | NME2; NME1-N | P22392; J3K | NDKB_HUMAN | Unmodified | 0 | 0 | 0 | 0 | 6325100 | 235640 | 2201500 | 0 | 0 | 154980 | 0 | 0 | 4204700 | 1610600 | 0 | 0 | 14545000 | 5996900 |
| FLRENPYYI | CLTC; CLTC | Q00610; A0A | CLH1_HUMAN | Unmodified | 27276000 | 14173000 | 15711000 | 13361000 | 929070 | 705810 | 0 | 0 | 7029100 | 359900 | 1209800 | 1.1E+08 | 852340 | 0 | 10774000 | 14286000 | 0 | 454680 |
| FLRENPYYI | CLTC; CLTC | Q00610; A0A | CLH1_HUMAN | Unmodified | 0 | 0 | 0 | 1878600 | 1937200 | 1212100 | 0 | 52349 | 0 | 0 | 0 | 0 | 4236700 | 2853500 | 0 | 0 | 0 | 3239100 |
| FLREWVES | ST13; ST13P5; S | P50502; Q8N | F10A1_HUMAN | Unmodified | 0 | 0 | 0 | 0 | 336110 | 0 | 0 | 0 | 2628000 | 0 | 0 | 2053900 | 0 | 0 | 0 | 451650 | 0 | 178310 |
| FLREWVES | ST13; ST13P5; S | P50502; Q8N | F10A1_HUMAN | Oxidation (M) | 0 | 0 | 0 | 0 | 0 | 0 | 0 | 0 | 340610 | 0 | 437680 | 2886200 | 656260 | 0 | 150900 | 128900 | 0 | 0 |
| FLRGYHQY | HLA-A; HLA-A; H | Q53Z42; A0A | Q53Z42_HUMA | Unmodified | 0 | 0 | 0 | 81744 | 0 | 0 | 0 | 0 | 0 | 0 | 651930 | 626860 | 0 | 0 | 0 | 0 | 0 | 672250 |
| FLRLEDFLD | PKN2; PKN2; PK | Q16513; B1A | PKN2_HUMAN | Unmodified | 140640 | 145820 | 207800 | 191110 | 106630 | 0 | 0 | 0 | 463500 | 0 | 0 | 940250 | 0 | 0 | 450530 | 201280 | 0 | 0 |
| FLRPGDDS | KLK3; KLK3; KLK | P07288; M0F | KLK3_HUMAN I | Oxidation (M) | 0 | 0 | 1101700 | 0 | 0 | 0 | 0 | 0 | 0 | 101460 | 0 | 0 | 179300 | 0 | 1251400 | 699580 | 0 | 0 |
| FLRYDCVIS | PLOD3; PLOD3 | O60568; H7C | PLOD3_HUMAI | Unmodified | 0 | 0 | 0 | 0 | 409380 | 0 | 61031 | 46553 | 482400 | 0 | 0 | 0 | 0 | 0 | 0 | 0 | 0 | 133860 |
| FLSAEEYQ( | SCYL1; SCYL1; | Q96KG9; E9I | SCYL1_HUMAN | Unmodified | 0 | 0 | 0 | 0 | 93022 | 0 | 0 | 0 | 0 | 0 | 0 | 631990 | 0 | 0 | 0 | 62412 | 273720 | 0 |
| FLSAIVSSV | IQGAP1; IQGAP | P46940; A0A | IQGA1_HUMAN | Oxidation (M) | 0 | 0 | 0 | 0 | 373610 | 0 | 141490 | 0 | 0 | 98248 | 0 | 0 | 0 | 0 | 0 | 0 | 160550 | 0 |
| FLSAVLAAC | OCRL | Q01968 | OCRL_HUMAN | Unmodified | 119330 | 163330 | 0 | 119640 | 1192400 | 0 | 0 | 103140 | 0 | 0 | 0 | 0 | 613230 | 498090 | 0 | 146370 | 624990 | 661000 |
| FLSDPQVH | DYNC1H1; DYN | Q14204; A0A | DYHC1_HUMA | Unmodified | 687430 | 304220 | 919700 | 688630 | 1247800 | 124580 | 105930 | 0 | 3968000 | 1151600 | 2213400 | 18551000 | 808550 | 282960 | 1217700 | 870450 | 0 | 275510 |
| FLSDRWMI | CBS; CBSL; CBS | P35520; P0D | CBS_HUMAN ( | Unmodified | 0 | 0 | 0 | 34698 | 283270 | 44677 | 211080 | 0 | 841420 | 231610 | 227840 | 5258700 | 301660 | 117050 | 50673 | 0 | 0 | 0 |
| FLSDRWMI | CBS; CBSL; CBS | P35520; P0D | CBS_HUMAN ( | Oxidation (M) | 29098 | 0 | 0 | 147050 | 223370 | 0 | 139170 | 58149 | 211340 | 166720 | 239250 | 3513100 | 0 | 165980 | 97245 | 0 | 0 | 0 |
| FLSEMLK | DSP | P15924 | DESP_HUMAN | Unmodified | 0 | 0 | 0 | 0 | 0 | 0 | 0 | 0 | 962540 | 61976 | 0 | 0 | 0 | 0 | 0 | 51477 | 0 | 0 |
| FLSESSSGI | KIDINS220; KID | Q9ULH0; E9 | KDIS_HUMAN I | Unmodified | 0 | 220610 | 132980 | 150560 | 0 | 15872 | 0 | 45230 | 218850 | 0 | 62591 | 570370 | 0 | 0 | 136160 | 197860 | 0 | 83642 |
| FLSFPTTK | HBA1; HBA2; H | P69905; G3\ | HBA_HUMAN H | Unmodified | 0 | 0 | 6072200 | 5501700 | 76181 | 645730 | 2515600 | 31354 | 0 | 0 | 0 | 429620 | 6734800 | 6498800 | 3394700 | 10338000 | 2303300 | 1943100 |
| FLSHSLVEI | CLUH; CLUH; C | O75153; I3L | CLU_HUMAN C | Unmodified | 0 | 0 | 0 | 0 | 0 | 0 | 0 | 0 | 0 | 0 | 245240 | 1546800 | 0 | 0 | 0 | 0 | 0 | 0 |
| FLSITEK | PLCB4; PLCB4 | Q15147; A0A | PLCB4_HUMAN | Unmodified | 0 | 101090 | 286190 | 417110 | 0 | 0 | 0 | 0 | 457590 | 168490 | 0 | 387410 | 875510 | 265910 | 515090 | 534390 | 747630 | 1152700 |
| FLSPQDTFI | RBM12 | Q9NTZ6 | RBM12_HUMAI | Unmodified | 0 | 0 | 0 | 0 | 0 | 0 | 0 | 0 | 0 | 0 | 122940 | 2155500 | 0 | 0 | 0 | 0 | 0 | 0 |
| FLSPVVPN' | EIF3L; EIF3L; EII | Q9Y262; B0( | EIF3L_HUMAN | Unmodified | 0 | 0 | 756050 | 375480 | 0 | 0 | 88290 | 0 | 0 | 0 | 864480 | 8885200 | 0 | 0 | 486000 | 655420 | 0 | 0 |
| FLSPVVPN' | EIF3L; EIF3L; EII | Q9Y262; B0( | EIF3L_HUMAN | Unmodified | 0 | 0 | 0 | 2199600 | 0 | 0 | 2717700 | 0 | 907780 | 605920 | 2228300 | 13792000 | 3700200 | 966390 | 0 | 0 | 0 | 0 |
| FLSQCAAR | IRAG2; IRAG2; I | Q12912; F5H | IRAG2_HUMAN | Unmodified | 65441 | 0 | 0 | 0 | 240790 | 0 | 0 | 0 | 0 | 0 | 116770 | 721670 | 0 | 0 | 0 | 0 | 0 | 0 |
| FLSQKQDV | MACF1; MACF1 | Q9UPN3; H3 | MACF1_HUMAI | Unmodified | 0 | 0 | 0 | 0 | 449680 | 0 | 0 | 0 | 0 | 0 | 223940 | 855490 | 0 | 0 | 0 | 0 | 684400 | 0 |
| FLSQLLER | ARHGEF2; ARH | Q92974; V9( | ARHG2_HUMA | Unmodified | 0 | 0 | 0 | 0 | 0 | 0 | 0 | 0 | 0 | 0 | 0 | 1303600 | 0 | 0 | 0 | 0 | 0 | 0 |
| FLSQPFQV. | ATP5F1B | P06576 | ATPB_HUMAN . | Oxidation (M) | 0 | 0 | 0 | 0 | 0 | 0 | 0 | 0 | 28803000 | 4688800 | 0 | 1040800 | 0 | 0 | 0 | 0 | 0 | 0 |
| FLSQPFQV. | ATP5F1B | P06576 | ATPB_HUMAN . | Unmodified | 0 | 0 | 0 | 0 | 0 | 0 | 0 | 0 | 23237000 | 1032300 | 0 | 0 | 0 | 0 | 0 | 0 | 0 | 0 |
| FLSQPFQV. | ATP5F1B | P06576 | ATPB_HUMAN . | Unmodified | 0 | 0 | 0 | 0 | 0 | 0 | 0 | 0 | 0 | 1254800 | 0 | 0 | 0 | 0 | 0 | 0 | 0 | 0 |
| FLSSAAAVS | CAD; CAD | P27708; F8V | PYR1_HUMAN | Unmodified | 0 | 192330 | 0 | 137260 | 0 | 0 | 0 | 0 | 1420300 | 0 | 0 | 1889200 | 0 | 0 | 399210 | 150980 | 0 | 0 |
| FLSSPTYR | RGS19 | P49795 | RGS19_HUMAI | Unmodified | 0 | 0 | 0 | 0 | 415070 | 0 | 0 | 0 | 193660 | 46815 | 0 | 0 | 0 | 0 | 115670 | 0 | 0 | 1957900 |
| FLSSPTYRA | RGS19 | P49795 | RGS19_HUMAI | Unmodified | 0 | 0 | 0 | 0 | 0 | 0 | 0 | 0 | 0 | 0 | 0 | 0 | 0 | 0 | 0 | 0 | 2553700 | 0 |
| FLSVISQQS | MYO10; MYO10 | Q9HD67; E9 | MYO10_HUMAI | Unmodified | 0 | 0 | 0 | 0 | 0 | 3300700 | 2857200 | 0 | 0 | 2476800 | 0 | 0 | 0 | 0 | 0 | 0 | 0 | 0 |
| FLSVLERQ | EPHX1 | P07099 | HYEP_HUMAN | Unmodified | 0 | 0 | 1444300 | 0 | 0 | 0 | 0 | 618550 | 4147800 | 1344900 | 0 | 0 | 0 | 917420 | 0 | 491240 | 0 | 0 |
| FLSVVQLNI | PROM2; PROM2 | Q8N271; F8\ | PROM2_HUMA | Unmodified | 0 | 0 | 59003 | 0 | 0 | 0 | 0 | 0 | 0 | 0 | 0 | 0 | 0 | 0 | 828740 | 0 | 0 | 0 |
| FLTAISVVK | PAXIP1 | Q6ZW49 | PAXI1_HUMAN | Unmodified | 0 | 0 | 0 | 0 | 0 | 0 | 0 | 188280 | 0 | 0 | 0 | 165080 | 0 | 0 | 0 | 0 | 0 | 0 |
| FLTALAQD( | PDCD6IP | Q8WUM4 | PDC6I_HUMAN | Unmodified | 0 | 2690300 | 0 | 390490 | 0 | 0 | 0 | 0 | 0 | 0 | 0 | 0 | 4832800 | 0 | 0 | 0 | 0 | 0 |
| FLTAVNLEI | GSTK1; GSTK1; | Q9Y2Q3; E9I | GSTK1_HUMAN | Oxidation (M) | 0 | 0 | 0 | 0 | 0 | 0 | 0 | 0 | 1454000 | 80208 | 0 | 0 | 0 | 0 | 0 | 0 | 0 | 0 |
| FLTAVNLEI | GSTK1; GSTK1; | Q9Y2Q3; E9I | GSTK1_HUMAN | Unmodified | 0 | 0 | 0 | 0 | 0 | 0 | 0 | 0 | 4714500 | 330200 | 0 | 0 | 0 | 0 | 0 | 0 | 0 | 305560 |
| FLTESHDR | HTRA1; HTRA1; | Q92743; A0A | HTRA1_HUMAN | Unmodified | 0 | 0 | 0 | 0 | 0 | 21125 | 0 | 0 | 3062600 | 64213 | 84640 | 0 | 0 | 10608 | 0 | 0 | 0 | 0 |
| FLTHTHGLI | SUSD2 | Q9UGT4 | SUSD2_HUMAI | Unmodified | 0 | 3489700 | 0 | 0 | 0 | 0 | 0 | 0 | 6934900 | 0 | 0 | 0 | 0 | 0 | 200660 | 91001 | 0 | 0 |
| FLTKGDNN | SEC11A; SEC11 | P67812; P0C | SC11A_HUMAN | Unmodified | 0 | 0 | 0 | 0 | 0 | 0 | 0 | 0 | 5664100 | 357660 | 381030 | 688170 | 0 | 0 | 0 | 0 | 0 | 0 |
| FLTKGDNN | SEC11A; SEC11 | P67812; P0C | SC11A_HUMAN | Unmodified | 0 | 0 | 0 | 0 | 105530 | 0 | 81216 | 0 | 36262000 | 3861900 | 3758400 | 77012 | 0 | 0 | 0 | 0 | 0 | 0 |
| FLTKGDNN | SEC11C; SEC11 | Q9BY50; K7E | SC11C_HUMAI | Unmodified | 0 | 0 | 0 | 0 | 0 | 0 | 0 | 0 | 2071500 | 266440 | 310310 | 185270 | 0 | 0 | 0 | 0 | 0 | 0 |
| FLTKGDNN | SEC11C; SEC11 | Q9BY50; K7E | SC11C_HUMAI | Unmodified | 0 | 0 | 0 | 0 | 0 | 124070 | 0 | 0 | 24313000 | 569740 | 2261600 | 1085500 | 0 | 0 | 0 | 0 | 0 | 0 |
| FLTLPPVLH | USP7; USP7; US | Q93009; A0A | UBP7_HUMAN | Oxidation (M) | 173590 | 0 | 68033 | 177090 | 279620 | 0 | 110390 | 0 | 366160 | 1078300 | 586610 | 0 | 277920 | 0 | 227490 | 0 | 132770 | 162410 |
| FLTLPPVLH | USP7; USP7; US | Q93009; A0A | UBP7_HUMAN | Unmodified | 0 | 0 | 0 | 0 | 0 | 0 | 0 | 0 | 159290 | 36837 | 18823 | 26451 | 0 | 0 | 0 | 0 | 0 | 0 |
| FLTPEEPFP | FAM199X | Q6PEV8 | F199X_HUMAN | Unmodified | 811030 | 1118400 | 497150 | 354460 | 18776 | 0 | 0 | 0 | 0 | 0 | 8726.9 | 0 | 32354 | 0 | 0 | 459490 | 0 | 0 |
| FLTPIYHPN | UBE2T | Q9NPD8 | UBE2T_HUMAN | Unmodified | 0 | 0 | 0 | 0 | 0 | 0 | 0 | 0 | 0 | 0 | 0 | 3851600 | 0 | 0 | 0 | 0 | 0 | 0 |
| FLTVLCSR | ANXA4; ANXA4 | P09525; Q6F | ANXA4_HUMAI | Unmodified | 2106300 | 706060 | 0 | 2459000 | 3570100 | 0 | 0 | 0 | 58328000 | 8181400 | 28297000 | 12662000 | 2586500 | 0 | 2059700 | 2804600 | 338940 | 992190 |
| FLTVMMK | RPS15A; N/A; R | P62244; H3E | RS15A_HUMAN | Oxidation (M) | 246930 | 206980 | 0 | 0 | 0 | 0 | 0 | 45573 | 0 | 0 | 294580 | 8824900 | 0 | 0 | 100050 | 411420 | 0 | 0 |
| FLTWILNR | NARS1 | O43776 | SYNC_HUMAN | Unmodified | 0 | 35437 | 0 | 0 | 0 | 0 | 0 | 0 | 0 | 0 | 0 | 1948400 | 0 | 0 | 0 | 156730 | 0 | 0 |
| FLVECFRNI | GNA13 | Q14344 | GNA13_HUMAI | Unmodified | 0 | 0 | 0 | 0 | 625330 | 0 | 608150 | 150810 | 0 | 0 | 0 | 0 | 0 | 0 | 0 | 0 | 1094400 | 689050 |
| FLVGFTNK | SSR1; SSR1; SS | P43307; C9J | SSRA_HUMAN | Unmodified | 0 | 0 | 794480 | 0 | 0 | 0 | 0 | 0 | 35882000 | 3903700 | 942770 | 522460 | 0 | 0 | 296450 | 0 | 0 | 0 |
| FLVGPDGV | GPX1; GPX1 | P07203; A0A | GPX1_HUMAN | Unmodified | 0 | 0 | 0 | 0 | 0 | 0 | 142720 | 96310 | 3215600 | 1900500 | 0 | 2062700 | 0 | 0 | 81646 | 111690 | 0 | 1129800 |
| FLVHNVKE | RPL32; RPL32; I | P62910; D3Y | RL32_HUMAN ( | Oxidation (M) | 0 | 0 | 0 | 0 | 0 | 0 | 2181000 | 0 | 0 | 0 | 0 | 0 | 1462900 | 516650 | 0 | 0 | 77118 | 0 |
| FLVIALK | TNIK; MINK1; M | Q9UKE5; Q8 | TNIK_HUMAN T | Unmodified | 50879 | 0 | 234840 | 0 | 287170 | 0 | 0 | 0 | 153800 | 551780 | 0 | 0 | 0 | 0 | 1172500 | 909510 | 0 | 0 |
| FLVIGNAGT | RAB4A | P20338 | RAB4A_HUMAN | Unmodified | 220850 | 63130 | 427760 | 1637100 | 599640 | 0 | 645580 | 1509500 | 635260 | 340900 | 0 | 3079900 | 2259300 | 0 | 1305400 | 1844700 | 0 | 0 |
| FLVIPLGSH | PACS2; PACS2 | Q86VP3; F8\ | PACS2_HUMAN | Unmodified | 0 | 0 | 0 | 0 | 0 | 0 | 0 | 0 | 0 | 0 | 0 | 1140000 | 0 | 0 | 0 | 0 | 0 | 0 |
| FLVLNYK | PTGR2; N/A; PT | Q8N8N7; G3 | PTGR2_HUMAN | Unmodified | 0 | 0 | 0 | 0 | 0 | 49794 | 59536 | 23859 | 0 | 374210 | 0 | 1937100 | 409960 | 342230 | 0 | 0 | 0 | 0 |
| FLVLPLTFN | XYLT2 | Q9H1B5 | XYLT2_HUMAN | Unmodified | 0 | 0 | 0 | 0 | 0 | 0 | 0 | 0 | 666840 | 0 | 0 | 143370 | 0 | 0 | 0 | 0 | 0 | 0 |
| FLVQNIHTI | BRIX1 | Q8TDN6 | BRX1_HUMAN | Unmodified | 0 | 0 | 0 | 0 | 0 | 0 | 0 | 354540 | 0 | 0 | 0 | 3234800 | 0 | 0 | 0 | 0 | 0 | 0 |
| FLVYVANFI | GDI2; GDI2; GD | P50395; Q5S | GDIB_HUMAN | Unmodified | 0 | 0 | 1188900 | 0 | 0 | 0 | 0 | 0 | 57608 | 0 | 0 | 2786500 | 0 | 0 | 0 | 0 | 0 | 0 |
| FLVYVANFI | GDI2; GDI2; GD | P50395; Q5S | GDIB_HUMAN | Unmodified | 7583200 | 2273200 | 3234500 | 5411300 | 0 | 0 | 0 | 0 | 7985400 | 1199400 | 2865400 | 18319000 | 0 | 0 | 14134000 | 6110800 | 0 | 0 |
| FLWESYR | EIF3A | Q14152 | EIF3A_HUMAN | Unmodified | 0 | 0 | 0 | 0 | 0 | 0 | 0 | 0 | 0 | 0 | 0 | 3121000 | 0 | 0 | 0 | 0 | 0 | 0 |
| FLYYEMGY | GPD2 | P43304 | GPDM_HUMAN | Unmodified | 0 | 0 | 0 | 0 | 0 | 0 | 0 | 0 | 1364400 | 188460 | 0 | 0 | 0 | 0 | 0 | 0 | 0 | 0 |
| FLYYEMGY | GPD2 | P43304 | GPDM_HUMAN | Oxidation (M) | 0 | 0 | 0 | 0 | 436700 | 0 | 0 | 0 | 495480 | 311850 | 0 | 0 | 0 | 0 | 0 | 0 | 0 | 0 |
| FMAASVAS | ATP8B1; ATP8B | O43520; A0A | AT8B1_HUMAN | Oxidation (M) | 0 | 0 | 0 | 0 | 0 | 0 | 0 | 0 | 0 | 0 | 0 | 488420 | 134250 | 0 | 0 | 0 | 0 | 0 |
| FMADCPHT | RAB14; RAB14 | P61106; X6R | RAB14_HUMAN | Oxidation (M) | 692460 | 681280 | 501780 | 651600 | 62993 | 0 | 0 | 0 | 664940 | 0 | 570450 | 2115400 | 0 | 0 | 591290 | 707560 | 0 | 0 |
| FMADCPHT | RAB14; RAB14 | P61106; X6R | RAB14_HUMAN | Unmodified | 0 | 153750 | 0 | 121570 | 122990 | 812550 | 0 | 0 | 629720 | 0 | 0 | 0 | 0 | 0 | 119220 | 124920 | 0 | 93313 |
| FMADTKIAI | FLOT2; FLOT2; I | Q14254; J3Q | FLOT2_HUMAN | Unmodified | 0 | 0 | 0 | 0 | 0 | 0 | 0 | 0 | 0 | 0 | 0 | 0 | 0 | 0 | 0 | 0 | 0 | 0 |
| FMCAQLPN | EHD1; EHD1; EI | Q9H4M9; C9 | EHD1_HUMAN | Oxidation (M) | 0 | 62209 | 0 | 77250 | 0 | 0 | 0 | 0 | 0 | 0 | 0 | 0 | 0 | 0 | 143360 | 0 | 0 | 48120 |
| FMCSQLPN | EHD4 | Q9H223 | EHD4_HUMAN | Oxidation (M) | 660870 | 426050 | 733550 | 636850 | 0 | 0 | 0 | 0 | 0 | 0 | 0 | 1481400 | 0 | 0 | 1044900 | 884430 | 0 | 0 |
| FMDQHPEI | NUDC | Q9Y266 | NUDC_HUMAN | Oxidation (M) | 0 | 0 | 0 | 0 | 0 | 0 | 0 | 0 | 0 | 0 | 0 | 0 | 274230 | 0 | 0 | 0 | 418110 | 93572 |
| FMDQHPEI | NUDC | Q9Y266 | NUDC_HUMAN | Unmodified | 0 | 0 | 0 | 0 | 0 | 0 | 0 | 0 | 0 | 188390 | 0 | 0 | 551470 | 543230 | 0 | 0 | 473950 | 579420 |
| FMDSAQVA | EPB41L5 | Q9HCM4 | E41L5_HUMAN | Unmodified | 0 | 0 | 171680 | 435310 | 0 | 0 | 0 | 0 | 3696000 | 251380 | 0 | 532390 | 40269 | 0 | 4491100 | 213870 | 32271 | 0 |
| FMDSAQVA | EPB41L5 | Q9HCM4 | E41L5_HUMAN | Oxidation (M) | 0 | 0 | 921840 | 2214800 | 0 | 0 | 0 | 0 | 1557600 | 0 | 0 | 0 | 0 | 0 | 1750000 | 2256900 | 0 | 0 |
| FMEDTAAE | ACY1; ABHD14A | Q03154; A0A | ACY1_HUMAN | Oxidation (M) | 61591 | 22000 | 81585 | 0 | 0 | 0 | 0 | 0 | 0 | 0 | 158090 | 627790 | 0 | 0 | 116170 | 0 | 0 | 0 |
| FMEELGED | ROR2; ROR2 | Q01974; B1A | ROR2_HUMAN | Unmodified | 0 | 0 | 0 | 0 | 0 | 0 | 0 | 0 | 1936300 | 0 | 0 | 0 | 0 | 0 | 0 | 0 | 0 | 0 |
| FMEENEGH | CD109 | Q6YHK3 | CD109_HUMAI | Unmodified | 0 | 0 | 0 | 0 | 0 | 0 | 0 | 0 | 1123500 | 0 | 0 | 0 | 0 | 0 | 0 | 0 | 0 | 0 |
| FMELNLAQ | VBP1 | P61758 | PFD3_HUMAN | Oxidation (M) | 152110 | 125610 | 350510 | 210200 | 462210 | 0 | 130920 | 0 | 0 | 117560 | 0 | 2019400 | 852280 | 0 | 303940 | 421400 | 954460 | 0 |
| FMELNLAQ | VBP1 | P61758 | PFD3_HUMAN | Unmodified | 0 | 0 | 0 | 0 | 0 | 0 | 0 | 0 | 0 | 270230 | 249540 | 1259500 | 1404000 | 0 | 0 | 0 | 1227900 | 1749500 |
| FMELNLAQ | VBP1 | P61758 | PFD3_HUMAN | Oxidation (M) | 0 | 0 | 0 | 0 | 446020 | 0 | 62973 | 115380 | 0 | 229150 | 0 | 886450 | 294500 | 120220 | 73740 | 107460 | 253000 | 0 |
| FMELNLAQ | VBP1 | P61758 | PFD3_HUMAN | Unmodified | 57922 | 0 | 0 | 0 | 918370 | 117770 | 827670 | 0 | 669670 | 681090 | 375220 | 2032900 | 1093500 | 0 | 0 | 0 | 947100 | 772990 |
| FMEPVTMQ | PLXNB2; PLXNE | O15031; H0Y | PLXB2_HUMAN | 2 Oxidation (M) | 0 | 0 | 238850 | 99487 | 0 | 0 | 0 | 0 | 0 | 0 | 0 | 0 | 0 | 0 | 0 | 371810 | 0 | 0 |
| FMEPVTMQ | PLXNB2; PLXNE | O15031; H0Y | PLXB2_HUMAN | 2 Oxidation (M) | 137890 | 0 | 354190 | 0 | 3733800 | 0 | 0 | 160280 | 0 | 0 | 0 | 0 | 1151000 | 2689100 | 0 | 0 | 1183900 | 2021100 |
| FMEPVTMQ | PLXNB2; PLXNE | O15031; H0Y | PLXB2_HUMAN | Oxidation (M) | 0 | 0 | 91250 | 0 | 7936500 | 0 | 0 | 0 | 492660 | 0 | 0 | 0 | 686160 | 0 | 0 | 0 | 1418600 | 6795400 |
| FMEPVTMQ | PLXNB2; PLXNE | O15031; H0Y | PLXB2_HUMAN | Unmodified | 0 | 0 | 0 | 514110 | 4118900 | 87106 | 306900 | 62176 | 656560 | 0 | 0 | 0 | 0 | 531000 | 0 | 0 | 447250 | 379180 |
| FMEQVIFK | ACTR3; ACTR3; | P61158; B4D | ARP3_HUMAN . | Oxidation (M) | 468990 | 463840 | 0 | 0 | 0 | 0 | 0 | 0 | 0 | 136880 | 0 | 7505200 | 0 | 0 | 477030 | 736240 | 0 | 0 |
| FMETSAK | RAP2C; RAP2A; | Q9Y3L5; P10 | RAP2C_HUMAI | Unmodified | 1924700 | 1571700 | 1503800 | 2658000 | 7386200 | 657240 | 2620800 | 23105 | 1436400 | 0 | 768970 | 4612400 | 4934900 | 2661000 | 1233500 | 75526 | 978030 | 3545100 |
| FMETVAEK | CLU; CLU | P10909; H0Y | CLUS_HUMAN | Unmodified | 0 | 284010 | 222820 | 220880 | 3490300 | 0 | 0 | 0 | 3756700 | 1056800 | 890150 | 0 | 0 | 0 | 192740 | 98312 | 296650 | 3817400 |
| FMEVMYGT | ECE1; ECE1; EC | P42892; B4D | ECE1_HUMAN | Unmodified | 0 | 0 | 0 | 0 | 399220 | 173660 | 0 | 0 | 2440300 | 204830 | 0 | 0 | 0 | 0 | 0 | 0 | 0 | 0 |
| FMEVMYGT | ECE1; ECE1; EC | P42892; B4D | ECE1_HUMAN | Unmodified | 0 | 0 | 0 | 0 | 0 | 0 | 0 | 0 | 2368300 | 0 | 0 | 0 | 0 | 0 | 0 | 0 | 0 | 0 |
| FMEVMYGT | ECE1; ECE1; EC | P42892; B4D | ECE1_HUMAN | Oxidation (M) | 0 | 0 | 0 | 0 | 0 | 44586 | 0 | 0 | 1817800 | 424560 | 0 | 0 | 0 | 0 | 0 | 0 | 0 | 0 |
| FMGTELNG | PHGDH; PHGD | O43175; A0A | SERA_HUMAN | Unmodified | 104200 | 0 | 477050 | 420270 | 1140000 | 4299900 | 214520 | 1343600 | 1148600 | 339540 | 1001300 | 12224000 | 7098100 | 2622500 | 558940 | 403330 | 1129600 | 644110 |
| FMGTELNG | PHGDH; PHGD | O43175; A0A | SERA_HUMAN | Oxidation (M) | 40283 | 0 | 248680 | 329900 | 524410 | 80735 | 379410 | 308350 | 0 | 119450 | 78624 | 2086400 | 3386600 | 1158900 | 615100 | 499410 | 25614 | 364110 |

| | | | | | | | | | | | | | | | | | | | | | | |
|---|---|---|---|---|---|---|---|---|---|---|---|---|---|---|---|---|---|---|---|---|---|---|
| FMLGKQEV | FKBP1A; FKBP1 | P62942; F6T | FKB1A_HUMAN | Unmodified | 1499300 | 1044800 | 0 | 205500 | 66241 | 0 | 0 | 0 | 1308500 | 0 | 0 | 2877300 | 120310 | 0 | 1620400 | 649110 | 0 | 0 |
| FMLGKQEV | FKBP1A; FKBP1 | P62942; F6T | FKB1A_HUMAN | Oxidation (M) | 13109000 | 3647200 | 395880 | 0 | 0 | 0 | 0 | 0 | 0 | 74333 | 0 | 1736400 | 113830 | 0 | 6196000 | 4509900 | 0 | 0 |
| FMLGKQEV | FKBP1A; FKBP1 | P62942; F6T | FKB1A_HUMAN | 2 Oxidation (M) | 198740 | 306060 | 0 | 7266400 | 485810 | 0 | 0 | 0 | 0 | 1016000 | 2143800 | 0 | 6276700 | 1744000 | 1322000 | 3220700 | 0 | 0 |
| FMLGKQEV | FKBP1A; FKBP1 | P62942; F6T | FKB1A_HUMAN | Oxidation (M) | 1560600 | 1384200 | 0 | 1442400 | 4430700 | 0 | 0 | 0 | 1307900 | 0 | 0 | 0 | 6193000 | 0 | 0 | 0 | 0 | 0 |
| FMNPFNMI | STIP1; STIP1 | P31948; F5H | STIP1_HUMAN | 2 Oxidation (M) | 79592 | 0 | 0 | 50216 | 0 | 0 | 0 | 0 | 108830 | 0 | 234230 | 1936000 | 0 | 0 | 98428 | 36493 | 0 | 0 |
| FMNPFNMI | STIP1; STIP1 | P31948; F5H | STIP1_HUMAN | Oxidation (M) | 27564 | 0 | 0 | 0 | 0 | 0 | 0 | 0 | 163600 | 0 | 54379 | 728550 | 0 | 0 | 0 | 0 | 0 | 0 |
| FMQASEDL | NME1; NME1-N | P15531; J3K | NDKA_HUMAN | Unmodified | 0 | 0 | 0 | 539960 | 9168000 | 0 | 959350 | 288230 | 3166900 | 797900 | 5643900 | 29034000 | 5472900 | 2690800 | 485100 | 439930 | 3477500 | 6765000 |
| FMQASEDL | NME1; NME1-N | P15531; J3K | NDKA_HUMAN | Oxidation (M) | 793670 | 0 | 1139200 | 0 | 6214900 | 2217300 | 1536800 | 644880 | 1277400 | 778620 | 5702600 | 25965000 | 7838600 | 6080700 | 2594100 | 2263400 | 4328600 | 6897800 |
| FMQASEDL | NME1; NME1-N | P15531; J3K | NDKA_HUMAN | Oxidation (M) | 0 | 0 | 0 | 0 | 4063400 | 0 | 919000 | 181870 | 0 | 194780 | 0 | 0 | 0 | 2476800 | 0 | 0 | 3421000 | 3792800 |
| FMQASEDL | NME1; NME1-N | P15531; J3K | NDKA_HUMAN | Unmodified | 0 | 0 | 0 | 0 | 2129700 | 185440 | 338550 | 0 | 0 | 0 | 0 | 0 | 849580 | 0 | 0 | 0 | 1767400 | 1730000 |
| FMQTFVLA | G3BP1; G3BP1; | Q13283; A0A | G3BP1_HUMAN | Oxidation (M) | 0 | 0 | 0 | 0 | 0 | 0 | 0 | 0 | 0 | 0 | 0 | 3573900 | 0 | 0 | 38560 | 0 | 0 | 0 |
| FMRDVEPE | MLF2; MLF2; MI | Q15773; F5H | MLF2_HUMAN | 3 Oxidation (M) | 1366900 | 1806400 | 362820 | 676850 | 0 | 0 | 0 | 0 | 0 | 0 | 0 | 0 | 0 | 0 | 317820 | 0 | 0 | 0 |
| FMTILCTR | ANXA6; ANXA6; | P08133; E7E | ANXA6_HUMAN | Unmodified | 0 | 0 | 0 | 379790 | 645160 | 0 | 119860 | 0 | 7997800 | 1020800 | 0 | 0 | 298660 | 0 | 245380 | 0 | 148970 | 1010600 |
| FMTILCTR | ANXA6; ANXA6; | P08133; E7E | ANXA6_HUMAN | Oxidation (M) | 31765 | 0 | 1412600 | 1276200 | 450960 | 165540 | 0 | 0 | 1267900 | 1542700 | 0 | 0 | 0 | 46139 | 1441600 | 1041900 | 353420 | 636210 |
| FMTPVIQDI | EIF3D; EIF3D; E | O15371; B0 | EIF3D_HUMAN | Oxidation (M) | 117460 | 11673000 | 0 | 0 | 0 | 0 | 0 | 0 | 0 | 0 | 0 | 1018000 | 0 | 0 | 0 | 0 | 0 | 0 |
| FMTTAMYD | UPF1 | Q92900 | RENT1_HUMAN | Oxidation (M) | 505970 | 464740 | 98846 | 410740 | 0 | 0 | 0 | 0 | 0 | 0 | 0 | 144590 | 0 | 0 | 42362 | 995150 | 0 | 0 |
| FMVQTIFAF | VAPA | Q9P0L0 | VAPA_HUMAN | 2 Oxidation (M) | 56469 | 0 | 0 | 0 | 1661300 | 131260 | 173990 | 22025 | 0 | 197590 | 202730 | 256960 | 817980 | 127110 | 123180 | 0 | 81543 | 287510 |
| FMVQTIFAF | VAPA | Q9P0L0 | VAPA_HUMAN | Oxidation (M) | 0 | 0 | 0 | 0 | 498380 | 38752 | 0 | 0 | 201190 | 49563 | 0 | 0 | 182570 | 0 | 0 | 0 | 28218 | 122770 |
| FMWGNLTI | NBR1; NBR1 | Q14596; B7Z | NBR1_HUMAN | Oxidation (M) | 0 | 0 | 0 | 0 | 0 | 0 | 3079700 | 0 | 0 | 0 | 0 | 0 | 0 | 84697 | 0 | 0 | 0 | 198680 |
| FNADEFED | RPL10; RPL10; | P27635; X1V | RL10_HUMAN | Oxidation (M) | 1186000 | 1488200 | 566900 | 481460 | 0 | 518740 | 0 | 0 | 133950 | 0 | 2259400 | 13139000 | 0 | 0 | 613240 | 1074500 | 0 | 0 |
| FNADEFED | RPL10; RPL10; | P27635; X1V | RL10_HUMAN | Unmodified | 86685 | 0 | 0 | 48706 | 469730 | 0 | 74001 | 0 | 350920 | 0 | 0 | 3548400 | 0 | 0 | 54919 | 18405 | 0 | 0 |
| FNADEFED | RPL10; RPL10; | P27635; X1V | RL10_HUMAN | Unmodified | 315790 | 304110 | 0 | 279650 | 0 | 0 | 0 | 0 | 293660 | 0 | 630190 | 2895400 | 0 | 0 | 130820 | 0 | 16039 | 0 |
| FNAESQGC | LOXL2; LOXL2 | Q9Y4K0; H0 | LOXL2_HUMAN | Unmodified | 0 | 0 | 0 | 0 | 0 | 38477 | 0 | 0 | 1748300 | 55125 | 54535 | 0 | 0 | 0 | 0 | 0 | 0 | 0 |
| FNAESQGC | LOXL2; LOXL2 | Q9Y4K0; H0 | LOXL2_HUMAN | Unmodified | 0 | 0 | 0 | 0 | 0 | 184810 | 0 | 0 | 320290 | 0 | 0 | 0 | 0 | 0 | 0 | 0 | 0 | 163540 |
| FNAKPGVII | DLG2; DLG2; D | Q15700; A0A | DLG2_HUMAN | Unmodified | 0 | 0 | 0 | 0 | 0 | 0 | 0 | 145580 | 0 | 0 | 0 | 0 | 3124100 | 3890700 | 0 | 0 | 0 | 0 |
| FNALFAQG | CLTC; CLTC | Q00610; A0A | CLH1_HUMAN | Unmodified | 8233400 | 10321000 | 4780600 | 2761000 | 1540300 | 0 | 693480 | 450140 | 1332500 | 0 | 1544900 | 25384000 | 2490500 | 0 | 8013000 | 8580100 | 465920 | 1056000 |
| FNALFAQG | CLTC; CLTC | Q00610; A0A | CLH1_HUMAN | Unmodified | 0 | 0 | 0 | 0 | 1058900 | 0 | 1152800 | 485110 | 0 | 0 | 0 | 0 | 885530 | 0 | 0 | 0 | 1394800 | 0 |
| FNASIAYDF | ASL; ASL; ASL | P04424; A0A | ARLY_HUMAN | Unmodified | 0 | 0 | 0 | 0 | 853160 | 171780 | 0 | 0 | 127470 | 120260 | 20282 | 0 | 0 | 245340 | 0 | 0 | 194180 | 498690 |
| FNASQLITC | PHB2; PHB2; PI | Q99623; F5C | PHB2_HUMAN | Unmodified | 0 | 0 | 0 | 0 | 0 | 0 | 0 | 0 | 1314800 | 529690 | 0 | 0 | 0 | 0 | 112580 | 0 | 0 | 0 |
| FNAVLCSR | ANXA11; ANXA1 | P50995; H0 | ANX11_HUMAN | Unmodified | 745050 | 505890 | 0 | 686310 | 3793200 | 1894700 | 443070 | 0 | 43267000 | 6677900 | 0 | 6903200 | 1888700 | 3257000 | 414940 | 675530 | 1617000 | 8754100 |
| FNDEHIPES | FLNB; FLNB | O75369; E7E | FLNB_HUMAN | Unmodified | 0 | 0 | 962380 | 1763200 | 4239100 | 509590 | 0 | 0 | 1572600 | 0 | 825270 | 10408000 | 0 | 0 | 1480000 | 700120 | 0 | 0 |
| FNDEHIPES | FLNB; FLNB | O75369; E7E | FLNB_HUMAN | Unmodified | 0 | 0 | 0 | 485760 | 0 | 0 | 350620 | 0 | 0 | 0 | 0 | 4678500 | 0 | 0 | 787040 | 0 | 0 | 0 |
| FNDLNYR | MANBA; MANB/ | O00462; A0A | MANBA_HUMA | Unmodified | 0 | 0 | 0 | 328740 | 0 | 0 | 0 | 0 | 872390 | 0 | 0 | 150270 | 0 | 0 | 0 | 0 | 0 | 0 |
| FNEEHIPDS | FLNA; FLNA; FL | P21333; A0A | FLNA_HUMAN | Unmodified | 6320300 | 7318900 | 9169000 | 14822000 | 3866000 | 1378600 | 1499500 | 1150700 | 3416400 | 396610 | 942940 | 14636000 | 3716200 | 2197100 | 10213000 | 9634900 | 1222900 | 547330 |
| FNEENYGV | ATXN2L; ATXN2 | Q8WWM7; H | ATX2L_HUMAN | Unmodified | 512590 | 211020 | 0 | 0 | 0 | 0 | 0 | 0 | 0 | 0 | 0 | 748830 | 0 | 0 | 0 | 0 | 0 | 0 |
| FNEIVAPGS | CDC42BPB; CD | Q9Y5S2; H0 | MRCKB_HUMA | Unmodified | 0 | 0 | 0 | 36617 | 0 | 0 | 0 | 0 | 82671 | 0 | 11249 | 207860 | 0 | 0 | 0 | 0 | 0 | 0 |
| FNEVAAQY | PIN4; PIN4; PIN | Q9Y237; H0 | PIN4_HUMAN | Unmodified | 0 | 0 | 76264 | 2563700 | 0 | 0 | 0 | 0 | 0 | 0 | 197850 | 2162000 | 196220 | 77012 | 2195600 | 177900 | 0 | 128070 |
| FNEVVSVLI | MYO6; MYO6; M | Q9UM54; A0 | MYO6_HUMAN | Unmodified | 0 | 0 | 3467700 | 4450400 | 467880 | 3829900 | 0 | 0 | 2264500 | 645240 | 1238400 | 4328600 | 1067700 | 875690 | 17164000 | 5820600 | 40948 | 0 |
| FNEYWRLI | S100A13 | Q99584 | S10AD_HUMAN | Unmodified | 0 | 0 | 0 | 0 | 54440000 | 0 | 0 | 0 | 0 | 0 | 0 | 0 | 813500 | 457890 | 0 | 0 | 21805000 | 49812000 |
| FNFFIHNL | PIK3C2A | O00443 | P3C2A_HUMAN | Unmodified | 0 | 0 | 0 | 0 | 0 | 0 | 0 | 0 | 319050 | 0 | 0 | 2037300 | 0 | 0 | 0 | 0 | 0 | 0 |
| FNFGEEEFI | DDX1; DDX1; D | Q92499; A0A | DDX1_HUMAN | Unmodified | 0 | 0 | 0 | 0 | 0 | 0 | 0 | 0 | 0 | 0 | 0 | 3679300 | 707400 | 0 | 0 | 0 | 0 | 0 |
| FNFGEEEFI | DDX1; DDX1; D | Q92499; A0A | DDX1_HUMAN | Unmodified | 0 | 0 | 0 | 0 | 0 | 0 | 61628 | 0 | 0 | 59484 | 0 | 0 | 293780 | 0 | 0 | 0 | 35860 | 191330 |
| FNFLEQAFI | KCTD12 | Q96CX2 | KCD12_HUMAN | Unmodified | 0 | 0 | 0 | 0 | 0 | 0 | 0 | 0 | 2124500 | 911080 | 0 | 1933200 | 0 | 0 | 0 | 0 | 0 | 0 |
| FNFLNPND | SF3A1 | Q15459 | SF3A1_HUMAN | Unmodified | 350070 | 255450 | 0 | 0 | 0 | 0 | 0 | 0 | 0 | 0 | 0 | 852200 | 0 | 0 | 0 | 172240 | 0 | 0 |
| FNGAHIPG | FLNC | Q14315 | FLNC_HUMAN | Unmodified | 0 | 0 | 0 | 0 | 664790 | 302500 | 0 | 0 | 0 | 0 | 0 | 0 | 0 | 0 | 0 | 0 | 356590 | 0 |
| FNGGGHIN | SOD2; SOD2; S | P04179; F5G | SODM_HUMAN | Unmodified | 0 | 0 | 0 | 0 | 0 | 0 | 0 | 0 | 1016000 | 0 | 0 | 0 | 0 | 0 | 0 | 0 | 0 | 0 |
| FNGGGHIN | SOD2; SOD2; S | P04179; F5G | SODM_HUMAN | Unmodified | 0 | 0 | 0 | 0 | 0 | 0 | 0 | 0 | 2565800 | 0 | 0 | 0 | 0 | 0 | 0 | 0 | 0 | 0 |
| FNGGHSPT | CD2AP | Q9Y5K6 | CD2AP_HUMAN | Unmodified | 1203500 | 1505500 | 1779100 | 1257900 | 0 | 0 | 9012.3 | 0 | 0 | 0 | 55705 | 645670 | 74540 | 0 | 2974000 | 2565300 | 0 | 0 |
| FNGIVEDFI | PBDC1 | Q9BVG4 | PBDC1_HUMAN | Unmodified | 21240 | 0 | 0 | 0 | 0 | 0 | 0 | 0 | 34652 | 0 | 34362 | 159060 | 237320 | 0 | 0 | 55146 | 643260 | 0 |
| FNGLLCHII | NOTCH1; NOTC | P46531; A0A | NOTC1_HUMA | Unmodified | 0 | 0 | 0 | 0 | 0 | 0 | 0 | 0 | 166250 | 0 | 0 | 180480 | 0 | 0 | 0 | 0 | 0 | 0 |
| FNGSHVVC | FLNB; FLNB | O75369; E7E | FLNB_HUMAN | Unmodified | 0 | 0 | 1219500 | 0 | 1671400 | 700020 | 0 | 0 | 247080 | 0 | 0 | 1780100 | 0 | 0 | 832670 | 664420 | 0 | 0 |
| FNGTHIPG | FLNA; FLNA; FL | P21333; A0A | FLNA_HUMAN | Unmodified | 175070 | 540830 | 236480 | 257120 | 0 | 0 | 264300 | 0 | 348160 | 62264 | 0 | 243740 | 0 | 0 | 2563100 | 564000 | 225270 | 0 |
| FNHEQHE | FNBP1; FNBP1; | Q96RU3; H0 | FNBP1_HUMAN | Unmodified | 0 | 0 | 0 | 0 | 2972500 | 0 | 0 | 0 | 0 | 0 | 0 | 0 | 0 | 0 | 0 | 0 | 0 | 0 |
| FNHNLLPS | TJP1; TJP1; TJP1 | Q07157; A0A | ZO1_HUMAN T | Unmodified | 0 | 0 | 0 | 0 | 0 | 0 | 0 | 0 | 187080 | 0 | 193740 | 0 | 0 | 0 | 0 | 657120 | 0 | 0 |
| FNHNLLPS | TJP1; TJP1; TJP1 | Q07157; A0A | ZO1_HUMAN T | Unmodified | 0 | 0 | 0 | 870100 | 7414800 | 1484500 | 0 | 70335 | 0 | 341480 | 0 | 0 | 5372700 | 8010500 | 0 | 0 | 696750 | 1555300 |
| FNILGTNTK | STAT3; STAT3; S | P40763; A0A | STAT3_HUMAN | Unmodified | 0 | 0 | 119340 | 0 | 487280 | 0 | 0 | 0 | 1521700 | 0 | 0 | 190170 | 0 | 0 | 120740 | 74716 | 0 | 0 |
| FNIPDLK | ATP6V1C1 | P21283 | VATC1_HUMAN | Unmodified | 73172 | 0 | 0 | 0 | 0 | 0 | 0 | 0 | 0 | 0 | 28844 | 1029200 | 59427 | 43308 | 150450 | 0 | 0 | 0 |
| FNITPEIK | CRAT; CRAT; CF | P43155; A0A | CACP_HUMAN | Unmodified | 0 | 0 | 0 | 0 | 0 | 0 | 0 | 0 | 747210 | 0 | 0 | 0 | 0 | 0 | 0 | 0 | 0 | 0 |
| FNKIPFYLQ | WDR48 | Q8TAF3 | WDR48_HUMA | Unmodified | 0 | 0 | 0 | 0 | 2549000 | 1150800 | 541660 | 280520 | 101010 | 338500 | 0 | 49261 | 1711000 | 1427300 | 0 | 0 | 3414200 | 738150 |
| FNLDATELS | STX6 | O43752 | STX6_HUMAN | Unmodified | 0 | 0 | 0 | 225700 | 267520 | 0 | 0 | 0 | 1476900 | 0 | 0 | 1688800 | 0 | 0 | 0 | 0 | 0 | 0 |
| FNLDLTHP | RPL22L1; RPL2 | Q6P5R6; H0 | RL22L_HUMAN | Unmodified | 0 | 711470 | 113830 | 1188700 | 0 | 0 | 663940 | 0 | 0 | 0 | 0 | 0 | 0 | 0 | 2555400 | 0 | 0 | 0 |
| FNLETEWK | OPA1; OPA1; OI | O60313; E5K | OPA1_HUMAN | Unmodified | 0 | 0 | 1464200 | 0 | 0 | 0 | 0 | 58529 | 0 | 0 | 0 | 4824800 | 0 | 0 | 0 | 0 | 0 | 0 |
| FNLILRQPN | TUFM | P49411 | EFTU_HUMAN | Oxidation (M) | 0 | 0 | 0 | 0 | 0 | 0 | 1518300 | 100760 | 8974500 | 3775900 | 409540 | 0 | 203780 | 0 | 0 | 20821 | 0 | 0 |
| FNLILRQPN | TUFM | P49411 | EFTU_HUMAN | Unmodified | 0 | 0 | 0 | 0 | 0 | 0 | 222050 | 0 | 6500500 | 466030 | 0 | 0 | 0 | 102150 | 0 | 0 | 0 | 129280 |
| FNLLAHLAI | MRPL50 | Q8N5N7 | RM50_HUMAN | Unmodified | 0 | 0 | 0 | 0 | 0 | 0 | 372110 | 0 | 810870 | 83839 | 0 | 0 | 81576 | 56625 | 0 | 0 | 35190 | 0 |
| FNLMSDAF | PCDH1 | Q08174 | PCDH1_HUMA | Oxidation (M) | 0 | 0 | 0 | 0 | 0 | 0 | 0 | 0 | 0 | 0 | 723020 | 1459000 | 0 | 0 | 0 | 0 | 0 | 0 |
| FNLMSDAF | PCDH1 | Q08174 | PCDH1_HUMA | Unmodified | 0 | 0 | 0 | 0 | 0 | 0 | 0 | 0 | 378520 | 0 | 1118600 | 1730700 | 252810 | 0 | 93859 | 0 | 170990 | 0 |
| FNLTGLNE | CSNK2B; CSNK | P67870; A0A | CSK2B_HUMAN | Unmodified | 0 | 0 | 0 | 0 | 0 | 0 | 0 | 0 | 754040 | 0 | 0 | 12790000 | 0 | 0 | 0 | 761710 | 0 | 0 |
| FNLTYVSHI | TARS1 | P26639 | SYTC_HUMAN | Unmodified | 0 | 0 | 0 | 0 | 0 | 0 | 0 | 0 | 0 | 0 | 0 | 2813600 | 0 | 0 | 180580 | 370840 | 0 | 0 |
| FNLTYVSHI | TARS1 | P26639 | SYTC_HUMAN | Unmodified | 0 | 0 | 0 | 0 | 0 | 0 | 0 | 0 | 0 | 0 | 0 | 945680 | 0 | 0 | 0 | 48067 | 0 | 0 |
| FNNGWMII | MITD1; MITD1; | Q8WV92; H7 | MITD1_HUMAN | Unmodified | 0 | 0 | 0 | 0 | 155770 | 0 | 0 | 0 | 732470 | 120670 | 0 | 1104100 | 0 | 0 | 0 | 0 | 0 | 0 |
| FNNPVFR | RNF31; RNF31; | Q96EP0; H0 | RNF31_HUMAN | Unmodified | 519300 | 0 | 0 | 0 | 0 | 0 | 0 | 0 | 0 | 0 | 0 | 0 | 0 | 76765 | 78653 | 0 | 0 | 0 |
| FNNWGGS | HADHB; HADHI | P55084; F5G | ECHB_HUMAN | Unmodified | 0 | 0 | 0 | 151090 | 0 | 0 | 0 | 435860 | 4783600 | 0 | 0 | 0 | 0 | 0 | 0 | 0 | 0 | 0 |
| FNPDIFSP | CLUH; CLUH; C | O75153; I3L | CLU_HUMAN C | Unmodified | 0 | 0 | 0 | 0 | 0 | 0 | 0 | 0 | 0 | 0 | 0 | 609090 | 0 | 0 | 0 | 0 | 201850 | 0 |
| FNPDIFSP | CLUH; CLUH; C | O75153; I3L | CLU_HUMAN C | Unmodified | 0 | 0 | 0 | 0 | 0 | 0 | 0 | 0 | 0 | 0 | 0 | 2913700 | 0 | 0 | 0 | 0 | 0 | 0 |
| FNPETDYL | ACO2; ACO2; A | Q99798; A2A | ACON_HUMAN | Unmodified | 0 | 0 | 0 | 0 | 0 | 0 | 0 | 0 | 2084700 | 0 | 0 | 0 | 0 | 0 | 0 | 0 | 0 | 0 |
| FNPETDYL | ACO2; ACO2; A | Q99798; A2A | ACON_HUMAN | Unmodified | 0 | 0 | 45442 | 0 | 0 | 0 | 0 | 0 | 3623900 | 724590 | 0 | 0 | 0 | 0 | 0 | 0 | 0 | 0 |
| FNPEWYK | ENPP1; ENPP1 | P22413; E9P | ENPP1_HUMAN | Unmodified | 0 | 0 | 0 | 0 | 0 | 0 | 0 | 0 | 0 | 0 | 0 | 0 | 547560 | 0 | 0 | 513500 | 535410 | 926780 |
| FNPFVTSD | RPL26; RPL26L | P61254; Q9 | RL26_HUMAN | Unmodified | 1812900 | 0 | 1077900 | 320690 | 533110 | 0 | 0 | 0 | 0 | 0 | 0 | 7967200 | 0 | 0 | 1697100 | 1340300 | 0 | 0 |
| FNPFVTSD | RPL26; RPL26L | P61254; Q9 | RL26_HUMAN | Unmodified | 0 | 390300 | 701980 | 416250 | 220120 | 58573 | 0 | 0 | 0 | 0 | 354930 | 1317200 | 409410 | 0 | 686440 | 384920 | 0 | 230790 |
| FNPHTSPH | NEDD1; NEDD1 | Q8NHV4; G3 | NEDD1_HUMA | Unmodified | 0 | 0 | 0 | 0 | 0 | 0 | 0 | 0 | 3208300 | 0 | 0 | 359280 | 0 | 0 | 0 | 0 | 0 | 0 |
| FNPLNAEV | LRRC20; LRRC2 | Q8TCA0; X6 | LRC20_HUMAN | Unmodified | 0 | 0 | 0 | 0 | 0 | 0 | 0 | 55915 | 0 | 0 | 0 | 1072900 | 0 | 0 | 0 | 0 | 0 | 0 |
| FNPNFYR | UBE2Z; UBE2Z | Q9H832; I3L | UBE2Z_HUMAN | Unmodified | 0 | 50470 | 67810 | 63450 | 0 | 0 | 0 | 0 | 175500 | 0 | 0 | 1063400 | 0 | 0 | 169860 | 117880 | 0 | 0 |
| FNQGMVD | CRELD2; CRELI | Q6UXH1; A6 | CREL2_HUMAN | Unmodified | 0 | 0 | 0 | 0 | 0 | 0 | 0 | 0 | 1148800 | 172120 | 258600 | 0 | 0 | 0 | 0 | 0 | 0 | 0 |
| FNQGMVD | CRELD2; CRELI | Q6UXH1; A6 | CREL2_HUMAN | Unmodified | 0 | 0 | 31626 | 0 | 0 | 0 | 0 | 0 | 5638000 | 613870 | 693300 | 0 | 3235400 | 2811500 | 0 | 0 | 73781 | 0 |
| FNQLPVELI | ABHD10 | Q9NUJ1 | ABHDA_HUMA | Unmodified | 0 | 0 | 0 | 0 | 0 | 0 | 61035 | 27568 | 5102100 | 2213100 | 110600 | 0 | 380550 | 342750 | 0 | 0 | 83305 | 151450 |
| FNQMVVK | ROCK1; ROCK1 | Q13464; A0A | ROCK1_HUMAN | Unmodified | 0 | 0 | 0 | 0 | 952960 | 0 | 0 | 0 | 208740 | 203880 | 0 | 0 | 0 | 280200 | 0 | 0 | 630420 | 0 |
| FNQTYQLA | TTLL12 | Q14166 | TTL12_HUMAN | Unmodified | 0 | 0 | 921290 | 501590 | 118560 | 0 | 0 | 83127 | 146340 | 0 | 0 | 14745000 | 0 | 0 | 1243900 | 1205100 | 50994 | 541790 |
| FNQVESDK | GALNT2 | Q10471 | GALT2_HUMAN | Unmodified | 0 | 0 | 0 | 0 | 133750 | 0 | 0 | 0 | 8021800 | 0 | 0 | 319710 | 0 | 0 | 0 | 0 | 0 | 0 |
| FNQVESDK | GALNT2 | Q10471 | GALT2_HUMAN | Unmodified | 0 | 0 | 0 | 0 | 0 | 0 | 88728 | 176480 | 12651000 | 0 | 0 | 0 | 0 | 0 | 0 | 0 | 0 | 0 |
| FNQVLDQF | PSMD3 | O43242 | PSMD3_HUMA | Unmodified | 0 | 0 | 0 | 0 | 0 | 0 | 0 | 0 | 0 | 0 | 293020 | 5082700 | 0 | 0 | 0 | 26884 | 0 | 0 |
| FNRLNRLT | SSB; SSB; SSB; | P05455; E9P | LA_HUMAN Lu | Unmodified | 0 | 0 | 0 | 0 | 80546 | 0 | 177750 | 0 | 0 | 271670 | 0 | 0 | 422560 | 91609 | 0 | 0 | 72109 | 50564 |
| FNSFYGDP | CASK; CASK; C | O14936; Q5 | CSKP_HUMAN | Unmodified | 0 | 0 | 0 | 0 | 0 | 0 | 0 | 0 | 506070 | 0 | 0 | 0 | 0 | 0 | 0 | 0 | 0 | 0 |
| FNSHESAA | TIA1; TIA1; TIA1 | P31483; F8V | TIA1_HUMAN N | Unmodified | 192280 | 0 | 0 | 138550 | 0 | 0 | 0 | 0 | 223520 | 0 | 0 | 1224000 | 0 | 0 | 0 | 0 | 0 | 0 |
| FNSHPFPG | KIRREL1; KIRRE | Q96J84; Q5V | KIRR1_HUMAN | Unmodified | 0 | 0 | 80627 | 0 | 6100500 | 1406800 | 0 | 0 | 0 | 108810 | 0 | 407940 | 504140 | 238370 | 80710 | 0 | 0 | 1642900 |
| FNSLNELVI | GRB2 | P62993 | GRB2_HUMAN | Unmodified | 7619900 | 6948400 | 6414700 | 919520 | 0 | 0 | 0 | 0 | 703420 | 192150 | 0 | 5556000 | 0 | 0 | 8572400 | 8953800 | 0 | 0 |
| FNSQSTSY | MRPL21; MRPL | Q7Z2W9; F5 | RM21_HUMAN | Unmodified | 0 | 0 | 0 | 0 | 530560 | 0 | 0 | 0 | 0 | 0 | 100330 | 0 | 342130 | 444630 | 0 | 99859 | 288150 | 0 |
| FNTPQQPK | OLA1; OLA1 | Q9NTK5; J3K | OLA1_HUMAN | Unmodified | 69748 | 356060 | 314130 | 498830 | 2761300 | 0 | 1967800 | 350850 | 1569400 | 600110 | 1186600 | 8151300 | 3174100 | 2428300 | 759290 | 474010 | 1925800 | 2613200 |
| FNTTSVIK | EFTUD2 | Q15029 | U5S1_HUMAN | Unmodified | 0 | 407780 | 0 | 0 | 836010 | 218560 | 421070 | 102150 | 101100 | 0 | 376260 | 1765800 | 0 | 0 | 0 | 0 | 210830 | 537880 |
| FNTVPGVQ | POMGNT1 | Q8WZA1 | PMGT1_HUMAN | Unmodified | 0 | 0 | 0 | 0 | 0 | 0 | 0 | 656610 | 3876800 | 435710 | 0 | 0 | 0 | 0 | 0 | 0 | 0 | 0 |
| FNTVPGVQ | POMGNT1 | Q8WZA1 | PMGT1_HUMAN | Unmodified | 0 | 0 | 0 | 0 | 0 | 0 | 0 | 0 | 0 | 770060 | 0 | 0 | 0 | 0 | 0 | 0 | 0 | 0 |
| FNVAVTR | MOV10; MOV10 | Q9HCE1; Q5 | MOV10_HUMA | Unmodified | 444740 | 1203700 | 0 | 45551 | 0 | 0 | 75245 | 346130 | 0 | 0 | 0 | 150840 | 0 | 0 | 361570 | 264210 | 0 | 0 |
| FNVEDGEI | EMC1 | Q8N766 | EMC1_HUMAN | Unmodified | 0 | 0 | 0 | 0 | 0 | 0 | 0 | 0 | 6592400 | 370280 | 182120 | 0 | 0 | 0 | 24269 | 24244 | 0 | 0 |
| FNVIQPGPI | DECR1; DECR1 | Q16698; H0 | DECR_HUMAN | Unmodified | 0 | 0 | 0 | 0 | 0 | 0 | 0 | 0 | 8729800 | 522700 | 0 | 0 | 0 | 0 | 0 | 0 | 0 | 0 |
| FNVLQYVV | EIF3A | Q14152 | EIF3A_HUMAN | Unmodified | 209490 | 0 | 519260 | 0 | 0 | 0 | 0 | 0 | 70236 | 43137 | 335700 | 12517000 | 0 | 0 | 124420 | 511720 | 0 | 0 |
| FNVLYDR | COPS6; COPS6 | Q7L5N1; E7 | CSN6_HUMAN | Unmodified | 0 | 0 | 0 | 0 | 0 | 0 | 0 | 0 | 0 | 0 | 0 | 3010700 | 0 | 0 | 0 | 0 | 0 | 0 |
| FNVLYDRQ | COPS6; COPS6 | Q7L5N1; E7 | CSN6_HUMAN | Unmodified | 0 | 0 | 0 | 0 | 129440 | 0 | 0 | 491180 | 164870 | 84763 | 243320 | 207280 | 414690 | 0 | 67729 | 0 | 0 | 248450 |
| FNVTTVDM | CLIC2 | O15247 | CLIC2_HUMAN | Unmodified | 0 | 0 | 0 | 0 | 945380 | 0 | 0 | 0 | 0 | 0 | 0 | 0 | 1219100 | 2933800 | 0 | 0 | 0 | 0 |
| FNVWDTA | RAN; RAN; RAN | P62826; H0 | RAN_HUMAN C | Unmodified | 1782200 | 1367100 | 2478400 | 1465900 | 11656000 | 2442200 | 6522100 | 3380500 | 4643600 | 2975000 | 8904200 | 1.1E+08 | 6487800 | 297960 | 4317300 | 2392400 | 3423700 | 5741200 |

| | | | | | | | | | | | | | | | | | | | | | | |
|---|---|---|---|---|---|---|---|---|---|---|---|---|---|---|---|---|---|---|---|---|---|---|
| FNVWDVG | ARF6 | P62330 | ARF6_HUMAN | Unmodified | 3194200 | 220460 | 4103800 | 3543400 | 7456200 | 1911400 | 0 | 253600 | 8278400 | 1952500 | 2341500 | 13756000 | 0 | 0 | 13543000 | 6787200 | 855260 | 1963800 |
| FNYGFEYL | DYNC1H1; DYN | Q14204; A0A | DYHC1_HUMA | Unmodified | 0 | 0 | 0 | 0 | 0 | 0 | 0 | 107960 | 1694200 | 0 | 717340 | 4335700 | 0 | 949200 | 0 | 0 | 0 | 1336000 |
| FNYSGSG | ILF3 | Q12906 | ILF3_HUMAN I | Unmodified | 964740 | 393850 | 263090 | 227390 | 387270 | 0 | 96608 | 44422 | 441800 | 0 | 0 | 2194400 | 0 | 0 | 0 | 137770 | 0 | 0 |
| FPAEDEFPI | CKB; CKB | P12277; G3V | KCRB_HUMAN | Unmodified | 0 | 0 | 0 | 0 | 0 | 0 | 376130 | 29169 | 166280 | 0 | 330710 | 374430 | 0 | 0 | 0 | 0 | 0 | 0 |
| FPAEDEFPI | CKB; CKB | P12277; G3V | KCRB_HUMAN | Oxidation (M) | 0 | 0 | 84315 | 123530 | 0 | 209640 | 311030 | 153180 | 0 | 0 | 819980 | 672970 | 0 | 0 | 130840 | 113650 | 0 | 0 |
| FPALTHYW | ANKH; ANKH | Q9HCJ1; D6 | ANKH_HUMAN | Unmodified | 0 | 0 | 105630 | 0 | 0 | 0 | 0 | 0 | 369250 | 0 | 29498 | 0 | 0 | 0 | 87162 | 122180 | 0 | 0 |
| FPDALRPF | EXT1 | Q16394 | EXT1_HUMAN | Unmodified | 0 | 0 | 0 | 0 | 0 | 0 | 0 | 0 | 5888700 | 508300 | 0 | 0 | 0 | 0 | 0 | 0 | 0 | 0 |
| FPDIYGPRI | SYT13; SYT13 | Q7L8C5; H0 | SYT13_HUMAN | Unmodified | 0 | 0 | 0 | 0 | 0 | 0 | 2326100 | 0 | 1512300 | 0 | 0 | 0 | 0 | 0 | 0 | 0 | 0 | 0 |
| FPEHELTFI | HYOU1; HYOU | Q9Y4L1; J3Q | HYOU1_HUMA | Unmodified | 306560 | 40002 | 645560 | 215350 | 0 | 0 | 0 | 0 | 9800200 | 227960 | 2543800 | 4289100 | 0 | 0 | 1138300 | 604990 | 0 | 0 |
| FPEVIINFPI | HDLBP; HDLBP | Q00341; A0A | VIGLN_HUMAN | Unmodified | 60371 | 36811 | 95867 | 47316 | 0 | 0 | 0 | 0 | 0 | 0 | 0 | 122210 | 23283 | 0 | 110360 | 100690 | 0 | 0 |
| FPFIRDQPT | MINK1; MINK1; | Q8N4C8; A0 | MINK1_HUMAN | Unmodified | 0 | 0 | 548110 | 740600 | 859500 | 0 | 0 | 0 | 1210400 | 168040 | 0 | 755850 | 1303500 | 197280 | 397470 | 345320 | 58061 | 393110 |
| FPGFAKPG | SNX9; SNX9; SN | Q9Y5X1; A0A | SNX9_HUMAN | Unmodified | 4101400 | 4472100 | 1942600 | 0 | 0 | 0 | 0 | 0 | 0 | 0 | 0 | 121920 | 0 | 0 | 1017000 | 1631200 | 0 | 0 |
| FPGINYPVL | HMGCL | P35914 | HMGCL_HUMA | Unmodified | 0 | 0 | 0 | 0 | 0 | 0 | 0 | 0 | 10828000 | 879640 | 0 | 0 | 0 | 0 | 0 | 0 | 0 | 0 |
| FPGYDSES | RPL5; RPL5 | P46777; A0A | RL5_HUMAN 6 | Unmodified | 0 | 0 | 0 | 0 | 0 | 0 | 0 | 0 | 0 | 0 | 1394300 | 206120 | 0 | 0 | 0 | 0 | 0 | 0 |
| FPHPIEISEI | NCR3LG1 | Q68D85 | NR3L1_HUMAN | Unmodified | 450940 | 333320 | 80600 | 139140 | 0 | 61966 | 0 | 0 | 1214400 | 0 | 44928 | 0 | 0 | 0 | 46192 | 28709 | 0 | 0 |
| FPILSANIK | NT5E; NT5E | P21589; Q9 | 5NTD_HUMAN | Unmodified | 389470 | 0 | 0 | 0 | 10527000 | 3252300 | 211430 | 181150 | 1331400 | 619520 | 0 | 0 | 0 | 0 | 476560 | 0 | 1462300 | 2382100 |
| FPLFGGWK | RPN1; RPN1 | P04843; B7Z | RPN1_HUMAN | Unmodified | 0 | 0 | 616610 | 0 | 0 | 0 | 0 | 0 | 2.41E+08 | 7052800 | 2450100 | 3991100 | 0 | 0 | 440210 | 0 | 0 | 0 |
| FPLKPGVPI | PLCB2 | Q00722 | PLCB2_HUMAN | Unmodified | 0 | 0 | 0 | 0 | 0 | 0 | 0 | 11551 | 0 | 0 | 921050 | 10700000 | 0 | 0 | 0 | 0 | 0 | 0 |
| FPLKPGVPI | PLCB2 | Q00722 | PLCB2_HUMAN | Unmodified | 0 | 0 | 0 | 0 | 0 | 0 | 0 | 0 | 0 | 0 | 1837100 | 2962700 | 0 | 0 | 0 | 0 | 0 | 0 |
| FPLSGLNM | USP43; USP43; | Q70EL4; V9 | UBP43_HUMAN | Oxidation (M) | 0 | 0 | 176190 | 0 | 0 | 0 | 0 | 0 | 0 | 0 | 0 | 0 | 83355 | 0 | 405830 | 331420 | 0 | 0 |
| FPLTTESAM | RPL23A; RPL23 | P62750; K7E | RL23A_HUMAN | Oxidation (M) | 1410300 | 703250 | 618850 | 0 | 0 | 0 | 80698 | 0 | 0 | 0 | 0 | 4870000 | 0 | 0 | 460150 | 452270 | 0 | 0 |
| FPLTTESAM | RPL23A; RPL23 | P62750; K7E | RL23A_HUMAN | Unmodified | 172170 | 0 | 0 | 0 | 0 | 0 | 0 | 0 | 250760 | 0 | 0 | 6442800 | 0 | 0 | 0 | 0 | 0 | 0 |
| FPNGVQLS | APMAP; APMAP | Q9HDC9; H | APMAP_HUMA | Oxidation (M) | 0 | 0 | 0 | 0 | 0 | 88473 | 0 | 0 | 1526300 | 0 | 141600 | 0 | 0 | 0 | 0 | 0 | 0 | 0 |
| FPNRLNLE | LGALS1 | P09382 | LEG1_HUMAN | Oxidation (M) | 148410 | 141850 | 0 | 0 | 3604800 | 521090 | 0 | 0 | 0 | 0 | 0 | 0 | 0 | 0 | 0 | 0 | 731040 | 1971600 |
| FPNRLNLE | LGALS1 | P09382 | LEG1_HUMAN | Unmodified | 0 | 0 | 0 | 0 | 1664500 | 0 | 0 | 0 | 1039400 | 0 | 0 | 0 | 0 | 0 | 0 | 0 | 288450 | 1642800 |
| FPPEETLK | CPE | P16870 | CBPE_HUMAN | Unmodified | 0 | 0 | 0 | 0 | 70681 | 0 | 0 | 0 | 1460300 | 592550 | 0 | 0 | 0 | 0 | 0 | 0 | 0 | 0 |
| FPPFFTLQF | VPS25; VPS25; | Q9BRG1; K7 | VPS25_HUMAN | Unmodified | 2134300 | 2985800 | 1513300 | 753200 | 0 | 0 | 0 | 0 | 138010 | 0 | 27338 | 1158800 | 0 | 0 | 4639900 | 3703700 | 0 | 0 |
| FPQLDSTSI | FASN; FASN | P49327; A0A | FAS_HUMAN F | Unmodified | 1654400 | 0 | 4121200 | 1897200 | 0 | 0 | 0 | 0 | 1064500 | 0 | 0 | 76197000 | 0 | 0 | 5577100 | 4209400 | 0 | 0 |
| FPQLDSTSI | FASN; FASN | P49327; A0A | FAS_HUMAN F | Unmodified | 0 | 0 | 713720 | 1047900 | 0 | 0 | 81434 | 0 | 1941000 | 0 | 2637600 | 10436000 | 0 | 1655200 | 2539600 | 1872300 | 169970 | 2221400 |
| FPRYAEIVH | ATP6V1B2; ATP | P21281; H0 | VATB2_HUMAN | Unmodified | 0 | 0 | 0 | 0 | 82211 | 0 | 0 | 0 | 0 | 0 | 0 | 0 | 88683 | 0 | 0 | 0 | 0 | 0 |
| FPSGNQG | VCP; VCP; VCP; | P55072; C9J | TERA_HUMAN | Unmodified | 0 | 0 | 0 | 0 | 0 | 0 | 0 | 0 | 0 | 0 | 0 | 1473200 | 0 | 0 | 1031800 | 384510 | 0 | 0 |
| FPSGSVLT | TBCE; TBCE; TB | Q15813; A0A | TBCE_HUMAN | Unmodified | 0 | 0 | 0 | 0 | 0 | 0 | 0 | 0 | 0 | 0 | 0 | 658620 | 0 | 0 | 0 | 0 | 0 | 0 |
| FPSLFINQF | CDKAL1 | Q5VV42 | CDKAL_HUMA | Unmodified | 0 | 0 | 0 | 0 | 0 | 0 | 0 | 0 | 36058 | 0 | 0 | 330150 | 0 | 0 | 0 | 0 | 0 | 0 |
| FPTLWSGA | HNRNPUL2; HN | Q1KMD3; H3 | HNRL2_HUMA | Unmodified | 0 | 0 | 0 | 0 | 0 | 0 | 0 | 0 | 0 | 0 | 496300 | 3538900 | 0 | 0 | 0 | 0 | 0 | 0 |
| FPVFNPATI | ALDH1A1; ALDH | P00352; Q5 | AL1A1_HUMAN | Unmodified | 0 | 0 | 0 | 0 | 0 | 0 | 0 | 0 | 0 | 0 | 342050 | 6017900 | 0 | 0 | 0 | 0 | 0 | 0 |
| FPVGPPKN | SCG2 | P13521 | SCG2_HUMAN | Oxidation (M) | 0 | 0 | 0 | 0 | 0 | 0 | 136030 | 260850 | 0 | 2455800 | 0 | 0 | 0 | 0 | 0 | 0 | 0 | 0 |
| FPVGPPKN | SCG2 | P13521 | SCG2_HUMAN | Unmodified | 0 | 0 | 0 | 0 | 0 | 111990 | 63602 | 0 | 245000 | 447270 | 0 | 0 | 0 | 0 | 0 | 0 | 0 | 0 |
| FPVISVPHL | SNX18 | Q96RF0 | SNX18_HUMAN | Unmodified | 2127700 | 0 | 14920000 | 9874700 | 438350 | 0 | 0 | 43476 | 0 | 0 | 0 | 0 | 4641100 | 1370100 | 22643000 | 14354000 | 0 | 0 |
| FQAALALSI | KIFBP; KIFBP; K | Q96EK5; A0A | KBP_HUMAN K | Unmodified | 0 | 0 | 31661 | 0 | 0 | 0 | 0 | 0 | 880290 | 564050 | 5251100 | 444090 | 43147 | 0 | 122050 | 163630 | 284590 | 0 |
| FQAKLEHE | MAPRE3; MAPR | Q9UPY8; Q1 | MARE3_HUMA | Unmodified | 0 | 0 | 0 | 0 | 460440 | 0 | 1155100 | 136120 | 275400 | 0 | 105660 | 0 | 0 | 0 | 0 | 0 | 0 | 0 |
| FQAKLEHE | MAPRE1 | Q15691 | MARE1_HUMA | Unmodified | 150490 | 288350 | 0 | 404590 | 2498200 | 375620 | 803270 | 98364 | 434170 | 0 | 2889100 | 3002400 | 1614600 | 954900 | 0 | 146090 | 1119600 | 993000 |
| FQAPARET | LIMA1; LIMA1; L | Q9UHB6; F8 | LIMA1_HUMAN | Unmodified | 0 | 0 | 0 | 0 | 1024400 | 276560 | 0 | 0 | 0 | 22357 | 0 | 97803 | 952400 | 0 | 0 | 0 | 408170 | 255230 |
| FQATSSGP | IQGAP2; IQGAP | Q13576; D6I | IQGA2_HUMAN | Unmodified | 0 | 0 | 422050 | 803280 | 0 | 0 | 0 | 0 | 0 | 0 | 0 | 45229 | 0 | 0 | 565360 | 454180 | 0 | 0 |
| FQATSSGP | IQGAP2; IQGAP | Q13576; D6I | IQGA2_HUMAN | Unmodified | 0 | 0 | 1782700 | 1146800 | 0 | 0 | 0 | 0 | 0 | 0 | 0 | 0 | 0 | 0 | 1879000 | 872460 | 0 | 0 |
| FQDADEKF | ECE1; ECE1; EC | P42892; B4D | ECE1_HUMAN | Unmodified | 0 | 0 | 0 | 0 | 0 | 0 | 0 | 41994 | 375050 | 0 | 0 | 0 | 0 | 227510 | 0 | 0 | 0 | 164940 |
| FQDDPDGI | SCG3 | Q8WXD2 | SCG3_HUMAN | Unmodified | 3704600 | 3934300 | 0 | 0 | 0 | 0 | 0 | 0 | 45048000 | 3600900 | 0 | 0 | 64684 | 0 | 332660 | 0 | 0 | 0 |
| FQDLGAAY | DNAJB11; DNA | Q9UBS4; A0 | DJB11_HUMAN | Unmodified | 0 | 0 | 0 | 0 | 0 | 23227 | 0 | 0 | 6987100 | 658330 | 545490 | 805720 | 95837 | 0 | 0 | 0 | 0 | 0 |
| FQDLGAAY | DNAJB11; DNA | Q9UBS4; A0 | DJB11_HUMAN | Unmodified | 0 | 0 | 0 | 0 | 0 | 0 | 1230100 | 0 | 3122500 | 0 | 0 | 0 | 0 | 0 | 0 | 0 | 0 | 0 |
| FQDLGVK | SEPTIN9; SEPTI | Q9UHD8; K7 | SEPT9_HUMAN | Unmodified | 0 | 0 | 74000 | 358490 | 3469700 | 299750 | 39188 | 0 | 10605000 | 3819900 | 1098200 | 1152700 | 256110 | 0 | 98972 | 114290 | 1083300 | 2852800 |
| FQDLGVKN | SEPTIN9; SEPTI | Q9UHD8; K7 | SEPT9_HUMAN | Unmodified | 0 | 0 | 0 | 0 | 465020 | 0 | 0 | 0 | 0 | 177720 | 0 | 0 | 0 | 0 | 0 | 0 | 493540 | 0 |
| FQDNFEFV | MAPRE1 | Q15691 | MARE1_HUMA | Unmodified | 0 | 0 | 0 | 0 | 0 | 0 | 0 | 0 | 0 | 0 | 115620 | 424360 | 0 | 0 | 0 | 0 | 0 | 0 |
| FQDTDGKI | PGD; PGD; PGD | P52209; K7E | 6PGD_HUMAN | Unmodified | 0 | 0 | 0 | 0 | 1627600 | 0 | 0 | 0 | 0 | 0 | 0 | 0 | 0 | 342230 | 0 | 0 | 428180 | 430660 |
| FQEFSPNL | EFNB2 | P52799 | EFNB2_HUMAN | Unmodified | 194420 | 190390 | 49850 | 164670 | 0 | 0 | 221410 | 0 | 183740 | 74689 | 0 | 0 | 0 | 0 | 180980 | 1895000 | 0 | 302600 |
| FQEHIIQAP | CHORDC1; CH | Q9UHD1; E9 | CHRD1_HUMA | Unmodified | 0 | 0 | 0 | 0 | 0 | 0 | 0 | 0 | 394480 | 0 | 589140 | 7042500 | 0 | 0 | 0 | 0 | 0 | 0 |
| FQELESETL | ACP3; ACP3 | P15309; E9P | PPAP_HUMAN | Unmodified | 0 | 0 | 0 | 0 | 0 | 0 | 0 | 0 | 10142000 | 2559500 | 0 | 0 | 0 | 0 | 0 | 0 | 0 | 0 |
| FQELESETL | ACP3; ACP3 | P15309; E9P | PPAP_HUMAN | Unmodified | 38113 | 0 | 0 | 0 | 287930 | 0 | 0 | 0 | 1202200 | 331700 | 0 | 0 | 0 | 0 | 0 | 0 | 0 | 0 |
| FQELIFEDF | NIBAN2 | Q96TA1 | NIBA2_HUMAN | Unmodified | 953960 | 1401500 | 1284600 | 0 | 0 | 0 | 0 | 0 | 4670000 | 0 | 0 | 2416000 | 0 | 0 | 2085200 | 1405300 | 0 | 0 |
| FQEQECPP | RRAS2 | P62070 | RRAS2_HUMAN | Unmodified | 1532600 | 420010 | 1132100 | 1061300 | 535010 | 0 | 0 | 0 | 337970 | 104350 | 0 | 0 | 0 | 237610 | 1444500 | 1222300 | 99799 | 148050 |
| FQETGFFK | STAMBP | O95630 | STABP_HUMAN | Unmodified | 2263800 | 1767400 | 685400 | 833720 | 754120 | 97828 | 0 | 55016 | 2701700 | 0 | 372920 | 2139000 | 490280 | 130600 | 1025800 | 1164000 | 0 | 98233 |
| FQETGFFKI | STAMBP | O95630 | STABP_HUMAN | Unmodified | 0 | 0 | 0 | 0 | 3592300 | 0 | 0 | 0 | 0 | 472160 | 0 | 0 | 3089900 | 1567600 | 0 | 0 | 689540 | 680580 |
| FQEYHIQQ | TIMM9 | Q9Y5J7 | TIM9_HUMAN | Unmodified | 0 | 0 | 0 | 0 | 0 | 0 | 0 | 0 | 986490 | 0 | 0 | 0 | 455010 | 0 | 0 | 0 | 0 | 0 |
| FQEYILALA | NCAM2; NCAM | O15394; H9I | NCAM2_HUMA | Unmodified | 182570 | 173880 | 430530 | 433180 | 0 | 0 | 0 | 0 | 0 | 0 | 0 | 0 | 0 | 349550 | 3673000 | 1735500 | 0 | 0 |
| FQFDGDK | PGD; PGD | P52209; K7E | 6PGD_HUMAN | Unmodified | 39484 | 144590 | 0 | 0 | 0 | 0 | 0 | 0 | 0 | 0 | 0 | 2999500 | 0 | 0 | 0 | 0 | 0 | 0 |
| FQFPSHVT | CDC42BPB | Q9Y5S2 | MRCKB_HUMA | Unmodified | 0 | 0 | 163130 | 168700 | 0 | 0 | 0 | 0 | 0 | 0 | 80508 | 1082900 | 0 | 0 | 274210 | 270680 | 0 | 0 |
| FQGEDTVV | KIF5B | P33176 | KINH_HUMAN | Unmodified | 0 | 0 | 0 | 0 | 0 | 0 | 0 | 0 | 1137200 | 0 | 0 | 5685800 | 0 | 0 | 0 | 0 | 0 | 0 |
| FQGEDTVV | KIF5B | P33176 | KINH_HUMAN | Unmodified | 0 | 0 | 0 | 0 | 0 | 0 | 0 | 0 | 0 | 0 | 0 | 0 | 1644800 | 0 | 0 | 0 | 1932400 | 0 |
| FQGIKHEC | DSTN | P60981 | DEST_HUMAN | Unmodified | 1432100 | 345890 | 265040 | 1067100 | 136690 | 0 | 0 | 0 | 296900 | 0 | 559840 | 13208000 | 369310 | 0 | 846840 | 1093400 | 0 | 107730 |
| FQGIKHEC | DSTN | P60981 | DEST_HUMAN | Unmodified | 61943 | 0 | 0 | 0 | 2208300 | 0 | 0 | 0 | 0 | 174120 | 0 | 0 | 0 | 3871500 | 0 | 0 | 2608800 | 814430 |
| FQGPDNG | TXNL1; TXNL1 | O43396; K7E | TXNL1_HUMAN | Unmodified | 90400 | 0 | 320210 | 283060 | 343790 | 0 | 0 | 0 | 257930 | 77296 | 355740 | 1201500 | 428780 | 433820 | 322920 | 174160 | 299690 | 0 |
| FQGPDNG | TXNL1; TXNL1 | O43396; K7E | TXNL1_HUMAN | Unmodified | 0 | 0 | 0 | 0 | 0 | 0 | 0 | 220460 | 0 | 0 | 0 | 0 | 959050 | 0 | 0 | 0 | 2567000 | 0 |
| FQGQCYK | VCAN; VCAN | P13611; E9P | CSPG2_HUMA | Unmodified | 0 | 0 | 0 | 0 | 0 | 0 | 0 | 74193 | 803640 | 111280 | 0 | 0 | 0 | 0 | 0 | 0 | 0 | 0 |
| FQGQDTILI | NUDT1 | P36639 | 8ODP_HUMAN | Unmodified | 0 | 1273600 | 0 | 0 | 114850 | 0 | 381240 | 0 | 0 | 63311 | 0 | 0 | 77379 | 0 | 0 | 0 | 0 | 0 |
| FQGQTCEI | ITGB1; ITGB1; I | P05556; H7 | ITB1_HUMAN I | Oxidation (M) | 413500 | 0 | 0 | 0 | 0 | 527130 | 0 | 0 | 692680 | 0 | 0 | 0 | 0 | 0 | 0 | 0 | 385060 | 1256600 |
| FQGQTCEI | ITGB1; ITGB1; I | P05556; H7 | ITB1_HUMAN I | Unmodified | 0 | 0 | 0 | 0 | 3109600 | 0 | 0 | 0 | 2736000 | 0 | 0 | 1922700 | 0 | 0 | 0 | 0 | 1067200 | 4292300 |
| FQGQTCEI | ITGB1; ITGB1; I | P05556; H7 | ITB1_HUMAN I | Unmodified | 0 | 0 | 0 | 0 | 0 | 0 | 0 | 0 | 0 | 0 | 0 | 0 | 0 | 0 | 0 | 0 | 727420 | 0 |
| FQHGVIAA | PSMB8; PSMB8 | P28062; Q5J | PSB8_HUMAN | Unmodified | 0 | 0 | 0 | 0 | 0 | 0 | 0 | 0 | 333250 | 0 | 0 | 3950700 | 0 | 0 | 0 | 0 | 0 | 0 |
| FQHPSHEL | LARP1; LARP1B | Q6PKG0; Q6 | LARP1_HUMAN | Unmodified | 346420 | 243620 | 0 | 0 | 0 | 0 | 0 | 0 | 0 | 0 | 0 | 1089800 | 0 | 272860 | 249000 | 1041200 | 0 | 0 |
| FQHVGTSV | SDF2L1 | Q9HCN8 | SDF2L_HUMAN | Unmodified | 0 | 0 | 0 | 0 | 0 | 0 | 0 | 0 | 792090 | 0 | 0 | 0 | 0 | 0 | 0 | 0 | 0 | 0 |
| FQIATVTEK | PSMB4 | P28070 | PSB4_HUMAN | Unmodified | 0 | 0 | 0 | 0 | 0 | 0 | 0 | 59135 | 0 | 0 | 923370 | 15291000 | 605530 | 0 | 539530 | 182000 | 0 | 0 |
| FQIINNTEG | YES1; YES1 | P07947; J3Q | YES_HUMAN T | Unmodified | 604490 | 3671700 | 1567100 | 802630 | 0 | 0 | 0 | 0 | 2508500 | 140860 | 287630 | 1871000 | 0 | 0 | 1584100 | 2813800 | 184800 | 569230 |
| FQIPPNAEI | FKBP4 | Q02790 | FKBP4_HUMAN | Unmodified | 0 | 0 | 0 | 1605700 | 0 | 0 | 0 | 0 | 0 | 0 | 943750 | 4458000 | 3124300 | 2099800 | 0 | 512800 | 1823700 | 0 |
| FQISNNGC | OCRL | Q01968 | OCRL_HUMAN | Unmodified | 451980 | 0 | 55670 | 0 | 0 | 0 | 0 | 18208 | 129460 | 40912 | 0 | 327380 | 0 | 0 | 0 | 0 | 0 | 115990 |
| FQLLEGPPI | GDI1; GDI1 | P31150; G5E | GDIA_HUMAN | Unmodified | 0 | 0 | 0 | 0 | 0 | 0 | 0 | 0 | 1042900 | 0 | 0 | 0 | 0 | 0 | 0 | 0 | 282630 | 0 |
| FQLQATTK | L1CAM | P32004 | L1CAM_HUMA | Unmodified | 3818700 | 2626000 | 130160 | 0 | 0 | 406130 | 150550 | 0 | 998810 | 1108800 | 0 | 478450 | 0 | 0 | 207980 | 120360 | 240040 | 1128200 |
| FQLTFPLR | NPTX1 | Q15818 | NPTX1_HUMAN | Unmodified | 0 | 0 | 28345 | 0 | 0 | 0 | 0 | 0 | 1.39E+08 | 1796200 | 0 | 4600300 | 0 | 0 | 0 | 0 | 0 | 0 |
| FQLTFPLRT | NPTX1 | Q15818 | NPTX1_HUMAN | Unmodified | 0 | 0 | 0 | 0 | 0 | 0 | 0 | 0 | 8677200 | 966670 | 0 | 0 | 0 | 0 | 0 | 0 | 0 | 0 |
| FQLTFPLRT | NPTX1 | Q15818 | NPTX1_HUMAN | Oxidation (M) | 0 | 0 | 0 | 0 | 0 | 0 | 0 | 0 | 1598700 | 4394300 | 0 | 166390 | 0 | 0 | 0 | 0 | 0 | 0 |
| FQMAYSNL | SRP14 | P37108 | SRP14_HUMAN | Oxidation (M) | 4255300 | 1135100 | 4967200 | 7126500 | 4179300 | 10411000 | 2847500 | 2769100 | 0 | 0 | 0 | 2923900 | 719790 | 3814100 | 21170000 | 17532000 | 2232000 | 10936000 |
| FQMAYSNL | SRP14 | P37108 | SRP14_HUMAN | Unmodified | 728710 | 541200 | 594880 | 1799800 | 0 | 0 | 0 | 0 | 0 | 0 | 0 | 1172400 | 0 | 0 | 3070700 | 1000300 | 0 | 0 |
| FQMTQEV | DNAJB11 | Q9UBS4 | DJB11_HUMAN | Unmodified | 0 | 0 | 0 | 0 | 0 | 0 | 0 | 0 | 3051600 | 0 | 0 | 0 | 0 | 0 | 0 | 0 | 0 | 0 |
| FQNDRTED | CAB39; CAB39 | Q9Y376; A0A | CAB39_HUMAN | Unmodified | 330180 | 285160 | 728350 | 123020 | 473290 | 428120 | 102320 | 0 | 661020 | 202140 | 444950 | 3975100 | 129700 | 0 | 889300 | 759670 | 251710 | 60969 |
| FQNPQTEG | AAK1; AAK1; AA | Q2M2I8; A0A | AAK1_HUMAN | Unmodified | 0 | 0 | 0 | 47384 | 0 | 0 | 0 | 0 | 0 | 0 | 0 | 0 | 121860 | 0 | 91224 | 81440 | 0 | 0 |
| FQNVADLH | GARS1; GARS1 | P41250; A0A | GARS_HUMAN | Unmodified | 0 | 0 | 0 | 0 | 255970 | 58379 | 136880 | 0 | 1003600 | 0 | 647840 | 1183800 | 0 | 0 | 0 | 0 | 0 | 484020 |
| FQNVAKEG | HSP90B1; HSP | P14625; A0A | ENPL_HUMAN | Unmodified | 0 | 0 | 0 | 0 | 121510 | 0 | 0 | 0 | 773720 | 2747800 | 0 | 0 | 0 | 0 | 0 | 0 | 75997 | 0 |
| FQPASAPA | CKAP5 | Q14008 | CKAP5_HUMAN | Unmodified | 0 | 0 | 0 | 40418 | 0 | 0 | 0 | 0 | 200480 | 0 | 0 | 883620 | 248910 | 99165 | 27179 | 50178 | 0 | 0 |
| FQPGETLTI | IQGAP1; IQGAP | P46940; H0 | IQGA1_HUMAN | Unmodified | 1227700 | 5986800 | 1574400 | 3516800 | 0 | 0 | 0 | 0 | 16919000 | 1163400 | 3618100 | 11856000 | 0 | 0 | 6342700 | 3567400 | 0 | 0 |
| FQPQSPDF | UFD1; UFD1; U | Q92890; C9 | UFD1_HUMAN | Unmodified | 0 | 0 | 0 | 0 | 782170 | 0 | 0 | 241650 | 0 | 128480 | 0 | 846770 | 845110 | 0 | 0 | 0 | 699610 | 0 |
| FQPVGTVE | IGSF3 | O75054 | IGSF3_HUMAN | Unmodified | 1354300 | 1710900 | 160710 | 186810 | 0 | 0 | 0 | 0 | 0 | 0 | 0 | 1035100 | 0 | 0 | 1798600 | 627930 | 0 | 0 |
| FQQIPASD | RGS14 | O43566 | RGS14_HUMA | Unmodified | 0 | 0 | 0 | 0 | 282260 | 0 | 0 | 0 | 0 | 34813 | 0 | 1754000 | 0 | 0 | 0 | 0 | 86063 | 321850 |
| FQQQILAQ | TAOK3; TAOK3 | Q9H2K8; G3 | TAOK3_HUMAN | Unmodified | 0 | 0 | 0 | 0 | 402940 | 0 | 512450 | 312180 | 0 | 0 | 0 | 971920 | 0 | 0 | 0 | 0 | 92735 | 332560 |
| FQQQQGLI | PFAS | O15067 | PUR4_HUMAN | Unmodified | 0 | 0 | 0 | 0 | 0 | 0 | 0 | 0 | 256110 | 0 | 0 | 1212900 | 0 | 0 | 0 | 0 | 0 | 0 |
| FQQVPTDA | LARP1; LARP1; | Q6PKG0; H0 | LARP1_HUMAN | Unmodified | 535940 | 674680 | 390250 | 470600 | 0 | 0 | 0 | 0 | 130060 | 0 | 378580 | 0 | 0 | 0 | 0 | 0 | 0 | 0 |
| FQRPGDP | SF1; SF1; SF1 | Q15637; H7 | SF01_HUMAN | Unmodified | 162100 | 0 | 0 | 0 | 738790 | 0 | 41923 | 0 | 419760 | 182980 | 643250 | 0 | 0 | 0 | 175460 | 198410 | 583730 | 73526 |
| FQSGDFH | CSE1L | P55060 | XPO2_HUMAN | Unmodified | 112350 | 86742 | 90707 | 67159 | 122990 | 0 | 0 | 0 | 103960 | 0 | 0 | 4856800 | 0 | 0 | 240560 | 277040 | 461850 | 0 |
| FQSIVIGCA | BAG2 | O95816 | BAG2_HUMAN | Unmodified | 0 | 0 | 0 | 59184 | 1564100 | 0 | 1338700 | 29514 | 0 | 99039 | 4633300 | 0 | 447510 | 436780 | 110920 | 0 | 0 | 313770 |
| FQSKYEPR | PDIA4; PDIA4 | P13667; A0A | PDIA4_HUMAN | Unmodified | 0 | 0 | 0 | 0 | 0 | 0 | 0 | 0 | 4380500 | 858540 | 403250 | 0 | 0 | 0 | 0 | 0 | 0 | 0 |
| FQSKYEPR | PDIA4 | P13667 | PDIA4_HUMAN | Unmodified | 0 | 0 | 0 | 0 | 0 | 0 | 0 | 0 | 0 | 467140 | 0 | 0 | 0 | 0 | 0 | 0 | 0 | 0 |
| FQSLGVAF | RAB7A; RAB7A; | P51149; A0A | RAB7A_HUMAN | Unmodified | 0 | 7951600 | 0 | 0 | 0 | 0 | 0 | 0 | 8661000 | 0 | 3266000 | 33575000 | 0 | 0 | 0 | 246740 | 0 | 0 |
| FQSMTSK | SERINC3; SERI | Q13530; Q5 | SERC3_HUMAN | Unmodified | 0 | 0 | 0 | 0 | 0 | 0 | 0 | 0 | 0 | 0 | 0 | 0 | 0 | 0 | 0 | 0 | 0 | 0 |

| | | | | | | | | | | | | | | | | | | | | | | |
|---|---|---|---|---|---|---|---|---|---|---|---|---|---|---|---|---|---|---|---|---|---|---|
| FQSPYEEQ | GOLIM4; GOLIN | O00461; F8V | GOLI4_HUMAN | Unmodified | 0 | 0 | 0 | 0 | 0 | 0 | 0 | 0 | 26207000 | 0 | 258450 | 0 | 0 | 0 | 0 | 0 | 0 | 0 |
| FQSPYEEQ | GOLIM4; GOLIN | O00461; F8V | GOLI4_HUMAN | Unmodified | 0 | 0 | 0 | 0 | 0 | 0 | 0 | 0 | 1942600 | 0 | 0 | 0 | 0 | 0 | 0 | 0 | 0 | 0 |
| FQSPYEEQ | GOLIM4; GOLIN | O00461; F8V | GOLI4_HUMAN | Unmodified | 0 | 0 | 0 | 0 | 0 | 0 | 0 | 0 | 6240800 | 1994300 | 0 | 0 | 0 | 0 | 0 | 0 | 0 | 0 |
| FQSSAVMA | H3C15 | Q71DI3 | H32_HUMAN H | Oxidation (M) | 0 | 246670 | 0 | 0 | 0 | 0 | 0 | 0 | 0 | 0 | 0 | 0 | 0 | 0 | 0 | 0 | 0 | 0 |
| FQSSHHPT | HSP90B1; HSP | P14625; A0A | ENPL_HUMAN | Unmodified | 2287800 | 675860 | 5235900 | 1228100 | 569760 | 144040 | 255870 | 173960 | 1.11E+09 | 14263000 | 61217000 | 1.01E+08 | 0 | 0 | 4606600 | 4608000 | 284730 | 240170 |
| FQSSHHPT | HSP90B1; HSP | P14625; A0A | ENPL_HUMAN | Oxidation (M) | 0 | 0 | 0 | 173740 | 0 | 0 | 0 | 0 | 18106000 | 7447000 | 13070000 | 603250 | 0 | 0 | 0 | 0 | 0 | 0 |
| FQSSHHPT | HSP90B1; HSP | P14625; A0A | ENPL_HUMAN | Unmodified | 0 | 0 | 0 | 0 | 834940 | 516380 | 0 | 0 | 1.04E+08 | 12650000 | 9305600 | 0 | 0 | 0 | 0 | 0 | 0 | 899210 |
| FQSSHHPT | HSP90B1; HSP | P14625; A0A | ENPL_HUMAN | Unmodified | 0 | 0 | 0 | 0 | 0 | 0 | 0 | 0 | 1679300 | 1919400 | 0 | 0 | 0 | 0 | 0 | 0 | 0 | 0 |
| FQSVESGA | THUMPD1; THU | Q9NXG2; H3 | THUM1_HUMA | Unmodified | 0 | 0 | 0 | 0 | 0 | 0 | 0 | 0 | 0 | 0 | 0 | 501100 | 0 | 0 | 0 | 0 | 0 | 0 |
| FQTEDDAR | SCRIB; SCRIB; S | Q14160; A0A | SCRIB_HUMAN | Unmodified | 0 | 0 | 0 | 0 | 0 | 0 | 0 | 0 | 408270 | 0 | 0 | 1609000 | 0 | 0 | 0 | 53055 | 0 | 0 |
| FQTEDDAR | SCRIB; SCRIB; S | Q14160; A0A | SCRIB_HUMAN | Unmodified | 0 | 0 | 0 | 76165 | 0 | 0 | 1255200 | 0 | 4671000 | 2143300 | 168260 | 59101 | 0 | 0 | 0 | 0 | 0 | 0 |
| FQTEIQTVN | DARS1; DARS1 | P14868; H7E | SYDC_HUMAN | Unmodified | 0 | 0 | 482080 | 0 | 0 | 162840 | 0 | 0 | 205410 | 1211100 | 589030 | 10343000 | 0 | 0 | 267290 | 281320 | 0 | 243440 |
| FQTLINDLD | DAAM1 | Q9Y4D1 | DAAM1_HUMA | Unmodified | 155550 | 0 | 0 | 228030 | 0 | 0 | 0 | 0 | 260310 | 0 | 156590 | 2148800 | 0 | 0 | 557000 | 379640 | 0 | 0 |
| FQTMEEK | RPL3; RPL3 | P39023; G5E | RL3_HUMAN 6( | Unmodified | 742260 | 1103900 | 558900 | 440310 | 0 | 0 | 0 | 0 | 311190 | 29708 | 486430 | 10912000 | 0 | 0 | 530470 | 390160 | 0 | 0 |
| FQTMEEKK | RPL3; RPL3 | P39023; G5E | RL3_HUMAN 6( | Unmodified | 0 | 292190 | 0 | 0 | 0 | 0 | 0 | 0 | 0 | 0 | 0 | 2056500 | 0 | 0 | 0 | 166250 | 0 | 0 |
| FQTTYGSR | VAMP7 | P51809 | VAMP7_HUMAI | Unmodified | 58144 | 39288 | 345260 | 0 | 0 | 0 | 0 | 0 | 0 | 0 | 0 | 887170 | 0 | 0 | 546290 | 329850 | 0 | 0 |
| FQVDLVSE | SLC12A2; SLC1 | P55011; G3X | S12A2_HUMAN | Unmodified | 325620 | 346160 | 764930 | 716970 | 271620 | 195940 | 0 | 0 | 574590 | 0 | 0 | 1039800 | 0 | 0 | 968220 | 865060 | 180490 | 321510 |
| FQVWDLG( | ARL1; ARL1; AR | P40616; B4D | ARL1_HUMAN | Unmodified | 725690 | 86159 | 0 | 199230 | 451870 | 146990 | 0 | 0 | 0 | 580700 | 823290 | 0 | 211610 | 86932 | 1693400 | 472960 | 0 | 451070 |
| FQYYEDSS | ITGB3; N/A | P05106; H3E | ITB3_HUMAN I | Unmodified | 2971100 | 3719700 | 2963400 | 5352700 | 0 | 0 | 0 | 0 | 154480 | 60817 | 0 | 0 | 99888 | 0 | 3954700 | 4864100 | 0 | 0 |
| FRAALRDYI | PPP5C; PPP5C; | P53041; A8N | PPP5_HUMAN | Unmodified | 0 | 0 | 0 | 0 | 2184400 | 686860 | 0 | 0 | 0 | 0 | 0 | 0 | 0 | 0 | 0 | 0 | 503360 | 1430500 |
| FRAPDEPQ | MISP | Q8IVT2 | MISP_HUMAN I | Unmodified | 0 | 0 | 0 | 0 | 2870300 | 432500 | 0 | 0 | 0 | 0 | 0 | 0 | 0 | 0 | 0 | 0 | 0 | 0 |
| FRAPPILPP | PRKAB1; PRKAE | Q9Y478; F5H | AAKB1_HUMAN | Unmodified | 0 | 0 | 0 | 0 | 0 | 0 | 0 | 0 | 546420 | 0 | 0 | 0 | 0 | 0 | 0 | 0 | 0 | 0 |
| FRCPEALF( | ACTG1; ACTB; A | P63261; P60 | ACTG_HUMAN | Oxidation (M) | 0 | 0 | 0 | 0 | 0 | 0 | 0 | 0 | 0 | 1147700 | 4116900 | 0 | 0 | 0 | 0 | 0 | 0 | 0 |
| FRCPEALF( | ACTG1; ACTB; A | P63261; P60 | ACTG_HUMAN | 2 Oxidation (M) | 19739000 | 3997900 | 0 | 22350000 | 57624000 | 25627000 | 9048600 | 0 | 21464000 | 5460000 | 9056600 | 34181000 | 32359000 | 23436000 | 52434000 | 14034000 | 16666000 | 31678000 |
| FRCPEALF( | ACTG1; ACTB; A | P63261; P60 | ACTG_HUMAN | Oxidation (M) | 0 | 1453200 | 240770 | 4025000 | 47805000 | 4674100 | 3261700 | 0 | 39573000 | 1775100 | 2156000 | 2337600 | 18838000 | 7515700 | 3171900 | 863190 | 8965600 | 2240200 |
| FRCPEALF( | ACTG1; ACTB; A | P63261; P60 | ACTG_HUMAN | Unmodified | 0 | 0 | 0 | 0 | 2135400 | 0 | 0 | 0 | 4776700 | 0 | 0 | 0 | 0 | 0 | 0 | 0 | 0 | 0 |
| FRDLTVEEL | UEVLD; UEVLD | Q8IX04; A8M | UEVLD_HUMAI | Unmodified | 45697 | 59370 | 101050 | 83845 | 0 | 0 | 0 | 0 | 707150 | 0 | 142810 | 1107100 | 128580 | 0 | 107170 | 78207 | 0 | 0 |
| FRDLTVEEL | UEVLD; UEVLD | Q8IX04; A8M | UEVLD_HUMAI | Unmodified | 0 | 0 | 0 | 0 | 0 | 275670 | 0 | 0 | 0 | 655850 | 109550 | 0 | 140720 | 0 | 0 | 0 | 0 | 0 |
| FREALGDA | DNAJC7; DNAJ( | Q99615; A0A | DNJC7_HUMAI | Unmodified | 177210 | 378670 | 460850 | 705880 | 0 | 0 | 0 | 0 | 0 | 0 | 366760 | 3141000 | 204290 | 0 | 336110 | 438180 | 0 | 0 |
| FREFFESNI | TSPAN14; TSPA | Q8NG11; A6 | TSN14_HUMAI | Unmodified | 0 | 267920 | 0 | 644910 | 0 | 0 | 110840 | 0 | 0 | 0 | 0 | 1268500 | 177120 | 158050 | 308500 | 263340 | 85403 | 0 |
| FRELIIEFSK | EGFR; EGFR; E( | P00533; Q5( | EGFR_HUMAN | Unmodified | 0 | 0 | 0 | 0 | 1868000 | 717440 | 0 | 0 | 0 | 0 | 0 | 0 | 0 | 136320 | 1164400 | 358660 | 549360 | 1019600 |
| FREMIPFAV | SEPTIN9; SEPTI | Q9UHD8; A( | SEPT9_HUMAN | Oxidation (M) | 0 | 0 | 0 | 0 | 4734400 | 0 | 655500 | 0 | 0 | 0 | 0 | 3055100 | 0 | 0 | 0 | 0 | 1102400 | 3008500 |
| FREMIPFAV | SEPTIN9; SEPTI | Q9UHD8; A( | SEPT9_HUMAN | Unmodified | 0 | 0 | 0 | 0 | 2108700 | 437610 | 0 | 0 | 405900 | 0 | 0 | 0 | 0 | 0 | 0 | 0 | 589580 | 0 |
| FRENLIYTY | MYO1C; MYO1( | O00159; I3L! | MYO1C_HUMA | Unmodified | 0 | 229050 | 0 | 0 | 0 | 0 | 0 | 0 | 0 | 0 | 0 | 0 | 0 | 0 | 648250 | 0 | 0 | 0 |
| FRGPFTDV | VAPB; VAPB | O95292; E5F | VAPB_HUMAN | Unmodified | 0 | 0 | 185250 | 85959 | 4775200 | 265400 | 1029800 | 355930 | 908720 | 545430 | 514000 | 1621600 | 923040 | 935440 | 63301 | 75372 | 548380 | 606600 |
| FRHFNALG | LTA4H | P09960 | LKHA4_HUMAI | Unmodified | 0 | 0 | 0 | 0 | 2113300 | 0 | 0 | 0 | 0 | 765530 | 0 | 0 | 0 | 0 | 0 | 0 | 1111700 | 0 |
| FRHHLGDI | GLB1; GLB1; GI | P16278; C9J | BGAL_HUMAN | Unmodified | 0 | 0 | 0 | 0 | 0 | 0 | 0 | 0 | 468370 | 0 | 0 | 0 | 0 | 0 | 0 | 0 | 0 | 0 |
| FRHPLLHIC | ERP44 | Q9BS26 | ERP44_HUMAN | Unmodified | 0 | 0 | 0 | 0 | 0 | 0 | 0 | 0 | 16696000 | 772370 | 201110 | 0 | 0 | 0 | 0 | 0 | 0 | 0 |
| FRHVFGQF | CORO1B; COR( | Q9BR76; F5H | COR1B_HUMA | Unmodified | 49743 | 0 | 102780 | 0 | 0 | 0 | 0 | 0 | 91628 | 0 | 270590 | 380080 | 305590 | 0 | 148240 | 105760 | 0 | 0 |
| FRIDELEPR | GANAB; GANAE | Q14697; F5H | GANAB_HUMA | Unmodified | 0 | 0 | 0 | 0 | 131940 | 0 | 233220 | 0 | 35303000 | 216580 | 0 | 0 | 0 | 0 | 0 | 0 | 0 | 0 |
| FRISLGLPV | RPL23; RPL23; I | P62829; J3K | RL23_HUMAN ( | Unmodified | 2143900 | 1325600 | 615590 | 698960 | 0 | 0 | 4703800 | 149580 | 0 | 435120 | 1387000 | 2685500 | 5878300 | 13788000 | 1256700 | 992710 | 0 | 1670800 |
| FRIVTLFSD | CD109 | Q6YHK3 | CD109_HUMAI | Unmodified | 0 | 0 | 0 | 0 | 0 | 0 | 0 | 0 | 793510 | 4127600 | 0 | 0 | 0 | 0 | 0 | 0 | 427540 | 0 |
| FRLADNSD | ADGRG6 | Q86SQ4 | AGRG6_HUMA | Unmodified | 0 | 0 | 0 | 849290 | 1051400 | 0 | 0 | 0 | 0 | 0 | 0 | 0 | 0 | 0 | 268400 | 523280 | 0 | 0 |
| FRLDCPLAI | VPS28; VPS28; | Q9UK41; E9I | VPS28_HUMAN | Unmodified | 3439600 | 1676100 | 0 | 2808300 | 0 | 0 | 0 | 0 | 2851700 | 0 | 0 | 601870 | 1575200 | 0 | 2910300 | 2360400 | 0 | 533200 |
| FRLDCPLAI | VPS28; VPS28; | Q9UK41; E9I | VPS28_HUMAN | Oxidation (M) | 14405000 | 8701400 | 9485900 | 8549400 | 523200 | 0 | 0 | 14002 | 903960 | 379400 | 470150 | 2067400 | 328640 | 122700 | 14558000 | 12424000 | 39560 | 364580 |
| FRLEAPDAI | ACO2; ACO2; A | Q99798; A2A | ACON_HUMAN | Unmodified | 0 | 0 | 0 | 0 | 0 | 0 | 391470 | 0 | 7478700 | 0 | 0 | 352130 | 0 | 0 | 0 | 0 | 0 | 0 |
| FRLPPGEY | CAPN1 | P07384 | CAN1_HUMAN | Unmodified | 371390 | 0 | 73835 | 857260 | 0 | 0 | 0 | 0 | 3478000 | 0 | 785920 | 3565300 | 0 | 0 | 0 | 309200 | 0 | 0 |
| FRLTGSSG | ATRN | O75882 | ATRN_HUMAN | Unmodified | 0 | 0 | 0 | 0 | 0 | 0 | 0 | 0 | 1863700 | 506050 | 0 | 0 | 0 | 0 | 0 | 0 | 0 | 46863 |
| FRLTSESTN | TMED4; TMED4 | Q7Z7H5; F8V | TMED4_HUMAI | Unmodified | 0 | 0 | 0 | 0 | 0 | 0 | 0 | 0 | 1581400 | 73321 | 0 | 0 | 0 | 0 | 38023 | 0 | 108190 | 310130 |
| FRNPPGGI | TMEM30A; TME | Q9NV96; A0 | CC50A_HUMAI | Unmodified | 325330 | 180630 | 0 | 1026600 | 0 | 124760 | 40966 | 0 | 16869000 | 1991800 | 955790 | 8912100 | 148300 | 57558 | 1193400 | 1131600 | 91560 | 0 |
| FRNPPGGI | TMEM30A; TME | Q9NV96; A0 | CC50A_HUMAI | Unmodified | 0 | 0 | 660750 | 0 | 4215700 | 543180 | 908010 | 0 | 15846000 | 1807200 | 3135300 | 2428800 | 578580 | 0 | 796990 | 665550 | 240620 | 678620 |
| FRPDMEEE | PSMB7 | Q99436 | PSB7_HUMAN | Unmodified | 0 | 0 | 0 | 0 | 195210 | 0 | 0 | 0 | 300090 | 93757 | 317410 | 2728300 | 0 | 0 | 0 | 0 | 0 | 0 |
| FRPENQAT | REPS2 | Q8NFH8 | REPS2_HUMAN | Unmodified | 0 | 0 | 0 | 691220 | 0 | 0 | 0 | 0 | 0 | 82168 | 0 | 0 | 542790 | 0 | 216880 | 175250 | 0 | 1014200 |
| FRPFYLSQI | RIPK1 | Q13546 | RIPK1_HUMAN | Unmodified | 0 | 0 | 0 | 0 | 365650 | 0 | 0 | 0 | 318630 | 300790 | 0 | 0 | 1087800 | 0 | 0 | 0 | 934230 | 864070 |
| FRPHQDAN | CEMIP2 | Q9UHN6 | CEIP2_HUMAN | Unmodified | 0 | 0 | 179900 | 0 | 66317 | 0 | 0 | 82241 | 407980 | 0 | 24963 | 70655 | 0 | 0 | 402890 | 68504 | 0 | 0 |
| FRPHQDAN | CEMIP2 | Q9UHN6 | CEIP2_HUMAN | Unmodified | 75571 | 0 | 32821 | 0 | 0 | 78271 | 24101 | 33857 | 978080 | 89050 | 31098 | 68913 | 0 | 0 | 0 | 47130 | 37020 | 38395 |
| FRPLNESE | KIF5B | P33176 | KINH_HUMAN | Unmodified | 0 | 0 | 0 | 818350 | 2016100 | 530580 | 57061 | 34456 | 2515500 | 165710 | 0 | 7673900 | 371630 | 352330 | 632080 | 646560 | 208890 | 758740 |
| FRPSDEHH | LHPP | Q9H008 | LHPP_HUMAN | Unmodified | 0 | 0 | 0 | 0 | 0 | 0 | 0 | 0 | 0 | 1494700 | 0 | 0 | 0 | 0 | 0 | 249630 | 442490 | 0 |
| FRPSEPHF | DPP4; DPP4; DI | P27487; A0A | DPP4_HUMAN | Unmodified | 1377200 | 1107300 | 5910400 | 7634200 | 0 | 0 | 0 | 0 | 0 | 0 | 0 | 0 | 0 | 0 | 11013000 | 7432900 | 337150 | 0 |
| FRQDLMNI | CCT2; CCT2; C( | P78371; F5G | TCPB_HUMAN | Unmodified | 69729 | 74301 | 0 | 183510 | 4400000 | 488850 | 666770 | 76414 | 2629200 | 677080 | 568230 | 6170000 | 3685400 | 1221800 | 166840 | 0 | 726420 | 1849800 |
| FRQIVSAV( | MARK3; MARK2 | P27448; Q7K | MARK3_HUMAI | Unmodified | 0 | 0 | 0 | 0 | 366970 | 0 | 370300 | 196390 | 153040 | 325950 | 0 | 0 | 847840 | 705340 | 118470 | 123760 | 218840 | 0 |
| FRQTSESTI | TMED9 | Q9BVK6 | TMED9_HUMAI | Unmodified | 17230 | 11049 | 0 | 0 | 0 | 0 | 0 | 0 | 1721300 | 0 | 0 | 0 | 0 | 0 | 0 | 120190 | 0 | 0 |
| FRSETITEEI | CBR1 | P16152 | CBR1_HUMAN | Oxidation (M) | 0 | 0 | 67775 | 621610 | 1081400 | 0 | 0 | 0 | 0 | 0 | 0 | 859160 | 0 | 0 | 459920 | 0 | 0 | 0 |
| FRSETITEEI | CBR1 | P16152 | CBR1_HUMAN | Unmodified | 0 | 0 | 0 | 0 | 1902700 | 0 | 0 | 0 | 0 | 88981 | 0 | 0 | 301920 | 1131000 | 0 | 0 | 719120 | 0 |
| FRSHEGET | SRSF9 | Q13242 | SRSF9_HUMAN | Unmodified | 562150 | 477730 | 186210 | 171090 | 0 | 0 | 0 | 0 | 0 | 0 | 25044 | 0 | 0 | 0 | 257650 | 228570 | 92494 | 85778 |
| FRTITSSYYI | RAB1A; RAB1B; | P62820; Q9H | RAB1A_HUMAN | Unmodified | 0 | 0 | 94049 | 497490 | 0 | 0 | 0 | 0 | 386020 | 0 | 358400 | 0 | 0 | 0 | 148950 | 84107 | 0 | 0 |
| FRVGDIIQII | CASK; CASK; C | O14936; A0A | CSKP_HUMAN | Unmodified | 68148 | 62917 | 481400 | 220190 | 0 | 0 | 0 | 0 | 1023900 | 177300 | 0 | 1607200 | 76519 | 52602 | 856880 | 430940 | 0 | 71831 |
| FRVGDIIQII | CASK; CASK; C | O14936; A0A | CSKP_HUMAN | Unmodified | 30060 | 0 | 0 | 323520 | 0 | 0 | 0 | 0 | 426780 | 101850 | 366550 | 0 | 1370000 | 0 | 313640 | 81206 | 0 | 449910 |
| FRYGGAPC | AP2A1 | O95782 | AP2A1_HUMAN | Unmodified | 0 | 0 | 0 | 0 | 374790 | 0 | 167710 | 0 | 0 | 308250 | 0 | 0 | 1015200 | 0 | 0 | 0 | 1021500 | 346280 |
| FRYSGKTE | EPB41L5 | Q9HCM4 | E41L5_HUMAN | Unmodified | 59794 | 73428 | 61809 | 550270 | 0 | 147390 | 0 | 0 | 452740 | 237660 | 0 | 0 | 586590 | 149570 | 483640 | 309140 | 0 | 134960 |
| FRYSYYDE | MEMO1 | Q9Y316 | MEMO1_HUMA | Unmodified | 381000 | 326650 | 0 | 0 | 0 | 0 | 0 | 0 | 0 | 0 | 0 | 308460 | 0 | 0 | 0 | 109470 | 0 | 0 |
| FSAFLDK | ALG13; ALG13 | A0A096LP10 | A0A096LP10_H | Unmodified | 0 | 0 | 0 | 0 | 0 | 0 | 0 | 0 | 0 | 0 | 0 | 0 | 943100 | 453870 | 249200 | 0 | 0 | 0 |
| FSAFLDKV | ALG13; ALG13 | A0A096LP10 | A0A096LP10_H | Unmodified | 0 | 0 | 0 | 0 | 0 | 0 | 0 | 0 | 0 | 0 | 0 | 0 | 0 | 0 | 0 | 0 | 150550 | 0 |
| FSAFLEK | PYGB | P11216 | PYGB_HUMAN | Unmodified | 0 | 70991 | 0 | 173030 | 0 | 0 | 347660 | 167870 | 0 | 0 | 0 | 1250900 | 0 | 617760 | 230740 | 234180 | 1449600 | 5198300 |
| FSAGADIR | EHHADH | Q08426 | ECHP_HUMAN | Unmodified | 0 | 132610 | 0 | 84571 | 0 | 0 | 0 | 0 | 2268000 | 28048 | 0 | 0 | 0 | 0 | 174030 | 91589 | 0 | 0 |
| FSAIASQLV | KIF20A | O95235 | KI20A_HUMAN | Oxidation (M) | 0 | 0 | 0 | 1001100 | 0 | 0 | 0 | 0 | 849610 | 966230 | 0 | 0 | 0 | 1625200 | 0 | 0 | 0 | 1275700 |
| FSALLSDC | NIBAN1 | Q9BZQ8 | NIBA1_HUMAN | Unmodified | 0 | 0 | 0 | 119810 | 0 | 0 | 0 | 0 | 626870 | 259040 | 0 | 0 | 0 | 0 | 0 | 0 | 0 | 0 |
| FSAMHEFC | GYS1 | P13807 | GYS1_HUMAN | Unmodified | 0 | 0 | 0 | 0 | 0 | 1134900 | 0 | 0 | 153440 | 0 | 0 | 452090 | 0 | 276440 | 0 | 0 | 93102 | 0 |
| FSAMKYEQ | PRKCSH; PRKC | P14314; K7E | GLU2B_HUMAI | Unmodified | 0 | 0 | 0 | 0 | 0 | 0 | 0 | 0 | 21623000 | 392000 | 0 | 0 | 0 | 0 | 0 | 0 | 0 | 0 |
| FSAMKYEQ | PRKCSH; PRKC | P14314; K7E | GLU2B_HUMAI | Oxidation (M) | 0 | 0 | 0 | 0 | 0 | 0 | 0 | 0 | 6392100 | 256140 | 0 | 0 | 0 | 0 | 0 | 0 | 91810 | 0 |
| FSAMKYEQ | PRKCSH; PRKC | P14314; K7E | GLU2B_HUMAI | Unmodified | 0 | 0 | 0 | 0 | 0 | 0 | 0 | 0 | 1430900 | 959480 | 0 | 0 | 0 | 0 | 0 | 0 | 0 | 0 |
| FSAMKYEQ | PRKCSH; PRKC | P14314; K7E | GLU2B_HUMAI | Oxidation (M) | 0 | 0 | 0 | 0 | 0 | 0 | 0 | 0 | 300490 | 596630 | 0 | 0 | 0 | 0 | 0 | 0 | 0 | 0 |
| FSAPVVPS | IPO7 | O95373 | IPO7_HUMAN I | Oxidation (M) | 82901 | 88212 | 61828 | 371770 | 2327200 | 126670 | 188430 | 62418 | 151290 | 236280 | 383600 | 1458900 | 917460 | 114690 | 263130 | 132470 | 467240 | 2097900 |
| FSARPSSIE | HACL1; HACL1 | Q9UJ83; C9J | HACL1_HUMAI | Unmodified | 0 | 0 | 0 | 0 | 0 | 0 | 192590 | 0 | 4788600 | 581240 | 0 | 0 | 0 | 0 | 0 | 0 | 32357 | 0 |
| FSASGELG | PCNA | P12004 | PCNA_HUMAN | Unmodified | 0 | 0 | 485250 | 0 | 217690 | 0 | 0 | 0 | 2386300 | 690690 | 1740000 | 7574600 | 3265900 | 851360 | 0 | 0 | 0 | 0 |
| FSASLAPHI | TXNDC15; TXNI | Q96J42; H0Y | TXD15_HUMAN | Unmodified | 0 | 0 | 0 | 0 | 0 | 0 | 0 | 0 | 2803600 | 236750 | 0 | 0 | 0 | 0 | 0 | 0 | 0 | 0 |
| FSATPRYM | PSMC2; PSMC2 | P35998; C9J | PRS7_HUMAN | Unmodified | 0 | 0 | 0 | 0 | 0 | 0 | 249100 | 142600 | 0 | 0 | 0 | 0 | 0 | 832980 | 0 | 0 | 202630 | 0 |
| FSAVALCK | UCHL1; UCHL1 | P09936; D6F | UCHL1_HUMAI | Unmodified | 0 | 0 | 0 | 0 | 4577500 | 976170 | 463540 | 0 | 8428600 | 1183300 | 0 | 27803000 | 0 | 0 | 0 | 208750 | 0 | 0 |
| FSAVQVLEI | DCLK1 | O15075 | DCLK1_HUMAI | Unmodified | 0 | 0 | 0 | 0 | 0 | 0 | 0 | 0 | 0 | 0 | 0 | 1633500 | 0 | 0 | 0 | 0 | 0 | 0 |
| FSAYIKNSN | CLIC1 | O00299 | CLIC1_HUMAN | Unmodified | 0 | 0 | 0 | 0 | 3678000 | 405140 | 1260900 | 881960 | 0 | 0 | 0 | 0 | 1047600 | 0 | 0 | 0 | 8005800 | 1379300 |
| FSAYIKNSN | CLIC1 | O00299 | CLIC1_HUMAN | Unmodified | 0 | 0 | 0 | 0 | 0 | 0 | 0 | 0 | 0 | 0 | 0 | 0 | 0 | 0 | 0 | 0 | 1440000 | 409540 |
| FSAYIKNSF | CLIC4 | Q9Y696 | CLIC4_HUMAN | Unmodified | 0 | 0 | 0 | 0 | 302490 | 0 | 0 | 0 | 0 | 0 | 0 | 0 | 0 | 0 | 0 | 0 | 269940 | 0 |
| FSCDIEQLI | SPATS2L; SPAT | Q9NUQ6; B8 | SPS2L_HUMAN | Unmodified | 0 | 0 | 0 | 0 | 0 | 0 | 169100 | 116090 | 0 | 0 | 0 | 3169600 | 284840 | 0 | 0 | 208360 | 0 | 0 |
| FSCMAWTI | PREP; PREP | P48147; A0A | PPCE_HUMAN | Unmodified | 0 | 0 | 0 | 0 | 0 | 0 | 0 | 0 | 182570 | 0 | 0 | 809530 | 0 | 0 | 0 | 0 | 92047 | 0 |
| FSCVPESM | MVB12B | Q9H7P6 | MB12B_HUMAI | Oxidation (M) | 217550 | 495260 | 151330 | 1311400 | 0 | 554170 | 12086 | 0 | 170150 | 807730 | 0 | 0 | 1062100 | 376600 | 771770 | 328530 | 51738 | 1673900 |
| FSCVPESM | MVB12B | Q9H7P6 | MB12B_HUMAI | Unmodified | 0 | 0 | 0 | 0 | 0 | 0 | 0 | 0 | 54312 | 68742 | 0 | 0 | 26780 | 24122 | 39569 | 0 | 0 | 0 |
| FSDDHYRV | NNT; NNT; NNT | Q13423; A0A | NNTM_HUMAN | Unmodified | 0 | 0 | 0 | 0 | 0 | 0 | 0 | 0 | 221190 | 101130 | 0 | 0 | 0 | 0 | 0 | 0 | 0 | 0 |
| FSDIQIR | MTHFD1; MTHF | P11586; V9G | C1TC_HUMAN | Unmodified | 0 | 0 | 0 | 0 | 0 | 0 | 0 | 0 | 1400500 | 690290 | 893120 | 4047000 | 0 | 0 | 0 | 0 | 67093 | 0 |
| FSDLGFEVI | CASP6; CASP6 | P55212; D6F | CASP6_HUMAI | Unmodified | 0 | 0 | 0 | 0 | 0 | 0 | 0 | 0 | 331490 | 0 | 0 | 1341400 | 0 | 0 | 0 | 0 | 0 | 0 |
| FSDLKLNG | CTNND1; CTNN | O60716; E9F | CTND1_HUMAI | Unmodified | 0 | 0 | 0 | 0 | 0 | 0 | 0 | 0 | 0 | 0 | 0 | 0 | 0 | 117310 | 0 | 0 | 548820 | 603660 |
| FSDLTEEEF | CTSF; CTSF; CT | Q9UBX1; A0 | CATF_HUMAN | Unmodified | 0 | 0 | 0 | 0 | 0 | 0 | 0 | 0 | 1135100 | 0 | 0 | 0 | 0 | 0 | 0 | 0 | 0 | 0 |
| FSDQAGPA | MARK2; MARK2 | E7ETY4; A0A | E7ETY4_HUMA | Unmodified | 171170 | 85643 | 201250 | 0 | 431180 | 183360 | 0 | 0 | 0 | 0 | 135920 | 650100 | 840720 | 414020 | 226300 | 272650 | 566810 | 255060 |
| FSDQAGPA | MARK2; MARK2 | E7ETY4; A0A | E7ETY4_HUMA | Unmodified | 0 | 0 | 0 | 241960 | 0 | 0 | 0 | 0 | 94294 | 0 | 0 | 0 | 445300 | 0 | 89057 | 149030 | 0 | 245690 |
| FSDRTIPTD | SCP2; SCP2; S( | P22307; H0Y | SCP2_HUMAN | Unmodified | 0 | 0 | 0 | 0 | 0 | 0 | 0 | 0 | 179880 | 363840 | 0 | 0 | 0 | 0 | 0 | 0 | 0 | 0 |
| FSDSYASVI | CTPS1; CTPS1; | P17812; B4E | PYRG1_HUMAI | Unmodified | 0 | 0 | 0 | 0 | 0 | 0 | 1095700 | 613130 | 0 | 0 | 0 | 6158600 | 0 | 0 | 0 | 1480700 | 0 | 0 |
| FSEAEHWL | LARS1; LARS1; | Q9P2J5; A0A | SYLC_HUMAN | Unmodified | 0 | 0 | 0 | 50826 | 0 | 0 | 0 | 0 | 0 | 0 | 109310 | 860790 | 153530 | 0 | 198460 | 43253 | 0 | 0 |
| FSEEEISFP | INPPL1; INPPL1 | O15357; A0A | SHIP2_HUMAN | Unmodified | 0 | 0 | 0 | 0 | 0 | 0 | 0 | 0 | 0 | 0 | 0 | 591430 | 0 | 0 | 0 | 0 | 0 | 0 |
| FSEEQRFN | XRCC5 | P13010 | XRCC5_HUMAI | Unmodified | 520970 | 213180 | 381510 | 0 | 0 | 0 | 148370 | 107670 | 790900 | 202870 | 2718000 | 3048700 | 218130 | 0 | 189470 | 0 | 0 | 0 |
| FSEFYSK | UBLCP1 | Q8WVY7 | UBCP1_HUMAI | Unmodified | 0 | 0 | 0 | 0 | 221080 | 0 | 0 | 0 | 0 | 0 | 0 | 900370 | 459820 | 126450 | 0 | 0 | 0 | 0 |

| | | | | | | | | | | | | | | | | | | | | | | |
|---|---|---|---|---|---|---|---|---|---|---|---|---|---|---|---|---|---|---|---|---|---|---|
| FSEGEATLF | QARS1; QARS1 | P47897; A0A | SYQ_HUMAN G | Unmodified | 0 | 0 | 0 | 0 | 0 | 0 | 685790 | 284800 | 360200 | 181620 | 253150 | 2082300 | 0 | 349020 | 0 | 0 | 0 | 0 |
| FSEGTSAD | PSMC6; PSMC6 | P62333; A0A | PRS10_HUMAN | Unmodified | 0 | 0 | 29144 | 0 | 32379 | 0 | 0 | 0 | 1180700 | 0 | 0 | 4012300 | 0 | 14168 | 0 | 0 | 73311 | 27484 |
| FSELQWR | POMT1; POMT1 | Q9Y6A1; A0A | POMT1_HUMAN | Unmodified | 0 | 0 | 0 | 0 | 0 | 0 | 0 | 0 | 1172500 | 0 | 0 | 0 | 0 | 0 | 0 | 0 | 0 | 0 |
| FSELTAEK | CCT5; CCT5; C | P48643; B7Z | TCPE_HUMAN | Unmodified | 277890 | 227480 | 0 | 710130 | 1704200 | 216670 | 0 | 266950 | 1574500 | 1767700 | 2756500 | 47244000 | 0 | 1841800 | 466890 | 403110 | 0 | 0 |
| FSEPDPSH | SMOC1 | Q9H4F8 | SMOC1_HUMA | Unmodified | 0 | 129840 | 0 | 0 | 0 | 0 | 0 | 0 | 2881400 | 374150 | 0 | 0 | 0 | 0 | 0 | 0 | 0 | 0 |
| FSEPHLRD | PGAP4 | Q9BRR3 | PGAP4_HUMA | Unmodified | 0 | 0 | 0 | 0 | 0 | 0 | 0 | 0 | 681530 | 0 | 0 | 0 | 0 | 0 | 0 | 0 | 35426 | 0 |
| FSEQVFK | FLOT1; FLOT1; | O75955; A0A | FLOT1_HUMAN | Unmodified | 119610 | 0 | 3770400 | 2653300 | 6381200 | 4429500 | 799430 | 179650 | 9736000 | 3110400 | 928710 | 7141100 | 1914400 | 1273700 | 6022200 | 0 | 44216000 | 3567000 |
| FSERLQDF | FOLH1B; FOLH | Q9HBA9; Q0 | FOH1B_HUMA | Unmodified | 2155300 | 2501900 | 25156000 | 14401000 | 0 | 0 | 0 | 0 | 0 | 0 | 0 | 0 | 177440 | 0 | 17165000 | 17863000 | 0 | 0 |
| FSERLQDF | FOLH1 | Q04609 | FOLH1_HUMAN | Unmodified | 0 | 0 | 0 | 2927100 | 0 | 0 | 0 | 0 | 0 | 0 | 0 | 0 | 0 | 0 | 0 | 0 | 0 | 0 |
| FSEVLKR | CKB; CKB; CKB | P12277; H0 | KCRB_HUMAN | Unmodified | 0 | 217520 | 745900 | 1068900 | 0 | 0 | 0 | 78391 | 507500 | 0 | 0 | 5500400 | 0 | 0 | 1816400 | 1550900 | 0 | 0 |
| FSFDLGK | FKBP4; FKBP4 | Q02790; F5 | FKBP4_HUMAN | Unmodified | 27026 | 0 | 388940 | 0 | 0 | 0 | 0 | 0 | 0 | 0 | 211060 | 2099400 | 0 | 0 | 414010 | 129450 | 0 | 0 |
| FSFDLGKG | FKBP4; FKBP4 | Q02790; F5 | FKBP4_HUMAN | Unmodified | 833710 | 0 | 0 | 0 | 0 | 0 | 232260 | 0 | 660770 | 0 | 1243100 | 8582400 | 725470 | 256070 | 727400 | 579240 | 0 | 0 |
| FSFMTCPF | UBE3A; UBE3A; | Q05086; A0 | UBE3A_HUMAN | Oxidation (M) | 0 | 0 | 12318 | 43868 | 0 | 0 | 0 | 0 | 0 | 0 | 30387 | 169480 | 0 | 0 | 20771 | 33292 | 0 | 0 |
| FSFPAPSSI | AHNAK2 | Q8IVF2 | AHNK2_HUMA | Unmodified | 0 | 36381 | 27324 | 0 | 938800 | 176210 | 0 | 0 | 0 | 0 | 0 | 0 | 0 | 0 | 0 | 0 | 0 | 375390 |
| FSFQCPGF | ILK; ILK | Q13418; A0 | ILK_HUMAN Int | Unmodified | 340360 | 368880 | 499860 | 0 | 194060 | 69150 | 113050 | 80039 | 0 | 0 | 0 | 382030 | 128190 | 0 | 446010 | 391640 | 82175 | 0 |
| FSFSGNTL | GANAB; GANAE | Q14697; F5 | GANAB_HUMA | Unmodified | 0 | 0 | 0 | 0 | 0 | 0 | 0 | 0 | 17778000 | 6912800 | 559600 | 0 | 0 | 0 | 0 | 0 | 0 | 0 |
| FSFSPEPTL | DCTPP1 | Q9H773 | DCTP1_HUMAI | Unmodified | 0 | 0 | 0 | 0 | 0 | 0 | 0 | 0 | 0 | 0 | 0 | 3984800 | 0 | 0 | 0 | 0 | 720420 | 0 |
| FSFSPEPTL | DCTPP1 | Q9H773 | DCTP1_HUMAI | Unmodified | 0 | 0 | 0 | 0 | 0 | 0 | 0 | 0 | 0 | 0 | 216120 | 1889800 | 0 | 0 | 0 | 0 | 0 | 0 |
| FSGEELDK | LRPAP1; LRPAP | P30533; A0A | AMRP_HUMAN | Unmodified | 0 | 0 | 0 | 0 | 0 | 0 | 0 | 0 | 7672000 | 654610 | 523570 | 47470 | 0 | 0 | 0 | 0 | 0 | 0 |
| FSGFEASH | PTPRJ; PTPRJ; P | Q12913; A0 | PTPRJ_HUMAN | Unmodified | 16452 | 569080 | 832500 | 1305900 | 6291400 | 7923500 | 2030200 | 690180 | 10402000 | 2417900 | 9067800 | 21602000 | 1714600 | 842730 | 1287000 | 927080 | 3817800 | 1848900 |
| FSGRTEYQ | EPB41L4B | Q9H329 | E41LB_HUMAN | Unmodified | 0 | 0 | 0 | 0 | 0 | 0 | 0 | 0 | 0 | 0 | 0 | 0 | 0 | 0 | 0 | 0 | 0 | 0 |
| FSGTTGQM | PI4KA | P42356 | PI4KA_HUMAN | Unmodified | 0 | 0 | 0 | 0 | 0 | 0 | 0 | 0 | 191800 | 0 | 6283600 | 0 | 0 | 0 | 220690 | 194220 | 0 | 0 |
| FSGWYDAI | PGAM1 | P18669 | PGAM1_HUMA | Unmodified | 1640800 | 1089200 | 2124900 | 1559500 | 558330 | 0 | 0 | 0 | 1986500 | 0 | 5207900 | 54911000 | 285330 | 0 | 4052800 | 2750500 | 0 | 0 |
| FSGWYDAI | PGAM1 | P18669 | PGAM1_HUMA | Unmodified | 0 | 0 | 0 | 0 | 0 | 0 | 0 | 0 | 0 | 0 | 0 | 22012000 | 0 | 0 | 3245300 | 3419400 | 0 | 0 |
| FSGYPLYH | FOLH1B; FOLH | Q9HBA9; Q0 | FOH1B_HUMA | Unmodified | 5320700 | 542830 | 6285200 | 19984000 | 0 | 0 | 0 | 0 | 249040 | 0 | 0 | 0 | 479480 | 0 | 45976000 | 8708000 | 0 | 0 |
| FSHDADPE | PSMD2; PSMD2 | Q13200; H7 | PSMD2_HUMA | 2 Oxidation (M) | 0 | 0 | 0 | 0 | 0 | 0 | 0 | 0 | 0 | 0 | 0 | 2923600 | 0 | 0 | 0 | 0 | 0 | 0 |
| FSHEEIAM | ALDOA; ALDOA | P04075; J3K | ALDOA_HUMAI | Oxidation (M) | 875230 | 322900 | 1119800 | 906120 | 6414800 | 1060500 | 525260 | 0 | 1096100 | 837000 | 3288600 | 10569000 | 1290700 | 440430 | 718590 | 1350900 | 1066800 | 850860 |
| FSHEEIAM | ALDOA; ALDOA | P04075; J3K | ALDOA_HUMAI | Unmodified | 0 | 138320 | 0 | 444530 | 12786000 | 1929400 | 0 | 0 | 5785200 | 773110 | 886100 | 5172400 | 779120 | 0 | 181720 | 109450 | 2168800 | 2763600 |
| FSHEEIAM | ALDOA; ALDOA | P04075; J3K | ALDOA_HUMAI | Oxidation (M) | 718490 | 219590 | 0 | 4977600 | 22266000 | 6663400 | 3880600 | 87759 | 4748500 | 8243700 | 25858000 | 77376000 | 8814200 | 2703900 | 3490600 | 3794500 | 4072200 | 2156100 |
| FSHEEIAM | ALDOA; ALDOA | P04075; J3K | ALDOA_HUMAI | Unmodified | 755670 | 509980 | 518250 | 2732700 | 1.28E+08 | 15117000 | 2994600 | 0 | 29798000 | 11681000 | 12006000 | 77085000 | 8269600 | 5669400 | 2124500 | 857430 | 23647000 | 28140000 |
| FSHLTSLP | LARP1; LARP1; | Q6PKG0; E5 | LARP1_HUMAN | Oxidation (M) | 0 | 0 | 697690 | 0 | 0 | 0 | 0 | 0 | 0 | 0 | 0 | 0 | 0 | 0 | 466140 | 629860 | 0 | 0 |
| FSHQGVQI | EIF3B; EIF3B | P55884; C9 | EIF3B_HUMAN | Unmodified | 738940 | 392990 | 485520 | 578080 | 0 | 0 | 0 | 61485 | 718620 | 0 | 339510 | 13264000 | 0 | 0 | 530830 | 657310 | 0 | 0 |
| FSIFGKDF | PRSS23 | O95084 | PRS23_HUMAN | Unmodified | 0 | 0 | 0 | 0 | 1299200 | 206000 | 0 | 0 | 0 | 0 | 0 | 0 | 201050 | 0 | 0 | 0 | 0 | 0 |
| FSIQDADR | PLSCR3; PLSCF | Q9NRY6; I3L | PLS3_HUMAN | Unmodified | 2961300 | 1774200 | 1039400 | 164280 | 553130 | 0 | 0 | 0 | 609060 | 191610 | 196430 | 1168100 | 0 | 66371 | 1821900 | 1671600 | 274750 | 0 |
| FSIQDADR | PLSCR3; PLSCF | Q9NRY6; I3L | PLS3_HUMAN | Unmodified | 8279000 | 4416300 | 1823600 | 756210 | 278430 | 0 | 0 | 0 | 2658700 | 885890 | 0 | 7602600 | 323140 | 0 | 2952500 | 2126900 | 0 | 0 |
| FSIQTMCPI | AIMP2; AIMP2 | Q13155; A8 | AIMP2_HUMAN | Oxidation (M) | 0 | 0 | 0 | 50895 | 0 | 0 | 0 | 0 | 0 | 0 | 0 | 1817800 | 145450 | 0 | 0 | 0 | 0 | 162330 |
| FSIQTMCPI | AIMP2; AIMP2 | Q13155; A8 | AIMP2_HUMAN | Unmodified | 0 | 0 | 0 | 0 | 0 | 0 | 0 | 0 | 348600 | 0 | 0 | 276990 | 0 | 0 | 0 | 0 | 0 | 0 |
| FSIRLVPN | CNDP2 | Q96KP4 | CNDP2_HUMA | Oxidation (M) | 0 | 0 | 0 | 0 | 0 | 0 | 0 | 0 | 0 | 0 | 0 | 0 | 3528200 | 5354600 | 0 | 0 | 0 | 0 |
| FSISPDEDS | SLC38A2; SLC3 | Q96QD8; F8 | S38A2_HUMAN | Unmodified | 3719500 | 1674900 | 681700 | 1601300 | 0 | 0 | 0 | 145370 | 3625800 | 796110 | 1287600 | 4329600 | 0 | 0 | 1113800 | 1038800 | 857980 | 1986200 |
| FSKGEILQ | PALS2; PALS2; | Q9NZW5; B | PALS2_HUMAN | Unmodified | 0 | 0 | 0 | 0 | 16293000 | 0 | 695720 | 0 | 0 | 0 | 2159600 | 1119700 | 0 | 1552400 | 0 | 0 | 5155300 | 15407000 |
| FSKVEDMA | MYH9; MYH10; | P35579; P35 | MYH9_HUMAN | Oxidation (M) | 0 | 0 | 0 | 2088300 | 0 | 5891300 | 0 | 0 | 0 | 1481400 | 522730 | 0 | 0 | 0 | 1591700 | 1710200 | 3080200 | 1942600 |
| FSKVEDMA | MYH9; MYH10; | P35579; P35 | MYH9_HUMAN | Unmodified | 0 | 0 | 0 | 892120 | 6965300 | 2320000 | 1494100 | 0 | 1108200 | 945620 | 0 | 1350300 | 1514400 | 997530 | 0 | 0 | 1393600 | 2747200 |
| FSLARQVD | TFRC; TFRC | P02786; F8 | TFR1_HUMAN | Unmodified | 0 | 0 | 0 | 0 | 3047200 | 126870 | 102890 | 0 | 0 | 0 | 0 | 0 | 659260 | 0 | 0 | 0 | 0 | 1241600 |
| FSLCSIR | ADAM10; ADAM | O14672; H0 | ADA10_HUMAN | Unmodified | 1127400 | 575120 | 4100600 | 1424000 | 0 | 0 | 0 | 0 | 848610 | 0 | 0 | 1695800 | 323990 | 0 | 5403200 | 4602700 | 0 | 0 |
| FSLDAVSL | DYNC1LI1; DYN | Q9Y6G9; C9 | DC1L1_HUMAN | Unmodified | 0 | 0 | 208530 | 131740 | 221050 | 50248 | 166540 | 120390 | 806860 | 275760 | 1078200 | 5499300 | 294760 | 103820 | 315530 | 303880 | 101470 | 98987 |
| FSLFAER | ITGA6; ITGA6 | P23229; H7 | ITA6_HUMAN In | Unmodified | 0 | 0 | 793210 | 491190 | 289860 | 0 | 0 | 0 | 0 | 0 | 0 | 1958400 | 0 | 0 | 1032300 | 1067100 | 568390 | 1219000 |
| FSLFAGGM | TMED9 | Q9BVK6 | TMED9_HUMAI | Oxidation (M) | 0 | 0 | 0 | 0 | 0 | 0 | 0 | 0 | 5925200 | 845720 | 423740 | 0 | 0 | 0 | 0 | 0 | 0 | 0 |
| FSLFAGGM | TMED9 | Q9BVK6 | TMED9_HUMAI | Unmodified | 0 | 0 | 0 | 0 | 0 | 0 | 0 | 0 | 17306000 | 0 | 0 | 0 | 0 | 0 | 0 | 0 | 0 | 0 |
| FSLGYCDF | MITD1; MITD1 | Q8WV92; H7 | MITD1_HUMAN | Unmodified | 0 | 0 | 0 | 0 | 2119300 | 805940 | 0 | 0 | 0 | 0 | 808340 | 0 | 0 | 0 | 0 | 0 | 0 | 0 |
| FSLIDLAGN | KIF2A; KIF2A; K | O00139; A0 | KIF2A_HUMAN | Unmodified | 0 | 0 | 0 | 0 | 0 | 0 | 0 | 0 | 0 | 0 | 0 | 1252300 | 0 | 0 | 0 | 0 | 0 | 0 |
| FSLQDPPN | UGDH | O60701 | UGDH_HUMAN | Unmodified | 0 | 0 | 104720 | 0 | 859190 | 0 | 0 | 173550 | 0 | 454970 | 3325800 | 0 | 0 | 0 | 0 | 0 | 116960 | 177690 |
| FSLVPHNY | NIBAN2 | Q96TA1 | NIBA2_HUMAN | Unmodified | 1584300 | 1656600 | 1058700 | 2292600 | 2282900 | 921710 | 0 | 0 | 4848200 | 746300 | 313720 | 3780800 | 199530 | 0 | 2868700 | 2193200 | 434610 | 443220 |
| FSLVPMEH | VPS13C; VPS13 | Q709C8; A0 | VP13C_HUMAN | Oxidation (M) | 0 | 0 | 0 | 0 | 0 | 0 | 0 | 0 | 0 | 0 | 0 | 641410 | 0 | 0 | 0 | 0 | 0 | 0 |
| FSLYFLAYE | GLO1 | Q04760 | LGUL_HUMAN | Unmodified | 0 | 0 | 0 | 1011400 | 4549500 | 499490 | 0 | 0 | 250690 | 0 | 689840 | 1070900 | 9697900 | 2880000 | 0 | 0 | 323790 | 2026800 |
| FSMPGFKG | AHNAK | Q09666 | AHNK_HUMAN | Oxidation (M) | 0 | 0 | 0 | 0 | 0 | 0 | 0 | 0 | 0 | 0 | 0 | 0 | 0 | 0 | 0 | 0 | 432410 | 777680 |
| FSMPGFKG | AHNAK | Q09666 | AHNK_HUMAN | Unmodified | 283360 | 0 | 0 | 0 | 5420400 | 563880 | 0 | 0 | 0 | 0 | 0 | 0 | 0 | 0 | 0 | 0 | 667550 | 1890500 |
| FSMPGFKG | AHNAK | Q09666 | AHNK_HUMAN | Unmodified | 0 | 0 | 0 | 0 | 3064500 | 329420 | 0 | 0 | 786050 | 70475 | 1842700 | 0 | 0 | 0 | 457640 | 0 | 0 | 1896900 |
| FSMPGFKG | AHNAK | Q09666 | AHNK_HUMAN | Unmodified | 0 | 0 | 0 | 0 | 4573800 | 50031 | 0 | 30187 | 0 | 0 | 0 | 86369 | 0 | 0 | 0 | 0 | 3731600 | 2827700 |
| FSMPGFKG | AHNAK | Q09666 | AHNK_HUMAN | Oxidation (M) | 0 | 0 | 0 | 0 | 2410200 | 0 | 0 | 0 | 0 | 0 | 0 | 0 | 76683 | 0 | 0 | 0 | 184150 | 539920 |
| FSMPGFKG | AHNAK | Q09666 | AHNK_HUMAN | Unmodified | 0 | 0 | 0 | 0 | 3001100 | 0 | 121390 | 0 | 0 | 0 | 0 | 0 | 605020 | 322850 | 0 | 23267 | 342310 | 2462200 |
| FSMPGFKG | AHNAK | Q09666 | AHNK_HUMAN | Oxidation (M) | 301440 | 0 | 0 | 30677 | 860400 | 0 | 0 | 0 | 0 | 0 | 0 | 0 | 0 | 0 | 0 | 178040 | 71363 | 0 |
| FSMPSLK | AHNAK | Q09666 | AHNK_HUMAN | Unmodified | 0 | 0 | 32689 | 0 | 810370 | 313860 | 0 | 0 | 0 | 0 | 0 | 0 | 0 | 0 | 0 | 0 | 323050 | 544370 |
| FSMPSLKG | AHNAK | Q09666 | AHNK_HUMAN | Oxidation (M) | 0 | 0 | 0 | 0 | 2772900 | 464630 | 0 | 0 | 0 | 0 | 0 | 0 | 0 | 0 | 0 | 0 | 1623400 | 1806700 |
| FSMPSLKG | AHNAK | Q09666 | AHNK_HUMAN | Unmodified | 0 | 0 | 0 | 0 | 15626000 | 1083700 | 0 | 0 | 0 | 0 | 0 | 0 | 227660 | 418430 | 0 | 0 | 4448900 | 7593000 |
| FSMPSLKG | AHNAK | Q09666 | AHNK_HUMAN | Unmodified | 0 | 0 | 0 | 0 | 2853800 | 0 | 0 | 0 | 0 | 0 | 0 | 0 | 0 | 0 | 0 | 0 | 575810 | 0 |
| FSMPSLKG | AHNAK | Q09666 | AHNK_HUMAN | Oxidation (M) | 0 | 0 | 0 | 0 | 555870 | 3513600 | 0 | 18649 | 0 | 0 | 0 | 0 | 0 | 110840 | 0 | 0 | 0 | 0 |
| FSMVVQD | PRDX5 | P30044 | PRDX5_HUMAN | Unmodified | 0 | 0 | 0 | 0 | 0 | 0 | 0 | 0 | 1304100 | 113800 | 104600 | 0 | 203180 | 0 | 0 | 0 | 0 | 0 |
| FSNISAAK | SERPINA7; CCT | P05543; P50 | THBG_HUMAN | Unmodified | 0 | 0 | 0 | 0 | 0 | 0 | 201540 | 115920 | 0 | 0 | 0 | 12354000 | 0 | 0 | 584950 | 0 | 0 | 0 |
| FSNNPALC | EGFR; EGFR | P00533; C9 | EGFR_HUMAN | Unmodified | 155270 | 0 | 0 | 763670 | 0 | 0 | 0 | 0 | 0 | 0 | 0 | 0 | 0 | 0 | 863300 | 1032200 | 0 | 0 |
| FSNTLELK | MYO1E | Q12965 | MYO1E_HUMAI | Unmodified | 358800 | 0 | 0 | 0 | 0 | 65452 | 0 | 296600 | 0 | 123110 | 44866 | 0 | 0 | 0 | 0 | 0 | 572590 | 6601700 |
| FSPAGPILS | PABPC1; PABP | P11940; Q5 | PABP1_HUMAN | Unmodified | 1.75E+08 | 1.65E+08 | 0 | 90448000 | 400240 | 0 | 0 | 107850 | 4932600 | 472940 | 1731000 | 17666000 | 1831400 | 761530 | 32016000 | 48326000 | 0 | 238350 |
| FSPAGPVL | PABPC4; PABP | Q13310; B1 | PABP4_HUMAN | Unmodified | 25902000 | 18927000 | 10242000 | 8548100 | 0 | 0 | 0 | 0 | 572950 | 0 | 0 | 2010200 | 0 | 0 | 6359800 | 11192000 | 0 | 0 |
| FSPDGELY | STRAP; STRAP | Q9Y3F4; H0 | STRAP_HUMAN | Unmodified | 0 | 0 | 0 | 0 | 0 | 0 | 0 | 0 | 0 | 0 | 0 | 6561200 | 0 | 34373 | 0 | 228420 | 0 | 0 |
| FSPDGNRF | WDR1; WDR1 | O75083; D6 | WDR1_HUMAN | Unmodified | 3006400 | 1120700 | 647250 | 0 | 9541200 | 1839900 | 0 | 83358 | 5563300 | 1505900 | 8453400 | 6365400 | 4810700 | 4648400 | 2673700 | 2172200 | 1089100 | 2267900 |
| FSPDGNRF | WDR1; WDR1 | O75083; D6 | WDR1_HUMAN | Unmodified | 0 | 0 | 0 | 0 | 6512200 | 0 | 167570 | 0 | 0 | 886760 | 0 | 0 | 4361500 | 1462700 | 0 | 0 | 1177300 | 825440 |
| FSPESVAAI | MFSD14A; N/A | Q96MC6; A0 | MF14A_HUMAN | Unmodified | 0 | 0 | 0 | 0 | 0 | 0 | 0 | 0 | 550700 | 0 | 194310 | 0 | 0 | 0 | 0 | 0 | 0 | 0 |
| FSPFSFK | ATP6V0A4; ATP | Q9HBG4; A0 | VPP4_HUMAN | Unmodified | 0 | 0 | 0 | 0 | 0 | 0 | 0 | 0 | 0 | 0 | 0 | 1150900 | 0 | 0 | 0 | 0 | 0 | 0 |
| FSPGAPGC | ZYX; ZYX; ZYX | Q15942; H0 | ZYX_HUMAN Z | Unmodified | 0 | 0 | 212970 | 0 | 2547800 | 944020 | 0 | 0 | 0 | 0 | 488560 | 32688 | 510560 | 336540 | 390340 | 365300 | 1053100 | 12556000 |
| FSPLTTNLI | ATP5PO; ATP5 | P48047; A0 | ATPO_HUMAN | Unmodified | 0 | 0 | 9482.3 | 16532 | 0 | 0 | 0 | 0 | 3978100 | 102350 | 44578 | 0 | 0 | 0 | 14574 | 0 | 0 | 0 |
| FSPNSSNP | RACK1; RACK1; | P63244; D6 | RACK1_HUMAI | Unmodified | 1164000 | 1171400 | 490600 | 0 | 0 | 0 | 0 | 119360 | 0 | 0 | 1139700 | 11524000 | 0 | 0 | 952210 | 975750 | 0 | 0 |
| FSPNTSQF | BUB3; BUB3; B | O43684; J3 | BUB3_HUMAN | Unmodified | 0 | 0 | 0 | 0 | 0 | 0 | 0 | 0 | 0 | 0 | 0 | 630930 | 43252 | 0 | 0 | 0 | 0 | 0 |
| FSPTDFLA | LRPPRC; LRPPF | P42704; B8 | LPPRC_HUMAN | Unmodified | 0 | 0 | 0 | 0 | 0 | 0 | 18543 | 0 | 1441800 | 0 | 0 | 489180 | 0 | 0 | 0 | 0 | 0 | 0 |
| FSPVTPKFT | ZYX; ZYX; ZYX | Q15942; H0 | ZYX_HUMAN Z | Unmodified | 0 | 0 | 0 | 0 | 0 | 0 | 0 | 89855 | 0 | 0 | 0 | 0 | 0 | 148820 | 0 | 0 | 648150 | 844060 |
| FSQAAKVA | DCTN2; DCTN2 | Q13561; A8 | DCTN2_HUMAI | Unmodified | 0 | 0 | 0 | 0 | 206170 | 0 | 0 | 0 | 0 | 0 | 0 | 0 | 141590 | 125760 | 0 | 0 | 121310 | 124890 |
| FSQELLSN | UBTF; UBTF; UE | P17480; E9 | UBF1_HUMAN | Oxidation (M) | 2185800 | 2023700 | 1899100 | 12230000 | 1.3E+08 | 0 | 0 | 0 | 43745000 | 19490000 | 16886000 | 34255000 | 31979000 | 12274000 | 7250000 | 5322500 | 10314000 | 35949000 |
| FSQEYESIE | PTPRF; PTPRF; | P10586; H0 | PTPRF_HUMAN | Unmodified | 2414600 | 3353800 | 0 | 0 | 16894000 | 5074500 | 0 | 0 | 8910500 | 0 | 0 | 0 | 2463000 | 0 | 6468700 | 4953200 | 0 | 0 |
| FSQFLETEY | PYGL; PYGL | P06737; E9 | PYGL_HUMAN | Unmodified | 0 | 0 | 0 | 0 | 875050 | 0 | 342990 | 0 | 0 | 0 | 0 | 0 | 0 | 0 | 0 | 0 | 314570 | 0 |
| FSQFLGDP | EFCAB14; EFC | O75071; A0 | EFC14_HUMAN | Unmodified | 0 | 195540 | 0 | 0 | 0 | 0 | 0 | 0 | 3181300 | 1478900 | 0 | 0 | 0 | 0 | 556890 | 0 | 0 | 0 |
| FSQIVNTLE | EPHB3 | P54753 | EPHB3_HUMAI | Unmodified | 1733900 | 808180 | 364480 | 230130 | 0 | 0 | 0 | 0 | 0 | 0 | 474650 | 7560700 | 0 | 267610 | 577000 | 325200 | 0 | 376180 |
| FSQLYPER | LSS; LSS | P48449; A0 | LSS_HUMAN L | Unmodified | 0 | 0 | 0 | 0 | 0 | 0 | 0 | 0 | 503030 | 0 | 0 | 0 | 0 | 0 | 0 | 0 | 0 | 0 |
| FSQLYPER | LSS; LSS | P48449; A0 | LSS_HUMAN L | Unmodified | 0 | 0 | 0 | 0 | 859190 | 232400 | 759080 | 208020 | 223300 | 726220 | 0 | 0 | 38477 | 51117 | 0 | 0 | 27318 | 226340 |
| FSQMLHPI | ERP44 | Q9BS26 | ERP44_HUMAN | Oxidation (M) | 0 | 0 | 0 | 0 | 0 | 0 | 0 | 0 | 539950 | 0 | 0 | 0 | 0 | 0 | 0 | 0 | 0 | 0 |
| FSRDDIIGE | SYT4; SYT4 | Q9H2B2; M | SYT4_HUMAN | Unmodified | 0 | 0 | 0 | 0 | 0 | 0 | 0 | 0 | 0 | 0 | 0 | 0 | 0 | 0 | 338500 | 0 | 0 | 0 |
| FSREEMHN | RFTN1; RFTN1; | Q14699; G3 | RFTN1_HUMAN | Unmodified | 0 | 0 | 0 | 0 | 0 | 0 | 0 | 0 | 0 | 0 | 0 | 432170 | 0 | 0 | 0 | 0 | 0 | 0 |
| FSSALQPG | VPS37C; VPS37 | A5D8V6; F5 | VP37C_HUMAN | Oxidation (M) | 214880 | 786440 | 210820 | 312930 | 0 | 0 | 0 | 0 | 155300 | 147470 | 0 | 15084 | 170470 | 0 | 1667600 | 0 | 0 | 0 |
| FSSASGAS | ANLN; ANLN; A | Q9NQW6; H | ANLN_HUMAN | Unmodified | 0 | 0 | 0 | 0 | 821990 | 0 | 35449 | 26984 | 0 | 0 | 0 | 0 | 52336 | 0 | 10406 | 0 | 0 | 191080 |
| FSSDASKV | FLNC | Q14315 | FLNC_HUMAN | Unmodified | 0 | 0 | 0 | 0 | 2155800 | 275040 | 0 | 0 | 0 | 0 | 0 | 0 | 0 | 0 | 0 | 0 | 5124000 | 0 |
| FSSDRTITE | PYGB; PYGB | P11216; H0 | PYGB_HUMAN | Unmodified | 0 | 0 | 114610 | 0 | 0 | 0 | 0 | 192160 | 714630 | 81415 | 0 | 812630 | 104890 | 0 | 233280 | 120440 | 135540 | 0 |
| FSSETWQN | ABHD14B; ABH | Q96IU4; F8 | ABHEB_HUMAI | Unmodified | 145260 | 0 | 0 | 69723 | 0 | 397760 | 0 | 0 | 0 | 0 | 662510 | 3331700 | 0 | 0 | 0 | 0 | 0 | 0 |
| FSSLLQALE | TRIM65; TRIM6 | Q6PJ69; H0 | TRI65_HUMAN | Unmodified | 0 | 0 | 68778 | 0 | 135420 | 0 | 0 | 0 | 0 | 0 | 150030 | 981290 | 0 | 0 | 372940 | 62879 | 0 | 0 |
| FSSLPLGR | NNMT | P40261 | NNMT_HUMAN | Unmodified | 0 | 0 | 0 | 0 | 14200000 | 1277800 | 0 | 0 | 1683800 | 0 | 0 | 0 | 0 | 0 | 0 | 0 | 0 | 0 |
| FSSPDEIDL | MIA3; MIA3 | Q5JRA6; A0 | TGO1_HUMAN | Unmodified | 0 | 0 | 0 | 36618 | 220120 | 0 | 0 | 0 | 6653400 | 2377900 | 377110 | 0 | 0 | 0 | 45535 | 0 | 0 | 0 |
| FSSQFVSK | SUN1; SUN1; S | O94901; H0 | SUN1_HUMAN | Unmodified | 0 | 0 | 0 | 0 | 0 | 0 | 0 | 0 | 3268100 | 0 | 40361 | 257720 | 0 | 0 | 0 | 0 | 0 | 0 |
| FSSQFVSK | SUN1; SUN1; S | O94901; H0 | SUN1_HUMAN | Unmodified | 0 | 0 | 0 | 0 | 0 | 0 | 0 | 0 | 1982800 | 0 | 0 | 0 | 0 | 0 | 0 | 0 | 0 | 0 |
| FSSQIHNN | LOXL2 | Q9Y4K0 | LOXL2_HUMAN | Unmodified | 0 | 0 | 0 | 278340 | 0 | 0 | 0 | 0 | 6441900 | 458050 | 42629 | 0 | 0 | 0 | 0 | 0 | 86220 | 350610 |
| FSSRAESF | MTAP; MTAP; M | Q13126; J3 | MTAP_HUMAN | Unmodified | 0 | 0 | 0 | 0 | 0 | 0 | 0 | 0 | 258660 | 41926 | 410090 | 0 | 0 | 0 | 0 | 0 | 0 | 0 |
| FSSSGDFL | RRP9 | O43818 | U3IP2_HUMAN | Unmodified | 0 | 0 | 0 | 0 | 0 | 0 | 0 | 0 | 0 | 0 | 0 | 3342800 | 0 | 0 | 0 | 0 | 0 | 0 |
| FSSSGGG | KRT9 | P35527 | K1C9_HUMAN | Unmodified | 73354 | 0 | 0 | 99982 | 409160 | 157350 | 0 | 0 | 760880 | 26930 | 0 | 0 | 359340 | 0 | 0 | 0 | 0 | 0 |
| FSSVSSPQ | KIAA1522 | Q9P206 | K1522_HUMAN | Unmodified | 0 | 0 | 754670 | 840780 | 0 | 0 | 0 | 0 | 0 | 36211 | 0 | 761490 | 0 | 0 | 1338200 | 1287600 | 0 | 0 |

| | | | | | | | | | | | | | | | | | | | | | | |
|---|---|---|---|---|---|---|---|---|---|---|---|---|---|---|---|---|---|---|---|---|---|---|
| FSTEGHEA | B4GALNT1 | Q00973 | B4GN1_HUMA | Unmodified | 0 | 0 | 0 | 0 | 0 | 0 | 0 | 0 | 2570900 | 127470 | 0 | 567440 | 0 | 0 | 0 | 0 | 0 | 0 |
| FSTEYELQ( | ANPEP | P15144 | AMPN_HUMAN | Unmodified | 0 | 0 | 0 | 0 | 0 | 0 | 0 | 0 | 0 | 205040 | 0 | 0 | 0 | 0 | 0 | 0 | 1674100 | 2834400 |
| FSTFFDDAI | DHX16; DHX16 | O60231; Q5! | DHX16_HUMAI | Unmodified | 29612 | 17895 | 0 | 0 | 0 | 0 | 0 | 0 | 0 | 0 | 44154 | 369240 | 0 | 0 | 0 | 0 | 0 | 0 |
| FSTPEQAAI | HNRNPL; HNRI | P14866; M0( | HNRPL_HUMAI | Unmodified | 0 | 20298 | 0 | 0 | 174560 | 0 | 143860 | 37746 | 0 | 77118 | 779190 | 6152300 | 253280 | 0 | 0 | 0 | 0 | 0 |
| FSTTQPPQI | ATP13A3; ATP1: | A0A2R8Y635 | A0A2R8Y635_H | Unmodified | 0 | 0 | 0 | 0 | 0 | 0 | 0 | 0 | 522390 | 0 | 0 | 0 | 0 | 0 | 0 | 0 | 0 | 0 |
| FSTWTNTE | EPHX1 | P07099 | HYEP_HUMAN | Unmodified | 0 | 0 | 358450 | 0 | 0 | 0 | 0 | 0 | 515530 | 182210 | 0 | 0 | 0 | 0 | 935580 | 0 | 0 | 0 |
| FSVCVLGD | RPL10A | P62906 | RL10A_HUMAN | Unmodified | 3412900 | 2087800 | 878500 | 1264800 | 1141800 | 457850 | 1616900 | 1240500 | 710320 | 564180 | 2458300 | 25903000 | 2034300 | 4183500 | 1088300 | 1211500 | 513510 | 1031300 |
| FSVGGMTD | LONP1; LONP1 | P36776; K7E | LONM_HUMAN | Unmodified | 0 | 0 | 0 | 0 | 0 | 0 | 0 | 0 | 1982100 | 218830 | 0 | 0 | 0 | 0 | 0 | 0 | 0 | 0 |
| FSVGGMTD | LONP1; LONP1 | P36776; K7E | LONM_HUMAN | Oxidation (M) | 0 | 0 | 0 | 0 | 0 | 0 | 0 | 0 | 933820 | 173060 | 2906900 | 0 | 0 | 0 | 0 | 0 | 0 | 0 |
| FSVHQQSE | ITGAV | P06756 | ITAV_HUMAN I | Unmodified | 0 | 0 | 0 | 0 | 0 | 0 | 0 | 0 | 131410 | 0 | 0 | 574490 | 0 | 0 | 1609700 | 0 | 18606 | 0 |
| FSVHQQSE | ITGAV | P06756 | ITAV_HUMAN I | Oxidation (M) | 0 | 0 | 476240 | 0 | 0 | 0 | 0 | 0 | 67278 | 0 | 31679 | 0 | 0 | 0 | 373670 | 207480 | 0 | 0 |
| FSVICPAVL | SLC39A8 | Q9C0K1 | S39A8_HUMAN | Unmodified | 660420 | 0 | 2702800 | 4089500 | 324380 | 0 | 0 | 0 | 1301800 | 0 | 0 | 684350 | 0 | 0 | 3649000 | 1165600 | 92055 | 0 |
| FSVIWQLVI | IGSF3 | O75054 | IGSF3_HUMAN | Unmodified | 757690 | 533160 | 73919 | 0 | 0 | 0 | 0 | 0 | 0 | 0 | 0 | 482120 | 0 | 0 | 524420 | 311850 | 0 | 0 |
| FSVPGFKAI | AHNAK | Q09666 | AHNK_HUMAN | Unmodified | 0 | 0 | 0 | 233150 | 5756200 | 880800 | 0 | 0 | 0 | 292350 | 0 | 0 | 0 | 0 | 0 | 131660 | 0 | 1940200 |
| FSVSPVVR | EEF2 | P13639 | EF2_HUMAN EI | Unmodified | 3451700 | 3603800 | 3096800 | 4064800 | 17212000 | 3127200 | 3222000 | 1431700 | 0 | 0 | 9600900 | 72063000 | 3878000 | 1817400 | 4358500 | 4270700 | 4815800 | 3764200 |
| FSVSPVVR\ | EEF2 | P13639 | EF2_HUMAN EI | Unmodified | 0 | 0 | 0 | 0 | 2092900 | 0 | 897780 | 0 | 0 | 207210 | 0 | 0 | 690730 | 569980 | 0 | 0 | 575370 | 0 |
| FSVSPVVR\ | EEF2 | P13639 | EF2_HUMAN EI | Unmodified | 0 | 0 | 0 | 0 | 29700000 | 6202800 | 7425300 | 2181900 | 0 | 2349300 | 270600 | 15717000 | 7628900 | 4194400 | 0 | 1871200 | 9720300 | 8103800 |
| FSVTPASC( | EPB41 | A0A2R8Y783 | A0A2R8Y783_H | Unmodified | 38158 | 0 | 47906 | 0 | 504080 | 0 | 176650 | 184770 | 467580 | 176800 | 0 | 0 | 238080 | 0 | 80176 | 0 | 0 | 0 |
| FSWNVWP | SEC23B; SEC23 | Q15437; Q1! | SC23B_HUMAI | Unmodified | 0 | 0 | 0 | 0 | 0 | 63864 | 0 | 0 | 0 | 25030 | 0 | 1724300 | 0 | 0 | 0 | 0 | 0 | 0 |
| FSWNVWP | SEC23B; SEC23 | Q15437; Q1! | SC23B_HUMAI | Unmodified | 13569 | 69000 | 0 | 0 | 109030 | 328470 | 0 | 0 | 873620 | 75524 | 0 | 1432800 | 143030 | 414730 | 0 | 0 | 0 | 999600 |
| FSYDLSQC | GOLM1 | Q8NBJ4 | GOLM1_HUMA | Unmodified | 0 | 0 | 0 | 0 | 0 | 0 | 0 | 0 | 9833000 | 311390 | 0 | 0 | 0 | 0 | 0 | 0 | 0 | 0 |
| FSYLAVIEG | PTPN1; PTPN1 | P18031; B4D | PTN1_HUMAN | Unmodified | 0 | 0 | 0 | 0 | 0 | 0 | 0 | 0 | 0 | 0 | 0 | 940890 | 0 | 0 | 0 | 0 | 0 | 0 |
| FSYLPIQK | ITGA6 | P23229 | ITA6_HUMAN I | Unmodified | 0 | 0 | 5600600 | 4528600 | 16167000 | 962710 | 0 | 0 | 407270 | 0 | 0 | 9533300 | 11477000 | 6259900 | 4841700 | 4094000 | 12277000 | 70497000 |
| FTAGDFST | ATP6V1D; ATP6 | Q9Y5K8; G3 | VATD_HUMAN | Unmodified | 80034 | 56725 | 0 | 178910 | 0 | 0 | 525780 | 436360 | 0 | 0 | 352700 | 1717200 | 0 | 711680 | 270740 | 234170 | 990040 | 67624 |
| FTAGDFST | ATP6V1D; ATP6 | Q9Y5K8; G3 | VATD_HUMAN | Unmodified | 0 | 0 | 0 | 0 | 0 | 0 | 0 | 220770 | 0 | 0 | 0 | 0 | 0 | 0 | 0 | 0 | 0 | 0 |
| FTAQGLPD | UPF1 | Q92900 | RENT1_HUMAI | Unmodified | 1721000 | 1382900 | 0 | 0 | 0 | 0 | 0 | 0 | 84148 | 0 | 0 | 869440 | 0 | 0 | 0 | 0 | 64918 | 0 |
| FTASAGIQ\ | ENO1; ENO1 | P06733; A0A | ENOA_HUMAN | Unmodified | 1027800 | 3087000 | 0 | 5202800 | 1.25E+08 | 5659600 | 6719500 | 1130200 | 10762000 | 15524000 | 4622800 | 10920000 | 50512000 | 19567000 | 3072000 | 2004600 | 40165000 | 52664000 |
| FTASAGIQ\ | ENO1; ENO1 | P06733; A0A | ENOA_HUMAN | Unmodified | 0 | 0 | 0 | 0 | 0 | 0 | 0 | 0 | 1324300 | 796020 | 3227900 | 970390 | 0 | 0 | 0 | 0 | 1262600 | 0 |
| FTASAGIQ\ | ENO1; ENO1 | P06733; A0A | ENOA_HUMAN | Unmodified | 0 | 0 | 0 | 0 | 1761600 | 0 | 845540 | 0 | 0 | 0 | 0 | 0 | 0 | 1060700 | 0 | 0 | 14030000 | 1425000 |
| FTASIQR | SMC4; SMC4 | Q9NTJ3; E9F | SMC4_HUMAN | Unmodified | 0 | 0 | 12952 | 0 | 729980 | 33662 | 56584 | 31951 | 577830 | 66775 | 86081 | 883670 | 101670 | 39312 | 19414 | 25065 | 0 | 0 |
| FTAVAQTDI | SERPINE2 | P07093 | GDN_HUMAN ( | Unmodified | 0 | 0 | 0 | 1021500 | 0 | 0 | 0 | 0 | 276970 | 0 | 398940 | 1843800 | 792600 | 761780 | 393330 | 442810 | 0 | 1225700 |
| FTCKEATD\ | ANPEP | P15144 | AMPN_HUMAN | Unmodified | 0 | 0 | 0 | 0 | 0 | 0 | 0 | 0 | 0 | 0 | 0 | 0 | 0 | 0 | 0 | 0 | 356260 | 465110 |
| FTCSAASS! | LRIG1 | Q96JA1 | LRIG1_HUMAN | Unmodified | 0 | 0 | 0 | 0 | 0 | 0 | 0 | 0 | 653650 | 0 | 0 | 0 | 0 | 0 | 0 | 0 | 0 | 0 |
| FTDEEVDEI | MYL12A; MYL12 | P19105; O14 | ML12A_HUMAI | Unmodified | 0 | 0 | 0 | 0 | 0 | 0 | 0 | 0 | 0 | 0 | 0 | 0 | 0 | 0 | 0 | 0 | 0 | 0 |
| FTDEEVDEI | MYL12A; MYL12 | P19105; O14 | ML12A_HUMAI | Unmodified | 0 | 0 | 0 | 0 | 0 | 0 | 0 | 0 | 0 | 0 | 0 | 445450 | 0 | 0 | 175450 | 0 | 0 | 0 |
| FTDEYQLFE | CAMK2D; CAMI | Q13557; E9F | KCC2D_HUMA | Unmodified | 0 | 0 | 0 | 0 | 0 | 0 | 0 | 0 | 0 | 169550 | 0 | 454790 | 0 | 0 | 0 | 0 | 0 | 0 |
| FTDFDEVR( | DNM2 | P50570 | DYN2_HUMAN | Unmodified | 1048300 | 0 | 0 | 1026700 | 0 | 0 | 0 | 0 | 0 | 0 | 774880 | 0 | 706300 | 457760 | 0 | 0 | 0 | 558690 |
| FTDLFEK | PDCD6IP; CSNI | Q8WUM4; Q | PDC6I_HUMAN | Unmodified | 80360000 | 47422000 | 29517000 | 25581000 | 17489000 | 6664900 | 1368200 | 411600 | 21282000 | 4057500 | 6817300 | 21113000 | 19939000 | 8645600 | 47723000 | 35023000 | 1182900 | 9940300 |
| FTDTRKDE( | EEF2 | P13639 | EF2_HUMAN EI | Unmodified | 859630 | 1198400 | 636020 | 2574900 | 0 | 0 | 0 | 0 | 1276500 | 0 | 924200 | 12821000 | 95262 | 0 | 1171300 | 1196000 | 0 | 0 |
| FTDYCDLN | SMOC1 | Q9H4F8 | SMOC1_HUMA | Unmodified | 0 | 0 | 0 | 0 | 0 | 0 | 0 | 0 | 543390 | 194750 | 0 | 0 | 0 | 0 | 0 | 0 | 0 | 176690 |
| FTEDLVGS\ | MEAK7; MEAK7 | Q6P9B6; H3 | MEAK7_HUMAI | Unmodified | 262840 | 353340 | 334760 | 406340 | 0 | 0 | 0 | 0 | 6769500 | 243700 | 216560 | 3405300 | 0 | 0 | 720770 | 699230 | 0 | 0 |
| FTFNKKET( | TM9SF2 | Q99805 | TM9S2_HUMAI | Unmodified | 17988 | 0 | 0 | 0 | 0 | 23340 | 0 | 0 | 734360 | 1078100 | 0 | 0 | 0 | 0 | 0 | 0 | 0 | 0 |
| FTFPDPPPI | ABCF1; ABCF1; | Q8NE71; H0 | ABCF1_HUMAI | Unmodified | 0 | 0 | 0 | 0 | 0 | 0 | 0 | 0 | 0 | 0 | 0 | 320730 | 0 | 0 | 0 | 0 | 0 | 0 |
| FTFSGREL\ | AHNAK | Q09666 | AHNK_HUMAN | Unmodified | 0 | 0 | 0 | 0 | 1328300 | 164840 | 0 | 0 | 0 | 0 | 0 | 0 | 0 | 0 | 0 | 0 | 141550 | 209840 |
| FTFYDGPPI | IARS1; IARS1; I/ | P41252; A0A | SYIC_HUMAN I | Unmodified | 0 | 0 | 0 | 0 | 0 | 0 | 0 | 0 | 599050 | 0 | 0 | 1720400 | 0 | 0 | 0 | 91889 | 0 | 0 |
| FTGDTLAM | PPFIBP2; PPFIB | Q8ND30; E9 | LIPB2_HUMAN | Oxidation (M) | 0 | 0 | 41795 | 0 | 0 | 0 | 0 | 0 | 0 | 0 | 0 | 0 | 0 | 0 | 0 | 139870 | 0 | 0 |
| FTGEGATAI | CHDH | Q8NE62 | CHDH_HUMAN | Unmodified | 0 | 0 | 0 | 0 | 0 | 239410 | 91699 | 0 | 3141500 | 1038900 | 0 | 0 | 0 | 0 | 0 | 0 | 0 | 0 |
| FTGIKHEW | CFL2 | Q9Y281 | COF2_HUMAN | Unmodified | 330240 | 0 | 0 | 0 | 6546200 | 0 | 0 | 0 | 0 | 0 | 0 | 492440 | 0 | 0 | 0 | 0 | 208230 | 0 |
| FTGIKHEW | CFL2 | Q9Y281 | COF2_HUMAN | Unmodified | 300370 | 425150 | 0 | 152690 | 231170 | 0 | 0 | 0 | 0 | 0 | 0 | 0 | 0 | 0 | 177270 | 0 | 0 | 153440 |
| FTGLSKEEI | SLC3A2; SLC3A | P08195; A0A | 4F2_HUMAN 4I | Unmodified | 8492600 | 5523000 | 3239400 | 4861900 | 2048400 | 455900 | 957000 | 0 | 7835500 | 413090 | 9227600 | 65668000 | 211640 | 0 | 3064400 | 7279100 | 0 | 234300 |
| FTGSEIR | UQCRC1; PMP( | P31930; O75 | QCR1_HUMAN | Unmodified | 0 | 131630 | 0 | 0 | 0 | 21560 | 0 | 0 | 16575000 | 359780 | 157990 | 145590 | 56172 | 0 | 0 | 0 | 33945 | 0 |
| FTGSQPFG | THBS1 | P07996 | TSP1_HUMAN | Unmodified | 0 | 0 | 519600 | 0 | 0 | 0 | 0 | 0 | 0 | 0 | 0 | 0 | 0 | 0 | 310730 | 131580 | 0 | 0 |
| FTGTLFWR | ATP8B1; ATP8B | O43520; A0/ | AT8B1_HUMAN | Unmodified | 0 | 0 | 0 | 0 | 0 | 0 | 0 | 0 | 0 | 0 | 0 | 2913500 | 0 | 0 | 153480 | 213990 | 0 | 0 |
| FTIGDHSR | WDR1; WDR1 | O75083; D6I | WDR1_HUMAN | Unmodified | 0 | 293760 | 62038 | 113910 | 0 | 0 | 0 | 0 | 132370 | 72405 | 0 | 1423200 | 0 | 0 | 38166 | 30963 | 0 | 0 |
| FTIINSK | EMB; EMB | Q6PCB8; D6 | EMB_HUMAN E | Unmodified | 100380 | 0 | 716610 | 0 | 123140 | 0 | 0 | 0 | 2079200 | 122900 | 0 | 0 | 0 | 0 | 1219800 | 1186600 | 87217 | 0 |
| FTISDHPQI | CAND1; CAND1 | Q86VP6; A0/ | CAND1_HUMA | Unmodified | 0 | 0 | 0 | 0 | 0 | 0 | 0 | 0 | 2122500 | 0 | 0 | 6403800 | 56002 | 0 | 0 | 1665600 | 0 | 0 |
| FTITPPTAQ | CAPZA1 | P52907 | CAZA1_HUMAI | Unmodified | 2129200 | 0 | 1788900 | 205010 | 8209300 | 2580200 | 894880 | 503220 | 3407300 | 4636100 | 6573100 | 11983000 | 9037300 | 0 | 4583600 | 5234500 | 832690 | 0 |
| FTITPSTTQ | CAPZA2 | P47755 | CAZA2_HUMAI | Unmodified | 0 | 0 | 202140 | 0 | 0 | 0 | 0 | 0 | 0 | 0 | 3781500 | 10004000 | 0 | 398070 | 0 | 2879000 | 0 | 0 |
| FTLNLQQ | GALNT2 | Q10471 | GALT2_HUMAI | Unmodified | 0 | 0 | 645660 | 1004400 | 0 | 0 | 0 | 77236 | 13662000 | 63764 | 376650 | 72837 | 0 | 0 | 1623300 | 944820 | 0 | 0 |
| FTLPTDHAI | STEAP3 | Q658P3 | STEA3_HUMAI | Unmodified | 0 | 0 | 1486400 | 1393200 | 877990 | 118030 | 0 | 0 | 1600800 | 0 | 897620 | 5093400 | 0 | 0 | 3271800 | 1389600 | 0 | 0 |
| FTLQDSR | PLEKHB2; PLEK | Q96CS7; A0/ | PKHB2_HUMAI | Unmodified | 1517000 | 1787600 | 2768900 | 0 | 0 | 0 | 0 | 0 | 0 | 0 | 84022 | 0 | 0 | 0 | 2740800 | 2398600 | 0 | 0 |
| FTLRDEGK | GSPT1; GSPT2; | P15170; Q8I | ERF3A_HUMAN | Unmodified | 1041400 | 0 | 2873400 | 824680 | 68073 | 0 | 61791 | 89557 | 3350600 | 0 | 736310 | 905780 | 0 | 0 | 21342 | 119930 | 62426 | 133680 |
| FTLTGLKPI | PTPRF; PTPRF; | P10586; H0\ | PTPRF_HUMAN | Unmodified | 584500 | 588710 | 166320 | 0 | 0 | 0 | 0 | 0 | 336170 | 0 | 0 | 0 | 0 | 0 | 386340 | 258620 | 0 | 0 |
| FTLTQSEAI | TTLL12 | Q14166 | TTL12_HUMAN | Unmodified | 0 | 0 | 0 | 449990 | 495100 | 0 | 0 | 0 | 0 | 0 | 0 | 352370 | 579130 | 0 | 1192400 | 0 | 80630 | 0 |
| FTMVQVWI | ATP6V1A; ATP6 | P38606; C9J | VATA_HUMAN | Oxidation (M) | 0 | 0 | 0 | 0 | 0 | 0 | 0 | 0 | 0 | 0 | 0 | 1258700 | 0 | 0 | 0 | 0 | 0 | 0 |
| FTNIGPDTI | FN1; FN1 | P02751; H0\ | FINC_HUMAN I | Oxidation (M) | 0 | 0 | 6030100 | 0 | 0 | 85069 | 0 | 0 | 402160 | 0 | 0 | 806770 | 0 | 0 | 440500 | 327370 | 0 | 0 |
| FTNIGPDTI | FN1; FN1 | P02751; H0\ | FINC_HUMAN I | Unmodified | 0 | 0 | 0 | 0 | 0 | 0 | 0 | 0 | 0 | 0 | 0 | 0 | 0 | 0 | 0 | 0 | 0 | 0 |
| FTNSDTPT! | TOR1AIP2 | Q8NFQ8 | TOIP2_HUMAN | Oxidation (M) | 0 | 0 | 0 | 0 | 34303 | 0 | 0 | 0 | 990100 | 46000 | 0 | 111690 | 0 | 0 | 0 | 0 | 0 | 0 |
| FTNSDTPT! | TOR1AIP2 | Q8NFQ8 | TOIP2_HUMAN | Unmodified | 0 | 0 | 0 | 0 | 0 | 0 | 0 | 0 | 6018400 | 241050 | 0 | 151110 | 0 | 0 | 0 | 0 | 0 | 0 |
| FTNSDTPT! | TOR1AIP2 | Q8NFQ8 | TOIP2_HUMAN | Oxidation (M) | 0 | 0 | 0 | 0 | 0 | 203750 | 0 | 0 | 2026100 | 2697600 | 0 | 0 | 0 | 0 | 0 | 0 | 51266 | 0 |
| FTNSDTPT! | TOR1AIP2 | Q8NFQ8 | TOIP2_HUMAN | Unmodified | 0 | 0 | 0 | 0 | 129470 | 0 | 0 | 0 | 1343000 | 1058100 | 0 | 0 | 0 | 0 | 0 | 0 | 0 | 0 |
| FTNSKLTSI | MSH2; MSH2; I | P43246; A0A | MSH2_HUMAN | Unmodified | 0 | 0 | 0 | 0 | 0 | 0 | 344480 | 0 | 0 | 0 | 0 | 0 | 0 | 0 | 0 | 0 | 0 | 0 |
| FTNSNTPN | TOR1AIP1; TOR | Q5JTV8; A0A | TOIP1_HUMAN | Oxidation (M) | 0 | 0 | 0 | 0 | 0 | 0 | 0 | 0 | 867950 | 0 | 148390 | 0 | 0 | 0 | 0 | 0 | 0 | 0 |
| FTNSNTPN | TOR1AIP1; TOR | Q5JTV8; A0A | TOIP1_HUMAN | Unmodified | 0 | 0 | 0 | 0 | 0 | 0 | 0 | 0 | 3080400 | 0 | 0 | 0 | 0 | 0 | 0 | 0 | 0 | 0 |
| FTNTQITEH | NCAPH; NCAPI | Q15003; E9F | CND2_HUMAN | Unmodified | 0 | 0 | 0 | 0 | 0 | 0 | 539320 | 387130 | 0 | 0 | 140890 | 797730 | 0 | 202790 | 0 | 163170 | 0 | 0 |
| FTPFTLGK | EFNA1 | P20827 | EFNA1_HUMAI | Unmodified | 1252600 | 888950 | 706030 | 698040 | 0 | 0 | 0 | 0 | 1451900 | 86708 | 125450 | 0 | 0 | 18265 | 517650 | 557730 | 37689 | 0 |
| FTPGTFTN( | RPSA; RPSA; RI | P08865; A0A | RSSA_HUMAN | Unmodified | 323300 | 336370 | 130000 | 170660 | 179450 | 0 | 362060 | 163950 | 0 | 0 | 166100 | 2873700 | 0 | 0 | 284630 | 0 | 0 | 91756 |
| FTPGTFTN( | RPSA; RPSA; RI | P08865; A0A | RSSA_HUMAN | Unmodified | 6857900 | 7863000 | 0 | 0 | 0 | 0 | 0 | 0 | 0 | 0 | 6679200 | 1.04E+08 | 683700 | 0 | 8152200 | 6751600 | 0 | 0 |
| FTPPLEDS( | EPHX1; EPHX1 | P07099; B1A | HYEP_HUMAN | Unmodified | 0 | 0 | 307910 | 336010 | 0 | 0 | 0 | 26766 | 706290 | 207410 | 0 | 0 | 0 | 0 | 380940 | 144410 | 0 | 0 |
| FTPPPQPQ | PTK7 | Q13308 | PTK7_HUMAN I | Unmodified | 0 | 0 | 0 | 0 | 216580 | 0 | 0 | 0 | 566060 | 0 | 309030 | 0 | 0 | 449860 | 0 | 0 | 0 | 0 |
| FTPVASKFS | ZYX; ZYX; ZYX | Q15942; H0\ | ZYX_HUMAN Z | Unmodified | 0 | 0 | 0 | 0 | 2666600 | 0 | 0 | 0 | 0 | 0 | 0 | 0 | 378890 | 0 | 0 | 0 | 1943800 | 0 |
| FTQAGSEV | ATP5F1B; ATP5I | P06576; H0\ | ATPB_HUMAN | Unmodified | 0 | 0 | 0 | 0 | 0 | 0 | 0 | 0 | 15714000 | 265390 | 298110 | 201670 | 0 | 0 | 0 | 0 | 72473 | 0 |
| FTQAGSEV | ATP5F1B; ATP5I | P06576; H0\ | ATPB_HUMAN | 2 Oxidation (M) | 0 | 0 | 0 | 0 | 0 | 0 | 0 | 0 | 3797400 | 0 | 0 | 0 | 0 | 0 | 0 | 0 | 0 | 0 |
| FTQDTQPH | DYNC1H1; DYN | Q14204; A0/ | DYHC1_HUMA | Unmodified | 130450 | 0 | 0 | 0 | 0 | 0 | 0 | 0 | 0 | 0 | 0 | 9063700 | 0 | 0 | 0 | 637030 | 0 | 0 |
| FTQGKSPA | CHST1 | O43916 | CHST1_HUMAI | Unmodified | 0 | 0 | 0 | 0 | 0 | 0 | 0 | 0 | 1346900 | 0 | 0 | 0 | 0 | 0 | 0 | 0 | 0 | 0 |
| FTQISPVWI | CHID1; CHID1; | Q9BWS9; E9 | CHID1_HUMAI | Unmodified | 0 | 0 | 0 | 0 | 0 | 0 | 0 | 0 | 35715000 | 6796000 | 1728500 | 898170 | 0 | 0 | 0 | 0 | 0 | 0 |
| FTQISPVWI | CHID1; CHID1; | Q9BWS9; E9 | CHID1_HUMAI | Unmodified | 0 | 0 | 0 | 0 | 0 | 0 | 0 | 0 | 4628600 | 0 | 0 | 0 | 0 | 0 | 0 | 0 | 0 | 0 |
| FTQLDLED | SMC4; SMC4; S | Q9NTJ3; E9F | SMC4_HUMAN | Unmodified | 0 | 0 | 0 | 0 | 1149300 | 364610 | 465220 | 473370 | 616380 | 215300 | 0 | 0 | 1271800 | 0 | 0 | 0 | 0 | 0 |
| FTQVTPTSI | FN1; FN1 | P02751; H0\ | FINC_HUMAN I | Unmodified | 111730 | 0 | 37044000 | 18957000 | 0 | 0 | 0 | 0 | 2859700 | 0 | 1671600 | 0 | 0 | 0 | 1012300 | 10048000 | 1696200 | 0 |
| FTQVTPTSI | FN1; FN1 | P02751; H0\ | FINC_HUMAN I | Unmodified | 0 | 0 | 822460 | 789870 | 0 | 0 | 0 | 0 | 0 | 794260 | 0 | 0 | 0 | 0 | 0 | 0 | 0 | 0 |
| FTRNEFNLI | RAB11A; RAB11 | P62491; Q1! | RB11A_HUMAI | Unmodified | 11449000 | 8226600 | 16238000 | 2383300 | 39132000 | 11050000 | 1971000 | 1329500 | 12790000 | 3842800 | 32468000 | 37363000 | 13113000 | 6576500 | 15018000 | 20973000 | 865000 | 4229400 |
| FTRVPSGG | MAN1B1 | Q9UKM7 | MA1B1_HUMAI | Unmodified | 0 | 0 | 0 | 0 | 0 | 0 | 0 | 0 | 12596000 | 0 | 0 | 0 | 0 | 0 | 0 | 0 | 0 | 0 |
| FTSPPGDK | PIK3C2B; PIK3( | O00750; F5( | P3C2B_HUMAI | Unmodified | 0 | 0 | 0 | 0 | 0 | 0 | 564300 | 0 | 0 | 0 | 0 | 336340 | 0 | 0 | 30183 | 342160 | 0 | 0 |
| FTSSPGLS | CASK; CASK; C/ | O14936; A0/ | CSKP_HUMAN | Unmodified | 0 | 0 | 0 | 0 | 0 | 0 | 0 | 0 | 293320 | 0 | 0 | 0 | 0 | 0 | 292030 | 285110 | 0 | 0 |
| FTSSPGLS | CASK; CASK; C/ | O14936; A0/ | CSKP_HUMAN | Unmodified | 379940 | 374350 | 2705700 | 2768500 | 0 | 0 | 0 | 0 | 5511100 | 736930 | 0 | 0 | 0 | 0 | 3374600 | 2925200 | 174360 | 0 |
| FTSVRGDK | SUPT16H | Q9Y5B9 | SP16H_HUMAI | Unmodified | 0 | 0 | 0 | 0 | 0 | 0 | 405090 | 62403 | 0 | 0 | 0 | 915530 | 0 | 0 | 0 | 0 | 0 | 0 |
| FTTEIHPSC | EPHA2 | P29317 | EPHA2_HUMAI | Unmodified | 0 | 0 | 0 | 148950 | 497320 | 0 | 0 | 0 | 0 | 0 | 0 | 0 | 89838 | 0 | 64761 | 0 | 2168900 | 6995000 |
| FTTEIHPSC | EPHA2 | P29317 | EPHA2_HUMAI | Unmodified | 0 | 266040 | 0 | 349620 | 9047800 | 3283100 | 0 | 0 | 0 | 0 | 0 | 0 | 590510 | 637390 | 0 | 0 | 33899000 | 62752000 |
| FTTEIHPSC | EPHA2 | P29317 | EPHA2_HUMAI | Unmodified | 0 | 0 | 0 | 0 | 0 | 0 | 0 | 0 | 0 | 0 | 0 | 0 | 0 | 0 | 0 | 0 | 3393500 | 365410 |
| FTTGDAMS | COTL1 | Q14019 | COTL1_HUMAI | Unmodified | 0 | 0 | 0 | 424500 | 20327 | 1029600 | 189940 | 128800 | 2833200 | 1243300 | 0 | 3620300 | 0 | 0 | 0 | 0 | 1328600 | 922350 |
| FTTLEGLLK | ZPR1; ZPR1; ZP | O75312; H7( | ZPR1_HUMAN | Unmodified | 0 | 0 | 0 | 0 | 0 | 0 | 0 | 0 | 219810 | 0 | 639840 | 1988700 | 0 | 0 | 0 | 0 | 0 | 0 |
| FTTPGFPD! | ST14 | Q9Y5Y6 | ST14_HUMAN ! | Unmodified | 5881200 | 5288800 | 12617000 | 16875000 | 0 | 0 | 0 | 0 | 6596500 | 0 | 780950 | 0 | 0 | 0 | 16652000 | 12446000 | 0 | 0 |
| FTTYEIR | SNX3 | O60493 | SNX3_HUMAN | Unmodified | 0 | 0 | 417630 | 608620 | 0 | 0 | 0 | 0 | 1011000 | 0 | 0 | 663650 | 294420 | 0 | 0 | 603880 | 0 | 0 |
| FTVDGIPYF | ITCH; ITCH; ITC | Q96J02; A0A | ITCH_HUMAN I | Unmodified | 4178000 | 4870900 | 2521800 | 493760 | 0 | 0 | 0 | 0 | 0 | 0 | 0 | 0 | 0 | 0 | 2649000 | 3636800 | 0 | 0 |
| FTVISVPHL | SNX33; SNX33 | Q8WV41; H3 | SNX33_HUMAI | Unmodified | 3754000 | 2431400 | 34821000 | 22880000 | 0 | 115670 | 0 | 10857 | 0 | 164530 | 72703 | 357740 | 276180 | 304000 | 69201000 | 34675000 | 0 | 0 |
| FTVLLMPN( | PA2G4 | Q9UQ80 | PA2G4_HUMAI | 2 Oxidation (M) | 0 | 0 | 0 | 0 | 0 | 0 | 0 | 0 | 0 | 0 | 0 | 0 | 0 | 0 | 0 | 0 | 0 | 0 |
| FTVSWYYR | PTGFRN | Q9P2B2 | FPRP_HUMAN | Unmodified | 2611700 | 0 | 4239500 | 4422400 | 0 | 0 | 0 | 0 | 0 | 0 | 0 | 5483800 | 0 | 0 | 8783500 | 8383400 | 0 | 0 |
| FTVVAISPP | MGAT2 | Q10469 | MGAT2_HUMAI | Unmodified | 0 | 0 | 0 | 0 | 0 | 0 | 0 | 0 | 810640 | 526470 | 0 | 0 | 0 | 0 | 0 | 0 | 643290 | 0 |
| FTVVAISPP | MGAT2 | Q10469 | MGAT2_HUMAI | Unmodified | 0 | 0 | 0 | 0 | 520220 | 129080 | 0 | 0 | 9716500 | 2112500 | 312300 | 687630 | 0 | 0 | 0 | 0 | 0 | 175050 |

| | | | | | | | | | | | | | | | | | | | | | | |
|---|---|---|---|---|---|---|---|---|---|---|---|---|---|---|---|---|---|---|---|---|---|---|
| FTVVVLR | PLA2G4A | P47712 | PA24A_HUMAN | Unmodified | 0 | 0 | 0 | 0 | 0 | 0 | 0 | 0 | 0 | 0 | 0 | 1134100 | 0 | 0 | 0 | 0 | 0 | 0 |
| FTWSSSHV | FAM83F | Q8NEG4 | FA83F_HUMAN | Unmodified | 28299 | 31055 | 72014 | 0 | 0 | 782890 | 36781 | 89543 | 0 | 0 | 0 | 203340 | 0 | 0 | 610840 | 579550 | 0 | 447460 |
| FTYGGCYG | SPINT1; SPINT1 | O43278; H3I | SPIT1_HUMAN | Unmodified | 541870 | 301920 | 265200 | 724480 | 0 | 0 | 0 | 0 | 929040 | 0 | 0 | 0 | 198620 | 0 | 542040 | 475590 | 0 | 0 |
| FTYLGSQD | ZC3HAV1; ZC3I | Q7Z2W4; C9 | ZCCHV_HUMA | Unmodified | 0 | 203420 | 504080 | 561570 | 186580 | 0 | 0 | 0 | 58062 | 0 | 0 | 0 | 0 | 0 | 206080 | 640180 | 0 | 0 |
| FTYRSDSFI | XRCC6; XRCC6 | P12956; B1A | XRCC6_HUMAI | Unmodified | 314650 | 155860 | 0 | 196370 | 0 | 0 | 0 | 0 | 0 | 0 | 473320 | 1714100 | 0 | 0 | 64144 | 0 | 0 | 0 |
| FVADGIFK | RPS3; RPS3; RF | P23396; E9P | RS3_HUMAN 4 | Unmodified | 1520100 | 0 | 2809000 | 427640 | 79822 | 0 | 290720 | 32344 | 260010 | 0 | 967740 | 15292000 | 30368 | 0 | 0 | 1253500 | 0 | 52796 |
| FVADGIFKA | RPS3; RPS3; RF | P23396; E9P | RS3_HUMAN 4 | Unmodified | 198090 | 1082700 | 11016 | 704510 | 0 | 0 | 0 | 0 | 0 | 0 | 0 | 346440 | 121460 | 45582 | 993200 | 50665 | 0 | 0 |
| FVAHSVLEI | TRAF5 | O00463 | TRAF5_HUMAN | Unmodified | 0 | 0 | 0 | 0 | 0 | 0 | 67388 | 53705 | 0 | 0 | 229340 | 797580 | 0 | 0 | 0 | 0 | 0 | 0 |
| FVALASNEI | DFFA; DFFA | O00273; K7E | DFFA_HUMAN | Unmodified | 0 | 0 | 0 | 0 | 0 | 0 | 0 | 0 | 0 | 0 | 0 | 0 | 1355500 | 1703400 | 0 | 0 | 0 | 0 |
| FVAPEEVLF | EIF2B2 | P49770 | EI2BB_HUMAN | Unmodified | 180880 | 0 | 0 | 143840 | 510870 | 0 | 0 | 0 | 0 | 107080 | 990270 | 1136700 | 377980 | 214480 | 192040 | 0 | 175110 | 694840 |
| FVASKNEQ | REPS1; REPS1; | Q96D71; H0 | REPS1_HUMAN | Unmodified | 0 | 0 | 577590 | 142400 | 79580 | 0 | 0 | 0 | 278020 | 30175 | 17652 | 139280 | 0 | 0 | 20621 | 37480 | 36637 | 0 |
| FVATMYHY | P3H4; CRTAP | Q92791; O7 | SC65_HUMAN | Unmodified | 0 | 3186400 | 0 | 0 | 0 | 0 | 0 | 0 | 1129700 | 343470 | 0 | 0 | 0 | 0 | 0 | 0 | 0 | 0 |
| FVATQAAVI | RABGGTA; RAB | Q92696; H0 | PGTA_HUMAN | Unmodified | 0 | 0 | 0 | 0 | 0 | 0 | 0 | 0 | 0 | 0 | 0 | 764080 | 0 | 0 | 0 | 14810000 | 0 | 0 |
| FVAVTSSN | DPYSL5 | Q9BPU6 | DPYL5_HUMAN | Unmodified | 45936 | 0 | 0 | 0 | 706310 | 0 | 0 | 0 | 0 | 0 | 342040 | 3685200 | 0 | 0 | 0 | 0 | 0 | 0 |
| FVAYNEEEI | AFDN | H0Y948 | H0Y948_HUMA | Unmodified | 0 | 0 | 0 | 0 | 0 | 0 | 0 | 0 | 0 | 0 | 0 | 0 | 550600 | 649150 | 0 | 0 | 0 | 0 |
| FVCAPPSR | NUP210 | Q8TEM1 | PO210_HUMAN | Unmodified | 0 | 0 | 0 | 0 | 0 | 0 | 0 | 0 | 1243900 | 127110 | 0 | 0 | 0 | 0 | 0 | 0 | 0 | 0 |
| FVCNDDV | IGF2R; IGF2R | P11717; A0A | MPRI_HUMAN | Unmodified | 0 | 0 | 0 | 635230 | 0 | 0 | 128780 | 0 | 23259000 | 257310 | 1563600 | 404290 | 0 | 0 | 0 | 0 | 0 | 0 |
| FVDDLFETI | PLXNA1; PLXNA | Q9UIW2; Q9 | PLXA1_HUMAN | Unmodified | 0 | 0 | 91999 | 0 | 0 | 0 | 0 | 0 | 0 | 0 | 14630 | 429670 | 0 | 0 | 481940 | 0 | 0 | 0 |
| FVDGEWYI | SND1 | Q7KZF4 | SND1_HUMAN | Unmodified | 425460 | 0 | 561330 | 757290 | 615220 | 0 | 0 | 0 | 0 | 88154 | 119680 | 7362200 | 976000 | 299960 | 831770 | 732440 | 186590 | 0 |
| FVDGLMIH | RPS3; RPS3; RF | P23396; E9P | RS3_HUMAN 4 | Oxidation (M) | 4941200 | 5049300 | 0 | 4025500 | 0 | 181750 | 1009100 | 75291 | 832450 | 0 | 2982200 | 2658200 | 1478400 | 0 | 4606700 | 0 | 0 | 1008300 |
| FVDGVSTV | SMC2 | O95347 | SMC2_HUMAN | Unmodified | 0 | 0 | 0 | 0 | 0 | 189790 | 51397 | 18768 | 222530 | 112090 | 0 | 663600 | 0 | 0 | 0 | 86437 | 0 | 0 |
| FVDHVFDE | RPN1; RPN1 | P04843; B7Z | RPN1_HUMAN | Unmodified | 226790 | 0 | 231230 | 0 | 439100 | 166310 | 347410 | 0 | 2.13E+08 | 23645000 | 2506600 | 2204600 | 0 | 0 | 394670 | 307660 | 338950 | 0 |
| FVDKVSVP | ITGB4; ITGB4 | P16144; J3Q | ITB4_HUMAN I | Oxidation (M) | 0 | 0 | 0 | 0 | 107050 | 0 | 0 | 0 | 0 | 0 | 0 | 0 | 0 | 0 | 0 | 0 | 997210 | 0 |
| FVDKVSVP | ITGB4; ITGB4 | P16144; J3Q | ITB4_HUMAN I | Unmodified | 0 | 0 | 0 | 0 | 1149500 | 0 | 0 | 0 | 0 | 0 | 0 | 0 | 0 | 0 | 0 | 0 | 3255900 | 3286800 |
| FVDLYGAQ | HADHA; HADH/ | P40939; A0A | ECHA_HUMAN | Unmodified | 0 | 0 | 0 | 0 | 0 | 0 | 0 | 0 | 2892500 | 0 | 0 | 0 | 0 | 0 | 0 | 0 | 83851 | 0 |
| FVDPGNHI | CLDND1; CLDN | Q9NY35; D6 | CLDN1_HUMAI | Unmodified | 1133100 | 536090 | 280820 | 60335 | 0 | 0 | 0 | 24748 | 0 | 0 | 0 | 6355000 | 0 | 0 | 773430 | 828040 | 0 | 0 |
| FVDPSQDI | AFDN; AFDN; A | P55196; Q5 | AFAD_HUMAN | Unmodified | 361210 | 0 | 0 | 369420 | 4367900 | 2792500 | 882180 | 386090 | 0 | 0 | 688000 | 1149700 | 3942700 | 4200700 | 0 | 29943 | 976710 | 1949800 |
| FVDVTEVVI | GGT1; GGT2; G | P19440; P36 | GGT1_HUMAN | Unmodified | 1339400 | 1770300 | 3828200 | 2157500 | 0 | 0 | 0 | 0 | 1682800 | 0 | 0 | 7438400 | 0 | 0 | 4620900 | 4502000 | 0 | 0 |
| FVEDEKDE | LRRC1 | Q9BTT6 | LRRC1_HUMAN | Unmodified | 124820 | 0 | 252140 | 247250 | 0 | 60526 | 0 | 0 | 987290 | 82483 | 440530 | 2673900 | 0 | 0 | 380040 | 582410 | 0 | 18569 |
| FVEDSFDP | RAB22A | Q9UL26 | RB22A_HUMAN | Unmodified | 22877 | 108020 | 0 | 50017 | 0 | 0 | 0 | 0 | 1292700 | 0 | 0 | 0 | 0 | 0 | 0 | 0 | 0 | 0 |
| FVEDSFDP | RAB22A | Q9UL26 | RB22A_HUMAN | Oxidation (M) | 2338900 | 4449900 | 0 | 0 | 0 | 0 | 0 | 0 | 0 | 1019100 | 425150 | 0 | 0 | 0 | 1865700 | 0 | 0 | 0 |
| FVEDTKKG | CBR1 | P16152 | CBR1_HUMAN | Unmodified | 0 | 0 | 0 | 0 | 885900 | 144940 | 0 | 0 | 0 | 47650 | 0 | 0 | 0 | 0 | 0 | 0 | 413740 | 491960 |
| FVEEIKSIAS | ITGA1 | P56199 | ITA1_HUMAN I | Unmodified | 0 | 0 | 0 | 20237 | 165190 | 0 | 0 | 0 | 0 | 0 | 0 | 0 | 0 | 0 | 0 | 0 | 3863000 | 5345900 |
| FVEFFGPG | ACO1 | P21399 | ACOC_HUMAN | Unmodified | 0 | 299990 | 103640 | 0 | 0 | 0 | 1023300 | 0 | 0 | 0 | 34053 | 732700 | 0 | 0 | 365200 | 70650 | 0 | 0 |
| FVEGLPINI | MDH1; MDH1; I | P40925; B8Z | MDHC_HUMAN | Unmodified | 557860 | 0 | 0 | 0 | 0 | 0 | 972130 | 0 | 5352200 | 658340 | 13607000 | 28833000 | 0 | 0 | 0 | 0 | 0 | 0 |
| FVEGLPINI | MDH1; MDH1; I | P40925; B8Z | MDHC_HUMAN | Unmodified | 0 | 0 | 0 | 0 | 21329000 | 3366800 | 1792500 | 1115900 | 6346800 | 2848400 | 5651000 | 24114000 | 15048000 | 6349700 | 0 | 151000 | 4562000 | 8924100 |
| FVEGLPINI | MDH1; MDH1; I | P40925; B8Z | MDHC_HUMAN | Oxidation (M) | 0 | 0 | 0 | 0 | 945860 | 439470 | 0 | 84018 | 0 | 0 | 0 | 0 | 1173000 | 477330 | 0 | 0 | 1173700 | 579020 |
| FVEIGAHKI | PLCG2; PLCG2 | P16885; A0A | PLCG2_HUMAI | Unmodified | 0 | 0 | 0 | 0 | 0 | 0 | 0 | 0 | 0 | 0 | 608230 | 5563500 | 0 | 0 | 0 | 0 | 0 | 0 |
| FVEIGAHKI | PLCG2; PLCG2 | P16885; A0A | PLCG2_HUMAI | Unmodified | 0 | 0 | 0 | 0 | 0 | 0 | 0 | 0 | 0 | 0 | 0 | 1270900 | 0 | 0 | 0 | 0 | 0 | 0 |
| FVEIHFNEK | MYO6; MYO6; I | Q9UM54; A0 | MYO6_HUMAN | Unmodified | 0 | 0 | 0 | 387660 | 0 | 0 | 0 | 0 | 0 | 0 | 0 | 249100 | 0 | 0 | 781770 | 1210000 | 519780 | 172790 |
| FVELPGAEI | EPRS1; EPRS1 | P07814; V9C | SYEP_HUMAN | Oxidation (M) | 0 | 0 | 310350 | 204830 | 0 | 0 | 380820 | 0 | 189000 | 0 | 768510 | 4386500 | 0 | 0 | 176170 | 0 | 0 | 0 |
| FVELPGAEI | EPRS1; EPRS1 | P07814; V9C | SYEP_HUMAN | Unmodified | 0 | 0 | 0 | 215510 | 0 | 0 | 0 | 0 | 429290 | 0 | 438560 | 1982100 | 0 | 0 | 0 | 0 | 0 | 0 |
| FVEQLGSY | MRPS16; MRPS | Q9Y3D3; A6I | RT16_HUMAN | Unmodified | 0 | 0 | 0 | 0 | 0 | 0 | 0 | 0 | 1040200 | 0 | 0 | 0 | 0 | 0 | 0 | 0 | 70725 | 0 |
| FVESILSNI | HUWE1 | Q7Z6Z7 | HUWE1_HUMA | Unmodified | 0 | 0 | 0 | 0 | 0 | 0 | 0 | 0 | 0 | 0 | 0 | 3431900 | 0 | 0 | 0 | 0 | 0 | 0 |
| FVEVGRVA | RPL14; RPL14 | P50914; E7E | RL14_HUMAN | Unmodified | 338070 | 0 | 0 | 2224000 | 2692300 | 244540 | 4219500 | 210960 | 0 | 751810 | 0 | 0 | 2539500 | 0 | 0 | 0 | 0 | 16260000 |
| FVFDIFQFA | UAP1 | Q16222 | UAP1_HUMAN | Unmodified | 0 | 0 | 140590 | 0 | 6711700 | 0 | 688530 | 0 | 0 | 372700 | 199840 | 0 | 2781500 | 1939100 | 0 | 0 | 1718800 | 2604900 |
| FVFDIFQFA | UAP1 | Q16222 | UAP1_HUMAN | Unmodified | 0 | 0 | 0 | 248010 | 1967400 | 470890 | 0 | 0 | 0 | 0 | 57546 | 378940 | 1206200 | 0 | 402640 | 82980 | 420550 | 818170 |
| FVFDRPLP | PSMA1; N/A; PS | P25786; B4D | PSA1_HUMAN | Unmodified | 0 | 0 | 1228900 | 1537400 | 0 | 0 | 0 | 41072 | 9806800 | 485600 | 1123700 | 20821000 | 461340 | 0 | 2119900 | 1874600 | 0 | 0 |
| FVFDVFRF | UAP1L1 | Q3KQV9 | UAP1L_HUMAN | Unmodified | 0 | 0 | 0 | 0 | 0 | 0 | 789390 | 0 | 0 | 0 | 0 | 0 | 0 | 0 | 0 | 0 | 0 | 0 |
| FVFSLVDAI | MDH2; MDH2 | P40926; G3 | MDHM_HUMAI | Oxidation (M) | 0 | 0 | 0 | 0 | 0 | 0 | 0 | 0 | 3569200 | 601860 | 0 | 1527100 | 0 | 0 | 0 | 117270 | 221070 | 0 |
| FVFSLVDAI | MDH2; MDH2 | P40926; G3 | MDHM_HUMAI | Unmodified | 0 | 0 | 0 | 0 | 0 | 0 | 0 | 0 | 13731000 | 0 | 0 | 156290 | 0 | 0 | 0 | 0 | 0 | 0 |
| FVGDADLE | CTNND1; CTNI | O60716; E9F | CTND1_HUMAI | Unmodified | 0 | 0 | 543550 | 345040 | 808310 | 0 | 0 | 0 | 0 | 0 | 403220 | 753980 | 294850 | 116780 | 782660 | 564630 | 338120 | 1767800 |
| FVGDADLE | CTNND1; CTNI | O60716; E9F | CTND1_HUMAI | Unmodified | 0 | 0 | 0 | 0 | 974560 | 0 | 0 | 0 | 0 | 0 | 0 | 0 | 0 | 0 | 0 | 0 | 662680 | 2263400 |
| FVGDPALV | DAG1 | Q14118 | DAG1_HUMAN | Unmodified | 1524500 | 2191700 | 0 | 0 | 1579300 | 0 | 0 | 154640 | 861380 | 271360 | 0 | 0 | 904120 | 0 | 0 | 291010 | 0 | 1056600 |
| FVGDPALV | DAG1 | Q14118 | DAG1_HUMAN | Unmodified | 4650300 | 2928800 | 0 | 875380 | 3348500 | 0 | 0 | 0 | 3017600 | 840260 | 0 | 0 | 1121900 | 0 | 711010 | 578230 | 327300 | 0 |
| FVGFPDK | HAPLN1 | P10915 | HPLN1_HUMAI | Unmodified | 0 | 0 | 0 | 0 | 0 | 256200 | 0 | 0 | 5896800 | 0 | 0 | 0 | 0 | 0 | 0 | 0 | 210860 | 85660 |
| FVGIKDED | TXN2; TXN2 | Q99757; M0 | THIOM_HUMAI | Unmodified | 0 | 0 | 0 | 0 | 527110 | 99586 | 0 | 0 | 6993600 | 1004100 | 616450 | 0 | 314230 | 117310 | 0 | 0 | 0 | 0 |
| FVGPGGYI | PNPT1 | Q8TCS8 | PNPT1_HUMAI | Unmodified | 0 | 0 | 0 | 0 | 0 | 0 | 248140 | 60213 | 566480 | 0 | 0 | 0 | 0 | 0 | 0 | 0 | 0 | 0 |
| FVGQDVEC | CTPS1; CTPS1; | P17812; B4E | PYRG1_HUMAI | Unmodified | 64487 | 0 | 0 | 0 | 0 | 0 | 0 | 0 | 0 | 0 | 0 | 1730300 | 0 | 0 | 108310 | 0 | 0 | 0 |
| FVHLLDQS | IQGAP1; IQGAP | P46940; A0A | IQGA1_HUMAN | Oxidation (M) | 99653 | 0 | 0 | 451670 | 12856000 | 2382900 | 0 | 0 | 0 | 1579200 | 0 | 0 | 1408900 | 0 | 1412800 | 0 | 371850 | 0 |
| FVHLLDQS | IQGAP1; IQGAP | P46940; A0A | IQGA1_HUMAN | Unmodified | 0 | 0 | 0 | 0 | 4732400 | 702650 | 0 | 0 | 0 | 0 | 0 | 0 | 0 | 0 | 0 | 517440 | 0 | 0 |
| FVHRDLAA | IGF1R; IGF1R | P08069; C9J | IGF1R_HUMAN | Unmodified | 0 | 0 | 0 | 0 | 404880 | 0 | 0 | 0 | 0 | 0 | 0 | 446630 | 0 | 0 | 0 | 0 | 0 | 0 |
| FVHSENQI | CSNK2A3; CSN | Q8NEV1; P6 | CSK23_HUMAN | Unmodified | 572820 | 237410 | 1011500 | 1602000 | 343390 | 1558600 | 112180 | 0 | 1958300 | 62660 | 1909200 | 12323000 | 0 | 0 | 1159700 | 674390 | 85879 | 131680 |
| FVIATSTK | RPL6 | Q02878 | RL6_HUMAN 6 | Unmodified | 1274000 | 2546500 | 2900700 | 671180 | 0 | 0 | 0 | 0 | 108070 | 0 | 2377100 | 66543000 | 0 | 0 | 3954700 | 2850700 | 0 | 0 |
| FVIATSTKI | RPL6 | Q02878 | RL6_HUMAN 6 | Unmodified | 116900 | 252330 | 0 | 0 | 0 | 0 | 0 | 0 | 0 | 0 | 0 | 920670 | 0 | 0 | 0 | 0 | 0 | 0 |
| FVIEENLH | NCBP1 | Q09161 | NCBP1_HUMAI | Unmodified | 0 | 0 | 0 | 0 | 0 | 0 | 0 | 0 | 0 | 0 | 0 | 2248800 | 0 | 0 | 0 | 0 | 0 | 0 |
| FVIGGPQG | MAT2A | P31153 | METK2_HUMAI | Unmodified | 187880 | 174570 | 0 | 0 | 0 | 0 | 0 | 0 | 310080 | 0 | 0 | 1352400 | 0 | 0 | 0 | 165490 | 191520 | 0 |
| FVIGGPQG | MAT2A | P31153 | METK2_HUMAI | Unmodified | 187210 | 0 | 0 | 0 | 393490 | 0 | 0 | 0 | 1042300 | 0 | 0 | 7982600 | 0 | 0 | 0 | 390980 | 0 | 901580 |
| FVIHCNSP | MACROH2A1; I | O75367; B4I | H2AY_HUMAN | Unmodified | 11742000 | 1871500 | 0 | 0 | 0 | 0 | 210800 | 112510 | 0 | 0 | 0 | 0 | 160160 | 185720 | 0 | 0 | 0 | 0 |
| FVIKPIDK | RDX; EZR; MSN | P35241; P15 | RADI_HUMAN I | Unmodified | 409610 | 420110 | 677310 | 918410 | 2616400 | 530340 | 427130 | 278290 | 1839600 | 708320 | 504780 | 6282300 | 585490 | 687060 | 1097700 | 755300 | 441300 | 241360 |
| FVINYDYPI | DDX5; DDX5; D | P17844; A0A | DDX5_HUMAN | Unmodified | 0 | 0 | 0 | 0 | 0 | 0 | 0 | 63303 | 989290 | 0 | 540650 | 4606100 | 0 | 0 | 0 | 0 | 0 | 0 |
| FVINYDYPI | DDX17; DDX17 | Q92841; A0/ | DDX17_HUMAI | Unmodified | 0 | 0 | 0 | 0 | 0 | 0 | 114120 | 0 | 736290 | 0 | 1008600 | 6177300 | 0 | 0 | 0 | 0 | 0 | 0 |
| FVKKPDDG | GLO1 | Q04760 | LGUL_HUMAN | Unmodified | 0 | 0 | 0 | 0 | 0 | 0 | 0 | 0 | 0 | 0 | 0 | 0 | 0 | 0 | 0 | 0 | 0 | 0 |
| FVKPGAEN | SCP2; SCP2 | P22307; E9P | SCP2_HUMAN | Unmodified | 0 | 0 | 0 | 0 | 0 | 0 | 0 | 0 | 1300100 | 0 | 0 | 0 | 0 | 0 | 0 | 0 | 0 | 0 |
| FVKQDQV | GSPT1; GSPT2; | P15170; Q8I | ERF3A_HUMAN | Unmodified | 376950 | 181910 | 0 | 928130 | 987310 | 368940 | 427390 | 351750 | 696710 | 609520 | 590710 | 3346800 | 2912400 | 912470 | 335350 | 223720 | 1005100 | 1471600 |
| FVKWDDD | PLCB1; PLCB1; | Q9NQ66; A0 | PLCB1_HUMAN | Unmodified | 0 | 0 | 0 | 0 | 0 | 0 | 0 | 0 | 769130 | 72726 | 0 | 0 | 0 | 0 | 0 | 0 | 0 | 24636 |
| FVLAALLK | HERC1 | Q15751 | HERC1_HUMAI | Unmodified | 0 | 0 | 0 | 0 | 0 | 0 | 0 | 0 | 0 | 0 | 9309 | 119750 | 0 | 0 | 0 | 9676.9 | 0 | 0 |
| FVLDKVPFI | ANLN; ANLN | Q9NQW6; H | ANLN_HUMAN | Unmodified | 0 | 0 | 0 | 0 | 1180000 | 0 | 202490 | 0 | 0 | 0 | 0 | 0 | 0 | 0 | 0 | 0 | 0 | 0 |
| FVLDYHFEI | CPNE2 | Q96FN4 | CPNE2_HUMAI | Unmodified | 0 | 0 | 0 | 0 | 0 | 0 | 450210 | 0 | 1366300 | 0 | 0 | 0 | 0 | 0 | 0 | 0 | 0 | 0 |
| FVLINWTG | DBNL; DBNL | Q9UJU6; F8 | DBNL_HUMAN | Unmodified | 0 | 0 | 0 | 0 | 0 | 0 | 0 | 0 | 0 | 0 | 0 | 0 | 0 | 0 | 0 | 0 | 0 | 0 |
| FVLINWTG | DBNL; DBNL | Q9UJU6; F8 | DBNL_HUMAN | Unmodified | 1260000 | 1623100 | 947950 | 1939000 | 7757900 | 1490700 | 820900 | 0 | 2133600 | 0 | 1868000 | 11066000 | 6025000 | 2914900 | 3209400 | 1875800 | 3168500 | 4401800 |
| FVLNLIK | BRIX1 | Q8TDN6 | BRX1_HUMAN | Unmodified | 73641 | 0 | 53350 | 0 | 0 | 0 | 0 | 40139 | 0 | 17175 | 0 | 3353900 | 0 | 0 | 36477 | 39953 | 0 | 0 |
| FVLSGANI | MCTS1; MCTS2 | Q9ULC4; A0 | MCTS1_HUMAI | Oxidation (M) | 429130 | 0 | 202320 | 418560 | 0 | 199010 | 364650 | 0 | 335180 | 334350 | 376180 | 2867500 | 910750 | 1015500 | 651370 | 618410 | 753150 | 1215100 |
| FVLSGANI | MCTS1; MCTS2 | Q9ULC4; A0 | MCTS1_HUMAI | Unmodified | 0 | 0 | 0 | 0 | 187020 | 0 | 0 | 19516 | 352590 | 0 | 87913 | 354380 | 377190 | 0 | 89625 | 0 | 0 | 417420 |
| FVLSGNLH | CPD | O75976 | CBPD_HUMAN | Unmodified | 0 | 0 | 0 | 0 | 1308000 | 0 | 0 | 0 | 98090000 | 9102700 | 1819200 | 0 | 0 | 0 | 0 | 0 | 0 | 0 |
| FVLSGNLH | CPD | O75976 | CBPD_HUMAN | Unmodified | 0 | 0 | 0 | 0 | 0 | 0 | 0 | 0 | 0 | 15123000 | 0 | 0 | 0 | 0 | 0 | 0 | 0 | 0 |
| FVLSGNLH | CPD | O75976 | CBPD_HUMAN | Unmodified | 0 | 0 | 0 | 0 | 0 | 0 | 0 | 0 | 0 | 1705000 | 0 | 0 | 0 | 0 | 0 | 0 | 0 | 0 |
| FVLSQAKD | TXNDC5 | Q8NBS9 | TXND5_HUMAI | Unmodified | 0 | 89704 | 175830 | 130820 | 0 | 0 | 51325 | 0 | 32852000 | 1701900 | 1197500 | 2310900 | 202450 | 120100 | 293120 | 143740 | 0 | 72940 |
| FVLSSGKF | CALR; CALR; C/ | P27797; A0A | CALR_HUMAN | Unmodified | 0 | 0 | 0 | 0 | 2195000 | 1131700 | 14812000 | 7158800 | 59767000 | 1.44E+08 | 0 | 0 | 2825100 | 2874500 | 0 | 0 | 876870 | 2803300 |
| FVLSSGKF | CALR; CALR; C/ | P27797; A0A | CALR_HUMAN | Unmodified | 0 | 0 | 0 | 160220 | 0 | 0 | 7469200 | 3823200 | 40718000 | 1.12E+08 | 0 | 0 | 0 | 174230 | 0 | 0 | 0 | 0 |
| FVLVKFDT | ERP29; ERP29 | P30040; F8V | ERP29_HUMAN | Unmodified | 0 | 0 | 0 | 0 | 0 | 0 | 0 | 0 | 30185000 | 4436200 | 1282800 | 165480 | 0 | 60344 | 0 | 0 | 0 | 0 |
| FVLVKFDT | ERP29; ERP29 | P30040; F8V | ERP29_HUMAN | Unmodified | 0 | 0 | 0 | 0 | 1071500 | 0 | 2256200 | 99424 | 37487000 | 43173000 | 3496000 | 0 | 0 | 0 | 0 | 0 | 300240 | 0 |
| FVMKPPQV | EIF2S2 | P20042 | IF2B_HUMAN E | Oxidation (M) | 79273 | 0 | 0 | 0 | 0 | 0 | 0 | 0 | 0 | 0 | 63415 | 826260 | 0 | 0 | 0 | 0 | 0 | 0 |
| FVMQEEFS | PDIA3 | P30101 | PDIA3_HUMAN | Oxidation (M) | 0 | 0 | 0 | 0 | 0 | 0 | 0 | 0 | 2820400 | 0 | 1008000 | 1475700 | 0 | 0 | 0 | 0 | 0 | 0 |
| FVMQEEFS | PDIA3 | P30101 | PDIA3_HUMAN | Unmodified | 0 | 0 | 0 | 0 | 0 | 0 | 115920 | 0 | 31206000 | 774700 | 2209300 | 2222500 | 0 | 0 | 344960 | 154740 | 0 | 0 |
| FVMQEEFS | PDIA3 | P30101 | PDIA3_HUMAN | Unmodified | 263380 | 0 | 1082100 | 1123900 | 6210400 | 0 | 1903800 | 1057300 | 2.05E+08 | 88012000 | 30093000 | 5677400 | 4348200 | 4842800 | 1352200 | 1311900 | 622860 | 4092200 |
| FVMQEEFS | PDIA3 | P30101 | PDIA3_HUMAN | Oxidation (M) | 0 | 0 | 417560 | 0 | 1064800 | 0 | 685960 | 559260 | 22581000 | 11711000 | 6304600 | 1475000 | 813710 | 747200 | 602080 | 497060 | 133580 | 404760 |
| FVMQEEFS | PDIA3 | P30101 | PDIA3_HUMAN | Unmodified | 0 | 0 | 0 | 0 | 0 | 0 | 127830 | 0 | 0 | 1684000 | 0 | 0 | 0 | 233190 | 0 | 0 | 0 | 0 |
| FVNCVRFS | WDR1; WDR1 | O75083; D6I | WDR1_HUMAN | Unmodified | 0 | 0 | 0 | 0 | 677480 | 307380 | 0 | 0 | 0 | 0 | 0 | 0 | 0 | 0 | 0 | 0 | 123900 | 0 |
| FVNDYDKD | RCN2; RCN2 | Q14257; A8 | RCN2_HUMAN | Unmodified | 0 | 0 | 0 | 0 | 0 | 0 | 0 | 0 | 302040 | 145820 | 91850 | 178470 | 0 | 0 | 0 | 0 | 0 | 0 |
| FVNFILKPQ | SIGIRR; SIGIRR | Q6IA17; C9J | SIGIR_HUMAN | Unmodified | 328330 | 0 | 1350900 | 586080 | 0 | 0 | 0 | 14210 | 852430 | 85358 | 0 | 1331600 | 0 | 0 | 1251700 | 1128300 | 0 | 0 |
| FVNLFPEV | AP2A2; AP2A2; | O94973; A0/ | AP2A2_HUMAN | Unmodified | 263180 | 409750 | 370420 | 190560 | 0 | 0 | 262910 | 0 | 609530 | 763740 | 535950 | 2657700 | 0 | 0 | 1230200 | 799920 | 21039 | 41352 |
| FVNLGIEPI | PSMC2; PSMC2 | P35998; C9J | PRS7_HUMAN | Unmodified | 0 | 0 | 443380 | 0 | 0 | 0 | 0 | 0 | 266850 | 0 | 0 | 2948000 | 0 | 0 | 245170 | 0 | 0 | 0 |
| FVNLGIEPI | PSMC2; PSMC2 | P35998; C9J | PRS7_HUMAN | Unmodified | 0 | 0 | 0 | 0 | 0 | 0 | 0 | 0 | 402650 | 0 | 362000 | 783540 | 0 | 0 | 0 | 0 | 0 | 0 |
| FVNRSPSP | DNPEP; DNPEP | Q9ULA0; E5I | DNPEP_HUMAI | Unmodified | 0 | 0 | 0 | 0 | 0 | 0 | 0 | 0 | 802290 | 0 | 0 | 446460 | 0 | 0 | 0 | 0 | 58094 | 0 |
| FVNVVPTF | FAU; FAU | P62861; E9P | RS30_HUMAN | Unmodified | 3267400 | 3003600 | 3191200 | 4591600 | 2227700 | 0 | 1551700 | 973770 | 0 | 391170 | 2673400 | 10383000 | 3613200 | 3542800 | 2020800 | 2231000 | 0 | 1640700 |
| FVNVVPTF | FAU; FAU | P62861; E9P | RS30_HUMAN | Unmodified | 833210 | 0 | 0 | 0 | 294480 | 0 | 0 | 201120 | 0 | 0 | 1877200 | 4527400 | 0 | 0 | 506760 | 416900 | 0 | 301320 |
| FVPAEMGT | FLNA; FLNA; FL | P21333; A0A | FLNA_HUMAN | Unmodified | 0 | 0 | 0 | 0 | 764530 | 482920 | 0 | 0 | 335350 | 174780 | 0 | 751490 | 0 | 0 | 64903 | 0 | 572720 | 0 |
| FVPDECPR | FKBP9 | O95302 | FKBP9_HUMAN | Unmodified | 0 | 0 | 58373 | 0 | 0 | 0 | 0 | 0 | 4877700 | 521820 | 469490 | 170650 | 0 | 0 | 0 | 0 | 0 | 0 |

| | | | | | | | | | | | | | | | | | | | | | | |
|---|---|---|---|---|---|---|---|---|---|---|---|---|---|---|---|---|---|---|---|---|---|---|
| FVPEKSDIE | DAAM1 | Q9Y4D1 | DAAM1_HUMA | Unmodified | 0 | 159400 | 0 | 0 | 0 | 0 | 0 | 0 | 880990 | 0 | 0 | 1122300 | 0 | 0 | 0 | 0 | 0 | 0 |
| FVPFSFSLL | ATP6V0A2; ATP6 | Q9Y487; F5H | VPP2_HUMAN | Unmodified | 0 | 0 | 117230 | 0 | 0 | 0 | 0 | 0 | 0 | 0 | 0 | 1050800 | 0 | 0 | 0 | 0 | 0 | 0 |
| FVPIQTEEM | EPB41L5 | Q9HCM4 | E41L5_HUMAN | Unmodified | 0 | 0 | 0 | 0 | 0 | 0 | 17892 | 0 | 352490 | 10171 | 0 | 0 | 368120 | 0 | 30096 | 0 | 0 | 24619 |
| FVPLTHLDI | SLITRK5; SLITRI | O94991; Q9I | SLIK5_HUMAN | Unmodified | 1072900 | 905110 | 0 | 0 | 0 | 0 | 0 | 0 | 711810 | 0 | 0 | 0 | 0 | 0 | 0 | 0 | 0 | 0 |
| FVPPALEK | NAXE; NAXE | Q8NCW5; A0 | NNRE_HUMAN | Unmodified | 355860 | 178090 | 340960 | 331010 | 0 | 0 | 0 | 0 | 2731700 | 484750 | 1635600 | 3052400 | 388900 | 0 | 348680 | 372880 | 0 | 0 |
| FVPQEMGV | FLNB; FLNB; FL | O75369; H7 | FLNB_HUMAN | Unmodified | 0 | 0 | 0 | 11311 | 1822400 | 0 | 0 | 0 | 258070 | 0 | 0 | 1249000 | 0 | 0 | 488330 | 222540 | 443210 | 0 |
| FVPQEMGV | FLNB; FLNB; FL | O75369; H7 | FLNB_HUMAN | Oxidation (M) | 18236 | 69404 | 77993 | 0 | 99668 | 289140 | 0 | 0 | 0 | 73887 | 0 | 594830 | 0 | 0 | 221460 | 266040 | 0 | 0 |
| FVPRYSLVL | APMAP; APMAP | Q9HDC9; H0 | APMAP_HUMAI | Unmodified | 0 | 0 | 0 | 0 | 0 | 0 | 0 | 0 | 0 | 734410 | 96027 | 0 | 0 | 0 | 0 | 0 | 0 | 0 |
| FVPYLIAGK | IMPDH2; IMPDH | P12268; A0A | IMDH2_HUMAI | Unmodified | 299540 | 0 | 304990 | 1129300 | 3027300 | 465920 | 4309300 | 338810 | 1528200 | 961890 | 1369600 | 1142400 | 2069900 | 0 | 696980 | 453250 | 2319200 | 2974000 |
| FVQAVIPTII | ATG3 | Q9NT62 | ATG3_HUMAN | Oxidation (M) | 0 | 0 | 0 | 0 | 0 | 0 | 0 | 0 | 0 | 0 | 0 | 495080 | 713760 | 931970 | 527060 | 507040 | 0 | 361570 |
| FVQCPDGE | RUVBL2; RUVBI | Q9Y230; M0 | RUVB2_HUMAI | Unmodified | 0 | 0 | 0 | 0 | 1813200 | 0 | 296600 | 0 | 0 | 0 | 0 | 0 | 0 | 1170800 | 0 | 0 | 358800 | 0 |
| FVQDHFDF | RAB31; RAB31 | Q13636; J3C | RAB31_HUMAN | Oxidation (M) | 0 | 0 | 0 | 0 | 4532100 | 690440 | 0 | 79096 | 0 | 0 | 0 | 0 | 0 | 0 | 205490 | 0 | 0 | 0 |
| FVQDHFDF | RAB31; RAB31 | Q13636; J3C | RAB31_HUMAN | Unmodified | 0 | 0 | 0 | 0 | 2126900 | 0 | 0 | 0 | 0 | 0 | 0 | 0 | 0 | 0 | 0 | 0 | 0 | 0 |
| FVQDTLKG | TRAPPC3; TRAF | O43617; A0A | TPPC3_HUMAN | Unmodified | 0 | 0 | 0 | 0 | 0 | 0 | 0 | 28187 | 0 | 0 | 0 | 4948700 | 0 | 0 | 0 | 1193100 | 0 | 0 |
| FVQEVVQS | ARHGEF1; ARH | Q92888; M0 | ARHG1_HUMA | Unmodified | 0 | 0 | 0 | 0 | 0 | 0 | 0 | 0 | 0 | 0 | 0 | 552660 | 0 | 0 | 0 | 0 | 0 | 0 |
| FVQGLDIGI | ITGA2; ITGA2; I | P17301; E9P | ITA2_HUMAN I | Unmodified | 2789800 | 2117300 | 2980200 | 3530500 | 13863000 | 3441800 | 0 | 902500 | 200140 | 451870 | 0 | 0 | 0 | 0 | 6267100 | 4084000 | 10179000 | 6587600 |
| FVQHDAVE | SHCBP1 | Q8NEM2 | SHCBP_HUMA | Unmodified | 0 | 0 | 0 | 0 | 1467000 | 0 | 0 | 0 | 0 | 0 | 0 | 0 | 0 | 0 | 0 | 0 | 0 | 0 |
| FVQVSEDS | SRGAP2; SRGA | O75044; P0I | SRGP2_HUMAI | Unmodified | 0 | 0 | 0 | 0 | 0 | 0 | 0 | 0 | 0 | 0 | 0 | 2086600 | 0 | 0 | 0 | 0 | 0 | 0 |
| FVQVSEDS | SRGAP2; SRGA | O75044; P0I | SRGP2_HUMAI | Unmodified | 0 | 0 | 0 | 0 | 0 | 0 | 0 | 0 | 0 | 0 | 0 | 0 | 0 | 0 | 0 | 0 | 0 | 0 |
| FVQVSEDS | SRGAP2; SRGA | O75044; P0I | SRGP2_HUMAI | Unmodified | 0 | 0 | 0 | 0 | 3681500 | 290800 | 292830 | 93618 | 0 | 199260 | 0 | 0 | 2978300 | 1055300 | 111860 | 0 | 1001000 | 403130 |
| FVRLPVANI | C1R; C1R; C1R; | P00736; B4I | C1R_HUMAN C | Unmodified | 0 | 0 | 0 | 0 | 0 | 0 | 0 | 0 | 828980 | 0 | 0 | 0 | 0 | 0 | 0 | 0 | 0 | 0 |
| FVRNPLGS | LRBA; LRBA; LR | P50851; A0A | LRBA_HUMAN | Unmodified | 0 | 0 | 0 | 241440 | 0 | 0 | 74752 | 0 | 327400 | 61112 | 0 | 1980000 | 271800 | 0 | 0 | 0 | 119280 | 259220 |
| FVSAIEGMI | SCARB2; SCARI | Q14108; A0A | SCRB2_HUMAI | Oxidation (M) | 745650 | 0 | 0 | 0 | 0 | 0 | 0 | 0 | 2727500 | 1763600 | 0 | 1346900 | 0 | 0 | 3251000 | 1486400 | 0 | 0 |
| FVSDEHYD | NECTIN3 | Q9NQS3 | NECT3_HUMAI | Unmodified | 683890 | 0 | 559330 | 1622500 | 0 | 0 | 0 | 0 | 3602600 | 228470 | 0 | 0 | 334680 | 0 | 1032400 | 740690 | 132200 | 679070 |
| FVSDEHYD | NECTIN3 | Q9NQS3 | NECT3_HUMAI | Unmodified | 0 | 0 | 0 | 0 | 6411300 | 0 | 711050 | 0 | 0 | 1538400 | 0 | 0 | 4407000 | 829180 | 0 | 0 | 1766100 | 9556700 |
| FVSDYETH | NRP1; NRP1; N | O14786; Q5 | NRP1_HUMAN | Unmodified | 0 | 0 | 214130 | 0 | 0 | 0 | 0 | 0 | 0 | 0 | 0 | 0 | 0 | 0 | 100330 | 278190 | 0 | 0 |
| FVSDYETH | NRP1; NRP1; N | O14786; Q5 | NRP1_HUMAN | Unmodified | 0 | 0 | 0 | 0 | 1176700 | 653090 | 0 | 0 | 0 | 0 | 0 | 0 | 0 | 0 | 0 | 0 | 409410 | 1781500 |
| FVSELWK | MYH9 | P35579 | MYH9_HUMAN | Unmodified | 350790 | 437530 | 2467500 | 842390 | 361510 | 0 | 0 | 0 | 1199000 | 207390 | 1816400 | 13201000 | 54006 | 0 | 3604100 | 2442400 | 69386 | 64878 |
| FVSELWKD | MYH9 | P35579 | MYH9_HUMAN | Unmodified | 0 | 0 | 0 | 0 | 0 | 0 | 0 | 0 | 0 | 0 | 0 | 782120 | 0 | 0 | 0 | 0 | 0 | 0 |
| FVSFEFSAC | PLCB2 | Q00722 | PLCB2_HUMAN | Unmodified | 0 | 0 | 0 | 0 | 0 | 0 | 0 | 0 | 0 | 0 | 0 | 3233600 | 0 | 0 | 0 | 0 | 0 | 0 |
| FVSFLGR | KPNA2; KPNA2; | P52292; J3Q | IMA1_HUMAN I | Unmodified | 472040 | 0 | 418460 | 0 | 0 | 0 | 0 | 9370.8 | 982900 | 0 | 0 | 2810100 | 85875 | 0 | 0 | 606180 | 0 | 0 |
| FVSFPTQVI | SLC35B2 | Q8TB61 | S35B2_HUMAN | Unmodified | 0 | 0 | 0 | 0 | 0 | 0 | 0 | 0 | 3114400 | 942350 | 0 | 1840200 | 0 | 0 | 0 | 0 | 0 | 0 |
| FVSISDLLV | GDI2 | P50395 | GDIB_HUMAN | Unmodified | 5841200 | 4721100 | 6719300 | 8163100 | 33717000 | 7549400 | 5054800 | 36642 | 37354000 | 12980000 | 33587000 | 82659000 | 12622000 | 9436900 | 13003000 | 10323000 | 6466900 | 11937000 |
| FVSRSEME | PALS2; PALS2 | Q9NZW5; B8 | PALS2_HUMAN | Unmodified | 32503 | 0 | 242010 | 429780 | 1279100 | 484780 | 0 | 0 | 960770 | 93458 | 963820 | 1206400 | 203780 | 164840 | 379960 | 384730 | 230080 | 709430 |
| FVTAVGTQ | NRP1; NRP1; N | O14786; Q5 | NRP1_HUMAN | Unmodified | 0 | 0 | 0 | 0 | 2158700 | 1097900 | 0 | 0 | 505350 | 0 | 0 | 0 | 576600 | 119280 | 0 | 0 | 495410 | 2947700 |
| FVTAVGTQ | NRP1; NRP1; N | O14786; Q5 | NRP1_HUMAN | Unmodified | 0 | 0 | 0 | 0 | 375750 | 160370 | 0 | 0 | 0 | 0 | 0 | 0 | 133640 | 0 | 0 | 0 | 208560 | 455940 |
| FVTDIDELG | CDK6 | Q00534 | CDK6_HUMAN | Unmodified | 0 | 0 | 0 | 0 | 671320 | 0 | 0 | 0 | 0 | 10301 | 0 | 0 | 0 | 0 | 0 | 0 | 1908400 | 648970 |
| FVTHVSDW | CREG1; CREG1 | O75629; A0A | CREG1_HUMAI | Unmodified | 0 | 0 | 0 | 0 | 0 | 0 | 0 | 0 | 7800600 | 1330800 | 677440 | 179350 | 0 | 0 | 0 | 0 | 121010 | 0 |
| FVTLLYR | ACP2; ACP2; AC | P11117; B7Z | PPAL_HUMAN I | Unmodified | 0 | 0 | 0 | 0 | 431350 | 114480 | 25305 | 11537 | 9512100 | 1840500 | 939900 | 639990 | 147200 | 57133 | 0 | 0 | 0 | 99875 |
| FVTLVFR | ACP3; ACP3 | P15309; E9P | PPAP_HUMAN I | Unmodified | 385530 | 0 | 448300 | 0 | 0 | 0 | 349570 | 0 | 5667700 | 1303100 | 0 | 0 | 0 | 0 | 0 | 972100 | 554930 | 0 |
| FVTNTTK | HDHD2; HDHD | Q9H0R4; K7 | HDHD2_HUMA | Unmodified | 0 | 0 | 0 | 61465 | 195600 | 12649 | 120320 | 87330 | 139740 | 31944 | 18129 | 397740 | 0 | 0 | 85859 | 0 | 39524 | 0 |
| FVTSCSRPI | UBE3C | Q15386 | UBE3C_HUMAI | Unmodified | 0 | 0 | 42226 | 0 | 0 | 0 | 0 | 0 | 0 | 0 | 62635 | 243340 | 0 | 0 | 0 | 0 | 0 | 0 |
| FVTSNTQEI | SRRT; SRRT | Q9BXP5; H7 | SRRT_HUMAN | Unmodified | 0 | 0 | 0 | 0 | 2841100 | 0 | 83403 | 36294 | 81182 | 142720 | 249960 | 1067600 | 354020 | 0 | 0 | 0 | 0 | 1016900 |
| FVVDGDTP | MYO1D; MYO1D | O94832; J3C | MYO1D_HUMA | Unmodified | 0 | 546910 | 355460 | 0 | 0 | 0 | 0 | 0 | 0 | 0 | 0 | 528700 | 0 | 0 | 541580 | 0 | 0 | 0 |
| FVVIQNEDI | ERBB2; ERBB2; | P04626; J3C | ERBB2_HUMAN | Unmodified | 715810 | 651570 | 547530 | 1173400 | 0 | 0 | 1060800 | 0 | 3768100 | 0 | 571950 | 0 | 0 | 0 | 4678900 | 1789800 | 0 | 0 |
| FVVLPTGD | FGFRL1 | Q8N441 | FGRL1_HUMAN | Unmodified | 2931500 | 3273500 | 3728400 | 3578400 | 967550 | 144120 | 0 | 0 | 307110 | 41618 | 0 | 0 | 1757600 | 1209100 | 3790400 | 2110700 | 0 | 0 |
| FVVQNVSA | HSPH1; HSPH1 | Q92598; A0A | HS105_HUMAN | Unmodified | 56671 | 35960 | 494060 | 1010900 | 76901 | 0 | 0 | 0 | 1196100 | 0 | 1147900 | 8009400 | 138240 | 0 | 828190 | 730220 | 54282 | 295050 |
| FVVQNVSA | HSPH1; HSPH1 | Q92598; A0A | HS105_HUMAN | Unmodified | 0 | 0 | 0 | 0 | 0 | 79862 | 328670 | 256120 | 0 | 227580 | 0 | 0 | 0 | 0 | 0 | 24906 | 393510 | 23894 |
| FVVTDGGI | CACNA2D1; CA | P54289; H0Y | CA2D1_HUMAI | Unmodified | 1293300 | 1201900 | 0 | 0 | 0 | 0 | 206410 | 135810 | 3673500 | 0 | 276390 | 0 | 0 | 0 | 0 | 0 | 0 | 0 |
| FVVTDGGI | CACNA2D1; CA | P54289; H0Y | CA2D1_HUMAI | Unmodified | 0 | 0 | 0 | 0 | 0 | 0 | 0 | 0 | 0 | 0 | 0 | 0 | 0 | 0 | 0 | 0 | 0 | 0 |
| FVVYEVLRE | UAP1 | Q16222 | UAP1_HUMAN | Unmodified | 0 | 0 | 113010 | 667770 | 3228800 | 487650 | 130110 | 0 | 103640 | 0 | 102480 | 0 | 2508900 | 1112300 | 382800 | 158750 | 963820 | 1447100 |
| FVYAGPAG | JMJD4; JMJD4 | Q9H9V9; A0 | JMJD4_HUMAN | Unmodified | 0 | 0 | 0 | 0 | 0 | 0 | 0 | 0 | 0 | 0 | 0 | 202910 | 342660 | 0 | 0 | 0 | 42984 | 51029 |
| FVYKEEHPI | GABARAP; GAB | O95166; I3L | GBRAP_HUMAI | Unmodified | 0 | 168450 | 0 | 0 | 0 | 0 | 376810 | 0 | 1142300 | 249430 | 140030 | 1256200 | 0 | 108980 | 0 | 206460 | 179610 | 353810 |
| FWAYEQYK | SLC25A24 | Q6NUK1 | SCMC1_HUMA | Unmodified | 0 | 0 | 0 | 0 | 0 | 0 | 0 | 0 | 0 | 413590 | 0 | 0 | 0 | 0 | 0 | 0 | 0 | 0 |
| FWAYEQYK | SLC25A24 | Q6NUK1 | SCMC1_HUMA | Unmodified | 0 | 0 | 0 | 0 | 0 | 0 | 0 | 0 | 2182500 | 544510 | 0 | 0 | 0 | 0 | 0 | 0 | 0 | 0 |
| FWCQQLAI | ITCH; ITCH; ITC | Q96J02; A0A | ITCH_HUMAN I | Oxidation (M) | 76995 | 0 | 0 | 0 | 0 | 0 | 0 | 0 | 1464400 | 0 | 364780 | 0 | 0 | 0 | 547870 | 587970 | 0 | 302330 |
| FWDASGV | LLGL1; LLGL1 | Q15334; A0A | L2GL1_HUMAN | Unmodified | 489380 | 0 | 0 | 351070 | 0 | 0 | 0 | 0 | 1052300 | 0 | 0 | 2319300 | 0 | 0 | 975460 | 1436200 | 0 | 0 |
| FWDYLHEI | FUT11 | Q495W5 | FUT11_HUMAN | Oxidation (M) | 0 | 0 | 0 | 0 | 0 | 0 | 0 | 0 | 1007300 | 369650 | 40377 | 0 | 0 | 0 | 0 | 0 | 0 | 0 |
| FWEEATPIV | ENPP5 | Q9UJA9 | ENPP5_HUMAI | Unmodified | 0 | 0 | 0 | 0 | 0 | 170330 | 0 | 0 | 949670 | 0 | 0 | 68501 | 0 | 0 | 0 | 0 | 310570 | 0 |
| FWEQSVR | GBA; GBA; GBA | P04062; A0A | GLCM_HUMAN | Unmodified | 0 | 0 | 0 | 0 | 0 | 0 | 0 | 0 | 2314000 | 31292 | 0 | 0 | 0 | 0 | 0 | 68763 | 0 | 0 |
| FWEVISDEI | TUBB3; TUBB3; | Q13509; G3 | TBB3_HUMAN | Unmodified | 0 | 0 | 0 | 0 | 1475200 | 0 | 0 | 0 | 2232200 | 0 | 0 | 0 | 0 | 0 | 0 | 0 | 0 | 907290 |
| FWEVISDEI | TUBB2A; TUBB2 | Q13885; Q9I | TBB2A_HUMAN | Unmodified | 0 | 0 | 0 | 0 | 0 | 0 | 0 | 0 | 15729000 | 1829000 | 5570300 | 20856000 | 0 | 0 | 0 | 0 | 444980 | 0 |
| FWEVISDEI | TUBB; TUBB | P07437; Q5J | TBB5_HUMAN | Unmodified | 0 | 0 | 2903800 | 875450 | 0 | 0 | 1416000 | 0 | 21763000 | 2774300 | 618090 | 34332000 | 1224200 | 0 | 0 | 880470 | 0 | 0 |
| FWEVISDEI | TUBB; TUBB | P07437; Q5J | TBB5_HUMAN | Unmodified | 0 | 0 | 0 | 0 | 5637700 | 0 | 1250100 | 0 | 0 | 0 | 0 | 0 | 0 | 0 | 0 | 0 | 0 | 5756500 |
| FWEVISDEI | TUBB4A; TUBB4 | P04350; P68 | TBB4A_HUMAN | Unmodified | 7735900 | 5097700 | 869880 | 5715700 | 4786400 | 613000 | 3616500 | 0 | 27283000 | 3331300 | 4497000 | 20276000 | 5524800 | 0 | 5995400 | 2279100 | 0 | 3486100 |
| FWHIADLH | SMPDL3B; SMP | Q92485; Q5 | ASM3B_HUMAI | Unmodified | 8097800 | 2803900 | 16624000 | 8399600 | 1328500 | 432440 | 0 | 0 | 9662500 | 682150 | 1001800 | 12006000 | 5680200 | 1535600 | 12909000 | 6624900 | 0 | 327770 |
| FWKGPSEA | DPP3; DPP3; DI | Q9NY33; G3 | DPP3_HUMAN | Unmodified | 0 | 0 | 0 | 0 | 0 | 0 | 0 | 0 | 406500 | 131360 | 513560 | 978890 | 494810 | 0 | 0 | 0 | 512050 | 0 |
| FWVNILK | PLXND1; PLXNI | Q9Y4D7; O1 | PLXD1_HUMAN | Unmodified | 0 | 0 | 724550 | 0 | 0 | 0 | 0 | 0 | 143850 | 0 | 0 | 1165800 | 0 | 0 | 1347100 | 1167400 | 0 | 0 |
| FWYFVSQL | RPL18A; RPL18 | Q02543; M0 | RL18A_HUMAN | Unmodified | 689770 | 0 | 0 | 0 | 0 | 0 | 0 | 0 | 0 | 0 | 0 | 3999800 | 0 | 0 | 0 | 0 | 0 | 0 |
| FWYFVSQL | RPL18A; RPL18 | Q02543; M0 | RL18A_HUMAN | Unmodified | 2085500 | 1154200 | 480590 | 243290 | 0 | 0 | 0 | 0 | 0 | 0 | 0 | 11790000 | 0 | 0 | 0 | 506800 | 0 | 0 |
| FWYQMILP | DPP4; DPP4; DI | P27487; A0A | DPP4_HUMAN | Oxidation (M) | 530210 | 414520 | 5609800 | 8107200 | 0 | 0 | 0 | 0 | 0 | 0 | 0 | 0 | 0 | 0 | 0 | 5078400 | 1303500 | 1732500 |
| FWYQMILP | DPP4; DPP4; DI | P27487; A0A | DPP4_HUMAN | Unmodified | 0 | 0 | 103610 | 629880 | 0 | 0 | 0 | 0 | 0 | 0 | 0 | 0 | 0 | 0 | 0 | 99894 | 0 | 0 |
| FYAFGRVF | EEF2 | P13639 | EF2_HUMAN El | Unmodified | 0 | 0 | 0 | 20805 | 144610 | 37244 | 75439 | 0 | 0 | 137220 | 0 | 178670 | 193370 | 109280 | 907750 | 0 | 77868 | 69407 |
| FYALSASFE | CALR; CALR; C/ | P27797; A0A | CALR_HUMAN | Unmodified | 807960 | 476340 | 432520 | 579110 | 315950 | 507440 | 748170 | 213350 | 3.56E+08 | 20812000 | 29963000 | 4439700 | 482620 | 346610 | 1047200 | 962630 | 0 | 518830 |
| FYAPWCGI | TMX3; PDIA4; P | Q96JJ7; P13 | TMX3_HUMAN | Unmodified | 0 | 0 | 0 | 0 | 0 | 0 | 320490 | 119580 | 3044900 | 2551600 | 132020 | 0 | 0 | 170690 | 0 | 0 | 0 | 0 |
| FYASSGEH | MYOF; MYOF | Q9NZM1; H0 | MYOF_HUMAN | Unmodified | 0 | 0 | 0 | 0 | 9966600 | 384360 | 0 | 9676.9 | 0 | 201930 | 0 | 0 | 0 | 0 | 0 | 0 | 0 | 0 |
| FYAYNPLA | AKR7L; AKR7A2 | Q8NHP1; O4 | ARK74_HUMAN | Unmodified | 0 | 176390 | 0 | 0 | 672500 | 0 | 0 | 0 | 4513900 | 2465300 | 0 | 3467600 | 0 | 0 | 0 | 0 | 0 | 0 |
| FYDLSSK | PSMD13; PSMD | Q9UNM6; A0 | PSD13_HUMAN | Unmodified | 0 | 0 | 30194 | 0 | 67933 | 167820 | 26841 | 36553 | 0 | 1289000 | 401520 | 5864800 | 0 | 0 | 0 | 0 | 0 | 27431 |
| FYDPMFK | FOLH1B; FOLH1 | Q9HBA9; Q0 | FOH1B_HUMAI | Oxidation (M) | 1240100 | 1077500 | 5621500 | 2141000 | 0 | 0 | 0 | 0 | 0 | 0 | 0 | 0 | 0 | 0 | 5230900 | 4878300 | 0 | 0 |
| FYDSNTVK | PAK2 | Q13177 | PAK2_HUMAN | Unmodified | 0 | 26592 | 0 | 0 | 71250 | 0 | 0 | 71663 | 0 | 28748 | 0 | 0 | 62422 | 0 | 0 | 0 | 499530 | 0 |
| FYEAFSK | HSP90AB1 | P08238 | HS90B_HUMAN | Unmodified | 2983100 | 1785900 | 4752000 | 4628900 | 17757000 | 4500100 | 3604500 | 1569400 | 8438300 | 10870000 | 10867000 | 1.09E+08 | 17546000 | 10641000 | 7072300 | 6426200 | 10485000 | 4428400 |
| FYEENGFL | PHYH; PHYH; P | O14832; B1A | PAHX_HUMAN | Unmodified | 0 | 0 | 0 | 0 | 0 | 0 | 0 | 0 | 14977000 | 1144000 | 0 | 206490 | 0 | 0 | 0 | 0 | 0 | 0 |
| FYEEVHDLI | NAP1L4; NAP1L | Q99733; P5 | NP1L4_HUMAN | Unmodified | 328310 | 469750 | 0 | 0 | 902930 | 0 | 366350 | 220670 | 712560 | 698050 | 4084400 | 2905000 | 1739400 | 815970 | 0 | 239260 | 288170 | 275480 |
| FYEEVHDLI | NAP1L4; NAP1L | Q99733; P5 | NP1L4_HUMAN | Unmodified | 5206200 | 5184400 | 6654500 | 5906300 | 11599000 | 2943700 | 2049300 | 884780 | 5569700 | 2402300 | 2076300 | 45360000 | 10664000 | 2968900 | 4527500 | 3850200 | 1762400 | 7479700 |
| FYEEVHDLI | NAP1L4; NAP1L | Q99733; C9J | NP1L4_HUMAN | Unmodified | 0 | 786810 | 0 | 475780 | 941550 | 0 | 755320 | 0 | 411090 | 0 | 0 | 0 | 1943800 | 942620 | 0 | 0 | 0 | 0 |
| FYEEVHDLI | NAP1L1; NAP1L | P55209; F8V | NP1L1_HUMAN | Unmodified | 0 | 0 | 0 | 0 | 7623800 | 403950 | 3787400 | 0 | 0 | 0 | 0 | 0 | 0 | 3073000 | 0 | 0 | 3324400 | 5808800 |
| FYELTQK | PLCB4; PLCB4 | Q15147; A0A | PLCB4_HUMAN | Unmodified | 416850 | 0 | 0 | 818840 | 0 | 0 | 0 | 0 | 0 | 939560 | 314560 | 0 | 0 | 0 | 415940 | 732460 | 1327800 | 0 |
| FYEQFSK | HSP90AA1 | P07900 | HS90A_HUMAI | Unmodified | 1470800 | 0 | 622100 | 631970 | 15301000 | 5640500 | 10484000 | 1875700 | 11825000 | 8755100 | 7600900 | 62826000 | 5629600 | 7252900 | 3248400 | 2450600 | 9429000 | 5213300 |
| FYEQMNGI | EEF1D; EEF1D; | P29692; E9P | EF1D_HUMAN | Unmodified | 0 | 0 | 0 | 0 | 0 | 0 | 0 | 0 | 204690 | 0 | 274350 | 3971500 | 0 | 0 | 0 | 0 | 0 | 0 |
| FYEQMNGI | EEF1D; EEF1D; | P29692; E9P | EF1D_HUMAN | Oxidation (M) | 0 | 0 | 0 | 0 | 0 | 0 | 0 | 0 | 0 | 0 | 186930 | 694950 | 0 | 0 | 0 | 0 | 0 | 0 |
| FYEQVVQA | PELO | Q9BRX2 | PELO_HUMAN | Unmodified | 532390 | 0 | 0 | 0 | 0 | 0 | 0 | 0 | 0 | 0 | 0 | 234440 | 0 | 0 | 358690 | 0 | 0 | 0 |
| FYESGNRA | MBLAC2 | Q68D91 | MBLC2_HUMAI | Unmodified | 0 | 0 | 746020 | 1048800 | 0 | 0 | 0 | 0 | 1133200 | 0 | 1768500 | 749100 | 0 | 0 | 198800 | 183960 | 0 | 0 |
| FYFNGEYA | PSPH; PSPH | P78330; C9J | SERB_HUMAN | Unmodified | 0 | 0 | 0 | 0 | 0 | 0 | 0 | 0 | 0 | 349720 | 606370 | 1581200 | 1334200 | 0 | 0 | 947800 | 0 | 0 |
| FYGAEIVSA | AKT1 | P31749 | AKT1_HUMAN I | Unmodified | 0 | 0 | 470460 | 1886100 | 0 | 0 | 0 | 0 | 0 | 0 | 1063000 | 1488500 | 0 | 0 | 4816500 | 2148400 | 0 | 0 |
| FYGAEIVSA | AKT2; AKT2; AK | P31751; J3Q | AKT2_HUMAN I | Unmodified | 0 | 0 | 0 | 0 | 0 | 0 | 0 | 0 | 0 | 0 | 0 | 0 | 0 | 0 | 801740 | 0 | 0 | 0 |
| FYGDEEK | CALR; CALR; C/ | P27797; A0A | CALR_HUMAN | Unmodified | 325800 | 0 | 500750 | 0 | 784670 | 147160 | 861180 | 461370 | 4.08E+08 | 48481000 | 20544000 | 6216400 | 416400 | 1065200 | 2936700 | 940550 | 65821 | 677930 |
| FYGDEEKD | CALR; CALR; C/ | P27797; A0A | CALR_HUMAN | Unmodified | 0 | 0 | 0 | 0 | 0 | 0 | 0 | 415440 | 30532000 | 12857000 | 2510200 | 0 | 0 | 0 | 0 | 0 | 0 | 0 |
| FYGDEEKD | CALR; CALR; C/ | P27797; A0A | CALR_HUMAN | Unmodified | 0 | 0 | 0 | 180470 | 593060 | 0 | 1457000 | 1604500 | 53042000 | 36363000 | 826010 | 0 | 890040 | 171410 | 0 | 0 | 185040 | 567540 |
| FYGEDGLP | CYB5D2; CYB5I | Q8WUJ1; I3L | NEUFC_HUMA | Unmodified | 0 | 0 | 0 | 0 | 0 | 0 | 0 | 0 | 563420 | 0 | 0 | 0 | 0 | 0 | 0 | 0 | 0 | 0 |
| FYGIPVTGV | MMP15 | P51511 | MMP15_HUMA | Unmodified | 0 | 1242000 | 1962100 | 1798000 | 0 | 0 | 376220 | 0 | 624180 | 0 | 0 | 0 | 0 | 0 | 4935000 | 2973800 | 0 | 0 |
| FYGLQVTG | MMP14; MMP14 | P50281; A0A | MMP14_HUMA | Oxidation (M) | 0 | 0 | 0 | 0 | 3719400 | 672680 | 0 | 0 | 0 | 0 | 0 | 0 | 0 | 0 | 0 | 0 | 1372400 | 2322500 |
| FYGLQVTG | MMP14; MMP14 | P50281; A0A | MMP14_HUMA | Unmodified | 0 | 0 | 0 | 0 | 4696700 | 762800 | 0 | 0 | 0 | 0 | 0 | 0 | 0 | 0 | 0 | 0 | 991330 | 1454600 |
| FYGPAGPY | PGRMC2 | O15173 | PGRC2_HUMAI | Unmodified | 0 | 0 | 0 | 0 | 537850 | 0 | 1018200 | 0 | 0 | 1027400 | 0 | 1801800 | 379250 | 0 | 0 | 0 | 0 | 0 |
| FYGPEGPY | PGRMC1 | O00264 | PGRC1_HUMAI | Unmodified | 0 | 0 | 0 | 0 | 0 | 0 | 0 | 0 | 654150 | 0 | 0 | 779150 | 0 | 0 | 0 | 0 | 0 | 0 |
| FYGPEGPY | PGRMC1 | O00264 | PGRC1_HUMAI | Unmodified | 0 | 0 | 0 | 0 | 0 | 0 | 1560800 | 66327 | 0 | 236660 | 0 | 0 | 0 | 0 | 0 | 0 | 0 | 0 |
| FYGPQVNN | WASL | O00401 | WASL_HUMAN | Unmodified | 981160 | 1137500 | 0 | 584910 | 0 | 0 | 0 | 0 | 0 | 284990 | 0 | 125760 | 0 | 0 | 1538400 | 863820 | 0 | 0 |
| FYGSPEEL | AOC1; AOC1 | P19801; C9J | AOC1_HUMAN | Unmodified | 0 | 0 | 0 | 0 | 0 | 0 | 0 | 47806 | 1980100 | 0 | 0 | 0 | 0 | 0 | 0 | 0 | 0 | 0 |
| FYGSRLDY | BIN3; BIN3 | Q9NQY0; H7 | BIN3_HUMAN I | Unmodified | 142220 | 215540 | 0 | 165050 | 0 | 0 | 0 | 0 | 0 | 0 | 0 | 0 | 0 | 0 | 128520 | 211600 | 0 | 0 |
| FYGVTVFK | P3H1; P3H1 | Q32P28; H7 | P3H1_HUMAN | Unmodified | 0 | 0 | 0 | 0 | 289040 | 0 | 246550 | 83890 | 4072900 | 2713100 | 0 | 0 | 0 | 0 | 0 | 0 | 0 | 0 |

| | | | | | | | | | | | | | | | | | | | | | | |
|---|---|---|---|---|---|---|---|---|---|---|---|---|---|---|---|---|---|---|---|---|---|---|
| FYHDWND | OLA1; OLA1; OL | Q9NTK5; J3K | OLA1_HUMAN | Unmodified | 298060 | 383760 | 221750 | 584500 | 0 | 0 | 134870 | 46587 | 736330 | 65764 | 1182500 | 11655000 | 346320 | 0 | 980810 | 902080 | 0 | 0 |
| FYHTGTEEI | NDST2; N/A | P52849; S4F | NDST2_HUMAI | Oxidation (M) | 0 | 37154 | 0 | 0 | 0 | 0 | 0 | 0 | 710140 | 79549 | 0 | 0 | 0 | 0 | 0 | 0 | 0 | 0 |
| FYHTGTEEI | NDST2; N/A | P52849; S4F | NDST2_HUMAI | Unmodified | 0 | 0 | 0 | 0 | 1140500 | 567110 | 0 | 0 | 1438500 | 121640 | 0 | 0 | 0 | 0 | 0 | 0 | 0 | 0 |
| FYIDPYK | GAR1 | Q9NY12 | GAR1_HUMAN | Unmodified | 0 | 0 | 232970 | 141940 | 0 | 0 | 799020 | 974800 | 0 | 0 | 136400 | 1423700 | 198640 | 352980 | 0 | 0 | 0 | 0 |
| FYLDLYPR | NLN; THOP1; N | Q9BYT8; P52 | NEUL_HUMAN | Unmodified | 0 | 0 | 0 | 0 | 0 | 0 | 0 | 0 | 0 | 0 | 0 | 2943100 | 0 | 0 | 0 | 0 | 0 | 0 |
| FYLETQAGI | CIT; CIT; CIT; CI | O14578; A0A | CTRO_HUMAN | Unmodified | 0 | 0 | 0 | 967090 | 1244900 | 838370 | 0 | 448570 | 0 | 0 | 0 | 0 | 64249 | 0 | 333770 | 166480 | 448910 | 0 |
| FYLLNIRLP | WLS; WLS; WLS | Q5T9L3; H0Y | WLS_HUMAN F | Unmodified | 0 | 0 | 0 | 0 | 160970 | 0 | 0 | 0 | 10383000 | 658680 | 0 | 0 | 67179 | 0 | 0 | 91400 | 0 | 0 |
| FYLPILVPS/ | GLT8D1; GLT8D | Q68CQ7; C9 | GL8D1_HUMAI | Unmodified | 0 | 0 | 0 | 0 | 49578 | 0 | 0 | 0 | 22618000 | 3106700 | 420700 | 936930 | 0 | 0 | 0 | 0 | 0 | 0 |
| FYLPILVPS/ | GLT8D1; GLT8D | Q68CQ7; C9 | GL8D1_HUMAI | Unmodified | 0 | 0 | 0 | 0 | 0 | 0 | 0 | 0 | 37461000 | 3094400 | 661200 | 819810 | 60839 | 0 | 0 | 0 | 0 | 0 |
| FYLYPDITR | SLC6A6; SLC6A | P31641; A0A | SC6A6_HUMAI | Unmodified | 0 | 0 | 0 | 0 | 394230 | 0 | 0 | 0 | 0 | 0 | 0 | 2654200 | 0 | 0 | 0 | 0 | 0 | 0 |
| FYNELTEIL' | PDCD6IP | Q8WUM4 | PDC6I_HUMAN | Unmodified | 77892000 | 89521000 | 22796000 | 34355000 | 22859000 | 5631800 | 691350 | 0 | 41947000 | 15957000 | 6477500 | 18667000 | 19807000 | 4436200 | 37590000 | 35790000 | 351360 | 3647000 |
| FYNELTEIL' | PDCD6IP | Q8WUM4 | PDC6I_HUMAN | Unmodified | 505840 | 0 | 312770 | 2370000 | 47828000 | 4931300 | 4695800 | 0 | 173070 | 18786000 | 309100 | 0 | 45576000 | 29317000 | 544590 | 158780 | 8282300 | 14649000 |
| FYNLVLLPF | BYSL; BYSL | Q13895; H7I | BYST_HUMAN | Unmodified | 0 | 0 | 0 | 0 | 0 | 0 | 0 | 0 | 0 | 0 | 0 | 1018600 | 0 | 0 | 0 | 0 | 0 | 0 |
| FYPEDVAEI | EZR; EZR | P15311; E7E | EZRI_HUMAN E | Unmodified | 0 | 31238 | 0 | 0 | 270920 | 15722 | 0 | 0 | 0 | 102050 | 0 | 0 | 179640 | 229270 | 73793 | 0 | 0 | 0 |
| FYPEDVSEI | MSN; MSN | P26038; V9G | MOES_HUMAN | Unmodified | 0 | 0 | 89050 | 519890 | 744590 | 256490 | 0 | 0 | 0 | 0 | 0 | 0 | 0 | 0 | 490140 | 62505 | 374330 | 158630 |
| FYPGQAPS | BCS1L; BCS1L | Q9Y276; H7( | BCS1_HUMAN | Unmodified | 0 | 0 | 0 | 0 | 0 | 0 | 145370 | 0 | 286370 | 13978 | 0 | 0 | 0 | 0 | 0 | 0 | 0 | 0 |
| FYPIPSLHS | SLK | Q9H2G2 | SLK_HUMAN ST | Unmodified | 0 | 0 | 0 | 505530 | 1790100 | 0 | 0 | 269550 | 1272600 | 959410 | 1101100 | 3040500 | 6175100 | 4357700 | 0 | 0 | 538450 | 971160 |
| FYPPKVELF | PAICS; PAICS; F | P22234; D6F | PUR6_HUMAN | Unmodified | 0 | 0 | 0 | 0 | 0 | 0 | 0 | 0 | 0 | 0 | 0 | 5145600 | 0 | 0 | 0 | 0 | 0 | 0 |
| FYPPVEPP/ | GALNT10 | Q86SR1 | GLT10_HUMAN | Unmodified | 0 | 64907 | 0 | 0 | 0 | 0 | 108960 | 0 | 3785500 | 0 | 0 | 0 | 0 | 0 | 0 | 0 | 0 | 0 |
| FYQDLKDR | MANF; MANF | P55145; H7( | MANF_HUMAN | Unmodified | 457820 | 0 | 0 | 0 | 293530 | 0 | 673220 | 0 | 0 | 22872000 | 0 | 0 | 861110 | 705320 | 0 | 0 | 0 | 343550 |
| FYQGFLFSI | PRKDC | P78527 | PRKDC_HUMAI | Unmodified | 0 | 96551 | 78884 | 0 | 0 | 0 | 0 | 802110 | 0 | 0 | 0 | 372220 | 0 | 1451100 | 0 | 0 | 0 | 0 |
| FYQPYSED | AP3S1; AP3S1 | Q92572; F5H | AP3S1_HUMAN | Unmodified | 91102 | 46960 | 0 | 0 | 0 | 0 | 196090 | 199840 | 0 | 0 | 0 | 1557700 | 0 | 0 | 0 | 74286 | 0 | 0 |
| FYQRFPEEI | AP3S2; AP3S2; | P59780; H0Y | AP3S2_HUMAN | Unmodified | 0 | 0 | 0 | 0 | 831340 | 0 | 354120 | 437420 | 0 | 320910 | 0 | 1594300 | 814830 | 1152800 | 0 | 0 | 2247600 | 0 |
| FYQVPVPLI | MRPS7; MRPS7 | Q9Y2R9; J3Q | RT07_HUMAN : | Unmodified | 0 | 0 | 0 | 0 | 0 | 0 | 0 | 0 | 1086300 | 0 | 0 | 0 | 0 | 0 | 0 | 0 | 0 | 0 |
| FYRDPTAF( | HAPLN1; HAPLI | P10915; D6F | HPLN1_HUMAI | Unmodified | 0 | 0 | 0 | 0 | 0 | 0 | 0 | 0 | 3304700 | 1902400 | 0 | 0 | 0 | 0 | 0 | 0 | 0 | 127240 |
| FYSFLVLK | RAD21L1; RAD2 | Q9H4I0; O6( | RD21L_HUMAN | Unmodified | 0 | 0 | 0 | 0 | 172190 | 0 | 241800 | 0 | 122210 | 209240 | 0 | 212020 | 257770 | 77970 | 0 | 13432 | 633850 | 0 |
| FYSLGHLSI | VANGL1 | Q8TAA9 | VANG1_HUMAI | Unmodified | 176400 | 0 | 447560 | 297440 | 0 | 0 | 0 | 0 | 670990 | 264030 | 0 | 0 | 0 | 0 | 754460 | 613810 | 0 | 0 |
| FYSLPQSP( | DARS2; DARS2; | Q6PI48; A0A | SYDM_HUMAN | Unmodified | 0 | 0 | 0 | 209110 | 0 | 0 | 0 | 0 | 1232900 | 0 | 0 | 0 | 0 | 0 | 0 | 0 | 0 | 0 |
| FYSLWDTG | STT3B | Q8TCJ2 | STT3B_HUMAN | Unmodified | 0 | 0 | 0 | 0 | 0 | 0 | 0 | 0 | 1863100 | 0 | 0 | 299640 | 0 | 0 | 0 | 0 | 0 | 0 |
| FYSQAIELN | PPP5C; PPP5C | P53041; H0Y | PPP5_HUMAN | Unmodified | 0 | 0 | 0 | 0 | 821230 | 0 | 0 | 90092 | 1091600 | 567010 | 86811 | 5377100 | 664180 | 0 | 18867 | 0 | 260520 | 239230 |
| FYSSEPNN | EPB41L5 | Q9HCM4 | E41L5_HUMAN | Unmodified | 855310 | 0 | 1769200 | 1327100 | 0 | 0 | 0 | 0 | 3958400 | 0 | 339480 | 1572900 | 0 | 0 | 5456200 | 4832300 | 0 | 0 |
| FYTDPSYFF | WASF2; WASF3 | Q9Y6W5; Q9 | WASF2_HUMAI | Unmodified | 0 | 0 | 0 | 0 | 1921300 | 0 | 1415000 | 0 | 0 | 324070 | 0 | 0 | 1221500 | 1429200 | 0 | 0 | 1844800 | 0 |
| FYTDPVEA\ | OXCT1 | P55809 | SCOT1_HUMAI | Unmodified | 0 | 0 | 0 | 0 | 0 | 0 | 0 | 0 | 4899800 | 0 | 0 | 860220 | 0 | 0 | 0 | 0 | 0 | 0 |
| FYTNPSYFF | WASF1; WASF1 | Q92558; Q5! | WASF1_HUMAI | Unmodified | 0 | 234870 | 0 | 0 | 0 | 9082400 | 387410 | 0 | 0 | 0 | 0 | 0 | 0 | 0 | 274920 | 0 | 0 | 0 |
| FYTVINHYI | NSG1 | P42857 | NSG1_HUMAN | Unmodified | 0 | 0 | 0 | 0 | 0 | 0 | 897180 | 420820 | 0 | 0 | 0 | 0 | 0 | 0 | 0 | 0 | 0 | 0 |
| FYVHNDIFI | G3BP1; G3BP1; | Q13283; A0A | G3BP1_HUMAI | Unmodified | 0 | 0 | 300900 | 0 | 0 | 0 | 0 | 0 | 448860 | 0 | 216250 | 6006100 | 0 | 0 | 156390 | 0 | 0 | 0 |
| FYWGHKEI | ACLY | P53396 | ACLY_HUMAN | Unmodified | 0 | 0 | 0 | 0 | 0 | 0 | 3492500 | 0 | 0 | 0 | 0 | 0 | 0 | 0 | 0 | 0 | 0 | 0 |
| FYYNSVIGI | TFPI | P10646 | TFPI1_HUMAN | Unmodified | 0 | 0 | 1531300 | 1436900 | 0 | 0 | 0 | 0 | 0 | 0 | 0 | 0 | 0 | 0 | 0 | 481260 | 0 | 0 |
| FYYVDHNT | WWP2 | O00308 | WWP2_HUMAN | Unmodified | 457380 | 928670 | 0 | 0 | 0 | 0 | 0 | 0 | 0 | 0 | 0 | 0 | 0 | 0 | 0 | 0 | 0 | 0 |
| GAAAAAAA | INTS3 | Q68E01 | INT3_HUMAN I | Unmodified | 0 | 0 | 0 | 0 | 0 | 0 | 0 | 0 | 0 | 0 | 81373 | 571410 | 0 | 91443 | 0 | 0 | 0 | 0 |
| GAAAERFF: | ADPGK; ADPGK | Q9BRR6; H3 | ADPGK_HUMAI | Unmodified | 0 | 0 | 0 | 0 | 0 | 0 | 0 | 0 | 4286000 | 2003100 | 0 | 0 | 0 | 0 | 0 | 0 | 0 | 0 |
| GAAAMIPG | EPB41L1; EPB4 | Q9H4G0; H7 | E41L1_HUMAN | Oxidation (M) | 0 | 0 | 153560 | 777380 | 0 | 0 | 0 | 0 | 0 | 0 | 296840 | 461830 | 0 | 0 | 319210 | 385000 | 0 | 0 |
| GAADAIAQ. | DPYSL4; DPYSL | O14531; Q5 | DPYL4_HUMAN | Unmodified | 0 | 0 | 0 | 0 | 0 | 0 | 0 | 0 | 0 | 0 | 0 | 0 | 0 | 0 | 0 | 0 | 157470 | 185780 |
| GAAGALLV | RAB2A; RAB2B; | P61019; Q8\ | RAB2A_HUMAN | Unmodified | 0 | 288040 | 0 | 0 | 0 | 0 | 0 | 0 | 1783700 | 0 | 0 | 2449900 | 0 | 0 | 0 | 445690 | 0 | 0 |
| GAAGALM\ | RAB14; RAB14 | P61106; X6R | RAB14_HUMAN | Unmodified | 213490 | 218270 | 0 | 198020 | 0 | 0 | 0 | 45592 | 990180 | 0 | 0 | 1247300 | 0 | 0 | 187770 | 160780 | 0 | 0 |
| GAAGALM\ | RAB14; RAB14 | P61106; X6R | RAB14_HUMAN | Oxidation (M) | 103150 | 434980 | 900870 | 1589900 | 0 | 0 | 0 | 0 | 0 | 0 | 0 | 5537500 | 0 | 0 | 409720 | 1700500 | 0 | 0 |
| GAAGGAE( | PCYT2; PCYT2; | Q99447; I3L | PCY2_HUMAN | Unmodified | 0 | 0 | 75938 | 2121000 | 794560 | 0 | 0 | 0 | 223440 | 120860 | 28873 | 428280 | 0 | 27886 | 6308800 | 0 | 0 | 155780 |
| GAAGLGDE | EPB41L4B | Q9H329 | E41LB_HUMAN | Unmodified | 884160 | 530270 | 1534800 | 1249500 | 0 | 0 | 0 | 0 | 705870 | 0 | 0 | 120510 | 0 | 1346900 | 2799300 | 1354500 | 0 | 37880 |
| GAAIFFEFK | AIDA | Q96BJ3 | AIDA_HUMAN / | Unmodified | 0 | 0 | 947120 | 0 | 1612600 | 0 | 1030300 | 465320 | 353200 | 486550 | 530790 | 815750 | 16086000 | 10206000 | 0 | 0 | 1281000 | 4876800 |
| GAALDVHE | CTBP2 | P56545 | CTBP2_HUMAN | Unmodified | 0 | 0 | 0 | 0 | 222540 | 0 | 0 | 0 | 0 | 67180 | 0 | 926710 | 0 | 0 | 0 | 0 | 90943 | 0 |
| GAAPNVVY | CPSF6; CPSF6; | Q16630; F8V | CPSF6_HUMAN | Unmodified | 2532500 | 1941100 | 797200 | 813390 | 0 | 0 | 0 | 58366 | 0 | 0 | 0 | 1302200 | 494640 | 0 | 396230 | 766910 | 0 | 0 |
| GAAPNVVY | CPSF6; CPSF6; | Q16630; F8V | CPSF6_HUMAN | Unmodified | 968200 | 981280 | 0 | 457360 | 0 | 0 | 67561 | 0 | 0 | 0 | 0 | 0 | 0 | 25717 | 0 | 0 | 0 | 0 |
| GAAPPDSP | LMCD1; LMCD1 | Q9NZU5; B4 | LMCD1_HUMA | Unmodified | 0 | 0 | 0 | 0 | 0 | 0 | 0 | 0 | 0 | 0 | 270330 | 813720 | 0 | 0 | 0 | 0 | 0 | 0 |
| GAASKPVD | TLN2; TLN2; TLI | Q9Y4G6; H0 | TLN2_HUMAN | Unmodified | 0 | 0 | 0 | 0 | 0 | 0 | 0 | 0 | 0 | 0 | 0 | 0 | 0 | 0 | 0 | 0 | 220180 | 416680 |
| GAASPSEP | N/A | N/A | N/A | Unmodified | 0 | 0 | 0 | 0 | 0 | 0 | 0 | 0 | 1161300 | 0 | 25774 | 606660 | 0 | 60567 | 0 | 0 | 0 | 1182500 |
| GAAVDEYF | CARM1; CARM1 | Q86X55; K7E | CARM1_HUMA | Unmodified | 0 | 0 | 0 | 0 | 0 | 0 | 0 | 0 | 0 | 0 | 0 | 1142900 | 0 | 0 | 0 | 0 | 0 | 0 |
| GAAVDEYF | CARM1; CARM1 | Q86X55; K7E | CARM1_HUMA | Unmodified | 665060 | 858540 | 128970 | 123470 | 0 | 0 | 0 | 0 | 0 | 0 | 46148 | 3976000 | 0 | 21507 | 221600 | 343490 | 0 | 0 |
| GAAVDGGI | BCAP31; BCAP3 | P51572; C9J | BAP31_HUMAN | Unmodified | 0 | 0 | 322950 | 0 | 579090 | 0 | 154940 | 69082 | 537570 | 0 | 0 | 0 | 134310 | 576870 | 1146200 | 1059600 | 0 | 191740 |
| GAAYAHAE | HACL1; HACL1 | Q9UJ83; C9J | HACL1_HUMAI | Unmodified | 0 | 0 | 0 | 0 | 0 | 0 | 0 | 0 | 962020 | 134210 | 3051500 | 0 | 0 | 41338 | 0 | 0 | 0 | 0 |
| GAAYAHAE | HACL1; HACL1 | Q9UJ83; C9J | HACL1_HUMAI | Unmodified | 0 | 0 | 0 | 0 | 0 | 0 | 0 | 0 | 894800 | 0 | 0 | 666270 | 703760 | 0 | 0 | 0 | 0 | 0 |
| GACASHVS | DBNL; DBNL; D | Q9UJU6; F8\ | DBNL_HUMAN | Unmodified | 0 | 0 | 0 | 547210 | 2759600 | 0 | 362580 | 150660 | 133270 | 91825 | 0 | 2555200 | 1758800 | 544500 | 0 | 0 | 1474000 | 1315600 |
| GACASHVS | DBNL; DBNL; D | Q9UJU6; F8\ | DBNL_HUMAN | Oxidation (M) | 183050 | 0 | 0 | 190570 | 296050 | 0 | 0 | 0 | 0 | 0 | 0 | 582490 | 553900 | 0 | 0 | 0 | 0 | 0 |
| GACGQGQ | HTRA1 | Q92743 | HTRA1_HUMAN | Unmodified | 0 | 0 | 0 | 0 | 0 | 0 | 0 | 0 | 447320 | 0 | 0 | 0 | 0 | 0 | 0 | 0 | 0 | 0 |
| GADAQEDI | PACSIN1; PACS | Q9BY11; F6U | PACN1_HUMAI | Unmodified | 3388100 | 3114000 | 531290 | 0 | 0 | 0 | 0 | 0 | 0 | 0 | 0 | 0 | 0 | 0 | 1127400 | 0 | 0 | 0 |
| GADAQEDI | PACSIN1; PACS | Q9BY11; F6U | PACN1_HUMAI | Unmodified | 2291200 | 644990 | 339170 | 518880 | 1231000 | 0 | 0 | 0 | 0 | 0 | 0 | 0 | 0 | 0 | 295690 | 183960 | 0 | 0 |
| GADCCVL\ | RAB7A; RAB7A; | P51149; A0A | RAB7A_HUMAN | Unmodified | 486790 | 464570 | 0 | 0 | 169930 | 788960 | 0 | 0 | 789290 | 0 | 0 | 1377800 | 0 | 0 | 0 | 531440 | 0 | 592140 |
| GADDAAD/ | CADM1; CADM: | Q9BY67; X5E | CADM1_HUMA | Unmodified | 0 | 0 | 0 | 0 | 2457000 | 295190 | 0 | 79612 | 3007400 | 0 | 2685200 | 1423800 | 0 | 0 | 0 | 155130 | 0 | 0 |
| GADDAAD/ | CADM1; CADM: | Q9BY67; X5E | CADM1_HUMA | Unmodified | 0 | 0 | 0 | 0 | 3824100 | 0 | 2631700 | 514940 | 0 | 470720 | 0 | 0 | 0 | 0 | 0 | 0 | 0 | 0 |
| GADFLVTE' | PKM; PKM; PKM | P14618; B4E | KPYM_HUMAN | Unmodified | 401790 | 310730 | 3987900 | 1998700 | 65094000 | 8331200 | 2339800 | 1824600 | 4177700 | 1367400 | 21531000 | 78855000 | 8409300 | 4793200 | 1980600 | 1489300 | 16251000 | 26490000 |
| GADFLVTE' | PKM; PKM; PKM | P14618; B4E | KPYM_HUMAN | Unmodified | 1275100 | 1010900 | 3888600 | 4154900 | 93490000 | 22986000 | 4955300 | 3412600 | 6994500 | 1537900 | 12624000 | 1.05E+08 | 15502000 | 5569300 | 6593500 | 6265500 | 21164000 | 38229000 |
| GADFLVTE' | PKM; PKM; PKM | P14618; B4E | KPYM_HUMAN | Unmodified | 0 | 0 | 0 | 0 | 15972000 | 1558500 | 1777600 | 0 | 0 | 0 | 0 | 0 | 0 | 0 | 0 | 0 | 13991000 | 7117200 |
| GADINAPD | MTPN | P58546 | MTPN_HUMAN | Unmodified | 0 | 0 | 0 | 1978300 | 1814200 | 0 | 8913500 | 204150 | 0 | 668690 | 831730 | 0 | 2486600 | 2539100 | 0 | 0 | 2216300 | 4724700 |
| GADKIEGAI | PLCG1 | P19174 | PLCG1_HUMAI | Unmodified | 1671700 | 0 | 0 | 1844400 | 0 | 0 | 0 | 0 | 0 | 0 | 0 | 933850 | 0 | 0 | 0 | 0 | 0 | 0 |
| GADKTVKG | MTPN; MTPN | P58546; C9J | MTPN_HUMAN | Unmodified | 0 | 0 | 0 | 76470 | 2015700 | 0 | 2898900 | 2823400 | 0 | 320640 | 0 | 0 | 1861800 | 1609200 | 0 | 0 | 5307900 | 1751400 |
| GADLNLVD | RAI14 | Q9P0K7 | RAI14_HUMAN | Unmodified | 0 | 0 | 0 | 0 | 1906100 | 0 | 0 | 0 | 0 | 0 | 0 | 1033300 | 0 | 0 | 0 | 0 | 633910 | 576850 |
| GADSAVFP | ATP11C; ATP11 | Q8NB49; A0 | AT11C_HUMAN | Unmodified | 0 | 0 | 0 | 267320 | 0 | 0 | 0 | 0 | 1115000 | 171080 | 0 | 899920 | 0 | 0 | 501550 | 204080 | 0 | 0 |
| GADSLEDF | DDX3X; DDX3Y; | O00571; O1! | DDX3X_HUMAI | Unmodified | 808610 | 0 | 0 | 0 | 0 | 0 | 0 | 0 | 0 | 0 | 10994000 | 0 | 0 | 0 | 0 | 0 | 0 | 0 |
| GADSVIME | ATP10D | Q9P241 | AT10D_HUMAN | Oxidation (M) | 0 | 0 | 0 | 0 | 104640 | 0 | 0 | 0 | 0 | 0 | 0 | 23342 | 0 | 0 | 0 | 0 | 0 | 0 |
| GAEAANVT | YBX1; YBX1; YB | P67809; C9J | YBOX1_HUMAN | Unmodified | 4080000 | 2887500 | 5595600 | 4421400 | 935240 | 285760 | 244470 | 262360 | 1261700 | 198330 | 4973400 | 22836000 | 858010 | 171900 | 2983100 | 4046200 | 76621 | 95789 |
| GAEAGSVV | FKRP | Q9H9S5 | FKRP_HUMAN | Unmodified | 0 | 37132 | 0 | 0 | 0 | 0 | 0 | 0 | 239780 | 0 | 0 | 0 | 0 | 0 | 104470 | 59688 | 0 | 285270 |
| GAEASAAS | MARCKSL1 | P49006 | MRP_HUMAN N | Unmodified | 9693700 | 0 | 20012000 | 63828000 | 37663000 | 6029800 | 21399000 | 41915000 | 1.36E+08 | 1.44E+08 | 1.42E+08 | 2.67E+08 | 2.23E+08 | 2.51E+08 | 61427000 | 55333000 | 3.52E+08 | 7.54E+08 |
| GAEDAPDA | CADM2 | Q8N3J6 | CADM2_HUMA | Unmodified | 288760 | 0 | 0 | 0 | 0 | 0 | 0 | 0 | 0 | 0 | 0 | 0 | 0 | 0 | 0 | 0 | 0 | 1648500 |
| GAEEEPPI | EIF2B5; EIF2B5; | Q13144; A0/ | EI2BE_HUMAN | Unmodified | 0 | 0 | 0 | 0 | 0 | 0 | 0 | 0 | 0 | 0 | 0 | 5197100 | 0 | 0 | 0 | 0 | 0 | 0 |
| GAEEMETV | PARK7; PARK7 | Q99497; K7E | PARK7_HUMAN | Unmodified | 0 | 0 | 0 | 0 | 0 | 0 | 0 | 0 | 1271500 | 0 | 0 | 0 | 0 | 0 | 0 | 0 | 0 | 0 |
| GAEEMETV | PARK7; PARK7 | Q99497; K7E | PARK7_HUMAN | 2 Oxidation (M) | 0 | 0 | 0 | 0 | 0 | 0 | 0 | 0 | 0 | 222770 | 599620 | 1136500 | 956940 | 284630 | 0 | 0 | 0 | 0 |
| GAEEMETV | PARK7; PARK7 | Q99497; K7E | PARK7_HUMAN | Oxidation (M) | 0 | 0 | 482410 | 2026100 | 7454100 | 0 | 0 | 63232 | 5593300 | 1675200 | 784760 | 5715300 | 15666000 | 8191300 | 462640 | 416870 | 2104900 | 3658300 |
| GAEEMETV | PARK7; PARK7 | Q99497; K7E | PARK7_HUMAN | 2 Oxidation (M) | 0 | 271650 | 848990 | 917090 | 0 | 0 | 0 | 295640 | 556600 | 869060 | 0 | 3674700 | 3865300 | 0 | 1153300 | 754150 | 492340 | 1241700 |
| GAEEMETV | PARK7; PARK7 | Q99497; K7E | PARK7_HUMAN | Unmodified | 0 | 0 | 0 | 341570 | 6760800 | 0 | 0 | 0 | 5957600 | 1296500 | 0 | 438140 | 3647400 | 3549500 | 0 | 0 | 0 | 1161900 |
| GAEEMETV | PARK7; PARK7 | Q99497; K7E | PARK7_HUMAN | Unmodified | 0 | 0 | 0 | 0 | 7519200 | 0 | 328330 | 0 | 0 | 600500 | 0 | 0 | 2649500 | 3628200 | 0 | 0 | 1459800 | 5398600 |
| GAEEMETV | PARK7; PARK7 | Q99497; K7E | PARK7_HUMAN | Oxidation (M) | 0 | 0 | 0 | 0 | 2770900 | 314950 | 1079700 | 93229 | 0 | 1233700 | 0 | 0 | 8763100 | 6946100 | 0 | 0 | 388200 | 3653500 |
| GAEHITTYT | CENPV | Q7Z7K6 | CENPV_HUMAI | Unmodified | 26702 | 0 | 0 | 0 | 61872 | 124230 | 1241900 | 537480 | 0 | 0 | 0 | 1672200 | 0 | 0 | 0 | 0 | 0 | 0 |
| GAEHLSLL' | SHB | Q15464 | SHB_HUMAN S | Unmodified | 0 | 0 | 0 | 0 | 0 | 0 | 0 | 0 | 0 | 116730 | 1969600 | 7818200 | 1634700 | 0 | 0 | 0 | 422770 | 0 |
| GAEIEYAM/ | MYDGF; MYDGI | Q969H8; M0 | MYDGF_HUMA | Oxidation (M) | 0 | 0 | 0 | 0 | 0 | 0 | 0 | 0 | 7949700 | 1066600 | 563680 | 845260 | 36486 | 0 | 0 | 0 | 0 | 0 |
| GAEIEYAM/ | MYDGF; MYDGI | Q969H8; M0 | MYDGF_HUMA | Unmodified | 0 | 0 | 0 | 0 | 0 | 0 | 0 | 0 | 35157000 | 3669200 | 1286700 | 490940 | 0 | 0 | 0 | 0 | 0 | 136460 |
| GAEILADTF | DDAH1; DDAH: | O94760; B4E | DDAH1_HUMA | Unmodified | 0 | 0 | 0 | 0 | 0 | 0 | 0 | 0 | 1393900 | 0 | 0 | 1243300 | 0 | 0 | 0 | 0 | 0 | 0 |
| GAEILADTF | DDAH1; DDAH: | O94760; B4E | DDAH1_HUMA | Unmodified | 0 | 0 | 0 | 0 | 0 | 0 | 0 | 0 | 0 | 602980 | 0 | 0 | 3195800 | 0 | 0 | 0 | 1351800 | 1330200 |
| GAEILEVLH | PSMD6; PSMD6 | Q15008; H7( | PSMD6_HUMA | Unmodified | 0 | 0 | 107880 | 0 | 0 | 0 | 0 | 0 | 0 | 0 | 0 | 2610500 | 0 | 0 | 303820 | 186560 | 0 | 0 |
| GAEILPRQI | MYO6; MYO6; M | Q9UM54; A0 | MYO6_HUMAN | Unmodified | 0 | 0 | 0 | 0 | 0 | 0 | 0 | 0 | 0 | 0 | 0 | 0 | 299480 | 118390 | 0 | 0 | 0 | 0 |
| GAELHLQP | ADAMTS4; ADA | O75173; Q5\ | ATS4_HUMAN / | Unmodified | 0 | 0 | 0 | 0 | 0 | 0 | 0 | 0 | 0 | 0 | 0 | 0 | 3058300 | 0 | 0 | 0 | 0 | 0 |
| GAELLRCIA | N/A | N/A | N/A | Unmodified | 0 | 0 | 0 | 0 | 0 | 0 | 0 | 0 | 131710 | 0 | 0 | 0 | 103730 | 0 | 0 | 0 | 119110 | 0 |
| GAEQAGV/ | AVPR1A; AVPR1 | P37288; F8V | V1AR_HUMAN | Unmodified | 124060 | 91163 | 0 | 0 | 0 | 0 | 0 | 0 | 853300 | 283730 | 0 | 0 | 247530 | 86526 | 0 | 0 | 0 | 0 |
| GAEVAHFN | NT5E; NT5E | P21589; Q9( | 5NTD_HUMAN | Oxidation (M) | 0 | 0 | 0 | 0 | 257770 | 0 | 0 | 0 | 0 | 0 | 0 | 0 | 0 | 0 | 0 | 0 | 0 | 256120 |
| GAEVNQVI | MYO1B; MYO1E | O43795; E9F | MYO1B_HUMA | Unmodified | 0 | 0 | 0 | 0 | 11456000 | 2897600 | 3207100 | 0 | 0 | 1132600 | 0 | 0 | 0 | 0 | 0 | 0 | 6519700 | 2807400 |
| GAEWHSH | CPD | O75976 | CBPD_HUMAN | Oxidation (M) | 0 | 0 | 0 | 0 | 0 | 0 | 0 | 0 | 3848900 | 250830 | 0 | 0 | 0 | 0 | 0 | 0 | 0 | 0 |
| GAEWHSH | CPD | O75976 | CBPD_HUMAN | Unmodified | 0 | 0 | 0 | 0 | 0 | 0 | 0 | 0 | 7299500 | 451870 | 100870 | 52082 | 0 | 0 | 0 | 0 | 0 | 0 |
| GAEYVSAR | SF3B1; SF3B1 | O75533; H7( | SF3B1_HUMAN | Unmodified | 0 | 0 | 0 | 0 | 0 | 0 | 0 | 0 | 0 | 0 | 0 | 33865000 | 0 | 0 | 0 | 0 | 0 | 0 |
| GAFGKPQ( | RPL10; RPL10L; | P27635; Q9( | RL10_HUMAN ( | Unmodified | 0 | 717700 | 552390 | 262290 | 0 | 0 | 38516 | 29491 | 0 | 0 | 291920 | 7403700 | 0 | 0 | 535100 | 415950 | 0 | 0 |
| GAFGVGQI | CNOT1 | A5YKK6 | CNOT1_HUMA | Unmodified | 0 | 0 | 0 | 0 | 0 | 0 | 0 | 0 | 0 | 0 | 0 | 936220 | 0 | 0 | 0 | 0 | 0 | 0 |
| GAFIDQSQ | RRM1; RRM1 | P23921; E9P | RIR1_HUMAN F | Unmodified | 0 | 0 | 0 | 0 | 0 | 0 | 0 | 0 | 0 | 0 | 429990 | 3615400 | 0 | 0 | 0 | 0 | 0 | 0 |

| | | | | | | | | | | | | | | | | | | | | | | |
|---|---|---|---|---|---|---|---|---|---|---|---|---|---|---|---|---|---|---|---|---|---|---|
| GAFLSNDA | MMP15 | P51511 | MMP15_HUMA | Unmodified | 0 | 0 | 0 | 0 | 0 | 0 | 0 | 0 | 0 | 0 | 0 | 0 | 142580 | 0 | 0 | 824370 | 0 | 0 |
| GAFQTGQ( | BIRC6; BIRC6 | Q9NR09; A0. | BIRC6_HUMAN | Unmodified | 0 | 0 | 0 | 0 | 0 | 0 | 0 | 0 | 0 | 0 | 0 | 0 | 0 | 0 | 0 | 0 | 0 | 0 |
| GAFSEEYK | CNP; CNP; CNI | P09543; K7E | CN37_HUMAN | Unmodified | 0 | 681620 | 1663700 | 1393900 | 2427300 | 1178700 | 0 | 0 | 3625800 | 27027 | 15409000 | 24599000 | 1765900 | 1210700 | 3867200 | 3123100 | 0 | 2080300 |
| GAFSEEYKI | CNP; CNP; CNI | P09543; K7E | CN37_HUMAN | Unmodified | 0 | 0 | 0 | 0 | 3911900 | 924430 | 913310 | 785160 | 0 | 0 | 0 | 0 | 3014100 | 0 | 0 | 0 | 2624200 | 1426300 |
| GAFSRLDP | DECR1 | Q16698 | DECR_HUMAN | Unmodified | 0 | 0 | 0 | 0 | 442120 | 0 | 267050 | 0 | 3789900 | 892610 | 0 | 0 | 0 | 0 | 0 | 0 | 92447 | 496330 |
| GAGAGHP( | SNTB1; SNTB1 | Q13884; A0/ | SNTB1_HUMAN | Unmodified | 0 | 0 | 0 | 0 | 0 | 0 | 0 | 0 | 1506900 | 172360 | 0 | 36895 | 0 | 0 | 0 | 0 | 0 | 0 |
| GAGATGKD | NPTX2 | P47972 | NPTX2_HUMAN | Oxidation (M) | 0 | 0 | 0 | 0 | 0 | 0 | 0 | 0 | 264540 | 147100 | 0 | 0 | 0 | 0 | 0 | 0 | 0 | 0 |
| GAGATGKD | NPTX2 | P47972 | NPTX2_HUMAN | Unmodified | 0 | 0 | 0 | 0 | 0 | 0 | 0 | 0 | 938460 | 0 | 0 | 0 | 0 | 0 | 0 | 0 | 0 | 0 |
| GAGAVGAL | PLEKHA2; PLEK | Q9HB19; A0. | PKHA2_HUMAN | Unmodified | 0 | 0 | 0 | 0 | 1511900 | 288590 | 8898600 | 10148000 | 0 | 143000 | 0 | 0 | 0 | 4768100 | 0 | 0 | 1870400 | 4964400 |
| GAGDGSD | MYH9 | P35579 | MYH9_HUMAN | Unmodified | 0 | 0 | 0 | 170180 | 3713100 | 412630 | 108700 | 52201 | 432650 | 0 | 0 | 380720 | 1108600 | 521500 | 212730 | 194410 | 1061100 | 1124900 |
| GAGGQGK | FLNB; FLNB | O75369; E7E | FLNB_HUMAN | Unmodified | 0 | 0 | 0 | 0 | 0 | 0 | 0 | 0 | 0 | 0 | 0 | 639610 | 0 | 0 | 86462 | 162320 | 0 | 0 |
| GAGGQSM | LTBP2; LTBP2; I | Q14767; G3 | LTBP2_HUMAN | Unmodified | 98267 | 0 | 0 | 269480 | 0 | 0 | 0 | 0 | 306300 | 73628 | 0 | 0 | 0 | 0 | 0 | 0 | 0 | 0 |
| GAGHPCYI | ABHD14B; ABH | Q96IU4; F8V | ABHEB_HUMAI | Unmodified | 0 | 0 | 0 | 0 | 2340500 | 0 | 0 | 0 | 0 | 217960 | 0 | 0 | 1364900 | 0 | 0 | 0 | 1704800 | 1642600 |
| GAGIEIYPG | SHCBP1 | Q8NEM2 | SHCBP_HUMA | Unmodified | 0 | 0 | 0 | 180900 | 3874800 | 0 | 0 | 0 | 0 | 0 | 0 | 0 | 0 | 0 | 0 | 0 | 346030 | 0 |
| GAGIGGLG | FLNB; FLNB | O75369; E7E | FLNB_HUMAN | Unmodified | 0 | 56100 | 871910 | 1098900 | 1690200 | 0 | 42812 | 22821 | 311700 | 0 | 1007600 | 6009700 | 269890 | 255370 | 693170 | 554390 | 108700 | 280350 |
| GAGKAPLN | FLNB; FLNB | O75369; E7E | FLNB_HUMAN | Unmodified | 0 | 0 | 0 | 0 | 2439400 | 450580 | 0 | 0 | 65759 | 114230 | 0 | 0 | 237940 | 0 | 121790 | 0 | 918580 | 139000 |
| GAGMPGQ | LMAN1 | P49257 | LMAN1_HUMAI | Oxidation (M) | 208270 | 0 | 0 | 309500 | 0 | 0 | 642790 | 0 | 97031000 | 4189100 | 2823500 | 357250 | 0 | 0 | 301630 | 0 | 0 | 194590 |
| GAGMPGQ | LMAN1 | P49257 | LMAN1_HUMAI | Unmodified | 0 | 0 | 0 | 0 | 0 | 0 | 0 | 0 | 5901400 | 0 | 0 | 0 | 0 | 0 | 0 | 0 | 0 | 0 |
| GAGNQYN | GLT8D1 | Q68CQ7 | GL8D1_HUMAI | Unmodified | 0 | 0 | 0 | 702710 | 0 | 0 | 0 | 0 | 11022000 | 568820 | 383960 | 531610 | 0 | 0 | 0 | 0 | 0 | 0 |
| GAGPGCYI | TPP2; TPP2; TPI | P29144; A0A | TPP2_HUMAN | Unmodified | 0 | 0 | 0 | 0 | 0 | 0 | 0 | 0 | 0 | 0 | 0 | 5439400 | 0 | 0 | 0 | 0 | 0 | 0 |
| GAGSIAGA | DSP | P15924 | DESP_HUMAN | Unmodified | 201810 | 174470 | 68670 | 0 | 0 | 0 | 0 | 0 | 324360 | 60289 | 63935 | 180550 | 130170 | 0 | 0 | 0 | 0 | 203990 |
| GAGTDDST | ANXA7 | P20073 | ANXA7_HUMAI | Unmodified | 1773200 | 1735100 | 1453600 | 3031600 | 0 | 0 | 0 | 0 | 62573000 | 8419500 | 902170 | 3556700 | 453680 | 200570 | 4773700 | 1641800 | 929540 | 0 |
| GAGTDEG( | ANXA4; ANXA4; | P09525; B4E | ANXA4_HUMAI | Unmodified | 474020 | 0 | 0 | 0 | 0 | 0 | 499850 | 0 | 11166000 | 0 | 0 | 3029200 | 0 | 0 | 0 | 0 | 0 | 0 |
| GAGTDEG( | ANXA4; ANXA4; | P09525; B4E | ANXA4_HUMAI | Unmodified | 0 | 0 | 0 | 0 | 0 | 0 | 0 | 27983 | 30278000 | 2193600 | 14373000 | 1422800 | 0 | 0 | 0 | 0 | 0 | 0 |
| GAGTDEG( | ANXA4; ANXA4; | P09525; B4E | ANXA4_HUMAI | Unmodified | 2593100 | 0 | 0 | 1782400 | 1486700 | 0 | 3790700 | 0 | 49810000 | 8767600 | 42777000 | 2124300 | 654900 | 77622 | 1599400 | 700580 | 0 | 0 |
| GAGTGGL( | FLNA; FLNA | P21333; Q6( | FLNA_HUMAN | Unmodified | 1637200 | 2305700 | 2808300 | 7696700 | 5692600 | 1677900 | 2332600 | 1769300 | 1731400 | 363250 | 0 | 3129400 | 5979100 | 3175500 | 3084200 | 2278100 | 1424800 | 1462500 |
| GAGTNED/ | ANXA3; ANXA3; | P12429; D6F | ANXA3_HUMAI | Unmodified | 0 | 0 | 0 | 0 | 0 | 717740 | 0 | 0 | 231890 | 0 | 0 | 0 | 0 | 0 | 0 | 0 | 209790 | 575640 |
| GAGTNED/ | ANXA3; ANXA3; | P12429; D6F | ANXA3_HUMAI | Unmodified | 0 | 0 | 0 | 0 | 20319000 | 5630100 | 0 | 0 | 0 | 0 | 0 | 0 | 0 | 0 | 0 | 0 | 0 | 4639400 |
| GAGTNED/ | ANXA3; ANXA3; | P12429; D6F | ANXA3_HUMAI | Oxidation (M) | 0 | 0 | 0 | 0 | 9425600 | 2863900 | 0 | 0 | 0 | 100180 | 0 | 0 | 0 | 0 | 0 | 0 | 2007200 | 1832100 |
| GAGTNED/ | ANXA3; ANXA3; | P12429; D6F | ANXA3_HUMAI | Unmodified | 0 | 0 | 0 | 0 | 3272800 | 320780 | 0 | 0 | 0 | 0 | 0 | 0 | 0 | 0 | 0 | 0 | 633090 | 642930 |
| GAGWLFG/ | PPP6C | O00743 | PPP6_HUMAN | Unmodified | 0 | 0 | 0 | 0 | 0 | 0 | 0 | 0 | 0 | 0 | 0 | 2547300 | 0 | 0 | 0 | 0 | 0 | 0 |
| GAGYTFG( | PPP2CB; PPP2( | P62714; P67 | PP2AB_HUMAI | Unmodified | 0 | 0 | 713630 | 0 | 0 | 0 | 0 | 0 | 5775900 | 0 | 2072500 | 9968900 | 0 | 0 | 0 | 2789400 | 0 | 0 |
| GAHGIIVV | RAB1A; RAB1A | P62820; E7E | RAB1A_HUMAI | Unmodified | 4959500 | 5548600 | 2905000 | 0 | 0 | 1634000 | 311320 | 0 | 15894000 | 1545000 | 4334200 | 15849000 | 0 | 0 | 7642900 | 5862200 | 0 | 4044700 |
| GAHGIIVV | RAB1B | Q9H0U4 | RAB1B_HUMAI | Unmodified | 4401000 | 4302700 | 2531400 | 7608100 | 9552900 | 0 | 0 | 13754 | 3486500 | 583260 | 4411100 | 8220600 | 730380 | 331030 | 11568000 | 8609400 | 935380 | 951080 |
| GAHLTALEI | MRPL53 | Q96EL3 | RM53_HUMAN | Oxidation (M) | 0 | 0 | 0 | 0 | 0 | 0 | 0 | 0 | 762660 | 0 | 0 | 0 | 0 | 0 | 0 | 0 | 0 | 0 |
| GAHNGQG | OLA1; OLA1 | Q9NTK5; J3K | OLA1_HUMAN | Unmodified | 186290 | 0 | 224270 | 832960 | 0 | 0 | 0 | 0 | 653310 | 109080 | 941520 | 11625000 | 1258900 | 306620 | 1536500 | 1077000 | 0 | 231320 |
| GAHSVGLN | RPSA; RPSA; RF | P08865; A0A | RSSA_HUMAN | 2 Oxidation (M) | 555920 | 711490 | 318510 | 391260 | 49056 | 0 | 0 | 0 | 205860 | 0 | 286640 | 2604600 | 407130 | 82760 | 883520 | 0 | 0 | 0 |
| GAHVTINAI | DBNL; DBNL; D | Q9UJU6; B4I | DBNL_HUMAN | Unmodified | 0 | 0 | 0 | 0 | 1965700 | 0 | 587280 | 0 | 0 | 0 | 0 | 0 | 997900 | 1020100 | 0 | 0 | 1263000 | 1385400 |
| GAHVTINAI | DBNL; DBNL; D | Q9UJU6; B4I | DBNL_HUMAN | Oxidation (M) | 0 | 0 | 0 | 0 | 0 | 0 | 109070 | 82684 | 0 | 0 | 0 | 0 | 397830 | 762740 | 0 | 0 | 312160 | 346240 |
| GAIFEEMLT | KCTD9 | Q7L273 | KCTD9_HUMAI | Oxidation (M) | 0 | 0 | 0 | 0 | 303670 | 0 | 0 | 0 | 0 | 0 | 0 | 0 | 130910 | 81217 | 0 | 0 | 346050 | 741660 |
| GAILTTMLA | CAMK2D; CAMI | Q13557; Q9I | KCC2D_HUMA | Unmodified | 0 | 0 | 0 | 0 | 0 | 0 | 0 | 0 | 365910 | 0 | 0 | 0 | 0 | 0 | 0 | 0 | 0 | 0 |
| GAILTTMLA | CAMK2D; CAMI | D6R938; H0 | D6R938_HUM/ | Unmodified | 0 | 0 | 0 | 0 | 0 | 0 | 0 | 0 | 0 | 0 | 0 | 0 | 0 | 0 | 0 | 0 | 2131100 | 0 |
| GAIQFVTH | SEC23B; SEC2 | Q15437; A0/ | SC23B_HUMAI | Unmodified | 104180 | 104640 | 0 | 0 | 0 | 0 | 0 | 0 | 0 | 45126 | 0 | 2460200 | 0 | 0 | 73280 | 212510 | 0 | 0 |
| GAIQFVTQ | SEC23A; SEC2 | Q15436; F5I | SC23A_HUMAI | Unmodified | 456880 | 3418700 | 0 | 71091 | 0 | 0 | 0 | 0 | 111810 | 0 | 0 | 2804000 | 0 | 0 | 245690 | 156180 | 0 | 0 |
| GAIQLGITH | PIP5K1A; PIP5K | Q99755; A2/ | PI51A_HUMAN | Unmodified | 414830 | 0 | 596040 | 102640 | 0 | 0 | 0 | 0 | 0 | 0 | 0 | 6142200 | 0 | 0 | 1272100 | 1510100 | 0 | 0 |
| GAIREYQT( | DYNC1H1; DYN | Q14204; A0/ | DYHC1_HUMA | Unmodified | 67953 | 72269 | 0 | 141190 | 0 | 0 | 0 | 0 | 409130 | 173880 | 264890 | 697850 | 189120 | 0 | 0 | 66356 | 39495 | 0 |
| GAISAEVYT | PRKAR1A; PRK/ | P10644; K7E | KAP0_HUMAN | Unmodified | 104080 | 0 | 0 | 0 | 1117300 | 0 | 612250 | 92344 | 1397300 | 599030 | 0 | 0 | 0 | 0 | 0 | 0 | 669510 | 1205300 |
| GAIVHYTILI | POGLUT3 | Q7Z4H8 | PLGT3_HUMAI | Unmodified | 0 | 0 | 0 | 0 | 0 | 0 | 0 | 0 | 1383900 | 0 | 0 | 0 | 0 | 0 | 0 | 0 | 0 | 0 |
| GAKAEEILE | RPL11; RPL11 | P62913; Q5 | RL11_HUMAN ( | Unmodified | 0 | 0 | 0 | 913070 | 113830 | 0 | 0 | 37916 | 199000 | 0 | 314110 | 2825700 | 0 | 129930 | 10467 | 474820 | 0 | 73604 |
| GAKLTPEEI | RPS8; RPS8 | P62241; Q5J | RS8_HUMAN 4 | Unmodified | 1877500 | 1182300 | 0 | 705700 | 1487000 | 979880 | 665160 | 1978900 | 274680 | 483710 | 168490 | 1064400 | 1723400 | 899630 | 67289 | 129610 | 45660 | 1218700 |
| GAKPQGHI | AHNAK | Q09666 | AHNK_HUMAN | Unmodified | 0 | 0 | 0 | 0 | 11406000 | 0 | 0 | 0 | 0 | 0 | 0 | 0 | 0 | 0 | 0 | 0 | 4096000 | 0 |
| GAKPQGHI | AHNAK | Q09666 | AHNK_HUMAN | Unmodified | 0 | 0 | 0 | 0 | 3909100 | 0 | 0 | 0 | 0 | 0 | 0 | 0 | 0 | 0 | 0 | 0 | 7723200 | 4815000 |
| GALAGEDT | PAK6 | Q9NQU5 | PAK6_HUMAN | Unmodified | 0 | 0 | 0 | 99627 | 401370 | 0 | 918920 | 157400 | 267330 | 127030 | 164860 | 301120 | 0 | 0 | 0 | 0 | 1489900 | 46854 |
| GALALAQA | MTHFD1; MTHF | P11586; V9C | C1TC_HUMAN | Unmodified | 205470 | 0 | 1001700 | 1271200 | 3024600 | 0 | 695890 | 485060 | 4368400 | 1680500 | 2524700 | 9360100 | 1469300 | 521240 | 881630 | 734190 | 1094400 | 2001000 |
| GALALAQA | MTHFD1; MTHF | P11586; V9C | C1TC_HUMAN | Unmodified | 0 | 0 | 0 | 0 | 0 | 0 | 77562 | 0 | 0 | 0 | 0 | 0 | 0 | 77244 | 0 | 0 | 240050 | 168730 |
| GALALEEK | MYH9 | P35579 | MYH9_HUMAN | Unmodified | 254380 | 0 | 929240 | 0 | 8394300 | 0 | 0 | 0 | 0 | 0 | 0 | 0 | 2071800 | 1385800 | 1387000 | 944290 | 989490 | 1531100 |
| GALEMVQN | TRAPPC3; TRAI | O43617; A0/ | TPPC3_HUMAI | Unmodified | 0 | 0 | 0 | 0 | 0 | 0 | 0 | 0 | 0 | 0 | 0 | 0 | 0 | 0 | 0 | 0 | 36320 | 0 |
| GALHTVSH | PPFIA1; PPFIA1 | Q13136; H0 | LIPA1_HUMAN | Unmodified | 0 | 0 | 0 | 0 | 756500 | 0 | 0 | 0 | 0 | 0 | 0 | 0 | 343070 | 0 | 0 | 0 | 330360 | 318930 |
| GALIHNVSI | PLCG2; PLCG2 | P16885; A0A | PLCG2_HUMAI | Unmodified | 0 | 0 | 0 | 0 | 0 | 123680 | 174960 | 0 | 0 | 0 | 255790 | 1356400 | 0 | 0 | 0 | 0 | 0 | 0 |
| GALIHNVSI | PLCG2; PLCG2 | P16885; A0A | PLCG2_HUMAI | Unmodified | 0 | 0 | 0 | 0 | 0 | 0 | 0 | 0 | 0 | 0 | 0 | 3780500 | 0 | 0 | 0 | 0 | 0 | 0 |
| GALIHNVSI | PLCG2; PLCG2 | P16885; A0A | PLCG2_HUMAI | Unmodified | 0 | 0 | 0 | 0 | 0 | 0 | 0 | 0 | 0 | 0 | 1691300 | 4786800 | 0 | 0 | 0 | 0 | 0 | 0 |
| GALINVPPI | VARS1; VARS1 | P26640; A0A | SYVC_HUMAN | Unmodified | 0 | 0 | 0 | 0 | 0 | 0 | 0 | 0 | 0 | 0 | 0 | 0 | 0 | 0 | 0 | 0 | 0 | 0 |
| GALPLDTV | ERP29 | P30040 | ERP29_HUMAI | Unmodified | 0 | 0 | 778570 | 0 | 1747700 | 0 | 7001800 | 5227100 | 5.24E+08 | 3.08E+08 | 13477000 | 5291500 | 0 | 287160 | 0 | 846440 | 0 | 0 |
| GALPLDTV | ERP29 | P30040 | ERP29_HUMAI | Unmodified | 0 | 0 | 0 | 0 | 0 | 0 | 8640200 | 2126900 | 4567800 | 70880000 | 200090 | 0 | 834130 | 4258300 | 0 | 0 | 179140 | 0 |
| GALQAVD( | ILVBL; ILVBL; IL | A1L0T0; E9P | HACL2_HUMAI | Unmodified | 0 | 0 | 0 | 0 | 0 | 0 | 2677400 | 0 | 0 | 0 | 0 | 769110 | 0 | 0 | 0 | 0 | 0 | 0 |
| GALQNIIPA | GAPDH; GAPDI | P04406; E7E | G3P_HUMAN ( | Unmodified | 41076000 | 19275000 | 45214000 | 53876000 | 73361000 | 24896000 | 9215100 | 8434200 | 68626000 | 10430000 | 2.17E+08 | 6.53E+08 | 41311000 | 15789000 | 67627000 | 48792000 | 28979000 | 63603000 |
| GALQNIIPA | GAPDH; GAPDI | P04406; E7E | G3P_HUMAN ( | Unmodified | 0 | 0 | 0 | 0 | 272610 | 0 | 0 | 217240 | 0 | 0 | 0 | 946430 | 0 | 0 | 0 | 0 | 3918000 | 465680 |
| GALQQGE/ | PBXIP1 | Q96AQ6 | PBIP1_HUMAN | Unmodified | 161900 | 0 | 0 | 0 | 460600 | 0 | 0 | 0 | 4047800 | 123400 | 298270 | 457550 | 0 | 0 | 0 | 0 | 0 | 0 |
| GALQYLVP | KPNB1 | Q14974 | IMB1_HUMAN I | Unmodified | 58125 | 73373 | 47483 | 58478 | 0 | 0 | 14944 | 0 | 51035 | 0 | 21358 | 198320 | 24248 | 0 | 443480 | 0 | 0 | 0 |
| GALSLQGS | SFTPA2; SFTPA | Q8IWL1; Q8I | SFPA2_HUMAN | Unmodified | 0 | 0 | 0 | 0 | 0 | 0 | 0 | 0 | 2247200 | 0 | 0 | 0 | 0 | 0 | 0 | 0 | 0 | 0 |
| GALSPGGF | ARVCF; ARVCF; | O00192; E9F | ARVC_HUMAN | Unmodified | 0 | 0 | 326260 | 135580 | 0 | 0 | 0 | 0 | 0 | 0 | 0 | 3406900 | 124760 | 116250 | 556040 | 485720 | 0 | 0 |
| GALSVAAS | NBR1 | Q14596 | NBR1_HUMAN | Unmodified | 0 | 0 | 0 | 0 | 0 | 0 | 2507600 | 0 | 1480500 | 310190 | 47935 | 0 | 0 | 0 | 0 | 0 | 433670 | 585620 |
| GALTTVEGI | ZPR1; ZPR1; ZP | O75312; H7 | ZPR1_HUMAN | Unmodified | 0 | 0 | 0 | 0 | 0 | 0 | 0 | 0 | 0 | 0 | 352860 | 1066400 | 424980 | 0 | 265120 | 0 | 0 | 0 |
| GALVLGSS | GYG1; GYG1; G | P46976; C9J | GLYG_HUMAN | Unmodified | 0 | 0 | 0 | 221410 | 671110 | 0 | 234630 | 257650 | 429090 | 438680 | 725600 | 8431100 | 1267300 | 841150 | 0 | 0 | 295190 | 429360 |
| GALVVVND | HSD17B4; HSD | P51659; E7E | DHB4_HUMAN | Unmodified | 0 | 1572700 | 0 | 28088 | 0 | 0 | 0 | 0 | 36938000 | 432930 | 0 | 1168800 | 75054 | 0 | 0 | 0 | 0 | 0 |
| GALVVVND | HSD17B4; HSD | P51659; E7E | DHB4_HUMAN | Unmodified | 0 | 0 | 0 | 0 | 0 | 0 | 0 | 0 | 10694000 | 0 | 0 | 0 | 0 | 0 | 0 | 0 | 0 | 0 |
| GALVVVND | HSD17B4; HSD | P51659; E7E | DHB4_HUMAN | Unmodified | 0 | 0 | 0 | 0 | 0 | 0 | 0 | 0 | 822590 | 123960 | 0 | 0 | 0 | 0 | 0 | 0 | 0 | 0 |
| GAMEELEK | SMAD1; SMAD! | Q15797; Q9 | SMAD1_HUMA | Unmodified | 0 | 0 | 0 | 0 | 0 | 0 | 0 | 0 | 0 | 0 | 0 | 0 | 0 | 0 | 172110 | 0 | 0 | 0 |
| GAMSTQQI | AARS1; AARS1; | P49588; A0A | SYAC_HUMAN | Unmodified | 75649 | 0 | 0 | 645340 | 2344500 | 621320 | 0 | 245640 | 1364900 | 639460 | 404610 | 4771800 | 2869300 | 1692600 | 635810 | 0 | 572130 | 1098400 |
| GAMSTQQI | AARS1; AARS1; | P49588; A0A | SYAC_HUMAN | Oxidation (M) | 0 | 0 | 0 | 0 | 1826400 | 0 | 0 | 0 | 0 | 0 | 0 | 0 | 2654400 | 4305500 | 0 | 0 | 1019300 | 0 |
| GANAGVQ | CES2 | O00748 | EST2_HUMAN ( | Unmodified | 0 | 0 | 0 | 0 | 0 | 0 | 0 | 0 | 2282800 | 147280 | 0 | 0 | 0 | 0 | 0 | 0 | 0 | 0 |
| GANDFMC | TCP1; TCP1; TC | P17987; F5G | TCPA_HUMAN | 2 Oxidation (M) | 0 | 0 | 0 | 0 | 0 | 0 | 0 | 0 | 0 | 0 | 0 | 340960 | 0 | 0 | 0 | 0 | 0 | 0 |
| GANDFMC | TCP1; TCP1; TC | P17987; F5G | TCPA_HUMAN | Oxidation (M) | 0 | 0 | 0 | 0 | 0 | 0 | 0 | 0 | 0 | 0 | 0 | 2400700 | 0 | 0 | 0 | 0 | 0 | 0 |
| GANDFMC | TCP1; TCP1; TC | P17987; F5G | TCPA_HUMAN | Unmodified | 0 | 0 | 0 | 0 | 0 | 0 | 0 | 0 | 279990 | 0 | 249810 | 2297700 | 0 | 0 | 0 | 0 | 0 | 0 |
| GANILLTDI | MAP4K5; MAP4 | Q9Y4K4; G3 | M4K5_HUMAN | Unmodified | 835500 | 0 | 0 | 0 | 1354500 | 0 | 0 | 0 | 0 | 0 | 319040 | 8500200 | 0 | 0 | 0 | 0 | 19151 | 0 |
| GANINAFD | ANKRD44; ANK | Q8N8A2; O1 | ANR44_HUMAI | Unmodified | 0 | 0 | 0 | 0 | 0 | 0 | 0 | 0 | 0 | 0 | 0 | 0 | 699560 | 0 | 0 | 0 | 0 | 0 |
| GANIQLLDI | DRG2; DRG2; D | P55039; J3Q | DRG2_HUMAN | Unmodified | 0 | 0 | 0 | 179730 | 0 | 0 | 0 | 0 | 129710 | 110930 | 80637 | 0 | 0 | 0 | 244690 | 89088 | 0 | 0 |
| GANLLLSA | CANT1 | Q8WVQ1 | CANT1_HUMAI | Unmodified | 0 | 0 | 0 | 0 | 0 | 0 | 0 | 0 | 17216000 | 5784500 | 807520 | 0 | 0 | 0 | 0 | 0 | 0 | 0 |
| GANPVEIR | HSPD1; HSPD1 | P10809; A0A | CH60_HUMAN | Unmodified | 79005 | 147660 | 0 | 189950 | 435760 | 130100 | 461030 | 221700 | 23003000 | 3579900 | 806330 | 549350 | 1068300 | 1481000 | 437370 | 323780 | 0 | 486690 |
| GANPVEIRI | HSPD1; HSPD1 | P10809; A0A | CH60_HUMAN | Oxidation (M) | 0 | 58915 | 0 | 0 | 0 | 0 | 354250 | 0 | 0 | 383400 | 0 | 0 | 185690 | 474880 | 48079 | 15520 | 0 | 0 |
| GANRTETV | PRKDC | P78527 | PRKDC_HUMAI | Unmodified | 0 | 0 | 0 | 0 | 0 | 0 | 0 | 0 | 0 | 0 | 0 | 926970 | 0 | 0 | 0 | 0 | 0 | 0 |
| GAPAPHCE | TAPBP; TAPBP; | O15533; A0/ | TPSN_HUMAN | Oxidation (M) | 0 | 0 | 0 | 0 | 0 | 0 | 0 | 0 | 452090 | 0 | 0 | 0 | 0 | 0 | 0 | 0 | 0 | 0 |
| GAPEGGQI | MST1R; MST1R | Q04912; J3K | RON_HUMAN I | Unmodified | 0 | 0 | 0 | 0 | 580510 | 0 | 0 | 0 | 490570 | 0 | 0 | 0 | 0 | 0 | 0 | 0 | 164240 | 270270 |
| GAPERILDI | ATP1A1 | P05023 | AT1A1_HUMAI | Unmodified | 0 | 0 | 0 | 0 | 2137400 | 0 | 1258300 | 0 | 0 | 0 | 0 | 0 | 2271200 | 1883400 | 0 | 0 | 0 | 628460 |
| GAPGVSVD | ACOT13 | Q9NPJ3 | ACO13_HUMAI | 2 Oxidation (M) | 0 | 0 | 0 | 0 | 0 | 0 | 0 | 0 | 0 | 0 | 0 | 0 | 38371000 | 17959000 | 0 | 0 | 0 | 0 |
| GAPLNTPV | ETFDH | Q16134 | ETFD_HUMAN | Unmodified | 0 | 0 | 0 | 0 | 0 | 0 | 0 | 0 | 867190 | 314760 | 39087 | 0 | 0 | 332320 | 0 | 0 | 0 | 0 |
| GAPLVKPLI | PTPN23 | Q9H3S7 | PTN23_HUMAI | Unmodified | 3511100 | 2909000 | 379270 | 0 | 15487000 | 2401000 | 3063600 | 815670 | 16314000 | 4972000 | 949080 | 3269200 | 1246100 | 1175500 | 0 | 0 | 1915400 | 8179400 |
| GAPLVVIC( | DPYSL3 | Q14195 | DPYL3_HUMAI | Unmodified | 0 | 0 | 0 | 0 | 0 | 0 | 0 | 68300 | 349160 | 0 | 16486000 | 43289000 | 0 | 0 | 0 | 0 | 0 | 0 |
| GAPPPGEP | BCAM | P50895 | BCAM_HUMAN | Unmodified | 858700 | 769260 | 0 | 0 | 961320 | 0 | 0 | 0 | 0 | 0 | 0 | 1555400 | 0 | 0 | 0 | 0 | 0 | 854230 |
| GAPPPGEP | BCAM | P50895 | BCAM_HUMAN | Oxidation (M) | 4690300 | 315500 | 0 | 0 | 856480 | 236520 | 0 | 0 | 891520 | 0 | 0 | 3078700 | 0 | 0 | 0 | 0 | 0 | 251190 |
| GAPPSSNII | MATR3; MATR3; | P43243; H0 | MATR3_HUMAI | Unmodified | 0 | 0 | 0 | 0 | 0 | 0 | 1235000 | 506990 | 0 | 0 | 0 | 771440 | 36044 | 0 | 0 | 0 | 0 | 0 |
| GAPTLPPA | BIRC5; BIRC5; I | O15392; H3I | BIRC5_HUMAN | Acetyl (Protein N | 0 | 0 | 223530 | 1493200 | 8179500 | 1171400 | 504200 | 0 | 765930 | 836520 | 0 | 46004 | 2274400 | 2678800 | 405960 | 335810 | 3868700 | 1324700 |
| GAPTTSLIS | ALDH7A1; ALDI | P49419; A0A | AL7A1_HUMAN | Unmodified | 0 | 0 | 0 | 0 | 333030 | 0 | 0 | 136850 | 0 | 0 | 0 | 1338800 | 0 | 0 | 0 | 0 | 0 | 447770 |
| GAPVGGHI | MYO1C; MYO1( | O00159; F5I | MYO1C_HUMA | Unmodified | 9386500 | 4757100 | 15728000 | 17347000 | 13010000 | 10721000 | 923090 | 0 | 1850700 | 0 | 0 | 3477400 | 0 | 2515400 | 32425000 | 23024000 | 0 | 0 |
| GAPVGKES | FLVCR1 | Q9Y5Y0 | FLVC1_HUMAN | Unmodified | 0 | 0 | 0 | 0 | 203350 | 0 | 0 | 0 | 0 | 0 | 0 | 0 | 1578600 | 714370 | 0 | 0 | 858900 | 0 |
| GAPVNISS | CSNK1D; CSNI | P48730; H7E | KC1D_HUMAN | Unmodified | 103910 | 163400 | 217760 | 251510 | 0 | 0 | 0 | 0 | 0 | 0 | 0 | 0 | 84216 | 0 | 203850 | 248480 | 0 | 0 |
| GAPYNALT | NENF | Q9UMX5 | NENF_HUMAN | Unmodified | 0 | 0 | 0 | 0 | 0 | 0 | 179120 | 172800 | 12278000 | 1272500 | 0 | 379880 | 195820 | 0 | 0 | 0 | 147880 | 0 |
| GAPYNALT | NENF | Q9UMX5 | NENF_HUMAN | Unmodified | 0 | 0 | 0 | 376640 | 93346 | 0 | 0 | 0 | 5700600 | 0 | 0 | 0 | 0 | 0 | 0 | 0 | 174560 | 0 |
| GAPYNALT | NENF | Q9UMX5 | NENF_HUMAN | Unmodified | 0 | 0 | 0 | 0 | 0 | 0 | 0 | 0 | 595930 | 342380 | 0 | 0 | 0 | 0 | 0 | 0 | 0 | 0 |

| | | | | | | | | | | | | | | | | | | | | | | |
|---|---|---|---|---|---|---|---|---|---|---|---|---|---|---|---|---|---|---|---|---|---|---|
| GAQAAIVV | RAB5A | P20339 | RAB5A_HUMAN | Unmodified | 1193700 | 1841600 | 517390 | 868460 | 0 | 0 | 0 | 0 | 3464300 | 0 | 679780 | 4398400 | 0 | 0 | 1551100 | 1108400 | 0 | 0 |
| GAQAAIVV | RAB5B; RAB5B | P61020; F8V | RAB5B_HUMAN | Unmodified | 4065100 | 3845700 | 0 | 5325100 | 887710 | 0 | 249170 | 0 | 3554200 | 964040 | 774680 | 2309700 | 413640 | 296540 | 0 | 1706800 | 0 | 0 |
| GAQAAIVV | RAB5C; N/A; RA | P51148; K7E | RAB5C_HUMAI | Unmodified | 0 | 400260 | 65831 | 0 | 0 | 0 | 0 | 0 | 0 | 0 | 1464800 | 5726500 | 0 | 0 | 61257 | 0 | 0 | 0 |
| GAQERLPT | UBFD1; UBFD1 | O14562; H3I | UBFD1_HUMAI | Oxidation (M) | 0 | 0 | 658950 | 0 | 0 | 0 | 0 | 0 | 0 | 0 | 365390 | 692350 | 0 | 0 | 0 | 0 | 0 | 0 |
| GAQGVILV | RAB18; RAB18; | Q9NP72; H0 | RAB18_HUMAN | Unmodified | 370570 | 298750 | 228100 | 305220 | 0 | 0 | 0 | 164360 | 1045500 | 0 | 438980 | 5028700 | 0 | 0 | 266090 | 321200 | 0 | 0 |
| GAQVEEIW | UCHL5; UCHL5 | Q9Y5K5; Q5I | UCHL5_HUMAI | Unmodified | 0 | 0 | 0 | 0 | 0 | 0 | 0 | 0 | 189590 | 89298 | 554620 | 1366200 | 0 | 0 | 0 | 0 | 0 | 0 |
| GAQVNAVI | PSMD10; PSMD | O75832; B1/ | PSD10_HUMAN | Unmodified | 0 | 0 | 0 | 0 | 0 | 0 | 225800 | 278690 | 948750 | 437490 | 0 | 2007900 | 2778900 | 2187700 | 298320 | 374120 | 2086500 | 1003500 |
| GARLIPEMI | HYOU1; HYOU1 | Q9Y4L1; A0/ | HYOU1_HUMA | Oxidation (M) | 0 | 0 | 0 | 0 | 0 | 487740 | 2835700 | 0 | 0 | 942660 | 0 | 0 | 0 | 0 | 0 | 0 | 0 | 0 |
| GARPPAAG | INF2; INF2; INF | Q27J81; A0A | INF2_HUMAN I | Unmodified | 0 | 0 | 0 | 0 | 0 | 0 | 0 | 0 | 2562000 | 577670 | 0 | 842180 | 0 | 0 | 0 | 0 | 0 | 0 |
| GARTDLRN | OSTF1 | Q92882 | OSTF1_HUMAN | Unmodified | 0 | 0 | 0 | 0 | 0 | 0 | 0 | 0 | 0 | 0 | 0 | 0 | 0 | 0 | 0 | 0 | 963720 | 0 |
| GASDTYVT | PTPRJ; PTPRJ; P | Q12913; A0/ | PTPRJ_HUMAN | Unmodified | 0 | 466420 | 0 | 0 | 3904100 | 517950 | 1038700 | 90615 | 3862400 | 1493000 | 6257300 | 3068700 | 2920800 | 1562900 | 0 | 348470 | 0 | 1006500 |
| GASEIILR | ATP2B4 | P23634 | AT2B4_HUMAN | Unmodified | 0 | 0 | 0 | 0 | 0 | 0 | 0 | 0 | 0 | 0 | 0 | 4034100 | 0 | 0 | 0 | 0 | 0 | 0 |
| GASEIILRK | ATP2B4 | P23634 | AT2B4_HUMAN | Unmodified | 0 | 0 | 379600 | 0 | 1634100 | 0 | 2254500 | 0 | 380170 | 142280 | 622470 | 2370300 | 67261 | 101180 | 211430 | 377940 | 218820 | 371580 |
| GASEILLKK | ATP2B3 | Q16720 | AT2B3_HUMAN | Unmodified | 184880 | 0 | 394980 | 456920 | 292340 | 0 | 234160 | 106980 | 322020 | 158110 | 218520 | 1342600 | 822070 | 617380 | 1247300 | 544690 | 92715 | 138340 |
| GASGSFVV | HP1BP3; HP1BI | Q5SSJ5; Q5S | HP1B3_HUMAI | Unmodified | 0 | 1500200 | 0 | 0 | 3465100 | 0 | 0 | 538790 | 255040 | 0 | 0 | 0 | 0 | 0 | 0 | 0 | 0 | 0 |
| GASGTFQL | HP1BP3; HP1BI | Q5SSJ5; B0C | HP1B3_HUMAI | Unmodified | 711560 | 473830 | 0 | 0 | 0 | 35274 | 96985 | 0 | 652680 | 0 | 0 | 0 | 0 | 0 | 0 | 0 | 27126 | 147240 |
| GASGTRED | COX5B | P10606 | COX5B_HUMAI | Unmodified | 0 | 170080 | 1239000 | 0 | 0 | 0 | 0 | 0 | 327650 | 1844900 | 0 | 0 | 0 | 0 | 427330 | 0 | 0 | 0 |
| GASLLLYQI | SRGAP2; SRGA | O75044; Q5\ | SRGP2_HUMAI | Unmodified | 0 | 0 | 0 | 0 | 0 | 0 | 0 | 0 | 260800 | 143490 | 0 | 561850 | 112280 | 0 | 123850 | 108410 | 335180 | 0 |
| GASQAGMI | CNN3 | Q15417 | CNN3_HUMAN | Unmodified | 0 | 0 | 0 | 0 | 2117600 | 0 | 0 | 0 | 0 | 0 | 0 | 0 | 0 | 0 | 0 | 0 | 0 | 471770 |
| GASQAGM | TAGLN3; TAGLI | Q9UI15; P37 | TAGL3_HUMAN | Unmodified | 704860 | 654870 | 463770 | 429410 | 836510 | 265300 | 0 | 0 | 828870 | 0 | 0 | 1244000 | 215810 | 0 | 248330 | 153470 | 119290 | 401090 |
| GASQAGM | TAGLN3; TAGLI | Q9UI15; P37 | TAGL3_HUMAN | Oxidation (M) | 1224500 | 648980 | 0 | 343400 | 437370 | 0 | 0 | 0 | 0 | 0 | 0 | 417440 | 0 | 0 | 0 | 150750 | 0 | 177670 |
| GASQAGM | TAGLN3; TAGLI | Q9UI15; P37 | TAGL3_HUMAN | 2 Oxidation (M) | 443410 | 280660 | 119790 | 68787 | 0 | 0 | 0 | 0 | 0 | 0 | 0 | 179660 | 0 | 0 | 266770 | 142340 | 0 | 0 |
| GASSAGEA | ARVCF; ARVCF; | O00192; E9F | ARVC_HUMAN | Unmodified | 235050 | 0 | 0 | 83027 | 91420 | 0 | 0 | 0 | 0 | 0 | 135160 | 75314 | 487740 | 0 | 0 | 2374000 | 171480 | 0 |
| GASSAGEA | ARVCF; ARVCF; | O00192; E9F | ARVC_HUMAN | Unmodified | 0 | 0 | 0 | 286160 | 0 | 0 | 0 | 0 | 0 | 0 | 0 | 0 | 239290 | 0 | 0 | 0 | 0 | 0 |
| GASSPLITV | NOMO1; NOMC | Q15155; P69 | NOMO1_HUMA | Unmodified | 0 | 0 | 0 | 0 | 0 | 0 | 0 | 0 | 4051900 | 0 | 1130200 | 0 | 0 | 0 | 0 | 0 | 0 | 0 |
| GASSSTND | CPD; CPD | O75976; J3Q | CBPD_HUMAN | Unmodified | 0 | 0 | 0 | 139350 | 519680 | 230240 | 41949 | 98155 | 6848000 | 7365400 | 0 | 0 | 144280 | 90033 | 0 | 0 | 220310 | 0 |
| GASWIDTA | SUMF2; SUMF2 | Q8NBJ7; J3Q | SUMF2_HUMAI | Unmodified | 0 | 0 | 0 | 30251 | 0 | 0 | 0 | 0 | 15992000 | 693510 | 321670 | 572350 | 408900 | 0 | 0 | 0 | 61883 | 0 |
| GATDIDKN | ITGAV | P06756 | ITAV_HUMAN I | Unmodified | 0 | 0 | 0 | 0 | 6044800 | 319170 | 0 | 0 | 0 | 1301400 | 0 | 0 | 0 | 0 | 0 | 0 | 1281700 | 769770 |
| GATGECNI | KCTD3 | Q9Y597 | KCTD3_HUMAI | Unmodified | 48185 | 0 | 0 | 0 | 0 | 0 | 0 | 0 | 0 | 116900 | 0 | 0 | 0 | 120790 | 0 | 0 | 102000 | 1449100 |
| GATLEPAQ | TAPBPL | Q9BX59 | TPSNR_HUMAI | Unmodified | 0 | 0 | 0 | 0 | 0 | 0 | 0 | 0 | 1778800 | 0 | 0 | 0 | 0 | 0 | 0 | 0 | 0 | 0 |
| GATLKITLD | PKM; PKM; PKM | P14618; B4D | KPYM_HUMAN | Unmodified | 0 | 0 | 0 | 0 | 1332600 | 0 | 0 | 0 | 0 | 370770 | 0 | 0 | 480850 | 0 | 0 | 0 | 1348800 | 867020 |
| GATLKITLD | PKM; PKM; PKM | P14618; B4D | KPYM_HUMAN | Oxidation (M) | 0 | 0 | 0 | 0 | 3561600 | 0 | 1029100 | 383910 | 0 | 0 | 0 | 0 | 340800 | 455070 | 0 | 0 | 3292400 | 1005800 |
| GATLSHYN | ACSF2; ACSF2 | Q96CM8; E9 | ACSF2_HUMAI | Unmodified | 0 | 0 | 0 | 0 | 0 | 0 | 810220 | 0 | 938190 | 0 | 0 | 0 | 0 | 0 | 0 | 0 | 0 | 0 |
| GATQQILDI | CCT2; CCT2; C | P78371; F5G | TCPB_HUMAN | Unmodified | 0 | 0 | 0 | 0 | 204280 | 158250 | 0 | 0 | 1851800 | 38941 | 1771100 | 19952000 | 0 | 0 | 0 | 0 | 0 | 0 |
| GATVVELFI | POMGNT2 | Q8NAT1 | PMGT2_HUMAI | Unmodified | 0 | 0 | 0 | 0 | 0 | 0 | 0 | 0 | 3067900 | 0 | 0 | 0 | 0 | 3971800 | 0 | 0 | 0 | 0 |
| GATYGKPV | RPL15; RPL15; | P61313; E7E | RL15_HUMAN | Unmodified | 0 | 3129900 | 4347500 | 3373800 | 0 | 0 | 0 | 58167 | 248280 | 0 | 0 | 29380000 | 1352600 | 0 | 1590600 | 0 | 0 | 117950 |
| GATYPSEIP | LMO7; LMO7; L | Q8WWI1; E9 | LMO7_HUMAN | Unmodified | 0 | 0 | 0 | 0 | 282850 | 0 | 0 | 0 | 0 | 0 | 0 | 0 | 0 | 0 | 0 | 0 | 0 | 0 |
| GAVADTRD | LAMB2 | P55268 | LAMB2_HUMAI | Unmodified | 0 | 0 | 0 | 0 | 0 | 0 | 0 | 136490 | 1467900 | 0 | 0 | 0 | 0 | 0 | 0 | 0 | 0 | 0 |
| GAVAEDGI | APEX1; APEX1; | P27695; A0A | APEX1_HUMAN | Unmodified | 46090 | 0 | 147460 | 0 | 1366600 | 161790 | 172360 | 0 | 486850 | 643690 | 3076200 | 6170500 | 1350800 | 229570 | 67659 | 0 | 107990 | 245290 |
| GAVAEDGI | APEX1; APEX1; | P27695; A0A | APEX1_HUMAN | Unmodified | 0 | 0 | 0 | 169720 | 1390300 | 131470 | 0 | 18356 | 1423600 | 100260 | 184950 | 3976900 | 1463800 | 109170 | 0 | 0 | 225480 | 479780 |
| GAVAIADAI | RANGAP1; RAN | P46060; H0Y | RAGP1_HUMAI | Unmodified | 0 | 0 | 0 | 0 | 0 | 0 | 0 | 0 | 0 | 0 | 0 | 2509300 | 0 | 0 | 0 | 0 | 0 | 0 |
| GAVAIADAI | RANGAP1; RAN | P46060; H0Y | RAGP1_HUMAI | Unmodified | 0 | 0 | 0 | 140060 | 2743500 | 308020 | 750620 | 432490 | 348590 | 471660 | 371400 | 455100 | 2637000 | 2001100 | 0 | 47085 | 2262200 | 1770300 |
| GAVAMAET | RANGAP1; RAN | P46060; H0Y | RAGP1_HUMAI | Unmodified | 92478 | 0 | 404900 | 0 | 1645100 | 0 | 0 | 285450 | 324410 | 438760 | 331180 | 7545700 | 4231200 | 0 | 0 | 0 | 541560 | 409780 |
| GAVASVTFI | MVP | Q14764 | MVP_HUMAN I | Unmodified | 1946500 | 1106600 | 0 | 0 | 1040400 | 207780 | 130900 | 0 | 886650 | 185350 | 0 | 14568000 | 0 | 0 | 153630 | 0 | 0 | 0 |
| GAVDAAVP | ATP6V1H; ATP6 | Q9UI12; E5R | VATH_HUMAN | Unmodified | 0 | 0 | 0 | 0 | 0 | 0 | 0 | 158460 | 474270 | 0 | 0 | 1272600 | 0 | 0 | 0 | 0 | 0 | 0 |
| GAVDGGLS | RPL5; RPL5; RP | P46777; Q5T | RL5_HUMAN 6 | Unmodified | 13556000 | 8918900 | 5680100 | 4179600 | 6524400 | 1424700 | 3120500 | 73143 | 2296300 | 1878500 | 7494600 | 38724000 | 8931400 | 3343800 | 6276300 | 4223600 | 2491900 | 4149600 |
| GAVEAAHC | ALDH16A1; ALD | Q8IZ83; F5H | A16A1_HUMAN | Unmodified | 0 | 0 | 0 | 0 | 0 | 0 | 233190 | 0 | 0 | 253000 | 0 | 1463700 | 0 | 0 | 0 | 0 | 0 | 0 |
| GAVEALAA | DDX21 | Q9NR30 | DDX21_HUMAI | Unmodified | 36334 | 24019 | 0 | 12159 | 0 | 0 | 73399 | 0 | 0 | 0 | 171830 | 179970 | 74901 | 0 | 0 | 0 | 0 | 0 |
| GAVEIIFK | PCCB; PCCB; P | P05166; C9J | PCCB_HUMAN | Unmodified | 306810 | 237640 | 0 | 0 | 0 | 0 | 0 | 1099400 | 1199300 | 105530 | 0 | 1070200 | 0 | 0 | 150670 | 212070 | 0 | 0 |
| GAVEKGEE | SFN | P31947 | 1433S_HUMAN | Unmodified | 586300 | 320890 | 0 | 0 | 548540 | 240470 | 0 | 0 | 2267800 | 237370 | 0 | 0 | 0 | 0 | 0 | 0 | 0 | 120690 |
| GAVEKGEE | SFN | P31947 | 1433S_HUMAN | Unmodified | 2480900 | 540190 | 0 | 0 | 81013000 | 13344000 | 0 | 0 | 6210300 | 8710700 | 0 | 631260 | 0 | 0 | 0 | 0 | 46757000 | 29155000 |
| GAVEPDRY | FOLH1B; FOLH | Q9HBA9; Q0 | FOH1B_HUMAI | Unmodified | 4497600 | 5490800 | 31637000 | 13820000 | 0 | 0 | 0 | 0 | 0 | 0 | 0 | 0 | 0 | 0 | 25954000 | 35288000 | 0 | 0 |
| GAVEPDRY | FOLH1 | Q04609 | FOLH1_HUMAI | Unmodified | 4876200 | 8620900 | 12407000 | 89208000 | 0 | 0 | 0 | 0 | 0 | 0 | 0 | 0 | 0 | 0 | 1.25E+08 | 9633300 | 0 | 0 |
| GAVGALLV | RAB11A; RAB11 | P62491; Q1 | RB11A_HUMAN | Unmodified | 0 | 0 | 586480 | 0 | 0 | 0 | 0 | 0 | 0 | 0 | 366140 | 1745200 | 0 | 0 | 0 | 0 | 0 | 0 |
| GAVHQLC | QDPR | P09417 | DHPR_HUMAN | Unmodified | 0 | 0 | 72602 | 78583 | 1212900 | 21160 | 0 | 405010 | 257150 | 0 | 271480 | 1772500 | 1825300 | 673750 | 0 | 0 | 609450 | 965770 |
| GAVIATELK | EIF3D | O15371 | EIF3D_HUMAN | Unmodified | 464530 | 0 | 499310 | 294170 | 0 | 0 | 0 | 0 | 0 | 0 | 735280 | 8626800 | 0 | 0 | 561350 | 1568000 | 0 | 0 |
| GAVLGRPT | PTPRS | Q13332 | PTPRS_HUMAN | Unmodified | 125870 | 0 | 117230 | 0 | 0 | 0 | 0 | 0 | 338000 | 0 | 0 | 0 | 0 | 0 | 77081 | 90455 | 0 | 683340 |
| GAVLPHFN | VAC14; VAC14 | Q08AM6; H3 | VAC14_HUMAI | Unmodified | 0 | 0 | 0 | 0 | 0 | 0 | 0 | 0 | 0 | 0 | 0 | 566320 | 0 | 0 | 0 | 0 | 0 | 0 |
| GAVLVCDI | PSAT1 | Q9Y617 | SERC_HUMAN | Unmodified | 0 | 0 | 0 | 0 | 0 | 0 | 2136000 | 0 | 0 | 0 | 680830 | 1715200 | 711500 | 0 | 0 | 0 | 0 | 0 |
| GAVQGGLI | PLIN4; PLIN4 | Q96Q06; A0 | PLIN4_HUMAN | Unmodified | 46239 | 0 | 0 | 0 | 0 | 0 | 0 | 103220 | 0 | 53861 | 0 | 331310 | 0 | 0 | 0 | 0 | 310600 | 660380 |
| GAVQTGVI | PLIN4; PLIN4 | Q96Q06; A0 | PLIN4_HUMAN | Unmodified | 0 | 0 | 0 | 0 | 0 | 0 | 0 | 55140 | 25970 | 0 | 0 | 41312 | 0 | 0 | 0 | 0 | 115760 | 248680 |
| GAVSAEQV | PFDN2 | Q9UHV9 | PFD2_HUMAN | Unmodified | 0 | 0 | 213700 | 0 | 494120 | 0 | 0 | 0 | 835630 | 120540 | 2743800 | 11077000 | 0 | 0 | 1008700 | 378780 | 0 | 0 |
| GAVSAEQV | PFDN2 | Q9UHV9 | PFD2_HUMAN | Unmodified | 0 | 0 | 0 | 0 | 4234700 | 0 | 445690 | 0 | 0 | 1632900 | 0 | 0 | 5320000 | 1578700 | 0 | 0 | 1406000 | 4648100 |
| GAVSSALYI | SLC12A2 | P55011 | S12A2_HUMAN | Oxidation (M) | 0 | 0 | 31988 | 0 | 359230 | 0 | 0 | 0 | 0 | 0 | 0 | 0 | 0 | 0 | 377890 | 0 | 395970 | 1607400 |
| GAVTDDEV | THOC7 | Q6I9Y2 | THOC7_HUMA | Acetyl (Protein N | 0 | 0 | 0 | 0 | 0 | 0 | 0 | 228720 | 430760 | 115890 | 0 | 355710 | 0 | 0 | 0 | 0 | 241220 | 0 |
| GAVVGN | TANC1 | Q9C0D5 | TANC1_HUMAI | Unmodified | 0 | 0 | 138910 | 0 | 0 | 0 | 0 | 0 | 0 | 0 | 0 | 0 | 367070 | 335730 | 629320 | 487660 | 0 | 0 |
| GAVWGATI | STRAP | Q9Y3F4 | STRAP_HUMAN | Unmodified | 222710 | 372620 | 251700 | 361390 | 2273700 | 0 | 583290 | 582460 | 119280 | 732120 | 1226800 | 6813400 | 2809300 | 2399700 | 368020 | 389100 | 668180 | 1120400 |
| GAVWGATI | STRAP | Q9Y3F4 | STRAP_HUMAN | Unmodified | 491450 | 0 | 452120 | 0 | 179540 | 0 | 294670 | 222820 | 0 | 24057 | 0 | 0 | 61711 | 0 | 221350 | 302280 | 308320 | 76531 |
| GAVWLNVI | MOGS; MOGS; | Q13724; A0/ | MOGS_HUMAN | Unmodified | 0 | 0 | 0 | 0 | 0 | 0 | 0 | 0 | 3353500 | 0 | 0 | 0 | 0 | 0 | 0 | 0 | 0 | 0 |
| GAVYSFDP | PSMB1 | P20618 | PSB1_HUMAN | Unmodified | 0 | 0 | 0 | 0 | 0 | 0 | 0 | 0 | 0 | 0 | 0 | 14724000 | 0 | 0 | 0 | 553670 | 0 | 0 |
| GAVYSFDP | PSMB1 | P20618 | PSB1_HUMAN | Unmodified | 0 | 0 | 237850 | 284140 | 0 | 156790 | 0 | 28285 | 0 | 0 | 733880 | 569860 | 341880 | 149360 | 0 | 0 | 82727 | 0 |
| GAWDWSI | GALNT7 | Q86SF2 | GALT7_HUMAN | Oxidation (M) | 0 | 0 | 0 | 0 | 0 | 0 | 0 | 0 | 556960 | 0 | 0 | 0 | 0 | 0 | 0 | 0 | 0 | 0 |
| GAWSNVLI | SLC25A4; SLC2 | P12235; P12 | ADT1_HUMAN | Unmodified | 0 | 0 | 0 | 0 | 0 | 0 | 0 | 0 | 8584300 | 1672300 | 0 | 0 | 0 | 0 | 0 | 0 | 0 | 0 |
| GAYDIFLN | LRPPRC | P42704 | LPPRC_HUMAI | Unmodified | 0 | 0 | 0 | 0 | 0 | 0 | 0 | 0 | 1191500 | 0 | 0 | 0 | 0 | 0 | 0 | 0 | 139230 | 0 |
| GAYEQVIK | ATP2C1; ATP2C | P98194; B4E | AT2C1_HUMAN | Unmodified | 0 | 0 | 0 | 0 | 0 | 0 | 0 | 0 | 831770 | 0 | 0 | 0 | 0 | 0 | 0 | 0 | 0 | 0 |
| GAYIYNALI | TARS1 | P26639 | SYTC_HUMAN | Unmodified | 0 | 0 | 13833 | 18727 | 0 | 27510 | 0 | 0 | 0 | 20730 | 0 | 274000 | 0 | 0 | 137640 | 31387 | 0 | 0 |
| GAYQDRDI | CCT4 | P50991 | TCPD_HUMAN | Unmodified | 0 | 0 | 0 | 0 | 0 | 0 | 0 | 0 | 0 | 27221 | 774530 | 8443900 | 0 | 0 | 31045 | 0 | 0 | 0 |
| GAYSLSIRI | FYN; FYN | P06241; E5R | FYN_HUMAN T | Unmodified | 0 | 0 | 0 | 0 | 0 | 0 | 0 | 0 | 1100400 | 0 | 0 | 0 | 0 | 0 | 0 | 0 | 0 | 0 |
| GAYSLSIRI | YES1; YES1 | P07947; J3Q | YES_HUMAN Ty | Unmodified | 0 | 0 | 0 | 0 | 1673900 | 0 | 0 | 0 | 2524300 | 0 | 0 | 0 | 0 | 0 | 2382000 | 985690 | 1800800 | 0 |
| GAYSVGPL | NOMO1; NOMC | Q15155; P69 | NOMO1_HUMA | Unmodified | 0 | 0 | 0 | 0 | 0 | 0 | 53955 | 0 | 12603000 | 0 | 984150 | 551110 | 0 | 283540 | 0 | 0 | 0 | 0 |
| GAYTQVIFL | GAA | P10253 | LYAG_HUMAN | Unmodified | 0 | 0 | 0 | 0 | 0 | 0 | 0 | 0 | 2496500 | 0 | 0 | 286980 | 0 | 0 | 0 | 0 | 0 | 0 |
| GCAEKSDY | MANF | P55145 | MANF_HUMAN | Unmodified | 0 | 0 | 0 | 0 | 0 | 0 | 0 | 0 | 0 | 741280 | 0 | 0 | 0 | 0 | 0 | 0 | 0 | 0 |
| GCALQCAI | HSPA4; HSPH1 | P34932; Q92 | HSP74_HUMAI | Unmodified | 0 | 0 | 0 | 0 | 482200 | 0 | 0 | 0 | 0 | 0 | 196930 | 7133800 | 1336300 | 540950 | 0 | 0 | 0 | 0 |
| GCCFVTFY | CELF1; CELF2; | Q92879; O9 | CELF1_HUMAN | Unmodified | 0 | 0 | 0 | 0 | 0 | 0 | 1465500 | 474900 | 0 | 0 | 0 | 0 | 0 | 0 | 0 | 0 | 0 | 0 |
| GCDVTFEW | IGF2R | P11717 | MPRI_HUMAN | Unmodified | 0 | 0 | 0 | 0 | 0 | 0 | 0 | 0 | 6758400 | 581040 | 218920 | 0 | 0 | 0 | 0 | 60758 | 0 | 0 |
| GCDVVVIP | MDH2 | P40926 | MDHM_HUMAI | Unmodified | 0 | 0 | 0 | 0 | 0 | 0 | 0 | 0 | 11888000 | 567130 | 0 | 0 | 0 | 51308 | 300460 | 0 | 0 | 0 |
| GCDVVVIP | MDH2 | P40926 | MDHM_HUMAI | Unmodified | 0 | 0 | 0 | 0 | 0 | 0 | 0 | 0 | 1468000 | 0 | 0 | 0 | 0 | 0 | 98613 | 0 | 0 | 0 |
| GCEVIQEIK | RO60; RO60 | P10155; H0Y | RO60_HUMAN | Unmodified | 1743300 | 991570 | 271850 | 0 | 0 | 0 | 0 | 0 | 0 | 0 | 0 | 276980 | 0 | 0 | 502530 | 650760 | 100640 | 0 |
| GCEVVPDV | FERMT3 | Q86UX7 | URP2_HUMAN | Unmodified | 59189 | 0 | 99519 | 681790 | 363600 | 0 | 752550 | 520460 | 0 | 0 | 0 | 0 | 598810 | 946710 | 151300 | 80472 | 0 | 0 |
| GCEVVPDV | FERMT3 | Q86UX7 | URP2_HUMAN | Unmodified | 0 | 0 | 0 | 0 | 839700 | 0 | 835420 | 247330 | 0 | 0 | 0 | 0 | 0 | 97463 | 0 | 0 | 0 | 153450 |
| GCEVVVSG | RPS3; RPS3; RP | P23396; E9P | RS3_HUMAN 4 | Unmodified | 0 | 0 | 1523600 | 1269100 | 0 | 0 | 1182300 | 419950 | 122730 | 70371 | 0 | 23327000 | 0 | 135770 | 736590 | 860900 | 0 | 0 |
| GCFAVPM | COLGALT1 | Q8NBJ5 | GT251_HUMAN | Unmodified | 4711800 | 0 | 0 | 0 | 0 | 0 | 0 | 0 | 270410 | 0 | 0 | 0 | 0 | 0 | 0 | 0 | 0 | 0 |
| GCFQAEIVI | ACAA1; ACAA1; | P09110; C9J | THIK_HUMAN | Unmodified | 0 | 0 | 0 | 0 | 0 | 0 | 0 | 0 | 0 | 0 | 0 | 0 | 0 | 0 | 0 | 0 | 0 | 0 |
| GCFQAEIVI | ACAA1; ACAA1; | P09110; C9J | THIK_HUMAN | Unmodified | 0 | 0 | 0 | 0 | 0 | 0 | 0 | 0 | 10029000 | 1693000 | 0 | 0 | 0 | 0 | 0 | 0 | 0 | 0 |
| GCFQAEIVI | ACAA1; ACAA1; | P09110; C9J | THIK_HUMAN | Unmodified | 0 | 0 | 0 | 0 | 1403400 | 0 | 0 | 0 | 0 | 602630 | 0 | 0 | 0 | 0 | 0 | 0 | 0 | 0 |
| GCGQRFDI | CHORDC1; CH | Q9UHD1; E9 | CHRD1_HUMA | Unmodified | 0 | 0 | 0 | 0 | 0 | 0 | 0 | 0 | 866650 | 0 | 0 | 1665700 | 0 | 0 | 0 | 0 | 0 | 0 |
| GCGTVLLS | RPL18; RPL18; | Q07020; F8V | RL18_HUMAN | Unmodified | 1539300 | 747100 | 174990 | 0 | 183190 | 0 | 476050 | 254690 | 0 | 0 | 0 | 4659500 | 477500 | 291050 | 0 | 402310 | 0 | 248300 |
| GCGTVLLS | RPL18; RPL18; | Q07020; F8V | RL18_HUMAN | Unmodified | 2537400 | 891150 | 1831400 | 200180 | 838000 | 0 | 0 | 0 | 0 | 0 | 336570 | 12001000 | 0 | 0 | 1537600 | 221950 | 89823 | 410640 |
| GCGVVKFE | HNRNPM; HNR | P52272; A0A | HNRPM_HUMA | Unmodified | 195150 | 102570 | 0 | 80205 | 317790 | 0 | 0 | 66878 | 267380 | 118260 | 0 | 3492700 | 0 | 0 | 30832 | 46848 | 0 | 141470 |
| GCILTLVER | EIF3CL; EIF3C | B5ME19; Q9 | EIFCL_HUMAN | Unmodified | 75487 | 62586 | 15649 | 96784 | 0 | 0 | 0 | 0 | 0 | 0 | 0 | 222520 | 94303 | 0 | 89352 | 0 | 0 | 0 |
| GCITIIGGG | PGK1 | P00558 | PGK1_HUMAN | Unmodified | 0 | 0 | 0 | 0 | 453990 | 0 | 199880 | 0 | 1313800 | 662860 | 0 | 10167000 | 995340 | 570810 | 0 | 0 | 962350 | 732960 |
| GCITIIGGG | PGK1 | P00558 | PGK1_HUMAN | Unmodified | 0 | 0 | 0 | 0 | 0 | 0 | 0 | 0 | 0 | 0 | 0 | 298250 | 0 | 0 | 0 | 0 | 502040 | 0 |
| GCLELIKET | MTHFD1; MTHF | P11586; V9G | C1TC_HUMAN | Unmodified | 0 | 0 | 0 | 0 | 906370 | 0 | 1299300 | 90394 | 0 | 1264000 | 0 | 0 | 1418000 | 453420 | 0 | 0 | 6034100 | 4729500 |
| GCLGNSK | GNAS; GNAS; G | P63092; Q5J | GNAS2_HUMAI | Unmodified | 0 | 139280 | 98822 | 1737700 | 3869300 | 3023600 | 1627700 | 2362400 | 0 | 0 | 181460 | 1953400 | 11087000 | 11770000 | 562640 | 1019000 | 11412000 | 34753000 |
| GCLLYGPP | PSMC6; PSMC6 | P62333; A0A | PRS10_HUMAN | Unmodified | 0 | 0 | 0 | 137570 | 0 | 0 | 0 | 222100 | 422500 | 140940 | 0 | 1302800 | 0 | 335760 | 0 | 0 | 1279300 | 0 |
| GCLTFVSK | ARPC1A; N/A | Q92747; A0/ | ARC1A_HUMAI | Unmodified | 0 | 0 | 0 | 0 | 0 | 0 | 0 | 0 | 0 | 0 | 0 | 665710 | 0 | 0 | 0 | 0 | 0 | 0 |
| GCLVTASA | PWP1; PWP1 | Q13610; B4I | PWP1_HUMAN | Unmodified | 0 | 79037 | 0 | 0 | 0 | 0 | 197680 | 348470 | 0 | 0 | 0 | 403070 | 0 | 0 | 0 | 0 | 0 | 0 |
| GCMTLEV | GAS6 | Q14393 | GAS6_HUMAN | Unmodified | 0 | 0 | 0 | 0 | 0 | 0 | 0 | 0 | 3036100 | 0 | 84240 | 0 | 0 | 0 | 120890 | 0 | 0 | 0 |
| GCPDNITV | PPM1L | Q5SGD2 | PPM1L_HUMAI | Oxidation (M) | 0 | 0 | 0 | 0 | 0 | 0 | 0 | 0 | 998710 | 0 | 0 | 0 | 0 | 0 | 0 | 0 | 0 | 0 |

| | | | | | | | | | | | | | | | | | | | | | | |
|---|---|---|---|---|---|---|---|---|---|---|---|---|---|---|---|---|---|---|---|---|---|---|
| GCPDNITV | PPM1L | Q5SGD2 | PPM1L_HUMAN | Unmodified | 0 | 0 | 0 | 0 | 0 | 2016600 | 0 | 0 | 4588300 | 0 | 0 | 0 | 0 | 0 | 0 | 0 | 0 | 0 |
| GCPEDAAV | IGF2R; IGF2R; I | P11717; A0A | MPRI_HUMAN | Unmodified | 0 | 0 | 136850 | 0 | 0 | 0 | 0 | 0 | 968100 | 556890 | 0 | 133110 | 0 | 0 | 0 | 0 | 0 | 0 |
| GCPPDDIE | ITGB1; ITGB1; I | P05556; E7E | ITB1_HUMAN I | Unmodified | 568680 | 206100 | 479200 | 400300 | 7533000 | 1298600 | 0 | 0 | 2535100 | 240550 | 236300 | 882660 | 1151500 | 600520 | 788790 | 632650 | 1071700 | 8927300 |
| GCPPDDIE | ITGB1; ITGB1; I | P05556; E7E | ITB1_HUMAN I | Unmodified | 0 | 0 | 0 | 0 | 2341500 | 0 | 0 | 0 | 0 | 0 | 0 | 0 | 0 | 0 | 0 | 0 | 1170500 | 2033400 |
| GCQLNFIE | ITGB6; ITGB6 | P18564; E9P | ITB6_HUMAN I | Unmodified | 0 | 0 | 0 | 0 | 0 | 0 | 0 | 0 | 0 | 0 | 1060900 | 1422100 | 0 | 0 | 0 | 0 | 0 | 0 |
| GCSFLPDP | PSAP; PSAP; PS | P07602; Q5E | SAP_HUMAN P | Unmodified | 0 | 0 | 0 | 0 | 1579000 | 4551300 | 1881600 | 1957600 | 35870000 | 16093000 | 1866000 | 507000 | 7800100 | 5218400 | 0 | 57906 | 0 | 1392800 |
| GCSGDERI | VIL1; VIL1; VIL1 | P09327; H7C | VILI_HUMAN Vi | Unmodified | 29706 | 1015900 | 37769 | 0 | 0 | 0 | 46878 | 0 | 87325 | 110810 | 0 | 824720 | 0 | 0 | 563360 | 0 | 0 | 0 |
| GCSNQYLV | B3GALT6 | Q96L58 | B3GT6_HUMAN | Unmodified | 24493 | 0 | 0 | 0 | 0 | 0 | 0 | 0 | 2617000 | 110660 | 0 | 223690 | 0 | 0 | 0 | 0 | 0 | 0 |
| GCTATLGN | RPS2; RPS2; RP | P15880; E9P | RS2_HUMAN 4 | Unmodified | 0 | 1852500 | 2278000 | 1912500 | 0 | 111340 | 158940 | 0 | 746260 | 183760 | 1817100 | 51750000 | 440180 | 229780 | 2083700 | 1717500 | 0 | 0 |
| GCTDNLTL | PDLIM1 | O00151 | PDLI1_HUMAN | Unmodified | 0 | 0 | 0 | 0 | 2605100 | 0 | 0 | 0 | 0 | 0 | 0 | 0 | 196100 | 0 | 0 | 48410 | 1540700 | 2663700 |
| GCTVWLTG | PAPSS2; PAPSS | O95340; O43 | PAPS2_HUMAN | Unmodified | 0 | 0 | 0 | 0 | 0 | 0 | 1126200 | 0 | 0 | 0 | 0 | 1199700 | 0 | 0 | 0 | 0 | 0 | 0 |
| GCVGEVSV | HSPG2 | P98160 | PGBM_HUMAN | Unmodified | 0 | 0 | 514210 | 0 | 0 | 0 | 0 | 0 | 0 | 0 | 0 | 0 | 0 | 0 | 0 | 0 | 0 | 0 |
| GCWDSIHV | CAPZB; CAPZB; | P47756; B1A | CAPZB_HUMAN | Unmodified | 3067400 | 968520 | 855240 | 952210 | 1829600 | 714620 | 505070 | 253390 | 1561200 | 833690 | 1867900 | 8759200 | 1669100 | 1300700 | 2518100 | 1440400 | 1452800 | 975580 |
| GCYLATGS | GEMIN5 | Q8TEQ6 | GEMI5_HUMAN | Unmodified | 0 | 0 | 0 | 0 | 0 | 58713 | 0 | 0 | 0 | 0 | 0 | 371330 | 0 | 0 | 0 | 0 | 16896 | 0 |
| GDAAVRDE | PSMD6; PSMD6 | Q15008; C9J | PSMD6_HUMA | Oxidation (M) | 0 | 0 | 0 | 0 | 0 | 0 | 0 | 12024 | 0 | 0 | 0 | 1154000 | 0 | 0 | 0 | 0 | 0 | 0 |
| GDAAVRDE | PSMD6; PSMD6 | Q15008; C9J | PSMD6_HUMA | Unmodified | 38303 | 0 | 0 | 0 | 0 | 0 | 0 | 0 | 0 | 0 | 0 | 1414500 | 0 | 0 | 0 | 0 | 0 | 0 |
| GDAMIMEE | PA2G4; PA2G4; | Q9UQ80; F8 | PA2G4_HUMAN | Unmodified | 192030 | 242990 | 0 | 0 | 0 | 0 | 0 | 0 | 855790 | 597810 | 544450 | 0 | 0 | 0 | 322380 | 0 | 0 | 0 |
| GDAMIMEE | PA2G4; PA2G4; | Q9UQ80; F8 | PA2G4_HUMAN | Oxidation (M) | 413020 | 380410 | 294310 | 0 | 0 | 0 | 0 | 0 | 248260 | 0 | 525850 | 3433600 | 0 | 0 | 0 | 0 | 0 | 0 |
| GDAMIMEE | PA2G4; PA2G4; | Q9UQ80; F8 | PA2G4_HUMAN | 2 Oxidation (M) | 208570 | 0 | 0 | 0 | 0 | 0 | 0 | 0 | 0 | 0 | 0 | 683070 | 0 | 0 | 394830 | 212120 | 0 | 0 |
| GDAPSPEE | YARS1; YARS1; | P54577; A0A | SYYC_HUMAN | Acetyl (Protein N | 0 | 0 | 0 | 0 | 329760 | 0 | 219940 | 378770 | 0 | 1084600 | 0 | 0 | 1726500 | 944520 | 0 | 0 | 430150 | 296080 |
| GDASKEDI | HADH; HADH; | Q16836; A0A | HCDH_HUMAN | Unmodified | 0 | 0 | 0 | 0 | 0 | 0 | 0 | 0 | 0 | 0 | 0 | 0 | 0 | 0 | 0 | 0 | 0 | 0 |
| GDASTGWI | CTHRC1; CTHR | Q96CG8; E5 | CTHR1_HUMAN | Unmodified | 0 | 0 | 0 | 0 | 0 | 0 | 0 | 0 | 1245700 | 92682 | 0 | 0 | 0 | 0 | 0 | 0 | 0 | 0 |
| GDATVSYE | EWSR1; EWSR1 | Q01844; H7I | EWS_HUMAN F | Unmodified | 0 | 0 | 0 | 0 | 0 | 0 | 212750 | 0 | 0 | 0 | 307470 | 1552600 | 0 | 0 | 51278 | 0 | 0 | 0 |
| GDAVRDVI | FLNA; FLNA; FL | P21333; A0A | FLNA_HUMAN | Unmodified | 764210 | 539630 | 0 | 1275300 | 854260 | 390490 | 848970 | 0 | 691100 | 0 | 404290 | 5721000 | 0 | 49687 | 2229700 | 2164500 | 182390 | 0 |
| GDEELDSL | H2AZ2; H2AZ1; | Q71UI9; P0C | H2AV_HUMAN | Unmodified | 0 | 0 | 0 | 0 | 0 | 0 | 0 | 212060 | 0 | 0 | 406780 | 1711000 | 0 | 292160 | 0 | 0 | 0 | 0 |
| GDERILSILI | CHGA; CHGA | P10645; G5E | CMGA_HUMAN | Unmodified | 0 | 0 | 0 | 0 | 0 | 0 | 0 | 0 | 0 | 155130 | 0 | 0 | 0 | 0 | 0 | 0 | 0 | 0 |
| GDERPHYY | SLC44A2; SLC4 | Q8IWA5; K7E | CTL2_HUMAN | Acetyl (Protein N | 5841300 | 3262100 | 3156600 | 3074200 | 342760 | 0 | 0 | 0 | 6005100 | 3221600 | 1879000 | 5568900 | 2167200 | 0 | 4294600 | 3750800 | 208990 | 607110 |
| GDFCIQVG | NME2P1; NME1 | O60361; P15 | NDK8_HUMAN | Unmodified | 619540 | 0 | 717760 | 441560 | 382130 | 0 | 682010 | 252190 | 3448100 | 197770 | 4222000 | 67161000 | 0 | 0 | 776470 | 1002300 | 0 | 295570 |
| GDFFYHSE | TMEM43 | Q9BTV4 | TMM43_HUMA | Unmodified | 0 | 0 | 0 | 0 | 0 | 841320 | 0 | 0 | 7799500 | 0 | 0 | 0 | 0 | 0 | 0 | 0 | 0 | 0 |
| GDFLFSSD | MYO1B; MYO1E | O43795; H7 | MYO1B_HUMA | Oxidation (M) | 0 | 0 | 0 | 0 | 0 | 0 | 0 | 0 | 920840 | 0 | 892600 | 4000400 | 0 | 0 | 0 | 0 | 0 | 201100 |
| GDFPSPIH | AOC1; AOC1 | P19801; C9J | AOC1_HUMAN | Unmodified | 0 | 0 | 0 | 0 | 0 | 0 | 0 | 0 | 4577200 | 0 | 0 | 0 | 0 | 0 | 0 | 41371 | 0 | 0 |
| GDFSSANN | FGA; FGA | P02671; A0A | FIBA_HUMAN F | Unmodified | 0 | 6097700 | 0 | 0 | 0 | 0 | 0 | 0 | 0 | 0 | 0 | 0 | 0 | 0 | 0 | 0 | 0 | 0 |
| GDGESQP | TNKS1BP1 | Q9C0C2 | TB182_HUMAN | Unmodified | 0 | 0 | 0 | 0 | 0 | 0 | 0 | 0 | 0 | 0 | 224740 | 0 | 0 | 0 | 0 | 0 | 0 | 0 |
| GDGPICLV | DDX5; DDX17; | P17844; Q92 | DDX5_HUMAN | Unmodified | 0 | 0 | 0 | 0 | 0 | 0 | 0 | 0 | 0 | 0 | 0 | 1991100 | 0 | 0 | 148400 | 234640 | 0 | 0 |
| GDGVVLVA | LSM3 | P62310 | LSM3_HUMAN | Unmodified | 0 | 0 | 0 | 0 | 0 | 0 | 0 | 0 | 0 | 0 | 0 | 11042000 | 0 | 0 | 0 | 375570 | 0 | 0 |
| GDHAWSF | SLC25A4; SLC2 | P12235; V9C | ADT1_HUMAN | Acetyl (Protein N | 0 | 0 | 0 | 0 | 0 | 0 | 319680 | 0 | 6929800 | 1353900 | 267150 | 0 | 0 | 0 | 0 | 0 | 0 | 0 |
| GDHLFALL | MOV10; MOV10 | Q9HCE1; Q5 | MOV10_HUMA | Unmodified | 7188300 | 8692600 | 1552600 | 1169100 | 0 | 0 | 0 | 210180 | 0 | 0 | 0 | 320650 | 0 | 0 | 4007500 | 4253400 | 0 | 0 |
| GDHRYPSS | UGGT1 | Q9NYU2 | UGGG1_HUMA | Unmodified | 0 | 0 | 0 | 0 | 0 | 0 | 0 | 0 | 4577700 | 0 | 0 | 0 | 0 | 0 | 0 | 0 | 0 | 0 |
| GDIFLVR | VCP; VCP; VCP; | P55072; C9J | TERA_HUMAN | Unmodified | 0 | 0 | 0 | 18640 | 0 | 0 | 0 | 0 | 1694800 | 0 | 492740 | 7874200 | 0 | 0 | 0 | 26859 | 44054 | 286960 |
| GDIIGVQGI | KARS1; KARS1; | Q15046; H3I | SYK_HUMAN Ly | Unmodified | 0 | 0 | 0 | 0 | 0 | 397530 | 0 | 0 | 0 | 0 | 430520 | 1664100 | 0 | 0 | 0 | 43856 | 0 | 0 |
| GDISISGPK | AHNAK | Q09666 | AHNK_HUMAN | Unmodified | 0 | 0 | 0 | 0 | 1223300 | 0 | 0 | 0 | 155700 | 0 | 0 | 0 | 0 | 0 | 0 | 0 | 0 | 466420 |
| GDIVIIHEKI | NOSTRIN; NOS | A0A0G2JPW | A0A0G2JPW3_I | Unmodified | 240390 | 255400 | 0 | 411920 | 0 | 0 | 0 | 0 | 0 | 0 | 0 | 1586100 | 0 | 140990 | 0 | 0 | 0 | 0 |
| GDLAYNYR | TMX3 | Q96JJ7 | TMX3_HUMAN | Unmodified | 0 | 0 | 0 | 0 | 47937 | 0 | 0 | 0 | 4939800 | 193590 | 456760 | 485180 | 20120 | 0 | 0 | 16322 | 28396 | 0 |
| GDLGIEIPA | PKM; PKLR; PKN | P14618; P30 | KPYM_HUMAN | Unmodified | 3562800 | 1935600 | 7912800 | 5291700 | 7956200 | 0 | 1945400 | 1816600 | 12690000 | 1250700 | 37519000 | 1.5E+08 | 1722800 | 964910 | 12315000 | 10564000 | 1685200 | 3232400 |
| GDLGIEIPA | PKM; PKLR; PKN | P14618; P30 | KPYM_HUMAN | Unmodified | 0 | 0 | 0 | 0 | 427810 | 0 | 1225600 | 0 | 0 | 0 | 0 | 127260 | 0 | 0 | 0 | 0 | 829110 | 673220 |
| GDLLEGAN | USP9X; USP9Y | Q93008; O0 | USP9X_HUMAN | Unmodified | 0 | 0 | 0 | 0 | 0 | 0 | 0 | 0 | 0 | 0 | 0 | 556780 | 0 | 0 | 0 | 0 | 0 | 0 |
| GDLLFLTNI | SEC11A; SEC11 | P67812; H0 | SC11A_HUMAN | Unmodified | 0 | 0 | 0 | 0 | 0 | 0 | 0 | 0 | 318270 | 134630 | 0 | 1332600 | 0 | 0 | 0 | 0 | 0 | 0 |
| GDLQTMLF | SUN1; SUN1; S | O94901; H0 | SUN1_HUMAN | Unmodified | 0 | 0 | 0 | 0 | 0 | 0 | 0 | 0 | 2080300 | 40926 | 0 | 0 | 0 | 0 | 0 | 0 | 0 | 0 |
| GDLSLRPE | TRIM26 | Q12899 | TRI26_HUMAN | Unmodified | 1453100 | 1183100 | 1932900 | 500810 | 0 | 0 | 0 | 0 | 0 | 0 | 0 | 0 | 0 | 0 | 3244700 | 1346400 | 0 | 0 |
| GDMANNS | ATP2B1; ATP2B | P20020; F8V | AT2B1_HUMAN | Acetyl (Protein N | 0 | 0 | 0 | 0 | 270780 | 0 | 0 | 0 | 0 | 0 | 0 | 0 | 1004500 | 1077700 | 0 | 0 | 0 | 479270 |
| GDMANNS | ATP2B1; ATP2B | P20020; F8V | AT2B1_HUMAN | Acetyl (Protein N | 0 | 0 | 0 | 0 | 0 | 0 | 0 | 0 | 0 | 0 | 0 | 1025500 | 1981300 | 2408700 | 0 | 0 | 0 | 0 |
| GDMDVSLF | AHNAK | Q09666 | AHNK_HUMAN | 2 Oxidation (M) | 877520 | 0 | 0 | 0 | 0 | 0 | 0 | 0 | 0 | 0 | 0 | 0 | 0 | 0 | 0 | 1325900 | 0 | 0 |
| GDMVTLPA | ADI1; ADI1 | Q9BV57; A0A | MTND_HUMAN | Oxidation (M) | 0 | 328610 | 149530 | 120390 | 0 | 0 | 0 | 0 | 0 | 0 | 0 | 2121800 | 0 | 0 | 340790 | 208540 | 0 | 0 |
| GDNITLLQ | SNRPE; SNRPE | P62304; A6N | RUXE_HUMAN | Unmodified | 952920 | 0 | 0 | 0 | 0 | 0 | 0 | 0 | 0 | 0 | 0 | 3822500 | 0 | 0 | 0 | 0 | 0 | 0 |
| GDNNAVD | SEC11A; SEC11 | P67812; P0C | SC11A_HUMAN | Unmodified | 0 | 0 | 112600 | 0 | 0 | 0 | 0 | 0 | 14173000 | 76593 | 1198900 | 1473400 | 0 | 0 | 0 | 69253 | 0 | 0 |
| GDNNEVDI | SEC11C; SEC11 | Q9BY50; K7E | SC11C_HUMAI | Unmodified | 0 | 0 | 0 | 0 | 0 | 0 | 0 | 0 | 3540900 | 285150 | 363310 | 1053700 | 0 | 0 | 0 | 0 | 0 | 0 |
| GDPAHAGI | PBXIP1 | Q96AQ6 | PBIP1_HUMAN | Unmodified | 0 | 0 | 0 | 0 | 0 | 0 | 0 | 0 | 88673 | 0 | 537290 | 0 | 0 | 0 | 0 | 0 | 0 | 0 |
| GDPECHLF | IGFBP2; IGFBP2 | P18065; C9J | IBP2_HUMAN I | Unmodified | 0 | 0 | 0 | 0 | 0 | 0 | 0 | 0 | 14505000 | 735220 | 0 | 0 | 0 | 0 | 0 | 0 | 0 | 0 |
| GDPGLGEI | TNKS1BP1; TNK | Q9C0C2; A0 | TB182_HUMAN | Unmodified | 0 | 64153 | 108580 | 272510 | 0 | 0 | 0 | 0 | 0 | 0 | 0 | 1469800 | 24629 | 0 | 235320 | 268260 | 0 | 0 |
| GDPIGGHI | MYO1D; MYO1D | O94832; J3Q | MYO1D_HUMA | Unmodified | 1935100 | 1469500 | 3367600 | 1659800 | 0 | 0 | 0 | 0 | 0 | 0 | 0 | 0 | 0 | 0 | 4926700 | 2438000 | 0 | 0 |
| GDPLGGVI | MYO1B; MYO1E | O43795; E9F | MYO1B_HUMA | Unmodified | 0 | 0 | 0 | 0 | 770630 | 0 | 0 | 0 | 0 | 0 | 351170 | 2662400 | 0 | 0 | 0 | 0 | 0 | 0 |
| GDPQVYEE | EIF3L; EIF3L; EI | Q9Y262; B0 | EIF3L_HUMAN | Unmodified | 0 | 0 | 0 | 0 | 0 | 0 | 0 | 0 | 0 | 0 | 365680 | 1776800 | 349630 | 0 | 0 | 0 | 0 | 0 |
| GDPRIEYQ | UGGT1 | Q9NYU2 | UGGG1_HUMA | Unmodified | 0 | 0 | 227750 | 0 | 0 | 0 | 23918 | 0 | 2910000 | 2913900 | 0 | 0 | 0 | 0 | 0 | 0 | 0 | 0 |
| GDQENVH | KPNB1; KPNB1 | Q14974; J3K | IMB1_HUMAN I | Oxidation (M) | 154490 | 0 | 463460 | 0 | 269040 | 0 | 0 | 0 | 0 | 115410 | 0 | 1655600 | 0 | 0 | 0 | 456710 | 0 | 0 |
| GDQLLSVN | LIN7A; LIN7C; L | O14910; Q9I | LIN7A_HUMAN | Unmodified | 0 | 0 | 0 | 0 | 0 | 0 | 0 | 0 | 0 | 0 | 0 | 0 | 0 | 0 | 529860 | 295650 | 0 | 0 |
| GDQLVPFS | TMEM43; N/A | Q9BTV4; V9C | TMM43_HUMA | Unmodified | 0 | 0 | 0 | 0 | 2394300 | 0 | 0 | 102960 | 1378900 | 0 | 0 | 258720 | 0 | 0 | 0 | 0 | 438620 | 1644000 |
| GDRAGVLA | GNB2; GNB4; G | P62879; Q9H | GBB2_HUMAN | Unmodified | 0 | 0 | 0 | 0 | 0 | 0 | 0 | 0 | 0 | 0 | 0 | 0 | 0 | 0 | 0 | 0 | 0 | 0 |
| GDRAGVLA | GNB2; GNB2 | P62879; E7E | GBB2_HUMAN | Oxidation (M) | 104470 | 88563 | 0 | 356810 | 462130 | 0 | 747920 | 0 | 123850 | 121070 | 851290 | 97826 | 860670 | 0 | 191800 | 0 | 0 | 1393900 |
| GDREQLLQ | YWHAH; YWHA | Q04917; F8V | 1433F_HUMAN | Acetyl (Protein N | 1067500 | 280210 | 355650 | 2245400 | 15228000 | 3629300 | 436140 | 0 | 6046100 | 3110700 | 5019800 | 15680000 | 6741500 | 5265300 | 0 | 2739600 | 6875500 | 8601400 |
| GDREQLLQ | YWHAH; YWHA | Q04917; F8V | 1433F_HUMAN | Acetyl (Protein N | 0 | 0 | 0 | 0 | 24088000 | 0 | 0 | 3962500 | 0 | 2515200 | 0 | 0 | 6310700 | 6883100 | 0 | 0 | 0 | 0 |
| GDRESSGK | TMED2; TMED2 | Q15363; E7E | TMED2_HUMAI | Unmodified | 0 | 0 | 0 | 0 | 0 | 0 | 0 | 0 | 4785000 | 108630 | 126900 | 0 | 0 | 0 | 0 | 0 | 0 | 0 |
| GDRESSGK | TMED2; TMED2 | Q15363; E7E | TMED2_HUMAI | Oxidation (M) | 0 | 0 | 0 | 0 | 0 | 0 | 0 | 0 | 975260 | 0 | 62118 | 0 | 0 | 0 | 0 | 0 | 0 | 0 |
| GDRFGNC | ADAM9; ADAM9 | Q13443; A0A | ADAM9_HUMA | Unmodified | 0 | 0 | 763900 | 2065000 | 267230 | 0 | 0 | 0 | 525840 | 0 | 32022 | 0 | 0 | 0 | 5323100 | 1941600 | 0 | 0 |
| GDRIVNTP | TM9SF4; TM9SF | Q92544; A0A | TM9S4_HUMAN | Unmodified | 0 | 0 | 0 | 0 | 0 | 366430 | 155100 | 0 | 4113800 | 0 | 0 | 0 | 0 | 0 | 0 | 0 | 0 | 0 |
| GDRSEDFG | ANXA1 | P04083 | ANXA1_HUMAN | Unmodified | 1930800 | 0 | 1999400 | 0 | 3341700 | 4702400 | 105660 | 293450 | 1340400 | 415220 | 3.29E+08 | 3.96E+09 | 0 | 324560 | 1046200 | 2317400 | 148950 | 612380 |
| GDRSEDFG | ANXA1 | P04083 | ANXA1_HUMAN | Unmodified | 0 | 0 | 0 | 0 | 790210 | 0 | 0 | 0 | 0 | 0 | 25783000 | 3004900 | 0 | 0 | 0 | 0 | 0 | 0 |
| GDSFTHTP | EGFR; EGFR; E | P00533; Q5C | EGFR_HUMAN | Unmodified | 195310 | 118690 | 971700 | 625650 | 0 | 92212 | 0 | 0 | 411860 | 153420 | 0 | 2810600 | 0 | 709670 | 1835800 | 1519100 | 0 | 0 |
| GDSKKDDI | KHDRBS1 | Q07666 | KHDR1_HUMAI | Unmodified | 0 | 0 | 0 | 0 | 0 | 0 | 0 | 0 | 529650 | 0 | 0 | 5111700 | 0 | 0 | 0 | 0 | 0 | 0 |
| GDSLKEPT | ZDHHC5; ZDHH | Q9C0B5; H0 | ZDHC5_HUMA | Unmodified | 0 | 0 | 0 | 0 | 0 | 0 | 0 | 0 | 0 | 281890 | 529460 | 230770 | 0 | 60866 | 0 | 0 | 0 | 0 |
| GDSPASSK | FN1; FN1 | P02751; H0 | FINC_HUMAN I | Unmodified | 0 | 0 | 314900 | 0 | 0 | 0 | 0 | 0 | 0 | 0 | 63934 | 249410 | 0 | 0 | 0 | 1290900 | 0 | 0 |
| GDSVIVVLI | SNRPD2; SNRP | P62316; K7E | SMD2_HUMAN | Unmodified | 673850 | 1443300 | 0 | 0 | 0 | 0 | 0 | 0 | 0 | 0 | 592600 | 3199800 | 0 | 0 | 0 | 0 | 0 | 0 |
| GDTFIGYV | EDIL3 | O43854 | EDIL3_HUMAN | Unmodified | 0 | 0 | 0 | 0 | 4159000 | 1126600 | 5476300 | 3061100 | 7253800 | 3011100 | 0 | 0 | 0 | 0 | 1039000 | 0 | 0 | 0 |
| GDTGGED | DCTPP1 | Q9H773 | DCTP1_HUMAI | Unmodified | 0 | 0 | 0 | 0 | 0 | 0 | 0 | 0 | 0 | 0 | 0 | 1728800 | 0 | 0 | 0 | 0 | 0 | 0 |
| GDTNIKPIL | RPA1; RPA1; RP | P27694; I3L5 | RFA1_HUMAN | Unmodified | 0 | 0 | 244280 | 0 | 0 | 0 | 0 | 0 | 0 | 0 | 0 | 2158800 | 0 | 0 | 0 | 0 | 0 | 0 |
| GDTVFFHP | PHYH; PHYH; P | O14832; B1A | PAHX_HUMAN | Unmodified | 0 | 0 | 0 | 0 | 0 | 0 | 0 | 0 | 26109000 | 1721800 | 0 | 322120 | 0 | 0 | 0 | 0 | 0 | 0 |
| GDVAEGDI | HNRNPA0 | Q13151 | ROA0_HUMAN | Unmodified | 0 | 0 | 0 | 0 | 0 | 0 | 261880 | 88610 | 801470 | 446850 | 0 | 1800900 | 0 | 0 | 0 | 0 | 0 | 0 |
| GDVDVNM | PLCG2; PLCG2 | P16885; A0A | PLCG2_HUMAI | Unmodified | 0 | 0 | 0 | 0 | 0 | 0 | 0 | 0 | 0 | 0 | 320850 | 515790 | 0 | 0 | 0 | 0 | 0 | 0 |
| GDVDVNM | PLCG2; PLCG2 | P16885; A0A | PLCG2_HUMAI | Unmodified | 0 | 0 | 0 | 0 | 0 | 0 | 0 | 0 | 0 | 0 | 754070 | 1944700 | 0 | 0 | 0 | 0 | 0 | 0 |
| GDVEEDET | TNPO1; TNPO1 | Q92973; A0A | TNPO1_HUMAI | Unmodified | 0 | 0 | 1301200 | 0 | 0 | 0 | 0 | 0 | 0 | 0 | 0 | 750780 | 0 | 0 | 0 | 0 | 267530 | 0 |
| GDVFPSYT | MFGE8; MFGE8 | Q08431; H0 | MFGM_HUMAN | Unmodified | 0 | 0 | 0 | 0 | 1530700 | 0 | 0 | 0 | 2513300 | 941630 | 240360 | 4340200 | 0 | 0 | 4354200 | 0 | 453890 | 0 |
| GDVLTIVAV | CSK; CSK; CSK | P41240; H3E | CSK_HUMAN T | Unmodified | 0 | 0 | 0 | 0 | 0 | 0 | 0 | 0 | 0 | 0 | 0 | 741440 | 0 | 0 | 0 | 0 | 0 | 0 |
| GDVQAQV | SEL1L | Q9UBV2 | SE1L1_HUMAN | Unmodified | 0 | 0 | 0 | 0 | 0 | 0 | 0 | 0 | 10158000 | 0 | 0 | 0 | 0 | 0 | 0 | 0 | 0 | 0 |
| GDVSNLDF | SLC7A5 | Q01650 | LAT1_HUMAN I | Unmodified | 162840 | 0 | 0 | 0 | 0 | 0 | 0 | 0 | 0 | 0 | 0 | 1032700 | 0 | 0 | 363960 | 0 | 0 | 0 |
| GDVTAEEA | MARCKSL1 | P49006 | MRP_HUMAN I | Unmodified | 1143500 | 1958900 | 11144000 | 9274900 | 0 | 0 | 0 | 146990 | 30084000 | 2171200 | 55187000 | 1.08E+08 | 0 | 493650 | 18459000 | 12543000 | 59288 | 0 |
| GDVTAQIAI | SOD2; SOD2; S | P04179; F5G | SODM_HUMAN | Unmodified | 0 | 0 | 0 | 0 | 0 | 0 | 0 | 0 | 2152900 | 162520 | 0 | 0 | 0 | 0 | 0 | 0 | 0 | 0 |
| GDVVLQSD | MYO1C; MYO1C | O00159; F5H | MYO1C_HUMA | Unmodified | 1148700 | 1154400 | 2837300 | 2852600 | 4321000 | 637160 | 0 | 0 | 373350 | 0 | 277870 | 671640 | 3079700 | 0 | 3752600 | 2273800 | 635740 | 1006600 |
| GDVVPKDV | TUBA3C; TUBA | P0DPH7; Q7 | TBA3C_HUMAN | Unmodified | 0 | 0 | 0 | 202240 | 1094400 | 205790 | 784350 | 725710 | 74643 | 0 | 0 | 728140 | 54533 | 0 | 195140 | 14990 | 0 | 0 |
| GDWSQFTI | ASPH | Q12797 | ASPH_HUMAN | Unmodified | 0 | 0 | 0 | 0 | 0 | 0 | 0 | 0 | 461470 | 0 | 0 | 0 | 0 | 0 | 0 | 0 | 0 | 0 |
| GDYAPILQI | DPP3; DPP3; DI | Q9NY33; G3 | DPP3_HUMAN | Unmodified | 0 | 0 | 0 | 0 | 0 | 0 | 130620 | 0 | 123560 | 0 | 0 | 2343600 | 0 | 0 | 0 | 0 | 345770 | 0 |
| GDYYRYLA | YWHAG | P61981 | 1433G_HUMAN | Unmodified | 0 | 0 | 96690 | 61036 | 540660 | 186830 | 0 | 0 | 0 | 0 | 62514 | 0 | 728340 | 504400 | 0 | 0 | 754670 | 402760 |
| GEAAAERP | MARCKS | P29966 | MARCS_HUMA | Unmodified | 480970 | 388800 | 3370500 | 1028200 | 0 | 0 | 0 | 47087 | 19278000 | 1330200 | 3171000 | 14760000 | 216380 | 86057 | 2115800 | 2057600 | 194100 | 1124400 |
| GEADILAAL | TRIM26; TRIM26 | Q12899; A0A | TRI26_HUMAN | Unmodified | 3233600 | 3915500 | 11831000 | 2715800 | 0 | 0 | 0 | 0 | 0 | 0 | 0 | 0 | 0 | 0 | 7887000 | 3692100 | 127450 | 0 |
| GEADILAAL | TRIM26; TRIM26 | Q12899; A0A | TRI26_HUMAN | Unmodified | 328230 | 0 | 0 | 0 | 0 | 0 | 0 | 0 | 0 | 0 | 0 | 0 | 0 | 0 | 0 | 0 | 0 | 0 |
| GEAEAFAIC | FLOT1; FLOT1; | O75955; A0A | FLOT1_HUMAN | Unmodified | 0 | 0 | 213520 | 220650 | 0 | 0 | 0 | 0 | 830540 | 79449 | 0 | 540560 | 0 | 0 | 301410 | 198480 | 0 | 0 |
| GEAEDMLN | PLCG2; PLCG2 | P16885; A0A | PLCG2_HUMAI | 2 Oxidation (M) | 0 | 0 | 0 | 0 | 0 | 0 | 0 | 0 | 0 | 0 | 0 | 414100 | 0 | 0 | 0 | 0 | 0 | 0 |
| GEAEDMLN | PLCG2; PLCG2 | P16885; A0A | PLCG2_HUMAI | Oxidation (M) | 0 | 0 | 0 | 0 | 0 | 0 | 0 | 0 | 0 | 0 | 0 | 2176000 | 0 | 0 | 0 | 0 | 0 | 0 |
| GEAEDMLN | PLCG2; PLCG2 | P16885; A0A | PLCG2_HUMAI | Unmodified | 0 | 0 | 0 | 0 | 0 | 0 | 0 | 0 | 0 | 0 | 0 | 4802200 | 0 | 0 | 0 | 0 | 0 | 0 |
| GEAGAGA | HGH1; HGH1 | Q9BTY7; E9F | HGH1_HUMAN | Acetyl (Protein N | 0 | 0 | 0 | 0 | 0 | 0 | 0 | 0 | 0 | 0 | 0 | 0 | 0 | 0 | 0 | 0 | 265500 | 187210 |
| GEAGAQGI | COL2A1 | P02458 | CO2A1_HUMAI | Unmodified | 0 | 0 | 407240 | 0 | 0 | 0 | 9580.7 | 0 | 0 | 0 | 502940 | 0 | 0 | 0 | 0 | 0 | 0 | 0 |

| | | | | | | | | | | | | | | | | | | | | | | |
|---|---|---|---|---|---|---|---|---|---|---|---|---|---|---|---|---|---|---|---|---|---|---|
| GEAGKFEA | ANK3; ANK3; AI | Q12955; H0Y | ANK3_HUMAN | Unmodified | 0 | 0 | 76757 | 545830 | 0 | 0 | 0 | 0 | 0 | 0 | 0 | 144700 | 114210 | 119570 | 902650 | 570230 | 0 | 0 |
| GEAGVPAE | FLNB; FLNB | O75369; E7E | FLNB_HUMAN | Unmodified | 0 | 0 | 0 | 0 | 0 | 0 | 0 | 0 | 0 | 0 | 0 | 2553300 | 0 | 0 | 0 | 0 | 0 | 0 |
| GEAVGVH( | DUSP23 | Q9BVJ7 | DUS23_HUMAI | Unmodified | 267150 | 123110 | 0 | 0 | 0 | 0 | 0 | 0 | 0 | 0 | 0 | 1901200 | 0 | 0 | 0 | 0 | 0 | 0 |
| GEAWALMI | SPON2; SPON2 | Q9BUD6; D6 | SPON2_HUMAI | Oxidation (M) | 0 | 0 | 427400 | 0 | 0 | 0 | 0 | 0 | 0 | 0 | 0 | 0 | 0 | 0 | 471630 | 311380 | 0 | 0 |
| GECLCGQ( | ITGB3; N/A | P05106; H3E | ITB3_HUMAN I | Unmodified | 0 | 0 | 0 | 519830 | 0 | 0 | 82587 | 0 | 0 | 0 | 0 | 0 | 0 | 47220 | 0 | 0 | 0 | 364930 |
| GECWCVN | IGFBP2; IGFBP2 | P18065; C9J | IBP2_HUMAN I | Unmodified | 0 | 0 | 0 | 0 | 0 | 0 | 0 | 0 | 1263100 | 189780 | 0 | 0 | 0 | 0 | 0 | 0 | 0 | 0 |
| GEDEEENN | RPN1; RPN1 | P04843; F8W | RPN1_HUMAN | Unmodified | 0 | 0 | 0 | 0 | 0 | 0 | 0 | 0 | 4691000 | 1539100 | 371960 | 263530 | 0 | 0 | 0 | 0 | 0 | 0 |
| GEDEEENN | RPN1; RPN1 | P04843; F8W | RPN1_HUMAN | Unmodified | 0 | 0 | 61035 | 0 | 0 | 0 | 0 | 0 | 4572500 | 0 | 0 | 0 | 0 | 0 | 0 | 0 | 0 | 0 |
| GEDFPANN | SSR1; SSR1; SS | P43307; C9J | SSRA_HUMAN | Unmodified | 0 | 0 | 0 | 0 | 0 | 0 | 0 | 0 | 5392600 | 0 | 0 | 0 | 0 | 0 | 215800 | 0 | 0 | 0 |
| GEDPLAGD | SRGAP2; SRGA | O75044; Q5\ | SRGP2_HUMAI | Oxidation (M) | 0 | 0 | 0 | 0 | 0 | 0 | 0 | 0 | 0 | 0 | 0 | 944290 | 0 | 0 | 0 | 0 | 0 | 0 |
| GEDQSELV | LEMD2; LEMD2 | Q8NC56; D6 | LEMD2_HUMAI | Unmodified | 0 | 0 | 0 | 0 | 0 | 0 | 18445 | 0 | 1976000 | 0 | 0 | 0 | 0 | 0 | 0 | 0 | 0 | 0 |
| GEDSVPDT | DDX1; DDX1; D | Q92499; A0A | DDX1_HUMAN | Unmodified | 0 | 0 | 0 | 0 | 0 | 0 | 0 | 0 | 0 | 0 | 0 | 2041600 | 0 | 0 | 0 | 0 | 0 | 0 |
| GEDVNRTL | MTPN | P58546 | MTPN_HUMAN | Unmodified | 173170 | 160790 | 0 | 290020 | 0 | 0 | 0 | 0 | 34107 | 0 | 0 | 244200 | 0 | 0 | 129310 | 0 | 0 | 0 |
| GEEDNSLS | CD82 | P27701 | CD82_HUMAN | Unmodified | 0 | 0 | 0 | 0 | 0 | 0 | 0 | 0 | 0 | 81803 | 0 | 1367800 | 0 | 0 | 543550 | 303980 | 0 | 0 |
| GEEDNSLS | CD82 | P27701 | CD82_HUMAN | Unmodified | 1317600 | 545730 | 3351400 | 0 | 0 | 0 | 0 | 0 | 0 | 0 | 0 | 2717600 | 0 | 0 | 3792400 | 349340 | 0 | 0 |
| GEELLSPLN | MYO1C; MYO1( | O00159; F5H | MYO1C_HUMA | Unmodified | 4747900 | 3059800 | 4520000 | 2250800 | 132170 | 0 | 0 | 0 | 395740 | 0 | 0 | 1321100 | 0 | 0 | 21117000 | 12525000 | 100790 | 242360 |
| GEELLSPLN | MYO1C; MYO1( | O00159; F5H | MYO1C_HUMA | Unmodified | 0 | 0 | 0 | 85524 | 0 | 0 | 0 | 0 | 0 | 0 | 0 | 0 | 0 | 0 | 0 | 0 | 0 | 0 |
| GEELSCEEI | SFN | P31947 | 1433S_HUMAN | Unmodified | 91263 | 0 | 0 | 0 | 0 | 0 | 0 | 0 | 414850 | 0 | 0 | 0 | 0 | 0 | 0 | 0 | 0 | 0 |
| GEELSCEEI | SFN | P31947 | 1433S_HUMAN | Unmodified | 293940 | 213960 | 0 | 234850 | 2325700 | 0 | 0 | 0 | 1808600 | 285820 | 0 | 0 | 0 | 0 | 0 | 159620 | 412630 | 0 |
| GEENLMDA | CCT8; CCT8 | P50990; H7( | TCPQ_HUMAN | Unmodified | 0 | 0 | 0 | 138750 | 0 | 0 | 0 | 0 | 332660 | 0 | 1340300 | 4046200 | 0 | 0 | 0 | 0 | 0 | 0 |
| GEFEGFVA | DPP3; DPP3; DI | Q9NY33; G3 | DPP3_HUMAN | Unmodified | 0 | 0 | 0 | 0 | 0 | 0 | 0 | 0 | 0 | 0 | 0 | 959510 | 0 | 0 | 0 | 0 | 0 | 0 |
| GEFTIETEG | GARS1; GARS1 | P41250; A0A | GARS_HUMAN | Unmodified | 0 | 0 | 0 | 0 | 0 | 0 | 0 | 0 | 0 | 62319 | 714770 | 4616100 | 124820 | 0 | 0 | 0 | 0 | 0 |
| GEFVTTVQ( | MDH1; MDH1; I | P40925; B8Z | MDHC_HUMAN | Unmodified | 0 | 0 | 0 | 0 | 1085300 | 0 | 0 | 142830 | 0 | 0 | 4279100 | 11686000 | 0 | 0 | 0 | 0 | 0 | 562300 |
| GEFVTTVQ( | MDH1; MDH1; I | P40925; B8Z | MDHC_HUMAN | Unmodified | 0 | 0 | 0 | 0 | 8625100 | 1516100 | 2198000 | 1059300 | 0 | 1695000 | 0 | 0 | 1663600 | 2945700 | 0 | 0 | 4185300 | 7527700 |
| GEGDIHEN | JPT1 | Q9UK76 | JUPI1_HUMAN | Unmodified | 0 | 0 | 754870 | 0 | 0 | 0 | 0 | 0 | 0 | 0 | 0 | 0 | 0 | 0 | 504780 | 329160 | 0 | 0 |
| GEGESPPV | MARCKSL1 | P49006 | MRP_HUMAN I | Unmodified | 345590 | 553170 | 7354100 | 2570300 | 0 | 0 | 371090 | 0 | 3225100 | 547610 | 35137000 | 57366000 | 0 | 0 | 7145300 | 5636500 | 0 | 0 |
| GEGESPPV | MARCKSL1 | P49006 | MRP_HUMAN I | Unmodified | 0 | 0 | 0 | 294480 | 0 | 0 | 0 | 0 | 0 | 0 | 0 | 11325000 | 0 | 0 | 2514600 | 0 | 0 | 0 |
| GEGESPPV | MARCKSL1 | P49006 | MRP_HUMAN I | Unmodified | 0 | 0 | 0 | 0 | 0 | 0 | 0 | 0 | 371340 | 0 | 1482400 | 4580000 | 0 | 0 | 0 | 859230 | 0 | 0 |
| GEGLPNFD | DNAJB11; N/A | Q9UBS4; H7 | DJB11_HUMAN | Unmodified | 20478000 | 693310 | 0 | 0 | 0 | 0 | 0 | 0 | 2183600 | 0 | 270030 | 0 | 0 | 0 | 272340 | 142050 | 0 | 0 |
| GEGMSQA | FAH; FAH | P16930; H3E | FAAA_HUMAN | Unmodified | 0 | 0 | 0 | 0 | 0 | 0 | 0 | 0 | 0 | 0 | 0 | 285640 | 0 | 0 | 0 | 0 | 41229 | 0 |
| GEGNETTN | L1CAM; L1CAM | P32004; H0Y | L1CAM_HUMAI | Oxidation (M) | 0 | 0 | 92529000 | 0 | 0 | 1.11E+08 | 6077100 | 0 | 1768200 | 5611900 | 5822400 | 10443000 | 0 | 0 | 26260000 | 0 | 0 | 0 |
| GEGPDVHI | AHNAK | Q09666 | AHNK_HUMAN | Unmodified | 0 | 0 | 0 | 0 | 362330 | 0 | 0 | 0 | 0 | 0 | 0 | 0 | 0 | 0 | 0 | 0 | 0 | 1928300 |
| GEGPEFDV | AHNAK | Q09666 | AHNK_HUMAN | Unmodified | 0 | 0 | 0 | 0 | 2105600 | 0 | 0 | 0 | 0 | 0 | 0 | 0 | 0 | 0 | 0 | 0 | 975640 | 1038700 |
| GEGQLGPA | EEF2 | P13639 | EF2_HUMAN EI | Unmodified | 1893200 | 1476200 | 1811800 | 3402600 | 483260 | 138130 | 0 | 37470 | 980870 | 200410 | 3771600 | 46974000 | 0 | 0 | 2499900 | 2666700 | 0 | 88247 |
| GEGVEIVDI | PPFIBP1; PPFIB | Q86W92; F5 | LIPB1_HUMAN | Unmodified | 0 | 0 | 0 | 0 | 0 | 0 | 0 | 0 | 0 | 0 | 0 | 0 | 0 | 0 | 0 | 0 | 0 | 0 |
| GEHPGLSI( | HMGB1P1; HM( | B2RPK0; P09 | HGB1A_HUMA | Unmodified | 0 | 0 | 0 | 0 | 0 | 0 | 52537 | 24071 | 0 | 68415 | 1909300 | 983730 | 0 | 0 | 0 | 0 | 0 | 0 |
| GEHSIVYLK | FKBP4 | Q02790 | FKBP4_HUMAN | Unmodified | 575820 | 282190 | 1149000 | 1934200 | 0 | 0 | 0 | 0 | 1187000 | 0 | 2882800 | 24475000 | 2516200 | 0 | 2621700 | 1892400 | 339850 | 0 |
| GEHSIVYLK | FKBP4 | Q02790 | FKBP4_HUMAN | Unmodified | 0 | 0 | 0 | 0 | 0 | 0 | 352100 | 51225 | 0 | 169370 | 182560 | 0 | 0 | 0 | 0 | 0 | 535590 | 0 |
| GEIEHHCS | GLG1; GLG1 | Q92896; H3I | GSLG1_HUMAI | Unmodified | 0 | 0 | 0 | 0 | 0 | 0 | 0 | 15797 | 674990 | 0 | 0 | 0 | 0 | 0 | 0 | 0 | 0 | 0 |
| GEIEQERID | ATP2B2; ATP2B | Q01814; P2( | AT2B2_HUMAN | Unmodified | 0 | 0 | 0 | 315510 | 449110 | 0 | 0 | 83078 | 0 | 0 | 0 | 0 | 1407900 | 1425900 | 0 | 0 | 202520 | 0 |
| GEIEQERLI | ATP2B3 | Q16720 | AT2B3_HUMAN | Unmodified | 0 | 0 | 0 | 0 | 0 | 0 | 0 | 0 | 0 | 0 | 0 | 1238900 | 0 | 0 | 0 | 0 | 0 | 0 |
| GEIEQRPTI | FARP1; FARP1; | Q9Y4F1; A0A | FARP1_HUMAN | Acetyl (Protein N | 0 | 0 | 0 | 0 | 0 | 55350 | 0 | 0 | 8087600 | 1186700 | 969090 | 3822400 | 3017700 | 0 | 219810 | 0 | 0 | 0 |
| GEIEQRPTI | FARP1; FARP1; | Q9Y4F1; A0A | FARP1_HUMAN | Acetyl (Protein N | 0 | 0 | 0 | 0 | 580210 | 0 | 0 | 0 | 0 | 1112900 | 0 | 0 | 0 | 0 | 0 | 0 | 0 | 804980 |
| GEILLRRTA | PTGFRN | Q9P2B2 | FPRP_HUMAN | Unmodified | 0 | 0 | 0 | 0 | 0 | 0 | 1190500 | 0 | 0 | 0 | 0 | 0 | 2008900 | 0 | 0 | 0 | 0 | 0 |
| GEILQIVNR | PALS2; PALS2; I | Q9NZW5; B8 | PALS2_HUMAN | Unmodified | 0 | 0 | 0 | 0 | 0 | 0 | 0 | 0 | 0 | 0 | 0 | 3050800 | 1973200 | 0 | 879590 | 541600 | 0 | 0 |
| GEISSHELN | SLC9A6; SLC9A | A0A6Q8PFS7 | A0A6Q8PFS7_I | Oxidation (M) | 0 | 0 | 82968 | 31985 | 0 | 0 | 0 | 0 | 1634100 | 127130 | 0 | 0 | 0 | 0 | 78366 | 0 | 0 | 0 |
| GEISSHELN | SLC9A6; SLC9A | A0A6Q8PFS7 | A0A6Q8PFS7_I | Unmodified | 0 | 0 | 63219 | 0 | 0 | 0 | 0 | 0 | 3558600 | 359940 | 0 | 368370 | 0 | 0 | 127790 | 0 | 0 | 0 |
| GEIVIGGQI | ACSL4; ACSL4 | O60488; H0Y | ACSL4_HUMAN | Oxidation (M) | 0 | 0 | 0 | 0 | 0 | 0 | 0 | 207220 | 0 | 0 | 400600 | 3133900 | 0 | 0 | 0 | 0 | 0 | 0 |
| GEKLDDLV | YKT6; YKT6 | O15498; A0A | YKT6_HUMAN S | Unmodified | 0 | 0 | 0 | 0 | 0 | 0 | 0 | 74380 | 181790 | 78662 | 0 | 0 | 244600 | 119480 | 0 | 0 | 132240 | 168450 |
| GELGCFLS | COLGALT1; CO | Q8NBJ5; M0 | GT251_HUMAN | Unmodified | 0 | 0 | 0 | 0 | 0 | 0 | 0 | 24136 | 4211000 | 306490 | 116960 | 0 | 0 | 0 | 0 | 0 | 0 | 0 |
| GELGCFLS | COLGALT1; CO | Q8NBJ5; M0 | GT251_HUMAN | Unmodified | 0 | 0 | 0 | 0 | 0 | 0 | 0 | 0 | 165990 | 0 | 0 | 0 | 0 | 0 | 0 | 0 | 0 | 0 |
| GELPAWAA | PROM2 | Q8N271 | PROM2_HUMA | Unmodified | 0 | 80545 | 1310300 | 0 | 0 | 0 | 0 | 0 | 0 | 0 | 0 | 154920 | 0 | 0 | 1667300 | 121410 | 0 | 0 |
| GELSGDFE | ANXA6 | P08133 | ANXA6_HUMAI | Unmodified | 89415 | 72790 | 1463700 | 0 | 0 | 0 | 894800 | 509420 | 235050 | 304430 | 0 | 505760 | 0 | 0 | 1071200 | 712930 | 0 | 0 |
| GELSTLLYN | PIGT; PIGT; PIG | Q969N2; A0A | PIGT_HUMAN ( | Unmodified | 0 | 0 | 0 | 0 | 0 | 36587 | 0 | 0 | 4390200 | 395700 | 0 | 2365600 | 0 | 0 | 0 | 0 | 0 | 0 |
| GELTTLIHQ | KTN1 | Q86UP2 | KTN1_HUMAN | Unmodified | 0 | 0 | 0 | 0 | 0 | 0 | 0 | 0 | 121010 | 0 | 0 | 969370 | 40330 | 0 | 0 | 0 | 0 | 0 |
| GELVGGLD | GLRX3 | O76003 | GLRX3_HUMAI | Unmodified | 0 | 0 | 0 | 0 | 3164400 | 0 | 105960 | 245480 | 748810 | 0 | 1476200 | 9815800 | 0 | 0 | 0 | 0 | 0 | 0 |
| GEMMDLQ | LDHB; LDHB; LI | P07195; A0A | LDHB_HUMAN | 2 Oxidation (M) | 0 | 0 | 427770 | 0 | 0 | 0 | 0 | 0 | 0 | 0 | 705290 | 7691100 | 0 | 0 | 0 | 404560 | 0 | 0 |
| GEMMDLQ | LDHB; LDHB; LI | P07195; A0A | LDHB_HUMAN | Oxidation (M) | 0 | 0 | 0 | 0 | 0 | 0 | 0 | 0 | 62040 | 0 | 125370 | 1230500 | 0 | 0 | 0 | 0 | 0 | 0 |
| GEMMDLQ | LDHA; LDHA; LI | P00338; F5G | LDHA_HUMAN | Oxidation (M) | 93933 | 53585 | 0 | 0 | 0 | 0 | 49744 | 28568 | 0 | 0 | 170160 | 11004000 | 183690 | 137320 | 267500 | 270580 | 0 | 0 |
| GEMMDLQ | LDHA; LDHA; LI | P00338; F5G | LDHA_HUMAN | 2 Oxidation (M) | 0 | 470630 | 1005300 | 649940 | 0 | 0 | 135720 | 0 | 0 | 0 | 240950 | 6930200 | 0 | 0 | 0 | 1575300 | 0 | 0 |
| GEMMDLQ | LDHA; LDHA; LI | P00338; F5G | LDHA_HUMAN | Unmodified | 39301 | 0 | 0 | 0 | 0 | 0 | 0 | 0 | 0 | 0 | 0 | 409780 | 0 | 0 | 0 | 0 | 0 | 0 |
| GENGTLSR | CHP1; CHP1; C | Q99653; F5( | CHP1_HUMAN | Unmodified | 0 | 0 | 61901 | 0 | 0 | 0 | 0 | 0 | 0 | 0 | 0 | 1406200 | 0 | 0 | 1616500 | 122440 | 0 | 0 |
| GENLEHLR | VAMP8; VAMP8 | Q9BV40; B8Z | VAMP8_HUMAI | Unmodified | 0 | 0 | 0 | 0 | 0 | 569630 | 0 | 0 | 742040 | 0 | 10041 | 365270 | 0 | 238010 | 0 | 0 | 0 | 0 |
| GENPLADD | SRGAP1; SRGA | Q7Z6B7; G5I | SRGP1_HUMAI | Unmodified | 105330 | 0 | 0 | 0 | 0 | 0 | 0 | 0 | 0 | 0 | 0 | 595810 | 0 | 0 | 0 | 0 | 0 | 0 |
| GENSWFS1 | SLC3A2; SLC3A | P08195; A0A | 4F2_HUMAN 4I | Unmodified | 4079600 | 1811700 | 1647500 | 669200 | 0 | 0 | 0 | 0 | 3632800 | 347330 | 1967800 | 48379000 | 0 | 0 | 3961400 | 3240300 | 0 | 0 |
| GEPAAAAA | MARCKS | P29966 | MARCS_HUMA | Unmodified | 4724500 | 3318000 | 14647000 | 17740000 | 21996 | 0 | 1570100 | 490630 | 2.27E+08 | 14680000 | 19140000 | 73573000 | 1955900 | 1408100 | 12480000 | 13233000 | 1660700 | 13643000 |
| GEPAAAAA | MARCKS | P29966 | MARCS_HUMA | Unmodified | 0 | 0 | 0 | 0 | 0 | 0 | 0 | 0 | 0 | 622480 | 0 | 0 | 0 | 0 | 0 | 0 | 0 | 0 |
| GEPGAAEP | KIAA2013 | Q8IYS2 | K2013_HUMAN | Unmodified | 0 | 0 | 0 | 0 | 0 | 0 | 0 | 0 | 4363100 | 577630 | 0 | 456150 | 0 | 0 | 0 | 0 | 0 | 0 |
| GEPGRPAL | DNPH1; DNPH1 | O43598; H0Y | DNPH1_HUMA | Unmodified | 99410 | 238060 | 0 | 0 | 0 | 0 | 0 | 0 | 0 | 0 | 0 | 4397300 | 0 | 0 | 461670 | 245740 | 0 | 0 |
| GEPHFIAV( | HLA-C; HLA-C; | Q53YP1; A0A | Q53YP1_HUMA | Unmodified | 0 | 0 | 0 | 36101 | 0 | 0 | 0 | 0 | 0 | 0 | 0 | 642490 | 0 | 0 | 0 | 223360 | 0 | 806520 |
| GEPLGVICI | DIP2B | Q9P265 | DIP2B_HUMAN | Unmodified | 0 | 0 | 0 | 0 | 0 | 0 | 0 | 0 | 376700 | 0 | 0 | 277860 | 0 | 0 | 0 | 0 | 0 | 0 |
| GEPLHIINS | VPS13C; VPS13 | Q709C8; A0A | VP13C_HUMAN | Unmodified | 0 | 0 | 0 | 0 | 0 | 0 | 0 | 0 | 0 | 239710 | 0 | 774220 | 0 | 0 | 0 | 0 | 0 | 289660 |
| GEPLTAGV | DIP2A; DIP2A | Q14689; A0A | DIP2A_HUMAN | Unmodified | 142430 | 4769700 | 1885100 | 2393000 | 0 | 0 | 0 | 0 | 656450 | 331580 | 43688 | 733620 | 0 | 0 | 531590 | 5267600 | 0 | 0 |
| GEPPHSEL | STK24; STK24; S | Q9Y6E0; H0Y | STK24_HUMAN | Unmodified | 0 | 0 | 0 | 0 | 0 | 0 | 0 | 0 | 0 | 0 | 88685 | 522360 | 46446 | 0 | 0 | 18946 | 0 | 0 |
| GEPPLPEEI | EIF3H; EIF3H; E | O15372; B3K | EIF3H_HUMAN | Unmodified | 0 | 0 | 0 | 0 | 0 | 0 | 0 | 0 | 0 | 0 | 346490 | 1783300 | 0 | 0 | 0 | 0 | 0 | 0 |
| GEPRAPW | HLA-C; HLA-C; | P10321; Q5( | HLAC_HUMAN | Unmodified | 0 | 0 | 0 | 0 | 1359400 | 0 | 0 | 0 | 3358200 | 1433900 | 319400 | 0 | 0 | 0 | 0 | 0 | 0 | 819480 |
| GEQGYGFI | SLC9A3R2 | Q15599 | NHRF2_HUMAI | Unmodified | 0 | 0 | 0 | 434550 | 0 | 0 | 0 | 0 | 0 | 0 | 0 | 0 | 226060 | 0 | 0 | 481410 | 127100 | 725540 |
| GEQMNSA | MSH2; MSH2; I | P43246; A0A | MSH2_HUMAN | Unmodified | 3706800 | 21893000 | 0 | 0 | 0 | 0 | 0 | 0 | 0 | 0 | 0 | 0 | 824470 | 0 | 869140 | 635750 | 0 | 0 |
| GEQPIFSTI | HOMER1 | Q86YM7 | HOME1_HUMA | Acetyl (Protein N | 0 | 0 | 0 | 0 | 0 | 0 | 0 | 32948 | 0 | 0 | 0 | 0 | 1371200 | 2678400 | 0 | 0 | 0 | 0 |
| GEQPIFTTI | HOMER2 | Q9NSB8 | HOME2_HUMA | Acetyl (Protein N | 0 | 0 | 0 | 0 | 0 | 0 | 393970 | 26301 | 0 | 0 | 0 | 0 | 6395000 | 4208000 | 0 | 0 | 0 | 0 |
| GEQPIFTTI | HOMER2 | Q9NSB8 | HOME2_HUMA | Acetyl (Protein N | 0 | 0 | 0 | 0 | 0 | 0 | 0 | 0 | 0 | 0 | 0 | 0 | 698080 | 1090300 | 0 | 0 | 0 | 0 |
| GEQQPISS | NHLRC2 | Q8NBF2 | NHLC2_HUMA | Unmodified | 0 | 0 | 0 | 0 | 0 | 0 | 0 | 0 | 0 | 0 | 0 | 427520 | 0 | 0 | 0 | 0 | 0 | 0 |
| GERDSQT( | GPS1; GPS1; GI | Q13098; A8K | CSN1_HUMAN | Unmodified | 0 | 195950 | 0 | 0 | 0 | 0 | 0 | 0 | 0 | 0 | 0 | 982780 | 0 | 0 | 0 | 0 | 0 | 0 |
| GERNFHVF | MYO1B; MYO1E | O43795; E9F | MYO1B_HUMA | Unmodified | 0 | 0 | 0 | 0 | 3391400 | 0 | 0 | 0 | 230990 | 162230 | 144590 | 0 | 35356 | 50556 | 0 | 0 | 854780 | 0 |
| GESDPAYQ | PDIA4; PDIA4 | P13667; A0A | PDIA4_HUMAN | Unmodified | 0 | 0 | 0 | 0 | 0 | 0 | 0 | 0 | 2926200 | 0 | 899820 | 0 | 0 | 0 | 0 | 0 | 0 | 755060 |
| GESFYQDF | RARS1 | P54136 | SYRC_HUMAN | Unmodified | 0 | 0 | 0 | 0 | 0 | 0 | 0 | 0 | 0 | 0 | 0 | 1043800 | 0 | 0 | 0 | 0 | 0 | 0 |
| GESGQSW | PTGES3; PTGES | Q15185; A0A | TEBP_HUMAN I | Unmodified | 0 | 0 | 812850 | 0 | 0 | 125950 | 0 | 0 | 3766900 | 152810 | 1222000 | 6891600 | 0 | 0 | 778620 | 787760 | 0 | 0 |
| GESGQSW | PTGES3; PTGES | Q15185; A0A | TEBP_HUMAN I | Unmodified | 0 | 0 | 0 | 0 | 2813600 | 0 | 0 | 0 | 588880 | 415170 | 186160 | 0 | 1019300 | 627780 | 0 | 0 | 324770 | 1581300 |
| GESLFHSK | EPCAM; EPCAM | P16422; B5N | EPCAM_HUMA | Unmodified | 0 | 0 | 0 | 0 | 0 | 0 | 0 | 0 | 0 | 0 | 0 | 2859800 | 0 | 0 | 0 | 0 | 0 | 0 |
| GESLSLPGI | NIBAN1 | Q9BZQ8 | NIBA1_HUMAN | Unmodified | 44978 | 0 | 176840 | 204990 | 0 | 0 | 43672 | 25486 | 4622700 | 671340 | 0 | 0 | 0 | 0 | 193210 | 162320 | 0 | 0 |
| GESPVDYD | PDIA6 | Q15084 | PDIA6_HUMAN | Unmodified | 0 | 0 | 0 | 0 | 0 | 0 | 0 | 0 | 8480100 | 191810 | 833740 | 1117700 | 0 | 0 | 0 | 0 | 0 | 0 |
| GESPVDYD | PDIA6 | Q15084 | PDIA6_HUMAN | Unmodified | 0 | 0 | 229500 | 0 | 0 | 0 | 0 | 0 | 20363000 | 134150 | 765300 | 883210 | 0 | 0 | 0 | 131150 | 0 | 0 |
| GESQTDIEI | CCT3; CCT3 | P49368; B4D | TCPG_HUMAN | Unmodified | 0 | 0 | 0 | 0 | 0 | 0 | 0 | 0 | 610240 | 0 | 456690 | 7555200 | 0 | 0 | 0 | 0 | 0 | 0 |
| GETGLFPA | SNX33 | Q8WV41 | SNX33_HUMAI | Unmodified | 2031200 | 10166000 | 6187600 | 2198400 | 0 | 0 | 0 | 0 | 0 | 0 | 11165 | 0 | 0 | 0 | 30834000 | 17493000 | 0 | 0 |
| GETPRVEE | SCARB2; SCARI | Q14108; A0A | SCRB2_HUMAI | Unmodified | 0 | 0 | 0 | 0 | 0 | 0 | 0 | 0 | 1049700 | 723080 | 0 | 531260 | 0 | 0 | 288880 | 112070 | 120940 | 0 |
| GEVERGP( | TMEM94 | Q12767 | TMM94_HUMAI | Unmodified | 0 | 0 | 0 | 47296 | 67586 | 0 | 29400 | 20065 | 134810 | 0 | 25538 | 289520 | 0 | 0 | 37086 | 0 | 0 | 74397 |
| GEVFRVVD | TJP2; TJP2; TJP2 | Q9UDY2; A0 | ZO2_HUMAN T | Unmodified | 0 | 0 | 0 | 60579 | 0 | 0 | 0 | 39746 | 668610 | 0 | 0 | 0 | 0 | 0 | 0 | 0 | 81274 | 0 |
| GEVGLLDN | PIPOX; PIPOX; I | Q9P0Z9; K7E | SOX_HUMAN P | Unmodified | 0 | 0 | 0 | 0 | 0 | 0 | 0 | 0 | 731780 | 0 | 0 | 0 | 0 | 0 | 0 | 0 | 0 | 0 |
| GEVLGDVL | NUMA1; NUMA | Q14980; F5H | NUMA1_HUMA | Unmodified | 0 | 0 | 0 | 0 | 0 | 12315 | 0 | 0 | 23265 | 0 | 23909 | 69163 | 0 | 0 | 0 | 0 | 0 | 0 |
| GEVPPKETI | MARCKSL1 | P49006 | MRP_HUMAN I | Unmodified | 0 | 0 | 0 | 0 | 0 | 0 | 0 | 0 | 914230 | 0 | 2070100 | 0 | 0 | 0 | 0 | 0 | 101210 | 1543200 |
| GEWTCKPI | FN1 | P02751 | FINC_HUMAN I | Unmodified | 0 | 0 | 3631100 | 0 | 146160 | 0 | 0 | 17215 | 0 | 0 | 0 | 0 | 0 | 0 | 0 | 31105 | 104380 | 277770 |
| GEYNVYST | PPP2R2A; PPP2 | P63151; Q6( | 2ABA_HUMAN | Unmodified | 0 | 0 | 0 | 0 | 0 | 0 | 0 | 0 | 0 | 0 | 0 | 1526300 | 0 | 0 | 945720 | 0 | 0 | 0 |
| GFAAMEET | MPRIP; MPRIP; | Q6WCQ1; H | MPRIP_HUMAN | Unmodified | 0 | 0 | 0 | 0 | 0 | 0 | 0 | 0 | 0 | 0 | 97708 | 661180 | 0 | 0 | 0 | 0 | 0 | 0 |
| GFAFVQYV | HNRNPC; HNR | P07910; G3\ | HNRPC_HUMA | Unmodified | 790720 | 398980 | 0 | 50226 | 699050 | 0 | 1324200 | 452430 | 1746900 | 661120 | 0 | 10225000 | 209360 | 0 | 0 | 305330 | 0 | 0 |
| GFAFVQYV | HNRNPC; HNR | P07910; G3\ | HNRPC_HUMA | Unmodified | 0 | 0 | 0 | 0 | 2216200 | 964640 | 4313500 | 912560 | 0 | 2268800 | 642910 | 0 | 2943300 | 1659800 | 0 | 0 | 0 | 2177400 |
| GFAFVTFDI | HNRNPA1L2; H | Q32P51; P09 | RA1L2_HUMAN | Unmodified | 815480 | 392300 | 721820 | 383350 | 638840 | 303560 | 235070 | 206630 | 2708800 | 843360 | 3491600 | 36812000 | 848590 | 0 | 873850 | 664990 | 204790 | 274820 |
| GFAFVTFDI | HNRNPA1L2; H | Q32P51; P09 | RA1L2_HUMAN | Unmodified | 0 | 0 | 0 | 0 | 1995700 | 432940 | 0 | 0 | 863720 | 2784900 | 0 | 3404000 | 2150500 | 0 | 0 | 0 | 0 | 562100 |
| GFAFVTFDI | HNRNPA3; HNI | P51991; A0A | ROA3_HUMAN | Unmodified | 92684 | 0 | 194810 | 247090 | 0 | 0 | 384820 | 203930 | 760850 | 254350 | 1169100 | 5104700 | 239210 | 177780 | 0 | 315610 | 76404 | 189800 |
| GFAFVTFDI | HNRNPA3; HNI | P51991; A0A | ROA3_HUMAN | Unmodified | 0 | 0 | 0 | 0 | 581330 | 0 | 2320300 | 146760 | 0 | 0 | 0 | 352940 | 1209800 | 0 | 0 | 0 | 327050 | 592540 |

| | | | | | | | | | | | | | | | | | | | | | | |
|---|---|---|---|---|---|---|---|---|---|---|---|---|---|---|---|---|---|---|---|---|---|---|
| GFAFVTFES | RBMXL3; RBMX | Q8N7X1; O7 | RMXL3_HUMAN | Unmodified | 0 | 0 | 0 | 0 | 0 | 0 | 0 | 0 | 0 | 0 | 0 | 6891700 | 0 | 0 | 0 | 0 | 0 | 0 |
| GFALLNFV | PLOD2; PLOD2 | O00469; E7E | PLOD2_HUMAN | Unmodified | 0 | 0 | 15371 | 17962 | 0 | 0 | 488800 | 0 | 762890 | 458880 | 0 | 106160 | 0 | 0 | 0 | 0 | 0 | 0 |
| GFALVGVG | CPSF6; CPSF6 | Q16630; F8V | CPSF6_HUMAN | Unmodified | 0 | 0 | 0 | 1545700 | 0 | 0 | 0 | 0 | 297510 | 182320 | 334900 | 1594300 | 1818700 | 0 | 0 | 0 | 0 | 0 |
| GFALVGVG | CPSF6; CPSF6 | Q16630; F8V | CPSF6_HUMAN | Unmodified | 1263800 | 85374 | 444520 | 498250 | 129680 | 0 | 0 | 0 | 0 | 88949 | 142220 | 0 | 860080 | 139420 | 0 | 487300 | 0 | 129010 |
| GFCFITFK | HNRNPD; HNR | Q14103; H0Y | HNRPD_HUMA | Unmodified | 0 | 0 | 0 | 0 | 0 | 0 | 0 | 0 | 374250 | 0 | 0 | 698780 | 0 | 0 | 0 | 0 | 0 | 0 |
| GFCFITFKE | HNRNPD; HNR | Q14103; H0Y | HNRPD_HUMA | Unmodified | 58415 | 0 | 0 | 0 | 141020 | 30628 | 188220 | 67317 | 579480 | 228240 | 0 | 4078500 | 177210 | 72237 | 0 | 0 | 0 | 143550 |
| GFCFITFKE | HNRNPD; HNR | Q14103; H0Y | HNRPD_HUMA | Unmodified | 0 | 0 | 0 | 0 | 0 | 0 | 0 | 0 | 0 | 0 | 0 | 6694000 | 0 | 0 | 0 | 0 | 0 | 0 |
| GFCFITYTD | HNRNPDL; HNI | O14979; A0A | HNRDL_HUMA | Unmodified | 0 | 0 | 180400 | 96504 | 0 | 2536900 | 0 | 0 | 303770 | 254670 | 0 | 1858900 | 84817 | 0 | 116730 | 70950 | 364010 | 0 |
| GFCQLVVS | SLC12A7; SLC1 | Q9Y666; A0A | S12A7_HUMAN | Unmodified | 0 | 0 | 0 | 0 | 0 | 0 | 0 | 0 | 0 | 0 | 0 | 1782300 | 0 | 0 | 0 | 0 | 0 | 0 |
| GFDEYMK | FABP5 | Q01469 | FABP5_HUMAN | Oxidation (M) | 130340 | 89093 | 0 | 0 | 1076500 | 0 | 0 | 0 | 0 | 0 | 0 | 1353500 | 0 | 0 | 0 | 0 | 1385800 | 3586400 |
| GFDEYMK | FABP5 | Q01469 | FABP5_HUMAN | Unmodified | 0 | 0 | 0 | 0 | 3844900 | 435980 | 0 | 0 | 148220 | 0 | 0 | 11629000 | 2759300 | 779880 | 0 | 0 | 2151900 | 5293100 |
| GFDILGIKP | RARS1 | P54136 | SYRC_HUMAN | Unmodified | 0 | 0 | 0 | 0 | 0 | 0 | 0 | 0 | 0 | 0 | 0 | 833000 | 0 | 0 | 0 | 0 | 0 | 0 |
| GFDPLLVEI | UBAP1; UBAP1 | Q9NZ09; A0A | UBAP1_HUMAN | Oxidation (M) | 2580000 | 2406200 | 165490 | 570360 | 0 | 0 | 0 | 0 | 0 | 0 | 0 | 0 | 210490 | 0 | 593870 | 411650 | 0 | 0 |
| GFDPNQLS | AGPS; AGPS; A( | O00116; A0A | ADAS_HUMAN | Unmodified | 0 | 0 | 0 | 0 | 0 | 0 | 0 | 0 | 1840000 | 655550 | 309630 | 0 | 0 | 0 | 0 | 0 | 0 | 0 |
| GFDQDMA | TXNRD1; TXNRI | Q16881; A0A | TRXR1_HUMAN | Unmodified | 0 | 65876 | 0 | 0 | 0 | 123850 | 0 | 0 | 128150 | 0 | 0 | 1974500 | 0 | 0 | 0 | 0 | 0 | 0 |
| GFDQTINLI | LSM8; LSM8; LS | O95777; C9J | LSM8_HUMAN | Unmodified | 0 | 0 | 0 | 0 | 0 | 0 | 0 | 14694 | 0 | 0 | 413730 | 1589400 | 0 | 0 | 0 | 0 | 143560 | 0 |
| GFDSFLGR | UTRN | P46939 | UTRN_HUMAN | Unmodified | 0 | 0 | 436500 | 283120 | 0 | 0 | 0 | 0 | 576580 | 0 | 0 | 105320 | 0 | 0 | 452060 | 491620 | 27747 | 0 |
| GFDTTAINK | ARHGEF7; ARH | Q14155; B1A | ARHG7_HUMA | Unmodified | 3558200 | 3232700 | 1359200 | 1011200 | 274180 | 0 | 85192 | 85962 | 947800 | 254380 | 0 | 443040 | 114760 | 0 | 1737000 | 1414700 | 142040 | 147160 |
| GFDVFNAL | NMT1; NMT2; N | P30419; O6( | NMT1_HUMAN | Unmodified | 0 | 0 | 0 | 0 | 0 | 0 | 182970 | 0 | 0 | 113220 | 0 | 0 | 0 | 0 | 0 | 0 | 0 | 0 |
| GFDWPLLV | GUSB | P08236 | BGLR_HUMAN | Unmodified | 0 | 0 | 475680 | 0 | 0 | 0 | 0 | 0 | 12640000 | 3716300 | 110920 | 164470 | 0 | 107850 | 473540 | 50522 | 0 | 0 |
| GFEAAVAA( | HMGCL | P35914 | HMGCL_HUMA | Unmodified | 0 | 0 | 0 | 0 | 259490 | 0 | 0 | 1131600 | 4832200 | 1650200 | 0 | 0 | 0 | 0 | 0 | 0 | 337470 | 0 |
| GFECKPHS | KLK11; KLK11; I | Q9UBX7; M0 | KLK11_HUMAN | Unmodified | 0 | 0 | 256030 | 0 | 233620 | 0 | 0 | 0 | 4094900 | 2024400 | 0 | 0 | 0 | 0 | 0 | 0 | 0 | 0 |
| GFEEQWYI | GUSB; GUSB; G | P08236; F8W | BGLR_HUMAN | Unmodified | 0 | 0 | 0 | 0 | 0 | 0 | 0 | 0 | 3435600 | 97802 | 0 | 0 | 0 | 0 | 0 | 0 | 0 | 0 |
| GFEKPTENS | ARPC5L | Q9BPX5 | ARP5L_HUMAN | Unmodified | 1087900 | 588550 | 1390400 | 1956300 | 9505800 | 1022200 | 2578000 | 1409600 | 1440000 | 1701300 | 1079500 | 8054300 | 5606100 | 2997700 | 2667100 | 2190100 | 4502300 | 4019200 |
| GFELLYQPI | ARVCF; ARVCF; | O00192; E9F | ARVC_HUMAN | Unmodified | 0 | 0 | 404900 | 0 | 0 | 0 | 17285 | 0 | 0 | 0 | 0 | 1715700 | 0 | 1754400 | 1452000 | 1010100 | 0 | 0 |
| GFELTFK | DDOST; DDOST | P39656; U3K | OST48_HUMAN | Unmodified | 0 | 0 | 79390 | 0 | 76211 | 0 | 633590 | 361850 | 9579200 | 2720300 | 0 | 987900 | 0 | 170550 | 0 | 0 | 0 | 122420 |
| GFESPSDN | ARPC5; ARPC5 | O15511; B1A | ARPC5_HUMAN | Oxidation (M) | 1170400 | 0 | 1750100 | 760510 | 2641100 | 0 | 732010 | 804230 | 0 | 0 | 902930 | 6435700 | 3439700 | 1655000 | 501480 | 577540 | 916640 | 546880 |
| GFESPSDN | ARPC5; ARPC5 | O15511; B1A | ARPC5_HUMAN | Unmodified | 0 | 0 | 0 | 0 | 3135900 | 309080 | 607260 | 0 | 1539700 | 176940 | 411270 | 1139000 | 2389900 | 1398100 | 0 | 0 | 609090 | 1138500 |
| GFETVSYLL | PIR | O00625 | PIR_HUMAN Pi | Oxidation (M) | 0 | 0 | 0 | 37944 | 0 | 0 | 0 | 0 | 0 | 0 | 0 | 2445000 | 0 | 0 | 0 | 0 | 0 | 0 |
| GFEVVYMT | HSP90AB1; HSI | P08238; Q58 | HS90B_HUMAN | Oxidation (M) | 0 | 0 | 0 | 0 | 0 | 1482900 | 0 | 0 | 1011900 | 695050 | 0 | 2431900 | 0 | 0 | 0 | 0 | 1397500 | 2368900 |
| GFFDPNTE | DSP | P15924 | DESP_HUMAN | Unmodified | 0 | 0 | 62023 | 0 | 413910 | 0 | 0 | 0 | 0 | 0 | 0 | 255020 | 123290 | 0 | 0 | 71275 | 1297600 | 273950 |
| GFFDPNTH | EPPK1; EPPK1 | P58107; A0A | EPIPL_HUMAN | Unmodified | 0 | 0 | 0 | 0 | 536640 | 0 | 0 | 0 | 2377100 | 0 | 0 | 0 | 1215600 | 0 | 0 | 0 | 0 | 0 |
| GFFEGTIAS | COPA; COPA; C | P53621; A0A | COPA_HUMAN | Unmodified | 0 | 0 | 628430 | 0 | 813890 | 679230 | 351220 | 336170 | 0 | 429710 | 440620 | 2961400 | 1307600 | 1185700 | 0 | 0 | 529100 | 1159200 |
| GFFICDQP | EPRS1; EPRS1 | P07814; V9C | SYEP_HUMAN | Unmodified | 0 | 0 | 97307 | 0 | 182000 | 0 | 196930 | 0 | 268630 | 92054 | 315950 | 2393200 | 3911400 | 914440 | 0 | 0 | 292920 | 506100 |
| GFFLVQTK | RP2 | O75695 | XRP2_HUMAN | Unmodified | 776600 | 805050 | 1092700 | 1226400 | 2896800 | 693570 | 584820 | 442650 | 3625000 | 3538500 | 1953500 | 13534000 | 4182300 | 2643800 | 2209000 | 2355600 | 3556700 | 6434700 |
| GFFSFFQR | COBLL1; COBLI | Q53SF7; A0A | COBL1_HUMAN | Unmodified | 3473400 | 0 | 404620 | 1004800 | 0 | 0 | 0 | 0 | 172920 | 0 | 0 | 0 | 2317600 | 1397600 | 0 | 0 | 0 | 0 |
| GFGDHIHV | TXNDC12; TXNI | O95881; V9( | TXD12_HUMAN | Unmodified | 0 | 0 | 0 | 0 | 0 | 272970 | 0 | 0 | 657850 | 208570 | 0 | 60593 | 0 | 0 | 0 | 0 | 0 | 0 |
| GFGDHIHV | TXNDC12; TXNI | O95881; V9( | TXD12_HUMAN | Unmodified | 0 | 0 | 0 | 0 | 910030 | 0 | 291370 | 325840 | 0 | 830330 | 0 | 0 | 0 | 0 | 0 | 0 | 0 | 194740 |
| GFGDLKSP | EEF1B2; EEF1B | P24534; F2Z | EF1B_HUMAN I | Unmodified | 0 | 0 | 0 | 0 | 17235000 | 1734800 | 3207100 | 135630 | 0 | 5284900 | 0 | 402810 | 7312000 | 3085000 | 0 | 0 | 13111000 | 5319500 |
| GFGFGLVK | VDAC2; VDAC2 | P45880; Q5J | VDAC2_HUMAI | Unmodified | 0 | 0 | 435430 | 196240 | 362160 | 0 | 0 | 289260 | 2.95E+08 | 36641000 | 1920400 | 0 | 297500 | 484490 | 0 | 0 | 0 | 0 |
| GFGFGLVK | VDAC2; VDAC2 | P45880; Q5J | VDAC2_HUMAI | Unmodified | 0 | 0 | 0 | 0 | 1269600 | 0 | 4635100 | 324320 | 68698000 | 1.94E+08 | 795850 | 0 | 0 | 1438700 | 0 | 0 | 760510 | 1917000 |
| GFGFGLVK | VDAC2; VDAC2 | P45880; Q5J | VDAC2_HUMAI | Unmodified | 0 | 0 | 0 | 0 | 0 | 0 | 1107600 | 0 | 295600 | 3305100 | 0 | 0 | 0 | 0 | 0 | 0 | 0 | 0 |
| GFGFGQG/ | CSRP1; CSRP1 | P21291; E9P | CSRP1_HUMAN | Unmodified | 1538600 | 1916700 | 1545500 | 1894900 | 18464000 | 3204600 | 6190300 | 3767100 | 5139100 | 3881500 | 683200 | 2970600 | 12087000 | 10849000 | 2930100 | 2420900 | 7924000 | 3786500 |
| GFGFILFK | HNRNPAB; HNI | Q99729; D6I | ROAA_HUMAN | Unmodified | 268650 | 126950 | 0 | 0 | 1072600 | 211500 | 273120 | 0 | 2705400 | 2265800 | 2834700 | 16699000 | 0 | 0 | 0 | 0 | 0 | 0 |
| GFGFITFTN | RBM3 | P98179 | RBM3_HUMAN | Oxidation (M) | 38525000 | 30191000 | 15662000 | 15460000 | 349280 | 125150 | 130070 | 43257 | 630870 | 397080 | 160540 | 861360 | 4228500 | 811860 | 17180000 | 20816000 | 76342 | 207780 |
| GFGFITFTN | RBM3 | P98179 | RBM3_HUMAN | Unmodified | 1019100 | 901880 | 0 | 495290 | 0 | 0 | 0 | 0 | 329970 | 59577 | 0 | 0 | 0 | 92383 | 482720 | 333640 | 0 | 0 |
| GFGFVCFS | PABPC4; PABP( | Q13310; P11 | PABP4_HUMAN | Unmodified | 88939000 | 33223000 | 17207000 | 28870000 | 1522500 | 0 | 1477900 | 712910 | 1294800 | 1353800 | 1313500 | 5062600 | 10274000 | 7837800 | 11006000 | 19028000 | 851040 | 0 |
| GFGFVDFN | NCL; NCL; NCL | P19338; C9J | NUCL_HUMAN | Unmodified | 0 | 0 | 0 | 0 | 0 | 0 | 0 | 240060 | 2104600 | 820160 | 3348200 | 7446600 | 922290 | 0 | 0 | 0 | 0 | 0 |
| GFGFVLFK | HNRNPDL; HNI | O14979; Q14 | HNRDL_HUMA | Unmodified | 646390 | 0 | 0 | 0 | 641850 | 166370 | 86319 | 0 | 2913400 | 1230500 | 3667300 | 34670000 | 0 | 0 | 0 | 0 | 0 | 260770 |
| GFGFVLFKI | HNRNPD; HNR | Q14103; H0Y | HNRPD_HUMA | Unmodified | 26604 | 0 | 0 | 0 | 0 | 0 | 102900 | 0 | 0 | 0 | 0 | 434370 | 356210 | 0 | 0 | 0 | 0 | 0 |
| GFGFVRFTI | TARDBP; TARDI | Q13148; A0A | TADBP_HUMAN | Unmodified | 0 | 0 | 0 | 0 | 0 | 0 | 7981000 | 810200 | 0 | 991710 | 0 | 0 | 4235100 | 1977900 | 0 | 0 | 0 | 4432100 |
| GFGFVSFEI | PABPC1; PABP( | P11940; Q9H | PABP1_HUMAN | Unmodified | 10506000 | 8587300 | 14302000 | 8608800 | 0 | 0 | 0 | 0 | 0 | 0 | 541720 | 1510600 | 0 | 0 | 5488500 | 5712300 | 0 | 0 |
| GFGFVSFEI | PABPC1; PABP( | P11940; Q9H | PABP1_HUMAN | Unmodified | 1.61E+08 | 1.03E+08 | 59765000 | 47194000 | 2441000 | 329610 | 0 | 0 | 3846000 | 2048600 | 2192400 | 16968000 | 3904900 | 1111800 | 28473000 | 40056000 | 768940 | 0 |
| GFGFVSFEI | PABPC1; PABP( | P11940; Q9H | PABP1_HUMAN | Unmodified | 2011100 | 857500 | 0 | 1550600 | 437360 | 0 | 0 | 0 | 0 | 0 | 0 | 0 | 1464100 | 1305200 | 0 | 0 | 0 | 0 |
| GFGFVSFEI | PABPC1; PABP( | P11940; Q9H | PABP1_HUMAN | Oxidation (M) | 3939000 | 167810 | 0 | 2429100 | 131260 | 0 | 0 | 0 | 0 | 243100 | 0 | 0 | 410610 | 0 | 0 | 0 | 467950 | 0 |
| GFGFVSYE | PABPC4; PABP( | Q13310; B1A | PABP4_HUMAN | Unmodified | 16120000 | 9997500 | 6511600 | 3379800 | 0 | 0 | 0 | 0 | 0 | 0 | 0 | 712270 | 427570 | 0 | 4355500 | 4099700 | 0 | 0 |
| GFGFVSYE | PABPC4; PABP( | Q13310; B1A | PABP4_HUMAN | Unmodified | 909030 | 557160 | 0 | 323370 | 0 | 0 | 67620 | 0 | 0 | 0 | 0 | 0 | 0 | 0 | 0 | 80581 | 0 | 0 |
| GFGFVSYE | PABPC4; PABP( | Q13310; B1A | PABP4_HUMAN | Unmodified | 212280 | 159970 | 0 | 209520 | 0 | 0 | 0 | 0 | 0 | 0 | 0 | 0 | 44502 | 0 | 0 | 0 | 0 | 0 |
| GFGFVTFD | HNRNPA2B1; H | P22626; A0A | ROA2_HUMAN | Unmodified | 1111700 | 537380 | 656200 | 403230 | 404260 | 218470 | 271150 | 0 | 3212900 | 422610 | 3725600 | 15867000 | 1328900 | 0 | 588960 | 470370 | 196880 | 237550 |
| GFGFVTFD | HNRNPA2B1; H | P22626; A0A | ROA2_HUMAN | Unmodified | 878260 | 0 | 0 | 0 | 2433300 | 347810 | 0 | 91085 | 2068500 | 2413900 | 616510 | 2115500 | 5179000 | 1348500 | 0 | 0 | 1970900 | 714520 |
| GFGFVTYA | HNRNPA1; HNI | P09651; A0A | ROA1_HUMAN | Unmodified | 0 | 0 | 0 | 0 | 0 | 0 | 0 | 0 | 0 | 0 | 0 | 432710 | 0 | 0 | 0 | 0 | 0 | 0 |
| GFGFVTYA | HNRNPA1; HNI | P09651; A0A | ROA1_HUMAN | Unmodified | 0 | 0 | 0 | 0 | 387400 | 0 | 0 | 0 | 3431700 | 761490 | 679690 | 4995300 | 1933600 | 0 | 0 | 0 | 0 | 0 |
| GFGFVTYA | HNRNPA1; HNI | P09651; A0A | ROA1_HUMAN | Oxidation (M) | 0 | 0 | 0 | 0 | 0 | 0 | 0 | 0 | 1151800 | 380090 | 913410 | 6501500 | 0 | 0 | 0 | 0 | 0 | 0 |
| GFGFVYFQ | HNRNPA0 | Q13151 | ROA0_HUMAN | Unmodified | 183760 | 0 | 0 | 187330 | 87338 | 0 | 102260 | 41417 | 382880 | 116680 | 0 | 657910 | 106130 | 85672 | 66499 | 81269 | 0 | 0 |
| GFGGAMTI | GBA; GBA; GBA | P04062; A0A | GLCM_HUMAN | Oxidation (M) | 0 | 0 | 0 | 0 | 298520 | 0 | 0 | 0 | 0 | 637160 | 0 | 0 | 369460 | 219030 | 0 | 0 | 0 | 0 |
| GFGGIGGII | ETF1; ETF1 | P62495; B7Z | ERF1_HUMAN I | Unmodified | 0 | 0 | 0 | 0 | 0 | 0 | 0 | 0 | 0 | 0 | 0 | 3711500 | 0 | 0 | 0 | 571130 | 0 | 0 |
| GFGGITHG | ACADVL | P49748 | ACADV_HUMAI | Unmodified | 0 | 0 | 0 | 0 | 0 | 0 | 0 | 0 | 1086900 | 0 | 0 | 0 | 0 | 0 | 0 | 0 | 0 | 0 |
| GFGGITHG | ACADVL | P49748 | ACADV_HUMAI | Unmodified | 0 | 0 | 0 | 0 | 0 | 0 | 0 | 0 | 809650 | 42793 | 0 | 0 | 0 | 0 | 0 | 0 | 0 | 0 |
| GFGGKFG\ | CTTN | Q14247 | SRC8_HUMAN | Unmodified | 0 | 0 | 0 | 0 | 7874600 | 353530 | 134370 | 0 | 0 | 208290 | 0 | 0 | 3444000 | 4718400 | 0 | 0 | 3593000 | 3026900 |
| GFGGKYG\ | CTTN | Q14247 | SRC8_HUMAN | Unmodified | 0 | 0 | 0 | 0 | 1477100 | 126910 | 0 | 46086 | 0 | 0 | 0 | 0 | 360980 | 629500 | 0 | 0 | 0 | 513590 |
| GFGHIGIA\ | GLO1 | Q04760 | LGUL_HUMAN | Unmodified | 1542000 | 1771100 | 1072000 | 1606700 | 0 | 0 | 0 | 0 | 0 | 0 | 1589500 | 11781000 | 2253200 | 0 | 2893000 | 3341900 | 0 | 0 |
| GFGPIFEE( | CNTN1; CNTN1 | Q12860; F8V | CNTN1_HUMA | Unmodified | 801870 | 1184100 | 0 | 0 | 0 | 0 | 0 | 0 | 4140800 | 0 | 0 | 0 | 0 | 0 | 0 | 0 | 0 | 0 |
| GFGRGGAI | CRIP1; N/A; N/A | P50238; K4D | CRIP1_HUMAN | Unmodified | 0 | 0 | 0 | 0 | 11492000 | 2116700 | 7809200 | 10860000 | 116360 | 1406400 | 0 | 0 | 0 | 230690 | 78152 | 0 | 737050 | 1645700 |
| GFGSDKEA | ANXA6; ANXA6; | P08133; E5R | ANXA6_HUMAN | Unmodified | 0 | 0 | 0 | 6526900 | 6573100 | 1881600 | 3860800 | 0 | 55456000 | 33301000 | 0 | 0 | 6075800 | 3130400 | 0 | 0 | 3012600 | 9861700 |
| GFGSDKEA | ANXA6; ANXA6; | P08133; E5R | ANXA6_HUMAN | Unmodified | 0 | 0 | 0 | 0 | 7051800 | 1259500 | 2369300 | 0 | 762360 | 17104000 | 0 | 0 | 8116300 | 3142000 | 0 | 0 | 2868900 | 13889000 |
| GFGSFRFP | VCP; VCP; VCP; | P55072; C9J | TERA_HUMAN T | Unmodified | 0 | 0 | 0 | 0 | 9798800 | 1988900 | 0 | 0 | 4595700 | 4003300 | 758940 | 5432800 | 13380000 | 2714000 | 0 | 0 | 1343000 | 1909000 |
| GFGTDEQA | ANXA11 | P50995 | ANX11_HUMAN | Unmodified | 1762200 | 905020 | 431250 | 2859400 | 1501800 | 640520 | 112000 | 47408 | 19692000 | 1153900 | 2983000 | 4911200 | 4250400 | 470200 | 0 | 1449900 | 820480 | 6056900 |
| GFGTDEQA | ANXA11 | P50995 | ANX11_HUMAN | Unmodified | 0 | 0 | 0 | 0 | 9644100 | 3979500 | 12040000 | 2813400 | 0 | 6605400 | 0 | 0 | 24503000 | 32427000 | 0 | 0 | 10188000 | 89487000 |
| GFGTDEQA | ANXA7 | P20073 | ANXA7_HUMAN | Unmodified | 1663500 | 2279200 | 2743600 | 5217800 | 2035500 | 606480 | 0 | 60496 | 3.95E+08 | 12538000 | 9057100 | 4585700 | 980990 | 778360 | 4563200 | 4649000 | 1530300 | 6274800 |
| GFGTDEQA | ANXA7 | P20073 | ANXA7_HUMAN | Unmodified | 0 | 0 | 0 | 0 | 0 | 0 | 0 | 0 | 49143000 | 1202400 | 0 | 0 | 0 | 0 | 0 | 0 | 0 | 1924100 |
| GFGTDEQA | ANXA7 | P20073 | ANXA7_HUMAN | Unmodified | 0 | 0 | 0 | 0 | 33927000 | 13775000 | 14075000 | 492590 | 19697000 | 1.63E+08 | 1531100 | 0 | 16018000 | 22220000 | 0 | 368660 | 40027000 | 99489000 |
| GFGTDFKK | HEXB; HEXB | P07686; Q5U | HEXB_HUMAN | Unmodified | 0 | 0 | 0 | 0 | 0 | 0 | 0 | 0 | 0 | 0 | 0 | 0 | 0 | 370610 | 0 | 0 | 0 | 0 |
| GFGYGQG/ | CSRP2; CSRP2 | Q16527; F8V | CSRP2_HUMAN | Unmodified | 331010 | 308420 | 0 | 0 | 0 | 0 | 484450 | 564170 | 235730 | 293550 | 522790 | 5788400 | 0 | 0 | 0 | 0 | 692610 | 731110 |
| GFHCESSA | EIF3B | P55884 | EIF3B_HUMAN | Unmodified | 0 | 0 | 0 | 0 | 1892700 | 4106400 | 58788 | 0 | 52243 | 0 | 0 | 948090 | 0 | 0 | 352050 | 71626 | 0 | 0 |
| GFHPHSKI | STAMBP | O95630 | STABP_HUMAN | Unmodified | 403220 | 0 | 0 | 0 | 0 | 0 | 0 | 0 | 308120 | 0 | 0 | 0 | 0 | 0 | 0 | 0 | 0 | 0 |
| GFHQEVFL | SDF4; SDF4; SD | Q9BRK5; H0 | CAB45_HUMAN | Unmodified | 0 | 0 | 0 | 171930 | 0 | 0 | 0 | 0 | 1379000 | 0 | 0 | 0 | 0 | 0 | 0 | 0 | 0 | 0 |
| GFHQVPFA | QARS1; QARS1; | P47897; H7( | SYQ_HUMAN G | Unmodified | 0 | 0 | 0 | 0 | 0 | 0 | 0 | 0 | 0 | 0 | 0 | 357050 | 0 | 0 | 0 | 0 | 0 | 0 |
| GFHQVPFA | QARS1; QARS1; | P47897; H7( | SYQ_HUMAN G | Unmodified | 0 | 0 | 0 | 0 | 0 | 0 | 0 | 0 | 0 | 0 | 0 | 1414500 | 0 | 0 | 1187800 | 0 | 0 | 0 |
| GFHQVPFA | QARS1; QARS1; | P47897; H7( | SYQ_HUMAN G | Unmodified | 0 | 0 | 0 | 0 | 1749700 | 427390 | 444700 | 0 | 671440 | 1492100 | 795260 | 0 | 856850 | 0 | 0 | 0 | 1534300 | 2822700 |
| GFHSGEDS | DST; DST; DST; | Q03001; A0A | DYST_HUMAN | Unmodified | 0 | 0 | 0 | 0 | 507050 | 0 | 166390 | 103700 | 218700 | 263240 | 0 | 0 | 0 | 0 | 0 | 0 | 252440 | 177960 |
| GFHTTIDIG | CNN3; CNN3 | Q15417; E9F | CNN3_HUMAN | Unmodified | 0 | 0 | 0 | 0 | 0 | 0 | 0 | 0 | 0 | 0 | 0 | 502940 | 0 | 0 | 0 | 0 | 0 | 0 |
| GFIMAEVM | OLA1; OLA1 | Q9NTK5; J3K | OLA1_HUMAN | 2 Oxidation (M) | 0 | 0 | 0 | 0 | 0 | 0 | 0 | 0 | 0 | 0 | 0 | 2293800 | 0 | 0 | 0 | 0 | 0 | 0 |
| GFIYPMLP( | MEGF8 | Q7Z7M0 | MEGF8_HUMAI | Oxidation (M) | 56964 | 0 | 0 | 0 | 0 | 0 | 0 | 0 | 657910 | 0 | 0 | 0 | 0 | 0 | 98405 | 143060 | 0 | 0 |
| GFKAEWLA | CANT1 | Q8WVQ1 | CANT1_HUMAN | Unmodified | 0 | 0 | 0 | 0 | 0 | 0 | 0 | 0 | 3486700 | 0 | 0 | 0 | 0 | 0 | 0 | 0 | 0 | 0 |
| GFKDQIYD | EIF4A1; EIF4A1; | P60842; J3K | IF4A1_HUMAN | Unmodified | 1742700 | 1361600 | 410730 | 4671200 | 0 | 306650 | 290050 | 46928 | 3377000 | 257690 | 1492800 | 20899000 | 0 | 209620 | 2104300 | 2724000 | 0 | 175880 |
| GFKDQIYEI | EIF4A2; EIF4A2 | Q14240; E7E | IF4A2_HUMAN | Unmodified | 0 | 0 | 0 | 0 | 0 | 0 | 0 | 0 | 0 | 0 | 0 | 1262600 | 0 | 0 | 0 | 0 | 0 | 0 |
| GFLAGYVV. | SLC35A4 | L0R6Q1 | S35U4_HUMAN | Unmodified | 0 | 0 | 0 | 0 | 0 | 0 | 1187900 | 251020 | 0 | 179440 | 775300 | 127710 | 0 | 0 | 0 | 0 | 110290 | 0 |
| GFLFGPSL/ | APRT; APRT; AP | P07741; H3E | APT_HUMAN A( | Unmodified | 0 | 0 | 0 | 0 | 402470 | 0 | 0 | 0 | 0 | 0 | 0 | 0 | 1157200 | 0 | 0 | 0 | 0 | 0 |
| GFLGCFPD | SNX9; SNX9; SN | Q9Y5X1; A0A | SNX9_HUMAN | Unmodified | 10983000 | 6012000 | 781030 | 716190 | 1150900 | 313670 | 16847 | 0 | 668270 | 175500 | 1106500 | 4689500 | 2661200 | 636500 | 2020900 | 1451400 | 157480 | 0 |
| GFLIDGYPI | AK5; AK1; AK1; / | Q9Y6K8; P0( | KAD5_HUMAN | Unmodified | 225990 | 237560 | 302400 | 0 | 360810 | 0 | 0 | 0 | 1918800 | 155710 | 0 | 4154900 | 241400 | 0 | 576520 | 385380 | 0 | 0 |
| GFLIDGYPI | AK1; AK1; AK1 | P00568; H0Y | KAD1_HUMAN | Unmodified | 0 | 0 | 0 | 0 | 1946200 | 151590 | 0 | 0 | 308490 | 1189300 | 65639 | 0 | 967160 | 589870 | 0 | 0 | 280100 | 968380 |
| GFLKEEDLT | CBS; CBSL; CBS | P35520; P0D | CBS_HUMAN C | Unmodified | 0 | 0 | 0 | 0 | 0 | 0 | 0 | 0 | 0 | 0 | 0 | 1631400 | 0 | 0 | 0 | 0 | 0 | 0 |
| GFLNSSELS | TSFM; TSFM; TS | P43897; F8V | EFTS_HUMAN I | Unmodified | 0 | 0 | 0 | 0 | 0 | 0 | 0 | 0 | 2480900 | 0 | 0 | 0 | 0 | 0 | 0 | 0 | 0 | 0 |
| GFLWDEGI | MOGS; MOGS; | Q13724; A0A | MOGS_HUMAN | Unmodified | 68677 | 0 | 0 | 0 | 1103200 | 253340 | 0 | 0 | 14057000 | 1839600 | 288960 | 1076800 | 0 | 0 | 59504 | 39208 | 232990 | 932080 |
| GFLWDEGI | MOGS; MOGS; | Q13724; A0A | MOGS_HUMAN | Unmodified | 0 | 0 | 0 | 0 | 37674 | 0 | 0 | 0 | 4917800 | 0 | 92691 | 0 | 0 | 0 | 0 | 0 | 0 | 0 |
| GFLYFNR | CPSF1 | Q10570 | CPSF1_HUMAN | Unmodified | 0 | 0 | 0 | 0 | 0 | 0 | 0 | 0 | 0 | 0 | 0 | 1217900 | 0 | 0 | 0 | 0 | 0 | 0 |
| GFMCQACI | EPHA3; EPHA3 | P29320; C9J | EPHA3_HUMAN | Oxidation (M) | 259980 | 0 | 0 | 0 | 0 | 0 | 0 | 0 | 178380 | 0 | 0 | 0 | 0 | 0 | 0 | 0 | 0 | 0 |
| GFMVQTGI | PPIL3; PPIL3; PI | Q9H2H8; C9 | PPIL3_HUMAN | Oxidation (M) | 87800 | 0 | 91300 | 340300 | 2420100 | 225850 | 0 | 0 | 0 | 164250 | 0 | 0 | 882380 | 241820 | 0 | 0 | 243830 | 399980 |
| GFMVQTGI | PPIL3; PPIL3; PI | Q9H2H8; C9 | PPIL3_HUMAN | Unmodified | 0 | 0 | 0 | 0 | 986660 | 0 | 317430 | 67284 | 221610 | 0 | 0 | 0 | 393270 | 332450 | 0 | 0 | 126010 | 342220 |

| | | | | | | | | | | | | | | | | | | | | | |
|---|---|---|---|---|---|---|---|---|---|---|---|---|---|---|---|---|---|---|---|---|---|
| GFNCESKF | FN1 | P02751 | FINC_HUMAN I | Unmodified | 32037 | 0 | 5524300 | 3691900 | 0 | 0 | 0 | 0 | 965170 | 0 | 430150 | 250600 | 158900 | 67051 | 88388 | 1410000 | 0 | 85907 |
| GFNGIHCC | EDIL3 | O43854 | EDIL3_HUMAN | Unmodified | 0 | 0 | 0 | 0 | 11053000 | 4645700 | 0 | 0 | 2496100 | 0 | 0 | 0 | 0 | 0 | 0 | 0 | 627260 | 12301000 |
| GFNKNDAI | GPC4 | O75487 | GPC4_HUMAN | Unmodified | 1245800 | 1907100 | 0 | 584190 | 0 | 0 | 123270 | 0 | 2790700 | 528930 | 0 | 0 | 10079000 | 5803200 | 0 | 0 | 1436400 | 5496900 |
| GFNLPYQD | ABCF1; ABCF1; | Q8NE71; H0 | ABCF1_HUMAN | Unmodified | 0 | 102470 | 171700 | 90012 | 0 | 0 | 417770 | 501230 | 0 | 0 | 0 | 1188400 | 305140 | 87193 | 160420 | 0 | 0 | 0 |
| GFNVVSAV | STEAP2; STEAP | Q8NFT2; C9. | STEA2_HUMAN | Unmodified | 0 | 0 | 302510 | 231060 | 0 | 0 | 0 | 0 | 0 | 0 | 0 | 0 | 0 | 0 | 1218000 | 0 | 0 | 0 |
| GFPEPSQA | NOXO1 | Q8NFA2 | NOXO1_HUMA | Unmodified | 0 | 0 | 0 | 0 | 0 | 0 | 0 | 0 | 0 | 0 | 1692500 | 1978900 | 0 | 0 | 0 | 0 | 0 | 0 |
| GFPFHSSE | APLP1; APLP1; | P51693; B7Z | APLP1_HUMAN | Unmodified | 0 | 0 | 0 | 0 | 0 | 0 | 0 | 0 | 2541900 | 0 | 0 | 0 | 0 | 0 | 0 | 0 | 0 | 0 |
| GFPTDATLI | SSB; SSB; SSB | P05455; E9P | LA_HUMAN Lu | Unmodified | 0 | 0 | 0 | 0 | 0 | 0 | 0 | 0 | 0 | 0 | 0 | 1491400 | 0 | 1198200 | 0 | 0 | 0 | 0 |
| GFPTIYFSP | PDIA3 | P30101 | PDIA3_HUMAN | Unmodified | 0 | 0 | 0 | 0 | 1065900 | 721510 | 985140 | 539870 | 1.67E+08 | 18641000 | 37521000 | 9997400 | 124070 | 1180100 | 165890 | 0 | 0 | 1960600 |
| GFPTIYFSP | PDIA3 | P30101 | PDIA3_HUMAN | Unmodified | 0 | 0 | 0 | 0 | 0 | 0 | 0 | 0 | 7644000 | 885400 | 2325200 | 886940 | 0 | 0 | 0 | 0 | 0 | 0 |
| GFQAVHST | TLL2 | Q9Y6L7 | TLL2_HUMAN T | Unmodified | 90501 | 0 | 0 | 0 | 0 | 0 | 0 | 0 | 351460 | 0 | 0 | 39992 | 0 | 0 | 0 | 0 | 0 | 0 |
| GFQFVSSS | CNBP; CNBP | P62633; D6F | CNBP_HUMAN | Unmodified | 103840 | 124480 | 109280 | 230880 | 0 | 17058 | 0 | 0 | 0 | 0 | 192070 | 104550 | 29115 | 0 | 868950 | 78637 | 16386 | 0 |
| GFQIYDGP | CEMIP2 | Q9UHN6 | CEIP2_HUMAN | Unmodified | 0 | 0 | 0 | 1372400 | 2904000 | 100670 | 0 | 0 | 6020200 | 253950 | 77730 | 23405 | 0 | 0 | 2417900 | 2004100 | 0 | 0 |
| GFQQISFVI | TOP2A | P11388 | TOP2A_HUMAN | Unmodified | 40885 | 0 | 0 | 0 | 901290 | 10170 | 259120 | 435010 | 49471 | 106110 | 0 | 639690 | 835400 | 296720 | 0 | 0 | 304770 | 355380 |
| GFQQISFVI | TOP2B; TOP2B | Q02880; E9F | TOP2B_HUMAN | Unmodified | 485180 | 67729 | 0 | 0 | 0 | 0 | 0 | 100900 | 0 | 156790 | 20810 | 298230 | 0 | 0 | 0 | 0 | 0 | 0 |
| GFRDEPYF | PATJ; PATJ; PAT | Q8NI35; A0A | INADL_HUMAN | Unmodified | 195720 | 0 | 0 | 809120 | 3725700 | 0 | 0 | 0 | 672900 | 290990 | 0 | 663260 | 1264600 | 1552500 | 1909900 | 0 | 1964400 | 2293400 |
| GFSAEQIAI | MAGT1; MAGT1 | Q9H0U3; A0 | MAGT1_HUMAI | Unmodified | 0 | 0 | 0 | 0 | 441440 | 0 | 0 | 0 | 13250000 | 1235400 | 0 | 0 | 452160 | 0 | 0 | 0 | 0 | 0 |
| GFSAISCT\ | MVB12A; MVB1 | Q96EY5; E9F | MB12A_HUMAI | Unmodified | 18522000 | 5953800 | 5456300 | 3805800 | 0 | 0 | 0 | 0 | 0 | 0 | 63094 | 3470600 | 0 | 0 | 18037000 | 14037000 | 0 | 0 |
| GFSEEHNT | CBX5; CBX5 | P45973; F8V | CBX5_HUMAN | Unmodified | 0 | 0 | 0 | 0 | 0 | 0 | 0 | 0 | 1074300 | 0 | 0 | 365420 | 0 | 0 | 0 | 0 | 0 | 0 |
| GFSEGLWE | HDGF | P51858 | HDGF_HUMAN | Unmodified | 0 | 0 | 0 | 0 | 3086600 | 0 | 0 | 14591 | 1825600 | 838220 | 2031500 | 8195300 | 2831200 | 0 | 0 | 69719 | 322200 | 192360 |
| GFSEGTAS | VPS13C | Q709C8 | VP13C_HUMAN | Unmodified | 0 | 0 | 0 | 0 | 0 | 0 | 0 | 0 | 0 | 0 | 0 | 1659600 | 0 | 0 | 0 | 0 | 0 | 0 |
| GFSFVATG | RPS6KA1; RPS6 | Q15418; Q5 | KS6A1_HUMAN | Oxidation (M) | 68540 | 3461100 | 2216500 | 5466400 | 0 | 0 | 0 | 0 | 0 | 0 | 0 | 0 | 0 | 0 | 8376800 | 6169100 | 0 | 0 |
| GFSIPECQI | CS; CS; CS; CS | O75390; F8V | CISY_HUMAN ( | Unmodified | 0 | 0 | 0 | 0 | 0 | 0 | 0 | 0 | 4776400 | 583260 | 0 | 0 | 0 | 0 | 0 | 0 | 0 | 0 |
| GFSLADIPY | GPC6 | Q9Y625 | GPC6_HUMAN | Unmodified | 0 | 812730 | 1648500 | 6905600 | 0 | 0 | 0 | 43066 | 2191100 | 246980 | 0 | 5005400 | 2697600 | 614470 | 0 | 2020600 | 0 | 573570 |
| GFSLEELR | RPL13; RPL13 | P26373; J3Q | RL13_HUMAN ( | Unmodified | 0 | 6389100 | 0 | 0 | 217400 | 0 | 0 | 0 | 688010 | 139180 | 2747100 | 23184000 | 0 | 0 | 6939200 | 0 | 204960 | 0 |
| GFSLLATEI | PARP1; PARP1; | P09874; A0A | PARP1_HUMAN | Unmodified | 1343000 | 643360 | 0 | 0 | 0 | 0 | 1238900 | 638130 | 0 | 0 | 871770 | 1482300 | 0 | 0 | 0 | 0 | 0 | 0 |
| GFSLLATEI | PARP1; PARP1; | P09874; A0A | PARP1_HUMAN | Unmodified | 0 | 0 | 0 | 0 | 424330 | 0 | 0 | 347650 | 0 | 0 | 0 | 0 | 0 | 0 | 0 | 0 | 0 | 0 |
| GFSLLATEI | PARP1; PARP1; | P09874; A0A | PARP1_HUMAN | Unmodified | 0 | 0 | 0 | 0 | 360890 | 0 | 336170 | 1724500 | 0 | 766920 | 0 | 0 | 1497100 | 553510 | 0 | 0 | 526750 | 656840 |
| GFSLSDVP | GPC1 | P35052 | GPC1_HUMAN | Unmodified | 1400600 | 2794800 | 1665300 | 5438500 | 2099500 | 1528000 | 0 | 0 | 1364900 | 0 | 0 | 71575 | 706720 | 0 | 2398900 | 1748100 | 0 | 0 |
| GFSNQIPL\ | SPPL2B; SPPL2 | Q8TCT7; A0/ | SPP2B_HUMAN | Unmodified | 0 | 0 | 0 | 0 | 0 | 0 | 0 | 0 | 2945500 | 177940 | 0 | 2211300 | 0 | 0 | 0 | 0 | 0 | 0 |
| GFSPQHKI | PPP3CA; PPP3C | Q08209; E7E | PP2BA_HUMAN | Unmodified | 0 | 0 | 0 | 0 | 0 | 0 | 347360 | 240420 | 0 | 0 | 0 | 0 | 1059400 | 1323700 | 0 | 0 | 381010 | 191140 |
| GFSVVADT | LASP1; LASP1 | Q14847; C9J | LASP1_HUMAN | Unmodified | 1235000 | 741440 | 1595700 | 1927600 | 3016700 | 0 | 0 | 957460 | 2239600 | 161380 | 471230 | 1698100 | 1807600 | 1015900 | 4889800 | 2474200 | 382050 | 1570900 |
| GFSVVADT | LASP1; LASP1 | Q14847; C9J | LASP1_HUMAN | Unmodified | 338860 | 0 | 0 | 1394800 | 13053000 | 2051800 | 1134600 | 393610 | 0 | 831530 | 0 | 0 | 5851600 | 4817200 | 232030 | 0 | 1773800 | 4422800 |
| GFSVVADT | LASP1; LASP1 | Q14847; C9J | LASP1_HUMAN | Unmodified | 0 | 0 | 0 | 0 | 6693500 | 570740 | 427040 | 576960 | 0 | 0 | 0 | 0 | 2354800 | 1879400 | 0 | 0 | 1199400 | 2252500 |
| GFTECEMT | ANLN | Q9NQW6 | ANLN_HUMAN | Oxidation (M) | 0 | 0 | 0 | 32457 | 382890 | 0 | 0 | 0 | 0 | 0 | 0 | 0 | 0 | 0 | 0 | 0 | 187810 | 135670 |
| GFTECEMT | ANLN | Q9NQW6 | ANLN_HUMAN | Unmodified | 0 | 0 | 0 | 0 | 1500000 | 161710 | 0 | 53045 | 0 | 0 | 0 | 0 | 0 | 0 | 0 | 0 | 603040 | 195530 |
| GFTGIDSD' | PAPSS2 | O95340 | PAPS2_HUMAN | Unmodified | 0 | 0 | 0 | 0 | 1193000 | 0 | 0 | 0 | 0 | 0 | 0 | 0 | 0 | 0 | 0 | 0 | 1062900 | 3613800 |
| GFTGIDSE\ | PAPSS1 | O43252 | PAPS1_HUMAN | Unmodified | 0 | 0 | 0 | 0 | 0 | 0 | 0 | 0 | 0 | 0 | 0 | 2681900 | 544990 | 0 | 0 | 0 | 0 | 0 |
| GFTIPEAFR | CLIC1 | O00299 | CLIC1_HUMAN | Unmodified | 2239900 | 2232100 | 1716900 | 1939200 | 632650 | 420890 | 0 | 100090 | 7267900 | 229150 | 1632200 | 21273000 | 440550 | 342770 | 4225200 | 2460600 | 241490 | 200840 |
| GFTIPEAFR | CLIC1 | O00299 | CLIC1_HUMAN | Unmodified | 0 | 0 | 0 | 588490 | 964230 | 0 | 0 | 0 | 729070 | 0 | 658980 | 0 | 810720 | 190530 | 0 | 0 | 471570 | 443240 |
| GFTLNDAA | LRPPRC | P42704 | LPPRC_HUMAN | Unmodified | 0 | 0 | 0 | 0 | 0 | 0 | 0 | 0 | 2025400 | 175210 | 0 | 0 | 0 | 0 | 0 | 0 | 0 | 0 |
| GFTMIGEH | CD55; CD55; C | P08174; B1A | DAF_HUMAN C | Oxidation (M) | 576790 | 0 | 0 | 0 | 0 | 0 | 0 | 0 | 0 | 0 | 344910 | 891870 | 0 | 0 | 0 | 0 | 0 | 0 |
| GFTMIGEH | CD55; CD55; C | P08174; B1A | DAF_HUMAN C | Oxidation (M) | 0 | 0 | 0 | 0 | 0 | 0 | 0 | 0 | 0 | 0 | 0 | 0 | 0 | 494880 | 0 | 0 | 2513500 | 4564400 |
| GFTMIGEH | CD55; CD55; C | P08174; B1A | DAF_HUMAN C | Unmodified | 0 | 0 | 0 | 0 | 0 | 444170 | 0 | 0 | 0 | 0 | 0 | 0 | 0 | 85259 | 0 | 0 | 1300000 | 3306800 |
| GFTPIPGTII | DSG2 | Q14126 | DSG2_HUMAN | Oxidation (M) | 0 | 0 | 2066300 | 0 | 3222900 | 135950 | 0 | 0 | 1259200 | 22708 | 3146500 | 37030000 | 978390 | 0 | 0 | 2907100 | 0 | 0 |
| GFTPIPGTII | DSG2 | Q14126 | DSG2_HUMAN | Unmodified | 0 | 0 | 0 | 0 | 1538800 | 1133200 | 0 | 0 | 3116900 | 0 | 0 | 2642800 | 0 | 0 | 0 | 0 | 0 | 0 |
| GFTPVAFAI | RHBDD2 | Q6NTF9 | RHBD2_HUMA | Unmodified | 0 | 0 | 0 | 0 | 0 | 0 | 0 | 0 | 581490 | 0 | 0 | 0 | 0 | 0 | 0 | 0 | 0 | 0 |
| GFVDDIIQF | PCCB; PCCB; P | P05166; C9J | PCCB_HUMAN | Unmodified | 0 | 0 | 0 | 0 | 0 | 0 | 0 | 0 | 965680 | 0 | 86377 | 0 | 0 | 0 | 0 | 0 | 0 | 0 |
| GFVEPDHY | TFRC; TFRC | P02786; G3\ | TFR1_HUMAN | Unmodified | 3727200 | 5579600 | 4817400 | 10037000 | 0 | 0 | 0 | 0 | 21562000 | 2380700 | 0 | 5760700 | 0 | 0 | 5892200 | 5662000 | 0 | 0 |
| GFVEPDHY | TFRC; TFRC | P02786; G3\ | TFR1_HUMAN | Unmodified | 0 | 0 | 0 | 0 | 3521400 | 1532500 | 992290 | 0 | 1528200 | 2892600 | 0 | 0 | 1480200 | 816440 | 0 | 0 | 0 | 1813100 |
| GFVFITFKE | HNRNPAB; HNI | Q99729; D6I | ROAA_HUMAN | Unmodified | 0 | 0 | 0 | 0 | 0 | 0 | 0 | 0 | 0 | 20882 | 0 | 2080400 | 0 | 0 | 0 | 0 | 0 | 0 |
| GFVFITFKE | HNRNPAB; HNI | Q99729; D6I | ROAA_HUMAN | Unmodified | 0 | 0 | 0 | 0 | 0 | 0 | 1099300 | 0 | 0 | 0 | 0 | 19059000 | 0 | 0 | 0 | 0 | 0 | 0 |
| GFVLDATD | CPD; CPD | O75976; J3C | CBPD_HUMAN | Unmodified | 0 | 0 | 0 | 0 | 0 | 0 | 0 | 0 | 9123000 | 1554700 | 0 | 0 | 0 | 0 | 0 | 0 | 0 | 0 |
| GFVLQDTV | MARS1; MARS1 | P56192; H0Y | SYMC_HUMAN | Unmodified | 0 | 0 | 0 | 0 | 0 | 0 | 0 | 0 | 33093 | 0 | 0 | 2927400 | 0 | 0 | 0 | 0 | 0 | 0 |
| GFVPSPTS( | VARS1; VARS1 | P26640; A0A | SYVC_HUMAN | Unmodified | 49315 | 35319 | 206840 | 0 | 0 | 0 | 0 | 0 | 0 | 0 | 0 | 7420000 | 0 | 0 | 242330 | 24498 | 0 | 0 |
| GFVQVDD( | RARS1 | P54136 | SYRC_HUMAN | Unmodified | 0 | 0 | 0 | 0 | 0 | 0 | 0 | 0 | 0 | 0 | 35624 | 790760 | 0 | 0 | 0 | 0 | 0 | 90117 |
| GFVVINQK | CCT6A; CCT6B | P40227; Q92 | TCPZ_HUMAN | Unmodified | 497200 | 357320 | 759540 | 838500 | 2406800 | 145520 | 294130 | 213910 | 2077800 | 578330 | 6326000 | 19987000 | 3781900 | 1162900 | 820520 | 916940 | 615470 | 1395200 |
| GFVVINQK | CCT6A | P40227 | TCPZ_HUMAN | Unmodified | 0 | 0 | 106710 | 0 | 143900 | 0 | 303520 | 0 | 162790 | 123160 | 0 | 23889 | 270880 | 216810 | 282410 | 0 | 2278700 | 116300 |
| GFVVINQK | CCT6B | Q92526 | TCPW_HUMAN | Unmodified | 0 | 0 | 0 | 0 | 0 | 0 | 0 | 0 | 0 | 0 | 0 | 0 | 0 | 0 | 0 | 0 | 309860 | 0 |
| GFYCGDD! | PLPP2; PLPP2 | O43688; K7E | PLPP2_HUMAN | Unmodified | 0 | 0 | 0 | 0 | 0 | 0 | 0 | 0 | 0 | 0 | 0 | 2243300 | 0 | 0 | 0 | 0 | 0 | 0 |
| GFYEGIML\ | LARS1; LARS1; | Q9P2J5; A0A | SYLC_HUMAN | Oxidation (M) | 0 | 0 | 0 | 0 | 0 | 0 | 0 | 0 | 0 | 0 | 0 | 1093800 | 0 | 0 | 0 | 0 | 0 | 0 |
| GFYEGIML\ | LARS1; LARS1; | Q9P2J5; A0A | SYLC_HUMAN | Oxidation (M) | 268630 | 0 | 0 | 0 | 1320400 | 0 | 910360 | 0 | 0 | 0 | 0 | 0 | 658720 | 0 | 0 | 0 | 183440 | 0 |
| GFYLNQST | CLUH; CLUH; C | O75153; I3L | CLU_HUMAN C | Unmodified | 0 | 0 | 0 | 0 | 0 | 0 | 0 | 0 | 0 | 0 | 0 | 1126800 | 0 | 0 | 20077 | 0 | 0 | 0 |
| GFYLNQST | CLUH; CLUH; C | O75153; I3L | CLU_HUMAN C | Unmodified | 0 | 0 | 0 | 0 | 0 | 0 | 0 | 0 | 0 | 0 | 0 | 2764000 | 0 | 0 | 0 | 0 | 0 | 0 |
| GGAAPEGI | NASP; NASP | P49321; E9P | NASP_HUMAN | Unmodified | 0 | 0 | 0 | 0 | 0 | 0 | 0 | 0 | 1719100 | 150540 | 708380 | 0 | 0 | 0 | 0 | 0 | 0 | 0 |
| GGAARPN( | RAB5C; N/A; RA | P51148; K7E | RAB5C_HUMAI | Unmodified | 617910 | 437960 | 0 | 0 | 0 | 0 | 0 | 0 | 0 | 0 | 0 | 0 | 42824 | 0 | 125470 | 66914 | 0 | 0 |
| GGAAVDPI | PARP1; PARP1 | P09874; A0A | PARP1_HUMAN | Unmodified | 1832700 | 126460 | 0 | 0 | 0 | 0 | 0 | 0 | 0 | 0 | 0 | 6739500 | 0 | 0 | 0 | 0 | 0 | 0 |
| GGAEQFMI | CCT7 | Q99832 | TCPH_HUMAN | Unmodified | 0 | 0 | 0 | 0 | 0 | 0 | 0 | 0 | 943790 | 0 | 1059400 | 7709900 | 0 | 0 | 550210 | 14686 | 0 | 0 |
| GGAEQFMI | CCT7 | Q99832 | TCPH_HUMAN | Oxidation (M) | 0 | 0 | 0 | 0 | 0 | 0 | 0 | 0 | 0 | 0 | 0 | 1806000 | 0 | 0 | 0 | 0 | 0 | 0 |
| GGAEVQIF | GATD3B; GATD | A0A0B4J2D5 | GAL3B_HUMAN | Unmodified | 0 | 0 | 0 | 0 | 413790 | 411590 | 56249 | 0 | 10619000 | 287400 | 0 | 0 | 478930 | 0 | 0 | 0 | 128310 | 0 |
| GGAFDGTI | EHD1; EHD1 | Q9H4M9; A0 | EHD1_HUMAN | Oxidation (M) | 0 | 0 | 0 | 1645300 | 2685400 | 0 | 0 | 0 | 0 | 0 | 520450 | 1101500 | 0 | 0 | 1590100 | 1377000 | 0 | 0 |
| GGAGVGSI | RPS19 | P39019 | RS19_HUMAN | Unmodified | 710110 | 657860 | 965150 | 323310 | 0 | 0 | 105600 | 95156 | 346070 | 22350 | 150800 | 1775000 | 0 | 0 | 0 | 774590 | 146630 | 158300 |
| GGALLTSP! | ERLIN1; ERLIN1 | O75477; B0( | ERLN1_HUMAN | Oxidation (M) | 15789 | 0 | 0 | 0 | 0 | 0 | 0 | 0 | 900900 | 167210 | 0 | 0 | 0 | 0 | 0 | 0 | 0 | 0 |
| GGALLTSP! | ERLIN1; ERLIN1 | O75477; B0( | ERLN1_HUMAN | Unmodified | 0 | 0 | 0 | 0 | 0 | 0 | 0 | 0 | 109390 | 0 | 0 | 0 | 0 | 0 | 0 | 0 | 0 | 0 |
| GGALLTST! | ERLIN2; ERLIN2 | O94905; E5F | ERLN2_HUMAN | Oxidation (M) | 29387 | 35693 | 0 | 0 | 0 | 0 | 0 | 0 | 1732600 | 369520 | 0 | 0 | 0 | 0 | 196880 | 84724 | 0 | 0 |
| GGALLTST! | ERLIN2; ERLIN2 | O94905; E5F | ERLN2_HUMAN | Unmodified | 0 | 0 | 0 | 0 | 0 | 0 | 0 | 0 | 143660 | 0 | 0 | 0 | 0 | 0 | 0 | 0 | 0 | 0 |
| GGARGHA( | SCRIB; SCRIB; ! | Q14160; A0/ | SCRIB_HUMAN | Unmodified | 0 | 0 | 0 | 0 | 199790 | 0 | 0 | 0 | 0 | 534610 | 0 | 0 | 143070 | 150800 | 0 | 0 | 254020 | 819830 |
| GGAVGGEI | B3GAT3; B3GAT | O94766; G3\ | B3GA3_HUMAI | Unmodified | 0 | 0 | 0 | 0 | 0 | 0 | 0 | 0 | 902360 | 0 | 0 | 0 | 0 | 0 | 0 | 0 | 0 | 0 |
| GGAVVDE( | SLC16A3; SLC1 | O15427; J3C | MOT4_HUMAN | Unmodified | 162220 | 0 | 1998700 | 0 | 0 | 0 | 0 | 0 | 0 | 0 | 0 | 2987900 | 0 | 0 | 0 | 0 | 0 | 0 |
| GGCSGLLF | ARSA; ARSA | P15289; A0A | ARSA_HUMAN | Unmodified | 0 | 0 | 0 | 0 | 0 | 0 | 0 | 0 | 2794100 | 159680 | 74354 | 0 | 0 | 0 | 0 | 0 | 0 | 0 |
| GGDGQINI | PEX14 | O75381 | PEX14_HUMAN | Unmodified | 0 | 0 | 0 | 0 | 0 | 0 | 0 | 0 | 662710 | 46679 | 0 | 271820 | 0 | 0 | 0 | 0 | 0 | 0 |
| GGDGTSM' | C1QL1; C1QL2; | O75973; Q7 | C1QRF_HUMAI | Unmodified | 0 | 0 | 0 | 0 | 0 | 0 | 0 | 0 | 1498800 | 0 | 0 | 0 | 0 | 0 | 0 | 0 | 0 | 0 |
| GGDHVPV! | SLC38A10 | Q9HBR0 | S38AA_HUMAN | Unmodified | 0 | 0 | 0 | 0 | 0 | 0 | 0 | 0 | 559750 | 115950 | 0 | 0 | 0 | 0 | 0 | 0 | 0 | 0 |
| GGDINFVF | NOMO1; NOMC | Q15155; P6 | NOMO1_HUMA | Unmodified | 0 | 0 | 0 | 0 | 0 | 0 | 0 | 0 | 1084500 | 1351600 | 0 | 0 | 0 | 0 | 0 | 0 | 0 | 0 |
| GGDLFDAI | DCLK1 | O15075 | DCLK1_HUMAI | Unmodified | 0 | 0 | 0 | 0 | 0 | 0 | 0 | 0 | 639200 | 57558 | 627410 | 365320 | 0 | 0 | 0 | 0 | 0 | 28864 |
| GGDPALIRI | SARS1; SARS1 | P49591; Q5T | SYSC_HUMAN | Unmodified | 0 | 0 | 644800 | 0 | 607340 | 182200 | 0 | 0 | 2211500 | 149080 | 134000 | 1237600 | 2388800 | 967690 | 725350 | 636550 | 0 | 510380 |
| GGDRIPAD | ATP1A1 | P05023 | AT1A1_HUMAN | Unmodified | 246640 | 160750 | 2942500 | 2281600 | 449330 | 0 | 0 | 0 | 2299500 | 280820 | 1614200 | 19025000 | 476130 | 159900 | 56055 | 3504600 | 0 | 0 |
| GGDSITAVI | NRP2 | O60462 | NRP2_HUMAN | Unmodified | 0 | 0 | 0 | 0 | 0 | 0 | 0 | 0 | 703220 | 0 | 0 | 0 | 0 | 0 | 0 | 0 | 0 | 0 |
| GGEARGEL | GAA | P10253 | LYAG_HUMAN | Unmodified | 0 | 0 | 0 | 0 | 0 | 0 | 0 | 0 | 2168700 | 233710 | 0 | 0 | 0 | 0 | 0 | 0 | 0 | 0 |
| GGEDAAV( | SLC38A10 | Q9HBR0 | S38AA_HUMAN | Unmodified | 561310 | 284140 | 194500 | 0 | 0 | 393670 | 0 | 0 | 534920 | 0 | 209800 | 328650 | 0 | 0 | 0 | 0 | 154030 | 204210 |
| GGEEPGRS | SLC4A2; SLC4A | P04920; C9J | B3A2_HUMAN | Unmodified | 291370 | 0 | 0 | 0 | 0 | 0 | 0 | 0 | 0 | 0 | 0 | 548330 | 0 | 0 | 0 | 630450 | 0 | 0 |
| GGEERVGE | VGF | O15240 | VGF_HUMAN N | Unmodified | 0 | 0 | 0 | 0 | 0 | 0 | 0 | 0 | 2848100 | 0 | 0 | 0 | 0 | 0 | 0 | 0 | 0 | 0 |
| GGEERVGE | VGF | O15240 | VGF_HUMAN N | Unmodified | 529900 | 814320 | 0 | 0 | 0 | 0 | 0 | 0 | 22614000 | 679330 | 0 | 0 | 0 | 0 | 0 | 0 | 0 | 0 |
| GGEIQPVS' | HSPE1; HSPE1; | P61604; B8Z | CH10_HUMAN | Unmodified | 0 | 0 | 0 | 0 | 0 | 55304 | 234750 | 40115 | 10183000 | 821120 | 454900 | 515100 | 0 | 0 | 0 | 0 | 0 | 0 |
| GGELLVHT | AGTRAP | Q6RW13 | ATRAP_HUMAN | Unmodified | 0 | 0 | 0 | 0 | 0 | 0 | 253160 | 214350 | 0 | 0 | 0 | 913930 | 0 | 0 | 0 | 172460 | 0 | 0 |
| GGELVYTD | PLEC; PLEC | Q15149; H0Y | PLEC_HUMAN | Unmodified | 0 | 0 | 0 | 0 | 0 | 105140 | 0 | 0 | 0 | 0 | 0 | 0 | 0 | 0 | 0 | 78432 | 974740 | 0 |
| GGENIYPAI | ACSF2; ACSF2 | Q96CM8; E9 | ACSF2_HUMAN | Unmodified | 0 | 0 | 0 | 0 | 0 | 0 | 0 | 0 | 1502600 | 0 | 0 | 0 | 0 | 0 | 0 | 0 | 0 | 0 |
| GGEPNQRI | DAG1 | Q14118 | DAG1_HUMAN | Unmodified | 1963700 | 2135900 | 785480 | 236390 | 0 | 0 | 0 | 0 | 0 | 0 | 0 | 752300 | 0 | 0 | 0 | 32921 | 771790 | 0 |
| GGFDWNL | GALNT2 | Q10471 | GALT2_HUMAN | Unmodified | 2449400 | 2080600 | 0 | 0 | 0 | 0 | 0 | 0 | 4411900 | 0 | 0 | 226560 | 0 | 0 | 1350900 | 0 | 0 | 0 |
| GGFSETRIE | EPB41L1; EPB4 | Q9H4G0; HC | E41L1_HUMAN | Unmodified | 0 | 0 | 0 | 0 | 793600 | 0 | 0 | 0 | 0 | 0 | 265430 | 0 | 211630 | 580150 | 0 | 0 | 172010 | 683950 |
| GGFVLLDG | SELENBP1; SEL | Q13228; A6F | SBP1_HUMAN | Unmodified | 0 | 0 | 302380 | 0 | 0 | 0 | 0 | 0 | 0 | 0 | 0 | 0 | 0 | 0 | 2930000 | 2289500 | 0 | 0 |
| GGGAFVQI | KIF5B | P33176 | KINH_HUMAN | Unmodified | 0 | 0 | 0 | 0 | 252460 | 0 | 0 | 0 | 261920 | 0 | 0 | 818700 | 0 | 0 | 0 | 0 | 0 | 0 |
| GGGALSAV | SRP54; SRP54; | P61011; G3\ | SRP54_HUMAN | Unmodified | 0 | 0 | 0 | 380540 | 153250 | 4393300 | 560640 | 406040 | 506630 | 234320 | 237050 | 1502100 | 0 | 0 | 0 | 398110 | 148960 | 0 |
| GGGAYYLI! | SLC12A1; SLC1 | Q13621; P5 | S12A1_HUMAN | Unmodified | 110260 | 0 | 1584500 | 161680 | 0 | 0 | 0 | 0 | 0 | 124000 | 0 | 1153300 | 0 | 0 | 464480 | 300330 | 0 | 0 |
| GGGDLPN! | DDAH2 | O95865 | DDAH2_HUMA | Unmodified | 356460 | 467700 | 466330 | 664140 | 260360 | 0 | 0 | 0 | 3880600 | 169570 | 417120 | 1034500 | 0 | 0 | 385700 | 319070 | 98298 | 456030 |
| GGGDPYSI | CNNM3 | Q8NE01 | CNNM3_HUMA | Unmodified | 0 | 0 | 0 | 0 | 0 | 0 | 0 | 0 | 0 | 0 | 0 | 0 | 0 | 0 | 0 | 0 | 0 | 0 |
| GGGEDFEI | SLC8A3 | P57103 | NAC3_HUMAN | Unmodified | 0 | 0 | 0 | 0 | 0 | 0 | 0 | 0 | 1500300 | 0 | 0 | 0 | 0 | 0 | 0 | 0 | 0 | 0 |
| GGGEQET( | PURB | Q96QR8 | PURB_HUMAN | Unmodified | 303460 | 336910 | 216570 | 232590 | 0 | 0 | 0 | 0 | 0 | 0 | 0 | 836170 | 0 | 0 | 229770 | 246250 | 0 | 0 |
| GGGEVIVR | RTCA | O00442 | RTCA_HUMAN | Unmodified | 0 | 0 | 0 | 0 | 0 | 0 | 201880 | 0 | 0 | 0 | 0 | 0 | 462390 | 637920 | 0 | 0 | 190620 | 413640 |

| | | | | | | | | | | | | | | | | | | | | | | |
|---|---|---|---|---|---|---|---|---|---|---|---|---|---|---|---|---|---|---|---|---|---|---|
| GGGGFGA | PURB | Q96QR8 | PURB_HUMAN | Unmodified | 786390 | 903550 | 0 | 0 | 0 | 0 | 0 | 0 | 0 | 0 | 0 | 455180 | 0 | 0 | 0 | 453200 | 0 | 0 |
| GGGGGGP | DRG1 | Q9Y295 | DRG1_HUMAN | Unmodified | 0 | 262300 | 307300 | 357780 | 513080 | 988120 | 129180 | 0 | 219120 | 334930 | 0 | 1553500 | 0 | 0 | 0 | 0 | 0 | 375050 |
| GGGGNFG | HNRNPA2B1; H | P22626; A0A | ROA2_HUMAN | Unmodified | 0 | 0 | 0 | 0 | 0 | 0 | 0 | 0 | 645910 | 0 | 2175700 | 1554900 | 0 | 0 | 0 | 0 | 0 | 0 |
| GGGGNFG | HNRNPA2B1; H | P22626; A0A | ROA2_HUMAN | Unmodified | 347440 | 207040 | 0 | 0 | 0 | 0 | 0 | 260280 | 662800 | 0 | 0 | 508480 | 0 | 0 | 0 | 0 | 0 | 0 |
| GGGGPGG | KHSRP; KHSRP | Q92945; M0 | FUBP2_HUMAN | Unmodified | 0 | 0 | 0 | 0 | 0 | 0 | 0 | 0 | 185130 | 43064 | 1676600 | 155730 | 0 | 0 | 0 | 0 | 0 | 0 |
| GGGGPGG | KHSRP; KHSRP | Q92945; M0 | FUBP2_HUMAN | Unmodified | 0 | 0 | 0 | 0 | 0 | 0 | 0 | 0 | 0 | 0 | 1101700 | 546990 | 0 | 0 | 0 | 0 | 0 | 0 |
| GGGGTGG | ZC3H10 | Q96K80 | ZC3HA_HUMAN | Unmodified | 1149800 | 996670 | 632230 | 1054300 | 0 | 0 | 0 | 0 | 0 | 0 | 0 | 0 | 0 | 0 | 451810 | 532110 | 0 | 0 |
| GGGHVAQ | RPS16; RPS16; | P62249; M0 | RS16_HUMAN | Unmodified | 2460300 | 2137200 | 1863100 | 499480 | 0 | 0 | 194930 | 113600 | 327730 | 113000 | 1217400 | 35410000 | 60078 | 0 | 2134000 | 1777900 | 83608 | 0 |
| GGGHVAQ | RPS16; RPS16; | P62249; M0 | RS16_HUMAN | Unmodified | 0 | 0 | 0 | 0 | 4890100 | 0 | 14003000 | 4115800 | 0 | 291940 | 121690 | 248970 | 307140 | 1869100 | 0 | 0 | 556920 | 1607000 |
| GGGLFLLA | AARSD1; PTGES | Q9BTE6; B3 | AASD1_HUMAN | Unmodified | 0 | 17020 | 94862 | 287760 | 0 | 0 | 0 | 0 | 0 | 0 | 667900 | 4737500 | 0 | 0 | 0 | 245190 | 0 | 0 |
| GGGLRLPL | SCRIB; SCRIB; | Q14160; A0A | SCRIB_HUMAN | Unmodified | 0 | 109520 | 322070 | 389370 | 204880 | 55795 | 0 | 0 | 4686200 | 257390 | 422060 | 1728700 | 154360 | 51157 | 489180 | 191780 | 627590 | 762360 |
| GGGPGPG | RRAS | P10301 | RRAS_HUMAN | Unmodified | 0 | 765040 | 2088100 | 0 | 2695800 | 448220 | 0 | 0 | 775780 | 0 | 0 | 177990 | 0 | 0 | 0 | 1997700 | 0 | 1110500 |
| GGGPGPG | RRAS | P10301 | RRAS_HUMAN | Unmodified | 129190 | 246750 | 0 | 271380 | 5456600 | 1547000 | 719720 | 348270 | 0 | 179660 | 0 | 0 | 1439600 | 860550 | 0 | 32560 | 3765100 | 5415100 |
| GGGPPREI | KIAA1522 | Q9P206 | K1522_HUMAN | Unmodified | 0 | 0 | 0 | 0 | 875300 | 0 | 0 | 0 | 550400 | 0 | 0 | 0 | 0 | 464050 | 0 | 0 | 0 | 725010 |
| GGGQIIPTA | EFTUD2; EEF2; | Q15029; P13 | U5S1_HUMAN | Unmodified | 1983200 | 1379300 | 1843200 | 194950 | 980660 | 0 | 234160 | 0 | 3298600 | 105400 | 4902200 | 83947000 | 267420 | 277120 | 2677200 | 1961600 | 304460 | 843940 |
| GGGTLCAC | DNASE2 | O00115 | DNS2A_HUMAN | Unmodified | 0 | 0 | 0 | 0 | 0 | 0 | 0 | 0 | 1869400 | 0 | 0 | 0 | 0 | 0 | 0 | 0 | 0 | 0 |
| GGHEIILSA | FMNL2 | Q96PY5 | FMNL2_HUMAN | Unmodified | 0 | 0 | 0 | 0 | 0 | 0 | 0 | 0 | 0 | 0 | 0 | 257620 | 0 | 0 | 0 | 0 | 0 | 0 |
| GGHFAAFE | EPHX1 | P07099 | HYEP_HUMAN | Unmodified | 0 | 0 | 730640 | 754280 | 0 | 0 | 0 | 0 | 2649300 | 344420 | 0 | 180550 | 0 | 0 | 893160 | 614000 | 0 | 0 |
| GGHFAAFE | EPHX1 | P07099 | HYEP_HUMAN | Unmodified | 0 | 0 | 0 | 0 | 0 | 0 | 0 | 0 | 5085900 | 1241600 | 0 | 0 | 0 | 0 | 369740 | 0 | 0 | 0 |
| GGHSGQL | LPP; LPP; LPP; L | Q93052; C9J | LPP_HUMAN Li | Unmodified | 0 | 0 | 0 | 1265400 | 10720000 | 0 | 268830 | 0 | 0 | 0 | 0 | 606710 | 7581400 | 5532400 | 0 | 804550 | 2380300 | 1680600 |
| GGHSSVST | CXCR4 | P61073 | CXCR4_HUMAN | Unmodified | 74952 | 0 | 0 | 0 | 76889 | 0 | 0 | 0 | 2933600 | 890830 | 0 | 0 | 0 | 0 | 0 | 0 | 0 | 0 |
| GGHVDLTN | OXCT1; OXCT1 | P55809; E9P | SCOT1_HUMAN | Unmodified | 0 | 0 | 0 | 0 | 0 | 0 | 0 | 0 | 1704100 | 0 | 0 | 0 | 0 | 0 | 0 | 0 | 0 | 0 |
| GGIMLPEK | HSPE1; HSPE1- | P61604; S4R | CH10_HUMAN | Oxidation (M) | 0 | 0 | 0 | 0 | 0 | 52144 | 0 | 0 | 16294000 | 1455900 | 0 | 0 | 0 | 1237800 | 0 | 71193 | 0 | 0 |
| GGIMLPEK | HSPE1; HSPE1- | P61604; S4R | CH10_HUMAN | Unmodified | 0 | 0 | 0 | 0 | 0 | 0 | 0 | 0 | 16755000 | 0 | 0 | 200480 | 2101500 | 0 | 0 | 0 | 0 | 0 |
| GGIMSLTE | VPS36; VPS36 | Q86VN1; A0 | VPS36_HUMAN | Oxidation (M) | 1057000 | 760450 | 251610 | 709120 | 0 | 0 | 0 | 0 | 0 | 0 | 0 | 0 | 0 | 0 | 1863700 | 761590 | 0 | 0 |
| GGINLTATC | DRG1 | Q9Y295 | DRG1_HUMAN | Unmodified | 0 | 0 | 0 | 184150 | 0 | 0 | 0 | 0 | 0 | 0 | 0 | 1768200 | 0 | 0 | 0 | 0 | 0 | 0 |
| GGISETRIE | EPB41L2; EPB4 | O43491; P11 | E41L2_HUMAN | Unmodified | 0 | 0 | 133340 | 348670 | 3733000 | 944550 | 469150 | 0 | 700890 | 0 | 0 | 0 | 1544300 | 1386900 | 0 | 0 | 1679000 | 7695100 |
| GGIVDEGA | PHGDH; PHGD | O43175; A0A | SERA_HUMAN | Unmodified | 0 | 0 | 9590000 | 7217700 | 0 | 0 | 0 | 0 | 6043100 | 471740 | 4126100 | 19126000 | 4309300 | 0 | 6692200 | 7551500 | 0 | 0 |
| GGIVGMTL | HSD17B10 | Q99714 | HCD2_HUMAN | Oxidation (M) | 0 | 0 | 0 | 0 | 0 | 0 | 0 | 0 | 2460100 | 274850 | 0 | 0 | 0 | 0 | 0 | 0 | 0 | 0 |
| GGKHHLGI | IGFBP2; IGFBP2 | P18065; C9J | IBP2_HUMAN I | Unmodified | 0 | 0 | 0 | 0 | 0 | 0 | 0 | 0 | 0 | 5141300 | 0 | 0 | 0 | 0 | 0 | 0 | 0 | 0 |
| GGKHHLGI | IGFBP2; IGFBP2 | P18065; C9J | IBP2_HUMAN I | Unmodified | 0 | 0 | 0 | 0 | 0 | 0 | 0 | 0 | 0 | 405220 | 0 | 0 | 0 | 0 | 0 | 0 | 0 | 0 |
| GGKPEPPA | RPS3; RPS3; RP | P23396; H0Y | RS3_HUMAN 4 | Unmodified | 379460 | 445730 | 0 | 0 | 290990 | 0 | 0 | 160180 | 0 | 0 | 969040 | 4000100 | 425610 | 0 | 0 | 429490 | 0 | 0 |
| GGKPGLTI | AHNAK | Q09666 | AHNK_HUMAN | Unmodified | 0 | 0 | 95286 | 0 | 4293200 | 0 | 0 | 0 | 0 | 0 | 0 | 0 | 0 | 0 | 0 | 0 | 0 | 0 |
| GGKSGELE | CHGA; CHGA | P10645; G5E | CMGA_HUMAN | Unmodified | 168980 | 0 | 0 | 0 | 86937 | 0 | 27092 | 0 | 415980 | 2162800 | 277790 | 173950 | 0 | 150940 | 0 | 0 | 0 | 122460 |
| GGLFQGG | WIPF2; WIPF2 | Q8TF74; J3Q | WIPF2_HUMAN | Unmodified | 2096800 | 1484700 | 0 | 1029400 | 0 | 0 | 0 | 0 | 0 | 0 | 0 | 848270 | 0 | 0 | 2945200 | 2000800 | 0 | 0 |
| GGLGVSSS | PAAF1; PAAF1; | Q9BRP4; F5H | PAAF1_HUMAN | Unmodified | 0 | 0 | 0 | 0 | 0 | 0 | 0 | 0 | 0 | 0 | 0 | 459220 | 0 | 0 | 0 | 0 | 0 | 0 |
| GGLPLEEV | ARSA; ARSA | P15289; A0A | ARSA_HUMAN | Unmodified | 0 | 0 | 0 | 0 | 0 | 0 | 0 | 0 | 2316000 | 0 | 0 | 0 | 0 | 0 | 0 | 0 | 0 | 0 |
| GGLQPPDS | APLP1; APLP1; | P51693; B7Z | APLP1_HUMAN | Oxidation (M) | 0 | 0 | 0 | 0 | 0 | 0 | 304070 | 110800 | 1035500 | 0 | 0 | 0 | 0 | 0 | 0 | 0 | 0 | 0 |
| GGLQPPDS | APLP1; APLP1; | P51693; B7Z | APLP1_HUMAN | Unmodified | 0 | 0 | 0 | 0 | 0 | 0 | 0 | 0 | 3382300 | 499830 | 0 | 0 | 0 | 0 | 0 | 0 | 0 | 0 |
| GGLSAVDA | DAG1 | Q14118 | DAG1_HUMAN | Unmodified | 0 | 0 | 0 | 0 | 0 | 0 | 0 | 0 | 0 | 0 | 0 | 0 | 0 | 0 | 0 | 0 | 0 | 0 |
| GGLSVAVP | GGT1; GGT2; G | P19440; P36 | GGT1_HUMAN | Unmodified | 0 | 5415800 | 9105000 | 6241300 | 561180 | 0 | 0 | 219930 | 8146200 | 590520 | 1863200 | 6733300 | 678180 | 0 | 15399000 | 10703000 | 149190 | 334530 |
| GGLSVAVP | GGT1; GGT2; G | P19440; P36 | GGT1_HUMAN | Unmodified | 106970 | 0 | 0 | 4537600 | 179280 | 0 | 0 | 36934 | 3871700 | 125470 | 3613300 | 0 | 0 | 0 | 142870 | 0 | 126590 | 0 |
| GGMALALA | RO60; RO60 | P10155; H0Y | RO60_HUMAN | Oxidation (M) | 2198500 | 2464500 | 748220 | 415830 | 0 | 0 | 0 | 17510 | 0 | 0 | 0 | 0 | 0 | 0 | 1509600 | 1529400 | 0 | 0 |
| GGMALALA | RO60; RO60 | P10155; H0Y | RO60_HUMAN | Unmodified | 0 | 0 | 0 | 0 | 0 | 0 | 0 | 0 | 0 | 0 | 0 | 0 | 0 | 0 | 817420 | 0 | 0 | 0 |
| GGMGSGG | KRT18; KRT18 | P05783; F8V | K1C18_HUMAN | 2 Oxidation (M) | 34268 | 0 | 0 | 0 | 0 | 0 | 0 | 0 | 0 | 0 | 1197900 | 2107200 | 0 | 0 | 0 | 0 | 0 | 0 |
| GGNCLLNI | ADAM9; ADAM9 | Q13443; A0A | ADAM9_HUMA | Unmodified | 2036500 | 430900 | 2026500 | 2323600 | 1354000 | 0 | 0 | 0 | 1600200 | 0 | 0 | 0 | 985360 | 0 | 4291800 | 2284600 | 0 | 0 |
| GGNFGGR | HNRNPA1L2; H | Q32P51; P09 | RA1L2_HUMAN | Unmodified | 0 | 0 | 0 | 0 | 0 | 0 | 96547 | 205910 | 0 | 303890 | 0 | 0 | 967380 | 371140 | 0 | 0 | 0 | 0 |
| GGNIGAKV | HSD17B4; HSD | P51659; E7E | DHB4_HUMAN | Unmodified | 0 | 0 | 0 | 0 | 0 | 0 | 0 | 0 | 1716400 | 986790 | 0 | 0 | 0 | 0 | 0 | 0 | 0 | 0 |
| GGNLRDKI | LRRC59; LRRC5 | Q96AG4; I3L | LRC59_HUMAN | Unmodified | 0 | 0 | 0 | 0 | 2983700 | 0 | 452590 | 0 | 468340 | 0 | 0 | 0 | 0 | 0 | 0 | 0 | 0 | 3178900 |
| GGNLVTGT | KHNYN | O15037 | KHNYN_HUMA | Unmodified | 0 | 0 | 0 | 0 | 0 | 0 | 0 | 0 | 0 | 0 | 0 | 0 | 0 | 0 | 0 | 0 | 0 | 0 |
| GGNLVTGT | KHNYN | O15037 | KHNYN_HUMA | Unmodified | 0 | 0 | 0 | 0 | 1016500 | 226080 | 0 | 0 | 0 | 0 | 0 | 386190 | 207780 | 55823 | 0 | 0 | 0 | 53923 |
| GGNMKEVI | CKB | P12277 | KCRB_HUMAN | Oxidation (M) | 0 | 0 | 0 | 0 | 1241000 | 0 | 11595000 | 0 | 0 | 0 | 0 | 0 | 0 | 3571600 | 0 | 0 | 0 | 0 |
| GGNMKEVI | CKB | P12277 | KCRB_HUMAN | Unmodified | 0 | 0 | 0 | 0 | 418750 | 0 | 2210000 | 0 | 0 | 56565 | 0 | 0 | 481090 | 1752000 | 0 | 0 | 0 | 0 |
| GGNRFEPY | HNRNPM; HNR | P52272; M0R | HNRPM_HUMA | Unmodified | 0 | 0 | 0 | 0 | 0 | 0 | 759870 | 0 | 0 | 0 | 0 | 0 | 0 | 0 | 0 | 0 | 0 | 0 |
| GGNVGINS | FASN; FASN | P49327; A0A | FAS_HUMAN Fa | Unmodified | 0 | 0 | 0 | 0 | 0 | 0 | 0 | 0 | 0 | 0 | 0 | 4911500 | 0 | 0 | 0 | 0 | 0 | 0 |
| GGPDFSNI | NUDCD2; NUD | Q8WVJ2; E5I | NUDC2_HUMA | Unmodified | 0 | 0 | 0 | 553540 | 1422500 | 139100 | 164310 | 130630 | 1843100 | 905800 | 299990 | 581030 | 1217200 | 708840 | 0 | 468770 | 878570 | 1232200 |
| GGPEETITC | SARG | Q9BW04 | SARG_HUMAN | Unmodified | 0 | 0 | 0 | 190370 | 0 | 0 | 0 | 0 | 0 | 0 | 0 | 0 | 1417300 | 0 | 0 | 0 | 0 | 0 |
| GGPEPTPL | GGCX | P38435 | VKGC_HUMAN | Unmodified | 0 | 0 | 0 | 0 | 0 | 0 | 0 | 0 | 2216800 | 0 | 0 | 0 | 0 | 0 | 0 | 0 | 0 | 0 |
| GGPEVQQ | FASN; FASN | P49327; A0A | FAS_HUMAN Fa | Oxidation (M) | 0 | 1021900 | 0 | 0 | 0 | 661170 | 0 | 0 | 0 | 0 | 0 | 17523000 | 0 | 0 | 0 | 0 | 171010 | 0 |
| GGPGSAVS | ANXA1; ANXA1; | P04083; Q5T | ANXA1_HUMAN | Unmodified | 760830 | 0 | 0 | 0 | 36307000 | 11257000 | 2003500 | 139670 | 294540 | 997890 | 3.58E+08 | 2.47E+09 | 0 | 0 | 0 | 441780 | 0 | 0 |
| GGPGSAVS | ANXA1; ANXA1; | P04083; Q5T | ANXA1_HUMAN | Oxidation (M) | 0 | 0 | 0 | 0 | 3825300 | 0 | 0 | 0 | 0 | 0 | 0 | 1507500 | 0 | 0 | 0 | 0 | 0 | 0 |
| GGPGSEC | MDK; MDK; MD | P21741; E9P | MK_HUMAN Mi | Unmodified | 0 | 0 | 0 | 0 | 0 | 0 | 0 | 0 | 8586700 | 763120 | 0 | 0 | 0 | 0 | 0 | 0 | 0 | 0 |
| GGPNIITLA | CSNK2A3; CSN | Q8NEV1; P68 | CSK23_HUMAN | Unmodified | 1935900 | 826760 | 1192900 | 1165000 | 0 | 0 | 20030 | 0 | 894960 | 42686 | 1554700 | 22446000 | 31720 | 0 | 1660000 | 1122900 | 0 | 0 |
| GGPNYQE | ACLY; ACLY | P53396; K7E | ACLY_HUMAN | Unmodified | 0 | 0 | 0 | 0 | 0 | 0 | 0 | 0 | 0 | 0 | 0 | 2964700 | 0 | 0 | 0 | 0 | 0 | 0 |
| GGPPFAFV | SRSF1; SRSF1 | Q07955; J3K | SRSF1_HUMAN | Unmodified | 2060700 | 1731300 | 0 | 0 | 0 | 0 | 0 | 0 | 0 | 0 | 0 | 0 | 0 | 0 | 1072100 | 360570 | 0 | 0 |
| GGPPPPPP | WASL | O00401 | WASL_HUMAN | Unmodified | 0 | 0 | 0 | 93920 | 0 | 0 | 0 | 0 | 0 | 122370 | 0 | 0 | 0 | 0 | 728810 | 392500 | 18910 | 0 |
| GGPQIAYE | WWP1; WWP1 | Q9H0M0; H0 | WWP1_HUMAN | Unmodified | 292820 | 404040 | 819220 | 597420 | 616590 | 0 | 0 | 0 | 411820 | 0 | 0 | 0 | 0 | 0 | 1169400 | 823590 | 0 | 440090 |
| GGPTPQEA | CHMP4B | Q9H444 | CHM4B_HUMA | Unmodified | 9249000 | 8194100 | 6137800 | 5305700 | 0 | 0 | 1249800 | 0 | 12642000 | 130340 | 3269800 | 4844200 | 988690 | 0 | 6605500 | 5912700 | 0 | 0 |
| GGPTVDPE | DNAJA3 | Q96EY1 | DNJA3_HUMAN | Unmodified | 0 | 0 | 0 | 0 | 0 | 0 | 0 | 0 | 1072000 | 0 | 0 | 0 | 0 | 0 | 0 | 0 | 0 | 0 |
| GGPVQVLE | SELENBP1 | Q13228 | SBP1_HUMAN | Unmodified | 0 | 0 | 1074100 | 1639900 | 0 | 0 | 0 | 0 | 0 | 0 | 0 | 0 | 898390 | 545320 | 1808500 | 1570900 | 0 | 771170 |
| GGPVQVLE | SELENBP1 | Q13228 | SBP1_HUMAN | Unmodified | 0 | 0 | 0 | 0 | 0 | 0 | 0 | 0 | 0 | 0 | 0 | 0 | 1379300 | 1360800 | 0 | 0 | 0 | 2072700 |
| GGQEPDRI | CNOT11 | Q9UKZ1 | CNO11_HUMA | Unmodified | 0 | 0 | 0 | 0 | 0 | 0 | 0 | 0 | 0 | 0 | 596360 | 1641900 | 0 | 0 | 0 | 715900 | 0 | 0 |
| GGQIGLQA | AHNAK | Q09666 | AHNK_HUMAN | Unmodified | 0 | 0 | 0 | 0 | 2352800 | 739240 | 0 | 0 | 0 | 0 | 0 | 0 | 0 | 0 | 122390 | 105100 | 0 | 1315200 |
| GGQIGLQA | AHNAK | Q09666 | AHNK_HUMAN | Unmodified | 0 | 0 | 0 | 0 | 12121000 | 0 | 0 | 0 | 0 | 0 | 0 | 0 | 0 | 0 | 0 | 0 | 2161500 | 7369700 |
| GGQQALL | CUL4A; CUL4A | Q13619; A0A | CUL4A_HUMAN | Unmodified | 0 | 0 | 0 | 0 | 0 | 216660 | 0 | 0 | 29311 | 107170 | 0 | 1692800 | 0 | 0 | 0 | 0 | 0 | 0 |
| GGQQLVFN | ANK3; ANK3; AN | Q12955; A0A | ANK3_HUMAN | Unmodified | 0 | 0 | 919630 | 924970 | 0 | 0 | 0 | 0 | 0 | 0 | 23340 | 147470 | 3945600 | 3694400 | 869770 | 1364900 | 0 | 0 |
| GGQQLVFN | ANK3; ANK3; AN | Q12955; A0A | ANK3_HUMAN | Unmodified | 0 | 0 | 0 | 0 | 803660 | 238270 | 0 | 0 | 0 | 0 | 0 | 0 | 769720 | 2291200 | 0 | 0 | 0 | 667130 |
| GGQTASFA | GTPBP1; GTPBP | O00178; B0 | GTPB1_HUMAN | Unmodified | 865900 | 601600 | 389890 | 1229600 | 0 | 0 | 0 | 0 | 0 | 258730 | 924060 | 68663000 | 0 | 151970 | 468540 | 578140 | 0 | 0 |
| GGQTREH | GSPT1; GSPT2; | P15170; Q8I | ERF3A_HUMAN | Unmodified | 0 | 0 | 478820 | 0 | 318730 | 56937 | 403010 | 346810 | 0 | 0 | 24566 | 4886500 | 0 | 107940 | 0 | 0 | 480330 | 487260 |
| GGQVIQIP | ATP2B1 | P20020 | AT2B1_HUMAN | Unmodified | 165310 | 410060 | 134660 | 748680 | 0 | 0 | 0 | 0 | 0 | 147540 | 127450 | 327580 | 0 | 0 | 2866200 | 631760 | 0 | 0 |
| GGQYFNDI | SV2A | Q7L0J3 | SV2A_HUMAN | Unmodified | 0 | 0 | 0 | 0 | 0 | 0 | 0 | 0 | 5093800 | 189140 | 0 | 345480 | 0 | 0 | 0 | 0 | 0 | 0 |
| GGQYFNDI | SV2A | Q7L0J3 | SV2A_HUMAN | Unmodified | 0 | 0 | 0 | 0 | 0 | 0 | 0 | 0 | 300270 | 1239500 | 0 | 0 | 0 | 0 | 0 | 0 | 0 | 0 |
| GGRGDVG | SRPRB; SRPRB | Q9Y5M8; H7 | SRPRB_HUMAN | Unmodified | 67313 | 0 | 102070 | 123720 | 0 | 0 | 140020 | 56151 | 288390 | 83309 | 275670 | 657900 | 382560 | 571680 | 0 | 54561 | 0 | 0 |
| GGRGGGR | NCL; OAZ2; EW | P19338; O95 | NUCL_HUMAN | Unmodified | 0 | 0 | 0 | 0 | 0 | 0 | 0 | 0 | 1213400 | 0 | 0 | 0 | 0 | 0 | 0 | 0 | 0 | 0 |
| GGRPQVAE | C1orf210 | Q8IVY1 | CA210_HUMAN | Unmodified | 3990000 | 7800900 | 0 | 3348200 | 0 | 0 | 0 | 0 | 0 | 733570 | 0 | 0 | 0 | 0 | 0 | 5027300 | 0 | 0 |
| GGRSPDEV | PACSIN3; PACS | Q9UKS6; H0 | PACN3_HUMAN | Unmodified | 5297400 | 5328000 | 2202100 | 1791100 | 0 | 0 | 0 | 0 | 118170 | 0 | 0 | 0 | 0 | 0 | 3506000 | 134250 | 0 | 0 |
| GGSASVW | ALDH16A1; ALD | Q8IZ83; F5H | A16A1_HUMAN | Unmodified | 135940 | 540980 | 0 | 0 | 0 | 0 | 0 | 0 | 0 | 0 | 0 | 922860 | 0 | 0 | 0 | 0 | 0 | 0 |
| GGSDDSSI | PARP1; PARP1 | P09874; A0A | PARP1_HUMAN | Unmodified | 673070 | 568270 | 175070 | 0 | 0 | 0 | 85998 | 69573 | 982260 | 168130 | 766040 | 4356200 | 0 | 0 | 261620 | 153070 | 0 | 0 |
| GGSDSDAS | HAPLN1; HAPLI | P10915; D6F | HPLN1_HUMAN | Unmodified | 0 | 0 | 0 | 0 | 0 | 0 | 0 | 0 | 880880 | 0 | 0 | 0 | 0 | 0 | 0 | 0 | 0 | 0 |
| GGSDSDAS | HAPLN1; HAPLI | P10915; D6F | HPLN1_HUMAN | Unmodified | 0 | 0 | 0 | 0 | 0 | 0 | 0 | 0 | 5997900 | 1412800 | 0 | 0 | 0 | 0 | 0 | 0 | 0 | 0 |
| GGSGSGP | HSPA1A; HSPA | P0DMV8; P0 | HS71A_HUMAN | Unmodified | 4958800 | 3934700 | 2482500 | 0 | 6161800 | 985440 | 1615000 | 633950 | 8314200 | 2143500 | 1028700 | 4743000 | 5002400 | 1848700 | 2887800 | 0 | 1208800 | 2413600 |
| GGSGTAGT | EFNB1 | P98172 | EFNB1_HUMAN | Unmodified | 0 | 0 | 0 | 0 | 0 | 0 | 0 | 0 | 0 | 0 | 0 | 791010 | 0 | 0 | 0 | 0 | 526910 | 1395800 |
| GGSGTAGT | EFNB1 | P98172 | EFNB1_HUMAN | Unmodified | 0 | 0 | 0 | 0 | 2286600 | 1461700 | 0 | 0 | 0 | 0 | 0 | 0 | 0 | 0 | 0 | 0 | 5768600 | 21965000 |
| GGSHDAPI | GDE1; GDE1 | Q9NZC3; H3 | GDE1_HUMAN | Unmodified | 0 | 0 | 0 | 0 | 0 | 0 | 0 | 0 | 2529100 | 174060 | 304450 | 1439400 | 0 | 0 | 0 | 0 | 0 | 0 |
| GGSLPKVE | NPM1; NPM1; N | P06748; A0A | NPM_HUMAN N | Unmodified | 37527 | 0 | 0 | 0 | 684630 | 36741 | 163920 | 193460 | 137820 | 288890 | 251350 | 1634000 | 1068900 | 570030 | 0 | 0 | 172190 | 242680 |
| GGSLPKVE | NPM1; NPM1; N | P06748; A0A | NPM_HUMAN N | Unmodified | 0 | 0 | 0 | 0 | 331360 | 0 | 213540 | 0 | 0 | 171480 | 0 | 0 | 283740 | 0 | 0 | 0 | 476760 | 245210 |
| GGSQMDV | UTRN | P46939 | UTRN_HUMAN | Oxidation (M) | 0 | 0 | 0 | 0 | 0 | 0 | 0 | 0 | 0 | 0 | 0 | 0 | 0 | 0 | 0 | 145950 | 0 | 0 |
| GGSRGEEV | CNP | P09543 | CN37_HUMAN | Unmodified | 553770 | 139590 | 3928200 | 4273600 | 9847.2 | 0 | 0 | 0 | 7063800 | 292400 | 14093000 | 65029000 | 0 | 0 | 5276900 | 5436600 | 108990 | 389870 |
| GGSRGEEV | CNP | P09543 | CN37_HUMAN | Unmodified | 0 | 0 | 0 | 0 | 3533600 | 161220 | 138160 | 111740 | 260130 | 738570 | 8237800 | 0 | 33768 | 1162600 | 0 | 0 | 657540 | 117260 |
| GGSTSYGT | FGA | P02671 | FIBA_HUMAN F | Unmodified | 0 | 634780 | 0 | 0 | 0 | 0 | 0 | 0 | 0 | 0 | 0 | 0 | 0 | 0 | 0 | 0 | 0 | 0 |
| GGSVLVTC | ICAM1; ICAM1 | P05362; K7E | ICAM1_HUMAN | Unmodified | 0 | 0 | 0 | 0 | 5509100 | 1050400 | 0 | 380460 | 48561 | 0 | 76478 | 0 | 0 | 0 | 85611 | 0 | 1137800 | 701280 |
| GGSVLVTC | ICAM1; ICAM1 | P05362; K7E | ICAM1_HUMAN | Unmodified | 0 | 0 | 0 | 0 | 815040 | 0 | 0 | 0 | 0 | 0 | 0 | 0 | 0 | 0 | 0 | 0 | 1529100 | 0 |
| GGSVPPQI | SNX18 | Q96RF0 | SNX18_HUMAN | Unmodified | 20924 | 112910 | 779310 | 465520 | 0 | 0 | 0 | 0 | 0 | 0 | 0 | 0 | 1197200 | 79792 | 1533500 | 700930 | 0 | 0 |
| GGSYLGRE | QSOX1 | O00391 | QSOX1_HUMAN | Unmodified | 0 | 0 | 0 | 0 | 0 | 0 | 706700 | 158900 | 0 | 879260 | 0 | 0 | 73779 | 1949400 | 86004 | 0 | 0 | 0 |
| GGSYSQA | HLA-A; HLA-A; | Q9GJ45; Q5I | Q9GJ45_HUMA | Unmodified | 0 | 0 | 0 | 0 | 1522900 | 0 | 0 | 0 | 0 | 0 | 0 | 0 | 0 | 0 | 0 | 0 | 0 | 0 |
| GGSYSQA | HLA-A; HLA-A; | Q53Z42; A0A | Q53Z42_HUMA | Unmodified | 0 | 0 | 0 | 348890 | 3333500 | 0 | 0 | 0 | 0 | 0 | 821520 | 17899000 | 0 | 0 | 89673 | 48632 | 0 | 0 |
| GGSYSQA | HLA-A; HLA-A; | Q53Z42; A0A | Q53Z42_HUMA | Unmodified | 0 | 0 | 0 | 0 | 0 | 238900 | 0 | 0 | 0 | 0 | 0 | 6029800 | 0 | 0 | 0 | 0 | 407370 | 1586400 |
| GGSYTQAA | HLA-A; HLA-A; | P04439; A0A | HLAA_HUMAN | Unmodified | 0 | 0 | 0 | 0 | 0 | 0 | 0 | 0 | 1566900 | 0 | 0 | 1913500 | 0 | 0 | 0 | 0 | 393790 | 0 |
| GGTFYVEP | ADAM10; ADAM | O14672; H3I | ADA10_HUMAN | Unmodified | 0 | 0 | 0 | 0 | 0 | 0 | 0 | 0 | 2420100 | 0 | 0 | 0 | 0 | 0 | 0 | 0 | 0 | 665360 |

| | | | | | | | | | | | | | | | | | | | | | | |
|---|---|---|---|---|---|---|---|---|---|---|---|---|---|---|---|---|---|---|---|---|---|---|
| GGTLKDDI | USP14 | P54578 | UBP14_HUMAN | Unmodified | 319850 | 133000 | 198670 | 0 | 34004 | 87973 | 0 | 74911 | 0 | 44091 | 0 | 5704700 | 200430 | 0 | 735120 | 81393 | 0 | 0 |
| GGTLTQYE | UGP2; UGP2; U | Q16851; E7E | UGPA_HUMAN | Unmodified | 66294 | 0 | 0 | 0 | 1176100 | 0 | 0 | 57304 | 1428600 | 451750 | 243680 | 2064200 | 555340 | 1218300 | 0 | 149510 | 137950 | 710420 |
| GGTWEFLC | SORL1 | Q92673 | SORL_HUMAN | Unmodified | 0 | 0 | 0 | 0 | 0 | 0 | 0 | 0 | 1577400 | 208600 | 105700 | 0 | 0 | 0 | 0 | 0 | 0 | 0 |
| GGVAEACF | TBCE; TBCE; TB | Q15813; A0A | TBCE_HUMAN | Unmodified | 0 | 0 | 0 | 0 | 0 | 0 | 0 | 0 | 0 | 0 | 0 | 0 | 0 | 0 | 0 | 0 | 0 | 0 |
| GGVAGGSI | EIF2S3; EIF2S3I | P41091; Q2V | IF2G_HUMAN E | Unmodified | 0 | 164580 | 511920 | 0 | 96160 | 0 | 12602 | 0 | 0 | 98554 | 0 | 221050 | 0 | 0 | 0 | 0 | 305740 | 198850 |
| GGVDVTLP | AHNAK | Q09666 | AHNK_HUMAN | Unmodified | 456070 | 0 | 0 | 0 | 3477600 | 953710 | 0 | 0 | 336190 | 0 | 0 | 2415800 | 0 | 0 | 0 | 0 | 1280500 | 1630700 |
| GGVEPAGF | LHFPL2 | Q6ZUX7 | LHPL2_HUMAN | Unmodified | 1367400 | 1430000 | 0 | 0 | 0 | 0 | 0 | 0 | 1975000 | 332010 | 0 | 0 | 0 | 0 | 227250 | 593900 | 137370 | 0 |
| GGVGAPGS | PPM1H | Q9ULR3 | PPM1H_HUMA | Unmodified | 0 | 0 | 0 | 0 | 0 | 0 | 0 | 0 | 0 | 0 | 0 | 1013600 | 0 | 0 | 0 | 0 | 0 | 0 |
| GGVGSRAD | PKP3; PKP3; PK | Q9Y446; E9F | PKP3_HUMAN | Unmodified | 164250 | 245490 | 821360 | 1200200 | 0 | 0 | 0 | 0 | 743610 | 0 | 957920 | 0 | 351790 | 166820 | 947020 | 805180 | 0 | 0 |
| GGVHLTKD | SUCLG2; SUCL | Q96I99; E9P | SUCB2_HUMAN | Unmodified | 0 | 0 | 0 | 0 | 0 | 0 | 239150 | 0 | 1240200 | 974720 | 0 | 0 | 0 | 0 | 0 | 0 | 0 | 0 |
| GGVLIQRN | ERBB2; ERBB2; | P04626; J3Q | ERBB2_HUMAN | Unmodified | 0 | 0 | 0 | 1290000 | 978320 | 0 | 0 | 0 | 0 | 0 | 0 | 0 | 286750 | 0 | 0 | 0 | 0 | 0 |
| GGVTEISAA | XPNPEP1; XPNI | Q9NQW7; Q | XPP1_HUMAN | Unmodified | 69256 | 0 | 0 | 123070 | 0 | 0 | 0 | 0 | 0 | 0 | 0 | 2211600 | 0 | 0 | 0 | 104680 | 0 | 0 |
| GGVTEISAA | XPNPEP1; XPNI | Q9NQW7; Q | XPP1_HUMAN | Unmodified | 188280 | 0 | 0 | 0 | 208910 | 0 | 49688 | 0 | 311670 | 0 | 423000 | 5932000 | 0 | 0 | 0 | 0 | 27804 | 0 |
| GGVTGSPE | AHNAK | Q09666 | AHNK_HUMAN | Unmodified | 0 | 0 | 0 | 0 | 2698600 | 106760 | 0 | 0 | 0 | 0 | 0 | 0 | 0 | 0 | 0 | 0 | 919910 | 984150 |
| GGVVLKED | BSG; BSG; BSG | P35613; A0A | BASI_HUMAN E | Unmodified | 824830 | 988020 | 2249000 | 5478800 | 9060100 | 1738300 | 1372400 | 844060 | 7122600 | 652200 | 985540 | 14840000 | 4572000 | 2641300 | 8197700 | 8477200 | 3521700 | 11459000 |
| GGVVLKED | BSG; BSG; BSG | P35613; A0A | BASI_HUMAN E | Unmodified | 0 | 0 | 0 | 0 | 1729000 | 0 | 0 | 0 | 0 | 0 | 0 | 1440200 | 1548500 | 1627200 | 0 | 2959100 | 524910 | 1662300 |
| GGVVTSNF | ACADVL; ACAD | P49748; G3V | ACADV_HUMAN | Unmodified | 0 | 0 | 0 | 0 | 0 | 0 | 0 | 0 | 1580500 | 116800 | 0 | 0 | 0 | 0 | 0 | 0 | 0 | 0 |
| GGWDHGD | KIAA1522 | Q9P206 | K1522_HUMAN | Unmodified | 0 | 0 | 396310 | 422970 | 0 | 0 | 0 | 0 | 0 | 0 | 180060 | 636030 | 0 | 0 | 706400 | 546960 | 0 | 0 |
| GGWSDNG | ADGRG2 | Q8IZP9 | AGRG2_HUMA | Unmodified | 0 | 0 | 0 | 0 | 0 | 0 | 0 | 0 | 756050 | 0 | 0 | 0 | 0 | 0 | 0 | 0 | 0 | 0 |
| GGYGGGM | CDC42SE2; CD | Q9NRR3; D6 | C42S2_HUMAN | Oxidation (M) | 0 | 0 | 0 | 152470 | 392450 | 0 | 0 | 0 | 0 | 0 | 0 | 0 | 0 | 0 | 302650 | 0 | 0 | 0 |
| GGYGGGM | CDC42SE2; CD | Q9NRR3; D6 | C42S2_HUMAN | 2 Oxidation (M) | 0 | 0 | 0 | 291760 | 0 | 0 | 0 | 48742 | 164980 | 174860 | 0 | 374780 | 149740 | 0 | 117490 | 188420 | 0 | 0 |
| GGYGGGM | CDC42SE2; CD | Q9NRR3; D6 | C42S2_HUMAN | Unmodified | 0 | 0 | 0 | 0 | 2685600 | 51813 | 0 | 0 | 0 | 0 | 0 | 0 | 0 | 468850 | 0 | 96190 | 0 | 1562200 |
| GGYGGGM | CDC42SE2; CD | Q9NRR3; D6 | C42S2_HUMAN | Oxidation (M) | 0 | 0 | 0 | 0 | 3923800 | 0 | 0 | 0 | 0 | 616990 | 0 | 0 | 1952900 | 310710 | 0 | 0 | 696330 | 1494700 |
| GGYIGSTYI | UBE2M | P61081 | UBC12_HUMAN | Unmodified | 571840 | 716450 | 1152500 | 956720 | 0 | 0 | 0 | 0 | 2397500 | 489550 | 993030 | 6071700 | 715010 | 0 | 0 | 0 | 230800 | 0 |
| GGYIGSTYI | UBE2M | P61081 | UBC12_HUMAN | Unmodified | 207400 | 0 | 0 | 0 | 0 | 0 | 0 | 0 | 0 | 76982 | 0 | 0 | 0 | 0 | 0 | 0 | 494140 | 439990 |
| GGYKFFAD | HPRT1 | P00492 | HPRT_HUMAN | Unmodified | 0 | 0 | 0 | 0 | 86930 | 0 | 266940 | 0 | 0 | 82000 | 0 | 0 | 0 | 9781.2 | 0 | 0 | 7742.3 | 0 |
| GGYLQGNY | SLC7A11 | Q9UPY5 | XCT_HUMAN C | Unmodified | 0 | 0 | 357990 | 497430 | 1431400 | 0 | 0 | 0 | 0 | 0 | 0 | 2289000 | 325410 | 0 | 844580 | 551240 | 0 | 0 |
| GGYVLHIG | AARS1; AARS1; | P49588; A0A | SYAC_HUMAN | Unmodified | 393450 | 0 | 1034800 | 747350 | 144970 | 0 | 184230 | 0 | 3319000 | 848750 | 2135200 | 9923500 | 2434100 | 1068400 | 1457300 | 980600 | 0 | 0 |
| GHAAFTSD | GSR | P00390 | GSHR_HUMAN | Unmodified | 244330 | 0 | 430490 | 508610 | 0 | 0 | 0 | 0 | 556800 | 134610 | 178490 | 2107300 | 0 | 0 | 596430 | 462000 | 0 | 0 |
| GHAGNPRI | SCRIB; SCRIB; S | Q14160; A0A | SCRIB_HUMAN | Unmodified | 0 | 0 | 0 | 840580 | 1110900 | 0 | 0 | 0 | 6647900 | 1677300 | 552250 | 2337700 | 765060 | 241570 | 811000 | 242270 | 1710900 | 2717400 |
| GHALYVEP | SLC39A3; SLC3 | Q9BRY0; F5H | S39A3_HUMAN | Unmodified | 0 | 0 | 366720 | 0 | 0 | 0 | 0 | 0 | 384410 | 54091 | 0 | 0 | 0 | 0 | 0 | 0 | 0 | 0 |
| GHAVGDIP | RPS23; RPS23 | P62266; D6F | RS23_HUMAN | Unmodified | 327500 | 364460 | 294790 | 136450 | 36525 | 0 | 0 | 0 | 0 | 0 | 242780 | 2073900 | 0 | 0 | 272590 | 245190 | 0 | 0 |
| GHAVGDIP | RPS23; RPS23 | P62266; D6F | RS23_HUMAN | Unmodified | 0 | 0 | 0 | 0 | 320630 | 56444 | 144640 | 152220 | 0 | 42982 | 0 | 0 | 0 | 0 | 0 | 0 | 0 | 0 |
| GHAVGDIP | RPS23 | P62266 | RS23_HUMAN | Unmodified | 0 | 0 | 0 | 0 | 0 | 0 | 311370 | 0 | 0 | 0 | 0 | 0 | 0 | 0 | 0 | 0 | 0 | 0 |
| GHAVLAGN | PDIA5 | Q14554 | PDIA5_HUMAN | Oxidation (M) | 0 | 0 | 0 | 0 | 0 | 0 | 0 | 0 | 1332800 | 0 | 0 | 0 | 0 | 0 | 0 | 0 | 0 | 0 |
| GHAVLAGN | PDIA5 | Q14554 | PDIA5_HUMAN | Unmodified | 0 | 0 | 0 | 0 | 0 | 0 | 0 | 0 | 7884500 | 0 | 0 | 0 | 0 | 0 | 0 | 0 | 0 | 679230 |
| GHCVDLPD | SPINT1; SPINT1 | O43278; H3I | SPIT1_HUMAN | Unmodified | 507870 | 252010 | 601070 | 308130 | 0 | 0 | 0 | 0 | 546160 | 0 | 0 | 0 | 0 | 0 | 322210 | 224580 | 0 | 0 |
| GHDLEFCL | SORT1 | Q99523 | SORT_HUMAN | Unmodified | 0 | 0 | 0 | 0 | 0 | 0 | 0 | 0 | 2802700 | 0 | 0 | 0 | 0 | 0 | 0 | 0 | 0 | 0 |
| GHDLNED( | CALU | O43852 | CALU_HUMAN | Unmodified | 0 | 0 | 0 | 0 | 0 | 0 | 134310 | 80705 | 8787300 | 0 | 1453300 | 821690 | 0 | 0 | 0 | 0 | 0 | 0 |
| GHDSTEEP | AOC1; AOC1 | P19801; C9J | AOC1_HUMAN | Unmodified | 0 | 0 | 1451400 | 0 | 0 | 0 | 0 | 57917 | 1070400 | 0 | 0 | 0 | 0 | 0 | 0 | 0 | 0 | 0 |
| GHEDGGIF | BTN2A1 | Q7KYR7 | BT2A1_HUMAN | Unmodified | 0 | 0 | 140120 | 0 | 0 | 0 | 0 | 0 | 0 | 0 | 0 | 1126800 | 0 | 0 | 0 | 194530 | 0 | 0 |
| GHEGDFLI | NCK1; NCK1 | P16333; C9J | NCK1_HUMAN | Unmodified | 158870 | 70240 | 56157 | 0 | 0 | 0 | 186470 | 0 | 0 | 0 | 0 | 336750 | 0 | 0 | 495990 | 286550 | 0 | 0 |
| GHEGSVAC | WDFY1; WDFY1 | Q8IWB7; C9 | WDFY1_HUMA | Unmodified | 0 | 0 | 0 | 0 | 0 | 0 | 0 | 0 | 0 | 0 | 0 | 345050 | 0 | 0 | 0 | 42298 | 0 | 0 |
| GHELVLAN | PTK7; PTK7; PTI | Q13308; Q8 | PTK7_HUMAN I | Unmodified | 0 | 0 | 0 | 889550 | 0 | 0 | 0 | 0 | 3111800 | 0 | 0 | 0 | 0 | 0 | 922360 | 727070 | 0 | 0 |
| GHENVEAA | PCCB; PCCB; P | P05166; C9J | PCCB_HUMAN | Unmodified | 0 | 0 | 0 | 0 | 0 | 0 | 0 | 0 | 719010 | 88147 | 86351 | 652380 | 0 | 0 | 0 | 0 | 28332 | 0 |
| GHETSALLI | ANKRD28 | O15084 | ANR28_HUMAN | Unmodified | 0 | 0 | 0 | 0 | 28780 | 0 | 0 | 0 | 0 | 100490 | 0 | 1124200 | 311750 | 0 | 0 | 0 | 0 | 20885 |
| GHFFVEDQ | PDLIM1 | O00151 | PDLI1_HUMAN | Unmodified | 0 | 0 | 0 | 95991 | 3855800 | 575960 | 1047000 | 174240 | 780550 | 200370 | 35585 | 0 | 395890 | 1124700 | 0 | 0 | 5359000 | 4084600 |
| GHFGPINS | EIF3I; EIF3I; EIF | Q13347; A0A | EIF3I_HUMAN I | Unmodified | 377580 | 101120 | 0 | 0 | 0 | 0 | 502070 | 439600 | 262900 | 0 | 838140 | 9854900 | 1740600 | 1439100 | 0 | 0 | 1780100 | 1554100 |
| GHFPAAYV | NOSTRIN; NOS | Q8IVI9; A0A | NOSTN_HUMA | Unmodified | 2768000 | 2806200 | 4361100 | 4643700 | 0 | 0 | 0 | 0 | 0 | 0 | 0 | 0 | 1551100 | 1041300 | 7168600 | 3814700 | 0 | 0 |
| GHGFITPAI | CARHSP1; CAR | Q9Y2V2; H3I | CHSP1_HUMAN | Unmodified | 0 | 0 | 0 | 0 | 1521300 | 0 | 675760 | 0 | 0 | 1779600 | 0 | 0 | 2419300 | 2033600 | 0 | 0 | 2706600 | 5108100 |
| GHIASVLNA | ME1 | P48163 | MAOX_HUMAN | Unmodified | 0 | 0 | 0 | 0 | 0 | 0 | 0 | 0 | 0 | 0 | 156920 | 2397300 | 0 | 0 | 0 | 0 | 0 | 0 |
| GHIAVSAAV | XPNPEP1; XPNI | Q9NQW7; Q | XPP1_HUMAN | Unmodified | 0 | 0 | 0 | 0 | 0 | 0 | 387150 | 246090 | 0 | 0 | 0 | 800550 | 0 | 132330 | 0 | 0 | 0 | 0 |
| GHIISDGG | GMPR2; GMPR2 | Q9P2T1; H0Y | GMPR2_HUMA | Unmodified | 0 | 0 | 0 | 0 | 0 | 100320 | 186540 | 84684 | 191320 | 110260 | 0 | 971590 | 424900 | 0 | 0 | 0 | 145350 | 355550 |
| GHIIVDEFQ | GSR | P00390 | GSHR_HUMAN | Unmodified | 309810 | 0 | 0 | 104710 | 1307700 | 1147700 | 238300 | 0 | 1922000 | 0 | 911930 | 8671100 | 1117600 | 661690 | 1727000 | 1413700 | 53863 | 526860 |
| GHLENNPA | RPLP0; RPLP0P | P05388; Q8N | RLA0_HUMAN | Unmodified | 20843000 | 19289000 | 16226000 | 11416000 | 0 | 172690 | 0 | 71066 | 2279500 | 0 | 5299200 | 64593000 | 1554100 | 250520 | 27084000 | 21887000 | 135730 | 304010 |
| GHLESSLLS | B3GAT3; B3GAT | O94766; G3V | B3GA3_HUMAN | Unmodified | 0 | 0 | 96798 | 76148 | 0 | 0 | 0 | 0 | 10781000 | 562500 | 894410 | 1264600 | 0 | 0 | 0 | 0 | 0 | 0 |
| GHLESSLLS | B3GAT3; B3GAT | O94766; G3V | B3GA3_HUMAN | Unmodified | 0 | 0 | 0 | 297150 | 0 | 0 | 0 | 0 | 8378900 | 456310 | 0 | 1058300 | 0 | 0 | 0 | 0 | 0 | 0 |
| GHLFGPAP | ROR2; ROR2 | Q01974; B1A | ROR2_HUMAN | Unmodified | 0 | 0 | 0 | 0 | 0 | 0 | 0 | 0 | 1985300 | 0 | 0 | 0 | 0 | 0 | 0 | 0 | 0 | 0 |
| GHLLLDSD | ABCC5 | O15440 | MRP5_HUMAN | Unmodified | 0 | 0 | 0 | 0 | 0 | 0 | 0 | 0 | 0 | 0 | 429180 | 2804900 | 0 | 0 | 0 | 0 | 0 | 0 |
| GHLPKVEM | AHNAK2 | Q8IVF2 | AHNK2_HUMA | Unmodified | 0 | 0 | 0 | 0 | 1224100 | 0 | 0 | 0 | 0 | 0 | 0 | 0 | 0 | 0 | 0 | 0 | 115210 | 1339900 |
| GHNGWVT | RACK1; RACK1; | P63244; D6F | RACK1_HUMAN | Oxidation (M) | 820410 | 0 | 0 | 0 | 1320300 | 0 | 0 | 0 | 0 | 0 | 230840 | 2904700 | 0 | 0 | 0 | 0 | 0 | 0 |
| GHNGWVT | RACK1; RACK1; | P63244; D6F | RACK1_HUMAN | Oxidation (M) | 0 | 5864300 | 824190 | 4941000 | 0 | 0 | 1357200 | 0 | 0 | 0 | 0 | 11246000 | 0 | 0 | 1447200 | 1160100 | 0 | 0 |
| GHPDLQGL | LMAN1 | P49257 | LMAN1_HUMA | Unmodified | 0 | 0 | 0 | 0 | 0 | 0 | 0 | 0 | 4630500 | 1201000 | 4778800 | 53507 | 0 | 0 | 0 | 0 | 0 | 0 |
| GHPDLQGL | LMAN1 | P49257 | LMAN1_HUMA | Unmodified | 237160 | 0 | 2709900 | 2682200 | 0 | 0 | 0 | 0 | 1.08E+08 | 4267100 | 7573900 | 3430400 | 829880 | 552110 | 2700700 | 1229800 | 0 | 0 |
| GHPEFSTN | USP5 | P45974 | UBP5_HUMAN | Unmodified | 0 | 0 | 0 | 0 | 0 | 0 | 0 | 0 | 0 | 0 | 0 | 0 | 0 | 0 | 0 | 0 | 0 | 0 |
| GHPLGATG | SCP2 | P22307 | SCP2_HUMAN | Unmodified | 0 | 0 | 0 | 0 | 0 | 0 | 61620 | 0 | 1119700 | 0 | 0 | 0 | 0 | 0 | 0 | 0 | 0 | 0 |
| GHQAFDV( | IGF2R; IGF2R; I | P11717; A0A | MPRI_HUMAN | Unmodified | 23009 | 45020 | 0 | 0 | 0 | 0 | 0 | 0 | 19253000 | 569390 | 1208200 | 2842300 | 0 | 0 | 0 | 27231 | 0 | 0 |
| GHQAFDV( | IGF2R; IGF2R; I | P11717; A0A | MPRI_HUMAN | Unmodified | 0 | 0 | 0 | 0 | 0 | 0 | 0 | 0 | 14203000 | 3450500 | 3175000 | 0 | 0 | 0 | 0 | 0 | 0 | 0 |
| GHQKLEEM | TSG101; TSG10 | Q99816; F5H | TS101_HUMAN | Unmodified | 662250 | 753590 | 205640 | 452910 | 35552 | 0 | 0 | 0 | 246420 | 0 | 99375 | 0 | 70747 | 0 | 941200 | 831620 | 0 | 0 |
| GHQKLEEM | TSG101; TSG10 | Q99816; F5H | TS101_HUMAN | Unmodified | 5313500 | 3498600 | 1326600 | 3116500 | 284750 | 0 | 0 | 0 | 3757800 | 324180 | 0 | 0 | 1317100 | 0 | 15233000 | 3937100 | 0 | 0 |
| GHQKLEEM | TSG101; TSG10 | Q99816; F5H | TS101_HUMAN | Oxidation (M) | 1750600 | 964750 | 800660 | 1539900 | 0 | 0 | 0 | 0 | 216920 | 0 | 264950 | 0 | 274530 | 0 | 3355200 | 2380500 | 0 | 0 |
| GHRIEEVPI | RPL4; RPL4 | P36578; H3E | RL4_HUMAN 6 | Unmodified | 0 | 0 | 100330 | 0 | 0 | 0 | 0 | 0 | 0 | 0 | 0 | 395320 | 0 | 0 | 0 | 0 | 0 | 0 |
| GHRIEEVPI | RPL4; RPL4 | P36578; H3E | RL4_HUMAN 6 | Unmodified | 0 | 0 | 317520 | 471800 | 0 | 0 | 0 | 0 | 0 | 0 | 717390 | 12271000 | 1172500 | 0 | 0 | 0 | 0 | 0 |
| GHRSVGGI | PSAT1 | Q9Y617 | SERC_HUMAN | Unmodified | 0 | 0 | 0 | 0 | 476140 | 0 | 0 | 0 | 0 | 0 | 0 | 0 | 1911100 | 0 | 0 | 0 | 476560 | 0 |
| GHSGRYW | SIGMAR1; SIGM | Q99720; B4D | SGMR1_HUMA | Unmodified | 0 | 0 | 0 | 0 | 0 | 0 | 0 | 0 | 1928300 | 49355 | 0 | 0 | 0 | 0 | 0 | 0 | 0 | 0 |
| GHTDSVQD | PAFAH1B1; PAF | P43034; A0A | LIS1_HUMAN P | Unmodified | 82341 | 0 | 580940 | 681980 | 1683300 | 540100 | 267770 | 170930 | 790720 | 554960 | 992670 | 4589800 | 0 | 322760 | 302740 | 208550 | 548030 | 596520 |
| GHTPTQPG | HSPG2 | P98160 | PGBM_HUMAN | Unmodified | 142460 | 0 | 5447900 | 2902700 | 0 | 0 | 0 | 0 | 110340 | 0 | 433810 | 386080 | 0 | 0 | 488530 | 1308400 | 0 | 0 |
| GHVDILAPT | DDX1; DDX1; D | Q92499; A0A | DDX1_HUMAN | Unmodified | 0 | 0 | 0 | 0 | 1079800 | 0 | 249580 | 28107 | 0 | 0 | 0 | 1514200 | 0 | 0 | 0 | 0 | 0 | 500100 |
| GHVFEESQ | EEF2 | P13639 | EF2_HUMAN EI | Unmodified | 0 | 0 | 0 | 131670 | 8610200 | 1328200 | 40415 | 314320 | 106830 | 1066700 | 1217500 | 0 | 2933800 | 1525500 | 0 | 0 | 2111400 | 2974700 |
| GHVFEESQ | EEF2 | P13639 | EF2_HUMAN EI | Oxidation (M) | 1600000 | 1919600 | 807010 | 1007300 | 3657800 | 1164400 | 499680 | 1150400 | 0 | 469000 | 1932500 | 20612000 | 2033900 | 654370 | 910290 | 1017500 | 1727500 | 1423400 |
| GHVGQDR | SEMA7A | O75326 | SEM7A_HUMAN | Unmodified | 0 | 0 | 0 | 0 | 2332200 | 0 | 0 | 0 | 0 | 0 | 1085900 | 0 | 614370 | 0 | 0 | 0 | 0 | 848020 |
| GHVLLHRT | DDAH1; DDAH1 | O94760; B4D | DDAH1_HUMA | Unmodified | 80865 | 0 | 692020 | 1073800 | 3462500 | 538700 | 170900 | 0 | 1581500 | 824690 | 422710 | 282290 | 3506000 | 1690400 | 0 | 54840 | 1058900 | 4597800 |
| GHVLLHRT | DDAH1; DDAH1 | O94760; B4D | DDAH1_HUMA | Unmodified | 0 | 0 | 0 | 0 | 1589300 | 87824 | 187610 | 0 | 0 | 139640 | 0 | 0 | 1456400 | 1323100 | 0 | 0 | 2992300 | 2165400 |
| GHVTQDAF | EFTUD2 | Q15029 | U5S1_HUMAN | Unmodified | 0 | 0 | 0 | 0 | 140640 | 0 | 51386 | 27457 | 0 | 0 | 161070 | 2145100 | 105990 | 0 | 0 | 0 | 73387 | 0 |
| GHYEVTGS | AHNAK | Q09666 | AHNK_HUMAN | Unmodified | 0 | 0 | 0 | 0 | 2512700 | 1567300 | 0 | 0 | 0 | 0 | 0 | 0 | 0 | 0 | 0 | 0 | 525370 | 2153000 |
| GHYEVTGS | AHNAK | Q09666 | AHNK_HUMAN | Unmodified | 0 | 0 | 0 | 0 | 3381400 | 209140 | 0 | 0 | 0 | 0 | 0 | 0 | 0 | 0 | 0 | 0 | 1041900 | 1434000 |
| GHYTEGAE | TUBB4A; TUBB | P04350; Q9E | TBB4A_HUMAN | Unmodified | 0 | 0 | 0 | 0 | 5294200 | 0 | 0 | 0 | 0 | 0 | 0 | 0 | 0 | 644260 | 0 | 0 | 528260 | 1224400 |
| GHYTEGAE | TUBB2A; TUBB; | Q13885; P07 | TBB2A_HUMAN | Unmodified | 553660 | 0 | 0 | 640420 | 2702700 | 0 | 1381300 | 0 | 7317100 | 2059800 | 4596200 | 5849400 | 0 | 0 | 469150 | 0 | 0 | 0 |
| GHYTEGAE | TUBB2A; TUBB; | Q13885; P07 | TBB2A_HUMAN | Unmodified | 22930000 | 12394000 | 17691000 | 24530000 | 1.96E+08 | 26152000 | 1.46E+08 | 0 | 1.09E+08 | 37836000 | 65211000 | 4.16E+08 | 2.02E+08 | 52763000 | 48572000 | 16085000 | 28220000 | 1.38E+08 |
| GHYTIGKEI | TUBA1A; TUBA1 | Q71U36; Q9 | TBA1A_HUMAN | Unmodified | 104910 | 0 | 0 | 0 | 0 | 43988 | 0 | 0 | 448590 | 182260 | 0 | 500170 | 0 | 0 | 0 | 0 | 0 | 0 |
| GHYTIGKEI | TUBA1A; TUBA1 | Q71U36; Q9 | TBA1A_HUMAN | Unmodified | 62001 | 0 | 0 | 0 | 0 | 0 | 707800 | 0 | 0 | 890030 | 0 | 878140 | 0 | 0 | 0 | 0 | 148280 | 63575 |
| GIAAPHAG | ZC2HC1A | Q96GY0 | ZC21A_HUMAN | Unmodified | 0 | 0 | 0 | 0 | 0 | 21139 | 0 | 0 | 111950 | 38768 | 0 | 0 | 93988 | 62270 | 0 | 0 | 0 | 64499 |
| GIADRLVLL | UEVLD | Q8IX04 | UEVLD_HUMAN | Unmodified | 37085 | 0 | 0 | 478100 | 5922600 | 3492400 | 0 | 0 | 183310 | 1452800 | 473610 | 507050 | 1478600 | 730420 | 0 | 0 | 0 | 0 |
| GIAEAVGLF | FOLH1 | Q04609 | FOLH1_HUMAN | Unmodified | 44873000 | 32678000 | 70130000 | 2.25E+08 | 0 | 0 | 0 | 0 | 582220 | 80335 | 0 | 0 | 15290000 | 5493800 | 1.89E+08 | 1.08E+08 | 0 | 0 |
| GIAEAVGLF | FOLH1 | Q04609 | FOLH1_HUMAN | Unmodified | 0 | 0 | 0 | 0 | 0 | 0 | 331480 | 30159 | 0 | 96736 | 0 | 0 | 4058100 | 1128700 | 0 | 0 | 0 | 0 |
| GIAFPTSIS | PA2G4; PA2G4; | Q9UQ80; F8 | PA2G4_HUMAN | Unmodified | 4140200 | 230340 | 154860 | 0 | 0 | 0 | 0 | 0 | 0 | 0 | 0 | 7957800 | 262800 | 0 | 128470 | 963270 | 0 | 0 |
| GIAGRQDII | TPPP3 | Q9BW30 | TPPP3_HUMAN | Unmodified | 209600 | 0 | 0 | 0 | 0 | 0 | 213590 | 163890 | 0 | 0 | 0 | 0 | 0 | 0 | 0 | 0 | 0 | 0 |
| GIALPETVD | SLC22A3 | O75751 | S22A3_HUMAN | Unmodified | 0 | 0 | 0 | 0 | 20163000 | 247940 | 0 | 0 | 0 | 0 | 0 | 0 | 0 | 0 | 0 | 1589700 | 357520 | 0 |
| GIALWGGE | EIF3B; EIF3B | P55884; C9J | EIF3B_HUMAN | Unmodified | 0 | 0 | 0 | 394040 | 0 | 0 | 0 | 32282 | 0 | 0 | 261670 | 6783100 | 166180 | 0 | 304430 | 441010 | 0 | 0 |
| GIAQMLEK | CSNK2B; CSNK | P67870; Q5S | CSK2B_HUMAN | Oxidation (M) | 67965 | 44210 | 0 | 116080 | 0 | 0 | 0 | 72174 | 0 | 0 | 808030 | 413710 | 92751 | 71595 | 100800 | 71271 | 0 | 0 |
| GIASTSDPF | TPR | P12270 | TPR_HUMAN N | Unmodified | 0 | 0 | 0 | 0 | 0 | 0 | 0 | 0 | 0 | 0 | 251670 | 582570 | 0 | 0 | 0 | 0 | 0 | 0 |
| GIAWWTDI | TMEM30A; TME | Q9NV96; A0A | CC50A_HUMAN | Unmodified | 0 | 0 | 650100 | 0 | 1907600 | 334130 | 625220 | 0 | 7084600 | 1433300 | 2461600 | 2841700 | 967170 | 148240 | 922750 | 777320 | 0 | 0 |
| GIAYIEFK | NCL; NCL; NCL | P19338; C9J | NUCL_HUMAN | Unmodified | 0 | 0 | 163000 | 252930 | 0 | 0 | 0 | 80438 | 0 | 0 | 0 | 490080 | 0 | 0 | 219690 | 194690 | 0 | 0 |
| GIAYIEFKTE | NCL; NCL; NCL | P19338; C9J | NUCL_HUMAN | Unmodified | 0 | 67638 | 0 | 80262 | 676670 | 78654 | 196650 | 37993 | 472050 | 0 | 868030 | 960630 | 592210 | 147150 | 178620 | 91385 | 124820 | 0 |
| GICEALEDS | MATN2; MATN2 | O00339; H0Y | MATN2_HUMA | Unmodified | 0 | 0 | 0 | 0 | 0 | 0 | 0 | 0 | 1077400 | 273300 | 0 | 0 | 0 | 0 | 0 | 0 | 0 | 0 |
| GICECGVC | ITGB1; ITGB1; I | P05556; H7C | ITB1_HUMAN I | Unmodified | 1656400 | 352860 | 0 | 770070 | 4741900 | 1282300 | 0 | 0 | 2691600 | 266510 | 169570 | 1652400 | 1093500 | 489620 | 697900 | 646000 | 2584500 | 8024900 |
| GICFIRTSR | TKT; TKT | P29401; A0A | TKT_HUMAN Tr | Unmodified | 0 | 0 | 172690 | 0 | 12687000 | 0 | 1698000 | 0 | 0 | 2197300 | 0 | 0 | 2408700 | 0 | 0 | 0 | 12564000 | 7348500 |
| GICNLVR | HEG1 | Q9ULI3 | HEG1_HUMAN | Unmodified | 0 | 0 | 0 | 641710 | 1173900 | 0 | 440240 | 0 | 0 | 0 | 0 | 0 | 503880 | 1370400 | 214380 | 221710 | 0 | 274550 |
| GICNLVRTI | HEG1 | Q9ULI3 | HEG1_HUMAN | Unmodified | 0 | 0 | 0 | 0 | 1732900 | 85204 | 3808400 | 0 | 0 | 0 | 0 | 0 | 383090 | 1007200 | 0 | 0 | 0 | 571230 |

| | | | | | | | | | | | | | | | | | | | | | | |
|---|---|---|---|---|---|---|---|---|---|---|---|---|---|---|---|---|---|---|---|---|---|---|
| GIDEGPEG | GPD1L; GPD1L | Q8N335; C9 | GPD1L_HUMAI | Unmodified | 0 | 0 | 0 | 0 | 0 | 0 | 0 | 0 | 379510 | 98240 | 226950 | 1106200 | 257510 | 229970 | 0 | 0 | 0 | 263220 |
| GIDEGPEG | GPD1L; GPD1L | Q8N335; C9 | GPD1L_HUMAI | Unmodified | 0 | 0 | 0 | 0 | 1339100 | 0 | 973170 | 0 | 0 | 578530 | 0 | 0 | 0 | 521930 | 0 | 0 | 928220 | 1631000 |
| GIDIHGVP\ | DDX1; DDX1; D | Q92499; A0/ | DDX1_HUMAN | Unmodified | 270690 | 289380 | 243260 | 453240 | 340760 | 131870 | 88735 | 18474 | 410080 | 101850 | 579190 | 1381900 | 780720 | 265350 | 436090 | 319720 | 95806 | 216730 |
| GIDIIRGPIL | RFC5; RFC5; RF | P40937; F5H | RFC5_HUMAN | Unmodified | 53114 | 56218 | 0 | 0 | 20095 | 0 | 0 | 0 | 72590 | 0 | 0 | 265250 | 0 | 0 | 56497 | 63095 | 42885 | 16114 |
| GIDLTQVK | DLAT; DLAT; DL | P10515; A0A | ODP2_HUMAN | Unmodified | 1214100 | 970690 | 28733 | 219040 | 274880 | 94312 | 0 | 0 | 1856800 | 330080 | 0 | 294980 | 728610 | 517440 | 76927 | 271100 | 157930 | 0 |
| GIDPFSLD/ | CCT6A | P40227 | TCPZ_HUMAN | Unmodified | 0 | 0 | 0 | 0 | 9947900 | 1864400 | 2845100 | 1404900 | 6133800 | 3781300 | 8133700 | 54037000 | 20312000 | 13484000 | 0 | 0 | 3782000 | 0 |
| GIDPFSLD/ | CCT6A | P40227 | TCPZ_HUMAN | Unmodified | 0 | 0 | 0 | 353090 | 0 | 0 | 0 | 0 | 0 | 0 | 0 | 0 | 12505000 | 0 | 0 | 0 | 0 | 0 |
| GIDPFSLD/ | CCT6A | P40227 | TCPZ_HUMAN | Unmodified | 0 | 0 | 0 | 0 | 0 | 0 | 561780 | 0 | 0 | 970410 | 0 | 0 | 2174600 | 2630000 | 0 | 0 | 2894100 | 1232500 |
| GIDRYNPEI | EIF3K; EIF3K; EI | Q9UBQ5; K7 | EIF3K_HUMAN | Unmodified | 0 | 0 | 0 | 0 | 0 | 0 | 0 | 0 | 0 | 0 | 0 | 1011700 | 0 | 0 | 0 | 0 | 0 | 0 |
| GIDRYNPEI | EIF3K; EIF3K; EI | Q9UBQ5; K7 | EIF3K_HUMAN | Unmodified | 0 | 0 | 0 | 876100 | 2386200 | 981230 | 946720 | 0 | 0 | 0 | 1188400 | 0 | 3618800 | 1366200 | 464060 | 429680 | 0 | 0 |
| GIDSDASYI | CTSS; CTSS; CT | P25774; A0A | CATS_HUMAN | Unmodified | 0 | 0 | 0 | 0 | 0 | 0 | 0 | 0 | 1311100 | 0 | 0 | 0 | 0 | 0 | 0 | 0 | 0 | 0 |
| GIDVQQVS | EIF4A1; EIF4A2; | P60842; Q14 | IF4A1_HUMAN | Unmodified | 0 | 1151700 | 0 | 1374600 | 0 | 0 | 0 | 0 | 0 | 236500 | 0 | 0 | 0 | 0 | 3252800 | 0 | 0 | 0 |
| GIEEEEILP( | GSPT1; GSPT1 | P15170; H3E | ERF3A_HUMAN | Unmodified | 0 | 123440 | 0 | 0 | 0 | 0 | 0 | 0 | 329850 | 0 | 131720 | 569260 | 0 | 0 | 0 | 109990 | 0 | 0 |
| GIEELFLDL | RAB21 | Q9UL25 | RAB21_HUMAN | Unmodified | 226490 | 0 | 0 | 507830 | 799540 | 150490 | 241840 | 0 | 339960 | 372450 | 210760 | 0 | 1062900 | 328220 | 0 | 97065 | 0 | 0 |
| GIEELFLDL | RAB21 | Q9UL25 | RAB21_HUMAN | Unmodified | 0 | 172030 | 0 | 255870 | 1103400 | 0 | 73951 | 0 | 647370 | 249200 | 192360 | 0 | 372540 | 0 | 0 | 93246 | 288500 | 0 |
| GIEILTDMS | CCT4 | P50991 | TCPD_HUMAN | Unmodified | 0 | 0 | 0 | 0 | 0 | 0 | 0 | 0 | 0 | 0 | 235350 | 419480 | 0 | 0 | 0 | 10605 | 0 | 0 |
| GIEILTDMS | CCT4 | P50991 | TCPD_HUMAN | Oxidation (M) | 0 | 0 | 0 | 947560 | 6061400 | 0 | 0 | 0 | 0 | 1831800 | 1837100 | 1465800 | 9767100 | 6211000 | 0 | 0 | 0 | 2991300 |
| GIELIPQVR | PLCB4; PLCB4 | Q15147; A0/ | PLCB4_HUMAI | Unmodified | 0 | 0 | 89597 | 0 | 103370 | 0 | 0 | 0 | 0 | 153320 | 0 | 0 | 365980 | 356240 | 0 | 0 | 0 | 2685200 |
| GIEMSEVR | DLD; DLD | P09622; E9P | DLDH_HUMAN | Unmodified | 0 | 0 | 0 | 0 | 0 | 0 | 0 | 0 | 0 | 0 | 0 | 0 | 0 | 0 | 0 | 0 | 0 | 0 |
| GIEMSEVRI | DLD; DLD | P09622; E9P | DLDH_HUMAN | Unmodified | 0 | 0 | 0 | 0 | 0 | 0 | 0 | 155460 | 6750800 | 840350 | 0 | 0 | 0 | 197470 | 0 | 0 | 0 | 0 |
| GIEMSEVRI | DLD; DLD | P09622; E9P | DLDH_HUMAN | Oxidation (M) | 15446 | 0 | 0 | 0 | 0 | 0 | 0 | 0 | 1146700 | 227490 | 179770 | 0 | 280390 | 0 | 0 | 0 | 0 | 0 |
| GIENYYDD | TSPAN15; TSPA | O95858; H7( | TSN15_HUMAI | Unmodified | 0 | 0 | 590400 | 131420 | 0 | 0 | 0 | 0 | 0 | 0 | 0 | 1081700 | 0 | 0 | 2362000 | 1339100 | 0 | 0 |
| GIEPTGNM | FLNB; FLNA; FL | O75369; P21 | FLNB_HUMAN | Unmodified | 0 | 0 | 493000 | 0 | 415460 | 1609000 | 558170 | 285860 | 0 | 69023 | 218240 | 4011200 | 0 | 796430 | 232190 | 162910 | 0 | 592550 |
| GIEQAVQS | IMMT; IMMT; IM | Q16891; B9/ | MIC60_HUMAN | Unmodified | 0 | 0 | 0 | 0 | 282230 | 0 | 0 | 0 | 1103800 | 145070 | 0 | 0 | 0 | 0 | 0 | 0 | 59067 | 0 |
| GIFEALRPL | DYNC1H1; DYN | Q14204; A0/ | DYHC1_HUMA | Unmodified | 0 | 0 | 0 | 0 | 0 | 0 | 0 | 0 | 0 | 0 | 0 | 161960 | 0 | 0 | 0 | 0 | 0 | 0 |
| GIFGFTDSI | WARS1; WARS1 | P23381; H0\ | SYWC_HUMAN | Unmodified | 0 | 0 | 0 | 0 | 0 | 0 | 0 | 0 | 0 | 0 | 0 | 668290 | 0 | 0 | 0 | 0 | 0 | 0 |
| GIFGSSAVI | KDM1A; KDM1A | O60341; R4( | KDM1A_HUMA | Unmodified | 0 | 0 | 171950 | 0 | 0 | 0 | 0 | 0 | 0 | 58211 | 0 | 1728400 | 0 | 0 | 0 | 0 | 0 | 0 |
| GIFLHLSN( | PLOD3; PLOD3 | O60568; H7( | PLOD3_HUMAI | Unmodified | 0 | 0 | 0 | 0 | 168860 | 0 | 0 | 0 | 26627000 | 1668300 | 1017600 | 194540 | 0 | 0 | 0 | 130530 | 85651 | 0 |
| GIFLVRESE | YES1; YES1 | P07947; J3Q | YES_HUMAN Ty | Unmodified | 0 | 0 | 0 | 1708100 | 2248700 | 1096200 | 0 | 0 | 1189200 | 1037000 | 0 | 0 | 0 | 0 | 0 | 0 | 1289800 | 2173000 |
| GIFNGFSV | SLC25A3; SLC2 | Q00325; F8\ | MPCP_HUMAN | Unmodified | 0 | 0 | 144870 | 0 | 0 | 0 | 0 | 0 | 6406000 | 552710 | 0 | 0 | 0 | 0 | 0 | 0 | 0 | 0 |
| GIFNGFSV | SLC25A3; SLC2 | Q00325; F8\ | MPCP_HUMAN | Unmodified | 0 | 0 | 0 | 0 | 0 | 0 | 789800 | 0 | 16882000 | 2126200 | 0 | 155760 | 0 | 0 | 0 | 0 | 0 | 0 |
| GIFNGFSV | SLC25A3; SLC2 | Q00325; F8\ | MPCP_HUMAN | Unmodified | 18697 | 0 | 0 | 0 | 0 | 0 | 0 | 0 | 6593700 | 2177200 | 0 | 0 | 0 | 0 | 0 | 0 | 0 | 0 |
| GIFPVLCKI | PKM; PKM; PKM | P14618; B4I | KPYM_HUMAN | Unmodified | 213910 | 0 | 0 | 572900 | 0 | 0 | 0 | 0 | 344250 | 272780 | 323160 | 2066600 | 0 | 0 | 1040300 | 273790 | 0 | 0 |
| GIFTSEIGTI | PRKDC | P78527 | PRKDC_HUMAI | Unmodified | 0 | 74126 | 372560 | 0 | 167390 | 178490 | 110670 | 0 | 257080 | 0 | 178110 | 1753700 | 0 | 0 | 70930 | 61154 | 0 | 0 |
| GIGFGDIFk | CD2AP | Q9Y5K6 | CD2AP_HUMAI | Unmodified | 779050 | 2201500 | 0 | 970570 | 3292100 | 0 | 757510 | 0 | 0 | 582750 | 0 | 0 | 12867000 | 9180700 | 128650 | 137370 | 209230 | 350130 |
| GIGGIFFDI | CPOX | P36551 | HEM6_HUMAN | Unmodified | 0 | 0 | 0 | 0 | 0 | 0 | 0 | 0 | 2017400 | 1401300 | 0 | 0 | 0 | 0 | 0 | 0 | 0 | 0 |
| GIGHVIKVI | UPF1 | Q92900 | RENT1_HUMAN | Unmodified | 324300 | 541950 | 0 | 551990 | 0 | 0 | 0 | 0 | 0 | 0 | 0 | 0 | 0 | 0 | 165060 | 79288 | 0 | 0 |
| GIGIENIHY | MCTS1; MCTS2 | Q9ULC4; A0. | MCTS1_HUMAI | Oxidation (M) | 0 | 0 | 965400 | 1405400 | 1298000 | 0 | 250000 | 0 | 503060 | 445440 | 0 | 2519000 | 1446900 | 1379100 | 0 | 903060 | 688220 | 2740700 |
| GIGLAIVRD | CBR1; CBR1; C | P16152; E9P | CBR1_HUMAN | Unmodified | 0 | 0 | 0 | 0 | 320670 | 0 | 0 | 0 | 0 | 0 | 0 | 0 | 0 | 0 | 0 | 0 | 592190 | 0 |
| GIGMGNIG | HNRNPM; HNR | P52272; M0( | HNRPM_HUMA | 3 Oxidation (M) | 0 | 317140 | 0 | 0 | 160840 | 0 | 0 | 170250 | 0 | 180010 | 0 | 0 | 1013900 | 0 | 355040 | 172940 | 355130 | 0 |
| GIGPVYSSI | ADSS2 | P30520 | PURA2_HUMAI | Unmodified | 0 | 0 | 0 | 88305 | 0 | 149210 | 0 | 596500 | 519350 | 35312 | 360560 | 1813500 | 0 | 0 | 41657 | 44239 | 0 | 0 |
| GIGTDEFTL | ANXA3; ANXA3 | P12429; D6F | ANXA3_HUMAI | Unmodified | 0 | 685850 | 0 | 0 | 350580 | 0 | 0 | 0 | 0 | 0 | 0 | 982700 | 0 | 0 | 0 | 0 | 0 | 0 |
| GIHEYDFEI | GNA13 | Q14344 | GNA13_HUMAI | Unmodified | 0 | 0 | 0 | 0 | 4881300 | 1012900 | 1068800 | 574450 | 0 | 1048100 | 1883400 | 0 | 0 | 465910 | 0 | 0 | 1494700 | 3875600 |
| GIHEYDFEI | GNA13 | Q14344 | GNA13_HUMAI | Unmodified | 0 | 0 | 0 | 0 | 1294600 | 0 | 508780 | 0 | 0 | 0 | 0 | 0 | 0 | 0 | 0 | 0 | 1203900 | 4699200 |
| GIHFQDYD | HPGD | P15428 | PGDH_HUMAN | Unmodified | 0 | 0 | 0 | 0 | 0 | 0 | 0 | 0 | 0 | 0 | 0 | 0 | 0 | 0 | 0 | 0 | 0 | 2289000 |
| GIHHGPSV | SPC24 | Q8NBT2 | SPC24_HUMAN | Unmodified | 0 | 0 | 0 | 0 | 867480 | 0 | 0 | 0 | 0 | 0 | 0 | 615070 | 867040 | 0 | 0 | 0 | 309400 | 475460 |
| GIHNSIGD' | DSP | P15924 | DESP_HUMAN | Unmodified | 0 | 0 | 0 | 0 | 0 | 0 | 0 | 0 | 628070 | 105910 | 0 | 0 | 73514 | 0 | 108950 | 0 | 0 | 0 |
| GIHPTIISES | CCT4 | P50991 | TCPD_HUMAN | Unmodified | 0 | 0 | 688780 | 0 | 8707000 | 239640 | 1112900 | 615470 | 5477900 | 3236600 | 5215600 | 31889000 | 27496000 | 16212000 | 127690 | 0 | 3917900 | 1103300 |
| GIHPTIISES | CCT4 | P50991 | TCPD_HUMAN | Unmodified | 0 | 0 | 0 | 0 | 0 | 0 | 529340 | 82044 | 0 | 0 | 0 | 0 | 399640 | 0 | 0 | 0 | 1454400 | 0 |
| GIHQSTIDL | TRIM25; TRIM25 | Q14258; A0/ | TRI25_HUMAN | Unmodified | 0 | 0 | 0 | 0 | 11097000 | 2217200 | 0 | 0 | 0 | 287900 | 0 | 0 | 813880 | 1203900 | 0 | 0 | 1711400 | 2266900 |
| GIHVIVLNC | POMGNT1; POI | Q8WZA1; Q6 | PMGT1_HUMAI | Unmodified | 0 | 0 | 0 | 0 | 0 | 0 | 0 | 0 | 2323500 | 0 | 0 | 0 | 0 | 0 | 0 | 0 | 0 | 0 |
| GIHYWEVE | TRIM25 | Q14258 | TRI25_HUMAN | Unmodified | 0 | 0 | 0 | 0 | 0 | 0 | 0 | 0 | 0 | 0 | 0 | 0 | 0 | 0 | 0 | 0 | 0 | 0 |
| GIIASSVGT | CTPS1; CTPS1; | P17812; A0A | PYRG1_HUMAI | Unmodified | 0 | 0 | 0 | 0 | 395620 | 0 | 270720 | 0 | 0 | 0 | 611680 | 2581200 | 0 | 0 | 0 | 0 | 431840 | 0 |
| GIIDSTHTE | ATP2B2; ATP2B | Q01814; A0/ | AT2B2_HUMAN | Unmodified | 0 | 0 | 0 | 0 | 0 | 0 | 933180 | 0 | 0 | 296530 | 0 | 0 | 0 | 0 | 0 | 0 | 0 | 0 |
| GIIDSTVGE | ATP2B4 | P23634 | AT2B4_HUMAN | Unmodified | 0 | 0 | 754380 | 1316500 | 48194000 | 4411400 | 9522300 | 1434100 | 1227500 | 1684900 | 1807000 | 0 | 1564000 | 979680 | 0 | 0 | 7042200 | 11937000 |
| GIIDSTVSD | ATP2B1; ATP2B | P20020; E7E | AT2B1_HUMAN | Unmodified | 0 | 0 | 4856200 | 4368400 | 0 | 412650 | 325530 | 0 | 3580400 | 464920 | 2686800 | 17990000 | 5475700 | 0 | 8308100 | 8309700 | 596050 | 766490 |
| GIIDSTVSD | ATP2B1; ATP2B | P20020; E7E | AT2B1_HUMAN | Unmodified | 0 | 0 | 0 | 0 | 2572100 | 0 | 0 | 0 | 0 | 498010 | 0 | 0 | 5406800 | 23209000 | 0 | 0 | 1263100 | 897960 |
| GIIDSTVSD | ATP2B1; ATP2B | P20020; E7E | AT2B1_HUMAN | Unmodified | 0 | 0 | 0 | 0 | 0 | 0 | 0 | 0 | 0 | 0 | 0 | 0 | 0 | 1885800 | 0 | 0 | 0 | 0 |
| GIILAGTEE( | ACAD9; ACAD9 | Q9H845; A0. | ACAD9_HUMAI | Unmodified | 0 | 0 | 0 | 607680 | 493000 | 0 | 109640 | 144560 | 1420100 | 673460 | 0 | 838420 | 403400 | 0 | 0 | 0 | 106600 | 0 |
| GIIRPGTAF | PLEC; PLEC | Q15149; H0\ | PLEC_HUMAN | Unmodified | 0 | 0 | 0 | 0 | 980010 | 0 | 0 | 0 | 0 | 0 | 0 | 0 | 0 | 0 | 0 | 0 | 3180400 | 509220 |
| GIISILDEE( | MYO1C; MYO1( | O00159; F5H | MYO1C_HUMA | Unmodified | 0 | 214860 | 0 | 324980 | 0 | 0 | 0 | 0 | 0 | 0 | 0 | 0 | 0 | 0 | 1949800 | 0 | 0 | 0 |
| GIKPSPAPI | CPNE8; CPNE8 | Q86YQ8; E7I | CPNE8_HUMAI | Unmodified | 0 | 0 | 0 | 0 | 4448600 | 1028800 | 562800 | 0 | 0 | 426080 | 0 | 0 | 0 | 0 | 0 | 0 | 0 | 833650 |
| GIKPVTLEL | ALDH9A1 | P49189 | AL9A1_HUMAN | Unmodified | 0 | 0 | 605290 | 284900 | 119460 | 0 | 119030 | 165140 | 1836200 | 345070 | 536930 | 2909200 | 1904800 | 0 | 464490 | 0 | 251440 | 543230 |
| GILAADES\ | ALDOC; ALDOC | P09972; J3Q | ALDOC_HUMA | Unmodified | 0 | 0 | 0 | 0 | 0 | 0 | 0 | 127770 | 0 | 0 | 0 | 4710400 | 947570 | 482920 | 0 | 0 | 1248000 | 1382200 |
| GILAADES\ | ALDOC; ALDOC | P09972; J3Q | ALDOC_HUMA | Oxidation (M) | 0 | 0 | 0 | 0 | 0 | 0 | 0 | 0 | 0 | 0 | 0 | 1757200 | 0 | 168010 | 0 | 0 | 279610 | 0 |
| GILADEDS! | FASN; FASN | P49327; A0A | FAS_HUMAN F | Unmodified | 826290 | 410860 | 3373400 | 2004000 | 1565600 | 816070 | 2836200 | 414940 | 1176900 | 481510 | 5418700 | 90458000 | 1237000 | 0 | 5039400 | 3464100 | 1704500 | 1243600 |
| GILDKEEAE | SCG3 | Q8WXD2 | SCG3_HUMAN | Unmodified | 0 | 0 | 0 | 0 | 48635 | 0 | 16701 | 109300 | 75966000 | 19640000 | 0 | 0 | 0 | 0 | 0 | 0 | 0 | 430050 |
| GILDKEEAE | SCG3 | Q8WXD2 | SCG3_HUMAN | Unmodified | 0 | 0 | 0 | 0 | 0 | 0 | 0 | 0 | 3949300 | 8486100 | 0 | 0 | 0 | 0 | 0 | 0 | 334770 | 0 |
| GILDLNTYI | PLCG2; PLCG2 | P16885; A0A | PLCG2_HUMAI | Unmodified | 0 | 0 | 283720 | 0 | 0 | 0 | 0 | 0 | 1714900 | 0 | 3765400 | 13947000 | 0 | 0 | 0 | 0 | 0 | 0 |
| GILDVQSV( | SVIP | Q8NHG7 | SVIP_HUMAN S | Unmodified | 4478800 | 3599400 | 2497900 | 2216800 | 1103200 | 231740 | 396200 | 81624 | 8210400 | 4575200 | 1569900 | 4582300 | 89213 | 2858200 | 2053600 | 2687800 | 698600 | 2808900 |
| GILEQGW( | COPS8; COPS8 | Q99627; H7( | CSN8_HUMAN | Unmodified | 0 | 0 | 0 | 0 | 0 | 0 | 0 | 0 | 0 | 0 | 371520 | 2897000 | 0 | 0 | 278910 | 0 | 0 | 0 |
| GILFVGSG\ | PGD; PGD; PGI | P52209; K7E | 6PGD_HUMAN | Unmodified | 944460 | 0 | 1547300 | 947650 | 677540 | 0 | 266600 | 0 | 10408000 | 445290 | 3098400 | 23173000 | 0 | 0 | 2200700 | 1861200 | 0 | 385700 |
| GILFVGSG\ | PGD; PGD; PGI | P52209; K7E | 6PGD_HUMAN | Oxidation (M) | 0 | 0 | 0 | 0 | 9093800 | 657390 | 537940 | 150790 | 0 | 2635800 | 888140 | 0 | 3987300 | 1898200 | 0 | 0 | 1960500 | 1615100 |
| GILFVGSG\ | PGD; PGD; PGI | P52209; K7E | 6PGD_HUMAN | Unmodified | 0 | 0 | 0 | 0 | 6456600 | 0 | 0 | 0 | 0 | 606760 | 0 | 0 | 2026700 | 0 | 0 | 0 | 0 | 2039500 |
| GILGNAIKV | GPX4; GPX4; GI | P36969; A0A | GPX4_HUMAN | Unmodified | 0 | 0 | 0 | 0 | 0 | 0 | 0 | 0 | 0 | 485100 | 0 | 0 | 0 | 819230 | 0 | 0 | 401180 | 280340 |
| GILGQHQF | ITGA2; ITGA2; I | P17301; E9P | ITA2_HUMAN I | Unmodified | 0 | 0 | 2368000 | 537520 | 0 | 0 | 0 | 0 | 0 | 0 | 0 | 353730 | 0 | 0 | 1855300 | 2875900 | 0 | 0 |
| GILGTLNK | SHTN1 | A0MZ66 | SHOT1_HUMAI | Unmodified | 0 | 0 | 0 | 0 | 500400 | 0 | 55552 | 41886 | 520030 | 114280 | 124590 | 211860 | 175310 | 519590 | 0 | 155780 | 0 | 0 |
| GILGYTEH( | GAPDH; GAPDH | P04406; E7E | G3P_HUMAN G | Unmodified | 0 | 0 | 0 | 0 | 2.58E+08 | 8877100 | 36392000 | 0 | 0 | 0 | 900900 | 0 | 75919000 | 56350000 | 0 | 0 | 1.47E+08 | 2.01E+08 |
| GILHEAAM( | DOCK9; DOCK9 | Q9BZ29; A0/ | DOCK9_HUMA | Unmodified | 0 | 0 | 0 | 0 | 408980 | 134130 | 58452 | 0 | 427900 | 42049 | 0 | 0 | 0 | 0 | 0 | 0 | 0 | 0 |
| GILIDTSRH | HEXB | P07686 | HEXB_HUMAN | Unmodified | 0 | 0 | 0 | 0 | 0 | 0 | 0 | 0 | 0 | 526880 | 0 | 0 | 270500 | 0 | 0 | 0 | 0 | 0 |
| GILLFGPP( | VPS4B; FIGNL1 | O75351; Q6I | VPS4B_HUMAN | Unmodified | 0 | 9587800 | 8439800 | 9550400 | 4391700 | 2159800 | 0 | 244070 | 17062000 | 5190600 | 11271000 | 14417000 | 13811000 | 3743300 | 6599500 | 8341000 | 0 | 2959000 |
| GILLFGTK | ACADVL; ACAD | P49748; K7E | ACADV_HUMAI | Unmodified | 169310 | 35276 | 0 | 0 | 0 | 0 | 0 | 12229 | 6135800 | 2318400 | 179770 | 1912100 | 0 | 0 | 58126 | 25432 | 0 | 0 |
| GILLYGPP( | NSF; NSF | P46459; I3L( | NSF_HUMAN V | Unmodified | 0 | 0 | 0 | 0 | 968820 | 615950 | 1073000 | 1292700 | 406670 | 259070 | 0 | 0 | 0 | 3920000 | 852820 | 631100 | 1335600 | 2156000 |
| GILQKNVPI | CTNNA1; CTNN | P35221; E5R | CTNA1_HUMAI | Unmodified | 0 | 0 | 0 | 0 | 0 | 0 | 0 | 0 | 0 | 0 | 0 | 0 | 327600 | 574630 | 0 | 0 | 0 | 0 |
| GILRQITLN | PRDX4; PRDX4 | Q13162; H7( | PRDX4_HUMAN | Unmodified | 0 | 0 | 0 | 0 | 0 | 0 | 0 | 0 | 716960 | 805960 | 618810 | 0 | 0 | 0 | 0 | 0 | 21750 | 0 |
| GILSGTSDI | VCL | P18206 | VINC_HUMAN | Unmodified | 0 | 545320 | 121370 | 543010 | 731070 | 0 | 0 | 0 | 401010 | 0 | 169850 | 383420 | 0 | 0 | 822670 | 84176 | 996260 | 0 |
| GILTLKYPIE | ACTC1; ACTA1; | P68032; P68 | ACTC_HUMAN | Oxidation (M) | 11941000 | 5640900 | 0 | 27675000 | 1.55E+08 | 32800000 | 36545000 | 0 | 12718000 | 23117000 | 14341000 | 10317000 | 1.39E+08 | 73273000 | 1069800 | 3602100 | 39363000 | 47123000 |
| GILTLKYPIE | ACTC1; ACTA1; | P68032; P68 | ACTC_HUMAN | Unmodified | 302510 | 0 | 0 | 1239900 | 34114000 | 6111800 | 6459200 | 0 | 10544000 | 3197900 | 0 | 321560 | 19535000 | 9819500 | 87876 | 69985 | 4439600 | 9602500 |
| GILVFQLK | AFDN; AFDN; A | P55196; Q5T | AFAD_HUMAN | Unmodified | 0 | 650780 | 143060 | 0 | 1342300 | 0 | 0 | 158300 | 777140 | 0 | 959570 | 2004200 | 1745100 | 1434600 | 0 | 0 | 0 | 282480 |
| GILYRTHSI | ATP5F1C | P36542 | ATPG_HUMAN | Unmodified | 0 | 0 | 0 | 0 | 0 | 0 | 0 | 0 | 0 | 710450 | 0 | 0 | 0 | 0 | 0 | 0 | 0 | 0 |
| GIMEEDEA( | JUP; JUP; JUP; J | P14923; C9J | PLAK_HUMAN . | Unmodified | 111240 | 44243 | 462780 | 1001700 | 620610 | 0 | 0 | 0 | 466090 | 0 | 532120 | 1585700 | 457800 | 308180 | 766180 | 841470 | 130910 | 188330 |
| GIMEEDEA( | JUP; JUP; JUP; J | P14923; C9J | PLAK_HUMAN . | Oxidation (M) | 0 | 0 | 0 | 0 | 318920 | 0 | 0 | 0 | 102990 | 0 | 311370 | 886150 | 0 | 0 | 0 | 0 | 0 | 0 |
| GIMEEDEA( | JUP; JUP; JUP; J | P14923; C9J | PLAK_HUMAN . | Unmodified | 0 | 0 | 0 | 0 | 14157000 | 1326700 | 0 | 0 | 0 | 775550 | 0 | 0 | 3012000 | 3154100 | 0 | 0 | 3031200 | 2865100 |
| GIMEEDEA( | JUP; JUP; JUP; J | P14923; C9J | PLAK_HUMAN . | Oxidation (M) | 96005 | 0 | 0 | 0 | 2422700 | 0 | 0 | 0 | 28788 | 0 | 0 | 0 | 1887500 | 1395100 | 0 | 0 | 758470 | 1554800 |
| GIMEEDEA( | JUP; JUP; JUP; J | P14923; C9J | PLAK_HUMAN . | Unmodified | 20588 | 45878 | 0 | 237940 | 11105000 | 1669800 | 433010 | 511230 | 100010 | 402510 | 0 | 124390 | 2485700 | 3427500 | 0 | 0 | 2972600 | 2485900 |
| GIMEEDEA( | JUP; JUP; JUP; J | P14923; C9J | PLAK_HUMAN . | Oxidation (M) | 0 | 0 | 0 | 0 | 816550 | 0 | 0 | 30918 | 0 | 0 | 0 | 0 | 1453400 | 761500 | 0 | 0 | 1186300 | 1394700 |
| GIMENPIVI | PALS2; PALS2; I | Q9NZW5; F8 | PALS2_HUMAN | Oxidation (M) | 0 | 0 | 234990 | 126890 | 1678700 | 692710 | 116850 | 249170 | 0 | 0 | 388010 | 0 | 888460 | 311590 | 0 | 0 | 42812 | 3093900 |
| GINSSNVEI | KPNA2; KPNA2; | P52292; J3Q | IMA1_HUMAN I | Unmodified | 0 | 0 | 0 | 376960 | 0 | 0 | 0 | 0 | 262850 | 413960 | 693920 | 0 | 0 | 0 | 202680 | 169260 | 300660 | 0 |
| GINSSNVEI | KPNA2; KPNA2; | P52292; J3Q | IMA1_HUMAN I | Unmodified | 733670 | 571280 | 1223400 | 1154500 | 3487700 | 1017200 | 406600 | 404860 | 1392000 | 0 | 584110 | 3051200 | 3326500 | 2060100 | 956800 | 463650 | 2000200 | 2460100 |
| GINTLVTYI | COX5A; COX5A | P20674; H3E | COX5A_HUMAI | Unmodified | 0 | 0 | 0 | 0 | 0 | 0 | 0 | 0 | 5055800 | 512060 | 0 | 0 | 0 | 0 | 0 | 0 | 0 | 0 |
| GINTLVTYI | COX5A; COX5A | P20674; H3E | COX5A_HUMAI | Oxidation (M) | 0 | 0 | 0 | 0 | 0 | 0 | 0 | 0 | 0 | 73074 | 0 | 0 | 0 | 0 | 0 | 0 | 0 | 0 |
| GINTLVTYI | COX5A; COX5A | P20674; H3E | COX5A_HUMAI | Oxidation (M) | 0 | 0 | 0 | 918470 | 0 | 0 | 0 | 0 | 0 | 409320 | 0 | 0 | 0 | 0 | 0 | 0 | 0 | 0 |
| GINVALAN( | FAM3C; FAM3C | Q92520; C9J | FAM3C_HUMAI | Unmodified | 0 | 1508600 | 213860 | 56695 | 306180 | 0 | 31096 | 0 | 18131000 | 1402300 | 1071200 | 1577700 | 0 | 0 | 0 | 3061200 | 0 | 0 |
| GINVALAN( | FAM3C; FAM3C | Q92520; C9J | FAM3C_HUMAI | Unmodified | 0 | 0 | 0 | 0 | 0 | 0 | 0 | 0 | 0 | 231420 | 0 | 0 | 0 | 0 | 0 | 0 | 0 | 0 |
| GIPAPEEEI | AHSA1; AHSA1; | O95433; H0' | AHSA1_HUMAN | Unmodified | 0 | 0 | 0 | 0 | 0 | 0 | 0 | 0 | 0 | 0 | 45688 | 602570 | 0 | 0 | 84987 | 0 | 0 | 0 |
| GIPDDRDG | USP9X; USP9Y; | Q93008; O0( | USP9X_HUMAN | Unmodified | 0 | 0 | 0 | 0 | 0 | 0 | 0 | 0 | 471600 | 331630 | 0 | 2596500 | 0 | 0 | 0 | 0 | 0 | 0 |
| GIPEDKNPI | GNG11; GNGT2 | P61952; O14 | GBG11_HUMAI | Unmodified | 1965200 | 907950 | 872020 | 0 | 687610 | 213470 | 4826400 | 6884500 | 0 | 0 | 0 | 0 | 2268100 | 3723500 | 72962 | 0 | 1017900 | 2938600 |
| GIPEFWLT\ | NAP1L1; NAP1L | P55209; F8V | NP1L1_HUMAN | Unmodified | 1982300 | 2573000 | 0 | 474060 | 733820 | 0 | 186190 | 0 | 1543800 | 538180 | 891890 | 2756400 | 0 | 0 | 0 | 0 | 0 | 28156 |
| GIPELEQYI | SCYL1; SCYL1; | Q96KG9; E9I | SCYL1_HUMAN | Unmodified | 0 | 0 | 0 | 0 | 0 | 895630 | 0 | 0 | 0 | 0 | 124290 | 1478000 | 0 | 0 | 196690 | 313290 | 0 | 0 |

| | | | | | | | | | | | | | | | | | | | | | | |
|---|---|---|---|---|---|---|---|---|---|---|---|---|---|---|---|---|---|---|---|---|---|---|
| GIPFAAPTK | CEL | P19835 | CEL_HUMAN B | Unmodified | 0 | 0 | 0 | 0 | 0 | 0 | 0 | 0 | 8538300 | 1454100 | 8740700 | 0 | 0 | 0 | 0 | 0 | 0 | 0 |
| GIPHLVTHI | RPS4Y2; RPS4X | Q8TD47; P62 | RS4Y2_HUMAN | Unmodified | 0 | 0 | 0 | 0 | 906680 | 0 | 876080 | 1358600 | 859000 | 394910 | 2319300 | 36610000 | 0 | 0 | 0 | 16776 | 102760 | 613840 |
| GIPIEVIPM | RPIA | P49247 | RPIA_HUMAN F | Oxidation (M) | 0 | 0 | 0 | 0 | 0 | 0 | 0 | 0 | 0 | 0 | 0 | 4652300 | 0 | 0 | 0 | 0 | 0 | 0 |
| GIPLATGD | EIF3G; EIF3G; E | O75821; K7E | EIF3G_HUMAN | Unmodified | 553440 | 860620 | 664370 | 0 | 2848700 | 0 | 0 | 0 | 1903200 | 0 | 1814000 | 9666300 | 3129000 | 1003100 | 0 | 966010 | 0 | 0 |
| GIPPLSLRP | EXOC2 | Q96KP1 | EXOC2_HUMA | Unmodified | 120390 | 0 | 230930 | 345780 | 4928000 | 774470 | 618580 | 21240 | 1731300 | 1210000 | 523380 | 1735700 | 970260 | 367410 | 677580 | 419520 | 687830 | 632590 |
| GIPRDVTEL | SLIT2; SLIT2; SL | O94813; X6F | SLIT2_HUMAN | Unmodified | 0 | 0 | 0 | 0 | 212600 | 0 | 0 | 0 | 0 | 0 | 0 | 0 | 0 | 0 | 0 | 0 | 0 | 4307700 |
| GIPVLVGLL | CTNND1; CTNN | O60716; E9F | CTND1_HUMA | Unmodified | 0 | 0 | 537000 | 0 | 621630 | 0 | 0 | 0 | 0 | 0 | 406060 | 729190 | 0 | 690960 | 0 | 517740 | 514200 | 663040 |
| GIPVLVGLL | CTNND1; CTNN | O60716; E9F | CTND1_HUMA | Unmodified | 0 | 0 | 2473200 | 3078200 | 4342000 | 1038700 | 0 | 0 | 0 | 0 | 1562700 | 3096400 | 2857600 | 1737400 | 2793500 | 3865900 | 1369700 | 3367500 |
| GIPVLVGLL | CTNND1; CTNN | O60716; E9F | CTND1_HUMA | Unmodified | 0 | 0 | 0 | 0 | 21189000 | 1371200 | 0 | 0 | 0 | 0 | 0 | 0 | 15286000 | 17854000 | 0 | 0 | 23715000 | 25976000 |
| GIPVVEHK | TOP2B; TOP2A; | Q02880; P11 | TOP2B_HUMAN | Unmodified | 0 | 128940 | 0 | 0 | 0 | 0 | 0 | 158260 | 73307 | 93409 | 0 | 420820 | 0 | 0 | 0 | 11879 | 0 | 0 |
| GIPVYPYPF | KDF1 | Q8NAX2 | KDF1_HUMAN | Unmodified | 0 | 0 | 619380 | 776770 | 0 | 0 | 0 | 0 | 0 | 0 | 0 | 0 | 0 | 0 | 858270 | 0 | 0 | 0 |
| GIQEAQVE | PDXDC1; PDXD | Q6P996; H3I | PDXD1_HUMA | Unmodified | 0 | 0 | 0 | 0 | 0 | 0 | 0 | 0 | 151080 | 43657 | 225210 | 2086000 | 332830 | 0 | 0 | 0 | 0 | 0 |
| GIQEEMEA | DPYSL2; DPYSL | Q16555; A0A | DPYL2_HUMAN | Unmodified | 0 | 0 | 2353800 | 617810 | 904670 | 0 | 0 | 50323 | 1678100 | 478920 | 0 | 3496500 | 0 | 0 | 2265400 | 581200 | 0 | 0 |
| GIQETEEMI | MUL1 | Q969V5 | MUL1_HUMAN | Unmodified | 0 | 0 | 0 | 0 | 0 | 0 | 0 | 0 | 1074400 | 0 | 0 | 0 | 0 | 0 | 0 | 0 | 0 | 35465 |
| GIQKFPGIN | HMGCL | P35914 | HMGCL_HUMA | Unmodified | 0 | 0 | 0 | 0 | 316760 | 0 | 161190 | 0 | 0 | 2208800 | 0 | 0 | 0 | 0 | 0 | 0 | 576140 | 0 |
| GIQNDLTK | CAVIN2 | O95810 | CAVN2_HUMA | Unmodified | 173280 | 0 | 89087 | 0 | 2695200 | 269610 | 617380 | 58752 | 561250 | 231580 | 0 | 596190 | 0 | 0 | 0 | 184080 | 693480 | 0 |
| GIQRGDLF | ANKFY1 | Q9P2R3 | ANFY1_HUMAN | Unmodified | 0 | 0 | 0 | 0 | 2890400 | 268000 | 1299500 | 0 | 518830 | 252030 | 91388 | 126350 | 0 | 0 | 0 | 0 | 212630 | 282430 |
| GIQTSDPN | HDHD2; HDHD | Q9H0R4; K7 | HDHD2_HUMA | Oxidation (M) | 0 | 0 | 0 | 0 | 0 | 0 | 0 | 0 | 1149400 | 0 | 0 | 2217900 | 0 | 0 | 1110400 | 0 | 0 | 0 |
| GIQVVGWT | GDE1 | Q9NZC3 | GDE1_HUMAN | Unmodified | 0 | 43456 | 0 | 0 | 275790 | 86961 | 752170 | 80264 | 350030 | 0 | 0 | 0 | 1329500 | 866860 | 0 | 0 | 761580 | 183810 |
| GIRHEVINI | GSTO1; GSTO1 | P78417; Q5T | GSTO1_HUMA | Unmodified | 0 | 0 | 0 | 389010 | 2680200 | 911030 | 1654400 | 322140 | 0 | 0 | 2945100 | 5881400 | 1512800 | 1261800 | 0 | 0 | 607370 | 469950 |
| GIRLTIAEEF | RPS4Y2; RPS4X | Q8TD47; P62 | RS4Y2_HUMAN | Unmodified | 0 | 0 | 0 | 0 | 545370 | 0 | 792180 | 0 | 0 | 315970 | 0 | 0 | 595440 | 291640 | 93806 | 0 | 0 | 401570 |
| GIRPAINVG | ATP5F1A | P25705 | ATPA_HUMAN | Unmodified | 0 | 0 | 0 | 0 | 0 | 0 | 0 | 0 | 625800 | 85317 | 0 | 0 | 0 | 0 | 0 | 0 | 0 | 0 |
| GIRVLPEFD | HEXB; HEXB | P07686; Q5U | HEXB_HUMAN | Unmodified | 0 | 0 | 528700 | 333630 | 3995200 | 603220 | 0 | 0 | 0 | 980420 | 34583 | 0 | 1631900 | 242220 | 0 | 0 | 223440 | 188550 |
| GISCMNTT | PPA1 | Q15181 | IPYR_HUMAN I | Unmodified | 0 | 0 | 0 | 0 | 415530 | 0 | 0 | 0 | 0 | 0 | 493000 | 2129900 | 0 | 537840 | 0 | 67052 | 0 | 762490 |
| GISCMNTT | PPA1 | Q15181 | IPYR_HUMAN I | Oxidation (M) | 0 | 0 | 141070 | 0 | 0 | 0 | 0 | 0 | 114220 | 0 | 0 | 4282700 | 0 | 0 | 367820 | 222380 | 0 | 0 |
| GISCMNTT | PPA1 | Q15181 | IPYR_HUMAN I | Oxidation (M) | 0 | 0 | 0 | 0 | 0 | 0 | 0 | 0 | 0 | 0 | 105150 | 2452800 | 429390 | 0 | 0 | 247740 | 0 | 0 |
| GISDLAQH | CCT3; CCT3 | P49368; B4D | TCPG_HUMAN | Oxidation (M) | 433380 | 726780 | 992740 | 1178700 | 2241200 | 458780 | 269240 | 71539 | 1583200 | 2229600 | 1888200 | 20402000 | 3617200 | 1297300 | 1348700 | 971810 | 351500 | 519770 |
| GISDLAQH | CCT3; CCT3 | P49368; B4D | TCPG_HUMAN | Unmodified | 0 | 0 | 0 | 0 | 0 | 183120 | 0 | 0 | 2176800 | 538360 | 661690 | 2025700 | 2016700 | 494030 | 0 | 0 | 280910 | 498110 |
| GISDPLTVF | AGPS; AGPS; A | O00116; B7Z | ADAS_HUMAN | Unmodified | 0 | 0 | 0 | 0 | 0 | 0 | 0 | 0 | 3630300 | 0 | 134230 | 0 | 0 | 0 | 0 | 0 | 0 | 0 |
| GISEETTTG | AHCY | P23526 | SAHH_HUMAN | Unmodified | 2201400 | 2758900 | 7657900 | 6187200 | 4829100 | 166070 | 816580 | 542540 | 11155000 | 2479800 | 14867000 | 83313000 | 3709400 | 0 | 6851300 | 5850300 | 2404600 | 7642500 |
| GISHVIVDE | DHX9 | Q08211 | DHX9_HUMAN | Unmodified | 0 | 0 | 0 | 0 | 0 | 10213000 | 0 | 0 | 0 | 296510 | 0 | 10468000 | 339110 | 0 | 0 | 0 | 0 | 0 |
| GISLNPEQ | SUB1 | P53999 | TCP4_HUMAN | Unmodified | 0 | 0 | 0 | 0 | 5330800 | 1272000 | 0 | 0 | 3087800 | 1715000 | 2575700 | 15995000 | 4877700 | 0 | 0 | 0 | 0 | 362640 |
| GISPVPINL | DNM2; DNM1; I | P50570; Q05 | DYN2_HUMAN | Unmodified | 606020 | 568530 | 632160 | 292790 | 0 | 0 | 0 | 219890 | 0 | 0 | 234850 | 922890 | 4480200 | 0 | 0 | 408000 | 0 | 0 |
| GISQEQMQ | ACTN4; ACTN4 | O43707; F5C | ACTN4_HUMA | Unmodified | 0 | 0 | 438600 | 319800 | 653540 | 0 | 0 | 0 | 0 | 186720 | 420100 | 6033700 | 0 | 0 | 0 | 0 | 139080 | 0 |
| GISQEQMQ | ACTN4; ACTN4 | O43707; F5C | ACTN4_HUMA | Oxidation (M) | 218380 | 96585 | 250500 | 0 | 302700 | 186670 | 0 | 0 | 407040 | 35969 | 330590 | 2583300 | 0 | 0 | 328000 | 451980 | 45917 | 0 |
| GISQEQMQ | ACTN4; ACTN4 | O43707; F5C | ACTN4_HUMA | Unmodified | 0 | 0 | 0 | 0 | 5856500 | 1819400 | 292220 | 0 | 0 | 905830 | 0 | 0 | 962490 | 0 | 0 | 0 | 1200100 | 438360 |
| GISQTNLIT | EPB41L3; EPB4 | Q9Y2J2; A0A | E41L3_HUMAN | Unmodified | 0 | 0 | 0 | 0 | 0 | 0 | 0 | 96624 | 0 | 0 | 0 | 0 | 0 | 0 | 0 | 0 | 0 | 3491300 |
| GISTKPVYI | TRIM25; TRIM25 | Q14258; A0A | TRI25_HUMAN | Unmodified | 0 | 0 | 0 | 0 | 627160 | 0 | 0 | 0 | 0 | 0 | 436690 | 1333600 | 0 | 0 | 160120 | 191080 | 206220 | 0 |
| GISVHISNA | TARDBP; TARDI | Q13148; A0A | TADBP_HUMAN | Unmodified | 521270 | 423470 | 414220 | 373170 | 412420 | 0 | 1513400 | 853410 | 837240 | 315030 | 0 | 4059700 | 1257800 | 12019 | 278380 | 273530 | 0 | 609070 |
| GITEPPFGI | DSG2; DSG2 | Q14126; J3K | DSG2_HUMAN | Unmodified | 0 | 0 | 37434 | 0 | 206420 | 0 | 0 | 0 | 1334900 | 137240 | 45003 | 142750 | 0 | 0 | 0 | 0 | 0 | 0 |
| GITEQQKE | XRCC5 | P13010 | XRCC5_HUMA | Unmodified | 0 | 0 | 0 | 0 | 389990 | 0 | 948050 | 0 | 0 | 390390 | 0 | 850370 | 0 | 0 | 0 | 0 | 0 | 504510 |
| GITFDSGG | LAP3; LAP3 | P28838; H0Y | AMPL_HUMAN | Unmodified | 0 | 0 | 0 | 408950 | 1621000 | 0 | 540270 | 529780 | 1312800 | 480280 | 553720 | 1648500 | 1146500 | 1222400 | 0 | 492950 | 0 | 1183100 |
| GITGVEDKI | TKT; TKT; TKT | P29401; A0A | TKT_HUMAN Tr | Unmodified | 0 | 0 | 0 | 0 | 2090200 | 226650 | 0 | 0 | 0 | 1104000 | 0 | 0 | 986020 | 0 | 0 | 0 | 1185800 | 0 |
| GITHIGYTD | NNT; NNT; NNT | Q13423; E9F | NNTM_HUMAN | Unmodified | 0 | 0 | 0 | 0 | 0 | 0 | 0 | 87522 | 13004000 | 1053700 | 0 | 0 | 0 | 0 | 65955 | 0 | 0 | 0 |
| GITINAAHV | TUFM | P49411 | EFTU_HUMAN | Unmodified | 0 | 0 | 0 | 0 | 667950 | 0 | 0 | 0 | 20867000 | 1002900 | 319670 | 80913 | 0 | 0 | 12336 | 0 | 392950 | 1003400 |
| GITNLCVIG | PFKP; PFKP; PF | Q01813; A0A | PFKAP_HUMAN | Unmodified | 0 | 0 | 0 | 0 | 0 | 0 | 0 | 0 | 0 | 0 | 0 | 4019200 | 0 | 0 | 0 | 0 | 0 | 0 |
| GITPNFMK | SLC25A24 | Q6NUK1 | SCMC1_HUMA | Unmodified | 0 | 0 | 0 | 0 | 0 | 0 | 0 | 0 | 1702500 | 0 | 0 | 0 | 42640 | 0 | 685630 | 0 | 0 | 0 |
| GIVDLIEER | GFM1; GFM1; G | Q96RP9; C9I | EFGM_HUMAN | Unmodified | 0 | 0 | 0 | 0 | 0 | 0 | 0 | 0 | 1144200 | 0 | 0 | 0 | 0 | 0 | 0 | 0 | 0 | 0 |
| GIVDQSQQ | YWHAZ; YWHA | P63104; B7Z | 1433Z_HUMAN | Unmodified | 0 | 0 | 317670 | 509370 | 338490 | 0 | 539660 | 92592 | 3562700 | 1497700 | 5350400 | 2867600 | 0 | 0 | 600410 | 592750 | 282970 | 0 |
| GIVDQSQQ | YWHAZ; YWHA | P63104; B7Z | 1433Z_HUMAN | Unmodified | 10822000 | 8086700 | 12310000 | 21813000 | 24009000 | 9394800 | 4439100 | 0 | 97421000 | 21037000 | 20954000 | 1.19E+08 | 9130300 | 8699200 | 22003000 | 31661000 | 20793000 | 16467000 |
| GIVEESVTG | AHCYL1; AHCY | O43865; Q9 | SAHH2_HUMA | Unmodified | 890730 | 831540 | 1773800 | 3684200 | 2001600 | 151850 | 102980 | 0 | 1638500 | 217940 | 0 | 7157300 | 0 | 102060 | 8462600 | 3097000 | 839250 | 0 |
| GIVEESVTG | AHCYL1; AHCY | O43865; Q9 | SAHH2_HUMA | Unmodified | 0 | 0 | 48856 | 145760 | 5117300 | 1594400 | 586650 | 0 | 0 | 244390 | 0 | 0 | 1671300 | 1097400 | 0 | 0 | 3400500 | 2190400 |
| GIVEFASKF | SFPQ | P23246 | SFPQ_HUMAN | Unmodified | 0 | 166140 | 0 | 0 | 0 | 0 | 84531 | 386730 | 246400 | 105550 | 328170 | 1615300 | 0 | 0 | 0 | 0 | 0 | 0 |
| GIVEFSGKI | NONO; NONO; | Q15233; C9I | NONO_HUMAN | Unmodified | 0 | 0 | 0 | 0 | 0 | 0 | 0 | 0 | 523850 | 137080 | 0 | 1521200 | 0 | 0 | 0 | 0 | 0 | 0 |
| GIVFEDVK | ACADM; ACADM | P11310; Q5T | ACADM_HUMA | Unmodified | 0 | 0 | 0 | 0 | 0 | 0 | 0 | 0 | 1811900 | 182710 | 185240 | 0 | 0 | 0 | 0 | 0 | 0 | 0 |
| GIVFVIQSQ | B3GLCT | Q6Y288 | B3GLT_HUMAN | Unmodified | 0 | 0 | 0 | 0 | 0 | 0 | 0 | 0 | 2368000 | 0 | 0 | 159480 | 0 | 0 | 0 | 0 | 0 | 0 |
| GIVGVENV | PYGL; PYGL | P06737; E9P | PYGL_HUMAN | Unmodified | 87678 | 143910 | 331710 | 182480 | 1187700 | 327620 | 159410 | 104890 | 0 | 0 | 0 | 0 | 202500 | 264530 | 247390 | 0 | 374220 | 947180 |
| GIVISFAR | EHHADH | Q08426 | ECHP_HUMAN | Unmodified | 0 | 0 | 0 | 0 | 0 | 0 | 0 | 0 | 1875200 | 236630 | 0 | 329470 | 0 | 0 | 0 | 0 | 0 | 0 |
| GIVKDIIHD | RPL8; RPL8; RP | P62917; E9P | RL8_HUMAN 6 | Unmodified | 0 | 0 | 0 | 0 | 2606900 | 265370 | 9631900 | 0 | 0 | 932680 | 0 | 0 | 6007000 | 3081400 | 0 | 0 | 394580 | 921210 |
| GIVLFVSK | OSMR | Q99650 | OSMR_HUMAN | Unmodified | 0 | 0 | 0 | 0 | 3033600 | 0 | 0 | 0 | 0 | 0 | 58061 | 0 | 0 | 0 | 0 | 0 | 0 | 0 |
| GIVNEQFLI | ACADVL; ACAD | P49748; J3Q | ACADV_HUMA | Unmodified | 0 | 0 | 0 | 0 | 373660 | 0 | 0 | 0 | 3859400 | 495680 | 0 | 0 | 0 | 0 | 0 | 0 | 0 | 0 |
| GIVPLAKVD | PDIA3; PDIA3 | P30101; H7E | PDIA3_HUMAN | Unmodified | 0 | 0 | 0 | 0 | 1112700 | 0 | 1551300 | 0 | 2913700 | 9784700 | 0 | 0 | 1445300 | 2224300 | 0 | 0 | 380060 | 1578700 |
| GIVPLAKVD | PDIA3; PDIA3 | P30101; H7E | PDIA3_HUMAN | Unmodified | 0 | 0 | 0 | 0 | 0 | 0 | 4400100 | 2255000 | 0 | 13883000 | 0 | 0 | 2013300 | 3187700 | 0 | 0 | 897130 | 733210 |
| GIVSKDEIT | ITGA6; ITGA6 | P23229; C9J | ITA6_HUMAN I | Unmodified | 985620 | 1668700 | 3126700 | 6680000 | 2774300 | 842640 | 218640 | 0 | 1174400 | 0 | 1788700 | 9685700 | 2896400 | 1814300 | 6360200 | 5296000 | 0 | 6280700 |
| GIVTFQFR | MGAT1 | P26572 | MGAT1_HUMA | Unmodified | 0 | 0 | 0 | 0 | 0 | 0 | 0 | 0 | 7857200 | 248830 | 0 | 0 | 0 | 0 | 0 | 0 | 0 | 177810 |
| GIVVTAYSP | AKR1B1; AKR1E | P15121; E9P | ALDR_HUMAN | Unmodified | 0 | 0 | 0 | 0 | 0 | 0 | 0 | 0 | 0 | 0 | 1695900 | 18682000 | 0 | 0 | 0 | 0 | 0 | 0 |
| GIVVYTGD | ATP1A1 | P05023 | AT1A1_HUMAN | Unmodified | 0 | 0 | 3442600 | 1303300 | 0 | 0 | 0 | 0 | 0 | 1713700 | 452600 | 45250000 | 0 | 0 | 7473900 | 7537800 | 0 | 0 |
| GIVVYTGD | ATP1A1 | P05023 | AT1A1_HUMAN | Oxidation (M) | 0 | 73119 | 12762000 | 8594300 | 948980 | 534400 | 0 | 0 | 0 | 0 | 932550 | 26910000 | 1960600 | 661260 | 10449000 | 8662300 | 680940 | 515730 |
| GIVVYTGD | ATP1A1 | P05023 | AT1A1_HUMAN | Unmodified | 740420 | 427300 | 6092300 | 7234100 | 1685300 | 0 | 0 | 255110 | 12500000 | 1554900 | 7646700 | 45596000 | 4525600 | 383690 | 13302000 | 5009000 | 0 | 0 |
| GIVYIFNGR | ITGAV | P06756 | ITAV_HUMAN I | Unmodified | 586750 | 0 | 595450 | 0 | 0 | 0 | 0 | 0 | 1492400 | 0 | 0 | 2610900 | 0 | 37357 | 979120 | 828830 | 0 | 0 |
| GIWHNNEI | CKMT1A; CKMT | P12532; F8V | KCRU_HUMAN | Unmodified | 153940 | 0 | 0 | 158830 | 0 | 0 | 0 | 0 | 438560 | 0 | 295660 | 1628200 | 0 | 54213 | 0 | 0 | 0 | 0 |
| GIWIPDGE | ERBB2; ERBB2; | P04626; J3Q | ERBB2_HUMAN | Unmodified | 0 | 0 | 0 | 0 | 5191800 | 0 | 0 | 0 | 0 | 0 | 0 | 0 | 0 | 0 | 0 | 1818600 | 0 | 0 |
| GIWLEAGK | TLN1 | Q9Y490 | TLN1_HUMAN | Unmodified | 263420 | 0 | 240970 | 0 | 967080 | 0 | 532360 | 352070 | 231410 | 0 | 103410 | 1433300 | 0 | 0 | 726150 | 539520 | 0 | 0 |
| GIYAVGDV | GSR | P00390 | GSHR_HUMAN | Unmodified | 0 | 0 | 0 | 0 | 2905500 | 539110 | 0 | 0 | 524830 | 429990 | 0 | 1699500 | 1929400 | 1116300 | 0 | 0 | 0 | 0 |
| GIYAYGFEK | EIF4A3; EIF4A1; | P38919; P60 | IF4A3_HUMAN | Unmodified | 0 | 0 | 3417700 | 892890 | 0 | 0 | 39476 | 0 | 776690 | 52401 | 0 | 6546000 | 0 | 0 | 1909600 | 1915400 | 0 | 0 |
| GIYAYGFEK | EIF4A3; EIF4A1; | P38919; P60 | IF4A3_HUMAN | Unmodified | 10801000 | 8981100 | 1450200 | 13317000 | 4840600 | 1865700 | 1110300 | 793190 | 9717000 | 1850600 | 7014000 | 41478000 | 11791000 | 1949900 | 8109700 | 8441900 | 2273200 | 3760700 |
| GIYFADMV | PARP1; PARP1 | P09874; A0A | PARP1_HUMAN | Oxidation (M) | 0 | 313040 | 0 | 0 | 0 | 0 | 769550 | 414290 | 0 | 182680 | 0 | 668380 | 0 | 0 | 0 | 0 | 0 | 0 |
| GIYFADMV | PARP1; PARP1 | P09874; A0A | PARP1_HUMAN | Unmodified | 0 | 0 | 0 | 0 | 0 | 0 | 1333200 | 163910 | 454990 | 199530 | 201870 | 0 | 0 | 0 | 0 | 0 | 0 | 0 |
| GIYKGDRE | TMED2; TMED2 | Q15363; E7E | TMED2_HUMA | Unmodified | 0 | 0 | 0 | 0 | 0 | 0 | 0 | 0 | 1054600 | 0 | 0 | 0 | 0 | 0 | 0 | 0 | 0 | 0 |
| GIYLRDPV | TPP2; TPP2; TPI | P29144; A0A | TPP2_HUMAN | Unmodified | 0 | 0 | 0 | 0 | 0 | 0 | 0 | 0 | 0 | 0 | 0 | 3150500 | 0 | 0 | 0 | 0 | 0 | 0 |
| GIYLWDVE | OAT | P04181 | OAT_HUMAN C | Unmodified | 0 | 0 | 0 | 0 | 0 | 0 | 0 | 0 | 0 | 0 | 0 | 0 | 0 | 0 | 0 | 0 | 0 | 0 |
| GKADAGPS | GPRIN1 | Q7Z2K8 | GRIN1_HUMAN | Unmodified | 0 | 0 | 0 | 0 | 0 | 0 | 0 | 56322 | 235700 | 65607 | 0 | 0 | 0 | 0 | 0 | 0 | 77042 | 259420 |
| GKADAGPS | GPRIN1 | Q7Z2K8 | GRIN1_HUMAN | Unmodified | 1287600 | 705340 | 0 | 0 | 204070 | 0 | 317700 | 434270 | 0 | 282310 | 0 | 0 | 0 | 0 | 0 | 0 | 823820 | 0 |
| GKADGGAI | CD46 | R4GN18 | R4GN18_HUM | Unmodified | 4024000 | 3950600 | 2056200 | 2195100 | 4007200 | 292980 | 161570 | 36222 | 915100 | 1611400 | 1184800 | 9443600 | 2474200 | 1540600 | 2996500 | 2832700 | 718330 | 2871300 |
| GKDGVGA | GFM1; GFM1; G | Q96RP9; C9 | EFGM_HUMAN | Unmodified | 0 | 0 | 0 | 0 | 0 | 0 | 0 | 0 | 442500 | 0 | 0 | 0 | 0 | 0 | 0 | 0 | 0 | 0 |
| GKDSLSDD | LYN; LYN | P07948; E5R | LYN_HUMAN T | Unmodified | 0 | 0 | 0 | 0 | 1460600 | 339340 | 0 | 0 | 13590000 | 914220 | 0 | 30542 | 0 | 0 | 0 | 0 | 0 | 0 |
| GKDTSCLA | CNPY3 | Q9BT09 | CNPY3_HUMA | Unmodified | 0 | 0 | 0 | 0 | 0 | 0 | 0 | 0 | 964040 | 0 | 0 | 0 | 0 | 0 | 0 | 0 | 0 | 0 |
| GKDYLDQL | MAPK13 | O15264 | MK13_HUMAN | Unmodified | 22244 | 0 | 0 | 0 | 45103 | 0 | 0 | 0 | 678160 | 554900 | 0 | 0 | 0 | 0 | 0 | 21593 | 0 | 0 |
| GKEGSALS | LSM12; LSM12 | Q3MHD2; K7 | LSM12_HUMAN | Unmodified | 0 | 0 | 0 | 0 | 259560 | 0 | 62936 | 47659 | 0 | 0 | 0 | 166560 | 480080 | 290940 | 0 | 0 | 183180 | 306980 |
| GKEGSALT | EPB41L3; EPB4 | Q9Y2J2; A0A | E41L3_HUMAN | Unmodified | 0 | 0 | 0 | 0 | 0 | 0 | 0 | 0 | 221280 | 173280 | 0 | 0 | 594780 | 644120 | 0 | 0 | 1252900 | 9620000 |
| GKELNEFR | PFDN2 | Q9UHV9 | PFD2_HUMAN | Unmodified | 0 | 0 | 0 | 0 | 1931500 | 234800 | 29766 | 692630 | 0 | 516600 | 0 | 32107 | 3266700 | 2372400 | 0 | 0 | 3105200 | 1540200 |
| GKENQLQL | USP14; USP14 | P54578; A6N | UBP14_HUMAN | Unmodified | 0 | 0 | 0 | 0 | 1748000 | 0 | 0 | 77247 | 0 | 0 | 0 | 0 | 930700 | 448960 | 0 | 0 | 430990 | 0 |
| GKENRLCY | MANF | P55145 | MANF_HUMAN | Unmodified | 0 | 0 | 0 | 0 | 0 | 0 | 0 | 0 | 0 | 2364600 | 0 | 0 | 0 | 0 | 0 | 0 | 0 | 0 |
| GKESEKEL | M6PR; M6PR; M | P20645; F5G | MPRD_HUMAN | Unmodified | 0 | 0 | 0 | 0 | 326380 | 0 | 1744800 | 986790 | 0 | 2046100 | 0 | 0 | 1360300 | 1249700 | 0 | 0 | 0 | 0 |
| GKEVENFE | L1RE1 | Q9UN81 | LORF1_HUMAN | Unmodified | 0 | 783940 | 0 | 252210 | 0 | 85724 | 0 | 0 | 0 | 0 | 0 | 0 | 0 | 0 | 0 | 268730 | 0 | 0 |
| GKFAQDVS | SNF8 | Q96H20 | SNF8_HUMAN | Unmodified | 361460 | 386290 | 0 | 174160 | 0 | 0 | 0 | 0 | 0 | 0 | 0 | 0 | 0 | 0 | 133010 | 98837 | 0 | 0 |
| GKFEDMAK | SP100; HMGB1 | P23497; B2F | SP100_HUMAN | Unmodified | 0 | 0 | 0 | 0 | 0 | 0 | 273770 | 185140 | 0 | 312880 | 0 | 93173 | 0 | 274560 | 0 | 0 | 0 | 0 |
| GKFGHEFL | MAGOH; MAGO | P61326; Q96 | MGN_HUMAN | Unmodified | 689920 | 1200900 | 308830 | 653020 | 0 | 0 | 0 | 0 | 0 | 0 | 0 | 0 | 1117000 | 0 | 601460 | 659220 | 0 | 0 |
| GKFPVQLE | NCSTN; NCSTN | Q92542; H0Y | NICA_HUMAN | Unmodified | 0 | 740450 | 209160 | 2405200 | 2062000 | 317560 | 0 | 0 | 19665000 | 3740400 | 123400 | 1095100 | 0 | 0 | 2938300 | 453660 | 219780 | 0 |
| GKFQDNFE | MAPRE1 | Q15691 | MARE1_HUMA | Unmodified | 77227 | 169220 | 0 | 72722 | 1875200 | 497270 | 931210 | 0 | 408010 | 718210 | 716530 | 1169900 | 618400 | 738150 | 511370 | 58607 | 1191100 | 1643300 |
| GKFQDNFE | MAPRE1 | Q15691 | MARE1_HUMA | Unmodified | 0 | 0 | 0 | 0 | 0 | 0 | 0 | 0 | 0 | 0 | 492640 | 1239300 | 1393100 | 0 | 0 | 0 | 980900 | 1014200 |
| GKFTPSPVI | VPS26C; VPS26 | O14972; A8N | VP26C_HUMAN | Unmodified | 0 | 0 | 0 | 68315 | 664650 | 140330 | 0 | 0 | 292480 | 49240 | 0 | 0 | 197840 | 245230 | 0 | 38997 | 0 | 0 |
| GKGASGSF | HP1BP3; HP1BI | Q5SSJ5; Q5S | HP1B3_HUMA | Unmodified | 169820 | 0 | 0 | 0 | 0 | 0 | 0 | 784480 | 0 | 0 | 0 | 0 | 0 | 0 | 0 | 0 | 0 | 0 |
| GKGASSST | CPD; CPD | O75976; J3Q | CBPD_HUMAN | Unmodified | 0 | 0 | 0 | 0 | 0 | 0 | 0 | 0 | 684670 | 590240 | 0 | 0 | 0 | 0 | 0 | 0 | 0 | 0 |
| GKGFDWPI | GUSB | P08236 | BGLR_HUMAN | Unmodified | 0 | 0 | 0 | 0 | 0 | 0 | 0 | 0 | 2293800 | 0 | 0 | 0 | 0 | 0 | 0 | 0 | 0 | 0 |
| GKGFSVVA | LASP1; LASP1 | Q14847; C9J | LASP1_HUMAN | Unmodified | 0 | 0 | 0 | 0 | 2746100 | 0 | 0 | 0 | 0 | 0 | 0 | 0 | 747020 | 396080 | 0 | 0 | 741570 | 603150 |
| GKGGEIQP | HSPE1; HSPE1 | P61604; B8Z | CH10_HUMAN | Unmodified | 0 | 0 | 0 | 0 | 0 | 0 | 147500 | 0 | 7875700 | 700780 | 270930 | 242240 | 1158100 | 0 | 0 | 0 | 0 | 0 |
| GKGGEIQP | HSPE1; HSPE1 | P61604; B8Z | CH10_HUMAN | Unmodified | 0 | 0 | 0 | 0 | 1027300 | 0 | 1619000 | 368510 | 1842700 | 3897800 | 0 | 0 | 3575300 | 3465300 | 0 | 0 | 470810 | 140950 |
| GKGGVTGS | AHNAK | Q09666 | AHNK_HUMAN | Unmodified | 0 | 111410 | 0 | 0 | 263600 | 0 | 0 | 0 | 263430 | 0 | 0 | 0 | 0 | 0 | 0 | 0 | 389690 | 171610 |

| | | | | | | | | | | | | | | | | | | | | | | |
|---|---|---|---|---|---|---|---|---|---|---|---|---|---|---|---|---|---|---|---|---|---|---|
| GKGGVTGS | AHNAK | Q09666 | AHNK_HUMAN | Unmodified | 0 | 0 | 0 | 0 | 13440000 | 0 | 0 | 0 | 0 | 0 | 0 | 0 | 0 | 0 | 0 | 0 | 4473800 | 0 |
| GKGIAGRQ | TPPP3 | Q9BW30 | TPPP3_HUMAN | Unmodified | 0 | 0 | 0 | 0 | 0 | 0 | 1064300 | 373660 | 0 | 0 | 0 | 0 | 0 | 0 | 0 | 0 | 0 | 0 |
| GKGILGNA | GPX4; GPX4; GI | P36969; A0A | GPX4_HUMAN | Unmodified | 0 | 0 | 0 | 0 | 0 | 0 | 0 | 20314 | 0 | 80283 | 0 | 0 | 318000 | 319160 | 0 | 0 | 0 | 0 |
| GKGILGNA | GPX4; GPX4; GI | P36969; A0A | GPX4_HUMAN | Unmodified | 0 | 0 | 0 | 0 | 0 | 0 | 108510 | 0 | 0 | 507750 | 0 | 0 | 256280 | 205850 | 0 | 0 | 655570 | 65575 |
| GKGPLPME | CC2D1A | Q6P1N0 | C2D1A_HUMAN | Unmodified | 0 | 0 | 0 | 164380 | 949070 | 104630 | 0 | 0 | 284980 | 0 | 0 | 0 | 421870 | 375990 | 0 | 0 | 154130 | 370580 |
| GKIEEDSE\ | CUTA; CUTA | O60888; C9I | CUTA_HUMAN | Unmodified | 0 | 0 | 0 | 0 | 997070 | 0 | 562610 | 0 | 2982200 | 194490 | 0 | 96366 | 1487200 | 0 | 0 | 0 | 2606700 | 2315100 |
| GKINSITVD | CAP1 | Q01518 | CAP1_HUMAN | Unmodified | 0 | 0 | 0 | 0 | 4156900 | 869720 | 2877000 | 1165300 | 574130 | 488550 | 0 | 0 | 3335300 | 1769600 | 0 | 0 | 2910300 | 1701400 |
| GKINSITVD | CAP1 | Q01518 | CAP1_HUMAN | Unmodified | 0 | 0 | 0 | 0 | 0 | 0 | 339440 | 348010 | 0 | 121780 | 0 | 0 | 102910 | 268520 | 0 | 0 | 188680 | 475250 |
| GKKPIEDP/ | CTNND1; CTNN | O60716; A0A | CTND1_HUMAN | Unmodified | 297670 | 406380 | 3530500 | 2166100 | 12772000 | 2578200 | 0 | 0 | 0 | 57827 | 2382400 | 6914400 | 5576000 | 6140800 | 6544300 | 5715000 | 3028600 | 10375000 |
| GKKQPSLH | IQGAP3; IQGAP | Q86VI3; F2Z | IQGA3_HUMAN | Unmodified | 0 | 0 | 0 | 0 | 1684100 | 231080 | 0 | 0 | 0 | 0 | 0 | 0 | 0 | 0 | 0 | 0 | 0 | 0 |
| GKLAPGFD | FKBP9; FKBP9P | O95302; Q7 | FKBP9_HUMAN | Unmodified | 0 | 0 | 0 | 0 | 0 | 0 | 133870 | 0 | 0 | 1639000 | 0 | 0 | 0 | 0 | 0 | 0 | 0 | 0 |
| GKLDGNQI | IDH2 | P48735 | IDHP_HUMAN | Unmodified | 0 | 0 | 0 | 0 | 0 | 0 | 0 | 0 | 19493000 | 293950 | 230670 | 439020 | 0 | 0 | 0 | 0 | 0 | 0 |
| GKLDGNQI | IDH2 | P48735 | IDHP_HUMAN | Oxidation (M) | 0 | 0 | 0 | 0 | 0 | 0 | 0 | 0 | 0 | 823110 | 0 | 0 | 0 | 0 | 0 | 0 | 0 | 0 |
| GKLDVQFS | FLNA; FLNA; FL | P21333; A0A | FLNA_HUMAN | Unmodified | 0 | 0 | 0 | 0 | 692710 | 0 | 0 | 0 | 0 | 0 | 0 | 1876200 | 0 | 0 | 0 | 0 | 122560 | 0 |
| GKLEAIITPI | MACROH2A1; N | O75367; D6I | H2AY_HUMAN | Unmodified | 3145300 | 2683200 | 0 | 0 | 0 | 0 | 254180 | 289100 | 0 | 0 | 132550 | 2127800 | 0 | 0 | 0 | 0 | 0 | 0 |
| GKLGQYIRI | SLC9A3R1; SLC | O14745; J3C | NHRF1_HUMAN | Unmodified | 0 | 0 | 0 | 0 | 9612900 | 1081400 | 3998500 | 200950 | 0 | 1206700 | 0 | 0 | 16131000 | 25633000 | 0 | 0 | 8564300 | 2548900 |
| GKLGVCFD | DDX21 | Q9NR30 | DDX21_HUMAN | Unmodified | 0 | 0 | 0 | 0 | 0 | 0 | 1442700 | 396180 | 0 | 0 | 0 | 0 | 0 | 0 | 0 | 0 | 0 | 548920 |
| GKLNHLC( | PGM3; PGM3; F | O95394; J3K | AGM1_HUMAN | Unmodified | 0 | 0 | 0 | 0 | 406690 | 0 | 0 | 0 | 0 | 0 | 0 | 0 | 1042100 | 816650 | 25954 | 0 | 0 | 946010 |
| GKLPGPTL | APRT; APRT; AP | P07741; H3E | APT_HUMAN A | Unmodified | 108970 | 0 | 538200 | 0 | 0 | 0 | 0 | 0 | 0 | 0 | 0 | 943590 | 0 | 0 | 1175200 | 279240 | 0 | 0 |
| GKLPIVNEI | IMPDH2; IMPDH | P12268; A0A | IMDH2_HUMAN | Unmodified | 0 | 0 | 0 | 806230 | 0 | 0 | 0 | 0 | 1297200 | 0 | 1142200 | 1751800 | 0 | 0 | 797870 | 541160 | 502810 | 367630 |
| GKLPIVNEI | IMPDH2; IMPDH | P12268; A0A | IMDH2_HUMAN | Unmodified | 0 | 0 | 0 | 0 | 0 | 0 | 0 | 0 | 0 | 0 | 0 | 0 | 0 | 0 | 0 | 0 | 2740400 | 0 |
| GKLPKQPV | RPL27A; RPL27 | P46776; E9P | RL27A_HUMAN | Unmodified | 0 | 0 | 0 | 661290 | 874110 | 0 | 1279800 | 1070100 | 128270 | 0 | 0 | 0 | 2584000 | 0 | 0 | 0 | 225350 | 1159800 |
| GKLSVQGE | MLEC; MLEC; M | Q14165; H0 | MLEC_HUMAN | Unmodified | 0 | 0 | 0 | 0 | 0 | 595510 | 0 | 0 | 2229600 | 709590 | 0 | 0 | 0 | 153810 | 0 | 0 | 76106 | 251840 |
| GKLTLEDQ | DAG1 | Q14118 | DAG1_HUMAN | Unmodified | 0 | 162500 | 0 | 292090 | 0 | 0 | 344310 | 0 | 0 | 0 | 0 | 0 | 2280800 | 1840500 | 393270 | 302340 | 777340 | 1418100 |
| GKLTLEDQ | DAG1 | Q14118 | DAG1_HUMAN | Unmodified | 166400 | 218460 | 178390 | 277970 | 2396200 | 333310 | 200290 | 56679 | 369780 | 240310 | 0 | 452580 | 1312500 | 985170 | 398660 | 260810 | 1239100 | 1155700 |
| GKLVDFEA | TTC13 | Q8NBP0 | TTC13_HUMAN | Unmodified | 0 | 0 | 0 | 0 | 281340 | 789300 | 0 | 0 | 2099500 | 0 | 0 | 0 | 0 | 0 | 0 | 0 | 0 | 0 |
| GKLVIFEG( | AVIL | O75366 | AVIL_HUMAN A | Unmodified | 465320 | 597030 | 112700 | 0 | 669720 | 0 | 0 | 0 | 593670 | 0 | 335660 | 3493600 | 0 | 0 | 1186000 | 829930 | 0 | 0 |
| GKNPEDLI | MMUT | P22033 | MUTA_HUMAN | Unmodified | 0 | 0 | 0 | 0 | 0 | 0 | 0 | 0 | 4812100 | 1320700 | 0 | 0 | 0 | 0 | 0 | 0 | 0 | 0 |
| GKNVIHPF | VARS1; VARS1 | P26640; A0A | SYVC_HUMAN | Unmodified | 0 | 0 | 0 | 0 | 306610 | 0 | 0 | 0 | 0 | 104230 | 0 | 0 | 0 | 53192 | 0 | 0 | 36900 | 0 |
| GKNVLINK | CALR; CALR; C/ | P27797; A0A | CALR_HUMAN | Unmodified | 70288 | 0 | 0 | 0 | 1089400 | 0 | 2083500 | 1720300 | 9906400 | 13878000 | 405740 | 0 | 0 | 64458 | 0 | 0 | 765070 | 938310 |
| GKNVLINKI | CALR; CALR; C/ | P27797; A0A | CALR_HUMAN | Unmodified | 0 | 0 | 0 | 0 | 233580 | 36971 | 1079100 | 41074 | 4784900 | 11950000 | 0 | 0 | 361980 | 218690 | 0 | 0 | 94027 | 303660 |
| GKPAVAAL | PMPCA | Q10713 | MPPA_HUMAN | Unmodified | 0 | 0 | 0 | 0 | 0 | 0 | 0 | 0 | 2357900 | 0 | 0 | 0 | 40053 | 0 | 0 | 0 | 0 | 0 |
| GKPNIGQE | OS9; OS9 | Q13438; B4E | OS9_HUMAN P | Unmodified | 0 | 0 | 0 | 0 | 0 | 0 | 0 | 0 | 15359000 | 787150 | 907780 | 461080 | 0 | 0 | 0 | 0 | 0 | 0 |
| GKPPEADG | TJP1; TJP1; TJP1 | Q07157; A0/ | ZO1_HUMAN T | Unmodified | 0 | 0 | 523960 | 861400 | 4559400 | 1054600 | 240070 | 117580 | 222960 | 337900 | 0 | 0 | 5704100 | 3428900 | 406790 | 314840 | 1084200 | 805430 |
| GKPYLTLE( | PLCB3 | Q01970 | PLCB3_HUMAN | Oxidation (M) | 0 | 199680 | 0 | 0 | 189380 | 52319 | 0 | 0 | 0 | 0 | 0 | 0 | 41549 | 0 | 48381 | 0 | 0 | 0 |
| GKQTMQVI | ACAA2; ACAA2; | P42765; A0A | THIM_HUMAN | Unmodified | 0 | 0 | 0 | 0 | 0 | 0 | 0 | 0 | 407520 | 0 | 0 | 0 | 0 | 0 | 0 | 0 | 0 | 0 |
| GKRGGHS | CXCR4 | P61073 | CXCR4_HUMAN | Unmodified | 0 | 0 | 0 | 0 | 142220 | 0 | 0 | 0 | 85871 | 1704400 | 0 | 0 | 0 | 0 | 0 | 0 | 0 | 0 |
| GKSEEELE/ | MICAL3; MICAL | Q7RTP6; A0/ | MICA3_HUMAN | Unmodified | 0 | 0 | 0 | 0 | 684190 | 0 | 0 | 0 | 3535100 | 0 | 0 | 0 | 0 | 0 | 0 | 0 | 0 | 0 |
| GKSESQMI | MME; MME; MM | P08473; Q3I | NEP_HUMAN N | Oxidation (M) | 0 | 0 | 346400 | 0 | 0 | 0 | 0 | 0 | 0 | 0 | 0 | 0 | 0 | 0 | 228700 | 140680 | 0 | 0 |
| GKSESQMI | MME; MME; MM | P08473; Q3I | NEP_HUMAN N | Unmodified | 0 | 0 | 486170 | 0 | 0 | 0 | 0 | 0 | 0 | 0 | 0 | 0 | 174860 | 0 | 610440 | 132600 | 0 | 0 |
| GKSESQMI | MME; MME; MM | P08473; Q3I | NEP_HUMAN N | Oxidation (M) | 0 | 0 | 1601500 | 981730 | 0 | 0 | 0 | 0 | 0 | 0 | 0 | 0 | 0 | 0 | 11134000 | 6867700 | 0 | 0 |
| GKSESQMI | MME; MME; MM | P08473; Q3I | NEP_HUMAN N | Unmodified | 0 | 350180 | 8864600 | 13237000 | 95560 | 0 | 0 | 0 | 701540 | 0 | 0 | 301940 | 2634500 | 0 | 21361000 | 14830000 | 0 | 0 |
| GKSPAQAE | EPB41L4B | Q9H329 | E41LB_HUMAN | Unmodified | 0 | 355070 | 0 | 2276500 | 0 | 0 | 971390 | 0 | 0 | 0 | 0 | 0 | 0 | 0 | 786330 | 1434100 | 0 | 0 |
| GKTEAYLE/ | SCG3 | Q8WXD2 | SCG3_HUMAN | Unmodified | 0 | 0 | 0 | 0 | 0 | 372430 | 0 | 147610 | 48591000 | 9401400 | 0 | 0 | 0 | 0 | 0 | 0 | 0 | 0 |
| GKTEQQTA | LAMC1 | P11047 | LAMC1_HUMAN | Unmodified | 390440 | 182780 | 348620 | 331450 | 465620 | 0 | 83055 | 0 | 786820 | 71524 | 337750 | 519800 | 0 | 84255 | 351580 | 234900 | 0 | 58925 |
| GKTEQQTA | LAMC1 | P11047 | LAMC1_HUMAN | Unmodified | 0 | 0 | 0 | 268650 | 737010 | 878890 | 123240 | 86458 | 290130 | 590770 | 0 | 0 | 1809900 | 1802100 | 0 | 0 | 0 | 4979400 |
| GKTMADSS | FERMT2; FERMT | Q96AC1; A0/ | FERM2_HUMAN | Oxidation (M) | 0 | 0 | 0 | 0 | 1533900 | 0 | 0 | 0 | 0 | 0 | 0 | 0 | 0 | 0 | 0 | 0 | 596790 | 1155000 |
| GKTPVEPE | RPS20; RPS20; | P60866; A0A | RS20_HUMAN | Unmodified | 504670 | 316980 | 0 | 152400 | 0 | 0 | 0 | 0 | 0 | 0 | 0 | 2240200 | 453500 | 0 | 162680 | 197720 | 0 | 0 |
| GKVAEIMQ | PPM1L | Q5SGD2 | PPM1L_HUMAN | Oxidation (M) | 0 | 0 | 0 | 0 | 0 | 0 | 0 | 0 | 1801700 | 993640 | 0 | 0 | 0 | 0 | 0 | 0 | 0 | 0 |
| GKVEEVELI | PRMT1; PRMT8; | Q99873; Q9I | ANM1_HUMAN | Unmodified | 139980 | 0 | 0 | 0 | 0 | 0 | 0 | 0 | 0 | 0 | 0 | 17703000 | 0 | 0 | 0 | 318430 | 0 | 0 |
| GKVKVGVN | GAPDH | P04406 | G3P_HUMAN G | Unmodified | 0 | 0 | 0 | 0 | 778660 | 0 | 221760 | 0 | 0 | 73993 | 0 | 0 | 0 | 226760 | 0 | 0 | 706410 | 352270 |
| GKVVDLLA | ADSS2 | P30520 | PURA2_HUMAN | Unmodified | 0 | 0 | 0 | 0 | 315840 | 45393 | 0 | 0 | 0 | 0 | 281980 | 1581900 | 156960 | 43794 | 0 | 0 | 41272 | 0 |
| GLAAAEPT/ | SLC1A5 | Q15758 | AAAT_HUMAN | Unmodified | 0 | 0 | 0 | 0 | 0 | 0 | 0 | 0 | 0 | 0 | 0 | 0 | 0 | 0 | 1584900 | 0 | 0 | 0 |
| GLAAALEN | GGT1; GGT1 | P19440; E7E | GGT1_HUMAN | Unmodified | 1690000 | 710980 | 3984200 | 326180 | 0 | 0 | 0 | 0 | 5093900 | 232410 | 1499800 | 6930300 | 0 | 0 | 5003700 | 0 | 0 | 0 |
| GLAADQTC | SNTB1; SNTB1 | Q13884; A0/ | SNTB1_HUMAN | Unmodified | 0 | 0 | 0 | 0 | 1592600 | 0 | 0 | 0 | 0 | 3106900 | 0 | 0 | 0 | 0 | 485890 | 0 | 0 | 2782900 |
| GLAAIKFEG | GART | P22102 | PUR2_HUMAN | Unmodified | 0 | 0 | 0 | 0 | 1647500 | 0 | 572440 | 0 | 0 | 0 | 0 | 465900 | 824050 | 0 | 0 | 0 | 724250 | 0 |
| GLAALPGD | LRIG1 | Q96JA1 | LRIG1_HUMAN | Unmodified | 0 | 0 | 0 | 0 | 0 | 0 | 0 | 0 | 1310800 | 0 | 0 | 0 | 0 | 0 | 0 | 0 | 0 | 0 |
| GLAEVQQI | AKAP12 | Q02952 | AKA12_HUMAN | Unmodified | 0 | 0 | 0 | 0 | 1905900 | 229190 | 0 | 0 | 0 | 0 | 0 | 0 | 0 | 0 | 0 | 0 | 451010 | 3641100 |
| GLAFIQDPI | GLO1 | Q04760 | LGUL_HUMAN | Unmodified | 47724 | 20003 | 0 | 68415 | 1281500 | 0 | 59663 | 0 | 158460 | 396340 | 99485 | 222620 | 1210300 | 913030 | 404380 | 0 | 597290 | 916070 |
| GLAFTDVD | FN1 | P02751 | FINC_HUMAN | Unmodified | 0 | 0 | 15705000 | 9163300 | 0 | 0 | 0 | 195840 | 399350 | 752100 | 1878300 | 407970 | 0 | 894760 | 0 | 104940 | 547680 | 3022900 |
| GLAFTDVD | FN1 | P02751 | FINC_HUMAN | Unmodified | 0 | 0 | 0 | 0 | 80267 | 0 | 0 | 0 | 0 | 33608 | 0 | 0 | 0 | 0 | 0 | 0 | 0 | 463900 |
| GLAGAVSE | TLN1 | Q9Y490 | TLN1_HUMAN | Unmodified | 617210 | 2078000 | 1751100 | 2134300 | 1571500 | 983210 | 903870 | 0 | 0 | 788330 | 709190 | 3465200 | 0 | 0 | 0 | 2663300 | 594870 | 0 |
| GLAGLGD\ | PYGB | P11216 | PYGB_HUMAN | Unmodified | 0 | 0 | 0 | 0 | 63497 | 0 | 0 | 0 | 264590 | 0 | 0 | 1616300 | 444370 | 0 | 0 | 430260 | 374400 | 2702000 |
| GLAGPPAS | KIAA1522 | Q9P206 | K1522_HUMAN | Unmodified | 0 | 0 | 631340 | 1297700 | 2478900 | 377920 | 0 | 22272 | 701690 | 0 | 0 | 978830 | 3969500 | 2123300 | 1068400 | 866850 | 870960 | 1344500 |
| GLAHRAKL | IDH1 | O75874 | IDHC_HUMAN | Unmodified | 0 | 0 | 0 | 0 | 583530 | 0 | 0 | 0 | 0 | 0 | 0 | 0 | 2397400 | 1733700 | 0 | 0 | 177410 | 41357 |
| GLAITFVSD | DDX39B; DDX3 | Q13838; O0 | DX39B_HUMAN | Unmodified | 1040300 | 502310 | 262210 | 201360 | 2013900 | 0 | 1075500 | 487850 | 1907000 | 1369600 | 1884800 | 9005100 | 140920 | 0 | 0 | 221750 | 0 | 0 |
| GLALSNSD | UCHL5; UCHL5 | Q9Y5K5; Q5I | UCHL5_HUMAN | Unmodified | 0 | 88375 | 0 | 208800 | 0 | 0 | 0 | 0 | 1303000 | 141130 | 0 | 6631100 | 97351 | 0 | 0 | 0 | 30583 | 0 |
| GLAPDLPEI | RPS13; RPS13 | P62277; J3K | RS13_HUMAN | Unmodified | 4538400 | 4708700 | 1341900 | 2215300 | 343040 | 0 | 97009 | 0 | 715850 | 393710 | 927190 | 40106000 | 0 | 0 | 2956300 | 4246000 | 0 | 0 |
| GLAPDLPEI | RPS13; RPS13 | P62277; J3K | RS13_HUMAN | Unmodified | 2589300 | 3581300 | 1033600 | 2400200 | 177550 | 0 | 415800 | 0 | 1149300 | 259370 | 924210 | 67742000 | 85018 | 0 | 2641600 | 1812800 | 0 | 154670 |
| GLAPEQPV | ACOT2; ACOT1 | P49753; Q8 | ACOT2_HUMAN | Unmodified | 0 | 0 | 0 | 0 | 0 | 0 | 0 | 0 | 2229900 | 0 | 0 | 148810 | 0 | 0 | 0 | 0 | 0 | 0 |
| GLAPTCAY | PGAP6 | Q9HCN3 | PGAP6_HUMAN | Unmodified | 324010 | 160420 | 74861 | 95482 | 0 | 0 | 0 | 0 | 947480 | 0 | 372650 | 1201800 | 0 | 0 | 367540 | 240650 | 0 | 0 |
| GLASWDE( | QSOX2 | Q6ZRP7 | QSOX2_HUMAN | Unmodified | 98082 | 39582 | 0 | 0 | 234830 | 0 | 0 | 11872 | 72675000 | 5719800 | 1752200 | 2741300 | 0 | 0 | 0 | 259040 | 0 | 0 |
| GLATFCLDI | PGRMC1; PGRM | O00264; O1 | PGRC1_HUMAN | Unmodified | 0 | 0 | 0 | 0 | 0 | 0 | 0 | 0 | 585270 | 0 | 0 | 1559700 | 0 | 0 | 0 | 0 | 0 | 0 |
| GLAVFISDI | AP2A2; AP2A1; | O94973; O9 | AP2A2_HUMAN | Unmodified | 4444400 | 5842200 | 5761700 | 8637800 | 928030 | 672420 | 0 | 0 | 1216500 | 83082 | 1038800 | 6619800 | 0 | 0 | 18792000 | 14847000 | 0 | 0 |
| GLAVFISDI | AP2A2; AP2A1; | O94973; O9 | AP2A2_HUMAN | Unmodified | 0 | 0 | 0 | 0 | 713920 | 0 | 0 | 0 | 147000 | 783830 | 114530 | 0 | 741600 | 0 | 0 | 0 | 196270 | 985370 |
| GLAYCETH | LIMS1; LIMS4; L | P48059; A0A | LIMS1_HUMAN | Unmodified | 0 | 0 | 0 | 0 | 0 | 0 | 2095200 | 0 | 709880 | 0 | 0 | 0 | 0 | 0 | 0 | 0 | 0 | 0 |
| GLCAIAQAI | RPS3; RPS3; RP | P23396; H0 | RS3_HUMAN 4 | Unmodified | 2331900 | 1972000 | 1042200 | 941380 | 847510 | 0 | 0 | 0 | 0 | 181030 | 685490 | 37148000 | 0 | 0 | 1752600 | 1969800 | 0 | 0 |
| GLCAIAQAI | RPS3; RPS3; RP | P23396; H0 | RS3_HUMAN 4 | Unmodified | 319760 | 0 | 171020 | 0 | 0 | 0 | 0 | 0 | 0 | 0 | 607360 | 1211700 | 0 | 0 | 0 | 0 | 0 | 0 |
| GLCGAIHS | ATP5F1C | P36542 | ATPG_HUMAN | Unmodified | 19499 | 0 | 0 | 0 | 213450 | 0 | 130290 | 54173 | 1454500 | 0 | 0 | 453110 | 0 | 0 | 0 | 0 | 0 | 126080 |
| GLCSPRPN | ITGA7; ITGA7 | Q13683; J3K | ITA7_HUMAN I | Unmodified | 0 | 0 | 0 | 0 | 0 | 0 | 1682500 | 0 | 0 | 0 | 0 | 0 | 0 | 0 | 0 | 0 | 0 | 0 |
| GLDARPEV | CDH1 | P12830 | CADH1_HUMA | Unmodified | 298770 | 207770 | 599890 | 2291100 | 509980 | 0 | 0 | 0 | 441230 | 0 | 0 | 413590 | 2905600 | 459190 | 2085300 | 1214600 | 0 | 0 |
| GLDARPEV | CDH1 | P12830 | CADH1_HUMA | Oxidation (M) | 0 | 393380 | 2645800 | 5286200 | 0 | 0 | 0 | 0 | 0 | 0 | 570220 | 1820900 | 0 | 0 | 6296000 | 3507100 | 0 | 0 |
| GLDCDTVT | PLXNA1 | Q9UIW2 | PLXA1_HUMAN | Unmodified | 0 | 231480 | 0 | 0 | 478630 | 0 | 0 | 0 | 269020 | 69997 | 0 | 291760 | 0 | 343490 | 0 | 88562 | 280010 | 655930 |
| GLDCDTVT | PLXNA1 | Q9UIW2 | PLXA1_HUMAN | Unmodified | 0 | 0 | 0 | 0 | 0 | 0 | 12807 | 0 | 0 | 0 | 0 | 327620 | 0 | 0 | 37635 | 0 | 68631 | 0 |
| GLDGARAE | SHMT2; SHMT2 | P34897; H0 | GLYM_HUMAN | Unmodified | 0 | 0 | 0 | 0 | 398350 | 0 | 280870 | 0 | 250780 | 6425900 | 0 | 0 | 382260 | 93933 | 0 | 0 | 89380 | 104040 |
| GLDGAVDI | ALDH16A1; ALD | Q8IZ83; F5H | A16A1_HUMAN | Unmodified | 0 | 0 | 0 | 0 | 0 | 0 | 0 | 215300 | 190740 | 0 | 0 | 984010 | 0 | 0 | 0 | 0 | 0 | 573830 |
| GLDGGGA | STMN3 | A0A0J9YW3 | A0A0J9YW36_H | Unmodified | 537800 | 529360 | 0 | 0 | 299600 | 0 | 407690 | 631340 | 645170 | 552000 | 3256200 | 2536000 | 42517 | 0 | 0 | 0 | 0 | 0 |
| GLDISNVK | DDX3X; DDX3X | O00571; A0/ | DDX3X_HUMAN | Unmodified | 2360900 | 1621100 | 0 | 972610 | 253230 | 29031 | 0 | 136250 | 1084300 | 353490 | 1248400 | 3267000 | 0 | 0 | 2086700 | 0 | 48860 | 0 |
| GLDLGGR( | AHNAK | Q09666 | AHNK_HUMAN | Unmodified | 37883 | 0 | 0 | 0 | 1548300 | 0 | 0 | 0 | 0 | 0 | 0 | 0 | 0 | 0 | 0 | 0 | 0 | 0 |
| GLDLGGR( | AHNAK | Q09666 | AHNK_HUMAN | Unmodified | 0 | 0 | 0 | 0 | 7240500 | 581250 | 0 | 0 | 0 | 0 | 0 | 0 | 0 | 0 | 0 | 0 | 1776300 | 6818500 |
| GLDMETDF | MOV10 | Q9HCE1 | MOV10_HUMA | Unmodified | 1702800 | 2278400 | 746890 | 428090 | 0 | 0 | 0 | 0 | 69562 | 0 | 66982 | 1160900 | 0 | 0 | 1027200 | 1119900 | 0 | 0 |
| GLDNLYK | EXOC1 | Q9NV70 | EXOC1_HUMAN | Unmodified | 0 | 0 | 0 | 0 | 798700 | 109860 | 1109300 | 119760 | 220890 | 0 | 0 | 753710 | 0 | 0 | 0 | 207550 | 185000 | 0 |
| GLDPARVN | MDH2; MDH2 | P40926; G3 | MDHM_HUMAN | Unmodified | 372500 | 0 | 677480 | 712730 | 4883000 | 1230900 | 7423700 | 1307600 | 57598000 | 23623000 | 1307400 | 0 | 1373900 | 5583400 | 0 | 0 | 1442200 | 4124500 |
| GLDPSTPA | LPIN1; LPIN1 | Q14693; A0/ | LPIN1_HUMAN | Unmodified | 0 | 0 | 0 | 0 | 0 | 0 | 0 | 0 | 0 | 0 | 159730 | 0 | 0 | 0 | 0 | 0 | 0 | 0 |
| GLDPYNVL | COX5B | P10606 | COX5B_HUMAN | Unmodified | 0 | 0 | 0 | 0 | 0 | 0 | 1154400 | 409910 | 3361000 | 1213600 | 242740 | 527730 | 278050 | 0 | 0 | 0 | 0 | 0 |
| GLDSNKNI | LRRC1 | Q9BTT6 | LRRC1_HUMAN | Unmodified | 0 | 0 | 0 | 0 | 3911300 | 0 | 110860 | 0 | 1562900 | 757870 | 0 | 0 | 1433400 | 844600 | 292460 | 141150 | 1511400 | 2749500 |
| GLDVDSLV | RPL17; RPL17; | P18621; A0A | RL17_HUMAN | Unmodified | 2618400 | 7079800 | 1202600 | 0 | 0 | 0 | 443260 | 0 | 748870 | 448120 | 4177200 | 14634000 | 352410 | 0 | 0 | 1255000 | 0 | 610930 |
| GLDVDSLV | RPL17; RPL17; | P18621; A0A | RL17_HUMAN | Unmodified | 5563300 | 6868900 | 0 | 3068500 | 8932900 | 1779900 | 7403600 | 139200 | 1703600 | 1559800 | 5373000 | 23647000 | 15005000 | 4545800 | 739320 | 947920 | 622430 | 3853400 |
| GLDWVKEI | APEX1; APEX1; | P27695; A0A | APEX1_HUMAN | Unmodified | 0 | 0 | 491480 | 0 | 0 | 0 | 0 | 0 | 0 | 0 | 720350 | 0 | 0 | 0 | 0 | 0 | 0 | 0 |
| GLDYEGG( | PLOD3; PLOD3 | O60568; H7 | PLOD3_HUMAN | Unmodified | 0 | 0 | 0 | 0 | 891510 | 11079 | 0 | 0 | 1825800 | 207560 | 0 | 0 | 0 | 10055 | 0 | 0 | 0 | 0 |
| GLDYGGV/ | NEDD4; SMURF | P46934; Q9 | NEDD4_HUMA | Unmodified | 0 | 1248500 | 387170 | 34215 | 104610 | 0 | 0 | 0 | 148730 | 0 | 1368800 | 2680500 | 0 | 686080 | 2077600 | 1555800 | 0 | 0 |
| GLDYGGV/ | NEDD4L; NEDD | Q96PU5; A0 | NED4L_HUMAN | Unmodified | 0 | 0 | 29387 | 31443 | 0 | 0 | 0 | 0 | 0 | 0 | 0 | 0 | 0 | 0 | 207030 | 30655 | 0 | 0 |
| GLEAGLDT | ELFN2 | Q5R3F8 | PPR29_HUMAN | Unmodified | 165360 | 0 | 0 | 0 | 1482400 | 1866500 | 946170 | 0 | 61970 | 280520 | 482400 | 289240 | 0 | 0 | 0 | 0 | 257240 | 269050 |
| GLEDSYEG | PLBD2 | Q8NHP8 | PLBL2_HUMAN | Unmodified | 0 | 0 | 243900 | 0 | 0 | 0 | 0 | 0 | 647150 | 139450 | 0 | 0 | 0 | 0 | 0 | 0 | 0 | 0 |
| GLEEENAQ | PBXIP1 | Q96AQ6 | PBIP1_HUMAN | Unmodified | 0 | 84535 | 0 | 0 | 0 | 0 | 0 | 0 | 3705900 | 182480 | 1342700 | 879150 | 0 | 0 | 0 | 0 | 0 | 0 |
| GLEEENAQ | PBXIP1 | Q96AQ6 | PBIP1_HUMAN | Unmodified | 0 | 0 | 0 | 0 | 0 | 0 | 0 | 0 | 793470 | 241830 | 0 | 0 | 0 | 0 | 0 | 0 | 172510 | 48459 |
| GLEEPEMD | HNRNPUL2; HN | Q1KMD3; H3 | HNRL2_HUMAN | Unmodified | 0 | 0 | 0 | 0 | 0 | 0 | 0 | 0 | 0 | 0 | 224650 | 996500 | 511630 | 237100 | 0 | 0 | 0 | 0 |
| GLEESLGM | OVCA2 | Q8WZ82 | OVCA2_HUMAN | Oxidation (M) | 22142 | 69740 | 0 | 0 | 0 | 0 | 0 | 0 | 0 | 0 | 0 | 237430 | 0 | 0 | 218840 | 37703 | 0 | 0 |
| GLEETVAK | KIF5B | P33176 | KINH_HUMAN | Unmodified | 0 | 0 | 860630 | 0 | 3644600 | 346480 | 463740 | 565160 | 733540 | 205970 | 0 | 4190500 | 1291600 | 371520 | 0 | 699550 | 1003700 | 2027400 |

| | | | | | | | | | | | | | | | | | | | | | | |
|---|---|---|---|---|---|---|---|---|---|---|---|---|---|---|---|---|---|---|---|---|---|---|
| GLEFPMTD | TOM1; TOM1; T | O60784; V9 | TOM1_HUMAN | 2 Oxidation (M) | 0 | 2357700 | 916070 | 3379700 | 0 | 0 | 0 | 0 | 0 | 0 | 0 | 328020 | 0 | 0 | 7206500 | 2258200 | 0 | 0 |
| GLEGTIRSD | NIBAN1 | Q9BZQ8 | NIBA1_HUMAN | Unmodified | 0 | 0 | 43184 | 321010 | 2774800 | 455750 | 2791400 | 41211 | 2444700 | 2844400 | 0 | 0 | 305280 | 639650 | 0 | 0 | 605600 | 740590 |
| GLEGTIRSD | NIBAN1 | Q9BZQ8 | NIBA1_HUMAN | Oxidation (M) | 0 | 0 | 0 | 0 | 0 | 0 | 128230 | 27371 | 0 | 228780 | 0 | 0 | 0 | 0 | 0 | 0 | 0 | 0 |
| GLEGWEK | ADSS2 | P30520 | PURA2_HUMAN | Unmodified | 0 | 0 | 0 | 0 | 0 | 0 | 0 | 0 | 0 | 84495 | 1325800 | 5998500 | 465170 | 0 | 0 | 0 | 500120 | 0 |
| GLEHRGKL | IDH2 | P48735 | IDHP_HUMAN | Unmodified | 0 | 0 | 0 | 0 | 0 | 0 | 0 | 0 | 0 | 720480 | 0 | 0 | 1371700 | 1655400 | 0 | 0 | 0 | 0 |
| GLEHVTVD | ENY2; ENY2 | Q9NPA8; E5 | ENY2_HUMAN | Unmodified | 0 | 0 | 0 | 0 | 2262800 | 0 | 458200 | 0 | 1871200 | 835860 | 1524000 | 86153 | 2276800 | 0 | 658230 | 0 | 1972700 | 2183000 |
| GLEISGTFT | AP2B1; AP2B1 | P63010; A0A | AP2B1_HUMAN | Unmodified | 360670 | 310770 | 0 | 283460 | 183480 | 0 | 0 | 0 | 425670 | 0 | 307230 | 2095100 | 176230 | 0 | 817540 | 153680 | 0 | 0 |
| GLEISGTFT | AP1B1 | Q10567 | AP1B1_HUMAN | Unmodified | 0 | 0 | 156470 | 363900 | 0 | 0 | 200400 | 0 | 618760 | 279470 | 0 | 1416900 | 0 | 0 | 0 | 288190 | 0 | 0 |
| GLELLENSI | GLMN; GLMN; | Q92990; M0 | GLMN_HUMAN | Unmodified | 0 | 0 | 0 | 0 | 0 | 0 | 0 | 0 | 0 | 0 | 0 | 692450 | 0 | 0 | 0 | 0 | 0 | 0 |
| GLELPSEIQ | FAM3B; FAM3B | P58499; A8N | FAM3B_HUMAN | Unmodified | 0 | 0 | 0 | 0 | 0 | 0 | 0 | 0 | 2977000 | 484520 | 820220 | 843590 | 0 | 0 | 0 | 0 | 0 | 0 |
| GLELQPQD | MYOF; MYOF | Q9NZM1; HC | MYOF_HUMAN | Unmodified | 0 | 0 | 0 | 0 | 3069100 | 1331100 | 0 | 63781 | 189470 | 0 | 0 | 0 | 79868 | 0 | 0 | 0 | 0 | 89458 |
| GLEPSPAA | DIP2B | Q9P265 | DIP2B_HUMAN | Unmodified | 0 | 0 | 0 | 0 | 0 | 0 | 0 | 0 | 0 | 0 | 0 | 3174800 | 0 | 0 | 0 | 0 | 0 | 0 |
| GLEQGIQD | MAN2A1 | Q16706 | MA2A1_HUMAN | Unmodified | 0 | 0 | 0 | 0 | 0 | 0 | 0 | 0 | 204920 | 42585 | 183730 | 598770 | 0 | 0 | 0 | 0 | 0 | 0 |
| GLEQGIQD | MAN2A1 | Q16706 | MA2A1_HUMAN | Unmodified | 0 | 0 | 0 | 0 | 0 | 0 | 0 | 0 | 3206700 | 352170 | 0 | 1430700 | 0 | 0 | 0 | 0 | 0 | 0 |
| GLEQLLVG | COL6A1; COL6 | P12109; A0A | CO6A1_HUMAN | Unmodified | 0 | 0 | 0 | 0 | 0 | 0 | 0 | 0 | 2450300 | 1375100 | 0 | 0 | 0 | 0 | 0 | 0 | 192830 | 3822700 |
| GLESLPPLF | MARS1 | P56192 | SYMC_HUMAN | Unmodified | 377390 | 0 | 1481300 | 489500 | 99231 | 0 | 0 | 56646 | 1763200 | 148180 | 1474800 | 16468000 | 137560 | 0 | 618200 | 480910 | 0 | 129940 |
| GLESTTLAD | CSRP1; CSRP1; | P21291; E9P | CSRP1_HUMAN | Unmodified | 1004200 | 720720 | 360490 | 1608900 | 13402000 | 3276200 | 2154000 | 1706800 | 3824000 | 4270000 | 486960 | 1149700 | 8364400 | 5926400 | 1107400 | 908430 | 6821600 | 3568400 |
| GLETFSQL | HEXA; HEXA; HI | P06865; H3E | HEXA_HUMAN | Unmodified | 0 | 0 | 0 | 0 | 0 | 0 | 0 | 0 | 515530 | 0 | 0 | 0 | 0 | 0 | 0 | 0 | 0 | 0 |
| GLETTATYD | ACOX1 | Q15067 | ACOX1_HUMAN | Unmodified | 0 | 0 | 0 | 0 | 0 | 0 | 0 | 0 | 2885200 | 0 | 517840 | 0 | 0 | 0 | 0 | 0 | 0 | 0 |
| GLEVTAYSF | AKR1A1 | P14550 | AK1A1_HUMAN | Unmodified | 0 | 0 | 0 | 719790 | 0 | 0 | 0 | 0 | 1706200 | 0 | 0 | 3040700 | 0 | 0 | 0 | 602240 | 0 | 0 |
| GLEVTDDS | AARS1; AARS1; | P49588; A0A | SYAC_HUMAN | Unmodified | 188760 | 316510 | 0 | 183460 | 402850 | 0 | 0 | 0 | 1790200 | 212150 | 944950 | 4891000 | 212480 | 46960 | 694220 | 933010 | 0 | 0 |
| GLEWLWSI | PKP3 | Q9Y446 | PKP3_HUMAN | Unmodified | 32498 | 66568 | 53870 | 155530 | 14756 | 0 | 0 | 0 | 34398 | 0 | 0 | 0 | 36783 | 0 | 571100 | 65228 | 0 | 0 |
| GLEYLYLN | DYNC1LI1; DYN | Q9Y6G9; C9 | DC1L1_HUMAN | Unmodified | 0 | 0 | 0 | 0 | 0 | 0 | 0 | 0 | 0 | 0 | 0 | 1956700 | 0 | 0 | 113610 | 60744 | 0 | 0 |
| GLFDEYGS | HSP90B2P; HSI | Q58FF3; P14 | ENPLL_HUMAN | Unmodified | 0 | 0 | 2008700 | 0 | 0 | 0 | 59978 | 0 | 1.81E+08 | 44944000 | 18584000 | 21777000 | 0 | 0 | 0 | 0 | 0 | 0 |
| GLFDEYGS | HSP90B2P; HSI | Q58FF3; P14 | ENPLL_HUMAN | Unmodified | 0 | 0 | 0 | 0 | 0 | 0 | 552560 | 195870 | 4064500 | 13374000 | 0 | 169610 | 0 | 0 | 0 | 0 | 0 | 0 |
| GLFEVNPW | APMAP; APMAP | Q9HDC9; H | APMAP_HUMAN | Unmodified | 0 | 0 | 0 | 0 | 0 | 0 | 0 | 0 | 9105600 | 938750 | 0 | 171840 | 0 | 0 | 0 | 0 | 0 | 0 |
| GLFEVNPW | APMAP; APMAP | Q9HDC9; H | APMAP_HUMAN | Unmodified | 0 | 0 | 0 | 302630 | 0 | 0 | 0 | 0 | 1.8E+08 | 7582900 | 1102000 | 510810 | 0 | 0 | 0 | 0 | 0 | 0 |
| GLFEVNPW | APMAP; APMAP | Q9HDC9; H | APMAP_HUMAN | Unmodified | 0 | 0 | 0 | 0 | 0 | 0 | 0 | 0 | 0 | 11301000 | 0 | 0 | 658470 | 362510 | 0 | 0 | 85670 | 0 |
| GLFFEHDL | CASC3; CASC3 | O15234; J3K | CASC3_HUMAN | Unmodified | 702010 | 480970 | 0 | 467760 | 449270 | 0 | 0 | 0 | 0 | 0 | 0 | 0 | 924810 | 691430 | 310780 | 168620 | 0 | 0 |
| GLFIIDDKG | PRDX4; PRDX4 | Q13162; H7 | PRDX4_HUMAN | Unmodified | 0 | 0 | 0 | 0 | 0 | 0 | 0 | 0 | 2189300 | 2725900 | 414500 | 0 | 0 | 0 | 0 | 0 | 0 | 0 |
| GLFIIDGK | PRDX2 | P32119 | PRDX2_HUMAN | Unmodified | 1680100 | 0 | 5510600 | 4772500 | 0 | 0 | 340560 | 607720 | 15166000 | 498320 | 15026000 | 1.4E+08 | 192950 | 0 | 6241600 | 3823900 | 0 | 0 |
| GLFIIDGKG | PRDX2 | P32119 | PRDX2_HUMAN | Unmodified | 0 | 0 | 0 | 687080 | 142180 | 0 | 0 | 0 | 2036800 | 616180 | 1066700 | 0 | 0 | 0 | 0 | 0 | 123300 | 132740 |
| GLFIIDGKG | PRDX2 | P32119 | PRDX2_HUMAN | Unmodified | 0 | 0 | 0 | 0 | 0 | 0 | 0 | 0 | 180250 | 2834800 | 220880 | 0 | 0 | 0 | 0 | 0 | 633580 | 636040 |
| GLFIIDPNG | PRDX3 | P30048 | PRDX3_HUMAN | Unmodified | 0 | 0 | 0 | 0 | 0 | 0 | 0 | 0 | 55339000 | 2652900 | 1411100 | 1892000 | 0 | 0 | 0 | 0 | 0 | 0 |
| GLFIKPTVF | ALDH1A3; ALDI | P47895; H0 | AL1A3_HUMAN | Oxidation (M) | 0 | 0 | 0 | 0 | 0 | 0 | 0 | 0 | 0 | 509680 | 541230 | 0 | 0 | 0 | 0 | 0 | 0 | 1221500 |
| GLFKENNE | ITGA2; ITGA2; I | P17301; E7E | ITA2_HUMAN I | Unmodified | 0 | 0 | 0 | 0 | 552720 | 100650 | 64219 | 0 | 0 | 0 | 0 | 0 | 0 | 148730 | 0 | 0 | 937400 | 1383300 |
| GLFLPEDEI | ASPH | Q12797 | ASPH_HUMAN | Unmodified | 0 | 0 | 0 | 0 | 0 | 0 | 0 | 0 | 8536000 | 731460 | 0 | 0 | 762670 | 544770 | 0 | 0 | 475020 | 0 |
| GLFLPEDEI | ASPH | Q12797 | ASPH_HUMAN | Unmodified | 0 | 0 | 0 | 0 | 0 | 0 | 0 | 0 | 1069400 | 550870 | 0 | 0 | 0 | 0 | 0 | 0 | 0 | 0 |
| GLFPASYV | SNX18 | Q96RF0 | SNX18_HUMAN | Unmodified | 0 | 0 | 4145800 | 4922400 | 428090 | 0 | 0 | 0 | 1067700 | 2112600 | 0 | 0 | 0 | 0 | 4056900 | 2698300 | 945490 | 0 |
| GLFRAAVP | ENO1; ENO1; E | P06733; K7E | ENOA_HUMAN | Unmodified | 0 | 0 | 0 | 0 | 80315000 | 5838200 | 6720100 | 0 | 0 | 10051000 | 0 | 0 | 27568000 | 0 | 0 | 0 | 35255000 | 78337000 |
| GLFTGLTPI | MTCH2; MTCH2 | Q9Y6C9; E9I | MTCH2_HUMA | Unmodified | 0 | 0 | 0 | 0 | 0 | 0 | 0 | 0 | 1770700 | 522370 | 0 | 41056 | 0 | 0 | 0 | 0 | 0 | 0 |
| GLGAGAG | PGRMC2 | O15173 | PGRC2_HUMAN | Unmodified | 0 | 0 | 0 | 0 | 0 | 0 | 0 | 0 | 0 | 0 | 0 | 0 | 0 | 0 | 0 | 0 | 0 | 0 |
| GLGDHDLI | PPM1H | Q9ULR3 | PPM1H_HUMA | Unmodified | 270480 | 0 | 0 | 0 | 286070 | 62713 | 95339 | 49668 | 76405 | 10450 | 15238 | 1418300 | 73462 | 0 | 179370 | 190570 | 132550 | 147270 |
| GLGDVDQI | LPL | P06858 | LIPL_HUMAN L | Unmodified | 0 | 0 | 0 | 0 | 0 | 0 | 28058 | 78159 | 1554500 | 122470 | 0 | 0 | 182120 | 0 | 0 | 0 | 0 | 0 |
| GLGEHEMI | LNPEP | Q9UIQ6 | LCAP_HUMAN | Unmodified | 0 | 0 | 420830 | 956500 | 442540 | 0 | 0 | 0 | 398960 | 0 | 385740 | 790000 | 284500 | 99003 | 381530 | 421020 | 0 | 0 |
| GLGEHEMI | LNPEP | Q9UIQ6 | LCAP_HUMAN | Oxidation (M) | 49167 | 0 | 255850 | 293750 | 141280 | 0 | 0 | 0 | 107270 | 0 | 127590 | 191230 | 0 | 69008 | 163280 | 167680 | 0 | 0 |
| GLGEISAAS | NCAM1; NCAM | P13591; A0A | NCAM1_HUMA | Unmodified | 0 | 0 | 170760 | 0 | 0 | 0 | 0 | 0 | 2780800 | 481740 | 0 | 0 | 0 | 0 | 0 | 0 | 0 | 0 |
| GLGFSIAG | SCRIB; SCRIB; | Q14160; A0 | SCRIB_HUMAN | Unmodified | 0 | 0 | 331360 | 0 | 0 | 0 | 0 | 67578 | 1528900 | 223250 | 0 | 1000500 | 401260 | 0 | 367350 | 276090 | 0 | 0 |
| GLGFSIAG | DLG2; DLG1; D | Q15700; Q1 | DLG2_HUMAN | Unmodified | 0 | 264750 | 150440 | 365380 | 6208300 | 1857600 | 110850 | 0 | 2409200 | 4572600 | 861930 | 252060 | 580470 | 552880 | 571780 | 444940 | 4212700 | 535200 |
| GLGGLIVV | ASRGL1 | Q7L266 | ASGL1_HUMAN | Unmodified | 0 | 0 | 0 | 0 | 0 | 0 | 99334 | 63827 | 2013200 | 1689900 | 104300 | 1724900 | 0 | 0 | 151060 | 0 | 107450 | 167340 |
| GLGHQVA | RTCB | Q9Y3I0 | RTCB_HUMAN | Unmodified | 0 | 0 | 0 | 310540 | 707970 | 0 | 0 | 0 | 797580 | 173630 | 402750 | 3043800 | 1234500 | 490060 | 0 | 0 | 0 | 406740 |
| GLGIKQDI | SEL1L | Q9UBV2 | SE1L1_HUMAN | Unmodified | 0 | 0 | 0 | 0 | 0 | 0 | 0 | 0 | 780670 | 44283 | 0 | 0 | 0 | 0 | 0 | 0 | 49191 | 0 |
| GLGKPGG | ARHGEF16 | Q5VV41 | ARHGG_HUMA | Unmodified | 124360 | 89505 | 0 | 0 | 0 | 0 | 0 | 0 | 704310 | 336990 | 0 | 0 | 847090 | 598380 | 238280 | 192160 | 0 | 0 |
| GLGLDDAL | RUVBL2; RUVBI | Q9Y230; A0 | RUVB2_HUMAN | Unmodified | 512640 | 0 | 0 | 0 | 0 | 0 | 0 | 0 | 0 | 0 | 0 | 5408300 | 0 | 0 | 0 | 0 | 0 | 0 |
| GLGLDESG | RUVBL1 | Q9Y265 | RUVB1_HUMAN | Unmodified | 430120 | 243630 | 389020 | 590820 | 4893100 | 820740 | 1752200 | 1376400 | 749430 | 565700 | 3714700 | 13112000 | 5206400 | 2784100 | 420330 | 398900 | 1807900 | 2314000 |
| GLGLGTEG | RNF31; RNF31 | Q96EP0; H0 | RNF31_HUMAN | Unmodified | 783010 | 805480 | 0 | 0 | 0 | 0 | 0 | 0 | 0 | 0 | 0 | 0 | 0 | 0 | 0 | 0 | 0 | 0 |
| GLGLSKAY | FLNA; FLNA; FL | P21333; A0A | FLNA_HUMAN | Unmodified | 0 | 0 | 0 | 0 | 0 | 0 | 0 | 0 | 78614 | 0 | 0 | 0 | 0 | 0 | 0 | 0 | 116910 | 28936 |
| GLGLSPDL | CTPS2 | Q9NRF8 | PYRG2_HUMAN | Unmodified | 0 | 0 | 0 | 0 | 0 | 0 | 0 | 0 | 0 | 0 | 0 | 1140400 | 0 | 0 | 0 | 0 | 0 | 0 |
| GLGLSPDL | CTPS1; CTPS1; | P17812; B4E | PYRG1_HUMAN | Unmodified | 0 | 0 | 0 | 0 | 0 | 0 | 0 | 0 | 693810 | 166540 | 699400 | 3186700 | 0 | 0 | 0 | 0 | 157250 | 0 |
| GLGMEVFF | CNOT1 | A5YKK6 | CNOT1_HUMA | Oxidation (M) | 0 | 0 | 0 | 0 | 0 | 0 | 0 | 0 | 0 | 0 | 0 | 276780 | 0 | 0 | 0 | 0 | 0 | 0 |
| GLGNQVE | DIP2B | Q9P265 | DIP2B_HUMAN | Unmodified | 298540 | 123070 | 296450 | 330840 | 0 | 0 | 0 | 0 | 0 | 246040 | 0 | 1413000 | 0 | 0 | 1095600 | 1128300 | 0 | 0 |
| GLGSTVQE | ISOC1; ISOC1 | Q96CN7; D6 | ISOC1_HUMAN | Unmodified | 0 | 0 | 0 | 0 | 840910 | 0 | 0 | 340350 | 1723000 | 839710 | 1789600 | 7979300 | 0 | 0 | 0 | 0 | 0 | 0 |
| GLGTDDN | ANXA4; ANXA4 | P09525; Q6 | ANXA4_HUMAN | Unmodified | 1573900 | 0 | 0 | 526440 | 894970 | 549260 | 0 | 0 | 1321100 | 0 | 8739900 | 18030000 | 500370 | 0 | 1152500 | 0 | 0 | 1003700 |
| GLGTDEDA | ANXA4; ANXA4; | P09525; B4E | ANXA4_HUMAN | Unmodified | 1183500 | 1012500 | 37057 | 1528100 | 3834500 | 1246100 | 0 | 0 | 53471000 | 7564500 | 10561000 | 3823900 | 2840400 | 0 | 2034800 | 187880 | 879030 | 1808500 |
| GLGTDEDA | ANXA4; ANXA4; | P09525; B4E | ANXA4_HUMAN | Unmodified | 0 | 0 | 0 | 0 | 0 | 1026200 | 0 | 0 | 0 | 3497400 | 3052600 | 34145 | 0 | 2965300 | 0 | 0 | 0 | 0 |
| GLGTDEDA | ANXA4; ANXA4; | P09525; B4E | ANXA4_HUMAN | Unmodified | 0 | 0 | 0 | 0 | 23808000 | 8993400 | 2287300 | 0 | 0 | 14476000 | 0 | 0 | 10884000 | 7035500 | 0 | 0 | 0 | 10162000 |
| GLGTDEDS | ANXA2P2; ANX | A6NMY6; P0 | AXA2L_HUMAN | Unmodified | 1459500 | 971600 | 0 | 934880 | 23294000 | 5769400 | 500870 | 53484 | 20911000 | 2439300 | 265790 | 0 | 333210 | 275930 | 649270 | 132100 | 4741000 | 7212300 |
| GLGTDEDS | ANXA2P2; ANX | A6NMY6; P0 | AXA2L_HUMAN | Unmodified | 0 | 0 | 0 | 0 | 11908000 | 0 | 49771 | 0 | 0 | 397070 | 0 | 0 | 0 | 0 | 0 | 0 | 4878200 | 6318800 |
| GLGTDEDT | ANXA6; ANXA6 | P08133; E5R | ANXA6_HUMAN | Unmodified | 4036900 | 1478600 | 9844300 | 17091000 | 16111000 | 4552600 | 6468100 | 124250 | 2.03E+08 | 31249000 | 260490 | 181360 | 11973000 | 5546100 | 20805000 | 6897600 | 4289800 | 25512000 |
| GLGTDEDT | ANXA6; ANXA6 | P08133; E5R | ANXA6_HUMAN | Unmodified | 0 | 0 | 0 | 647100 | 40998000 | 918490 | 8870300 | 66059 | 1726000 | 64839000 | 0 | 0 | 30254000 | 13461000 | 0 | 0 | 10159000 | 61643000 |
| GLGTDEDT | ANXA1; ANXA1 | P04083; Q5 | ANXA1_HUMAN | Unmodified | 0 | 0 | 0 | 0 | 1518800 | 595090 | 0 | 0 | 0 | 0 | 33287000 | 3.91E+08 | 191610 | 0 | 0 | 0 | 0 | 0 |
| GLGTDEDT | ANXA1; ANXA1 | P04083; Q5 | ANXA1_HUMAN | Unmodified | 0 | 0 | 0 | 0 | 1.68E+08 | 55555000 | 6257100 | 0 | 0 | 1739000 | 2.98E+08 | 25357000 | 0 | 96551 | 0 | 0 | 6054700 | 19021000 |
| GLGTDEDT | ANXA1; ANXA1 | P04083; Q5 | ANXA1_HUMAN | Unmodified | 0 | 0 | 0 | 0 | 23076000 | 4873300 | 0 | 0 | 0 | 0 | 25704000 | 1649800 | 0 | 0 | 0 | 0 | 2039800 | 3459400 |
| GLGTGTLY | CLNS1A; CLNS | P54105; E9P | ICLN_HUMAN I | Unmodified | 0 | 0 | 0 | 219940 | 481290 | 1918900 | 0 | 0 | 658500 | 85131 | 2951100 | 6544700 | 601020 | 0 | 0 | 0 | 0 | 0 |
| GLGTRDN | ANXA6 | P08133 | ANXA6_HUMAN | Unmodified | 93500 | 29885 | 30914 | 863790 | 991020 | 0 | 425630 | 413460 | 7997200 | 9495700 | 0 | 0 | 5949900 | 1856100 | 37545 | 43249 | 1239600 | 3271100 |
| GLGWVQF | SLIRP; SLIRP; S | Q9GZT3; G3 | SLIRP_HUMAN | Unmodified | 0 | 0 | 0 | 0 | 0 | 0 | 0 | 0 | 2310100 | 0 | 0 | 0 | 0 | 520090 | 0 | 0 | 0 | 0 |
| GLHGICMT | STC2; STC2 | O76061; H0 | STC2_HUMAN | Unmodified | 99797 | 0 | 0 | 0 | 0 | 0 | 0 | 0 | 790970 | 0 | 0 | 0 | 0 | 0 | 0 | 0 | 36040 | 0 |
| GLHLYGTQ | TSPAN4; TSPAN | O14817; A8 | TSN4_HUMAN | Unmodified | 0 | 924780 | 0 | 522200 | 0 | 0 | 0 | 0 | 0 | 0 | 1859000 | 0 | 0 | 0 | 5733000 | 0 | 0 | 822620 |
| GLHVVEVT | FLNB; FLNB | O75369; E7E | FLNB_HUMAN | Unmodified | 73933 | 78372 | 1613600 | 2825900 | 69923000 | 10899000 | 0 | 92155 | 1631900 | 1759800 | 1889300 | 4616100 | 0 | 0 | 5018000 | 1586200 | 13888000 | 11263000 |
| GLHVVEVT | FLNB; FLNB | O75369; E7E | FLNB_HUMAN | Unmodified | 0 | 0 | 0 | 0 | 3106100 | 0 | 0 | 0 | 0 | 0 | 0 | 0 | 0 | 0 | 0 | 0 | 7054800 | 0 |
| GLIAAICAG | PARK7; PARK7; | Q99497; K7E | PARK7_HUMAN | Unmodified | 708780 | 0 | 300780 | 1342100 | 6648300 | 808330 | 0 | 0 | 1991200 | 2778200 | 452190 | 486730 | 17166000 | 7040900 | 2053000 | 437850 | 2729700 | 5923500 |
| GLIAAICAG | PARK7; PARK7; | Q99497; K7E | PARK7_HUMAN | Unmodified | 0 | 0 | 0 | 0 | 7207000 | 0 | 1127000 | 0 | 0 | 3687900 | 0 | 0 | 26283000 | 20087000 | 0 | 0 | 9143500 | 11670000 |
| GLIAAICAG | PARK7; PARK7; | Q99497; K7E | PARK7_HUMAN | Unmodified | 0 | 0 | 0 | 0 | 0 | 0 | 619600 | 0 | 0 | 201980 | 0 | 0 | 658430 | 0 | 0 | 0 | 1470500 | 470210 |
| GLIDEVNQ | FGA; FGA | P02671; A0A | FIBA_HUMAN F | Unmodified | 0 | 949050 | 0 | 0 | 0 | 0 | 0 | 0 | 0 | 0 | 0 | 0 | 0 | 0 | 0 | 0 | 0 | 0 |
| GLIDEVNQ | FGA; FGA | P02671; A0A | FIBA_HUMAN F | Unmodified | 0 | 11545000 | 0 | 0 | 0 | 0 | 0 | 0 | 0 | 0 | 0 | 0 | 368670 | 4036800 | 0 | 0 | 440230 | 0 |
| GLIDGAES | PTPRJ; PTPRJ; P | Q12913; A0 | PTPRJ_HUMAN | Unmodified | 0 | 0 | 0 | 0 | 0 | 0 | 0 | 0 | 3399600 | 0 | 967490 | 2344300 | 0 | 0 | 0 | 0 | 0 | 0 |
| GLIENPALL | NDUFB4 | O95168 | NDUB4_HUMA | Unmodified | 0 | 0 | 0 | 0 | 0 | 0 | 0 | 0 | 2042500 | 202110 | 0 | 0 | 0 | 0 | 0 | 0 | 0 | 0 |
| GLIEPISMI | CORO2B | Q9UQ03 | COR2B_HUMA | Oxidation (M) | 0 | 0 | 0 | 0 | 0 | 0 | 0 | 0 | 0 | 0 | 487710 | 4387100 | 0 | 0 | 0 | 0 | 0 | 0 |
| GLIEVKTDE | ACAT2 | Q9BWD1 | THIC_HUMAN | Unmodified | 0 | 0 | 0 | 0 | 0 | 0 | 0 | 0 | 975460 | 189450 | 0 | 456410 | 218990 | 0 | 0 | 311920 | 63766 | 0 |
| GLIEVKTDE | ACAT2 | Q9BWD1 | THIC_HUMAN | Oxidation (M) | 0 | 0 | 0 | 0 | 2933700 | 0 | 821640 | 473540 | 0 | 2804800 | 0 | 0 | 1447100 | 0 | 0 | 0 | 386880 | 0 |
| GLIEVKTDE | ACAT2 | Q9BWD1 | THIC_HUMAN | Unmodified | 0 | 0 | 0 | 0 | 13083000 | 435640 | 1351600 | 0 | 599720 | 1749600 | 0 | 0 | 1480600 | 0 | 0 | 0 | 410280 | 905460 |
| GLIGNLYL | GLB1L2 | Q8IW92 | GLBL2_HUMAN | Unmodified | 0 | 0 | 0 | 0 | 0 | 0 | 0 | 0 | 2617700 | 0 | 0 | 0 | 0 | 0 | 0 | 0 | 0 | 0 |
| GLINAWTA | EDIL3 | O43854 | EDIL3_HUMAN | Unmodified | 0 | 0 | 0 | 0 | 94370000 | 4562100 | 8894500 | 0 | 843040 | 4211400 | 0 | 0 | 0 | 23613 | 0 | 0 | 4162200 | 30204000 |
| GLIPREQGI | PLEC; PLEC | Q15149; H0 | PLEC_HUMAN | Unmodified | 0 | 0 | 0 | 0 | 0 | 0 | 0 | 0 | 0 | 0 | 0 | 0 | 0 | 0 | 0 | 0 | 8604200 | 0 |
| GLIPVFALG | CPSF3; CPSF3 | Q9UKF6; G5 | CPSF3_HUMAN | Unmodified | 0 | 0 | 0 | 0 | 0 | 0 | 0 | 0 | 0 | 0 | 66475 | 662510 | 0 | 0 | 0 | 0 | 0 | 0 |
| GLIVYQLEN | RAE1; RAE1; RA | P78406; E9P | RAE1L_HUMAN | Unmodified | 178060 | 287710 | 0 | 1865200 | 0 | 0 | 109720 | 0 | 0 | 752090 | 0 | 1976100 | 0 | 0 | 0 | 0 | 0 | 0 |
| GLKEGIPAL | EEF2 | P13639 | EF2_HUMAN EI | Unmodified | 0 | 0 | 0 | 0 | 2908600 | 461510 | 4054300 | 0 | 39028 | 996780 | 0 | 0 | 2102300 | 1591400 | 0 | 0 | 5886200 | 1644600 |
| GLKLTFDS | VDAC1; VDAC1 | P21796; C9J | VDAC1_HUMAN | Unmodified | 0 | 0 | 0 | 0 | 0 | 0 | 0 | 23496 | 249780 | 975200 | 0 | 0 | 0 | 0 | 0 | 0 | 0 | 0 |
| GLKLTFDS | VDAC1; VDAC1 | P21796; C9J | VDAC1_HUMAN | Unmodified | 0 | 0 | 0 | 0 | 0 | 0 | 0 | 0 | 5207800 | 10658000 | 0 | 0 | 89257 | 0 | 0 | 0 | 0 | 0 |
| GLKPGVVY | FN1 | P02751 | FINC_HUMAN I | Unmodified | 0 | 0 | 716150 | 0 | 0 | 0 | 0 | 0 | 0 | 0 | 0 | 0 | 0 | 0 | 0 | 3586500 | 0 | 0 |
| GLKPSCTII | FASN; FASN | P49327; A0A | FAS_HUMAN F | Oxidation (M) | 0 | 0 | 0 | 0 | 0 | 0 | 0 | 0 | 0 | 0 | 0 | 2551900 | 0 | 0 | 0 | 0 | 0 | 0 |
| GLKYELISE | STRBP; ILF3 | Q96SI9; Q12 | STRBP_HUMAN | Unmodified | 3852600 | 911850 | 0 | 0 | 0 | 0 | 0 | 0 | 0 | 0 | 0 | 1177700 | 0 | 0 | 0 | 0 | 0 | 0 |
| GLLAQGLR | NIBAN2 | Q96TA1 | NIBA2_HUMAN | Unmodified | 1191000 | 0 | 0 | 0 | 0 | 0 | 0 | 0 | 0 | 0 | 0 | 0 | 0 | 0 | 1996100 | 2212400 | 0 | 0 |
| GLLATASEI | WDR6; WDR6; | Q9NNW5; E | WDR6_HUMAN | Unmodified | 349680 | 1108800 | 217680 | 0 | 170870 | 0 | 158850 | 0 | 0 | 44384 | 215700 | 2333400 | 0 | 0 | 0 | 784130 | 131340 | 0 |
| GLLCDPNF | MELTF | P08582 | TRFM_HUMAN | Unmodified | 0 | 0 | 0 | 0 | 0 | 0 | 0 | 0 | 2337100 | 0 | 150410 | 0 | 0 | 0 | 0 | 0 | 0 | 0 |
| GLLDVTCK | SKP1; SKP1; SK | P63208; E7E | SKP1_HUMAN | Unmodified | 0 | 0 | 0 | 420030 | 399190 | 409610 | 339810 | 0 | 0 | 370740 | 453520 | 3737300 | 0 | 1102100 | 0 | 617540 | 0 | 0 |
| GLLEFEHQ | GANAB; GANAE | Q14697; F5I | GANAB_HUMA | Unmodified | 0 | 0 | 0 | 0 | 255860 | 0 | 0 | 0 | 18026000 | 0 | 1045000 | 0 | 0 | 0 | 0 | 0 | 0 | 183100 |
| GLLEFEHQ | GANAB; GANAE | Q14697; F5I | GANAB_HUMA | Unmodified | 0 | 0 | 1349600 | 0 | 66878 | 0 | 0 | 0 | 1.53E+08 | 12203000 | 2479700 | 3778300 | 0 | 0 | 926760 | 655790 | 0 | 101770 |
| GLLENELK | LMCD1; LMCD1 | Q9NZU5; H7 | LMCD1_HUMA | Unmodified | 0 | 0 | 307410 | 166010 | 0 | 0 | 26329 | 27265 | 87460 | 0 | 697010 | 3629900 | 33201 | 0 | 185930 | 0 | 0 | 0 |

| | | | | | | | | | | | | | | | | | | | | | | |
|---|---|---|---|---|---|---|---|---|---|---|---|---|---|---|---|---|---|---|---|---|---|---|
| GLLFPVCTI | DKK3; DKK3 | Q9UBP4; F6 | DKK3_HUMAN | Unmodified | 0 | 0 | 0 | 0 | 0 | 0 | 718660 | 0 | 0 | 0 | 2237400 | 0 | 0 | 0 | 0 | 0 | 0 | 0 |
| GLLFVTNID | ITGA7; ITGA7; I | Q13683; J3K | ITA7_HUMAN I | Unmodified | 0 | 53926 | 0 | 0 | 1400100 | 0 | 3042600 | 233290 | 466720 | 1196700 | 0 | 1041400 | 642620 | 0 | 0 | 0 | 33826 | 644650 |
| GLLGALTS | CYRIB; CYRIB | Q9NUQ9; E5 | CYRIB_HUMAN | Unmodified | 0 | 0 | 0 | 0 | 152370 | 0 | 152830 | 0 | 0 | 0 | 0 | 2133800 | 0 | 0 | 79185 | 0 | 41046 | 0 |
| GLLGCNIIF | PSMD9; PSMD9 | O00233; J3K | PSMD9_HUMA | Unmodified | 0 | 0 | 0 | 0 | 1571900 | 0 | 637320 | 323520 | 0 | 0 | 295450 | 0 | 0 | 0 | 0 | 0 | 3932800 | 2198500 |
| GLLGMCV( | FKBP9 | O95302 | FKBP9_HUMAN | Oxidation (M) | 0 | 0 | 0 | 0 | 0 | 0 | 0 | 0 | 1406800 | 1152000 | 0 | 0 | 0 | 0 | 0 | 0 | 0 | 0 |
| GLLGMCV( | FKBP9 | O95302 | FKBP9_HUMAN | Unmodified | 0 | 0 | 0 | 0 | 0 | 0 | 24000 | 0 | 4222300 | 1271900 | 550970 | 0 | 0 | 0 | 0 | 0 | 0 | 0 |
| GLLHSFDL | EIF2A; EIF2A; EI | Q9BY44; H7 | EIF2A_HUMAN | Unmodified | 0 | 0 | 0 | 0 | 0 | 0 | 0 | 0 | 0 | 0 | 413200 | 1862900 | 0 | 0 | 0 | 246260 | 420520 | 355230 |
| GLLLDTSR | HEXA; HEXA; HI | P06865; H3E | HEXA_HUMAN | Unmodified | 105190 | 0 | 567860 | 0 | 72731 | 0 | 0 | 89133 | 1529600 | 106140 | 251690 | 84278 | 0 | 0 | 198730 | 0 | 0 | 0 |
| GLLLGDFK | CCER2 | I3L3R5 | CCER2_HUMAI | Unmodified | 0 | 0 | 0 | 0 | 510530 | 0 | 1003300 | 1237400 | 0 | 0 | 0 | 0 | 0 | 0 | 0 | 0 | 226620 | 416700 |
| GLLLSRENI | ILVBL; ILVBL | A1L0T0; M0F | HACL2_HUMAI | Unmodified | 52113 | 0 | 140680 | 297310 | 620040 | 80949 | 1338600 | 600900 | 263750 | 0 | 160320 | 310240 | 296930 | 322820 | 57186 | 116800 | 0 | 542460 |
| GLLLTDPVI | GOLGA7B; GOL | Q2TAP0; M0I | GOG7B_HUMA | Unmodified | 0 | 0 | 0 | 0 | 0 | 0 | 0 | 0 | 3437000 | 688870 | 1748200 | 257860 | 0 | 0 | 0 | 0 | 0 | 0 |
| GLLLTGHEI | LLGL1; LLGL1 | Q15334; A0/ | L2GL1_HUMAN | Unmodified | 0 | 0 | 0 | 0 | 0 | 0 | 0 | 0 | 0 | 0 | 0 | 0 | 0 | 0 | 0 | 0 | 0 | 0 |
| GLLLYGPP( | KATNAL2; KATN | Q8IYT4; K7E | KATL2_HUMAN | Unmodified | 1767900 | 1252400 | 0 | 2595600 | 1250700 | 0 | 0 | 0 | 11399000 | 573880 | 0 | 30434000 | 1677800 | 0 | 1685500 | 2329100 | 0 | 0 |
| GLLNAIVIK | OAT | P04181 | OAT_HUMAN C | Unmodified | 0 | 0 | 0 | 0 | 0 | 0 | 0 | 0 | 5248300 | 725580 | 0 | 354360 | 0 | 0 | 0 | 0 | 0 | 0 |
| GLLNAIVIKI | OAT | P04181 | OAT_HUMAN C | Unmodified | 0 | 235380 | 0 | 0 | 0 | 0 | 0 | 0 | 778090 | 1282000 | 0 | 0 | 0 | 0 | 178140 | 155500 | 0 | 0 |
| GLLNGASC | SF3B3 | Q15393 | SF3B3_HUMAN | Unmodified | 1348300 | 2495800 | 953030 | 475650 | 0 | 0 | 72147 | 57249 | 831680 | 0 | 66784 | 1020800 | 0 | 0 | 623950 | 3298500 | 0 | 0 |
| GLLPEELTF | ETFA; ETFA; ETF | P13804; H0Y | ETFA_HUMAN I | Unmodified | 0 | 0 | 0 | 0 | 376900 | 0 | 652230 | 0 | 3907800 | 1047000 | 0 | 0 | 421920 | 0 | 0 | 0 | 0 | 0 |
| GLLPGCVY | NOMO1; NOMC | Q15155; P69 | NOMO1_HUM/ | Unmodified | 0 | 0 | 0 | 0 | 0 | 0 | 0 | 0 | 5227100 | 259150 | 462560 | 0 | 0 | 0 | 0 | 89810 | 0 | 0 |
| GLLPQLLG | SLC25A13 | Q9UJS0 | CMC2_HUMAN | Unmodified | 0 | 0 | 0 | 0 | 0 | 0 | 0 | 0 | 2277700 | 541570 | 0 | 0 | 0 | 0 | 0 | 0 | 0 | 0 |
| GLLQLQWI | TOMM70 | O94826 | TOM70_HUMAI | Unmodified | 0 | 0 | 0 | 0 | 198950 | 0 | 433780 | 0 | 0 | 324400 | 0 | 0 | 816590 | 420260 | 0 | 0 | 169030 | 431790 |
| GLLQQIGD | MGAT1; MGAT1 | P26572; D6F | MGAT1_HUMAI | Unmodified | 0 | 0 | 0 | 0 | 0 | 0 | 0 | 0 | 13629000 | 435750 | 0 | 0 | 0 | 0 | 0 | 0 | 0 | 0 |
| GLLRQPQC | ITGA3; ITGA3; I | P26006; K7E | ITA3_HUMAN I | Unmodified | 0 | 0 | 0 | 0 | 2.02E+08 | 62727000 | 462400 | 542520 | 1243400 | 1744100 | 1401000 | 6870200 | 0 | 0 | 0 | 0 | 1.3E+08 | 1.48E+08 |
| GLLSDSMT | TPD52L2; TPD5 | O43399; A0/ | TPD54_HUMAN | Unmodified | 0 | 0 | 0 | 0 | 0 | 0 | 0 | 0 | 1570700 | 0 | 0 | 123950 | 0 | 0 | 1507500 | 934350 | 0 | 0 |
| GLLSIRYPN | ACTR1A; ACTR1 | P61163; A0A | ACTZ_HUMAN . | Unmodified | 0 | 0 | 0 | 0 | 102300 | 0 | 0 | 0 | 498490 | 0 | 0 | 0 | 0 | 0 | 0 | 0 | 0 | 76837 |
| GLLSLLNDI | CCDC28A; CCI | Q8IWP9; B4I | CC28A_HUMAI | Unmodified | 0 | 0 | 0 | 0 | 0 | 0 | 0 | 0 | 0 | 0 | 0 | 615830 | 0 | 0 | 0 | 0 | 0 | 0 |
| GLLSQGSP | GCLC; GCLC; G | P48506; A0A | GSH1_HUMAN | Unmodified | 32115 | 0 | 0 | 0 | 774630 | 179920 | 0 | 0 | 39146 | 0 | 0 | 0 | 0 | 63896 | 0 | 0 | 0 | 0 |
| GLLSQGSP | GCLC; GCLC; G | P48506; A0A | GSH1_HUMAN | Unmodified | 0 | 0 | 0 | 0 | 1497100 | 0 | 0 | 0 | 383690 | 0 | 87890 | 1690700 | 0 | 0 | 373460 | 0 | 0 | 0 |
| GLLSSLDH | CYC1 | P08574 | CY1_HUMAN C | Unmodified | 0 | 0 | 0 | 0 | 0 | 0 | 0 | 0 | 10972000 | 363120 | 0 | 0 | 0 | 0 | 0 | 0 | 0 | 0 |
| GLLSSLDH | CYC1 | P08574 | CY1_HUMAN C | Unmodified | 0 | 0 | 0 | 0 | 0 | 0 | 0 | 0 | 13819000 | 432930 | 0 | 0 | 141490 | 0 | 0 | 0 | 0 | 0 |
| GLMPDGTS | ADH5; ADH5 | P11766; H0Y | ADHX_HUMAN | Oxidation (M) | 0 | 0 | 72771 | 45613 | 15645 | 0 | 0 | 0 | 43560 | 0 | 183850 | 1764500 | 0 | 0 | 0 | 67438 | 41689 | 0 |
| GLMPDGTS | ADH5; ADH5 | P11766; H0Y | ADHX_HUMAN | Unmodified | 0 | 63276 | 0 | 0 | 402660 | 0 | 0 | 0 | 409950 | 41746 | 1292300 | 17292000 | 0 | 0 | 0 | 286350 | 308890 | 0 |
| GLMPDGTS | ADH5; ADH5 | P11766; H0Y | ADHX_HUMAN | Unmodified | 0 | 0 | 0 | 0 | 2728900 | 411610 | 2269900 | 475520 | 0 | 381620 | 259830 | 0 | 453130 | 944620 | 0 | 0 | 2764600 | 1037100 |
| GLMSYRIIT | MANSC1; MANS | Q9H8J5; F5H | MANS1_HUMA | Oxidation (M) | 508610 | 0 | 0 | 0 | 0 | 0 | 0 | 0 | 0 | 385060 | 0 | 0 | 0 | 0 | 0 | 0 | 0 | 0 |
| GLNSRLEA | ANLN; ANLN | Q9NQW6; C | ANLN_HUMAN | Unmodified | 0 | 0 | 0 | 222440 | 4146400 | 884960 | 0 | 0 | 0 | 0 | 0 | 0 | 389790 | 0 | 0 | 0 | 3614300 | 3362100 |
| GLNTDELG | SPART | Q8N0X7 | SPART_HUMAN | Unmodified | 0 | 0 | 131040 | 208190 | 0 | 0 | 0 | 0 | 0 | 0 | 145500 | 807890 | 0 | 0 | 0 | 234540 | 0 | 0 |
| GLNTPEDS | ADAMTS4; ADA | O75173; Q5 | ATS4_HUMAN / | Unmodified | 0 | 0 | 0 | 0 | 0 | 0 | 0 | 0 | 0 | 0 | 0 | 633660 | 0 | 0 | 0 | 0 | 0 | 0 |
| GLPAMDEC | SYT4 | Q9H2B2 | SYT4_HUMAN S | Oxidation (M) | 0 | 0 | 0 | 295090 | 0 | 0 | 0 | 0 | 0 | 0 | 0 | 531170 | 0 | 0 | 2955700 | 640540 | 0 | 0 |
| GLPAMDEC | SYT4 | Q9H2B2 | SYT4_HUMAN S | 2 Oxidation (M) | 0 | 0 | 0 | 144440 | 0 | 0 | 0 | 0 | 0 | 0 | 0 | 0 | 0 | 0 | 272570 | 909980 | 0 | 0 |
| GLPDNISS | UGP2; UGP2; U | Q16851; A0/ | UGPA_HUMAN | Unmodified | 0 | 0 | 0 | 0 | 0 | 0 | 0 | 0 | 0 | 0 | 0 | 1989100 | 0 | 0 | 0 | 0 | 0 | 0 |
| GLPDNISS | UGP2; UGP2; U | Q16851; A0/ | UGPA_HUMAN | Unmodified | 0 | 0 | 0 | 0 | 143780 | 0 | 181570 | 0 | 0 | 138260 | 0 | 0 | 0 | 0 | 0 | 0 | 248990 | 0 |
| GLPEPSVW | PTK7; PTK7; PTI | Q13308; Q8 | PTK7_HUMAN I | Unmodified | 0 | 0 | 0 | 296590 | 0 | 0 | 0 | 0 | 567510 | 0 | 0 | 0 | 0 | 0 | 227910 | 309480 | 0 | 0 |
| GLPFEAEN | RBM12 | Q9NTZ6 | RBM12_HUMAI | Unmodified | 0 | 0 | 0 | 0 | 0 | 0 | 0 | 0 | 0 | 227000 | 950900 | 3542600 | 0 | 0 | 0 | 0 | 0 | 0 |
| GLPGDISA/ | MMP15 | P51511 | MMP15_HUMA | Unmodified | 0 | 0 | 1034000 | 618160 | 0 | 0 | 0 | 0 | 0 | 0 | 0 | 0 | 0 | 0 | 1667400 | 730910 | 0 | 0 |
| GLPGDRGI | COL4A2 | P08572 | CO4A2_HUMAI | Unmodified | 0 | 0 | 0 | 0 | 0 | 0 | 0 | 0 | 1036700 | 0 | 0 | 0 | 0 | 0 | 0 | 0 | 0 | 0 |
| GLPLLVALL | ARVCF; ARVCF; | O00192; E9F | ARVC_HUMAN | Unmodified | 110670 | 124720 | 610610 | 696330 | 0 | 0 | 0 | 0 | 297880 | 0 | 0 | 1759600 | 0 | 0 | 1787800 | 1156000 | 0 | 0 |
| GLPLLVALL | ARVCF; ARVCF; | O00192; E9F | ARVC_HUMAN | Unmodified | 0 | 0 | 1549500 | 4379800 | 0 | 0 | 0 | 0 | 331710 | 0 | 145470 | 377170 | 0 | 0 | 2124500 | 8377400 | 0 | 0 |
| GLPLYPDP | N/A; AAK1; AAK | Q6ZSR9; A0/ | YJ005_HUMAN | Unmodified | 106230 | 47202 | 144940 | 258090 | 330370 | 0 | 73507 | 0 | 127490 | 73794 | 0 | 0 | 1242700 | 995950 | 175210 | 166410 | 158910 | 387290 |
| GLPNVQRS | IDH1 | O75874 | IDHC_HUMAN | Unmodified | 0 | 0 | 0 | 315040 | 6994900 | 360500 | 503180 | 0 | 1252400 | 1803600 | 166350 | 0 | 19913000 | 11150000 | 697330 | 0 | 2606900 | 4431200 |
| GLPNVQRS | IDH1 | O75874 | IDHC_HUMAN | Oxidation (M) | 0 | 1164000 | 0 | 0 | 3687200 | 0 | 2417300 | 0 | 0 | 834990 | 0 | 0 | 12134000 | 9672800 | 0 | 0 | 1284900 | 0 |
| GLPNVQRS | IDH1 | O75874 | IDHC_HUMAN | Unmodified | 0 | 0 | 0 | 0 | 6194900 | 0 | 3108200 | 0 | 0 | 3051900 | 0 | 0 | 16454000 | 0 | 0 | 0 | 0 | 0 |
| GLPNVQRS | IDH1 | O75874 | IDHC_HUMAN | Oxidation (M) | 0 | 0 | 0 | 0 | 0 | 0 | 0 | 0 | 0 | 0 | 0 | 0 | 5724700 | 2597200 | 0 | 0 | 0 | 0 |
| GLPPVDFV | MAN1A1 | P33908 | MA1A1_HUMAI | Unmodified | 103380 | 0 | 0 | 0 | 0 | 0 | 0 | 0 | 2195600 | 0 | 367980 | 2468900 | 0 | 0 | 0 | 0 | 0 | 0 |
| GLPPVDFV | MAN1A1 | P33908 | MA1A1_HUMAI | Unmodified | 0 | 0 | 0 | 0 | 0 | 0 | 0 | 0 | 4511900 | 243450 | 1329600 | 2697100 | 0 | 0 | 0 | 0 | 0 | 0 |
| GLPQTAVC | ANK3; ANK3; AI | Q12955; H0Y | ANK3_HUMAN | Unmodified | 0 | 0 | 1217600 | 2440900 | 0 | 0 | 0 | 0 | 0 | 0 | 0 | 1552000 | 3398600 | 4319300 | 3980700 | 1887100 | 0 | 0 |
| GLPTGDSP | PNPO; PNPO | Q9NVS9; J3( | PNPO_HUMAN | Oxidation (M) | 0 | 0 | 0 | 0 | 0 | 0 | 202920 | 0 | 0 | 0 | 0 | 0 | 7101700 | 1916200 | 0 | 0 | 2454400 | 0 |
| GLPTVRPC | GPC6 | Q9Y625 | GPC6_HUMAN | Unmodified | 633910 | 343920 | 0 | 185380 | 0 | 0 | 0 | 0 | 257660 | 0 | 0 | 0 | 0 | 0 | 0 | 26708 | 0 | 0 |
| GLPTVRPC | GPC6 | Q9Y625 | GPC6_HUMAN | Oxidation (M) | 1400300 | 450750 | 0 | 950100 | 0 | 0 | 0 | 0 | 0 | 0 | 0 | 0 | 0 | 0 | 0 | 282720 | 0 | 0 |
| GLPVGSPP | UTRN; UTRN | P46939; Q5 | UTRN_HUMAN | Unmodified | 0 | 0 | 1385100 | 0 | 1427200 | 0 | 0 | 0 | 2588600 | 0 | 0 | 890380 | 989670 | 761380 | 2125700 | 1599500 | 723380 | 2859700 |
| GLPVLVELL | ARVCF; ARVCF; | O00192; E9F | ARVC_HUMAN | Unmodified | 23397 | 40469 | 154130 | 140590 | 0 | 0 | 0 | 0 | 20674 | 0 | 19248 | 141160 | 0 | 0 | 1071200 | 693630 | 0 | 0 |
| GLQDVLRT | CCT6A; CCT6B | P40227; Q9 | TCPZ_HUMAN | Unmodified | 0 | 0 | 0 | 0 | 1097500 | 206010 | 188270 | 0 | 0 | 1607000 | 0 | 0 | 4920800 | 1304000 | 0 | 0 | 227650 | 1466200 |
| GLQDVLRT | CCT6A; CCT6B | P40227; Q9 | TCPZ_HUMAN | Oxidation (M) | 0 | 0 | 0 | 0 | 0 | 0 | 0 | 0 | 0 | 0 | 0 | 0 | 1284600 | 0 | 0 | 0 | 374510 | 0 |
| GLQDVLRT | CCT6A; CCT6B | P40227; Q9 | TCPZ_HUMAN | Unmodified | 0 | 0 | 0 | 216230 | 0 | 0 | 648240 | 0 | 0 | 417300 | 216250 | 0 | 1388800 | 0 | 0 | 0 | 189150 | 0 |
| GLQEAAEE | VGF | O15240 | VGF_HUMAN N | Unmodified | 0 | 1653700 | 40372 | 606770 | 0 | 19462000 | 1788800 | 0 | 438030 | 0 | 67546 | 797770 | 0 | 0 | 1763000 | 125700 | 0 | 372750 |
| GLQEGYEN | UBE2O; UBE2O | Q9C0C9; K7 | UBE2O_HUMAI | Unmodified | 0 | 0 | 0 | 0 | 0 | 0 | 0 | 0 | 0 | 0 | 42917 | 1209900 | 0 | 0 | 0 | 0 | 0 | 0 |
| GLQEIEESI | MITD1; MITD1; I | Q8WV92; H7 | MITD1_HUMAN | Unmodified | 0 | 0 | 0 | 0 | 0 | 0 | 0 | 0 | 0 | 0 | 0 | 3934600 | 0 | 0 | 0 | 0 | 0 | 0 |
| GLQEVGLP | UMPS | P11172 | UMPS_HUMAN | Unmodified | 107030 | 140530 | 0 | 187580 | 0 | 0 | 13345 | 0 | 687430 | 145330 | 295770 | 2979800 | 66354 | 363270 | 214990 | 144830 | 0 | 504470 |
| GLQKEEVV | GMPS | P49915 | GUAA_HUMAN | Unmodified | 0 | 0 | 0 | 0 | 0 | 0 | 127910 | 299510 | 288140 | 0 | 240170 | 660140 | 0 | 0 | 0 | 0 | 0 | 0 |
| GLQLSALAI | CNNM3 | Q8NE01 | CNNM3_HUM/ | Unmodified | 0 | 0 | 0 | 0 | 0 | 0 | 0 | 0 | 207450 | 187480 | 0 | 498110 | 0 | 0 | 0 | 0 | 0 | 0 |
| GLQNSASD | SRCIN1; SRCIN | Q9C0H9; A0 | SRCN1_HUMAI | Unmodified | 0 | 0 | 0 | 0 | 1761400 | 0 | 84785 | 0 | 0 | 0 | 0 | 714870 | 0 | 0 | 0 | 0 | 0 | 671670 |
| GLQPGEEE | USP13; USP13; | Q92995; A0/ | UBP13_HUMAN | Unmodified | 1613600 | 489380 | 0 | 580650 | 0 | 0 | 0 | 0 | 357600 | 0 | 0 | 214060 | 0 | 0 | 0 | 1317600 | 0 | 0 |
| GLQQQNSI | IQGAP1; IQGAP | P46940; A0A | IQGA1_HUMAN | Unmodified | 0 | 0 | 0 | 0 | 744850 | 340310 | 0 | 0 | 0 | 0 | 0 | 0 | 0 | 0 | 0 | 131100 | 0 | 0 |
| GLQRLEYR | GFPT1; GFPT1 | Q06210; A0/ | GFPT1_HUMAN | Unmodified | 0 | 0 | 0 | 0 | 241110 | 0 | 0 | 0 | 0 | 0 | 0 | 0 | 0 | 595020 | 0 | 0 | 0 | 0 |
| GLQRLLLIN | SPR | P35270 | SPRE_HUMAN | Unmodified | 0 | 0 | 0 | 0 | 0 | 0 | 382360 | 0 | 0 | 0 | 0 | 0 | 141100 | 0 | 0 | 0 | 0 | 0 |
| GLQSGVDI | CNN2; CNN2; ( | Q99439; H3I | CNN2_HUMAN | Unmodified | 0 | 0 | 0 | 116180 | 9102800 | 0 | 650620 | 711950 | 325830 | 61167 | 447220 | 155940 | 0 | 0 | 0 | 0 | 449970 | 241040 |
| GLQSGVDI | CNN2; CNN2; ( | Q99439; H3I | CNN2_HUMAN | Unmodified | 0 | 112560 | 0 | 280160 | 2427600 | 285660 | 0 | 0 | 0 | 0 | 0 | 0 | 204980 | 351550 | 0 | 0 | 270390 | 228610 |
| GLQSIFQE( | SMS; SMS | P52788; H7( | SPSY_HUMAN | Oxidation (M) | 0 | 0 | 0 | 0 | 2827000 | 0 | 0 | 0 | 0 | 0 | 0 | 3121300 | 10684000 | 0 | 0 | 0 | 1225000 | 3310200 |
| GLQSIFQE( | SMS; SMS | P52788; H7( | SPSY_HUMAN | Unmodified | 0 | 0 | 0 | 0 | 1672300 | 0 | 530720 | 0 | 1038600 | 0 | 0 | 931940 | 5198000 | 2696800 | 0 | 0 | 2636700 | 2118700 |
| GLQSLPTH | ERBB2; ERBB2 | P04626; B4D | ERBB2_HUMAN | Unmodified | 413320 | 513600 | 1199400 | 1370400 | 1051500 | 105780 | 0 | 0 | 0 | 0 | 0 | 0 | 0 | 301140 | 1790700 | 2054700 | 0 | 0 |
| GLQTSQD/ | CALR; CALR; C/ | P27797; A0A | CALR_HUMAN | Unmodified | 0 | 0 | 0 | 0 | 0 | 0 | 0 | 0 | 0 | 9226900 | 0 | 0 | 203860 | 0 | 0 | 0 | 0 | 0 |
| GLQVRPAP | MVD | P53602 | MVD1_HUMAN | Oxidation (M) | 0 | 4618300 | 0 | 0 | 0 | 0 | 0 | 0 | 0 | 0 | 0 | 2922700 | 0 | 0 | 0 | 0 | 0 | 0 |
| GLRPEVICI | DIP2A; DIP2A | Q14689; A0/ | DIP2A_HUMAN | Unmodified | 92655 | 0 | 0 | 391460 | 0 | 0 | 0 | 0 | 791860 | 0 | 0 | 496970 | 0 | 0 | 108150 | 80031 | 0 | 0 |
| GLSAEPGV | CHGA | P10645 | CMGA_HUMAN | Unmodified | 0 | 0 | 0 | 0 | 0 | 0 | 96798 | 162870 | 9680400 | 461880 | 0 | 0 | 0 | 0 | 0 | 0 | 232600 | 0 |
| GLSEDTTEI | NCL; NCL; NCL | P19338; C9J | NUCL_HUMAN | Unmodified | 0 | 0 | 246840 | 0 | 0 | 0 | 0 | 0 | 1744300 | 319540 | 6531500 | 11434000 | 294520 | 0 | 0 | 0 | 0 | 0 |
| GLSEDTTEI | NCL; NCL; NCL | P19338; C9J | NUCL_HUMAN | Unmodified | 204850 | 248170 | 0 | 177560 | 2422300 | 0 | 0 | 693700 | 2268600 | 0 | 1894600 | 2008000 | 2501000 | 583570 | 0 | 0 | 0 | 0 |
| GLSEDTTEI | NCL; NCL; NCL | P19338; C9J | NUCL_HUMAN | Unmodified | 0 | 0 | 0 | 0 | 3048600 | 578840 | 0 | 421820 | 0 | 1714200 | 0 | 0 | 1231600 | 1250100 | 0 | 0 | 1529600 | 550630 |
| GLSEDVSIS | EFTUD2; EFTUD | Q15029; K7E | U5S1_HUMAN | Unmodified | 0 | 160100 | 0 | 0 | 0 | 0 | 0 | 0 | 0 | 0 | 0 | 1313800 | 282500 | 0 | 0 | 0 | 0 | 0 |
| GLSFFAFEI | TCF25; TCF25; | Q9BQ70; H3 | TCF25_HUMAN | Unmodified | 0 | 1355100 | 337760 | 0 | 0 | 0 | 0 | 0 | 0 | 0 | 0 | 0 | 0 | 0 | 525440 | 458800 | 0 | 0 |
| GLSFLFPLL | EIF4G2; EIF4G2 | P78344; H0Y | IF4G2_HUMAN | Unmodified | 46170 | 0 | 0 | 0 | 83334 | 29469 | 40304 | 0 | 92299 | 120110 | 22832 | 248790 | 162980 | 96919 | 0 | 45581 | 67765 | 64131 |
| GLSFSEAT/ | ADD3 | Q9UEY8 | ADDG_HUMAN | Unmodified | 0 | 0 | 0 | 0 | 0 | 0 | 0 | 0 | 0 | 0 | 0 | 0 | 0 | 0 | 0 | 0 | 0 | 0 |
| GLSGTAVLI | RNPEP; RNPEP; | Q9H4A4; A6I | AMPB_HUMAN | Unmodified | 0 | 0 | 0 | 0 | 0 | 0 | 0 | 0 | 458290 | 0 | 502790 | 2003200 | 0 | 0 | 0 | 0 | 0 | 0 |
| GLSHSPW/ | PBK; PBK | Q96KB5; E5I | TOPK_HUMAN | Unmodified | 0 | 0 | 800810 | 0 | 1268400 | 0 | 181410 | 150260 | 891450 | 510710 | 620110 | 0 | 0 | 194670 | 0 | 0 | 572840 | 78679 |
| GLSHVAM( | TTC13 | Q8NBP0 | TTC13_HUMAN | Unmodified | 0 | 0 | 0 | 0 | 0 | 0 | 0 | 0 | 0 | 0 | 0 | 0 | 0 | 0 | 0 | 0 | 0 | 0 |
| GLSKIERAI | ATP2A2; ATP2A | P16615; H7( | AT2A2_HUMAN | Unmodified | 0 | 0 | 0 | 0 | 224300 | 0 | 525810 | 0 | 0 | 0 | 0 | 0 | 0 | 132020 | 0 | 0 | 0 | 0 |
| GLSLGGSD | OGFOD3 | Q6PK18 | OGFD3_HUMA | Unmodified | 0 | 0 | 103550 | 0 | 324320 | 149420 | 0 | 0 | 2612300 | 798480 | 0 | 0 | 0 | 0 | 0 | 0 | 413560 | 1271800 |
| GLSLNAAR | SLC12A7; SLC1 | Q9Y666; A0/ | S12A7_HUMAN | Unmodified | 81904 | 100280 | 0 | 192730 | 0 | 30134 | 81663 | 0 | 0 | 54444 | 0 | 51569 | 0 | 0 | 600050 | 514810 | 0 | 0 |
| GLSLQHIG | ERI3; ERI3 | O43414; H0Y | ERI3_HUMAN E | Unmodified | 1949400 | 255380 | 816520 | 1407900 | 297160 | 0 | 0 | 88638 | 0 | 0 | 0 | 0 | 1927300 | 1331600 | 181150 | 1853100 | 0 | 420190 |
| GLSLVDKEI | RRM2; RRM2 | P31350; A0A | RIR2_HUMAN F | Unmodified | 0 | 0 | 0 | 0 | 839900 | 0 | 0 | 166440 | 0 | 0 | 0 | 0 | 863530 | 0 | 0 | 0 | 0 | 1185600 |
| GLSMDREL | GPRASP2 | Q96D09 | GASP2_HUMAI | Oxidation (M) | 0 | 0 | 0 | 0 | 0 | 0 | 0 | 0 | 0 | 0 | 0 | 0 | 0 | 0 | 394330 | 576040 | 0 | 0 |
| GLSNHFQ\ | TOMM40; TOMI | O96008; K7E | TOM40_HUMAI | Unmodified | 0 | 0 | 0 | 0 | 0 | 0 | 0 | 0 | 2547500 | 1073200 | 0 | 0 | 0 | 0 | 0 | 0 | 0 | 0 |
| GLSPAQAD | EPB41L2; EPB4 | O43491; A0/ | E41L2_HUMAN | Unmodified | 0 | 0 | 0 | 0 | 809980 | 0 | 0 | 0 | 0 | 0 | 0 | 1095900 | 0 | 0 | 696840 | 0 | 0 | 224560 |
| GLSPAQAD | EPB41L2; EPB4 | O43491; A0/ | E41L2_HUMAN | Unmodified | 0 | 0 | 0 | 0 | 0 | 0 | 0 | 0 | 0 | 0 | 0 | 0 | 0 | 0 | 0 | 0 | 570500 | 0 |
| GLSQSALP | RPS13; RPS13; | P62277; J3K | RS13_HUMAN . | Unmodified | 165860 | 175290 | 215660 | 243140 | 0 | 0 | 0 | 0 | 0 | 0 | 0 | 478710 | 0 | 0 | 78582 | 482700 | 0 | 0 |
| GLSQSALP | RPS13; RPS13; | P62277; J3K | RS13_HUMAN . | Unmodified | 3420300 | 2607400 | 2129600 | 1917200 | 611700 | 104110 | 208430 | 411600 | 662590 | 241500 | 312310 | 3825100 | 2497600 | 552370 | 1578200 | 1682000 | 152010 | 0 |
| GLSRDMQ( | MVB12A; MVB1 | Q96EY5; M0 | MB12A_HUMAI | Unmodified | 0 | 0 | 0 | 928790 | 1360200 | 0 | 0 | 71860 | 0 | 305740 | 0 | 0 | 4677900 | 1369400 | 0 | 0 | 0 | 2269500 |
| GLSRDMQ( | MVB12A; MVB1 | Q96EY5; M0 | MB12A_HUMAI | Unmodified | 0 | 0 | 0 | 0 | 0 | 0 | 0 | 0 | 0 | 479880 | 0 | 0 | 11642000 | 0 | 0 | 0 | 0 | 347890 |
| GLSRDMQ( | MVB12A; MVB1 | Q96EY5; M0 | MB12A_HUMAI | Oxidation (M) | 0 | 0 | 0 | 0 | 0 | 0 | 0 | 0 | 0 | 0 | 0 | 0 | 4304100 | 487640 | 0 | 0 | 0 | 0 |
| GLSRQPAL | APEH; APEH; AI | P13798; C9J | ACPH_HUMAN | Unmodified | 0 | 0 | 0 | 1070300 | 0 | 0 | 0 | 0 | 0 | 0 | 915290 | 0 | 0 | 0 | 0 | 129910 | 0 | 0 |
| GLSSDLEG | AKAP12 | Q02952 | AKA12_HUMAN | Unmodified | 0 | 0 | 0 | 0 | 1410500 | 0 | 0 | 13698 | 0 | 0 | 150120 | 0 | 0 | 0 | 0 | 0 | 98613 | 2238100 |
| GLSSDLEG | AKAP12 | Q02952 | AKA12_HUMAN | Unmodified | 0 | 0 | 0 | 0 | 0 | 0 | 81561 | 41363 | 0 | 0 | 0 | 424670 | 0 | 0 | 0 | 0 | 2257700 | 0 |

| | | | | | | | | | | | | | | | | | | | | | | |
|---|---|---|---|---|---|---|---|---|---|---|---|---|---|---|---|---|---|---|---|---|---|---|
| GLSSILDAA | ARHGEF1; ARH | Q92888; M0 | ARHG1_HUMA | Unmodified | 10591000 | 0 | 3570400 | 0 | 444920 | 0 | 0 | 0 | 0 | 942940 | 0 | 3591500 | 0 | 0 | 11468000 | 8753700 | 0 | 0 |
| GLSTEDATS | ACBD7 | Q8N6N7 | ACBD7_HUMAI | Unmodified | 0 | 0 | 0 | 0 | 0 | 0 | 0 | 291240 | 0 | 0 | 0 | 441530 | 0 | 0 | 0 | 0 | 0 | 0 |
| GLSVREILA | PDHA1; PDHA1 | P08559; Q5J | ODPA_HUMAN | Unmodified | 0 | 0 | 0 | 0 | 0 | 0 | 0 | 0 | 202990 | 219250 | 0 | 0 | 0 | 0 | 0 | 0 | 0 | 0 |
| GLTASNQF | FARP1; FARP1; | Q9Y4F1; C9J | FARP1_HUMAN | Unmodified | 0 | 0 | 0 | 0 | 0 | 0 | 0 | 0 | 639430 | 0 | 0 | 2031600 | 0 | 0 | 0 | 0 | 0 | 0 |
| GLTAVSNN | OXCT1 | P55809 | SCOT1_HUMAI | Unmodified | 0 | 0 | 0 | 0 | 0 | 0 | 0 | 0 | 627500 | 839790 | 0 | 0 | 0 | 0 | 0 | 0 | 0 | 0 |
| GLTAVSNN | OXCT1 | P55809 | SCOT1_HUMAI | Unmodified | 0 | 0 | 0 | 0 | 0 | 0 | 1886300 | 0 | 0 | 1235100 | 0 | 0 | 0 | 0 | 0 | 0 | 0 | 0 |
| GLTAYHDIS | ITM2C; ITM2C; I | Q9NQX7; E7 | ITM2C_HUMAN | Unmodified | 0 | 0 | 89534 | 0 | 0 | 0 | 0 | 103420 | 2772400 | 0 | 107580 | 0 | 0 | 0 | 0 | 0 | 0 | 0 |
| GLTDLSAC | NOP56; NOP56 | O00567; Q5J | NOP56_HUMAI | Unmodified | 0 | 0 | 0 | 0 | 1077800 | 0 | 10189 | 162940 | 107380 | 0 | 142130 | 0 | 175020 | 0 | 0 | 0 | 646590 | 1034000 |
| GLTEQEVE | CUL3 | Q13618 | CUL3_HUMAN | Unmodified | 0 | 0 | 97994 | 0 | 1664700 | 0 | 0 | 0 | 0 | 246720 | 827620 | 3156700 | 0 | 0 | 0 | 229690 | 0 | 0 |
| GLTETGLYF | RACGAP1; RAC | Q9H0H5; H0 | RGAP1_HUMAI | Unmodified | 0 | 0 | 750280 | 0 | 0 | 484190 | 0 | 178630 | 513620 | 0 | 859680 | 4868700 | 0 | 0 | 698070 | 0 | 0 | 0 |
| GLTETGLYF | RACGAP1; RAC | Q9H0H5; H0 | RGAP1_HUMAI | Unmodified | 0 | 0 | 0 | 0 | 1457500 | 336140 | 0 | 0 | 0 | 317100 | 0 | 0 | 0 | 0 | 0 | 0 | 0 | 0 |
| GLTEVTQSI | TAX1BP1; TAX1 | Q86VP1; B8 | TAXB1_HUMAN | Unmodified | 0 | 0 | 0 | 0 | 285590 | 0 | 0 | 67861 | 596180 | 0 | 0 | 0 | 0 | 165080 | 0 | 0 | 0 | 297930 |
| GLTEVTQSI | TAX1BP1; TAX1 | Q86VP1; B8 | TAXB1_HUMAN | Unmodified | 0 | 0 | 0 | 0 | 567700 | 0 | 853540 | 0 | 859600 | 0 | 0 | 0 | 233740 | 0 | 0 | 0 | 319820 | 1533500 |
| GLTGQAEA | YOD1 | Q5VVQ6 | OTU1_HUMAN | Unmodified | 0 | 0 | 0 | 0 | 442860 | 63388 | 119600 | 0 | 0 | 0 | 0 | 0 | 2229000 | 1625200 | 0 | 0 | 1141600 | 346210 |
| GLTIVGSVL | SLC12A7; SLC1 | Q9Y666; A0 | S12A7_HUMAN | Unmodified | 0 | 0 | 0 | 0 | 0 | 0 | 181490 | 0 | 2391100 | 611440 | 0 | 420890 | 131950 | 64890 | 232820 | 0 | 0 | 0 |
| GLTLNCLG | NAA16; NAA15; | Q6N069; Q9 | NAA16_HUMAN | Unmodified | 0 | 0 | 0 | 0 | 0 | 0 | 0 | 0 | 0 | 0 | 0 | 0 | 0 | 0 | 0 | 0 | 0 | 0 |
| GLTMLDHE | SAE1; SAE1; SA | Q9UBE0; B3 | SAE1_HUMAN | Oxidation (M) | 0 | 0 | 0 | 0 | 375270 | 0 | 0 | 0 | 324180 | 310080 | 292690 | 1250800 | 0 | 0 | 0 | 0 | 0 | 0 |
| GLTPSQIG | RPS13; RPS13 | P62277; J3K | RS13_HUMAN | Unmodified | 0 | 0 | 0 | 0 | 0 | 0 | 0 | 0 | 659130 | 0 | 1834300 | 13404000 | 0 | 0 | 0 | 0 | 0 | 0 |
| GLTPSQIG | RPS13; RPS13 | P62277; J3K | RS13_HUMAN | Unmodified | 298120 | 537100 | 724240 | 1422500 | 0 | 0 | 390590 | 0 | 0 | 0 | 0 | 0 | 1803600 | 342250 | 0 | 0 | 0 | 0 |
| GLTPTGML | PPP3CA; PPP3C | Q08209; E7E | PP2BA_HUMAN | Oxidation (M) | 0 | 0 | 0 | 0 | 0 | 0 | 0 | 0 | 0 | 0 | 545170 | 1608200 | 0 | 0 | 0 | 0 | 0 | 0 |
| GLTSARAAI | ATP1A1; ATP1A | P05023; Q5T | AT1A1_HUMAN | Unmodified | 0 | 0 | 0 | 0 | 230060 | 114630 | 0 | 0 | 0 | 0 | 0 | 171010 | 471650 | 0 | 0 | 0 | 128420 | 160710 |
| GLTSVINQ | LDHB; LDHB | P07195; A0A | LDHB_HUMAN | Unmodified | 1311300 | 1177300 | 3047700 | 2326000 | 17376000 | 0 | 0 | 253660 | 480310 | 0 | 15283000 | 1.61E+08 | 7455300 | 3437500 | 2339300 | 1917700 | 5535400 | 6512400 |
| GLTSVINQ | LDHB; LDHB | P07195; A0A | LDHB_HUMAN | Unmodified | 0 | 0 | 0 | 0 | 952360 | 0 | 0 | 0 | 0 | 0 | 0 | 0 | 646650 | 625780 | 0 | 0 | 2155300 | 343030 |
| GLTTLTPVS | SLC6A8 | P48029 | SC6A8_HUMAN | Unmodified | 0 | 0 | 857720 | 892530 | 0 | 0 | 0 | 0 | 0 | 0 | 533770 | 5272400 | 0 | 0 | 0 | 1149100 | 0 | 0 |
| GLTTVFSGI | ESPN; ESPN; ES | B1AK53; A0 | ESPN_HUMAN | Unmodified | 0 | 0 | 0 | 0 | 0 | 0 | 0 | 0 | 0 | 0 | 0 | 583910 | 0 | 0 | 0 | 0 | 0 | 0 |
| GLTVPIASI | NAXE | Q8NCW5 | NNRE_HUMAN | Unmodified | 263570 | 0 | 57040 | 99314 | 172750 | 71823 | 15390 | 0 | 384800 | 412750 | 225120 | 511020 | 221920 | 202560 | 328870 | 178330 | 428480 | 0 |
| GLTVQQWI | PGM3; PGM3; F | O95394; J3K | AGM1_HUMAN | Unmodified | 0 | 0 | 0 | 0 | 1416900 | 2336000 | 0 | 0 | 0 | 0 | 235260 | 0 | 3208000 | 4132000 | 0 | 0 | 23290 | 0 |
| GLTWDAA | MARVELD3; MA | Q96A59; J3K | MALD3_HUMAI | Unmodified | 4295600 | 0 | 0 | 0 | 0 | 0 | 0 | 0 | 0 | 0 | 1389000 | 0 | 0 | 0 | 0 | 872790 | 0 | 0 |
| GLTWDAA | MARVELD3; MA | Q96A59; J3K | MALD3_HUMAI | Unmodified | 1526000 | 0 | 2438700 | 0 | 0 | 0 | 0 | 0 | 0 | 0 | 2554200 | 3127700 | 0 | 0 | 2781200 | 2036900 | 0 | 0 |
| GLVASLISV | SMC2 | O95347 | SMC2_HUMAN | Unmodified | 0 | 0 | 0 | 0 | 0 | 0 | 328250 | 51929 | 291730 | 181320 | 0 | 855920 | 337460 | 0 | 0 | 0 | 0 | 0 |
| GLVAVITG | HSD17B10; HS | Q99714; Q5I | HCD2_HUMAN | Unmodified | 0 | 0 | 0 | 0 | 0 | 0 | 0 | 0 | 3583500 | 560130 | 0 | 0 | 0 | 0 | 0 | 0 | 0 | 0 |
| GLVDGGE | CGN | Q9P2M7 | CING_HUMAN | Unmodified | 0 | 0 | 0 | 0 | 0 | 0 | 0 | 0 | 0 | 0 | 0 | 453870 | 0 | 0 | 0 | 0 | 0 | 0 |
| GLVDKFNC | CRELD2; CRELI | Q6UXH1; A6 | CREL2_HUMAN | Unmodified | 0 | 0 | 0 | 0 | 0 | 0 | 0 | 0 | 0 | 193730 | 0 | 748710 | 0 | 0 | 0 | 0 | 0 | 0 |
| GLVDKLQA | CIAPIN1; CIAPII | Q6FI81; H3B | CPIN1_HUMAN | Unmodified | 0 | 0 | 0 | 0 | 0 | 0 | 0 | 0 | 0 | 0 | 0 | 700520 | 0 | 0 | 0 | 0 | 117370 | 0 |
| GLVDREPP | ADAM17; ADAM | P78536; A6H | ADA17_HUMAN | Unmodified | 0 | 0 | 0 | 0 | 0 | 0 | 0 | 0 | 626210 | 0 | 0 | 0 | 0 | 0 | 0 | 0 | 0 | 601620 |
| GLVDSFNK | CRELD1; CRELI | Q96HD1; A0 | CREL1_HUMAN | Unmodified | 0 | 0 | 0 | 0 | 0 | 0 | 0 | 0 | 4076500 | 0 | 5169400 | 0 | 0 | 0 | 0 | 0 | 0 | 0 |
| GLVEAVES | TRIM14 | Q14142 | TRI14_HUMAN | Unmodified | 0 | 0 | 28368 | 267190 | 231320 | 0 | 0 | 0 | 0 | 0 | 0 | 0 | 0 | 0 | 215470 | 0 | 0 | 0 |
| GLVEDSGS | GPC2; GPC2 | Q8N158; A0 | GPC2_HUMAN | Unmodified | 0 | 0 | 0 | 0 | 0 | 0 | 166210 | 0 | 2202100 | 0 | 0 | 193980 | 0 | 0 | 45469 | 0 | 0 | 0 |
| GLVEPVDV | FLNA; FLNA | P21333; Q60 | FLNA_HUMAN | Unmodified | 0 | 0 | 0 | 0 | 0 | 0 | 910300 | 383450 | 590210 | 0 | 0 | 1242400 | 0 | 0 | 0 | 0 | 0 | 0 |
| GLVGEIIK | NME1; NME2; N | P15531; P22 | NDKA_HUMAN | Unmodified | 52708 | 49414 | 0 | 0 | 15891000 | 3937900 | 57664 | 140710 | 5303900 | 0 | 30418000 | 51790000 | 6588900 | 232700 | 3382900 | 0 | 2937300 | 10210000 |
| GLVGEIIKR | NME1; NME2; N | P15531; P22 | NDKA_HUMAN | Unmodified | 0 | 37336 | 0 | 0 | 0 | 0 | 0 | 0 | 0 | 0 | 407280 | 1454500 | 0 | 0 | 0 | 0 | 0 | 0 |
| GLVGEIIKR | NME1; NME2; N | P15531; P22 | NDKA_HUMAN | Unmodified | 0 | 0 | 0 | 0 | 6385200 | 492420 | 3576300 | 0 | 0 | 1156900 | 0 | 0 | 7683400 | 5124700 | 0 | 0 | 7185800 | 5159100 |
| GLVLDHGA | CCT6A; CCT6B; | P40227; Q92 | TCPZ_HUMAN | Unmodified | 26882 | 0 | 0 | 0 | 0 | 0 | 0 | 0 | 0 | 0 | 0 | 1570900 | 557310 | 0 | 379640 | 0 | 0 | 0 |
| GLVLDHGA | CCT6A; CCT6B; | P40227; Q92 | TCPZ_HUMAN | Unmodified | 0 | 0 | 223170 | 0 | 0 | 0 | 119370 | 0 | 719160 | 0 | 368920 | 5159400 | 2046500 | 448330 | 209510 | 0 | 0 | 0 |
| GLVLGPIHI | SLC3A2; SLC3A | P08195; A0A | 4F2_HUMAN 4I | Unmodified | 0 | 0 | 0 | 3342300 | 16155000 | 2874700 | 862780 | 261100 | 0 | 1672200 | 0 | 47098000 | 1317100 | 764040 | 6198600 | 3349700 | 478560 | 2805700 |
| GLVNLQSC | AP3M1 | Q9Y2T2 | AP3M1_HUMAI | Unmodified | 0 | 316850 | 0 | 0 | 3989200 | 0 | 122630 | 486810 | 0 | 594940 | 145180 | 4332400 | 1115300 | 295880 | 0 | 0 | 0 | 0 |
| GLVQALGL | AKR1A1 | P14550 | AK1A1_HUMAN | Unmodified | 0 | 0 | 806330 | 65996 | 291430 | 0 | 0 | 0 | 3426600 | 370990 | 340470 | 5454900 | 264070 | 0 | 108290 | 71435 | 107470 | 153840 |
| GLVQALQT | FASN; FASN | P49327; A0A | FAS_HUMAN F | Unmodified | 2985500 | 2580100 | 6413000 | 4107100 | 9346600 | 1651900 | 5613500 | 1601700 | 3628900 | 1819700 | 9887600 | 1.18E+08 | 13380000 | 7339400 | 6250000 | 5462900 | 4134200 | 4463600 |
| GLVQALQT | FASN; FASN | P49327; A0A | FAS_HUMAN F | Unmodified | 0 | 0 | 0 | 0 | 861600 | 0 | 4923900 | 283000 | 0 | 721930 | 0 | 0 | 1498000 | 2023700 | 0 | 0 | 3455200 | 1291800 |
| GLVRAEEL | ITGA7; ITGA7; I | Q13683; J3K | ITA7_HUMAN I | Unmodified | 0 | 0 | 0 | 0 | 0 | 0 | 296220 | 0 | 0 | 538980 | 0 | 0 | 0 | 0 | 0 | 0 | 0 | 0 |
| GLVRAEEL | ITGA7; ITGA7; I | Q13683; J3K | ITA7_HUMAN I | Unmodified | 0 | 0 | 0 | 0 | 0 | 0 | 1282200 | 0 | 0 | 0 | 0 | 0 | 0 | 0 | 0 | 0 | 0 | 0 |
| GLVRLGEK | TRAF2; TRAF2; T | Q12933; B1 | TRAF2_HUMAN | Unmodified | 0 | 0 | 0 | 0 | 2774400 | 0 | 20871 | 0 | 0 | 68826 | 20313 | 57984 | 0 | 0 | 0 | 0 | 0 | 115270 |
| GLVRQEEA | PLK1 | P53350 | PLK1_HUMAN | Unmodified | 48773 | 35925 | 0 | 90261 | 0 | 0 | 0 | 0 | 476370 | 57390 | 0 | 206250 | 0 | 0 | 69643 | 36381 | 0 | 0 |
| GLVTVEDE | PLCB4; PLCB4 | Q15147; A0 | PLCB4_HUMAN | Oxidation (M) | 0 | 0 | 0 | 0 | 107600 | 0 | 0 | 0 | 0 | 0 | 0 | 0 | 0 | 0 | 271720 | 0 | 655730 | 0 |
| GLVTVKPC | GPC4 | O75487 | GPC4_HUMAN | Oxidation (M) | 664300 | 586060 | 0 | 45945 | 0 | 0 | 0 | 0 | 0 | 0 | 0 | 151920 | 0 | 0 | 0 | 74751 | 0 | 0 |
| GLVVDMD | AARS1; AARS1; | P49588; A0A | SYAC_HUMAN | Unmodified | 0 | 0 | 0 | 0 | 1567200 | 380230 | 0 | 0 | 2950900 | 464910 | 566800 | 3122300 | 0 | 2176800 | 0 | 0 | 591450 | 1025700 |
| GLVVDMD | AARS1; AARS1; | P49588; A0A | SYAC_HUMAN | Oxidation (M) | 0 | 0 | 650040 | 0 | 0 | 133330 | 30373 | 0 | 0 | 0 | 0 | 1532700 | 390520 | 0 | 0 | 92701 | 0 | 158600 |
| GLVVDMD | AARS1; AARS1; | P49588; A0A | SYAC_HUMAN | Oxidation (M) | 0 | 0 | 0 | 0 | 1617000 | 0 | 0 | 37928 | 0 | 801590 | 0 | 0 | 3807500 | 0 | 0 | 0 | 1156500 | 902810 |
| GLVVDMD | AARS1; AARS1; | P49588; A0A | SYAC_HUMAN | Unmodified | 0 | 0 | 0 | 0 | 5210800 | 0 | 524320 | 0 | 0 | 0 | 0 | 0 | 2781600 | 3079100 | 0 | 0 | 982230 | 0 |
| GLVVLGFP | GPX1 | P07203 | GPX1_HUMAN | Unmodified | 0 | 0 | 0 | 0 | 0 | 0 | 0 | 0 | 643280 | 201220 | 541720 | 1190400 | 0 | 362190 | 0 | 0 | 0 | 0 |
| GLVVLGFP | GPX1 | P07203 | GPX1_HUMAN | Unmodified | 0 | 0 | 0 | 0 | 0 | 0 | 0 | 0 | 2056800 | 777580 | 0 | 1014000 | 0 | 277680 | 0 | 0 | 2104400 | 1254100 |
| GLVVLTPEF | ADAM15 | Q13444 | ADA15_HUMAN | Unmodified | 0 | 0 | 0 | 0 | 0 | 0 | 0 | 0 | 457240 | 0 | 0 | 0 | 0 | 0 | 0 | 0 | 0 | 0 |
| GLVVPVIR | DLST | P36957 | ODO2_HUMAN | Unmodified | 0 | 0 | 0 | 0 | 0 | 0 | 0 | 0 | 8024800 | 732870 | 106830 | 292180 | 0 | 0 | 404500 | 0 | 0 | 0 |
| GLVVSGSK | WDR33; WDR3 | Q9C0J8; C9J | WDR33_HUMA | Unmodified | 0 | 0 | 0 | 0 | 0 | 0 | 97679 | 0 | 0 | 0 | 0 | 0 | 0 | 0 | 0 | 0 | 0 | 0 |
| GLVVTDLK | CHID1; CHID1; | Q9BWS9; E9 | CHID1_HUMAN | Unmodified | 38346 | 0 | 0 | 0 | 0 | 0 | 1452500 | 988660 | 6040400 | 254700 | 0 | 0 | 0 | 338620 | 0 | 0 | 258310 | 0 |
| GLVVTDLK | CHID1; CHID1; | Q9BWS9; E9 | CHID1_HUMAN | Unmodified | 0 | 0 | 0 | 0 | 0 | 0 | 0 | 0 | 24196000 | 473910 | 1562500 | 456110 | 0 | 0 | 0 | 0 | 0 | 265350 |
| GLVYETSVI | CS; CS; CS; CS | O75390; F8V | CISY_HUMAN ( | Unmodified | 0 | 0 | 0 | 0 | 0 | 0 | 0 | 0 | 3153500 | 0 | 0 | 0 | 0 | 0 | 0 | 0 | 0 | 0 |
| GLVYETSVI | CS; CS; CS; CS | O75390; F8V | CISY_HUMAN ( | Unmodified | 0 | 0 | 144840 | 0 | 0 | 0 | 0 | 0 | 64274000 | 1790200 | 0 | 425390 | 0 | 0 | 0 | 161400 | 0 | 0 |
| GLVYETSVI | CS; CS; CS; CS | O75390; F8V | CISY_HUMAN ( | Unmodified | 0 | 0 | 0 | 0 | 705000 | 0 | 0 | 0 | 0 | 10972000 | 0 | 0 | 0 | 1991900 | 0 | 0 | 121120 | 2638600 |
| GLWIPEGE | EGFR; EGFR; E( | P00533; Q50 | EGFR_HUMAN | Unmodified | 0 | 0 | 0 | 0 | 1745700 | 303380 | 0 | 0 | 0 | 0 | 0 | 0 | 0 | 0 | 0 | 0 | 0 | 763640 |
| GLWVPQH | MICU2 | Q8IYU8 | MICU2_HUMAN | Unmodified | 0 | 0 | 0 | 0 | 0 | 0 | 0 | 0 | 612910 | 286180 | 0 | 0 | 0 | 0 | 0 | 0 | 0 | 0 |
| GLYGIKDD | LDHA | P00338 | LDHA_HUMAN | Unmodified | 0 | 0 | 0 | 0 | 0 | 0 | 0 | 0 | 0 | 0 | 0 | 0 | 0 | 0 | 0 | 0 | 0 | 107940 |
| GLYNEHTV | ALKBH5 | Q6P6C2 | ALKB5_HUMAN | Unmodified | 0 | 0 | 0 | 0 | 0 | 0 | 0 | 0 | 0 | 0 | 89377 | 766840 | 0 | 0 | 0 | 0 | 0 | 0 |
| GMAAAGN | RTCB | Q9Y3I0 | RTCB_HUMAN | Oxidation (M) | 0 | 0 | 0 | 0 | 0 | 0 | 0 | 0 | 0 | 0 | 297430 | 896740 | 0 | 0 | 0 | 0 | 0 | 0 |
| GMAAAGN | RTCB | Q9Y3I0 | RTCB_HUMAN | Unmodified | 0 | 0 | 0 | 99438 | 0 | 0 | 0 | 0 | 0 | 0 | 268470 | 2318600 | 0 | 0 | 67801 | 0 | 0 | 0 |
| GMAFLASK | KIT; CSF1R | P10721; P07 | KIT_HUMAN M | Unmodified | 0 | 480070 | 0 | 0 | 1197600 | 0 | 888540 | 0 | 1814800 | 1412700 | 323200 | 1117000 | 0 | 0 | 3834700 | 0 | 784480 | 0 |
| GMAFLHTL | ILK; ILK | Q13418; A0 | ILK_HUMAN Int | Oxidation (M) | 0 | 0 | 764900 | 662000 | 15643 | 0 | 0 | 0 | 337760 | 0 | 0 | 493290 | 0 | 0 | 0 | 0 | 0 | 0 |
| GMAFTLQE | ME2; ME2; ME2 | P23368; A0A | MAOM_HUMAN | Unmodified | 0 | 0 | 0 | 0 | 0 | 0 | 0 | 0 | 1100900 | 0 | 0 | 0 | 0 | 0 | 0 | 0 | 0 | 0 |
| GMDEQLTS | UTRN | P46939 | UTRN_HUMAN | Oxidation (M) | 0 | 0 | 0 | 387310 | 746950 | 0 | 0 | 0 | 0 | 68953 | 0 | 161020 | 415770 | 67979 | 484140 | 401980 | 715680 | 0 |
| GMDEQLTS | UTRN | P46939 | UTRN_HUMAN | Unmodified | 0 | 11009 | 255560 | 344290 | 1717400 | 0 | 0 | 0 | 456410 | 332710 | 0 | 403890 | 479310 | 644950 | 534290 | 0 | 733970 | 805230 |
| GMDLQPG | LRRC47 | Q8N1G4 | LRC47_HUMAN | Unmodified | 0 | 0 | 0 | 0 | 0 | 0 | 0 | 0 | 373690 | 0 | 0 | 575710 | 238010 | 0 | 0 | 0 | 0 | 0 |
| GMFYNSYF | PREP; PREP | P48147; A0A | PPCE_HUMAN | Unmodified | 0 | 0 | 0 | 0 | 0 | 0 | 0 | 0 | 0 | 0 | 1082600 | 1116800 | 174140 | 0 | 0 | 0 | 0 | 0 |
| GMGGAMD | OXCT1; OXCT1 | P55809; E9P | SCOT1_HUMAI | Oxidation (M) | 0 | 0 | 0 | 0 | 0 | 0 | 0 | 0 | 633650 | 0 | 0 | 0 | 0 | 0 | 0 | 0 | 0 | 0 |
| GMGGAMD | OXCT1; OXCT1 | P55809; E9P | SCOT1_HUMAI | Unmodified | 0 | 95015 | 0 | 0 | 0 | 0 | 332830 | 0 | 2080600 | 359710 | 0 | 0 | 0 | 0 | 0 | 0 | 0 | 0 |
| GMGLSLMI | FASN; FASN | P49327; A0A | FAS_HUMAN F | Unmodified | 0 | 0 | 0 | 0 | 0 | 0 | 0 | 0 | 0 | 0 | 0 | 5146400 | 0 | 0 | 0 | 0 | 0 | 0 |
| GMGLVNH | PPAT; PPAT | Q06203; D6I | PUR1_HUMAN | Oxidation (M) | 0 | 41739 | 0 | 0 | 0 | 0 | 0 | 14577 | 111140 | 0 | 269210 | 1527900 | 278850 | 218010 | 0 | 0 | 121910 | 2826800 |
| GMITVTDPI | UBA6; UBA6 | A0AVT1; H0Y | UBA6_HUMAN | Oxidation (M) | 0 | 0 | 0 | 0 | 0 | 0 | 0 | 0 | 0 | 0 | 414140 | 1657600 | 46373 | 0 | 0 | 130500 | 0 | 2180400 |
| GMLHEEKF | MIA3; MIA3 | Q5JRA6; A0A | TGO1_HUMAN | Unmodified | 0 | 0 | 0 | 0 | 0 | 0 | 190680 | 0 | 8223600 | 1595700 | 1142100 | 0 | 0 | 270420 | 0 | 0 | 0 | 116060 |
| GMLHEEKF | MIA3; MIA3 | Q5JRA6; A0A | TGO1_HUMAN | Oxidation (M) | 0 | 0 | 0 | 0 | 0 | 0 | 0 | 0 | 2116900 | 757890 | 0 | 0 | 0 | 0 | 0 | 0 | 0 | 0 |
| GMLREDAV | MSN | P26038 | MOES_HUMAN | Oxidation (M) | 935170 | 1214700 | 1682500 | 3378200 | 1896900 | 891700 | 0 | 25917 | 0 | 0 | 0 | 0 | 458860 | 192390 | 3040400 | 2207800 | 804210 | 303670 |
| GMLREDSN | RDX; RDX; RDX | P35241; A0A | RADI_HUMAN I | Oxidation (M) | 31472 | 0 | 0 | 0 | 0 | 0 | 0 | 0 | 0 | 0 | 0 | 1435100 | 0 | 0 | 165220 | 122230 | 0 | 0 |
| GMLTSTTEI | CDS2 | O95674 | CDS2_HUMAN | Unmodified | 0 | 0 | 0 | 0 | 0 | 0 | 0 | 0 | 1228800 | 117880 | 0 | 0 | 0 | 0 | 0 | 0 | 0 | 0 |
| GMNMYLTI | NPEPPSL1; NPI | A6NEC2; P5 | PSAL_HUMAN I | Oxidation (M) | 0 | 0 | 0 | 0 | 996200 | 0 | 0 | 0 | 244210 | 75174 | 0 | 976260 | 0 | 0 | 0 | 0 | 772320 | 76584 |
| GMNMYLTI | NPEPPSL1; NPI | A6NEC2; P5 | PSAL_HUMAN I | Unmodified | 0 | 0 | 0 | 0 | 0 | 120570 | 0 | 31169 | 289410 | 0 | 170160 | 1809800 | 0 | 206520 | 0 | 104600 | 889050 | 1169800 |
| GMSISRPN | B4GALT1 | P15291 | B4GT1_HUMAN | Unmodified | 0 | 0 | 0 | 0 | 0 | 0 | 0 | 0 | 735490 | 0 | 0 | 0 | 0 | 0 | 0 | 0 | 0 | 0 |
| GMSLNLEP | ATP5F1A; ATP5 | P25705; K7E | ATPA_HUMAN | Oxidation (M) | 0 | 0 | 0 | 0 | 0 | 0 | 0 | 0 | 0 | 934390 | 0 | 0 | 0 | 0 | 0 | 0 | 0 | 0 |
| GMSLNLEP | ATP5F1A; ATP5 | P25705; K7E | ATPA_HUMAN | Oxidation (M) | 0 | 0 | 0 | 143530 | 0 | 0 | 0 | 0 | 0 | 1230800 | 0 | 0 | 252140 | 0 | 0 | 0 | 0 | 0 |
| GMSVSDL | IMMT; IMMT; IM | Q16891; B9 | MIC60_HUMAN | Oxidation (M) | 0 | 0 | 0 | 0 | 0 | 0 | 0 | 0 | 412390 | 0 | 0 | 0 | 0 | 0 | 0 | 0 | 0 | 0 |
| GMTELEPS | SLC39A10; SLC | Q9ULF5; E7 | S39AA_HUMAN | Oxidation (M) | 0 | 0 | 0 | 0 | 541710 | 0 | 154740 | 150180 | 191790 | 0 | 0 | 0 | 0 | 248380 | 0 | 0 | 506190 | 0 |
| GMTLVTPL | DHX9 | Q08211 | DHX9_HUMAN | Oxidation (M) | 118740 | 0 | 138020 | 0 | 0 | 0 | 714530 | 0 | 44702 | 0 | 256240 | 2536900 | 0 | 0 | 0 | 0 | 0 | 0 |
| GMTPGEAE | EPB41L1; EPB4 | Q9H4G0; Q4 | E41L1_HUMAN | Oxidation (M) | 0 | 0 | 431220 | 0 | 0 | 0 | 0 | 0 | 0 | 0 | 391430 | 1990500 | 0 | 0 | 12539000 | 3604000 | 0 | 0 |
| GMTPGEAE | EPB41L1; EPB4 | Q9H4G0; Q4 | E41L1_HUMAN | Oxidation (M) | 0 | 0 | 0 | 0 | 0 | 0 | 0 | 0 | 0 | 0 | 144960 | 800550 | 0 | 0 | 285810 | 193000 | 0 | 0 |
| GMTTVDDF | DPYSL3 | Q14195 | DPYL3_HUMAN | Oxidation (M) | 0 | 0 | 0 | 0 | 0 | 0 | 1034100 | 859090 | 0 | 112590 | 5796200 | 34321000 | 0 | 0 | 0 | 0 | 0 | 0 |
| GMTTVDDF | DPYSL3 | Q14195 | DPYL3_HUMAN | Unmodified | 0 | 0 | 0 | 0 | 0 | 0 | 762840 | 74168 | 0 | 0 | 1992000 | 10404000 | 0 | 0 | 0 | 0 | 0 | 0 |
| GMTYDALH | ABHD12; ABHD | Q8N2K0; A0 | ABD12_HUMAN | Oxidation (M) | 28310 | 17123 | 0 | 33495 | 0 | 0 | 0 | 0 | 8547000 | 2293500 | 144600 | 312840 | 0 | 0 | 0 | 0 | 0 | 23535 |
| GMTYDALH | ABHD12; ABHD | Q8N2K0; A0 | ABD12_HUMAN | Unmodified | 0 | 0 | 0 | 0 | 0 | 0 | 0 | 0 | 1846000 | 120830 | 0 | 0 | 0 | 0 | 0 | 0 | 0 | 0 |
| GMVFGIPD | HSPG2 | P98160 | PGBM_HUMAN | Oxidation (M) | 0 | 0 | 69498 | 0 | 0 | 29644 | 0 | 0 | 0 | 0 | 0 | 0 | 0 | 0 | 0 | 0 | 0 | 0 |
| GMVLLCDF | CNTN1 | Q12860 | CNTN1_HUMA | Unmodified | 0 | 0 | 0 | 42990 | 0 | 0 | 0 | 0 | 297630 | 0 | 0 | 0 | 0 | 0 | 0 | 0 | 0 | 0 |
| GMVLLCDF | CNTN1 | Q12860 | CNTN1_HUMA | Oxidation (M) | 374240 | 269540 | 0 | 0 | 0 | 0 | 0 | 0 | 0 | 0 | 0 | 0 | 0 | 0 | 0 | 0 | 0 | 0 |

| | | | | | | | | | | | | | | | | | | | | | | |
|---|---|---|---|---|---|---|---|---|---|---|---|---|---|---|---|---|---|---|---|---|---|---|
| GMYDGPVI | DPYSL3; DPYSL | Q14195; H0Y | DPYL3_HUMAN | Oxidation (M) | 0 | 0 | 0 | 0 | 0 | 0 | 0 | 0 | 0 | 0 | 4029700 | 6738500 | 0 | 0 | 132140 | 0 | 0 | 0 |
| GMYDVGLI | UAP1 | Q16222 | UAP1_HUMAN | Oxidation (M) | 221500 | 87330 | 0 | 0 | 870260 | 0 | 0 | 24413 | 0 | 0 | 0 | 797520 | 594450 | 0 | 0 | 0 | 83834 | 185600 |
| GNAAWQE | EPRS1; EPRS1; | P07814; V9G | SYEP_HUMAN | Unmodified | 0 | 0 | 0 | 0 | 608900 | 0 | 0 | 0 | 254250 | 131440 | 335470 | 4603000 | 0 | 0 | 0 | 211100 | 0 | 0 |
| GNAEGSSI | HDGF | P51858 | HDGF_HUMAN | Unmodified | 0 | 0 | 0 | 0 | 0 | 0 | 0 | 0 | 0 | 0 | 658570 | 0 | 0 | 0 | 0 | 0 | 0 | 0 |
| GNAEPDPF | AVIL | O75366 | AVIL_HUMAN A | Unmodified | 0 | 282340 | 0 | 0 | 0 | 0 | 0 | 0 | 0 | 0 | 330890 | 6048500 | 0 | 0 | 0 | 0 | 0 | 0 |
| GNAFGSC( | ADAM15 | Q13444 | ADA15_HUMAN | Unmodified | 883900 | 0 | 0 | 0 | 0 | 0 | 0 | 0 | 84543 | 0 | 0 | 1218000 | 0 | 0 | 281980 | 0 | 0 | 0 |
| GNAFGSC( | ADAM15 | Q13444 | ADA15_HUMAN | Unmodified | 2008100 | 455100 | 516760 | 846310 | 0 | 0 | 0 | 0 | 1514500 | 0 | 0 | 2655200 | 51991 | 0 | 708250 | 552290 | 33092 | 0 |
| GNAGGIQF | NAXE | Q8NCW5 | NNRE_HUMAN | Unmodified | 0 | 0 | 472030 | 0 | 493630 | 0 | 19680 | 64585 | 2295100 | 973250 | 1400300 | 2287700 | 423900 | 139750 | 0 | 1073300 | 308620 | 299760 |
| GNAGGIQF | NAXE | Q8NCW5 | NNRE_HUMAN | Unmodified | 803910 | 0 | 587880 | 785340 | 1611600 | 0 | 563710 | 215280 | 200420 | 2487400 | 2997800 | 5467100 | 2591200 | 1153800 | 0 | 1053000 | 1965700 | 2256900 |
| GNAGQSN | FASN; FASN | P49327; A0A | FAS_HUMAN F | Unmodified | 0 | 0 | 325230 | 509990 | 0 | 0 | 0 | 0 | 332380 | 0 | 372440 | 14528000 | 0 | 0 | 533720 | 416460 | 0 | 0 |
| GNAGQSN | FASN; FASN | P49327; A0A | FAS_HUMAN F | Oxidation (M) | 0 | 0 | 0 | 0 | 0 | 0 | 0 | 0 | 0 | 0 | 0 | 5201200 | 0 | 0 | 1136800 | 512030 | 0 | 0 |
| GNAIYNLLI | NCAPD2 | Q15021 | CND1_HUMAN | Unmodified | 0 | 0 | 0 | 0 | 0 | 0 | 0 | 0 | 0 | 0 | 0 | 79003 | 0 | 0 | 0 | 0 | 0 | 0 |
| GNAMVEE( | SPTAN1; SPTAN | Q13813; A0A | SPTN1_HUMAN | Oxidation (M) | 0 | 0 | 0 | 0 | 0 | 0 | 0 | 0 | 264100 | 198470 | 587030 | 1373300 | 0 | 0 | 0 | 0 | 0 | 0 |
| GNAMVEE( | SPTAN1; SPTAN | Q13813; A0A | SPTN1_HUMAN | Unmodified | 0 | 0 | 0 | 0 | 0 | 0 | 0 | 0 | 0 | 0 | 0 | 2524800 | 0 | 0 | 0 | 0 | 0 | 0 |
| GNANDIGN | PRSS23 | O95084 | PRS23_HUMAN | Unmodified | 0 | 0 | 0 | 0 | 1724100 | 0 | 0 | 0 | 0 | 0 | 0 | 0 | 0 | 0 | 0 | 0 | 0 | 0 |
| GNDIDPEA | MET; MET | P08581; H7C | MET_HUMAN H | Unmodified | 0 | 0 | 0 | 0 | 2078600 | 0 | 0 | 0 | 315790 | 0 | 0 | 0 | 0 | 0 | 0 | 0 | 0 | 0 |
| GNDIIAAAK | VCL | P18206 | VINC_HUMAN ' | Unmodified | 0 | 0 | 636960 | 0 | 1324400 | 0 | 713260 | 913710 | 637190 | 569440 | 768440 | 3771600 | 902970 | 575770 | 327510 | 0 | 3035500 | 1166100 |
| GNDIIAAAK | VCL | P18206 | VINC_HUMAN ' | Unmodified | 54268 | 67175 | 0 | 130070 | 76874 | 0 | 0 | 0 | 0 | 0 | 115950 | 1183700 | 0 | 0 | 160340 | 159220 | 32737 | 84386 |
| GNDISSGT | PEBP1 | P30086 | PEBP1_HUMAN | Unmodified | 2953100 | 3123000 | 2175800 | 2475300 | 4666500 | 858550 | 0 | 60261 | 15606000 | 7540000 | 15790000 | 42057000 | 7348000 | 6031700 | 5659300 | 13434000 | 2335400 | 7032400 |
| GNDISSGT | PEBP1 | P30086 | PEBP1_HUMAN | Unmodified | 911630 | 0 | 0 | 0 | 2990200 | 0 | 0 | 0 | 0 | 1535000 | 0 | 0 | 8859500 | 2469700 | 0 | 0 | 1360500 | 2827800 |
| GNDQVRFI | ASS1; ASS1 | P00966; Q5T | ASSY_HUMAN . | Unmodified | 751500 | 0 | 0 | 0 | 0 | 0 | 240070 | 43873 | 0 | 0 | 0 | 0 | 238480 | 0 | 65525 | 0 | 0 | 0 |
| GNDTPLAL | HUWE1 | Q7Z6Z7 | HUWE1_HUMA | Unmodified | 0 | 0 | 0 | 0 | 0 | 0 | 0 | 0 | 0 | 0 | 0 | 818560 | 0 | 0 | 0 | 0 | 0 | 0 |
| GNEAVASR | GOLM1 | Q8NBJ4 | GOLM1_HUMA | Unmodified | 0 | 0 | 0 | 0 | 0 | 0 | 0 | 0 | 770960 | 0 | 0 | 0 | 0 | 0 | 0 | 0 | 0 | 0 |
| GNEAVASR | GOLM1 | Q8NBJ4 | GOLM1_HUMA | Unmodified | 0 | 0 | 0 | 0 | 0 | 0 | 0 | 0 | 1815500 | 217440 | 0 | 0 | 0 | 0 | 0 | 0 | 0 | 0 |
| GNECFQK( | STIP1 | P31948 | STIP1_HUMAN | Unmodified | 0 | 0 | 0 | 0 | 0 | 0 | 0 | 0 | 352610 | 0 | 270700 | 959790 | 1093600 | 0 | 0 | 0 | 0 | 0 |
| GNECFQK( | STIP1 | P31948 | STIP1_HUMAN | Oxidation (M) | 0 | 0 | 0 | 0 | 0 | 0 | 0 | 0 | 0 | 0 | 62780 | 409720 | 203200 | 99615 | 0 | 0 | 0 | 0 |
| GNEEAQIFI | EFTUD2 | Q15029 | U5S1_HUMAN | Unmodified | 4732000 | 3317600 | 642600 | 1232000 | 180710 | 0 | 0 | 30672 | 121050 | 0 | 0 | 2452000 | 2903100 | 536690 | 430630 | 807610 | 67921 | 71922 |
| GNEENPHI | B4GALT3 | O60512 | B4GT3_HUMAN | Unmodified | 0 | 0 | 0 | 0 | 0 | 0 | 0 | 0 | 2444900 | 0 | 0 | 0 | 0 | 0 | 0 | 0 | 0 | 0 |
| GNEFEDYC | DDX6; DDX6 | P26196; Q8I | DDX6_HUMAN | Unmodified | 0 | 0 | 0 | 0 | 0 | 0 | 0 | 0 | 0 | 0 | 0 | 1011300 | 0 | 0 | 165280 | 0 | 0 | 0 |
| GNEFEDYC | DDX6; DDX6 | P26196; Q8I | DDX6_HUMAN | Unmodified | 0 | 116060 | 157260 | 202020 | 0 | 0 | 0 | 0 | 252130 | 0 | 183810 | 1405700 | 0 | 0 | 158890 | 179240 | 0 | 0 |
| GNFGGSF/ | HNRNPM; HNR | P52272; M0I | HNRPM_HUMA | Unmodified | 0 | 0 | 355160 | 0 | 0 | 0 | 0 | 322920 | 195910 | 0 | 0 | 0 | 1210300 | 582880 | 0 | 0 | 0 | 0 |
| GNFGGSF/ | HNRNPM; HNR | P52272; M0I | HNRPM_HUMA | Unmodified | 28997 | 0 | 0 | 0 | 0 | 375020 | 1874500 | 1770200 | 314810 | 0 | 857570 | 1103700 | 200820 | 365590 | 77578 | 0 | 0 | 0 |
| GNFIPYANI | ABCA2; ABCA2; | Q9BZC7; A0. | ABCA2_HUMAI | Unmodified | 639150 | 0 | 0 | 0 | 0 | 114140 | 0 | 0 | 976380 | 0 | 90510 | 0 | 0 | 0 | 0 | 0 | 0 | 0 |
| GNFNYIEFI | MYL12A; MYL12 | P19105; O14 | ML12A_HUMAI | Unmodified | 0 | 0 | 0 | 0 | 2518300 | 705880 | 0 | 0 | 819790 | 254200 | 542620 | 2605800 | 971760 | 0 | 852160 | 597510 | 342840 | 502340 |
| GNFNYIEFI | MYL12A; MYL12 | P19105; O14 | ML12A_HUMAI | Unmodified | 0 | 0 | 0 | 0 | 5302700 | 463660 | 499640 | 0 | 0 | 0 | 0 | 0 | 984420 | 736600 | 0 | 0 | 2425700 | 1130400 |
| GNFSAFM( | GFPT2 | O94808 | GFPT2_HUMAN | Oxidation (M) | 0 | 0 | 0 | 0 | 0 | 0 | 0 | 0 | 0 | 0 | 0 | 0 | 0 | 0 | 0 | 0 | 0 | 812820 |
| GNFSPLELI | ARHGAP29; ARI | Q52LW3; F8' | RHG29_HUMAI | Unmodified | 0 | 0 | 0 | 0 | 433970 | 0 | 0 | 0 | 0 | 90252 | 0 | 0 | 471010 | 0 | 0 | 134940 | 740300 | 1118600 |
| GNFSSFM( | GFPT1; GFPT1; | Q06210; A0A | GFPT1_HUMAN | Oxidation (M) | 0 | 0 | 31015 | 0 | 175930 | 0 | 84301 | 0 | 0 | 0 | 0 | 163440 | 335830 | 0 | 46031 | 45443 | 73619 | 0 |
| GNGYFQVI | WDR48 | Q8TAF3 | WDR48_HUMA | Unmodified | 0 | 0 | 0 | 0 | 0 | 0 | 0 | 169710 | 0 | 0 | 102990 | 536420 | 0 | 0 | 209530 | 54310 | 0 | 0 |
| GNHETDNI | PPP5C; PPP5C; | P53041; A8M | PPP5_HUMAN . | Oxidation (M) | 0 | 30157 | 331830 | 0 | 0 | 0 | 0 | 0 | 0 | 0 | 0 | 1553500 | 0 | 0 | 0 | 69643 | 0 | 0 |
| GNHETDNI | PPP5C; PPP5C; | P53041; A8M | PPP5_HUMAN . | Unmodified | 0 | 0 | 0 | 0 | 0 | 0 | 0 | 0 | 310060 | 0 | 0 | 2236800 | 0 | 0 | 0 | 0 | 0 | 0 |
| GNIDNNTP | EDIL3 | O43854 | EDIL3_HUMAN | Unmodified | 1015500 | 415970 | 0 | 0 | 7315300 | 0 | 0 | 0 | 33560000 | 0 | 0 | 0 | 0 | 113080 | 0 | 0 | 0 | 2005800 |
| GNIELASSI | LIMCH1; LIMCH | Q9UPQ0; H0 | LIMC1_HUMAN | Unmodified | 0 | 0 | 0 | 303100 | 0 | 0 | 0 | 0 | 0 | 0 | 221200 | 651880 | 0 | 0 | 389180 | 215290 | 0 | 0 |
| GNIFLTSY( | PLXNB2 | O15031 | PLXB2_HUMAN | Unmodified | 139180 | 57610 | 43973 | 164270 | 0 | 0 | 0 | 0 | 0 | 0 | 0 | 0 | 100190 | 0 | 988520 | 880510 | 0 | 0 |
| GNIGTGLL( | SLC36A1; SLC3 | Q7Z2H8; E7I | S36A1_HUMAN | Unmodified | 0 | 0 | 0 | 0 | 0 | 0 | 0 | 0 | 1021000 | 56320 | 0 | 0 | 0 | 0 | 0 | 0 | 0 | 0 |
| GNIHLKPSI | ITGB4 | P16144 | ITB4_HUMAN I | Unmodified | 0 | 0 | 0 | 0 | 279440 | 375280 | 28643 | 0 | 0 | 0 | 0 | 150100 | 0 | 0 | 0 | 0 | 224270 | 573780 |
| GNIIISTPEK | SNRNP200 | O75643 | U520_HUMAN | Unmodified | 364680 | 185570 | 0 | 0 | 0 | 0 | 0 | 144750 | 0 | 146050 | 0 | 1150200 | 0 | 0 | 51578 | 91859 | 0 | 0 |
| GNIIISTPEK | SNRNP200 | O75643 | U520_HUMAN | Unmodified | 125870 | 0 | 0 | 0 | 0 | 0 | 0 | 0 | 62031 | 0 | 0 | 2396400 | 0 | 0 | 18539 | 0 | 0 | 0 |
| GNILIPGIN | CNDP2 | Q96KP4 | CNDP2_HUMA | Unmodified | 0 | 0 | 217580 | 0 | 0 | 0 | 115330 | 0 | 412590 | 2273400 | 2187300 | 3196400 | 44154 | 0 | 1270600 | 1924400 | 0 | 24972 |
| GNILNLNG | FOLH1 | Q04609 | FOLH1_HUMAI | Unmodified | 861850 | 1004200 | 2053100 | 4334400 | 0 | 0 | 0 | 0 | 0 | 0 | 0 | 0 | 337420 | 0 | 4101600 | 3200000 | 0 | 0 |
| GNILNLNG | FOLH1 | Q04609 | FOLH1_HUMAI | Unmodified | 7650100 | 6729600 | 12067000 | 79328000 | 0 | 0 | 0 | 0 | 0 | 0 | 0 | 0 | 0 | 41656 | 42352000 | 41141000 | 0 | 0 |
| GNIQLSYSI | IGF2R; IGF2R; I | P11717; A0A | MPRI_HUMAN ( | Unmodified | 0 | 0 | 0 | 0 | 0 | 0 | 0 | 0 | 18237000 | 2773300 | 1501300 | 2246100 | 772700 | 0 | 0 | 278530 | 0 | 0 |
| GNIQLSYSI | IGF2R; IGF2R; I | P11717; A0A | MPRI_HUMAN ( | Unmodified | 0 | 0 | 76428 | 0 | 0 | 0 | 204020 | 0 | 9932400 | 1401200 | 256810 | 340120 | 226570 | 118630 | 0 | 0 | 0 | 0 |
| GNIWSAEK | ANLN | Q9NQW6 | ANLN_HUMAN | Unmodified | 0 | 0 | 0 | 0 | 5949700 | 0 | 0 | 0 | 0 | 0 | 0 | 207700 | 125990 | 0 | 0 | 0 | 1346100 | 25451 |
| GNKEDYDI | SCG3 | Q8WXD2 | SCG3_HUMAN | Unmodified | 0 | 0 | 0 | 0 | 0 | 0 | 0 | 0 | 4138700 | 751940 | 0 | 0 | 0 | 0 | 0 | 0 | 0 | 0 |
| GNLAGLTLI | CLSTN1; CLSTN | O94985; Q5! | CSTN1_HUMAI | Unmodified | 0 | 0 | 444430 | 0 | 0 | 165630 | 0 | 0 | 3231100 | 0 | 0 | 0 | 1973000 | 0 | 0 | 0 | 0 | 0 |
| GNLANVIR | SLC25A4; SLC2 | P12235; P12 | ADT1_HUMAN . | Unmodified | 0 | 0 | 244080 | 0 | 0 | 0 | 0 | 0 | 7566900 | 1054600 | 853260 | 1076300 | 0 | 0 | 0 | 0 | 0 | 0 |
| GNLEWLDI | VPS25; VPS25 | Q9BRG1; K7 | VPS25_HUMAN | Unmodified | 2758300 | 3211200 | 2105100 | 1068000 | 0 | 0 | 0 | 0 | 0 | 0 | 0 | 1057500 | 420510 | 198330 | 5068800 | 4436100 | 0 | 0 |
| GNLGMNC | GLG1; GLG1 | Q92896; H3I | GSLG1_HUMAI | Oxidation (M) | 0 | 0 | 0 | 0 | 0 | 0 | 0 | 0 | 3613100 | 0 | 0 | 0 | 0 | 0 | 0 | 0 | 0 | 0 |
| GNLGMNC | GLG1; GLG1 | Q92896; H3I | GSLG1_HUMAI | Unmodified | 0 | 0 | 0 | 0 | 0 | 0 | 0 | 0 | 5143200 | 0 | 0 | 0 | 0 | 0 | 0 | 0 | 0 | 0 |
| GNLLQCIC | FN1 | P02751 | FINC_HUMAN I | Unmodified | 0 | 0 | 2571700 | 1281600 | 0 | 0 | 0 | 0 | 0 | 0 | 0 | 0 | 0 | 522400 | 0 | 850090 | 0 | 0 |
| GNLYSFGC | RCC2 | Q9P258 | RCC2_HUMAN | Unmodified | 0 | 0 | 0 | 0 | 0 | 0 | 0 | 0 | 0 | 0 | 1349300 | 7434000 | 0 | 341700 | 0 | 0 | 0 | 0 |
| GNLYWTD\ | NID1 | P14543 | NID1_HUMAN I | Unmodified | 0 | 0 | 511470 | 0 | 0 | 0 | 0 | 0 | 2594300 | 955910 | 573000 | 0 | 0 | 0 | 0 | 0 | 0 | 0 |
| GNMEVARI | CHMP1B | Q7LBR1 | CHM1B_HUMA | Unmodified | 0 | 0 | 0 | 0 | 1222500 | 240220 | 0 | 0 | 0 | 649650 | 0 | 0 | 340800 | 263060 | 0 | 0 | 0 | 0 |
| GNMEVARI | CHMP1B | Q7LBR1 | CHM1B_HUMA | Unmodified | 0 | 0 | 0 | 0 | 50579000 | 4355700 | 1435500 | 219940 | 0 | 10677000 | 0 | 0 | 7630500 | 6971700 | 0 | 0 | 1252100 | 5634300 |
| GNMEVARI | CHMP1B | Q7LBR1 | CHM1B_HUMA | Oxidation (M) | 0 | 0 | 0 | 0 | 14457000 | 1437900 | 449460 | 258130 | 0 | 2801400 | 59045 | 0 | 2927900 | 2594200 | 0 | 0 | 388110 | 1015400 |
| GNNFLYTN | WDR1; WDR1 | O75083; D6I | WDR1_HUMAN | Unmodified | 720590 | 745560 | 489050 | 547280 | 4162000 | 1092300 | 0 | 0 | 861090 | 293560 | 1061900 | 5800500 | 3678300 | 1738500 | 1040600 | 647330 | 617340 | 1452300 |
| GNNLQDTI | MTOR; MTOR | P42345; B1A | MTOR_HUMAN | Unmodified | 0 | 0 | 0 | 0 | 0 | 0 | 0 | 0 | 0 | 0 | 0 | 506590 | 0 | 0 | 0 | 0 | 0 | 0 |
| GNNVLYIS | LSM6 | P62312 | LSM6_HUMAN | Unmodified | 0 | 0 | 0 | 0 | 0 | 0 | 0 | 0 | 0 | 0 | 0 | 1545200 | 0 | 0 | 0 | 0 | 0 | 0 |
| GNNVYCLI | COPA; COPA; C | P53621; A0A | COPA_HUMAN | Unmodified | 0 | 0 | 0 | 0 | 0 | 141290 | 0 | 0 | 0 | 0 | 0 | 945910 | 0 | 485700 | 0 | 0 | 0 | 0 |
| GNPADVTE | PLPP2; PLPP2; I | O43688; K7E | PLPP2_HUMAN | Unmodified | 0 | 0 | 0 | 0 | 0 | 45715 | 26216 | 17725 | 13821000 | 1403100 | 0 | 12044000 | 0 | 0 | 102720 | 444330 | 0 | 0 |
| GNPECDGI | ST14 | Q9Y5Y6 | ST14_HUMAN S | Unmodified | 511310 | 135280 | 701390 | 972730 | 0 | 0 | 0 | 0 | 183410 | 0 | 0 | 22216 | 198180 | 39335 | 773390 | 679890 | 0 | 0 |
| GNPENIED | KRIT1; KRIT1; KI | O00522; C9J | KRIT1_HUMAN | Unmodified | 0 | 0 | 0 | 241470 | 1397000 | 382800 | 754280 | 0 | 0 | 183140 | 0 | 258220 | 0 | 973830 | 325980 | 188950 | 885930 | 948700 |
| GNPICSLH | PFAS; PFAS | O15067; J3K | PUR4_HUMAN | Unmodified | 0 | 0 | 0 | 0 | 0 | 0 | 0 | 272830 | 0 | 0 | 482600 | 1279600 | 0 | 0 | 0 | 0 | 0 | 0 |
| GNPTVEVD | ENO1; ENO1; E | P06733; K7E | ENOA_HUMAN | Unmodified | 3618300 | 0 | 2839100 | 3295500 | 3296200 | 0 | 0 | 1151700 | 13846000 | 2349300 | 3208400 | 53005000 | 1626100 | 1495400 | 4498900 | 6901200 | 0 | 1670900 |
| GNPTVEVD | ENO2; ENO2; E | P09104; F5H | ENOG_HUMAN | Unmodified | 0 | 0 | 0 | 0 | 0 | 0 | 0 | 0 | 0 | 0 | 0 | 1044500 | 0 | 0 | 0 | 0 | 0 | 0 |
| GNQLQTLF | SLITRK6 | Q9H5Y7 | SLIK6_HUMAN | Unmodified | 1735000 | 1433700 | 0 | 0 | 0 | 0 | 0 | 0 | 1066200 | 0 | 0 | 0 | 0 | 0 | 0 | 0 | 0 | 0 |
| GNQLWEYI | GALNT1 | Q10472 | GALT1_HUMAN | Unmodified | 0 | 0 | 0 | 0 | 306870 | 0 | 0 | 0 | 4570300 | 730270 | 1813500 | 1388600 | 0 | 0 | 0 | 0 | 0 | 0 |
| GNQLWEYI | GALNT1 | Q10472 | GALT1_HUMAN | Unmodified | 182490 | 0 | 0 | 0 | 0 | 0 | 0 | 0 | 0 | 372880 | 0 | 0 | 0 | 84063 | 0 | 0 | 0 | 81407 |
| GNRPVILTY | NDRG3; NDRG: | Q9UGV2; Q5 | NDRG3_HUMA | Unmodified | 0 | 156290 | 431990 | 1669500 | 3806900 | 1090800 | 0 | 0 | 139100 | 464930 | 4329100 | 45301000 | 4731000 | 2039700 | 1591600 | 1305600 | 1630200 | 816850 |
| GNRPVILTY | NDRG1; NDRG: | Q92597; A0A | NDRG1_HUMA | Oxidation (M) | 324080 | 116360 | 1300800 | 0 | 4744400 | 2349300 | 0 | 47526 | 4627200 | 3312900 | 452820 | 3800200 | 1296400 | 1265600 | 618120 | 1709800 | 6386200 | 9485500 |
| GNRPVILTY | NDRG1; NDRG: | Q92597; A0A | NDRG1_HUMA | Unmodified | 0 | 0 | 0 | 0 | 14213000 | 172610 | 0 | 0 | 9075300 | 5428300 | 0 | 6372200 | 1469400 | 995060 | 0 | 0 | 9103100 | 9649100 |
| GNSLQEILI | HDLBP; HDLBP | Q00341; A0A | VIGLN_HUMAN | Unmodified | 0 | 0 | 0 | 0 | 263930 | 0 | 0 | 0 | 0 | 0 | 0 | 0 | 0 | 0 | 0 | 619930 | 0 | 0 |
| GNSPPSSG | NAA10 | P41227 | NAA10_HUMAI | Unmodified | 0 | 0 | 58926 | 0 | 0 | 0 | 37691 | 0 | 0 | 0 | 61808 | 915030 | 0 | 0 | 0 | 0 | 22943 | 80206 |
| GNSYMIQR | ITGA3 | P26006 | ITA3_HUMAN I | Unmodified | 62921 | 42355 | 0 | 0 | 231060 | 0 | 0 | 0 | 171010 | 0 | 398170 | 886160 | 0 | 0 | 97203 | 33542 | 0 | 168190 |
| GNSYMIQR | ITGA3 | P26006 | ITA3_HUMAN I | Unmodified | 0 | 0 | 0 | 0 | 5373800 | 3226700 | 0 | 0 | 188800 | 20039 | 0 | 63624 | 0 | 68821 | 0 | 0 | 0 | 4541200 |
| GNTAAQM/ | SERPINB6; SER | P35237; A0A | SPB6_HUMAN | Unmodified | 24409 | 0 | 0 | 138060 | 306460 | 147140 | 186690 | 0 | 192280 | 0 | 216260 | 1512300 | 250840 | 458870 | 118510 | 66682 | 540860 | 344150 |
| GNTALHIAS | ANK3; ANK2; AI | Q12955; Q0 | ANK3_HUMAN | Unmodified | 0 | 0 | 1594900 | 860000 | 0 | 0 | 533910 | 158360 | 0 | 0 | 0 | 0 | 1387800 | 0 | 2104200 | 1487200 | 432270 | 0 |
| GNTLKEDV | MAP4K4; MAP4 | O95819; C9J | M4K4_HUMAN | Unmodified | 0 | 0 | 65249 | 222940 | 0 | 0 | 0 | 0 | 639770 | 90973 | 492970 | 2644000 | 0 | 0 | 162260 | 126940 | 0 | 0 |
| GNTSGSHI | UPF1 | Q92900 | RENT1_HUMAI | Unmodified | 1742200 | 1902100 | 1566300 | 1992900 | 0 | 0 | 0 | 0 | 369710 | 0 | 0 | 1182400 | 0 | 0 | 661070 | 683370 | 0 | 265590 |
| GNVFSSPT | CALD1; CALD1; | Q05682; E7E | CALD1_HUMAI | Unmodified | 0 | 0 | 477550 | 0 | 173550 | 0 | 0 | 0 | 127440 | 0 | 0 | 184140 | 0 | 0 | 1101900 | 608020 | 0 | 0 |
| GNVFSSPT | CALD1; CALD1; | Q05682; E7E | CALD1_HUMAI | Unmodified | 0 | 0 | 0 | 2104000 | 524960 | 0 | 0 | 0 | 0 | 0 | 0 | 0 | 6436800 | 4825100 | 515190 | 256740 | 1078500 | 684710 |
| GNVGFVFT | RPLP0; RPLP0P | P05388; Q8I | RLA0_HUMAN ( | Unmodified | 2798700 | 2874100 | 6850400 | 969320 | 0 | 0 | 0 | 96957 | 0 | 0 | 556590 | 13468000 | 0 | 232350 | 8109900 | 5047000 | 0 | 331680 |
| GNVGFVFT | RPLP0; RPLP0P | P05388; Q8I | RLA0_HUMAN ( | Unmodified | 2110800 | 2699200 | 0 | 0 | 0 | 0 | 0 | 0 | 0 | 0 | 0 | 7173400 | 0 | 0 | 3511400 | 1807700 | 0 | 0 |
| GNVGFVFT | RPLP0; RPLP0P | P05388; Q8I | RLA0_HUMAN ( | Oxidation (M) | 560850 | 3240300 | 178120 | 692420 | 0 | 0 | 0 | 0 | 0 | 0 | 0 | 0 | 0 | 0 | 12207000 | 1240600 | 0 | 0 |
| GNVGFVFT | RPLP0; RPLP0P | P05388; Q8I | RLA0_HUMAN ( | Unmodified | 100090 | 160110 | 0 | 0 | 0 | 0 | 0 | 0 | 0 | 0 | 0 | 0 | 0 | 0 | 4512400 | 0 | 0 | 0 |
| GNVGVVLF | DUT; DUT; DUT | P33316; H0Y | DUT_HUMAN D | Unmodified | 0 | 0 | 0 | 0 | 0 | 0 | 0 | 0 | 0 | 0 | 0 | 2241000 | 0 | 0 | 0 | 0 | 0 | 0 |
| GNVLLEDG | SEMA4B | Q9NPR2 | SEM4B_HUMAI | Unmodified | 0 | 0 | 113180 | 0 | 0 | 0 | 0 | 17789 | 333300 | 0 | 0 | 0 | 0 | 0 | 0 | 0 | 0 | 0 |
| GNVQVVIP | UBA1 | P22314 | UBA1_HUMAN | Unmodified | 71961 | 0 | 51234 | 114120 | 0 | 0 | 374170 | 0 | 1539700 | 469070 | 491590 | 1604700 | 560420 | 1602200 | 144610 | 265520 | 264290 | 749840 |
| GNVVFGEF | DPYSL3; DPYSL | Q14195; H0Y | DPYL3_HUMAN | Unmodified | 0 | 0 | 0 | 0 | 0 | 0 | 0 | 0 | 0 | 0 | 15766000 | 14291000 | 0 | 756810 | 0 | 0 | 0 | 0 |
| GNWDEQF | SERPINB6; SER | P35237; A0A | SPB6_HUMAN | Unmodified | 0 | 0 | 0 | 0 | 0 | 0 | 0 | 0 | 0 | 0 | 179100 | 2357100 | 0 | 0 | 0 | 0 | 0 | 0 |
| GNWDEQF | SERPINB6; SER | P35237; A0A | SPB6_HUMAN | Unmodified | 84940 | 76776 | 0 | 0 | 0 | 279660 | 0 | 0 | 177980 | 0 | 0 | 5323200 | 0 | 0 | 262550 | 142700 | 0 | 0 |
| GNWEQPQ | UBAP2L; UBAP: | Q14157; Q5' | UBP2L_HUMAN | Unmodified | 241320 | 0 | 46377 | 0 | 158630 | 0 | 0 | 0 | 101100 | 96285 | 554630 | 1502800 | 0 | 0 | 0 | 33699 | 0 | 157590 |
| GNWLLVG! | ITGA2; ITGA2; I | P17301; D6F | ITA2_HUMAN I | Unmodified | 1640700 | 1404300 | 585860 | 1555200 | 3266800 | 1560900 | 0 | 0 | 29305 | 40030 | 0 | 178060 | 735490 | 245130 | 2783900 | 1092800 | 898700 | 5909000 |
| GNWLLVG! | ITGA2; ITGA2; I | P17301; D6F | ITA2_HUMAN I | Oxidation (M) | 0 | 0 | 0 | 355850 | 6548900 | 0 | 0 | 0 | 0 | 0 | 0 | 0 | 0 | 0 | 0 | 0 | 763810 | 6086800 |
| GNWLLVG! | ITGA2; ITGA2; I | P17301; D6F | ITA2_HUMAN I | Unmodified | 0 | 0 | 0 | 0 | 638530 | 0 | 0 | 0 | 0 | 0 | 0 | 0 | 0 | 0 | 0 | 0 | 0 | 531940 |
| GNWLLVG! | ITGA2; ITGA2; I | P17301; D6F | ITA2_HUMAN I | Oxidation (M) | 0 | 0 | 0 | 0 | 4843900 | 0 | 0 | 0 | 0 | 0 | 0 | 0 | 0 | 0 | 0 | 0 | 2394200 | 10266000 |
| GPAAAQGS | UBQLN2 | Q9UHD9 | UBQL2_HUMAI | Unmodified | 293220 | 374300 | 0 | 0 | 0 | 0 | 0 | 0 | 0 | 0 | 0 | 124790 | 0 | 0 | 0 | 317580 | 0 | 0 |
| GPAFNMAS | AHNAK | Q09666 | AHNK_HUMAN | Unmodified | 0 | 540060 | 0 | 0 | 306690 | 0 | 0 | 20886 | 0 | 245040 | 0 | 0 | 0 | 0 | 0 | 0 | 83730 | 228640 |

| | | | | | | | | | | | | | | | | | | | | | | |
|---|---|---|---|---|---|---|---|---|---|---|---|---|---|---|---|---|---|---|---|---|---|---|
| GPAGDATV | SLC1A5; SLC1A | Q15758; M0 | AAAT_HUMAN | Unmodified | 0 | 21547 | 0 | 0 | 1187700 | 0 | 0 | 0 | 1690100 | 232840 | 4770200 | 8865500 | 0 | 0 | 0 | 0 | 0 | 0 |
| GPAGDATV | SLC1A5; SLC1A | Q15758; M0 | AAAT_HUMAN | Unmodified | 0 | 0 | 0 | 0 | 510800 | 0 | 0 | 0 | 0 | 0 | 0 | 0 | 0 | 0 | 0 | 0 | 505490 | 0 |
| GPAHCLLL | QPRT | Q15274 | NADC_HUMAN | Unmodified | 0 | 0 | 0 | 0 | 0 | 0 | 0 | 0 | 0 | 0 | 0 | 1423100 | 0 | 0 | 0 | 0 | 39147 | 0 |
| GPALTPIRD | LSR; LSR | Q86X29; S4F | LSR_HUMAN Li | Unmodified | 1313600 | 777930 | 2432700 | 675480 | 0 | 0 | 0 | 0 | 4505900 | 284430 | 350530 | 5389100 | 0 | 0 | 3075600 | 4780500 | 0 | 0 |
| GPAPRETK | ARPC3; ARPC3; | O15145; F8\ | ARPC3_HUMAN | Unmodified | 0 | 0 | 0 | 110480 | 5214800 | 336210 | 456930 | 29147 | 0 | 794810 | 149460 | 0 | 4082000 | 1098200 | 0 | 0 | 3791300 | 445080 |
| GPASTPTYS | CC2D1A; CC2D | Q6P1N0; K7 | C2D1A_HUMAI | Unmodified | 592920 | 0 | 0 | 627740 | 302530 | 0 | 0 | 0 | 570980 | 115530 | 0 | 0 | 0 | 0 | 881110 | 0 | 0 | 0 |
| GPATVEDL | GIPC1; GIPC1 | O14908; K7E | GIPC1_HUMAN | Unmodified | 0 | 0 | 0 | 0 | 0 | 0 | 0 | 0 | 0 | 0 | 0 | 0 | 0 | 0 | 0 | 0 | 0 | 0 |
| GPAVGIDL | HSPA8; HSPA8; | P11142; E9P | HSP7C_HUMAI | Unmodified | 69514 | 0 | 0 | 0 | 0 | 0 | 0 | 0 | 0 | 0 | 0 | 6923100 | 0 | 0 | 0 | 0 | 0 | 0 |
| GPAVGIDL | HSPA8; HSPA8; | P11142; E9P | HSP7C_HUMAI | Unmodified | 0 | 1786800 | 0 | 2472400 | 0 | 0 | 1185200 | 0 | 5483400 | 3633500 | 0 | 0 | 2478400 | 0 | 0 | 0 | 5049900 | 0 |
| GPAYGLSR | TAGLN2; TAGLN | P37802; X6R | TAGL2_HUMAN | Unmodified | 163480 | 0 | 275270 | 166440 | 0 | 0 | 87847 | 101960 | 286190 | 63935 | 716290 | 1480100 | 0 | 43052 | 160370 | 227320 | 0 | 0 |
| GPCIIYNEI | RPL4; RPL4 | P36578; H3E | RL4_HUMAN 6( | Unmodified | 1677300 | 658520 | 0 | 0 | 0 | 0 | 0 | 90500 | 0 | 0 | 249800 | 12057000 | 0 | 0 | 320540 | 0 | 0 | 0 |
| GPDAASKL | PSMC1 | P62191 | PRS4_HUMAN | Unmodified | 0 | 0 | 0 | 0 | 0 | 0 | 0 | 0 | 0 | 0 | 0 | 841750 | 0 | 0 | 0 | 0 | 0 | 0 |
| GPDEDGEE | TTYH3; TTYH3 | Q9C0H2; H7 | TTYH3_HUMAN | Unmodified | 0 | 0 | 0 | 0 | 0 | 0 | 0 | 0 | 60807 | 297280 | 54118 | 824990 | 0 | 0 | 0 | 0 | 0 | 0 |
| GPDGLTAF | MTPN; MTPN | P58546; C9J | MTPN_HUMAN | Unmodified | 1117400 | 439430 | 2259300 | 760450 | 0 | 0 | 0 | 0 | 850570 | 1000600 | 3835800 | 10031000 | 71778 | 0 | 2472100 | 2210200 | 0 | 0 |
| GPDGYDP( | MOV10; MOV1C | Q9HCE1; Q8 | MOV10_HUMA | Unmodified | 439500 | 0 | 129070 | 0 | 0 | 0 | 0 | 159100 | 0 | 0 | 0 | 0 | 320110 | 0 | 0 | 0 | 0 | 0 |
| GPEDTAQL | STXBP2; STXBP | Q15833; M0 | STXB2_HUMAN | Unmodified | 300760 | 458200 | 1145200 | 906210 | 829240 | 165990 | 168020 | 91437 | 1096400 | 195470 | 244540 | 2190900 | 434360 | 447670 | 1031100 | 708790 | 276050 | 624150 |
| GPELLTMV | VCP; VCP; VCP; | P55072; C9J | TERA_HUMAN | Oxidation (M) | 0 | 0 | 0 | 0 | 0 | 0 | 0 | 0 | 315180 | 0 | 0 | 246640 | 0 | 0 | 0 | 0 | 0 | 0 |
| GPEPVVSQ | GPC3; GPC3 | P51654; H0Y | GPC3_HUMAN | Unmodified | 0 | 0 | 0 | 0 | 0 | 0 | 0 | 0 | 3443200 | 782240 | 0 | 0 | 0 | 0 | 0 | 0 | 0 | 0 |
| GPEQVPVP | SLC38A10; SLC | Q9HBR0; H0 | S38AA_HUMAN | Unmodified | 0 | 0 | 0 | 0 | 0 | 0 | 0 | 0 | 3283700 | 0 | 0 | 0 | 0 | 0 | 0 | 0 | 0 | 0 |
| GPFLLGIK | ACO1 | P21399 | ACOC_HUMAN | Unmodified | 0 | 0 | 0 | 0 | 0 | 0 | 0 | 0 | 0 | 0 | 0 | 2339100 | 0 | 0 | 0 | 0 | 0 | 0 |
| GPFLVALG | PDLIM5; PDLIM | Q96HC4; HC | PDLI5_HUMAN | Unmodified | 146300 | 0 | 288410 | 0 | 0 | 0 | 0 | 0 | 0 | 0 | 277740 | 3061100 | 0 | 0 | 419370 | 385560 | 0 | 0 |
| GPFSRYDY | RPN1; RPN1 | P04843; B7Z | RPN1_HUMAN | Unmodified | 0 | 0 | 0 | 0 | 0 | 0 | 0 | 0 | 13825000 | 587750 | 279370 | 308830 | 0 | 0 | 0 | 0 | 0 | 0 |
| GPFSRYDY | RPN1; RPN1 | P04843; B7Z | RPN1_HUMAN | Unmodified | 0 | 0 | 0 | 0 | 372930 | 0 | 334500 | 0 | 1.04E+08 | 11084000 | 3201600 | 0 | 0 | 0 | 0 | 0 | 0 | 0 |
| GPFSVEAD | LFNG; LFNG | Q8NES3; A0 | LFNG_HUMAN | Unmodified | 0 | 0 | 98809 | 0 | 201060 | 0 | 0 | 84463 | 2483100 | 70067 | 0 | 0 | 84241 | 42674 | 41618 | 25766 | 0 | 0 |
| GPFTDVVT | VAPA; VAPB; VA | Q9P0L0; Q9 | VAPA_HUMAN | Unmodified | 0 | 0 | 0 | 0 | 0 | 0 | 0 | 98594 | 513730 | 0 | 994070 | 2197000 | 0 | 0 | 0 | 0 | 0 | 275860 |
| GPGLGSTC | PURA | Q00577 | PURA_HUMAN | Unmodified | 0 | 0 | 0 | 0 | 0 | 0 | 0 | 0 | 726570 | 0 | 219030 | 1983400 | 0 | 0 | 0 | 794600 | 0 | 0 |
| GPGLYYVD | PSMB5 | P28074 | PSB5_HUMAN | Unmodified | 0 | 0 | 0 | 0 | 0 | 0 | 0 | 0 | 0 | 0 | 0 | 1184400 | 0 | 0 | 0 | 0 | 0 | 0 |
| GPGPTVKP | BROX; BROX | Q5VW32; F5 | BROX_HUMAN | Unmodified | 2123300 | 1318200 | 0 | 305420 | 0 | 0 | 0 | 0 | 0 | 0 | 105120 | 0 | 0 | 0 | 3134600 | 2882900 | 0 | 0 |
| GPGPTVKP | BROX; BROX | Q5VW32; F5 | BROX_HUMAN | Unmodified | 0 | 635030 | 548170 | 639620 | 0 | 0 | 0 | 0 | 558390 | 54352 | 182260 | 744920 | 0 | 0 | 575850 | 0 | 0 | 0 |
| GPGYPQAE | SH3GL2 | Q99962 | SH3G2_HUMAI | Unmodified | 0 | 0 | 0 | 0 | 0 | 0 | 0 | 0 | 1397300 | 549880 | 0 | 0 | 0 | 0 | 0 | 0 | 0 | 0 |
| GPGYPQAE | SH3GL2 | Q99962 | SH3G2_HUMAI | Oxidation (M) | 0 | 0 | 0 | 0 | 88605 | 0 | 0 | 0 | 1034200 | 855520 | 0 | 0 | 77138 | 65733 | 0 | 0 | 39746 | 53600 |
| GPHLVQSE | LMAN1 | P49257 | LMAN1_HUMAI | Unmodified | 0 | 0 | 2145900 | 1454900 | 0 | 0 | 0 | 0 | 3.57E+08 | 7425400 | 8114100 | 8581000 | 0 | 0 | 0 | 0 | 0 | 0 |
| GPHLVQSE | LMAN1 | P49257 | LMAN1_HUMAI | Unmodified | 0 | 0 | 0 | 0 | 0 | 0 | 784070 | 0 | 4680900 | 14005000 | 850220 | 0 | 0 | 0 | 0 | 0 | 0 | 4644700 |
| GPHVDVS( | AHNAK | Q09666 | AHNK_HUMAN | Unmodified | 0 | 0 | 0 | 121810 | 5105800 | 920240 | 0 | 18107 | 118130 | 0 | 328380 | 0 | 112800 | 0 | 468500 | 269000 | 724160 | 2071100 |
| GPIDAITGE | PLXNA1; PLXNA | Q9UIW2; O7 | PLXA1_HUMAN | Unmodified | 0 | 616650 | 0 | 0 | 0 | 0 | 0 | 0 | 587140 | 0 | 0 | 1190500 | 0 | 0 | 196240 | 0 | 0 | 0 |
| GPIKFNVW | RAN; RAN; RAN | P62826; H0Y | RAN_HUMAN ( | Unmodified | 282840 | 461150 | 0 | 216860 | 201480 | 0 | 0 | 0 | 0 | 172330 | 0 | 2120000 | 86611 | 0 | 0 | 0 | 146660 | 0 |
| GPILMELQ | PDHA1; PDHA2 | P08559; P29 | ODPA_HUMAN | Unmodified | 0 | 0 | 0 | 0 | 0 | 0 | 0 | 0 | 760870 | 0 | 0 | 0 | 0 | 0 | 0 | 0 | 0 | 0 |
| GPIVPLNV | SLC12A2; SLC1 | P55011; G3 | S12A2_HUMAN | Unmodified | 0 | 0 | 0 | 2161400 | 1265700 | 0 | 358100 | 98368 | 499460 | 0 | 579640 | 1724700 | 0 | 0 | 0 | 3090700 | 493380 | 6024500 |
| GPKAEVTAI | AHNAK2 | Q8IVF2 | AHNK2_HUMA | Unmodified | 0 | 0 | 0 | 0 | 1455100 | 0 | 0 | 0 | 0 | 0 | 0 | 0 | 0 | 0 | 0 | 512050 | 139460 | 3977400 |
| GPKPTAPS | NCMAP | Q5T1S8 | NCMAP_HUMA | Unmodified | 0 | 0 | 0 | 0 | 0 | 0 | 0 | 0 | 0 | 0 | 768220 | 5163500 | 0 | 0 | 0 | 0 | 0 | 0 |
| GPKVDIDA | AHNAK | Q09666 | AHNK_HUMAN | Unmodified | 0 | 0 | 0 | 184420 | 16658000 | 1503200 | 455850 | 0 | 0 | 0 | 0 | 0 | 0 | 0 | 364810 | 0 | 1080400 | 4944500 |
| GPKVDIDTI | AHNAK | Q09666 | AHNK_HUMAN | Unmodified | 0 | 0 | 0 | 0 | 909260 | 0 | 0 | 0 | 0 | 0 | 122480 | 0 | 983540 | 0 | 0 | 0 | 65843 | 522750 |
| GPKVDINA | AHNAK | Q09666 | AHNK_HUMAN | Unmodified | 0 | 0 | 0 | 0 | 2374600 | 0 | 0 | 0 | 0 | 0 | 0 | 0 | 0 | 0 | 0 | 0 | 306290 | 0 |
| GPKVDINA | AHNAK | Q09666 | AHNK_HUMAN | Unmodified | 0 | 0 | 0 | 0 | 1553800 | 0 | 0 | 0 | 0 | 0 | 0 | 0 | 0 | 0 | 0 | 0 | 0 | 0 |
| GPKVDISAI | AHNAK | Q09666 | AHNK_HUMAN | Unmodified | 0 | 0 | 0 | 0 | 756070 | 0 | 0 | 0 | 0 | 0 | 0 | 0 | 0 | 0 | 0 | 0 | 0 | 0 |
| GPKVDVSA | AHNAK | Q09666 | AHNK_HUMAN | Oxidation (M) | 0 | 0 | 0 | 0 | 4688500 | 660910 | 0 | 0 | 0 | 0 | 0 | 0 | 0 | 0 | 0 | 0 | 600070 | 1098100 |
| GPKVDVSA | AHNAK | Q09666 | AHNK_HUMAN | Unmodified | 221540 | 0 | 0 | 0 | 5763100 | 282770 | 0 | 0 | 0 | 0 | 0 | 0 | 0 | 0 | 0 | 0 | 736370 | 2132600 |
| GPKVQANL | AHNAK | Q09666 | AHNK_HUMAN | Unmodified | 0 | 0 | 0 | 0 | 0 | 0 | 0 | 0 | 0 | 0 | 0 | 0 | 0 | 0 | 0 | 0 | 265790 | 0 |
| GPLAELPS | TNKS1BP1; TNI | Q9C0C2; E9 | TB182_HUMAN | Unmodified | 553230 | 166260 | 271460 | 129530 | 0 | 0 | 0 | 0 | 0 | 0 | 82938 | 702170 | 0 | 0 | 454610 | 580920 | 162780 | 0 |
| GPLASQIS( | NT5E; NT5E | P21589; Q9 | 5NTD_HUMAN | Unmodified | 0 | 0 | 0 | 0 | 19301000 | 9661100 | 0 | 0 | 549410 | 13269 | 0 | 0 | 0 | 0 | 0 | 0 | 3904100 | 6999000 |
| GPLATGGII | TMA7; hCG_20 | Q9Y2S6; A0 | TMA7_HUMAN | Unmodified | 440630 | 447390 | 684840 | 496450 | 793630 | 0 | 0 | 0 | 622380 | 61015 | 83577 | 405120 | 246250 | 0 | 509190 | 429390 | 0 | 0 |
| GPLATGGII | TMA7; hCG_20 | Q9Y2S6; A0 | TMA7_HUMAN | Unmodified | 514920 | 0 | 0 | 792470 | 176130 | 0 | 0 | 0 | 237740 | 0 | 135250 | 233160 | 286540 | 96697 | 904130 | 1016300 | 58716 | 233190 |
| GPLEHLYS | IGFBP2; IGFBP2 | P18065; C9J | IBP2_HUMAN | Unmodified | 0 | 0 | 0 | 0 | 52433 | 0 | 0 | 0 | 5874200 | 0 | 0 | 0 | 0 | 332100 | 0 | 0 | 93129 | 0 |
| GPLEYVPS | MVP | Q14764 | MVP_HUMAN | Unmodified | 0 | 0 | 0 | 0 | 0 | 0 | 83963 | 0 | 0 | 109960 | 0 | 7898800 | 0 | 0 | 0 | 0 | 0 | 0 |
| GPLGCSEF | IGF2R | P11717 | MPRI_HUMAN | Unmodified | 0 | 0 | 0 | 0 | 0 | 0 | 0 | 0 | 3683600 | 875930 | 196050 | 1017600 | 160050 | 0 | 0 | 0 | 0 | 0 |
| GPLGIDPQ | CDC42BPA; CD | Q5VT25; A0 | MRCKA_HUMA | Unmodified | 0 | 0 | 0 | 0 | 0 | 0 | 0 | 0 | 0 | 0 | 0 | 0 | 0 | 0 | 0 | 0 | 0 | 0 |
| GPLGPNW | PITPNA; PITPNA | O00169; I3L | PIPNA_HUMAN | Unmodified | 186260 | 155880 | 239750 | 212310 | 0 | 0 | 0 | 0 | 0 | 0 | 0 | 1982900 | 0 | 0 | 800240 | 813870 | 0 | 0 |
| GPLIAPGP | SLC6A8 | P48029 | SC6A8_HUMAN | Unmodified | 0 | 29826 | 176050 | 497920 | 0 | 0 | 0 | 0 | 0 | 0 | 0 | 3256200 | 0 | 0 | 671970 | 524450 | 0 | 0 |
| GPLINSEFY | GLB1; GLB1; GI | P16278; C9J | BGAL_HUMAN | Unmodified | 0 | 0 | 0 | 0 | 0 | 0 | 0 | 0 | 2787500 | 0 | 0 | 0 | 0 | 0 | 635550 | 0 | 0 | 0 |
| GPLKNTSD | CTNNA1; CTNN | P35221; E5R | CTNA1_HUMAI | Unmodified | 0 | 0 | 0 | 0 | 859820 | 0 | 76973 | 0 | 0 | 0 | 0 | 0 | 563640 | 0 | 0 | 0 | 51031 | 223280 |
| GPLLEEQA | FAM98B | Q52LJ0 | FA98B_HUMAN | Unmodified | 0 | 0 | 0 | 0 | 0 | 0 | 0 | 0 | 0 | 0 | 0 | 620410 | 0 | 0 | 0 | 0 | 0 | 0 |
| GPLLGHTD | STRN | O43815 | STRN_HUMAN | Unmodified | 0 | 0 | 0 | 0 | 0 | 0 | 0 | 0 | 0 | 0 | 0 | 825980 | 0 | 0 | 0 | 0 | 0 | 0 |
| GPLLYCAR | LRRC47 | Q8N1G4 | LRC47_HUMAN | Unmodified | 237080 | 0 | 188460 | 133450 | 461870 | 0 | 103060 | 0 | 802660 | 0 | 0 | 1586600 | 890750 | 428980 | 158990 | 150400 | 353810 | 324010 |
| GPLMMYIS | EEF2 | P13639 | EF2_HUMAN El | 2 Oxidation (M) | 456140 | 271400 | 0 | 0 | 668350 | 205750 | 0 | 0 | 178850 | 0 | 358540 | 6997000 | 0 | 0 | 0 | 315610 | 0 | 82231 |
| GPLMMYIS | EEF2 | P13639 | EF2_HUMAN El | Oxidation (M) | 0 | 0 | 0 | 0 | 0 | 0 | 0 | 0 | 480260 | 0 | 1139000 | 3929700 | 0 | 0 | 0 | 0 | 0 | 0 |
| GPLPAAPP | RTN4; RTN4 | Q9NQC3; F8 | RTN4_HUMAN | Unmodified | 0 | 0 | 0 | 0 | 0 | 0 | 0 | 0 | 0 | 0 | 1223000 | 0 | 0 | 0 | 0 | 0 | 0 | 0 |
| GPLQSVQV | RPS16; RPS16; | P62249; M0 | RS16_HUMAN | Unmodified | 3749600 | 2882900 | 750700 | 3170300 | 384350 | 0 | 150230 | 87044 | 1109400 | 0 | 3104200 | 19485000 | 2044100 | 792700 | 3087400 | 3308700 | 0 | 1206300 |
| GPLQSVQV | RPS16; RPS16; | P62249; M0 | RS16_HUMAN | Unmodified | 0 | 0 | 0 | 0 | 249390 | 0 | 0 | 102600 | 0 | 0 | 0 | 1699500 | 0 | 0 | 0 | 0 | 0 | 0 |
| GPLREEKIS | KIAA0319L; KIA | Q8IZA0; B1A | K319L_HUMAN | Unmodified | 0 | 0 | 0 | 0 | 108710 | 0 | 0 | 0 | 0 | 534510 | 0 | 0 | 0 | 0 | 0 | 0 | 0 | 0 |
| GPLVNASL | PRMT5; PRMT5; | O14744; H0 | ANM5_HUMAN | Unmodified | 40745 | 48799 | 0 | 83090 | 0 | 0 | 0 | 0 | 263170 | 0 | 245070 | 10084000 | 0 | 0 | 192280 | 143530 | 0 | 0 |
| GPNAVQLV | HUWE1 | Q7Z6Z7 | HUWE1_HUMA | Unmodified | 0 | 0 | 0 | 0 | 721670 | 0 | 0 | 26051 | 0 | 0 | 0 | 1107400 | 388920 | 117160 | 0 | 0 | 0 | 95713 |
| GPNGYGFI | SLC9A3R1; SLC | O14745; J3Q | NHRF1_HUMAI | Unmodified | 0 | 63974 | 407810 | 580760 | 1071700 | 258590 | 448610 | 300330 | 142200 | 100390 | 236140 | 1653400 | 6478600 | 6658700 | 705500 | 653620 | 83512 | 435490 |
| GPNSYPGS | AFDN; AFDN; A | P55196; H0Y | AFAD_HUMAN | Unmodified | 0 | 0 | 88121 | 0 | 0 | 0 | 0 | 0 | 0 | 0 | 0 | 0 | 524270 | 175210 | 0 | 0 | 0 | 0 |
| GPPDFSSD | TMPO; TMPO; T | P42167; P42 | LAP2B_HUMAN | Unmodified | 0 | 0 | 0 | 0 | 0 | 0 | 0 | 0 | 892220 | 0 | 1073000 | 1709300 | 0 | 0 | 0 | 0 | 0 | 0 |
| GPPGPQGI | COL2A1 | P02458 | CO2A1_HUMAI | Unmodified | 0 | 0 | 0 | 0 | 0 | 0 | 0 | 0 | 0 | 0 | 1042900 | 0 | 0 | 0 | 0 | 0 | 0 | 0 |
| GPPHLQIRI | MAN1B1; MAN1 | Q9UKM7; HC | MA1B1_HUMAI | Unmodified | 0 | 0 | 0 | 0 | 0 | 0 | 0 | 0 | 3935000 | 0 | 0 | 0 | 0 | 0 | 0 | 0 | 0 | 0 |
| GPPLWEEC | PDIA5; PDIA5 | Q14554; C9J | PDIA5_HUMAN | Unmodified | 0 | 0 | 0 | 0 | 0 | 0 | 0 | 0 | 3891200 | 0 | 0 | 0 | 0 | 0 | 0 | 0 | 0 | 0 |
| GPPLWEEC | PDIA5; PDIA5 | Q14554; C9J | PDIA5_HUMAN | Unmodified | 0 | 0 | 0 | 0 | 527210 | 0 | 0 | 0 | 18981000 | 955880 | 0 | 0 | 0 | 0 | 0 | 0 | 0 | 0 |
| GPPPPPGE | SF3B2; SF3B2; | Q13435; E9F | SF3B2_HUMAN | Unmodified | 97351 | 0 | 0 | 0 | 0 | 0 | 0 | 0 | 0 | 0 | 0 | 634110 | 0 | 0 | 0 | 0 | 0 | 0 |
| GPPPPPGE | SF3B2; SF3B2; | Q13435; E9F | SF3B2_HUMAN | Oxidation (M) | 589200 | 547610 | 0 | 261620 | 0 | 0 | 0 | 0 | 0 | 0 | 0 | 1013100 | 0 | 0 | 0 | 431200 | 0 | 0 |
| GPPPSGIA | DCTN1 | E7EX90 | E7EX90_HUMA | Unmodified | 0 | 0 | 0 | 0 | 680920 | 80896 | 0 | 0 | 217730 | 74493 | 0 | 50520 | 527800 | 0 | 0 | 0 | 144460 | 224870 |
| GPPPTDPY | CPSF6; CPSF6 | Q16630; F8 | CPSF6_HUMAN | Unmodified | 1194500 | 465690 | 0 | 0 | 0 | 0 | 0 | 0 | 0 | 0 | 0 | 0 | 0 | 0 | 181780 | 0 | 0 | 0 |
| GPPQEEEE | HDGF | P51858 | HDGF_HUMAN | Unmodified | 0 | 0 | 0 | 0 | 0 | 0 | 0 | 0 | 0 | 0 | 1311000 | 172030 | 0 | 0 | 0 | 0 | 0 | 0 |
| GPPQQHG | FNBP1L; FNBP1 | Q5T0N5; A0 | FBP1L_HUMAN | Unmodified | 1325300 | 949480 | 0 | 0 | 0 | 0 | 0 | 0 | 0 | 0 | 0 | 0 | 0 | 0 | 0 | 0 | 0 | 0 |
| GPPQSPVF | ATXN2L; ATXN2 | Q8WWM7; F | ATX2L_HUMAN | Unmodified | 0 | 0 | 0 | 0 | 0 | 0 | 0 | 0 | 0 | 0 | 0 | 0 | 0 | 0 | 0 | 0 | 0 | 0 |
| GPPSTPVP | COBL; COBL; C | O75128; J3K | COBL_HUMAN | Unmodified | 263310 | 252600 | 0 | 0 | 0 | 0 | 0 | 89535 | 0 | 88315 | 0 | 300970 | 0 | 0 | 0 | 0 | 0 | 0 |
| GPPVRLFTI | NENF | Q9UMX5 | NENF_HUMAN | Unmodified | 0 | 0 | 0 | 0 | 0 | 0 | 0 | 0 | 0 | 2007200 | 0 | 0 | 0 | 0 | 0 | 0 | 0 | 0 |
| GPQGYGFI | SLC9A3R2; SLC | Q15599; H3 | NHRF2_HUMAI | Unmodified | 360520 | 374000 | 922800 | 1180100 | 2489200 | 770220 | 93661 | 0 | 977270 | 180960 | 405600 | 2630600 | 3724300 | 1855800 | 1207900 | 821170 | 1363600 | 2231400 |
| GPQITGPS | AHNAK | Q09666 | AHNK_HUMAN | Unmodified | 0 | 0 | 0 | 0 | 2830200 | 102820 | 0 | 0 | 0 | 0 | 0 | 1432200 | 0 | 0 | 0 | 16801 | 1726700 | 287900 |
| GPQLFHMI | PSMA5 | P28066 | PSA5_HUMAN | Unmodified | 51340 | 0 | 131840 | 0 | 0 | 153060 | 0 | 0 | 920920 | 0 | 0 | 1468000 | 0 | 0 | 0 | 74642 | 0 | 0 |
| GPQLFHMI | PSMA5 | P28066 | PSA5_HUMAN | Oxidation (M) | 176790 | 78662 | 755170 | 360700 | 0 | 0 | 0 | 0 | 0 | 0 | 0 | 7952000 | 95621 | 0 | 669520 | 794180 | 0 | 0 |
| GPQVSGEI | AHNAK | Q09666 | AHNK_HUMAN | Unmodified | 0 | 0 | 0 | 0 | 976670 | 0 | 0 | 0 | 0 | 0 | 0 | 0 | 139150 | 0 | 0 | 0 | 0 | 435020 |
| GPQYGSLE | PACSIN1; PACS | Q9BY11; F6 | PACN1_HUMAI | Unmodified | 13718000 | 7358100 | 0 | 0 | 0 | 86011 | 0 | 35247 | 1612600 | 1359900 | 91347 | 94373 | 428640 | 0 | 3052400 | 1063600 | 112190 | 176560 |
| GPQYGTLE | PACSIN3; PACS | Q9UKS6; E9 | PACN3_HUMAI | Unmodified | 13358000 | 10593000 | 7074100 | 4043000 | 2874700 | 0 | 0 | 0 | 577280 | 44966 | 3505600 | 214130 | 5226000 | 2427400 | 9864500 | 5957800 | 583700 | 1709800 |
| GPQYGTVE | PACSIN2; PACS | Q9UNF0; B0 | PACN2_HUMAI | Unmodified | 28505000 | 24917000 | 20738000 | 22443000 | 3292400 | 89641 | 0 | 65629 | 255820 | 77134 | 546970 | 0 | 34796000 | 5944600 | 38851000 | 35702000 | 159270 | 967940 |
| GPRISAPN | AHNAK | Q09666 | AHNK_HUMAN | Unmodified | 0 | 0 | 0 | 0 | 4605600 | 565250 | 475460 | 280740 | 0 | 0 | 0 | 0 | 0 | 0 | 0 | 0 | 3891500 | 3065900 |
| GPRISAPN | AHNAK | Q09666 | AHNK_HUMAN | Unmodified | 0 | 0 | 0 | 0 | 2382500 | 0 | 0 | 0 | 0 | 0 | 0 | 0 | 0 | 0 | 0 | 0 | 2724400 | 0 |
| GPRLDFEG | AHNAK | Q09666 | AHNK_HUMAN | Unmodified | 0 | 0 | 0 | 0 | 1051200 | 0 | 0 | 0 | 0 | 0 | 0 | 0 | 0 | 0 | 0 | 0 | 130080 | 614490 |
| GPRLDFEG | AHNAK | Q09666 | AHNK_HUMAN | Unmodified | 0 | 0 | 0 | 0 | 1312300 | 400430 | 0 | 0 | 0 | 0 | 0 | 0 | 0 | 0 | 0 | 0 | 283610 | 373480 |
| GPSAAALA | SLC22A3 | O75751 | S22A3_HUMAN | Unmodified | 0 | 0 | 0 | 0 | 0 | 0 | 0 | 0 | 0 | 0 | 0 | 0 | 0 | 0 | 0 | 1519100 | 0 | 0 |
| GPSAAGE | METAP2; METAI | P50579; F8V | MAP2_HUMAN | Unmodified | 0 | 0 | 0 | 0 | 0 | 0 | 0 | 0 | 0 | 0 | 627850 | 434040 | 0 | 0 | 0 | 0 | 0 | 0 |
| GPSASAPS | NCAM1 | A0A0D9SF3( | A0A0D9SF30_I | Unmodified | 0 | 0 | 0 | 0 | 0 | 0 | 0 | 0 | 583170 | 0 | 0 | 186750 | 0 | 0 | 0 | 0 | 0 | 0 |
| GPSDFNGI | IST1 | P53990 | IST1_HUMAN IS | Oxidation (M) | 0 | 0 | 0 | 3082200 | 14984000 | 0 | 0 | 0 | 6069200 | 784030 | 0 | 2234200 | 0 | 0 | 0 | 0 | 0 | 1820900 |
| GPSDFNGI | IST1 | P53990 | IST1_HUMAN IS | Unmodified | 0 | 0 | 0 | 0 | 0 | 0 | 0 | 0 | 1981900 | 0 | 185280 | 0 | 0 | 0 | 0 | 0 | 0 | 0 |
| GPSFDVQV | DPP3; DPP3; DI | Q9NY33; G3 | DPP3_HUMAN | Unmodified | 0 | 0 | 0 | 395700 | 0 | 0 | 0 | 0 | 306980 | 123070 | 0 | 1752300 | 0 | 0 | 302870 | 124980 | 0 | 0 |
| GPSFNQEF | PRRC2C; PRRC | Q9Y520; A0 | PRC2C_HUMAI | Unmodified | 112270 | 0 | 67293 | 0 | 0 | 0 | 0 | 0 | 0 | 55632 | 0 | 241260 | 0 | 0 | 0 | 49858 | 0 | 0 |
| GPSGCVES | QARS1; QARS1 | P47897; H7 | SYQ_HUMAN ( | Unmodified | 0 | 0 | 0 | 0 | 0 | 0 | 0 | 0 | 0 | 0 | 0 | 849050 | 0 | 0 | 0 | 0 | 0 | 0 |
| GPSGLLYI | AGRN; AGRN; A | O00468; A0 | AGRIN_HUMAN | Unmodified | 0 | 0 | 683780 | 0 | 0 | 0 | 0 | 0 | 0 | 0 | 0 | 0 | 0 | 0 | 0 | 0 | 0 | 0 |
| GPSGLLVY | CYB5R3; CYB5I | P00387; B1A | NB5R3_HUMAI | Unmodified | 0 | 0 | 0 | 0 | 0 | 0 | 0 | 0 | 0 | 0 | 0 | 0 | 0 | 0 | 0 | 0 | 0 | 0 |

| | | | | | | | | | | | | | | | | | | | | | | |
|---|---|---|---|---|---|---|---|---|---|---|---|---|---|---|---|---|---|---|---|---|---|---|
| GPSGQGQ | MOGS; MOGS; | Q13724; A0/ | MOGS_HUMAN | Unmodified | 0 | 0 | 0 | 0 | 0 | 0 | 0 | 0 | 821940 | 0 | 0 | 0 | 0 | 0 | 0 | 0 | 0 | 0 |
| GPSGYGFN | SLC9A3R1 | O14745 | NHRF1_HUMAI | Unmodified | 0 | 0 | 282320 | 323050 | 1627800 | 555720 | 384240 | 371590 | 364860 | 286330 | 193660 | 1397500 | 5333000 | 5248900 | 415810 | 467170 | 955540 | 559360 |
| GPSHPLDL | SORBS3; SORB | O60504; H0\ | VINEX_HUMAN | Unmodified | 2982900 | 523840 | 0 | 0 | 0 | 0 | 0 | 0 | 0 | 0 | 0 | 0 | 0 | 0 | 168400 | 0 | 0 | 0 |
| GPSLKGDL | AHNAK | Q09666 | AHNK_HUMAN | Oxidation (M) | 0 | 0 | 0 | 0 | 1482900 | 1033300 | 0 | 160430 | 0 | 0 | 0 | 0 | 0 | 0 | 0 | 0 | 198430 | 881130 |
| GPSLKGDL | AHNAK | Q09666 | AHNK_HUMAN | Unmodified | 44122 | 0 | 0 | 0 | 4496800 | 0 | 0 | 0 | 275160 | 0 | 0 | 0 | 124860 | 0 | 55852 | 0 | 938300 | 6762700 |
| GPSRASSV | CRIP2; CRIP2 | P52943; H0\ | CRIP2_HUMAN | Unmodified | 0 | 0 | 0 | 0 | 0 | 0 | 0 | 0 | 0 | 156390 | 0 | 0 | 0 | 0 | 0 | 0 | 0 | 378140 |
| GPSRASSV | CRIP2; CRIP2 | P52943; H0\ | CRIP2_HUMAN | Unmodified | 0 | 0 | 0 | 0 | 0 | 0 | 253100 | 0 | 0 | 0 | 0 | 0 | 0 | 0 | 0 | 0 | 597000 | 0 |
| GPSSVEDII | NPM1; NPM1; N | P06748; A0A | NPM_HUMAN I | Unmodified | 0 | 0 | 784620 | 799110 | 933050 | 0 | 0 | 17997 | 3983500 | 792750 | 9642500 | 11723000 | 0 | 0 | 713230 | 453650 | 107660 | 373990 |
| GPSTEYGF | TARS2; TARS2; | Q9BW92; F6 | SYTM_HUMAN | Unmodified | 0 | 0 | 0 | 0 | 0 | 0 | 0 | 0 | 318070 | 0 | 0 | 0 | 0 | 0 | 0 | 0 | 0 | 1406300 |
| GPSTNPDS | EPB41 | A0A2R8Y78 | A0A2R8Y783_H | Unmodified | 0 | 0 | 0 | 0 | 0 | 0 | 0 | 0 | 1380500 | 0 | 0 | 0 | 0 | 0 | 0 | 0 | 250060 | 0 |
| GPSTVTDLI | AP2A2; AP2A2; | O94973; A0/ | AP2A2_HUMAN | Unmodified | 29059 | 0 | 0 | 45961 | 0 | 0 | 0 | 0 | 0 | 0 | 0 | 703250 | 0 | 0 | 103290 | 59480 | 0 | 0 |
| GPSWDPFI | HSPB1; HSPB1; | P04792; A0A | HSPB1_HUMAI | Unmodified | 2160900 | 961870 | 3245000 | 3162300 | 0 | 0 | 0 | 0 | 5143300 | 344840 | 0 | 0 | 586790 | 0 | 5666500 | 3422200 | 0 | 0 |
| GPSYGLSA | CNN2; CNN3; ( | Q99439; Q1! | CNN2_HUMAN | Unmodified | 0 | 0 | 0 | 0 | 4873000 | 0 | 0 | 0 | 0 | 0 | 0 | 0 | 0 | 0 | 0 | 0 | 0 | 0 |
| GPTEADELI | G6PD | P11413 | G6PD_HUMAN | Unmodified | 0 | 0 | 0 | 0 | 89917 | 0 | 0 | 0 | 0 | 147300 | 0 | 1327700 | 0 | 0 | 0 | 0 | 0 | 0 |
| GPTEQLVS | PPM1A | P35813 | PPM1A_HUMAI | Unmodified | 205600 | 119600 | 0 | 0 | 0 | 0 | 0 | 0 | 0 | 0 | 85152 | 0 | 117360 | 0 | 1683900 | 1269100 | 216160 | 0 |
| GPTISLTQI | ATP6AP1; ATP6. | Q15904; A0/ | VAS1_HUMAN | Unmodified | 0 | 0 | 127240 | 0 | 0 | 32924 | 0 | 0 | 0 | 0 | 76103 | 394870 | 0 | 0 | 0 | 0 | 0 | 0 |
| GPTMPDEL | GPSM1; GPSM1 | Q86YR5; A0/ | GPSM1_HUMA | Oxidation (M) | 28264 | 0 | 0 | 0 | 0 | 0 | 0 | 0 | 174050 | 0 | 0 | 105370 | 0 | 0 | 68551 | 26948 | 0 | 0 |
| GPVAGWIN | CNDP2; CNDP2 | Q96KP4; A0/ | CNDP2_HUMA | Unmodified | 24241 | 0 | 847270 | 97051 | 0 | 0 | 0 | 0 | 736190 | 0 | 693550 | 4829900 | 264100 | 0 | 4245300 | 1331400 | 0 | 0 |
| GPVDAVQK | PLXNB2 | O15031 | PLXB2_HUMAN | Unmodified | 98061 | 70343 | 644460 | 0 | 0 | 0 | 0 | 0 | 290510 | 11702 | 0 | 861230 | 0 | 0 | 685930 | 441690 | 0 | 0 |
| GPVEAKPE | NCAM1; NCAM | P13591; A0A | NCAM1_HUMA | Unmodified | 0 | 0 | 0 | 0 | 0 | 0 | 80403 | 96180 | 8626200 | 1402400 | 208630 | 5040200 | 0 | 0 | 0 | 0 | 0 | 0 |
| GPVEGYEE | TRMT112; TRM1 | Q9UI30; F5G | TR112_HUMAN | Unmodified | 279350 | 0 | 338370 | 133710 | 0 | 0 | 0 | 0 | 0 | 0 | 0 | 1532200 | 0 | 0 | 462140 | 348830 | 0 | 0 |
| GPVGDSHI | DIP2B | Q9P265 | DIP2B_HUMAN | Unmodified | 0 | 0 | 0 | 0 | 0 | 2613300 | 1637000 | 0 | 740320 | 0 | 0 | 1466900 | 0 | 0 | 0 | 0 | 0 | 0 |
| GPVGLAIDI | LRP1 | Q07954 | LRP1_HUMAN | Unmodified | 0 | 0 | 0 | 0 | 0 | 0 | 0 | 0 | 1945300 | 0 | 0 | 0 | 0 | 0 | 0 | 0 | 0 | 0 |
| GPVGLEGL | ALDH18A1 | P54886 | P5CS_HUMAN | Unmodified | 0 | 0 | 0 | 0 | 0 | 0 | 453640 | 0 | 1251800 | 127950 | 0 | 0 | 0 | 0 | 0 | 0 | 0 | 0 |
| GPVGTVSE | MYOF | Q9NZM1 | MYOF_HUMAN | Unmodified | 0 | 0 | 0 | 0 | 17909000 | 3847800 | 0 | 0 | 109570 | 123220 | 0 | 326060 | 0 | 0 | 0 | 0 | 167180 | 2182300 |
| GPVGTVSE | MYOF | Q9NZM1 | MYOF_HUMAN | Unmodified | 0 | 60176 | 0 | 0 | 2836500 | 446290 | 0 | 0 | 0 | 0 | 0 | 0 | 0 | 0 | 0 | 0 | 0 | 221480 |
| GPVGTVSE | MYOF | Q9NZM1 | MYOF_HUMAN | Unmodified | 0 | 0 | 0 | 0 | 6219300 | 274490 | 0 | 0 | 0 | 0 | 0 | 0 | 0 | 0 | 0 | 0 | 0 | 506420 |
| GPVHIVHS | EMC1; EMC1 | Q8N766; H7 | EMC1_HUMAN | Unmodified | 0 | 0 | 0 | 0 | 392310 | 0 | 0 | 0 | 3748400 | 653130 | 381880 | 0 | 0 | 0 | 0 | 0 | 0 | 0 |
| GPVKPTGG | DDX6; DDX6 | P26196; Q8I | DDX6_HUMAN | Unmodified | 0 | 0 | 0 | 0 | 0 | 0 | 0 | 0 | 295560 | 0 | 608300 | 1615800 | 0 | 0 | 0 | 0 | 0 | 0 |
| GPVKPTGG | DDX6; DDX6 | P26196; Q8I | DDX6_HUMAN | Oxidation (M) | 125250 | 155250 | 314240 | 0 | 0 | 0 | 0 | 0 | 0 | 0 | 162600 | 344440 | 0 | 0 | 264070 | 220330 | 0 | 0 |
| GPVREGD\ | RPS28 | P62857 | RS28_HUMAN | Unmodified | 0 | 0 | 0 | 0 | 0 | 0 | 0 | 0 | 0 | 0 | 1271400 | 2157000 | 0 | 0 | 0 | 202320 | 0 | 0 |
| GPVREGD\ | RPS28 | P62857 | RS28_HUMAN | Unmodified | 9715700 | 6825600 | 4581700 | 8223800 | 3701400 | 1044900 | 321710 | 89238 | 5136200 | 218690 | 2500500 | 22473000 | 6991400 | 1465900 | 5483200 | 4554100 | 1504900 | 3842300 |
| GPVTDVAY | WDR1 | O75083 | WDR1_HUMAN | Unmodified | 1177600 | 260100 | 0 | 746100 | 836320 | 0 | 0 | 36814 | 0 | 0 | 1576200 | 5282700 | 0 | 0 | 687770 | 579360 | 0 | 0 |
| GPVTPLHLI | PLEKHN1 | Q494U1 | PKHN1_HUMAI | Unmodified | 0 | 0 | 0 | 0 | 0 | 0 | 0 | 0 | 0 | 0 | 0 | 0 | 0 | 0 | 538110 | 755220 | 0 | 0 |
| GPVYHEYC | POGLUT3 | Q7Z4H8 | PLGT3_HUMAN | Unmodified | 0 | 0 | 0 | 0 | 0 | 0 | 0 | 0 | 1781200 | 834320 | 0 | 0 | 0 | 0 | 0 | 0 | 0 | 0 |
| GPVYIGELF | TOLLIP; TOLLIP; | Q9H0E2; F2 | TOLIP_HUMAN | Unmodified | 19490000 | 6953200 | 8812600 | 13554000 | 0 | 0 | 0 | 0 | 4821400 | 180710 | 877070 | 3759800 | 0 | 0 | 29217000 | 39044000 | 0 | 31275 |
| GPVYTEQP | DLL3; DLL3 | Q9NYJ7; M0I | DLL3_HUMAN | Unmodified | 0 | 0 | 0 | 0 | 0 | 0 | 0 | 0 | 940960 | 0 | 0 | 0 | 0 | 0 | 0 | 0 | 0 | 0 |
| GPWTCVGI | DNASE2 | O00115 | DNS2A_HUMAI | Unmodified | 0 | 0 | 0 | 130130 | 0 | 0 | 231460 | 88447 | 1415100 | 109670 | 217750 | 0 | 0 | 0 | 0 | 0 | 0 | 0 |
| GPWTCVGI | DNASE2 | O00115 | DNS2A_HUMAI | Unmodified | 0 | 0 | 0 | 0 | 0 | 0 | 0 | 0 | 2311600 | 0 | 0 | 95638 | 0 | 0 | 0 | 0 | 0 | 0 |
| GPYPLGIDI | ATP6V0A2 | Q9Y487 | VPP2_HUMAN | Unmodified | 0 | 0 | 0 | 0 | 0 | 0 | 0 | 0 | 111820 | 63329 | 0 | 130070 | 0 | 0 | 28852 | 0 | 0 | 0 |
| GQAAVQQI | CBR3; CBR1; C | O75828; P16 | CBR3_HUMAN | Unmodified | 432530 | 0 | 255880 | 1064300 | 115770 | 0 | 0 | 0 | 0 | 0 | 0 | 4312000 | 0 | 0 | 1179000 | 1193100 | 224980 | 0 |
| GQATDIAIC | CLPP; CLPP | Q16740; M0I | CLPP_HUMAN | Oxidation (M) | 0 | 189300 | 0 | 0 | 0 | 562930 | 0 | 0 | 728190 | 0 | 69571 | 0 | 0 | 0 | 0 | 0 | 238290 | 0 |
| GQATDIAIC | CLPP; CLPP | Q16740; M0I | CLPP_HUMAN | Unmodified | 0 | 0 | 0 | 0 | 0 | 0 | 0 | 0 | 780200 | 0 | 8593.5 | 0 | 0 | 0 | 0 | 0 | 0 | 0 |
| GQAVDYEC | PDIA4; PDIA4 | P13667; A0A | PDIA4_HUMAN | Unmodified | 193280 | 183660 | 421540 | 370090 | 1200100 | 173250 | 4164200 | 3931400 | 22643000 | 11960000 | 3395000 | 2153500 | 901870 | 1136400 | 640130 | 0 | 311780 | 732070 |
| GQCAKVSS | MYO1C; MYO1( | O00159; F5I | MYO1C_HUMA | Unmodified | 0 | 0 | 0 | 0 | 544980 | 0 | 0 | 0 | 0 | 0 | 0 | 0 | 469350 | 0 | 0 | 0 | 289470 | 0 |
| GQCDLELII | CLTC; CLTC | Q00610; A0/ | CLH1_HUMAN | Unmodified | 200740 | 0 | 98947 | 0 | 0 | 0 | 0 | 0 | 373480 | 0 | 0 | 8401600 | 0 | 0 | 0 | 0 | 0 | 0 |
| GQDCIWG | SEZ6; SEZ6; SE | Q53EL9; C9J | SEZ6_HUMAN | Unmodified | 0 | 0 | 0 | 0 | 0 | 0 | 0 | 0 | 1296400 | 0 | 0 | 0 | 0 | 0 | 0 | 0 | 0 | 0 |
| GQDIFQR | DNAJC13 | O75165 | DJC13_HUMAN | Unmodified | 0 | 0 | 0 | 0 | 0 | 0 | 0 | 0 | 0 | 184180 | 0 | 1315000 | 0 | 0 | 0 | 0 | 0 | 0 |
| GQDIRIPTP | CKMT1A | P12532 | KCRU_HUMAN | Unmodified | 0 | 59580 | 0 | 17354 | 0 | 0 | 0 | 0 | 0 | 0 | 0 | 1446200 | 0 | 0 | 63181 | 67669 | 0 | 0 |
| GQDMETE/ | TMED2; TMED2 | Q15363; F5( | TMED2_HUMAI | Unmodified | 0 | 0 | 0 | 0 | 0 | 0 | 0 | 0 | 208340 | 0 | 0 | 0 | 0 | 0 | 0 | 0 | 0 | 0 |
| GQDPLADI | MVP | Q14764 | MVP_HUMAN I | Unmodified | 245770 | 89334 | 0 | 0 | 0 | 0 | 0 | 0 | 265170 | 0 | 0 | 7226800 | 0 | 0 | 0 | 0 | 97768 | 0 |
| GQDSIEPVI | EIF2B3 | Q9NR50 | EI2BG_HUMAN | Unmodified | 0 | 0 | 0 | 0 | 0 | 0 | 0 | 0 | 0 | 0 | 0 | 714720 | 0 | 0 | 0 | 0 | 0 | 0 |
| GQDVGRY( | SSR4 | P51571 | SSRD_HUMAN | Unmodified | 0 | 0 | 0 | 0 | 0 | 0 | 0 | 0 | 23846000 | 1614900 | 1214200 | 445990 | 0 | 0 | 0 | 0 | 0 | 0 |
| GQEAFVPG | CA12; CA12 | O43570; B3I | CAH12_HUMAI | Unmodified | 276980 | 100560 | 0 | 60594 | 0 | 0 | 0 | 0 | 0 | 0 | 0 | 0 | 11143 | 0 | 70015 | 147470 | 0 | 0 |
| GQEDAILS\ | DLG3; DLG3 | Q92796; Q5. | DLG3_HUMAN | Unmodified | 0 | 0 | 0 | 0 | 0 | 0 | 0 | 0 | 0 | 0 | 0 | 820260 | 0 | 0 | 0 | 0 | 0 | 0 |
| GQEGSVCI | DKK1 | O94907 | DKK1_HUMAN | Unmodified | 0 | 0 | 0 | 0 | 0 | 0 | 0 | 0 | 843740 | 0 | 0 | 0 | 0 | 0 | 0 | 0 | 221160 | 41176 |
| GQENNVFI | MGAM2 | Q2M2H8 | MGAL_HUMAN | Unmodified | 0 | 0 | 0 | 0 | 0 | 0 | 0 | 0 | 0 | 0 | 0 | 1255700 | 0 | 3986700 | 0 | 0 | 0 | 0 |
| GQEPLGPC | TCF25; TCF25; | Q9BQ70; H3 | TCF25_HUMAN | Unmodified | 0 | 0 | 95563 | 0 | 0 | 0 | 0 | 0 | 0 | 0 | 0 | 0 | 0 | 0 | 348370 | 73253 | 0 | 0 |
| GQEPLGPC | TCF25; TCF25; | Q9BQ70; H3 | TCF25_HUMAN | Unmodified | 399120 | 504250 | 329650 | 157670 | 0 | 0 | 0 | 0 | 0 | 0 | 0 | 0 | 0 | 0 | 370850 | 273620 | 0 | 0 |
| GQESAGIV | PPAT; PPAT; PP. | Q06203; D6I | PUR1_HUMAN | Unmodified | 0 | 0 | 0 | 0 | 0 | 0 | 0 | 0 | 0 | 0 | 762770 | 5437900 | 0 | 0 | 0 | 213530 | 0 | 0 |
| GQETSTNP | IDH1 | O75874 | IDHC_HUMAN | Unmodified | 285210 | 0 | 647860 | 1652100 | 4013000 | 480620 | 446950 | 0 | 2637200 | 1969900 | 879400 | 2886800 | 12680000 | 8983000 | 1010000 | 2294000 | 1049900 | 934870 |
| GQETSTNP | IDH1 | O75874 | IDHC_HUMAN | Unmodified | 0 | 0 | 0 | 0 | 453560 | 0 | 292070 | 0 | 0 | 441670 | 0 | 0 | 2036200 | 2515600 | 0 | 0 | 314860 | 0 |
| GQFAALPG | AVIL | O75366 | AVIL_HUMAN A | Unmodified | 0 | 0 | 0 | 0 | 0 | 0 | 0 | 0 | 0 | 0 | 2588000 | 31374000 | 0 | 0 | 0 | 0 | 0 | 0 |
| GQFAEHLL | DHX36; DHX36 | Q9H2U1; A0 | DHX36_HUMAI | Unmodified | 171690 | 179350 | 53916 | 0 | 0 | 47804 | 0 | 0 | 0 | 0 | 120670 | 1685500 | 0 | 0 | 286020 | 384620 | 0 | 0 |
| GQFETYLR | ASAH1; ASAH1; | Q13510; A0/ | ASAH1_HUMAI | Unmodified | 0 | 0 | 505810 | 297020 | 0 | 0 | 0 | 0 | 4409000 | 298560 | 508970 | 1391500 | 0 | 0 | 336410 | 296480 | 0 | 0 |
| GQFNFDHI | UCK2; UCK2 | Q9BZX2; A0/ | UCK2_HUMAN | Unmodified | 0 | 0 | 0 | 0 | 757810 | 0 | 236240 | 0 | 0 | 0 | 0 | 364880 | 762510 | 0 | 0 | 104030 | 1234500 | 0 |
| GQFSTDEL | CLTCL1; CLTC; | P53675; Q0( | CLH2_HUMAN | Unmodified | 4925200 | 12263000 | 1854800 | 1812600 | 0 | 0 | 0 | 0 | 0 | 0 | 0 | 3214300 | 0 | 0 | 5850700 | 5850500 | 0 | 0 |
| GQFSTDEL | CLTCL1; CLTC; | P53675; Q0( | CLH2_HUMAN | Unmodified | 10475000 | 3964800 | 4875500 | 2134000 | 0 | 0 | 0 | 0 | 0 | 511400 | 423020 | 40101000 | 0 | 0 | 2471500 | 6066800 | 0 | 0 |
| GQGAPARI | CFAP298; CFAF | P57076; H7( | CF298_HUMAN | Unmodified | 0 | 0 | 0 | 0 | 123250 | 0 | 0 | 0 | 90226 | 0 | 0 | 335610 | 200880 | 0 | 0 | 108980 | 0 | 85606 |
| GQGLTGPT | EHHADH | Q08426 | ECHP_HUMAN | Unmodified | 0 | 0 | 0 | 0 | 0 | 0 | 0 | 0 | 979260 | 0 | 0 | 0 | 0 | 0 | 0 | 0 | 0 | 0 |
| GQGLTGPT | EHHADH | Q08426 | ECHP_HUMAN | Unmodified | 0 | 0 | 0 | 0 | 0 | 0 | 0 | 0 | 1287300 | 52859 | 0 | 0 | 0 | 0 | 0 | 0 | 0 | 0 |
| GQGSSPV/ | VCL | P18206 | VINC_HUMAN | Unmodified | 0 | 0 | 0 | 0 | 306500 | 0 | 0 | 0 | 1235300 | 146550 | 0 | 1981900 | 0 | 0 | 0 | 0 | 869050 | 783760 |
| GQGSVSAS | SEC13 | P55735 | SEC13_HUMAN | Unmodified | 0 | 0 | 0 | 0 | 0 | 0 | 0 | 0 | 0 | 0 | 28361 | 1812600 | 0 | 0 | 0 | 44330 | 0 | 0 |
| GQGVYLGI | ERP29; ERP29 | P30040; F8V | ERP29_HUMAN | Oxidation (M) | 0 | 0 | 0 | 0 | 0 | 0 | 0 | 0 | 3066900 | 1031900 | 0 | 0 | 0 | 0 | 0 | 0 | 0 | 0 |
| GQGVYLGI | ERP29; ERP29 | P30040; F8V | ERP29_HUMAN | Oxidation (M) | 0 | 0 | 0 | 0 | 0 | 0 | 0 | 0 | 0 | 949840 | 0 | 0 | 0 | 0 | 0 | 0 | 0 | 0 |
| GQHEMSEI | DSG2 | Q14126 | DSG2_HUMAN | Unmodified | 0 | 0 | 0 | 0 | 0 | 0 | 0 | 0 | 0 | 0 | 265840 | 0 | 0 | 0 | 0 | 0 | 0 | 0 |
| GQHEMSEI | DSG2 | Q14126 | DSG2_HUMAN | Unmodified | 0 | 0 | 0 | 496650 | 2794400 | 687960 | 0 | 0 | 1438400 | 87272 | 828010 | 1299900 | 0 | 0 | 0 | 0 | 0 | 0 |
| GQHEVHG | SDF2L1 | Q9HCN8 | SDF2L_HUMAN | Unmodified | 0 | 0 | 0 | 0 | 0 | 0 | 0 | 0 | 661610 | 0 | 128090 | 0 | 0 | 0 | 0 | 0 | 0 | 0 |
| GQHSISYTI | PELI2; PELI3; PI | Q9HAT8; Q8 | PELI2_HUMAN | Unmodified | 0 | 0 | 0 | 0 | 0 | 0 | 0 | 0 | 440780 | 0 | 0 | 308050 | 0 | 0 | 0 | 0 | 0 | 0 |
| GQHVPGSI | FLNC; FLNA; FL | Q14315; P21 | FLNC_HUMAN | Unmodified | 0 | 2391500 | 0 | 0 | 0 | 0 | 0 | 0 | 0 | 0 | 0 | 0 | 0 | 0 | 1977400 | 945270 | 0 | 0 |
| GQIELTGK | AGPS; AGPS; A( | O00116; A0/ | ADAS_HUMAN | Unmodified | 87246 | 83171 | 166030 | 456580 | 494030 | 283260 | 278880 | 202490 | 5669000 | 1292700 | 2273500 | 0 | 123410 | 0 | 155100 | 121540 | 0 | 0 |
| GQIIGNFQ. | DSG2 | Q14126 | DSG2_HUMAN | Unmodified | 1183600 | 294250 | 356260 | 1672100 | 0 | 0 | 0 | 0 | 10466000 | 1327400 | 605010 | 2133100 | 0 | 0 | 202050 | 518770 | 0 | 0 |
| GQIKEIGW | PFKP; PFKP; PF | Q01813; A0/ | PFKAP_HUMAN | Unmodified | 0 | 0 | 0 | 0 | 1463200 | 0 | 258170 | 0 | 0 | 0 | 0 | 0 | 0 | 0 | 0 | 0 | 2680500 | 1237800 |
| GQILMPNI( | RPL32; RPL32; I | P62910; D3Y | RL32_HUMAN ( | Oxidation (M) | 0 | 0 | 0 | 0 | 0 | 0 | 0 | 0 | 0 | 0 | 0 | 845820 | 0 | 0 | 0 | 0 | 0 | 0 |
| GQILMPNI( | RPL32; RPL32; I | P62910; D3Y | RL32_HUMAN ( | Oxidation (M) | 1811900 | 0 | 0 | 0 | 393460 | 0 | 0 | 61626 | 0 | 0 | 131010 | 2230300 | 369760 | 0 | 0 | 0 | 0 | 0 |
| GQILMPNI( | RPL32; RPL32; I | P62910; D3Y | RL32_HUMAN ( | Unmodified | 1124000 | 1158800 | 642190 | 656080 | 0 | 0 | 0 | 0 | 601070 | 0 | 316190 | 7211200 | 0 | 0 | 867200 | 420440 | 0 | 0 |
| GQILNLTQ. | PI4K2A; N/A | Q9BTU6; E9I | P4K2A_HUMAN | Unmodified | 0 | 0 | 0 | 0 | 0 | 0 | 0 | 0 | 388300 | 0 | 0 | 685280 | 0 | 0 | 0 | 0 | 0 | 0 |
| GQILWFR | ATP2B2; ATP2B | Q01814; Q1( | AT2B2_HUMAN | Unmodified | 0 | 0 | 329530 | 372120 | 0 | 0 | 0 | 0 | 0 | 0 | 0 | 0 | 0 | 0 | 1144100 | 1538800 | 0 | 0 |
| GQINNSIKI | SERPINB5 | P36952 | SPB5_HUMAN | Unmodified | 0 | 0 | 0 | 0 | 0 | 627630 | 494280 | 0 | 131930 | 0 | 0 | 0 | 2122200 | 0 | 0 | 0 | 1583200 | 2669600 |
| GQIPEIKQT | VBP1 | P61758 | PFD3_HUMAN | Unmodified | 0 | 0 | 0 | 0 | 3908500 | 0 | 556460 | 222680 | 0 | 2751900 | 104790 | 0 | 4132700 | 3251500 | 0 | 0 | 2356400 | 3490400 |
| GQIQAVAS | TCP11L1; TCP1 | Q9NUJ3; E9I | T11L1_HUMAN | Unmodified | 0 | 0 | 0 | 0 | 0 | 0 | 0 | 0 | 248640 | 0 | 0 | 478010 | 0 | 0 | 0 | 0 | 0 | 0 |
| GQIQEIVLV | HSPA2 | P54652 | HSP72_HUMAI | Unmodified | 533640 | 585680 | 0 | 0 | 0 | 0 | 0 | 0 | 0 | 0 | 0 | 1295800 | 0 | 0 | 0 | 0 | 0 | 0 |
| GQIQEIVLV | HSPA2 | P54652 | HSP72_HUMAI | Unmodified | 245680 | 408620 | 0 | 0 | 2814500 | 0 | 0 | 0 | 0 | 0 | 0 | 0 | 0 | 0 | 0 | 0 | 252160 | 877580 |
| GQKCEFQI | HSPD1; HSPD1 | P10809; A0A | CH60_HUMAN | Unmodified | 0 | 0 | 0 | 0 | 0 | 0 | 0 | 0 | 340450 | 1125300 | 0 | 0 | 0 | 1526600 | 0 | 0 | 0 | 0 |
| GQKSPGAL | AKAP2; PALM2/ | Q9Y2D5; C9. | AKAP2_HUMAN | Unmodified | 0 | 0 | 0 | 0 | 492950 | 0 | 0 | 0 | 0 | 0 | 0 | 0 | 0 | 0 | 0 | 0 | 353020 | 0 |
| GQKSPGAL | AKAP2; PALM2/ | Q9Y2D5; C9. | AKAP2_HUMAN | Unmodified | 0 | 0 | 0 | 0 | 567380 | 0 | 0 | 0 | 0 | 0 | 0 | 0 | 0 | 0 | 0 | 0 | 380270 | 339260 |
| GQLEAARP | TNFAIP2 | Q03169 | TNAP2_HUMAN | Unmodified | 0 | 0 | 0 | 0 | 0 | 0 | 0 | 0 | 0 | 80685 | 62823 | 1582400 | 0 | 0 | 0 | 0 | 0 | 0 |
| GQLHHNP | ALPL | P05186 | PPBT_HUMAN | Unmodified | 339990 | 427950 | 0 | 0 | 0 | 0 | 0 | 0 | 2746700 | 694930 | 0 | 0 | 0 | 0 | 0 | 0 | 0 | 0 |
| GQLITGMD | FKBP9 | O95302 | FKBP9_HUMAN | Unmodified | 0 | 56850000 | 0 | 1100500 | 0 | 0 | 0 | 55432 | 5628100 | 0 | 0 | 249180 | 0 | 0 | 0 | 0 | 1054000 | 0 |
| GQLLAEQL | RAB3B | P20337 | RAB3B_HUMAN | Unmodified | 0 | 54824 | 0 | 69984 | 271740 | 0 | 0 | 0 | 26163 | 102230 | 0 | 0 | 0 | 0 | 0 | 0 | 0 | 0 |
| GQLLCPTC | LMCD1; LMCD1 | Q9NZU5; B4 | LMCD1_HUMA | Unmodified | 0 | 0 | 447160 | 0 | 224370 | 0 | 0 | 0 | 0 | 0 | 0 | 826870 | 32842 | 0 | 0 | 0 | 0 | 0 |
| GQLLIFGAI | RCC2 | Q9P258 | RCC2_HUMAN | Unmodified | 0 | 0 | 0 | 0 | 0 | 0 | 276320 | 0 | 545860 | 409500 | 2542800 | 3117000 | 0 | 0 | 0 | 0 | 0 | 0 |
| GQLNHIPS | PATJ; PATJ; PAT. | Q8NI35; A0A | INADL_HUMAN | Unmodified | 0 | 0 | 0 | 0 | 2219400 | 0 | 0 | 0 | 204420 | 0 | 0 | 0 | 1906600 | 4939000 | 0 | 92863 | 855630 | 1555000 |
| GQLPDYTS | DRG1; DRG1 | Q9Y295; H0\ | DRG1_HUMAN | Unmodified | 0 | 0 | 0 | 0 | 0 | 0 | 0 | 37641 | 0 | 0 | 0 | 1430000 | 0 | 0 | 0 | 0 | 0 | 0 |
| GQLPKPFA | ALG9; N/A; ALG | Q9H6U8; A0 | ALG9_HUMAN | Unmodified | 0 | 0 | 0 | 160320 | 1674800 | 0 | 0 | 139110 | 851630 | 353670 | 0 | 0 | 0 | 544390 | 0 | 0 | 0 | 375880 |
| GQLQELST | MACF1; MACF1 | Q9UPN3; H3 | MACF1_HUMAI | Unmodified | 0 | 0 | 0 | 0 | 377480 | 0 | 0 | 0 | 410000 | 171360 | 456080 | 0 | 486530 | 119340 | 0 | 0 | 629870 | 184680 |
| GQLQGLVS | KLK14; KLK14 | Q9P0G3; A0. | KLK14_HUMAN | Unmodified | 4624000 | 0 | 0 | 11995 | 444130 | 0 | 0 | 0 | 5614500 | 0 | 0 | 0 | 0 | 4784200 | 0 | 117000 | 0 | 0 |
| GQLTTDQV | ACADVL; ACAD | P49748; K7E | ACADV_HUMAI | Unmodified | 0 | 0 | 0 | 0 | 734880 | 0 | 1197400 | 0 | 20178000 | 5283600 | 0 | 0 | 0 | 0 | 0 | 0 | 8112500 | 679360 |
| GQLVLFHA | NAGPA; NAGPA | Q9UK23; K7I | NAGPA_HUMA | Unmodified | 0 | 0 | 0 | 0 | 0 | 0 | 381130 | 0 | 576410 | 0 | 143190 | 0 | 0 | 0 | 0 | 0 | 43777 | 0 |

| | | | | | | | | | | | | | | | | | | | | | | |
|---|---|---|---|---|---|---|---|---|---|---|---|---|---|---|---|---|---|---|---|---|---|---|
| GQLVPLET | AK1; AK1; AK1 | P00568; H0Y | KAD1_HUMAN | 2 Oxidation (M) | 0 | 0 | 0 | 0 | 0 | 0 | 0 | 0 | 0 | 292410 | 0 | 0 | 1051900 | 0 | 0 | 0 | 319940 | 0 |
| GQMQKPFI | PIN1; PIN1 | Q13526; K7E | PIN1_HUMAN | Oxidation (M) | 16704000 | 16835000 | 6021600 | 8090600 | 74102 | 0 | 0 | 0 | 1255700 | 0 | 678530 | 1988700 | 121890 | 46298 | 9756400 | 10156000 | 45072 | 0 |
| GQMQKPFI | PIN1; PIN1 | Q13526; K7E | PIN1_HUMAN | Unmodified | 3972400 | 3707800 | 528420 | 886870 | 0 | 0 | 0 | 0 | 0 | 0 | 27130 | 0 | 0 | 0 | 1292000 | 607680 | 0 | 0 |
| GQNALVEL | HBS1L; HBS1L; | Q9Y450; J3Q | HBS1L_HUMAN | Unmodified | 1310900 | 1403200 | 121450 | 845100 | 1901400 | 166900 | 430310 | 0 | 1419400 | 622310 | 356410 | 1968500 | 1039500 | 560360 | 703560 | 705180 | 0 | 713540 |
| GQNLTPYV | FAM234B; FAM | A2RU67; Q6 | F234B_HUMAN | Unmodified | 0 | 0 | 0 | 0 | 0 | 0 | 0 | 0 | 3023800 | 0 | 0 | 262620 | 0 | 0 | 0 | 0 | 0 | 0 |
| GQNSLALH | RALA; RALA; RA | P11233; H7C | RALA_HUMAN | Unmodified | 357110 | 1441100 | 1251500 | 1763300 | 1054600 | 249070 | 484510 | 343480 | 1679000 | 841970 | 499610 | 4113300 | 3592200 | 0 | 0 | 1899200 | 551230 | 1125000 |
| GQNSLALH | RALA; RALA; RA | P11233; H7C | RALA_HUMAN | Unmodified | 0 | 0 | 0 | 0 | 291100 | 0 | 336910 | 39827 | 0 | 0 | 0 | 0 | 650290 | 595590 | 0 | 0 | 713980 | 665570 |
| GQNVLLTE | TNIK; MINK1; M | Q9UKE5; Q8 | TNIK_HUMAN | Unmodified | 238200 | 0 | 765280 | 660080 | 319520 | 0 | 0 | 0 | 1640500 | 253990 | 1560200 | 3519000 | 240180 | 170200 | 1365200 | 578640 | 0 | 287090 |
| GQPDTVQI | PTPN23 | Q9H3S7 | PTN23_HUMAN | Unmodified | 959130 | 560880 | 0 | 0 | 0 | 0 | 0 | 0 | 0 | 0 | 0 | 0 | 0 | 0 | 0 | 0 | 0 | 0 |
| GQPDVVV | ACAT1; ACAT1; | P24752; A0A | THIL_HUMAN | Unmodified | 0 | 0 | 117230 | 76847 | 207020 | 0 | 0 | 0 | 4086500 | 0 | 117510 | 133070 | 279520 | 0 | 0 | 0 | 0 | 113380 |
| GQPGDIYH | GBA; GBA; GBA | P04062; A0A | GLCM_HUMAN | Unmodified | 0 | 0 | 0 | 0 | 0 | 0 | 0 | 0 | 3599800 | 436720 | 88273 | 0 | 0 | 0 | 0 | 37947 | 0 | 0 |
| GQPGHAYI | UTRN | P46939 | UTRN_HUMAN | Unmodified | 0 | 0 | 0 | 59126 | 1091900 | 0 | 0 | 0 | 88385 | 0 | 0 | 0 | 0 | 0 | 200060 | 150810 | 0 | 99729 |
| GQPIYIQFS | PTBP1; PTBP1; | P26599; A6N | PTBP1_HUMAN | Unmodified | 230400 | 0 | 155640 | 0 | 0 | 0 | 0 | 0 | 0 | 0 | 0 | 5233500 | 0 | 0 | 0 | 127350 | 0 | 0 |
| GQPLGPAC | NOMO1; NOMC | Q15155; P69 | NOMO1_HUMA | Unmodified | 0 | 0 | 0 | 0 | 0 | 0 | 0 | 0 | 5405800 | 251550 | 624340 | 0 | 0 | 0 | 0 | 0 | 0 | 0 |
| GQPLGPAC | NOMO1; NOMC | Q15155; P69 | NOMO1_HUMA | Unmodified | 0 | 0 | 0 | 0 | 382000 | 0 | 366870 | 0 | 1962700 | 3446900 | 700930 | 0 | 0 | 0 | 0 | 0 | 0 | 0 |
| GQPLQAAV | ERICH5 | Q6P6B1 | ERIC5_HUMAN | Unmodified | 0 | 0 | 0 | 438180 | 0 | 0 | 0 | 0 | 213510 | 0 | 0 | 0 | 0 | 0 | 0 | 0 | 0 | 870060 |
| GQPSEGES | GATD3B; GATD | A0A0B4J2D5 | GAL3B_HUMAN | Unmodified | 0 | 0 | 0 | 25791 | 0 | 62126 | 94008 | 17741 | 1051400 | 330600 | 0 | 0 | 711860 | 55770 | 0 | 0 | 147610 | 268480 |
| GQQIGKVV | RPL26; RPL26L | P61254; Q9L | RL26_HUMAN | Unmodified | 0 | 0 | 0 | 0 | 0 | 0 | 580970 | 536760 | 0 | 0 | 0 | 0 | 0 | 79914 | 0 | 0 | 0 | 0 |
| GQQLVMM | GSPT1; GSPT2; | P15170; Q8I | ERF3A_HUMAN | Unmodified | 0 | 0 | 0 | 0 | 1033800 | 0 | 0 | 0 | 0 | 0 | 0 | 1057700 | 645930 | 0 | 0 | 0 | 0 | 0 |
| GQQLVMM | GSPT1; GSPT2; | P15170; Q8I | ERF3A_HUMAN | Oxidation (M) | 0 | 59958 | 0 | 0 | 0 | 0 | 0 | 0 | 0 | 0 | 108750 | 0 | 303830 | 0 | 111980 | 88107 | 0 | 0 |
| GQQPEAIR | EHD2 | Q9NZN4 | EHD2_HUMAN | Unmodified | 0 | 0 | 0 | 0 | 2507400 | 910460 | 0 | 43335 | 0 | 62144 | 0 | 0 | 470900 | 237700 | 0 | 0 | 200690 | 180090 |
| GQRLPQAL | VPS37B; VPS37 | Q9H9H4; F5 | VP37B_HUMAN | Unmodified | 43642000 | 21337000 | 42553000 | 47686000 | 20068000 | 1080900 | 0 | 0 | 7176400 | 0 | 6073600 | 6948200 | 44072000 | 28553000 | 1.26E+08 | 38247000 | 1692100 | 28066000 |
| GQRPETLH | TMEM97; TMEM | Q5BJF2; J3K | SGMR2_HUMA | Unmodified | 0 | 0 | 0 | 0 | 0 | 0 | 0 | 0 | 1286700 | 0 | 9882.7 | 627400 | 0 | 20446 | 0 | 0 | 0 | 0 |
| GQRSWSV | UEVLD | Q8IX04 | UEVLD_HUMAN | Oxidation (M) | 0 | 0 | 0 | 0 | 246810 | 0 | 0 | 0 | 0 | 448300 | 0 | 0 | 1150600 | 2230900 | 0 | 0 | 1903400 | 2978100 |
| GQSEDPG | SLC3A2; SLC3A | P08195; A0A | 4F2_HUMAN 4I | Unmodified | 1993200 | 2886100 | 1158100 | 2745000 | 3059600 | 1351000 | 0 | 0 | 8298000 | 2757800 | 10286000 | 34844000 | 715350 | 0 | 3745000 | 3183600 | 0 | 0 |
| GQSEDPG | SLC3A2; SLC3A | P08195; A0A | 4F2_HUMAN 4I | Unmodified | 18827000 | 0 | 6852200 | 12145000 | 15037000 | 8583400 | 372360 | 0 | 55419000 | 0 | 29089000 | 2.1E+08 | 0 | 0 | 16343000 | 10788000 | 274570 | 4356700 |
| GQSEVSAA | GAS6 | Q14393 | GAS6_HUMAN | Unmodified | 0 | 0 | 0 | 0 | 0 | 0 | 0 | 0 | 792220 | 0 | 0 | 0 | 0 | 0 | 0 | 0 | 0 | 0 |
| GQSFFIDAI | NUBP1 | P53384 | NUBP1_HUMAN | Unmodified | 5778100 | 0 | 0 | 0 | 108600 | 0 | 0 | 0 | 185630 | 0 | 62624 | 67723 | 0 | 0 | 0 | 470570 | 1170200 | 0 |
| GQSGTYG | BCAM; BCAM | P50895; A0A | BCAM_HUMAN | Unmodified | 3323300 | 1502900 | 0 | 0 | 0 | 0 | 0 | 0 | 1074500 | 0 | 0 | 0 | 0 | 0 | 0 | 111240 | 0 | 0 |
| GQSMFVEN | GPRC5C; GPRC | Q9NQ84; A8 | GPC5C_HUMA | Unmodified | 0 | 124930 | 744730 | 1140800 | 1445000 | 0 | 0 | 0 | 4160400 | 841800 | 0 | 0 | 1592700 | 1336200 | 608790 | 27035 | 30972 | 1079700 |
| GQSMFVEN | GPRC5C; GPRC | Q9NQ84; A8 | GPC5C_HUMA | Oxidation (M) | 0 | 0 | 0 | 481790 | 895460 | 0 | 0 | 0 | 0 | 0 | 826000 | 0 | 404480 | 0 | 0 | 0 | 0 | 0 |
| GQSPLHIL | ANKFY1 | Q9P2R3 | ANFY1_HUMAN | Unmodified | 0 | 0 | 408240 | 0 | 0 | 0 | 15952 | 0 | 0 | 0 | 0 | 670220 | 0 | 0 | 125560 | 0 | 0 | 0 |
| GQSSLALH | RALB; RALB; RA | P11234; C9J | RALB_HUMAN | Unmodified | 0 | 0 | 0 | 0 | 457530 | 170760 | 0 | 0 | 0 | 27705 | 118960 | 1743000 | 0 | 0 | 177130 | 158920 | 188710 | 204040 |
| GQTHTLED | HSD17B10; HS | Q99714; Q5I | HCD2_HUMAN | Unmodified | 0 | 0 | 0 | 0 | 0 | 0 | 54943 | 0 | 4380900 | 237910 | 194380 | 164550 | 0 | 0 | 0 | 0 | 0 | 0 |
| GQTLTLTQ | ATP2C1; ATP2C | P98194; B4E | AT2C1_HUMAN | Unmodified | 66864 | 0 | 0 | 0 | 0 | 0 | 0 | 0 | 2151000 | 142620 | 0 | 875150 | 0 | 223440 | 0 | 0 | 0 | 0 |
| GQTLVVQF | CALR; CALR; C/ | P27797; A0A | CALR_HUMAN | Unmodified | 0 | 0 | 0 | 0 | 0 | 0 | 0 | 0 | 39938000 | 2171500 | 8614700 | 1020400 | 0 | 0 | 0 | 127830 | 0 | 0 |
| GQTLVVQF | CALR; CALR; C/ | P27797; A0A | CALR_HUMAN | Unmodified | 0 | 0 | 0 | 0 | 0 | 0 | 7166000 | 0 | 57672000 | 55572000 | 4920800 | 0 | 1359300 | 1199500 | 0 | 0 | 0 | 838440 |
| GQTPAQAE | EPB41L5 | Q9HCM4 | E41L5_HUMAN | Unmodified | 0 | 0 | 621310 | 542280 | 0 | 0 | 0 | 0 | 933850 | 0 | 0 | 442340 | 0 | 0 | 760510 | 682390 | 0 | 0 |
| GQTQEEEV | CASC3; CASC3 | O15234; J3K | CASC3_HUMAN | Unmodified | 0 | 284210 | 0 | 0 | 0 | 0 | 0 | 0 | 23402 | 0 | 0 | 45761 | 0 | 0 | 161670 | 402390 | 0 | 0 |
| GQTVEDLL | INTS1 | Q8N201 | INT1_HUMAN I | Oxidation (M) | 21026000 | 14742000 | 0 | 10935000 | 0 | 0 | 0 | 0 | 13553000 | 0 | 0 | 0 | 0 | 0 | 9256100 | 0 | 0 | 0 |
| GQVAKKED | SBDS; SBDS | Q9Y3A5; A0/ | SBDS_HUMAN | Unmodified | 406900 | 0 | 0 | 0 | 649910 | 0 | 0 | 0 | 0 | 0 | 0 | 880690 | 585810 | 643300 | 0 | 0 | 0 | 0 |
| GQVCLPVI | UBE2L5; UBE2L | A0A1B0GUS | UB2L5_HUMAN | Unmodified | 307740 | 144470 | 285760 | 0 | 0 | 0 | 376210 | 0 | 0 | 0 | 850710 | 3509300 | 0 | 377450 | 586290 | 565870 | 0 | 0 |
| GQVCVMIH | RTCB | Q9Y3I0 | RTCB_HUMAN | Oxidation (M) | 0 | 0 | 7662 | 0 | 0 | 0 | 0 | 16755 | 0 | 0 | 66612 | 993550 | 0 | 0 | 0 | 0 | 0 | 25814 |
| GQVFDQN | CPM; CPM | P14384; H0Y | CBPM_HUMAN | Unmodified | 0 | 0 | 0 | 0 | 1526500 | 603830 | 0 | 0 | 8164600 | 598760 | 0 | 0 | 0 | 0 | 0 | 0 | 0 | 0 |
| GQVINLDQ | FAM91A1; FAM | Q658Y4; H0Y | F91A1_HUMAN | Unmodified | 0 | 0 | 0 | 78535 | 90519 | 0 | 62554 | 0 | 0 | 203430 | 0 | 394450 | 56287 | 85321 | 0 | 99847 | 92424 | 0 |
| GQVKEEGI | PARP1; PARP1 | P09874; A0A | PARP1_HUMAN | Unmodified | 0 | 0 | 0 | 0 | 0 | 0 | 0 | 227190 | 20074 | 95679 | 0 | 0 | 0 | 0 | 0 | 0 | 0 | 0 |
| GQVKNPG | SH3GL1; SH3G | Q99961; M0I | SH3G1_HUMAN | Oxidation (M) | 0 | 0 | 0 | 0 | 0 | 0 | 0 | 0 | 0 | 0 | 0 | 0 | 0 | 0 | 0 | 0 | 0 | 0 |
| GQVLFDKV | EPB41L2; EPB4 | O43491; Q9 | E41L2_HUMAN | Unmodified | 0 | 0 | 0 | 0 | 2116600 | 0 | 260820 | 0 | 0 | 0 | 0 | 0 | 150650 | 0 | 0 | 0 | 2739200 | 1509700 |
| GQVLFDLV | EPB41L1; EPB4 | Q9H4G0; Q4 | E41L1_HUMAN | Unmodified | 0 | 0 | 0 | 186320 | 0 | 0 | 0 | 0 | 0 | 60394 | 0 | 0 | 0 | 0 | 783560 | 0 | 0 | 0 |
| GQVLNIQN | SSB; SSB; SSB | P05455; E9P | LA_HUMAN Lu | Unmodified | 0 | 0 | 0 | 0 | 0 | 0 | 0 | 0 | 0 | 0 | 0 | 603380 | 0 | 0 | 0 | 0 | 0 | 0 |
| GQVPENEA | CDH1; CDH1; ( | P12830; H3E | CADH1_HUMA | Unmodified | 1164700 | 0 | 0 | 1473400 | 0 | 0 | 0 | 0 | 3331800 | 0 | 819610 | 0 | 0 | 0 | 769790 | 1394000 | 0 | 0 |
| GQVPPLVT | SEC24C; SEC2 | P53992; G5E | SC24C_HUMAN | Unmodified | 0 | 0 | 0 | 0 | 0 | 0 | 0 | 0 | 0 | 0 | 0 | 0 | 0 | 0 | 0 | 0 | 0 | 0 |
| GQVPSHLH | ENTPD3 | O75355 | ENTP3_HUMAN | Unmodified | 0 | 0 | 0 | 0 | 0 | 0 | 0 | 0 | 2292300 | 188900 | 0 | 0 | 0 | 0 | 0 | 0 | 0 | 142880 |
| GQVQEVG | PFKL | P17858 | PFKAL_HUMAN | Unmodified | 295310 | 117620 | 307660 | 316280 | 0 | 0 | 55491 | 0 | 1094900 | 430050 | 0 | 900110 | 0 | 0 | 176810 | 476790 | 69691 | 0 |
| GQVQEVG | PFKL | P17858 | PFKAL_HUMAN | Unmodified | 0 | 0 | 100390 | 0 | 376710 | 0 | 220560 | 0 | 0 | 0 | 0 | 0 | 0 | 0 | 0 | 0 | 266260 | 2528500 |
| GQVVSLIR | CAPN1; CAPN1 | P07384; E9P | CAN1_HUMAN | Unmodified | 3.61E+08 | 3.09E+08 | 58888000 | 1.99E+08 | 0 | 0 | 6318300 | 0 | 119800 | 14460 | 0 | 1824700 | 49185000 | 21161000 | 1.02E+08 | 1.38E+08 | 0 | 0 |
| GQYFGELA | PRKAR2A; PRK | P13861; P31 | KAP2_HUMAN | Unmodified | 198910 | 0 | 983300 | 1493900 | 2031900 | 715020 | 405050 | 0 | 2762000 | 1178900 | 2837600 | 3276000 | 4225200 | 1985400 | 2433800 | 1623800 | 1043400 | 1101000 |
| GQYFGELA | PRKAR2A; PRK | P13861; H7C | KAP2_HUMAN | Unmodified | 0 | 0 | 0 | 0 | 2096100 | 0 | 514120 | 0 | 0 | 0 | 0 | 0 | 5587400 | 5277500 | 0 | 0 | 1690800 | 2650800 |
| GQYHTLQ | PKP3; PKP3; PK | Q9Y446; E9F | PKP3_HUMAN | Unmodified | 76651 | 65924 | 0 | 0 | 0 | 0 | 0 | 0 | 0 | 0 | 56056 | 212520 | 0 | 0 | 234640 | 203720 | 0 | 0 |
| GQYISPFHI | PPA1; PPA1 | Q15181; Q5 | IPYR_HUMAN I | Unmodified | 0 | 0 | 151610 | 222050 | 583190 | 149820 | 0 | 0 | 0 | 0 | 1928400 | 8322200 | 595470 | 373950 | 214820 | 419570 | 0 | 853740 |
| GQYISPFHI | PPA1; PPA1 | Q15181; Q5 | IPYR_HUMAN I | Oxidation (M) | 0 | 0 | 0 | 0 | 1675500 | 0 | 0 | 0 | 0 | 105990 | 0 | 0 | 3300300 | 907770 | 0 | 0 | 1179400 | 0 |
| GQYQGLS | UGGT1 | Q9NYU2 | UGGG1_HUMA | Unmodified | 0 | 0 | 0 | 0 | 0 | 0 | 0 | 0 | 23062000 | 0 | 0 | 0 | 0 | 0 | 0 | 0 | 0 | 0 |
| GRAEGEWI | BCLAF1; BCLAF | Q9NYF8; E9I | BCLF1_HUMAN | Unmodified | 0 | 0 | 0 | 0 | 0 | 0 | 0 | 0 | 0 | 0 | 0 | 447300 | 0 | 0 | 0 | 0 | 0 | 0 |
| GRAENALL | PEAK1; PEAK1 | Q9H792; H0 | PEAK1_HUMAN | Unmodified | 0 | 0 | 0 | 0 | 0 | 0 | 0 | 0 | 0 | 0 | 206390 | 167930 | 0 | 0 | 0 | 0 | 0 | 0 |
| GRAQLGV | CCT6A | P40227 | TCPZ_HUMAN | Unmodified | 0 | 0 | 0 | 0 | 2039100 | 83474 | 1369100 | 0 | 153800 | 435640 | 0 | 0 | 5219700 | 1678500 | 0 | 0 | 2261300 | 1306500 |
| GRCETSAS | SLC19A1; SLC1 | P41440; H3E | S19A1_HUMAN | Unmodified | 157990 | 69626 | 413070 | 466660 | 0 | 0 | 0 | 0 | 0 | 0 | 0 | 0 | 0 | 80970 | 783970 | 736450 | 8938.2 | 0 |
| GRDDITVH | MANEAL; MANE | Q5VSG8; E9I | MANEL_HUMAN | Unmodified | 0 | 0 | 0 | 0 | 54851 | 0 | 0 | 0 | 861460 | 27007 | 128790 | 0 | 0 | 0 | 0 | 29572 | 306250 | 0 |
| GRDIVVLG | PSMA7 | O14818 | PSA7_HUMAN | Unmodified | 0 | 0 | 0 | 0 | 451820 | 0 | 0 | 0 | 0 | 0 | 0 | 2264700 | 651500 | 0 | 0 | 96108 | 103630 | 287420 |
| GRDIVVLG | PSMA7 | O14818 | PSA7_HUMAN | Unmodified | 0 | 0 | 0 | 0 | 1456600 | 0 | 55248 | 0 | 766650 | 382900 | 702920 | 2574800 | 1031600 | 2734900 | 0 | 0 | 400630 | 0 |
| GRDLESRL | BROX; BROX; B | Q5VW32; F5 | BROX_HUMAN | Unmodified | 0 | 0 | 0 | 0 | 0 | 0 | 0 | 0 | 0 | 1402600 | 0 | 0 | 663790 | 0 | 0 | 0 | 0 | 0 |
| GRDPALPT | MOGS; MOGS; | Q13724; A0/ | MOGS_HUMAN | Unmodified | 0 | 0 | 0 | 0 | 0 | 0 | 0 | 0 | 17189000 | 1572700 | 315900 | 0 | 0 | 0 | 0 | 0 | 0 | 0 |
| GRDTLQEA | LAMC1 | P11047 | LAMC1_HUMAN | Unmodified | 249800 | 0 | 0 | 0 | 0 | 867390 | 0 | 0 | 841530 | 0 | 0 | 0 | 0 | 71981 | 180920 | 104320 | 0 | 1640200 |
| GRDVIAQS | EIF4A3; EIF4A3 | P38919; I3L | IF4A3_HUMAN | Unmodified | 730620 | 0 | 430510 | 0 | 0 | 0 | 1312400 | 33025 | 0 | 38314 | 0 | 1494300 | 0 | 0 | 394250 | 320950 | 0 | 0 |
| GRDWNVD | GDI1; GDI2; GD | P31150; P50 | GDIA_HUMAN | Unmodified | 0 | 0 | 0 | 0 | 1220400 | 0 | 0 | 0 | 7299400 | 2339600 | 4392600 | 22219000 | 0 | 1653900 | 0 | 0 | 301670 | 1084900 |
| GRDYCSEE | C1orf21 | Q9H246 | CA021_HUMAN | Unmodified | 44039 | 0 | 0 | 291450 | 0 | 0 | 0 | 60577 | 0 | 0 | 0 | 0 | 1058000 | 894880 | 0 | 0 | 0 | 0 |
| GREDGDAI | DIS3L2 | Q8IYB7 | DI3L2_HUMAN | Unmodified | 95277 | 0 | 0 | 0 | 0 | 0 | 0 | 0 | 0 | 0 | 63142 | 1031800 | 0 | 0 | 0 | 138540 | 0 | 0 |
| GREEHYEE | GOLIM4; GOLIN | O00461; F8V | GOLI4_HUMAN | Unmodified | 0 | 0 | 0 | 0 | 0 | 518260 | 167580 | 0 | 5073700 | 2412100 | 479600 | 0 | 0 | 0 | 0 | 0 | 0 | 271100 |
| GREFGVTT | ADSS2 | P30520 | PURA2_HUMAN | Unmodified | 0 | 0 | 0 | 0 | 0 | 0 | 0 | 0 | 0 | 0 | 71907 | 1007900 | 0 | 0 | 0 | 0 | 0 | 0 |
| GRELYERPI | MYO1D; MYO1I | O94832; J3Q | MYO1D_HUMA | Unmodified | 14742000 | 20755000 | 4017600 | 4505800 | 2077500 | 1327600 | 0 | 0 | 3014300 | 708760 | 0 | 881930 | 1825600 | 1234200 | 34727000 | 16011000 | 1128400 | 3525200 |
| GREMFEVT | CHID1 | Q9BWS9 | CHID1_HUMAN | 2 Oxidation (M) | 0 | 0 | 0 | 0 | 0 | 0 | 0 | 0 | 1289800 | 0 | 0 | 0 | 0 | 0 | 0 | 0 | 0 | 0 |
| GREMFEVT | CHID1 | Q9BWS9 | CHID1_HUMAN | Oxidation (M) | 0 | 0 | 0 | 0 | 0 | 0 | 0 | 0 | 1951700 | 0 | 0 | 0 | 0 | 0 | 0 | 0 | 0 | 0 |
| GREMFEVT | CHID1 | Q9BWS9 | CHID1_HUMAN | Unmodified | 0 | 0 | 0 | 0 | 0 | 0 | 0 | 0 | 3093500 | 0 | 0 | 580750 | 0 | 0 | 0 | 0 | 0 | 0 |
| GREQDLIL | CASKIN2; CASK | Q8WXE0; J3 | CSKI2_HUMAN | Unmodified | 0 | 0 | 0 | 163800 | 389750 | 0 | 0 | 0 | 0 | 0 | 0 | 271680 | 0 | 0 | 104750 | 134250 | 453080 | 389340 |
| GRESVLIIS | GET3 | O43681 | GET3_HUMAN | Unmodified | 78408 | 53811 | 0 | 724920 | 2416600 | 64789 | 215640 | 18026 | 0 | 121300 | 748310 | 3474900 | 1151400 | 796190 | 213450 | 130040 | 2878600 | 8843000 |
| GRESYIETE | CLTCL1; CLTCL | P53675; A0A | CLH2_HUMAN | Unmodified | 0 | 0 | 0 | 0 | 0 | 0 | 4426500 | 0 | 0 | 0 | 0 | 0 | 0 | 0 | 0 | 0 | 0 | 0 |
| GREVLDGF | ROBO1; ROBO1 | Q9Y6N7; A0 | ROBO1_HUMA | Unmodified | 0 | 0 | 0 | 0 | 0 | 0 | 0 | 0 | 0 | 47622 | 0 | 0 | 0 | 0 | 0 | 0 | 0 | 603290 |
| GRFAVHRI | RPS4X | P62701 | RS4X_HUMAN | Unmodified | 0 | 0 | 0 | 0 | 488150 | 114200 | 2033700 | 548440 | 0 | 297700 | 122850 | 0 | 191400 | 277950 | 0 | 0 | 0 | 375400 |
| GRFQDNLI | MAPRE2; MAPR | Q15555; K7E | MARE2_HUMAN | Unmodified | 108700 | 134430 | 128420 | 372000 | 1004900 | 66850 | 1860800 | 0 | 327230 | 644490 | 13755 | 39027 | 922140 | 1341000 | 535770 | 146290 | 57166 | 1321000 |
| GRFQDNLI | MAPRE2; MAPR | Q15555; K7E | MARE2_HUMAN | Unmodified | 0 | 0 | 0 | 0 | 0 | 0 | 1089100 | 0 | 0 | 273260 | 0 | 0 | 0 | 0 | 0 | 0 | 0 | 442500 |
| GRFYTDSN | MAN2B1 | O00754 | MA2B1_HUMAN | Unmodified | 84089 | 0 | 137430 | 0 | 0 | 0 | 0 | 0 | 2956000 | 583320 | 0 | 0 | 0 | 0 | 0 | 0 | 0 | 0 |
| GRFYYVDH | WWP2 | O00308 | WWP2_HUMAN | Unmodified | 821600 | 114380 | 298110 | 455590 | 0 | 0 | 0 | 0 | 0 | 0 | 0 | 0 | 0 | 0 | 755040 | 386280 | 0 | 0 |
| GRGDIQP | MTSS1; MTSS1; | O43312; H0Y | MTSS1_HUMAN | Unmodified | 0 | 0 | 0 | 0 | 0 | 0 | 0 | 0 | 0 | 0 | 0 | 0 | 0 | 0 | 0 | 0 | 0 | 0 |
| GRGGAVG | B3GAT3; B3GA | O94766; G3 | B3GA3_HUMAN | Unmodified | 0 | 0 | 0 | 0 | 0 | 0 | 0 | 0 | 2406600 | 0 | 0 | 0 | 0 | 0 | 0 | 0 | 0 | 0 |
| GRGITGVEI | TKT; TKT; TKT | P29401; A0A | TKT_HUMAN Tr | Unmodified | 66802 | 0 | 0 | 0 | 0 | 0 | 0 | 0 | 579100 | 324630 | 0 | 3767500 | 0 | 0 | 0 | 182770 | 0 | 0 |
| GRGLEYLY | DYNC1LI1; DYN | Q9Y6G9; C9 | DC1L1_HUMAN | Unmodified | 85904 | 0 | 189860 | 0 | 0 | 0 | 0 | 0 | 0 | 375510 | 0 | 3307700 | 0 | 0 | 0 | 93825 | 0 | 0 |
| GRGMELSI | CANT1; CANT1 | Q8WVQ1; K | CANT1_HUMAN | Oxidation (M) | 0 | 0 | 0 | 0 | 0 | 0 | 0 | 0 | 8614800 | 0 | 0 | 0 | 0 | 0 | 0 | 0 | 0 | 0 |
| GRGMELSI | CANT1; CANT1 | Q8WVQ1; K | CANT1_HUMAN | Unmodified | 0 | 0 | 0 | 0 | 0 | 0 | 0 | 0 | 2895600 | 0 | 52813 | 0 | 79423 | 0 | 0 | 378610 | 0 | 0 |
| GRGVGSPI | ZDHHC5 | Q9C0B5 | ZDHC5_HUMA | Unmodified | 0 | 0 | 0 | 0 | 601570 | 0 | 0 | 112530 | 0 | 0 | 0 | 0 | 787020 | 0 | 0 | 0 | 358720 | 0 |
| GRHMGIGI | RPL19; RPL19; | P84098; J3K | RL19_HUMAN | Unmodified | 144280 | 14763 | 0 | 0 | 0 | 0 | 0 | 0 | 0 | 0 | 0 | 271000 | 0 | 0 | 92343 | 41047 | 0 | 0 |
| GRHVVLGA | NAA10 | P41227 | NAA10_HUMAN | Unmodified | 0 | 0 | 0 | 0 | 753790 | 184650 | 0 | 0 | 203920 | 0 | 175580 | 2208900 | 661060 | 0 | 0 | 0 | 0 | 0 |
| GRIAVGSD | DPYSL2; DPYSL | Q16555; A0/ | DPYL2_HUMAN | Unmodified | 0 | 0 | 0 | 0 | 6439700 | 0 | 1141800 | 0 | 0 | 818540 | 0 | 0 | 0 | 0 | 0 | 0 | 644740 | 1082700 |
| GRIAVGSD | DPYSL2; DPYSL | Q16555; A0/ | DPYL2_HUMAN | Unmodified | 0 | 0 | 0 | 0 | 396060 | 0 | 1007600 | 0 | 0 | 0 | 0 | 0 | 0 | 0 | 0 | 0 | 0 | 0 |
| GRIAVGSD | CRMP1; CRMP1 | Q14194; E9F | DPYL1_HUMAN | Unmodified | 0 | 0 | 0 | 0 | 0 | 0 | 1353800 | 0 | 0 | 845330 | 0 | 0 | 0 | 0 | 0 | 0 | 0 | 0 |
| GRILDDDTI | DYNC1H1; DYN | Q14204; A0/ | DYHC1_HUMA | Unmodified | 0 | 0 | 107600 | 403700 | 0 | 0 | 0 | 0 | 1155700 | 0 | 0 | 945680 | 0 | 0 | 283050 | 0 | 0 | 0 |
| GRILTVEDI | TKT; TKT | P29401; A0A | TKT_HUMAN Tr | Unmodified | 0 | 0 | 0 | 0 | 26918000 | 2634900 | 0 | 0 | 0 | 5831600 | 0 | 0 | 6886700 | 0 | 0 | 0 | 2565200 | 4423500 |
| GRISCTIAN | FN1 | P02751 | FINC_HUMAN I | Unmodified | 0 | 0 | 0 | 0 | 0 | 284280 | 0 | 0 | 0 | 0 | 0 | 0 | 0 | 609610 | 0 | 0 | 0 | 0 |
| GRISVGSD | DPYSL3; DPYSL | Q14195; H0 | DPYL3_HUMAN | Unmodified | 0 | 0 | 0 | 0 | 1660400 | 0 | 1162400 | 96774 | 0 | 0 | 4488000 | 12784000 | 0 | 0 | 0 | 0 | 101130 | 367540 |
| GRLDECQI | ST6GALNAC4; | Q9H4F1; A6I | SIA7D_HUMAN | Oxidation (M) | 0 | 0 | 0 | 0 | 0 | 0 | 0 | 0 | 1303000 | 0 | 0 | 0 | 0 | 581480 | 0 | 0 | 0 | 0 |
| GRLDECQI | ST6GALNAC4; | Q9H4F1; A6I | SIA7D_HUMAN | Unmodified | 0 | 0 | 0 | 0 | 0 | 0 | 0 | 0 | 1911800 | 202630 | 0 | 413020 | 0 | 142610 | 0 | 0 | 0 | 0 |

| | | | | | | | | | | | | | | | | | | | | | | |
|---|---|---|---|---|---|---|---|---|---|---|---|---|---|---|---|---|---|---|---|---|---|---|
| GRLDNGQ | PACSIN2 | Q9UNF0 | PACN2_HUMAI | Unmodified | 47171000 | 61738000 | 24591000 | 44920000 | 4405300 | 1091700 | 1206000 | 349250 | 2151200 | 1008700 | 1100900 | 3090700 | 71893000 | 17451000 | 47630000 | 34154000 | 1578800 | 2162000 |
| GRLDSGQI | PACSIN1; PACS | Q9BY11; F6U | PACN1_HUMAI | Unmodified | 8875500 | 10985000 | 448640 | 1703400 | 0 | 0 | 0 | 0 | 0 | 0 | 0 | 134120 | 0 | 0 | 3745800 | 806810 | 0 | 0 |
| GRLDYLSS | SLC3A2; SLC3A | P08195; A0A | 4F2_HUMAN 4I | Unmodified | 4616000 | 4319900 | 3802400 | 5249300 | 31799000 | 3464200 | 2829300 | 715940 | 16259000 | 2932800 | 20176000 | 62062000 | 4735600 | 2752900 | 7948900 | 0 | 1081900 | 4069000 |
| GRLHLESP | IGSF3 | O75054 | IGSF3_HUMAN | Unmodified | 545730 | 593670 | 664850 | 549310 | 559630 | 107410 | 0 | 0 | 332550 | 0 | 0 | 1008300 | 446210 | 0 | 1281200 | 442850 | 95203 | 0 |
| GRLLPYLAI | GP1BB | P13224 | GP1BB_HUMAI | Unmodified | 0 | 3338800 | 0 | 3732100 | 0 | 0 | 0 | 0 | 0 | 0 | 0 | 27648 | 0 | 0 | 3528400 | 3646700 | 0 | 0 |
| GRLMPGQI | FKBP10; FKBP1 | Q96AY3; H0 | FKB10_HUMAN | Unmodified | 0 | 0 | 0 | 0 | 0 | 0 | 0 | 0 | 44054 | 0 | 477820 | 355290 | 0 | 0 | 0 | 0 | 0 | 0 |
| GRLSKEDIE | HSPA8; HSPA8; | P11142; E9P | HSP7C_HUMAI | Unmodified | 0 | 0 | 0 | 0 | 0 | 0 | 0 | 0 | 1690700 | 0 | 0 | 0 | 0 | 0 | 0 | 0 | 0 | 0 |
| GRLSKEEIE | HSPA1L; HSPA1 | P34931; P0D | HS71L_HUMAN | Unmodified | 4779300 | 16647000 | 3253600 | 1737500 | 167660 | 0 | 0 | 0 | 7344700 | 0 | 597280 | 9056500 | 0 | 572400 | 14087000 | 7534500 | 0 | 0 |
| GRLTECLE | AP2A2; AP2A2; | O94973; A0A | AP2A2_HUMAN | Unmodified | 0 | 0 | 0 | 0 | 0 | 0 | 0 | 0 | 0 | 0 | 0 | 735440 | 0 | 0 | 0 | 0 | 0 | 0 |
| GRLTVMTD | LMAN2; LMAN2 | Q12907; D6I | LMAN2_HUMAI | Unmodified | 0 | 0 | 0 | 0 | 0 | 0 | 0 | 0 | 775390 | 0 | 245680 | 0 | 0 | 0 | 0 | 47057 | 0 | 0 |
| GRLTVMTD | LMAN2; LMAN2 | Q12907; D6I | LMAN2_HUMAI | Oxidation (M) | 0 | 0 | 0 | 0 | 0 | 0 | 0 | 0 | 6281200 | 1072200 | 0 | 0 | 0 | 0 | 0 | 0 | 0 | 0 |
| GRLTVMTD | LMAN2; LMAN2 | Q12907; D6I | LMAN2_HUMAI | Unmodified | 0 | 0 | 0 | 0 | 142390 | 0 | 0 | 0 | 68787000 | 6754100 | 88897 | 0 | 354900 | 209700 | 0 | 0 | 36287 | 0 |
| GRLVECLE | AP2A1 | O95782 | AP2A1_HUMAN | Unmodified | 0 | 0 | 78089 | 0 | 0 | 0 | 0 | 0 | 0 | 0 | 0 | 2179700 | 0 | 0 | 0 | 0 | 0 | 307280 |
| GRLYPWG | SEPTIN2; SEPTI | Q15019; H7 | SEPT2_HUMAN | Unmodified | 0 | 0 | 0 | 0 | 8526800 | 499930 | 739980 | 0 | 647930 | 0 | 0 | 389600 | 1040800 | 0 | 0 | 0 | 6220700 | 4881600 |
| GRMSMKE | TUBB2A; TUBB; | Q13885; P07 | TBB2A_HUMAN | Unmodified | 0 | 0 | 0 | 0 | 3038100 | 0 | 303130 | 20801 | 0 | 685940 | 0 | 0 | 0 | 0 | 0 | 0 | 225290 | 1276400 |
| GRMSMKE | TUBB2A; TUBB; | Q13885; P07 | TBB2A_HUMAN | Oxidation (M) | 0 | 0 | 0 | 0 | 529170 | 0 | 161090 | 25150 | 0 | 100190 | 0 | 0 | 188350 | 0 | 0 | 0 | 18180 | 0 |
| GRNSATSA | MARK2; MARK2 | Q7KZI7; E7E | MARK2_HUMAI | Unmodified | 0 | 0 | 0 | 0 | 0 | 0 | 0 | 0 | 0 | 0 | 0 | 0 | 750980 | 0 | 225090 | 0 | 0 | 0 |
| GRPCKIVEI | EIF5A; EIF5A2; | P63241; Q9 | IF5A1_HUMAN | Unmodified | 0 | 0 | 0 | 0 | 0 | 0 | 0 | 0 | 0 | 0 | 0 | 0 | 0 | 0 | 0 | 0 | 403670 | 0 |
| GRPPPTPLI | FKBP15 | Q5T1M5 | FKB15_HUMAN | Unmodified | 0 | 0 | 0 | 0 | 353790 | 0 | 0 | 0 | 0 | 0 | 0 | 0 | 0 | 0 | 0 | 0 | 0 | 145490 |
| GRPYDYN | PDIA4; PDIA4 | P13667; A0A | PDIA4_HUMAN | Unmodified | 0 | 0 | 178510 | 0 | 0 | 457320 | 1428800 | 1051900 | 8340900 | 8690800 | 1789800 | 497270 | 0 | 0 | 0 | 0 | 0 | 0 |
| GRQDSNSI | GOLIM4; GOLIM | O00461; F8 | GOLI4_HUMAN | Unmodified | 0 | 0 | 0 | 0 | 915890 | 69153 | 608340 | 266220 | 2142300 | 7185000 | 0 | 0 | 302840 | 268520 | 0 | 0 | 41994 | 581170 |
| GRQDSNSI | GOLIM4; GOLIM | O00461; F8 | GOLI4_HUMAN | Oxidation (M) | 0 | 0 | 0 | 0 | 167820 | 0 | 0 | 147010 | 65941 | 617160 | 0 | 0 | 0 | 0 | 0 | 0 | 0 | 0 |
| GRQMSCLI | GLG1 | Q92896 | GSLG1_HUMAI | Unmodified | 0 | 0 | 0 | 0 | 0 | 0 | 0 | 0 | 0 | 0 | 0 | 0 | 0 | 0 | 0 | 0 | 0 | 0 |
| GRQPGEVI | TRIM26; TRIM26 | Q12899; A2A | TRI26_HUMAN | Unmodified | 4358900 | 3364000 | 4716500 | 6147400 | 0 | 0 | 0 | 50881 | 88261 | 160820 | 23132 | 515230 | 1565700 | 990660 | 4739900 | 4420400 | 0 | 48293 |
| GRQVSSEC | GLG1; GLG1 | Q92896; H3I | GSLG1_HUMAI | Unmodified | 0 | 0 | 0 | 0 | 0 | 0 | 0 | 0 | 2627600 | 0 | 432880 | 0 | 0 | 0 | 0 | 0 | 0 | 0 |
| GRQYPWG | SEPTIN7; SEPTI | Q16181; E7E | SEPT7_HUMAN | Unmodified | 53797 | 0 | 0 | 992030 | 2710400 | 0 | 0 | 0 | 0 | 0 | 0 | 626120 | 0 | 0 | 178620 | 256220 | 0 | 0 |
| GRRAEDGS | ANXA2; ANXA2; | P07355; H0 | ANXA2_HUMAI | Unmodified | 455450 | 959410 | 0 | 0 | 7408200 | 3689000 | 0 | 0 | 13316000 | 803600 | 0 | 0 | 0 | 0 | 107800 | 0 | 371220 | 2232300 |
| GRSAINEV | RPL31; RPL31; | P62899; B8Z | RL31_HUMAN | Unmodified | 512450 | 0 | 189260 | 254670 | 0 | 0 | 0 | 0 | 0 | 0 | 0 | 318140 | 243990 | 0 | 0 | 0 | 0 | 0 |
| GRSEDIYRI | UGGT1 | Q9NYU2 | UGGG1_HUMA | Unmodified | 0 | 0 | 0 | 0 | 0 | 0 | 0 | 0 | 0 | 10696000 | 0 | 0 | 0 | 0 | 0 | 0 | 0 | 0 |
| GRSELIVVE | SCIN; SCIN; SC | Q9Y6U3; E5I | ADSV_HUMAN | Unmodified | 0 | 0 | 0 | 0 | 0 | 0 | 0 | 0 | 1933900 | 1307100 | 0 | 1023900 | 0 | 0 | 0 | 0 | 0 | 0 |
| GRSPDELP | DIAPH1; DIAPH | O60610; A0A | DIAP1_HUMAN | Unmodified | 0 | 0 | 33840 | 0 | 189990 | 0 | 0 | 0 | 0 | 0 | 0 | 0 | 0 | 0 | 0 | 28164 | 94707 | 115280 |
| GRSYLAPG | PECR; PECR | Q9BY49; B4I | PECR_HUMAN | Unmodified | 0 | 0 | 0 | 0 | 0 | 0 | 0 | 0 | 0 | 1320500 | 0 | 0 | 0 | 0 | 0 | 0 | 0 | 0 |
| GRTFDEIAS | SLC2A1 | P11166 | GTR1_HUMAN | Unmodified | 0 | 176480 | 372230 | 308070 | 0 | 0 | 0 | 0 | 290650 | 0 | 198280 | 2165800 | 0 | 0 | 643150 | 639080 | 0 | 0 |
| GRTFDEIAS | SLC2A1 | P11166 | GTR1_HUMAN | Unmodified | 0 | 0 | 0 | 0 | 0 | 110820 | 752710 | 0 | 0 | 37850 | 0 | 0 | 0 | 0 | 0 | 0 | 0 | 0 |
| GRTFNLTA | PDXDC1; PDXD | Q6P996; H3I | PDXD1_HUMAI | Unmodified | 0 | 0 | 0 | 0 | 0 | 0 | 0 | 0 | 0 | 0 | 0 | 0 | 1022000 | 0 | 0 | 0 | 0 | 0 |
| GRTGDLLE | OPLAH | O14841 | OPLA_HUMAN | Unmodified | 0 | 0 | 0 | 0 | 0 | 0 | 0 | 0 | 0 | 371760 | 0 | 608710 | 0 | 0 | 0 | 0 | 0 | 0 |
| GRTGLIPSI | OSTF1 | Q92882 | OSTF1_HUMAN | Unmodified | 0 | 0 | 0 | 672870 | 1400600 | 0 | 0 | 0 | 1038200 | 698970 | 982880 | 3056400 | 0 | 934590 | 0 | 0 | 0 | 1053600 |
| GRTLGEID | FAM126A | H7C0W7 | H7C0W7_HUM | Unmodified | 0 | 0 | 0 | 0 | 4652900 | 0 | 0 | 326630 | 0 | 0 | 0 | 0 | 1797300 | 0 | 0 | 0 | 212100 | 8114500 |
| GRTVIIEQS | HSPD1; HSPD1 | P10809; A0A | CH60_HUMAN | Unmodified | 0 | 0 | 0 | 0 | 1208000 | 0 | 4038700 | 542850 | 23569000 | 5402300 | 0 | 0 | 5882300 | 4708900 | 0 | 189400 | 412120 | 1575700 |
| GRTVIIEQS | HSPD1; HSPD1 | P10809; A0A | CH60_HUMAN | Unmodified | 0 | 0 | 0 | 0 | 0 | 0 | 2479200 | 293930 | 0 | 0 | 0 | 0 | 0 | 0 | 0 | 0 | 305630 | 0 |
| GRVALLSG | TKFC; TKFC; TK | Q3LXA3; H0 | TKFC_HUMAN | Unmodified | 0 | 0 | 0 | 0 | 0 | 0 | 0 | 0 | 0 | 0 | 168650 | 0 | 369460 | 245290 | 0 | 0 | 0 | 417720 |
| GRVAVGSD | DPYSL4; DPYSL | O14531; Q5 | DPYL4_HUMAN | Unmodified | 0 | 0 | 0 | 0 | 0 | 0 | 0 | 0 | 0 | 333810 | 0 | 0 | 0 | 0 | 0 | 0 | 193270 | 507820 |
| GRVFANAP | PFKL | P17858 | PFKAL_HUMAN | Unmodified | 0 | 0 | 0 | 0 | 1091400 | 0 | 938410 | 839090 | 134770 | 280210 | 0 | 0 | 866410 | 446280 | 0 | 0 | 768320 | 1769500 |
| GRVFEVSL | RPS3A; RPS3A; | P61247; D6F | RS3A_HUMAN | Unmodified | 11018000 | 0 | 0 | 0 | 9210700 | 0 | 11909000 | 41604 | 2154600 | 2224000 | 2612600 | 3036900 | 6121500 | 5486900 | 0 | 4358700 | 0 | 6097700 |
| GRVHFTSN | CXADR | P78310 | CXAR_HUMAN | Unmodified | 1902600 | 1123900 | 1091500 | 853140 | 33161000 | 2537900 | 1198800 | 400990 | 2310200 | 43180 | 881860 | 11440000 | 1348600 | 709350 | 0 | 1261800 | 926800 | 65284 |
| GRVNFDD | GFPT1; GFPT1; | Q06210; A0A | GFPT1_HUMAN | Unmodified | 0 | 73959 | 0 | 610640 | 0 | 0 | 0 | 0 | 1567900 | 0 | 0 | 186950 | 1714100 | 0 | 535380 | 457970 | 0 | 2403400 |
| GRVNFDD | GFPT1; GFPT1; | Q06210; A0A | GFPT1_HUMAN | Unmodified | 0 | 0 | 0 | 0 | 388190 | 0 | 0 | 0 | 0 | 0 | 0 | 0 | 1438300 | 913930 | 0 | 0 | 0 | 0 |
| GRVPYPGN | YES1; SRC; YES | P07947; P12 | YES_HUMAN Ty | Unmodified | 273360 | 260160 | 154260 | 504330 | 0 | 122280 | 42404 | 0 | 321010 | 47862 | 0 | 881130 | 0 | 0 | 726290 | 370300 | 151520 | 0 |
| GRVQLFEIS | ARRDC1; ARRD | Q8N5I2; Q5 | ARRD1_HUMAI | Unmodified | 2084800 | 1979500 | 1084500 | 1919000 | 68020 | 0 | 0 | 0 | 1819600 | 0 | 0 | 911160 | 0 | 0 | 2877900 | 2959900 | 0 | 102830 |
| GRVQLFEIS | ARRDC1; ARRD | Q8N5I2; Q5 | ARRD1_HUMAI | Unmodified | 0 | 0 | 313800 | 6159000 | 0 | 0 | 0 | 0 | 685300 | 0 | 0 | 0 | 2896700 | 0 | 0 | 0 | 0 | 982830 |
| GRVSVPTH | VCAN; VCAN; V | P13611; Q8 | CSPG2_HUMAI | Unmodified | 0 | 0 | 0 | 0 | 0 | 0 | 0 | 0 | 3921300 | 957360 | 0 | 0 | 0 | 0 | 0 | 0 | 0 | 0 |
| GRVVDIYSI | PRKAG1; PRKA | P54619; F8V | AAKG1_HUMAI | Unmodified | 0 | 0 | 0 | 50967 | 476880 | 0 | 49810 | 0 | 188170 | 0 | 0 | 0 | 0 | 176740 | 0 | 0 | 87298 | 769990 |
| GRWQGNI | ILK; ILK | Q13418; A0A | ILK_HUMAN Int | Unmodified | 0 | 414700 | 617070 | 974520 | 2368000 | 0 | 2483600 | 663680 | 522380 | 166440 | 0 | 0 | 684350 | 1330000 | 472230 | 584160 | 507090 | 1220300 |
| GRYEAEGC | BICD2 | Q8TD16 | BICD2_HUMAN | Unmodified | 0 | 0 | 0 | 0 | 779460 | 0 | 0 | 0 | 78799 | 0 | 0 | 0 | 0 | 0 | 0 | 0 | 0 | 0 |
| GRYGLFPA | CTTN | Q14247 | SRC8_HUMAN | Unmodified | 0 | 0 | 0 | 1134300 | 7265700 | 2341000 | 0 | 0 | 0 | 290340 | 0 | 1118600 | 0 | 5819500 | 0 | 0 | 2600400 | 2141900 |
| GSAFSTSIS | FARSA; FARSA; | Q9Y285; K7E | SYFA_HUMAN | Unmodified | 0 | 28207 | 169520 | 0 | 0 | 0 | 49348 | 0 | 1025100 | 0 | 0 | 1856000 | 0 | 0 | 0 | 539870 | 0 | 428420 |
| GSAGAPSV | PPM1A; PPM1A | P35813; E9P | PPM1A_HUMAI | Unmodified | 372010 | 345990 | 339050 | 197690 | 0 | 0 | 0 | 0 | 94508 | 76941 | 156890 | 1844300 | 0 | 0 | 865920 | 436790 | 0 | 0 |
| GSAIDEDIV | TOR1B | O14657 | TOR1B_HUMAI | Unmodified | 0 | 0 | 0 | 0 | 0 | 0 | 0 | 0 | 2196000 | 0 | 129250 | 0 | 0 | 0 | 0 | 0 | 0 | 0 |
| GSAIDEDIV | TOR1B | O14657 | TOR1B_HUMAI | Unmodified | 0 | 0 | 0 | 0 | 0 | 0 | 0 | 0 | 1023700 | 506510 | 0 | 0 | 0 | 0 | 0 | 0 | 0 | 0 |
| GSAITGPV | RPL23; RPL23 | P62829; J3K | RL23_HUMAN | Unmodified | 8285500 | 4961900 | 2899100 | 278300 | 3589300 | 764780 | 3287600 | 2714300 | 697240 | 413450 | 2315600 | 25698000 | 293970 | 2452500 | 0 | 3569600 | 852040 | 2381900 |
| GSAPPGPV | GSTO1 | P78417 | GSTO1_HUMAI | Unmodified | 119170 | 103880 | 246580 | 0 | 2220600 | 709560 | 0 | 0 | 210160 | 0 | 3296600 | 8278100 | 0 | 0 | 371620 | 446810 | 0 | 0 |
| GSAVVGQ | H6PD | O95479 | G6PE_HUMAN | Unmodified | 0 | 0 | 0 | 0 | 0 | 0 | 0 | 0 | 3585700 | 234050 | 0 | 0 | 0 | 0 | 0 | 0 | 0 | 0 |
| GSAVWCQ | PSAP; PSAP; PS | P07602; Q5E | SAP_HUMAN P | Unmodified | 0 | 0 | 0 | 0 | 0 | 0 | 0 | 0 | 9186500 | 673850 | 0 | 0 | 0 | 0 | 319730 | 0 | 0 | 0 |
| GSCDSLCI | SLITRK6 | Q9H5Y7 | SLIK6_HUMAN | Unmodified | 261290 | 0 | 0 | 0 | 0 | 0 | 0 | 0 | 0 | 0 | 0 | 0 | 0 | 0 | 0 | 0 | 0 | 0 |
| GSCNLSRV | GFPT1; GFPT1; | Q06210; A0A | GFPT1_HUMAN | Unmodified | 0 | 0 | 0 | 0 | 0 | 0 | 0 | 0 | 0 | 311800 | 0 | 0 | 594580 | 0 | 0 | 0 | 0 | 0 |
| GSCPFGGI | MKRN1 | Q9UHC7 | MKRN1_HUMA | Unmodified | 1324800 | 573020 | 0 | 0 | 0 | 0 | 0 | 0 | 0 | 0 | 0 | 0 | 0 | 0 | 212590 | 381130 | 0 | 0 |
| GSDCGIVN | ALDH7A1; ALDH | P49419; A0A | AL7A1_HUMAN | Unmodified | 0 | 0 | 0 | 0 | 0 | 339640 | 0 | 0 | 1354400 | 0 | 0 | 0 | 0 | 0 | 0 | 0 | 0 | 2280300 |
| GSDEPPVF | LAP3; LAP3 | P28838; H0 | AMPL_HUMAN | Unmodified | 0 | 0 | 0 | 0 | 1824100 | 0 | 462600 | 122060 | 2543900 | 230970 | 1604400 | 2830800 | 0 | 1447900 | 0 | 0 | 0 | 835600 |
| GSDFDCEL | HNRNPK; HNRI | P61978; Q5 | HNRPK_HUMA | Unmodified | 668470 | 359060 | 0 | 595290 | 452730 | 0 | 65041 | 68700 | 880230 | 100070 | 2377500 | 14723000 | 365370 | 0 | 538970 | 217250 | 0 | 0 |
| GSDGAPD | ERAP1 | Q9NZ08 | ERAP1_HUMAN | Unmodified | 0 | 0 | 18107 | 16169 | 0 | 20816 | 0 | 0 | 2175200 | 1080100 | 1852100 | 270420 | 0 | 533490 | 74819 | 29801 | 0 | 31360 |
| GSDHSASI | NANS; NANS; N | Q9NR45; Q5 | SIAS_HUMAN S | Unmodified | 0 | 0 | 681300 | 325200 | 936030 | 0 | 519320 | 0 | 0 | 165890 | 0 | 1330300 | 1825200 | 0 | 960790 | 416450 | 1176200 | 1199000 |
| GSDQAIITL | BSG; BSG; BSG | P35613; A0A | BASI_HUMAN E | Unmodified | 0 | 0 | 5793900 | 2046600 | 0 | 218540 | 0 | 0 | 776650 | 0 | 0 | 16898000 | 151520 | 0 | 10786000 | 6798200 | 0 | 200380 |
| GSDVIIMLV | RAB6A; RAB6B; | P20340; Q9I | RAB6A_HUMAN | Oxidation (M) | 0 | 0 | 0 | 0 | 380890 | 0 | 0 | 0 | 0 | 0 | 0 | 680420 | 0 | 0 | 0 | 0 | 0 | 994000 |
| GSDVIIMLV | RAB6A; RAB6B; | P20340; Q9I | RAB6A_HUMAN | Oxidation (M) | 138960 | 94209 | 366400 | 25730 | 0 | 0 | 0 | 0 | 5190500 | 0 | 466600 | 3971400 | 0 | 0 | 775570 | 819490 | 248150 | 0 |
| GSDVIIMLV | RAB6A; RAB6B; | P20340; Q9I | RAB6A_HUMAN | Unmodified | 0 | 0 | 0 | 0 | 0 | 0 | 0 | 0 | 3279800 | 0 | 0 | 0 | 0 | 0 | 0 | 0 | 0 | 0 |
| GSDVIIMLV | RAB6A; RAB6B; | P20340; Q9I | RAB6A_HUMAN | Oxidation (M) | 1049700 | 1081700 | 174350 | 1702300 | 0 | 0 | 0 | 0 | 672600 | 0 | 1247300 | 14720000 | 0 | 247840 | 653910 | 1094300 | 0 | 0 |
| GSEAAQLL | HDDC3; HDDC | Q8N4P3; H0 | MESH1_HUMA | Acetyl (Protein N | 0 | 0 | 0 | 0 | 0 | 0 | 0 | 0 | 0 | 189890 | 101680 | 654990 | 0 | 0 | 0 | 0 | 0 | 0 |
| GSEGLYM | EIF2A; EIF2A; EI | Q9BY44; H7 | EIF2A_HUMAN | Oxidation (M) | 498490 | 0 | 190530 | 289780 | 0 | 0 | 0 | 0 | 245280 | 0 | 0 | 5717600 | 0 | 0 | 609700 | 875810 | 0 | 0 |
| GSELEITLTI | IARS1; IARS1; I | P41252; A0A | SYIC_HUMAN I | Unmodified | 98894 | 0 | 0 | 400670 | 0 | 0 | 0 | 327330 | 0 | 0 | 372730 | 1671000 | 243340 | 0 | 0 | 0 | 0 | 0 |
| GSESGIFT | FGA | P02671 | FIBA_HUMAN F | Unmodified | 0 | 1067000 | 0 | 0 | 0 | 0 | 0 | 0 | 0 | 0 | 0 | 0 | 0 | 0 | 0 | 0 | 0 | 343210 |
| GSEVGFHC | AHNAK | Q09666 | AHNK_HUMAN | Unmodified | 0 | 81615 | 0 | 0 | 4093800 | 0 | 0 | 0 | 0 | 0 | 0 | 0 | 0 | 0 | 179970 | 0 | 667680 | 2274200 |
| GSFIQHSV | GALNT16 | Q8N428 | GLT16_HUMAN | Unmodified | 0 | 0 | 0 | 0 | 0 | 0 | 0 | 0 | 7840400 | 2884100 | 0 | 0 | 0 | 0 | 0 | 0 | 0 | 0 |
| GSFSEQGII | PDIA6 | Q15084 | PDIA6_HUMAN | Unmodified | 0 | 0 | 142240 | 0 | 0 | 0 | 0 | 0 | 1.07E+08 | 3608900 | 4708000 | 5012000 | 68822 | 0 | 805940 | 565160 | 0 | 0 |
| GSFSEQGII | PDIA6 | Q15084 | PDIA6_HUMAN | Unmodified | 0 | 0 | 0 | 0 | 10827 | 0 | 0 | 0 | 5044900 | 6296800 | 125410 | 0 | 0 | 0 | 0 | 0 | 0 | 28068 |
| GSFSLSVR | LCK; LYN; LCK; | P06239; P07 | LCK_HUMAN Ty | Unmodified | 348990 | 0 | 0 | 344300 | 0 | 0 | 0 | 0 | 731850 | 0 | 0 | 0 | 0 | 0 | 781090 | 860260 | 0 | 0 |
| GSFSLSVRI | LYN | P07948 | LYN_HUMAN Ty | Unmodified | 0 | 0 | 0 | 537910 | 5523100 | 0 | 0 | 0 | 1999900 | 1742600 | 0 | 0 | 657380 | 0 | 0 | 0 | 0 | 0 |
| GSFSSTAA | GPR180 | Q86V85 | GP180_HUMAN | Unmodified | 0 | 0 | 17801 | 0 | 0 | 0 | 25075 | 0 | 9236000 | 833530 | 109450 | 109090 | 0 | 0 | 0 | 0 | 0 | 0 |
| GSGEMMTI | OPA1; OPA1; O | O60313; E5K | OPA1_HUMAN | Unmodified | 0 | 0 | 0 | 0 | 0 | 0 | 0 | 0 | 0 | 0 | 30441 | 990870 | 0 | 0 | 7879.9 | 0 | 0 | 0 |
| GSGGEWK | AHNAK | Q09666 | AHNK_HUMAN | Unmodified | 58329 | 0 | 0 | 0 | 4603300 | 730250 | 0 | 0 | 0 | 107870 | 0 | 0 | 0 | 0 | 0 | 0 | 468270 | 2749500 |
| GSGGWTL | DDO | Q99489 | OXDD_HUMAN | Unmodified | 0 | 0 | 0 | 0 | 0 | 0 | 0 | 0 | 559710 | 0 | 0 | 156270 | 0 | 0 | 0 | 97465 | 0 | 0 |
| GSGIFDES | JPT2; JPT2; JPT2 | Q9H910; H3 | JUPI2_HUMAN | Unmodified | 450900 | 428760 | 450190 | 686210 | 0 | 0 | 0 | 0 | 477270 | 167880 | 820880 | 1677500 | 0 | 0 | 591710 | 742570 | 0 | 0 |
| GSGIVTRCI | MX2; MX1; MX1 | P20592; P20 | MX2_HUMAN I | Unmodified | 0 | 0 | 0 | 0 | 0 | 0 | 0 | 0 | 0 | 0 | 0 | 0 | 69162 | 0 | 0 | 0 | 0 | 2268400 |
| GSGNLEAII | CCT2; CCT2; C | P78371; F5G | TCPB_HUMAN | Unmodified | 0 | 0 | 718980 | 480930 | 14346000 | 517880 | 4338200 | 4219700 | 2067800 | 1455900 | 4850200 | 18060000 | 15114000 | 6104900 | 287190 | 0 | 4719400 | 6546300 |
| GSGNLEAII | CCT2; CCT2; C | P78371; F5G | TCPB_HUMAN | Unmodified | 23449 | 27235 | 210050 | 135660 | 1623000 | 0 | 516650 | 0 | 631550 | 416340 | 778080 | 5103300 | 1652800 | 852400 | 132880 | 151470 | 595250 | 1003500 |
| GSGPLSPS | PTPRF; PTPRF; | P10586; H0 | PTPRF_HUMAN | Unmodified | 0 | 0 | 0 | 0 | 162770 | 0 | 0 | 0 | 1066800 | 0 | 0 | 0 | 0 | 0 | 0 | 0 | 0 | 0 |
| GSGSQSS | SMC3 | Q9UQE7 | SMC3_HUMAN | Unmodified | 0 | 0 | 0 | 0 | 0 | 0 | 0 | 0 | 0 | 0 | 0 | 1354700 | 0 | 0 | 0 | 0 | 0 | 0 |
| GSGTAEVE | PKM; PKM; PKM | P14618; H3E | KPYM_HUMAN | Unmodified | 469150 | 972100 | 2370000 | 2240800 | 33908000 | 3985000 | 0 | 1174400 | 2301100 | 1331300 | 4454900 | 27267000 | 5478300 | 2111900 | 2428800 | 1529900 | 7075100 | 13935000 |
| GSGTAEVE | PKM; PKM; PKM | P14618; H3E | KPYM_HUMAN | Unmodified | 0 | 0 | 0 | 0 | 605300 | 47934 | 1230600 | 144440 | 0 | 51298 | 0 | 0 | 242930 | 199360 | 0 | 0 | 1949300 | 370210 |
| GSGTASDE | HUWE1 | Q7Z6Z7 | HUWE1_HUMA | Unmodified | 0 | 0 | 0 | 0 | 0 | 0 | 0 | 0 | 192590 | 0 | 0 | 1283100 | 0 | 0 | 0 | 0 | 0 | 0 |
| GSGVHGG | PPFIBP2; PPFIB | Q8ND30; E9 | LIPB2_HUMAN | Unmodified | 140520 | 69630 | 521780 | 416060 | 0 | 0 | 0 | 0 | 0 | 0 | 0 | 541210 | 0 | 0 | 502160 | 564990 | 0 | 0 |
| GSHQISLD | EGFR; EGFR | P00533; E9P | EGFR_HUMAN | Unmodified | 0 | 0 | 649310 | 1159300 | 68852000 | 8997100 | 0 | 66801 | 216680 | 118490 | 0 | 0 | 13085000 | 10602000 | 2380400 | 0 | 9923600 | 22984000 |
| GSIFVVFDS | SSB; SSB; SSB | P05455; E9P | LA_HUMAN Lu | Unmodified | 0 | 0 | 0 | 0 | 30840 | 0 | 0 | 0 | 0 | 126160 | 223530 | 402950 | 0 | 0 | 0 | 25692 | 0 | 0 |
| GSIFVVFDS | SSB; SSB; SSB | P05455; E9P | LA_HUMAN Lu | Unmodified | 0 | 0 | 146450 | 108770 | 0 | 0 | 0 | 0 | 451950 | 336070 | 0 | 5183700 | 0 | 0 | 1623100 | 0 | 0 | 0 |
| GSIHDFPG | ANXA6; ANXA6; | P08133; E5R | ANXA6_HUMAN | Unmodified | 0 | 0 | 0 | 0 | 540790 | 32283 | 442500 | 0 | 24837000 | 2176300 | 0 | 0 | 70097 | 0 | 197060 | 0 | 121410 | 222150 |
| GSIHDFPG | ANXA6; ANXA6; | P08133; E5R | ANXA6_HUMAN | Oxidation (M) | 0 | 0 | 0 | 0 | 0 | 0 | 0 | 0 | 4894700 | 0 | 0 | 0 | 0 | 0 | 838060 | 0 | 0 | 0 |
| GSIHKFVP | IMPDH2; IMPDH | P12268; A0A | IMDH2_HUMAI | Unmodified | 0 | 0 | 0 | 0 | 0 | 0 | 1829900 | 0 | 0 | 0 | 0 | 0 | 0 | 0 | 0 | 0 | 810440 | 0 |
| GSIMLDRE | SLC4A4 | Q9Y6R1 | S4A4_HUMAN | 2 Oxidation (M) | 0 | 0 | 0 | 0 | 1127400 | 0 | 0 | 0 | 0 | 0 | 0 | 0 | 384080 | 281130 | 0 | 0 | 0 | 0 |
| GSIQVDGE | HSPG2 | P98160 | PGBM_HUMAN | Unmodified | 0 | 0 | 0 | 352690 | 0 | 0 | 0 | 0 | 0 | 0 | 0 | 0 | 0 | 0 | 0 | 0 | 0 | 932420 |
| GSISPMNA | N/A | N/A | N/A | Oxidation (M) | 891850 | 1164600 | 670700 | 0 | 0 | 0 | 0 | 0 | 0 | 0 | 0 | 0 | 0 | 0 | 61522 | 0 | 0 | 0 |

| | | | | | | | | | | | | | | | | | | | | | | |
|---|---|---|---|---|---|---|---|---|---|---|---|---|---|---|---|---|---|---|---|---|---|---|
| GSITISAEEI | CPNE3; CPNE3 | O75131; H0Y | CPNE3_HUMAI | Unmodified | 386170 | 222540 | 3409700 | 2423600 | 0 | 0 | 305330 | 92095 | 22627000 | 1070500 | 0 | 1422000 | 0 | 0 | 2204800 | 2099700 | 0 | 0 |
| GSITISAEEI | CPNE3; CPNE3 | O75131; H0Y | CPNE3_HUMAI | Unmodified | 879970 | 0 | 338700 | 1702300 | 0 | 0 | 376480 | 92842 | 24093000 | 1352100 | 973740 | 1495400 | 0 | 0 | 738650 | 1324300 | 0 | 0 |
| GSIVGGHE | RET; RET | P07949; C9J | RET_HUMAN P | Unmodified | 0 | 0 | 0 | 0 | 0 | 0 | 0 | 0 | 6046500 | 0 | 0 | 0 | 256520 | 0 | 0 | 0 | 255170 | 0 |
| GSIVWQEV | HEXB; HEXB; H | P07686; H0Y | HEXB_HUMAN | Unmodified | 0 | 0 | 0 | 0 | 0 | 0 | 0 | 0 | 3193100 | 393820 | 335320 | 167250 | 0 | 0 | 0 | 620570 | 0 | 0 |
| GSKGVYQY | MYO1E; MYO1E | Q12965; H0Y | MYO1E_HUMAI | Acetyl (Protein N | 0 | 0 | 0 | 0 | 0 | 0 | 0 | 0 | 0 | 101110 | 0 | 0 | 0 | 0 | 0 | 0 | 1157600 | 335000 |
| GSKPGGV( | OTUD7B | Q6GQQ9 | OTU7B_HUMAI | Unmodified | 0 | 276660 | 215740 | 353250 | 1824800 | 0 | 0 | 0 | 0 | 137100 | 0 | 296810 | 1252500 | 728920 | 169870 | 346440 | 181070 | 1300800 |
| GSLAADKV | HSD17B4; HSD | P51659; E7E | DHB4_HUMAN | Unmodified | 0 | 0 | 0 | 24975 | 0 | 0 | 0 | 0 | 3479400 | 0 | 0 | 0 | 0 | 0 | 0 | 0 | 0 | 0 |
| GSLAADKV | HSD17B4; HSD | P51659; E7E | DHB4_HUMAN | Unmodified | 0 | 0 | 0 | 0 | 0 | 0 | 0 | 0 | 0 | 597640 | 0 | 0 | 0 | 0 | 0 | 0 | 0 | 0 |
| GSLAAVAH | OGDH; OGDH; | Q02218; E9F | ODO1_HUMAN | Unmodified | 0 | 0 | 0 | 0 | 319470 | 0 | 522050 | 0 | 1266400 | 0 | 0 | 0 | 0 | 0 | 0 | 0 | 0 | 178590 |
| GSLAEAVG | EPN1 | Q9Y6I3 | EPN1_HUMAN | Unmodified | 0 | 896790 | 0 | 0 | 0 | 0 | 0 | 0 | 0 | 0 | 0 | 0 | 0 | 0 | 0 | 0 | 0 | 0 |
| GSLAEAVG | EPN1 | Q9Y6I3 | EPN1_HUMAN | Unmodified | 396130 | 0 | 433800 | 0 | 0 | 0 | 0 | 0 | 0 | 0 | 0 | 0 | 0 | 0 | 342010 | 374220 | 0 | 0 |
| GSLASLDSI | CTNND1; CTNN | O60716; E9F | CTND1_HUMAI | Unmodified | 108790 | 0 | 486060 | 731620 | 0 | 0 | 0 | 0 | 0 | 102190 | 98939 | 288450 | 98092 | 0 | 1786200 | 1638600 | 0 | 187000 |
| GSLASLDSI | CTNND1; CTNN | O60716; E9F | CTND1_HUMAI | Unmodified | 0 | 0 | 0 | 0 | 303310 | 0 | 0 | 0 | 0 | 0 | 269000 | 2395500 | 340600 | 0 | 0 | 0 | 124600 | 286240 |
| GSLCSGC( | PXN; PXN; PXN | P49023; A0A | PAXI_HUMAN F | Unmodified | 0 | 0 | 0 | 0 | 0 | 0 | 0 | 0 | 0 | 0 | 0 | 0 | 0 | 0 | 0 | 0 | 347780 | 337980 |
| GSLEEAEAI | ANLN; ANLN | Q9NQW6; H | ANLN_HUMAN | Unmodified | 0 | 0 | 0 | 111420 | 1686600 | 0 | 0 | 0 | 0 | 0 | 0 | 0 | 0 | 0 | 0 | 0 | 144040 | 0 |
| GSLEGPSA | ESPN; ESPN; ES | B1AK53; A0A | ESPN_HUMAN | Unmodified | 0 | 0 | 0 | 0 | 0 | 0 | 0 | 0 | 0 | 0 | 0 | 1177200 | 0 | 0 | 0 | 0 | 0 | 0 |
| GSLEITESQ | ZDHHC5; ZDHH | Q9C0B5; H0 | ZDHC5_HUMA | Unmodified | 273880 | 0 | 1290400 | 1518900 | 343430 | 0 | 0 | 0 | 1151800 | 191150 | 2120000 | 3730400 | 805820 | 0 | 1436700 | 1861900 | 0 | 0 |
| GSLESPATI | PSMD12 | O00232 | PSD12_HUMAN | Unmodified | 0 | 0 | 130850 | 0 | 112720 | 0 | 46158 | 0 | 0 | 0 | 0 | 6647300 | 0 | 0 | 121930 | 0 | 0 | 0 |
| GSLESPATI | PSMD12 | O00232 | PSD12_HUMAN | Unmodified | 0 | 0 | 0 | 0 | 0 | 0 | 0 | 0 | 0 | 0 | 432280 | 3053400 | 0 | 0 | 0 | 0 | 0 | 0 |
| GSLEVLNLI | SBDS; SBDS | Q9Y3A5; A0A | SBDS_HUMAN | Unmodified | 211770 | 0 | 0 | 0 | 0 | 0 | 0 | 0 | 0 | 0 | 219260 | 520690 | 0 | 0 | 446230 | 382840 | 0 | 0 |
| GSLEVLNLI | SBDS; SBDS | Q9Y3A5; A0A | SBDS_HUMAN | Unmodified | 0 | 690920 | 0 | 0 | 0 | 136090 | 0 | 0 | 0 | 0 | 217660 | 604860 | 792190 | 362750 | 486010 | 532990 | 597350 | 344800 |
| GSLFGGSV | PDCD6IP; PDCI | Q8WUM4; A0 | PDC6I_HUMAN | Unmodified | 13647000 | 26728000 | 12765000 | 10688000 | 5559200 | 606990 | 0 | 0 | 125900 | 4254700 | 0 | 4103100 | 3233400 | 3174000 | 23375000 | 18286000 | 555190 | 4270100 |
| GSLGATGE | AHNAK | Q09666 | AHNK_HUMAN | Oxidation (M) | 0 | 0 | 0 | 0 | 4253300 | 0 | 0 | 0 | 0 | 0 | 0 | 0 | 0 | 0 | 0 | 0 | 0 | 1774100 |
| GSLGVDGE | FAM234A; FAM2 | Q9H0X4; B4I | F234A_HUMAN | Unmodified | 1116600 | 621940 | 0 | 1152600 | 0 | 0 | 0 | 0 | 1671700 | 886380 | 764750 | 2685300 | 0 | 0 | 0 | 1645300 | 0 | 0 |
| GSLKEELLK | SWAP70; SWAF | Q9UH65; E9 | SWP70_HUMAI | Acetyl (Protein N | 0 | 317490 | 0 | 0 | 3257700 | 590220 | 385080 | 0 | 445460 | 151640 | 0 | 0 | 0 | 0 | 0 | 0 | 217480 | 524970 |
| GSLKPHDY | PHYH; PHYH; P | O14832; B1A | PAHX_HUMAN | Unmodified | 0 | 0 | 0 | 0 | 24471 | 0 | 0 | 25542 | 6576700 | 1092500 | 0 | 0 | 0 | 19165 | 0 | 0 | 0 | 0 |
| GSLLDFLK | HCK; YES1; SR( | P08631; P07 | HCK_HUMAN T | Unmodified | 1748200 | 2643200 | 3114100 | 1742300 | 6766300 | 1664200 | 1013500 | 0 | 11762000 | 4529700 | 0 | 14291000 | 1249900 | 1231100 | 4258300 | 4042600 | 3400100 | 1923700 |
| GSLLDFLKI | YES1; YES1 | P07947; J3Q | YES_HUMAN Ty | Unmodified | 19919 | 0 | 0 | 0 | 609690 | 220130 | 0 | 25025 | 289270 | 734700 | 0 | 76903 | 379480 | 460280 | 0 | 81322 | 1791100 | 1025900 |
| GSLLDFLK( | SRC | P12931 | SRC_HUMAN P | Unmodified | 0 | 0 | 0 | 0 | 0 | 0 | 0 | 0 | 0 | 146120 | 0 | 204990 | 0 | 0 | 0 | 48026 | 1694500 | 0 |
| GSLLDFLKS | LYN | P07948 | LYN_HUMAN Ty | Unmodified | 35576 | 46579 | 0 | 81591 | 1917000 | 471060 | 24927 | 0 | 311210 | 1078300 | 0 | 0 | 85840 | 0 | 40545 | 46386 | 453800 | 0 |
| GSLLEDVVI | CLGN | O14967 | CLGN_HUMAN | Unmodified | 0 | 0 | 0 | 0 | 0 | 0 | 0 | 0 | 2266500 | 27919 | 0 | 0 | 0 | 0 | 0 | 0 | 0 | 0 |
| GSLLFTAGI | BLVRA | P53004 | BIEA_HUMAN E | Unmodified | 0 | 0 | 0 | 203190 | 392290 | 0 | 104630 | 0 | 0 | 154540 | 0 | 0 | 1118900 | 412470 | 236370 | 0 | 309980 | 187450 |
| GSLLTLQR | ASPH | Q12797 | ASPH_HUMAN | Unmodified | 608880 | 67706 | 0 | 0 | 0 | 0 | 0 | 0 | 6082800 | 0 | 0 | 0 | 0 | 0 | 0 | 0 | 0 | 0 |
| GSLLWNQ( | EMC10; EMC10 | Q5UCC4; M( | EMC10_HUMAI | Unmodified | 0 | 0 | 0 | 0 | 0 | 0 | 0 | 0 | 4653800 | 0 | 0 | 132090 | 0 | 0 | 0 | 0 | 0 | 0 |
| GSLNITTPG | VIL1; VIL1 | P09327; C9J | VILI_HUMAN Vi | Unmodified | 68175 | 0 | 0 | 0 | 0 | 0 | 0 | 0 | 80625 | 17055 | 146580 | 2250400 | 0 | 0 | 0 | 0 | 0 | 0 |
| GSLPAGEH | ARRDC1; ARRD | Q8N5I2; Q5T | ARRD1_HUMAI | Unmodified | 9368400 | 5936600 | 1947700 | 5401000 | 0 | 0 | 213080 | 0 | 2736900 | 370870 | 0 | 1151200 | 0 | 0 | 27362000 | 8802400 | 0 | 0 |
| GSLQILDQI | MRI1 | Q9BV20 | MTNA_HUMAN | Unmodified | 0 | 24819 | 0 | 19379 | 0 | 0 | 0 | 0 | 0 | 84168 | 39842 | 183240 | 12821 | 0 | 29866 | 0 | 0 | 47870 |
| GSLREDDL | PALD1 | Q9ULE6 | PALD_HUMAN | Unmodified | 0 | 0 | 0 | 0 | 0 | 0 | 0 | 0 | 257200 | 181130 | 325520 | 902620 | 269660 | 181060 | 0 | 0 | 408050 | 657490 |
| GSLTFEPLT | XPNPEP1; XPNI | Q9NQW7; Q | XPP1_HUMAN | Unmodified | 0 | 0 | 222230 | 187780 | 0 | 0 | 0 | 0 | 447580 | 0 | 270140 | 3307300 | 0 | 0 | 433080 | 724140 | 0 | 0 |
| GSLTSEEFA | VPS36 | Q86VN1 | VPS36_HUMAN | Unmodified | 96555 | 2091000 | 2510900 | 622560 | 383470 | 0 | 112940 | 0 | 280470 | 157660 | 265330 | 464400 | 640580 | 0 | 5132500 | 1183100 | 75302 | 0 |
| GSLVAPDR | AARS1; AARS1; | P49588; A0A | SYAC_HUMAN | Unmodified | 0 | 0 | 0 | 0 | 0 | 0 | 0 | 0 | 0 | 0 | 0 | 0 | 0 | 1452400 | 0 | 0 | 0 | 0 |
| GSLVDYLR | CSK | P41240 | CSK_HUMAN T | Unmodified | 0 | 0 | 0 | 0 | 0 | 0 | 0 | 0 | 0 | 0 | 134580 | 2581500 | 318760 | 0 | 45491 | 0 | 0 | 156410 |
| GSNKLVIEE | CCT4 | P50991 | TCPD_HUMAN | Unmodified | 0 | 0 | 0 | 46333 | 0 | 0 | 0 | 0 | 99530 | 332470 | 95380 | 0 | 0 | 293420 | 0 | 0 | 0 | 193730 |
| GSNLPSPV | FMNL2 | Q96PY5 | FMNL2_HUMAI | Unmodified | 0 | 0 | 0 | 0 | 0 | 0 | 0 | 0 | 0 | 0 | 0 | 1201600 | 0 | 0 | 0 | 0 | 0 | 0 |
| GSNMDFRI | EIF4H; EIF4H; E | Q15056; A0A | IF4H_HUMAN E | Unmodified | 177780 | 40339 | 0 | 0 | 0 | 0 | 22923 | 0 | 0 | 0 | 116890 | 208790 | 0 | 0 | 388400 | 306900 | 0 | 0 |
| GSNNVAL( | COPB2; COPB2 | P35606; H0Y | COPB2_HUMAI | Unmodified | 0 | 0 | 0 | 0 | 0 | 0 | 0 | 0 | 0 | 0 | 0 | 1674300 | 0 | 0 | 0 | 0 | 0 | 0 |
| GSNRSSLN | DIP2B | Q9P265 | DIP2B_HUMAN | Unmodified | 0 | 0 | 0 | 0 | 550160 | 0 | 0 | 0 | 806980 | 0 | 0 | 0 | 0 | 0 | 0 | 0 | 450760 | 1737200 |
| GSNTELTVI | NCAM2; NCAM | O15394; H9I | NCAM2_HUMA | Unmodified | 330390 | 1457600 | 2838300 | 3131000 | 0 | 0 | 0 | 72459 | 0 | 0 | 160470 | 139990 | 0 | 0 | 4024400 | 2627800 | 1070700 | 125370 |
| GSNTTSHL | UQCRC2; UQCI | P22695; H3E | QCR2_HUMAN | Unmodified | 0 | 0 | 0 | 0 | 0 | 0 | 0 | 13080 | 895930 | 293110 | 109360 | 0 | 0 | 0 | 0 | 0 | 0 | 0 |
| GSPESRLSI | GANAB; GANAE | Q14697; F5H | GANAB_HUMA | Unmodified | 0 | 0 | 0 | 118570 | 0 | 0 | 0 | 0 | 0 | 2066000 | 0 | 0 | 0 | 0 | 0 | 0 | 0 | 0 |
| GSPGADGI | COL1A1 | P02452 | CO1A1_HUMAI | Unmodified | 0 | 0 | 0 | 0 | 0 | 0 | 0 | 0 | 0 | 0 | 0 | 0 | 0 | 0 | 0 | 0 | 0 | 0 |
| GSPGCPLC | VPS37D; VPS37 | Q86XT2; C9J | VP37D_HUMAN | Unmodified | 2104800 | 911120 | 589290 | 1773100 | 0 | 0 | 494140 | 0 | 0 | 425430 | 0 | 0 | 1442700 | 0 | 941790 | 564820 | 0 | 459590 |
| GSPLLIGVF | GFPT1; GFPT1; | Q06210; A0A | GFPT1_HUMAN | Unmodified | 0 | 0 | 0 | 0 | 0 | 0 | 0 | 0 | 0 | 0 | 0 | 0 | 388680 | 207590 | 0 | 0 | 0 | 0 |
| GSPLVVISC | CRMP1; DPYSL | Q14194; Q1( | DPYL1_HUMAN | Unmodified | 1461300 | 344180 | 0 | 2268300 | 0 | 0 | 1079700 | 90400 | 2007000 | 509810 | 0 | 5243100 | 0 | 537320 | 0 | 0 | 0 | 0 |
| GSPMEISLI | LTA4H | P09960 | LKHA4_HUMAN | Oxidation (M) | 0 | 0 | 0 | 0 | 0 | 0 | 42129 | 0 | 0 | 0 | 1024700 | 1578100 | 0 | 0 | 0 | 0 | 0 | 0 |
| GSPMEISLI | LTA4H | P09960 | LKHA4_HUMAN | Unmodified | 0 | 0 | 0 | 0 | 0 | 0 | 0 | 0 | 0 | 0 | 173750 | 551600 | 0 | 0 | 0 | 0 | 0 | 0 |
| GSPNANEF | LAP3; LAP3 | P28838; H0Y | AMPL_HUMAN | Unmodified | 0 | 0 | 0 | 0 | 0 | 0 | 0 | 0 | 0 | 0 | 246980 | 1393400 | 0 | 0 | 0 | 0 | 0 | 274450 |
| GSPQAGVI | SNTB1 | Q13884 | SNTB1_HUMAN | Unmodified | 0 | 0 | 0 | 0 | 0 | 0 | 0 | 0 | 2166600 | 288780 | 0 | 0 | 0 | 0 | 0 | 0 | 0 | 0 |
| GSQDHSG | HSH2D | Q96JZ2 | HSH2D_HUMA | Unmodified | 0 | 0 | 1197600 | 939140 | 0 | 0 | 0 | 0 | 0 | 0 | 0 | 0 | 0 | 0 | 1717100 | 1214600 | 0 | 0 |
| GSQFGQS( | EPRS1; EPRS1 | P07814; V9C | SYEP_HUMAN | Unmodified | 0 | 0 | 0 | 0 | 0 | 0 | 0 | 0 | 0 | 0 | 0 | 1796300 | 472510 | 0 | 0 | 0 | 0 | 0 |
| GSQFGQS( | EPRS1; EPRS1 | P07814; V9C | SYEP_HUMAN | Unmodified | 0 | 0 | 0 | 0 | 143120 | 0 | 271890 | 138240 | 0 | 88371 | 0 | 0 | 280800 | 390650 | 0 | 0 | 173480 | 247150 |
| GSQGYLVV | SUN1; SUN1; S | O94901; H0Y | SUN1_HUMAN | Unmodified | 135960 | 61159 | 0 | 0 | 0 | 0 | 0 | 0 | 1272900 | 0 | 0 | 555450 | 0 | 0 | 227220 | 235810 | 0 | 0 |
| GSQPPPAA | EPB41L2; EPB4 | O43491; E9F | E41L2_HUMAN | Unmodified | 0 | 0 | 222040 | 182350 | 1543000 | 467060 | 26279 | 0 | 0 | 0 | 63488 | 519760 | 430310 | 0 | 295180 | 1185900 | 213120 | 722280 |
| GSQQYRAI | TUBB; TUBB; TU | P07437; Q5J | TBB5_HUMAN | Unmodified | 0 | 0 | 0 | 0 | 6089300 | 0 | 5265300 | 362200 | 82458 | 3953500 | 0 | 0 | 3683600 | 56413 | 0 | 0 | 0 | 0 |
| GSQSSKAP | MARCKSL1 | P49006 | MRP_HUMAN N | Unmodified | 0 | 0 | 0 | 0 | 0 | 0 | 0 | 0 | 0 | 0 | 0 | 0 | 0 | 722240 | 0 | 0 | 0 | 145950 |
| GSQTNLKD | KIF23; KIF23; KI | Q02241; A0A | KIF23_HUMAN | Unmodified | 0 | 0 | 0 | 0 | 4426100 | 0 | 0 | 0 | 0 | 1188700 | 0 | 0 | 116870 | 0 | 0 | 14585 | 867420 | 0 |
| GSRCWGE | ERBB2; ERBB2; | P04626; J3Q | ERBB2_HUMAN | Unmodified | 0 | 0 | 0 | 0 | 228110 | 120120 | 0 | 0 | 0 | 0 | 0 | 0 | 106680 | 0 | 0 | 0 | 0 | 0 |
| GSRIELGD | NAA50; NAA50; | Q9GZZ1; A0A | NAA50_HUMAN | Unmodified | 0 | 0 | 0 | 111210 | 0 | 0 | 0 | 0 | 280160 | 172820 | 0 | 1293200 | 368370 | 516390 | 0 | 0 | 242890 | 0 |
| GSRLLESSI | ITGA5 | P08648 | ITA5_HUMAN I | Unmodified | 0 | 0 | 0 | 0 | 15038000 | 5844300 | 0 | 0 | 0 | 0 | 0 | 0 | 1059300 | 0 | 0 | 0 | 4181200 | 5150300 |
| GSRNEVIG | SYT4 | Q9H2B2 | SYT4_HUMAN | Unmodified | 0 | 0 | 1884200 | 0 | 0 | 0 | 0 | 0 | 526100 | 0 | 0 | 0 | 0 | 0 | 9027500 | 3201600 | 0 | 0 |
| GSRVDIETF | AHNAK | Q09666 | AHNK_HUMAN | Unmodified | 0 | 0 | 0 | 397170 | 5730000 | 1042300 | 0 | 0 | 128560 | 0 | 0 | 0 | 258450 | 212860 | 0 | 0 | 1439500 | 3525000 |
| GSRVDIETF | AHNAK | Q09666 | AHNK_HUMAN | Unmodified | 0 | 0 | 0 | 0 | 8689700 | 0 | 0 | 0 | 0 | 0 | 0 | 0 | 0 | 0 | 0 | 0 | 0 | 4762700 |
| GSSAAGLT | MCM5; MCM5 | P33992; B1A | MCM5_HUMAN | Unmodified | 0 | 0 | 0 | 30845 | 87091 | 0 | 0 | 126490 | 268530 | 59681 | 158160 | 565030 | 88377 | 0 | 0 | 0 | 0 | 100070 |
| GSSAGFDF | PSMA6; PSMA6 | P60900; G3V | PSA6_HUMAN | Unmodified | 0 | 0 | 0 | 0 | 0 | 0 | 72709 | 0 | 0 | 664200 | 1216600 | 0 | 384140 | 100590 | 0 | 0 | 0 | 0 |
| GSSAGFDF | PSMA6; PSMA6 | P60900; G3V | PSA6_HUMAN | Unmodified | 0 | 0 | 0 | 0 | 361680 | 0 | 0 | 0 | 0 | 2919600 | 0 | 0 | 0 | 0 | 0 | 0 | 0 | 0 |
| GSSAGGL( | PTPRF; PTPRF; | P10586; H0Y | PTPRF_HUMAN | Unmodified | 753060 | 615280 | 0 | 0 | 0 | 0 | 0 | 39916 | 1068500 | 0 | 0 | 0 | 0 | 0 | 209990 | 182460 | 0 | 0 |
| GSSAGQLV | TMEM219; TMEI | Q86XT9; I3L | TM219_HUMAN | Unmodified | 0 | 0 | 0 | 0 | 0 | 0 | 0 | 0 | 913030 | 133430 | 0 | 0 | 0 | 0 | 0 | 0 | 0 | 0 |
| GSSCFECT | COMT; COMT | P21964; A0A | COMT_HUMAN | Unmodified | 1222100 | 0 | 0 | 0 | 0 | 0 | 0 | 0 | 0 | 0 | 0 | 1142400 | 0 | 0 | 830000 | 903880 | 0 | 0 |
| GSSGPTQII | STRN4; STRN4; | Q9NRL3; M0 | STRN4_HUMAN | Unmodified | 0 | 0 | 0 | 0 | 0 | 0 | 0 | 0 | 0 | 0 | 0 | 2042600 | 0 | 0 | 0 | 0 | 0 | 0 |
| GSSLHVGS | PLXNB2; PLXNE | O15031; H0Y | PLXB2_HUMAN | Unmodified | 69815 | 0 | 1232600 | 970000 | 3678400 | 0 | 0 | 0 | 0 | 0 | 0 | 1006800 | 2247000 | 632750 | 1314500 | 1902100 | 450920 | 2989400 |
| GSSLKIEER | CALD1; CALD1; | Q05682; C9J | CALD1_HUMAI | Unmodified | 31456 | 207080 | 0 | 0 | 208480 | 0 | 510760 | 8777.7 | 0 | 0 | 0 | 0 | 1815700 | 1551500 | 0 | 0 | 925950 | 786740 |
| GSSNFCVK | AHCYL1 | O43865 | SAHH2_HUMAI | Unmodified | 0 | 0 | 0 | 0 | 3101100 | 0 | 187080 | 0 | 0 | 78844 | 0 | 0 | 1427500 | 0 | 0 | 0 | 1268700 | 1743400 |
| GSSQPNLS | EPN2; EPN2; EF | O95208; F6F | EPN2_HUMAN | Unmodified | 849050 | 666760 | 564310 | 789580 | 0 | 0 | 0 | 0 | 220390 | 0 | 0 | 398050 | 199600 | 28164 | 996940 | 859100 | 0 | 0 |
| GSSSASIVK | DSG2 | Q14126 | DSG2_HUMAN | Unmodified | 0 | 0 | 0 | 0 | 2410300 | 0 | 0 | 0 | 0 | 0 | 0 | 146500 | 341490 | 229200 | 0 | 34473 | 0 | 0 |
| GSSTVRFT( | ANPEP | P15144 | AMPN_HUMAN | Unmodified | 0 | 0 | 0 | 0 | 0 | 0 | 0 | 0 | 0 | 0 | 0 | 0 | 0 | 0 | 0 | 0 | 2442600 | 1204400 |
| GSTAENAE | EGFR; EGFR | P00533; E9P | EGFR_HUMAN | Unmodified | 0 | 0 | 598970 | 832890 | 2243700 | 352920 | 0 | 0 | 0 | 0 | 0 | 0 | 264870 | 0 | 595490 | 651770 | 104540 | 684880 |
| GSTAENAE | EGFR; EGFR | P00533; E9P | EGFR_HUMAN | Unmodified | 0 | 0 | 0 | 0 | 9635100 | 0 | 0 | 53909 | 0 | 0 | 0 | 0 | 0 | 688300 | 0 | 0 | 560870 | 1974400 |
| GSTAPVGG | PDIA6 | Q15084 | PDIA6_HUMAN | Unmodified | 0 | 0 | 0 | 0 | 0 | 0 | 0 | 0 | 18159000 | 467850 | 681260 | 2723800 | 0 | 0 | 247400 | 847500 | 0 | 0 |
| GSTAVGVR | PSMA7 | O14818 | PSA7_HUMAN | Unmodified | 0 | 0 | 0 | 0 | 1025000 | 0 | 230560 | 0 | 0 | 431860 | 0 | 0 | 807720 | 550200 | 0 | 0 | 818190 | 402250 |
| GSTDDKGF | CNDP2; CNDP2 | Q96KP4; A0A | CNDP2_HUMA | Unmodified | 85716 | 0 | 0 | 290370 | 0 | 0 | 0 | 0 | 1619600 | 54890 | 2335600 | 7947700 | 215550 | 0 | 896370 | 350210 | 0 | 0 |
| GSTDNLMI | CCT8 | P50990 | TCPQ_HUMAN | Unmodified | 0 | 0 | 0 | 0 | 0 | 0 | 0 | 0 | 243740 | 0 | 0 | 2571300 | 0 | 0 | 0 | 60576 | 0 | 0 |
| GSTDNLMI | CCT8 | P50990 | TCPQ_HUMAN | Oxidation (M) | 0 | 0 | 0 | 0 | 0 | 0 | 49637 | 0 | 0 | 0 | 158390 | 691410 | 0 | 0 | 0 | 0 | 0 | 0 |
| GSTEQDAA | APLP1; APLP1; | P51693; B7Z | APLP1_HUMAN | Unmodified | 0 | 0 | 0 | 1356000 | 0 | 0 | 0 | 0 | 253980 | 0 | 0 | 0 | 0 | 0 | 0 | 0 | 0 | 0 |
| GSTFRPHD | KIAA1522 | Q9P206 | K1522_HUMAN | Unmodified | 0 | 0 | 1169400 | 595740 | 1379700 | 0 | 62756 | 0 | 0 | 240680 | 239580 | 1002700 | 2185400 | 2362700 | 389720 | 478820 | 414290 | 870030 |
| GSTHPQP( | GLS | O94925 | GLSK_HUMAN | Unmodified | 0 | 0 | 0 | 0 | 0 | 0 | 0 | 0 | 306280 | 0 | 0 | 0 | 0 | 0 | 0 | 0 | 0 | 0 |
| GSTHPQP( | GLS | O94925 | GLSK_HUMAN | Unmodified | 0 | 0 | 0 | 0 | 867180 | 454840 | 0 | 0 | 988710 | 0 | 0 | 0 | 0 | 0 | 0 | 0 | 0 | 0 |
| GSTIETEQK | ENAH; ENAH | Q8N8S7; A0 | ENAH_HUMAN | Unmodified | 0 | 0 | 0 | 0 | 8020700 | 0 | 0 | 1498100 | 3071400 | 1519700 | 929630 | 9431000 | 2880300 | 1967800 | 649620 | 1030600 | 2301500 | 3748600 |
| GSTIFLGFS | CEMIP2; CEMIP | Q9UHN6; B4 | CEIP2_HUMAN | Unmodified | 0 | 29087 | 0 | 0 | 4346400 | 0 | 0 | 0 | 738440 | 524950 | 0 | 0 | 354370 | 0 | 0 | 0 | 93850 | 0 |
| GSTISQSLA | PLXNB3; PLXNE | Q9ULL4; H7( | PLXB3_HUMAN | Unmodified | 0 | 0 | 87453 | 61666 | 0 | 0 | 0 | 0 | 0 | 0 | 0 | 0 | 0 | 0 | 169280 | 136300 | 0 | 0 |
| GSTPYGGV | ACAT1; ACAT1; | P24752; H0Y | THIL_HUMAN A | Unmodified | 0 | 0 | 0 | 0 | 0 | 0 | 0 | 0 | 4176300 | 0 | 112900 | 0 | 0 | 0 | 196150 | 0 | 29859 | 0 |
| GSTPYKGD | SCRIB; SCRIB; | Q14160; H0Y | SCRIB_HUMAN | Unmodified | 0 | 0 | 325530 | 0 | 0 | 0 | 0 | 0 | 4440200 | 267700 | 298870 | 1976200 | 0 | 0 | 498870 | 755860 | 0 | 0 |
| GSTPYKGD | SCRIB; SCRIB; | Q14160; H0Y | SCRIB_HUMAN | Unmodified | 0 | 0 | 0 | 0 | 0 | 0 | 0 | 0 | 2344100 | 82285 | 0 | 0 | 0 | 0 | 0 | 0 | 0 | 0 |
| GSTSLLVHI | FZD6 | O60353 | FZD6_HUMAN | Unmodified | 0 | 0 | 0 | 0 | 2873300 | 0 | 0 | 0 | 880310 | 0 | 0 | 0 | 0 | 0 | 0 | 0 | 0 | 696760 |
| GSVAIWSG | CD46 | P15529 | MCP_HUMAN N | Unmodified | 10624000 | 4897500 | 2441000 | 3448600 | 4521700 | 749540 | 0 | 0 | 4783300 | 1007600 | 698930 | 8278300 | 2728100 | 2141200 | 4442500 | 3351000 | 526750 | 1653100 |
| GSVAYVPQ | ABCC3 | O15438 | MRP3_HUMAN | Unmodified | 0 | 0 | 0 | 0 | 2151100 | 0 | 0 | 0 | 0 | 0 | 0 | 0 | 0 | 0 | 0 | 0 | 0 | 0 |
| GSVAYVPQ | ABCC1; ABCC1 | P33527; I3L | MRP1_HUMAN | Unmodified | 0 | 0 | 0 | 0 | 0 | 0 | 0 | 0 | 0 | 0 | 0 | 0 | 0 | 0 | 0 | 0 | 0 | 0 |
| GSVDHEN\ | CANT1 | Q8WVQ1 | CANT1_HUMAI | Unmodified | 42129 | 153270 | 149430 | 113630 | 0 | 0 | 0 | 0 | 21714000 | 734320 | 0 | 148300 | 0 | 0 | 354640 | 142050 | 0 | 0 |
| GSVDSNW | TOMM40 | O96008 | TOM40_HUMAI | Unmodified | 0 | 0 | 0 | 0 | 0 | 0 | 0 | 0 | 5755200 | 1606800 | 0 | 0 | 0 | 0 | 0 | 0 | 0 | 0 |
| GSVLEPEG | ACACA; ACACA | Q13085; Q5 | ACACA_HUMAI | Unmodified | 0 | 0 | 153470 | 0 | 0 | 0 | 0 | 0 | 0 | 0 | 181600 | 943940 | 0 | 0 | 0 | 0 | 0 | 0 |

| | | | | | | | | | | | | | | | | | | | | | | |
|---|---|---|---|---|---|---|---|---|---|---|---|---|---|---|---|---|---|---|---|---|---|---|
| GSVLPNSD | SCP2; SCP2 | P22307; H0Y | SCP2_HUMAN | Unmodified | 0 | 0 | 0 | 0 | 175750 | 0 | 0 | 0 | 23034000 | 46818 | 63380 | 0 | 0 | 0 | 0 | 0 | 196280 | 109470 |
| GSVLPNSD | SCP2; SCP2 | P22307; H0Y | SCP2_HUMAN | Unmodified | 0 | 0 | 0 | 0 | 0 | 0 | 0 | 0 | 652520 | 0 | 0 | 0 | 0 | 0 | 0 | 0 | 0 | 0 |
| GSVLQEAK | SCARB1 | B7ZKQ9 | B7ZKQ9_HUMA | Unmodified | 2191300 | 2101500 | 1615300 | 1435100 | 11869000 | 0 | 0 | 0 | 567680 | 0 | 2541500 | 6075400 | 0 | 0 | 2408100 | 1609600 | 0 | 0 |
| GSVSSEAS | DTX3L | Q8TDB6 | DTX3L_HUMAN | Unmodified | 0 | 0 | 0 | 0 | 0 | 0 | 0 | 0 | 212090 | 0 | 46381 | 0 | 0 | 0 | 0 | 189920 | 0 | 499720 |
| GSVTLSEIR | ANLN; ANLN; A | Q9NQW6; H | ANLN_HUMAN | Unmodified | 75994 | 0 | 283200 | 265800 | 4186300 | 0 | 77825 | 0 | 264240 | 69054 | 257410 | 440630 | 233180 | 0 | 238410 | 140600 | 279340 | 358360 |
| GSVTVLSG | ALKBH5 | Q6P6C2 | ALKB5_HUMAN | Unmodified | 0 | 0 | 0 | 0 | 0 | 0 | 0 | 0 | 0 | 209400 | 0 | 729300 | 0 | 0 | 0 | 0 | 0 | 0 |
| GSVVNVNF | MINK1; MAP4K | Q8N4C8; O9 | MINK1_HUMAN | Unmodified | 0 | 0 | 454120 | 0 | 0 | 0 | 0 | 0 | 543700 | 98208 | 228040 | 831100 | 0 | 0 | 1212800 | 431680 | 0 | 83394 |
| GSVVNVNF | MAP4K4; MAP4 | O95819; C9J | M4K4_HUMAN | Unmodified | 133120 | 0 | 768550 | 789030 | 0 | 0 | 0 | 0 | 5951400 | 266640 | 1059400 | 8541800 | 0 | 0 | 716940 | 1169900 | 161670 | 252610 |
| GSVVNVNF | MAP4K4; MAP4 | O95819; C9J | M4K4_HUMAN | Unmodified | 0 | 0 | 0 | 507150 | 407950 | 222580 | 64537 | 64863 | 2105500 | 462930 | 968560 | 4197200 | 0 | 0 | 0 | 0 | 477470 | 1177500 |
| GSVYIGGA | HSPG2 | P98160 | PGBM_HUMAN | Unmodified | 0 | 0 | 307180 | 252720 | 0 | 0 | 0 | 0 | 77763 | 0 | 0 | 0 | 0 | 0 | 106230 | 0 | 0 | 81397 |
| GSYGSDAE | TJP2; TJP2; TJP2 | Q9UDY2; A0 | ZO2_HUMAN T | Unmodified | 0 | 0 | 0 | 0 | 109630 | 0 | 0 | 0 | 2550500 | 0 | 220960 | 720230 | 0 | 0 | 0 | 173480 | 0 | 0 |
| GSYLTHEA | CADM4 | Q8NFZ8 | CADM4_HUMA | Unmodified | 99950 | 0 | 149120 | 0 | 0 | 0 | 0 | 0 | 0 | 0 | 0 | 6093300 | 0 | 0 | 430400 | 170830 | 0 | 20721 |
| GSYNPVTH | HEXA; HEXA; HI | P06865; H3E | HEXA_HUMAN | Unmodified | 0 | 0 | 0 | 0 | 1747300 | 0 | 0 | 30709 | 1540600 | 0 | 0 | 187350 | 0 | 0 | 0 | 0 | 769580 | 0 |
| GSYSLSHV | HEXB; HEXB | P07686; Q5L | HEXB_HUMAN | Unmodified | 322630 | 0 | 826130 | 0 | 0 | 0 | 0 | 0 | 2855200 | 390080 | 0 | 0 | 0 | 0 | 904110 | 0 | 0 | 0 |
| GSYSLSVR | SLA2; HCK; HC | Q9H6Q3; P0 | SLAP2_HUMAN | Unmodified | 0 | 188020 | 0 | 284340 | 0 | 0 | 0 | 0 | 0 | 0 | 872790 | 4166500 | 0 | 0 | 149560 | 185260 | 0 | 0 |
| GSYVSIHS | DDX39B; DDX3 | Q13838; O0 | DX39B_HUMAN | Unmodified | 49570 | 39610 | 0 | 0 | 0 | 180810 | 11192 | 0 | 0 | 0 | 446200 | 1752500 | 0 | 0 | 33380 | 0 | 0 | 0 |
| GSYVSIHS | DDX39B; DDX3 | Q13838; O0 | DX39B_HUMAN | Unmodified | 268100 | 519740 | 0 | 0 | 701650 | 0 | 787980 | 0 | 485060 | 0 | 848990 | 3568600 | 508490 | 198700 | 0 | 0 | 0 | 408050 |
| GTAAAAAA | RPL14 | P50914 | RL14_HUMAN ( | Unmodified | 0 | 0 | 0 | 0 | 0 | 0 | 0 | 0 | 357350 | 191610 | 0 | 0 | 0 | 0 | 0 | 0 | 166960 | 486530 |
| GTAAILEYL | LRRC40 | Q9H9A6 | LRC40_HUMAN | Unmodified | 0 | 0 | 0 | 0 | 0 | 0 | 0 | 0 | 0 | 0 | 0 | 2873000 | 0 | 0 | 0 | 0 | 0 | 0 |
| GTADAFIVF | IGF2R; IGF2R | P11717; A0A | MPRI_HUMAN ( | Unmodified | 151500 | 0 | 705200 | 436310 | 0 | 0 | 0 | 0 | 37607000 | 2057800 | 1827900 | 5563600 | 798870 | 0 | 403570 | 507710 | 133640 | 151960 |
| GTADAFIVF | IGF2R; IGF2R | P11717; A0A | MPRI_HUMAN ( | Unmodified | 0 | 0 | 0 | 65809 | 0 | 0 | 0 | 0 | 0 | 2367700 | 0 | 0 | 0 | 0 | 0 | 0 | 0 | 0 |
| GTADVTHD | SLC2A1; SLC2A | P11166; A0A | GTR1_HUMAN | Unmodified | 0 | 0 | 285160 | 0 | 0 | 0 | 0 | 0 | 0 | 0 | 0 | 388740 | 0 | 0 | 96450 | 0 | 0 | 0 |
| GTADVTHD | SLC2A1; SLC2A | P11166; A0A | GTR1_HUMAN | Unmodified | 413270 | 0 | 991780 | 780140 | 0 | 0 | 0 | 0 | 6804900 | 702120 | 840160 | 24732000 | 0 | 0 | 1313200 | 2751800 | 592230 | 0 |
| GTADVTHD | SLC2A1; SLC2A | P11166; A0A | GTR1_HUMAN | Oxidation (M) | 0 | 0 | 931490 | 490140 | 0 | 0 | 0 | 0 | 304930 | 0 | 208670 | 3427600 | 0 | 0 | 1371300 | 632270 | 0 | 0 |
| GTAEKLKPI | ITGB1; ITGB1; I | P05556; E7E | ITB1_HUMAN I | Unmodified | 0 | 0 | 0 | 0 | 8825500 | 0 | 0 | 0 | 0 | 0 | 0 | 0 | 0 | 0 | 0 | 0 | 10153000 | 9969000 |
| GTAFGGWI | ADH5 | P11766 | ADHX_HUMAN | Unmodified | 0 | 0 | 185490 | 0 | 0 | 0 | 0 | 0 | 526180 | 76279 | 1682700 | 11852000 | 0 | 0 | 0 | 0 | 0 | 89420 |
| GTAVVNGE | PRDX3 | P30048 | PRDX3_HUMAN | Unmodified | 119700 | 84159 | 183900 | 186970 | 0 | 0 | 0 | 0 | 10979000 | 863170 | 321720 | 904560 | 0 | 0 | 0 | 298420 | 44173 | 0 |
| GTAVVNGE | PRDX3 | P30048 | PRDX3_HUMAN | Unmodified | 0 | 0 | 0 | 0 | 262150 | 224590 | 223720 | 0 | 1608200 | 2075700 | 0 | 0 | 0 | 0 | 0 | 0 | 0 | 0 |
| GTAYTFFTF | DDX17; DDX17 | Q92841; A0A | DDX17_HUMAN | Unmodified | 0 | 0 | 0 | 0 | 161010 | 0 | 0 | 0 | 0 | 0 | 0 | 3950300 | 0 | 0 | 0 | 0 | 0 | 0 |
| GTAYVVYEI | SF3B6 | Q9Y3B4 | SF3B6_HUMAN | Unmodified | 504560 | 437220 | 310630 | 236210 | 0 | 0 | 0 | 0 | 665670 | 0 | 242470 | 1182000 | 0 | 0 | 0 | 215640 | 0 | 0 |
| GTCEQGPH | GRN; GRN; GRN | P28799; K7E | GRN_HUMAN F | Unmodified | 0 | 0 | 0 | 0 | 0 | 0 | 0 | 0 | 588760 | 0 | 0 | 0 | 0 | 0 | 0 | 0 | 40193 | 56524 |
| GTCLRFAG | NHLRC2 | Q8NBF2 | NHLC2_HUMA | Unmodified | 0 | 0 | 0 | 0 | 0 | 0 | 297900 | 0 | 0 | 0 | 0 | 0 | 501230 | 237250 | 0 | 0 | 160540 | 249070 |
| GTCSGFEP | LMCD1; LMCD1 | Q9NZU5; H7 | LMCD1_HUMA | Unmodified | 0 | 0 | 0 | 0 | 0 | 0 | 0 | 0 | 0 | 0 | 0 | 1259800 | 0 | 0 | 0 | 0 | 0 | 0 |
| GTCVEGTIF | USP7; USP7; US | Q93009; A0A | UBP7_HUMAN | Unmodified | 0 | 0 | 0 | 0 | 0 | 0 | 25589 | 0 | 0 | 0 | 60847 | 738910 | 0 | 0 | 185900 | 0 | 78961 | 0 |
| GTDECAIES | CTSC; CTSC; C | P53634; A0A | CATC_HUMAN | Unmodified | 0 | 0 | 0 | 0 | 0 | 0 | 0 | 0 | 0 | 0 | 915070 | 1382400 | 0 | 0 | 0 | 0 | 0 | 0 |
| GTDECAIES | CTSC; CTSC; C | P53634; A0A | CATC_HUMAN | Unmodified | 0 | 0 | 0 | 0 | 376190 | 0 | 0 | 0 | 914720 | 0 | 2693900 | 1162100 | 0 | 0 | 0 | 0 | 0 | 0 |
| GTDELYAVI | PRKCB; PRKCB | P05771; H3E | KPCB_HUMAN | Unmodified | 0 | 0 | 91784 | 0 | 0 | 0 | 0 | 0 | 0 | 0 | 0 | 0 | 617020 | 0 | 0 | 156440 | 0 | 0 |
| GTDERQPV | PLA2G3 | Q9NZ20 | PA2G3_HUMAN | Oxidation (M) | 204900 | 0 | 0 | 0 | 0 | 0 | 0 | 0 | 0 | 0 | 0 | 0 | 0 | 0 | 0 | 0 | 0 | 0 |
| GTDIMYTG | SLC25A5 | P05141 | ADT2_HUMAN | Oxidation (M) | 0 | 0 | 0 | 0 | 0 | 0 | 0 | 0 | 2372700 | 827040 | 3407700 | 0 | 0 | 0 | 0 | 0 | 0 | 0 |
| GTDIMYTG | SLC25A5 | P05141 | ADT2_HUMAN | Unmodified | 0 | 0 | 0 | 0 | 0 | 0 | 0 | 0 | 3832000 | 0 | 0 | 0 | 0 | 0 | 0 | 0 | 0 | 0 |
| GTDQSTMF | MVB12B | Q9H7P6 | MB12B_HUMAN | Unmodified | 0 | 0 | 238780 | 0 | 0 | 0 | 0 | 0 | 1317500 | 168550 | 0 | 131940 | 0 | 0 | 0 | 323800 | 0 | 0 |
| GTDSQAMI | FKBP5 | Q13451 | FKBP5_HUMAN | Unmodified | 157760 | 0 | 169800 | 129880 | 2219400 | 0 | 77827 | 75988 | 258620 | 0 | 0 | 703080 | 1411800 | 464800 | 222640 | 55442 | 0 | 0 |
| GTDTVAGL | NAMPT; NAMPT | P43490; A0A | NAMPT_HUMA | Unmodified | 0 | 0 | 0 | 0 | 0 | 0 | 0 | 0 | 0 | 0 | 0 | 1446300 | 0 | 0 | 0 | 0 | 0 | 0 |
| GTDTVAGL | NAMPT; NAMPT | P43490; A0A | NAMPT_HUMA | Unmodified | 0 | 0 | 109700 | 290100 | 1003200 | 0 | 0 | 0 | 0 | 0 | 290770 | 1922100 | 0 | 0 | 448380 | 144610 | 0 | 0 |
| GTDVNVFN | ANXA1 | P04083 | ANXA1_HUMAN | Unmodified | 0 | 7819200 | 0 | 12448000 | 42827000 | 29393000 | 2.72E+08 | 31347 | 1171000 | 832450 | 21382000 | 5.8E+08 | 46205000 | 99265000 | 4138900 | 29857 | 22568000 | 1.2E+08 |
| GTDVNVFN | ANXA1 | P04083 | ANXA1_HUMAN | Unmodified | 0 | 0 | 0 | 0 | 11795000 | 1989600 | 0 | 0 | 0 | 0 | 4078700 | 1192700 | 0 | 0 | 0 | 0 | 0 | 0 |
| GTDVPKWI | ANXA2P2; ANXA | A6NMY6; P0 | AXA2L_HUMAN | Unmodified | 0 | 0 | 0 | 0 | 420450 | 0 | 17926 | 0 | 0 | 230660 | 0 | 0 | 0 | 0 | 0 | 0 | 104090 | 1123000 |
| GTDVPKWI | ANXA2P2; ANXA | A6NMY6; P0 | AXA2L_HUMAN | Unmodified | 0 | 0 | 0 | 0 | 8625500 | 0 | 0 | 0 | 0 | 0 | 0 | 0 | 0 | 174170 | 0 | 0 | 2123200 | 2596300 |
| GTEAPAVV | PAK2 | Q13177 | PAK2_HUMAN | Unmodified | 698950 | 0 | 0 | 1565400 | 8924400 | 0 | 531890 | 0 | 5983300 | 1918700 | 2598500 | 5609500 | 6743400 | 2984300 | 905420 | 1134700 | 2636900 | 5276800 |
| GTEDELDK | TPM1; TPM3; TP | P09493; P06 | TPM1_HUMAN | Unmodified | 0 | 0 | 0 | 0 | 2525500 | 0 | 0 | 0 | 0 | 144820 | 0 | 0 | 0 | 0 | 0 | 0 | 49508 | 0 |
| GTEDITSPH | RUVBL1; RUVBL | Q9Y265; E7E | RUVB1_HUMAN | Unmodified | 259110 | 0 | 171890 | 89861 | 0 | 0 | 33576 | 32153 | 0 | 71947 | 413410 | 9875000 | 0 | 0 | 326220 | 409780 | 0 | 0 |
| GTEDITSPH | RUVBL1; RUVBL | Q9Y265; E7E | RUVB1_HUMAN | Oxidation (M) | 57455 | 0 | 0 | 0 | 0 | 0 | 0 | 0 | 0 | 0 | 238990 | 4802600 | 0 | 0 | 454920 | 0 | 0 | 0 |
| GTEELYAIK | PRKCA | P17252 | KPCA_HUMAN | Unmodified | 0 | 0 | 0 | 0 | 895200 | 0 | 0 | 0 | 0 | 0 | 0 | 371930 | 0 | 23506 | 149330 | 0 | 6029500 | 0 |
| GTEHSGST | ATP6V0A1; ATP | Q93050; B7Z | VPP1_HUMAN | Unmodified | 379520 | 225600 | 1664200 | 458570 | 0 | 0 | 0 | 103880 | 2385200 | 199040 | 312080 | 2189600 | 0 | 0 | 671430 | 579770 | 0 | 0 |
| GTEITHAVV | EIF3CL; EIF3C | B5ME19; Q9 | EIFCL_HUMAN | Unmodified | 0 | 0 | 0 | 0 | 0 | 0 | 0 | 0 | 0 | 0 | 0 | 1924300 | 0 | 0 | 0 | 39173 | 0 | 0 |
| GTEITHAVV | EIF3CL; EIF3C | B5ME19; Q9 | EIFCL_HUMAN | Unmodified | 113290 | 0 | 195670 | 0 | 0 | 0 | 0 | 0 | 0 | 27782 | 165850 | 1603100 | 0 | 0 | 0 | 142310 | 0 | 0 |
| GTEVQVDD | RUVBL2 | Q9Y230 | RUVB2_HUMAN | Unmodified | 0 | 0 | 0 | 0 | 357140 | 76590 | 131440 | 62357 | 155230 | 0 | 334300 | 2505100 | 0 | 0 | 0 | 0 | 0 | 178790 |
| GTEVQVDD | RUVBL2 | Q9Y230 | RUVB2_HUMAN | Unmodified | 0 | 0 | 0 | 41726 | 0 | 0 | 0 | 0 | 0 | 0 | 594420 | 2370500 | 0 | 0 | 0 | 0 | 0 | 0 |
| GTFCSFDT | ABAT; ABAT; AB | P80404; H3E | GABT_HUMAN | Unmodified | 0 | 0 | 0 | 0 | 0 | 0 | 0 | 0 | 932700 | 63019 | 0 | 0 | 0 | 0 | 0 | 0 | 0 | 0 |
| GTFKEQND | PXN; PXN; PXN | P49023; A0A | PAXI_HUMAN F | Unmodified | 0 | 0 | 671390 | 42626 | 0 | 0 | 0 | 0 | 0 | 0 | 0 | 0 | 372210 | 0 | 0 | 0 | 1150300 | 0 |
| GTFLDDTG | MUC5AC | P98088 | MUC5A_HUMA | Unmodified | 0 | 0 | 0 | 0 | 0 | 0 | 0 | 0 | 0 | 0 | 0 | 0 | 0 | 0 | 0 | 0 | 0 | 407830 |
| GTGARPYT | AARS1; AARS1; | P49588; A0A | SYAC_HUMAN | Unmodified | 0 | 0 | 0 | 0 | 56192 | 0 | 0 | 12718 | 0 | 0 | 0 | 202940 | 48270 | 0 | 0 | 0 | 0 | 0 |
| GTGASGSF | H1-3; H1-4; H1 | P16402; P10 | H13_HUMAN H | Unmodified | 0 | 0 | 0 | 0 | 0 | 0 | 0 | 5208500 | 0 | 0 | 0 | 0 | 0 | 0 | 0 | 0 | 0 | 0 |
| GTGASGSF | H1-3; H1-4; H1 | P16402; P10 | H13_HUMAN H | Unmodified | 0 | 0 | 0 | 0 | 0 | 0 | 0 | 3620700 | 0 | 0 | 0 | 0 | 0 | 0 | 0 | 0 | 0 | 0 |
| GTGASGSF | H1-3; H1-4; H1 | P16402; P10 | H13_HUMAN H | Unmodified | 0 | 60173 | 0 | 0 | 855230 | 0 | 3128400 | 2.53E+08 | 0 | 669530 | 0 | 0 | 0 | 163480 | 38239 | 0 | 18989 | 671440 |
| GTGATADF | NETO1 | Q8TDF5 | NETO1_HUMAN | Unmodified | 0 | 0 | 0 | 0 | 0 | 0 | 0 | 0 | 0 | 0 | 0 | 0 | 0 | 0 | 0 | 767590 | 0 | 0 |
| GTGGSTGE | CTAG1A; CTAG | P78358; O75 | CTG1B_HUMAN | Unmodified | 757720 | 0 | 169260 | 319900 | 0 | 0 | 0 | 0 | 0 | 0 | 0 | 0 | 0 | 0 | 0 | 0 | 0 | 0 |
| GTGGVDTA | CKMT1A | P12532 | KCRU_HUMAN | Unmodified | 0 | 0 | 0 | 0 | 0 | 0 | 0 | 0 | 0 | 0 | 0 | 0 | 0 | 0 | 0 | 0 | 0 | 0 |
| GTGGVDTA | CKMT1A | P12532 | KCRU_HUMAN | Unmodified | 0 | 37781 | 3547600 | 0 | 0 | 0 | 0 | 0 | 0 | 0 | 0 | 3260800 | 0 | 0 | 0 | 0 | 0 | 0 |
| GTGGVDTA | CKB; CKB | P12277; H0Y | KCRB_HUMAN | Unmodified | 0 | 0 | 0 | 0 | 0 | 0 | 0 | 0 | 0 | 0 | 0 | 0 | 0 | 0 | 0 | 0 | 0 | 0 |
| GTGIVSAP | RPS2; RPS2; RP | P15880; E9P | RS2_HUMAN 4 | Unmodified | 162860 | 1147300 | 1268300 | 804180 | 354360 | 119720 | 764610 | 42127 | 334970 | 206260 | 1383700 | 30706000 | 557050 | 0 | 1058900 | 1166200 | 0 | 732130 |
| GTGLFCST | MAN2A1 | Q16706 | MA2A1_HUMAN | Unmodified | 0 | 0 | 240730 | 0 | 158660 | 0 | 0 | 0 | 1487200 | 464490 | 0 | 2340300 | 0 | 0 | 0 | 83795 | 0 | 0 |
| GTGLLSSD | ZC3HAV1; ZC3H | Q7Z2W4; C9 | ZCCHV_HUMA | Unmodified | 0 | 379970 | 0 | 1001900 | 974450 | 2013600 | 0 | 0 | 531690 | 0 | 190930 | 0 | 220020 | 0 | 604270 | 0 | 0 | 0 |
| GTGLLSSD | ZC3HAV1; ZC3H | Q7Z2W4; C9 | ZCCHV_HUMA | Unmodified | 0 | 0 | 0 | 0 | 3193400 | 1167400 | 0 | 0 | 0 | 0 | 0 | 0 | 614050 | 369370 | 0 | 0 | 0 | 0 |
| GTGLQPGE | USP5 | P45974 | UBP5_HUMAN | Unmodified | 0 | 0 | 291800 | 0 | 0 | 0 | 0 | 0 | 298810 | 0 | 0 | 1224900 | 0 | 0 | 0 | 391350 | 0 | 0 |
| GTGLTAVA | SUSD2 | Q9UGT4 | SUSD2_HUMAN | Unmodified | 2370100 | 935020 | 0 | 0 | 0 | 0 | 0 | 0 | 8774700 | 552070 | 0 | 0 | 0 | 0 | 927140 | 0 | 0 | 0 |
| GTGREQQI | HSPA9; HSPA9; | P38646; H0Y | GRP75_HUMAN | Unmodified | 0 | 0 | 0 | 0 | 117900 | 0 | 0 | 0 | 1936400 | 0 | 0 | 0 | 2635600 | 0 | 0 | 0 | 0 | 1530500 |
| GTGRIPDQ | TMED10; TMED | P49755; G3V | TMEDA_HUMAN | Oxidation (M) | 31258 | 453510 | 1161300 | 932110 | 877790 | 257900 | 422310 | 70239 | 90106000 | 6088200 | 6378900 | 3646800 | 886680 | 472410 | 951880 | 644720 | 0 | 680030 |
| GTGRIPDQ | TMED10; TMED | P49755; G3V | TMEDA_HUMAN | Unmodified | 0 | 0 | 0 | 0 | 0 | 0 | 0 | 0 | 11167000 | 384310 | 0 | 0 | 0 | 0 | 0 | 0 | 0 | 70767 |
| GTGRIPDQ | TMED10; TMED | P49755; G3V | TMEDA_HUMAN | Unmodified | 0 | 0 | 0 | 0 | 768480 | 0 | 0 | 0 | 2384300 | 7704600 | 0 | 0 | 0 | 3138600 | 0 | 0 | 0 | 1757000 |
| GTGRIPDQ | TMED10; TMED | P49755; G3V | TMEDA_HUMAN | Oxidation (M) | 0 | 0 | 0 | 0 | 0 | 0 | 0 | 0 | 0 | 3110500 | 0 | 0 | 0 | 0 | 0 | 0 | 0 | 857640 |
| GTGVIQLYE | DNAAF10 | Q96MX6 | DAA10_HUMAN | Unmodified | 247940 | 274800 | 0 | 0 | 0 | 0 | 0 | 30220 | 0 | 0 | 0 | 2608100 | 13489 | 0 | 0 | 0 | 0 | 0 |
| GTGVVTSV | LARS1; LARS1; | Q9P2J5; A0A | SYLC_HUMAN | Unmodified | 381390 | 300790 | 156470 | 0 | 0 | 0 | 0 | 0 | 0 | 0 | 582640 | 3148900 | 0 | 0 | 598620 | 0 | 0 | 0 |
| GTGVVTSV | LARS1; LARS1; | Q9P2J5; A0A | SYLC_HUMAN | Unmodified | 0 | 0 | 901230 | 0 | 0 | 0 | 0 | 0 | 643760 | 0 | 0 | 4536000 | 522780 | 0 | 447230 | 0 | 0 | 0 |
| GTHGVIVV | RAB35 | Q15286 | RAB35_HUMAN | Unmodified | 3181300 | 3967500 | 1005900 | 1695500 | 0 | 0 | 0 | 19420 | 676630 | 0 | 868480 | 0 | 0 | 0 | 6149600 | 3361400 | 0 | 0 |
| GTHGVIVV | RAB35 | Q15286 | RAB35_HUMAN | Unmodified | 8127700 | 7452400 | 1533000 | 2439100 | 0 | 0 | 0 | 0 | 794840 | 0 | 0 | 1167300 | 0 | 0 | 7146500 | 6405400 | 0 | 0 |
| GTHMQHA | HMGCS1 | Q01581 | HMCS1_HUMA | 2 Oxidation (M) | 0 | 0 | 0 | 0 | 0 | 0 | 0 | 0 | 0 | 0 | 0 | 9469100 | 0 | 0 | 0 | 0 | 0 | 0 |
| GTHMQHA | HMGCS1 | Q01581 | HMCS1_HUMA | Oxidation (M) | 0 | 0 | 171840 | 0 | 0 | 0 | 0 | 0 | 0 | 0 | 0 | 2113600 | 0 | 0 | 0 | 0 | 0 | 0 |
| GTHSPPLTI | CAPN15 | O75808 | CAN15_HUMAN | Unmodified | 1209200 | 886700 | 0 | 0 | 0 | 0 | 0 | 0 | 0 | 216350 | 0 | 0 | 0 | 0 | 0 | 0 | 0 | 0 |
| GTHTGVW | FASN; FASN; FA | P49327; A0A | FAS_HUMAN Fa | Unmodified | 349520 | 0 | 646390 | 0 | 0 | 0 | 0 | 0 | 0 | 0 | 374670 | 10824000 | 0 | 0 | 0 | 195030 | 0 | 0 |
| GTIADYQPI | PKP2 | Q99959 | PKP2_HUMAN | Unmodified | 0 | 0 | 0 | 0 | 0 | 0 | 0 | 0 | 0 | 0 | 0 | 542510 | 0 | 0 | 94096 | 0 | 0 | 0 |
| GTIEILSDV | RPLP0; RPLP0P | P05388; Q8N | RLA0_HUMAN ( | Unmodified | 0 | 517550 | 0 | 0 | 0 | 0 | 0 | 0 | 0 | 0 | 0 | 0 | 0 | 0 | 483160 | 38067 | 0 | 0 |
| GTIEINPR | CRYZ | Q08257 | QOR_HUMAN ( | Unmodified | 0 | 0 | 0 | 0 | 0 | 0 | 0 | 0 | 1124700 | 0 | 0 | 828500 | 0 | 0 | 0 | 0 | 0 | 0 |
| GTINFLHAI | ERP44 | Q9BS26 | ERP44_HUMAN | Unmodified | 0 | 0 | 0 | 108140 | 0 | 0 | 0 | 164400 | 54814000 | 4408200 | 0 | 0 | 0 | 0 | 0 | 0 | 0 | 0 |
| GTIPDMIAI | NAE1 | Q13564 | ULA1_HUMAN | Unmodified | 0 | 0 | 0 | 0 | 0 | 1413300 | 0 | 189890 | 0 | 0 | 0 | 824980 | 0 | 0 | 0 | 0 | 0 | 0 |
| GTIQTGVD | PLIN4; PLIN4 | Q96Q06; A0A | PLIN4_HUMAN | Unmodified | 0 | 0 | 310240 | 99259 | 0 | 0 | 0 | 60622 | 65159 | 136010 | 0 | 0 | 296440 | 0 | 0 | 0 | 88806 | 201380 |
| GTIQVITQG | PHGDH; PHGD | O43175; A0A | SERA_HUMAN | Unmodified | 285890 | 176360 | 1186800 | 2026500 | 1305700 | 0 | 2278000 | 2699700 | 1844400 | 1819500 | 1480500 | 13823000 | 18613000 | 17725000 | 2806400 | 2432600 | 1316100 | 2277000 |
| GTITDAPGI | ANXA11 | P50995 | ANX11_HUMAN | Unmodified | 0 | 0 | 0 | 0 | 0 | 0 | 0 | 0 | 5927700 | 354910 | 0 | 0 | 0 | 0 | 0 | 0 | 0 | 0 |
| GTITDAPGI | ANXA11 | P50995 | ANX11_HUMAN | Unmodified | 0 | 0 | 160530 | 500690 | 0 | 53288 | 0 | 0 | 46452000 | 2395700 | 3001800 | 2254100 | 0 | 0 | 0 | 447890 | 0 | 1269700 |
| GTITDAPGI | ANXA11 | P50995 | ANX11_HUMAN | Unmodified | 0 | 0 | 1477800 | 1611800 | 5642500 | 1721500 | 0 | 0 | 3.48E+08 | 28978000 | 14120000 | 48863000 | 1130100 | 988660 | 1910400 | 1012600 | 1227500 | 16416000 |
| GTITVSAQE | CPNE1; CPNE1 | Q99829; Q5J | CPNE1_HUMAN | Unmodified | 0 | 0 | 1140600 | 936450 | 0 | 0 | 0 | 0 | 3519000 | 0 | 787990 | 3773200 | 0 | 0 | 1082800 | 887420 | 0 | 0 |
| GTIVLASPG | PTGR1; PTGR1; | Q14914; F2Z | PTGR1_HUMAN | Unmodified | 99362 | 50676 | 74693 | 0 | 1657900 | 0 | 0 | 0 | 1610200 | 285330 | 0 | 0 | 0 | 16317 | 90614 | 229540 | 0 | 0 |
| GTIVLASPG | PTGR1; PTGR1; | Q14914; F2Z | PTGR1_HUMAN | Unmodified | 0 | 0 | 0 | 0 | 9854000 | 1891700 | 0 | 0 | 0 | 2331800 | 0 | 369640 | 1363800 | 435980 | 0 | 0 | 0 | 0 |
| GTKAPQAA | OLA1; OLA1 | Q9NTK5; J3K | OLA1_HUMAN | Unmodified | 0 | 0 | 0 | 0 | 441780 | 30787 | 309080 | 201940 | 0 | 149410 | 0 | 0 | 0 | 318390 | 0 | 0 | 408340 | 317350 |
| GTKAQVER | F11R; F11R | Q9Y624; A0A | JAM1_HUMAN | Acetyl (Protein N | 0 | 0 | 0 | 0 | 0 | 0 | 0 | 0 | 0 | 0 | 0 | 0 | 0 | 0 | 0 | 0 | 0 | 0 |
| GTKNENKF | SLC44A2 | Q8IWA5 | CTL2_HUMAN ( | Unmodified | 0 | 0 | 0 | 1849400 | 273360 | 116030 | 0 | 0 | 5501400 | 485720 | 461200 | 530030 | 414820 | 0 | 0 | 0 | 110920 | 381230 |
| GTKQPLLEI | NSG1 | P42857 | NSG1_HUMAN | Oxidation (M) | 0 | 0 | 0 | 0 | 0 | 0 | 3727500 | 0 | 0 | 0 | 0 | 0 | 0 | 0 | 0 | 0 | 0 | 0 |
| GTLDPVEK | HSPA8; HSPA8; | P11142; E9P | HSP7C_HUMAN | Unmodified | 23444000 | 23751000 | 16894000 | 24105000 | 2449500 | 0 | 344750 | 120760 | 26734000 | 1277400 | 5470200 | 84500000 | 3130800 | 949800 | 22131000 | 19545000 | 323920 | 0 |
| GTLLGESLI | TIPRL | O75663 | TIPRL_HUMAN | Unmodified | 0 | 0 | 0 | 0 | 0 | 0 | 0 | 0 | 0 | 0 | 0 | 973940 | 0 | 0 | 0 | 0 | 0 | 0 |

| | | | | | | | | | | | | | | | | | | | | | | |
|---|---|---|---|---|---|---|---|---|---|---|---|---|---|---|---|---|---|---|---|---|---|---|
| GTLRLPSQ | EPHA1 | P21709 | EPHA1_HUMAN | Unmodified | 0 | 0 | 115410 | 0 | 0 | 0 | 0 | 0 | 0 | 0 | 163710 | 564490 | 0 | 0 | 296850 | 170770 | 17330 | 0 |
| GTLSGWIL | CANX; CANX; C | P27824; A0A | CALX_HUMAN | Unmodified | 0 | 0 | 0 | 0 | 0 | 0 | 0 | 0 | 18872000 | 2261800 | 0 | 1426900 | 0 | 0 | 0 | 0 | 0 | 0 |
| GTLSVMED | BCAM; BCAM; E | P50895; A0A | BCAM_HUMAN | Unmodified | 884690 | 450960 | 0 | 0 | 0 | 0 | 0 | 0 | 0 | 0 | 0 | 0 | 0 | 0 | 0 | 0 | 0 | 0 |
| GTLTEAFP | TPP2; TPP2; TPI | P29144; A0A | TPP2_HUMAN | Unmodified | 0 | 0 | 0 | 0 | 0 | 0 | 0 | 0 | 468750 | 0 | 564150 | 2796300 | 0 | 0 | 0 | 105820 | 222110 | 0 |
| GTLTLCPY | PSMD2 | Q13200 | PSMD2_HUMA | Unmodified | 0 | 0 | 0 | 0 | 0 | 0 | 0 | 0 | 0 | 0 | 0 | 1720200 | 0 | 0 | 0 | 0 | 0 | 0 |
| GTLVQTKG | H1-3; H1-4; H1 | P16402; P10 | H13_HUMAN H | Unmodified | 0 | 79583 | 0 | 0 | 154470 | 0 | 321200 | 11414000 | 108930 | 37368 | 0 | 0 | 0 | 0 | 0 | 0 | 19653 | 79908 |
| GTLVQTKG | H1-3; H1-4; H1 | P16402; P10 | H13_HUMAN H | Unmodified | 0 | 0 | 0 | 0 | 684420 | 0 | 1132100 | 36877000 | 0 | 0 | 0 | 0 | 0 | 0 | 0 | 0 | 0 | 0 |
| GTMTTGHN | ILF2; ILF2; ILF2; | Q12905; B4I | ILF2_HUMAN Ir | Oxidation (M) | 0 | 1449100 | 0 | 0 | 0 | 0 | 0 | 0 | 0 | 0 | 0 | 3605100 | 0 | 0 | 0 | 0 | 0 | 0 |
| GTMVTIEGI | MTAP; MTAP; M | Q13126; J3Q | MTAP_HUMAN | Unmodified | 0 | 0 | 0 | 0 | 0 | 0 | 0 | 0 | 491310 | 94032 | 338340 | 1730100 | 0 | 0 | 0 | 0 | 45827 | 0 |
| GTNEDMVI | EDIL3 | O43854 | EDIL3_HUMAN | Unmodified | 0 | 0 | 0 | 0 | 623950 | 1161900 | 0 | 0 | 4481600 | 142030 | 0 | 0 | 0 | 0 | 0 | 0 | 0 | 302550 |
| GTNEDMVI | EDIL3 | O43854 | EDIL3_HUMAN | Oxidation (M) | 0 | 0 | 0 | 128170 | 1.38E+08 | 19122000 | 3369200 | 1959300 | 7038100 | 2923700 | 0 | 0 | 0 | 0 | 87387 | 0 | 3680100 | 18940000 |
| GTNEDMVI | EDIL3 | O43854 | EDIL3_HUMAN | Unmodified | 0 | 0 | 0 | 0 | 23568000 | 3340100 | 801080 | 0 | 4867400 | 0 | 0 | 0 | 0 | 0 | 0 | 0 | 752180 | 5854500 |
| GTNIQENE | PELO | Q9BRX2 | PELO_HUMAN | Unmodified | 224790 | 0 | 0 | 152160 | 466130 | 686570 | 0 | 85944 | 0 | 0 | 0 | 327100 | 315770 | 211650 | 0 | 338370 | 0 | 534350 |
| GTPEPGPL | DNPEP; DNPEP | Q9ULA0; E5I | DNPEP_HUMAI | Unmodified | 0 | 0 | 0 | 0 | 0 | 0 | 0 | 0 | 0 | 0 | 0 | 1248600 | 0 | 0 | 0 | 0 | 250190 | 0 |
| GTPEQPQC | GLRX5 | Q86SX6 | GLRX5_HUMAI | Unmodified | 0 | 0 | 765420 | 0 | 0 | 712960 | 105510 | 0 | 10001000 | 0 | 84624 | 82808 | 0 | 0 | 0 | 0 | 402630 | 802640 |
| GTPGEGLG | DDAH2; DDAH2 | O95865; Q5! | DDAH2_HUMA | Acetyl (Protein N | 0 | 0 | 0 | 0 | 5015200 | 0 | 0 | 0 | 0 | 708820 | 0 | 0 | 0 | 0 | 532130 | 0 | 28059000 | 7065800 |
| GTPLISPLIK | FASN; FASN | P49327; A0A | FAS_HUMAN Fa | Unmodified | 420540 | 0 | 3581600 | 546600 | 0 | 0 | 0 | 0 | 0 | 0 | 7829900 | 52955000 | 0 | 0 | 0 | 1959900 | 0 | 0 |
| GTPMKPNT | ICAM1; ICAM1 | P05362; E7E | ICAM1_HUMAI | Unmodified | 0 | 0 | 0 | 0 | 12135000 | 2686900 | 0 | 64578 | 0 | 0 | 0 | 0 | 0 | 0 | 0 | 0 | 0 | 2657800 |
| GTPQEPRC | GLRX3 | O76003 | GLRX3_HUMAI | Unmodified | 0 | 0 | 0 | 0 | 634560 | 198170 | 359310 | 227590 | 0 | 95525 | 97895 | 0 | 464460 | 330850 | 0 | 0 | 790940 | 512240 |
| GTPQQIDY | FUBP1; FUBP1 | Q96AE4; E9I | FUBP1_HUMAI | Unmodified | 163810 | 0 | 0 | 0 | 0 | 0 | 0 | 0 | 0 | 0 | 396970 | 1391500 | 0 | 18028 | 0 | 0 | 0 | 0 |
| GTQCVEQI | BCCIP | Q9P287 | BCCIP_HUMAI | Unmodified | 0 | 0 | 0 | 0 | 0 | 0 | 0 | 0 | 0 | 0 | 0 | 1501200 | 0 | 0 | 0 | 0 | 0 | 0 |
| GTQGVVTN | EIF3B | P55884 | EIF3B_HUMAN | Unmodified | 0 | 167940 | 99996 | 127840 | 67974 | 0 | 0 | 0 | 102580 | 0 | 250520 | 2557300 | 34915 | 0 | 74096 | 159050 | 0 | 22889 |
| GTQVYSPE | SNRNP200; SN | O75643; A0A | U520_HUMAN | Unmodified | 0 | 0 | 163740 | 0 | 93660 | 0 | 0 | 0 | 0 | 0 | 0 | 2214500 | 17817 | 0 | 0 | 0 | 0 | 0 |
| GTRDDEYD | RAB11A; RAB11 | P62491; Q15 | RB11A_HUMAI | Acetyl (Protein N | 10383000 | 14347000 | 15539000 | 41047000 | 85904000 | 15714000 | 20629000 | 9918300 | 12831000 | 9964100 | 15464000 | 43376000 | 26408000 | 25176000 | 14139000 | 17371000 | 5502800 | 10168000 |
| GTRPEDSS | ASDURF; N/A; A | L0R819; U3K | ASURF_HUMAI | Unmodified | 0 | 0 | 0 | 0 | 0 | 0 | 0 | 0 | 0 | 0 | 157920 | 1431800 | 0 | 0 | 0 | 0 | 0 | 0 |
| GTSEAQPL | SLC16A10; SLC | Q8TF71; A0A | MOT10_HUMAI | Unmodified | 0 | 0 | 1059800 | 0 | 0 | 0 | 0 | 0 | 0 | 0 | 0 | 2667500 | 0 | 0 | 1731400 | 754690 | 0 | 0 |
| GTSENLPE | SLC43A1 | O75387 | LAT3_HUMAN L | Unmodified | 172180 | 102620 | 5288400 | 3435800 | 110900 | 195980 | 11330 | 0 | 1663600 | 65285 | 35267 | 1002900 | 0 | 1004500 | 8868600 | 6097700 | 23948 | 1107900 |
| GTSFDAAA | ETFA; ETFA; ETF | P13804; H0Y | ETFA_HUMAN I | Unmodified | 0 | 0 | 0 | 0 | 0 | 0 | 0 | 0 | 1820800 | 0 | 0 | 0 | 0 | 0 | 0 | 0 | 0 | 0 |
| GTSGGLLH | DIPK2A; DIPK2A | Q8NDZ4; E7 | DIK2A_HUMAN | Unmodified | 0 | 0 | 152670 | 0 | 0 | 0 | 0 | 0 | 30338000 | 998470 | 0 | 887510 | 0 | 0 | 0 | 0 | 0 | 0 |
| GTSGGLLH | DIPK2A; DIPK2A | Q8NDZ4; E7 | DIK2A_HUMAN | Unmodified | 0 | 0 | 0 | 0 | 0 | 0 | 0 | 0 | 7572700 | 328140 | 0 | 0 | 0 | 0 | 0 | 0 | 0 | 0 |
| GTSGSLAD | DIP2B | Q9P265 | DIP2B_HUMAN | Unmodified | 0 | 0 | 0 | 0 | 0 | 0 | 0 | 0 | 0 | 0 | 138480 | 1459700 | 0 | 207920 | 0 | 0 | 0 | 0 |
| GTSISTKPP | RFC4; RFC4; RF | P35249; C9J | RFC4_HUMAN | Unmodified | 0 | 0 | 0 | 0 | 554890 | 0 | 0 | 255390 | 174290 | 133240 | 0 | 0 | 523530 | 182980 | 0 | 0 | 248130 | 215500 |
| GTSLENLIM | MACF1; MACF1 | Q9UPN3; H3 | MACF1_HUMAI | Oxidation (M) | 0 | 0 | 0 | 13413 | 0 | 0 | 0 | 0 | 174990 | 47920 | 0 | 0 | 21080 | 0 | 29045 | 0 | 0 | 0 |
| GTSYQSPH | RUVBL2; RUVBI | Q9Y230; M0I | RUVB2_HUMAI | Unmodified | 29552 | 0 | 30612 | 0 | 0 | 0 | 0 | 26114 | 0 | 0 | 118880 | 1732000 | 0 | 0 | 0 | 0 | 28349 | 74603 |
| GTSYQSPH | RUVBL2; RUVBI | Q9Y230; M0I | RUVB2_HUMAI | Unmodified | 0 | 0 | 0 | 0 | 25625 | 0 | 0 | 0 | 0 | 0 | 230270 | 689270 | 0 | 0 | 0 | 0 | 0 | 0 |
| GTTAPGPII | INCENP; INCEN | Q9NQS7; E9 | INCE_HUMAN I | Acetyl (Protein N | 0 | 0 | 0 | 20957 | 2258700 | 2636700 | 0 | 0 | 297250 | 98350 | 54103 | 153190 | 0 | 945150 | 0 | 0 | 0 | 0 |
| GTTAVLTEK | PSMB7 | Q99436 | PSB7_HUMAN | Unmodified | 0 | 0 | 304960 | 0 | 3194600 | 718640 | 283070 | 0 | 1465200 | 158490 | 674020 | 3922600 | 1500300 | 0 | 185860 | 81174 | 1379100 | 2790800 |
| GTTGTQAS | ADSL; ADSL; AD | P30566; A0A | PUR8_HUMAN | Unmodified | 259760 | 237660 | 0 | 989900 | 1273500 | 850000 | 238380 | 0 | 876190 | 215830 | 5548900 | 32044000 | 0 | 266090 | 910340 | 749300 | 162900 | 489320 |
| GTTITLVLK | HSP90B1; HSP9 | P14625; A0A | ENPL_HUMAN | Unmodified | 0 | 0 | 0 | 0 | 0 | 0 | 0 | 0 | 15783000 | 355670 | 9503100 | 2521300 | 0 | 0 | 756370 | 0 | 0 | 0 |
| GTTPLMQK | CTNND1; CTNN | A0A669KB62 | A0A669KB62_H | Unmodified | 235540 | 177520 | 392280 | 0 | 6180000 | 907140 | 181500 | 0 | 436760 | 0 | 613200 | 3928300 | 0 | 0 | 637840 | 0 | 2631300 | 9206900 |
| GTVEGFEP | SRP14; SRP14 | P37108; H0Y | SRP14_HUMAI | Unmodified | 556690 | 876230 | 1920300 | 1259800 | 0 | 0 | 0 | 0 | 0 | 0 | 242510 | 866030 | 0 | 0 | 4120500 | 3536500 | 0 | 0 |
| GTVEKGPQ | PACSIN3; PACS | Q9UKS6; E9I | PACN3_HUMAI | Unmodified | 1241500 | 2196000 | 282480 | 1669300 | 0 | 0 | 0 | 0 | 0 | 0 | 71332 | 94639 | 1717100 | 176920 | 140070 | 365790 | 0 | 57169 |
| GTVEPQLE | FLNA; FLNA; FL | P21333; A0A | FLNA_HUMAN | Unmodified | 1060100 | 1822400 | 0 | 1005700 | 129840 | 45277 | 0 | 0 | 122100 | 30953 | 0 | 2993400 | 220660 | 0 | 921260 | 598440 | 132960 | 73693 |
| GTVEPQLE | FLNA; FLNA; FL | P21333; A0A | FLNA_HUMAN | Unmodified | 267130 | 295580 | 271330 | 1568500 | 2655800 | 0 | 356530 | 0 | 365920 | 0 | 0 | 0 | 923970 | 696280 | 350470 | 187060 | 0 | 0 |
| GTVGEPTY | DYNC1H1; DYN | Q14204; A0A | DYHC1_HUMA | Unmodified | 0 | 0 | 0 | 0 | 0 | 0 | 0 | 0 | 0 | 0 | 0 | 3423900 | 0 | 0 | 0 | 249040 | 0 | 0 |
| GTVGGRLA | PCYOX1L; PCY( | Q8NBM8; D6 | PCYXL_HUMAN | Unmodified | 0 | 0 | 0 | 0 | 0 | 0 | 44850 | 53871 | 0 | 497380 | 0 | 0 | 0 | 0 | 0 | 0 | 0 | 0 |
| GTVGSILDI | ABCE1; ABCE1; | P61221; H0Y | ABCE1_HUMAI | Unmodified | 184770 | 192480 | 238820 | 0 | 1631400 | 343670 | 157090 | 69408 | 0 | 0 | 377060 | 4403100 | 0 | 255070 | 233380 | 247090 | 0 | 0 |
| GTVIKVPLK | MYO6; MYO6; M | Q9UM54; A0 | MYO6_HUMAN | Unmodified | 0 | 0 | 0 | 557960 | 0 | 0 | 0 | 0 | 0 | 0 | 0 | 0 | 0 | 0 | 0 | 0 | 0 | 0 |
| GTVLPQGP | LPIN1 | Q14693 | LPIN1_HUMAN | Unmodified | 0 | 0 | 0 | 0 | 0 | 0 | 0 | 0 | 0 | 0 | 22075 | 346200 | 0 | 0 | 0 | 0 | 0 | 0 |
| GTVQALHA | DCXR; DCXR; D | Q7Z4W1; J3I | DCXR_HUMAN | Unmodified | 0 | 0 | 68480 | 0 | 0 | 0 | 0 | 0 | 0 | 0 | 0 | 0 | 0 | 0 | 124970 | 104550 | 0 | 0 |
| GTVRDYPD | ANXA3; ANXA3; | P12429; D6I | ANXA3_HUMAI | Unmodified | 0 | 0 | 0 | 0 | 970040 | 0 | 0 | 0 | 0 | 0 | 0 | 0 | 0 | 0 | 0 | 0 | 0 | 0 |
| GTVRPAND | ANXA6; ANXA6 | P08133; E5R | ANXA6_HUMAI | Unmodified | 2908500 | 3278600 | 7042500 | 7805200 | 13812000 | 3912600 | 3791100 | 3229100 | 21369000 | 5043900 | 3530500 | 13226000 | 7797200 | 4633500 | 10396000 | 6052200 | 5201700 | 14776000 |
| GTVSFHSH | CSDE1; CSDE1 | O75534; E9I | CSDE1_HUMAI | Unmodified | 65949 | 0 | 64474 | 0 | 0 | 0 | 0 | 0 | 48352 | 0 | 34809 | 773850 | 0 | 0 | 0 | 140710 | 0 | 0 |
| GTVSFHSH | CSDE1; CSDE1 | O75534; E9I | CSDE1_HUMAI | Unmodified | 0 | 0 | 73371 | 310030 | 299700 | 0 | 0 | 0 | 0 | 0 | 510640 | 0 | 657190 | 402450 | 0 | 0 | 109220 | 0 |
| GTVTSTAT( | DIP2A; DIP2A | Q14689; A0A | DIP2A_HUMAN | Unmodified | 0 | 0 | 0 | 0 | 387400 | 0 | 0 | 0 | 450510 | 103730 | 0 | 0 | 0 | 0 | 218360 | 0 | 0 | 58903 |
| GTVVTGTLI | TUFM | P49411 | EFTU_HUMAN | Unmodified | 97057 | 58990 | 65256 | 903240 | 0 | 0 | 34291 | 44667 | 3384500 | 0 | 0 | 0 | 0 | 0 | 501770 | 25659 | 0 | 0 |
| GTVVYGEP | DPYSL2; DPYSL | Q16555; A0A | DPYL2_HUMAI | Unmodified | 0 | 0 | 0 | 0 | 27930000 | 2936300 | 3458500 | 685350 | 3617300 | 1924000 | 0 | 6554900 | 0 | 1318100 | 0 | 0 | 1733500 | 3016000 |
| GTVYLTPYI | WBP2; WBP2; V | Q969T9; K7E | WBP2_HUMAN | Unmodified | 0 | 0 | 179800 | 0 | 672010 | 0 | 350140 | 0 | 0 | 581760 | 0 | 0 | 0 | 0 | 169520 | 116440 | 1159700 | 0 |
| GTWERPG( | SELENBP1; SEL | Q13228; A6I | SBP1_HUMAN | Unmodified | 0 | 0 | 0 | 0 | 0 | 0 | 0 | 0 | 0 | 0 | 0 | 0 | 0 | 0 | 1671300 | 1166000 | 0 | 0 |
| GTYLTDETI | CD46 | P15529 | MCP_HUMAN I | Unmodified | 0 | 0 | 0 | 0 | 0 | 0 | 0 | 0 | 966900 | 151990 | 59981 | 830680 | 0 | 0 | 0 | 16914 | 0 | 0 |
| GTYLTDETI | CD46 | P15529 | MCP_HUMAN I | Unmodified | 171230 | 155230 | 0 | 88603 | 481920 | 86586 | 185400 | 9583.6 | 94982 | 0 | 0 | 313410 | 0 | 0 | 84501 | 80921 | 0 | 0 |
| GTYTEFLK | ABCC4; ABCC4 | O15439; A0A | MRP4_HUMAN | Unmodified | 0 | 0 | 1649900 | 413330 | 0 | 0 | 0 | 0 | 0 | 0 | 0 | 0 | 0 | 0 | 0 | 1319200 | 0 | 0 |
| GVAALTSD | TLN1 | Q9Y490 | TLN1_HUMAN | Unmodified | 0 | 1058800 | 0 | 180200 | 814370 | 0 | 0 | 0 | 0 | 243090 | 0 | 0 | 212360 | 158170 | 1236600 | 0 | 0 | 0 |
| GVAAMQSI | TMCO3 | Q6UWJ1 | TMCO3_HUMA | Unmodified | 68256 | 0 | 0 | 0 | 0 | 0 | 30293 | 0 | 599310 | 0 | 57815 | 36349 | 0 | 0 | 0 | 0 | 106520 | 190670 |
| GVACLGCI | LMCD1; LMCD1 | Q9NZU5; H7 | LMCD1_HUMA | Unmodified | 0 | 0 | 0 | 0 | 83580 | 0 | 0 | 0 | 0 | 0 | 0 | 820370 | 0 | 0 | 0 | 0 | 45493 | 240160 |
| GVAEGEVD | MGAT4B; MGAT | Q9UQ53; H0 | MGT4B_HUMAI | Unmodified | 0 | 0 | 613090 | 1807900 | 0 | 132720 | 0 | 54640 | 2049100 | 823290 | 0 | 0 | 0 | 0 | 74597 | 479850 | 149100 | 0 |
| GVAEVLQD | SLC44A2; SLC4 | Q8IWA5; K7I | CTL2_HUMAN | Unmodified | 9006000 | 2966400 | 0 | 0 | 0 | 0 | 0 | 0 | 4481200 | 2113900 | 0 | 4823300 | 0 | 0 | 0 | 0 | 0 | 0 |
| GVAEVLQD | SLC44A2; SLC4 | Q8IWA5; K7I | CTL2_HUMAN | Unmodified | 0 | 0 | 0 | 0 | 0 | 0 | 0 | 0 | 1390800 | 1971300 | 396740 | 0 | 0 | 0 | 0 | 0 | 0 | 0 |
| GVAGALRP | UQCRFS1P1; U | P0C7P4; P47 | UCRIL_HUMAN | Unmodified | 0 | 0 | 0 | 0 | 0 | 0 | 0 | 0 | 15375000 | 0 | 1120800 | 0 | 0 | 0 | 0 | 0 | 0 | 0 |
| GVAIGQTSI | NUP210 | Q8TEM1 | PO210_HUMAI | Unmodified | 0 | 1244900 | 0 | 0 | 0 | 0 | 0 | 0 | 2527000 | 0 | 0 | 0 | 70369 | 0 | 0 | 0 | 58893 | 0 |
| GVAINFVK | EIF4A3; EIF4A3 | P38919; I3L3 | IF4A3_HUMAN | Unmodified | 522970 | 336580 | 488330 | 97317 | 0 | 0 | 0 | 0 | 0 | 0 | 0 | 1135300 | 0 | 0 | 258430 | 252270 | 0 | 0 |
| GVAINMVT | EIF4A1 | P60842 | IF4A1_HUMAN | Oxidation (M) | 0 | 0 | 183820 | 0 | 0 | 0 | 0 | 0 | 0 | 0 | 0 | 852220 | 0 | 0 | 1066800 | 0 | 0 | 0 |
| GVAINMVT | EIF4A1 | P60842 | IF4A1_HUMAN | Unmodified | 0 | 0 | 422950 | 158360 | 0 | 0 | 0 | 0 | 0 | 0 | 0 | 541840 | 0 | 0 | 0 | 218830 | 0 | 0 |
| GVAINMVT | EIF4A1 | P60842 | IF4A1_HUMAN | Unmodified | 1474500 | 1638300 | 2039200 | 2051500 | 0 | 496030 | 623910 | 111130 | 2704400 | 249900 | 2760300 | 8254400 | 2317300 | 596700 | 3020100 | 2057700 | 290490 | 1044500 |
| GVAINMVT | EIF4A1 | P60842 | IF4A1_HUMAN | Oxidation (M) | 0 | 684620 | 372150 | 949060 | 290030 | 0 | 0 | 0 | 62160 | 173570 | 398570 | 3908300 | 702600 | 0 | 1006000 | 549290 | 0 | 0 |
| GVALSNVII | SERPINB5 | P36952 | SPB5_HUMAN | Unmodified | 0 | 80875 | 0 | 389460 | 0 | 2331100 | 0 | 87198 | 0 | 0 | 0 | 0 | 0 | 0 | 123630 | 0 | 0 | 340660 |
| GVAPLWMI | SLC25A3; SLC2 | Q00325; F8V | MPCP_HUMAN | Oxidation (M) | 0 | 0 | 0 | 53699 | 0 | 0 | 0 | 0 | 11616000 | 1482400 | 0 | 0 | 0 | 0 | 0 | 0 | 0 | 0 |
| GVAPLWMI | SLC25A3; SLC2 | Q00325; F8V | MPCP_HUMAN | Unmodified | 0 | 0 | 0 | 0 | 0 | 0 | 0 | 0 | 26168000 | 1596300 | 0 | 0 | 0 | 0 | 0 | 0 | 0 | 0 |
| GVAVDYLP | P4HA1 | P13674 | P4HA1_HUMAI | Unmodified | 0 | 0 | 0 | 0 | 0 | 0 | 0 | 0 | 2794800 | 0 | 0 | 0 | 0 | 0 | 0 | 0 | 0 | 0 |
| GVCGACKI | PXN; PXN; PXN | P49023; A0A | PAXI_HUMAN F | Unmodified | 0 | 0 | 0 | 103600 | 0 | 0 | 0 | 0 | 0 | 0 | 0 | 0 | 0 | 0 | 0 | 0 | 3151000 | 3770100 |
| GVCLNEK | IGFBP5; IGFBP5 | P24593; C9J | IBP5_HUMAN I | Unmodified | 0 | 0 | 0 | 0 | 348290 | 0 | 0 | 105880 | 3852800 | 340300 | 0 | 0 | 330590 | 237260 | 0 | 11003 | 0 | 0 |
| GVDDLDFF | ACTR3; ACTR3; | P61158; B4D | ARP3_HUMAN | Unmodified | 2381100 | 602420 | 0 | 0 | 0 | 0 | 0 | 0 | 0 | 1595500 | 839390 | 10055000 | 0 | 0 | 7086300 | 1848300 | 0 | 0 |
| GVDDSTGF | PDE2A; PDE2A; | O00408; H0Y | PDE2A_HUMAI | Unmodified | 0 | 0 | 0 | 0 | 0 | 0 | 0 | 0 | 0 | 0 | 0 | 2935300 | 0 | 0 | 0 | 0 | 0 | 0 |
| GVDEATIID | ANXA1; ANXA1; | P04083; Q5T | ANXA1_HUMAI | Unmodified | 539030 | 0 | 144930 | 0 | 18490000 | 4463200 | 1006600 | 0 | 1078300 | 605000 | 1.79E+08 | 8.71E+08 | 0 | 0 | 0 | 846010 | 1736700 | 2796600 |
| GVDEATIID | ANXA1; ANXA1; | P04083; Q5T | ANXA1_HUMAI | Unmodified | 0 | 0 | 0 | 0 | 0 | 0 | 0 | 0 | 0 | 0 | 34031000 | 5E+08 | 0 | 0 | 0 | 87095 | 0 | 0 |
| GVDEATIID | ANXA1; ANXA1; | P04083; Q5T | ANXA1_HUMAI | Unmodified | 0 | 0 | 0 | 0 | 3804400 | 444560 | 0 | 0 | 0 | 79933 | 185060 | 0 | 0 | 0 | 0 | 0 | 470270 | 0 |
| GVDEVTIVI | ANXA2; ANXA2; | P07355; H0Y | ANXA2_HUMAI | Unmodified | 2427700 | 2348400 | 665620 | 1275400 | 13584000 | 4748400 | 1022800 | 420020 | 47161000 | 7819200 | 770410 | 445940 | 197300 | 208440 | 1140200 | 724830 | 2658700 | 4072000 |
| GVDEVTIVI | ANXA2; ANXA2; | P07355; H0Y | ANXA2_HUMAI | Unmodified | 0 | 0 | 0 | 0 | 5399700 | 0 | 520230 | 0 | 0 | 0 | 0 | 0 | 0 | 0 | 0 | 0 | 304500 | 0 |
| GVDIVMDP | VAT1; VAT1; VA | Q99536; K7E | VAT1_HUMAN ! | Oxidation (M) | 0 | 0 | 0 | 0 | 0 | 0 | 0 | 0 | 1871600 | 505720 | 1247300 | 2068300 | 0 | 0 | 232740 | 0 | 0 | 5514500 |
| GVDIVMDP | VAT1; VAT1; VA | Q99536; K7E | VAT1_HUMAN ! | Unmodified | 0 | 485340 | 0 | 0 | 8104900 | 0 | 183550 | 0 | 6174900 | 439410 | 0 | 1005100 | 935800 | 0 | 0 | 0 | 0 | 3314600 |
| GVDIYPENI | SYT4 | Q9H2B2 | SYT4_HUMAN ! | Unmodified | 277910 | 119010 | 3409200 | 2964800 | 691690 | 0 | 431800 | 48510 | 0 | 117920 | 0 | 567340 | 80293 | 129550 | 9586800 | 4882700 | 0 | 0 |
| GVDIYTR | ACY1; ABHD14 | Q03154; A0A | ACY1_HUMAN | Unmodified | 0 | 48610 | 0 | 0 | 617970 | 0 | 0 | 13390 | 129260 | 0 | 278720 | 1301200 | 0 | 0 | 29714 | 0 | 23080 | 0 |
| GVDLFESFI | QTRT2 | Q9H974 | QTRT2_HUMAI | Unmodified | 104690 | 101660 | 0 | 17529 | 0 | 0 | 0 | 0 | 93212 | 0 | 28484 | 70273 | 0 | 0 | 13525 | 0 | 0 | 0 |
| GVDLHEQS | RAB5B | P61020 | RAB5B_HUMAI | Unmodified | 793600 | 0 | 0 | 0 | 0 | 0 | 0 | 37389 | 293140 | 238830 | 0 | 64979 | 0 | 0 | 0 | 0 | 0 | 0 |
| GVDLQENI | RAB5C; N/A | P51148; K7E | RAB5C_HUMAI | Unmodified | 2726200 | 1025500 | 2425600 | 0 | 0 | 0 | 0 | 0 | 1730300 | 0 | 226820 | 9674100 | 0 | 0 | 3319500 | 422270 | 0 | 0 |
| GVDLTEPT( | RAB5A | P20339 | RAB5A_HUMAI | Unmodified | 1885200 | 0 | 0 | 0 | 0 | 0 | 0 | 0 | 334390 | 0 | 0 | 1401000 | 0 | 0 | 913990 | 62079 | 0 | 0 |
| GVDLVLNS | FASN; FASN | P49327; A0A | FAS_HUMAN Fa | Unmodified | 710270 | 0 | 1920500 | 2192500 | 556830 | 0 | 719760 | 212490 | 933670 | 0 | 7082900 | 52048000 | 0 | 0 | 6151100 | 3433500 | 0 | 0 |
| GVDLYLR | XPOT; XPOT | O43592; F8V | XPOT_HUMAN | Unmodified | 105880 | 122890 | 1041300 | 0 | 0 | 0 | 0 | 0 | 0 | 0 | 0 | 1627300 | 0 | 0 | 669650 | 283130 | 0 | 0 |
| GVDRIDEN | SEC23IP | Q9Y6Y8 | S23IP_HUMAN | Unmodified | 0 | 0 | 0 | 71487 | 0 | 295830 | 0 | 16755 | 0 | 0 | 0 | 1107800 | 0 | 0 | 44875 | 0 | 0 | 0 |
| GVEEEEED | SNRPF | P62306 | RUXF_HUMAN | Unmodified | 1631600 | 1937700 | 628290 | 0 | 0 | 309360 | 0 | 0 | 357630 | 0 | 245120 | 987990 | 0 | 0 | 0 | 538950 | 0 | 0 |
| GVEEEEED | SNRPF | P62306 | RUXF_HUMAN | Oxidation (M) | 657110 | 1137000 | 197790 | 0 | 0 | 0 | 0 | 0 | 102870 | 0 | 82350 | 740300 | 0 | 0 | 387680 | 246830 | 0 | 0 |
| GVEGAYSD | GNPTAB | Q3T906 | GNPTA_HUMAI | Unmodified | 0 | 0 | 0 | 0 | 0 | 0 | 0 | 0 | 0 | 0 | 0 | 0 | 0 | 0 | 0 | 0 | 0 | 0 |
| GVEGLIDIE | PDAP1 | Q13442 | HAP28_HUMAI | Unmodified | 1823000 | 1667600 | 2672700 | 3220700 | 2731800 | 763400 | 579370 | 260120 | 906870 | 258160 | 650610 | 1845500 | 1911300 | 649710 | 2283100 | 2135000 | 1077600 | 4519100 |
| GVEGTWVI | ACSL4 | O60488 | ACSL4_HUMAI | Oxidation (M) | 0 | 0 | 0 | 0 | 1319900 | 0 | 0 | 0 | 0 | 0 | 1266200 | 1242600 | 0 | 0 | 0 | 0 | 0 | 0 |
| GVEICIATP | DDX5; DDX17; I | P17844; Q92 | DDX5_HUMAN | Unmodified | 0 | 0 | 0 | 0 | 0 | 0 | 0 | 0 | 0 | 0 | 320960 | 2159100 | 0 | 0 | 0 | 0 | 0 | 0 |
| GVEIENLGI | NCOA4; NCOA4 | Q13772; B4I | NCOA4_HUMA | Unmodified | 0 | 0 | 0 | 0 | 246690 | 0 | 908440 | 189170 | 957340 | 224340 | 0 | 0 | 667180 | 0 | 0 | 0 | 393110 | 939440 |
| GVEITGFPE | GART | P22102 | PUR2_HUMAN | Unmodified | 0 | 0 | 0 | 993880 | 4860300 | 0 | 333720 | 0 | 51094 | 597790 | 0 | 1027800 | 729620 | 6636700 | 1110900 | 178370 | 3113200 | 2848900 |
| GVELEPTD | GRK6; GRK6 | P43250; F8V | GRK6_HUMAN | Unmodified | 74350 | 0 | 558770 | 523630 | 0 | 0 | 185620 | 378490 | 0 | 0 | 512130 | 506060 | 1321000 | 915080 | 801090 | 202110 | 62851 | 110950 |
| GVEPSPSP | UFD1; UFD1 | Q92890; C9J | UFD1_HUMAN | Unmodified | 136010 | 0 | 0 | 0 | 539310 | 0 | 0 | 0 | 133080 | 70071 | 0 | 0 | 0 | 0 | 0 | 0 | 92130 | 0 |

| | | | | | | | | | | | | | | | | | | | | | | |
|---|---|---|---|---|---|---|---|---|---|---|---|---|---|---|---|---|---|---|---|---|---|---|
| GVETIANDV | DLAT; DLAT; DL | P10515; A0A | ODP2_HUMAN | Unmodified | 0 | 0 | 0 | 64725 | 0 | 0 | 0 | 20222 | 7819800 | 1612900 | 70125 | 0 | 205070 | 0 | 45705 | 18814 | 279990 | 0 |
| GVEVDPSL | OPA1; OPA1; O | O60313; E5K | OPA1_HUMAN | Unmodified | 0 | 0 | 0 | 0 | 644400 | 0 | 0 | 0 | 216000 | 0 | 970620 | 8276500 | 241740 | 0 | 116130 | 0 | 0 | 0 |
| GVEVTVGH | GATD3B; GATD | A0A0B4J2DE | GAL3B_HUMAN | Unmodified | 0 | 0 | 0 | 0 | 0 | 0 | 0 | 0 | 5429000 | 0 | 0 | 0 | 100110 | 0 | 0 | 0 | 0 | 0 |
| GVFLANQK | SORCS2; SORC | Q96PQ0; B5 | SORC2_HUMA | Unmodified | 0 | 0 | 328460 | 0 | 726920 | 0 | 1528400 | 0 | 2454100 | 0 | 0 | 137910 | 0 | 2774400 | 381530 | 0 | 360820 | 1886800 |
| GVFLANQK | SORCS2; SORC | Q96PQ0; B5 | SORC2_HUMA | Unmodified | 0 | 0 | 0 | 0 | 0 | 0 | 0 | 0 | 1951700 | 223780 | 0 | 0 | 0 | 0 | 0 | 0 | 0 | 0 |
| GVFLDTILP | TLL2 | Q9Y6L7 | TLL2_HUMAN T | Unmodified | 0 | 0 | 0 | 0 | 0 | 32791 | 0 | 0 | 1390800 | 0 | 276680 | 0 | 0 | 0 | 0 | 99958 | 0 | 0 |
| GVFVQSVL | HSD17B12; HS | Q53GQ0; A0 | DHB12_HUMA | Unmodified | 0 | 0 | 0 | 0 | 0 | 0 | 116000 | 0 | 0 | 0 | 45696 | 801100 | 0 | 0 | 0 | 0 | 0 | 0 |
| GVGAAATA | TLN1 | Q9Y490 | TLN1_HUMAN | Unmodified | 0 | 38963 | 0 | 0 | 258270 | 62403 | 0 | 0 | 0 | 0 | 0 | 0 | 0 | 0 | 101320 | 0 | 0 | 0 |
| GVGASGSF | H1-0 | P07305 | H10_HUMAN H | Unmodified | 2856800 | 1538700 | 26619 | 0 | 236000 | 40214 | 72980 | 141060 | 0 | 34675 | 0 | 0 | 0 | 0 | 0 | 0 | 0 | 0 |
| GVGDSFKK | HYOU1; HYOU1 | Q9Y4L1; E9F | HYOU1_HUMA | Unmodified | 0 | 0 | 263130 | 0 | 0 | 0 | 1350000 | 888970 | 0 | 7430900 | 0 | 0 | 473170 | 1744100 | 0 | 0 | 91259 | 428840 |
| GVGFDRTL | HYOU1; HYOU1 | Q9Y4L1; E9F | HYOU1_HUMA | Oxidation (M) | 0 | 24189 | 65590 | 565650 | 0 | 1361900 | 302150 | 46858 | 12951000 | 4868900 | 3941700 | 0 | 140720 | 0 | 92592 | 42159 | 276490 | 0 |
| GVGFDRTL | HYOU1; HYOU1 | Q9Y4L1; E9F | HYOU1_HUMA | Unmodified | 0 | 0 | 0 | 0 | 0 | 0 | 0 | 0 | 17589000 | 1049900 | 252230 | 0 | 119510 | 0 | 0 | 0 | 0 | 0 |
| GVGFGDIF | SH3KBP1; SH3I | Q96B97; Q5' | SH3K1_HUMAN | Unmodified | 0 | 0 | 0 | 0 | 0 | 0 | 48312 | 9753.4 | 0 | 0 | 0 | 0 | 1001600 | 760140 | 0 | 0 | 69045 | 0 |
| GVGGAPPT | LRP1 | Q07954 | LRP1_HUMAN | Unmodified | 0 | 0 | 0 | 0 | 0 | 0 | 0 | 0 | 1546700 | 0 | 0 | 0 | 0 | 0 | 0 | 0 | 0 | 0 |
| GVGGAVP( | PKP3; PKP3; PK | Q9Y446; E9F | PKP3_HUMAN | Unmodified | 124790 | 0 | 833140 | 618780 | 0 | 0 | 0 | 0 | 549740 | 0 | 1050300 | 2309500 | 0 | 0 | 0 | 0 | 0 | 0 |
| GVGGAVP( | PKP3; PKP3; PK | Q9Y446; E9F | PKP3_HUMAN | Unmodified | 0 | 0 | 554370 | 1014100 | 483760 | 0 | 0 | 0 | 0 | 0 | 0 | 0 | 2611500 | 726870 | 0 | 0 | 687330 | 515480 |
| GVGGKLPN | G3BP2; G3BP2; | Q9UN86; A0 | G3BP2_HUMAI | Unmodified | 0 | 0 | 0 | 0 | 0 | 0 | 0 | 0 | 0 | 0 | 0 | 1607900 | 0 | 0 | 134950 | 0 | 0 | 0 |
| GVGGSQPI | ANPEP | P15144 | AMPN_HUMAN | Unmodified | 0 | 0 | 0 | 0 | 0 | 0 | 59804 | 0 | 0 | 148650 | 0 | 0 | 1805000 | 0 | 0 | 0 | 697230 | 5808800 |
| GVGIISEGN | ATP1A1; ATP1A | P05023; P50 | AT1A1_HUMAN | Unmodified | 4137500 | 2371000 | 39142000 | 33158000 | 31426000 | 0 | 760720 | 594630 | 30947000 | 5846500 | 32838000 | 2.08E+08 | 13611000 | 4088000 | 74100000 | 67652000 | 1873000 | 1039800 |
| GVGIISEGN | ATP1A1; ATP1A | P05023; P13 | AT1A1_HUMAN | Unmodified | 0 | 0 | 262080 | 17480000 | 33780000 | 4415300 | 4242600 | 0 | 591140 | 10627000 | 2962900 | 0 | 80888000 | 36148000 | 10826000 | 0 | 14280000 | 10121000 |
| GVGIISEGN | ATP1A1; ATP1A | P05023; P13 | AT1A1_HUMAN | Unmodified | 0 | 0 | 0 | 0 | 1.2E+08 | 241670 | 45581000 | 0 | 0 | 18848000 | 638670 | 0 | 1.45E+08 | 1.06E+08 | 0 | 0 | 19132000 | 28517000 |
| GVGISVLEN | PSAT1 | Q9Y617 | SERC_HUMAN | Unmodified | 0 | 0 | 151760 | 317290 | 260770 | 0 | 0 | 0 | 571780 | 0 | 602070 | 2385900 | 0 | 0 | 0 | 452670 | 0 | 0 |
| GVGISVLEN | PSAT1 | Q9Y617 | SERC_HUMAN | Oxidation (M) | 0 | 0 | 306400 | 0 | 0 | 601920 | 0 | 0 | 208430 | 0 | 1422000 | 2868400 | 0 | 0 | 0 | 578820 | 0 | 0 |
| GVGISVLEN | PSAT1 | Q9Y617 | SERC_HUMAN | Unmodified | 0 | 0 | 0 | 0 | 538910 | 0 | 110430 | 0 | 927100 | 0 | 0 | 0 | 724600 | 329510 | 0 | 0 | 0 | 0 |
| GVGMVADI | DDOST; DDOST | P39656; A0A | OST48_HUMAN | Oxidation (M) | 0 | 0 | 0 | 0 | 0 | 0 | 0 | 0 | 0 | 1190200 | 0 | 0 | 0 | 0 | 0 | 0 | 0 | 0 |
| GVGQADW | TNKS1BP1 | Q9C0C2 | TB182_HUMAN | Unmodified | 0 | 0 | 392240 | 348410 | 0 | 0 | 0 | 0 | 0 | 0 | 0 | 703840 | 617600 | 0 | 587280 | 422510 | 0 | 0 |
| GVGSPEPC | ZDHHC5 | Q9C0B5 | ZDHC5_HUMA | Unmodified | 294150 | 313950 | 1228300 | 1307500 | 1575700 | 0 | 0 | 0 | 981310 | 216070 | 485390 | 1214200 | 1668800 | 993220 | 0 | 1199300 | 894290 | 3695800 |
| GVGTDEAC | ANXA11 | P50995 | ANX11_HUMAN | Unmodified | 0 | 0 | 0 | 291210 | 203280 | 56171 | 0 | 0 | 41204000 | 1379700 | 1904700 | 1809800 | 181450 | 80822 | 437570 | 156620 | 0 | 1278700 |
| GVGTDEAC | ANXA11 | P50995 | ANX11_HUMAN | Unmodified | 0 | 0 | 0 | 767590 | 1199100 | 734480 | 0 | 0 | 4454000 | 1365200 | 501300 | 0 | 511930 | 0 | 0 | 0 | 0 | 360030 |
| GVIALCIED | SDHA; SDHA; N | P31040; D6F | SDHA_HUMAN | Unmodified | 0 | 0 | 0 | 0 | 0 | 0 | 0 | 0 | 1922300 | 0 | 0 | 0 | 0 | 0 | 0 | 0 | 0 | 0 |
| GVIDLDVFI | TSG101; TSG10 | Q99816; J3Q | TS101_HUMAN | Unmodified | 0 | 1376200 | 0 | 0 | 0 | 0 | 0 | 0 | 0 | 0 | 0 | 0 | 232080 | 0 | 2786900 | 325270 | 0 | 0 |
| GVIDLIFEK | EIF4G1; EIF4G1 | Q04637; C9J | IF4G1_HUMAN | Unmodified | 0 | 0 | 0 | 0 | 0 | 0 | 0 | 0 | 0 | 0 | 0 | 1846400 | 0 | 0 | 0 | 0 | 0 | 0 |
| GVIDMGNS | SPTAN1; SPTAN | Q13813; A0A | SPTN1_HUMAN | Oxidation (M) | 0 | 0 | 0 | 0 | 0 | 0 | 0 | 0 | 0 | 0 | 0 | 1238700 | 0 | 0 | 0 | 0 | 0 | 0 |
| GVIDMGNS | SPTAN1; SPTAN | Q13813; A0A | SPTN1_HUMAN | Unmodified | 0 | 0 | 0 | 0 | 0 | 0 | 0 | 0 | 0 | 0 | 0 | 877910 | 0 | 0 | 0 | 0 | 0 | 379070 |
| GVIDVFLH/ | MAN1B1; MAN1 | Q9UKM7; HC | MA1B1_HUMAN | Unmodified | 0 | 0 | 0 | 0 | 0 | 0 | 0 | 0 | 2897700 | 937100 | 50499 | 89941 | 0 | 0 | 74548 | 0 | 0 | 0 |
| GVIINTASV | HSD17B10 | Q99714 | HCD2_HUMAN | Unmodified | 0 | 0 | 0 | 0 | 0 | 0 | 0 | 0 | 2715500 | 0 | 0 | 0 | 0 | 0 | 0 | 0 | 0 | 0 |
| GVIIQGARC | NRP2 | O60462 | NRP2_HUMAN | Unmodified | 0 | 0 | 0 | 0 | 0 | 0 | 444540 | 0 | 0 | 306740 | 0 | 0 | 0 | 0 | 0 | 0 | 0 | 0 |
| GVILYSDPA | FOLH1 | Q04609 | FOLH1_HUMAI | Unmodified | 7618200 | 7380100 | 19100000 | 62331000 | 0 | 0 | 0 | 0 | 208500 | 76435 | 0 | 0 | 7547300 | 4666100 | 23323000 | 25637000 | 0 | 0 |
| GVINIIPGS | ALDH1L2; ALDH | Q3SY69; A0A | AL1L2_HUMAN | Unmodified | 0 | 0 | 0 | 0 | 0 | 0 | 0 | 0 | 787270 | 0 | 0 | 0 | 0 | 0 | 0 | 0 | 0 | 0 |
| GVIVDKDF | CCT5; CCT5; C | P48643; B7Z | TCPE_HUMAN | Unmodified | 0 | 0 | 0 | 0 | 2605100 | 591470 | 1322700 | 144280 | 0 | 1316900 | 0 | 0 | 5095600 | 2835700 | 0 | 0 | 1621000 | 3197600 |
| GVIVDKDF | CCT5; CCT5; C | P48643; B7Z | TCPE_HUMAN | Oxidation (M) | 0 | 0 | 0 | 0 | 212760 | 0 | 39642 | 0 | 0 | 0 | 0 | 0 | 569100 | 208310 | 0 | 0 | 0 | 0 |
| GVKGDEC( | ATRN | O75882 | ATRN_HUMAN | Unmodified | 41218 | 0 | 0 | 0 | 0 | 0 | 0 | 0 | 249960 | 0 | 0 | 0 | 0 | 0 | 0 | 0 | 0 | 0 |
| GVKIDKTD | ARHGDIA; ARH | P52565; J3K | GDIR1_HUMAN | Oxidation (M) | 54487 | 63629 | 0 | 0 | 0 | 0 | 62931 | 0 | 0 | 0 | 0 | 942660 | 0 | 0 | 0 | 0 | 0 | 236520 |
| GVLCDKD( | PPM1L | Q5SGD2 | PPM1L_HUMAN | Unmodified | 0 | 0 | 0 | 0 | 0 | 0 | 0 | 0 | 3234200 | 157450 | 0 | 0 | 0 | 0 | 0 | 0 | 0 | 0 |
| GVLDVPAC | PLCG1; PLCG1 | P19174; A0A | PLCG1_HUMAI | Unmodified | 0 | 0 | 0 | 0 | 0 | 160080 | 0 | 0 | 153940 | 0 | 320360 | 1794700 | 0 | 0 | 0 | 115660 | 0 | 0 |
| GVLENEFG | IL1RAP | Q9NPH3 | IL1AP_HUMAN | Unmodified | 0 | 0 | 0 | 0 | 0 | 0 | 0 | 0 | 2729700 | 303360 | 0 | 0 | 452210 | 0 | 518880 | 442850 | 0 | 0 |
| GVLEQEEK | CLGN | O14967 | CLGN_HUMAN | Unmodified | 0 | 79438 | 0 | 95302 | 0 | 0 | 0 | 0 | 357480 | 317280 | 54354 | 185930 | 413760 | 208420 | 35209 | 39328 | 0 | 66064 |
| GVLETFSG | AK3 | Q9UIJ7 | KAD3_HUMAN | Unmodified | 0 | 0 | 0 | 0 | 0 | 0 | 0 | 0 | 0 | 0 | 0 | 0 | 0 | 0 | 0 | 0 | 0 | 0 |
| GVLFGVPG | PRDX5 | P30044 | PRDX5_HUMAI | Unmodified | 86578 | 0 | 811660 | 304820 | 4895300 | 424030 | 2146500 | 754380 | 12538000 | 4111900 | 2004700 | 7411800 | 6876500 | 6256700 | 603970 | 277810 | 2929500 | 5312600 |
| GVLFGVPG | PRDX5 | P30044 | PRDX5_HUMAI | Unmodified | 0 | 0 | 0 | 0 | 0 | 0 | 995270 | 0 | 0 | 403810 | 0 | 0 | 0 | 1494300 | 0 | 0 | 4238500 | 0 |
| GVLFHSGC | PAFAH1B1; PAF | P43034; I3L | LIS1_HUMAN | Unmodified | 0 | 0 | 182700 | 0 | 0 | 0 | 0 | 0 | 173340 | 0 | 98291 | 1188400 | 0 | 0 | 0 | 0 | 0 | 0 |
| GVLFYGPP | VCP; VCP; VCP; | P55072; C9J | TERA_HUMAN | Unmodified | 0 | 0 | 0 | 104570 | 5627600 | 0 | 1754900 | 1311200 | 4344300 | 222360 | 0 | 17939000 | 10277000 | 6160300 | 0 | 0 | 1532300 | 2339100 |
| GVLLDIDDI | IQGAP2; IQGAP | Q13576; F5H | IQGA2_HUMAN | Unmodified | 0 | 0 | 1049800 | 0 | 348590 | 0 | 788200 | 0 | 0 | 0 | 0 | 0 | 0 | 0 | 25844 | 0 | 0 | 0 |
| GVLLEIEDL | IQGAP1; IQGAP | P46940; H0Y | IQGA1_HUMAN | Unmodified | 519790 | 934480 | 83919 | 347100 | 893990 | 420710 | 474720 | 0 | 345850 | 302510 | 0 | 498310 | 0 | 0 | 1366800 | 388430 | 0 | 78799 |
| GVLLFGPP | PSMC2; PSMC2 | P35998; C9J | PRS7_HUMAN | Unmodified | 0 | 0 | 630430 | 0 | 0 | 0 | 0 | 0 | 0 | 0 | 0 | 3730900 | 0 | 0 | 238840 | 259390 | 0 | 0 |
| GVLLYGPP | PSMC5; SPATA | P62195; Q8 | PRS8_HUMAN | Unmodified | 549840 | 176560 | 1193100 | 919800 | 855380 | 180150 | 230390 | 1967500 | 0 | 368490 | 1141300 | 7320500 | 3655000 | 3092700 | 793360 | 806070 | 3412600 | 216780 |
| GVLMYGPP | PSMC3; PSMC3 | P17980; E9P | PRS6A_HUMAN | Unmodified | 99396 | 0 | 0 | 0 | 0 | 0 | 0 | 162950 | 0 | 0 | 326380 | 0 | 319710 | 220740 | 0 | 0 | 190110 | 346670 |
| GVLMYGPP | PSMC3; PSMC3 | P17980; E9P | PRS6A_HUMAN | Oxidation (M) | 0 | 0 | 0 | 0 | 183290 | 0 | 0 | 0 | 0 | 0 | 0 | 935400 | 0 | 0 | 0 | 0 | 166820 | 0 |
| GVLQTAAE | GEMIN5 | Q8TEQ6 | GEMI5_HUMAN | Unmodified | 0 | 0 | 0 | 0 | 0 | 0 | 0 | 0 | 0 | 0 | 0 | 669110 | 0 | 0 | 0 | 0 | 0 | 0 |
| GVLRQITVI | PRDX2 | P32119 | PRDX2_HUMAI | Unmodified | 0 | 0 | 14930 | 0 | 566450 | 388000 | 356600 | 0 | 282250 | 1752300 | 673510 | 0 | 1265500 | 542400 | 0 | 0 | 0 | 72958 |
| GVLSDLTK | TSPAN14; TSPA | Q8NG11; H7 | TSN14_HUMAN | Unmodified | 904530 | 0 | 0 | 0 | 19276000 | 407170 | 15036000 | 9346600 | 10467000 | 32782000 | 2.27E+08 | 6460200 | 5731700 | 37744000 | 13276000 | 1785100 | 3899000 | 40403000 |
| GVMGGQS | USO1 | O60763 | USO1_HUMAN | Unmodified | 0 | 0 | 0 | 0 | 0 | 0 | 0 | 0 | 0 | 0 | 0 | 1165700 | 0 | 0 | 0 | 0 | 0 | 0 |
| GVMGGQS | USO1 | O60763 | USO1_HUMAN | Oxidation (M) | 0 | 0 | 54802 | 0 | 0 | 0 | 80991 | 0 | 0 | 0 | 0 | 1300100 | 154790 | 0 | 81927 | 56188 | 0 | 0 |
| GVMLAVDA | HSPD1; HSPD1 | P10809; A0A | CH60_HUMAN | Unmodified | 0 | 0 | 0 | 0 | 0 | 0 | 0 | 0 | 424470 | 0 | 0 | 0 | 0 | 0 | 0 | 0 | 0 | 0 |
| GVMTTGSD | NUP210 | Q8TEM1 | PO210_HUMAN | 2 Oxidation (M) | 0 | 0 | 0 | 0 | 0 | 0 | 0 | 0 | 9892400 | 8023200 | 1027700 | 0 | 0 | 0 | 0 | 0 | 0 | 0 |
| GVMTTGSD | NUP210 | Q8TEM1 | PO210_HUMAN | Oxidation (M) | 0 | 0 | 0 | 0 | 0 | 0 | 0 | 0 | 16140000 | 3015200 | 0 | 0 | 0 | 0 | 0 | 0 | 0 | 0 |
| GVMTTGSD | NUP210 | Q8TEM1 | PO210_HUMAN | Unmodified | 0 | 0 | 0 | 0 | 0 | 0 | 0 | 0 | 2803700 | 843240 | 0 | 0 | 0 | 0 | 0 | 0 | 0 | 0 |
| GVNCIDYY | COPB2; COPB2 | P35606; H0Y | COPB2_HUMAI | Unmodified | 0 | 0 | 0 | 0 | 0 | 0 | 0 | 0 | 0 | 454840 | 1149900 | 2961700 | 0 | 0 | 168220 | 0 | 2955100 | 937660 |
| GVNIGGAC | CRIP2; CRIP2 | P52943; H0Y | CRIP2_HUMAN | Unmodified | 283070 | 193590 | 0 | 0 | 0 | 0 | 0 | 290730 | 3812700 | 259220 | 0 | 0 | 428560 | 0 | 213480 | 0 | 0 | 597190 |
| GVNIGGAC | CRIP2; CRIP2 | P52943; H0Y | CRIP2_HUMAN | Unmodified | 0 | 0 | 0 | 0 | 0 | 0 | 0 | 1929200 | 19680000 | 2253300 | 0 | 0 | 0 | 0 | 0 | 0 | 2374800 | 7201100 |
| GVNIGGAC | CRIP2; CRIP2 | P52943; H0Y | CRIP2_HUMAN | Unmodified | 0 | 0 | 0 | 0 | 0 | 0 | 0 | 701870 | 0 | 1811400 | 0 | 0 | 0 | 0 | 0 | 0 | 5172900 | 5598700 |
| GVNLDHTE | GRK5 | P34947 | GRK5_HUMAN | Unmodified | 0 | 0 | 0 | 0 | 151220 | 0 | 38163 | 48788 | 70456 | 27140 | 1660900 | 1320900 | 80138 | 0 | 0 | 0 | 265880 | 282880 |
| GVNLPGAA | PKM; PKM; PKM | P14618; B4D | KPYM_HUMAN | Unmodified | 2249600 | 655050 | 3221200 | 2866400 | 25843000 | 4742100 | 0 | 1030600 | 6362800 | 0 | 13392000 | 67583000 | 1728000 | 0 | 5290000 | 6171000 | 522030 | 2287700 |
| GVNLPGAA | PKM; PKM; PKM | P14618; B4D | KPYM_HUMAN | Unmodified | 0 | 0 | 0 | 0 | 39036000 | 5526400 | 1743800 | 363970 | 0 | 1993500 | 529820 | 539890 | 0 | 3210700 | 0 | 0 | 8457900 | 13991000 |
| GVNLYVKN | PABPC1; PABP( | P11940; A0A | PABP1_HUMAN | Unmodified | 0 | 0 | 0 | 0 | 0 | 0 | 0 | 0 | 0 | 0 | 0 | 0 | 4610900 | 0 | 0 | 0 | 0 | 0 |
| GVNSFMV | DPYSL3 | Q14195 | DPYL3_HUMAN | 2 Oxidation (M) | 0 | 0 | 0 | 0 | 0 | 0 | 0 | 0 | 0 | 0 | 875230 | 2611100 | 0 | 0 | 0 | 0 | 0 | 0 |
| GVNSYPSL | DNAJC10; DNA | Q8IXB1; A0A | DJC10_HUMAN | Unmodified | 0 | 0 | 0 | 0 | 0 | 0 | 0 | 0 | 18197000 | 2910200 | 852160 | 7244500 | 0 | 0 | 0 | 0 | 0 | 0 |
| GVNTFSPE | PSMA5 | P28066 | PSA5_HUMAN | Unmodified | 0 | 1286400 | 0 | 0 | 0 | 0 | 87346 | 0 | 272120 | 194960 | 0 | 4533400 | 0 | 119520 | 373410 | 261540 | 0 | 83468 |
| GVNTGAVC | CRIP2; CRIP2; | P52943; H0Y | CRIP2_HUMAN | Unmodified | 1296900 | 542130 | 0 | 59578 | 666540 | 146010 | 1018700 | 677210 | 5403900 | 73455 | 0 | 0 | 0 | 0 | 545310 | 0 | 779650 | 1661500 |
| GVNTGAVC | CRIP2; CRIP2; | P52943; H0Y | CRIP2_HUMAN | Unmodified | 555660 | 456170 | 0 | 0 | 3683900 | 0 | 3994000 | 6441300 | 4346600 | 7619200 | 0 | 0 | 0 | 0 | 0 | 0 | 20298000 | 29905000 |
| GVNTGGRI | MIA3; MIA3 | Q5JRA6; A0A | TGO1_HUMAN | Unmodified | 0 | 0 | 0 | 0 | 0 | 0 | 0 | 0 | 779610 | 156600 | 0 | 0 | 0 | 0 | 0 | 0 | 98735 | 0 |
| GVNTVFHC | NSDHL; NSDHL | Q15738; C9J | NSDHL_HUMAN | Unmodified | 0 | 0 | 0 | 0 | 66905 | 0 | 262860 | 84245 | 97901 | 0 | 0 | 0 | 78394 | 0 | 0 | 0 | 39320 | 0 |
| GVNVSALS | ATP6V1A; ATP6 | P38606; C9J | VATA_HUMAN | Unmodified | 289780 | 271580 | 104520 | 0 | 0 | 0 | 0 | 0 | 880190 | 201260 | 491650 | 4750400 | 139640 | 0 | 138580 | 535150 | 0 | 0 |
| GVNWAAFI | COPA; COPA; C | P53621; A0A | COPA_HUMAN | Oxidation (M) | 0 | 0 | 0 | 0 | 0 | 0 | 0 | 84346 | 0 | 0 | 0 | 2908600 | 0 | 0 | 0 | 437120 | 0 | 0 |
| GVPALVAL | ARVCF; ARVCF; | O00192; E9F | ARVC_HUMAN | Unmodified | 41641 | 0 | 608810 | 740660 | 0 | 0 | 0 | 0 | 159470 | 0 | 119670 | 1018100 | 0 | 0 | 1708200 | 1259900 | 0 | 0 |
| GVPELVLK | ITGA6 | P23229 | ITA6_HUMAN I | Unmodified | 0 | 0 | 0 | 4678800 | 9368000 | 0 | 567040 | 381410 | 249820 | 0 | 671880 | 4950700 | 4395000 | 1830500 | 0 | 0 | 3807000 | 27105000 |
| GVPELVLKI | ITGA6 | P23229 | ITA6_HUMAN I | Unmodified | 0 | 0 | 0 | 0 | 272630 | 122520 | 0 | 61606 | 0 | 0 | 0 | 0 | 506640 | 275920 | 0 | 0 | 1065100 | 2293800 |
| GVPFTLTTV | CLIC3 | O95833 | CLIC3_HUMAN | Unmodified | 0 | 0 | 0 | 0 | 1625200 | 0 | 0 | 0 | 0 | 0 | 490670 | 0 | 0 | 0 | 0 | 0 | 0 | 0 |
| GVPIIFADE | DAG1 | Q14118 | DAG1_HUMAN | Oxidation (M) | 0 | 0 | 0 | 0 | 0 | 0 | 0 | 0 | 7362100 | 0 | 0 | 2327000 | 1615700 | 0 | 0 | 0 | 0 | 0 |
| GVPIIFADE | DAG1 | Q14118 | DAG1_HUMAN | Unmodified | 0 | 0 | 0 | 0 | 0 | 0 | 0 | 0 | 2840200 | 346320 | 0 | 0 | 797210 | 0 | 0 | 0 | 0 | 0 |
| GVPLVYEAI | PCK2; PCK2 | Q16822; H0Y | PCKGM_HUMA | Unmodified | 0 | 0 | 0 | 0 | 0 | 0 | 63918 | 0 | 3452900 | 218750 | 109260 | 0 | 188630 | 87337 | 0 | 0 | 0 | 0 |
| GVPLYRHI | ENO1; ENO1; E | P06733; K7E | ENOA_HUMAN | Unmodified | 0 | 0 | 0 | 0 | 3.21E+08 | 18103000 | 7816400 | 0 | 0 | 26996000 | 0 | 0 | 1.25E+08 | 79211000 | 0 | 0 | 74695000 | 1.46E+08 |
| GVPRGLYD | DPYSL2; DPYSL | Q16555; A0A | DPYL2_HUMAN | Unmodified | 0 | 0 | 0 | 0 | 3532700 | 536800 | 379660 | 481230 | 0 | 0 | 0 | 0 | 0 | 0 | 0 | 0 | 0 | 445350 |
| GVPSLMPC | GPC2 | Q8N158 | GPC2_HUMAN | Unmodified | 0 | 0 | 0 | 0 | 2124300 | 101210 | 148460 | 0 | 0 | 0 | 0 | 0 | 289350 | 711770 | 0 | 0 | 173430 | 155250 |
| GVPSVPVA | SLC12A7; SLC1 | Q9Y666; A0A | S12A7_HUMAN | Unmodified | 249330 | 0 | 0 | 0 | 185110 | 0 | 0 | 0 | 3704900 | 0 | 0 | 1467600 | 0 | 0 | 745160 | 0 | 0 | 0 |
| GVPTGFILF | MTHFD1; MTHF | P11586; V9C | C1TC_HUMAN | Unmodified | 0 | 0 | 224570 | 0 | 0 | 0 | 0 | 0 | 2339100 | 0 | 616980 | 14189000 | 0 | 0 | 714430 | 570540 | 0 | 0 |
| GVPTGFILF | MTHFD1; MTHF | P11586; V9C | C1TC_HUMAN | Unmodified | 258070 | 0 | 411400 | 1727500 | 5104100 | 606960 | 677340 | 90318 | 7949500 | 1038600 | 2952500 | 6880800 | 3278200 | 0 | 0 | 536820 | 1861500 | 4272100 |
| GVPWNFTI | EPB41; EPB41; | P11171; A0A | EPB41_HUMAN | Unmodified | 0 | 526750 | 726700 | 856440 | 0 | 0 | 0 | 0 | 0 | 0 | 0 | 0 | 0 | 0 | 1880200 | 1810900 | 0 | 0 |
| GVPYRPTD | SBF1 | O95248 | MTMR5_HUMA | Unmodified | 147710 | 0 | 0 | 169670 | 0 | 0 | 0 | 0 | 0 | 0 | 230350 | 808570 | 0 | 0 | 0 | 163210 | 0 | 0 |
| GVQFAPFS | AGPS; AGPS; A( | O00116; B7Z | ADAS_HUMAN | Unmodified | 0 | 0 | 0 | 0 | 0 | 0 | 0 | 0 | 3508100 | 0 | 0 | 261060 | 0 | 0 | 0 | 0 | 0 | 0 |
| GVQGFQD | FAM120A | Q9NZB2 | F120A_HUMAN | Unmodified | 138770 | 0 | 132530 | 168350 | 0 | 0 | 1138500 | 232010 | 100020 | 0 | 0 | 252770 | 304900 | 0 | 0 | 0 | 0 | 284940 |
| GVQLQTHF | ARCN1; ARCN1 | P48444; B0Y | COPD_HUMAN | Unmodified | 0 | 0 | 0 | 0 | 0 | 0 | 0 | 0 | 0 | 0 | 281430 | 0 | 0 | 0 | 0 | 0 | 0 | 0 |
| GVQVETISF | FKBP1A; FKBP1 | P62942; Q5V | FKB1A_HUMAN | Unmodified | 7308800 | 10412000 | 6308300 | 4361500 | 0 | 93807 | 117420 | 0 | 2673600 | 199560 | 1233400 | 8898700 | 0 | 227290 | 11679000 | 9010900 | 462570 | 535770 |
| GVQVETISF | FKBP1A; FKBP1 | P62942; Q5V | FKB1A_HUMAN | Unmodified | 5553300 | 2311600 | 7009700 | 5877100 | 27373000 | 10762000 | 15117000 | 6679500 | 3266600 | 6014900 | 0 | 3924000 | 38950000 | 32487000 | 7462400 | 121590 | 41853000 | 39250000 |
| GVQVNYDI | SEL1L | Q9UBV2 | SE1L1_HUMAN | Unmodified | 0 | 0 | 0 | 0 | 0 | 0 | 0 | 0 | 1229400 | 299650 | 0 | 0 | 0 | 0 | 0 | 0 | 0 | 0 |
| GVQYLNEII | EEF2 | P13639 | EF2_HUMAN EI | Unmodified | 0 | 1126100 | 3286300 | 1207100 | 2737900 | 438970 | 0 | 0 | 2148400 | 0 | 5355200 | 40376000 | 859840 | 0 | 2734200 | 2342800 | 297400 | 848150 |
| GVQYLNEII | EEF2 | P13639 | EF2_HUMAN EI | Unmodified | 0 | 139740 | 0 | 302600 | 2630900 | 188650 | 475490 | 0 | 0 | 343830 | 57862 | 121550 | 1234600 | 911810 | 229570 | 0 | 79205 | 1276500 |
| GVRVKETAI | FLNA; FLNA; FL | P21333; F8W | FLNA_HUMAN | Unmodified | 0 | 0 | 0 | 0 | 2065300 | 381870 | 2210700 | 78717 | 0 | 0 | 0 | 0 | 1083600 | 1969600 | 0 | 0 | 2160800 | 0 |
| GVSEHDFA | PLXNB1 | O43157 | PLXB1_HUMAN | Unmodified | 0 | 0 | 0 | 0 | 0 | 0 | 0 | 0 | 514350 | 0 | 0 | 0 | 0 | 0 | 0 | 0 | 0 | 0 |

| | | | | | | | | | | | | | | | | | | | | | | |
|---|---|---|---|---|---|---|---|---|---|---|---|---|---|---|---|---|---|---|---|---|---|---|
| GVSEIVQN | FABP1; FABP1 | P07148; A8N | FABPL_HUMAN | Unmodified | 29119 | 0 | 333940 | 0 | 0 | 0 | 353550 | 0 | 0 | 0 | 103030 | 486690 | 0 | 0 | 32717 | 0 | 0 | 0 |
| GVSFYEVPI | MYO1C; MYO1C | O00159; I3L | MYO1C_HUMA | Unmodified | 374590 | 256490 | 145390 | 641190 | 0 | 0 | 0 | 0 | 0 | 0 | 0 | 0 | 48199 | 0 | 1742600 | 368060 | 0 | 0 |
| GVSHGQFI | MGAT1 | P26572 | MGAT1_HUMAI | Unmodified | 0 | 0 | 0 | 0 | 37072 | 0 | 0 | 0 | 1448300 | 268880 | 0 | 0 | 0 | 0 | 0 | 0 | 0 | 0 |
| GVSKAVEH | ENO1; ENO1; E | P06733; K7E | ENOA_HUMAN | Unmodified | 0 | 0 | 0 | 0 | 123640 | 0 | 0 | 55249 | 0 | 0 | 0 | 0 | 0 | 0 | 0 | 0 | 988040 | 201540 |
| GVSLDLSN | CARMIL1 | Q5VZK9 | CARL1_HUMAN | Unmodified | 0 | 0 | 0 | 0 | 0 | 0 | 0 | 52838 | 0 | 0 | 0 | 931980 | 0 | 0 | 0 | 0 | 0 | 0 |
| GVSLTNHH | PRKCSH; PRKC | P14314; K7E | GLU2B_HUMAI | Unmodified | 28845 | 0 | 151060 | 0 | 0 | 0 | 0 | 0 | 27972000 | 590980 | 375540 | 603820 | 0 | 0 | 73975 | 92140 | 0 | 0 |
| GVSLTNHH | PRKCSH; PRKC | P14314; K7E | GLU2B_HUMAI | Unmodified | 0 | 0 | 0 | 0 | 0 | 0 | 0 | 0 | 6409400 | 0 | 0 | 0 | 0 | 0 | 0 | 0 | 0 | 0 |
| GVSQMPLF | ATIC; ATIC | P31939; C9J | PUR9_HUMAN | Unmodified | 0 | 0 | 0 | 0 | 0 | 0 | 0 | 0 | 664110 | 0 | 158470 | 4280000 | 0 | 0 | 0 | 0 | 0 | 0 |
| GVSSLSIQF | CARMIL1 | Q5VZK9 | CARL1_HUMAN | Unmodified | 0 | 0 | 0 | 0 | 0 | 0 | 0 | 0 | 0 | 0 | 77487 | 1921700 | 0 | 0 | 0 | 0 | 0 | 0 |
| GVSSQETA | NAMPT; NAMPT | P43490; A0A | NAMPT_HUMAI | Unmodified | 45769 | 0 | 1649900 | 131960 | 0 | 0 | 0 | 0 | 0 | 56155 | 0 | 10767000 | 0 | 0 | 1302600 | 1222900 | 0 | 0 |
| GVSVWPV | B3GAT3; B3GAT | O94766; G3 | B3GA3_HUMAI | Unmodified | 0 | 0 | 0 | 0 | 0 | 0 | 0 | 0 | 2573500 | 264500 | 282950 | 724840 | 0 | 0 | 0 | 0 | 0 | 0 |
| GVSVWPV | B3GAT3; B3GAT | O94766; G3 | B3GA3_HUMAI | Unmodified | 0 | 0 | 0 | 0 | 0 | 0 | 0 | 0 | 15203000 | 477070 | 132550 | 150980 | 0 | 0 | 0 | 0 | 0 | 0 |
| GVTDLDAV | NOS1AP; NOS1 | O75052; E9F | CAPON_HUMA | Unmodified | 0 | 0 | 0 | 0 | 116120 | 27995 | 0 | 0 | 295230 | 0 | 0 | 0 | 205430 | 76438 | 0 | 0 | 18897 | 55864 |
| GVTDVVITF | OGFOD3 | Q6PK18 | OGFD3_HUMA | Unmodified | 0 | 0 | 0 | 0 | 0 | 0 | 0 | 0 | 2371500 | 0 | 0 | 0 | 0 | 0 | 0 | 0 | 31350 | 0 |
| GVTFLFPIQ | DDX21 | Q9NR30 | DDX21_HUMAI | Unmodified | 0 | 0 | 0 | 0 | 0 | 0 | 0 | 0 | 0 | 0 | 184830 | 1116100 | 0 | 0 | 0 | 0 | 0 | 0 |
| GVTFNVTT | CLIC1 | O00299 | CLIC1_HUMAN | Unmodified | 3599200 | 4227800 | 2891600 | 6531500 | 75327000 | 14868000 | 6813000 | 5643500 | 13857000 | 8271300 | 6496800 | 20187000 | 46763000 | 0 | 7109600 | 4252600 | 32120000 | 25401000 |
| GVTFNVTT | CLIC1 | O00299 | CLIC1_HUMAN | Unmodified | 0 | 0 | 0 | 0 | 629020 | 0 | 0 | 0 | 1362800 | 0 | 0 | 0 | 0 | 0 | 0 | 0 | 0 | 0 |
| GVTIASGG | MACROH2A1; N | O75367; D6I | H2AY_HUMAN | Unmodified | 15831000 | 2057200 | 0 | 0 | 0 | 0 | 0 | 268350 | 0 | 0 | 0 | 48609000 | 0 | 0 | 0 | 0 | 0 | 0 |
| GVTIASGG | MACROH2A1; N | O75367; D6I | H2AY_HUMAN | Unmodified | 1354500 | 782980 | 0 | 0 | 0 | 0 | 0 | 0 | 0 | 0 | 0 | 2419800 | 0 | 0 | 0 | 0 | 0 | 0 |
| GVTIASGG | MACROH2A2; N | Q9P0M6; A0. | H2AW_HUMAN | Unmodified | 505720 | 697420 | 0 | 0 | 1128300 | 101670 | 0 | 0 | 0 | 0 | 198280 | 18183000 | 144510 | 0 | 0 | 0 | 0 | 495430 |
| GVTIIGPAT | ACLY | P53396 | ACLY_HUMAN | Unmodified | 417270 | 0 | 0 | 1052300 | 2760800 | 257210 | 4081300 | 1169600 | 1334500 | 521130 | 2137300 | 16040000 | 1161500 | 0 | 1191400 | 705210 | 46881 | 1175500 |
| GVTIKPTVD | EIF2S3; EIF2S3I | P41091; Q2 | IF2G_HUMAN E | Unmodified | 270630 | 138320 | 413050 | 692410 | 1308400 | 313530 | 139580 | 79813 | 554510 | 252420 | 1511400 | 6564500 | 1015800 | 297590 | 380910 | 455330 | 208600 | 1077900 |
| GVTIPYRPK | PCBP2 | F8VRH0 | F8VRH0_HUM/ | Unmodified | 32823000 | 16511000 | 0 | 61715000 | 0 | 0 | 16455000 | 110440 | 1483000 | 0 | 0 | 2203300 | 70638000 | 19922000 | 0 | 15767000 | 0 | 3479400 |
| GVTIPYRPK | PCBP2; PCBP2 | Q15366; H3I | PCBP2_HUMAI | Unmodified | 34208000 | 18276000 | 280300 | 3071200 | 0 | 0 | 0 | 416270 | 731890 | 0 | 0 | 9431700 | 0 | 0 | 4197700 | 6653600 | 0 | 0 |
| GVTIPYRPK | PCBP2; PCBP2 | F8VZX2; H3E | F8VZX2_HUMA | Unmodified | 5625000 | 8180900 | 0 | 4683000 | 0 | 0 | 812550 | 329030 | 0 | 0 | 0 | 1931600 | 4343700 | 1140700 | 948260 | 1171900 | 347940 | 0 |
| GVTQFGNK | NUDT21 | O43809 | CPSF5_HUMAI | Unmodified | 0 | 2551000 | 1181200 | 1548900 | 262900 | 174090 | 161150 | 12100 | 166210 | 222570 | 94431 | 496960 | 547490 | 155930 | 649530 | 1226400 | 36966 | 74646 |
| GVTQIDND | ATP6V1C1; ATP | P21283; E7E | VATC1_HUMAI | Unmodified | 0 | 0 | 0 | 0 | 506090 | 0 | 0 | 0 | 247330 | 0 | 0 | 1230100 | 0 | 0 | 0 | 0 | 0 | 0 |
| GVTQRLDL | SPECC1; SPEC | Q5M775; A0. | CYTSB_HUMAI | Unmodified | 0 | 0 | 27594 | 109900 | 336950 | 55151 | 0 | 0 | 0 | 35649 | 0 | 0 | 239540 | 174650 | 71466 | 47674 | 60462 | 98248 |
| GVTQYYAY | DDX6 | P26196 | DDX6_HUMAN | Unmodified | 574310 | 408550 | 213450 | 220810 | 0 | 0 | 0 | 0 | 0 | 119640 | 0 | 1521000 | 0 | 0 | 658710 | 374200 | 0 | 0 |
| GVVDSDDI | HSP90B1; HSP | P14625; A0A | ENPL_HUMAN | Unmodified | 280190 | 0 | 3643100 | 1529100 | 0 | 0 | 0 | 462160 | 2.8E+08 | 11635000 | 78297000 | 45198000 | 0 | 0 | 4443300 | 2091100 | 0 | 0 |
| GVVDSDDI | HSP90B1; HSP | P14625; A0A | ENPL_HUMAN | Unmodified | 0 | 0 | 1559100 | 941340 | 5645800 | 0 | 24334000 | 13717000 | 2.09E+08 | 4.19E+08 | 30387000 | 7873400 | 4208400 | 6224500 | 0 | 0 | 613600 | 535260 |
| GVVDSDDI | HSP90B1; HSP | P14625; A0A | ENPL_HUMAN | Unmodified | 0 | 0 | 0 | 0 | 0 | 0 | 3372900 | 51497 | 1520100 | 9637500 | 0 | 0 | 0 | 0 | 0 | 0 | 66575 | 0 |
| GVVDSEDL | HSP90AB1; HSI | P08238; Q5 | HS90B_HUMAI | Unmodified | 6428700 | 2002900 | 16478000 | 3792400 | 0 | 0 | 0 | 116170 | 39023000 | 2471800 | 43594000 | 4.18E+08 | 0 | 0 | 5388600 | 14954000 | 233920 | 0 |
| GVVDSEDL | HSP90AB1; HSI | P08238; Q5 | HS90B_HUMAI | Unmodified | 0 | 0 | 0 | 0 | 4516500 | 1120400 | 0 | 0 | 20298000 | 3659900 | 3042400 | 12295000 | 0 | 0 | 0 | 262930 | 1643500 | 1003300 |
| GVVDSEDL | HSP90AB1; HSI | P08238; Q5 | HS90B_HUMAI | Oxidation (M) | 428240 | 286740 | 280790 | 2485800 | 0 | 0 | 0 | 0 | 1186100 | 1480500 | 2667300 | 9993000 | 0 | 0 | 611290 | 526350 | 479700 | 0 |
| GVVEEWLS | FAM126A; FAM | Q9BYI3; A0A | HYCCI_HUMAI | Unmodified | 0 | 0 | 0 | 0 | 3547000 | 276720 | 0 | 0 | 30890 | 0 | 44035 | 48187 | 0 | 0 | 0 | 0 | 0 | 3586900 |
| GVVEVTHD | SERPINH1; SER | P50454; E9P | SERPH_HUMAI | Unmodified | 0 | 0 | 0 | 0 | 1759200 | 0 | 2558900 | 1416900 | 70360000 | 20712000 | 260550 | 182970 | 0 | 0 | 0 | 52241 | 97168 | 813290 |
| GVVEVTHD | SERPINH1; SER | P50454; E9P | SERPH_HUMAI | Unmodified | 0 | 0 | 0 | 0 | 239460 | 0 | 693450 | 0 | 0 | 3207000 | 0 | 0 | 0 | 141360 | 0 | 0 | 0 | 119200 |
| GVVFDVTS | NENF | Q9UMX5 | NENF_HUMAN | Unmodified | 0 | 0 | 0 | 0 | 156960 | 430480 | 0 | 0 | 24338000 | 6002400 | 0 | 0 | 0 | 0 | 0 | 0 | 71227 | 338250 |
| GVVFDVTS | NENF | Q9UMX5 | NENF_HUMAN | Unmodified | 0 | 0 | 0 | 0 | 0 | 0 | 184590 | 0 | 3257400 | 2179600 | 0 | 0 | 0 | 44658 | 0 | 0 | 0 | 0 |
| GVVFDVTS | NENF | Q9UMX5 | NENF_HUMAN | Unmodified | 0 | 0 | 0 | 0 | 0 | 0 | 0 | 0 | 0 | 7889300 | 0 | 0 | 0 | 0 | 0 | 0 | 0 | 0 |
| GVVFNVTT | CLIC5; CLIC5; ( | Q9NZA1; A0. | CLIC5_HUMAN | Unmodified | 0 | 0 | 0 | 0 | 0 | 0 | 0 | 164740 | 0 | 0 | 0 | 1940600 | 0 | 0 | 0 | 0 | 0 | 0 |
| GVVFSVTT | CLIC4 | Q9Y696 | CLIC4_HUMAN | Unmodified | 0 | 0 | 112960 | 0 | 5573300 | 1647100 | 0 | 958890 | 775920 | 1394100 | 16091 | 1042100 | 2993700 | 3491600 | 0 | 431500 | 3726200 | 3871000 |
| GVVGPGP/ | OSBP | P22059 | OSBP1_HUMAI | Unmodified | 0 | 0 | 0 | 0 | 21810 | 0 | 0 | 0 | 0 | 0 | 909780 | 3643700 | 0 | 0 | 0 | 856700 | 0 | 0 |
| GVVIGTGEI | ATP2C1; ATP2C | P98194; B4E | AT2C1_HUMAN | Unmodified | 0 | 0 | 0 | 0 | 0 | 0 | 0 | 0 | 833060 | 0 | 0 | 415180 | 0 | 0 | 0 | 0 | 74459 | 0 |
| GVVKDEPE | IGF1R; IGF1R | P08069; C9J | IGF1R_HUMAN | Unmodified | 0 | 0 | 0 | 0 | 2580000 | 631690 | 350620 | 0 | 0 | 0 | 0 | 0 | 0 | 0 | 0 | 0 | 0 | 334630 |
| GVVPDNHI | HACL1; HACL1 | Q9UJ83; C9J | HACL1_HUMAI | Unmodified | 0 | 0 | 0 | 0 | 0 | 0 | 0 | 0 | 1446900 | 73760 | 0 | 0 | 0 | 0 | 0 | 0 | 0 | 84026 |
| GVVPFQAG | ZC3HAV1; ZC3I | Q7Z2W4; C9 | ZCCHV_HUMA | Unmodified | 0 | 0 | 1282600 | 1747200 | 0 | 0 | 0 | 0 | 0 | 0 | 0 | 0 | 0 | 0 | 0 | 0 | 0 | 0 |
| GVVPLAGT | ALDOC; ALDOC | P09972; J3Q | ALDOC_HUMA | Unmodified | 3135300 | 0 | 0 | 4277000 | 7613800 | 0 | 0 | 191860 | 14725000 | 1306300 | 28490000 | 73875000 | 0 | 0 | 2394200 | 0 | 1339500 | 2572100 |
| GVVPLAGT | ALDOC; ALDOC | P09972; J3Q | ALDOC_HUMA | Unmodified | 0 | 0 | 0 | 0 | 31529000 | 7996700 | 3265000 | 1809600 | 0 | 2693700 | 2000100 | 0 | 5345100 | 1985300 | 0 | 0 | 9906800 | 13845000 |
| GVVQELQC | EEF1D; EEF1D; | P29692; E9P | EF1D_HUMAN | Unmodified | 22182 | 45761 | 98143 | 0 | 0 | 0 | 0 | 0 | 334070 | 0 | 3038200 | 16132000 | 0 | 0 | 0 | 192520 | 0 | 0 |
| GVVQELQC | EEF1D; EEF1D; | P29692; E9P | EF1D_HUMAN | Unmodified | 65643 | 200470 | 0 | 408640 | 589460 | 0 | 0 | 0 | 1033600 | 101560 | 1125300 | 2486500 | 0 | 83347 | 173300 | 0 | 197440 | 468340 |
| GVVTIEQIV | ALOX5 | P09917 | LOX5_HUMAN | Unmodified | 0 | 0 | 0 | 0 | 0 | 0 | 0 | 0 | 1460000 | 0 | 333250 | 1775300 | 0 | 125370 | 0 | 0 | 0 | 0 |
| GVVVATGD | ATP1A3; ATP1A | P13637; M0 | AT1A3_HUMAN | Unmodified | 0 | 0 | 0 | 0 | 0 | 0 | 0 | 0 | 743650 | 0 | 0 | 0 | 0 | 0 | 0 | 0 | 0 | 0 |
| GVWRPPL | CLGN | O14967 | CLGN_HUMAN | Unmodified | 0 | 0 | 0 | 0 | 0 | 0 | 0 | 0 | 56229000 | 2910600 | 2887400 | 0 | 0 | 0 | 0 | 0 | 112150 | 0 |
| GVYHDDK | GXYLT1 | Q4G148 | GXLT1_HUMAN | Unmodified | 0 | 0 | 0 | 0 | 0 | 0 | 28046 | 0 | 419640 | 184690 | 0 | 0 | 0 | 0 | 0 | 0 | 0 | 0 |
| GVYQFCFS | TMED3; TMED3 | Q9Y3Q3; G3 | TMED3_HUMAI | Unmodified | 0 | 0 | 0 | 0 | 0 | 0 | 0 | 0 | 1885900 | 1441700 | 0 | 0 | 0 | 0 | 0 | 0 | 0 | 0 |
| GWAAAVTF | PFAS; PFAS | O15067; H0 | PUR4_HUMAN | Unmodified | 0 | 0 | 0 | 0 | 0 | 0 | 0 | 0 | 0 | 0 | 150410 | 1113900 | 0 | 0 | 0 | 0 | 0 | 0 |
| GWALLHP | PLOD3; PLOD3 | O60568; H7 | PLOD3_HUMAI | Unmodified | 1657700 | 875020 | 0 | 0 | 69193 | 0 | 0 | 31597 | 476350 | 0 | 0 | 110610 | 0 | 0 | 0 | 40260 | 0 | 0 |
| GWDLTVK | ISG15 | P05161 | ISG15_HUMAN | Unmodified | 1293300 | 1375900 | 0 | 0 | 450410 | 196110 | 0 | 0 | 0 | 0 | 0 | 1857300 | 840840 | 283080 | 504870 | 491510 | 674570 | 1008200 |
| GWDQEPA | LSR; LSR | Q86X29; S4F | LSR_HUMAN Li | Unmodified | 127180 | 101260 | 0 | 0 | 0 | 0 | 0 | 0 | 0 | 0 | 713280 | 0 | 362240 | 0 | 0 | 404940 | 0 | 0 |
| GWDQEPA | LSR; LSR | Q86X29; S4F | LSR_HUMAN Li | Unmodified | 358410 | 0 | 445270 | 617050 | 683340 | 0 | 0 | 0 | 1128100 | 339170 | 200840 | 1426000 | 420700 | 0 | 473270 | 311910 | 0 | 0 |
| GWDQEPA | LSR; LSR | Q86X29; S4F | LSR_HUMAN Li | Unmodified | 0 | 59258 | 0 | 0 | 0 | 0 | 0 | 0 | 0 | 322730 | 0 | 0 | 671500 | 0 | 0 | 0 | 0 | 0 |
| GWDQGLL | FKBP2; FKBP2; | P26885; A0A | FKBP2_HUMAN | Oxidation (M) | 0 | 23166 | 0 | 0 | 0 | 0 | 0 | 35483 | 5708900 | 2414000 | 0 | 0 | 0 | 0 | 0 | 0 | 0 | 0 |
| GWDQGLL | FKBP2; FKBP2; | P26885; A0A | FKBP2_HUMAN | Unmodified | 0 | 0 | 0 | 0 | 0 | 315830 | 0 | 0 | 11755000 | 2212300 | 0 | 0 | 0 | 0 | 0 | 0 | 0 | 0 |
| GWDQGLL | FKBP2; FKBP2; | P26885; A0A | FKBP2_HUMAN | Unmodified | 0 | 0 | 0 | 0 | 0 | 0 | 0 | 0 | 4542300 | 0 | 0 | 0 | 0 | 0 | 11059 | 0 | 0 | 0 |
| GWDQGLL | FKBP2; FKBP2; | P26885; A0A | FKBP2_HUMAN | Oxidation (M) | 0 | 0 | 0 | 0 | 0 | 0 | 0 | 0 | 2709200 | 0 | 220800 | 0 | 0 | 286690 | 0 | 0 | 0 | 0 |
| GWEEGVA | FKBP1A; FKBP1 | P62942; F6T | FKB1A_HUMAN | Unmodified | 1994700 | 956470 | 0 | 0 | 0 | 0 | 0 | 0 | 0 | 0 | 0 | 951520 | 0 | 0 | 1176600 | 763490 | 0 | 0 |
| GWEEGVA | FKBP1A; FKBP1 | P62942; F6T | FKB1A_HUMAN | Oxidation (M) | 439600 | 368210 | 392540 | 0 | 0 | 0 | 0 | 0 | 0 | 0 | 0 | 0 | 0 | 0 | 679120 | 485680 | 0 | 0 |
| GWEFMWN | CKMT1A; CKMT | P12532; P17 | KCRU_HUMAN | Oxidation (M) | 0 | 0 | 0 | 0 | 0 | 0 | 0 | 0 | 0 | 0 | 0 | 976560 | 0 | 0 | 0 | 0 | 0 | 0 |
| GWENHVE | DYNC1H1; DYN | Q14204; A0/ | DYHC1_HUMA | Unmodified | 0 | 0 | 0 | 0 | 0 | 140970 | 0 | 0 | 604260 | 144810 | 0 | 428700 | 0 | 194710 | 0 | 0 | 0 | 0 |
| GWEWGKD | OTUD7B; OTUD | Q6GQQ9; Q | OTU7B_HUMAI | Unmodified | 0 | 0 | 0 | 0 | 0 | 0 | 0 | 0 | 0 | 0 | 0 | 0 | 0 | 0 | 0 | 0 | 0 | 21803 |
| GWGLLGPI | DHX30; DHX30 | Q7L2E3; H7I | DHX30_HUMAI | Unmodified | 40308 | 0 | 0 | 59134 | 0 | 0 | 0 | 0 | 232920 | 0 | 0 | 108260 | 0 | 0 | 0 | 24449 | 0 | 0 |
| GWIPGNEE | MYOF | Q9NZM1 | MYOF_HUMAN | Unmodified | 0 | 298000 | 0 | 0 | 9871400 | 0 | 0 | 0 | 508970 | 0 | 0 | 330800 | 0 | 0 | 0 | 0 | 199960 | 886360 |
| GWIPGNEE | MYOF | Q9NZM1 | MYOF_HUMAN | Unmodified | 0 | 0 | 0 | 0 | 1881200 | 63427 | 0 | 0 | 0 | 0 | 0 | 0 | 0 | 0 | 0 | 0 | 0 | 432390 |
| GWKEVHEI | RO60; RO60; R | P10155; H0 | RO60_HUMAN | Unmodified | 2391300 | 3522200 | 0 | 968240 | 0 | 0 | 0 | 0 | 0 | 0 | 182320 | 0 | 678420 | 100120 | 603650 | 456820 | 0 | 0 |
| GWLHPND | SHC1; SHC1; S | P29353; Q5 | SHC1_HUMAN | Unmodified | 0 | 0 | 0 | 0 | 765680 | 0 | 99048 | 113460 | 410360 | 14084 | 274040 | 884810 | 0 | 0 | 0 | 0 | 411940 | 0 |
| GWLRDPS/ | VCL | P18206 | VINC_HUMAN | Unmodified | 382190 | 282700 | 0 | 0 | 224620 | 0 | 0 | 0 | 0 | 0 | 0 | 7512800 | 0 | 0 | 1141100 | 1203100 | 524400 | 0 |
| GWNYILEK | NAMPT; NAMPT | P43490; A0A | NAMPT_HUMAI | Unmodified | 0 | 0 | 379400 | 0 | 0 | 404600 | 0 | 0 | 0 | 0 | 281530 | 6975400 | 0 | 0 | 336680 | 506050 | 0 | 0 |
| GWNYILEK | NAMPT; NAMPT | P43490; A0A | NAMPT_HUMAI | Unmodified | 0 | 0 | 0 | 0 | 30802000 | 1653400 | 0 | 0 | 0 | 446490 | 0 | 0 | 1576900 | 1029900 | 0 | 0 | 937860 | 0 |
| GWPEVWA | MIA3 | Q5JRA6 | TGO1_HUMAN | Unmodified | 0 | 0 | 32746 | 0 | 0 | 0 | 0 | 0 | 4520600 | 389940 | 841940 | 0 | 0 | 0 | 0 | 0 | 0 | 0 |
| GWPLYLST | IDH1 | O75874 | IDHC_HUMAN | Unmodified | 1289100 | 356390 | 1576600 | 1410200 | 0 | 0 | 1613900 | 164890 | 2258100 | 1324400 | 0 | 7155900 | 9359800 | 0 | 2887900 | 1764400 | 0 | 0 |
| GWPLYLST | IDH1 | O75874 | IDHC_HUMAN | Unmodified | 0 | 0 | 0 | 0 | 776220 | 0 | 456110 | 31141 | 0 | 531840 | 0 | 0 | 3024500 | 2675400 | 0 | 0 | 523440 | 139680 |
| GWQDVTA | TPD52; N/A | P55327; H0 | TPD52_HUMAN | Unmodified | 0 | 0 | 0 | 0 | 653700 | 0 | 202110 | 0 | 963570 | 738190 | 0 | 0 | 478590 | 0 | 0 | 0 | 349220 | 385780 |
| GWQDVTA | TPD52; N/A | P55327; H0 | TPD52_HUMAN | Unmodified | 0 | 0 | 0 | 0 | 3316200 | 0 | 1081800 | 596760 | 2543600 | 0 | 0 | 0 | 2287500 | 2670900 | 0 | 129940 | 1577800 | 1662300 |
| GWQDVTA | TPD52; N/A | P55327; H0 | TPD52_HUMAN | Unmodified | 0 | 0 | 0 | 0 | 297810 | 59537 | 1035300 | 147190 | 0 | 898540 | 0 | 0 | 440580 | 650740 | 0 | 0 | 898540 | 0 |
| GWRPSSRI | CHGA; CHGA | P10645; G5E | CMGA_HUMAN | Unmodified | 0 | 0 | 0 | 0 | 0 | 0 | 0 | 0 | 6643900 | 961630 | 0 | 0 | 0 | 0 | 0 | 0 | 0 | 0 |
| GWRVPGP | TMEM132A; TM | Q24JP5; H0 | T132A_HUMAN | Unmodified | 116480 | 0 | 0 | 0 | 0 | 0 | 0 | 0 | 1799300 | 0 | 0 | 0 | 0 | 0 | 0 | 0 | 0 | 0 |
| GWSFLLGK | LNPEP | Q9UIQ6 | LCAP_HUMAN | Unmodified | 0 | 0 | 386800 | 0 | 0 | 0 | 0 | 0 | 301580 | 0 | 0 | 1270200 | 0 | 0 | 0 | 218840 | 0 | 0 |
| GWTQWYD | ESYT2; ESYT2; E | A0FGR8; A0/ | ESYT2_HUMAN | Oxidation (M) | 42282 | 0 | 0 | 0 | 951010 | 0 | 236500 | 0 | 0 | 0 | 0 | 0 | 0 | 0 | 0 | 0 | 128750 | 264090 |
| GWVLGPN | PSMD8; PSMD8 | P48556; R4 | PSMD8_HUMA | Unmodified | 0 | 0 | 0 | 0 | 0 | 0 | 0 | 0 | 0 | 0 | 0 | 2766700 | 0 | 0 | 0 | 0 | 0 | 0 |
| GWWPCYA | MYOF | Q9NZM1 | MYOF_HUMAN | Unmodified | 0 | 0 | 0 | 0 | 4130900 | 567070 | 0 | 0 | 0 | 0 | 0 | 0 | 0 | 0 | 0 | 0 | 0 | 535430 |
| GWWYAHF | MYO6; MYO6; N | Q9UM54; A0 | MYO6_HUMAN | Unmodified | 0 | 0 | 397550 | 1650900 | 0 | 0 | 0 | 0 | 0 | 150790 | 81732 | 0 | 1681300 | 0 | 1798200 | 787740 | 0 | 0 |
| GYADIVQLI | OSTF1 | Q92882 | OSTF1_HUMAN | Unmodified | 373160 | 0 | 255390 | 557680 | 2324700 | 231130 | 412510 | 0 | 1746300 | 649270 | 806490 | 1604900 | 3014700 | 2281100 | 1465200 | 594430 | 717700 | 960890 |
| GYAEVVVA | CPSF7; CPSF7; | Q8N684; C9 | CPSF7_HUMAN | Unmodified | 3149100 | 0 | 7886600 | 3891400 | 231050 | 0 | 0 | 83519 | 0 | 0 | 174570 | 2201100 | 1024200 | 487060 | 0 | 0 | 0 | 0 |
| GYAFIEFAS | NCL; NCL; NCL | P19338; C9J | NUCL_HUMAN | Unmodified | 0 | 0 | 0 | 0 | 643390 | 0 | 0 | 0 | 32219 | 20498 | 805340 | 3685900 | 1168500 | 0 | 0 | 0 | 0 | 0 |
| GYAFITFCG | HNRNPR; HNRI | O43390; B4I | HNRPR_HUMA | Unmodified | 0 | 0 | 0 | 660640 | 0 | 0 | 0 | 0 | 0 | 0 | 0 | 1805800 | 19508 | 0 | 0 | 0 | 0 | 0 |
| GYAFNHSA | ACTR2; ACTR2 | P61160; F5H | ARP2_HUMAN | Unmodified | 1596400 | 1627200 | 1290900 | 593840 | 0 | 0 | 0 | 0 | 0 | 143730 | 0 | 20298000 | 0 | 0 | 2724400 | 3632700 | 0 | 0 |
| GYAFVHFE | PABPC4; PABP | Q13310; B1/ | PABP4_HUMAN | Unmodified | 1664600 | 1138400 | 1471900 | 0 | 0 | 0 | 0 | 0 | 0 | 0 | 0 | 0 | 0 | 0 | 623730 | 500850 | 0 | 0 |
| GYAFVHFE | PABPC4; PABP | Q13310; B1/ | PABP4_HUMAN | Unmodified | 5771300 | 14190000 | 0 | 5468500 | 337830 | 0 | 230900 | 169020 | 101000 | 0 | 0 | 0 | 1589600 | 613880 | 0 | 0 | 0 | 0 |
| GYAFVTFC | SYNCRIP; SYN | O60506; B7 | HNRPQ_HUMA | Unmodified | 115200 | 0 | 0 | 0 | 0 | 0 | 0 | 0 | 856870 | 0 | 0 | 5298000 | 0 | 0 | 0 | 0 | 0 | 0 |
| GYAVLGGE | FASN; FASN | P49327; A0A | FAS_HUMAN F | Unmodified | 234610 | 104700 | 1578200 | 667970 | 0 | 152660 | 0 | 0 | 133320 | 45872 | 0 | 36330000 | 324040 | 0 | 3075900 | 998320 | 92322 | 0 |
| GYAVNVFD | NSDHL; NSDHL | Q15738; C9J | NSDHL_HUMA | Unmodified | 0 | 0 | 0 | 0 | 106890 | 0 | 0 | 0 | 8606100 | 0 | 0 | 2412400 | 42677 | 45120 | 0 | 0 | 0 | 0 |
| GYAWGLN | MAN1A1 | P33908 | MA1A1_HUMAI | Unmodified | 0 | 0 | 0 | 0 | 0 | 0 | 0 | 0 | 1386800 | 0 | 0 | 898710 | 0 | 0 | 0 | 0 | 0 | 0 |
| GYCAYGTR | MKRN2; MKRN | Q9H000; C9 | MKRN2_HUMA | Unmodified | 390770 | 198010 | 56414 | 107310 | 0 | 0 | 0 | 0 | 244570 | 0 | 0 | 0 | 835350 | 0 | 48570 | 96630 | 52650 | 53773 |
| GYCIYGDR | MKRN1; MKRN | Q9UHC7; C9 | MKRN1_HUMA | Unmodified | 4055500 | 2526100 | 592420 | 1956200 | 0 | 0 | 0 | 0 | 0 | 0 | 0 | 0 | 0 | 0 | 1302000 | 592430 | 0 | 0 |
| GYDAPLCN | ACSL4; ACSL4 | O60488; H0 | ACSL4_HUMAN | Unmodified | 0 | 0 | 0 | 0 | 546780 | 122050 | 513640 | 0 | 37502 | 0 | 741440 | 1925400 | 0 | 0 | 0 | 0 | 694190 | 575250 |
| GYDAPLCN | ACSL4; ACSL4 | O60488; H0 | ACSL4_HUMAN | Unmodified | 0 | 0 | 0 | 0 | 1333600 | 0 | 37047 | 33785 | 0 | 0 | 635330 | 4004400 | 22248 | 0 | 0 | 0 | 1213200 | 567220 |
| GYDELVPH | AGL | P35573 | GDE_HUMAN ( | Unmodified | 0 | 0 | 0 | 0 | 0 | 0 | 0 | 0 | 0 | 0 | 0 | 3384900 | 0 | 0 | 0 | 0 | 0 | 0 |

| | | | | | | | | | | | | | | | | | | | | | | |
|---|---|---|---|---|---|---|---|---|---|---|---|---|---|---|---|---|---|---|---|---|---|---|
| GYDFESETI | GFPT1; GFPT1; | Q06210; A0A | GFPT1_HUMAN | Unmodified | 0 | 0 | 657880 | 0 | 0 | 0 | 0 | 0 | 870430 | 286260 | 268810 | 2090200 | 0 | 0 | 688860 | 387450 | 0 | 0 |
| GYDQDAYD | HLA-B; HLA-B; | D5H3J5; A0A | D5H3J5_HUMA | Unmodified | 0 | 0 | 0 | 0 | 0 | 0 | 0 | 0 | 567550 | 0 | 0 | 0 | 0 | 0 | 0 | 0 | 0 | 0 |
| GYDQIIPK | POGLUT1 | Q8NBL1 | PGLT1_HUMAN | Unmodified | 0 | 0 | 1811600 | 1431900 | 0 | 1605500 | 82344 | 0 | 1672700 | 0 | 0 | 0 | 0 | 368010 | 2280300 | 0 | 0 | 357260 |
| GYDQSAYD | HLA-C; HLA-C; | P10321; Q5 | HLAC_HUMAN | Unmodified | 0 | 0 | 0 | 0 | 0 | 0 | 0 | 0 | 1936600 | 0 | 0 | 374400 | 0 | 0 | 0 | 0 | 0 | 0 |
| GYDQYAYD | HLA-C; HLA-C; | Q53YP1; A0A | Q53YP1_HUMA | Unmodified | 0 | 0 | 0 | 0 | 0 | 0 | 0 | 0 | 0 | 0 | 316800 | 3082400 | 0 | 0 | 0 | 0 | 0 | 0 |
| GYDSAGV | GFPT1; GFPT1 | Q06210; A0A | GFPT1_HUMAN | Unmodified | 0 | 0 | 781180 | 0 | 0 | 0 | 0 | 0 | 0 | 0 | 0 | 615150 | 0 | 0 | 218410 | 154890 | 0 | 0 |
| GYDSAGV | GFPT1; GFPT1 | Q06210; A0A | GFPT1_HUMAN | Unmodified | 243160 | 0 | 0 | 0 | 0 | 0 | 0 | 0 | 0 | 0 | 0 | 0 | 905780 | 439760 | 0 | 0 | 0 | 0 |
| GYDVIAQA | EIF4A1; EIF4A2; | P60842; Q14 | IF4A1_HUMAN | Unmodified | 5824000 | 0 | 3523600 | 2920200 | 4339400 | 1213800 | 2268000 | 2857300 | 1963800 | 2345000 | 3349900 | 13684000 | 3845800 | 3101500 | 431080 | 5995600 | 2173100 | 2224700 |
| GYDYGPHI | FASN; FASN | P49327; A0A | FAS_HUMAN F | Unmodified | 0 | 0 | 0 | 0 | 0 | 0 | 0 | 0 | 0 | 0 | 0 | 12315000 | 0 | 0 | 3713300 | 1743600 | 0 | 0 |
| GYEEVHVP | SNRNP200; SN | O75643; A0A | U520_HUMAN | Unmodified | 0 | 0 | 0 | 0 | 0 | 0 | 0 | 0 | 0 | 130870 | 0 | 3802900 | 0 | 0 | 0 | 0 | 0 | 0 |
| GYEEWLLN | ACTN2; ACTN4; | P35609; O43 | ACTN2_HUMAI | Unmodified | 330100 | 0 | 0 | 0 | 0 | 0 | 0 | 0 | 1330800 | 359160 | 0 | 5506300 | 0 | 0 | 1622700 | 832700 | 0 | 0 |
| GYEIDEDIV | TOR1A | O14656 | TOR1A_HUMAI | Unmodified | 0 | 211440 | 0 | 0 | 0 | 169250 | 0 | 0 | 5381500 | 152140 | 0 | 0 | 0 | 0 | 0 | 0 | 265490 | 382660 |
| GYELAHQR | GGT1; GGT2; G | P19440; P36 | GGT1_HUMAN | Unmodified | 394940 | 363160 | 957640 | 0 | 0 | 0 | 0 | 0 | 1166000 | 108010 | 0 | 1375100 | 0 | 0 | 1233600 | 910360 | 0 | 0 |
| GYELLFQPI | CTNND1; CTNN | O60716; A0A | CTND1_HUMAI | Unmodified | 0 | 141150 | 2502600 | 260650 | 0 | 0 | 0 | 0 | 0 | 0 | 2899400 | 17481000 | 100620 | 0 | 19056000 | 7340000 | 0 | 0 |
| GYEVIYLTE | HSP90B1; HSP9 | P14625; A0A | ENPL_HUMAN | Unmodified | 0 | 0 | 0 | 0 | 0 | 0 | 0 | 0 | 481200 | 768270 | 0 | 0 | 0 | 0 | 0 | 0 | 0 | 0 |
| GYEVIYLTE | HSP90B1; HSP9 | P14625; A0A | ENPL_HUMAN | Unmodified | 0 | 0 | 0 | 0 | 0 | 0 | 0 | 0 | 2877900 | 574000 | 0 | 0 | 0 | 0 | 0 | 0 | 0 | 0 |
| GYEVIYLTE | HSP90B1; HSP9 | P14625; A0A | ENPL_HUMAN | Unmodified | 0 | 0 | 0 | 0 | 0 | 0 | 0 | 0 | 0 | 2643400 | 0 | 0 | 0 | 0 | 0 | 0 | 0 | 0 |
| GYFAVTEK | FLII; FLII; FLII | Q13045; J3Q | FLII_HUMAN Pr | Unmodified | 0 | 0 | 0 | 0 | 700060 | 0 | 333130 | 383440 | 0 | 237900 | 0 | 425110 | 278110 | 181920 | 0 | 0 | 298660 | 141590 |
| GYFDEEMN | PLEC; EPPK1; P | Q15149; P58 | PLEC_HUMAN | Unmodified | 0 | 0 | 0 | 0 | 298240 | 0 | 0 | 0 | 824930 | 207210 | 0 | 0 | 594080 | 535140 | 0 | 0 | 736030 | 873590 |
| GYFEYIEEN | HNRNPU; HNR | Q00839; A0A | HNRPU_HUMA | Unmodified | 621340 | 204330 | 0 | 0 | 0 | 0 | 0 | 0 | 501710 | 0 | 0 | 1203500 | 0 | 0 | 0 | 0 | 0 | 0 |
| GYFSEEMN | PLEC; PLEC | Q15149; H0Y | PLEC_HUMAN | Unmodified | 0 | 0 | 0 | 0 | 262570 | 0 | 0 | 0 | 158550 | 25710 | 0 | 0 | 0 | 0 | 0 | 0 | 322460 | 0 |
| GYFVQNTY | GUSB | P08236 | BGLR_HUMAN | Unmodified | 0 | 0 | 0 | 0 | 0 | 0 | 0 | 0 | 0 | 385400 | 0 | 22628 | 0 | 0 | 0 | 0 | 0 | 0 |
| GYFVQPTV | ALDH1A1 | P00352 | AL1A1_HUMAN | Unmodified | 0 | 0 | 0 | 0 | 975690 | 164480 | 0 | 0 | 1274400 | 68299 | 1020500 | 28919000 | 0 | 0 | 0 | 0 | 159500 | 134410 |
| GYFVQPTV | ALDH1A1 | P00352 | AL1A1_HUMAN | Oxidation (M) | 89422 | 85577 | 0 | 0 | 0 | 0 | 0 | 0 | 0 | 149540 | 8092000 | 2.85E+08 | 0 | 0 | 457100 | 0 | 0 | 0 |
| GYFVQTPE | HACL1; HACL1 | Q9UJ83; B3K | HACL1_HUMAI | Unmodified | 0 | 0 | 0 | 0 | 0 | 0 | 0 | 0 | 0 | 0 | 0 | 0 | 0 | 0 | 0 | 0 | 0 | 0 |
| GYGFGLIKI | VDAC1; VDAC1 | P21796; C9J | VDAC1_HUMAI | Unmodified | 0 | 0 | 0 | 0 | 730900 | 0 | 2232500 | 59324 | 29075000 | 59238000 | 510440 | 0 | 0 | 2287900 | 0 | 0 | 0 | 0 |
| GYGFVHFE | PABPC1; PABP( | P11940; A0A | PABP1_HUMAN | Unmodified | 5548200 | 6174700 | 563410 | 467660 | 0 | 0 | 0 | 0 | 0 | 0 | 0 | 939990 | 0 | 207770 | 2401100 | 2411200 | 0 | 0 |
| GYGFVHFE | PABPC1; PABP( | P11940; A0A | PABP1_HUMAN | Unmodified | 59759000 | 41564000 | 30131000 | 48513000 | 5864400 | 927250 | 2057900 | 190380 | 1253700 | 2958000 | 1529300 | 2260300 | 20982000 | 10632000 | 11288000 | 10506000 | 1417200 | 1784700 |
| GYGIQIEQI | ADAM17 | P78536 | ADA17_HUMAN | Unmodified | 0 | 838080 | 127560 | 199860 | 480530 | 0 | 0 | 985950 | 1149300 | 941230 | 723740 | 0 | 0 | 749700 | 0 | 147490 | 0 | 0 |
| GYGIQIEQI | ADAM17 | P78536 | ADA17_HUMAN | Unmodified | 0 | 0 | 0 | 0 | 2386200 | 0 | 0 | 0 | 0 | 0 | 0 | 0 | 0 | 0 | 0 | 0 | 622600 | 1740300 |
| GYGPDGH | DBNL; DBNL; D | Q9UJU6; H7 | DBNL_HUMAN | Oxidation (M) | 108150 | 203890 | 28370 | 200970 | 204650 | 0 | 0 | 0 | 203350 | 0 | 0 | 0 | 0 | 0 | 276680 | 59208 | 0 | 0 |
| GYGYGQG | CSRP2; CSRP2; | Q16527; F8V | CSRP2_HUMAI | Unmodified | 474230 | 0 | 0 | 0 | 193080 | 0 | 0 | 0 | 0 | 0 | 226400 | 3060400 | 116090 | 0 | 0 | 0 | 0 | 125360 |
| GYGYGQG | CSRP1; CSRP1; | P21291; E9P | CSRP1_HUMAI | Unmodified | 730070 | 0 | 949410 | 0 | 643570 | 0 | 0 | 0 | 0 | 0 | 406610 | 660290 | 0 | 0 | 1544200 | 868390 | 0 | 0 |
| GYHGPLDI | ALG9; N/A; ALG | Q9H6U8; A0 | ALG9_HUMAN | Unmodified | 0 | 0 | 0 | 0 | 861860 | 0 | 266850 | 0 | 1919400 | 209210 | 0 | 190100 | 0 | 0 | 0 | 0 | 0 | 266490 |
| GYHGSTVV | POMT1; POMT1 | Q9Y6A1; A0A | POMT1_HUMAI | Unmodified | 0 | 0 | 0 | 0 | 0 | 0 | 0 | 0 | 1034500 | 59450 | 0 | 0 | 0 | 0 | 0 | 0 | 0 | 0 |
| GYHLNEEG | FBLN1; FBLN1 | P23142; B1A | FBLN1_HUMAN | Unmodified | 0 | 0 | 0 | 0 | 0 | 0 | 0 | 0 | 1315200 | 0 | 0 | 0 | 0 | 0 | 0 | 0 | 0 | 0 |
| GYHQYAYD | HLA-A; HLA-A; I | Q53Z42; A0A | Q53Z42_HUMA | Unmodified | 0 | 0 | 0 | 0 | 0 | 0 | 78730 | 0 | 0 | 16665 | 284680 | 6545200 | 0 | 0 | 501230 | 318130 | 167970 | 167080 |
| GYHQYAYD | HLA-A; HLA-A; I | Q53Z42; A0A | Q53Z42_HUMA | Unmodified | 0 | 166120 | 0 | 0 | 0 | 0 | 0 | 91566 | 0 | 0 | 0 | 1422100 | 0 | 0 | 0 | 0 | 0 | 0 |
| GYHYATCL | GFPT1; GFPT1 | Q06210; A0A | GFPT1_HUMAN | Unmodified | 42544 | 0 | 0 | 72234 | 0 | 0 | 0 | 0 | 0 | 152380 | 0 | 771950 | 0 | 0 | 312450 | 0 | 0 | 0 |
| GYIFLEYAS | EIF3B; EIF3B | P55884; C9J | EIF3B_HUMAN | Unmodified | 0 | 1886000 | 1945200 | 3149900 | 2668400 | 0 | 0 | 0 | 1988100 | 1560000 | 0 | 12555000 | 2340300 | 0 | 0 | 2568100 | 1090700 | 0 |
| GYIGEGDM | ACOT8; ACOT8; | O14734; E9F | ACOT8_HUMAI | Unmodified | 0 | 0 | 0 | 0 | 0 | 0 | 0 | 0 | 920230 | 64033 | 0 | 0 | 0 | 0 | 0 | 0 | 0 | 0 |
| GYIGVVNR | DNM2; DNM1; I | P50570; Q05 | DYN2_HUMAN | Unmodified | 826100 | 73337 | 481610 | 482100 | 0 | 0 | 0 | 0 | 0 | 0 | 109270 | 1045600 | 552110 | 0 | 545840 | 558470 | 0 | 0 |
| GYIGVVNR | DNM2; DNM1; I | P50570; Q05 | DYN2_HUMAN | Unmodified | 0 | 0 | 399260 | 0 | 1312600 | 179670 | 67891 | 32295 | 0 | 308470 | 0 | 0 | 1738200 | 346670 | 0 | 0 | 263140 | 427170 |
| GYIPIYTPF | SARS1; SARS1 | P49591; Q5T | SYSC_HUMAN | Oxidation (M) | 0 | 0 | 0 | 0 | 0 | 0 | 0 | 0 | 0 | 0 | 0 | 961370 | 0 | 0 | 230950 | 0 | 0 | 0 |
| GYIPIYTPF | SARS1; SARS1 | P49591; Q5T | SYSC_HUMAN | Oxidation (M) | 0 | 328160 | 229070 | 0 | 0 | 0 | 0 | 0 | 0 | 0 | 0 | 1065400 | 0 | 0 | 0 | 0 | 0 | 0 |
| GYIQMTPLI | ABCC1; ABCC1 | P33527; A0A | MRP1_HUMAN | Oxidation (M) | 0 | 0 | 0 | 0 | 0 | 0 | 148960 | 0 | 0 | 0 | 0 | 1130800 | 0 | 0 | 0 | 0 | 0 | 0 |
| GYISPYFIN | HSPD1; HSPD1 | P10809; A0A | CH60_HUMAN | Unmodified | 0 | 0 | 0 | 0 | 0 | 0 | 0 | 302800 | 10098000 | 1445800 | 269800 | 707090 | 0 | 0 | 0 | 0 | 0 | 0 |
| GYIWNYGA | PPA1; PPA1 | Q15181; Q5 | IPYR_HUMAN I | Unmodified | 1010600 | 0 | 902240 | 1401200 | 3167200 | 0 | 0 | 0 | 928820 | 0 | 0 | 36024000 | 2976300 | 579720 | 2324700 | 1766700 | 0 | 3379500 |
| GYIWNYGA | PPA1; PPA1 | Q15181; Q5 | IPYR_HUMAN I | Unmodified | 0 | 0 | 0 | 0 | 1440300 | 0 | 0 | 0 | 0 | 0 | 0 | 0 | 2760300 | 3440500 | 0 | 0 | 0 | 3084600 |
| GYIWNYGT | PPA2; PPA2; PP | Q9H2U2; D6 | IPYR2_HUMAN | Unmodified | 0 | 0 | 0 | 0 | 0 | 0 | 0 | 194920 | 4472500 | 1039100 | 0 | 508070 | 0 | 0 | 0 | 0 | 0 | 0 |
| GYKPPDEG | COPS5; COPS5 | Q92905; E5F | CSN5_HUMAN | Unmodified | 0 | 0 | 159890 | 0 | 0 | 0 | 527430 | 0 | 0 | 0 | 103330 | 1130600 | 0 | 0 | 141090 | 129060 | 408050 | 0 |
| GYLDDPTV | G6PD; G6PD | P11413; E7E | G6PD_HUMAN | Unmodified | 0 | 0 | 0 | 0 | 0 | 0 | 0 | 0 | 0 | 0 | 0 | 5009300 | 0 | 0 | 0 | 0 | 0 | 0 |
| GYLDKLEP | ATP5F1A | P25705 | ATPA_HUMAN | Unmodified | 0 | 134770 | 0 | 0 | 1501100 | 295360 | 1435200 | 0 | 3019100 | 0 | 0 | 1943000 | 962400 | 686400 | 0 | 0 | 454700 | 873530 |
| GYLETPQP | TOR3A; TOR3A | Q9H497; Q5 | TOR3A_HUMAI | Unmodified | 0 | 0 | 0 | 0 | 0 | 0 | 0 | 0 | 419810 | 0 | 0 | 0 | 0 | 0 | 0 | 0 | 0 | 0 |
| GYLGPEQL | MDH2 | P40926 | MDHM_HUMAI | Unmodified | 0 | 0 | 0 | 0 | 231800 | 0 | 285680 | 208940 | 11357000 | 2451900 | 308380 | 0 | 0 | 758540 | 0 | 0 | 0 | 0 |
| GYLLPGAP | DIPK1B | Q5VUD6 | DIK1B_HUMAN | Unmodified | 0 | 0 | 0 | 0 | 0 | 0 | 202780 | 86223 | 7735500 | 0 | 0 | 0 | 0 | 0 | 0 | 0 | 0 | 0 |
| GYLPLPR | FLVCR1 | Q9Y5Y0 | FLVC1_HUMAN | Unmodified | 185110 | 132390 | 478070 | 346960 | 954410 | 0 | 0 | 0 | 909940 | 400660 | 200930 | 1561000 | 823560 | 345560 | 320080 | 346490 | 405030 | 365720 |
| GYLPLPRG | FLVCR1 | Q9Y5Y0 | FLVC1_HUMAN | Unmodified | 0 | 0 | 0 | 0 | 2508600 | 0 | 0 | 0 | 0 | 0 | 0 | 0 | 0 | 769300 | 0 | 0 | 778220 | 0 |
| GYLTGMAC | ARSA; ARSA | P15289; A0A | ARSA_HUMAN | Unmodified | 0 | 0 | 0 | 53164 | 0 | 0 | 0 | 0 | 1265300 | 0 | 0 | 0 | 0 | 0 | 0 | 0 | 337490 | 0 |
| GYLTKEDLI | S100A10 | P60903 | S10AA_HUMAN | Unmodified | 0 | 0 | 0 | 0 | 12753000 | 1754400 | 0 | 0 | 0 | 0 | 0 | 0 | 0 | 0 | 0 | 0 | 2248800 | 3217900 |
| GYLTKEDLI | S100A10 | P60903 | S10AA_HUMAN | Oxidation (M) | 0 | 0 | 0 | 0 | 6512500 | 1235600 | 0 | 0 | 0 | 251440 | 0 | 0 | 0 | 0 | 0 | 0 | 169360 | 1052700 |
| GYLTLSDS | CANT1; CANT1 | Q8WVQ1; K7 | CANT1_HUMAI | Unmodified | 0 | 0 | 0 | 0 | 0 | 0 | 0 | 44505 | 3593900 | 0 | 0 | 0 | 0 | 0 | 0 | 0 | 0 | 0 |
| GYNQFAYD | HLA-C; HLA-C; | A0A0G2JIF0; | A0A0G2JIF0_HI | Unmodified | 0 | 0 | 0 | 0 | 0 | 0 | 0 | 0 | 0 | 0 | 224060 | 3621100 | 0 | 0 | 0 | 0 | 0 | 0 |
| GYNVNDEI | BASP1; BASP1 | P80723; U3K | BASP1_HUMAN | Unmodified | 0 | 0 | 0 | 126630 | 0 | 0 | 316510 | 553170 | 471110 | 101220 | 0 | 201630 | 0 | 0 | 194770 | 146150 | 642550 | 2484600 |
| GYNVNDEI | BASP1; BASP1 | P80723; U3K | BASP1_HUMAN | Unmodified | 0 | 0 | 0 | 0 | 1370600 | 249200 | 53030 | 811380 | 0 | 104400 | 0 | 0 | 0 | 30841 | 0 | 0 | 739790 | 597000 |
| GYPHWPAI | HDGF | P51858 | HDGF_HUMAN | Unmodified | 0 | 0 | 0 | 0 | 1499600 | 0 | 0 | 43584 | 520730 | 394960 | 642560 | 1716200 | 1393900 | 194460 | 0 | 0 | 0 | 0 |
| GYPPEDIF | NANS | Q9NR45 | SIAS_HUMAN S | Unmodified | 0 | 0 | 18204 | 0 | 170430 | 48027 | 20088 | 0 | 0 | 0 | 46517 | 133480 | 156000 | 74232 | 0 | 17746 | 32609 | 60384 |
| GYPPEDIF | NANS | Q9NR45 | SIAS_HUMAN S | Unmodified | 0 | 0 | 1787000 | 597580 | 3330100 | 0 | 622490 | 92385 | 488930 | 0 | 0 | 3709100 | 2222700 | 1442100 | 319330 | 523420 | 403120 | 775150 |
| GYPTFHYY | PDIA5; PDIA5 | Q14554; H7 | PDIA5_HUMAN | Unmodified | 0 | 0 | 0 | 0 | 0 | 0 | 0 | 0 | 2794100 | 942340 | 0 | 0 | 0 | 0 | 0 | 0 | 0 | 0 |
| GYPTLLLFR | TXNDC5 | Q8NBS9 | TXND5_HUMAI | Unmodified | 0 | 0 | 161680 | 166480 | 0 | 0 | 0 | 0 | 59253000 | 1539300 | 2360600 | 1201400 | 0 | 0 | 268340 | 332430 | 0 | 0 |
| GYPTLLLFR | TXNDC5 | Q8NBS9 | TXND5_HUMAI | Unmodified | 0 | 0 | 0 | 0 | 0 | 0 | 0 | 0 | 819270 | 1251200 | 0 | 0 | 0 | 0 | 0 | 0 | 0 | 0 |
| GYPTLLWF | TXNDC5 | Q8NBS9 | TXND5_HUMAI | Unmodified | 0 | 0 | 0 | 0 | 0 | 0 | 0 | 0 | 25165000 | 0 | 0 | 0 | 0 | 0 | 0 | 0 | 31243 | 0 |
| GYPTLLWF | TXNDC5 | Q8NBS9 | TXND5_HUMAI | Unmodified | 0 | 0 | 0 | 0 | 186800 | 0 | 0 | 0 | 22270000 | 5173500 | 1086100 | 0 | 0 | 0 | 0 | 0 | 0 | 0 |
| GYQQDAYI | HLA-A; HLA-A; I | Q9GJ45; A0A | Q9GJ45_HUMA | Unmodified | 0 | 0 | 0 | 0 | 361580 | 190560 | 0 | 0 | 0 | 0 | 0 | 0 | 0 | 0 | 0 | 0 | 0 | 0 |
| GYQVTYVR | PTPRF; PTPRF; I | P10586; H0Y | PTPRF_HUMAN | Unmodified | 417660 | 216380 | 0 | 0 | 0 | 0 | 0 | 0 | 0 | 0 | 0 | 0 | 0 | 0 | 143700 | 124630 | 0 | 0 |
| GYRMVVNI | HINT1 | P49773 | HINT1_HUMAN | Unmodified | 0 | 0 | 0 | 0 | 3154200 | 0 | 1102700 | 0 | 0 | 2068500 | 0 | 0 | 406360 | 393130 | 0 | 0 | 775620 | 1498900 |
| GYRMVVNI | HINT1 | P49773 | HINT1_HUMAN | Oxidation (M) | 0 | 0 | 0 | 0 | 2276300 | 0 | 0 | 0 | 0 | 1933300 | 0 | 0 | 3710100 | 1582700 | 0 | 0 | 1395500 | 0 |
| GYRQDAYD | HLA-A; HLA-A; I | P04439; A0A | HLAA_HUMAN | Unmodified | 0 | 0 | 0 | 0 | 0 | 0 | 0 | 0 | 8559300 | 0 | 0 | 0 | 0 | 0 | 0 | 0 | 0 | 0 |
| GYRQDAYD | HLA-A; HLA-A; I | P04439; A0A | HLAA_HUMAN | Unmodified | 0 | 0 | 0 | 0 | 0 | 0 | 0 | 0 | 5430700 | 1102700 | 0 | 2549000 | 0 | 0 | 0 | 0 | 1001800 | 0 |
| GYRQHWV | FERMT3 | Q86UX7 | URP2_HUMAN | Unmodified | 0 | 0 | 0 | 0 | 0 | 0 | 1726800 | 0 | 0 | 0 | 0 | 0 | 0 | 731670 | 0 | 0 | 1012900 | 0 |
| GYSDEIYV | ARHGAP5; ARH | Q13017; A0A | RHG05_HUMAI | Unmodified | 0 | 0 | 0 | 0 | 1874500 | 364200 | 0 | 256600 | 0 | 0 | 0 | 0 | 0 | 0 | 0 | 0 | 58230 | 0 |
| GYSFSLTTF | PSMA2; N/A; PS | P25787; A0A | PSA2_HUMAN | Unmodified | 0 | 0 | 0 | 0 | 0 | 0 | 0 | 0 | 1632900 | 0 | 1291100 | 8111700 | 0 | 0 | 0 | 0 | 0 | 0 |
| GYSFTTTAE | ACTG1; ACTB; A | P63261; P60 | ACTG_HUMAN | Unmodified | 35665000 | 37340000 | 53863000 | 8119300 | 1132800 | 525110 | 0 | 0 | 9326000 | 1660800 | 7760800 | 5.9E+08 | 456820 | 427090 | 1.19E+08 | 1.12E+08 | 302870 | 554860 |
| GYSFTTTAE | ACTG1; ACTB; A | P63261; P60 | ACTG_HUMAN | Unmodified | 42758000 | 95925000 | 51606000 | 29151000 | 2336500 | 693260 | 0 | 0 | 26754000 | 497690 | 12286000 | 2.87E+08 | 462060 | 74348 | 1.33E+08 | 77285000 | 496090 | 584760 |
| GYSFTTTAE | ACTG1; ACTB; A | P63261; P60 | ACTG_HUMAN | Unmodified | 3424500 | 15156000 | 6534700 | 13795000 | 30096000 | 10652000 | 3237700 | 204750 | 25486000 | 3078900 | 19636000 | 16509000 | 3825500 | 0 | 22302000 | 9183100 | 4141300 | 10031000 |
| GYSLNLIPF | GPC2; GPC2 | Q8N158; A0 | GPC2_HUMAN | Unmodified | 0 | 0 | 0 | 0 | 0 | 0 | 0 | 0 | 1924500 | 0 | 0 | 0 | 0 | 0 | 0 | 0 | 0 | 0 |
| GYSLVSGG | SHMT2; SHMT2 | P34897; H0Y | GLYM_HUMAN | Unmodified | 0 | 0 | 0 | 0 | 0 | 0 | 0 | 0 | 19300000 | 1858600 | 223770 | 1073700 | 0 | 0 | 0 | 0 | 0 | 0 |
| GYTIHWNG | CEMIP2; CEMIP | Q9UHN6; H3 | CEIP2_HUMAN | Unmodified | 594740 | 100010 | 423390 | 242910 | 752240 | 292500 | 0 | 0 | 2150800 | 600070 | 214930 | 438840 | 0 | 0 | 1078800 | 580900 | 0 | 0 |
| GYTLDPNG | MATN2; MATN2 | O00339; H0Y | MATN2_HUMAI | Unmodified | 0 | 0 | 0 | 0 | 0 | 0 | 0 | 0 | 408370 | 0 | 0 | 0 | 0 | 0 | 0 | 0 | 0 | 0 |
| GYTLVEYET | RBM8A; RBM8A | Q9Y5S9; A0A | RBM8A_HUMAI | Unmodified | 422390 | 385040 | 620970 | 0 | 133860 | 0 | 0 | 0 | 395730 | 0 | 0 | 2557700 | 0 | 0 | 423550 | 425640 | 0 | 0 |
| GYTPEGTVI | NOA1 | Q8NC60 | NOA1_HUMAN | Unmodified | 0 | 0 | 0 | 0 | 0 | 0 | 0 | 0 | 283540 | 0 | 0 | 0 | 0 | 0 | 0 | 0 | 0 | 0 |
| GYVSCALG | HMGCL | P35914 | HMGCL_HUMA | Unmodified | 0 | 0 | 0 | 0 | 0 | 0 | 0 | 0 | 1232400 | 0 | 0 | 0 | 0 | 0 | 0 | 0 | 0 | 0 |
| GYVSDIDC | GCLC; GCLC; G | P48506; A0A | GSH1_HUMAN | Unmodified | 0 | 0 | 0 | 0 | 0 | 0 | 0 | 0 | 413490 | 0 | 0 | 793330 | 0 | 0 | 0 | 0 | 0 | 0 |
| GYVVGQAL | ARRDC1 | Q8N5I2 | ARRD1_HUMAI | Unmodified | 1989400 | 545100 | 2822600 | 393330 | 0 | 0 | 129180 | 0 | 0 | 0 | 0 | 1831200 | 0 | 0 | 4060200 | 3302500 | 0 | 0 |
| GYVVRISG | RPS6; RPS6 | P62753; A2A | RS6_HUMAN 4 | Unmodified | 0 | 0 | 0 | 0 | 0 | 0 | 156510 | 101270 | 0 | 0 | 0 | 0 | 0 | 193650 | 0 | 0 | 0 | 0 |
| GYVVWQE | HEXA; HEXA; HI | P06865; H3E | HEXA_HUMAN | Unmodified | 0 | 0 | 0 | 0 | 0 | 0 | 0 | 0 | 2753600 | 189010 | 0 | 0 | 0 | 0 | 0 | 0 | 0 | 0 |
| GYYFEIPSI | CACNA2D2; CA | Q9NY47; C9 | CA2D2_HUMAI | Unmodified | 0 | 0 | 0 | 0 | 0 | 0 | 0 | 0 | 6091400 | 0 | 239350 | 768300 | 0 | 0 | 0 | 0 | 0 | 0 |
| GYYFGTSSI | LMAN2L | Q9H0V9 | LMA2L_HUMAN | Unmodified | 0 | 0 | 0 | 0 | 0 | 0 | 0 | 0 | 37004000 | 2492100 | 0 | 0 | 0 | 0 | 0 | 0 | 0 | 0 |
| GYYSPYSV | PLEC; PLEC | Q15149; H0Y | PLEC_HUMAN | Unmodified | 0 | 0 | 0 | 0 | 405140 | 0 | 47670 | 0 | 0 | 0 | 0 | 0 | 0 | 0 | 0 | 0 | 716020 | 902870 |
| GYYWPSIQ | ADGRG6; ADGI | Q86SQ4; H0 | AGRG6_HUMA | Unmodified | 0 | 0 | 0 | 271050 | 0 | 49007 | 0 | 0 | 0 | 0 | 47846 | 0 | 0 | 0 | 0 | 156390 | 0 | 0 |
| GYYYEIPSI | CACNA2D1 | P54289 | CA2D1_HUMAI | Unmodified | 0 | 0 | 0 | 0 | 0 | 0 | 0 | 0 | 4112400 | 0 | 0 | 0 | 0 | 0 | 0 | 0 | 0 | 0 |
| HAACPVLV | P4HA1 | P13674 | P4HA1_HUMAI | Unmodified | 0 | 0 | 0 | 0 | 0 | 0 | 0 | 0 | 761360 | 15254 | 49756 | 0 | 0 | 0 | 0 | 0 | 21704 | 0 |
| HAAENPGK | HNRNPU; HNR | Q00839; A0A | HNRPU_HUMA | Oxidation (M) | 0 | 0 | 0 | 0 | 359890 | 0 | 2917400 | 3061200 | 0 | 0 | 0 | 0 | 0 | 139200 | 0 | 0 | 0 | 0 |
| HAAENPGK | HNRNPU; HNR | Q00839; A0A | HNRPU_HUMA | Unmodified | 0 | 47161 | 0 | 0 | 0 | 0 | 14295000 | 4371000 | 0 | 485060 | 0 | 0 | 0 | 0 | 0 | 0 | 226020 | 901440 |
| HAAGVLAC | HYI; HYI; HYI; H | Q5T013; F6L | HYI_HUMAN Pu | Unmodified | 0 | 0 | 0 | 0 | 0 | 0 | 0 | 0 | 295270 | 0 | 0 | 0 | 0 | 0 | 0 | 0 | 152010 | 0 |
| HAALSLSK | ILKAP; ILKAP; IL | Q9H0C8; H7 | ILKAP_HUMAN | Unmodified | 0 | 0 | 0 | 0 | 0 | 0 | 104770 | 82053 | 0 | 0 | 0 | 434330 | 0 | 32372 | 32711 | 0 | 0 | 0 |
| HAASTVQII | NOP58 | Q9Y2X3 | NOP58_HUMAI | Unmodified | 0 | 0 | 0 | 0 | 0 | 0 | 379710 | 218140 | 0 | 34851 | 0 | 0 | 0 | 0 | 39380 | 0 | 0 | 0 |
| HAAVLVETI | CLASP2; CLASF | O75122; B3K | CLAP2_HUMAN | Unmodified | 0 | 0 | 0 | 0 | 228610 | 0 | 0 | 57321 | 128160 | 0 | 92787 | 736560 | 107470 | 0 | 0 | 0 | 0 | 0 |
| HAAYILLTG | FAM234A; FAM2 | Q9H0X4; H7 | F234A_HUMAN | Unmodified | 0 | 0 | 0 | 0 | 0 | 3854600 | 1918900 | 0 | 0 | 0 | 0 | 1286500 | 0 | 871800 | 0 | 0 | 839100 | 0 |

| | | | | | | | | | | | | | | | | | | | | | | |
|---|---|---|---|---|---|---|---|---|---|---|---|---|---|---|---|---|---|---|---|---|---|---|
| HAAYSVGV | IGSF8 | Q969P0 | IGSF8_HUMAN | Unmodified | 3295900 | 4809700 | 0 | 262520 | 0 | 0 | 0 | 0 | 1271500 | 0 | 0 | 1682000 | 0 | 0 | 4241700 | 1826200 | 0 | 0 |
| HAAYSVGV | IGSF8 | Q969P0 | IGSF8_HUMAN | Oxidation (M) | 2871400 | 2333900 | 1993400 | 1376200 | 0 | 0 | 0 | 0 | 0 | 0 | 0 | 0 | 0 | 0 | 1991700 | 1919000 | 0 | 136240 |
| HADDHTPK | TJP1; TJP1; TJP1 | Q07157; A0A | ZO1_HUMAN T | Unmodified | 0 | 0 | 0 | 0 | 1042600 | 0 | 0 | 0 | 0 | 0 | 0 | 0 | 693710 | 725980 | 0 | 0 | 113400 | 0 |
| HADEFLLD | FUT8; FUT8 | Q9BYC5; A0A | FUT8_HUMAN | Unmodified | 0 | 0 | 0 | 0 | 78150 | 0 | 0 | 0 | 14434000 | 652410 | 0 | 244130 | 0 | 0 | 0 | 0 | 0 | 0 |
| HADHSSLT | HUWE1 | Q7Z6Z7 | HUWE1_HUMA | Unmodified | 0 | 0 | 64249 | 0 | 0 | 0 | 0 | 0 | 179780 | 0 | 118380 | 980710 | 0 | 0 | 100770 | 90428 | 0 | 0 |
| HADIVTTTT | SHMT2; SHMT2 | P34897; H0Y | GLYM_HUMAN | Unmodified | 0 | 0 | 0 | 0 | 0 | 0 | 0 | 0 | 26651000 | 2384400 | 0 | 0 | 166780 | 0 | 0 | 0 | 0 | 0 |
| HADLEIR | MVP; MVP; MVP | Q14764; H3I | MVP_HUMAN N | Unmodified | 0 | 6282500 | 6472100 | 0 | 75161000 | 23861000 | 6682900 | 3842000 | 474900 | 0 | 89918 | 6488100 | 0 | 188030 | 1710700 | 0 | 117520 | 341470 |
| HADPAHQI | KLRG2 | A4D1S0 | KLRG2_HUMAN | Unmodified | 0 | 0 | 142840 | 237490 | 0 | 0 | 0 | 0 | 0 | 0 | 0 | 0 | 0 | 0 | 0 | 0 | 0 | 0 |
| HAEAQAAA | SELENON; SELE | Q9NZV5; H9 | SELN_HUMAN | Unmodified | 0 | 0 | 0 | 0 | 0 | 0 | 0 | 0 | 668630 | 0 | 127090 | 289880 | 0 | 0 | 0 | 0 | 0 | 0 |
| HAEATLGS | MOB1B; MOB1A | Q7L9L4; Q9H | MOB1B_HUMA | Unmodified | 227760 | 293540 | 75009 | 156220 | 1088700 | 263820 | 0 | 0 | 190670 | 45675 | 902230 | 2264400 | 0 | 0 | 163590 | 189760 | 290700 | 1047800 |
| HAEELERL | SRP68 | Q9UHB9 | SRP68_HUMAN | Unmodified | 0 | 0 | 0 | 0 | 0 | 0 | 0 | 0 | 0 | 0 | 0 | 803130 | 0 | 0 | 0 | 0 | 0 | 0 |
| HAEFESLC | LRBA; LRBA; LR | P50851; A0A | LRBA_HUMAN | Unmodified | 0 | 0 | 0 | 0 | 0 | 0 | 40721 | 0 | 0 | 0 | 0 | 1707900 | 0 | 0 | 0 | 106400 | 0 | 0 |
| HAELIASTF | ATXN10; ATXN1 | Q9UBB4; B1 | ATX10_HUMAN | Unmodified | 0 | 0 | 0 | 0 | 0 | 0 | 0 | 0 | 0 | 0 | 0 | 1623600 | 0 | 0 | 0 | 0 | 89841 | 29012 |
| HAEMVHTC | CAP1; CAP1; CA | Q01518; Q5T | CAP1_HUMAN | Unmodified | 0 | 0 | 0 | 0 | 0 | 162950 | 0 | 0 | 486790 | 0 | 0 | 2334500 | 0 | 0 | 0 | 0 | 12395 | 0 |
| HAEPEQNV | IGF2R; IGF2R; I | P11717; A0A | MPRI_HUMAN | Unmodified | 0 | 0 | 507450 | 0 | 0 | 0 | 0 | 0 | 26262000 | 5877300 | 3593300 | 3502800 | 671360 | 382190 | 383110 | 0 | 0 | 0 |
| HAEPEQNV | IGF2R; IGF2R; I | P11717; A0A | MPRI_HUMAN | Unmodified | 0 | 0 | 742970 | 172440 | 321690 | 0 | 0 | 0 | 35175000 | 4891500 | 5079200 | 1003500 | 1099700 | 219390 | 266130 | 405490 | 0 | 137080 |
| HAEPSSSP | USP6NL; USP6I | Q92738; X6F | US6NL_HUMAI | Unmodified | 0 | 0 | 0 | 0 | 0 | 0 | 0 | 0 | 0 | 0 | 0 | 0 | 0 | 0 | 0 | 0 | 241050 | 722950 |
| HAEQERDE | MYH10 | P35580 | MYH10_HUMAI | Unmodified | 0 | 0 | 696520 | 1008500 | 1289300 | 433190 | 75830 | 0 | 265430 | 0 | 550170 | 9906200 | 741050 | 0 | 811050 | 828470 | 108500 | 122740 |
| HAESVGDC | LRPAP1 | P30533 | AMRP_HUMAN | Unmodified | 0 | 0 | 0 | 0 | 0 | 0 | 0 | 36571 | 1279100 | 0 | 0 | 89531 | 0 | 0 | 0 | 0 | 0 | 0 |
| HAEVVASIK | SLC9A3R2; SLC | Q15599; H3I | NHRF2_HUMAI | Unmodified | 37945 | 0 | 306540 | 603270 | 132630 | 0 | 0 | 0 | 345410 | 0 | 59752 | 914470 | 0 | 0 | 1232300 | 525700 | 59940 | 184620 |
| HAGAYHCI | SEMA4C | Q9C0C4 | SEM4C_HUMAI | Unmodified | 0 | 0 | 0 | 0 | 0 | 0 | 0 | 0 | 259530 | 0 | 0 | 0 | 0 | 0 | 0 | 0 | 0 | 0 |
| HAHGDQY | IDH2 | P48735 | IDHP_HUMAN | Unmodified | 77295 | 0 | 0 | 0 | 0 | 0 | 246780 | 0 | 281000 | 1919000 | 0 | 0 | 0 | 0 | 0 | 0 | 0 | 0 |
| HAHSSSLQ | SVIL; SVIL | O95425; A0A | SVIL_HUMAN S | Unmodified | 0 | 0 | 0 | 0 | 0 | 0 | 0 | 0 | 0 | 0 | 49759 | 106130 | 0 | 0 | 11106 | 0 | 0 | 0 |
| HAIPNLVK | CTNND1; CTNN | O60716; A0A | CTND1_HUMAI | Unmodified | 0 | 872350 | 0 | 0 | 16507000 | 3126300 | 0 | 0 | 0 | 0 | 1708200 | 7080000 | 22378000 | 16991000 | 10721000 | 8011900 | 11626000 | 17623000 |
| HAIVICNR | COPA; COPA; C | P53621; A0A | COPA_HUMAN | Unmodified | 1573000 | 0 | 843720 | 0 | 261860 | 0 | 0 | 0 | 0 | 64762 | 23792 | 684040 | 0 | 0 | 0 | 0 | 170650 | 0 |
| HAKPPDVD | CLGN | O14967 | CLGN_HUMAN | Unmodified | 0 | 0 | 0 | 0 | 0 | 0 | 203650 | 50905 | 4686500 | 1507400 | 489010 | 0 | 119200 | 142940 | 0 | 0 | 0 | 0 |
| HAKRPDAD | CANX; CANX; C | P27824; A0A | CALX_HUMAN | Unmodified | 0 | 0 | 0 | 0 | 633910 | 0 | 0 | 0 | 8473100 | 30794000 | 388450 | 0 | 0 | 0 | 0 | 0 | 0 | 65614 |
| HALDAHQ | ATP6V1F | Q16864 | VATF_HUMAN | Unmodified | 0 | 0 | 0 | 0 | 0 | 0 | 0 | 0 | 1647900 | 0 | 598000 | 4756600 | 0 | 0 | 0 | 0 | 0 | 0 |
| HALGAVAQ | IQGAP3; IQGAP | Q86VI3; F2Z | IQGA3_HUMAN | Unmodified | 0 | 0 | 0 | 0 | 0 | 0 | 25972 | 0 | 0 | 0 | 0 | 1778400 | 0 | 0 | 0 | 0 | 0 | 0 |
| HALHCTILA | COPS4; COPS4 | Q9BT78; D6I | CSN4_HUMAN | Unmodified | 0 | 0 | 0 | 0 | 242610 | 47156 | 1513400 | 795630 | 0 | 0 | 108940 | 1305900 | 0 | 1177600 | 0 | 0 | 1009900 | 223690 |
| HALIIYDDL | ATP5F1A | P25705 | ATPA_HUMAN | Unmodified | 107290 | 0 | 219380 | 0 | 0 | 0 | 401320 | 370130 | 24893000 | 3701400 | 866180 | 445060 | 126350 | 0 | 107900 | 56023 | 0 | 0 |
| HALLDVTPI | TJP2; TJP2; TJP2 | Q9UDY2; A0 | ZO2_HUMAN T | Unmodified | 0 | 0 | 0 | 0 | 673130 | 0 | 0 | 34194 | 268590 | 476690 | 0 | 689740 | 0 | 0 | 0 | 0 | 0 | 0 |
| HALLDVTPI | TJP1; TJP1; TJP1 | Q07157; A0A | ZO1_HUMAN T | Unmodified | 0 | 0 | 0 | 0 | 5747900 | 0 | 0 | 0 | 0 | 0 | 0 | 0 | 2123500 | 2331300 | 0 | 0 | 0 | 0 |
| HALLEEENI | PLXNB2; PLXNE | O15031; A6( | PLXB2_HUMAN | Unmodified | 0 | 0 | 2673600 | 2693800 | 0 | 0 | 0 | 0 | 4885700 | 0 | 0 | 1627700 | 0 | 0 | 15179000 | 6976500 | 0 | 0 |
| HALQRDIF | EXOC1 | Q9NV70 | EXOC1_HUMAI | Unmodified | 0 | 0 | 0 | 0 | 5163100 | 226870 | 0 | 0 | 0 | 0 | 0 | 0 | 3427800 | 1786400 | 0 | 0 | 0 | 2727500 |
| HAMNLEAV | GCDH | Q92947 | GCDH_HUMAN | Unmodified | 0 | 0 | 0 | 0 | 0 | 0 | 0 | 0 | 2354800 | 0 | 0 | 0 | 0 | 0 | 0 | 0 | 0 | 0 |
| HANAKPFE | NUDT5; NUDT5 | Q9UKK9; A6I | NUDT5_HUMAI | Unmodified | 0 | 0 | 0 | 0 | 0 | 0 | 61472 | 0 | 0 | 0 | 587970 | 0 | 0 | 0 | 0 | 0 | 0 | 0 |
| HANAKPFE | NUDT5; NUDT5 | Q9UKK9; A6I | NUDT5_HUMAI | Unmodified | 168570 | 271450 | 0 | 539410 | 13972000 | 3202900 | 5837200 | 154900 | 4005600 | 5024200 | 3299100 | 9683900 | 3908300 | 4057900 | 268340 | 264060 | 5155200 | 0 |
| HANSVDTS | DOCK10; DOCK | Q96BY6; A0A | DOC10_HUMA | Unmodified | 0 | 0 | 0 | 0 | 0 | 0 | 0 | 0 | 0 | 0 | 0 | 0 | 0 | 0 | 0 | 0 | 0 | 0 |
| HAPDVVLD | SRGAP1; SRGA | Q7Z6B7; G5I | SRGP1_HUMAI | Unmodified | 0 | 0 | 0 | 0 | 1773500 | 314460 | 0 | 0 | 0 | 0 | 130820 | 459710 | 375290 | 0 | 0 | 0 | 562570 | 772290 |
| HAPGLIAAI | BROX; BROX | Q5VW32; F5 | BROX_HUMAN | Unmodified | 4910800 | 3114800 | 706340 | 2338300 | 1249900 | 0 | 872460 | 0 | 2446900 | 1755200 | 284540 | 1405200 | 44433 | 147330 | 4435700 | 3065900 | 999570 | 10668 |
| HAPVLELE | CAP2; CAP2; CA | P40123; B7Z | CAP2_HUMAN | Unmodified | 0 | 0 | 0 | 0 | 885030 | 0 | 426120 | 391380 | 0 | 0 | 0 | 0 | 507750 | 0 | 0 | 0 | 458280 | 539700 |
| HAPVLELE | CAP2; CAP2; CA | P40123; B7Z | CAP2_HUMAN | Unmodified | 0 | 0 | 0 | 0 | 526560 | 0 | 86948 | 48341 | 0 | 0 | 0 | 100120 | 131140 | 78183 | 49900 | 0 | 259640 | 197230 |
| HAQDSDPI | CDCA3; CDCA3 | Q99618; F8V | CDCA3_HUMA | Unmodified | 301110 | 214860 | 263090 | 520780 | 529590 | 130320 | 0 | 0 | 335570 | 158340 | 0 | 103510 | 518790 | 103850 | 0 | 0 | 0 | 0 |
| HAQDSDPI | CDCA3; CDCA3 | Q99618; F8V | CDCA3_HUMA | Oxidation (M) | 0 | 0 | 0 | 0 | 1858300 | 0 | 0 | 0 | 0 | 36762 | 0 | 0 | 0 | 0 | 0 | 0 | 0 | 0 |
| HAQDSDPI | CDCA3; CDCA3 | Q99618; F8V | CDCA3_HUMA | Unmodified | 0 | 0 | 0 | 0 | 3836800 | 0 | 116550 | 0 | 0 | 0 | 0 | 0 | 0 | 764420 | 0 | 0 | 238150 | 0 |
| HAQPTCPG | FASN; FASN | P49327; A0A | FAS_HUMAN F | Unmodified | 0 | 0 | 0 | 0 | 0 | 0 | 0 | 0 | 0 | 0 | 0 | 3178800 | 0 | 0 | 0 | 95884 | 0 | 0 |
| HARERVTPI | PDLIM1 | O00151 | PDLI1_HUMAN | Unmodified | 0 | 0 | 0 | 0 | 2750200 | 183420 | 317020 | 173420 | 0 | 264410 | 0 | 0 | 0 | 393990 | 0 | 0 | 9006200 | 5511700 |
| HARNQVD | RACGAP1; RAC | Q9H0H5; F8 | RGAP1_HUMAI | Unmodified | 0 | 0 | 0 | 0 | 32433000 | 218200 | 0 | 0 | 0 | 2469400 | 274500 | 94982 | 1900700 | 1043200 | 0 | 0 | 3754600 | 3863200 |
| HASAPSHV | TBC1D4; TBC1I | O60343; A0A | TBCD4_HUMAI | Unmodified | 0 | 0 | 0 | 0 | 0 | 0 | 0 | 0 | 0 | 0 | 0 | 211780 | 0 | 0 | 0 | 20874 | 0 | 0 |
| HASGGSTV | HUWE1; HUWE | Q7Z6Z7; H0Y | HUWE1_HUMA | Unmodified | 0 | 0 | 0 | 0 | 0 | 0 | 0 | 0 | 0 | 0 | 0 | 2455800 | 0 | 0 | 0 | 115040 | 0 | 0 |
| HASNMLGI | VPS35 | Q96QK1 | VPS35_HUMAN | Unmodified | 0 | 0 | 0 | 0 | 0 | 0 | 0 | 0 | 0 | 0 | 0 | 880440 | 0 | 0 | 0 | 0 | 0 | 0 |
| HASPEDIKI | DNAJB6; DNAJE | O75190; A0A | DNJB6_HUMAN | Unmodified | 0 | 189820 | 148090 | 0 | 211440 | 80693 | 0 | 0 | 48450 | 0 | 72466 | 120410 | 0 | 0 | 166860 | 71100 | 0 | 0 |
| HASQDELA | AVIL | O75366 | AVIL_HUMAN A | Unmodified | 0 | 0 | 0 | 0 | 0 | 0 | 0 | 0 | 0 | 0 | 0 | 18161000 | 0 | 0 | 0 | 0 | 0 | 0 |
| HASQDELA | AVIL | O75366 | AVIL_HUMAN A | Unmodified | 0 | 0 | 0 | 0 | 0 | 0 | 0 | 0 | 0 | 0 | 0 | 1627200 | 0 | 0 | 0 | 0 | 0 | 0 |
| HASSGSFL | GLRX3 | O76003 | GLRX3_HUMAN | Unmodified | 0 | 0 | 0 | 0 | 0 | 0 | 0 | 19453 | 0 | 0 | 386910 | 6760000 | 0 | 0 | 0 | 0 | 103410 | 0 |
| HASSGSFL | GLRX3 | O76003 | GLRX3_HUMAN | Unmodified | 0 | 0 | 0 | 0 | 1278200 | 0 | 0 | 0 | 0 | 0 | 0 | 0 | 1685200 | 0 | 0 | 0 | 921480 | 802560 |
| HATAEEVEI | BRIX1 | Q8TDN6 | BRX1_HUMAN | Unmodified | 0 | 0 | 0 | 0 | 0 | 784870 | 644630 | 0 | 80399 | 0 | 0 | 550940 | 0 | 0 | 0 | 0 | 0 | 202380 |
| HATFVDGL | CEMIP2; CEMIP | Q9UHN6; F5 | CEIP2_HUMAN | Unmodified | 881200 | 950240 | 1321300 | 480810 | 153560 | 0 | 0 | 0 | 4903500 | 325670 | 0 | 516740 | 0 | 0 | 729440 | 2201300 | 0 | 0 |
| HATRHSPC | IRF6; IRF6; IRF6 | O14896; B1A | IRF6_HUMAN I | Unmodified | 0 | 0 | 0 | 0 | 0 | 0 | 0 | 0 | 0 | 72011 | 0 | 0 | 0 | 5304200 | 0 | 0 | 0 | 0 |
| HATRIIDEV | BAG2 | O95816 | BAG2_HUMAN | Unmodified | 0 | 0 | 0 | 0 | 0 | 0 | 0 | 0 | 0 | 0 | 0 | 0 | 0 | 0 | 0 | 0 | 280820 | 132110 |
| HATRIIDEV | BAG2 | O95816 | BAG2_HUMAN | Unmodified | 0 | 628480 | 10240 | 0 | 2224400 | 0 | 52574 | 0 | 0 | 352300 | 0 | 354500 | 936080 | 614020 | 416270 | 0 | 2036300 | 614960 |
| HATWRGTS | DIPK2A; DIPK2A | Q8NDZ4; E7 | DIK2A_HUMAN | Unmodified | 0 | 0 | 0 | 0 | 0 | 0 | 0 | 0 | 1162600 | 997960 | 0 | 0 | 0 | 0 | 0 | 0 | 0 | 0 |
| HAVDGRQI | GLT8D1; GLT8D | Q68CQ7; C9 | GL8D1_HUMAI | Unmodified | 0 | 0 | 0 | 0 | 0 | 0 | 0 | 0 | 0 | 243630 | 0 | 375160 | 0 | 0 | 0 | 0 | 0 | 0 |
| HAVDGRQI | GLT8D1; GLT8D | Q68CQ7; C9 | GL8D1_HUMAI | Unmodified | 0 | 0 | 0 | 0 | 0 | 0 | 0 | 0 | 47093000 | 1239600 | 0 | 0 | 0 | 0 | 0 | 0 | 0 | 0 |
| HAVSEGTK | H2BE1; H2BC4 | A0A2R8Y619 | H2BE1_HUMAN | Unmodified | 373400 | 0 | 0 | 0 | 2558900 | 675800 | 9013900 | 54696000 | 0 | 209660 | 0 | 900530 | 475150 | 0 | 0 | 0 | 130250 | 3800200 |
| HAVSEGTK | H2BC11; H2BC | P06899; O60 | H2B1J_HUMAN | Unmodified | 0 | 0 | 0 | 0 | 374350 | 710440 | 10932000 | 1.03E+08 | 0 | 25265 | 0 | 0 | 174760 | 0 | 0 | 0 | 532350 | 4323700 |
| HAVSEGTK | H2BE1; H2BC4 | A0A2R8Y619 | H2BE1_HUMAN | Unmodified | 754490 | 0 | 0 | 0 | 3141800 | 1453000 | 42916000 | 2.84E+08 | 394530 | 185420 | 0 | 2020100 | 7478700 | 1908500 | 0 | 0 | 87186 | 8139200 |
| HAVTEAEIC | PPP1R9B | Q96SB3 | NEB2_HUMAN | Unmodified | 0 | 0 | 0 | 0 | 0 | 0 | 750140 | 696360 | 0 | 0 | 0 | 0 | 0 | 1330900 | 0 | 0 | 1573700 | 505380 |
| HAVVMFQT | LAMP3; LAMP3 | Q9UQV4; E7 | LAMP3_HUMAI | Oxidation (M) | 0 | 38237 | 0 | 155530 | 67304 | 0 | 41372 | 26574 | 755170 | 514130 | 0 | 0 | 39656 | 62845 | 0 | 0 | 0 | 0 |
| HAVVMFQT | LAMP3; LAMP3 | Q9UQV4; E7 | LAMP3_HUMAI | Unmodified | 0 | 0 | 0 | 0 | 0 | 0 | 151650 | 0 | 7414200 | 1361100 | 0 | 0 | 0 | 0 | 0 | 0 | 53056 | 0 |
| HAVVNLIN | CTNNB1; CTNN | P35222; A0A | CTNB1_HUMAI | Unmodified | 7855200 | 4481300 | 12714000 | 18480000 | 36597000 | 11098000 | 0 | 0 | 5059400 | 1439100 | 7041200 | 26346000 | 17609000 | 16724000 | 23795000 | 21136000 | 25986000 | 30573000 |
| HAVVNLIN | CTNNB1; CTNN | P35222; A0A | CTNB1_HUMAI | Unmodified | 0 | 0 | 0 | 306800 | 78239000 | 15015000 | 832020 | 0 | 0 | 1889400 | 0 | 0 | 57102000 | 55583000 | 0 | 0 | 1E+08 | 79755000 |
| HAVVNLIN | CTNNB1; CTNN | P35222; A0A | CTNB1_HUMAI | Unmodified | 0 | 0 | 0 | 0 | 2619100 | 0 | 0 | 0 | 0 | 0 | 0 | 0 | 1313900 | 2769100 | 0 | 0 | 6266700 | 407370 |
| HAWIWHD | LOXL2 | Q9Y4K0 | LOXL2_HUMAN | Unmodified | 0 | 0 | 0 | 0 | 0 | 0 | 0 | 0 | 333830 | 0 | 84790 | 0 | 0 | 0 | 0 | 0 | 0 | 0 |
| HAWNNYK | MAN1A1 | P33908 | MA1A1_HUMAI | Unmodified | 0 | 0 | 0 | 0 | 0 | 0 | 0 | 0 | 0 | 0 | 0 | 0 | 0 | 0 | 0 | 0 | 0 | 0 |
| HAYGDQYI | IDH1 | O75874 | IDHC_HUMAN | Unmodified | 0 | 0 | 0 | 0 | 1423300 | 0 | 0 | 0 | 836390 | 0 | 1806700 | 0 | 7971100 | 0 | 0 | 0 | 0 | 0 |
| HAYGLGEH | OTUD6B; OTUD | Q8N6M0; A0 | OTU6B_HUMAI | Unmodified | 0 | 0 | 0 | 0 | 0 | 36545 | 22194 | 0 | 205550 | 0 | 59558 | 672260 | 0 | 0 | 0 | 0 | 176560 | 0 |
| HAYSFHQS | BTBD17 | A6NE02 | BTBDH_HUMAI | Unmodified | 0 | 0 | 0 | 0 | 0 | 0 | 808160 | 0 | 3472700 | 187600 | 0 | 0 | 0 | 0 | 0 | 0 | 0 | 0 |
| HCDKAEFF | C1GALT1C1 | Q96EU7 | C1GLC_HUMAI | Unmodified | 0 | 0 | 0 | 0 | 0 | 0 | 0 | 0 | 0 | 0 | 0 | 0 | 0 | 0 | 0 | 0 | 0 | 0 |
| HCFTSNEA | THUMPD3; THU | Q9BV44; H7 | THUM3_HUMA | Unmodified | 0 | 0 | 0 | 0 | 128750 | 0 | 0 | 0 | 0 | 97539 | 0 | 23957000 | 349380 | 245710 | 67142 | 0 | 255100 | 0 |
| HCGYTQLS | USP54; USP54; | Q70EL1; A0A | UBP54_HUMAI | Unmodified | 0 | 0 | 0 | 0 | 0 | 0 | 0 | 0 | 0 | 0 | 0 | 0 | 0 | 0 | 0 | 0 | 0 | 0 |
| HCILDVSG | DLG2; DLG3; D | Q15700; Q9 | DLG2_HUMAN | Unmodified | 54869 | 0 | 1343400 | 797950 | 907470 | 888380 | 0 | 0 | 2455800 | 1230500 | 532000 | 3697800 | 2154900 | 5133100 | 961160 | 313620 | 585110 | 667210 |
| HCIMQANA | PDCD6IP; PDCI | Q8WUM4; F | PDC6I_HUMAN | Oxidation (M) | 14421000 | 8800600 | 6702600 | 5927800 | 5582100 | 1193000 | 468740 | 242190 | 3587800 | 2680100 | 2066900 | 3218900 | 9792300 | 3801400 | 6864300 | 7267200 | 2378100 | 2600500 |
| HCIMQANA | PDCD6IP; PDCI | Q8WUM4; F | PDC6I_HUMAN | Unmodified | 66772000 | 50105000 | 12151000 | 17330000 | 20994000 | 3435000 | 3362900 | 76245 | 19522000 | 14492000 | 2059300 | 0 | 43712000 | 15863000 | 19265000 | 10663000 | 4665400 | 15255000 |
| HCIMQANA | PDCD6IP; PDCI | Q8WUM4; F | PDC6I_HUMAN | Unmodified | 0 | 0 | 0 | 0 | 745640 | 0 | 0 | 0 | 0 | 0 | 0 | 0 | 1007800 | 0 | 0 | 0 | 490740 | 339560 |
| HCLQTVWI | PSAP; PSAP; PS | P07602; Q5E | SAP_HUMAN P | Unmodified | 0 | 0 | 0 | 0 | 0 | 1520600 | 1469800 | 378610 | 2946400 | 3573000 | 0 | 96056 | 434750 | 0 | 0 | 0 | 0 | 0 |
| HCNMVLEI | SNRPD2 | P62316 | SMD2_HUMAN | Unmodified | 0 | 26577 | 0 | 0 | 0 | 41460 | 0 | 0 | 0 | 0 | 0 | 586550 | 0 | 0 | 0 | 0 | 78413 | 0 |
| HCNNGIPE | NIBAN2 | Q96TA1 | NIBA2_HUMAN | Unmodified | 1731500 | 901780 | 187520 | 1003600 | 0 | 0 | 0 | 0 | 693390 | 0 | 0 | 1685200 | 0 | 0 | 1116600 | 1095900 | 0 | 105790 |
| HCRAPCC | TRAF2; TRAF2 | Q12933; B1A | TRAF2_HUMAN | Unmodified | 0 | 0 | 0 | 0 | 323340 | 322210 | 0 | 0 | 0 | 0 | 0 | 0 | 0 | 0 | 0 | 0 | 0 | 0 |
| HCSQVDS | CTTN | Q14247 | SRC8_HUMAN | Unmodified | 0 | 0 | 0 | 550760 | 1291200 | 0 | 0 | 0 | 0 | 0 | 0 | 0 | 720000 | 509450 | 0 | 347140 | 353280 | 717170 |
| HCTLRMEL | EDIL3 | O43854 | EDIL3_HUMAN | Unmodified | 0 | 0 | 0 | 0 | 2018700 | 609810 | 0 | 0 | 0 | 0 | 1740600 | 297370 | 0 | 0 | 0 | 0 | 0 | 1268600 |
| HCVKEVVE | GATD3B; GATD | A0A0B4J2D5 | GAL3B_HUMAN | Unmodified | 0 | 0 | 0 | 0 | 0 | 0 | 232530 | 0 | 1066000 | 314510 | 261070 | 0 | 0 | 0 | 0 | 0 | 99278 | 0 |
| HCVLNELA | TOR2A | Q5JU69 | TOR2A_HUMAN | Unmodified | 0 | 0 | 0 | 0 | 0 | 0 | 0 | 0 | 2305400 | 187520 | 0 | 0 | 0 | 0 | 0 | 0 | 0 | 0 |
| HCYCEEG | ADAM15 | Q13444 | ADA15_HUMAN | Unmodified | 509270 | 0 | 0 | 0 | 0 | 0 | 0 | 0 | 0 | 0 | 0 | 0 | 0 | 0 | 0 | 0 | 0 | 0 |
| HDAATCFV | NAPA; NAPA; N | P54920; M0I | SNAA_HUMAN | Unmodified | 812060 | 341390 | 0 | 0 | 0 | 0 | 0 | 0 | 0 | 0 | 0 | 420190 | 0 | 0 | 907300 | 544890 | 0 | 0 |
| HDADGQA | PSMD3 | O43242 | PSMD3_HUMA | Unmodified | 157150 | 0 | 0 | 0 | 0 | 0 | 0 | 0 | 0 | 0 | 58003 | 3014000 | 0 | 0 | 444700 | 523230 | 0 | 0 |
| HDDPALTL | PDXDC1; PDXD | Q6P996; H3I | PDXD1_HUMAI | Unmodified | 0 | 0 | 0 | 0 | 0 | 0 | 0 | 0 | 0 | 0 | 0 | 803830 | 0 | 0 | 0 | 0 | 0 | 0 |
| HDFSTVLT | EXOC7; EXOC7 | Q9UPT5; B4I | EXOC7_HUMAI | Unmodified | 0 | 0 | 0 | 0 | 0 | 0 | 0 | 0 | 316440 | 0 | 0 | 382790 | 0 | 0 | 0 | 0 | 0 | 0 |
| HDLCAAAC | STC2; STC2 | O76061; H0Y | STC2_HUMAN | Unmodified | 0 | 0 | 0 | 0 | 0 | 0 | 0 | 0 | 1598800 | 0 | 0 | 1260900 | 0 | 0 | 124840 | 0 | 17523 | 168570 |
| HDLLVGAP | ITGA2B; ITGA2E | P08514; A0A | ITA2B_HUMAN | Oxidation (M) | 997550 | 0 | 1598500 | 0 | 0 | 0 | 117270 | 0 | 488930 | 0 | 0 | 0 | 0 | 0 | 7444800 | 3001600 | 0 | 0 |
| HDLNPLIK | IGF2R; IGF2R; I | P11717; A0A | MPRI_HUMAN | Unmodified | 0 | 0 | 299610 | 0 | 943790 | 8472400 | 210990 | 1118000 | 5971200 | 1277500 | 678080 | 1480200 | 158630 | 10741000 | 0 | 0 | 1009500 | 8188000 |
| HDPTFEPLI | SUSD2 | Q9UGT4 | SUSD2_HUMAI | Unmodified | 7313000 | 6553900 | 0 | 0 | 2797900 | 986450 | 0 | 0 | 77062000 | 6245100 | 0 | 0 | 0 | 0 | 10294000 | 725920 | 0 | 0 |
| HDVVFLITK | CLTC; CLTC | Q00610; A0A | CLH1_HUMAN | Unmodified | 1830100 | 1464700 | 0 | 724750 | 1378200 | 4410100 | 2566000 | 1342800 | 171500 | 0 | 0 | 3108600 | 148820 | 2821900 | 415780 | 0 | 196110 | 2067400 |
| HEADANPF | MPP7; MPP7 | Q5T2T1; S4F | MPP7_HUMAN | Unmodified | 0 | 0 | 0 | 0 | 1221000 | 138400 | 0 | 0 | 0 | 55526 | 0 | 0 | 169670 | 164500 | 0 | 0 | 33681 | 331960 |
| HEALFQLVI | NOSTRIN; NOS | Q8IVI9; A0A | NOSTN_HUMA | Unmodified | 0 | 0 | 287140 | 220480 | 0 | 0 | 0 | 0 | 0 | 0 | 0 | 0 | 0 | 0 | 0 | 0 | 0 | 0 |
| HEANNPQI | STIP1; STIP1; ST | P31948; F5H | STIP1_HUMAN | Unmodified | 0 | 0 | 0 | 0 | 199670 | 0 | 0 | 0 | 433660 | 70706 | 7920.7 | 366070 | 0 | 12187 | 120110 | 0 | 102600 | 61782 |
| HEAQRLEQ | MVP | Q14764 | MVP_HUMAN N | Unmodified | 0 | 0 | 0 | 0 | 865270 | 0 | 15175 | 0 | 1955800 | 94798 | 0 | 10576000 | 0 | 0 | 0 | 0 | 0 | 0 |
| HECQANG | DSTN | P60981 | DEST_HUMAN | Unmodified | 0 | 0 | 0 | 0 | 0 | 0 | 0 | 0 | 0 | 0 | 0 | 0 | 0 | 0 | 0 | 0 | 0 | 0 |

| | | | | | | | | | | | | | | | | | | | | | | |
|---|---|---|---|---|---|---|---|---|---|---|---|---|---|---|---|---|---|---|---|---|---|---|
| HEDANKAV | PABPC4; PABP( | Q13310; B1/ | PABP4_HUMAN | Unmodified | 2020300 | 1621200 | 0 | 747720 | 0 | 0 | 0 | 0 | 0 | 0 | 0 | 642510 | 0 | 0 | 137590 | 519450 | 0 | 0 |
| HEDANKAV | PABPC4; PABP( | Q13310; B1/ | PABP4_HUMAN | Unmodified | 3944100 | 1743800 | 0 | 2117000 | 0 | 0 | 39810 | 0 | 0 | 156020 | 0 | 0 | 0 | 0 | 0 | 60462 | 159560 | 0 |
| HEEAPGHF | CSRP1; CSRP1; | P21291; E9P | CSRP1_HUMAI | Unmodified | 0 | 0 | 1093900 | 401780 | 0 | 0 | 0 | 0 | 564670 | 0 | 0 | 0 | 0 | 0 | 166920 | 0 | 47449 | 50224 |
| HEERQDEH | HSPB1; HSPB1; | P04792; A0A | HSPB1_HUMAI | Unmodified | 45375 | 0 | 699730 | 0 | 0 | 0 | 0 | 0 | 998350 | 0 | 0 | 0 | 0 | 0 | 341580 | 367010 | 0 | 0 |
| HEGALETLI | PDCD6IP; PDCI | Q8WUM4; F8 | PDC6I_HUMAN | Unmodified | 3087500 | 4714600 | 4009100 | 1095900 | 160580 | 0 | 0 | 99458 | 768600 | 0 | 0 | 2333300 | 65405 | 0 | 4035500 | 2828500 | 0 | 0 |
| HEIEGTGLI | NIBAN2 | Q96TA1 | NIBA2_HUMAN | Unmodified | 1412700 | 0 | 0 | 0 | 0 | 0 | 0 | 0 | 628770 | 0 | 0 | 2796900 | 0 | 0 | 2358300 | 815710 | 0 | 0 |
| HEILQWVL | LRRC59 | Q96AG4 | LRC59_HUMAN | Unmodified | 0 | 0 | 0 | 0 | 0 | 193690 | 836200 | 22676 | 1978300 | 147890 | 74688 | 165930 | 0 | 0 | 0 | 0 | 0 | 0 |
| HEIVQTLSL | SELENBP1; SEL | Q13228; A6F | SBP1_HUMAN | Unmodified | 0 | 0 | 393880 | 0 | 0 | 0 | 0 | 0 | 0 | 0 | 0 | 0 | 0 | 0 | 298930 | 154060 | 0 | 0 |
| HEKEDGAI | CCT8 | P50990 | TCPQ_HUMAN | Unmodified | 0 | 0 | 0 | 0 | 0 | 0 | 0 | 0 | 154480 | 0 | 188590 | 1691100 | 0 | 0 | 113530 | 58628 | 0 | 0 |
| HELIEFR | CLTC; CLTC; CI | Q00610; J3K | CLH1_HUMAN | Unmodified | 1319400 | 1444800 | 0 | 472440 | 458840 | 168040 | 574470 | 332190 | 395720 | 53605 | 65301 | 5764800 | 0 | 0 | 1525600 | 1627800 | 0 | 0 |
| HELLPSVNI | PON2; PON2; P | Q15165; G3 | PON2_HUMAN | Unmodified | 0 | 0 | 0 | 0 | 0 | 116380 | 0 | 0 | 1301200 | 0 | 0 | 0 | 0 | 0 | 0 | 0 | 0 | 1094600 |
| HELLQPFN | PA2G4 | Q9UQ80 | PA2G4_HUMAI | Unmodified | 1948200 | 3215300 | 2224200 | 897990 | 639820 | 179200 | 304530 | 139280 | 1531700 | 288910 | 3116900 | 35700000 | 0 | 149520 | 3112500 | 2769100 | 0 | 0 |
| HELQANCY | CFL1; CFL1; CF | P23528; G3 | COF1_HUMAN | Unmodified | 3203400 | 1209700 | 4703300 | 1856100 | 3528000 | 702680 | 0 | 97146 | 2503400 | 160460 | 1755200 | 23806000 | 847740 | 0 | 4943500 | 2771800 | 620970 | 1034900 |
| HELQANCY | CFL1; CFL1; CF | P23528; G3 | COF1_HUMAN | Unmodified | 0 | 0 | 0 | 0 | 3333700 | 1159100 | 276740 | 0 | 0 | 138100 | 88340 | 0 | 0 | 225950 | 0 | 0 | 1150000 | 1626200 |
| HELQVEMK | DIAPH1; DIAPH | O60610; A0/ | DIAP1_HUMAN | Unmodified | 0 | 0 | 0 | 0 | 0 | 0 | 0 | 0 | 269070 | 0 | 109430 | 733120 | 0 | 0 | 0 | 0 | 0 | 0 |
| HELTEISNV | AHNAK | Q09666 | AHNK_HUMAN | Unmodified | 0 | 0 | 0 | 0 | 1052900 | 0 | 0 | 0 | 0 | 0 | 0 | 0 | 0 | 0 | 0 | 0 | 238400 | 441140 |
| HEMLPEFY | CAND1; CAND1 | Q86VP6; A0/ | CAND1_HUMA | Oxidation (M) | 0 | 0 | 0 | 0 | 0 | 0 | 0 | 0 | 149740 | 12346 | 0 | 818610 | 0 | 0 | 0 | 0 | 0 | 0 |
| HENVIGLLI | MAPK14; MAPK | Q16539; B5 | MK14_HUMAN | Unmodified | 0 | 0 | 95920 | 0 | 0 | 0 | 16428 | 0 | 161540 | 0 | 87867 | 0 | 0 | 0 | 370110 | 0 | 72170 | 0 |
| HEQEYMEV | TMED2; TMED2 | Q15363; E7E | TMED2_HUMAI | Unmodified | 28829 | 0 | 0 | 0 | 0 | 0 | 0 | 0 | 4409600 | 445060 | 0 | 832370 | 0 | 0 | 163150 | 71186 | 0 | 0 |
| HEQEYMEV | TMED2; TMED2 | Q15363; E7E | TMED2_HUMAI | Unmodified | 0 | 0 | 0 | 0 | 0 | 0 | 0 | 0 | 3849300 | 0 | 152350 | 0 | 0 | 0 | 0 | 0 | 0 | 0 |
| HEQILVLDI | VAPA; VAPA | Q9P0L0; J3Q | VAPA_HUMAN | Unmodified | 0 | 0 | 0 | 0 | 0 | 0 | 433650 | 34239 | 1447000 | 256400 | 1219300 | 4203700 | 668770 | 0 | 0 | 0 | 0 | 0 |
| HEQNIDCC | CALR; CALR; C/ | P27797; A0A | CALR_HUMAN | Unmodified | 0 | 0 | 0 | 0 | 0 | 261840 | 1672400 | 808180 | 2.3E+08 | 25131000 | 15651000 | 8511900 | 0 | 0 | 376600 | 734070 | 272230 | 1099400 |
| HESQMDS | REEP5; REEP5 | Q00765; E2( | REEP5_HUMAN | Unmodified | 45763 | 0 | 168060 | 80025 | 0 | 0 | 0 | 0 | 306900 | 33502 | 377460 | 194300 | 0 | 0 | 135150 | 66043 | 0 | 0 |
| HETDMFN( | GAS7; GAS7 | O60861; A0/ | GAS7_HUMAN | Oxidation (M) | 0 | 0 | 0 | 0 | 0 | 0 | 0 | 0 | 0 | 0 | 837630 | 2865800 | 0 | 0 | 0 | 0 | 0 | 0 |
| HETLPYGR | ENPP1 | P22413 | ENPP1_HUMAN | Unmodified | 0 | 0 | 642690 | 0 | 0 | 0 | 0 | 0 | 0 | 0 | 0 | 0 | 0 | 0 | 309180 | 297890 | 0 | 0 |
| HEVININLK | GSTO1; GSTO1 | P78417; Q5 | GSTO1_HUMAI | Unmodified | 33782 | 0 | 33245 | 28783 | 0 | 348170 | 0 | 0 | 0 | 0 | 0 | 2044300 | 20714 | 113590 | 170320 | 65479 | 0 | 1057700 |
| HEVSGETV | ECPAS; ECPAS | Q5VYK3; J3K | ECM29_HUMAI | Unmodified | 0 | 0 | 0 | 0 | 0 | 0 | 0 | 0 | 0 | 0 | 703640 | 782640 | 0 | 0 | 0 | 124920 | 0 | 0 |
| HEVTILGGI | UBE2H; UBE2H | P62256; C9J | UBE2H_HUMAI | Unmodified | 0 | 430720 | 116630 | 0 | 0 | 0 | 0 | 0 | 0 | 0 | 21364 | 659030 | 0 | 0 | 0 | 275340 | 0 | 0 |
| HEWQVNG | CFL2 | Q9Y281 | COF2_HUMAN | Unmodified | 0 | 0 | 0 | 0 | 2741500 | 0 | 0 | 166360 | 0 | 0 | 0 | 0 | 0 | 0 | 0 | 0 | 0 | 0 |
| HEYGSPGI | CYFIP1; CYFIP2 | Q7L576; Q9( | CYFP1_HUMAN | Unmodified | 0 | 0 | 0 | 0 | 0 | 0 | 0 | 0 | 0 | 0 | 0 | 3159300 | 0 | 0 | 3902600 | 0 | 0 | 0 |
| HFAGAVCY | MYO6; MYO6; M | Q9UM54; A0 | MYO6_HUMAN | Unmodified | 330950 | 0 | 0 | 2562100 | 0 | 0 | 0 | 0 | 1848800 | 329580 | 0 | 0 | 0 | 0 | 2109300 | 1507000 | 0 | 0 |
| HFAGDVLG | CHID1; CHID1; | Q9BWS9; E9 | CHID1_HUMAN | Unmodified | 0 | 0 | 0 | 0 | 0 | 0 | 0 | 0 | 27055000 | 810260 | 1190500 | 338640 | 0 | 0 | 0 | 0 | 0 | 0 |
| HFCPNVPII | RHOB | P62745 | RHOB_HUMAN | Unmodified | 0 | 0 | 0 | 0 | 1260000 | 0 | 821310 | 354240 | 505770 | 370610 | 501660 | 2872300 | 1695900 | 840140 | 1420200 | 1030300 | 0 | 0 |
| HFCPNVPII | RHOA; RHOC; F | P61586; P08 | RHOA_HUMAN | Unmodified | 284360 | 0 | 307070 | 0 | 1324600 | 0 | 1321500 | 41880 | 1574400 | 0 | 1065200 | 1728000 | 274670 | 345370 | 1019800 | 407800 | 940160 | 1292400 |
| HFCPNVPII | RHOA; RHOC; F | P61586; P08 | RHOA_HUMAN | Unmodified | 25991000 | 19152000 | 10750000 | 26533000 | 1.23E+08 | 21528000 | 14889000 | 11394000 | 45845000 | 26028000 | 43859000 | 89628000 | 87183000 | 50508000 | 24881000 | 20074000 | 55537000 | 1.1E+08 |
| HFCPNVPII | RHOA; RHOC; F | P61586; P08 | RHOA_HUMAN | Unmodified | 0 | 0 | 0 | 0 | 3654500 | 0 | 0 | 0 | 0 | 0 | 0 | 0 | 242340 | 0 | 0 | 0 | 2459200 | 1770900 |
| HFDWLYAF | SNX18 | Q96RF0 | SNX18_HUMAN | Unmodified | 0 | 0 | 0 | 0 | 0 | 0 | 0 | 0 | 0 | 0 | 0 | 0 | 0 | 5050600 | 0 | 0 | 0 | 0 |
| HFDWLYEF | SNX9; SNX9; SI | Q9Y5X1; A0/ | SNX9_HUMAN | Unmodified | 1771200 | 1161200 | 214560 | 0 | 0 | 0 | 0 | 0 | 313480 | 0 | 0 | 782420 | 0 | 0 | 282590 | 0 | 0 | 0 |
| HFDWLYNI | SNX33; SNX33 | Q8WV41; H3 | SNX33_HUMAN | Unmodified | 0 | 0 | 2437800 | 795640 | 0 | 0 | 0 | 0 | 0 | 0 | 0 | 0 | 0 | 0 | 5031800 | 2361600 | 0 | 0 |
| HFEAKIEK | LRPAP1 | P30533 | AMRP_HUMAN | Unmodified | 0 | 0 | 0 | 0 | 0 | 0 | 0 | 0 | 0 | 353410 | 0 | 0 | 0 | 0 | 0 | 0 | 0 | 0 |
| HFEANNGI | LARS1; LARS1; | Q9P2J5; A0A | SYLC_HUMAN | Unmodified | 0 | 0 | 114540 | 0 | 0 | 0 | 0 | 0 | 0 | 0 | 0 | 3132900 | 0 | 0 | 73745 | 60302 | 0 | 0 |
| HFEATDILV | EIF5B; EIF5B | O60841; A0/ | IF2P_HUMAN E | Unmodified | 101750 | 0 | 93937 | 109550 | 0 | 0 | 0 | 0 | 115290 | 0 | 396200 | 3696400 | 0 | 0 | 0 | 95145 | 445960 | 0 |
| HFELGGDI | RPL36A; RPL36 | P83881; Q9( | RL36A_HUMAN | Unmodified | 0 | 2498600 | 0 | 909370 | 627760 | 149130 | 501980 | 0 | 354490 | 121300 | 777520 | 6895200 | 1189300 | 667540 | 0 | 0 | 157900 | 588570 |
| HFELGGDI | RPL36A; RPL36 | P83881; Q9( | RL36A_HUMAN | Unmodified | 311330 | 0 | 596980 | 0 | 26676 | 0 | 0 | 0 | 0 | 0 | 109250 | 1846200 | 0 | 0 | 0 | 0 | 0 | 0 |
| HFEQAIER | AFG3L2 | Q9Y4W6 | AFG32_HUMAN | Unmodified | 0 | 0 | 0 | 0 | 0 | 0 | 554990 | 0 | 0 | 390300 | 0 | 366780 | 0 | 0 | 0 | 0 | 0 | 0 |
| HFFINICHI | IGF2R; IGF2R; I | P11717; A0A | MPRI_HUMAN | Unmodified | 0 | 0 | 596130 | 229660 | 0 | 0 | 0 | 0 | 10810000 | 1209200 | 902480 | 1323700 | 176170 | 0 | 190700 | 161380 | 0 | 0 |
| HFFINICHI | IGF2R; IGF2R; I | P11717; A0A | MPRI_HUMAN | Unmodified | 0 | 0 | 0 | 0 | 0 | 116520 | 783050 | 0 | 0 | 2509300 | 0 | 0 | 0 | 1958500 | 0 | 0 | 0 | 0 |
| HFHNTLEH | VPS35 | Q96QK1 | VPS35_HUMAN | Unmodified | 59850 | 4089000 | 0 | 0 | 0 | 0 | 0 | 0 | 0 | 54477 | 104800 | 1396700 | 141560 | 0 | 0 | 0 | 0 | 0 |
| HFHPLFEYI | VPS35 | Q96QK1 | VPS35_HUMAN | Unmodified | 53398 | 0 | 95164 | 0 | 0 | 0 | 0 | 0 | 0 | 0 | 0 | 1318800 | 0 | 0 | 0 | 0 | 0 | 0 |
| HFHPLFEYI | VPS35 | Q96QK1 | VPS35_HUMAN | Unmodified | 0 | 0 | 419600 | 201090 | 323760 | 0 | 197800 | 0 | 218700 | 255300 | 0 | 1633000 | 0 | 0 | 0 | 0 | 1827700 | 0 |
| HFILDECDI | DDX39B; DDX3 | Q13838; F6T | DX39B_HUMAN | Unmodified | 0 | 0 | 0 | 0 | 139090 | 0 | 62201 | 0 | 0 | 0 | 0 | 4395300 | 0 | 0 | 0 | 0 | 0 | 0 |
| HFIPYNAK | GULP1; GULP1 | Q9UBP9; B8 | GULP1_HUMAI | Unmodified | 42624 | 48783 | 114160 | 311520 | 109510 | 0 | 0 | 0 | 0 | 0 | 0 | 58488 | 361270 | 0 | 0 | 0 | 0 | 145090 |
| HFLLEEDKI | FASN; FASN | P49327; A0A | FAS_HUMAN F | Unmodified | 2236400 | 983420 | 4067600 | 3966900 | 0 | 0 | 0 | 0 | 1802400 | 0 | 2307900 | 2.25E+08 | 0 | 0 | 12068000 | 8339500 | 0 | 0 |
| HFLLEEDKI | FASN; FASN | P49327; A0A | FAS_HUMAN F | Unmodified | 0 | 0 | 0 | 0 | 0 | 0 | 58217 | 0 | 0 | 0 | 0 | 5423800 | 0 | 0 | 0 | 0 | 0 | 0 |
| HFLVEDK | SEC24C | P53992 | SC24C_HUMAI | Unmodified | 0 | 0 | 0 | 0 | 0 | 0 | 61755 | 0 | 48249 | 106050 | 0 | 245560 | 0 | 0 | 0 | 0 | 0 | 936000 |
| HFNALGG | LTA4H | P09960 | LKHA4_HUMAN | Unmodified | 0 | 0 | 0 | 0 | 1192400 | 949630 | 0 | 0 | 466880 | 0 | 289130 | 4711800 | 0 | 677350 | 0 | 0 | 0 | 0 |
| HFNLLSSK | HTATIP2 | Q9BUP3 | HTAI2_HUMAN | Unmodified | 0 | 0 | 0 | 0 | 0 | 0 | 151310 | 0 | 3306600 | 912230 | 0 | 0 | 0 | 0 | 0 | 0 | 0 | 0 |
| HFNLLSSK | HTATIP2 | Q9BUP3 | HTAI2_HUMAN | Unmodified | 0 | 0 | 0 | 0 | 0 | 0 | 0 | 0 | 0 | 0 | 0 | 0 | 0 | 0 | 0 | 0 | 0 | 0 |
| HFNYDVC( | UTRN; UTRN; U | P46939; H0 | UTRN_HUMAN | Unmodified | 0 | 609240 | 0 | 698160 | 0 | 0 | 0 | 0 | 257650 | 246230 | 0 | 0 | 79185 | 0 | 219640 | 160280 | 73968 | 60931 |
| HFPEFPATI | CCDC50 | Q8IVM0 | CCD50_HUMA | Unmodified | 1101900 | 2244000 | 1606100 | 2587400 | 0 | 0 | 0 | 0 | 0 | 0 | 0 | 0 | 0 | 0 | 1602400 | 0 | 0 | 0 |
| HFPEFPATI | CCDC50 | Q8IVM0 | CCD50_HUMA | Oxidation (M) | 0 | 0 | 0 | 0 | 0 | 0 | 0 | 0 | 0 | 0 | 0 | 0 | 4347600 | 2148900 | 0 | 0 | 0 | 0 |
| HFPEFPATI | CCDC50 | Q8IVM0 | CCD50_HUMA | Unmodified | 0 | 0 | 0 | 0 | 0 | 0 | 0 | 0 | 0 | 0 | 0 | 0 | 997660 | 0 | 0 | 0 | 0 | 0 |
| HFPSVNWI | ATP6V1A | P38606 | VATA_HUMAN | Unmodified | 441220 | 0 | 292160 | 526540 | 828140 | 0 | 1382500 | 0 | 1291300 | 542360 | 398640 | 1743800 | 2370500 | 1038900 | 544280 | 415810 | 610060 | 918680 |
| HFSGLEEA | CCT8; CCT8 | P50990; H7( | TCPQ_HUMAN | Unmodified | 272130 | 188640 | 401530 | 823220 | 643830 | 174640 | 207350 | 0 | 2041300 | 662030 | 2122500 | 16901000 | 2399000 | 492330 | 495150 | 704740 | 340910 | 586780 |
| HFSGLEEA | CCT8; CCT8 | P50990; H7( | TCPQ_HUMAN | Unmodified | 0 | 397960 | 0 | 1227700 | 3551500 | 299480 | 2643400 | 0 | 0 | 1061600 | 0 | 0 | 5442600 | 1217600 | 66336 | 0 | 4945300 | 3612100 |
| HFSVEGQL | HSP90AA2P; H | Q14568; P0 | HS902_HUMAN | Unmodified | 2340400 | 1759200 | 2332800 | 5229800 | 13841000 | 1715600 | 1051700 | 23434 | 15980000 | 10149000 | 16809000 | 93210000 | 5578800 | 3063600 | 5191900 | 4421300 | 3716600 | 4432800 |
| HFSVTAEG | ATP2A3 | Q93084 | AT2A3_HUMAN | Unmodified | 0 | 0 | 0 | 0 | 0 | 0 | 0 | 0 | 0 | 0 | 0 | 1254500 | 0 | 0 | 0 | 0 | 0 | 0 |
| HFTEDIQTI | SRPK1; SRPK2; | Q96SB4; P7 | SRPK1_HUMAN | Unmodified | 0 | 41039 | 0 | 0 | 0 | 18659 | 0 | 0 | 0 | 0 | 78698 | 823410 | 0 | 0 | 0 | 0 | 0 | 0 |
| HFTEFVPLI | ATP6V1A | P38606 | VATA_HUMAN | Unmodified | 54368 | 54025 | 66138 | 50714 | 0 | 0 | 0 | 0 | 119710 | 0 | 152130 | 1529800 | 0 | 0 | 148100 | 53257 | 0 | 0 |
| HFTEFVPLI | ATP6V1A | P38606 | VATA_HUMAN | Unmodified | 67546 | 0 | 0 | 0 | 0 | 0 | 97571 | 0 | 1208000 | 560360 | 0 | 753770 | 0 | 795210 | 0 | 0 | 0 | 0 |
| HFTGYRIEL | INSR | P06213 | INSR_HUMAN I | Unmodified | 104430 | 41202 | 338510 | 235950 | 663600 | 0 | 690400 | 0 | 2483200 | 0 | 0 | 0 | 0 | 0 | 0 | 189100 | 347730 | 0 |
| HFTIDDFEI | AURKB; AURKB | Q96GD4; J3 | AURKB_HUMAI | Unmodified | 0 | 0 | 0 | 1043100 | 0 | 0 | 0 | 0 | 586300 | 0 | 0 | 0 | 0 | 0 | 487100 | 0 | 0 | 0 |
| HFTILDAPC | GSPT1; GSPT2; | P15170; Q8 | ERF3A_HUMAN | Unmodified | 0 | 0 | 0 | 0 | 1588500 | 0 | 378450 | 175400 | 0 | 74731 | 495730 | 5020900 | 3113000 | 933730 | 359840 | 420890 | 990610 | 657300 |
| HFVALSTN | GPI; GPI; GPI; C | P06744; A0A | G6PI_HUMAN | Unmodified | 704870 | 208710 | 867780 | 1032400 | 2177000 | 0 | 301190 | 0 | 3876200 | 1174200 | 3153000 | 30481000 | 2005300 | 708320 | 802450 | 1063300 | 828220 | 503230 |
| HFVALSTN | GPI; GPI; GPI; C | P06744; A0A | G6PI_HUMAN | Unmodified | 0 | 0 | 0 | 0 | 417070 | 0 | 113170 | 93286 | 0 | 0 | 0 | 0 | 104040 | 193210 | 0 | 0 | 243470 | 153390 |
| HFVCEGCI | LMCD1; LMCD1 | Q9NZU5; B4 | LMCD1_HUMA | Unmodified | 46038 | 370810 | 0 | 0 | 1368900 | 3861000 | 854080 | 569130 | 0 | 307260 | 0 | 1048800 | 0 | 581870 | 0 | 0 | 0 | 0 |
| HFVGMLPE | DSTN; DSTN | P60981; F6R | DEST_HUMAN | Unmodified | 518790 | 585400 | 0 | 856480 | 4404300 | 579160 | 408280 | 0 | 1217800 | 399720 | 801650 | 8442500 | 2906800 | 1734100 | 577180 | 322850 | 925790 | 1882600 |
| HFVGMLPE | DSTN; DSTN | P60981; F6R | DEST_HUMAN | Oxidation (M) | 558330 | 395960 | 0 | 352500 | 1815900 | 228150 | 204620 | 0 | 438940 | 182710 | 1668600 | 3381400 | 1724300 | 812770 | 743750 | 0 | 302520 | 589120 |
| HFVGMLPE | DSTN; DSTN | P60981; F6R | DEST_HUMAN | Unmodified | 0 | 0 | 0 | 0 | 0 | 0 | 221630 | 0 | 0 | 0 | 0 | 0 | 1655300 | 0 | 0 | 0 | 1995600 | 557370 |
| HFVGMLPE | DSTN; DSTN | P60981; F6R | DEST_HUMAN | Oxidation (M) | 0 | 0 | 0 | 0 | 1049500 | 682230 | 376570 | 0 | 0 | 0 | 0 | 0 | 879720 | 2049400 | 0 | 102630 | 0 | 0 |
| HFVGYPTN | PTGR1; PTGR1; | Q14914; F6 | PTGR1_HUMAN | Unmodified | 0 | 0 | 0 | 363070 | 8384700 | 713620 | 561240 | 0 | 593220 | 688600 | 0 | 320100 | 0 | 869830 | 0 | 0 | 0 | 0 |
| HFVLDECD | DDX39A; DDX3 | O00148; A0/ | DX39A_HUMAN | Unmodified | 54646 | 0 | 0 | 0 | 0 | 0 | 397060 | 232990 | 0 | 0 | 0 | 264310 | 0 | 0 | 0 | 0 | 0 | 167890 |
| HFYQPIWT | SQOR; SQOR; S | Q9Y6N5; H3 | SQOR_HUMAN | Unmodified | 0 | 0 | 0 | 0 | 0 | 0 | 0 | 0 | 8666000 | 297280 | 0 | 0 | 0 | 0 | 0 | 0 | 0 | 0 |
| HFYSLVK | DHTKD1 | Q96HY7 | DHTK1_HUMAI | Unmodified | 0 | 0 | 0 | 0 | 0 | 0 | 0 | 56249 | 594760 | 0 | 0 | 0 | 0 | 0 | 0 | 0 | 0 | 0 |
| HFYVVIPQI | FNBP1L; FNBP1 | Q5T0N5; A0/ | FBP1L_HUMAN | Unmodified | 1359800 | 2093300 | 0 | 171790 | 665220 | 103150 | 239370 | 0 | 266080 | 132300 | 0 | 0 | 932720 | 518160 | 274460 | 508810 | 293890 | 564540 |
| HFYWYLTN | RPS10; RPS10-I | P46783; A0A | RS10_HUMAN | Unmodified | 0 | 0 | 0 | 0 | 0 | 0 | 0 | 0 | 0 | 0 | 0 | 1639900 | 0 | 0 | 0 | 0 | 0 | 0 |
| HGAGAEIS | PIP4K2C | Q8TBX8 | PI42C_HUMAN | Unmodified | 186140 | 131160 | 214370 | 535500 | 3732700 | 459770 | 752790 | 691190 | 1840000 | 1394600 | 167570 | 996110 | 1418600 | 1664900 | 190190 | 229660 | 2802200 | 4768800 |
| HGAGAEIS | PIP4K2B; PIP4K | P78356; P48 | PI42B_HUMAN | Unmodified | 386110 | 386970 | 568720 | 1281800 | 5666900 | 1078800 | 7074700 | 7140100 | 0 | 496120 | 975050 | 4568400 | 7385000 | 8041100 | 940230 | 760390 | 4363100 | 9334800 |
| HGANPDLI | HECTD1; HECT | Q9ULT8; G3 | HECD1_HUMA | Unmodified | 0 | 0 | 0 | 0 | 705500 | 67131 | 21541 | 0 | 46062 | 24293 | 57165 | 320220 | 0 | 0 | 0 | 152470 | 43016 | 182650 |
| HGANTVPII | JAG1 | P78504 | JAG1_HUMAN I | Unmodified | 0 | 0 | 0 | 0 | 0 | 0 | 0 | 62816 | 0 | 69404 | 0 | 35615 | 1003200 | 884450 | 0 | 0 | 623390 | 1134000 |
| HGATHVFA | SMARCA5 | O60264 | SMCA5_HUMA | Unmodified | 398580 | 0 | 0 | 0 | 0 | 0 | 0 | 0 | 0 | 0 | 0 | 0 | 0 | 0 | 0 | 0 | 0 | 0 |
| HGCAFLVD | ABAT; ABAT; AB | P80404; H3E | GABT_HUMAN | Unmodified | 0 | 0 | 0 | 0 | 0 | 0 | 0 | 0 | 1011200 | 0 | 0 | 0 | 0 | 0 | 0 | 0 | 0 | 0 |
| HGDEIYIAP | APIP | Q96GX9 | MTNB_HUMAN | Unmodified | 0 | 0 | 0 | 0 | 0 | 0 | 262120 | 129560 | 0 | 0 | 549720 | 689910 | 113180 | 0 | 0 | 0 | 0 | 0 |
| HGDGTTLD | NPEPPS; NPEPI | P55786; A0A | PSA_HUMAN P | Unmodified | 0 | 0 | 0 | 0 | 183840 | 0 | 0 | 0 | 301850 | 0 | 0 | 794770 | 0 | 0 | 0 | 0 | 308520 | 87170 |
| HGDLHSFI | AXL; AXL | P30530; M0I | UFO_HUMAN T | Unmodified | 0 | 0 | 0 | 0 | 347970 | 159430 | 0 | 0 | 421500 | 0 | 0 | 0 | 0 | 0 | 0 | 119980 | 0 | 0 |
| HGDLRNFI | MET | P08581 | MET_HUMAN H | Unmodified | 0 | 0 | 0 | 0 | 1330500 | 0 | 0 | 0 | 0 | 165930 | 0 | 0 | 0 | 0 | 0 | 0 | 748020 | 1552500 |
| HGDMVQL | POMT1; POMT1 | Q9Y6A1; A0/ | POMT1_HUMAI | Unmodified | 0 | 0 | 0 | 0 | 0 | 0 | 0 | 0 | 2528600 | 0 | 0 | 0 | 0 | 0 | 0 | 0 | 0 | 0 |
| HGDPGDA | PDCD5; PDCD5 | O14737; X6F | PDCD5_HUMA | Unmodified | 0 | 0 | 436550 | 0 | 223830 | 71173 | 36523 | 0 | 84519 | 0 | 31542 | 319520 | 0 | 0 | 0 | 472140 | 22744 | 132460 |
| HGDRSPID | ACP3; ACP3 | P15309; E9P | PPAP_HUMAN | Unmodified | 685330 | 136510 | 505940 | 70650 | 0 | 0 | 0 | 0 | 5672100 | 0 | 0 | 0 | 0 | 0 | 203470 | 373580 | 22875 | 0 |
| HGDRVILG | KIF14 | Q15058 | KIF14_HUMAN | Unmodified | 0 | 0 | 0 | 0 | 719170 | 0 | 0 | 0 | 0 | 0 | 0 | 0 | 0 | 0 | 0 | 0 | 0 | 0 |
| HGEIDYEAI | PSMC6; PSMC6 | P62333; A0A | PRS10_HUMAN | Unmodified | 0 | 0 | 0 | 597930 | 1659800 | 0 | 122590 | 198280 | 0 | 0 | 529950 | 4371700 | 924060 | 0 | 0 | 397630 | 0 | 851310 |
| HGEMWAIS | ERBIN | Q96RT1 | ERBIN_HUMAN | Oxidation (M) | 0 | 0 | 0 | 0 | 1046900 | 0 | 0 | 0 | 0 | 190300 | 0 | 0 | 0 | 0 | 0 | 0 | 233560 | 0 |
| HGESAWN | PGAM1 | P18669 | PGAM1_HUMA | Unmodified | 719080 | 537650 | 1495900 | 1635400 | 0 | 0 | 0 | 0 | 1099100 | 0 | 1339000 | 48465000 | 0 | 0 | 2676600 | 2735200 | 0 | 0 |
| HGESAWN | PGAM1 | P18669 | PGAM1_HUMA | Unmodified | 0 | 0 | 0 | 880080 | 6382100 | 1659500 | 87177 | 28019 | 1368800 | 0 | 3097100 | 0 | 1304000 | 967270 | 0 | 0 | 477580 | 1532500 |
| HGESGSM | PVR; PVR; PVR | P15151; A0A | PVR_HUMAN P | Oxidation (M) | 1842100 | 4972700 | 30494 | 761970 | 627790 | 71387 | 0 | 0 | 128160 | 34962 | 762570 | 5073200 | 0 | 0 | 824280 | 207140 | 0 | 0 |
| HGESGSM | PVR; PVR; PVR | P15151; A0A | PVR_HUMAN P | Unmodified | 3113700 | 2713600 | 1836700 | 1402400 | 1164700 | 426340 | 0 | 0 | 786530 | 147930 | 4222600 | 20823000 | 343150 | 108220 | 1665700 | 1148600 | 66702 | 33863 |
| HGESVLVL | EXT2 | Q93063 | EXT2_HUMAN I | Unmodified | 0 | 0 | 0 | 0 | 0 | 0 | 0 | 0 | 1466100 | 101460 | 79732 | 0 | 0 | 0 | 0 | 0 | 0 | 0 |
| HGEVCPA( | PRDX4 | Q13162 | PRDX4_HUMAN | Unmodified | 0 | 0 | 0 | 0 | 0 | 0 | 0 | 0 | 31201000 | 5079700 | 5061800 | 801780 | 0 | 0 | 0 | 0 | 0 | 558360 |
| HGEVCPA( | PRDX4 | Q13162 | PRDX4_HUMAN | Unmodified | 0 | 0 | 0 | 0 | 0 | 0 | 0 | 0 | 0 | 1066200 | 0 | 0 | 0 | 0 | 0 | 0 | 0 | 0 |

| | | | | | | | | | | | | | | | | | | | | | | |
|---|---|---|---|---|---|---|---|---|---|---|---|---|---|---|---|---|---|---|---|---|---|---|
| HGGEDYVI | CYC1 | P08574 | CY1_HUMAN C | Unmodified | 0 | 0 | 0 | 0 | 0 | 0 | 0 | 0 | 6476800 | 329510 | 0 | 0 | 0 | 0 | 0 | 0 | 0 | 0 |
| HGGKAPLF | FLNA; FLNA; FL | P21333; A0A | FLNA_HUMAN | Unmodified | 0 | 0 | 0 | 0 | 13263000 | 2108700 | 0 | 0 | 0 | 676100 | 0 | 0 | 4672400 | 0 | 0 | 0 | 6969200 | 1150900 |
| HGGPGPG | PPP1R13L | Q8WUF5 | IASPP_HUMAN | Unmodified | 417880 | 0 | 0 | 0 | 0 | 0 | 0 | 0 | 0 | 0 | 0 | 0 | 0 | 0 | 0 | 32079 | 0 | 0 |
| HGGSLGL( | PSMD1; PSMD1 | Q99460; A0A | PSMD1_HUMA | Unmodified | 239460 | 382740 | 0 | 0 | 0 | 0 | 474240 | 264520 | 0 | 0 | 160480 | 907520 | 711040 | 0 | 0 | 173460 | 340950 | 0 |
| HGGTCHN | NOTCH1; NOT( | P46531; A0A | NOTC1_HUMA | Unmodified | 44838 | 30757 | 0 | 135270 | 0 | 0 | 0 | 0 | 0 | 0 | 21143 | 62815 | 0 | 0 | 91168 | 54009 | 0 | 0 |
| HGGYKPSI | CKB; CKB; CKB | P12277; G3\ | KCRB_HUMAN | Unmodified | 6427000 | 4273000 | 1982000 | 69414000 | 0 | 0 | 2651200 | 0 | 59337000 | 0 | 0 | 1.38E+08 | 5598900 | 0 | 51711000 | 30722000 | 0 | 0 |
| HGILQFLHI | GCLC; GCLC; G | P48506; A0A | GSH1_HUMAN | Unmodified | 0 | 0 | 0 | 0 | 2440800 | 718070 | 0 | 0 | 0 | 17632 | 160630 | 1379900 | 306090 | 127130 | 0 | 0 | 55061 | 140680 |
| HGINCFINI | CCT2; CCT2; C( | P78371; F5G | TCPB_HUMAN | Unmodified | 0 | 0 | 268830 | 0 | 0 | 0 | 0 | 152160 | 0 | 0 | 365210 | 4200100 | 0 | 0 | 0 | 0 | 1574400 | 2068000 |
| HGIPTAQW | GART; GART; G/ | P22102; C9J | PUR2_HUMAN | Unmodified | 0 | 0 | 0 | 0 | 0 | 0 | 0 | 0 | 0 | 0 | 0 | 1491900 | 0 | 0 | 0 | 0 | 0 | 0 |
| HGIVEDWI | ACTR3; ACTR3; | P61158; B4D | ARP3_HUMAN | Oxidation (M) | 99585 | 324090 | 0 | 0 | 0 | 0 | 44831 | 48829 | 2365800 | 1486100 | 286520 | 6060500 | 40441 | 0 | 515050 | 583810 | 0 | 0 |
| HGIVEDWI | ACTR3; ACTR3; | P61158; B4D | ARP3_HUMAN | Unmodified | 127340 | 0 | 0 | 0 | 0 | 0 | 0 | 0 | 0 | 0 | 66550 | 1350600 | 0 | 0 | 241460 | 0 | 0 | 0 |
| HGKLQDAI | LRPPRC | P42704 | LPPRC_HUMAN | Unmodified | 0 | 0 | 0 | 0 | 0 | 0 | 0 | 0 | 0 | 0 | 0 | 0 | 0 | 0 | 0 | 0 | 0 | 0 |
| HGKNPVMI | ILF3 | Q12906 | ILF3_HUMAN Ir | Oxidation (M) | 398560 | 0 | 0 | 0 | 0 | 0 | 0 | 0 | 340050 | 19090 | 0 | 0 | 0 | 0 | 0 | 126070 | 0 | 0 |
| HGKYMAC( | TUBA1A; TUBA4 | Q71U36; P68 | TBA1A_HUMAN | Unmodified | 0 | 0 | 0 | 133270 | 1103000 | 225140 | 2407500 | 0 | 1013000 | 672760 | 0 | 0 | 798530 | 1091700 | 0 | 0 | 1071700 | 1117900 |
| HGKYMAC( | TUBA1A; TUBA4 | Q71U36; P68 | TBA1A_HUMAN | Oxidation (M) | 0 | 0 | 0 | 0 | 0 | 0 | 2175300 | 0 | 0 | 0 | 0 | 0 | 4414200 | 1179500 | 0 | 0 | 0 | 0 |
| HGLANLTA | ANKFY1 | Q9P2R3 | ANFY1_HUMAN | Unmodified | 0 | 0 | 0 | 0 | 2374700 | 0 | 0 | 0 | 269690 | 0 | 0 | 0 | 0 | 0 | 0 | 0 | 0 | 0 |
| HGLEAISIM | VARS1; VARS1 | P26640; A0A | SYVC_HUMAN | Unmodified | 0 | 141050 | 0 | 0 | 228330 | 0 | 122520 | 0 | 267370 | 0 | 0 | 2707200 | 0 | 0 | 149820 | 144150 | 0 | 0 |
| HGLEVIYMI | HSP90AA1 | P07900 | HS90A_HUMAN | Oxidation (M) | 20099000 | 0 | 0 | 0 | 0 | 0 | 0 | 0 | 1117000 | 4640300 | 1547200 | 2320300 | 0 | 0 | 0 | 0 | 4015300 | 2927200 |
| HGLEVIYMI | HSP90AA1 | P07900 | HS90A_HUMAN | Unmodified | 0 | 0 | 0 | 0 | 2670000 | 0 | 0 | 0 | 987160 | 1174200 | 0 | 0 | 0 | 0 | 0 | 0 | 0 | 1134000 |
| HGLEVSEM | SF3B3; SF3B3 | Q15393; J3Q | SF3B3_HUMAN | Oxidation (M) | 0 | 0 | 0 | 0 | 222330 | 0 | 0 | 0 | 0 | 0 | 0 | 3330600 | 0 | 0 | 0 | 0 | 0 | 164850 |
| HGLGYSLL | MOV10; MOV10 | Q9HCE1; Q5 | MOV10_HUMA | Unmodified | 1549900 | 1212500 | 0 | 1507200 | 0 | 0 | 0 | 0 | 491060 | 0 | 0 | 0 | 0 | 0 | 769420 | 1240900 | 0 | 0 |
| HGLIPNLL( | AGL | P35573 | GDE_HUMAN ( | Unmodified | 0 | 0 | 0 | 0 | 0 | 0 | 0 | 0 | 0 | 0 | 0 | 572690 | 0 | 0 | 0 | 0 | 0 | 0 |
| HGLLVPNN | EIF6; EIF6 | P56537; B7Z | IF6_HUMAN Eu | Unmodified | 3383600 | 4081400 | 1759600 | 1505300 | 0 | 0 | 0 | 26994 | 1131200 | 324680 | 682010 | 10614000 | 0 | 0 | 2804900 | 2327000 | 0 | 0 |
| HGLRALRE | PSMA1; PSMA1 | P25786; F5G | PSA1_HUMAN | Unmodified | 0 | 0 | 0 | 0 | 5203500 | 164800 | 1503200 | 0 | 0 | 3259500 | 0 | 0 | 3248700 | 3623900 | 0 | 0 | 3787600 | 3173500 |
| HGLSNSIY | KIT | P10721 | KIT_HUMAN Ma | Unmodified | 0 | 0 | 0 | 0 | 0 | 0 | 478820 | 27032 | 1019800 | 0 | 0 | 0 | 0 | 0 | 0 | 0 | 0 | 0 |
| HGLSPLEV | RGS14; RGS14 | O43566; H0Y | RGS14_HUMAN | Unmodified | 0 | 0 | 0 | 0 | 0 | 0 | 0 | 0 | 0 | 0 | 215850 | 2047500 | 0 | 0 | 0 | 0 | 0 | 0 |
| HGLYLPTR | FASN; FASN | P49327; A0A | FAS_HUMAN F | Unmodified | 109480 | 0 | 563520 | 227920 | 0 | 0 | 257900 | 0 | 179500 | 0 | 286900 | 12691000 | 0 | 0 | 919170 | 794400 | 0 | 0 |
| HGLYLPTR | FASN; FASN | P49327; A0A | FAS_HUMAN F | Unmodified | 119910 | 0 | 0 | 1363700 | 556900 | 184930 | 413030 | 51268 | 374580 | 0 | 2974900 | 7349600 | 0 | 0 | 985480 | 305960 | 0 | 0 |
| HGLYNLK | IGFBP2; IGFBP2 | P18065; C9J | IBP2_HUMAN I | Unmodified | 0 | 0 | 0 | 0 | 0 | 787530 | 0 | 0 | 44538000 | 21584 | 0 | 0 | 0 | 0 | 0 | 0 | 0 | 0 |
| HGNQYIQ\ | MARS1 | P56192 | SYMC_HUMAN | Unmodified | 0 | 0 | 0 | 0 | 0 | 0 | 0 | 0 | 109990 | 0 | 0 | 2391200 | 0 | 0 | 0 | 0 | 0 | 0 |
| HGNRGET( | COL1A2; COL1/ | P08123; A0A | CO1A2_HUMAI | Unmodified | 0 | 0 | 0 | 0 | 0 | 0 | 0 | 0 | 975310 | 0 | 0 | 0 | 0 | 0 | 0 | 0 | 0 | 0 |
| HGNVASL\ | CD2AP | Q9Y5K6 | CD2AP_HUMAI | Unmodified | 3257300 | 5261200 | 0 | 4030200 | 412830 | 0 | 0 | 0 | 0 | 0 | 0 | 0 | 0 | 0 | 4958100 | 3354800 | 0 | 0 |
| HGNVASL\ | CD2AP | Q9Y5K6 | CD2AP_HUMAI | Unmodified | 0 | 0 | 0 | 0 | 2150800 | 0 | 0 | 0 | 0 | 0 | 0 | 0 | 2919200 | 1719200 | 0 | 0 | 0 | 0 |
| HGPAVADL | BTBD17 | A6NE02 | BTBDH_HUMAI | Unmodified | 0 | 0 | 0 | 0 | 0 | 0 | 0 | 0 | 5721900 | 0 | 0 | 0 | 0 | 0 | 0 | 0 | 0 | 0 |
| HGPINVNI | NINJ1 | Q92982 | NINJ1_HUMAN | Unmodified | 221430 | 0 | 1600500 | 971510 | 371890 | 0 | 0 | 0 | 437520 | 0 | 0 | 396550 | 246020 | 0 | 4117200 | 2098900 | 66329 | 0 |
| HGPVYEDI | APEH; APEH; AI | P13798; C9J | ACPH_HUMAN | Unmodified | 0 | 0 | 0 | 0 | 0 | 0 | 0 | 0 | 0 | 0 | 0 | 0 | 963380 | 0 | 0 | 0 | 948090 | 0 |
| HGRGQVLI | EPB41L1; EPB4 | Q9H4G0; Q4 | E41L1_HUMAN | Unmodified | 0 | 0 | 107470 | 407850 | 2791800 | 431020 | 0 | 0 | 117330 | 0 | 899430 | 0 | 700140 | 0 | 616080 | 0 | 0 | 643130 |
| HGSLGFLP | RPL3; RPL3; RP | P39023; H7( | RL3_HUMAN 6( | Unmodified | 517460 | 164910 | 160140 | 175710 | 0 | 0 | 0 | 0 | 0 | 0 | 0 | 2904100 | 0 | 0 | 550140 | 416510 | 0 | 0 |
| HGSNIEAM | ACAT2 | Q9BWD1 | THIC_HUMAN / | Unmodified | 0 | 0 | 93250 | 88741 | 0 | 0 | 0 | 0 | 1191800 | 68526 | 208960 | 7792000 | 0 | 0 | 0 | 0 | 0 | 0 |
| HGSYEDA\ | TCP1; TCP1; TC | P17987; F5H | TCPA_HUMAN | Unmodified | 338570 | 0 | 1592400 | 493080 | 772790 | 217380 | 0 | 0 | 2534900 | 185210 | 4319100 | 20079000 | 1570500 | 156510 | 937590 | 1185100 | 274290 | 518200 |
| HGTALQRS | OSBP | P22059 | OSBP1_HUMAI | Unmodified | 0 | 0 | 0 | 0 | 4061900 | 0 | 1684900 | 0 | 0 | 130470 | 0 | 0 | 2764100 | 3283100 | 0 | 0 | 90773 | 650530 |
| HGTCAAQ\ | RNASET2; N/A; | O00584; H0Y | RNT2_HUMAN | Unmodified | 0 | 0 | 92126 | 0 | 0 | 0 | 0 | 0 | 939910 | 313070 | 391350 | 87886 | 531620 | 590740 | 72218 | 0 | 26385 | 48385 |
| HGTCAAQ\ | RNASET2; N/A; | O00584; H0Y | RNT2_HUMAN | Unmodified | 0 | 0 | 0 | 0 | 0 | 0 | 0 | 0 | 1656100 | 774480 | 416430 | 0 | 2175700 | 0 | 83701 | 0 | 0 | 282040 |
| HGTDDGV | FGB; FGB | P02675; D6F | FIBB_HUMAN F | Oxidation (M) | 0 | 1211000 | 0 | 0 | 0 | 0 | 0 | 0 | 0 | 0 | 0 | 0 | 0 | 395350 | 0 | 0 | 0 | 0 |
| HGTEVATA | RIPOR2; RIPOR | Q9Y4F9; F5C | RIPR2_HUMAN | Unmodified | 0 | 0 | 0 | 0 | 0 | 0 | 0 | 0 | 0 | 0 | 325840 | 1042100 | 83045 | 38108 | 0 | 24718 | 0 | 0 |
| HGTPQKYI | SLC44A2; SLC4 | Q8IWA5; K7I | CTL2_HUMAN | Unmodified | 710480 | 585290 | 400630 | 172810 | 0 | 12973 | 36023 | 0 | 187100 | 50355 | 0 | 1526600 | 0 | 0 | 972390 | 672360 | 13469 | 46472 |
| HGVDTHLF | SNTA1 | Q13424 | SNTA1_HUMAN | Unmodified | 0 | 0 | 0 | 0 | 0 | 0 | 0 | 0 | 0 | 0 | 0 | 0 | 0 | 0 | 4039500 | 826400 | 0 | 0 |
| HGVDVEV( | MOV10; MOV10 | Q9HCE1; Q5 | MOV10_HUMA | Unmodified | 2463200 | 2746500 | 0 | 0 | 0 | 0 | 0 | 0 | 0 | 0 | 0 | 195230 | 0 | 0 | 2123500 | 2004900 | 0 | 0 |
| HGVDVEV( | MOV10; MOV10 | Q9HCE1; Q5 | MOV10_HUMA | Unmodified | 2985300 | 2174300 | 0 | 0 | 0 | 0 | 0 | 0 | 0 | 0 | 0 | 0 | 174560 | 0 | 0 | 896190 | 0 | 0 |
| HGVEAKNY | TMED10 | P49755 | TMEDA_HUMAI | Unmodified | 0 | 0 | 0 | 0 | 0 | 0 | 82727 | 15861 | 279360 | 7176000 | 0 | 0 | 0 | 0 | 0 | 0 | 0 | 0 |
| HGVFVGS/ | PCK2; PCK2 | Q16822; H0Y | PCKGM_HUMA | Unmodified | 0 | 0 | 0 | 0 | 0 | 0 | 0 | 0 | 2288200 | 0 | 0 | 0 | 0 | 0 | 0 | 0 | 0 | 0 |
| HGVGRIDI\ | ASS1 | P00966 | ASSY_HUMAN | Unmodified | 0 | 0 | 212110 | 237990 | 0 | 17960 | 0 | 0 | 412710 | 0 | 22571 | 0 | 0 | 199590 | 273080 | 0 | 75256 | 0 |
| HGVGRIDI\ | ASS1 | P00966 | ASSY_HUMAN | Unmodified | 0 | 0 | 0 | 0 | 289150 | 234260 | 855740 | 0 | 173260 | 317420 | 0 | 0 | 780100 | 638450 | 0 | 0 | 69427 | 742050 |
| HGVGRIDI\ | ASS1 | P00966 | ASSY_HUMAN | Oxidation (M) | 0 | 0 | 0 | 0 | 0 | 0 | 522100 | 49601 | 0 | 950020 | 0 | 0 | 687590 | 466410 | 0 | 0 | 0 | 0 |
| HGVIVAAD | PSMB5; PSMB5 | P28074; H0Y | PSB5_HUMAN | Unmodified | 65808 | 0 | 76936 | 81929 | 0 | 0 | 0 | 0 | 0 | 0 | 0 | 830840 | 0 | 0 | 92657 | 118390 | 0 | 0 |
| HGVPISVT( | FLOT1; FLOT1; | O75955; A2/ | FLOT1_HUMAN | Unmodified | 0 | 0 | 2659900 | 2205300 | 3521300 | 0 | 242690 | 0 | 6387600 | 1896100 | 1500500 | 8238500 | 727230 | 0 | 5275100 | 3104900 | 0 | 1959600 |
| HGVPPPCI | PTPN23 | Q9H3S7 | PTN23_HUMAN | Unmodified | 287640 | 170020 | 0 | 0 | 213340 | 156540 | 0 | 0 | 0 | 0 | 0 | 0 | 0 | 0 | 0 | 52430 | 0 | 0 |
| HGVQECIL | ARHGEF2; ARH | Q92974; V9( | ARHG2_HUMA | Unmodified | 0 | 0 | 0 | 0 | 0 | 0 | 0 | 7157.9 | 334000 | 0 | 0 | 1874100 | 0 | 0 | 0 | 0 | 0 | 0 |
| HGVSHKVI | GARS1; GARS1 | P41250; A0A | GARS_HUMAN | Unmodified | 55442 | 34451 | 49698 | 0 | 0 | 0 | 0 | 0 | 0 | 100310 | 204800 | 1876600 | 451810 | 130090 | 116570 | 116000 | 0 | 0 |
| HGVSHKVI | GARS1; GARS1 | P41250; A0A | GARS_HUMAN | Unmodified | 0 | 0 | 0 | 748170 | 0 | 64183 | 108380 | 0 | 1673200 | 637680 | 975610 | 9366300 | 876220 | 0 | 388780 | 663370 | 0 | 77850 |
| HGVVPLAT | RPL21; RPL21; | P46778; G3\ | RL21_HUMAN ( | Oxidation (M) | 473330 | 403010 | 295720 | 237230 | 0 | 0 | 0 | 0 | 0 | 0 | 169580 | 1616500 | 0 | 0 | 219900 | 327320 | 0 | 0 |
| HGVVPLAT | RPL21; RPL21; | P46778; G3\ | RL21_HUMAN ( | Unmodified | 713330 | 778380 | 213390 | 652910 | 362860 | 0 | 0 | 0 | 0 | 0 | 309920 | 7333600 | 170000 | 0 | 574480 | 381920 | 0 | 367310 |
| HGVYNPNI | MDH2; MDH2 | P40926; G3X | MDHM_HUMAN | Unmodified | 0 | 1582700 | 0 | 0 | 0 | 92300 | 0 | 192190 | 2404100 | 370550 | 0 | 0 | 399380 | 0 | 3855900 | 0 | 34842 | 49993 |
| HGVYNPNI | MDH2; MDH2 | P40926; G3X | MDHM_HUMAN | Unmodified | 0 | 0 | 0 | 0 | 0 | 0 | 3551200 | 0 | 0 | 0 | 0 | 0 | 0 | 0 | 192880 | 0 | 0 | 0 |
| HGWEDVTI | STEAP1 | Q9UHE8 | STEA1_HUMAN | Unmodified | 519400 | 1326600 | 7404800 | 3910600 | 0 | 0 | 0 | 0 | 504220 | 0 | 0 | 0 | 0 | 0 | 16248000 | 5179300 | 0 | 0 |
| HGYENPTY | APLP1; APLP1; | P51693; B7Z | APLP1_HUMAN | Unmodified | 0 | 0 | 0 | 0 | 0 | 0 | 0 | 0 | 1789800 | 66207 | 0 | 0 | 0 | 0 | 0 | 0 | 0 | 26780 |
| HGYFCFHE | NIBAN1 | Q9BZQ8 | NIBA1_HUMAN | Unmodified | 0 | 0 | 0 | 274950 | 116070 | 102310 | 0 | 0 | 1538700 | 429610 | 0 | 0 | 0 | 0 | 0 | 164440 | 0 | 0 |
| HGYFCFHE | NIBAN1 | Q9BZQ8 | NIBA1_HUMAN | Unmodified | 0 | 0 | 0 | 372460 | 0 | 0 | 0 | 0 | 915620 | 0 | 0 | 0 | 0 | 0 | 0 | 0 | 0 | 0 |
| HGYIGEFEI | RPS15A; N/A; R | P62244; H3E | RS15A_HUMAN | Unmodified | 2882000 | 898060 | 465950 | 0 | 110490 | 0 | 360920 | 0 | 0 | 0 | 0 | 47426000 | 0 | 0 | 1096000 | 1010200 | 0 | 0 |
| HGYIGEFEI | RPS15A; N/A; R | P62244; H3E | RS15A_HUMAN | Unmodified | 2482400 | 1944400 | 1242800 | 1027200 | 960290 | 0 | 4032900 | 324510 | 825020 | 0 | 2970100 | 37299000 | 407160 | 727420 | 2086800 | 2361600 | 0 | 0 |
| HGYIPGKN | CORO1C; COR( | Q9ULV4; H0 | COR1C_HUMA | Unmodified | 0 | 0 | 0 | 44301 | 0 | 0 | 0 | 0 | 0 | 0 | 0 | 0 | 0 | 202130 | 0 | 0 | 1046300 | 0 |
| HGYTSSLE | SEMA4C | Q9C0C4 | SEM4C_HUMAI | Unmodified | 149130 | 486020 | 0 | 0 | 0 | 0 | 0 | 0 | 4403700 | 0 | 0 | 43354 | 0 | 0 | 284040 | 189400 | 0 | 259300 |
| HHAAYVNI | SOD2; SOD2; S | P04179; G8J | SODM_HUMAN | Unmodified | 66109 | 81688 | 40698 | 0 | 339250 | 75828 | 78646 | 46455 | 15137000 | 1341000 | 528650 | 0 | 237760 | 240290 | 43587 | 0 | 151560 | 114030 |
| HHCPNTPI | RAC1 | P63000 | RAC1_HUMAN | Unmodified | 6783100 | 6616900 | 10883000 | 3610100 | 350820 | 0 | 391890 | 56873 | 964420 | 143970 | 934730 | 18872000 | 620070 | 409560 | 8880300 | 7450400 | 322490 | 0 |
| HHEASIRLI | NUP210 | Q8TEM1 | PO210_HUMAN | Unmodified | 0 | 0 | 0 | 0 | 0 | 0 | 0 | 0 | 8140900 | 0 | 0 | 0 | 0 | 3653200 | 0 | 0 | 0 | 397660 |
| HHEASIRLI | NUP210 | Q8TEM1 | PO210_HUMAN | Oxidation (M) | 0 | 0 | 0 | 0 | 0 | 0 | 0 | 0 | 3709900 | 48672 | 0 | 0 | 0 | 35160 | 0 | 0 | 0 | 0 |
| HHEGFTN\ | FUCA1 | P04066 | FUCO_HUMAN | Unmodified | 0 | 0 | 0 | 0 | 716610 | 0 | 0 | 0 | 1221400 | 0 | 0 | 0 | 0 | 0 | 0 | 0 | 0 | 0 |
| HHGPQTLY | GANAB; GANAE | Q14697; F5H | GANAB_HUMA | Unmodified | 0 | 0 | 1368100 | 1117200 | 0 | 0 | 675680 | 0 | 5.62E+08 | 82287000 | 6102200 | 7715100 | 0 | 0 | 2101900 | 1113800 | 0 | 0 |
| HHGPQTLY | GANAB; GANAE | Q14697; F5H | GANAB_HUMA | Unmodified | 0 | 0 | 0 | 0 | 0 | 0 | 0 | 0 | 1170100 | 4225500 | 0 | 0 | 0 | 0 | 0 | 0 | 0 | 0 |
| HHGQEGS | GMPPB; GMPPI | Q9Y5P6; A0/ | GMPPB_HUMA | Unmodified | 50697 | 0 | 135290 | 0 | 0 | 0 | 0 | 0 | 0 | 0 | 0 | 943380 | 0 | 0 | 0 | 104890 | 0 | 0 |
| HHHTQIAS | GGT1; GGT2; G | P19440; P36 | GGT1_HUMAN | Unmodified | 0 | 0 | 0 | 396550 | 0 | 0 | 0 | 0 | 293820 | 0 | 0 | 0 | 0 | 0 | 2233600 | 655180 | 0 | 0 |
| HHISIAEIYI | NAPA; NAPA; N. | P54920; M0I | SNAA_HUMAN | Unmodified | 0 | 0 | 0 | 0 | 0 | 0 | 1084200 | 0 | 572280 | 724440 | 0 | 477430 | 0 | 2432100 | 0 | 0 | 729450 | 1057700 |
| HHITPLLSA | MTPN | P58546 | MTPN_HUMAN | Unmodified | 0 | 0 | 0 | 809380 | 1072700 | 254980 | 511230 | 0 | 753670 | 118400 | 0 | 19729000 | 2696800 | 0 | 3851200 | 1220900 | 503550 | 1483800 |
| HHLGDDV | GLB1; GLB1; GI | P16278; C9J | BGAL_HUMAN | Unmodified | 0 | 0 | 0 | 0 | 0 | 0 | 0 | 0 | 1124500 | 0 | 0 | 527060 | 0 | 0 | 0 | 0 | 0 | 0 |
| HHLGLEEP | IGFBP2; IGFBP2 | P18065; C9J | IBP2_HUMAN I | Unmodified | 0 | 0 | 0 | 0 | 0 | 0 | 61409 | 51198 | 1070100 | 381680 | 0 | 0 | 0 | 0 | 0 | 0 | 0 | 0 |
| HHLGLEEP | IGFBP2; IGFBP2 | P18065; C9J | IBP2_HUMAN I | Unmodified | 0 | 0 | 0 | 0 | 0 | 0 | 0 | 0 | 5408400 | 1671200 | 0 | 0 | 0 | 69977 | 56000 | 0 | 0 | 0 |
| HHNQSTAI | EIF3A | Q14152 | EIF3A_HUMAN | Oxidation (M) | 302290 | 351480 | 0 | 0 | 0 | 0 | 0 | 0 | 0 | 0 | 0 | 6390600 | 0 | 0 | 209480 | 361590 | 0 | 0 |
| HHNQSTAI | EIF3A | Q14152 | EIF3A_HUMAN | Unmodified | 0 | 0 | 0 | 0 | 100650 | 1359700 | 0 | 0 | 0 | 0 | 0 | 8845800 | 0 | 0 | 0 | 0 | 0 | 0 |
| HHPHFLTH | MYO1C; MYO1( | O00159; F5H | MYO1C_HUMA | Unmodified | 0 | 0 | 0 | 259110 | 1454000 | 168090 | 0 | 0 | 0 | 0 | 0 | 0 | 0 | 0 | 369840 | 0 | 0 | 83123 |
| HHPTTLAV( | CD70 | P32970 | CD70_HUMAN | Unmodified | 0 | 0 | 0 | 0 | 2186600 | 0 | 0 | 0 | 0 | 0 | 0 | 0 | 0 | 0 | 0 | 0 | 0 | 0 |
| HHQASETH | SLC39A9 | Q9NUM3 | S39A9_HUMAN | Unmodified | 0 | 0 | 0 | 0 | 0 | 0 | 0 | 0 | 4931300 | 294570 | 161270 | 692260 | 0 | 0 | 0 | 0 | 0 | 0 |
| HHQASETH | SLC39A9 | Q9NUM3 | S39A9_HUMAN | Unmodified | 0 | 0 | 0 | 0 | 0 | 0 | 0 | 0 | 224300 | 307150 | 0 | 0 | 0 | 0 | 0 | 0 | 0 | 0 |
| HHSDGSV | TAPBP; TAPBP; | O15533; A0/ | TPSN_HUMAN | Unmodified | 0 | 0 | 0 | 0 | 0 | 0 | 0 | 0 | 33028000 | 2030200 | 2395400 | 1049800 | 0 | 0 | 0 | 0 | 0 | 0 |
| HHSLGGQ | PDIA6 | Q15084 | PDIA6_HUMAN | Unmodified | 0 | 0 | 1103400 | 0 | 811350 | 0 | 565010 | 0 | 1.1E+08 | 12871000 | 13399000 | 5825900 | 0 | 317790 | 1666300 | 0 | 0 | 0 |
| HHVLGTITI | NOMO1; NOMC | Q15155; P69 | NOMO1_HUM/ | Unmodified | 0 | 0 | 0 | 0 | 0 | 0 | 0 | 0 | 2148500 | 0 | 56736 | 44170 | 0 | 0 | 0 | 0 | 0 | 0 |
| HHVLHDQI | OGDH; OGDH; | Q02218; E9F | ODO1_HUMAN | Unmodified | 0 | 0 | 0 | 0 | 0 | 0 | 0 | 0 | 620970 | 0 | 0 | 0 | 0 | 0 | 0 | 0 | 0 | 0 |
| HHVLQND | DST; DST; DST; | Q03001; A0/ | DYST_HUMAN | Unmodified | 0 | 0 | 0 | 0 | 500290 | 271420 | 0 | 0 | 221460 | 148560 | 0 | 731920 | 0 | 0 | 0 | 0 | 112140 | 0 |
| HIADLAGN | ENO3; ENO3; E | P13929; K7E | ENOB_HUMAN | Unmodified | 0 | 0 | 0 | 0 | 0 | 0 | 0 | 0 | 1747500 | 623040 | 295650 | 2696300 | 0 | 0 | 0 | 0 | 0 | 275940 |
| HIADLAGN | ENO1; ENO1; E | P06733; K7E | ENOA_HUMAN | Unmodified | 8261800 | 0 | 4130900 | 32251000 | 1.57E+08 | 30136000 | 2430800 | 0 | 1.37E+08 | 39774000 | 46936000 | 2.6E+08 | 91048000 | 19776000 | 20713000 | 17000000 | 12575000 | 54578000 |
| HIAEDADRI | TPM1; TPM1; TF | P09493; H0Y | TPM1_HUMAN | Unmodified | 0 | 0 | 0 | 0 | 6068700 | 1109700 | 52487 | 0 | 0 | 267940 | 0 | 0 | 518260 | 719520 | 0 | 0 | 103660 | 0 |
| HIAEEADRI | TPM4; TPM3; TF | P67936; P06 | TPM4_HUMAN | Unmodified | 899180 | 942530 | 0 | 2341900 | 61795000 | 25429000 | 1450300 | 941270 | 2333200 | 2005400 | 4105500 | 7107200 | 25026000 | 8669300 | 0 | 0 | 5976200 | 6489300 |
| HIAEVSQE\ | STXBP1; STXBP | P61764; A0A | STXB1_HUMAN | Unmodified | 524760 | 532640 | 281620 | 474450 | 1285700 | 228150 | 1048200 | 650200 | 2829100 | 1358100 | 760680 | 3599300 | 1117400 | 606510 | 561080 | 269350 | 232320 | 928770 |
| HIAEVSQE\ | STXBP1; STXBP | P61764; A0A | STXB1_HUMAN | Unmodified | 0 | 0 | 0 | 0 | 1003400 | 0 | 3434300 | 1971300 | 0 | 247550 | 0 | 0 | 214670 | 0 | 0 | 0 | 351380 | 408480 |
| HIAEVSQE\ | STXBP1; STXBP | P61764; A0A | STXB1_HUMAN | Unmodified | 0 | 0 | 0 | 0 | 0 | 0 | 1070900 | 176490 | 0 | 0 | 0 | 0 | 0 | 0 | 0 | 0 | 233010 | 0 |
| HIAFIGDSI | CASD1; CASD1 | Q96PB1; C9. | CASD1_HUMAI | Unmodified | 0 | 0 | 644680 | 0 | 0 | 0 | 0 | 0 | 1303900 | 64152 | 0 | 0 | 0 | 0 | 0 | 1078800 | 0 | 0 |
| HIAFSGNK | MRPL13; MRPL | Q9BYD1; E5I | RM13_HUMAN | Unmodified | 0 | 0 | 0 | 0 | 0 | 0 | 230320 | 0 | 836170 | 254030 | 0 | 0 | 2031600 | 238820 | 0 | 15359 | 0 | 0 |
| HIANYISGI | SF3B3 | Q15393 | SF3B3_HUMAN | Unmodified | 0 | 0 | 0 | 0 | 0 | 0 | 0 | 0 | 107750 | 45902 | 0 | 1503600 | 0 | 0 | 0 | 0 | 0 | 0 |
| HIAVVLEEI | STXBP3 | O00186 | STXB3_HUMAN | Unmodified | 398140 | 0 | 467840 | 0 | 1744100 | 504440 | 0 | 0 | 0 | 570170 | 739680 | 4601000 | 1786100 | 1413400 | 841570 | 1028900 | 298220 | 1327200 |
| HICYFQIDI | DPP4; DPP4; DI | P27487; A0A | DPP4_HUMAN | Unmodified | 0 | 0 | 583420 | 882360 | 0 | 0 | 0 | 0 | 0 | 0 | 0 | 0 | 0 | 0 | 2063400 | 2195200 | 0 | 0 |

| | | | | | | | | | | | | | | | | | | | | | | |
|---|---|---|---|---|---|---|---|---|---|---|---|---|---|---|---|---|---|---|---|---|---|---|
| HIDCAAIYC | AKR1A1; AKR1A | P14550; V9C | AK1A1_HUMAN | Unmodified | 0 | 0 | 0 | 0 | 552570 | 0 | 0 | 0 | 2236800 | 0 | 0 | 0 | 0 | 0 | 0 | 1013100 | 0 | 757680 |
| HIDCAHVY | AKR1B1; AKR1E | P15121; E9P | ALDR_HUMAN | Unmodified | 0 | 0 | 0 | 0 | 1310700 | 0 | 0 | 0 | 0 | 0 | 3081600 | 25137000 | 0 | 0 | 0 | 0 | 186220 | 0 |
| HIDFSLR | RPS9 | P46781 | RS9_HUMAN 4 | Unmodified | 2107700 | 823070 | 1102100 | 0 | 84383 | 117660 | 0 | 0 | 321570 | 801090 | 0 | 16233000 | 0 | 0 | 601780 | 802340 | 0 | 0 |
| HIDRNYEPI | EPS8; EPS8; EP | Q12929; A0A | EPS8_HUMAN | Unmodified | 173790 | 167410 | 0 | 412660 | 416700 | 0 | 0 | 0 | 0 | 9938.6 | 0 | 125710 | 0 | 0 | 0 | 0 | 0 | 665440 |
| HIDSAHLYI | AKR1C3; AKR1C | P42330; Q0A | AK1C3_HUMAN | Unmodified | 354390 | 0 | 0 | 0 | 1669300 | 0 | 0 | 0 | 0 | 0 | 0 | 0 | 0 | 0 | 0 | 0 | 0 | 0 |
| HIDYFNNQ | MYO1D; MYO1D | O94832; J3Q | MYO1D_HUMA | Unmodified | 13931000 | 20079000 | 2811800 | 0 | 707000 | 0 | 0 | 0 | 0 | 606010 | 382440 | 1919100 | 197180 | 0 | 0 | 0 | 0 | 562620 |
| HIEEDEVRS | STX3; STX3; ST | Q13277; A0A | STX3_HUMAN S | Unmodified | 0 | 0 | 908500 | 309440 | 226540 | 0 | 0 | 0 | 190220 | 103370 | 74273 | 401000 | 0 | 45881 | 1109700 | 148690 | 195090 | 0 |
| HIEEENLID | MUC13 | Q9H3R2 | MUC13_HUMA | Unmodified | 3471000 | 0 | 0 | 0 | 0 | 0 | 0 | 0 | 0 | 180120 | 0 | 0 | 0 | 0 | 0 | 0 | 0 | 0 |
| HIEPELAGF | CCZ1; CCZ1B | P86791; P86 | CCZ1_HUMAN | Unmodified | 0 | 0 | 0 | 0 | 0 | 0 | 0 | 0 | 0 | 0 | 0 | 1011700 | 0 | 0 | 0 | 0 | 0 | 0 |
| HIEVQVAQ | SPAG9; SPAG9 | O60271; A0A | JIP4_HUMAN C | Unmodified | 0 | 0 | 0 | 0 | 0 | 0 | 0 | 0 | 45088 | 0 | 132140 | 564130 | 23833 | 0 | 0 | 0 | 0 | 104570 |
| HIEWESVL | CTNND1; CTNN | O60716; A0A | CTND1_HUMA | Unmodified | 4358700 | 1821200 | 2464400 | 11673000 | 1436800 | 2526800 | 0 | 0 | 0 | 0 | 4860500 | 9434500 | 2325100 | 3714800 | 11637000 | 9830200 | 5828700 | 8868900 |
| HIEWESVL | CTNND1; CTNN | O60716; A0A | CTND1_HUMA | Unmodified | 0 | 0 | 0 | 0 | 1109800 | 163890 | 0 | 0 | 0 | 0 | 0 | 0 | 0 | 282540 | 0 | 0 | 337990 | 2445600 |
| HIEWESVL | CTNND1; CTNN | O60716; A0A | CTND1_HUMA | Unmodified | 0 | 0 | 0 | 0 | 1923400 | 0 | 0 | 0 | 0 | 0 | 0 | 0 | 2640600 | 1311300 | 0 | 0 | 2406800 | 3735100 |
| HIFALFNTE | DNM3; DNM1; I | Q9UQ16; Q0 | DYN3_HUMAN | Unmodified | 30354 | 61163 | 27181 | 22733 | 0 | 0 | 0 | 0 | 418330 | 70104 | 0 | 0 | 0 | 0 | 0 | 0 | 0 | 0 |
| HIFDHVIGF | ABCC3 | O15438 | MRP3_HUMAN | Unmodified | 0 | 0 | 0 | 0 | 4832900 | 621750 | 0 | 0 | 0 | 0 | 0 | 0 | 0 | 0 | 0 | 0 | 226700 | 0 |
| HIFENVIGF | ABCC1; ABCC1 | P33527; I3L4 | MRP1_HUMAN | Unmodified | 0 | 0 | 0 | 0 | 1279700 | 0 | 0 | 0 | 0 | 0 | 0 | 4003500 | 0 | 0 | 0 | 309800 | 0 | 0 |
| HIFSEDTSD | SERPINB5 | P36952 | SPB5_HUMAN | Oxidation (M) | 0 | 0 | 0 | 0 | 919150 | 0 | 0 | 0 | 0 | 0 | 289710 | 563480 | 429080 | 0 | 0 | 96238 | 777930 | 867920 |
| HIFSEDTSD | SERPINB5 | P36952 | SPB5_HUMAN | Unmodified | 0 | 0 | 0 | 23727 | 0 | 0 | 0 | 0 | 0 | 0 | 0 | 0 | 1153400 | 1247200 | 0 | 0 | 1486400 | 2820800 |
| HIFSLEQLE | MYO10; MYO10 | Q9HD67; D6 | MYO10_HUMA | Unmodified | 0 | 0 | 0 | 0 | 0 | 0 | 0 | 0 | 0 | 0 | 0 | 0 | 0 | 0 | 631260 | 0 | 0 | 0 |
| HIFVLRERP | TAOK1 | Q7L7X3 | TAOK1_HUMAN | Unmodified | 0 | 0 | 0 | 0 | 0 | 0 | 0 | 0 | 0 | 236350 | 0 | 0 | 241680 | 113410 | 0 | 0 | 0 | 104550 |
| HIGELYSNI | PTPRG | P23470 | PTPRG_HUMAN | Unmodified | 0 | 0 | 0 | 445670 | 0 | 0 | 0 | 0 | 0 | 0 | 0 | 786020 | 0 | 0 | 0 | 847750 | 0 | 0 |
| HIGKVVVQ | FASN; FASN | P49327; A0A | FAS_HUMAN Fa | Unmodified | 0 | 0 | 0 | 0 | 334130 | 0 | 368890 | 0 | 0 | 59268 | 0 | 0 | 0 | 136060 | 0 | 0 | 166950 | 61946 |
| HIGLVYSGI | PSMA2; N/A; PS | P25787; A0A | PSA2_HUMAN | Oxidation (M) | 0 | 0 | 475950 | 507430 | 0 | 0 | 0 | 0 | 883760 | 106080 | 233740 | 7922200 | 106560 | 0 | 0 | 752170 | 0 | 0 |
| HIGLVYSGI | PSMA2; N/A; PS | P25787; A0A | PSA2_HUMAN | Unmodified | 0 | 0 | 0 | 0 | 0 | 72913 | 0 | 0 | 0 | 0 | 0 | 4049000 | 0 | 0 | 0 | 0 | 61406 | 0 |
| HIGLVYSGI | PSMA2; N/A; PS | P25787; A0A | PSA2_HUMAN | Oxidation (M) | 0 | 0 | 0 | 0 | 484930 | 0 | 457040 | 0 | 0 | 607480 | 0 | 0 | 0 | 0 | 0 | 30465 | 0 | 0 |
| HIGYDDSS | MAT2A | P31153 | METK2_HUMAN | Unmodified | 0 | 0 | 0 | 0 | 605760 | 0 | 0 | 122240 | 0 | 144970 | 0 | 0 | 628770 | 160090 | 0 | 0 | 491000 | 554710 |
| HIIYVDNK | ACSL4 | O60488 | ACSL4_HUMAN | Unmodified | 0 | 0 | 0 | 0 | 0 | 0 | 0 | 0 | 0 | 0 | 0 | 1955500 | 0 | 0 | 0 | 0 | 0 | 0 |
| HIKENDYY | STT3A | P46977 | STT3A_HUMAN | Unmodified | 0 | 0 | 0 | 0 | 0 | 0 | 0 | 0 | 356470 | 0 | 0 | 0 | 0 | 142760 | 0 | 59015 | 0 | 0 |
| HIKENDYY | STT3A | P46977 | STT3A_HUMAN | Unmodified | 0 | 0 | 0 | 0 | 43850 | 0 | 0 | 0 | 1180700 | 637410 | 0 | 0 | 0 | 0 | 0 | 0 | 0 | 0 |
| HILANFK | N/A; TPT1; TPT1 | Q56UQ5; P1 | TPT1L_HUMAN | Unmodified | 0 | 229960 | 1092800 | 906450 | 1226400 | 0 | 282480 | 174090 | 475860 | 155250 | 839530 | 1878800 | 1402200 | 1053700 | 1172300 | 949390 | 608370 | 627730 |
| HILILQNK | EIF2S3; EIF2S3 | P41091; Q2 | IF2G_HUMAN E | Unmodified | 0 | 0 | 629630 | 0 | 0 | 0 | 0 | 254940 | 14241000 | 2688300 | 2933800 | 1246300 | 133140 | 0 | 0 | 161900 | 0 | 0 |
| HILILQNKI | EIF2S3; EIF2S3 | P41091; Q2 | IF2G_HUMAN E | Unmodified | 0 | 142270 | 0 | 176390 | 1022000 | 0 | 139760 | 0 | 0 | 261780 | 591550 | 2803200 | 450040 | 355930 | 0 | 370510 | 150630 | 0 |
| HILLAVAND | MACROH2A2; M | Q9P0M6; O7 | H2AW_HUMAN | Unmodified | 7706700 | 5969800 | 0 | 0 | 844880 | 1162900 | 0 | 0 | 0 | 0 | 205440 | 16986000 | 0 | 0 | 0 | 0 | 0 | 2143600 |
| HILPLTND | ITGB6; ITGB6; I | P18564; A0A | ITB6_HUMAN I | Unmodified | 0 | 0 | 0 | 0 | 0 | 0 | 0 | 0 | 0 | 0 | 0 | 1311300 | 0 | 0 | 0 | 0 | 0 | 0 |
| HIMGQNV | RPL5; RPL5 | P46777; A0A | RL5_HUMAN 6 | Unmodified | 1224700 | 669030 | 142370 | 0 | 330830 | 0 | 0 | 0 | 520020 | 0 | 428310 | 5672500 | 0 | 0 | 397270 | 182240 | 0 | 152520 |
| HIMGQNV | RPL5; RPL5 | P46777; A0A | RL5_HUMAN 6 | Oxidation (M) | 1398900 | 1626100 | 201910 | 274240 | 0 | 0 | 0 | 0 | 0 | 0 | 0 | 8681300 | 384420 | 0 | 971250 | 694160 | 0 | 0 |
| HIMGQNV | RPL5; RPL5 | P46777; A0A | RL5_HUMAN 6 | 2 Oxidation (M) | 507870 | 0 | 0 | 0 | 0 | 0 | 0 | 0 | 0 | 0 | 0 | 165730 | 0 | 0 | 0 | 60817 | 0 | 0 |
| HINDSAPE | CKAP5 | Q14008 | CKAP5_HUMAN | Unmodified | 730420 | 0 | 0 | 0 | 453140 | 249430 | 425590 | 25401 | 1424400 | 541580 | 571480 | 0 | 2215600 | 906960 | 0 | 0 | 513330 | 278710 |
| HINPSQDE | ADGRG2 | Q8IZP9 | AGRG2_HUMA | Unmodified | 0 | 0 | 0 | 0 | 4005100 | 2034000 | 0 | 0 | 9680500 | 2043200 | 0 | 0 | 65916 | 0 | 0 | 0 | 0 | 0 |
| HINQLLR | GPC3; GPC3 | P51654; H0 | GPC3_HUMAN | Unmodified | 0 | 0 | 0 | 0 | 0 | 0 | 0 | 0 | 5098800 | 1924800 | 0 | 0 | 0 | 0 | 0 | 0 | 0 | 0 |
| HINTWVAE | SERPINB6; SER | P35237; A0A | SPB6_HUMAN | Unmodified | 0 | 0 | 45540 | 51702 | 0 | 0 | 0 | 0 | 0 | 0 | 80999 | 3154400 | 0 | 163640 | 303270 | 84246 | 0 | 0 |
| HINVTING | ABCC1; ABCC1 | P33527; I3L4 | MRP1_HUMAN | Unmodified | 0 | 0 | 42077 | 0 | 0 | 0 | 0 | 0 | 0 | 0 | 0 | 401300 | 0 | 0 | 0 | 0 | 0 | 0 |
| HIQEHYGG | GALK1; GALK1 | P51570; K7E | GALK1_HUMAN | Unmodified | 0 | 0 | 0 | 1140700 | 553220 | 0 | 0 | 0 | 0 | 0 | 0 | 0 | 0 | 0 | 1752500 | 0 | 259730 | 0 |
| HIQELSGD | UROS; UROS; U | P10746; A0A | HEM4_HUMAN | Unmodified | 0 | 0 | 0 | 0 | 218490 | 0 | 0 | 0 | 0 | 0 | 468560 | 1319700 | 0 | 0 | 0 | 0 | 0 | 239180 |
| HIQQYHME | OS9; OS9 | Q13438; F8V | OS9_HUMAN P | Unmodified | 0 | 0 | 0 | 0 | 0 | 0 | 0 | 0 | 255080 | 0 | 122300 | 0 | 0 | 0 | 0 | 0 | 0 | 0 |
| HIRILPWSV | EDIL3 | O43854 | EDIL3_HUMAN | Unmodified | 0 | 0 | 0 | 0 | 562560 | 148420 | 0 | 0 | 440900 | 67979 | 0 | 0 | 0 | 0 | 0 | 0 | 0 | 121440 |
| HISCPLLILI | ABHD12; ABHD | Q8N2K0; A0A | ABD12_HUMAN | Unmodified | 0 | 0 | 0 | 0 | 0 | 0 | 0 | 0 | 18129000 | 1862700 | 0 | 0 | 0 | 0 | 0 | 0 | 0 | 0 |
| HISCPLLILI | ABHD12; ABHD | Q8N2K0; A0A | ABD12_HUMAN | Unmodified | 0 | 0 | 0 | 0 | 0 | 0 | 0 | 0 | 31910000 | 3907400 | 0 | 0 | 0 | 0 | 0 | 0 | 0 | 0 |
| HISEGCVFI | TXNDC17; TXN | Q9BRA2; I3L | TXD17_HUMAN | Unmodified | 214870 | 0 | 0 | 0 | 0 | 0 | 96914 | 34133 | 0 | 0 | 0 | 1032500 | 284000 | 0 | 231740 | 199470 | 0 | 0 |
| HISLTLPAT | MVB12B | Q9H7P6 | MB12B_HUMA | Unmodified | 4162000 | 0 | 1251400 | 3486700 | 0 | 0 | 178000 | 780220 | 3037500 | 200110 | 66901 | 0 | 0 | 0 | 2973600 | 5274300 | 0 | 0 |
| HISQISVAE | HINT1 | P49773 | HINT1_HUMAN | Oxidation (M) | 0 | 0 | 0 | 0 | 8443200 | 1186400 | 2096400 | 0 | 4138600 | 2312200 | 3272300 | 4583800 | 22951000 | 4206000 | 2084800 | 0 | 1426800 | 2315600 |
| HISQISVAE | HINT1 | P49773 | HINT1_HUMAN | Unmodified | 0 | 0 | 0 | 133790 | 4754500 | 1883600 | 827830 | 0 | 404180 | 3329700 | 0 | 0 | 4396600 | 6512300 | 0 | 0 | 1229100 | 1402000 |
| HISVAMDK | B4GALT1 | P15291 | B4GT1_HUMAN | Unmodified | 0 | 0 | 0 | 0 | 0 | 0 | 0 | 0 | 0 | 0 | 0 | 0 | 0 | 0 | 0 | 0 | 0 | 0 |
| HITIFSPEG | PSMA6; PSMA6 | P60900; G3 | PSA6_HUMAN | Unmodified | 0 | 0 | 0 | 0 | 350640 | 0 | 60253 | 0 | 0 | 93805 | 0 | 0 | 157730 | 81502 | 0 | 0 | 189190 | 112510 |
| HITTLQASF | RAB21 | Q9UL25 | RAB21_HUMAN | Unmodified | 266310 | 646120 | 260890 | 313710 | 0 | 0 | 0 | 0 | 1335700 | 280550 | 887400 | 2710100 | 381580 | 0 | 593370 | 1227000 | 0 | 0 |
| HITTLQASF | RAB21 | Q9UL25 | RAB21_HUMAN | Unmodified | 0 | 0 | 292410 | 259740 | 517660 | 83859 | 157200 | 61791 | 299470 | 0 | 284670 | 419930 | 498800 | 208920 | 0 | 0 | 0 | 0 |
| HITTPREPL | NOTCH2 | Q04721 | NOTC2_HUMA | Unmodified | 158510 | 0 | 0 | 401860 | 5953300 | 597820 | 942440 | 0 | 576300 | 635800 | 0 | 0 | 1060200 | 2856600 | 315000 | 198830 | 4820800 | 23595000 |
| HIVASLQEF | ZDHHC5 | Q9C0B5 | ZDHC5_HUMA | Unmodified | 0 | 186540 | 794630 | 1636500 | 4313800 | 506600 | 38570 | 24302 | 865230 | 666300 | 653010 | 1294800 | 2917200 | 1533400 | 1161200 | 1220500 | 1339000 | 2672600 |
| HIVAVLPEI | NAGK; NAGK; N | Q9UJ70; C9J | NAGK_HUMAN | Unmodified | 0 | 0 | 0 | 0 | 0 | 0 | 0 | 0 | 0 | 0 | 0 | 1644600 | 0 | 0 | 163650 | 0 | 0 | 0 |
| HIVENAVQ | IPO5; IPO5 | O00410; H0 | IPO5_HUMAN I | Unmodified | 0 | 0 | 215550 | 0 | 0 | 0 | 0 | 0 | 0 | 0 | 175620 | 6951200 | 0 | 0 | 193220 | 188970 | 0 | 0 |
| HIVIDGSN | N4BP1 | O75113 | N4BP1_HUMAN | Unmodified | 335140 | 303610 | 0 | 0 | 70067 | 0 | 0 | 0 | 0 | 0 | 0 | 0 | 0 | 0 | 0 | 0 | 0 | 0 |
| HIVLSGGS | ACTR2; ACTR2 | P61160; F5H | ARP2_HUMAN | Oxidation (M) | 0 | 0 | 0 | 33841 | 0 | 0 | 0 | 340300 | 0 | 0 | 0 | 2642400 | 0 | 0 | 0 | 0 | 0 | 0 |
| HIVLSGGS | ACTR2; ACTR2 | P61160; F5H | ARP2_HUMAN | Oxidation (M) | 1387200 | 0 | 0 | 0 | 0 | 0 | 0 | 0 | 0 | 0 | 0 | 8052100 | 0 | 0 | 0 | 1599000 | 0 | 0 |
| HIVLSGGS | ACTR2; ACTR2 | P61160; F5H | ARP2_HUMAN | Unmodified | 0 | 0 | 0 | 0 | 0 | 0 | 0 | 0 | 0 | 0 | 0 | 1457300 | 0 | 0 | 0 | 0 | 0 | 0 |
| HIVLSGGS | ACTR2; ACTR2 | P61160; F5H | ARP2_HUMAN | Oxidation (M) | 0 | 0 | 0 | 947580 | 3513100 | 5722800 | 628080 | 0 | 140650 | 790430 | 0 | 2188000 | 1743100 | 1223400 | 0 | 0 | 874370 | 1713000 |
| HIVNHDDV | ERBIN | Q96RT1 | ERBIN_HUMAN | Unmodified | 0 | 0 | 148440 | 0 | 1645900 | 0 | 0 | 0 | 0 | 0 | 179830 | 0 | 0 | 0 | 0 | 0 | 0 | 0 |
| HIVNHDDV | ERBIN | Q96RT1 | ERBIN_HUMAN | Oxidation (M) | 0 | 0 | 0 | 0 | 1604500 | 0 | 0 | 0 | 0 | 0 | 0 | 0 | 0 | 0 | 0 | 0 | 317920 | 0 |
| HIVSNDSS | PLA2G4A | P47712 | PA24A_HUMAN | Unmodified | 0 | 0 | 0 | 0 | 0 | 0 | 0 | 0 | 0 | 0 | 0 | 0 | 0 | 0 | 0 | 0 | 0 | 0 |
| HIVTGTTRF | MOV10; MOV10 | Q9HCE1; Q8 | MOV10_HUMA | Unmodified | 8548300 | 4250500 | 3964100 | 2848500 | 0 | 0 | 0 | 0 | 0 | 0 | 0 | 675890 | 0 | 0 | 7192000 | 6947100 | 0 | 0 |
| HIVVSCAA | PYCR2; PYCR2; | Q96C36; A0 | P5CR2_HUMAN | Unmodified | 0 | 0 | 0 | 0 | 0 | 0 | 0 | 0 | 1389300 | 791130 | 0 | 0 | 0 | 0 | 0 | 0 | 0 | 620450 |
| HIYLLPSGF | GOT1 | P17174 | AATC_HUMAN | Unmodified | 0 | 0 | 0 | 0 | 1621500 | 0 | 0 | 0 | 0 | 921230 | 0 | 0 | 0 | 0 | 0 | 0 | 1013400 | 0 |
| HIYYITGETI | HSP90AA5P; HS | Q58FG0; P0 | HS905_HUMAN | Unmodified | 559820 | 427800 | 1188300 | 1022000 | 1277700 | 231090 | 367660 | 0 | 11620000 | 1350400 | 4001000 | 43094000 | 0 | 0 | 1614100 | 913650 | 89868 | 0 |
| HIYYITGETI | HSP90AA1 | P07900 | HS90A_HUMAN | Unmodified | 1497800 | 764040 | 0 | 1876700 | 13229000 | 4731700 | 875980 | 0 | 27343000 | 12456000 | 14408000 | 38546000 | 3098500 | 2246600 | 872190 | 792220 | 4429100 | 7865100 |
| HIYYITGETI | HSP90AA1 | P07900 | HS90A_HUMAN | Unmodified | 0 | 0 | 0 | 0 | 0 | 0 | 0 | 0 | 1162000 | 2008900 | 1290600 | 1320800 | 1043000 | 1657100 | 0 | 0 | 0 | 0 |
| HKAGFLDL | TLN1 | Q9Y490 | TLN1_HUMAN | Unmodified | 0 | 0 | 0 | 0 | 0 | 0 | 1795900 | 0 | 0 | 0 | 0 | 0 | 0 | 1519600 | 0 | 0 | 252850 | 0 |
| HKEFVGNV | MFGE8; MFGE8 | Q08431; H0 | MFGM_HUMAN | Unmodified | 0 | 0 | 0 | 1857500 | 45568000 | 9140300 | 738580 | 0 | 271400 | 0 | 0 | 76736 | 2934600 | 1952000 | 0 | 0 | 5674800 | 7929300 |
| HKELAPYD | RPS19; RPS19; | P39019; M0 | RS19_HUMAN | Unmodified | 152550 | 267420 | 0 | 149150 | 107960 | 0 | 166220 | 0 | 0 | 213860 | 0 | 0 | 659720 | 0 | 0 | 0 | 0 | 0 |
| HKELAPYD | RPS19 | P39019 | RS19_HUMAN | Unmodified | 0 | 0 | 0 | 0 | 903100 | 0 | 826830 | 0 | 0 | 0 | 0 | 0 | 2221400 | 1118400 | 0 | 0 | 318060 | 862120 |
| HKELFDEL | ADK; ADK; ADK | P55263; A0A | ADK_HUMAN A | Unmodified | 0 | 0 | 0 | 0 | 1188600 | 822830 | 511940 | 0 | 0 | 0 | 0 | 1093100 | 0 | 634050 | 0 | 0 | 0 | 1590700 |
| HKENLQHT | TAOK3; TAOK3 | Q9H2K8; G3 | TAOK3_HUMAN | Unmodified | 0 | 0 | 0 | 0 | 5653500 | 0 | 1930500 | 0 | 0 | 0 | 0 | 0 | 1897600 | 3338800 | 0 | 0 | 0 | 2247500 |
| HKFNTVPG | POMGNT1 | Q8WZA1 | PMGT1_HUMA | Unmodified | 0 | 0 | 0 | 0 | 0 | 0 | 0 | 0 | 1046400 | 264630 | 0 | 0 | 0 | 0 | 0 | 0 | 0 | 0 |
| HKGPPVFT | PCYT2; PCYT2; | Q99447; I3L | PCY2_HUMAN | Unmodified | 0 | 0 | 0 | 86028 | 0 | 0 | 0 | 0 | 0 | 0 | 0 | 265230 | 592410 | 0 | 45088 | 0 | 0 | 0 |
| HKIDDLTA | ANKRD26; ANK | Q9UPS8; A0 | ANR26_HUMAN | Unmodified | 0 | 0 | 0 | 0 | 217800 | 0 | 0 | 0 | 0 | 0 | 0 | 0 | 0 | 0 | 0 | 0 | 162440 | 314050 |
| HKISFLCDI | CDCP1 | Q9H5V8 | CDCP1_HUMA | Oxidation (M) | 0 | 0 | 0 | 105780 | 3586300 | 385960 | 0 | 0 | 0 | 0 | 228410 | 0 | 0 | 0 | 2992100 | 0 | 0 | 6205800 |
| HKISFLCDI | CDCP1 | Q9H5V8 | CDCP1_HUMA | Unmodified | 0 | 0 | 0 | 0 | 0 | 0 | 0 | 0 | 0 | 0 | 0 | 0 | 0 | 0 | 0 | 0 | 539980 | 738290 |
| HKLDVTSV | CCT5; CCT5; C | P48643; B7Z | TCPE_HUMAN | Unmodified | 0 | 212650 | 744280 | 825920 | 2536500 | 709030 | 1031800 | 698460 | 3830900 | 1644700 | 3302000 | 1956000 | 9015200 | 4154100 | 671640 | 652990 | 1064000 | 2975800 |
| HKLDVTSV | CCT5; CCT5; C | P48643; B7Z | TCPE_HUMAN | Unmodified | 0 | 0 | 0 | 0 | 0 | 0 | 0 | 0 | 0 | 0 | 0 | 0 | 669130 | 813420 | 0 | 0 | 0 | 0 |
| HKLIQEEQ | TTC17; TTC17 | Q96AE7; A0 | TTC17_HUMAN | Unmodified | 0 | 0 | 0 | 0 | 0 | 0 | 0 | 0 | 1130900 | 168020 | 0 | 0 | 0 | 0 | 0 | 0 | 0 | 0 |
| HKLTPEYLE | ERLIN1 | O75477 | ERLN1_HUMAN | Unmodified | 0 | 0 | 0 | 0 | 936710 | 147510 | 707540 | 249150 | 6234300 | 1937000 | 849690 | 0 | 0 | 0 | 0 | 0 | 0 | 0 |
| HKLTPEYLE | ERLIN1 | O75477 | ERLN1_HUMAN | Unmodified | 0 | 0 | 0 | 0 | 0 | 0 | 0 | 0 | 817980 | 0 | 0 | 0 | 0 | 0 | 0 | 0 | 0 | 0 |
| HKNPKTGI | CANX; CANX; C | P27824; A0A | CALX_HUMAN | Unmodified | 0 | 0 | 32896 | 0 | 0 | 74506 | 0 | 0 | 52460 | 1387800 | 0 | 0 | 0 | 0 | 0 | 0 | 0 | 0 |
| HKPAFTEE | CLUH; CLUH; C | O75153; I3L | CLU_HUMAN C | Unmodified | 0 | 0 | 0 | 0 | 0 | 0 | 0 | 0 | 0 | 0 | 521500 | 5341700 | 0 | 0 | 0 | 0 | 0 | 0 |
| HKPGGGD | MAP4; MAP4; M | P27816; H7 | MAP4_HUMAN | Unmodified | 0 | 0 | 0 | 0 | 192170 | 41664 | 0 | 0 | 26140 | 0 | 0 | 0 | 345460 | 157340 | 0 | 0 | 124190 | 501950 |
| HKPGIVQE | AFDN; AFDN; A | P55196; Q5 | AFAD_HUMAN | Unmodified | 0 | 0 | 0 | 0 | 4043200 | 736770 | 0 | 0 | 0 | 127860 | 971470 | 1472200 | 8016200 | 6013900 | 0 | 0 | 244770 | 537240 |
| HKPWLEPT | CLSTN1; CLSTN | O94985; A0 | CSTN1_HUMAN | Unmodified | 0 | 0 | 0 | 0 | 0 | 0 | 0 | 0 | 5853200 | 251340 | 0 | 0 | 0 | 0 | 0 | 0 | 0 | 0 |
| HKRTEEEEI | RGS10 | O43665 | RGS10_HUMAN | Unmodified | 0 | 0 | 0 | 0 | 1484500 | 0 | 275790 | 0 | 0 | 0 | 0 | 0 | 0 | 0 | 0 | 0 | 233880 | 0 |
| HKSDFGKF | CALR; CALR; C | P27797; A0A | CALR_HUMAN | Unmodified | 0 | 0 | 0 | 0 | 264890 | 0 | 965870 | 546000 | 3033600 | 15065000 | 58314 | 0 | 233750 | 0 | 0 | 0 | 456840 | 221930 |
| HKSETDTS | CCT6A | P40227 | TCPZ_HUMAN | Unmodified | 0 | 155010 | 0 | 0 | 234350 | 80721 | 0 | 0 | 1665300 | 384580 | 1985200 | 10149000 | 1274900 | 386080 | 0 | 0 | 0 | 293810 |
| HKSEVEIPA | ADD2 | P35612 | ADDB_HUMAN | Unmodified | 0 | 0 | 0 | 0 | 2557700 | 0 | 0 | 0 | 0 | 0 | 0 | 0 | 0 | 0 | 0 | 0 | 0 | 0 |
| HKWEAAH | HLA-A; HLA-A; | Q53Z42; A0 | Q53Z42_HUMA | Unmodified | 10454 | 0 | 0 | 0 | 0 | 0 | 0 | 0 | 0 | 0 | 0 | 3972700 | 161340 | 75600 | 0 | 0 | 0 | 0 |
| HKWNDFA | OPA1; OPA1; O | O60313; E5 | OPA1_HUMAN | Unmodified | 0 | 0 | 0 | 0 | 0 | 0 | 0 | 0 | 0 | 0 | 64137 | 862260 | 0 | 0 | 0 | 0 | 0 | 0 |
| HLADHGQ | MRPS14 | O60783 | RT14_HUMAN | Unmodified | 0 | 0 | 0 | 0 | 115920 | 0 | 0 | 33894 | 616390 | 0 | 0 | 0 | 0 | 12904 | 0 | 0 | 0 | 56414 |
| HLAEELR | MYH15; CCNDI | Q9Y2K3; Q9 | MYH15_HUMA | Unmodified | 1082300 | 1167100 | 604560 | 646510 | 0 | 0 | 0 | 7639.5 | 13530 | 0 | 0 | 0 | 0 | 0 | 1473400 | 1029300 | 0 | 0 |
| HLAELNHQ | PSAP; PSAP | P07602; C9J | SAP_HUMAN P | Unmodified | 13771 | 0 | 0 | 0 | 0 | 0 | 168170 | 52288 | 1132300 | 0 | 32659 | 0 | 0 | 45112 | 0 | 0 | 0 | 0 |
| HLAGLGLT | SERPINH1; SER | P50454; E9P | SERPH_HUMA | Unmodified | 0 | 0 | 0 | 0 | 50191 | 0 | 1196200 | 280080 | 44227000 | 12144000 | 0 | 0 | 0 | 0 | 0 | 0 | 0 | 0 |
| HLAGLGLT | SERPINH1; SER | P50454; E9P | SERPH_HUMA | Unmodified | 0 | 0 | 0 | 0 | 0 | 0 | 0 | 0 | 2392600 | 500730 | 0 | 0 | 0 | 0 | 79789 | 0 | 0 | 0 |
| HLAGLGLT | SERPINH1; SER | P50454; E9P | SERPH_HUMA | Unmodified | 0 | 0 | 0 | 0 | 0 | 0 | 149820 | 0 | 0 | 5624800 | 0 | 0 | 0 | 0 | 0 | 0 | 0 | 0 |
| HLASLYAN | CPD | O75976 | CBPD_HUMAN | Unmodified | 0 | 0 | 0 | 0 | 0 | 0 | 0 | 0 | 1464100 | 0 | 0 | 0 | 0 | 0 | 0 | 0 | 0 | 0 |
| HLATSDML | ERMP1; ERMP1 | Q7Z2K6; A0 | ERMP1_HUMAN | Unmodified | 0 | 0 | 0 | 0 | 0 | 0 | 0 | 0 | 611310 | 0 | 0 | 0 | 0 | 0 | 0 | 0 | 0 | 0 |

| | | | | | | | | | | | | | | | | | | | | | | |
|---|---|---|---|---|---|---|---|---|---|---|---|---|---|---|---|---|---|---|---|---|---|---|
| HLAVEWGI | DECR2; DECR2 | Q9NUI1; Q4' | DECR2_HUMAI | Unmodified | 0 | 0 | 0 | 0 | 760660 | 0 | 0 | 0 | 2818000 | 0 | 0 | 0 | 0 | 0 | 0 | 0 | 402490 | 0 |
| HLCQQLQ/ | NUMA1; NUMA | Q14980; H0' | NUMA1_HUMA | Unmodified | 0 | 0 | 0 | 0 | 0 | 0 | 0 | 0 | 78756 | 162950 | 0 | 185440 | 285250 | 130350 | 0 | 0 | 182890 | 149060 |
| HLDACETM | ATP2B2; ATP2B | Q01814; Q1( | AT2B2_HUMAN | Oxidation (M) | 158760 | 0 | 0 | 89303 | 0 | 0 | 0 | 0 | 0 | 0 | 0 | 682480 | 0 | 0 | 65713 | 92316 | 0 | 0 |
| HLDACETM | ATP2B2; ATP2B | Q01814; Q1( | AT2B2_HUMAN | Unmodified | 0 | 0 | 0 | 0 | 0 | 0 | 0 | 0 | 0 | 0 | 389350 | 0 | 0 | 0 | 1145400 | 0 | 0 | 0 |
| HLDFLKQP | MAP1B; MAP1B | P46821; D6F | MAP1B_HUMAI | Unmodified | 36728 | 0 | 0 | 0 | 0 | 0 | 805260 | 145070 | 0 | 404840 | 0 | 0 | 901090 | 269440 | 0 | 0 | 497170 | 0 |
| HLDGEED( | MAGED2; MAGI | Q9UNF1; Q5 | MAGD2_HUMA | Unmodified | 0 | 0 | 0 | 0 | 0 | 0 | 0 | 0 | 129460 | 0 | 0 | 65831 | 0 | 0 | 18934 | 0 | 0 | 0 |
| HLDGRQIV | DNAJA2 | O60884 | DNJA2_HUMAN | Unmodified | 0 | 0 | 0 | 0 | 1347900 | 0 | 0 | 116850 | 0 | 0 | 0 | 0 | 1924000 | 545330 | 0 | 0 | 129050 | 178560 |
| HLDHVAAL | PYGB | P11216 | PYGB_HUMAN | Unmodified | 75315 | 0 | 0 | 0 | 0 | 0 | 0 | 0 | 721350 | 478950 | 0 | 915830 | 0 | 0 | 245110 | 136910 | 0 | 403480 |
| HLDIEDK | CHCHD3 | C9JRZ6 | C9JRZ6_HUMA | Unmodified | 0 | 0 | 0 | 82017 | 0 | 30594 | 39337 | 0 | 175100 | 0 | 0 | 177920 | 0 | 21923 | 0 | 30683 | 119260 | 0 |
| HLDLSNVA | PACSIN2; PACS | Q9UNF0; A0 | PACN2_HUMAI | Unmodified | 45826000 | 44626000 | 30120000 | 36582000 | 7406200 | 1307600 | 1504900 | 1290800 | 1740000 | 1441400 | 2022700 | 7457300 | 76967000 | 25253000 | 58655000 | 41523000 | 1809200 | 2813000 |
| HLDLSNVA | PACSIN2; PACS | Q9UNF0; A0 | PACN2_HUMAI | Unmodified | 0 | 0 | 0 | 0 | 170260 | 648440 | 0 | 0 | 0 | 0 | 0 | 0 | 1163800 | 0 | 74456 | 0 | 47561 | 1202400 |
| HLDQCEH( | PLA2G3 | Q9NZ20 | PA2G3_HUMAN | Unmodified | 0 | 0 | 0 | 0 | 0 | 0 | 0 | 0 | 0 | 0 | 0 | 0 | 0 | 0 | 0 | 0 | 0 | 0 |
| HLDSVLQC | IGFBP6; IGFBP( | P24592; A0A | IBP6_HUMAN I | Unmodified | 0 | 0 | 0 | 0 | 0 | 0 | 0 | 19956 | 2054800 | 0 | 0 | 287130 | 0 | 0 | 0 | 0 | 0 | 0 |
| HLEEAFTSE | STT3B | Q8TCJ2 | STT3B_HUMAN | Unmodified | 70957 | 0 | 0 | 0 | 0 | 0 | 0 | 0 | 31595000 | 1835700 | 2951800 | 0 | 0 | 0 | 64455 | 0 | 0 | 0 |
| HLEEAFTSE | STT3B | Q8TCJ2 | STT3B_HUMAN | Unmodified | 0 | 0 | 0 | 0 | 97032 | 0 | 233890 | 0 | 0 | 1305200 | 0 | 0 | 0 | 0 | 0 | 0 | 0 | 0 |
| HLEEEEERI | SLITRK6 | Q9H5Y7 | SLIK6_HUMAN | Unmodified | 2045000 | 957070 | 0 | 0 | 0 | 0 | 0 | 0 | 2922700 | 0 | 0 | 0 | 0 | 0 | 0 | 213700 | 0 | 215250 |
| HLEEEEERI | SLITRK6 | Q9H5Y7 | SLIK6_HUMAN | Unmodified | 917480 | 0 | 0 | 0 | 0 | 0 | 0 | 0 | 2948200 | 5190300 | 0 | 0 | 178950 | 203200 | 0 | 29125 | 56329 | 820480 |
| HLEEGRLYI | ME1 | P48163 | MAOX_HUMAN | Unmodified | 0 | 0 | 0 | 0 | 0 | 0 | 0 | 0 | 0 | 0 | 0 | 612880 | 0 | 0 | 0 | 0 | 0 | 0 |
| HLEEGRLYI | ME1 | P48163 | MAOX_HUMAN | Unmodified | 0 | 0 | 0 | 212880 | 2379700 | 1671300 | 0 | 0 | 0 | 0 | 0 | 0 | 0 | 0 | 0 | 0 | 0 | 778540 |
| HLEEIVHVE | AMPD2; AMPD2 | Q01433; H0' | AMPD2_HUMA | Unmodified | 0 | 0 | 0 | 0 | 0 | 0 | 0 | 0 | 0 | 0 | 0 | 958820 | 0 | 0 | 89161 | 93808 | 0 | 0 |
| HLEEPGET( | CHGB; CHGB | P05060; A0A | SCG1_HUMAN | Unmodified | 0 | 0 | 0 | 0 | 0 | 0 | 0 | 0 | 2987000 | 1009700 | 0 | 0 | 0 | 0 | 0 | 0 | 0 | 0 |
| HLEFSHDC | NANS | Q9NR45 | SIAS_HUMAN S | Unmodified | 0 | 0 | 0 | 0 | 0 | 0 | 0 | 0 | 0 | 0 | 0 | 666030 | 0 | 0 | 0 | 0 | 0 | 0 |
| HLEGLSEE/ | PGAM1 | P18669 | PGAM1_HUMA | Oxidation (M) | 0 | 0 | 0 | 0 | 935460 | 0 | 0 | 0 | 0 | 0 | 0 | 0 | 0 | 0 | 0 | 0 | 1085100 | 1194600 |
| HLEHSTIIYI | DCAF7; DCAF7 | P61962; A0A | DCAF7_HUMAI | Unmodified | 120180 | 0 | 0 | 0 | 0 | 0 | 0 | 0 | 0 | 0 | 252730 | 3323700 | 257550 | 318520 | 115490 | 0 | 0 | 0 |
| HLEIFHTEF | CUL1; CUL1; CI | Q13616; A0/ | CUL1_HUMAN | Unmodified | 0 | 0 | 0 | 0 | 0 | 0 | 0 | 0 | 0 | 0 | 0 | 2081300 | 862540 | 530880 | 0 | 0 | 0 | 0 |
| HLEIIYAIN( | PYGB | P11216 | PYGB_HUMAN | Unmodified | 0 | 0 | 0 | 0 | 293740 | 55331 | 0 | 0 | 1519900 | 185470 | 0 | 0 | 0 | 0 | 0 | 0 | 40688 | 125170 |
| HLEINPDH | HSP90AB1 | P08238 | HS90B_HUMAI | Unmodified | 2689000 | 2249300 | 6612900 | 1444700 | 4776300 | 0 | 394200 | 26565 | 19001000 | 3238300 | 22331000 | 1.38E+08 | 3734000 | 975870 | 9713100 | 8611400 | 1923800 | 1612400 |
| HLEINPDH | HSP90AB1 | P08238 | HS90B_HUMAI | Unmodified | 2984100 | 984700 | 7706600 | 8197900 | 23340000 | 4606700 | 1357300 | 138060 | 11565000 | 3946600 | 11113000 | 24844000 | 15925000 | 7262400 | 5060700 | 3767600 | 6635700 | 15849000 |
| HLEINPDH | HSP90AA1 | P07900 | HS90A_HUMAI | Unmodified | 0 | 449780 | 7062900 | 23988000 | 2872900 | 616470 | 313990 | 154070 | 32487000 | 3132000 | 23711000 | 1.71E+08 | 1255400 | 455610 | 0 | 6084100 | 2074700 | 1569900 |
| HLEINPDH | HSP90AA1 | P07900 | HS90A_HUMAI | Unmodified | 875130 | 0 | 5177100 | 4135000 | 17068000 | 4455000 | 474820 | 0 | 10578000 | 3750200 | 10413000 | 23634000 | 3827300 | 1133800 | 2983400 | 1889400 | 4441600 | 9954300 |
| HLELYNQF | ARHGEF39 | Q8N4T4 | ARG39_HUMAI | Unmodified | 784750 | 259690 | 860970 | 1118900 | 0 | 0 | 49830 | 0 | 0 | 0 | 0 | 0 | 0 | 0 | 1679000 | 524960 | 0 | 0 |
| HLEMNPHI | DHX9 | Q08211 | DHX9_HUMAN | Unmodified | 0 | 32146 | 0 | 0 | 56806 | 0 | 0 | 65258 | 100240 | 21515 | 71729 | 1797300 | 0 | 0 | 0 | 0 | 0 | 0 |
| HLERQIEEI | GPRIN1 | Q7Z2K8 | GRIN1_HUMAN | Unmodified | 0 | 0 | 0 | 0 | 206560 | 0 | 170220 | 0 | 0 | 0 | 0 | 0 | 0 | 0 | 0 | 0 | 556460 | 1289700 |
| HLESRVEEI | PPM1B; N/A; PF | O75688; A0/ | PPM1B_HUMAI | Unmodified | 187150 | 0 | 0 | 0 | 109000 | 0 | 0 | 0 | 377250 | 0 | 0 | 850460 | 349020 | 0 | 135410 | 51844 | 0 | 0 |
| HLFASVHD | SLC26A6; SLC2 | Q9BXS9; A0/ | S26A6_HUMAN | Unmodified | 313610 | 370280 | 0 | 176560 | 3517800 | 131120 | 0 | 0 | 0 | 0 | 146540 | 598580 | 1676000 | 664950 | 0 | 0 | 625480 | 661560 |
| HLFCPDLLI | DECR2; DECR2 | Q9NUI1; Q4' | DECR2_HUMAI | Unmodified | 0 | 0 | 0 | 0 | 0 | 0 | 0 | 0 | 815020 | 0 | 0 | 0 | 500740 | 0 | 0 | 0 | 0 | 0 |
| HLFETDVQ | MPP7; MPP7 | Q5T2T1; U5( | MPP7_HUMAN | Unmodified | 0 | 0 | 0 | 119330 | 5853200 | 347870 | 0 | 0 | 113330 | 224910 | 0 | 0 | 1218800 | 761490 | 0 | 0 | 172430 | 309670 |
| HLFLTSKPI | OLA1; OLA1; OL | Q9NTK5; J3K | OLA1_HUMAN | Oxidation (M) | 0 | 0 | 0 | 373850 | 0 | 0 | 272600 | 0 | 0 | 0 | 549350 | 4144000 | 260430 | 248410 | 260810 | 197110 | 0 | 0 |
| HLFLTSKPI | OLA1; OLA1; OL | Q9NTK5; J3K | OLA1_HUMAN | Unmodified | 0 | 0 | 0 | 0 | 0 | 0 | 138540 | 0 | 0 | 58215 | 0 | 501150 | 0 | 0 | 51188 | 32053 | 97495 | 0 |
| HLFSSTENI | RAB11FIP1; RAI | Q6WKZ4; E7 | RFIP1_HUMAN | Unmodified | 0 | 0 | 0 | 0 | 1919200 | 0 | 0 | 0 | 0 | 0 | 0 | 0 | 0 | 0 | 0 | 0 | 0 | 0 |
| HLFTGPVL! | FAH; FAH | P16930; H0' | FAAA_HUMAN | Unmodified | 0 | 0 | 0 | 0 | 0 | 0 | 0 | 0 | 0 | 0 | 0 | 763510 | 0 | 0 | 0 | 65100 | 84726 | 0 |
| HLGESVPD | BTN2A1 | Q7KYR7 | BT2A1_HUMAN | Unmodified | 137070 | 0 | 0 | 0 | 0 | 0 | 0 | 0 | 0 | 0 | 0 | 2073600 | 0 | 0 | 0 | 404820 | 0 | 0 |
| HLGGIPWT | UQCRC1 | P31930 | QCR1_HUMAN | Unmodified | 0 | 0 | 0 | 0 | 0 | 0 | 0 | 0 | 4946300 | 0 | 0 | 0 | 0 | 0 | 0 | 0 | 0 | 0 |
| HLGGIPWT | UQCRC1 | P31930 | QCR1_HUMAN | Unmodified | 0 | 0 | 0 | 0 | 0 | 0 | 0 | 0 | 6718500 | 0 | 0 | 0 | 0 | 499270 | 0 | 0 | 0 | 0 |
| HLGPATAFI | FAT1; FAT1 | Q14517; A0/ | FAT1_HUMAN I | Unmodified | 0 | 0 | 0 | 0 | 0 | 0 | 0 | 0 | 0 | 0 | 0 | 1568000 | 0 | 165420 | 0 | 0 | 0 | 0 |
| HLGTEIQN' | TBC1D2 | Q9BYX2 | TBD2A_HUMAN | Unmodified | 0 | 0 | 0 | 0 | 889490 | 0 | 0 | 0 | 0 | 0 | 0 | 0 | 0 | 0 | 0 | 0 | 0 | 0 |
| HLGTSEVD | SLC38A9; SLC3 | Q8NBW4; E7 | S38A9_HUMAN | Unmodified | 0 | 0 | 0 | 0 | 0 | 0 | 0 | 0 | 520450 | 0 | 0 | 0 | 0 | 0 | 0 | 0 | 0 | 287360 |
| HLGVCISV/ | HNRNPR; HNRI | O43390; B4I | HNRPR_HUMA | Unmodified | 0 | 0 | 0 | 0 | 0 | 0 | 0 | 0 | 0 | 173470 | 393800 | 6198000 | 0 | 0 | 0 | 0 | 0 | 0 |
| HLGYKPEE' | MYO1C; MYO1( | O00159; F5I | MYO1C_HUMA | Unmodified | 208880 | 259510 | 984730 | 1001800 | 0 | 0 | 0 | 0 | 58287 | 0 | 0 | 242170 | 189480 | 0 | 1855100 | 1708200 | 0 | 0 |
| HLHPIQTSI | NUP210 | Q8TEM1 | PO210_HUMAI | Unmodified | 0 | 0 | 0 | 0 | 0 | 0 | 0 | 0 | 15672000 | 786020 | 194610 | 0 | 0 | 0 | 0 | 0 | 0 | 0 |
| HLHQAVSI | DST; DST; DST; | Q03001; A0/ | DYST_HUMAN | Unmodified | 0 | 0 | 0 | 0 | 607840 | 0 | 0 | 0 | 334260 | 291610 | 0 | 0 | 0 | 0 | 0 | 0 | 392790 | 0 |
| HLHTAMFT | DNAJC13; DNA | O75165; A0/ | DJC13_HUMAN | Unmodified | 0 | 0 | 0 | 0 | 0 | 0 | 0 | 0 | 0 | 31490 | 0 | 258280 | 0 | 0 | 0 | 0 | 0 | 0 |
| HLIATQLLS | ADGRE5 | P48960 | AGRE5_HUMAI | Oxidation (M) | 109580 | 142650 | 152400 | 235040 | 0 | 0 | 0 | 28156 | 0 | 0 | 0 | 0 | 0 | 0 | 360250 | 456500 | 0 | 0 |
| HLIEQDFP( | EHD1; EHD1; EI | Q9H4M9; C9 | EHD1_HUMAN | Oxidation (M) | 205600 | 196690 | 189330 | 214180 | 204210 | 100750 | 0 | 0 | 0 | 0 | 350080 | 2164400 | 76539 | 0 | 301230 | 440970 | 83777 | 89773 |
| HLIEQDFP( | EHD1; EHD1; EI | Q9H4M9; C9 | EHD1_HUMAN | Unmodified | 0 | 0 | 0 | 0 | 0 | 0 | 0 | 0 | 282480 | 0 | 757050 | 4533600 | 0 | 89760 | 0 | 65658 | 179170 | 411590 |
| HLILPEKYP | SNRNP200; SN | O75643; A0/ | U520_HUMAN | Unmodified | 0 | 279620 | 0 | 0 | 0 | 0 | 0 | 0 | 0 | 0 | 0 | 544940 | 0 | 0 | 0 | 0 | 0 | 0 |
| HLIPAANT( | YWHAE | P62258 | 1433E_HUMAN | Unmodified | 1754000 | 943110 | 15252000 | 10153000 | 5535200 | 5275700 | 2272900 | 880530 | 7302800 | 5843400 | 7827500 | 59757000 | 3638300 | 5561800 | 11419000 | 14645000 | 4912200 | 336640 |
| HLIPAANT( | YWHAE | P62258 | 1433E_HUMAN | Unmodified | 0 | 0 | 0 | 0 | 1469200 | 0 | 3448100 | 1029000 | 0 | 2735000 | 0 | 0 | 5252400 | 10873000 | 0 | 0 | 6677000 | 0 |
| HLIPMNPN | PLS1; PLS1 | Q14651; C9J | PLSI_HUMAN F | Oxidation (M) | 0 | 0 | 0 | 0 | 1665500 | 0 | 0 | 0 | 0 | 0 | 0 | 2116100 | 0 | 0 | 0 | 0 | 19224 | 0 |
| HLISTHFAF | HSPG2 | P98160 | PGBM_HUMAN | Unmodified | 0 | 0 | 8774000 | 3838400 | 0 | 0 | 0 | 163610 | 2224100 | 0 | 0 | 0 | 0 | 0 | 0 | 3747600 | 0 | 0 |
| HLIVLINK | GSPT1; GSPT2; | P15170; Q8I | ERF3A_HUMAN | Unmodified | 47623 | 64672 | 121380 | 0 | 84197 | 0 | 18103 | 0 | 92235 | 26985 | 156810 | 981390 | 0 | 62859 | 95815 | 135410 | 32862 | 103140 |
| HLKALTSEI | MSN | P26038 | MOES_HUMAN | Unmodified | 0 | 0 | 0 | 0 | 479710 | 0 | 116810 | 31056 | 0 | 0 | 0 | 0 | 0 | 0 | 0 | 0 | 2186900 | 0 |
| HLKEETIQII | USP25 | Q9UHP3 | UBP25_HUMAI | Unmodified | 0 | 0 | 0 | 0 | 1065100 | 129230 | 54392 | 31533 | 0 | 0 | 0 | 0 | 0 | 0 | 0 | 0 | 73859 | 163490 |
| HLKPGDRV | SORD; SORD | Q00796; A0/ | DHSO_HUMAN | Unmodified | 0 | 0 | 0 | 0 | 0 | 0 | 0 | 0 | 0 | 0 | 0 | 1113300 | 0 | 0 | 828270 | 540530 | 0 | 0 |
| HLLFEVQP( | RNF141; RNF14 | Q8WVD5; E9 | RN141_HUMAI | Unmodified | 748930 | 1079400 | 0 | 0 | 182630 | 0 | 0 | 0 | 0 | 0 | 0 | 0 | 0 | 0 | 453850 | 473650 | 0 | 0 |
| HLLFQSHN | SEPTIN10; SEP1 | Q9P0V9; B5I | SEP10_HUMAN | Unmodified | 0 | 0 | 0 | 0 | 480580 | 0 | 0 | 0 | 0 | 0 | 0 | 0 | 251390 | 0 | 0 | 0 | 119940 | 460290 |
| HLLGFSAK | ATF6; ATF6; ATI | P18850; A0A | ATF6A_HUMAN | Unmodified | 0 | 0 | 0 | 0 | 0 | 0 | 0 | 0 | 0 | 0 | 0 | 0 | 0 | 0 | 0 | 0 | 0 | 0 |
| HLLIGLPS( | EMC1; EMC1 | Q8N766; H7 | EMC1_HUMAN | Unmodified | 0 | 0 | 0 | 0 | 0 | 0 | 0 | 0 | 39969000 | 3436500 | 1254500 | 1168000 | 4867000 | 0 | 0 | 0 | 0 | 0 |
| HLLIGVSSI | ATP5F1C | P36542 | ATPG_HUMAN | Unmodified | 0 | 0 | 0 | 0 | 0 | 0 | 762880 | 0 | 12373000 | 3902100 | 76117 | 55708 | 0 | 90803 | 0 | 0 | 0 | 58529 |
| HLLPLTDR' | ITGB5; ITGB5; I | P18084; H7( | ITB5_HUMAN I | Unmodified | 2156900 | 1266300 | 3074000 | 2413600 | 749770 | 564890 | 0 | 0 | 2415900 | 290230 | 803330 | 8191000 | 977850 | 207170 | 4394900 | 2830000 | 312270 | 466490 |
| HLLPVETQI | DYNC1H1; DYN | Q14204; A0/ | DYHC1_HUMA | Unmodified | 0 | 0 | 0 | 0 | 0 | 0 | 0 | 0 | 212170 | 0 | 0 | 4768100 | 0 | 0 | 0 | 0 | 0 | 0 |
| HLLQAPLD | SEC23B; SEC2: | Q15437; A0/ | SC23B_HUMAI | Unmodified | 0 | 0 | 0 | 0 | 1039300 | 0 | 0 | 0 | 3034300 | 0 | 422070 | 9490600 | 0 | 0 | 0 | 0 | 0 | 0 |
| HLLQLKEEI | KIF20B | Q96Q89 | KI20B_HUMAN | Unmodified | 0 | 0 | 0 | 0 | 1645100 | 0 | 0 | 0 | 0 | 0 | 0 | 0 | 0 | 0 | 0 | 0 | 375210 | 671490 |
| HLLRETFEI | ASNS | P08243 | ASNS_HUMAN | Unmodified | 89758 | 0 | 535800 | 985740 | 1052300 | 653600 | 291740 | 0 | 402710 | 1196500 | 1403200 | 3168000 | 2812300 | 2088700 | 259420 | 951970 | 622220 | 790220 |
| HLLVHAPEI | ACLY | P53396 | ACLY_HUMAN | Unmodified | 102330 | 0 | 0 | 165500 | 0 | 0 | 0 | 0 | 0 | 0 | 0 | 5350900 | 0 | 0 | 489730 | 32446 | 0 | 0 |
| HLLVRRESI | MICALL1; MICA | Q8N3F8; B0( | MILK1_HUMAN | Unmodified | 0 | 0 | 0 | 0 | 838880 | 0 | 0 | 0 | 0 | 0 | 0 | 0 | 413670 | 313560 | 0 | 0 | 0 | 0 |
| HLMEKASR | MGLL | H7C4E0 | H7C4E0_HUM/ | 2 Oxidation (M) | 0 | 0 | 0 | 0 | 0 | 0 | 0 | 1710500 | 0 | 0 | 577230 | 1895900 | 2101000 | 0 | 0 | 0 | 0 | 0 |
| HLMELNAL | DHX36; DHX36 | Q9H2U1; E7 | DHX36_HUMAI | Oxidation (M) | 0 | 0 | 0 | 0 | 0 | 0 | 0 | 0 | 0 | 0 | 0 | 1022000 | 0 | 0 | 238260 | 0 | 0 | 0 |
| HLMLLRQE | ANKFY1; ANKF\ | Q9P2R3; I3L | ANFY1_HUMAN | Unmodified | 81575 | 0 | 0 | 68657 | 834270 | 0 | 0 | 0 | 0 | 0 | 0 | 0 | 0 | 932520 | 0 | 0 | 0 | 0 |
| HLMLPDFD | XRCC5; XRCC5 | P13010; C9J | XRCC5_HUMAI | Oxidation (M) | 0 | 0 | 0 | 0 | 0 | 0 | 0 | 0 | 0 | 0 | 0 | 295530 | 0 | 0 | 212560 | 0 | 0 | 0 |
| HLNDETTS | CYRIB | Q9NUQ9 | CYRIB_HUMAN | Unmodified | 0 | 0 | 0 | 0 | 93347 | 136330 | 0 | 0 | 0 | 143390 | 308690 | 0 | 268880 | 0 | 0 | 0 | 0 | 990960 |
| HLNEIDLFI | PTGES3; PTGES | Q15185; A0/ | TEBP_HUMAN I | Unmodified | 2238300 | 0 | 0 | 0 | 6577100 | 1203300 | 1689500 | 0 | 14553000 | 5739400 | 3672600 | 28542000 | 7063400 | 5888800 | 0 | 0 | 2894000 | 4147000 |
| HLNEIDLFI | PTGES3; PTGES | Q15185; A0/ | TEBP_HUMAN I | Unmodified | 0 | 0 | 0 | 0 | 779120 | 129380 | 178220 | 0 | 0 | 371170 | 0 | 0 | 578780 | 385370 | 0 | 0 | 582300 | 0 |
| HLNLAENS | PACSIN1; PACS | Q9BY11; F6l | PACN1_HUMAI | Unmodified | 1131200 | 1043700 | 0 | 104150 | 0 | 0 | 0 | 0 | 0 | 0 | 0 | 0 | 0 | 0 | 331000 | 210450 | 0 | 0 |
| HLNLAENS | PACSIN1; PACS | Q9BY11; F6l | PACN1_HUMAI | Unmodified | 5687800 | 7258100 | 97944 | 529460 | 0 | 0 | 0 | 0 | 0 | 0 | 0 | 0 | 221850 | 0 | 0 | 0 | 88378 | 0 |
| HLNPDTEL | TCF25; TCF25; | Q9BQ70; H3 | TCF25_HUMAN | Unmodified | 0 | 0 | 0 | 0 | 0 | 0 | 0 | 38029 | 0 | 0 | 0 | 0 | 0 | 0 | 191860 | 149480 | 0 | 0 |
| HLNPDTEL | TCF25; TCF25; | Q9BQ70; H3 | TCF25_HUMAN | Unmodified | 225910 | 257710 | 0 | 921020 | 0 | 0 | 0 | 0 | 0 | 0 | 0 | 0 | 0 | 0 | 35059 | 264050 | 0 | 0 |
| HLPGAFFL( | ZCCHC3 | Q9NUD5 | ZCHC3_HUMA | Unmodified | 0 | 0 | 0 | 0 | 0 | 0 | 0 | 0 | 0 | 0 | 0 | 0 | 0 | 0 | 0 | 0 | 0 | 0 |
| HLPNDPMI | CS; CS; CS; CS | O75390; A0/ | CISY_HUMAN ( | Oxidation (M) | 0 | 0 | 0 | 0 | 0 | 0 | 0 | 0 | 709040 | 0 | 0 | 0 | 0 | 0 | 0 | 0 | 0 | 0 |
| HLPNDPMI | CS; CS; CS; CS | O75390; A0/ | CISY_HUMAN ( | Unmodified | 0 | 0 | 0 | 0 | 0 | 0 | 0 | 0 | 1402000 | 142490 | 0 | 0 | 0 | 0 | 0 | 0 | 0 | 0 |
| HLPSSGGA | POU2F3 | Q9UKI9 | PO2F3_HUMAN | Unmodified | 0 | 0 | 0 | 0 | 0 | 0 | 0 | 0 | 0 | 0 | 0 | 516850 | 0 | 0 | 0 | 0 | 0 | 0 |
| HLQAVDYA | SV2A | Q7L0J3 | SV2A_HUMAN | Unmodified | 0 | 0 | 0 | 0 | 296620 | 0 | 373580 | 0 | 2328500 | 431050 | 0 | 264680 | 0 | 0 | 0 | 0 | 122560 | 1420200 |
| HLQDLME( | TOP1; TOP1 | P11387; A0A | TOP1_HUMAN | Unmodified | 0 | 0 | 41539 | 0 | 827450 | 0 | 0 | 85371 | 248610 | 0 | 0 | 9348000 | 0 | 0 | 120920 | 20068 | 0 | 0 |
| HLQDLSGF | LRPAP1 | P30533 | AMRP_HUMAN | Unmodified | 0 | 0 | 0 | 0 | 0 | 0 | 0 | 0 | 1751100 | 0 | 0 | 0 | 0 | 0 | 0 | 0 | 0 | 0 |
| HLQDLSGF | LRPAP1 | P30533 | AMRP_HUMAN | Unmodified | 1445500 | 2047900 | 0 | 0 | 2870500 | 479230 | 72151 | 8468.8 | 1866800 | 4634800 | 0 | 0 | 24914 | 0 | 0 | 0 | 60396 | 491270 |
| HLQENTQS | TRAF6 | Q9Y4K3 | TRAF6_HUMAN | Unmodified | 0 | 0 | 0 | 0 | 0 | 0 | 499140 | 384180 | 0 | 0 | 43487 | 141100 | 30508 | 15575 | 0 | 0 | 0 | 0 |
| HLQEQLNE | NF2; NF2 | P35240; A0A | MERL_HUMAN | Unmodified | 0 | 0 | 0 | 240030 | 3515400 | 692970 | 0 | 0 | 179760 | 0 | 0 | 255070 | 0 | 965300 | 0 | 0 | 1539000 | 5856400 |
| HLQEQLNE | NF2; NF2 | P35240; A0A | MERL_HUMAN | Unmodified | 0 | 0 | 0 | 0 | 0 | 0 | 0 | 0 | 0 | 0 | 0 | 0 | 266880 | 385730 | 0 | 0 | 2069700 | 1335900 |
| HLQKEAQA | AIDA | Q96BJ3 | AIDA_HUMAN / | Unmodified | 0 | 0 | 0 | 0 | 0 | 0 | 0 | 37177 | 0 | 0 | 0 | 0 | 0 | 0 | 0 | 0 | 0 | 129970 |
| HLQLAIR | H2AX; H2AC6; I | P16104; Q9: | H2AX_HUMAN | Unmodified | 1848900 | 1626900 | 0 | 0 | 0 | 0 | 0 | 0 | 0 | 0 | 0 | 10246000 | 0 | 0 | 0 | 0 | 0 | 0 |
| HLQLAIRGI | H2AZ2; H2AZ1; | Q71UI9; P0C | H2AV_HUMAN | Unmodified | 2789800 | 4633000 | 0 | 0 | 855210 | 0 | 6926400 | 197220 | 0 | 0 | 0 | 5132200 | 0 | 0 | 0 | 0 | 0 | 4320500 |
| HLQLAIRNI | H2AX; H2AC6; I | P16104; Q9: | H2AX_HUMAN | Unmodified | 5.06E+08 | 3.52E+08 | 8319300 | 2300400 | 79454000 | 29211000 | 53150000 | 1.16E+08 | 5929700 | 4606900 | 9347300 | 3.69E+09 | 6757500 | 5506400 | 0 | 2036300 | 2447400 | 33484000 |
| HLQLAIRNI | H2AC18; H2AC | Q6FI13; Q99 | H2A2A_HUMAI | Unmodified | 17040000 | 18272000 | 0 | 0 | 15564000 | 4829500 | 17609000 | 325910 | 0 | 880240 | 0 | 8029200 | 3574300 | 1258900 | 0 | 0 | 0 | 16173000 |
| HLQLAIRNI | H2AC6; H2AW; | Q93077; Q7I | H2A1C_HUMAI | Unmodified | 1183300 | 1168100 | 0 | 0 | 0 | 0 | 162740 | 0 | 0 | 0 | 0 | 916200 | 0 | 0 | 0 | 0 | 0 | 663490 |
| HLQLAVRN | H2AC21 | Q8IUE6 | H2A2B_HUMAI | Unmodified | 19902000 | 2190800 | 0 | 0 | 2700400 | 1117900 | 3046300 | 6660000 | 0 | 0 | 0 | 74462000 | 0 | 0 | 0 | 0 | 0 | 0 |
| HLQRSLLE( | SLITRK6 | Q9H5Y7 | SLIK6_HUMAN | Unmodified | 0 | 0 | 0 | 0 | 0 | 0 | 0 | 0 | 0 | 189430 | 0 | 0 | 0 | 66093 | 0 | 0 | 183400 | 202720 |
| HLQVNVTN | MYO1D; MYO1I | O94832; K7E | MYO1D_HUMA | Unmodified | 1757600 | 845500 | 265610 | 511250 | 0 | 0 | 0 | 0 | 0 | 0 | 0 | 186230 | 0 | 0 | 559130 | 518100 | 0 | 0 |
| HLQVNVTN | MYO1D; MYO1I | O94832; K7E | MYO1D_HUMA | Unmodified | 3374300 | 1571200 | 0 | 898660 | 0 | 0 | 176620 | 22762 | 272420 | 0 | 0 | 465650 | 0 | 182050 | 1245000 | 1043900 | 260540 | 0 |
| HLRAVPTD | RAB11A; RAB11 | P62491; Q1! | RB11A_HUMAN | Unmodified | 0 | 0 | 0 | 0 | 3210600 | 1286500 | 260420 | 0 | 0 | 425970 | 0 | 949290 | 1167300 | 0 | 0 | 0 | 182310 | 307300 |

| | | | | | | | | | | | | | | | | | | | | | | |
|---|---|---|---|---|---|---|---|---|---|---|---|---|---|---|---|---|---|---|---|---|---|---|
| HLRSPVLTI | GAS6 | Q14393 | GAS6_HUMAN | Unmodified | 0 | 0 | 0 | 0 | 138590 | 0 | 0 | 758030 | 28545000 | 0 | 0 | 0 | 564650 | 2172400 | 0 | 0 | 294270 | 715290 |
| HLSAGDVA | GALNT10 | Q86SR1 | GLT10_HUMAN | Unmodified | 0 | 0 | 0 | 0 | 143880 | 0 | 39107 | 35999 | 678350 | 0 | 0 | 0 | 0 | 0 | 0 | 0 | 133630 | 0 |
| HLSAGDVA | GALNT10 | Q86SR1 | GLT10_HUMAN | Unmodified | 0 | 0 | 0 | 0 | 0 | 0 | 83471 | 29477 | 893850 | 34044 | 0 | 22134 | 0 | 0 | 0 | 0 | 122870 | 33242 |
| HLSLAHNN | TLR9; N/A | Q9NR96; H0 | TLR9_HUMAN | Unmodified | 0 | 0 | 0 | 0 | 0 | 0 | 0 | 0 | 0 | 0 | 219410 | 711770 | 0 | 0 | 0 | 0 | 0 | 0 |
| HLSMFILLP | SERPINB5 | P36952 | SPB5_HUMAN | Oxidation (M) | 0 | 0 | 0 | 0 | 0 | 0 | 0 | 0 | 0 | 0 | 0 | 0 | 0 | 710840 | 0 | 0 | 692940 | 2244600 |
| HLSMFILLP | SERPINB5 | P36952 | SPB5_HUMAN | Oxidation (M) | 0 | 0 | 0 | 0 | 0 | 0 | 0 | 0 | 0 | 0 | 0 | 0 | 0 | 0 | 0 | 0 | 1215000 | 0 |
| HLSPYATLT | ESYT1 | Q9BSJ8 | ESYT1_HUMAN | Unmodified | 0 | 0 | 0 | 0 | 2557100 | 0 | 0 | 0 | 0 | 220820 | 0 | 0 | 0 | 0 | 0 | 0 | 0 | 811770 |
| HLSSQNRY | IMPDH2; IMPDH | P12268; A0A | IMDH2_HUMAN | Unmodified | 0 | 0 | 0 | 0 | 320290 | 90779 | 94766 | 42325 | 0 | 69092 | 58434 | 0 | 226150 | 0 | 0 | 0 | 127570 | 296790 |
| HLSSQNRY | IMPDH2; IMPDH | P12268; A0A | IMDH2_HUMAN | Unmodified | 0 | 0 | 0 | 0 | 2811800 | 183450 | 2265600 | 688700 | 0 | 712650 | 83892 | 0 | 1164600 | 627480 | 0 | 0 | 1875800 | 1693100 |
| HLSVNDLP | PRDX3 | P30048 | PRDX3_HUMAN | Unmodified | 0 | 0 | 0 | 0 | 0 | 0 | 0 | 0 | 25851000 | 3392800 | 566160 | 0 | 1045000 | 0 | 0 | 0 | 0 | 0 |
| HLSVNDLP | PRDX3 | P30048 | PRDX3_HUMAN | Unmodified | 0 | 0 | 0 | 0 | 0 | 0 | 0 | 0 | 0 | 3473100 | 0 | 0 | 0 | 105980 | 0 | 0 | 0 | 0 |
| HLTAALDR | TTC7B | Q86TV6 | TTC7B_HUMAN | Unmodified | 0 | 0 | 0 | 250940 | 396020 | 273540 | 137380 | 0 | 0 | 0 | 0 | 232880 | 239170 | 104930 | 442550 | 302570 | 118270 | 626740 |
| HLTAALDR | TTC7B | Q86TV6 | TTC7B_HUMAN | Oxidation (M) | 0 | 0 | 0 | 66528 | 1079900 | 0 | 239270 | 0 | 0 | 46327 | 0 | 0 | 348540 | 222540 | 0 | 0 | 850220 | 1240500 |
| HLTAALDR | TTC7B | Q86TV6 | TTC7B_HUMAN | Unmodified | 0 | 0 | 0 | 0 | 0 | 0 | 160610 | 0 | 0 | 0 | 0 | 0 | 326360 | 0 | 0 | 0 | 0 | 1118500 |
| HLTDAYFK | RPL6 | Q02878 | RL6_HUMAN 6 | Unmodified | 2306900 | 0 | 1462100 | 1133300 | 233390 | 0 | 200080 | 0 | 0 | 130050 | 0 | 9437900 | 370990 | 0 | 845180 | 791900 | 0 | 0 |
| HLTDAYFKI | RPL6 | Q02878 | RL6_HUMAN 6 | Unmodified | 0 | 0 | 0 | 0 | 0 | 0 | 0 | 0 | 0 | 0 | 0 | 0 | 0 | 0 | 0 | 0 | 0 | 0 |
| HLTGEFEK | RAN; RAN; RAN | P62826; H0 | RAN_HUMAN G | Unmodified | 0 | 396040 | 284720 | 0 | 0 | 0 | 929260 | 391070 | 0 | 0 | 0 | 3910600 | 14601000 | 0 | 0 | 0 | 0 | 0 |
| HLTHAQST | EBP | Q15125 | EBP_HUMAN 3 | Unmodified | 0 | 0 | 0 | 0 | 0 | 0 | 0 | 0 | 6162500 | 774220 | 0 | 0 | 0 | 0 | 0 | 0 | 0 | 13524 |
| HLTPTGREI | LSAMP; LSAMP; | Q13449; F5H | LSAMP_HUMAN | Unmodified | 0 | 0 | 0 | 0 | 0 | 0 | 0 | 0 | 1799400 | 0 | 0 | 0 | 0 | 0 | 0 | 0 | 0 | 0 |
| HLTPVTLEL | ALDH3A1; ALDH | P30838; P43 | AL3A1_HUMAN | Unmodified | 68235 | 70148 | 0 | 0 | 0 | 0 | 37153 | 0 | 2754600 | 519740 | 0 | 0 | 0 | 0 | 67007 | 0 | 0 | 57196 |
| HLTSRHPE | JUP | P14923 | PLAK_HUMAN | Unmodified | 319410 | 271950 | 2535200 | 4261900 | 514660 | 0 | 0 | 277450 | 0 | 0 | 1386900 | 7334800 | 0 | 0 | 4283500 | 4813400 | 0 | 0 |
| HLTSRHQE | CTNNB1; CTNN | P35222; A0A | CTNB1_HUMAN | Unmodified | 0 | 0 | 0 | 284320 | 0 | 124130 | 0 | 0 | 0 | 0 | 0 | 0 | 0 | 0 | 0 | 0 | 0 | 0 |
| HLTYENVEI | RAB11A; RAB11 | P62491; Q15 | RB11A_HUMAN | Unmodified | 0 | 0 | 0 | 787020 | 1764800 | 484510 | 963910 | 0 | 0 | 816110 | 0 | 0 | 618500 | 377120 | 0 | 51266 | 314980 | 385580 |
| HLVCSSGG | SDSL; SDSL | Q96GA7; F8 | SDSL_HUMAN | Unmodified | 0 | 98478 | 0 | 0 | 0 | 0 | 0 | 0 | 129160 | 0 | 0 | 0 | 360050 | 0 | 0 | 0 | 176780 | 54338 |
| HLVDPIDD | DGLUCY; DGLU | Q7Z3D6; H0 | GLUCM_HUMA | Unmodified | 0 | 162430 | 0 | 0 | 0 | 0 | 9117.3 | 0 | 968100 | 342720 | 0 | 0 | 0 | 0 | 17199 | 0 | 0 | 0 |
| HLVESTNE | UGGT1 | Q9NYU2 | UGGG1_HUMA | Oxidation (M) | 0 | 0 | 0 | 0 | 0 | 0 | 0 | 0 | 1570300 | 434240 | 206150 | 0 | 0 | 0 | 0 | 0 | 0 | 0 |
| HLVESTNE | UGGT1 | Q9NYU2 | UGGG1_HUMA | Unmodified | 0 | 0 | 0 | 0 | 0 | 0 | 102550 | 0 | 7476800 | 2556000 | 566910 | 0 | 0 | 0 | 0 | 0 | 0 | 0 |
| HLVGVCYT | CYC1 | P08574 | CY1_HUMAN C | Unmodified | 0 | 0 | 0 | 0 | 0 | 0 | 0 | 0 | 5005400 | 48934 | 0 | 0 | 0 | 0 | 0 | 0 | 0 | 0 |
| HLVLLDTA | MAN2B1; N/A | O00754; M0 | MA2B1_HUMAN | Unmodified | 199990 | 0 | 325630 | 470930 | 0 | 0 | 0 | 0 | 4785400 | 323570 | 712330 | 660860 | 123510 | 0 | 276030 | 216210 | 0 | 0 |
| HLVMGDIP | NASP; NASP; N | P49321; E9P | NASP_HUMAN | Oxidation (M) | 0 | 0 | 0 | 0 | 568700 | 0 | 629880 | 0 | 0 | 1970300 | 222940 | 193620 | 0 | 378480 | 199490 | 0 | 394500 | 0 |
| HLVMGDIP | NASP; NASP; N | P49321; E9P | NASP_HUMAN | Unmodified | 0 | 0 | 0 | 0 | 0 | 0 | 0 | 0 | 80458 | 168790 | 0 | 0 | 0 | 0 | 0 | 0 | 0 | 0 |
| HLVQALQA | GOLGA3 | Q08378 | GOGA3_HUMA | Unmodified | 0 | 0 | 0 | 37293 | 0 | 0 | 0 | 0 | 0 | 0 | 0 | 0 | 293680 | 0 | 0 | 0 | 0 | 0 |
| HLVSEEQF | CHAT | P28329 | CLAT_HUMAN | Unmodified | 0 | 3158900 | 39894 | 99265 | 0 | 0 | 0 | 258140 | 0 | 0 | 228200 | 1070100 | 0 | 0 | 0 | 0 | 338660 | 0 |
| HLVSPEALI | CSNK2A2; CSN | P19784; H3E | CSK22_HUMAN | Unmodified | 0 | 0 | 0 | 0 | 0 | 0 | 0 | 0 | 0 | 0 | 0 | 1900500 | 0 | 0 | 0 | 0 | 0 | 0 |
| HLVVKETE | FREM1; FREM1 | Q5H8C1; F8 | FREM1_HUMAN | Unmodified | 0 | 0 | 0 | 0 | 13199000 | 0 | 0 | 0 | 0 | 0 | 0 | 0 | 0 | 0 | 0 | 0 | 0 | 0 |
| HLVYESDK | RCN1 | Q15293 | RCN1_HUMAN | Unmodified | 0 | 0 | 261310 | 0 | 35107 | 0 | 0 | 0 | 8795500 | 382450 | 625900 | 677840 | 72494 | 0 | 222990 | 227300 | 0 | 0 |
| HLVYESDK | RCN1 | Q15293 | RCN1_HUMAN | Unmodified | 0 | 0 | 27096 | 0 | 0 | 0 | 0 | 0 | 8158600 | 2225400 | 687670 | 68034 | 87958 | 175000 | 0 | 0 | 0 | 0 |
| HLVYESDQ | CALU; CALU | O43852; H0 | CALU_HUMAN | Unmodified | 0 | 0 | 188900 | 0 | 37431 | 0 | 0 | 15208 | 19180000 | 580100 | 1198800 | 408610 | 0 | 0 | 308890 | 69320 | 0 | 0 |
| HLWDYTF | ACTR2; ACTR2 | P61160; F5H | ARP2_HUMAN | Unmodified | 444740 | 241950 | 400410 | 1209800 | 2924500 | 223610 | 0 | 35287 | 949820 | 0 | 513140 | 7506200 | 1214900 | 603260 | 1293100 | 343590 | 569280 | 2013100 |
| HLWDYTF | ACTR2; ACTR2 | P61160; F5H | ARP2_HUMAN | Unmodified | 0 | 0 | 0 | 0 | 395630 | 0 | 992720 | 0 | 0 | 0 | 0 | 0 | 0 | 0 | 0 | 0 | 462770 | 0 |
| HLYTLDGG | RACK1; RACK1; | P63244; D6F | RACK1_HUMAN | Unmodified | 0 | 0 | 0 | 0 | 0 | 0 | 0 | 0 | 0 | 0 | 0 | 3313900 | 0 | 0 | 0 | 44310 | 0 | 0 |
| HMALDMG | SNAP25 | P60880 | SNP25_HUMAN | Unmodified | 0 | 0 | 0 | 0 | 0 | 0 | 0 | 0 | 1827700 | 0 | 0 | 0 | 658250 | 0 | 0 | 0 | 0 | 0 |
| HMALDMG | SNAP25 | P60880 | SNP25_HUMAN | Oxidation (M) | 0 | 0 | 0 | 0 | 0 | 0 | 0 | 0 | 288390 | 0 | 0 | 0 | 0 | 0 | 0 | 0 | 29496 | 0 |
| HMDLCLTV | GALNT2 | Q10471 | GALT2_HUMAN | Oxidation (M) | 0 | 0 | 0 | 0 | 0 | 0 | 0 | 0 | 3717800 | 1443500 | 0 | 0 | 0 | 0 | 0 | 0 | 0 | 0 |
| HMDLCLTV | GALNT2 | Q10471 | GALT2_HUMAN | Unmodified | 0 | 0 | 0 | 0 | 0 | 0 | 0 | 0 | 3598600 | 425420 | 0 | 569760 | 0 | 0 | 0 | 0 | 0 | 0 |
| HMEASLQE | IGFBP5 | P24593 | IBP5_HUMAN | Unmodified | 631640 | 1002400 | 0 | 0 | 0 | 0 | 0 | 0 | 3108400 | 543570 | 0 | 0 | 0 | 0 | 0 | 0 | 0 | 0 |
| HMGAVFLL | ITGA3 | P26006 | ITA3_HUMAN | Oxidation (M) | 0 | 0 | 0 | 0 | 907180 | 0 | 0 | 0 | 475920 | 0 | 90112 | 1365200 | 0 | 0 | 0 | 0 | 182490 | 0 |
| HMGNRVYI | FLNC | Q14315 | FLNC_HUMAN | Unmodified | 0 | 0 | 0 | 0 | 4789200 | 170500 | 0 | 0 | 853750 | 0 | 0 | 0 | 0 | 0 | 271400 | 0 | 2200700 | 1245100 |
| HMGNRVYI | FLNC | Q14315 | FLNC_HUMAN | Oxidation (M) | 0 | 0 | 0 | 0 | 1087700 | 0 | 0 | 0 | 0 | 0 | 0 | 0 | 0 | 0 | 0 | 0 | 713780 | 285900 |
| HMMQNYL | SNX33; SNX33 | Q8WV41; H3 | SNX33_HUMAN | Unmodified | 0 | 0 | 203870 | 0 | 0 | 0 | 224200 | 0 | 0 | 0 | 0 | 0 | 0 | 0 | 425990 | 149890 | 0 | 0 |
| HMQHENV | MAPK13 | O15264 | MK13_HUMAN | Oxidation (M) | 0 | 5009800 | 0 | 303830 | 0 | 0 | 31956 | 0 | 0 | 0 | 0 | 4047000 | 0 | 0 | 0 | 0 | 0 | 0 |
| HMSEFME | PSMA1; PSMA1 | P25786; F5C | PSA1_HUMAN | Oxidation (M) | 0 | 0 | 0 | 0 | 0 | 0 | 0 | 0 | 605200 | 0 | 0 | 1068000 | 0 | 0 | 0 | 0 | 0 | 0 |
| HMYHSLYL | RPL19; RPL19; | P84098; J3K | RL19_HUMAN | Oxidation (M) | 0 | 0 | 0 | 0 | 0 | 0 | 0 | 0 | 0 | 0 | 0 | 1858100 | 0 | 0 | 0 | 0 | 0 | 0 |
| HMYVFGDI | ERP44 | Q9BS26 | ERP44_HUMAN | Unmodified | 0 | 0 | 0 | 0 | 0 | 0 | 0 | 0 | 7573400 | 79397 | 0 | 0 | 0 | 0 | 0 | 126750 | 0 | 0 |
| HMYVFGDI | ERP44 | Q9BS26 | ERP44_HUMAN | Oxidation (M) | 0 | 0 | 0 | 0 | 0 | 0 | 0 | 0 | 22654000 | 5053600 | 889620 | 0 | 0 | 0 | 0 | 0 | 126400 | 0 |
| HMYVFGDI | ERP44 | Q9BS26 | ERP44_HUMAN | Unmodified | 0 | 0 | 0 | 0 | 0 | 0 | 0 | 0 | 14538000 | 631250 | 0 | 0 | 0 | 0 | 0 | 0 | 0 | 0 |
| HMYVFGDI | ERP44 | Q9BS26 | ERP44_HUMAN | Oxidation (M) | 0 | 0 | 0 | 0 | 0 | 0 | 96666 | 0 | 596630 | 3037100 | 0 | 0 | 81164 | 0 | 0 | 0 | 0 | 0 |
| HNAAFCYK | SORD; SORD | Q00796; A0A | DHSO_HUMAN | Unmodified | 0 | 0 | 245310 | 67796 | 0 | 0 | 0 | 0 | 0 | 0 | 0 | 461980 | 0 | 0 | 228750 | 170520 | 0 | 0 |
| HNADDIVA | DIP2B | Q9P265 | DIP2B_HUMAN | Unmodified | 0 | 0 | 0 | 0 | 0 | 0 | 0 | 0 | 0 | 0 | 0 | 675980 | 0 | 0 | 0 | 242620 | 0 | 0 |
| HNAVFDQI | CASK; CASK; C | O14936; A0 | CSKP_HUMAN | Unmodified | 0 | 633850 | 0 | 644800 | 0 | 481090 | 422930 | 0 | 708500 | 203160 | 0 | 3006100 | 31055 | 557800 | 33962 | 1318700 | 429730 | 251060 |
| HNAVFDQI | CASK; CASK; C | O14936; A0 | CSKP_HUMAN | Unmodified | 0 | 0 | 0 | 1651600 | 2293600 | 92079 | 0 | 0 | 248550 | 3239900 | 0 | 0 | 0 | 2219300 | 2167700 | 0 | 536760 | 221810 |
| HNAVFDQI | CASK; CASK; C | O14936; A0 | CSKP_HUMAN | Unmodified | 0 | 0 | 0 | 0 | 5649100 | 582860 | 0 | 0 | 269040 | 0 | 0 | 0 | 1332100 | 0 | 167060 | 0 | 2340000 | 3348900 |
| HNDDEQY | HSP90AA2P; HS | Q14568; P0 | HS902_HUMAN | Unmodified | 3660100 | 2703200 | 3204700 | 4713800 | 3379500 | 0 | 583080 | 0 | 15327000 | 1584000 | 11029000 | 91741000 | 1561100 | 0 | 8995000 | 5320100 | 1024400 | 0 |
| HNDDEQY | HSP90AB1 | P08238 | HS90B_HUMAN | Unmodified | 0 | 0 | 471090 | 1164900 | 0 | 0 | 0 | 0 | 5566700 | 1055200 | 6319400 | 1537000 | 1107100 | 0 | 567590 | 0 | 0 | 432450 |
| HNDDEQY | HSP90AB1 | P08238 | HS90B_HUMAN | Unmodified | 0 | 0 | 0 | 0 | 44642000 | 24579000 | 4426700 | 235970 | 2848900 | 17925000 | 15582000 | 0 | 32800000 | 10759000 | 0 | 0 | 23616000 | 30642000 |
| HNDDEQY | HSP90AA1 | P07900 | HS90A_HUMAN | Unmodified | 0 | 0 | 0 | 0 | 0 | 0 | 0 | 0 | 2029800 | 466770 | 708030 | 1032200 | 0 | 0 | 0 | 0 | 0 | 0 |
| HNDDEQY | HSP90AA1 | P07900 | HS90A_HUMAN | Oxidation (M) | 217830 | 0 | 0 | 1018800 | 0 | 344070 | 0 | 0 | 1955300 | 516180 | 2529100 | 2702100 | 0 | 0 | 0 | 0 | 0 | 0 |
| HNDDEQY | HSP90AA1 | P07900 | HS90A_HUMAN | Oxidation (M) | 0 | 0 | 0 | 406270 | 8439600 | 3222400 | 1691100 | 265570 | 357350 | 5174700 | 2349200 | 0 | 1912000 | 936530 | 0 | 0 | 2835100 | 0 |
| HNDDEQY | HSP90AA1 | P07900 | HS90A_HUMAN | Unmodified | 0 | 0 | 0 | 0 | 12193000 | 2789500 | 0 | 0 | 753900 | 4809200 | 959750 | 0 | 887360 | 0 | 0 | 0 | 3109300 | 8567700 |
| HNELTGDN | CISD2; CISD2 | Q8N5K1; D6 | CISD2_HUMAN | Unmodified | 0 | 0 | 0 | 0 | 741870 | 0 | 532970 | 253000 | 19859000 | 1862100 | 4331900 | 1158600 | 398110 | 447060 | 0 | 0 | 0 | 0 |
| HNFLQAHN | PTPN23 | Q9H3S7 | PTN23_HUMAN | Unmodified | 350410 | 298830 | 95333 | 0 | 166580 | 89129 | 0 | 0 | 44136 | 0 | 0 | 0 | 216860 | 0 | 0 | 0 | 0 | 0 |
| HNFTPLAR | ACAA2; ACAA2; | P42765; A0A | THIM_HUMAN | Unmodified | 0 | 0 | 0 | 0 | 0 | 439960 | 0 | 45732 | 1133700 | 331130 | 0 | 0 | 0 | 0 | 0 | 0 | 0 | 17754 |
| HNGAAEPE | PKP3; PKP3 | Q9Y446; E9F | PKP3_HUMAN | Unmodified | 0 | 0 | 0 | 1014000 | 0 | 0 | 0 | 0 | 0 | 120150 | 259580 | 1175000 | 0 | 0 | 1250300 | 1537200 | 0 | 0 |
| HNIMDFA | CLTC; CLTC; CI | Q00610; J3K | CLH1_HUMAN | 3 Oxidation (M) | 0 | 460660 | 0 | 0 | 0 | 0 | 0 | 0 | 0 | 0 | 0 | 3305700 | 0 | 0 | 0 | 0 | 0 | 0 |
| HNINIGITN | FLVCR1; FLVCR | Q9Y5Y0; H7 | FLVC1_HUMAN | Unmodified | 0 | 255410 | 1212400 | 1384400 | 412960 | 0 | 0 | 0 | 1719300 | 472230 | 0 | 1535800 | 0 | 0 | 1467600 | 1236700 | 0 | 0 |
| HNIQALLK | PRKAR1A; PRK | P10644; X6R | KAP0_HUMAN | Unmodified | 0 | 0 | 0 | 0 | 2222800 | 547050 | 1339800 | 217580 | 974930 | 473450 | 83741 | 0 | 261980 | 0 | 0 | 0 | 1027000 | 1753400 |
| HNIQDEDT | PLXNB2; PLXNE | O15031; A6 | PLXB2_HUMAN | Unmodified | 0 | 0 | 0 | 0 | 0 | 0 | 0 | 0 | 0 | 0 | 0 | 3767700 | 0 | 0 | 5400100 | 3863500 | 0 | 0 |
| HNIQFSSFI | GLRX3 | O76003 | GLRX3_HUMAN | Unmodified | 0 | 0 | 208800 | 1244200 | 13941000 | 3259300 | 3852800 | 0 | 1107600 | 2198400 | 4567500 | 4481000 | 5078900 | 1911100 | 742170 | 389550 | 3158500 | 7315500 |
| HNIRLMGE | PFDN2 | Q9UHV9 | PFD2_HUMAN | Unmodified | 0 | 161650 | 0 | 0 | 1033900 | 108370 | 30029 | 0 | 0 | 214730 | 0 | 0 | 0 | 389020 | 0 | 0 | 493370 | 923120 |
| HNKPEVEV | ARPC4; ARPC4- | P59998; A0A | ARPC4_HUMAN | Unmodified | 0 | 0 | 0 | 0 | 0 | 0 | 0 | 0 | 0 | 0 | 0 | 0 | 0 | 0 | 0 | 0 | 0 | 0 |
| HNLEQDLV | TAOK1 | Q7L7X3 | TAOK1_HUMAN | Unmodified | 488160 | 0 | 0 | 645640 | 0 | 0 | 0 | 0 | 0 | 0 | 121530 | 2073300 | 0 | 0 | 483870 | 388070 | 0 | 0 |
| HNLLRPET | MAN1B1; MAN1 | Q9UKM7; H0 | MA1B1_HUMAN | Unmodified | 0 | 0 | 0 | 0 | 0 | 0 | 0 | 0 | 263060 | 1698600 | 0 | 0 | 1503400 | 0 | 0 | 0 | 2550300 | 0 |
| HNMDIGTV | YTHDF2 | Q9Y5A9 | YTHD2_HUMAN | Unmodified | 0 | 101530 | 0 | 0 | 0 | 0 | 0 | 0 | 0 | 0 | 0 | 444130 | 788190 | 0 | 0 | 0 | 0 | 0 |
| HNMDIGTV | YTHDF2 | Q9Y5A9 | YTHD2_HUMAN | Oxidation (M) | 62839 | 88282 | 0 | 0 | 0 | 0 | 0 | 0 | 0 | 0 | 0 | 0 | 366090 | 0 | 0 | 0 | 0 | 0 |
| HNNCMAS | CKMT1A; CKMT | P12532; C9 | KCRU_HUMAN | Unmodified | 0 | 0 | 0 | 24278 | 0 | 0 | 0 | 0 | 0 | 0 | 0 | 1547600 | 0 | 0 | 0 | 0 | 0 | 0 |
| HNNDTQH | HSP90B1; HSP | P14625; A0A | ENPL_HUMAN | Unmodified | 0 | 0 | 0 | 0 | 0 | 0 | 0 | 0 | 6055200 | 0 | 0 | 1003800 | 0 | 0 | 0 | 0 | 0 | 0 |
| HNNLDLVI | IDH3B; IDH3B; | O43837; A0 | IDH3B_HUMAN | Unmodified | 0 | 0 | 0 | 0 | 482430 | 0 | 0 | 0 | 2338100 | 0 | 0 | 0 | 0 | 0 | 564090 | 0 | 77852 | 0 |
| HNPPDDIG | MANEA; MANEA | Q5SRI9; X6R | MANEA_HUMA | Oxidation (M) | 0 | 0 | 0 | 0 | 0 | 0 | 0 | 0 | 3738600 | 0 | 0 | 0 | 0 | 0 | 0 | 0 | 0 | 0 |
| HNPPDDIG | MANEA; MANEA | Q5SRI9; X6R | MANEA_HUMA | Unmodified | 0 | 0 | 0 | 0 | 0 | 0 | 0 | 0 | 5295900 | 0 | 0 | 0 | 0 | 0 | 0 | 0 | 0 | 0 |
| HNPVFGVN | ACTR3B; ACTR | Q9P1U1; P6 | ARP3B_HUMAN | Oxidation (M) | 124760 | 266660 | 0 | 0 | 0 | 0 | 0 | 0 | 0 | 0 | 0 | 3536200 | 0 | 0 | 1014300 | 612850 | 0 | 0 |
| HNQLPLVII | P4HB; P4HB; P | P07237; A0A | PDIA1_HUMAN | Unmodified | 517920 | 0 | 917370 | 0 | 2864000 | 0 | 2117200 | 298680 | 58371000 | 50984000 | 9656800 | 2148000 | 0 | 0 | 0 | 1456700 | 742710 | 867770 |
| HNQLPLVII | P4HB; P4HB; P | P07237; A0A | PDIA1_HUMAN | Unmodified | 0 | 0 | 0 | 0 | 0 | 0 | 0 | 0 | 0 | 1315500 | 0 | 0 | 0 | 0 | 0 | 0 | 0 | 0 |
| HNSILQLD | ATP6V0A2 | Q9Y487 | VPP2_HUMAN | Unmodified | 0 | 0 | 0 | 0 | 94090 | 0 | 0 | 0 | 1890700 | 714140 | 0 | 690380 | 0 | 0 | 0 | 51595 | 0 | 0 |
| HNTAVSQL | SMC4; SMC4 | Q9NTJ3; E9F | SMC4_HUMAN | Unmodified | 13065000 | 5169800 | 16625000 | 0 | 0 | 0 | 0 | 141120 | 0 | 0 | 1042800 | 1127000 | 49922 | 39391 | 8762400 | 3541500 | 0 | 0 |
| HNTNSDPI | POMT2 | Q9UKY4 | POMT2_HUMA | Unmodified | 0 | 0 | 0 | 0 | 0 | 0 | 0 | 0 | 3277700 | 111530 | 0 | 0 | 0 | 0 | 0 | 0 | 0 | 0 |
| HNTVLMA | SLC12A7; SLC1 | Q9Y666; A0 | S12A7_HUMAN | Oxidation (M) | 0 | 0 | 0 | 817100 | 0 | 0 | 1320200 | 0 | 0 | 0 | 0 | 1294600 | 0 | 0 | 0 | 0 | 0 | 0 |
| HNVEVLGII | GSPT1; GSPT1 | P15170; H3E | ERF3A_HUMAN | Unmodified | 0 | 0 | 295740 | 1744500 | 4583200 | 0 | 0 | 0 | 0 | 0 | 1552400 | 5632100 | 0 | 3996800 | 0 | 1300000 | 1982300 | 3219200 |
| HNVMISTE | SELENBP1; SEL | Q13228; A6 | SBP1_HUMAN | Unmodified | 0 | 0 | 0 | 0 | 0 | 0 | 0 | 0 | 0 | 0 | 0 | 0 | 0 | 0 | 0 | 853310 | 0 | 0 |
| HNVQTLDI | PPAT | Q06203 | PUR1_HUMAN | Unmodified | 0 | 0 | 0 | 0 | 0 | 412850 | 109890 | 0 | 0 | 0 | 0 | 2753800 | 0 | 0 | 0 | 0 | 0 | 0 |
| HNYCRNP | PLAT; PLAT; PL | P00750; E7E | TPA_HUMAN T | Unmodified | 0 | 213160 | 0 | 0 | 0 | 0 | 0 | 0 | 0 | 0 | 0 | 0 | 0 | 0 | 0 | 0 | 181660 | 2842700 |
| HNYLPFIMI | UCHL5; UCHL5 | Q9Y5K5; H0 | UCHL5_HUMA | Oxidation (M) | 0 | 0 | 0 | 0 | 0 | 0 | 0 | 0 | 0 | 0 | 0 | 0 | 278050 | 0 | 0 | 0 | 0 | 0 |
| HNYQLELR | HLA-DQB1; HL | P01920; B4 | DQB1_HUMAN | Unmodified | 0 | 47386 | 0 | 0 | 0 | 0 | 0 | 0 | 1215000 | 0 | 0 | 0 | 0 | 0 | 0 | 0 | 0 | 0 |
| HPAKPDPS | RBBP7; RBBP7; | Q16576; E9 | RBBP7_HUMAN | Unmodified | 0 | 27104 | 614690 | 0 | 0 | 0 | 0 | 0 | 471840 | 0 | 660830 | 3550100 | 97034 | 0 | 150070 | 757270 | 0 | 32995 |
| HPASEAEI | PTK7; PTK7; PT | Q13308; Q8 | PTK7_HUMAN | Unmodified | 0 | 0 | 491170 | 846610 | 0 | 0 | 0 | 20902 | 6548700 | 335720 | 454540 | 401690 | 512940 | 0 | 1950700 | 1312300 | 0 | 411080 |
| HPDADSLY | YARS1; YARS1 | P54577; A0 | SYYC_HUMAN | Unmodified | 0 | 0 | 0 | 0 | 0 | 0 | 0 | 0 | 641870 | 0 | 245690 | 2070700 | 0 | 0 | 108880 | 139390 | 85398 | 0 |
| HPDSHQLF | G3BP1; G3BP1; | Q13283; A0 | G3BP1_HUMAN | Unmodified | 0 | 0 | 0 | 0 | 0 | 0 | 0 | 0 | 0 | 0 | 0 | 2672800 | 0 | 714630 | 0 | 0 | 0 | 0 |
| HPDYAILA | RRM1 | P23921 | RIR1_HUMAN | Unmodified | 0 | 0 | 0 | 0 | 0 | 0 | 152780 | 0 | 0 | 0 | 0 | 660030 | 0 | 0 | 0 | 161730 | 359680 | 0 |
| HPEAEMAC | JUP | P14923 | PLAK_HUMAN | Unmodified | 233260 | 127800 | 1702700 | 0 | 0 | 35901 | 56082 | 0 | 1191200 | 193380 | 830040 | 8679700 | 0 | 0 | 2701600 | 2489600 | 0 | 45473 |
| HPEAEMAC | JUP | P14923 | PLAK_HUMAN | Oxidation (M) | 53708 | 35044 | 791610 | 0 | 0 | 0 | 0 | 0 | 0 | 0 | 204900 | 1246400 | 0 | 0 | 1919500 | 1302700 | 0 | 0 |

| | | | | | | | | | | | | | | | | | | | | | | |
|---|---|---|---|---|---|---|---|---|---|---|---|---|---|---|---|---|---|---|---|---|---|---|
| HPELFEAL( | PSMC5; PSMC5 | P62195; J3Q | PRS8_HUMAN | Unmodified | 0 | 0 | 0 | 0 | 0 | 0 | 0 | 0 | 0 | 0 | 0 | 2158400 | 0 | 0 | 0 | 0 | 0 | 0 |
| HPESNTAG | CLIC4 | Q9Y696 | CLIC4_HUMAN | Unmodified | 153150 | 0 | 278080 | 364410 | 2881100 | 429750 | 202220 | 195500 | 2482900 | 209490 | 578030 | 3886000 | 565250 | 542090 | 656780 | 355580 | 655890 | 1999100 |
| HPESNTAG | CLIC4 | Q9Y696 | CLIC4_HUMAN | Oxidation (M) | 341640 | 120700 | 157160 | 197740 | 965140 | 258260 | 0 | 0 | 0 | 146350 | 640190 | 958910 | 317310 | 266520 | 0 | 507640 | 199740 | 0 |
| HPEVDVLII | ACLY | P53396 | ACLY_HUMAN | Unmodified | 0 | 0 | 0 | 25796 | 0 | 0 | 0 | 0 | 361690 | 184550 | 56991 | 403340 | 17541 | 0 | 54110 | 0 | 0 | 0 |
| HPGFEATF | ST14 | Q9Y5Y6 | ST14_HUMAN S | Unmodified | 0 | 0 | 3059500 | 280460 | 0 | 0 | 0 | 0 | 0 | 0 | 0 | 0 | 0 | 0 | 0 | 1993000 | 0 | 0 |
| HPGSFDVV | RPS4X; RPS4Y1 | P62701; P22 | RS4X_HUMAN | Unmodified | 0 | 0 | 0 | 0 | 0 | 0 | 0 | 0 | 0 | 0 | 0 | 3310800 | 0 | 0 | 0 | 0 | 0 | 0 |
| HPHDIIDDI | RPL12 | P30050 | RL12_HUMAN ( | Unmodified | 962770 | 1107500 | 0 | 440620 | 0 | 0 | 0 | 0 | 0 | 0 | 3281400 | 7649600 | 0 | 0 | 477940 | 473500 | 0 | 0 |
| HPIGVLLN/ | MEMO1 | Q9Y316 | MEMO1_HUMA | Unmodified | 337610 | 0 | 48150 | 0 | 0 | 0 | 0 | 0 | 0 | 0 | 0 | 539900 | 0 | 0 | 497710 | 140050 | 0 | 0 |
| HPLDHDC: | ZFAND2B; ZFAN | Q8WV99; B4 | ZFN2B_HUMAN | Unmodified | 0 | 0 | 0 | 0 | 0 | 0 | 0 | 0 | 6561600 | 0 | 0 | 0 | 0 | 0 | 0 | 0 | 0 | 0 |
| HPQAHVDI | FDXR; FDXR; FD | P22570; A0A | ADRO_HUMAN | Unmodified | 0 | 0 | 0 | 0 | 0 | 212670 | 0 | 0 | 628520 | 176700 | 0 | 178980 | 0 | 0 | 0 | 0 | 0 | 0 |
| HPQLDYVF | UTRN | P46939 | UTRN_HUMAN | Unmodified | 0 | 0 | 0 | 0 | 320170 | 0 | 0 | 0 | 0 | 0 | 0 | 258010 | 0 | 0 | 1111800 | 598170 | 45120 | 0 |
| HPQLLYESI | CSNK1A1L; CSI | Q8N752; P4 | KC1AL_HUMAN | Unmodified | 0 | 0 | 0 | 0 | 0 | 0 | 0 | 0 | 0 | 0 | 0 | 494170 | 0 | 0 | 266660 | 173430 | 0 | 0 |
| HPQPGAVE | CAD; CAD; CAD | P27708; F8V | PYR1_HUMAN | Unmodified | 0 | 0 | 0 | 0 | 0 | 0 | 0 | 0 | 0 | 0 | 0 | 284530 | 0 | 0 | 0 | 0 | 0 | 0 |
| HPSKPDPS | RBBP4 | Q09028 | RBBP4_HUMAN | Unmodified | 0 | 0 | 0 | 0 | 0 | 0 | 0 | 0 | 1045400 | 0 | 0 | 1435600 | 0 | 0 | 0 | 0 | 0 | 0 |
| HPSQQDTF | EPB41L1; EPB4 | Q9H4G0; Q4 | E41L1_HUMAN | Unmodified | 0 | 0 | 84643 | 0 | 0 | 0 | 0 | 0 | 0 | 0 | 771880 | 682140 | 0 | 0 | 549540 | 465740 | 0 | 0 |
| HPSVNENF | PLAA; PLAA | Q9Y263; H0' | PLAP_HUMAN | Unmodified | 0 | 0 | 0 | 0 | 0 | 0 | 0 | 0 | 904180 | 0 | 0 | 418190 | 0 | 0 | 0 | 0 | 0 | 0 |
| HPVCSGTC | SPINT1 | O43278 | SPIT1_HUMAN | Unmodified | 437100 | 130680 | 0 | 0 | 0 | 31975 | 0 | 0 | 93852 | 0 | 0 | 0 | 0 | 0 | 83463 | 64416 | 0 | 0 |
| HPVISESEV | SNX9; SNX9; SN | Q9Y5X1; A0A | SNX9_HUMAN | Unmodified | 11279000 | 15774000 | 576020 | 805780 | 2316800 | 0 | 84708 | 0 | 0 | 0 | 0 | 3140500 | 488320 | 0 | 1732500 | 2950300 | 0 | 1041900 |
| HPVSEVGP | SLC20A1 | Q8WUM9 | S20A1_HUMAN | Unmodified | 0 | 0 | 0 | 0 | 0 | 0 | 0 | 0 | 0 | 0 | 508730 | 956060 | 0 | 0 | 0 | 0 | 0 | 0 |
| HPVSLEQY | PSMD8; PSMD8 | P48556; K7E | PSMD8_HUMA | Oxidation (M) | 0 | 0 | 97967 | 0 | 0 | 0 | 0 | 0 | 0 | 0 | 512910 | 3550000 | 242860 | 0 | 0 | 0 | 0 | 0 |
| HPVTGQFL | TFRC; TFRC | P02786; G3V | TFR1_HUMAN | Unmodified | 711810 | 564630 | 754880 | 866460 | 250190 | 0 | 0 | 100740 | 8648100 | 494900 | 0 | 3829600 | 0 | 0 | 1230800 | 885640 | 0 | 0 |
| HPWDDISY | ACLY | P53396 | ACLY_HUMAN | 2 Oxidation (M) | 0 | 0 | 0 | 0 | 0 | 0 | 0 | 0 | 0 | 0 | 0 | 1657800 | 0 | 0 | 86590 | 0 | 0 | 0 |
| HPYNFPEG | GALNT16 | Q8N428 | GLT16_HUMAN | Unmodified | 0 | 0 | 0 | 0 | 0 | 0 | 0 | 0 | 1774700 | 0 | 0 | 0 | 0 | 0 | 0 | 0 | 0 | 0 |
| HQAFEAEL | SPTAN1; SPTAN | Q13813; A0A | SPTN1_HUMAN | Unmodified | 0 | 0 | 0 | 0 | 0 | 0 | 0 | 0 | 36944 | 0 | 0 | 739290 | 0 | 0 | 81005 | 0 | 0 | 0 |
| HQAFEAEL | SPTAN1; SPTAN | Q13813; A0A | SPTN1_HUMAN | Unmodified | 0 | 0 | 412660 | 214440 | 894370 | 384540 | 45838 | 0 | 589170 | 226410 | 701400 | 9391900 | 379600 | 121390 | 592800 | 373290 | 136220 | 0 |
| HQAFEAEL | SPTAN1; SPTAN | Q13813; A0A | SPTN1_HUMAN | Unmodified | 0 | 0 | 40037 | 0 | 0 | 1641900 | 0 | 0 | 0 | 0 | 0 | 1450900 | 0 | 0 | 0 | 51732 | 674310 | 0 |
| HQAFMAEL | SPTBN1; SPTBN | Q01082; A0A | SPTB2_HUMAN | Unmodified | 0 | 0 | 0 | 0 | 0 | 0 | 0 | 0 | 0 | 0 | 0 | 0 | 0 | 0 | 0 | 0 | 0 | 0 |
| HQALQAEI | SPTAN1; SPTAN | Q13813; A0A | SPTN1_HUMAN | Unmodified | 0 | 0 | 89139 | 0 | 0 | 0 | 0 | 0 | 0 | 0 | 0 | 890900 | 0 | 0 | 0 | 0 | 0 | 0 |
| HQAQIDHY | RTN4; RTN4; RT | Q9NQC3; F8 | RTN4_HUMAN | Unmodified | 0 | 0 | 566010 | 310270 | 774590 | 0 | 268350 | 177910 | 305100 | 0 | 0 | 2290200 | 0 | 0 | 505010 | 814770 | 142830 | 155710 |
| HQASINELI | EPB41L1; EPB4 | Q9H4G0; H7 | E41L1_HUMAN | Unmodified | 0 | 0 | 0 | 0 | 839550 | 45210 | 0 | 0 | 0 | 0 | 87445 | 300580 | 301080 | 288850 | 0 | 0 | 413150 | 1066500 |
| HQASISELI | EPB41L2; EPB4 | O43491; Q6 | E41L2_HUMAN | Unmodified | 0 | 0 | 0 | 0 | 23989 | 483490 | 72988 | 0 | 0 | 193190 | 0 | 0 | 0 | 0 | 0 | 0 | 225160 | 635380 |
| HQDGLVV\ | MUC5AC | P98088 | MUC5A_HUMA | Unmodified | 0 | 0 | 0 | 0 | 0 | 0 | 0 | 0 | 0 | 0 | 0 | 0 | 0 | 0 | 0 | 0 | 0 | 196920 |
| HQDGLVV\ | MUC5AC | P98088 | MUC5A_HUMA | Unmodified | 0 | 0 | 0 | 0 | 0 | 0 | 0 | 0 | 0 | 0 | 0 | 0 | 0 | 0 | 0 | 0 | 0 | 450730 |
| HQDVPSQI | AASDHPPT | Q9NRN7 | ADPPT_HUMAN | Unmodified | 53962 | 70293 | 0 | 2591900 | 17058 | 0 | 0 | 0 | 0 | 117620 | 370520 | 2652100 | 684140 | 0 | 0 | 83743 | 0 | 0 |
| HQEAEMA( | CTNNB1; CTNN | P35222; A0A | CTNB1_HUMAI | Unmodified | 2758900 | 1928700 | 5809800 | 5741700 | 117950 | 0 | 0 | 0 | 356560 | 35942 | 1099800 | 10447000 | 0 | 0 | 10946000 | 10114000 | 359180 | 63752 |
| HQEAEMA( | CTNNB1; CTNN | P35222; A0A | CTNB1_HUMAI | Oxidation (M) | 850640 | 616420 | 0 | 1253100 | 0 | 127400 | 0 | 0 | 621420 | 237450 | 0 | 3855100 | 0 | 340210 | 1560900 | 4805800 | 0 | 364740 |
| HQEAEMA( | CTNNB1; CTNN | P35222; A0A | CTNB1_HUMAI | Oxidation (M) | 0 | 0 | 0 | 841940 | 0 | 0 | 0 | 0 | 0 | 0 | 278150 | 0 | 0 | 0 | 0 | 0 | 0 | 0 |
| HQEGEIFD | RPL6 | Q02878 | RL6_HUMAN 6( | Unmodified | 2559600 | 1261400 | 1602600 | 666420 | 0 | 0 | 0 | 0 | 0 | 0 | 2657200 | 20214000 | 0 | 0 | 1998300 | 1907000 | 0 | 0 |
| HQEGEIFD | RPL6 | Q02878 | RL6_HUMAN 6( | Unmodified | 3038800 | 3469800 | 0 | 2850100 | 460060 | 427660 | 68191 | 49934 | 477660 | 247660 | 0 | 4018400 | 1372300 | 0 | 0 | 0 | 100810 | 235900 |
| HQEHILRF/ | TRIM28 | Q13263 | TIF1B_HUMAN | Unmodified | 0 | 0 | 0 | 0 | 0 | 0 | 4519100 | 0 | 0 | 0 | 0 | 0 | 0 | 0 | 0 | 0 | 0 | 0 |
| HQEPVYSV | TBL1XR1; TBL1) | Q9BZK7; A0/ | TBL1R_HUMAN | Unmodified | 0 | 0 | 0 | 0 | 101870 | 0 | 0 | 29537 | 489910 | 0 | 389690 | 2574800 | 0 | 0 | 0 | 0 | 0 | 0 |
| HQFVLSRP | SH3BP4 | Q9P0V3 | SH3B4_HUMAI | Unmodified | 0 | 0 | 0 | 0 | 390560 | 0 | 0 | 0 | 1112000 | 531150 | 0 | 0 | 0 | 0 | 154890 | 251280 | 151930 | 164870 |
| HQGGWTD | FAM98A | Q8NCA5 | FA98A_HUMAN | Unmodified | 0 | 0 | 146540 | 105280 | 0 | 0 | 0 | 0 | 156050 | 0 | 0 | 0 | 0 | 0 | 0 | 0 | 0 | 0 |
| HQGSELHF | HSPG2 | P98160 | PGBM_HUMAN | Unmodified | 0 | 0 | 781370 | 757090 | 0 | 0 | 0 | 0 | 0 | 0 | 0 | 0 | 0 | 0 | 0 | 516870 | 0 | 0 |
| HQGVMVG | ACTG1; ACTC1; | P63261; P68 | ACTG_HUMAN | 2 Oxidation (M) | 15108000 | 13549000 | 4916300 | 7258700 | 21370000 | 10081000 | 737440 | 974140 | 4710500 | 906020 | 9772900 | 57312000 | 21334000 | 3024400 | 7659000 | 16122000 | 1955100 | 7745000 |
| HQGVMVG | ACTG1; ACTC1; | P63261; P68 | ACTG_HUMAN | Oxidation (M) | 13061000 | 16729000 | 29645000 | 26523000 | 57715000 | 38548000 | 1719400 | 1392500 | 26252000 | 5634800 | 7543900 | 93519000 | 40319000 | 16761000 | 33718000 | 31509000 | 5061500 | 23901000 |
| HQGVMVG | ACTG1; ACTC1; | P63261; P68 | ACTG_HUMAN | Unmodified | 9572800 | 11551000 | 14684000 | 13897000 | 51874000 | 19281000 | 1858100 | 865670 | 37934000 | 3402800 | 9623500 | 1.23E+08 | 26481000 | 12136000 | 20734000 | 10678000 | 3212500 | 13700000 |
| HQGVMVG | ACTG1; ACTC1; | P63261; P68 | ACTG_HUMAN | 2 Oxidation (M) | 0 | 94398 | 0 | 898940 | 1.29E+08 | 28184000 | 5249300 | 5980500 | 1573300 | 8529100 | 36391 | 0 | 68615000 | 15649000 | 0 | 0 | 12840000 | 15591000 |
| HQGVMVG | ACTG1; ACTC1; | P63261; P68 | ACTG_HUMAN | Oxidation (M) | 351730 | 677410 | 0 | 2496100 | 2.57E+08 | 44765000 | 18540000 | 12563000 | 1640000 | 21436000 | 0 | 251760 | 1.67E+08 | 58290000 | 0 | 0 | 43564000 | 78080000 |
| HQGVMVG | ACTG1; ACTC1; | P63261; P68 | ACTG_HUMAN | Unmodified | 0 | 602840 | 0 | 1265000 | 3.73E+08 | 55964000 | 19269000 | 2692500 | 5986800 | 24863000 | 638850 | 841380 | 1.6E+08 | 55711000 | 0 | 0 | 55378000 | 1.06E+08 |
| HQGVMVG | ACTG1; ACTC1; | P63261; P68 | ACTG_HUMAN | 2 Oxidation (M) | 255910 | 0 | 0 | 7761700 | 16173000 | 11227000 | 369800 | 2551200 | 0 | 390900 | 0 | 0 | 3630700 | 17136000 | 0 | 0 | 7772700 | 20586000 |
| HQGVMVG | ACTG1; ACTC1; | P63261; P68 | ACTG_HUMAN | Oxidation (M) | 0 | 0 | 0 | 4615500 | 24115000 | 3620900 | 6202100 | 1753900 | 1405400 | 2321900 | 0 | 0 | 9672700 | 2483700 | 0 | 0 | 11383000 | 11724000 |
| HQGVMVG | ACTG1; ACTC1; | P63261; P68 | ACTG_HUMAN | Unmodified | 387940 | 0 | 0 | 4205800 | 28010000 | 7479600 | 2007600 | 567850 | 942340 | 923700 | 168950 | 0 | 15094000 | 5890300 | 0 | 0 | 9672400 | 4153100 |
| HQHDRVC | ITGA7; ITGA7 | Q13683; J3K | ITA7_HUMAN I | Unmodified | 0 | 0 | 0 | 0 | 299960 | 0 | 820020 | 0 | 127620 | 0 | 0 | 0 | 0 | 0 | 0 | 0 | 0 | 0 |
| HQILEQAVI | SPTBN1; SPTBN | Q01082; A0/ | SPTB2_HUMAN | Unmodified | 0 | 0 | 0 | 0 | 943580 | 283630 | 0 | 0 | 698860 | 354470 | 466820 | 0 | 0 | 0 | 0 | 0 | 458110 | 209330 |
| HQILPKEQ | UQCRB | P14927 | QCR7_HUMAN | Unmodified | 0 | 0 | 0 | 0 | 0 | 0 | 0 | 0 | 0 | 5127000 | 0 | 0 | 0 | 0 | 0 | 0 | 685750 | 692460 |
| HQLAEQKF | RNF41; RNF41; | Q9H4P4; F8 | RNF41_HUMAN | Unmodified | 0 | 0 | 0 | 0 | 2100900 | 575320 | 90880 | 0 | 0 | 48613 | 0 | 0 | 282300 | 756310 | 0 | 0 | 0 | 0 |
| HQLQVQM | EEA1 | Q15075 | EEA1_HUMAN | Unmodified | 0 | 0 | 0 | 0 | 1962200 | 87781 | 107360 | 0 | 0 | 0 | 0 | 0 | 130440 | 0 | 0 | 0 | 35032 | 128510 |
| HQLYIDET\ | FGB | P02675 | FIBB_HUMAN F | Unmodified | 0 | 7810900 | 0 | 0 | 0 | 0 | 0 | 0 | 0 | 0 | 0 | 0 | 252630 | 0 | 0 | 0 | 0 | 0 |
| HQNGRFV( | CTNND1; CTNN | O60716; E9F | CTND1_HUMA | Unmodified | 0 | 0 | 0 | 0 | 0 | 109580 | 0 | 0 | 185230 | 27418 | 0 | 186100 | 295740 | 109100 | 0 | 54235 | 112340 | 697590 |
| HQNLLKEL | CHGA; CHGA | P10645; G5E | CMGA_HUMAN | Unmodified | 0 | 0 | 0 | 0 | 0 | 0 | 0 | 0 | 0 | 634950 | 0 | 0 | 0 | 0 | 0 | 0 | 0 | 0 |
| HQNVQLPI | PTBP1; PTBP1; | P26599; A0A | PTBP1_HUMAN | Unmodified | 0 | 1105200 | 121900 | 165410 | 0 | 0 | 0 | 0 | 149100 | 0 | 340690 | 1033900 | 0 | 0 | 0 | 35147 | 0 | 0 |
| HQNVQLPI | PTBP1; PTBP1; | P26599; A0A | PTBP1_HUMAN | Unmodified | 30811 | 0 | 0 | 38810 | 291800 | 91348 | 205860 | 58949 | 0 | 715760 | 0 | 196350 | 309270 | 138510 | 0 | 0 | 0 | 0 |
| HQPPSTN( | WWP2 | O00308 | WWP2_HUMAN | Unmodified | 702190 | 581660 | 226800 | 359190 | 0 | 0 | 0 | 93714 | 44606 | 0 | 0 | 0 | 37397 | 0 | 503930 | 550770 | 0 | 0 |
| HQPTAIIAK | TKT; TKT; TKT | P29401; A0/ | TKT_HUMAN Tr | Unmodified | 0 | 0 | 0 | 0 | 223220 | 46431 | 0 | 0 | 0 | 98991 | 39225 | 135470 | 0 | 0 | 0 | 0 | 0 | 0 |
| HQQALEV\ | SCRN3 | Q0VDG4 | SCRN3_HUMA | Unmodified | 0 | 0 | 0 | 0 | 600250 | 0 | 195500 | 179740 | 270270 | 0 | 67973 | 0 | 358400 | 348690 | 0 | 0 | 184760 | 0 |
| HQQLLEEK | MYH10 | P35580 | MYH10_HUMA | Unmodified | 0 | 0 | 388670 | 68436 | 0 | 0 | 0 | 0 | 261440 | 0 | 0 | 1050000 | 141100 | 120130 | 0 | 237160 | 0 | 0 |
| HQQLLGE\ | CUL1; CUL1; CI | Q13616; A0/ | CUL1_HUMAN | Unmodified | 0 | 0 | 0 | 0 | 0 | 0 | 0 | 0 | 0 | 0 | 0 | 964520 | 0 | 2792100 | 0 | 0 | 0 | 0 |
| HQQQLLA: | XPO4 | Q9C0E2 | XPO4_HUMAN | Unmodified | 0 | 0 | 0 | 0 | 0 | 0 | 0 | 98810 | 0 | 46393 | 0 | 2091700 | 0 | 0 | 0 | 0 | 0 | 0 |
| HQQSSTLC | DOCK9; DOCK9 | Q9BZ29; A0/ | DOCK9_HUMA | Unmodified | 0 | 15512 | 91629 | 0 | 0 | 0 | 0 | 0 | 16204 | 102400 | 0 | 0 | 0 | 0 | 79676 | 119780 | 0 | 0 |
| HQSFVLVG | DHX15 | O43143 | DHX15_HUMAI | Unmodified | 45793 | 0 | 0 | 0 | 0 | 0 | 54341 | 62494 | 497830 | 115070 | 0 | 6270500 | 0 | 52342 | 89137 | 0 | 15219 | 0 |
| HQTLQGLA | TWF2; TWF2; T\ | Q6IBS0; D6F | TWF2_HUMAN | Unmodified | 0 | 0 | 0 | 0 | 0 | 0 | 0 | 0 | 1770600 | 266460 | 1312400 | 12050000 | 0 | 0 | 482740 | 0 | 0 | 0 |
| HQTLQGLA | TWF2; TWF2; T\ | Q6IBS0; D6F | TWF2_HUMAN | Unmodified | 0 | 0 | 0 | 0 | 0 | 0 | 0 | 0 | 0 | 0 | 0 | 0 | 0 | 0 | 0 | 0 | 1020100 | 0 |
| HQTNISELI | EPB41L3; EPB4 | Q9Y2J2; A0A | E41L3_HUMAN | Unmodified | 0 | 51759 | 0 | 0 | 0 | 0 | 61528 | 11598 | 0 | 0 | 0 | 0 | 157470 | 217240 | 0 | 0 | 546240 | 5954700 |
| HQTNISELI | EPB41L3; EPB4 | Q9Y2J2; A0A | E41L3_HUMAN | Unmodified | 0 | 0 | 0 | 0 | 0 | 0 | 0 | 74991 | 0 | 0 | 0 | 0 | 0 | 0 | 0 | 0 | 1556000 | 1534900 |
| HQVDKGP\ | PLXNB1 | O43157 | PLXB1_HUMAN | Unmodified | 0 | 0 | 0 | 45369 | 0 | 0 | 0 | 0 | 0 | 78243 | 100230 | 378190 | 0 | 0 | 113580 | 0 | 0 | 0 |
| HQVEQLSS | NUMA1; NUMA | Q14980; F5H | NUMA1_HUMA | Unmodified | 40364 | 0 | 0 | 0 | 0 | 0 | 0 | 0 | 527490 | 161390 | 0 | 271870 | 0 | 0 | 0 | 0 | 0 | 0 |
| HQVEYLGL | MYO1D; MYO1D | O94832; J3Q | MYO1D_HUMA | Unmodified | 7968300 | 15787000 | 2028700 | 2471200 | 0 | 0 | 0 | 0 | 1829900 | 0 | 0 | 1361700 | 0 | 0 | 6867400 | 4356500 | 0 | 0 |
| HRAAEAAIN | PRKRA; PRKRA; | O75569; A0/ | PRKRA_HUMAI | Unmodified | 0 | 0 | 0 | 0 | 0 | 0 | 1459200 | 332770 | 0 | 30269 | 0 | 0 | 151910 | 0 | 0 | 0 | 0 | 0 |
| HRADHPPA | LIMA1; LIMA1; L | Q9UHB6; F8 | LIMA1_HUMAN | Unmodified | 109360 | 136960 | 205630 | 0 | 897160 | 84280 | 36171 | 0 | 9408.3 | 82804 | 0 | 0 | 232780 | 112670 | 293440 | 86625 | 20670 | 0 |
| HRAEAPPL | PRKRA; PRKRA; | O75569; A0/ | PRKRA_HUMAI | Unmodified | 562020 | 808460 | 718960 | 943150 | 0 | 0 | 1243400 | 385180 | 0 | 0 | 0 | 0 | 1328700 | 392620 | 615530 | 665840 | 0 | 298760 |
| HRAFEDEM | SPTBN1; SPTBN | Q01082; A0/ | SPTB2_HUMAN | Unmodified | 0 | 0 | 0 | 0 | 0 | 0 | 0 | 0 | 0 | 0 | 0 | 934050 | 0 | 0 | 0 | 0 | 0 | 0 |
| HRALSIENS | LDOC1 | O95751 | LDOC1_HUMAI | Oxidation (M) | 0 | 0 | 0 | 0 | 165480 | 0 | 59611 | 0 | 0 | 0 | 0 | 0 | 0 | 0 | 339190 | 271630 | 152060 | 188090 |
| HRALSIENS | LDOC1 | O95751 | LDOC1_HUMAI | Unmodified | 0 | 0 | 117770 | 284660 | 0 | 0 | 0 | 0 | 0 | 0 | 0 | 0 | 0 | 0 | 431640 | 262390 | 0 | 0 |
| HRDFVAEP | BANF1; BANF1 | O75531; E9F | BAF_HUMAN B | Unmodified | 0 | 0 | 0 | 0 | 0 | 0 | 0 | 12678 | 172320 | 28757 | 0 | 4069300 | 0 | 0 | 0 | 0 | 0 | 0 |
| HRDFVAEP | BANF1; BANF1 | O75531; E9F | BAF_HUMAN B | Oxidation (M) | 74801 | 80166 | 129970 | 0 | 0 | 0 | 0 | 0 | 0 | 0 | 71463 | 1079500 | 831670 | 0 | 94946 | 60533 | 0 | 0 |
| HRDFVAEP | BANF1 | O75531 | BAF_HUMAN B | Oxidation (M) | 12210000 | 7440000 | 3172700 | 5428200 | 58531000 | 16249000 | 12241000 | 2877200 | 9202500 | 8841000 | 8951200 | 2.53E+08 | 25826000 | 21431000 | 5237800 | 5597700 | 9764700 | 40769000 |
| HRDFVAEP | BANF1 | O75531 | BAF_HUMAN B | Unmodified | 521610 | 888210 | 0 | 0 | 9793700 | 0 | 3851700 | 0 | 6849300 | 3599400 | 0 | 5452800 | 6186200 | 5054200 | 596350 | 0 | 3262800 | 18326000 |
| HRDGGEAI | AHCYL2 | Q96HN2 | SAHH3_HUMAI | Unmodified | 0 | 0 | 222800 | 312190 | 156350 | 0 | 0 | 0 | 0 | 0 | 0 | 0 | 0 | 0 | 435220 | 0 | 0 | 0 |
| HRDLLHNE | CPSF7 | Q8N684 | CPSF7_HUMAN | Unmodified | 770470 | 462930 | 184110 | 326220 | 0 | 39126 | 0 | 0 | 0 | 0 | 0 | 0 | 17149 | 0 | 376570 | 0 | 0 | 0 |
| HRDTGILD: | MBP; MBP; MBF | P02686; H7E | MBP_HUMAN M | Unmodified | 0 | 0 | 0 | 0 | 700420 | 0 | 0 | 0 | 0 | 0 | 0 | 0 | 171590 | 0 | 0 | 0 | 116920 | 169420 |
| HRDYETAT | ACTN4; ACTN4 | O43707; F5( | ACTN4_HUMAI | Unmodified | 0 | 0 | 0 | 0 | 3755100 | 728580 | 0 | 0 | 439030 | 395880 | 0 | 1303800 | 170420 | 0 | 0 | 0 | 1062400 | 425560 |
| HREAQVSA | CIT; CIT; CIT; CI | O14578; A0/ | CTRO_HUMAN | Unmodified | 0 | 0 | 0 | 0 | 2933600 | 0 | 545440 | 0 | 0 | 0 | 0 | 0 | 0 | 28967 | 0 | 0 | 695830 | 0 |
| HREYMEQ( | MARVELD2; MA | Q8N4S9; A1 | MALD2_HUMAI | Oxidation (M) | 0 | 32917 | 0 | 0 | 0 | 0 | 0 | 0 | 0 | 0 | 0 | 187130 | 0 | 0 | 341260 | 154230 | 0 | 0 |
| HREYMEQ( | MARVELD2; MA | Q8N4S9; A1 | MALD2_HUMAI | Unmodified | 0 | 108750 | 0 | 112450 | 255110 | 0 | 0 | 0 | 0 | 0 | 0 | 0 | 0 | 0 | 0 | 164270 | 213370 | 0 |
| HRFDPVET | ERO1B; ERO1B | Q86YB8; Q5' | ERO1B_HUMAI | Unmodified | 0 | 0 | 0 | 539260 | 0 | 0 | 0 | 0 | 359030 | 0 | 0 | 0 | 57039 | 2270700 | 0 | 0 | 0 | 0 |
| HRHIAVVLI | STXBP3 | O00186 | STXB3_HUMAN | Unmodified | 79913 | 0 | 0 | 147000 | 0 | 0 | 0 | 0 | 404190 | 108920 | 0 | 0 | 786250 | 420160 | 95255 | 0 | 164880 | 539010 |
| HRHMGAV | ITGA3 | P26006 | ITA3_HUMAN I | Oxidation (M) | 0 | 0 | 0 | 0 | 3098400 | 2005800 | 0 | 0 | 0 | 0 | 0 | 0 | 0 | 0 | 0 | 0 | 2074100 | 453020 |
| HRHMGAV | ITGA3 | P26006 | ITA3_HUMAN I | Unmodified | 0 | 0 | 0 | 0 | 1395600 | 222430 | 0 | 0 | 64389 | 0 | 0 | 0 | 0 | 0 | 0 | 0 | 94625 | 487480 |
| HRISKEEAN | RPL11; RPL11 | P62913; Q5\ | RL11_HUMAN ( | Unmodified | 1534300 | 0 | 301220 | 0 | 345580 | 251480 | 0 | 0 | 2786100 | 591850 | 885510 | 5634800 | 615800 | 479820 | 0 | 382540 | 31723 | 279080 |
| HRLVNACL | BOLA2; BOLA2E | Q9H3K6; A0. | BOLA2_HUMAN | Unmodified | 0 | 289760 | 0 | 0 | 8151100 | 1152700 | 0 | 0 | 0 | 0 | 1332200 | 3058400 | 4236400 | 1670300 | 238900 | 0 | 1328100 | 1235000 |
| HRNGGLIT | SNF8; SNF8; SN | Q96H20; D6 | SNF8_HUMAN | Unmodified | 2605500 | 3078100 | 2009600 | 1601700 | 0 | 0 | 0 | 0 | 0 | 0 | 0 | 403640 | 1681500 | 0 | 5195200 | 3363800 | 0 | 0 |
| HRPDGKY\ | MTHFD1; MTHF | P11586; V9C | C1TC_HUMAN | Unmodified | 0 | 0 | 0 | 0 | 2445700 | 0 | 650720 | 40002 | 123290 | 599620 | 0 | 0 | 275890 | 0 | 0 | 0 | 2264800 | 1954700 |
| HRPELIDY( | ACTN1; ACTN1; | P12814; G3\ | ACTN1_HUMAI | Unmodified | 491880 | 625770 | 620360 | 1726800 | 3760800 | 128890 | 0 | 0 | 908960 | 0 | 2502900 | 16736000 | 1629600 | 0 | 152270 | 1737700 | 0 | 2873500 |
| HRPELIEYD | ACTN4; ACTN4 | O43707; F5( | ACTN4_HUMAI | Unmodified | 0 | 0 | 0 | 0 | 9929000 | 4336400 | 232130 | 186870 | 6968100 | 2040700 | 2037300 | 20369000 | 0 | 0 | 0 | 0 | 1578200 | 1808500 |
| HRPQVAII( | PNP; PNP; PNP | P00491; G3\ | PNPH_HUMAN | Unmodified | 0 | 0 | 54795 | 0 | 580810 | 0 | 0 | 0 | 0 | 0 | 159120 | 6352400 | 0 | 0 | 331740 | 0 | 0 | 0 |
| HRPSEADE | AP3D1; AP3D1 | O14617; A0/ | AP3D1_HUMAN | Unmodified | 78434 | 0 | 0 | 104890 | 0 | 0 | 0 | 0 | 367730 | 41588 | 260530 | 1008900 | 0 | 0 | 0 | 66768 | 0 | 0 |
| HRSLTAHS | IQGAP3; IQGAP | Q86VI3; F2Z | IQGA3_HUMAN | Unmodified | 0 | 0 | 0 | 0 | 971180 | 145160 | 0 | 0 | 204360 | 0 | 0 | 0 | 0 | 0 | 0 | 0 | 69017 | 0 |
| HRSNSFSC | AHNAK | Q09666 | AHNK_HUMAN | Unmodified | 0 | 0 | 0 | 0 | 7158000 | 984430 | 0 | 0 | 0 | 0 | 0 | 0 | 0 | 0 | 0 | 0 | 793200 | 3877200 |

| | | | | | | | | | | | | | | | | | | | | | | |
|---|---|---|---|---|---|---|---|---|---|---|---|---|---|---|---|---|---|---|---|---|---|---|
| HRSPSGAG | GPRIN1 | Q7Z2K8 | GRIN1_HUMAN | Unmodified | 0 | 0 | 0 | 0 | 0 | 0 | 0 | 0 | 0 | 0 | 0 | 0 | 0 | 0 | 0 | 0 | 0 | 0 |
| HRTEALMD | LAMB2 | P55268 | LAMB2_HUMAN | Unmodified | 0 | 0 | 0 | 0 | 0 | 288980 | 0 | 0 | 1202200 | 255900 | 155710 | 0 | 0 | 401460 | 0 | 0 | 0 | 310870 |
| HRVIGSGC | LDHB; LDHB; LI | P07195; A0A | LDHB_HUMAN | Unmodified | 815100 | 440720 | 739100 | 1048600 | 1100200 | 438460 | 45095 | 19576 | 0 | 0 | 2305500 | 28969000 | 933740 | 586870 | 1044100 | 1191100 | 375770 | 0 |
| HRVQELGH | TLN1 | Q9Y490 | TLN1_HUMAN | Unmodified | 0 | 0 | 0 | 2790900 | 5522800 | 1504900 | 3071100 | 1351700 | 459520 | 465560 | 230130 | 1812900 | 5258000 | 7795700 | 0 | 0 | 1707500 | 3400100 |
| HRVTMNEF | AKT1 | P31749 | AKT1_HUMAN | Unmodified | 0 | 0 | 0 | 0 | 0 | 0 | 0 | 0 | 1028200 | 0 | 0 | 697650 | 349040 | 852760 | 0 | 0 | 0 | 0 |
| HRVVQPTL | AOC1 | P19801 | AOC1_HUMAN | Unmodified | 0 | 0 | 0 | 0 | 0 | 0 | 0 | 0 | 3756500 | 190170 | 0 | 0 | 0 | 0 | 0 | 0 | 0 | 0 |
| HSAESQILK | NNMT | P40261 | NNMT_HUMAN | Unmodified | 0 | 0 | 0 | 0 | 257160 | 0 | 0 | 0 | 0 | 0 | 0 | 0 | 0 | 0 | 0 | 0 | 0 | 0 |
| HSAGELQC | CPT2; CPT2; CF | P23786; A0A | CPT2_HUMAN | Unmodified | 0 | 0 | 0 | 0 | 0 | 0 | 0 | 0 | 619620 | 0 | 0 | 0 | 0 | 0 | 0 | 0 | 0 | 0 |
| HSANMNFS | ERBIN | Q96RT1 | ERBIN_HUMAN | Unmodified | 0 | 0 | 0 | 0 | 1441100 | 0 | 0 | 0 | 0 | 0 | 0 | 0 | 0 | 0 | 0 | 0 | 0 | 0 |
| HSANMNFS | ERBIN | Q96RT1 | ERBIN_HUMAN | Oxidation (M) | 0 | 0 | 32794 | 21921 | 613000 | 86423 | 20173 | 0 | 0 | 0 | 0 | 39349 | 190320 | 114610 | 62013 | 23408 | 124520 | 240340 |
| HSANMNFS | ERBIN | Q96RT1 | ERBIN_HUMAN | Unmodified | 0 | 0 | 0 | 0 | 1832000 | 399270 | 0 | 0 | 0 | 0 | 0 | 0 | 300150 | 274720 | 0 | 0 | 118640 | 0 |
| HSATPFLK | ASNS; ASNS; AS | P08243; F8W | ASNS_HUMAN | Unmodified | 0 | 0 | 0 | 0 | 0 | 0 | 0 | 0 | 0 | 150820 | 135220 | 710210 | 0 | 0 | 0 | 0 | 0 | 0 |
| HSATPFLKV | ASNS; ASNS; AS | P08243; F8W | ASNS_HUMAN | Unmodified | 0 | 0 | 0 | 2963000 | 0 | 0 | 0 | 0 | 0 | 0 | 0 | 1609000 | 863080 | 722530 | 0 | 0 | 0 | 0 |
| HSCIFPGQ | HEG1; HEG1 | Q9ULI3; H7C | HEG1_HUMAN | Unmodified | 266750 | 0 | 0 | 586700 | 179050 | 0 | 0 | 0 | 0 | 0 | 0 | 0 | 0 | 0 | 374240 | 221460 | 0 | 0 |
| HSDITAHS | GAS6 | Q14393 | GAS6_HUMAN | Unmodified | 0 | 0 | 0 | 0 | 0 | 0 | 0 | 0 | 3637000 | 157120 | 0 | 132820 | 0 | 0 | 0 | 0 | 0 | 0 |
| HSDLPPLN | SPNS1; SPNS1; | Q9H2V7; H3 | SPNS1_HUMAN | Unmodified | 102140 | 0 | 0 | 0 | 0 | 0 | 0 | 0 | 300900 | 0 | 0 | 64396 | 0 | 0 | 166690 | 117540 | 0 | 0 |
| HSDLQEVE | HEG1 | Q9ULI3 | HEG1_HUMAN | Unmodified | 153260 | 0 | 0 | 0 | 1777700 | 0 | 0 | 0 | 0 | 0 | 0 | 0 | 0 | 0 | 0 | 0 | 0 | 0 |
| HSDPHLLE | SLC4A7 | C9JRP1 | C9JRP1_HUMA | Unmodified | 0 | 0 | 0 | 0 | 5337200 | 0 | 0 | 0 | 0 | 0 | 0 | 0 | 0 | 0 | 0 | 0 | 1866900 | 2295800 |
| HSDSKEDE | CD276; CD276; | Q5ZPR3; A0A | CD276_HUMAN | Unmodified | 0 | 0 | 2740400 | 0 | 3457400 | 0 | 0 | 0 | 927080 | 0 | 0 | 0 | 0 | 0 | 0 | 2300400 | 0 | 0 |
| HSDYINAN | PTPRG | P23470 | PTPRG_HUMAN | Unmodified | 0 | 0 | 605560 | 630480 | 6680200 | 0 | 0 | 0 | 0 | 0 | 288430 | 923060 | 312490 | 0 | 671070 | 917970 | 0 | 0 |
| HSDYPLSL | FUT11 | Q495W5 | FUT11_HUMAN | Unmodified | 15172 | 0 | 10788 | 0 | 0 | 0 | 0 | 0 | 1873800 | 0 | 0 | 0 | 0 | 0 | 0 | 0 | 0 | 0 |
| HSEAATAQ | HNRNPD; HNR | Q14103; D6I | HNRPD_HUMA | Unmodified | 0 | 0 | 0 | 0 | 87081 | 53991 | 0 | 167900 | 329420 | 209780 | 2416400 | 10586000 | 565680 | 87352 | 0 | 0 | 165090 | 681950 |
| HSEAFEAL | CKAP4 | Q07065 | CKAP4_HUMAN | Unmodified | 0 | 0 | 0 | 0 | 0 | 0 | 0 | 0 | 517000 | 0 | 0 | 0 | 0 | 0 | 0 | 0 | 0 | 0 |
| HSEALEALE | FARP1; FARP1; | Q9Y4F1; C9J | FARP1_HUMAN | Unmodified | 0 | 0 | 106420 | 189970 | 303570 | 0 | 0 | 0 | 490120 | 369790 | 1057400 | 1850500 | 0 | 0 | 0 | 142810 | 0 | 0 |
| HSEGLLPV | SLC38A3; SLC3 | Q99624; A0A | S38A3_HUMAN | Unmodified | 0 | 0 | 0 | 0 | 0 | 0 | 62203 | 0 | 1207100 | 0 | 0 | 571720 | 0 | 0 | 0 | 0 | 0 | 0 |
| HSEGLLPV | SLC38A3; SLC3 | Q99624; A0A | S38A3_HUMAN | Unmodified | 0 | 0 | 0 | 0 | 0 | 0 | 0 | 0 | 1188900 | 0 | 0 | 0 | 0 | 0 | 0 | 0 | 0 | 0 |
| HSEHAFDH | ENTPD3 | O75355 | ENTP3_HUMAN | Unmodified | 0 | 0 | 52357 | 0 | 0 | 0 | 0 | 0 | 515640 | 125330 | 0 | 0 | 0 | 0 | 117720 | 81272 | 29499 | 0 |
| HSEIQQLEI | STX4 | Q12846 | STX4_HUMAN | Unmodified | 148290 | 282130 | 0 | 0 | 0 | 0 | 0 | 0 | 707600 | 135570 | 378330 | 2478500 | 0 | 0 | 397550 | 674240 | 0 | 0 |
| HSELSELN | STAM2; STAM | O75886; Q9 | STAM2_HUMAN | Unmodified | 2237800 | 2169900 | 9051700 | 3093300 | 4400300 | 49910000 | 2062200 | 604610 | 550570 | 62655 | 0 | 0 | 1872800 | 2997800 | 5430700 | 785230 | 673920 | 20829000 |
| HSELVHVH | HIP1R | O75146 | HIP1R_HUMAN | Unmodified | 0 | 0 | 0 | 0 | 364980 | 0 | 0 | 0 | 1265000 | 0 | 0 | 261460 | 955710 | 1309000 | 0 | 0 | 47201 | 95154 |
| HSEPSKLTI | MAN1B1; MAN1 | Q9UKM7; H0 | MA1B1_HUMAN | Unmodified | 0 | 0 | 0 | 0 | 0 | 0 | 86578 | 46737 | 11369000 | 1082400 | 0 | 0 | 51997 | 0 | 0 | 0 | 0 | 0 |
| HSERDRVC | P3H1 | Q32P28 | P3H1_HUMAN | Unmodified | 0 | 0 | 0 | 0 | 0 | 0 | 0 | 0 | 5563500 | 492040 | 171130 | 0 | 0 | 0 | 0 | 0 | 0 | 0 |
| HSEYNPQH | SMAD1 | Q15797 | SMAD1_HUMA | Unmodified | 116080 | 97484 | 180290 | 167240 | 0 | 0 | 0 | 0 | 131380 | 24717 | 0 | 239040 | 78236 | 0 | 177970 | 214330 | 43825 | 0 |
| HSFGPLDY | GLS; GLS | B8ZZA8; B8Z | B8ZZA8_HUMA | Unmodified | 0 | 0 | 0 | 0 | 0 | 0 | 0 | 56527 | 4984600 | 0 | 0 | 0 | 0 | 0 | 0 | 0 | 0 | 0 |
| HSFYMLDK | ITGA1 | P56199 | ITA1_HUMAN | Oxidation (M) | 791380 | 0 | 0 | 0 | 0 | 0 | 0 | 0 | 0 | 0 | 0 | 0 | 0 | 0 | 0 | 0 | 0 | 0 |
| HSGFEDEL | CHGA; CHGA | P10645; G5E | CMGA_HUMAN | Unmodified | 0 | 124490 | 0 | 0 | 0 | 0 | 0 | 0 | 1.36E+08 | 40686000 | 0 | 0 | 0 | 0 | 0 | 112590 | 334320 | 0 |
| HSGFEDEL | CHGA | P10645 | CMGA_HUMAN | Unmodified | 0 | 0 | 0 | 0 | 0 | 0 | 0 | 0 | 0 | 1677000 | 0 | 0 | 0 | 0 | 0 | 0 | 0 | 0 |
| HSGLWHS | TOR1B | O14657 | TOR1B_HUMAN | Unmodified | 0 | 101660 | 0 | 0 | 0 | 0 | 35107 | 0 | 2304600 | 210900 | 0 | 0 | 0 | 0 | 0 | 0 | 0 | 0 |
| HSGNITFD | RPL12 | P30050 | RL12_HUMAN | Unmodified | 11639000 | 11813000 | 0 | 4573000 | 1783000 | 0 | 0 | 0 | 1449500 | 856870 | 5129300 | 46653000 | 6641200 | 1628500 | 5210300 | 4287000 | 0 | 658290 |
| HSGPNSAI | HNRNPF | P52597 | HNRPF_HUMA | Unmodified | 275920 | 284650 | 101220 | 229700 | 227810 | 0 | 112730 | 0 | 1301000 | 328030 | 1148400 | 5994400 | 188290 | 48476 | 245990 | 326240 | 58970 | 73162 |
| HSHPSDEK | SLC4A2; SLC4A | P04920; A0A | B3A2_HUMAN | Unmodified | 0 | 0 | 0 | 0 | 760970 | 122930 | 67526 | 0 | 0 | 0 | 0 | 0 | 399260 | 338900 | 0 | 0 | 132110 | 213870 |
| HSIAQLDPI | CDCA8 | Q53HL2 | BOREA_HUMA | Unmodified | 0 | 0 | 837320 | 1225600 | 1471400 | 0 | 0 | 0 | 1105300 | 0 | 523890 | 1097200 | 1698500 | 703050 | 0 | 0 | 0 | 0 |
| HSIAQLDPI | CDCA8 | Q53HL2 | BOREA_HUMA | Unmodified | 0 | 0 | 0 | 431400 | 0 | 0 | 0 | 0 | 0 | 0 | 0 | 0 | 0 | 0 | 0 | 0 | 0 | 0 |
| HSLEGHLR | SLC7A2; SLC7A | P52569; A0A | CTR2_HUMAN | Unmodified | 0 | 0 | 0 | 0 | 0 | 0 | 0 | 0 | 0 | 0 | 0 | 1042900 | 0 | 0 | 0 | 0 | 0 | 0 |
| HSLFHPED | KLK3; KLK3; KLI | P07288; Q8I | KLK3_HUMAN | Oxidation (M) | 0 | 0 | 2019600 | 3262300 | 0 | 0 | 0 | 0 | 0 | 0 | 0 | 0 | 799560 | 0 | 9723600 | 1939300 | 0 | 0 |
| HSLFIRHPE | CREG1; CREG1 | O75629; A0A | CREG1_HUMA | Unmodified | 0 | 0 | 0 | 0 | 0 | 0 | 65252 | 0 | 2320400 | 0 | 0 | 0 | 0 | 0 | 0 | 0 | 0 | 0 |
| HSLNSSSA | LRPPRC | P42704 | LPPRC_HUMAN | Unmodified | 0 | 0 | 0 | 0 | 0 | 0 | 0 | 0 | 2469300 | 169440 | 0 | 0 | 0 | 0 | 0 | 0 | 0 | 0 |
| HSLPSGLG | ARFIP1 | P53367 | ARFP1_HUMAN | Unmodified | 0 | 0 | 915170 | 787860 | 0 | 0 | 0 | 0 | 0 | 0 | 0 | 0 | 517580 | 0 | 0 | 0 | 0 | 0 |
| HSLVSFVVI | SPON2 | Q9BUD6 | SPON2_HUMA | Unmodified | 0 | 96834 | 7256800 | 4575000 | 0 | 0 | 41139 | 0 | 0 | 57528 | 269940 | 0 | 720590 | 1307300 | 8876300 | 3317900 | 0 | 0 |
| HSMAYQDI | MUC13; MUC1 | Q9H3R2; C9 | MUC13_HUMA | Oxidation (M) | 1297000 | 913890 | 0 | 0 | 0 | 0 | 0 | 0 | 2539600 | 728310 | 0 | 0 | 0 | 0 | 0 | 0 | 232070 | 106630 |
| HSMLFIEAS | RAB18; RAB18; | Q9NP72; H0 | RAB18_HUMAN | Oxidation (M) | 498380 | 66991 | 0 | 0 | 0 | 0 | 143910 | 101250 | 0 | 128400 | 210640 | 505410 | 0 | 0 | 548620 | 89071 | 0 | 57318 |
| HSMNILNR | DDX19B; DDX1 | Q9UMR2; Q | DD19B_HUMA | Unmodified | 0 | 0 | 0 | 0 | 0 | 0 | 0 | 380230 | 0 | 0 | 0 | 0 | 803790 | 0 | 0 | 117230 | 0 | 0 |
| HSNLVQLL | CSK | P41240 | CSK_HUMAN | Unmodified | 91670 | 0 | 58643 | 242000 | 0 | 0 | 0 | 0 | 0 | 0 | 932040 | 2220000 | 304390 | 0 | 418150 | 186290 | 0 | 0 |
| HSPESSRG | GLIPR2 | Q9H4G4 | GAPR1_HUMA | Unmodified | 931170 | 892030 | 996880 | 2017900 | 4424400 | 0 | 2503000 | 644380 | 534080 | 191710 | 0 | 0 | 1739900 | 2693700 | 3677600 | 2830400 | 0 | 1267200 |
| HSPIAPSSF | INCENP | Q9NQS7 | INCE_HUMAN | Unmodified | 0 | 0 | 0 | 1496900 | 1666700 | 208830 | 0 | 0 | 0 | 0 | 0 | 0 | 612980 | 0 | 1100400 | 0 | 340990 | 228930 |
| HSPPDDLG | MANEAL; MANE | Q5VSG8; H0 | MANEL_HUMA | Unmodified | 0 | 0 | 0 | 0 | 0 | 0 | 0 | 0 | 980160 | 0 | 0 | 0 | 0 | 0 | 0 | 265270 | 0 | 0 |
| HSPQEAPH | CNTFR; CNTFR | P26992; Q5 | CNTFR_HUMA | Unmodified | 231990 | 0 | 0 | 0 | 0 | 0 | 0 | 0 | 311740 | 0 | 0 | 0 | 0 | 0 | 0 | 0 | 0 | 0 |
| HSQAVEEL | MYH9 | P35579 | MYH9_HUMAN | Unmodified | 1429100 | 1859400 | 2338500 | 5372500 | 40873000 | 8168200 | 4505300 | 1328200 | 4765100 | 6755200 | 1610500 | 9995400 | 9090000 | 12576000 | 7247300 | 4960300 | 13213000 | 14909000 |
| HSQAVEEL | MYH9 | P35579 | MYH9_HUMAN | Unmodified | 0 | 0 | 164040 | 381330 | 692460 | 0 | 0 | 0 | 273920 | 506790 | 2212700 | 1198500 | 0 | 0 | 0 | 111490 | 346080 | 0 |
| HSQAVEEL | MYH9 | P35579 | MYH9_HUMAN | Unmodified | 0 | 0 | 0 | 0 | 509900 | 0 | 1362100 | 0 | 0 | 194430 | 0 | 0 | 0 | 295410 | 0 | 0 | 6347700 | 1416400 |
| HSQFIGYPI | HSP90AA5P; HS | Q58FG0; P0 | HS905_HUMAN | Unmodified | 4077800 | 2905300 | 3420000 | 4329000 | 21779000 | 7097000 | 3512800 | 0 | 34649000 | 15391000 | 16209000 | 83284000 | 11837000 | 3738500 | 9033600 | 3769000 | 12745000 | 11806000 |
| HSQFLGYP | HSP90AB2P | Q58FF8 | H90B2_HUMAN | Unmodified | 6067700 | 6357700 | 6847300 | 18066000 | 58465000 | 14378000 | 5131700 | 0 | 35196000 | 16141000 | 29124000 | 1.51E+08 | 25303000 | 14449000 | 9792600 | 14871000 | 19873000 | 39561000 |
| HSQNGFIV | SNX1 | Q13596 | SNX1_HUMAN | Unmodified | 0 | 0 | 0 | 0 | 530550 | 190540 | 112550 | 151170 | 0 | 0 | 0 | 0 | 0 | 231940 | 0 | 0 | 0 | 258740 |
| HSQPWQV | KLK3; KLK3; KLI | P07288; Q8I | KLK3_HUMAN | Unmodified | 0 | 0 | 0 | 896920 | 0 | 0 | 0 | 46921 | 0 | 0 | 0 | 0 | 0 | 0 | 5178200 | 1663100 | 0 | 0 |
| HSQYHVD | PGAM5 | Q96HS1 | PGAM5_HUMA | Unmodified | 0 | 0 | 0 | 0 | 0 | 0 | 0 | 0 | 965560 | 400200 | 0 | 558720 | 0 | 0 | 0 | 0 | 0 | 0 |
| HSSDINHL | FNBP1L; FNBP1 | Q5T0N5; A0 | FBP1L_HUMAN | Unmodified | 414790 | 323940 | 0 | 84668 | 0 | 0 | 0 | 0 | 0 | 0 | 0 | 0 | 0 | 0 | 159980 | 191960 | 0 | 0 |
| HSSGCAFL | BIRC5; BIRC5; | O15392; H3I | BIRC5_HUMAN | Unmodified | 0 | 0 | 108040 | 0 | 295730 | 33287 | 0 | 0 | 0 | 0 | 0 | 140450 | 119600 | 109330 | 0 | 0 | 242890 | 0 |
| HSSIFDQLI | MPP1; MPP1; M | Q00013; C9J | EM55_HUMAN | Unmodified | 0 | 0 | 0 | 156410 | 739820 | 0 | 180590 | 0 | 0 | 0 | 0 | 0 | 0 | 191810 | 0 | 0 | 0 | 431510 |
| HSSLAGCC | CLTC; CLTC | Q00610; A0A | CLH1_HUMAN | Unmodified | 12318000 | 6130700 | 1136400 | 0 | 0 | 0 | 139440 | 65027 | 0 | 0 | 0 | 1641000 | 0 | 0 | 2532300 | 4366700 | 0 | 0 |
| HSSLAGCC | CLTC; CLTC | Q00610; A0A | CLH1_HUMAN | Unmodified | 1797300 | 1293700 | 243990 | 641480 | 0 | 69568 | 207540 | 145920 | 0 | 161760 | 0 | 1143400 | 76954 | 127870 | 1149200 | 360260 | 145070 | 0 |
| HSSLITPLQ | PRKDC | P78527 | PRKDC_HUMA | Unmodified | 283890 | 406100 | 921980 | 95220 | 0 | 0 | 1135500 | 103980 | 169820 | 202840 | 48662 | 3980500 | 0 | 0 | 285790 | 218700 | 60011 | 238860 |
| HSSTEVLLS | SDF2 | Q99470 | SDF2_HUMAN | Unmodified | 0 | 0 | 0 | 0 | 1710500 | 0 | 1019900 | 0 | 50384000 | 11105000 | 2802800 | 979470 | 0 | 526780 | 0 | 0 | 0 | 540330 |
| HSSVYPTQ | ILF3; ILF3; ILF3; | Q12906; K7E | ILF3_HUMAN | Unmodified | 0 | 0 | 0 | 0 | 0 | 0 | 0 | 0 | 862780 | 196420 | 575910 | 1459000 | 656660 | 0 | 0 | 0 | 0 | 0 |
| HSSVYPTQ | ILF3; ILF3; ILF3; | Q12906; K7E | ILF3_HUMAN | Oxidation (M) | 3597600 | 887260 | 95903 | 573030 | 0 | 0 | 0 | 0 | 0 | 0 | 0 | 493180 | 0 | 0 | 208030 | 417920 | 0 | 0 |
| HSSYTCIC | PEPD; PEPD; PE | P12955; A0A | PEPD_HUMAN | Unmodified | 0 | 0 | 0 | 0 | 0 | 0 | 0 | 0 | 0 | 0 | 0 | 1542500 | 0 | 0 | 0 | 0 | 0 | 0 |
| HSTFHGED | STIM1; STIM1; S | Q13586; E9F | STIM1_HUMAN | Unmodified | 745880 | 0 | 0 | 0 | 0 | 0 | 0 | 0 | 2496200 | 726380 | 0 | 0 | 0 | 0 | 0 | 0 | 0 | 0 |
| HSTGIDIGI | SLC25A24; SLC | Q6NUK1; A0 | SCMC1_HUMA | Unmodified | 0 | 0 | 0 | 0 | 0 | 0 | 0 | 0 | 1613500 | 745650 | 0 | 0 | 0 | 0 | 0 | 0 | 0 | 0 |
| HSTGIDIGI | SLC25A24; SLC | Q6NUK1; A0 | SCMC1_HUMA | Unmodified | 0 | 0 | 0 | 0 | 0 | 0 | 0 | 0 | 6126100 | 1669400 | 0 | 0 | 0 | 0 | 0 | 0 | 0 | 0 |
| HSTLDFML | SMS | P52788 | SPSY_HUMAN | Oxidation (M) | 0 | 0 | 0 | 0 | 0 | 0 | 0 | 0 | 0 | 113430 | 0 | 1011700 | 320520 | 0 | 0 | 0 | 0 | 0 |
| HSTLDFML | SMS | P52788 | SPSY_HUMAN | Unmodified | 0 | 0 | 0 | 0 | 0 | 17073 | 0 | 0 | 0 | 0 | 0 | 334080 | 0 | 0 | 0 | 0 | 0 | 0 |
| HSTVLENTI | MELTF | P08582 | TRFM_HUMAN | Unmodified | 1446000 | 0 | 0 | 0 | 4610100 | 1006700 | 0 | 715380 | 3182400 | 856050 | 0 | 15478000 | 0 | 0 | 0 | 0 | 0 | 645680 |
| HSTVQHSY | DSG2 | Q14126 | DSG2_HUMAN | Unmodified | 0 | 0 | 0 | 0 | 2178200 | 0 | 0 | 0 | 0 | 0 | 0 | 0 | 0 | 207030 | 0 | 0 | 182340 | 310760 |
| HSVLGAAV | DNAJC11 | Q9NVH1 | DJC11_HUMAN | Unmodified | 0 | 0 | 0 | 0 | 0 | 0 | 0 | 0 | 2499400 | 211180 | 0 | 0 | 0 | 0 | 0 | 0 | 0 | 0 |
| HSVNNPYS | SCP2; SCP2 | P22307; E9P | SCP2_HUMAN | Unmodified | 0 | 0 | 0 | 0 | 0 | 0 | 0 | 0 | 3320800 | 0 | 0 | 0 | 0 | 0 | 0 | 0 | 0 | 0 |
| HSVVCPAS | LY6D | Q14210 | LY6D_HUMAN | Unmodified | 0 | 0 | 0 | 0 | 12726 | 0 | 1793800 | 610870 | 0 | 0 | 0 | 0 | 408920 | 0 | 0 | 0 | 0 | 0 |
| HSVYDTTN | DST; DST; DST; | Q03001; A0A | DYST_HUMAN | Unmodified | 0 | 0 | 0 | 0 | 23698 | 270480 | 0 | 0 | 0 | 245430 | 0 | 0 | 0 | 0 | 0 | 192860 | 176700 | 0 |
| HSYTASYD | DPP4; DPP4; DI | P27487; A0A | DPP4_HUMAN | Unmodified | 170850 | 381080 | 2233900 | 1911700 | 0 | 0 | 0 | 0 | 0 | 0 | 0 | 0 | 0 | 0 | 6639700 | 4317400 | 0 | 0 |
| HSYTASYD | DPP4; DPP4; DI | P27487; A0A | DPP4_HUMAN | Unmodified | 0 | 0 | 5793600 | 6864400 | 0 | 0 | 0 | 0 | 65046 | 0 | 0 | 0 | 0 | 0 | 10357000 | 8765100 | 0 | 0 |
| HTAAPTDP | PLP2 | Q04941 | PLP2_HUMAN | Unmodified | 723860 | 473910 | 7354300 | 0 | 10717000 | 0 | 2275600 | 3096500 | 29672000 | 5871600 | 3109200 | 18051000 | 3802900 | 1278300 | 10327000 | 5080700 | 8683700 | 9689200 |
| HTADRWR | HSPB1; HSPB1; | P04792; A0A | HSPB1_HUMA | Unmodified | 0 | 0 | 0 | 0 | 1149400 | 0 | 0 | 0 | 0 | 2713400 | 0 | 0 | 1396800 | 0 | 0 | 0 | 965400 | 717990 |
| HTAMVSW | FLNA; FLNA; FL | P21333; A0A | FLNA_HUMAN | Oxidation (M) | 1238500 | 956390 | 606920 | 3796700 | 0 | 0 | 0 | 0 | 0 | 0 | 0 | 2274300 | 545040 | 0 | 1946200 | 1654500 | 762640 | 0 |
| HTAMVSW | FLNA; FLNA; FL | P21333; A0A | FLNA_HUMAN | Oxidation (M) | 153150 | 0 | 329040 | 2751500 | 2969100 | 897370 | 1279900 | 0 | 0 | 330570 | 0 | 0 | 0 | 1626200 | 0 | 0 | 1182000 | 714540 |
| HTATHLLN | AARS2 | Q5JTZ9 | SYAM_HUMAN | Unmodified | 0 | 0 | 0 | 0 | 0 | 0 | 39877 | 59927 | 305450 | 0 | 0 | 81242 | 0 | 0 | 0 | 0 | 92781 | 0 |
| HTCEQGD | MOGS; MOGS; | Q13724; A0A | MOGS_HUMAN | Unmodified | 0 | 0 | 0 | 0 | 0 | 0 | 0 | 0 | 6828000 | 0 | 0 | 0 | 0 | 0 | 0 | 0 | 0 | 0 |
| HTDLDHEC | NLGN1; NLGN1 | Q8N2Q7; J3I | NLGN1_HUMA | Unmodified | 307890 | 137400 | 0 | 0 | 0 | 0 | 0 | 0 | 0 | 0 | 0 | 0 | 0 | 0 | 0 | 0 | 0 | 0 |
| HTDNVIQW | IQGAP1; IQGAP | P46940; A0A | IQGA1_HUMAN | Unmodified | 0 | 0 | 0 | 0 | 2680000 | 0 | 0 | 0 | 585930 | 238430 | 59917 | 0 | 361590 | 0 | 581680 | 0 | 1484400 | 276980 |
| HTDPVPDP | ATP2A2; ATP2A | P16615; H7C | AT2A2_HUMAN | Unmodified | 34323 | 0 | 69379 | 79160 | 0 | 0 | 0 | 0 | 0 | 0 | 86557 | 296130 | 0 | 0 | 83409 | 34356 | 0 | 0 |
| HTEAAPGD | SARG | Q9BW04 | SARG_HUMAN | Unmodified | 0 | 440580 | 371850 | 786510 | 0 | 435250 | 0 | 0 | 0 | 0 | 0 | 0 | 1287000 | 1104200 | 344470 | 582890 | 0 | 0 |
| HTEAPLYV | PTK7 | Q13308 | PTK7_HUMAN | Unmodified | 0 | 0 | 0 | 0 | 0 | 0 | 0 | 0 | 1489300 | 0 | 0 | 0 | 0 | 3005300 | 868160 | 427550 | 371420 | 0 |
| HTEDVGVV | LOXL2; LOXL2; | Q9Y4K0; E5F | LOXL2_HUMAN | Unmodified | 0 | 0 | 0 | 0 | 0 | 0 | 0 | 0 | 723210 | 0 | 183120 | 26635 | 0 | 0 | 0 | 0 | 0 | 0 |
| HTEPNETTI | NCAM1; NCAM | P13591; A0A | NCAM1_HUMA | Unmodified | 0 | 0 | 0 | 0 | 0 | 0 | 91663 | 0 | 3130500 | 230790 | 0 | 1733000 | 0 | 0 | 0 | 0 | 0 | 0 |
| HTFKGDYS | NECTIN1 | Q15223 | NECT1_HUMA | Unmodified | 0 | 18074 | 11296 | 0 | 0 | 0 | 0 | 0 | 0 | 0 | 0 | 77446 | 0 | 0 | 22962 | 15817 | 0 | 0 |
| HTGCCGD | PPA1; PPA1 | Q15181; Q5 | IPYR_HUMAN | Unmodified | 191620 | 0 | 168340 | 214560 | 0 | 0 | 0 | 0 | 0 | 0 | 1318900 | 11027000 | 568230 | 0 | 209230 | 278860 | 0 | 0 |
| HTGFLEISC | TRMT10C | Q7L0Y3 | TM10C_HUMA | Unmodified | 0 | 0 | 0 | 0 | 0 | 0 | 0 | 0 | 1142100 | 0 | 0 | 0 | 0 | 0 | 0 | 0 | 0 | 0 |
| HTGPITCL | WDR82 | Q6UXN9 | WDR82_HUMA | Unmodified | 239420 | 131330 | 276340 | 0 | 0 | 0 | 0 | 35672 | 0 | 0 | 0 | 3461100 | 0 | 0 | 0 | 0 | 0 | 0 |
| HTGPNSPD | HNRNPH1; HNI | P31943; P55 | HNRH1_HUMA | Unmodified | 1573800 | 0 | 879930 | 0 | 0 | 0 | 227910 | 151500 | 235430 | 358920 | 1100000 | 8057400 | 0 | 0 | 0 | 568750 | 0 | 434850 |
| HTGPNSPD | HNRNPH1; HNI | P31943; P55 | HNRH1_HUMA | Unmodified | 0 | 0 | 0 | 0 | 0 | 0 | 0 | 115750 | 0 | 0 | 0 | 0 | 0 | 0 | 0 | 0 | 0 | 0 |
| HTGYVIELC | QARS1; QARS1 | P47897; H7C | SYQ_HUMAN | Unmodified | 0 | 227320 | 158300 | 166690 | 142660 | 0 | 0 | 0 | 234410 | 0 | 485090 | 6625400 | 0 | 0 | 301780 | 0 | 0 | 0 |
| HTHEGDPL | TWF2; TWF2; TV | Q6IBS0; A0A | TWF2_HUMAN | Oxidation (M) | 0 | 0 | 0 | 0 | 0 | 0 | 0 | 0 | 0 | 0 | 0 | 5470600 | 0 | 0 | 0 | 0 | 495210 | 0 |

| | | | | | | | | | | | | | | | | | | | | | | |
|---|---|---|---|---|---|---|---|---|---|---|---|---|---|---|---|---|---|---|---|---|---|---|
| HTHIDKPD | TM9SF2 | Q99805 | TM9S2_HUMAN | Oxidation (M) | 0 | 0 | 271660 | 0 | 0 | 0 | 0 | 0 | 6422700 | 785110 | 1956800 | 2055900 | 0 | 0 | 0 | 0 | 0 | 0 |
| HTHIDKPD | TM9SF2 | Q99805 | TM9S2_HUMAN | Unmodified | 602550 | 217240 | 211060 | 0 | 79862 | 0 | 149630 | 0 | 71148000 | 4369200 | 1914200 | 14576000 | 0 | 0 | 237180 | 119470 | 0 | 0 |
| HTHIDKPD | TM9SF2 | Q99805 | TM9S2_HUMAN | Unmodified | 0 | 0 | 0 | 0 | 0 | 0 | 0 | 0 | 1259500 | 278910 | 0 | 0 | 0 | 0 | 0 | 0 | 0 | 0 |
| HTHLAVSF | IGSF8; IGSF8 | Q969P0; C9J | IGSF8_HUMAN | Unmodified | 1053500 | 1340600 | 1435600 | 621680 | 325520 | 0 | 0 | 0 | 0 | 0 | 0 | 0 | 2876500 | 1312800 | 2189900 | 1952200 | 0 | 0 |
| HTHQLYNF | MITD1; MITD1 | Q8WV92; H7 | MITD1_HUMAN | Unmodified | 0 | 0 | 0 | 0 | 0 | 0 | 0 | 0 | 2034800 | 0 | 820620 | 6034400 | 0 | 0 | 0 | 0 | 0 | 0 |
| HTIAVVWG | FLNB; FLNB | O75369; E7E | FLNB_HUMAN | Unmodified | 0 | 0 | 0 | 0 | 0 | 0 | 0 | 0 | 330370 | 0 | 0 | 3591700 | 0 | 0 | 0 | 0 | 0 | 0 |
| HTIAVVWG | FLNB; FLNB | O75369; E7E | FLNB_HUMAN | Unmodified | 0 | 0 | 0 | 716800 | 6616400 | 954230 | 0 | 0 | 0 | 0 | 0 | 403990 | 703550 | 0 | 0 | 0 | 1489500 | 822130 |
| HTLADNFN | M6PR; M6PR | P20645; H0Y | MPRD_HUMAN | Unmodified | 3552700 | 1874700 | 3473200 | 1570300 | 0 | 0 | 0 | 648350 | 35949000 | 4958000 | 844440 | 55037000 | 0 | 3860200 | 2511400 | 2841000 | 0 | 0 |
| HTLADNFN | M6PR; M6PR | P20645; H0Y | MPRD_HUMAN | Unmodified | 183730 | 339140 | 784380 | 0 | 0 | 0 | 0 | 0 | 6742800 | 325710 | 619020 | 5392200 | 159930 | 0 | 420850 | 258500 | 0 | 0 |
| HTLDGAAC | MOB4; HSPE1-I | Q9Y3A3; S4F | PHOCN_HUMA | Unmodified | 0 | 0 | 0 | 0 | 0 | 0 | 0 | 0 | 0 | 0 | 0 | 1796700 | 0 | 0 | 0 | 0 | 0 | 0 |
| HTLGENIAI | ECE1; ECE1; EC | P42892; B4E | ECE1_HUMAN | Unmodified | 161880 | 0 | 0 | 0 | 135730 | 0 | 0 | 0 | 12482000 | 313520 | 0 | 520030 | 0 | 0 | 191080 | 0 | 61669 | 0 |
| HTLGHLLS | PLOD1 | Q02809 | PLOD1_HUMAN | Unmodified | 0 | 0 | 0 | 0 | 0 | 0 | 0 | 0 | 3082700 | 112670 | 47652 | 576940 | 0 | 0 | 0 | 80351 | 0 | 0 |
| HTLSYVDV | SDHA; SDHA; S | P31040; D6F | SDHA_HUMAN | Unmodified | 0 | 0 | 0 | 0 | 0 | 0 | 0 | 0 | 1622400 | 0 | 0 | 0 | 0 | 0 | 0 | 0 | 0 | 0 |
| HTLSYVDV | SDHA; SDHA; S | P31040; D6F | SDHA_HUMAN | Unmodified | 0 | 0 | 0 | 0 | 0 | 0 | 359400 | 0 | 153050 | 2679700 | 0 | 0 | 0 | 0 | 0 | 0 | 0 | 780730 |
| HTLTQIK | CCT6A | P40227 | TCPZ_HUMAN | Unmodified | 0 | 509730 | 1121000 | 915920 | 0 | 0 | 0 | 145920 | 1855700 | 901980 | 1377800 | 10029000 | 435410 | 1553000 | 891610 | 837870 | 0 | 0 |
| HTMDGPR | HS2ST1 | Q7LGA3 | HS2ST_HUMAN | Oxidation (M) | 0 | 0 | 0 | 0 | 0 | 0 | 0 | 0 | 10413000 | 0 | 0 | 0 | 0 | 0 | 0 | 0 | 0 | 0 |
| HTMDGPR | HS2ST1 | Q7LGA3 | HS2ST_HUMAN | Unmodified | 0 | 0 | 0 | 0 | 292550 | 0 | 0 | 0 | 9053000 | 0 | 0 | 0 | 0 | 0 | 0 | 0 | 0 | 0 |
| HTNYTMEH | ACTN2; ACTN4 | P35609; O43 | ACTN2_HUMAN | Unmodified | 26628 | 721520 | 364200 | 0 | 0 | 43711 | 0 | 0 | 190740 | 0 | 0 | 9100400 | 0 | 0 | 806330 | 634360 | 0 | 0 |
| HTNYTMEH | ACTN2; ACTN4 | P35609; O43 | ACTN2_HUMAN | Oxidation (M) | 0 | 0 | 0 | 0 | 0 | 0 | 0 | 0 | 0 | 0 | 0 | 8576900 | 0 | 0 | 0 | 0 | 0 | 0 |
| HTNYTMEH | ACTN4; ACTN1 | O43707; P12 | ACTN4_HUMAN | Oxidation (M) | 84535 | 0 | 0 | 332730 | 643510 | 298910 | 530730 | 0 | 0 | 0 | 136970 | 1420200 | 774250 | 0 | 776070 | 0 | 204880 | 0 |
| HTPGGPW | GMPPA; GMPPA | Q96IJ6; A0A | GMPPA_HUMA | Unmodified | 0 | 0 | 0 | 0 | 496600 | 0 | 0 | 0 | 0 | 434100 | 0 | 0 | 550610 | 476670 | 0 | 0 | 441900 | 1175400 |
| HTQENCET | CCT3; CCT3 | P49368; B4E | TCPG_HUMAN | Oxidation (M) | 0 | 0 | 0 | 0 | 0 | 0 | 0 | 0 | 0 | 0 | 468610 | 3765700 | 0 | 0 | 0 | 0 | 0 | 0 |
| HTQQRAAA | EFNB1 | P98172 | EFNB1_HUMAN | Unmodified | 0 | 0 | 0 | 0 | 0 | 0 | 0 | 0 | 0 | 0 | 0 | 0 | 0 | 0 | 0 | 0 | 104690 | 703290 |
| HTQVCING | ADAM10 | O14672 | ADA10_HUMAN | Unmodified | 0 | 0 | 0 | 1801700 | 0 | 0 | 0 | 0 | 0 | 0 | 0 | 3743800 | 0 | 0 | 3819800 | 0 | 0 | 0 |
| HTQWGIDF | FNBP1L | Q5T0N5 | FBP1L_HUMAN | Unmodified | 1192800 | 930610 | 245890 | 0 | 0 | 0 | 0 | 0 | 141820 | 0 | 0 | 0 | 0 | 0 | 654980 | 0 | 0 | 0 |
| HTRDSDIIE | ENDOD1 | O94919 | ENDD1_HUMA | Unmodified | 0 | 0 | 0 | 0 | 0 | 0 | 0 | 0 | 1191400 | 197620 | 0 | 0 | 92806 | 0 | 0 | 0 | 0 | 0 |
| HTRTQEVH | ADGRG6 | Q86SQ4 | AGRG6_HUMA | Unmodified | 0 | 0 | 289930 | 274130 | 2501900 | 601570 | 0 | 0 | 0 | 0 | 0 | 0 | 74924 | 0 | 0 | 0 | 0 | 0 |
| HTSALCNS | TLN1 | Q9Y490 | TLN1_HUMAN | Unmodified | 55842 | 23745 | 137550 | 0 | 190940 | 86717 | 0 | 77325 | 41277 | 26348 | 0 | 142540 | 0 | 181560 | 0 | 82012 | 92534 | 153620 |
| HTSISESSF | UTRN | P46939 | UTRN_HUMAN | Unmodified | 0 | 0 | 0 | 0 | 1487700 | 218820 | 15609 | 0 | 0 | 598420 | 0 | 0 | 958490 | 840720 | 0 | 0 | 265510 | 1668200 |
| HTSVQTTS | FN1 | P02751 | FINC_HUMAN | Unmodified | 0 | 0 | 1183500 | 336290 | 0 | 0 | 0 | 0 | 0 | 0 | 0 | 0 | 0 | 0 | 0 | 2352400 | 0 | 0 |
| HTTEAAAG | PKP3 | Q9Y446 | PKP3_HUMAN | Unmodified | 1273400 | 0 | 3470900 | 3590500 | 0 | 0 | 0 | 0 | 541050 | 56257 | 1188600 | 11615000 | 0 | 0 | 6283700 | 6864600 | 0 | 0 |
| HTTSIFDDF | YTHDF1; YTHDI | Q9BYJ9; Q7Z | YTHD1_HUMAN | Unmodified | 0 | 680940 | 0 | 0 | 0 | 0 | 0 | 0 | 0 | 0 | 0 | 677970 | 224450 | 0 | 0 | 1046500 | 0 | 0 |
| HTTSIFDDF | YTHDF2 | Q9Y5A9 | YTHD2_HUMAN | Unmodified | 887650 | 815650 | 651340 | 673150 | 0 | 0 | 0 | 0 | 0 | 0 | 0 | 1069900 | 0 | 0 | 971690 | 501970 | 0 | 0 |
| HTVDDGLC | CAND1; CAND1 | Q86VP6; A0A | CAND1_HUMA | Unmodified | 0 | 0 | 0 | 0 | 0 | 0 | 0 | 65004 | 589160 | 84107 | 0 | 1229700 | 0 | 0 | 0 | 68782 | 0 | 0 |
| HTVDDGLC | CAND1; CAND1 | Q86VP6; A0A | CAND1_HUMA | Unmodified | 0 | 0 | 0 | 0 | 0 | 0 | 0 | 0 | 0 | 0 | 0 | 0 | 0 | 0 | 1104500 | 0 | 0 | 0 |
| HTVYAALQ | CYFIP1; CYFIP1 | Q7L576; A0A | CYFP1_HUMAN | Unmodified | 0 | 152610 | 102590 | 37805 | 0 | 0 | 0 | 0 | 0 | 0 | 0 | 878640 | 0 | 0 | 395560 | 300380 | 0 | 0 |
| HTYRCLNG | ST14 | Q9Y5Y6 | ST14_HUMAN | Unmodified | 0 | 0 | 959140 | 0 | 0 | 0 | 0 | 125800 | 87745 | 0 | 0 | 0 | 0 | 351000 | 41630 | 0 | 0 | 91940 |
| HVAAGTQC | JUP | P14923 | PLAK_HUMAN | Unmodified | 557910 | 0 | 2414300 | 1085900 | 0 | 34520 | 0 | 0 | 41493 | 0 | 0 | 9082200 | 0 | 0 | 5935200 | 6256000 | 0 | 0 |
| HVAEVLEY | EIF2S1; EIF2S1 | P05198; H0Y | IF2A_HUMAN | Unmodified | 0 | 0 | 0 | 0 | 0 | 0 | 0 | 0 | 1128400 | 0 | 1725700 | 9274800 | 251710 | 0 | 4163700 | 838820 | 0 | 0 |
| HVALSVGC | NUDCD2; NUD | Q8WVJ2; E5I | NUDC2_HUMA | Unmodified | 0 | 56189 | 76716 | 193870 | 1647500 | 0 | 164170 | 0 | 0 | 198680 | 0 | 0 | 1206500 | 924430 | 0 | 0 | 539380 | 1361100 |
| HVATEYQE | KIF4A | O95239 | KIF4A_HUMAN | Unmodified | 0 | 0 | 0 | 0 | 124720 | 0 | 0 | 21657 | 58359 | 85113 | 0 | 25819 | 336300 | 34166 | 0 | 0 | 277660 | 135630 |
| HVATVQNE | C1orf21; C1orf2 | Q9H246; A0A | CA021_HUMAN | Unmodified | 0 | 0 | 0 | 0 | 76639 | 0 | 135670 | 117530 | 0 | 43193 | 0 | 0 | 509280 | 694760 | 0 | 0 | 110400 | 131820 |
| HVCDEEEV | SLC4A4; SLC4A | Q9Y6R1; A0A | S4A4_HUMAN | Unmodified | 0 | 0 | 0 | 0 | 6386200 | 664690 | 0 | 0 | 0 | 0 | 0 | 0 | 1601600 | 489740 | 209930 | 0 | 0 | 0 |
| HVDKGTAE | EMC1 | Q8N766 | EMC1_HUMAN | Unmodified | 0 | 0 | 0 | 0 | 0 | 0 | 0 | 0 | 9744000 | 0 | 0 | 0 | 0 | 0 | 0 | 0 | 0 | 708810 |
| HVDLLINK | SCP2; SCP2; SC | P22307; H0Y | SCP2_HUMAN | Unmodified | 0 | 0 | 0 | 0 | 748300 | 0 | 0 | 0 | 3144300 | 512230 | 0 | 160000 | 0 | 0 | 0 | 0 | 265470 | 12084000 |
| HVDRSDFF | PAPOLA; PAPOI | P51003; A0A | PAPOA_HUMAI | Unmodified | 0 | 0 | 0 | 0 | 0 | 0 | 0 | 121570 | 0 | 0 | 0 | 941550 | 0 | 0 | 0 | 0 | 0 | 0 |
| HVDSLSQF | SEPTIN9; SEPTI | Q9UHD8; K7 | SEPT9_HUMAN | Unmodified | 0 | 0 | 0 | 0 | 567170 | 0 | 0 | 0 | 0 | 0 | 0 | 0 | 112010 | 52593 | 0 | 0 | 217300 | 757790 |
| HVEAVAYY | SGTA; SGTA; SC | O43765; A0A | SGTA_HUMAN | Unmodified | 0 | 0 | 0 | 0 | 0 | 0 | 431960 | 77215 | 123640 | 129340 | 0 | 454540 | 1032900 | 0 | 0 | 47192 | 0 | 0 |
| HVEAVAYY | SGTA; SGTA; SC | O43765; A0A | SGTA_HUMAN | Unmodified | 0 | 0 | 0 | 0 | 932170 | 0 | 640200 | 0 | 0 | 739550 | 233360 | 1960500 | 974230 | 0 | 0 | 0 | 362250 | 1318000 |
| HVEAVYIDI | PITPNA; PITPNA | Q00169; I3L | PIPNA_HUMAN | Unmodified | 0 | 0 | 0 | 74790 | 615750 | 0 | 0 | 0 | 132540 | 0 | 0 | 530650 | 315420 | 0 | 0 | 47851 | 99446 | 183670 |
| HVEDVPAF | AZGP1; AZGP1 | P25311; H7E | ZA2G_HUMAN | Unmodified | 0 | 0 | 0 | 0 | 0 | 663980 | 0 | 0 | 0 | 0 | 433760 | 0 | 0 | 0 | 183470 | 0 | 0 | 0 |
| HVEDVPAF | AZGP1; AZGP1 | P25311; H7E | ZA2G_HUMAN | Unmodified | 0 | 0 | 0 | 495260 | 353690 | 0 | 0 | 0 | 0 | 0 | 11424000 | 0 | 0 | 262410 | 0 | 0 | 0 | 0 |
| HVEEFSPR | MCM6 | Q14566 | MCM6_HUMAN | Unmodified | 0 | 0 | 0 | 0 | 332150 | 0 | 0 | 44232 | 0 | 122710 | 0 | 0 | 0 | 0 | 0 | 0 | 54003 | 0 |
| HVEESVKP | TRAF4; TRAF4 | Q9BUZ4; F6S | TRAF4_HUMAN | Oxidation (M) | 0 | 0 | 0 | 0 | 0 | 0 | 0 | 0 | 0 | 0 | 0 | 0 | 0 | 0 | 426640 | 0 | 0 | 0 |
| HVEMYQW | TMEM43 | Q9BTV4 | TMM43_HUMAI | Unmodified | 0 | 0 | 0 | 0 | 0 | 0 | 0 | 0 | 4500600 | 212260 | 0 | 0 | 0 | 0 | 0 | 0 | 0 | 0 |
| HVEMYQW | TMEM43 | Q9BTV4 | TMM43_HUMAI | Unmodified | 0 | 0 | 0 | 0 | 0 | 0 | 0 | 0 | 3825400 | 363980 | 0 | 0 | 0 | 0 | 0 | 0 | 0 | 0 |
| HVETNSYD | VIL1; VIL1 | P09327; C9J | VILI_HUMAN Vi | Unmodified | 176760 | 0 | 0 | 0 | 0 | 0 | 0 | 136420 | 1149500 | 165400 | 1488500 | 8271600 | 0 | 0 | 0 | 0 | 0 | 0 |
| HVETNSYD | VIL1; VIL1 | P09327; C9J | VILI_HUMAN Vi | Unmodified | 0 | 0 | 0 | 0 | 0 | 0 | 0 | 0 | 0 | 1589900 | 0 | 299230 | 0 | 0 | 0 | 0 | 0 | 0 |
| HVETNTYD | AVIL; AVIL | O75366; F8V | AVIL_HUMAN A | Unmodified | 0 | 0 | 0 | 0 | 0 | 0 | 170840 | 86603 | 0 | 0 | 1092900 | 17658000 | 0 | 0 | 0 | 0 | 0 | 0 |
| HVETNTYD | AVIL; AVIL | O75366; F8V | AVIL_HUMAN A | Unmodified | 0 | 0 | 34654 | 0 | 0 | 0 | 0 | 0 | 0 | 0 | 0 | 7812100 | 0 | 0 | 25029 | 26759 | 0 | 0 |
| HVFAIFNTE | DNM2 | P50570 | DYN2_HUMAN | Unmodified | 661770 | 953480 | 417760 | 379700 | 0 | 0 | 0 | 0 | 848560 | 0 | 23724 | 0 | 383200 | 0 | 354560 | 389690 | 0 | 0 |
| HVFAIFNTE | DNM2 | P50570 | DYN2_HUMAN | Unmodified | 0 | 708490 | 0 | 0 | 1578100 | 326470 | 0 | 0 | 237500 | 399150 | 0 | 0 | 1352000 | 1183200 | 0 | 0 | 157040 | 739260 |
| HVFGESDE | TPI1; TPI1; TPI1 | P60174; U3K | TPIS_HUMAN | Unmodified | 9675600 | 3478600 | 2021300 | 9086300 | 21853000 | 839740 | 876230 | 1052500 | 14453000 | 3034600 | 23287000 | 73106000 | 60437000 | 12288000 | 11069000 | 14444000 | 5748300 | 15540000 |
| HVFGQAVK | CORO1C; COR | Q9ULV4; F8V | COR1C_HUMA | Unmodified | 0 | 0 | 0 | 0 | 0 | 0 | 0 | 0 | 0 | 0 | 0 | 3447100 | 0 | 0 | 0 | 0 | 175850 | 1195100 |
| HVFLTGPP | NTPCR; NTPCR | Q9BSD7; Q5 | NTPCR_HUMAI | Unmodified | 147870 | 192700 | 0 | 145170 | 0 | 0 | 411410 | 334800 | 0 | 0 | 0 | 0 | 458010 | 0 | 243370 | 200140 | 87812 | 0 |
| HVFWGSG | TFRC; TFRC | P02786; G3V | TFR1_HUMAN | Unmodified | 5554500 | 6116800 | 6968600 | 11218000 | 687330 | 219860 | 0 | 0 | 45023000 | 2034900 | 426600 | 2500900 | 565380 | 0 | 6835900 | 6672900 | 51352 | 0 |
| HVGMAVA | PSMA3 | P25788 | PSA3_HUMAN | Oxidation (M) | 942110 | 0 | 0 | 78938 | 0 | 0 | 0 | 0 | 0 | 0 | 0 | 3076000 | 0 | 0 | 0 | 489320 | 0 | 0 |
| HVGMAVA | PSMA3 | P25788 | PSA3_HUMAN | Unmodified | 0 | 0 | 0 | 0 | 0 | 0 | 0 | 0 | 100880 | 0 | 0 | 1408700 | 0 | 0 | 91880 | 0 | 0 | 0 |
| HVGNKTEC | ATP2B1; ATP2B | P20020; E7E | AT2B1_HUMAN | Unmodified | 0 | 0 | 422030 | 3007300 | 0 | 0 | 0 | 0 | 901680 | 383770 | 319550 | 1872500 | 0 | 0 | 4177000 | 1692000 | 0 | 0 |
| HVGNQQY | FLNB; FLNB; FL | O75369; A0A | FLNB_HUMAN | Unmodified | 0 | 180070 | 171110 | 307640 | 37219000 | 1353000 | 0 | 0 | 1323700 | 0 | 0 | 1846100 | 553940 | 580990 | 4673100 | 2224500 | 9682100 | 1257400 |
| HVGSNLCL | GALNT2 | Q10471 | GALT2_HUMAN | Unmodified | 0 | 0 | 0 | 0 | 0 | 0 | 0 | 0 | 3040200 | 95701 | 0 | 107830 | 0 | 0 | 0 | 0 | 0 | 0 |
| HVGSNLCL | GALNT2 | Q10471 | GALT2_HUMAN | Unmodified | 0 | 0 | 0 | 0 | 0 | 0 | 31096 | 9139.1 | 0 | 498840 | 0 | 0 | 0 | 0 | 0 | 0 | 0 | 0 |
| HVGSRLYS | FLNA; FLNA; FL | P21333; A0A | FLNA_HUMAN | Unmodified | 0 | 0 | 0 | 0 | 13060000 | 1264100 | 33408000 | 517310 | 0 | 929950 | 0 | 0 | 5274900 | 14201000 | 0 | 0 | 16340000 | 5961500 |
| HVGWEGA | SLC7A1 | P30825 | SL7A1_HUMAN | Unmodified | 0 | 0 | 601540 | 0 | 0 | 0 | 0 | 0 | 0 | 0 | 0 | 824680 | 366550 | 0 | 833830 | 0 | 0 | 0 |
| HVHLENAT | C11orf52 | Q96A22 | CK052_HUMAN | Unmodified | 10281000 | 6500200 | 1911700 | 1772400 | 0 | 0 | 605870 | 0 | 4852200 | 643760 | 0 | 0 | 0 | 1363200 | 3964800 | 3327800 | 0 | 0 |
| HVHLENAT | C11orf52 | Q96A22 | CK052_HUMAN | Unmodified | 0 | 0 | 0 | 0 | 0 | 0 | 0 | 0 | 0 | 1197300 | 0 | 0 | 6468300 | 2603000 | 0 | 0 | 0 | 698410 |
| HVHPTTAP | MPRIP; MPRIP; | Q6WCQ1; J3 | MPRIP_HUMAN | Unmodified | 0 | 0 | 0 | 0 | 0 | 0 | 0 | 0 | 0 | 0 | 0 | 0 | 0 | 0 | 0 | 0 | 380980 | 0 |
| HVILSEIK | TCF25; TCF25; | Q9BQ70; H3 | TCF25_HUMAN | Unmodified | 231380 | 483260 | 193910 | 0 | 0 | 0 | 0 | 0 | 0 | 0 | 0 | 0 | 0 | 0 | 361100 | 335370 | 11901 | 0 |
| HVIPMNPN | PLS3; PLS3; PLS | P13797; U3K | PLST_HUMAN | Oxidation (M) | 792340 | 0 | 0 | 0 | 0 | 0 | 0 | 0 | 0 | 0 | 0 | 0 | 0 | 0 | 0 | 0 | 0 | 0 |
| HVISYSLSF | UQCRQ | O14949 | QCR8_HUMAN | Unmodified | 0 | 127700 | 0 | 0 | 0 | 0 | 0 | 0 | 5622700 | 170450 | 0 | 0 | 0 | 0 | 63241 | 54330 | 0 | 0 |
| HVIYAPSSH | FOLH1B; FOLH | Q9HBA9; Q0 | FOH1B_HUMAI | Unmodified | 0 | 12433000 | 53828000 | 2.14E+08 | 0 | 0 | 0 | 0 | 11470 | 0 | 0 | 16459 | 0 | 0 | 75816000 | 61295000 | 0 | 0 |
| HVKGILLYC | NSF; NSF | P46459; I3L | NSF_HUMAN V | Unmodified | 0 | 0 | 0 | 0 | 0 | 0 | 0 | 0 | 0 | 0 | 0 | 0 | 0 | 0 | 0 | 0 | 151040 | 0 |
| HVLDTLIQL | HUWE1; HUWE | Q7Z6Z7; H0Y | HUWE1_HUMA | Unmodified | 0 | 0 | 0 | 0 | 0 | 0 | 0 | 0 | 0 | 0 | 0 | 1028600 | 0 | 0 | 0 | 0 | 0 | 0 |
| HVLFPLKSI | YARS1; YARS1 | P54577; A0A | SYYC_HUMAN | Unmodified | 0 | 0 | 0 | 0 | 908490 | 0 | 422940 | 0 | 0 | 314260 | 0 | 0 | 3605900 | 1724300 | 0 | 0 | 597420 | 644610 |
| HVLHLPTV | MGAT4B; MGAT | Q9UQ53; E5 | MGT4B_HUMA | Unmodified | 0 | 0 | 0 | 0 | 0 | 0 | 441020 | 0 | 1419000 | 290530 | 0 | 0 | 0 | 0 | 0 | 0 | 0 | 0 |
| HVLHVQLN | ECH1; N/A; ECH | Q13011; M0 | ECH1_HUMAN | Unmodified | 0 | 0 | 0 | 0 | 0 | 0 | 55186 | 0 | 549880 | 255250 | 0 | 0 | 0 | 0 | 0 | 0 | 0 | 0 |
| HVLLQEEV | GOLM2 | Q6P4E1 | GOLM2_HUMA | Unmodified | 0 | 0 | 0 | 0 | 0 | 0 | 0 | 110520 | 1449100 | 262550 | 0 | 932200 | 1267700 | 591480 | 0 | 0 | 0 | 0 |
| HVLLQEEV | GOLM2 | Q6P4E1 | GOLM2_HUMA | Unmodified | 0 | 0 | 0 | 86407 | 0 | 0 | 0 | 0 | 683470 | 0 | 0 | 0 | 0 | 0 | 0 | 0 | 0 | 0 |
| HVLNLLDE | EXOC4 | Q96A65 | EXOC4_HUMAI | Unmodified | 0 | 0 | 159940 | 0 | 0 | 0 | 0 | 0 | 0 | 0 | 340970 | 873950 | 0 | 0 | 0 | 359990 | 0 | 0 |
| HVLSVLSK | GSS; GSS; GSS | P48637; A0A | GSHB_HUMAN | Unmodified | 0 | 0 | 0 | 0 | 0 | 0 | 0 | 0 | 0 | 0 | 0 | 1929000 | 0 | 0 | 0 | 0 | 0 | 0 |
| HVLTAAHC | PRSS23 | O95084 | PRS23_HUMAN | Unmodified | 0 | 0 | 21725 | 0 | 957250 | 334350 | 0 | 0 | 0 | 0 | 0 | 0 | 104040 | 0 | 0 | 0 | 0 | 316620 |
| HVLTGSAD | EIF3I; EIF3I; EIF | Q13347; A0A | EIF3I_HUMAN | Unmodified | 105730 | 0 | 0 | 41113 | 0 | 0 | 64698 | 0 | 0 | 0 | 0 | 0 | 162510 | 129950 | 0 | 0 | 165130 | 227170 |
| HVLTLTDQ | ITGB3; N/A; ITG | P05106; H3E | ITB3_HUMAN I | Unmodified | 4332400 | 6020300 | 4005600 | 10295000 | 304700 | 0 | 338630 | 232410 | 387870 | 0 | 0 | 0 | 1174700 | 0 | 8851200 | 8597700 | 23509 | 120090 |
| HVLTLTDQ | ITGB3; N/A; ITG | P05106; H3E | ITB3_HUMAN I | Unmodified | 1909300 | 1552600 | 1675500 | 7375200 | 8680800 | 758060 | 13193000 | 1815600 | 0 | 0 | 0 | 0 | 7794600 | 8599900 | 2955900 | 2322200 | 265220 | 2436000 |
| HVLTLTDQ | ITGB3; N/A; ITG | P05106; H3E | ITB3_HUMAN I | Unmodified | 7024400 | 6640100 | 9341300 | 28072000 | 30187000 | 1993800 | 44006000 | 4785500 | 0 | 0 | 0 | 0 | 28407000 | 33624000 | 9239400 | 4493400 | 1573600 | 7382900 |
| HVLTTLGEI | MYL6B | P14649 | MYL6B_HUMAI | Unmodified | 0 | 1353900 | 0 | 0 | 114660 | 0 | 39956 | 37020 | 304310 | 0 | 93173 | 366990 | 0 | 0 | 1219900 | 0 | 21231 | 0 |
| HVLVTLGEI | MYL6; MYL6; MY | P60660; G3V | MYL6_HUMAN | Unmodified | 1164500 | 1392800 | 1302500 | 1749000 | 10633000 | 1228200 | 59498 | 39392 | 4013300 | 285480 | 1893700 | 9687000 | 803400 | 342890 | 2788100 | 1909700 | 289660 | 966780 |
| HVMELRQ | TAOK3; TAOK3 | Q9H2K8; G3 | TAOK3_HUMAN | Unmodified | 0 | 0 | 0 | 0 | 1659700 | 0 | 0 | 595720 | 0 | 0 | 0 | 0 | 0 | 1343000 | 0 | 0 | 0 | 0 |
| HVMTNLGE | CALM2; CALM3 | P0DP24; P0I | CALM2_HUMAI | Unmodified | 0 | 0 | 61567 | 0 | 379620 | 85019 | 84500 | 0 | 72640 | 0 | 0 | 0 | 0 | 0 | 135110 | 142660 | 0 | 71987 |
| HVNPVQAL | CTNNA1; CTNN | P35221; E5R | CTNA1_HUMAI | Unmodified | 0 | 0 | 2474500 | 4323200 | 15016000 | 1321900 | 0 | 0 | 269650 | 354680 | 1108000 | 1507700 | 9343400 | 9864400 | 1817100 | 2040500 | 0 | 0 |
| HVNSNQC | GALNT13; GALN | Q8IUC8; H7 | GLT13_HUMAN | Unmodified | 0 | 0 | 0 | 0 | 0 | 0 | 0 | 0 | 276880 | 0 | 0 | 0 | 0 | 0 | 0 | 0 | 0 | 0 |
| HVPGGGN | MAP4; MAP4; M | P27816; H0Y | MAP4_HUMAN | Unmodified | 87601 | 62838 | 0 | 0 | 774170 | 125970 | 76455 | 37391 | 0 | 465060 | 227460 | 1129000 | 754210 | 215540 | 0 | 0 | 199520 | 1351000 |
| HVPRAVFV | TUBA4A; TUBA4 | P68366; C9J | TBA4A_HUMAN | Unmodified | 0 | 0 | 0 | 0 | 179970 | 0 | 406320 | 0 | 0 | 0 | 0 | 0 | 176940 | 311810 | 0 | 0 | 62591 | 43757 |
| HVPRAVFV | TUBA1A; TUBA1 | Q71U36; Q9 | TBA1A_HUMAN | Unmodified | 0 | 0 | 0 | 0 | 7843100 | 0 | 1976800 | 25168 | 0 | 3182200 | 0 | 0 | 870650 | 0 | 0 | 0 | 2306300 | 1424300 |
| HVPRAVFV | TUBA1A; TUBA1 | Q71U36; Q9 | TBA1A_HUMAN | Unmodified | 0 | 0 | 0 | 0 | 2790900 | 0 | 0 | 0 | 0 | 0 | 0 | 0 | 0 | 0 | 0 | 0 | 324770 | 1304800 |
| HVPVVYVD | DYNC1H1; DYN | Q14204; A0A | DYHC1_HUMA | Unmodified | 0 | 0 | 0 | 0 | 0 | 0 | 0 | 0 | 0 | 0 | 0 | 1818300 | 0 | 0 | 0 | 0 | 0 | 0 |
| HVQDIVFQ | PICK1 | Q9NRD5 | PICK1_HUMAN | Unmodified | 0 | 0 | 0 | 53476 | 0 | 0 | 0 | 0 | 0 | 0 | 0 | 424320 | 0 | 0 | 88778 | 64676 | 0 | 0 |
| HVQLLGR | CYFIP1; CYFIP2 | Q7L576; Q9 | CYFP1_HUMAN | Unmodified | 144100 | 0 | 0 | 146690 | 0 | 0 | 0 | 0 | 0 | 205430 | 0 | 1133500 | 0 | 0 | 271640 | 219570 | 32108 | 22928 |
| HVQLSHEF | DDC; DDC; DD | P20711; A0A | DDC_HUMAN | Unmodified | 0 | 54426 | 21698 | 0 | 0 | 0 | 0 | 0 | 278890 | 82472 | 0 | 0 | 0 | 0 | 0 | 0 | 0 | 0 |

| | | | | | | | | | | | | | | | | | | | | | | |
|---|---|---|---|---|---|---|---|---|---|---|---|---|---|---|---|---|---|---|---|---|---|---|
| HVQTHFD( | TAX1BP1; TAX1 | Q86VP1; B8Z | TAXB1_HUMAN | Unmodified | 0 | 0 | 0 | 0 | 113930 | 0 | 0 | 0 | 980440 | 117650 | 0 | 0 | 0 | 0 | 0 | 0 | 37075 | 228880 |
| HVRGQNVI | ROR1; ROR1 | Q01973; A0A | ROR1_HUMAN | Unmodified | 0 | 0 | 0 | 0 | 1113100 | 1190400 | 227380 | 0 | 0 | 0 | 0 | 0 | 0 | 0 | 0 | 0 | 0 | 160480 |
| HVSGGLFS | IGSF3 | O75054 | IGSF3_HUMAN | Unmodified | 0 | 752410 | 0 | 193110 | 0 | 0 | 0 | 0 | 0 | 0 | 0 | 1909400 | 0 | 0 | 1223200 | 0 | 0 | 0 |
| HVSILDLP( | DTD2 | Q96FN9 | DTD2_HUMAN | Unmodified | 0 | 0 | 0 | 0 | 0 | 0 | 0 | 0 | 0 | 0 | 0 | 0 | 0 | 0 | 176640 | 0 | 0 | 0 |
| HVSLYQPQ | SEMA4B; SEMA | Q9NPR2; H0 | SEM4B_HUMAI | Unmodified | 0 | 0 | 0 | 0 | 0 | 0 | 0 | 0 | 1270400 | 217120 | 0 | 0 | 0 | 0 | 0 | 0 | 0 | 0 |
| HVSPAGAA | ATIC | P31939 | PUR9_HUMAN | Unmodified | 1182400 | 0 | 1608900 | 1888600 | 10400000 | 1259700 | 752770 | 559350 | 3157300 | 1634400 | 2088400 | 9435300 | 6025700 | 5010000 | 1476700 | 2191500 | 2673700 | 4265200 |
| HVSPVTPPI | ADAM9; ADAMS | Q13443; A0A | ADAM9_HUMA | Unmodified | 441940 | 0 | 668720 | 913650 | 551220 | 199370 | 0 | 0 | 0 | 0 | 0 | 0 | 0 | 0 | 905740 | 852200 | 72525 | 0 |
| HVSSTSSSI | ATP2C1; ATP2C | P98194; B4E | AT2C1_HUMAN | Unmodified | 0 | 0 | 0 | 0 | 0 | 0 | 0 | 68696 | 2677000 | 592620 | 0 | 198530 | 0 | 0 | 0 | 0 | 0 | 0 |
| HVSTSSDE | LRRC8D; LRRC | Q7L1W4; E9 | LRC8D_HUMAI | Unmodified | 0 | 0 | 0 | 0 | 595720 | 0 | 0 | 0 | 445760 | 143310 | 0 | 0 | 407000 | 156400 | 0 | 0 | 0 | 564790 |
| HVTQEFVS | ITGB4 | P16144 | ITB4_HUMAN I | Unmodified | 0 | 0 | 0 | 0 | 2020400 | 0 | 0 | 0 | 0 | 0 | 0 | 0 | 0 | 0 | 0 | 0 | 4310500 | 5699100 |
| HVVDGISR | PTPRF; PTPRF | P10586; H0Y | PTPRF_HUMAN | Unmodified | 137450 | 0 | 0 | 0 | 0 | 0 | 0 | 0 | 0 | 0 | 0 | 0 | 0 | 0 | 0 | 0 | 0 | 0 |
| HVVEDLIA( | MME; MME; MM | P08473; A0A | NEP_HUMAN N | Unmodified | 0 | 0 | 10740000 | 19483000 | 0 | 0 | 0 | 0 | 12654000 | 250520 | 0 | 437300 | 103020 | 0 | 34027000 | 24690000 | 0 | 0 |
| HVVFGHVk | PPIF | P30405 | PPIF_HUMAN F | Unmodified | 0 | 0 | 0 | 0 | 0 | 0 | 46282 | 0 | 1391400 | 253030 | 28909 | 0 | 0 | 0 | 0 | 0 | 26960 | 89357 |
| HVVFGHVk | PPIF | P30405 | PPIF_HUMAN F | Unmodified | 0 | 0 | 0 | 0 | 0 | 0 | 316650 | 0 | 0 | 450280 | 0 | 0 | 0 | 0 | 0 | 0 | 0 | 0 |
| HVVFGKVL | PPIB; PPIB; PPII | P23284; A0A | PPIB_HUMAN F | Unmodified | 0 | 0 | 0 | 0 | 0 | 0 | 0 | 0 | 196410 | 349950 | 0 | 0 | 0 | 0 | 0 | 0 | 0 | 0 |
| HVVFGQVI | PPID | Q08752 | PPID_HUMAN I | Unmodified | 26199 | 0 | 0 | 0 | 194820 | 0 | 0 | 0 | 116690 | 0 | 0 | 3443600 | 0 | 0 | 0 | 0 | 301870 | 318380 |
| HVVFIAQR | RPS7; RPS7; RP | P62081; A0A | RS7_HUMAN 4 | Unmodified | 487080 | 388530 | 235500 | 239230 | 133660 | 0 | 80564 | 0 | 285510 | 0 | 0 | 6674700 | 81451 | 28346 | 403440 | 350080 | 0 | 49737 |
| HVVFTAETI | PFAS | O15067 | PUR4_HUMAN | Unmodified | 0 | 0 | 0 | 0 | 611520 | 0 | 0 | 0 | 0 | 0 | 0 | 1987500 | 0 | 0 | 0 | 0 | 0 | 0 |
| HVVFYPTLI | CHID1; CHID1; | Q9BWS9; H( | CHID1_HUMAN | Unmodified | 0 | 0 | 29906 | 0 | 0 | 0 | 0 | 0 | 5570000 | 772160 | 0 | 0 | 0 | 0 | 0 | 0 | 0 | 0 |
| HVVLGAIEI | NAA10 | P41227 | NAA10_HUMAI | Unmodified | 0 | 0 | 0 | 0 | 0 | 0 | 0 | 0 | 0 | 49487 | 298180 | 4454100 | 0 | 0 | 0 | 0 | 80477 | 85759 |
| HVVQSIST( | ATP5PB; ATP5P | P24539; Q5( | AT5F1_HUMAN | Unmodified | 0 | 0 | 0 | 0 | 2606500 | 117920 | 87111 | 0 | 7948300 | 1324300 | 116210 | 0 | 0 | 656280 | 0 | 174650 | 111620 | 260920 |
| HVVQSIST( | ATP5PB; ATP5P | P24539; Q5( | AT5F1_HUMAN | Unmodified | 0 | 0 | 0 | 0 | 0 | 0 | 848510 | 0 | 466010 | 1138700 | 0 | 0 | 0 | 0 | 0 | 0 | 0 | 0 |
| HVYGNGYS | NECTIN1 | Q15223 | NECT1_HUMAI | Unmodified | 0 | 39888 | 0 | 0 | 95397 | 0 | 0 | 0 | 0 | 0 | 29084 | 253420 | 0 | 0 | 0 | 0 | 23886 | 0 |
| HWAAAQS( | H6PD | O95479 | G6PE_HUMAN | Unmodified | 0 | 0 | 0 | 0 | 0 | 312570 | 0 | 0 | 436120 | 33649 | 0 | 104490 | 0 | 0 | 0 | 0 | 0 | 0 |
| HWDQNER | SLC3A2; SLC3A | P08195; A0A | 4F2_HUMAN 4I | Unmodified | 0 | 658560 | 0 | 597900 | 60483 | 41573 | 0 | 0 | 3267500 | 168120 | 0 | 13875000 | 0 | 0 | 1140100 | 717980 | 0 | 0 |
| HWEEILQQ | OPA1; OPA1; OI | O60313; E5K | OPA1_HUMAN | Unmodified | 0 | 0 | 0 | 0 | 0 | 0 | 0 | 0 | 0 | 46121 | 0 | 2971600 | 0 | 0 | 0 | 0 | 0 | 0 |
| HWELTAEG | FARSA; FARSA; | Q9Y285; K7E | SYFA_HUMAN I | Unmodified | 0 | 0 | 0 | 0 | 0 | 0 | 0 | 0 | 0 | 118290 | 0 | 2645600 | 0 | 0 | 0 | 0 | 15193 | 0 |
| HWELTAEG | FARSA; FARSA; | Q9Y285; K7E | SYFA_HUMAN I | Unmodified | 0 | 0 | 0 | 0 | 0 | 0 | 0 | 0 | 0 | 0 | 540790 | 3047000 | 195060 | 0 | 0 | 0 | 0 | 0 |
| HWGGNVL | RPL7A; RPL7A | P62424; Q5T | RL7A_HUMAN ( | Unmodified | 3338900 | 4712400 | 3451000 | 850980 | 0 | 0 | 0 | 0 | 0 | 0 | 757720 | 3383900 | 0 | 0 | 1986500 | 2201200 | 0 | 0 |
| HWILPQDY | RCN1 | Q15293 | RCN1_HUMAN | Unmodified | 55151 | 0 | 2632500 | 0 | 0 | 0 | 0 | 0 | 6696600 | 0 | 624990 | 1163300 | 0 | 0 | 371490 | 168030 | 0 | 0 |
| HWKPLGQ | ROBO1; ROBO1 | Q9Y6N7; A0A | ROBO1_HUMA | Unmodified | 0 | 0 | 0 | 29240 | 0 | 0 | 0 | 0 | 0 | 432200 | 0 | 0 | 0 | 0 | 0 | 0 | 446970 | 4723200 |
| HWLDSPW | OGDH; OGDH; | Q02218; E9F | ODO1_HUMAN | Unmodified | 0 | 0 | 0 | 0 | 0 | 0 | 177210 | 0 | 69280 | 441380 | 16740 | 0 | 101770 | 870720 | 0 | 0 | 0 | 0 |
| HWMLDKL' | RPS4Y2; RPS4X | Q8TD47; P62 | RS4Y2_HUMAN | Oxidation (M) | 0 | 0 | 0 | 65725 | 0 | 0 | 296690 | 0 | 0 | 0 | 0 | 0 | 0 | 0 | 0 | 0 | 0 | 0 |
| HWNIDPVk | B3GLCT | Q6Y288 | B3GLT_HUMAN | Unmodified | 0 | 0 | 0 | 0 | 0 | 0 | 0 | 19062 | 445740 | 136140 | 0 | 0 | 0 | 0 | 62441 | 0 | 0 | 0 |
| HWPFMVVI | HSPA8; HSPA8; | P11142; E9P | HSP7C_HUMAI | Oxidation (M) | 15707000 | 18129000 | 20157000 | 29054000 | 59062000 | 14540000 | 8342000 | 989700 | 20992000 | 15798000 | 10021000 | 85664000 | 63517000 | 38456000 | 14789000 | 15292000 | 6316600 | 20831000 |
| HWPFMVVI | HSPA8; HSPA8; | P11142; E9P | HSP7C_HUMAI | Unmodified | 5626300 | 773810 | 3560800 | 5520000 | 50650000 | 6833600 | 4851700 | 130850 | 27819000 | 7432300 | 1760300 | 17931000 | 24005000 | 14447000 | 4652000 | 2602700 | 2890800 | 14448000 |
| HWPFQVIN | HSPA1A; HSPA1 | P0DMV8; P0 | HS71A_HUMAI | Unmodified | 8991700 | 8274400 | 6710400 | 3481300 | 12035000 | 2674400 | 868480 | 0 | 24977000 | 8238600 | 2426600 | 18128000 | 15518000 | 3390200 | 6970100 | 3854000 | 1598000 | 7041600 |
| HWPTFISD | XPO1; XPO1; XF | O14980; A0A | XPO1_HUMAN | Unmodified | 0 | 0 | 0 | 0 | 0 | 0 | 0 | 0 | 0 | 0 | 171100 | 1571300 | 0 | 0 | 0 | 0 | 0 | 0 |
| HWSDSRYI | COLGALT1; CO | Q8NBJ5; M0 | GT251_HUMAN | Unmodified | 23342 | 0 | 0 | 0 | 0 | 0 | 0 | 0 | 1809900 | 149460 | 48804 | 0 | 0 | 0 | 0 | 0 | 0 | 0 |
| HWTGILR | PLSCR1; PLSCF | O15162; C9J | PLS1_HUMAN I | Unmodified | 0 | 0 | 0 | 0 | 0 | 0 | 0 | 0 | 1065100 | 586650 | 0 | 0 | 0 | 0 | 0 | 0 | 0 | 0 |
| HYAGDVVY | MYO1D; MYO1I | O94832; J3Q | MYO1D_HUMA | Unmodified | 0 | 5283600 | 409640 | 1078900 | 304250 | 0 | 31975 | 0 | 674640 | 311450 | 0 | 385830 | 0 | 0 | 6730400 | 994830 | 0 | 0 |
| HYAGEVQY | MYO10; MYO10 | Q9HD67; D6 | MYO10_HUMAI | Unmodified | 0 | 0 | 0 | 317980 | 0 | 0 | 0 | 0 | 276310 | 86256 | 0 | 73128 | 0 | 19593 | 265900 | 413490 | 0 | 0 |
| HYAGEVQY | MYO10; MYO10 | Q9HD67; D6 | MYO10_HUMAI | Unmodified | 0 | 0 | 0 | 0 | 231990 | 45496 | 0 | 0 | 0 | 0 | 0 | 0 | 0 | 0 | 0 | 0 | 0 | 76351 |
| HYAHTDCF | TUFM | P49411 | EFTU_HUMAN | Unmodified | 0 | 0 | 0 | 0 | 0 | 0 | 0 | 0 | 9903800 | 0 | 0 | 0 | 0 | 0 | 0 | 0 | 0 | 0 |
| HYALNVPL' | DIS3L2 | Q8IYB7 | DI3L2_HUMAN | Unmodified | 0 | 0 | 0 | 0 | 0 | 0 | 0 | 0 | 0 | 0 | 0 | 1739500 | 0 | 0 | 0 | 159560 | 0 | 0 |
| HYASEEIKE | SPTBN1; SPTBN | Q01082; A0A | SPTB2_HUMAN | Unmodified | 0 | 0 | 0 | 400930 | 117280 | 0 | 0 | 32048 | 0 | 0 | 68622 | 712060 | 0 | 0 | 0 | 0 | 0 | 0 |
| HYCAIADAI | PLCG2; PLCG2 | P16885; A0A | PLCG2_HUMAI | Unmodified | 0 | 0 | 0 | 0 | 228400 | 86122 | 0 | 0 | 0 | 0 | 370550 | 3955100 | 0 | 0 | 0 | 0 | 0 | 0 |
| HYCTVANP | MVP; MVP; MVP | Q14764; H3I | MVP_HUMAN N | Unmodified | 323830 | 17682000 | 0 | 0 | 0 | 0 | 0 | 0 | 607070 | 0 | 0 | 13165000 | 0 | 0 | 0 | 0 | 0 | 0 |
| HYCYPHFT | GNAS; GNAS; G | Q5JWF2; P63 | GNAS1_HUMAI | Unmodified | 199550 | 0 | 0 | 0 | 0 | 0 | 0 | 0 | 0 | 0 | 0 | 1476100 | 0 | 0 | 315200 | 389530 | 0 | 0 |
| HYCYPHFT | GNAS; GNAS; G | Q5JWF2; P63 | GNAS1_HUMAI | Unmodified | 152780 | 0 | 0 | 0 | 0 | 0 | 0 | 0 | 0 | 0 | 0 | 3176700 | 0 | 0 | 1075900 | 475240 | 0 | 0 |
| HYDLESVP | MOV10; MOV10 | Q9HCE1; Q5 | MOV10_HUMA | Oxidation (M) | 3501000 | 1320700 | 0 | 0 | 0 | 0 | 0 | 0 | 0 | 0 | 0 | 0 | 0 | 0 | 1083900 | 2060500 | 0 | 0 |
| HYEDGYP( | CTNND1; CTNN | O60716; A0A | CTND1_HUMAI | Unmodified | 978800 | 1068100 | 3310700 | 4217500 | 425800 | 203910 | 0 | 0 | 0 | 0 | 960290 | 4660500 | 463110 | 236550 | 6101500 | 0 | 296850 | 619910 |
| HYEISLQLD | TMTC4 | Q5T4D3 | TMTC4_HUMAI | Unmodified | 0 | 0 | 0 | 0 | 0 | 0 | 0 | 0 | 320500 | 36641 | 0 | 0 | 0 | 0 | 0 | 0 | 0 | 0 |
| HYFQNTQ( | ARF4; ARF5; AR | P18085; P84 | ARF4_HUMAN , | Unmodified | 0 | 884760 | 2173900 | 0 | 0 | 0 | 0 | 0 | 3442400 | 321020 | 0 | 6459100 | 0 | 0 | 4080100 | 2121900 | 0 | 0 |
| HYFQNTQ( | ARF4; ARF5; AR | P18085; P84 | ARF4_HUMAN , | Unmodified | 7141900 | 6656200 | 5006500 | 3163400 | 0 | 0 | 0 | 0 | 3456800 | 72138 | 0 | 32326000 | 0 | 0 | 11757000 | 9933700 | 0 | 0 |
| HYFQNTQ( | ARF5; ARF5 | P84085; C9J | ARF5_HUMAN , | Unmodified | 647180 | 945430 | 0 | 995330 | 0 | 0 | 0 | 0 | 0 | 0 | 288020 | 0 | 0 | 0 | 0 | 0 | 0 | 0 |
| HYGFQLYQ | PSMA4; PSMA4 | P25789; H0Y | PSA4_HUMAN | Unmodified | 212580 | 295410 | 385120 | 718900 | 2608600 | 0 | 299440 | 0 | 2384600 | 1322700 | 1171300 | 7597000 | 1532000 | 1050100 | 736190 | 545460 | 1332000 | 0 |
| HYGGLTGL | PGAM1; PGAM4 | P18669; Q8N | PGAM1_HUMA | Unmodified | 0 | 0 | 1222400 | 0 | 72645 | 0 | 0 | 0 | 0 | 0 | 829240 | 17560000 | 0 | 0 | 1407100 | 1514000 | 0 | 0 |
| HYGGLTGL | PGAM1; PGAM4 | P18669; Q8N | PGAM1_HUMA | Unmodified | 257090 | 0 | 0 | 1710600 | 1278300 | 0 | 0 | 0 | 200740 | 67861 | 0 | 4997500 | 0 | 0 | 0 | 0 | 1508100 | 4020500 |
| HYGGLTGL | PGAM1; PGAM4 | P18669; Q8N | PGAM1_HUMA | Unmodified | 0 | 0 | 0 | 0 | 0 | 77072 | 42974 | 0 | 0 | 0 | 0 | 0 | 0 | 0 | 0 | 0 | 204410 | 0 |
| HYGNKPIG | C11orf54; C11( | Q9H0W9; A( | CK054_HUMAN | Unmodified | 0 | 0 | 34674 | 51492 | 290520 | 0 | 0 | 0 | 351970 | 149710 | 0 | 734480 | 562270 | 255870 | 0 | 0 | 1073000 | 0 |
| HYGPGWV | PPIB; PPIB; PPII | P23284; A0A | PPIB_HUMAN F | Oxidation (M) | 0 | 78788 | 0 | 0 | 0 | 0 | 347530 | 72463 | 19040000 | 575150 | 8415500 | 2100700 | 0 | 0 | 739150 | 447180 | 0 | 0 |
| HYGPGWV | PPIB; PPIB; PPII | P23284; A0A | PPIB_HUMAN F | Unmodified | 0 | 0 | 664750 | 246920 | 0 | 0 | 180870 | 42935 | 1.01E+08 | 4698600 | 6051900 | 14950000 | 179780 | 630610 | 1483900 | 353110 | 0 | 0 |
| HYGPGWV | PPIB; PPIB; PPII | P23284; A0A | PPIB_HUMAN F | Oxidation (M) | 0 | 0 | 0 | 0 | 0 | 0 | 0 | 0 | 11852000 | 25071000 | 8737000 | 1062100 | 1622200 | 5851200 | 0 | 0 | 0 | 0 |
| HYGPGWV | PPIB; PPIB; PPII | P23284; A0A | PPIB_HUMAN F | Unmodified | 0 | 0 | 0 | 0 | 0 | 0 | 0 | 0 | 12825000 | 4389500 | 0 | 370810 | 0 | 0 | 338230 | 0 | 0 | 0 |
| HYGYNSYS | CTSB; CTSB; C1 | P07858; E9P | CATB_HUMAN | Unmodified | 0 | 0 | 0 | 0 | 1397400 | 751270 | 0 | 24330 | 823350 | 0 | 0 | 0 | 0 | 0 | 0 | 0 | 0 | 0 |
| HYIFNPRKI | UGGT1 | Q9NYU2 | UGGG1_HUMA | Unmodified | 106750 | 123470 | 89435 | 197700 | 0 | 0 | 0 | 0 | 3308300 | 1964400 | 0 | 180030 | 0 | 0 | 133220 | 128900 | 0 | 0 |
| HYLDFINH' | DYNC1H1; DYN | Q14204; A0A | DYHC1_HUMA | Unmodified | 0 | 0 | 0 | 0 | 0 | 0 | 0 | 0 | 0 | 1229300 | 0 | 3992800 | 1471000 | 0 | 0 | 904790 | 0 | 0 |
| HYLFYDGE | VPS26A; VPS26 | O75436; S4F | VP26A_HUMAN | Unmodified | 0 | 0 | 247140 | 202090 | 352520 | 0 | 0 | 21713 | 739120 | 0 | 246760 | 3258700 | 200530 | 128090 | 211770 | 158210 | 0 | 114450 |
| HYLLTLFSV | ADAMTS1; ADA | Q9UHI8; E5F | ATS1_HUMAN , | Unmodified | 0 | 0 | 87935 | 0 | 0 | 0 | 0 | 0 | 46748 | 0 | 0 | 0 | 0 | 0 | 0 | 130770 | 0 | 0 |
| HYNGEAYE | DNAJA1 | P31689 | DNJA1_HUMAN | Unmodified | 0 | 0 | 0 | 0 | 0 | 0 | 0 | 0 | 0 | 0 | 0 | 0 | 0 | 0 | 0 | 0 | 0 | 0 |
| HYQFASGA | PDCD6IP; PDCI | Q8WUM4; C | PDC6I_HUMAN | Unmodified | 20994000 | 27462000 | 9777800 | 6677700 | 4331000 | 1614500 | 0 | 0 | 3659400 | 2229500 | 1324300 | 6696200 | 3683300 | 2399000 | 12182000 | 15257000 | 0 | 2537300 |
| HYQFASGA | PDCD6IP; PDCI | Q8WUM4; C | PDC6I_HUMAN | Unmodified | 198570 | 692270 | 8504.7 | 0 | 0 | 0 | 54948 | 0 | 0 | 151540 | 0 | 0 | 0 | 0 | 387480 | 0 | 46765 | 0 |
| HYQHVAA\ | APLP1; APLP1; , | P51693; B7Z | APLP1_HUMAN | Unmodified | 0 | 0 | 0 | 0 | 0 | 0 | 0 | 0 | 1053800 | 0 | 0 | 0 | 0 | 0 | 0 | 0 | 0 | 0 |
| HYQINQQV | FN1 | P02751 | FINC_HUMAN I | Unmodified | 609000 | 3250000 | 34516000 | 11019000 | 0 | 3716400 | 0 | 76606 | 3770200 | 1781200 | 940080 | 275190 | 3545900 | 3577600 | 1667800 | 4955400 | 392640 | 4429800 |
| HYRGPAGI | SLC1A5; SLC1A | Q15758; M0 | AAAT_HUMAN I | Unmodified | 0 | 460050 | 0 | 0 | 28175000 | 0 | 1168800 | 223950 | 4217200 | 0 | 4222900 | 30775000 | 8500500 | 3212300 | 1664700 | 0 | 875090 | 2551400 |
| HYRGPAGI | SLC1A5; SLC1A | Q15758; M0 | AAAT_HUMAN I | Oxidation (M) | 0 | 0 | 0 | 0 | 14059000 | 599540 | 5296600 | 614920 | 0 | 1376300 | 0 | 0 | 5150900 | 2720800 | 0 | 0 | 5412200 | 1365000 |
| HYRGPAGI | SLC1A5; SLC1A | Q15758; M0 | AAAT_HUMAN I | Unmodified | 0 | 0 | 0 | 0 | 32993000 | 3890200 | 8458000 | 1325900 | 0 | 4353000 | 0 | 0 | 15803000 | 9466800 | 0 | 0 | 12621000 | 6284600 |
| HYRNIPGP | SMARCA1; SMA | P28370; O6( | SMCA1_HUMA | Unmodified | 0 | 0 | 0 | 0 | 0 | 0 | 253750 | 0 | 0 | 48839 | 0 | 0 | 0 | 0 | 0 | 0 | 0 | 0 |
| HYSPEDEP | N/A; AAK1 | Q6ZSR9; A0A | YJ005_HUMAN | Unmodified | 0 | 0 | 0 | 280670 | 0 | 0 | 0 | 0 | 149050 | 0 | 0 | 0 | 1374500 | 659810 | 200770 | 821740 | 0 | 565740 |
| HYTEAIKRI | STIP1 | P31948 | STIP1_HUMAN | Unmodified | 0 | 0 | 0 | 0 | 713580 | 0 | 0 | 214310 | 0 | 909020 | 0 | 0 | 2844100 | 0 | 0 | 0 | 1721300 | 1008500 |
| HYTEHADC | YES1; YES1 | P07947; J3Q | YES_HUMAN Ty | Unmodified | 929150 | 344650 | 854230 | 313190 | 845430 | 272030 | 0 | 66911 | 780430 | 376710 | 235970 | 1222300 | 571170 | 625980 | 1186800 | 1005100 | 846940 | 1515900 |
| HYTEHADC | YES1; YES1 | P07947; J3Q | YES_HUMAN Ty | Unmodified | 366920 | 0 | 0 | 0 | 1500700 | 404760 | 0 | 0 | 0 | 152310 | 0 | 259910 | 953460 | 0 | 0 | 0 | 2478200 | 5685000 |
| HYYEVSCH | DDX1; DDX1; D | Q92499; A0A | DDX1_HUMAN | Unmodified | 0 | 0 | 0 | 0 | 0 | 0 | 0 | 0 | 0 | 0 | 0 | 1691400 | 0 | 0 | 0 | 0 | 0 | 0 |
| HYYTGTQG | ARF6 | P62330 | ARF6_HUMAN , | Unmodified | 0 | 0 | 0 | 0 | 0 | 0 | 594990 | 0 | 0 | 295360 | 473440 | 642090 | 0 | 0 | 1654600 | 1125000 | 0 | 0 |
| HYYTGTQG | ARF6 | P62330 | ARF6_HUMAN , | Unmodified | 0 | 0 | 609740 | 0 | 0 | 0 | 0 | 0 | 0 | 0 | 0 | 1689600 | 0 | 0 | 0 | 0 | 0 | 0 |
| IAAADYIPT\ | GNAZ | P19086 | GNAZ_HUMAN | Unmodified | 0 | 0 | 0 | 0 | 0 | 0 | 0 | 0 | 0 | 0 | 0 | 309780 | 0 | 0 | 0 | 0 | 0 | 0 |
| IAAAGLDVI | GRHPR; GRHPF | Q9UBQ7; U3 | GRHPR_HUMA | Unmodified | 1331500 | 0 | 1194000 | 0 | 20162000 | 2721600 | 1349500 | 255670 | 6075000 | 3640000 | 3093400 | 12239000 | 9353100 | 5266900 | 5291200 | 2435100 | 2083100 | 3669100 |
| IAAAILNTPI | GOT2 | P00505 | AATM_HUMAN | Unmodified | 0 | 49847 | 202540 | 97261 | 543640 | 92457 | 146610 | 0 | 17917000 | 7676000 | 443760 | 1293200 | 237290 | 183810 | 192170 | 175440 | 0 | 0 |
| IAAANGIGF | PGM1; PGM1 | P36871; A0A | PGM1_HUMAN | Unmodified | 0 | 0 | 0 | 0 | 0 | 0 | 0 | 0 | 0 | 325010 | 0 | 413940 | 182670 | 71188 | 44575 | 60626 | 0 | 0 |
| IAADFRNKI | HSPH1; HSPH1 | Q92598; A0A | HS105_HUMAN | Unmodified | 0 | 0 | 0 | 168070 | 0 | 0 | 753050 | 652500 | 0 | 0 | 740420 | 0 | 0 | 0 | 0 | 0 | 1197700 | 1668800 |
| IAAEHCIPLI | XPO4 | Q9C0E2 | XPO4_HUMAN | Unmodified | 0 | 0 | 0 | 0 | 0 | 0 | 0 | 0 | 0 | 0 | 0 | 163140 | 0 | 0 | 0 | 0 | 0 | 0 |
| IAAENELN( | IPO8 | O15397 | IPO8_HUMAN I | Unmodified | 0 | 0 | 176360 | 319700 | 5121800 | 1348300 | 243890 | 0 | 57449 | 238470 | 121300 | 574730 | 913330 | 567120 | 133490 | 285650 | 1137200 | 1138300 |
| IAAEYYDNI | DEGS1 | O15121 | DEGS1_HUMAI | Unmodified | 0 | 0 | 0 | 0 | 470980 | 0 | 0 | 0 | 3838700 | 0 | 0 | 0 | 0 | 0 | 0 | 0 | 325520 | 0 |
| IAAGDRLR( | UGGT1 | Q9NYU2 | UGGG1_HUMA | Unmodified | 0 | 0 | 0 | 0 | 0 | 0 | 0 | 0 | 0 | 4856800 | 0 | 0 | 0 | 0 | 0 | 0 | 0 | 0 |
| IAAGEKIPLS | MCCC1; MCCC | Q96RQ3; E9I | MCCA_HUMAN | Unmodified | 0 | 0 | 0 | 0 | 0 | 0 | 0 | 0 | 2136100 | 0 | 0 | 0 | 0 | 0 | 0 | 0 | 0 | 0 |
| IAAHLADFT | DNAJC13 | O75165 | DJC13_HUMAN | Unmodified | 0 | 0 | 0 | 0 | 0 | 0 | 0 | 0 | 0 | 0 | 0 | 3031700 | 0 | 0 | 0 | 0 | 0 | 0 |
| IAALDPENI | EIF2AK3; EIF2AI | Q9NZJ5; A0A | E2AK3_HUMAN | Unmodified | 0 | 0 | 0 | 0 | 0 | 0 | 0 | 0 | 0 | 0 | 0 | 0 | 0 | 736920 | 0 | 0 | 0 | 0 |
| IAALQAFAD | SPTAN1; SPTAN | Q13813; A0A | SPTN1_HUMAN | Unmodified | 43207 | 0 | 501340 | 296850 | 1720400 | 667110 | 0 | 0 | 1565400 | 929120 | 1075000 | 8056600 | 347710 | 192270 | 1333000 | 501230 | 0 | 0 |
| IAAPLTKVD | FLOT2; FLOT2; I | Q14254; J3Q | FLOT2_HUMAN | Unmodified | 0 | 0 | 0 | 9694800 | 20639000 | 2998200 | 4712800 | 1013800 | 20644000 | 19506000 | 6928900 | 11348000 | 7124000 | 9719600 | 0 | 0 | 878840 | 2892000 |
| IAAQEVPIEI | ACOT8; ACOT8; | O14734; E9F | ACOT8_HUMAI | Unmodified | 0 | 0 | 0 | 0 | 0 | 0 | 0 | 0 | 47983000 | 736780 | 0 | 0 | 0 | 0 | 0 | 0 | 0 | 0 |
| IAAQTLLGT | PYCR3; PYCR3; | Q53H96; B5I | P5CR3_HUMAN | Unmodified | 0 | 0 | 0 | 0 | 334370 | 0 | 0 | 41036 | 0 | 518120 | 158580 | 3397600 | 1740000 | 0 | 0 | 0 | 297790 | 401570 |
| IAARIYEENI | SCG3 | Q8WXD2 | SCG3_HUMAN | Unmodified | 0 | 0 | 0 | 0 | 0 | 0 | 0 | 0 | 0 | 1296300 | 0 | 0 | 0 | 0 | 0 | 0 | 0 | 0 |
| IAASATCGE | LAMA5 | O15230 | LAMA5_HUMAN | Unmodified | 695240 | 0 | 0 | 41606 | 0 | 0 | 0 | 0 | 0 | 0 | 0 | 0 | 331430 | 147040 | 0 | 0 | 0 | 1736200 |
| IAASEEELK | CHST1 | O43916 | CHST1_HUMAI | Unmodified | 0 | 0 | 0 | 0 | 0 | 0 | 0 | 0 | 636030 | 0 | 0 | 0 | 0 | 0 | 0 | 0 | 0 | 0 |
| IAASEEELKI | CHST1 | O43916 | CHST1_HUMAI | Unmodified | 0 | 0 | 0 | 0 | 0 | 0 | 32522 | 0 | 12866000 | 805950 | 0 | 0 | 0 | 0 | 0 | 0 | 0 | 0 |
| IAASSSFR | SH3GL1 | Q99961 | SH3G1_HUMAI | Unmodified | 0 | 0 | 0 | 0 | 0 | 0 | 87837 | 104240 | 1245100 | 27278 | 0 | 492380 | 0 | 0 | 0 | 121280 | 55634 | 578020 |
| IAAVHNVPI | SRXN1 | Q9BYN0 | SRXN1_HUMAI | Unmodified | 274660 | 0 | 324890 | 547070 | 257350 | 131210 | 0 | 0 | 0 | 0 | 33164 | 0 | 0 | 0 | 543010 | 559630 | 0 | 0 |
| IAAYLQSD( | GSTM3 | P21266 | GSTM3_HUMAI | Unmodified | 62720 | 0 | 195690 | 0 | 0 | 0 | 0 | 0 | 0 | 0 | 0 | 0 | 0 | 554590 | 498500 | 479850 | 0 | 234510 |

| | | | | | | | | | | | | | | | | | | | | | | |
|---|---|---|---|---|---|---|---|---|---|---|---|---|---|---|---|---|---|---|---|---|---|---|
| IACAAVDC | PDIA5; PDIA5 | Q14554; H7 | PDIA5_HUMAN | Unmodified | 0 | 0 | 0 | 0 | 0 | 0 | 0 | 0 | 2947700 | 786200 | 92636 | 0 | 0 | 0 | 0 | 0 | 0 | 0 |
| IACAAVDC | PDIA5; PDIA5 | Q14554; H7 | PDIA5_HUMAN | Unmodified | 0 | 0 | 0 | 0 | 0 | 0 | 0 | 0 | 195070 | 158350 | 0 | 0 | 0 | 62291 | 0 | 0 | 0 | 0 |
| IACAAVDC | PDIA5; PDIA5 | Q14554; H7 | PDIA5_HUMAN | Unmodified | 0 | 0 | 0 | 0 | 0 | 0 | 0 | 0 | 0 | 966160 | 0 | 0 | 0 | 0 | 0 | 0 | 53491 | 0 |
| IACLDFSLC | TCP1; TCP1; TC | P17987; F5C | TCPA_HUMAN | Unmodified | 0 | 0 | 0 | 199430 | 373330 | 0 | 856740 | 474320 | 1029300 | 1217200 | 692230 | 8982900 | 1511600 | 885290 | 0 | 0 | 0 | 0 |
| IADAFSPAS | LSR; LSR; LSR | Q86X29; M0 | LSR_HUMAN Li | Unmodified | 0 | 0 | 0 | 0 | 0 | 0 | 0 | 0 | 1575600 | 839940 | 0 | 0 | 0 | 0 | 0 | 0 | 0 | 0 |
| IADDKYND | HSP90B1; HSP | P14625; A0A | ENPL_HUMAN | Unmodified | 0 | 171690 | 0 | 313870 | 4097500 | 454210 | 2468000 | 1124600 | 83982000 | 46974000 | 14418000 | 13522000 | 1895700 | 898970 | 833870 | 0 | 1536400 | 2406400 |
| IADDKYND | HSP90B1; HSP | P14625; A0A | ENPL_HUMAN | Unmodified | 0 | 0 | 0 | 0 | 319610 | 88242 | 3749300 | 86266 | 8131600 | 33138000 | 1064700 | 0 | 298240 | 937820 | 0 | 0 | 0 | 194750 |
| IADFGLARL | YES1; FYN; LCK | P07947; P06 | YES_HUMAN Ty | Unmodified | 0 | 0 | 0 | 0 | 0 | 0 | 0 | 0 | 0 | 0 | 0 | 0 | 0 | 0 | 0 | 0 | 2058600 | 2203000 |
| IADFGLAR | HCK; LYN; HCK | P08631; P07 | HCK_HUMAN T | Unmodified | 0 | 0 | 0 | 0 | 914660 | 181010 | 584500 | 0 | 0 | 120780 | 0 | 0 | 179480 | 81119 | 0 | 0 | 0 | 0 |
| IADFGMCK | PRKCB | P05771 | KPCB_HUMAN | Oxidation (M) | 0 | 0 | 0 | 0 | 0 | 0 | 167230 | 100600 | 0 | 0 | 0 | 0 | 0 | 0 | 0 | 0 | 0 | 0 |
| IADFGWSV | AURKB; AURKB | Q96GD4; J3 | AURKB_HUMA | Unmodified | 391720 | 768590 | 718250 | 2850400 | 3290900 | 568510 | 149460 | 0 | 0 | 329200 | 122710 | 0 | 0 | 282330 | 650080 | 529740 | 406340 | 0 |
| IADFGWSV | AURKA; AURKA | O14965; A3 | AURKA_HUMA | Unmodified | 276170 | 280250 | 275600 | 473200 | 0 | 0 | 0 | 0 | 0 | 0 | 400220 | 0 | 0 | 0 | 639190 | 197950 | 0 | 0 |
| IADFLNSFD | BRK1 | Q8WUW1 | BRK1_HUMAN | Oxidation (M) | 60260 | 24880 | 47170 | 163670 | 47643 | 103290 | 83705 | 18666 | 156810 | 120440 | 89357 | 1109300 | 206290 | 126590 | 330650 | 259850 | 174330 | 0 |
| IADFLNSFD | BRK1 | Q8WUW1 | BRK1_HUMAN | Unmodified | 0 | 0 | 0 | 0 | 728630 | 0 | 150430 | 0 | 0 | 0 | 0 | 0 | 372280 | 550880 | 0 | 0 | 734170 | 518570 |
| IADFLNSFD | BRK1 | Q8WUW1 | BRK1_HUMAN | Oxidation (M) | 0 | 0 | 0 | 0 | 1303900 | 0 | 331220 | 142720 | 0 | 517710 | 0 | 0 | 1361300 | 864460 | 0 | 0 | 1546700 | 1057600 |
| IADFQDVL | TBC1D13 | Q9NVG8 | TBC13_HUMA | Unmodified | 0 | 0 | 35424 | 34857 | 0 | 0 | 0 | 21722 | 0 | 0 | 143220 | 1093800 | 0 | 0 | 69715 | 69324 | 0 | 0 |
| IADGSVKR | AKR1C4; AKR1 | P17516; P52 | AK1C4_HUMA | Unmodified | 898700 | 301630 | 0 | 0 | 51364000 | 3547500 | 0 | 0 | 0 | 255010 | 0 | 0 | 431200 | 281350 | 0 | 0 | 2463800 | 1500100 |
| IADGYEQA | CCT5; CCT5; C | P48643; B7Z | TCPE_HUMAN | Unmodified | 40175 | 62651 | 0 | 2119200 | 0 | 0 | 67044 | 40990 | 2931600 | 80633 | 2243800 | 14902000 | 1311500 | 846280 | 0 | 1909000 | 0 | 0 |
| IADLGNAC | SRPK1; SRPK2; | Q96SB4; P7 | SRPK1_HUMA | Unmodified | 0 | 0 | 0 | 0 | 0 | 0 | 0 | 0 | 0 | 0 | 0 | 2597900 | 0 | 0 | 0 | 0 | 0 | 0 |
| IADMGHLK | PCNA | P12004 | PCNA_HUMAN | Unmodified | 0 | 0 | 0 | 0 | 1189400 | 0 | 0 | 0 | 851260 | 417240 | 1437500 | 0 | 0 | 0 | 0 | 0 | 218330 | 18841 |
| IADPEHDH | MAPK3; MAPK3 | P27361; E9P | MK03_HUMAN | Unmodified | 0 | 0 | 0 | 252130 | 0 | 0 | 0 | 0 | 0 | 0 | 0 | 534850 | 0 | 0 | 204910 | 107110 | 0 | 0 |
| IADQLEEL | BAG1; BAG1; B | Q99933; C9 | BAG1_HUMAN | Unmodified | 0 | 0 | 0 | 0 | 600330 | 645040 | 0 | 165460 | 0 | 0 | 0 | 1021100 | 0 | 0 | 0 | 0 | 0 | 172140 |
| IADRGAEY | SF3B1; SF3B1 | O75533; H7 | SF3B1_HUMA | Unmodified | 0 | 0 | 0 | 0 | 0 | 0 | 0 | 0 | 0 | 0 | 0 | 990270 | 0 | 0 | 0 | 0 | 0 | 0 |
| IADRLGLEL | PRPS1; PRPS1L | P60891; P21 | PRPS1_HUMA | Unmodified | 0 | 0 | 0 | 0 | 975800 | 0 | 3351600 | 153490 | 0 | 321690 | 0 | 0 | 1087500 | 585460 | 0 | 0 | 3659000 | 384060 |
| IADSIGCST | ENOPH1; ENO | Q9UHY7; D6 | ENOPH_HUMA | Unmodified | 0 | 31735 | 0 | 0 | 0 | 0 | 0 | 0 | 0 | 0 | 170730 | 982320 | 0 | 0 | 0 | 107990 | 67359 | 0 |
| IADSKDHV | ANTXR1; ANTX | Q9H6X2; A0 | ANTR1_HUMA | Unmodified | 0 | 0 | 0 | 87390 | 382350 | 257530 | 0 | 0 | 97129 | 0 | 0 | 0 | 0 | 0 | 0 | 0 | 0 | 80292 |
| IADSKRAFE | FLOT2; FLOT2; | Q14254; J3C | FLOT2_HUMA | Unmodified | 0 | 518400 | 0 | 0 | 1423900 | 0 | 2069300 | 790220 | 0 | 1536800 | 0 | 0 | 654650 | 1505500 | 125160 | 142180 | 0 | 0 |
| IAEAFRLPT | GIPC1 | O14908 | GIPC1_HUMA | Oxidation (M) | 0 | 0 | 0 | 181500 | 14865000 | 0 | 575540 | 0 | 634040 | 413770 | 180260 | 0 | 2223800 | 3065300 | 751930 | 0 | 8499100 | 10457000 |
| IAEAFRLPT | GIPC1 | O14908 | GIPC1_HUMA | 2 Oxidation (M) | 0 | 0 | 0 | 435500 | 3500400 | 912120 | 0 | 0 | 0 | 387200 | 0 | 552530 | 0 | 1402100 | 519780 | 278600 | 3492900 | 3127900 |
| IAEANKLKL | ERLIN2; ERLIN | O94905; E5 | ERLN2_HUMA | Unmodified | 0 | 0 | 0 | 0 | 0 | 0 | 0 | 0 | 0 | 455820 | 0 | 0 | 0 | 0 | 0 | 0 | 0 | 0 |
| IAECSSQL | MYH10 | P35580 | MYH10_HUMA | Unmodified | 0 | 0 | 0 | 0 | 0 | 0 | 0 | 0 | 0 | 0 | 0 | 2166900 | 0 | 0 | 225950 | 91324 | 0 | 0 |
| IAECTFTK | CTHRC1 | Q96CG8 | CTHR1_HUMA | Unmodified | 0 | 0 | 0 | 0 | 0 | 36436 | 0 | 23428 | 4661800 | 2275300 | 0 | 398290 | 192540 | 0 | 0 | 0 | 722520 | 488760 |
| IAEDDGKP | DAG1 | Q14118 | DAG1_HUMAN | Unmodified | 5400200 | 4909100 | 1214500 | 803960 | 2040200 | 347910 | 54081 | 0 | 2289000 | 171160 | 146470 | 90930 | 977100 | 304470 | 979910 | 718390 | 385780 | 385430 |
| IAEESNFPF | NSF; NSF | P46459; I3L | NSF_HUMAN V | Unmodified | 384270 | 0 | 658180 | 0 | 3082000 | 0 | 3326200 | 1819900 | 1218100 | 760640 | 1027800 | 3303100 | 9772300 | 7809300 | 887740 | 840640 | 2175200 | 8962000 |
| IAEFSRTPE | TUBGCP2; TUB | Q9BSJ2; F2Z | GCP2_HUMAN | Unmodified | 47511 | 0 | 0 | 94800 | 410270 | 0 | 141900 | 100770 | 0 | 158090 | 0 | 0 | 340280 | 168820 | 0 | 0 | 88147 | 211890 |
| IAEFTTNLT | MYH9 | P35579 | MYH9_HUMAN | Unmodified | 82915 | 126460 | 619500 | 316020 | 0 | 2630000 | 2473600 | 1742400 | 690770 | 55716 | 435790 | 3493700 | 0 | 6318700 | 828550 | 532390 | 340270 | 4266700 |
| IAEFTTNLT | MYH9 | P35579 | MYH9_HUMAN | Unmodified | 0 | 0 | 0 | 0 | 950060 | 0 | 0 | 0 | 0 | 0 | 0 | 0 | 409530 | 0 | 0 | 0 | 0 | 456520 |
| IAEFTTNLT | MYH9 | P35579 | MYH9_HUMAN | Unmodified | 0 | 0 | 0 | 0 | 246910 | 0 | 0 | 0 | 0 | 0 | 0 | 0 | 166780 | 0 | 0 | 0 | 253490 | 147650 |
| IAEGAQQG | NAGK; NAGK; N | Q9UJ70; C9J | NAGK_HUMAN | Unmodified | 0 | 0 | 0 | 0 | 0 | 0 | 0 | 30976 | 0 | 0 | 0 | 418030 | 0 | 0 | 80135 | 0 | 0 | 0 |
| IAEIEAEMA | DRG1; DRG1 | Q9Y295; F8 | DRG1_HUMAN | Unmodified | 0 | 0 | 0 | 0 | 0 | 0 | 0 | 441520 | 0 | 0 | 400980 | 1229200 | 116760 | 0 | 0 | 0 | 0 | 0 |
| IAEIEAEMA | DRG1; DRG1 | Q9Y295; F8 | DRG1_HUMAN | Unmodified | 0 | 0 | 0 | 0 | 1120300 | 0 | 0 | 0 | 240170 | 0 | 0 | 0 | 875670 | 0 | 0 | 0 | 72839 | 395360 |
| IAEIFMR | DECR2; DECR2 | Q9NUI1; Q4 | DECR2_HUMA | Unmodified | 1322000 | 2465900 | 0 | 0 | 0 | 0 | 0 | 0 | 862670 | 79911 | 0 | 0 | 0 | 0 | 0 | 0 | 254110 | 0 |
| IAELIYK | PDE2A; PDE2A | O00408; E9 | PDE2A_HUMA | Unmodified | 0 | 0 | 0 | 0 | 0 | 0 | 0 | 38809 | 0 | 0 | 0 | 4641600 | 0 | 0 | 0 | 0 | 24384 | 0 |
| IAELLENVT | USP10; USP10 | Q14694; J3K | UBP10_HUMA | Unmodified | 0 | 0 | 0 | 0 | 0 | 0 | 0 | 0 | 299970 | 0 | 587880 | 5193600 | 0 | 0 | 0 | 0 | 0 | 0 |
| IAELLSPGS | SERPINB6; SER | P35237; A0A | SPB6_HUMAN | Unmodified | 112420 | 0 | 99070 | 267690 | 46868 | 0 | 0 | 0 | 232280 | 0 | 329740 | 4877700 | 0 | 0 | 458360 | 284270 | 0 | 0 |
| IAELMPGA | PSMC5 | P62195 | PRS8_HUMAN | Unmodified | 0 | 0 | 0 | 128740 | 712230 | 0 | 201820 | 96554 | 0 | 0 | 226540 | 1045100 | 820880 | 557180 | 0 | 0 | 441280 | 0 |
| IAELMPGA | PSMC5 | P62195 | PRS8_HUMAN | Oxidation (M) | 0 | 0 | 0 | 0 | 0 | 0 | 0 | 0 | 0 | 0 | 0 | 438130 | 252370 | 0 | 46732 | 0 | 0 | 0 |
| IAENPANP | WASHC2A; WA | Q641Q2; Q9 | WAC2A_HUMA | Unmodified | 0 | 0 | 0 | 0 | 684410 | 0 | 0 | 0 | 0 | 0 | 0 | 155990 | 0 | 83274 | 0 | 0 | 90336 | 270430 |
| IAENVEEV | STX1A; STX1A; | Q16623; A8 | STX1A_HUMA | Unmodified | 0 | 0 | 0 | 0 | 0 | 0 | 0 | 0 | 938820 | 0 | 0 | 0 | 0 | 0 | 0 | 0 | 0 | 0 |
| IAENVKLEL | TAX1BP1; TAX1 | Q86VP1; B8 | TAXB1_HUMA | Unmodified | 0 | 623790 | 248970 | 328380 | 1867900 | 1576800 | 184810 | 334240 | 4817100 | 1316600 | 0 | 254260 | 98773 | 0 | 326530 | 0 | 1438700 | 4256700 |
| IAEPDARFS | CRIM1 | Q9NZV1 | CRIM1_HUMA | Unmodified | 0 | 0 | 0 | 0 | 4482700 | 838340 | 331420 | 0 | 0 | 0 | 0 | 0 | 0 | 0 | 0 | 0 | 64553 | 673850 |
| IAEPDARFS | CRIM1 | Q9NZV1 | CRIM1_HUMA | Oxidation (M) | 0 | 0 | 0 | 0 | 4479300 | 636390 | 398640 | 0 | 0 | 0 | 0 | 0 | 0 | 0 | 0 | 0 | 0 | 374820 |
| IAEPSVCG | MCM4; MCM4; | P33991; A0A | MCM4_HUMAN | Unmodified | 0 | 0 | 0 | 0 | 88748 | 0 | 22099 | 0 | 248180 | 46437 | 273890 | 267260 | 89462 | 0 | 0 | 14940 | 0 | 0 |
| IAEQEHTQ | MACF1; MACF1 | Q9UPN3; H | MACF1_HUMA | Unmodified | 0 | 0 | 0 | 0 | 0 | 0 | 0 | 0 | 243520 | 0 | 273130 | 962690 | 0 | 0 | 0 | 0 | 0 | 0 |
| IAEQQKAQ | PLEC; PLEC | Q15149; H0 | PLEC_HUMAN | Unmodified | 0 | 0 | 0 | 0 | 0 | 0 | 0 | 0 | 0 | 0 | 0 | 0 | 0 | 0 | 0 | 0 | 304870 | 566110 |
| IAEQVASF | CTNNA1; CTNN | P35221; G3 | CTNA1_HUMA | Unmodified | 6227300 | 2452900 | 8623700 | 12435000 | 24387000 | 5364800 | 0 | 0 | 3175800 | 951030 | 3975100 | 16325000 | 15449000 | 2971700 | 20470000 | 16710000 | 0 | 0 |
| IAESHLQS | PPP1R1B; PPP1 | Q9UD71; J3 | PPR1B_HUMA | Unmodified | 0 | 0 | 0 | 0 | 0 | 0 | 0 | 0 | 0 | 0 | 552460 | 8496400 | 0 | 0 | 0 | 0 | 0 | 0 |
| IAESIPYLK | ASPH | Q12797 | ASPH_HUMAN | Unmodified | 0 | 0 | 0 | 0 | 405960 | 0 | 723130 | 20462 | 7667900 | 773910 | 500610 | 0 | 165680 | 248120 | 0 | 0 | 0 | 59945 |
| IAESLGGS | EFR3A; EFR3A | Q14156; A0 | EFR3A_HUMA | Unmodified | 0 | 0 | 443280 | 362960 | 0 | 0 | 0 | 0 | 813160 | 304260 | 0 | 1354600 | 0 | 0 | 876810 | 0 | 617130 | 0 |
| IAESLGGS | EFR3A; EFR3A | Q14156; A0 | EFR3A_HUMA | Unmodified | 0 | 0 | 0 | 0 | 1746200 | 0 | 0 | 0 | 0 | 0 | 0 | 0 | 0 | 0 | 0 | 0 | 0 | 0 |
| IAESYQNIL | EXT1 | Q16394 | EXT1_HUMAN | Unmodified | 0 | 0 | 0 | 0 | 888450 | 3472400 | 1936400 | 25099 | 7814100 | 15197000 | 0 | 0 | 13625000 | 5824500 | 0 | 39386 | 19573000 | 45613000 |
| IAETDFEK | DNAJC7; DNAJ | Q99615; A0 | DNJC7_HUMA | Unmodified | 0 | 0 | 0 | 0 | 1583300 | 186160 | 416400 | 0 | 807630 | 501780 | 241110 | 1349400 | 0 | 0 | 0 | 0 | 0 | 199370 |
| IAETYSIEM | SNX33 | Q8WV41 | SNX33_HUMA | Oxidation (M) | 0 | 653740 | 1879500 | 1110800 | 0 | 0 | 0 | 0 | 0 | 0 | 0 | 0 | 0 | 0 | 3517500 | 2127600 | 0 | 0 |
| IAETYSIEM | SNX33 | Q8WV41 | SNX33_HUMA | Unmodified | 0 | 0 | 567340 | 552010 | 0 | 0 | 0 | 0 | 5035500 | 419110 | 427300 | 0 | 458440 | 0 | 2507800 | 712260 | 0 | 0 |
| IAETYSIEM | SNX33 | Q8WV41 | SNX33_HUMA | Oxidation (M) | 261010 | 264780 | 1540100 | 2202600 | 181430 | 0 | 0 | 130190 | 0 | 0 | 0 | 0 | 354460 | 274630 | 3672000 | 1828500 | 0 | 0 |
| IAETYSIEM | SNX33 | Q8WV41 | SNX33_HUMA | Unmodified | 0 | 0 | 0 | 4513000 | 0 | 0 | 0 | 0 | 0 | 0 | 0 | 0 | 416370 | 0 | 0 | 0 | 0 | 0 |
| IAEVDASV | UQCRC1 | P31930 | QCR1_HUMAN | Unmodified | 71982 | 0 | 0 | 0 | 654440 | 0 | 0 | 0 | 39966000 | 638850 | 580730 | 443030 | 0 | 0 | 0 | 0 | 0 | 747340 |
| IAEVDASV | UQCRC1 | P31930 | QCR1_HUMAN | Unmodified | 0 | 0 | 0 | 0 | 0 | 0 | 0 | 0 | 17440000 | 2540800 | 0 | 0 | 0 | 0 | 0 | 0 | 0 | 0 |
| IAEVDCTAE | TXNDC5 | Q8NBS9 | TXND5_HUMA | Unmodified | 0 | 0 | 0 | 0 | 0 | 0 | 28429 | 0 | 7772500 | 530870 | 0 | 68559 | 0 | 0 | 0 | 0 | 0 | 0 |
| IAEVDCTAE | TXNDC5 | Q8NBS9 | TXND5_HUMA | Unmodified | 0 | 0 | 0 | 0 | 731290 | 241900 | 366780 | 402430 | 13169000 | 13250000 | 0 | 0 | 463850 | 646520 | 163390 | 0 | 0 | 0 |
| IAEVGAGG | PURB | Q96QR8 | PURB_HUMAN | Unmodified | 0 | 0 | 0 | 0 | 0 | 0 | 0 | 125780 | 0 | 0 | 0 | 0 | 0 | 0 | 0 | 0 | 0 | 0 |
| IAEVGGVP | C11orf54; C11 | Q9H0W9; A | CK054_HUMA | Unmodified | 45487 | 0 | 0 | 0 | 0 | 0 | 0 | 0 | 257070 | 40718 | 65160 | 532980 | 0 | 0 | 0 | 0 | 223460 | 81588 |
| IAFAITAIK | RPS18; RPS18; | P62269; A0A | RS18_HUMAN | Unmodified | 2390800 | 1565200 | 2139400 | 867030 | 2759600 | 237810 | 4071500 | 3148500 | 850810 | 1451000 | 2485400 | 37997000 | 1516500 | 741500 | 2247600 | 2271000 | 832600 | 1897300 |
| IAFGGETD | ALDH3A2; ALD | P51648; A0A | AL3A2_HUMA | Unmodified | 340130 | 0 | 0 | 0 | 0 | 0 | 0 | 0 | 1939500 | 358540 | 0 | 0 | 0 | 0 | 159800 | 0 | 0 | 0 |
| IAFGGETD | ALDH3A2; ALD | P51648; A0A | AL3A2_HUMA | Unmodified | 0 | 0 | 0 | 0 | 0 | 0 | 0 | 0 | 7284500 | 2361600 | 0 | 0 | 0 | 0 | 0 | 0 | 0 | 0 |
| IAFGGETD | ALDH3A2; ALD | P51648; A0A | AL3A2_HUMA | Unmodified | 0 | 0 | 0 | 0 | 733810 | 0 | 0 | 0 | 448620 | 1039100 | 0 | 0 | 0 | 0 | 0 | 0 | 0 | 0 |
| IAFTEWIEP | SMARCA5 | O60264 | SMCA5_HUMA | Unmodified | 400470 | 0 | 266870 | 290750 | 173850 | 0 | 901780 | 0 | 0 | 0 | 0 | 167420 | 1634900 | 374490 | 204890 | 206720 | 0 | 102930 |
| IAFTGSTEV | ALDH1A3; ALD | P47895; O9 | AL1A3_HUMA | Unmodified | 0 | 0 | 0 | 0 | 0 | 0 | 0 | 0 | 0 | 0 | 0 | 0 | 0 | 0 | 0 | 0 | 1264000 | 4255100 |
| IAFTGSTEV | ALDH1A3; ALD | P47895; H0 | AL1A3_HUMA | Unmodified | 0 | 0 | 0 | 0 | 1082300 | 0 | 0 | 0 | 834840 | 0 | 0 | 0 | 0 | 0 | 0 | 0 | 17094000 | 10291000 |
| IAGAIGPCV | SEC23B; SEC2 | Q15437; A0 | SC23B_HUMA | Unmodified | 0 | 0 | 0 | 0 | 0 | 0 | 0 | 0 | 0 | 0 | 0 | 1453900 | 0 | 0 | 0 | 0 | 0 | 0 |
| IAGDQSTL | SMPDL3B; SMP | Q92485; F8 | ASM3B_HUMA | Unmodified | 3301200 | 2535100 | 11436000 | 5105400 | 253060 | 52190 | 0 | 0 | 2747400 | 425220 | 714140 | 7740200 | 219970 | 87197 | 989220 | 9907000 | 0 | 0 |
| IAGEASRL | H2BC4; H2BC1 | P62807; Q6 | H2B1C_HUMA | Unmodified | 0 | 0 | 0 | 0 | 670570 | 258430 | 684970 | 2534000 | 314320 | 195500 | 0 | 0 | 0 | 0 | 0 | 0 | 50138 | 148890 |
| IAGGTSNE | ESRP1; ESRP1; | Q6NXG1; H | ESRP1_HUMA | Unmodified | 0 | 0 | 0 | 0 | 0 | 0 | 0 | 0 | 318920 | 0 | 0 | 1303000 | 0 | 0 | 0 | 0 | 0 | 0 |
| IAGKVPGL | ATXN2L; ATXN2 | Q8WWM7; | ATX2L_HUMA | Unmodified | 1049600 | 0 | 0 | 0 | 0 | 0 | 0 | 0 | 0 | 0 | 110730 | 0 | 0 | 0 | 0 | 0 | 0 | 0 |
| IAGLAVSLT | NCSTN; NCSTN | Q92542; Q5 | NICA_HUMAN | Unmodified | 0 | 0 | 497520 | 0 | 0 | 0 | 0 | 11254 | 1010700 | 0 | 0 | 897010 | 0 | 0 | 0 | 0 | 0 | 0 |
| IAGLAVSLT | NCSTN; NCSTN | Q92542; Q5 | NICA_HUMAN | Unmodified | 0 | 0 | 0 | 0 | 0 | 0 | 0 | 0 | 25871000 | 0 | 0 | 0 | 0 | 0 | 0 | 586790 | 0 | 0 |
| IAGLCNR | ATP1A1; ATP1A | P05023; P50 | AT1A1_HUMA | Unmodified | 0 | 0 | 7485200 | 9462800 | 5158500 | 1211900 | 242880 | 0 | 6471700 | 2212600 | 5038900 | 45929000 | 10008000 | 1637900 | 10010000 | 8293200 | 1517100 | 0 |
| IAGLCNRA | ATP1A1 | P05023 | AT1A1_HUMA | Unmodified | 0 | 0 | 0 | 0 | 13969000 | 1485400 | 2430000 | 53696 | 0 | 3208600 | 430850 | 0 | 21587000 | 10209000 | 0 | 0 | 5172000 | 4062200 |
| IAGMDPVV | PANK4; PANK4 | Q9NVE7; E9 | PANK4_HUMA | Unmodified | 0 | 0 | 0 | 247000 | 0 | 0 | 0 | 0 | 1347200 | 0 | 0 | 345270 | 0 | 0 | 74969 | 83805 | 0 | 0 |
| IAGQVAAA | RPS19; RPS19 | P39019; A0A | RS19_HUMAN | Unmodified | 1945100 | 851810 | 864310 | 820970 | 0 | 0 | 0 | 0 | 160240 | 67791 | 510010 | 14960000 | 0 | 0 | 877330 | 782690 | 0 | 0 |
| IAGRAVLIA | RUVBL2; RUVB | Q9Y230; M0 | RUVB2_HUMA | Unmodified | 0 | 0 | 0 | 0 | 1642600 | 0 | 572310 | 238800 | 0 | 218090 | 0 | 0 | 1358500 | 933180 | 0 | 0 | 513230 | 460380 |
| IAGYVTHL | RPS17; RPS17; | P08708; H0 | RS17_HUMAN | Oxidation (M) | 62000 | 63629 | 0 | 0 | 0 | 0 | 168230 | 274750 | 0 | 87756 | 0 | 995660 | 0 | 0 | 64298 | 65822 | 0 | 0 |
| IAGYVTHL | RPS17; RPS17; | P08708; H0 | RS17_HUMAN | Unmodified | 84085 | 0 | 106930 | 205500 | 0 | 0 | 0 | 35976 | 0 | 0 | 0 | 2766100 | 155570 | 0 | 0 | 0 | 0 | 0 |
| IAHFLDR | EIF3E; EIF3E; E | P60228; A0A | EIF3E_HUMAN | Unmodified | 0 | 183950 | 0 | 0 | 0 | 0 | 0 | 21730 | 0 | 0 | 0 | 1432000 | 262430 | 0 | 196700 | 0 | 0 | 0 |
| IAHIMGPP | KHSRP; KHSRP | Q92945; M0 | FUBP2_HUMA | Unmodified | 0 | 0 | 0 | 0 | 451790 | 0 | 38991 | 0 | 641110 | 229550 | 304680 | 1426600 | 0 | 144580 | 0 | 9914.5 | 34110 | 199280 |
| IAHVELAD | HSPG2 | P98160 | PGBM_HUMAN | Unmodified | 91551 | 0 | 5482600 | 2695600 | 161600 | 190830 | 0 | 0 | 1668000 | 220190 | 532250 | 1272700 | 156330 | 0 | 680460 | 2115100 | 46863 | 97948 |
| IAIDHNQM | ERGIC2; ERGIC | Q96RQ1; A0 | ERGI2_HUMA | Unmodified | 0 | 0 | 0 | 0 | 0 | 15337 | 0 | 0 | 1551000 | 181060 | 0 | 0 | 0 | 0 | 0 | 0 | 25855 | 0 |
| IAIIGAGIG | PCYOX1; PCYO | Q9UHG3; F8 | PCYOX_HUMA | Unmodified | 0 | 223630 | 0 | 0 | 0 | 0 | 0 | 0 | 24269000 | 2044500 | 585460 | 292360 | 0 | 0 | 939370 | 44779 | 0 | 0 |
| IAIIGAGIG | PCYOX1; PCYO | Q9UHG3; F8 | PCYOX_HUMA | Unmodified | 0 | 0 | 0 | 0 | 0 | 0 | 0 | 0 | 0 | 8831400 | 0 | 0 | 0 | 0 | 0 | 0 | 0 | 0 |
| IAIKNCDG | CTNND1; CTNN | O60716; A0 | CTND1_HUMA | Unmodified | 0 | 0 | 0 | 0 | 0 | 0 | 0 | 0 | 0 | 0 | 0 | 0 | 534480 | 3370200 | 0 | 0 | 2189400 | 306770 |
| IAILGYR | RHEB; RHEB | Q15382; F8 | RHEB_HUMAN | Unmodified | 815120 | 1437000 | 658190 | 726320 | 3017200 | 648950 | 435580 | 364430 | 578770 | 970790 | 315010 | 1804000 | 763580 | 535560 | 1187900 | 865630 | 0 | 1262800 |
| IAILTCPFE | CCT5; CCT5; C | P48643; B7Z | TCPE_HUMAN | Unmodified | 203020 | 0 | 1105500 | 697140 | 4073700 | 738040 | 2083600 | 939690 | 2137800 | 1606600 | 2510200 | 16577000 | 8504400 | 4492700 | 0 | 0 | 3382300 | 5514800 |
| IAIPGLAGA | PTBP1; PTBP1; | P26599; A6 | PTBP1_HUMA | Unmodified | 455930 | 521880 | 249210 | 0 | 53045 | 196570 | 0 | 0 | 721450 | 337030 | 309120 | 4971200 | 0 | 0 | 0 | 475800 | 0 | 0 |
| IAIVNHDK | ABCE1; ABCE1; | P61221; D6 | ABCE1_HUMA | Unmodified | 48767 | 73433 | 0 | 100060 | 1777200 | 0 | 134380 | 53688 | 273750 | 0 | 0 | 2127900 | 1030400 | 0 | 68498 | 99521 | 636880 | 558420 |
| IAIWTTECE | EIF4E; EIF4E | P06730; D6 | IF4E_HUMAN E | Unmodified | 183000 | 0 | 0 | 0 | 0 | 0 | 0 | 0 | 0 | 0 | 0 | 3338800 | 322890 | 0 | 0 | 0 | 0 | 0 |
| IAIWTTECE | EIF4E; EIF4E | P06730; D6 | IF4E_HUMAN E | Unmodified | 255570 | 164130 | 0 | 506460 | 5174200 | 1508400 | 0 | 0 | 3999300 | 563400 | 0 | 0 | 0 | 0 | 0 | 253800 | 0 | 746600 |
| IAIWTTECE | EIF4E; EIF4E | P06730; D6 | IF4E_HUMAN E | Unmodified | 0 | 0 | 0 | 0 | 0 | 0 | 0 | 0 | 116070 | 0 | 0 | 0 | 3497400 | 1364700 | 0 | 0 | 0 | 0 |
| IAIYELLFK | RPS10; RPS10- | P46783; A0A | RS10_HUMAN | Unmodified | 11816000 | 11520000 | 4163900 | 7747900 | 8028200 | 1041900 | 2803400 | 88778 | 7156600 | 3059700 | 10063000 | 58675000 | 4942100 | 2670700 | 9222000 | 3859900 | 0 | 6546800 |
| IAKEEIFGP | ALDH1A3; ALD | P47895; H0 | AL1A3_HUMA | Unmodified | 0 | 0 | 0 | 0 | 1316500 | 0 | 0 | 0 | 0 | 0 | 0 | 0 | 0 | 0 | 0 | 0 | 6681200 | 5970300 |

| | | | | | | | | | | | | | | | | | | | | | | |
|---|---|---|---|---|---|---|---|---|---|---|---|---|---|---|---|---|---|---|---|---|---|---|
| IAKEEIFGP\ | ALDH1A1 | P00352 | AL1A1_HUMAN | Unmodified | 0 | 0 | 0 | 0 | 0 | 0 | 0 | 0 | 1292300 | 0 | 3646800 | 68051000 | 0 | 0 | 0 | 0 | 0 | 0 |
| IAKEEIFGP\ | ALDH1A1 | P00352 | AL1A1_HUMAN | Oxidation (M) | 0 | 0 | 0 | 0 | 0 | 0 | 0 | 0 | 215120 | 248760 | 7876500 | 1.4E+08 | 0 | 0 | 0 | 0 | 0 | 0 |
| IAKSEPPTL | SLK | Q9H2G2 | SLK_HUMAN S | Unmodified | 0 | 0 | 0 | 0 | 480510 | 0 | 713960 | 0 | 0 | 261120 | 0 | 0 | 1361900 | 0 | 0 | 0 | 643820 | 0 |
| IALALASR | GGT7 | Q9UJ14 | GGT7_HUMAN | Unmodified | 0 | 0 | 0 | 0 | 0 | 0 | 0 | 0 | 560350 | 238160 | 0 | 0 | 295610 | 0 | 0 | 0 | 0 | 19354 |
| IALDFQR | LGALS3; LGALS | P17931; G3\ | LEG3_HUMAN | Unmodified | 0 | 0 | 0 | 0 | 0 | 0 | 0 | 0 | 0 | 0 | 0 | 4165500 | 0 | 0 | 0 | 0 | 0 | 0 |
| IALDKNPET | STK10 | O94804 | STK10_HUMAN | Unmodified | 0 | 0 | 0 | 0 | 10100000 | 0 | 0 | 0 | 0 | 1266100 | 910730 | 0 | 1318700 | 1189000 | 0 | 0 | 7826500 | 6389100 |
| IALENQLEC | REPS2 | Q8NFH8 | REPS2_HUMAN | Unmodified | 0 | 0 | 169880 | 543000 | 0 | 0 | 0 | 0 | 1186000 | 0 | 0 | 0 | 122060 | 0 | 380550 | 473880 | 0 | 0 |
| IALESEGRP | SQSTM1 | Q13501 | SQSTM_HUMA | Oxidation (M) | 677700 | 0 | 594130 | 0 | 418130 | 0 | 0 | 0 | 0 | 0 | 0 | 0 | 0 | 0 | 0 | 0 | 295810 | 0 |
| IALESEGRP | SQSTM1 | Q13501 | SQSTM_HUMA | Unmodified | 0 | 0 | 0 | 0 | 1332600 | 0 | 251540 | 22418 | 0 | 0 | 0 | 0 | 805840 | 322500 | 0 | 0 | 303220 | 1378200 |
| IALGNEEGI | CDC42BPA; CD | Q5VT25; A0 | MRCKA_HUMA | Unmodified | 0 | 0 | 0 | 0 | 0 | 0 | 0 | 0 | 311650 | 0 | 0 | 1271400 | 0 | 0 | 593380 | 0 | 0 | 0 |
| IALLEEAK | RDX; RDX; RDX | P35241; A0 | RADI_HUMAN | Unmodified | 0 | 0 | 2331700 | 6221500 | 31209000 | 1404900 | 295080 | 1980200 | 2997200 | 2265600 | 1916200 | 13423000 | 37890000 | 30377000 | 3899400 | 0 | 4781900 | 10431000 |
| IALLEEAKK | RDX; RDX; RDX | P35241; A0 | RADI_HUMAN | Unmodified | 0 | 0 | 0 | 0 | 0 | 0 | 0 | 0 | 0 | 0 | 0 | 0 | 162200 | 3413100 | 0 | 0 | 0 | 0 |
| IALLEEARR | EZR; EZR | P15311; E7E | EZRI_HUMAN E | Unmodified | 22020 | 0 | 3172400 | 0 | 1195600 | 464370 | 35157 | 25150 | 1218000 | 536500 | 372840 | 1719200 | 0 | 23298 | 269640 | 84235 | 0 | 214240 |
| IALRNDLD( | RABGAP1L; RAB | B7ZAP0; E9F | RBG10_HUMAI | Unmodified | 0 | 0 | 0 | 0 | 552250 | 0 | 162790 | 0 | 466190 | 303340 | 592180 | 569570 | 218970 | 0 | 0 | 0 | 0 | 0 |
| IALTDNALI | RPL7; RPL7 | P18124; A8 | RL7_HUMAN 6 | Unmodified | 7372600 | 7189300 | 2702400 | 2662500 | 413920 | 136250 | 622120 | 414630 | 1388800 | 325960 | 4044500 | 64477000 | 0 | 479240 | 3383200 | 3081400 | 0 | 203120 |
| IALTDNALI | RPL7; RPL7 | P18124; A8 | RL7_HUMAN 6 | Unmodified | 0 | 0 | 0 | 0 | 243780 | 0 | 2932600 | 407150 | 0 | 0 | 0 | 0 | 84957 | 172270 | 0 | 0 | 0 | 244080 |
| IALTQSAAN | HOMER2; HOM | Q9NSB8; H0 | HOME2_HUMA | Unmodified | 0 | 0 | 78587 | 82470 | 0 | 0 | 0 | 0 | 184510 | 0 | 0 | 0 | 1441700 | 2097900 | 60436 | 141080 | 0 | 0 |
| IALTQSFSK | DIP2A; DIP2C; | Q14689; Q9 | DIP2A_HUMAN | Unmodified | 129580 | 107990 | 349030 | 306180 | 0 | 0 | 0 | 0 | 0 | 0 | 0 | 723880 | 0 | 0 | 869140 | 755220 | 0 | 0 |
| IALYCHQL | CTNNA1; CTNN | P35221; P26 | CTNA1_HUMAI | Unmodified | 1219000 | 494830 | 1134300 | 5234400 | 6869800 | 1135000 | 30825 | 0 | 399010 | 259590 | 978340 | 7931800 | 8199500 | 7285800 | 5443600 | 6618800 | 0 | 0 |
| IALYETPTG | PGM1; PGM1 | P36871; A0A | PGM1_HUMAN | Unmodified | 0 | 0 | 160080 | 1069000 | 1164800 | 1040500 | 0 | 193250 | 660550 | 272460 | 0 | 1171400 | 1991700 | 1471400 | 127400 | 169250 | 1409700 | 7341600 |
| IAMENDDG | ATXN2L; ATXN2 | Q8WWM7; H | AX2L_HUMAN | Unmodified | 0 | 0 | 0 | 0 | 134320 | 1097100 | 0 | 0 | 0 | 0 | 0 | 376930 | 0 | 0 | 0 | 0 | 0 | 0 |
| IAMHSDSQ | CAPN7; CAPN7 | Q9Y6W3; H7 | CAN7_HUMAN | Unmodified | 0 | 0 | 0 | 0 | 0 | 0 | 0 | 29222 | 966530 | 127730 | 0 | 2306700 | 0 | 0 | 0 | 0 | 0 | 0 |
| IAMTCEQI | CNOT1; CNOT1 | A5YKK6; H3 | CNOT1_HUMA | Unmodified | 0 | 0 | 0 | 0 | 285770 | 117410 | 0 | 0 | 0 | 42786 | 0 | 0 | 0 | 0 | 0 | 0 | 0 | 104370 |
| IANDNSLN | GOT1 | P17174 | AATC_HUMAN | Unmodified | 324140 | 1238700 | 0 | 910730 | 887650 | 89938 | 0 | 0 | 872680 | 262620 | 1385400 | 3766200 | 1007200 | 0 | 190080 | 0 | 86559 | 158100 |
| IANDNSLN | GOT1 | P17174 | AATC_HUMAN | Unmodified | 0 | 0 | 0 | 0 | 0 | 0 | 0 | 0 | 0 | 0 | 0 | 0 | 4665900 | 0 | 0 | 0 | 0 | 0 |
| IANDNSLN | GOT1 | P17174 | AATC_HUMAN | Unmodified | 0 | 0 | 0 | 0 | 1921100 | 0 | 1350700 | 0 | 0 | 0 | 0 | 0 | 9770400 | 6431200 | 0 | 0 | 1183900 | 916680 |
| IANEQISAS | NRP2 | O60462 | NRP2_HUMAN | Unmodified | 0 | 0 | 0 | 0 | 0 | 0 | 0 | 0 | 1818900 | 0 | 0 | 0 | 0 | 0 | 0 | 0 | 0 | 0 |
| IANFKIEPP | TOP1 | P11387 | TOP1_HUMAN | Unmodified | 0 | 0 | 0 | 0 | 0 | 0 | 0 | 0 | 0 | 28934 | 0 | 1120300 | 49050 | 33558 | 0 | 0 | 99946 | 52572 |
| IANILKDKD | PDIA4; PDIA4 | P13667; A0 | PDIA4_HUMAN | Unmodified | 0 | 0 | 0 | 0 | 0 | 0 | 1261400 | 581300 | 0 | 2754800 | 0 | 0 | 0 | 0 | 0 | 0 | 0 | 0 |
| IANLAYVTK | MOV10; MOV1C | Q9HCE1; Q | MOV10_HUMA | Unmodified | 7692300 | 10024000 | 1147800 | 2585600 | 1918900 | 0 | 0 | 24640 | 7530200 | 3161700 | 0 | 998890 | 0 | 0 | 6301700 | 7223000 | 0 | 0 |
| IANLGSCN | S100A14 | Q9HCY8 | S10AE_HUMAN | Unmodified | 0 | 57890 | 0 | 0 | 0 | 0 | 0 | 38640 | 3746800 | 617440 | 0 | 0 | 0 | 0 | 0 | 0 | 417360 | 0 |
| IANLQTDLS | FLNA; FLNA; FL | P21333; A0 | FLNA_HUMAN | Unmodified | 182310 | 1813400 | 193300 | 223430 | 0 | 0 | 0 | 0 | 135620 | 0 | 0 | 2199200 | 0 | 0 | 2162600 | 399510 | 241650 | 0 |
| IANPDQLL | ABCD3; ABCD3 | P28288; A0 | ABCD3_HUMAI | Unmodified | 0 | 0 | 0 | 0 | 0 | 0 | 0 | 0 | 1937500 | 0 | 0 | 0 | 0 | 0 | 0 | 0 | 0 | 0 |
| IANPVEGSS | PCBP1 | Q15365 | PCBP1_HUMAI | Unmodified | 18021000 | 12901000 | 3889500 | 5727500 | 0 | 0 | 0 | 0 | 2383000 | 0 | 0 | 5468300 | 2277300 | 0 | 4082300 | 3594800 | 0 | 0 |
| IANPVEGSS | PCBP1 | Q15365 | PCBP1_HUMAI | Unmodified | 0 | 0 | 0 | 558020 | 0 | 0 | 1025300 | 0 | 0 | 1369300 | 0 | 0 | 10587000 | 2594500 | 0 | 0 | 0 | 0 |
| IANPVEGST | PCBP2; PCBP2; | Q15366; F8 | PCBP2_HUMAI | Unmodified | 25785000 | 23621000 | 10300000 | 12468000 | 0 | 0 | 364330 | 572120 | 1853000 | 557600 | 774970 | 10659000 | 1773800 | 344050 | 10989000 | 13682000 | 0 | 0 |
| IANPVEGST | PCBP2; PCBP2; | Q15366; F8 | PCBP2_HUMAI | Unmodified | 0 | 0 | 189750 | 0 | 0 | 0 | 0 | 0 | 0 | 651230 | 0 | 0 | 13973000 | 2827300 | 3169100 | 99424 | 0 | 0 |
| IANTHLNRI | KIF23; KIF23; K | Q02241; A0 | KIF23_HUMAN | Unmodified | 23842 | 0 | 0 | 0 | 1554000 | 594550 | 0 | 0 | 542770 | 276030 | 236690 | 55584 | 0 | 53678 | 42042 | 17048 | 238850 | 87083 |
| IANTHLNRI | KIF23; KIF23; K | Q02241; A0 | KIF23_HUMAN | Unmodified | 0 | 0 | 0 | 0 | 4894700 | 1119800 | 158680 | 0 | 0 | 315860 | 0 | 0 | 424580 | 114590 | 0 | 0 | 615600 | 0 |
| IAPAEAPDA | IGF2BP3 | O00425 | IF2B3_HUMAN | Unmodified | 1511000 | 549300 | 109770 | 392550 | 0 | 0 | 28190 | 0 | 0 | 146400 | 66433 | 2756900 | 0 | 0 | 75558 | 0 | 0 | 0 |
| IAPAVVHIE | HTRA1; HTRA1 | Q92743; A0 | HTRA1_HUMAN | Unmodified | 0 | 0 | 0 | 0 | 461420 | 0 | 0 | 0 | 2396000 | 1225400 | 0 | 0 | 0 | 0 | 0 | 0 | 352550 | 2161000 |
| IAPELYHK | KARS1 | Q15046 | SYK_HUMAN L | Unmodified | 0 | 0 | 0 | 0 | 0 | 0 | 0 | 0 | 0 | 0 | 0 | 1152100 | 0 | 0 | 0 | 0 | 0 | 0 |
| IAPNGRPF | NEDD4L; NEDD | Q96PU5; A0 | NED4L_HUMAN | Unmodified | 4535900 | 4789400 | 4529800 | 2546700 | 643700 | 0 | 0 | 0 | 0 | 0 | 0 | 699000 | 0 | 0 | 1827100 | 5817000 | 0 | 0 |
| IAPNTPQLA | NOMO2 | Q5JPE7 | NOMO2_HUMA | Unmodified | 0 | 0 | 0 | 218370 | 0 | 0 | 0 | 0 | 3466100 | 1517200 | 402990 | 107580 | 0 | 0 | 0 | 0 | 0 | 0 |
| IAPPEAPVT | PARP1; PARP1 | P09874; A0 | PARP1_HUMAN | Oxidation (M) | 223410 | 292170 | 0 | 0 | 0 | 0 | 0 | 375280 | 266090 | 0 | 191190 | 2299600 | 0 | 0 | 0 | 0 | 0 | 0 |
| IAPPPVVSS | NRP1; NRP1; N | O14786; Q5 | NRP1_HUMAN | Unmodified | 0 | 321030 | 476630 | 0 | 2621000 | 1658200 | 0 | 0 | 347330 | 0 | 0 | 0 | 0 | 0 | 611740 | 1806000 | 2387700 | 3069100 |
| IAPQYYDM | DHX15 | O43143 | DHX15_HUMAN | Unmodified | 0 | 0 | 0 | 0 | 0 | 0 | 52270 | 24427 | 0 | 80992 | 0 | 552760 | 0 | 0 | 0 | 0 | 0 | 0 |
| IAPVENDN | PSME4 | Q14997 | PSME4_HUMAI | Unmodified | 0 | 0 | 0 | 0 | 0 | 0 | 0 | 0 | 0 | 0 | 0 | 1782000 | 0 | 0 | 0 | 0 | 0 | 0 |
| IAPVHIDSE | PDS5A | Q29RF7 | PDS5A_HUMAI | Unmodified | 282820 | 204710 | 0 | 0 | 164370 | 0 | 45754 | 0 | 0 | 43731 | 0 | 861600 | 62417 | 0 | 0 | 0 | 0 | 161600 |
| IAPVHIDTE | PDS5B | Q9NTI5 | PDS5B_HUMAN | Unmodified | 0 | 0 | 154620 | 0 | 565520 | 0 | 0 | 285360 | 119850 | 0 | 173060 | 1942600 | 0 | 0 | 0 | 0 | 117430 | 0 |
| IAQDAFDD | FMNL2; FMNL2 | Q96PY5; C9 | FMNL2_HUMAI | Unmodified | 0 | 0 | 0 | 0 | 2054200 | 0 | 0 | 0 | 0 | 446000 | 0 | 0 | 108170 | 0 | 0 | 0 | 0 | 0 |
| IAQDLEMY | EZR; EZR | P15311; E7E | EZRI_HUMAN E | Oxidation (M) | 727300 | 5106900 | 144740 | 1364600 | 8296100 | 2079600 | 1298500 | 0 | 6345300 | 5034400 | 777960 | 3190800 | 2318000 | 2528300 | 1613000 | 656460 | 1903500 | 0 |
| IAQDLEMY | EZR; EZR | P15311; E7E | EZRI_HUMAN E | Unmodified | 0 | 0 | 0 | 0 | 1017400 | 0 | 0 | 0 | 750840 | 672550 | 0 | 0 | 0 | 0 | 0 | 0 | 345740 | 0 |
| IAQDLEMY | EZR; EZR | P15311; E7E | EZRI_HUMAN E | Oxidation (M) | 0 | 0 | 0 | 0 | 1522000 | 0 | 0 | 0 | 0 | 631850 | 0 | 0 | 0 | 0 | 0 | 0 | 0 | 0 |
| IAQDLEMY | RDX; RDX; RDX | P35241; A0 | RADI_HUMAN I | Oxidation (M) | 0 | 0 | 221820 | 917010 | 2298900 | 1801100 | 0 | 0 | 0 | 576710 | 48129 | 1139600 | 0 | 0 | 4818700 | 962450 | 1073500 | 0 |
| IAQDLEMY | MSN | P26038 | MOES_HUMAN | Oxidation (M) | 1524300 | 263430 | 0 | 2583500 | 35501000 | 7817700 | 6975500 | 87432 | 1326300 | 0 | 246010 | 0 | 3861900 | 2729900 | 2162300 | 1442800 | 16938000 | 4382000 |
| IAQDLEMY | MSN | P26038 | MOES_HUMAN | Unmodified | 0 | 0 | 0 | 0 | 2321700 | 0 | 0 | 0 | 0 | 0 | 0 | 0 | 152540 | 156440 | 0 | 0 | 2017600 | 0 |
| IAQDLGLEI | PCDHA6; PCDH | Q9UN73; Q9 | PCDA6_HUMAI | Unmodified | 0 | 0 | 0 | 0 | 0 | 0 | 0 | 0 | 239860 | 47325 | 0 | 0 | 0 | 0 | 0 | 0 | 0 | 0 |
| IAQFLSDIP | ACAA1; ACAA1; | P09110; H7C | THIK_HUMAN 3 | Unmodified | 0 | 0 | 0 | 0 | 0 | 0 | 0 | 0 | 27790000 | 721430 | 488250 | 0 | 0 | 0 | 0 | 0 | 0 | 0 |
| IAQGVSGS | IMPDH1; IMPDH | P20839; C9J | IMDH1_HUMAI | Unmodified | 0 | 0 | 0 | 0 | 210400 | 59822 | 58490 | 93888 | 0 | 0 | 89845 | 292540 | 0 | 0 | 56058 | 0 | 0 | 115340 |
| IAQGVSGS | IMPDH1; IMPDH | P20839; C9J | IMDH1_HUMAI | Unmodified | 0 | 0 | 0 | 0 | 176890 | 0 | 184870 | 70861 | 86905 | 0 | 0 | 0 | 298110 | 168380 | 0 | 0 | 168270 | 0 |
| IAQITGPPD | FUBP1; FUBP1 | Q96AE4; E9F | FUBP1_HUMAN | Unmodified | 0 | 0 | 0 | 0 | 0 | 0 | 0 | 0 | 4066800 | 0 | 1578400 | 4879800 | 0 | 0 | 0 | 0 | 0 | 0 |
| IAQLEEELE | MYH9 | P35579 | MYH9_HUMAN | Unmodified | 0 | 0 | 0 | 0 | 0 | 0 | 0 | 0 | 1495900 | 0 | 0 | 4722400 | 0 | 0 | 1500900 | 1002800 | 0 | 0 |
| IAQLEEELE | MYH9 | P35579 | MYH9_HUMAN | Unmodified | 0 | 0 | 0 | 0 | 0 | 2942100 | 0 | 0 | 1335500 | 0 | 0 | 2065100 | 84716 | 567320 | 0 | 0 | 5793700 | 1029100 |
| IAQLEEELE | MYH9 | P35579 | MYH9_HUMAN | Unmodified | 450670 | 0 | 897260 | 3291500 | 38756000 | 8634000 | 924570 | 0 | 0 | 1682100 | 2748400 | 3263700 | 15374000 | 589600 | 5857500 | 0 | 9991700 | 22558000 |
| IAQLEEELE | MYH10 | P35580 | MYH10_HUMAI | Oxidation (M) | 0 | 0 | 0 | 0 | 2169200 | 0 | 0 | 0 | 0 | 0 | 0 | 768230 | 1077300 | 0 | 589040 | 0 | 0 | 0 |
| IAQLEEQLD | MYH9 | P35579 | MYH9_HUMAN | Unmodified | 449480 | 499840 | 4432400 | 2455200 | 30038000 | 8605700 | 0 | 1639800 | 3197500 | 2788800 | 2767800 | 20914000 | 11795000 | 9475000 | 5316300 | 3235400 | 4960400 | 12042000 |
| IAQLEEQLD | MYH9 | P35579 | MYH9_HUMAN | Unmodified | 0 | 0 | 0 | 0 | 3550200 | 0 | 231870 | 59056 | 0 | 1394900 | 0 | 142130 | 427950 | 580860 | 0 | 0 | 1555200 | 2204200 |
| IAQLEEQLD | MYH9 | P35579 | MYH9_HUMAN | Unmodified | 0 | 0 | 0 | 0 | 26003000 | 2051900 | 9711800 | 3048700 | 0 | 1392800 | 0 | 0 | 10042000 | 14638000 | 0 | 0 | 23389000 | 15609000 |
| IAQLFSISP( | TFCP2; TFCP2; | Q12800; F8 | TFCP2_HUMAN | Unmodified | 0 | 0 | 0 | 0 | 35289 | 0 | 0 | 0 | 0 | 0 | 80928 | 294840 | 0 | 0 | 0 | 0 | 0 | 12802 |
| IAQLICER | DUT; DUT; DUT | P33316; H0 | DUT_HUMAN D | Unmodified | 0 | 0 | 48625 | 29697 | 0 | 0 | 0 | 0 | 408580 | 0 | 50316 | 4490000 | 0 | 0 | 0 | 0 | 0 | 42001 |
| IAQPGDHV | MCM7; MCM7 | P33993; C9 | MCM7_HUMAN | Unmodified | 0 | 0 | 0 | 0 | 0 | 0 | 0 | 0 | 295860 | 109130 | 192790 | 849470 | 0 | 0 | 0 | 0 | 0 | 0 |
| IAQPNYIPT | GNAI1; GNAI1 | P63096; A0 | GNAI1_HUMAN | Unmodified | 0 | 0 | 232560 | 0 | 0 | 0 | 60957 | 0 | 580410 | 0 | 3730200 | 13141000 | 0 | 63289 | 112350 | 258380 | 0 | 0 |
| IAQQLER | CHMP6; CHMP | Q96FZ7; I3L | CHMP6_HUMA | Unmodified | 360590 | 447760 | 0 | 0 | 216370 | 0 | 0 | 6283.4 | 487310 | 50627 | 64159 | 1148000 | 0 | 0 | 939590 | 965620 | 0 | 80247 |
| IAQSAELAD | MACF1; MACF1 | Q9UPN3; H3 | MACF1_HUMAI | Unmodified | 0 | 0 | 0 | 0 | 0 | 0 | 0 | 0 | 419630 | 0 | 0 | 0 | 0 | 0 | 0 | 0 | 0 | 0 |
| IAQSDYIPT | GNAI2 | P04899 | GNAI2_HUMAN | Unmodified | 4712000 | 6594900 | 1055100 | 2765800 | 0 | 836080 | 226590 | 213710 | 11059000 | 1388300 | 10755000 | 90690000 | 120440 | 57559 | 2377900 | 3201600 | 519220 | 502230 |
| IAQSDYIPT | GNAI2 | P04899 | GNAI2_HUMAN | Unmodified | 149150 | 0 | 0 | 0 | 684260 | 0 | 0 | 137020 | 190750 | 593590 | 0 | 766240 | 0 | 0 | 0 | 0 | 650510 | 1223200 |
| IAQSDYIPT | GNAI2 | P04899 | GNAI2_HUMAN | Unmodified | 0 | 0 | 0 | 0 | 1964400 | 0 | 0 | 0 | 0 | 333450 | 0 | 1054000 | 0 | 0 | 0 | 0 | 0 | 0 |
| IAQSEAELS | MYO6; MYO6; N | Q9UM54; A0 | MYO6_HUMAN | Unmodified | 0 | 0 | 628640 | 1621500 | 0 | 0 | 0 | 0 | 0 | 0 | 0 | 0 | 0 | 0 | 1519400 | 1060200 | 0 | 0 |
| IARNTQIIIQ | MMUT | P22033 | MUTA_HUMAN | Unmodified | 0 | 0 | 0 | 0 | 0 | 0 | 0 | 0 | 0 | 788460 | 0 | 0 | 0 | 0 | 0 | 0 | 0 | 0 |
| IARQESTSV | DSG2 | Q14126 | DSG2_HUMAN | Unmodified | 0 | 0 | 0 | 0 | 3439900 | 1304400 | 0 | 0 | 3557200 | 1153100 | 585910 | 604970 | 0 | 0 | 0 | 0 | 0 | 0 |
| IARQESTSV | DSG2 | Q14126 | DSG2_HUMAN | Unmodified | 0 | 0 | 0 | 0 | 640510 | 0 | 0 | 0 | 1020800 | 0 | 0 | 0 | 0 | 0 | 0 | 0 | 0 | 0 |
| IASANMDG | LRP2 | P98164 | LRP2_HUMAN | Unmodified | 0 | 0 | 0 | 0 | 0 | 0 | 0 | 0 | 529430 | 354080 | 0 | 0 | 0 | 0 | 0 | 0 | 0 | 0 |
| IASAPEPR | CC2D1A; CC2D | Q6P1N0; K7 | C2D1A_HUMAI | Unmodified | 558630 | 722510 | 464310 | 753520 | 1235900 | 503130 | 0 | 0 | 1004600 | 198460 | 171520 | 0 | 1174900 | 258980 | 95512 | 400460 | 292640 | 461330 |
| IASLEVENC | EEF1D; EEF1D; | P29692; E9P | EF1D_HUMAN | Unmodified | 0 | 83522 | 707390 | 402980 | 0 | 0 | 0 | 0 | 2204000 | 0 | 3270000 | 24722000 | 0 | 0 | 804830 | 712390 | 0 | 0 |
| IASLPQEVC | RPRD1B; RPRD | Q9NQG5; A2 | RPR1B_HUMAN | Unmodified | 0 | 0 | 30715 | 0 | 951560 | 0 | 0 | 0 | 1151200 | 2071700 | 0 | 161550 | 704220 | 0 | 147780 | 0 | 0 | 0 |
| IASPEGQD | RTCB | Q9Y3I0 | RTCB_HUMAN | Unmodified | 0 | 0 | 0 | 0 | 1878100 | 200000 | 0 | 0 | 223930 | 0 | 978760 | 5169000 | 1719500 | 1441800 | 0 | 0 | 592090 | 1113900 |
| IASQMITEG | COPS4; COPS4 | Q9BT78; D6 | CSN4_HUMAN | Oxidation (M) | 0 | 0 | 0 | 0 | 0 | 94163 | 0 | 0 | 0 | 0 | 0 | 615160 | 0 | 76000 | 0 | 0 | 0 | 0 |
| IASQMITEG | COPS4; COPS4 | Q9BT78; D6 | CSN4_HUMAN | Unmodified | 0 | 0 | 0 | 0 | 0 | 0 | 0 | 0 | 0 | 0 | 0 | 2311100 | 0 | 0 | 0 | 0 | 0 | 0 |
| IASVLGIST | FAM177A1; FAN | Q8N128; G3 | F177A_HUMAN | Unmodified | 0 | 0 | 0 | 0 | 0 | 0 | 0 | 633430 | 0 | 0 | 0 | 0 | 0 | 411600 | 0 | 2326900 | 0 | 0 |
| IASWADLVI | UMPS | P11172 | UMPS_HUMAN | Unmodified | 0 | 0 | 0 | 0 | 0 | 0 | 0 | 0 | 934380 | 587230 | 689660 | 4552100 | 0 | 0 | 0 | 0 | 0 | 0 |
| IATCLELR | AIDA | Q96BJ3 | AIDA_HUMAN | Unmodified | 0 | 0 | 0 | 0 | 0 | 0 | 0 | 48328 | 583010 | 293730 | 1145200 | 763220 | 3882600 | 2126000 | 0 | 0 | 865720 | 2680800 |
| IATCLELRS | AIDA | Q96BJ3 | AIDA_HUMAN | Unmodified | 0 | 0 | 0 | 0 | 0 | 0 | 0 | 0 | 0 | 0 | 0 | 0 | 3434100 | 3057700 | 0 | 0 | 1299800 | 2821800 |
| IATCQPLS( | PROM2 | Q8N271 | PROM2_HUMA | Unmodified | 1048500 | 445920 | 152570 | 190500 | 0 | 0 | 0 | 0 | 0 | 0 | 0 | 523970 | 0 | 0 | 547320 | 707870 | 0 | 0 |
| IATDPTIHT\ | ALG9; N/A; ALG | Q9H6U8; A0 | ALG9_HUMAN | Unmodified | 0 | 0 | 0 | 0 | 0 | 0 | 313300 | 0 | 6163200 | 0 | 0 | 0 | 0 | 0 | 0 | 0 | 0 | 0 |
| IATEKDFLE | PSMC2; PSMC2 | P35998; C9J | PRS7_HUMAN | Unmodified | 0 | 32136 | 35721 | 0 | 256630 | 0 | 261750 | 257450 | 0 | 0 | 54332 | 0 | 0 | 0 | 0 | 0 | 0 | 216230 |
| IATFEALEA | N/A | N/A | N/A | Unmodified | 169070 | 66302 | 0 | 139560 | 4583700 | 747600 | 126710 | 59279 | 1554700 | 0 | 0 | 283090 | 363940 | 185740 | 0 | 23163 | 148100 | 182480 |
| IATGHNIID | SYPL1; SYPL1; | Q16563; C9J | SYPL1_HUMAN | Unmodified | 0 | 0 | 447270 | 156150 | 0 | 0 | 0 | 0 | 945290 | 0 | 0 | 1287000 | 0 | 0 | 1048500 | 850960 | 0 | 0 |
| IATGSFLK | PAICS; PAICS; | P22234; D6F | PUR6_HUMAN | Unmodified | 62721 | 150110 | 116750 | 248750 | 5917500 | 301430 | 60181 | 689780 | 2563400 | 1882000 | 5108700 | 59228000 | 8238800 | 4786100 | 0 | 54933 | 4965600 | 4803300 |
| IATITTHTP | MAGI1; MAGI1; | Q96QZ7; H7 | MAGI1_HUMAN | Unmodified | 0 | 0 | 89418 | 189680 | 75104 | 0 | 0 | 0 | 463660 | 72611 | 251220 | 543610 | 322330 | 131500 | 215610 | 262950 | 41107 | 101630 |
| IATLGYLPT( | GNA11; GNA11 | P29992; A0A | GNA11_HUMAI | Unmodified | 61725 | 96580 | 0 | 0 | 0 | 0 | 0 | 0 | 507400 | 69225 | 86308 | 3177100 | 0 | 0 | 0 | 0 | 0 | 0 |
| IATLISSFLK | PGM3; PGM3; F | O95394; J3K | AGM1_HUMAN | Unmodified | 117050 | 112630 | 127790 | 310070 | 0 | 70416 | 0 | 0 | 1078500 | 346210 | 452980 | 2776100 | 1405500 | 451950 | 0 | 338430 | 76314 | 579580 |
| IAVAAASKP | CRABP2; CRAB | P29373; Q5 | RABP2_HUMAN | Unmodified | 163780 | 147550 | 0 | 0 | 16956000 | 0 | 12354000 | 3352900 | 6521700 | 12154000 | 0 | 366510 | 0 | 0 | 0 | 0 | 1.37E+08 | 1.17E+08 |
| IAVAAQNC' | TPI1 | P60174 | TPIS_HUMAN T | Unmodified | 3962000 | 1442400 | 3090900 | 4708900 | 24982000 | 3952000 | 2361600 | 935460 | 2935400 | 1965600 | 5330100 | 33915000 | 20033000 | 0 | 4734500 | 3465300 | 14685000 | 19871000 |
| IAVAGAQH | PACS2; PACS2 | Q86VP3; F8\ | PACS2_HUMAI | Unmodified | 0 | 0 | 0 | 0 | 0 | 0 | 0 | 0 | 0 | 0 | 0 | 611760 | 0 | 0 | 0 | 0 | 0 | 0 |
| IAVAGDLF( | GAS6 | Q14393 | GAS6_HUMAN | Unmodified | 0 | 0 | 0 | 0 | 0 | 0 | 0 | 0 | 4749100 | 19801 | 0 | 0 | 0 | 0 | 0 | 0 | 0 | 0 |
| IAVAHNGE | PPAT; PPAT | Q06203; D6I | PUR1_HUMAN | Unmodified | 0 | 0 | 0 | 0 | 530800 | 493670 | 0 | 0 | 564210 | 93100 | 1296100 | 8827400 | 1303600 | 457250 | 0 | 0 | 336130 | 277120 |
| IAVDRVNA/ | SEMA3C | Q99985 | SEM3C_HUMAI | Unmodified | 16680000 | 298000 | 1650300 | 0 | 0 | 0 | 60808 | 0 | 0 | 0 | 0 | 135300 | 0 | 0 | 16346000 | 706690 | 27186 | 0 |
| IAVEPVNPS | EFTUD2 | Q15029 | U5S1_HUMAN | Unmodified | 798130 | 0 | 0 | 0 | 0 | 288900 | 738160 | 0 | 0 | 0 | 1010000 | 2306000 | 1965100 | 0 | 0 | 0 | 684580 | 2150700 |

| | | | | | | | | | | | | | | | | | | | | | | |
|---|---|---|---|---|---|---|---|---|---|---|---|---|---|---|---|---|---|---|---|---|---|---|
| IAVHCTVR | RPL11; RPL11 | P62913; Q5 | RL11_HUMAN | Unmodified | 1583700 | 539780 | 318560 | 118230 | 0 | 0 | 0 | 0 | 0 | 0 | 591100 | 3902800 | 0 | 0 | 414610 | 505030 | 0 | 0 |
| IAVHCTVR | RPL11; RPL11 | P62913; Q5 | RL11_HUMAN | Unmodified | 0 | 0 | 0 | 0 | 4481700 | 0 | 10308000 | 839110 | 0 | 713420 | 0 | 0 | 2493500 | 5449200 | 0 | 0 | 582410 | 2390000 |
| IAVIADLDTI | CANT1; CANT1 | Q8WVQ1; K | CANT1_HUMAI | Unmodified | 0 | 0 | 0 | 0 | 0 | 0 | 0 | 0 | 2933600 | 179440 | 0 | 0 | 0 | 0 | 0 | 0 | 0 | 0 |
| IAVIADLDTI | CANT1; CANT1 | Q8WVQ1; K | CANT1_HUMAI | Unmodified | 0 | 0 | 0 | 0 | 0 | 0 | 0 | 0 | 1643200 | 1027700 | 0 | 0 | 0 | 0 | 0 | 0 | 0 | 0 |
| IAVPLIQGY | GLB1L2 | Q8IW92 | GLBL2_HUMAN | Unmodified | 0 | 0 | 0 | 0 | 0 | 0 | 0 | 0 | 751980 | 593480 | 62353 | 281520 | 0 | 0 | 0 | 0 | 0 | 205800 |
| IAVPRNPEE | SFTPA2; SFTPA | Q8IWL1; Q8I | SFPA2_HUMAN | Unmodified | 0 | 0 | 0 | 0 | 0 | 0 | 516650 | 0 | 4900100 | 2921100 | 0 | 0 | 627510 | 0 | 0 | 0 | 651720 | 690990 |
| IAVQPGTV | COL4A2 | P08572 | CO4A2_HUMAI | Unmodified | 0 | 0 | 0 | 743600 | 0 | 455190 | 0 | 0 | 224540 | 0 | 143080 | 0 | 1204800 | 0 | 0 | 0 | 0 | 236490 |
| IAVSIEDEAI | FAM120A; FAM | Q9NZB2; A0 | F120A_HUMAN | Unmodified | 0 | 0 | 0 | 0 | 0 | 1277200 | 0 | 0 | 0 | 0 | 0 | 687220 | 0 | 0 | 0 | 276470 | 0 | 0 |
| IAVSKPSGF | PLD3; PLD3 | Q8IV08; M0F | PLD3_HUMAN | Unmodified | 0 | 0 | 0 | 0 | 830550 | 0 | 0 | 0 | 3336400 | 631240 | 1630300 | 880420 | 0 | 0 | 0 | 0 | 0 | 0 |
| IAVSNIVTQ | TRAPPC8; TRAF | Q9Y2L5; J3Q | TPPC8_HUMAN | Unmodified | 0 | 0 | 0 | 0 | 612380 | 103950 | 584950 | 420320 | 0 | 203470 | 111180 | 0 | 435230 | 220950 | 57952 | 0 | 103070 | 161030 |
| IAVSNLHK | RRM1 | P23921 | RIR1_HUMAN | Unmodified | 0 | 0 | 91008 | 0 | 0 | 111970 | 192840 | 123520 | 0 | 0 | 43409 | 497130 | 75728 | 130010 | 0 | 0 | 14600 | 365420 |
| IAVVEEDGF | PPFIA2; PPFIA2 | O75334; G3 | LIPA2_HUMAN | Unmodified | 0 | 321000 | 482180 | 1182200 | 0 | 0 | 0 | 0 | 390710 | 0 | 255320 | 263400 | 190610 | 102880 | 406740 | 474230 | 202670 | 0 |
| IAVVGAGIG | PCYOX1L; PCY | Q8NBM8; D6 | PCYXL_HUMAN | Unmodified | 0 | 0 | 0 | 0 | 0 | 0 | 0 | 0 | 30314000 | 1329200 | 1115700 | 308490 | 0 | 0 | 0 | 0 | 31516 | 0 |
| IAVVLTDGK | COL21A1; COL | Q96P44; F5 | COLA1_HUMAI | Unmodified | 0 | 26919 | 18321 | 0 | 0 | 0 | 0 | 214050 | 2025500 | 355410 | 113210 | 236290 | 93115 | 0 | 59217 | 94859 | 91864 | 0 |
| IAVVLTDGK | COL21A1; COL | Q96P44; F5 | COLA1_HUMAI | Unmodified | 0 | 0 | 0 | 0 | 632060 | 4118500 | 806120 | 0 | 2239900 | 771280 | 0 | 0 | 0 | 0 | 0 | 0 | 195570 | 956460 |
| IAVYDDNN | PLCB4; PLCB4 | Q15147; A0 | PLCB4_HUMAN | Unmodified | 0 | 0 | 0 | 783340 | 0 | 0 | 0 | 0 | 0 | 0 | 52337 | 0 | 0 | 0 | 0 | 0 | 0 | 96918 |
| IAVYHLQEL | ZNF512; ZNF51 | Q96ME7; G3 | ZN512_HUMAN | Unmodified | 791980 | 0 | 0 | 0 | 0 | 0 | 0 | 0 | 0 | 0 | 0 | 0 | 0 | 0 | 0 | 0 | 0 | 0 |
| IAVYSCPFD | CCT8 | P50990 | TCPQ_HUMAN | Oxidation (M) | 0 | 0 | 85381 | 0 | 1003400 | 0 | 0 | 0 | 407040 | 231720 | 472740 | 3345000 | 629420 | 424350 | 52289 | 196240 | 231340 | 1381300 |
| IAVYSCPFD | CCT8 | P50990 | TCPQ_HUMAN | Oxidation (M) | 0 | 0 | 0 | 0 | 0 | 0 | 0 | 0 | 0 | 0 | 0 | 0 | 1456600 | 689290 | 0 | 0 | 1724100 | 0 |
| IAWALSR | GLO1 | Q04760 | LGUL_HUMAN | Unmodified | 774470 | 495810 | 242000 | 258850 | 2943700 | 498640 | 292670 | 207610 | 1966700 | 1948900 | 527500 | 6364000 | 5654400 | 2969000 | 1389400 | 1979400 | 1616100 | 2641500 |
| IAWESPQG | FN1 | P02751 | FINC_HUMAN | Unmodified | 0 | 0 | 23588000 | 13797000 | 0 | 0 | 0 | 0 | 646940 | 263970 | 914970 | 0 | 532840 | 527410 | 0 | 0 | 0 | 1071900 |
| IAWTHITIPE | TOLLIP; TOLLIP | Q9H0E2; E9 | TOLIP_HUMAN | Unmodified | 19835000 | 11552000 | 4025000 | 3317100 | 0 | 0 | 0 | 0 | 1880500 | 0 | 0 | 865220 | 0 | 0 | 17554000 | 15252000 | 0 | 0 |
| IAYEEIFVK | TFPI | P10646 | TFPI1_HUMAN | Unmodified | 0 | 0 | 1772600 | 2611200 | 254400 | 0 | 0 | 0 | 0 | 0 | 0 | 0 | 0 | 0 | 0 | 559620 | 568270 | 0 |
| IAYHSANAI | PKHD1L1 | Q86WI1 | PKHL1_HUMAN | Unmodified | 0 | 0 | 0 | 0 | 0 | 0 | 0 | 0 | 386270 | 0 | 0 | 0 | 0 | 0 | 0 | 0 | 0 | 0 |
| IAYSLLGLK | EPHA2 | P29317 | EPHA2_HUMAI | Unmodified | 0 | 0 | 0 | 0 | 309980 | 0 | 256220 | 0 | 0 | 0 | 0 | 3442100 | 0 | 0 | 0 | 7727.6 | 580720 | 0 |
| IAYSLLGLK | EPHA2 | P29317 | EPHA2_HUMAI | Unmodified | 22616 | 126100 | 0 | 89826 | 3400400 | 388400 | 0 | 0 | 0 | 0 | 0 | 0 | 327010 | 0 | 0 | 0 | 16537000 | 16585000 |
| IAYSNDGK | EDIL3 | O43854 | EDIL3_HUMAN | Unmodified | 0 | 0 | 0 | 0 | 6283200 | 3387700 | 201250 | 219650 | 4581200 | 244860 | 0 | 1039700 | 0 | 0 | 0 | 0 | 185090 | 692920 |
| IAYSNDGK | EDIL3 | O43854 | EDIL3_HUMAN | Oxidation (M) | 0 | 0 | 0 | 0 | 19006000 | 6473100 | 2871600 | 1887500 | 0 | 963310 | 0 | 0 | 0 | 0 | 0 | 0 | 1637900 | 9735400 |
| IAYSNDGK | EDIL3 | O43854 | EDIL3_HUMAN | Unmodified | 0 | 0 | 0 | 0 | 48610000 | 6768400 | 3726400 | 0 | 0 | 0 | 0 | 0 | 0 | 0 | 0 | 0 | 3161900 | 14715000 |
| IAYTFAR | LONP1; LONP1 | P36776; K7E | LONM_HUMAN | Unmodified | 0 | 0 | 0 | 0 | 0 | 0 | 0 | 0 | 3071500 | 1193900 | 0 | 139800 | 0 | 0 | 0 | 0 | 0 | 0 |
| ICANHYITP | RANBP1; RANB | P43487; F6 | RANG_HUMAN | Oxidation (M) | 0 | 221420 | 0 | 709810 | 0 | 324430 | 1131300 | 166200 | 0 | 0 | 0 | 4310600 | 0 | 0 | 189610 | 0 | 0 | 0 |
| ICCQFDFK | MAN2A2; MAN2 | P49641; Q1 | MA2A2_HUMAI | Unmodified | 0 | 0 | 0 | 0 | 0 | 0 | 0 | 0 | 1467800 | 130930 | 0 | 751700 | 0 | 0 | 0 | 0 | 0 | 0 |
| ICDDELILIK | TCP1; TCP1; TC | P17987; F5G | TCPA_HUMAN | Unmodified | 0 | 155770 | 0 | 255690 | 0 | 0 | 0 | 0 | 226620 | 0 | 637390 | 17848000 | 0 | 0 | 719720 | 0 | 0 | 0 |
| ICDFENASI | HSD17B4; HSD | P51659; A0A | DHB4_HUMAN | Unmodified | 231800 | 0 | 0 | 0 | 617590 | 0 | 0 | 0 | 16362000 | 0 | 994380 | 0 | 0 | 0 | 0 | 0 | 0 | 0 |
| ICDFGLARI | KIT | P10721 | KIT_HUMAN M | Unmodified | 0 | 0 | 0 | 0 | 0 | 0 | 118140 | 0 | 0 | 445140 | 0 | 0 | 0 | 0 | 0 | 0 | 0 | 0 |
| ICDQISDA | MAT2A | P31153 | METK2_HUMAN | Unmodified | 0 | 0 | 0 | 0 | 0 | 0 | 1718600 | 0 | 688690 | 0 | 0 | 3709600 | 0 | 0 | 0 | 0 | 0 | 537380 |
| ICDQWDAI | ACTN4; ACTN4 | O43707; F5 | ACTN4_HUMAI | Unmodified | 0 | 0 | 0 | 0 | 0 | 0 | 0 | 0 | 309180 | 0 | 0 | 2413700 | 0 | 0 | 0 | 194450 | 0 | 0 |
| ICDQWDAI | ACTN4; ACTN4 | O43707; F5 | ACTN4_HUMAI | Unmodified | 1369400 | 411580 | 563710 | 493650 | 1551300 | 811840 | 0 | 0 | 2796200 | 491530 | 696720 | 26301000 | 162580 | 0 | 948980 | 1019100 | 762920 | 0 |
| ICDQWDNI | ACTN1; ACTN1 | P12814; A0A | ACTN1_HUMAI | Unmodified | 0 | 0 | 0 | 0 | 2203000 | 1395600 | 615570 | 60549 | 0 | 0 | 0 | 5800100 | 0 | 0 | 0 | 0 | 2339900 | 0 |
| ICEEAFAR | OGDH; OGDH; | Q02218; E9F | ODO1_HUMAN | Unmodified | 0 | 0 | 0 | 0 | 0 | 0 | 0 | 0 | 1033900 | 128520 | 0 | 0 | 0 | 0 | 0 | 0 | 0 | 0 |
| ICEPGYSP1 | CTSB; CTSB; C1 | P07858; E9P | CATB_HUMAN | Unmodified | 0 | 0 | 0 | 0 | 1580800 | 987280 | 0 | 0 | 0 | 88958 | 0 | 0 | 136920 | 689520 | 0 | 0 | 0 | 0 |
| ICEVINEAV | TPP2; TPP2; TPI | P29144; A0A | TPP2_HUMAN | Unmodified | 0 | 0 | 0 | 0 | 0 | 0 | 0 | 0 | 0 | 0 | 0 | 6409900 | 0 | 0 | 0 | 0 | 0 | 0 |
| ICGDIHGQ | PPP1CB; PPP1 | P62140; E7E | PP1B_HUMAN | Unmodified | 0 | 0 | 0 | 0 | 0 | 0 | 0 | 0 | 667260 | 71538 | 232300 | 9390500 | 0 | 0 | 0 | 0 | 0 | 0 |
| ICGDIHGQ | PPP1CA; PPP1 | P62136; P36 | PP1A_HUMAN | Unmodified | 0 | 90793 | 0 | 0 | 0 | 0 | 0 | 0 | 241000 | 29259 | 0 | 2178300 | 0 | 0 | 0 | 0 | 0 | 0 |
| ICGEEQLD | PTPRB; PTPRB | P23467; F8V | PTPRB_HUMAN | Unmodified | 442250 | 174150 | 0 | 0 | 0 | 0 | 0 | 0 | 0 | 0 | 0 | 0 | 0 | 0 | 0 | 0 | 0 | 0 |
| ICGLEEDLK | ROCK2; ROCK2 | O75116; E9F | ROCK2_HUMAI | Unmodified | 0 | 0 | 0 | 0 | 162020 | 5210600 | 358890 | 107770 | 37759000 | 18913000 | 26312000 | 0 | 513060 | 0 | 0 | 0 | 0 | 217910 |
| ICGVEDAV | PSPH; PSPH | P78330; C9J | SERB_HUMAN | Unmodified | 0 | 0 | 0 | 0 | 765140 | 0 | 0 | 0 | 515730 | 297980 | 397420 | 620590 | 615790 | 302400 | 73999 | 112190 | 0 | 0 |
| ICHQIEYYF | SSB; SSB; SSB | P05455; E9P | LA_HUMAN Lu | Unmodified | 0 | 0 | 0 | 0 | 0 | 0 | 0 | 0 | 152220 | 0 | 116780 | 751710 | 0 | 0 | 0 | 0 | 0 | 0 |
| ICHQIEYYF | SSB; SSB; SSB | P05455; E9P | LA_HUMAN Lu | Unmodified | 0 | 0 | 0 | 0 | 335790 | 0 | 0 | 0 | 0 | 505890 | 0 | 1115000 | 0 | 0 | 0 | 0 | 227570 | 0 |
| ICKICEQG | N/A | N/A | N/A | Unmodified | 0 | 0 | 0 | 0 | 3368700 | 690680 | 177790 | 159450 | 0 | 0 | 0 | 10297 | 0 | 2131000 | 0 | 0 | 127570 | 1975800 |
| ICKPVLKEG | DKK1 | O94907 | DKK1_HUMAN | Unmodified | 0 | 0 | 0 | 0 | 0 | 0 | 0 | 0 | 0 | 0 | 0 | 0 | 0 | 0 | 0 | 0 | 2161200 | 0 |
| ICLDILK | UBE2E3; UBE2 | Q969T4; P61 | UB2E3_HUMAN | Unmodified | 1713600 | 579640 | 480440 | 0 | 0 | 0 | 0 | 6831.7 | 476880 | 0 | 0 | 5592600 | 0 | 0 | 1152500 | 598610 | 0 | 0 |
| ICLEDNVLI | FAM3C; FAM3C | Q92520; C9 | FAM3C_HUMAI | Unmodified | 0 | 0 | 0 | 0 | 0 | 0 | 169930 | 0 | 9139700 | 1086200 | 0 | 0 | 0 | 0 | 0 | 0 | 0 | 0 |
| ICLEDNVLI | FAM3C; FAM3C | Q92520; C9 | FAM3C_HUMAI | Oxidation (M) | 0 | 0 | 0 | 0 | 0 | 0 | 0 | 0 | 750520 | 580010 | 0 | 0 | 0 | 26782 | 0 | 0 | 0 | 0 |
| ICLEQLRAF | TRIM56; TRIM5 | Q9BRZ2; C9 | TRI56_HUMAN | Unmodified | 0 | 0 | 0 | 31722 | 633830 | 0 | 74204 | 33890 | 0 | 29003 | 0 | 840240 | 116340 | 0 | 0 | 0 | 62415 | 75687 |
| ICLLSLRTQ | PALS1 | Q8N3R9 | PALS1_HUMAN | Unmodified | 0 | 0 | 0 | 0 | 524920 | 178650 | 89859 | 0 | 0 | 0 | 0 | 0 | 186990 | 160500 | 0 | 0 | 264410 | 279820 |
| ICLQPPPTS | VARS1; VARS1; | P26640; A2A | SYVC_HUMAN | Unmodified | 0 | 0 | 0 | 0 | 0 | 0 | 0 | 0 | 0 | 0 | 231620 | 675550 | 0 | 0 | 0 | 0 | 0 | 0 |
| ICLVNDPRI | BLMH; BLMH | Q13867; K7E | BLMH_HUMAN | Unmodified | 0 | 0 | 55537 | 0 | 26753 | 0 | 0 | 0 | 607460 | 63018 | 172240 | 568840 | 0 | 0 | 122960 | 116050 | 0 | 0 |
| ICMNEDPA | ILK; ILK | Q13418; A0 | ILK_HUMAN Int | Unmodified | 271570 | 284290 | 394810 | 512320 | 531320 | 233620 | 331490 | 173660 | 68132 | 26949 | 0 | 198170 | 312340 | 711090 | 463000 | 385090 | 93588 | 512220 |
| ICNAVSPDI | MTHFD2; MTHF | P13995; A0A | MTDC_HUMAN | Unmodified | 0 | 0 | 0 | 0 | 0 | 0 | 0 | 0 | 2113700 | 255660 | 134220 | 1138000 | 0 | 0 | 0 | 0 | 0 | 0 |
| ICPNDTGG | SEMA3C | Q99985 | SEM3C_HUMAI | Unmodified | 0 | 0 | 0 | 327270 | 0 | 0 | 0 | 0 | 0 | 0 | 0 | 0 | 0 | 0 | 0 | 0 | 0 | 0 |
| ICPQGSTC | GPC4 | O75487 | GPC4_HUMAN | Unmodified | 362980 | 131210 | 55071 | 130930 | 0 | 0 | 0 | 0 | 541910 | 46117 | 0 | 0 | 282620 | 423340 | 0 | 57375 | 70411 | 293670 |
| ICPVETLVE | ECHS1 | P30084 | ECHM_HUMAN | Unmodified | 0 | 0 | 0 | 0 | 0 | 0 | 726020 | 0 | 305890 | 0 | 0 | 0 | 0 | 0 | 0 | 0 | 0 | 0 |
| ICRDLSHIG | PCNA | P12004 | PCNA_HUMAN | Unmodified | 0 | 0 | 0 | 0 | 658020 | 165590 | 121290 | 0 | 0 | 642970 | 177240 | 106030 | 2079300 | 0 | 0 | 0 | 64173 | 1358800 |
| ICREVPTTL | UTRN | P46939 | UTRN_HUMAN | Unmodified | 0 | 0 | 0 | 0 | 0 | 0 | 0 | 0 | 0 | 0 | 0 | 0 | 0 | 0 | 0 | 0 | 293020 | 0 |
| ICRNFQCV | ADAM9; ADAM | Q13443; A0 | ADAM9_HUMA | Unmodified | 0 | 0 | 0 | 0 | 4790700 | 628580 | 0 | 0 | 0 | 0 | 0 | 0 | 1638700 | 1453400 | 0 | 0 | 517560 | 0 |
| ICSPDKMI | NSF; NSF | P46459; I3L | NSF_HUMAN V | Oxidation (M) | 0 | 0 | 0 | 0 | 0 | 0 | 188630 | 203570 | 0 | 0 | 0 | 0 | 514400 | 441990 | 0 | 0 | 264020 | 203280 |
| ICSVDGIPA | RRAS | P10301 | RRAS_HUMAN | Unmodified | 78673 | 0 | 76919 | 0 | 0 | 0 | 0 | 0 | 0 | 0 | 0 | 0 | 0 | 0 | 138380 | 330140 | 0 | 0 |
| ICSWNVD | APEX1; APEX1; | P27695; A0A | APEX1_HUMAN | Unmodified | 0 | 0 | 0 | 0 | 0 | 0 | 0 | 0 | 618400 | 0 | 1226900 | 6469600 | 0 | 0 | 96168 | 70669 | 0 | 0 |
| ICSWNVD | APEX1; APEX1; | P27695; A0A | APEX1_HUMAN | Unmodified | 0 | 0 | 0 | 0 | 2264900 | 396230 | 1413800 | 0 | 0 | 1572000 | 0 | 0 | 0 | 819890 | 0 | 0 | 721930 | 606110 |
| ICTVSENFL | GPM6A; GPM6 | P51674; D6F | GPM6A_HUMA | Unmodified | 6902600 | 2043000 | 0 | 0 | 0 | 0 | 0 | 0 | 348390 | 0 | 0 | 371860 | 0 | 0 | 0 | 274050 | 0 | 0 |
| ICVNGDDA | GPX4; GPX4; GI | P36969; A0A | GPX4_HUMAN | Unmodified | 1642300 | 936470 | 484730 | 752390 | 173460 | 0 | 612900 | 234700 | 1552600 | 694890 | 54461 | 1678100 | 1700600 | 2657400 | 871000 | 1078300 | 1043800 | 0 |
| ICYIFHETF | DNM1L | O00429 | DNM1L_HUMA | Unmodified | 0 | 0 | 0 | 0 | 0 | 0 | 0 | 0 | 0 | 0 | 0 | 1388500 | 0 | 0 | 0 | 0 | 0 | 0 |
| IDAASPLEK | ADH1B; ADH1C | P00325; P00 | ADH1B_HUMAI | Unmodified | 0 | 160110 | 65481 | 190810 | 0 | 0 | 0 | 0 | 312210 | 62443 | 1376700 | 783870 | 0 | 0 | 0 | 164160 | 158440 | 0 |
| IDAFHYIQL | NT5DC3 | Q86UY8 | NT5D3_HUMAI | Unmodified | 0 | 0 | 0 | 0 | 0 | 26016 | 0 | 104980 | 890520 | 0 | 0 | 0 | 0 | 0 | 0 | 0 | 0 | 0 |
| IDAFHYVQI | NT5DC2; NT5D | Q9H857; C9 | NT5D2_HUMAI | Unmodified | 0 | 0 | 0 | 0 | 0 | 0 | 0 | 0 | 879320 | 92928 | 0 | 0 | 0 | 0 | 0 | 0 | 0 | 0 |
| IDASKNEEI | HNRNPD; HNR | Q14103; D6I | HNRPD_HUMA | Unmodified | 0 | 0 | 0 | 0 | 0 | 0 | 0 | 0 | 189330 | 21726 | 1924600 | 6376800 | 0 | 0 | 0 | 0 | 0 | 0 |
| IDATSASVL | PDIA4; PDIA4 | P13667; A0A | PDIA4_HUMAN | Unmodified | 0 | 0 | 0 | 0 | 0 | 0 | 0 | 0 | 12282000 | 0 | 1269000 | 1952400 | 0 | 0 | 0 | 0 | 0 | 0 |
| IDATSASVL | PDIA4; PDIA4 | P13667; A0A | PDIA4_HUMAN | Unmodified | 184400 | 0 | 0 | 0 | 642520 | 0 | 1913200 | 207540 | 68475000 | 12105000 | 1890800 | 1393700 | 675280 | 0 | 0 | 239340 | 0 | 0 |
| IDAVNAETI | PMPCB; PMPCE | O75439; G3 | MPPB_HUMAN | Unmodified | 0 | 0 | 0 | 0 | 0 | 0 | 0 | 0 | 1110800 | 194170 | 0 | 73404 | 0 | 0 | 0 | 0 | 0 | 0 |
| IDCDNLEQ | DDX1; DDX1; D | Q92499; A0 | DDX1_HUMAN | Unmodified | 0 | 0 | 0 | 0 | 0 | 0 | 0 | 0 | 0 | 0 | 0 | 2268600 | 0 | 0 | 0 | 0 | 0 | 0 |
| IDCPTLAR | PDE2A; PDE2A; | O00408; C9 | PDE2A_HUMAI | Unmodified | 79004 | 51224 | 0 | 0 | 0 | 1011000 | 56995 | 0 | 0 | 0 | 0 | 2241900 | 0 | 0 | 0 | 0 | 0 | 0 |
| IDCVSKNEI | ATP1B3 | P54709 | AT1B3_HUMAN | Unmodified | 0 | 0 | 0 | 0 | 0 | 0 | 0 | 0 | 2867600 | 0 | 0 | 3953800 | 0 | 0 | 722360 | 549310 | 0 | 0 |
| IDCVSKNEI | ATP1B3 | P54709 | AT1B3_HUMAN | Oxidation (M) | 0 | 0 | 0 | 0 | 1215200 | 0 | 0 | 0 | 557490 | 0 | 0 | 19237000 | 0 | 0 | 0 | 0 | 0 | 0 |
| IDDIRKPGE | CCT5; CCT5; C | P48643; B7Z | TCPE_HUMAN | Unmodified | 0 | 0 | 0 | 0 | 0 | 0 | 0 | 0 | 0 | 0 | 708290 | 2826700 | 0 | 320540 | 0 | 0 | 0 | 0 |
| IDDIVSGHI | CCT3; CCT3 | P49368; B4D | TCPG_HUMAN | Unmodified | 29794 | 0 | 56332 | 0 | 792000 | 4783400 | 0 | 0 | 177270 | 60873 | 390700 | 2072100 | 0 | 0 | 71155 | 0 | 29716 | 45692 |
| IDDIVSGHI | CCT3; CCT3 | P49368; B4D | TCPG_HUMAN | Unmodified | 146000 | 0 | 53454 | 0 | 0 | 61237 | 0 | 0 | 0 | 0 | 305910 | 1074000 | 0 | 0 | 144430 | 181230 | 0 | 0 |
| IDDLFSGK | CD81; CD81; C | P60033; A6 | CD81_HUMAN | Unmodified | 1823100 | 365300 | 0 | 60469 | 0 | 0 | 0 | 0 | 41743 | 0 | 36248 | 1385200 | 72267 | 0 | 0 | 35183 | 0 | 0 |
| IDDLIGDLV | ENPP4 | Q9Y6X5 | ENPP4_HUMAI | Unmodified | 0 | 0 | 0 | 0 | 0 | 0 | 0 | 0 | 0 | 425520 | 81906 | 0 | 0 | 0 | 0 | 0 | 0 | 0 |
| IDDLQMVL | ATP6V0A1; ATP | Q93050; B7 | VPP1_HUMAN | Unmodified | 0 | 0 | 0 | 0 | 0 | 0 | 0 | 0 | 902310 | 86218 | 0 | 0 | 0 | 0 | 0 | 0 | 0 | 0 |
| IDDLQMVL | ATP6V0A1; ATP | Q93050; B7 | VPP1_HUMAN | Oxidation (M) | 0 | 0 | 0 | 0 | 0 | 0 | 0 | 0 | 747410 | 0 | 0 | 778060 | 0 | 0 | 0 | 0 | 0 | 0 |
| IDDPTDSKI | CALR; CALR; C | P27797; A0A | CALR_HUMAN | Unmodified | 0 | 0 | 0 | 0 | 0 | 0 | 1655300 | 2164900 | 59500000 | 8877600 | 1835700 | 0 | 0 | 0 | 0 | 0 | 0 | 0 |
| IDDVVNTR | CCT4 | P50991 | TCPD_HUMAN | Unmodified | 83490 | 48185 | 0 | 723330 | 0 | 19184 | 0 | 0 | 898150 | 1079600 | 3078400 | 5263600 | 2861300 | 0 | 0 | 0 | 0 | 0 |
| IDEARQELI | ARF6 | P62330 | ARF6_HUMAN | Unmodified | 0 | 174660 | 144910 | 0 | 0 | 601020 | 0 | 0 | 0 | 0 | 0 | 0 | 0 | 0 | 0 | 647420 | 0 | 112610 |
| IDEIKNDN | FAS | P25445 | TNR6_HUMAN | Unmodified | 433680 | 244800 | 481470 | 180500 | 192120 | 72542 | 0 | 0 | 265430 | 35461 | 0 | 888800 | 159300 | 58179 | 775570 | 646800 | 0 | 0 |
| IDELEPR | GANAB; GANAE | Q14697; F5 | GANAB_HUMA | Unmodified | 8709100 | 0 | 171300 | 3338400 | 0 | 0 | 0 | 165300 | 24529000 | 2557200 | 3453100 | 780690 | 0 | 0 | 70730 | 108280 | 66038 | 0 |
| IDELERQVL | NPTX1 | Q15818 | NPTX1_HUMAN | Unmodified | 0 | 0 | 0 | 0 | 0 | 0 | 14273 | 0 | 20927000 | 195840 | 0 | 0 | 0 | 0 | 0 | 0 | 0 | 0 |
| IDEMPEAA | HDGF | P51858 | HDGF_HUMAN | Unmodified | 0 | 0 | 404660 | 0 | 253900 | 0 | 104040 | 0 | 286260 | 0 | 1751700 | 313060 | 221430 | 0 | 866950 | 889760 | 0 | 181300 |
| IDEMPEAA | HDGF | P51858 | HDGF_HUMAN | Oxidation (M) | 0 | 0 | 0 | 556170 | 1310000 | 153840 | 0 | 0 | 0 | 108490 | 277070 | 2203500 | 0 | 148560 | 0 | 0 | 312290 | 527830 |
| IDESIHLQL | EIF3L; EIF3L; EI | Q9Y262; B0 | EIF3L_HUMAN | Unmodified | 37414 | 0 | 0 | 0 | 0 | 0 | 0 | 0 | 0 | 0 | 182970 | 718290 | 0 | 0 | 0 | 0 | 0 | 0 |
| IDESSLTGE | ATP2B4; ATP2B | P23634; P20 | AT2B4_HUMAN | Unmodified | 184620 | 0 | 2363000 | 1050000 | 870360 | 0 | 984870 | 108870 | 1250600 | 426560 | 3051900 | 17332000 | 471120 | 0 | 4265700 | 1502300 | 31236 | 0 |
| IDEYDYSKI | SDHA; SDHA; S | P31040; D6F | SDHA_HUMAN | Unmodified | 0 | 0 | 0 | 0 | 0 | 0 | 0 | 0 | 945430 | 0 | 139670 | 0 | 0 | 0 | 0 | 0 | 0 | 0 |
| IDFEKEPLG | ACO1 | P21399 | ACOC_HUMAN | Unmodified | 0 | 0 | 0 | 0 | 0 | 0 | 0 | 0 | 0 | 0 | 0 | 4371800 | 0 | 0 | 0 | 0 | 0 | 0 |
| IDFGEAAR | GNAI3 | P08754 | GNAI3_HUMAN | Unmodified | 112710 | 865720 | 982550 | 0 | 262690 | 0 | 31576 | 0 | 204540 | 86982 | 880870 | 13774000 | 0 | 0 | 1049600 | 1150400 | 57690 | 0 |
| IDFVGELNI | NPEPPSL1; NPI | A6NEC2; P5 | PSAL_HUMAN | Unmodified | 0 | 0 | 0 | 0 | 0 | 0 | 0 | 0 | 0 | 176100 | 984940 | 4283000 | 0 | 0 | 746080 | 551600 | 122950 | 370470 |
| IDGALHREI | TRMT10C; TRM | Q7L0Y3; C9 | TM10C_HUMAI | Unmodified | 0 | 0 | 0 | 0 | 0 | 0 | 22172 | 0 | 0 | 132720 | 0 | 122030 | 74655 | 0 | 0 | 0 | 0 | 22444 |
| IDGDKDGF | CALU | O43852 | CALU_HUMAN | Unmodified | 0 | 0 | 0 | 268010 | 0 | 0 | 122000 | 0 | 18628000 | 1030700 | 5545400 | 1529600 | 0 | 0 | 388890 | 348040 | 0 | 0 |
| IDGNLVVRI | CYB5R3; CYB5I | P00387; B1A | NB5R3_HUMAI | Unmodified | 0 | 0 | 278450 | 0 | 0 | 0 | 0 | 0 | 0 | 0 | 0 | 0 | 0 | 0 | 0 | 0 | 0 | 0 |
| IDGSANLK | ATP1B3; ATP1B | P54709; C9J | AT1B3_HUMAN | Unmodified | 147640 | 1049900 | 2705900 | 1764600 | 1753900 | 261710 | 124770 | 0 | 1902600 | 147530 | 339510 | 15777000 | 1276900 | 849820 | 4289400 | 1850300 | 187640 | 934530 |

| | | | | | | | | | | | | | | | | | | | | | | |
|---|---|---|---|---|---|---|---|---|---|---|---|---|---|---|---|---|---|---|---|---|---|---|
| IDHILDAL | RPN1; RPN1 | P04843; B7Z | RPN1_HUMAN | Unmodified | 0 | 0 | 0 | 0 | 0 | 0 | 0 | 0 | 0 | 0 | 0 | 880540 | 0 | 0 | 0 | 0 | 0 | 0 |
| IDHLSFGEI | ERGIC2; ERGIC | Q96RQ1; A0. | ERGI2_HUMAN | Unmodified | 0 | 0 | 0 | 0 | 0 | 0 | 0 | 0 | 681950 | 177650 | 0 | 0 | 0 | 0 | 0 | 0 | 0 | 0 |
| IDHQGKLP | HTRA1; HTRA1 | Q92743; A0/ | HTRA1_HUMAN | Unmodified | 0 | 0 | 0 | 0 | 670530 | 0 | 0 | 0 | 1179000 | 127640 | 0 | 0 | 0 | 0 | 0 | 0 | 827550 | 101690 |
| IDIIPNPQEI | HSP90AB1; HSI | P08238; Q5‹ | HS90B_HUMAN | Unmodified | 2043300 | 2247800 | 9238100 | 3438300 | 0 | 0 | 0 | 0 | 10837000 | 712360 | 13138000 | 63838000 | 828460 | 0 | 6700800 | 10482000 | 0 | 0 |
| IDIIPNTQEI | HSP90AB2P | Q58FF8 | H90B2_HUMAN | Oxidation (M) | 7719800 | 2456100 | 0 | 0 | 0 | 0 | 0 | 0 | 0 | 0 | 0 | 0 | 0 | 0 | 0 | 0 | 0 | 0 |
| IDINKEIEQI | DSG2 | Q14126 | DSG2_HUMAN | Unmodified | 575100 | 234160 | 0 | 757830 | 2004800 | 531300 | 0 | 0 | 1132200 | 0 | 890540 | 845880 | 953090 | 287190 | 300500 | 346250 | 79213 | 193600 |
| IDIPSFDWI | ABAT; ABAT; AB | P80404; H3E | GABT_HUMAN | Unmodified | 0 | 0 | 0 | 0 | 0 | 0 | 0 | 0 | 484270 | 113300 | 0 | 0 | 0 | 0 | 0 | 0 | 0 | 0 |
| IDISPVLLQI | MRE11; MRE11; | P49959; F5G | MRE11_HUMAN | Unmodified | 0 | 0 | 0 | 0 | 556400 | 499080 | 24697 | 0 | 0 | 0 | 26561 | 1346900 | 0 | 0 | 0 | 0 | 193350 | 0 |
| IDIVENR | ASS1 | P00966 | ASSY_HUMAN . | Unmodified | 313660 | 0 | 435530 | 558890 | 0 | 0 | 163880 | 0 | 1158400 | 155640 | 0 | 0 | 0 | 0 | 828720 | 2028200 | 0 | 0 |
| IDKLMIEMI | ENO1; ENO1; E | P06733; A0A | ENOA_HUMAN 2 | Oxidation (M) | 0 | 0 | 0 | 0 | 0 | 58331 | 0 | 0 | 0 | 246340 | 0 | 0 | 591540 | 246240 | 0 | 0 | 440560 | 1230800 |
| IDKLMIEMI | ENO1; ENO1; E | P06733; A0A | ENOA_HUMAN | Oxidation (M) | 0 | 0 | 0 | 0 | 9812700 | 1731300 | 444430 | 492130 | 147980 | 2291100 | 0 | 0 | 3257000 | 1777200 | 0 | 0 | 2304200 | 4778200 |
| IDKLMIEMI | ENO1; ENO1; E | P06733; A0A | ENOA_HUMAN | Unmodified | 0 | 0 | 0 | 0 | 20907000 | 1302300 | 641730 | 80397 | 462090 | 2156600 | 538660 | 0 | 4110800 | 2068900 | 0 | 0 | 3530000 | 9398900 |
| IDKLMIEMI | ENO1; ENO1; E | P06733; A0A | ENOA_HUMAN | Unmodified | 0 | 0 | 0 | 646810 | 389380 | 0 | 0 | 0 | 0 | 0 | 0 | 0 | 0 | 0 | 0 | 0 | 434610 | 251190 |
| IDKPILK | RPL8; RPL8; RP | P62917; E9P | RL8_HUMAN 6( | Unmodified | 0 | 597880 | 253820 | 0 | 0 | 0 | 0 | 0 | 66931 | 0 | 1439400 | 2609400 | 0 | 0 | 410620 | 343490 | 0 | 0 |
| IDKTDYMV( | ARHGDIA; ARH( | P52565; J3K | GDIR1_HUMAN | Unmodified | 354380 | 424360 | 741570 | 238300 | 0 | 0 | 0 | 0 | 361120 | 0 | 174580 | 11454000 | 0 | 0 | 1223600 | 673610 | 0 | 0 |
| IDKTDYMV( | ARHGDIA; ARH( | P52565; J3K | GDIR1_HUMAN | Oxidation (M) | 326820 | 461660 | 0 | 0 | 0 | 0 | 0 | 0 | 0 | 0 | 152120 | 5292400 | 0 | 0 | 1671500 | 1271300 | 0 | 0 |
| IDKTDYMV( | ARHGDIA; ARH( | P52565; J3K | GDIR1_HUMAN | Oxidation (M) | 0 | 0 | 0 | 0 | 0 | 0 | 0 | 0 | 0 | 0 | 0 | 6570500 | 0 | 0 | 2157000 | 0 | 0 | 0 |
| IDKWDGSA | SPCS2; SPCS2; | Q15005; E9F | SPCS2_HUMAN | Unmodified | 0 | 0 | 0 | 0 | 508370 | 0 | 142410 | 182360 | 459150 | 114400 | 53437 | 0 | 0 | 0 | 0 | 0 | 80321 | 185640 |
| IDLAVLLGK | UBAP2L; UBAP2 | Q14157; F8V | UBP2L_HUMAN | Unmodified | 20102 | 20326 | 23415 | 12623 | 0 | 0 | 0 | 12977 | 81188 | 0 | 87143 | 888170 | 0 | 0 | 23325 | 69254 | 0 | 0 |
| IDLENRQV/ | RAB7A; RAB7A; | P51149; A0A | RAB7A_HUMAN | Unmodified | 0 | 0 | 0 | 0 | 557240 | 0 | 0 | 0 | 0 | 195240 | 260820 | 0 | 0 | 410970 | 0 | 0 | 12198 | 374670 |
| IDLENTLEQ | CCDC6 | Q16204 | CCDC6_HUMA | Unmodified | 0 | 0 | 0 | 0 | 0 | 0 | 0 | 0 | 0 | 0 | 0 | 705070 | 0 | 0 | 0 | 0 | 0 | 0 |
| IDLFEREEV | PCYOX1; PCYO | Q9UHG3; F8 | PCYOX_HUMAI | Unmodified | 0 | 0 | 0 | 0 | 0 | 0 | 0 | 0 | 26225000 | 851130 | 303370 | 0 | 0 | 0 | 0 | 0 | 0 | 0 |
| IDLLDDSCI | RAB23 | Q9ULC3 | RAB23_HUMAN | Unmodified | 0 | 0 | 0 | 217630 | 0 | 0 | 0 | 0 | 0 | 0 | 0 | 0 | 0 | 0 | 269340 | 414900 | 0 | 0 |
| IDLMPHDLI | TUBGCP2; TUB | Q9BSJ2; F2Z | GCP2_HUMAN | Oxidation (M) | 0 | 0 | 0 | 0 | 0 | 0 | 0 | 0 | 0 | 0 | 0 | 196890 | 0 | 0 | 0 | 0 | 0 | 0 |
| IDLNYLR | NIT1 | Q86X76 | NIT1_HUMAN [ | Unmodified | 0 | 0 | 0 | 0 | 0 | 0 | 0 | 0 | 0 | 0 | 0 | 1361500 | 0 | 0 | 0 | 0 | 0 | 0 |
| IDLPEVQIEI | ZNF638 | Q14966 | ZN638_HUMAN | Unmodified | 0 | 0 | 0 | 0 | 0 | 0 | 0 | 0 | 0 | 0 | 0 | 504330 | 1825400 | 0 | 0 | 0 | 0 | 0 |
| IDLPEYQGI | ITPA | Q9BY32 | ITPA_HUMAN I | Unmodified | 0 | 0 | 0 | 0 | 0 | 0 | 0 | 0 | 0 | 0 | 0 | 3156300 | 0 | 0 | 0 | 0 | 87350 | 0 |
| IDLTLLNR | PRPF8; PRPF8 | Q6P2Q9; I3L | PRP8_HUMAN | Unmodified | 0 | 0 | 0 | 0 | 0 | 0 | 0 | 0 | 0 | 0 | 49954 | 3157100 | 0 | 0 | 0 | 0 | 0 | 0 |
| IDMSSNNG | EPRS1; EPRS1 | P07814; V9G | SYEP_HUMAN | Oxidation (M) | 0 | 0 | 31289 | 0 | 0 | 0 | 0 | 0 | 0 | 0 | 0 | 1027900 | 0 | 0 | 0 | 31598 | 0 | 0 |
| IDMSSNNG | EPRS1; EPRS1 | P07814; V9G | SYEP_HUMAN | Unmodified | 0 | 0 | 0 | 0 | 0 | 0 | 0 | 0 | 0 | 0 | 0 | 814810 | 130950 | 0 | 0 | 0 | 0 | 0 |
| IDNDGDGI | RCN1 | Q15293 | RCN1_HUMAN | Unmodified | 0 | 0 | 0 | 0 | 0 | 0 | 0 | 0 | 0 | 141870 | 664880 | 0 | 0 | 0 | 0 | 0 | 0 | 0 |
| IDNLDVNR | PSIP1 | O75475 | PSIP1_HUMAN | Unmodified | 44991 | 0 | 135660 | 226730 | 0 | 0 | 0 | 0 | 0 | 0 | 278350 | 4113300 | 0 | 0 | 325420 | 225910 | 0 | 0 |
| IDNQYATV/ | N/A | N/A | N/A | Unmodified | 1.99E+09 | 1.54E+09 | 1.33E+09 | 1.46E+09 | 2.78E+08 | 4.07E+08 | 73073000 | 15733000 | 7.07E+08 | 2.17E+08 | 2.74E+08 | 5.02E+08 | 6.98E+08 | 2.09E+08 | 1.94E+09 | 1.64E+09 | 94808000 | 1.66E+08 |
| IDNSQVES( | CALR; CALR; C/ | P27797; A0A | CALR_HUMAN | Unmodified | 0 | 0 | 0 | 0 | 0 | 0 | 0 | 0 | 67146000 | 8274300 | 0 | 0 | 0 | 0 | 0 | 0 | 0 | 0 |
| IDNSQVES( | CALR; CALR; C/ | P27797; A0A | CALR_HUMAN | Unmodified | 0 | 0 | 1973900 | 0 | 0 | 0 | 0 | 0 | 3.09E+08 | 31823000 | 27647000 | 4215700 | 0 | 0 | 0 | 2754000 | 0 | 0 |
| IDPLAPLDK | ADH5; ADH5 | P11766; H0Y | ADHX_HUMAN | Unmodified | 0 | 0 | 0 | 0 | 0 | 162710 | 0 | 81154 | 0 | 234640 | 3986000 | 23335000 | 0 | 0 | 1090700 | 65821 | 0 | 1048700 |
| IDPSSLSFN | SCARB1; SCARI | Q8WTV0; A0 | SCRB1_HUMAI | Oxidation (M) | 408580 | 229930 | 436990 | 0 | 0 | 0 | 0 | 0 | 0 | 0 | 0 | 3819100 | 0 | 0 | 997210 | 313930 | 0 | 0 |
| IDPTLCR | EIF2S3; EIF2S3I | P41091; Q2\ | IF2G_HUMAN E | Unmodified | 0 | 0 | 0 | 0 | 85277 | 267250 | 0 | 0 | 0 | 0 | 0 | 1160300 | 0 | 91561 | 0 | 0 | 0 | 78445 |
| IDPVNGQII | CDH2; CDH2; ( | P19022; C9J | CADH2_HUMA | Unmodified | 0 | 0 | 0 | 0 | 0 | 0 | 0 | 0 | 3634000 | 485490 | 2486400 | 2142100 | 0 | 0 | 0 | 0 | 0 | 2106400 |
| IDPYGFERI | GRTP1; GRTP1 | Q5TC63; A0/ | GRTP1_HUMAN | Unmodified | 587770 | 537690 | 1111600 | 6332400 | 0 | 0 | 0 | 0 | 0 | 63437 | 492720 | 0 | 0 | 0 | 2555900 | 2463400 | 0 | 0 |
| IDPYVFDRI | PLA2G4A | P47712 | PA24A_HUMAN | Unmodified | 0 | 0 | 0 | 0 | 0 | 0 | 0 | 0 | 0 | 0 | 1557300 | 6784300 | 0 | 0 | 0 | 41355 | 0 | 0 |
| IDQLEGDH | ACTN1; ACTN1; | P12814; H0Y | ACTN1_HUMAI | Unmodified | 4439800 | 0 | 0 | 0 | 0 | 0 | 0 | 0 | 0 | 0 | 534270 | 9231500 | 0 | 0 | 0 | 1226600 | 0 | 4717600 |
| IDQLEGDH | ACTN1; ACTN1; | P12814; H0Y | ACTN1_HUMAI | Oxidation (M) | 0 | 0 | 0 | 0 | 0 | 0 | 0 | 0 | 1437600 | 0 | 116150 | 6408900 | 0 | 0 | 0 | 0 | 351520 | 0 |
| IDQLIRPIN/ | PREX1 | Q8TCU6 | PREX1_HUMAN | Unmodified | 0 | 0 | 0 | 0 | 0 | 0 | 77309 | 0 | 0 | 0 | 0 | 319160 | 0 | 0 | 0 | 0 | 395900 | 0 |
| IDQLNREL/ | IMMT; IMMT; IM | Q16891; B9/ | MIC60_HUMAN | Unmodified | 0 | 0 | 0 | 0 | 0 | 0 | 60342 | 0 | 843270 | 164070 | 54948 | 76604 | 60159 | 0 | 0 | 0 | 0 | 0 |
| IDQLQEELI | OPA1; OPA1; O | O60313; E5K | OPA1_HUMAN | Unmodified | 0 | 0 | 0 | 0 | 0 | 0 | 0 | 0 | 0 | 59929 | 767890 | 870130 | 0 | 0 | 0 | 0 | 0 | 0 |
| IDQLQEELI | OPA1 | A0A2R8YDM | A0A2R8YDM2_ | Unmodified | 0 | 0 | 0 | 0 | 0 | 0 | 0 | 0 | 5650000 | 0 | 0 | 0 | 0 | 0 | 0 | 0 | 0 | 0 |
| IDQVNQLLI | COPS2; COPS2 | P61201; B4D | CSN2_HUMAN | Unmodified | 0 | 0 | 0 | 0 | 0 | 0 | 0 | 0 | 0 | 0 | 0 | 7559600 | 0 | 0 | 712230 | 335290 | 0 | 0 |
| IDRFGEDA( | YOD1 | Q5VVQ6 | OTU1_HUMAN | Unmodified | 651370 | 1009400 | 306670 | 0 | 201460 | 0 | 44012 | 7995.6 | 0 | 0 | 0 | 0 | 0 | 0 | 417250 | 409210 | 0 | 138440 |
| IDRPEAISEI | SAR1B; SAR1B; | Q9Y6B6; D6I | SAR1B_HUMAN | Unmodified | 0 | 0 | 0 | 0 | 75402 | 0 | 0 | 77803 | 0 | 0 | 172530 | 0 | 237810 | 0 | 0 | 0 | 0 | 0 |
| IDRQQFEE | GOLGA7 | Q7Z5G4 | GOGA7_HUMA | Unmodified | 425210 | 0 | 419360 | 0 | 0 | 0 | 0 | 0 | 0 | 206280 | 0 | 6866200 | 0 | 0 | 539090 | 681640 | 0 | 0 |
| IDRYTQQG | MTHFD1L; MTH | Q6UB35; A0. | C1TM_HUMAN | Unmodified | 0 | 0 | 119300 | 494770 | 109930 | 0 | 0 | 0 | 526130 | 0 | 139460 | 0 | 881140 | 575460 | 4377500 | 51234 | 0 | 0 |
| IDSDISGTL | CNPY2; N/A; CI | Q9Y2B0; F8\ | CNPY2_HUMAI | Unmodified | 456880 | 0 | 0 | 0 | 0 | 93636 | 0 | 0 | 4544800 | 0 | 1976600 | 489510 | 0 | 0 | 0 | 89014 | 0 | 0 |
| IDSGSEVIV | MMUT | P22033 | MUTA_HUMAN | Unmodified | 0 | 0 | 0 | 0 | 0 | 0 | 0 | 56540 | 1955800 | 0 | 0 | 0 | 0 | 0 | 0 | 0 | 0 | 0 |
| IDSIADHVN | STX17 | P56962 | STX17_HUMAN | Unmodified | 0 | 0 | 0 | 0 | 0 | 0 | 0 | 92547 | 0 | 0 | 0 | 1432500 | 0 | 0 | 0 | 0 | 0 | 0 |
| IDSILEVVQ | PALS2; PALS2 | Q9NZW5; B8 | PALS2_HUMAN | Unmodified | 0 | 0 | 713760 | 217780 | 0 | 0 | 0 | 0 | 628900 | 0 | 3641700 | 1588600 | 485440 | 0 | 816150 | 404660 | 0 | 113050 |
| IDSNLDELS | SNAP29; SNAP2 | O95721; C9J | SNP29_HUMAN | Oxidation (M) | 42803 | 0 | 53990 | 0 | 0 | 0 | 0 | 0 | 129670 | 0 | 34986 | 81775 | 0 | 0 | 93327 | 94927 | 0 | 0 |
| IDSPGFKP/ | TJP1; TJP1; TJP1 | Q07157; A0/ | ZO1_HUMAN T | Unmodified | 0 | 0 | 201320 | 158980 | 0 | 0 | 0 | 0 | 0 | 0 | 0 | 0 | 0 | 0 | 226440 | 310950 | 0 | 0 |
| IDSQTNSSI | PTPRK; PTPRK; | Q15262; Q5 | PTPRK_HUMAN | Unmodified | 687480 | 666150 | 0 | 0 | 0 | 0 | 0 | 0 | 1080900 | 382320 | 0 | 2693700 | 0 | 0 | 0 | 0 | 0 | 0 |
| IDSRLEQYT | CALD1; CALD1; | Q05682; C9J | CALD1_HUMAI | Unmodified | 130160 | 194140 | 1147800 | 571920 | 0 | 0 | 0 | 0 | 0 | 0 | 0 | 0 | 1049800 | 963100 | 381080 | 175480 | 228470 | 756120 |
| IDTGWLDR | ACACA | Q13085 | ACACA_HUMAI | Unmodified | 0 | 11838 | 0 | 28101 | 0 | 0 | 28734 | 0 | 39516 | 0 | 51429 | 453820 | 0 | 0 | 173700 | 53641 | 0 | 0 |
| IDTIAADESI | EPHA10; EPHA | Q5JZY3; Q15 | EPHAA_HUMAI | Unmodified | 0 | 0 | 0 | 0 | 0 | 0 | 0 | 0 | 159370 | 0 | 233500 | 0 | 0 | 0 | 0 | 0 | 40309 | 0 |
| IDTIAADESI | EPHA10; EPHA | Q5JZY3; Q15 | EPHAA_HUMAI | Unmodified | 316080 | 0 | 0 | 0 | 0 | 0 | 936200 | 0 | 1436200 | 708610 | 521740 | 0 | 0 | 1106000 | 0 | 0 | 0 | 0 |
| IDTIAADESI | EPHA3; EPHA6; | P29320; Q9L | EPHA3_HUMAI | Oxidation (M) | 617710 | 914130 | 0 | 0 | 0 | 0 | 0 | 0 | 0 | 0 | 0 | 0 | 0 | 0 | 0 | 0 | 0 | 0 |
| IDTIAADESI | EPHA3; EPHA6; | P29320; Q9L | EPHA3_HUMAI | Unmodified | 0 | 153040 | 0 | 0 | 0 | 0 | 181910 | 0 | 0 | 71788 | 331500 | 0 | 0 | 0 | 0 | 35059 | 0 | 0 |
| IDTIAPDEIT | EPHA2 | P29317 | EPHA2_HUMAI | Unmodified | 0 | 0 | 0 | 0 | 774950 | 0 | 0 | 0 | 0 | 173760 | 0 | 0 | 0 | 0 | 0 | 0 | 433410 | 1817700 |
| IDTIEIITDR | HNRNPA2B1; H | P22626; A0A | ROA2_HUMAN | Unmodified | 0 | 0 | 369610 | 175030 | 0 | 0 | 216280 | 0 | 763850 | 311620 | 1443900 | 7007400 | 0 | 0 | 0 | 0 | 196910 | 0 |
| IDTIEIITDR( | HNRNPA2B1; H | P22626; A0A | ROA2_HUMAN | Unmodified | 84232 | 0 | 140820 | 0 | 0 | 0 | 60566 | 34765 | 1278800 | 94892 | 537740 | 3860600 | 337010 | 0 | 0 | 73436 | 0 | 0 |
| IDTIEIITDR( | HNRNPA2B1; H | P22626; A0A | ROA2_HUMAN | Unmodified | 0 | 0 | 0 | 0 | 256120 | 0 | 52857 | 25164 | 524950 | 103040 | 0 | 858470 | 489280 | 0 | 0 | 0 | 0 | 33623 |
| IDTNLNK | SDHB; SDHB; S | P21912; A0A | SDHB_HUMAN | Unmodified | 0 | 0 | 0 | 0 | 0 | 0 | 0 | 0 | 0 | 0 | 0 | 11483000 | 0 | 0 | 0 | 0 | 0 | 0 |
| IDTRNELES | HSPA5; HSPA5; | P11021; A0A | BIP_HUMAN Er | Unmodified | 0 | 123500 | 1690400 | 533440 | 128000 | 505470 | 788390 | 42798 | 1.57E+08 | 35017000 | 8725100 | 9332300 | 0 | 131440 | 4187500 | 1430600 | 0 | 0 |
| IDVGEAEPI | YARS1; YARS1 | P54577; A0A | SYYC_HUMAN | Unmodified | 0 | 0 | 0 | 0 | 0 | 0 | 0 | 0 | 0 | 0 | 0 | 0 | 0 | 0 | 0 | 0 | 0 | 0 |
| IDVPANRYI | FARSB | Q9NSD9 | SYFB_HUMAN | Unmodified | 0 | 660420 | 46335 | 0 | 0 | 0 | 0 | 0 | 0 | 0 | 225670 | 1213400 | 0 | 0 | 109420 | 45646 | 0 | 0 |
| IDVTNKVV/ | SH3GL3; SH3G | Q99963; H3I | SH3G3_HUMAI | Unmodified | 0 | 0 | 0 | 0 | 0 | 0 | 0 | 0 | 0 | 0 | 0 | 0 | 119620 | 165930 | 0 | 0 | 0 | 0 |
| IDVVVNNA( | HSD17B4; HSD | P51659; E7E | DHB4_HUMAN | Unmodified | 0 | 0 | 0 | 0 | 0 | 0 | 0 | 0 | 1231600 | 0 | 420670 | 0 | 0 | 0 | 0 | 0 | 0 | 0 |
| IDYGEYMD | SSBP1; SSBP1; | Q04837; A0/ | SSBP_HUMAN | Unmodified | 0 | 0 | 0 | 0 | 0 | 0 | 0 | 0 | 1918200 | 0 | 0 | 0 | 0 | 0 | 0 | 0 | 0 | 0 |
| IDYIAGLDS | APRT; APRT; AP | P07741; H3E | APT_HUMAN A( | Unmodified | 520470 | 327560 | 556500 | 199840 | 0 | 0 | 0 | 0 | 0 | 0 | 0 | 6391800 | 0 | 0 | 1515600 | 1190700 | 0 | 0 |
| IEAGYIQTG | HBS1L; HBS1L; | Q9Y450; H0Y | HBS1L_HUMAN | Unmodified | 0 | 0 | 0 | 0 | 0 | 0 | 0 | 0 | 0 | 0 | 0 | 490970 | 0 | 0 | 0 | 0 | 0 | 0 |
| IEAINQAIAI | FAM98A; FAM98 | Q8NCA5; E9 | FA98A_HUMAN | Unmodified | 199330 | 0 | 335210 | 0 | 0 | 0 | 0 | 159650 | 0 | 0 | 1224800 | 3603300 | 0 | 484440 | 397260 | 293590 | 0 | 0 |
| IEALQNHEI | KPNA2; KPNA2; | P52292; A0A | IMA1_HUMAN I | Unmodified | 0 | 0 | 221730 | 135790 | 0 | 0 | 0 | 0 | 93902 | 0 | 0 | 468810 | 0 | 0 | 205220 | 0 | 23304 | 0 |
| IEANEALVK | ATP5F1D | P30049 | ATPD_HUMAN | Unmodified | 0 | 0 | 228340 | 0 | 631570 | 0 | 324510 | 129190 | 8676000 | 254410 | 454810 | 2390700 | 845000 | 520900 | 360360 | 0 | 262490 | 0 |
| IEAPALAFLI | LRRC8A | Q8IWT6 | LRC8A_HUMAN | Unmodified | 65661 | 56353 | 0 | 0 | 0 | 0 | 0 | 0 | 0 | 0 | 0 | 3382300 | 0 | 0 | 0 | 171190 | 0 | 0 |
| IECGPKYPI | UBE2V1; UBE2V | Q13404; G3Y | UB2V1_HUMAN | Unmodified | 1186100 | 454620 | 150210 | 620050 | 0 | 0 | 0 | 0 | 1736500 | 0 | 0 | 2306900 | 95659 | 0 | 0 | 811420 | 0 | 0 |
| IEDDMDG( | ITGA6; ITGA6 | P23229; C9J | ITA6_HUMAN I | Oxidation (M) | 0 | 0 | 0 | 0 | 0 | 0 | 0 | 0 | 0 | 0 | 0 | 463090 | 0 | 0 | 385620 | 359180 | 0 | 0 |
| IEDFLER | RPS9; RPS9; RP | P46781; B5N | RS9_HUMAN 4 | Unmodified | 0 | 437310 | 777290 | 0 | 0 | 0 | 0 | 0 | 0 | 0 | 2850200 | 11502000 | 0 | 0 | 518370 | 0 | 0 | 0 |
| IEDGNDFG | PSME2; PSME2; | Q9UL46; A0/ | PSME2_HUMAI | Unmodified | 0 | 0 | 0 | 0 | 0 | 0 | 0 | 0 | 0 | 0 | 0 | 1885600 | 0 | 0 | 0 | 0 | 0 | 0 |
| IEDGNNFG | PSME1; PSME1 | Q06323; H0Y | PSME1_HUMAI | Unmodified | 0 | 0 | 0 | 0 | 702640 | 0 | 0 | 0 | 368600 | 0 | 1242300 | 4709700 | 0 | 0 | 55921 | 61149 | 0 | 62273 |
| IEDILEVIEK | KYNU; KYNU; K | Q16719; A0/ | KYNU_HUMAN | Unmodified | 0 | 0 | 385110 | 0 | 0 | 88282 | 0 | 138570 | 0 | 94848 | 134770 | 171770 | 0 | 0 | 0 | 0 | 151100 | 123230 |
| IEDLSQQA( | NACA; NACA; N | E9PAV3; Q13 | NACAM_HUMA | Unmodified | 1591200 | 1630600 | 0 | 840770 | 605050 | 0 | 268900 | 0 | 353200 | 0 | 0 | 0 | 260540 | 0 | 0 | 0 | 49222 | 123880 |
| IEDSEPHIP | ATP2B1; ATP2B | P20020; E7E | AT2B1_HUMAN | Unmodified | 0 | 0 | 1226600 | 0 | 0 | 0 | 0 | 45060 | 0 | 0 | 277790 | 0 | 0 | 1020500 | 5038200 | 5570900 | 0 | 0 |
| IEDVGSDEI | HSP90AB1; HSI | P08238; Q5‹ | HS90B_HUMAI | Unmodified | 0 | 0 | 0 | 0 | 0 | 0 | 0 | 0 | 606150 | 143950 | 1347700 | 1402100 | 0 | 0 | 287290 | 262530 | 0 | 0 |
| IEDVGSDEI | HSP90AB1; HSI | P08238; Q5‹ | HS90B_HUMAI | Unmodified | 0 | 0 | 0 | 0 | 0 | 0 | 0 | 0 | 321240 | 0 | 1660100 | 3362500 | 0 | 0 | 261640 | 146700 | 0 | 0 |
| IEDVTPIPSI | RPS14; RPS14 | P62263; A0A | RS14_HUMAN . | Unmodified | 0 | 0 | 0 | 0 | 0 | 0 | 0 | 0 | 0 | 0 | 0 | 4768800 | 0 | 0 | 73832 | 196200 | 0 | 0 |
| IEDVTPIPSI | RPS14; RPS14 | P62263; A0A | RS14_HUMAN . | Unmodified | 265370 | 0 | 83236 | 48970 | 0 | 0 | 0 | 0 | 189480 | 54315 | 4796400 | 8016100 | 0 | 0 | 92513 | 0 | 0 | 54037 |
| IEDYFPEFA | GNAS; GNAS; G | Q5JWF2; P6 | GNAS1_HUMAI | Unmodified | 0 | 0 | 0 | 0 | 0 | 0 | 0 | 0 | 248970 | 0 | 2057000 | 3395100 | 0 | 0 | 0 | 0 | 0 | 0 |
| IEEALGDK | ENO3 | P13929 | ENOB_HUMAN | Unmodified | 0 | 0 | 0 | 0 | 0 | 0 | 0 | 0 | 0 | 0 | 0 | 791460 | 0 | 0 | 0 | 0 | 0 | 0 |
| IEEDTENM/ | VPS37B; VPS37 | Q9H9H4; F5 | VP37B_HUMAN | Unmodified | 711020 | 299120 | 2156700 | 80472 | 0 | 0 | 0 | 0 | 55020 | 0 | 137090 | 350320 | 0 | 0 | 2389300 | 1650700 | 81090 | 0 |
| IEEELGDEA | ENO2; ENO2 | P09104; F5H | ENOG_HUMAN | Unmodified | 13364000 | 901530 | 21434000 | 2527400 | 0 | 0 | 81221 | 88228 | 1832800 | 184050 | 83103 | 2046700 | 0 | 0 | 0 | 17076000 | 0 | 0 |
| IEEELGDEA | ENO2; ENO2 | P09104; F5H | ENOG_HUMAN | Unmodified | 0 | 0 | 0 | 0 | 0 | 0 | 0 | 0 | 0 | 0 | 0 | 0 | 0 | 0 | 0 | 0 | 1151700 | 2548100 |
| IEEELGSK | ENO1; ENO1 | P06733; A0A | ENOA_HUMAN | Unmodified | 0 | 1716600 | 5381400 | 4530300 | 6690500 | 1331800 | 0 | 7177.7 | 39932000 | 4275800 | 14608000 | 1.57E+08 | 2717900 | 0 | 71692 | 8915200 | 1614700 | 0 |
| IEEESEAM/ | VPS37C; VPS37 | A5D8V6; F5H | VP37C_HUMAN | Unmodified | 336910 | 277270 | 631420 | 0 | 0 | 156420 | 0 | 0 | 254040 | 53224 | 0 | 0 | 0 | 0 | 474760 | 774040 | 0 | 0 |
| IEEGLDQIN | SNAP23; SNAP2 | O00161; H3I | SNP23_HUMAN | Unmodified | 149790 | 0 | 0 | 0 | 303400 | 143500 | 0 | 0 | 0 | 734500 | 1143600 | 3971300 | 0 | 0 | 436870 | 190850 | 0 | 342080 |
| IEELQLIVNI | PSMC5; PSMC5 | P62195; J3Q | PRS8_HUMAN | Unmodified | 0 | 0 | 0 | 0 | 0 | 0 | 0 | 0 | 0 | 0 | 0 | 803030 | 0 | 0 | 0 | 0 | 0 | 234800 |
| IEELTGLLN | PPFIBP2; PPFIB | Q8ND30; E9 | LIPB2_HUMAN | Unmodified | 0 | 0 | 0 | 0 | 0 | 0 | 0 | 0 | 0 | 0 | 0 | 0 | 0 | 0 | 594930 | 579910 | 0 | 0 |
| IEEMVEPLR | UXS1 | Q8NBZ7 | UXS1_HUMAN | Unmodified | 0 | 0 | 0 | 376310 | 0 | 0 | 0 | 0 | 1805900 | 0 | 0 | 0 | 0 | 0 | 240580 | 242570 | 609870 | 0 |
| IEENATGFS | MITD1; MITD1; I | Q8WV92; H7 | MITD1_HUMAN | Unmodified | 185920 | 0 | 0 | 114210 | 0 | 0 | 0 | 0 | 0 | 0 | 0 | 8544200 | 0 | 0 | 149800 | 77368 | 0 | 0 |
| IEEVPELPL\ | RPL4; RPL4 | P36578; H3E | RL4_HUMAN 6( | Unmodified | 41681 | 64508 | 0 | 0 | 0 | 52208 | 190800 | 68492 | 0 | 0 | 939950 | 4264600 | 0 | 0 | 0 | 0 | 0 | 0 |
| IEEVPELPL\ | RPL4; RPL4 | P36578; H3E | RL4_HUMAN 6( | Unmodified | 0 | 0 | 0 | 0 | 0 | 0 | 0 | 0 | 0 | 0 | 0 | 5255100 | 0 | 0 | 0 | 0 | 0 | 0 |
| IEEYEPVHS | CEMIP2; CEMIP | Q9UHN6; H3 | CEIP2_HUMAN | Unmodified | 597460 | 374130 | 1047600 | 211440 | 0 | 0 | 0 | 0 | 1100800 | 448270 | 0 | 0 | 0 | 0 | 1096600 | 766390 | 0 | 0 |

| | | | | | | | | | | | | | | | | | | | | | | |
|---|---|---|---|---|---|---|---|---|---|---|---|---|---|---|---|---|---|---|---|---|---|---|
| IEFDNDAD | ITGA6 | P23229 | ITA6_HUMAN I | Oxidation (M) | 0 | 0 | 0 | 0 | 0 | 0 | 0 | 0 | 0 | 0 | 0 | 2743600 | 0 | 0 | 1108600 | 1019900 | 0 | 0 |
| IEFGVDVTT | PAICS; PAICS; F | P22234; D6F | PUR6_HUMAN | Unmodified | 0 | 0 | 66669 | 0 | 637140 | 0 | 112450 | 35927 | 1042400 | 223930 | 0 | 2655000 | 0 | 0 | 88957 | 288300 | 0 | 0 |
| IEFIGQIELY | VPS26B | Q4G0F5 | VP26B_HUMAN | Unmodified | 0 | 0 | 0 | 0 | 2947100 | 20268000 | 16729000 | 7331400 | 0 | 0 | 0 | 0 | 1474700 | 32294000 | 0 | 0 | 17015000 | 21689000 |
| IEFLQSHEN | KPNA6; KPNA1; | O60684; P52 | IMA7_HUMAN I | Unmodified | 0 | 0 | 799750 | 0 | 0 | 0 | 0 | 0 | 484460 | 0 | 164140 | 0 | 0 | 0 | 401730 | 453340 | 0 | 0 |
| IEFSEFIQAL | NCS1 | P62166 | NCS1_HUMAN | Unmodified | 0 | 20456 | 0 | 0 | 302750 | 0 | 0 | 0 | 0 | 0 | 0 | 0 | 0 | 0 | 62176 | 15443 | 0 | 0 |
| IEFSLPDLE | PSMC2; PSMC2 | P35998; C9J | PRS7_HUMAN | Unmodified | 0 | 181540 | 0 | 0 | 0 | 0 | 0 | 175670 | 918180 | 457240 | 0 | 733210 | 0 | 0 | 0 | 0 | 0 | 0 |
| IEGDMIVC/ | RPL5; RPL5; RP | P46777; Q5T | RL5_HUMAN 6 | Oxidation (M) | 3331200 | 1615700 | 335880 | 840510 | 0 | 0 | 0 | 166090 | 0 | 0 | 0 | 14194000 | 0 | 0 | 1884400 | 685770 | 0 | 0 |
| IEGDMIVC/ | RPL5; RPL5; RP | P46777; Q5T | RL5_HUMAN 6 | Unmodified | 756990 | 480810 | 0 | 37453 | 0 | 140300 | 0 | 0 | 156950 | 138410 | 0 | 3333400 | 0 | 0 | 201570 | 0 | 0 | 0 |
| IEGDPQGV | HDLBP; HDLBP | Q00341; A0/ | VIGLN_HUMAN | Unmodified | 0 | 80285 | 0 | 0 | 0 | 0 | 0 | 0 | 0 | 0 | 0 | 313490 | 0 | 0 | 0 | 77389 | 0 | 0 |
| IEGEGSVL( | MVP | Q14764 | MVP_HUMAN M | Unmodified | 427120 | 659350 | 0 | 0 | 0 | 0 | 0 | 0 | 337170 | 0 | 0 | 7302400 | 0 | 0 | 0 | 0 | 0 | 0 |
| IEGFTNVK | GIPC1 | O14908 | GIPC1_HUMAN | Unmodified | 0 | 0 | 0 | 0 | 0 | 0 | 0 | 0 | 0 | 0 | 0 | 0 | 0 | 0 | 934230 | 1208900 | 0 | 0 |
| IEGHQDAV | WDFY1; N/A; W | Q8IWB7; A0/ | WDFY1_HUMA | Unmodified | 103100 | 0 | 0 | 0 | 0 | 0 | 0 | 0 | 0 | 0 | 0 | 972920 | 0 | 0 | 0 | 0 | 0 | 0 |
| IEGLDIHFII | EPHX1 | P07099 | HYEP_HUMAN | Unmodified | 0 | 0 | 1302400 | 367750 | 0 | 0 | 0 | 0 | 0 | 0 | 0 | 0 | 0 | 0 | 0 | 0 | 0 | 0 |
| IEGLLAAFP | ARCN1; ARCN1 | P48444; E9F | COPD_HUMAN | Unmodified | 0 | 0 | 0 | 0 | 0 | 0 | 0 | 0 | 0 | 0 | 358000 | 544240 | 0 | 0 | 0 | 0 | 0 | 0 |
| IEGLQNLVI | PPP1R7; PPP1R | Q15435; C9J | PP1R7_HUMAN | Unmodified | 0 | 0 | 0 | 0 | 0 | 0 | 0 | 0 | 211940 | 0 | 289520 | 4365300 | 0 | 0 | 0 | 0 | 0 | 0 |
| IEGSGDQV | DNM2; DNM2 | P50570; K7E | DYN2_HUMAN | Unmodified | 245640 | 74735 | 0 | 270470 | 0 | 0 | 0 | 0 | 0 | 0 | 0 | 0 | 0 | 0 | 0 | 287290 | 0 | 0 |
| IEGTPLETI( | NDUFB3; NDUF | O43676; C9J | NDUB3_HUMA | Unmodified | 0 | 0 | 0 | 0 | 0 | 0 | 0 | 0 | 494920 | 0 | 0 | 0 | 0 | 0 | 0 | 0 | 0 | 0 |
| IEGVYAR | RPL35A; RPL35 | P18077; F8V | RL35A_HUMAN | Unmodified | 896290 | 822490 | 1320900 | 133920 | 214930 | 39103 | 0 | 23316 | 202480 | 0 | 213130 | 5775000 | 0 | 0 | 0 | 693720 | 144930 | 224160 |
| IEGVYARDI | RPL35A; RPL35 | P18077; F8V | RL35A_HUMAN | Unmodified | 5791300 | 4238900 | 1681200 | 643330 | 2506700 | 425480 | 4218400 | 0 | 1899300 | 462270 | 3099000 | 26880000 | 758410 | 1102400 | 1528300 | 1648500 | 342110 | 2640300 |
| IEGVYARDI | RPL35A; RPL35 | P18077; F8V | RL35A_HUMAN | Unmodified | 8876400 | 8020100 | 6949300 | 6371100 | 382690 | 0 | 1778600 | 114200 | 794980 | 51412 | 1408900 | 49391000 | 0 | 0 | 6733300 | 5109600 | 0 | 0 |
| IEGYEDQVI | CAPZA2; CAPZ/ | P47755; A0A | CAZA2_HUMAN | Unmodified | 0 | 0 | 364140 | 0 | 0 | 0 | 0 | 0 | 0 | 0 | 282310 | 1916000 | 0 | 0 | 0 | 324120 | 0 | 0 |
| IEGYEDQVI | CAPZA1 | P52907 | CAZA1_HUMAN | Unmodified | 639790 | 418480 | 819260 | 253910 | 0 | 0 | 0 | 0 | 767760 | 0 | 924490 | 5023800 | 0 | 0 | 804660 | 511970 | 0 | 0 |
| IEHFMPLL\ | AP1M2; AP1M2 | Q9Y6Q5; K7I | AP1M2_HUMAI | Unmodified | 0 | 0 | 0 | 0 | 0 | 0 | 0 | 0 | 0 | 0 | 0 | 447000 | 0 | 0 | 0 | 0 | 0 | 0 |
| IEHTETDTV | TMEM63B; TME | Q5T3F8; H3I | CSCL2_HUMAI | Unmodified | 0 | 0 | 0 | 0 | 0 | 0 | 0 | 0 | 0 | 0 | 0 | 0 | 0 | 0 | 211660 | 126870 | 0 | 0 |
| IEHVEQNTI | PARP4 | Q9UKK3 | PARP4_HUMAN | Unmodified | 35351 | 0 | 0 | 12292 | 0 | 0 | 0 | 0 | 0 | 0 | 0 | 3232100 | 0 | 0 | 0 | 0 | 0 | 0 |
| IEIAQKHPD | TJP2; TJP2; TJP2 | Q9UDY2; A0 | ZO2_HUMAN T | Unmodified | 0 | 0 | 0 | 0 | 0 | 0 | 0 | 0 | 1663600 | 245500 | 0 | 0 | 0 | 0 | 0 | 0 | 0 | 0 |
| IEIESFYEGI | HSPA5; HSPA5; | P11021; A0A | BIP_HUMAN Er | Unmodified | 0 | 0 | 0 | 0 | 0 | 0 | 0 | 0 | 7455900 | 1823600 | 410010 | 127420 | 0 | 0 | 0 | 0 | 0 | 0 |
| IEIESFYEGI | HSPA5; HSPA5; | P11021; A0A | BIP_HUMAN Er | Unmodified | 0 | 0 | 0 | 0 | 0 | 0 | 0 | 0 | 0 | 2275900 | 0 | 0 | 541860 | 0 | 0 | 0 | 0 | 0 |
| IEIFQTLPVI | NCAM2; NCAM; | O15394; H9I | NCAM2_HUMA | Unmodified | 0 | 1206600 | 10678000 | 2907900 | 0 | 0 | 0 | 0 | 0 | 0 | 0 | 0 | 81187 | 0 | 20695000 | 13989000 | 0 | 0 |
| IEIKLSDIPE | UQCRFS1P1; U | P0C7P4; P4; | UCRIL_HUMAN | Unmodified | 0 | 0 | 0 | 0 | 0 | 0 | 0 | 0 | 1091500 | 2067700 | 0 | 0 | 0 | 0 | 0 | 0 | 0 | 0 |
| IEILAPPNG | AIMP1 | Q12904 | AIMP1_HUMAN | Unmodified | 0 | 0 | 0 | 0 | 0 | 0 | 0 | 0 | 0 | 0 | 668940 | 1378600 | 0 | 0 | 0 | 0 | 0 | 0 |
| IEINFPAEYI | UBE2L3 | P68036 | UB2L3_HUMAN | Unmodified | 0 | 36714 | 488590 | 0 | 254890 | 0 | 0 | 39355 | 146730 | 0 | 1559800 | 7401300 | 0 | 0 | 1446200 | 764160 | 0 | 0 |
| IEIPIHIVPH | BRCC3; BRCC3 | P46736; A0A | BRCC3_HUMAI | Unmodified | 0 | 0 | 0 | 0 | 0 | 0 | 0 | 0 | 0 | 0 | 0 | 371580 | 0 | 0 | 0 | 0 | 0 | 0 |
| IEIPLFHVLI | ANKRD13A; AN | Q8IZ07; Q6Z | AN13A_HUMAI | Unmodified | 911720 | 509240 | 676600 | 0 | 0 | 0 | 0 | 0 | 0 | 0 | 0 | 456060 | 0 | 0 | 941130 | 973430 | 0 | 0 |
| IEIQTLTGLI | CASP6; CASP6 | P55212; D6F | CASP6_HUMAI | Unmodified | 0 | 0 | 0 | 0 | 0 | 0 | 0 | 0 | 0 | 0 | 92993 | 838670 | 0 | 0 | 0 | 0 | 0 | 0 |
| IEIQTLTGLI | CASP6; CASP6 | P55212; D6F | CASP6_HUMAI | Unmodified | 0 | 0 | 0 | 0 | 0 | 0 | 0 | 0 | 0 | 0 | 406070 | 6827400 | 0 | 0 | 0 | 0 | 0 | 0 |
| IEKELETVC | YWHAH; YWHA | Q04917; A0/ | 1433F_HUMAN | Unmodified | 0 | 0 | 0 | 0 | 551790 | 0 | 0 | 0 | 0 | 0 | 0 | 0 | 135740 | 0 | 0 | 0 | 189630 | 3453600 |
| IEKVDLPLII | SAAL1 | Q96ER3 | SAAL1_HUMAN | Unmodified | 0 | 4241500 | 31702 | 588450 | 0 | 0 | 88831 | 0 | 61879 | 43831 | 37214 | 259040 | 324130 | 0 | 0 | 128430 | 0 | 0 |
| IELETYK | CCDC6 | Q16204 | CCDC6_HUMA | Unmodified | 0 | 0 | 394040 | 0 | 0 | 0 | 0 | 0 | 244570 | 0 | 249640 | 1623900 | 0 | 0 | 0 | 0 | 0 | 0 |
| IELHGKPIE | IGF2BP3; IGF2E | O00425; F8V | IF2B3_HUMAN | Unmodified | 464700 | 0 | 361250 | 0 | 0 | 0 | 0 | 0 | 0 | 0 | 0 | 1095300 | 0 | 0 | 0 | 0 | 0 | 0 |
| IELLHNIDP | ARSB | P15848 | ARSB_HUMAN | Unmodified | 0 | 33941 | 0 | 0 | 0 | 0 | 0 | 0 | 820370 | 42039 | 0 | 280510 | 0 | 0 | 0 | 0 | 0 | 0 |
| IELQRNPPI | FUBP1; FUBP1 | Q96AE4; E9F | FUBP1_HUMAN | Oxidation (M) | 0 | 0 | 0 | 0 | 0 | 0 | 0 | 0 | 0 | 0 | 0 | 2069100 | 0 | 0 | 0 | 0 | 0 | 0 |
| IELQRNPPI | FUBP1; FUBP1 | Q96AE4; E9F | FUBP1_HUMAN | Unmodified | 39359 | 0 | 0 | 87963 | 0 | 0 | 0 | 0 | 0 | 0 | 0 | 1586900 | 0 | 0 | 0 | 0 | 0 | 0 |
| IENESDMIF | STXBP3 | O00186 | STXB3_HUMAN | Oxidation (M) | 0 | 0 | 0 | 0 | 122530 | 0 | 0 | 0 | 145030 | 0 | 1511500 | 668340 | 93093 | 0 | 501770 | 282930 | 0 | 84616 |
| IENESDMIF | STXBP3 | O00186 | STXB3_HUMAN | Unmodified | 138510 | 104970 | 0 | 0 | 0 | 0 | 0 | 0 | 117710 | 0 | 300060 | 2191200 | 0 | 0 | 505460 | 318540 | 0 | 0 |
| IENIDHLGF | ITM2B; ITM2B | Q9Y287; A0/ | ITM2B_HUMAN | Unmodified | 0 | 0 | 0 | 0 | 0 | 0 | 0 | 0 | 234720 | 0 | 0 | 0 | 0 | 0 | 0 | 77961 | 0 | 0 |
| IENLELVPV | VIL1; VIL1 | P09327; B4E | VILI_HUMAN Vi | Unmodified | 0 | 0 | 0 | 0 | 0 | 0 | 0 | 0 | 0 | 0 | 407250 | 10389000 | 0 | 0 | 0 | 0 | 0 | 0 |
| IENLSNLH( | PPP1R7; PPP1R | Q15435; C9J | PP1R7_HUMAN | Oxidation (M) | 0 | 0 | 0 | 0 | 0 | 0 | 0 | 0 | 0 | 0 | 0 | 2888700 | 0 | 0 | 0 | 0 | 0 | 0 |
| IENSELLDK | ZC3HAV1; ZC3I | Q7Z2W4; C9 | ZCCHV_HUMA | Unmodified | 67585 | 60311 | 272780 | 0 | 0 | 0 | 0 | 158530 | 206380 | 419770 | 0 | 104670 | 0 | 0 | 0 | 486180 | 0 | 0 |
| IENVPTGPI | PPIH; PPIH; PPI | O43447; A0/ | PPIH_HUMAN I | Unmodified | 160080 | 143610 | 250020 | 0 | 0 | 0 | 0 | 0 | 0 | 0 | 141390 | 317980 | 68482 | 0 | 501900 | 194430 | 0 | 0 |
| IENVVLVVF | XPNPEP1; XPNI | Q9NQW7; Q | XPP1_HUMAN | Unmodified | 0 | 0 | 0 | 0 | 0 | 0 | 0 | 0 | 0 | 0 | 0 | 3739200 | 0 | 0 | 175950 | 107340 | 0 | 0 |
| IEPADAHVI | NAA50; NAA50; | Q9GZZ1; A0/ | NAA50_HUMAI | Unmodified | 0 | 0 | 0 | 0 | 0 | 0 | 0 | 27782 | 0 | 0 | 280400 | 338110 | 0 | 0 | 0 | 0 | 0 | 0 |
| IEPEPFENC | GSS; GSS; GSS | P48637; A0A | GSHB_HUMAN | Unmodified | 256980 | 0 | 266920 | 469180 | 590180 | 154310 | 0 | 0 | 1093700 | 302050 | 1278000 | 3593400 | 0 | 0 | 566710 | 740420 | 0 | 0 |
| IEPGVDPDI | RNPEP; RNPEP | Q9H4A4; A6I | AMPB_HUMAN | Unmodified | 0 | 0 | 0 | 0 | 0 | 0 | 0 | 0 | 0 | 0 | 0 | 4642000 | 0 | 0 | 266810 | 68852 | 0 | 0 |
| IEPGVSVSL | LARS1; LARS1; | Q9P2J5; A0A | SYLC_HUMAN | Unmodified | 0 | 0 | 0 | 0 | 0 | 0 | 0 | 0 | 0 | 0 | 0 | 14256000 | 0 | 0 | 140620 | 82644 | 0 | 0 |
| IEPHHTAVI | PIR | O00625 | PIR_HUMAN Pi | Unmodified | 0 | 0 | 427430 | 0 | 0 | 0 | 0 | 0 | 1184800 | 0 | 597030 | 13941000 | 0 | 0 | 71628 | 428770 | 0 | 0 |
| IEPHHTAVI | PIR | O00625 | PIR_HUMAN Pi | Unmodified | 0 | 0 | 0 | 0 | 0 | 0 | 0 | 0 | 0 | 0 | 745540 | 571040 | 0 | 0 | 0 | 0 | 0 | 0 |
| IEPHTGLLL | DSP | P15924 | DESP_HUMAN | Unmodified | 0 | 0 | 0 | 0 | 0 | 0 | 0 | 0 | 304950 | 0 | 0 | 0 | 0 | 0 | 0 | 0 | 15525 | 0 |
| IEPNLPSYR | DPP4; DPP4; DI | P27487; A0A | DPP4_HUMAN | Unmodified | 0 | 345350 | 9411600 | 2935500 | 0 | 0 | 0 | 0 | 0 | 0 | 0 | 0 | 0 | 0 | 7897200 | 4953800 | 0 | 0 |
| IEPNYPK | CORO2B | Q9UQ03 | COR2B_HUMA | Unmodified | 0 | 0 | 0 | 0 | 0 | 0 | 0 | 0 | 0 | 0 | 0 | 1679200 | 0 | 0 | 0 | 0 | 0 | 0 |
| IEPSPYK | TM9SF2 | Q99805 | TM9S2_HUMAN | Unmodified | 0 | 0 | 0 | 0 | 0 | 0 | 0 | 0 | 476970 | 0 | 0 | 109830 | 0 | 31727 | 0 | 0 | 0 | 0 |
| IEPSVNFLK | LMAN2; LMAN2 | Q12907; D6I | LMAN2_HUMAI | Unmodified | 0 | 0 | 1406800 | 0 | 0 | 0 | 0 | 0 | 72339000 | 2485000 | 1204800 | 1926600 | 0 | 0 | 0 | 0 | 0 | 0 |
| IEPVGMVAI | N/A | N/A | N/A | Oxidation (M) | 0 | 111080 | 0 | 0 | 0 | 0 | 0 | 0 | 0 | 531640 | 0 | 0 | 0 | 0 | 0 | 0 | 308310 | 0 |
| IEQELTAAK | CHMP4B | Q9H444 | CHM4B_HUMA | Unmodified | 0 | 0 | 0 | 0 | 0 | 0 | 0 | 0 | 482060 | 0 | 0 | 0 | 0 | 0 | 0 | 0 | 0 | 0 |
| IEQEQKFTV | ATP2B3; ATP2B | Q16720; P2( | AT2B3_HUMAN | Unmodified | 0 | 0 | 0 | 790630 | 0 | 0 | 0 | 0 | 0 | 0 | 0 | 0 | 0 | 0 | 86152 | 112590 | 0 | 0 |
| IEQFVYSSP | SARS1; SARS1 | P49591; Q5T | SYSC_HUMAN | Unmodified | 0 | 0 | 0 | 537070 | 0 | 345680 | 0 | 0 | 1162900 | 0 | 289100 | 8526100 | 270920 | 0 | 811070 | 724590 | 0 | 0 |
| IEQKYDADI | TAGLN3; TAGLI | Q9UI15; C9J | TAGL3_HUMAN | Unmodified | 0 | 0 | 0 | 0 | 0 | 0 | 0 | 0 | 456490 | 30464 | 0 | 1639800 | 0 | 0 | 0 | 0 | 0 | 0 |
| IEQLQNHE | KPNA4; KPNA4; | O00629; H7( | IMA3_HUMAN I | Unmodified | 6131800 | 2895400 | 724310 | 3198500 | 190050 | 0 | 0 | 0 | 480500 | 0 | 294990 | 1214000 | 0 | 0 | 5429600 | 8159200 | 0 | 0 |
| IEQSQHLF( | ARFIP1 | P53367 | ARFP1_HUMAN | Unmodified | 0 | 0 | 0 | 445690 | 0 | 0 | 0 | 0 | 0 | 0 | 0 | 677090 | 0 | 0 | 275090 | 207020 | 362530 | 0 |
| IEQVDKEDI | NOP58 | Q9Y2X3 | NOP58_HUMAI | Unmodified | 0 | 0 | 0 | 0 | 0 | 0 | 0 | 0 | 0 | 0 | 78848 | 0 | 0 | 0 | 0 | 0 | 0 | 0 |
| IEQYATR | UTRN; UTRN; U | P46939; H0\ | UTRN_HUMAN | Unmodified | 0 | 0 | 0 | 0 | 0 | 0 | 0 | 0 | 0 | 0 | 0 | 0 | 0 | 0 | 308710 | 241010 | 0 | 0 |
| IERDSREHI | IGFBP5 | P24593 | IBP5_HUMAN I | Unmodified | 0 | 0 | 0 | 0 | 0 | 0 | 0 | 0 | 4814100 | 0 | 0 | 0 | 0 | 0 | 0 | 0 | 0 | 0 |
| IEREHQISP | EHD3 | Q9NZN3 | EHD3_HUMAN | Unmodified | 0 | 0 | 0 | 93102 | 0 | 0 | 266750 | 180580 | 0 | 75576 | 0 | 0 | 69412 | 0 | 1257800 | 0 | 0 | 0 |
| IEREHQISP | EHD1; EHD1; E | Q9H4M9; C9 | EHD1_HUMAN | Unmodified | 2088700 | 2196100 | 3042200 | 2997300 | 13062000 | 2911300 | 339200 | 0 | 2012700 | 327140 | 3137500 | 21632000 | 4892700 | 2542800 | 3644100 | 3432200 | 2368800 | 13494000 |
| IERTVQIEA | SEPTIN2; SEPTI | Q15019; C9I | SEPT2_HUMAN | Unmodified | 0 | 0 | 0 | 0 | 4781100 | 0 | 352280 | 0 | 0 | 0 | 0 | 0 | 0 | 0 | 0 | 0 | 602800 | 2150800 |
| IESGELDPE | RAB4A; RAB4A | P20338; A0A | RAB4A_HUMAN | Unmodified | 0 | 0 | 168550 | 133200 | 0 | 0 | 0 | 0 | 224450 | 0 | 53004 | 268450 | 0 | 0 | 141470 | 57756 | 0 | 0 |
| IESGELDPE | RAB4A; RAB4A | P20338; A0A | RAB4A_HUMAN | Oxidation (M) | 0 | 47541 | 375470 | 278600 | 0 | 0 | 0 | 0 | 0 | 0 | 0 | 368060 | 0 | 0 | 0 | 338540 | 0 | 0 |
| IESGEMGLI | EFCAB14; EFC/ | O75071; A0/ | EFC14_HUMAN | Oxidation (M) | 0 | 0 | 0 | 0 | 0 | 0 | 0 | 0 | 906770 | 193300 | 0 | 53899 | 0 | 0 | 0 | 0 | 0 | 0 |
| IESGEMGLI | EFCAB14; EFC/ | O75071; A0/ | EFC14_HUMAN | Unmodified | 118100 | 0 | 250710 | 0 | 0 | 0 | 169530 | 37912 | 7382500 | 1069200 | 543830 | 0 | 0 | 0 | 0 | 0 | 0 | 0 |
| IESILMSLPI | CPOX; N/A | P36551; H0\ | HEM6_HUMAN | Oxidation (M) | 0 | 0 | 0 | 0 | 0 | 0 | 0 | 0 | 0 | 0 | 0 | 592480 | 0 | 0 | 0 | 0 | 0 | 0 |
| IESKADPSII | CCDC93; CCD( | Q567U6; F8\ | CCD93_HUMA | Unmodified | 0 | 0 | 0 | 0 | 255640 | 46185 | 0 | 0 | 0 | 0 | 0 | 0 | 128470 | 57997 | 0 | 0 | 0 | 122040 |
| IESKIEEMV | UXS1 | Q8NBZ7 | UXS1_HUMAN | Unmodified | 0 | 0 | 0 | 0 | 0 | 0 | 0 | 0 | 1215200 | 0 | 0 | 0 | 0 | 0 | 0 | 0 | 0 | 0 |
| IESLSSQLS | LMNB1; LMNB1 | P20700; A0A | LMNB1_HUMAI | Unmodified | 0 | 0 | 0 | 0 | 147670 | 0 | 0 | 0 | 0 | 0 | 683670 | 0 | 168460 | 0 | 0 | 0 | 0 | 0 |
| IESPGYLTS | NRP1; NRP1; N | O14786; Q5. | NRP1_HUMAN | Unmodified | 0 | 0 | 804140 | 0 | 0 | 0 | 0 | 0 | 0 | 0 | 0 | 0 | 0 | 0 | 0 | 0 | 0 | 0 |
| IESPHPVD\ | DHX36; DHX36 | Q9H2U1; A0 | DHX36_HUMAI | Unmodified | 239390 | 669710 | 0 | 0 | 135100 | 0 | 23142 | 0 | 0 | 28433 | 150060 | 1874700 | 0 | 0 | 650060 | 419050 | 0 | 183220 |
| IESVLSSSG | GM2A | P17900 | SAP3_HUMAN | Unmodified | 0 | 0 | 149570 | 57930 | 0 | 0 | 74550 | 0 | 1627200 | 0 | 302950 | 0 | 0 | 0 | 473040 | 346960 | 0 | 0 |
| IETALTSLH | NPTX1 | Q15818 | NPTX1_HUMAN | Unmodified | 149420 | 448310 | 0 | 291730 | 99217 | 0 | 0 | 0 | 29389000 | 718510 | 346460 | 558100 | 74966 | 36421 | 257730 | 386050 | 29154 | 0 |
| IETALTSLH( | NPTX1 | Q15818 | NPTX1_HUMAN | Unmodified | 0 | 0 | 0 | 0 | 0 | 0 | 0 | 0 | 0 | 2367600 | 0 | 0 | 0 | 148910 | 0 | 0 | 0 | 0 |
| IETATTLK | IPO11; IPO11 | Q9UI26; F8V | IPO11_HUMAN | Unmodified | 0 | 0 | 0 | 22856 | 0 | 0 | 0 | 0 | 0 | 0 | 0 | 899280 | 0 | 0 | 34951 | 46353 | 0 | 0 |
| IETELRDICI | YWHAZ; YWHAZ | P63104; B0A | 1433Z_HUMAN | Unmodified | 0 | 91815 | 0 | 684000 | 3988000 | 54869 | 295560 | 0 | 61689 | 434960 | 26914 | 0 | 4856400 | 2469900 | 1044800 | 0 | 5449900 | 7506800 |
| IETMLGDV/ | VARS1; VARS1 | P26640; A0A | SYVC_HUMAN | Oxidation (M) | 66923 | 0 | 0 | 0 | 0 | 0 | 0 | 0 | 0 | 0 | 135350 | 1604000 | 0 | 0 | 102490 | 98500 | 34760 | 0 |
| IETMLGDV/ | VARS1; VARS1 | P26640; A0A | SYVC_HUMAN | Unmodified | 0 | 0 | 0 | 0 | 0 | 0 | 0 | 0 | 0 | 0 | 0 | 0 | 0 | 0 | 0 | 0 | 0 | 0 |
| IETNENNLE | CKAP4 | Q07065 | CKAP4_HUMAN | Unmodified | 0 | 0 | 0 | 0 | 0 | 0 | 55487 | 27889 | 2760500 | 168830 | 184740 | 0 | 0 | 199010 | 0 | 0 | 0 | 135060 |
| IETVKYPTYI | FUT8; FUT8 | Q9BYC5; G3 | FUT8_HUMAN | Unmodified | 0 | 0 | 0 | 0 | 0 | 0 | 0 | 0 | 7719700 | 1172500 | 0 | 0 | 0 | 0 | 0 | 0 | 0 | 0 |
| IETVNESWI | ACO1 | P21399 | ACOC_HUMAN | Unmodified | 0 | 161640 | 77829 | 0 | 0 | 0 | 0 | 0 | 0 | 0 | 281940 | 1174100 | 0 | 0 | 99778 | 77373 | 23788 | 0 |
| IETWHKPD | PITPNB; PITPNE | P48739; A0A | PIPNB_HUMAN | Unmodified | 690520 | 430560 | 635370 | 632000 | 914020 | 0 | 0 | 0 | 1213800 | 108480 | 1800200 | 2878300 | 370310 | 0 | 1339300 | 796360 | 0 | 405210 |
| IETWHKPD | PITPNA; PITPN/ | Q00169; I3L | PIPNA_HUMAN | Unmodified | 416560 | 0 | 594710 | 430780 | 0 | 0 | 0 | 0 | 0 | 0 | 0 | 1359000 | 0 | 0 | 472970 | 0 | 0 | 0 |
| IEVEKPFAI/ | PPIB; PPIB; PPII | P23284; A0A | PPIB_HUMAN F | Unmodified | 0 | 0 | 1801600 | 0 | 0 | 0 | 0 | 0 | 15758000 | 937010 | 1346800 | 2897300 | 0 | 0 | 0 | 0 | 0 | 0 |
| IEVIEIMTDI | HNRNPA1L2; H | Q32P51; P0 | RA1L2_HUMAN | Unmodified | 0 | 0 | 0 | 0 | 0 | 0 | 0 | 0 | 0 | 0 | 1537500 | 904530 | 0 | 0 | 0 | 0 | 0 | 0 |
| IEVIEIMTDI | HNRNPA1L2; H | Q32P51; P0 | RA1L2_HUMAN | Unmodified | 0 | 0 | 0 | 0 | 0 | 0 | 0 | 0 | 987270 | 193180 | 273190 | 5386700 | 0 | 0 | 0 | 0 | 0 | 0 |
| IEVIEIMTDI | HNRNPA1L2; H | Q32P51; P0 | RA1L2_HUMAN | Oxidation (M) | 0 | 0 | 0 | 0 | 0 | 0 | 0 | 0 | 0 | 0 | 0 | 2160100 | 103270 | 0 | 0 | 0 | 0 | 0 |
| IEVIEIMTDI | HNRNPA1L2; H | Q32P51; P0 | RA1L2_HUMAN | Unmodified | 0 | 0 | 0 | 0 | 0 | 0 | 0 | 0 | 97445 | 0 | 0 | 988310 | 0 | 0 | 0 | 0 | 0 | 0 |
| IEVIKPGDL | ETFB; ETFB | P38117; M0 | ETFB_HUMAN | Unmodified | 0 | 0 | 0 | 93983 | 46135 | 39758 | 49717 | 23556 | 5291800 | 256900 | 0 | 0 | 166420 | 47831 | 0 | 0 | 0 | 48186 |
| IEVLQQHEI | KPNA3; KPNA3 | O00505; H0 | IMA4_HUMAN I | Unmodified | 185270 | 128450 | 0 | 159290 | 0 | 0 | 129620 | 0 | 676400 | 0 | 263660 | 2450800 | 0 | 0 | 235330 | 181320 | 0 | 0 |
| IEVLYGDEI | POGLUT3 | Q7Z4H8 | PLGT3_HUMAN | Unmodified | 0 | 0 | 0 | 0 | 0 | 0 | 0 | 0 | 2083400 | 0 | 0 | 0 | 0 | 0 | 0 | 0 | 0 | 0 |
| IEVMSLGR | TMEFF2 | Q9UIK5 | TEFF2_HUMAN | Oxidation (M) | 247560 | 0 | 0 | 0 | 0 | 0 | 0 | 0 | 0 | 0 | 0 | 0 | 0 | 0 | 0 | 0 | 0 | 0 |
| IEVPLYSLLI | HSPH1; HSPH1 | Q92598; A0/ | HS105_HUMAN | Unmodified | 0 | 0 | 82512 | 0 | 0 | 0 | 0 | 0 | 0 | 0 | 0 | 880900 | 0 | 0 | 0 | 0 | 0 | 0 |
| IEVSQLLK | CD47; CD47 | Q08722; A0/ | CD47_HUMAN | Unmodified | 4487400 | 0 | 15294000 | 4125000 | 95264 | 0 | 840070 | 0 | 72297000 | 8043200 | 31722000 | 1.9E+08 | 1632400 | 224820 | 11615000 | 17042000 | 0 | 998190 |
| IEVSQLLKG | CD47; CD47 | Q08722; A0/ | CD47_HUMAN | Unmodified | 0 | 0 | 0 | 0 | 0 | 0 | 0 | 0 | 7201500 | 1069800 | 2409500 | 2881100 | 0 | 0 | 0 | 0 | 127460 | 735050 |

| | | | | | | | | | | | | | | | | | | | | | | |
|---|---|---|---|---|---|---|---|---|---|---|---|---|---|---|---|---|---|---|---|---|---|---|
| IEVSREEIFE | SMURF1; SMUF | Q9HCE7; Q9 | SMUF1_HUMAI | Unmodified | 662310 | 0 | 0 | 146000 | 0 | 0 | 0 | 0 | 0 | 156290 | 0 | 0 | 0 | 0 | 0 | 0 | 0 | 0 |
| IEVVCNDRI | UBL5; UBL5 | Q9BZL1; K7E | UBL5_HUMAN | Unmodified | 0 | 0 | 64572 | 99859 | 0 | 0 | 0 | 0 | 0 | 0 | 0 | 0 | 86597 | 0 | 99727 | 94272 | 0 | 0 |
| IEWLESHQ | HSPA5; HSPA5; | P11021; A0A | BIP_HUMAN Er | Unmodified | 269430 | 0 | 6328500 | 148300 | 0 | 0 | 641260 | 0 | 52112000 | 3873900 | 6280900 | 2788100 | 0 | 0 | 4988100 | 688850 | 0 | 0 |
| IEWLESHQ | HSPA5; HSPA5; | P11021; A0A | BIP_HUMAN Er | Unmodified | 0 | 0 | 0 | 135500 | 0 | 0 | 0 | 142700 | 16673000 | 5094300 | 1406300 | 838430 | 0 | 0 | 217690 | 105770 | 235530 | 0 |
| IEYDTFGEL | FH | P07954 | FUMH_HUMAN | Unmodified | 0 | 0 | 0 | 0 | 0 | 0 | 0 | 0 | 922040 | 0 | 0 | 638430 | 0 | 0 | 0 | 0 | 0 | 0 |
| IEYNDQND | FLNB; FLNB | O75369; E7E | FLNB_HUMAN | Unmodified | 0 | 0 | 231780 | 86405 | 181820 | 0 | 0 | 0 | 0 | 28451 | 0 | 884680 | 0 | 0 | 62129 | 108870 | 0 | 0 |
| IEYQFFEDF | UGGT1 | Q9NYU2 | UGGG1_HUMA | Unmodified | 0 | 0 | 0 | 0 | 0 | 0 | 257080 | 0 | 2130200 | 195320 | 0 | 0 | 1349600 | 0 | 2174700 | 887640 | 0 | 0 |
| IEYQFFEDF | UGGT1 | Q9NYU2 | UGGG1_HUMA | Unmodified | 0 | 0 | 0 | 0 | 0 | 0 | 0 | 0 | 13940000 | 308750 | 535890 | 206750 | 0 | 0 | 0 | 0 | 0 | 0 |
| IFAAGFDSS | PELI2; PELI2; PE | Q9HAT8; H0 | PELI2_HUMAN | Unmodified | 0 | 0 | 0 | 0 | 0 | 0 | 0 | 0 | 2133400 | 1177600 | 0 | 0 | 0 | 0 | 0 | 0 | 0 | 0 |
| IFAAGFDSS | PELI2; PELI2; PE | Q9HAT8; H0 | PELI2_HUMAN | Unmodified | 0 | 0 | 0 | 0 | 0 | 0 | 0 | 9433.5 | 0 | 114600 | 0 | 1679400 | 0 | 0 | 0 | 0 | 0 | 0 |
| IFALAFVK | DOCK1; DOCK1 | Q14185; A0A | DOCK1_HUMA | Unmodified | 0 | 0 | 0 | 392140 | 305290 | 56439 | 226930 | 0 | 0 | 0 | 0 | 296130 | 0 | 190720 | 1074100 | 0 | 102430 | 74375 |
| IFANTESYL | NCBP1 | Q09161 | NCBP1_HUMAI | Unmodified | 0 | 0 | 1709200 | 575870 | 0 | 0 | 0 | 0 | 0 | 0 | 0 | 938530 | 0 | 0 | 0 | 120180 | 0 | 0 |
| IFAPNHVVA | RPL18A; RPL18 | Q02543; M0 | RL18A_HUMAN | Unmodified | 0 | 0 | 0 | 0 | 0 | 0 | 0 | 0 | 0 | 0 | 73403 | 2383900 | 0 | 0 | 0 | 0 | 0 | 0 |
| IFAQDGEG | FLNB; FLNB | O75369; E7E | FLNB_HUMAN | Unmodified | 0 | 0 | 0 | 0 | 0 | 0 | 0 | 0 | 113970 | 0 | 692410 | 999270 | 0 | 0 | 0 | 0 | 27482 | 153690 |
| IFAQDGEG | FLNB; FLNB | O75369; E7E | FLNB_HUMAN | Oxidation (M) | 0 | 0 | 0 | 1983400 | 14485000 | 0 | 486920 | 171240 | 0 | 372310 | 0 | 3800000 | 1577800 | 632580 | 0 | 330160 | 4316000 | 0 |
| IFAQDGEG | FLNB; FLNB | O75369; E7E | FLNB_HUMAN | Unmodified | 0 | 0 | 170630 | 0 | 55712000 | 3877000 | 0 | 0 | 0 | 0 | 0 | 0 | 3716300 | 0 | 0 | 0 | 8935900 | 10578000 |
| IFAQYLVPH | PDS5A; PDS5A | Q29RF7; H0 | PDS5A_HUMAN | Unmodified | 244760 | 0 | 0 | 0 | 0 | 0 | 2853000 | 0 | 0 | 0 | 376550 | 581230 | 0 | 11707000 | 0 | 0 | 0 | 0 |
| IFAREILDSI | ENO3; ENO3; E | P13929; E5R | ENOB_HUMAN | Unmodified | 0 | 0 | 0 | 0 | 0 | 0 | 0 | 0 | 0 | 0 | 0 | 0 | 0 | 0 | 0 | 0 | 0 | 938290 |
| IFASIAPSIY | MCM2; MCM2 | P49736; H0 | MCM2_HUMAN | Unmodified | 0 | 0 | 0 | 0 | 459720 | 0 | 0 | 0 | 0 | 130230 | 280180 | 1404200 | 331840 | 0 | 0 | 0 | 0 | 0 |
| IFCCHGGL | PPP1CA; PPP1C | P62136; P62 | PP1A_HUMAN | Unmodified | 0 | 86179 | 0 | 169160 | 268040 | 0 | 0 | 0 | 1570500 | 0 | 172100 | 1276700 | 0 | 0 | 494560 | 273360 | 215030 | 0 |
| IFCCHGGL | PPP1CA; PPP1C | P62136; P62 | PP1A_HUMAN | Oxidation (M) | 176910 | 24811 | 199510 | 149830 | 0 | 0 | 86217 | 34277 | 445190 | 70035 | 0 | 1173400 | 69597 | 0 | 88318 | 169630 | 0 | 0 |
| IFCCHGGL | PPP1CA; PPP1C | P62136; P62 | PP1A_HUMAN | Unmodified | 304370 | 0 | 0 | 315940 | 0 | 0 | 0 | 0 | 1877800 | 0 | 0 | 0 | 0 | 0 | 471160 | 0 | 0 | 0 |
| IFCVHGGL | PPP4C; PPP4C; | P60510; H3E | PP4C_HUMAN | Unmodified | 0 | 0 | 0 | 0 | 0 | 0 | 0 | 0 | 0 | 0 | 0 | 853150 | 0 | 0 | 0 | 0 | 0 | 0 |
| IFDANTKPN | UGDH | O60701 | UGDH_HUMAN | Unmodified | 619140 | 0 | 0 | 301180 | 39316000 | 7194500 | 0 | 0 | 0 | 0 | 132260 | 346900 | 9435500 | 10493000 | 0 | 0 | 0 | 2924600 |
| IFDAVGFTF | NCDN; NCDN | Q9UBB6; C9 | NCDN_HUMAN | Unmodified | 0 | 0 | 0 | 0 | 0 | 143630 | 0 | 0 | 0 | 0 | 0 | 801530 | 0 | 0 | 0 | 0 | 0 | 0 |
| IFDAVGFTF | NCDN; NCDN | Q9UBB6; C9 | NCDN_HUMAN | Unmodified | 69117 | 0 | 94983 | 376230 | 0 | 0 | 0 | 0 | 0 | 0 | 172460 | 348620 | 687560 | 330610 | 168130 | 91583 | 1780200 | 700810 |
| IFDDSPCPI | SLC44A2 | Q8IWA5 | CTL2_HUMAN | Unmodified | 6561500 | 3772000 | 6609000 | 1798200 | 0 | 508260 | 0 | 0 | 7026400 | 1527700 | 1454000 | 17888000 | 147270 | 0 | 9225500 | 8986900 | 0 | 0 |
| IFDEILVNA | TOP2B; TOP2A; | Q02880; P11 | TOP2B_HUMAN | Unmodified | 168600 | 85393 | 0 | 0 | 0 | 0 | 287750 | 0 | 175890 | 144430 | 144470 | 1017100 | 53299 | 56388 | 0 | 33638 | 0 | 0 |
| IFDFDDDG | CIB1 | Q99828 | CIB1_HUMAN | Unmodified | 336380 | 353930 | 0 | 186430 | 138240 | 102580 | 0 | 0 | 378420 | 0 | 179600 | 643470 | 0 | 0 | 318230 | 231840 | 0 | 0 |
| IFDIDEAEE | EIF2S2 | P20042 | IF2B_HUMAN E | Unmodified | 0 | 0 | 423030 | 0 | 812240 | 0 | 0 | 0 | 802600 | 0 | 1786900 | 0 | 0 | 0 | 662040 | 0 | 0 | 0 |
| IFDPQNPD | CRKL | P46109 | CRKL_HUMAN | Unmodified | 675330 | 0 | 438200 | 244320 | 3421600 | 691290 | 236370 | 184470 | 243680 | 0 | 648020 | 1038100 | 581360 | 368730 | 0 | 312850 | 705130 | 845220 |
| IFDPRVDIV | GGCX | P38435 | VKGC_HUMAN | Unmodified | 0 | 0 | 0 | 0 | 0 | 66531 | 0 | 0 | 3884900 | 0 | 0 | 0 | 0 | 0 | 0 | 0 | 25312 | 0 |
| IFDSEEILA | NARS1 | O43776 | SYNC_HUMAN | Unmodified | 0 | 0 | 0 | 0 | 0 | 0 | 0 | 0 | 0 | 0 | 0 | 1638500 | 0 | 0 | 0 | 125900 | 0 | 0 |
| IFDSEEILA | NARS1 | O43776 | SYNC_HUMAN | Unmodified | 659320 | 0 | 0 | 2740300 | 0 | 0 | 0 | 0 | 3737800 | 0 | 0 | 19420000 | 0 | 0 | 2782900 | 1430400 | 258300 | 0 |
| IFDTSSGHI | WDR45B | Q5MNZ6 | WIPI3_HUMAN | Unmodified | 212750 | 160310 | 93994 | 0 | 0 | 0 | 241510 | 0 | 360890 | 0 | 129330 | 678910 | 0 | 0 | 487210 | 548850 | 0 | 0 |
| IFDTSSGHI | WDR45B | Q5MNZ6 | WIPI3_HUMAN | Unmodified | 171050 | 166780 | 0 | 236950 | 0 | 45518 | 0 | 0 | 0 | 0 | 167000 | 616140 | 0 | 0 | 627470 | 251640 | 0 | 0 |
| IFDYSPLDF | USP13; USP13; | Q92995; A0A | UBP13_HUMAN | Oxidation (M) | 1135400 | 1138000 | 0 | 946660 | 0 | 0 | 0 | 0 | 0 | 0 | 0 | 239470 | 578180 | 0 | 0 | 0 | 0 | 0 |
| IFEDPTDSD | RASAL2; RASAL | Q9UJF2; H0 | NGAP_HUMAN | Unmodified | 0 | 0 | 0 | 0 | 2186900 | 353990 | 0 | 47745 | 266420 | 258460 | 169040 | 467880 | 0 | 232290 | 0 | 0 | 191200 | 227400 |
| IFEEQGQT | PIK3R2 | O00459 | P85B_HUMAN | Unmodified | 0 | 0 | 96435 | 68261 | 40443 | 72490 | 0 | 0 | 0 | 0 | 0 | 160320 | 0 | 0 | 55017 | 82827 | 0 | 0 |
| IFEGTNDIL | ACADVL; ACAD | P49748; J3Q | ACADV_HUMAI | Unmodified | 0 | 0 | 0 | 0 | 0 | 0 | 0 | 0 | 3486900 | 779070 | 383160 | 0 | 0 | 0 | 0 | 0 | 252990 | 0 |
| IFEIFGPVAI | NAF1 | Q96HR8 | NAF1_HUMAN | Unmodified | 0 | 0 | 0 | 0 | 0 | 0 | 0 | 0 | 18106 | 0 | 0 | 91249 | 0 | 0 | 0 | 0 | 0 | 0 |
| IFFAGDTIPI | FLNB; FLNB | O75369; E7E | FLNB_HUMAN | Unmodified | 0 | 0 | 0 | 0 | 13590000 | 0 | 0 | 0 | 0 | 21846 | 549920 | 6387800 | 227030 | 0 | 0 | 466000 | 5912400 | 3976500 |
| IFFAGTETA | MAOA | P21397 | AOFA_HUMAN | Unmodified | 0 | 0 | 0 | 0 | 0 | 0 | 0 | 0 | 0 | 0 | 0 | 0 | 0 | 0 | 1635300 | 401030 | 0 | 0 |
| IFFENHPYF | MX1; MX1; MX1 | P20591; A0A | MX1_HUMAN I | Unmodified | 0 | 0 | 0 | 0 | 0 | 0 | 0 | 0 | 0 | 0 | 0 | 0 | 0 | 0 | 0 | 0 | 0 | 1035900 |
| IFFVNPAPP | AGRN | O00468 | AGRIN_HUMAN | Unmodified | 1460900 | 0 | 2046600 | 0 | 0 | 0 | 0 | 0 | 258710 | 0 | 0 | 0 | 0 | 442080 | 950020 | 486360 | 0 | 0 |
| IFGGEIKTH | P4HB; P4HB; P4 | P07237; A0A | PDIA1_HUMAN | Unmodified | 0 | 0 | 0 | 0 | 0 | 0 | 578000 | 0 | 0 | 2718100 | 0 | 0 | 0 | 0 | 0 | 0 | 0 | 0 |
| IFGGLDML | ANP32B | Q92688 | AN32B_HUMAI | Oxidation (M) | 0 | 0 | 0 | 0 | 0 | 0 | 0 | 0 | 0 | 0 | 266220 | 9135200 | 0 | 0 | 0 | 0 | 0 | 0 |
| IFGGLDML | ANP32B | Q92688 | AN32B_HUMAI | Oxidation (M) | 0 | 35937 | 0 | 341910 | 116790 | 0 | 0 | 0 | 392210 | 164340 | 356500 | 1087200 | 0 | 0 | 0 | 0 | 0 | 0 |
| IFGGLDML | ANP32B | Q92688 | AN32B_HUMAI | Unmodified | 0 | 0 | 0 | 0 | 20801 | 0 | 0 | 0 | 141420 | 0 | 0 | 0 | 0 | 0 | 0 | 219820 | 0 | 0 |
| IFGGVQQL | CACNA2D2; CA | Q9NY47; C9 | CA2D2_HUMAI | Unmodified | 332310 | 0 | 0 | 0 | 0 | 0 | 0 | 0 | 12446000 | 0 | 605500 | 1330200 | 0 | 0 | 0 | 0 | 48442 | 0 |
| IFGSEAAW | ACADVL; ACAD | P49748; J3Q | ACADV_HUMAI | Unmodified | 0 | 0 | 0 | 0 | 0 | 0 | 0 | 0 | 5714500 | 197890 | 0 | 0 | 0 | 0 | 0 | 0 | 0 | 0 |
| IFGSNKWT | ARL6IP1; N/A; A | Q15041; H3I | AR6P1_HUMAN | Unmodified | 0 | 0 | 0 | 0 | 0 | 0 | 137050 | 0 | 371470 | 309820 | 0 | 817860 | 0 | 0 | 0 | 0 | 0 | 0 |
| IFGVTTLDI | MDH2; MDH2 | P40926; G3X | MDHM_HUMAN | Unmodified | 0 | 0 | 0 | 0 | 877770 | 0 | 1487700 | 0 | 91654000 | 13888000 | 5836400 | 3791500 | 1653800 | 0 | 0 | 0 | 254810 | 0 |
| IFGVTTLDI | MDH2; MDH2 | P40926; G3X | MDHM_HUMAN | Unmodified | 0 | 0 | 0 | 0 | 896800 | 416610 | 5281200 | 0 | 0 | 9650200 | 0 | 0 | 8068500 | 5298400 | 0 | 0 | 1289900 | 1786000 |
| IFHAGDCG | MGAT2 | Q10469 | MGAT2_HUMAI | Unmodified | 0 | 0 | 0 | 0 | 121360 | 0 | 0 | 0 | 288360 | 47788 | 0 | 0 | 0 | 0 | 0 | 0 | 0 | 0 |
| IFHELTQTD | MTHFD1; MTHF | P11586; V9G | C1TC_HUMAN | Unmodified | 360970 | 295390 | 899100 | 700970 | 1893800 | 419310 | 652060 | 355000 | 4380200 | 1777700 | 1462500 | 23555000 | 613310 | 198540 | 831600 | 789040 | 595260 | 578660 |
| IFHLEVCSF | STAM | Q92783 | STAM1_HUMAN | Unmodified | 72669 | 0 | 0 | 207120 | 1602400 | 0 | 581790 | 0 | 80249 | 1174000 | 0 | 0 | 3123500 | 570030 | 0 | 0 | 184630 | 904180 |
| IFHLEVCSF | STAM | Q92783 | STAM1_HUMAN | Unmodified | 0 | 0 | 0 | 0 | 1440700 | 0 | 561990 | 0 | 0 | 1295100 | 0 | 0 | 680660 | 0 | 0 | 0 | 301080 | 578550 |
| IFIAQPAW | DNAJC13 | O75165 | DJC13_HUMAN | Unmodified | 0 | 0 | 0 | 0 | 0 | 0 | 0 | 0 | 0 | 0 | 0 | 1155900 | 0 | 0 | 0 | 0 | 0 | 0 |
| IFIGTFK | SNRPB; SNRPN | P14678; P63 | RSMB_HUMAN | Unmodified | 2080400 | 1701200 | 1073000 | 1629400 | 845950 | 133220 | 198870 | 124710 | 1669800 | 1003000 | 5776100 | 11238000 | 1732200 | 435040 | 797020 | 904250 | 183410 | 504030 |
| IFILGPSHH | MEMO1 | Q9Y316 | MEMO1_HUMA | Unmodified | 0 | 404110 | 0 | 0 | 212690 | 0 | 0 | 0 | 0 | 0 | 182160 | 1153700 | 0 | 0 | 0 | 0 | 0 | 0 |
| IFINLPR | TXNL1; TXNL1 | O43396; K7E | TXNL1_HUMAN | Unmodified | 0 | 0 | 532330 | 0 | 351230 | 0 | 0 | 99870 | 1407600 | 545460 | 712020 | 1377200 | 0 | 352350 | 852440 | 784700 | 1163900 | 564220 |
| IFINNEWHI | ALDH1A1; ALDH | P00352; Q5S | AL1A1_HUMAN | Unmodified | 0 | 78864 | 0 | 0 | 0 | 0 | 501840 | 41216 | 5767800 | 1448900 | 52128000 | 9.55E+08 | 0 | 0 | 0 | 165950 | 0 | 0 |
| IFINNEWHI | ALDH1A1; ALDH | P00352; Q5S | AL1A1_HUMAN | Unmodified | 0 | 383050 | 0 | 0 | 0 | 0 | 0 | 0 | 0 | 0 | 622920 | 19150000 | 0 | 0 | 0 | 0 | 0 | 0 |
| IFINNEWHI | ALDH1A3; ALDH | P47895; H0Y | AL1A3_HUMAN | Unmodified | 0 | 0 | 0 | 0 | 1166300 | 0 | 0 | 0 | 249250 | 0 | 0 | 0 | 0 | 0 | 0 | 0 | 11422000 | 8742000 |
| IFINNEWHI | ALDH1A3; ALDH | P47895; H0Y | AL1A3_HUMAN | Unmodified | 0 | 0 | 0 | 0 | 147440 | 0 | 0 | 12994 | 0 | 25874 | 0 | 0 | 0 | 0 | 0 | 0 | 1127100 | 446950 |
| IFINNEWHI | ALDH1A3; ALDH | P47895; H0Y | AL1A3_HUMAN | Unmodified | 0 | 0 | 0 | 0 | 0 | 0 | 0 | 0 | 0 | 0 | 0 | 0 | 0 | 0 | 0 | 0 | 914110 | 437110 |
| IFIQKKEEAI | PARD6B | Q9BYG5 | PAR6B_HUMAN | Unmodified | 0 | 0 | 0 | 0 | 3948900 | 389420 | 1120700 | 0 | 0 | 0 | 0 | 0 | 0 | 0 | 0 | 0 | 0 | 0 |
| IFIYDGRGD | PPWD1; PPWD1 | Q96BP3; D6I | PPWD1_HUMA | Unmodified | 0 | 0 | 32439 | 0 | 294300 | 70695 | 185100 | 0 | 0 | 82045 | 130790 | 1482800 | 0 | 0 | 0 | 0 | 20950 | 17942 |
| IFKSQGAAI | MDH1; MDH1; I | P40925; C9J | MDHC_HUMAN | Unmodified | 0 | 0 | 0 | 0 | 0 | 0 | 138100 | 86946 | 0 | 0 | 0 | 0 | 0 | 0 | 0 | 258940 | 0 | 0 |
| IFLAQKPAP | AP3B1 | O00203 | AP3B1_HUMAN | Unmodified | 71035 | 58840 | 134400 | 91054 | 510720 | 66159 | 197460 | 28842 | 424830 | 0 | 336520 | 3491100 | 93681 | 0 | 179090 | 114440 | 42845 | 125230 |
| IFLEFERVES | RBM17 | Q96I25 | SPF45_HUMAN | Unmodified | 0 | 0 | 0 | 0 | 0 | 0 | 0 | 0 | 0 | 0 | 148670 | 3205700 | 0 | 0 | 0 | 0 | 0 | 0 |
| IFLGGFELK | MUC5AC | P98088 | MUC5A_HUMA | Unmodified | 0 | 0 | 0 | 94979 | 0 | 0 | 0 | 0 | 0 | 0 | 0 | 0 | 0 | 0 | 0 | 0 | 0 | 0 |
| IFLITLDNSI | UBE4A | Q14139 | UBE4A_HUMAN | Unmodified | 0 | 0 | 0 | 0 | 0 | 0 | 0 | 0 | 0 | 0 | 0 | 1473500 | 0 | 0 | 0 | 0 | 0 | 0 |
| IFLLTNNNL | MYO1B; MYO1E | O43795; H7 | MYO1B_HUMA | Unmodified | 297920 | 0 | 182060 | 374780 | 21197000 | 2715600 | 2089800 | 157400 | 2531400 | 4977400 | 5946500 | 15991000 | 1170100 | 0 | 813720 | 415460 | 2842000 | 4471300 |
| IFLQDIK | FTH1; FTH1; FTI | P02794; G3V | FRIH_HUMAN I | Unmodified | 18495 | 0 | 818260 | 173010 | 0 | 0 | 0 | 0 | 17770000 | 522420 | 0 | 2381500 | 0 | 0 | 1139100 | 482590 | 0 | 0 |
| IFLQDIKKPI | FTH1; FTH1; FTI | P02794; G3V | FRIH_HUMAN I | Unmodified | 0 | 0 | 0 | 0 | 0 | 0 | 0 | 0 | 5702500 | 5662000 | 0 | 0 | 0 | 0 | 0 | 0 | 0 | 926490 |
| IFLSHEELEI | ABCC1; ABCC1 | P33527; I3L4 | MRP1_HUMAN | Unmodified | 0 | 0 | 546270 | 0 | 0 | 0 | 0 | 0 | 177360 | 0 | 574490 | 7159300 | 0 | 0 | 0 | 465670 | 0 | 0 |
| IFLSHEELEI | ABCC1; ABCC1 | P33527; I3L4 | MRP1_HUMAN | Unmodified | 0 | 0 | 0 | 0 | 0 | 0 | 0 | 0 | 0 | 0 | 0 | 1555600 | 353660 | 0 | 0 | 327150 | 0 | 0 |
| IFLSTFQSE | GPR39 | O43194 | GPR39_HUMAI | Unmodified | 0 | 0 | 0 | 0 | 763650 | 168270 | 92516 | 0 | 0 | 0 | 0 | 0 | 136290 | 0 | 0 | 0 | 372610 | 2531000 |
| IFNLYPR | CRMP1; DPYS; | Q14194; Q14 | DPYL1_HUMAN | Unmodified | 0 | 0 | 0 | 0 | 0 | 0 | 0 | 408850 | 0 | 35135 | 0 | 52060000 | 0 | 0 | 0 | 0 | 0 | 0 |
| IFNLYPRK | CRMP1; DPYS; | Q14194; Q14 | DPYL1_HUMAN | Unmodified | 0 | 0 | 0 | 0 | 0 | 0 | 275860 | 0 | 0 | 0 | 0 | 1979400 | 0 | 0 | 0 | 0 | 0 | 0 |
| IFNTFGPR | UXS1; UXS1; U | Q8NBZ7; C9 | UXS1_HUMAN | Unmodified | 0 | 0 | 0 | 0 | 0 | 0 | 0 | 0 | 3868000 | 778010 | 0 | 0 | 0 | 0 | 0 | 0 | 0 | 0 |
| IFNTNNLW | UGP2; UGP2; U | Q16851; E7E | UGPA_HUMAN | Unmodified | 0 | 0 | 0 | 0 | 1338100 | 0 | 0 | 0 | 0 | 1106600 | 0 | 0 | 496350 | 583140 | 0 | 0 | 334700 | 375080 |
| IFPATEK | CALCOCO2 | Q13137 | CACO2_HUMA | Unmodified | 0 | 0 | 0 | 0 | 0 | 0 | 0 | 0 | 350960 | 61225 | 0 | 0 | 236120 | 0 | 0 | 0 | 216660 | 1384300 |
| IFPGLAAD | SNTB2; SNTB2; | Q13425; A0A | SNTB2_HUMAN | Unmodified | 132980 | 120130 | 1809500 | 0 | 0 | 0 | 0 | 0 | 0 | 0 | 706760 | 602380 | 2526300 | 1403400 | 240400 | 145710 | 0 | 0 |
| IFPGNWDN | MFGE8 | Q08431 | MFGM_HUMAN | Unmodified | 0 | 145270 | 1453700 | 0 | 2796000 | 0 | 69385 | 0 | 0 | 0 | 0 | 0 | 0 | 0 | 0 | 0 | 0 | 0 |
| IFPGNWDN | MFGE8 | Q08431 | MFGM_HUMAN | Unmodified | 283010 | 255000 | 9083000 | 2018900 | 15700000 | 2796200 | 0 | 0 | 910410 | 328060 | 0 | 855600 | 931350 | 0 | 1187700 | 764410 | 439150 | 3744800 |
| IFPLLGLHE | COG4; COG4; C | Q9H9E3; A0A | COG4_HUMAN | Unmodified | 0 | 0 | 93260 | 0 | 0 | 0 | 0 | 0 | 0 | 0 | 0 | 335740 | 0 | 0 | 0 | 0 | 0 | 0 |
| IFQAENAK | EXOC8 | Q8IYI6 | EXOC8_HUMAI | Unmodified | 0 | 0 | 0 | 0 | 381990 | 263250 | 75849 | 21309 | 350280 | 443610 | 0 | 146450 | 0 | 74860 | 0 | 0 | 292000 | 21890 |
| IFQAGVVS | ST14 | Q9Y5Y6 | ST14_HUMAN | Unmodified | 527210 | 866490 | 444240 | 1915900 | 0 | 0 | 0 | 0 | 0 | 0 | 0 | 0 | 0 | 0 | 3026100 | 2504500 | 0 | 0 |
| IFQAHKNC | TLN1 | Q9Y490 | TLN1_HUMAN | Unmodified | 0 | 0 | 0 | 0 | 131390 | 0 | 333240 | 527530 | 0 | 0 | 0 | 0 | 0 | 70613 | 0 | 0 | 94366 | 21573 |
| IFQEALVQI | NIBAN1; NIBAN | Q9BZQ8; H0 | NIBA1_HUMAN | Unmodified | 0 | 0 | 0 | 92614 | 506750 | 0 | 0 | 0 | 168730 | 1569200 | 0 | 68315 | 0 | 0 | 0 | 0 | 0 | 51587 |
| IFQGNVHN | SNRNP40; SNR | Q96DI7; Q9 | SNR40_HUMAI | Unmodified | 186740 | 0 | 0 | 0 | 0 | 0 | 0 | 0 | 0 | 2360200 | 0 | 1131000 | 0 | 0 | 0 | 141610 | 0 | 0 |
| IFQKGESP | PDIA6 | Q15084 | PDIA6_HUMAN | Unmodified | 142670 | 0 | 0 | 58682 | 27844 | 0 | 61788 | 0 | 39969000 | 62685 | 2756700 | 1901200 | 0 | 0 | 151030 | 0 | 0 | 0 |
| IFQKGESP | PDIA6 | Q15084 | PDIA6_HUMAN | Unmodified | 102200 | 96334 | 0 | 0 | 0 | 0 | 0 | 0 | 51030000 | 6083000 | 4910900 | 1045700 | 0 | 0 | 196200 | 124350 | 0 | 0 |
| IFQNAPTDI | USP5 | P45974 | UBP5_HUMAN | Unmodified | 5328300 | 8797300 | 0 | 3266100 | 4586200 | 0 | 2405200 | 538310 | 3153400 | 1333300 | 1035600 | 5424100 | 16040000 | 6048000 | 2350000 | 0 | 3323400 | 8711000 |
| IFQNLDGA | PLOD1 | Q02809 | PLOD1_HUMAI | Unmodified | 0 | 0 | 32759 | 66146 | 0 | 17917 | 0 | 26745 | 8523700 | 685310 | 177570 | 165330 | 0 | 0 | 0 | 41692 | 0 | 0 |
| IFQNLNGA | PLOD3; PLOD3 | O60568; H7 | PLOD3_HUMAI | Unmodified | 0 | 0 | 0 | 0 | 1457200 | 242120 | 0 | 0 | 7767200 | 0 | 0 | 0 | 0 | 0 | 0 | 0 | 1931000 | 0 |
| IFQNLNGA | PLOD3; PLOD3 | O60568; H7 | PLOD3_HUMAI | Unmodified | 0 | 0 | 0 | 0 | 0 | 0 | 0 | 0 | 1262700 | 113420 | 0 | 0 | 0 | 0 | 0 | 0 | 0 | 0 |
| IFQTLNGA | PLOD2; PLOD2; | O00469; C9J | PLOD2_HUMAI | Unmodified | 0 | 0 | 0 | 0 | 0 | 0 | 0 | 0 | 4015400 | 166900 | 176110 | 140940 | 0 | 0 | 0 | 0 | 0 | 0 |
| IFQTLNGA | PLOD2; PLOD2; | O00469; C9J | PLOD2_HUMAI | Unmodified | 0 | 0 | 0 | 0 | 522200 | 106390 | 0 | 0 | 244040 | 0 | 0 | 0 | 0 | 0 | 0 | 0 | 0 | 0 |
| IFQVAQELS | TSNAX; TSNAX; | Q99598; Q5 | TSNAX_HUMAN | Unmodified | 0 | 1421500 | 0 | 0 | 0 | 0 | 0 | 0 | 922910 | 425480 | 0 | 1625700 | 0 | 0 | 2975400 | 0 | 0 | 0 |
| IFRDGEEA | PDIA3; PDIA3 | P30101; H7E | PDIA3_HUMAN | Unmodified | 93181 | 0 | 2212400 | 1641700 | 609240 | 96477 | 252510 | 0 | 3.51E+08 | 17259000 | 43598000 | 15414000 | 1780000 | 937940 | 3044300 | 2367300 | 61378 | 1176500 |
| IFRDGEEA | PDIA3 | P30101 | PDIA3_HUMAN | Unmodified | 0 | 0 | 0 | 0 | 10700000 | 1077000 | 15775000 | 1725200 | 13341000 | 84477000 | 11539000 | 0 | 6192900 | 7182100 | 0 | 0 | 3493100 | 9506600 |
| IFRIFEGTNI | ACADVL; ACAD | P49748; J3Q | ACADV_HUMAI | Unmodified | 0 | 0 | 1134400 | 0 | 0 | 0 | 0 | 0 | 311600 | 601760 | 0 | 0 | 0 | 0 | 1098600 | 0 | 0 | 0 |
| IFRKQENAI | PNMA2 | Q9UL42 | PNMA2_HUMA | Unmodified | 0 | 1041600 | 0 | 0 | 0 | 0 | 0 | 0 | 0 | 0 | 0 | 0 | 0 | 0 | 0 | 0 | 0 | 0 |
| IFRQMDTN | HPCA; HPCAL1 | P84074; P37 | HPCA_HUMAN | Unmodified | 0 | 0 | 0 | 0 | 762420 | 165940 | 49002 | 12712 | 531980 | 0 | 84580 | 0 | 0 | 0 | 12972 | 0 | 83630 | 489180 |
| IFSAYIK | ARPC1B; ARPC | O15143; C9J | ARC1B_HUMAI | Unmodified | 1643400 | 988800 | 0 | 642120 | 152100 | 112190 | 420190 | 0 | 1232600 | 225850 | 0 | 10552000 | 548770 | 0 | 1293600 | 1307800 | 0 | 0 |

| | | | | | | | | | | | | | | | | | | | | | | |
|---|---|---|---|---|---|---|---|---|---|---|---|---|---|---|---|---|---|---|---|---|---|---|
| IFSEPGSLS | GRTP1 | Q5TC63 | GRTP1_HUMAN | Oxidation (M) | 0 | 0 | 295180 | 0 | 0 | 0 | 0 | 0 | 0 | 0 | 0 | 0 | 0 | 0 | 620220 | 455770 | 0 | 0 |
| IFSFAHQTI | VPS35 | Q96QK1 | VPS35_HUMAN | Unmodified | 0 | 0 | 34071 | 0 | 0 | 0 | 0 | 0 | 106650 | 75663 | 37921 | 2260200 | 0 | 0 | 179370 | 0 | 0 | 0 |
| IFSFDGK | COL18A1; COL | P39060; H7C | COIA1_HUMAN | Unmodified | 0 | 0 | 113950 | 58441 | 0 | 417370 | 0 | 0 | 777460 | 66432 | 80663 | 0 | 0 | 0 | 39034 | 211690 | 0 | 0 |
| IFSFDGKD | COL18A1; COL | P39060; H7C | COIA1_HUMAN | Unmodified | 0 | 0 | 0 | 0 | 0 | 4326900 | 241740 | 0 | 0 | 906960 | 0 | 0 | 0 | 0 | 0 | 0 | 0 | 2209100 |
| IFSGPSSEC | ITGA2; ITGA2; I | P17301; D6F | ITA2_HUMAN I | Unmodified | 5950500 | 121670 | 2419700 | 9075000 | 97011000 | 12025000 | 8298400 | 0 | 891540 | 0 | 0 | 842660 | 10754000 | 10927000 | 10922000 | 5854800 | 48263000 | 1.15E+08 |
| IFSGSSHQ | PRPS1; PRPS1L | P60891; P21 | PRPS1_HUMAN | Unmodified | 300660 | 213390 | 732650 | 1071300 | 4245500 | 863960 | 4157100 | 2364600 | 0 | 1233200 | 1842800 | 9091000 | 12789000 | 3677600 | 1114600 | 1044900 | 3292500 | 4380200 |
| IFSGSSHQ | PRPS1; PRPS1L | P60891; P21 | PRPS1_HUMAN | Unmodified | 0 | 0 | 0 | 0 | 0 | 0 | 3862400 | 0 | 0 | 131590 | 0 | 0 | 696660 | 1783100 | 0 | 0 | 2015800 | 394960 |
| IFSIIAEGEN | IPO5; IPO5 | O00410; H0 | IPO5_HUMAN I | Unmodified | 0 | 0 | 0 | 0 | 1228800 | 0 | 0 | 0 | 0 | 0 | 0 | 1216600 | 570290 | 0 | 0 | 0 | 1055400 | 0 |
| IFSNTDPDI | LACTB2 | Q53H82 | LACB2_HUMAN | Unmodified | 0 | 0 | 0 | 0 | 138320 | 0 | 0 | 0 | 2703600 | 61719 | 0 | 101700 | 0 | 0 | 0 | 34750 | 0 | 0 |
| IFSNWGHF | SUN1; SUN1; S | O94901; H0 | SUN1_HUMAN | Unmodified | 0 | 0 | 0 | 0 | 0 | 0 | 0 | 0 | 4347000 | 0 | 0 | 0 | 0 | 0 | 0 | 0 | 0 | 0 |
| IFSPNVVNI | DNM1L; DNM1I | O00429; B4I | DNM1L_HUMA | Oxidation (M) | 0 | 0 | 0 | 0 | 0 | 0 | 0 | 0 | 0 | 0 | 71195 | 432450 | 0 | 0 | 251310 | 0 | 0 | 0 |
| IFSQETLTK | PCYOX1 | Q9UHG3 | PCYOX_HUMAI | Unmodified | 0 | 0 | 0 | 0 | 0 | 0 | 324420 | 0 | 15538000 | 8454800 | 0 | 0 | 0 | 283550 | 0 | 0 | 72401 | 0 |
| IFSSVQWL | UBE2O; UBE2O | Q9C0C9; K7 | UBE2O_HUMAI | Unmodified | 0 | 0 | 213160 | 325920 | 1745500 | 281280 | 0 | 0 | 500250 | 0 | 493740 | 6926200 | 1146600 | 611010 | 461050 | 230960 | 2207100 | 0 |
| IFSVGVSVL | KHNYN; KHNYN | O15037; G3 | KHNYN_HUMA | Unmodified | 467630 | 600560 | 136770 | 0 | 0 | 0 | 0 | 0 | 0 | 0 | 0 | 0 | 0 | 0 | 0 | 364000 | 0 | 0 |
| IFSYIAR | NOS1AP; NOS1 | O75052; E9F | CAPON_HUMA | Unmodified | 0 | 0 | 143220 | 97965 | 0 | 360300 | 0 | 0 | 669890 | 0 | 0 | 0 | 0 | 390060 | 0 | 233540 | 0 | 0 |
| IFTAEELEAI | HSPA13 | P48723 | HSP13_HUMAI | Unmodified | 179140 | 0 | 0 | 0 | 1125400 | 0 | 0 | 0 | 13543000 | 4610900 | 575580 | 0 | 0 | 0 | 0 | 185970 | 0 | 642780 |
| IFTFAEK | YARS1; YARS1; | P54577; A0A | SYYC_HUMAN | Unmodified | 0 | 0 | 0 | 584620 | 4361300 | 372280 | 0 | 1039800 | 2772600 | 708750 | 0 | 3250400 | 6213100 | 6536200 | 1027200 | 1037600 | 2589900 | 3711300 |
| IFTFICK | GULP1 | Q9UBP9 | GULP1_HUMAI | Unmodified | 0 | 0 | 0 | 297370 | 0 | 0 | 0 | 0 | 0 | 0 | 186300 | 0 | 540350 | 0 | 0 | 202380 | 0 | 0 |
| IFTIESTR | DYNC1H1; DYN | Q14204; A0A | DYHC1_HUMA | Unmodified | 132470 | 0 | 0 | 0 | 0 | 0 | 0 | 0 | 0 | 0 | 0 | 5011800 | 0 | 0 | 0 | 377480 | 0 | 0 |
| IFTSIGEDYI | PHB; PHB; PHB | P35232; C9J | PHB_HUMAN F | Unmodified | 0 | 0 | 0 | 0 | 0 | 0 | 0 | 0 | 409550 | 0 | 0 | 0 | 0 | 673680 | 0 | 0 | 0 | 0 |
| IFTTGDVNI | SLC25A24; SLC | Q6NUK1; J3I | SCMC1_HUMA | Oxidation (M) | 0 | 0 | 0 | 0 | 0 | 0 | 42094 | 0 | 2414000 | 1247300 | 0 | 0 | 0 | 0 | 0 | 0 | 0 | 0 |
| IFTVAQVDI | SLC12A7; SLC1 | Q9Y666; A0A | S12A7_HUMAN | Oxidation (M) | 0 | 0 | 166470 | 149220 | 0 | 0 | 286240 | 0 | 349500 | 121890 | 0 | 0 | 0 | 0 | 497970 | 585210 | 0 | 0 |
| IFVAELKDP | HELZ2 | Q9BYK8 | HELZ2_HUMAN | Unmodified | 0 | 0 | 61580 | 229790 | 0 | 0 | 0 | 0 | 0 | 0 | 0 | 0 | 0 | 0 | 207650 | 153720 | 1022600 | 0 |
| IFVEESIYDI | ALDH1A1 | P00352 | AL1A1_HUMAN | Unmodified | 0 | 0 | 0 | 0 | 0 | 0 | 0 | 0 | 0 | 0 | 75117 | 384690 | 0 | 0 | 68623 | 0 | 0 | 0 |
| IFVEESIYDI | ALDH1A1 | P00352 | AL1A1_HUMAN | Unmodified | 0 | 0 | 0 | 0 | 0 | 0 | 29897 | 19978 | 3236800 | 1268800 | 27257000 | 4.26E+08 | 0 | 0 | 0 | 0 | 0 | 0 |
| IFVFLEHQT | DSC2 | Q02487 | DSC2_HUMAN | Unmodified | 0 | 0 | 0 | 0 | 0 | 0 | 0 | 0 | 3183200 | 648030 | 0 | 0 | 0 | 0 | 0 | 0 | 0 | 0 |
| IFVGGIKED | HNRNPA1L2; H | Q32P51; P0 | RA1L2_HUMAN | Unmodified | 0 | 0 | 0 | 0 | 0 | 0 | 2315900 | 0 | 0 | 0 | 0 | 5212100 | 0 | 0 | 0 | 0 | 0 | 0 |
| IFVGGIKED | HNRNPA3; HNI | P51991; A0A | ROA3_HUMAN | Unmodified | 0 | 0 | 0 | 0 | 0 | 0 | 0 | 0 | 0 | 68191 | 0 | 3481300 | 0 | 0 | 0 | 29514 | 191400 | 0 |
| IFVGGIKED | HNRNPA3; HNI | P51991; A0A | ROA3_HUMAN | Unmodified | 0 | 0 | 0 | 0 | 5186600 | 0 | 19009000 | 261380 | 2225500 | 4678000 | 7265300 | 7813200 | 11974000 | 5119900 | 0 | 349940 | 1359700 | 10031000 |
| IFVGGIPHN | DAZAP1; DAZAF | Q96EP5; K7E | DAZP1_HUMAI | Unmodified | 0 | 147960 | 47789 | 0 | 0 | 0 | 0 | 0 | 351670 | 0 | 186430 | 4619100 | 0 | 605710 | 63319 | 0 | 0 | 0 |
| IFVGGIPHN | DAZAP1; DAZAF | Q96EP5; K7E | DAZP1_HUMAI | Unmodified | 0 | 0 | 0 | 0 | 590600 | 0 | 1975900 | 0 | 1067000 | 961020 | 1328000 | 3632000 | 2529700 | 789650 | 0 | 0 | 309610 | 949310 |
| IFVGGLNPI | HNRNPAB; HNI | D6R9P3; D6I | D6R9P3_HUM/ | Unmodified | 432390 | 3197100 | 0 | 0 | 1718500 | 0 | 109300 | 718980 | 4173600 | 2061700 | 5810700 | 31149000 | 2027700 | 0 | 0 | 2831800 | 0 | 0 |
| IFVGGLPYI | RBM24; RBM38 | Q9BX46; Q9I | RBM24_HUMAI | Unmodified | 0 | 0 | 0 | 0 | 0 | 0 | 0 | 0 | 0 | 0 | 0 | 1203100 | 0 | 0 | 0 | 0 | 0 | 0 |
| IFVGGLPYI | RBM24; RBM38 | Q9BX46; Q9I | RBM24_HUMAI | Unmodified | 0 | 0 | 0 | 0 | 0 | 0 | 0 | 0 | 83807 | 0 | 274040 | 643410 | 0 | 0 | 0 | 0 | 0 | 0 |
| IFVGGLSPI | HNRNPD; HNR | Q14103; H0 | HNRPD_HUMA | Unmodified | 554000 | 0 | 155300 | 0 | 3047900 | 711280 | 0 | 1152500 | 3796300 | 1934300 | 11743000 | 42327000 | 3148800 | 988580 | 0 | 168180 | 0 | 941540 |
| IFVGTKGIP | RPS4X | P62701 | RS4X_HUMAN | Unmodified | 0 | 0 | 0 | 0 | 0 | 0 | 793720 | 0 | 0 | 0 | 0 | 0 | 830760 | 0 | 0 | 0 | 40183 | 114160 |
| IFVLGPKYS | EFL1 | Q7Z2Z2 | EFL1_HUMAN I | Unmodified | 0 | 0 | 0 | 0 | 50895 | 0 | 0 | 0 | 0 | 57528 | 0 | 0 | 0 | 0 | 0 | 0 | 51219 | 0 |
| IFVMATHGI | PRPSAP2; PRPS | O60256; I3L | KPRB_HUMAN | Unmodified | 0 | 0 | 0 | 0 | 410910 | 0 | 0 | 0 | 668190 | 0 | 193270 | 1297200 | 0 | 0 | 0 | 0 | 399280 | 0 |
| IFVMATHGI | PRPSAP2; PRPS | O60256; I3L | KPRB_HUMAN | Oxidation (M) | 57205 | 0 | 0 | 143060 | 0 | 0 | 0 | 0 | 425840 | 126580 | 0 | 2364100 | 233350 | 0 | 131650 | 202010 | 0 | 0 |
| IFVNDDR | ILF3; ILF3; ILF3; | Q12906; K7E | ILF3_HUMAN I | Unmodified | 369890 | 288630 | 0 | 0 | 72540 | 0 | 142710 | 28802 | 0 | 107620 | 203680 | 413930 | 114340 | 0 | 0 | 0 | 125260 | 0 |
| IFVNDDRH | ILF3; ILF3; ILF3; | Q12906; K7E | ILF3_HUMAN I | Unmodified | 1983800 | 0 | 0 | 972240 | 0 | 0 | 0 | 0 | 0 | 0 | 624260 | 2366200 | 0 | 120440 | 0 | 0 | 43390 | 0 |
| IFVTTKNSK | VCL | P18206 | VINC_HUMAN | Unmodified | 0 | 0 | 350420 | 0 | 0 | 0 | 851190 | 509760 | 0 | 89898 | 0 | 2172000 | 0 | 0 | 0 | 594060 | 3590500 | 0 |
| IFYDRAEYL | MOV10; MOV10 | Q9HCE1; Q5 | MOV10_HUMA | Unmodified | 1885500 | 1801000 | 0 | 702900 | 0 | 0 | 0 | 0 | 0 | 0 | 0 | 0 | 0 | 0 | 843640 | 1245300 | 0 | 0 |
| IFYFSVPPF | H6PD | O95479 | G6PE_HUMAN | Unmodified | 0 | 0 | 0 | 0 | 0 | 0 | 0 | 0 | 128630 | 13011 | 0 | 0 | 0 | 0 | 0 | 0 | 0 | 0 |
| IFYPEIEEV | DUT; DUT; DUT | P33316; H0 | DUT_HUMAN D | Unmodified | 0 | 0 | 2455500 | 0 | 0 | 0 | 0 | 0 | 676210 | 138640 | 2041400 | 777330 | 0 | 0 | 0 | 0 | 0 | 0 |
| IFYPEIEEV | DUT; DUT; DUT | P33316; H0 | DUT_HUMAN D | Unmodified | 0 | 0 | 0 | 0 | 6254800 | 0 | 0 | 0 | 4669800 | 0 | 3000200 | 16494000 | 0 | 0 | 0 | 0 | 0 | 0 |
| IFYPETTDI | IQGAP1; IQGAP | P46940; A0A | IQGA1_HUMAN | Unmodified | 2101700 | 1281000 | 1316600 | 3013100 | 50308000 | 11934000 | 2843500 | 1677200 | 5661500 | 3439900 | 1687400 | 15078000 | 5406700 | 5976100 | 2696000 | 2444000 | 5354800 | 2969900 |
| IFYPETTDV | IQGAP2; IQGAP | Q13576; D6I | IQGA2_HUMAN | Unmodified | 613210 | 249560 | 2145500 | 2800700 | 0 | 0 | 0 | 0 | 0 | 0 | 323910 | 0 | 1765500 | 0 | 2297100 | 1430500 | 0 | 0 |
| IFYSITGPG | CDH3; CDH3 | P22223; J3Q | CADH3_HUMA | Unmodified | 0 | 0 | 0 | 0 | 12785000 | 3378300 | 0 | 0 | 0 | 194430 | 0 | 0 | 0 | 0 | 0 | 0 | 0 | 673450 |
| IFYVSHDS | NOS1AP; NOS1 | O75052; E9F | CAPON_HUMA | Unmodified | 21491 | 0 | 106750 | 138650 | 0 | 0 | 0 | 0 | 594310 | 0 | 0 | 0 | 0 | 0 | 220790 | 214660 | 0 | 0 |
| IGAADYQP | GNAO1; GNAO | P09471; H3E | GNAO_HUMAN | Unmodified | 0 | 0 | 9067.6 | 0 | 0 | 0 | 0 | 38312 | 1729300 | 114510 | 31226 | 0 | 0 | 0 | 0 | 0 | 0 | 0 |
| IGACPSAH | PSAP; PSAP; PS | P07602; Q5E | SAP_HUMAN P | Unmodified | 0 | 0 | 0 | 0 | 313530 | 685870 | 1625800 | 1122400 | 9979100 | 6775900 | 0 | 0 | 879390 | 1521600 | 0 | 0 | 213370 | 1686800 |
| IGADFLAR | GOT1 | P17174 | AATC_HUMAN | Unmodified | 1075700 | 536670 | 1068500 | 1665600 | 739550 | 0 | 0 | 0 | 2793300 | 634410 | 5019700 | 29843000 | 1611600 | 222260 | 2023600 | 2468200 | 0 | 0 |
| IGAEVYHNI | ENO1; ENO1; E | P06733; K7E | ENOA_HUMAN | Unmodified | 599450 | 636520 | 1229800 | 2686200 | 23431000 | 4052200 | 619620 | 466210 | 14624000 | 4163200 | 3161300 | 37317000 | 10181000 | 0 | 2308400 | 1872800 | 4201900 | 10348000 |
| IGAEVYHNI | ENO1; ENO1 | P06733; A0A | ENOA_HUMAN | Unmodified | 0 | 0 | 0 | 0 | 281150 | 0 | 210300 | 0 | 118060 | 0 | 0 | 277660 | 0 | 296600 | 0 | 0 | 1099900 | 387410 |
| IGAEVYHNI | ENO1; ENO1 | P06733; A0A | ENOA_HUMAN | Unmodified | 0 | 0 | 0 | 0 | 0 | 0 | 0 | 0 | 0 | 135210 | 0 | 0 | 0 | 0 | 0 | 0 | 1494800 | 468960 |
| IGAFGYME | RHOA; RHOA | P61586; A0A | RHOA_HUMAN | Unmodified | 1662100 | 1733200 | 841080 | 0 | 1753400 | 342400 | 238930 | 77388 | 10361000 | 769970 | 4047900 | 17386000 | 599500 | 460020 | 1660300 | 1272400 | 574700 | 1036900 |
| IGAFGYME | RHOA; RHOA | P61586; A0A | RHOA_HUMAN | Oxidation (M) | 1811500 | 1458100 | 1251700 | 1096200 | 325290 | 266540 | 0 | 0 | 2476900 | 601100 | 1370200 | 7194200 | 270510 | 129820 | 1440200 | 1815000 | 94169 | 221220 |
| IGALEGYR | NARS1; NARS1 | O43776; K7E | SYNC_HUMAN | Unmodified | 555210 | 691970 | 252090 | 168490 | 0 | 108700 | 0 | 172600 | 781940 | 312850 | 0 | 7504500 | 81219 | 289030 | 505350 | 346910 | 333490 | 74504 |
| IGALEGYR | NARS1; NARS1 | O43776; K7E | SYNC_HUMAN | Unmodified | 0 | 119340 | 302190 | 0 | 91744 | 47168 | 133120 | 0 | 608110 | 287960 | 720570 | 2838700 | 172460 | 220330 | 507420 | 327700 | 0 | 53457 |
| IGALGFSG | JAK2 | O60674 | JAK2_HUMAN | Unmodified | 0 | 0 | 0 | 163790 | 0 | 0 | 0 | 496930 | 0 | 0 | 0 | 142180 | 0 | 0 | 0 | 167880 | 0 | 0 |
| IGARVYALF | PLS3; PLS3 | P13797; A0A | PLST_HUMAN I | Unmodified | 0 | 0 | 0 | 0 | 5462300 | 0 | 0 | 0 | 0 | 0 | 0 | 0 | 438030 | 0 | 0 | 0 | 7803700 | 0 |
| IGARVYALF | LCP1 | P13796 | PLSL_HUMAN I | Unmodified | 0 | 0 | 0 | 0 | 0 | 0 | 347360 | 24504 | 0 | 0 | 0 | 0 | 0 | 0 | 0 | 0 | 587280 | 0 |
| IGASTLLSD | DERA; DERA; D | Q9Y315; G3 | DEOC_HUMAN | Unmodified | 0 | 0 | 0 | 0 | 0 | 0 | 0 | 0 | 0 | 0 | 0 | 1646400 | 0 | 0 | 0 | 0 | 0 | 0 |
| IGAVNCGD | DNAJC10; DNA | Q8IXB1; A0A | DJC10_HUMAN | Unmodified | 0 | 0 | 0 | 0 | 0 | 0 | 0 | 0 | 5944000 | 71193 | 85322 | 0 | 0 | 0 | 0 | 0 | 0 | 0 |
| IGAVNCGD | DNAJC10; DNA | Q8IXB1; A0A | DJC10_HUMAN | Unmodified | 0 | 0 | 0 | 0 | 0 | 0 | 0 | 0 | 375420 | 706420 | 0 | 0 | 0 | 0 | 0 | 0 | 0 | 0 |
| IGCGEWKF | CLGN | O14967 | CLGN_HUMAN | Oxidation (M) | 0 | 0 | 0 | 0 | 0 | 347360 | 0 | 0 | 3030300 | 1361100 | 979030 | 0 | 0 | 0 | 0 | 0 | 0 | 0 |
| IGCGEWKF | CLGN | O14967 | CLGN_HUMAN | Unmodified | 0 | 378280 | 0 | 546970 | 0 | 0 | 77217 | 0 | 7789200 | 420060 | 93787 | 0 | 0 | 0 | 0 | 0 | 0 | 0 |
| IGDEDVGR | PPIB; PPIB; PPI | P23284; A0A | PPIB_HUMAN F | Unmodified | 0 | 0 | 558400 | 47779 | 17199 | 0 | 378360 | 14804 | 5944400 | 0 | 0 | 2919500 | 0 | 0 | 79665 | 0 | 0 | 0 |
| IGDEDVGR | PPIB; PPIB; PPI | P23284; A0A | PPIB_HUMAN F | Unmodified | 394760 | 0 | 538160 | 32195 | 239330 | 24565 | 1303700 | 18321 | 40393000 | 48453000 | 10237000 | 1530200 | 1456100 | 1462700 | 2301800 | 677400 | 48826 | 158890 |
| IGDEDVGR | PPIB; PPIB; PPI | P23284; A0A | PPIB_HUMAN F | Unmodified | 0 | 0 | 0 | 0 | 0 | 0 | 228360 | 0 | 159060 | 1198300 | 0 | 0 | 0 | 0 | 0 | 0 | 0 | 0 |
| IGDEYFTFI | CCT3; CCT3 | P49368; B4D | TCPG_HUMAN | Unmodified | 0 | 0 | 0 | 0 | 0 | 0 | 0 | 0 | 0 | 0 | 0 | 1634100 | 0 | 0 | 0 | 0 | 0 | 0 |
| IGDFGLAK | TYK2; TYK2 | P29597; E9P | TYK2_HUMAN I | Unmodified | 70541 | 107030 | 0 | 79796 | 1209800 | 395460 | 81112 | 77239 | 110330 | 62364 | 0 | 391810 | 67080 | 82468 | 223110 | 269360 | 348840 | 1519700 |
| IGDFGLVTS | EIF2AK2; EIF2A | P19525; A0A | E2AK2_HUMAN | Unmodified | 0 | 0 | 0 | 0 | 0 | 0 | 0 | 0 | 0 | 0 | 0 | 1824500 | 0 | 0 | 0 | 0 | 0 | 0 |
| IGDFGMTR | INSR; IGF1R; IG | P06213; P08 | INSR_HUMAN I | Unmodified | 0 | 0 | 0 | 0 | 2973500 | 0 | 312680 | 0 | 0 | 318840 | 0 | 0 | 0 | 632530 | 0 | 0 | 0 | 851520 |
| IGDFIDVSE | MRPL39; MRPL | Q9NYK5; C9 | RM39_HUMAN | Unmodified | 0 | 0 | 0 | 0 | 0 | 0 | 0 | 0 | 1543300 | 0 | 0 | 0 | 0 | 4027200 | 0 | 0 | 0 | 0 |
| IGDKDVGR | PPIC | P45877 | PPIC_HUMAN I | Unmodified | 0 | 0 | 0 | 0 | 0 | 0 | 0 | 0 | 0 | 247020 | 0 | 0 | 0 | 0 | 0 | 0 | 0 | 0 |
| IGDKLDIIC | EFNB1 | P98172 | EFNB1_HUMAI | Unmodified | 0 | 63088 | 0 | 0 | 0 | 0 | 0 | 0 | 0 | 0 | 0 | 0 | 0 | 0 | 0 | 439760 | 291480 | 0 |
| IGDLFNGR | SRPK1; SRPK2; | Q96SB4; P78 | SRPK1_HUMAN | Unmodified | 0 | 0 | 0 | 0 | 0 | 0 | 0 | 0 | 0 | 0 | 0 | 1835500 | 0 | 0 | 0 | 0 | 0 | 0 |
| IGDLQAFQ | SLC3A2; SLC3A | A0A7P0T9F7 | A0A7P0T9F7_H | Unmodified | 256880 | 255050 | 0 | 0 | 0 | 0 | 0 | 0 | 637600 | 109440 | 325330 | 1350300 | 0 | 0 | 169840 | 174920 | 0 | 0 |
| IGDLQAFQ | SLC3A2; SLC3A | P08195; A0A | 4F2_HUMAN 4I | Unmodified | 0 | 905140 | 0 | 278950 | 632190 | 0 | 0 | 0 | 1279100 | 156330 | 3327000 | 7551300 | 0 | 0 | 1255200 | 147520 | 80600 | 0 |
| IGDMGGFA | MVB12A; MVB1 | Q96EY5; M0 | MB12A_HUMAI | Oxidation (M) | 16346000 | 5803600 | 8846400 | 0 | 0 | 0 | 0 | 0 | 785410 | 418060 | 0 | 1799300 | 5338500 | 0 | 12919000 | 10698000 | 0 | 0 |
| IGDMGGFA | MVB12A; MVB1 | Q96EY5; M0 | MB12A_HUMAI | Oxidation (M) | 4505100 | 2355600 | 0 | 4112100 | 0 | 0 | 0 | 0 | 0 | 213210 | 0 | 0 | 4135600 | 0 | 3347700 | 3194600 | 0 | 0 |
| IGDMGGFA | MVB12A; MVB1 | Q96EY5; M0 | MB12A_HUMAI | Unmodified | 0 | 0 | 0 | 1000200 | 0 | 0 | 0 | 0 | 0 | 0 | 0 | 0 | 715520 | 0 | 0 | 0 | 0 | 0 |
| IGDMRIMEI | DOCK7; DOCK7 | Q96N67; A0 | DOCK7_HUMA | 2 Oxidation (M) | 0 | 153690 | 247290 | 5067300 | 2724700 | 9080500 | 2871000 | 326150 | 172300 | 1711400 | 8053200 | 362780 | 1593600 | 5442500 | 0 | 0 | 1503400 | 14679000 |
| IGDQWDK | FN1 | P02751 | FINC_HUMAN I | Unmodified | 904320 | 6663200 | 25942000 | 8777400 | 413690 | 0 | 234960 | 0 | 1621800 | 1593700 | 1156000 | 3364900 | 6662800 | 2991000 | 16597000 | 15055000 | 183500 | 2054800 |
| IGDTPMVR | CBS; CBSL; CB | P35520; P0D | CBS_HUMAN C | Unmodified | 0 | 0 | 117570 | 0 | 636520 | 0 | 0 | 0 | 0 | 27649 | 417870 | 2503000 | 0 | 0 | 0 | 81117 | 0 | 0 |
| IGDTSVSYI | LMNB1 | P20700 | LMNB1_HUMAI | Unmodified | 0 | 0 | 0 | 0 | 0 | 0 | 361210 | 107020 | 0 | 0 | 0 | 0 | 63260 | 36142 | 0 | 0 | 0 | 0 |
| IGDTVSFSI | ITGB3; N/A | P05106; H3E | ITB3_HUMAN I | Unmodified | 2214400 | 3654200 | 1042200 | 4191800 | 6369600 | 0 | 13463000 | 1722000 | 0 | 0 | 0 | 0 | 4793300 | 5831900 | 0 | 0 | 13646 | 2158800 |
| IGDTVTCV | TMEFF2 | Q9UIK5 | TEFF2_HUMAN | Unmodified | 673840 | 0 | 0 | 0 | 0 | 0 | 0 | 0 | 0 | 0 | 0 | 0 | 0 | 0 | 0 | 0 | 0 | 0 |
| IGDTWSK | FN1 | P02751 | FINC_HUMAN I | Unmodified | 0 | 0 | 7424000 | 2961900 | 0 | 303160 | 0 | 0 | 1053300 | 167480 | 447330 | 154370 | 382830 | 385270 | 0 | 2348400 | 0 | 888510 |
| IGDTWSKK | FN1 | P02751 | FINC_HUMAN I | Unmodified | 0 | 1405400 | 3848800 | 440480 | 0 | 255860 | 0 | 0 | 298260 | 75673 | 0 | 129630 | 82194 | 0 | 4563000 | 714890 | 0 | 0 |
| IGDVSSSA | YTHDF1 | Q9BYJ9 | YTHD1_HUMAI | Unmodified | 250840 | 122340 | 65846 | 0 | 0 | 11675 | 28802 | 86941 | 424740 | 39396 | 0 | 122080 | 1064700 | 207720 | 66452 | 91870 | 0 | 27836 |
| IGEAEAAVI | FLOT2; FLOT2; | Q14254; J3C | FLOT2_HUMAN | Unmodified | 0 | 0 | 0 | 0 | 0 | 0 | 275680 | 0 | 5809200 | 0 | 0 | 2951900 | 1753300 | 0 | 0 | 0 | 500600 | 877530 |
| IGEAEAAVI | FLOT2; FLOT2; | Q14254; J3C | FLOT2_HUMAN | Oxidation (M) | 0 | 0 | 423050 | 0 | 0 | 0 | 0 | 0 | 351970 | 0 | 0 | 931730 | 552170 | 0 | 716610 | 275970 | 0 | 0 |
| IGEAEAAVI | FLOT2; FLOT2; | Q14254; J3C | FLOT2_HUMAN | Unmodified | 0 | 0 | 258480 | 0 | 627130 | 0 | 0 | 0 | 12930000 | 523870 | 0 | 1329500 | 0 | 0 | 0 | 16391 | 0 | 0 |
| IGEAEAAVI | FLOT2; FLOT2; | Q14254; J3C | FLOT2_HUMAN | Oxidation (M) | 1139900 | 0 | 0 | 0 | 4382100 | 586330 | 0 | 0 | 0 | 1009600 | 0 | 0 | 1735000 | 725860 | 0 | 0 | 0 | 0 |
| IGEAEAAVI | FLOT2; FLOT2; | Q14254; J3C | FLOT2_HUMAN | Unmodified | 0 | 0 | 0 | 0 | 0 | 0 | 0 | 0 | 1809800 | 517180 | 0 | 108490 | 0 | 436530 | 0 | 0 | 0 | 0 |
| IGEEEIQKP | SF3A1 | Q15459 | SF3A1_HUMAN | Unmodified | 0 | 0 | 0 | 0 | 0 | 0 | 0 | 0 | 46858 | 0 | 625920 | 0 | 0 | 0 | 0 | 0 | 0 | 0 |
| IGEEFLTDL | PYGB | P11216 | PYGB_HUMAN | Unmodified | 0 | 0 | 18277 | 30254 | 219770 | 42940 | 48097 | 0 | 1035500 | 582430 | 144290 | 865510 | 254770 | 139070 | 0 | 29201 | 646380 | 1566800 |
| IGEEFLTDL | PYGB | P11216 | PYGB_HUMAN | Unmodified | 82996 | 90768 | 184210 | 366700 | 1852100 | 230660 | 403750 | 0 | 1885300 | 984130 | 717220 | 3265400 | 1117900 | 769760 | 396860 | 279230 | 1519100 | 5296300 |
| IGEEQSAEI | RBBP7; RBBP7; | Q16576; Q5 | RBBP7_HUMAI | Unmodified | 172070 | 0 | 256900 | 881950 | 0 | 0 | 0 | 0 | 0 | 0 | 731530 | 3417000 | 0 | 0 | 293870 | 234270 | 0 | 0 |
| IGEEQSPEI | RBBP4; RBBP4; | Q09028; H0 | RBBP4_HUMAI | Unmodified | 0 | 0 | 0 | 0 | 0 | 431250 | 0 | 291440 | 0 | 3348900 | 2700400 | 6857600 | 197910 | 1282600 | 0 | 0 | 0 | 0 |
| IGEGSTGIV | PAK4 | O96013 | PAK4_HUMAN | Unmodified | 333980 | 430590 | 149100 | 398930 | 0 | 0 | 0 | 0 | 0 | 0 | 0 | 490460 | 0 | 0 | 0 | 473580 | 0 | 0 |
| IGEGTYGT | CDK5 | Q00535 | CDK5_HUMAN | Unmodified | 1235900 | 0 | 470020 | 0 | 2045300 | 311590 | 1438100 | 1024000 | 1271700 | 599050 | 329120 | 0 | 1473900 | 917450 | 547570 | 0 | 1872000 | 2389300 |
| IGEGTYGV | CDK1; CDK3; C | P06493; Q0 | CDK1_HUMAN | Unmodified | 844630 | 866380 | 1088200 | 2622200 | 5487800 | 1300100 | 1488600 | 1600800 | 2868800 | 1988800 | 962730 | 2359800 | 6367900 | 3430000 | 1467100 | 1243900 | 1588300 | 3726400 |
| IGEGTYGV | CDK1; CDK1; C | P06493; A0A | CDK1_HUMAN | Unmodified | 0 | 237250 | 0 | 0 | 123090 | 0 | 137750 | 0 | 0 | 0 | 0 | 0 | 0 | 0 | 0 | 0 | 430020 | 0 |
| IGEHMEEH | TXNRD1; TXNRI | Q16881; A0 | TRXR1_HUMAN | Unmodified | 0 | 25146 | 0 | 166870 | 605980 | 0 | 0 | 95050 | 771130 | 129090 | 228330 | 11586000 | 53429 | 0 | 452550 | 208430 | 126740 | 0 |
| IGEHMEEH | TXNRD1; TXNRI | Q16881; A0 | TRXR1_HUMAN | Oxidation (M) | 3475800 | 0 | 190830 | 0 | 158190 | 0 | 0 | 0 | 34126 | 0 | 57312 | 855900 | 0 | 0 | 0 | 81088 | 0 | 0 |

| | | | | | | | | | | | | | | | | | | | | | | |
|---|---|---|---|---|---|---|---|---|---|---|---|---|---|---|---|---|---|---|---|---|---|---|
| IGEIATGSP | BAIAP2L1 | Q9UHR4 | BI2L1_HUMAN | Unmodified | 0 | 358940 | 405420 | 2799100 | 59583000 | 5299500 | 0 | 0 | 2925200 | 2372500 | 720500 | 1078600 | 24531000 | 19214000 | 2269700 | 772590 | 1616100 | 0 |
| IGEIATGSP | BAIAP2L1 | Q9UHR4 | BI2L1_HUMAN | Unmodified | 44273 | 0 | 0 | 588240 | 7935000 | 1226400 | 0 | 0 | 1266700 | 498170 | 167200 | 242950 | 2243500 | 2035200 | 287680 | 0 | 0 | 242500 |
| IGELVGVLV | MYO1D; MYO1D | O94832; J3Q | MYO1D_HUMA | Unmodified | 26573000 | 28863000 | 5478100 | 9185100 | 0 | 0 | 0 | 0 | 4265400 | 455650 | 1332700 | 3822200 | 0 | 0 | 27978000 | 13970000 | 0 | 0 |
| IGELVGVLV | MYO1D; MYO1D | O94832; K7E | MYO1D_HUMA | Unmodified | 4532000 | 5585200 | 0 | 1685900 | 727280 | 0 | 0 | 0 | 273890 | 994970 | 0 | 129840 | 2064700 | 323130 | 367620 | 929390 | 454540 | 2379300 |
| IGELVGVLV | MYO1D; MYO1D | O94832; K7E | MYO1D_HUMA | Unmodified | 1470800 | 1361800 | 0 | 979370 | 0 | 0 | 0 | 0 | 245330 | 0 | 0 | 0 | 311550 | 0 | 0 | 0 | 0 | 0 |
| IGETVVDLE | DYSF | O75923 | DYSF_HUMAN | Unmodified | 0 | 0 | 0 | 0 | 630300 | 0 | 26819 | 0 | 0 | 171910 | 0 | 0 | 101090 | 0 | 0 | 0 | 0 | 0 |
| IGEVDVEQI | CCNB1; CCNB1 | P14635; H0Y | CCNB1_HUMA | Unmodified | 100360 | 0 | 99093 | 0 | 0 | 0 | 91859 | 75157 | 947410 | 69244 | 323190 | 1329300 | 0 | 35148 | 241190 | 155450 | 208410 | 0 |
| IGEWELIQE | USP5 | P45974 | UBP5_HUMAN | Unmodified | 1102100 | 1991900 | 135350 | 722300 | 0 | 0 | 0 | 0 | 0 | 0 | 0 | 298470 | 0 | 0 | 1354100 | 0 | 0 | 0 |
| IGFAAEQLA | TUSC3; TUSC3; | Q13454; D6I | TUSC3_HUMAI | Unmodified | 0 | 0 | 0 | 0 | 0 | 0 | 0 | 0 | 3220000 | 553200 | 0 | 610200 | 0 | 0 | 0 | 0 | 0 | 0 |
| IGFDVVTLS | NTPCR; NTPCR | Q9BSD7; Q5 | NTPCR_HUMAI | Unmodified | 0 | 0 | 134970 | 323710 | 0 | 0 | 0 | 0 | 495010 | 0 | 0 | 0 | 1082600 | 429130 | 314870 | 137530 | 0 | 0 |
| IGFGPDPTI | LTBP1; LTBP1; L | Q14766; E7E | LTBP1_HUMAN | Unmodified | 0 | 0 | 0 | 0 | 0 | 0 | 0 | 0 | 0 | 0 | 0 | 0 | 0 | 0 | 0 | 0 | 198490 | 2512000 |
| IGFPWSEIF | RDX; EZR; MSN | P35241; P15 | RADI_HUMAN I | Unmodified | 0 | 0 | 15210000 | 37455000 | 38437000 | 0 | 1545100 | 322470 | 59351000 | 8594500 | 11557000 | 98428000 | 14500000 | 8527300 | 30123000 | 0 | 4525800 | 3750300 |
| IGFPWSEIF | RDX; EZR; MSN | P35241; P15 | RADI_HUMAN I | Unmodified | 0 | 0 | 50934 | 425550 | 2.11E+08 | 34353000 | 34254000 | 100170 | 3236600 | 53824000 | 2232800 | 50749 | 20740000 | 49264000 | 0 | 0 | 36088000 | 10928000 |
| IGFPWSEIF | RDX; EZR; MSN | P35241; P15 | RADI_HUMAN I | Unmodified | 0 | 0 | 0 | 0 | 18829000 | 0 | 2897300 | 0 | 1678500 | 6241800 | 557020 | 0 | 5142800 | 8258800 | 0 | 0 | 6056900 | 1273600 |
| IGFSGPKLE | AHNAK | Q09666 | AHNK_HUMAN | Unmodified | 0 | 0 | 0 | 0 | 3010500 | 0 | 0 | 0 | 0 | 0 | 0 | 0 | 0 | 0 | 0 | 0 | 516090 | 1509700 |
| IGFVGFPS\ | DRG1 | Q9Y295 | DRG1_HUMAN | Unmodified | 0 | 0 | 0 | 0 | 0 | 0 | 0 | 0 | 514460 | 0 | 1226300 | 3643900 | 0 | 0 | 0 | 114180 | 0 | 0 |
| IGGAQNRS | MIF | P14174 | MIF_HUMAN M | Unmodified | 0 | 0 | 0 | 0 | 5915500 | 861930 | 0 | 1818500 | 0 | 704370 | 0 | 0 | 2959000 | 2544300 | 0 | 0 | 14153000 | 6164200 |
| IGGDLTAA\ | YTHDF3; YTHDI | Q7Z739; A0A | YTHD3_HUMAI | Unmodified | 1037700 | 951560 | 0 | 0 | 1214000 | 0 | 0 | 3044300 | 150480 | 0 | 0 | 362800 | 337010 | 0 | 0 | 255520 | 0 | 0 |
| IGGGIDVP\ | KHSRP; KHSRP | Q92945; M0I | FUBP2_HUMAI | Unmodified | 568540 | 0 | 0 | 0 | 0 | 0 | 0 | 58233 | 2381300 | 745110 | 2539600 | 4966000 | 0 | 0 | 0 | 0 | 0 | 0 |
| IGGIGTVPV | EEF1A2; EEF1A | Q05639; Q5\ | EF1A2_HUMAN | Unmodified | 2.99E+08 | 2.56E+08 | 1.31E+08 | 69554000 | 1830800 | 884850 | 0 | 0 | 16251000 | 1581900 | 19401000 | 2.06E+08 | 2198400 | 1795700 | 2.61E+08 | 3.21E+08 | 784670 | 2488000 |
| IGGIGTVPV | EEF1A2; EEF1A | Q05639; A0A | EF1A2_HUMAN | Oxidation (M) | 0 | 0 | 227480 | 0 | 14182000 | 136560 | 0 | 0 | 0 | 0 | 0 | 0 | 31084000 | 4040600 | 0 | 202150 | 3131000 | 0 |
| IGGIGTVPV | EEF1A2; EEF1A | Q05639; A0A | EF1A2_HUMAN | Unmodified | 0 | 0 | 0 | 0 | 0 | 0 | 0 | 0 | 0 | 0 | 0 | 0 | 2430200 | 971910 | 831200 | 0 | 0 | 1645600 |
| IGGIGTVPV | EEF1A1P5; EEF | Q5VTE0; P68 | EF1A3_HUMAN | Unmodified | 3185800 | 0 | 10800000 | 1653200 | 649900 | 406050 | 206450 | 54913 | 804210 | 289780 | 241920 | 3035700 | 496110 | 0 | 4868400 | 2721500 | 524830 | 233370 |
| IGGIGTVPV | EEF1A1P5; EEF | Q5VTE0; P68 | EF1A3_HUMAN | Oxidation (M) | 35498000 | 1.82E+08 | 17430000 | 2.16E+08 | 58098000 | 21904000 | 12085000 | 0 | 7084500 | 7901500 | 22101000 | 8524100 | 1.14E+08 | 23958000 | 1.09E+08 | 11765000 | 26256000 | 74866000 |
| IGGIGTVPV | EEF1A1P5; EEF | Q5VTE0; P68 | EF1A3_HUMAN | Unmodified | 2254100 | 14709000 | 459690 | 12212000 | 35839000 | 0 | 3747800 | 0 | 6525000 | 1666500 | 0 | 0 | 34180000 | 6817600 | 5460900 | 0 | 10261000 | 26457000 |
| IGGNEGID\ | FUBP1; FUBP1 | Q96AE4; E9F | FUBP1_HUMAI | Unmodified | 0 | 0 | 0 | 0 | 0 | 0 | 0 | 0 | 0 | 0 | 1182600 | 3919600 | 0 | 0 | 0 | 0 | 0 | 0 |
| IGGSVHLIV | DHTKD1; DHTK | Q96HY7; H7 | DHTK1_HUMAI | Unmodified | 0 | 0 | 0 | 0 | 0 | 0 | 0 | 0 | 2711000 | 0 | 0 | 0 | 0 | 0 | 0 | 0 | 0 | 0 |
| IGGVQQDT | PHB2; PHB2; PI | Q99623; F5H | PHB2_HUMAN | Unmodified | 0 | 0 | 0 | 0 | 235490 | 0 | 175050 | 139340 | 1264100 | 284650 | 0 | 0 | 0 | 319150 | 0 | 0 | 338180 | 0 |
| IGGVTERM | FH | P07954 | FUMH_HUMAN | Unmodified | 0 | 33892 | 56471 | 120450 | 1926300 | 330710 | 3802800 | 399730 | 5050800 | 3122900 | 0 | 287950 | 3150700 | 2551800 | 39494 | 57189 | 3629600 | 2387800 |
| IGHFEFHG | GPR180 | Q86V85 | GP180_HUMAI | Unmodified | 0 | 0 | 0 | 0 | 0 | 0 | 0 | 0 | 2203100 | 162240 | 0 | 0 | 0 | 0 | 0 | 0 | 0 | 0 |
| IGHGGTLD | TRUB1 | Q8WWH5 | TRUB1_HUMAI | Unmodified | 0 | 0 | 0 | 0 | 0 | 0 | 540150 | 0 | 0 | 116130 | 0 | 0 | 265670 | 0 | 0 | 0 | 0 | 0 |
| IGHKVESES | ENOPH1; ENOF | Q9UHY7; D6 | ENOPH_HUMA | Unmodified | 0 | 0 | 0 | 0 | 0 | 0 | 0 | 0 | 0 | 11749 | 0 | 736530 | 0 | 40099 | 0 | 73852 | 0 | 0 |
| IGHKVESES | ENOPH1; ENOF | Q9UHY7; D6 | ENOPH_HUMA | Unmodified | 0 | 0 | 15417 | 0 | 734230 | 85794 | 0 | 0 | 497760 | 452020 | 468540 | 2072100 | 50940 | 0 | 0 | 0 | 368180 | 274610 |
| IGHRSVDS | PIP5K1A; PIP5K | Q99755; A2A | PI51A_HUMAN | Unmodified | 0 | 18449 | 0 | 0 | 729560 | 0 | 0 | 101320 | 0 | 44724 | 0 | 0 | 1073600 | 3.28E+08 | 0 | 0 | 335940 | 773200 |
| IGHYILGDT | PRKAA1; PRKAA | Q13131; Q9 | AAPK1_HUMAI | Unmodified | 1833200 | 836810 | 569540 | 545970 | 2286800 | 632600 | 350220 | 34954 | 1895800 | 746620 | 300760 | 976200 | 2321900 | 1195200 | 2480000 | 1336500 | 700410 | 4639800 |
| IGHYVLGD | PRKAA2; PRKAA | P54646; A0A | AAPK2_HUMAI | Unmodified | 0 | 0 | 0 | 0 | 0 | 0 | 328890 | 87693 | 0 | 123010 | 107680 | 0 | 252690 | 205920 | 0 | 0 | 199570 | 1406800 |
| IGIEIIKR | HSPD1; HSPD1 | P10809; A0A | CH60_HUMAN | Unmodified | 0 | 0 | 0 | 0 | 0 | 0 | 0 | 0 | 1121100 | 508050 | 562180 | 0 | 246620 | 204810 | 0 | 0 | 0 | 0 |
| IGIGELITR | CLUH; CLUH; C | O75153; I3L | CLU_HUMAN C | Unmodified | 45423 | 0 | 0 | 0 | 4324600 | 771870 | 259690 | 101080 | 1648000 | 730600 | 593760 | 2286000 | 0 | 0 | 0 | 0 | 516260 | 1143200 |
| IGIIGGTGL | MTAP; MTAP; N | Q13126; F8V | MTAP_HUMAN | Unmodified | 0 | 0 | 5164400 | 0 | 6926100 | 2859600 | 1587400 | 470830 | 0 | 2220300 | 4839600 | 11377000 | 5197800 | 3619700 | 0 | 0 | 2736300 | 6031200 |
| IGIMPGHII | SUCLG1; SUCL | P53597; A0A | SUCA_HUMAN | Unmodified | 0 | 0 | 0 | 0 | 0 | 0 | 0 | 0 | 710530 | 187250 | 0 | 0 | 0 | 0 | 0 | 0 | 0 | 86770 |
| IGIPEFK | CPM; CPM; CPN | P14384; F8V | CBPM_HUMAN | Unmodified | 0 | 0 | 0 | 0 | 0 | 0 | 0 | 0 | 0 | 0 | 0 | 0 | 0 | 0 | 0 | 0 | 0 | 0 |
| IGIVGLPNV | OLA1; OLA1; OL | Q9NTK5; J3K | OLA1_HUMAN | Unmodified | 728550 | 0 | 1916300 | 1818300 | 3757400 | 870230 | 4880800 | 3033800 | 2107500 | 1725000 | 6461200 | 15364000 | 7701700 | 4089600 | 2131800 | 2620700 | 4713200 | 2067500 |
| IGKFENGV | MGAT4A | Q9UM21 | MGT4A_HUMAI | Oxidation (M) | 0 | 0 | 0 | 0 | 0 | 0 | 0 | 0 | 4148500 | 165310 | 0 | 0 | 0 | 0 | 0 | 0 | 0 | 0 |
| IGKFENGV | MGAT4A | Q9UM21 | MGT4A_HUMAI | Unmodified | 0 | 0 | 0 | 0 | 0 | 0 | 0 | 0 | 3448700 | 0 | 0 | 0 | 0 | 0 | 0 | 0 | 0 | 0 |
| IGKPAPDFI | PRDX2; PRDX2 | P32119; A6N | PRDX2_HUMAI | Unmodified | 0 | 0 | 0 | 914430 | 3798300 | 277570 | 0 | 718520 | 3006700 | 6858300 | 1773200 | 299420 | 6001500 | 2159700 | 0 | 0 | 5635500 | 6638500 |
| IGKPHTVP( | RPS2; RPS2; RP | P15880; E9P | RS2_HUMAN 4 | Unmodified | 0 | 124870 | 109050 | 0 | 165840 | 0 | 67326 | 33172 | 0 | 0 | 72096 | 1872500 | 0 | 0 | 0 | 0 | 437880 | 0 |
| IGKVDCTQ | TXNDC5 | Q8NBS9 | TXND5_HUMAI | Unmodified | 0 | 0 | 0 | 0 | 0 | 0 | 0 | 0 | 29797000 | 5639900 | 1295000 | 0 | 0 | 0 | 0 | 0 | 0 | 0 |
| IGKVTSEEL | UQCRC2; UQCI | P22695; H3E | QCR2_HUMAN | Unmodified | 0 | 0 | 0 | 0 | 0 | 0 | 0 | 0 | 9000500 | 1892500 | 0 | 0 | 0 | 0 | 0 | 0 | 0 | 0 |
| IGLADYALE | SUN2; SUN2 | Q9UH99; B0 | SUN2_HUMAN | Unmodified | 0 | 0 | 0 | 0 | 0 | 0 | 0 | 0 | 2452100 | 0 | 0 | 0 | 0 | 0 | 0 | 0 | 0 | 0 |
| IGLAEEIR | ACACA; ACACA | Q13085; Q5 | ACACA_HUMAI | Unmodified | 0 | 0 | 0 | 0 | 23935 | 0 | 0 | 0 | 2047900 | 30329 | 168840 | 793880 | 43012 | 0 | 0 | 0 | 0 | 0 |
| IGLDDEEKI | MYO6; MYO6; N | Q9UM54; A0 | MYO6_HUMAN | Unmodified | 0 | 0 | 0 | 0 | 0 | 0 | 0 | 0 | 0 | 0 | 0 | 0 | 0 | 0 | 0 | 0 | 0 | 0 |
| IGLETSLR | RUVBL2 | Q9Y230 | RUVB2_HUMAI | Unmodified | 216120 | 0 | 0 | 0 | 0 | 0 | 0 | 0 | 0 | 0 | 0 | 11919000 | 0 | 0 | 0 | 304590 | 0 | 0 |
| IGLETVGVI | TXNRD1; TXNRI | Q16881; A0A | TRXR1_HUMAN | Unmodified | 0 | 0 | 164940 | 54410 | 0 | 0 | 414590 | 351810 | 2813000 | 3078700 | 4387500 | 17247000 | 833380 | 0 | 490080 | 1448700 | 0 | 817780 |
| IGLETVGVI | TXNRD1; TXNRI | Q16881; A0A | TRXR1_HUMAN | Unmodified | 0 | 0 | 0 | 0 | 14263000 | 1367400 | 1560100 | 1102100 | 0 | 1940000 | 0 | 0 | 3442700 | 1623900 | 0 | 0 | 5380000 | 3500300 |
| IGLFFWPK | EPB41L5; EPB4 | Q9HCM4; Q | E41L5_HUMAN | Unmodified | 2949400 | 2458300 | 4073800 | 2169300 | 517630 | 435210 | 0 | 0 | 3610500 | 2154600 | 317140 | 1744000 | 0 | 370040 | 14112000 | 9749500 | 381690 | 0 |
| IGLFNSSAI | GYS1 | P13807 | GYS1_HUMAN | Unmodified | 136680 | 0 | 0 | 0 | 0 | 0 | 234150 | 108890 | 0 | 177920 | 0 | 4716300 | 0 | 0 | 0 | 0 | 0 | 0 |
| IGLIHGHQ | VPS29; VPS29 | Q9UBQ0; F8 | VPS29_HUMAN | Unmodified | 0 | 0 | 0 | 0 | 0 | 0 | 0 | 0 | 690540 | 0 | 117580 | 548520 | 0 | 0 | 0 | 0 | 0 | 0 |
| IGLIQFCLS | CCT4 | P50991 | TCPD_HUMAN | Unmodified | 16453 | 0 | 0 | 0 | 0 | 0 | 0 | 0 | 25979 | 0 | 1173500 | 8233800 | 0 | 0 | 0 | 12790 | 0 | 0 |
| IGLIQGNR | PFKM; PFKM; PI | P08237; A0A | PFKAM_HUMAI | Unmodified | 0 | 0 | 0 | 0 | 0 | 0 | 0 | 0 | 0 | 0 | 864310 | 6138800 | 0 | 0 | 0 | 0 | 0 | 0 |
| IGLIQGNR\ | PFKM; PFKM; PI | P08237; A0A | PFKAM_HUMAI | Unmodified | 0 | 0 | 0 | 0 | 1046000 | 0 | 0 | 0 | 0 | 0 | 845350 | 0 | 4155100 | 802750 | 0 | 0 | 714850 | 1465500 |
| IGLMSLSK | UTRN; UTRN; U | P46939; H0Y | UTRN_HUMAN | Unmodified | 0 | 0 | 85894 | 0 | 0 | 0 | 0 | 0 | 881260 | 437910 | 0 | 0 | 0 | 139600 | 315250 | 173700 | 339570 | 175830 |
| IGLMSLSK | UTRN; UTRN; U | P46939; H0Y | UTRN_HUMAN | Oxidation (M) | 0 | 0 | 84676 | 0 | 0 | 0 | 0 | 0 | 0 | 133980 | 0 | 0 | 0 | 0 | 211750 | 454430 | 0 | 0 |
| IGLVEALC( | DNAJA2 | O60884 | DNJA2_HUMAN | Unmodified | 0 | 121860 | 0 | 0 | 1011500 | 42835 | 0 | 0 | 0 | 135800 | 0 | 0 | 0 | 355590 | 172610 | 0 | 100810 | 0 |
| IGLYPANY\ | PACSIN3 | Q9UKS6 | PACN3_HUMAI | Unmodified | 0 | 0 | 1647700 | 0 | 0 | 0 | 0 | 0 | 0 | 0 | 0 | 0 | 0 | 0 | 686680 | 0 | 0 | 0 |
| IGMDGSSF | LRP1 | Q07954 | LRP1_HUMAN | Unmodified | 0 | 0 | 0 | 0 | 0 | 0 | 0 | 0 | 566790 | 25584 | 0 | 0 | 0 | 0 | 0 | 48450 | 0 | 0 |
| IGNAKGDD | PSAT1 | Q9Y617 | SERC_HUMAN | Unmodified | 1047900 | 0 | 0 | 383920 | 885940 | 0 | 0 | 0 | 1706600 | 0 | 3690500 | 7490600 | 1295100 | 570480 | 382660 | 1423500 | 0 | 0 |
| IGNCPFSQ | CLIC1; CLIC5; ( | O00299; Q9I | CLIC1_HUMAN | Unmodified | 1543200 | 1068500 | 975600 | 2829400 | 14909000 | 4616800 | 516050 | 517390 | 3910200 | 1804400 | 1697500 | 10370000 | 9600000 | 12069000 | 2430000 | 1955700 | 7432500 | 8125200 |
| IGNELEK | TJP2; TJP2; TJP2 | Q9UDY2; A0 | ZO2_HUMAN T | Unmodified | 0 | 178270 | 28619 | 252800 | 0 | 0 | 12154 | 15216 | 2366000 | 488600 | 0 | 916640 | 0 | 0 | 178040 | 157190 | 119190 | 0 |
| IGNFSTDIK | LCP1 | P13796 | PLSL_HUMAN I | Unmodified | 0 | 0 | 0 | 103320 | 0 | 0 | 0 | 0 | 405200 | 316870 | 0 | 0 | 0 | 60829 | 146600 | 108880 | 773430 | 0 |
| IGNLQTDL | FLNB | O75369 | FLNB_HUMAN | Unmodified | 0 | 0 | 0 | 0 | 0 | 0 | 0 | 0 | 0 | 0 | 0 | 3678200 | 0 | 0 | 0 | 0 | 0 | 0 |
| IGNSYFKEI | STIP1 | P31948 | STIP1_HUMAN | Unmodified | 0 | 0 | 0 | 1859400 | 5757000 | 2413800 | 960800 | 764010 | 5174800 | 4786300 | 3434300 | 18285000 | 12073000 | 5266200 | 9032300 | 0 | 5087800 | 5871100 |
| IGNTEDGA | UBE2O | Q9C0C9 | UBE2O_HUMAI | Unmodified | 0 | 0 | 0 | 0 | 0 | 0 | 0 | 0 | 0 | 0 | 76530 | 4293200 | 0 | 0 | 0 | 0 | 0 | 0 |
| IGNTGGMI | ACLY | P53396 | ACLY_HUMAN | Unmodified | 0 | 0 | 0 | 0 | 2113100 | 372810 | 2983700 | 0 | 0 | 0 | 730160 | 3971400 | 506190 | 0 | 51071 | 0 | 0 | 800710 |
| IGNTGGMI | ACLY | P53396 | ACLY_HUMAN | Oxidation (M) | 0 | 0 | 0 | 0 | 278670 | 201270 | 0 | 1021300 | 0 | 437740 | 0 | 12933000 | 0 | 0 | 0 | 0 | 137070 | 0 |
| IGNVEISQ\ | RPA2; RPA2 | P15927; Q5T | RFA2_HUMAN | Unmodified | 0 | 0 | 0 | 0 | 0 | 0 | 45556 | 0 | 0 | 93658 | 42330 | 0 | 157810 | 52651 | 0 | 0 | 68479 | 61926 |
| IGPALSCGI | ALDH1A1; ALDI | P00352; Q5 | AL1A1_HUMAN | Unmodified | 0 | 0 | 561050 | 364270 | 0 | 0 | 0 | 0 | 150980 | 0 | 469590 | 5366500 | 0 | 0 | 686850 | 0 | 0 | 0 |
| IGPALSCGI | ALDH1A1; ALDI | P00352; Q5 | AL1A1_HUMAN | Unmodified | 0 | 0 | 0 | 0 | 0 | 0 | 0 | 0 | 0 | 0 | 0 | 5194900 | 0 | 0 | 0 | 0 | 0 | 0 |
| IGPALSCGI | ALDH1A1; ALDI | P00352; Q5 | AL1A1_HUMAN | Unmodified | 0 | 0 | 0 | 0 | 0 | 0 | 0 | 0 | 2585400 | 3966700 | 21088000 | 6.47E+08 | 0 | 0 | 0 | 0 | 0 | 0 |
| IGPAPAFT( | FBLN2 | P98095 | FBLN2_HUMAN | Unmodified | 0 | 0 | 20273 | 0 | 0 | 0 | 0 | 0 | 456600 | 0 | 166020 | 39127 | 0 | 0 | 0 | 0 | 0 | 0 |
| IGPEPTTDS | EHD1; EHD1; E | Q9H4M9; C9 | EHD1_HUMAN | Unmodified | 0 | 0 | 0 | 1251000 | 1202900 | 0 | 0 | 0 | 1940600 | 0 | 979310 | 1822200 | 0 | 0 | 1147000 | 328650 | 0 | 0 |
| IGPEPTTDS | EHD1; EHD1; E | Q9H4M9; C9 | EHD1_HUMAN | Oxidation (M) | 0 | 2963400 | 1070000 | 3039000 | 0 | 912140 | 0 | 0 | 0 | 0 | 3719500 | 13164000 | 0 | 0 | 3940500 | 1946700 | 968580 | 2184400 |
| IGPGDVLTI | SEZ6; SEZ6; SEZ | Q53EL9; C9J | SEZ6_HUMAN | Unmodified | 0 | 0 | 0 | 0 | 0 | 0 | 0 | 0 | 0 | 0 | 0 | 0 | 0 | 0 | 0 | 0 | 0 | 0 |
| IGPIATPDYI | ACSS2 | Q9NR19 | ACSA_HUMAN | Unmodified | 0 | 0 | 0 | 0 | 732270 | 0 | 0 | 251470 | 0 | 0 | 112770 | 0 | 365580 | 0 | 0 | 235880 | 806070 | 0 |
| IGPILDNST | PPP2R1A; PPP2 | P30153; B3K | 2AAA_HUMAN | Unmodified | 2186500 | 2402200 | 11972000 | 12287000 | 33400000 | 6950900 | 7644800 | 1780500 | 33510000 | 12935000 | 14286000 | 1.29E+08 | 24642000 | 18011000 | 11123000 | 8408000 | 14101000 | 35252000 |
| IGPILDNST | PPP2R1A; PPP2 | P30153; B3K | 2AAA_HUMAN | Unmodified | 0 | 0 | 0 | 0 | 6768800 | 0 | 1312600 | 0 | 0 | 1323100 | 0 | 0 | 0 | 0 | 0 | 0 | 6808400 | 3107800 |
| IGPILDTNA | PPP2R1B; PPP2 | P30154; H0Y | 2AAB_HUMAN | Unmodified | 0 | 0 | 280160 | 0 | 2183500 | 313950 | 137850 | 0 | 1063600 | 0 | 1243400 | 4055000 | 2960000 | 2548000 | 819910 | 538590 | 178660 | 511140 |
| IGPITPLEFY | BLMH; BLMH; B | Q13867; K7E | BLMH_HUMAN | Unmodified | 0 | 0 | 2178100 | 0 | 0 | 0 | 0 | 0 | 5023600 | 0 | 1119900 | 7513700 | 0 | 0 | 0 | 0 | 0 | 0 |
| IGPKKPLPD | RPS3; RPS3; RP | P23396; E9P | RS3_HUMAN 4 | Unmodified | 0 | 147010 | 0 | 0 | 0 | 0 | 0 | 0 | 0 | 0 | 0 | 895500 | 0 | 0 | 0 | 0 | 0 | 0 |
| IGPLGLSPI | RPL12 | P30050 | RL12_HUMAN ( | Unmodified | 5727100 | 6533300 | 2893000 | 3245300 | 4563600 | 0 | 1660000 | 914820 | 1333100 | 1120500 | 5426300 | 49914000 | 0 | 0 | 3697500 | 3780700 | 621760 | 3446300 |
| IGPQQSIST | RAB2B | Q8WUD1 | RAB2B_HUMAI | Unmodified | 0 | 89427 | 0 | 0 | 0 | 0 | 298470 | 0 | 738940 | 294920 | 451530 | 1935300 | 0 | 0 | 123100 | 284410 | 0 | 0 |
| IGPSILNSD | RPE; RPE; RPE; | Q96AT9; F8V | RPE_HUMAN R | Unmodified | 0 | 74205 | 0 | 259590 | 0 | 0 | 437670 | 0 | 622800 | 155020 | 737410 | 2325000 | 35734 | 0 | 338650 | 50923 | 178250 | 459050 |
| IGPYPDYS | BMPR2 | Q13873 | BMPR2_HUMAI | Unmodified | 0 | 0 | 0 | 0 | 4040000 | 0 | 2066200 | 0 | 968110 | 786690 | 0 | 0 | 0 | 917340 | 0 | 373640 | 521280 | 2344700 |
| IGPYQPNV | MATR3; MATR3; | P43243; B3K | MATR3_HUMAI | Unmodified | 0 | 0 | 152960 | 0 | 0 | 0 | 0 | 0 | 1343000 | 0 | 1751200 | 12757000 | 0 | 0 | 0 | 0 | 0 | 0 |
| IGQGASGT | PAK2 | Q13177 | PAK2_HUMAN | Unmodified | 125230 | 142400 | 0 | 366530 | 5578300 | 0 | 0 | 281760 | 1010500 | 928520 | 571290 | 1049000 | 3125900 | 2038400 | 298870 | 184860 | 1290100 | 3627300 |
| IGQGYLIK | RPL3; RPL3; RP | P39023; B5N | RL3_HUMAN 6 | Unmodified | 5319400 | 4678300 | 5833700 | 2661900 | 1232500 | 0 | 825420 | 760890 | 0 | 238100 | 3408800 | 39238000 | 615540 | 0 | 3571500 | 2818900 | 0 | 154610 |
| IGQLEEQLE | MYH10 | P35580 | MYH10_HUMAI | Unmodified | 0 | 0 | 315700 | 458290 | 2071900 | 339710 | 0 | 0 | 0 | 0 | 0 | 2151100 | 447450 | 175260 | 253780 | 434870 | 0 | 0 |
| IGQLEEQLE | MYH10 | P35580 | MYH10_HUMAI | Unmodified | 0 | 0 | 0 | 0 | 6105800 | 606080 | 704380 | 0 | 0 | 0 | 0 | 0 | 1553300 | 1978400 | 0 | 0 | 342240 | 820230 |
| IGQQPQQF | KHSRP; KHSRP | Q92945; M0 | FUBP2_HUMAI | Unmodified | 0 | 32267 | 0 | 25800 | 290650 | 106880 | 146720 | 0 | 1539200 | 969650 | 987670 | 2959900 | 0 | 276220 | 24872 | 0 | 0 | 297830 |
| IGQTSSSVS | ITGA2 | P17301 | ITA2_HUMAN I | Unmodified | 5062700 | 1977900 | 0 | 4739900 | 3278900 | 696110 | 0 | 0 | 175960 | 0 | 0 | 805140 | 0 | 0 | 9873500 | 0 | 516110 | 3507400 |
| IGQTSSSVS | ITGA2 | P17301 | ITA2_HUMAN I | Unmodified | 1236600 | 1128100 | 0 | 2242700 | 3266500 | 699670 | 0 | 0 | 0 | 0 | 0 | 0 | 494480 | 0 | 99026 | 311520 | 944090 | 3469700 |
| IGQTSSSVS | ITGA2 | P17301 | ITA2_HUMAN I | Unmodified | 0 | 0 | 0 | 261970 | 17352000 | 3218200 | 0 | 0 | 0 | 360500 | 0 | 0 | 2464600 | 289030 | 0 | 0 | 4683600 | 18643000 |
| IGQVHYALI | LNPEP | Q9UIQ6 | LCAP_HUMAN | Unmodified | 94544 | 0 | 163570 | 0 | 0 | 0 | 302580 | 0 | 173190 | 87522 | 0 | 1098700 | 164530 | 163660 | 121230 | 81901 | 0 | 0 |
| IGRDQELYI | TARS1 | P26639 | SYTC_HUMAN | Unmodified | 164640 | 0 | 0 | 369520 | 0 | 507340 | 508310 | 0 | 0 | 0 | 0 | 1827600 | 0 | 362840 | 0 | 198150 | 0 | 0 |
| IGRFVVQN | HSPH1; HSPH1 | Q92598; A0A | HS105_HUMAI | Unmodified | 0 | 0 | 0 | 0 | 1705700 | 0 | 6305500 | 3734600 | 0 | 427100 | 0 | 0 | 1542400 | 2574000 | 0 | 0 | 4593100 | 3128100 |
| IGRIEDVTP | RPS14; RPS14 | P62263; A0A | RS14_HUMAN | Unmodified | 5814700 | 4454800 | 2817800 | 3887300 | 3221900 | 431560 | 0 | 201910 | 0 | 0 | 1530300 | 21000000 | 2537800 | 942880 | 2152900 | 2579000 | 0 | 0 |
| IGRIEDVTP | RPS14; RPS14 | P62263; A0A | RS14_HUMAN | Unmodified | 4180200 | 3285500 | 481810 | 2427200 | 2555300 | 361630 | 587580 | 399930 | 993130 | 824690 | 780810 | 20869000 | 3499300 | 1247900 | 1132900 | 1721600 | 148040 | 323490 |
| IGRNECVV | EIF2S1; EIF2S1; | P05198; H0Y | IF2A_HUMAN E | Unmodified | 0 | 0 | 0 | 0 | 0 | 0 | 0 | 0 | 0 | 139870 | 0 | 1444900 | 0 | 0 | 0 | 0 | 0 | 0 |
| IGRNECVV | EIF2S1; EIF2S1; | P05198; H0Y | IF2A_HUMAN E | Unmodified | 117470 | 0 | 204140 | 0 | 0 | 0 | 0 | 0 | 0 | 0 | 0 | 2405000 | 0 | 0 | 302750 | 485810 | 0 | 0 |
| IGRPSETGI | DDB1; DDB1; D | Q16531; A0A | DDB1_HUMAN | Unmodified | 0 | 2428300 | 0 | 0 | 0 | 0 | 0 | 0 | 52680 | 0 | 0 | 315400 | 0 | 0 | 0 | 0 | 59398 | 0 |

| | | | | | | | | | | | | | | | | | | | | | | |
|---|---|---|---|---|---|---|---|---|---|---|---|---|---|---|---|---|---|---|---|---|---|---|
| IGRTHTQD. | FH | P07954 | FUMH_HUMAN | Unmodified | 0 | 0 | 0 | 0 | 942300 | 0 | 2703000 | 0 | 0 | 2111400 | 0 | 0 | 4695900 | 2250300 | 0 | 0 | 1435300 | 1156700 |
| IGSCTQQD | DARS1; DARS1; | P14868; H7( | SYDC_HUMAN | Unmodified | 84090 | 39457 | 323500 | 301330 | 3059300 | 569500 | 388470 | 350260 | 1987500 | 1197800 | 2372200 | 13500000 | 3316200 | 4747100 | 158300 | 1391700 | 1060600 | 2253000 |
| IGSDPLAYE | SLC9A1 | P19634 | SL9A1_HUMAN | Unmodified | 0 | 0 | 0 | 0 | 1654100 | 0 | 0 | 0 | 0 | 0 | 0 | 0 | 2545200 | 3230800 | 1531100 | 0 | 0 | 888150 |
| IGSDPLAYE | SLC9A1 | P19634 | SL9A1_HUMAN | Unmodified | 0 | 0 | 0 | 0 | 0 | 0 | 0 | 0 | 0 | 0 | 0 | 0 | 0 | 5917400 | 0 | 0 | 0 | 1372600 |
| IGSDVELLL | LYRM7 | Q5U5X0 | LYRM7_HUMAN | Unmodified | 0 | 0 | 0 | 0 | 0 | 0 | 0 | 0 | 593960 | 0 | 0 | 0 | 0 | 0 | 0 | 0 | 0 | 0 |
| IGSENPSD' | USP13; USP13; | Q92995; A0/ | UBP13_HUMAN | Unmodified | 445280 | 832680 | 0 | 0 | 0 | 0 | 424780 | 0 | 0 | 0 | 0 | 0 | 0 | 0 | 304240 | 357390 | 0 | 0 |
| IGSFGPQEI | ACACA; ACACA | Q13085; Q5! | ACACA_HUMAI | Unmodified | 0 | 0 | 0 | 0 | 0 | 0 | 21440 | 0 | 0 | 0 | 0 | 303470 | 0 | 0 | 0 | 0 | 0 | 0 |
| IGSFTIQNV | HSPA4L; HSPA4 | O95757; E9F | HS74L_HUMAN | Unmodified | 0 | 0 | 0 | 0 | 478980 | 0 | 0 | 62683 | 532360 | 0 | 1388100 | 2884900 | 0 | 231890 | 0 | 0 | 0 | 363970 |
| IGSGFDNII | CTBP1; CTBP1; | Q13363; E9F | CTBP1_HUMAN | Unmodified | 0 | 0 | 0 | 0 | 0 | 0 | 0 | 0 | 0 | 0 | 0 | 1020600 | 165340 | 0 | 0 | 0 | 0 | 0 |
| IGSGKFTLS | RPS6KA1; RPS6 | Q15418; E9F | KS6A1_HUMAN | Unmodified | 55908 | 0 | 0 | 159770 | 333300 | 0 | 0 | 0 | 533580 | 0 | 0 | 203090 | 315610 | 0 | 0 | 58074 | 161100 | 261260 |
| IGSLIDVNC | MAGOH; MAGC | P61326; Q96 | MGN_HUMAN I | Unmodified | 683630 | 214990 | 240690 | 1218400 | 6754600 | 1478400 | 337620 | 0 | 130020 | 2889700 | 360330 | 3941000 | 2827600 | 2485200 | 157110 | 139510 | 769060 | 2582900 |
| IGSLIDVNC | MAGOH; MAGC | P61326; Q96 | MGN_HUMAN I | Unmodified | 0 | 0 | 78580 | 0 | 0 | 0 | 0 | 0 | 0 | 0 | 0 | 481580 | 0 | 0 | 0 | 0 | 0 | 0 |
| IGSMTGAE | SLC4A2; SLC4A | P04920; A0A | B3A2_HUMAN . | Oxidation (M) | 0 | 0 | 243820 | 0 | 0 | 0 | 0 | 0 | 0 | 0 | 0 | 81107 | 0 | 0 | 106750 | 0 | 0 | 0 |
| IGSPEYIK | EDIL3 | O43854 | EDIL3_HUMAN | Unmodified | 0 | 0 | 0 | 0 | 0 | 14558000 | 0 | 0 | 0 | 553450 | 0 | 166000 | 0 | 0 | 0 | 0 | 0 | 0 |
| IGSSELQEF | SLC39A14; SLC | Q15043; E5F | S39AE_HUMAN | Unmodified | 2034500 | 1058400 | 462470 | 1675700 | 0 | 0 | 0 | 0 | 1699000 | 58623 | 328130 | 2168000 | 0 | 0 | 1778000 | 1068400 | 0 | 0 |
| IGSSLPQDI | TACC2; TACC2; | O95359; Q4' | TACC2_HUMAN | Unmodified | 510630 | 0 | 0 | 44264 | 218920 | 24799 | 0 | 0 | 40266 | 21233 | 0 | 396130 | 290050 | 207280 | 0 | 28488 | 44120 | 275980 |
| IGSTIFGER | PLPBP | O94903 | PLPHP_HUMAN | Unmodified | 140450 | 131310 | 0 | 87490 | 306030 | 0 | 0 | 0 | 699490 | 0 | 167510 | 980130 | 0 | 0 | 0 | 88609 | 0 | 0 |
| IGSTTNPFL | PSD3; PSD3; PS | Q9NYI0; E5F | PSD3_HUMAN | Unmodified | 0 | 0 | 0 | 236920 | 1141300 | 241200 | 0 | 29834 | 774840 | 249850 | 0 | 1529900 | 0 | 146920 | 120780 | 144190 | 394960 | 744980 |
| IGSVKYEGI | CANT1 | Q8WVQ1 | CANT1_HUMAI | Unmodified | 0 | 0 | 0 | 0 | 509490 | 0 | 0 | 0 | 31921000 | 6251400 | 1026600 | 0 | 1006400 | 0 | 0 | 0 | 0 | 776160 |
| IGSYIER | KIT | P10721 | KIT_HUMAN Ma | Unmodified | 0 | 0 | 0 | 0 | 0 | 0 | 0 | 63422 | 1227600 | 514500 | 281340 | 5092000 | 0 | 0 | 0 | 0 | 134050 | 0 |
| IGSYLAER | EFR3A; EFR3A; | Q14156; E5F | EFR3A_HUMAN | Unmodified | 0 | 54946 | 0 | 0 | 0 | 515260 | 0 | 0 | 197130 | 0 | 0 | 951340 | 0 | 0 | 682890 | 539830 | 0 | 0 |
| IGSYLSAST | SCYL1; SCYL1; | Q96KG9; H0 | SCYL1_HUMAN | Unmodified | 0 | 0 | 236550 | 0 | 0 | 758230 | 0 | 0 | 120750 | 0 | 0 | 470740 | 0 | 0 | 62313 | 0 | 0 | 21515 |
| IGTDIQDNI | FDPS; FDPS | P14324; A0A | FPPS_HUMAN | Unmodified | 387350 | 0 | 355340 | 473380 | 0 | 0 | 28715 | 0 | 2726800 | 515640 | 693530 | 10623000 | 0 | 0 | 448420 | 226570 | 0 | 197930 |
| IGTLELGAF | LRIG1 | Q96JA1 | LRIG1_HUMAN | Unmodified | 0 | 0 | 0 | 0 | 0 | 0 | 0 | 0 | 955560 | 0 | 0 | 0 | 0 | 0 | 0 | 0 | 0 | 0 |
| IGTNQMAV | EIF2B1 | Q14232 | EI2BA_HUMAN | Unmodified | 301440 | 299530 | 0 | 0 | 0 | 0 | 0 | 0 | 517090 | 0 | 605980 | 745800 | 0 | 0 | 0 | 0 | 0 | 0 |
| IGVFNTSG( | RAB23 | Q9ULC3 | RAB23_HUMAN | Unmodified | 0 | 0 | 1005800 | 616500 | 0 | 0 | 0 | 0 | 0 | 0 | 0 | 663320 | 0 | 0 | 1561200 | 552890 | 0 | 0 |
| IGVIRLLDPI | DST; DST; DST; | Q03001; A0/ | DYST_HUMAN | Unmodified | 0 | 0 | 0 | 0 | 3067500 | 0 | 105690 | 28235 | 385100 | 0 | 0 | 475320 | 0 | 0 | 0 | 0 | 0 | 0 |
| IGVLDEGK | RPS9; RPS9; RP | P46781; B5N | RS9_HUMAN 4 | Unmodified | 7911.6 | 2420100 | 0 | 0 | 91650 | 0 | 0 | 0 | 92024 | 194450 | 1718900 | 59439000 | 0 | 0 | 117020 | 0 | 241110 | 0 |
| IGWRNDAS | ITGB3; N/A; ITG | P05106; H3E | ITB3_HUMAN Ii | Unmodified | 6578700 | 9408800 | 7777400 | 29191000 | 48200000 | 6872200 | 93937000 | 1070800 | 0 | 0 | 0 | 0 | 48683000 | 47634000 | 8390400 | 6175900 | 1647000 | 15740000 |
| IGYDYKLEC | ACSL4; ACSL4 | O60488; H0' | ACSL4_HUMAN | Unmodified | 0 | 0 | 0 | 93428 | 869910 | 0 | 506260 | 319120 | 0 | 109920 | 1346600 | 3927600 | 0 | 0 | 0 | 0 | 565670 | 381770 |
| IGYDYKLEC | ACSL4; ACSL4 | O60488; H0' | ACSL4_HUMAN | Unmodified | 0 | 0 | 0 | 0 | 597790 | 255250 | 219500 | 69519 | 0 | 0 | 613740 | 3112100 | 0 | 126740 | 0 | 0 | 480650 | 206680 |
| IGYNPATVF | EEF1A2; EEF1A: | Q05639; A0/ | EF1A2_HUMAN | 2 Oxidation (M) | 4514800 | 4567200 | 2013100 | 5251400 | 7159300 | 681230 | 2721600 | 0 | 0 | 0 | 0 | 0 | 11890000 | 1964500 | 19297000 | 1106700 | 2575100 | 737350 |
| IGYNPDTV/ | EEF1A1P5; EEF | Q5VTE0; P68 | EF1A3_HUMAN | 2 Oxidation (M) | 2616600 | 5960800 | 0 | 4013200 | 4779400 | 0 | 1170400 | 0 | 0 | 0 | 0 | 325960 | 6379700 | 2364100 | 187430 | 1358100 | 4727000 | 3734600 |
| IGYPDDIVS | MME; MME; MM | P08473; A0A | NEP_HUMAN N | Unmodified | 0 | 0 | 4418700 | 2292500 | 0 | 0 | 385010 | 0 | 595950 | 0 | 0 | 0 | 495320 | 160830 | 3795600 | 2412300 | 414700 | 0 |
| IGYPKPALL | WARS1; WARS1 | P23381; H0\ | SYWC_HUMAN | Unmodified | 577620 | 0 | 0 | 1199700 | 15688000 | 3016000 | 1244400 | 0 | 8971100 | 5994500 | 1374000 | 1251800 | 11298000 | 3395200 | 0 | 441590 | 3298900 | 13528000 |
| IGYSSPLTL | ACSL4 | O60488 | ACSL4_HUMAN | Unmodified | 0 | 0 | 0 | 0 | 234380 | 0 | 117400 | 0 | 0 | 0 | 861790 | 6137300 | 0 | 0 | 0 | 0 | 0 | 0 |
| IHAENAIR | CHMP1B | Q7LBR1 | CHM1B_HUMA | Unmodified | 0 | 0 | 0 | 0 | 8359.2 | 0 | 0 | 0 | 9926500 | 51775 | 759160 | 13912000 | 0 | 0 | 0 | 0 | 13377 | 0 |
| IHAGDEVIC | CNK3/IPCEF1; | G9CGD6; Q6 | CNIPF_HUMAN | Unmodified | 0 | 0 | 0 | 0 | 0 | 0 | 0 | 0 | 560730 | 251530 | 0 | 93369 | 0 | 0 | 0 | 0 | 0 | 0 |
| IHAGPITKH | PSMC6; PSMC6 | P62333; A0A | PRS10_HUMAN | Unmodified | 0 | 0 | 0 | 0 | 914270 | 0 | 512390 | 0 | 0 | 785730 | 0 | 0 | 753250 | 590230 | 0 | 0 | 929800 | 735130 |
| IHAQWQK | HMGCS1 | Q01581 | HMCS1_HUMA | Unmodified | 0 | 265910 | 0 | 0 | 0 | 0 | 0 | 97907 | 193900 | 44770 | 24032 | 1068900 | 0 | 0 | 71544 | 34991 | 0 | 0 |
| IHAREIFDS | ENO1; ENO1; E | P06733; K7E | ENOA_HUMAN | Unmodified | 0 | 0 | 0 | 109310 | 1.78E+08 | 9469800 | 11811000 | 29776 | 0 | 32844000 | 191060 | 0 | 71898000 | 57494000 | 0 | 0 | 1.88E+08 | 1.54E+08 |
| IHAYSQLLE | PSMD6 | Q15008 | PSMD6_HUMA | Unmodified | 0 | 0 | 83056 | 0 | 0 | 0 | 0 | 0 | 415300 | 239770 | 151870 | 1518500 | 0 | 0 | 284060 | 0 | 0 | 0 |
| IHCTQTLLE | CRABP1 | P29762 | RABP1_HUMAI | Unmodified | 0 | 0 | 0 | 0 | 0 | 0 | 0 | 0 | 1356700 | 343640 | 0 | 0 | 0 | 0 | 0 | 0 | 382250 | 2697500 |
| IHDLREQM | DLG1; DLG1; D | Q12959; A0/ | DLG1_HUMAN | Unmodified | 0 | 0 | 0 | 0 | 0 | 0 | 0 | 0 | 856620 | 0 | 0 | 0 | 0 | 0 | 0 | 0 | 0 | 0 |
| IHDLREQM | DLG1; DLG1; D | Q12959; A0/ | DLG1_HUMAN | Unmodified | 0 | 0 | 0 | 0 | 587130 | 0 | 0 | 0 | 0 | 0 | 0 | 0 | 0 | 0 | 0 | 0 | 202820 | 336290 |
| IHDPQLLTE | DARS1; DARS1 | P14868; H7E | SYDC_HUMAN | Unmodified | 0 | 0 | 0 | 0 | 0 | 0 | 0 | 0 | 211180 | 0 | 230760 | 5462500 | 0 | 0 | 84078 | 0 | 0 | 0 |
| IHEADINAD | SERINC3 | Q13530 | SERC3_HUMAI | Unmodified | 340140 | 220040 | 989210 | 474650 | 0 | 0 | 0 | 0 | 0 | 0 | 0 | 0 | 0 | 0 | 269730 | 168440 | 0 | 125340 |
| IHEAIIQSPI | ARMT1; ARMT1 | Q9H993; F5( | ARMT1_HUMAI | Unmodified | 81130 | 0 | 0 | 0 | 0 | 0 | 0 | 39600 | 0 | 0 | 135740 | 231470 | 0 | 0 | 127130 | 0 | 0 | 0 |
| IHEEREILGI | POFUT2 | Q9Y2G5 | OFUT2_HUMAI | Unmodified | 0 | 0 | 0 | 0 | 0 | 0 | 0 | 0 | 1021300 | 0 | 0 | 0 | 0 | 0 | 0 | 0 | 0 | 0 |
| IHEGCEEP/ | CLTC; CLTC | Q00610; A0/ | CLH1_HUMAN | Unmodified | 36178000 | 13260000 | 13435000 | 21875000 | 1174700 | 137690 | 0 | 0 | 3608300 | 619230 | 39406 | 1.21E+08 | 3593200 | 303990 | 17865000 | 14817000 | 435200 | 913130 |
| IHEQGLIAII | CASK; CASK; C/ | O14936; A0/ | CSKP_HUMAN | Unmodified | 0 | 0 | 0 | 671020 | 0 | 0 | 0 | 0 | 1082700 | 3189000 | 0 | 323040 | 0 | 0 | 0 | 0 | 0 | 0 |
| IHFIEAQDL | ESYT2; ESYT2; E | A0FGR8; A0/ | ESYT2_HUMAN | Unmodified | 0 | 0 | 162220 | 0 | 0 | 0 | 0 | 125580 | 0 | 0 | 0 | 146650 | 0 | 0 | 99127 | 108950 | 0 | 0 |
| IHFNWLPP | ITGB4 | P16144 | ITB4_HUMAN Ii | Oxidation (M) | 0 | 0 | 0 | 0 | 238550 | 0 | 0 | 0 | 0 | 0 | 0 | 0 | 0 | 0 | 0 | 0 | 1362400 | 0 |
| IHFNWLPP | ITGB4 | P16144 | ITB4_HUMAN Ii | Unmodified | 0 | 0 | 0 | 0 | 0 | 0 | 0 | 0 | 0 | 0 | 0 | 0 | 0 | 0 | 0 | 0 | 1536700 | 0 |
| IHFPLATYA | TUBA3C; TUBA: | P0DPH7; Q7 | TBA3C_HUMAI | Unmodified | 8584700 | 12766000 | 27108000 | 14701000 | 21881000 | 17229000 | 46078000 | 8318100 | 1.58E+08 | 61676000 | 27070000 | 7.95E+08 | 13080000 | 10369000 | 26695000 | 21398000 | 12975000 | 18568000 |
| IHFPSDK | SPINT1; SPINT1 | O43278; H3I | SPIT1_HUMAN | Unmodified | 218040 | 172760 | 137990 | 154780 | 0 | 0 | 0 | 0 | 761130 | 9454.2 | 171490 | 72366 | 0 | 0 | 266250 | 223510 | 843670 | 0 |
| IHFRPELPA | MAOA | P21397 | AOFA_HUMAN | Unmodified | 0 | 0 | 974480 | 0 | 181080 | 0 | 0 | 0 | 0 | 0 | 0 | 0 | 0 | 0 | 433680 | 328770 | 0 | 0 |
| IHFSGYDN | PRSS23 | O95084 | PRS23_HUMAI | Unmodified | 717120 | 0 | 0 | 0 | 0 | 0 | 0 | 0 | 0 | 0 | 0 | 0 | 0 | 0 | 0 | 170950 | 0 | 0 |
| IHFVPGWD | IARS2 | Q9NSE4 | SYIM_HUMAN I | Unmodified | 0 | 0 | 0 | 0 | 0 | 0 | 0 | 0 | 2018400 | 529640 | 0 | 0 | 0 | 0 | 0 | 0 | 0 | 0 |
| IHGAVQAS | DUSP13; DUSP | Q9UII6; U3K | DS13B_HUMAI | Unmodified | 0 | 0 | 0 | 0 | 0 | 0 | 0 | 0 | 20056000 | 147320 | 0 | 0 | 0 | 0 | 0 | 0 | 0 | 0 |
| IHGFTVNQ | PITRM1; PITRM: | Q5JRX3; A0A | PREP_HUMAN | Unmodified | 0 | 0 | 0 | 0 | 535450 | 0 | 0 | 0 | 1020700 | 124350 | 65529 | 0 | 0 | 0 | 0 | 0 | 0 | 0 |
| IHGLDWHI | WDR59; WDR5! | Q6PJI9; B7Z | WDR59_HUMA | Unmodified | 0 | 0 | 0 | 0 | 0 | 0 | 0 | 0 | 0 | 0 | 0 | 353850 | 0 | 0 | 0 | 0 | 0 | 0 |
| IHGQNVPF | CLSTN1; CLSTN | O94985; A0/ | CSTN1_HUMAI | Unmodified | 0 | 0 | 0 | 0 | 127910 | 0 | 0 | 33426 | 3479100 | 739430 | 0 | 0 | 0 | 90730 | 0 | 0 | 0 | 0 |
| IHGRVGDS | AGA | P20933 | ASPG_HUMAN | Oxidation (M) | 0 | 0 | 0 | 0 | 0 | 0 | 0 | 0 | 0 | 1656700 | 0 | 0 | 0 | 0 | 0 | 0 | 0 | 0 |
| IHGSGHVE | ANK3; ANK3; AI | Q12955; H0' | ANK3_HUMAN | Unmodified | 0 | 0 | 692050 | 661200 | 0 | 0 | 0 | 0 | 0 | 0 | 0 | 267500 | 5295300 | 5513300 | 1207700 | 789690 | 214420 | 384110 |
| IHHDVNEL | TUBGCP2; TUB | Q9BSJ2; F2Z | GCP2_HUMAN | Unmodified | 0 | 0 | 0 | 65422 | 0 | 182930 | 0 | 0 | 99005 | 0 | 113000 | 557300 | 0 | 173690 | 0 | 0 | 0 | 418460 |
| IHIDLPNEC | PSMC6; PSMC6 | P62333; A0A | PRS10_HUMAN | Unmodified | 0 | 0 | 0 | 0 | 2353400 | 0 | 0 | 0 | 700480 | 1228200 | 0 | 253990 | 5478500 | 812710 | 0 | 0 | 1851600 | 92301 |
| IHIEENEDR | TTC17 | Q96AE7 | TTC17_HUMAN | Unmodified | 0 | 0 | 0 | 0 | 0 | 0 | 0 | 0 | 1069500 | 0 | 0 | 50586 | 0 | 0 | 0 | 0 | 0 | 0 |
| IHIFAQQNI | ELP2; ELP2 | Q6IA86; F5G | ELP2_HUMAN I | Unmodified | 0 | 0 | 54507 | 0 | 0 | 1496400 | 0 | 0 | 0 | 0 | 0 | 3046300 | 54176 | 0 | 158790 | 158030 | 0 | 0 |
| IHIMPSLNF | CPE; CPE; CPE; | P16870; C9J | CBPE_HUMAN | Oxidation (M) | 0 | 0 | 0 | 0 | 0 | 0 | 0 | 0 | 14476000 | 2389200 | 0 | 0 | 0 | 0 | 1501800 | 0 | 0 | 0 |
| IHIMPSLNF | CPE; CPE; CPE; | P16870; C9J | CBPE_HUMAN | Unmodified | 0 | 99426 | 0 | 0 | 0 | 0 | 0 | 0 | 40890000 | 2707100 | 0 | 0 | 0 | 0 | 0 | 0 | 0 | 0 |
| IHKPDPWL | EHHADH | Q08426 | ECHP_HUMAN | Unmodified | 0 | 0 | 0 | 0 | 0 | 0 | 0 | 0 | 1492000 | 0 | 0 | 0 | 0 | 0 | 0 | 0 | 0 | 0 |
| IHLAQSLHI | CSE1L | P55060 | XPO2_HUMAN | Unmodified | 0 | 0 | 0 | 0 | 153780 | 0 | 0 | 0 | 70109 | 75153 | 0 | 604670 | 0 | 0 | 0 | 0 | 66142 | 137180 |
| IHLFDIDVP | NIT2; NIT2 | Q9NQR4; H7 | NIT2_HUMAN ( | Unmodified | 0 | 0 | 0 | 0 | 0 | 0 | 0 | 0 | 318950 | 0 | 299650 | 508780 | 0 | 0 | 0 | 0 | 0 | 0 |
| IHLFDIDVP | NIT2; NIT2 | Q9NQR4; H7 | NIT2_HUMAN ( | Unmodified | 0 | 0 | 0 | 0 | 6056100 | 0 | 212370 | 0 | 0 | 0 | 83851 | 0 | 1471400 | 1278000 | 0 | 0 | 2314200 | 1323400 |
| IHLMPSMN | CPD | O75976 | CBPD_HUMAN | Unmodified | 0 | 0 | 0 | 0 | 0 | 0 | 359220 | 288730 | 12639000 | 709090 | 0 | 0 | 0 | 174560 | 0 | 141060 | 72886 | 0 |
| IHLMPSMN | CPD | O75976 | CBPD_HUMAN | Oxidation (M) | 0 | 103280 | 0 | 0 | 0 | 0 | 0 | 0 | 16750000 | 1015800 | 0 | 0 | 0 | 0 | 0 | 0 | 0 | 0 |
| IHLMPSMN | CPD | O75976 | CBPD_HUMAN | 2 Oxidation (M) | 0 | 0 | 0 | 0 | 0 | 0 | 0 | 0 | 0 | 0 | 0 | 0 | 0 | 0 | 0 | 0 | 0 | 0 |
| IHLPETVEQ | CASK; CASK; C/ | O14936; A0/ | CSKP_HUMAN | Unmodified | 0 | 0 | 411110 | 894450 | 0 | 0 | 0 | 67430 | 1374600 | 0 | 0 | 0 | 1671900 | 0 | 0 | 656500 | 0 | 0 |
| IHLPETVEQ | CASK; CASK; C/ | O14936; A0/ | CSKP_HUMAN | Unmodified | 0 | 0 | 0 | 0 | 401490 | 0 | 0 | 0 | 2938600 | 414240 | 0 | 441660 | 651720 | 0 | 1510200 | 303130 | 90277 | 0 |
| IHLPLNYPF | PCDH1 | Q08174 | PCDH1_HUMA | Unmodified | 0 | 0 | 0 | 0 | 0 | 0 | 0 | 0 | 0 | 0 | 587850 | 1792900 | 0 | 0 | 0 | 0 | 0 | 0 |
| IHLQLWDT. | RAB27A; RAB27 | P51159; H3E | RB27A_HUMAN | Unmodified | 422890 | 0 | 667550 | 1096000 | 0 | 0 | 0 | 0 | 0 | 0 | 0 | 13509000 | 0 | 0 | 723260 | 4579400 | 0 | 0 |
| IHLVDKWN | SCARB1; SCARI | Q8WTV0; A0 | SCRB1_HUMAI | Unmodified | 583960 | 488080 | 0 | 224750 | 0 | 0 | 0 | 0 | 104900 | 56573 | 49368 | 1343200 | 0 | 0 | 201270 | 0 | 0 | 0 |
| IHLVVLVKP | CXADR | P78310 | CXAR_HUMAN | Unmodified | 0 | 0 | 0 | 0 | 1463300 | 0 | 0 | 0 | 0 | 0 | 0 | 792780 | 0 | 0 | 0 | 0 | 0 | 0 |
| IHLVVLVKP | CXADR | P78310 | CXAR_HUMAN | Unmodified | 0 | 0 | 0 | 0 | 2901600 | 0 | 0 | 0 | 0 | 0 | 0 | 0 | 0 | 0 | 0 | 0 | 0 | 0 |
| IHMGSCAE | ACAT1; ACAT1; | P24752; A0A | THIL_HUMAN A | Unmodified | 0 | 0 | 0 | 0 | 0 | 0 | 0 | 260860 | 1777200 | 493630 | 0 | 0 | 252610 | 562560 | 0 | 0 | 151460 | 217120 |
| IHNAENIQI | EIF4G1; EIF4G1 | Q04637; C9J | IF4G1_HUMAN | Unmodified | 0 | 0 | 0 | 0 | 0 | 0 | 0 | 0 | 0 | 0 | 0 | 434360 | 0 | 0 | 46338 | 58827 | 0 | 0 |
| IHNAITVHI | PALS1; PALS1 | Q8N3R9; G3 | PALS1_HUMAN | Unmodified | 0 | 0 | 75080 | 0 | 1832900 | 419990 | 0 | 0 | 144430 | 171200 | 0 | 0 | 851800 | 754860 | 46798 | 36014 | 541060 | 1080900 |
| IHNFGLIQE | ACADVL; ACAD | P49748; J3K | ACADV_HUMAI | Unmodified | 0 | 0 | 0 | 0 | 0 | 0 | 0 | 0 | 1080700 | 395990 | 0 | 284680 | 0 | 0 | 0 | 0 | 0 | 0 |
| IHNLGLNC | SEC24D; SEC2- | O94855; E9F | SC24D_HUMAI | Unmodified | 0 | 0 | 0 | 0 | 0 | 0 | 0 | 0 | 0 | 0 | 0 | 940290 | 0 | 31762 | 0 | 0 | 0 | 0 |
| IHPELLAK | MACROH2A2; I | Q9P0M6; O7 | H2AW_HUMAN | Unmodified | 119390 | 230960 | 0 | 0 | 0 | 0 | 0 | 0 | 0 | 0 | 0 | 3028500 | 0 | 0 | 0 | 0 | 0 | 0 |
| IHPFHILIAL | RO60; RO60 | P10155; G5E | RO60_HUMAN | Unmodified | 1254200 | 1424200 | 0 | 792010 | 0 | 0 | 0 | 0 | 0 | 0 | 0 | 181740 | 172520 | 0 | 1503400 | 588060 | 0 | 0 |
| IHPMAYQL | CSNK2B; CSNK | P67870; Q5! | CSK2B_HUMAI | Oxidation (M) | 1507800 | 0 | 0 | 0 | 2413600 | 0 | 0 | 0 | 167410 | 82736 | 0 | 2325000 | 1517100 | 0 | 0 | 412360 | 0 | 0 |
| IHPMAYQL | CSNK2B; CSNK | P67870; Q5! | CSK2B_HUMAI | Oxidation (M) | 0 | 0 | 0 | 0 | 2312200 | 0 | 1038700 | 35844 | 0 | 481410 | 0 | 0 | 682650 | 897500 | 0 | 0 | 2085400 | 808190 |
| IHPQTIIAG\ | CCT2; CCT2; C( | P78371; F5G | TCPB_HUMAN | Unmodified | 68260 | 14967 | 0 | 0 | 0 | 2726500 | 0 | 0 | 778710 | 0 | 0 | 7034800 | 0 | 0 | 201260 | 183070 | 118240 | 0 |
| IHPQTIIAG\ | CCT2; CCT2; C( | P78371; F5G | TCPB_HUMAN | Unmodified | 0 | 0 | 0 | 0 | 2707100 | 454740 | 0 | 68700 | 0 | 0 | 260230 | 0 | 6855400 | 0 | 0 | 0 | 1813500 | 2464700 |
| IHPTSVISG | TCP1; TCP1; TC | P17987; F5H | TCPA_HUMAN | Unmodified | 106080 | 0 | 482000 | 127900 | 625340 | 47292 | 31415 | 48324 | 2232300 | 346080 | 3225500 | 14456000 | 935800 | 0 | 680580 | 649260 | 121000 | 295650 |
| IHPTSVISG | TCP1; TCP1; TC | P17987; F5H | TCPA_HUMAN | Unmodified | 0 | 0 | 0 | 0 | 0 | 0 | 537380 | 0 | 0 | 324290 | 0 | 0 | 1093300 | 0 | 0 | 0 | 1001000 | 790470 |
| IHPVDKLTI | ACO2; ACO2; A | Q99798; A2/ | ACON_HUMAN | Unmodified | 0 | 0 | 0 | 0 | 0 | 0 | 265590 | 0 | 1179600 | 714280 | 0 | 0 | 0 | 0 | 0 | 0 | 0 | 0 |
| IHPVSTMVI | LDHB; LDHB; N | P07195; A0A | LDHB_HUMAN | Unmodified | 0 | 0 | 0 | 0 | 285170 | 0 | 0 | 0 | 0 | 87498 | 1343900 | 13231000 | 0 | 0 | 0 | 0 | 62013 | 210970 |
| IHQALKEDI | CUL5; CUL5 | Q93034; H0' | CUL5_HUMAN | Unmodified | 68159 | 50223 | 0 | 113140 | 122600 | 54827 | 47766 | 0 | 60773 | 91085 | 167420 | 800960 | 110320 | 154250 | 201700 | 79374 | 129670 | 165200 |
| IHQIELIPNI | CDC42BPA; CD | Q5VT25; A0/ | MRCKA_HUMA | Unmodified | 92301 | 148170 | 275190 | 884370 | 0 | 423240 | 0 | 0 | 2304500 | 661340 | 610950 | 1270400 | 587230 | 742270 | 845600 | 692550 | 365840 | 1061200 |
| IHQIEYAME | PSMA1; N/A | P25786; B4D | PSA1_HUMAN | Unmodified | 240670 | 0 | 510260 | 0 | 0 | 0 | 0 | 0 | 2574700 | 1179800 | 2133500 | 9216800 | 934160 | 0 | 161400 | 0 | 0 | 0 |
| IHQIEYAME | PSMA1; N/A | P25786; B4D | PSA1_HUMAN | Oxidation (M) | 0 | 0 | 0 | 268870 | 0 | 0 | 0 | 0 | 0 | 0 | 0 | 1048400 | 239510 | 0 | 310620 | 0 | 0 | 0 |
| IHQIGPGM | DNAJA1 | P31689 | DNJA1_HUMAN | Unmodified | 0 | 251630 | 0 | 141490 | 0 | 0 | 0 | 0 | 6774100 | 0 | 0 | 0 | 0 | 2570200 | 0 | 0 | 0 | 0 |
| IHQIGPGM | DNAJA1 | P31689 | DNJA1_HUMAN | Oxidation (M) | 1219100 | 642120 | 0 | 0 | 0 | 0 | 0 | 0 | 2178400 | 213090 | 0 | 691680 | 0 | 0 | 1044400 | 1338600 | 0 | 0 |
| IHQQELEV( | UTRN | P46939 | UTRN_HUMAN | Unmodified | 0 | 0 | 0 | 0 | 0 | 0 | 0 | 0 | 2765100 | 331840 | 0 | 0 | 0 | 0 | 725450 | 1904600 | 0 | 1186400 |
| IHQSTNQTI | FOCAD | Q5VW36 | FOCAD_HUMA | Unmodified | 0 | 0 | 45987 | 0 | 0 | 0 | 0 | 0 | 0 | 0 | 141830 | 837310 | 0 | 0 | 0 | 72522 | 0 | 0 |
| IHSDLAEEF | DSG2; DSG2 | Q14126; J3K | DSG2_HUMAN | Unmodified | 0 | 0 | 0 | 0 | 0 | 841410 | 0 | 0 | 1053300 | 0 | 35776 | 0 | 0 | 0 | 0 | 0 | 49684 | 0 |

| | | | | | | | | | | | | | | | | | | | | | | |
|---|---|---|---|---|---|---|---|---|---|---|---|---|---|---|---|---|---|---|---|---|---|---|
| IHSDLAEEF | DSG2; DSG2 | Q14126; J3K | DSG2_HUMAN | Unmodified | 109010 | 0 | 721020 | 0 | 0 | 957800 | 14944 | 17028 | 0 | 0 | 0 | 0 | 281110 | 225740 | 683650 | 760810 | 0 | 127050 |
| IHSGSVTAL | TEP1; TEP1 | Q99973; G3 | TEP1_HUMAN T | Unmodified | 44253 | 0 | 0 | 0 | 0 | 0 | 0 | 0 | 0 | 0 | 0 | 310230 | 0 | 0 | 0 | 0 | 0 | 0 |
| IHSLCYND | ELAPOR1; ELAP | Q6UXG2; E9 | ELAP1_HUMAN | Unmodified | 295600 | 0 | 0 | 72108 | 0 | 28022 | 0 | 0 | 577960 | 0 | 0 | 0 | 0 | 0 | 0 | 0 | 72151 | 0 |
| IHSLTHLDS | BRCC3; BRCC3 | P46736; A0A | BRCC3_HUMA | Unmodified | 0 | 0 | 0 | 0 | 0 | 0 | 0 | 0 | 98865 | 0 | 163950 | 929580 | 0 | 0 | 0 | 0 | 0 | 125890 |
| IHVFYIDYG | SND1 | Q7KZF4 | SND1_HUMAN | Unmodified | 0 | 0 | 2571600 | 1594700 | 316460 | 0 | 0 | 0 | 544940 | 101480 | 3141800 | 3972500 | 2690200 | 0 | 1285700 | 6046400 | 962250 | 807550 |
| IHVFYIDYG | SND1 | Q7KZF4 | SND1_HUMAN | Unmodified | 0 | 0 | 0 | 0 | 0 | 0 | 0 | 0 | 0 | 427990 | 0 | 89262 | 4072500 | 1954100 | 0 | 0 | 0 | 0 |
| IHVLEAQDI | ESYT1 | Q9BSJ8 | ESYT1_HUMAN | Unmodified | 625320 | 275240 | 653830 | 466830 | 1476100 | 125810 | 62309 | 17715 | 1308400 | 348280 | 410060 | 1681200 | 123260 | 112240 | 386290 | 219620 | 99699 | 182650 |
| IHVNPPAE | MGAT4A | Q9UM21 | MGT4A_HUMA | Unmodified | 0 | 754940 | 0 | 567340 | 322330 | 356290 | 819070 | 278740 | 3742100 | 653590 | 0 | 0 | 0 | 0 | 0 | 1244900 | 138260 | 0 |
| IHVPKPLDF | POGLUT3 | Q7Z4H8 | PLGT3_HUMAN | Unmodified | 0 | 0 | 0 | 0 | 0 | 0 | 0 | 0 | 6864100 | 813080 | 0 | 0 | 0 | 0 | 0 | 0 | 0 | 0 |
| IHVSDQEL | UBA1; UBA1 | P22314; Q5J | UBA1_HUMAN | Unmodified | 0 | 0 | 0 | 0 | 0 | 716450 | 0 | 21000 | 75370 | 163580 | 551260 | 0 | 0 | 0 | 0 | 0 | 0 | 0 |
| IHVSDQEL | UBA1; UBA1 | P22314; Q5J | UBA1_HUMAN | Unmodified | 1041800 | 0 | 0 | 3885400 | 7797100 | 1778300 | 5088500 | 0 | 16223000 | 5145900 | 8212000 | 25323000 | 7174500 | 3974700 | 0 | 837640 | 6150200 | 1337600 |
| IHVSYQET | KTN1 | Q86UP2 | KTN1_HUMAN | Unmodified | 0 | 0 | 0 | 0 | 0 | 0 | 0 | 0 | 0 | 0 | 0 | 0 | 0 | 0 | 0 | 0 | 0 | 0 |
| IHVWNGES | WDR82 | Q6UXN9 | WDR82_HUMA | Unmodified | 0 | 0 | 0 | 0 | 0 | 0 | 0 | 0 | 0 | 0 | 0 | 2589000 | 0 | 0 | 0 | 40454 | 0 | 0 |
| IHVYGYSM | PHPT1 | Q9NRX4 | PHP14_HUMAN | Unmodified | 0 | 0 | 0 | 0 | 1092700 | 0 | 0 | 0 | 1152500 | 232030 | 0 | 1965800 | 0 | 0 | 0 | 0 | 278630 | 2180600 |
| IHVYGYSM | PHPT1 | Q9NRX4 | PHP14_HUMAN | Oxidation (M) | 374590 | 0 | 263020 | 292660 | 0 | 0 | 233000 | 0 | 618850 | 0 | 0 | 2135200 | 393160 | 390160 | 599530 | 572430 | 0 | 0 |
| IHYIGQNEF | HMOX2; HMOX | P30519; I3L1 | HMOX2_HUMA | Unmodified | 0 | 0 | 0 | 0 | 0 | 0 | 0 | 0 | 0 | 0 | 0 | 7768300 | 0 | 0 | 0 | 0 | 28229 | 0 |
| IHYLDTTTL | CRKL | P46109 | CRKL_HUMAN | Unmodified | 468600 | 343050 | 0 | 0 | 0 | 0 | 0 | 0 | 0 | 69390 | 0 | 2716300 | 0 | 0 | 291780 | 291090 | 0 | 0 |
| IHYLDTTTL | CRK; CRK | P46108; I3L2 | CRK_HUMAN A | Unmodified | 1409700 | 3450100 | 0 | 0 | 0 | 0 | 0 | 0 | 0 | 0 | 0 | 0 | 0 | 0 | 0 | 0 | 0 | 0 |
| IHYPDFLV | PROM2 | Q8N271 | PROM2_HUMA | Unmodified | 3603400 | 2450900 | 2712500 | 2879700 | 0 | 0 | 0 | 0 | 0 | 0 | 1653700 | 4824400 | 872790 | 0 | 3609200 | 3116600 | 0 | 0 |
| IIADIFEYTA | MMUT | P22033 | MUTA_HUMAN | Unmodified | 0 | 0 | 0 | 0 | 0 | 0 | 0 | 0 | 2664700 | 274170 | 0 | 0 | 229900 | 0 | 0 | 0 | 0 | 0 |
| IIADLEEEQ | UTRN; UTRN | P46939; Q5 | UTRN_HUMAN | Unmodified | 0 | 0 | 279130 | 161660 | 0 | 0 | 0 | 0 | 454270 | 0 | 0 | 0 | 0 | 0 | 0 | 0 | 0 | 0 |
| IIADLEEEQ | UTRN; UTRN | P46939; Q5 | UTRN_HUMAN | Unmodified | 0 | 0 | 0 | 2399200 | 0 | 0 | 395440 | 0 | 973190 | 0 | 0 | 0 | 2284300 | 0 | 0 | 0 | 0 | 0 |
| IIAEALTR | NCLN; NCLN; N | Q969V3; K7E | NCLN_HUMAN | Unmodified | 0 | 124020 | 119740 | 0 | 992800 | 99021 | 0 | 111970 | 21298000 | 1594600 | 1241000 | 1079800 | 0 | 0 | 0 | 0 | 0 | 0 |
| IIAFVGSPV | PSMD4; PSMD4 | P55036; A6P | PSMD4_HUMA | Unmodified | 0 | 0 | 0 | 0 | 0 | 0 | 0 | 0 | 0 | 0 | 0 | 7696000 | 0 | 0 | 0 | 0 | 0 | 0 |
| IIAGDKEVV | SEC31A; SEC31 | O94979; D6I | SC31A_HUMAN | Unmodified | 0 | 0 | 0 | 0 | 0 | 0 | 0 | 0 | 0 | 0 | 249940 | 626760 | 0 | 0 | 0 | 0 | 0 | 0 |
| IIAKGEELLS | MYO1C; MYO1C | O00159; F5H | MYO1C_HUMA | Unmodified | 2305400 | 4405600 | 125500 | 6307500 | 760880 | 765710 | 0 | 0 | 0 | 0 | 0 | 462020 | 0 | 2822200 | 7044200 | 4523700 | 0 | 0 |
| IIAKGEELLS | MYO1C; MYO1C | O00159; F5H | MYO1C_HUMA | Unmodified | 0 | 0 | 0 | 6069300 | 34819000 | 16371000 | 3395400 | 0 | 0 | 727600 | 0 | 0 | 5087300 | 3864000 | 0 | 0 | 12779000 | 3794300 |
| IIALDGDTK | TKT; TKT | P29401; A0A | TKT_HUMAN Tr | Unmodified | 0 | 0 | 358350 | 247960 | 412210 | 0 | 44748 | 62419 | 2399200 | 1130700 | 4797800 | 18109000 | 196600 | 0 | 0 | 440880 | 0 | 0 |
| IIALGHSGF | NT5E; NT5E | P21589; Q96 | 5NTD_HUMAN | Oxidation (M) | 157200 | 0 | 0 | 0 | 12094000 | 5364300 | 79601 | 0 | 0 | 0 | 213320 | 86766 | 0 | 0 | 0 | 0 | 1790200 | 3803000 |
| IIALGHSGF | NT5E; NT5E | P21589; Q96 | 5NTD_HUMAN | Unmodified | 0 | 0 | 0 | 0 | 35290000 | 9596600 | 0 | 0 | 0 | 0 | 0 | 0 | 0 | 0 | 0 | 0 | 2282700 | 13102000 |
| IIALGHSGF | NT5E; NT5E | P21589; Q96 | 5NTD_HUMAN | Unmodified | 0 | 0 | 0 | 0 | 26729000 | 5158500 | 0 | 26425 | 0 | 0 | 0 | 0 | 0 | 0 | 0 | 0 | 9522600 | 14980000 |
| IIALGHSGF | NT5E; NT5E | P21589; Q96 | 5NTD_HUMAN | Oxidation (M) | 0 | 0 | 0 | 0 | 4856100 | 1370400 | 0 | 0 | 0 | 0 | 0 | 0 | 0 | 0 | 0 | 0 | 1277000 | 1385500 |
| IIANALSSEI | PRKDC | P78527 | PRKDC_HUMA | Unmodified | 0 | 0 | 0 | 0 | 0 | 0 | 0 | 0 | 930400 | 0 | 0 | 2014200 | 0 | 0 | 0 | 0 | 0 | 0 |
| IIAPANLERI | ABI2; ABI2; ABI2 | Q9NYB9; E7 | ABI2_HUMAN A | Unmodified | 0 | 0 | 0 | 0 | 1126000 | 407260 | 86949 | 77793 | 520500 | 0 | 1043600 | 3996800 | 235300 | 0 | 0 | 0 | 661290 | 1715000 |
| IIAPANMER | ABI1; ABI1; ABI1 | Q8IZP0; A0A | ABI1_HUMAN A | Unmodified | 330600 | 0 | 695200 | 1274400 | 3512000 | 0 | 0 | 0 | 0 | 0 | 0 | 3018100 | 1664500 | 690810 | 1145600 | 762510 | 722540 | 1106500 |
| IIAPLVTR | PLOD2; PLOD2 | O00469; E7E | PLOD2_HUMA | Unmodified | 0 | 0 | 0 | 0 | 0 | 0 | 0 | 0 | 5489800 | 1577000 | 0 | 0 | 0 | 0 | 0 | 0 | 0 | 0 |
| IIAQCLNK | NDUFS2; NDUF | O75306; A0A | NDUS2_HUMA | Unmodified | 0 | 0 | 0 | 0 | 0 | 0 | 137240 | 27510 | 620190 | 219670 | 0 | 0 | 0 | 0 | 0 | 0 | 0 | 0 |
| IIASSPEMN | PYCR2; PYCR2; | Q96C36; A0 | P5CR2_HUMA | Unmodified | 0 | 0 | 0 | 0 | 0 | 0 | 0 | 83693 | 3103300 | 693250 | 0 | 0 | 0 | 0 | 0 | 0 | 0 | 0 |
| IIATSQGLLI | SEMA3C | Q99985 | SEM3C_HUMA | Unmodified | 102740 | 0 | 951510 | 743570 | 0 | 0 | 0 | 0 | 0 | 0 | 0 | 0 | 0 | 0 | 1748300 | 1490000 | 0 | 0 |
| IIAVDINK | ADH1B; ADH1C | P00325; P00 | ADH1B_HUMA | Unmodified | 88355 | 374410 | 0 | 0 | 185160 | 0 | 163810 | 0 | 1258800 | 94377 | 4622100 | 1553400 | 0 | 87094 | 0 | 0 | 50880 | 0 |
| IICAQQCS | EGFR; EGFR; E | P00533; Q5 | EGFR_HUMAN | Unmodified | 0 | 0 | 0 | 690760 | 5713200 | 1919900 | 65844 | 0 | 7967.6 | 86140 | 0 | 0 | 1899700 | 1116000 | 0 | 0 | 1584200 | 1388800 |
| IICDLVEEK | MYO1C; MYO1C | O00159; F5H | MYO1C_HUMA | Unmodified | 1776400 | 1595800 | 0 | 2967600 | 5834100 | 0 | 154800 | 0 | 0 | 0 | 0 | 0 | 0 | 0 | 6557900 | 4245000 | 3149000 | 0 |
| IICDLVEEK | MYO1C; MYO1C | O00159; F5H | MYO1C_HUMA | Unmodified | 0 | 0 | 0 | 0 | 513220 | 0 | 0 | 0 | 0 | 0 | 0 | 0 | 0 | 0 | 0 | 0 | 695480 | 0 |
| IICMGAKE | GGCT; N/A | O75223; M0 | GGCT_HUMAN | Unmodified | 0 | 0 | 0 | 0 | 147430 | 0 | 73996 | 35567 | 330010 | 0 | 0 | 0 | 591040 | 96525 | 0 | 0 | 452430 | 128800 |
| IICNTKDNE | TXNRD1; TXNR | Q16881; A0 | TRXR1_HUMAN | Unmodified | 0 | 0 | 0 | 0 | 10353000 | 1100000 | 1146700 | 0 | 0 | 3211900 | 0 | 0 | 4990500 | 3699300 | 0 | 0 | 19616000 | 13892000 |
| IICQGFTGK | SUCLG1; SUCL | P53597; A0A | SUCA_HUMAN | Unmodified | 0 | 0 | 0 | 0 | 0 | 0 | 0 | 83181 | 3537000 | 872120 | 0 | 0 | 290100 | 0 | 0 | 0 | 102410 | 291400 |
| IICQGFTGK | SUCLG1; SUCL | P53597; A0A | SUCA_HUMAN | Unmodified | 0 | 0 | 0 | 0 | 0 | 0 | 648170 | 125040 | 0 | 267940 | 0 | 0 | 0 | 0 | 0 | 0 | 146250 | 0 |
| IIDEVVNK | BAG2 | O95816 | BAG2_HUMAN | Unmodified | 63457 | 0 | 326410 | 305220 | 1281700 | 335390 | 0 | 11921 | 838430 | 200420 | 0 | 308580 | 713910 | 224040 | 197750 | 215120 | 0 | 0 |
| IIDEVVNKF | BAG2 | O95816 | BAG2_HUMAN | Unmodified | 0 | 0 | 0 | 0 | 695360 | 125770 | 0 | 0 | 175660 | 0 | 0 | 0 | 0 | 0 | 15592 | 0 | 385810 | 0 |
| IIDGLLVMR | PPIH; PPIH; PPI | O43447; A0 | PPIH_HUMAN | Unmodified | 0 | 106090 | 0 | 0 | 0 | 0 | 0 | 0 | 0 | 0 | 0 | 0 | 632030 | 0 | 0 | 0 | 0 | 0 |
| IIDGRDLMF | STS; STS; STS; | P08842; A0A | STS_HUMAN S | Unmodified | 0 | 0 | 0 | 0 | 288710 | 0 | 0 | 0 | 1071800 | 0 | 0 | 0 | 0 | 0 | 0 | 0 | 0 | 0 |
| IIDISDVFR | DARS2; DARS2; | Q6PI48; A0A | SYDM_HUMAN | Unmodified | 19434 | 0 | 20090 | 0 | 0 | 1478400 | 0 | 0 | 6851700 | 206950 | 279070 | 138620 | 0 | 0 | 0 | 11655 | 0 | 0 |
| IIDISDVFR | DARS2; DARS2; | Q6PI48; A0A | SYDM_HUMAN | Unmodified | 0 | 0 | 0 | 0 | 0 | 0 | 0 | 0 | 1502800 | 2383200 | 0 | 0 | 217990 | 0 | 0 | 0 | 0 | 0 |
| IIDKEGDD | SMC4; SMC4; S | Q9NTJ3; C9 | SMC4_HUMAN | Unmodified | 79499 | 0 | 59449 | 104120 | 347940 | 0 | 0 | 0 | 767680 | 309440 | 161410 | 964150 | 497600 | 0 | 74370 | 103540 | 0 | 0 |
| IIDKTQLNP | MARK3; MARK1 | P27448; Q9 | MARK3_HUMA | Unmodified | 0 | 0 | 0 | 0 | 0 | 0 | 0 | 255050 | 0 | 0 | 0 | 0 | 0 | 0 | 0 | 0 | 0 | 0 |
| IIDPLPPID | DDX42; DDX42 | Q86XP3; A0 | DDX42_HUMA | Unmodified | 0 | 0 | 0 | 0 | 0 | 0 | 0 | 0 | 0 | 0 | 1627500 | 1805200 | 0 | 0 | 0 | 157500 | 0 | 0 |
| IIDQVYR | PLXNB2 | O15031 | PLXB2_HUMAN | Unmodified | 586360 | 465400 | 2276500 | 822780 | 0 | 0 | 0 | 0 | 1931000 | 274550 | 300390 | 7337600 | 0 | 0 | 3283600 | 2619100 | 0 | 0 |
| IIDRIHGTE | NUMA1; NUMA | Q14980; F5 | NUMA1_HUMA | Unmodified | 0 | 0 | 0 | 0 | 0 | 0 | 0 | 0 | 0 | 0 | 0 | 427070 | 0 | 0 | 0 | 0 | 0 | 0 |
| IIDSARSAY | SFN | P31947 | 1433S_HUMAN | Unmodified | 0 | 0 | 0 | 0 | 1071000 | 674250 | 0 | 0 | 0 | 623830 | 0 | 0 | 516880 | 0 | 0 | 0 | 661070 | 0 |
| IIDSGPQLS | FAM91A1; FAM | Q658Y4; E7E | F91A1_HUMAN | Unmodified | 0 | 0 | 0 | 0 | 0 | 0 | 0 | 0 | 1295100 | 0 | 0 | 1391000 | 0 | 0 | 0 | 0 | 1004100 | 0 |
| IIDSKNYAG | SUSD2 | Q9UGT4 | SUSD2_HUMA | Unmodified | 0 | 0 | 186000 | 0 | 1313200 | 58554 | 0 | 0 | 0 | 0 | 0 | 0 | 0 | 0 | 0 | 0 | 0 | 0 |
| IIDSLFNTV | UGDH | O60701 | UGDH_HUMAN | Unmodified | 0 | 0 | 0 | 0 | 916410 | 0 | 0 | 0 | 0 | 0 | 0 | 688810 | 322550 | 0 | 0 | 0 | 233450 | 0 |
| IIDSLFNTV | UGDH | O60701 | UGDH_HUMAN | Unmodified | 0 | 0 | 0 | 0 | 0 | 0 | 0 | 0 | 0 | 0 | 0 | 0 | 0 | 0 | 0 | 0 | 0 | 0 |
| IIDTSLTRD | FKBP11; FKBP1 | Q9NYL4; F8 | FKB11_HUMAN | Unmodified | 0 | 0 | 0 | 421340 | 0 | 0 | 0 | 0 | 1634500 | 0 | 1312500 | 0 | 0 | 0 | 320940 | 0 | 0 | 0 |
| IIDVSTVK | REXO2; REXO2; | Q9Y3B8; F5 | ORN_HUMAN | Unmodified | 0 | 217800 | 494490 | 605240 | 3089200 | 797850 | 146470 | 79585 | 1999200 | 609720 | 1306300 | 5798600 | 8684500 | 2098000 | 501750 | 637070 | 653100 | 2326700 |
| IIDVVYNAS | RPS8; RPS8 | P62241; Q5J | RS8_HUMAN 4 | Unmodified | 7910800 | 7965400 | 3452400 | 3192500 | 97060 | 1638200 | 307550 | 371440 | 1688400 | 0 | 2208200 | 56629000 | 975230 | 0 | 2372300 | 3902700 | 254010 | 736370 |
| IIDVVYNAS | RPS8; RPS8 | P62241; Q5J | RS8_HUMAN 4 | Unmodified | 0 | 0 | 0 | 0 | 301450 | 0 | 0 | 4816000 | 0 | 0 | 0 | 0 | 3396600 | 1000800 | 0 | 0 | 0 | 0 |
| IIEDLRAQIF | KRT18; KRT18 | P05783; F8V | K1C18_HUMAN | Unmodified | 0 | 0 | 0 | 0 | 1250600 | 0 | 0 | 18579 | 1930300 | 1504400 | 4084700 | 324070 | 2314300 | 0 | 0 | 0 | 662100 | 0 |
| IIEDQQESL | SSB | P05455 | LA_HUMAN Lu | Unmodified | 0 | 0 | 0 | 0 | 0 | 0 | 0 | 0 | 0 | 0 | 0 | 2481500 | 0 | 0 | 0 | 0 | 0 | 0 |
| IIEEAPAPG | MCCC1; MCCC | Q96RQ3; E9 | MCCA_HUMAN | Unmodified | 0 | 0 | 0 | 0 | 0 | 0 | 0 | 0 | 1124500 | 78710 | 0 | 0 | 0 | 0 | 0 | 0 | 0 | 0 |
| IIEELQDLG | OS9 | Q13438 | OS9_HUMAN P | Unmodified | 0 | 0 | 0 | 0 | 0 | 0 | 0 | 0 | 10273000 | 521430 | 626860 | 0 | 0 | 36856 | 0 | 0 | 0 | 0 |
| IIEETLALK | ARPC2; ARPC2; | O15144; G5 | ARPC2_HUMA | Unmodified | 1081200 | 959810 | 1932900 | 501540 | 0 | 0 | 0 | 0 | 686630 | 0 | 1540800 | 12987000 | 52816 | 0 | 2597500 | 1374900 | 0 | 0 |
| IIEGGAAHK | DLG1; DLG1; D | Q12959; H7 | DLG1_HUMAN | Unmodified | 0 | 0 | 0 | 0 | 1619400 | 344920 | 0 | 21387 | 378870 | 953840 | 136770 | 292830 | 227120 | 0 | 0 | 0 | 658870 | 294820 |
| IIEIEDEAEK | TRIM29; TRIM2 | Q14134; E9 | TRI29_HUMAN | Unmodified | 0 | 0 | 0 | 0 | 0 | 0 | 0 | 0 | 0 | 0 | 448470 | 2674800 | 0 | 0 | 0 | 0 | 0 | 0 |
| IIELEQK | SPECC1; SPEC | Q5M775; J3 | CYTSB_HUMA | Unmodified | 0 | 0 | 0 | 0 | 0 | 0 | 0 | 0 | 2421500 | 0 | 0 | 0 | 0 | 0 | 0 | 0 | 0 | 0 |
| IIELPFQNK | SERPINB5 | P36952 | SPB5_HUMAN | Unmodified | 0 | 0 | 0 | 0 | 2433600 | 0 | 0 | 0 | 0 | 0 | 0 | 0 | 2081800 | 1781300 | 0 | 0 | 4720300 | 6477900 |
| IIELSQTTA | NCAM2; NCAM | O15394; H9 | NCAM2_HUMA | Unmodified | 791830 | 984410 | 3800200 | 1927500 | 0 | 0 | 0 | 0 | 0 | 0 | 0 | 0 | 0 | 0 | 6966700 | 7448700 | 0 | 0 |
| IIENAENEY | CAPZA1 | P52907 | CAZA1_HUMA | Unmodified | 0 | 0 | 0 | 0 | 16849000 | 0 | 0 | 0 | 3065000 | 1046800 | 948970 | 599990 | 4559700 | 2904300 | 0 | 0 | 2933300 | 639210 |
| IIENAENEY | CAPZA1 | P52907 | CAZA1_HUMA | Oxidation (M) | 2175300 | 1792600 | 0 | 0 | 8776600 | 0 | 187190 | 0 | 0 | 2953700 | 0 | 4222600 | 8026400 | 709780 | 2176200 | 1488900 | 4022800 | 4130100 |
| IIETLTQQL | PFDN2 | Q9UHV9 | PFD2_HUMAN | Unmodified | 433850 | 190910 | 233290 | 1095300 | 2257700 | 0 | 1101600 | 564310 | 1383700 | 2977600 | 9529000 | 13060000 | 6825000 | 2298300 | 713950 | 516600 | 4729500 | 2239900 |
| IIETNEEIID | N/A | N/A | N/A | Oxidation (M) | 0 | 0 | 0 | 0 | 3217000 | 0 | 0 | 0 | 0 | 0 | 1761400 | 0 | 0 | 0 | 0 | 0 | 13661000 | 2502200 |
| IIETPGIR | LRPPRC | P42704 | LPPRC_HUMA | Unmodified | 0 | 0 | 0 | 0 | 0 | 0 | 0 | 0 | 1951200 | 95736 | 193180 | 0 | 0 | 0 | 0 | 0 | 0 | 0 |
| IIEVAPQVA | SEPHS1 | P49903 | SPS1_HUMAN | Unmodified | 0 | 0 | 0 | 0 | 0 | 0 | 0 | 97966 | 685310 | 0 | 854220 | 1443000 | 3482100 | 0 | 0 | 0 | 0 | 0 |
| IIEVEEEQE | FBLN1; FBLN1; | P23142; B1A | FBLN1_HUMA | Unmodified | 0 | 0 | 0 | 0 | 0 | 0 | 0 | 0 | 2370200 | 0 | 0 | 0 | 0 | 0 | 0 | 0 | 0 | 0 |
| IIEVGDTPK | SEC61A1; SEC6 | P61619; Q9 | S61A1_HUMA | Unmodified | 81055 | 139490 | 0 | 0 | 0 | 0 | 274790 | 0 | 5179900 | 1147000 | 437090 | 1003200 | 0 | 0 | 0 | 0 | 0 | 0 |
| IIEVGDTPK | SEC61A1; SEC6 | P61619; Q9 | S61A1_HUMA | Unmodified | 0 | 0 | 0 | 0 | 0 | 0 | 0 | 0 | 563280 | 0 | 0 | 0 | 0 | 0 | 0 | 0 | 0 | 0 |
| IIEVSGQK | RAB14; RAB14 | P61106; X6R | RAB14_HUMA | Unmodified | 2245600 | 2765700 | 0 | 0 | 1561400 | 784980 | 29158 | 61540 | 3859100 | 1070300 | 0 | 12904000 | 1276200 | 0 | 0 | 3383500 | 201870 | 278790 |
| IIEVVDAIM | PFKP; PFKP; PF | Q01813; A0 | PFKAP_HUMA | Unmodified | 0 | 0 | 0 | 0 | 0 | 0 | 0 | 0 | 0 | 0 | 0 | 2585400 | 0 | 0 | 0 | 0 | 502990 | 279290 |
| IIEVVDAIM | PFKP; PFKP; PF | Q01813; A0 | PFKAP_HUMA | Oxidation (M) | 0 | 0 | 0 | 0 | 0 | 0 | 0 | 0 | 0 | 0 | 0 | 2734100 | 0 | 0 | 127190 | 81873 | 0 | 0 |
| IIFDDFR | STT3A | P46977 | STT3A_HUMAN | Unmodified | 0 | 0 | 0 | 0 | 0 | 0 | 0 | 0 | 19361000 | 2280600 | 339180 | 1727400 | 0 | 667670 | 0 | 0 | 0 | 0 |
| IIFDDFREA | STT3A | P46977 | STT3A_HUMAN | Unmodified | 0 | 0 | 0 | 0 | 0 | 0 | 0 | 0 | 3154100 | 879090 | 0 | 0 | 0 | 0 | 0 | 0 | 0 | 483940 |
| IIFDDFREA | STT3A | P46977 | STT3A_HUMAN | Unmodified | 0 | 0 | 0 | 0 | 0 | 0 | 0 | 0 | 3234700 | 266900 | 0 | 0 | 1511700 | 0 | 0 | 0 | 0 | 0 |
| IIFEDDRCL | HINT1; HINT1; | P49773; H0 | HINT1_HUMAN | Unmodified | 1489200 | 0 | 0 | 3235300 | 19447000 | 7237200 | 1471800 | 0 | 15067000 | 19027000 | 4125500 | 4356700 | 53588000 | 9302200 | 7433200 | 0 | 5824300 | 11042000 |
| IIFEDDRCL | HINT1; HINT1; | P49773; H0 | HINT1_HUMAN | Unmodified | 562960 | 0 | 273160 | 1270000 | 6475600 | 0 | 0 | 0 | 0 | 3596100 | 0 | 3288500 | 29348000 | 6747000 | 0 | 0 | 794620 | 4551200 |
| IIFVVGGPG | AK1; AK1 | P00568; Q5 | KAD1_HUMAN | Unmodified | 44433000 | 31413000 | 795600 | 33319000 | 1535600 | 0 | 0 | 211500 | 3959400 | 983080 | 1749200 | 6880400 | 1088800 | 346980 | 1798000 | 27905000 | 237860 | 520400 |
| IIFVVGGPG | AK1; AK1 | P00568; Q5 | KAD1_HUMAN | Unmodified | 0 | 0 | 0 | 0 | 6026800 | 872970 | 2251800 | 0 | 1730100 | 2808000 | 0 | 0 | 0 | 7165600 | 0 | 0 | 3918100 | 1903500 |
| IIGATDSSG | CBX3; CBX1; C | Q13185; P8 | CBX3_HUMAN | 2 Oxidation (M) | 582990 | 38850 | 36658 | 0 | 0 | 0 | 0 | 0 | 0 | 0 | 0 | 1484700 | 80163 | 0 | 64460 | 0 | 0 | 0 |
| IIGFGSALL | AP2A2; AP2A2 | O94973; A0 | AP2A2_HUMA | Unmodified | 0 | 0 | 0 | 57691 | 2397300 | 0 | 195590 | 0 | 81248 | 1401100 | 0 | 0 | 3899300 | 1649100 | 0 | 0 | 2161300 | 2302000 |
| IIGGDPKG | WDR1; WDR1 | O75083; D6 | WDR1_HUMAN | Unmodified | 186790 | 0 | 0 | 0 | 11791000 | 1622800 | 2095900 | 2056800 | 0 | 1242500 | 0 | 0 | 6339100 | 7513700 | 0 | 0 | 6907500 | 3181200 |
| IIGIPSPSSL | FRYL; FRYL; FR | O94915; F2 | FRYL_HUMAN | Unmodified | 0 | 0 | 87515 | 0 | 776670 | 167650 | 0 | 0 | 566750 | 167610 | 176830 | 2793800 | 0 | 0 | 0 | 140460 | 32343 | 29963 |
| IIGLDQVAG | MYH9 | P35579 | MYH9_HUMAN | Oxidation (M) | 24442 | 0 | 0 | 0 | 177380 | 98089 | 294200 | 86229 | 0 | 0 | 547770 | 1306100 | 0 | 0 | 0 | 15410 | 359150 | 68077 |
| IIGLKPEGV | ATP1B3; ATP1B | P54709; H7 | AT1B3_HUMA | Unmodified | 0 | 0 | 2294400 | 852790 | 0 | 0 | 0 | 0 | 0 | 0 | 505990 | 8145500 | 0 | 0 | 0 | 140650 | 0 | 0 |
| IIGLKPEGV | ATP1B3; ATP1B | P54709; H7 | AT1B3_HUMA | Unmodified | 0 | 0 | 268100 | 176740 | 94292 | 0 | 98587 | 0 | 233310 | 0 | 388520 | 0 | 0 | 0 | 195890 | 89997 | 0 | 0 |
| IIGNLLYYR | IQGAP1; IQGAP | P46940; H0 | IQGA1_HUMA | Unmodified | 0 | 929140 | 0 | 0 | 634690 | 813640 | 0 | 0 | 0 | 680950 | 0 | 6234700 | 0 | 0 | 887990 | 1067600 | 0 | 0 |
| IIGTLQNSA | MME; MME; MM | P08473; A0A | NEP_HUMAN N | Unmodified | 162570 | 0 | 1794400 | 4751100 | 0 | 0 | 0 | 0 | 1463500 | 511870 | 0 | 0 | 0 | 0 | 4096200 | 2908200 | 0 | 0 |
| IIGTLQNSA | MME; MME; MM | P08473; A0A | NEP_HUMAN N | Unmodified | 48850 | 0 | 29395000 | 70055000 | 169660 | 0 | 0 | 0 | 22632000 | 4071400 | 2385500 | 0 | 17835000 | 4167900 | 74685000 | 47738000 | 51791 | 0 |
| IIHDFPQFY | RPA3; RPA3 | P35244; B5 | RFA3_HUMAN | Unmodified | 525440 | 440710 | 125030 | 189870 | 3876700 | 354300 | 2749100 | 0 | 731240 | 1798200 | 1501600 | 7110900 | 6782900 | 1926700 | 195370 | 133060 | 2640600 | 2260200 |
| IIHEAGYSE | GNAI1; GNAI1 | P63096; A0A | GNAI1_HUMA | Unmodified | 0 | 0 | 154310 | 437900 | 714200 | 525560 | 368460 | 159930 | 687010 | 471430 | 2767400 | 13945000 | 1972700 | 3069000 | 0 | 438010 | 220680 | 639850 |
| IIHEAGYSE | GNAI1; GNAI1 | P63096; A0A | GNAI1_HUMA | Unmodified | 0 | 0 | 0 | 0 | 0 | 0 | 0 | 0 | 0 | 0 | 0 | 0 | 322240 | 1044800 | 0 | 0 | 85001 | 71230 |

| | | | | | | | | | | | | | | | | | | | | | | |
|---|---|---|---|---|---|---|---|---|---|---|---|---|---|---|---|---|---|---|---|---|---|---|
| IIHEDGFSG | GNAO1; GNAO1 | P09471; A0A | GNAO_HUMAN | Unmodified | 0 | 0 | 0 | 0 | 199280 | 0 | 3481400 | 195550 | 4952000 | 5488100 | 0 | 0 | 0 | 0 | 0 | 0 | 0 | 0 |
| IIHEDGYSE | GNAI3 | P08754 | GNAI3_HUMAN | Unmodified | 813360 | 508690 | 667880 | 785060 | 903800 | 414170 | 523100 | 401990 | 797480 | 581050 | 1364700 | 6418300 | 904200 | 841560 | 1731300 | 1462000 | 818680 | 1592000 |
| IIHEDGYSE | GNAI3 | P08754 | GNAI3_HUMAN | Unmodified | 0 | 0 | 0 | 0 | 0 | 76437 | 234840 | 184260 | 244270 | 190350 | 0 | 0 | 217270 | 317990 | 0 | 0 | 722810 | 559590 |
| IIHEDGYSE | GNAI2 | P04899 | GNAI2_HUMAN | Unmodified | 0 | 0 | 0 | 0 | 148270 | 63979 | 135300 | 47447 | 2217300 | 393510 | 2228800 | 43586000 | 0 | 0 | 50591 | 1297300 | 299080 | 663050 |
| IIHEDGYSE | GNAI2 | P04899 | GNAI2_HUMAN | Unmodified | 1173700 | 0 | 0 | 2641600 | 3158800 | 1845900 | 786010 | 68717 | 5352500 | 1306100 | 7939500 | 14227000 | 0 | 791410 | 334860 | 183330 | 3582600 | 5314300 |
| IIHGAGYSE | GNA11 | P29992 | GNA11_HUMAN | Unmodified | 0 | 0 | 0 | 0 | 0 | 38836 | 0 | 0 | 0 | 0 | 0 | 3353900 | 0 | 0 | 0 | 0 | 0 | 0 |
| IIHGAGYSE | GNA11 | P29992 | GNA11_HUMAN | Unmodified | 0 | 41777 | 0 | 0 | 707900 | 0 | 0 | 104060 | 80354 | 242020 | 0 | 0 | 0 | 26052 | 0 | 0 | 0 | 422290 |
| IIHGQDFD( | GNA13 | Q14344 | GNA13_HUMAN | Unmodified | 123610 | 106630 | 0 | 1050300 | 523330 | 229260 | 335380 | 193900 | 3549200 | 527240 | 1507500 | 10250000 | 0 | 0 | 39249 | 152540 | 509220 | 706770 |
| IIHGSGYSD | GNAQ; GNAQ | P50148; B1A | GNAQ_HUMAN | Unmodified | 0 | 200680 | 0 | 0 | 0 | 267910 | 501390 | 0 | 0 | 0 | 0 | 813440 | 0 | 0 | 0 | 0 | 0 | 0 |
| IIHGSGYSD | GNA14 | O95837 | GNA14_HUMAN | Unmodified | 0 | 0 | 0 | 265570 | 1155400 | 0 | 0 | 499830 | 0 | 1159900 | 888590 | 0 | 374520 | 531060 | 0 | 0 | 35344 | 96413 |
| IIIAEVVNK | MYO1B; MYO1B | O43795; H7( | MYO1B_HUMA | Unmodified | 100110 | 0 | 448960 | 715370 | 35305000 | 5657500 | 2907900 | 2449700 | 3796800 | 1504300 | 10246000 | 42008000 | 114510 | 1410800 | 1715700 | 832580 | 4053300 | 9199600 |
| IIIAEVVNKII | MYO1B; MYO1B | O43795; H7( | MYO1B_HUMA | Unmodified | 0 | 0 | 0 | 0 | 1283600 | 0 | 199190 | 0 | 0 | 62965 | 0 | 0 | 0 | 0 | 0 | 0 | 1811300 | 0 |
| IIIEELPK | CTHRC1; CTHR | Q96CG8; E5 | CTHR1_HUMAN | Unmodified | 0 | 0 | 0 | 0 | 0 | 0 | 0 | 0 | 14018000 | 3796100 | 0 | 0 | 0 | 0 | 0 | 0 | 0 | 0 |
| IIIPEIQK | CSE1L | P55060 | XPO2_HUMAN | Unmodified | 0 | 0 | 0 | 0 | 568400 | 0 | 585940 | 212000 | 1395900 | 789750 | 2138100 | 29311000 | 0 | 15202 | 0 | 0 | 0 | 451680 |
| IIIPPFLAYG | FKBP10 | Q96AY3 | FKB10_HUMAN | Unmodified | 0 | 0 | 0 | 0 | 0 | 0 | 0 | 0 | 0 | 74024 | 1577600 | 1050000 | 0 | 0 | 0 | 0 | 0 | 0 |
| IIIQESALDY | GLG1 | Q92896 | GSLG1_HUMAN | Unmodified | 0 | 0 | 0 | 0 | 0 | 0 | 0 | 0 | 4159800 | 382580 | 0 | 5375300 | 0 | 0 | 0 | 0 | 0 | 0 |
| IIKDGEQHE | AIFM1; AIFM1; A | O95831; E9F | AIFM1_HUMAN | Unmodified | 0 | 0 | 0 | 0 | 0 | 0 | 0 | 0 | 381760 | 65171 | 0 | 1154200 | 0 | 0 | 0 | 0 | 0 | 0 |
| IIKPFPAPQ | ADAM17 | P78536 | ADA17_HUMAN | Unmodified | 0 | 0 | 0 | 0 | 0 | 0 | 0 | 0 | 1642700 | 0 | 0 | 0 | 0 | 0 | 0 | 0 | 154890 | 0 |
| IIKPVLTQE | MCM3; MCM3; | P25205; Q7Z | MCM3_HUMAN | Unmodified | 0 | 0 | 0 | 0 | 0 | 0 | 339070 | 0 | 625000 | 0 | 371200 | 2224700 | 0 | 0 | 0 | 0 | 0 | 0 |
| IILAAFR | ATP6V1H; ATP6 | Q9UI12; G3\ | VATH_HUMAN | Unmodified | 0 | 0 | 0 | 0 | 0 | 0 | 0 | 0 | 71836 | 0 | 0 | 889450 | 0 | 0 | 0 | 0 | 0 | 71185 |
| IILDLISESP | HNRNPK; HNRI | P61978; Q5T | HNRPK_HUMA | Unmodified | 4979100 | 3667200 | 1874300 | 3494700 | 12359000 | 2497700 | 9324400 | 1496700 | 11583000 | 9456700 | 15893000 | 1.01E+08 | 13998000 | 4734300 | 2367400 | 3096900 | 1782700 | 4215300 |
| IILDLISESP | HNRNPK; HNRI | P61978; Q5T | HNRPK_HUMA | Unmodified | 203180 | 0 | 0 | 484110 | 1158200 | 0 | 2217500 | 413120 | 0 | 2091300 | 0 | 97164 | 0 | 821680 | 0 | 0 | 446840 | 1684500 |
| IILEAEKMD | RHEB; RHEB | Q15382; C9J | RHEB_HUMAN | Oxidation (M) | 140780 | 125710 | 0 | 143880 | 559450 | 185820 | 46078 | 48715 | 156260 | 97550 | 0 | 118640 | 155390 | 90762 | 0 | 79791 | 120330 | 226360 |
| IILEAEKMD | RHEB; RHEB | Q15382; C9J | RHEB_HUMAN | Unmodified | 418660 | 334400 | 0 | 562590 | 2627800 | 846690 | 191790 | 71253 | 624570 | 561580 | 0 | 208270 | 1540700 | 570380 | 0 | 0 | 465580 | 1685600 |
| IILEALR | SCG2; SCG2; S | P13521; C9J | SCG2_HUMAN | Unmodified | 62729 | 86498 | 0 | 98813 | 1813800 | 250790 | 1111900 | 936980 | 3378700 | 396700 | 751810 | 2655600 | 1439300 | 836150 | 79717 | 82438 | 2329500 | 1961000 |
| IILEALRQAE | SCG2; SCG2 | P13521; C9J | SCG2_HUMAN | Unmodified | 0 | 0 | 0 | 0 | 0 | 0 | 0 | 0 | 1287700 | 324610 | 0 | 0 | 0 | 0 | 0 | 0 | 0 | 0 |
| IILHNTMES | YKT6; YKT6 | O15498; A0A | YKT6_HUMAN S | Unmodified | 0 | 0 | 0 | 0 | 0 | 0 | 0 | 0 | 0 | 0 | 333450 | 164330 | 0 | 0 | 0 | 0 | 0 | 0 |
| IILHNTMES | YKT6; YKT6 | O15498; A0A | YKT6_HUMAN S | Unmodified | 0 | 0 | 0 | 0 | 0 | 0 | 0 | 110790 | 4389700 | 0 | 152350 | 406250 | 0 | 0 | 0 | 0 | 0 | 0 |
| IILLAEGR | AHCY | P23526 | SAHH_HUMAN | Unmodified | 0 | 745690 | 986360 | 1116400 | 2579600 | 0 | 53096 | 174300 | 2658800 | 1496000 | 2935000 | 21573000 | 2041500 | 1034300 | 2824800 | 2084000 | 1560400 | 0 |
| IILNAVFGFI | TOR1A; TOR1A | O14656; A0A | TOR1A_HUMAN | Unmodified | 0 | 0 | 0 | 0 | 0 | 0 | 0 | 0 | 982890 | 0 | 0 | 0 | 0 | 0 | 0 | 0 | 0 | 0 |
| IILNSHSPA | USP24 | Q9UPU5 | UBP24_HUMAN | Unmodified | 0 | 0 | 0 | 0 | 0 | 0 | 0 | 0 | 0 | 0 | 0 | 1824300 | 0 | 0 | 0 | 0 | 0 | 0 |
| IILNTAVAEF | CNOT10; CNOT | Q9H9A5; H7 | CNO10_HUMA | Unmodified | 0 | 0 | 0 | 0 | 0 | 0 | 0 | 0 | 0 | 0 | 129320 | 462470 | 0 | 0 | 0 | 0 | 0 | 0 |
| IILPEGAK | RPN1; RPN1 | P04843; B7Z | RPN1_HUMAN | Unmodified | 0 | 161450 | 0 | 0 | 0 | 0 | 540630 | 0 | 63868000 | 8699300 | 1860900 | 1528900 | 0 | 0 | 0 | 0 | 0 | 695940 |
| IILPEGAKN | RPN1; RPN1 | P04843; B7Z | RPN1_HUMAN | Unmodified | 0 | 0 | 0 | 0 | 0 | 0 | 1272500 | 0 | 730750 | 633140 | 0 | 0 | 0 | 0 | 0 | 0 | 0 | 0 |
| IILQDEDVT | PLXNA1 | Q9UIW2 | PLXA1_HUMAN | Unmodified | 133640 | 0 | 0 | 542510 | 3442900 | 717920 | 0 | 0 | 568310 | 235380 | 220320 | 1661300 | 2292700 | 1590600 | 431910 | 286390 | 958660 | 2771100 |
| IILQDEDVT | PLXNA1 | Q9UIW2 | PLXA1_HUMAN | Unmodified | 0 | 0 | 0 | 128610 | 773760 | 139050 | 0 | 0 | 0 | 0 | 0 | 0 | 0 | 459560 | 0 | 0 | 1347300 | 769830 |
| IILSPFATTT | SH3BP4 | Q9P0V3 | SH3B4_HUMAN | Unmodified | 0 | 0 | 0 | 0 | 0 | 0 | 0 | 0 | 2583100 | 767960 | 432800 | 1001200 | 0 | 354180 | 0 | 0 | 0 | 0 |
| IILTARPFRL | GANAB; GANAE | Q14697; F5H | GANAB_HUMA | Unmodified | 70742 | 66867 | 431730 | 373390 | 0 | 0 | 45273 | 15840 | 2.2E+08 | 18749000 | 5803000 | 1365500 | 48945 | 0 | 1365100 | 303120 | 0 | 0 |
| IIMLFTDGG | CACNA2D1 | P54289 | CA2D1_HUMAN | Unmodified | 0 | 0 | 0 | 0 | 0 | 0 | 422950 | 0 | 3236800 | 0 | 0 | 0 | 0 | 0 | 0 | 0 | 0 | 0 |
| IIMLFTDGG | CACNA2D1 | P54289 | CA2D1_HUMAN | Oxidation (M) | 1037400 | 3119100 | 0 | 0 | 0 | 0 | 1247500 | 77395 | 5813300 | 642700 | 0 | 0 | 0 | 0 | 0 | 0 | 0 | 0 |
| IIMTSSASG | HSD17B4; HSD | P51659; A0A | DHB4_HUMAN | Oxidation (M) | 0 | 0 | 0 | 0 | 0 | 0 | 0 | 0 | 8263600 | 1626300 | 0 | 200700 | 0 | 104920 | 0 | 0 | 0 | 0 |
| IIMTSSASG | HSD17B4; HSD | P51659; A0A | DHB4_HUMAN | Unmodified | 0 | 0 | 0 | 0 | 0 | 0 | 0 | 73858 | 25824000 | 954460 | 190580 | 0 | 0 | 0 | 0 | 0 | 0 | 156140 |
| IINADSEDP | PSMC2; PSMC2 | P35998; C9J | PRS7_HUMAN | Unmodified | 95651 | 0 | 519700 | 202740 | 410340 | 0 | 89263 | 30728 | 667390 | 61407 | 281350 | 2498900 | 555370 | 0 | 259280 | 172340 | 0 | 250000 |
| IINADSEDP | PSMC2; PSMC2 | P35998; C9J | PRS7_HUMAN | Unmodified | 0 | 0 | 0 | 666500 | 5776300 | 0 | 3289900 | 0 | 939300 | 1407900 | 0 | 3474700 | 8362500 | 3921800 | 0 | 184080 | 5671200 | 5989800 |
| IINAFFPEG | CHP1; CHP1; C | Q99653; F5C | CHP1_HUMAN | Unmodified | 472020 | 783330 | 887990 | 423070 | 0 | 0 | 0 | 9660.1 | 576740 | 0 | 76791 | 7116700 | 0 | 81309 | 3620700 | 2705100 | 0 | 0 |
| IINDNATYC | NOP56; NOP56 | O00567; Q5J | NOP56_HUMAN | Unmodified | 0 | 0 | 0 | 0 | 0 | 0 | 0 | 0 | 0 | 0 | 0 | 319700 | 0 | 0 | 0 | 19106 | 0 | 0 |
| IINEGAAILF | PPP3CB; PPP3C | P16298; Q5F | PP2BB_HUMAN | Unmodified | 0 | 0 | 0 | 0 | 0 | 0 | 0 | 0 | 308450 | 0 | 0 | 1798400 | 0 | 0 | 0 | 0 | 106090 | 0 |
| IINEPTAAAI | HSPA1L; HSPA6 | P34931; P17 | HS71L_HUMAN | Unmodified | 0 | 0 | 0 | 0 | 0 | 0 | 0 | 373850 | 40624000 | 17063000 | 10122000 | 7652400 | 0 | 0 | 0 | 0 | 171310 | 0 |
| IINEPTAAAI | HSPA1L; HSPA5 | P34931; P11 | HS71L_HUMAN | Unmodified | 43523000 | 35906000 | 56445000 | 5120500 | 1.08E+08 | 20182000 | 0 | 217640 | 1.72E+08 | 92433000 | 45833000 | 1.35E+08 | 1.81E+08 | 1823900 | 35994000 | 37754000 | 9807200 | 35210000 |
| IINEPTAAAI | HSPA8; HSPA2; | P11142; P54 | HSP7C_HUMAN | Unmodified | 37727000 | 35083000 | 14426000 | 29915000 | 12696000 | 3826500 | 1713700 | 724210 | 20862000 | 5875800 | 3362700 | 35932000 | 26934000 | 5195200 | 30084000 | 16241000 | 2262800 | 5592200 |
| IINEPTAAAI | HSPA8; HSPA8; | P11142; E9P | HSP7C_HUMAN | Unmodified | 0 | 0 | 0 | 0 | 2197000 | 0 | 1996500 | 0 | 0 | 1039600 | 0 | 0 | 5633200 | 3043400 | 0 | 0 | 1257700 | 0 |
| IINEPTAAAI | HSPA5; HSPA5; | P11021; A0A | BIP_HUMAN Er | Unmodified | 9997700 | 1377800 | 5339000 | 2951700 | 0 | 0 | 0 | 568530 | 4.94E+08 | 11023000 | 26336000 | 17617000 | 1828200 | 63675 | 10736000 | 7206900 | 0 | 0 |
| IINEPTAAAI | HSPA5; HSPA5; | P11021; A0A | BIP_HUMAN Er | Unmodified | 0 | 0 | 0 | 4712600 | 23738000 | 4104300 | 29603000 | 9846000 | 2.62E+08 | 4.34E+08 | 37376000 | 0 | 14639000 | 15597000 | 2558200 | 0 | 1889300 | 18787000 |
| IINEPTAAAI | HSPA6; HSPA1A | P17066; P0D | HSP76_HUMAN | Unmodified | 14006000 | 7277200 | 4286800 | 5657800 | 0 | 0 | 0 | 0 | 12087000 | 2062000 | 0 | 13212000 | 875400 | 0 | 0 | 6159200 | 0 | 0 |
| IINEPTAAAI | HSPA1A; HSPA1 | P0DMV8; P0 | HS71A_HUMAN | Unmodified | 2064400 | 0 | 2818500 | 0 | 1116500 | 0 | 0 | 0 | 2506500 | 308220 | 295960 | 2235300 | 0 | 0 | 0 | 1622000 | 0 | 0 |
| IINEPTAAAI | HSPA1A; HSPA1 | P0DMV8; P0 | HS71A_HUMAN | Unmodified | 0 | 0 | 0 | 24442000 | 1046100 | 0 | 0 | 0 | 14526000 | 2049600 | 941770 | 1858400 | 3537000 | 794760 | 0 | 0 | 0 | 0 |
| IINEVSKPL/ | MANF | P55145 | MANF_HUMAN | Unmodified | 365590 | 270310 | 592270 | 570430 | 2802100 | 361640 | 1778600 | 944780 | 1.62E+08 | 41111000 | 22841000 | 6014900 | 2116600 | 2468300 | 722900 | 666620 | 393240 | 3256300 |
| IINEVSKPL/ | MANF | P55145 | MANF_HUMAN | Unmodified | 0 | 0 | 0 | 0 | 0 | 0 | 430000 | 0 | 0 | 2080500 | 0 | 0 | 0 | 0 | 0 | 0 | 154660 | 0 |
| IINFAPSLLF | IPO8; IPO8 | O15397; F5H | IPO8_HUMAN I | Unmodified | 69739 | 84574 | 124400 | 104010 | 0 | 0 | 0 | 0 | 0 | 0 | 0 | 1702300 | 0 | 0 | 211300 | 122940 | 0 | 0 |
| IINILTDK | LRBA; LRBA; LR | P50851; A0A | LRBA_HUMAN | Unmodified | 0 | 0 | 0 | 0 | 0 | 0 | 0 | 0 | 0 | 0 | 0 | 3200800 | 0 | 0 | 0 | 0 | 0 | 0 |
| IINKREGIT/ | PLS1; PLS1 | Q14651; C9J | PLSI_HUMAN F | Unmodified | 0 | 0 | 0 | 0 | 6547500 | 1749400 | 0 | 0 | 0 | 619470 | 0 | 0 | 2459400 | 3845300 | 0 | 0 | 0 | 865840 |
| IINLGPVHP | PIP4P1 | Q86T03 | PP4P1_HUMAN | Oxidation (M) | 0 | 0 | 285160 | 0 | 0 | 1502200 | 0 | 60118 | 2464700 | 1054300 | 0 | 2411200 | 0 | 0 | 0 | 0 | 0 | 0 |
| IINLGPVHP | PIP4P1 | Q86T03 | PP4P1_HUMAN | Unmodified | 0 | 0 | 0 | 0 | 0 | 0 | 0 | 0 | 1356900 | 0 | 0 | 87678 | 0 | 0 | 0 | 0 | 0 | 0 |
| IINNTENLV | PSAT1 | Q9Y617 | SERC_HUMAN | Unmodified | 155390 | 0 | 1369000 | 2876700 | 1307500 | 198850 | 0 | 0 | 2168100 | 204030 | 5076100 | 25599000 | 2559400 | 473470 | 3784700 | 2223200 | 0 | 0 |
| IINNTENLV | PSAT1 | Q9Y617 | SERC_HUMAN | Unmodified | 0 | 0 | 0 | 0 | 4857900 | 617470 | 496840 | 0 | 0 | 575760 | 117010 | 0 | 8505300 | 4831400 | 0 | 0 | 0 | 0 |
| IINPQFKEE | CASD1; CASD1 | Q96PB1; C9. | CASD1_HUMAN | Unmodified | 0 | 0 | 0 | 0 | 0 | 0 | 0 | 0 | 363770 | 0 | 0 | 0 | 0 | 0 | 0 | 0 | 0 | 0 |
| IINQISTNEI | ANGPTL2 | Q9UKU9 | ANGL2_HUMAN | Unmodified | 0 | 0 | 0 | 0 | 77273 | 0 | 161070 | 0 | 928310 | 0 | 0 | 0 | 0 | 0 | 0 | 0 | 0 | 0 |
| IINTPEVVR | EHD1; EHD1; El | Q9H4M9; C9 | EHD1_HUMAN | Unmodified | 725600 | 600100 | 955660 | 211890 | 0 | 0 | 0 | 26394 | 0 | 0 | 1231900 | 16841000 | 0 | 0 | 1710500 | 1530600 | 242570 | 0 |
| IINTPEVVR' | EHD1; EHD1; El | Q9H4M9; C9 | EHD1_HUMAN | Unmodified | 0 | 0 | 0 | 0 | 181180 | 37075 | 0 | 0 | 0 | 0 | 0 | 0 | 0 | 0 | 0 | 0 | 405620 | 132180 |
| IIPAIWFR | ERGIC1 | Q969X5 | ERGI1_HUMAN | Unmodified | 0 | 0 | 1208400 | 0 | 0 | 0 | 0 | 0 | 27053000 | 2592200 | 89434 | 807270 | 0 | 0 | 1465400 | 712670 | 0 | 0 |
| IIPFNRLTIG | FGG; FGG; FGG | P02679; C9J | FIBG_HUMAN I | Unmodified | 0 | 0 | 0 | 0 | 0 | 0 | 0 | 0 | 0 | 0 | 0 | 0 | 0 | 1123300 | 0 | 0 | 0 | 0 |
| IIPGADADV | DPYSL5 | Q9BPU6 | DPYL5_HUMAN | Unmodified | 0 | 0 | 0 | 0 | 0 | 0 | 36270 | 49828 | 0 | 1302400 | 0 | 2281900 | 0 | 0 | 0 | 0 | 0 | 26551 |
| IIPGGVAER | LIN7A; LIN7A; L | O14910; H0Y | LIN7A_HUMAN | Unmodified | 0 | 0 | 111980 | 83994 | 0 | 0 | 19696 | 0 | 1440500 | 410010 | 0 | 0 | 0 | 0 | 0 | 0 | 0 | 0 |
| IIPGSKEETI | MPP7; MPP7 | Q5T2T1; S4F | MPP7_HUMAN | Unmodified | 57454 | 36175 | 0 | 275490 | 373880 | 191720 | 0 | 0 | 669680 | 0 | 115280 | 635570 | 668900 | 242540 | 307030 | 0 | 0 | 0 |
| IIPKPEFPR/ | MRPL24 | Q96A35 | RM24_HUMAN | Unmodified | 0 | 0 | 0 | 0 | 0 | 0 | 0 | 0 | 2034800 | 0 | 0 | 0 | 0 | 0 | 0 | 0 | 0 | 0 |
| IIPLLTDPK | EPHX1 | P07099 | HYEP_HUMAN | Unmodified | 0 | 0 | 3947900 | 0 | 0 | 0 | 441470 | 0 | 1395500 | 575210 | 0 | 1026600 | 0 | 0 | 1415400 | 574560 | 0 | 0 |
| IIPLYSTLPP | DHX15 | O43143 | DHX15_HUMAN | Unmodified | 968040 | 4249800 | 337620 | 69736 | 509510 | 54253 | 69876 | 0 | 1122200 | 250560 | 497540 | 1811000 | 663870 | 0 | 663910 | 667630 | 0 | 0 |
| IIPPGSGIIH | ACO1 | P21399 | ACOC_HUMAN | Unmodified | 753600 | 0 | 0 | 761930 | 0 | 0 | 0 | 0 | 777940 | 0 | 1496400 | 11267000 | 0 | 0 | 1956900 | 996630 | 0 | 0 |
| IIPRHLQLA | H2AX; H2AC6; I | P16104; Q93 | H2AX_HUMAN | Unmodified | 235770 | 113220 | 0 | 0 | 2334500 | 1295600 | 8456300 | 582840 | 0 | 182730 | 0 | 0 | 714750 | 382590 | 0 | 0 | 0 | 2082600 |
| IIPTLEEYQI | HNRNPK | Q5T6W2 | Q5T6W2_HUM. | Unmodified | 0 | 0 | 0 | 0 | 0 | 0 | 0 | 0 | 0 | 0 | 0 | 4663600 | 0 | 0 | 0 | 0 | 0 | 0 |
| IIQANGYSF | ERBIN; ERBIN | Q96RT1; B4I | ERBIN_HUMAN | Unmodified | 0 | 0 | 0 | 0 | 2483800 | 0 | 0 | 0 | 0 | 0 | 0 | 0 | 0 | 0 | 0 | 437590 | 0 | 0 |
| IIQEFLSK | MSH2 | P43246 | MSH2_HUMAN | Unmodified | 0 | 0 | 0 | 0 | 998270 | 0 | 4099700 | 1893300 | 284310 | 446150 | 216700 | 206850 | 0 | 3052000 | 0 | 0 | 192140 | 2775400 |
| IIQEQDAGI | STX8; STX8 | Q9UNK0; K7 | STX8_HUMAN S | Unmodified | 0 | 0 | 98273 | 0 | 0 | 0 | 0 | 0 | 1002700 | 266980 | 456210 | 1531700 | 0 | 0 | 494440 | 115340 | 0 | 0 |
| IIQFNPGPE | GNPDA1; GNPI | P46926; D6F | GNPI1_HUMAN | Unmodified | 0 | 0 | 0 | 0 | 406070 | 0 | 185600 | 112480 | 0 | 178940 | 707670 | 1556100 | 774440 | 743800 | 0 | 0 | 478040 | 0 |
| IIQGQGVDI | TIGAR | Q9NQ88 | TIGAR_HUMAN | Unmodified | 0 | 0 | 0 | 0 | 472860 | 0 | 0 | 0 | 0 | 158520 | 328540 | 2682600 | 522030 | 0 | 0 | 101140 | 162360 | 0 |
| IIQHCSNYS | DNM1L | O00429 | DNM1L_HUMA | Unmodified | 16472 | 0 | 0 | 0 | 0 | 0 | 0 | 0 | 0 | 0 | 0 | 645520 | 0 | 0 | 0 | 0 | 0 | 0 |
| IIQLIEGK | BPNT1; BPNT1 | O95861; A6N | BPNT1_HUMAN | Unmodified | 430530 | 196920 | 1283400 | 726820 | 1526500 | 606270 | 939280 | 406220 | 1613100 | 521400 | 2149500 | 11383000 | 2701100 | 1097900 | 2286800 | 979320 | 760300 | 2600600 |
| IIQLLDDYP | RPLP0; RPLP0P | P05388; Q8N | RLA0_HUMAN ( | Unmodified | 18940000 | 19433000 | 10132000 | 12596000 | 4897100 | 0 | 3438400 | 2457200 | 1404800 | 1418000 | 8610400 | 69952000 | 6313400 | 4653700 | 28437000 | 15897000 | 527090 | 5893700 |
| IIQQIVIQNI | DAAM1 | Q9Y4D1 | DAAM1_HUMA | Unmodified | 0 | 0 | 0 | 0 | 0 | 0 | 0 | 0 | 0 | 0 | 0 | 8863900 | 0 | 0 | 0 | 0 | 0 | 0 |
| IIQSLLDEF\ | VPS37A; VPS37 | Q8NEZ2; E5I | VP37A_HUMAN | Unmodified | 285770 | 474180 | 35580 | 0 | 0 | 0 | 0 | 0 | 0 | 0 | 0 | 159420 | 0 | 0 | 0 | 0 | 0 | 0 |
| IIQTPGLWE | RAB18; RAB18; | Q9NP72; A0. | RAB18_HUMAN | Unmodified | 0 | 0 | 0 | 0 | 0 | 0 | 0 | 0 | 0 | 0 | 796040 | 3726700 | 0 | 0 | 0 | 0 | 0 | 0 |
| IIRDEQYTA | LNPEP | Q9UIQ6 | LCAP_HUMAN | Unmodified | 0 | 0 | 0 | 0 | 0 | 0 | 0 | 0 | 1131900 | 0 | 0 | 0 | 0 | 0 | 0 | 0 | 0 | 0 |
| IIRDEQYTA | LNPEP | Q9UIQ6 | LCAP_HUMAN | Unmodified | 0 | 0 | 0 | 0 | 186040 | 0 | 0 | 0 | 763490 | 0 | 0 | 0 | 0 | 0 | 215790 | 192740 | 0 | 0 |
| IIRFHDIISD | P4HA1 | P13674 | P4HA1_HUMAN | Unmodified | 0 | 0 | 0 | 608890 | 88442 | 0 | 0 | 0 | 2360900 | 3906400 | 1027800 | 0 | 74539 | 0 | 0 | 232990 | 54075 | 0 |
| IIRSENFEEI | CRABP2; CRAB | P29373; Q59 | RABP2_HUMAN | Unmodified | 0 | 0 | 0 | 0 | 4292300 | 1075800 | 2379900 | 413060 | 2491700 | 4001600 | 0 | 0 | 0 | 0 | 0 | 0 | 27411000 | 9919100 |
| IIRVLFPGN | MAP1B; MAP1B | P46821; D6F | MAP1B_HUMAN | Unmodified | 0 | 0 | 0 | 0 | 222440 | 0 | 4613200 | 0 | 0 | 2093200 | 91498 | 0 | 2224400 | 1392500 | 0 | 0 | 551880 | 433060 |
| IIRYPDSHC | G3BP2; G3BP2; | Q9UN86; A0 | G3BP2_HUMAN | Unmodified | 0 | 0 | 0 | 0 | 0 | 0 | 0 | 0 | 0 | 0 | 0 | 2406000 | 0 | 0 | 0 | 0 | 0 | 0 |
| IISAASEGG | DARS1 | P14868 | SYDC_HUMAN | Unmodified | 0 | 0 | 124950 | 2521600 | 1671100 | 406610 | 1038700 | 273770 | 1523900 | 857420 | 614080 | 7286600 | 1404100 | 503540 | 0 | 0 | 907960 | 0 |
| IISANGCK | ATP1A1 | P05023 | AT1A1_HUMAN | Unmodified | 478130 | 0 | 1099600 | 0 | 0 | 0 | 348060 | 238220 | 0 | 0 | 0 | 2797600 | 232250 | 300070 | 1978200 | 479370 | 0 | 0 |
| IISANGCKV | ATP1A1 | P05023 | AT1A1_HUMAN | Unmodified | 0 | 0 | 745350 | 5738100 | 239180 | 0 | 0 | 0 | 2830200 | 244920 | 2363900 | 20087000 | 584470 | 0 | 5764300 | 8842000 | 495020 | 149850 |
| IISANGCKV | ATP1A1 | P05023 | AT1A1_HUMAN | Unmodified | 0 | 0 | 0 | 4724400 | 0 | 0 | 0 | 0 | 0 | 0 | 2519400 | 0 | 2717300 | 0 | 0 | 244580 | 1369500 | 0 |
| IISAPEMR | IGFBP5; IGFBP5 | P24593; C9J | IBP5_HUMAN I | Unmodified | 0 | 0 | 0 | 0 | 19545 | 0 | 0 | 0 | 9249500 | 0 | 0 | 0 | 0 | 0 | 0 | 0 | 0 | 0 |
| IISAPEMRQ | IGFBP5 | P24593 | IBP5_HUMAN I | Unmodified | 0 | 0 | 0 | 0 | 0 | 0 | 0 | 0 | 1527300 | 323260 | 0 | 0 | 0 | 0 | 0 | 0 | 0 | 0 |
| IISAPEMRQ | IGFBP5 | P24593 | IBP5_HUMAN I | Unmodified | 0 | 0 | 0 | 0 | 0 | 0 | 70435 | 75496 | 7210900 | 3633100 | 0 | 0 | 0 | 0 | 0 | 0 | 147480 | 129460 |
| IISDNLTYC | NOP58; NOP58 | Q9Y2X3; H7I | NOP58_HUMAN | Unmodified | 0 | 0 | 0 | 0 | 0 | 0 | 0 | 223640 | 0 | 0 | 0 | 544600 | 0 | 0 | 0 | 0 | 0 | 93185 |
| IISGTLQATI | PSD3; PSD3 | Q9NYI0; E5F | PSD3_HUMAN | Unmodified | 0 | 0 | 0 | 0 | 1166700 | 0 | 0 | 0 | 110790 | 74722 | 0 | 223620 | 410820 | 247090 | 0 | 0 | 581690 | 1181100 |
| IISHVWENI | PIK3CB | P42338 | PK3CB_HUMAN | Unmodified | 0 | 0 | 0 | 0 | 0 | 0 | 0 | 0 | 0 | 0 | 0 | 404390 | 0 | 1877900 | 217700 | 161760 | 0 | 0 |
| IISIFSGTEK | PAK2 | Q13177 | PAK2_HUMAN | Unmodified | 0 | 0 | 0 | 0 | 5528600 | 2151900 | 828570 | 578380 | 0 | 0 | 623650 | 5725500 | 6476900 | 2541500 | 0 | 0 | 5285600 | 5481600 |
| IISKIENHE( | PKM; PKM; PKM | P14618; B4D | KPYM_HUMAN | Unmodified | 703360 | 572140 | 34326 | 4789800 | 16251000 | 2200500 | 518750 | 46113 | 6053400 | 1000100 | 5544500 | 23280000 | 3280900 | 1040900 | 1085700 | 1066000 | 3650500 | 5696700 |

| | | | | | | | | | | | | | | | | | | | | | | |
|---|---|---|---|---|---|---|---|---|---|---|---|---|---|---|---|---|---|---|---|---|---|---|
| IISKQDDLM | MICU2; MICU2 | Q8IYU8; A0A | MICU2_HUMAN | Unmodified | 0 | 0 | 0 | 0 | 0 | 0 | 0 | 0 | 1214600 | 178690 | 0 | 0 | 0 | 0 | 0 | 0 | 0 | 0 |
| IISLDLPVAE | UBR4; UBR4 | Q5T4S7; X6F | UBR4_HUMAN | Unmodified | 0 | 0 | 0 | 0 | 0 | 0 | 0 | 16253 | 0 | 0 | 0 | 430400 | 0 | 0 | 0 | 0 | 0 | 0 |
| IISLDLPVAE | UBR4; UBR4 | Q5T4S7; X6F | UBR4_HUMAN | Unmodified | 0 | 0 | 0 | 0 | 209550 | 0 | 0 | 0 | 0 | 427630 | 289790 | 3296700 | 0 | 0 | 0 | 0 | 0 | 137770 |
| IISLSSSVPE | COBL; COBL; C | O75128; J3K | COBL_HUMAN | Unmodified | 717910 | 649770 | 323960 | 341920 | 0 | 0 | 0 | 0 | 0 | 0 | 0 | 231100 | 117660 | 0 | 177790 | 59309 | 0 | 0 |
| IISNASCTT | GAPDH; GAPDH | P04406; E7E | G3P_HUMAN G | Unmodified | 10691000 | 1491300 | 3141400 | 7545000 | 70794000 | 11961000 | 30281000 | 21849000 | 8082400 | 8148600 | 0 | 1.04E+08 | 61370000 | 39989000 | 4932500 | 7045100 | 1.79E+08 | 1.02E+08 |
| IISNEEGYR | DPP4; DPP4; DI | P27487; A0A | DPP4_HUMAN | Unmodified | 701000 | 1017700 | 7025800 | 7471600 | 0 | 0 | 70615 | 0 | 56005 | 0 | 0 | 0 | 0 | 0 | 16345000 | 9728100 | 0 | 0 |
| IISNLFLGR | MYO18A; MYO1 | Q92614; A0A | MY18A_HUMAN | Unmodified | 0 | 0 | 0 | 0 | 0 | 0 | 0 | 0 | 0 | 0 | 0 | 746820 | 0 | 0 | 0 | 0 | 0 | 0 |
| IISPTLLAEY | METTL1; METTL | Q9UBP6; H0 | TRMB_HUMAN | Unmodified | 0 | 0 | 0 | 0 | 0 | 0 | 0 | 0 | 0 | 0 | 0 | 228280 | 0 | 0 | 0 | 0 | 0 | 0 |
| IISREVSDG | ATIC | P31939 | PUR9_HUMAN | Unmodified | 0 | 0 | 0 | 0 | 6023400 | 0 | 2526500 | 0 | 475890 | 1746200 | 0 | 0 | 12838000 | 5608900 | 0 | 0 | 7541000 | 3418400 |
| IISSDRDLL | XRCC6; XRCC6 | P12956; B1A | XRCC6_HUMAN | Unmodified | 1823500 | 807600 | 0 | 644970 | 2885500 | 727280 | 2823700 | 0 | 1832600 | 2137900 | 2835900 | 17797000 | 2251400 | 536820 | 412040 | 0 | 798660 | 1264100 |
| IISSIEQK | YWHAE; YWHAE | P62258; I3L3 | 1433E_HUMAN | Unmodified | 0 | 0 | 8837500 | 0 | 0 | 0 | 58338 | 296290 | 1992900 | 746280 | 1378300 | 18928000 | 2389300 | 0 | 11415000 | 9504500 | 0 | 0 |
| IISSIEQKEE | YWHAE; YWHAE | P62258; I3L3 | 1433E_HUMAN | Unmodified | 1925900 | 977210 | 7031600 | 5874400 | 4330800 | 0 | 828920 | 537700 | 3570200 | 878800 | 4499600 | 42006000 | 4027000 | 2861500 | 13664000 | 14730000 | 271770 | 2628900 |
| IISSIEQKEE | YWHAE; YWHAE | P62258; I3L3 | 1433E_HUMAN | Unmodified | 8725700 | 3660700 | 18863000 | 26562000 | 11933000 | 10272000 | 2031300 | 961680 | 21310000 | 11240000 | 30935000 | 2.17E+08 | 28842000 | 11495000 | 36487000 | 43835000 | 7005000 | 3476000 |
| IISSRDDQ | MPP7; MPP7 | Q5T2T1; U5C | MPP7_HUMAN | Unmodified | 0 | 0 | 0 | 0 | 9047700 | 0 | 0 | 0 | 1144800 | 606400 | 0 | 0 | 1420200 | 1704900 | 0 | 0 | 848790 | 0 |
| IISTITR | CLSTN1; CLSTN | O94985; Q5 | CSTN1_HUMAN | Unmodified | 0 | 0 | 0 | 241530 | 0 | 0 | 0 | 0 | 8285100 | 221200 | 0 | 87054 | 0 | 0 | 186870 | 0 | 0 | 0 |
| IISTLANKN | FSD1L; FSD1L; | Q9BXM9; A0 | FSD1L_HUMAN | Unmodified | 0 | 0 | 0 | 0 | 0 | 0 | 452210 | 0 | 974470 | 0 | 234290 | 1127300 | 0 | 0 | 0 | 0 | 0 | 0 |
| IISTQRVVV | NCAM2; NCAM | O15394; H9I | NCAM2_HUMA | Unmodified | 0 | 0 | 137020 | 1014300 | 0 | 0 | 331800 | 85633 | 0 | 0 | 0 | 0 | 0 | 0 | 0 | 0 | 164380 | 115210 |
| IISTTASKTE | VAPB | O95292 | VAPB_HUMAN | Unmodified | 135230 | 27432 | 0 | 164090 | 550840 | 35331 | 106550 | 33750 | 389270 | 0 | 0 | 0 | 343070 | 465510 | 0 | 0 | 67249 | 254020 |
| IISVNSVDL | DLG1; DLG1; D | Q12959; A0A | DLG1_HUMAN | Unmodified | 0 | 0 | 369300 | 0 | 0 | 5332500 | 0 | 0 | 629680 | 100800 | 0 | 790380 | 0 | 0 | 2137100 | 878530 | 0 | 0 |
| IISYAQGFM | PGD; PGD | P52209; K7E | 6PGD_HUMAN | Oxidation (M) | 0 | 0 | 0 | 0 | 0 | 0 | 0 | 0 | 0 | 0 | 0 | 1137600 | 0 | 0 | 381190 | 112650 | 0 | 0 |
| IISYTVVFR | PTPRF; PTPRF; | P10586; H0Y | PTPRF_HUMAN | Unmodified | 482840 | 835000 | 0 | 0 | 0 | 0 | 0 | 0 | 390120 | 0 | 0 | 0 | 0 | 0 | 294590 | 148130 | 0 | 0 |
| IITEGASILR | PPP3CA; PPP3C | Q08209; E9F | PP2BA_HUMAN | Unmodified | 0 | 0 | 0 | 1597700 | 0 | 0 | 0 | 0 | 0 | 0 | 709550 | 3432000 | 1373700 | 0 | 0 | 0 | 0 | 0 |
| IITEGFEAAK | CCT6A | P40227 | TCPZ_HUMAN | Unmodified | 584620 | 2224500 | 76576 | 43704 | 7596200 | 22240000 | 4424600 | 2817300 | 1703600 | 3460600 | 11106000 | 40013000 | 2079000 | 1434800 | 1343600 | 1153400 | 2767100 | 6854800 |
| IITEGFEAAK | CCT6A | P40227 | TCPZ_HUMAN | Unmodified | 0 | 152060 | 0 | 0 | 706290 | 543710 | 595370 | 293240 | 0 | 611940 | 0 | 0 | 2088700 | 1800200 | 0 | 0 | 1142700 | 1692300 |
| IITENDHVL | SORL1; SORL1; | Q92673; E9F | SORL_HUMAN | Unmodified | 0 | 0 | 0 | 0 | 0 | 0 | 0 | 0 | 4015600 | 0 | 0 | 0 | 0 | 0 | 0 | 0 | 0 | 0 |
| IITESFFEVP | VCAN; VCAN | P13611; D6F | CSPG2_HUMAN | Unmodified | 0 | 0 | 0 | 0 | 0 | 0 | 0 | 0 | 1087700 | 0 | 0 | 0 | 0 | 0 | 0 | 0 | 0 | 0 |
| IITGGAAAQ | DLG1; DLG1; D | Q12959; H7( | DLG1_HUMAN | Unmodified | 41943 | 182580 | 0 | 0 | 297850 | 422190 | 0 | 0 | 631510 | 250680 | 0 | 756520 | 0 | 0 | 0 | 0 | 189090 | 0 |
| IITGGAAAQ | DLG1; DLG1; D | Q12959; H7( | DLG1_HUMAN | Unmodified | 0 | 0 | 0 | 0 | 0 | 454940 | 0 | 0 | 0 | 664820 | 135180 | 0 | 0 | 0 | 0 | 0 | 0 | 0 |
| IITKDFPQY | ANK3; ANK3; AN | Q12955; A0A | ANK3_HUMAN | Unmodified | 0 | 0 | 0 | 0 | 0 | 0 | 0 | 0 | 0 | 0 | 0 | 0 | 478850 | 828710 | 0 | 0 | 0 | 0 |
| IITLAGPTN | PCBP2; PCBP2; | Q15366; F8V | PCBP2_HUMAN | Unmodified | 70437000 | 69258000 | 47616000 | 55047000 | 1642800 | 461330 | 1714500 | 0 | 8421900 | 1252800 | 7416100 | 50551000 | 14310000 | 2821500 | 32904000 | 44246000 | 0 | 1070200 |
| IITLEEGDIL | FAHD1 | Q6P587 | FAHD1_HUMAN | Unmodified | 0 | 0 | 0 | 0 | 0 | 0 | 0 | 0 | 1254600 | 0 | 0 | 0 | 0 | 0 | 0 | 0 | 0 | 0 |
| IITLTGPTN | PCBP1 | Q15365 | PCBP1_HUMAN | Unmodified | 27496000 | 24302000 | 13119000 | 9252700 | 1124600 | 0 | 138580 | 53923 | 9428400 | 1136500 | 10869000 | 1.05E+08 | 4169600 | 2473100 | 11158000 | 9478400 | 726230 | 0 |
| IITNRLEYE | PCYT2; PCYT2; | Q99447; I3L | PCY2_HUMAN | Unmodified | 200950 | 142190 | 335460 | 0 | 3379400 | 8066700 | 3780200 | 2300800 | 1295900 | 0 | 786980 | 2804500 | 458040 | 5157700 | 590960 | 261500 | 247000 | 5271700 |
| IITRDFPQY | ANK2; ANK2; AN | Q01484; A0A | ANK2_HUMAN | Unmodified | 0 | 0 | 0 | 0 | 0 | 0 | 0 | 0 | 103940 | 267560 | 14002 | 0 | 0 | 0 | 0 | 0 | 0 | 0 |
| IITSELYR | EIF2B5; EIF2B5 | Q13144; A0A | EI2BE_HUMAN | Unmodified | 0 | 0 | 0 | 0 | 0 | 0 | 0 | 41344 | 1200000 | 236210 | 0 | 1715600 | 0 | 0 | 0 | 0 | 0 | 25178 |
| IITVEKHPD | YARS1; YARS1 | P54577; A0A | SYYC_HUMAN | Unmodified | 0 | 0 | 0 | 0 | 1324500 | 163580 | 375410 | 0 | 1492400 | 41556 | 182800 | 228640 | 2016800 | 1012400 | 0 | 0 | 352040 | 876800 |
| IITVEKHPD | YARS1; YARS1 | P54577; A0A | SYYC_HUMAN | Unmodified | 0 | 0 | 0 | 0 | 0 | 0 | 207200 | 0 | 129070 | 571940 | 0 | 0 | 1901000 | 719980 | 0 | 0 | 0 | 0 |
| IITVLTNPAI | NDST2; N/A | P52849; S4F | NDST2_HUMAN | Unmodified | 0 | 0 | 0 | 0 | 0 | 0 | 0 | 0 | 1223600 | 151980 | 0 | 0 | 0 | 0 | 0 | 0 | 0 | 0 |
| IITVLTNPAI | NDST2; N/A | P52849; S4F | NDST2_HUMAN | Unmodified | 0 | 0 | 0 | 0 | 0 | 0 | 0 | 0 | 6008100 | 143370 | 0 | 0 | 0 | 0 | 0 | 0 | 0 | 0 |
| IIVDELK | RACK1; RACK1; | P63244; D6F | RACK1_HUMAN | Unmodified | 250180 | 0 | 0 | 51784 | 1570200 | 80933 | 0 | 0 | 98644 | 1659900 | 906890 | 14042000 | 0 | 0 | 1697300 | 733020 | 458590 | 701350 |
| IIVDELKQE | RACK1; RACK1; | P63244; D6F | RACK1_HUMAN | Unmodified | 8101600 | 5639900 | 7186400 | 2900900 | 270020 | 0 | 133580 | 79643 | 3472300 | 271150 | 11228000 | 1.4E+08 | 203770 | 0 | 6464900 | 6412200 | 32819 | 275380 |
| IIVLGLLPR | PAFAH1B2; PAF | P68402; J3K | PA1B2_HUMAN | Unmodified | 0 | 0 | 773850 | 0 | 0 | 0 | 0 | 0 | 1086500 | 153290 | 2475700 | 23904000 | 0 | 0 | 1985400 | 1734500 | 0 | 0 |
| IIVNDRQLA | RTCB | Q9Y3I0 | RTCB_HUMAN | Unmodified | 0 | 0 | 0 | 0 | 647630 | 0 | 419600 | 0 | 0 | 0 | 219340 | 0 | 1868300 | 1903300 | 0 | 0 | 0 | 923600 |
| IIVSDHVEF | IPO8; IPO8 | O15397; F5H | IPO8_HUMAN I | Unmodified | 0 | 0 | 0 | 0 | 0 | 0 | 0 | 0 | 0 | 0 | 0 | 41738000 | 0 | 0 | 0 | 0 | 0 | 0 |
| IIWELIK | IDH1; IDH1; IDH | O75874; C9J | IDHC_HUMAN | Unmodified | 1775600 | 1310400 | 1655900 | 3423600 | 1646800 | 239750 | 197300 | 0 | 4515500 | 511090 | 3356000 | 14641000 | 4562400 | 1592300 | 5286800 | 3973800 | 125930 | 254820 |
| IIWQFIK | IDH2; IDH2 | P48735; H0Y | IDHP_HUMAN | Unmodified | 0 | 0 | 0 | 0 | 0 | 0 | 0 | 0 | 25238000 | 941730 | 401590 | 0 | 0 | 0 | 0 | 0 | 0 | 0 |
| IIYGGSVTG | TPI1 | P60174 | TPIS_HUMAN T | Unmodified | 6121300 | 2006200 | 4165900 | 5805200 | 21848000 | 3637100 | 1147400 | 531290 | 6050500 | 2636000 | 7439100 | 80478000 | 21232000 | 8375700 | 6685200 | 4051100 | 9145300 | 13318000 |
| IIYIVHDEVK | PSMA3; PSMA3 | P25788; G3V | PSA3_HUMAN | Unmodified | 0 | 0 | 989350 | 0 | 893740 | 295690 | 0 | 0 | 1444500 | 843560 | 1634800 | 6212200 | 430350 | 0 | 0 | 0 | 172280 | 544370 |
| IIYIVHDEVK | PSMA3; PSMA3 | P25788; G3V | PSA3_HUMAN | Unmodified | 84904 | 197690 | 155570 | 372470 | 1565800 | 445240 | 166160 | 75561 | 1849000 | 1322500 | 1030000 | 6844900 | 1302600 | 929880 | 163280 | 227510 | 542080 | 645730 |
| IIYSHFTCA | GNAQ; GNA11; | P50148; P29 | GNAQ_HUMAN | Unmodified | 153050 | 179310 | 0 | 0 | 0 | 0 | 0 | 0 | 0 | 0 | 0 | 2406800 | 0 | 0 | 0 | 0 | 0 | 0 |
| IKAAVEDPF | DLST | P36957 | ODO2_HUMAN | Unmodified | 0 | 0 | 0 | 0 | 286550 | 0 | 2946300 | 37198 | 0 | 2058800 | 0 | 0 | 0 | 370020 | 0 | 0 | 0 | 0 |
| IKAAVPSIKI | PGK1 | P00558 | PGK1_HUMAN | Unmodified | 0 | 0 | 0 | 0 | 0 | 0 | 0 | 0 | 522620 | 0 | 0 | 0 | 0 | 0 | 0 | 0 | 1606300 | 189330 |
| IKADIIYPGI | LACTB2 | Q53H82 | LACB2_HUMAN | Unmodified | 0 | 251270 | 0 | 68905 | 3663500 | 0 | 0 | 0 | 19504000 | 5388400 | 433630 | 615400 | 397670 | 0 | 128450 | 0 | 1122600 | 1246300 |
| IKAEYEGD | PAICS; PAICS; F | P22234; D6F | PUR6_HUMAN | Unmodified | 0 | 0 | 0 | 0 | 0 | 2080500 | 0 | 0 | 215770 | 0 | 193410 | 5368500 | 0 | 0 | 0 | 0 | 0 | 0 |
| IKAIPQLQG | RPL6 | Q02878 | RL6_HUMAN 60 | Unmodified | 0 | 0 | 0 | 0 | 0 | 0 | 0 | 0 | 0 | 0 | 0 | 0 | 1060500 | 497910 | 0 | 0 | 0 | 0 |
| IKAPGFAHI | NDUFS2; NDUF | O75306; A0A | NDUS2_HUMA | Unmodified | 0 | 0 | 0 | 0 | 43627 | 0 | 196330 | 25923 | 888100 | 422790 | 0 | 0 | 0 | 0 | 0 | 0 | 0 | 0 |
| IKAQDEAFA | MRPL11; MRPL | Q9Y3B7; H0Y | RM11_HUMAN | Unmodified | 0 | 97875 | 0 | 0 | 1652700 | 179860 | 0 | 0 | 1222200 | 366970 | 0 | 0 | 0 | 0 | 0 | 0 | 0 | 0 |
| IKCEPKEGS | CXADR | P78310 | CXAR_HUMAN | Unmodified | 80796 | 0 | 0 | 0 | 4077000 | 0 | 53002 | 0 | 0 | 173090 | 0 | 0 | 327420 | 846410 | 0 | 0 | 48878 | 262670 |
| IKDAFDRNI | PGD | P52209 | 6PGD_HUMAN | Unmodified | 0 | 0 | 0 | 0 | 760560 | 459160 | 1304200 | 0 | 760510 | 1793900 | 308800 | 1622200 | 0 | 0 | 665440 | 239380 | 792520 | 0 |
| IKDCVDAP | SLC43A1 | O75387 | LAT3_HUMAN L | Unmodified | 275610 | 0 | 2832600 | 1250200 | 0 | 0 | 0 | 0 | 0 | 0 | 0 | 810120 | 0 | 0 | 6154100 | 4624400 | 0 | 0 |
| IKDDLEDLI | ECT2; ECT2 | Q9H8V3; H7 | ECT2_HUMAN | Unmodified | 0 | 0 | 0 | 0 | 767550 | 181170 | 0 | 0 | 102400 | 234080 | 0 | 0 | 0 | 0 | 0 | 0 | 10358 | 0 |
| IKDIAWTED | WDR1; WDR1 | O75083; D6I | WDR1_HUMAN | Unmodified | 1551000 | 229070 | 439670 | 1285800 | 14604000 | 2927800 | 1465200 | 660680 | 2274400 | 928570 | 1119100 | 5791000 | 6272300 | 2671000 | 1770500 | 367210 | 0 | 3943000 |
| IKDIAWTED | WDR1; WDR1 | O75083; D6I | WDR1_HUMAN | Unmodified | 0 | 0 | 0 | 196490 | 2344100 | 747080 | 74328 | 144480 | 368290 | 0 | 0 | 1776100 | 1745300 | 799860 | 0 | 0 | 0 | 560090 |
| IKDLSTVEA | ANP32E; ANP32 | Q9BTT0; E9F | AN32E_HUMAN | Unmodified | 0 | 0 | 0 | 0 | 2580800 | 0 | 0 | 0 | 0 | 1313600 | 0 | 0 | 1913900 | 0 | 0 | 0 | 1159300 | 0 |
| IKDPDASKI | CALR; CALR; CA | P27797; A0A | CALR_HUMAN | Unmodified | 316060 | 0 | 2378300 | 0 | 0 | 0 | 631590 | 782220 | 4.03E+08 | 9201100 | 8728200 | 23131000 | 0 | 0 | 3396000 | 754670 | 0 | 0 |
| IKDRLPLAV | SEPTIN7; SEPTI | Q16181; E7E | SEPT7_HUMAN | Unmodified | 0 | 0 | 0 | 0 | 2514800 | 104970 | 0 | 0 | 0 | 0 | 0 | 0 | 251260 | 127280 | 0 | 0 | 802960 | 1206100 |
| IKDVDEKAI | HTRA1; HTRA1 | Q92743; A0A | HTRA1_HUMAN | Unmodified | 0 | 0 | 0 | 0 | 260130 | 0 | 0 | 0 | 834930 | 234820 | 0 | 0 | 0 | 0 | 0 | 0 | 0 | 951100 |
| IKEDRPITIK | VPS28; VPS28; | Q9UK41; E9I | VPS28_HUMAN | Unmodified | 0 | 0 | 1480500 | 966820 | 0 | 0 | 0 | 0 | 0 | 0 | 0 | 0 | 0 | 0 | 3059500 | 1931800 | 0 | 0 |
| IKEELEDLN | STX2; STX2 | P32856; J3K | STX2_HUMAN S | Unmodified | 105390 | 99106 | 754320 | 0 | 0 | 0 | 0 | 0 | 825520 | 109820 | 0 | 7927900 | 0 | 0 | 740660 | 555380 | 0 | 0 |
| IKEEMNEDI | TAOK3; TAOK3 | Q9H2K8; G3 | TAOK3_HUMAN | Unmodified | 0 | 0 | 0 | 636640 | 529700 | 0 | 85851 | 150570 | 0 | 0 | 27701 | 212300 | 378470 | 273350 | 0 | 0 | 75464 | 265810 |
| IKEETEIIEG | RUVBL2; RUVBL | Q9Y230; M0I | RUVB2_HUMAN | Unmodified | 0 | 0 | 0 | 160860 | 0 | 0 | 0 | 0 | 0 | 0 | 641810 | 2504300 | 0 | 0 | 146400 | 0 | 0 | 0 |
| IKEIRESTG | PCBP1; PCBP3; | Q15365; P57 | PCBP1_HUMAN | Unmodified | 0 | 0 | 0 | 0 | 1236700 | 0 | 1358900 | 349470 | 0 | 439010 | 0 | 0 | 4522800 | 0 | 0 | 0 | 545150 | 733270 |
| IKEIRESTG | PCBP1; PCBP3; | Q15365; P57 | PCBP1_HUMAN | Oxidation (M) | 0 | 0 | 0 | 0 | 0 | 0 | 0 | 0 | 0 | 192470 | 0 | 0 | 1602400 | 0 | 0 | 0 | 0 | 0 |
| IKEKYIDQE | HSP90AB1; HSP | P08238; Q58 | HS90B_HUMAN | Unmodified | 0 | 0 | 0 | 0 | 1614400 | 0 | 1340800 | 514800 | 0 | 483720 | 0 | 0 | 1042200 | 886950 | 0 | 0 | 2894300 | 940410 |
| IKELRENTQ | HNRNPK; HNRN | P61978; Q51 | HNRPK_HUMA | Unmodified | 0 | 0 | 0 | 0 | 1961400 | 244010 | 5810100 | 11196 | 0 | 837110 | 0 | 0 | 3400900 | 1564900 | 0 | 0 | 859840 | 749280 |
| IKEPLLLPE | CDH3; CDH3 | P22223; J3Q | CADH3_HUMA | Unmodified | 0 | 0 | 0 | 0 | 7371200 | 1766900 | 0 | 0 | 0 | 0 | 0 | 0 | 0 | 0 | 0 | 0 | 0 | 0 |
| IKETAQAM | EXT2 | Q93063 | EXT2_HUMAN E | Unmodified | 0 | 0 | 47881 | 0 | 0 | 401000 | 0 | 0 | 673810 | 35632 | 0 | 0 | 276350 | 0 | 0 | 0 | 0 | 0 |
| IKETVDQVE | DYNC1H1; DYN | Q14204; A0A | DYHC1_HUMA | Unmodified | 105290 | 60784 | 691970 | 251820 | 1271000 | 0 | 358780 | 0 | 2812500 | 1290000 | 478380 | 10640000 | 0 | 0 | 958260 | 1017700 | 351980 | 765560 |
| IKFDAGTLL | VPS36; VPS36 | Q86VN1; A0 | VPS36_HUMAN | Unmodified | 2477100 | 2365300 | 1076700 | 1002600 | 0 | 0 | 0 | 0 | 0 | 0 | 0 | 0 | 0 | 0 | 4871200 | 2276300 | 0 | 0 |
| IKFDDVVQ | PLCG2; PLCG2 | P16885; A0A | PLCG2_HUMAN | Unmodified | 0 | 0 | 0 | 0 | 0 | 0 | 0 | 0 | 0 | 0 | 3670400 | 22203000 | 0 | 0 | 0 | 0 | 0 | 0 |
| IKGCWDSI | CAPZB; CAPZB; | P47756; B1A | CAPZB_HUMAN | Unmodified | 0 | 0 | 0 | 0 | 893100 | 0 | 0 | 0 | 0 | 501490 | 0 | 0 | 309400 | 627430 | 0 | 0 | 422090 | 456830 |
| IKGDMDIS | AHNAK | Q09666 | AHNK_HUMAN | Oxidation (M) | 0 | 0 | 0 | 0 | 1878500 | 273350 | 0 | 0 | 0 | 0 | 0 | 0 | 0 | 0 | 0 | 0 | 1175900 | 954300 |
| IKGDMDIS | AHNAK | Q09666 | AHNK_HUMAN | Unmodified | 0 | 0 | 0 | 0 | 1776200 | 0 | 0 | 0 | 0 | 0 | 0 | 0 | 0 | 0 | 0 | 0 | 791960 | 713980 |
| IKGDVPSV | AHNAK | Q09666 | AHNK_HUMAN | Unmodified | 0 | 361860 | 243390 | 405940 | 40707000 | 4770900 | 0 | 0 | 0 | 0 | 0 | 0 | 1073600 | 783520 | 262440 | 0 | 9382600 | 16171000 |
| IKGEEFPLT | RTN4IP1; RTN4 | Q8WWV3; G | RT4I1_HUMAN | Unmodified | 0 | 0 | 0 | 235300 | 0 | 0 | 0 | 0 | 1139400 | 0 | 0 | 0 | 0 | 0 | 298530 | 0 | 0 | 0 |
| IKGEELSEA | DCTN1; DCTN1 | Q14203; A0A | DCTN1_HUMAN | Unmodified | 0 | 0 | 262660 | 0 | 302710 | 0 | 0 | 0 | 618260 | 53036 | 380720 | 2142200 | 0 | 0 | 406720 | 22134 | 176980 | 0 |
| IKGEELSEA | DCTN1; DCTN1 | Q14203; A0A | DCTN1_HUMAN | Unmodified | 0 | 0 | 0 | 0 | 222200 | 0 | 1993500 | 0 | 0 | 2095000 | 0 | 0 | 506600 | 779310 | 0 | 0 | 864730 | 0 |
| IKGEHPGL | HMGB1P1; HM | B2RPK0; P09 | HGB1A_HUMA | Unmodified | 0 | 0 | 1160600 | 646790 | 0 | 350420 | 43082 | 396580 | 0 | 126000 | 0 | 1193900 | 168530 | 0 | 497910 | 0 | 30114 | 35990 |
| IKGEHPGL | HMGB1P1; HM | B2RPK0; P09 | HGB1A_HUMA | Unmodified | 0 | 0 | 419680 | 0 | 0 | 0 | 1033500 | 299200 | 767190 | 384840 | 1289700 | 8851100 | 688480 | 0 | 328060 | 0 | 0 | 0 |
| IKGGADVS | AHNAK | Q09666 | AHNK_HUMAN | Unmodified | 87686 | 0 | 0 | 0 | 477430 | 0 | 0 | 0 | 0 | 0 | 0 | 0 | 0 | 0 | 0 | 0 | 1999000 | 5153300 |
| IKGGLFADI | PLAT; PLAT; PLA | P00750; E7E | TPA_HUMAN Ti | Unmodified | 0 | 0 | 0 | 0 | 0 | 0 | 0 | 0 | 0 | 0 | 0 | 0 | 0 | 0 | 0 | 0 | 0 | 3584000 |
| IKGIVEESV | AHCYL2; AHCY | Q96HN2; H0 | SAHH3_HUMAN | Unmodified | 0 | 0 | 0 | 0 | 1236700 | 0 | 0 | 0 | 0 | 0 | 0 | 0 | 0 | 0 | 0 | 0 | 1021100 | 0 |
| IKHTRTQE | ADGRG6 | Q86SQ4 | AGRG6_HUMA | Unmodified | 0 | 0 | 0 | 0 | 801930 | 0 | 0 | 0 | 0 | 0 | 0 | 0 | 0 | 0 | 0 | 0 | 0 | 0 |
| IKIGVVGGS | PMM2; PMM2; F | O15305; H3I | PMM2_HUMAN | Unmodified | 0 | 0 | 0 | 0 | 0 | 0 | 0 | 0 | 0 | 0 | 0 | 0 | 318870 | 0 | 0 | 0 | 0 | 0 |
| IKIPRHELT | AHNAK | Q09666 | AHNK_HUMAN | Unmodified | 0 | 0 | 0 | 0 | 23297000 | 2031800 | 0 | 0 | 0 | 0 | 0 | 0 | 1483500 | 0 | 0 | 0 | 8635900 | 9917600 |
| IKIYEFPETI | SEPTIN7; SEPTI | Q16181; E7E | SEPT7_HUMAN | Unmodified | 48746 | 49215 | 0 | 293480 | 1242900 | 0 | 0 | 154120 | 0 | 0 | 0 | 0 | 525520 | 541500 | 0 | 0 | 0 | 1385300 |
| IKIYEFPETI | SEPTIN7; SEPTI | Q16181; E7E | SEPT7_HUMAN | Unmodified | 0 | 0 | 0 | 0 | 2382800 | 0 | 348340 | 0 | 0 | 0 | 0 | 0 | 0 | 0 | 0 | 0 | 1063300 | 1098100 |
| IKLDGYELP | EHD4 | Q9H223 | EHD4_HUMAN | Unmodified | 1601900 | 1105400 | 987370 | 3045300 | 17318000 | 8284500 | 0 | 0 | 843520 | 0 | 0 | 2533900 | 0 | 4883200 | 9804700 | 3225600 | 14599000 | 15679000 |
| IKLQIWDTA | RAB10 | P61026 | RAB10_HUMAN | Unmodified | 0 | 0 | 0 | 139500 | 0 | 0 | 0 | 0 | 0 | 0 | 52737 | 0 | 0 | 0 | 0 | 0 | 0 | 0 |
| IKLQLWDT | RAB39A | Q14964 | RB39A_HUMAN | Unmodified | 13055000 | 711470 | 77237 | 3159800 | 665640 | 0 | 384370 | 88685 | 3425800 | 1033000 | 441670 | 4907800 | 304960 | 0 | 3873400 | 6454900 | 620490 | 526350 |
| IKLQVWDT | RAB13 | P51153 | RAB13_HUMAN | Unmodified | 3669200 | 1871300 | 0 | 332320 | 0 | 0 | 206640 | 27078 | 0 | 0 | 0 | 556880 | 0 | 0 | 310360 | 328700 | 0 | 0 |
| IKLQVWDT | RAB13 | P51153 | RAB13_HUMAN | Unmodified | 0 | 0 | 0 | 0 | 1394600 | 235950 | 0 | 0 | 0 | 216560 | 0 | 0 | 690870 | 298290 | 0 | 0 | 188000 | 639850 |
| IKLTIYDWD | MYOF | Q9NZM1 | MYOF_HUMAN | Unmodified | 0 | 0 | 0 | 0 | 30393000 | 4144400 | 0 | 0 | 0 | 0 | 0 | 0 | 0 | 0 | 0 | 0 | 228700 | 3496300 |
| IKLYQDQL | TOR1B | O14657 | TOR1B_HUMAN | Unmodified | 0 | 0 | 0 | 0 | 0 | 0 | 0 | 0 | 0 | 341750 | 0 | 0 | 0 | 0 | 0 | 0 | 0 | 0 |
| IKNCLNPQ | CPNE3; CPNE3 | O75131; A0A | CPNE3_HUMAN | Unmodified | 0 | 0 | 0 | 0 | 120220 | 0 | 378580 | 288270 | 254900 | 381870 | 0 | 52477 | 0 | 0 | 0 | 0 | 328430 | 285870 |
| IKNDFQNL | COLEC12 | Q5KU26 | COL12_HUMAN | Unmodified | 138210 | 0 | 368170 | 736410 | 0 | 0 | 0 | 0 | 229090 | 0 | 0 | 0 | 0 | 0 | 368830 | 0 | 0 | 0 |
| IKNENTEGS | TOP2A | P11388 | TOP2A_HUMAN | Unmodified | 0 | 0 | 0 | 0 | 0 | 0 | 0 | 287280 | 0 | 197970 | 0 | 0 | 0 | 0 | 0 | 0 | 0 | 0 |
| IKNEVDSTL | SDHB; SDHB; S | P21912; A0A | SDHB_HUMAN | Unmodified | 0 | 0 | 0 | 0 | 0 | 93647 | 0 | 0 | 1489500 | 247300 | 0 | 0 | 0 | 0 | 0 | 0 | 0 | 0 |
| IKNEVDSTL | SDHB; SDHB; S | P21912; A0A | SDHB_HUMAN | Unmodified | 0 | 0 | 0 | 0 | 231220 | 0 | 0 | 0 | 4241400 | 726520 | 0 | 0 | 0 | 0 | 0 | 0 | 0 | 0 |

| | | | | | | | | | | | | | | | | | | | | | | |
|---|---|---|---|---|---|---|---|---|---|---|---|---|---|---|---|---|---|---|---|---|---|---|
| IKPGISEFAT | ENTPD4; ENTPI | Q9Y227; E5F | ENTP4_HUMAN | Unmodified | 0 | 0 | 0 | 0 | 0 | 0 | 0 | 0 | 5770500 | 533570 | 0 | 0 | 0 | 0 | 0 | 0 | 0 | 0 |
| IKPHLMSQ | P4HB; P4HB; P4 | P07237; A0A | PDIA1_HUMAN | Oxidation (M) | 0 | 0 | 82313 | 0 | 0 | 0 | 965450 | 0 | 7088500 | 1689000 | 520510 | 0 | 0 | 0 | 150780 | 0 | 0 | 0 |
| IKPHLMSQ | P4HB; P4HB; P4 | P07237; A0A | PDIA1_HUMAN | Unmodified | 0 | 0 | 0 | 0 | 0 | 0 | 0 | 0 | 15345000 | 683130 | 871990 | 874640 | 0 | 0 | 0 | 214680 | 0 | 0 |
| IKPHLMSQ | P4HB; P4HB; P4 | P07237; A0A | PDIA1_HUMAN | Oxidation (M) | 0 | 0 | 215660 | 0 | 2317300 | 237130 | 6418100 | 1247500 | 57278000 | 59507000 | 7781300 | 1856600 | 3508400 | 4833500 | 1153200 | 1069400 | 689240 | 1850300 |
| IKPHLMSQ | P4HB; P4HB; P4 | P07237; A0A | PDIA1_HUMAN | Unmodified | 0 | 0 | 0 | 0 | 0 | 0 | 3746400 | 179530 | 83878000 | 27600000 | 3958600 | 1011600 | 1662600 | 2642300 | 0 | 0 | 285540 | 0 |
| IKPHQGQH | VEGFA; VEGFA; | P15692; A0A | VEGFA_HUMAN | Unmodified | 0 | 0 | 0 | 0 | 248920 | 0 | 0 | 0 | 987780 | 73060 | 150970 | 0 | 0 | 0 | 0 | 0 | 131970 | 137700 |
| IKPLGPMLL | ECPAS; ECPAS; | Q5VYK3; R4 | ECM29_HUMAI | Oxidation (M) | 0 | 0 | 0 | 0 | 0 | 0 | 0 | 0 | 0 | 0 | 0 | 349050 | 0 | 0 | 0 | 0 | 0 | 0 |
| IKPSLGSGS | PLBD2 | Q8NHP8 | PLBL2_HUMAN | Unmodified | 0 | 0 | 0 | 0 | 0 | 0 | 85822 | 0 | 1024900 | 0 | 0 | 0 | 0 | 0 | 0 | 0 | 0 | 0 |
| IKPSQNTEI | MTPAP | Q9NVV4 | PAPD1_HUMAI | Unmodified | 0 | 0 | 0 | 0 | 0 | 0 | 0 | 0 | 491680 | 139980 | 521540 | 0 | 0 | 0 | 0 | 0 | 0 | 0 |
| IKPSTLQVE | CNRIP1; CNRIP | Q96F85; B8Z | CNRP1_HUMAI | Unmodified | 0 | 0 | 30868 | 156770 | 0 | 0 | 673020 | 0 | 0 | 48480 | 0 | 0 | 207720 | 347080 | 329990 | 0 | 482530 | 0 |
| IKPVFIEDAI | CACNA2D1; CA | P54289; E7E | CA2D1_HUMAI | Unmodified | 3449800 | 2438600 | 0 | 0 | 0 | 0 | 315630 | 248580 | 7508200 | 0 | 0 | 0 | 0 | 0 | 0 | 0 | 0 | 0 |
| IKQESNQIC | ANK3; ANK3; AI | Q12955; A0A | ANK3_HUMAN | Unmodified | 0 | 0 | 0 | 0 | 0 | 0 | 0 | 0 | 0 | 0 | 0 | 423440 | 6323100 | 0 | 2151000 | 0 | 0 | 0 |
| IKQHLEND | TXNL1; TXNL1; | O43396; K7E | TXNL1_HUMAN | Unmodified | 311640 | 0 | 0 | 1361300 | 926900 | 370220 | 0 | 0 | 2368600 | 259260 | 578620 | 797890 | 595940 | 659830 | 301750 | 522390 | 837290 | 2533200 |
| IKQIESDRC | CDCA8 | Q53HL2 | BOREA_HUMAI | Unmodified | 0 | 0 | 0 | 147680 | 260450 | 41245 | 28725 | 17145 | 98540 | 33925 | 0 | 0 | 98906 | 115100 | 0 | 0 | 36478 | 30351 |
| IKQIIEDQS | DLG3; DLG3 | Q92796; Q5J | DLG3_HUMAN | Unmodified | 0 | 0 | 0 | 0 | 0 | 0 | 0 | 0 | 0 | 1132000 | 0 | 0 | 0 | 424980 | 0 | 215330 | 995650 | 0 |
| IKSDYAQLL | ATP6V1A | P38606 | VATA_HUMAN | Unmodified | 190040 | 0 | 16816 | 0 | 0 | 0 | 0 | 0 | 708820 | 64451 | 103580 | 194140 | 74312 | 40198 | 148740 | 49256 | 0 | 0 |
| IKSEHPGLS | HMGB2; HMGB | P26583; D6F | HMGB2_HUMA | Unmodified | 0 | 0 | 0 | 0 | 0 | 0 | 0 | 65128 | 0 | 0 | 0 | 875000 | 320830 | 81965 | 0 | 0 | 0 | 0 |
| IKSGEEDFE | PIN1; PIN1; PIN | Q13526; K7E | PIN1_HUMAN | Unmodified | 19947000 | 7610900 | 0 | 9488400 | 4190000 | 1403500 | 4243600 | 995680 | 1059600 | 1827700 | 0 | 0 | 7099100 | 7096100 | 5172200 | 6520700 | 5366300 | 5793600 |
| IKSQGQDV | UCHL3; UCHL3 | P15374; Q5T | UCHL3_HUMAI | Unmodified | 0 | 0 | 0 | 0 | 0 | 0 | 101910 | 0 | 0 | 72317 | 0 | 221000 | 427750 | 1106000 | 0 | 0 | 0 | 0 |
| IKSTNPGIS | HMGB3; HMGB | O15347; E7E | HMGB3_HUMA | Unmodified | 0 | 0 | 0 | 0 | 0 | 0 | 0 | 0 | 0 | 706210 | 0 | 0 | 0 | 74036 | 0 | 0 | 0 | 0 |
| IKVAEDEAE | SLC3A2; SLC3A | P08195; A0A | 4F2_HUMAN 4I | Unmodified | 12166000 | 8594400 | 3299100 | 12143000 | 46155000 | 9404300 | 2156200 | 969100 | 10920000 | 2309800 | 15351000 | 43080000 | 2973300 | 0 | 11689000 | 10741000 | 3463500 | 7013300 |
| IKVAEDEAE | SLC3A2; SLC3A | P08195; A0A | 4F2_HUMAN 4I | Unmodified | 0 | 0 | 0 | 0 | 2133200 | 0 | 0 | 0 | 0 | 0 | 0 | 0 | 0 | 0 | 0 | 0 | 1125200 | 0 |
| IKVDEFVTH | ADH5 | P11766 | ADHX_HUMAN | Unmodified | 231950 | 0 | 0 | 0 | 5468600 | 661040 | 292680 | 0 | 1236400 | 0 | 9029400 | 41551000 | 2991600 | 477220 | 0 | 0 | 599660 | 3704400 |
| IKVDNHLFI | PIP4K2B; PIP4K | P78356; P48 | PI42B_HUMAN | Unmodified | 210200 | 0 | 322780 | 905450 | 2675500 | 0 | 2798500 | 1052700 | 0 | 0 | 0 | 3004700 | 414030 | 1896700 | 102530 | 608700 | 567020 | 2914100 |
| IKVEELENE | PPFIBP2; PPFIB | Q8ND30; E9 | LIPB2_HUMAN | Unmodified | 0 | 0 | 0 | 1644400 | 0 | 0 | 0 | 0 | 0 | 0 | 0 | 0 | 1837900 | 0 | 0 | 0 | 0 | 0 |
| IKVLGSGAI | EGFR; EGFR; EC | P00533; Q5C | EGFR_HUMAN | Unmodified | 0 | 0 | 0 | 0 | 2881100 | 0 | 0 | 0 | 0 | 0 | 0 | 0 | 0 | 0 | 0 | 0 | 203150 | 452430 |
| IKVNNHLFI | PIP4K2C; PIP4K | Q8TBX8; F8V | PI42C_HUMAN | Unmodified | 77619 | 0 | 0 | 0 | 1055000 | 0 | 0 | 0 | 0 | 0 | 0 | 0 | 719140 | 1309300 | 0 | 0 | 114160 | 376220 |
| IKWGDAGA | GAPDH; GAPDH | P04406; E7E | G3P_HUMAN G | Oxidation (M) | 16365000 | 0 | 2104600 | 25713000 | 2.43E+08 | 22195000 | 52130000 | 490140 | 11915000 | 28257000 | 29760000 | 76504000 | 1.72E+08 | 43301000 | 19055000 | 9873700 | 92279000 | 1.37E+08 |
| IKWGDAGA | GAPDH; GAPDH | P04406; E7E | G3P_HUMAN G | Unmodified | 0 | 0 | 0 | 0 | 68628000 | 3984600 | 11938000 | 0 | 11776000 | 3932600 | 1032400 | 3538700 | 29462000 | 14320000 | 0 | 0 | 22285000 | 59730000 |
| IKWGDAGA | GAPDH; GAPDH | P04406; E7E | G3P_HUMAN G | Oxidation (M) | 0 | 0 | 0 | 0 | 0 | 0 | 991620 | 0 | 0 | 0 | 0 | 0 | 0 | 0 | 0 | 0 | 1724000 | 398440 |
| IKWGDAGA | GAPDH; GAPDH | P04406; E7E | G3P_HUMAN G | Unmodified | 0 | 0 | 0 | 0 | 2470500 | 0 | 3010200 | 0 | 0 | 0 | 0 | 0 | 1061900 | 0 | 0 | 0 | 7403900 | 2823300 |
| IKYPENFFL | PPP1CA; PPP1C | P62136; P62 | PP1A_HUMAN | Unmodified | 1214700 | 0 | 2350000 | 577450 | 395720 | 337490 | 0 | 0 | 1824200 | 982780 | 766990 | 20645000 | 0 | 0 | 4128600 | 2618000 | 252730 | 0 |
| ILAAALTEC | RANGAP1 | P46060 | RAGP1_HUMAI | Unmodified | 0 | 0 | 0 | 0 | 1836200 | 262050 | 142670 | 169310 | 335340 | 215710 | 228910 | 1738000 | 810920 | 459800 | 0 | 0 | 287170 | 875710 |
| ILAAILHLGI | MYO1D; MYO1D | O94832; J3Q | MYO1D_HUMA | Unmodified | 2460500 | 4469900 | 904500 | 0 | 0 | 0 | 0 | 0 | 198480 | 305270 | 0 | 713990 | 0 | 0 | 3365800 | 1820800 | 0 | 0 |
| ILAALEAVG | PLCB4; PLCB4; | Q15147; A0A | PLCB4_HUMAN | Unmodified | 0 | 0 | 188610 | 61282 | 0 | 34084 | 0 | 0 | 0 | 0 | 0 | 359740 | 0 | 0 | 209180 | 245070 | 0 | 0 |
| ILAALEAVG | PLCB4; PLCB4; | Q15147; A0A | PLCB4_HUMAN | Unmodified | 276130 | 191950 | 403140 | 1207100 | 0 | 0 | 0 | 0 | 0 | 0 | 0 | 0 | 0 | 0 | 1220200 | 1445200 | 0 | 0 |
| ILAANPEAK | SUCO; SUCO | Q9UBS9; B4 | SUCO_HUMAN | Unmodified | 0 | 0 | 69307 | 0 | 75931 | 536540 | 0 | 0 | 1525400 | 249290 | 115980 | 581040 | 0 | 0 | 0 | 29714 | 0 | 0 |
| ILACAPLYH | ITGAV | P06756 | ITAV_HUMAN I | Unmodified | 516800 | 0 | 0 | 271190 | 0 | 0 | 0 | 0 | 793240 | 108750 | 0 | 2161500 | 0 | 0 | 460610 | 227230 | 0 | 0 |
| ILACDDLDI | SUCLA2; SUCL | Q9P2R7; A0A | SUCB1_HUMAI | Unmodified | 0 | 0 | 0 | 0 | 0 | 0 | 0 | 0 | 660380 | 0 | 0 | 0 | 0 | 0 | 0 | 0 | 0 | 0 |
| ILADATAKM | TLN1 | Q9Y490 | TLN1_HUMAN | Unmodified | 0 | 0 | 0 | 0 | 1047800 | 3493600 | 158760 | 201670 | 0 | 0 | 0 | 0 | 0 | 0 | 0 | 0 | 0 | 3357300 |
| ILADEALKL | EHHADH | Q08426 | ECHP_HUMAN | Unmodified | 0 | 0 | 0 | 0 | 0 | 0 | 0 | 0 | 0 | 187510 | 0 | 0 | 0 | 0 | 0 | 0 | 0 | 0 |
| ILADQQDK | RUVBL1; RUVBI | Q9Y265; H7 | RUVB1_HUMAI | Unmodified | 349430 | 0 | 247570 | 328860 | 4346200 | 317850 | 748900 | 896660 | 1601400 | 114350 | 136970 | 7081500 | 2127800 | 1227100 | 232630 | 184120 | 862980 | 1309000 |
| ILADQQDK | RUVBL1; RUVBI | Q9Y265; H7 | RUVB1_HUMAI | Unmodified | 0 | 0 | 0 | 0 | 0 | 0 | 284540 | 428050 | 0 | 238020 | 0 | 789890 | 0 | 746490 | 0 | 68665 | 710850 | 0 |
| ILADQQDK | RUVBL1; RUVBI | Q9Y265; H7 | RUVB1_HUMAI | Oxidation (M) | 0 | 0 | 0 | 0 | 0 | 0 | 0 | 79814 | 9846.6 | 0 | 0 | 39181 | 0 | 0 | 0 | 0 | 0 | 75973 |
| ILAEADGLS | NAGK; NAGK; N | Q9UJ70; H7 | NAGK_HUMAN | Unmodified | 177550 | 168610 | 247120 | 224990 | 1691900 | 0 | 0 | 0 | 1428000 | 0 | 519660 | 1040100 | 1229400 | 771110 | 347720 | 229430 | 252530 | 2396600 |
| ILAEGGGAI | TUFM | P49411 | EFTU_HUMAN | Unmodified | 0 | 0 | 35701 | 75026 | 1951600 | 115540 | 1775200 | 105140 | 9229800 | 0 | 114540 | 104410 | 324920 | 501360 | 28653 | 16464 | 281260 | 400220 |
| ILAFPCNQI | GPX4; GPX4; GI | P36969; A0A | GPX4_HUMAN | Unmodified | 1870200 | 0 | 0 | 1154100 | 0 | 0 | 0 | 0 | 0 | 1082500 | 546680 | 1862000 | 1797200 | 0 | 1550500 | 1388300 | 0 | 0 |
| ILAGLGFDI | ABCF1; ABCF1; | Q8NE71; H0 | ABCF1_HUMAN | Oxidation (M) | 291840 | 390860 | 0 | 276330 | 0 | 0 | 0 | 0 | 0 | 0 | 0 | 4625600 | 0 | 0 | 281530 | 43607 | 0 | 0 |
| ILAGSADSE | PCSK1N | Q9UHG2 | PCS1N_HUMAI | Unmodified | 0 | 0 | 0 | 0 | 0 | 172160 | 0 | 0 | 7148100 | 6099100 | 80575 | 0 | 0 | 0 | 0 | 0 | 549670 | 543180 |
| ILAIELENLK | SHTN1 | A0MZ66 | SHOT1_HUMAI | Unmodified | 0 | 0 | 159970 | 0 | 0 | 0 | 450080 | 0 | 657420 | 543610 | 0 | 930810 | 876950 | 914080 | 0 | 101920 | 0 | 0 |
| ILAIGLINEA | IQGAP1; IQGAP | P46940; A0A | IQGA1_HUMAN | Unmodified | 1103200 | 2393100 | 68910 | 0 | 1453400 | 0 | 0 | 32297 | 922380 | 0 | 114200 | 1958800 | 0 | 0 | 1369300 | 555950 | 219310 | 99183 |
| ILAIVGTAES | ATP7A | Q04656 | ATP7A_HUMAN | Unmodified | 131130 | 98104 | 890650 | 2574700 | 599690 | 0 | 925840 | 160620 | 0 | 129950 | 794530 | 2414100 | 3128100 | 2798400 | 2007700 | 1473300 | 100960 | 696110 |
| ILALCMGN | RDX; MSN; RDX | P35241; P26 | RADI_HUMAN I | Oxidation (M) | 0 | 0 | 0 | 594580 | 4920700 | 2750300 | 0 | 0 | 1727500 | 43066 | 0 | 1549700 | 1329100 | 2549300 | 643440 | 0 | 2310000 | 2310500 |
| ILALCMGN | RDX; MSN; RDX | P35241; P26 | RADI_HUMAN I | Unmodified | 0 | 0 | 0 | 0 | 3819200 | 0 | 0 | 0 | 0 | 0 | 0 | 0 | 0 | 188230 | 0 | 0 | 0 | 939420 |
| ILALCMGN | RDX; MSN; RDX | P35241; P26 | RADI_HUMAN I | 2 Oxidation (M) | 488130 | 269300 | 357820 | 1039100 | 1948400 | 349890 | 415420 | 0 | 0 | 166030 | 0 | 1072500 | 1154700 | 989220 | 737060 | 893090 | 101700 | 47938 |
| ILALCMGN | RDX; MSN; RDX | P35241; P26 | RADI_HUMAN I | Unmodified | 0 | 0 | 0 | 0 | 2363600 | 0 | 0 | 0 | 0 | 0 | 0 | 0 | 400740 | 0 | 0 | 0 | 734110 | 0 |
| ILALVPPWT | ELP3; ELP3 | Q9H9T3; B4I | ELP3_HUMAN I | Unmodified | 0 | 0 | 0 | 0 | 0 | 0 | 0 | 0 | 0 | 0 | 0 | 2683800 | 0 | 0 | 0 | 0 | 0 | 0 |
| ILAPAYFILG | ASAH1; ASAH1; | Q13510; A0A | ASAH1_HUMAI | Unmodified | 0 | 0 | 0 | 0 | 0 | 0 | 53650 | 0 | 1279000 | 17803 | 0 | 0 | 0 | 0 | 0 | 0 | 0 | 0 |
| ILAPGEEEN | AIDA | Q96BJ3 | AIDA_HUMAN | Unmodified | 0 | 0 | 0 | 0 | 0 | 0 | 0 | 0 | 0 | 0 | 0 | 0 | 0 | 0 | 0 | 579610 | 0 | 0 |
| ILAPGEEEN | AIDA | Q96BJ3 | AIDA_HUMAN | Unmodified | 0 | 0 | 0 | 17549000 | 0 | 0 | 0 | 0 | 0 | 0 | 0 | 0 | 0 | 0 | 4482400 | 2231500 | 0 | 0 |
| ILAPSPDSL | ABCA3; ABCA3 | Q99758; H0 | ABCA3_HUMAI | Unmodified | 166200 | 82311 | 0 | 0 | 0 | 0 | 0 | 0 | 2638000 | 0 | 0 | 0 | 0 | 0 | 0 | 0 | 0 | 0 |
| ILAPYTDFH | RICTOR | Q6R327 | RICTR_HUMAN | Unmodified | 0 | 32302 | 0 | 107870 | 0 | 140850 | 0 | 0 | 0 | 0 | 0 | 1206700 | 0 | 0 | 85621 | 104600 | 0 | 122870 |
| ILAQATSDL | TLN1 | Q9Y490 | TLN1_HUMAN | Unmodified | 0 | 0 | 0 | 0 | 0 | 0 | 0 | 0 | 0 | 0 | 96096 | 269570 | 0 | 0 | 0 | 0 | 0 | 0 |
| ILAQATSDL | TLN1 | Q9Y490 | TLN1_HUMAN | Unmodified | 2635600 | 0 | 0 | 2217400 | 10743000 | 0 | 0 | 0 | 0 | 2288200 | 0 | 2257500 | 9704600 | 5581100 | 2765800 | 3086700 | 1828600 | 0 |
| ILAQITGTEI | PDLIM5; PDLIM | Q96HC4; HC | PDLI5_HUMAN | Unmodified | 0 | 0 | 0 | 0 | 0 | 0 | 0 | 0 | 0 | 0 | 1545800 | 2746900 | 0 | 0 | 0 | 0 | 0 | 661320 |
| ILAQITGTEI | PDLIM5; PDLIM | Q96HC4; HC | PDLI5_HUMAN | Unmodified | 0 | 0 | 0 | 0 | 661790 | 158950 | 0 | 0 | 0 | 0 | 0 | 0 | 0 | 0 | 0 | 0 | 0 | 494780 |
| ILAQQTGR | ABHD11; ABHD | Q8NFV4; H7 | ABHDB_HUMA | Unmodified | 737700 | 1376300 | 45573 | 286340 | 92416 | 0 | 25581 | 261230 | 1054400 | 253910 | 83239 | 691780 | 51297 | 23190 | 370260 | 237010 | 290990 | 576420 |
| ILAQVVGD | EFTUD2 | Q15029 | U5S1_HUMAN | Unmodified | 0 | 0 | 0 | 0 | 0 | 0 | 0 | 18438 | 0 | 0 | 0 | 385820 | 0 | 0 | 0 | 23134 | 0 | 0 |
| ILARGENLE | VAMP8; VAMP8 | Q9BV40; B8 | VAMP8_HUMAI | Unmodified | 0 | 0 | 0 | 0 | 0 | 0 | 0 | 0 | 568730 | 100310 | 399780 | 0 | 0 | 0 | 1112800 | 0 | 0 | 0 |
| ILASGSVQL | SDK2 | Q58EX2 | SDK2_HUMAN | Unmodified | 0 | 0 | 0 | 0 | 0 | 0 | 0 | 0 | 538840 | 0 | 0 | 0 | 0 | 0 | 0 | 0 | 0 | 0 |
| ILASPVELAI | UGGT1 | Q9NYU2 | UGGG1_HUMA | Unmodified | 0 | 0 | 0 | 0 | 0 | 0 | 0 | 0 | 414660 | 141480 | 0 | 0 | 0 | 0 | 0 | 0 | 0 | 0 |
| ILASTQFEP | ERAP1 | Q9NZ08 | ERAP1_HUMAN | Unmodified | 0 | 0 | 0 | 0 | 0 | 0 | 0 | 0 | 535140 | 0 | 747140 | 0 | 0 | 0 | 0 | 0 | 0 | 0 |
| ILATPPQED | TKT; TKT; TKT | P29401; A0A | TKT_HUMAN Tr | Unmodified | 0 | 0 | 0 | 0 | 4281000 | 1306400 | 0 | 0 | 12022000 | 2178000 | 14527000 | 40435000 | 0 | 0 | 0 | 1219200 | 789670 | 1628900 |
| ILATPPQED | TKT; TKT; TKT | P29401; A0A | TKT_HUMAN Tr | Unmodified | 0 | 0 | 0 | 0 | 5254800 | 0 | 0 | 0 | 0 | 476560 | 0 | 0 | 0 | 0 | 0 | 0 | 0 | 1099700 |
| ILATPPQED | TKT; TKT; TKT | P29401; A0A | TKT_HUMAN Tr | Oxidation (M) | 0 | 0 | 0 | 0 | 4739900 | 655420 | 0 | 0 | 0 | 0 | 0 | 0 | 0 | 0 | 0 | 0 | 631350 | 793800 |
| ILATPPQED | TKT; TKT; TKT | P29401; A0A | TKT_HUMAN Tr | Oxidation (M) | 0 | 0 | 0 | 0 | 8462400 | 0 | 0 | 0 | 0 | 1540700 | 0 | 0 | 0 | 0 | 0 | 0 | 0 | 2688600 |
| ILATPPQED | TKT; TKT; TKT | P29401; A0A | TKT_HUMAN Tr | Unmodified | 0 | 0 | 0 | 0 | 14603000 | 0 | 0 | 0 | 0 | 2164300 | 0 | 0 | 0 | 0 | 0 | 0 | 3692700 | 8393400 |
| ILAVNGQD | SCRIB; SCRIB; | Q14160; A0A | SCRIB_HUMAN | Unmodified | 0 | 0 | 0 | 0 | 0 | 0 | 0 | 0 | 1070600 | 0 | 0 | 0 | 0 | 0 | 555330 | 334740 | 0 | 0 |
| ILCFYGPPC | LONP1; LONP1 | P36776; K7E | LONM_HUMAN | Unmodified | 0 | 0 | 0 | 0 | 0 | 0 | 0 | 0 | 3014400 | 1565500 | 0 | 0 | 0 | 0 | 0 | 0 | 0 | 0 |
| ILCFYGPPC | LONP1; LONP1 | P36776; K7E | LONM_HUMAN | Unmodified | 0 | 0 | 0 | 0 | 0 | 0 | 162100 | 0 | 0 | 225250 | 0 | 0 | 0 | 224210 | 0 | 0 | 0 | 0 |
| ILCHMQLS | TM9SF1; TM9SF | O15321; E9F | TM9S1_HUMAN | Unmodified | 0 | 0 | 0 | 0 | 0 | 0 | 0 | 0 | 585690 | 0 | 0 | 0 | 0 | 0 | 0 | 0 | 0 | 0 |
| ILCQEEQD | BRCC3; BRCC3 | P46736; A0A | BRCC3_HUMAI | Unmodified | 0 | 0 | 0 | 0 | 0 | 0 | 0 | 0 | 181340 | 0 | 0 | 705620 | 0 | 0 | 0 | 0 | 52176 | 0 |
| ILCWVMTG | C1GALT1; C1G | Q9NS00; C9 | C1GLT_HUMAN | Unmodified | 0 | 0 | 0 | 0 | 0 | 0 | 0 | 0 | 0 | 0 | 0 | 10566000 | 0 | 0 | 0 | 0 | 0 | 0 |
| ILDAAGANI | GRHPR; GRHPF | Q9UBQ7; U3 | GRHPR_HUMA | Unmodified | 0 | 0 | 79859 | 0 | 0 | 72996 | 0 | 0 | 840020 | 282560 | 0 | 0 | 416880 | 0 | 0 | 380950 | 0 | 0 |
| ILDATDQES | KIAA0319L; KIA | Q8IZA0; C9J | K319L_HUMAN | Unmodified | 130240 | 0 | 1341100 | 0 | 765080 | 0 | 0 | 0 | 1100700 | 0 | 228800 | 0 | 0 | 0 | 855810 | 1116700 | 0 | 0 |
| ILDAVVAQE | EFTUD2; EFTUD | Q15029; K7E | U5S1_HUMAN | Unmodified | 222580 | 313530 | 6.64E+08 | 383730 | 0 | 0 | 0 | 1157600 | 0 | 0 | 0 | 2872400 | 0 | 0 | 289180 | 0 | 0 | 0 |
| ILDDDTIITT | DYNC1H1; DYN | Q14204; A0A | DYHC1_HUMA | Unmodified | 249690 | 0 | 175540 | 620020 | 0 | 0 | 0 | 0 | 2443100 | 683130 | 1204600 | 5870700 | 0 | 0 | 1130500 | 482480 | 0 | 0 |
| ILDDICVAK | KARS1 | Q15046 | SYK_HUMAN Ly | Unmodified | 0 | 0 | 0 | 0 | 825990 | 0 | 1382200 | 141950 | 151080 | 54279 | 762390 | 9724900 | 160340 | 0 | 28095 | 34982 | 0 | 0 |
| ILDDWGET | MANF | P55145 | MANF_HUMAN | Unmodified | 0 | 0 | 0 | 0 | 0 | 0 | 0 | 122900 | 14518000 | 4156400 | 1449500 | 460160 | 0 | 0 | 0 | 101340 | 0 | 0 |
| ILDDWGET | MANF | P55145 | MANF_HUMAN | Unmodified | 0 | 0 | 0 | 0 | 0 | 1029200 | 0 | 0 | 0 | 1604100 | 0 | 0 | 0 | 0 | 0 | 0 | 108950 | 0 |
| ILDEIEEHN | SEPTIN2; SEPTI | Q15019; H7 | SEPT2_HUMAN | Unmodified | 322580 | 0 | 1101400 | 497790 | 1763600 | 342980 | 68665 | 104250 | 0 | 619920 | 5167800 | 7883600 | 521610 | 0 | 514830 | 1170400 | 1180000 | 1598800 |
| ILDENDNK | FAT1; FAT1 | Q14517; A0A | FAT1_HUMAN I | Unmodified | 228630 | 101750 | 383130 | 424120 | 0 | 0 | 0 | 0 | 0 | 176190 | 220010 | 0 | 593600 | 613470 | 330960 | 98139 | 373130 | 1100000 |
| ILDETQEAV | CHMP6; CHMP | Q96FZ7; I3L | CHMP6_HUMA | Unmodified | 0 | 0 | 159720 | 0 | 0 | 0 | 0 | 0 | 158580 | 0 | 0 | 712620 | 0 | 237880 | 398390 | 431060 | 0 | 0 |
| ILDGGNPFI | PSD2 | Q9BQI7 | PSD2_HUMAN | Unmodified | 0 | 0 | 0 | 0 | 0 | 0 | 226080 | 0 | 0 | 28451 | 4570900 | 1328200 | 0 | 0 | 412070 | 0 | 0 | 0 |
| ILDHQQQN | TLN2; TLN2; TLI | Q9Y4G6; H0 | TLN2_HUMAN | Unmodified | 0 | 0 | 0 | 0 | 0 | 0 | 0 | 24594 | 0 | 0 | 0 | 0 | 0 | 0 | 0 | 0 | 0 | 473910 |
| ILDIVPPTFS | NCBP1 | Q09161 | NCBP1_HUMAI | Unmodified | 0 | 0 | 0 | 0 | 0 | 0 | 0 | 0 | 0 | 0 | 0 | 256190 | 0 | 0 | 0 | 0 | 0 | 0 |
| ILDKCNEIII | HSPA8; HSPA8; | P11142; E9P | HSP7C_HUMAI | Unmodified | 51322 | 146800 | 0 | 0 | 15426000 | 2345400 | 9840600 | 105760 | 0 | 7682800 | 0 | 0 | 22306000 | 14939000 | 0 | 0 | 7150200 | 10057000 |
| ILDLEDLER | DAAM1 | Q9Y4D1 | DAAM1_HUMA | Unmodified | 0 | 0 | 0 | 160160 | 173760 | 0 | 242320 | 0 | 1655900 | 0 | 672030 | 2732800 | 0 | 0 | 0 | 299700 | 674990 | 0 |
| ILDLEDLER | DAAM1 | Q9Y4D1 | DAAM1_HUMA | Unmodified | 238700 | 163270 | 187280 | 952540 | 1058800 | 47090 | 159430 | 0 | 1183700 | 0 | 307380 | 317830 | 1196000 | 526040 | 921490 | 390030 | 0 | 0 |
| ILDLGITGP | DPYSL2; DPYSL | Q16555; A0A | DPYL2_HUMAN | Unmodified | 1639300 | 1028200 | 924590 | 1336000 | 1865900 | 2547700 | 0 | 0 | 14847000 | 832670 | 1312600 | 18204000 | 0 | 284320 | 1270400 | 931120 | 0 | 0 |
| ILDLIESGK | ALDH1A1 | P00352 | AL1A1_HUMAN | Unmodified | 0 | 0 | 0 | 0 | 0 | 0 | 0 | 0 | 0 | 0 | 0 | 17246000 | 0 | 0 | 0 | 0 | 0 | 0 |
| ILDLIESGKI | ALDH1A1 | P00352 | AL1A1_HUMAN | Unmodified | 0 | 0 | 0 | 0 | 0 | 0 | 0 | 0 | 319230 | 0 | 3176900 | 1.08E+08 | 0 | 0 | 0 | 0 | 0 | 0 |
| ILDLIESGKI | ALDH1A1 | P00352 | AL1A1_HUMAN | Unmodified | 0 | 0 | 0 | 0 | 0 | 0 | 0 | 0 | 0 | 0 | 382440 | 1566700 | 0 | 0 | 0 | 0 | 0 | 0 |
| ILDLSNFEII | BIRC6; BIRC6 | Q9NR09; A0 | BIRC6_HUMAN | Unmodified | 0 | 0 | 0 | 0 | 0 | 0 | 0 | 0 | 0 | 0 | 0 | 554160 | 0 | 0 | 0 | 0 | 0 | 0 |
| ILDPNTGEI | RANGAP1 | P46060 | RAGP1_HUMAI | Unmodified | 0 | 0 | 0 | 0 | 549540 | 0 | 0 | 0 | 369720 | 0 | 114640 | 855810 | 1171600 | 1009800 | 0 | 101730 | 0 | 926090 |
| ILDPTASVP | KIAA1549 | Q9HCM3 | K1549_HUMAN | Unmodified | 0 | 0 | 0 | 0 | 6710200 | 0 | 0 | 65973 | 0 | 0 | 0 | 0 | 952590 | 0 | 0 | 0 | 0 | 917900 |
| ILDQGEDFI | ERP29; ERP29 | P30040; F8V | ERP29_HUMAN | Oxidation (M) | 0 | 0 | 149320 | 0 | 0 | 0 | 31333 | 0 | 24641000 | 9449000 | 3234800 | 430270 | 0 | 0 | 0 | 0 | 0 | 350200 |
| ILDQGEDFI | ERP29; ERP29 | P30040; F8V | ERP29_HUMAN | Unmodified | 0 | 0 | 0 | 0 | 0 | 0 | 0 | 0 | 1.26E+08 | 19690000 | 1133400 | 340340 | 0 | 0 | 0 | 0 | 0 | 0 |
| ILDQGEDFI | ERP29; ERP29 | P30040; F8V | ERP29_HUMAN | Unmodified | 0 | 0 | 0 | 0 | 0 | 0 | 0 | 0 | 0 | 11146000 | 357810 | 0 | 0 | 0 | 0 | 0 | 0 | 0 |

| | | | | | | | | | | | | | | | | | | | | | | |
|---|---|---|---|---|---|---|---|---|---|---|---|---|---|---|---|---|---|---|---|---|---|---|
| ILDQGEDFI | ERP29; ERP29 | P30040; F8V | ERP29_HUMAN | Oxidation (M) | 0 | 0 | 0 | 0 | 0 | 0 | 0 | 0 | 0 | 4367600 | 0 | 0 | 0 | 0 | 0 | 0 | 0 | 0 |
| ILDQSEAEK | MVP | Q14764 | MVP_HUMAN I | Unmodified | 2151000 | 998090 | 0 | 0 | 2982700 | 0 | 189850 | 0 | 2310400 | 589870 | 0 | 27804000 | 0 | 110940 | 28895 | 0 | 0 | 0 |
| ILDRCSSILI | ATP1A1 | P05023 | AT1A1_HUMAN | Unmodified | 0 | 0 | 0 | 501910 | 0 | 0 | 0 | 0 | 0 | 0 | 1038800 | 0 | 0 | 436550 | 0 | 0 | 0 | 0 |
| ILDRCSSILI | ATP1A1 | P05023 | AT1A1_HUMAN | Unmodified | 0 | 0 | 0 | 2504200 | 1168900 | 0 | 0 | 0 | 908150 | 966890 | 5191200 | 0 | 4807700 | 1821300 | 0 | 72643 | 0 | 0 |
| ILDRHLAILS | TXNDC9; TXND | O14530; B8Z | TXND9_HUMAN | Unmodified | 0 | 0 | 0 | 0 | 0 | 0 | 93077 | 0 | 0 | 0 | 0 | 0 | 0 | 0 | 0 | 0 | 244880 | 332640 |
| ILDRHLAILS | TXNDC9; TXND | O14530; B8Z | TXND9_HUMAN | Unmodified | 0 | 0 | 0 | 0 | 665390 | 0 | 0 | 0 | 0 | 0 | 0 | 0 | 0 | 0 | 0 | 0 | 200560 | 247050 |
| ILDSVGIEAI | RPLP2 | P05387 | RLA2_HUMAN | Unmodified | 0 | 0 | 0 | 0 | 0 | 0 | 0 | 0 | 0 | 0 | 1281500 | 433230 | 0 | 0 | 0 | 0 | 0 | 0 |
| ILDSVGIEAI | RPLP2 | P05387 | RLA2_HUMAN | Unmodified | 63072000 | 1.07E+08 | 64858000 | 77974000 | 19947000 | 2374700 | 3447000 | 1347300 | 11986000 | 3857500 | 23955000 | 2.85E+08 | 90589000 | 33749000 | 1.31E+08 | 1.33E+08 | 2802900 | 6943800 |
| ILDSVGIEAI | RPLP2 | P05387 | RLA2_HUMAN | Unmodified | 0 | 0 | 0 | 0 | 0 | 0 | 0 | 0 | 0 | 0 | 0 | 0 | 11718000 | 5601200 | 0 | 0 | 0 | 0 |
| ILDTADENC | ERLEC1 | Q96DZ1 | ERLEC_HUMAN | Unmodified | 0 | 0 | 0 | 0 | 31700 | 0 | 0 | 0 | 643550 | 251510 | 0 | 0 | 50931 | 72920 | 0 | 0 | 0 | 21406 |
| ILDTALSK | HMBS; HMBS; I | P08397; A0A | HEM3_HUMAN | Unmodified | 0 | 120530 | 0 | 138270 | 632650 | 243280 | 938860 | 210540 | 248810 | 72424 | 477320 | 2724300 | 0 | 203450 | 0 | 0 | 0 | 434450 |
| ILDVAQDLK | CNOT1 | A5YKK6 | CNOT1_HUMA | Unmodified | 0 | 0 | 0 | 0 | 0 | 0 | 0 | 0 | 0 | 0 | 0 | 1044500 | 0 | 0 | 0 | 0 | 0 | 0 |
| ILDVASLEV | GCN1 | Q92616 | GCN1_HUMAN | Unmodified | 0 | 0 | 303580 | 0 | 1396800 | 0 | 0 | 0 | 0 | 0 | 0 | 2395800 | 631610 | 0 | 0 | 377770 | 0 | 745660 |
| ILDVNDNIF | DSG2 | Q14126 | DSG2_HUMAN | Unmodified | 0 | 0 | 0 | 0 | 1018900 | 457520 | 0 | 0 | 2575500 | 585100 | 374710 | 655500 | 354130 | 0 | 0 | 0 | 0 | 0 |
| ILDWHVAN | TALDO1; TALDO | P37837; F2Z | TALDO_HUMAN | Unmodified | 0 | 0 | 0 | 0 | 0 | 0 | 0 | 0 | 0 | 0 | 0 | 2538200 | 0 | 0 | 0 | 44087 | 0 | 0 |
| ILDWHVAN | TALDO1; TALDO | P37837; F2Z | TALDO_HUMAN | Unmodified | 0 | 0 | 0 | 0 | 0 | 0 | 0 | 0 | 1231900 | 0 | 0 | 0 | 560410 | 0 | 0 | 255340 | 0 | 0 |
| ILDYSCSQI | USP9X | Q93008 | USP9X_HUMAN | Unmodified | 0 | 0 | 0 | 0 | 0 | 0 | 0 | 0 | 0 | 0 | 0 | 403830 | 0 | 0 | 0 | 0 | 0 | 0 |
| ILDYSVHAE | CNPY4; CNPY4 | Q8N129; A0A | CNPY4_HUMAI | Unmodified | 0 | 0 | 0 | 0 | 0 | 0 | 0 | 0 | 0 | 0 | 0 | 0 | 0 | 0 | 0 | 0 | 0 | 0 |
| ILDYSVHAE | CNPY4; CNPY4 | Q8N129; A0A | CNPY4_HUMAI | Unmodified | 0 | 0 | 0 | 0 | 0 | 0 | 153190 | 87192 | 4610800 | 0 | 0 | 0 | 0 | 0 | 0 | 0 | 0 | 0 |
| ILEAHQNVA | FERMT1; FERMT | Q9BQL6; G3 | FERM1_HUMAN | Unmodified | 0 | 0 | 0 | 0 | 0 | 0 | 0 | 0 | 0 | 0 | 0 | 0 | 0 | 0 | 0 | 0 | 350660 | 537340 |
| ILEAHQNVA | FERMT2; FERMT | Q96AC1; H0 | FERM2_HUMAN | Oxidation (M) | 0 | 303940 | 0 | 119990 | 818390 | 627270 | 76273 | 0 | 114520 | 0 | 0 | 0 | 0 | 0 | 175470 | 197110 | 623060 | 454470 |
| ILEAHQNVA | FERMT2; FERMT | Q96AC1; H0 | FERM2_HUMAN | Unmodified | 127670 | 0 | 0 | 98904 | 2646400 | 930430 | 0 | 0 | 290340 | 0 | 0 | 0 | 0 | 0 | 139000 | 151170 | 1202800 | 1008300 |
| ILEALETEK | KIF14 | Q15058 | KIF14_HUMAN | Unmodified | 212350 | 462080 | 0 | 285460 | 4898800 | 244720 | 0 | 0 | 278250 | 0 | 245970 | 664880 | 191920 | 95001 | 595570 | 329680 | 515620 | 0 |
| ILEATESIK | RIPOR2; RIPOR | Q9Y4F9; F5C | RIPR2_HUMAN | Unmodified | 0 | 0 | 0 | 0 | 0 | 0 | 82581 | 100190 | 0 | 105480 | 899320 | 789330 | 0 | 0 | 0 | 92529 | 0 | 0 |
| ILEAVGNIA | DOP1B | Q9Y3R5 | DOP2_HUMAN | Unmodified | 0 | 0 | 0 | 0 | 0 | 0 | 0 | 0 | 1372100 | 250240 | 0 | 172720 | 0 | 0 | 0 | 0 | 0 | 0 |
| ILEDDLK | ABCE1; ABCE1; | P61221; H0Y | ABCE1_HUMAN | Unmodified | 210060 | 224080 | 398010 | 358880 | 2317500 | 698820 | 472010 | 208300 | 368090 | 244110 | 1017900 | 6415000 | 649150 | 522770 | 446280 | 406600 | 810030 | 918250 |
| ILEDNSIPQ | PDHB | P11177 | ODPB_HUMAN | Unmodified | 0 | 0 | 0 | 0 | 0 | 0 | 600270 | 0 | 2622500 | 544920 | 188740 | 0 | 0 | 0 | 0 | 0 | 0 | 0 |
| ILEDNSIPQ | PDHB | P11177 | ODPB_HUMAN | Unmodified | 0 | 0 | 0 | 0 | 0 | 0 | 1433200 | 0 | 0 | 1314800 | 0 | 0 | 561350 | 0 | 0 | 0 | 469650 | 179100 |
| ILEDQEENI | PGLS; PGLS | O95336; M0 | 6PGL_HUMAN | Unmodified | 421520 | 0 | 0 | 0 | 0 | 0 | 0 | 0 | 0 | 281910 | 386990 | 0 | 0 | 0 | 0 | 1205600 | 0 | 0 |
| ILEDTQFVD | GAPVD1; GAPV | Q14C86; C9 | GAPD1_HUMAI | Unmodified | 0 | 0 | 0 | 0 | 0 | 0 | 0 | 0 | 0 | 0 | 0 | 1216500 | 0 | 0 | 0 | 85251 | 259650 | 0 |
| ILEFFGLK | P4HB; P4HB; P4 | P07237; A0A | PDIA1_HUMAN | Unmodified | 228910 | 186130 | 45592 | 0 | 58578 | 30166 | 0 | 103920 | 48691000 | 1855800 | 4187500 | 3794300 | 0 | 0 | 1220600 | 696200 | 14917 | 0 |
| ILEFFGLKK | P4HB; P4HB; P4 | P07237; A0A | PDIA1_HUMAN | Unmodified | 0 | 0 | 0 | 0 | 0 | 0 | 0 | 0 | 2606300 | 0 | 0 | 206310 | 0 | 0 | 0 | 0 | 0 | 0 |
| ILEFFGLKK | P4HB; P4HB; P4 | P07237; A0A | PDIA1_HUMAN | Unmodified | 0 | 0 | 0 | 0 | 166280 | 0 | 0 | 0 | 42045000 | 3414600 | 4171800 | 1586600 | 0 | 0 | 0 | 0 | 887230 | 0 |
| ILEFIAVSQI | LONP1; LONP1 | P36776; K7E | LONM_HUMAN | Unmodified | 0 | 0 | 0 | 0 | 0 | 0 | 0 | 0 | 2729500 | 0 | 0 | 0 | 0 | 0 | 0 | 0 | 0 | 0 |
| ILEFIAVSQI | LONP1; LONP1 | P36776; K7E | LONM_HUMAN | Unmodified | 0 | 0 | 0 | 0 | 0 | 0 | 340200 | 0 | 263920 | 748940 | 0 | 0 | 0 | 0 | 0 | 0 | 0 | 0 |
| ILEGSGVDA | TCAF1; TCAF1; | Q9Y4C2; C9 | TCAF1_HUMAN | Unmodified | 501430 | 583270 | 0 | 0 | 0 | 0 | 0 | 0 | 0 | 0 | 0 | 0 | 0 | 0 | 0 | 353540 | 0 | 0 |
| ILEGTETTK | MRPS22; MRPS | P82650; A0A | RT22_HUMAN | Unmodified | 0 | 0 | 0 | 0 | 0 | 0 | 0 | 0 | 639170 | 0 | 0 | 0 | 0 | 0 | 0 | 0 | 0 | 0 |
| ILEIEDLFSS | PALS1; PALS1 | Q8N3R9; G3 | PALS1_HUMAN | Unmodified | 0 | 0 | 179010 | 625730 | 8920100 | 829360 | 555910 | 0 | 906920 | 1106500 | 234540 | 182690 | 3109700 | 2180200 | 588700 | 0 | 5289200 | 4252600 |
| ILELDQFK | XRCC6; XRCC6 | P12956; B1A | XRCC6_HUMAI | Unmodified | 0 | 0 | 0 | 0 | 0 | 0 | 0 | 0 | 521140 | 0 | 1178600 | 3536100 | 0 | 0 | 0 | 0 | 0 | 0 |
| ILELDQFKG | XRCC6; XRCC6 | P12956; B1A | XRCC6_HUMAI | Unmodified | 0 | 0 | 0 | 0 | 452710 | 174150 | 174650 | 253680 | 0 | 483440 | 277690 | 423220 | 330150 | 126340 | 0 | 0 | 114550 | 107740 |
| ILELEEK | TRIM38; ITGA4; | O00635; P13 | TRI38_HUMAN | Unmodified | 0 | 0 | 0 | 0 | 2025500 | 395670 | 62192 | 0 | 154180 | 0 | 359160 | 461520 | 133020 | 0 | 0 | 0 | 229630 | 0 |
| ILELEEKHR | CIT; CIT; CIT; CI | A0A7I2V4H7 | A0A7I2V4H7_H | Unmodified | 0 | 0 | 0 | 0 | 3896900 | 0 | 0 | 0 | 0 | 427190 | 0 | 0 | 127420 | 0 | 0 | 0 | 1514100 | 0 |
| ILELPYSIHA | TTC17 | Q96AE7 | TTC17_HUMAN | Unmodified | 2321600 | 0 | 0 | 0 | 0 | 30318 | 0 | 0 | 4122800 | 1116100 | 0 | 150630 | 0 | 0 | 0 | 0 | 573680 | 0 |
| ILELSGSSS | PRKDC | P78527 | PRKDC_HUMAI | Unmodified | 0 | 0 | 0 | 0 | 0 | 0 | 0 | 0 | 264630 | 0 | 0 | 1355400 | 0 | 0 | 0 | 0 | 0 | 0 |
| ILEMDPSN | TMTC3 | Q6ZXV5 | TMTC3_HUMAI | Unmodified | 0 | 0 | 0 | 0 | 0 | 0 | 0 | 0 | 4885600 | 0 | 0 | 0 | 0 | 0 | 0 | 0 | 0 | 0 |
| ILENAQRFN | STAT1; STAT1; S | P42224; A0A | STAT1_HUMAN | Oxidation (M) | 0 | 0 | 0 | 0 | 0 | 301930 | 0 | 0 | 0 | 0 | 0 | 0 | 0 | 0 | 0 | 0 | 241770 | 712420 |
| ILENEKDLE | TBCA; TBCA; TB | O75347; E5F | TBCA_HUMAN | Unmodified | 1765500 | 1726600 | 0 | 720400 | 0 | 0 | 0 | 0 | 5308100 | 371180 | 3672800 | 9470100 | 739100 | 0 | 0 | 1026500 | 0 | 0 |
| ILENEKDLE | TBCA; TBCA; TB | O75347; E5F | TBCA_HUMAN | Unmodified | 1084600 | 899750 | 0 | 1492800 | 0 | 0 | 0 | 0 | 1976900 | 241060 | 0 | 1849000 | 1293500 | 0 | 0 | 0 | 0 | 0 |
| ILENPQSLS | XPO7; XPO7 | Q9UIA9; E7E | XPO7_HUMAN | Unmodified | 0 | 0 | 0 | 0 | 0 | 0 | 0 | 0 | 279350 | 0 | 0 | 1131800 | 0 | 0 | 0 | 0 | 0 | 0 |
| ILENVEVKG | PPID; PPID | Q08752; H0Y | PPID_HUMAN I | Unmodified | 0 | 0 | 0 | 0 | 0 | 0 | 0 | 0 | 165060 | 0 | 0 | 577470 | 307300 | 0 | 0 | 0 | 0 | 0 |
| ILEPGLNILI | STOML2 | Q9UJZ1 | STML2_HUMAN | Unmodified | 0 | 0 | 0 | 0 | 0 | 0 | 0 | 0 | 306920 | 0 | 0 | 0 | 0 | 0 | 0 | 0 | 0 | 0 |
| ILEPPEGQI | MOB4; HSPE1-I | Q9Y3A3; S4F | PHOCN_HUMA | Unmodified | 0 | 0 | 219510 | 0 | 671890 | 0 | 0 | 0 | 439530 | 0 | 94954 | 1266800 | 0 | 0 | 150760 | 0 | 250970 | 195050 |
| ILEQEEEEE | SRRT | Q9BXP5 | SRRT_HUMAN | Unmodified | 0 | 0 | 0 | 0 | 0 | 0 | 0 | 0 | 0 | 0 | 341680 | 0 | 0 | 0 | 0 | 0 | 0 | 0 |
| ILEQQNSSI | SEPTIN7; SEPTI | Q16181; E7E | SEPT7_HUMAN | Unmodified | 293140 | 0 | 764370 | 1203000 | 1234300 | 320430 | 0 | 0 | 604390 | 232030 | 126880 | 3351300 | 0 | 0 | 999500 | 597790 | 12033 | 535270 |
| ILEQQNSSI | SEPTIN7; SEPTI | Q16181; E7E | SEPT7_HUMAN | Unmodified | 0 | 0 | 0 | 0 | 10926000 | 2216600 | 414830 | 376440 | 0 | 655510 | 0 | 0 | 3217300 | 1901500 | 0 | 0 | 2119800 | 3254500 |
| ILESASSNS | MAN2A1 | Q16706 | MA2A1_HUMAI | Unmodified | 0 | 0 | 0 | 0 | 126820 | 0 | 45085 | 0 | 7408100 | 4605700 | 1134400 | 4209800 | 182680 | 0 | 0 | 0 | 0 | 0 |
| ILESELLR | PSD2 | Q9BQI7 | PSD2_HUMAN | Unmodified | 0 | 0 | 0 | 0 | 0 | 0 | 0 | 0 | 0 | 0 | 2050100 | 0 | 0 | 0 | 0 | 0 | 0 | 0 |
| ILESSQTLLS | SSNA1; SSNA1 | O43805; A0A | SSNA1_HUMAI | Unmodified | 0 | 0 | 0 | 0 | 0 | 0 | 294410 | 36298 | 0 | 0 | 0 | 519590 | 0 | 0 | 0 | 0 | 0 | 0 |
| ILESSQTLLS | SSNA1; SSNA1 | O43805; A0A | SSNA1_HUMAI | Unmodified | 0 | 0 | 0 | 0 | 394420 | 74304 | 170970 | 0 | 0 | 106760 | 0 | 0 | 379930 | 0 | 0 | 0 | 518630 | 413990 |
| ILETTTFFQI | UGGT1 | Q9NYU2 | UGGG1_HUMA | Unmodified | 0 | 0 | 0 | 0 | 0 | 0 | 0 | 0 | 22445000 | 8116300 | 2593400 | 536170 | 386630 | 0 | 0 | 0 | 0 | 0 |
| ILETWGELL | RABGAP1 | Q9Y3P9 | RBGP1_HUMAI | Unmodified | 0 | 0 | 0 | 0 | 0 | 0 | 0 | 0 | 0 | 0 | 328400 | 994350 | 0 | 0 | 0 | 0 | 0 | 0 |
| ILEVCVWD | DNAJC3 | Q13217 | DNJC3_HUMAI | Unmodified | 0 | 0 | 0 | 0 | 0 | 0 | 0 | 0 | 0 | 98863 | 0 | 0 | 0 | 0 | 0 | 0 | 0 | 0 |
| ILEVHIDK | DNAJA1 | P31689 | DNJA1_HUMAN | Unmodified | 0 | 0 | 0 | 0 | 0 | 0 | 0 | 0 | 1629000 | 725330 | 0 | 480860 | 955850 | 0 | 0 | 0 | 0 | 0 |
| ILEVHVDK | DNAJA2 | O60884 | DNJA2_HUMAN | Unmodified | 267790 | 0 | 0 | 0 | 4247300 | 634050 | 264060 | 0 | 890840 | 730200 | 0 | 1544000 | 2067400 | 1272400 | 0 | 0 | 219190 | 399370 |
| ILEVNHVN | SNX27 | Q96L92 | SNX27_HUMAN | Unmodified | 0 | 317350 | 0 | 0 | 166390 | 0 | 0 | 0 | 0 | 0 | 0 | 1047600 | 0 | 0 | 0 | 324630 | 0 | 0 |
| ILEVSGVDL | PATJ; PATJ; PAT | Q8NI35; A0A | INADL_HUMAN | Unmodified | 0 | 0 | 0 | 0 | 1752800 | 507580 | 0 | 0 | 717200 | 0 | 0 | 0 | 0 | 0 | 0 | 0 | 0 | 841890 |
| ILEVVKDK | COX5A; COX5A | P20674; H3E | COX5A_HUMAI | Unmodified | 0 | 0 | 0 | 0 | 0 | 0 | 0 | 0 | 637850 | 187690 | 0 | 0 | 0 | 0 | 0 | 0 | 0 | 0 |
| ILEVVNQIQ | ENDOD1 | O94919 | ENDD1_HUMA | Unmodified | 0 | 0 | 276390 | 126900 | 646860 | 0 | 0 | 0 | 5122300 | 690440 | 0 | 0 | 0 | 0 | 674190 | 846240 | 135270 | 0 |
| ILEVVNQIQ | ENDOD1 | O94919 | ENDD1_HUMA | Oxidation (M) | 0 | 0 | 0 | 0 | 0 | 0 | 0 | 0 | 5608100 | 2355300 | 0 | 0 | 1783100 | 0 | 0 | 0 | 0 | 0 |
| ILEVVNQIQ | ENDOD1 | O94919 | ENDD1_HUMA | Unmodified | 0 | 0 | 0 | 1712600 | 0 | 0 | 0 | 23276 | 4738700 | 491290 | 0 | 0 | 0 | 0 | 0 | 0 | 0 | 0 |
| ILEYKRQVC | DSP | P15924 | DESP_HUMAN | Unmodified | 0 | 0 | 0 | 0 | 118210 | 0 | 0 | 0 | 0 | 273080 | 0 | 0 | 90494 | 88077 | 0 | 0 | 209010 | 103610 |
| ILFDGIPLEI | MMUT | P22033 | MUTA_HUMAN | Unmodified | 0 | 0 | 0 | 0 | 0 | 0 | 0 | 0 | 1212300 | 940740 | 0 | 0 | 0 | 0 | 0 | 0 | 0 | 0 |
| ILFFNTPK | PSMD8; PSMD8 | P48556; K7E | PSMD8_HUMA | Unmodified | 0 | 0 | 0 | 0 | 0 | 0 | 0 | 0 | 156400 | 0 | 0 | 6056600 | 0 | 0 | 0 | 0 | 0 | 0 |
| ILFIFIDSDI | P4HB; P4HB; P4 | P07237; A0A | PDIA1_HUMAN | Unmodified | 0 | 290710 | 545310 | 1428800 | 0 | 0 | 516500 | 182410 | 74430000 | 18016000 | 6789600 | 3618900 | 462460 | 0 | 1815600 | 903420 | 0 | 0 |
| ILFLDPSGK | TXNDC12 | O95881 | TXD12_HUMAN | Unmodified | 0 | 0 | 0 | 0 | 0 | 0 | 0 | 12227 | 16494000 | 1271100 | 2024900 | 0 | 0 | 0 | 0 | 0 | 0 | 0 |
| ILFLDPSGK | TXNDC12 | O95881 | TXD12_HUMAN | Unmodified | 0 | 0 | 0 | 0 | 0 | 0 | 476970 | 0 | 26004000 | 10268000 | 0 | 0 | 2407700 | 1879900 | 0 | 0 | 0 | 0 |
| ILFNNAVK | ASNS; ASNS | P08243; F8W | ASNS_HUMAN | Unmodified | 193960 | 265990 | 307510 | 159250 | 174460 | 72533 | 902600 | 567990 | 877740 | 0 | 1198700 | 13222000 | 224810 | 302670 | 1254600 | 868930 | 27820 | 430530 |
| ILFQPQTN | SEC24D; SEC24 | O94855; E9F | SC24D_HUMAI | Unmodified | 0 | 0 | 0 | 0 | 0 | 0 | 0 | 0 | 0 | 0 | 0 | 1098700 | 0 | 0 | 0 | 0 | 0 | 0 |
| ILFRDVVET | PLCH1; PLCH1 | Q4KWH8; A0 | PLCH1_HUMAI | Unmodified | 0 | 0 | 0 | 275300 | 89293 | 0 | 0 | 0 | 216530 | 202310 | 0 | 165680 | 0 | 0 | 0 | 0 | 697230 | 0 |
| ILFSSREEQ | GAK | O14976 | GAK_HUMAN | Unmodified | 70964 | 44034 | 68691 | 56238 | 0 | 0 | 123200 | 329240 | 79752 | 0 | 0 | 298800 | 735540 | 231280 | 87935 | 79398 | 0 | 424850 |
| ILFYHPNEV | CCZ1; CCZ1B; | P86791; P86 | CCZ1_HUMAN | Unmodified | 40570 | 0 | 0 | 183480 | 960490 | 148120 | 94753 | 0 | 0 | 109870 | 0 | 0 | 0 | 613710 | 0 | 0 | 0 | 455530 |
| ILGADTSVD | ATP5F1A; ATP5I | P25705; K7E | ATPA_HUMAN | Unmodified | 0 | 0 | 0 | 94178 | 0 | 0 | 91154 | 230760 | 3119900 | 0 | 1923800 | 0 | 0 | 0 | 0 | 0 | 0 | 0 |
| ILGEGIAAS | EHHADH | Q08426 | ECHP_HUMAN | Unmodified | 0 | 0 | 0 | 0 | 0 | 808820 | 454030 | 0 | 8185600 | 0 | 0 | 0 | 0 | 0 | 0 | 0 | 55846 | 0 |
| ILGEGSFST | PDPK2P; PDPK | Q6A1A2; O1 | PDPK2_HUMAI | Unmodified | 0 | 204380 | 567610 | 709130 | 0 | 0 | 0 | 0 | 0 | 0 | 0 | 0 | 563490 | 0 | 1019700 | 1099000 | 0 | 0 |
| ILGEGSFST | PDPK2P; PDPK | Q6A1A2; O1 | PDPK2_HUMAI | Unmodified | 0 | 0 | 0 | 111470 | 1200600 | 0 | 0 | 0 | 0 | 218070 | 0 | 0 | 6894200 | 6080300 | 0 | 0 | 0 | 1112000 |
| ILGGLLSAY | MAN1B1; MAN1 | Q9UKM7; H0 | MA1B1_HUMAI | Unmodified | 0 | 0 | 0 | 0 | 0 | 0 | 0 | 0 | 848920 | 315730 | 0 | 0 | 0 | 0 | 40595 | 0 | 0 | 0 |
| ILGGVISAIS | CAPNS1; CAPN | P04632; A0A | CPNS1_HUMAI | Unmodified | 0 | 0 | 0 | 0 | 0 | 0 | 0 | 0 | 447730 | 0 | 66586 | 63710 | 0 | 0 | 0 | 0 | 0 | 0 |
| ILGLAIESQI | SNAP23; SNAP2 | O00161; H3I | SNP23_HUMAI | Unmodified | 838250 | 1730400 | 1388300 | 3196400 | 5149500 | 901450 | 0 | 16634 | 761770 | 452910 | 3983200 | 7715400 | 5394100 | 1480900 | 4290800 | 3139500 | 1216500 | 3588500 |
| ILGLAPSR | TMCO1; TMCO1 | Q9UM00; J3 | TMCO1_HUMA | Unmodified | 0 | 0 | 76707 | 0 | 0 | 0 | 0 | 0 | 1717100 | 135920 | 0 | 483970 | 0 | 0 | 0 | 0 | 0 | 0 |
| ILGLGDLG | ME2; ME2; ME2 | P23368; A0A | MAOM_HUMAN | Oxidation (M) | 532930 | 0 | 0 | 0 | 0 | 0 | 0 | 0 | 533910 | 144070 | 0 | 0 | 0 | 0 | 0 | 0 | 0 | 0 |
| ILGLGDLG | ME2; ME2; ME2 | P23368; A0A | MAOM_HUMAN | Unmodified | 0 | 0 | 0 | 0 | 0 | 0 | 0 | 0 | 90308 | 0 | 29451 | 0 | 0 | 0 | 0 | 0 | 0 | 0 |
| ILGLLDAYL | EEF1G | P26641 | EF1G_HUMAN | Unmodified | 463650 | 281720 | 287150 | 0 | 0 | 0 | 0 | 0 | 822280 | 0 | 2049500 | 20510000 | 0 | 0 | 1319000 | 872260 | 0 | 0 |
| ILGNKATFS | ANK3; ANK3; A | Q12955; A0A | ANK3_HUMAN | Unmodified | 0 | 0 | 0 | 0 | 0 | 0 | 0 | 0 | 0 | 0 | 0 | 0 | 2105000 | 1292700 | 0 | 0 | 0 | 0 |
| ILGPAESDE | UBR4; UBR4 | Q5T4S7; A0A | UBR4_HUMAN | Unmodified | 128390 | 0 | 0 | 0 | 0 | 0 | 0 | 0 | 0 | 0 | 126410 | 2716200 | 0 | 0 | 0 | 0 | 0 | 0 |
| ILGPGLNK | RPL10A | P62906 | RL10A_HUMAN | Unmodified | 3075400 | 4157800 | 0 | 0 | 2544700 | 265500 | 0 | 394910 | 1503300 | 893660 | 2493400 | 83049000 | 2287300 | 734490 | 2193700 | 0 | 260250 | 508750 |
| ILGPQGNTI | KHDRBS1 | Q07666 | KHDR1_HUMAI | Unmodified | 0 | 0 | 10487000 | 0 | 1330400 | 994570 | 0 | 73591 | 1107100 | 1519100 | 0 | 95628 | 0 | 0 | 0 | 0 | 220720 | 389370 |
| ILGSDGAFI | ITGA2; ITGA2; I | P17301; E7E | ITA2_HUMAN I | Unmodified | 26713 | 0 | 2067800 | 40942 | 0 | 2204200 | 0 | 0 | 0 | 516050 | 57154 | 90234 | 9642300 | 0 | 2881600 | 2902900 | 3773200 | 10846000 |
| ILGSDGAFI | ITGA2; ITGA2; I | P17301; E7E | ITA2_HUMAN I | Unmodified | 0 | 0 | 0 | 2141200 | 8128900 | 4179300 | 0 | 0 | 0 | 0 | 0 | 462430 | 932570 | 88022 | 0 | 0 | 6332100 | 3649300 |
| ILGSGISSS | CCT8 | P50990 | TCPQ_HUMAN | Oxidation (M) | 204550 | 0 | 166690 | 169410 | 98727 | 0 | 182630 | 99714 | 240330 | 108690 | 483360 | 4011100 | 166300 | 123750 | 320150 | 266070 | 99432 | 0 |
| ILGSGISSS | CCT8 | P50990 | TCPQ_HUMAN | Unmodified | 0 | 0 | 0 | 0 | 0 | 270880 | 147160 | 0 | 527170 | 155680 | 0 | 603180 | 0 | 0 | 0 | 0 | 0 | 0 |
| ILGSGISSS | CCT8 | P50990 | TCPQ_HUMAN | Unmodified | 0 | 0 | 81089 | 272980 | 1115600 | 0 | 0 | 0 | 1708700 | 52486 | 0 | 4761500 | 1348500 | 0 | 0 | 0 | 0 | 0 |
| ILGSGISSS | CCT8 | P50990 | TCPQ_HUMAN | Oxidation (M) | 244700 | 0 | 0 | 0 | 1100000 | 0 | 0 | 0 | 0 | 0 | 877170 | 6967200 | 633200 | 0 | 654450 | 529790 | 0 | 0 |
| ILGSGISSS | CCT8 | P50990 | TCPQ_HUMAN | Unmodified | 0 | 0 | 0 | 0 | 4275700 | 0 | 0 | 0 | 0 | 0 | 0 | 656270 | 3790500 | 1989500 | 0 | 0 | 1907500 | 3045000 |
| ILGSGISSS | CCT8 | P50990 | TCPQ_HUMAN | Oxidation (M) | 0 | 0 | 0 | 0 | 1861400 | 372640 | 336190 | 0 | 0 | 366530 | 0 | 2066300 | 3869200 | 0 | 0 | 0 | 0 | 2100400 |
| ILGTAGTEE | CRYZ; CRYZ; CI | Q08257; A6 | QOR_HUMAN | Unmodified | 0 | 0 | 0 | 0 | 176750 | 0 | 0 | 0 | 1133200 | 401160 | 0 | 0 | 0 | 0 | 0 | 0 | 220690 | 0 |
| ILGTQEITAI | EHBP1L1 | Q8N3D4 | EH1L1_HUMAN | Unmodified | 0 | 0 | 0 | 0 | 734460 | 0 | 0 | 0 | 848270 | 0 | 0 | 132630 | 0 | 0 | 0 | 0 | 0 | 0 |
| ILGVDKNA | DNAJC7; DNAJ | Q99615; A0A | DNJC7_HUMAI | Unmodified | 43922 | 50628 | 0 | 350050 | 289590 | 0 | 293460 | 92056 | 210270 | 495370 | 0 | 0 | 660910 | 171410 | 0 | 0 | 41797 | 0 |
| ILGVGPDD | LSS; LSS | P48449; C9J | LSS_HUMAN L | Unmodified | 0 | 0 | 0 | 0 | 0 | 0 | 0 | 0 | 0 | 0 | 0 | 1892100 | 0 | 0 | 0 | 0 | 0 | 0 |
| ILGYDRCD | HAPLN1 | P10915 | HPLN1_HUMAI | Unmodified | 0 | 0 | 0 | 0 | 0 | 0 | 0 | 54238 | 691960 | 0 | 0 | 0 | 110100 | 0 | 0 | 0 | 0 | 0 |

| | | | | | | | | | | | | | | | | | | | | | | |
|---|---|---|---|---|---|---|---|---|---|---|---|---|---|---|---|---|---|---|---|---|---|---|
| ILHDADRIH | PI4KA | P42356 | PI4KA_HUMAN | Unmodified | 41268 | 0 | 465390 | 0 | 488100 | 0 | 141920 | 0 | 654940 | 0 | 0 | 1223300 | 0 | 0 | 544230 | 788910 | 0 | 0 |
| ILHLLGQEC | COPG1 | Q9Y678 | COPG1_HUMA | Unmodified | 0 | 0 | 182140 | 0 | 0 | 0 | 0 | 0 | 0 | 0 | 0 | 1129300 | 0 | 0 | 0 | 0 | 0 | 0 |
| ILHLLTQEA | ALDH18A1 | P54886 | P5CS_HUMAN | Unmodified | 0 | 0 | 0 | 0 | 0 | 0 | 0 | 0 | 743150 | 0 | 0 | 0 | 0 | 0 | 0 | 0 | 0 | 0 |
| ILHLPTSWI | CTSC; CTSC; C1 | P53634; A0A | CATC_HUMAN | Unmodified | 0 | 0 | 1067900 | 0 | 529750 | 335170 | 0 | 0 | 3444700 | 0 | 6424900 | 8445300 | 340820 | 0 | 0 | 726550 | 0 | 0 |
| ILHLQGEGI | KIF20A | O95235 | KI20A_HUMAN | Unmodified | 40449 | 0 | 1270200 | 2132900 | 1924000 | 0 | 0 | 0 | 907380 | 334910 | 0 | 200040 | 795790 | 0 | 527530 | 0 | 227520 | 0 |
| ILHLQRVID | TRAF3; TRAF3; 1 | Q13114; A6N | TRAF3_HUMAN | Unmodified | 0 | 85721 | 0 | 0 | 3067100 | 0 | 0 | 0 | 926170 | 91384 | 0 | 566690 | 0 | 0 | 388290 | 0 | 0 | 0 |
| ILHQEAMN | PLCG2; PLCG2 | P16885; A0A | PLCG2_HUMAI | Oxidation (M) | 0 | 0 | 0 | 0 | 0 | 0 | 0 | 0 | 0 | 0 | 3335500 | 26565000 | 0 | 0 | 0 | 0 | 0 | 0 |
| ILHQEAMN | PLCG2; PLCG2 | P16885; A0A | PLCG2_HUMAI | Oxidation (M) | 0 | 0 | 0 | 0 | 0 | 0 | 0 | 0 | 0 | 171720 | 3460600 | 787150 | 0 | 0 | 0 | 0 | 41454 | 0 |
| ILHQHLGAI | MYOF; MYOF | Q9NZM1; H0 | MYOF_HUMAN | Unmodified | 0 | 0 | 0 | 0 | 1144300 | 0 | 0 | 0 | 195410 | 258180 | 0 | 0 | 0 | 156760 | 0 | 0 | 0 | 122410 |
| ILHQHLGAI | MYOF; MYOF | Q9NZM1; H0 | MYOF_HUMAN | Unmodified | 0 | 0 | 0 | 0 | 2934400 | 0 | 0 | 0 | 0 | 0 | 0 | 0 | 0 | 0 | 0 | 0 | 0 | 0 |
| ILHQRAMD | GLG1; GLG1 | Q92896; H3I | GSLG1_HUMAI | Unmodified | 0 | 0 | 0 | 0 | 0 | 0 | 0 | 177030 | 0 | 456310 | 0 | 0 | 0 | 0 | 0 | 0 | 0 | 0 |
| ILHQRAMD | GLG1; GLG1 | Q92896; H3I | GSLG1_HUMAI | Oxidation (M) | 0 | 0 | 0 | 0 | 165590 | 0 | 0 | 0 | 0 | 525460 | 0 | 0 | 38293 | 29125 | 0 | 0 | 0 | 0 |
| ILHVNGFN | GNAS; GNAS; G | A0A590UJG5 | A0A590UJG5_H | Unmodified | 127590 | 97001 | 266470 | 433410 | 1335600 | 309410 | 0 | 55527 | 0 | 0 | 515970 | 3828700 | 851980 | 645820 | 657790 | 660520 | 704890 | 1826900 |
| ILHVNGFN | GNAS; GNAS; G | A0A590UJJ0 | A0A590UJJ0_H | Unmodified | 252850 | 279650 | 0 | 680620 | 1640900 | 588350 | 96394 | 53429 | 465460 | 75352 | 572130 | 4655100 | 1514200 | 843030 | 1309200 | 1171800 | 979240 | 1956200 |
| ILHVNGFN | GNAS; GNAS; G | Q5JWF2; P63 | GNAS1_HUMAI | Unmodified | 0 | 0 | 737310 | 502030 | 532440 | 0 | 79229 | 43272 | 348620 | 0 | 2657000 | 11501000 | 271340 | 0 | 1608300 | 1131600 | 368460 | 964580 |
| ILHVNGFN | GNAS; GNAS; G | Q5JWF2; P63 | GNAS1_HUMAI | Unmodified | 0 | 0 | 0 | 0 | 9291700 | 0 | 0 | 0 | 0 | 0 | 431070 | 0 | 3696300 | 3152700 | 0 | 0 | 69595 | 11637000 |
| ILIANTGMD | CCT2; CCT2; C( | P78371; F5G | TCPB_HUMAN | Unmodified | 0 | 0 | 603280 | 302410 | 1836800 | 286570 | 463420 | 0 | 2867000 | 440610 | 2433500 | 13166000 | 1717400 | 1139700 | 778480 | 0 | 648880 | 1102300 |
| ILIANTGMD | CCT2; CCT2; C( | P78371; F5G | TCPB_HUMAN | Oxidation (M) | 207290 | 0 | 0 | 270830 | 1117400 | 0 | 209220 | 0 | 960620 | 0 | 1554300 | 10132000 | 1374300 | 517010 | 578250 | 539760 | 551910 | 1063900 |
| ILIANTGMD | CCT2; CCT2; C( | P78371; F5G | TCPB_HUMAN | Oxidation (M) | 0 | 0 | 0 | 0 | 0 | 266420 | 1225700 | 1199900 | 0 | 697550 | 0 | 0 | 6853700 | 5980800 | 0 | 0 | 1629700 | 1057900 |
| ILIANTGMD | CCT2; CCT2; C( | P78371; F5G | TCPB_HUMAN | Unmodified | 22976 | 0 | 0 | 0 | 11940000 | 0 | 4348600 | 1590200 | 602640 | 2583200 | 405650 | 0 | 15034000 | 8593300 | 0 | 0 | 6260300 | 6928300 |
| ILIDSIYK | PLCH1; PLCH1 | Q4KWH8; A0 | PLCH1_HUMAI | Unmodified | 0 | 0 | 0 | 0 | 0 | 0 | 1292900 | 751320 | 11592000 | 6796000 | 102780 | 1894000 | 0 | 348660 | 0 | 0 | 0 | 0 |
| ILIDTGEPAI | LACTB2 | Q53H82 | LACB2_HUMAN | Unmodified | 0 | 0 | 0 | 0 | 484250 | 0 | 0 | 0 | 2556400 | 878910 | 0 | 0 | 0 | 0 | 0 | 0 | 462090 | 0 |
| ILIDVSNNN | STEAP2; STEAP | Q8NFT2; C9 | STEA2_HUMAN | Oxidation (M) | 198780 | 283880 | 6409200 | 958400 | 0 | 0 | 0 | 0 | 0 | 0 | 0 | 237230 | 0 | 0 | 4315700 | 2891600 | 0 | 0 |
| ILIEDSDQN | AP3D1; AP3D1; | O14617; A0A | AP3D1_HUMAN | Unmodified | 0 | 0 | 0 | 0 | 0 | 0 | 0 | 0 | 0 | 0 | 0 | 1109000 | 0 | 0 | 0 | 0 | 0 | 0 |
| ILIEDWK | PRKCSH; PRKC | P14314; K7E | GLU2B_HUMAI | Unmodified | 0 | 0 | 0 | 0 | 153850 | 0 | 306010 | 154730 | 34924000 | 10407000 | 2533500 | 805970 | 0 | 0 | 0 | 0 | 31282 | 69525 |
| ILIEDWKK | PRKCSH; PRKC | P14314; K7E | GLU2B_HUMAI | Unmodified | 0 | 0 | 0 | 0 | 0 | 0 | 337730 | 0 | 7601700 | 4366700 | 0 | 150130 | 0 | 0 | 0 | 0 | 0 | 0 |
| ILIIGESGV( | RAB18; RAB18; | Q9NP72; A0 | RAB18_HUMAN | Unmodified | 843550 | 1426400 | 0 | 2137700 | 3330300 | 0 | 2218000 | 922780 | 2271300 | 1243000 | 5470000 | 17335000 | 3555900 | 2765400 | 393020 | 0 | 275810 | 0 |
| ILILGLDGA | ARL1; ARL1; AR | P40616; F8V | ARL1_HUMAN | Unmodified | 0 | 0 | 0 | 0 | 0 | 0 | 78577 | 36220 | 1482700 | 147190 | 4302800 | 1397300 | 0 | 0 | 0 | 0 | 0 | 0 |
| ILIRFFPLES | DDT; DDT; DDT | P30046; H7( | DOPD_HUMAN | Unmodified | 0 | 0 | 0 | 0 | 343750 | 0 | 1131900 | 0 | 33700 | 396660 | 0 | 0 | 487610 | 400300 | 162150 | 89138 | 522570 | 296740 |
| ILISLATGHI | ANXA6 | E5RK69 | E5RK69_HUMA | Unmodified | 0 | 0 | 2245100 | 3252100 | 4085200 | 0 | 0 | 0 | 54366000 | 14725000 | 0 | 0 | 0 | 3811200 | 8916300 | 0 | 875500 | 4419400 |
| ILITGGAGF | UXS1; UXS1; U) | Q8NBZ7; C9 | UXS1_HUMAN | Unmodified | 0 | 0 | 0 | 145450 | 122990 | 0 | 0 | 0 | 3944500 | 499170 | 0 | 0 | 0 | 0 | 0 | 0 | 0 | 264290 |
| ILITTVPPNI | ILF2; ILF2 | Q12905; B4I | ILF2_HUMAN Ir | Unmodified | 1092700 | 1102700 | 0 | 0 | 0 | 0 | 0 | 0 | 0 | 0 | 0 | 2668500 | 0 | 0 | 0 | 554730 | 0 | 0 |
| ILIVGGGVA | NNT; NNT; NNT | Q13423; E9F | NNTM_HUMAN | Unmodified | 0 | 0 | 0 | 0 | 0 | 0 | 641390 | 0 | 4201500 | 1717800 | 0 | 0 | 0 | 0 | 0 | 0 | 0 | 0 |
| ILKEDILNYI | DBT; DBT; DBT; | P11182; A0A | ODB2_HUMAN | Unmodified | 0 | 0 | 0 | 55533 | 0 | 0 | 0 | 0 | 2938800 | 124270 | 0 | 0 | 0 | 0 | 0 | 0 | 0 | 0 |
| ILKEEQELY | MYO6; MYO6; M | Q9UM54; A0 | MYO6_HUMAN | Unmodified | 84552 | 0 | 0 | 3983600 | 85476 | 0 | 0 | 0 | 1652800 | 255530 | 367550 | 2887900 | 226450 | 0 | 4837200 | 3364600 | 0 | 0 |
| ILKEQERLA | STIP1 | P31948 | STIP1_HUMAN | Unmodified | 0 | 0 | 0 | 0 | 3327200 | 487590 | 0 | 186330 | 0 | 1233600 | 0 | 0 | 13090000 | 9525800 | 0 | 0 | 9455800 | 11101000 |
| ILKKDVVIQ | PRKCB; PRKCA | P05771; P17 | KPCB_HUMAN | Unmodified | 0 | 0 | 0 | 0 | 0 | 0 | 2795600 | 530680 | 0 | 0 | 0 | 0 | 205720 | 544000 | 0 | 0 | 0 | 0 |
| ILKLPSFELI | DDB1; DDB1; D | Q16531; A0A | DDB1_HUMAN | Unmodified | 0 | 0 | 0 | 0 | 0 | 0 | 0 | 0 | 0 | 0 | 0 | 732140 | 0 | 0 | 0 | 0 | 0 | 0 |
| ILKPTDENL | MRPL15; MRPL | Q9P015; E5F | RM15_HUMAN | Unmodified | 0 | 0 | 0 | 0 | 0 | 0 | 0 | 0 | 868070 | 182410 | 0 | 0 | 0 | 0 | 0 | 0 | 0 | 0 |
| ILKYNLLQN | N/A | N/A | N/A | Unmodified | 0 | 0 | 6936200 | 1546300 | 3942000 | 7020800 | 0 | 567170 | 1088600 | 1031800 | 10939000 | 672830 | 0 | 6498600 | 857810 | 0 | 455470 | 5473500 |
| ILLAELEQLI | VIM; VIM; VIM | P08670; B0Y | VIME_HUMAN | Unmodified | 0 | 0 | 0 | 0 | 32760000 | 7214600 | 0 | 0 | 632810 | 0 | 0 | 0 | 0 | 0 | 0 | 587540 | 5847900 | 26850000 |
| ILLAELEQLI | VIM; VIM; VIM | P08670; B0Y | VIME_HUMAN | Unmodified | 0 | 0 | 0 | 0 | 42517000 | 6051900 | 5594100 | 763380 | 0 | 67662 | 0 | 0 | 1460000 | 984130 | 0 | 0 | 22477000 | 37779000 |
| ILLAVNGK | PGRMC2 | O15173 | PGRC2_HUMAI | Unmodified | 65285 | 0 | 233210 | 54752 | 0 | 0 | 0 | 173440 | 201540 | 0 | 187800 | 2093000 | 0 | 0 | 0 | 0 | 0 | 0 |
| ILLDAQHES | BNIP3; BNIP3; E | Q12983; F6F | BNIP3_HUMAN | Unmodified | 0 | 0 | 0 | 0 | 2751900 | 30791 | 839340 | 40720 | 235910 | 87753 | 0 | 266750 | 0 | 324180 | 0 | 0 | 371610 | 2118400 |
| ILLDNDHY/ | LONP2; LONP2 | Q86WA8; H3 | LONP2_HUMAI | Oxidation (M) | 0 | 0 | 0 | 0 | 0 | 0 | 0 | 0 | 557440 | 173500 | 0 | 0 | 0 | 0 | 0 | 0 | 0 | 0 |
| ILLDNDHY/ | LONP2; LONP2 | Q86WA8; H3 | LONP2_HUMAI | Oxidation (M) | 0 | 34519 | 0 | 0 | 0 | 0 | 0 | 0 | 0 | 0 | 0 | 1238700 | 0 | 0 | 0 | 0 | 0 | 0 |
| ILLDSGLEK | KIF14 | Q15058 | KIF14_HUMAN | Unmodified | 0 | 0 | 0 | 0 | 2755600 | 739100 | 328610 | 96428 | 0 | 0 | 0 | 461450 | 293060 | 311170 | 0 | 0 | 702800 | 301420 |
| ILLENLGEA | VPS13C | Q709C8 | VP13C_HUMAN | Unmodified | 0 | 0 | 0 | 0 | 0 | 0 | 0 | 0 | 0 | 0 | 0 | 2989400 | 0 | 0 | 181110 | 0 | 0 | 0 |
| ILLFRHDPT | PRKD1; PRKD1; | Q15139; A0A | KPCD1_HUMAI | Unmodified | 0 | 0 | 112010 | 0 | 0 | 0 | 0 | 0 | 53030 | 0 | 0 | 0 | 0 | 230550 | 311170 | 158510 | 115460 | 0 |
| ILLGLNFYG | CHID1; CHID1; | Q9BWS9; H0 | CHID1_HUMAN | Oxidation (M) | 0 | 0 | 0 | 0 | 0 | 0 | 0 | 0 | 0 | 195690 | 0 | 0 | 17636 | 1192600 | 36863 | 0 | 0 | 0 |
| ILLITEDLK | PPID | Q08752 | PPID_HUMAN | Unmodified | 0 | 0 | 0 | 0 | 0 | 0 | 0 | 0 | 0 | 0 | 1273800 | 5594700 | 333960 | 0 | 0 | 0 | 0 | 0 |
| ILLLEQFFLI | PIPOX | Q9P0Z9 | SOX_HUMAN P | Unmodified | 0 | 0 | 0 | 0 | 0 | 0 | 0 | 0 | 8177200 | 2475300 | 0 | 0 | 33166 | 0 | 0 | 0 | 0 | 0 |
| ILLLGAGES | GNA13; GNA12 | Q14344; Q0 | GNA13_HUMAI | Unmodified | 0 | 0 | 0 | 0 | 12929000 | 2125000 | 10611000 | 4325400 | 0 | 2690900 | 0 | 0 | 809320 | 1987600 | 0 | 0 | 10422000 | 10470000 |
| ILLLGLDNA | ARL3 | P36405 | ARL3_HUMAN | Unmodified | 0 | 0 | 0 | 0 | 0 | 0 | 0 | 0 | 0 | 0 | 3219300 | 11628000 | 1450800 | 0 | 0 | 0 | 0 | 0 |
| ILLLGLDNA | ARL3 | P36405 | ARL3_HUMAN | Unmodified | 0 | 0 | 0 | 0 | 0 | 229090 | 767060 | 95759 | 0 | 534780 | 0 | 0 | 5297900 | 5442400 | 0 | 0 | 730350 | 893430 |
| ILLMDEDQI | SEMA3C | Q99985 | SEM3C_HUMAI | Oxidation (M) | 0 | 0 | 1040900 | 446220 | 0 | 0 | 0 | 0 | 0 | 0 | 0 | 0 | 0 | 0 | 540730 | 218750 | 0 | 0 |
| ILLNACCP( | CBR1 | P16152 | CBR1_HUMAN | Unmodified | 0 | 0 | 0 | 0 | 3514100 | 0 | 0 | 0 | 0 | 0 | 0 | 0 | 0 | 0 | 0 | 0 | 667310 | 1997400 |
| ILLRNPGN( | VCL | P18206 | VINC_HUMAN | Oxidation (M) | 169240 | 373860 | 0 | 0 | 1670700 | 843480 | 287470 | 0 | 0 | 476650 | 0 | 4390800 | 0 | 0 | 0 | 409270 | 3316800 | 0 |
| ILLRNPGN( | VCL | P18206 | VINC_HUMAN | Unmodified | 0 | 0 | 0 | 0 | 3953500 | 1173300 | 0 | 0 | 0 | 0 | 0 | 0 | 3984800 | 1476600 | 0 | 0 | 5557400 | 0 |
| ILLRNPGN( | VCL | P18206 | VINC_HUMAN | Oxidation (M) | 0 | 0 | 0 | 0 | 460910 | 474490 | 1040600 | 0 | 0 | 468500 | 0 | 0 | 232490 | 2381000 | 0 | 0 | 7833800 | 0 |
| ILLRNPGN( | VCL | P18206 | VINC_HUMAN | Unmodified | 0 | 0 | 0 | 0 | 0 | 860970 | 522750 | 0 | 0 | 0 | 0 | 0 | 0 | 294900 | 0 | 0 | 3013000 | 0 |
| ILLSSGRD/ | CCT2; CCT2; C( | P78371; F5G | TCPB_HUMAN | Oxidation (M) | 0 | 0 | 0 | 0 | 0 | 0 | 0 | 89939 | 0 | 706880 | 0 | 0 | 2592800 | 1367700 | 0 | 0 | 0 | 0 |
| ILLSWVR | UTRN; DMD; DI | P46939; P11 | UTRN_HUMAN | Unmodified | 0 | 0 | 535560 | 216360 | 0 | 0 | 0 | 0 | 304720 | 213790 | 0 | 0 | 0 | 0 | 1381900 | 1075600 | 82553 | 0 |
| ILLTCSLND | BSG; BSG; BSG | P35613; A0A | BASI_HUMAN E | Unmodified | 0 | 0 | 0 | 0 | 0 | 0 | 0 | 0 | 0 | 0 | 0 | 0 | 0 | 0 | 1086600 | 0 | 0 | 0 |
| ILLTCSLND | BSG; BSG; BSG | P35613; A0A | BASI_HUMAN E | Unmodified | 0 | 0 | 0 | 0 | 0 | 0 | 1619500 | 0 | 0 | 0 | 0 | 0 | 0 | 758610 | 0 | 0 | 1515600 | 5635700 |
| ILLTEPPMN | ACTR2; ACTR2 | P61160; F5H | ARP2_HUMAN | Unmodified | 0 | 477630 | 0 | 797260 | 7942600 | 781710 | 1166500 | 596510 | 1779600 | 384780 | 1621500 | 7323600 | 8796100 | 1376500 | 895890 | 753140 | 1294800 | 2863000 |
| ILLTEPPMN | ACTR2; ACTR2 | P61160; F5H | ARP2_HUMAN | Oxidation (M) | 0 | 0 | 0 | 0 | 0 | 0 | 0 | 246830 | 0 | 0 | 0 | 0 | 666720 | 668120 | 0 | 667720 | 0 | 0 |
| ILLTQENPF | NSUN2 | Q08J23 | NSUN2_HUMA | Unmodified | 0 | 0 | 0 | 0 | 0 | 0 | 0 | 0 | 0 | 0 | 160800 | 1406800 | 0 | 0 | 0 | 0 | 0 | 0 |
| ILLTQENPF | NSUN2 | Q08J23 | NSUN2_HUMA | Unmodified | 0 | 0 | 0 | 0 | 343090 | 61488 | 500970 | 142920 | 55566 | 0 | 222040 | 792330 | 0 | 218440 | 0 | 0 | 0 | 207580 |
| ILLWDTR | WDR77; DPH7; | Q9BQA1; Q9 | MEP50_HUMAN | Unmodified | 0 | 0 | 167650 | 0 | 0 | 0 | 0 | 0 | 257360 | 0 | 0 | 8296600 | 0 | 0 | 0 | 212010 | 0 | 0 |
| ILMAIDSEL | XPOT; XPOT | O43592; F8V | XPOT_HUMAN | Oxidation (M) | 0 | 0 | 0 | 0 | 0 | 0 | 0 | 0 | 0 | 0 | 0 | 999160 | 0 | 0 | 0 | 0 | 0 | 0 |
| ILMDKPEM | NPEPPS; NPEPI | P55786; A0A | PSA_HUMAN P | 2 Oxidation (M) | 29675 | 0 | 0 | 122440 | 697780 | 161270 | 0 | 0 | 231550 | 247380 | 170900 | 1199600 | 211520 | 0 | 108010 | 177960 | 555540 | 652540 |
| ILMDKPEM | NPEPPS; NPEPI | P55786; A0A | PSA_HUMAN P | Oxidation (M) | 0 | 0 | 0 | 52807 | 2062300 | 292650 | 260930 | 0 | 507750 | 319300 | 506790 | 2924700 | 445350 | 429380 | 342820 | 0 | 769210 | 1093600 |
| ILMEHIHK | RPL19; RPL19; | P84098; J3K | RL19_HUMAN | Unmodified | 152590 | 0 | 106340 | 0 | 0 | 0 | 0 | 0 | 57816 | 0 | 102300 | 905220 | 0 | 0 | 0 | 72757 | 0 | 0 |
| ILMELDGPI | SPAG1 | Q07617 | SPAG1_HUMAI | Unmodified | 0 | 0 | 0 | 0 | 1414600 | 205180 | 0 | 0 | 0 | 0 | 0 | 0 | 0 | 0 | 0 | 0 | 401240 | 612460 |
| ILMLGLDA/ | ARF6 | P62330 | ARF6_HUMAN | Oxidation (M) | 867150 | 798960 | 1397000 | 265500 | 0 | 0 | 0 | 0 | 0 | 0 | 0 | 3818200 | 0 | 0 | 3543400 | 3105600 | 0 | 0 |
| ILMPELASL | PLCB2 | Q00722 | PLCB2_HUMAN | Unmodified | 0 | 0 | 0 | 0 | 0 | 0 | 0 | 0 | 0 | 0 | 0 | 773920 | 0 | 0 | 0 | 0 | 0 | 0 |
| ILMPELASL | PLCB2 | Q00722 | PLCB2_HUMAN | Oxidation (M) | 0 | 0 | 0 | 0 | 0 | 0 | 0 | 0 | 0 | 0 | 67127 | 4414500 | 0 | 0 | 0 | 0 | 0 | 0 |
| ILMSPSAK | INPP5A; INPP5/ | Q14642; B1/ | I5P1_HUMAN I | Unmodified | 0 | 0 | 0 | 0 | 0 | 0 | 0 | 0 | 287120 | 16005 | 0 | 688190 | 22560 | 0 | 188740 | 0 | 24222 | 0 |
| ILMTLTVK | PLCG2; PLCG2 | P16885; A0A | PLCG2_HUMAI | Oxidation (M) | 0 | 0 | 0 | 0 | 0 | 0 | 68710 | 0 | 0 | 0 | 963880 | 3332000 | 0 | 0 | 0 | 0 | 0 | 0 |
| ILMTLTVK | PLCG2; PLCG2 | P16885; A0A | PLCG2_HUMAI | Unmodified | 0 | 0 | 0 | 0 | 0 | 0 | 0 | 0 | 0 | 0 | 563340 | 2106200 | 0 | 0 | 0 | 0 | 0 | 0 |
| ILMVGLDA/ | ARF4; ARF5; AR | P18085; P84 | ARF4_HUMAN | Oxidation (M) | 0 | 425520 | 0 | 443920 | 0 | 0 | 0 | 71162 | 930300 | 0 | 816530 | 7544900 | 0 | 0 | 0 | 0 | 0 | 0 |
| ILMVGLDA/ | ARF4; ARF5; AR | P18085; P84 | ARF4_HUMAN | Unmodified | 1797800 | 0 | 0 | 0 | 211890 | 0 | 0 | 0 | 1235600 | 193560 | 400560 | 7825300 | 0 | 0 | 0 | 0 | 0 | 0 |
| ILNAGAWS | CUL5 | Q93034 | CUL5_HUMAN | Unmodified | 317210 | 0 | 0 | 1303700 | 863460 | 264110 | 1073200 | 602090 | 0 | 250090 | 0 | 0 | 376770 | 0 | 210990 | 140800 | 391440 | 444280 |
| ILNDDTALK | RAD23B; RAD2 | P54727; Q5V | RD23B_HUMAN | Unmodified | 2731600 | 3457200 | 2167000 | 101090 | 0 | 0 | 166450 | 483180 | 1468600 | 513090 | 984790 | 1561800 | 0 | 137360 | 4023300 | 4850500 | 0 | 0 |
| ILNEDGSPI | NENF | Q9UMX5 | NENF_HUMAN | Unmodified | 0 | 0 | 0 | 0 | 0 | 0 | 0 | 0 | 74424000 | 60904000 | 0 | 0 | 0 | 0 | 0 | 0 | 0 | 0 |
| ILNEKPTTD | SLK | Q9H2G2 | SLK_HUMAN S | Unmodified | 0 | 32869 | 259700 | 229870 | 402920 | 0 | 0 | 0 | 786810 | 62247 | 1293600 | 2323300 | 0 | 0 | 588750 | 339170 | 0 | 90433 |
| ILNETTGPK | ERBB4; ERBB4 | Q15303; A0A | ERBB4_HUMAN | Unmodified | 0 | 0 | 0 | 0 | 0 | 0 | 0 | 0 | 0 | 85340 | 108540 | 733680 | 377380 | 403920 | 0 | 101510 | 0 | 0 |
| ILNHVLQH | COMT; COMT; ( | P21964; F8V | COMT_HUMAN | Unmodified | 0 | 0 | 0 | 0 | 0 | 0 | 0 | 0 | 0 | 0 | 0 | 0 | 4603200 | 0 | 0 | 0 | 0 | 0 |
| ILNIFGVIK | TFRC; TFRC | P02786; G3 | TFR1_HUMAN | Unmodified | 5435700 | 6028800 | 5925400 | 6152000 | 4871000 | 1920300 | 2513800 | 427260 | 33849000 | 9585300 | 2079800 | 5399000 | 1019800 | 692630 | 10913000 | 7974300 | 1127700 | 1554200 |
| ILNLSGNEI | NXF1; NXF1; N) | Q9UBU9; E9 | NXF1_HUMAN | Unmodified | 0 | 0 | 0 | 0 | 0 | 0 | 0 | 0 | 0 | 0 | 0 | 847550 | 0 | 0 | 0 | 0 | 0 | 0 |
| ILNMSSGR | HIBADH; HIBAD | P31937; H7E | 3HIDH_HUMAN | Unmodified | 0 | 0 | 0 | 0 | 0 | 0 | 144060 | 0 | 426530 | 203710 | 0 | 0 | 0 | 0 | 0 | 0 | 0 | 0 |
| ILNPEEIEK | PSMA7; PSMA7 | O14818; H0 | PSA7_HUMAN | Unmodified | 572100 | 0 | 441410 | 480300 | 3062000 | 854640 | 232770 | 0 | 8112300 | 2521700 | 5004800 | 23707000 | 2299600 | 1364400 | 390960 | 216300 | 176720 | 1192400 |
| ILNPEEIEK | PSMA7; PSMA7 | O14818; H0 | PSA7_HUMAN | Unmodified | 0 | 0 | 0 | 499850 | 5974900 | 635080 | 508360 | 0 | 337160 | 1990600 | 197180 | 0 | 3562900 | 1365100 | 0 | 0 | 893640 | 1686100 |
| ILNPFFMEI | ST3GAL4; ST3G | Q11206; E9F | SIA4C_HUMAN | Unmodified | 0 | 99542 | 0 | 0 | 0 | 0 | 0 | 0 | 703810 | 0 | 0 | 0 | 0 | 0 | 0 | 0 | 0 | 0 |
| ILNPFFMEI | ST3GAL4; ST3G | Q11206; E9F | SIA4C_HUMAN | Oxidation (M) | 0 | 0 | 0 | 0 | 0 | 0 | 0 | 0 | 157020 | 16396 | 0 | 268280 | 0 | 0 | 0 | 0 | 0 | 0 |
| ILNPLLDR | PKP3 | Q9Y446 | PKP3_HUMAN | Unmodified | 161970 | 147050 | 435860 | 315280 | 0 | 0 | 0 | 0 | 0 | 0 | 738150 | 3355700 | 0 | 0 | 1248300 | 1121100 | 0 | 0 |
| ILNPLLDRV | PKP3 | Q9Y446 | PKP3_HUMAN | Unmodified | 0 | 2236800 | 3391300 | 4519100 | 0 | 0 | 0 | 0 | 1034700 | 0 | 404190 | 6271800 | 227140 | 0 | 4974900 | 6720800 | 0 | 0 |
| ILNSDEHA | DGKH; DGKH | Q86XP1; A0/ | DGKH_HUMAN | Unmodified | 0 | 0 | 0 | 110310 | 0 | 0 | 55594 | 0 | 0 | 0 | 0 | 0 | 89085 | 0 | 0 | 0 | 473450 | 345710 |
| ILNVDNPH | GRIK3 | Q13003 | GRIK3_HUMAN | Unmodified | 0 | 0 | 0 | 702310 | 0 | 0 | 0 | 0 | 778330 | 0 | 0 | 845620 | 0 | 0 | 0 | 0 | 0 | 0 |
| ILPDGEDFI | INPPL1 | O15357 | SHIP2_HUMAN | Unmodified | 0 | 0 | 0 | 0 | 0 | 0 | 0 | 0 | 0 | 0 | 0 | 3163900 | 0 | 0 | 0 | 0 | 0 | 0 |
| ILPDILK | CBS; CBSL; CB! | P35520; P0D | CBS_HUMAN C | Unmodified | 0 | 0 | 0 | 0 | 505360 | 0 | 386460 | 226750 | 681930 | 544150 | 703460 | 6946100 | 0 | 0 | 0 | 0 | 0 | 0 |
| ILPEEITNLF | LRRC40 | Q9H9A6 | LRC40_HUMAN | Unmodified | 0 | 0 | 75778 | 0 | 0 | 0 | 145100 | 80456 | 288770 | 0 | 460020 | 2198000 | 0 | 0 | 86601 | 95338 | 0 | 0 |
| ILPEQGLMI | PLAA; PLAA; PL/ | Q9Y263; H0 | PLAP_HUMAN | Oxidation (M) | 0 | 0 | 0 | 0 | 0 | 0 | 100750 | 0 | 0 | 0 | 323240 | 819470 | 0 | 0 | 0 | 0 | 0 | 0 |
| ILPFQYVLC | TFCP2; TFCP2 | Q12800; F8V | TFCP2_HUMAN | Unmodified | 0 | 0 | 0 | 0 | 112540 | 30211 | 0 | 0 | 0 | 0 | 143810 | 229320 | 0 | 0 | 0 | 0 | 0 | 0 |
| ILPLTGIPVL | PGAP1; PGAP1 | Q75T13; B4D | PGAP1_HUMAI | Unmodified | 0 | 0 | 0 | 0 | 0 | 0 | 0 | 0 | 2049400 | 55979 | 0 | 0 | 0 | 0 | 0 | 0 | 0 | 0 |
| ILPSYQEPH | MPP2; MPP2; M | Q14168; A0/ | MPP2_HUMAN | Unmodified | 0 | 0 | 434770 | 0 | 0 | 0 | 0 | 0 | 0 | 0 | 0 | 0 | 0 | 0 | 611010 | 308970 | 0 | 0 |
| ILPSYRDTII | PALS2; PALS2; | Q9NZW5; B8 | PALS2_HUMAN | Unmodified | 0 | 0 | 1046100 | 3776600 | 9338500 | 2909000 | 0 | 1865400 | 2609500 | 2222400 | 8761700 | 452960 | 0 | 875530 | 2521200 | 1536400 | 6329800 | 5907000 |
| ILPTLEAVA/ | ILF2; ILF2; ILF2 | Q12905; B4I | ILF2_HUMAN Ir | Unmodified | 830660 | 801980 | 320890 | 0 | 0 | 0 | 4492.8 | 0 | 0 | 0 | 573590 | 3212500 | 0 | 0 | 394060 | 379100 | 0 | 0 |

| | | | | | | | | | | | | | | | | | | | | | | |
|---|---|---|---|---|---|---|---|---|---|---|---|---|---|---|---|---|---|---|---|---|---|---|
| ILPVFDEPP | TMOD3; TMOD3 | Q9NYL9; H0 | TMOD3_HUMA | Unmodified | 0 | 0 | 342880 | 279560 | 10366000 | 2804000 | 746550 | 73968 | 566340 | 852080 | 463890 | 2335500 | 1319300 | 680830 | 362990 | 316250 | 727910 | 465530 |
| ILPVFDEPP | TMOD3; TMOD3 | Q9NYL9; H0 | TMOD3_HUMA | Unmodified | 0 | 0 | 0 | 0 | 0 | 0 | 0 | 0 | 0 | 0 | 0 | 2931600 | 1372800 | 0 | 0 | 1578600 | 0 | 0 |
| ILPVFDEPP | TMOD3; TMOD3 | Q9NYL9; H0 | TMOD3_HUMA | Unmodified | 0 | 0 | 0 | 434090 | 3130300 | 0 | 0 | 58059 | 0 | 777410 | 0 | 0 | 1187400 | 0 | 0 | 0 | 545030 | 0 |
| ILPWSWYG | EDIL3 | O43854 | EDIL3_HUMAN | Unmodified | 0 | 0 | 0 | 0 | 64353000 | 29350000 | 1025400 | 0 | 23181000 | 5497300 | 0 | 0 | 0 | 0 | 0 | 0 | 2200600 | 31053000 |
| ILQCYREN | CHCHD3; CHC | Q9NX63; C9 | MIC19_HUMAN | Unmodified | 0 | 0 | 0 | 0 | 0 | 0 | 0 | 0 | 620530 | 726500 | 0 | 0 | 0 | 0 | 0 | 0 | 0 | 0 |
| ILQDGGLQ | PHGDH; PHGD | O43175; A0A | SERA_HUMAN | Unmodified | 739040 | 435030 | 7035900 | 8199700 | 7953100 | 1336700 | 14616000 | 8860800 | 6607900 | 4663700 | 3560200 | 20117000 | 50230000 | 52991000 | 7267800 | 729290 | 3330400 | 9473400 |
| ILQDIASGS | NDUFS1; NDUF | P28331; B4D | NDUS1_HUMA | Unmodified | 0 | 0 | 0 | 0 | 0 | 0 | 0 | 0 | 3168400 | 819160 | 275780 | 0 | 0 | 0 | 0 | 0 | 35934 | 0 |
| ILQDSLGG | KIF5B; KIF5C; K | P33176; O60 | KINH_HUMAN | Unmodified | 0 | 0 | 0 | 0 | 197870 | 0 | 0 | 48498 | 496360 | 433800 | 296290 | 895950 | 279380 | 0 | 126060 | 121090 | 98453 | 0 |
| ILQDVPFN | ZYG11B; ZYG11 | Q9C0D3; A8 | ZY11B_HUMAN | Unmodified | 0 | 0 | 0 | 0 | 1479400 | 0 | 0 | 0 | 0 | 961630 | 0 | 0 | 0 | 0 | 0 | 0 | 0 | 621190 |
| ILQERAVLG | NSDHL; NSDHL | Q15738; C9J | NSDHL_HUMA | Unmodified | 0 | 0 | 0 | 0 | 0 | 0 | 630380 | 162150 | 0 | 237310 | 0 | 0 | 0 | 179960 | 0 | 0 | 0 | 0 |
| ILQEYITQE | AP1M1; AP1M1 | Q9BXS5; K7I | AP1M1_HUMAI | Unmodified | 0 | 0 | 0 | 585340 | 0 | 0 | 0 | 0 | 1148500 | 69828 | 0 | 880160 | 995790 | 0 | 0 | 237840 | 0 | 0 |
| ILQEYITQQ | AP1M2; AP1M2 | Q9Y6Q5; K7I | AP1M2_HUMAI | Unmodified | 0 | 0 | 518110 | 0 | 0 | 0 | 0 | 0 | 0 | 0 | 0 | 3959000 | 0 | 0 | 731940 | 1723700 | 0 | 0 |
| ILQEYVEH | ASNS | P08243 | ASNS_HUMAN | Oxidation (M) | 0 | 0 | 0 | 0 | 0 | 0 | 0 | 0 | 0 | 0 | 68222 | 423950 | 0 | 0 | 93077 | 56879 | 0 | 0 |
| ILQGGVGI | CSNK1A1; CSN | P48729; A0A | KC1A_HUMAN | Unmodified | 1045800 | 568900 | 1646700 | 979190 | 0 | 0 | 0 | 0 | 0 | 0 | 85789 | 4270500 | 0 | 0 | 2444200 | 1635000 | 0 | 0 |
| ILQGGVGI | CSNK1A1; CSN | P48729; A0A | KC1A_HUMAN | Unmodified | 0 | 0 | 0 | 121120 | 0 | 0 | 113890 | 0 | 0 | 475030 | 0 | 0 | 41548 | 0 | 0 | 0 | 189240 | 108220 |
| ILQHKDEL | SERPINB5 | P36952 | SPB5_HUMAN | Unmodified | 0 | 0 | 0 | 0 | 1124000 | 0 | 0 | 0 | 0 | 0 | 0 | 0 | 2752500 | 1141400 | 0 | 0 | 4650000 | 9037700 |
| ILQIITELIK | DHX29; DHX29 | Q7Z478; A0A | DHX29_HUMAI | Unmodified | 0 | 0 | 0 | 0 | 0 | 0 | 0 | 0 | 0 | 0 | 0 | 270610 | 0 | 0 | 0 | 0 | 0 | 0 |
| ILQIITELIKT | DHX29; DHX29 | Q7Z478; A0A | DHX29_HUMAI | Unmodified | 0 | 0 | 10799 | 32628 | 0 | 16482 | 19383 | 0 | 46977 | 23375 | 0 | 171330 | 65758 | 43677 | 52277 | 24345 | 0 | 21774 |
| ILQINKDD | SRP72; SRP72; | O76094; D6I | SRP72_HUMAN | Unmodified | 216780 | 0 | 0 | 0 | 700140 | 0 | 916210 | 712020 | 0 | 134190 | 487760 | 2399600 | 706910 | 0 | 0 | 0 | 1451100 | 0 |
| ILQLCMGN | EZR; EZR | P15311; E7E | EZRI_HUMAN E | Oxidation (M) | 632230 | 685510 | 0 | 79304 | 6758600 | 446960 | 319660 | 0 | 5564400 | 1130200 | 0 | 5650200 | 1380100 | 2045500 | 355940 | 281300 | 347640 | 0 |
| ILQLCMGN | EZR; EZR | P15311; E7E | EZRI_HUMAN E | Unmodified | 0 | 0 | 0 | 0 | 1904100 | 0 | 0 | 10258 | 2078700 | 594820 | 0 | 0 | 0 | 246930 | 0 | 0 | 178790 | 0 |
| ILQLCMGN | EZR; EZR | P15311; E7E | EZRI_HUMAN E | 2 Oxidation (M) | 550850 | 0 | 0 | 802020 | 1735500 | 0 | 0 | 0 | 1095600 | 623680 | 0 | 3981500 | 926350 | 950090 | 708590 | 529910 | 0 | 0 |
| ILQLCMGN | EZR; EZR | P15311; E7E | EZRI_HUMAN E | Unmodified | 0 | 0 | 0 | 0 | 3081000 | 434620 | 0 | 0 | 1438400 | 1074900 | 0 | 0 | 585180 | 1077300 | 0 | 0 | 300290 | 329120 |
| ILQNYDNK | RGPD3; RGPD2 | A6NKT7; P0D | RGPD3_HUMAI | Unmodified | 0 | 0 | 0 | 0 | 0 | 0 | 0 | 0 | 53983 | 343480 | 43726 | 202430 | 0 | 0 | 0 | 0 | 0 | 0 |
| ILQRSQAQ | LSM12 | Q3MHD2 | LSM12_HUMAN | Unmodified | 0 | 0 | 0 | 0 | 81786 | 0 | 37267 | 316260 | 0 | 0 | 0 | 0 | 0 | 0 | 0 | 0 | 762830 | 0 |
| ILQVSFK | ELAVL1 | Q15717 | ELAV1_HUMAN | Unmodified | 0 | 158420 | 669990 | 0 | 0 | 0 | 149440 | 100530 | 0 | 171400 | 0 | 8396800 | 94182 | 61442 | 342770 | 454200 | 0 | 0 |
| ILQYSEPYK | AHNAK2 | Q8IVF2 | AHNK2_HUMAI | Unmodified | 0 | 0 | 0 | 0 | 0 | 385760 | 0 | 0 | 0 | 0 | 0 | 0 | 0 | 0 | 0 | 0 | 183220 | 1815200 |
| ILQYSEPYK | AHNAK2 | Q8IVF2 | AHNK2_HUMAI | Unmodified | 0 | 0 | 0 | 77146 | 4788200 | 516960 | 0 | 0 | 0 | 0 | 0 | 0 | 0 | 0 | 0 | 0 | 1726600 | 5597100 |
| ILREFLGKS | ABHD12; ABHD | Q8N2K0; A0A | ABD12_HUMAN | Unmodified | 0 | 0 | 0 | 0 | 0 | 0 | 0 | 0 | 8470800 | 351020 | 0 | 0 | 0 | 0 | 0 | 0 | 0 | 0 |
| ILRPWQSS | SF1; SF1 | Q15637; A0A | SF01_HUMAN S | Unmodified | 854170 | 4932700 | 332660 | 343220 | 0 | 0 | 0 | 0 | 350770 | 83142 | 185570 | 803010 | 107630 | 0 | 218570 | 159200 | 0 | 0 |
| ILRQTEVLL | SH3GLB2; SH3 | Q9NR46; B7 | SHLB2_HUMAN | Unmodified | 0 | 0 | 0 | 0 | 3227900 | 0 | 0 | 0 | 0 | 1058800 | 0 | 0 | 1386700 | 0 | 0 | 0 | 654790 | 1037800 |
| ILSACEK | COPA; COPA; C | P53621; A0A | COPA_HUMAN | Unmodified | 0 | 1137700 | 223830 | 0 | 406550 | 1048200 | 1079600 | 41192 | 280960 | 1439700 | 0 | 1995000 | 516300 | 0 | 0 | 37120 | 971700 | 1154700 |
| ILSANGEA | ATP2B1; ATP2B | P20020; E7E | AT2B1_HUMAN | Unmodified | 8488100 | 3153000 | 1155900 | 1316700 | 591850 | 496640 | 368780 | 345840 | 1499700 | 965600 | 653120 | 14852000 | 2457300 | 1265300 | 7087000 | 5424100 | 582250 | 254940 |
| ILSCGEVIH | PPA2; PPA2; PP | Q9H2U2; D6 | IPYR2_HUMAN | Unmodified | 0 | 0 | 0 | 0 | 139250 | 0 | 311930 | 126690 | 2183700 | 1235500 | 0 | 2916400 | 175400 | 0 | 0 | 0 | 174650 | 232710 |
| ILSDDVPIR | RAD23A; RAD2 | P54725; A0A | RD23A_HUMAI | Unmodified | 341440 | 970460 | 137990 | 0 | 0 | 0 | 265250 | 0 | 0 | 0 | 0 | 182220 | 0 | 0 | 104530 | 102140 | 0 | 0 |
| ILSDDVPIR | RAD23A; RAD2 | P54725; A0A | RD23A_HUMAI | Unmodified | 5735200 | 5165700 | 750330 | 491960 | 0 | 0 | 113800 | 59694 | 0 | 0 | 0 | 0 | 147300 | 100040 | 970240 | 2095700 | 0 | 0 |
| ILSDDVPIR | RAD23A; RAD2 | P54725; A0A | RD23A_HUMAI | Unmodified | 6182800 | 6196100 | 1054500 | 3793800 | 585160 | 0 | 734880 | 258380 | 0 | 133950 | 0 | 0 | 2514000 | 897840 | 1206800 | 1612800 | 0 | 0 |
| ILSDLLPFC | ARHGEF1; ARH | Q92888; M0 | ARHG1_HUMA | Unmodified | 0 | 0 | 0 | 0 | 817820 | 1749400 | 0 | 0 | 0 | 120390 | 0 | 465480 | 0 | 0 | 0 | 0 | 571940 | 0 |
| ILSGPFVQ | GART; GART | P22102; A0A | PUR2_HUMAN | Unmodified | 0 | 0 | 0 | 0 | 0 | 0 | 0 | 150400 | 470330 | 313920 | 0 | 2043400 | 596000 | 340300 | 0 | 0 | 0 | 0 |
| ILSGVVTK | RPS11; RPS11 | P62280; M0 | RS11_HUMAN | Unmodified | 311060 | 216560 | 317290 | 403990 | 143820 | 0 | 280520 | 384020 | 0 | 0 | 0 | 3367700 | 0 | 0 | 227100 | 184300 | 0 | 150660 |
| ILSILRHQN | CHGA; CHGA | P10645; G5E | CMGA_HUMAN | Unmodified | 0 | 0 | 0 | 0 | 0 | 0 | 0 | 0 | 153490 | 681050 | 0 | 0 | 0 | 0 | 0 | 0 | 0 | 0 |
| ILSILRHQN | CHGA; CHGA | P10645; G5E | CMGA_HUMAN | Unmodified | 0 | 0 | 0 | 0 | 0 | 0 | 0 | 0 | 0 | 219290 | 0 | 0 | 0 | 0 | 0 | 0 | 0 | 0 |
| ILSISADIET | HNRNPK; HNRI | P61978; Q5 | HNRPK_HUMA | Unmodified | 0 | 235810 | 0 | 57079 | 72813 | 0 | 181360 | 0 | 0 | 131000 | 26218 | 0 | 482380 | 0 | 290480 | 0 | 0 | 0 |
| ILSKPIEVQ | DDX46; DDX46 | Q7L014; A0A | DDX46_HUMAI | Unmodified | 0 | 0 | 0 | 0 | 0 | 0 | 0 | 0 | 0 | 0 | 0 | 1214500 | 0 | 0 | 0 | 0 | 0 | 0 |
| ILSLLEGQK | ALDH3A2; ALDI | P51648; A0A | AL3A2_HUMAN | Unmodified | 0 | 256150 | 4661500 | 314620 | 0 | 0 | 0 | 35122 | 4201400 | 418640 | 0 | 7306300 | 0 | 0 | 200400 | 1269700 | 0 | 0 |
| ILSNNPSK | GSS; GSS; GSS | P48637; A0A | GSHB_HUMAN | Unmodified | 77374 | 0 | 163330 | 270570 | 1170100 | 0 | 0 | 81311 | 0 | 230620 | 0 | 2217200 | 0 | 0 | 0 | 0 | 448340 | 464740 |
| ILSSDDYGI | SPTBN1; SPTBI | Q01082; A0A | SPTB2_HUMAN | Unmodified | 1311200 | 316860 | 0 | 0 | 152710 | 0 | 14371 | 0 | 1015800 | 174660 | 0 | 2205200 | 109080 | 0 | 0 | 0 | 116840 | 0 |
| ILSSIQDMF | EPHB3 | P54753 | EPHB3_HUMAI | Unmodified | 185840 | 0 | 0 | 0 | 0 | 0 | 0 | 64288 | 0 | 181270 | 0 | 1337400 | 162160 | 0 | 182770 | 0 | 203850 | 0 |
| ILSSLGSTG | N/A | N/A | N/A | Unmodified | 0 | 0 | 0 | 0 | 0 | 0 | 153820 | 261800 | 0 | 0 | 0 | 0 | 0 | 0 | 0 | 0 | 240570 | 524580 |
| ILSTIDRELL | PSMC4 | P43686 | PRS6B_HUMAN | Unmodified | 0 | 129790 | 568390 | 960930 | 3475800 | 0 | 0 | 0 | 2572100 | 1722400 | 2328200 | 10058000 | 5317100 | 2164000 | 311010 | 335290 | 1781600 | 2972900 |
| ILSTSTIGPE | PALM2; PALM2 | Q8IXS6; B1A | PALM2_HUMAN | Unmodified | 0 | 0 | 0 | 0 | 1436800 | 0 | 440070 | 434140 | 1041500 | 0 | 0 | 0 | 0 | 0 | 0 | 0 | 145410 | 556570 |
| ILSVADSNI | PLS1; PLS1; PLS | Q14651; C9J | PLSI_HUMAN F | Unmodified | 0 | 0 | 0 | 0 | 0 | 0 | 0 | 0 | 250440 | 0 | 494170 | 926650 | 0 | 0 | 0 | 30425 | 0 | 0 |
| ILSYKIGKE | CAPZA2; CAPZ | P47755; P52 | CAZA2_HUMAN | Unmodified | 0 | 0 | 0 | 0 | 758010 | 0 | 0 | 657730 | 0 | 0 | 0 | 0 | 0 | 0 | 0 | 0 | 2777200 | 603890 |
| ILTCVLLPQ | LRRC1 | Q9BTT6 | LRRC1_HUMAN | Unmodified | 0 | 0 | 0 | 0 | 0 | 0 | 0 | 0 | 1505600 | 361310 | 258170 | 797620 | 0 | 0 | 0 | 0 | 0 | 0 |
| ILTDYGFEG | NDUFS3 | O75489 | NDUS3_HUMA | Unmodified | 0 | 0 | 0 | 0 | 0 | 0 | 0 | 0 | 576280 | 0 | 0 | 0 | 0 | 0 | 0 | 0 | 0 | 0 |
| ILTERGYSF | ACTG1; ACTB; | P63261; P60 | ACTG_HUMAN | Unmodified | 7892800 | 4036700 | 9635000 | 11140000 | 2591600 | 1567600 | 1399100 | 655740 | 4213700 | 1380900 | 1436700 | 16005000 | 786330 | 0 | 7560300 | 12911000 | 0 | 418740 |
| ILTERGYSF | ACTG1; ACTB; | P63261; P60 | ACTG_HUMAN | Unmodified | 1069200 | 522530 | 2307700 | 24496000 | 19667000 | 15149000 | 4136200 | 149420 | 13581000 | 8166600 | 8040100 | 0 | 8261500 | 1829300 | 170930 | 136830 | 10273000 | 1441600 |
| ILTFDQLAL | RPL18; RPL18; | Q07020; F8V | RL18_HUMAN | Unmodified | 1258300 | 1277500 | 807180 | 698340 | 0 | 0 | 915200 | 388200 | 0 | 0 | 1165300 | 14285000 | 976030 | 0 | 1064400 | 851940 | 0 | 0 |
| ILTFDQLAL | RPL18; RPL18; | Q07020; F8V | RL18_HUMAN | Unmodified | 0 | 0 | 0 | 0 | 0 | 0 | 2444800 | 0 | 0 | 0 | 0 | 0 | 2127700 | 0 | 0 | 0 | 0 | 0 |
| ILTFLESTYI | NUP210 | Q8TEM1 | PO210_HUMAI | Oxidation (M) | 0 | 0 | 0 | 0 | 0 | 0 | 0 | 0 | 2594000 | 366960 | 0 | 0 | 0 | 0 | 0 | 0 | 0 | 0 |
| ILTFLESTYI | NUP210 | Q8TEM1 | PO210_HUMAI | Unmodified | 17768 | 0 | 0 | 0 | 0 | 0 | 0 | 0 | 686690 | 0 | 0 | 0 | 0 | 0 | 0 | 0 | 0 | 0 |
| ILTGGADKI | PRPF19 | Q9UMS4 | PRP19_HUMAN | Unmodified | 0 | 0 | 0 | 0 | 953570 | 0 | 888770 | 475780 | 0 | 0 | 0 | 0 | 1194000 | 0 | 0 | 0 | 1020700 | 271090 |
| ILTGHTQS | NLE1; NLE1; NL | Q9NVX2; A0 | NLE1_HUMAN | Unmodified | 72791 | 20459 | 0 | 0 | 0 | 0 | 0 | 0 | 0 | 0 | 0 | 910530 | 513770 | 453980 | 0 | 0 | 0 | 0 |
| ILTHVAEM | SUN2 | Q9UH99 | SUN2_HUMAN | Unmodified | 0 | 0 | 0 | 0 | 0 | 0 | 0 | 37735 | 3145000 | 75604 | 0 | 0 | 0 | 0 | 0 | 0 | 0 | 0 |
| ILTHVAEM | SUN2 | Q9UH99 | SUN2_HUMAN | Oxidation (M) | 0 | 0 | 0 | 990590 | 0 | 0 | 0 | 0 | 3201800 | 194340 | 0 | 0 | 0 | 0 | 1007700 | 0 | 0 | 0 |
| ILTIEVK | DNAJB1; DNAJE | P25685; M0 | DNJB1_HUMAN | Unmodified | 41300 | 124870 | 0 | 330770 | 0 | 0 | 0 | 0 | 0 | 0 | 0 | 0 | 1039100 | 527620 | 0 | 0 | 0 | 907760 |
| ILTLEPTALT | BTN3A3 | O00478 | BT3A3_HUMAN | Unmodified | 0 | 0 | 0 | 0 | 0 | 0 | 0 | 0 | 0 | 0 | 144490 | 316690 | 0 | 0 | 0 | 0 | 0 | 419580 |
| ILTNFTGV | CNOT1 | A5YKK6 | CNOT1_HUMA | Oxidation (M) | 0 | 0 | 36011 | 187780 | 0 | 0 | 103780 | 0 | 0 | 0 | 0 | 721120 | 0 | 0 | 0 | 59628 | 0 | 0 |
| ILTPEVR | GDF15; GDF15 | Q99988; A0A | GDF15_HUMAI | Unmodified | 0 | 0 | 0 | 0 | 0 | 0 | 0 | 0 | 3968200 | 851490 | 0 | 0 | 0 | 0 | 0 | 0 | 0 | 0 |
| ILTPIIFKEE | MRPS22; MRPS | P82650; A0A | RT22_HUMAN | Unmodified | 0 | 0 | 0 | 0 | 0 | 0 | 0 | 0 | 4234500 | 372630 | 0 | 116540 | 0 | 0 | 0 | 0 | 0 | 0 |
| ILTPLVSLD | GLOD4; GLOD4 | Q9HC38; I3L | GLOD4_HUMA | Unmodified | 436120 | 0 | 782550 | 1115300 | 4449700 | 1101300 | 580060 | 383810 | 2305300 | 1420300 | 4784400 | 9370500 | 3919600 | 1519000 | 1693800 | 1704300 | 996110 | 1384300 |
| ILTSVDQYL | NIBAN2 | Q96TA1 | NIBA2_HUMAN | Unmodified | 0 | 614310 | 0 | 0 | 8994600 | 0 | 0 | 0 | 175850 | 1290800 | 0 | 0 | 842150 | 618350 | 1303600 | 0 | 1034800 | 1385700 |
| ILTTASSHE | UGP2; UGP2; U | Q16851; C9J | UGPA_HUMAN | Unmodified | 0 | 0 | 128040 | 0 | 4918000 | 0 | 1668000 | 703110 | 4036200 | 3371400 | 698520 | 3811300 | 2563200 | 1255100 | 0 | 241920 | 1516800 | 2610800 |
| ILTTEGRNA | DHX9 | Q08211 | DHX9_HUMAN | Unmodified | 0 | 0 | 0 | 0 | 193120 | 0 | 363850 | 540500 | 0 | 192570 | 0 | 0 | 281100 | 0 | 0 | 0 | 0 | 0 |
| ILTTEIGLHI | ABCC4 | O15439 | MRP4_HUMAN | Unmodified | 0 | 0 | 10655000 | 4101200 | 0 | 0 | 0 | 0 | 0 | 0 | 0 | 790520 | 0 | 0 | 0 | 0 | 0 | 0 |
| ILTTEIGLHI | ABCC4 | O15439 | MRP4_HUMAN | Unmodified | 0 | 0 | 491430 | 379420 | 277380 | 0 | 184730 | 0 | 0 | 0 | 0 | 183330 | 0 | 0 | 971960 | 810480 | 0 | 0 |
| ILTTGFSR | CORO1A; COR | P31146; H3E | COR1A_HUMAI | Unmodified | 459480 | 0 | 582870 | 605430 | 0 | 0 | 320050 | 0 | 967860 | 722070 | 3668700 | 17300000 | 0 | 1267000 | 0 | 664120 | 0 | 853410 |
| ILTTIEDLD | SMC2; SMC2 | O95347; A0A | SMC2_HUMAN | Unmodified | 0 | 0 | 83209 | 0 | 153070 | 0 | 225390 | 0 | 307460 | 251350 | 0 | 674440 | 159520 | 240810 | 52062 | 0 | 0 | 209200 |
| ILTTNTWSS | UGDH | O60701 | UGDH_HUMAN | Unmodified | 0 | 0 | 0 | 0 | 33121000 | 7556300 | 1738300 | 0 | 0 | 0 | 0 | 3175200 | 7324300 | 9459600 | 0 | 0 | 2355000 | 3051300 |
| ILTVPESQK | ABCA2; ABCA2; | Q9BZC7; A0 | ABCA2_HUMAI | Unmodified | 0 | 0 | 0 | 0 | 212530 | 0 | 0 | 0 | 2244800 | 0 | 0 | 0 | 0 | 0 | 0 | 0 | 0 | 0 |
| ILVALCGGI | ANXA1 | P04083 | ANXA1_HUMAN | Unmodified | 0 | 0 | 0 | 0 | 5238000 | 1139000 | 682660 | 175720 | 0 | 144000 | 12642000 | 1.38E+08 | 0 | 0 | 0 | 40810 | 899430 | 1486700 |
| ILVATNLFG | DDX39B; DDX3 | Q13838; O0 | DX39B_HUMAN | Unmodified | 0 | 0 | 0 | 0 | 0 | 0 | 0 | 0 | 4095000 | 2534700 | 2606700 | 27230000 | 0 | 0 | 0 | 530850 | 0 | 0 |
| ILVDALQK | ATP6AP2; ATP6 | O75787; A0A | RENR_HUMAN | Unmodified | 0 | 0 | 315250 | 144270 | 54939 | 156480 | 0 | 0 | 13034000 | 1837200 | 2773000 | 4891600 | 78893 | 0 | 333750 | 332840 | 260790 | 18336 |
| ILVDVEKVD | PYGB | P11216 | PYGB_HUMAN | Unmodified | 87979 | 0 | 0 | 255590 | 0 | 0 | 0 | 0 | 842010 | 382940 | 0 | 1394000 | 0 | 0 | 310930 | 264860 | 0 | 0 |
| ILVDVSNPT | STEAP3; STEAP | Q658P3; B8 | STEA3_HUMAN | Unmodified | 0 | 0 | 1253300 | 592780 | 0 | 0 | 0 | 0 | 1920200 | 546330 | 1953700 | 9121700 | 270140 | 0 | 1436300 | 1163800 | 0 | 0 |
| ILVELATFLE | MYOF | Q9NZM1 | MYOF_HUMAN | Unmodified | 0 | 0 | 0 | 0 | 642690 | 0 | 0 | 0 | 444080 | 0 | 0 | 0 | 0 | 0 | 0 | 0 | 0 | 619890 |
| ILVERYLIE | IST1; IST1; IST1 | P53990; H3E | IST1_HUMAN IS | Unmodified | 0 | 0 | 0 | 0 | 1.19E+08 | 13429000 | 4868700 | 325740 | 361490 | 26054000 | 3525800 | 0 | 28797000 | 24305000 | 0 | 0 | 5189500 | 12591000 |
| ILVGNKND | RAB35; RAB35 | Q15286; F5H | RAB35_HUMAN | Unmodified | 6790000 | 5112500 | 1674200 | 4148000 | 0 | 0 | 0 | 803650 | 2004900 | 385830 | 1472600 | 5886700 | 534650 | 0 | 2793900 | 3550700 | 0 | 34090 |
| ILVGNKND | RAB35; RAB35 | Q15286; F5H | RAB35_HUMAN | Unmodified | 6348500 | 3968400 | 0 | 3702600 | 0 | 0 | 0 | 0 | 230810 | 0 | 1024700 | 5426500 | 0 | 0 | 3953600 | 5368300 | 0 | 0 |
| ILVGPSEIG | FLNC | Q14315 | FLNC_HUMAN | Unmodified | 0 | 0 | 0 | 0 | 784580 | 0 | 0 | 0 | 0 | 0 | 0 | 0 | 0 | 0 | 0 | 0 | 1300200 | 200130 |
| ILVGTNLVR | VPS35 | Q96QK1 | VPS35_HUMAN | Unmodified | 358110 | 347280 | 131350 | 0 | 0 | 0 | 0 | 0 | 831800 | 203120 | 730140 | 0 | 0 | 0 | 997670 | 0 | 0 | 0 |
| ILVHGFQN | IGSF21 | Q96ID5 | IGS21_HUMAN | Oxidation (M) | 0 | 0 | 0 | 0 | 0 | 0 | 0 | 0 | 4144100 | 294850 | 174150 | 0 | 0 | 0 | 0 | 0 | 477520 | 0 |
| ILVHGFQN | IGSF21 | Q96ID5 | IGS21_HUMAN | Unmodified | 0 | 0 | 0 | 0 | 0 | 0 | 0 | 0 | 726470 | 0 | 0 | 0 | 0 | 0 | 0 | 0 | 0 | 0 |
| ILVKKEELT | EIF4A1; EIF4A1; | P60842; J3K | IF4A1_HUMAN | Unmodified | 294560 | 1215200 | 0 | 278120 | 0 | 0 | 0 | 0 | 0 | 0 | 0 | 4027600 | 0 | 0 | 0 | 0 | 0 | 0 |
| ILVKRDELT | EIF4A3; EIF4A3 | P38919; I3L | IF4A3_HUMAN | Unmodified | 252590 | 213700 | 0 | 0 | 0 | 0 | 0 | 0 | 0 | 0 | 0 | 806790 | 0 | 0 | 0 | 0 | 0 | 0 |
| ILVLDDTN | CNP; CNP; CNI | P09543; K7E | CN37_HUMAN | Unmodified | 209270 | 0 | 80431 | 705580 | 0 | 0 | 174030 | 0 | 776020 | 185040 | 4939400 | 24753000 | 0 | 339260 | 1547600 | 1623300 | 0 | 0 |
| ILVLDDTN | CNP; CNP; CNI | P09543; K7E | CN37_HUMAN | Unmodified | 0 | 0 | 371060 | 0 | 0 | 0 | 0 | 0 | 0 | 0 | 1117600 | 14085000 | 0 | 0 | 0 | 820660 | 0 | 0 |
| ILVNHQSFI | RAB32 | Q13637 | RAB32_HUMAN | Unmodified | 0 | 0 | 606760 | 0 | 0 | 0 | 0 | 0 | 0 | 0 | 0 | 0 | 0 | 0 | 164780 | 182730 | 0 | 0 |
| ILVPEGTRD | DLAT; DLAT; DL | P10515; A0A | ODP2_HUMAN | Unmodified | 0 | 0 | 0 | 88659 | 0 | 0 | 373640 | 0 | 5653400 | 1781800 | 0 | 0 | 0 | 0 | 0 | 0 | 0 | 0 |
| ILVPIQQVL | HSPA13 | P48723 | HSP13_HUMAI | Unmodified | 0 | 0 | 0 | 663770 | 0 | 666110 | 0 | 0 | 13137000 | 2317600 | 785130 | 530900 | 0 | 0 | 421740 | 0 | 0 | 0 |
| ILVQERLTQ | RTRAF; RTRAF; | Q9Y224; G3 | RTRAF_HUMAN | Unmodified | 0 | 0 | 0 | 240110 | 3176500 | 620000 | 446940 | 0 | 0 | 635590 | 637930 | 0 | 5980200 | 1495000 | 0 | 0 | 1081700 | 0 |
| ILVQGCHA | SNTB2 | Q13425 | SNTB2_HUMAI | Unmodified | 0 | 0 | 0 | 0 | 0 | 0 | 0 | 0 | 0 | 0 | 0 | 0 | 0 | 0 | 1171400 | 827830 | 0 | 0 |
| ILVTGGSGI | GFUS; GFUS; G | Q13630; E9F | FCL_HUMAN G | Unmodified | 0 | 0 | 0 | 307820 | 3227600 | 0 | 255750 | 98521 | 1698600 | 949530 | 0 | 1647500 | 1357600 | 0 | 0 | 737180 | 3052900 | 4182400 |
| ILVVNAAYF | SERPINB5 | P36952 | SPB5_HUMAN | Unmodified | 0 | 0 | 0 | 0 | 0 | 0 | 0 | 44050 | 0 | 0 | 0 | 0 | 2803600 | 1619600 | 0 | 0 | 3153900 | 6702400 |
| ILVVNAAYF | SERPINB5 | P36952 | SPB5_HUMAN | Oxidation (M) | 0 | 0 | 0 | 0 | 0 | 0 | 0 | 0 | 0 | 0 | 0 | 0 | 0 | 195040 | 0 | 0 | 907810 | 276130 |
| ILVVNAAYF | SERPINB5 | P36952 | SPB5_HUMAN | Unmodified | 0 | 1689000 | 0 | 0 | 0 | 595010 | 20483 | 0 | 0 | 0 | 0 | 0 | 13901 | 33044 | 0 | 0 | 284780 | 0 |
| ILWLTTR | LLGL2; LLGL2; | Q6P1M3; J3 | L2GL2_HUMAN | Unmodified | 554660 | 0 | 468610 | 0 | 0 | 0 | 0 | 0 | 1184600 | 176710 | 0 | 2191200 | 0 | 0 | 992330 | 728130 | 0 | 0 |
| ILYAPAGSE | ABCA2; ABCA2; | Q9BZC7; A0 | ABCA2_HUMAI | Unmodified | 0 | 0 | 0 | 0 | 0 | 0 | 0 | 0 | 673740 | 171910 | 0 | 0 | 0 | 0 | 0 | 0 | 0 | 0 |
| ILYDILAFA | DNAJC10; DNA | Q8IXB1; A0A | DJC10_HUMAN | Unmodified | 0 | 0 | 0 | 0 | 0 | 0 | 498700 | 0 | 18701000 | 9587300 | 1029500 | 308590 | 0 | 0 | 0 | 0 | 0 | 0 |

| | | | | | | | | | | | | | | | | | | | | | | |
|---|---|---|---|---|---|---|---|---|---|---|---|---|---|---|---|---|---|---|---|---|---|---|
| ILYLIQAWA | HGS; HGS; HG! | O14964; A0/ | HGS_HUMAN H | Unmodified | 117460 | 88227 | 0 | 0 | 0 | 0 | 0 | 0 | 0 | 24714 | 0 | 0 | 0 | 0 | 70165 | 47024 | 0 | 0 |
| ILYLIQAWA | HGS; HGS; HG! | O14964; A0/ | HGS_HUMAN H | Unmodified | 823100 | 1001400 | 339660 | 564490 | 860650 | 335310 | 5207800 | 0 | 985850 | 3552300 | 124070 | 307410 | 0 | 0 | 359780 | 198790 | 857320 | 270420 |
| ILYLTPEQE | ETF1; ETF1 | P62495; B7Z | ERF1_HUMAN I | Unmodified | 521670 | 356100 | 537750 | 0 | 476260 | 0 | 0 | 0 | 0 | 0 | 829490 | 5379100 | 141660 | 0 | 974310 | 580910 | 0 | 0 |
| ILYSHLVDY | GNA11; GNA11 | P29992; K7E | GNA11_HUMAI | Unmodified | 910780 | 903180 | 0 | 0 | 0 | 0 | 0 | 0 | 0 | 140320 | 0 | 4541500 | 0 | 0 | 0 | 0 | 50558 | 0 |
| ILYSHLVDY | GNA11; GNA11 | P29992; K7E | GNA11_HUMAI | Unmodified | 256440 | 0 | 59036 | 36181 | 0 | 0 | 0 | 0 | 0 | 0 | 0 | 1244500 | 0 | 0 | 0 | 0 | 0 | 0 |
| ILYSQCGD' | MYL6B; MYL6; N | P14649; P60 | MYL6B_HUMAI | Oxidation (M) | 598190 | 256100 | 641600 | 989900 | 4548800 | 1100100 | 17201 | 54010 | 1174500 | 502720 | 751150 | 4575600 | 1287900 | 414960 | 840430 | 741670 | 507780 | 694020 |
| ILYSQCGD' | MYL6B; MYL6; N | P14649; P60 | MYL6B_HUMAI | Unmodified | 0 | 0 | 0 | 1169300 | 10803000 | 2915000 | 0 | 0 | 3904500 | 1270600 | 578920 | 8175000 | 3311600 | 68401 | 1171400 | 0 | 1616200 | 2801200 |
| ILYSQCGD' | MYL6B; MYL6; N | P14649; P60 | MYL6B_HUMAI | Unmodified | 0 | 0 | 0 | 0 | 53363000 | 2099300 | 324640 | 0 | 1353900 | 0 | 0 | 0 | 5700100 | 7855500 | 0 | 0 | 9424100 | 13898000 |
| ILYSQCGD' | MYL6B; MYL6; N | P14649; P60 | MYL6B_HUMAI | Oxidation (M) | 0 | 0 | 0 | 0 | 14874000 | 1724500 | 0 | 0 | 0 | 0 | 0 | 0 | 5592600 | 6342000 | 0 | 0 | 6432700 | 2361800 |
| ILYTWTPGT | SYNJ1; SYNJ1; S | O43426; C9J | SYNJ1_HUMAN | Unmodified | 0 | 0 | 0 | 0 | 140860 | 0 | 276260 | 0 | 0 | 0 | 0 | 0 | 0 | 0 | 0 | 0 | 0 | 0 |
| IMADIRAQY | KRT18; KRT18 | P05783; F8V | K1C18_HUMAN | Unmodified | 0 | 0 | 0 | 0 | 1541500 | 163560 | 0 | 0 | 0 | 0 | 0 | 0 | 771200 | 261760 | 0 | 0 | 412950 | 0 |
| IMAFREEIAI | UTRN; UTRN | P46939; Q5T | UTRN_HUMAN | Oxidation (M) | 0 | 0 | 374170 | 0 | 588140 | 0 | 0 | 0 | 562720 | 675150 | 0 | 0 | 0 | 0 | 2411000 | 1963400 | 686040 | 0 |
| IMAISAGDA | PKP2 | Q99959 | PKP2_HUMAN | Unmodified | 0 | 0 | 0 | 0 | 994970 | 0 | 54171 | 83651 | 96386 | 0 | 0 | 0 | 261530 | 0 | 0 | 0 | 773580 | 0 |
| IMASLNLLF | GLMN; GLMN; ( | Q92990; M0 | GLMN_HUMAN | Oxidation (M) | 0 | 0 | 0 | 0 | 0 | 0 | 0 | 0 | 0 | 0 | 0 | 204460 | 0 | 0 | 0 | 0 | 0 | 0 |
| IMATPEQV( | XRCC6; XRCC6 | P12956; B1A | XRCC6_HUMAI | Unmodified | 0 | 0 | 0 | 400530 | 2013300 | 0 | 1909700 | 1042600 | 831570 | 628230 | 1346500 | 8431700 | 0 | 0 | 0 | 0 | 326350 | 521610 |
| IMATPEQV( | XRCC6; XRCC6 | P12956; B1A | XRCC6_HUMAI | Oxidation (M) | 0 | 0 | 0 | 0 | 0 | 0 | 0 | 319620 | 0 | 0 | 547660 | 2788000 | 0 | 252430 | 0 | 0 | 0 | 0 |
| IMDPNIVG! | ATP5F1B; ATP5I | P06576; H0Y | ATPB_HUMAN , | Unmodified | 0 | 0 | 0 | 0 | 0 | 0 | 0 | 0 | 41612000 | 374300 | 0 | 0 | 0 | 0 | 0 | 0 | 0 | 0 |
| IMDPNIVG! | ATP5F1B; ATP5I | P06576; H0Y | ATPB_HUMAN , | Oxidation (M) | 68883 | 2.44E+09 | 579520 | 0 | 0 | 0 | 0 | 21328 | 14574000 | 635080 | 382970 | 772710 | 0 | 0 | 49384 | 0 | 0 | 0 |
| IMDPNIVG! | ATP5F1B | P06576 | ATPB_HUMAN , | Unmodified | 0 | 0 | 0 | 0 | 0 | 0 | 0 | 0 | 3277400 | 1970300 | 0 | 0 | 0 | 0 | 0 | 0 | 0 | 0 |
| IMDPNIVG! | ATP5F1B | P06576 | ATPB_HUMAN , | Oxidation (M) | 0 | 0 | 0 | 0 | 0 | 0 | 0 | 0 | 1047800 | 1871000 | 0 | 0 | 0 | 0 | 0 | 0 | 0 | 0 |
| IMDVGWPE | TNS3; TNS3; TN | Q68CZ2; C9 | TENS3_HUMAN | Unmodified | 0 | 0 | 0 | 0 | 985810 | 0 | 0 | 0 | 19889 | 0 | 0 | 0 | 0 | 0 | 0 | 0 | 0 | 487630 |
| IMEAMEK | PIN4; PIN4; PIN | Q9Y237; H0Y | PIN4_HUMAN I | Unmodified | 0 | 0 | 0 | 0 | 0 | 0 | 0 | 0 | 0 | 0 | 0 | 471840 | 0 | 0 | 0 | 0 | 0 | 0 |
| IMEETTPTT | SLC24A1; SLC2 | O60721; F5H | NCKX1_HUMAI | Oxidation (M) | 0 | 0 | 0 | 0 | 1135200 | 699330 | 0 | 0 | 844850 | 947060 | 1617700 | 11348000 | 0 | 0 | 0 | 0 | 550320 | 602120 |
| IMEGPAFN | PDHB | P11177 | ODPB_HUMAN | Oxidation (M) | 0 | 0 | 0 | 0 | 0 | 0 | 0 | 0 | 9388800 | 836290 | 0 | 0 | 0 | 37181 | 67715 | 0 | 0 | 0 |
| IMEGPAFN | PDHB | P11177 | ODPB_HUMAN | Unmodified | 0 | 0 | 0 | 34002 | 16153 | 0 | 0 | 0 | 3226900 | 0 | 0 | 0 | 30667 | 41394 | 0 | 0 | 0 | 0 |
| IMEIVDAITT | PFKM; PFKM; PI | P08237; A0A | PFKAM_HUMAI | Oxidation (M) | 0 | 0 | 0 | 0 | 0 | 0 | 0 | 0 | 0 | 0 | 0 | 2161200 | 0 | 0 | 497140 | 186420 | 0 | 0 |
| IMELEGR | GOLM1; GOLM1 | Q8NBJ4; F8V | GOLM1_HUMA | Oxidation (M) | 0 | 0 | 0 | 0 | 0 | 0 | 0 | 0 | 2990000 | 0 | 0 | 0 | 0 | 0 | 0 | 0 | 0 | 0 |
| IMEMSAK | ALDH9A1 | P49189 | AL9A1_HUMAN | Unmodified | 21730 | 0 | 43251 | 0 | 0 | 0 | 0 | 0 | 56938 | 30680 | 0 | 0 | 0 | 0 | 32966 | 66612 | 777290 | 0 |
| IMEYYEKKE | ATP6V1E2; ATP( | Q96A05; P36 | VATE2_HUMAN | Unmodified | 0 | 0 | 231780 | 0 | 558700 | 0 | 558330 | 316680 | 0 | 237800 | 0 | 0 | 0 | 179000 | 0 | 0 | 9300.3 | 0 |
| IMFEVQDLI | DYNC1H1; DYN | Q14204; A0/ | DYHC1_HUMA | Oxidation (M) | 0 | 0 | 0 | 0 | 0 | 0 | 0 | 0 | 0 | 0 | 0 | 1919600 | 0 | 0 | 0 | 0 | 0 | 0 |
| IMGGNSILI | STXBP6 | Q8NFX7 | STXB6_HUMAN | Oxidation (M) | 0 | 0 | 151720 | 143460 | 0 | 0 | 0 | 0 | 0 | 0 | 0 | 0 | 0 | 0 | 243400 | 249240 | 0 | 0 |
| IMGIPEEEQ | MYH9 | P35579 | MYH9_HUMAN | 2 Oxidation (M) | 0 | 0 | 0 | 0 | 0 | 0 | 0 | 0 | 0 | 0 | 0 | 4322900 | 0 | 0 | 0 | 0 | 0 | 0 |
| IMGKILDQ( | ERP29; ERP29 | P30040; F8V | ERP29_HUMAN | Oxidation (M) | 0 | 0 | 0 | 0 | 0 | 0 | 0 | 0 | 0 | 5236700 | 0 | 0 | 0 | 0 | 0 | 0 | 0 | 0 |
| IMGKILDQ( | ERP29; ERP29 | P30040; F8V | ERP29_HUMAN | Unmodified | 0 | 0 | 0 | 0 | 0 | 0 | 0 | 0 | 0 | 1313700 | 0 | 0 | 0 | 0 | 0 | 0 | 0 | 0 |
| IMGKILDQ( | ERP29; ERP29 | P30040; F8V | ERP29_HUMAN | 2 Oxidation (M) | 0 | 0 | 0 | 0 | 0 | 0 | 0 | 48744 | 0 | 1608900 | 0 | 0 | 0 | 0 | 0 | 0 | 0 | 0 |
| IMGKILDQ( | ERP29; ERP29 | P30040; F8V | ERP29_HUMAN | Oxidation (M) | 0 | 0 | 0 | 0 | 0 | 0 | 0 | 0 | 0 | 1519900 | 0 | 0 | 0 | 0 | 0 | 0 | 0 | 0 |
| IMGLDLPD | SHMT2; SHMT2 | P34897; G3V | GLYM_HUMAN | Oxidation (M) | 0 | 0 | 0 | 0 | 0 | 0 | 0 | 0 | 5350900 | 454420 | 0 | 0 | 0 | 0 | 0 | 0 | 0 | 0 |
| IMGLDLPD | SHMT2; SHMT2 | P34897; G3V | GLYM_HUMAN | Unmodified | 0 | 0 | 0 | 0 | 0 | 0 | 97971 | 0 | 5775800 | 0 | 191710 | 0 | 0 | 0 | 0 | 0 | 0 | 0 |
| IMGLDLPD | SHMT2; SHMT2 | P34897; G3V | GLYM_HUMAN | 2 Oxidation (M) | 0 | 0 | 0 | 0 | 0 | 0 | 0 | 0 | 2508300 | 423400 | 0 | 0 | 0 | 0 | 0 | 0 | 0 | 0 |
| IMGPNYTP | EEF2 | P13639 | EF2_HUMAN El | Unmodified | 0 | 0 | 0 | 0 | 0 | 0 | 0 | 0 | 0 | 0 | 0 | 2941200 | 0 | 0 | 0 | 0 | 0 | 0 |
| IMGPNYTP | EEF2 | P13639 | EF2_HUMAN El | Unmodified | 0 | 0 | 279650 | 0 | 0 | 0 | 0 | 0 | 156860 | 0 | 272410 | 7236100 | 0 | 0 | 0 | 344770 | 0 | 0 |
| IMGPNYTP | EEF2 | P13639 | EF2_HUMAN El | Oxidation (M) | 0 | 0 | 0 | 0 | 0 | 0 | 38416 | 18841 | 0 | 0 | 143530 | 3374200 | 0 | 0 | 468190 | 426200 | 0 | 0 |
| IMHAINR | DPP4; DPP4; DI | P27487; A0A | DPP4_HUMAN | Unmodified | 0 | 0 | 0 | 440240 | 0 | 0 | 0 | 0 | 0 | 0 | 0 | 0 | 0 | 0 | 438810 | 187390 | 499040 | 811310 |
| IMHFPTWV | DDX1; DDX1; D | Q92499; A0/ | DDX1_HUMAN | Oxidation (M) | 0 | 0 | 0 | 0 | 0 | 0 | 0 | 0 | 0 | 0 | 0 | 5374600 | 0 | 0 | 0 | 294210 | 0 | 0 |
| IMIMTDQD | TOP2B; TOP2A; | Q02880; P11 | TOP2B_HUMAI | Oxidation (M) | 0 | 0 | 0 | 0 | 0 | 0 | 147280 | 89575 | 0 | 0 | 0 | 210850 | 0 | 0 | 0 | 0 | 0 | 0 |
| IMIMTDQD | TOP2B; TOP2A; | Q02880; P11 | TOP2B_HUMAI | Unmodified | 0 | 0 | 0 | 0 | 0 | 0 | 651650 | 0 | 0 | 0 | 0 | 798470 | 0 | 0 | 0 | 0 | 75469 | 84622 |
| IMKDILEK | HSP90AA1 | P07900 | HS90A_HUMAI | Unmodified | 0 | 0 | 0 | 0 | 17677000 | 448960 | 313720 | 681180 | 0 | 1813200 | 0 | 0 | 1224100 | 954820 | 0 | 0 | 2326500 | 2757400 |
| IMLEDGNL | DPYSL3 | Q14195 | DPYL3_HUMAN | Oxidation (M) | 248900 | 0 | 0 | 0 | 938780 | 0 | 149090 | 149930 | 231400 | 0 | 20433000 | 80370000 | 182780 | 0 | 0 | 0 | 0 | 0 |
| IMLEDGNL | DPYSL3 | Q14195 | DPYL3_HUMAN | Unmodified | 41182 | 554000 | 0 | 0 | 1272700 | 0 | 909290 | 0 | 1494700 | 0 | 25233000 | 88381000 | 0 | 0 | 0 | 0 | 80654 | 0 |
| IMLEDGNL | DPYSL3 | Q14195 | DPYL3_HUMAN | Oxidation (M) | 0 | 0 | 0 | 0 | 3145800 | 0 | 2995600 | 0 | 0 | 0 | 0 | 0 | 797300 | 0 | 0 | 0 | 1014900 | 0 |
| IMLFTGGP | SEC23B; SEC23 | Q15437; A0/ | SC23B_HUMAI | 2 Oxidation (M) | 0 | 0 | 0 | 0 | 0 | 0 | 0 | 0 | 0 | 0 | 0 | 2040100 | 0 | 0 | 0 | 0 | 0 | 0 |
| IMLFTNED | XRCC6; XRCC6 | P12956; B1A | XRCC6_HUMAI | Oxidation (M) | 132000 | 0 | 0 | 0 | 0 | 0 | 0 | 113990 | 149680 | 0 | 0 | 1594100 | 0 | 0 | 36526 | 50152 | 0 | 0 |
| IMLFTNED | XRCC6; XRCC6 | P12956; B1A | XRCC6_HUMAI | Unmodified | 128510 | 0 | 0 | 0 | 0 | 0 | 0 | 0 | 0 | 0 | 0 | 2279400 | 0 | 0 | 573440 | 0 | 0 | 0 |
| IMLKGDNI | SNRPE; SNRPE | P62304; A6N | RUXE_HUMAN | Oxidation (M) | 345270 | 385380 | 0 | 0 | 0 | 242050 | 0 | 0 | 0 | 1329400 | 0 | 100060 | 88230 | 0 | 0 | 20042 | 1792200 | 0 |
| IMLKGDNI | SNRPE; SNRPE | P62304; A6N | RUXE_HUMAN | Unmodified | 0 | 0 | 0 | 0 | 292200 | 35677 | 0 | 0 | 63552 | 84959 | 0 | 0 | 504270 | 0 | 0 | 0 | 0 | 478260 |
| IMLLDEFTT | STXBP3 | O00186 | STXB3_HUMAN | Oxidation (M) | 1031700 | 529280 | 2207500 | 4761200 | 9606100 | 2866600 | 1721700 | 615980 | 1914800 | 2623400 | 3628500 | 8252200 | 9094100 | 7730900 | 5631600 | 5192000 | 3118900 | 11862000 |
| IMLNTPED\ | PSMD11 | O00231 | PSD11_HUMAN | Oxidation (M) | 0 | 0 | 0 | 281860 | 0 | 0 | 28368 | 0 | 0 | 0 | 0 | 3149500 | 674820 | 0 | 0 | 274040 | 0 | 0 |
| IMLPGVLR | DKC1; DKC1 | O60832; H7 | DKC1_HUMAN | Oxidation (M) | 0 | 0 | 0 | 0 | 0 | 0 | 0 | 0 | 327560 | 0 | 98160 | 942370 | 0 | 0 | 0 | 0 | 0 | 0 |
| IMLPWDPT | RPS3; RPS3; RP | P23396; E9P | RS3_HUMAN 4 | Oxidation (M) | 4203100 | 1920400 | 2260300 | 4120000 | 2183800 | 0 | 1563800 | 1391300 | 736560 | 323590 | 6240100 | 55060000 | 0 | 0 | 0 | 4077300 | 0 | 328080 |
| IMLPWDPT | RPS3; RPS3; RP | P23396; E9P | RS3_HUMAN 4 | Unmodified | 0 | 0 | 0 | 0 | 0 | 0 | 0 | 0 | 0 | 122020 | 0 | 1354100 | 0 | 0 | 0 | 0 | 0 | 222800 |
| IMMEFER | CHMP2A; CHM | O43633; M0 | CHM2A_HUMA | Unmodified | 0 | 0 | 0 | 0 | 541500 | 0 | 0 | 0 | 6565900 | 161530 | 807100 | 3063500 | 0 | 0 | 0 | 0 | 0 | 0 |
| IMNTFSVM | TUBB2A; TUBB2 | Q13885; Q9 | TBB2A_HUMAN | 2 Oxidation (M) | 0 | 0 | 0 | 0 | 0 | 0 | 0 | 0 | 899870 | 0 | 0 | 637960 | 0 | 0 | 0 | 0 | 0 | 0 |
| IMNTFSVM | TUBB2A; TUBB2 | Q13885; Q9 | TBB2A_HUMAN | Oxidation (M) | 0 | 0 | 0 | 0 | 0 | 0 | 0 | 0 | 0 | 0 | 0 | 1877700 | 0 | 0 | 0 | 0 | 47472 | 0 |
| IMNTFSVVI | TUBB4A; TUBB; | P04350; P07 | TBB4A_HUMAN | Oxidation (M) | 1191500 | 0 | 3122900 | 0 | 0 | 0 | 0 | 0 | 1384500 | 0 | 0 | 25188000 | 0 | 0 | 2330100 | 2742500 | 0 | 0 |
| IMNTFSVVI | TUBB4A; TUBB; | P04350; P07 | TBB4A_HUMAN | Unmodified | 0 | 0 | 0 | 0 | 0 | 0 | 0 | 0 | 1026400 | 0 | 0 | 1739700 | 0 | 0 | 0 | 0 | 0 | 0 |
| IMNVIGEPI | ATP5F1B; ATP5I | P06576; H0Y | ATPB_HUMAN , | Unmodified | 0 | 0 | 0 | 0 | 0 | 0 | 0 | 0 | 2666200 | 0 | 0 | 0 | 0 | 0 | 0 | 0 | 193270 | 0 |
| IMNVIGEPI | ATP5F1B; ATP5I | P06576; H0Y | ATPB_HUMAN , | Unmodified | 0 | 0 | 42704 | 0 | 421680 | 0 | 295060 | 59852 | 78082000 | 5823500 | 144390 | 0 | 252620 | 201620 | 0 | 0 | 0 | 209710 |
| IMNVIGEPI | ATP5F1B; ATP5I | P06576; H0Y | ATPB_HUMAN , | Oxidation (M) | 0 | 0 | 0 | 0 | 0 | 0 | 303170 | 113730 | 6657700 | 2146200 | 132730 | 203530 | 205970 | 0 | 0 | 0 | 0 | 0 |
| IMPEDIIINC | TOP1 | P11387 | TOP1_HUMAN | Oxidation (M) | 0 | 0 | 0 | 0 | 0 | 0 | 0 | 0 | 0 | 0 | 0 | 1370300 | 0 | 0 | 0 | 0 | 0 | 0 |
| IMQDFESD | DHFR | P00374 | DYR_HUMAN D | Oxidation (M) | 0 | 75893 | 0 | 399760 | 514600 | 27440 | 0 | 0 | 0 | 0 | 0 | 0 | 497730 | 0 | 259810 | 0 | 0 | 0 |
| IMQDFESD | DHFR | P00374 | DYR_HUMAN D | Unmodified | 0 | 0 | 0 | 436330 | 1948300 | 534850 | 159700 | 0 | 743230 | 315380 | 303110 | 588930 | 0 | 401140 | 237710 | 80945 | 0 | 1228200 |
| IMQNTDPH | EIF3CL; EIF3C | B5ME19; Q9 | EIFCL_HUMAN | Oxidation (M) | 0 | 0 | 0 | 0 | 0 | 0 | 0 | 0 | 1011400 | 21267 | 0 | 841740 | 0 | 0 | 0 | 0 | 0 | 0 |
| IMQQMSDI | DDAH1; DDAH1 | O94760; B4I | DDAH1_HUMA | Unmodified | 0 | 156070 | 142160 | 0 | 802510 | 182710 | 36692 | 0 | 1050700 | 241360 | 163340 | 0 | 672900 | 683450 | 336410 | 215630 | 0 | 1261000 |
| IMQSSSEV( | HSPD1; HSPD1 | P10809; A0A | CH60_HUMAN | 2 Oxidation (M) | 0 | 0 | 0 | 0 | 0 | 0 | 0 | 0 | 3780900 | 1264200 | 0 | 0 | 0 | 0 | 0 | 0 | 0 | 0 |
| IMQSSSEV( | HSPD1; HSPD1 | P10809; A0A | CH60_HUMAN | Oxidation (M) | 0 | 0 | 0 | 0 | 0 | 0 | 0 | 0 | 2992500 | 0 | 0 | 0 | 0 | 0 | 0 | 0 | 0 | 0 |
| IMQSSSEV( | HSPD1; HSPD1 | P10809; A0A | CH60_HUMAN | 3 Oxidation (M) | 0 | 0 | 0 | 0 | 0 | 0 | 0 | 0 | 0 | 1478600 | 151810 | 0 | 0 | 0 | 0 | 0 | 0 | 0 |
| IMRPDDAN | ACOT7 | O00154 | BACH_HUMAN | Unmodified | 0 | 0 | 494950 | 169580 | 6710000 | 0 | 653070 | 1026500 | 6327700 | 3332700 | 1555600 | 9673900 | 2340300 | 652370 | 0 | 83275 | 5192600 | 8615600 |
| IMSKIWVQI | PIK3C2B; PIK3C | O00750; F5C | P3C2B_HUMAI | 3 Oxidation (M) | 0 | 0 | 0 | 0 | 0 | 0 | 3310600 | 4019100 | 0 | 5562000 | 0 | 0 | 652700 | 1309900 | 0 | 0 | 8444000 | 9694800 |
| IMSVDEQK | PNMA2 | Q9UL42 | PNMA2_HUMA | Unmodified | 4481200 | 5274500 | 0 | 0 | 0 | 0 | 0 | 0 | 0 | 0 | 0 | 0 | 0 | 0 | 0 | 0 | 0 | 0 |
| IMVANIEEV | SEC22B | O75396 | SC22B_HUMAI | Oxidation (M) | 0 | 0 | 0 | 0 | 0 | 0 | 0 | 0 | 993370 | 474600 | 0 | 0 | 0 | 0 | 0 | 0 | 0 | 0 |
| IMVCEKDP. | CNTFR; CNTFR | P26992; Q5T | CNTFR_HUMAI | Oxidation (M) | 508000 | 0 | 0 | 0 | 0 | 0 | 0 | 0 | 0 | 0 | 0 | 795730 | 0 | 0 | 0 | 0 | 0 | 0 |
| IMVDRINLA | PLEC; PLEC | Q15149; H0Y | PLEC_HUMAN | Oxidation (M) | 0 | 0 | 0 | 0 | 466190 | 150490 | 73330 | 121460 | 0 | 80368 | 0 | 417600 | 0 | 263120 | 0 | 0 | 1427300 | 1216100 |
| IMVDRINLA | PLEC; PLEC | Q15149; H0Y | PLEC_HUMAN | Unmodified | 0 | 0 | 0 | 0 | 776360 | 0 | 0 | 0 | 0 | 0 | 0 | 0 | 0 | 0 | 0 | 0 | 1886500 | 2194300 |
| IMVLGANW | SEC14L2; N/A; | O76054; H7 | S14L2_HUMAN | Oxidation (M) | 0 | 0 | 0 | 0 | 75462 | 0 | 95148 | 0 | 0 | 23807 | 0 | 0 | 0 | 0 | 0 | 0 | 137170 | 0 |
| IMVSRSEID | ANXA3; ANXA3 | P12429; D6F | ANXA3_HUMAI | Oxidation (M) | 0 | 0 | 0 | 0 | 0 | 0 | 0 | 0 | 0 | 0 | 0 | 0 | 0 | 0 | 0 | 0 | 0 | 0 |
| IMVSRSEID | ANXA6; ANXA6; | P08133; E5R | ANXA6_HUMAI | Oxidation (M) | 0 | 49678 | 0 | 795570 | 880030 | 0 | 0 | 0 | 0 | 0 | 0 | 0 | 278460 | 0 | 89123 | 81840 | 0 | 119200 |
| IMVSRSEID | ANXA6; ANXA6; | P08133; E5R | ANXA6_HUMAI | Unmodified | 0 | 0 | 0 | 492340 | 1475300 | 0 | 0 | 0 | 7208700 | 1896700 | 0 | 0 | 0 | 582370 | 0 | 0 | 292940 | 1422500 |
| IMVSRSEID | ANXA1 | P04083 | ANXA1_HUMAI | Oxidation (M) | 324920 | 344280 | 0 | 1937500 | 25781000 | 5776600 | 1797200 | 958000 | 0 | 407220 | 51538000 | 1.84E+08 | 2773700 | 2455100 | 357580 | 81808 | 433560 | 2826600 |
| IMVSRSEID | ANXA1 | P04083 | ANXA1_HUMAI | Unmodified | 0 | 0 | 0 | 508160 | 31915000 | 1972000 | 1069800 | 223610 | 47750 | 580460 | 10612000 | 86731000 | 2373800 | 1924500 | 0 | 46025 | 958660 | 4648600 |
| IMVSRSEID | ANXA1 | P04083 | ANXA1_HUMAI | 2 Oxidation (M) | 0 | 0 | 0 | 660530 | 0 | 1466000 | 29260 | 0 | 0 | 0 | 4381000 | 45193000 | 0 | 0 | 600380 | 0 | 0 | 0 |
| IMVSRSEID | ANXA1 | P04083 | ANXA1_HUMAI | Unmodified | 0 | 0 | 0 | 0 | 1901200 | 0 | 0 | 0 | 0 | 0 | 0 | 0 | 0 | 0 | 0 | 0 | 707140 | 405360 |
| IMVSRSELI | ANXA6 | P08133 | ANXA6_HUMAI | Oxidation (M) | 0 | 0 | 0 | 789670 | 0 | 0 | 0 | 0 | 3044800 | 0 | 0 | 0 | 93600 | 0 | 537210 | 0 | 0 | 0 |
| IMVSRSETI | ANXA11; ANXA1 | P50995; H0Y | ANX11_HUMAI | Oxidation (M) | 0 | 0 | 0 | 0 | 0 | 0 | 0 | 0 | 8517800 | 0 | 1087100 | 2998100 | 0 | 0 | 462650 | 0 | 103960 | 0 |
| IMVSRSETI | ANXA11; ANXA1 | P50995; H0Y | ANX11_HUMAI | Unmodified | 27069 | 0 | 0 | 36782 | 118160 | 0 | 0 | 0 | 12271000 | 0 | 60107 | 272950 | 28544 | 0 | 0 | 0 | 0 | 0 |
| IMVSRSETI | ANXA11; ANXA1 | P50995; H0Y | ANX11_HUMAI | Oxidation (M) | 670290 | 610880 | 331880 | 892310 | 17123000 | 8694400 | 0 | 0 | 29394000 | 36329000 | 7305800 | 0 | 14685000 | 15196000 | 481260 | 413130 | 5200200 | 59565000 |
| IMVSRSETI | ANXA11; ANXA1 | P50995; H0Y | ANX11_HUMAI | Unmodified | 0 | 0 | 0 | 0 | 31852000 | 231480 | 474200 | 0 | 56926000 | 19474000 | 358730 | 0 | 10345000 | 945710 | 0 | 0 | 288030 | 46918000 |
| IMVSRSEVI | ANXA2P2; ANX/ | A6NMY6; P0 | AXA2L_HUMAN | Oxidation (M) | 0 | 0 | 0 | 230150 | 17250000 | 2406200 | 480290 | 273930 | 6521600 | 2928000 | 0 | 0 | 599100 | 189980 | 0 | 0 | 1269800 | 2440100 |
| IMVSRSEVI | ANXA2P2; ANX/ | A6NMY6; P0 | AXA2L_HUMAN | Unmodified | 0 | 0 | 0 | 0 | 28448000 | 6343700 | 1404800 | 0 | 13133000 | 17968 | 0 | 0 | 0 | 0 | 0 | 0 | 932650 | 5870500 |
| IMVVGSTIN | UBFD1; UBFD1 | O14562; H3I | UBFD1_HUMAI | Oxidation (M) | 0 | 0 | 0 | 0 | 0 | 0 | 0 | 65080 | 0 | 253070 | 329220 | 434330 | 0 | 606570 | 0 | 32518 | 418070 | 1232700 |
| IMWSQRDI | PABPC4; PABP( | Q13310; P11 | PABP4_HUMAN | Unmodified | 17009000 | 8596500 | 5616000 | 8253900 | 0 | 0 | 0 | 0 | 765930 | 0 | 0 | 3242700 | 0 | 0 | 1218500 | 4958300 | 0 | 0 |
| IMWSQRDI | PABPC4; PABP( | Q13310; P11 | PABP4_HUMAN | Oxidation (M) | 25271000 | 7713900 | 11892000 | 7706300 | 0 | 0 | 0 | 0 | 0 | 0 | 0 | 1288800 | 122750 | 0 | 4766500 | 10116000 | 0 | 0 |
| IMYCTDPG | SEZ6L; SEZ6L; S | Q9BYH1; B0 | SE6L1_HUMAN | Unmodified | 0 | 0 | 0 | 0 | 0 | 0 | 0 | 0 | 1815100 | 0 | 0 | 0 | 0 | 0 | 0 | 0 | 0 | 0 |
| IMYCTDPG | SEZ6L; SEZ6L; S | Q9BYH1; B0 | SE6L1_HUMAN | Oxidation (M) | 0 | 0 | 0 | 0 | 0 | 0 | 0 | 0 | 359060 | 0 | 0 | 0 | 0 | 0 | 0 | 0 | 0 | 0 |
| IMYSHLVD' | GNAQ | P50148 | GNAQ_HUMAN | Oxidation (M) | 329030 | 982320 | 0 | 0 | 0 | 305410 | 0 | 0 | 547780 | 0 | 0 | 2499100 | 0 | 0 | 298910 | 325710 | 0 | 0 |
| IMYTVFEHT | GARS1; GARS1; | P41250; A0A | GARS_HUMAN | Oxidation (M) | 0 | 0 | 0 | 1161800 | 0 | 0 | 0 | 0 | 1710300 | 0 | 0 | 18704000 | 0 | 0 | 419380 | 0 | 0 | 0 |
| IMYTVFEHT | GARS1; GARS1; | P41250; A0A | GARS_HUMAN | Unmodified | 0 | 0 | 0 | 0 | 0 | 0 | 0 | 0 | 1189900 | 0 | 0 | 2267500 | 0 | 0 | 0 | 0 | 0 | 0 |
| INAAHGFS | MAPKAP1; MAP | Q9BPZ7; B1/ | SIN1_HUMAN T | Unmodified | 0 | 0 | 119650 | 94648 | 180750 | 0 | 0 | 0 | 0 | 0 | 123960 | 1532900 | 28858 | 0 | 480400 | 380110 | 0 | 0 |
| INAANYAS\ | ABHD14B; ABH | Q96IU4; F8V | ABHEB_HUMAI | Unmodified | 0 | 262980 | 219200 | 0 | 0 | 1156900 | 0 | 63091 | 963380 | 102160 | 475760 | 3650300 | 1594600 | 521540 | 0 | 284620 | 712070 | 2217800 |
| INAEQQLK | CIT; CIT; CIT; CI | O14578; A0/ | CTRO_HUMAN | Unmodified | 0 | 0 | 0 | 0 | 11110000 | 899070 | 0 | 0 | 0 | 0 | 0 | 0 | 0 | 0 | 0 | 0 | 0 | 0 |

| | | | | | | | | | | | | | | | | | | | | | | |
|---|---|---|---|---|---|---|---|---|---|---|---|---|---|---|---|---|---|---|---|---|---|---|
| INAEQQLKI | CIT; CIT; CIT; CI | O14578; A0/ | CTRO_HUMAN | Unmodified | 0 | 0 | 0 | 0 | 0 | 0 | 0 | 0 | 0 | 0 | 0 | 0 | 0 | 0 | 0 | 0 | 315970 | 0 |
| INAEQQLKI | CIT; CIT; CIT; CI | O14578; A0/ | CTRO_HUMAN | Unmodified | 0 | 0 | 0 | 0 | 4825800 | 184560 | 654060 | 0 | 0 | 0 | 0 | 0 | 0 | 233160 | 0 | 0 | 2910000 | 499450 |
| INAERSRGI | C1orf198 | Q9H425 | CA198_HUMAN | Unmodified | 0 | 0 | 0 | 0 | 1120000 | 0 | 1161800 | 341260 | 0 | 221340 | 0 | 0 | 676470 | 691960 | 0 | 0 | 268470 | 865150 |
| INAFNWAK | FARP1; FARP1; | Q9Y4F1; A0/ | FARP1_HUMAN | Unmodified | 75550 | 0 | 530270 | 207830 | 0 | 0 | 0 | 0 | 2096900 | 290900 | 703240 | 939450 | 0 | 0 | 1295000 | 984480 | 0 | 0 |
| INAGMLAQ | RPA3 | P35244 | RFA3_HUMAN | Oxidation (M) | 0 | 106740 | 0 | 0 | 380960 | 316070 | 257370 | 0 | 0 | 262470 | 0 | 0 | 2980300 | 444400 | 0 | 0 | 1087600 | 458410 |
| INALTAASE | CCT7 | Q99832 | TCPH_HUMAN | Unmodified | 0 | 0 | 0 | 19501 | 50317 | 0 | 28550 | 0 | 0 | 55491 | 0 | 0 | 122710 | 34532 | 0 | 0 | 47351 | 42842 |
| INCELSQG | SORL1 | Q92673 | SORL_HUMAN | Unmodified | 0 | 0 | 0 | 0 | 0 | 0 | 0 | 0 | 989760 | 171090 | 0 | 0 | 0 | 0 | 0 | 0 | 0 | 0 |
| INCVAAVFS | PSD3; PSD3; PS | Q9NYI0; E5F | PSD3_HUMAN | Unmodified | 0 | 0 | 0 | 0 | 531410 | 0 | 0 | 0 | 337710 | 0 | 0 | 159970 | 0 | 0 | 0 | 0 | 499580 | 0 |
| INDFVLSPC | EIF2A; EIF2A; EI | Q9BY44; C9I | EIF2A_HUMAN | Unmodified | 0 | 0 | 0 | 66624 | 0 | 0 | 0 | 152310 | 0 | 0 | 433050 | 2636600 | 23989 | 0 | 111750 | 112400 | 0 | 0 |
| INDIIEVK | MYO1D; MYO1D | O94832; J3Q | MYO1D_HUMA | Unmodified | 2902200 | 3803600 | 0 | 0 | 0 | 0 | 0 | 91379 | 33414 | 0 | 209400 | 539490 | 0 | 0 | 1538500 | 176000 | 0 | 0 |
| INDRFEFPE | USP7; USP7; US | Q93009; A0/ | UBP7_HUMAN | Unmodified | 0 | 0 | 0 | 0 | 0 | 0 | 0 | 0 | 0 | 0 | 175040 | 606230 | 0 | 0 | 0 | 0 | 0 | 0 |
| INEAIVAVQ | RTRAF | Q9Y224 | RTRAF_HUMAN | Unmodified | 0 | 0 | 53760 | 0 | 0 | 0 | 61839 | 0 | 0 | 0 | 255310 | 965030 | 63386 | 0 | 0 | 134460 | 0 | 0 |
| INEAIVAVQ | RTRAF | Q9Y224 | RTRAF_HUMAN | Unmodified | 0 | 0 | 0 | 346170 | 0 | 0 | 0 | 0 | 0 | 0 | 796460 | 1460300 | 0 | 0 | 0 | 142830 | 0 | 0 |
| INEAIVAVQ | RTRAF | Q9Y224 | RTRAF_HUMAN | Unmodified | 0 | 0 | 0 | 352240 | 3408000 | 526280 | 0 | 0 | 112120 | 370730 | 0 | 0 | 4122000 | 1232800 | 0 | 0 | 944590 | 2120400 |
| INEALITTILI | RICTOR; RICTO | Q6R327; D6I | RICTR_HUMAN | Unmodified | 0 | 0 | 0 | 0 | 0 | 0 | 0 | 0 | 0 | 0 | 0 | 61277 | 0 | 0 | 120340 | 16884 | 0 | 0 |
| INEANQLM | GALNT7; GALNT | Q86SF2; H0Y | GALT7_HUMAN | Unmodified | 0 | 0 | 0 | 0 | 0 | 0 | 0 | 0 | 2637800 | 93139 | 0 | 0 | 0 | 0 | 0 | 0 | 0 | 0 |
| INEANQLM | GALNT7; GALNT | Q86SF2; H0Y | GALT7_HUMAN | Oxidation (M) | 0 | 203650 | 66491 | 0 | 0 | 0 | 0 | 0 | 955190 | 0 | 0 | 273160 | 0 | 0 | 72549 | 0 | 0 | 0 |
| INEAVECLL | EPRS1; EPRS1 | P07814; V9G | SYEP_HUMAN | Unmodified | 0 | 0 | 0 | 0 | 131750 | 0 | 0 | 0 | 261370 | 108400 | 472250 | 8192400 | 417770 | 0 | 111250 | 62095 | 0 | 0 |
| INEAVNDLT | TTC13; TTC13; T | Q8NBP0; E5 | TTC13_HUMAN | Unmodified | 207470 | 0 | 0 | 0 | 188220 | 0 | 0 | 38148 | 2004900 | 226010 | 352120 | 0 | 0 | 0 | 0 | 0 | 0 | 0 |
| INEGFDLLF | ADH1C | P00326 | ADH1G_HUMA | Unmodified | 0 | 0 | 0 | 0 | 0 | 0 | 0 | 0 | 0 | 0 | 2684000 | 0 | 0 | 0 | 0 | 0 | 0 | 0 |
| INEGIGQGI | ACOX1 | Q15067 | ACOX1_HUMAI | Unmodified | 0 | 0 | 0 | 0 | 0 | 0 | 0 | 0 | 3819200 | 603530 | 967950 | 337180 | 0 | 0 | 0 | 0 | 0 | 0 |
| INEGLEHLA | NAPG; NAPG; N | Q99747; J3Q | SNAG_HUMAN | Unmodified | 0 | 0 | 0 | 0 | 1748300 | 0 | 466190 | 420310 | 0 | 169970 | 0 | 0 | 720070 | 1786500 | 0 | 0 | 918960 | 1585500 |
| INEILSNALI | HNRNPM; HNR | P52272; M0F | HNRPM_HUMA | Unmodified | 664040 | 357070 | 0 | 0 | 376840 | 0 | 0 | 384730 | 2220100 | 511940 | 382180 | 1345100 | 0 | 0 | 267240 | 192550 | 789330 | 0 |
| INEIRQMS( | PCBP1; PCBP3; | Q15365; P57 | PCBP1_HUMAI | Unmodified | 114660 | 44145 | 0 | 837690 | 3548900 | 704280 | 4945100 | 1413600 | 0 | 1226800 | 0 | 0 | 14542000 | 7633000 | 0 | 0 | 992250 | 1904200 |
| INEIRQMS( | PCBP1; PCBP3; | Q15365; P57 | PCBP1_HUMAI | Oxidation (M) | 196850 | 0 | 0 | 445620 | 1685800 | 270410 | 1653600 | 29364 | 0 | 463430 | 0 | 0 | 5502800 | 1542400 | 0 | 0 | 0 | 765210 |
| INEKPQVIA | EDF1 | O60869 | EDF1_HUMAN | Unmodified | 2244000 | 4387000 | 5461900 | 4116400 | 0 | 0 | 0 | 0 | 4042400 | 0 | 2613800 | 4971900 | 226260 | 0 | 7228700 | 6372600 | 778180 | 0 |
| INELMPK | MANF | P55145 | MANF_HUMAN | Unmodified | 0 | 0 | 0 | 0 | 426180 | 0 | 384970 | 0 | 33648000 | 11724000 | 3099600 | 0 | 357840 | 0 | 267590 | 0 | 146230 | 941240 |
| INELMPK | MANF | P55145 | MANF_HUMAN | Oxidation (M) | 0 | 0 | 0 | 0 | 14630 | 0 | 0 | 0 | 3567200 | 1651700 | 650030 | 63585 | 0 | 0 | 0 | 0 | 0 | 172480 |
| INELTGIK | PSMC2; PSMC2 | P35998; C9J | PRS7_HUMAN | Unmodified | 0 | 0 | 0 | 0 | 0 | 0 | 212960 | 219210 | 0 | 157120 | 1072200 | 1785000 | 0 | 0 | 0 | 0 | 0 | 0 |
| INEMVNR | NAGK; NAGK; N | Q9UJ70; H7( | NAGK_HUMAN | Unmodified | 40451 | 206150 | 0 | 76068 | 24369 | 0 | 0 | 0 | 909670 | 0 | 52331 | 247710 | 104240 | 0 | 0 | 0 | 0 | 0 |
| INEVQTDV( | TWF1; TWF1 | Q12792; F8V | TWF1_HUMAN | Unmodified | 983100 | 654750 | 736590 | 707340 | 387570 | 0 | 0 | 0 | 237360 | 0 | 0 | 656100 | 392550 | 408290 | 1120100 | 609720 | 0 | 327350 |
| INEVQTDV( | TWF1; TWF1 | Q12792; F8V | TWF1_HUMAN | Unmodified | 218170 | 195920 | 0 | 0 | 0 | 0 | 0 | 0 | 0 | 0 | 0 | 0 | 0 | 71877 | 0 | 0 | 0 | 0 |
| INEWLTLVE | DYNC1H1; DYN | Q14204; A0/ | DYHC1_HUMA | Unmodified | 0 | 0 | 0 | 0 | 0 | 0 | 0 | 0 | 1060800 | 489050 | 565910 | 4343200 | 0 | 0 | 0 | 0 | 0 | 0 |
| INFDDNAE | SUCLG2; SUCL | Q96I99; E9P | SUCB2_HUMAI | Unmodified | 0 | 0 | 0 | 0 | 0 | 0 | 0 | 0 | 949600 | 0 | 0 | 0 | 0 | 0 | 0 | 0 | 0 | 0 |
| INFDDNAE | SUCLG2; SUCL | Q96I99; E9P | SUCB2_HUMAI | Unmodified | 0 | 0 | 0 | 0 | 0 | 0 | 0 | 0 | 1404300 | 284880 | 0 | 0 | 0 | 0 | 0 | 0 | 0 | 0 |
| INFDSNSA | SUCLA2; SUCL/ | Q9P2R7; A0/ | SUCB1_HUMAI | Unmodified | 0 | 0 | 0 | 0 | 149670 | 0 | 0 | 43443 | 883980 | 0 | 0 | 0 | 152210 | 0 | 0 | 0 | 0 | 0 |
| INFVIDNNI | CSDE1; CSDE1 | O75534; E9F | CSDE1_HUMAI | Unmodified | 0 | 0 | 0 | 0 | 0 | 0 | 0 | 0 | 0 | 0 | 0 | 0 | 0 | 0 | 0 | 320850 | 0 | 0 |
| INGEVSSIS | CD2AP | Q9Y5K6 | CD2AP_HUMAI | Unmodified | 0 | 0 | 1032300 | 0 | 0 | 0 | 0 | 0 | 0 | 0 | 0 | 0 | 0 | 0 | 1121100 | 515030 | 0 | 0 |
| INGEVSSIS | CD2AP | Q9Y5K6 | CD2AP_HUMAI | Unmodified | 2087200 | 1646100 | 2006300 | 2547400 | 816610 | 2320400 | 549550 | 0 | 2107700 | 0 | 0 | 202520 | 3783600 | 2033200 | 4121200 | 1963800 | 374230 | 0 |
| INGIEYVPF | CAPN7 | Q9Y6W3 | CAN7_HUMAN | Oxidation (M) | 63377 | 0 | 0 | 49310 | 0 | 0 | 0 | 0 | 0 | 0 | 0 | 510290 | 0 | 0 | 0 | 0 | 0 | 0 |
| INGIPQQH | CEACAM5; CEA | P06731; A0A | CEAM5_HUMAI | Unmodified | 0 | 0 | 0 | 0 | 0 | 0 | 0 | 0 | 2E+08 | 18914000 | 0 | 0 | 0 | 0 | 0 | 0 | 0 | 0 |
| INGIPVEEL | L1CAM | P32004 | L1CAM_HUMAI | Unmodified | 0 | 0 | 0 | 0 | 0 | 0 | 0 | 0 | 692990 | 0 | 1336800 | 0 | 0 | 0 | 0 | 0 | 0 | 0 |
| INGKDSIEK | RUVBL1; RUVBI | Q9Y265; H7( | RUVB1_HUMAI | Unmodified | 0 | 0 | 0 | 0 | 3143800 | 0 | 697310 | 0 | 0 | 0 | 0 | 0 | 1117200 | 817010 | 0 | 0 | 927230 | 845270 |
| INGTVTENI | TJP1; TJP1; TJP1 | Q07157; A0/ | ZO1_HUMAN T | Unmodified | 0 | 0 | 0 | 0 | 433560 | 0 | 0 | 0 | 0 | 0 | 0 | 0 | 246860 | 131340 | 0 | 0 | 112910 | 59001 |
| INGTVTENI | TJP2; TJP2; TJP2 | Q9UDY2; A0 | ZO2_HUMAN T | Unmodified | 0 | 0 | 0 | 0 | 1846700 | 0 | 0 | 0 | 3105900 | 714610 | 0 | 747250 | 272560 | 0 | 0 | 0 | 0 | 0 |
| INHEGEVN | RBBP7; RBBP4; | Q16576; Q0! | RBBP7_HUMAN | Unmodified | 0 | 0 | 0 | 0 | 0 | 0 | 0 | 0 | 0 | 0 | 0 | 0 | 1037100 | 133840 | 0 | 0 | 153400 | 306940 |
| INIHEYYLG | ERLEC1 | Q96DZ1 | ERLEC_HUMAN | Oxidation (M) | 0 | 0 | 0 | 0 | 0 | 0 | 0 | 0 | 2698300 | 594740 | 0 | 0 | 0 | 0 | 0 | 0 | 0 | 0 |
| INIHEYYLG | ERLEC1 | Q96DZ1 | ERLEC_HUMAN | Unmodified | 0 | 0 | 0 | 0 | 0 | 0 | 63746 | 0 | 8776700 | 776610 | 0 | 0 | 0 | 0 | 0 | 0 | 0 | 0 |
| INIRVLDPF | TKT; TKT | P29401; A0A | TKT_HUMAN Tr | Unmodified | 0 | 0 | 0 | 0 | 5117100 | 933580 | 134470 | 0 | 0 | 3254300 | 0 | 0 | 380710 | 0 | 0 | 0 | 1287400 | 1465800 |
| INISEGNCI | PCBP1; PCBP3; | Q15365; P57 | PCBP1_HUMAI | Unmodified | 0 | 0 | 11353000 | 9371300 | 0 | 0 | 0 | 0 | 1969600 | 367180 | 2811700 | 16716000 | 475860 | 0 | 9747700 | 10449000 | 0 | 0 |
| INISEGNCI | PCBP2; PCBP2; | Q15366; F8V | PCBP2_HUMAI | Unmodified | 492460 | 0 | 195480 | 2725000 | 2463900 | 195920 | 4416500 | 0 | 0 | 878180 | 464060 | 0 | 20907000 | 4557500 | 0 | 0 | 0 | 0 |
| INISEGNCI | PCBP1 | Q15365 | PCBP1_HUMAI | Unmodified | 747090 | 0 | 0 | 1083500 | 0 | 0 | 1297400 | 0 | 565020 | 381080 | 1446600 | 0 | 1885100 | 743180 | 0 | 0 | 348220 | 0 |
| INISQVIAV | POLR2A; POLR2 | P24928; A0A | RPB1_HUMAN | Unmodified | 310010 | 0 | 0 | 0 | 0 | 0 | 0 | 0 | 201730 | 244850 | 340780 | 2027000 | 0 | 0 | 0 | 0 | 0 | 0 |
| INISQVIAV | POLR2A; POLR2 | P24928; A0A | RPB1_HUMAN | Unmodified | 0 | 0 | 0 | 0 | 0 | 0 | 361120 | 0 | 0 | 303450 | 0 | 0 | 92907 | 89165 | 0 | 0 | 139190 | 117150 |
| INIYHNTAS | EVL; EVL; EVL; E | Q9UI08; G3V | EVL_HUMAN Er | Unmodified | 0 | 0 | 0 | 0 | 0 | 0 | 0 | 0 | 627440 | 0 | 0 | 1106700 | 0 | 0 | 0 | 277980 | 194060 | 0 |
| INIYSNSLE | ATP11C; ATP11 | Q8NB49; A0. | AT11C_HUMAN | Unmodified | 0 | 0 | 157200 | 156850 | 200690 | 0 | 0 | 0 | 203740 | 114270 | 64044 | 2652400 | 0 | 0 | 382130 | 328950 | 122240 | 0 |
| INKESLLPV | UGP2; UGP2; U | Q16851; E7E | UGPA_HUMAN | Unmodified | 0 | 0 | 0 | 0 | 466380 | 0 | 63041 | 0 | 0 | 266210 | 0 | 0 | 191500 | 102280 | 0 | 0 | 0 | 0 |
| INKMEWW: | NUDC | Q9Y266 | NUDC_HUMAN | Unmodified | 0 | 0 | 0 | 0 | 0 | 0 | 0 | 0 | 1362700 | 654120 | 0 | 1572500 | 0 | 0 | 0 | 0 | 213440 | 0 |
| INKMEWW: | NUDC | Q9Y266 | NUDC_HUMAN | Unmodified | 0 | 0 | 0 | 0 | 113270 | 0 | 99683 | 0 | 0 | 135410 | 0 | 0 | 101470 | 0 | 0 | 0 | 0 | 118160 |
| INKPRAEDS | NPTN; NPTN | Q9Y639; Q9I | NPTN_HUMAN | Unmodified | 0 | 0 | 0 | 0 | 1049500 | 0 | 0 | 0 | 0 | 0 | 1016300 | 0 | 670970 | 0 | 0 | 0 | 0 | 795610 |
| INKSESVVY | MPZL1 | O95297 | MPZL1_HUMAN | Unmodified | 0 | 1880100 | 0 | 1215700 | 211130 | 37213 | 0 | 0 | 1183800 | 0 | 71551 | 1968100 | 0 | 0 | 298860 | 500330 | 79474 | 691190 |
| INKTEICSQ | STEAP1 | Q9UHE8 | STEA1_HUMAN | Unmodified | 2143400 | 0 | 0 | 8968500 | 0 | 0 | 0 | 0 | 0 | 0 | 0 | 0 | 4886300 | 3594600 | 10087000 | 8739800 | 0 | 0 |
| INLAEDVLA | NCLN; NCLN; N | Q969V3; K7E | NCLN_HUMAN | Unmodified | 0 | 0 | 0 | 0 | 0 | 0 | 0 | 0 | 9479900 | 1666400 | 0 | 0 | 0 | 0 | 0 | 0 | 0 | 0 |
| INLAEDVLA | NCLN; NCLN; N | Q969V3; K7E | NCLN_HUMAN | Unmodified | 0 | 0 | 0 | 0 | 0 | 0 | 0 | 0 | 2000700 | 0 | 0 | 0 | 0 | 0 | 0 | 0 | 0 | 0 |
| INLAVANT( | FARP1; FARP1; | Q9Y4F1; A0/ | FARP1_HUMAN | Unmodified | 0 | 75632 | 0 | 107270 | 47753 | 0 | 0 | 0 | 0 | 95620 | 0 | 0 | 0 | 0 | 719810 | 0 | 0 | 0 |
| INLIAPPR | EIF2S1; EIF2S1; | P05198; H0Y | IF2A_HUMAN E | Unmodified | 634370 | 549410 | 869130 | 1051000 | 276100 | 0 | 0 | 0 | 1097500 | 0 | 0 | 0 | 0 | 0 | 739010 | 627200 | 0 | 0 |
| INLQGDIVE | ADD1; ADD1; A | P35611; E7E | ADDA_HUMAN | Unmodified | 253210 | 66425 | 0 | 0 | 0 | 0 | 0 | 0 | 210590 | 0 | 0 | 0 | 0 | 0 | 632350 | 412540 | 0 | 0 |
| INLVLSR | DHFR2; DHFR; | Q86XF0; P0( | DYR2_HUMAN | Unmodified | 253780 | 350030 | 477480 | 382670 | 0 | 0 | 0 | 0 | 315430 | 122200 | 921150 | 334730 | 482240 | 0 | 604600 | 778420 | 0 | 0 |
| INLWHLEIT | PPP2R2A; PPP2 | P63151; Q6( | 2ABA_HUMAN | Unmodified | 0 | 0 | 132010 | 0 | 0 | 10903 | 0 | 0 | 0 | 0 | 0 | 1403300 | 0 | 0 | 0 | 0 | 0 | 0 |
| INMNGVNS | UBE2V1; UBE2V | Q13404; G3V | UB2V1_HUMAI | Unmodified | 117240 | 0 | 0 | 0 | 0 | 0 | 0 | 0 | 0 | 0 | 0 | 806290 | 0 | 0 | 0 | 0 | 0 | 0 |
| INMVEELEI | KARS1 | Q15046 | SYK_HUMAN Ly | Unmodified | 0 | 0 | 0 | 0 | 0 | 0 | 91472 | 0 | 184680 | 0 | 493980 | 2325300 | 105510 | 0 | 0 | 0 | 0 | 0 |
| INNAAFEA\ | PRPS2; PRPS2 | P11908; H7( | PRPS2_HUMAN | Unmodified | 456340 | 1395400 | 0 | 0 | 0 | 171800 | 1182900 | 352910 | 2488100 | 633510 | 0 | 3883000 | 0 | 0 | 387010 | 440560 | 0 | 0 |
| INNACFEA\ | PRPS1; PRPS1; | P60891; A0A | PRPS1_HUMAN | Unmodified | 23196 | 0 | 0 | 0 | 0 | 0 | 0 | 0 | 0 | 0 | 0 | 1903200 | 0 | 0 | 0 | 0 | 0 | 0 |
| INNFSADIK | PLS3 | P13797 | PLST_HUMAN I | Unmodified | 0 | 953730 | 0 | 0 | 3026900 | 1772300 | 0 | 296370 | 0 | 431970 | 0 | 0 | 0 | 0 | 0 | 0 | 5266000 | 508770 |
| INNIAVAVG | GPR180 | Q86V85 | GP180_HUMAN | Unmodified | 0 | 0 | 0 | 0 | 1275800 | 0 | 0 | 0 | 8324600 | 953330 | 168350 | 0 | 0 | 0 | 0 | 0 | 0 | 0 |
| INNVPAEGI | CYRIB; CYRIB | Q9NUQ9; A( | CYRIB_HUMAN | Unmodified | 0 | 0 | 959280 | 813600 | 924610 | 0 | 0 | 0 | 4647200 | 1032000 | 2734000 | 27329000 | 0 | 0 | 536190 | 694800 | 629060 | 459690 |
| INNYLTVPA | SLC9A1 | P19634 | SL9A1_HUMAN | Oxidation (M) | 390800 | 0 | 0 | 0 | 0 | 305540 | 0 | 0 | 68299 | 0 | 0 | 708640 | 1823000 | 0 | 746620 | 534230 | 0 | 0 |
| INNYLTVPA | SLC9A1 | P19634 | SL9A1_HUMAN | Unmodified | 100990 | 74595 | 0 | 240780 | 0 | 0 | 0 | 0 | 0 | 0 | 0 | 344060 | 1259800 | 0 | 225680 | 99065 | 0 | 0 |
| INPDEREEI | GCLM | P48507 | GSH0_HUMAN | Unmodified | 0 | 0 | 0 | 0 | 0 | 0 | 171090 | 118020 | 0 | 0 | 0 | 0 | 964690 | 407700 | 0 | 0 | 0 | 0 |
| INPDGSQS | CNPY2; N/A; CI | Q9Y2B0; F8V | CNPY2_HUMAI | Unmodified | 0 | 0 | 0 | 0 | 49220 | 0 | 0 | 0 | 55863000 | 480740 | 3105500 | 0 | 0 | 0 | 0 | 0 | 0 | 127790 |
| INPICNDH | PBK; PBK | Q96KB5; E5I | TOPK_HUMAN | Unmodified | 0 | 0 | 0 | 0 | 379070 | 0 | 0 | 106420 | 0 | 0 | 0 | 0 | 0 | 273470 | 0 | 0 | 353350 | 0 |
| INPSSMFD | PYGB | P11216 | PYGB_HUMAN | Unmodified | 0 | 0 | 0 | 0 | 263540 | 0 | 0 | 0 | 0 | 0 | 0 | 0 | 212450 | 0 | 0 | 0 | 410330 | 1746300 |
| INPYMSSP( | RPL17; RPL17; I | P18621; A0A | RL17_HUMAN ( | 2 Oxidation (M) | 724510 | 457760 | 264670 | 799390 | 0 | 76783 | 0 | 44661 | 0 | 0 | 457950 | 6884300 | 0 | 0 | 216830 | 410240 | 0 | 0 |
| INPYMSSP( | RPL17; RPL17; I | P18621; A0A | RL17_HUMAN ( | Oxidation (M) | 994770 | 0 | 0 | 0 | 0 | 0 | 0 | 0 | 0 | 0 | 0 | 2178400 | 0 | 0 | 484680 | 0 | 0 | 0 |
| INQDPLGI( | NAGA | P17050 | NAGAB_HUMA | Unmodified | 0 | 0 | 198460 | 0 | 0 | 0 | 0 | 62667 | 1460100 | 427900 | 0 | 0 | 0 | 0 | 116380 | 0 | 0 | 0 |
| INQDPQVI | PGM1 | P36871 | PGM1_HUMAN | Oxidation (M) | 0 | 0 | 0 | 0 | 0 | 0 | 0 | 0 | 0 | 250540 | 80118 | 0 | 0 | 0 | 0 | 0 | 0 | 675180 |
| INQEELAS( | FYB1 | O15117 | FYB1_HUMAN I | Unmodified | 0 | 0 | 1480100 | 0 | 0 | 0 | 0 | 0 | 0 | 0 | 0 | 2073100 | 0 | 0 | 0 | 0 | 0 | 0 |
| INQIQMK | CUL4B; CUL4A; | Q13620; Q1: | CUL4B_HUMAI | Unmodified | 0 | 0 | 0 | 0 | 0 | 0 | 0 | 0 | 108810 | 0 | 38532 | 925270 | 0 | 0 | 0 | 0 | 0 | 0 |
| INQMATAPI | SMC3 | Q9UQE7 | SMC3_HUMAN | Unmodified | 0 | 0 | 0 | 0 | 1272100 | 174040 | 294350 | 0 | 0 | 0 | 0 | 0 | 0 | 0 | 0 | 0 | 0 | 0 |
| INQVFHGS | PRKDC | P78527 | PRKDC_HUMAI | Unmodified | 0 | 0 | 0 | 0 | 0 | 0 | 407620 | 601590 | 0 | 0 | 0 | 0 | 0 | 0 | 0 | 0 | 0 | 0 |
| INQYPESN. | STEAP2; STEAP: | Q8NFT2; C9. | STEA2_HUMAN | Unmodified | 0 | 0 | 169390 | 162460 | 0 | 36636 | 0 | 0 | 0 | 0 | 0 | 0 | 0 | 0 | 800860 | 142500 | 0 | 0 |
| INREQFWE | DBN1; DBN1 | Q16643; D6I | DREB_HUMAN | Unmodified | 0 | 0 | 0 | 0 | 0 | 0 | 0 | 0 | 0 | 0 | 440080 | 1995500 | 0 | 0 | 0 | 0 | 0 | 0 |
| INRLSTPIA( | LMO7; LMO7; LI | Q8WWI1; E9 | LMO7_HUMAN | Unmodified | 0 | 0 | 0 | 0 | 0 | 0 | 1825600 | 0 | 0 | 228350 | 0 | 0 | 0 | 0 | 0 | 0 | 6226600 | 5597600 |
| INRSMQNV | CNN2; CNN2; ( | Q99439; H3I | CNN2_HUMAN | Oxidation (M) | 0 | 0 | 0 | 0 | 7330500 | 1207000 | 852560 | 0 | 0 | 0 | 0 | 0 | 0 | 1299600 | 0 | 0 | 1062000 | 0 |
| INSANGLEI | LSAMP; LSAMP; | Q13449; F5I | LSAMP_HUMAI | Unmodified | 65135 | 96631 | 71499 | 0 | 0 | 0 | 0 | 0 | 965400 | 0 | 0 | 0 | 0 | 0 | 179000 | 178170 | 0 | 0 |
| INSAPQQIE | NUP210 | Q8TEM1 | PO210_HUMAN | Unmodified | 0 | 0 | 0 | 0 | 0 | 0 | 0 | 0 | 3407100 | 705730 | 72229 | 0 | 0 | 0 | 0 | 0 | 0 | 0 |
| INSDDSNL | RAB34; RAB34 | Q9BZG1; C9 | RAB34_HUMAN | Unmodified | 0 | 0 | 0 | 0 | 0 | 0 | 0 | 0 | 296080 | 374780 | 0 | 1194200 | 0 | 0 | 0 | 0 | 0 | 0 |
| INSENADAI | UBR1 | Q8IWV7 | UBR1_HUMAN | Unmodified | 0 | 0 | 0 | 0 | 0 | 0 | 0 | 0 | 0 | 20721 | 0 | 158900 | 0 | 0 | 0 | 0 | 0 | 0 |
| INSGGKLPI | G3BP1; G3BP1; | Q13283; A0/ | G3BP1_HUMAI | Unmodified | 0 | 0 | 0 | 0 | 0 | 0 | 0 | 0 | 1760100 | 591590 | 2648100 | 8712200 | 0 | 0 | 0 | 0 | 0 | 0 |
| INSITVDNC | CAP1 | Q01518 | CAP1_HUMAN | Unmodified | 171130 | 88412 | 0 | 338760 | 2666100 | 423970 | 980200 | 0 | 1260700 | 632870 | 741050 | 4649200 | 1162100 | 910180 | 258720 | 374680 | 1196300 | 1788800 |
| INSITVDNC | CAP1 | Q01518 | CAP1_HUMAN | Unmodified | 0 | 0 | 0 | 0 | 0 | 0 | 392640 | 163220 | 1139700 | 221130 | 239390 | 1455300 | 927040 | 387600 | 0 | 209450 | 0 | 0 |
| INSQIKIDAI | UBA6; UBA6 | A0AVT1; H0Y | UBA6_HUMAN | Unmodified | 0 | 0 | 0 | 0 | 1206600 | 0 | 0 | 0 | 0 | 106050 | 0 | 0 | 0 | 93379 | 0 | 0 | 526460 | 457530 |
| INSSGESGI | EIF5B; EIF5B | O60841; A0/ | IF2P_HUMAN E | Unmodified | 0 | 0 | 0 | 0 | 0 | 0 | 0 | 0 | 0 | 64347 | 0 | 662290 | 0 | 0 | 0 | 0 | 0 | 0 |
| INSSLQLPI | SEMA4B; SEMA | Q9NPR2; H0 | SEM4B_HUMAI | Unmodified | 0 | 0 | 28138 | 0 | 0 | 88316 | 0 | 0 | 1128900 | 43175 | 0 | 0 | 0 | 0 | 0 | 0 | 0 | 0 |
| INSSSVKQI | ANLN; ANLN; A | Q9NQW6; H | ANLN_HUMAN | Unmodified | 0 | 0 | 0 | 478550 | 13776000 | 1741100 | 188300 | 78240 | 0 | 0 | 0 | 0 | 435410 | 1011200 | 0 | 0 | 6949100 | 4682200 |
| INSTATPDC | SLK | Q9H2G2 | SLK_HUMAN S | Unmodified | 126250 | 0 | 307940 | 61078 | 883150 | 0 | 0 | 0 | 930030 | 17977 | 186380 | 1794400 | 0 | 382680 | 487670 | 399760 | 0 | 0 |
| INSVEVYD( | PTK7; PTK7 | Q13308; H7( | PTK7_HUMAN I | Unmodified | 0 | 0 | 0 | 290630 | 0 | 0 | 0 | 0 | 2465900 | 0 | 0 | 0 | 0 | 0 | 426590 | 295820 | 0 | 0 |
| INSVGSTAS | KIT | P10721 | KIT_HUMAN Ma | Unmodified | 0 | 0 | 0 | 0 | 646150 | 0 | 0 | 0 | 2911400 | 275090 | 1896100 | 6092900 | 0 | 0 | 0 | 0 | 645180 | 0 |
| INSYGYGD | DNAJA3 | Q96EY1 | DNJA3_HUMAN | Unmodified | 0 | 0 | 0 | 0 | 0 | 0 | 0 | 0 | 5009700 | 233070 | 0 | 0 | 0 | 0 | 210440 | 0 | 0 | 0 |

| | | | | | | | | | | | | | | | | | | | | | | |
|---|---|---|---|---|---|---|---|---|---|---|---|---|---|---|---|---|---|---|---|---|---|---|
| INTFYSEK | XPR1 | Q9UBH6 | XPR1_HUMAN | Unmodified | 211030 | 0 | 309500 | 240620 | 1262700 | 0 | 271400 | 0 | 119720 | 75462 | 391710 | 0 | 301880 | 319690 | 156630 | 178010 | 57815 | 547640 |
| INTQEYLDV | CACNA2D1 | P54289 | CA2D1_HUMAI | Oxidation (M) | 3723600 | 2525700 | 0 | 0 | 0 | 0 | 0 | 0 | 4386200 | 261770 | 0 | 0 | 8441100 | 4354300 | 0 | 0 | 0 | 0 |
| INTQEYLDV | CACNA2D1 | P54289 | CA2D1_HUMAI | Unmodified | 0 | 0 | 0 | 0 | 0 | 0 | 0 | 0 | 5749900 | 0 | 0 | 0 | 0 | 596870 | 0 | 0 | 0 | 0 |
| INTQEYLDV | CACNA2D2; CA | Q9NY47; C9 | CA2D2_HUMAI | Oxidation (M) | 0 | 0 | 0 | 0 | 0 | 0 | 0 | 0 | 5708800 | 1426100 | 0 | 2030900 | 0 | 0 | 0 | 0 | 0 | 0 |
| INTQEYLDV | CACNA2D2; CA | Q9NY47; C9 | CA2D2_HUMAI | Unmodified | 0 | 0 | 0 | 0 | 0 | 0 | 142000 | 0 | 6165100 | 125730 | 0 | 0 | 0 | 0 | 0 | 0 | 0 | 0 |
| INVADIING | MGAT5 | Q09328 | MGT5A_HUMAI | Unmodified | 0 | 0 | 0 | 0 | 0 | 0 | 0 | 0 | 2953500 | 72788 | 173110 | 950910 | 0 | 0 | 0 | 0 | 0 | 0 |
| INVGGLNH | KCNB2 | Q92953 | KCNB2_HUMAI | Unmodified | 0 | 0 | 0 | 0 | 0 | 226440 | 19378 | 24018 | 1491300 | 627670 | 634110 | 3407000 | 151960 | 0 | 0 | 0 | 0 | 0 |
| INVGIGEIK | WLS; WLS; WLS | Q5T9L3; H0\ | WLS_HUMAN F | Unmodified | 0 | 0 | 0 | 0 | 0 | 0 | 119490 | 50320 | 556380 | 0 | 64713 | 336030 | 0 | 0 | 0 | 0 | 0 | 0 |
| INVGIGEIK | WLS; WLS; WLS | Q5T9L3; H0\ | WLS_HUMAN F | Unmodified | 0 | 0 | 0 | 0 | 0 | 0 | 0 | 0 | 1159000 | 140280 | 0 | 0 | 0 | 0 | 0 | 0 | 0 | 0 |
| INVMNRGI | ILK; ILK; ILK | Q13418; A0/ | ILK_HUMAN Int | Unmodified | 0 | 109830 | 0 | 362200 | 7254600 | 628670 | 5820700 | 0 | 0 | 87238 | 0 | 0 | 3022100 | 7116700 | 44079 | 0 | 1370000 | 6912900 |
| INVMNRGI | ILK; ILK; ILK | Q13418; A0/ | ILK_HUMAN Int | Oxidation (M) | 0 | 0 | 0 | 0 | 0 | 0 | 0 | 0 | 0 | 252290 | 0 | 0 | 1342000 | 0 | 0 | 0 | 172550 | 0 |
| INVNEIFYD | RAP1B; RAP1A; | P61224; P62 | RAP1B_HUMAN | Unmodified | 4555000 | 12757000 | 2392300 | 13765000 | 4201400 | 598310 | 1759700 | 0 | 1479300 | 1219400 | 3314900 | 4138300 | 8890600 | 4635500 | 22848000 | 3696500 | 933480 | 2681500 |
| INVNEIFYD | RAP1B; RAP1A; | P61224; P62 | RAP1B_HUMAN | Unmodified | 0 | 0 | 0 | 0 | 56480 | 0 | 39897 | 0 | 0 | 0 | 0 | 0 | 179790 | 155970 | 0 | 0 | 0 | 0 |
| INVNEIFYD | RAP1B; RAP1A; | P61224; P62 | RAP1B_HUMAN | Unmodified | 0 | 0 | 0 | 0 | 3092200 | 361840 | 4854900 | 0 | 0 | 0 | 27815 | 0 | 3663300 | 3875300 | 0 | 0 | 438690 | 907150 |
| INVPELPTP | ZW10 | O43264 | ZW10_HUMAN | Unmodified | 0 | 204270 | 0 | 0 | 0 | 0 | 0 | 0 | 0 | 0 | 169780 | 0 | 0 | 0 | 0 | 58973 | 0 | 0 |
| INVPVPQF | ITM2C; ITM2C; | Q9NQX7; E7 | ITM2C_HUMAN | Unmodified | 0 | 0 | 244420 | 0 | 0 | 0 | 0 | 0 | 8550000 | 1036100 | 388970 | 0 | 0 | 0 | 0 | 0 | 0 | 766110 |
| INVSGLTTK | GOT1 | P17174 | AATC_HUMAN | Unmodified | 1075900 | 329200 | 679860 | 2188300 | 1873700 | 0 | 0 | 0 | 3095800 | 892290 | 7145500 | 25622000 | 2813400 | 514040 | 2702000 | 2044800 | 283070 | 137460 |
| INVYSDLNI | ATP11A; ATP11 | P98196; H0 | AT11A_HUMAN | Unmodified | 0 | 0 | 0 | 305310 | 0 | 0 | 0 | 0 | 0 | 0 | 574050 | 679820 | 0 | 0 | 0 | 326190 | 0 | 0 |
| INVYYNEA | TUBB2A | Q13885 | TBB2A_HUMAN | Unmodified | 116140 | 0 | 156760 | 0 | 0 | 0 | 0 | 0 | 660000 | 0 | 0 | 3074900 | 0 | 0 | 114050 | 44826 | 60404 | 0 |
| INVYYNEA | TUBB4B | P68371 | TBB4B_HUMAN | Unmodified | 0 | 1309500 | 7467500 | 2060600 | 567470 | 0 | 192930 | 274600 | 0 | 0 | 5444800 | 46312000 | 0 | 0 | 4713100 | 2665500 | 0 | 177130 |
| INVYYNEA | TUBB4B | P68371 | TBB4B_HUMAN | Unmodified | 2091200 | 1276500 | 0 | 3688100 | 0 | 857890 | 0 | 422220 | 6968400 | 1546900 | 1775100 | 6777300 | 2323000 | 0 | 857490 | 655570 | 0 | 569530 |
| INVYYNEA | TUBB2B | Q9BVA1 | TBB2B_HUMAN | Unmodified | 0 | 0 | 0 | 0 | 0 | 0 | 0 | 0 | 6984700 | 149250 | 2366400 | 74050000 | 0 | 0 | 0 | 0 | 0 | 0 |
| INVYYNEA | TUBB2B | Q9BVA1 | TBB2B_HUMAN | Unmodified | 0 | 0 | 0 | 0 | 0 | 0 | 0 | 0 | 3995600 | 1014200 | 1183000 | 4416800 | 0 | 0 | 0 | 0 | 0 | 0 |
| INVYYNES | TUBB6; TUBB6; | Q9BUF5; K7 | TBB6_HUMAN | Unmodified | 0 | 0 | 0 | 0 | 0 | 0 | 0 | 0 | 311760 | 0 | 0 | 0 | 0 | 0 | 0 | 0 | 0 | 0 |
| INYTEGR | GPI; GPI; GPI; G | P06744; K7E | G6PI_HUMAN | Unmodified | 648460 | 499690 | 2125700 | 2249200 | 5645600 | 1151000 | 0 | 186300 | 7316400 | 1979900 | 875350 | 36393000 | 2402000 | 757480 | 1991100 | 1416200 | 553150 | 1367100 |
| INYTEGRA | GPI; GPI; GPI; G | P06744; K7E | G6PI_HUMAN | Unmodified | 0 | 0 | 0 | 0 | 8253100 | 250050 | 335110 | 0 | 0 | 4210500 | 0 | 0 | 3331200 | 1620800 | 0 | 0 | 1613100 | 1296100 |
| IPADQGIAC | PDE2A; PDE2A; | O00408; H0 | PDE2A_HUMAN | Unmodified | 0 | 0 | 0 | 0 | 0 | 0 | 0 | 0 | 0 | 0 | 0 | 2088100 | 0 | 0 | 0 | 0 | 0 | 0 |
| IPADTFAAL | IQGAP1; IQGAP | P46940; A0/ | IQGA1_HUMAN | Unmodified | 0 | 0 | 63951 | 39243 | 107070 | 0 | 165950 | 19679 | 0 | 1477200 | 614170 | 4097700 | 0 | 0 | 169750 | 195340 | 0 | 0 |
| IPAEVSQAT | LRRC1 | Q9BTT6 | LRRC1_HUMAN | Unmodified | 253260 | 0 | 500950 | 436920 | 0 | 0 | 1820400 | 0 | 3268200 | 0 | 1026700 | 5562100 | 0 | 0 | 605820 | 713550 | 0 | 0 |
| IPAEVSQAT | LRRC1 | Q9BTT6 | LRRC1_HUMAN | Unmodified | 0 | 0 | 0 | 67701 | 298250 | 0 | 0 | 0 | 0 | 139030 | 0 | 0 | 324250 | 0 | 0 | 0 | 99876 | 146710 |
| IPAHQVLYS | MYOF; MYOF | Q9NZM1; H | MYOF_HUMAN | Unmodified | 0 | 0 | 0 | 0 | 1430100 | 813900 | 0 | 0 | 211560 | 0 | 0 | 0 | 0 | 0 | 0 | 16646 | 0 | 0 |
| IPAHQVLYS | MYOF; MYOF | Q9NZM1; H | MYOF_HUMAN | Unmodified | 0 | 0 | 0 | 0 | 789110 | 0 | 0 | 0 | 0 | 0 | 0 | 0 | 0 | 0 | 0 | 0 | 0 | 64080 |
| IPALDLLIK | PRKDC; PRKDC | P78527; F5C | PRKDC_HUMAI | Unmodified | 71728 | 81293 | 0 | 0 | 62913 | 0 | 63814 | 81909 | 155900 | 107780 | 126710 | 139340 | 0 | 0 | 106110 | 93522 | 59507 | 55365 |
| IPAMTIAK | HSPD1; HSPD1 | P10809; A0/ | CH60_HUMAN | Unmodified | 145030 | 68476 | 160860 | 264040 | 0 | 0 | 987250 | 0 | 28920000 | 2630500 | 917900 | 0 | 2044500 | 1486200 | 167640 | 92127 | 192260 | 570200 |
| IPASWTNP | TPP2; TPP2; TPI | P29144; A0/ | TPP2_HUMAN | Unmodified | 0 | 0 | 0 | 0 | 0 | 0 | 0 | 51536 | 0 | 0 | 609530 | 6235700 | 0 | 0 | 0 | 0 | 0 | 0 |
| IPAYFVTVS | VARS1; VARS1; | P26640; H0 | SYVC_HUMAN | Unmodified | 38697 | 0 | 0 | 0 | 0 | 0 | 0 | 0 | 0 | 0 | 0 | 1420600 | 0 | 0 | 180120 | 0 | 0 | 0 |
| IPDEELR | SH3GL2; SH3G | Q99962; Q9 | SH3G2_HUMAI | Unmodified | 0 | 9508100 | 14570000 | 0 | 0 | 39119 | 0 | 7534.2 | 1225900 | 0 | 0 | 509520 | 0 | 0 | 11469000 | 13071000 | 1397500 | 0 |
| IPDEELRQ | SH3GL2; SH3G | Q99962; Q9 | SH3G2_HUMAI | Unmodified | 3557200 | 1668800 | 9559500 | 0 | 0 | 0 | 0 | 0 | 0 | 0 | 0 | 0 | 0 | 0 | 5863100 | 3032100 | 96496 | 0 |
| IPDEELRQ | SH3GL1 | Q99961 | SH3G1_HUMAI | Unmodified | 2408800 | 1869500 | 0 | 3383200 | 0 | 0 | 0 | 0 | 0 | 0 | 0 | 0 | 0 | 0 | 0 | 0 | 0 | 0 |
| IPDISSFLL | POMK; POMK; F | Q9H5K3; A0 | SG196_HUMAN | Oxidation (M) | 0 | 0 | 0 | 0 | 0 | 0 | 0 | 0 | 745650 | 117620 | 28394 | 42553 | 0 | 0 | 0 | 0 | 0 | 0 |
| IPDISSFLL | POMK; POMK; F | Q9H5K3; A0 | SG196_HUMAN | Unmodified | 162080 | 675900 | 0 | 0 | 26022 | 0 | 0 | 0 | 279740 | 0 | 0 | 0 | 32385 | 0 | 27962 | 0 | 0 | 0 |
| IPDIVLWPT | AGPS; AGPS; A | O00116; A0/ | ADAS_HUMAN | Unmodified | 0 | 127290 | 0 | 0 | 0 | 0 | 198630 | 83939 | 5930100 | 983850 | 423980 | 0 | 0 | 0 | 0 | 0 | 0 | 0 |
| IPDPEAVKF | CANX; CANX; C | P27824; A0/ | CALX_HUMAN | Unmodified | 0 | 0 | 0 | 0 | 0 | 0 | 0 | 0 | 1479400 | 169010 | 0 | 0 | 0 | 0 | 0 | 0 | 0 | 0 |
| IPDPSAVKF | CLGN | O14967 | CLGN_HUMAN | Unmodified | 0 | 0 | 0 | 0 | 0 | 0 | 0 | 0 | 4739600 | 670660 | 845240 | 0 | 0 | 0 | 0 | 0 | 0 | 0 |
| IPDQLVILD | TMED10; TMED | P49755; G3 | TMEDA_HUMAI | Oxidation (M) | 0 | 0 | 0 | 0 | 0 | 0 | 0 | 0 | 15577000 | 1616200 | 1966100 | 1994900 | 0 | 0 | 272310 | 376200 | 0 | 0 |
| IPDQLVILD | TMED10; TMED | P49755; G3 | TMEDA_HUMAI | Unmodified | 0 | 0 | 0 | 0 | 0 | 0 | 0 | 0 | 72659000 | 1199100 | 700570 | 0 | 0 | 0 | 0 | 0 | 0 | 0 |
| IPDWFLNR | RPS18; RPS18 | P62269; J3J | RS18_HUMAN | Unmodified | 4073000 | 0 | 0 | 0 | 0 | 0 | 0 | 0 | 0 | 0 | 0 | 51004000 | 0 | 0 | 0 | 0 | 0 | 0 |
| IPDWFLNR | RPS18; RPS18 | P62269; J3J | RS18_HUMAN | Unmodified | 665780 | 707830 | 919270 | 1858000 | 1599000 | 0 | 0 | 0 | 0 | 0 | 0 | 2034600 | 0 | 0 | 450940 | 624790 | 132780 | 0 |
| IPEFIAFR | NAA25 | Q14CX7 | NAA25_HUMAI | Unmodified | 0 | 44638 | 34797 | 0 | 0 | 0 | 0 | 0 | 0 | 0 | 87855 | 2213300 | 0 | 0 | 29582 | 63144 | 0 | 248990 |
| IPEFSSFVV | DARS2; DARS2; | Q6PI48; A0/ | SYDM_HUMAN | Unmodified | 0 | 0 | 0 | 0 | 0 | 0 | 0 | 0 | 1860300 | 0 | 0 | 0 | 0 | 0 | 0 | 0 | 0 | 0 |
| IPEGLFDPS | ACTL6A; ACTL6 | O96019; O9 | ACL6A_HUMAN | Unmodified | 0 | 0 | 0 | 0 | 0 | 0 | 0 | 0 | 326360 | 158970 | 900400 | 5294400 | 0 | 0 | 0 | 0 | 0 | 0 |
| IPEINSSDN | FLNB; FLNB | O75369; E7E | FLNB_HUMAN | Oxidation (M) | 0 | 0 | 96206 | 0 | 0 | 0 | 0 | 0 | 0 | 0 | 0 | 1224400 | 0 | 0 | 0 | 0 | 0 | 0 |
| IPEIQATMR | CHMP3 | Q9Y3E7 | CHMP3_HUMA | Unmodified | 0 | 0 | 0 | 0 | 0 | 0 | 0 | 0 | 2651100 | 0 | 0 | 0 | 0 | 0 | 0 | 0 | 0 | 0 |
| IPEQILGK | MAP2K1 | Q02750 | MP2K1_HUMAI | Unmodified | 99532 | 8364.6 | 0 | 100060 | 1353800 | 166490 | 0 | 17861 | 239370 | 0 | 289020 | 767550 | 1242400 | 510130 | 186460 | 217250 | 0 | 0 |
| IPFRVGLYT | PCDHA2; PCDH | Q9Y5H9; Q9 | PCDA2_HUMAI | Unmodified | 0 | 0 | 0 | 0 | 0 | 0 | 0 | 0 | 1380500 | 267210 | 0 | 0 | 0 | 0 | 0 | 0 | 0 | 0 |
| IPFTVSLDN | SUSD2 | Q9UGT4 | SUSD2_HUMAI | Unmodified | 0 | 0 | 0 | 0 | 0 | 0 | 0 | 0 | 1914700 | 0 | 0 | 393250 | 0 | 0 | 0 | 0 | 0 | 0 |
| IPGDQVVS | HSPG2; HSPG2 | P98160; A0/ | PGBM_HUMAN | Unmodified | 0 | 0 | 0 | 0 | 0 | 0 | 0 | 0 | 0 | 0 | 0 | 0 | 0 | 0 | 0 | 0 | 0 | 0 |
| IPGEKDSVI | CD55; CD55; C | P08174; B1/ | DAF_HUMAN C | Unmodified | 0 | 0 | 0 | 0 | 987600 | 0 | 64516 | 0 | 0 | 0 | 0 | 0 | 0 | 0 | 0 | 0 | 858430 | 3758700 |
| IPGGIIEDS | CCT3; CCT3; C | P49368; E9P | TCPG_HUMAN | Unmodified | 0 | 0 | 107600 | 0 | 0 | 0 | 0 | 0 | 1568200 | 0 | 3735700 | 16370000 | 0 | 0 | 155630 | 122040 | 0 | 0 |
| IPGIIIAASA | MCM5; MCM5; | P33992; B1/ | MCM5_HUMAN | Unmodified | 67712 | 0 | 99563 | 27044 | 107020 | 25347 | 0 | 48957 | 0 | 141090 | 297880 | 634260 | 71201 | 0 | 0 | 33801 | 0 | 0 |
| IPGLLGVF | CSE1L | P55060 | XPO2_HUMAN | Unmodified | 0 | 0 | 0 | 0 | 0 | 0 | 0 | 0 | 0 | 0 | 0 | 965100 | 0 | 0 | 0 | 0 | 0 | 0 |
| IPGLLSPHF | FASN; FASN | P49327; A0/ | FAS_HUMAN F | Unmodified | 1335400 | 302040 | 1345000 | 151760 | 0 | 353440 | 0 | 0 | 0 | 0 | 53504 | 64966000 | 0 | 0 | 4745700 | 2021000 | 0 | 0 |
| IPGMLIIDT | EIF5B; EIF5B | O60841; A0/ | IF2P_HUMAN E | Oxidation (M) | 0 | 0 | 0 | 0 | 0 | 0 | 0 | 0 | 52097 | 0 | 0 | 2231700 | 264780 | 341130 | 0 | 0 | 0 | 0 |
| IPGNIMSAI | TMEM68; TMEM | Q96MH6; Q | TMM68_HUMAI | Unmodified | 0 | 0 | 0 | 0 | 243240 | 55336 | 16122 | 0 | 181380 | 32920 | 0 | 0 | 0 | 0 | 0 | 0 | 101300 | 304480 |
| IPGPGTPV | SIT1; SIT1 | Q9Y3P8; A0/ | SIT1_HUMAN S | Unmodified | 0 | 0 | 0 | 0 | 0 | 0 | 0 | 0 | 0 | 1780800 | 0 | 0 | 0 | 0 | 0 | 0 | 0 | 0 |
| IPGSNTRLE | ARHGAP39 | Q9C0H5 | RHG39_HUMAI | Unmodified | 117210 | 120700 | 0 | 0 | 0 | 0 | 0 | 0 | 519060 | 0 | 58422 | 89882 | 0 | 0 | 0 | 0 | 0 | 0 |
| IPGSPPESN | GDI2; GDI2; GD | P50395; V9 | GDIB_HUMAN | Oxidation (M) | 190130 | 0 | 665250 | 634590 | 0 | 0 | 0 | 0 | 0 | 0 | 0 | 2186000 | 0 | 0 | 962940 | 66678 | 0 | 0 |
| IPGVPWCF | TFF3 | Q07654 | TFF3_HUMAN | Unmodified | 0 | 0 | 0 | 0 | 0 | 20405 | 0 | 0 | 3255400 | 35408 | 0 | 0 | 0 | 0 | 0 | 0 | 0 | 0 |
| IPHGVSGV | DPYSL5 | Q9BPU6 | DPYL5_HUMAN | Unmodified | 0 | 0 | 0 | 0 | 0 | 0 | 0 | 0 | 206830 | 0 | 97099 | 1322300 | 0 | 0 | 0 | 0 | 0 | 0 |
| IPHSIFSLH | LRRC8A | Q8IWT6 | LRC8A_HUMAN | Unmodified | 0 | 0 | 0 | 0 | 0 | 0 | 0 | 0 | 0 | 0 | 0 | 293680 | 0 | 0 | 0 | 0 | 80534 | 92126 |
| IPIEIVNK | STEAP2; STEAP | Q8NFT2; B5 | STEA2_HUMAN | Unmodified | 108000 | 0 | 0 | 0 | 249050 | 0 | 0 | 0 | 0 | 0 | 0 | 0 | 0 | 0 | 1569400 | 1012600 | 389460 | 164650 |
| IPIFSAAGL | ATP6V1B2; ATP | P21281; P15 | VATB2_HUMAN | Unmodified | 0 | 0 | 0 | 0 | 83022 | 0 | 0 | 0 | 1275700 | 43770 | 493640 | 3772600 | 0 | 0 | 0 | 0 | 0 | 0 |
| IPILFFANK | ARL6; ARL6; AR | Q9H0F7; H7 | ARL6_HUMAN | Unmodified | 0 | 0 | 71128 | 0 | 0 | 0 | 0 | 0 | 0 | 0 | 0 | 750630 | 21073 | 0 | 0 | 148790 | 0 | 0 |
| IPILVINK | STEAP1B; STEA | Q6NZ63; Q9 | STEAL_HUMAN | Unmodified | 0 | 11620 | 1108200 | 0 | 2259100 | 0 | 42098 | 107970 | 0 | 77041 | 0 | 0 | 56350 | 0 | 3169500 | 2021300 | 557140 | 239450 |
| IPISLVLR | OXSR1; OXSR1 | O95747; C9 | OXSR1_HUMAN | Unmodified | 0 | 0 | 0 | 0 | 0 | 0 | 0 | 0 | 0 | 0 | 0 | 925390 | 0 | 0 | 0 | 0 | 0 | 0 |
| IPKEEQEET | SEC24D; SEC2 | O94855; E9 | SC24D_HUMAI | Unmodified | 0 | 0 | 0 | 0 | 807590 | 0 | 0 | 0 | 0 | 0 | 0 | 522490 | 0 | 0 | 0 | 0 | 48484 | 0 |
| IPLAEWESI | UQCRC1 | P31930 | QCR1_HUMAN | Unmodified | 0 | 0 | 0 | 0 | 0 | 0 | 0 | 0 | 4187600 | 0 | 603400 | 0 | 0 | 0 | 0 | 0 | 0 | 0 |
| IPLLSDLTH | PRDX4; PRDX4; | Q13162; A6 | PRDX4_HUMAI | Unmodified | 40884 | 0 | 36441 | 0 | 0 | 26947 | 0 | 27762 | 3507000 | 502960 | 380560 | 1118600 | 0 | 0 | 94923 | 72050 | 41662 | 0 |
| IPLLVVGN | RALB; RALB; RA | P11234; C9 | RALB_HUMAN | Unmodified | 45119 | 0 | 0 | 0 | 0 | 0 | 0 | 0 | 0 | 0 | 318630 | 835820 | 0 | 0 | 0 | 0 | 0 | 0 |
| IPLLVVGN | RALB; RALB; RA | P11234; C9 | RALB_HUMAN | Unmodified | 0 | 0 | 0 | 0 | 1174300 | 0 | 0 | 0 | 0 | 110880 | 248440 | 3158100 | 0 | 614630 | 537240 | 458200 | 378440 | 774450 |
| IPLLVVGN | RALB; RALB; RA | P11234; C9 | RALB_HUMAN | Unmodified | 0 | 0 | 0 | 0 | 0 | 0 | 0 | 0 | 0 | 0 | 0 | 1189900 | 0 | 0 | 330520 | 343920 | 0 | 0 |
| IPLLYDAEI | ITGA2; ITGA2 | P17301; E7E | ITA2_HUMAN I | Unmodified | 9386500 | 0 | 16777000 | 675700 | 13010000 | 0 | 0 | 0 | 0 | 311750 | 0 | 0 | 2211700 | 0 | 0 | 23024000 | 4186400 | 3878100 |
| IPLNDLFR | LAMP2 | P13473 | LAMP2_HUMAN | Unmodified | 317190 | 867200 | 1068000 | 1400900 | 0 | 0 | 0 | 0 | 12564000 | 1036800 | 58296 | 349900 | 45607 | 0 | 870990 | 1170000 | 0 | 0 |
| IPLNKFPGF | SNX9; SNX9; SI | Q9Y5X1; A0/ | SNX9_HUMAN | Unmodified | 253150 | 0 | 0 | 0 | 0 | 0 | 0 | 0 | 0 | 0 | 0 | 0 | 0 | 0 | 0 | 0 | 0 | 0 |
| IPLNKFPGF | SNX9; SNX9; SI | Q9Y5X1; A0/ | SNX9_HUMAN | Unmodified | 76678000 | 39574000 | 0 | 799130 | 2787600 | 0 | 0 | 0 | 2007800 | 268670 | 880440 | 10604000 | 974270 | 0 | 688260 | 768440 | 0 | 0 |
| IPLNNGAG | ERGIC1; ERGIC | Q969X5; F2 | ERGI1_HUMAN | Unmodified | 0 | 0 | 289890 | 0 | 0 | 0 | 0 | 0 | 3579600 | 0 | 0 | 0 | 0 | 0 | 189810 | 81682 | 0 | 0 |
| IPLPAGYRE | LSS; LSS | P48449; C9 | LSS_HUMAN La | Unmodified | 178150 | 154730 | 0 | 0 | 0 | 0 | 0 | 0 | 0 | 115240 | 48100 | 2367100 | 0 | 0 | 0 | 0 | 0 | 0 |
| IPLSQEEIT | MCCC1; MCCC | Q96RQ3; E9 | MCCA_HUMAN | Unmodified | 0 | 0 | 0 | 0 | 0 | 0 | 0 | 43072 | 1886400 | 0 | 0 | 0 | 0 | 0 | 0 | 0 | 0 | 0 |
| IPLVNISLLF | GNPTAB | Q3T906 | GNPTA_HUMAI | Unmodified | 0 | 0 | 0 | 0 | 0 | 0 | 0 | 0 | 333670 | 0 | 37679 | 0 | 0 | 0 | 0 | 0 | 0 | 0 |
| IPMPDFDLI | AHNAK | Q09666 | AHNK_HUMAN | Unmodified | 0 | 0 | 0 | 0 | 1274000 | 113830 | 0 | 0 | 0 | 0 | 0 | 0 | 0 | 0 | 0 | 0 | 0 | 784030 |
| IPNIYAIGD | DLD; DLD | P09622; E9P | DLDH_HUMAN | Oxidation (M) | 0 | 0 | 0 | 0 | 0 | 0 | 0 | 0 | 3023400 | 854370 | 851810 | 0 | 1953900 | 737990 | 0 | 0 | 0 | 156990 |
| IPNIYAIGD | DLD; DLD | P09622; E9P | DLDH_HUMAN | Unmodified | 0 | 0 | 0 | 0 | 0 | 0 | 99558 | 0 | 2017200 | 203890 | 0 | 0 | 0 | 0 | 0 | 927890 | 0 | 0 |
| IPNPDFFEI | CANX; CANX; C | P27824; A0/ | CALX_HUMAN | Unmodified | 0 | 0 | 0 | 0 | 0 | 0 | 229140 | 0 | 5884100 | 1303400 | 1079000 | 537000 | 0 | 0 | 0 | 23864 | 0 | 183110 |
| IPPPVIMVQ | ABCF2 | Q9UG63 | ABCF2_HUMAN | Oxidation (M) | 47838 | 0 | 100890 | 33968 | 0 | 0 | 0 | 0 | 0 | 0 | 17475 | 1748500 | 0 | 0 | 187480 | 263630 | 0 | 0 |
| IPPYHYHV | MVP; MVP; MVF | Q14764; H3 | MVP_HUMAN M | Unmodified | 384660 | 477390 | 0 | 0 | 1327300 | 0 | 42036 | 0 | 360530 | 0 | 0 | 18169000 | 0 | 0 | 0 | 0 | 310440 | 0 |
| IPPYHYHV | MVP; MVP; MVF | Q14764; H3 | MVP_HUMAN M | Unmodified | 641390 | 2439500 | 0 | 0 | 0 | 0 | 0 | 0 | 1546700 | 656600 | 0 | 38728000 | 0 | 0 | 0 | 0 | 0 | 0 |
| IPQAIAQLS | NDUFA9; N/A | Q16795; A0/ | NDUA9_HUMA | Unmodified | 0 | 0 | 0 | 0 | 0 | 0 | 0 | 0 | 2978000 | 150750 | 387800 | 2704600 | 0 | 0 | 0 | 0 | 0 | 0 |
| IPQDRPTSI | TAOK1 | Q7L7X3 | TAOK1_HUMAN | Unmodified | 200240 | 0 | 0 | 0 | 367340 | 0 | 0 | 0 | 701810 | 195940 | 499530 | 2257200 | 201360 | 0 | 638760 | 0 | 242270 | 0 |
| IPRDLFEDE | SYNCRIP; SYNC | O60506; B7 | HNRPQ_HUMA | Unmodified | 0 | 0 | 119590 | 0 | 0 | 0 | 195270 | 0 | 361840 | 238860 | 462940 | 541790 | 0 | 0 | 229480 | 89977 | 0 | 0 |
| IPRDLYEDE | HNRNPR; HNRI | O43390; B4 | HNRPR_HUMA | Unmodified | 0 | 0 | 0 | 0 | 0 | 0 | 496060 | 0 | 330850 | 0 | 552070 | 882530 | 0 | 0 | 0 | 0 | 0 | 0 |
| IPRHELTEI | AHNAK | Q09666 | AHNK_HUMAN | Unmodified | 0 | 0 | 0 | 0 | 5693000 | 896010 | 0 | 0 | 0 | 0 | 0 | 0 | 0 | 0 | 0 | 0 | 1005300 | 1840800 |
| IPSAVGYQ | ATP5F1B; ATP5 | P06576; H0 | ATPB_HUMAN | 2 Oxidation (M) | 0 | 0 | 0 | 0 | 0 | 0 | 0 | 0 | 1188300 | 0 | 209490 | 0 | 0 | 0 | 0 | 0 | 0 | 0 |
| IPSAVGYQ | ATP5F1B; ATP5 | P06576; H0 | ATPB_HUMAN | Unmodified | 0 | 0 | 0 | 0 | 0 | 0 | 0 | 0 | 2820500 | 0 | 0 | 0 | 0 | 0 | 0 | 0 | 0 | 0 |
| IPSAVSTVS | SLC9A1 | P19634 | SL9A1_HUMAN | Unmodified | 85258 | 0 | 468390 | 118630 | 0 | 0 | 0 | 0 | 0 | 181720 | 138150 | 528430 | 108680 | 0 | 486790 | 306300 | 0 | 0 |
| IPSAVSTVS | SLC9A1 | P19634 | SL9A1_HUMAN | Oxidation (M) | 158040 | 95855 | 644260 | 135710 | 0 | 0 | 0 | 0 | 0 | 0 | 204360 | 0 | 0 | 0 | 1269400 | 700250 | 0 | 0 |
| IPSDTFYDI | DAG1 | Q14118 | DAG1_HUMAN | Unmodified | 680150 | 0 | 0 | 0 | 279250 | 0 | 0 | 0 | 0 | 0 | 0 | 0 | 0 | 0 | 0 | 0 | 0 | 0 |
| IPSEQEQLI | UPF1 | Q92900 | RENT1_HUMAN | Unmodified | 196230 | 149390 | 147410 | 550960 | 592570 | 108110 | 0 | 0 | 0 | 159990 | 0 | 0 | 923530 | 341250 | 95242 | 0 | 125330 | 309050 |
| IPSISDLK | GFM1; GFM1; G | Q96RP9; C9 | EFGM_HUMAN | Unmodified | 0 | 0 | 0 | 0 | 0 | 0 | 0 | 0 | 838150 | 0 | 0 | 0 | 0 | 0 | 0 | 0 | 0 | 0 |
| IPSTPDSFN | PSMB4 | P28070 | PSB4_HUMAN | Oxidation (M) | 0 | 0 | 0 | 152880 | 0 | 0 | 0 | 0 | 0 | 0 | 0 | 0 | 2821000 | 0 | 0 | 3954600 | 0 | 0 |

| | | | | | | | | | | | | | | | | | | | | | | |
|---|---|---|---|---|---|---|---|---|---|---|---|---|---|---|---|---|---|---|---|---|---|---|
| IPTEAPQLE | BROX | Q5VW32 | BROX_HUMAN | Unmodified | 13160000 | 12580000 | 4158000 | 5559900 | 13152000 | 2880400 | 622910 | 502420 | 10810000 | 3543600 | 3383400 | 29094000 | 6765200 | 2187500 | 6840800 | 4673900 | 291700 | 2692500 |
| IPTGQEYA | CAMK2D; CAMI | Q13557; E9F | KCC2D_HUMA | Unmodified | 0 | 0 | 0 | 0 | 0 | 0 | 53639 | 29796 | 604370 | 83621 | 61849 | 286500 | 67707 | 0 | 0 | 0 | 0 | 495440 |
| IPTHLFTFIC | RO60; RO60 | P10155; H0Y | RO60_HUMAN | Unmodified | 32264000 | 6094600 | 0 | 0 | 0 | 0 | 0 | 0 | 0 | 0 | 0 | 0 | 0 | 0 | 22704000 | 7728600 | 0 | 0 |
| IPTHLFTFIC | RO60; RO60 | P10155; H0Y | RO60_HUMAN | Unmodified | 7644700 | 14033000 | 4334200 | 3091400 | 0 | 0 | 0 | 0 | 0 | 0 | 0 | 880800 | 0 | 0 | 11874000 | 4808200 | 0 | 0 |
| IPTISTQLK | VCL | P18206 | VINC_HUMAN | Unmodified | 0 | 0 | 0 | 0 | 0 | 0 | 0 | 0 | 0 | 0 | 479220 | 3570700 | 0 | 0 | 0 | 64735 | 220110 | 0 |
| IPTTFENGR | APOD; APOD; A | P05090; C9J | APOD_HUMAN | Unmodified | 35731 | 0 | 0 | 0 | 0 | 0 | 0 | 0 | 62326000 | 742120 | 0 | 0 | 0 | 0 | 0 | 0 | 0 | 0 |
| IPTYNTPLK | KIF23; KIF23; KI | Q02241; H7I | KIF23_HUMAN | Unmodified | 0 | 0 | 329860 | 182110 | 1486100 | 275470 | 811920 | 144580 | 1149100 | 96688 | 850600 | 2006600 | 101500 | 46489 | 437490 | 149450 | 106810 | 172320 |
| IPVDEEAFV | PAM | P19021 | AMD_HUMAN | Unmodified | 0 | 0 | 0 | 0 | 0 | 0 | 0 | 0 | 1673100 | 0 | 0 | 354930 | 0 | 0 | 0 | 0 | 0 | 0 |
| IPVDTYNNI | VPS35 | Q96QK1 | VPS35_HUMAN | Unmodified | 0 | 0 | 549590 | 0 | 0 | 0 | 0 | 0 | 638630 | 279670 | 1002400 | 18917000 | 0 | 0 | 1369500 | 782820 | 0 | 0 |
| IPVFEDEQS | CNNM4 | Q6P4Q7 | CNNM4_HUMA | Unmodified | 0 | 0 | 48605 | 0 | 0 | 0 | 0 | 0 | 0 | 0 | 0 | 0 | 0 | 0 | 290060 | 79466 | 0 | 0 |
| IPVFHNGS | SLC4A7; SLC4A | Q9Y6M7; C9 | S4A7_HUMAN | Unmodified | 0 | 0 | 0 | 0 | 0 | 0 | 0 | 0 | 0 | 0 | 0 | 1067300 | 0 | 0 | 0 | 0 | 44931 | 201570 |
| IPVGPETLG | ATP5F1B; ATP5 | P06576; H0Y | ATPB_HUMAN | Unmodified | 0 | 0 | 330720 | 0 | 0 | 0 | 0 | 0 | 9276900 | 172120 | 0 | 383330 | 0 | 0 | 0 | 0 | 0 | 0 |
| IPVHPNDH | FH | P07954 | FUMH_HUMAN | Unmodified | 0 | 0 | 46555 | 0 | 0 | 0 | 0 | 0 | 1137700 | 0 | 664660 | 19155 | 464260 | 0 | 0 | 0 | 0 | 0 |
| IPVIAVDSD | EHHADH | Q08426 | ECHP_HUMAN | Unmodified | 0 | 0 | 0 | 0 | 0 | 0 | 0 | 0 | 991230 | 220550 | 0 | 0 | 0 | 0 | 0 | 0 | 0 | 0 |
| IPVPAERITL | DVL2; DVL2; DV | O14641; I3L | DVL2_HUMAN | Unmodified | 0 | 0 | 208500 | 0 | 455290 | 135930 | 0 | 57544 | 197700 | 217540 | 48096 | 0 | 1585600 | 1615700 | 0 | 0 | 119260 | 382710 |
| IPVQLVFK | CD109 | Q6YHK3 | CD109_HUMAN | Unmodified | 0 | 0 | 0 | 0 | 0 | 0 | 0 | 0 | 2172700 | 0 | 0 | 0 | 0 | 0 | 0 | 0 | 0 | 0 |
| IPVSGPFLV | IGF2BP3; IGF2E | O00425; F8V | IF2B3_HUMAN | Unmodified | 869220 | 439890 | 300820 | 324460 | 0 | 0 | 53062 | 39592 | 54346 | 0 | 335040 | 1737200 | 217240 | 85750 | 198740 | 96816 | 0 | 30505 |
| IPVTIITGYL | CBWD2; CBWD | Q8IUF1; Q4V | CBWD2_HUMA | Unmodified | 0 | 0 | 0 | 0 | 0 | 0 | 0 | 0 | 0 | 0 | 0 | 936360 | 0 | 0 | 0 | 0 | 0 | 0 |
| IPVTLNMK | LYPLA2; LYPLA2 | O95372; Q5 | LYPA2_HUMAN | Unmodified | 218690 | 166030 | 85509 | 0 | 0 | 0 | 35048 | 0 | 210030 | 23590 | 0 | 4468700 | 0 | 0 | 180090 | 108920 | 0 | 0 |
| IPVYEEERS | CNNM3 | Q8NE01 | CNNM3_HUMA | Oxidation (M) | 48218 | 0 | 0 | 33070 | 0 | 0 | 0 | 0 | 0 | 0 | 0 | 486660 | 0 | 0 | 643530 | 190430 | 0 | 0 |
| IPWSEFFDI | POFUT2; POFU | Q9Y2G5; S6 | OFUT2_HUMAN | Unmodified | 0 | 0 | 0 | 0 | 0 | 0 | 0 | 0 | 238730 | 20392 | 0 | 0 | 0 | 0 | 0 | 0 | 0 | 0 |
| IPYLPITNFN | FLNB | O75369 | FLNB_HUMAN | Unmodified | 139540 | 0 | 1339700 | 689550 | 1244700 | 513570 | 0 | 0 | 0 | 45853 | 620970 | 6385200 | 206630 | 0 | 1417800 | 875520 | 424520 | 700220 |
| IPYYNIHPM | H6PD | O95479 | G6PE_HUMAN | Unmodified | 0 | 0 | 0 | 0 | 0 | 0 | 0 | 0 | 1199300 | 0 | 0 | 0 | 0 | 0 | 0 | 0 | 0 | 0 |
| IQAATDSKI | CEMIP2; CEMIP | Q9UHN6; B4 | CEIP2_HUMAN | Unmodified | 0 | 0 | 0 | 76745 | 0 | 622100 | 0 | 69622 | 330330 | 648640 | 0 | 0 | 241710 | 211530 | 0 | 0 | 88218 | 122550 |
| IQAEFEK | SF3A3 | Q12874 | SF3A3_HUMAN | Unmodified | 0 | 0 | 0 | 0 | 0 | 0 | 0 | 0 | 0 | 0 | 0 | 1156000 | 0 | 0 | 0 | 0 | 0 | 0 |
| IQAEISAHE | UTRN | P46939 | UTRN_HUMAN | Unmodified | 0 | 0 | 1130300 | 3719700 | 0 | 0 | 0 | 0 | 1334500 | 0 | 0 | 362820 | 0 | 0 | 902380 | 2391700 | 0 | 0 |
| IQAGEGET | EEA1 | Q15075 | EEA1_HUMAN | Unmodified | 0 | 0 | 0 | 0 | 1826300 | 0 | 0 | 0 | 0 | 0 | 0 | 0 | 0 | 0 | 0 | 0 | 208650 | 0 |
| IQAGEVSQ | PYM1 | Q9BRP8 | PYM1_HUMAN | Unmodified | 0 | 0 | 0 | 0 | 0 | 0 | 0 | 0 | 0 | 0 | 0 | 0 | 0 | 0 | 0 | 183730 | 0 | 0 |
| IQAGEVSQ | PYM1 | Q9BRP8 | PYM1_HUMAN | Unmodified | 449880 | 429970 | 0 | 0 | 97736 | 0 | 0 | 0 | 221470 | 137980 | 0 | 0 | 0 | 0 | 0 | 0 | 0 | 0 |
| IQAIAWAR | CTPS1; CTPS1; | P17812; B4E | PYRG1_HUMAN | Unmodified | 107910 | 0 | 176690 | 172670 | 73960 | 0 | 39991 | 25798 | 552300 | 156270 | 310560 | 4329600 | 0 | 0 | 307760 | 196340 | 47699 | 0 |
| IQAKIPGLK | RTN4; RTN4; RT | Q9NQC3; F8 | RTN4_HUMAN | Unmodified | 0 | 0 | 0 | 0 | 225130 | 0 | 118770 | 0 | 0 | 0 | 0 | 0 | 133020 | 121840 | 0 | 0 | 15465 | 262510 |
| IQALESNLE | TBC1D4; TBC1D | O60343; A0A | TBCD4_HUMAN | Unmodified | 0 | 0 | 0 | 0 | 0 | 0 | 0 | 0 | 0 | 0 | 0 | 851900 | 0 | 0 | 0 | 0 | 0 | 0 |
| IQALGLGEI | PLOD1; PLOD1; | Q02809; Q5J | PLOD1_HUMAN | Unmodified | 0 | 0 | 0 | 0 | 0 | 0 | 0 | 0 | 6398400 | 2683700 | 0 | 0 | 0 | 0 | 0 | 0 | 0 | 2118900 |
| IQALQQQA | TPM4; TPM4; TF | P67936; A0A | TPM4_HUMAN | Unmodified | 165820 | 183430 | 0 | 0 | 988860 | 37372 | 0 | 0 | 0 | 0 | 490280 | 2882200 | 0 | 0 | 313970 | 357450 | 0 | 0 |
| IQASQQDD | LSR; LSR | Q86X29; S4F | LSR_HUMAN Li | Oxidation (M) | 384160 | 93271 | 814470 | 254120 | 0 | 0 | 0 | 0 | 699550 | 127580 | 52226 | 775300 | 0 | 0 | 311010 | 281420 | 0 | 0 |
| IQASQQDD | LSR; LSR | Q86X29; S4F | LSR_HUMAN Li | Unmodified | 1413800 | 1467300 | 3494900 | 2717500 | 0 | 0 | 0 | 0 | 17019000 | 2136100 | 756460 | 6425100 | 0 | 0 | 307120 | 1917000 | 0 | 0 |
| IQASQQDD | LSR; LSR | Q86X29; S4F | LSR_HUMAN Li | Oxidation (M) | 0 | 0 | 0 | 175900 | 448120 | 0 | 0 | 0 | 1777600 | 1379500 | 268150 | 0 | 320440 | 150640 | 71613 | 56353 | 0 | 0 |
| IQASQQDD | LSR; LSR | Q86X29; S4F | LSR_HUMAN Li | Unmodified | 0 | 0 | 0 | 0 | 1132300 | 0 | 0 | 0 | 6082200 | 1309600 | 0 | 0 | 0 | 0 | 0 | 0 | 0 | 0 |
| IQASTMAFI | TAGLN2; TAGLN | P37802; X6R | TAGL2_HUMAN | Oxidation (M) | 2696100 | 1707500 | 902340 | 983480 | 7637300 | 3324000 | 1138000 | 796660 | 962960 | 1138600 | 918800 | 4114200 | 597490 | 1428900 | 1203300 | 1284800 | 4966600 | 4478800 |
| IQAYDYLEC | RHOB | P62745 | RHOB_HUMAN | Unmodified | 438860 | 330790 | 0 | 0 | 491460 | 0 | 0 | 0 | 766860 | 0 | 0 | 3790500 | 0 | 0 | 1223600 | 683550 | 0 | 0 |
| IQDAALSCI | ANKRD26; ANK | Q9UPS8; E7I | ANR26_HUMAN | Unmodified | 0 | 0 | 0 | 0 | 0 | 0 | 0 | 0 | 0 | 0 | 0 | 0 | 0 | 0 | 0 | 0 | 141930 | 398290 |
| IQDAGPASI | ZC3HAV1; ZC3H | Q7Z2W4; C9 | ZCCHV_HUMA | Unmodified | 437090 | 237480 | 522980 | 2440100 | 2525000 | 1353700 | 0 | 0 | 334050 | 37335 | 0 | 0 | 1272900 | 345760 | 379680 | 314350 | 0 | 0 |
| IQDAHRLH | WDR1 | O75083 | WDR1_HUMAN | Unmodified | 3243600 | 1802500 | 455660 | 4291200 | 1.04E+08 | 19366000 | 4897300 | 0 | 1242700 | 5730000 | 3751600 | 447300 | 13447000 | 10083000 | 643120 | 0 | 10520000 | 18176000 |
| IQDAHRLH | WDR1 | O75083 | WDR1_HUMAN | Unmodified | 0 | 0 | 0 | 0 | 1499200 | 0 | 0 | 0 | 0 | 0 | 0 | 0 | 0 | 798980 | 0 | 0 | 2242400 | 0 |
| IQDDEFVVI | KNTC1; KNTC1 | P50748; E7E | KNTC1_HUMAN | Unmodified | 0 | 0 | 0 | 0 | 0 | 0 | 0 | 0 | 861500 | 0 | 0 | 0 | 0 | 0 | 0 | 0 | 0 | 0 |
| IQDEQTTY | JAM3 | Q9BX67 | JAM3_HUMAN | Unmodified | 0 | 0 | 0 | 0 | 0 | 0 | 0 | 0 | 2432200 | 0 | 0 | 1666600 | 0 | 0 | 0 | 0 | 0 | 0 |
| IQDGLGELI | CUL1; CUL1; CI | Q13616; A0A | CUL1_HUMAN | Unmodified | 0 | 0 | 0 | 0 | 0 | 0 | 0 | 0 | 0 | 0 | 0 | 811510 | 0 | 0 | 0 | 0 | 0 | 0 |
| IQDHQVVII | ENAH; ENAH; E | Q8N8S7; A0 | ENAH_HUMAN | Unmodified | 0 | 0 | 0 | 0 | 0 | 0 | 92501 | 44970 | 191780 | 200190 | 238630 | 862570 | 91401 | 152570 | 0 | 0 | 124740 | 0 |
| IQDKQYEQ | ARHGAP29 | Q52LW3 | RHG29_HUMAN | Unmodified | 0 | 0 | 0 | 0 | 9956700 | 0 | 31447 | 81621 | 0 | 0 | 0 | 0 | 0 | 0 | 0 | 0 | 988900 | 1728400 |
| IQDLEHHL | RABGAP1 | Q9Y3P9 | RBGP1_HUMAN | Unmodified | 0 | 0 | 0 | 0 | 197790 | 0 | 0 | 0 | 158400 | 0 | 186760 | 398050 | 0 | 0 | 0 | 0 | 0 | 0 |
| IQDLHLPGI | ADAM15 | Q13444 | ADA15_HUMAN | Unmodified | 0 | 0 | 0 | 0 | 0 | 0 | 0 | 0 | 1266000 | 0 | 0 | 0 | 0 | 0 | 0 | 0 | 0 | 0 |
| IQDVLFLYK | FAM234A; FAM2 | Q9H0X4; C9 | F234A_HUMAN | Unmodified | 81592 | 56635 | 61738 | 0 | 0 | 0 | 0 | 0 | 0 | 0 | 0 | 571870 | 0 | 0 | 215820 | 278170 | 0 | 0 |
| IQDYDVSLI | USP7; USP7; US | Q93009; A0A | UBP7_HUMAN | Unmodified | 0 | 0 | 0 | 0 | 125150 | 0 | 0 | 0 | 208790 | 337260 | 0 | 1662200 | 0 | 0 | 0 | 0 | 0 | 319350 |
| IQEAASQG | RFTN1; RFTN1; | Q14699; G3 | RFTN1_HUMAN | Unmodified | 0 | 0 | 0 | 0 | 1396500 | 0 | 0 | 0 | 1214200 | 0 | 0 | 317170 | 0 | 0 | 0 | 0 | 0 | 212070 |
| IQEAGTEV | MDH2; MDH2 | P40926; G3 | MDHM_HUMAN | Unmodified | 458680 | 297210 | 1119700 | 268130 | 119200 | 0 | 256690 | 77922 | 81555000 | 4881800 | 3676500 | 2950400 | 962260 | 352220 | 885460 | 745320 | 0 | 0 |
| IQEAKDVY | NAMPT; NAMPT | P43490; A0A | NAMPT_HUMAN | Unmodified | 0 | 0 | 0 | 0 | 4275800 | 319280 | 0 | 0 | 1919100 | 597380 | 0 | 0 | 0 | 0 | 0 | 0 | 0 | 0 |
| IQEELDRFL | R3HCC1; R3HC | Q9Y3T6; E5F | R3HC1_HUMAN | Unmodified | 218010 | 397150 | 0 | 207370 | 0 | 347890 | 0 | 0 | 0 | 0 | 0 | 74568 | 0 | 0 | 316280 | 307770 | 0 | 0 |
| IQEELSALR | SUN2 | Q9UH99 | SUN2_HUMAN | Unmodified | 8368200 | 9053900 | 0 | 4775400 | 469790 | 0 | 0 | 23948 | 890530 | 0 | 0 | 2714900 | 583220 | 0 | 8762200 | 6294900 | 0 | 0 |
| IQEELSALR | SUN2 | Q9UH99 | SUN2_HUMAN | Unmodified | 0 | 384790 | 0 | 0 | 0 | 0 | 0 | 0 | 8759800 | 0 | 0 | 0 | 0 | 0 | 0 | 0 | 0 | 0 |
| IQEELSALR | SUN2 | Q9UH99 | SUN2_HUMAN | Unmodified | 0 | 0 | 0 | 0 | 0 | 0 | 366240 | 80189 | 65427000 | 0 | 0 | 0 | 0 | 0 | 0 | 0 | 0 | 0 |
| IQEENVIPR | LRPPRC | P42704 | LPPRC_HUMAN | Unmodified | 0 | 0 | 0 | 0 | 501800 | 0 | 0 | 0 | 2353800 | 0 | 183260 | 158010 | 0 | 0 | 0 | 0 | 0 | 0 |
| IQEENVIPR | LRPPRC | P42704 | LPPRC_HUMAN | Unmodified | 0 | 0 | 0 | 0 | 0 | 0 | 0 | 0 | 973330 | 0 | 0 | 0 | 0 | 0 | 0 | 0 | 0 | 0 |
| IQEGVFDIN | RAB2A | P61019 | RAB2A_HUMAN | Unmodified | 457820 | 1383600 | 1627800 | 1515500 | 1486100 | 236920 | 0 | 74938 | 10810000 | 2429200 | 1060100 | 9041900 | 710720 | 428540 | 1267800 | 1105200 | 0 | 0 |
| IQEILTQVK | IGF2BP3 | O00425 | IF2B3_HUMAN | Unmodified | 965630 | 949320 | 0 | 0 | 195530 | 0 | 173960 | 102260 | 1058400 | 0 | 153280 | 2517300 | 935610 | 322180 | 0 | 63047 | 0 | 0 |
| IQELEDLLA | LMNB1; LMNB1 | P20700; E9P | LMNB1_HUMAN | Unmodified | 306390 | 0 | 457200 | 730580 | 0 | 0 | 0 | 0 | 2197000 | 256740 | 2552100 | 1422600 | 0 | 0 | 0 | 150380 | 0 | 0 |
| IQELEEPEE | CES2 | O00748 | EST2_HUMAN | Unmodified | 0 | 0 | 0 | 0 | 0 | 0 | 0 | 0 | 2517300 | 732470 | 0 | 0 | 0 | 0 | 0 | 0 | 0 | 0 |
| IQELQEK | CIT; CIT; CIT; CI | O14578; A0A | CTRO_HUMAN | Unmodified | 0 | 0 | 94705 | 144410 | 1175700 | 72361 | 0 | 48095 | 458290 | 0 | 261510 | 1984700 | 0 | 0 | 144680 | 0 | 215470 | 163680 |
| IQELQEKLE | CIT; CIT; CIT; CI | O14578; A0A | CTRO_HUMAN | Unmodified | 0 | 0 | 0 | 0 | 0 | 0 | 0 | 0 | 0 | 0 | 0 | 0 | 0 | 0 | 0 | 0 | 0 | 307510 |
| IQEPLFK | PPP2R5B; PPP2 | Q15173; Q1 | 2A5B_HUMAN | Unmodified | 0 | 0 | 172900 | 205630 | 0 | 0 | 0 | 0 | 468960 | 154650 | 202590 | 1414800 | 277530 | 264810 | 287520 | 272030 | 0 | 0 |
| IQEPNTFPA | GNS; GNS; GNS | P15586; H7 | GNS_HUMAN | Unmodified | 0 | 0 | 482510 | 0 | 0 | 0 | 0 | 0 | 17926000 | 7009600 | 2101700 | 862980 | 1913800 | 0 | 0 | 0 | 0 | 0 |
| IQEQESSGI | PPP1R2B; PPP1 | Q6NXS1; P4 | IPP2B_HUMAN | Unmodified | 0 | 0 | 0 | 0 | 0 | 0 | 0 | 0 | 0 | 0 | 0 | 1519600 | 0 | 0 | 0 | 0 | 0 | 0 |
| IQEQVQQT | PKP2; PKP2 | Q99959; A0A | PKP2_HUMAN | Unmodified | 0 | 0 | 0 | 0 | 0 | 0 | 0 | 0 | 0 | 0 | 0 | 519440 | 0 | 0 | 0 | 0 | 0 | 0 |
| IQEQVQQT | PKP2; PKP2 | Q99959; A0A | PKP2_HUMAN | Unmodified | 0 | 0 | 0 | 0 | 169370 | 0 | 0 | 0 | 0 | 0 | 0 | 2272200 | 0 | 0 | 0 | 0 | 166920 | 615980 |
| IQESHPELF | SFXN1 | Q9H9B4 | SFXN1_HUMAN | Unmodified | 0 | 0 | 444100 | 0 | 0 | 0 | 0 | 0 | 396950 | 0 | 0 | 124510 | 0 | 0 | 0 | 0 | 0 | 0 |
| IQESYGDV | ATP2B1; ATP2B | P20020; F8V | AT2B1_HUMAN | Unmodified | 252050 | 0 | 453010 | 1779600 | 1519200 | 0 | 129310 | 123400 | 298650 | 213360 | 997960 | 5684900 | 9530400 | 4827600 | 3672200 | 2612800 | 0 | 1040400 |
| IQEVADELC | ARF4; ARF4; AR | P18085; C9J | ARF4_HUMAN | Unmodified | 959980 | 956440 | 2037100 | 680590 | 827930 | 0 | 0 | 0 | 1194100 | 491380 | 702390 | 7656000 | 0 | 0 | 3187600 | 2294000 | 156790 | 7452000 |
| IQEVGEPSI | SEC62; SEC62; | Q99442; F8V | SEC62_HUMAN | Unmodified | 0 | 0 | 0 | 0 | 0 | 0 | 0 | 0 | 0 | 0 | 0 | 0 | 0 | 0 | 0 | 0 | 0 | 0 |
| IQEVGEPSI | SEC62; SEC62; | Q99442; F8V | SEC62_HUMAN | Unmodified | 0 | 0 | 0 | 941500 | 0 | 0 | 130820 | 51585 | 1002600 | 2978500 | 1283900 | 0 | 0 | 0 | 0 | 0 | 0 | 0 |
| IQEVGEPSI | SEC62; SEC62; | Q99442; F8V | SEC62_HUMAN | Unmodified | 0 | 0 | 0 | 0 | 0 | 0 | 486720 | 338770 | 0 | 503970 | 0 | 0 | 0 | 0 | 0 | 0 | 0 | 0 |
| IQEVWRQF | PDPK1; PDPK1 | O15530; E9F | PDPK1_HUMAN | Unmodified | 0 | 0 | 0 | 0 | 76243 | 0 | 107100 | 0 | 0 | 0 | 0 | 0 | 705290 | 1197600 | 0 | 0 | 0 | 259330 |
| IQFFAIEIAR | PBDC1 | Q9BVG4 | PBDC1_HUMAN | Unmodified | 53729 | 68197 | 48614 | 136930 | 0 | 0 | 0 | 0 | 251720 | 187780 | 221440 | 973200 | 0 | 0 | 152490 | 218070 | 0 | 0 |
| IQFFAIEIAR | PBDC1 | Q9BVG4 | PBDC1_HUMAN | Unmodified | 0 | 0 | 0 | 0 | 135030 | 0 | 0 | 0 | 0 | 317550 | 0 | 0 | 348960 | 358780 | 0 | 0 | 1446900 | 0 |
| IQFGTLSDI | MAN2A1 | Q16706 | MA2A1_HUMAN | Unmodified | 0 | 0 | 0 | 0 | 0 | 0 | 0 | 0 | 90482 | 0 | 0 | 186020 | 0 | 0 | 0 | 0 | 0 | 0 |
| IQFHNVKPI | NIPSNAP1; NIP | Q9BPW8; F8 | NIPS1_HUMAN | Unmodified | 0 | 0 | 0 | 0 | 0 | 0 | 0 | 0 | 1389700 | 0 | 0 | 0 | 0 | 0 | 0 | 0 | 0 | 0 |
| IQFSVPAPF | PPP1R1B; PPP1 | Q9UD71; J3 | PPR1B_HUMAN | Unmodified | 0 | 0 | 0 | 0 | 0 | 0 | 0 | 0 | 0 | 0 | 184430 | 1733500 | 0 | 0 | 0 | 0 | 0 | 0 |
| IQFVEHEEL | CHSY1 | Q86X52 | CHSS1_HUMAN | Unmodified | 0 | 0 | 0 | 0 | 620540 | 0 | 0 | 0 | 3628500 | 0 | 0 | 0 | 0 | 0 | 0 | 0 | 0 | 0 |
| IQFVGACN | DYNC1H1; DYN | Q14204; A0A | DYHC1_HUMA | Unmodified | 0 | 0 | 0 | 0 | 0 | 0 | 0 | 0 | 0 | 0 | 0 | 945730 | 0 | 0 | 106090 | 56829 | 0 | 0 |
| IQGDLAGR | JAM3 | Q9BX67 | JAM3_HUMAN | Unmodified | 0 | 0 | 0 | 0 | 0 | 130450 | 0 | 0 | 1036200 | 250330 | 0 | 1146300 | 0 | 0 | 0 | 838190 | 30497 | 40250 |
| IQGDLAGR | JAM3 | Q9BX67 | JAM3_HUMAN | Unmodified | 0 | 0 | 0 | 0 | 595260 | 0 | 0 | 0 | 17406000 | 2180200 | 0 | 12723000 | 0 | 0 | 0 | 0 | 0 | 0 |
| IQGFQAKG | TRIM26; TRIM2 | Q12899; A0A | TRI26_HUMAN | Unmodified | 2823600 | 4628200 | 0 | 3814300 | 0 | 0 | 0 | 0 | 0 | 0 | 0 | 0 | 401500 | 0 | 1000300 | 836840 | 0 | 0 |
| IQGFQAKG | TRIM26; TRIM2 | Q12899; A0A | TRI26_HUMAN | Unmodified | 686140 | 1184300 | 0 | 1337800 | 0 | 0 | 0 | 0 | 0 | 0 | 0 | 0 | 0 | 0 | 272570 | 282040 | 0 | 0 |
| IQGGYENV | PLOD1 | Q02809 | PLOD1_HUMAN | Unmodified | 0 | 0 | 0 | 0 | 0 | 0 | 0 | 0 | 1481000 | 0 | 0 | 0 | 0 | 0 | 0 | 0 | 0 | 0 |
| IQGLTVEQ | DYNC1H1; DYN | Q14204; A0A | DYHC1_HUMA | Unmodified | 118410 | 555540 | 91121 | 0 | 0 | 0 | 0 | 0 | 478640 | 67479 | 187180 | 3044900 | 0 | 0 | 154390 | 152050 | 159220 | 148430 |
| IQHEELNN | EEA1 | Q15075 | EEA1_HUMAN | Unmodified | 0 | 0 | 0 | 0 | 521460 | 0 | 0 | 0 | 0 | 0 | 0 | 0 | 0 | 0 | 0 | 0 | 0 | 0 |
| IQHGSGFG | TIPRL | O75663 | TIPRL_HUMAN | Unmodified | 262230 | 272640 | 0 | 1178100 | 0 | 0 | 0 | 0 | 1234700 | 189090 | 884120 | 2051700 | 295570 | 0 | 0 | 638160 | 0 | 105300 |
| IQHLSTIDY | FRMD8 | Q9BZ67 | FRMD8_HUMAN | Unmodified | 0 | 0 | 416000 | 231220 | 0 | 194910 | 0 | 0 | 608930 | 0 | 197800 | 0 | 456140 | 176180 | 290140 | 391400 | 290100 | 1220300 |
| IQIAPDSG | FUBP1; FUBP1; | Q96AE4; C9 | FUBP1_HUMAN | Unmodified | 0 | 0 | 0 | 0 | 0 | 0 | 0 | 0 | 1127500 | 0 | 1893700 | 3836200 | 0 | 0 | 0 | 0 | 0 | 0 |
| IQIGTQAIE | UBR4; UBR4 | Q5T4S7; A0A | UBR4_HUMAN | Unmodified | 0 | 0 | 0 | 0 | 207630 | 0 | 44676 | 0 | 0 | 60699 | 0 | 711630 | 0 | 0 | 0 | 0 | 320420 | 64867 |
| IQIIGLGTQ | VPS13C | Q709C8 | VP13C_HUMAN | Unmodified | 0 | 0 | 0 | 0 | 0 | 0 | 0 | 0 | 0 | 0 | 205550 | 1444100 | 0 | 0 | 75583 | 0 | 0 | 0 |
| IQIPRPDDF | HDLBP; HDLBP | Q00341; A0A | VIGLN_HUMAN | Unmodified | 1129300 | 0 | 0 | 0 | 808800 | 0 | 0 | 0 | 1230300 | 374150 | 878010 | 2949800 | 3379700 | 0 | 685800 | 546290 | 293220 | 577620 |
| IQKLESDVS | FGB | P02675 | FIBB_HUMAN | Oxidation (M) | 0 | 978790 | 0 | 0 | 0 | 0 | 0 | 0 | 0 | 0 | 0 | 0 | 0 | 0 | 0 | 0 | 0 | 0 |
| IQKLLQDFF | HSPA8; HSPA2; | P11142; P54 | HSP7C_HUMAN | Unmodified | 0 | 0 | 0 | 0 | 0 | 0 | 1080800 | 0 | 0 | 0 | 0 | 0 | 0 | 0 | 0 | 0 | 1367600 | 326610 |
| IQKVDVYDI | LSAMP; LSAMP; | Q13449; F5H | LSAMP_HUMAN | Unmodified | 0 | 0 | 0 | 0 | 0 | 0 | 0 | 0 | 1961500 | 262560 | 0 | 0 | 0 | 0 | 0 | 0 | 0 | 0 |
| IQLEQVQE | DCTN1; DCTN1 | Q14203; Q6 | DCTN1_HUMAN | Unmodified | 0 | 0 | 0 | 0 | 0 | 0 | 0 | 0 | 212500 | 0 | 354590 | 1081000 | 189260 | 0 | 0 | 0 | 0 | 0 |
| IQLFLAR | THOC1; THOC1 | Q96FV9; A0A | THOC1_HUMA | Unmodified | 68472 | 0 | 0 | 0 | 0 | 0 | 0 | 24632 | 0 | 0 | 0 | 250570 | 0 | 51019 | 0 | 0 | 0 | 0 |
| IQLINNMLI | PGK1 | P00558 | PGK1_HUMAN | Unmodified | 0 | 0 | 0 | 0 | 11849000 | 0 | 0 | 0 | 6701200 | 0 | 2902500 | 21358000 | 765280 | 301440 | 0 | 0 | 863960 | 2432600 |
| IQLINNMLI | PGK1 | P00558 | PGK1_HUMAN | Oxidation (M) | 444910 | 358130 | 1811900 | 40870 | 652230 | 670450 | 0 | 111130 | 2321600 | 315990 | 3743900 | 23783000 | 798520 | 258350 | 2986900 | 2669400 | 541190 | 961930 |
| IQLLDLPGI | DRG1 | Q9Y295 | DRG1_HUMAN | Unmodified | 0 | 0 | 0 | 0 | 0 | 0 | 0 | 0 | 139160 | 0 | 174120 | 0 | 0 | 0 | 0 | 30268 | 0 | 0 |
| IQLLDLPGI | DRG1 | Q9Y295 | DRG1_HUMAN | Unmodified | 0 | 0 | 0 | 0 | 0 | 0 | 0 | 0 | 0 | 0 | 81142 | 783230 | 0 | 0 | 0 | 0 | 0 | 0 |

| | | | | | | | | | | | | | | | | | | | | | | |
|---|---|---|---|---|---|---|---|---|---|---|---|---|---|---|---|---|---|---|---|---|---|---|
| IQLLVFK | EIF3L; EIF3L; EI | Q9Y262; B0( | EIF3L_HUMAN | Unmodified | 0 | 0 | 141270 | 0 | 0 | 0 | 0 | 42774 | 0 | 0 | 149250 | 4134200 | 0 | 0 | 0 | 0 | 0 | 0 |
| IQLNNPTEI | KIAA2013 | Q8IYS2 | K2013_HUMAN | Unmodified | 316920 | 1044900 | 0 | 0 | 0 | 0 | 1466800 | 0 | 42242000 | 2692100 | 449400 | 618620 | 95726 | 0 | 0 | 0 | 0 | 0 |
| IQLQPTSIEI | GMPPB; GMPPI | Q9Y5P6; A0/ | GMPPB_HUMA | Unmodified | 0 | 127690 | 0 | 0 | 0 | 0 | 0 | 0 | 1061500 | 262790 | 0 | 2411400 | 0 | 0 | 89000 | 0 | 0 | 0 |
| IQLSDYTK | DPP4; DPP4; DI | P27487; A0A | DPP4_HUMAN | Unmodified | 961830 | 1464900 | 10111000 | 18952000 | 0 | 0 | 0 | 0 | 0 | 183880 | 0 | 0 | 0 | 0 | 16244000 | 9780600 | 3502400 | 1916900 |
| IQLVAVNYI | DCTN4 | Q9UJW0 | DCTN4_HUMAI | Unmodified | 0 | 0 | 0 | 0 | 0 | 0 | 0 | 0 | 266620 | 47918 | 163540 | 706830 | 110950 | 0 | 0 | 0 | 0 | 0 |
| IQMLDDTQ | FARP1; FARP1; | Q9Y4F1; A0/ | FARP1_HUMAN | Unmodified | 0 | 0 | 0 | 0 | 1411900 | 0 | 0 | 0 | 5987900 | 888350 | 0 | 1208700 | 167430 | 0 | 791210 | 350210 | 0 | 717090 |
| IQMLDDTQ | FARP1; FARP1; | Q9Y4F1; A0/ | FARP1_HUMAN | Oxidation (M) | 0 | 0 | 0 | 0 | 0 | 0 | 0 | 0 | 1263800 | 632970 | 696540 | 3097000 | 0 | 0 | 386920 | 1390700 | 0 | 272300 |
| IQMSYSVK | TM9SF3; TM9SF | Q9HD45; Q5 | TM9S3_HUMAN | Oxidation (M) | 0 | 0 | 0 | 0 | 0 | 0 | 0 | 0 | 2882800 | 369560 | 60406 | 0 | 0 | 0 | 50546 | 43682 | 25440 | 0 |
| IQMSYSVK | TM9SF3; TM9SF | Q9HD45; Q5 | TM9S3_HUMAN | Unmodified | 0 | 0 | 0 | 0 | 0 | 0 | 209510 | 0 | 34955000 | 4386800 | 933520 | 1880700 | 0 | 0 | 0 | 0 | 0 | 0 |
| IQNDAGVR | KHSRP; KHSRP | Q92945; M0I | FUBP2_HUMAN | Unmodified | 0 | 0 | 0 | 0 | 640400 | 0 | 578300 | 369950 | 104400 | 1074900 | 0 | 0 | 288740 | 0 | 0 | 0 | 341890 | 740440 |
| IQNGAQGI | HNRNPH1; HNI | P31943; E9P | HNRH1_HUMA | Unmodified | 392620 | 1160700 | 0 | 0 | 154720 | 184970 | 0 | 132090 | 610530 | 36865 | 11795 | 2832800 | 0 | 0 | 1847600 | 0 | 169260 | 120800 |
| IQNPQILK | LRBA; LRBA; LR | P50851; A0A | LRBA_HUMAN | Unmodified | 55939 | 0 | 0 | 0 | 0 | 0 | 20618 | 0 | 98306 | 0 | 0 | 4228000 | 0 | 0 | 0 | 122380 | 0 | 0 |
| IQNSGDFY | PTPN6; PTPN6; | P29350; F5G | PTN6_HUMAN | Unmodified | 0 | 0 | 0 | 0 | 0 | 0 | 0 | 0 | 0 | 0 | 0 | 0 | 0 | 0 | 0 | 0 | 0 | 0 |
| IQNSSGTD | KNTC1 | P50748 | KNTC1_HUMAI | Unmodified | 0 | 0 | 0 | 0 | 0 | 0 | 0 | 0 | 0 | 0 | 0 | 738230 | 0 | 0 | 0 | 0 | 0 | 0 |
| IQNTGDYY | PTPN11; PTPN1 | Q06124; A0/ | PTN11_HUMAN | Unmodified | 180120 | 120580 | 99175 | 0 | 0 | 0 | 0 | 0 | 0 | 0 | 0 | 1796700 | 0 | 0 | 209320 | 151330 | 0 | 0 |
| IQNTLHCC | TSPAN6; TSPAN | O43657; A0/ | TSN6_HUMAN | Unmodified | 1958000 | 552250 | 2167100 | 3845100 | 1336100 | 0 | 0 | 0 | 20198000 | 1926100 | 228720 | 9647900 | 1176400 | 0 | 4680400 | 2917500 | 0 | 0 |
| IQPALQEK | PFDN5; PFDN5 | Q99471; A0/ | PFD5_HUMAN | Unmodified | 317290 | 181570 | 282300 | 649950 | 3273200 | 0 | 1100200 | 739620 | 2080100 | 2213200 | 1023300 | 9047400 | 5302000 | 1718100 | 744710 | 443880 | 4294700 | 3238700 |
| IQPDTIIQV | HEXA; HEXA; HI | P06865; H3I | HEXA_HUMAN | Unmodified | 0 | 0 | 0 | 0 | 0 | 0 | 0 | 0 | 690130 | 0 | 0 | 0 | 0 | 0 | 0 | 0 | 284700 | 0 |
| IQPETGPL( | PLXNB2 | O15031 | PLXB2_HUMAN | Unmodified | 0 | 0 | 290430 | 221090 | 0 | 0 | 0 | 0 | 227310 | 55262 | 0 | 0 | 0 | 695900 | 434450 | 345570 | 0 | 0 |
| IQPGNPNY | AP2B1; AP2B1 | P63010; A0A | AP2B1_HUMAN | Unmodified | 0 | 0 | 0 | 0 | 872420 | 0 | 0 | 0 | 2291200 | 399880 | 0 | 5464900 | 1169900 | 0 | 0 | 0 | 578030 | 0 |
| IQPLRVQRI | PTPRF; PTPRF | P10586; A2A | PTPRF_HUMAN | Unmodified | 0 | 0 | 0 | 0 | 0 | 0 | 0 | 0 | 1647700 | 0 | 0 | 0 | 0 | 0 | 0 | 0 | 0 | 0 |
| IQPQMDM\ | FN3KRP | Q9HA64 | KT3K_HUMAN I | Oxidation (M) | 0 | 0 | 0 | 0 | 0 | 0 | 264300 | 0 | 0 | 0 | 0 | 0 | 0 | 0 | 0 | 0 | 0 | 0 |
| IQPQSLQG | DNAAF2 | Q9NVR5 | KTU_HUMAN P | Unmodified | 0 | 0 | 0 | 0 | 0 | 0 | 0 | 0 | 0 | 0 | 0 | 358620 | 0 | 0 | 0 | 0 | 0 | 0 |
| IQPYTEQST | KIAA0319L; KIA | Q8IZA0; C9J | K319L_HUMAN | Unmodified | 0 | 46849 | 0 | 0 | 0 | 46483 | 29224 | 0 | 3500800 | 575030 | 0 | 390320 | 0 | 0 | 301660 | 400370 | 0 | 0 |
| IQQDADSV | ACBD3 | Q9H3P7 | GCP60_HUMAI | Unmodified | 0 | 0 | 0 | 0 | 0 | 0 | 440440 | 0 | 0 | 0 | 0 | 1388000 | 0 | 0 | 0 | 0 | 0 | 0 |
| IQQEIAVQN | OTUB1; OTUB1 | Q96FW1; J3I | OTUB1_HUMAI | Unmodified | 0 | 0 | 0 | 0 | 0 | 0 | 0 | 0 | 0 | 0 | 439900 | 739740 | 0 | 0 | 0 | 0 | 0 | 0 |
| IQQELQTAI | CHMP4A; CHM | Q9BY43; E9I | CHM4A_HUMA | Unmodified | 0 | 27932 | 0 | 0 | 125520 | 203080 | 55912 | 0 | 3472500 | 983970 | 331120 | 1379600 | 0 | 0 | 0 | 0 | 107960 | 394860 |
| IQQGLFDV | RAB2B; RAB2B | Q8WUD1; A( | RAB2B_HUMAN | Unmodified | 122520 | 0 | 0 | 0 | 0 | 0 | 0 | 0 | 375500 | 0 | 0 | 3693200 | 0 | 0 | 189040 | 242090 | 87530 | 0 |
| IQQIAEGEK | FLOT2; FLOT2; I | Q14254; J3Q | FLOT2_HUMAN | Unmodified | 134780 | 0 | 1735500 | 701040 | 0 | 0 | 0 | 122690 | 959000 | 889540 | 740040 | 7152300 | 0 | 0 | 2138800 | 2101000 | 0 | 0 |
| IQQIAEGEK | FLOT2; FLOT2; I | Q14254; J3Q | FLOT2_HUMAN | Unmodified | 0 | 0 | 0 | 0 | 0 | 0 | 36563 | 0 | 0 | 253440 | 0 | 0 | 0 | 0 | 0 | 0 | 0 | 0 |
| IQQIIEGK | OPA1; OPA1; OI | O60313; E5I | OPA1_HUMAN | Unmodified | 0 | 0 | 0 | 0 | 0 | 0 | 0 | 0 | 0 | 0 | 0 | 4279400 | 0 | 0 | 0 | 0 | 0 | 0 |
| IQQLAETIK | TRAF5 | O00463 | TRAF5_HUMAN | Unmodified | 0 | 0 | 0 | 0 | 116720 | 0 | 33835 | 67816 | 0 | 0 | 50877 | 758490 | 0 | 0 | 0 | 0 | 123360 | 0 |
| IQQLAISGL | AP3M2; AP3M1; | P53677; Q9\ | AP3M2_HUMAI | Unmodified | 107560 | 269150 | 0 | 151090 | 750410 | 0 | 0 | 193710 | 0 | 0 | 0 | 3408000 | 353950 | 421190 | 29119 | 0 | 161040 | 365410 |
| IQQLLNTLK | DIP2A; DIP2B; D | Q14689; Q9I | DIP2A_HUMAN | Unmodified | 315530 | 343620 | 1852700 | 2554600 | 2813600 | 0 | 573290 | 111440 | 5720100 | 1792700 | 2641200 | 14101000 | 982260 | 1593900 | 2693200 | 2032600 | 609010 | 2999200 |
| IQQLQEILQ | RAI14 | Q9P0K7 | RAI14_HUMAN | Unmodified | 10329 | 11580 | 102320 | 50297 | 0 | 0 | 0 | 0 | 0 | 0 | 90538 | 464950 | 0 | 0 | 171720 | 136400 | 0 | 0 |
| IQQLTALG/ | PFDN5; PFDN5 | Q99471; A0/ | PFD5_HUMAN | Unmodified | 0 | 0 | 0 | 0 | 2180800 | 0 | 0 | 17391 | 1109500 | 1052100 | 0 | 5392300 | 2669400 | 24384 | 0 | 0 | 1376300 | 0 |
| IQQLVNTLI | DIP2C; DIP2C; I | Q9Y2E4; A0/ | DIP2C_HUMAN | Unmodified | 0 | 0 | 0 | 0 | 0 | 0 | 0 | 0 | 640590 | 223300 | 0 | 0 | 0 | 0 | 357360 | 240860 | 818430 | 0 |
| IQQNTFTR | FLNC; FLNB; FL | Q14315; O7! | FLNC_HUMAN | Unmodified | 1665400 | 1839400 | 2249700 | 4646500 | 2739300 | 711260 | 0 | 171360 | 802750 | 263690 | 279490 | 3046200 | 1096700 | 726240 | 3558500 | 2320400 | 0 | 775890 |
| IQQNTFTR\ | FLNC; FLNB; FL | Q14315; O7! | FLNC_HUMAN | Unmodified | 0 | 0 | 0 | 0 | 17450000 | 6627100 | 9086700 | 1922600 | 0 | 644730 | 0 | 0 | 8884200 | 8610000 | 0 | 0 | 5496400 | 7621200 |
| IQQQQPPP | VARS1; VARS1; | P26640; A0A | SYVC_HUMAN | Unmodified | 0 | 0 | 0 | 0 | 264400 | 0 | 0 | 38630 | 0 | 0 | 0 | 1701200 | 0 | 0 | 0 | 0 | 0 | 338490 |
| IQQRGQG/ | CFAP298; CFAF | P57076; H7( | CF298_HUMAN | Unmodified | 0 | 0 | 0 | 0 | 697270 | 0 | 0 | 0 | 0 | 186400 | 0 | 0 | 932730 | 718040 | 0 | 0 | 707480 | 754010 |
| IQQYFPSN\ | PLCG2; PLCG2 | P16885; A0A | PLCG2_HUMAI | Unmodified | 0 | 0 | 0 | 0 | 0 | 0 | 0 | 0 | 0 | 0 | 295330 | 508700 | 0 | 0 | 0 | 0 | 0 | 0 |
| IQRFLSQPI | ATP5F1B | P06576 | ATPB_HUMAN , | Oxidation (M) | 0 | 0 | 0 | 0 | 0 | 0 | 0 | 0 | 0 | 2351600 | 42954 | 0 | 0 | 0 | 0 | 0 | 0 | 0 |
| IQRPPEDSI | UGP2; UGP2; U | Q16851; A0/ | UGPA_HUMAN | Unmodified | 167940 | 0 | 222350 | 661480 | 8388900 | 0 | 2130800 | 1053500 | 3996700 | 3713200 | 1358200 | 6882400 | 3718000 | 2390500 | 406050 | 272410 | 2431900 | 2963100 |
| IQRTTEEDF | PTGFRN | Q9P2B2 | FPRP_HUMAN | Unmodified | 0 | 0 | 0 | 1027900 | 0 | 0 | 0 | 0 | 0 | 0 | 0 | 0 | 908700 | 0 | 0 | 0 | 0 | 0 |
| IQSGHSLE( | UBA6 | A0AVT1 | UBA6_HUMAN | Unmodified | 72151 | 203140 | 0 | 0 | 131700 | 0 | 312300 | 135030 | 364910 | 59908 | 95433 | 1319800 | 0 | 104100 | 40026 | 67160 | 0 | 242800 |
| IQSHSRGE | PPP2R2A; PPP2 | P63151; E5R | 2ABA_HUMAN | Unmodified | 0 | 0 | 0 | 0 | 0 | 0 | 0 | 143400 | 5158100 | 0 | 0 | 3276100 | 442680 | 0 | 470820 | 0 | 357030 | 0 |
| IQSIKKEDS | EFCAB14; EFC/ | O75071; A0/ | EFC14_HUMAN | Unmodified | 0 | 0 | 0 | 0 | 0 | 0 | 0 | 0 | 6136200 | 1289600 | 0 | 0 | 0 | 0 | 0 | 0 | 0 | 0 |
| IQSLGDSK | AARS1; AARS1; | P49588; A0A | SYAC_HUMAN | Unmodified | 0 | 0 | 96576 | 163080 | 703240 | 350370 | 0 | 18247 | 75695 | 73987 | 0 | 1485300 | 707090 | 244320 | 144760 | 111020 | 269970 | 818570 |
| IQSQFTDA( | DSP | P15924 | DESP_HUMAN | Unmodified | 0 | 0 | 0 | 0 | 0 | 0 | 0 | 0 | 592420 | 326910 | 0 | 175500 | 0 | 0 | 0 | 124030 | 0 | 0 |
| IQSSHNFQ | GOLM1; GOLM1 | Q8NBJ4; F8\ | GOLM1_HUMA | Unmodified | 137450 | 0 | 206740 | 0 | 1275800 | 1017100 | 0 | 0 | 64591000 | 4063000 | 0 | 781910 | 614760 | 0 | 191750 | 0 | 0 | 1025000 |
| IQSTVTQP( | NOMO1; NOMC | Q15155; P6! | NOMO1_HUM/ | Unmodified | 0 | 302660 | 0 | 0 | 679650 | 0 | 0 | 0 | 16894000 | 2166400 | 1652500 | 1151500 | 0 | 30916 | 0 | 108450 | 0 | 0 |
| IQSTVTQP( | NOMO1; NOMC | Q15155; P6! | NOMO1_HUM/ | Unmodified | 0 | 0 | 0 | 0 | 322840 | 0 | 854910 | 226110 | 266870 | 3168900 | 0 | 0 | 0 | 309760 | 0 | 0 | 0 | 90906 |
| IQTDSVVAT | HMBS; HMBS; I | P08397; A0A | HEM3_HUMAN | Unmodified | 47219 | 0 | 0 | 0 | 850310 | 0 | 0 | 112560 | 0 | 0 | 568950 | 2566600 | 481360 | 0 | 0 | 85243 | 285400 | 431040 |
| IQTFLLLDE | ABCC4; ABCC4 | O15439; A0/ | MRP4_HUMAN | Unmodified | 0 | 0 | 218190 | 154890 | 0 | 0 | 0 | 0 | 0 | 0 | 0 | 0 | 0 | 0 | 528280 | 257100 | 0 | 0 |
| IQTLPNMR | TP53BP2 | Q13625 | ASPP2_HUMAN | Unmodified | 0 | 0 | 0 | 0 | 277270 | 0 | 0 | 0 | 0 | 0 | 0 | 0 | 0 | 0 | 0 | 0 | 161930 | 470210 |
| IQTLTSSVR | POR; POR; POR | P16435; H0\ | NCPR_HUMAN | Unmodified | 0 | 19508 | 121010 | 76524 | 0 | 0 | 0 | 0 | 638570 | 0 | 0 | 1037500 | 0 | 0 | 114970 | 81899 | 0 | 0 |
| IQTNNNPF | RELA; RELA; RE | Q04206; A0/ | TF65_HUMAN 1 | Unmodified | 0 | 0 | 219600 | 0 | 0 | 0 | 0 | 0 | 0 | 131920 | 0 | 264260 | 0 | 0 | 78318 | 55439 | 0 | 0 |
| IQTNNNPF | RELA; RELA; RE | Q04206; A0/ | TF65_HUMAN 1 | Unmodified | 124490 | 0 | 0 | 0 | 0 | 0 | 0 | 0 | 580190 | 0 | 0 | 1732300 | 0 | 0 | 216140 | 0 | 0 | 0 |
| IQTPELGEV | MIA3; MIA3 | Q5JRA6; A0/ | TGO1_HUMAN | Unmodified | 0 | 0 | 0 | 0 | 0 | 0 | 0 | 0 | 2279400 | 0 | 179910 | 0 | 0 | 0 | 0 | 0 | 0 | 0 |
| IQTQLQRD | ANPEP | P15144 | AMPN_HUMAN | Unmodified | 0 | 0 | 0 | 0 | 0 | 0 | 0 | 0 | 0 | 0 | 0 | 0 | 0 | 0 | 0 | 0 | 783920 | 679740 |
| IQTQPGYAI | SLC25A3; SLC2 | Q00325; F8\ | MPCP_HUMAN | Unmodified | 0 | 0 | 194850 | 89852 | 0 | 0 | 0 | 0 | 53156000 | 3413800 | 329630 | 662840 | 0 | 0 | 89798 | 67048 | 0 | 0 |
| IQTQPGYAI | SLC25A3; SLC2 | Q00325; F8\ | MPCP_HUMAN | Unmodified | 0 | 0 | 0 | 0 | 0 | 0 | 0 | 0 | 14389000 | 1319500 | 300090 | 2426900 | 0 | 0 | 403620 | 0 | 374110 | 0 |
| IQVEHGVT( | PIK3C2B; PIK3C | O00750; F5C | P3C2B_HUMAI | Unmodified | 0 | 0 | 0 | 0 | 0 | 0 | 0 | 0 | 0 | 80278 | 0 | 215280 | 0 | 0 | 519010 | 376580 | 0 | 0 |
| IQVHYYED( | CAPZA2; CAPZ/ | P47755; P52 | CAZA2_HUMAN | Unmodified | 2445000 | 522450 | 755640 | 527630 | 318030 | 0 | 0 | 0 | 261310 | 0 | 2228600 | 10366000 | 200790 | 431730 | 2074200 | 1452300 | 251510 | 0 |
| IQVHYYED( | CAPZA1 | P52907 | CAZA1_HUMAN | Unmodified | 0 | 245810 | 0 | 0 | 9019600 | 0 | 600070 | 0 | 861520 | 371860 | 232740 | 1812200 | 8925300 | 5203800 | 0 | 0 | 5531500 | 3428300 |
| IQVLQQQA | TPM3; TPM3; TI | D6R904; A0/ | D6R904_HUM/ | Unmodified | 0 | 0 | 111790 | 0 | 0 | 0 | 0 | 0 | 0 | 0 | 1133900 | 1044400 | 0 | 0 | 0 | 100810 | 0 | 0 |
| IQVLQQQA | TPM3; TPM3; TI | D6R904; A0/ | D6R904_HUM/ | Unmodified | 2449700 | 1923500 | 5782600 | 3343000 | 3233400 | 0 | 0 | 0 | 9481700 | 424220 | 5555700 | 62200000 | 1440800 | 501620 | 6315000 | 4825100 | 0 | 0 |
| IQVLQQQA | TPM3; TPM3; TI | D6R904; A0/ | D6R904_HUM/ | Unmodified | 0 | 0 | 1906900 | 0 | 4338600 | 0 | 0 | 0 | 4190800 | 0 | 0 | 0 | 0 | 0 | 0 | 1119100 | 1633300 | 493220 |
| IQVLVEPDI | LGALS3 | P17931 | LEG3_HUMAN | Unmodified | 0 | 0 | 0 | 0 | 0 | 0 | 0 | 0 | 0 | 0 | 0 | 2124000 | 1350300 | 0 | 0 | 0 | 0 | 0 |
| IQVQDLMQ | GOLGA4; GOLG | Q13439; C9J | GOGA4_HUMA | Unmodified | 0 | 0 | 0 | 0 | 0 | 4582300 | 958650 | 381880 | 0 | 0 | 495880 | 0 | 0 | 1774500 | 229140 | 0 | 0 | 4190200 |
| IQVRLPDG | UBXN1; UBXN1 | Q04323; E9I | UBXN1_HUMAI | Unmodified | 419200 | 694010 | 158420 | 965750 | 219110 | 0 | 0 | 0 | 644700 | 0 | 884150 | 3401700 | 0 | 0 | 354060 | 64137 | 0 | 0 |
| IQVSELCK | GLG1; GLG1 | Q92896; H3I | GSLG1_HUMAI | Unmodified | 0 | 0 | 701680 | 0 | 785520 | 0 | 418860 | 63428 | 5719300 | 582200 | 0 | 1392800 | 0 | 0 | 0 | 0 | 0 | 0 |
| IQVSKLESS | KIF4A | O95239 | KIF4A_HUMAN | Unmodified | 0 | 0 | 0 | 0 | 1388400 | 282210 | 2162700 | 5143200 | 0 | 0 | 0 | 0 | 1053100 | 1051200 | 0 | 0 | 1166600 | 692560 |
| IQVTPPGF( | XRCC6; XRCC6 | P12956; B1/ | XRCC6_HUMAI | Unmodified | 431940 | 0 | 0 | 0 | 0 | 0 | 0 | 0 | 144880 | 0 | 305140 | 1397700 | 0 | 0 | 0 | 0 | 0 | 0 |
| IQVTPPGF( | XRCC6; XRCC6 | P12956; B1/ | XRCC6_HUMAI | Unmodified | 0 | 0 | 0 | 0 | 0 | 0 | 0 | 0 | 0 | 0 | 0 | 370760 | 0 | 0 | 0 | 0 | 0 | 0 |
| IQVVVLSAS | HSPG2 | P98160 | PGBM_HUMAN | Unmodified | 0 | 0 | 1722300 | 437630 | 0 | 0 | 0 | 0 | 248460 | 0 | 0 | 0 | 0 | 0 | 0 | 627910 | 0 | 0 |
| IQVWHEEI | MSN | P26038 | MOES_HUMAN | Unmodified | 0 | 0 | 327480 | 0 | 0 | 0 | 0 | 0 | 0 | 0 | 0 | 0 | 0 | 0 | 0 | 0 | 0 | 0 |
| IQYQLVDIS | SH3BGRL3; SH | Q9H299; Q5 | SH3L3_HUMAN | Unmodified | 0 | 0 | 0 | 0 | 0 | 0 | 0 | 0 | 0 | 0 | 0 | 0 | 0 | 0 | 0 | 0 | 2613400 | 2331200 |
| IRAALNAVF | SLC9A3R1; SLC | O14745; J3Q | NHRF1_HUMAI | Unmodified | 0 | 0 | 0 | 0 | 8811100 | 0 | 0 | 0 | 0 | 109610 | 0 | 0 | 18875000 | 27520000 | 0 | 0 | 3373900 | 1098600 |
| IRADQDTIR | NPTXR; NPTXR | O95502; A0/ | NPTXR_HUMAN | Unmodified | 0 | 0 | 0 | 0 | 0 | 0 | 0 | 0 | 14406000 | 1489200 | 170240 | 0 | 0 | 0 | 0 | 0 | 0 | 0 |
| IRAQTEGIN | RUVBL1; RUVBI | Q9Y265; H7( | RUVB1_HUMAI | Unmodified | 858980 | 0 | 773920 | 1160700 | 8804900 | 1854200 | 2936000 | 67433 | 1553300 | 950810 | 2338600 | 23388000 | 7895100 | 5198300 | 1159200 | 1002200 | 1605300 | 2692300 |
| IRDAGNDS | OXR1; OXR1 | Q8N573; H0 | OXR1_HUMAN | Unmodified | 38665 | 52645 | 39596 | 93437 | 0 | 94870 | 0 | 0 | 1337800 | 262030 | 72119 | 257820 | 0 | 0 | 0 | 16710 | 21107 | 0 |
| IRDEMVATI | ATP2A2; ATP2A | P16615; H7( | AT2A2_HUMAN | Unmodified | 0 | 0 | 0 | 0 | 0 | 0 | 0 | 0 | 0 | 0 | 0 | 576630 | 0 | 0 | 0 | 0 | 0 | 0 |
| IRDIEEAIPI | WLS; WLS; WLS | Q5T9L3; H0\ | WLS_HUMAN F | Unmodified | 0 | 0 | 0 | 44122 | 0 | 0 | 0 | 0 | 4416700 | 236550 | 0 | 0 | 0 | 0 | 0 | 0 | 0 | 0 |
| IRDKPEEQ\ | CRK | P46108 | CRK_HUMAN A | Unmodified | 0 | 0 | 93663 | 0 | 0 | 0 | 0 | 0 | 185380 | 0 | 0 | 329270 | 0 | 0 | 269470 | 250730 | 0 | 0 |
| IRDPNQGG | EIF4G1; EIF4G1 | Q04637; C9J | IF4G1_HUMAN | Unmodified | 97001 | 141610 | 0 | 202450 | 245180 | 0 | 0 | 0 | 594900 | 123420 | 0 | 719410 | 527950 | 0 | 0 | 85946 | 0 | 0 |
| IRDTSQEP( | ANK3; ANK3; AI | Q12955; A0/ | ANK3_HUMAN | Unmodified | 16027 | 0 | 241850 | 290580 | 0 | 0 | 0 | 0 | 0 | 0 | 18565 | 0 | 145620 | 136820 | 533280 | 501860 | 0 | 0 |
| IRDWGSGS | ATP2A3 | Q93084 | AT2A3_HUMAN | Unmodified | 0 | 0 | 0 | 520830 | 0 | 0 | 0 | 0 | 279570 | 247470 | 451910 | 1566400 | 0 | 0 | 0 | 0 | 73172 | 0 |
| IREDLPNLE | VPS35 | Q96QK1 | VPS35_HUMAN | Unmodified | 878230 | 0 | 1082800 | 1160000 | 3391400 | 0 | 0 | 0 | 2742200 | 467380 | 3307300 | 14461000 | 1993800 | 0 | 1588600 | 1106000 | 0 | 0 |
| IREEFPDRII | TUBB6; TUBB6; | Q9BUF5; K7I | TBB6_HUMAN | Oxidation (M) | 0 | 0 | 0 | 0 | 1338400 | 0 | 0 | 0 | 0 | 0 | 0 | 1513300 | 0 | 0 | 0 | 0 | 0 | 0 |
| IREEFPDRII | TUBB4A; TUBB4 | P04350; M0I | TBB4A_HUMAN | Oxidation (M) | 0 | 0 | 0 | 0 | 0 | 0 | 0 | 1923800 | 613700 | 7400200 | 4770500 | 0 | 22295000 | 7736800 | 0 | 0 | 0 | 0 |
| IREEWNNF | JAK1; JAK1; JAK | P23458; A0A | JAK1_HUMAN 1 | Unmodified | 0 | 0 | 0 | 0 | 1643100 | 687790 | 0 | 0 | 488330 | 0 | 0 | 0 | 0 | 0 | 0 | 0 | 126580 | 190820 |
| IREEYPDRII | TUBB2A; TUBB2 | Q13885; Q9I | TBB2A_HUMAN | Oxidation (M) | 0 | 0 | 0 | 0 | 0 | 465680 | 0 | 0 | 6395800 | 3063300 | 435990 | 24471000 | 0 | 0 | 0 | 0 | 0 | 0 |
| IREEYPDRII | TUBB2A; TUBB2 | Q13885; Q9I | TBB2A_HUMAN | Unmodified | 0 | 0 | 0 | 0 | 472610 | 0 | 121860 | 0 | 4791500 | 956120 | 106080 | 0 | 0 | 0 | 0 | 0 | 669750 | 770630 |
| IREEYPDRII | TUBB2A; TUBB2 | Q13885; Q9I | TBB2A_HUMAN | 2 Oxidation (M) | 0 | 0 | 0 | 0 | 0 | 0 | 0 | 0 | 0 | 1319900 | 1878200 | 9106000 | 0 | 0 | 0 | 0 | 0 | 128120 |
| IREEYPDRII | TUBB; TUBB4B; | P07437; P68 | TBB5_HUMAN | Oxidation (M) | 5619000 | 1665000 | 10147000 | 10660000 | 42926000 | 15045000 | 62704000 | 14037000 | 17128000 | 12792000 | 17481000 | 1.25E+08 | 54113000 | 19260000 | 10635000 | 9362200 | 13041000 | 38776000 |
| IREEYPDRII | TUBB; TUBB4B; | P07437; P68 | TBB5_HUMAN | Unmodified | 986490 | 0 | 1165000 | 2391500 | 75445000 | 4375600 | 66922000 | 2693000 | 45534000 | 15627000 | 2939300 | 26120000 | 28490000 | 13946000 | 8396600 | 1216600 | 10056000 | 43346000 |
| IREFDSSTL | YIPF6 | Q96EC8 | YIPF6_HUMAN | Unmodified | 0 | 0 | 3272500 | 698880 | 0 | 0 | 0 | 0 | 0 | 0 | 0 | 1844300 | 0 | 0 | 519710 | 420580 | 0 | 0 |
| IREGMAAL( | ACADVL; ACAD | P49748; G3\ | ACADV_HUMAI | Oxidation (M) | 0 | 0 | 0 | 0 | 0 | 0 | 0 | 0 | 2198600 | 223330 | 0 | 0 | 0 | 0 | 0 | 0 | 0 | 0 |
| IREIADGLC | TPT1; TPT1; TPT | P13693; E9P | TCTP_HUMAN | Unmodified | 64309 | 0 | 0 | 146770 | 2658000 | 743560 | 847800 | 0 | 785290 | 769780 | 2002200 | 5599700 | 4059000 | 3399900 | 264160 | 0 | 1601100 | 1701400 |
| IREILSDDK | CLASP2; CLASF | O75122; B3I | CLAP2_HUMAN | Unmodified | 0 | 0 | 0 | 0 | 0 | 0 | 0 | 0 | 0 | 0 | 0 | 666890 | 0 | 611100 | 0 | 0 | 0 | 0 |
| IRELESQISI | MYH9 | P35579 | MYH9_HUMAN | Unmodified | 339410 | 0 | 0 | 0 | 0 | 0 | 0 | 0 | 0 | 0 | 0 | 648490 | 0 | 0 | 852110 | 546350 | 0 | 0 |
| IRELESQISI | MYH9 | P35579 | MYH9_HUMAN | Unmodified | 313080 | 0 | 213150 | 2195400 | 3768600 | 880470 | 0 | 0 | 2508600 | 352230 | 1032200 | 1034300 | 570380 | 372930 | 3602300 | 2734600 | 565380 | 408390 |
| IRELGSLPC | SMC3 | Q9UQE7 | SMC3_HUMAN | Unmodified | 0 | 19455 | 124770 | 0 | 860550 | 138880 | 271280 | 111480 | 0 | 409360 | 0 | 280900 | 139090 | 0 | 89814 | 0 | 0 | 0 |
| IRELLQQMI | LANCL1; LANCI | O43813; E9I | LANC1_HUMAI | Unmodified | 0 | 0 | 0 | 0 | 194080 | 0 | 0 | 0 | 726470 | 0 | 0 | 263830 | 0 | 0 | 1172600 | 0 | 0 | 167160 |
| IRELTAVVQ | RPS3; RPS3; RP | P23396; H0\ | RS3_HUMAN 4 | Unmodified | 765110 | 670410 | 225840 | 0 | 0 | 0 | 139550 | 0 | 0 | 0 | 241350 | 7340100 | 0 | 0 | 444120 | 250510 | 23865 | 0 |
| IREPLTAQC | CC2D1A; CC2D | Q6P1N0; A0, | C2D1A_HUMAI | Unmodified | 2201200 | 2500100 | 904820 | 710480 | 1262100 | 134150 | 0 | 0 | 0 | 0 | 0 | 1267300 | 0 | 0 | 587990 | 609280 | 0 | 93340 |
| IREQNLQD | ADD1; ADD1; A | P35611; E7E | ADDA_HUMAN | Unmodified | 84014 | 0 | 0 | 0 | 2912400 | 0 | 52258 | 84487 | 401700 | 148270 | 74614 | 202550 | 0 | 135860 | 0 | 0 | 434360 | 0 |

| | | | | | | | | | | | | | | | | | | | | | | |
|---|---|---|---|---|---|---|---|---|---|---|---|---|---|---|---|---|---|---|---|---|---|---|
| IRETTDFKV | FLNB; FLNB | O75369; E7E | FLNB_HUMAN | Unmodified | 0 | 0 | 0 | 47533 | 11572000 | 5429300 | 0 | 0 | 357890 | 0 | 0 | 2362600 | 1260700 | 992640 | 0 | 293140 | 149470 | 2737800 |
| IREVLGFG( | PLCB3 | Q01970 | PLCB3_HUMAN | Unmodified | 0 | 0 | 0 | 0 | 1283900 | 0 | 0 | 0 | 0 | 0 | 0 | 0 | 0 | 0 | 0 | 0 | 0 | 0 |
| IREYFGGF( | HNRNPD; HNR | Q14103; H0Y | HNRPD_HUMA | Oxidation (M) | 0 | 0 | 0 | 0 | 862740 | 0 | 359110 | 0 | 0 | 787140 | 0 | 3979400 | 1193800 | 926660 | 0 | 0 | 0 | 0 |
| IRFHDFLGI | PRDX6 | P30041 | PRDX6_HUMAN | Unmodified | 0 | 0 | 2569500 | 2394300 | 1203400 | 0 | 0 | 0 | 855850 | 0 | 428010 | 551460 | 5608000 | 641540 | 0 | 0 | 727880 | 0 |
| IRFHDFLGI | PRDX6 | P30041 | PRDX6_HUMAN | Unmodified | 0 | 0 | 0 | 0 | 834620 | 0 | 0 | 0 | 0 | 570650 | 0 | 0 | 6322500 | 1615100 | 0 | 0 | 1542000 | 5782300 |
| IRFQESEEF | HSPA4 | P34932 | HSP74_HUMAN | Unmodified | 2.46E+08 | 17205000 | 14893000 | 34133000 | 4646200 | 339430 | 717660 | 296930 | 648690 | 878200 | 865280 | 8404000 | 1046900 | 539890 | 28025000 | 13949000 | 1113500 | 2020400 |
| IRGAALDVI | CTBP2 | P56545 | CTBP2_HUMAN | Unmodified | 0 | 0 | 0 | 0 | 146900 | 0 | 0 | 0 | 0 | 0 | 0 | 349150 | 152820 | 0 | 0 | 0 | 0 | 0 |
| IRGAALDVI | CTBP1; CTBP1; | Q13363; H0Y | CTBP1_HUMAN | Unmodified | 0 | 0 | 0 | 0 | 0 | 0 | 331830 | 0 | 0 | 0 | 36510 | 719280 | 1631700 | 923350 | 0 | 0 | 0 | 0 |
| IRGETLGII( | CTBP1; CTBP1 | Q13363; D6I | CTBP1_HUMAN | Unmodified | 0 | 0 | 0 | 0 | 0 | 0 | 0 | 0 | 0 | 0 | 0 | 1965200 | 33221 | 0 | 0 | 0 | 0 | 0 |
| IRGIGFGDI | CD2AP | Q9Y5K6 | CD2AP_HUMAI | Unmodified | 0 | 0 | 4295100 | 0 | 591260 | 0 | 0 | 0 | 326840 | 0 | 0 | 180660 | 2916200 | 0 | 0 | 0 | 0 | 0 |
| IRGIGFGDI | CD2AP | Q9Y5K6 | CD2AP_HUMAI | Unmodified | 0 | 204430 | 0 | 0 | 3784700 | 243600 | 0 | 0 | 0 | 162050 | 0 | 0 | 13253000 | 4418500 | 0 | 0 | 0 | 0 |
| IRGIVEESV | AHCYL1 | O43865 | SAHH2_HUMAI | Unmodified | 0 | 0 | 909580 | 0 | 570860 | 0 | 0 | 0 | 0 | 0 | 0 | 0 | 0 | 0 | 0 | 0 | 0 | 0 |
| IRGIVEESV | AHCYL1 | O43865 | SAHH2_HUMAI | Unmodified | 0 | 0 | 0 | 0 | 5758000 | 1370100 | 1852000 | 0 | 0 | 239140 | 0 | 0 | 1715600 | 2098400 | 0 | 0 | 5514700 | 2255700 |
| IRGTSYQSI | RUVBL2; RUVBI | Q9Y230; M0I | RUVB2_HUMAI | Unmodified | 0 | 0 | 0 | 0 | 311890 | 1259300 | 0 | 0 | 0 | 0 | 0 | 2323300 | 0 | 0 | 0 | 0 | 0 | 2271300 |
| IRGTSYQSI | RUVBL2; RUVBI | Q9Y230; M0I | RUVB2_HUMAI | Unmodified | 0 | 0 | 0 | 0 | 0 | 0 | 0 | 0 | 0 | 0 | 668830 | 865120 | 0 | 0 | 0 | 0 | 0 | 0 |
| IRHGWED\ | STEAP1 | Q9UHE8 | STEA1_HUMAN | Unmodified | 437170 | 441500 | 7320900 | 8631800 | 0 | 0 | 0 | 0 | 0 | 0 | 0 | 0 | 1971400 | 916170 | 13770000 | 6558900 | 0 | 0 |
| IRHYAGDV | MYO1D; MYO1I | O94832; J3Q | MYO1D_HUMA | Unmodified | 230560 | 245870 | 0 | 149940 | 0 | 0 | 0 | 0 | 171250 | 0 | 0 | 0 | 0 | 0 | 284860 | 201000 | 0 | 0 |
| IRIWGTEGI | CIAO1 | O76071 | CIAO1_HUMAN | Unmodified | 56560 | 0 | 0 | 99380 | 237190 | 0 | 44015 | 45171 | 193650 | 104440 | 516630 | 789130 | 0 | 0 | 0 | 0 | 296750 | 145540 |
| IRKEPLEFE | DNAJC13; DNA | O75165; A0A | DJC13_HUMAN | Unmodified | 75539 | 0 | 0 | 0 | 13150000 | 1306800 | 0 | 0 | 0 | 0 | 0 | 1750200 | 1896300 | 0 | 331130 | 150110 | 2402300 | 1221300 |
| IRKFQILND | CYFIP1 | Q7L576 | CYFP1_HUMAN | Unmodified | 0 | 0 | 0 | 63393 | 244220 | 35785 | 0 | 0 | 0 | 49984 | 0 | 0 | 148660 | 0 | 0 | 0 | 81782 | 118080 |
| IRKLDNGG | YES1; FYN; FYN | P07947; P06 | YES_HUMAN Ty | Unmodified | 0 | 0 | 0 | 147630 | 527610 | 211560 | 0 | 0 | 518150 | 325260 | 92252 | 0 | 668930 | 253230 | 0 | 0 | 381480 | 486190 |
| IRLEETLEQ | OSBP | P22059 | OSBP1_HUMAI | Unmodified | 0 | 0 | 0 | 600130 | 371930 | 62323 | 0 | 0 | 1389300 | 78779 | 735700 | 2957800 | 731550 | 160280 | 313190 | 208710 | 79460 | 163170 |
| IRLESEEEG | CDK1; CDK1; C | P06493; A0A | CDK1_HUMAN | Unmodified | 0 | 0 | 433230 | 533030 | 0 | 0 | 0 | 0 | 666960 | 0 | 0 | 0 | 0 | 0 | 547130 | 493940 | 0 | 0 |
| IRLESEEEG | CDK1; CDK1; C | P06493; A0A | CDK1_HUMAN | Unmodified | 0 | 0 | 0 | 802950 | 5451000 | 900870 | 1385900 | 48123 | 153820 | 3160700 | 0 | 0 | 6501500 | 1991800 | 0 | 0 | 5158200 | 4937000 |
| IRLHVENEI | ARCN1; ARCN1 | P48444; B0Y | COPD_HUMAN | Unmodified | 0 | 0 | 0 | 0 | 0 | 0 | 0 | 13635 | 0 | 0 | 172800 | 468970 | 141830 | 0 | 0 | 0 | 0 | 0 |
| IRLISLTDEI | HSP90B1; HSP | P14625; A0A | ENPL_HUMAN | Unmodified | 0 | 0 | 0 | 0 | 1267400 | 0 | 3306000 | 0 | 14210000 | 31245000 | 2673700 | 830800 | 1423800 | 1472300 | 0 | 0 | 0 | 1191400 |
| IRLQAYHT( | AK2; AK2; AK2; | P54819; A0A | KAD2_HUMAN | Unmodified | 0 | 0 | 0 | 0 | 0 | 0 | 0 | 0 | 347850 | 0 | 0 | 0 | 0 | 0 | 0 | 0 | 0 | 0 |
| IRLQAYHT( | AK2; AK2; AK2; | P54819; A0A | KAD2_HUMAN | Unmodified | 0 | 0 | 0 | 0 | 1858300 | 0 | 1200500 | 0 | 2024300 | 4263500 | 901840 | 2919900 | 4241200 | 1599700 | 0 | 0 | 367050 | 356810 |
| IRLSEGEAK | MIA3; MIA3 | Q5JRA6; A0A | TGO1_HUMAN | Unmodified | 0 | 0 | 0 | 0 | 0 | 0 | 0 | 0 | 9368700 | 4442700 | 0 | 0 | 0 | 0 | 0 | 0 | 0 | 0 |
| IRLSPEPWI | ACSL3; ACSL3; | O95573; A0A | ACSL3_HUMAN | Unmodified | 0 | 0 | 0 | 142550 | 1318400 | 0 | 299290 | 0 | 0 | 0 | 0 | 217260 | 1906500 | 513850 | 0 | 0 | 0 | 265190 |
| IRMEPDET\ | RAD23A; RAD2 | P54725; A0A | RD23A_HUMAN | Unmodified | 1567300 | 735600 | 427020 | 30149 | 85767 | 0 | 27515 | 0 | 109210 | 25757 | 0 | 280960 | 257340 | 0 | 0 | 0 | 161570 | 0 |
| IRMNVDQA | RRAS2; RRAS2 | P62070; E9P | RRAS2_HUMAN | Unmodified | 0 | 0 | 809270 | 1571200 | 1496000 | 0 | 0 | 0 | 3807300 | 0 | 0 | 0 | 0 | 474990 | 1842700 | 0 | 0 | 427670 |
| IRNEDLIFN | JAK2; JAK2 | O60674; A0A | JAK2_HUMAN T | Unmodified | 0 | 0 | 0 | 0 | 0 | 0 | 0 | 0 | 0 | 341540 | 0 | 0 | 292490 | 0 | 0 | 0 | 438150 | 1762300 |
| IRNIGISAHI | GFM1; GFM1; G | Q96RP9; C9I | EFGM_HUMAN | Unmodified | 0 | 0 | 0 | 0 | 0 | 0 | 0 | 0 | 702120 | 0 | 0 | 0 | 0 | 0 | 0 | 0 | 49836 | 0 |
| IRNIIQQAII | DNAJC9 | Q8WXX5 | DNJC9_HUMAI | Unmodified | 0 | 0 | 0 | 0 | 0 | 0 | 0 | 0 | 0 | 340910 | 0 | 0 | 0 | 0 | 0 | 0 | 0 | 0 |
| IRNSFVNN | ARHGAP5 | Q13017 | RHG05_HUMAI | Unmodified | 0 | 0 | 0 | 0 | 1517200 | 0 | 0 | 0 | 0 | 0 | 0 | 0 | 0 | 0 | 0 | 0 | 107410 | 0 |
| IRNVAYDTI | PLOD3; PLOD3 | O60568; H7( | PLOD3_HUMAI | Unmodified | 0 | 0 | 0 | 0 | 0 | 0 | 0 | 0 | 9952700 | 2967000 | 0 | 0 | 0 | 0 | 0 | 0 | 0 | 0 |
| IRNVGLCE/ | CCZ1; CCZ1B; ( | P86791; P86 | CCZ1_HUMAN | Unmodified | 0 | 0 | 0 | 0 | 320910 | 0 | 0 | 0 | 79358 | 43218 | 0 | 0 | 223050 | 138190 | 0 | 0 | 23387 | 0 |
| IRPFFPQQ | FGB; FGB | P02675; D6F | FIBB_HUMAN F | Unmodified | 0 | 9883700 | 0 | 0 | 0 | 0 | 0 | 0 | 106680 | 0 | 0 | 0 | 0 | 3709900 | 0 | 0 | 0 | 0 |
| IRPGEQEQ | EPB41; EPB41; | P11171; A0A | EPB41_HUMAN | Unmodified | 854460 | 0 | 448270 | 1612900 | 0 | 0 | 0 | 0 | 0 | 0 | 0 | 1840700 | 0 | 0 | 1367800 | 1859500 | 0 | 0 |
| IRPGEYEQI | EPB41L1; EPB4 | Q9H4G0; A0 | E41L1_HUMAN | Unmodified | 445040 | 169190 | 573900 | 907750 | 0 | 0 | 0 | 0 | 522630 | 0 | 461920 | 3019700 | 0 | 0 | 2378500 | 1656200 | 0 | 0 |
| IRPTAVTLEI | SUN2; SUN2 | Q9UH99; B0 | SUN2_HUMAN | Unmodified | 0 | 0 | 0 | 0 | 0 | 0 | 0 | 0 | 7551100 | 417540 | 0 | 0 | 0 | 0 | 0 | 0 | 0 | 0 |
| IRQEDENFI | ERBIN | Q96RT1 | ERBIN_HUMAN | Unmodified | 0 | 611760 | 0 | 2651600 | 31368000 | 0 | 495020 | 1062800 | 0 | 321960 | 905390 | 0 | 0 | 0 | 0 | 257980 | 0 | 0 |
| IRQEEIEIEV | FLOT2; FLOT2; I | Q14254; J3C | FLOT2_HUMAN | Unmodified | 0 | 0 | 1930300 | 13778000 | 5528700 | 75359 | 640690 | 0 | 7868300 | 5625700 | 2569600 | 4505900 | 504790 | 347130 | 5555200 | 2296400 | 0 | 173530 |
| IRQEEIEIEV | FLOT2; FLOT2; I | Q14254; J3C | FLOT2_HUMAN | Unmodified | 195140 | 87472 | 0 | 555190 | 4971100 | 274480 | 0 | 157290 | 2278300 | 1221900 | 348020 | 0 | 430820 | 0 | 215060 | 233060 | 113510 | 468900 |
| IRQLEEEK | MYH10 | P35580 | MYH10_HUMAI | Unmodified | 0 | 0 | 0 | 0 | 0 | 0 | 0 | 114880 | 0 | 0 | 14617 | 1585500 | 0 | 0 | 0 | 0 | 0 | 0 |
| IRRPHDYQ | U2AF2; U2AF2 | P26368; K7E | U2AF2_HUMAN | Oxidation (M) | 1004200 | 1084300 | 0 | 0 | 0 | 0 | 0 | 0 | 0 | 0 | 0 | 951390 | 0 | 0 | 0 | 730760 | 0 | 0 |
| IRRQLPAP( | AHNAK2 | Q8IVF2 | AHNK2_HUMAI | Unmodified | 0 | 0 | 0 | 0 | 2933300 | 0 | 0 | 0 | 0 | 0 | 0 | 0 | 0 | 0 | 0 | 0 | 478730 | 4051900 |
| IRSHVATC! | RNF114; RNF11 | Q9Y508; A0A | RN114_HUMAN | Unmodified | 0 | 0 | 105580 | 0 | 0 | 0 | 0 | 0 | 172940 | 0 | 0 | 0 | 0 | 427880 | 0 | 0 | 357240 | 0 |
| IRSLDNGG | LYN | P07948 | LYN_HUMAN T | Unmodified | 0 | 0 | 115140 | 401030 | 2883500 | 1199100 | 448220 | 199860 | 0 | 821090 | 0 | 0 | 1228100 | 2429500 | 123060 | 0 | 0 | 0 |
| IRSLQEQAI | TPM1; TPM1; TF | H0YKJ4; H0Y | H0YKJ4_HUMA | Unmodified | 186790 | 0 | 211580 | 292540 | 56406 | 0 | 0 | 0 | 361910 | 72078 | 44652 | 1236900 | 375470 | 84446 | 455520 | 0 | 0 | 14716 |
| IRSSSGILSI | NRP1; NRP1; N | O14786; Q5. | NRP1_HUMAN | Oxidation (M) | 0 | 0 | 0 | 0 | 0 | 183680 | 0 | 0 | 0 | 0 | 0 | 0 | 0 | 0 | 0 | 0 | 0 | 1143300 |
| IRSTLNEIYI | CAPZB; CAPZB; | P47756; B1A | CAPZB_HUMAN | Unmodified | 2931000 | 2926200 | 1666000 | 2430200 | 8760400 | 2364700 | 1260100 | 715300 | 3851000 | 2158200 | 0 | 11409000 | 4488700 | 3167400 | 3206400 | 2498200 | 2479700 | 3781700 |
| IRTDAGFTL | VCL | P18206 | VINC_HUMAN | Unmodified | 0 | 0 | 0 | 0 | 0 | 0 | 133670 | 146100 | 216810 | 155230 | 0 | 679910 | 494550 | 0 | 0 | 186710 | 475920 | 0 |
| IRTFLAQH/ | ENY2 | Q9NPA8 | ENY2_HUMAN | Unmodified | 0 | 0 | 0 | 0 | 1411900 | 0 | 107890 | 0 | 336680 | 304450 | 242870 | 0 | 610230 | 311660 | 0 | 0 | 882090 | 781790 |
| IRTHNSEVI | JAG1 | P78504 | JAG1_HUMAN I | Unmodified | 0 | 0 | 0 | 0 | 160690 | 0 | 137400 | 20793 | 268310 | 279870 | 571690 | 480390 | 2976700 | 4196400 | 18277 | 74349 | 1133800 | 2672300 |
| IRTLDNGGI | HCK; HCK; HCI | P08631; J3KI | HCK_HUMAN T | Unmodified | 42246 | 112760 | 0 | 0 | 0 | 0 | 0 | 0 | 0 | 0 | 0 | 1211000 | 0 | 0 | 0 | 0 | 0 | 0 |
| IRTLQEQLE | RRBP1; RRBP1; | Q9P2E9; A0A | RRBP1_HUMAN | Unmodified | 0 | 0 | 0 | 0 | 0 | 0 | 0 | 83108 | 0 | 0 | 0 | 270210 | 582150 | 381760 | 0 | 37095 | 0 | 0 |
| IRTVEINGE | RAB35; RAB35 | Q15286; F5I | RAB35_HUMAN | Unmodified | 228480 | 0 | 0 | 0 | 0 | 0 | 50442 | 0 | 0 | 0 | 0 | 0 | 0 | 0 | 479000 | 203880 | 0 | 92411 |
| IRTVQDALE | USP10; USP10; | Q14694; H3I | UBP10_HUMAN | Unmodified | 0 | 0 | 0 | 0 | 0 | 0 | 0 | 0 | 81590 | 0 | 98734 | 622010 | 0 | 0 | 0 | 40838 | 0 | 0 |
| IRVDVADQ. | EIF4B; EIF4B; EI | P23588; F8V | IF4B_HUMAN E | Unmodified | 606850 | 242380 | 451240 | 119860 | 3179900 | 897810 | 0 | 0 | 0 | 0 | 1659300 | 4712600 | 0 | 0 | 879210 | 670380 | 0 | 0 |
| IRVESLLVT/ | DHX15 | O43143 | DHX15_HUMAI | Unmodified | 0 | 0 | 0 | 0 | 611750 | 0 | 672520 | 0 | 650790 | 487280 | 809770 | 3542100 | 433440 | 0 | 0 | 0 | 0 | 366720 |
| IRVLEAEKE | CEP55 | Q53EZ4 | CEP55_HUMAN | Unmodified | 0 | 0 | 0 | 73154 | 12093000 | 1422100 | 210920 | 67761 | 0 | 3374100 | 0 | 0 | 548010 | 590640 | 0 | 0 | 562230 | 493600 |
| IRVVLNKAI | EHD4 | Q9H223 | EHD4_HUMAN | Unmodified | 68629 | 118800 | 0 | 0 | 0 | 0 | 0 | 0 | 0 | 0 | 0 | 0 | 0 | 0 | 0 | 116560 | 55942 | 0 |
| IRVVLNKAI | EHD4 | Q9H223 | EHD4_HUMAN | Oxidation (M) | 0 | 0 | 0 | 0 | 0 | 0 | 0 | 0 | 0 | 0 | 0 | 0 | 0 | 0 | 0 | 305330 | 0 | 0 |
| IRWNDGLI | PAPSS2 | O95340 | PAPS2_HUMAN | Unmodified | 0 | 0 | 0 | 0 | 1605900 | 199520 | 24212 | 0 | 0 | 0 | 0 | 0 | 0 | 0 | 0 | 0 | 1650600 | 1367300 |
| IRYESGDH | POR; POR; POR | P16435; H0Y | NCPR_HUMAN | Unmodified | 0 | 0 | 0 | 0 | 0 | 0 | 0 | 0 | 555030 | 0 | 0 | 1919200 | 0 | 0 | 0 | 0 | 0 | 0 |
| IRYESLTDP | HSP90AB4P; H: | Q58FF6; P0I | H90B4_HUMAN | Unmodified | 0 | 0 | 1548100 | 0 | 0 | 0 | 0 | 0 | 755140 | 0 | 766550 | 10339000 | 0 | 0 | 1699500 | 0 | 0 | 0 |
| IRYESLTDP | HSP90AB1; HSI | P08238; Q5I | HS90B_HUMAN | Unmodified | 5373800 | 8451800 | 8352600 | 19527000 | 44388000 | 16132000 | 2333900 | 2091400 | 56569000 | 15754000 | 52291000 | 3.03E+08 | 20568000 | 5964100 | 15040000 | 14280000 | 16594000 | 31380000 |
| IRYGGRDY | MYOF; MYOF | Q9NZM1; H( | MYOF_HUMAN | Unmodified | 0 | 0 | 0 | 40628 | 1662300 | 395530 | 0 | 0 | 0 | 0 | 0 | 0 | 0 | 216520 | 0 | 0 | 0 | 0 |
| ISAAASDS( | EPS8; EPS8; EP | Q12929; A0A | EPS8_HUMAN | Unmodified | 191170 | 335050 | 0 | 104180 | 3957500 | 962590 | 20367 | 234950 | 250150 | 0 | 0 | 783570 | 306170 | 781420 | 0 | 0 | 668050 | 390530 |
| ISAAATFIQI | PKP2 | Q99959 | PKP2_HUMAN | Unmodified | 92789 | 0 | 128480 | 230120 | 0 | 0 | 0 | 0 | 121550 | 0 | 431110 | 3555900 | 0 | 0 | 381530 | 195180 | 0 | 76311 |
| ISAASSVPR | SH3GL3 | Q99963 | SH3G3_HUMAI | Unmodified | 0 | 3406800 | 3033800 | 2112800 | 0 | 0 | 0 | 0 | 0 | 0 | 0 | 0 | 1448500 | 1089200 | 881370 | 143540 | 0 | 0 |
| ISADTHQF! | ERGIC2; ERGIC | Q96RQ1; A0. | ERGI2_HUMAN | Unmodified | 0 | 0 | 0 | 0 | 0 | 0 | 0 | 0 | 3970700 | 730840 | 121660 | 0 | 0 | 0 | 0 | 0 | 0 | 0 |
| ISADTHQF! | ERGIC2; ERGIC | Q96RQ1; A0. | ERGI2_HUMAN | Unmodified | 0 | 0 | 0 | 0 | 37222 | 0 | 0 | 0 | 15488000 | 0 | 362380 | 0 | 54599 | 0 | 0 | 0 | 0 | 130820 |
| ISAESPTGL | SYTL4 | Q96C24 | SYTL4_HUMAN | Unmodified | 0 | 0 | 0 | 0 | 2374300 | 187990 | 146100 | 178830 | 1276800 | 465640 | 0 | 0 | 145580 | 0 | 0 | 148090 | 330560 | 0 |
| ISAFGYLEC | RHOC; RHOC; I | P08134; E9P | RHOC_HUMAN | Unmodified | 3455600 | 2778300 | 0 | 4482300 | 1415100 | 0 | 0 | 0 | 0 | 0 | 0 | 1771100 | 1647200 | 40406 | 7471300 | 5579800 | 584560 | 2133900 |
| ISAILHPST\ | WDR36; WDR3( | Q8NI36; A0A | WDR36_HUMA | Unmodified | 0 | 0 | 0 | 0 | 0 | 0 | 0 | 0 | 0 | 0 | 0 | 719450 | 0 | 0 | 0 | 0 | 0 | 0 |
| ISAKAALAH | CDK2; CDK2; C | P24941; G3\ | CDK2_HUMAN | Unmodified | 0 | 0 | 0 | 0 | 129650 | 0 | 0 | 0 | 0 | 0 | 0 | 0 | 46625 | 0 | 0 | 0 | 36099 | 0 |
| ISALALLK | COPS5 | Q92905 | CSN5_HUMAN | Unmodified | 69494 | 0 | 0 | 0 | 497790 | 0 | 0 | 0 | 1110500 | 0 | 94914 | 1592900 | 0 | 0 | 0 | 188830 | 804640 | 786930 |
| ISALNIVGD | NDEL1; NDE1; I | Q9GZM8; Q! | NDEL1_HUMAI | Unmodified | 50644 | 0 | 0 | 0 | 831640 | 105120 | 0 | 0 | 446830 | 230150 | 152410 | 0 | 952450 | 0 | 0 | 0 | 238530 | 532040 |
| ISANENSL/ | LIMA1; LIMA1; L | Q9UHB6; F8 | LIMA1_HUMAN | Unmodified | 702960 | 0 | 270960 | 0 | 2035600 | 0 | 0 | 0 | 86844 | 0 | 0 | 866620 | 0 | 0 | 899780 | 498370 | 0 | 106740 |
| ISANENSL/ | LIMA1; LIMA1; L | Q9UHB6; F8 | LIMA1_HUMAN | Unmodified | 0 | 0 | 0 | 0 | 1480800 | 0 | 0 | 0 | 0 | 0 | 0 | 0 | 671160 | 750030 | 0 | 0 | 0 | 401870 |
| ISAPNVDFI | AHNAK | Q09666 | AHNK_HUMAN | Unmodified | 0 | 0 | 91818 | 0 | 10502000 | 1256500 | 0 | 0 | 0 | 0 | 0 | 1453600 | 0 | 0 | 0 | 0 | 1059600 | 4040900 |
| ISAPNVDFI | AHNAK | Q09666 | AHNK_HUMAN | Unmodified | 0 | 0 | 0 | 0 | 2045900 | 160060 | 0 | 0 | 0 | 0 | 0 | 0 | 207680 | 157810 | 0 | 0 | 939470 | 1734000 |
| ISAQVVDY\ | PTPN23 | Q9H3S7 | PTN23_HUMAN | Unmodified | 0 | 1560600 | 0 | 0 | 0 | 0 | 0 | 0 | 0 | 0 | 0 | 0 | 0 | 0 | 0 | 0 | 0 | 0 |
| ISASLAELK | MGAT5 | Q09328 | MGT5A_HUMAI | Unmodified | 0 | 0 | 0 | 0 | 0 | 0 | 0 | 0 | 527610 | 0 | 20509 | 834570 | 0 | 0 | 0 | 0 | 0 | 0 |
| ISATEDTPE | KIFBP; KIFBP; K | Q96EK5; A0A | KBP_HUMAN K | Unmodified | 0 | 0 | 0 | 0 | 0 | 0 | 0 | 0 | 0 | 0 | 0 | 1353600 | 0 | 0 | 0 | 0 | 0 | 0 |
| ISATLKREF( | NUDT9; NUDT9 | Q9BW91; D6 | NUDT9_HUMAI | Unmodified | 0 | 0 | 0 | 0 | 364790 | 0 | 0 | 0 | 0 | 244520 | 0 | 0 | 677440 | 0 | 0 | 0 | 0 | 0 |
| ISATSIFFES | SHMT2; SHMT1 | P34897; P34 | GLYM_HUMAN | Unmodified | 0 | 0 | 0 | 0 | 0 | 0 | 0 | 0 | 2857300 | 71921 | 0 | 247880 | 0 | 0 | 0 | 0 | 0 | 0 |
| ISATSIFFES | SHMT2; SHMT2 | P34897; G3\ | GLYM_HUMAN | Unmodified | 0 | 2100400 | 0 | 0 | 0 | 0 | 0 | 0 | 2509200 | 871630 | 0 | 0 | 0 | 0 | 0 | 0 | 18905 | 392850 |
| ISAVSVAER | DHX9 | Q08211 | DHX9_HUMAN | Unmodified | 0 | 0 | 0 | 0 | 0 | 0 | 0 | 0 | 1027000 | 161160 | 178100 | 1695500 | 0 | 0 | 0 | 0 | 57770 | 0 |
| ISCTIANR | FN1 | P02751 | FINC_HUMAN I | Unmodified | 530860 | 1350700 | 11642000 | 4909200 | 314620 | 1451300 | 0 | 36316 | 3648800 | 834250 | 636520 | 3652400 | 1737300 | 2392800 | 0 | 2655600 | 198110 | 2165300 |
| ISCTIANRC | FN1 | P02751 | FINC_HUMAN I | Unmodified | 0 | 0 | 0 | 0 | 0 | 2461200 | 0 | 0 | 0 | 415330 | 0 | 0 | 801650 | 2362100 | 0 | 0 | 97877 | 1102400 |
| ISDAIMNM( | GPC4 | O75487 | GPC4_HUMAN | Unmodified | 425330 | 417250 | 0 | 0 | 0 | 0 | 0 | 0 | 3836500 | 0 | 0 | 0 | 729300 | 0 | 0 | 0 | 40255 | 81157 |
| ISDAIMNM( | GPC4 | O75487 | GPC4_HUMAN | 2 Oxidation (M) | 821670 | 470940 | 0 | 0 | 0 | 0 | 0 | 0 | 0 | 0 | 0 | 0 | 0 | 0 | 0 | 0 | 0 | 0 |
| ISDAIMNM( | GPC4 | O75487 | GPC4_HUMAN | Oxidation (M) | 659860 | 489090 | 0 | 212120 | 0 | 0 | 0 | 0 | 661940 | 0 | 0 | 0 | 0 | 0 | 0 | 0 | 0 | 0 |
| ISDDLMQK | PSMD3; PSMD3 | O43242; H0Y | PSMD3_HUMA | Unmodified | 61997 | 0 | 47474 | 0 | 71861 | 36643 | 106310 | 0 | 509780 | 76354 | 178960 | 2434000 | 0 | 0 | 62007 | 505320 | 0 | 0 |
| ISDEDWDII | HSD17B4; HSD | P51659; E7E | DHB4_HUMAN | Unmodified | 160760 | 0 | 0 | 0 | 0 | 0 | 0 | 0 | 12717000 | 0 | 1792300 | 0 | 0 | 0 | 0 | 172830 | 0 | 0 |
| ISDEDWDII | HSD17B4; HSD | P51659; E7E | DHB4_HUMAN | Unmodified | 0 | 0 | 0 | 0 | 0 | 0 | 0 | 0 | 20574000 | 1174800 | 0 | 0 | 0 | 0 | 0 | 0 | 0 | 0 |
| ISDEGIAYL' | RFC4; RFC4 | P35249; C9J | RFC4_HUMAN | Unmodified | 0 | 0 | 0 | 0 | 0 | 0 | 0 | 0 | 516840 | 0 | 0 | 0 | 0 | 0 | 0 | 0 | 0 | 0 |
| ISDFGLSK | CAMK1; ZAP70; | Q14012; P4 | KCC1A_HUMAI | Unmodified | 0 | 0 | 0 | 0 | 0 | 0 | 237010 | 48867 | 0 | 75857 | 273580 | 1020400 | 0 | 0 | 0 | 72652 | 0 | 0 |
| ISDFGLSRI | RET | P07949 | RET_HUMAN P | Unmodified | 0 | 0 | 0 | 0 | 0 | 0 | 0 | 0 | 2219100 | 352560 | 0 | 0 | 0 | 0 | 0 | 0 | 0 | 0 |
| ISDFGLSRI | RET | P07949 | RET_HUMAN P | Unmodified | 0 | 0 | 0 | 0 | 0 | 0 | 0 | 0 | 1043500 | 0 | 0 | 0 | 0 | 0 | 0 | 0 | 0 | 0 |
| ISDIGLADL | ABCC5 | O15440 | MRP5_HUMAN | Unmodified | 0 | 0 | 0 | 0 | 0 | 0 | 0 | 0 | 0 | 0 | 0 | 4972300 | 0 | 0 | 0 | 0 | 0 | 0 |
| ISDLGGGV | PDK3 | Q15120 | PDK3_HUMAN | Unmodified | 244090 | 270610 | 0 | 0 | 0 | 799430 | 169810 | 102690 | 1069200 | 241320 | 0 | 446130 | 0 | 0 | 0 | 114010 | 179190 | 589530 |
| ISDLGLAVI | GRK6; GRK6; G | P43250; D6F | GRK6_HUMAN | Unmodified | 0 | 464080 | 1294200 | 1052200 | 0 | 0 | 0 | 0 | 1640000 | 0 | 0 | 1156900 | 0 | 0 | 1483800 | 750580 | 0 | 0 |
| ISDLGLAVI | GRK5 | P34947 | GRK5_HUMAN | Unmodified | 90051 | 179050 | 0 | 147390 | 0 | 0 | 0 | 0 | 118380 | 0 | 0 | 1735500 | 0 | 0 | 886430 | 20233 | 0 | 262190 |
| ISDLGLAVI | GRK5 | P34947 | GRK5_HUMAN | Unmodified | 0 | 0 | 0 | 0 | 0 | 0 | 0 | 0 | 0 | 0 | 142560 | 766930 | 0 | 0 | 0 | 0 | 0 | 0 |

| | | | | | | | | | | | | | | | | | | | | | | |
|---|---|---|---|---|---|---|---|---|---|---|---|---|---|---|---|---|---|---|---|---|---|---|
| ISDLGLSR | ROR1; ROR1 | Q01973; A0/ | ROR1_HUMAN | Unmodified | 35601 | 34504 | 187980 | 0 | 0 | 0 | 0 | 0 | 0 | 0 | 0 | 1002400 | 0 | 0 | 228800 | 133060 | 0 | 0 |
| ISDLGLSRE | ROR1; ROR1 | Q01973; A0/ | ROR1_HUMAN | Unmodified | 0 | 0 | 0 | 166640 | 2090900 | 1551400 | 413920 | 0 | 0 | 0 | 205500 | 0 | 0 | 90936 | 0 | 0 | 0 | 0 |
| ISDLGVAEA | STK11; STK11; S | Q15831; K7E | STK11_HUMAN | Unmodified | 126670 | 0 | 82582 | 207970 | 38888 | 0 | 0 | 0 | 228790 | 0 | 0 | 517790 | 0 | 0 | 217900 | 193110 | 52310 | 0 |
| ISDLISSITQ | KDF1 | Q8NAX2 | KDF1_HUMAN | Unmodified | 0 | 0 | 524070 | 761020 | 96179 | 0 | 0 | 0 | 2262800 | 381230 | 619660 | 1383100 | 0 | 0 | 2899000 | 1753800 | 0 | 0 |
| ISDNGPYE | IGSF21 | Q96ID5 | IGS21_HUMAN | Unmodified | 0 | 0 | 0 | 0 | 0 | 0 | 0 | 0 | 1351800 | 0 | 0 | 576890 | 0 | 0 | 0 | 0 | 0 | 0 |
| ISDPNFVK | PI4K2A; N/A | Q9BTU6; E9I | P4K2A_HUMAN | Unmodified | 0 | 0 | 0 | 0 | 0 | 0 | 0 | 0 | 0 | 0 | 0 | 404110 | 0 | 0 | 0 | 107850 | 0 | 0 |
| ISDSEGFK | SFPQ | P23246 | SFPQ_HUMAN | Unmodified | 0 | 0 | 0 | 0 | 0 | 0 | 0 | 0 | 601390 | 352470 | 37574 | 4385600 | 1947300 | 0 | 0 | 0 | 0 | 0 |
| ISDYVGTCI | MBLAC2 | Q68D91 | MBLC2_HUMAI | Unmodified | 0 | 0 | 213740 | 0 | 1196700 | 652740 | 0 | 0 | 0 | 0 | 115200 | 985060 | 0 | 0 | 217120 | 198440 | 423220 | 0 |
| ISEDERMEI | C1GALT1C1 | Q96EU7 | C1GLC_HUMAI | Unmodified | 0 | 0 | 79492 | 330870 | 0 | 0 | 0 | 0 | 582610 | 0 | 0 | 59541 | 0 | 0 | 0 | 0 | 0 | 0 |
| ISEDETERN | ACIN1; ACIN1; | Q9UKV3; G3 | ACINU_HUMAN | Unmodified | 0 | 0 | 0 | 0 | 0 | 0 | 0 | 0 | 0 | 0 | 23612 | 156190 | 0 | 0 | 0 | 0 | 0 | 0 |
| ISEDKNPD' | JUP | P14923 | PLAK_HUMAN . | Unmodified | 0 | 0 | 485250 | 0 | 749270 | 0 | 0 | 0 | 870350 | 279020 | 267670 | 2587500 | 183030 | 0 | 1203500 | 1466100 | 346710 | 165070 |
| ISEDTAILK | KIAA0319L; KIA | Q8IZA0; B1A | K319L_HUMAN | Unmodified | 237600 | 692630 | 542150 | 467970 | 0 | 125840 | 190350 | 145550 | 10274000 | 3298400 | 0 | 552900 | 0 | 1737900 | 479250 | 587340 | 0 | 553700 |
| ISEECIAQW | YARS1; YARS1 | P54577; A0A | SYYC_HUMAN | Unmodified | 0 | 0 | 0 | 0 | 0 | 0 | 487730 | 0 | 0 | 0 | 0 | 2138600 | 0 | 0 | 0 | 0 | 0 | 0 |
| ISEEDELDT | PEA15; PEA15 | Q15121; B1/ | PEA15_HUMAN | Unmodified | 1188700 | 5775700 | 855770 | 0 | 0 | 0 | 0 | 0 | 685550 | 0 | 0 | 0 | 0 | 208130 | 0 | 229340 | 0 | 0 |
| ISEEDELDT | PEA15; PEA15 | Q15121; B1/ | PEA15_HUMAN | Unmodified | 0 | 0 | 0 | 0 | 954390 | 0 | 228830 | 0 | 0 | 0 | 0 | 0 | 1328500 | 1199800 | 0 | 0 | 0 | 0 |
| ISEIEDAAFI | ERLIN2; ERLIN1 | O94905; O7! | ERLN2_HUMAN | Unmodified | 0 | 0 | 0 | 0 | 0 | 0 | 0 | 0 | 5161100 | 363960 | 0 | 0 | 0 | 0 | 0 | 0 | 0 | 0 |
| ISEIEDAAFI | ERLIN2; ERLIN1 | O94905; O7! | ERLN2_HUMAN | Unmodified | 0 | 0 | 0 | 0 | 0 | 0 | 0 | 0 | 2849500 | 2058700 | 165720 | 0 | 370040 | 0 | 0 | 0 | 0 | 0 |
| ISEILLDHG | ANK1 | P16157 | ANK1_HUMAN | Unmodified | 245370 | 0 | 163980 | 672460 | 0 | 0 | 1498400 | 329810 | 0 | 0 | 0 | 0 | 558470 | 0 | 278260 | 270960 | 0 | 0 |
| ISEIQMTAP | DST; DST; DST; | Q03001; A0/ | DYST_HUMAN | Oxidation (M) | 482850 | 996990 | 2975700 | 2108400 | 0 | 0 | 0 | 0 | 0 | 0 | 0 | 0 | 0 | 0 | 0 | 0 | 0 | 0 |
| ISEKEEVTTI | LMNB2 | Q03252 | LMNB2_HUMAI | Unmodified | 0 | 0 | 0 | 0 | 1235000 | 0 | 0 | 0 | 0 | 117210 | 0 | 0 | 0 | 0 | 0 | 0 | 79253 | 0 |
| ISELDAFLK | PSME1 | Q06323 | PSME1_HUMAI | Unmodified | 0 | 0 | 0 | 0 | 0 | 0 | 0 | 27939 | 926580 | 0 | 580120 | 1455000 | 0 | 0 | 0 | 159420 | 0 | 0 |
| ISELDAFLK | PSME1 | Q06323 | PSME1_HUMAI | Unmodified | 0 | 0 | 0 | 0 | 146580 | 0 | 575170 | 0 | 0 | 77081 | 0 | 0 | 109530 | 0 | 0 | 0 | 46091 | 285240 |
| ISELERNVE | LAMB1; LAMB1; | P07942; G3) | LAMB1_HUMAI | Unmodified | 370330 | 0 | 312050 | 310410 | 0 | 0 | 0 | 0 | 0 | 0 | 0 | 0 | 0 | 0 | 0 | 0 | 0 | 56772 |
| ISELGAGN | MAP2K1 | Q02750 | MP2K1_HUMAI | Unmodified | 0 | 0 | 0 | 0 | 729470 | 0 | 0 | 0 | 0 | 0 | 381500 | 2167400 | 0 | 0 | 510350 | 379880 | 0 | 0 |
| ISELGAGN | MAP2K2 | P36507 | MP2K2_HUMAI | Unmodified | 0 | 0 | 0 | 0 | 0 | 0 | 0 | 40242 | 0 | 0 | 125890 | 420470 | 0 | 0 | 0 | 0 | 0 | 0 |
| ISELKAEAV | IGFBP5; IGFBP! | P24593; C9J | IBP5_HUMAN I | Unmodified | 0 | 0 | 0 | 0 | 0 | 0 | 0 | 57316 | 1794300 | 2569900 | 0 | 0 | 15407 | 0 | 0 | 0 | 0 | 0 |
| ISEMGSYQ | ABCC1; ABCC1 | P33527; I3L4 | MRP1_HUMAN | Oxidation (M) | 0 | 0 | 224710 | 136990 | 0 | 466780 | 0 | 0 | 0 | 614890 | 1268300 | 2359100 | 0 | 0 | 264800 | 310490 | 0 | 0 |
| ISEMGSYQ | ABCC1; ABCC1 | P33527; I3L4 | MRP1_HUMAN | Unmodified | 0 | 0 | 0 | 0 | 0 | 0 | 0 | 0 | 0 | 0 | 40483 | 225490 | 0 | 0 | 0 | 0 | 0 | 0 |
| ISEPEAEISI | ERBIN | Q96RT1 | ERBIN_HUMAN | Oxidation (M) | 0 | 0 | 0 | 0 | 5272000 | 1454800 | 0 | 0 | 0 | 0 | 0 | 363870 | 0 | 0 | 0 | 326800 | 452990 | 0 |
| ISEPEAEISI | ERBIN | Q96RT1 | ERBIN_HUMAN | Unmodified | 0 | 0 | 0 | 0 | 5279500 | 0 | 0 | 23132 | 0 | 0 | 0 | 0 | 0 | 0 | 0 | 278670 | 0 | 0 |
| ISEQFTAMI | TUBB2A; TUBB4 | Q13885; P04 | TBB2A_HUMAN | Unmodified | 0 | 0 | 0 | 0 | 493020 | 0 | 0 | 55993 | 1335100 | 115220 | 0 | 9028100 | 0 | 0 | 0 | 0 | 460280 | 0 |
| ISEQFTAMI | TUBB2A; TUBB4 | Q13885; P04 | TBB2A_HUMAN | Oxidation (M) | 1144700 | 561620 | 0 | 503820 | 1244300 | 0 | 981130 | 520940 | 2093900 | 649170 | 1552400 | 38394000 | 489820 | 30262 | 655090 | 755290 | 523250 | 252430 |
| ISEQFTAMI | TUBB2A; TUBB4 | Q13885; P04 | TBB2A_HUMAN | Unmodified | 0 | 0 | 0 | 0 | 0 | 0 | 0 | 0 | 1370400 | 0 | 0 | 0 | 0 | 0 | 0 | 0 | 0 | 0 |
| ISEQTYQLS | STXBP1; STXBP | P61764; A0A | STXB1_HUMAN | Unmodified | 96663 | 126290 | 0 | 0 | 0 | 0 | 0 | 0 | 0 | 21328 | 0 | 428810 | 0 | 0 | 0 | 0 | 0 | 0 |
| ISERNELA | N/A | N/A | N/A | Unmodified | 67486000 | 47916000 | 6846500 | 3413300 | 79189000 | 22000000 | 7320500 | 885120 | 41756000 | 23682000 | 11325000 | 25366000 | 1.03E+08 | 18356000 | 13816000 | 7217900 | 7045800 | 39794000 |
| ISERTPSAL | ALDOC; ALDOC | P09972; J3Q | ALDOC_HUMA | Unmodified | 0 | 0 | 0 | 389190 | 0 | 0 | 0 | 0 | 0 | 0 | 1593600 | 1791600 | 3067700 | 923700 | 0 | 370460 | 1601800 | 1321000 |
| ISESAKQEL | SNX6; SNX6 | Q9UNH7; A0 | SNX6_HUMAN | Unmodified | 127190 | 0 | 299580 | 259770 | 9289100 | 1548700 | 1441500 | 954780 | 1008600 | 586770 | 627300 | 4725500 | 3546400 | 1991200 | 333160 | 309700 | 0 | 0 |
| ISEVLQK | CKAP4 | Q07065 | CKAP4_HUMAI | Unmodified | 0 | 0 | 45827 | 72214 | 0 | 0 | 0 | 64046 | 3864900 | 234530 | 320820 | 0 | 1403400 | 1017100 | 96259 | 78572 | 111560 | 82870 |
| ISFCADDK | GULP1 | Q9UBP9 | GULP1_HUMAI | Unmodified | 49612 | 0 | 174730 | 309500 | 0 | 0 | 0 | 0 | 0 | 0 | 0 | 0 | 161790 | 0 | 71697 | 187270 | 0 | 0 |
| ISFEDEPR | SLC4A7; SLC4A | Q9Y6M7; A0. | S4A7_HUMAN | Unmodified | 0 | 0 | 0 | 0 | 135660 | 0 | 0 | 0 | 0 | 0 | 1552300 | 1356300 | 0 | 0 | 0 | 111070 | 0 | 501750 |
| ISFEDEPRK | SLC4A7; SLC4A | Q9Y6M7; A0. | S4A7_HUMAN | Unmodified | 0 | 0 | 0 | 68621 | 3578000 | 134420 | 649280 | 0 | 204370 | 0 | 97993 | 58170 | 274740 | 182020 | 0 | 15128 | 846060 | 3815100 |
| ISFEEFCAV | PPP3R1; PPP3R | P63098; F6U | CANB1_HUMAI | Unmodified | 0 | 0 | 0 | 67808 | 0 | 0 | 0 | 0 | 0 | 0 | 50944 | 1096400 | 0 | 0 | 0 | 117780 | 0 | 0 |
| ISFGTPAPG | MOV10 | Q9HCE1 | MOV10_HUMA | 2 Oxidation (M) | 9290100 | 2998600 | 234600 | 0 | 0 | 0 | 0 | 58409 | 0 | 0 | 0 | 0 | 0 | 0 | 1697300 | 0 | 340170 | 906800 |
| ISFKFDHLF | CAPZA1 | P52907 | CAZA1_HUMAN | Unmodified | 0 | 300300 | 0 | 0 | 4133200 | 445350 | 210480 | 0 | 0 | 0 | 0 | 0 | 2221400 | 1037200 | 0 | 0 | 610880 | 1077100 |
| ISFLCDDLT | CDCP1 | Q9H5V8 | CDCP1_HUMA | Unmodified | 0 | 0 | 0 | 0 | 0 | 0 | 0 | 0 | 154090 | 0 | 0 | 6326800 | 0 | 0 | 751610 | 745410 | 0 | 0 |
| ISFLCDDLT | CDCP1 | Q9H5V8 | CDCP1_HUMA | Oxidation (M) | 0 | 0 | 0 | 351220 | 1164400 | 0 | 0 | 0 | 0 | 0 | 0 | 0 | 909290 | 348920 | 0 | 0 | 0 | 1678000 |
| ISFLENNLE | KIF5B; KIF5C; K | P33176; O60 | KINH_HUMAN | Unmodified | 122990 | 0 | 132450 | 152070 | 5974800 | 892990 | 1492800 | 485300 | 6184900 | 1891400 | 2268300 | 11101000 | 819510 | 383630 | 243550 | 235910 | 1910600 | 925410 |
| ISFPAIQAAI | WARS1; WARS1 | P23381; H0Y | SYWC_HUMAN | Unmodified | 0 | 0 | 77745 | 361780 | 0 | 0 | 0 | 0 | 1633100 | 0 | 27133 | 502600 | 0 | 0 | 1095900 | 0 | 0 | 0 |
| ISFPWNEIF | NF2 | P35240 | MERL_HUMAN | Unmodified | 0 | 0 | 0 | 0 | 819340 | 0 | 0 | 0 | 0 | 188410 | 0 | 0 | 668070 | 0 | 0 | 0 | 1295700 | 995450 |
| ISFQPAVAC | ITM2C; ITM2C | Q9NQX7; C9 | ITM2C_HUMAN | Unmodified | 0 | 0 | 0 | 0 | 0 | 0 | 0 | 0 | 1929100 | 368570 | 0 | 0 | 0 | 0 | 0 | 0 | 0 | 0 |
| ISFQPAVAC | ITM2C; ITM2C | Q9NQX7; C9 | ITM2C_HUMAN | Unmodified | 0 | 0 | 0 | 0 | 0 | 512920 | 250680 | 0 | 2669000 | 77044 | 0 | 0 | 150580 | 0 | 0 | 0 | 0 | 0 |
| ISFQPAVAC | ITM2C; ITM2C | Q9NQX7; C9 | ITM2C_HUMAN | Unmodified | 0 | 85868 | 0 | 235700 | 4928600 | 1001200 | 0 | 14433 | 14159000 | 827200 | 0 | 1033100 | 0 | 0 | 78827 | 0 | 0 | 495710 |
| ISGADINSI | PSMC4 | P43686 | PRS6B_HUMAN | Oxidation (M) | 0 | 651630 | 0 | 0 | 0 | 0 | 0 | 0 | 0 | 0 | 29928 | 325090 | 0 | 0 | 0 | 0 | 0 | 0 |
| ISGAIGPCV | SEC23A; SEC23 | Q15436; F5H | SC23A_HUMAN | Unmodified | 0 | 0 | 0 | 0 | 0 | 0 | 0 | 0 | 0 | 0 | 0 | 651040 | 0 | 0 | 0 | 0 | 0 | 0 |
| ISGCLHDIC | KIF14 | Q15058 | KIF14_HUMAN | Unmodified | 0 | 0 | 0 | 0 | 1800200 | 370190 | 195500 | 0 | 0 | 94163 | 172320 | 385480 | 685170 | 0 | 0 | 0 | 754010 | 445250 |
| ISGDTCSG | SORL1 | Q92673 | SORL_HUMAN | Unmodified | 0 | 0 | 0 | 0 | 0 | 0 | 0 | 0 | 357100 | 0 | 0 | 0 | 0 | 0 | 0 | 0 | 0 | 0 |
| ISGETIFVT/ | CLTC; CLTC | Q00610; A0/ | CLH1_HUMAN | Unmodified | 4562300 | 2064500 | 566440 | 419280 | 0 | 0 | 0 | 0 | 0 | 0 | 0 | 6760600 | 0 | 0 | 2638200 | 3202900 | 0 | 0 |
| ISGETIFVT/ | CLTC; CLTC | Q00610; A0/ | CLH1_HUMAN | Unmodified | 29107000 | 47968000 | 10319000 | 7356900 | 306890 | 0 | 0 | 0 | 3391000 | 0 | 664880 | 77357000 | 0 | 0 | 19396000 | 27209000 | 0 | 0 |
| ISGFELDPE | SLC5A6 | Q9Y289 | SC5A6_HUMAI | Unmodified | 0 | 0 | 0 | 0 | 0 | 0 | 0 | 0 | 0 | 0 | 255570 | 5043200 | 0 | 0 | 0 | 0 | 0 | 0 |
| ISGGNDKQ | RPS6; RPS6 | P62753; A2A | RS6_HUMAN 4 | Unmodified | 34015 | 0 | 0 | 0 | 299950 | 0 | 46478 | 0 | 0 | 181400 | 0 | 0 | 554160 | 222190 | 0 | 0 | 0 | 790640 |
| ISGGSVVEI | IDH1; IDH1; IDI | O75874; C9J | IDHC_HUMAN | Unmodified | 0 | 0 | 0 | 0 | 221710 | 43451 | 0 | 0 | 345280 | 0 | 0 | 1047700 | 137650 | 0 | 0 | 0 | 0 | 0 |
| ISGGSVVEI | IDH1; IDH1; IDI | O75874; C9J | IDHC_HUMAN | Oxidation (M) | 0 | 0 | 68523 | 0 | 0 | 0 | 0 | 0 | 78993 | 0 | 329280 | 1055800 | 0 | 0 | 332930 | 0 | 0 | 0 |
| ISGGYENVI | PLOD2; PLOD2 | O00469; E7E | PLOD2_HUMAI | Oxidation (M) | 0 | 0 | 0 | 0 | 0 | 0 | 0 | 0 | 907820 | 164140 | 0 | 0 | 0 | 0 | 0 | 0 | 0 | 0 |
| ISGGYENVI | PLOD2; PLOD2 | O00469; E7E | PLOD2_HUMAI | Unmodified | 0 | 0 | 0 | 0 | 0 | 0 | 0 | 0 | 6926400 | 325520 | 0 | 204370 | 0 | 0 | 0 | 0 | 0 | 0 |
| ISGIFLTNH | QSOX2; QSOX2 | Q6ZRP7; H0 | QSOX2_HUMAI | Unmodified | 0 | 0 | 139920 | 0 | 0 | 0 | 151270 | 17806 | 1719300 | 279940 | 664220 | 0 | 0 | 0 | 0 | 0 | 0 | 0 |
| ISGLIYEETI | H4C1 | P62805 | H4_HUMAN Hi | Unmodified | 2.62E+08 | 1.25E+08 | 1340500 | 509620 | 799160 | 0 | 0 | 834170 | 4118800 | 838980 | 10407000 | 2.22E+09 | 477360 | 529180 | 648090 | 1045900 | 0 | 0 |
| ISGLIYEETI | H4C1 | P62805 | H4_HUMAN Hi | Unmodified | 4.14E+08 | 1.52E+08 | 1014800 | 739210 | 3240300 | 2567800 | 8888000 | 7748200 | 13579000 | 13732000 | 7653500 | 8.9E+08 | 0 | 0 | 428860 | 574090 | 0 | 3135600 |
| ISGTSIAFK | MARK3; MARK1 | P27448; Q9I | MARK3_HUMAI | Unmodified | 0 | 0 | 120910 | 254150 | 0 | 0 | 60298 | 82692 | 348870 | 92069 | 0 | 262540 | 52948 | 0 | 216220 | 242180 | 0 | 0 |
| ISGTVNIR | TARS1 | P26639 | SYTC_HUMAN | Unmodified | 41072 | 214270 | 0 | 0 | 0 | 0 | 82346 | 97088 | 0 | 364290 | 0 | 10976000 | 51928 | 0 | 0 | 0 | 0 | 0 |
| ISHLPLVEE | MCM2; MCM2 | P49736; H0Y | MCM2_HUMAN | Unmodified | 0 | 0 | 0 | 0 | 0 | 0 | 425350 | 35004 | 0 | 0 | 0 | 1060500 | 0 | 0 | 0 | 0 | 0 | 0 |
| ISHLVLPVC | TOR1AIP1; TOR | Q5JTV8; A0A | TOIP1_HUMAN | Unmodified | 0 | 0 | 0 | 0 | 0 | 0 | 0 | 22485 | 11447000 | 0 | 443560 | 0 | 0 | 0 | 0 | 0 | 0 | 0 |
| ISHLVLPVC | TOR1AIP1; TOR | Q5JTV8; A0A | TOIP1_HUMAN | Unmodified | 0 | 0 | 0 | 0 | 0 | 0 | 0 | 0 | 1392800 | 434900 | 872370 | 0 | 0 | 0 | 0 | 0 | 0 | 0 |
| ISHLVLPVC | TOR1AIP2 | Q8NFQ8 | TOIP2_HUMAN | Unmodified | 0 | 0 | 0 | 0 | 0 | 0 | 0 | 0 | 2091400 | 1324900 | 0 | 0 | 0 | 0 | 0 | 0 | 0 | 0 |
| ISIENSEEAI | ADGRG1; ADGI | Q9Y653; H3I | AGRG1_HUMA | Unmodified | 0 | 0 | 616090 | 579870 | 0 | 0 | 0 | 0 | 5165400 | 0 | 483320 | 0 | 0 | 0 | 796850 | 0 | 0 | 0 |
| ISIIRFPDTV | NOMO1; NOMC | Q15155; P69 | NOMO1_HUMA | Unmodified | 0 | 0 | 0 | 0 | 0 | 0 | 226830 | 32888 | 9574700 | 1901200 | 0 | 0 | 430000 | 0 | 0 | 0 | 0 | 0 |
| ISILINR | TSPYL2 | Q9H2G4 | TSYL2_HUMAN | Unmodified | 0 | 61148 | 0 | 36411 | 494040 | 27067000 | 641460 | 91405 | 3192000 | 2443600 | 3087100 | 1386500 | 0 | 0 | 39153 | 124720 | 629640 | 0 |
| ISIMHEDW | NOMO1; NOMC | Q15155; P69 | NOMO1_HUMA | Oxidation (M) | 0 | 0 | 0 | 0 | 0 | 0 | 0 | 0 | 882780 | 0 | 0 | 0 | 0 | 0 | 0 | 0 | 0 | 0 |
| ISIMHEDW | NOMO1; NOMC | Q15155; P69 | NOMO1_HUMA | Unmodified | 0 | 0 | 0 | 0 | 0 | 0 | 0 | 0 | 1212700 | 0 | 0 | 0 | 0 | 0 | 0 | 0 | 0 | 0 |
| ISIPDVDLD | AHNAK | Q09666 | AHNK_HUMAN | Unmodified | 0 | 0 | 0 | 0 | 3955300 | 0 | 0 | 0 | 0 | 0 | 0 | 0 | 0 | 0 | 0 | 0 | 0 | 2610600 |
| ISIPDVDLD | AHNAK | Q09666 | AHNK_HUMAN | Unmodified | 0 | 0 | 0 | 0 | 12748000 | 2609100 | 0 | 0 | 0 | 0 | 0 | 0 | 441970 | 0 | 0 | 0 | 4917800 | 6046200 |
| ISIPDVDLD | AHNAK | Q09666 | AHNK_HUMAN | Unmodified | 0 | 0 | 0 | 0 | 0 | 0 | 0 | 0 | 0 | 0 | 0 | 0 | 0 | 0 | 0 | 0 | 1091300 | 0 |
| ISIPDVGLH | AHNAK | Q09666 | AHNK_HUMAN | Unmodified | 0 | 0 | 0 | 0 | 832570 | 0 | 0 | 0 | 0 | 0 | 0 | 0 | 0 | 0 | 0 | 0 | 549430 | 568290 |
| ISIPIRNNY | NPTXR; NPTXR | O95502; A0/ | NPTXR_HUMAN | Oxidation (M) | 0 | 0 | 0 | 0 | 0 | 0 | 0 | 43732 | 4017100 | 3814000 | 0 | 0 | 0 | 1043400 | 0 | 0 | 0 | 0 |
| ISIPIRNNY | NPTXR; NPTXR | O95502; A0/ | NPTXR_HUMAN | Unmodified | 0 | 0 | 0 | 0 | 0 | 0 | 36141 | 0 | 3167600 | 1737400 | 0 | 0 | 0 | 0 | 0 | 0 | 0 | 0 |
| ISIVEALTLL | NDUFAF4 | Q9P032 | NDUF4_HUMA | Unmodified | 0 | 0 | 0 | 0 | 0 | 0 | 0 | 0 | 2402500 | 129920 | 56063 | 40658 | 0 | 0 | 0 | 0 | 0 | 0 |
| ISKEIGGDV | CAP1; CAP1; C/ | Q01518; Q5 | CAP1_HUMAN | Unmodified | 0 | 0 | 0 | 0 | 0 | 0 | 413690 | 18363 | 0 | 67910 | 0 | 0 | 0 | 0 | 0 | 0 | 175370 | 97179 |
| ISKESEDFI | MCM6 | Q14566 | MCM6_HUMAN | Unmodified | 0 | 0 | 0 | 0 | 0 | 0 | 295900 | 223560 | 0 | 359140 | 0 | 0 | 594370 | 258680 | 0 | 0 | 190660 | 123520 |
| ISKEVLAGI | HNRNPU; HNR | Q00839; A0/ | HNRPU_HUMA | Unmodified | 0 | 0 | 0 | 0 | 0 | 0 | 2209700 | 0 | 0 | 0 | 0 | 0 | 0 | 0 | 0 | 0 | 0 | 0 |
| ISKLGSGNI | FOLH1B; FOLH | Q9HBA9; Q0 | FOH1B_HUMAI | Unmodified | 164680 | 369350 | 0 | 3371600 | 1523200 | 29205 | 0 | 0 | 0 | 0 | 0 | 0 | 735030 | 260800 | 0 | 168690 | 0 | 31856 |
| ISKLGSGNI | FOLH1B; FOLH | Q9HBA9; Q0 | FOH1B_HUMAI | Unmodified | 0 | 0 | 0 | 1071900 | 0 | 0 | 0 | 0 | 124110 | 0 | 0 | 0 | 2810200 | 1418800 | 0 | 0 | 0 | 58159 |
| ISKPAPYWI | PRDX4; PRDX4 | Q13162; A6I | PRDX4_HUMAN | Unmodified | 0 | 0 | 0 | 0 | 0 | 0 | 0 | 0 | 67545000 | 34353000 | 24396000 | 8628300 | 0 | 0 | 674330 | 52601 | 0 | 0 |
| ISKPAPYWI | PRDX4; PRDX4 | Q13162; A6I | PRDX4_HUMAN | Unmodified | 0 | 0 | 0 | 0 | 0 | 0 | 0 | 0 | 4289600 | 2416200 | 0 | 0 | 0 | 0 | 0 | 0 | 0 | 0 |
| ISKSSFRE | S100A16 | Q96FQ6 | S10AG_HUMAI | Unmodified | 0 | 0 | 0 | 0 | 0 | 0 | 0 | 0 | 0 | 0 | 0 | 0 | 0 | 0 | 0 | 0 | 369150 | 130180 |
| ISKYPTLK | ERP44 | Q9BS26 | ERP44_HUMAN | Unmodified | 0 | 0 | 0 | 0 | 3845100 | 671190 | 0 | 0 | 7042100 | 1294000 | 0 | 0 | 0 | 0 | 0 | 0 | 281050 | 359670 |
| ISLADIAQK | PSMD3 | O43242 | PSMD3_HUMA | Unmodified | 0 | 0 | 104400 | 0 | 0 | 0 | 0 | 0 | 140010 | 0 | 0 | 8012800 | 0 | 0 | 0 | 215220 | 0 | 0 |
| ISLAIPNLGI | ABRAXAS2 | Q15018 | ABRX2_HUMAN | Unmodified | 23360 | 0 | 0 | 0 | 0 | 0 | 0 | 0 | 0 | 0 | 0 | 925370 | 0 | 0 | 0 | 0 | 0 | 0 |
| ISLAPTDVK | ATXN2L; ATXN2 | Q8WWM7; I | ATX2L_HUMAN | Unmodified | 329310 | 235410 | 91567 | 0 | 0 | 0 | 0 | 0 | 0 | 144810 | 0 | 0 | 535760 | 0 | 0 | 0 | 0 | 228840 |
| ISLEVQLEQ | REPS1; REPS1; | Q96D71; H0 | REPS1_HUMAN | Unmodified | 0 | 0 | 0 | 0 | 0 | 0 | 0 | 0 | 0 | 0 | 0 | 0 | 0 | 0 | 135230 | 0 | 0 | 0 |
| ISLGLPVGA | RPL23; RPL23; | P62829; J3K | RL23_HUMAN ( | Unmodified | 2618800 | 2929200 | 0 | 0 | 54482 | 0 | 3243800 | 2651500 | 1182200 | 442740 | 3320600 | 30490000 | 0 | 0 | 1855700 | 1177500 | 72459 | 1982300 |
| ISLIQIFR | PRPF8; PRPF8 | Q6P2Q9; I3L | PRP8_HUMAN | Unmodified | 0 | 0 | 155530 | 0 | 0 | 0 | 0 | 0 | 0 | 0 | 0 | 487940 | 0 | 0 | 110620 | 0 | 0 | 0 |
| ISLLPNDED | ZFYVE16 | Q7Z3T8 | ZFY16_HUMAN | Unmodified | 0 | 0 | 0 | 0 | 1167900 | 0 | 0 | 0 | 227480 | 47769 | 0 | 0 | 485320 | 0 | 0 | 0 | 0 | 0 |
| ISLNDVSK | PARP4 | Q9UKK3 | PARP4_HUMAN | Unmodified | 595980 | 1232300 | 0 | 382150 | 0 | 0 | 74708 | 0 | 0 | 0 | 0 | 546850 | 0 | 0 | 587460 | 295500 | 0 | 0 |
| ISLNSLCY | CPNE3; CPNE3 | O75131; H0 | CPNE3_HUMAI | Oxidation (M) | 2100900 | 1080800 | 1189900 | 2340800 | 747000 | 1207300 | 826280 | 1409700 | 2628000 | 880530 | 722520 | 1646900 | 986090 | 1491700 | 1599000 | 1370000 | 2197800 | 2487000 |
| ISLNSLCY | CPNE3; CPNE3 | O75131; H0 | CPNE3_HUMAI | Unmodified | 0 | 0 | 0 | 0 | 0 | 0 | 0 | 0 | 3999500 | 393360 | 0 | 0 | 0 | 0 | 0 | 0 | 0 | 0 |
| ISLNTLTLN | FLOT1; FLOT1; I | O75955; A2/ | FLOT1_HUMAN | Unmodified | 0 | 0 | 1432300 | 1365700 | 393180 | 0 | 0 | 0 | 3006600 | 537770 | 475270 | 4461700 | 0 | 0 | 3744900 | 2696900 | 0 | 0 |
| ISLNTLTLN | FLOT1; FLOT1; I | O75955; A2/ | FLOT1_HUMAN | Unmodified | 0 | 0 | 0 | 0 | 1261100 | 0 | 0 | 0 | 1128300 | 1380000 | 0 | 622820 | 0 | 0 | 0 | 0 | 393560 | 0 |
| ISLPLGSEE | SEMA4B; SEMA | Q9NPR2; H0 | SEM4B_HUMAI | Unmodified | 0 | 0 | 0 | 0 | 0 | 0 | 0 | 0 | 1124000 | 0 | 0 | 0 | 0 | 0 | 0 | 0 | 0 | 0 |
| ISLQWLR | DPP4; DPP4; DI | P27487; A0A | DPP4_HUMAN | Unmodified | 646620 | 519120 | 6277000 | 4549900 | 0 | 0 | 0 | 0 | 0 | 0 | 178550 | 54588 | 0 | 361400 | 18969000 | 12304000 | 0 | 0 |
| ISLSGVHHI | CLSTN1; CLSTN | O94985; Q5 | CSTN1_HUMAI | Unmodified | 0 | 0 | 0 | 0 | 134540 | 267720 | 0 | 0 | 4310700 | 274490 | 0 | 0 | 252250 | 171320 | 0 | 0 | 258540 | 0 |

| | | | | | | | | | | | | | | | | | | | | | | |
|---|---|---|---|---|---|---|---|---|---|---|---|---|---|---|---|---|---|---|---|---|---|---|
| ISLSNESNK | PYGL | P06737 | PYGL_HUMAN | Unmodified | 0 | 34974 | 0 | 0 | 599940 | 0 | 0 | 0 | 0 | 252980 | 0 | 0 | 0 | 0 | 0 | 0 | 328610 | 1088200 |
| ISMENSK | PLCB4; PLCB4 | Q15147; A0A | PLCB4_HUMAN | Unmodified | 0 | 55915 | 87227 | 0 | 0 | 0 | 0 | 0 | 0 | 0 | 0 | 0 | 0 | 0 | 33024 | 0 | 161880 | 754370 |
| ISMENSKAI | PLCB4; PLCB4 | Q15147; A0A | PLCB4_HUMAN | Unmodified | 0 | 0 | 0 | 0 | 0 | 0 | 0 | 0 | 0 | 0 | 0 | 0 | 74108 | 67975 | 0 | 0 | 16427 | 362260 |
| ISMINNPAK | UGGT1 | Q9NYU2 | UGGG1_HUMA | Unmodified | 0 | 0 | 0 | 48358 | 0 | 0 | 0 | 0 | 18745000 | 4006700 | 1034100 | 0 | 0 | 0 | 0 | 0 | 467200 | 0 |
| ISMINNPAK | UGGT1 | Q9NYU2 | UGGG1_HUMA | Oxidation (M) | 0 | 0 | 0 | 13293 | 102890 | 0 | 0 | 0 | 971010 | 351540 | 0 | 38093 | 0 | 0 | 0 | 0 | 0 | 0 |
| ISMPDFDLI | AHNAK | Q09666 | AHNK_HUMAN | Unmodified | 0 | 0 | 0 | 0 | 4633200 | 0 | 0 | 0 | 0 | 0 | 0 | 0 | 0 | 0 | 0 | 0 | 1104700 | 2283200 |
| ISMPDFDLI | AHNAK | Q09666 | AHNK_HUMAN | Oxidation (M) | 0 | 0 | 0 | 0 | 11468000 | 1964300 | 0 | 0 | 0 | 0 | 0 | 0 | 0 | 0 | 0 | 0 | 4583600 | 3250500 |
| ISMPDFDLI | AHNAK | Q09666 | AHNK_HUMAN | Oxidation (M) | 0 | 0 | 0 | 0 | 0 | 515300 | 0 | 0 | 160000 | 60107 | 0 | 0 | 0 | 235520 | 0 | 0 | 1018500 | 146620 |
| ISMPDFDLI | AHNAK | Q09666 | AHNK_HUMAN | Unmodified | 0 | 2279100 | 0 | 0 | 0 | 0 | 0 | 0 | 31252 | 0 | 0 | 0 | 0 | 0 | 0 | 0 | 2021300 | 1250800 |
| ISMPDFDLI | AHNAK | Q09666 | AHNK_HUMAN | Unmodified | 0 | 0 | 0 | 0 | 5591600 | 553160 | 0 | 0 | 0 | 0 | 0 | 0 | 0 | 0 | 375770 | 0 | 1188500 | 2397800 |
| ISMPDFDLI | AHNAK | Q09666 | AHNK_HUMAN | Unmodified | 0 | 0 | 0 | 0 | 318630 | 0 | 0 | 0 | 0 | 0 | 0 | 0 | 0 | 0 | 0 | 0 | 577060 | 204450 |
| ISMPDIDFN | AHNAK | Q09666 | AHNK_HUMAN | Oxidation (M) | 0 | 0 | 0 | 0 | 1848000 | 724270 | 0 | 0 | 0 | 0 | 0 | 0 | 504660 | 0 | 0 | 0 | 0 | 770430 |
| ISMPDIDFN | AHNAK | Q09666 | AHNK_HUMAN | Unmodified | 0 | 0 | 0 | 0 | 4308900 | 874540 | 108660 | 0 | 0 | 0 | 0 | 0 | 0 | 0 | 731790 | 765200 | 759420 | 2266900 |
| ISMPDIDLN | AHNAK | Q09666 | AHNK_HUMAN | Unmodified | 0 | 0 | 0 | 0 | 5945100 | 504480 | 2987100 | 0 | 0 | 0 | 0 | 0 | 40813 | 42788 | 0 | 0 | 1821300 | 2025600 |
| ISMPDIDLN | AHNAK | Q09666 | AHNK_HUMAN | Oxidation (M) | 0 | 0 | 0 | 0 | 3125400 | 339600 | 0 | 0 | 0 | 0 | 0 | 0 | 0 | 0 | 0 | 0 | 1588200 | 1676000 |
| ISMPDLDLI | AHNAK | Q09666 | AHNK_HUMAN | Oxidation (M) | 0 | 0 | 0 | 0 | 1643900 | 958650 | 0 | 0 | 0 | 0 | 0 | 0 | 0 | 0 | 0 | 0 | 1081600 | 981620 |
| ISMPDLDLI | AHNAK | Q09666 | AHNK_HUMAN | Unmodified | 0 | 0 | 0 | 0 | 5870400 | 1064700 | 0 | 0 | 0 | 0 | 0 | 0 | 174110 | 0 | 0 | 0 | 2676900 | 5143600 |
| ISMPDLDLI | AHNAK | Q09666 | AHNK_HUMAN | Unmodified | 0 | 0 | 0 | 0 | 22037000 | 4008900 | 0 | 0 | 0 | 0 | 0 | 0 | 0 | 0 | 0 | 0 | 0 | 8859000 |
| ISMPDLDLI | AHNAK | Q09666 | AHNK_HUMAN | Unmodified | 0 | 0 | 0 | 0 | 8314100 | 0 | 0 | 0 | 0 | 0 | 0 | 0 | 0 | 405030 | 0 | 0 | 4433700 | 1170500 |
| ISMPDVDFI | AHNAK | Q09666 | AHNK_HUMAN | Unmodified | 0 | 0 | 0 | 0 | 4381500 | 18869 | 0 | 0 | 0 | 0 | 0 | 0 | 0 | 0 | 0 | 0 | 0 | 0 |
| ISMPDVDFI | AHNAK | Q09666 | AHNK_HUMAN | Oxidation (M) | 0 | 115830 | 0 | 0 | 906750 | 0 | 0 | 0 | 0 | 0 | 0 | 0 | 0 | 0 | 0 | 0 | 874570 | 596610 |
| ISMPDVDLI | AHNAK | Q09666 | AHNK_HUMAN | Oxidation (M) | 0 | 0 | 0 | 0 | 21823000 | 5669200 | 0 | 0 | 0 | 0 | 0 | 0 | 109930 | 0 | 0 | 83089 | 8583500 | 12012000 |
| ISMPDVDLI | AHNAK | Q09666 | AHNK_HUMAN | Unmodified | 0 | 0 | 0 | 0 | 2307600 | 207070 | 0 | 0 | 0 | 0 | 0 | 0 | 0 | 0 | 0 | 0 | 661470 | 1001400 |
| ISMPDVDLI | AHNAK | Q09666 | AHNK_HUMAN | Oxidation (M) | 0 | 0 | 0 | 0 | 3167000 | 0 | 0 | 0 | 0 | 0 | 0 | 0 | 0 | 0 | 0 | 0 | 2171700 | 486280 |
| ISMPDVDLI | AHNAK | Q09666 | AHNK_HUMAN | Unmodified | 0 | 0 | 0 | 0 | 6819100 | 0 | 0 | 0 | 0 | 0 | 0 | 0 | 0 | 246140 | 0 | 0 | 8189100 | 15816000 |
| ISMPDVDLI | AHNAK | Q09666 | AHNK_HUMAN | Unmodified | 0 | 0 | 0 | 0 | 0 | 0 | 0 | 0 | 0 | 0 | 0 | 0 | 0 | 0 | 0 | 0 | 508730 | 0 |
| ISMPDVDLI | AHNAK | Q09666 | AHNK_HUMAN | Oxidation (M) | 0 | 0 | 0 | 0 | 5119400 | 787390 | 41800 | 0 | 0 | 0 | 0 | 0 | 0 | 0 | 0 | 0 | 1963300 | 1058600 |
| ISMPDVDLI | AHNAK | Q09666 | AHNK_HUMAN | Unmodified | 0 | 0 | 0 | 0 | 757620 | 0 | 20597 | 0 | 0 | 12451 | 0 | 0 | 0 | 0 | 0 | 0 | 1028000 | 927280 |
| ISMPDVDLI | AHNAK | Q09666 | AHNK_HUMAN | Unmodified | 0 | 0 | 0 | 0 | 0 | 0 | 57258 | 0 | 0 | 16243 | 0 | 0 | 0 | 0 | 0 | 0 | 941650 | 2689700 |
| ISMPDVDLI | AHNAK | Q09666 | AHNK_HUMAN | Unmodified | 0 | 0 | 0 | 0 | 2269900 | 0 | 0 | 0 | 0 | 0 | 0 | 0 | 0 | 0 | 0 | 0 | 843270 | 983740 |
| ISMPDVDLI | AHNAK | Q09666 | AHNK_HUMAN | Oxidation (M) | 0 | 0 | 0 | 0 | 1270900 | 0 | 0 | 0 | 0 | 0 | 0 | 0 | 0 | 0 | 0 | 0 | 775330 | 584080 |
| ISMPDVGLI | AHNAK | Q09666 | AHNK_HUMAN | Unmodified | 0 | 0 | 0 | 0 | 1826300 | 0 | 0 | 0 | 0 | 0 | 0 | 0 | 0 | 0 | 0 | 0 | 936060 | 1275200 |
| ISMPDVNLI | AHNAK | Q09666 | AHNK_HUMAN | Unmodified | 0 | 0 | 0 | 0 | 3163100 | 0 | 0 | 0 | 0 | 0 | 0 | 0 | 0 | 0 | 0 | 0 | 1138000 | 1654400 |
| ISMPDVNLI | AHNAK | Q09666 | AHNK_HUMAN | Oxidation (M) | 0 | 0 | 0 | 0 | 488450 | 0 | 0 | 0 | 0 | 0 | 0 | 891770 | 0 | 0 | 0 | 0 | 480240 | 243160 |
| ISMPEIDLN | AHNAK | Q09666 | AHNK_HUMAN | Unmodified | 0 | 0 | 0 | 0 | 3238400 | 215790 | 0 | 0 | 0 | 0 | 0 | 0 | 0 | 0 | 0 | 249930 | 2877800 | 0 |
| ISMPEIDLN | AHNAK | Q09666 | AHNK_HUMAN | Oxidation (M) | 0 | 0 | 0 | 0 | 1486500 | 0 | 0 | 0 | 2274800 | 551750 | 0 | 201430 | 0 | 0 | 0 | 0 | 0 | 582500 |
| ISMPEIDLN | AHNAK | Q09666 | AHNK_HUMAN | Unmodified | 0 | 0 | 0 | 0 | 0 | 0 | 0 | 0 | 0 | 0 | 0 | 0 | 0 | 0 | 0 | 0 | 1000600 | 0 |
| ISMPEVDLI | AHNAK | Q09666 | AHNK_HUMAN | Unmodified | 0 | 0 | 0 | 0 | 339420 | 616530 | 0 | 376400 | 0 | 0 | 0 | 0 | 0 | 0 | 0 | 0 | 870020 | 4652800 |
| ISMPEVDLI | AHNAK | Q09666 | AHNK_HUMAN | Unmodified | 0 | 0 | 0 | 0 | 8187200 | 1098800 | 0 | 0 | 0 | 0 | 0 | 0 | 0 | 0 | 0 | 0 | 4695100 | 4046100 |
| ISMPEVDLI | AHNAK | Q09666 | AHNK_HUMAN | Oxidation (M) | 0 | 0 | 0 | 0 | 2361600 | 416200 | 0 | 0 | 568260 | 0 | 0 | 0 | 0 | 0 | 0 | 0 | 0 | 0 |
| ISMPGFKG | AHNAK | Q09666 | AHNK_HUMAN | Unmodified | 0 | 0 | 0 | 0 | 2190800 | 0 | 0 | 0 | 0 | 0 | 0 | 0 | 0 | 0 | 0 | 0 | 144640 | 3259200 |
| ISMPGFKG | AHNAK | Q09666 | AHNK_HUMAN | Unmodified | 0 | 0 | 0 | 0 | 3579600 | 531960 | 0 | 0 | 0 | 0 | 0 | 0 | 0 | 0 | 0 | 0 | 853130 | 1005500 |
| ISMQDVDL | AHNAK | Q09666 | AHNK_HUMAN | Unmodified | 0 | 0 | 0 | 42134 | 1869300 | 0 | 0 | 0 | 506590 | 33450 | 0 | 0 | 0 | 0 | 0 | 0 | 337720 | 1433300 |
| ISMSHEEEI | GMPPB; GMPPI | Q9Y5P6; A0A | GMPPB_HUMA | Oxidation (M) | 0 | 0 | 0 | 0 | 0 | 0 | 0 | 0 | 0 | 494200 | 0 | 594540 | 0 | 0 | 0 | 0 | 0 | 0 |
| ISMVNNIIH | SHCBP1 | Q8NEM2 | SHCBP_HUMA | Oxidation (M) | 0 | 0 | 0 | 0 | 14650000 | 0 | 0 | 0 | 0 | 0 | 0 | 0 | 0 | 0 | 0 | 0 | 0 | 0 |
| ISNDRLGS | PPM1H | Q9ULR3 | PPM1H_HUMA | Unmodified | 0 | 0 | 0 | 0 | 0 | 0 | 0 | 0 | 706500 | 0 | 0 | 2756700 | 0 | 0 | 0 | 0 | 0 | 0 |
| ISNDRLGS | PPM1H | Q9ULR3 | PPM1H_HUMA | Unmodified | 0 | 0 | 0 | 0 | 0 | 0 | 218610 | 0 | 745710 | 485870 | 817720 | 3122100 | 478550 | 0 | 0 | 0 | 0 | 0 |
| ISNIPSLIFL | NXN | Q6DKJ4 | NXN_HUMAN I | Unmodified | 0 | 0 | 0 | 0 | 0 | 0 | 0 | 0 | 0 | 0 | 18882 | 182790 | 0 | 0 | 0 | 0 | 0 | 0 |
| ISNIVIKDD | PIGS | Q96S52 | PIGS_HUMAN ( | Unmodified | 0 | 0 | 0 | 0 | 0 | 0 | 0 | 0 | 2974800 | 0 | 0 | 0 | 0 | 0 | 0 | 0 | 0 | 0 |
| ISNLLSDYC | POLR2B; POLR | P30876; C9J | RPB2_HUMAN | Unmodified | 136440 | 177670 | 65207 | 28750 | 0 | 0 | 0 | 11362 | 1057600 | 18834 | 106430 | 1212700 | 0 | 0 | 103600 | 51959 | 0 | 0 |
| ISNSATYSC | PDLIM5; PDLIM | Q96HC4; HC | PDLI5_HUMAN | Unmodified | 0 | 0 | 488080 | 658620 | 0 | 0 | 0 | 0 | 0 | 0 | 345030 | 1295400 | 0 | 0 | 686940 | 479710 | 0 | 0 |
| ISNTAISISD | GART; GART; G | P22102; C9J | PUR2_HUMAN | Unmodified | 318270 | 204830 | 0 | 0 | 0 | 0 | 1403100 | 658350 | 1050200 | 436040 | 525720 | 2346900 | 2367200 | 1401200 | 0 | 0 | 222090 | 0 |
| ISNYGWDC | CACYBP | Q9HB71 | CYBP_HUMAN | Unmodified | 127510 | 139160 | 0 | 161290 | 422510 | 0 | 312060 | 0 | 846090 | 710620 | 1921500 | 9616000 | 57958 | 372380 | 0 | 0 | 122830 | 103200 |
| ISNYGWDC | CACYBP | Q9HB71 | CYBP_HUMAN | Unmodified | 119380 | 0 | 0 | 93334 | 2407700 | 270750 | 2962400 | 1834300 | 720700 | 2181800 | 499010 | 0 | 5168000 | 3181900 | 0 | 48623 | 1696000 | 917850 |
| ISPDAWQV | XPOT; XPOT; XP | O43592; F8V | XPOT_HUMAN | Unmodified | 95615 | 0 | 0 | 0 | 94681 | 32457 | 0 | 0 | 69329 | 0 | 0 | 1910700 | 109760 | 109160 | 0 | 37943 | 0 | 60584 |
| ISPDRVYIN | MIF | P14174 | MIF_HUMAN M | Oxidation (M) | 0 | 0 | 0 | 0 | 19213000 | 1480700 | 0 | 0 | 0 | 399370 | 2152700 | 0 | 2625500 | 12985000 | 652860 | 0 | 20301000 | 24921000 |
| ISPDRVYIN | MIF | P14174 | MIF_HUMAN M | Unmodified | 0 | 0 | 0 | 0 | 1507700 | 0 | 0 | 0 | 0 | 368650 | 0 | 0 | 0 | 0 | 0 | 0 | 1346100 | 1788500 |
| ISPLCHNLI | JAK1; JAK1; JAK | P23458; A0A | JAK1_HUMAN T | Unmodified | 0 | 0 | 0 | 0 | 293350 | 131270 | 121890 | 0 | 387660 | 98889 | 0 | 323820 | 0 | 0 | 0 | 12147 | 0 | 0 |
| ISPLQFR | COPA; COPA; C | P53621; A0A | COPA_HUMAN | Unmodified | 78064 | 0 | 99612 | 169560 | 0 | 165020 | 0 | 0 | 428360 | 227090 | 280740 | 3529300 | 619830 | 0 | 220160 | 323170 | 84200 | 205540 |
| ISPMLSPFI | BABAM2; BABA | Q9NXR7; F8 | BABA2_HUMAN | Oxidation (M) | 92116 | 0 | 0 | 0 | 0 | 0 | 0 | 0 | 0 | 0 | 0 | 890940 | 0 | 0 | 0 | 0 | 0 | 0 |
| ISPQLLLAA | CNNM4 | Q6P4Q7 | CNNM4_HUMA | Unmodified | 0 | 0 | 331690 | 380800 | 0 | 0 | 0 | 0 | 463130 | 0 | 0 | 12093000 | 0 | 0 | 855010 | 806260 | 0 | 0 |
| ISPQLLLAA | CNNM4 | Q6P4Q7 | CNNM4_HUMA | Unmodified | 0 | 0 | 0 | 19938 | 0 | 0 | 0 | 0 | 0 | 312040 | 0 | 0 | 254890 | 202350 | 0 | 0 | 0 | 0 |
| ISPVEESED | SDC4 | P31431 | SDC4_HUMAN | Unmodified | 379300 | 0 | 394230 | 176150 | 0 | 0 | 0 | 0 | 0 | 0 | 0 | 0 | 0 | 0 | 188580 | 242660 | 0 | 0 |
| ISQAEEED | HINT2 | Q9BX68 | HINT2_HUMAN | Unmodified | 0 | 0 | 0 | 0 | 0 | 0 | 0 | 0 | 43798000 | 11747000 | 0 | 0 | 0 | 0 | 0 | 0 | 0 | 0 |
| ISQATAQIK | STX12; STX12 | Q86Y82; B1A | STX12_HUMAN | Unmodified | 812710 | 840960 | 2289000 | 2630000 | 570900 | 0 | 306150 | 42527 | 3475600 | 1081800 | 252290 | 861740 | 1475500 | 1116400 | 3822900 | 4093900 | 25863 | 333460 |
| ISQDADLK | RAI14 | Q9P0K7 | RAI14_HUMAN | Unmodified | 0 | 0 | 0 | 0 | 0 | 0 | 0 | 0 | 0 | 0 | 0 | 762720 | 0 | 0 | 0 | 0 | 0 | 0 |
| ISQDADLKT | RAI14 | Q9P0K7 | RAI14_HUMAN | Unmodified | 0 | 0 | 0 | 0 | 375600 | 108980 | 0 | 29419 | 0 | 34922 | 0 | 0 | 0 | 99206 | 0 | 0 | 350340 | 334610 |
| ISQEIGNLN | P3H1; P3H1 | Q32P28; H7 | P3H1_HUMAN | Oxidation (M) | 0 | 0 | 0 | 0 | 1148100 | 0 | 0 | 0 | 411390 | 0 | 0 | 0 | 0 | 0 | 34445 | 0 | 0 | 0 |
| ISQEIGNLN | P3H1; P3H1 | Q32P28; H7 | P3H1_HUMAN | Unmodified | 0 | 0 | 0 | 66216 | 0 | 0 | 0 | 0 | 3026400 | 243690 | 0 | 0 | 0 | 0 | 0 | 0 | 0 | 0 |
| ISQKDIEQS | ANXA4; ANXA4 | P09525; Q6 | ANXA4_HUMAN | Unmodified | 0 | 0 | 0 | 0 | 0 | 0 | 0 | 0 | 0 | 1537000 | 0 | 0 | 0 | 0 | 0 | 0 | 0 | 0 |
| ISQLEMAR | MSN | P26038 | MOES_HUMAN | Oxidation (M) | 0 | 167930 | 0 | 584460 | 2477600 | 0 | 0 | 0 | 0 | 0 | 0 | 0 | 609840 | 879520 | 336820 | 329860 | 752720 | 0 |
| ISQLEMAR | MSN | P26038 | MOES_HUMAN | Unmodified | 0 | 1811700 | 524490 | 1735100 | 9839600 | 4748000 | 0 | 0 | 196900 | 0 | 0 | 0 | 2988300 | 0 | 854350 | 0 | 2343800 | 5439800 |
| ISQLEMAR | MSN | P26038 | MOES_HUMAN | Unmodified | 0 | 0 | 0 | 0 | 1183200 | 295770 | 0 | 0 | 0 | 0 | 0 | 139890 | 0 | 113490 | 0 | 0 | 0 | 596730 |
| ISQLEMAR | MSN | P26038 | MOES_HUMAN | Unmodified | 0 | 0 | 0 | 0 | 3217100 | 0 | 0 | 0 | 0 | 0 | 0 | 0 | 359680 | 79966 | 0 | 0 | 564440 | 1324600 |
| ISQLQAELF | B3GAT3; B3GA | O94766; E9F | B3GA3_HUMAN | Unmodified | 0 | 491270 | 251170 | 589460 | 720740 | 0 | 0 | 43468 | 12325000 | 1162300 | 126180 | 1889400 | 0 | 474050 | 0 | 67225 | 2358000 | 1601200 |
| ISQLQAELF | B3GAT3; B3GA | O94766; E9F | B3GA3_HUMAN | Unmodified | 0 | 0 | 0 | 0 | 0 | 0 | 0 | 0 | 9314400 | 0 | 0 | 0 | 0 | 0 | 0 | 0 | 0 | 0 |
| ISQMPVILT | GMPS | P49915 | GUAA_HUMAN | Oxidation (M) | 0 | 109210 | 281440 | 339610 | 346570 | 101460 | 106400 | 0 | 560470 | 205050 | 1420600 | 9715300 | 344590 | 133790 | 373430 | 322270 | 0 | 0 |
| ISQPSDINT | PIK3C2B; PIK3C | O00750; F5C | P3C2B_HUMAN | Unmodified | 0 | 0 | 349020 | 493570 | 0 | 30889 | 0 | 0 | 0 | 0 | 0 | 397110 | 1312600 | 0 | 0 | 727200 | 0 | 0 |
| ISQSGDFLI | FKBP9; FKBP9 | O95302; C9J | FKBP9_HUMAN | Unmodified | 0 | 0 | 0 | 0 | 0 | 0 | 0 | 0 | 0 | 0 | 0 | 1418900 | 0 | 0 | 0 | 0 | 0 | 0 |
| ISQSNYIPT | GNAI3 | P08754 | GNAI3_HUMAN | Unmodified | 760490 | 76480 | 759080 | 295300 | 0 | 0 | 230510 | 150260 | 531990 | 438010 | 0 | 10019000 | 181380 | 0 | 94547 | 85928 | 0 | 0 |
| ISQTVEILK | SEPTIN5; SEPTI | Q99719; F8V | SEPT5_HUMAN | Unmodified | 0 | 0 | 0 | 0 | 0 | 0 | 0 | 0 | 0 | 0 | 0 | 170620 | 0 | 0 | 0 | 173100 | 0 | 0 |
| ISQTYQQQ | ANXA4; ANXA4; | P09525; B4E | ANXA4_HUMAN | Unmodified | 3453700 | 3184000 | 3821100 | 663160 | 241930 | 223060 | 28013 | 0 | 28573000 | 2307200 | 16124000 | 6437700 | 79021 | 0 | 633940 | 384870 | 39820 | 0 |
| ISRDADCR | ECH1; N/A; ECH | Q13011; M0 | ECH1_HUMAN | Unmodified | 0 | 0 | 0 | 0 | 0 | 0 | 174380 | 0 | 0 | 510460 | 0 | 0 | 0 | 87554 | 0 | 0 | 0 | 0 |
| ISRDELLQV | CHP1 | Q99653 | CHP1_HUMAN | Unmodified | 652300 | 936220 | 0 | 3528800 | 766300 | 315960 | 0 | 0 | 2684400 | 220280 | 1455500 | 6390100 | 2631700 | 0 | 6382200 | 2788000 | 0 | 323480 |
| ISRGTIEILS | RPLP0; RPLP0P | P05388; Q8 | RLA0_HUMAN ( | Unmodified | 134780 | 0 | 67782 | 0 | 0 | 0 | 179000 | 0 | 0 | 0 | 94959 | 0 | 243620 | 161190 | 323150 | 0 | 0 | 0 |
| ISRPGDSDI | HNRNPL; HNRI | P14866; B4D | HNRPL_HUMAI | Unmodified | 0 | 0 | 0 | 0 | 0 | 0 | 0 | 0 | 0 | 0 | 0 | 849590 | 0 | 0 | 33412 | 21858 | 0 | 0 |
| ISRPLPEVG | B4GALT4; B4G | O60513; C9J | B4GT4_HUMAN | Oxidation (M) | 0 | 0 | 0 | 0 | 0 | 0 | 0 | 0 | 1800700 | 90690 | 0 | 0 | 0 | 0 | 0 | 0 | 0 | 0 |
| ISRPLPEVG | B4GALT4; B4G | O60513; C9J | B4GT4_HUMAN | Unmodified | 0 | 0 | 0 | 0 | 0 | 0 | 0 | 0 | 4439700 | 348290 | 0 | 0 | 0 | 0 | 0 | 0 | 0 | 0 |
| ISRPPTSVG | B4GALT3 | O60512 | B4GT3_HUMAN | Unmodified | 0 | 0 | 68481 | 0 | 0 | 0 | 0 | 38323 | 1860400 | 0 | 235940 | 0 | 0 | 0 | 0 | 0 | 0 | 0 |
| ISRVEFTFP | AURKA | O14965 | AURKA_HUMAI | Unmodified | 118140 | 0 | 0 | 0 | 0 | 0 | 0 | 0 | 0 | 0 | 0 | 0 | 0 | 0 | 198050 | 127950 | 0 | 0 |
| ISRYQGVN | PABPC4; PABP | Q13310; B1A | PABP4_HUMAN | Unmodified | 471940 | 2541700 | 558690 | 2875700 | 0 | 0 | 0 | 0 | 0 | 0 | 0 | 0 | 729120 | 0 | 0 | 955590 | 0 | 0 |
| ISSDLDGH | TKT; TKT; TKT | P29401; F8V | TKT_HUMAN Tr | Unmodified | 0 | 0 | 0 | 0 | 15641000 | 1878100 | 303160 | 90445 | 1510400 | 4127800 | 2812300 | 3788500 | 1809500 | 614540 | 0 | 0 | 1478100 | 2834300 |
| ISSEAHREV | PEPD; PEPD; PE | P12955; A0A | PEPD_HUMAN | Unmodified | 0 | 0 | 0 | 0 | 937940 | 0 | 215960 | 87426 | 0 | 461700 | 5758.6 | 0 | 0 | 0 | 0 | 0 | 0 | 381090 |
| ISSIQSIVPA | HSPD1; HSPD1 | P10809; A0A | CH60_HUMAN | Unmodified | 0 | 0 | 0 | 0 | 0 | 0 | 0 | 0 | 6496300 | 0 | 1978000 | 0 | 0 | 0 | 0 | 0 | 0 | 0 |
| ISSLLEEQF | RPS8; RPS8 | P62241; Q5J | RS8_HUMAN 4 | Unmodified | 0 | 0 | 6153400 | 7085600 | 6682500 | 1653500 | 11602000 | 8844100 | 1163600 | 905050 | 8349100 | 63334000 | 10282000 | 3355200 | 0 | 0 | 1410100 | 6262900 |
| ISSNPNPV | ESYT2; ESYT2; | A0FGR8; A0A | ESYT2_HUMAN | Unmodified | 0 | 0 | 0 | 0 | 0 | 689090 | 0 | 0 | 0 | 0 | 0 | 105160 | 0 | 0 | 0 | 0 | 0 | 0 |
| ISSPTETER | S100A11 | P31949 | S10AB_HUMAN | Unmodified | 0 | 2106900 | 1630600 | 1753600 | 4988900 | 1128500 | 0 | 69110 | 14724000 | 1090100 | 47932 | 0 | 2094000 | 457960 | 2324900 | 0 | 643180 | 1708000 |
| ISSPTETER | S100A11 | P31949 | S10AB_HUMAN | Unmodified | 622420 | 0 | 0 | 2033400 | 1.29E+08 | 25102000 | 0 | 0 | 5060800 | 19771000 | 0 | 0 | 32846000 | 29364000 | 1130300 | 0 | 36085000 | 83145000 |
| ISSPTETER | S100A11 | P31949 | S10AB_HUMAN | Unmodified | 0 | 0 | 0 | 0 | 1669800 | 0 | 0 | 0 | 0 | 287150 | 0 | 0 | 335630 | 0 | 0 | 0 | 1342700 | 772080 |
| ISSSSFIEPI | TMTC3 | Q6ZXV5 | TMTC3_HUMAI | Unmodified | 0 | 0 | 0 | 0 | 0 | 0 | 0 | 14202 | 5569400 | 683900 | 391640 | 0 | 0 | 0 | 0 | 0 | 0 | 0 |
| ISSTPAKLE | HSPG2 | P98160 | PGBM_HUMAN | Unmodified | 0 | 93489 | 784630 | 2386000 | 962360 | 1850900 | 93053 | 31362 | 445520 | 276290 | 0 | 0 | 2716700 | 1468300 | 370810 | 696870 | 174750 | 2293900 |
| ISSVEGTLA | TLL2 | Q9Y6L7 | TLL2_HUMAN T | Unmodified | 0 | 0 | 0 | 0 | 0 | 0 | 140720 | 0 | 1942800 | 0 | 0 | 0 | 0 | 0 | 0 | 0 | 0 | 0 |
| ISSYDDRFI | KHNYN | O15037 | KHNYN_HUMA | Unmodified | 219880 | 165170 | 0 | 138730 | 327870 | 0 | 0 | 0 | 83067 | 0 | 0 | 0 | 0 | 0 | 89534 | 58256 | 0 | 0 |
| ISTEVGITN | RPN2 | P04844 | RPN2_HUMAN | Unmodified | 0 | 344510 | 111130 | 0 | 0 | 0 | 0 | 0 | 2703000 | 234140 | 0 | 0 | 0 | 0 | 1485600 | 0 | 0 | 0 |
| ISTEVGITN | RPN2 | P04844 | RPN2_HUMAN | Unmodified | 0 | 0 | 0 | 0 | 0 | 0 | 883320 | 121160 | 71401000 | 27582000 | 5576400 | 0 | 1099400 | 0 | 0 | 0 | 961040 | 372600 |
| ISTLPQLNS | CCNB1 | P14635 | CCNB1_HUMA | Unmodified | 449880 | 0 | 0 | 567750 | 1061500 | 287920 | 194260 | 66376 | 677050 | 667620 | 281000 | 681480 | 986170 | 913520 | 0 | 207030 | 224550 | 382750 |
| ISTLSCENK | IL18 | Q14116 | IL18_HUMAN I | Unmodified | 0 | 0 | 204640 | 0 | 314730 | 0 | 0 | 0 | 0 | 60972 | 0 | 6179900 | 0 | 0 | 44074 | 0 | 855420 | 465560 |
| ISTLYYQLG | DNAJC3; DNAJ | Q13217; X6F | DNJC3_HUMAI | Unmodified | 0 | 0 | 0 | 0 | 0 | 0 | 0 | 0 | 9397300 | 0 | 265490 | 0 | 0 | 0 | 0 | 0 | 0 | 0 |
| ISTLYYQLG | DNAJC3; DNAJ | Q13217; X6F | DNJC3_HUMAI | Unmodified | 0 | 0 | 0 | 0 | 0 | 0 | 0 | 0 | 1675800 | 0 | 0 | 0 | 0 | 0 | 0 | 0 | 249210 | 0 |

| | | | | | | | | | | | | | | | | | | | | | | |
|---|---|---|---|---|---|---|---|---|---|---|---|---|---|---|---|---|---|---|---|---|---|---|
| ISTMRLPDE | IGFBP2; IGFBP2 | P18065; C9J | IBP2_HUMAN I | Unmodified | 0 | 0 | 0 | 0 | 0 | 284490 | 0 | 0 | 4508500 | 130880 | 0 | 0 | 0 | 0 | 0 | 0 | 0 | 0 |
| ISTMRLPDE | IGFBP2; IGFBP2 | P18065; C9J | IBP2_HUMAN I | Oxidation (M) | 0 | 0 | 0 | 404000 | 0 | 0 | 2566000 | 0 | 20883000 | 3047200 | 0 | 0 | 507250 | 0 | 180730 | 0 | 3808800 | 213400 |
| ISTMRLPDE | IGFBP2; IGFBP2 | P18065; C9J | IBP2_HUMAN I | Unmodified | 0 | 0 | 0 | 0 | 1433000 | 0 | 0 | 0 | 21943000 | 0 | 0 | 0 | 0 | 0 | 0 | 0 | 0 | 0 |
| ISTPPEAYW | B3GNT2 | Q9NY97 | B3GN2_HUMAI | Unmodified | 0 | 0 | 0 | 0 | 0 | 0 | 0 | 0 | 3321500 | 1578900 | 0 | 0 | 0 | 0 | 0 | 0 | 0 | 0 |
| ISTSAWSS | VPS13C; VPS13 | Q709C8; A0 | VP13C_HUMAN | Unmodified | 0 | 0 | 0 | 0 | 1191900 | 376540 | 459600 | 154550 | 0 | 65317 | 0 | 0 | 1855300 | 815770 | 0 | 0 | 0 | 0 |
| ISTVVSSK | SRP14; SRP14 | P37108; H0Y | SRP14_HUMAN | Unmodified | 4488800 | 0 | 2305200 | 7851900 | 1479700 | 168140 | 509210 | 311920 | 595670 | 164150 | 315580 | 5265200 | 307000 | 167990 | 15940000 | 7513800 | 127910 | 528400 |
| ISTYGLPAG | CD2AP | Q9Y5K6 | CD2AP_HUMAI | Unmodified | 26645000 | 20146000 | 26922000 | 27619000 | 6156200 | 1208500 | 2492900 | 0 | 1174400 | 614810 | 792010 | 3536300 | 36776000 | 13368000 | 29006000 | 20113000 | 0 | 972170 |
| ISVAGVTSS | GOT2 | P00505 | AATM_HUMAN | Unmodified | 868830 | 184150 | 0 | 554260 | 157810 | 0 | 0 | 0 | 20098000 | 1851800 | 599160 | 862230 | 113420 | 0 | 0 | 0 | 0 | 0 |
| ISVFIQTDK | CD109 | Q6YHK3 | CD109_HUMAI | Unmodified | 81462 | 0 | 0 | 0 | 0 | 0 | 0 | 361320 | 1864300 | 175190 | 0 | 148050 | 0 | 0 | 0 | 0 | 0 | 0 |
| ISVIVETVYT | RPN1 | P04843 | RPN1_HUMAN | Unmodified | 0 | 0 | 0 | 0 | 0 | 0 | 3730900 | 0 | 4.11E+08 | 1.21E+08 | 16641000 | 7963100 | 0 | 0 | 0 | 0 | 0 | 0 |
| ISVLQNNYI | EEA1 | Q15075 | EEA1_HUMAN I | Unmodified | 0 | 0 | 0 | 0 | 1089300 | 0 | 0 | 0 | 0 | 0 | 0 | 0 | 0 | 0 | 0 | 0 | 0 | 0 |
| ISVNDFIIK | DLAT; DLAT; DL | P10515; A0A | ODP2_HUMAN | Unmodified | 102820 | 0 | 0 | 0 | 0 | 0 | 0 | 444330 | 4410200 | 1361400 | 0 | 0 | 0 | 0 | 0 | 0 | 41914 | 0 |
| ISVREPMQ | ATP5F1A; ATP5I | P25705; K7E | ATPA_HUMAN | Unmodified | 0 | 0 | 0 | 0 | 0 | 0 | 1001700 | 0 | 17778000 | 3473100 | 0 | 219110 | 403100 | 541970 | 0 | 0 | 0 | 0 |
| ISVTQPDSI | CD109 | Q6YHK3 | CD109_HUMAI | Unmodified | 398770 | 0 | 0 | 0 | 0 | 0 | 0 | 0 | 10402000 | 2210900 | 0 | 0 | 0 | 0 | 0 | 0 | 784730 | 499660 |
| ISVVNVNP | TNIK | Q9UKE5 | TNIK_HUMAN T | Unmodified | 0 | 0 | 0 | 0 | 0 | 0 | 0 | 0 | 0 | 0 | 0 | 1045300 | 0 | 0 | 0 | 0 | 0 | 0 |
| ISVVPTQP | WDR4 | P57081 | WDR4_HUMAN | Unmodified | 0 | 0 | 0 | 0 | 0 | 0 | 0 | 0 | 0 | 0 | 0 | 1254000 | 0 | 0 | 0 | 0 | 0 | 0 |
| ISVYDYDTF | MYOF; MYOF | Q9NZM1; H0 | MYOF_HUMAN | Unmodified | 0 | 0 | 0 | 0 | 17134000 | 5996300 | 0 | 0 | 0 | 0 | 0 | 0 | 0 | 0 | 0 | 0 | 363670 | 0 |
| ISVYYNEAS | TUBB3; TUBB3; | Q13509; G3 | TBB3_HUMAN | Unmodified | 348860 | 163540 | 0 | 0 | 546500 | 0 | 0 | 149540 | 6054200 | 179010 | 268600 | 4350300 | 0 | 0 | 0 | 0 | 0 | 185510 |
| ISVYYNEAS | TUBB3; TUBB3; | Q13509; G3 | TBB3_HUMAN | Unmodified | 0 | 0 | 0 | 0 | 1782500 | 0 | 0 | 0 | 0 | 197640 | 0 | 0 | 0 | 0 | 0 | 0 | 0 | 0 |
| ISVYYNEAT | TUBB; TUBB | P07437; Q5J | TBB5_HUMAN | Unmodified | 1152600 | 1118000 | 2776300 | 997590 | 343490 | 0 | 0 | 0 | 16043000 | 274660 | 2145800 | 1.62E+08 | 0 | 75994 | 4893800 | 4431600 | 0 | 0 |
| ISVYYNEAT | TUBB; TUBB | P07437; Q5J | TBB5_HUMAN | Unmodified | 724850 | 0 | 0 | 782960 | 281050 | 0 | 0 | 0 | 5507800 | 300960 | 1294900 | 14466000 | 389580 | 0 | 0 | 216170 | 0 | 0 |
| ISWEEYK | RCN1 | Q15293 | RCN1_HUMAN | Unmodified | 0 | 0 | 0 | 19193000 | 0 | 0 | 0 | 0 | 4267100 | 520400 | 0 | 0 | 0 | 0 | 0 | 0 | 0 | 0 |
| ISWQAAIFK | PI4KA; PI4KAP2 | P42356; A4C | PI4KA_HUMAN | Unmodified | 49383 | 0 | 435050 | 177270 | 0 | 0 | 39465 | 0 | 203260 | 82594 | 0 | 712080 | 0 | 0 | 1197600 | 1010700 | 0 | 0 |
| ISWYLRDII | ECH1; N/A; ECH | Q13011; M0 | ECH1_HUMAN | Unmodified | 0 | 0 | 0 | 0 | 0 | 735040 | 0 | 0 | 2841700 | 301010 | 0 | 0 | 0 | 0 | 0 | 0 | 0 | 0 |
| ISYFTFIGTF | TXNL1; TXNL1; | O43396; K7E | TXNL1_HUMAN | Oxidation (M) | 351630 | 517600 | 212020 | 1040600 | 0 | 0 | 0 | 0 | 949670 | 198040 | 156140 | 3466500 | 142160 | 0 | 1087100 | 779890 | 164310 | 0 |
| ISYFTFIGTF | TXNL1; TXNL1; | O43396; K7E | TXNL1_HUMAN | Unmodified | 0 | 0 | 104510 | 0 | 0 | 0 | 0 | 0 | 395110 | 0 | 0 | 0 | 0 | 0 | 0 | 0 | 0 | 0 |
| ISYHTQHK | SULF2; SULF2 | Q8IWU5; H0 | SULF2_HUMAN | Unmodified | 0 | 0 | 0 | 0 | 0 | 0 | 0 | 0 | 0 | 0 | 0 | 96847 | 24801 | 0 | 0 | 0 | 0 | 164660 |
| ISYLLTDK | ATP9A; ATP9A | O75110; A0A | ATP9A_HUMAN | Unmodified | 0 | 0 | 0 | 0 | 0 | 0 | 0 | 0 | 1138800 | 443900 | 1255800 | 0 | 0 | 0 | 0 | 0 | 0 | 0 |
| ISYRIVSLEF | DSG2; DSG2 | Q14126; J3K | DSG2_HUMAN | Unmodified | 109660 | 0 | 0 | 0 | 7739500 | 924900 | 0 | 0 | 0 | 1654800 | 0 | 0 | 2144000 | 0 | 419800 | 0 | 160340 | 565850 |
| ISYVQDENI | INCENP; INCEN | Q9NQS7; E9 | INCE_HUMAN I | Unmodified | 0 | 0 | 206050 | 814600 | 0 | 0 | 0 | 0 | 0 | 461960 | 33539 | 0 | 127150 | 0 | 0 | 0 | 52078 | 0 |
| ITADLLSNG | SEPTIN9; SEPTI | Q9UHD8; A0 | SEPT9_HUMAN | Unmodified | 0 | 0 | 0 | 0 | 43835 | 21815 | 265060 | 0 | 192910 | 0 | 903480 | 1805400 | 0 | 0 | 0 | 0 | 29995 | 0 |
| ITAELSWPS | TMEM131 | Q92545 | TM131_HUMAN | Unmodified | 0 | 0 | 0 | 0 | 68378 | 184590 | 0 | 0 | 492740 | 0 | 0 | 0 | 1225900 | 0 | 0 | 0 | 0 | 0 |
| ITAFVVER | ACADVL | P49748 | ACADV_HUMAI | Unmodified | 0 | 0 | 0 | 0 | 0 | 0 | 0 | 0 | 16991000 | 0 | 0 | 0 | 0 | 0 | 0 | 0 | 0 | 0 |
| ITAFVVERG | ACADVL | P49748 | ACADV_HUMAI | Unmodified | 0 | 0 | 0 | 0 | 702570 | 0 | 0 | 54177 | 1400700 | 4356200 | 0 | 0 | 0 | 301390 | 0 | 0 | 0 | 0 |
| ITAHLVHEL | ACAA2; ACAA2; | P42765; A0A | THIM_HUMAN | Unmodified | 0 | 0 | 0 | 0 | 0 | 0 | 0 | 0 | 979130 | 67675 | 0 | 0 | 0 | 0 | 0 | 0 | 0 | 0 |
| ITALDEFATI | SPTAN1; SPTAN | Q13813; A0A | SPTN1_HUMAN | Unmodified | 0 | 0 | 0 | 0 | 0 | 0 | 0 | 0 | 0 | 0 | 0 | 2965800 | 0 | 0 | 0 | 0 | 0 | 0 |
| ITAMDTED | RANBP3; RANB | Q9H6Z4; B7 | RANB3_HUMAI | Unmodified | 0 | 0 | 0 | 0 | 0 | 0 | 0 | 0 | 0 | 0 | 0 | 0 | 0 | 0 | 0 | 0 | 0 | 0 |
| ITANLFR | MAN2A1 | Q16706 | MA2A1_HUMAI | Unmodified | 0 | 0 | 0 | 0 | 0 | 0 | 346950 | 231820 | 3961400 | 430140 | 87049 | 509830 | 0 | 0 | 0 | 0 | 0 | 0 |
| ITASEALK | CAMK2D; CAMK | Q13557; E9F | KCC2D_HUMA | Unmodified | 0 | 26094 | 0 | 183960 | 216060 | 0 | 0 | 14551 | 213600 | 62390 | 59892 | 0 | 0 | 0 | 211160 | 0 | 273960 | 99329 |
| ITASLCDLK | HMGCS1 | Q01581 | HMCS1_HUMA | Unmodified | 108190 | 0 | 0 | 208420 | 0 | 452900 | 410140 | 208930 | 0 | 0 | 0 | 11654000 | 0 | 0 | 0 | 0 | 0 | 0 |
| ITASYEDR | F11R | Q9Y624 | JAM1_HUMAN | Unmodified | 722310 | 659200 | 825070 | 0 | 0 | 0 | 0 | 0 | 272960 | 55936 | 0 | 253750 | 0 | 0 | 662220 | 551230 | 21714 | 0 |
| ITASYEDRV | F11R | Q9Y624 | JAM1_HUMAN | Unmodified | 8118900 | 15475000 | 5874400 | 11085000 | 17442000 | 2953500 | 0 | 0 | 15157000 | 83769 | 2480200 | 2616700 | 25995000 | 7837800 | 23510000 | 6436500 | 5368900 | 12576000 |
| ITCLCQVP | AARS1; AARS1; | P49588; A0A | SYAC_HUMAN | Unmodified | 0 | 0 | 279790 | 90332 | 0 | 0 | 0 | 0 | 528170 | 0 | 270650 | 2624800 | 54351 | 0 | 258870 | 155410 | 0 | 0 |
| ITDFGLAK | EGFR; EGFR; E | P00533; Q5 | EGFR_HUMAN | Unmodified | 217530 | 821090 | 3416800 | 2700000 | 9769900 | 3536700 | 214370 | 155150 | 823970 | 420250 | 0 | 1933300 | 1572200 | 1762300 | 3509000 | 14688000 | 1299600 | 3161100 |
| ITDFGLAR | MAP3K11; ERB | Q16584; P0 | M3K11_HUMAI | Unmodified | 935060 | 306760 | 498680 | 942700 | 756350 | 261650 | 0 | 26678 | 0 | 0 | 1279400 | 3552200 | 57320 | 412070 | 11017000 | 5143800 | 1241400 | 1670600 |
| ITDIGEVSQ | MET | P08581 | MET_HUMAN H | Oxidation (M) | 0 | 0 | 0 | 0 | 0 | 0 | 0 | 0 | 307780 | 81843 | 38887 | 0 | 0 | 0 | 76009 | 0 | 0 | 0 |
| ITDLANLSA | GATD3B; GATD | A0A0B4J2D5 | GAL3B_HUMAN | Unmodified | 24813 | 0 | 0 | 0 | 0 | 0 | 1228000 | 103770 | 14196000 | 1088500 | 0 | 0 | 514320 | 0 | 0 | 0 | 0 | 0 |
| ITDNELELY | SLC12A2; SLC1 | P55011; G3 | S12A2_HUMAN | Unmodified | 358980 | 0 | 0 | 169610 | 0 | 0 | 0 | 0 | 0 | 102920 | 144710 | 0 | 0 | 0 | 780370 | 670630 | 0 | 0 |
| ITDPEILESF | OPLAH | O14841 | OPLA_HUMAN | Unmodified | 0 | 0 | 0 | 0 | 0 | 0 | 0 | 20987 | 0 | 0 | 0 | 1210800 | 0 | 0 | 0 | 0 | 0 | 0 |
| ITDRNLINA | ANKRD28 | O15084 | ANR28_HUMAI | Unmodified | 0 | 0 | 0 | 0 | 0 | 0 | 123770 | 0 | 0 | 0 | 0 | 1419000 | 0 | 0 | 0 | 0 | 0 | 1334400 |
| ITDSAGHIL | TMED10; TMED | P49755; G3 | TMEDA_HUMAI | Unmodified | 566160 | 318950 | 2494700 | 1717900 | 2521900 | 436450 | 1653800 | 1087700 | 6.08E+08 | 34521000 | 14731000 | 11442000 | 3720900 | 3583200 | 2905300 | 2505300 | 248810 | 3753500 |
| ITDSAGHIL | TMED10; TMED | P49755; G3 | TMEDA_HUMAI | Unmodified | 0 | 172790 | 0 | 218900 | 6325400 | 0 | 5955700 | 2179300 | 1.61E+08 | 93005000 | 3730500 | 216240 | 6313900 | 8120700 | 0 | 32886 | 248000 | 10468000 |
| ITDSAGHIL | TMED10; TMED | P49755; G3 | TMEDA_HUMAI | Unmodified | 0 | 0 | 0 | 0 | 0 | 0 | 491520 | 210610 | 596250 | 4140300 | 0 | 0 | 0 | 93233 | 0 | 0 | 91163 | 0 |
| ITDVEHITG | ENPP1 | P22413 | ENPP1_HUMAI | Unmodified | 0 | 0 | 0 | 0 | 0 | 0 | 0 | 0 | 0 | 0 | 0 | 338230 | 0 | 0 | 0 | 74572 | 0 | 0 |
| ITDVEHITG | ENPP1 | P22413 | ENPP1_HUMAI | Unmodified | 0 | 0 | 0 | 406720 | 0 | 0 | 0 | 0 | 0 | 243290 | 0 | 0 | 0 | 0 | 0 | 0 | 466820 | 768900 |
| ITEDYYVHL | TM9SF4; TM9SF | Q92544; A0A | TM9S4_HUMAI | Unmodified | 0 | 0 | 0 | 31375 | 0 | 0 | 177590 | 0 | 16674000 | 7447300 | 169880 | 534100 | 0 | 0 | 144100 | 30600 | 0 | 0 |
| ITEFCHR | P4HB; P4HB; P4 | P07237; A0A | PDIA1_HUMAN | Unmodified | 0 | 0 | 57589 | 0 | 0 | 0 | 0 | 14708 | 4569300 | 0 | 256130 | 0 | 0 | 0 | 0 | 0 | 0 | 0 |
| ITEFCHRFL | P4HB; P4HB; P4 | P07237; A0A | PDIA1_HUMAN | Unmodified | 0 | 0 | 0 | 0 | 0 | 0 | 217580 | 0 | 1306100 | 1425000 | 327860 | 337200 | 0 | 0 | 0 | 0 | 0 | 0 |
| ITEGDVVV | GART | P22102 | PUR2_HUMAN | Unmodified | 0 | 0 | 0 | 0 | 0 | 0 | 0 | 0 | 0 | 0 | 0 | 1520000 | 0 | 0 | 0 | 0 | 0 | 0 |
| ITEGVSDTN | UEVLD | Q8IX04 | UEVLD_HUMAI | Unmodified | 0 | 0 | 0 | 0 | 0 | 0 | 0 | 0 | 0 | 0 | 0 | 0 | 0 | 0 | 0 | 0 | 0 | 0 |
| ITELKEEIEV | PFDN4; PFDN4 | Q9NQP4; E9 | PFD4_HUMAN | Unmodified | 222560 | 357260 | 227800 | 437450 | 919850 | 0 | 28135 | 0 | 3090400 | 600250 | 3104900 | 5899300 | 333720 | 72714 | 287050 | 320890 | 90761 | 175330 |
| ITELTDENV | POLR2C; POLR | P19387; H3E | RPB3_HUMAN | Unmodified | 0 | 0 | 0 | 0 | 0 | 0 | 0 | 0 | 2043200 | 0 | 0 | 512420 | 0 | 0 | 0 | 150060 | 0 | 0 |
| ITENIGCVM | PSMA6; PSMA6 | P60900; G3 | PSA6_HUMAN | Unmodified | 0 | 0 | 0 | 0 | 0 | 0 | 0 | 0 | 1047600 | 0 | 0 | 0 | 0 | 0 | 0 | 0 | 0 | 0 |
| ITENIGCVM | PSMA6; PSMA6 | P60900; G3 | PSA6_HUMAN | Unmodified | 0 | 0 | 0 | 0 | 1692800 | 0 | 0 | 0 | 0 | 504520 | 0 | 0 | 678010 | 0 | 0 | 0 | 339870 | 0 |
| ITESEEVVS | LMNA; LMNA; L | P02545; A0A | LMNA_HUMAN | Unmodified | 0 | 0 | 0 | 0 | 3018200 | 639490 | 0 | 0 | 0 | 0 | 0 | 0 | 1160800 | 230870 | 0 | 0 | 890950 | 827570 |
| ITESRQVVE | CIT; CIT; CIT; CI | O14578; A0A | CTRO_HUMAN | Unmodified | 0 | 0 | 0 | 0 | 2776200 | 448060 | 78824 | 0 | 405530 | 320550 | 0 | 0 | 0 | 0 | 0 | 0 | 458000 | 0 |
| ITETEDPVK | CTPS2 | Q9NRF8 | PYRG2_HUMAI | Unmodified | 0 | 0 | 0 | 0 | 0 | 0 | 0 | 0 | 64873 | 0 | 77759 | 1146800 | 0 | 0 | 0 | 97938 | 0 | 0 |
| ITEYLCEPLI | AP1B1; AP2B1; | Q10567; P63 | AP1B1_HUMAN | Unmodified | 150950 | 93578 | 0 | 0 | 0 | 0 | 0 | 0 | 0 | 0 | 0 | 2451100 | 0 | 0 | 360800 | 489110 | 0 | 0 |
| ITEYLER | CAPN7 | Q9Y6W3 | CAN7_HUMAN | Unmodified | 0 | 0 | 0 | 0 | 0 | 0 | 0 | 0 | 0 | 0 | 0 | 0 | 0 | 0 | 0 | 0 | 0 | 0 |
| ITEYLERVQ | CAPN7 | Q9Y6W3 | CAN7_HUMAN | Unmodified | 0 | 0 | 0 | 0 | 0 | 0 | 0 | 0 | 0 | 1884000 | 0 | 3734600 | 0 | 0 | 0 | 0 | 0 | 0 |
| ITFCTGIR | PSMD4; PSMD4 | P55036; A6P | PSMD4_HUMA | Unmodified | 0 | 0 | 0 | 141920 | 0 | 0 | 0 | 0 | 113420 | 0 | 0 | 1600700 | 0 | 0 | 96246 | 0 | 0 | 0 |
| ITFDDYIAC | SRI; SRI; SRI | P30626; B4D | SORCN_HUMA | Unmodified | 628010 | 0 | 0 | 142830 | 0 | 0 | 0 | 23872 | 5160400 | 162750 | 0 | 252160 | 0 | 0 | 620380 | 347890 | 0 | 0 |
| ITFHGEGD | DNAJA1 | P31689 | DNJA1_HUMAN | Unmodified | 366740 | 1397500 | 98776 | 1591300 | 4846300 | 617820 | 0 | 0 | 2385300 | 1576600 | 271790 | 403730 | 8494200 | 0 | 1351700 | 418650 | 0 | 0 |
| ITFHGEGD | DNAJA1 | P31689 | DNJA1_HUMAN | Unmodified | 0 | 0 | 0 | 2894500 | 5259700 | 0 | 0 | 0 | 0 | 4461800 | 0 | 0 | 29220 | 10118000 | 0 | 0 | 0 | 0 |
| ITFLLQAIR | IPO5; IPO5; IPO | O00410; E7E | IPO5_HUMAN I | Unmodified | 288990 | 129040 | 0 | 0 | 0 | 0 | 0 | 0 | 140860 | 97606 | 0 | 8765700 | 0 | 0 | 0 | 255260 | 0 | 0 |
| ITFPKEGD | DNAJB1; DNAJE | P25685; M0 | DNJB1_HUMAN | Unmodified | 0 | 0 | 0 | 0 | 3684200 | 0 | 1384100 | 0 | 0 | 1249400 | 0 | 0 | 2207200 | 1458900 | 0 | 0 | 2396200 | 2118100 |
| ITFPREGDE | DNAJB4 | Q9UDY4 | DNJB4_HUMAN | Unmodified | 237960 | 343770 | 17141 | 0 | 3235700 | 901050 | 47111 | 0 | 192290 | 95529 | 0 | 24769 | 0 | 0 | 0 | 34839 | 686170 | 0 |
| ITFTGEADG | DNAJA2 | O60884 | DNJA2_HUMAN | Unmodified | 0 | 611650 | 0 | 516400 | 0 | 0 | 0 | 0 | 0 | 0 | 0 | 0 | 0 | 0 | 737750 | 0 | 0 | 0 |
| ITFVPADSD | BSND | Q8WZ55 | BSND_HUMAN | Unmodified | 0 | 0 | 0 | 0 | 0 | 0 | 0 | 123640 | 0 | 0 | 0 | 512560 | 0 | 0 | 0 | 59259 | 0 | 0 |
| ITGEAFVQF | HNRNPF | P52597 | HNRPF_HUMAI | Unmodified | 368360 | 0 | 136870 | 171410 | 0 | 0 | 0 | 0 | 1147300 | 0 | 0 | 3014400 | 0 | 0 | 385930 | 261620 | 183270 | 58110 |
| ITGHFYAC | IGF2BP3 | O00425 | IF2B3_HUMAN | Unmodified | 553460 | 189920 | 0 | 90823 | 0 | 0 | 0 | 0 | 0 | 0 | 0 | 837830 | 0 | 94801 | 0 | 0 | 0 | 0 |
| ITGHFYAC | IGF2BP3 | O00425 | IF2B3_HUMAN | Unmodified | 0 | 0 | 0 | 0 | 0 | 0 | 89030 | 0 | 0 | 0 | 0 | 1138400 | 1138100 | 0 | 0 | 0 | 0 | 0 |
| ITGKYCECI | ITGB3; N/A | P05106; H3E | ITB3_HUMAN I | Unmodified | 0 | 28119 | 0 | 324670 | 0 | 669280 | 0 | 33543 | 0 | 0 | 564220 | 0 | 148990 | 1084100 | 0 | 0 | 0 | 416110 |
| ITGLDPAGI | LPL | P06858 | LIPL_HUMAN L | Unmodified | 0 | 0 | 0 | 0 | 0 | 0 | 0 | 0 | 961700 | 0 | 0 | 0 | 0 | 0 | 0 | 0 | 0 | 0 |
| ITGMLLEID | PABPC4; PABP | Q13310; P11 | PABP4_HUMAN | 2 Oxidation (M) | 0 | 2249500 | 0 | 0 | 0 | 0 | 0 | 0 | 0 | 0 | 0 | 0 | 0 | 0 | 244240 | 0 | 535890 | 0 |
| ITGNMGLA | SCP2 | P22307 | SCP2_HUMAN | Oxidation (M) | 0 | 0 | 0 | 0 | 56367 | 0 | 0 | 0 | 10745000 | 2491500 | 0 | 169590 | 0 | 0 | 591470 | 0 | 24566 | 0 |
| ITGNMGLA | SCP2 | P22307 | SCP2_HUMAN | Unmodified | 0 | 0 | 0 | 0 | 0 | 0 | 629300 | 0 | 34740000 | 2875600 | 339080 | 0 | 0 | 0 | 0 | 0 | 0 | 0 |
| ITGNMGLA | SCP2 | P22307 | SCP2_HUMAN | 2 Oxidation (M) | 0 | 0 | 0 | 0 | 67946 | 0 | 0 | 0 | 2923200 | 726390 | 206590 | 0 | 0 | 0 | 0 | 0 | 122060 | 0 |
| ITGNMGLA | SCP2 | P22307 | SCP2_HUMAN | 2 Oxidation (M) | 0 | 0 | 0 | 0 | 0 | 0 | 0 | 0 | 0 | 512110 | 0 | 0 | 0 | 0 | 0 | 0 | 0 | 0 |
| ITGNPVVT | MOV10; MOV10 | Q9HCE1; Q5 | MOV10_HUMA | Unmodified | 3054300 | 1661300 | 764630 | 0 | 0 | 0 | 92170 | 0 | 0 | 0 | 51903 | 117120 | 0 | 0 | 0 | 1527900 | 0 | 0 |
| ITGNPVVT | MOV10; MOV10 | Q9HCE1; Q5 | MOV10_HUMA | Unmodified | 494630 | 6016500 | 1438700 | 1675100 | 0 | 0 | 0 | 0 | 0 | 0 | 0 | 0 | 0 | 0 | 3799600 | 4164600 | 0 | 0 |
| ITGQLESLE | MACF1; MACF1 | Q9UPN3; H3 | MACF1_HUMAI | Unmodified | 0 | 0 | 0 | 97742 | 1167100 | 0 | 0 | 0 | 880350 | 1671600 | 426440 | 0 | 0 | 0 | 0 | 0 | 1034000 | 0 |
| ITGYIIK | FN1; FN1; FN1 | P02751; H0Y | FINC_HUMAN I | Unmodified | 313290 | 1484600 | 12706000 | 0 | 114820 | 750890 | 0 | 0 | 0 | 149550 | 91633 | 1074400 | 0 | 73449 | 0 | 0 | 75148 | 0 |
| ITGYIIKYEK | FN1; FN1; FN1 | P02751; H0Y | FINC_HUMAN I | Unmodified | 275810 | 980700 | 88570 | 465260 | 0 | 0 | 0 | 0 | 0 | 0 | 0 | 108960 | 0 | 0 | 139140 | 714160 | 0 | 0 |
| ITHLSTLQV | HAGH; HAGH; I | Q16775; H3 | GLO2_HUMAN | Unmodified | 0 | 936100 | 320600 | 0 | 1488300 | 0 | 0 | 0 | 2981600 | 1016200 | 570460 | 1065900 | 1918000 | 0 | 0 | 631110 | 1127500 | 0 |
| ITHSPLTIC | GNAI2 | P04899 | GNAI2_HUMAN | Unmodified | 0 | 0 | 0 | 0 | 0 | 0 | 0 | 0 | 0 | 0 | 0 | 635810 | 0 | 0 | 0 | 0 | 0 | 0 |
| ITHSPLTIC | GNAI2 | P04899 | GNAI2_HUMAN | Unmodified | 0 | 0 | 0 | 0 | 0 | 0 | 0 | 0 | 0 | 0 | 1108400 | 18428000 | 0 | 0 | 0 | 0 | 0 | 0 |
| ITHSPLTIC | GNAI2 | P04899 | GNAI2_HUMAN | Unmodified | 0 | 0 | 0 | 0 | 8248900 | 2049200 | 0 | 0 | 1895700 | 1523600 | 0 | 1796200 | 0 | 0 | 0 | 0 | 8956700 | 19893000 |
| ITHVVSST | SHCBP1; SHCB | Q8NEM2; I3 | SHCBP_HUMA | Oxidation (M) | 0 | 0 | 0 | 0 | 561180 | 0 | 0 | 0 | 51212 | 0 | 0 | 79998 | 0 | 0 | 0 | 0 | 0 | 0 |
| ITHVVSST | SHCBP1; SHCB | Q8NEM2; I3 | SHCBP_HUMA | Unmodified | 0 | 0 | 0 | 0 | 1576300 | 214870 | 0 | 0 | 144140 | 188020 | 0 | 423960 | 0 | 0 | 0 | 0 | 0 | 0 |
| ITIAAYLPLK | NUP210 | Q8TEM1 | PO210_HUMAN | Unmodified | 0 | 0 | 0 | 0 | 0 | 0 | 0 | 0 | 2366900 | 3315900 | 484240 | 0 | 0 | 0 | 0 | 0 | 0 | 0 |
| ITIGQAPTE | MTHFD1; MTHF | P11586; V9 | C1TC_HUMAN | Unmodified | 27573000 | 2364800 | 0 | 0 | 9312000 | 1567400 | 2370800 | 2163300 | 3716900 | 2230300 | 7947000 | 8372700 | 2912900 | 1728800 | 87133 | 65943 | 6240500 | 6230400 |
| ITIIPQDPVL | ABCC1; ABCC1 | P33527; I3L | MRP1_HUMAN | Unmodified | 0 | 51559 | 455630 | 348190 | 0 | 0 | 0 | 0 | 97470 | 0 | 362530 | 4064800 | 0 | 0 | 1011900 | 860710 | 0 | 0 |
| ITILVTHQL | ABCC4; ABCC4 | O15439; A0A | MRP4_HUMAN | Unmodified | 0 | 362530 | 2687600 | 4433900 | 926740 | 0 | 2709200 | 39525 | 0 | 0 | 0 | 0 | 512610 | 934700 | 2576300 | 2198300 | 0 | 0 |
| ITIPNAFLG | DVL2; DVL2; DV | O14641; I3L | DVL2_HUMAN | Unmodified | 0 | 0 | 0 | 40691 | 60081 | 0 | 0 | 0 | 0 | 0 | 0 | 0 | 244470 | 111750 | 33715 | 0 | 0 | 0 |
| ITISPLQELT | IGF2BP3 | O00425 | IF2B3_HUMAN | Unmodified | 700510 | 465220 | 95638 | 120880 | 0 | 0 | 0 | 0 | 0 | 101860 | 153900 | 1121900 | 0 | 0 | 151680 | 47432 | 0 | 0 |
| ITITGYR | NOMO1; NOMO | Q15155; P69 | NOMO1_HUMA | Unmodified | 0 | 0 | 364680 | 0 | 0 | 0 | 0 | 0 | 11071000 | 625200 | 0 | 491730 | 0 | 0 | 0 | 0 | 0 | 0 |
| ITITNDK | HSPA1L; HSPA | P34931; P17 | HS71L_HUMAN | Unmodified | 15515000 | 24746000 | 28565000 | 4942400 | 6854100 | 6225500 | 3307700 | 504570 | 22018000 | 3366100 | 371320 | 14408000 | 2405700 | 2527700 | 17752000 | 8806300 | 3309900 | 2409800 |
| ITITNDQNF | HSPA5; HSPA5; | P11021; A0A | BIP_HUMAN Er | Unmodified | 259120 | 221170 | 359880 | 0 | 1027000 | 1164500 | 0 | 64051 | 20822000 | 644320 | 642680 | 4291800 | 0 | 0 | 1655500 | 708610 | 597320 | 325580 |

| | | | | | | | | | | | | | | | | | | | | | | |
|---|---|---|---|---|---|---|---|---|---|---|---|---|---|---|---|---|---|---|---|---|---|---|
| ITITNDQNF | HSPA5; HSPA5; | P11021; A0A | BIP_HUMAN Er | Unmodified | 188700 | 189020 | 1028800 | 144040 | 0 | 0 | 1185700 | 0 | 31886000 | 1682000 | 1275600 | 1641800 | 0 | 0 | 2030700 | 530100 | 192120 | 0 |
| ITITNDQNF | HSPA5; HSPA5; | P11021; A0A | BIP_HUMAN Er | Unmodified | 0 | 0 | 0 | 0 | 0 | 0 | 0 | 0 | 16209000 | 3445000 | 350350 | 0 | 0 | 0 | 0 | 0 | 0 | 0 |
| ITKADLEQF | UTRN | P46939 | UTRN_HUMAN | Unmodified | 0 | 0 | 0 | 420200 | 7510300 | 0 | 350880 | 0 | 0 | 1963400 | 0 | 0 | 2608600 | 1375200 | 0 | 0 | 3791000 | 11234000 |
| ITKFENAFL | ATP5F1A | P25705 | ATPA_HUMAN , | Unmodified | 0 | 0 | 0 | 0 | 2870200 | 0 | 211810 | 0 | 14261000 | 489420 | 0 | 0 | 0 | 0 | 0 | 0 | 67202 | 124100 |
| ITKFENAFL | ATP5F1A | P25705 | ATPA_HUMAN , | Unmodified | 0 | 0 | 0 | 0 | 0 | 0 | 0 | 0 | 20119000 | 3244500 | 458210 | 0 | 0 | 0 | 0 | 0 | 0 | 0 |
| ITKFIEENKE | SMC4; SMC4; S | Q9NTJ3; E9F | SMC4_HUMAN | Unmodified | 0 | 0 | 0 | 0 | 977840 | 0 | 0 | 0 | 0 | 0 | 0 | 0 | 0 | 0 | 0 | 0 | 314370 | 0 |
| ITKPGSIDS | EIF4G1; EIF4G1 | Q04637; E7E | IF4G1_HUMAN | Unmodified | 368790 | 241610 | 0 | 1023300 | 193070 | 173520 | 0 | 0 | 313760 | 124930 | 0 | 0 | 689800 | 0 | 420760 | 422300 | 118280 | 0 |
| ITLDNAYMI | PKM; PKM; PKM | P14618; B4D | KPYM_HUMAN | Oxidation (M) | 594340 | 0 | 1694700 | 1006500 | 51649000 | 402640 | 2426300 | 3113700 | 3063200 | 1802700 | 14448000 | 38378000 | 3318700 | 2290200 | 1266700 | 1256200 | 7886000 | 16374000 |
| ITLDNAYMI | PKM; PKM; PKM | P14618; B4D | KPYM_HUMAN | Unmodified | 514630 | 747940 | 742810 | 1178900 | 77282000 | 7473900 | 4021100 | 2026600 | 7602600 | 3038500 | 8814500 | 61389000 | 7376700 | 5079700 | 920230 | 690480 | 8356200 | 27432000 |
| ITLDNAYMI | PKM; PKM; PKM | P14618; B4D | KPYM_HUMAN | Oxidation (M) | 0 | 0 | 0 | 0 | 39135000 | 4511000 | 13003000 | 125870 | 0 | 2693500 | 0 | 0 | 6579300 | 5129500 | 0 | 0 | 27608000 | 22487000 |
| ITLDNAYMI | PKM; PKM; PKM | P14618; B4D | KPYM_HUMAN | Unmodified | 0 | 0 | 0 | 0 | 7167200 | 378240 | 0 | 0 | 0 | 0 | 0 | 0 | 0 | 506150 | 0 | 0 | 3241100 | 3937500 |
| ITLDNAYMI | PKM; PKM; PKM | P14618; B4D | KPYM_HUMAN | Oxidation (M) | 0 | 0 | 0 | 0 | 0 | 0 | 2559600 | 0 | 0 | 0 | 0 | 0 | 0 | 833040 | 0 | 0 | 8997700 | 2238800 |
| ITLDNAYMI | PKM; PKM; PKM | P14618; B4D | KPYM_HUMAN | Unmodified | 0 | 0 | 0 | 0 | 0 | 0 | 339190 | 184710 | 0 | 0 | 0 | 0 | 0 | 0 | 0 | 0 | 2146000 | 0 |
| ITLEDVISH | IQGAP2; IQGAP | Q13576; D6I | IQGA2_HUMAN | Unmodified | 0 | 0 | 0 | 0 | 0 | 0 | 0 | 0 | 0 | 0 | 0 | 0 | 0 | 0 | 0 | 0 | 0 | 0 |
| ITLEGPTED | HDLBP; HDLBP | Q00341; A0/ | VIGLN_HUMAN | Unmodified | 0 | 0 | 0 | 0 | 0 | 0 | 4814500 | 2223800 | 0 | 0 | 0 | 0 | 0 | 1288800 | 0 | 0 | 3360400 | 0 |
| ITLEHLGIC | ZNF185; ZNF18 | O15231; H0 | ZN185_HUMAN | Unmodified | 0 | 0 | 0 | 0 | 0 | 0 | 0 | 0 | 0 | 0 | 0 | 0 | 0 | 0 | 0 | 0 | 818550 | 0 |
| ITLESFLAW | ZC3H15 | Q8WU90 | ZC3HF_HUMAI | Unmodified | 25481 | 0 | 0 | 0 | 980640 | 0 | 0 | 0 | 0 | 0 | 416730 | 1750300 | 1091900 | 297490 | 0 | 0 | 250250 | 806960 |
| ITLFLEEDK | ANLN | Q9NQW6 | ANLN_HUMAN | Unmodified | 0 | 0 | 0 | 0 | 9857000 | 2021400 | 231410 | 30221 | 0 | 0 | 0 | 0 | 618220 | 0 | 0 | 0 | 1901200 | 1474300 |
| ITLHITDASI | ZPR1; ZPR1; ZP | O75312; H7 | ZPR1_HUMAN | Unmodified | 0 | 0 | 0 | 0 | 0 | 0 | 0 | 0 | 0 | 0 | 0 | 0 | 0 | 0 | 0 | 0 | 0 | 0 |
| ITLPVDFVT | PGK1 | P00558 | PGK1_HUMAN | Unmodified | 0 | 0 | 1348500 | 0 | 0 | 0 | 0 | 0 | 298970 | 0 | 1893100 | 5828900 | 0 | 0 | 0 | 0 | 0 | 0 |
| ITLPVDFVT | PGK1 | P00558 | PGK1_HUMAN | Unmodified | 0 | 0 | 17380000 | 30550000 | 71379000 | 14635000 | 0 | 4958100 | 85157000 | 133990 | 71035000 | 2.81E+08 | 2.22E+08 | 63299000 | 33699000 | 27110000 | 1.01E+08 | 2.04E+08 |
| ITLPVDFVT | PGK1 | P00558 | PGK1_HUMAN | Oxidation (M) | 0 | 0 | 0 | 0 | 0 | 0 | 0 | 0 | 0 | 0 | 0 | 0 | 0 | 0 | 0 | 0 | 2725300 | 0 |
| ITLQDVVSH | IQGAP1; IQGAP | P46940; A0A | IQGA1_HUMAN | Unmodified | 174350 | 78981 | 1221500 | 716440 | 12491000 | 2125100 | 0 | 229810 | 2780900 | 586600 | 1123300 | 6245000 | 1718400 | 172080 | 1756300 | 1141700 | 0 | 489600 |
| ITLRSELLG | EDIL3 | O43854 | EDIL3_HUMAN | Unmodified | 280700 | 0 | 0 | 0 | 76904000 | 21251000 | 2070100 | 707680 | 10480000 | 5169600 | 0 | 0 | 0 | 0 | 0 | 0 | 5869900 | 41496000 |
| ITLSSSSEE | PARP12 | Q9H0J9 | PAR12_HUMAN | Unmodified | 158480 | 412900 | 715020 | 469630 | 0 | 106820 | 0 | 0 | 0 | 0 | 0 | 0 | 0 | 0 | 431760 | 469350 | 0 | 0 |
| ITLTSDPRL | UFM1; UFM1 | P61960; H0 | UFM1_HUMAN | Unmodified | 691290 | 622310 | 2816200 | 2238800 | 940540 | 153160 | 0 | 0 | 1086000 | 399140 | 1595800 | 1107500 | 1591800 | 756430 | 9462500 | 5647500 | 885740 | 1448400 |
| ITLVSAAPG | ACOT13 | Q9NPJ3 | ACO13_HUMAI | Unmodified | 89638 | 115790 | 102270 | 73740 | 875470 | 92215 | 0 | 0 | 1013400 | 391200 | 255800 | 0 | 165250 | 137540 | 95532 | 64222 | 0 | 132490 |
| ITLVSSGSG | FLOT1; FLOT1; I | O75955; A0/ | FLOT1_HUMAN | Oxidation (M) | 0 | 0 | 721090 | 1060700 | 1738400 | 718390 | 126770 | 0 | 1621900 | 304450 | 1053900 | 1824700 | 0 | 0 | 962050 | 764040 | 174760 | 1635500 |
| ITLVSSGSG | FLOT1; FLOT1; I | O75955; A0/ | FLOT1_HUMAN | Unmodified | 0 | 0 | 1056500 | 920220 | 7550100 | 1557700 | 289350 | 0 | 10079000 | 1324300 | 2226300 | 4430600 | 933550 | 0 | 1887000 | 622340 | 389470 | 1268400 |
| ITMPVIFNE | OSBPL1A | Q9BXW6 | OSBL1_HUMAN | Oxidation (M) | 0 | 0 | 0 | 0 | 0 | 0 | 0 | 0 | 0 | 0 | 32029 | 257000 | 0 | 0 | 6778.4 | 0 | 0 | 0 |
| ITNDFYPEE | EIF3B; EIF3B | P55884; C9J | EIF3B_HUMAN | Unmodified | 0 | 165500 | 711110 | 0 | 0 | 0 | 0 | 0 | 0 | 0 | 906600 | 5700000 | 0 | 0 | 390720 | 487310 | 0 | 0 |
| ITNDFYPEE | EIF3B; EIF3B | P55884; C9J | EIF3B_HUMAN | Unmodified | 0 | 325510 | 0 | 0 | 0 | 0 | 0 | 0 | 0 | 0 | 321460 | 2248000 | 0 | 830250 | 0 | 0 | 0 | 0 |
| ITNLTQQLE | DSP | P15924 | DESP_HUMAN | Unmodified | 0 | 0 | 0 | 0 | 115290 | 0 | 0 | 0 | 573740 | 0 | 0 | 117370 | 0 | 0 | 36429 | 35645 | 70048 | 0 |
| ITNNIDPVG | GNB1; GNB1; G | P62873; F6U | GBB1_HUMAN | Unmodified | 0 | 0 | 0 | 0 | 0 | 0 | 0 | 0 | 204520 | 176140 | 677420 | 698270 | 0 | 0 | 0 | 0 | 0 | 0 |
| ITNQVHGL | PSMD7 | P51665 | PSMD7_HUMA | Unmodified | 0 | 0 | 0 | 0 | 57822 | 0 | 343890 | 0 | 0 | 0 | 0 | 1337800 | 0 | 0 | 0 | 0 | 14280 | 0 |
| ITNQVIYLN | DYNC1H1; DYN | Q14204; A0/ | DYHC1_HUMA | Unmodified | 0 | 0 | 0 | 0 | 0 | 0 | 0 | 0 | 0 | 0 | 0 | 669740 | 0 | 0 | 0 | 0 | 0 | 0 |
| ITNQVIYLN | DYNC1H1; DYN | Q14204; A0/ | DYHC1_HUMA | Unmodified | 0 | 0 | 0 | 0 | 0 | 0 | 0 | 0 | 0 | 0 | 0 | 2168600 | 0 | 0 | 0 | 0 | 0 | 0 |
| ITNSLTVLC | EIF3J; EIF3J; EIF | O75822; H0 | EIF3J_HUMAN I | Unmodified | 0 | 0 | 0 | 0 | 0 | 0 | 0 | 0 | 0 | 0 | 321910 | 2456300 | 0 | 0 | 0 | 0 | 0 | 0 |
| ITNSLTVLC | EIF3J; EIF3J; EIF | O75822; H0 | EIF3J_HUMAN I | Unmodified | 0 | 0 | 0 | 0 | 2598400 | 212280 | 972700 | 0 | 0 | 0 | 0 | 0 | 1911500 | 0 | 0 | 0 | 0 | 45731 |
| ITNVSAIIKP | PI4KA; PI4KA | P42356; A8N | PI4KA_HUMAN | Unmodified | 0 | 0 | 374070 | 234910 | 0 | 0 | 0 | 18963 | 0 | 0 | 0 | 345220 | 0 | 0 | 609590 | 441240 | 0 | 0 |
| ITPAHDQN | VARS1; VARS1 | P26640; A0A | SYVC_HUMAN | Unmodified | 0 | 0 | 0 | 0 | 0 | 0 | 0 | 0 | 0 | 0 | 61822 | 399270 | 0 | 0 | 0 | 0 | 0 | 0 |
| ITPENLPQI | MATR3; MATR3; | P43243; D6F | MATR3_HUMAI | Unmodified | 73023 | 33056 | 47000 | 26293 | 0 | 0 | 73586 | 18731 | 155400 | 182070 | 74567 | 267280 | 49027 | 25853 | 0 | 48043 | 51239 | 94398 |
| ITPGARGAI | CNP; CNP; CNI | P09543; K7E | CN37_HUMAN | Unmodified | 0 | 0 | 0 | 0 | 802220 | 113750 | 0 | 152910 | 0 | 0 | 0 | 0 | 0 | 0 | 0 | 0 | 203870 | 946750 |
| ITPSYVAFTI | HSPA5; HSPA5; | P11021; A0A | BIP_HUMAN Er | Unmodified | 0 | 0 | 0 | 0 | 0 | 457990 | 0 | 40872 | 25645000 | 482720 | 2147000 | 964690 | 0 | 0 | 707270 | 0 | 0 | 0 |
| ITPSYVAFTI | HSPA5; HSPA5; | P11021; A0A | BIP_HUMAN Er | Unmodified | 0 | 0 | 8200400 | 710740 | 38687 | 0 | 26580 | 263070 | 7.07E+08 | 7976500 | 21573000 | 16816000 | 0 | 1522400 | 15504000 | 3589100 | 165670 | 107830 |
| ITPSYVAFTI | HSPA5; HSPA5; | P11021; A0A | BIP_HUMAN Er | Unmodified | 0 | 279910 | 0 | 580110 | 0 | 0 | 1454500 | 0 | 0 | 4081100 | 0 | 0 | 0 | 0 | 0 | 1349500 | 0 | 0 |
| ITQCSVEIQ | STX7 | O15400 | STX7_HUMAN | Unmodified | 0 | 0 | 280950 | 0 | 0 | 0 | 0 | 0 | 0 | 0 | 228370 | 783180 | 0 | 0 | 235100 | 350360 | 0 | 0 |
| ITQDIFQQL | MARS1; MARS1 | P56192; H0 | SYMC_HUMAN | Unmodified | 16194 | 14908 | 85978 | 0 | 86724 | 31868 | 0 | 17183 | 107830 | 57012 | 147340 | 945220 | 0 | 0 | 43415 | 53610 | 25847 | 72744 |
| ITQDIFQQL | MARS1; MARS1 | P56192; H0 | SYMC_HUMAN | Unmodified | 0 | 0 | 0 | 0 | 187230 | 0 | 0 | 0 | 598660 | 0 | 0 | 5719300 | 0 | 0 | 0 | 0 | 0 | 0 |
| ITQLPVR | LRIG1 | Q96JA1 | LRIG1_HUMAN | Unmodified | 0 | 0 | 580100 | 0 | 0 | 670000 | 0 | 189790 | 304760 | 0 | 0 | 0 | 0 | 440770 | 0 | 0 | 97116 | 452970 |
| ITQQMPK | HDLBP; HDLBP | Q00341; A0/ | VIGLN_HUMAN | Unmodified | 106570 | 0 | 244820 | 117590 | 317040 | 0 | 0 | 45050 | 0 | 0 | 0 | 667620 | 642370 | 339920 | 109540 | 125720 | 0 | 203400 |
| ITQSEFDR | SH3GLB1; SH3 | Q9Y371; A0/ | SHLB1_HUMAN | Unmodified | 78463 | 89904 | 101870 | 137670 | 0 | 0 | 0 | 0 | 570550 | 0 | 0 | 1157900 | 0 | 0 | 574000 | 276370 | 0 | 0 |
| ITQSEFDR | SH3GLB1; SH3 | Q9Y371; A0/ | SHLB1_HUMAN | Unmodified | 46471 | 0 | 0 | 248100 | 0 | 0 | 0 | 0 | 550970 | 106170 | 146040 | 0 | 285930 | 0 | 126000 | 45289 | 0 | 0 |
| ITQVDFPPI | DYNC1I2; DYN | Q13409; E7E | DC1I2_HUMAN | Unmodified | 0 | 0 | 0 | 285300 | 0 | 0 | 451060 | 253760 | 328560 | 40895 | 259070 | 927100 | 2101900 | 1311400 | 69359 | 0 | 1064600 | 1890100 |
| ITQVTWQK | NECTIN1 | Q15223 | NECT1_HUMAI | Unmodified | 743190 | 864710 | 211700 | 203120 | 1632700 | 2397800 | 333380 | 0 | 267970 | 80165 | 184210 | 359610 | 974920 | 300650 | 157660 | 100120 | 861230 | 259340 |
| ITQWEDPR | ITCH; NEDD4L; | Q96J02; Q96 | ITCH_HUMAN I | Unmodified | 1955900 | 2309900 | 0 | 0 | 0 | 0 | 0 | 0 | 0 | 0 | 0 | 353270 | 0 | 0 | 2209200 | 2505400 | 0 | 0 |
| ITQWEDPR | NEDD4L; NEDD | Q96PU5; A0 | NED4L_HUMAI | Unmodified | 2136800 | 4830800 | 2424200 | 3859000 | 0 | 0 | 0 | 0 | 1041700 | 0 | 618530 | 1031300 | 0 | 0 | 7129100 | 3063900 | 0 | 0 |
| ITQWEDPR | ITCH; ITCH; ITC | Q96J02; A0A | ITCH_HUMAN I | Unmodified | 0 | 0 | 225950 | 0 | 0 | 0 | 0 | 0 | 0 | 0 | 0 | 0 | 0 | 0 | 113200 | 0 | 0 | 0 |
| ITQYVEEVK | STX2; STX2 | P32856; J3K | STX2_HUMAN | Unmodified | 0 | 0 | 0 | 0 | 0 | 0 | 0 | 0 | 0 | 0 | 0 | 1369200 | 0 | 0 | 0 | 0 | 0 | 0 |
| ITQYVEEVK | STX2; STX2 | P32856; J3K | STX2_HUMAN | Unmodified | 0 | 0 | 0 | 0 | 0 | 0 | 0 | 0 | 335860 | 0 | 448690 | 5109300 | 0 | 0 | 0 | 0 | 28553 | 0 |
| ITRITSLTQL | TRIM32; TRIM3 | Q13049; Q5 | TRI32_HUMAN | Unmodified | 0 | 0 | 0 | 0 | 637600 | 0 | 928540 | 0 | 0 | 274060 | 0 | 0 | 0 | 0 | 0 | 0 | 45322 | 0 |
| ITRLTYEIED | DSP | P15924 | DESP_HUMAN | Unmodified | 0 | 0 | 0 | 0 | 253100 | 0 | 78885 | 0 | 475600 | 0 | 0 | 314130 | 0 | 0 | 0 | 0 | 286470 | 0 |
| ITRPLLLLNI | CPNE2; CPNE2 | Q96FN4; H3 | CPNE2_HUMAI | Unmodified | 0 | 0 | 0 | 0 | 1284600 | 128840 | 307390 | 201970 | 707350 | 381050 | 0 | 0 | 0 | 0 | 0 | 0 | 0 | 0 |
| ITRTKEEALI | PIN1; PIN1; PIN | Q13526; K7E | PIN1_HUMAN I | Unmodified | 130510 | 0 | 0 | 1357200 | 4464500 | 1897100 | 3964400 | 166610 | 0 | 2680200 | 0 | 0 | 16225000 | 5964500 | 0 | 0 | 10910000 | 7273100 |
| ITRYQGVNI | PABPC1; PABP | P11940; A0A | PABP1_HUMAN | Unmodified | 10804000 | 7032600 | 10498000 | 17705000 | 0 | 0 | 0 | 0 | 0 | 144460 | 0 | 0 | 1762100 | 928420 | 2316700 | 2720600 | 0 | 0 |
| ITSAVDGKI | ERBIN | Q96RT1 | ERBIN_HUMAN | Unmodified | 0 | 0 | 0 | 0 | 2352000 | 546370 | 0 | 2038400 | 0 | 190720 | 0 | 0 | 313960 | 346830 | 0 | 0 | 201870 | 412900 |
| ITSAVDGKI | ERBIN | Q96RT1 | ERBIN_HUMAN | Unmodified | 0 | 0 | 0 | 0 | 4270400 | 0 | 0 | 0 | 0 | 0 | 0 | 0 | 0 | 0 | 0 | 0 | 804180 | 0 |
| ITSAVWGPI | EIF3I; EIF3I; EIF | Q13347; A0/ | EIF3I_HUMAN I | Unmodified | 0 | 0 | 0 | 0 | 0 | 0 | 0 | 0 | 0 | 0 | 0 | 3158600 | 0 | 0 | 0 | 0 | 0 | 0 |
| ITSAYLQDII | NDUFA10; NDU | O95299; A0/ | NDUAA_HUMA | Unmodified | 0 | 0 | 0 | 0 | 0 | 0 | 0 | 0 | 1786900 | 752080 | 114940 | 0 | 0 | 0 | 0 | 0 | 0 | 0 |
| ITSCIFQLL | PAICS; PAICS | P22234; E9P | PUR6_HUMAN | Unmodified | 23197 | 73469 | 27785 | 743550 | 956960 | 135140 | 321600 | 0 | 332060 | 665170 | 343640 | 723760 | 3600700 | 1186700 | 867110 | 131070 | 1081600 | 1062900 |
| ITSDEPLTK | ATXN10; ATXN1 | Q9UBB4; B1 | ATX10_HUMAN | Unmodified | 0 | 0 | 187280 | 0 | 0 | 244760 | 0 | 0 | 768340 | 1385100 | 84204 | 1059600 | 0 | 635450 | 0 | 89542 | 0 | 0 |
| ITSDEPLTK | ATXN10; ATXN1 | Q9UBB4; B1 | ATX10_HUMAN | Unmodified | 180860 | 206800 | 579610 | 173580 | 0 | 0 | 0 | 0 | 0 | 60144 | 548300 | 6003500 | 45189 | 0 | 1444500 | 1602500 | 0 | 0 |
| ITSDFRIGF | ITGB1; ITGB1; I | P05556; H7 | ITB1_HUMAN I | Unmodified | 0 | 0 | 0 | 277330 | 1.28E+08 | 7378600 | 2344200 | 49570 | 1577900 | 2715100 | 0 | 0 | 4289300 | 3731100 | 38990 | 0 | 24487000 | 70549000 |
| ITSEAEDLV | PSME3; PSME3 | P61289; A0A | PSME3_HUMAI | Unmodified | 0 | 0 | 0 | 0 | 0 | 0 | 0 | 0 | 0 | 0 | 617880 | 0 | 0 | 0 | 0 | 0 | 0 | 0 |
| ITSEALLVT | CAND1; CAND | Q86VP6; H0 | CAND1_HUMA | Unmodified | 0 | 0 | 0 | 0 | 442700 | 0 | 219800 | 44395 | 187740 | 0 | 555450 | 7893700 | 513930 | 290970 | 625930 | 366970 | 768930 | 426310 |
| ITSEIPQTE | SARNP; SARNP | P82979; F8V | SARNP_HUMAI | Unmodified | 0 | 0 | 0 | 0 | 0 | 0 | 0 | 0 | 0 | 250170 | 243840 | 704950 | 0 | 0 | 0 | 0 | 0 | 0 |
| ITSENPDE | ACACB; ACACA | O00763; Q1 | ACACB_HUMAI | Unmodified | 0 | 0 | 443380 | 290460 | 0 | 0 | 0 | 0 | 0 | 0 | 0 | 2433300 | 0 | 0 | 426210 | 497480 | 0 | 0 |
| ITSFEEAK | PPP3CA; PPP3 | Q08209; E7E | PP2BA_HUMAN | Unmodified | 0 | 0 | 341540 | 0 | 0 | 0 | 232170 | 0 | 360750 | 251270 | 0 | 1964600 | 4138500 | 2127300 | 792930 | 0 | 414830 | 1128400 |
| ITSGPFEPD | PA2G4 | Q9UQ80 | PA2G4_HUMAI | Unmodified | 0 | 0 | 2161900 | 0 | 687150 | 0 | 0 | 0 | 0 | 0 | 0 | 8940500 | 0 | 0 | 2572500 | 1027300 | 0 | 0 |
| ITSGPFEPD | PA2G4 | Q9UQ80 | PA2G4_HUMAI | Unmodified | 0 | 0 | 0 | 0 | 0 | 0 | 0 | 0 | 1524100 | 0 | 0 | 0 | 0 | 0 | 0 | 0 | 0 | 0 |
| ITSGPFEPD | PA2G4 | Q9UQ80 | PA2G4_HUMAI | Oxidation (M) | 2640000 | 5833100 | 0 | 1819100 | 0 | 0 | 0 | 0 | 1055100 | 0 | 0 | 13096000 | 0 | 0 | 379420 | 1771000 | 0 | 0 |
| ITSIVKDSS | SDCBP; SDCBP | O00560; G5I | SDCB1_HUMAI | Unmodified | 0 | 0 | 0 | 254300 | 69265 | 0 | 0 | 0 | 629330 | 570210 | 0 | 0 | 294250 | 334130 | 171170 | 0 | 32993 | 126770 |
| ITSLQEEVK | ROCK1; ROCK1 | Q13464; A0/ | ROCK1_HUMAI | Unmodified | 0 | 0 | 0 | 247410 | 0 | 0 | 356150 | 266380 | 1790300 | 461200 | 356220 | 926660 | 0 | 0 | 185390 | 0 | 0 | 441650 |
| ITSLTQLTD | TRIM32; TRIM3 | Q13049; Q5 | TRI32_HUMAN | Unmodified | 0 | 94497 | 30006 | 0 | 86925 | 0 | 0 | 0 | 0 | 0 | 0 | 0 | 0 | 0 | 0 | 0 | 251150 | 0 |
| ITSNFVIPSI | MKRN1; MKRN | Q9UHC7; Q | MKRN1_HUMA | Unmodified | 2647900 | 2286100 | 0 | 849150 | 0 | 0 | 0 | 0 | 0 | 0 | 0 | 0 | 0 | 0 | 2440100 | 1526200 | 0 | 0 |
| ITSNFVIPSI | MKRN1; MKRN | Q9UHC7; Q | MKRN1_HUMA | Unmodified | 11914000 | 11091000 | 0 | 4477200 | 0 | 0 | 0 | 0 | 0 | 0 | 0 | 0 | 2594300 | 0 | 1899500 | 2706800 | 0 | 0 |
| ITSPVLIIHG | ABHD17B | Q5VST6 | AB17B_HUMAN | Unmodified | 0 | 227420 | 0 | 86985 | 42670 | 0 | 0 | 0 | 0 | 0 | 0 | 0 | 0 | 0 | 248550 | 0 | 0 | 0 |
| ITSVAWSPI | GEMIN5 | Q8TEQ6 | GEMI5_HUMAN | Unmodified | 0 | 0 | 0 | 0 | 1270200 | 0 | 135170 | 47450 | 0 | 0 | 169640 | 748080 | 0 | 0 | 0 | 0 | 0 | 0 |
| ITTGAQDDI | AFG3L2 | Q9Y4W6 | AFG32_HUMAN | Unmodified | 2029400 | 263570 | 0 | 0 | 0 | 0 | 0 | 0 | 235100 | 0 | 0 | 0 | 0 | 0 | 0 | 0 | 412350 | 0 |
| ITTGAQDDI | AFG3L2 | Q9Y4W6 | AFG32_HUMAN | Unmodified | 107810 | 0 | 0 | 0 | 433340 | 0 | 0 | 25690 | 3551100 | 0 | 0 | 0 | 0 | 0 | 0 | 0 | 0 | 0 |
| ITTGSSSAG | PPP1R12A; PPP | O14974; A0/ | MYPT1_HUMAN | Unmodified | 0 | 0 | 0 | 0 | 2734900 | 0 | 0 | 0 | 0 | 0 | 0 | 0 | 319340 | 0 | 0 | 0 | 0 | 0 |
| ITTHFELK | AK3 | Q9UIJ7 | KAD3_HUMAN | Unmodified | 0 | 0 | 0 | 0 | 0 | 0 | 0 | 0 | 773500 | 221000 | 0 | 0 | 0 | 0 | 0 | 0 | 0 | 0 |
| ITTQITAGAI | NRXN1; NRXN1 | Q9ULB1; A0/ | NRX1A_HUMAI | Unmodified | 42518 | 0 | 0 | 0 | 170100 | 0 | 0 | 0 | 537080 | 72339 | 0 | 0 | 4887000 | 0 | 0 | 0 | 0 | 0 |
| ITTQLDQVT | EEA1 | Q15075 | EEA1_HUMAN I | Unmodified | 0 | 0 | 0 | 0 | 966870 | 184840 | 0 | 0 | 150120 | 0 | 1142400 | 341920 | 392720 | 0 | 0 | 0 | 0 | 0 |
| ITTRKTPCG | RPS20; RPS20; | P60866; A0A | RS20_HUMAN | Unmodified | 0 | 0 | 0 | 0 | 0 | 0 | 0 | 186010 | 0 | 0 | 0 | 0 | 0 | 297360 | 0 | 0 | 0 | 0 |
| ITTVSLSAPI | SORL1; SORL1; | Q92673; E9F | SORL_HUMAN | Unmodified | 0 | 479900 | 0 | 0 | 0 | 0 | 0 | 0 | 4320100 | 0 | 0 | 0 | 0 | 0 | 1979900 | 0 | 0 | 0 |
| ITVHFINRD | FDX1 | P10109 | ADX_HUMAN A | Unmodified | 0 | 0 | 0 | 0 | 0 | 0 | 0 | 0 | 979880 | 68593 | 0 | 0 | 0 | 0 | 0 | 0 | 0 | 0 |
| ITVLEALRH | ATAD3A; ATAD3 | Q9NVI7; Q5 | ATD3A_HUMAN | Unmodified | 0 | 0 | 0 | 0 | 0 | 0 | 0 | 0 | 2708900 | 140910 | 0 | 0 | 0 | 0 | 0 | 0 | 0 | 0 |
| ITVPDTYEA | PI4KA | P42356 | PI4KA_HUMAN | Unmodified | 0 | 0 | 365920 | 221180 | 0 | 0 | 0 | 0 | 691950 | 148710 | 0 | 3203700 | 19922 | 0 | 851200 | 779200 | 0 | 0 |
| ITVPDTYEA | PI4KA | P42356 | PI4KA_HUMAN | Unmodified | 0 | 0 | 176040 | 173540 | 0 | 0 | 0 | 0 | 0 | 0 | 0 | 0 | 0 | 0 | 155720 | 106200 | 0 | 0 |
| ITVPFLEQC | PREP | P48147 | PPCE_HUMAN | Unmodified | 0 | 0 | 0 | 0 | 0 | 0 | 0 | 0 | 142820 | 0 | 112140 | 928290 | 0 | 0 | 0 | 0 | 0 | 0 |
| ITVPFLEQC | PREP | P48147 | PPCE_HUMAN | Unmodified | 49800 | 0 | 0 | 253750 | 1411400 | 225720 | 287400 | 0 | 497350 | 633050 | 781630 | 171530 | 1318400 | 978600 | 0 | 0 | 1055100 | 1140000 |
| ITVTSEVPF | RPL22; RPL22; I | P35268; K7E | RL22_HUMAN | Unmodified | 642120 | 749320 | 0 | 0 | 0 | 0 | 384200 | 676660 | 390590 | 249390 | 2570600 | 17993000 | 0 | 0 | 0 | 0 | 0 | 0 |
| ITVTSEVPF | RPL22; RPL22; I | P35268; K7E | RL22_HUMAN | Unmodified | 3433400 | 4175200 | 0 | 0 | 0 | 0 | 0 | 0 | 0 | 587370 | 603210 | 20482000 | 0 | 0 | 0 | 0 | 0 | 0 |
| ITVVPSAKD | GTPBP4 | Q9BZE4 | GTPB4_HUMAI | Unmodified | 0 | 0 | 0 | 0 | 0 | 0 | 102910 | 0 | 0 | 0 | 0 | 0 | 26789 | 16865 | 0 | 0 | 0 | 0 |
| ITVWLDLLK | FARP1; FARP1; | Q9Y4F1; A0/ | FARP1_HUMAN | Unmodified | 0 | 0 | 100680 | 0 | 0 | 0 | 50139 | 0 | 122430 | 733470 | 236100 | 0 | 261210 | 296140 | 477670 | 159340 | 30523 | 249820 |
| ITVYEALNH | CASK; CASK; C/ | O14936; A0/ | CSKP_HUMAN | Unmodified | 889130 | 900540 | 4773300 | 2198100 | 427980 | 141300 | 0 | 0 | 5049400 | 1573500 | 0 | 3779700 | 347840 | 175160 | 5896200 | 5325600 | 0 | 101120 |
| ITWSIIR | ATP6V1A | P38606 | VATA_HUMAN | Unmodified | 0 | 0 | 0 | 0 | 0 | 0 | 0 | 0 | 0 | 0 | 0 | 3270900 | 0 | 0 | 186900 | 0 | 0 | 0 |
| ITWSNPPA | GOT1 | P17174 | AATC_HUMAN | Unmodified | 138050 | 0 | 0 | 312740 | 0 | 0 | 332890 | 0 | 681700 | 0 | 1259700 | 10313000 | 0 | 0 | 1244500 | 783710 | 0 | 0 |

| | | | | | | | | | | | | | | | | | | | | | | |
|---|---|---|---|---|---|---|---|---|---|---|---|---|---|---|---|---|---|---|---|---|---|---|
| ITYGETGGI | FN1; FN1 | P02751; H0Y | FINC_HUMAN | Unmodified | 0 | 71385 | 1068100 | 313590 | 0 | 0 | 0 | 0 | 0 | 360140 | 84874 | 217780 | 0 | 0 | 0 | 1201800 | 0 | 0 |
| ITYKAENLG | TM9SF4; TM9SF | Q92544; A0A | TM9S4_HUMAN | Unmodified | 0 | 0 | 0 | 67179 | 0 | 0 | 0 | 16238 | 22403000 | 681720 | 730230 | 901140 | 0 | 0 | 0 | 0 | 0 | 0 |
| ITYLQNPK | FUT8; FUT8; FU | Q9BYC5; A0A | FUT8_HUMAN | Unmodified | 0 | 0 | 0 | 146930 | 0 | 0 | 0 | 0 | 4629900 | 647680 | 0 | 0 | 0 | 0 | 60124 | 80703 | 0 | 0 |
| ITYLQNPKD | FUT8; FUT8; FU | Q9BYC5; A0A | FUT8_HUMAN | Unmodified | 0 | 0 | 0 | 0 | 0 | 0 | 0 | 0 | 0 | 123210 | 0 | 0 | 0 | 0 | 0 | 0 | 0 | 0 |
| ITYYHLSFP | FBLN1 | B1AHL2 | B1AHL2_HUMA | Unmodified | 76922 | 0 | 0 | 1320600 | 0 | 0 | 0 | 0 | 43200000 | 1063700 | 0 | 0 | 0 | 0 | 0 | 0 | 0 | 0 |
| IVADFSASV | GMPS | P49915 | GUAA_HUMAN | Unmodified | 0 | 0 | 0 | 0 | 0 | 0 | 189690 | 0 | 0 | 811770 | 0 | 0 | 658030 | 0 | 0 | 0 | 0 | 162020 |
| IVADKDYS | LDHB; LDHB; LI | P07195; A0A | LDHB_HUMAN | Unmodified | 0 | 0 | 0 | 0 | 18598000 | 53882 | 1606000 | 738640 | 290680 | 103240 | 713560 | 3833100 | 10339000 | 5644100 | 0 | 61198 | 8231900 | 9755800 |
| IVADQLCAI | IST1; IST1; IST1 | P53990; H3E | IST1_HUMAN I | Unmodified | 0 | 0 | 1906400 | 0 | 6750500 | 3090800 | 0 | 103170 | 10934000 | 5272900 | 6132700 | 21300000 | 2356000 | 0 | 879890 | 685950 | 494440 | 1080500 |
| IVAECNAVI | CTNNA1; CTNN | P35221; P26 | CTNA1_HUMAI | Unmodified | 1974000 | 808620 | 0 | 7038900 | 0 | 0 | 0 | 15347 | 238450 | 0 | 0 | 9601900 | 1920500 | 0 | 7899100 | 7580900 | 0 | 0 |
| IVAECNAVI | CTNNA1; CTNN | P35221; G3X | CTNA1_HUMAI | Oxidation (M) | 0 | 0 | 0 | 245050 | 3617200 | 1033100 | 0 | 0 | 0 | 0 | 0 | 0 | 5619300 | 2437100 | 0 | 0 | 0 | 0 |
| IVAECNAVI | CTNNA1; CTNN | P35221; G3X | CTNA1_HUMAI | Unmodified | 0 | 0 | 0 | 0 | 1419100 | 0 | 0 | 0 | 0 | 51448 | 0 | 0 | 2004300 | 1509800 | 0 | 0 | 0 | 0 |
| IVAERPGTN | TRIM28; TRIM28 | Q13263; M0I | TIF1B_HUMAN | Oxidation (M) | 114210 | 0 | 0 | 0 | 0 | 0 | 137040 | 123870 | 0 | 0 | 0 | 125610 | 0 | 0 | 0 | 0 | 0 | 128580 |
| IVAERPGTN | TRIM28; TRIM28 | Q13263; M0I | TIF1B_HUMAN | Unmodified | 77580 | 55014 | 0 | 0 | 0 | 0 | 645940 | 301200 | 155510 | 0 | 357170 | 687230 | 0 | 0 | 0 | 0 | 0 | 157360 |
| IVAERPGTN | TRIM28; TRIM28 | Q13263; M0I | TIF1B_HUMAN | Unmodified | 0 | 0 | 0 | 0 | 676130 | 0 | 0 | 447360 | 0 | 911290 | 0 | 2499500 | 840980 | 0 | 0 | 0 | 645710 | 1223000 |
| IVAFENAFE | USO1 | O60763 | USO1_HUMAN | Unmodified | 0 | 0 | 0 | 0 | 0 | 0 | 0 | 0 | 0 | 0 | 0 | 1234900 | 0 | 0 | 0 | 0 | 0 | 0 |
| IVAGYASR | RPIA | P49247 | RPIA_HUMAN F | Unmodified | 0 | 0 | 0 | 0 | 511760 | 45896 | 91601 | 0 | 347560 | 644420 | 523680 | 3459300 | 450620 | 64833 | 36608 | 17562 | 62869 | 68866 |
| IVAIAVGDS | CEMIP2; CEMIP | Q9UHN6; F5 | CEIP2_HUMAN | Unmodified | 132720 | 456260 | 860020 | 148860 | 0 | 0 | 659070 | 460820 | 4100400 | 392190 | 0 | 898890 | 0 | 0 | 804340 | 709030 | 0 | 0 |
| IVAISEDYPI | DSG2 | Q14126 | DSG2_HUMAN | Unmodified | 0 | 0 | 0 | 0 | 0 | 0 | 0 | 0 | 354200 | 365340 | 0 | 0 | 0 | 0 | 0 | 0 | 0 | 0 |
| IVAISEDYPI | DSG2 | Q14126 | DSG2_HUMAN | Unmodified | 109730 | 592550 | 0 | 300210 | 1635700 | 544420 | 0 | 0 | 4398100 | 726120 | 276740 | 1933000 | 297250 | 0 | 173590 | 262500 | 0 | 0 |
| IVALNAHTF | FBLL1; FBL; FBL | A6NHQ2; P2 | FBLL1_HUMAN | Unmodified | 0 | 19926 | 75522 | 0 | 0 | 0 | 47003 | 10993 | 0 | 0 | 0 | 1946600 | 0 | 0 | 53249 | 0 | 0 | 60657 |
| IVALSSSLS | SNRNP200 | O75643 | U520_HUMAN | Unmodified | 185690 | 0 | 0 | 0 | 0 | 0 | 0 | 0 | 0 | 0 | 0 | 5584700 | 0 | 0 | 289910 | 0 | 0 | 0 |
| IVAPISDSPI | PGLS; PGLS | O95336; M0I | 6PGL_HUMAN | Unmodified | 760340 | 366810 | 0 | 0 | 0 | 0 | 0 | 28913 | 2150200 | 0 | 578090 | 4911600 | 0 | 0 | 0 | 0 | 0 | 0 |
| IVAPISDSPI | PGLS; PGLS | O95336; M0I | 6PGL_HUMAN | Unmodified | 0 | 0 | 0 | 0 | 4259400 | 0 | 2039200 | 294880 | 3998000 | 6700000 | 132920 | 760280 | 3031900 | 6246100 | 0 | 0 | 6528000 | 15732000 |
| IVAPPGGR | DPYSL2; DPYSL | Q16555; A0A | DPYL2_HUMAN | Unmodified | 0 | 0 | 0 | 0 | 1281800 | 223720 | 0 | 0 | 0 | 233680 | 0 | 0 | 0 | 0 | 0 | 0 | 0 | 519450 |
| IVASTLSNP | GOT1 | P17174 | AATC_HUMAN | Unmodified | 0 | 0 | 0 | 1381400 | 3843500 | 1296100 | 158880 | 14270 | 2426400 | 3227900 | 5065700 | 15289000 | 4089900 | 0 | 3233400 | 2868300 | 0 | 0 |
| IVATKPLYV | PABPC1; PABP | P11940; Q9H | PABP1_HUMAN | Unmodified | 1.61E+08 | 1.77E+08 | 79598000 | 68508000 | 582310 | 0 | 0 | 0 | 3528500 | 236670 | 0 | 17084000 | 4357600 | 536560 | 48553000 | 1.2E+08 | 0 | 0 |
| IVATKPLYV | PABPC1; PABP | P11940; Q9H | PABP1_HUMAN | Unmodified | 6353200 | 9089900 | 3803900 | 2495100 | 0 | 0 | 0 | 0 | 0 | 0 | 0 | 1549600 | 1252800 | 0 | 0 | 0 | 0 | 0 |
| IVAVTGAEA | AARS1; AARS1; | P49588; A0A | SYAC_HUMAN | Unmodified | 0 | 0 | 403990 | 293660 | 4343800 | 1673300 | 603490 | 275870 | 2864100 | 2814400 | 2240400 | 8266600 | 6609200 | 3708300 | 458180 | 767050 | 1145300 | 1990800 |
| IVCDLIENK | MYO1E; MYO1E | Q12965; H0Y | MYO1E_HUMAI | Unmodified | 0 | 0 | 0 | 0 | 0 | 0 | 0 | 0 | 0 | 0 | 0 | 1064400 | 0 | 0 | 0 | 0 | 0 | 0 |
| IVCDRNEP | PELI2; PELI2; PE | Q9HAT8; H0 | PELI2_HUMAN | Unmodified | 569000 | 0 | 0 | 0 | 0 | 0 | 0 | 0 | 1162300 | 363090 | 0 | 359010 | 0 | 59171 | 0 | 44475 | 0 | 0 |
| IVDDLKDE | SF3B1 | O75533 | SF3B1_HUMAN | Unmodified | 0 | 157830 | 0 | 0 | 0 | 0 | 0 | 0 | 0 | 0 | 0 | 3995200 | 0 | 0 | 0 | 0 | 0 | 0 |
| IVDDWANI | CANX; CANX; C | P27824; A0A | CALX_HUMAN | Unmodified | 0 | 0 | 0 | 0 | 0 | 0 | 256730 | 0 | 2.14E+08 | 12168000 | 7956400 | 4633400 | 0 | 0 | 0 | 0 | 348760 | 0 |
| IVDDWANI | CANX; CANX; C | P27824; A0A | CALX_HUMAN | Unmodified | 0 | 0 | 0 | 175530 | 299240 | 0 | 510870 | 119280 | 3.43E+08 | 28745000 | 7505000 | 5358200 | 0 | 0 | 0 | 0 | 0 | 0 |
| IVDFYPEDI | NR2C2AP | Q86WQ0 | NR2CA_HUMAI | Unmodified | 0 | 0 | 0 | 0 | 0 | 0 | 0 | 0 | 0 | 0 | 0 | 0 | 0 | 0 | 1260500 | 0 | 0 | 0 |
| IVDGKVVSI | KRT18 | P05783 | K1C18_HUMAN | Unmodified | 0 | 0 | 0 | 27722 | 0 | 0 | 0 | 0 | 0 | 0 | 0 | 0 | 0 | 0 | 0 | 0 | 1018500 | 208120 |
| IVDGPDPEI | DNAJC13 | O75165 | DJC13_HUMAN | Unmodified | 0 | 0 | 0 | 0 | 0 | 0 | 0 | 0 | 0 | 0 | 0 | 0 | 0 | 0 | 0 | 0 | 0 | 0 |
| IVDIIDTTGS | TPP2; TPP2; TPI | P29144; A0A | TPP2_HUMAN | Unmodified | 0 | 0 | 0 | 0 | 0 | 0 | 0 | 0 | 0 | 0 | 1031800 | 0 | 0 | 0 | 0 | 0 | 0 | 0 |
| IVDIIDTTGS | TPP2; TPP2; TPI | P29144; A0A | TPP2_HUMAN | Unmodified | 0 | 0 | 0 | 0 | 0 | 0 | 0 | 0 | 1139700 | 514850 | 1051900 | 1565000 | 0 | 0 | 0 | 0 | 0 | 0 |
| IVDKPGDH | TRNT1 | Q96Q11 | TRNT1_HUMAN | Unmodified | 175820 | 2992800 | 0 | 61749 | 0 | 0 | 0 | 0 | 0 | 0 | 366330 | 5072300 | 217450 | 178030 | 146440 | 58106 | 0 | 0 |
| IVDNGYGY | CARS1; CARS1 | P49589; B4D | SYCC_HUMAN | Unmodified | 0 | 0 | 0 | 0 | 0 | 0 | 0 | 0 | 0 | 0 | 0 | 2338800 | 0 | 0 | 0 | 0 | 0 | 0 |
| IVDPEIALK | CSTF2; CSTF2; | P33240; E7E | CSTF2_HUMAN | Unmodified | 0 | 0 | 37325 | 0 | 175490 | 59980 | 0 | 62398 | 322740 | 364940 | 0 | 1843500 | 676240 | 166890 | 0 | 0 | 156870 | 0 |
| IVDQNTKL | CPE; CPE; CPE; | P16870; C9J | CBPE_HUMAN | Unmodified | 0 | 0 | 0 | 0 | 0 | 65710 | 492660 | 566000 | 30697 | 2353400 | 0 | 0 | 0 | 0 | 0 | 0 | 121130 | 536790 |
| IVDRIDND | RCN1 | Q15293 | RCN1_HUMAN | Unmodified | 97953 | 0 | 0 | 331240 | 1989200 | 773880 | 1206000 | 809430 | 76510000 | 23068000 | 10186000 | 5837400 | 2509000 | 3243400 | 334220 | 393810 | 0 | 2291100 |
| IVDVAKFTS | CASK; CASK; C | O14936; A0A | CSKP_HUMAN | Unmodified | 0 | 0 | 0 | 0 | 0 | 0 | 0 | 68193 | 0 | 0 | 0 | 0 | 890530 | 419680 | 0 | 0 | 339300 | 0 |
| IVDVAKFTS | CASK; CASK; C | Q5JS79; A0A | Q5JS79_HUMA | Unmodified | 0 | 0 | 0 | 0 | 296990 | 0 | 366650 | 0 | 0 | 399120 | 0 | 0 | 366690 | 476710 | 0 | 0 | 61083 | 267650 |
| IVDVIGEK | PRKAR2A; PRKA | P13861; H7C | KAP2_HUMAN | Unmodified | 184770 | 444190 | 2957200 | 1846200 | 4699200 | 616730 | 1734400 | 841610 | 4590700 | 837230 | 674770 | 4823200 | 16909000 | 11140000 | 3228000 | 2408900 | 59420 | 4366900 |
| IVDVYENLY | MAPKAPK2 | P49137 | MAPK2_HUMAI | Unmodified | 0 | 0 | 0 | 0 | 0 | 0 | 0 | 0 | 0 | 0 | 0 | 0 | 0 | 0 | 0 | 0 | 0 | 0 |
| IVDWKEDC | DIS3L2 | Q8IYB7 | DI3L2_HUMAN | Unmodified | 0 | 0 | 0 | 0 | 0 | 0 | 0 | 0 | 0 | 0 | 0 | 165500 | 0 | 0 | 53017 | 26547 | 0 | 0 |
| IVDYLLEK | MYO10; MYO10 | Q9HD67; D6 | MYO10_HUMAI | Unmodified | 0 | 0 | 139530 | 175150 | 44321 | 0 | 0 | 0 | 481000 | 0 | 0 | 0 | 0 | 0 | 372850 | 219970 | 27093 | 0 |
| IVEAAENEY | CAPZA2 | P47755 | CAZA2_HUMAN | Oxidation (M) | 0 | 0 | 0 | 0 | 3203200 | 0 | 0 | 0 | 0 | 458270 | 0 | 2571100 | 0 | 0 | 0 | 0 | 0 | 0 |
| IVEAAENEY | CAPZA2 | P47755 | CAZA2_HUMAN | Unmodified | 0 | 0 | 0 | 0 | 3300800 | 0 | 188610 | 72151 | 1088900 | 0 | 1925000 | 322730 | 1854400 | 0 | 0 | 0 | 0 | 1900200 |
| IVEANPLLE | MYO6; MYO6; M | Q9UM54; A0 | MYO6_HUMAN | Unmodified | 0 | 0 | 1004900 | 0 | 0 | 0 | 0 | 0 | 27896 | 0 | 0 | 1293400 | 0 | 0 | 4057900 | 0 | 0 | 0 |
| IVEDEPNK | CSE1L | P55060 | XPO2_HUMAN | Unmodified | 0 | 0 | 0 | 26855 | 0 | 0 | 0 | 0 | 0 | 0 | 0 | 7141300 | 0 | 0 | 0 | 0 | 7591.4 | 28281 |
| IVEDEPNKI | CSE1L | P55060 | XPO2_HUMAN | Unmodified | 0 | 0 | 0 | 0 | 167820 | 969520 | 347390 | 360550 | 0 | 0 | 0 | 762020 | 0 | 0 | 0 | 0 | 0 | 205420 |
| IVEEPQSNI | OXSR1 | O95747 | OXSR1_HUMAI | Unmodified | 0 | 0 | 71803 | 0 | 239580 | 0 | 0 | 30152 | 0 | 333520 | 196030 | 210220 | 1082500 | 249350 | 0 | 0 | 202010 | 187620 |
| IVEFHHNH | FARP1; FARP1; | Q9Y4F1; A0A | FARP1_HUMAN | Unmodified | 0 | 0 | 0 | 2019600 | 0 | 0 | 0 | 0 | 4325700 | 1204700 | 604290 | 3884200 | 2032100 | 0 | 0 | 0 | 0 | 0 |
| IVEGISQPI | PMVK | Q15126 | PMVK_HUMAN | Unmodified | 81200 | 63321 | 0 | 54425 | 0 | 0 | 0 | 45149 | 271250 | 0 | 0 | 519240 | 37002 | 0 | 0 | 89069 | 0 | 21759 |
| IVEGISQPI | PMVK | Q15126 | PMVK_HUMAN | Unmodified | 0 | 0 | 0 | 0 | 0 | 0 | 0 | 0 | 0 | 279060 | 0 | 984230 | 0 | 0 | 0 | 0 | 326490 | 0 |
| IVEIPFNSTI | ATP1A1 | P05023 | AT1A1_HUMAN | Unmodified | 0 | 0 | 94701000 | 78507000 | 22016000 | 3503300 | 5302700 | 958570 | 22744000 | 3908300 | 45736000 | 3.34E+08 | 20409000 | 14365000 | 84861000 | 1.05E+08 | 1962400 | 3333700 |
| IVEIPFNSTI | ATP1A1 | P05023 | AT1A1_HUMAN | Unmodified | 56232 | 0 | 0 | 0 | 6358600 | 0 | 631710 | 131590 | 2481000 | 613720 | 0 | 2052600 | 16238000 | 2472900 | 0 | 0 | 0 | 0 |
| IVEIPFNSTI | ATP1A1 | P05023 | AT1A1_HUMAN | Unmodified | 0 | 0 | 0 | 457320 | 80326000 | 4612600 | 5279600 | 0 | 1629300 | 7730300 | 0 | 0 | 57170000 | 43560000 | 0 | 0 | 11964000 | 12357000 |
| IVEIPFNSTI | ATP1A1 | P05023 | AT1A1_HUMAN | Oxidation (M) | 0 | 0 | 0 | 11527000 | 88119000 | 0 | 9597600 | 226520 | 458780 | 23076000 | 0 | 0 | 1.28E+08 | 1.27E+08 | 0 | 0 | 17659000 | 8613500 |
| IVELISR | TOM1L2; TOM1 | Q6ZVM7; F5I | TM1L2_HUMAN | Unmodified | 5326800 | 4969500 | 3508700 | 4186400 | 10033000 | 2735500 | 18214000 | 11326000 | 959980 | 1122800 | 0 | 3620100 | 5189400 | 6272800 | 6259800 | 4402300 | 112210 | 2884100 |
| IVELMGR | GPD2 | P43304 | GPDM_HUMAN | Unmodified | 0 | 0 | 0 | 0 | 0 | 0 | 59507 | 0 | 2913600 | 127800 | 0 | 451260 | 91220 | 0 | 0 | 0 | 0 | 0 |
| IVELSEANK | CIT; CIT; CIT; CI | O14578; A0A | CTRO_HUMAN | Unmodified | 0 | 0 | 3038300 | 530150 | 2803800 | 1408900 | 583870 | 113460 | 943720 | 561800 | 0 | 978020 | 109270 | 0 | 0 | 0 | 200980 | 474360 |
| IVEMSTSK | EIF5A; EIF5A2; E | P63241; Q9C | IF5A1_HUMAN | Oxidation (M) | 1070200 | 748460 | 135410 | 1421200 | 712230 | 711450 | 503530 | 254100 | 350430 | 613560 | 1082400 | 3699500 | 2559900 | 1320300 | 963910 | 791090 | 1266300 | 5475800 |
| IVEMSTSK | EIF5A; EIF5A2; E | P63241; Q9C | IF5A1_HUMAN | Unmodified | 6434100 | 5017000 | 4219800 | 5815100 | 66880000 | 8183100 | 2687100 | 1848700 | 9361800 | 3606000 | 5329600 | 50661000 | 17859000 | 8272500 | 5755400 | 5309100 | 13287000 | 31420000 |
| IVENGQER | DNAJB6; DNAJE | O75190; Q8I | DNJB6_HUMAN | Unmodified | 5204500 | 7571300 | 1985000 | 1727600 | 0 | 0 | 0 | 0 | 0 | 0 | 0 | 535650 | 0 | 0 | 2918100 | 2015400 | 110690 | 0 |
| IVENLGILT | EPS8L2; EPS8L | Q9H6S3; A0A | ES8L2_HUMAN | Unmodified | 0 | 0 | 104130 | 0 | 0 | 0 | 0 | 535120 | 173460 | 66713 | 111670 | 348090 | 0 | 0 | 522010 | 352730 | 0 | 0 |
| IVENLGILT | EPS8L2; EPS8L | Q9H6S3; A0A | ES8L2_HUMAN | Unmodified | 0 | 0 | 0 | 906720 | 1091000 | 381750 | 0 | 0 | 463720 | 0 | 298510 | 0 | 1460400 | 0 | 514150 | 0 | 0 | 0 |
| IVENLGILT | EPS8L2; EPS8L | Q9H6S3; A0A | ES8L2_HUMAN | Unmodified | 0 | 0 | 0 | 0 | 0 | 0 | 0 | 0 | 1300900 | 523050 | 0 | 868660 | 3861100 | 1986400 | 0 | 445320 | 0 | 0 |
| IVEPPENIQ | CNOT1; CNOT1 | A5YKK6; H3E | CNOT1_HUMA | Unmodified | 0 | 0 | 0 | 0 | 0 | 0 | 167360 | 49823 | 166340 | 0 | 0 | 705180 | 0 | 0 | 0 | 0 | 0 | 0 |
| IVEPYIAWG | RPL7; RPL7 | P18124; A8M | RL7_HUMAN 6 | Unmodified | 2916700 | 3655600 | 946370 | 1455200 | 0 | 0 | 0 | 0 | 0 | 0 | 0 | 21954000 | 0 | 0 | 2141500 | 2029200 | 0 | 0 |
| IVEQDTMPI | DDX3X; DDX3Y | O00571; O15 | DDX3X_HUMAI | Oxidation (M) | 129540 | 142130 | 186330 | 0 | 0 | 0 | 0 | 0 | 0 | 0 | 0 | 0 | 0 | 0 | 0 | 0 | 0 | 0 |
| IVERDGLSE | COASY | Q13057 | COASY_HUMAI | Unmodified | 447490 | 208420 | 355850 | 369640 | 0 | 0 | 0 | 0 | 420600 | 30751 | 0 | 249700 | 0 | 0 | 596510 | 378320 | 24593 | 299960 |
| IVESYQIR | CNTN1 | Q12860 | CNTN1_HUMA | Unmodified | 553430 | 526760 | 0 | 212520 | 0 | 274990 | 0 | 0 | 1047100 | 435420 | 360730 | 1164100 | 0 | 0 | 136750 | 89339 | 0 | 0 |
| IVETHQPIV | COG4; COG4; ( | Q9H9E3; A0A | COG4_HUMAN | Unmodified | 0 | 0 | 0 | 0 | 0 | 0 | 0 | 0 | 0 | 505080 | 0 | 483620 | 0 | 0 | 0 | 0 | 0 | 0 |
| IVETQNEEY | FGD4; FGD4; F( | Q96M96; B7 | FGD4_HUMAN | Unmodified | 0 | 0 | 0 | 0 | 193160 | 0 | 0 | 0 | 0 | 0 | 0 | 1202900 | 134830 | 87343 | 0 | 0 | 0 | 0 |
| IVEVGPRD | HMGCLL1; HM( | Q8TB92; P35 | HMGC2_HUMA | Unmodified | 0 | 0 | 0 | 0 | 65005 | 0 | 571010 | 131230 | 1429900 | 2164600 | 0 | 0 | 125670 | 83159 | 0 | 0 | 104430 | 262310 |
| IVEVLLMK | RBMXL2; RBMX | O75526; P38 | RMXL2_HUMAN | Unmodified | 0 | 0 | 0 | 0 | 42333 | 0 | 0 | 0 | 1336900 | 124710 | 435580 | 1203400 | 104510 | 0 | 0 | 0 | 0 | 0 |
| IVEVLLMK | RBMXL2; RBMX | O75526; P38 | RMXL2_HUMAN | Oxidation (M) | 0 | 0 | 0 | 0 | 0 | 0 | 0 | 0 | 0 | 0 | 501910 | 1754700 | 0 | 0 | 0 | 0 | 0 | 0 |
| IVEVNGVC | SLC9A3R1 | O14745 | NHRF1_HUMAI | Unmodified | 0 | 0 | 0 | 0 | 0 | 0 | 0 | 0 | 0 | 0 | 0 | 1702900 | 0 | 0 | 403760 | 187410 | 0 | 0 |
| IVEWNGK | SCARB2; SCARI | Q14108; A0A | SCRB2_HUMAI | Unmodified | 165160 | 134530 | 582660 | 127870 | 241440 | 139280 | 279040 | 0 | 4609400 | 1730000 | 0 | 1104500 | 167330 | 87441 | 756900 | 346800 | 0 | 24023 |
| IVFAAGNF | OAT | P04181 | OAT_HUMAN C | Unmodified | 0 | 0 | 73123 | 0 | 0 | 0 | 308610 | 0 | 2603500 | 494100 | 191650 | 254490 | 0 | 0 | 0 | 0 | 0 | 0 |
| IVFDSIDNL | NAGK; NAGK; N | Q9UJ70; C9J | NAGK_HUMAN | Unmodified | 0 | 0 | 0 | 0 | 0 | 0 | 0 | 0 | 1547900 | 370750 | 280590 | 1347800 | 0 | 0 | 0 | 0 | 0 | 0 |
| IVFEDGNIN | CRMP1; CRMP1 | Q14194; E9F | DPYL1_HUMAN | Unmodified | 0 | 0 | 0 | 0 | 0 | 0 | 1754800 | 1042700 | 2450800 | 1102400 | 619000 | 0 | 0 | 0 | 0 | 0 | 0 | 0 |
| IVFNLCPLC | GEMIN5 | Q8TEQ6 | GEMI5_HUMAN | Unmodified | 0 | 0 | 0 | 0 | 0 | 0 | 0 | 0 | 0 | 0 | 0 | 877840 | 61675 | 74885 | 0 | 0 | 0 | 0 |
| IVFPHGITLI | LRP1; LRP1 | Q07954; Q6I | LRP1_HUMAN | Unmodified | 0 | 0 | 232180 | 106210 | 0 | 0 | 0 | 0 | 1254900 | 218620 | 0 | 0 | 0 | 0 | 134040 | 84405 | 0 | 0 |
| IVFSGNLF | NIBAN2 | Q96TA1 | NIBA2_HUMAN | Unmodified | 0 | 0 | 0 | 0 | 500370 | 0 | 168550 | 0 | 6725900 | 167470 | 1279200 | 7378400 | 0 | 0 | 0 | 0 | 0 | 0 |
| IVFSGNLF | NIBAN2 | Q96TA1 | NIBA2_HUMAN | Unmodified | 1075500 | 940990 | 609750 | 553120 | 0 | 96993 | 0 | 0 | 1751100 | 0 | 0 | 1599300 | 0 | 0 | 1069300 | 980850 | 124190 | 53178 |
| IVFVPGCSI | RARS1 | P54136 | SYRC_HUMAN | Unmodified | 0 | 0 | 121160 | 249120 | 286670 | 0 | 0 | 0 | 333010 | 178880 | 653760 | 6247600 | 0 | 0 | 0 | 188790 | 0 | 0 |
| IVFVTGNA | ITPA | Q9BY32 | ITPA_HUMAN I | Unmodified | 142160 | 0 | 0 | 0 | 1188900 | 95431 | 0 | 0 | 0 | 0 | 614840 | 4490600 | 0 | 0 | 0 | 0 | 914720 | 1589800 |
| IVGAPMHD | MTHFD1; MTHF | P11586; V9G | C1TC_HUMAN | Oxidation (M) | 0 | 0 | 0 | 149710 | 865500 | 3222000 | 936720 | 0 | 0 | 0 | 390440 | 1933500 | 1071000 | 999870 | 0 | 0 | 3064700 | 1448100 |
| IVGAPMHD | MTHFD1; MTHF | P11586; V9G | C1TC_HUMAN | Unmodified | 0 | 0 | 0 | 0 | 0 | 0 | 797540 | 0 | 402310 | 0 | 0 | 0 | 0 | 0 | 0 | 0 | 1566200 | 3147900 |
| IVGNGSEQ | RABGAP1; RAB( | Q9Y3P9; B5I | RBGP1_HUMAI | Unmodified | 0 | 0 | 0 | 0 | 175830 | 0 | 0 | 118130 | 0 | 0 | 0 | 0 | 0 | 0 | 0 | 0 | 0 | 0 |
| IVGNLLYYF | IQGAP2; IQGAP | Q13576; E7E | IQGA2_HUMAN | Unmodified | 0 | 0 | 1064800 | 120800 | 0 | 0 | 0 | 0 | 0 | 0 | 0 | 0 | 0 | 0 | 746500 | 470290 | 0 | 0 |
| IVGPEENLS | PLOD2; PLOD2 | O00469; E7E | PLOD2_HUMAI | Unmodified | 0 | 0 | 0 | 0 | 220860 | 0 | 0 | 0 | 11252000 | 2399800 | 623000 | 0 | 0 | 0 | 0 | 0 | 0 | 0 |
| IVGPSGAA | FLNA; FLNA; FL | P21333; A0A | FLNA_HUMAN | Unmodified | 651990 | 698380 | 0 | 3183100 | 3814400 | 0 | 925770 | 541030 | 0 | 0 | 0 | 2530100 | 2567800 | 2148200 | 780920 | 526220 | 1396600 | 1100000 |
| IVGSKPLYV | PABPC4; PABP( | Q13310; B1A | PABP4_HUMAN | Unmodified | 51284000 | 29092000 | 14776000 | 12758000 | 0 | 0 | 0 | 0 | 377240 | 0 | 0 | 1581600 | 427590 | 0 | 9721200 | 9348100 | 0 | 0 |
| IVGSKPLYV | PABPC4; PABP( | Q13310; B1A | PABP4_HUMAN | Unmodified | 147740 | 2484100 | 0 | 1021400 | 0 | 306730 | 0 | 0 | 0 | 0 | 0 | 137740 | 0 | 0 | 0 | 0 | 0 | 0 |
| IVGYFVSG | ACAA2; ACAA2; | P42765; A0A | THIM_HUMAN | Oxidation (M) | 0 | 0 | 0 | 0 | 0 | 0 | 0 | 0 | 0 | 245040 | 0 | 0 | 0 | 0 | 0 | 0 | 0 | 0 |
| IVGYVGTS | PATJ; PATJ; PAT | Q8NI35; A0A | INADL_HUMAN | Unmodified | 0 | 0 | 0 | 0 | 0 | 0 | 0 | 0 | 547860 | 135140 | 0 | 262790 | 0 | 0 | 0 | 0 | 178320 | 0 |
| IVGYVLAK | NAA11; NAA10; | Q9BSU3; P4 | NAA11_HUMAN | Unmodified | 70728 | 45375 | 139740 | 36864 | 0 | 0 | 0 | 0 | 81449 | 13195 | 357140 | 4535600 | 0 | 0 | 229320 | 132920 | 0 | 0 |
| IVHLQETC | NLN; NLN | Q9BYT8; E9F | NEUL_HUMAN | Unmodified | 0 | 0 | 0 | 0 | 0 | 151500 | 0 | 0 | 523200 | 0 | 139320 | 1046600 | 0 | 0 | 0 | 0 | 0 | 0 |
| IVHQSLNI | TMEM131 | Q92545 | TM131_HUMAN | Unmodified | 0 | 0 | 0 | 0 | 0 | 0 | 0 | 0 | 1291300 | 205400 | 532660 | 0 | 0 | 0 | 0 | 0 | 0 | 0 |
| IVIDFDSIN | DCTD | P32321 | DCTD_HUMAN | Unmodified | 0 | 0 | 0 | 0 | 0 | 0 | 0 | 285770 | 0 | 0 | 0 | 1771300 | 212860 | 0 | 350110 | 0 | 0 | 0 |
| IVIDFDSIN | DCTD | P32321 | DCTD_HUMAN | Unmodified | 0 | 0 | 0 | 43590 | 1393500 | 257850 | 410070 | 278260 | 0 | 410650 | 0 | 0 | 2389100 | 708880 | 0 | 0 | 2278500 | 1689600 |
| IVIGDASET | ATP4A | P20648 | ATP4A_HUMAN | Unmodified | 0 | 0 | 0 | 0 | 0 | 0 | 0 | 0 | 0 | 0 | 1521000 | 0 | 0 | 0 | 0 | 0 | 0 | 0 |
| IVIGYQSHA | EIF4E; EIF4E | P06730; D6F | IF4E_HUMAN E | Unmodified | 696620 | 742370 | 860100 | 1007600 | 2981000 | 5323500 | 1280000 | 457250 | 2172400 | 975070 | 4123700 | 18634000 | 5171100 | 6486300 | 769070 | 0 | 661340 | 1627300 |

| | | | | | | | | | | | | | | | | | | | | | | |
|---|---|---|---|---|---|---|---|---|---|---|---|---|---|---|---|---|---|---|---|---|---|---|
| IVILEYQPSI | SF3B3; SF3B3; | Q15393; J3Q | SF3B3_HUMAN | Unmodified | 0 | 0 | 0 | 0 | 0 | 0 | 0 | 0 | 0 | 0 | 0 | 6494300 | 0 | 0 | 0 | 0 | 0 | 0 |
| IVILPDYLEI | USP5 | P45974 | UBP5_HUMAN | Unmodified | 301870 | 0 | 0 | 0 | 0 | 0 | 0 | 0 | 66489 | 0 | 77341 | 431740 | 0 | 0 | 0 | 0 | 0 | 0 |
| IVITDCGQL | PPIF | P30405 | PPIF_HUMAN F | Unmodified | 0 | 0 | 0 | 0 | 0 | 0 | 327210 | 61902 | 911410 | 592540 | 0 | 0 | 0 | 0 | 0 | 0 | 434200 | 0 |
| IVIVGDGG( | RHOF; RHOF | Q9HBH0; F5 | RHOF_HUMAN | Unmodified | 1200800 | 1164100 | 170850 | 0 | 0 | 134950 | 0 | 0 | 581590 | 0 | 0 | 510350 | 0 | 0 | 881500 | 890410 | 568130 | 771380 |
| IVIVPSLNPI | CPD | O75976 | CBPD_HUMAN | Unmodified | 0 | 0 | 0 | 0 | 0 | 0 | 0 | 0 | 24597000 | 5981600 | 0 | 539180 | 0 | 0 | 0 | 0 | 0 | 0 |
| IVIVPSLNPI | CPD | O75976 | CBPD_HUMAN | Unmodified | 0 | 0 | 2193800 | 1064000 | 0 | 0 | 0 | 0 | 53998000 | 1385800 | 739330 | 710340 | 0 | 0 | 0 | 124900 | 141400 | 0 |
| IVIVPSLNPI | CPD | O75976 | CBPD_HUMAN | Unmodified | 0 | 0 | 0 | 0 | 0 | 0 | 0 | 0 | 1227100 | 2176100 | 0 | 0 | 0 | 0 | 0 | 63411 | 0 | 0 |
| IVKADEHVI | PRKAR2A; PRKA | P13861; H7( | KAP2_HUMAN | Unmodified | 0 | 0 | 0 | 339290 | 0 | 0 | 0 | 0 | 896370 | 0 | 0 | 996270 | 0 | 0 | 594310 | 0 | 0 | 0 |
| IVKDLMSK/ | PGK1 | P00558 | PGK1_HUMAN | Unmodified | 0 | 0 | 0 | 0 | 608150 | 0 | 29260 | 200780 | 0 | 0 | 0 | 0 | 81059 | 135550 | 0 | 0 | 1204000 | 372170 |
| IVKDSLSDI | HSDL2 | Q6YN16 | HSDL2_HUMAI | Unmodified | 0 | 0 | 0 | 0 | 0 | 213170 | 0 | 0 | 857790 | 157350 | 0 | 0 | 0 | 0 | 0 | 0 | 168700 | 344600 |
| IVKPNGEKI | RPS7; RPS7; RP | P62081; A0A | RS7_HUMAN 4 | Oxidation (M) | 0 | 5423300 | 0 | 523260 | 0 | 0 | 0 | 0 | 0 | 0 | 685240 | 2213000 | 365140 | 0 | 1228800 | 0 | 0 | 0 |
| IVKSESGY( | SNX27; SNX27 | Q96L92; A0/ | SNX27_HUMAN | Unmodified | 0 | 0 | 0 | 0 | 866760 | 0 | 367900 | 0 | 0 | 0 | 0 | 0 | 0 | 0 | 0 | 0 | 0 | 0 |
| IVLAAAGG\ | PMPCB; PMPCE | O75439; G3\ | MPPB_HUMAN | Unmodified | 0 | 0 | 0 | 0 | 0 | 0 | 0 | 0 | 2024300 | 0 | 0 | 0 | 0 | 0 | 0 | 0 | 0 | 0 |
| IVLANDPD/ | PGM2 | Q96G03 | PGM2_HUMAN | Unmodified | 0 | 0 | 0 | 0 | 0 | 0 | 0 | 0 | 119420 | 0 | 1906500 | 3502700 | 0 | 0 | 492580 | 401080 | 0 | 0 |
| IVLDNSVFS | CLTCL1; CLTC; | P53675; Q0( | CLH2_HUMAN | Unmodified | 16756000 | 14441000 | 6069200 | 163460 | 0 | 0 | 0 | 52465 | 735650 | 0 | 0 | 1.04E+08 | 0 | 0 | 9339400 | 17975000 | 0 | 0 |
| IVLDNSVFS | CLTCL1; CLTC; | P53675; Q0( | CLH2_HUMAN | Unmodified | 0 | 2582400 | 488530 | 708640 | 0 | 304080 | 0 | 0 | 58225 | 0 | 0 | 862290 | 392930 | 0 | 2169900 | 241680 | 0 | 0 |
| IVLDSDAAE | GBE1; GBE1 | Q04446; E9F | GLGB_HUMAN | Unmodified | 0 | 0 | 0 | 0 | 0 | 0 | 0 | 0 | 265240 | 85640 | 0 | 572260 | 0 | 0 | 160580 | 135060 | 0 | 0 |
| IVLEDGTLH | DPYSL2; DPYSL | Q16555; A0/ | DPYL2_HUMAN | Unmodified | 984110 | 773960 | 783700 | 574390 | 11716000 | 2882000 | 777350 | 454030 | 11267000 | 1953300 | 1391200 | 22489000 | 1077400 | 627400 | 770130 | 493720 | 803950 | 1930300 |
| IVLGGTTVH | VPS45; VPS45; | Q9NRW7; A( | VPS45_HUMAN | Unmodified | 43638 | 1504200 | 642580 | 0 | 0 | 0 | 0 | 0 | 587170 | 163990 | 209620 | 0 | 77383 | 0 | 0 | 194880 | 0 | 0 |
| IVLLDSSLE | CCT3; CCT3; C( | P49368; Q5S | TCPG_HUMAN | Unmodified | 299440 | 0 | 0 | 0 | 0 | 0 | 0 | 0 | 2272100 | 308900 | 3274600 | 14798000 | 691380 | 0 | 1730000 | 1159500 | 269700 | 0 |
| IVLLDSSLE | CCT3; CCT3; C( | P49368; Q5S | TCPG_HUMAN | Unmodified | 0 | 0 | 0 | 0 | 0 | 0 | 0 | 0 | 0 | 0 | 0 | 1683500 | 0 | 0 | 0 | 0 | 0 | 0 |
| IVLLGDMN | RAB20 | Q9NX57 | RAB20_HUMAN | Unmodified | 0 | 0 | 0 | 0 | 0 | 627170 | 0 | 0 | 1776100 | 195090 | 0 | 0 | 0 | 0 | 0 | 0 | 0 | 0 |
| IVLNFNPHI | NRP2; NRP2 | O60462; C9J | NRP2_HUMAN | Unmodified | 0 | 0 | 85791 | 0 | 0 | 0 | 265760 | 0 | 1490900 | 172530 | 0 | 0 | 891460 | 0 | 0 | 0 | 0 | 0 |
| IVLNPNKP\ | ADPGK | Q9BRR6 | ADPGK_HUMAI | Unmodified | 0 | 0 | 0 | 0 | 0 | 0 | 0 | 0 | 8426800 | 0 | 0 | 928860 | 0 | 0 | 0 | 0 | 0 | 0 |
| IVLQNGAH | CRYZ; CRYZ | Q08257; A6I | QOR_HUMAN ( | Unmodified | 0 | 0 | 0 | 0 | 0 | 0 | 0 | 0 | 680240 | 309240 | 0 | 0 | 468510 | 186290 | 0 | 0 | 0 | 0 |
| IVLRDNVPI | LRP2 | P98164 | LRP2_HUMAN I | Unmodified | 0 | 0 | 0 | 0 | 0 | 0 | 0 | 0 | 1207900 | 0 | 0 | 0 | 0 | 0 | 0 | 0 | 0 | 0 |
| IVLTNPVCT | EIF2S3; EIF2S3I | P41091; Q2\ | IF2G_HUMAN E | Unmodified | 357130 | 172500 | 466650 | 1222800 | 1582600 | 0 | 419100 | 289010 | 0 | 843200 | 1087500 | 6995100 | 1188400 | 1165600 | 535180 | 638010 | 437200 | 1229200 |
| IVLVDDSIV | PPAT | Q06203 | PUR1_HUMAN | Unmodified | 0 | 0 | 0 | 183880 | 0 | 0 | 220600 | 0 | 496500 | 519540 | 881420 | 2723300 | 2766000 | 967390 | 0 | 0 | 0 | 384130 |
| IVMTPSR | IGSF21 | Q96ID5 | IGS21_HUMAN | Unmodified | 0 | 0 | 0 | 0 | 0 | 0 | 0 | 0 | 1049500 | 56515 | 0 | 2632000 | 0 | 0 | 436750 | 0 | 0 | 0 |
| IVNFDPK | GLT8D1; GLT8D | Q68CQ7; C9 | GL8D1_HUMAI | Unmodified | 0 | 0 | 0 | 0 | 0 | 0 | 0 | 0 | 17519000 | 1291100 | 140970 | 367580 | 0 | 0 | 0 | 0 | 0 | 0 |
| IVNFDPKLL | GLT8D1; GLT8D | Q68CQ7; C9 | GL8D1_HUMAI | Unmodified | 41926 | 53247 | 0 | 0 | 0 | 0 | 0 | 0 | 0 | 261740 | 0 | 0 | 0 | 0 | 0 | 0 | 0 | 0 |
| IVNFLLQHS | ANK3 | Q12955 | ANK3_HUMAN | Unmodified | 61830 | 0 | 118030 | 3434100 | 0 | 0 | 0 | 0 | 0 | 0 | 0 | 182460 | 4323700 | 7992300 | 5095600 | 3559800 | 0 | 0 |
| IVNLGSSKT | PHF5A | Q7RTV0 | PHF5A_HUMAI | Unmodified | 1941900 | 922100 | 0 | 484010 | 0 | 0 | 3615200 | 0 | 0 | 111060 | 0 | 0 | 280660 | 0 | 0 | 189340 | 0 | 0 |
| IVNLGSSKT | PHF5A | Q7RTV0 | PHF5A_HUMAI | Unmodified | 433220 | 0 | 0 | 0 | 0 | 0 | 0 | 0 | 140690 | 0 | 0 | 483130 | 1080800 | 0 | 0 | 154200 | 0 | 0 |
| IVNPKGEEI | PSME2; PSME2; | Q9UL46; A0/ | PSME2_HUMAI | Unmodified | 0 | 0 | 0 | 0 | 2169500 | 0 | 0 | 0 | 1402000 | 1161300 | 222140 | 1694600 | 2033200 | 0 | 0 | 0 | 2200900 | 0 |
| IVNPKGEEI | PSME2; PSME2; | Q9UL46; A0/ | PSME2_HUMAI | Oxidation (M) | 0 | 0 | 0 | 0 | 576020 | 0 | 193420 | 159000 | 262850 | 377510 | 170790 | 305910 | 648080 | 196700 | 0 | 0 | 890080 | 693770 |
| IVNQPSSLI | HUWE1 | Q7Z6Z7 | HUWE1_HUMA | Unmodified | 0 | 0 | 0 | 0 | 0 | 218590 | 0 | 189840 | 0 | 0 | 0 | 1628800 | 402280 | 0 | 0 | 0 | 243400 | 242970 |
| IVNSAQTG | ATP5MG; N/A; / | O75964; A0/ | ATP5L_HUMAN | Unmodified | 263830 | 0 | 196710 | 64261 | 232240 | 72582 | 189180 | 246750 | 9163700 | 2535900 | 0 | 269250 | 0 | 92539 | 136290 | 70378 | 0 | 0 |
| IVNTPEVIR | EHD3 | Q9NZN3 | EHD3_HUMAN | Unmodified | 0 | 0 | 955660 | 0 | 0 | 0 | 0 | 0 | 0 | 0 | 0 | 0 | 0 | 0 | 0 | 0 | 0 | 0 |
| IVNTPFQVL | TM9SF4; TM9SF | Q92544; A0/ | TM9S4_HUMAN | Unmodified | 0 | 0 | 0 | 0 | 0 | 0 | 0 | 0 | 13813000 | 231780 | 0 | 286980 | 0 | 0 | 0 | 0 | 0 | 0 |
| IVNTPFQVL | TM9SF4; TM9SF | Q92544; A0/ | TM9S4_HUMAN | Oxidation (M) | 0 | 0 | 0 | 0 | 0 | 41910 | 0 | 0 | 5425200 | 881440 | 959560 | 755030 | 0 | 17536 | 0 | 0 | 0 | 0 |
| IVNTPFQVL | TM9SF4; TM9SF | Q92544; A0/ | TM9S4_HUMAN | Unmodified | 0 | 0 | 0 | 0 | 0 | 0 | 0 | 0 | 3104600 | 0 | 0 | 0 | 0 | 0 | 0 | 0 | 0 | 0 |
| IVPETNLK | DIPK1A; DIPK1/ | Q5T7M9; A0 | DIK1A_HUMAN | Unmodified | 0 | 0 | 0 | 0 | 0 | 0 | 0 | 0 | 1739700 | 124190 | 0 | 0 | 132750 | 0 | 0 | 0 | 0 | 0 |
| IVPEWQDY | UGGT1; UGGT1 | Q9NYU2; H7 | UGGG1_HUMA | Unmodified | 0 | 0 | 0 | 0 | 0 | 0 | 73698 | 247630 | 22824000 | 4765700 | 1916100 | 0 | 0 | 0 | 0 | 0 | 0 | 0 |
| IVPEWQDY | UGGT1; UGGT1 | Q9NYU2; H7 | UGGG1_HUMA | Unmodified | 0 | 0 | 0 | 0 | 0 | 0 | 0 | 0 | 0 | 1505900 | 0 | 0 | 0 | 592780 | 0 | 0 | 0 | 0 |
| IVPGQFLA\ | SF3B3 | Q15393 | SF3B3_HUMAN | Unmodified | 694660 | 0 | 0 | 460770 | 0 | 0 | 0 | 0 | 0 | 0 | 791170 | 3109100 | 0 | 0 | 0 | 589110 | 0 | 0 |
| IVPHICNLL | CLASP1; CLASF | Q7Z460; F8\ | CLAP1_HUMAN | Unmodified | 0 | 0 | 0 | 0 | 0 | 0 | 0 | 0 | 0 | 0 | 0 | 500160 | 0 | 0 | 899220 | 0 | 0 | 0 |
| IVPNVLLEC | CS; CS; CS; CS | O75390; A0/ | CISY_HUMAN ( | Unmodified | 0 | 0 | 57746 | 0 | 276570 | 0 | 0 | 89272 | 34874000 | 2607400 | 0 | 2362300 | 0 | 0 | 0 | 137140 | 0 | 458290 |
| IVPSLLFNI | EFR3A; EFR3A | Q14156; A0/ | EFR3A_HUMAN | Unmodified | 0 | 0 | 0 | 0 | 0 | 0 | 207400 | 0 | 1040200 | 306280 | 0 | 0 | 0 | 0 | 0 | 0 | 0 | 0 |
| IVPSPDWF | SPON2 | Q9BUD6 | SPON2_HUMAI | Unmodified | 0 | 0 | 0 | 0 | 0 | 0 | 0 | 0 | 0 | 0 | 0 | 0 | 0 | 0 | 432000 | 0 | 0 | 0 |
| IVPSPDWF | SPON2 | Q9BUD6 | SPON2_HUMAI | Unmodified | 0 | 0 | 0 | 0 | 0 | 0 | 0 | 0 | 0 | 0 | 0 | 0 | 0 | 0 | 5469300 | 0 | 0 | 0 |
| IVPSYRTQS | CASK; CASK; C/ | A0A2U3TZM | A0A2U3TZM4_I | Unmodified | 0 | 0 | 0 | 0 | 1337100 | 548340 | 0 | 91567 | 0 | 0 | 0 | 0 | 0 | 0 | 0 | 0 | 0 | 0 |
| IVPSYRTQS | CASK; CASK | A0A2U3TZN( | A0A2U3TZN6_I | Unmodified | 0 | 0 | 0 | 0 | 0 | 0 | 0 | 0 | 176200 | 428220 | 0 | 0 | 0 | 0 | 0 | 0 | 0 | 0 |
| IVPTDMND | ALG9; N/A; ALG | Q9H6U8; A0 | ALG9_HUMAN | Unmodified | 0 | 0 | 0 | 0 | 0 | 0 | 0 | 0 | 434840 | 0 | 0 | 0 | 0 | 0 | 0 | 0 | 39956 | 0 |
| IVPTDMND | ALG9; N/A; ALG | Q9H6U8; A0 | ALG9_HUMAN | Unmodified | 0 | 0 | 0 | 0 | 0 | 0 | 0 | 0 | 14364000 | 78036 | 0 | 0 | 0 | 0 | 0 | 0 | 0 | 0 |
| IVPTDMND | ALG9; N/A; ALG | Q9H6U8; A0 | ALG9_HUMAN | Oxidation (M) | 0 | 0 | 0 | 0 | 0 | 0 | 0 | 0 | 6643800 | 490000 | 660700 | 191090 | 0 | 0 | 0 | 0 | 0 | 0 |
| IVPTVYEDK | ERGIC1 | Q969X5 | ERGI1_HUMAN | Unmodified | 264360 | 53502 | 634620 | 477670 | 307840 | 0 | 0 | 0 | 56235000 | 4192700 | 890450 | 860840 | 0 | 0 | 0 | 566160 | 0 | 0 |
| IVPTVYEDK | ERGIC1 | Q969X5 | ERGI1_HUMAN | Unmodified | 0 | 0 | 0 | 0 | 0 | 0 | 0 | 0 | 3809300 | 1647900 | 0 | 0 | 0 | 0 | 0 | 0 | 0 | 0 |
| IVPTVYEDK | ERGIC1 | Q969X5 | ERGI1_HUMAN | Unmodified | 0 | 0 | 0 | 0 | 0 | 0 | 0 | 0 | 0 | 231160 | 0 | 0 | 0 | 0 | 0 | 0 | 0 | 0 |
| IVPVEITISLI | CMPK1; CMPK1 | P30085; Q5T | KCY_HUMAN U | Oxidation (M) | 0 | 0 | 0 | 0 | 1808500 | 65060 | 124000 | 0 | 0 | 905140 | 0 | 0 | 2879700 | 1095700 | 0 | 0 | 1577900 | 2095800 |
| IVPVEITISLI | CMPK1; CMPK1 | P30085; Q5T | KCY_HUMAN U | 2 Oxidation (M) | 0 | 0 | 0 | 0 | 2177700 | 51601 | 0 | 0 | 0 | 199030 | 0 | 0 | 3141900 | 1640600 | 0 | 0 | 1065900 | 1836100 |
| IVQAEGEAI | PHB2; PHB2; PI | Q99623; F5H | PHB2_HUMAN | Unmodified | 0 | 0 | 0 | 0 | 512340 | 44125 | 844120 | 449900 | 2089400 | 1115900 | 544760 | 468790 | 431070 | 430620 | 0 | 0 | 0 | 0 |
| IVQEDRAVI | DYNC1H1; DYN | Q14204; A0/ | DYHC1_HUMA | Unmodified | 50072 | 0 | 171460 | 381830 | 787630 | 0 | 1335700 | 2256200 | 1056900 | 347320 | 785900 | 544420 | 2479800 | 701380 | 0 | 118680 | 0 | 0 |
| IVQEDRAVI | DYNC1H1; DYN | Q14204; A0/ | DYHC1_HUMA | Unmodified | 0 | 0 | 0 | 0 | 2800800 | 440960 | 1946700 | 51152 | 0 | 1514300 | 0 | 0 | 2327700 | 0 | 0 | 0 | 1035500 | 0 |
| IVQELPQLL | NCEH1; NCEH1 | Q6PIU2; A0A | NCEH1_HUMA | Unmodified | 0 | 0 | 0 | 0 | 1074300 | 0 | 0 | 0 | 3456100 | 952300 | 0 | 0 | 0 | 0 | 0 | 0 | 0 | 0 |
| IVQFSPSGI | PAM; PAM | P19021; H7E | AMD_HUMAN F | Unmodified | 0 | 0 | 0 | 0 | 0 | 0 | 0 | 0 | 2008200 | 0 | 2269900 | 0 | 0 | 0 | 0 | 0 | 0 | 0 |
| IVQIPFCGE | EPRS1 | P07814 | SYEP_HUMAN | Unmodified | 0 | 0 | 0 | 0 | 0 | 0 | 0 | 0 | 0 | 0 | 0 | 0 | 0 | 0 | 0 | 0 | 0 | 0 |
| IVQIPFCGE | EPRS1 | P07814 | SYEP_HUMAN | Unmodified | 0 | 0 | 0 | 0 | 0 | 0 | 0 | 0 | 650450 | 215980 | 539770 | 1065000 | 674620 | 0 | 0 | 0 | 0 | 0 |
| IVQLLGQN | UCK2; UCK2 | Q9BZX2; A0/ | UCK2_HUMAN | Unmodified | 0 | 0 | 0 | 0 | 1997800 | 220940 | 843420 | 740520 | 0 | 0 | 224550 | 286930 | 1589800 | 725220 | 0 | 0 | 1056300 | 642640 |
| IVQMTEAE\ | PPP1CB; PPP1( | P62140; C9J | PP1B_HUMAN | Unmodified | 0 | 0 | 0 | 995110 | 0 | 0 | 0 | 55962 | 2652200 | 262760 | 1839700 | 9483000 | 0 | 0 | 1276800 | 512720 | 0 | 0 |
| IVQMTEAE\ | PPP1CB; PPP1( | P62140; C9J | PP1B_HUMAN | Oxidation (M) | 129340 | 150660 | 221350 | 0 | 0 | 71535 | 0 | 0 | 464090 | 0 | 69919 | 1365400 | 0 | 0 | 484040 | 447450 | 0 | 0 |
| IVQMTEAE\ | PPP1CB; PPP1( | P62140; C9J | PP1B_HUMAN | Unmodified | 0 | 0 | 0 | 0 | 2450600 | 363050 | 483530 | 31130 | 18782000 | 1694800 | 0 | 0 | 708560 | 814650 | 0 | 0 | 1768800 | 720030 |
| IVQMTEAE\ | PPP1CB; PPP1( | P62140; C9J | PP1B_HUMAN | Oxidation (M) | 0 | 0 | 0 | 0 | 498750 | 275850 | 197590 | 0 | 0 | 311310 | 0 | 0 | 352580 | 0 | 0 | 0 | 0 | 0 |
| IVQNSNGY | UGDH; UGDH; | O60701; E7E | UGDH_HUMAN | Unmodified | 0 | 0 | 0 | 0 | 4085900 | 0 | 0 | 0 | 0 | 0 | 0 | 0 | 0 | 504320 | 0 | 0 | 0 | 0 |
| IVQQWNL( | PLOD2; PLOD2; | O00469; C9J | PLOD2_HUMAI | Unmodified | 0 | 0 | 0 | 0 | 0 | 0 | 0 | 0 | 12126000 | 2034500 | 0 | 0 | 0 | 0 | 0 | 0 | 0 | 0 |
| IVQTYHVNI | ACTN1; ACTN1; | P12814; A0A | ACTN1_HUMAI | Oxidation (M) | 0 | 0 | 0 | 0 | 883400 | 0 | 0 | 0 | 0 | 0 | 0 | 7149300 | 0 | 0 | 1163500 | 114820 | 0 | 0 |
| IVQVALNGI | KPNA6; KPNA1; | O60684; P52 | IMA7_HUMAN I | Unmodified | 98290 | 138050 | 24475 | 0 | 0 | 0 | 0 | 0 | 0 | 0 | 0 | 54451 | 65224 | 0 | 236960 | 102900 | 0 | 0 |
| IVRASNGD. | HSPA9; HSPA9; | P38646; H0\ | GRP75_HUMAI | Unmodified | 0 | 0 | 0 | 0 | 1048400 | 946760 | 873630 | 134590 | 0 | 2052900 | 0 | 0 | 0 | 525900 | 0 | 0 | 226510 | 0 |
| IVRASQESE | SUN2 | Q9UH99 | SUN2_HUMAN | Unmodified | 0 | 0 | 0 | 0 | 0 | 0 | 115440 | 0 | 0 | 468870 | 0 | 0 | 0 | 0 | 0 | 0 | 0 | 0 |
| IVRGDQPA. | PGRMC1 | O00264 | PGRC1_HUMAI | Unmodified | 0 | 289010 | 0 | 0 | 0 | 0 | 0 | 0 | 1047700 | 1432700 | 0 | 0 | 0 | 0 | 0 | 0 | 0 | 0 |
| IVRPWAEG | OS9; OS9 | Q13438; B4E | OS9_HUMAN P | Unmodified | 26152 | 0 | 84292 | 0 | 0 | 83033 | 110990 | 87344 | 30720000 | 445460 | 837310 | 342930 | 77482 | 0 | 42243 | 0 | 0 | 0 |
| IVRPWAEG | OS9; OS9 | Q13438; B4E | OS9_HUMAN P | Unmodified | 0 | 0 | 0 | 0 | 0 | 0 | 0 | 0 | 2969600 | 395310 | 0 | 0 | 376740 | 0 | 0 | 0 | 0 | 0 |
| IVRSEQEF( | MCCC1; MCCC | Q96RQ3; E9I | MCCA_HUMAN | Unmodified | 0 | 0 | 290900 | 235900 | 0 | 324540 | 378940 | 0 | 673520 | 0 | 0 | 0 | 0 | 0 | 0 | 0 | 0 | 0 |
| IVRTAELSP | MPP1 | Q00013 | EM55_HUMAN | Unmodified | 0 | 0 | 0 | 0 | 2522100 | 0 | 0 | 0 | 0 | 0 | 0 | 0 | 0 | 0 | 0 | 0 | 842700 | 0 |
| IVSAGEQQ | LLGL1; LLGL1 | Q15334; A0/ | L2GL1_HUMAN | Unmodified | 0 | 0 | 271370 | 0 | 0 | 0 | 0 | 0 | 1601100 | 0 | 0 | 1683700 | 0 | 0 | 970600 | 445330 | 0 | 0 |
| IVSGEAESV | LONP1; LONP1 | P36776; K7E | LONM_HUMAN | Unmodified | 0 | 0 | 0 | 0 | 0 | 0 | 0 | 0 | 4161200 | 0 | 0 | 0 | 0 | 0 | 0 | 0 | 0 | 0 |
| IVSGIITPIH | PSMF1; PSMF1; | Q92530; Q5( | PSMF1_HUMAI | Unmodified | 0 | 0 | 0 | 0 | 0 | 0 | 0 | 0 | 0 | 1105200 | 0 | 620960 | 0 | 0 | 0 | 0 | 980030 | 0 |
| IVSGKDYN | LDHA; LDHA; LI | P00338; F5G | LDHA_HUMAN | Unmodified | 0 | 291770 | 0 | 2476800 | 15211000 | 2435100 | 986910 | 501370 | 1139500 | 3379000 | 743970 | 3507500 | 5374900 | 3041600 | 0 | 0 | 17927000 | 18175000 |
| IVSGNLRIL | UGP2; UGP2; U | Q16851; E7E | UGPA_HUMAN | Unmodified | 0 | 0 | 0 | 0 | 1936900 | 142840 | 1101700 | 437660 | 0 | 870810 | 0 | 320350 | 362830 | 375860 | 0 | 0 | 1735400 | 798280 |
| IVSKIDGDK | CALU | O43852 | CALU_HUMAN | Unmodified | 0 | 0 | 0 | 86564 | 7455000 | 4122900 | 1275100 | 541680 | 32738000 | 8671700 | 1966200 | 962670 | 992630 | 819790 | 156210 | 24208 | 484370 | 1921000 |
| IVSLEPAYP | DSG2; DSG2 | Q14126; J3K | DSG2_HUMAN | Unmodified | 303750 | 135770 | 381510 | 506420 | 2153200 | 806530 | 0 | 0 | 7639200 | 911490 | 306530 | 484170 | 552910 | 912280 | 200500 | 263680 | 0 | 204240 |
| IVSLFAEHN | EIF2S3; EIF2S3I | P41091; Q2\ | IF2G_HUMAN E | Unmodified | 0 | 0 | 0 | 0 | 473640 | 0 | 0 | 0 | 745350 | 115020 | 66749 | 896530 | 0 | 0 | 0 | 0 | 0 | 0 |
| IVSLPECFN | NIT2; NIT2; NIT | Q9NQR4; F8 | NIT2_HUMAN ( | Unmodified | 0 | 0 | 164370 | 0 | 581870 | 146500 | 205080 | 106950 | 0 | 284950 | 312020 | 957900 | 1147900 | 833800 | 0 | 0 | 2185700 | 787260 |
| IVSQLLTLM | VCP; VCP; VCP; | P55072; C9J | TERA_HUMAN | Unmodified | 108040 | 0 | 0 | 142010 | 596420 | 114000 | 49305 | 0 | 417250 | 287820 | 42344 | 269110 | 1333900 | 450410 | 128520 | 40929 | 135580 | 180160 |
| IVSRPEELR | CCT3; CCT3 | P49368; B4D | TCPG_HUMAN | Unmodified | 2455300 | 1585100 | 2842900 | 7034500 | 2564100 | 3309300 | 1740500 | 0 | 17054000 | 4189900 | 26705000 | 1.21E+08 | 31732000 | 7239800 | 5723900 | 3889200 | 3530200 | 10046000 |
| IVSSSDVGI | SPTBN1; SPTBN | Q01082; A0/ | SPTB2_HUMAN | Unmodified | 0 | 0 | 0 | 0 | 0 | 0 | 0 | 0 | 1248600 | 289400 | 0 | 4550800 | 0 | 0 | 0 | 0 | 322800 | 0 |
| IVSWLEYES | FKBP4 | Q02790 | FKBP4_HUMAN | Unmodified | 265920 | 0 | 266310 | 732710 | 751010 | 0 | 806800 | 77319 | 722790 | 0 | 2434600 | 8247900 | 4501900 | 3182000 | 893550 | 731260 | 736660 | 0 |
| IVSWSQVG | ACAT2 | Q9BWD1 | THIC_HUMAN / | Oxidation (M) | 0 | 0 | 0 | 0 | 3829500 | 349340 | 0 | 0 | 738480 | 1702900 | 295210 | 985530 | 1907100 | 0 | 0 | 0 | 1298100 | 679760 |
| IVTCGADRI | ARPC1A; ARPC | Q92747; E9F | ARC1A_HUMAI | Unmodified | 317870 | 135640 | 834050 | 257580 | 0 | 0 | 0 | 0 | 0 | 0 | 0 | 4684700 | 0 | 0 | 624240 | 900530 | 0 | 0 |
| IVTCGTDRI | ARPC1B; ARPC | O15143; C9J | ARC1B_HUMAI | Unmodified | 0 | 0 | 0 | 0 | 0 | 0 | 0 | 0 | 0 | 0 | 0 | 6518800 | 0 | 0 | 0 | 0 | 0 | 0 |
| IVTDSDSKT | SLK | Q9H2G2 | SLK_HUMAN S | Unmodified | 317300 | 270110 | 236500 | 651300 | 0 | 0 | 49714 | 30595 | 1813400 | 177110 | 1633300 | 4565600 | 1631800 | 934310 | 417900 | 437580 | 0 | 0 |
| IVTEETMTT | ANK3 | Q12955 | ANK3_HUMAN | Oxidation (M) | 0 | 0 | 698570 | 1422600 | 0 | 0 | 0 | 0 | 0 | 480650 | 0 | 0 | 1456200 | 223470 | 1321200 | 1272500 | 345900 | 0 |
| IVTEETMTT | ANK3 | Q12955 | ANK3_HUMAN | Unmodified | 0 | 0 | 581110 | 1389500 | 0 | 0 | 0 | 0 | 0 | 0 | 0 | 0 | 2861400 | 1515800 | 1620500 | 2681700 | 0 | 0 |
| IVTGGSDN | SHMT1 | P34896 | GLYC_HUMAN | Unmodified | 0 | 0 | 447160 | 1179000 | 0 | 0 | 0 | 0 | 0 | 261580 | 869030 | 6528700 | 0 | 0 | 1087900 | 0 | 0 | 0 |
| IVTGGSDN | SHMT1 | P34896 | GLYC_HUMAN | Unmodified | 0 | 0 | 0 | 0 | 0 | 188010 | 0 | 0 | 0 | 330950 | 0 | 0 | 256690 | 0 | 0 | 0 | 0 | 0 |
| IVTIHECDS | CBARP; CBARP | Q8N350; A0 | CBARP_HUMAI | Unmodified | 0 | 0 | 0 | 0 | 0 | 0 | 0 | 0 | 0 | 0 | 0 | 0 | 488170 | 735520 | 212320 | 0 | 516740 | 1985100 |
| IVTLEEEPIK | BIRC6; BIRC6 | Q9NR09; A0 | BIRC6_HUMAN | Unmodified | 0 | 0 | 0 | 0 | 0 | 0 | 0 | 0 | 0 | 0 | 0 | 775210 | 0 | 0 | 0 | 0 | 0 | 0 |
| IVTLFSDFK | CD109 | Q6YHK3 | CD109_HUMAI | Unmodified | 348560 | 0 | 0 | 0 | 0 | 0 | 0 | 0 | 3686600 | 523560 | 0 | 0 | 0 | 0 | 0 | 0 | 229270 | 269410 |
| IVTNNLK | TPM2; TPM2; TF | P07951; A0A | TPM2_HUMAN | Unmodified | 0 | 0 | 0 | 0 | 0 | 0 | 0 | 0 | 0 | 0 | 361240 | 0 | 0 | 0 | 0 | 0 | 0 | 0 |

| | | | | | | | | | | | | | | | | | | | | | | |
|---|---|---|---|---|---|---|---|---|---|---|---|---|---|---|---|---|---|---|---|---|---|---|
| IVTSEEVIIR | NPTN; NPTN; N | Q9Y639; Q9I | NPTN_HUMAN | Unmodified | 185350 | 370930 | 223810 | 864470 | 0 | 0 | 0 | 26414 | 1396000 | 129360 | 276670 | 10131000 | 0 | 37141 | 2054000 | 1247700 | 0 | 47456 |
| IVTTYIR | DNM2; DNM2 | P50570; K7E | DYN2_HUMAN | Unmodified | 1658900 | 2695500 | 3914000 | 738170 | 0 | 0 | 27027 | 11856 | 862950 | 0 | 61989 | 2585800 | 0 | 0 | 1862400 | 988010 | 0 | 0 |
| IVTVVPQDT | ELP1; ELP1; ELF | O95163; A0A | ELP1_HUMAN I | Unmodified | 0 | 0 | 0 | 0 | 0 | 0 | 0 | 0 | 241740 | 167430 | 577460 | 1029200 | 0 | 0 | 0 | 0 | 0 | 0 |
| IVVDGSGQ | SMURF1 | Q9HCE7 | SMUF1_HUMAI | Unmodified | 612770 | 166430 | 0 | 81472 | 0 | 0 | 0 | 0 | 0 | 141010 | 15874 | 67239 | 194710 | 63011 | 148640 | 93861 | 0 | 119680 |
| IVVFQYSD( | DDB1; DDB1; D | Q16531; A0/ | DDB1_HUMAN | Unmodified | 0 | 0 | 183780 | 0 | 0 | 0 | 0 | 0 | 339820 | 106080 | 351820 | 3051100 | 0 | 0 | 416290 | 0 | 112920 | 0 |
| IVVFQYSD( | DDB1; DDB1; D | Q16531; A0/ | DDB1_HUMAN | Unmodified | 235960 | 0 | 0 | 0 | 190120 | 0 | 57113 | 0 | 155480 | 456970 | 0 | 336370 | 512870 | 0 | 0 | 0 | 1278100 | 170810 |
| IVVHLHPAF | VPS36; VPS36 | Q86VN1; A0, | VPS36_HUMAN | Unmodified | 15288000 | 12966000 | 7527600 | 6664900 | 0 | 0 | 0 | 0 | 503930 | 187880 | 189980 | 4283700 | 1405800 | 0 | 21446000 | 15643000 | 70423 | 0 |
| IVVLLQR | PSME1; PSME1 | Q06323; H0' | PSME1_HUMAI | Unmodified | 0 | 0 | 0 | 0 | 0 | 0 | 0 | 0 | 0 | 1629300 | 795020 | 3130400 | 0 | 0 | 0 | 0 | 463870 | 0 |
| IVVMNNLLI | PIP5K1A; PIPSL | Q99755; A2/ | PI51A_HUMAN | Oxidation (M) | 0 | 0 | 118800 | 0 | 0 | 0 | 0 | 0 | 0 | 0 | 0 | 714380 | 0 | 0 | 111560 | 266490 | 0 | 0 |
| IVVNLTGR | RPS15A; N/A; R | P62244; H3E | RS15A_HUMAN | Unmodified | 3809200 | 2488900 | 1767400 | 1250900 | 0 | 0 | 0 | 1290700 | 0 | 0 | 823150 | 97546000 | 0 | 0 | 3785400 | 3234100 | 0 | 0 |
| IVVNLTGRL | RPS15A; N/A; R | P62244; H3E | RS15A_HUMAN | Unmodified | 0 | 473970 | 226630 | 458070 | 586870 | 0 | 1375800 | 700180 | 230580 | 277940 | 958540 | 5094600 | 1360700 | 351260 | 0 | 158410 | 124310 | 381370 |
| IVVPPHLGY | FKBP9; FKBP9 | O95302; C9J | FKBP9_HUMAN | Unmodified | 0 | 0 | 0 | 89838 | 0 | 0 | 0 | 0 | 1022800 | 0 | 1085800 | 0 | 0 | 0 | 0 | 0 | 0 | 0 |
| IVVSFQEEN | NAV2; NAV2; N/ | Q8IVL1; A0A | NAV2_HUMAN | Oxidation (M) | 0 | 0 | 0 | 0 | 0 | 365880 | 0 | 0 | 0 | 0 | 0 | 205910 | 0 | 0 | 0 | 0 | 0 | 146270 |
| IVVTRSEIDI | ANXA7 | P20073 | ANXA7_HUMAI | Unmodified | 0 | 0 | 0 | 0 | 0 | 0 | 783620 | 232170 | 1.67E+08 | 20051000 | 3584900 | 0 | 0 | 2097200 | 0 | 0 | 1995500 | 4626000 |
| IVVVGDSQ | RND3; RND3 | P61587; Q53 | RND3_HUMAN | Unmodified | 0 | 39714 | 0 | 0 | 0 | 0 | 0 | 0 | 0 | 0 | 0 | 0 | 0 | 0 | 0 | 0 | 1340200 | 0 |
| IVVVGDSQ | RND3; RND3 | P61587; Q53 | RND3_HUMAN | Unmodified | 0 | 0 | 0 | 0 | 37775 | 0 | 73369 | 26469 | 0 | 0 | 0 | 0 | 0 | 0 | 0 | 0 | 424210 | 218970 |
| IVVVTAGVF | LDHB; LDHB; LI | P07195; A0A | LDHB_HUMAN | Unmodified | 2818600 | 2782100 | 1794100 | 2704300 | 395960 | 206480 | 59514 | 94420 | 82834 | 73560 | 18992000 | 2.3E+08 | 214250 | 78896 | 3174700 | 3290900 | 227290 | 221230 |
| IVVVTAGVF | LDHB; LDHB; LI | P07195; A0A | LDHB_HUMAN | Unmodified | 1723800 | 915450 | 1728600 | 3071500 | 1039600 | 480110 | 39006 | 31995 | 74736 | 89097 | 3742800 | 39132000 | 449740 | 297430 | 2619100 | 1637100 | 375320 | 274170 |
| IVVVWVGT | PAFAH1B3; PAF | Q15102; M0I | PA1B3_HUMAN | Unmodified | 0 | 0 | 237890 | 0 | 1187900 | 0 | 1283800 | 60256 | 1273000 | 2592700 | 1916600 | 16103000 | 724420 | 363290 | 0 | 482300 | 0 | 0 |
| IVWLQEK | THOP1; THOP1 | P52888; K7E | THOP1_HUMAI | Unmodified | 0 | 0 | 72632 | 0 | 0 | 0 | 0 | 0 | 201840 | 238120 | 268690 | 1825300 | 0 | 70097 | 110910 | 100550 | 0 | 0 |
| IVYGHLDDI | ACO2; ACO2; A | Q99798; A2/ | ACON_HUMAN | Unmodified | 0 | 0 | 0 | 0 | 0 | 0 | 0 | 0 | 0 | 0 | 0 | 0 | 0 | 0 | 0 | 0 | 0 | 0 |
| IVYGHLDDI | ACO2; ACO2; A | Q99798; A2/ | ACON_HUMAN | Unmodified | 0 | 0 | 0 | 0 | 0 | 0 | 1025200 | 484790 | 6937500 | 1504100 | 0 | 0 | 215410 | 0 | 0 | 0 | 0 | 241990 |
| IVYLYTK | RPL34 | P49207 | RL34_HUMAN ( | Unmodified | 5247200 | 4493400 | 4181300 | 3047600 | 0 | 0 | 0 | 0 | 555640 | 0 | 851800 | 18253000 | 1386300 | 0 | 4900800 | 4369600 | 0 | 240980 |
| IVYMHMER | RGS13 | O14921 | RGS13_HUMAI | Unmodified | 0 | 0 | 0 | 0 | 0 | 39752 | 0 | 0 | 0 | 0 | 409640 | 1229200 | 0 | 0 | 0 | 0 | 0 | 0 |
| IVYPTEK | LASP1; LASP1; I | Q14847; K7E | LASP1_HUMAN | Unmodified | 1210000 | 0 | 0 | 0 | 7780600 | 2045100 | 990360 | 678810 | 2677800 | 0 | 0 | 1372300 | 2918900 | 2800000 | 0 | 0 | 2297900 | 3194600 |
| IVYPTEKVN | LASP1; LASP1; I | Q14847; K7E | LASP1_HUMAN | Unmodified | 0 | 0 | 0 | 157840 | 0 | 0 | 0 | 563210 | 0 | 0 | 0 | 0 | 0 | 786920 | 0 | 0 | 2597100 | 0 |
| IWCFGPDG | EEF2 | P13639 | EF2_HUMAN EI | Unmodified | 2676300 | 1324800 | 99617 | 2686000 | 18869000 | 3838200 | 2823400 | 52305 | 4221200 | 2975700 | 6225100 | 7268700 | 11193000 | 4863400 | 2373900 | 1535100 | 7376900 | 12163000 |
| IWDLADPE | EPS15L1; EPS1 | Q9UBC2; M( | EP15R_HUMAN | Unmodified | 191650 | 0 | 0 | 0 | 919380 | 0 | 187970 | 55315 | 0 | 0 | 0 | 685180 | 0 | 0 | 0 | 0 | 0 | 0 |
| IWDLEGK | RACK1; RACK1; | P63244; D6F | RACK1_HUMAI | Unmodified | 2637200 | 1943300 | 1816800 | 1046700 | 451810 | 172620 | 0 | 297550 | 0 | 486810 | 2838000 | 35244000 | 490720 | 0 | 1637800 | 1376400 | 0 | 0 |
| IWDLNMEN | PPP2R2A; PPP2 | P63151; A0A | 2ABA_HUMAN | Oxidation (M) | 2086100 | 75343 | 374570 | 0 | 0 | 164190 | 0 | 0 | 526460 | 0 | 350300 | 4066100 | 0 | 0 | 1699300 | 1124500 | 0 | 0 |
| IWDLQAGA | CORO7; CORO | P57737; A0A | CORO7_HUMA | Unmodified | 0 | 0 | 0 | 0 | 0 | 0 | 0 | 0 | 246500 | 0 | 642730 | 2724700 | 0 | 0 | 388280 | 148560 | 0 | 0 |
| IWDPTPSH | SF3B1 | O75533 | SF3B1_HUMAN | Unmodified | 0 | 0 | 0 | 0 | 0 | 0 | 0 | 0 | 0 | 0 | 371530 | 0 | 143580 | 0 | 0 | 0 | 0 | 0 |
| IWDPTPSH | SF3B1 | O75533 | SF3B1_HUMAN | Unmodified | 421030 | 0 | 0 | 0 | 0 | 0 | 0 | 0 | 0 | 0 | 389600 | 0 | 0 | 0 | 0 | 227530 | 0 | 0 |
| IWDTTQK | WDR1; WDR1 | O75083; D6I | WDR1_HUMAN | Unmodified | 0 | 0 | 0 | 0 | 1497500 | 9762.6 | 151590 | 70626 | 0 | 175620 | 506780 | 0 | 22634 | 0 | 0 | 0 | 239160 | 708050 |
| IWEGSSK | PRKCD | Q05655 | KPCD_HUMAN | Unmodified | 0 | 0 | 0 | 0 | 0 | 0 | 0 | 0 | 0 | 0 | 0 | 0 | 0 | 0 | 0 | 0 | 505500 | 0 |
| IWEIPEGGI | CORO2B | Q9UQ03 | COR2B_HUMA | Unmodified | 0 | 0 | 0 | 0 | 0 | 0 | 0 | 0 | 0 | 0 | 0 | 3245400 | 0 | 0 | 0 | 0 | 0 | 0 |
| IWEIPEGGI | CORO2B | Q9UQ03 | COR2B_HUMA | Unmodified | 0 | 0 | 0 | 0 | 0 | 0 | 0 | 0 | 0 | 0 | 0 | 5886300 | 0 | 0 | 0 | 0 | 0 | 0 |
| IWEQIQPDI | AIMP1 | Q12904 | AIMP1_HUMAN | Unmodified | 0 | 0 | 0 | 0 | 0 | 0 | 0 | 0 | 0 | 0 | 0 | 2737200 | 1104900 | 1790700 | 0 | 0 | 595970 | 612650 |
| IWGDCTVR | SARS1; SARS1 | P49591; Q5T | SYSC_HUMAN | Unmodified | 0 | 0 | 0 | 0 | 0 | 0 | 0 | 0 | 0 | 0 | 0 | 1017300 | 0 | 0 | 0 | 0 | 0 | 0 |
| IWGEDLRH | NAMPT; NAMPT | P43490; A0A | NAMPT_HUMAI | Unmodified | 0 | 0 | 376940 | 0 | 2007800 | 737840 | 0 | 0 | 0 | 0 | 0 | 0 | 395540 | 0 | 0 | 0 | 0 | 0 |
| IWGLGFLP( | DNAJC10; DNA | Q8IXB1; A0A | DJC10_HUMAN | Unmodified | 0 | 457770 | 0 | 0 | 0 | 16912 | 63988 | 0 | 338600 | 113730 | 0 | 0 | 0 | 0 | 0 | 0 | 0 | 0 |
| IWHHTFYN | POTEJ; POTEE; I | P0CG39; Q6 | POTEJ_HUMAN | Unmodified | 82228000 | 88935000 | 64204000 | 77588000 | 63786000 | 33905000 | 5191000 | 0 | 1.06E+08 | 6390900 | 41244000 | 5.12E+08 | 36858000 | 11935000 | 1.26E+08 | 1.02E+08 | 6427000 | 22439000 |
| IWHHTFYN | POTEJ; POTEE; I | P0CG39; Q6 | POTEJ_HUMAN | Unmodified | 0 | 0 | 0 | 0 | 6330000 | 190720 | 0 | 0 | 0 | 80120 | 0 | 599450 | 0 | 0 | 0 | 0 | 1996200 | 0 |
| IWHHTFYN | ACTC1; ACTA1; | P68032; P68 | ACTC_HUMAN | Unmodified | 0 | 0 | 0 | 0 | 0 | 0 | 0 | 0 | 0 | 0 | 0 | 0 | 1550400 | 0 | 0 | 0 | 1240300 | 0 |
| IWHHTFYN | ACTG1; ACTB; / | P63261; P60 | ACTG_HUMAN | Unmodified | 2340300 | 1073500 | 6270100 | 1.01E+08 | 8.1E+09 | 1.07E+09 | 8.82E+08 | 1206900 | 39175000 | 2.56E+08 | 98514000 | 1702300 | 2.42E+09 | 1.79E+09 | 895860 | 4249100 | 1.09E+09 | 1.95E+09 |
| IWKLADCD | EHD4 | Q9H223 | EHD4_HUMAN | Oxidation (M) | 0 | 0 | 0 | 0 | 611710 | 0 | 0 | 0 | 0 | 0 | 0 | 0 | 509330 | 828330 | 0 | 0 | 3680300 | 2093700 |
| IWKLADVD | EHD1; EHD1 | Q9H4M9; A0 | EHD1_HUMAN | Unmodified | 0 | 0 | 0 | 0 | 26130000 | 2359800 | 1472000 | 0 | 0 | 1016900 | 0 | 0 | 12184000 | 6605000 | 0 | 0 | 63580000 | 35109000 |
| IWLVDSK | ME1 | P48163 | MAOX_HUMAN | Unmodified | 0 | 0 | 0 | 175760 | 1421800 | 1936000 | 114320 | 0 | 60035 | 51955 | 214320 | 2327800 | 0 | 466290 | 319210 | 111620 | 889340 | 406970 |
| IWLVDSKG | ME1 | P48163 | MAOX_HUMAN | Unmodified | 0 | 0 | 0 | 0 | 0 | 0 | 0 | 0 | 0 | 0 | 0 | 0 | 0 | 0 | 0 | 0 | 563020 | 0 |
| IWMYNCHI | LOXL2 | Q9Y4K0 | LOXL2_HUMAN | Oxidation (M) | 0 | 0 | 0 | 0 | 0 | 0 | 0 | 0 | 616510 | 0 | 0 | 0 | 0 | 0 | 0 | 0 | 0 | 0 |
| IWMYNCHI | LOXL2 | Q9Y4K0 | LOXL2_HUMAN | Unmodified | 0 | 0 | 0 | 0 | 0 | 0 | 0 | 0 | 1356000 | 0 | 0 | 0 | 0 | 0 | 0 | 0 | 0 | 619660 |
| IWNLSENS | TBL1XR1; TBL1) | Q9BZK7; A0/ | TBL1R_HUMAN | Unmodified | 0 | 0 | 0 | 0 | 0 | 0 | 0 | 0 | 0 | 0 | 0 | 1537100 | 0 | 0 | 0 | 0 | 115240 | 0 |
| IWNSSLQT | CRAT; CRAT; CI | P43155; A0A | CACP_HUMAN | Unmodified | 0 | 0 | 0 | 0 | 828840 | 0 | 0 | 0 | 1758100 | 434320 | 0 | 0 | 0 | 0 | 0 | 0 | 0 | 0 |
| IWNTMMG | SLC38A9; SLC3 | Q8NBW4; D( | S38A9_HUMAN | 2 Oxidation (M) | 0 | 0 | 0 | 0 | 0 | 0 | 0 | 0 | 0 | 0 | 0 | 0 | 1321700 | 13383000 | 0 | 0 | 0 | 0 |
| IWNVHSVL | EIF3L; EIF3L; EI | Q9Y262; B0( | EIF3L_HUMAN | Unmodified | 147310 | 392300 | 73788 | 302730 | 0 | 0 | 37275 | 0 | 278720 | 0 | 50371 | 659600 | 78669 | 0 | 734930 | 72681 | 0 | 0 |
| IWNVIYEEN | ERO1A; ERO1A | Q96HE7; G3 | ERO1A_HUMAI | Unmodified | 0 | 0 | 0 | 0 | 0 | 0 | 0 | 0 | 7781800 | 0 | 831710 | 0 | 0 | 0 | 0 | 0 | 0 | 0 |
| IWQYVYSK | ACTR1A; ACTR1 | P61163; P42 | ACTZ_HUMAN . | Unmodified | 0 | 0 | 0 | 0 | 0 | 0 | 0 | 0 | 0 | 0 | 0 | 0 | 0 | 0 | 0 | 0 | 0 | 0 |
| IWRDTANV | CDH1 | P12830 | CADH1_HUMA | Unmodified | 2313200 | 0 | 407960 | 1786400 | 859230 | 0 | 0 | 0 | 8029900 | 32065 | 236700 | 1485700 | 0 | 0 | 0 | 786750 | 0 | 0 |
| IWRDYVAP | SND1 | Q7KZF4 | SND1_HUMAN | Unmodified | 163100 | 168850 | 284200 | 322160 | 0 | 0 | 0 | 0 | 0 | 0 | 0 | 0 | 311510 | 467670 | 351960 | 262700 | 0 | 0 |
| IWRRSYDV | PGAM1; PGAM4 | P18669; Q8I | PGAM1_HUMA | Oxidation (M) | 0 | 0 | 0 | 0 | 1048400 | 0 | 0 | 0 | 0 | 0 | 0 | 0 | 0 | 831470 | 0 | 0 | 2149400 | 929040 |
| IWRRSYDV | PGAM1; PGAM4 | P18669; Q8I | PGAM1_HUMA | Unmodified | 0 | 0 | 0 | 0 | 1497600 | 0 | 701360 | 0 | 0 | 0 | 0 | 0 | 0 | 0 | 0 | 0 | 1236600 | 0 |
| IWSVPNAS | PRPF19 | Q9UMS4 | PRP19_HUMAN | Unmodified | 0 | 0 | 0 | 0 | 0 | 0 | 0 | 0 | 365400 | 0 | 148250 | 2619100 | 0 | 0 | 0 | 0 | 0 | 0 |
| IWYEHRLIC | IDH1 | O75874 | IDHC_HUMAN | Unmodified | 0 | 0 | 1647300 | 0 | 0 | 0 | 0 | 0 | 0 | 0 | 0 | 0 | 921690 | 0 | 0 | 0 | 64572 | 167890 |
| IWYEHRLIC | IDH2; IDH2 | P48735; H0Y | IDHP_HUMAN | Unmodified | 0 | 0 | 0 | 0 | 0 | 0 | 0 | 0 | 0 | 2471500 | 0 | 0 | 0 | 0 | 0 | 0 | 156920 | 0 |
| IWYLLDGK | MRPL13; MRPL | Q9BYD1; E5I | RM13_HUMAN | Unmodified | 0 | 0 | 0 | 0 | 0 | 40492 | 0 | 0 | 581500 | 0 | 0 | 0 | 0 | 0 | 0 | 0 | 101410 | 186210 |
| IYADSLKPN | AFDN; AFDN; A | P55196; Q5I | AFAD_HUMAN | Unmodified | 0 | 812340 | 0 | 0 | 828100 | 469270 | 0 | 118640 | 721650 | 0 | 1049200 | 3381700 | 1043800 | 434690 | 0 | 511770 | 0 | 0 |
| IYALPDDLV | PLS1 | Q14651 | PLSI_HUMAN F | Unmodified | 0 | 0 | 0 | 0 | 2766100 | 0 | 0 | 0 | 0 | 568630 | 520260 | 3154100 | 1341800 | 1525000 | 0 | 0 | 0 | 422950 |
| IYAMHWGT | GNB2; GNB1; G | P62879; P62 | GBB2_HUMAN | Unmodified | 4338900 | 4964700 | 1614000 | 3092600 | 1197100 | 481320 | 540250 | 290730 | 11382000 | 248090 | 8997400 | 1.22E+08 | 563300 | 1018700 | 5711100 | 4643200 | 1205500 | 694570 |
| IYAMHWGT | GNB2; GNB1; G | P62879; P62 | GBB2_HUMAN | Oxidation (M) | 1816100 | 423030 | 919140 | 741270 | 379660 | 496280 | 0 | 98329 | 2007200 | 57710 | 1610700 | 31089000 | 163670 | 209250 | 2762200 | 2643800 | 207690 | 0 |
| IYAMHWGT | GNB2; GNB1; G | P62879; P62 | GBB2_HUMAN | Unmodified | 0 | 0 | 0 | 327900 | 7870800 | 1426100 | 6458600 | 0 | 1239000 | 4027200 | 1324700 | 192790 | 622900 | 3471900 | 0 | 0 | 4960000 | 3143500 |
| IYAMHWGT | GNB2; GNB1; G | P62879; P62 | GBB2_HUMAN | Oxidation (M) | 53329 | 0 | 283700 | 240390 | 1698000 | 1056400 | 2318300 | 392490 | 0 | 1553800 | 1293800 | 633180 | 544650 | 2602100 | 0 | 0 | 4082600 | 1549500 |
| IYANFFPYG | HPCA; HPCAL1 | P84074; P37 | HPCA_HUMAN | Unmodified | 0 | 0 | 0 | 0 | 0 | 0 | 0 | 0 | 0 | 0 | 0 | 1275000 | 0 | 0 | 0 | 0 | 0 | 0 |
| IYAPQGLLL | GOLGA7 | Q7Z5G4 | GOGA7_HUMA | Unmodified | 1332600 | 2300600 | 2980800 | 2736800 | 1131400 | 0 | 0 | 16780 | 0 | 297360 | 2085500 | 20021000 | 0 | 0 | 4194800 | 4366500 | 0 | 0 |
| IYDAQNELS | APPL1; APPL1; / | Q9UKG1; C9 | DP13A_HUMAN | Unmodified | 916650 | 605570 | 0 | 974720 | 2158500 | 0 | 0 | 0 | 2676600 | 807590 | 529100 | 2130800 | 3066300 | 0 | 410400 | 537460 | 0 | 0 |
| IYDDYYPDI | CXADR | P78310 | CXAR_HUMAN | Unmodified | 0 | 0 | 0 | 0 | 13007000 | 2713000 | 1033100 | 0 | 1604400 | 191760 | 0 | 0 | 0 | 0 | 0 | 0 | 0 | 0 |
| IYDILSDNG | ERBIN | Q96RT1 | ERBIN_HUMAN | Unmodified | 0 | 0 | 650710 | 1053600 | 24013000 | 4304100 | 180400 | 161440 | 660180 | 455680 | 1311300 | 1105300 | 2037400 | 1739100 | 878400 | 897480 | 1398300 | 3706200 |
| IYDILSDNG | ERBIN | Q96RT1 | ERBIN_HUMAN | Unmodified | 0 | 0 | 0 | 0 | 2192700 | 0 | 0 | 0 | 0 | 0 | 0 | 0 | 0 | 0 | 0 | 0 | 0 | 0 |
| IYDLNKPEA | STRAP | Q9Y3F4 | STRAP_HUMAN | Unmodified | 0 | 137370 | 979200 | 999300 | 2159200 | 788520 | 0 | 0 | 199210 | 0 | 1069700 | 13249000 | 2012700 | 0 | 0 | 0 | 349990 | 1671500 |
| IYDREQTRY | IQGAP1; IQGAP | P46940; A0A | IQGA1_HUMAN | Unmodified | 0 | 0 | 0 | 0 | 13620000 | 0 | 0 | 0 | 0 | 0 | 0 | 0 | 0 | 0 | 0 | 0 | 0 | 0 |
| IYDSLLALSI | CLDN8 | P56748 | CLD8_HUMAN | Unmodified | 0 | 0 | 0 | 0 | 0 | 0 | 0 | 0 | 0 | 0 | 0 | 51990 | 0 | 0 | 365310 | 411930 | 0 | 0 |
| IYDYDPLYK | APOBEC3B; AP( | Q9UH17; P3 | ABC3B_HUMAI | Unmodified | 1189000 | 1394000 | 1103500 | 290790 | 0 | 0 | 0 | 0 | 0 | 0 | 0 | 0 | 0 | 0 | 2504700 | 1050700 | 0 | 0 |
| IYDYNSVIR | SNX9; SNX9; SI | Q9Y5X1; A0/ | SNX9_HUMAN | Unmodified | 5737100 | 0 | 0 | 0 | 403490 | 309230 | 0 | 0 | 0 | 0 | 0 | 0 | 0 | 0 | 0 | 0 | 0 | 0 |
| IYEDGDDD | CACYBP | Q9HB71 | CYBP_HUMAN | Unmodified | 0 | 0 | 0 | 48882 | 163660 | 0 | 0 | 14500 | 276980 | 48711 | 0 | 960210 | 0 | 0 | 54624 | 0 | 32432 | 0 |
| IYEDGDDD | CACYBP | Q9HB71 | CYBP_HUMAN | Oxidation (M) | 0 | 0 | 0 | 0 | 0 | 0 | 0 | 0 | 0 | 0 | 0 | 801070 | 0 | 0 | 0 | 0 | 0 | 0 |
| IYEDGDDD | CACYBP | Q9HB71 | CYBP_HUMAN | Unmodified | 0 | 0 | 427070 | 0 | 0 | 0 | 0 | 0 | 225630 | 227010 | 1946300 | 1684200 | 611090 | 0 | 0 | 761410 | 0 | 0 |
| IYEDGDDD | CACYBP | Q9HB71 | CYBP_HUMAN | Unmodified | 0 | 0 | 0 | 0 | 1450400 | 238190 | 742150 | 908610 | 0 | 419230 | 0 | 0 | 2255300 | 0 | 0 | 0 | 1868200 | 392480 |
| IYEEAWTK | NAA15; NAA15 | Q9BXJ9; A0A | NAA15_HUMAI | Unmodified | 0 | 0 | 0 | 0 | 353830 | 229060 | 37008 | 0 | 0 | 0 | 169470 | 4812000 | 0 | 0 | 0 | 0 | 698200 | 0 |
| IYEENDRA\ | SCG3 | Q8WXD2 | SCG3_HUMAN | Unmodified | 0 | 0 | 0 | 0 | 0 | 0 | 23064 | 0 | 63786000 | 12748000 | 0 | 0 | 0 | 0 | 0 | 0 | 0 | 0 |
| IYEENDRA\ | SCG3 | Q8WXD2 | SCG3_HUMAN | Unmodified | 0 | 0 | 0 | 0 | 0 | 0 | 0 | 84677 | 1216200 | 5352800 | 0 | 0 | 0 | 0 | 0 | 0 | 0 | 0 |
| IYEFPETDD | SEPTIN7; SEPTI | Q16181; E7E | SEPT7_HUMAN | Unmodified | 45061 | 0 | 784510 | 494000 | 1966000 | 384140 | 0 | 19479 | 763390 | 38323 | 1359900 | 6444000 | 390900 | 0 | 563010 | 463720 | 125330 | 688830 |
| IYEFPETDD | SEPTIN7; SEPTI | Q16181; E7E | SEPT7_HUMAN | Unmodified | 0 | 0 | 0 | 0 | 12830000 | 2466000 | 1663800 | 763810 | 0 | 674230 | 709020 | 1046500 | 0 | 0 | 0 | 0 | 5244700 | 0 |
| IYEFPETDD | SEPTIN7; SEPTI | Q16181; E7E | SEPT7_HUMAN | Unmodified | 21834 | 0 | 225690 | 148020 | 10650000 | 0 | 1026700 | 747060 | 45035000 | 2702400 | 2568400 | 1759600 | 1545900 | 0 | 119630 | 153030 | 2688000 | 2457900 |
| IYEFTLQR | MYO1B; MYO1E | O43795; E9F | MYO1B_HUMA | Unmodified | 125180 | 219410 | 0 | 0 | 503930 | 0 | 0 | 68780 | 0 | 0 | 550950 | 5837800 | 0 | 0 | 392270 | 0 | 0 | 0 |
| IYELAAGGT | FH | P07954 | FUMH_HUMAN | Unmodified | 0 | 0 | 0 | 0 | 0 | 0 | 0 | 0 | 1159100 | 0 | 23284 | 0 | 0 | 0 | 94931 | 62153 | 0 | 0 |
| IYESHVGIS | GBE1; GBE1 | Q04446; E9F | GLGB_HUMAN | Unmodified | 0 | 0 | 0 | 0 | 0 | 0 | 0 | 0 | 0 | 0 | 0 | 687630 | 0 | 0 | 0 | 0 | 0 | 0 |
| IYFDDFSC( | ANAPC5 | F5H0N1 | F5H0N1_HUM/ | Unmodified | 0 | 0 | 41558 | 0 | 0 | 0 | 0 | 0 | 0 | 0 | 0 | 0 | 0 | 0 | 1069500 | 580050 | 0 | 0 |
| IYFGKDIPN | ERLIN2; ERLIN2 | O94905; E5F | ERLN2_HUMAN | Oxidation (M) | 0 | 285080 | 0 | 822290 | 2241900 | 20258000 | 2975000 | 178090 | 451580 | 6579200 | 0 | 0 | 1189800 | 5505400 | 0 | 0 | 479380 | 7198300 |
| IYFGKDIPN | ERLIN2; ERLIN2 | O94905; E5F | ERLN2_HUMAN | 2 Oxidation (M) | 0 | 198690 | 0 | 0 | 0 | 0 | 0 | 0 | 0 | 3912200 | 0 | 0 | 461270 | 0 | 0 | 0 | 0 | 550580 |
| IYFGSNIPN | ERLIN1 | O75477 | ERLN1_HUMAN | Unmodified | 0 | 0 | 0 | 0 | 1585200 | 0 | 0 | 0 | 2536300 | 577150 | 0 | 115660 | 0 | 0 | 0 | 0 | 0 | 0 |
| IYFGSNIPN | ERLIN1 | O75477 | ERLN1_HUMAN | Oxidation (M) | 0 | 0 | 0 | 0 | 0 | 0 | 0 | 29920 | 1802600 | 1974000 | 171330 | 0 | 0 | 0 | 0 | 0 | 0 | 0 |
| IYFMAGSSI | HSP90B1; HSP9 | P14625; A0A | ENPL_HUMAN | Unmodified | 74179 | 0 | 445090 | 0 | 0 | 0 | 0 | 0 | 44149000 | 3167000 | 2758900 | 5481600 | 0 | 179010 | 271540 | 0 | 0 | 0 |
| IYFMAGSSI | HSP90B1; HSP9 | P14625; A0A | ENPL_HUMAN | Unmodified | 0 | 0 | 0 | 0 | 0 | 0 | 0 | 0 | 18638000 | 11167000 | 0 | 0 | 0 | 0 | 0 | 0 | 0 | 0 |
| IYFTDSSSK | APMAP; APMAP | Q9HDC9; H( | APMAP_HUMAI | Unmodified | 0 | 0 | 0 | 0 | 0 | 0 | 0 | 0 | 43048000 | 3915900 | 303760 | 79403 | 0 | 0 | 72915 | 179020 | 0 | 0 |
| IYFTDSSSK | APMAP; APMAP | Q9HDC9; H( | APMAP_HUMAI | Unmodified | 0 | 0 | 0 | 0 | 0 | 0 | 119360 | 0 | 9866200 | 2125100 | 0 | 0 | 0 | 0 | 0 | 0 | 0 | 0 |
| IYFTDSSSK | APMAP; APMAP | Q9HDC9; H( | APMAP_HUMAI | Unmodified | 0 | 0 | 0 | 0 | 0 | 0 | 0 | 0 | 4723400 | 2806100 | 50727 | 0 | 0 | 0 | 0 | 0 | 0 | 0 |
| IYGADDIEL | MTHFD1; MTHF | P11586; V9G | C1TC_HUMAN | Unmodified | 0 | 0 | 850120 | 1400100 | 1894300 | 413100 | 1048500 | 369130 | 5799900 | 890200 | 2384800 | 15063000 | 0 | 0 | 1469300 | 983180 | 1089200 | 950890 |
| IYGADDIEL | MTHFD1; MTHF | P11586; V9G | C1TC_HUMAN | Unmodified | 265390 | 0 | 0 | 1314200 | 23621000 | 1981500 | 14103000 | 1042900 | 6744400 | 11136000 | 1415000 | 4335000 | 9664100 | 3989700 | 0 | 356800 | 17848000 | 15454000 |
| IYGAPGVEI | LLGL1; LLGL1 | Q15334; A0/ | L2GL1_HUMAN | Unmodified | 0 | 0 | 0 | 0 | 0 | 0 | 411640 | 0 | 530370 | 0 | 0 | 5757000 | 0 | 0 | 0 | 0 | 0 | 0 |
| IYGFYDECI | PPP1CA; PPP1( | P62136; P62 | PP1A_HUMAN | Unmodified | 645010 | 446170 | 638970 | 0 | 0 | 0 | 0 | 0 | 776830 | 0 | 0 | 4303300 | 0 | 0 | 897020 | 561880 | 0 | 0 |

| | | | | | | | | | | | | | | | | | | | | | | |
|---|---|---|---|---|---|---|---|---|---|---|---|---|---|---|---|---|---|---|---|---|---|---|
| IYGFYDECI | PPP1CA; PPP1C | P62136; P62 | PP1A_HUMAN | Unmodified | 532920 | 571450 | 0 | 0 | 0 | 0 | 0 | 0 | 263530 | 0 | 0 | 2966000 | 0 | 0 | 0 | 750300 | 0 | 0 |
| IYGISFPDP | TARS1 | P26639 | SYTC_HUMAN | Unmodified | 0 | 0 | 565170 | 821600 | 984910 | 441690 | 0 | 0 | 1574800 | 194160 | 3308300 | 8781200 | 646960 | 250810 | 920380 | 738250 | 199990 | 1224000 |
| IYHLPDAES | SEPTIN2; SEPTI | Q15019; H7 | SEPT2_HUMAN | Unmodified | 0 | 28807 | 0 | 0 | 0 | 0 | 0 | 0 | 0 | 0 | 157140 | 1045800 | 0 | 0 | 32952 | 0 | 0 | 0 |
| IYHLPDAES | SEPTIN2; SEPTI | Q15019; H7 | SEPT2_HUMAN | Unmodified | 0 | 0 | 0 | 123930 | 0 | 0 | 0 | 0 | 565650 | 0 | 893690 | 6102600 | 0 | 0 | 344160 | 83792 | 0 | 0 |
| IYHLPDAES | SEPTIN2; SEPTI | Q15019; H7 | SEPT2_HUMAN | Unmodified | 0 | 0 | 0 | 0 | 4062100 | 229970 | 0 | 0 | 0 | 526900 | 0 | 0 | 1536600 | 0 | 0 | 0 | 1165600 | 1005000 |
| IYHLQSKVF | PITPNA; PITPNA | Q00169; I3L | PIPNA_HUMAN | Unmodified | 0 | 0 | 0 | 0 | 0 | 0 | 0 | 0 | 0 | 0 | 0 | 0 | 853620 | 585730 | 0 | 0 | 418610 | 591490 |
| IYHLRQEDS | IGSF3 | O75054 | IGSF3_HUMAN | Unmodified | 0 | 0 | 0 | 0 | 0 | 0 | 49096 | 134420 | 0 | 0 | 225410 | 1639000 | 0 | 0 | 0 | 0 | 0 | 0 |
| IYHPNIDEK | UBE2L5; UBE2L | A0A1B0GUS | UB2L5_HUMAN | Unmodified | 0 | 0 | 0 | 0 | 966030 | 0 | 0 | 0 | 0 | 0 | 0 | 2976700 | 0 | 0 | 0 | 0 | 103100 | 55928 |
| IYHPNINSN | UBE2D3; UBE2I | P61077; P51 | UB2D3_HUMAI | Unmodified | 149620 | 670270 | 0 | 131220 | 0 | 0 | 0 | 0 | 0 | 0 | 0 | 735520 | 0 | 0 | 1857600 | 223340 | 0 | 0 |
| IYHPNVDK | UBE2NL; UBE2I | Q5JXB2; P61 | UE2NL_HUMAI | Unmodified | 0 | 0 | 0 | 0 | 0 | 0 | 0 | 0 | 0 | 0 | 0 | 2080500 | 0 | 0 | 0 | 0 | 0 | 0 |
| IYIDDNNKI | PIP4K2A; PIP4K | P48426; S4F | PI42A_HUMAN | Unmodified | 0 | 0 | 0 | 0 | 0 | 0 | 700980 | 402180 | 0 | 0 | 0 | 0 | 181510 | 320170 | 0 | 0 | 796190 | 560060 |
| IYIDKKYET | SUPT16H | Q9Y5B9 | SP16H_HUMAI | Oxidation (M) | 0 | 0 | 0 | 0 | 0 | 0 | 706400 | 0 | 0 | 0 | 0 | 0 | 0 | 0 | 0 | 0 | 85354 | 0 |
| IYIDSNNNF | CLTC; CLTC | Q00610; A0A | CLH1_HUMAN | Unmodified | 18686000 | 14824000 | 9499000 | 15152000 | 4489000 | 1594700 | 1008300 | 899970 | 3611800 | 539000 | 1291200 | 1.09E+08 | 8703200 | 4721400 | 8323600 | 11571000 | 1280100 | 4390900 |
| IYIGDDNPL | ITGAV | P06756 | ITAV_HUMAN I | Unmodified | 1472700 | 623460 | 1101300 | 1734900 | 4504000 | 1040700 | 421640 | 0 | 2815900 | 1032000 | 315420 | 3953200 | 1000500 | 85861 | 2913000 | 1882400 | 2139900 | 762980 |
| IYIPLPEEAA | VPS4A; N/A | Q9UN37; I3L | VPS4A_HUMAN | Unmodified | 73216 | 0 | 0 | 0 | 0 | 0 | 0 | 0 | 2458200 | 122480 | 1251100 | 4868800 | 0 | 0 | 0 | 0 | 368560 | 0 |
| IYIPLPEPH | VPS4B | O75351 | VPS4B_HUMAN | Unmodified | 695050 | 486410 | 844740 | 1489700 | 356470 | 0 | 0 | 0 | 0 | 418880 | 492370 | 2608500 | 398060 | 329060 | 638210 | 499680 | 145920 | 0 |
| IYISLLK | CTNND1; CTNN | O60716; A0A | CTND1_HUMAI | Unmodified | 0 | 0 | 2801300 | 0 | 4697300 | 622400 | 0 | 12797 | 0 | 0 | 3457800 | 12782000 | 4869200 | 1007300 | 3837900 | 2682300 | 1898300 | 2551800 |
| IYITLTGVH | CACYBP | Q9HB71 | CYBP_HUMAN | Unmodified | 0 | 0 | 0 | 0 | 0 | 0 | 0 | 0 | 0 | 0 | 0 | 2563100 | 2505200 | 0 | 252750 | 0 | 0 | 0 |
| IYIVRGLEL | MYOF; MYOF | Q9NZM1; H0 | MYOF_HUMAN | Unmodified | 0 | 0 | 0 | 0 | 5690100 | 1235300 | 0 | 0 | 0 | 0 | 0 | 0 | 0 | 0 | 0 | 0 | 0 | 0 |
| IYKDGERII | PRKAR2A; PRKA | P13861; H7 | KAP2_HUMAN | Unmodified | 0 | 0 | 0 | 0 | 982600 | 0 | 34666 | 31990 | 0 | 94633 | 0 | 0 | 1303200 | 503050 | 0 | 0 | 114960 | 543600 |
| IYKGPLGC | IGF2R | P11717 | MPRI_HUMAN | Unmodified | 0 | 0 | 0 | 0 | 0 | 0 | 0 | 0 | 845290 | 669750 | 0 | 0 | 0 | 0 | 0 | 0 | 0 | 0 |
| IYKLDDPS | EIF2S3; EIF2S3 | P41091; H7E | IF2G_HUMAN E | Unmodified | 0 | 0 | 0 | 263740 | 0 | 0 | 0 | 0 | 0 | 0 | 0 | 2405000 | 0 | 0 | 0 | 0 | 0 | 0 |
| IYKVPSTEA | GDI2; GDI2; GD | P50395; Q5 | GDIB_HUMAN | Oxidation (M) | 0 | 0 | 0 | 455510 | 0 | 0 | 0 | 0 | 0 | 541800 | 0 | 0 | 661560 | 0 | 727980 | 452800 | 0 | 0 |
| IYLDENYEF | ITM2C; ITM2C; | Q9NQX7; E7 | ITM2C_HUMAN | Unmodified | 0 | 0 | 0 | 0 | 0 | 0 | 0 | 0 | 970270 | 0 | 0 | 0 | 0 | 0 | 0 | 0 | 0 | 0 |
| IYLEGKIDY | SSBP1; SSBP1; | Q04837; A0A | SSBP_HUMAN | Oxidation (M) | 0 | 0 | 0 | 0 | 0 | 0 | 0 | 0 | 1656400 | 600850 | 328890 | 437380 | 220920 | 0 | 0 | 0 | 0 | 0 |
| IYLEGKIDY | SSBP1; SSBP1; | Q04837; A0A | SSBP_HUMAN | Unmodified | 0 | 0 | 0 | 0 | 279570 | 88677 | 79483 | 0 | 3635100 | 691960 | 125290 | 0 | 287900 | 141970 | 73626 | 0 | 0 | 0 |
| IYLPYLHEW | TSG101; TSG10 | Q99816; F5H | TS101_HUMAN | Unmodified | 6363900 | 4180900 | 2297800 | 404370 | 0 | 0 | 0 | 0 | 0 | 0 | 45405 | 2700100 | 0 | 35261 | 15425000 | 4717800 | 0 | 0 |
| IYLYLTK | GARS1; GARS1; | P41250; A0A | GARS_HUMAN | Unmodified | 0 | 0 | 0 | 0 | 0 | 0 | 0 | 0 | 0 | 0 | 0 | 6846300 | 0 | 0 | 0 | 0 | 0 | 0 |
| IYLYTLNDN | FN1; FN1 | P02751; H0Y | FINC_HUMAN I | Unmodified | 0 | 0 | 18429000 | 2131700 | 0 | 264270 | 0 | 0 | 969590 | 973530 | 0 | 0 | 0 | 0 | 427690 | 6759000 | 0 | 0 |
| IYNEIPDINI | PDCD4 | Q53EL6 | PDCD4_HUMA | Unmodified | 0 | 0 | 78774 | 212610 | 0 | 394550 | 0 | 0 | 368450 | 1270300 | 667220 | 2843700 | 0 | 0 | 792500 | 231240 | 0 | 0 |
| IYNKIPGTS | PTPRA | P18433 | PTPRA_HUMAN | Unmodified | 203350 | 106030 | 0 | 376040 | 708960 | 0 | 167360 | 0 | 216290 | 0 | 278450 | 1236800 | 535520 | 271160 | 242010 | 252890 | 279800 | 799950 |
| IYNSIYIGS | SF3B1 | O75533 | SF3B1_HUMAN | Unmodified | 351050 | 0 | 0 | 0 | 0 | 0 | 0 | 0 | 0 | 0 | 0 | 3247800 | 0 | 0 | 0 | 0 | 0 | 0 |
| IYNVIGTLR | FOLH1B; FOLH | Q9HBA9; Q0 | FOH1B_HUMAI | Unmodified | 9124100 | 6607200 | 57814000 | 53176000 | 0 | 0 | 0 | 0 | 0 | 0 | 0 | 0 | 0 | 0 | 46803000 | 43863000 | 0 | 0 |
| IYPDPELEV | TRAF4; TRAF4; | Q9BUZ4; A0 | TRAF4_HUMAN | Unmodified | 103090 | 260750 | 67411 | 244830 | 0 | 0 | 0 | 0 | 0 | 0 | 0 | 52659 | 0 | 0 | 531740 | 106340 | 0 | 0 |
| IYPLPDDPS | MYOF; MYOF | Q9NZM1; H0 | MYOF_HUMAN | Unmodified | 0 | 0 | 0 | 0 | 0 | 512980 | 0 | 0 | 417440 | 45151 | 0 | 0 | 0 | 0 | 0 | 0 | 0 | 0 |
| IYPLPDDPS | MYOF; MYOF | Q9NZM1; H0 | MYOF_HUMAN | Unmodified | 0 | 0 | 0 | 0 | 5069500 | 1091600 | 0 | 0 | 0 | 0 | 0 | 0 | 0 | 0 | 0 | 0 | 0 | 476500 |
| IYPLPHMY | SDHB; SDHB; S | P21912; A0A | SDHB_HUMAN | Oxidation (M) | 0 | 0 | 0 | 0 | 0 | 0 | 0 | 0 | 1193500 | 474320 | 0 | 0 | 0 | 0 | 0 | 0 | 0 | 0 |
| IYPLPHMY | SDHB; SDHB; S | P21912; A0A | SDHB_HUMAN | Unmodified | 0 | 0 | 0 | 0 | 0 | 0 | 0 | 0 | 2540600 | 527480 | 0 | 0 | 0 | 0 | 0 | 0 | 0 | 0 |
| IYPTFLHLH | SSRP1; SSRP1 | Q08945; E9F | SSRP1_HUMAN | Unmodified | 0 | 0 | 0 | 0 | 0 | 0 | 0 | 0 | 0 | 0 | 0 | 1172800 | 0 | 0 | 0 | 0 | 0 | 0 |
| IYQDAYRAI | TFAM; TFAM | Q00059; H7I | TFAM_HUMAN | Unmodified | 0 | 0 | 0 | 0 | 250700 | 0 | 0 | 0 | 837530 | 0 | 0 | 0 | 0 | 0 | 0 | 0 | 0 | 0 |
| IYQGEDLPI | ATL3; ATL3 | Q6DD88; F5 | ATLA3_HUMAN | Unmodified | 0 | 0 | 413740 | 401520 | 3501400 | 168360 | 2466100 | 847450 | 0 | 490030 | 360190 | 296160 | 1854300 | 788630 | 597120 | 321590 | 437610 | 1703500 |
| IYQGEELPI | ATL1; ATL2; ATL | Q8WXF7; Q8 | ATLA1_HUMAN | Unmodified | 0 | 0 | 0 | 0 | 475320 | 0 | 1299000 | 670680 | 0 | 114960 | 430650 | 0 | 184850 | 0 | 0 | 0 | 0 | 441370 |
| IYQIYEGTS | ACADM; ACADN | P11310; Q51 | ACADM_HUMA | Unmodified | 0 | 0 | 0 | 0 | 0 | 0 | 0 | 0 | 4008700 | 624580 | 0 | 0 | 0 | 0 | 0 | 0 | 0 | 0 |
| IYQNIQDG | RAB14 | P61106 | RAB14_HUMAN | Unmodified | 6653000 | 9111200 | 11365000 | 19709000 | 5648500 | 0 | 2328500 | 1366100 | 28849000 | 3240100 | 19127000 | 91354000 | 5013100 | 553110 | 9358500 | 8200200 | 0 | 1426300 |
| IYQRAFGG | SDHA; SDHA; N | P31040; D6F | SDHA_HUMAN | Unmodified | 0 | 0 | 0 | 0 | 0 | 0 | 0 | 0 | 0 | 234300 | 0 | 0 | 0 | 0 | 0 | 0 | 0 | 0 |
| IYQTTTERP | ATG5; ATG5; AT | Q9H1Y0; A0 | ATG5_HUMAN | Unmodified | 101000 | 507250 | 0 | 0 | 0 | 89348 | 202820 | 68577 | 189920 | 0 | 0 | 1440900 | 43994 | 0 | 134350 | 58652 | 1142900 | 235800 |
| IYQYVINK | CTPS1; CTPS1; | P17812; A0A | PYRG1_HUMAI | Unmodified | 258790 | 0 | 0 | 242170 | 865690 | 136500 | 659220 | 374940 | 0 | 1149500 | 597910 | 3050400 | 568550 | 309730 | 0 | 406250 | 895250 | 909040 |
| IYRAEDIPQ | MYOF | Q9NZM1 | MYOF_HUMAN | Oxidation (M) | 0 | 0 | 0 | 0 | 5823000 | 0 | 0 | 0 | 0 | 0 | 0 | 0 | 0 | 0 | 0 | 0 | 0 | 5526800 |
| IYRAEDIPQ | MYOF | Q9NZM1 | MYOF_HUMAN | Unmodified | 0 | 0 | 0 | 0 | 11374000 | 2438600 | 0 | 0 | 0 | 0 | 0 | 0 | 0 | 0 | 0 | 0 | 0 | 678870 |
| IYRQGNLFI | NT5DC2; NT5D | Q9H857; H7 | NT5D2_HUMAI | Unmodified | 0 | 0 | 0 | 0 | 0 | 0 | 0 | 0 | 523320 | 0 | 0 | 0 | 0 | 0 | 0 | 0 | 0 | 0 |
| IYSAERFLIA | TXNRD1; TXNRI | Q16881; A0A | TRXR1_HUMAN | Unmodified | 0 | 0 | 70308 | 0 | 2783600 | 0 | 0 | 0 | 0 | 0 | 0 | 0 | 0 | 0 | 0 | 0 | 482400 | 945770 |
| IYSEVTCFF | MAN2A1 | Q16706 | MA2A1_HUMAI | Unmodified | 0 | 0 | 0 | 0 | 0 | 0 | 0 | 0 | 212980 | 0 | 0 | 303920 | 0 | 0 | 0 | 0 | 0 | 0 |
| IYSFGLGGI | SORT1 | Q99523 | SORT_HUMAN | Unmodified | 0 | 44695 | 0 | 0 | 0 | 0 | 0 | 111620 | 8935500 | 986350 | 0 | 1169200 | 0 | 0 | 0 | 58391 | 0 | 0 |
| IYSHDGTD | UGGT1 | Q9NYU2 | UGGG1_HUMA | Unmodified | 0 | 0 | 0 | 0 | 0 | 0 | 0 | 0 | 14434000 | 0 | 1179900 | 0 | 0 | 0 | 0 | 0 | 0 | 0 |
| IYSISMK | FOLH1 | Q04609 | FOLH1_HUMAI | Oxidation (M) | 0 | 414660 | 0 | 0 | 163090 | 0 | 0 | 82166 | 0 | 78859 | 0 | 378020 | 84396 | 73103 | 4945400 | 2689300 | 0 | 0 |
| IYSISMK | FOLH1 | Q04609 | FOLH1_HUMAI | Unmodified | 0 | 0 | 10773000 | 0 | 130380 | 0 | 268510 | 0 | 0 | 135730 | 0 | 543160 | 0 | 0 | 9294800 | 6848200 | 0 | 0 |
| IYSISMKHP | FOLH1 | Q04609 | FOLH1_HUMAI | Oxidation (M) | 60895 | 0 | 0 | 226120 | 0 | 0 | 38197 | 0 | 0 | 0 | 0 | 0 | 0 | 0 | 40933 | 59519 | 0 | 0 |
| IYSITATNA | ELAPOR2; ELAP | A8MWY0; H7 | ELAP2_HUMAN | Unmodified | 0 | 0 | 0 | 0 | 0 | 0 | 0 | 0 | 4467300 | 392410 | 0 | 0 | 0 | 0 | 0 | 0 | 0 | 0 |
| IYSLSSQPI | PACS1; PACS1; | Q6VY07; E9F | PACS1_HUMAI | Unmodified | 117160 | 0 | 171940 | 0 | 721370 | 198840 | 0 | 62327 | 0 | 321890 | 0 | 0 | 472210 | 562360 | 411230 | 452390 | 162880 | 398010 |
| IYSSEAIK | MYO6; MYO6; M | Q9UM54; A0 | MYO6_HUMAN | Unmodified | 0 | 0 | 6497800 | 1365700 | 515330 | 0 | 0 | 0 | 899850 | 0 | 0 | 1766600 | 3387800 | 902760 | 9689100 | 4466400 | 433450 | 0 |
| IYTFNQSR | UGP2; UGP2; U | Q16851; E7E | UGPA_HUMAN | Unmodified | 0 | 156090 | 0 | 0 | 0 | 0 | 142420 | 0 | 204500 | 0 | 0 | 1896600 | 0 | 0 | 51206 | 0 | 0 | 0 |
| IYTSPTWSA | LGALS3BP | Q08380 | LG3BP_HUMAI | Unmodified | 0 | 0 | 0 | 0 | 0 | 0 | 0 | 0 | 3846100 | 0 | 7041800 | 928590 | 0 | 0 | 0 | 0 | 0 | 0 |
| IYTSPTWSA | LGALS3BP | Q08380 | LG3BP_HUMAI | Unmodified | 0 | 0 | 0 | 0 | 0 | 1574700 | 0 | 0 | 12388000 | 3379100 | 9948400 | 876640 | 4922100 | 4616100 | 0 | 0 | 960970 | 14844000 |
| IYVASVHQI | PUF60; PUF60; | Q9UHX1; A0 | PUF60_HUMAI | Unmodified | 0 | 97225 | 0 | 0 | 100990 | 1316800 | 483040 | 1053800 | 2554300 | 354270 | 944500 | 3805300 | 499110 | 619140 | 0 | 0 | 452180 | 3918500 |
| IYVDDGLIS | PKM; PKM; PKM | P14618; B4D | KPYM_HUMAN | Unmodified | 0 | 0 | 0 | 3087600 | 14812000 | 2005200 | 1964100 | 443120 | 776440 | 1632400 | 2595000 | 4315400 | 6728600 | 4881600 | 2155000 | 874050 | 6025900 | 8531500 |
| IYVDDGLIS | PKM; PKM; PKM | P14618; B4D | KPYM_HUMAN | Unmodified | 0 | 0 | 0 | 0 | 2954700 | 0 | 3499400 | 729360 | 0 | 126240 | 0 | 0 | 443940 | 895360 | 0 | 0 | 13546000 | 2516300 |
| IYVGNLPPI | SRSF1; SRSF1; | Q07955; J3K | SRSF1_HUMAI | Unmodified | 669270 | 606710 | 0 | 0 | 0 | 0 | 0 | 0 | 0 | 0 | 0 | 1413600 | 0 | 0 | 0 | 609650 | 0 | 0 |
| IYVGNLPTI | SRSF9; SRSF9; | Q13242; S4F | SRSF9_HUMAI | Unmodified | 0 | 646140 | 0 | 0 | 0 | 0 | 0 | 0 | 0 | 0 | 0 | 218360 | 0 | 0 | 0 | 0 | 0 | 0 |
| IYVGNLPTI | SRSF9; SRSF9; | Q13242; S4F | SRSF9_HUMAI | Unmodified | 533540 | 404970 | 0 | 123340 | 0 | 0 | 182520 | 151700 | 0 | 354550 | 1278200 | 0 | 0 | 0 | 221880 | 339530 | 257970 | 476410 |
| IYVGNLPTI | SRSF9; SRSF9; | Q13242; S4F | SRSF9_HUMAI | Unmodified | 344280 | 536190 | 0 | 472880 | 0 | 0 | 321850 | 0 | 0 | 0 | 0 | 0 | 0 | 0 | 0 | 0 | 1448400 | 0 |
| IYVSANSG | ACACA; ACACA | Q13085; Q5 | ACACA_HUMAI | Unmodified | 0 | 65310 | 0 | 0 | 0 | 0 | 0 | 0 | 0 | 0 | 0 | 822570 | 0 | 0 | 128450 | 0 | 0 | 0 |
| IYVVDLSNI | CDCP1 | Q9H5V8 | CDCP1_HUMA | Unmodified | 909260 | 428990 | 2938700 | 2256200 | 3430000 | 440340 | 0 | 3629900 | 114410 | 0 | 3652000 | 19473000 | 1768000 | 4725800 | 3933700 | 4013300 | 1520100 | 16353000 |
| IYVVDLSNI | CDCP1 | Q9H5V8 | CDCP1_HUMA | Oxidation (M) | 0 | 0 | 105310 | 1319300 | 1830600 | 1981100 | 0 | 0 | 0 | 521280 | 1481100 | 0 | 1386600 | 329360 | 116970 | 71602 | 5988500 | 6271700 |
| IYVVDLSNI | CDCP1 | Q9H5V8 | CDCP1_HUMA | Unmodified | 0 | 0 | 0 | 245940 | 3158300 | 722260 | 0 | 0 | 0 | 198580 | 0 | 0 | 762930 | 577720 | 50657 | 0 | 3276000 | 7332500 |
| IYVVDVGSI | SELENBP1; SEL | Q13228; A6F | SBP1_HUMAN | Unmodified | 0 | 529390 | 649320 | 0 | 0 | 360110 | 227330 | 0 | 0 | 0 | 0 | 0 | 0 | 0 | 772180 | 705860 | 63974 | 810570 |
| IYVVDVGSI | SELENBP1; SEL | Q13228; A6F | SBP1_HUMAN | Unmodified | 0 | 0 | 0 | 0 | 0 | 0 | 0 | 0 | 0 | 0 | 0 | 0 | 1297100 | 1039400 | 0 | 0 | 204890 | 1976000 |
| IYWGTATT | YARS1; YARS1; | P54577; A0A | SYYC_HUMAN | Oxidation (M) | 0 | 0 | 0 | 0 | 206210 | 0 | 118490 | 0 | 0 | 0 | 0 | 660150 | 226760 | 192960 | 0 | 60626 | 52458 | 0 |
| IYWSDLSQ | LDLR; LDLR; LD | P01130; J3K | LDLR_HUMAN | Unmodified | 139940 | 0 | 0 | 0 | 0 | 0 | 0 | 0 | 2397100 | 68164 | 0 | 0 | 0 | 0 | 0 | 0 | 0 | 0 |
| IYYHVVETG | TM9SF2 | Q99805 | TM9S2_HUMAI | Oxidation (M) | 261480 | 173800 | 294780 | 224450 | 60433 | 0 | 66336 | 0 | 18449000 | 1487300 | 0 | 5168100 | 0 | 0 | 205610 | 258710 | 0 | 0 |
| IYYHVVETG | TM9SF2 | Q99805 | TM9S2_HUMAI | Unmodified | 466930 | 0 | 0 | 337470 | 0 | 0 | 0 | 0 | 29080000 | 1954700 | 1162400 | 2297400 | 0 | 0 | 512770 | 0 | 0 | 0 |
| IYYSEETSS | PLCG1 | P19174 | PLCG1_HUMAI | Unmodified | 0 | 0 | 0 | 0 | 0 | 146660 | 0 | 0 | 0 | 0 | 0 | 2498900 | 0 | 0 | 0 | 0 | 1513500 | 632580 |
| IYYVDHFTF | ITCH; ITCH; ITC | Q96J02; A0A | ITCH_HUMAN I | Unmodified | 5607800 | 6747900 | 4948100 | 2032000 | 0 | 0 | 0 | 0 | 861270 | 0 | 0 | 1001100 | 0 | 0 | 5946200 | 3876700 | 0 | 86954 |
| KAAAEDVN | PFDN4; PFDN4 | Q9NQP4; E9 | PFD4_HUMAN | Unmodified | 0 | 0 | 0 | 0 | 1559500 | 0 | 137950 | 252380 | 983200 | 793780 | 552790 | 4109300 | 1959500 | 857600 | 0 | 0 | 936650 | 1133600 |
| KAAALEFLN | STIP1; STIP1; S | P31948; F5H | STIP1_HUMAN | Unmodified | 0 | 0 | 0 | 0 | 0 | 0 | 0 | 0 | 0 | 0 | 325970 | 2138700 | 303180 | 0 | 0 | 0 | 0 | 0 |
| KAAAPAPEI | EEF1G | P26641 | EF1G_HUMAN | Unmodified | 0 | 0 | 0 | 0 | 1580300 | 462200 | 0 | 0 | 1397500 | 500560 | 1438300 | 8208500 | 0 | 0 | 0 | 0 | 1008500 | 1540600 |
| KAAAPAPEI | EEF1G | P26641 | EF1G_HUMAN | Oxidation (M) | 0 | 0 | 0 | 0 | 411770 | 149540 | 0 | 0 | 0 | 90767 | 954400 | 3503800 | 485150 | 0 | 0 | 0 | 606580 | 0 |
| KAAAQLLQ | SEPTIN11; SEP | Q9NVA2; H0 | SEP11_HUMAN | Unmodified | 0 | 0 | 0 | 0 | 318510 | 0 | 0 | 0 | 0 | 0 | 0 | 250190 | 0 | 0 | 0 | 0 | 183390 | 212030 |
| KAAAQLLQ | SEPTIN11; SEP | Q9NVA2; H0 | SEP11_HUMAN | Unmodified | 0 | 0 | 0 | 0 | 1740000 | 325420 | 156540 | 272940 | 269560 | 132590 | 0 | 910430 | 59297 | 0 | 0 | 0 | 412010 | 440410 |
| KAAATTAQI | VIL1; VIL1 | P09327; B4D | VILI_HUMAN Vi | Unmodified | 0 | 0 | 0 | 0 | 0 | 158150 | 1269300 | 347320 | 625450 | 524510 | 797050 | 3454200 | 0 | 889920 | 0 | 0 | 10059 | 564920 |
| KAACIAEQ | KIF20A | O95235 | KI20A_HUMAN | Unmodified | 0 | 0 | 0 | 465560 | 1816900 | 104300 | 168120 | 0 | 284650 | 288230 | 0 | 0 | 1425100 | 1206600 | 0 | 0 | 806590 | 91459 |
| KAADMGNI | SEL1L | Q9UBV2 | SE1L1_HUMAN | Oxidation (M) | 289050 | 0 | 0 | 240760 | 0 | 0 | 0 | 0 | 1219200 | 103920 | 0 | 0 | 0 | 0 | 133080 | 87895 | 0 | 0 |
| KAAGHPGI | TLN1 | Q9Y490 | TLN1_HUMAN | Unmodified | 0 | 0 | 75421 | 0 | 3757400 | 1414500 | 3139700 | 1127800 | 0 | 291790 | 0 | 0 | 714670 | 1863200 | 23022 | 0 | 1672200 | 281060 |
| KAAGIDEQI | LAP3 | P28838 | AMPL_HUMAN | Unmodified | 0 | 0 | 0 | 0 | 0 | 0 | 0 | 0 | 0 | 0 | 0 | 465330 | 0 | 81608 | 0 | 0 | 0 | 0 |
| KAALEAQN | CELF1; CELF1; | Q92879; G5I | CELF1_HUMAN | Unmodified | 705200 | 0 | 0 | 0 | 0 | 0 | 0 | 0 | 0 | 0 | 0 | 154910 | 0 | 0 | 0 | 0 | 0 | 0 |
| KAAYDIEV | FLOT1; FLOT1; | O75955; A2A | FLOT1_HUMAN | Unmodified | 0 | 0 | 0 | 216890 | 764310 | 0 | 56373 | 288130 | 897710 | 172610 | 0 | 0 | 239440 | 249160 | 0 | 0 | 0 | 0 |
| KACADATL | GNB1; GNB1; G | P62873; F6U | GBB1_HUMAN | Unmodified | 1520500 | 0 | 0 | 2765100 | 1880300 | 2201300 | 748860 | 265020 | 8868200 | 0 | 9538000 | 9141200 | 1057200 | 521920 | 790040 | 1005900 | 2367400 | 2353000 |
| KACADATL | GNB1; GNB1; G | P62873; F6U | GBB1_HUMAN | Oxidation (M) | 0 | 0 | 0 | 0 | 2113900 | 0 | 0 | 0 | 0 | 0 | 0 | 0 | 295450 | 0 | 0 | 0 | 4329900 | 2950600 |
| KACADATL | GNB1; GNB1; G | P62873; F6U | GBB1_HUMAN | Unmodified | 0 | 0 | 0 | 0 | 1660300 | 63291 | 0 | 0 | 0 | 0 | 0 | 802160 | 75196 | 109390 | 0 | 0 | 1857400 | 4496700 |
| KACGDSTL | GNB2; GNB2; G | P62879; C9J | GBB2_HUMAN | Unmodified | 2942900 | 2023200 | 0 | 4356400 | 0 | 905460 | 455600 | 147660 | 1625500 | 0 | 6384500 | 14211000 | 1338500 | 0 | 0 | 2046600 | 1242700 | 0 |
| KADCTITM | SCP2; SCP2 | P22307; H0Y | SCP2_HUMAN | Oxidation (M) | 0 | 0 | 0 | 0 | 0 | 0 | 0 | 0 | 16392000 | 170990 | 0 | 0 | 0 | 0 | 0 | 0 | 0 | 0 |
| KADCTITM | SCP2; SCP2 | P22307; H0Y | SCP2_HUMAN | Unmodified | 0 | 0 | 0 | 0 | 0 | 0 | 0 | 0 | 3843700 | 0 | 0 | 0 | 0 | 0 | 0 | 0 | 0 | 0 |
| KADGVHTV | TMED1 | Q13445 | TMED1_HUMAI | Unmodified | 0 | 0 | 0 | 0 | 0 | 0 | 0 | 0 | 19962000 | 216710 | 155130 | 204590 | 0 | 0 | 0 | 243230 | 0 | 0 |
| KADGVHTV | TMED1 | Q13445 | TMED1_HUMAI | Unmodified | 0 | 0 | 0 | 0 | 0 | 257600 | 0 | 0 | 14093000 | 9201500 | 0 | 0 | 0 | 0 | 0 | 0 | 0 | 0 |
| KADIGVAM | ATP1A1; ATP1A | P05023; P13 | AT1A1_HUMAN | Unmodified | 0 | 1431900 | 10316000 | 9861000 | 2617600 | 0 | 2194300 | 39851 | 23045000 | 2573000 | 7921400 | 75506000 | 12142000 | 4160600 | 13758000 | 11184000 | 1139000 | 283360 |
| KADIGVAM | ATP1A1; ATP1A | P05023; P13 | AT1A1_HUMAN | Oxidation (M) | 732690 | 784020 | 7566700 | 5409800 | 2411500 | 558590 | 905310 | 28253 | 3111900 | 2493500 | 4258600 | 16926000 | 3296200 | 1096300 | 6177200 | 9664300 | 295980 | 612730 |
| KADLNTITE | UTRN | P46939 | UTRN_HUMAN | Unmodified | 0 | 0 | 0 | 506710 | 0 | 0 | 0 | 0 | 0 | 0 | 0 | 0 | 0 | 0 | 0 | 0 | 0 | 0 |
| KADPQEAI | NAPA; NAPA; N | P54920; M0 | SNAA_HUMAN | Unmodified | 327750 | 0 | 0 | 159080 | 267060 | 0 | 0 | 0 | 488750 | 261160 | 0 | 1040100 | 0 | 387360 | 364550 | 120710 | 0 | 390490 |
| KADSVANC | RRBP1; RRBP1; | Q9P2E9; V9 | RRBP1_HUMAN | Unmodified | 0 | 0 | 0 | 0 | 0 | 0 | 217000 | 0 | 0 | 72633 | 28794 | 242730 | 1532900 | 207260 | 0 | 0 | 0 | 91211 |
| KADVGFAM | ATP2B2; ATP2B | Q01814; Q1 | AT2B2_HUMAN | Unmodified | 0 | 0 | 0 | 953070 | 1738200 | 0 | 1606600 | 0 | 2332200 | 727670 | 1296200 | 8259500 | 1205500 | 0 | 2386500 | 0 | 0 | 0 |

| | | | | | | | | | | | | | | | | | | | | | | |
|---|---|---|---|---|---|---|---|---|---|---|---|---|---|---|---|---|---|---|---|---|---|---|
| KADVGFAM | ATP2B2; ATP2B | Q01814; Q1 | AT2B2_HUMAN | Oxidation (M) | 437990 | 357060 | 2751000 | 2618300 | 0 | 0 | 0 | 0 | 0 | 0 | 0 | 14730000 | 599550 | 704760 | 3805800 | 3056100 | 0 | 0 |
| KADVIPVHY | TPP2; TPP2; TPI | P29144; A0A | TPP2_HUMAN | Unmodified | 0 | 480720 | 144720 | 2512300 | 475220 | 0 | 0 | 0 | 719450 | 242860 | 0 | 9055600 | 2356700 | 0 | 130670 | 229360 | 0 | 367610 |
| KAEAAASAI | CHMP2A; CHM | O43633; M0 | CHM2A_HUMA | Unmodified | 0 | 228740 | 0 | 0 | 0 | 0 | 0 | 0 | 1503200 | 0 | 0 | 2427500 | 0 | 0 | 233420 | 776420 | 0 | 0 |
| KAEAAASAI | CHMP2A; CHM | O43633; M0 | CHM2A_HUMA | Unmodified | 6212800 | 5854500 | 2335700 | 9146600 | 34604000 | 6336100 | 0 | 18032 | 31981000 | 12565000 | 4461600 | 14055000 | 11284000 | 5316000 | 5059000 | 1680500 | 1751800 | 3137000 |
| KAEAATEA | DCUN1D3 | Q8IWE4 | DCNL3_HUMA | Unmodified | 0 | 0 | 0 | 0 | 0 | 0 | 0 | 0 | 0 | 0 | 0 | 0 | 0 | 272960 | 0 | 0 | 466500 | 1631500 |
| KAEAFQLY | FLOT1; FLOT1; | O75955; A0 | FLOT1_HUMAN | Oxidation (M) | 75982 | 0 | 243820 | 258540 | 2489500 | 195240 | 87762 | 0 | 2099200 | 311810 | 248020 | 438290 | 1186300 | 179900 | 3257200 | 89850 | 0 | 112730 |
| KAEAGAGS | RPS10; RPS10P | P46783; Q9 | RS10_HUMAN | Unmodified | 4446600 | 3122900 | 2096900 | 1957400 | 237880 | 0 | 0 | 0 | 421720 | 0 | 0 | 7642700 | 0 | 0 | 1935300 | 2368300 | 0 | 0 |
| KAEALEAAI | ANAPC5; ANAP | Q9UJX4; F5 | APC5_HUMAN | Unmodified | 0 | 35456 | 0 | 0 | 0 | 0 | 0 | 0 | 0 | 0 | 1794400 | 1516900 | 0 | 0 | 0 | 0 | 0 | 0 |
| KAECPRAE | PLXNB2 | O15031 | PLXB2_HUMAN | Unmodified | 0 | 0 | 0 | 931680 | 1945700 | 422990 | 0 | 0 | 0 | 0 | 0 | 445980 | 1574000 | 584510 | 0 | 304060 | 1284100 | 1242400 |
| KAEEIANEN | AARS1; AARS1; | P49588; A0A | SYAC_HUMAN | Unmodified | 0 | 1160800 | 0 | 357960 | 7390800 | 1106000 | 0 | 158560 | 591760 | 1322200 | 1219000 | 6796000 | 5222300 | 2806000 | 0 | 1185600 | 583940 | 1698900 |
| KAEEIANEN | AARS1; AARS1; | P49588; A0A | SYAC_HUMAN | Oxidation (M) | 0 | 0 | 0 | 0 | 753730 | 122560 | 222310 | 54694 | 0 | 102570 | 173870 | 752890 | 760220 | 188750 | 0 | 0 | 239830 | 493080 |
| KAEEQVQA | TAX1BP1; TAX1 | Q86VP1; B8 | TAXB1_HUMAN | Unmodified | 0 | 0 | 0 | 0 | 0 | 0 | 259640 | 0 | 1271500 | 0 | 0 | 0 | 164670 | 0 | 0 | 0 | 157060 | 252060 |
| KAEEVQAW | MRPS26 | Q9BYN8 | RT26_HUMAN | Unmodified | 0 | 0 | 0 | 0 | 0 | 0 | 0 | 0 | 0 | 0 | 0 | 0 | 0 | 0 | 0 | 0 | 0 | 0 |
| KAEFLDSIK | N/A | N/A | N/A | Unmodified | 0 | 0 | 0 | 0 | 0 | 16175000 | 0 | 163920 | 0 | 0 | 0 | 0 | 0 | 0 | 0 | 0 | 31935 | 18147 |
| KAEGAATEI | BASP1; BASP1 | P80723; U3 | BASP1_HUMAN | Unmodified | 165200 | 15598 | 0 | 0 | 2891600 | 763650 | 48989 | 126370 | 8913000 | 534960 | 0 | 0 | 0 | 36045 | 0 | 0 | 32048 | 1277800 |
| KAEGAATEI | BASP1 | P80723 | BASP1_HUMAN | Unmodified | 0 | 337030 | 0 | 0 | 12877000 | 4295600 | 275680 | 279420 | 26336000 | 1754200 | 0 | 0 | 243250 | 0 | 0 | 0 | 656900 | 6565300 |
| KAEGVALTI | PACSIN1; PACS | Q9BY11; F6 | PACN1_HUMA | Unmodified | 1617500 | 1628400 | 0 | 0 | 0 | 0 | 0 | 0 | 0 | 0 | 0 | 0 | 0 | 0 | 1322800 | 925620 | 0 | 0 |
| KAEIGIAMG | ATP2A1; ATP2A | O14983; P1 | AT2A1_HUMAN | Unmodified | 0 | 0 | 0 | 0 | 475610 | 0 | 973320 | 0 | 0 | 109350 | 319900 | 0 | 0 | 0 | 0 | 0 | 0 | 0 |
| KAELLDNEI | CACYBP | Q9HB71 | CYBP_HUMAN | Unmodified | 0 | 0 | 0 | 814040 | 0 | 302390 | 0 | 0 | 94378 | 47469 | 0 | 484200 | 0 | 67194 | 0 | 0 | 88517 | 96234 |
| KAELLDNEI | CACYBP | Q9HB71 | CYBP_HUMAN | Unmodified | 656880 | 0 | 202930 | 444760 | 10303000 | 1804200 | 7537800 | 5197100 | 12389000 | 7364600 | 21812000 | 69634000 | 18745000 | 13354000 | 0 | 1328200 | 3389900 | 2514600 |
| KAELVKLQ | IQGAP1; IQGAP | P46940; H0 | IQGA1_HUMAN | Unmodified | 0 | 0 | 0 | 0 | 1714800 | 0 | 0 | 0 | 0 | 0 | 0 | 0 | 0 | 0 | 0 | 0 | 958150 | 0 |
| KAEMGRFS | SLC38A2; SLC3 | Q96QD8; F8 | S38A2_HUMAN | Oxidation (M) | 0 | 0 | 0 | 0 | 11223000 | 1385700 | 4420200 | 201870 | 0 | 724390 | 0 | 0 | 4077400 | 3460900 | 0 | 0 | 6511100 | 15291000 |
| KAEMGRFS | SLC38A2; SLC3 | Q96QD8; F8 | S38A2_HUMAN | Unmodified | 0 | 0 | 0 | 0 | 11246000 | 0 | 1747100 | 0 | 0 | 1581700 | 0 | 0 | 1558600 | 1570300 | 0 | 0 | 2445100 | 21405000 |
| KAEQEVEQ | FMNL1; FMNL1 | O95466; K7 | FMNL1_HUMA | Unmodified | 0 | 0 | 0 | 0 | 2944900 | 790490 | 0 | 0 | 0 | 0 | 0 | 0 | 0 | 0 | 0 | 0 | 0 | 0 |
| KAERELSE | EZR; EZR | P15311; E7E | EZRI_HUMAN E | Unmodified | 1659100 | 2636800 | 344790 | 1830600 | 6815300 | 1982900 | 0 | 346300 | 10226000 | 2254900 | 1616400 | 4798100 | 5524600 | 809830 | 0 | 27902 | 658290 | 188550 |
| KAESFAQEI | SHTN1 | A0MZ66 | SHOT1_HUMA | Unmodified | 0 | 0 | 0 | 140020 | 185720 | 0 | 0 | 0 | 0 | 0 | 0 | 0 | 775350 | 0 | 0 | 81148 | 0 | 98553 |
| KAETLPEVE | PPFIA2; PPFIA1 | O75334; Q1 | LIPA2_HUMAN | Unmodified | 0 | 0 | 0 | 82013 | 0 | 0 | 0 | 263900 | 984510 | 415160 | 0 | 0 | 0 | 0 | 778230 | 315710 | 0 | 0 |
| KAEVEGKD | HNRNPU; HNR | Q00839; A0 | HNRPU_HUMA | Unmodified | 0 | 230240 | 0 | 138850 | 3768000 | 631360 | 9791200 | 5131700 | 399650 | 177450 | 152440 | 86897 | 6552700 | 2728200 | 0 | 0 | 493850 | 1656900 |
| KAEVRLQFI | G6PD | P11413 | G6PD_HUMAN | Unmodified | 0 | 0 | 0 | 0 | 2189000 | 0 | 0 | 0 | 0 | 0 | 0 | 0 | 0 | 0 | 0 | 0 | 0 | 0 |
| KAEYPFSSE | ATP2C1; ATP2C | P98194; B4E | AT2C1_HUMAN | Unmodified | 93839 | 0 | 0 | 35351 | 0 | 0 | 0 | 0 | 1628800 | 141780 | 0 | 0 | 0 | 0 | 161370 | 0 | 0 | 0 |
| KAFCEPGN | YARS1; YARS1 | P54577; A0A | SYYC_HUMAN | Unmodified | 0 | 1115200 | 0 | 47878 | 38957 | 0 | 741520 | 166190 | 916780 | 227180 | 0 | 344690 | 1717200 | 1698700 | 19826 | 97470 | 0 | 733970 |
| KAFEESLST | ECE1; ECE1; EC | P42892; B4D | ECE1_HUMAN | Unmodified | 0 | 0 | 0 | 0 | 150950 | 0 | 0 | 0 | 1591100 | 683610 | 0 | 0 | 0 | 0 | 0 | 0 | 162890 | 0 |
| KAFSGYLG | ESD; ESD; ESD; | P10768; X6R | ESTD_HUMAN | Unmodified | 0 | 0 | 218170 | 0 | 0 | 0 | 315910 | 337450 | 103960 | 236730 | 219180 | 1338200 | 256610 | 286770 | 0 | 0 | 277220 | 0 |
| KAFTELQA | PFDN1; PFDN1 | O60925; E5F | PFD1_HUMAN | Unmodified | 0 | 0 | 0 | 0 | 684460 | 0 | 0 | 0 | 68304 | 117890 | 33531 | 565190 | 0 | 0 | 0 | 0 | 313060 | 170560 |
| KAGGANYD | COTL1; COTL1 | Q14019; H3 | COTL1_HUMAN | Unmodified | 0 | 0 | 0 | 0 | 3969500 | 0 | 0 | 0 | 0 | 457470 | 0 | 0 | 0 | 0 | 0 | 0 | 10697000 | 49917 |
| KAGKLDPH | MYH9 | P35579 | MYH9_HUMAN | Unmodified | 559240 | 118870 | 374190 | 526430 | 293700 | 342870 | 0 | 0 | 954480 | 309290 | 828030 | 5172100 | 0 | 0 | 882980 | 734600 | 68878 | 0 |
| KAGNFYVP | RPL7; RPL7; RP | P18124; C9J | RL7_HUMAN 6 | Unmodified | 0 | 0 | 0 | 0 | 0 | 0 | 0 | 0 | 0 | 0 | 0 | 784420 | 0 | 209050 | 0 | 0 | 0 | 0 |
| KAGPLGGS | NECTIN1 | Q15223 | NECT1_HUMA | Unmodified | 0 | 0 | 0 | 0 | 0 | 0 | 0 | 0 | 0 | 0 | 957700 | 1195300 | 0 | 0 | 0 | 0 | 0 | 0 |
| KAGPLGGS | NECTIN1 | Q15223 | NECT1_HUMA | Unmodified | 277730 | 0 | 0 | 0 | 368650 | 0 | 0 | 0 | 521270 | 0 | 0 | 6147200 | 0 | 0 | 0 | 250280 | 0 | 0 |
| KAGQVFLE | CNP | P09543 | CN37_HUMAN | Unmodified | 0 | 0 | 350610 | 867200 | 667220 | 0 | 0 | 0 | 0 | 0 | 813970 | 5759200 | 771740 | 0 | 0 | 0 | 0 | 547960 |
| KAGTQIENI | ACTN4; ACTN4 | O43707; K7 | ACTN4_HUMA | Unmodified | 396370 | 275030 | 595310 | 1758200 | 551270 | 774360 | 80508 | 0 | 1434500 | 270870 | 466690 | 6656800 | 302670 | 156520 | 1004400 | 1478300 | 353700 | 0 |
| KAGTQIENI | ACTN1; ACTN1 | P12814; G3 | ACTN1_HUMA | Unmodified | 122960 | 1474200 | 70767 | 3103200 | 351190 | 377820 | 117900 | 0 | 162940 | 0 | 394800 | 3169500 | 0 | 0 | 372870 | 0 | 0 | 182610 |
| KAGVFSDL | AOC1; AOC1; A | P19801; C9J | AOC1_HUMAN | Unmodified | 0 | 0 | 0 | 160050 | 10502000 | 3246900 | 271210 | 230400 | 622640 | 316550 | 0 | 0 | 332640 | 2635100 | 0 | 0 | 6016200 | 10023000 |
| KAHAAIREN | RPL5; RPL5; RP | P46777; A0A | RL5_HUMAN 6 | Unmodified | 0 | 760660 | 0 | 0 | 1789300 | 385280 | 407520 | 511750 | 0 | 351790 | 195430 | 2081000 | 616280 | 0 | 0 | 0 | 0 | 0 |
| KAHIRQDLI | CARMIL1; CARM | Q5VZK9; A0 | CARL1_HUMAN | Unmodified | 0 | 0 | 0 | 0 | 688100 | 0 | 0 | 0 | 0 | 0 | 0 | 107340 | 72129 | 0 | 0 | 0 | 124850 | 231690 |
| KAHLMEIQ | RPL3; RPL3; RP | P39023; B5 | RL3_HUMAN 6 | Oxidation (M) | 241690 | 233630 | 0 | 52945 | 0 | 0 | 0 | 0 | 0 | 0 | 46943 | 1866800 | 0 | 0 | 158690 | 137950 | 0 | 0 |
| KAHLMEIQ | RPL3; RPL3; RP | P39023; B5 | RL3_HUMAN 6 | Unmodified | 0 | 150270 | 0 | 0 | 0 | 0 | 0 | 0 | 65459 | 0 | 0 | 4545500 | 0 | 0 | 89442 | 0 | 0 | 0 |
| KAHLTNQY | PABPC4; PABP | Q13310; B1 | PABP4_HUMAN | Unmodified | 13704000 | 1485800 | 2248600 | 8449500 | 0 | 0 | 0 | 0 | 0 | 0 | 0 | 0 | 100940 | 0 | 2510200 | 1968600 | 0 | 0 |
| KAHLTNQY | PABPC4; PABP | Q13310; B1 | PABP4_HUMAN | Oxidation (M) | 1273800 | 430470 | 384200 | 687090 | 0 | 0 | 0 | 0 | 0 | 0 | 0 | 0 | 0 | 0 | 357420 | 452210 | 0 | 0 |
| KAHQDIHT | GOLIM4; GOLI | O00461; F8 | GOLI4_HUMAN | Unmodified | 0 | 0 | 0 | 0 | 1693000 | 0 | 0 | 65966 | 7107200 | 848340 | 270720 | 0 | 0 | 0 | 0 | 0 | 0 | 0 |
| KAHVLAAS | CTNNA1; CTNN | P35221; E5R | CTNA1_HUMA | Unmodified | 694630 | 757300 | 0 | 4252200 | 11839000 | 1863400 | 0 | 35462 | 520490 | 197610 | 2903800 | 7653300 | 6032000 | 3124800 | 6526200 | 3235100 | 1979700 | 1083200 |
| KAHVLAAS | CTNNA1; CTNN | P35221; E5R | CTNA1_HUMA | Unmodified | 0 | 0 | 0 | 5438100 | 5821800 | 0 | 0 | 0 | 0 | 0 | 0 | 0 | 7584700 | 0 | 0 | 994930 | 0 | 0 |
| KAIADWYN | RO60; RO60 | P10155; H0 | RO60_HUMAN | Unmodified | 1966600 | 2258100 | 0 | 279670 | 0 | 0 | 0 | 0 | 253850 | 49842 | 0 | 772350 | 0 | 0 | 717040 | 484920 | 0 | 0 |
| KAIDTSGMI | EFR3A; EFR3A | Q14156; A0 | EFR3A_HUMAN | Unmodified | 0 | 0 | 0 | 106440 | 726170 | 0 | 86583 | 68774 | 225070 | 0 | 0 | 67376 | 188050 | 145490 | 0 | 0 | 0 | 570350 |
| KAIEDEGGI | SAFB; SAFB | Q15424; K7 | SAFB1_HUMAN | Unmodified | 0 | 0 | 0 | 0 | 0 | 0 | 0 | 0 | 0 | 0 | 0 | 732000 | 0 | 0 | 0 | 0 | 0 | 0 |
| KAIIIFVPVP | RPS7; RPS7 | P62081; B5 | RS7_HUMAN 4 | Unmodified | 1246300 | 838530 | 0 | 0 | 0 | 0 | 0 | 0 | 0 | 0 | 0 | 10965000 | 0 | 0 | 0 | 385680 | 0 | 0 |
| KAKSETVLT | ITGA3 | P26006 | ITA3_HUMAN I | Unmodified | 0 | 0 | 0 | 0 | 758530 | 133710 | 119190 | 0 | 0 | 0 | 0 | 0 | 114080 | 0 | 0 | 0 | 724330 | 1190400 |
| KALAAAGY | H1-3; H1-4; H1 | P16402; P10 | H13_HUMAN H | Unmodified | 5619700 | 4626800 | 0 | 0 | 4119800 | 708500 | 2513200 | 1.42E+08 | 0 | 98679 | 0 | 165960 | 235880 | 0 | 0 | 0 | 0 | 830230 |
| KALAAAGY | H1-3; H1-4; H1 | P16402; P10 | H13_HUMAN H | Unmodified | 0 | 0 | 0 | 0 | 141400 | 0 | 164610 | 8685500 | 0 | 0 | 0 | 0 | 0 | 0 | 0 | 0 | 0 | 86515 |
| KALAAGGY | H1-5 | P16401 | H15_HUMAN H | Unmodified | 8786000 | 4801700 | 0 | 0 | 2037600 | 0 | 340720 | 17286000 | 371490 | 61171 | 1356200 | 3339300 | 0 | 0 | 0 | 0 | 0 | 31479 |
| KALAAGGY | H1-5 | P16401 | H15_HUMAN H | Unmodified | 15343 | 17209 | 0 | 0 | 0 | 0 | 0 | 526120 | 0 | 0 | 0 | 0 | 0 | 16185 | 0 | 0 | 0 | 0 |
| KALDQFVN | SMC3 | Q9UQE7 | SMC3_HUMAN | Unmodified | 0 | 0 | 0 | 0 | 512900 | 101860 | 158510 | 59521 | 0 | 43337 | 0 | 0 | 150290 | 44982 | 0 | 0 | 0 | 0 |
| KALIEAEK | ERLIN2; ERLIN2 | O94905; E5F | ERLN2_HUMAN | Unmodified | 0 | 0 | 0 | 0 | 0 | 0 | 0 | 0 | 561060 | 123990 | 0 | 0 | 0 | 0 | 0 | 0 | 0 | 0 |
| KALIEVLQP | WARS1 | P23381 | SYWC_HUMAN | Unmodified | 0 | 0 | 0 | 0 | 1188200 | 0 | 0 | 0 | 744610 | 393350 | 0 | 391810 | 0 | 0 | 0 | 0 | 0 | 779470 |
| KALLIQGSN | COL2A1 | P02458 | CO2A1_HUMA | Unmodified | 0 | 0 | 0 | 0 | 0 | 0 | 0 | 0 | 0 | 0 | 1700000 | 0 | 0 | 0 | 0 | 0 | 0 | 0 |
| KALTGHLEI | ANXA1; ANXA1; | P04083; Q5 | ANXA1_HUMA | Unmodified | 0 | 0 | 0 | 0 | 3278700 | 469550 | 56300 | 299750 | 0 | 66300 | 13670000 | 47731000 | 0 | 0 | 0 | 0 | 266640 | 1397300 |
| KALYGTTSE | MME; MME; MM | P08473; A0A | NEP_HUMAN N | Unmodified | 0 | 0 | 0 | 0 | 0 | 0 | 0 | 0 | 0 | 0 | 0 | 242580 | 0 | 0 | 0 | 2112400 | 0 | 0 |
| KALYGTTSE | MME; MME; MM | P08473; A0A | NEP_HUMAN N | Unmodified | 0 | 0 | 0 | 1105200 | 0 | 0 | 0 | 0 | 181540 | 0 | 0 | 0 | 177850 | 0 | 329020 | 737770 | 0 | 31623 |
| KALYPSSV | MYO1B; MYO1B | O43795; H7 | MYO1B_HUMA | Unmodified | 0 | 0 | 0 | 0 | 716680 | 109120 | 0 | 0 | 205870 | 0 | 137030 | 0 | 0 | 0 | 0 | 0 | 100270 | 438960 |
| KALYPSSV | MYO1B; MYO1B | O43795; H7 | MYO1B_HUMA | Unmodified | 0 | 0 | 0 | 0 | 17757000 | 0 | 0 | 0 | 0 | 0 | 0 | 0 | 5479200 | 0 | 0 | 0 | 4314100 | 0 |
| KAMEGAGT | ANXA6; ANXA6; | P08133; E7E | ANXA6_HUMA | Oxidation (M) | 0 | 0 | 0 | 0 | 58065 | 0 | 0 | 0 | 0 | 101680 | 0 | 0 | 0 | 0 | 0 | 0 | 83856 | 92645 |
| KANELLLST | SH2D4A | Q9H788 | SH24A_HUMA | Oxidation (M) | 33406 | 0 | 0 | 0 | 165060 | 29149 | 0 | 0 | 0 | 0 | 0 | 0 | 0 | 0 | 143590 | 0 | 0 | 0 |
| KANGVLEA | SLC44A2; SLC4 | Q8IWA5; K7 | CTL2_HUMAN | Unmodified | 790990 | 1322200 | 566150 | 0 | 0 | 0 | 0 | 27841 | 0 | 0 | 0 | 1532600 | 0 | 158290 | 0 | 666780 | 0 | 0 |
| KANIRNMS | EEF2 | P13639 | EF2_HUMAN E | Unmodified | 0 | 0 | 0 | 0 | 1814900 | 3253200 | 2783100 | 1017400 | 0 | 1341800 | 0 | 0 | 4183200 | 328930 | 0 | 0 | 687810 | 8636700 |
| KANLQIDQ | MYH9 | P35579 | MYH9_HUMAN | Unmodified | 413270 | 442760 | 842260 | 1939200 | 2493400 | 341570 | 0 | 21519 | 3376600 | 518910 | 401160 | 6895600 | 758110 | 316490 | 2025200 | 1286700 | 356290 | 418310 |
| KANLQIDQ | MYH9 | P35579 | MYH9_HUMAN | Unmodified | 0 | 0 | 0 | 592030 | 13555000 | 3365700 | 0 | 0 | 242640 | 468300 | 583580 | 0 | 3147300 | 0 | 0 | 0 | 2236900 | 3066200 |
| KAPDFVFY | RDX; EZR; MSN | P35241; P15 | RADI_HUMAN | Unmodified | 2586500 | 3348000 | 3048100 | 6641800 | 44572000 | 24052000 | 2022800 | 1280100 | 20979000 | 9497400 | 5951700 | 24916000 | 8989900 | 12129000 | 5854100 | 5455600 | 4774600 | 4737200 |
| KAPNEFLTS | CNK3/IPCEF1; | G9CGD6; Q6 | CNIPF_HUMAN | Unmodified | 0 | 0 | 0 | 0 | 0 | 0 | 0 | 0 | 0 | 0 | 0 | 189170 | 0 | 0 | 0 | 0 | 0 | 0 |
| KAPPDGWI | BUD31; BUD31 | P41223; F8 | BUD31_HUMA | Unmodified | 0 | 0 | 0 | 345180 | 37157 | 256070 | 76788 | 0 | 0 | 0 | 556170 | 0 | 0 | 0 | 0 | 0 | 0 | 0 |
| KAPSDLYQ | CAPN1; CAPN1 | P07384; E9P | CAN1_HUMAN | Unmodified | 0 | 0 | 69514 | 0 | 0 | 0 | 77362 | 50385 | 149100 | 32825 | 0 | 2429500 | 0 | 0 | 0 | 0 | 0 | 0 |
| KAQCPIVEI | RPS5; RPS5; RP | P46782; M0 | RS5_HUMAN 4 | Unmodified | 167050 | 260090 | 46676 | 0 | 0 | 0 | 0 | 0 | 0 | 0 | 0 | 1193100 | 273350 | 315990 | 69464 | 25420 | 0 | 0 |
| KAQEEAPQ | EPB41L1; EPB4 | Q9H4G0; Q4 | E41L1_HUMAN | Unmodified | 0 | 0 | 939250 | 451480 | 0 | 0 | 0 | 0 | 0 | 0 | 2065900 | 2879600 | 0 | 0 | 299980 | 1391900 | 0 | 0 |
| KAQEGGG | FAM98A; FAM9 | Q8NCA5; E9 | FA98A_HUMAN | Unmodified | 0 | 0 | 0 | 0 | 225130 | 0 | 0 | 0 | 0 | 0 | 0 | 2150900 | 406430 | 293310 | 0 | 263760 | 0 | 0 |
| KAQGSTSS | SIRPA | P78324 | SHPS1_HUMA | Unmodified | 0 | 0 | 0 | 0 | 1808300 | 263130 | 111050 | 169270 | 0 | 0 | 0 | 0 | 121780 | 0 | 0 | 0 | 1135800 | 1091700 |
| KAQQELEE | MSN | P26038 | MOES_HUMAN | Unmodified | 220070 | 21397 | 0 | 313660 | 4035500 | 0 | 92534 | 0 | 0 | 0 | 0 | 0 | 0 | 0 | 167340 | 162460 | 106070 | 341380 |
| KAQQELEE | MSN | P26038 | MOES_HUMAN | Unmodified | 296710 | 0 | 432250 | 0 | 13879000 | 3328000 | 153070 | 36757 | 0 | 0 | 0 | 97484 | 135620 | 184600 | 0 | 821170 | 1345400 | 4206300 |
| KARAYGPG | FLNA; FLNA; FL | P21333; A0A | FLNA_HUMAN | Unmodified | 0 | 0 | 0 | 0 | 803910 | 94735 | 0 | 331710 | 0 | 112830 | 0 | 0 | 0 | 0 | 0 | 0 | 677070 | 0 |
| KARESYVE | CLTC; CLTC; CI | Q00610; J3K | CLH1_HUMAN | Unmodified | 0 | 1625700 | 0 | 0 | 1021800 | 0 | 1200300 | 96702 | 0 | 410350 | 0 | 1165400 | 1437200 | 0 | 0 | 571240 | 457410 | 736960 |
| KASAAQLR | UBE2M | P61081 | UBC12_HUMA | Unmodified | 0 | 0 | 0 | 0 | 0 | 0 | 254520 | 0 | 0 | 367350 | 0 | 0 | 0 | 0 | 0 | 0 | 792980 | 0 |
| KASELPVSE | ATP2C1; ATP2C | P98194; B4E | AT2C1_HUMAN | Unmodified | 0 | 0 | 0 | 0 | 0 | 0 | 0 | 0 | 1175000 | 0 | 0 | 328130 | 0 | 0 | 0 | 0 | 0 | 0 |
| KASGPPVS | H1-3; H1-4; H1 | P16402; P10 | H13_HUMAN H | Unmodified | 12292000 | 16412000 | 0 | 0 | 3184800 | 1130300 | 8150700 | 2.44E+08 | 370730 | 316640 | 1634700 | 1003600 | 386470 | 0 | 0 | 0 | 0 | 2473000 |
| KASGPPVS | H1-3; H1-4; H1 | P16402; P10 | H13_HUMAN H | Unmodified | 0 | 0 | 0 | 0 | 0 | 0 | 411280 | 8864400 | 0 | 0 | 104470 | 0 | 0 | 0 | 0 | 0 | 0 | 0 |
| KASSEGGT | ARPC1B; ARPC | O15143; C9 | ARC1B_HUMA | Unmodified | 176800 | 429980 | 0 | 68472 | 4606000 | 16422 | 1343900 | 1353000 | 0 | 2761000 | 0 | 5332800 | 967890 | 0 | 1604400 | 275300 | 9975100 | 10093000 |
| KASVVTLP | DYNC1H1; DYN | Q14204; A0 | DYHC1_HUMA | Unmodified | 0 | 0 | 0 | 0 | 0 | 0 | 0 | 0 | 0 | 93918 | 0 | 1862000 | 0 | 0 | 257020 | 173240 | 0 | 0 |
| KATAAVIFL | LYPLA1; LYPLA | O75608; B4 | LYPA1_HUMAN | Unmodified | 0 | 0 | 0 | 135570 | 0 | 0 | 0 | 0 | 0 | 0 | 0 | 177010 | 0 | 0 | 192240 | 0 | 0 | 0 |
| KATDGVTL | PACSIN2; PACS | Q9UNF0; A0 | PACN2_HUMA | Unmodified | 492740 | 0 | 2958600 | 0 | 0 | 0 | 0 | 0 | 0 | 0 | 0 | 0 | 0 | 0 | 2652100 | 632690 | 0 | 0 |
| KATEGKTSI | STXBP3 | O00186 | STXB3_HUMAN | Unmodified | 0 | 0 | 0 | 0 | 1013100 | 0 | 425270 | 84622 | 0 | 0 | 0 | 0 | 394350 | 741290 | 0 | 0 | 528950 | 898180 |
| KATGPPVSI | H1-5 | P16401 | H15_HUMAN H | Unmodified | 9038700 | 13404000 | 0 | 0 | 595750 | 0 | 506750 | 26405000 | 0 | 173820 | 612360 | 541860 | 252440 | 110580 | 0 | 0 | 56930 | 0 |
| KATGPPVSI | H1-5 | P16401 | H15_HUMAN H | Unmodified | 0 | 0 | 0 | 0 | 0 | 0 | 0 | 1583200 | 0 | 0 | 0 | 0 | 0 | 0 | 0 | 0 | 0 | 0 |
| KATPYTFP | GALNT1; GALN | Q10472; K7 | GALT1_HUMAN | Unmodified | 0 | 0 | 35947 | 0 | 0 | 0 | 0 | 0 | 1566600 | 82560 | 124080 | 1580600 | 0 | 0 | 0 | 0 | 0 | 0 |
| KATVNLLGI | REEP5 | Q00765 | REEP5_HUMAN | Unmodified | 0 | 0 | 185590 | 0 | 968430 | 0 | 0 | 0 | 0 | 0 | 0 | 0 | 0 | 0 | 0 | 0 | 0 | 0 |
| KATVNLLGI | REEP5 | Q00765 | REEP5_HUMAN | Unmodified | 0 | 0 | 0 | 0 | 0 | 0 | 1642300 | 0 | 0 | 0 | 402710 | 0 | 0 | 0 | 0 | 0 | 0 | 0 |
| KAVAGQQ | TNIP1; TNIP1; T | Q15025; A0 | TNIP1_HUMAN | Unmodified | 0 | 0 | 0 | 0 | 0 | 0 | 0 | 0 | 0 | 0 | 0 | 0 | 0 | 0 | 0 | 0 | 530540 | 538570 |
| KAVDVFFPI | CLTC; CLTC | Q00610; A0 | CLH1_HUMAN | Oxidation (M) | 4155200 | 3103200 | 0 | 0 | 142310 | 0 | 0 | 0 | 0 | 498890 | 0 | 3964300 | 0 | 283210 | 0 | 0 | 0 | 0 |
| KAVEEAERI | USP8; USP8; U | P40818; H0 | UBP8_HUMAN | Unmodified | 0 | 0 | 0 | 0 | 2192200 | 556110 | 0 | 19968 | 376720 | 84305 | 0 | 110170 | 96808 | 37337 | 0 | 0 | 41431 | 127090 |
| KAVEEPLN | CD47; CD47 | Q08722; H7 | CD47_HUMAN | Unmodified | 361390 | 0 | 1121600 | 0 | 0 | 166850 | 289910 | 0 | 1416200 | 1181700 | 2482300 | 4714100 | 0 | 0 | 0 | 276810 | 170860 | 0 |
| KAVEEPLN | CD47; CD47 | Q08722; H7 | CD47_HUMAN | Unmodified | 0 | 0 | 0 | 0 | 0 | 0 | 0 | 0 | 0 | 377660 | 0 | 0 | 0 | 0 | 0 | 0 | 0 | 0 |
| KAVIFCLSA | DSTN; DSTN | P60981; F6R | DEST_HUMAN | Unmodified | 0 | 0 | 0 | 0 | 0 | 0 | 0 | 0 | 0 | 0 | 0 | 1230000 | 0 | 0 | 0 | 0 | 0 | 0 |
| KAVIFCLSA | DSTN; DSTN | P60981; F6R | DEST_HUMAN | Unmodified | 407810 | 263150 | 0 | 284460 | 1230100 | 0 | 330940 | 178840 | 217430 | 244460 | 0 | 2772000 | 1750300 | 0 | 257660 | 0 | 713640 | 792680 |

| | | | | | | | | | | | | | | | | | | | | | | |
|---|---|---|---|---|---|---|---|---|---|---|---|---|---|---|---|---|---|---|---|---|---|---|
| KAVIQHFQ | APP; APP; APP; | P05067; H7 | A4_HUMAN Am | Unmodified | 0 | 0 | 0 | 0 | 0 | 0 | 0 | 98029 | 1413600 | 216290 | 0 | 0 | 0 | 0 | 0 | 0 | 0 | 0 |
| KAVLFCLS | CFL2; CFL2 | Q9Y281; F8 | COF2_HUMAN | Unmodified | 0 | 0 | 0 | 0 | 532420 | 0 | 0 | 0 | 45053 | 0 | 0 | 0 | 0 | 0 | 722540 | 0 | 0 | 0 |
| KAVLFCLS | CFL1; CFL1; CF | P23528; G3 | COF1_HUMAN | Unmodified | 880750 | 900200 | 769400 | 750640 | 10340000 | 1981700 | 526970 | 309570 | 2490900 | 591980 | 495770 | 7465300 | 3709100 | 2563100 | 714420 | 801190 | 1964500 | 3892600 |
| KAVLFCLS | CFL1; CFL1; CF | P23528; G3 | COF1_HUMAN | Unmodified | 0 | 1805600 | 1681200 | 2863700 | 13475000 | 3150700 | 1272400 | 602040 | 1708600 | 808510 | 1986200 | 13103000 | 9229400 | 0 | 1752600 | 1609200 | 394430 | 381110 |
| KAVLVALK | VARS1; VARS1 | P26640; A0A | SYVC_HUMAN | Unmodified | 0 | 0 | 0 | 0 | 0 | 0 | 0 | 0 | 0 | 0 | 0 | 435760 | 0 | 0 | 0 | 27585 | 0 | 0 |
| KAVQFGTG | KIFBP; KIFBP; K | Q96EK5; A0 | KBP_HUMAN K | Unmodified | 0 | 0 | 0 | 0 | 0 | 0 | 0 | 0 | 253380 | 0 | 0 | 196780 | 149240 | 100890 | 0 | 0 | 85722 | 85367 |
| KAYVRLAP | RPL23A; RPL23 | P62750; K7E | RL23A_HUMAN | Unmodified | 0 | 0 | 0 | 0 | 429120 | 0 | 180490 | 65532 | 0 | 0 | 0 | 0 | 434670 | 220790 | 0 | 0 | 146860 | 227420 |
| KCAADLGL | HINT1 | P49773 | HINT1_HUMAN | Unmodified | 0 | 0 | 0 | 84878 | 950450 | 172340 | 779020 | 0 | 152450 | 441790 | 152700 | 719200 | 2932900 | 1325200 | 0 | 57514 | 1235100 | 690430 |
| KCAGCTNF | FHL2 | Q14192 | FHL2_HUMAN | Unmodified | 189720 | 0 | 0 | 0 | 208720 | 285290 | 0 | 0 | 0 | 0 | 0 | 0 | 0 | 0 | 0 | 0 | 518510 | 289080 |
| KCATVTEN | SPINT2; SPINT2 | O43291; K7E | SPIT2_HUMAN | Unmodified | 0 | 0 | 0 | 47865 | 0 | 0 | 0 | 0 | 4591800 | 429200 | 0 | 0 | 0 | 0 | 0 | 0 | 0 | 0 |
| KCCILTEEG | PHLDA1; PHLD | Q8WV24; R4 | PHLA1_HUMAN | Unmodified | 0 | 0 | 0 | 0 | 0 | 0 | 0 | 0 | 0 | 0 | 0 | 0 | 0 | 0 | 0 | 0 | 3571500 | 2320900 |
| KCEPIIMTV | CORO1C; COR | Q9ULV4; Q6 | COR1C_HUMA | Unmodified | 0 | 0 | 0 | 0 | 743290 | 0 | 0 | 0 | 0 | 0 | 153520 | 0 | 0 | 0 | 0 | 0 | 0 | 567780 |
| KCEVLCSQ | TM9SF4; TM9SF | Q92544; A0 | TM9S4_HUMAN | Unmodified | 0 | 0 | 0 | 0 | 0 | 0 | 0 | 0 | 31420000 | 2798100 | 1152100 | 0 | 0 | 0 | 0 | 0 | 0 | 0 |
| KCFIEEIPD | TMED9 | Q9BVK6 | TMED9_HUMA | Unmodified | 0 | 0 | 0 | 0 | 0 | 0 | 603360 | 836740 | 2671000 | 0 | 0 | 268140 | 0 | 0 | 0 | 142110 | 0 | 0 |
| KCFIEEIPD | TMED9 | Q9BVK6 | TMED9_HUMA | Oxidation (M) | 0 | 0 | 0 | 0 | 648080 | 0 | 601700 | 0 | 2025200 | 16199000 | 972060 | 0 | 2316200 | 1996900 | 0 | 0 | 0 | 1530400 |
| KCLELFSEL | HSP90AB1 | P08238 | HS90B_HUMAN | Unmodified | 0 | 0 | 0 | 0 | 0 | 0 | 0 | 0 | 0 | 0 | 0 | 1366900 | 0 | 593960 | 0 | 0 | 0 | 0 |
| KCLELFTEL | HSP90AA1 | P07900 | HS90A_HUMAN | Unmodified | 0 | 0 | 0 | 0 | 0 | 0 | 0 | 0 | 274270 | 109220 | 279260 | 1979300 | 161710 | 89650 | 0 | 0 | 0 | 0 |
| KCLELFTEL | HSP90AA1 | P07900 | HS90A_HUMAN | Unmodified | 0 | 0 | 0 | 0 | 2293400 | 512870 | 0 | 0 | 0 | 160280 | 0 | 0 | 0 | 0 | 0 | 0 | 1652000 | 1208700 |
| KCPFTGNV | RPS11; RPS11 | P62280; M0 | RS11_HUMAN | Unmodified | 1702900 | 641100 | 1185200 | 668870 | 0 | 0 | 0 | 131450 | 437310 | 0 | 1324700 | 31403000 | 0 | 0 | 1277400 | 895030 | 157990 | 142970 |
| KCQDVIVA | ZW10 | O43264 | ZW10_HUMAN | Oxidation (M) | 0 | 0 | 0 | 0 | 0 | 0 | 0 | 0 | 0 | 0 | 0 | 0 | 7713600 | 0 | 0 | 0 | 0 | 0 |
| KCTVSVET | MYO1D; MYO1D | O94832; K7E | MYO1D_HUMA | Unmodified | 0 | 0 | 0 | 0 | 2799300 | 242030 | 0 | 0 | 0 | 554870 | 0 | 0 | 4998100 | 3692000 | 0 | 0 | 3160600 | 16568000 |
| KDDESNLV | KLC1; KLC1; KL | Q07866; G3 | KLC1_HUMAN | Unmodified | 0 | 0 | 0 | 0 | 72404 | 0 | 330220 | 0 | 0 | 0 | 38846 | 669690 | 0 | 0 | 0 | 0 | 0 | 169580 |
| KDDEVQV | RPL26; RPL26L | P61254; Q9 | RL26_HUMAN | Unmodified | 430210 | 368370 | 0 | 0 | 0 | 0 | 0 | 0 | 0 | 0 | 316950 | 3312700 | 0 | 0 | 0 | 0 | 0 | 0 |
| KDDPLTNL | ACTN1; ACTN1 | P12814; G3 | ACTN1_HUMA | Unmodified | 0 | 0 | 132450 | 366570 | 0 | 0 | 0 | 0 | 0 | 0 | 0 | 2193300 | 0 | 0 | 272970 | 244010 | 206540 | 0 |
| KDDPTLLS | PCMT1; PCMT1 | P22061; F6S | PIMT_HUMAN | Unmodified | 0 | 0 | 0 | 0 | 0 | 0 | 62370 | 0 | 107040 | 47534 | 46501 | 765140 | 0 | 0 | 0 | 0 | 0 | 0 |
| KDDPVTNL | ACTN4; ACTN4 | O43707; F5 | ACTN4_HUMA | Unmodified | 404140 | 218210 | 0 | 0 | 0 | 0 | 0 | 0 | 2403700 | 0 | 0 | 4944600 | 0 | 0 | 839160 | 1118900 | 0 | 202360 |
| KDDRDASF | BCAM; BCAM; B | P50895; A0A | BCAM_HUMAN | Unmodified | 1334300 | 747680 | 0 | 0 | 0 | 0 | 0 | 0 | 257780 | 0 | 0 | 0 | 0 | 0 | 74445 | 0 | 0 | 0 |
| KDEDIARLL | CCDC50 | Q8IVM0 | CCD50_HUMA | Unmodified | 163140 | 0 | 0 | 1939000 | 243660 | 0 | 0 | 71685 | 0 | 0 | 0 | 0 | 420380 | 0 | 0 | 147850 | 0 | 0 |
| KDEDLFRE | BANF1 | O75531 | BAF_HUMAN B | Unmodified | 0 | 2179900 | 0 | 0 | 2021600 | 348490 | 0 | 49535 | 2365100 | 0 | 1756000 | 3589600 | 0 | 0 | 0 | 0 | 156530 | 979690 |
| KDEELSSAI | TSG101; TSG10 | Q99816; F5 | TS101_HUMAN | Unmodified | 1252500 | 514750 | 837140 | 0 | 0 | 0 | 0 | 0 | 56987 | 0 | 0 | 146110 | 0 | 0 | 948290 | 891390 | 0 | 0 |
| KDEESGG | COPG1; COPG | Q9Y678; D6 | COPG1_HUMA | Unmodified | 0 | 0 | 0 | 0 | 0 | 0 | 0 | 0 | 0 | 0 | 43835 | 488910 | 0 | 0 | 73242 | 0 | 0 | 0 |
| KDEGEAWI | PAAF1 | Q9BRP4 | PAAF1_HUMAN | Unmodified | 0 | 0 | 0 | 0 | 327370 | 0 | 0 | 406360 | 451280 | 0 | 0 | 0 | 97636 | 348300 | 0 | 0 | 0 | 1491400 |
| KDEKTDTL | XRCC5 | P13010 | XRCC5_HUMA | Unmodified | 0 | 0 | 0 | 0 | 0 | 0 | 0 | 0 | 0 | 0 | 304800 | 1858000 | 0 | 0 | 0 | 0 | 0 | 0 |
| KDFDELNV | SIL1; SIL1 | Q9H173; D6 | SIL1_HUMAN | Unmodified | 0 | 384780 | 328210 | 433990 | 923400 | 6454600 | 0 | 210950 | 0 | 0 | 645190 | 0 | 0 | 1397300 | 750380 | 0 | 0 | 5683300 |
| KDFIATQG | PTPRJ; PTPRJ; P | Q12913; A0 | PTPRJ_HUMAN | Unmodified | 0 | 0 | 0 | 0 | 0 | 0 | 0 | 0 | 0 | 0 | 1402300 | 1130000 | 0 | 0 | 0 | 0 | 0 | 0 |
| KDGNASG | EEF1A1P5; EEF | Q5VTE0; P6 | EF1A3_HUMAN | Unmodified | 2376600 | 709100 | 1834200 | 729540 | 0 | 0 | 0 | 0 | 418600 | 0 | 394980 | 1955100 | 93499 | 0 | 0 | 1075200 | 0 | 0 |
| KDGQLIYT | PSEN1; PSEN1; | P49768; G3 | PSN1_HUMAN | Unmodified | 0 | 0 | 0 | 0 | 0 | 0 | 0 | 0 | 0 | 0 | 0 | 479290 | 0 | 0 | 0 | 0 | 0 | 0 |
| KDHRDNL | FASN; FASN | P49327; A0A | FAS_HUMAN F | Unmodified | 0 | 0 | 0 | 0 | 0 | 0 | 0 | 0 | 290580 | 0 | 0 | 2132200 | 0 | 0 | 244450 | 117430 | 0 | 0 |
| KDHVLDL | GALNT6; GALN | Q8NCL4; S4 | GALT6_HUMAN | Oxidation (M) | 0 | 0 | 0 | 0 | 465540 | 0 | 0 | 0 | 0 | 0 | 0 | 0 | 0 | 1254100 | 0 | 0 | 325400 | 2021900 |
| KDITNQEE | STEAP1 | Q9UHE8 | STEA1_HUMAN | Unmodified | 546810 | 88289 | 317490 | 1352600 | 0 | 0 | 0 | 0 | 0 | 0 | 0 | 0 | 1422200 | 675830 | 215170 | 274810 | 0 | 0 |
| KDLHDAN | SLC16A1; SLC1 | P53985; Q5 | MOT1_HUMAN | Unmodified | 0 | 0 | 429680 | 441680 | 604480 | 162480 | 30901 | 0 | 0 | 165420 | 418290 | 1059200 | 327650 | 79868 | 460220 | 358110 | 0 | 0 |
| KDLYANTV | ACTG1; ACTB; | P63261; P60 | ACTG_HUMAN | Oxidation (M) | 7320700 | 8755700 | 4665900 | 5880400 | 1368200 | 0 | 0 | 211980 | 3193100 | 1015600 | 7690000 | 34794000 | 558210 | 0 | 10203000 | 12695000 | 662540 | 0 |
| KDLYANTV | ACTG1; ACTB; | P63261; P60 | ACTG_HUMAN | Unmodified | 1068900 | 1621200 | 0 | 0 | 972680 | 0 | 449340 | 517520 | 2803100 | 307990 | 1124800 | 4300600 | 0 | 0 | 0 | 0 | 287960 | 0 |
| KDLYANTV | ACTG1; ACTB; | P63261; P60 | ACTG_HUMAN | 2 Oxidation (M) | 0 | 0 | 0 | 0 | 1291600 | 0 | 336080 | 0 | 0 | 0 | 0 | 0 | 873810 | 0 | 0 | 0 | 502320 | 0 |
| KDLYANTV | ACTG1; ACTB; | P63261; P60 | ACTG_HUMAN | Oxidation (M) | 0 | 0 | 0 | 0 | 3086200 | 1875100 | 0 | 0 | 0 | 0 | 0 | 0 | 883760 | 0 | 177950 | 0 | 2597200 | 929380 |
| KDLYANTV | ACTG1; ACTB; | P63261; P60 | ACTG_HUMAN | Unmodified | 0 | 0 | 0 | 0 | 2484600 | 0 | 0 | 0 | 0 | 0 | 437870 | 0 | 337130 | 0 | 0 | 0 | 796170 | 0 |
| KDNDFIYH | PDCD6IP | Q8WUM4 | PDC6I_HUMAN | Unmodified | 9172200 | 5143800 | 943090 | 702630 | 0 | 0 | 0 | 0 | 1008600 | 131430 | 377180 | 502400 | 597990 | 90119 | 1431600 | 3829300 | 0 | 0 |
| KDPELWGS | CLTC; CLTC | Q00610; A0 | CLH1_HUMAN | Unmodified | 1256900 | 0 | 0 | 0 | 0 | 0 | 0 | 0 | 0 | 0 | 0 | 1647000 | 0 | 0 | 0 | 388670 | 0 | 0 |
| KDPELWGS | CLTC; CLTC | Q00610; A0 | CLH1_HUMAN | Unmodified | 0 | 1067300 | 0 | 0 | 0 | 0 | 0 | 0 | 0 | 0 | 0 | 2750400 | 0 | 0 | 1102200 | 0 | 0 | 0 |
| KDPTGMD | SPCS2; SPCS2; | Q15005; E9 | SPCS2_HUMA | Oxidation (M) | 0 | 0 | 0 | 0 | 0 | 0 | 0 | 0 | 716710 | 350390 | 0 | 473130 | 0 | 0 | 0 | 0 | 0 | 0 |
| KDPTGMD | SPCS2; SPCS2; | Q15005; E9 | SPCS2_HUMA | Unmodified | 0 | 0 | 0 | 737420 | 0 | 15523 | 0 | 0 | 3693500 | 287000 | 152840 | 0 | 0 | 0 | 0 | 0 | 0 | 0 |
| KDQGELER | MYH9 | P35579 | MYH9_HUMAN | Unmodified | 0 | 0 | 0 | 0 | 1481900 | 273200 | 503330 | 0 | 0 | 839540 | 502960 | 0 | 0 | 488910 | 0 | 0 | 480190 | 0 |
| KDRVTDAL | HSPD1; HSPD1 | P10809; A0A | CH60_HUMAN | Unmodified | 83718 | 163190 | 0 | 160260 | 0 | 0 | 0 | 0 | 8674700 | 531230 | 114600 | 0 | 0 | 190440 | 0 | 0 | 0 | 0 |
| KDSETGEN | HSPA9; HSPA9; | P38646; A0A | GRP75_HUMA | Unmodified | 0 | 0 | 0 | 0 | 0 | 0 | 0 | 0 | 1699300 | 682050 | 0 | 0 | 0 | 0 | 0 | 0 | 0 | 0 |
| KDSQICEL | MANF | P55145 | MANF_HUMAN | Unmodified | 0 | 0 | 0 | 0 | 0 | 0 | 0 | 0 | 295560 | 316470 | 82094 | 0 | 0 | 0 | 0 | 0 | 0 | 0 |
| KDSSQDLL | NAALADL2 | Q58DX5 | NADL2_HUMA | Unmodified | 0 | 892680 | 0 | 234430 | 0 | 0 | 0 | 0 | 0 | 0 | 406950 | 0 | 813010 | 0 | 406370 | 195640 | 0 | 0 |
| KDVIELTDE | PDIA6 | Q15084 | PDIA6_HUMAN | Unmodified | 0 | 0 | 0 | 0 | 0 | 0 | 0 | 0 | 4102700 | 0 | 2669500 | 0 | 0 | 0 | 0 | 0 | 0 | 0 |
| KDVLETFT | CD9; CD9; CD9 | P21926; A0A | CD9_HUMAN C | Unmodified | 44624000 | 53081000 | 39001000 | 1.01E+08 | 0 | 568570 | 1179300 | 0 | 26864000 | 13254000 | 0 | 18036000 | 5249300 | 0 | 84588000 | 48667000 | 0 | 1586600 |
| KDWDDDQ | CCT8; CCT8 | P50990; H7 | TCPQ_HUMAN | Unmodified | 0 | 0 | 0 | 0 | 0 | 0 | 0 | 0 | 88248 | 0 | 152300 | 695240 | 0 | 0 | 0 | 0 | 0 | 0 |
| KDYEEIGPS | ACTR3; ACTR3 | P61158; B4 | ARP3_HUMAN | Unmodified | 96297 | 52174 | 74604 | 127160 | 0 | 0 | 0 | 0 | 0 | 0 | 160030 | 798380 | 0 | 0 | 121470 | 107430 | 0 | 0 |
| KEAAENSL | YWHAE; YWHA | P62258; K7E | 1433E_HUMAN | Unmodified | 0 | 0 | 0 | 11635 | 0 | 0 | 0 | 130700 | 97171 | 44237 | 0 | 1605000 | 0 | 301280 | 57636 | 79069 | 0 | 0 |
| KEAESSPF | HSP90B1; HSP | P14625; A0A | ENPL_HUMAN | Unmodified | 0 | 0 | 69243 | 0 | 0 | 0 | 0 | 18561 | 11859000 | 1327200 | 1545700 | 2216600 | 0 | 299960 | 584730 | 26410 | 0 | 130830 |
| KEAESSPF | HSP90B1; HSP | P14625; A0A | ENPL_HUMAN | Unmodified | 0 | 0 | 0 | 0 | 0 | 72805 | 0 | 0 | 1032600 | 2008600 | 438640 | 0 | 0 | 0 | 0 | 0 | 0 | 0 |
| KEAPAPPK | RPL23A | P62750 | RL23A_HUMAN | Unmodified | 0 | 7698 | 0 | 0 | 0 | 0 | 0 | 0 | 15350 | 0 | 55737 | 311460 | 23316 | 51831 | 0 | 12999 | 0 | 0 |
| KEAPPMEK | RPS15; RPS15; | P62841; K7E | RS15_HUMAN | Unmodified | 0 | 0 | 0 | 0 | 0 | 0 | 0 | 0 | 0 | 0 | 195500 | 893210 | 0 | 0 | 0 | 0 | 0 | 0 |
| KEASDPQP | CAPZA1 | P52907 | CAZA1_HUMA | Unmodified | 0 | 0 | 0 | 0 | 0 | 0 | 76211 | 79026 | 110480 | 133730 | 293680 | 0 | 0 | 86569 | 0 | 0 | 0 | 0 |
| KEAVTFDQ | PLAA; PLAA; PL | Q9Y263; H0 | PLAP_HUMAN | Unmodified | 0 | 0 | 0 | 0 | 63554 | 0 | 0 | 35461 | 93669 | 60822 | 0 | 999700 | 0 | 95594 | 0 | 0 | 0 | 0 |
| KEDEVEEW | EZR; EZR | P15311; E7E | EZRI_HUMAN E | Unmodified | 0 | 0 | 0 | 0 | 1809000 | 120030 | 0 | 0 | 47011 | 0 | 0 | 0 | 389090 | 83795 | 0 | 0 | 0 | 81236 |
| KEDLVFIFW | CFL1; CFL2; CF | P23528; Q9 | COF1_HUMAN | Unmodified | 3063100 | 5677000 | 965320 | 912040 | 3184200 | 0 | 463460 | 0 | 1563200 | 562170 | 42060 | 5858300 | 2722500 | 659420 | 5230700 | 2829600 | 1163900 | 1111400 |
| KEDLYLKPI | EEF2 | P13639 | EF2_HUMAN E | Unmodified | 0 | 0 | 538250 | 0 | 570400 | 104840 | 19094 | 11120 | 161960 | 63793 | 518060 | 13637000 | 0 | 0 | 0 | 0 | 178620 | 67072 |
| KEDWNEID | GALNT2 | Q10471 | GALT2_HUMAN | Unmodified | 0 | 0 | 0 | 0 | 0 | 0 | 0 | 0 | 13468000 | 877720 | 0 | 0 | 0 | 0 | 0 | 0 | 0 | 0 |
| KEEDSHTT | PLPP1 | O14494 | PLPP1_HUMAN | Unmodified | 0 | 0 | 10949000 | 10949000 | 0 | 0 | 0 | 0 | 0 | 0 | 0 | 6285200 | 0 | 0 | 10049000 | 5533100 | 0 | 0 |
| KEEEEEEE | FAM177A1; FA | Q8N128; G3 | F177A_HUMAN | Unmodified | 0 | 0 | 923370 | 317850 | 0 | 0 | 0 | 0 | 0 | 0 | 0 | 149960 | 0 | 0 | 1195000 | 486070 | 0 | 0 |
| KEEEEEEE | MLEC | Q14165 | MLEC_HUMAN | Unmodified | 0 | 0 | 96830 | 0 | 0 | 0 | 0 | 0 | 2012100 | 151980 | 155430 | 0 | 0 | 0 | 107130 | 42941 | 0 | 0 |
| KEEEEEEE | MLEC | Q14165 | MLEC_HUMAN | Unmodified | 0 | 0 | 0 | 0 | 0 | 0 | 49430 | 0 | 2900300 | 0 | 852310 | 172640 | 0 | 0 | 0 | 0 | 0 | 0 |
| KEEEKEEL | CPE | P16870 | CBPE_HUMAN | Unmodified | 0 | 0 | 0 | 0 | 0 | 0 | 0 | 0 | 671370 | 175950 | 565460 | 0 | 688840 | 0 | 0 | 0 | 0 | 0 |
| KEEELQAA | MYH9 | P35579 | MYH9_HUMAN | Unmodified | 0 | 0 | 0 | 0 | 488340 | 0 | 0 | 0 | 2962800 | 0 | 0 | 0 | 0 | 0 | 3645900 | 0 | 0 | 0 |
| KEEELQGA | MYH10 | P35580 | MYH10_HUMA | Unmodified | 0 | 0 | 1176200 | 172440 | 0 | 447340 | 0 | 0 | 0 | 0 | 460020 | 5167300 | 0 | 0 | 0 | 444570 | 0 | 0 |
| KEEENASV | PRKDC | P78527 | PRKDC_HUMA | Unmodified | 0 | 0 | 142720 | 0 | 0 | 0 | 0 | 0 | 0 | 0 | 0 | 3206900 | 0 | 0 | 0 | 0 | 0 | 0 |
| KEEGWWF | NOSTRIN; NOS | A0A0G2JPW | A0A0G2JPW3_ | Unmodified | 1125500 | 782900 | 2434400 | 450010 | 0 | 0 | 0 | 0 | 894510 | 0 | 690970 | 8891400 | 0 | 0 | 3333800 | 1541500 | 0 | 0 |
| KEEGWWF | NOSTRIN; NOS | A0A0G2JPW | A0A0G2JPW3_ | Unmodified | 1035400 | 588900 | 1267200 | 245840 | 54547 | 0 | 0 | 0 | 0 | 62936 | 0 | 176610 | 0 | 0 | 2041600 | 897870 | 0 | 0 |
| KEELMFFL | DSTN; DSTN | P60981; F6R | DEST_HUMAN | Oxidation (M) | 0 | 138140 | 0 | 0 | 0 | 0 | 30899 | 0 | 0 | 0 | 0 | 0 | 99937 | 0 | 113010 | 0 | 0 | 0 |
| KEELTLEGI | EIF4A1; EIF4A1; | P60842; J3K | IF4A1_HUMAN | Unmodified | 326870 | 327000 | 311180 | 65769 | 115480 | 0 | 0 | 0 | 0 | 0 | 0 | 3959600 | 0 | 0 | 726560 | 543710 | 0 | 0 |
| KEENLADW | EPRS1 | P07814 | SYEP_HUMAN | Unmodified | 0 | 0 | 0 | 0 | 0 | 0 | 0 | 0 | 657710 | 0 | 0 | 4225300 | 0 | 0 | 255850 | 90592 | 0 | 0 |
| KEESEESDI | RPLP1; RPLP2; | P05386; P05 | RLA1_HUMAN | Oxidation (M) | 0 | 0 | 0 | 0 | 0 | 0 | 0 | 17277 | 0 | 0 | 1547900 | 7026300 | 0 | 0 | 0 | 20534 | 0 | 0 |
| KEFLEPDS | NUCB2 | H0YD18 | H0YD18_HUM | Unmodified | 0 | 0 | 0 | 191980 | 1425100 | 0 | 204610 | 83783 | 0 | 0 | 0 | 395500 | 2832700 | 2674100 | 0 | 132550 | 0 | 158300 |
| KEFSPFGSI | PABPC4; PABP | Q13310; B1 | PABP4_HUMAN | Unmodified | 922120 | 608650 | 0 | 0 | 0 | 0 | 0 | 0 | 0 | 0 | 0 | 0 | 0 | 0 | 232690 | 213730 | 99188 | 0 |
| KEFSPFGTI | PABPC1; PABP | P11940; A0A | PABP1_HUMAN | Unmodified | 23610000 | 13386000 | 1816900 | 3950900 | 0 | 0 | 0 | 0 | 0 | 0 | 0 | 0 | 0 | 0 | 0 | 5204700 | 0 | 0 |
| KEGGGDS | MARCKSL1 | P49006 | MRP_HUMAN | Unmodified | 0 | 0 | 0 | 0 | 0 | 0 | 0 | 0 | 3752700 | 1657100 | 17187000 | 6796900 | 2161300 | 1153900 | 0 | 0 | 1144000 | 2550800 |
| KEGGLGPL | PRDX2; PRDX2 | P32119; A6 | PRDX2_HUMA | Unmodified | 1709700 | 1100400 | 899940 | 2621800 | 261100 | 293960 | 965130 | 0 | 15319000 | 4566800 | 32832000 | 62960000 | 1010100 | 178860 | 2438200 | 1994900 | 830910 | 624150 |
| KEGGLGPL | PRDX2; PRDX2 | P32119; A6 | PRDX2_HUMA | Unmodified | 2300600 | 1166900 | 530380 | 1810600 | 0 | 0 | 0 | 0 | 8216800 | 316280 | 6409000 | 56212000 | 264080 | 0 | 1578700 | 1172100 | 0 | 0 |
| KEGNASG | EEF1A2; EEF1A | Q05639; A0 | EF1A2_HUMAN | Unmodified | 0 | 0 | 935370 | 255210 | 0 | 0 | 0 | 0 | 0 | 0 | 0 | 0 | 0 | 0 | 694960 | 400660 | 0 | 0 |
| KEGPYDVV | PARK7; PARK7 | Q99497; K7E | PARK7_HUMAN | Unmodified | 0 | 0 | 0 | 0 | 0 | 0 | 0 | 0 | 763540 | 0 | 0 | 1472100 | 0 | 0 | 0 | 0 | 0 | 0 |
| KEGSYSSL | JMJD1C; JMJD1 | Q15652; A0 | JHD2C_HUMA | Oxidation (M) | 0 | 10361000 | 0 | 0 | 0 | 0 | 0 | 0 | 0 | 0 | 0 | 0 | 0 | 0 | 0 | 0 | 314640 | 0 |
| KEGYDFHS | ACTR1A; ACTR1 | P61163; A0A | ACTZ_HUMAN | Unmodified | 0 | 0 | 0 | 0 | 0 | 0 | 0 | 0 | 0 | 0 | 0 | 572020 | 0 | 0 | 0 | 0 | 0 | 0 |
| KEIADYLAA | IST1; IST1; IST1 | P53990; H3 | IST1_HUMAN I | Unmodified | 0 | 162560 | 0 | 610150 | 0 | 0 | 41224 | 0 | 1346100 | 0 | 436630 | 6623500 | 122650 | 0 | 0 | 336910 | 0 | 0 |
| KEIEMHSIN | KDSR; KDSR; K | Q06136; K7 | KDSR_HUMAN | Unmodified | 0 | 0 | 0 | 0 | 0 | 0 | 0 | 0 | 539450 | 0 | 0 | 0 | 0 | 0 | 0 | 0 | 0 | 0 |
| KELGGLEG | CNPY3 | Q9BT09 | CNPY3_HUMA | Unmodified | 0 | 0 | 0 | 0 | 0 | 0 | 0 | 25096 | 4810400 | 2908600 | 1793000 | 0 | 0 | 0 | 0 | 0 | 0 | 345860 |
| KELNYFAK | PGK1 | P00558 | PGK1_HUMAN | Unmodified | 0 | 0 | 0 | 0 | 0 | 0 | 0 | 0 | 203550 | 0 | 0 | 0 | 315980 | 204580 | 0 | 0 | 857470 | 443030 |
| KELQAAGK | GPI; GPI | P06744; A0A | G6PI_HUMAN | Unmodified | 0 | 0 | 0 | 154930 | 0 | 0 | 0 | 0 | 461240 | 226100 | 514720 | 1565800 | 0 | 0 | 0 | 24483 | 0 | 144970 |
| KEMSGDV | ANXA6; ANXA6; | P08133; E7E | ANXA6_HUMA | Oxidation (M) | 0 | 0 | 0 | 0 | 756900 | 0 | 829190 | 0 | 963500 | 0 | 0 | 0 | 486320 | 26670 | 0 | 0 | 341980 | 1493000 |
| KENALVQM | PTBP1; PTBP1; | P26599; A0A | PTBP1_HUMAN | 2 Oxidation (M) | 0 | 0 | 0 | 0 | 1190500 | 0 | 0 | 0 | 761720 | 137840 | 0 | 234670 | 0 | 0 | 0 | 0 | 0 | 0 |
| KENKEYAS | ANKRD26 | E7ESJ3 | E7ESJ3_HUMA | Unmodified | 0 | 0 | 0 | 0 | 2482300 | 0 | 0 | 0 | 0 | 524230 | 0 | 0 | 3956200 | 1347800 | 0 | 0 | 1142000 | 6588300 |
| KENPLQFK | RDX; EZR; RDX; | P35241; P15 | RADI_HUMAN | Unmodified | 934760 | 0 | 0 | 1031200 | 817720 | 0 | 108180 | 72887 | 3486500 | 1026300 | 649840 | 7018200 | 106350 | 0 | 1029000 | 1538400 | 606800 | 0 |
| KEPAVLELE | CAP1 | Q01518 | CAP1_HUMAN | Unmodified | 0 | 0 | 0 | 0 | 0 | 0 | 0 | 0 | 1637200 | 149980 | 1705400 | 6007500 | 0 | 0 | 0 | 0 | 0 | 0 |
| KEPELFQT | EHD1; EHD1; E | Q9H4M9; C9 | EHD1_HUMAN | Unmodified | 78285 | 0 | 243290 | 79493 | 0 | 0 | 0 | 0 | 0 | 0 | 0 | 1317100 | 0 | 0 | 181750 | 263250 | 0 | 0 |
| KEPELFQT | EHD1; EHD1; E | Q9H4M9; C9 | EHD1_HUMAN | Unmodified | 0 | 0 | 0 | 0 | 723250 | 87562 | 0 | 0 | 0 | 96064 | 0 | 0 | 154970 | 43296 | 0 | 0 | 312780 | 606960 |
| KEPFELEAF | HSPA4L; HSPA | O95757; E9 | HS74L_HUMAN | Unmodified | 0 | 0 | 0 | 0 | 0 | 0 | 0 | 0 | 0 | 0 | 0 | 1103000 | 0 | 0 | 0 | 0 | 44169 | 0 |
| KEPFFHGH | CSNK2A1; CSN | P68400; A0A | CSK21_HUMAN | Unmodified | 0 | 0 | 211000 | 0 | 0 | 514870 | 0 | 0 | 0 | 0 | 0 | 3063000 | 18292 | 1377800 | 285750 | 0 | 0 | 0 |

| | | | | | | | | | | | | | | | | | | | | | | |
|---|---|---|---|---|---|---|---|---|---|---|---|---|---|---|---|---|---|---|---|---|---|---|
| KEPVYLSG | UGGT1 | Q9NYU2 | UGGG1_HUMA | Unmodified | 0 | 0 | 0 | 0 | 0 | 0 | 0 | 0 | 3321600 | 224100 | 0 | 0 | 0 | 0 | 0 | 0 | 0 | 414230 |
| KEQTADGV | NUDT5; NUDT5 | Q9UKK9; H0 | NUDT5_HUMA | Unmodified | 0 | 0 | 0 | 0 | 354800 | 0 | 0 | 44605 | 1108700 | 467190 | 2303800 | 2216200 | 136580 | 0 | 0 | 0 | 0 | 0 |
| KESEAVEW | MSN | P26038 | MOES_HUMAN | Unmodified | 69258 | 73724 | 189330 | 375080 | 11340000 | 2476600 | 571120 | 409320 | 961600 | 232020 | 79414 | 0 | 1168300 | 2480400 | 263440 | 211120 | 1973500 | 8110900 |
| KESKGPIVF | SLC12A2; SLC1 | P55011; G3 | S12A2_HUMAN | Unmodified | 0 | 0 | 0 | 0 | 325210 | 0 | 0 | 0 | 0 | 0 | 0 | 0 | 0 | 0 | 0 | 0 | 112300 | 2644600 |
| KESYSIYVY | H2BC19P; H2B | Q6DRA6; P0 | H2B2D_HUMA | Unmodified | 21964000 | 14263000 | 0 | 0 | 0 | 0 | 296280 | 1145500 | 65581 | 0 | 0 | 1.17E+08 | 0 | 0 | 0 | 0 | 64245 | 0 |
| KESYSVYV | H2BC4; H2BC1 | P62807; O6 | H2B1C_HUMA | Unmodified | 66788000 | 41654000 | 662720 | 0 | 0 | 0 | 881890 | 5665200 | 362640 | 0 | 140620 | 6.43E+08 | 0 | 0 | 0 | 265190 | 0 | 0 |
| KEVEVVEII | MVP; MVP; MVP | Q14764; H3 | MVP_HUMAN | Unmodified | 371310 | 154220 | 0 | 0 | 0 | 0 | 0 | 0 | 0 | 0 | 0 | 2032300 | 0 | 0 | 0 | 0 | 0 | 0 |
| KEVEVVEII | MVP; MVP; MVP | Q14764; H3 | MVP_HUMAN | Unmodified | 524780 | 0 | 0 | 0 | 0 | 0 | 0 | 0 | 565810 | 0 | 0 | 386550 | 0 | 0 | 0 | 0 | 0 | 0 |
| KEVIPVNVF | EPRS1; EPRS1 | P07814; V9 | SYEP_HUMAN | Oxidation (M) | 0 | 0 | 0 | 0 | 0 | 0 | 0 | 0 | 0 | 0 | 0 | 0 | 0 | 0 | 0 | 0 | 0 | 0 |
| KEVVHTVSI | RUVBL2; RUVBI | Q9Y230; M0 | RUVB2_HUMA | Unmodified | 0 | 0 | 0 | 0 | 0 | 0 | 0 | 0 | 0 | 0 | 0 | 3787500 | 0 | 0 | 0 | 0 | 0 | 0 |
| KEWEEAEL | APLP2 | Q06481 | APLP2_HUMAN | Unmodified | 0 | 0 | 0 | 0 | 0 | 0 | 0 | 0 | 802480 | 157020 | 0 | 0 | 0 | 0 | 0 | 0 | 96985 | 300320 |
| KEWSGLLE | PFKP; PFKP; PF | Q01813; A0 | PFKAP_HUMAN | Unmodified | 0 | 0 | 0 | 0 | 0 | 0 | 0 | 0 | 0 | 0 | 0 | 2295200 | 0 | 0 | 0 | 0 | 0 | 0 |
| KFACNGTV | EIF1B; EIF1; EIF | O60739; P41 | EIF1B_HUMAN | Unmodified | 0 | 0 | 0 | 382310 | 877080 | 0 | 0 | 0 | 0 | 0 | 0 | 0 | 797800 | 913250 | 0 | 315040 | 0 | 0 |
| KFAEAFEAI | CCT8 | P50990 | TCPQ_HUMAN | Unmodified | 0 | 0 | 290780 | 0 | 1143000 | 0 | 0 | 0 | 1041400 | 370460 | 1188900 | 12259000 | 1381900 | 0 | 279130 | 291220 | 225510 | 1072700 |
| KFAEAFEAI | CCT8 | P50990 | TCPQ_HUMAN | Unmodified | 0 | 0 | 0 | 0 | 5585400 | 132610 | 1295800 | 0 | 0 | 1020800 | 0 | 0 | 7158200 | 2087900 | 0 | 0 | 3304400 | 4802400 |
| KFAEQDAK | FKBP5 | Q13451 | FKBP5_HUMAN | Unmodified | 29169 | 45111 | 16221 | 0 | 359390 | 0 | 0 | 54600 | 0 | 37918 | 0 | 197740 | 1224700 | 0 | 31836 | 52084 | 49397 | 34185 |
| KFAEQTLR | PLEC | Q15149 | PLEC_HUMAN | Unmodified | 0 | 0 | 0 | 0 | 0 | 0 | 0 | 0 | 0 | 0 | 0 | 0 | 0 | 0 | 0 | 0 | 0 | 0 |
| KFAETQPK | EEF1G | P26641 | EF1G_HUMAN | Unmodified | 1886000 | 1587500 | 2311400 | 0 | 30220 | 561510 | 0 | 121140 | 14762 | 82303 | 0 | 13282000 | 134160 | 0 | 0 | 1007600 | 0 | 0 |
| KFAETQPKI | EEF1G | P26641 | EF1G_HUMAN | Unmodified | 0 | 0 | 0 | 0 | 0 | 0 | 0 | 0 | 0 | 0 | 0 | 718720 | 0 | 0 | 0 | 0 | 0 | 14322 |
| KFAMEPEEI | PDIA4; PDIA4 | P13667; A0A | PDIA4_HUMAN | Unmodified | 0 | 0 | 0 | 0 | 0 | 0 | 0 | 0 | 1665900 | 0 | 0 | 0 | 101590 | 0 | 0 | 0 | 0 | 73981 |
| KFAMEPEEI | PDIA4; PDIA4 | P13667; A0A | PDIA4_HUMAN | Oxidation (M) | 0 | 0 | 0 | 0 | 0 | 0 | 294910 | 0 | 5032800 | 1775800 | 555810 | 0 | 0 | 0 | 644660 | 0 | 0 | 0 |
| KFAMEPEEI | PDIA4; PDIA4 | P13667; A0A | PDIA4_HUMAN | Unmodified | 0 | 0 | 0 | 0 | 0 | 0 | 0 | 0 | 3874700 | 0 | 0 | 0 | 0 | 0 | 0 | 0 | 0 | 0 |
| KFANSLVG | SPTBN1; SPTBN | Q01082; A0 | SPTB2_HUMAN | Unmodified | 0 | 0 | 0 | 0 | 0 | 0 | 0 | 0 | 0 | 0 | 0 | 2302500 | 0 | 0 | 0 | 0 | 0 | 0 |
| KFCFSIQQ | CD99L2 | Q8TCZ2 | C99L2_HUMAN | Unmodified | 0 | 0 | 0 | 0 | 955250 | 0 | 513330 | 0 | 0 | 308580 | 0 | 0 | 0 | 0 | 0 | 0 | 346530 | 782220 |
| KFDDGEDF | DNAJC3; DNAJ | Q13217; X6 | DNJC3_HUMA | Unmodified | 0 | 0 | 0 | 0 | 0 | 0 | 0 | 0 | 8314400 | 360510 | 369860 | 0 | 0 | 0 | 0 | 0 | 0 | 241110 |
| KFDEGRNN | P4HB; P4HB; P4 | P07237; A0A | PDIA1_HUMAN | Unmodified | 0 | 0 | 126390 | 0 | 142560 | 345050 | 132060 | 15039 | 18206000 | 6585100 | 3539700 | 326720 | 0 | 183650 | 0 | 0 | 29717 | 0 |
| KFDEKENV | MCTS1 | Q9ULC4 | MCTS1_HUMA | Unmodified | 0 | 31435 | 0 | 0 | 535990 | 0 | 176980 | 157800 | 0 | 116940 | 0 | 0 | 446530 | 744250 | 0 | 0 | 1158800 | 0 |
| KFDEVLVN | HSPA4; HSPA4; | P34932; A0A | HSP74_HUMAN | Unmodified | 112280 | 0 | 0 | 0 | 55974 | 0 | 0 | 0 | 441160 | 0 | 272620 | 5894200 | 0 | 0 | 0 | 0 | 0 | 0 |
| KFDEVLVN | HSPA4; HSPA4; | P34932; A0A | HSP74_HUMAN | Unmodified | 0 | 0 | 0 | 0 | 0 | 0 | 0 | 0 | 0 | 0 | 0 | 6411700 | 0 | 0 | 0 | 0 | 0 | 0 |
| KFDLGQDV | GDI1; GDI2; GD | P31150; P50 | GDIA_HUMAN | Unmodified | 25716 | 524280 | 125680 | 1159700 | 0 | 0 | 0 | 22689 | 0 | 0 | 0 | 914450 | 0 | 0 | 7105300 | 51307 | 0 | 0 |
| KFDQLLAEI | MYH14; MYH9; | Q7Z406; P3 | MYH14_HUMA | Unmodified | 0 | 0 | 2078700 | 1589200 | 11996000 | 2141700 | 3280900 | 729860 | 1044300 | 1656800 | 698030 | 6858500 | 4741300 | 3298300 | 2227100 | 0 | 4397100 | 3196400 |
| KFDQLLAEI | MYH9 | P35579 | MYH9_HUMAN | Unmodified | 0 | 0 | 0 | 0 | 1218300 | 0 | 342910 | 99066 | 0 | 1045700 | 0 | 0 | 515200 | 661770 | 0 | 0 | 3161300 | 619120 |
| KFDSSHDF | FKBP5 | Q13451 | FKBP5_HUMAN | Unmodified | 347740 | 0 | 146510 | 0 | 2431000 | 0 | 0 | 0 | 0 | 0 | 405370 | 2121000 | 2969600 | 866540 | 0 | 0 | 0 | 0 |
| KFDSSYDR | FKBP10; FKBP1 | Q96AY3; C9 | FKB10_HUMAN | Unmodified | 474720 | 0 | 0 | 0 | 265420 | 0 | 0 | 0 | 0 | 0 | 364070 | 0 | 0 | 0 | 0 | 0 | 202840 | 0 |
| KFDVNTSA | CLTC; CLTC | Q00610; A0 | CLH1_HUMAN | Unmodified | 21961000 | 18883000 | 1638400 | 2971700 | 0 | 0 | 46804 | 0 | 433530 | 858970 | 170190 | 14179000 | 423820 | 0 | 26880000 | 7687700 | 276200 | 153030 |
| KFDVNTSA | CLTC; CLTC | Q00610; A0 | CLH1_HUMAN | Unmodified | 230950 | 4012400 | 0 | 2554800 | 0 | 0 | 0 | 0 | 0 | 0 | 0 | 157010 | 0 | 0 | 2325400 | 0 | 0 | 0 |
| KFEDATVQ | HSPA2 | P54652 | HSP72_HUMAN | Unmodified | 0 | 0 | 0 | 0 | 0 | 0 | 0 | 0 | 0 | 0 | 0 | 0 | 0 | 0 | 0 | 0 | 0 | 0 |
| KFEHFSGL | LOXL2 | Q9Y4K0 | LOXL2_HUMAN | Unmodified | 0 | 0 | 0 | 0 | 0 | 2789800 | 0 | 0 | 4320200 | 1088200 | 0 | 0 | 0 | 0 | 0 | 0 | 0 | 857310 |
| KFEQGFITD | IMPDH1; IMPDH | P20839; C9 | IMDH1_HUMAN | Unmodified | 0 | 0 | 173980 | 1182400 | 0 | 0 | 0 | 0 | 0 | 0 | 0 | 0 | 2891300 | 0 | 0 | 0 | 0 | 0 |
| KFEQLKQE | PABPC4; PABP | Q13310; B1 | PABP4_HUMAN | Unmodified | 0 | 1265400 | 0 | 1589800 | 36443 | 0 | 0 | 0 | 0 | 0 | 119160 | 0 | 0 | 19768 | 15458 | 13950 | 157640 | 207190 |
| KFESEILEA | DHX9 | Q08211 | DHX9_HUMAN | Unmodified | 0 | 0 | 0 | 0 | 548800 | 0 | 878450 | 0 | 0 | 300680 | 0 | 0 | 293040 | 238650 | 0 | 0 | 0 | 0 |
| KFFSFIEGC | MAP4K4; MAP4 | O95819; A0 | M4K4_HUMAN | Unmodified | 0 | 0 | 0 | 0 | 311440 | 0 | 251500 | 0 | 0 | 437890 | 0 | 0 | 406370 | 0 | 0 | 0 | 1349900 | 1144900 |
| KFGADDKN | SYT4 | Q9H2B2 | SYT4_HUMAN | Unmodified | 0 | 0 | 1094500 | 0 | 147720 | 118680 | 15949 | 0 | 0 | 71115 | 23609 | 104250 | 0 | 0 | 2217600 | 0 | 0 | 0 |
| KFGATLEIV | ETF1; ETF1 | P62495; B7Z | ERF1_HUMAN | Unmodified | 119300 | 289790 | 0 | 173500 | 0 | 0 | 0 | 0 | 0 | 0 | 0 | 1037400 | 110430 | 0 | 0 | 246840 | 98319 | 0 |
| KFGATLEIV | ETF1; ETF1 | P62495; B7Z | ERF1_HUMAN | Unmodified | 0 | 0 | 0 | 0 | 1408000 | 0 | 0 | 0 | 0 | 0 | 0 | 0 | 0 | 1682200 | 0 | 0 | 1315100 | 675340 |
| KFGDPVVQ | HSPA1A; HSPA | P0DMV8; P0 | HS71A_HUMAN | Unmodified | 2220200 | 2882800 | 162350 | 1609400 | 16015000 | 1731400 | 183170 | 0 | 10516000 | 6301400 | 0 | 3725300 | 7622400 | 3693700 | 0 | 244220 | 1123600 | 2542600 |
| KFGDPVVQ | HSPA1A; HSPA | P0DMV8; P0 | HS71A_HUMAN | Oxidation (M) | 2067000 | 1799800 | 0 | 1072900 | 0 | 0 | 0 | 0 | 0 | 2714500 | 0 | 1200200 | 5068400 | 1392100 | 647960 | 573000 | 0 | 0 |
| KFGEEIARL | PDCD6IP | Q8WUM4 | PDC6I_HUMAN | Unmodified | 2143500 | 1165500 | 2555400 | 8168100 | 48821000 | 9123000 | 9979700 | 0 | 0 | 48259000 | 2308800 | 0 | 71323000 | 37602000 | 617900 | 513950 | 7767700 | 29333000 |
| KFGGQHCI | PLAU; PLAU; PL | P00749; E7E | UROK_HUMAN | Unmodified | 0 | 0 | 0 | 0 | 0 | 0 | 0 | 0 | 35400 | 5800 | 0 | 0 | 0 | 0 | 0 | 0 | 0 | 2597100 |
| KFGGQHCI | PLAU; PLAU; PL | P00749; E7E | UROK_HUMAN | Unmodified | 0 | 2170200 | 0 | 758540 | 113000 | 0 | 0 | 0 | 0 | 0 | 0 | 0 | 0 | 0 | 0 | 0 | 304600 | 253940 |
| KFGTINIVH | PSAT1 | Q9Y617 | SERC_HUMAN | Unmodified | 0 | 218000 | 908090 | 1268600 | 2538500 | 0 | 0 | 0 | 678820 | 980840 | 1866000 | 5986300 | 4124800 | 2769600 | 758550 | 1015500 | 253090 | 455220 |
| KFGVLSDN | PPAT | Q06203 | PUR1_HUMAN | Unmodified | 0 | 0 | 0 | 0 | 0 | 0 | 0 | 0 | 0 | 0 | 0 | 1123200 | 0 | 0 | 0 | 0 | 0 | 0 |
| KFGVTDFP | QSOX1 | O00391 | QSOX1_HUMA | Unmodified | 0 | 0 | 0 | 0 | 0 | 0 | 0 | 0 | 905470 | 0 | 116550 | 0 | 0 | 0 | 0 | 0 | 0 | 0 |
| KFGVVLDE | COPE; COPE; C | O14579; M0 | COPE_HUMAN | Unmodified | 0 | 0 | 0 | 0 | 0 | 0 | 0 | 0 | 1567800 | 205920 | 186870 | 6725200 | 0 | 0 | 0 | 0 | 0 | 0 |
| KFGYVDFE | NCL; NCL; NCL | P19338; C9 | NUCL_HUMAN | Unmodified | 253830 | 590850 | 0 | 137630 | 0 | 0 | 0 | 0 | 2632100 | 956710 | 4097100 | 15742000 | 395970 | 0 | 309050 | 154940 | 0 | 0 |
| KFHEAFIPS | DIS3L2; DIS3L2 | Q8IYB7; C9 | DI3L2_HUMAN | Unmodified | 77565 | 0 | 68856 | 350450 | 101150 | 0 | 0 | 0 | 0 | 65576 | 80176 | 774160 | 0 | 0 | 337830 | 85183 | 0 | 0 |
| KFHGILDQ | PSMD11 | O00231 | PSD11_HUMAN | Unmodified | 0 | 0 | 115510 | 0 | 0 | 0 | 0 | 0 | 0 | 0 | 298640 | 852200 | 954580 | 0 | 0 | 0 | 0 | 0 |
| KFHPEHFV | PXN; PXN; PXN | P49023; A0A | PAXI_HUMAN | Unmodified | 0 | 0 | 0 | 0 | 1009300 | 0 | 27718 | 0 | 0 | 63770 | 0 | 0 | 0 | 565910 | 0 | 0 | 3978000 | 1974000 |
| KFHVPRQN | EPS8; EPS8; EP | Q12929; A0 | EPS8_HUMAN | Unmodified | 0 | 0 | 0 | 0 | 1307000 | 0 | 0 | 0 | 0 | 0 | 0 | 0 | 0 | 288490 | 0 | 0 | 395890 | 0 |
| KFIDIAAAK | DNAJC3; DNAJ | Q13217; X6 | DNJC3_HUMA | Unmodified | 0 | 0 | 0 | 0 | 0 | 0 | 0 | 0 | 1407100 | 305270 | 0 | 0 | 0 | 0 | 0 | 0 | 295260 | 0 |
| KFIGVEAG | CEMIP2; CEMIP | Q9UHN6; F5 | CEIP2_HUMAN | Unmodified | 270980 | 0 | 0 | 0 | 2214200 | 242760 | 0 | 0 | 537870 | 406970 | 0 | 0 | 496660 | 236310 | 0 | 241280 | 454970 | 279800 |
| KFIQENIFG | PDIA3 | P30101 | PDIA3_HUMAN | Unmodified | 0 | 0 | 0 | 0 | 100330 | 0 | 0 | 0 | 44254000 | 1898900 | 979230 | 0 | 0 | 0 | 0 | 0 | 92165 | 0 |
| KFIQENIFG | PDIA3 | P30101 | PDIA3_HUMAN | Oxidation (M) | 0 | 0 | 268820 | 138400 | 0 | 1103700 | 183980 | 0 | 8768800 | 2229500 | 4353700 | 4728400 | 197600 | 0 | 354820 | 222340 | 0 | 0 |
| KFIQENIFG | PDIA3 | P30101 | PDIA3_HUMAN | Unmodified | 0 | 0 | 0 | 0 | 0 | 0 | 0 | 0 | 12840000 | 7288400 | 0 | 0 | 0 | 0 | 0 | 0 | 0 | 390770 |
| KFIQENIFG | PDIA3 | P30101 | PDIA3_HUMAN | Oxidation (M) | 0 | 0 | 0 | 0 | 0 | 0 | 0 | 0 | 6140400 | 6445300 | 2313900 | 0 | 0 | 0 | 0 | 0 | 0 | 0 |
| KFISDKDA | PDIA3 | P30101 | PDIA3_HUMAN | Unmodified | 0 | 0 | 0 | 0 | 4658400 | 687430 | 6018100 | 0 | 26462000 | 70453000 | 2227900 | 0 | 3493500 | 5939900 | 0 | 0 | 734830 | 4365200 |
| KFKDDSNI | RAB4A | P20338 | RAB4A_HUMAN | Unmodified | 0 | 0 | 0 | 0 | 216700 | 0 | 0 | 0 | 0 | 214100 | 0 | 412740 | 0 | 0 | 0 | 0 | 0 | 0 |
| KFKPGYLE | PPA2; PPA2; PP | Q9H2U2; D6 | IPYR2_HUMAN | Unmodified | 0 | 0 | 0 | 0 | 0 | 0 | 0 | 0 | 688780 | 0 | 56711 | 245750 | 0 | 0 | 0 | 0 | 0 | 0 |
| KFKVEYHA | ADK; ADK; ADK; | P55263; A0A | ADK_HUMAN A | Unmodified | 0 | 0 | 0 | 0 | 3926300 | 799910 | 0 | 0 | 352850 | 876750 | 485040 | 1279200 | 1683400 | 1018100 | 0 | 0 | 2039600 | 2662800 |
| KFLDAGHK | PDIA3 | P30101 | PDIA3_HUMAN | Unmodified | 0 | 0 | 0 | 0 | 0 | 0 | 0 | 0 | 0 | 0 | 0 | 0 | 0 | 0 | 0 | 0 | 0 | 0 |
| KFLDAGHK | PDIA3 | P30101 | PDIA3_HUMAN | Unmodified | 0 | 0 | 0 | 0 | 0 | 0 | 368240 | 0 | 4930100 | 5338500 | 0 | 0 | 284930 | 0 | 0 | 0 | 0 | 0 |
| KFLDGNEL | CLIC1 | O00299 | CLIC1_HUMAN | Unmodified | 0 | 0 | 0 | 377010 | 2709100 | 108350 | 1031200 | 0 | 0 | 576570 | 0 | 1285700 | 2038700 | 0 | 0 | 0 | 2239800 | 0 |
| KFLEESVSN | UCHL3; UCHL3 | P15374; Q5 | UCHL3_HUMA | Oxidation (M) | 0 | 0 | 0 | 0 | 0 | 0 | 18876 | 0 | 0 | 0 | 0 | 999320 | 0 | 0 | 0 | 0 | 0 | 0 |
| KFLESGGQ | P4HB; P4HB; P4 | P07237; A0A | PDIA1_HUMAN | Oxidation (M) | 0 | 0 | 0 | 0 | 0 | 0 | 0 | 0 | 12867000 | 832150 | 0 | 0 | 0 | 0 | 0 | 0 | 0 | 0 |
| KFLESGGQ | P4HB; P4HB; P4 | P07237; A0A | PDIA1_HUMAN | Unmodified | 0 | 0 | 0 | 0 | 0 | 0 | 0 | 0 | 20157000 | 366080 | 0 | 0 | 0 | 0 | 0 | 0 | 0 | 0 |
| KFLKPYTLE | ATP1B3; ATP1B | P54709; C9 | AT1B3_HUMAN | Unmodified | 0 | 0 | 874040 | 524050 | 171000 | 0 | 0 | 0 | 1264500 | 0 | 982570 | 10224000 | 0 | 0 | 0 | 0 | 0 | 0 |
| KFLTESHDI | HTRA1; HTRA1; | Q92743; A0 | HTRA1_HUMAN | Unmodified | 0 | 1233500 | 0 | 0 | 133340 | 42478 | 0 | 0 | 110790 | 168010 | 0 | 0 | 0 | 0 | 0 | 915100 | 360680 | 2690200 |
| KFLVYVANI | GDI2; GDI2; GD | P50395; Q5 | GDIB_HUMAN | Unmodified | 286270 | 215910 | 0 | 550300 | 0 | 0 | 0 | 0 | 1142000 | 0 | 108910 | 4425700 | 0 | 0 | 0 | 786200 | 0 | 0 |
| KFMADCPI | RAB14; RAB14 | P61106; X6R | RAB14_HUMAN | Oxidation (M) | 0 | 0 | 0 | 681600 | 0 | 0 | 0 | 0 | 432880 | 85838 | 1363200 | 1610400 | 0 | 0 | 0 | 494300 | 0 | 0 |
| KFMADCPI | RAB14; RAB14 | P61106; X6R | RAB14_HUMAN | Oxidation (M) | 0 | 0 | 0 | 0 | 1067000 | 0 | 1009700 | 0 | 0 | 0 | 0 | 0 | 936820 | 895230 | 0 | 0 | 913800 | 0 |
| KFMGTELN | PHGDH; PHGD | O43175; A0 | SERA_HUMAN | Unmodified | 0 | 0 | 0 | 0 | 104470 | 0 | 89673 | 58746 | 0 | 99563 | 0 | 439010 | 0 | 332990 | 0 | 0 | 0 | 0 |
| KFMLFREAI | SPTAN1; SPTAN | Q13813; A0 | SPTN1_HUMAN | Oxidation (M) | 0 | 0 | 0 | 0 | 779990 | 0 | 0 | 0 | 0 | 69325 | 0 | 0 | 382630 | 0 | 0 | 333350 | 1552200 | 273130 |
| KFNALFAQ | CLTC; CLTC | Q00610; A0 | CLH1_HUMAN | Unmodified | 17278000 | 1546000 | 1266000 | 1624900 | 646680 | 429120 | 540820 | 307300 | 2800500 | 522150 | 835830 | 34624000 | 868220 | 0 | 979190 | 1648600 | 567110 | 340380 |
| KFNALFAQ | CLTC; CLTC | Q00610; A0 | CLH1_HUMAN | Unmodified | 0 | 0 | 0 | 0 | 435700 | 86778 | 378120 | 0 | 0 | 0 | 0 | 0 | 1576900 | 202370 | 0 | 0 | 2090200 | 241930 |
| KFPQYVFG | DDO | Q99489 | OXDD_HUMAN | Unmodified | 0 | 0 | 0 | 0 | 0 | 0 | 0 | 0 | 1519900 | 627150 | 0 | 0 | 0 | 0 | 0 | 0 | 0 | 0 |
| KFPVFNPA | ALDH1A1; ALDI | P00352; Q5 | AL1A1_HUMAN | Unmodified | 0 | 0 | 0 | 255600 | 0 | 0 | 0 | 0 | 0 | 0 | 0 | 802950 | 0 | 0 | 0 | 0 | 0 | 0 |
| KFPVFNPA | ALDH1A1; ALDI | P00352; Q5 | AL1A1_HUMAN | Unmodified | 0 | 0 | 0 | 0 | 0 | 102950 | 0 | 0 | 256000 | 0 | 23969000 | 4.17E+08 | 0 | 0 | 0 | 0 | 0 | 0 |
| KFQAPARE | LIMA1; LIMA1; L | Q9UHB6; F8 | LIMA1_HUMAN | Unmodified | 31166 | 0 | 0 | 0 | 0 | 0 | 0 | 0 | 0 | 0 | 0 | 0 | 1637200 | 464050 | 0 | 0 | 303650 | 126960 |
| KFQEAVEK | MAN1B1; MAN1 | Q9UKM7; H | MA1B1_HUMA | Unmodified | 0 | 0 | 0 | 0 | 0 | 0 | 0 | 0 | 2227700 | 0 | 0 | 0 | 0 | 0 | 0 | 0 | 0 | 0 |
| KFQGIKHE | DSTN | P60981 | DEST_HUMAN | Unmodified | 1457000 | 955270 | 0 | 179930 | 202670 | 0 | 0 | 0 | 701000 | 0 | 344640 | 9079000 | 318380 | 124030 | 271010 | 783660 | 0 | 0 |
| KFQGLQLE | VPS37D | Q86XT2 | VP37D_HUMAN | Unmodified | 0 | 0 | 27081 | 236090 | 0 | 0 | 0 | 0 | 0 | 255820 | 0 | 0 | 0 | 267950 | 157870 | 72638 | 175580 | 0 |
| KFQILNDEI | CYFIP1 | Q7L576 | CYFP1_HUMAN | Unmodified | 492950 | 410990 | 126060 | 808880 | 1011100 | 218960 | 138590 | 0 | 277390 | 206260 | 0 | 405300 | 439220 | 231850 | 703950 | 321810 | 192210 | 399760 |
| KFQQHIQA | TAOK1 | Q7L7X3 | TAOK1_HUMAN | Unmodified | 32248 | 37409 | 16803 | 0 | 0 | 0 | 0 | 0 | 111760 | 16212 | 11069 | 173450 | 0 | 0 | 0 | 88959 | 0 | 0 |
| KFQQHIQA | TAOK1 | Q7L7X3 | TAOK1_HUMAN | Unmodified | 0 | 100710 | 47239 | 0 | 560620 | 199470 | 0 | 175540 | 944310 | 0 | 0 | 348820 | 100860 | 132390 | 0 | 29431 | 0 | 72937 |
| KFQQQILA | TAOK3; TAOK3 | Q9H2K8; G3 | TAOK3_HUMAN | Unmodified | 0 | 0 | 0 | 0 | 1512100 | 0 | 580380 | 353250 | 66195 | 40454 | 110590 | 0 | 1507800 | 1085300 | 0 | 392940 | 0 | 0 |
| KFQSFIESC | TNIK; TNIK | Q9UKE5; C9 | TNIK_HUMAN | Unmodified | 0 | 0 | 0 | 0 | 243010 | 55086 | 552340 | 0 | 171120 | 241950 | 355520 | 0 | 76486 | 295740 | 0 | 0 | 63004 | 270450 |
| KFRLDCPL | VPS28; VPS28; | Q9UK41; E9 | VPS28_HUMAN | Unmodified | 0 | 0 | 0 | 628500 | 0 | 0 | 0 | 0 | 0 | 0 | 0 | 0 | 0 | 0 | 234800 | 0 | 0 | 0 |
| KFSEDFGQ | VPS4A | Q9UN37 | VPS4A_HUMAN | Unmodified | 1164300 | 781210 | 536870 | 1299100 | 71209 | 0 | 423500 | 363810 | 2005000 | 2461600 | 1139200 | 1987400 | 4918700 | 4062200 | 552370 | 581740 | 0 | 0 |
| KFSNQYKA | RAB7A; RAB7A; | P51149; C9 | RAB7A_HUMAN | Unmodified | 0 | 0 | 0 | 0 | 281850 | 0 | 1403600 | 362630 | 0 | 0 | 0 | 0 | 1520800 | 0 | 0 | 0 | 1073400 | 208840 |
| KFTCQGPV | SLIT2; SLIT2; SL | O94813; H0 | SLIT2_HUMAN | Unmodified | 0 | 0 | 0 | 0 | 0 | 0 | 0 | 52586 | 0 | 0 | 0 | 0 | 0 | 0 | 0 | 0 | 0 | 5836100 |
| KFTEDFGQ | VPS4B | O75351 | VPS4B_HUMAN | Unmodified | 1745300 | 0 | 1541100 | 2732200 | 8869800 | 1492200 | 942270 | 0 | 4555400 | 3603000 | 0 | 4613600 | 6258600 | 3339400 | 0 | 1845800 | 0 | 3768600 |
| KFTFGLDW | USP13; USP5; U | Q92995; P4 | UBP13_HUMAN | Unmodified | 0 | 0 | 0 | 0 | 0 | 0 | 0 | 19754 | 0 | 0 | 342310 | 1933500 | 0 | 0 | 0 | 0 | 1592600 | 0 |
| KFTGIKHEV | CFL2 | Q9Y281 | COF2_HUMAN | Unmodified | 525740 | 360770 | 162460 | 306500 | 25865000 | 0 | 0 | 0 | 0 | 0 | 80894 | 1072700 | 969160 | 0 | 456260 | 297890 | 0 | 2088300 |
| KFTTDDSIC | PFKP; PFKP; PF | Q01813; A0 | PFKAP_HUMAN | Unmodified | 0 | 0 | 0 | 0 | 2572800 | 2636700 | 362000 | 41345 | 900480 | 404210 | 669050 | 2755700 | 696990 | 586230 | 125790 | 118920 | 1391900 | 2220400 |
| KFTYLGSQI | ZC3HAV1; ZC3H | Q7Z2W4; C9 | ZCCHV_HUMA | Unmodified | 0 | 0 | 0 | 0 | 0 | 0 | 0 | 0 | 0 | 0 | 0 | 0 | 0 | 0 | 0 | 0 | 0 | 0 |
| KFVEAMAE | STX2 | P32856 | STX2_HUMAN | Oxidation (M) | 0 | 0 | 0 | 0 | 0 | 0 | 31412 | 0 | 0 | 0 | 196410 | 2968100 | 0 | 0 | 380980 | 0 | 0 | 0 |
| KFVEAMAE | STX2 | P32856 | STX2_HUMAN | Unmodified | 0 | 0 | 0 | 0 | 0 | 0 | 0 | 0 | 0 | 0 | 0 | 643730 | 0 | 0 | 0 | 0 | 0 | 0 |
| KFVETPGQ | SSB; SSB | P05455; E7E | LA_HUMAN Lu | Unmodified | 0 | 0 | 0 | 0 | 490360 | 0 | 0 | 0 | 75287 | 0 | 274250 | 912160 | 0 | 205210 | 0 | 0 | 62004 | 283290 |
| KFVHLLDQ | IQGAP1; IQGAP | P46940; A0A | IQGA1_HUMAN | Oxidation (M) | 0 | 0 | 0 | 342530 | 1578300 | 0 | 0 | 0 | 0 | 321860 | 0 | 0 | 0 | 0 | 0 | 0 | 396350 | 0 |

| | | | | | | | | | | | | | | | | | | | | | | |
|---|---|---|---|---|---|---|---|---|---|---|---|---|---|---|---|---|---|---|---|---|---|---|
| KFVIHPESN | SF3B3 | Q15393 | SF3B3_HUMAN | Unmodified | 0 | 0 | 0 | 424460 | 0 | 0 | 0 | 0 | 0 | 0 | 0 | 16330000 | 353860 | 0 | 0 | 0 | 0 | 0 |
| KFVIKPIDK | RDX; EZR; MSN | P35241; P15 | RADI_HUMAN I | Unmodified | 118690 | 140110 | 100690 | 0 | 7197300 | 1298900 | 393540 | 174160 | 369470 | 811850 | 332300 | 876980 | 2209100 | 2017600 | 115860 | 113190 | 963320 | 513030 |
| KFVLCPECI | EIF5; EIF5; EIF5 | P55010; H0Y | IF5_HUMAN Eu | Unmodified | 0 | 0 | 0 | 0 | 0 | 0 | 0 | 0 | 0 | 0 | 0 | 789170 | 0 | 448930 | 0 | 0 | 652730 | 686900 |
| KFVLDYHFI | CPNE2 | Q96FN4 | CPNE2_HUMAI | Unmodified | 0 | 0 | 0 | 0 | 1714000 | 0 | 1401200 | 145020 | 1500800 | 0 | 0 | 0 | 0 | 0 | 0 | 0 | 0 | 990850 |
| KFVLDYHFI | CPNE2 | Q96FN4 | CPNE2_HUMAI | Unmodified | 0 | 0 | 0 | 0 | 1850900 | 168890 | 3021000 | 0 | 0 | 678830 | 0 | 0 | 199660 | 294960 | 0 | 0 | 0 | 0 |
| KFVNFILKP | SIGIRR; SIGIRR | Q6IA17; C9J | SIGIR_HUMAN | Unmodified | 0 | 340970 | 0 | 687100 | 0 | 0 | 0 | 0 | 376950 | 0 | 0 | 753580 | 0 | 0 | 551260 | 1202000 | 0 | 0 |
| KFVPGCFV | CDCP1 | Q9H5V8 | CDCP1_HUMA | Unmodified | 114560 | 0 | 136150 | 367320 | 121420 | 84794 | 0 | 0 | 116350 | 0 | 235640 | 1956300 | 331400 | 0 | 708500 | 421350 | 1224600 | 463950 |
| KFVTDSNA | CKAP5; CKAP5 | Q14008; E9F | CKAP5_HUMAN | Unmodified | 0 | 0 | 0 | 0 | 0 | 0 | 394470 | 138200 | 0 | 407530 | 0 | 1340800 | 0 | 0 | 0 | 0 | 0 | 0 |
| KFVVYEVLF | UAP1 | Q16222 | UAP1_HUMAN | Unmodified | 0 | 0 | 0 | 0 | 11535000 | 1638200 | 568370 | 0 | 212310 | 589770 | 0 | 1168700 | 13473000 | 5954500 | 0 | 0 | 2382000 | 3098300 |
| KFWSSLPS | GPC4 | O75487 | GPC4_HUMAN | Unmodified | 1249300 | 902470 | 51751 | 0 | 270880 | 0 | 0 | 0 | 1057400 | 0 | 0 | 525460 | 0 | 0 | 0 | 0 | 0 | 0 |
| KFYEAFSK | HSP90AB1 | P08238 | HS90B_HUMAN | Unmodified | 16901 | 0 | 326260 | 656030 | 1807400 | 576850 | 909240 | 384120 | 731480 | 1175500 | 1352100 | 12255000 | 1404200 | 1068200 | 569680 | 521470 | 968750 | 779320 |
| KFYEQFSK | HSP90AA1 | P07900 | HS90A_HUMAN | Unmodified | 0 | 231790 | 153740 | 0 | 2150800 | 403920 | 542930 | 49381 | 646190 | 914940 | 728370 | 5579300 | 500280 | 214860 | 294290 | 175760 | 754880 | 1124900 |
| KFYGPEGP | PGRMC1 | O00264 | PGRC1_HUMAI | Unmodified | 0 | 0 | 0 | 0 | 0 | 0 | 0 | 0 | 781470 | 0 | 0 | 1499500 | 0 | 0 | 0 | 0 | 0 | 0 |
| KFYPLEIDY | PARP1 | P09874 | PARP1_HUMAN | Unmodified | 0 | 0 | 0 | 0 | 0 | 0 | 0 | 0 | 535970 | 420040 | 649110 | 1688800 | 0 | 0 | 0 | 0 | 0 | 0 |
| KGADIMYT | SLC25A4; SLC2 | P12235; P12 | ADT1_HUMAN | Unmodified | 0 | 0 | 0 | 0 | 0 | 0 | 0 | 0 | 1030300 | 0 | 0 | 0 | 0 | 0 | 0 | 0 | 0 | 0 |
| KGADLNLV | RAI14 | Q9P0K7 | RAI14_HUMAN | Unmodified | 0 | 0 | 0 | 0 | 0 | 0 | 0 | 0 | 0 | 0 | 0 | 0 | 0 | 0 | 0 | 0 | 0 | 0 |
| KGADSLED | DDX3X; DDX3Y | O00571; O1 | DDX3X_HUMAI | Unmodified | 2655600 | 0 | 0 | 0 | 0 | 0 | 0 | 0 | 0 | 0 | 0 | 0 | 0 | 0 | 0 | 0 | 0 | 0 |
| KGATLKITLI | PKM; PKM; PKM | P14618; B4D | KPYM_HUMAN | Oxidation (M) | 0 | 0 | 0 | 0 | 1512100 | 0 | 0 | 0 | 0 | 0 | 0 | 0 | 0 | 0 | 0 | 0 | 2092800 | 821630 |
| KGATLKITLI | PKM; PKM; PKM | P14618; B4D | KPYM_HUMAN | Unmodified | 0 | 0 | 0 | 0 | 0 | 0 | 566890 | 109220 | 0 | 0 | 0 | 0 | 0 | 0 | 0 | 0 | 861980 | 0 |
| KGAVAEDG | APEX1; APEX1; | P27695; A0A | APEX1_HUMAN | Unmodified | 0 | 0 | 0 | 0 | 0 | 0 | 205850 | 205940 | 0 | 929820 | 1619300 | 1932900 | 439470 | 495410 | 80313 | 0 | 0 | 0 |
| KGAVSSAL | SLC12A2 | P55011 | S12A2_HUMAN | Oxidation (M) | 170650 | 200080 | 111350 | 610610 | 540830 | 0 | 188700 | 0 | 54760 | 0 | 80414 | 323300 | 225250 | 0 | 1203900 | 631610 | 0 | 976330 |
| KGCPPDDI | ITGB1; ITGB1; I | P05556; E7E | ITB1_HUMAN I | Unmodified | 747380 | 1840200 | 423150 | 28896 | 24630000 | 5920300 | 346810 | 41273 | 2715400 | 692580 | 0 | 1122400 | 2417000 | 1151700 | 0 | 1215600 | 5470500 | 42458000 |
| KGDDQSR | CCT3; CCT3 | P49368; B4D | TCPG_HUMAN | Unmodified | 0 | 0 | 0 | 0 | 602010 | 58557 | 99200 | 136240 | 0 | 58378 | 0 | 0 | 695600 | 282900 | 0 | 0 | 517890 | 457550 |
| KGDECELL | TUFM | P49411 | EFTU_HUMAN | Unmodified | 0 | 0 | 0 | 0 | 659820 | 0 | 415780 | 125360 | 5119900 | 1353800 | 158540 | 178140 | 3174200 | 184420 | 0 | 0 | 0 | 255580 |
| KGDEYIING | ACADM; ACADM | P11310; Q5T | ACADM_HUMA | Unmodified | 0 | 0 | 0 | 0 | 0 | 0 | 0 | 0 | 261460 | 0 | 0 | 0 | 0 | 0 | 0 | 0 | 0 | 0 |
| KGDIFLVR | VCP; VCP; VCP; | P55072; C9J | TERA_HUMAN | Unmodified | 382750 | 0 | 0 | 163870 | 0 | 0 | 0 | 0 | 0 | 0 | 0 | 5234400 | 0 | 0 | 170440 | 230360 | 0 | 0 |
| KGDIIQLQF | EPRS1; EPRS1 | P07814; V9G | SYEP_HUMAN | Unmodified | 0 | 0 | 0 | 76256 | 0 | 0 | 0 | 0 | 0 | 0 | 281280 | 2572500 | 0 | 0 | 88350 | 87283 | 0 | 0 |
| KGDILTLLN | SPTAN1; SPTAN | Q13813; A0A | SPTN1_HUMAN | Unmodified | 0 | 0 | 283810 | 0 | 0 | 0 | 0 | 0 | 344210 | 26713 | 551870 | 3786300 | 0 | 0 | 0 | 0 | 0 | 0 |
| KGDIVDIK | RPL21; RPL21; I | P46778; G3V | RL21_HUMAN | Unmodified | 2525900 | 2444800 | 1527500 | 973500 | 476640 | 79354 | 191540 | 0 | 330920 | 0 | 0 | 9082500 | 164460 | 102270 | 1316800 | 1528900 | 139500 | 195480 |
| KGDIVDIKG | RPL21; RPL21; I | P46778; G3V | RL21_HUMAN | Unmodified | 0 | 68769 | 0 | 7743800 | 4289900 | 159280 | 483220 | 2842400 | 0 | 0 | 2400000 | 0 | 1646100 | 9418400 | 0 | 0 | 447290 | 10912000 |
| KGDLLLNR | PSMD6 | Q15008 | PSMD6_HUMA | Unmodified | 0 | 0 | 159920 | 276040 | 591220 | 0 | 89992 | 0 | 0 | 56670 | 247700 | 2290000 | 446430 | 222910 | 211560 | 112110 | 0 | 0 |
| KGDLLNAH | FKBP7; FKBP7; | Q9Y680; F8W | FKBP7_HUMAN | Unmodified | 0 | 0 | 0 | 0 | 560720 | 0 | 32574 | 0 | 2882400 | 788590 | 779040 | 444110 | 209090 | 374550 | 0 | 0 | 0 | 0 |
| KGDLSLRP | TRIM26 | Q12899 | TRI26_HUMAN | Unmodified | 3049000 | 3000900 | 1423900 | 1596900 | 0 | 0 | 0 | 0 | 0 | 0 | 0 | 0 | 0 | 0 | 1968700 | 1651000 | 0 | 0 |
| KGDPVYVY | MIA3 | Q5JRA6 | TGO1_HUMAN | Unmodified | 0 | 0 | 0 | 0 | 0 | 0 | 0 | 0 | 2325800 | 0 | 201000 | 0 | 0 | 0 | 0 | 0 | 0 | 0 |
| KGDVEGS | SMC3 | Q9UQE7 | SMC3_HUMAN | Unmodified | 0 | 0 | 0 | 0 | 0 | 0 | 0 | 0 | 0 | 14630 | 75687 | 204590 | 0 | 0 | 0 | 0 | 0 | 0 |
| KGDVLHMI | FKBP2; FKBP2; | P26885; A0A | FKBP2_HUMAN | Oxidation (M) | 0 | 0 | 0 | 0 | 0 | 0 | 0 | 0 | 5510700 | 1606300 | 0 | 0 | 0 | 0 | 0 | 0 | 0 | 0 |
| KGDVVIVLT | PKM; PKM; PKM | P14618; B4D | KPYM_HUMAN | Unmodified | 0 | 0 | 0 | 61836 | 106480 | 0 | 0 | 0 | 1202500 | 0 | 0 | 408110 | 122680 | 0 | 77560 | 0 | 112130 | 0 |
| KGDVVIVLT | PKM; PKM; PKM | P14618; B4D | KPYM_HUMAN | Oxidation (M) | 628530 | 528830 | 1145600 | 4180300 | 1234100 | 0 | 0 | 0 | 1359200 | 534310 | 815670 | 15914000 | 712320 | 0 | 1749300 | 1502800 | 0 | 0 |
| KGEAGHMI | FAXDC2; FAXD | Q96IV6; E5R | FXDC2_HUMAI | Acetyl (Protein N | 0 | 0 | 0 | 0 | 0 | 0 | 0 | 0 | 0 | 0 | 0 | 0 | 0 | 0 | 0 | 0 | 0 | 0 |
| KGEDEKLD | IFNAR2; IFNAR2 | F8WDE1; H7 | F8WDE1_HUM | Unmodified | 597230 | 0 | 0 | 809770 | 457580 | 0 | 0 | 0 | 234890 | 353550 | 162520 | 621500 | 2836200 | 1121000 | 59073 | 0 | 137080 | 246560 |
| KGEDVLGS | MARS1; MARS1 | P56192; F5H | SYMC_HUMAN | Unmodified | 0 | 0 | 0 | 0 | 352750 | 0 | 291990 | 318810 | 0 | 0 | 0 | 944190 | 110720 | 0 | 0 | 0 | 150140 | 128350 |
| KGEFETGFI | GSPT1; GSPT2; | P15170; Q8I | ERF3A_HUMAN | Unmodified | 0 | 0 | 0 | 0 | 250490 | 0 | 0 | 0 | 215490 | 79690 | 252120 | 1739300 | 685120 | 464040 | 0 | 0 | 111120 | 282880 |
| KGEGDAAT | GAP43 | P17677 | NEUM_HUMAN | Unmodified | 0 | 0 | 0 | 0 | 0 | 0 | 0 | 2763200 | 310770 | 0 | 0 | 0 | 0 | 0 | 0 | 0 | 0 | 0 |
| KGEGDAAT | GAP43 | P17677 | NEUM_HUMAN | Unmodified | 0 | 0 | 0 | 0 | 0 | 0 | 894680 | 1192100 | 0 | 0 | 0 | 0 | 0 | 0 | 0 | 0 | 0 | 0 |
| KGEGLPNF | DNAJB11; N/A | Q9UBS4; H7 | DJB11_HUMAN | Unmodified | 42893 | 54006 | 250980 | 398020 | 428940 | 459430 | 577910 | 196250 | 56314000 | 9817500 | 6158700 | 1906800 | 331280 | 285910 | 238180 | 95885 | 0 | 410910 |
| KGEIFELK | AP1B1; AP2B1; | Q10567; P63 | AP1B1_HUMAN | Unmodified | 132130 | 173200 | 559480 | 206080 | 169510 | 0 | 0 | 0 | 0 | 0 | 375100 | 3103000 | 327150 | 93848 | 640230 | 388500 | 62311 | 0 |
| KGEITGEVI | FLNB; FLNB | O75369; E7E | FLNB_HUMAN | Unmodified | 0 | 0 | 69741 | 0 | 0 | 0 | 0 | 0 | 171500 | 0 | 0 | 386690 | 0 | 0 | 0 | 74442 | 0 | 0 |
| KGEKDIPGI | RPS6; RPS6 | P62753; A2A | RS6_HUMAN 4 | Unmodified | 0 | 0 | 0 | 0 | 373690 | 159940 | 2583600 | 1532200 | 619490 | 0 | 0 | 0 | 1426500 | 432340 | 0 | 0 | 72570 | 295670 |
| KGELSMED | ABCC5; ABCC5 | O15440; C9J | MRP5_HUMAN | Oxidation (M) | 0 | 47395 | 0 | 0 | 0 | 0 | 0 | 0 | 0 | 0 | 0 | 487100 | 0 | 0 | 0 | 0 | 0 | 0 |
| KGEPVSAE | MTHFD1; MTHF | P11586; V9G | C1TC_HUMAN | Oxidation (M) | 0 | 0 | 0 | 0 | 0 | 0 | 0 | 0 | 0 | 0 | 0 | 566890 | 0 | 0 | 0 | 0 | 0 | 0 |
| KGEQELAE | SLC1A4 | P43007 | SATT_HUMAN I | Unmodified | 155690 | 0 | 0 | 0 | 0 | 0 | 0 | 0 | 628780 | 366690 | 39578 | 618260 | 0 | 0 | 0 | 0 | 57659 | 0 |
| KGEQELAE | SLC1A4 | P43007 | SATT_HUMAN I | Unmodified | 0 | 0 | 0 | 0 | 165090 | 0 | 74989 | 0 | 5059300 | 205650 | 166190 | 3174800 | 0 | 0 | 0 | 0 | 0 | 0 |
| KGESGQSV | PTGES3; PTGES | Q15185; A0A | TEBP_HUMAN I | Unmodified | 0 | 1630500 | 0 | 573570 | 164470 | 0 | 49055 | 22399 | 1777100 | 139550 | 411840 | 1930100 | 0 | 0 | 1348200 | 0 | 0 | 89298 |
| KGESGQSV | PTGES3; PTGES | Q15185; A0A | TEBP_HUMAN I | Unmodified | 0 | 0 | 0 | 0 | 1531900 | 412060 | 128300 | 182320 | 137870 | 993980 | 0 | 0 | 1434500 | 606250 | 0 | 0 | 747580 | 906830 |
| KGESQTDII | CCT3; CCT3 | P49368; B4D | TCPG_HUMAN | Unmodified | 0 | 0 | 0 | 0 | 0 | 0 | 78828 | 0 | 0 | 0 | 1342300 | 260560 | 0 | 0 | 0 | 0 | 41982 | 24714 |
| KGESQTDII | CCT3; CCT3 | P49368; B4D | TCPG_HUMAN | Unmodified | 292440 | 211810 | 942760 | 162520 | 0 | 0 | 0 | 53636 | 2815400 | 0 | 5787800 | 29011000 | 0 | 0 | 1283900 | 3058200 | 111910 | 0 |
| KGEYVFPAI | MOV10; MOV10 | Q9HCE1; Q5 | MOV10_HUMA | Unmodified | 501240 | 380190 | 0 | 0 | 0 | 136230 | 0 | 0 | 0 | 0 | 0 | 0 | 0 | 0 | 244280 | 320920 | 0 | 0 |
| KGFSEGLW | HDGF | P51858 | HDGF_HUMAN | Unmodified | 0 | 0 | 0 | 0 | 0 | 0 | 0 | 0 | 0 | 0 | 856710 | 2367100 | 0 | 0 | 0 | 0 | 4267100 | 1979800 |
| KGGDGIKP | OLA1; OLA1; OL | Q9NTK5; J3K | OLA1_HUMAN | Unmodified | 0 | 0 | 0 | 0 | 0 | 0 | 0 | 59140 | 0 | 0 | 802790 | 0 | 459600 | 0 | 0 | 0 | 0 | 0 |
| KGGEMEILI | IQGAP2; IQGAP | Q13576; E9F | IQGA2_HUMAN | Unmodified | 0 | 0 | 285170 | 338970 | 0 | 0 | 0 | 0 | 0 | 0 | 0 | 297370 | 0 | 0 | 162570 | 114040 | 0 | 0 |
| KGGETSEM | FGB; FGB | P02675; D6F | FIBB_HUMAN F | Oxidation (M) | 0 | 581490 | 0 | 0 | 0 | 0 | 0 | 0 | 0 | 0 | 0 | 0 | 0 | 0 | 0 | 0 | 0 | 0 |
| KGGPGRG | IST1 | P53990 | IST1_HUMAN I | 4 Oxidation (M) | 0 | 0 | 0 | 0 | 1843700 | 0 | 0 | 0 | 0 | 0 | 1543800 | 0 | 0 | 0 | 0 | 0 | 0 | 0 |
| KGGPGSE | MDK; MDK; MDI | P21741; E9P | MK_HUMAN Mi | Unmodified | 54190 | 0 | 0 | 0 | 0 | 0 | 0 | 0 | 24044000 | 2994000 | 0 | 0 | 0 | 0 | 0 | 0 | 0 | 0 |
| KGGSWIQE | PRKDC | P78527 | PRKDC_HUMAI | Unmodified | 0 | 0 | 285310 | 185420 | 0 | 0 | 1261900 | 333110 | 274490 | 389360 | 0 | 1437200 | 249680 | 0 | 74820 | 98242 | 0 | 724100 |
| KGGSYSQ | HLA-A; HLA-A; I | Q53Z42; A0A | Q53Z42_HUMA | Unmodified | 0 | 0 | 0 | 0 | 0 | 0 | 0 | 0 | 1368600 | 234400 | 994570 | 6058900 | 0 | 0 | 0 | 0 | 196060 | 850370 |
| KGGSYTQA | HLA-A; HLA-A; I | P04439; A0A | HLAA_HUMAN | Unmodified | 0 | 0 | 0 | 0 | 0 | 0 | 0 | 0 | 1601600 | 327830 | 0 | 3051300 | 0 | 0 | 0 | 0 | 276380 | 0 |
| KGHFPAAY | NOSTRIN; NOS | Q8IVI9; A0A | NOSTN_HUMA | Unmodified | 2214700 | 1701000 | 0 | 1525300 | 87198 | 0 | 0 | 0 | 0 | 0 | 0 | 0 | 0 | 0 | 1905900 | 3211800 | 0 | 0 |
| KGIELIPQV | PLCB4; PLCB4 | Q15147; A0A | PLCB4_HUMAN | Unmodified | 0 | 0 | 0 | 0 | 461780 | 0 | 0 | 0 | 0 | 42909 | 0 | 0 | 652860 | 486020 | 0 | 0 | 608590 | 3063800 |
| KGIGPVYS | ADSS2 | P30520 | PURA2_HUMAI | Unmodified | 0 | 0 | 0 | 0 | 347640 | 0 | 0 | 103630 | 103960 | 0 | 0 | 1037100 | 162050 | 0 | 0 | 0 | 365980 | 0 |
| KGIIRPGTA | PLEC; PLEC | Q15149; H0Y | PLEC_HUMAN | Unmodified | 0 | 0 | 0 | 0 | 0 | 0 | 0 | 0 | 0 | 0 | 0 | 0 | 0 | 0 | 0 | 0 | 1599200 | 1438500 |
| KGILEIESAI | FGD4; FGD4; F | Q96M96; B7 | FGD4_HUMAN | Unmodified | 0 | 4925500 | 0 | 0 | 0 | 0 | 0 | 0 | 0 | 0 | 0 | 0 | 0 | 0 | 0 | 0 | 0 | 0 |
| KGILGQHC | ITGA2 | P17301 | ITA2_HUMAN I | Unmodified | 0 | 0 | 804490 | 0 | 0 | 0 | 0 | 0 | 0 | 0 | 0 | 0 | 0 | 549050 | 951260 | 407390 | 0 | 232040 |
| KGIVDQSQ | YWHAZ; YWHA | P63104; B7Z | 1433Z_HUMAN | Unmodified | 440130 | 275640 | 689170 | 338750 | 0 | 253690 | 0 | 0 | 2253400 | 813150 | 527260 | 4007700 | 120540 | 0 | 756130 | 685800 | 366500 | 396180 |
| KGIVDQSQ | YWHAZ; YWHA | P63104; B7Z | 1433Z_HUMAN | Unmodified | 569510 | 2226700 | 4636000 | 4240800 | 4976300 | 3117300 | 2061200 | 367280 | 16856000 | 4359100 | 6893300 | 34325000 | 2123500 | 1144700 | 6939500 | 6333000 | 5883100 | 3700400 |
| KGKEDEGE | DYNC1H1; DYN | Q14204; A0A | DYHC1_HUMA | Unmodified | 0 | 0 | 0 | 0 | 0 | 0 | 0 | 0 | 210460 | 0 | 0 | 2153100 | 0 | 0 | 966630 | 0 | 0 | 0 |
| KGLSFFAFI | TCF25; TCF25; | Q9BQ70; H3 | TCF25_HUMAN | Unmodified | 409820 | 0 | 0 | 403740 | 0 | 0 | 0 | 0 | 0 | 0 | 0 | 0 | 0 | 0 | 900730 | 391970 | 0 | 0 |
| KGLSSDLE | AKAP12 | Q02952 | AKA12_HUMAN | Unmodified | 0 | 0 | 0 | 0 | 148840 | 0 | 13105 | 0 | 0 | 0 | 0 | 0 | 0 | 0 | 0 | 0 | 569370 | 714320 |
| KGLSTEDA | ACBD7 | Q8N6N7 | ACBD7_HUMAI | Unmodified | 0 | 0 | 0 | 0 | 0 | 0 | 0 | 134090 | 0 | 0 | 0 | 0 | 0 | 0 | 0 | 0 | 0 | 0 |
| KGLTPSQI | RPS13; RPS13 | P62277; J3K | RS13_HUMAN | Unmodified | 3639900 | 2901400 | 1105000 | 423180 | 0 | 0 | 0 | 0 | 0 | 0 | 1316000 | 23164000 | 0 | 0 | 3204800 | 2170000 | 0 | 0 |
| KGNAEGSS | HDGF | P51858 | HDGF_HUMAN | Unmodified | 0 | 0 | 0 | 0 | 0 | 0 | 0 | 0 | 0 | 0 | 572140 | 2454600 | 0 | 0 | 0 | 0 | 0 | 0 |
| KGNAEPDF | AVIL | O75366 | AVIL_HUMAN A | Unmodified | 0 | 0 | 0 | 0 | 959720 | 0 | 0 | 0 | 0 | 0 | 0 | 1387300 | 0 | 0 | 0 | 0 | 0 | 0 |
| KGNEAVAS | GOLM1 | Q8NBJ4 | GOLM1_HUMA | Unmodified | 0 | 0 | 0 | 0 | 368000 | 0 | 0 | 0 | 2919600 | 126880 | 0 | 0 | 481250 | 471120 | 0 | 0 | 192270 | 162640 |
| KGNFNYIEI | MYL12A; MYL12 | P19105; O1 | ML12A_HUMAN | Unmodified | 322880 | 93209 | 0 | 845050 | 6909900 | 1121900 | 0 | 0 | 744800 | 396700 | 721230 | 9302900 | 1823900 | 864870 | 730830 | 576880 | 721770 | 1758500 |
| KGNFNYIEI | MYL12A; MYL12 | P19105; O1 | ML12A_HUMAN | Unmodified | 0 | 0 | 0 | 0 | 10175000 | 16329 | 273610 | 0 | 0 | 0 | 0 | 0 | 2637800 | 1856900 | 0 | 0 | 5009800 | 2210500 |
| KGNKEDYD | SCG3 | Q8WXD2 | SCG3_HUMAN | Unmodified | 0 | 0 | 0 | 0 | 0 | 0 | 0 | 0 | 0 | 0 | 0 | 0 | 0 | 0 | 0 | 0 | 0 | 0 |
| KGNLEWLD | VPS25; VPS25 | Q9BRG1; K7 | VPS25_HUMAN | Unmodified | 343560 | 361590 | 294560 | 0 | 0 | 0 | 197910 | 0 | 0 | 0 | 0 | 0 | 0 | 0 | 823120 | 76120 | 338780 | 211750 |
| KGNTALHI | ANK3; ANK2; A | Q12955; Q0 | ANK3_HUMAN | Unmodified | 0 | 0 | 1968800 | 4484900 | 545020 | 0 | 0 | 0 | 269550 | 0 | 0 | 176870 | 1415000 | 9067800 | 9154600 | 7644900 | 0 | 0 |
| KGNVVFGE | DPYSL3; DPYSL | Q14195; H0 | DPYL3_HUMAN | Unmodified | 0 | 0 | 0 | 0 | 0 | 0 | 0 | 0 | 0 | 0 | 0 | 6634100 | 0 | 0 | 0 | 0 | 490000 | 0 |
| KGPDGYDI | MOV10; MOV10 | Q9HCE1; Q5 | MOV10_HUMA | Unmodified | 3844800 | 7954300 | 2286100 | 3043900 | 0 | 0 | 71427 | 0 | 0 | 0 | 0 | 253630 | 327620 | 58595 | 4380800 | 5776600 | 0 | 0 |
| KGPEDTAQ | STXBP2; STXBP | Q15833; M0 | STXB2_HUMAN | Unmodified | 612880 | 730540 | 0 | 1856500 | 815690 | 0 | 0 | 356770 | 1083500 | 0 | 0 | 2944400 | 1557400 | 0 | 0 | 2095000 | 0 | 694420 |
| KGPGLYYV | PSMB8; PSMB8 | P28062; Q5J | PSB8_HUMAN | Unmodified | 0 | 0 | 0 | 0 | 0 | 0 | 0 | 0 | 0 | 0 | 108250 | 3403300 | 0 | 0 | 0 | 0 | 0 | 0 |
| KGPGPGGI | NRGN | Q92686 | NEUG_HUMAN | Unmodified | 0 | 0 | 0 | 0 | 0 | 0 | 0 | 0 | 0 | 99536 | 0 | 0 | 0 | 0 | 0 | 0 | 208090 | 1324000 |
| KGPLEWQI | GEMIN6; GEMIN | Q8WXD5; B8 | GEMI6_HUMAN | Unmodified | 0 | 0 | 0 | 0 | 173460 | 0 | 54755 | 0 | 0 | 49489 | 0 | 345570 | 113590 | 154670 | 0 | 0 | 39104 | 0 |
| KGPLIAPGF | SLC6A8 | P48029 | SC6A8_HUMAN | Unmodified | 35704 | 0 | 0 | 0 | 0 | 0 | 0 | 0 | 0 | 0 | 0 | 484420 | 0 | 0 | 0 | 0 | 0 | 0 |
| KGPLIAPGF | SLC6A8 | P48029 | SC6A8_HUMAN | Unmodified | 0 | 0 | 999370 | 1900900 | 0 | 0 | 0 | 0 | 0 | 243590 | 3031500 | 9973800 | 0 | 0 | 2398600 | 3832700 | 0 | 829810 |
| KGPQGYGI | SLC9A3R2; SLC | Q15599; H3I | NHRF2_HUMAI | Unmodified | 156960 | 0 | 238650 | 753290 | 0 | 0 | 0 | 11634 | 309810 | 202980 | 0 | 579620 | 1073900 | 836830 | 521400 | 263180 | 430340 | 26148 |
| KGPQVRD | KRT18; KRT18 | P05783; F8V | K1C18_HUMAN | Unmodified | 0 | 0 | 0 | 0 | 6086000 | 1386600 | 0 | 0 | 623710 | 4629700 | 2070600 | 0 | 3944500 | 1043800 | 0 | 0 | 995270 | 852980 |
| KGPRQDP | TNFRSF21 | O75509 | TNR21_HUMAN | Unmodified | 0 | 0 | 0 | 0 | 219840 | 0 | 0 | 0 | 0 | 0 | 56838 | 97204 | 0 | 95979 | 0 | 0 | 0 | 1128800 |
| KGPSGYGF | SLC9A3R1 | O14745 | NHRF1_HUMAI | Unmodified | 0 | 0 | 739650 | 1656500 | 7545500 | 2104700 | 1664600 | 1246900 | 451450 | 338020 | 863510 | 6728100 | 24299000 | 39990000 | 2771700 | 2204700 | 2800200 | 790140 |
| KGQAVDYE | PDIA4; PDIA4 | P13667; A0A | PDIA4_HUMAN | Unmodified | 0 | 0 | 0 | 0 | 4825800 | 970710 | 13005000 | 7568400 | 75271000 | 38968000 | 11391000 | 3410100 | 0 | 3137300 | 0 | 0 | 909920 | 2917700 |
| KGQGLTGF | EHHADH | Q08426 | ECHP_HUMAN | Unmodified | 0 | 0 | 0 | 0 | 0 | 0 | 0 | 0 | 1701600 | 0 | 0 | 0 | 0 | 0 | 0 | 0 | 0 | 0 |
| KGQIELTG | AGPS; AGPS; A | O00116; A0A | ADAS_HUMAN | Unmodified | 0 | 0 | 0 | 0 | 0 | 0 | 0 | 0 | 1620700 | 575130 | 0 | 0 | 0 | 0 | 0 | 0 | 0 | 0 |
| KGQTHTLE | HSD17B10; HS | Q99714; Q5I | HCD2_HUMAN | Unmodified | 0 | 0 | 0 | 0 | 0 | 0 | 78460 | 0 | 67219000 | 810640 | 1285800 | 314310 | 0 | 0 | 398930 | 0 | 0 | 0 |
| KGRDTLQE | LAMC1 | P11047 | LAMC1_HUMAI | Unmodified | 0 | 0 | 0 | 0 | 86838 | 187700 | 0 | 0 | 0 | 0 | 0 | 512860 | 0 | 139400 | 0 | 0 | 0 | 635090 |
| KGSCNLSF | GFPT1; GFPT1; | Q06210; A0A | GFPT1_HUMAN | Unmodified | 0 | 0 | 0 | 0 | 91980 | 0 | 0 | 0 | 0 | 0 | 0 | 0 | 877710 | 571310 | 0 | 0 | 128660 | 0 |
| KGSDSSIPI | JMJD1C; JMJD1 | Q15652; A0A | JHD2C_HUMAN | Unmodified | 0 | 0 | 0 | 0 | 0 | 0 | 0 | 18692000 | 0 | 0 | 0 | 0 | 0 | 0 | 0 | 0 | 0 | 0 |
| KGSGVGE | APLP2 | Q06481 | APLP2_HUMAN | Unmodified | 0 | 0 | 0 | 0 | 251770 | 0 | 0 | 0 | 1707700 | 651000 | 0 | 132050 | 0 | 0 | 0 | 35168 | 0 | 0 |
| KGSITSVQ | ATP5F1B; ATP5 | P06576; H0Y | ATPB_HUMAN | Unmodified | 0 | 162410 | 0 | 0 | 0 | 0 | 0 | 0 | 0 | 4126300 | 0 | 0 | 0 | 0 | 0 | 0 | 0 | 0 |
| KGSLESPA | PSMD12 | O00232 | PSD12_HUMAN | Unmodified | 0 | 0 | 0 | 0 | 0 | 0 | 0 | 0 | 0 | 0 | 0 | 1698000 | 0 | 0 | 0 | 114590 | 0 | 0 |

| | | | | | | | | | | | | | | | | | | | | | | |
|---|---|---|---|---|---|---|---|---|---|---|---|---|---|---|---|---|---|---|---|---|---|---|
| KGSPVSEI( | SNTB1; SNTB1 | Q13884; A0A | SNTB1_HUMAN | Unmodified | 0 | 0 | 0 | 0 | 0 | 0 | 0 | 0 | 1204600 | 0 | 0 | 0 | 0 | 0 | 0 | 0 | 0 | 0 |
| KGSQTNLK | KIF23; KIF23; KI | Q02241; A0A | KIF23_HUMAN | Unmodified | 0 | 0 | 0 | 416000 | 8602500 | 1795700 | 30729 | 84024 | 1165400 | 734620 | 311640 | 1069400 | 346540 | 758390 | 0 | 0 | 577040 | 0 |
| KGSVVNVN | MAP4K4; MAP4 | O95819; C9J | M4K4_HUMAN | Unmodified | 0 | 0 | 362580 | 0 | 0 | 0 | 0 | 0 | 1294500 | 0 | 0 | 2094400 | 0 | 0 | 240520 | 0 | 0 | 0 |
| KGTAVFWY | P4HA1 | P13674 | P4HA1_HUMAN | Unmodified | 0 | 53880 | 0 | 0 | 0 | 0 | 0 | 0 | 3274900 | 4730100 | 808840 | 0 | 0 | 0 | 0 | 40570 | 0 | 0 |
| KGTAVFWY | P4HA2 | O15460 | P4HA2_HUMAN | Unmodified | 0 | 0 | 28850 | 0 | 0 | 0 | 0 | 0 | 4084300 | 1049700 | 0 | 0 | 0 | 0 | 0 | 0 | 0 | 0 |
| KGTDIMYT( | SLC25A5 | P05141 | ADT2_HUMAN | Oxidation (M) | 0 | 0 | 0 | 0 | 0 | 0 | 0 | 0 | 2166100 | 0 | 0 | 0 | 0 | 0 | 0 | 0 | 0 | 0 |
| KGTDIMYT( | SLC25A5 | P05141 | ADT2_HUMAN | Unmodified | 0 | 0 | 0 | 0 | 0 | 0 | 0 | 0 | 2418100 | 173580 | 0 | 0 | 0 | 0 | 0 | 0 | 0 | 0 |
| KGTDIMYT( | SLC25A5 | P05141 | ADT2_HUMAN | Oxidation (M) | 0 | 44918 | 0 | 0 | 0 | 0 | 0 | 0 | 940820 | 278270 | 0 | 0 | 0 | 0 | 0 | 0 | 0 | 176210 |
| KGTDVNVF | ANXA1 | P04083 | ANXA1_HUMAN | Unmodified | 94638 | 0 | 0 | 0 | 4135100 | 1216700 | 37766 | 0 | 301780 | 616060 | 89074000 | 8.01E+08 | 0 | 0 | 0 | 113060 | 295100 | 448290 |
| KGTDVNVF | ANXA1 | P04083 | ANXA1_HUMAN | Unmodified | 0 | 0 | 0 | 0 | 2018900 | 318060 | 0 | 0 | 0 | 0 | 3580700 | 512040 | 0 | 0 | 0 | 0 | 0 | 834650 |
| KGTDVPKW | ANXA2P2; ANX | A6NMY6; P0 | AXA2L_HUMAN | Unmodified | 0 | 0 | 0 | 0 | 248620 | 0 | 0 | 0 | 0 | 145650 | 0 | 0 | 0 | 0 | 0 | 0 | 131570 | 141350 |
| KGTEVQVD | RUVBL2 | Q9Y230 | RUVB2_HUMAN | Unmodified | 163030 | 0 | 0 | 0 | 0 | 0 | 0 | 20802 | 0 | 0 | 0 | 3811200 | 0 | 0 | 0 | 21739 | 0 | 0 |
| KGTGLLSSI | ZC3HAV1; ZC3H | Q7Z2W4; C9 | ZCCHV_HUMA | Unmodified | 0 | 0 | 0 | 0 | 1019200 | 315340 | 0 | 0 | 0 | 0 | 0 | 0 | 274320 | 0 | 0 | 0 | 0 | 161140 |
| KGTMTTGH | ILF2; ILF2; ILF2; | Q12905; B4D | ILF2_HUMAN Ir | Oxidation (M) | 0 | 0 | 0 | 0 | 0 | 0 | 0 | 0 | 0 | 0 | 93375 | 560590 | 0 | 0 | 0 | 0 | 0 | 0 |
| KGTVACGC | VCAN; VCAN | P13611; E9P | CSPG2_HUMAN | Unmodified | 0 | 0 | 0 | 0 | 0 | 0 | 0 | 0 | 460900 | 241020 | 0 | 0 | 0 | 0 | 0 | 0 | 0 | 208680 |
| KGTVEGFE | SRP14; SRP14 | P37108; H0Y | SRP14_HUMAN | Unmodified | 5452700 | 5346900 | 6454400 | 17331000 | 2482300 | 0 | 2521300 | 852100 | 790490 | 339070 | 3126100 | 10416000 | 1938400 | 849410 | 22713000 | 13586000 | 424610 | 2054200 |
| KGTYLTDE | CD46 | P15529 | MCP_HUMAN I | Unmodified | 0 | 815850 | 0 | 0 | 42647 | 0 | 0 | 0 | 0 | 158760 | 194270 | 0 | 0 | 0 | 268720 | 0 | 0 | 0 |
| KGVAINFVI | EIF4A3; EIF4A3 | P38919; I3L3 | IF4A3_HUMAN | Unmodified | 0 | 0 | 0 | 0 | 0 | 0 | 0 | 0 | 1572000 | 58661 | 0 | 1270300 | 0 | 0 | 0 | 0 | 0 | 0 |
| KGVAINFVI | EIF4A3; EIF4A3 | P38919; I3L3 | IF4A3_HUMAN | Unmodified | 345150 | 339800 | 0 | 200180 | 0 | 0 | 0 | 0 | 0 | 0 | 0 | 206710 | 0 | 157340 | 0 | 0 | 0 | 0 |
| KGVAINFVI | EIF4A2 | Q14240 | IF4A2_HUMAN | Unmodified | 0 | 0 | 0 | 0 | 0 | 0 | 0 | 0 | 0 | 0 | 0 | 0 | 0 | 0 | 0 | 0 | 0 | 0 |
| KGVAINMV | EIF4A1 | P60842 | IF4A1_HUMAN | Unmodified | 374130 | 376520 | 493150 | 643860 | 0 | 0 | 0 | 0 | 598380 | 119360 | 1036500 | 1878500 | 0 | 0 | 936270 | 314070 | 261850 | 0 |
| KGVEFPMA | TOM1L2 | Q6ZVM7 | TM1L2_HUMAN | Oxidation (M) | 1048000 | 899010 | 0 | 207540 | 0 | 0 | 0 | 0 | 0 | 0 | 0 | 0 | 0 | 0 | 671720 | 647450 | 0 | 0 |
| KGVLETFS( | AK3 | Q9UIJ7 | KAD3_HUMAN | Unmodified | 0 | 0 | 0 | 0 | 0 | 0 | 661470 | 0 | 1831000 | 253130 | 0 | 0 | 0 | 0 | 0 | 0 | 0 | 0 |
| KGVNLPGA | PKM; PKM; PKM | P14618; B4D | KPYM_HUMAN | Unmodified | 1890500 | 0 | 5116500 | 5943300 | 45373000 | 7679100 | 1897200 | 1496600 | 9605600 | 2319400 | 30935000 | 86353000 | 8543300 | 3362000 | 11392000 | 8357900 | 4727900 | 16306000 |
| KGVNLPGA | PKM; PKM; PKM | P14618; B4D | KPYM_HUMAN | Unmodified | 0 | 0 | 0 | 750730 | 28370000 | 4762400 | 0 | 206020 | 0 | 2241800 | 0 | 0 | 0 | 2961900 | 0 | 0 | 4702900 | 11643000 |
| KGVPIIFAD | DAG1 | Q14118 | DAG1_HUMAN | Oxidation (M) | 0 | 0 | 0 | 0 | 3921600 | 1752000 | 0 | 0 | 5615900 | 0 | 0 | 0 | 2760900 | 2731500 | 0 | 0 | 2430800 | 0 |
| KGVPSVPV | SLC12A7; SLC1 | Q9Y666; A0A | S12A7_HUMAN | Unmodified | 0 | 0 | 0 | 0 | 0 | 0 | 0 | 0 | 949530 | 0 | 0 | 1812500 | 0 | 0 | 318200 | 610090 | 0 | 0 |
| KGVQFPPS | TOM1L1; TOM1 | O75674; J3K | TM1L1_HUMAN | Unmodified | 5861800 | 2782900 | 0 | 3374300 | 0 | 0 | 0 | 0 | 602150 | 0 | 0 | 383150 | 1377200 | 0 | 4397400 | 4672100 | 0 | 0 |
| KGVQLQTH | ARCN1; ARCN1 | P48444; B0Y | COPD_HUMAN | Unmodified | 47944 | 0 | 41378 | 45721 | 159890 | 28313 | 0 | 0 | 218670 | 341500 | 129070 | 692630 | 189320 | 41000 | 153020 | 48541 | 0 | 74726 |
| KGVSHGQI | MGAT1 | P26572 | MGAT1_HUMAN | Unmodified | 0 | 0 | 0 | 0 | 0 | 0 | 0 | 0 | 3294100 | 265060 | 0 | 0 | 0 | 0 | 0 | 0 | 0 | 0 |
| KGYDAPLC | ACSL4; ACSL4 | O60488; H0Y | ACSL4_HUMAN | Unmodified | 0 | 0 | 0 | 0 | 475950 | 195290 | 246740 | 17921 | 0 | 47098 | 0 | 1453300 | 61468 | 27699 | 0 | 0 | 1214500 | 354140 |
| KGYEVIYLT | HSP90B1; HSP9 | P14625; A0A | ENPL_HUMAN | Unmodified | 0 | 0 | 0 | 0 | 0 | 0 | 1255700 | 0 | 35188000 | 18644000 | 2210700 | 231410 | 0 | 0 | 0 | 0 | 0 | 0 |
| KGYEVIYLT | HSP90B1; HSP9 | P14625; A0A | ENPL_HUMAN | Unmodified | 0 | 0 | 0 | 0 | 0 | 0 | 0 | 0 | 15746000 | 1980500 | 873110 | 0 | 0 | 0 | 0 | 0 | 0 | 0 |
| KGYLTLSDS | CANT1; CANT1 | Q8WVQ1; K7 | CANT1_HUMAN | Unmodified | 0 | 0 | 0 | 0 | 0 | 0 | 0 | 0 | 2349800 | 215030 | 0 | 381960 | 0 | 0 | 0 | 0 | 0 | 0 |
| KGYNVNDI | BASP1; BASP1 | P80723; U3K | BASP1_HUMAN | Unmodified | 0 | 0 | 376760 | 0 | 224280 | 69427 | 0 | 91114 | 22268 | 97100 | 0 | 0 | 0 | 0 | 0 | 0 | 34216 | 105420 |
| KGYPEVALI | COPA; COPA; C | P53621; A0A | COPA_HUMAN | Unmodified | 0 | 0 | 0 | 0 | 0 | 0 | 0 | 0 | 0 | 0 | 0 | 936760 | 0 | 0 | 0 | 184770 | 0 | 0 |
| KHDAQFTV | EPHA3; EPHA3 | P29320; C9J | EPHA3_HUMAN | Oxidation (M) | 132000 | 215990 | 0 | 0 | 0 | 0 | 0 | 0 | 0 | 0 | 0 | 0 | 0 | 0 | 0 | 0 | 0 | 0 |
| KHEAFETDI | SPTAN1; SPTAN | Q13813; A0A | SPTN1_HUMAN | Unmodified | 0 | 0 | 0 | 0 | 0 | 0 | 0 | 0 | 0 | 0 | 55412 | 983830 | 117240 | 0 | 0 | 0 | 0 | 0 |
| KHEAIETDI | SPTBN1; SPTBN | Q01082; A0A | SPTB2_HUMAN | Unmodified | 0 | 0 | 0 | 0 | 0 | 0 | 0 | 0 | 196820 | 0 | 139700 | 1095100 | 0 | 0 | 0 | 0 | 0 | 0 |
| KHEAIETDI | SPTBN2; SPTBN | O15020; A4C | SPTN2_HUMAN | Unmodified | 0 | 0 | 0 | 0 | 0 | 0 | 0 | 0 | 0 | 0 | 0 | 0 | 0 | 0 | 0 | 0 | 0 | 0 |
| KHEALFQL | NOSTRIN; NOS | Q8IVI9; A0A0 | NOSTN_HUMA | Unmodified | 4381800 | 2599300 | 6056700 | 6033000 | 0 | 0 | 0 | 0 | 0 | 0 | 0 | 0 | 0 | 1792200 | 11229000 | 5548000 | 0 | 0 |
| KHEYWFAV | OXR1; OXR1 | Q8N573; H0 | OXR1_HUMAN | Unmodified | 0 | 593080 | 0 | 0 | 0 | 0 | 0 | 69692 | 0 | 0 | 0 | 1016800 | 0 | 0 | 0 | 0 | 0 | 0 |
| KHFFINICH | IGF2R; IGF2R; I | P11717; A0A | MPRI_HUMAN | Unmodified | 0 | 35368 | 62420 | 0 | 0 | 0 | 0 | 0 | 1828400 | 206470 | 0 | 0 | 0 | 0 | 0 | 0 | 839800 | 0 |
| KHFPEFPA | CCDC50 | Q8IVM0 | CCD50_HUMA | Unmodified | 0 | 0 | 2593500 | 3464900 | 0 | 0 | 0 | 0 | 0 | 0 | 0 | 0 | 2289900 | 0 | 3259500 | 1594600 | 0 | 0 |
| KHFPSVNV | ATP6V1A | P38606 | VATA_HUMAN | Unmodified | 89694 | 0 | 0 | 0 | 0 | 0 | 0 | 0 | 0 | 0 | 0 | 1242800 | 0 | 0 | 151790 | 0 | 0 | 0 |
| KHFTILDAP | GSPT1; GSPT2; | P15170; Q8I | ERF3A_HUMAN | Unmodified | 0 | 0 | 0 | 0 | 734610 | 104090 | 306820 | 130330 | 349940 | 126670 | 203360 | 1039200 | 2087600 | 641160 | 0 | 0 | 484880 | 0 |
| KHFVGYPT | PTGR1; PTGR1; | Q14914; F6X | PTGR1_HUMAN | Unmodified | 0 | 0 | 0 | 325970 | 12465000 | 2832700 | 83661 | 39229 | 903810 | 1188500 | 0 | 0 | 1150600 | 0 | 0 | 0 | 836100 | 1575200 |
| KHGIDVRV | AAMDC; AAMD( | Q9H7C9; E9 | AAMDC_HUMA | Unmodified | 0 | 0 | 0 | 0 | 0 | 0 | 0 | 0 | 0 | 0 | 0 | 0 | 403390 | 0 | 0 | 0 | 493150 | 810390 |
| KHINTWVA | SERPINB6; SER | P35237; A0A | SPB6_HUMAN | Unmodified | 0 | 0 | 0 | 0 | 0 | 0 | 0 | 0 | 0 | 0 | 0 | 1435800 | 0 | 0 | 0 | 0 | 0 | 0 |
| KHISQISVA | HINT1 | P49773 | HINT1_HUMAN | Oxidation (M) | 0 | 0 | 0 | 0 | 6350000 | 0 | 5199600 | 0 | 875800 | 2582900 | 0 | 1885600 | 10990000 | 2789600 | 0 | 2184800 | 3980800 | 0 |
| KHISQISVA | HINT1 | P49773 | HINT1_HUMAN | Unmodified | 0 | 0 | 0 | 0 | 0 | 0 | 2784600 | 0 | 1705100 | 0 | 0 | 0 | 2362400 | 0 | 0 | 0 | 1271400 | 1224300 |
| KHISQISVA | HINT1 | P49773 | HINT1_HUMAN | Oxidation (M) | 0 | 689510 | 0 | 1476600 | 17463000 | 961630 | 17074000 | 0 | 4421200 | 7468600 | 721850 | 8884900 | 32069000 | 10980000 | 0 | 454680 | 10278000 | 3858500 |
| KHISQISVA | HINT1 | P49773 | HINT1_HUMAN | Unmodified | 0 | 0 | 0 | 0 | 16125000 | 0 | 5793900 | 0 | 11186000 | 4496100 | 0 | 1432000 | 15413000 | 7625500 | 0 | 0 | 6669100 | 7307000 |
| KHLEINPDI | HSP90AB1 | P08238 | HS90B_HUMAN | Unmodified | 2647000 | 173700 | 156230 | 671410 | 4178000 | 886060 | 0 | 0 | 13679000 | 3743700 | 19570000 | 1.41E+08 | 6456200 | 1542200 | 5682900 | 5478100 | 257970 | 2237600 |
| KHLEINPDI | HSP90AB1 | P08238 | HS90B_HUMAN | Unmodified | 1962800 | 690960 | 0 | 3800900 | 24135000 | 6686900 | 609630 | 0 | 7157400 | 1136200 | 5168400 | 18313000 | 13286000 | 6038400 | 652810 | 0 | 6776500 | 20473000 |
| KHLEINPDI | HSP90AA1 | P07900 | HS90A_HUMAN | Unmodified | 1189500 | 1005000 | 477580 | 4080500 | 2493800 | 1784100 | 494190 | 0 | 16300000 | 2403600 | 14296000 | 52719000 | 1611400 | 0 | 1142600 | 1405400 | 793140 | 4160500 |
| KHLEINPDI | HSP90AA1 | P07900 | HS90A_HUMAN | Unmodified | 0 | 26229 | 210570 | 3858500 | 15560000 | 2856500 | 1135600 | 0 | 7240500 | 3097200 | 5446600 | 6731200 | 3490900 | 1088300 | 947400 | 831990 | 4582100 | 8831100 |
| KHPDADSL | AIMP1; AIMP1 | Q12904; A0A | AIMP1_HUMAN | Unmodified | 0 | 0 | 0 | 309000 | 0 | 0 | 0 | 0 | 0 | 0 | 1248800 | 1109500 | 862240 | 0 | 0 | 0 | 0 | 0 |
| KHPDASVN | HMGB1; HMGB | P09429; Q5T | HMGB1_HUMA | Unmodified | 0 | 0 | 0 | 222840 | 0 | 0 | 340720 | 483870 | 0 | 564290 | 2326100 | 5110300 | 1260400 | 0 | 299980 | 120460 | 0 | 0 |
| KHPDASVN | HMGB1; HMGB | P09429; Q5T | HMGB1_HUMA | Unmodified | 0 | 0 | 0 | 0 | 965150 | 0 | 491260 | 175490 | 0 | 516910 | 815600 | 3425700 | 815020 | 0 | 0 | 114740 | 0 | 0 |
| KHPDSSVN | HMGB2; HMGB | P26583; D6F | HMGB2_HUMA | Unmodified | 0 | 0 | 0 | 239650 | 0 | 141730 | 71561 | 0 | 865510 | 242020 | 0 | 1371200 | 0 | 0 | 0 | 0 | 149570 | 365280 |
| KHPEVDVL | ACLY | P53396 | ACLY_HUMAN | Unmodified | 62553 | 34441 | 103710 | 270860 | 0 | 0 | 74203 | 0 | 341940 | 114630 | 147900 | 5314400 | 66338 | 0 | 190640 | 167720 | 0 | 0 |
| KHQALQAE | SPTAN1; SPTAN | Q13813; A0A | SPTN1_HUMAN | Unmodified | 0 | 0 | 0 | 0 | 133270 | 0 | 0 | 0 | 205340 | 184160 | 974810 | 3337200 | 0 | 0 | 0 | 0 | 54823 | 0 |
| KHQFLAEIL | DIP2B | Q9P265 | DIP2B_HUMAN | Unmodified | 0 | 0 | 0 | 0 | 0 | 0 | 0 | 80655 | 335860 | 0 | 0 | 1481700 | 0 | 0 | 0 | 0 | 0 | 0 |
| KHQLLEAD | SPTAN1; SPTAN | Q13813; A0A | SPTN1_HUMAN | Unmodified | 0 | 0 | 0 | 0 | 0 | 0 | 554580 | 0 | 0 | 165010 | 0 | 2334200 | 0 | 0 | 0 | 0 | 460570 | 0 |
| KHSGFEDE | CHGA; CHGA | P10645; G5E | CMGA_HUMAN | Unmodified | 0 | 0 | 0 | 0 | 0 | 0 | 0 | 0 | 52641000 | 30425000 | 0 | 0 | 0 | 0 | 0 | 0 | 0 | 0 |
| KHSGFEDE | CHGA | P10645 | CMGA_HUMAN | Unmodified | 0 | 0 | 0 | 0 | 0 | 0 | 0 | 0 | 0 | 648410 | 0 | 0 | 0 | 0 | 0 | 0 | 0 | 0 |
| KHSQFIGYI | HSP90AA5P; H | Q58FG0; P0 | HS905_HUMAN | Unmodified | 1140800 | 6993700 | 337070 | 3080100 | 14088000 | 4522200 | 1890700 | 24578 | 11857000 | 8157400 | 8253000 | 40258000 | 6037500 | 799180 | 966330 | 812210 | 4517100 | 8851500 |
| KHSQFIGYI | HSP90AB1; HSI | P08238; Q58 | HS90B_HUMAN | Unmodified | 710600 | 1156300 | 1195500 | 2459200 | 23500000 | 4519700 | 1573100 | 31957 | 8659400 | 5674000 | 6068000 | 27339000 | 16794000 | 6590600 | 1172000 | 1126200 | 8485800 | 7090300 |
| KHTADENP | ECT2 | Q9H8V3 | ECT2_HUMAN | Unmodified | 11062 | 0 | 0 | 636070 | 215080 | 32352 | 0 | 0 | 0 | 59352 | 0 | 0 | 0 | 50379 | 84580 | 0 | 0 | 0 |
| KHTLGEVW | NMD3; NMD3 | Q96D46; C9 | NMD3_HUMAN | Unmodified | 0 | 0 | 0 | 0 | 0 | 0 | 0 | 0 | 0 | 0 | 0 | 570560 | 0 | 0 | 0 | 0 | 0 | 0 |
| KHVLFGTV | THY1; THY1; TH | P04216; J3Q | THY1_HUMAN | Unmodified | 0 | 0 | 0 | 0 | 0 | 0 | 0 | 0 | 0 | 0 | 0 | 1404400 | 0 | 0 | 0 | 0 | 0 | 0 |
| KHVNPVQ | CTNNA1; CTNN | P35221; E5R | CTNA1_HUMAN | Unmodified | 742280 | 773350 | 1549800 | 3305300 | 13041000 | 1955200 | 0 | 0 | 445580 | 410720 | 835040 | 3066200 | 12709000 | 8243800 | 2942000 | 2791400 | 0 | 0 |
| KIAAEYYDN | DEGS1 | O15121 | DEGS1_HUMAN | Unmodified | 0 | 0 | 0 | 187710 | 0 | 0 | 0 | 0 | 1573100 | 0 | 0 | 0 | 0 | 0 | 0 | 0 | 0 | 0 |
| KIADDKYNI | HSP90B1; HSP9 | P14625; A0A | ENPL_HUMAN | Unmodified | 213730 | 0 | 0 | 0 | 0 | 0 | 218920 | 531430 | 49568000 | 44909000 | 6202400 | 8848600 | 0 | 0 | 324370 | 292920 | 0 | 285800 |
| KIAEAGSRI | CTNNA1; CTNN | P35221; P26 | CTNA1_HUMAN | Unmodified | 136810 | 0 | 477770 | 1471000 | 5404100 | 2168600 | 37509 | 0 | 0 | 244290 | 375950 | 256570 | 4380900 | 4219600 | 437620 | 482180 | 25076 | 20593 |
| KIAFAITAIK | RPS18; RPS18; | P62269; A0A | RS18_HUMAN | Unmodified | 0 | 0 | 0 | 0 | 0 | 0 | 0 | 0 | 0 | 0 | 0 | 2196000 | 0 | 0 | 0 | 0 | 0 | 0 |
| KICDGIFYT | PRKAA1 | Q13131 | AAPK1_HUMAN | Unmodified | 74445 | 0 | 0 | 0 | 356860 | 54566 | 198760 | 0 | 276750 | 0 | 0 | 28595 | 189750 | 237310 | 112230 | 55766 | 588660 | 486190 |
| KICTVSENI | GPM6A; GPM6A | P51674; D6F | GPM6A_HUMA | Unmodified | 3506400 | 912520 | 0 | 0 | 0 | 0 | 396890 | 262750 | 0 | 0 | 0 | 0 | 794280 | 0 | 0 | 0 | 0 | 0 |
| KIDAAQNW | VCL | P18206 | VINC_HUMAN | Unmodified | 0 | 0 | 0 | 0 | 0 | 0 | 0 | 0 | 0 | 0 | 0 | 0 | 0 | 0 | 447720 | 339340 | 0 | 0 |
| KIDAAQNW | VCL | P18206 | VINC_HUMAN | Unmodified | 0 | 0 | 0 | 0 | 0 | 1824800 | 643720 | 0 | 0 | 476500 | 0 | 493900 | 0 | 0 | 1952300 | 1826900 | 6572000 | 0 |
| KIDDLIGDL | ENPP4 | Q9Y6X5 | ENPP4_HUMAN | Unmodified | 0 | 0 | 607110 | 0 | 0 | 0 | 0 | 0 | 5539200 | 528800 | 1108900 | 1933800 | 0 | 0 | 0 | 359170 | 0 | 0 |
| KIDDLIGDL | ENPP4 | Q9Y6X5 | ENPP4_HUMAN | Unmodified | 0 | 0 | 0 | 0 | 0 | 0 | 0 | 0 | 2721700 | 2936400 | 1029700 | 0 | 0 | 0 | 0 | 0 | 0 | 0 |
| KIDGDTIIFS | NRCAM; NRCAI | Q92823; A0A | NRCAM_HUMA | Unmodified | 0 | 0 | 0 | 0 | 844160 | 0 | 0 | 0 | 1195300 | 0 | 0 | 0 | 0 | 0 | 0 | 0 | 0 | 0 |
| KIDGQQTIL | CAPZA2; CAPZA | P47755; A0A | CAZA2_HUMAN | Unmodified | 0 | 699760 | 456460 | 607570 | 2702700 | 1060800 | 858440 | 90608 | 745330 | 215450 | 871340 | 4434900 | 2011500 | 1420400 | 710880 | 565290 | 1698600 | 4043200 |
| KIDKYTEVL | RPL13A; RPL13 | P40429; A0A | RL13A_HUMAN | Unmodified | 0 | 631770 | 0 | 0 | 800720 | 0 | 0 | 324260 | 0 | 0 | 0 | 557940 | 1440500 | 387290 | 0 | 0 | 103900 | 431210 |
| KIDQEGKIF | ACAD9; ACAD9 | Q9H845; A0 | ACAD9_HUMAN | Unmodified | 0 | 0 | 0 | 0 | 196440 | 0 | 0 | 0 | 2099900 | 0 | 0 | 0 | 0 | 0 | 0 | 0 | 368990 | 0 |
| KIEDNNTL | RPL23A; RPL23 | P62750; K7E | RL23A_HUMAN | Unmodified | 0 | 0 | 0 | 0 | 349280 | 0 | 0 | 0 | 0 | 113050 | 257430 | 1439300 | 0 | 0 | 0 | 0 | 0 | 0 |
| KIEEQLTLE | SERPINB1 | P30740 | ILEU_HUMAN L | Unmodified | 61985 | 93124 | 0 | 0 | 0 | 0 | 0 | 0 | 445360 | 0 | 0 | 0 | 0 | 0 | 0 | 0 | 0 | 0 |
| KIEESETIEI | OXR1; OXR1 | Q8N573; H0 | OXR1_HUMAN | Unmodified | 0 | 0 | 0 | 0 | 0 | 0 | 0 | 0 | 129730 | 29493 | 147820 | 1075800 | 0 | 0 | 0 | 0 | 0 | 0 |
| KIEFSLPDL | PSMC2; PSMC2 | P35998; C9J | PRS7_HUMAN | Unmodified | 105770 | 0 | 205790 | 194180 | 122010 | 0 | 0 | 0 | 289670 | 0 | 222060 | 3033900 | 0 | 0 | 255700 | 156820 | 0 | 0 |
| KIEFSLPDL | PSMC2; PSMC2 | P35998; C9J | PRS7_HUMAN | Unmodified | 0 | 0 | 0 | 0 | 1131400 | 69376 | 698590 | 0 | 434220 | 248520 | 0 | 0 | 912780 | 0 | 0 | 0 | 352670 | 0 |
| KIEIGDGAE | TWF2; TWF2 | Q6IBS0; A0A | TWF2_HUMAN | Unmodified | 0 | 0 | 0 | 0 | 0 | 0 | 0 | 0 | 0 | 0 | 0 | 1639500 | 0 | 0 | 0 | 0 | 0 | 1890200 |
| KIEISQHAK | RPL37A; RPL37 | P61513; M0F | RL37A_HUMAN | Unmodified | 0 | 0 | 0 | 0 | 0 | 0 | 406660 | 185130 | 0 | 0 | 143500 | 0 | 116070 | 0 | 0 | 0 | 0 | 0 |
| KIELDVKYN | BAIAP2L1 | Q9UHR4 | BI2L1_HUMAN | Unmodified | 0 | 0 | 0 | 0 | 382110 | 0 | 0 | 0 | 0 | 0 | 0 | 0 | 0 | 0 | 0 | 0 | 180390 | 0 |
| KIENSELLD | ZC3HAV1; ZC3H | Q7Z2W4; C9 | ZCCHV_HUMA | Unmodified | 0 | 0 | 287560 | 0 | 0 | 0 | 0 | 0 | 0 | 0 | 0 | 0 | 0 | 0 | 0 | 0 | 0 | 0 |
| KIENSELLD | ZC3HAV1; ZC3H | Q7Z2W4; C9 | ZCCHV_HUMA | Unmodified | 0 | 0 | 0 | 0 | 464270 | 0 | 0 | 0 | 0 | 0 | 0 | 0 | 0 | 0 | 0 | 0 | 27746 | 0 |
| KIEPEAVLQ | EPB41L1; EPB4 | Q9H4G0; H7 | E41L1_HUMAN | Unmodified | 38234 | 0 | 0 | 561850 | 350700 | 0 | 0 | 0 | 0 | 0 | 570850 | 1142900 | 806020 | 0 | 250850 | 596650 | 141860 | 370420 |
| KIEPELDGS | GPI; GPI; GPI | P06744; K7E | G6PI_HUMAN | Unmodified | 0 | 1745900 | 2480700 | 10220000 | 12116000 | 10462000 | 12274000 | 308860 | 22246000 | 15596000 | 15278000 | 60354000 | 21564000 | 11929000 | 4047600 | 3892500 | 17977000 | 18667000 |
| KIEPELDGS | GPI; GPI; GPI | P06744; K7E | G6PI_HUMAN | Unmodified | 0 | 0 | 0 | 0 | 0 | 0 | 0 | 0 | 0 | 0 | 0 | 0 | 0 | 0 | 0 | 0 | 766600 | 0 |
| KIEQELTAA | CHMP4B | Q9H444 | CHM4B_HUMA | Unmodified | 3810200 | 3872100 | 2552400 | 2488300 | 14210000 | 2880900 | 891430 | 0 | 7675400 | 3304900 | 870990 | 2834000 | 8933400 | 5355800 | 3229600 | 2767400 | 835000 | 0 |
| KIEQELTAA | CHMP4B | Q9H444 | CHM4B_HUMA | Unmodified | 7156800 | 7707200 | 3759300 | 6856400 | 32891000 | 2315500 | 1548300 | 489320 | 14864000 | 6825800 | 2822900 | 4550400 | 22384000 | 10877000 | 4599900 | 5158700 | 1349900 | 7876100 |
| KIERLDTDI | DDX19B; DDX1 | Q9UMR2; Q9 | DD19B_HUMAN | Unmodified | 208830 | 127450 | 105550 | 150690 | 958200 | 146410 | 83772 | 44297 | 110710 | 82442 | 121470 | 794990 | 468120 | 236650 | 0 | 109190 | 108630 | 0 |
| KIESFGSK | PPIF | P30405 | PPIF_HUMAN F | Unmodified | 0 | 0 | 113720 | 0 | 0 | 0 | 376680 | 57568 | 0 | 1541500 | 60079 | 0 | 139210 | 95735 | 0 | 0 | 0 | 0 |
| KIFDIDEAE | EIF2S2 | P20042 | IF2B_HUMAN E | Unmodified | 0 | 155360 | 0 | 521210 | 806300 | 0 | 0 | 0 | 924440 | 0 | 0 | 1999600 | 0 | 0 | 0 | 446120 | 206580 | 834030 |
| KIFQEALVC | NIBAN1; NIBAN | Q9BZQ8; H0 | NIBA1_HUMAN | Unmodified | 0 | 0 | 0 | 0 | 341310 | 0 | 0 | 0 | 427480 | 456560 | 0 | 0 | 0 | 0 | 0 | 0 | 0 | 0 |
| KIFVGGIKE | HNRNPA3; HNI | P51991; A0A | ROA3_HUMAN | Unmodified | 0 | 0 | 0 | 0 | 0 | 0 | 0 | 0 | 0 | 0 | 0 | 365960 | 0 | 0 | 0 | 0 | 0 | 0 |
| KIFVGGLNI | HNRNPAB; HNI | D6R9P3; D6I | D6R9P3_HUMA | Unmodified | 225530 | 0 | 0 | 0 | 587720 | 0 | 0 | 0 | 0 | 1029900 | 0 | 8446300 | 0 | 0 | 0 | 0 | 0 | 0 |
| KIGCGNFG | CSNK1G1; CSN | Q9HCP0; P7 | KC1G1_HUMAN | Unmodified | 0 | 0 | 0 | 0 | 422930 | 144330 | 137390 | 0 | 0 | 0 | 0 | 0 | 244010 | 170500 | 0 | 0 | 269880 | 153090 |

| | | | | | | | | | | | | | | | | | | | | | | |
|---|---|---|---|---|---|---|---|---|---|---|---|---|---|---|---|---|---|---|---|---|---|---|
| KIGDEYFTF | CCT3; CCT3 | P49368; B4D | TCPG_HUMAN | Unmodified | 0 | 0 | 216210 | 0 | 0 | 0 | 0 | 0 | 325010 | 0 | 0 | 1386100 | 0 | 0 | 0 | 0 | 0 | 0 |
| KIGDEYFTF | CCT3; CCT3 | P49368; B4D | TCPG_HUMAN | Unmodified | 0 | 73984 | 665010 | 284890 | 1272600 | 164420 | 0 | 0 | 2548300 | 548860 | 629670 | 8881600 | 620790 | 0 | 848210 | 0 | 78454 | 0 |
| KIGEAEAAV | FLOT2; FLOT2; I | Q14254; J3Q | FLOT2_HUMAN | Unmodified | 0 | 0 | 0 | 0 | 0 | 0 | 0 | 0 | 0 | 0 | 402680 | 384390 | 0 | 0 | 0 | 0 | 0 | 0 |
| KIGEEEIQK | SF3A1 | Q15459 | SF3A1_HUMAN | Unmodified | 378900 | 259760 | 0 | 0 | 0 | 0 | 0 | 0 | 0 | 0 | 0 | 210650 | 0 | 0 | 0 | 43109 | 0 | 0 |
| KIGGIFAFK | SCP2; SCP2 | P22307; H0Y | SCP2_HUMAN | Unmodified | 0 | 0 | 0 | 0 | 0 | 0 | 0 | 0 | 22059000 | 6012800 | 0 | 0 | 0 | 0 | 0 | 0 | 0 | 0 |
| KIGLDDEEI | MYO6; MYO6; M | Q9UM54; A0 | MYO6_HUMAN | Unmodified | 230250 | 202330 | 344120 | 768370 | 0 | 0 | 0 | 0 | 404620 | 0 | 0 | 0 | 0 | 0 | 667140 | 512190 | 0 | 0 |
| KIGQQPQC | KHSRP; KHSRP | Q92945; M0 | FUBP2_HUMAN | Unmodified | 0 | 0 | 0 | 0 | 567900 | 0 | 131680 | 316980 | 1438800 | 977110 | 425750 | 3715500 | 0 | 0 | 0 | 0 | 279810 | 518340 |
| KIGYNPAT\ | EEF1A2; EEF1A | Q05639; A0A | EF1A2_HUMAN | 2 Oxidation (M) | 23011000 | 28868000 | 686530 | 26482000 | 10542000 | 0 | 5474300 | 0 | 0 | 0 | 0 | 0 | 21584000 | 20963000 | 47874000 | 195850 | 0 | 0 |
| KIGYNPDT\ | EEF1A1P5; EEF | Q5VTE0; P68 | EF1A3_HUMAN | 2 Oxidation (M) | 17538000 | 4685800 | 3404500 | 21312000 | 54503000 | 1945000 | 1205400 | 0 | 1946400 | 2059300 | 178150 | 2755500 | 939140 | 22872000 | 31027000 | 4109600 | 26600000 | 34349000 |
| KIGYNPDT\ | EEF1A1P5; EEF | Q5VTE0; P68 | EF1A3_HUMAN | Oxidation (M) | 433250 | 1118400 | 0 | 0 | 0 | 0 | 0 | 0 | 232060 | 0 | 0 | 0 | 1725200 | 758370 | 0 | 0 | 918740 | 2354300 |
| KIHEDELNI | CCPG1; CCPG | Q9ULG6; H3 | CCPG1_HUMA | Unmodified | 0 | 0 | 223340 | 189630 | 0 | 0 | 0 | 14426 | 205480 | 0 | 0 | 0 | 0 | 0 | 0 | 0 | 0 | 0 |
| KIHIDLPNE | PSMC6; PSMC6 | P62333; A0A | PRS10_HUMAN | Unmodified | 0 | 0 | 0 | 236260 | 1803700 | 289590 | 867410 | 0 | 446400 | 892960 | 0 | 0 | 2620600 | 1169600 | 0 | 0 | 1257000 | 1373300 |
| KIHPQTIIAC | CCT2; CCT2; C | P78371; F5G | TCPB_HUMAN | Unmodified | 0 | 0 | 0 | 604890 | 10247000 | 869510 | 7284600 | 0 | 1045300 | 3494900 | 851510 | 0 | 25573000 | 16265000 | 0 | 0 | 10749000 | 5919900 |
| KIHQIELIPN | CDC42BPA; CD | Q5VT25; A0A | MRCKA_HUMA | Unmodified | 0 | 0 | 0 | 0 | 0 | 0 | 0 | 0 | 0 | 0 | 0 | 210110 | 0 | 0 | 0 | 52911 | 31611 | 84003 |
| KIHTEPQLS | QARS1; QARS1 | P47897; A0A | SYQ_HUMAN G | Unmodified | 0 | 0 | 0 | 0 | 0 | 0 | 0 | 0 | 0 | 0 | 144860 | 1676100 | 0 | 0 | 26961 | 24158 | 0 | 0 |
| KIHVYGYSI | PHPT1 | Q9NRX4 | PHP14_HUMAN | Oxidation (M) | 336130 | 0 | 0 | 0 | 185960 | 0 | 0 | 137310 | 508600 | 1224200 | 271820 | 2163800 | 3367400 | 1588400 | 1710800 | 732470 | 781100 | 683870 |
| KIHVYGYSI | PHPT1 | Q9NRX4 | PHP14_HUMAN | Unmodified | 501250 | 0 | 0 | 772480 | 3825900 | 0 | 1321800 | 0 | 2286600 | 1288400 | 530510 | 2292100 | 3423800 | 2086100 | 765050 | 323550 | 1547000 | 3194300 |
| KIIADIKK | CHMP2A; CHM | O43633; M0 | CHM2A_HUMA | Unmodified | 0 | 220750 | 0 | 0 | 806550 | 141800 | 0 | 0 | 1646200 | 269190 | 177620 | 2613800 | 0 | 138310 | 0 | 103930 | 0 | 0 |
| KIIAKGEELI | MYO1C; MYO1C | O00159; F5H | MYO1C_HUMA | Unmodified | 1946100 | 1876200 | 0 | 3851600 | 0 | 0 | 0 | 0 | 139450 | 0 | 0 | 0 | 0 | 0 | 1009100 | 1059100 | 0 | 0 |
| KIIEDQQES | SSB | P05455 | LA_HUMAN Lup | Unmodified | 0 | 0 | 82889 | 0 | 0 | 0 | 0 | 0 | 0 | 0 | 496000 | 2089000 | 328700 | 18135 | 0 | 0 | 0 | 0 |
| KIIENELEGI | DRG1 | Q9Y295 | DRG1_HUMAN | Unmodified | 0 | 0 | 0 | 0 | 0 | 0 | 0 | 0 | 0 | 0 | 0 | 868450 | 0 | 0 | 0 | 0 | 0 | 0 |
| KIIETPGIR | LRPPRC | P42704 | LPPRC_HUMAN | Unmodified | 0 | 0 | 0 | 0 | 0 | 0 | 0 | 0 | 486870 | 53900 | 0 | 0 | 0 | 0 | 0 | 0 | 27054 | 481370 |
| KIIPTLEEYC | HNRNPK | Q5T6W2 | Q5T6W2_HUM | Unmodified | 0 | 0 | 0 | 0 | 0 | 0 | 0 | 0 | 0 | 0 | 0 | 1428400 | 0 | 0 | 0 | 0 | 0 | 0 |
| KIKVDEFVT | ADH5 | P11766 | ADHX_HUMAN | Unmodified | 0 | 0 | 0 | 966340 | 10788000 | 1792100 | 0 | 30710 | 753390 | 0 | 4086800 | 11640000 | 2928600 | 1968500 | 0 | 0 | 3042000 | 4926800 |
| KILAIELENI | SHTN1 | A0MZ66 | SHOT1_HUMAI | Unmodified | 0 | 0 | 0 | 0 | 0 | 0 | 1096300 | 644980 | 0 | 979990 | 0 | 0 | 0 | 0 | 0 | 0 | 0 | 0 |
| KILATPPQE | TKT; TKT; TKT | P29401; A0A | TKT_HUMAN Tr | Unmodified | 0 | 0 | 0 | 0 | 608200 | 0 | 0 | 0 | 823220 | 193940 | 556860 | 1652700 | 0 | 0 | 0 | 0 | 106860 | 0 |
| KILDDWGE | MANF | P55145 | MANF_HUMAN | Unmodified | 0 | 0 | 0 | 0 | 0 | 0 | 0 | 0 | 8799100 | 2909200 | 0 | 0 | 0 | 0 | 0 | 0 | 0 | 0 |
| KILDDWGE | MANF | P55145 | MANF_HUMAN | Unmodified | 0 | 0 | 0 | 0 | 0 | 0 | 110520 | 45243 | 0 | 560210 | 0 | 0 | 0 | 0 | 0 | 0 | 0 | 0 |
| KILDSVGIE | RPLP2 | P05387 | RLA2_HUMAN ( | Unmodified | 8886300 | 9543000 | 5036400 | 11225000 | 0 | 0 | 0 | 0 | 260060 | 0 | 1446300 | 23779000 | 4967400 | 519550 | 8707300 | 8557400 | 0 | 0 |
| KILEVHIDK | DNAJA1 | P31689 | DNJA1_HUMAN | Unmodified | 0 | 0 | 0 | 147300 | 1171300 | 0 | 0 | 0 | 1104000 | 1039400 | 0 | 167200 | 1482300 | 0 | 77840 | 77423 | 107720 | 207260 |
| KILEVVNQI | ENDOD1 | O94919 | ENDD1_HUMA | Unmodified | 0 | 0 | 0 | 0 | 0 | 0 | 0 | 0 | 691860 | 55673 | 0 | 0 | 0 | 0 | 136420 | 0 | 113830 | 0 |
| KILGNKATF | ANK3; ANK3; AI | Q12955; A0A | ANK3_HUMAN | Unmodified | 0 | 0 | 0 | 0 | 0 | 0 | 0 | 0 | 0 | 0 | 0 | 0 | 694010 | 0 | 0 | 0 | 0 | 0 |
| KILIEDWK | PRKCSH; PRKC | P14314; K7E | GLU2B_HUMAI | Unmodified | 0 | 0 | 0 | 0 | 0 | 0 | 0 | 0 | 5274400 | 3142200 | 0 | 0 | 0 | 0 | 0 | 0 | 0 | 0 |
| KILIEDWKK | PRKCSH; PRKC | P14314; K7E | GLU2B_HUMAI | Unmodified | 0 | 0 | 0 | 0 | 0 | 0 | 209310 | 57706 | 2517500 | 2222900 | 0 | 0 | 0 | 0 | 0 | 0 | 0 | 0 |
| KILQDGGL | PHGDH; PHGD | O43175; A0A | SERA_HUMAN | Unmodified | 0 | 0 | 0 | 175810 | 0 | 0 | 527900 | 130160 | 0 | 0 | 0 | 0 | 1805500 | 312340 | 0 | 220910 | 0 | 0 |
| KILYDILAFA | DNAJC10; DNA | Q8IXB1; A0A | DJC10_HUMAN | Unmodified | 0 | 0 | 0 | 0 | 0 | 0 | 0 | 0 | 5394600 | 1551200 | 0 | 0 | 0 | 0 | 0 | 0 | 0 | 0 |
| KINEELESC | SMAP; C11orf5 | O00193; E9F | SMAP_HUMAN | Oxidation (M) | 0 | 0 | 0 | 0 | 0 | 0 | 0 | 0 | 0 | 0 | 0 | 905980 | 0 | 0 | 0 | 0 | 0 | 0 |
| KINELMPK | MANF | P55145 | MANF_HUMAN | Unmodified | 0 | 0 | 0 | 0 | 0 | 0 | 0 | 0 | 311720 | 407980 | 0 | 0 | 0 | 0 | 0 | 0 | 0 | 0 |
| KINELTGIK | PSMC2; PSMC2 | P35998; C9J | PRS7_HUMAN | Unmodified | 0 | 0 | 0 | 0 | 0 | 0 | 0 | 168410 | 0 | 0 | 0 | 1032200 | 0 | 0 | 0 | 0 | 0 | 0 |
| KINELTGIKI | PSMC2; PSMC2 | P35998; C9J | PRS7_HUMAN | Unmodified | 0 | 0 | 0 | 0 | 1514900 | 0 | 0 | 0 | 0 | 0 | 0 | 0 | 0 | 0 | 0 | 0 | 465000 | 0 |
| KINPENSKI | NUDC | Q9Y266 | NUDC_HUMAN | Unmodified | 0 | 0 | 0 | 0 | 0 | 0 | 0 | 0 | 200500 | 149120 | 138020 | 309710 | 288580 | 0 | 0 | 0 | 0 | 0 |
| KINPICNDI | PBK; PBK | Q96KB5; E5F | TOPK_HUMAN | Unmodified | 180200 | 0 | 0 | 112390 | 0 | 0 | 0 | 0 | 0 | 0 | 0 | 0 | 983780 | 0 | 0 | 0 | 67398 | 0 |
| KINPICNDI | PBK; PBK | Q96KB5; E5F | TOPK_HUMAN | Unmodified | 0 | 0 | 0 | 0 | 0 | 0 | 0 | 70003 | 0 | 0 | 0 | 0 | 0 | 0 | 0 | 0 | 975290 | 0 |
| KINQLISETI | XPR1 | Q9UBH6 | XPR1_HUMAN | Unmodified | 0 | 0 | 0 | 0 | 2018600 | 159270 | 499190 | 0 | 0 | 402510 | 208070 | 495170 | 739050 | 0 | 0 | 0 | 0 | 0 |
| KINRLSTPI | LMO7; LMO7; LI | Q8WWI1; E9 | LMO7_HUMAN | Unmodified | 0 | 0 | 0 | 0 | 0 | 0 | 0 | 0 | 0 | 0 | 0 | 0 | 0 | 0 | 0 | 0 | 1169800 | 1200900 |
| KINSDSEEI | UTRN | P46939 | UTRN_HUMAN | Unmodified | 10218 | 1212100 | 765470 | 1996700 | 0 | 0 | 0 | 0 | 0 | 0 | 404650 | 0 | 0 | 0 | 165690 | 1348600 | 0 | 0 |
| KIPGFCEG | SERINC3 | Q13530 | SERC3_HUMAN | Unmodified | 1560100 | 498200 | 761420 | 254260 | 0 | 0 | 0 | 0 | 538900 | 328920 | 197750 | 1672200 | 0 | 0 | 0 | 704630 | 0 | 0 |
| KIPLAQSEA | CNPY1; CNPY1 | A0A1B0GVE | A0A1B0GVE0_I | Unmodified | 0 | 0 | 0 | 0 | 0 | 0 | 0 | 0 | 0 | 0 | 1111200 | 0 | 0 | 0 | 0 | 0 | 0 | 0 |
| KIPNPDFFE | CANX; CANX; C | P27824; A0A | CALX_HUMAN | Unmodified | 72185 | 0 | 113410 | 0 | 0 | 0 | 0 | 18100 | 9.86E+08 | 39391000 | 14763000 | 10435000 | 0 | 0 | 1142700 | 77578 | 0 | 0 |
| KIPVFHNG | SLC4A7; SLC4A | Q9Y6M7; C9 | S4A7_HUMAN | Unmodified | 0 | 0 | 0 | 0 | 0 | 0 | 0 | 0 | 0 | 0 | 255050 | 1372800 | 0 | 194570 | 0 | 0 | 0 | 0 |
| KIPVLVQSK | GTPBP1 | O00178 | GTPB1_HUMAN | Unmodified | 0 | 134560 | 67439 | 114460 | 0 | 0 | 0 | 0 | 0 | 0 | 0 | 0 | 0 | 0 | 183280 | 156900 | 83585 | 0 |
| KIQALQQQ | TPM4; TPM4; TF | P67936; A0A | TPM4_HUMAN | Unmodified | 847210 | 661780 | 0 | 2793500 | 0 | 2104900 | 0 | 48714 | 331390 | 819200 | 388780 | 339710 | 2242500 | 495430 | 2559900 | 0 | 0 | 0 |
| KIQASTMAI | TAGLN2; TAGLI | P37802; X6R | TAGL2_HUMAN | Unmodified | 653000 | 643900 | 0 | 432050 | 11792000 | 1325200 | 1690000 | 287630 | 918200 | 1052000 | 940920 | 1995400 | 2002700 | 1248400 | 104210 | 168770 | 3231700 | 4559000 |
| KIQDEQTT\ | JAM3 | Q9BX67 | JAM3_HUMAN | Unmodified | 0 | 0 | 0 | 0 | 0 | 0 | 0 | 0 | 2460600 | 196480 | 0 | 2942600 | 0 | 0 | 0 | 0 | 0 | 0 |
| KIQDHQVV | ENAH; ENAH; E | Q8N8S7; A0 | ENAH_HUMAN | Unmodified | 0 | 0 | 271240 | 199240 | 162060 | 0 | 99572 | 338050 | 474330 | 359190 | 325250 | 4131900 | 301140 | 191550 | 210430 | 318320 | 0 | 0 |
| KIQESYGD | ATP2B1; ATP2B | P20020; F8V | AT2B1_HUMAN | Unmodified | 0 | 0 | 69859 | 887720 | 916320 | 0 | 0 | 59309 | 1042400 | 0 | 0 | 577410 | 2551200 | 1682700 | 491520 | 401040 | 365090 | 264130 |
| KIQMSNLM | ATP6V1E1; ATP | P36543; C9J | VATE1_HUMAN | Unmodified | 0 | 0 | 0 | 0 | 406450 | 0 | 140580 | 0 | 0 | 0 | 0 | 1197100 | 274150 | 162530 | 0 | 0 | 0 | 0 |
| KIQMSNLM | ATP6V1E1; ATP | P36543; C9J | VATE1_HUMAN | Unmodified | 0 | 0 | 0 | 0 | 0 | 0 | 194980 | 0 | 0 | 59497 | 0 | 0 | 0 | 0 | 0 | 0 | 54756 | 0 |
| KIQQNTFTI | FLNC; FLNB; FL | Q14315; O7 | FLNC_HUMAN | Unmodified | 0 | 0 | 0 | 0 | 5668900 | 758220 | 0 | 0 | 0 | 0 | 0 | 0 | 0 | 0 | 0 | 0 | 990680 | 1232000 |
| KIQVLQQQ | TPM3; TPM3; TF | D6R904; A0 | D6R904_HUMA | Unmodified | 0 | 0 | 0 | 0 | 0 | 0 | 0 | 0 | 0 | 34973 | 1559200 | 369850 | 0 | 0 | 0 | 0 | 0 | 0 |
| KIQVLQQQ | TPM3; TPM3; TF | D6R904; A0 | D6R904_HUMA | Unmodified | 14545 | 0 | 0 | 0 | 0 | 0 | 406400 | 0 | 0 | 39705 | 3857300 | 669390 | 0 | 17780 | 0 | 0 | 0 | 0 |
| KIREGQVE | CAVIN1 | Q6NZI2 | CAVN1_HUMAI | Unmodified | 0 | 0 | 0 | 0 | 3714300 | 0 | 0 | 0 | 0 | 0 | 0 | 0 | 0 | 0 | 0 | 0 | 0 | 0 |
| KIRNAQLA | TARS1; TARS3 | P26639; A2F | SYTC_HUMAN | Unmodified | 0 | 0 | 0 | 0 | 712990 | 84012 | 129750 | 0 | 0 | 76424 | 0 | 0 | 130740 | 128670 | 0 | 0 | 116120 | 124190 |
| KISEIEDAAI | ERLIN2; ERLIN2 | O94905; E5F | ERLN2_HUMAN | Unmodified | 0 | 1902200 | 0 | 0 | 0 | 0 | 0 | 44163 | 37948000 | 1557300 | 178320 | 2599700 | 0 | 0 | 0 | 0 | 0 | 0 |
| KISEIEDAAI | ERLIN2; ERLIN2 | O94905; E5F | ERLN2_HUMAN | Unmodified | 0 | 0 | 0 | 0 | 254630 | 0 | 0 | 0 | 11546000 | 1609700 | 328380 | 0 | 0 | 0 | 0 | 0 | 0 | 0 |
| KISELDAFL | PSME1 | Q06323 | PSME1_HUMAI | Unmodified | 0 | 0 | 0 | 0 | 0 | 711280 | 0 | 0 | 2103200 | 2606700 | 463080 | 0 | 2881300 | 1118700 | 0 | 0 | 4442700 | 11861000 |
| KISGGSVVI | IDH1; IDH1; IDI | O75874; C9J | IDHC_HUMAN | Unmodified | 133910 | 162790 | 190090 | 382800 | 262980 | 0 | 0 | 0 | 1937000 | 0 | 429600 | 2848300 | 487420 | 305210 | 1641300 | 1191500 | 0 | 84723 |
| KISGGSVVI | IDH1; IDH1; IDI | O75874; C9J | IDHC_HUMAN | 2 Oxidation (M) | 36366 | 140580 | 0 | 54132 | 0 | 0 | 82023 | 0 | 82852 | 0 | 118990 | 521970 | 1786400 | 70565 | 0 | 0 | 0 | 0 |
| KISGGSVVI | IDH1; IDH1; IDI | O75874; C9J | IDHC_HUMAN | Oxidation (M) | 0 | 0 | 0 | 0 | 0 | 0 | 0 | 0 | 140680 | 0 | 54706 | 263190 | 671700 | 70442 | 1024200 | 651190 | 0 | 0 |
| KISGGSVVI | IDH1; IDH1; IDI | O75874; C9J | IDHC_HUMAN | Oxidation (M) | 0 | 0 | 0 | 0 | 0 | 0 | 0 | 0 | 0 | 36489 | 0 | 0 | 1004200 | 458180 | 0 | 0 | 0 | 120780 |
| KISGGSVVI | IDH1; IDH1; IDI | O75874; C9J | IDHC_HUMAN | 2 Oxidation (M) | 0 | 0 | 0 | 0 | 0 | 0 | 0 | 0 | 0 | 0 | 0 | 0 | 1635600 | 1687800 | 0 | 0 | 0 | 0 |
| KISLEDIQA | DNAJC9 | Q8WXX5 | DNJC9_HUMAI | Unmodified | 0 | 0 | 0 | 0 | 0 | 0 | 128920 | 0 | 943580 | 784320 | 472450 | 867220 | 330640 | 0 | 0 | 0 | 0 | 287240 |
| KISRLDAEL | CHMP5 | Q9NZZ3 | CHMP5_HUMA | Unmodified | 0 | 0 | 0 | 0 | 27123000 | 0 | 0 | 0 | 10557000 | 9048200 | 4830200 | 3863800 | 7085400 | 5647600 | 447520 | 0 | 483130 | 2311300 |
| KISSDLDGI | TKT; TKT; TKT | P29401; F8V | TKT_HUMAN Tr | Unmodified | 0 | 0 | 0 | 0 | 481420 | 0 | 64393 | 0 | 0 | 461800 | 1088000 | 1088400 | 52200 | 0 | 42697 | 0 | 240780 | 0 |
| KISSIQSIVF | HSPD1; HSPD1 | P10809; A0A | CH60_HUMAN | Unmodified | 0 | 0 | 0 | 0 | 0 | 0 | 0 | 0 | 9392300 | 3768300 | 0 | 0 | 0 | 0 | 0 | 0 | 0 | 0 |
| KISSIQSIVF | HSPD1; HSPD1 | P10809; A0A | CH60_HUMAN | Unmodified | 0 | 0 | 0 | 0 | 0 | 0 | 0 | 0 | 1232500 | 0 | 0 | 0 | 0 | 0 | 0 | 0 | 0 | 0 |
| KISTVVSSK | SRP14; SRP14 | P37108; H0Y | SRP14_HUMAN | Unmodified | 1622500 | 0 | 2329600 | 2434700 | 942250 | 56779 | 498630 | 169920 | 0 | 80786 | 0 | 2558000 | 371510 | 151360 | 6449400 | 5227200 | 0 | 0 |
| KITAFVPND | RPS23; RPS23 | P62266; D6F | RS23_HUMAN | Unmodified | 696350 | 1734300 | 0 | 1521400 | 1092600 | 0 | 0 | 0 | 0 | 0 | 0 | 262760 | 2276800 | 1132900 | 0 | 0 | 0 | 1064700 |
| KITESVAET | SYAP1 | Q96A49 | SYAP1_HUMAN | Unmodified | 0 | 0 | 0 | 0 | 608360 | 48754 | 259960 | 54361 | 216070 | 271590 | 135490 | 653190 | 1130100 | 653300 | 1488000 | 0 | 917800 | 425380 |
| KITIKNDPS | NARS1; NARS1 | O43776; K7E | SYNC_HUMAN | Unmodified | 0 | 0 | 0 | 0 | 934370 | 132930 | 493840 | 231430 | 415900 | 247670 | 0 | 542570 | 2612100 | 1591300 | 0 | 0 | 0 | 758160 |
| KITNSLTVL | EIF3J; EIF3J; EIF | O75822; H0Y | EIF3J_HUMAN | Unmodified | 0 | 0 | 0 | 0 | 320800 | 0 | 0 | 0 | 0 | 0 | 0 | 734080 | 404930 | 0 | 0 | 0 | 0 | 0 |
| KIVDQNTKI | CPE; CPE; CPE; | P16870; C9J | CBPE_HUMAN | Unmodified | 0 | 0 | 0 | 0 | 0 | 0 | 0 | 258900 | 0 | 920560 | 0 | 0 | 0 | 0 | 0 | 0 | 0 | 0 |
| KIVEGISQP | PMVK | Q15126 | PMVK_HUMAN | Unmodified | 203030 | 196430 | 73106 | 212520 | 0 | 0 | 20939 | 0 | 0 | 0 | 38438 | 242720 | 83697 | 0 | 135200 | 164020 | 42083 | 18027 |
| KIVGAEIQH | PXDN | Q92626 | PXDN_HUMAN | Unmodified | 0 | 0 | 0 | 0 | 0 | 0 | 0 | 0 | 614630 | 0 | 0 | 0 | 0 | 0 | 0 | 0 | 0 | 0 |
| KIVIYVQQT | RABGAP1 | Q9Y3P9 | RBGP1_HUMAI | Unmodified | 0 | 0 | 0 | 44964 | 647350 | 0 | 219660 | 48686 | 118260 | 0 | 0 | 496820 | 147750 | 125580 | 42661 | 55894 | 29699 | 86105 |
| KIVNSAQT | ATP5MG; N/A; | O75964; A0A | ATP5L_HUMAN | Unmodified | 0 | 0 | 0 | 0 | 0 | 0 | 0 | 0 | 1538900 | 378530 | 0 | 0 | 0 | 0 | 0 | 0 | 0 | 0 |
| KIVSWLEYE | FKBP4 | Q02790 | FKBP4_HUMAN | Unmodified | 0 | 0 | 0 | 0 | 0 | 0 | 0 | 0 | 0 | 0 | 276430 | 974810 | 1040400 | 0 | 2158300 | 0 | 0 | 0 |
| KIWCFGPD | EEF2 | P13639 | EF2_HUMAN El | Unmodified | 690650 | 308550 | 0 | 3172200 | 7508700 | 839170 | 779890 | 0 | 3336800 | 1666400 | 2508300 | 18900000 | 5885100 | 1425200 | 1463700 | 1804900 | 4025800 | 3178700 |
| KIYFTDSSS | APMAP; APMAP | Q9HDC9; H | APMAP_HUMAI | Unmodified | 0 | 0 | 0 | 0 | 0 | 0 | 0 | 0 | 3597400 | 212890 | 0 | 0 | 0 | 0 | 0 | 0 | 0 | 0 |
| KIYIGDDNI | ITGAV | P06756 | ITAV_HUMAN I | Unmodified | 0 | 0 | 465290 | 870220 | 1758400 | 661080 | 244910 | 0 | 1725700 | 301610 | 495180 | 1943900 | 864200 | 389770 | 1425900 | 1193600 | 656740 | 39400 |
| KIYQNIQD | RAB14 | P61106 | RAB14_HUMAN | Unmodified | 1491100 | 870610 | 0 | 2931700 | 1227800 | 229240 | 0 | 0 | 7696600 | 682880 | 5444100 | 23010000 | 1596100 | 0 | 566160 | 1440800 | 0 | 0 |
| KKATDGVT | PACSIN2; PACS | Q9UNF0; A0 | PACN2_HUMAI | Unmodified | 0 | 0 | 0 | 0 | 0 | 0 | 0 | 0 | 0 | 0 | 0 | 0 | 0 | 0 | 1138100 | 665460 | 0 | 0 |
| KKDDETVD | CDC23 | Q9UJX2 | CDC23_HUMA | Unmodified | 0 | 0 | 0 | 39871 | 76554 | 24635 | 59006 | 37476 | 0 | 41021 | 0 | 801720 | 0 | 0 | 0 | 54819 | 23155 | 81763 |
| KKDEELSS | TSG101; TSG10 | Q99816; F5H | TS101_HUMAN | Unmodified | 64904000 | 46425000 | 63317000 | 38412000 | 3195400 | 1066000 | 0 | 185260 | 8337200 | 3371100 | 2145000 | 7320600 | 10669000 | 4687100 | 93681000 | 65259000 | 285800 | 2117700 |
| KKDEGSYS | SDC1 | P18827 | SDC1_HUMAN | Unmodified | 802270 | 860860 | 0 | 0 | 19723000 | 3465100 | 0 | 0 | 2405100 | 227930 | 0 | 0 | 0 | 565590 | 0 | 0 | 2330100 | 8345900 |
| KKDLSLEEI | STMN1; STMN2 | P16949; Q9 | STMN1_HUMAI | Unmodified | 0 | 0 | 0 | 0 | 3877600 | 0 | 3747700 | 2197100 | 3300300 | 10910000 | 115210 | 268380 | 1932700 | 524700 | 0 | 0 | 1260800 | 3067700 |
| KKDQVTAQ | XRCC5 | P13010 | XRCC5_HUMAI | Unmodified | 0 | 0 | 0 | 0 | 1450100 | 0 | 641040 | 1306100 | 313800 | 509650 | 1398100 | 7499500 | 626830 | 0 | 0 | 0 | 461120 | 769180 |
| KKDTSLEEI | STMN3; STMN3 | Q9NZ72; A0 | STMN3_HUMAI | Unmodified | 0 | 0 | 0 | 0 | 0 | 0 | 418580 | 232920 | 0 | 0 | 0 | 1184400 | 0 | 42699 | 0 | 0 | 0 | 0 |
| KKEEEATEV | RDX; RDX; RDX | P35241; A0A | RADI_HUMAN I | Unmodified | 174100 | 127030 | 511600 | 498050 | 4873300 | 1207200 | 199070 | 199060 | 1044700 | 381290 | 346370 | 1713300 | 4005000 | 3076100 | 641000 | 477070 | 569450 | 1410000 |
| KKELEEEV | SEPTIN11; SEP | Q9NVA2; H0 | SEP11_HUMAN | Unmodified | 0 | 0 | 0 | 301390 | 4336900 | 1549600 | 96056 | 220710 | 194280 | 388540 | 162210 | 840010 | 330240 | 88763 | 0 | 0 | 958830 | 974510 |
| KKELEEIVQ | HSPA5; HSPA5 | P11021; A0A | BIP_HUMAN Er | Unmodified | 756610 | 0 | 0 | 0 | 5628500 | 0 | 13703000 | 2273200 | 2.27E+08 | 1.05E+08 | 7047900 | 9333600 | 2758300 | 4985100 | 4362000 | 1282200 | 705140 | 488720 |
| KKEPELFQ | EHD1; EHD1; E | Q9H4M9; C9 | EHD1_HUMAN | Unmodified | 937210 | 634890 | 489300 | 261460 | 138990 | 0 | 0 | 0 | 399960 | 54423 | 847700 | 14122000 | 82088 | 0 | 939080 | 1073300 | 0 | 0 |
| KKEPEPNF | PSMD1; PSMD1 | Q99460; A0 | PSMD1_HUMA | Unmodified | 70269 | 49478 | 238090 | 132900 | 273580 | 0 | 0 | 0 | 372120 | 91597 | 311720 | 1589800 | 371190 | 0 | 270550 | 0 | 51799 | 172670 |
| KKEPVGDD | SARS1; SARS1 | P49591; Q5 | SYSC_HUMAN | Unmodified | 0 | 0 | 0 | 0 | 1825200 | 0 | 0 | 0 | 544580 | 677600 | 386500 | 450490 | 0 | 1587700 | 0 | 0 | 1408900 | 1347200 |
| KKEVPAVPI | RPL7 | P18124 | RL7_HUMAN 6 | Unmodified | 0 | 0 | 0 | 0 | 1068600 | 306890 | 0 | 0 | 437260 | 0 | 0 | 0 | 930570 | 0 | 671230 | 0 | 0 | 0 |
| KKFDAISAE | UTRN | P46939 | UTRN_HUMAN | Unmodified | 0 | 0 | 0 | 0 | 198710 | 0 | 0 | 0 | 0 | 134660 | 0 | 0 | 0 | 0 | 150000 | 11992 | 112100 | 125710 |
| KKFDDGEI | DNAJC3; DNAJ | Q13217; X6F | DNJC3_HUMAI | Unmodified | 0 | 0 | 0 | 0 | 0 | 0 | 0 | 0 | 13668000 | 0 | 0 | 0 | 0 | 0 | 0 | 0 | 0 | 0 |
| KKGENIEQ | UBAP1; UBAP1 | Q9NZ09; A0 | UBAP1_HUMAI | Unmodified | 385040 | 783760 | 54365 | 220040 | 0 | 0 | 0 | 0 | 0 | 0 | 0 | 0 | 0 | 0 | 183260 | 0 | 0 | 0 |
| KKGGEMEI | IQGAP2; IQGAP | Q13576; E9F | IQGA2_HUMAN | Unmodified | 0 | 0 | 374070 | 0 | 376560 | 0 | 0 | 0 | 0 | 0 | 0 | 0 | 0 | 0 | 170790 | 180780 | 0 | 135660 |

| | | | | | | | | | | | | | | | | | | | | | | |
|---|---|---|---|---|---|---|---|---|---|---|---|---|---|---|---|---|---|---|---|---|---|---|
| KKGIAWWT | TMEM30A; TME | Q9NV96; A0/ | CC50A_HUMAI | Unmodified | 0 | 0 | 0 | 92326 | 1279700 | 168400 | 0 | 0 | 728680 | 959900 | 288290 | 0 | 921570 | 54616 | 0 | 0 | 325230 | 361960 |
| KKGTVVYG | DPYSL2; DPYSL | Q16555; A0/ | DPYL2_HUMAN | Unmodified | 0 | 0 | 0 | 0 | 2832100 | 0 | 0 | 0 | 0 | 613290 | 0 | 0 | 0 | 0 | 0 | 0 | 457160 | 0 |
| KKHPDASV | HMGB1; HMGB | P09429; Q5T | HMGB1_HUMA | Unmodified | 0 | 0 | 0 | 0 | 1368400 | 0 | 143490 | 0 | 75433 | 0 | 467930 | 7235000 | 303440 | 161600 | 25602 | 0 | 0 | 0 |
| KKHPDSSV | HMGB2; HMGB | P26583; D6F | HMGB2_HUMA | Unmodified | 0 | 0 | 0 | 0 | 0 | 0 | 0 | 0 | 0 | 333890 | 0 | 0 | 0 | 0 | 0 | 0 | 0 | 0 |
| KKLDDQSE | FERMT2; FERMT | Q96AC1; G3 | FERM2_HUMAI | Unmodified | 0 | 0 | 0 | 0 | 11159000 | 330340 | 111450 | 0 | 0 | 66387 | 0 | 0 | 0 | 0 | 0 | 0 | 3819900 | 15147000 |
| KKLYCIYVA | ATP5F1A; ATP5I | P25705; K7E | ATPA_HUMAN , | Unmodified | 0 | 0 | 0 | 0 | 0 | 0 | 165980 | 0 | 275270 | 570910 | 0 | 0 | 0 | 0 | 0 | 0 | 0 | 0 |
| KKMEDSVG | MYH9 | P35579 | MYH9_HUMAN | Unmodified | 0 | 0 | 155260 | 153860 | 1553800 | 250450 | 0 | 0 | 155780 | 0 | 0 | 0 | 253350 | 103430 | 247410 | 138870 | 253980 | 370230 |
| KKMEDSVG | MYH9 | P35579 | MYH9_HUMAN | Oxidation (M) | 0 | 0 | 0 | 0 | 2592500 | 0 | 0 | 0 | 63016 | 1221200 | 0 | 277700 | 0 | 0 | 0 | 0 | 0 | 0 |
| KKNPEVPV | HMGB3; HMGB | O15347; E7E | HMGB3_HUMA | Unmodified | 0 | 158200 | 0 | 0 | 0 | 0 | 0 | 0 | 0 | 288850 | 236870 | 0 | 0 | 151060 | 0 | 0 | 64011 | 355650 |
| KKPEDSPS | RANBP2 | P49792 | RBP2_HUMAN | Unmodified | 0 | 0 | 0 | 0 | 0 | 0 | 0 | 0 | 0 | 0 | 0 | 1843800 | 0 | 0 | 0 | 0 | 0 | 0 |
| KKPFMLDE | EIF2S2 | P20042 | IF2B_HUMAN E | Unmodified | 0 | 0 | 0 | 0 | 0 | 0 | 0 | 0 | 893560 | 0 | 374170 | 2066500 | 0 | 0 | 0 | 0 | 0 | 0 |
| KKPFVYTQ | RNPEP; RNPEP | Q9H4A4; A6 | AMPB_HUMAN | Unmodified | 0 | 0 | 0 | 0 | 0 | 0 | 0 | 0 | 523800 | 0 | 292150 | 5274400 | 0 | 0 | 344670 | 233590 | 0 | 0 |
| KKPGDASS | MYO9B; MYO9E | Q13459; M0 | MYO9B_HUMA | Unmodified | 0 | 0 | 0 | 0 | 1539900 | 0 | 0 | 0 | 0 | 0 | 0 | 0 | 0 | 0 | 0 | 0 | 0 | 0 |
| KKQIEALQ | EEA1 | Q15075 | EEA1_HUMAN I | Unmodified | 0 | 0 | 0 | 0 | 621460 | 79742 | 68926 | 0 | 0 | 38328 | 0 | 0 | 166510 | 125220 | 0 | 0 | 76697 | 99441 |
| KKTDELPQ | FN1; FN1 | P02751; H0Y | FINC_HUMAN | Unmodified | 0 | 3217500 | 0 | 888620 | 0 | 1846700 | 0 | 0 | 0 | 709950 | 0 | 0 | 0 | 0 | 0 | 1278000 | 0 | 0 |
| KKVDDDLC | MYH10 | P35580 | MYH10_HUMA | Unmodified | 0 | 0 | 341690 | 350470 | 4846400 | 756980 | 181930 | 168470 | 163720 | 55760 | 0 | 4391300 | 1025000 | 0 | 388590 | 453100 | 104420 | 325090 |
| KKVEEVLEE | CBX1; CBX1; CI | P83916; C9J | CBX1_HUMAN | Unmodified | 0 | 0 | 0 | 0 | 0 | 0 | 0 | 0 | 719010 | 204130 | 0 | 729710 | 143160 | 0 | 0 | 0 | 118680 | 117990 |
| KKVPLQYN | CIT; CIT; CIT; CI | A0A7I2V4A6 | A0A7I2V4A6_H | Unmodified | 0 | 0 | 0 | 0 | 1049600 | 151460 | 69258 | 0 | 0 | 0 | 0 | 0 | 0 | 0 | 0 | 0 | 0 | 116620 |
| KKVWLDPN | RPL19; RPL19; | P84098; J3K | RL19_HUMAN | Unmodified | 202250 | 1170000 | 0 | 543720 | 120720 | 0 | 0 | 0 | 0 | 0 | 0 | 0 | 1748100 | 0 | 0 | 0 | 0 | 114450 |
| KKYDVDTLI | COMT; COMT; | P21964; E7E | COMT_HUMAN | Oxidation (M) | 563170 | 584950 | 0 | 0 | 26307 | 0 | 0 | 0 | 0 | 0 | 165820 | 1655000 | 0 | 0 | 0 | 826660 | 0 | 0 |
| KKYPYWPH | ATP5PD | O75947 | ATP5H_HUMAN | Unmodified | 0 | 0 | 0 | 0 | 0 | 0 | 236230 | 0 | 2382300 | 853330 | 0 | 0 | 0 | 0 | 0 | 0 | 0 | 0 |
| KLAAGHEL | GJA1 | P17302 | CXA1_HUMAN | Unmodified | 0 | 0 | 0 | 0 | 2999600 | 0 | 0 | 0 | 0 | 0 | 0 | 0 | 0 | 0 | 0 | 0 | 0 | 777610 |
| KLAAQSCA | DHX9 | Q08211 | DHX9_HUMAN | Unmodified | 0 | 0 | 0 | 0 | 0 | 0 | 904790 | 46513 | 0 | 108340 | 245240 | 1553000 | 0 | 0 | 0 | 0 | 0 | 0 |
| KLAAVTYN | SEPTIN7; SEPTI | Q16181; E7E | SEPT7_HUMAN | Unmodified | 0 | 0 | 0 | 0 | 2184900 | 141560 | 402160 | 900940 | 0 | 4300700 | 0 | 99141 | 577410 | 349560 | 0 | 0 | 2482600 | 597600 |
| KLADEQLS | ABHD11 | Q8NFV4 | ABHDB_HUMA | Unmodified | 0 | 0 | 0 | 0 | 0 | 0 | 0 | 0 | 698890 | 0 | 0 | 0 | 0 | 0 | 0 | 0 | 0 | 0 |
| KLADMYGC | CDH2; CDH2 | P19022; C9J | CADH2_HUMA | Unmodified | 0 | 0 | 0 | 0 | 0 | 0 | 0 | 0 | 0 | 358060 | 640040 | 680950 | 0 | 0 | 0 | 0 | 9699800 | 9210400 |
| KLADMYGC | CDH2; CDH2 | P19022; C9J | CADH2_HUMA | Oxidation (M) | 0 | 0 | 0 | 0 | 0 | 0 | 0 | 0 | 130060 | 0 | 40901 | 0 | 0 | 0 | 0 | 0 | 2410700 | 1043800 |
| KLADMYGC | CDH1; CDH3 | P12830; P22 | CADH1_HUMA | Unmodified | 0 | 0 | 3632500 | 1624900 | 13156000 | 2927900 | 0 | 20846 | 288940 | 206650 | 735370 | 2594200 | 13722000 | 11695000 | 4261900 | 0 | 953160 | 1049000 |
| KLADMYGC | CDH1; CDH3 | P12830; P22 | CADH1_HUMA | Oxidation (M) | 0 | 0 | 0 | 769530 | 3698200 | 1191800 | 0 | 0 | 64054 | 0 | 156630 | 737490 | 1420500 | 1563000 | 0 | 1002200 | 238260 | 0 |
| KLADQFLEI | GOLM2; GOLM | Q6P4E1; H0Y | GOLM2_HUMA | Unmodified | 0 | 0 | 0 | 0 | 333620 | 0 | 0 | 0 | 5031700 | 438890 | 249240 | 0 | 0 | 0 | 0 | 0 | 0 | 662250 |
| KLAFDMAT | OSTF1 | Q92882 | OSTF1_HUMA | Oxidation (M) | 0 | 0 | 0 | 0 | 0 | 0 | 201270 | 179580 | 0 | 0 | 138110 | 1547900 | 0 | 0 | 0 | 0 | 270410 | 0 |
| KLAFDMAT | OSTF1 | Q92882 | OSTF1_HUMA | Unmodified | 0 | 0 | 0 | 0 | 0 | 0 | 0 | 41446 | 509200 | 40623 | 0 | 205770 | 821990 | 1148800 | 0 | 0 | 0 | 360810 |
| KLAGQRFT | FHL1; FHL1; FH | Q13642; A0/ | FHL1_HUMAN | Unmodified | 0 | 0 | 0 | 0 | 0 | 0 | 2228200 | 0 | 0 | 0 | 0 | 0 | 0 | 1649200 | 0 | 0 | 0 | 0 |
| KLALQLHPI | DNAJB11; DNA | Q9UBS4; A0 | DJB11_HUMAN | Unmodified | 0 | 0 | 0 | 48328 | 0 | 0 | 0 | 0 | 11273000 | 336870 | 515970 | 3231100 | 0 | 0 | 0 | 0 | 0 | 0 |
| KLALQWHF | DNAJC3; DNAJ | Q13217; X6F | DNJC3_HUMA | Unmodified | 0 | 0 | 0 | 0 | 0 | 0 | 0 | 0 | 2153400 | 0 | 0 | 160750 | 0 | 0 | 0 | 0 | 0 | 0 |
| KLANSEPV | MACF1; MACF1 | Q9UPN3; H3 | MACF1_HUMA | Unmodified | 0 | 0 | 0 | 0 | 414070 | 964500 | 0 | 0 | 0 | 0 | 0 | 0 | 0 | 131130 | 0 | 0 | 835570 | 0 |
| KLAPEYEK | PDIA4; PDIA4 | P13667; A0A | PDIA4_HUMA | Unmodified | 0 | 0 | 0 | 0 | 0 | 0 | 0 | 1508500 | 0 | 0 | 1391800 | 467730 | 0 | 0 | 0 | 0 | 0 | 0 |
| KLAPITYPQ | RAC2; RAC2; R/ | P15153; B1A | RAC2_HUMAN | Unmodified | 0 | 0 | 0 | 0 | 1712600 | 0 | 393820 | 328230 | 0 | 0 | 1040600 | 2600000 | 0 | 0 | 0 | 0 | 1169500 | 1577000 |
| KLAPITYPQ | RAC2; RAC2; R/ | P15153; B1A | RAC2_HUMAN | Unmodified | 0 | 0 | 0 | 0 | 0 | 0 | 470260 | 0 | 0 | 0 | 0 | 0 | 0 | 0 | 0 | 0 | 992240 | 0 |
| KLAPPLVTL | AP2B1; AP2B1; | P63010; A0A | AP2B1_HUMA | Unmodified | 196010 | 1841200 | 139350 | 310330 | 0 | 0 | 0 | 0 | 195950 | 63826 | 0 | 1029400 | 0 | 0 | 1135100 | 346170 | 0 | 0 |
| KLAQQYYL | PSMA2; N/A; N/ | P25787; A0A | PSA2_HUMAN | Unmodified | 0 | 1315300 | 0 | 0 | 0 | 0 | 0 | 0 | 2949100 | 0 | 0 | 15063000 | 0 | 0 | 0 | 1529600 | 0 | 0 |
| KLASAAYP | YARS1; YARS1 | P54577; A0A | SYYC_HUMAN | Unmodified | 0 | 0 | 0 | 0 | 197540 | 0 | 0 | 0 | 0 | 0 | 0 | 0 | 1364500 | 0 | 0 | 0 | 121460 | 156500 |
| KLAVGDNN | IQGAP1; IQGAP | P46940; A0A | IQGA1_HUMA | Unmodified | 0 | 0 | 0 | 0 | 1713600 | 0 | 0 | 0 | 0 | 253490 | 0 | 0 | 0 | 0 | 0 | 0 | 0 | 0 |
| KLAVNNIA | BTF3L4; BTF3L4 | Q96K17; E9F | BT3L4_HUMAN | Oxidation (M) | 0 | 0 | 0 | 0 | 979880 | 0 | 0 | 0 | 392660 | 0 | 0 | 2277900 | 0 | 0 | 0 | 0 | 0 | 0 |
| KLCQGLFF | HYOU1; HYOU1 | Q9Y4L1; A0/ | HYOU1_HUMA | Unmodified | 0 | 0 | 0 | 0 | 0 | 0 | 0 | 0 | 2666500 | 3642000 | 1024200 | 0 | 0 | 0 | 0 | 0 | 0 | 0 |
| KLDALCNII | COPA; COPA; C | P53621; A0A | COPA_HUMAN | Unmodified | 0 | 0 | 0 | 0 | 0 | 0 | 0 | 0 | 0 | 0 | 0 | 3111100 | 0 | 0 | 0 | 135040 | 0 | 0 |
| KLDAQVQE | MYH10 | P35580 | MYH10_HUMA | Unmodified | 0 | 0 | 0 | 0 | 3812400 | 0 | 730840 | 95871 | 8046600 | 723190 | 0 | 189240 | 1182900 | 1967300 | 0 | 0 | 494780 | 0 |
| KLDDFVET | SCARB2; SCARI | Q14108; A0/ | SCRB2_HUMAI | Unmodified | 0 | 0 | 0 | 0 | 0 | 0 | 0 | 0 | 2156800 | 523700 | 0 | 0 | 0 | 0 | 0 | 0 | 0 | 0 |
| KLDDQSED | FERMT2; FERM | Q96AC1; G3 | FERM2_HUMAI | Unmodified | 0 | 0 | 0 | 0 | 9003300 | 3182500 | 0 | 0 | 0 | 0 | 0 | 0 | 0 | 0 | 0 | 0 | 3998900 | 16203000 |
| KLDDYLNT | CLIC5; CLIC5; | Q9NZA1; A0 | CLIC5_HUMAN | Unmodified | 0 | 0 | 0 | 0 | 0 | 0 | 0 | 0 | 0 | 0 | 0 | 930180 | 0 | 0 | 975290 | 0 | 0 | 0 |
| KLDEAVAE | AHCY | P23526 | SAHH_HUMAN | Unmodified | 886230 | 733240 | 4028400 | 2125400 | 7911100 | 2542200 | 0 | 2403000 | 4175600 | 2265800 | 8405300 | 40749000 | 6612800 | 4363600 | 3500200 | 2537600 | 2502100 | 7375500 |
| KLDEAVAE | AHCY | P23526 | SAHH_HUMAN | Unmodified | 0 | 0 | 0 | 0 | 3953000 | 451100 | 0 | 247270 | 0 | 942290 | 0 | 0 | 1989400 | 1866400 | 0 | 0 | 3437400 | 3603500 |
| KLDEDSLTI | STK4; STK4; STI | Q13043; F5H | STK4_HUMAN | Unmodified | 0 | 0 | 0 | 0 | 236370 | 0 | 103820 | 0 | 0 | 0 | 57211 | 311040 | 58509 | 0 | 0 | 0 | 118440 | 347950 |
| KLDELYGT | RPL4; RPL4 | P36578; H3E | RL4_HUMAN 6 | Unmodified | 550520 | 609610 | 0 | 0 | 0 | 0 | 0 | 0 | 0 | 0 | 0 | 6207800 | 0 | 0 | 117860 | 342260 | 0 | 0 |
| KLDELYGT | RPL4; RPL4 | P36578; H3E | RL4_HUMAN 6 | Unmodified | 2896800 | 0 | 2052300 | 0 | 0 | 97350 | 86266 | 0 | 0 | 0 | 0 | 12499000 | 0 | 0 | 1594400 | 1878700 | 0 | 0 |
| KLDESIYD | STRN3 | Q13033 | STRN3_HUMA | Unmodified | 0 | 0 | 0 | 0 | 0 | 0 | 0 | 0 | 470770 | 0 | 0 | 2173900 | 0 | 0 | 0 | 0 | 0 | 0 |
| KLDLEAWF | SARS1; SARS1 | P49591; Q5T | SYSC_HUMAN | Unmodified | 0 | 0 | 192040 | 330360 | 0 | 0 | 0 | 0 | 1791200 | 149440 | 0 | 3984800 | 0 | 0 | 595040 | 534840 | 0 | 0 |
| KLDNGGY | YES1; FYN; FYN | P07947; P06 | YES_HUMAN Ty | Unmodified | 392070 | 460110 | 1267300 | 363430 | 0 | 114260 | 0 | 0 | 1287100 | 0 | 228530 | 2279500 | 0 | 0 | 1183500 | 380150 | 271480 | 113730 |
| KLDNGGY | YES1; YES1 | P07947; J3Q | YES_HUMAN Ty | Unmodified | 0 | 0 | 0 | 0 | 446940 | 0 | 0 | 0 | 0 | 240490 | 0 | 0 | 435720 | 293040 | 0 | 0 | 870190 | 581290 |
| KLDPDSIPS | SEC24D | O94855 | SC24D_HUMAN | Unmodified | 0 | 0 | 0 | 165640 | 2110500 | 0 | 0 | 0 | 0 | 0 | 87104 | 361960 | 0 | 0 | 0 | 0 | 0 | 0 |
| KLDPELHLI | ILF2; ILF2 | Q12905; B4E | ILF2_HUMAN I | Unmodified | 1364000 | 1338300 | 641840 | 340200 | 0 | 0 | 21698 | 21147 | 137640 | 86513 | 399280 | 8095500 | 23883 | 0 | 0 | 742150 | 0 | 0 |
| KLDPGSEE | EEF1G | P26641 | EF1G_HUMAN | Unmodified | 0 | 0 | 963980 | 0 | 0 | 0 | 0 | 0 | 1218900 | 796470 | 856310 | 54828000 | 0 | 0 | 1601400 | 1542200 | 0 | 0 |
| KLDQEMEC | RAD50; N/A; RA | Q92878; A0/ | RAD50_HUMAI | Unmodified | 0 | 0 | 0 | 0 | 0 | 0 | 0 | 0 | 0 | 0 | 0 | 896720 | 0 | 0 | 0 | 0 | 0 | 0 |
| KLDSELTAF | DIAPH1; DIAPH | O60610; A0/ | DIAP1_HUMAN | Unmodified | 0 | 0 | 0 | 301600 | 4439900 | 272750 | 0 | 0 | 668460 | 942060 | 0 | 0 | 260870 | 1399400 | 0 | 112750 | 1505100 | 2129300 |
| KLDSKVNII | SEPTIN8; SEPTI | Q92599; Q1 | SEPT8_HUMAN | Unmodified | 0 | 0 | 0 | 0 | 2899100 | 0 | 613660 | 268740 | 0 | 417220 | 0 | 0 | 0 | 0 | 0 | 0 | 1326400 | 1078900 |
| KLDVLSND | FBP1; FBP1 | P09467; A0A | F16P1_HUMAN | Oxidation (M) | 0 | 0 | 0 | 0 | 0 | 0 | 0 | 0 | 721290 | 515320 | 0 | 0 | 0 | 0 | 0 | 0 | 0 | 0 |
| KLDYGQH | EIF4A3; EIF4A3 | P38919; I3L | IF4A3_HUMAN | Unmodified | 0 | 0 | 0 | 0 | 0 | 0 | 15760 | 0 | 0 | 0 | 0 | 2237000 | 0 | 0 | 511360 | 285980 | 0 | 0 |
| KLEAAEDIA | PHB; PHB | P35232; C9J | PHB_HUMAN F | Unmodified | 0 | 0 | 0 | 0 | 0 | 0 | 0 | 0 | 4307500 | 452410 | 0 | 477580 | 438440 | 0 | 0 | 0 | 351950 | 81053 |
| KLEAAEER | STMN1; STMN2 | P16949; Q9 | STMN1_HUMAI | Unmodified | 0 | 0 | 0 | 0 | 0 | 0 | 0 | 0 | 1241400 | 57935 | 23928 | 429080 | 37142 | 0 | 0 | 0 | 0 | 0 |
| KLEAMTNS | VCL | P18206 | VINC_HUMAN | Unmodified | 0 | 0 | 0 | 0 | 0 | 0 | 0 | 0 | 0 | 0 | 0 | 0 | 0 | 0 | 0 | 0 | 374060 | 0 |
| KLEAQETLN | BAZ2A; BAZ2A; | Q9UIF9; F8V | BAZ2A_HUMAN | Unmodified | 126190 | 0 | 0 | 0 | 0 | 0 | 511890 | 34969 | 0 | 0 | 0 | 0 | 0 | 0 | 0 | 0 | 0 | 0 |
| KLEATIIGE | DPP7; DPP7 | Q9UHL4; R4 | DPP2_HUMAN | Unmodified | 0 | 0 | 0 | 0 | 0 | 0 | 0 | 0 | 1191800 | 0 | 0 | 192420 | 0 | 0 | 0 | 0 | 0 | 0 |
| KLEDGPKF | EEF1A1P5; EEF | Q5VTE0; P68 | EF1A3_HUMAN | Unmodified | 0 | 0 | 0 | 0 | 446760 | 0 | 742430 | 1044000 | 0 | 0 | 0 | 0 | 422400 | 1156400 | 0 | 0 | 1935300 | 438630 |
| KLEDPQSS | SLC19A2; SLC1 | O60779; A0/ | S19A2_HUMAN | Unmodified | 664920 | 246510 | 0 | 309970 | 0 | 0 | 0 | 0 | 490110 | 169040 | 0 | 807940 | 0 | 0 | 312740 | 0 | 0 | 0 |
| KLEDVKNS | TPD52; N/A | P55327; H0Y | TPD52_HUMAN | Unmodified | 0 | 0 | 0 | 0 | 807930 | 0 | 43475 | 43453 | 0 | 1961000 | 0 | 0 | 774120 | 1029600 | 0 | 0 | 635640 | 243340 |
| KLEDYYKID | STXBP3 | O00186 | STXB3_HUMAN | Unmodified | 0 | 0 | 0 | 431460 | 6666400 | 0 | 1292500 | 959810 | 314090 | 1473100 | 347130 | 0 | 8050400 | 9246800 | 0 | 0 | 4178600 | 6158200 |
| KLEEEGEQ | SCP2; SCP2 | P22307; H0Y | SCP2_HUMAN | Unmodified | 0 | 0 | 95035 | 0 | 0 | 0 | 533410 | 237840 | 41775000 | 13707000 | 662370 | 0 | 31821 | 373220 | 92861 | 120190 | 235340 | 257970 |
| KLEEEGEQ | SCP2; SCP2 | P22307; H0Y | SCP2_HUMAN | Unmodified | 0 | 0 | 0 | 0 | 0 | 0 | 594890 | 195550 | 30623000 | 7312500 | 537140 | 0 | 0 | 367690 | 0 | 0 | 0 | 0 |
| KLEEEQIILE | MYH9 | P35579 | MYH9_HUMAN | Unmodified | 0 | 0 | 0 | 1232400 | 8769800 | 2023700 | 759540 | 0 | 1757500 | 1382300 | 2525600 | 2556200 | 3771800 | 1721100 | 1658800 | 1036900 | 4257200 | 2717700 |
| KLEEEQIILE | MYH9 | P35579 | MYH9_HUMAN | Unmodified | 0 | 0 | 0 | 0 | 3165700 | 0 | 0 | 0 | 0 | 0 | 0 | 0 | 122350 | 375030 | 0 | 0 | 3978900 | 1761100 |
| KLEEIIHQIT | RAB3GAP1 | Q15042 | RB3GP_HUMAI | Unmodified | 0 | 0 | 17019 | 0 | 157630 | 0 | 0 | 0 | 0 | 0 | 36800 | 94788 | 95825 | 38724 | 156720 | 0 | 0 | 77330 |
| KLEELELDE | MAP2K2; N/A | P36507; A0A | MP2K2_HUMAI | Unmodified | 0 | 0 | 0 | 0 | 1541800 | 182750 | 749180 | 0 | 479080 | 424000 | 0 | 0 | 0 | 26324 | 144740 | 0 | 180090 | 706070 |
| KLEELELDE | MAP2K1 | Q02750 | MP2K1_HUMAI | Unmodified | 66411 | 0 | 0 | 473830 | 0 | 0 | 0 | 58996 | 438730 | 0 | 0 | 978830 | 0 | 372000 | 328350 | 515870 | 0 | 0 |
| KLEELEQK | STAT1; STAT1; S | P42224; A0A | STAT1_HUMAN | Unmodified | 0 | 0 | 0 | 58758 | 204820 | 0 | 0 | 0 | 9558.5 | 0 | 0 | 280440 | 0 | 0 | 0 | 0 | 29862 | 0 |
| KLEELEQK | STAT1; STAT1; S | P42224; A0A | STAT1_HUMAN | Unmodified | 70297 | 38111 | 0 | 0 | 1341300 | 145480 | 0 | 31687 | 146150 | 74137 | 101200 | 0 | 218860 | 0 | 78442 | 0 | 391950 | 1058600 |
| KLEENEVIP | SDC4 | P31431 | SDC4_HUMAN | Unmodified | 0 | 0 | 1663100 | 1769600 | 1048200 | 707690 | 0 | 0 | 779390 | 146370 | 0 | 0 | 430710 | 274440 | 1008400 | 1684900 | 0 | 208360 |
| KLEERGFD | BANF1 | O75531 | BAF_HUMAN B | Unmodified | 0 | 0 | 0 | 0 | 0 | 0 | 0 | 2817500 | 196330 | 0 | 97128 | 0 | 1703600 | 0 | 0 | 0 | 1630200 | 2638400 |
| KLEEVVNK | ZC3H15 | Q8WU90 | ZC3HF_HUMAI | Unmodified | 0 | 0 | 0 | 0 | 0 | 0 | 15659 | 75840 | 43165 | 0 | 49699 | 704810 | 123490 | 48030 | 82658 | 0 | 25451 | 12627 |
| KLEEVVQIL | ITPA | Q9BY32 | ITPA_HUMAN I | Unmodified | 0 | 0 | 43755 | 0 | 1531500 | 4975600 | 6213800 | 797420 | 0 | 149520 | 0 | 708030 | 0 | 4349000 | 0 | 0 | 0 | 3098800 |
| KLEEVVQIL | ITPA | Q9BY32 | ITPA_HUMAN I | Unmodified | 0 | 0 | 0 | 0 | 0 | 0 | 0 | 0 | 0 | 0 | 995000 | 7079200 | 0 | 0 | 0 | 0 | 107970 | 0 |
| KLEGDSTD | MYH9 | P35579 | MYH9_HUMAN | Unmodified | 0 | 0 | 0 | 1280300 | 0 | 0 | 0 | 0 | 0 | 0 | 0 | 0 | 0 | 0 | 0 | 0 | 0 | 0 |
| KLEGNSPQ | RAB8A | P61006 | RAB8A_HUMAI | Unmodified | 3290900 | 1486700 | 62198 | 0 | 10780000 | 0 | 0 | 0 | 0 | 1569400 | 0 | 16361000 | 0 | 0 | 2965500 | 2173400 | 0 | 0 |
| KLEGNSPQ | RAB8A | P61006 | RAB8A_HUMAI | Unmodified | 0 | 0 | 0 | 0 | 0 | 0 | 84387 | 62172 | 0 | 103380 | 0 | 0 | 945480 | 647480 | 0 | 0 | 826000 | 0 |
| KLEHLITEL | DYNC1H1; DYN | Q14204; A0/ | DYHC1_HUMA | Unmodified | 89037 | 0 | 0 | 0 | 0 | 0 | 0 | 0 | 543360 | 144290 | 0 | 5833800 | 0 | 0 | 318770 | 0 | 0 | 0 |
| KLEIGFNGI | TM9SF3; TM9SF | Q9HD45; Q5 | TM9S3_HUMAN | Unmodified | 0 | 0 | 0 | 0 | 0 | 0 | 0 | 0 | 2456800 | 199220 | 0 | 528520 | 0 | 0 | 0 | 0 | 0 | 0 |
| KLEKEEEE | HMGA1 | P17096 | HMGA1_HUMA | Unmodified | 0 | 0 | 0 | 0 | 6434800 | 1753700 | 57628 | 28577 | 277340 | 162080 | 6055100 | 5819100 | 1082100 | 517900 | 0 | 0 | 2467400 | 7302000 |
| KLELDGEKI | ADD2 | P35612 | ADDB_HUMAN | Unmodified | 0 | 0 | 0 | 0 | 2452000 | 629270 | 0 | 0 | 0 | 0 | 0 | 0 | 0 | 0 | 0 | 0 | 0 | 0 |
| KLELSDNR | ANP32A; ANP3 | P39687; H0Y | AN32A_HUMAI | Unmodified | 0 | 0 | 0 | 0 | 9752100 | 0 | 186330 | 0 | 1581800 | 2437700 | 23579000 | 8378700 | 11546000 | 1138100 | 210740 | 97587 | 0 | 0 |
| KLEMDLKD | MYH9 | P35579 | MYH9_HUMAN | Unmodified | 53405 | 0 | 0 | 0 | 9149200 | 0 | 1059800 | 0 | 0 | 292870 | 0 | 0 | 1164500 | 1495500 | 0 | 0 | 2288800 | 2057200 |
| KLEMDLKD | MYH9 | P35579 | MYH9_HUMAN | Oxidation (M) | 0 | 0 | 0 | 0 | 959820 | 0 | 226010 | 0 | 0 | 983620 | 0 | 0 | 338850 | 328700 | 0 | 0 | 528800 | 691260 |
| KLENCNYA | LCP1; PLS3; PL | P13796; P13 | PLSL_HUMAN | Unmodified | 0 | 0 | 0 | 0 | 255430 | 0 | 0 | 16142 | 143930 | 0 | 0 | 665840 | 331120 | 336470 | 0 | 0 | 128390 | 0 |
| KLENCNYA | PLS3; PLS3 | P13797; A0A | PLST_HUMAN | Unmodified | 44836 | 0 | 0 | 0 | 6776300 | 1347700 | 43279 | 0 | 0 | 0 | 0 | 0 | 333190 | 1677500 | 0 | 0 | 3119200 | 2101600 |
| KLENSPLG | OCIAD1; OCIAD | Q9NX40; D6 | OCAD1_HUMA | Unmodified | 0 | 0 | 0 | 0 | 0 | 0 | 0 | 0 | 511650 | 0 | 0 | 0 | 0 | 0 | 0 | 0 | 0 | 0 |
| KLEPISNDI | MYOF | Q9NZM1 | MYOF_HUMAN | Unmodified | 190560 | 70544 | 0 | 0 | 17972000 | 4867200 | 235720 | 0 | 0 | 129840 | 0 | 0 | 0 | 0 | 0 | 0 | 612030 | 1480000 |
| KLEPIWNE | TMX3; TMX3; TM | Q96JJ7; B4D | TMX3_HUMAN | Oxidation (M) | 0 | 0 | 0 | 0 | 0 | 0 | 0 | 33607 | 8467100 | 3319600 | 1387500 | 0 | 0 | 0 | 0 | 0 | 0 | 0 |
| KLEPIWNE | TMX3; TMX3; TM | Q96JJ7; B4D | TMX3_HUMAN | Unmodified | 0 | 0 | 0 | 0 | 0 | 0 | 0 | 0 | 14419000 | 1730400 | 460560 | 0 | 0 | 0 | 0 | 0 | 0 | 0 |
| KLEQTVEQ | CALCOCO2; CA | Q13137; D6I | CACO2_HUMA | Unmodified | 307640 | 61390 | 0 | 0 | 269430 | 0 | 389790 | 35963 | 400120 | 0 | 0 | 0 | 0 | 0 | 0 | 0 | 148000 | 967560 |
| KLEQTVEQ | CALCOCO2; CA | Q13137; D6I | CACO2_HUMA | Unmodified | 0 | 0 | 0 | 0 | 0 | 0 | 90832 | 0 | 0 | 103110 | 0 | 0 | 0 | 0 | 0 | 0 | 0 | 841820 |
| KLESFYIQK | HEXB; HEXB; HI | P07686; H0Y | HEXB_HUMAN | Unmodified | 0 | 0 | 0 | 0 | 0 | 0 | 0 | 0 | 641010 | 381920 | 0 | 0 | 0 | 0 | 135250 | 0 | 0 | 0 |

| | | | | | | | | | | | | | | | | | | | | | | |
|---|---|---|---|---|---|---|---|---|---|---|---|---|---|---|---|---|---|---|---|---|---|---|
| KLETAVNLA | VDAC1 | P21796 | VDAC1_HUMAI | Unmodified | 0 | 0 | 0 | 0 | 0 | 0 | 218710 | 0 | 63994000 | 5963800 | 1223900 | 0 | 0 | 0 | 0 | 0 | 296150 | 0 |
| KLETAVNLA | VDAC1 | P21796 | VDAC1_HUMAI | Unmodified | 0 | 0 | 0 | 0 | 0 | 0 | 0 | 0 | 3251300 | 9756500 | 0 | 0 | 0 | 0 | 0 | 0 | 0 | 0 |
| KLETLSLNN | LRRC57; N/A; L | Q8N9N7; A0 | LRC57_HUMAN | Unmodified | 441190 | 117010 | 293070 | 190660 | 0 | 232820 | 0 | 0 | 444320 | 0 | 385480 | 1420400 | 0 | 0 | 1002600 | 431020 | 250010 | 0 |
| KLETLSLNN | LRRC57; N/A; L | Q8N9N7; A0 | LRC57_HUMAN | Unmodified | 0 | 0 | 0 | 103190 | 16610000 | 3408100 | 2441100 | 0 | 262380 | 1983000 | 425740 | 0 | 3855400 | 1645800 | 0 | 0 | 8025200 | 21731000 |
| KLEVEANN | CAPZB; CAPZB; | P47756; A0A | CAPZB_HUMAI | Unmodified | 3062300 | 6458800 | 559490 | 0 | 0 | 0 | 0 | 49009 | 979350 | 0 | 0 | 23395000 | 0 | 0 | 2252600 | 1269100 | 0 | 0 |
| KLEVNEAEI | CAVIN1 | Q6NZI2 | CAVN1_HUMAI | Unmodified | 0 | 0 | 0 | 0 | 8757200 | 1579500 | 0 | 0 | 0 | 0 | 0 | 151460 | 0 | 0 | 0 | 0 | 0 | 0 |
| KLFDTLNEI | HSPA13 | P48723 | HSP13_HUMAN | Unmodified | 0 | 0 | 0 | 0 | 0 | 0 | 0 | 0 | 762980 | 546170 | 132260 | 0 | 0 | 550140 | 252780 | 0 | 0 | 69669 |
| KLFEAEEQI | EHD1; EHD1; EI | Q9H4M9; C9 | EHD1_HUMAN | Unmodified | 0 | 0 | 0 | 0 | 469070 | 0 | 0 | 86848 | 0 | 0 | 0 | 8171100 | 0 | 0 | 0 | 749380 | 0 | 0 |
| KLFIGGLSF | HNRNPA3; HNI | P51991; A0A | ROA3_HUMAN | Unmodified | 0 | 0 | 0 | 0 | 457840 | 0 | 3726500 | 3569100 | 0 | 0 | 0 | 0 | 0 | 0 | 0 | 0 | 0 | 0 |
| KLFIGGLSF | HNRNPA1L2; H | Q32P51; P09 | RA1L2_HUMAN | Unmodified | 485190 | 205110 | 0 | 0 | 801980 | 0 | 32629 | 19588 | 0 | 0 | 377040 | 3188700 | 243590 | 52194 | 0 | 212160 | 1659800 | 4628100 |
| KLFLVQLQI | NUDT21; NUDT | O43809; H3I | CPSF5_HUMAN | Unmodified | 0 | 0 | 0 | 0 | 112650 | 0 | 244180 | 121290 | 0 | 247350 | 0 | 5224000 | 0 | 0 | 0 | 0 | 0 | 0 |
| KLFNDYGG | ANPEP | P15144 | AMPN_HUMAN | Unmodified | 0 | 0 | 0 | 0 | 0 | 0 | 0 | 0 | 0 | 0 | 0 | 0 | 0 | 0 | 0 | 0 | 572440 | 0 |
| KLFTAESLI | ARCN1; ARCN1 | P48444; B0Y | COPD_HUMAN | Unmodified | 0 | 0 | 0 | 0 | 0 | 0 | 0 | 0 | 0 | 0 | 0 | 2393800 | 0 | 0 | 0 | 0 | 0 | 0 |
| KLFVGGIKE | HNRNPA2B1; H | P22626; A0A | ROA2_HUMAN | Unmodified | 221390 | 0 | 0 | 0 | 0 | 0 | 0 | 0 | 0 | 0 | 0 | 1116300 | 0 | 0 | 0 | 0 | 0 | 0 |
| KLFYSTFAT | ECHS1 | P30084 | ECHM_HUMAN | Unmodified | 0 | 0 | 0 | 0 | 0 | 0 | 479120 | 721130 | 1398100 | 0 | 0 | 0 | 395710 | 0 | 0 | 0 | 11820 | 0 |
| KLGADFHG | UBAP1 | Q9NZ09 | UBAP1_HUMAI | Unmodified | 6994200 | 2258200 | 0 | 0 | 5518700 | 620950 | 0 | 0 | 0 | 0 | 0 | 0 | 0 | 0 | 0 | 630920 | 0 | 0 |
| KLGADFHG | UBAP1 | Q9NZ09 | UBAP1_HUMAI | Unmodified | 577660 | 1574800 | 0 | 420070 | 5667700 | 1549000 | 0 | 0 | 0 | 0 | 0 | 0 | 2691300 | 0 | 0 | 0 | 0 | 576300 |
| KLGAGQFG | HCK; HCK; HCK | P08631; J3KI | HCK_HUMAN T | Oxidation (M) | 0 | 0 | 0 | 0 | 188300 | 0 | 192830 | 0 | 0 | 0 | 2067800 | 2620900 | 504360 | 0 | 0 | 0 | 327990 | 0 |
| KLGDTYPSI | RNPEP; RNPEP | Q9H4A4; A6I | AMPB_HUMAN | Unmodified | 0 | 0 | 0 | 122050 | 330050 | 0 | 0 | 0 | 0 | 0 | 360360 | 3548300 | 310350 | 97763 | 0 | 0 | 103820 | 0 |
| KLGEMWN | HMGB3; HMGB | O15347; E7E | HMGB3_HUMA | Oxidation (M) | 0 | 0 | 0 | 0 | 480760 | 0 | 0 | 29408 | 986710 | 745840 | 0 | 0 | 0 | 0 | 0 | 0 | 0 | 0 |
| KLGEMWN | HMGB1; HMGB | P09429; Q5T | HMGB1_HUMA | Oxidation (M) | 0 | 0 | 517750 | 0 | 643870 | 0 | 309280 | 333350 | 0 | 838130 | 1538800 | 3956200 | 1040300 | 50277 | 394930 | 317990 | 299590 | 0 |
| KLGEMWN | HMGB1; HMGB | P09429; Q5T | HMGB1_HUMA | Unmodified | 0 | 0 | 0 | 62576 | 1335400 | 0 | 695530 | 353190 | 574210 | 326150 | 2548100 | 7513100 | 1412900 | 0 | 0 | 0 | 0 | 0 |
| KLGEMWSI | HMGB2 | P26583 | HMGB2_HUMA | Unmodified | 0 | 32897 | 0 | 0 | 0 | 0 | 9798.3 | 214790 | 335000 | 280820 | 341270 | 1310200 | 344130 | 18216 | 0 | 0 | 0 | 22340 |
| KLGEWVGI | RPS12 | P25398 | RS12_HUMAN | Unmodified | 0 | 0 | 0 | 0 | 0 | 0 | 0 | 0 | 0 | 0 | 0 | 2115600 | 444950 | 0 | 0 | 0 | 0 | 234650 |
| KLGGSLAD | CCT2; CCT2; C | P78371; F5G | TCPB_HUMAN | Unmodified | 389450 | 198310 | 605480 | 474710 | 0 | 379730 | 6713400 | 47478 | 4655300 | 6032900 | 1914200 | 17145000 | 25068000 | 8335200 | 1997800 | 1448300 | 7179000 | 9402000 |
| KLGGSLAD | CCT2; CCT2; C | P78371; F5G | TCPB_HUMAN | Unmodified | 0 | 0 | 0 | 0 | 983620 | 0 | 568960 | 0 | 881860 | 562540 | 2179900 | 7000500 | 2640100 | 445490 | 0 | 0 | 1703300 | 1180100 |
| KLGGSQED | TALDO1; TALDO | P37837; F2Z | TALDO_HUMAI | Unmodified | 0 | 0 | 0 | 0 | 0 | 0 | 0 | 0 | 153530 | 0 | 543810 | 0 | 0 | 55425 | 0 | 0 | 0 | 112020 |
| KLGGTIDD | CCT4 | P50991 | TCPD_HUMAN | Unmodified | 0 | 0 | 588360 | 812870 | 4219700 | 3049500 | 1559500 | 746360 | 0 | 1843100 | 1099000 | 5260400 | 0 | 5715600 | 0 | 0 | 2526200 | 7375700 |
| KLGINSLQI | TPD52; TPD52; | P55327; H0Y | TPD52_HUMAN | Unmodified | 0 | 0 | 0 | 0 | 0 | 0 | 0 | 0 | 0 | 0 | 0 | 0 | 0 | 0 | 0 | 0 | 0 | 0 |
| KLGLLGDS | CEL | P19835 | CEL_HUMAN B | Unmodified | 0 | 0 | 0 | 0 | 0 | 0 | 0 | 0 | 44064000 | 6859300 | 29076000 | 2656800 | 0 | 0 | 0 | 0 | 0 | 0 |
| KLGLLGDS | CEL | P19835 | CEL_HUMAN B | Unmodified | 0 | 0 | 0 | 0 | 0 | 0 | 0 | 0 | 129050 | 7917800 | 97029 | 0 | 0 | 0 | 0 | 0 | 0 | 0 |
| KLGLVFDD | CAP1 | Q01518 | CAP1_HUMAN | Unmodified | 0 | 0 | 0 | 18207 | 771580 | 196500 | 25051 | 0 | 0 | 0 | 0 | 0 | 383450 | 82928 | 0 | 0 | 310300 | 151030 |
| KLGLVFDD | CAP1 | Q01518 | CAP1_HUMAN | Unmodified | 0 | 0 | 0 | 0 | 1625500 | 0 | 668870 | 0 | 0 | 261300 | 0 | 0 | 1936500 | 907880 | 0 | 0 | 636040 | 628240 |
| KLGNDAVS | CARMIL1; CARI | Q5VZK9; A0A | CARL1_HUMAN | Unmodified | 0 | 0 | 0 | 0 | 0 | 0 | 0 | 0 | 0 | 0 | 0 | 0 | 0 | 0 | 0 | 0 | 448120 | 0 |
| KLGNNCVI | HSD17B10; HS | Q99714; Q5I | HCD2_HUMAN | Unmodified | 0 | 0 | 0 | 0 | 0 | 0 | 1139800 | 764430 | 5115300 | 1475600 | 0 | 0 | 0 | 0 | 0 | 0 | 0 | 916810 |
| KLGNNCVI | HSD17B10; HS | Q99714; Q5I | HCD2_HUMAN | Unmodified | 0 | 0 | 0 | 0 | 0 | 0 | 2136600 | 0 | 0 | 950690 | 0 | 0 | 0 | 0 | 0 | 0 | 0 | 0 |
| KLGPEGELI | SLC1A5; SLC1A | Q15758; M0I | AAAT_HUMAN | Unmodified | 0 | 0 | 173550 | 200650 | 0 | 0 | 149630 | 93958 | 0 | 0 | 2738500 | 4113500 | 0 | 0 | 425310 | 400860 | 0 | 0 |
| KLGQSESQ | PSMD9; PSMD9 | O00233; F5I | PSMD9_HUMA | Unmodified | 0 | 0 | 0 | 0 | 718520 | 215400 | 122730 | 65644 | 0 | 177910 | 22062 | 0 | 1435400 | 709300 | 0 | 0 | 814070 | 1206200 |
| KLGRSEIQI | DSG2 | Q14126 | DSG2_HUMAN | Unmodified | 0 | 0 | 0 | 0 | 1406000 | 460160 | 0 | 0 | 0 | 1440800 | 0 | 0 | 1232700 | 0 | 0 | 0 | 0 | 0 |
| KLGVNNIS | BTF3; BTF3; BTI | P20290; A0A | BTF3_HUMAN | Oxidation (M) | 0 | 0 | 0 | 0 | 4189100 | 525590 | 1110700 | 0 | 0 | 731200 | 1290500 | 11249000 | 6998000 | 1208900 | 0 | 0 | 0 | 3915700 |
| KLHEYNTQ | PIK3R1; PIK3R1 | P27986; H0Y | P85A_HUMAN | Unmodified | 0 | 0 | 72996 | 156090 | 0 | 0 | 0 | 0 | 0 | 0 | 57456 | 98025 | 374630 | 0 | 102590 | 141690 | 0 | 0 |
| KLHGLQAII | NUP210 | Q8TEM1 | PO210_HUMAN | Unmodified | 0 | 0 | 0 | 0 | 0 | 0 | 0 | 0 | 17929000 | 1184500 | 0 | 0 | 0 | 0 | 0 | 0 | 0 | 0 |
| KLHVGYLQ | ATP1B3 | P54709 | AT1B3_HUMAN | Unmodified | 0 | 0 | 0 | 0 | 0 | 0 | 0 | 0 | 0 | 0 | 1121000 | 0 | 0 | 0 | 1465200 | 0 | 0 | 0 |
| KLIDDYDSI | SCG3 | Q8WXD2 | SCG3_HUMAN | Unmodified | 0 | 0 | 0 | 0 | 0 | 0 | 0 | 0 | 1668000 | 680380 | 0 | 0 | 0 | 0 | 0 | 0 | 0 | 0 |
| KLIELQAGK | PRKCSH; PRKC | P14314; K7E | GLU2B_HUMAI | Unmodified | 0 | 0 | 0 | 0 | 0 | 0 | 14288 | 13643 | 5324900 | 1837600 | 0 | 0 | 0 | 0 | 0 | 0 | 0 | 0 |
| KLIELQAGK | PRKCSH; PRKC | P14314; K7E | GLU2B_HUMAI | Unmodified | 0 | 0 | 0 | 0 | 88995 | 0 | 0 | 57401 | 6503900 | 3156900 | 256960 | 94240 | 0 | 0 | 0 | 0 | 0 | 0 |
| KLILDSARA | TKT; TKT | P29401; A0A | TKT_HUMAN Tr | Unmodified | 0 | 0 | 0 | 0 | 3138900 | 0 | 375410 | 215490 | 0 | 656370 | 0 | 0 | 0 | 468930 | 0 | 0 | 579110 | 0 |
| KLITDALNK | PRKG2 | Q13237 | KGP2_HUMAN | Unmodified | 0 | 0 | 0 | 0 | 0 | 0 | 130590 | 37359 | 0 | 0 | 0 | 0 | 0 | 0 | 0 | 0 | 0 | 0 |
| KLIYFQLHR | TRIM28 | Q13263 | TIF1B_HUMAN | Unmodified | 0 | 0 | 0 | 0 | 151760 | 0 | 632910 | 0 | 0 | 1021400 | 0 | 1605800 | 0 | 0 | 0 | 0 | 153700 | 384420 |
| KLKLDGLD | LRPAP1; LRPAP | P30533; A0A | AMRP_HUMAN | Unmodified | 0 | 0 | 0 | 0 | 499140 | 0 | 0 | 0 | 252720 | 97562 | 0 | 0 | 0 | 0 | 0 | 0 | 19365 | 0 |
| KLKPEYDIN | OLA1; OLA1; OL | Q9NTK5; J3K | OLA1_HUMAN | Unmodified | 0 | 0 | 0 | 0 | 0 | 0 | 0 | 0 | 0 | 0 | 0 | 0 | 0 | 0 | 0 | 0 | 465290 | 275370 |
| KLLAAEPLS | COLGALT1 | Q8NBJ5 | GT251_HUMAN | Unmodified | 0 | 0 | 0 | 0 | 0 | 0 | 0 | 0 | 748580 | 163970 | 0 | 0 | 0 | 0 | 0 | 0 | 0 | 0 |
| KLLAAGQL | DNAJC3; DNAJ | Q13217; X6F | DNJC3_HUMAI | Unmodified | 0 | 0 | 0 | 0 | 0 | 0 | 0 | 0 | 0 | 4101800 | 0 | 0 | 0 | 0 | 0 | 0 | 0 | 0 |
| KLLADQAE | RPL19; RPL19; I | P84098; J3K | RL19_HUMAN | Unmodified | 43104 | 0 | 101050 | 0 | 611280 | 0 | 0 | 51315 | 1004900 | 80501 | 167480 | 1453300 | 0 | 0 | 236450 | 90863 | 17253 | 90132 |
| KLLADQAE | RPL19; RPL19; I | P84098; J3K | RL19_HUMAN | Unmodified | 0 | 0 | 0 | 0 | 0 | 0 | 0 | 0 | 0 | 0 | 0 | 3955700 | 0 | 0 | 0 | 0 | 0 | 0 |
| KLLDLVQQ | SNU13; SNU13 | P55769; A0A | NH2L1_HUMAI | Unmodified | 285640 | 0 | 0 | 0 | 114460 | 0 | 1282600 | 650050 | 0 | 0 | 0 | 1784600 | 857910 | 815120 | 0 | 92193 | 691330 | 428710 |
| KLLEAHEE | NAPA; NAPA; N | P54920; M0I | SNAA_HUMAN | Unmodified | 2468300 | 1244800 | 411980 | 463710 | 3765200 | 212140 | 1458100 | 753650 | 3386600 | 614130 | 2095300 | 8058600 | 3361400 | 528880 | 433280 | 937540 | 1162200 | 2479100 |
| KLLEDRIAE | MYH9 | P35579 | MYH9_HUMAN | Unmodified | 0 | 0 | 0 | 0 | 38270000 | 3479700 | 1283900 | 0 | 341060 | 2422000 | 189710 | 0 | 9331300 | 7184500 | 0 | 0 | 7979200 | 7853100 |
| KLLELDPEI | P4HA1 | P13674 | P4HA1_HUMAI | Unmodified | 0 | 0 | 0 | 0 | 0 | 0 | 0 | 0 | 169640 | 1076300 | 148290 | 0 | 0 | 0 | 0 | 0 | 0 | 0 |
| KLLERLPEV | NUCB1 | Q02818 | NUCB1_HUMA | Unmodified | 0 | 0 | 0 | 0 | 516520 | 173540 | 0 | 30641 | 3424900 | 1147200 | 541460 | 342050 | 175780 | 359560 | 0 | 0 | 98172 | 972000 |
| KLLEVQPQ | EHBP1; EHBP1 | Q8NDI1; H7I | EHBP1_HUMAI | Unmodified | 0 | 0 | 0 | 0 | 195920 | 0 | 0 | 0 | 0 | 58556 | 0 | 0 | 0 | 0 | 0 | 0 | 172650 | 210260 |
| KLLFQEAHI | TAOK1 | Q7L7X3 | TAOK1_HUMAI | Unmodified | 0 | 0 | 0 | 294390 | 0 | 0 | 0 | 0 | 0 | 0 | 0 | 358710 | 0 | 0 | 0 | 0 | 0 | 0 |
| KLLGLGQK | NASP; NASP; N | P49321; E9P | NASP_HUMAN | Unmodified | 0 | 0 | 0 | 0 | 117410 | 0 | 139180 | 0 | 272850 | 283520 | 104760 | 359270 | 261190 | 0 | 0 | 0 | 0 | 0 |
| KLLINAHEC | PLCB4; PLCB4 | Q15147; A0A | PLCB4_HUMAI | Unmodified | 33763 | 0 | 0 | 311560 | 0 | 0 | 0 | 23545 | 0 | 51104 | 0 | 0 | 0 | 0 | 511230 | 288450 | 1171200 | 2408800 |
| KLLLDPEFT | WDR11 | Q9BZH6 | WDR11_HUMA | Unmodified | 0 | 0 | 0 | 0 | 0 | 0 | 0 | 0 | 0 | 0 | 0 | 260200 | 0 | 0 | 0 | 69610 | 0 | 0 |
| KLLMMAGI | RPS2; RPS2; RP | P15880; E9P | RS2_HUMAN 4 | Oxidation (M) | 0 | 0 | 0 | 0 | 0 | 0 | 0 | 0 | 0 | 0 | 0 | 3471000 | 0 | 0 | 0 | 0 | 0 | 0 |
| KLLMMAGI | RPS2; RPS2; RP | P15880; E9P | RS2_HUMAN 4 | Unmodified | 0 | 0 | 606510 | 0 | 0 | 0 | 0 | 0 | 0 | 0 | 0 | 701250 | 0 | 0 | 0 | 0 | 0 | 0 |
| KLLPDDPYI | GSTO1; GSTO1 | P78417; Q5T | GSTO1_HUMAI | Unmodified | 0 | 0 | 0 | 0 | 895060 | 0 | 0 | 79952 | 0 | 0 | 848880 | 4377100 | 0 | 0 | 0 | 0 | 359820 | 0 |
| KLLPLVSDE | PYGB | P11216 | PYGB_HUMAN | Unmodified | 0 | 0 | 0 | 0 | 476910 | 0 | 0 | 0 | 81253 | 1218300 | 0 | 605290 | 1508400 | 0 | 0 | 0 | 1345100 | 5940500 |
| KLLSESAQI | TMX1 | Q9H3N1 | TMX1_HUMAN | Unmodified | 0 | 0 | 0 | 0 | 684260 | 0 | 0 | 0 | 0 | 0 | 0 | 0 | 0 | 77775 | 0 | 0 | 0 | 0 |
| KLLTEEGQI | SLC25A24 | Q6NUK1 | SCMC1_HUMA | Unmodified | 0 | 0 | 0 | 0 | 0 | 0 | 0 | 0 | 1091200 | 192060 | 104260 | 0 | 0 | 0 | 0 | 0 | 0 | 0 |
| KLLVPGKIC | VPS29; VPS29 | Q9UBQ0; F8 | VPS29_HUMAN | Unmodified | 0 | 0 | 0 | 0 | 624410 | 0 | 185990 | 0 | 0 | 201410 | 0 | 0 | 510390 | 1071500 | 0 | 0 | 550360 | 110270 |
| KLMEDRIAE | MYH10 | P35580 | MYH10_HUMAI | Unmodified | 0 | 0 | 0 | 0 | 966860 | 183010 | 0 | 0 | 0 | 0 | 0 | 0 | 403830 | 283550 | 0 | 103270 | 156360 | 0 |
| KLMFEQQK | PLCG2; PLCG2 | P16885; A0A | PLCG2_HUMAI | Unmodified | 20866 | 0 | 0 | 0 | 1044700 | 0 | 0 | 0 | 0 | 0 | 192430 | 1719700 | 0 | 225820 | 0 | 0 | 0 | 0 |
| KLMHQAAL | TXNRD1; TXNRI | Q16881; A0A | TRXR1_HUMAN | Oxidation (M) | 0 | 0 | 0 | 0 | 0 | 0 | 0 | 0 | 0 | 0 | 0 | 1061200 | 0 | 0 | 0 | 0 | 0 | 0 |
| KLMHQAAL | TXNRD1; TXNRI | Q16881; A0A | TRXR1_HUMAN | Unmodified | 0 | 625070 | 0 | 0 | 0 | 0 | 0 | 0 | 0 | 0 | 0 | 2200100 | 0 | 0 | 0 | 0 | 0 | 0 |
| KLMLDTWI | CUL5 | Q93034 | CUL5_HUMAN | Oxidation (M) | 0 | 0 | 0 | 0 | 0 | 0 | 43683 | 0 | 973180 | 0 | 0 | 458940 | 0 | 206170 | 0 | 0 | 0 | 0 |
| KLMWLFG | PFAS; PFAS; PF | O15067; J3Q | PUR4_HUMAN | Oxidation (M) | 0 | 188670 | 0 | 0 | 0 | 0 | 0 | 0 | 0 | 0 | 0 | 97269 | 0 | 0 | 65865 | 13619 | 0 | 0 |
| KLNAFGNA | EHD1; EHD3; EI | Q9H4M9; Q9 | EHD1_HUMAN | Unmodified | 262280 | 495480 | 0 | 1486300 | 0 | 0 | 0 | 0 | 0 | 129970 | 0 | 3772700 | 0 | 0 | 1030100 | 585650 | 0 | 0 |
| KLNDQANT | KCNN2; KCNN2 | Q9H2S1; A0 | KCNN2_HUMA | Unmodified | 0 | 0 | 389130 | 402260 | 0 | 0 | 0 | 0 | 0 | 0 | 0 | 212160 | 0 | 0 | 284910 | 0 | 138030 | 205630 |
| KLNEEIERL | CDC42BPB | Q9Y5S2 | MRCKB_HUMA | Unmodified | 0 | 0 | 0 | 0 | 1071400 | 0 | 0 | 0 | 169250 | 153560 | 0 | 0 | 0 | 0 | 0 | 0 | 337350 | 1258500 |
| KLNEILQAF | EIF3CL; EIF3C | B5ME19; Q9 | EIFCL_HUMAN | Unmodified | 0 | 188770 | 570310 | 0 | 0 | 0 | 0 | 0 | 0 | 0 | 0 | 2108400 | 0 | 0 | 0 | 0 | 0 | 0 |
| KLNEQVTQ | PDCD11; PDCD | Q14690; A0A | RRP5_HUMAN | Unmodified | 0 | 0 | 0 | 0 | 74918 | 0 | 82340 | 74316 | 0 | 0 | 0 | 0 | 0 | 0 | 0 | 0 | 0 | 0 |
| KLNESLDEI | BAIAP2L1 | Q9UHR4 | BI2L1_HUMAN | Unmodified | 0 | 0 | 0 | 0 | 3006800 | 700240 | 0 | 0 | 250330 | 269870 | 0 | 0 | 1116400 | 1653400 | 0 | 269060 | 0 | 0 |
| KLNESLDEI | BAIAP2L1 | Q9UHR4 | BI2L1_HUMAN | Unmodified | 0 | 0 | 0 | 0 | 3420300 | 1236600 | 0 | 0 | 567460 | 335170 | 0 | 0 | 2094400 | 1456000 | 0 | 0 | 0 | 0 |
| KLNIARNE | ACAT1; ACAT1; | P24752; A0A | THIL_HUMAN A | Unmodified | 0 | 0 | 0 | 0 | 159360 | 0 | 0 | 0 | 1813300 | 1557600 | 0 | 0 | 751910 | 0 | 0 | 0 | 0 | 0 |
| KLNKTDEEI | STXBP1; STXBP | P61764; A0A | STXB1_HUMAN | Unmodified | 0 | 0 | 0 | 0 | 0 | 0 | 1540900 | 890810 | 0 | 407430 | 10807 | 752570 | 0 | 0 | 0 | 0 | 0 | 0 |
| KLNQALLD | FTL | P02792 | FRIL_HUMAN F | Unmodified | 0 | 0 | 0 | 0 | 0 | 0 | 0 | 0 | 18941000 | 648340 | 0 | 699900 | 0 | 0 | 0 | 0 | 0 | 0 |
| KLNQNSRS | EIF3E; EIF3E; EI | P60228; B3K | EIF3E_HUMAN | Unmodified | 0 | 0 | 0 | 0 | 1402200 | 0 | 0 | 239330 | 0 | 272370 | 0 | 0 | 0 | 0 | 0 | 0 | 0 | 1023600 |
| KLNVTEQEI | ENO1; ENO1; E | P06733; A0A | ENOA_HUMAN | Unmodified | 0 | 767570 | 4417300 | 0 | 0 | 0 | 0 | 0 | 21230000 | 5332200 | 0 | 0 | 18136000 | 5030000 | 4042800 | 2941000 | 10648000 | 13659000 |
| KLNVTEQEI | ENO1; ENO1; E | P06733; A0A | ENOA_HUMAN | Unmodified | 601210 | 419850 | 0 | 3374800 | 1.46E+08 | 16070000 | 1740200 | 1098500 | 15125000 | 17623000 | 5252800 | 8103600 | 48295000 | 22533000 | 490110 | 608330 | 27132000 | 67884000 |
| KLPEEEAE | PNPO; PNPO; P | Q9NVS9; J3 | PNPO_HUMAN | Unmodified | 0 | 0 | 0 | 0 | 970640 | 0 | 132390 | 75264 | 0 | 0 | 0 | 939340 | 2061000 | 1698600 | 0 | 0 | 867890 | 2103100 |
| KLPEELGRI | SPTBN1; SPTBN | Q01082; A0A | SPTB2_HUMAN | Unmodified | 97780 | 0 | 0 | 0 | 175680 | 111010 | 0 | 0 | 239400 | 42610 | 462940 | 2725800 | 0 | 0 | 0 | 0 | 0 | 0 |
| KLPEMDID | HSPA4 | P34932 | HSP74_HUMAN | Unmodified | 0 | 0 | 0 | 207840 | 1367200 | 0 | 155530 | 53277 | 875050 | 149370 | 734300 | 1600700 | 595080 | 283260 | 0 | 188220 | 308930 | 686810 |
| KLPFPDETI | GPAA1; GPAA1; | O43292; E9I | GPAA1_HUMAI | Unmodified | 0 | 0 | 0 | 0 | 0 | 0 | 0 | 0 | 5487800 | 694780 | 119880 | 0 | 0 | 0 | 0 | 0 | 0 | 0 |
| KLPFPDETI | GPAA1; GPAA1; | O43292; E9I | GPAA1_HUMAI | Unmodified | 0 | 0 | 0 | 0 | 0 | 0 | 0 | 0 | 578820 | 0 | 0 | 0 | 0 | 0 | 0 | 0 | 0 | 0 |
| KLPIQEFHL | FEN1; FEN1 | P39748; I3L3 | FEN1_HUMAN | Unmodified | 0 | 0 | 0 | 0 | 0 | 0 | 0 | 0 | 0 | 144810 | 0 | 1348200 | 0 | 0 | 0 | 0 | 0 | 0 |
| KLPIVETLG | ATP2C1; ATP2C | P98194; B4E | AT2C1_HUMAN | Unmodified | 0 | 0 | 0 | 0 | 0 | 0 | 0 | 0 | 240220 | 0 | 0 | 147060 | 0 | 0 | 0 | 0 | 0 | 0 |
| KLPNNPAF | SOGA1; SOGA1 | O94964; X6I | SOGA1_HUMAI | Unmodified | 0 | 0 | 0 | 140890 | 0 | 0 | 0 | 0 | 0 | 0 | 0 | 0 | 520080 | 511910 | 0 | 0 | 0 | 0 |
| KLPPLPLTL | TOMM40 | O96008 | TOM40_HUMAI | Unmodified | 0 | 0 | 0 | 0 | 0 | 0 | 0 | 0 | 168840 | 0 | 0 | 0 | 0 | 0 | 0 | 0 | 0 | 0 |
| KLPPLPSLT | LAMTOR1 | Q6IAA8 | LTOR1_HUMAN | Unmodified | 0 | 0 | 0 | 905780 | 1743100 | 0 | 0 | 0 | 2743100 | 1440500 | 491390 | 1523800 | 0 | 0 | 2363700 | 0 | 0 | 0 |
| KLPPVLAIQ | USP9X | Q93008 | USP9X_HUMAN | Unmodified | 0 | 0 | 0 | 0 | 0 | 0 | 0 | 0 | 0 | 0 | 0 | 531710 | 0 | 0 | 0 | 0 | 0 | 0 |
| KLPQVEHV | MTREX | P42285 | MTREX_HUMAN | Unmodified | 0 | 0 | 0 | 0 | 0 | 0 | 0 | 0 | 0 | 0 | 0 | 984490 | 0 | 0 | 0 | 0 | 0 | 0 |
| KLPRTFCQI | EEF2 | P13639 | EF2_HUMAN EI | Unmodified | 0 | 0 | 0 | 0 | 585980 | 89768 | 959250 | 0 | 22945 | 145010 | 113360 | 0 | 239540 | 399740 | 0 | 0 | 410900 | 265110 |
| KLPSVEGLI | LAMTOR3 | Q9UHA4 | LTOR3_HUMAN | Unmodified | 896870 | 955050 | 1302800 | 0 | 13582000 | 1376300 | 3283400 | 71403 | 10432000 | 5414300 | 7426100 | 11934000 | 15414000 | 8134900 | 4951300 | 1744800 | 1231200 | 7693100 |
| KLPVESIQI | VPS25; VPS25; | Q9BRG1; K7 | VPS25_HUMAN | Unmodified | 1212500 | 3351800 | 424710 | 921610 | 0 | 0 | 0 | 0 | 0 | 0 | 21914 | 250870 | 57306 | 0 | 7088700 | 846410 | 0 | 0 |
| KLPVESIQI | VPS25; VPS25; | Q9BRG1; K7 | VPS25_HUMAN | Unmodified | 3114500 | 3808800 | 1168800 | 1411900 | 0 | 0 | 0 | 0 | 646010 | 0 | 838780 | 888490 | 1079000 | 0 | 8785800 | 2546200 | 0 | 0 |
| KLQATVQE | CORO1A; COR | P31146; H3E | COR1A_HUMAI | Unmodified | 0 | 182210 | 0 | 87412 | 0 | 0 | 0 | 0 | 0 | 48260 | 395860 | 1237500 | 123540 | 72769 | 0 | 0 | 0 | 0 |
| KLQCVDLH | KLK3; KLK3; KLI | P07288; Q8I | KLK3_HUMAN | Unmodified | 663930 | 0 | 992130 | 605360 | 0 | 0 | 0 | 0 | 0 | 0 | 0 | 0 | 0 | 0 | 609910 | 400920 | 0 | 0 |

| | | | | | | | | | | | | | | | | | | | | | | |
|---|---|---|---|---|---|---|---|---|---|---|---|---|---|---|---|---|---|---|---|---|---|---|
| KLQDKIEK | PACSIN2; PACS | Q9UNF0; A0 | PACN2_HUMAI | Unmodified | 3043500 | 2414300 | 0 | 0 | 0 | 0 | 0 | 300610 | 0 | 0 | 0 | 303430 | 0 | 0 | 1026100 | 2024600 | 0 | 0 |
| KLQDLYMA | DHX29; DHX29 | Q7Z478; A0A | DHX29_HUMAI | Oxidation (M) | 0 | 0 | 0 | 89012 | 314370 | 0 | 0 | 0 | 0 | 0 | 48371 | 197830 | 474760 | 0 | 0 | 0 | 0 | 0 |
| KLQEEELLA | CCDC50 | Q8IVM0 | CCD50_HUMA | Oxidation (M) | 3169900 | 4830200 | 2942200 | 4530500 | 0 | 0 | 0 | 23048 | 0 | 0 | 0 | 0 | 0 | 0 | 4337800 | 5557800 | 0 | 0 |
| KLQEEELLA | CCDC50 | Q8IVM0 | CCD50_HUMA | Unmodified | 2450900 | 2783500 | 0 | 0 | 0 | 0 | 0 | 0 | 0 | 0 | 0 | 0 | 0 | 0 | 1391200 | 837510 | 0 | 0 |
| KLQEEGWI | CD2AP | Q9Y5K6 | CD2AP_HUMAI | Unmodified | 2583700 | 1833500 | 2278300 | 684290 | 0 | 0 | 0 | 0 | 0 | 0 | 0 | 1393700 | 0 | 0 | 4535100 | 2808600 | 850240 | 0 |
| KLQEEGWI | CD2AP | Q9Y5K6 | CD2AP_HUMAI | Unmodified | 3173600 | 2369100 | 1922300 | 0 | 0 | 0 | 0 | 0 | 0 | 0 | 0 | 0 | 0 | 0 | 4716700 | 2516800 | 0 | 0 |
| KLQEESDLI | EIF3J | O75822 | EIF3J_HUMAN I | Unmodified | 0 | 0 | 0 | 269820 | 0 | 0 | 0 | 10130 | 369770 | 0 | 327930 | 6000500 | 455310 | 0 | 182680 | 169590 | 0 | 212910 |
| KLQELSSK/ | FASN; FASN | P49327; A0A | FAS_HUMAN F | Unmodified | 0 | 0 | 0 | 0 | 0 | 0 | 0 | 0 | 0 | 0 | 0 | 4649300 | 0 | 0 | 0 | 0 | 0 | 0 |
| KLQFSLK | BTF3; BTF3; BTI | P20290; A0A | BTF3_HUMAN | Unmodified | 0 | 0 | 0 | 0 | 161950 | 0 | 0 | 0 | 0 | 0 | 0 | 2524800 | 0 | 0 | 0 | 0 | 0 | 0 |
| KLQGEVEK | PFDN6; PFDN6 | O15212; A2A | PFD6_HUMAN | Unmodified | 0 | 0 | 0 | 0 | 2816800 | 135340 | 1919600 | 845890 | 0 | 612330 | 0 | 0 | 1962300 | 2684200 | 0 | 0 | 5373700 | 1691800 |
| KLQIQCVVI | EEF1D; EEF1B2 | P29692; P24 | EF1D_HUMAN | Unmodified | 0 | 0 | 134710 | 0 | 0 | 0 | 49911 | 0 | 0 | 0 | 137690 | 613610 | 0 | 0 | 75774 | 26958 | 0 | 0 |
| KLQLELNQ | ROCK1; ROCK1 | Q13464; A0A | ROCK1_HUMAI | Unmodified | 0 | 0 | 0 | 0 | 878090 | 70973 | 0 | 0 | 0 | 171140 | 0 | 0 | 0 | 102460 | 0 | 0 | 0 | 214720 |
| KLQLERPV\ | KIAA1522 | Q9P206 | K1522_HUMAN | Unmodified | 0 | 0 | 0 | 0 | 563460 | 0 | 0 | 0 | 0 | 129640 | 269250 | 169660 | 1136100 | 239840 | 0 | 0 | 154920 | 0 |
| KLQLSGHA | STIM1; STIM1; S | Q13586; E9F | STIM1_HUMAN | Unmodified | 0 | 0 | 95147 | 0 | 0 | 0 | 0 | 0 | 1007900 | 0 | 48215 | 0 | 0 | 0 | 0 | 0 | 0 | 0 |
| KLQNQLQ( | VPS4B | O75351 | VPS4B_HUMAN | Unmodified | 0 | 0 | 0 | 2535700 | 596690 | 0 | 0 | 0 | 8835100 | 799540 | 1523300 | 5778500 | 1126400 | 453600 | 637620 | 560900 | 0 | 570330 |
| KLQQELEFI | PSMC4 | P43686 | PRS6B_HUMAN | Unmodified | 0 | 0 | 0 | 562580 | 2707800 | 438760 | 0 | 0 | 0 | 1127500 | 0 | 1687700 | 2512300 | 2127500 | 0 | 0 | 1423800 | 2596200 |
| KLQQGIPP! | NUCB2; NUCB. | P80303; Q2L | NUCB2_HUMA | Unmodified | 0 | 0 | 0 | 0 | 158530 | 0 | 0 | 0 | 810210 | 0 | 0 | 0 | 44516 | 0 | 44846 | 0 | 94098 | 0 |
| KLQQGIPP! | NUCB2; NUCB. | P80303; Q2L | NUCB2_HUMA | Unmodified | 0 | 0 | 0 | 0 | 0 | 0 | 1815600 | 991260 | 9382700 | 3350000 | 0 | 0 | 438400 | 511600 | 0 | 0 | 0 | 0 |
| KLQTLQSEI | GIT2; GIT2; GIT. | Q14161; R4( | GIT2_HUMAN / | Unmodified | 0 | 0 | 18255 | 36344 | 250950 | 32307 | 84560 | 155570 | 125920 | 87578 | 0 | 0 | 100500 | 111270 | 38620 | 29502 | 185560 | 258420 |
| KLSALALLS | IPO11; IPO11 | Q9UI26; H0Y | IPO11_HUMAN | Unmodified | 0 | 0 | 0 | 0 | 0 | 0 | 0 | 460780 | 0 | 0 | 0 | 232250 | 0 | 0 | 0 | 28783 | 328550 | 0 |
| KLSCSLEDI | PPFIBP2; PPFIB | Q8ND30; HC | LIPB2_HUMAN | Unmodified | 0 | 0 | 0 | 197830 | 0 | 0 | 0 | 0 | 0 | 0 | 0 | 0 | 181800 | 221550 | 90602 | 72432 | 0 | 0 |
| KLSDDNTI( | SPTAN1; SPTAN | Q13813; A0A | SPTN1_HUMAN | Unmodified | 0 | 0 | 0 | 0 | 0 | 0 | 0 | 0 | 0 | 49874 | 0 | 1927700 | 0 | 0 | 0 | 0 | 0 | 0 |
| KLSEDSLTI | STK3; STK3; STI | Q13188; Q8I | STK3_HUMAN ! | Unmodified | 0 | 0 | 0 | 0 | 2548700 | 58116 | 0 | 0 | 0 | 126940 | 0 | 0 | 164270 | 0 | 0 | 0 | 0 | 152040 |
| KLSEEEILEI | RCN2; RCN2 | Q14257; A8N | RCN2_HUMAN | Unmodified | 0 | 0 | 0 | 0 | 0 | 0 | 0 | 0 | 10985000 | 0 | 0 | 445090 | 0 | 0 | 0 | 0 | 0 | 0 |
| KLSEKLDSI | TFRC; TFRC | P02786; G3\ | TFR1_HUMAN | Unmodified | 142420 | 0 | 0 | 272140 | 0 | 0 | 0 | 0 | 3383900 | 0 | 0 | 0 | 0 | 0 | 0 | 0 | 0 | 0 |
| KLSELLR | HSP90AA1 | P07900 | HS90A_HUMAN | Unmodified | 669330 | 0 | 0 | 982390 | 0 | 645400 | 85987 | 0 | 0 | 1932900 | 2350200 | 27839000 | 1261200 | 663000 | 563720 | 0 | 0 | 0 |
| KLSELLRYY | HSP90AA1 | P07900 | HS90A_HUMAN | Unmodified | 0 | 0 | 0 | 0 | 1330100 | 0 | 0 | 0 | 0 | 0 | 0 | 0 | 0 | 0 | 0 | 0 | 346950 | 0 |
| KLSELLRYY | HSP90AA1 | P07900 | HS90A_HUMAN | Oxidation (M) | 0 | 0 | 0 | 0 | 0 | 0 | 0 | 0 | 0 | 1842000 | 0 | 0 | 1566700 | 0 | 0 | 0 | 0 | 0 |
| KLSGLNAFI | MBD3; MBD3; I | O95983; A0A | MBD3_HUMAN | Unmodified | 130850 | 0 | 0 | 0 | 402900 | 0 | 304840 | 0 | 119670 | 499030 | 0 | 0 | 474490 | 81400 | 0 | 0 | 119100 | 0 |
| KLSLGQYD | TNS3; TNS3 | Q68CZ2; E9I | TENS3_HUMAN | Unmodified | 0 | 0 | 0 | 0 | 0 | 0 | 0 | 0 | 0 | 0 | 0 | 0 | 0 | 0 | 0 | 0 | 1455300 | 0 |
| KLSLLNGG | KIF20A | O95235 | KI20A_HUMAN | Unmodified | 21090 | 0 | 96752 | 451020 | 1330600 | 0 | 0 | 0 | 0 | 0 | 0 | 0 | 719570 | 525270 | 172360 | 114530 | 621030 | 317340 |
| KLSMYGVD | EPB41; EPB41; | P11171; A0A | EPB41_HUMAN | Unmodified | 134310 | 285530 | 364720 | 0 | 866450 | 0 | 674580 | 0 | 0 | 0 | 0 | 823480 | 0 | 0 | 0 | 373800 | 185700 | 374210 |
| KLSMYGVD | EPB41; EPB41; | P11171; A0A | EPB41_HUMAN | Oxidation (M) | 211700 | 0 | 215200 | 628870 | 0 | 0 | 354500 | 277600 | 207090 | 0 | 25471 | 313980 | 110250 | 256420 | 375980 | 626680 | 0 | 0 |
| KLSNLNAD | LRRC1 | Q9BTT6 | LRRC1_HUMAN | Unmodified | 22701 | 0 | 87024 | 0 | 1343300 | 460700 | 0 | 0 | 416220 | 227540 | 69163 | 697390 | 611180 | 291220 | 36016 | 59093 | 42791 | 469560 |
| KLSQPITV( | CAPN5; CAPN5 | O15484; E7E | CAN5_HUMAN | Unmodified | 0 | 0 | 191150 | 0 | 518300 | 0 | 0 | 0 | 368930 | 251210 | 0 | 283010 | 1038200 | 1207300 | 0 | 291630 | 418930 | 1205500 |
| KLSRADLTE | UQCRC1 | P31930 | QCR1_HUMAN | Unmodified | 0 | 0 | 0 | 0 | 0 | 0 | 0 | 0 | 0 | 755090 | 0 | 0 | 0 | 0 | 0 | 0 | 0 | 0 |
| KLSSAMSA | MDH1; MDH1; I | P40925; B8Z | MDHC_HUMAN | Unmodified | 0 | 0 | 0 | 0 | 229360 | 0 | 307660 | 347400 | 0 | 357530 | 1687900 | 0 | 164920 | 0 | 0 | 0 | 744000 | 0 |
| KLSSWDQ/ | SF3B1 | O75533 | SF3B1_HUMAN | Unmodified | 289780 | 466720 | 122270 | 0 | 273550 | 0 | 0 | 0 | 0 | 193570 | 224670 | 0 | 0 | 0 | 0 | 244420 | 17230 | 0 |
| KLSTIALAL( | PPP2R1A; PPP2 | P30153; P30 | 2AAA_HUMAN | Unmodified | 0 | 0 | 0 | 1298200 | 142680 | 0 | 0 | 0 | 477140 | 171290 | 599690 | 824820 | 403930 | 0 | 0 | 0 | 16246 | 0 |
| KLSVACFY( | DDX21 | Q9NR30 | DDX21_HUMAI | Unmodified | 0 | 0 | 0 | 0 | 0 | 0 | 220470 | 33560 | 0 | 0 | 0 | 702590 | 0 | 382590 | 0 | 0 | 0 | 0 |
| KLTAGEAC. | ECI2; ECI2 | O75521; A0A | ECI2_HUMAN E | Unmodified | 0 | 0 | 0 | 0 | 0 | 0 | 0 | 0 | 8930600 | 1739200 | 0 | 0 | 0 | 0 | 0 | 0 | 0 | 0 |
| KLTAYLDLN | ITM2B; ITM2B | Q9Y287; A0A | ITM2B_HUMAN | Unmodified | 0 | 0 | 0 | 0 | 0 | 0 | 0 | 0 | 0 | 0 | 0 | 0 | 0 | 0 | 0 | 0 | 0 | 0 |
| KLTDIINND | GPD1L; GPD1L | Q8N335; C9 | GPD1L_HUMAI | Unmodified | 0 | 0 | 48196 | 44370 | 102610 | 0 | 0 | 0 | 366080 | 28323 | 159990 | 766410 | 140360 | 0 | 0 | 83253 | 27654 | 136660 |
| KLTDPTGW | DSC2; DSC2 | Q02487; A0A | DSC2_HUMAN | Unmodified | 0 | 0 | 0 | 0 | 386310 | 117960 | 0 | 0 | 2235500 | 737790 | 0 | 0 | 456970 | 346540 | 0 | 0 | 0 | 661070 |
| KLTEDDLIL | PLAA; PLAA; PL | Q9Y263; H0Y | PLAP_HUMAN I | Unmodified | 0 | 0 | 0 | 0 | 0 | 0 | 0 | 0 | 0 | 0 | 258730 | 797480 | 0 | 0 | 0 | 113290 | 0 | 0 |
| KLTELENEL | AFDN; AFDN; A | P55196; H0Y | AFAD_HUMAN | Unmodified | 20843 | 0 | 0 | 0 | 614930 | 0 | 0 | 0 | 484540 | 127540 | 295810 | 0 | 1905500 | 1186500 | 0 | 51253 | 188510 | 194310 |
| KLTELFSVI | KDF1 | Q8NAX2 | KDF1_HUMAN | Unmodified | 0 | 0 | 3458500 | 607520 | 0 | 0 | 0 | 0 | 2941400 | 912420 | 690990 | 672930 | 585290 | 0 | 890910 | 700070 | 0 | 0 |
| KLTELGTVI | IQGAP1; IQGAP | P46940; H0Y | IQGA1_HUMAN | Unmodified | 368430 | 465930 | 667320 | 616820 | 5572100 | 1958000 | 316400 | 163350 | 895330 | 983970 | 0 | 1859500 | 0 | 886570 | 340950 | 476750 | 0 | 0 |
| KLTFDEEAY | HTATIP2; HTATI | Q9BUP3; E9I | HTAI2_HUMAN | Unmodified | 0 | 0 | 0 | 0 | 237230 | 191770 | 0 | 96955 | 1210000 | 397110 | 0 | 0 | 0 | 144890 | 0 | 0 | 45998 | 0 |
| KLTGGSNV | PICALM; PICALI | Q13492; H0Y | PICAL_HUMAN | Unmodified | 394390 | 117890 | 0 | 0 | 194780 | 0 | 57081 | 33605 | 59783 | 243200 | 0 | 74894 | 1879100 | 979900 | 0 | 0 | 57895 | 202060 |
| KLTGIKHEL | CFL1; CFL1; CF | P23528; G3V | COF1_HUMAN | Unmodified | 20031000 | 8249900 | 4045700 | 13995000 | 2.12E+08 | 28907000 | 7107700 | 2339800 | 11342000 | 7123900 | 14664000 | 78951000 | 49569000 | 26442000 | 9554800 | 7093500 | 32050000 | 55360000 |
| KLTGKDVN | RPS7 | P62081 | RS7_HUMAN 4 | Unmodified | 810990 | 966570 | 0 | 0 | 6510900 | 1219100 | 5943600 | 227470 | 407480 | 2256400 | 0 | 0 | 19563000 | 8523300 | 0 | 0 | 3404300 | 5724000 |
| KLTPGEVYI | NPTX1 | Q15818 | NPTX1_HUMAN | Unmodified | 0 | 0 | 278020 | 0 | 0 | 0 | 0 | 0 | 15996000 | 939510 | 0 | 0 | 0 | 347140 | 0 | 0 | 0 | 0 |
| KLTPITYPQ | RAC1 | P63000 | RAC1_HUMAN | Oxidation (M) | 2509500 | 1783300 | 0 | 1138500 | 3876200 | 0 | 831000 | 770160 | 0 | 383200 | 0 | 2437100 | 2670300 | 1358400 | 2338300 | 2345700 | 1267300 | 2904800 |
| KLTPITYPQ | RAC1 | P63000 | RAC1_HUMAN | Unmodified | 971820 | 1264200 | 466700 | 1183900 | 11168000 | 0 | 870430 | 409600 | 1450200 | 0 | 811470 | 2085300 | 3484000 | 2454700 | 864630 | 864200 | 2566600 | 8663600 |
| KLTSIKIQNI | PTPRA | P18433 | PTPRA_HUMAN | Unmodified | 0 | 0 | 0 | 0 | 290470 | 15776 | 74315 | 0 | 102430 | 56866 | 122020 | 0 | 122060 | 85317 | 0 | 0 | 351680 | 523420 |
| KLTTDPDLI | LARP4; LARP4; | Q71RC2; A0. | LARP4_HUMAN | Unmodified | 411310 | 564230 | 255670 | 296350 | 0 | 0 | 0 | 0 | 0 | 0 | 51237 | 238320 | 0 | 0 | 297580 | 179290 | 0 | 0 |
| KLTWTYAAI | VPS13C; VPS13 | Q709C8; A0A | VP13C_HUMAN | Unmodified | 0 | 0 | 0 | 0 | 0 | 0 | 0 | 0 | 0 | 0 | 0 | 0 | 0 | 0 | 0 | 0 | 0 | 0 |
| KLVAIVDPI | GANAB; GANAE | Q14697; F5I | GANAB_HUMA | Unmodified | 0 | 0 | 0 | 0 | 0 | 0 | 0 | 0 | 13123000 | 708920 | 900170 | 498160 | 0 | 0 | 0 | 0 | 0 | 0 |
| KLVAIVDPI | GANAB; GANAE | Q14697; F5I | GANAB_HUMA | Unmodified | 0 | 0 | 0 | 0 | 0 | 0 | 0 | 0 | 9090100 | 129370 | 0 | 0 | 0 | 0 | 0 | 0 | 0 | 0 |
| KLVDYDSA | BIN1 | O00499 | BIN1_HUMAN I | Unmodified | 0 | 0 | 0 | 0 | 0 | 0 | 0 | 0 | 0 | 0 | 0 | 0 | 646950 | 314640 | 0 | 0 | 475770 | 213990 |
| KLVEAEELE | CIT; CIT; CIT | O14578; A0A | CTRO_HUMAN | Unmodified | 0 | 0 | 0 | 261230 | 0 | 0 | 0 | 401910 | 0 | 0 | 0 | 23340 | 0 | 0 | 0 | 143770 | 0 | 0 |
| KLVESLPQI | MRPL12; N/A | P52815; B4D | RM12_HUMAN | Unmodified | 0 | 0 | 0 | 0 | 0 | 0 | 0 | 0 | 1534000 | 582690 | 0 | 186550 | 0 | 52922 | 0 | 0 | 0 | 0 |
| KLVGFISAII | NMT1; NMT1; N | P30419; A0A | NMT1_HUMAN | Unmodified | 0 | 0 | 0 | 0 | 0 | 0 | 0 | 0 | 0 | 0 | 0 | 1755100 | 0 | 0 | 0 | 0 | 0 | 0 |
| KLVGYLDRI | PSAP; PSAP; PS | P07602; Q5E | SAP_HUMAN P | Unmodified | 318350 | 353980 | 1672200 | 2274500 | 7659600 | 3653000 | 674000 | 934810 | 1.16E+08 | 73148000 | 12532000 | 549100 | 17008000 | 13447000 | 1120400 | 422400 | 198140 | 3324800 |
| KLVHEGPL | ARHGEF1; ARH | Q92888; M0I | ARHG1_HUMA | Unmodified | 0 | 0 | 0 | 0 | 1214700 | 0 | 125060 | 0 | 0 | 0 | 0 | 0 | 0 | 0 | 0 | 0 | 0 | 0 |
| KLVIIEGDLI | TPM3; TPM3; TF | P06753; J3KI | TPM3_HUMAN | Unmodified | 0 | 0 | 0 | 0 | 0 | 0 | 0 | 0 | 0 | 0 | 1475700 | 0 | 0 | 0 | 0 | 0 | 0 | 0 |
| KLVIPSELG | FKBP2; FKBP2 | P26885; A0A | FKBP2_HUMAN | Unmodified | 0 | 0 | 0 | 131210 | 0 | 0 | 0 | 157250 | 21244000 | 0 | 0 | 0 | 0 | 0 | 0 | 39346 | 0 | 0 |
| KLVIVGDG/ | RHOA; RHOC; I | P61586; P08 | RHOA_HUMAN | Unmodified | 389770 | 282600 | 574570 | 1644600 | 10663000 | 2554100 | 1186200 | 292220 | 5658000 | 4695500 | 2880200 | 12596000 | 3349700 | 5105900 | 1105600 | 2130800 | 4965100 | 9651200 |
| KLVNTINP( | UPF1 | Q92900 | RENT1_HUMAN | Unmodified | 506480 | 663220 | 245540 | 0 | 0 | 0 | 0 | 0 | 42031 | 0 | 0 | 334940 | 0 | 0 | 194910 | 123790 | 0 | 0 |
| KLVVATEK | EMC1; EMC1 | Q8N766; Q5 | EMC1_HUMAN | Unmodified | 0 | 0 | 0 | 0 | 0 | 0 | 0 | 0 | 3184200 | 67371 | 0 | 91561 | 0 | 0 | 0 | 0 | 0 | 0 |
| KLVVENVD | NCKAP1 | Q9Y2A7 | NCKP1_HUMAI | Oxidation (M) | 430510 | 283160 | 291980 | 116800 | 0 | 0 | 0 | 0 | 55110 | 0 | 0 | 1298900 | 95977 | 0 | 764530 | 688630 | 0 | 0 |
| KLVVVGDG | RHOB | P62745 | RHOB_HUMAN | Unmodified | 0 | 0 | 0 | 0 | 0 | 0 | 0 | 41863 | 0 | 0 | 0 | 360030 | 0 | 0 | 0 | 0 | 0 | 0 |
| KLVWVPSD | MYH9; MYH9; I | P35579; Q5E | MYH9_HUMAN | Unmodified | 0 | 0 | 0 | 0 | 6823900 | 629840 | 271910 | 0 | 361680 | 0 | 0 | 2619300 | 1856900 | 718050 | 0 | 0 | 573040 | 3188900 |
| KLVYLYLMI | AP1B1; AP2B1; | Q10567; P6: | AP1B1_HUMAN | Oxidation (M) | 0 | 74716 | 0 | 0 | 716020 | 914410 | 0 | 177050 | 0 | 255620 | 0 | 2677900 | 844320 | 0 | 0 | 199140 | 436320 | 520850 |
| KLWVGQLI | MICAL1 | Q8TDZ2 | MICA1_HUMAN | Unmodified | 0 | 0 | 0 | 43210 | 75106 | 0 | 0 | 0 | 69090 | 0 | 0 | 0 | 0 | 0 | 0 | 0 | 0 | 17712 |
| KLYDIDVAI | RPL23A; RPL23 | P62750; K7E | RL23A_HUMAN | Unmodified | 466940 | 416780 | 181080 | 0 | 539350 | 69584 | 126850 | 72167 | 122770 | 55809 | 516540 | 5272600 | 339750 | 123460 | 0 | 185140 | 55957 | 165090 |
| KLYGAQFH | GMPS | P49915 | GUAA_HUMAN | Unmodified | 0 | 0 | 0 | 0 | 0 | 0 | 0 | 0 | 271870 | 0 | 0 | 2114100 | 0 | 0 | 0 | 0 | 0 | 0 |
| KLYGDADY | CTPS1; CTPS1; | P17812; B4E | PYRG1_HUMAN | Unmodified | 0 | 0 | 0 | 0 | 0 | 0 | 0 | 0 | 0 | 0 | 0 | 794400 | 0 | 0 | 0 | 0 | 0 | 0 |
| KLYSIAAPA | ABHD12; ABHD | Q8N2K0; A0. | ABD12_HUMAN | Unmodified | 0 | 21827 | 0 | 0 | 0 | 0 | 0 | 0 | 1312700 | 197880 | 0 | 0 | 0 | 0 | 0 | 0 | 0 | 0 |
| KLYTLEEHL | ROCK2; ROCK2 | O75116; E9F | ROCK2_HUMAI | Unmodified | 0 | 0 | 0 | 0 | 2331000 | 0 | 0 | 0 | 0 | 0 | 0 | 0 | 580710 | 0 | 0 | 0 | 2391100 | 2288100 |
| KLYVSNLGI | PPAT; PPAT | Q06203; D6I | PUR1_HUMAN | Unmodified | 750510 | 194860 | 0 | 0 | 0 | 0 | 0 | 232260 | 0 | 0 | 0 | 3802700 | 0 | 0 | 0 | 0 | 0 | 0 |
| KMDDSAL( | SNX18 | Q96RF0 | SNX18_HUMAN | Oxidation (M) | 0 | 0 | 1515500 | 1244200 | 139870 | 0 | 0 | 63489 | 0 | 0 | 0 | 0 | 2133700 | 409240 | 1908900 | 1182900 | 0 | 0 |
| KMDDSAL( | SNX18 | Q96RF0 | SNX18_HUMAN | Unmodified | 816380 | 1024200 | 1601100 | 1972700 | 1506500 | 0 | 0 | 11377 | 0 | 0 | 0 | 0 | 9349400 | 3247400 | 2738500 | 1765700 | 0 | 0 |
| KMDDSAL( | SNX18 | Q96RF0 | SNX18_HUMAN | Oxidation (M) | 1439400 | 1328800 | 3815200 | 4425200 | 0 | 0 | 0 | 0 | 0 | 0 | 0 | 0 | 6043200 | 2292700 | 3729700 | 4384600 | 0 | 0 |
| KMDDSVL( | SNX33 | Q8WV41 | SNX33_HUMAN | Oxidation (M) | 0 | 26546 | 31310 | 1679000 | 467870 | 0 | 0 | 0 | 36815 | 0 | 0 | 0 | 0 | 985020 | 2912700 | 1071400 | 185540 | 0 |
| KMDDSVL( | SNX33 | Q8WV41 | SNX33_HUMAN | Oxidation (M) | 2514900 | 4818800 | 0 | 12281000 | 0 | 0 | 0 | 0 | 0 | 0 | 0 | 0 | 1548400 | 0 | 27069000 | 13603000 | 0 | 0 |
| KMDEYVAS | AHCYL1 | O43865 | SAHH2_HUMAI | Oxidation (M) | 192450 | 230580 | 0 | 2888300 | 15768000 | 1059500 | 1793900 | 0 | 160860 | 690380 | 2222000 | 248080 | 5997100 | 1101100 | 4407000 | 271230 | 6994100 | 2503600 |
| KMDEYVAS | AHCYL1 | O43865 | SAHH2_HUMAI | Unmodified | 0 | 0 | 0 | 0 | 0 | 0 | 0 | 0 | 0 | 0 | 0 | 0 | 0 | 0 | 0 | 0 | 153670 | 457930 |
| KMDEYVAS | AHCYL1 | O43865 | SAHH2_HUMAI | Oxidation (M) | 0 | 0 | 0 | 0 | 2758800 | 0 | 556040 | 0 | 0 | 0 | 0 | 0 | 0 | 0 | 0 | 0 | 2959800 | 1285700 |
| KMDEYVAS | AHCYL2; AHCY | Q96HN2; HC | SAHH3_HUMAI | Oxidation (M) | 0 | 738450 | 0 | 0 | 0 | 405420 | 3562700 | 0 | 0 | 0 | 0 | 0 | 691120 | 537700 | 1299100 | 0 | 1330300 | 0 |
| KMDPAEED | SDC2; SDC2; S | P34741; E9P | SDC2_HUMAN | Oxidation (M) | 356950 | 303790 | 0 | 0 | 0 | 0 | 0 | 0 | 0 | 0 | 0 | 106860 | 0 | 0 | 0 | 141370 | 0 | 0 |
| KMDPAEED | SDC2; SDC2; S | P34741; E9P | SDC2_HUMAN | Unmodified | 0 | 526440 | 0 | 0 | 0 | 0 | 0 | 0 | 0 | 0 | 0 | 0 | 0 | 0 | 0 | 0 | 0 | 0 |
| KMDPSTCA | ADAM10 | O14672 | ADA10_HUMAN | Oxidation (M) | 1583000 | 869920 | 2442200 | 2767200 | 671650 | 300480 | 0 | 0 | 495350 | 0 | 0 | 3591300 | 1065800 | 512060 | 3423900 | 3422100 | 255970 | 282050 |
| KMDPSTCA | ADAM10 | O14672 | ADA10_HUMAN | Unmodified | 761770 | 353620 | 1430600 | 2780700 | 0 | 0 | 0 | 0 | 0 | 0 | 47679 | 4526200 | 0 | 75059 | 2475700 | 1480100 | 0 | 0 |
| KMDPSTCA | ADAM10 | O14672 | ADA10_HUMAN | Unmodified | 0 | 0 | 0 | 0 | 1912400 | 0 | 241020 | 0 | 0 | 37621 | 0 | 0 | 1331200 | 1030300 | 0 | 0 | 279830 | 725950 |
| KMDPSTCA | ADAM10 | O14672 | ADA10_HUMAN | Oxidation (M) | 0 | 0 | 0 | 0 | 0 | 0 | 0 | 0 | 0 | 0 | 0 | 0 | 0 | 911860 | 0 | 0 | 0 | 0 |
| KMDSPGAI | TRAF3; TRAF3; | Q13114; A6I | TRAF3_HUMAN | Oxidation (M) | 0 | 0 | 0 | 0 | 1768400 | 0 | 0 | 0 | 185610 | 0 | 0 | 0 | 0 | 0 | 0 | 0 | 0 | 0 |
| KMDSPGAI | TRAF3; TRAF3; | Q13114; A6I | TRAF3_HUMAN | Unmodified | 0 | 0 | 0 | 0 | 0 | 0 | 0 | 0 | 0 | 0 | 0 | 0 | 0 | 0 | 0 | 0 | 0 | 0 |
| KMDVGGL! | SYT2; SYT1; SY | Q8N9I0; P21 | SYT2_HUMAN ! | Unmodified | 0 | 0 | 0 | 0 | 0 | 0 | 763670 | 196880 | 343960 | 340470 | 0 | 0 | 0 | 128800 | 0 | 0 | 0 | 0 |
| KMEDSVG( | MYH9 | P35579 | MYH9_HUMAN | Unmodified | 46728 | 183260 | 0 | 0 | 3555200 | 916370 | 0 | 0 | 0 | 0 | 0 | 1139500 | 647760 | 0 | 0 | 0 | 795470 | 1042800 |
| KMEDSVG( | MYH9 | P35579 | MYH9_HUMAN | Oxidation (M) | 62828 | 0 | 276920 | 246590 | 1573100 | 322110 | 0 | 27988 | 0 | 98626 | 0 | 1146900 | 509130 | 490500 | 476420 | 325260 | 440650 | 669430 |
| KMEEEILLL | MYH10 | P35580 | MYH10_HUMAI | Unmodified | 0 | 0 | 0 | 0 | 2015000 | 0 | 194110 | 40801 | 2240900 | 0 | 0 | 538620 | 0 | 0 | 0 | 0 | 0 | 0 |
| KMEMEME( | SEPTIN7; SEPTI | Q16181; E7E | SEPT7_HUMAN | Unmodified | 0 | 0 | 0 | 0 | 780650 | 0 | 0 | 0 | 177670 | 0 | 0 | 0 | 0 | 0 | 0 | 0 | 104390 | 336450 |
| KMEMEME( | SEPTIN7; SEPTI | Q16181; E7E | SEPT7_HUMAN | 2 Oxidation (M) | 0 | 0 | 0 | 0 | 789650 | 0 | 0 | 0 | 0 | 0 | 0 | 87540 | 185320 | 0 | 0 | 11629 | 0 | 343560 |
| KMEMEME( | SEPTIN7; SEPTI | Q16181; E7E | SEPT7_HUMAN | Oxidation (M) | 0 | 0 | 0 | 0 | 1645400 | 0 | 0 | 0 | 32967 | 0 | 0 | 0 | 0 | 71116 | 0 | 0 | 122920 | 151200 |
| KMESDFEC | DIAPH1; DIAPH | O60610; A0A | DIAP1_HUMAN | Unmodified | 0 | 0 | 0 | 0 | 656560 | 252360 | 0 | 0 | 0 | 0 | 555550 | 1614700 | 703770 | 0 | 0 | 0 | 600340 | 1826700 |
| KMESDFEC | DIAPH1; DIAPH | O60610; A0A | DIAP1_HUMAN | Oxidation (M) | 295720 | 0 | 0 | 0 | 477810 | 63759 | 44514 | 0 | 0 | 185220 | 0 | 0 | 422850 | 265740 | 0 | 79619 | 246950 | 517970 |
| KMFESFIES | PRKAR2A; PRKA | P13861; H7( | KAP2_HUMAN | Oxidation (M) | 0 | 0 | 0 | 0 | 0 | 0 | 0 | 0 | 0 | 93863 | 140280 | 352300 | 517010 | 354850 | 333380 | 0 | 0 | 0 |

| | | | | | | | | | | | | | | | | | | | | | |
|---|---|---|---|---|---|---|---|---|---|---|---|---|---|---|---|---|---|---|---|---|---|
| KMGAPESC | FTH1; FTH1; FTI | P02794; G3\ | FRIH_HUMAN I | Oxidation (M) | 0 | 0 | 0 | 0 | 0 | 0 | 253870 | 0 | 5533600 | 2705200 | 0 | 0 | 209750 | 509320 | 0 | 0 | 766170 | 0 |
| KMGAPESC | FTH1; FTH1; FTI | P02794; G3\ | FRIH_HUMAN I | Unmodified | 0 | 0 | 0 | 0 | 0 | 0 | 0 | 0 | 4282800 | 559400 | 0 | 0 | 0 | 0 | 0 | 0 | 0 | 0 |
| KMGDHLTN | FTL | P02792 | FRIL_HUMAN F | Unmodified | 0 | 0 | 0 | 0 | 0 | 0 | 0 | 0 | 1936600 | 204890 | 0 | 0 | 0 | 0 | 0 | 0 | 0 | 0 |
| KMGQNLAI | ITGB4; ITGB4 | P16144; J3Q | ITB4_HUMAN I | Oxidation (M) | 0 | 0 | 0 | 845960 | 425650 | 0 | 0 | 0 | 0 | 0 | 0 | 0 | 0 | 0 | 0 | 0 | 665420 | 402720 |
| KMISTQQLI | DZIP1 | Q86YF9 | DZIP1_HUMAN | Oxidation (M) | 0 | 0 | 0 | 0 | 0 | 0 | 0 | 0 | 0 | 0 | 0 | 1120600 | 0 | 0 | 0 | 0 | 0 | 0 |
| KMMADEAI | ECH1 | Q13011 | ECH1_HUMAN | Oxidation (M) | 0 | 0 | 0 | 0 | 0 | 0 | 0 | 0 | 354880 | 0 | 0 | 0 | 0 | 0 | 0 | 0 | 0 | 0 |
| KMPSASCV | SORL1; SORL1; | Q92673; E9F | SORL_HUMAN | Unmodified | 0 | 0 | 0 | 0 | 0 | 0 | 0 | 0 | 494740 | 0 | 0 | 0 | 0 | 0 | 0 | 0 | 0 | 0 |
| KMQELLQT | EHD1; EHD1; EI | Q9H4M9; C9 | EHD1_HUMAN | Unmodified | 0 | 0 | 0 | 120260 | 501990 | 0 | 145940 | 98013 | 0 | 0 | 0 | 749590 | 0 | 0 | 0 | 0 | 571550 | 702160 |
| KMQPDQQ | VASP; VASP | P50552; K7E | VASP_HUMAN | Unmodified | 31583 | 0 | 0 | 122450 | 160610 | 0 | 0 | 0 | 180840 | 81833 | 0 | 1228200 | 0 | 189770 | 53339 | 29651 | 96602 | 0 |
| KMQQNIQE | MYH9 | P35579 | MYH9_HUMAN | Oxidation (M) | 0 | 0 | 0 | 0 | 0 | 2086700 | 0 | 0 | 589100 | 264580 | 2640300 | 4353700 | 0 | 0 | 0 | 674360 | 0 | 1597600 |
| KMQQNIQE | MYH9 | P35579 | MYH9_HUMAN | Unmodified | 0 | 0 | 0 | 0 | 7181000 | 0 | 2107900 | 0 | 0 | 0 | 0 | 1604700 | 0 | 0 | 0 | 0 | 0 | 2651700 |
| KMTTSTDH | ITGA1 | P56199 | ITA1_HUMAN I | Unmodified | 0 | 25396 | 0 | 0 | 0 | 0 | 0 | 0 | 279450 | 0 | 242680 | 0 | 20680 | 0 | 155900 | 0 | 165490 | 620520 |
| KMTVTDQV | APOD; APOD | P05090; C9J | APOD_HUMAN | Oxidation (M) | 0 | 0 | 307900 | 87658 | 0 | 48153 | 0 | 0 | 5102000 | 851610 | 0 | 0 | 485430 | 238060 | 0 | 0 | 0 | 0 |
| KMTVTDQV | APOD; APOD | P05090; C9J | APOD_HUMAN | Unmodified | 0 | 483410 | 0 | 187460 | 0 | 0 | 0 | 0 | 3942500 | 475680 | 0 | 0 | 0 | 0 | 214270 | 101220 | 0 | 0 |
| KMTVTDQV | APOD | P05090 | APOD_HUMAN | Oxidation (M) | 0 | 27018 | 76867 | 0 | 0 | 0 | 53887 | 0 | 230250 | 640180 | 0 | 228780 | 0 | 0 | 0 | 0 | 0 | 0 |
| KMTVTDQV | APOD | P05090 | APOD_HUMAN | Unmodified | 0 | 0 | 0 | 0 | 0 | 0 | 0 | 0 | 40273 | 2836200 | 0 | 0 | 0 | 0 | 0 | 274470 | 0 | 0 |
| KMYDDAIE | MAN1A2; MAN1 | O60476; H0 | MA1A2_HUMAI | Unmodified | 0 | 0 | 0 | 0 | 0 | 0 | 0 | 0 | 957400 | 0 | 0 | 0 | 0 | 0 | 0 | 0 | 0 | 0 |
| KNCPHIVV | DDX39B; DDX3 | Q13838; F6T | DX39B_HUMAN | Unmodified | 0 | 0 | 0 | 0 | 0 | 0 | 0 | 0 | 0 | 0 | 0 | 0 | 0 | 0 | 0 | 0 | 0 | 0 |
| KNDDGDT | STX2 | P32856 | STX2_HUMAN S | Unmodified | 0 | 0 | 0 | 0 | 379840 | 0 | 109810 | 61858 | 171160 | 76200 | 0 | 1294400 | 0 | 37664 | 0 | 0 | 571940 | 60965 |
| KNDDGWY | ABI1; ABI1; ABI1 | Q8IZP0; A0A | ABI1_HUMAN A | Unmodified | 232580 | 50238 | 213940 | 152410 | 0 | 0 | 0 | 0 | 629700 | 0 | 126710 | 1421400 | 0 | 0 | 1082300 | 678060 | 0 | 0 |
| KNDHGDG | FKBP7 | Q9Y680 | FKBP7_HUMAN | Unmodified | 0 | 0 | 0 | 0 | 27284 | 15923 | 0 | 0 | 813970 | 48574 | 0 | 0 | 0 | 0 | 188250 | 0 | 0 | 0 |
| KNDIYGED | GDI2 | P50395 | GDIB_HUMAN | Unmodified | 0 | 0 | 0 | 0 | 2916000 | 0 | 0 | 0 | 0 | 0 | 0 | 331190 | 1177800 | 0 | 49344 | 0 | 81361 | 50549 |
| KNEALAPPI | MGAT2 | Q10469 | MGAT2_HUMAI | Unmodified | 0 | 0 | 0 | 0 | 0 | 0 | 0 | 0 | 8385100 | 0 | 0 | 0 | 0 | 0 | 0 | 0 | 0 | 0 |
| KNEALAPPI | MGAT2 | Q10469 | MGAT2_HUMAI | Unmodified | 0 | 0 | 0 | 0 | 0 | 0 | 0 | 0 | 1085300 | 0 | 0 | 0 | 0 | 0 | 0 | 0 | 0 | 0 |
| KNEELEQL | PNPO; PNPO; P | Q9NVS9; J3C | PNPO_HUMAN | Unmodified | 424950 | 403490 | 555850 | 1739800 | 1506600 | 396070 | 0 | 0 | 741760 | 186670 | 1060700 | 4762200 | 1754400 | 1363600 | 1562700 | 1362600 | 256280 | 2119400 |
| KNEFQGEL | GOLM1; GOLM1 | Q8NBJ4; F8V | GOLM1_HUMA | Unmodified | 0 | 32205 | 276820 | 0 | 596800 | 0 | 0 | 0 | 29509000 | 1353500 | 0 | 258940 | 125440 | 0 | 0 | 0 | 0 | 0 |
| KNEGLTWF | NUDCD1; NUD | Q96RS6; A0A | NUDC1_HUMA | Unmodified | 18272 | 69473 | 0 | 0 | 238460 | 0 | 0 | 31707 | 69942 | 97858 | 0 | 402630 | 46082 | 0 | 0 | 0 | 0 | 0 |
| KNEGVNW | CHMP1A; CHM | Q9HD42; F8 | CHM1A_HUMA | Unmodified | 351510 | 0 | 0 | 0 | 1048000 | 0 | 202980 | 0 | 0 | 0 | 314990 | 0 | 0 | 0 | 0 | 148960 | 0 | 211390 |
| KNELFSNL | SLC9A3R1 | O14745 | NHRF1_HUMAI | Unmodified | 501590 | 0 | 2061300 | 2498100 | 10182000 | 0 | 4643800 | 3074100 | 0 | 0 | 626920 | 2893400 | 28327000 | 25181000 | 2885300 | 2209700 | 5158900 | 3084900 |
| KNFESLSE | RPN2; RPN2; RI | P04844; F2Z | RPN2_HUMAN | Unmodified | 0 | 0 | 0 | 0 | 0 | 0 | 0 | 0 | 8397300 | 7907200 | 0 | 0 | 0 | 0 | 0 | 0 | 0 | 0 |
| KNFEWVAE | PAICS; PAICS; F | P22234; D6F | PUR6_HUMAN | Unmodified | 0 | 0 | 0 | 0 | 0 | 114890 | 0 | 0 | 0 | 511430 | 0 | 0 | 4963600 | 2526100 | 0 | 0 | 1398100 | 887280 |
| KNGFVVLK | EIF5A; EIF5AL1; | P63241; Q6I | IF5A1_HUMAN | Unmodified | 0 | 361130 | 15725 | 22136 | 849680 | 0 | 483670 | 265310 | 51369 | 0 | 74921 | 362480 | 0 | 2439700 | 1018200 | 337400 | 176060 | 0 |
| KNGPLEVA | MACROH2A1; N | O75367; B4I | H2AY_HUMAN | Unmodified | 4162100 | 1225500 | 0 | 0 | 0 | 0 | 0 | 900170 | 0 | 0 | 0 | 0 | 0 | 118410 | 0 | 0 | 0 | 0 |
| KNGQLAAS | FSCN1; FSCN1 | Q16658; C9J | FSCN1_HUMAI | Oxidation (M) | 0 | 0 | 0 | 0 | 0 | 0 | 0 | 0 | 0 | 0 | 0 | 1664300 | 0 | 0 | 0 | 0 | 0 | 0 |
| KNIILEEGK | CFL1; CFL1; CF | P23528; G3\ | COF1_HUMAN | Unmodified | 1830800 | 1155600 | 2130100 | 1145400 | 2429000 | 221910 | 0 | 0 | 1129000 | 61562 | 977090 | 14429000 | 1060600 | 0 | 2936600 | 2071000 | 0 | 0 |
| KNIILEEGK | CFL1; CFL1; CF | P23528; G3\ | COF1_HUMAN | Unmodified | 0 | 4842100 | 0 | 4463300 | 3.35E+08 | 31167000 | 18089000 | 0 | 9890300 | 12513000 | 0 | 5660000 | 1.07E+08 | 53754000 | 0 | 0 | 76337000 | 1.64E+08 |
| KNILDSKPT | CORO1C; COR | Q9ULV4; H0 | COR1C_HUMA | Unmodified | 73159 | 0 | 0 | 144060 | 3777600 | 847020 | 110510 | 187720 | 312280 | 261660 | 707960 | 3714700 | 493170 | 504990 | 81508 | 99859 | 459010 | 958300 |
| KNINCSIEE | HMGCL | P35914 | HMGCL_HUMA | Unmodified | 0 | 0 | 0 | 0 | 0 | 0 | 0 | 0 | 526450 | 0 | 0 | 0 | 0 | 250310 | 0 | 0 | 0 | 0 |
| KNIPEGSH | MOB1B; MOB1A | Q7L9L4; Q9 | MOB1B_HUMA | Unmodified | 212080 | 515410 | 0 | 305160 | 9452400 | 1907200 | 459000 | 209830 | 432380 | 820140 | 1233900 | 2668900 | 4118600 | 3107600 | 602340 | 247310 | 3843700 | 9036300 |
| KNIYGNIED | AGPS; AGPS; A( | O00116; A0A | ADAS_HUMAN | Unmodified | 0 | 0 | 0 | 0 | 0 | 0 | 1069200 | 0 | 10229000 | 2576900 | 209300 | 0 | 0 | 0 | 0 | 0 | 397570 | 0 |
| KNIYLNSGI | ADAM17 | P78536 | ADA17_HUMAN | Unmodified | 0 | 0 | 0 | 0 | 363840 | 67507 | 36870 | 0 | 121990 | 0 | 0 | 0 | 71468 | 85622 | 0 | 0 | 171480 | 470800 |
| KNKEEAAE | RPS6; RPS6 | P62753; A2A | RS6_HUMAN 4 | Unmodified | 741100 | 0 | 0 | 0 | 522140 | 0 | 0 | 645860 | 0 | 0 | 0 | 3190600 | 0 | 0 | 0 | 22408 | 0 | 54492 |
| KNLALSRE | LSR; LSR | Q86X29; S4F | LSR_HUMAN Li | Unmodified | 0 | 0 | 0 | 0 | 0 | 0 | 0 | 0 | 0 | 1325100 | 0 | 0 | 382380 | 625890 | 0 | 0 | 0 | 0 |
| KNLDSTTV | CSRP1; CSRP1; | P21291; E9P | CSRP1_HUMAI | Unmodified | 83444 | 0 | 0 | 107290 | 8161100 | 0 | 650800 | 1888300 | 486870 | 1888600 | 0 | 0 | 2972700 | 3030100 | 490820 | 88670 | 847180 | 0 |
| KNLEAQHK | SEPTIN7; SEPTI | Q16181; E7E | SEPT7_HUMAN | Unmodified | 0 | 0 | 0 | 0 | 1319500 | 172070 | 103440 | 0 | 0 | 42708 | 0 | 0 | 132060 | 0 | 0 | 0 | 497470 | 765100 |
| KNLFETPIL | MFGE8 | Q08431 | MFGM_HUMAN | Unmodified | 0 | 0 | 0 | 824090 | 0 | 729720 | 0 | 0 | 0 | 186830 | 54735 | 853310 | 1713000 | 0 | 1817600 | 645770 | 344890 | 893970 |
| KNLHGDGI | LMAN2; LMAN2 | Q12907; D6I | LMAN2_HUMAI | Unmodified | 0 | 0 | 0 | 76847 | 0 | 1466900 | 0 | 0 | 8717500 | 983020 | 0 | 0 | 0 | 0 | 0 | 0 | 0 | 0 |
| KNLQDFAA | PROM1 | O43490 | PROM1_HUMA | Unmodified | 0 | 0 | 0 | 0 | 0 | 0 | 0 | 0 | 0 | 0 | 0 | 0 | 0 | 0 | 0 | 0 | 0 | 0 |
| KNLQYYDI | RAN; RAN; RAN | P62826; J3K | RAN_HUMAN G | Unmodified | 0 | 0 | 863420 | 399140 | 5551700 | 993340 | 1383500 | 1574300 | 571520 | 743180 | 985750 | 10203000 | 3187500 | 1402600 | 0 | 0 | 1292600 | 3376500 |
| KNLTQDEM | EIF3A | Q14152 | EIF3A_HUMAN | Oxidation (M) | 18671 | 0 | 0 | 40420 | 505610 | 0 | 0 | 0 | 0 | 0 | 33326 | 191640 | 124600 | 68624 | 227000 | 38405 | 45928 | 81144 |
| KNLTQDEM | EIF3A | Q14152 | EIF3A_HUMAN | Unmodified | 0 | 0 | 0 | 0 | 369310 | 0 | 79415 | 37956 | 46685 | 0 | 79937 | 785060 | 0 | 0 | 0 | 0 | 38012 | 0 |
| KNLVDPFV | DYSF; MYOF | O75923; Q9I | DYSF_HUMAN | Unmodified | 0 | 0 | 0 | 0 | 1608300 | 0 | 0 | 0 | 0 | 0 | 0 | 0 | 0 | 0 | 0 | 0 | 0 | 0 |
| KNLVDPFV | MYOF | Q9NZM1 | MYOF_HUMAN | Unmodified | 0 | 0 | 0 | 0 | 990930 | 0 | 0 | 0 | 0 | 0 | 0 | 0 | 0 | 0 | 0 | 0 | 0 | 0 |
| KNLVEQHI | RPN1; RPN1 | P04843; B7Z | RPN1_HUMAN | Unmodified | 0 | 0 | 0 | 0 | 0 | 0 | 0 | 0 | 4162900 | 445870 | 0 | 0 | 0 | 0 | 0 | 0 | 0 | 0 |
| KNNPAIVII | LMAN1 | P49257 | LMAN1_HUMAI | Unmodified | 0 | 0 | 0 | 0 | 0 | 0 | 0 | 0 | 13212000 | 172720 | 0 | 0 | 0 | 0 | 0 | 0 | 0 | 0 |
| KNNRQPYA | RPL4; RPL4; RP | P36578; H3E | RL4_HUMAN 6 | Unmodified | 2643100 | 2693200 | 233370 | 917260 | 87352 | 0 | 0 | 154900 | 0 | 0 | 0 | 809470 | 2402500 | 118810 | 0 | 451410 | 0 | 0 |
| KNPAVTQL | CPD | O75976 | CBPD_HUMAN | Unmodified | 0 | 0 | 0 | 0 | 0 | 0 | 0 | 0 | 6098000 | 162660 | 73551 | 0 | 0 | 0 | 0 | 0 | 0 | 0 |
| KNPQAVLD | PAK2; PAK3; PA | Q13177; O7! | PAK2_HUMAN | Unmodified | 0 | 0 | 0 | 0 | 2877600 | 448380 | 0 | 0 | 715940 | 291330 | 0 | 2870200 | 1129500 | 674970 | 237630 | 0 | 669260 | 1799600 |
| KNPVTSST | NEDD1 | Q8NHV4 | NEDD1_HUMA | Unmodified | 0 | 0 | 0 | 0 | 252910 | 598970 | 1315000 | 216210 | 0 | 0 | 0 | 0 | 0 | 1414200 | 315350 | 358130 | 0 | 2709400 |
| KNQILIIDE | ABCC4 | O15439 | MRP4_HUMAN | Unmodified | 0 | 0 | 319050 | 140360 | 468690 | 0 | 0 | 0 | 0 | 0 | 0 | 0 | 179140 | 236890 | 0 | 487830 | 0 | 0 |
| KNSLRVEG | EPB41L2; EPB4 | O43491; Q6 | E41L2_HUMAN | Unmodified | 0 | 0 | 0 | 0 | 571940 | 79592 | 0 | 0 | 0 | 0 | 0 | 0 | 0 | 0 | 0 | 0 | 103790 | 194330 |
| KNSVVEAS | YWHAH; YWHA | Q04917; A2I | 1433F_HUMAN | Unmodified | 625920 | 999950 | 549040 | 0 | 4513700 | 0 | 357260 | 0 | 2226600 | 635200 | 1504500 | 9384600 | 0 | 1023300 | 1620800 | 1220800 | 0 | 0 |
| KNSVVEAS | YWHAH; YWHA | Q04917; A2I | 1433F_HUMAN | Unmodified | 0 | 0 | 0 | 0 | 2335800 | 0 | 0 | 0 | 0 | 340520 | 0 | 0 | 0 | 642960 | 0 | 0 | 588200 | 588560 |
| KNSYMNPE | MME; MME; MM | P08473; A0A | NEP_HUMAN N | Unmodified | 0 | 6647 | 283400 | 0 | 0 | 0 | 0 | 10409 | 0 | 0 | 0 | 0 | 0 | 0 | 0 | 0 | 0 | 0 |
| KNVEPFTS | MRPL1; MRPL1 | Q9BYD6; H0 | RM01_HUMAN | Unmodified | 0 | 0 | 0 | 0 | 0 | 0 | 0 | 0 | 518870 | 0 | 0 | 0 | 0 | 0 | 0 | 0 | 0 | 0 |
| KNVIDPPIY | EDIL3 | O43854 | EDIL3_HUMAN | Unmodified | 0 | 83817 | 0 | 0 | 655560 | 0 | 0 | 0 | 172640 | 0 | 0 | 0 | 0 | 0 | 0 | 0 | 0 | 529590 |
| KNVSINTV | TES; TES | Q9UGI8; F8V | TES_HUMAN Te | Unmodified | 0 | 0 | 0 | 0 | 2039800 | 0 | 0 | 0 | 0 | 902570 | 196490 | 1056200 | 1520800 | 1475800 | 506690 | 367230 | 1599600 | 5859800 |
| KNWYLPAF | ERMP1; ERMP1 | Q7Z2K6; E7E | ERMP1_HUMAI | Unmodified | 0 | 0 | 0 | 128300 | 0 | 0 | 0 | 6938.3 | 2878000 | 0 | 296900 | 256650 | 0 | 0 | 0 | 0 | 0 | 0 |
| KNYPDPVS | IARS1; IARS1; I | P41252; A0A | SYIC_HUMAN I | Unmodified | 0 | 0 | 0 | 0 | 0 | 0 | 0 | 0 | 0 | 0 | 0 | 779960 | 0 | 0 | 0 | 0 | 0 | 0 |
| KPAAVLKP | XRN2 | Q9H0D6 | XRN2_HUMAN | Unmodified | 0 | 0 | 0 | 0 | 0 | 0 | 0 | 0 | 0 | 0 | 0 | 639640 | 0 | 0 | 0 | 0 | 0 | 0 |
| KPADLQNL | CLIC4 | Q9Y696 | CLIC4_HUMAN | Unmodified | 0 | 0 | 0 | 0 | 0 | 0 | 0 | 0 | 0 | 0 | 0 | 2975900 | 0 | 0 | 287680 | 0 | 0 | 0 |
| KPALTAIEG | SEL1L | Q9UBV2 | SE1L1_HUMAN | Unmodified | 0 | 0 | 0 | 0 | 0 | 0 | 0 | 0 | 5777400 | 155730 | 618650 | 0 | 0 | 0 | 0 | 720380 | 0 | 0 |
| KPANDITS | SERBP1 | Q8NC51 | PAIRB_HUMAN | Unmodified | 0 | 0 | 73792 | 49735 | 583860 | 0 | 0 | 0 | 0 | 0 | 150660 | 212910 | 0 | 0 | 694860 | 0 | 0 | 690740 |
| KPASFMTSI | ACLY | P53396 | ACLY_HUMAN | Unmodified | 0 | 0 | 0 | 0 | 365740 | 123210 | 0 | 0 | 0 | 145370 | 0 | 3881900 | 349160 | 0 | 69154 | 0 | 153380 | 0 |
| KPATPAEDI | EEF1D; EEF1D; | P29692; E9P | EF1D_HUMAN | Unmodified | 0 | 0 | 0 | 0 | 0 | 0 | 0 | 0 | 0 | 0 | 0 | 2451900 | 0 | 0 | 0 | 0 | 0 | 0 |
| KPATSYVR | RPL28; RPL28 | P46779; H0 | RL28_HUMAN | Unmodified | 0 | 0 | 0 | 0 | 0 | 0 | 158200 | 150830 | 0 | 0 | 0 | 0 | 114000 | 0 | 0 | 0 | 55461 | 0 |
| KPDAYTEQ | TMEM9; TMEM9 | Q9P0T7; B1A | TMEM9_HUMAI | Unmodified | 0 | 0 | 0 | 0 | 0 | 0 | 0 | 0 | 0 | 0 | 597040 | 0 | 0 | 0 | 149060 | 0 | 0 | 0 |
| KPDDLPAF | SLC44A2 | Q8IWA5 | CTL2_HUMAN | Unmodified | 1453100 | 1724500 | 201410 | 0 | 0 | 0 | 0 | 0 | 434550 | 311710 | 12747 | 2344100 | 0 | 0 | 974150 | 801780 | 498710 | 0 |
| KPDTIEVQ | RDX; EZR; MSN | P35241; P15 | RADI_HUMAN I | Unmodified | 123820 | 0 | 0 | 233580 | 1874800 | 1248900 | 82563 | 43125 | 1861300 | 574560 | 444650 | 2811600 | 520480 | 155390 | 277820 | 0 | 830070 | 241730 |
| KPDTIEVQ | RDX; EZR; MSN | P35241; P15 | RADI_HUMAN I | Oxidation (M) | 0 | 0 | 0 | 0 | 0 | 86934 | 40570 | 0 | 0 | 0 | 0 | 1325000 | 0 | 0 | 0 | 61031 | 24419 | 136080 |
| KPEAQLPA | IQGAP2; IQGAP | Q13576; D6I | IQGA2_HUMAN | Oxidation (M) | 0 | 0 | 641660 | 0 | 0 | 0 | 0 | 0 | 0 | 0 | 0 | 0 | 0 | 0 | 0 | 0 | 0 | 0 |
| KPEDVLDD | BSG; BSG; BSG | P35613; A0A | BASI_HUMAN E | Unmodified | 0 | 0 | 1580500 | 371240 | 0 | 102000 | 292190 | 0 | 1675100 | 0 | 0 | 3988300 | 0 | 0 | 1456300 | 2199100 | 0 | 0 |
| KPEDWDEI | CALR; CALR; CA | P27797; A0A | CALR_HUMAN | Unmodified | 0 | 0 | 0 | 0 | 0 | 0 | 0 | 0 | 7771600 | 0 | 0 | 0 | 0 | 0 | 0 | 0 | 0 | 0 |
| KPEDWDEI | CALR; CALR; CA | P27797; A0A | CALR_HUMAN | Oxidation (M) | 0 | 0 | 0 | 0 | 0 | 0 | 0 | 0 | 1481800 | 1937100 | 588880 | 0 | 0 | 0 | 0 | 0 | 0 | 0 |
| KPEDWDEI | CANX; CANX; C | P27824; A0A | CALX_HUMAN | Unmodified | 0 | 0 | 0 | 0 | 0 | 0 | 0 | 0 | 0 | 1007100 | 359640 | 431510 | 0 | 0 | 0 | 0 | 0 | 0 |
| KPEKPLILQ | CD2AP | Q9Y5K6 | CD2AP_HUMAI | Unmodified | 5923400 | 4183200 | 2040700 | 4457100 | 0 | 0 | 0 | 0 | 431620 | 0 | 292310 | 1082600 | 935000 | 425640 | 7600100 | 4218300 | 0 | 0 |
| KPENTTRAI | CMTM6 | Q9NX76 | CKLF6_HUMAN | Unmodified | 0 | 406550 | 0 | 0 | 205340 | 0 | 0 | 0 | 2119700 | 0 | 0 | 2249200 | 0 | 0 | 0 | 0 | 0 | 0 |
| KPESEGYL | CEP55; CEP55 | Q53EZ4; H0 | CEP55_HUMAN | Unmodified | 616700 | 565200 | 1885500 | 1594900 | 2282600 | 581180 | 32459 | 0 | 4930900 | 352080 | 529130 | 1922600 | 0 | 63269 | 3158100 | 1273500 | 0 | 0 |
| KPEVPFFGI | RALGAPA1; RAL | Q6GYQ0; A0 | RGPA1_HUMAI | Unmodified | 0 | 0 | 0 | 0 | 0 | 0 | 0 | 0 | 0 | 0 | 0 | 152880 | 0 | 0 | 23562 | 29920 | 0 | 0 |
| KPFKLSGL | MARCKSL1 | P49006 | MRP_HUMAN N | Unmodified | 0 | 0 | 0 | 0 | 0 | 0 | 0 | 25883 | 0 | 328530 | 0 | 0 | 1539100 | 1781200 | 0 | 0 | 0 | 5912000 |
| KPFLLLAIK | B3GNT2 | Q9NY97 | B3GN2_HUMAI | Unmodified | 0 | 0 | 0 | 0 | 0 | 0 | 0 | 0 | 594040 | 33937 | 0 | 0 | 0 | 0 | 0 | 0 | 0 | 0 |
| KPFPINHGI | UCHL3; UCHL3 | P15374; Q5 | UCHL3_HUMAI | Unmodified | 0 | 0 | 0 | 0 | 0 | 0 | 0 | 0 | 0 | 0 | 0 | 1352700 | 0 | 0 | 0 | 0 | 0 | 0 |
| KPFSIEEVE | ADH1C; ADH1A | P00326; P07 | ADH1G_HUMA | Unmodified | 0 | 0 | 0 | 0 | 0 | 0 | 0 | 0 | 0 | 0 | 3465400 | 1454000 | 0 | 0 | 0 | 0 | 0 | 0 |
| KPGGFDISI | XRCC6; XRCC6 | P12956; B1A | XRCC6_HUMAI | Unmodified | 0 | 0 | 0 | 0 | 0 | 0 | 0 | 0 | 415810 | 0 | 0 | 9597900 | 0 | 0 | 0 | 0 | 0 | 0 |
| KPGINVASI | GANAB; GANAE | Q14697; F5H | GANAB_HUMA | Unmodified | 0 | 0 | 0 | 0 | 0 | 0 | 0 | 0 | 28584000 | 2660500 | 0 | 0 | 0 | 0 | 0 | 0 | 0 | 0 |
| KPGMFFNF | G6PD | P11413 | G6PD_HUMAN | Oxidation (M) | 0 | 0 | 0 | 0 | 7132800 | 0 | 0 | 0 | 1074600 | 0 | 0 | 1701900 | 640520 | 0 | 0 | 0 | 0 | 0 |
| KPGMFFNF | G6PD | P11413 | G6PD_HUMAN | Unmodified | 0 | 0 | 0 | 0 | 8854700 | 0 | 355930 | 121010 | 1014300 | 0 | 0 | 0 | 2391400 | 0 | 0 | 0 | 0 | 0 |
| KPIAGQVV | PXN; PXN; PXN | P49023; A0A | PAXI_HUMAN F | Unmodified | 0 | 0 | 0 | 0 | 1122200 | 0 | 0 | 0 | 0 | 0 | 0 | 0 | 0 | 0 | 56806 | 0 | 232960 | 973290 |
| KPIDYTVLD | ABI1; ABI1; ABI1 | Q8IZP0; A0A | ABI1_HUMAN A | Unmodified | 137270 | 256460 | 0 | 0 | 544380 | 0 | 0 | 0 | 756330 | 0 | 0 | 2554100 | 911660 | 0 | 852330 | 821610 | 0 | 0 |
| KPIEDPANI | CTNND1; CTNN | O60716; A0A | CTND1_HUMAI | Unmodified | 217230 | 134880 | 931860 | 1414100 | 494080 | 0 | 0 | 0 | 0 | 0 | 427010 | 2416700 | 193350 | 1074800 | 1509300 | 1602300 | 0 | 260240 |
| KPIIGILMQI | GGH; GGH; GG | Q92820; A0A | GGH_HUMAN | Unmodified | 0 | 0 | 0 | 0 | 0 | 0 | 0 | 0 | 8336700 | 0 | 0 | 0 | 0 | 0 | 0 | 0 | 0 | 0 |
| KPIPDEHLI | SEC31A; SEC31 | O94979; H0 | SC31A_HUMAN | Unmodified | 0 | 0 | 0 | 0 | 0 | 0 | 0 | 0 | 157420 | 0 | 0 | 656190 | 97783 | 0 | 0 | 0 | 0 | 0 |
| KPITTGGVT | FHL2 | Q14192 | FHL2_HUMAN | Unmodified | 0 | 0 | 0 | 0 | 1114500 | 0 | 0 | 0 | 0 | 0 | 78233 | 0 | 0 | 0 | 0 | 0 | 736770 | 1689400 |
| KPLEALYG | BPHL | Q86WA6 | BPHL_HUMAN | Unmodified | 0 | 0 | 0 | 0 | 0 | 0 | 0 | 0 | 871910 | 0 | 0 | 0 | 0 | 0 | 0 | 0 | 492070 | 365170 |
| KPLFHGDS | CDK1; CDK1; C | P06493; A0A | CDK1_HUMAN | Unmodified | 289400 | 300450 | 537980 | 201430 | 0 | 0 | 0 | 0 | 315530 | 0 | 0 | 0 | 0 | 0 | 664300 | 604600 | 0 | 0 |
| KPLLESGTI | UBA1 | P22314 | UBA1_HUMAN | Unmodified | 0 | 0 | 0 | 132740 | 0 | 0 | 0 | 0 | 579060 | 0 | 427370 | 5738600 | 0 | 0 | 639780 | 474760 | 0 | 0 |
| KPLPDHVS | RPS3; RPS3; RP | P23396; E9P | RS3_HUMAN 4 | Unmodified | 0 | 0 | 0 | 0 | 0 | 0 | 0 | 0 | 0 | 0 | 0 | 5117800 | 0 | 0 | 0 | 0 | 0 | 0 |
| KPLPDHVS | RPS3; RPS3; RP | P23396; E9P | RS3_HUMAN 4 | Unmodified | 5464300 | 0 | 4180400 | 2312900 | 0 | 0 | 0 | 0 | 0 | 0 | 6101800 | 52816000 | 0 | 0 | 3659000 | 3393900 | 0 | 0 |
| KPLTLSLH | TOR1B; TOR1B | O14657; H0 | TOR1B_HUMAN | Unmodified | 0 | 0 | 0 | 0 | 58710 | 0 | 0 | 0 | 3232900 | 130990 | 0 | 0 | 0 | 0 | 29785 | 0 | 0 | 0 |

| | | | | | | | | | | | | | | | | | | | | | | |
|---|---|---|---|---|---|---|---|---|---|---|---|---|---|---|---|---|---|---|---|---|---|---|
| KPLTLSLHC | TOR1A; TOR1A | O14656; A0A | TOR1A_HUMAN | Unmodified | 0 | 51140 | 188610 | 0 | 1172200 | 4440200 | 1838200 | 1388100 | 3943900 | 0 | 655080 | 0 | 276190 | 2103000 | 130390 | 0 | 474790 | 2504700 |
| KPLVLCGD | APEX1; APEX1; | P27695; G3V | APEX1_HUMAN | Unmodified | 0 | 0 | 0 | 0 | 0 | 0 | 0 | 0 | 0 | 0 | 0 | 3138700 | 0 | 0 | 0 | 0 | 0 | 0 |
| KPLYLHLH | AIDA | Q96BJ3 | AIDA_HUMAN | Unmodified | 0 | 0 | 0 | 1068000 | 2271200 | 138500 | 83926 | 0 | 0 | 0 | 0 | 0 | 4905400 | 4167600 | 0 | 190910 | 608410 | 6419600 |
| KPMVLGHE | SORD; SORD; S | Q00796; H0Y | DHSO_HUMAN | Unmodified | 0 | 0 | 1168500 | 1469300 | 572060 | 143410 | 0 | 0 | 404590 | 160380 | 898930 | 3991600 | 2892400 | 1240400 | 525040 | 1242700 | 101730 | 70537 |
| KPMVLGHE | SORD; SORD; S | Q00796; H0Y | DHSO_HUMAN | Oxidation (M) | 0 | 62384 | 776620 | 1466300 | 348200 | 87984 | 0 | 0 | 132990 | 67417 | 356180 | 3695000 | 1215900 | 931180 | 1196600 | 1047500 | 67229 | 107630 |
| KPMVMPDI | N/A | N/A | N/A | 2 Oxidation (M) | 63033000 | 27207000 | 62868000 | 55819000 | 0 | 0 | 0 | 0 | 0 | 0 | 0 | 0 | 0 | 246270 | 1.17E+08 | 1.01E+08 | 0 | 0 |
| KPNEGADC | SERBP1 | Q8NC51 | PAIRB_HUMAN | Unmodified | 0 | 0 | 0 | 0 | 0 | 0 | 0 | 0 | 87198 | 21904 | 1575600 | 0 | 0 | 45895 | 0 | 0 | 152150 | 0 |
| KPNEGADC | SERBP1 | Q8NC51 | PAIRB_HUMAN | Unmodified | 0 | 156200 | 0 | 0 | 0 | 0 | 0 | 0 | 0 | 0 | 0 | 17970 | 0 | 0 | 0 | 0 | 0 | 0 |
| KPNVGCQ | UBA6 | A0AVT1 | UBA6_HUMAN | Unmodified | 0 | 0 | 0 | 0 | 0 | 0 | 0 | 17872 | 0 | 0 | 0 | 1007300 | 0 | 0 | 0 | 0 | 0 | 111440 |
| KPPKPQLM | EIF3A | Q14152 | EIF3A_HUMAN | Unmodified | 129650 | 82931 | 64043 | 115530 | 108830 | 0 | 0 | 1425900 | 0 | 0 | 0 | 4820100 | 0 | 1022400 | 180630 | 251050 | 0 | 0 |
| KPPKPQLM | EIF3A | Q14152 | EIF3A_HUMAN | Oxidation (M) | 0 | 0 | 28765 | 0 | 0 | 0 | 0 | 0 | 0 | 0 | 0 | 922860 | 0 | 0 | 69355 | 72175 | 0 | 0 |
| KPPLLNNA | PARP1; PARP1 | P09874; A0A | PARP1_HUMAN | Unmodified | 383090 | 339760 | 0 | 0 | 0 | 0 | 0 | 0 | 412300 | 0 | 718750 | 3610300 | 0 | 0 | 0 | 0 | 0 | 0 |
| KPPPSPQP | OS9 | B4E321 | B4E321_HUMA | Unmodified | 0 | 0 | 0 | 0 | 0 | 0 | 0 | 0 | 1325200 | 166080 | 0 | 98832 | 0 | 0 | 0 | 0 | 0 | 0 |
| KPPPVPALI | TRIM25; TRIM25 | Q14258; A0A | TRI25_HUMAN | Unmodified | 0 | 0 | 0 | 0 | 0 | 0 | 0 | 0 | 0 | 0 | 1014100 | 0 | 0 | 0 | 0 | 0 | 0 | 0 |
| KPPSVTPIF | POMGNT1 | Q8WZA1 | PMGT1_HUMA | Unmodified | 0 | 0 | 0 | 0 | 0 | 0 | 0 | 0 | 6805300 | 0 | 0 | 0 | 0 | 0 | 0 | 0 | 0 | 0 |
| KPQEMIDIV | B3GNT2 | Q9NY97 | B3GN2_HUMA | Unmodified | 0 | 0 | 0 | 0 | 0 | 0 | 0 | 0 | 1267800 | 0 | 0 | 0 | 0 | 0 | 0 | 0 | 0 | 0 |
| KPQYDIWG | ADCY6 | O43306 | ADCY6_HUMA | Unmodified | 0 | 195290 | 0 | 584890 | 0 | 0 | 0 | 0 | 0 | 0 | 0 | 0 | 0 | 0 | 0 | 0 | 0 | 0 |
| KPRPPPAL | SET | Q01105 | SET_HUMAN P | Unmodified | 0 | 0 | 0 | 0 | 0 | 0 | 0 | 0 | 0 | 0 | 329410 | 351840 | 0 | 0 | 0 | 0 | 0 | 0 |
| KPSAGDDF | CD99; CD99; C | P14209; A8N | CD99_HUMAN | Unmodified | 0 | 0 | 2749100 | 0 | 0 | 0 | 0 | 0 | 0 | 0 | 0 | 0 | 0 | 0 | 2883500 | 1605600 | 0 | 0 |
| KPSGGYPQ | N/A | N/A | N/A | Oxidation (M) | 0 | 95034 | 0 | 527480 | 0 | 0 | 0 | 109770 | 0 | 0 | 0 | 194820 | 1818500 | 29293 | 0 | 0 | 23551 | 400850 |
| KPSLSLTLT | PLPP2 | O43688 | PLPP2_HUMAN | Unmodified | 0 | 0 | 0 | 0 | 0 | 0 | 0 | 0 | 880740 | 0 | 426420 | 7582900 | 0 | 0 | 0 | 0 | 0 | 0 |
| KPSLVSDLI | SNTB2; SNTB2; | Q13425; H7I | SNTB2_HUMAN | Unmodified | 0 | 0 | 5517100 | 0 | 1283500 | 0 | 0 | 0 | 0 | 0 | 0 | 1261000 | 0 | 0 | 0 | 2037100 | 0 | 0 |
| KPSVPETEA | UMAD1; UMAD1 | C9J7I0; A0A1 | UMAD1_HUMA | Unmodified | 0 | 0 | 0 | 0 | 0 | 0 | 0 | 0 | 0 | 0 | 0 | 0 | 0 | 0 | 0 | 0 | 0 | 0 |
| KPTSALDEI | CLPTM1L; CLPT | Q96KA5; G5I | CLP1L_HUMAN | Unmodified | 0 | 0 | 0 | 0 | 0 | 0 | 0 | 0 | 17331000 | 133020 | 439600 | 0 | 0 | 0 | 0 | 0 | 0 | 0 |
| KPTYDPVSI | TNK2; TNK2; TN | Q07912; A0A | ACK1_HUMAN | Unmodified | 56134 | 107610 | 285580 | 220820 | 0 | 0 | 0 | 0 | 0 | 0 | 0 | 411200 | 0 | 240940 | 300450 | 190780 | 0 | 0 |
| KPVAGALD | COPS8; COPS8 | Q99627; H7C | CSN8_HUMAN | Unmodified | 0 | 0 | 0 | 0 | 0 | 0 | 0 | 0 | 0 | 0 | 0 | 491680 | 0 | 0 | 0 | 0 | 0 | 0 |
| KPVEGYDIS | ARPC4; ARPC4; | P59998; R4C | ARPC4_HUMAN | Oxidation (M) | 998580 | 962530 | 0 | 0 | 0 | 0 | 0 | 0 | 0 | 0 | 0 | 1940100 | 0 | 0 | 0 | 756520 | 0 | 0 |
| KPVILEVTP | DNAJC13; DNA | O75165; A0A | DJC13_HUMAN | Unmodified | 0 | 0 | 0 | 0 | 0 | 0 | 0 | 0 | 0 | 0 | 0 | 5697600 | 0 | 0 | 103700 | 109130 | 0 | 0 |
| KPVTDCVIS | HSPH1 | Q92598 | HS105_HUMAN | Unmodified | 19246 | 0 | 0 | 0 | 0 | 0 | 146310 | 20302 | 0 | 0 | 0 | 407470 | 0 | 0 | 66268 | 48611 | 0 | 0 |
| KPVTDCVIS | HSPH1 | Q92598 | HS105_HUMAN | Unmodified | 0 | 0 | 0 | 0 | 0 | 0 | 0 | 0 | 0 | 0 | 0 | 1521200 | 0 | 0 | 0 | 0 | 0 | 0 |
| KPVVDCVV | HSPA4; HSPA4; | P34932; A0A | HSP74_HUMAN | Unmodified | 45560 | 0 | 0 | 0 | 0 | 0 | 0 | 32451 | 0 | 0 | 0 | 880640 | 0 | 0 | 0 | 0 | 0 | 0 |
| KPVVDCVV | HSPA4; HSPA4; | P34932; A0A | HSP74_HUMAN | Unmodified | 82615 | 0 | 0 | 0 | 0 | 0 | 0 | 0 | 0 | 0 | 0 | 2410500 | 0 | 0 | 0 | 0 | 0 | 0 |
| KPVVSTISK | SLC7A11 | Q9UPY5 | XCT_HUMAN C | Unmodified | 122190 | 103760 | 0 | 242740 | 470270 | 40202 | 0 | 0 | 0 | 0 | 73245 | 443470 | 77806 | 48043 | 566590 | 758000 | 65870 | 0 |
| KPWLAYPH | PCYOX1 | Q9UHG3 | PCYOX_HUMA | Unmodified | 0 | 0 | 804920 | 0 | 0 | 0 | 321000 | 0 | 5676700 | 1870800 | 319950 | 0 | 0 | 0 | 0 | 0 | 0 | 0 |
| KPWTAVDT | IGF2R; IGF2R | P11717; A0A | MPRI_HUMAN | Unmodified | 0 | 0 | 197130 | 0 | 25518 | 0 | 0 | 0 | 9318100 | 1157500 | 203500 | 532920 | 0 | 0 | 0 | 0 | 0 | 0 |
| KPYCNAHY | LASP1; LASP1; | Q14847; C9J | LASP1_HUMAN | Unmodified | 1936600 | 0 | 0 | 0 | 0 | 1588800 | 0 | 3335900 | 0 | 1030700 | 0 | 3290300 | 0 | 0 | 0 | 0 | 2019800 | 11406000 |
| KPYIWEYSI | EPRS1; EPRS1 | P07814; V9C | SYEP_HUMAN | Unmodified | 0 | 0 | 0 | 0 | 0 | 0 | 0 | 0 | 206850 | 0 | 0 | 2565300 | 0 | 0 | 0 | 0 | 0 | 373290 |
| KQAEETYEI | IPO5; IPO5; IPO | O00410; E7E | IPO5_HUMAN I | Unmodified | 335870 | 0 | 0 | 0 | 0 | 0 | 0 | 0 | 105070 | 47152 | 127060 | 2829100 | 738520 | 0 | 255010 | 0 | 0 | 0 |
| KQAEILQES | TBCA; TBCA; TB | O75347; E5F | TBCA_HUMAN | Unmodified | 400590 | 224910 | 0 | 362000 | 738900 | 0 | 14271 | 0 | 924490 | 235990 | 657090 | 1627400 | 1789400 | 321350 | 363770 | 282720 | 0 | 0 |
| KQAELEEIY | ADGRE5 | P48960 | AGRE5_HUMA | Unmodified | 826330 | 383350 | 839390 | 0 | 2778100 | 0 | 0 | 0 | 738940 | 24115 | 186420 | 0 | 0 | 0 | 668500 | 715590 | 177170 | 1894400 |
| KQAELEEIY | ADGRE5 | P48960 | AGRE5_HUMA | Unmodified | 0 | 0 | 0 | 0 | 24948000 | 3523600 | 0 | 0 | 0 | 43338 | 0 | 0 | 0 | 0 | 0 | 0 | 0 | 31441000 |
| KQAGPASV | PDIA3 | P30101 | PDIA3_HUMAN | Unmodified | 587280 | 160600 | 0 | 0 | 2527700 | 0 | 1066800 | 402600 | 12254000 | 4579300 | 2211900 | 781050 | 604830 | 656100 | 0 | 0 | 466610 | 3602900 |
| KQDEEINQ | SH2D4A; SH2D | Q9H788; H0 | SH24A_HUMA | Unmodified | 0 | 0 | 0 | 0 | 940840 | 125720 | 0 | 0 | 179710 | 0 | 0 | 0 | 367810 | 285460 | 0 | 0 | 0 | 0 |
| KQDHTIVD | ITGB4 | P16144 | ITB4_HUMAN I | Oxidation (M) | 0 | 0 | 0 | 0 | 1317100 | 240350 | 109350 | 61545 | 0 | 78714 | 0 | 1309700 | 1422300 | 127060 | 0 | 38321 | 688760 | 1510700 |
| KQDHTIVD | ITGB4 | P16144 | ITB4_HUMAN I | Unmodified | 0 | 0 | 0 | 0 | 842820 | 0 | 0 | 0 | 0 | 0 | 0 | 515720 | 0 | 0 | 0 | 0 | 321040 | 2282300 |
| KQDIENILH | ACO1 | P21399 | ACOC_HUMAN | Unmodified | 67446 | 45327 | 0 | 488930 | 1999200 | 481490 | 1075900 | 0 | 550750 | 818990 | 961620 | 6691800 | 1383700 | 0 | 306710 | 399660 | 798810 | 992090 |
| KQDPDTWI | USP13; USP13; | Q92995; A0A | UBP13_HUMA | Unmodified | 1048300 | 1032800 | 494040 | 0 | 0 | 0 | 0 | 0 | 0 | 0 | 0 | 0 | 0 | 0 | 681500 | 343320 | 0 | 0 |
| KQEAEWK | CLTA; CLTA; CL | P09496; C9J | CLCA_HUMAN | Unmodified | 0 | 28242 | 0 | 0 | 0 | 0 | 0 | 0 | 0 | 0 | 0 | 10271000 | 0 | 0 | 0 | 0 | 0 | 0 |
| KQEEFDVA | TPP2; TPP2; TPI | P29144; A0A | TPP2_HUMAN | Unmodified | 0 | 0 | 0 | 0 | 0 | 0 | 0 | 0 | 0 | 0 | 237650 | 4613800 | 0 | 0 | 0 | 84294 | 0 | 0 |
| KQEELETLC | CALCOCO2; CA | Q13137; D6I | CACO2_HUMA | Unmodified | 549450 | 2348300 | 0 | 97855 | 483300 | 217310 | 773270 | 211020 | 1704800 | 329690 | 43709 | 297950 | 745190 | 460430 | 0 | 77187 | 364840 | 3197800 |
| KQEFLEK | CHMP4BP1; CH | P59074; Q9I | CHM4P_HUMA | Unmodified | 1937700 | 2327500 | 0 | 0 | 732120 | 721350 | 195180 | 91894 | 2848100 | 1666300 | 595490 | 1548800 | 4230800 | 0 | 2209500 | 0 | 96220 | 103010 |
| KQEFLEQK | CHMP4A; CHM | Q9BY43; E9I | CHM4A_HUMA | Unmodified | 1400700 | 0 | 106840 | 0 | 1404600 | 0 | 77594 | 20404 | 686480 | 137420 | 458980 | 1090600 | 409630 | 123390 | 132830 | 80535 | 153220 | 113060 |
| KQEFLEQK | CHMP4A; CHM | Q9BY43; E9I | CHM4A_HUMA | Unmodified | 74416 | 162510 | 0 | 105910 | 1086300 | 0 | 0 | 0 | 0 | 1728800 | 0 | 0 | 535190 | 399510 | 0 | 0 | 405300 | 0 |
| KQEGTPEG | PSMC1 | P62191 | PRS4_HUMAN | Unmodified | 0 | 0 | 703480 | 0 | 1546800 | 584210 | 0 | 333920 | 74958 | 122570 | 10709000 | 0 | 796100 | 0 | 0 | 0 | 0 | 830860 |
| KQEIVAEKE | CORO1C; COR | Q9ULV4; F8V | COR1C_HUMA | Unmodified | 0 | 0 | 0 | 0 | 697870 | 101320 | 39281 | 0 | 26503 | 28496 | 0 | 0 | 0 | 0 | 0 | 0 | 0 | 0 |
| KQELEEICI | MYH9 | P35579 | MYH9_HUMAN | Unmodified | 54492 | 85500 | 0 | 62237 | 0 | 0 | 0 | 0 | 0 | 138680 | 877460 | 1508100 | 0 | 0 | 308350 | 179900 | 0 | 0 |
| KQELEEICI | MYH9 | P35579 | MYH9_HUMAN | Unmodified | 180510 | 0 | 112070 | 75464 | 0 | 0 | 0 | 0 | 0 | 0 | 0 | 556130 | 0 | 0 | 274350 | 182940 | 0 | 0 |
| KQELEEILH | MYH10 | P35580 | MYH10_HUMA | Unmodified | 0 | 0 | 0 | 0 | 0 | 0 | 0 | 0 | 0 | 0 | 0 | 536140 | 0 | 0 | 0 | 277230 | 0 | 0 |
| KQELIEDLC | HOOK1 | Q9UJC3 | HOOK1_HUMA | Unmodified | 88912 | 62779 | 0 | 335300 | 0 | 0 | 0 | 0 | 1597100 | 163690 | 0 | 544010 | 497200 | 762220 | 280380 | 165550 | 0 | 0 |
| KQENANA | PNMA2 | Q9UL42 | PNMA2_HUMA | Unmodified | 195660 | 11167000 | 0 | 0 | 0 | 0 | 0 | 0 | 0 | 0 | 0 | 0 | 0 | 0 | 0 | 0 | 0 | 0 |
| KQEVQAWI | USP5 | P45974 | UBP5_HUMAN | Unmodified | 2152600 | 756450 | 839340 | 429280 | 0 | 0 | 0 | 0 | 0 | 0 | 614940 | 1542900 | 0 | 0 | 0 | 529180 | 0 | 0 |
| KQEVQAWI | USP5 | P45974 | UBP5_HUMAN | Unmodified | 0 | 0 | 0 | 0 | 0 | 97838 | 0 | 0 | 0 | 77840 | 0 | 0 | 600330 | 256550 | 0 | 0 | 0 | 0 |
| KQEWIQAII | SWAP70; SWAF | Q9UH65; E7 | SWP70_HUMA | Unmodified | 0 | 0 | 0 | 0 | 628570 | 0 | 0 | 0 | 0 | 0 | 86638 | 354170 | 0 | 0 | 0 | 0 | 0 | 201010 |
| KQEWTAIIP | ITFG1; ITFG1; IT | Q8TB96; F5C | TIP_HUMAN T- | Unmodified | 0 | 0 | 0 | 0 | 0 | 0 | 0 | 0 | 3879700 | 1386000 | 0 | 0 | 0 | 0 | 0 | 0 | 0 | 0 |
| KQEYLEVQ | HGS; HGS; HGS | O14964; A0A | HGS_HUMAN I | Unmodified | 378400 | 426330 | 1916000 | 0 | 82057 | 0 | 0 | 0 | 828620 | 896410 | 0 | 428670 | 0 | 0 | 188770 | 0 | 0 | 67031 |
| KQFGAQAN | ACTN1; ACTN1; | P12814; H0Y | ACTN1_HUMA | Unmodified | 124520 | 106860 | 46012 | 35217 | 0 | 0 | 0 | 0 | 67434 | 0 | 0 | 3239100 | 260770 | 0 | 254730 | 339480 | 0 | 0 |
| KQFSLENV | N4BP1 | O75113 | N4BP1_HUMA | Unmodified | 179110 | 280950 | 0 | 0 | 0 | 0 | 0 | 0 | 0 | 0 | 0 | 0 | 0 | 0 | 0 | 0 | 0 | 0 |
| KQGEDNST | MTDH; MTDH | Q86UE4; E5I | LYRIC_HUMAN | Unmodified | 0 | 0 | 0 | 0 | 0 | 0 | 0 | 0 | 0 | 0 | 0 | 634380 | 186390 | 0 | 0 | 0 | 0 | 0 |
| KQGEVSKA | MACROH2A1 | O75367 | H2AY_HUMAN | Unmodified | 101230 | 0 | 0 | 0 | 0 | 0 | 0 | 2153800 | 0 | 0 | 0 | 0 | 0 | 0 | 0 | 0 | 0 | 0 |
| KQGLLPLTI | ACO2; ACO2; A | Q99798; A2A | ACON_HUMAN | Unmodified | 0 | 0 | 0 | 0 | 191730 | 0 | 1426100 | 0 | 4506900 | 543610 | 302190 | 894960 | 0 | 0 | 0 | 0 | 79490 | 163500 |
| KQHPYTFP | GALNT2 | Q10471 | GALT2_HUMAN | Unmodified | 0 | 0 | 37304 | 35116 | 0 | 0 | 0 | 0 | 7764100 | 834870 | 41333 | 0 | 0 | 0 | 29550 | 24728 | 0 | 0 |
| KQIAVEAQI | FLOT2; FLOT2; | Q14254; J3Q | FLOT2_HUMAN | Unmodified | 0 | 0 | 0 | 0 | 4375700 | 0 | 214230 | 0 | 1046000 | 286770 | 615400 | 919770 | 376250 | 0 | 575450 | 316860 | 0 | 0 |
| KQIEELKGC | UCHL1; UCHL1 | P09936; D6F | UCHL1_HUMA | Unmodified | 597950 | 245880 | 0 | 0 | 32427000 | 2741900 | 2347100 | 2149100 | 2023200 | 4820800 | 1346100 | 4059700 | 0 | 0 | 0 | 0 | 0 | 0 |
| KQIPAITCIC | IQGAP1; IQGAP | P46940; A0A | IQGA1_HUMAN | Unmodified | 0 | 0 | 0 | 0 | 3987900 | 0 | 1176100 | 239520 | 0 | 253070 | 0 | 953730 | 709540 | 372160 | 0 | 0 | 52392 | 0 |
| KQIQFADD | AHCYL1 | O43865 | SAHH2_HUMA | Unmodified | 0 | 0 | 0 | 459110 | 18682000 | 3610200 | 593260 | 29630 | 298360 | 0 | 0 | 0 | 112010 | 53051 | 0 | 0 | 3222600 | 5424100 |
| KQIQFADD | AHCYL1 | O43865 | SAHH2_HUMA | Oxidation (M) | 0 | 0 | 0 | 0 | 2994500 | 586930 | 0 | 0 | 0 | 0 | 0 | 0 | 0 | 0 | 0 | 0 | 369570 | 0 |
| KQISDVERI | DIAPH1; DIAPH | O60610; A0A | DIAP1_HUMAN | Unmodified | 0 | 0 | 101530 | 242500 | 195360 | 0 | 0 | 0 | 397690 | 0 | 0 | 2942000 | 0 | 1017600 | 177610 | 117330 | 420030 | 0 |
| KQIWTLEQ | FSCN1 | Q16658 | FSCN1_HUMA | Unmodified | 632640 | 0 | 0 | 471640 | 555390 | 180000 | 0 | 0 | 0 | 158850 | 899620 | 3665600 | 0 | 0 | 0 | 0 | 376390 | 571250 |
| KQIYSVDQ | PLCG2; PLCG2; | P16885; A0A | PLCG2_HUMA | Unmodified | 0 | 0 | 0 | 0 | 0 | 0 | 0 | 0 | 0 | 0 | 55712 | 715320 | 0 | 0 | 0 | 30426 | 93907 | 0 |
| KQKVEGTE | NCL; NCL; NCL | P19338; C9J | NUCL_HUMAN | Unmodified | 0 | 0 | 0 | 0 | 0 | 6335800 | 6345500 | 0 | 0 | 2405900 | 1095800 | 452930 | 726160 | 11182000 | 0 | 0 | 3306200 | 5245600 |
| KQLCPYAA | MKRN1; MKRN1 | Q9UHC7; C9 | MKRN1_HUMA | Unmodified | 1012400 | 0 | 0 | 0 | 0 | 44817 | 0 | 0 | 0 | 0 | 0 | 0 | 178180 | 0 | 0 | 0 | 0 | 0 |
| KQLDSTIGI | TXNRD1; TXNRI | Q16881; A0A | TRXR1_HUMAN | Unmodified | 0 | 0 | 0 | 0 | 2725500 | 0 | 0 | 0 | 0 | 0 | 0 | 0 | 2023600 | 0 | 0 | 0 | 1854300 | 1589200 |
| KQLGDNEA | VTA1; VTA1; VTA | Q9NP79; Q5 | VTA1_HUMAN | Unmodified | 6007200 | 0 | 0 | 0 | 0 | 0 | 0 | 0 | 0 | 0 | 0 | 1262600 | 320840 | 0 | 0 | 0 | 0 | 0 |
| KQLQALSS | RDX; RDX; RDX | P35241; A0A | RADI_HUMAN I | Unmodified | 94945 | 585040 | 0 | 2147100 | 6624600 | 2715100 | 0 | 0 | 1699100 | 494140 | 0 | 2715400 | 9402400 | 6551100 | 0 | 42987 | 3792700 | 5825200 |
| KQMVIDVLI | RPS24; RPS24; | P62847; A0A | RS24_HUMAN | Unmodified | 57378 | 38663 | 0 | 29996 | 0 | 0 | 35403 | 0 | 128150 | 0 | 259060 | 2239700 | 0 | 0 | 21992 | 0 | 0 | 0 |
| KQNDVFGI | GDI1 | P31150 | GDIA_HUMAN | Unmodified | 0 | 0 | 0 | 0 | 0 | 0 | 0 | 89447 | 0 | 0 | 403960 | 0 | 0 | 0 | 0 | 0 | 0 | 0 |
| KQPPVSPG | HMGA1 | P17096 | HMGA1_HUMA | Unmodified | 0 | 0 | 0 | 0 | 0 | 0 | 0 | 0 | 0 | 0 | 698580 | 466340 | 0 | 0 | 0 | 0 | 0 | 0 |
| KQPPVSPG | HMGA1 | P17096 | HMGA1_HUMA | Unmodified | 0 | 0 | 0 | 0 | 1046900 | 359280 | 85140 | 0 | 0 | 0 | 8677700 | 17397000 | 0 | 0 | 0 | 0 | 168220 | 826080 |
| KQQADSN | PTPRJ; PTPRJ; P | Q12913; A0A | PTPRJ_HUMAN | Unmodified | 2144800 | 1233200 | 0 | 797440 | 1054200 | 0 | 47233 | 27807 | 62230 | 0 | 1003300 | 4889600 | 758380 | 0 | 1008500 | 81747 | 0 | 97500 |
| KQQAIELTC | RAD21; RAD21; | O60216; E5F | RAD21_HUMAN | Unmodified | 0 | 0 | 0 | 0 | 468960 | 0 | 0 | 0 | 0 | 658670 | 0 | 0 | 670110 | 0 | 0 | 0 | 0 | 0 |
| KQQELMDI | CALCOCO2; CA | Q13137; D6I | CACO2_HUMA | Unmodified | 0 | 63397 | 0 | 45389 | 793020 | 0 | 846350 | 0 | 927330 | 300060 | 164810 | 0 | 387550 | 0 | 0 | 0 | 455350 | 3858600 |
| KQQELMDI | CALCOCO2; CA | Q13137; D6I | CACO2_HUMA | Oxidation (M) | 132550 | 0 | 0 | 0 | 0 | 0 | 177520 | 0 | 0 | 0 | 0 | 0 | 0 | 0 | 0 | 0 | 0 | 315100 |
| KQQISLATC | CCT5; CCT5; CC | P48643; B7Z | TCPE_HUMAN | Unmodified | 0 | 0 | 50887 | 0 | 0 | 0 | 0 | 99763 | 0 | 188440 | 3749300 | 730410 | 3226400 | 0 | 0 | 0 | 0 | 3433700 |
| KQQLYVSS | SEMA3C | Q99985 | SEM3C_HUMA | Unmodified | 130730 | 0 | 0 | 951270 | 0 | 0 | 0 | 0 | 0 | 0 | 0 | 0 | 0 | 58077 | 637940 | 337450 | 199330 | 206390 |
| KQQNVDQ | DHRS4L2; DHR | Q6PKH6; Q9 | DR4L2_HUMAN | Unmodified | 0 | 0 | 0 | 0 | 0 | 0 | 0 | 0 | 783240 | 0 | 0 | 0 | 0 | 0 | 0 | 0 | 0 | 0 |
| KQQPNPGI | TTLL12 | Q14166 | TTL12_HUMAN | Unmodified | 0 | 0 | 0 | 0 | 0 | 0 | 114910 | 56177 | 0 | 0 | 116090 | 767420 | 0 | 0 | 0 | 0 | 0 | 0 |
| KQQQQQQ | NUCB1 | Q02818 | NUCB1_HUMA | Unmodified | 0 | 0 | 0 | 0 | 0 | 0 | 0 | 0 | 44601 | 131860 | 198330 | 0 | 0 | 0 | 0 | 0 | 0 | 0 |
| KQQSIAGS | AIMP1; AIMP1 | Q12904; A0A | AIMP1_HUMAN | Unmodified | 0 | 0 | 0 | 0 | 0 | 0 | 0 | 0 | 0 | 0 | 157870 | 2599100 | 0 | 0 | 0 | 0 | 0 | 0 |
| KQSKPVTTI | HSPD1; HSPD1 | P10809; A0A | CH60_HUMAN | Unmodified | 0 | 0 | 0 | 0 | 0 | 0 | 0 | 0 | 7673100 | 0 | 0 | 0 | 355260 | 0 | 0 | 0 | 0 | 0 |
| KQSLGELIC | TPI1 | P60174 | TPIS_HUMAN T | Unmodified | 0 | 0 | 0 | 0 | 0 | 0 | 0 | 0 | 385940 | 0 | 27087000 | 0 | 0 | 0 | 403520 | 0 | 0 | 0 |
| KQTDFEHR | PFKP; PFKP; PF | Q01813; A0A | PFKAP_HUMAN | Unmodified | 0 | 0 | 0 | 0 | 0 | 352110 | 0 | 0 | 371730 | 0 | 284840 | 861190 | 148250 | 0 | 0 | 0 | 0 | 817720 |
| KQTIDNSQ | YWHAQ | P27348 | 1433T_HUMAN | Unmodified | 53413 | 0 | 0 | 563680 | 0 | 0 | 0 | 0 | 229390 | 0 | 490020 | 0 | 0 | 0 | 281310 | 162010 | 0 | 0 |
| KQTIDNSQ | YWHAQ | P27348 | 1433T_HUMAN | Unmodified | 0 | 181480 | 0 | 0 | 0 | 0 | 0 | 0 | 0 | 33611 | 1492600 | 22163000 | 211730 | 0 | 0 | 0 | 0 | 0 |
| KQVAIIEK | TAOK3; TAOK3 | Q9H2K8; G3 | TAOK3_HUMAN | Unmodified | 0 | 0 | 0 | 172960 | 407150 | 111070 | 359660 | 249290 | 0 | 0 | 0 | 57502 | 530580 | 564460 | 0 | 0 | 245740 | 165410 |
| KQVEIAQRI | ASS1; ASS1; AS | P00966; Q5T | ASSY_HUMAN | Unmodified | 0 | 0 | 0 | 0 | 0 | 0 | 579690 | 0 | 0 | 22662 | 0 | 0 | 360390 | 412930 | 0 | 18007 | 0 | 0 |
| KQVSNLLA | ZC3H10 | Q96K80 | ZC3HA_HUMA | Unmodified | 351850 | 356480 | 0 | 1342200 | 0 | 0 | 782560 | 0 | 0 | 0 | 0 | 0 | 3357700 | 1693500 | 0 | 0 | 1787400 | 0 |
| KQVVNIPSI | RPS9 | P46781 | RS9_HUMAN 4 | Unmodified | 513540 | 242830 | 0 | 156450 | 0 | 0 | 0 | 0 | 0 | 0 | 0 | 11088000 | 0 | 0 | 498370 | 124690 | 0 | 0 |
| KQVVNIPSI | RPS9 | P46781 | RS9_HUMAN 4 | Unmodified | 258320 | 336300 | 353700 | 1180300 | 199090 | 134610 | 448560 | 0 | 248570 | 63474 | 424660 | 1746100 | 345300 | 113240 | 649570 | 198220 | 0 | 228070 |
| KQWITKWI | ITGA6; ITGA6 | P23229; H7E | ITA6_HUMAN I | Unmodified | 0 | 0 | 0 | 0 | 422430 | 0 | 0 | 0 | 0 | 0 | 0 | 0 | 1225100 | 0 | 0 | 0 | 660650 | 4491400 |

| | | | | | | | | | | | | | | | | | | | | | | |
|---|---|---|---|---|---|---|---|---|---|---|---|---|---|---|---|---|---|---|---|---|---|---|
| KQWKPAL | SNTB1; SNTB1 | Q13884; A0 | SNTB1_HUMAN | Unmodified | 0 | 0 | 0 | 0 | 43767 | 0 | 0 | 0 | 470440 | 313370 | 0 | 0 | 0 | 0 | 0 | 0 | 0 | 73293 |
| KQYDSFTF | TMED7; TMED7 | Q9Y3B3; A0 | TMED7_HUMAI | Unmodified | 0 | 0 | 0 | 0 | 0 | 0 | 0 | 0 | 3247600 | 0 | 463260 | 325180 | 0 | 0 | 97169 | 152360 | 0 | 0 |
| KQYDTYGE | DNAJB11; DNA | Q9UBS4; A0 | DJB11_HUMAN | Unmodified | 0 | 0 | 0 | 0 | 0 | 0 | 0 | 0 | 234170 | 0 | 0 | 217350 | 0 | 0 | 0 | 0 | 196570 | 0 |
| KRAAEMM | KIF5B | P33176 | KINH_HUMAN | Unmodified | 0 | 0 | 0 | 0 | 987200 | 0 | 0 | 0 | 337040 | 184340 | 0 | 0 | 0 | 165420 | 0 | 0 | 171320 | 0 |
| KRAEFTVE | FLNA; FLNA; FL | P21333; A0A | FLNA_HUMAN | Unmodified | 516800 | 1104400 | 425710 | 1472600 | 1024000 | 456290 | 435410 | 242250 | 120590 | 213550 | 38744 | 732440 | 721480 | 462570 | 1150500 | 0 | 614670 | 243810 |
| KRATEILNI | KIF5C; KIF5C; K | O60282; A0 | KIF5C_HUMAN | Unmodified | 0 | 0 | 0 | 0 | 83711 | 0 | 87447 | 0 | 0 | 64056 | 0 | 0 | 341000 | 216010 | 0 | 0 | 0 | 0 |
| KRDLEQAIS | RELA; RELA; RE | Q04206; A0 | TF65_HUMAN | Unmodified | 16055000 | 7352400 | 7930300 | 13179000 | 0 | 63677 | 114820 | 0 | 574240 | 0 | 0 | 656650 | 4172300 | 677100 | 7722200 | 7001300 | 0 | 0 |
| KRDNTNEI | ITGB1; ITGB1; I | P05556; H7C | ITB1_HUMAN | Unmodified | 0 | 0 | 682800 | 0 | 3527200 | 55171000 | 0 | 0 | 0 | 0 | 0 | 0 | 0 | 0 | 0 | 0 | 0 | 1E+08 |
| KRDQTASA | COBLL1; COBL | Q53SF7; A0 | COBL1_HUMAI | Unmodified | 372090 | 0 | 294420 | 204880 | 0 | 0 | 0 | 0 | 0 | 0 | 0 | 0 | 743390 | 655980 | 157140 | 282760 | 0 | 0 |
| KRDSLGTA | BLOC1S5; BLO | Q8TDH9; G5 | BL1S5_HUMAN | Unmodified | 0 | 0 | 0 | 62794 | 566010 | 0 | 0 | 54852 | 0 | 149090 | 0 | 237910 | 504880 | 0 | 0 | 0 | 0 | 243520 |
| KRDVVETQ | DOCK9; DOCK | Q9BZ29; A0 | DOCK9_HUMA | Unmodified | 41833 | 0 | 0 | 0 | 600300 | 0 | 0 | 0 | 253750 | 178490 | 0 | 171650 | 120670 | 97154 | 0 | 50367 | 114480 | 75202 |
| KREGSDSY | PLCG2; PLCG2 | P16885; A0 | PLCG2_HUMAN | Unmodified | 0 | 0 | 0 | 0 | 0 | 0 | 0 | 0 | 0 | 0 | 0 | 2072900 | 0 | 0 | 0 | 0 | 0 | 0 |
| KREGVTPV | AKAP12 | Q02952 | AKA12_HUMAN | Unmodified | 0 | 0 | 0 | 0 | 770880 | 0 | 0 | 0 | 0 | 70474 | 0 | 0 | 0 | 0 | 0 | 0 | 606870 | 933370 |
| KREPADVP | PTPRM; PTPRM | P28827; E7E | PTPRM_HUMAI | Unmodified | 0 | 0 | 0 | 0 | 1648100 | 0 | 0 | 0 | 0 | 0 | 0 | 0 | 0 | 0 | 248490 | 333390 | 152720 | 301630 |
| KRFDTEEEI | RARS1 | P54136 | SYRC_HUMAN | Unmodified | 0 | 0 | 0 | 0 | 90680 | 0 | 0 | 0 | 115720 | 0 | 169830 | 1394200 | 0 | 0 | 0 | 0 | 0 | 0 |
| KRGAPPSS | MATR3; MATR3 | P43243; H0 | MATR3_HUMAI | Unmodified | 0 | 103400 | 0 | 0 | 0 | 0 | 0 | 650500 | 0 | 580960 | 0 | 2716900 | 3413800 | 1520800 | 0 | 0 | 201430 | 1253900 |
| KRGFAFVTI | HNRNPA1L2; H | Q32P51; P0 | RA1L2_HUMAN | Unmodified | 438990 | 194090 | 628810 | 353390 | 2748500 | 786620 | 0 | 0 | 7464400 | 3366500 | 8315500 | 24543000 | 1738700 | 5250200 | 170720 | 0 | 179530 | 1043700 |
| KRGFAFVTI | HNRNPA3; HNI | P51991; A0A | ROA3_HUMAN | Unmodified | 0 | 0 | 0 | 0 | 1559300 | 391360 | 2872400 | 73435 | 168880 | 706850 | 1193800 | 4847400 | 0 | 1570300 | 0 | 0 | 0 | 884620 |
| KRGFEVVY | HSP90AB1 | P08238 | HS90B_HUMAI | Unmodified | 0 | 0 | 0 | 0 | 1814400 | 0 | 0 | 0 | 1083400 | 791150 | 0 | 751680 | 1820600 | 0 | 0 | 0 | 1120900 | 2584500 |
| KRGFEVVY | HSP90AB1 | P08238 | HS90B_HUMAI | Oxidation (M) | 0 | 0 | 0 | 0 | 5330700 | 0 | 0 | 0 | 0 | 1659600 | 0 | 5499300 | 6176300 | 0 | 0 | 0 | 0 | 1007500 |
| KRGFGFVT | HNRNPA2B1; H | P22626; A0A | ROA2_HUMAN | Unmodified | 0 | 0 | 0 | 0 | 581710 | 0 | 1153300 | 0 | 4505700 | 1850500 | 5600300 | 12438000 | 5098200 | 780230 | 0 | 0 | 0 | 0 |
| KRGFGFVY | HNRNPA0 | Q13151 | ROA0_HUMAN | Unmodified | 0 | 0 | 0 | 0 | 767210 | 0 | 0 | 0 | 274930 | 904530 | 0 | 0 | 1229500 | 0 | 0 | 0 | 0 | 0 |
| KRGHVFEE | EEF2 | P13639 | EF2_HUMAN EI | Unmodified | 0 | 0 | 0 | 0 | 1773700 | 0 | 0 | 0 | 0 | 0 | 0 | 0 | 0 | 0 | 0 | 0 | 342080 | 0 |
| KRHEMPPI | MYH9; MYH9 | P35579; Q5E | MYH9_HUMAN | Oxidation (M) | 0 | 0 | 0 | 210750 | 165000 | 55178 | 0 | 0 | 0 | 46808 | 0 | 0 | 204100 | 0 | 0 | 0 | 0 | 30965 |
| KRHPYNFP | GALNT16 | Q8N428 | GLT16_HUMAN | Unmodified | 0 | 0 | 0 | 0 | 0 | 0 | 0 | 0 | 681210 | 0 | 0 | 0 | 0 | 0 | 0 | 0 | 0 | 0 |
| KRILDEIEEI | SEPTIN2; SEPT | Q15019; H7 | SEPT2_HUMAN | Unmodified | 0 | 0 | 0 | 268860 | 6759000 | 1644900 | 159330 | 0 | 262550 | 216350 | 0 | 385750 | 555650 | 647180 | 0 | 0 | 1636800 | 1176300 |
| KRILHCLGI | BLVRA | P53004 | BIEA_HUMAN E | Unmodified | 0 | 0 | 0 | 0 | 2296000 | 189660 | 123830 | 0 | 0 | 56506 | 0 | 0 | 1216700 | 1326200 | 0 | 0 | 830820 | 2627800 |
| KRIQVISQS | DOCK10; DOC | Q96BY6; H0 | DOC10_HUMA | Unmodified | 0 | 0 | 341780 | 0 | 1072000 | 0 | 0 | 0 | 0 | 0 | 0 | 0 | 0 | 0 | 0 | 0 | 610610 | 0 |
| KRLEMETL | KIF14 | Q15058 | KIF14_HUMAN | Unmodified | 0 | 0 | 0 | 0 | 2024000 | 311930 | 431430 | 0 | 0 | 0 | 0 | 144830 | 0 | 0 | 0 | 0 | 622710 | 284280 |
| KRLIDLHTI | SCFD1; SCFD1 | Q8WVM8; A | SCFD1_HUMAI | Unmodified | 0 | 0 | 0 | 0 | 6971600 | 1437000 | 1505500 | 0 | 0 | 659840 | 0 | 0 | 0 | 0 | 0 | 0 | 0 | 496740 |
| KRLVQSPN | RPS27L; RPS2 | Q71UM5; P4 | RS27L_HUMAN | Oxidation (M) | 0 | 0 | 0 | 0 | 0 | 0 | 71625 | 0 | 99860 | 0 | 0 | 0 | 180010 | 97650 | 0 | 0 | 0 | 0 |
| KRMGPGA | RAB1A; RAB1A | P62820; E7E | RAB1A_HUMAN | Unmodified | 308170 | 354100 | 74635 | 89189 | 7866400 | 930240 | 1824600 | 1100400 | 1146200 | 2044800 | 711140 | 917060 | 4231200 | 3121400 | 125190 | 151470 | 1134300 | 4126300 |
| KRPDEVPD | NPTN; NPTN | Q9Y639; H3I | NPTN_HUMAN | Unmodified | 360190 | 565620 | 351860 | 92110 | 435310 | 0 | 127970 | 0 | 0 | 0 | 132040 | 1821400 | 0 | 0 | 375920 | 182420 | 0 | 226270 |
| KRPDEVPD | NPTN; NPTN | Q9Y639; H3I | NPTN_HUMAN | Oxidation (M) | 0 | 0 | 0 | 0 | 0 | 0 | 0 | 0 | 0 | 0 | 0 | 957370 | 0 | 0 | 271280 | 146180 | 0 | 0 |
| KRPETVAT | MYO1C; MYO1 | O00159; F5F | MYO1C_HUMA | Unmodified | 3661600 | 2559600 | 6369100 | 7304300 | 2464500 | 1812500 | 305300 | 149930 | 599470 | 83132 | 237610 | 1351900 | 0 | 17285 | 9382100 | 10994000 | 658420 | 282100 |
| KRQLEESVI | KIF5B | P33176 | KINH_HUMAN | Unmodified | 0 | 0 | 0 | 0 | 0 | 0 | 0 | 0 | 0 | 15242 | 0 | 1440700 | 0 | 0 | 25026 | 0 | 0 | 0 |
| KRSEELLS | CEP55; CEP55 | Q53EZ4; H0 | CEP55_HUMAN | Unmodified | 40931 | 689150 | 0 | 3768400 | 28534000 | 3134500 | 91192 | 0 | 2786000 | 6641700 | 0 | 0 | 2873000 | 1645900 | 1519200 | 27834 | 376870 | 724960 |
| KRSSITEPE | TP53BP2 | Q13625 | ASPP2_HUMAI | Unmodified | 0 | 0 | 0 | 0 | 169380 | 0 | 0 | 24196 | 0 | 0 | 0 | 0 | 138160 | 95470 | 0 | 0 | 0 | 206440 |
| KRTELFIAA | RPL8; RPL8; RP | P62917; E9P | RL8_HUMAN 6 | Unmodified | 0 | 0 | 0 | 0 | 1291300 | 0 | 2619700 | 0 | 0 | 277540 | 0 | 7321400 | 2226200 | 0 | 0 | 0 | 659870 | 1058500 |
| KRTGPAAT | P4HB; P4HB; P | P07237; A0A | PDIA1_HUMAN | Unmodified | 0 | 0 | 0 | 0 | 0 | 0 | 0 | 0 | 116550 | 1562300 | 0 | 0 | 0 | 0 | 0 | 0 | 0 | 647290 |
| KRVEDAYIL | CCT6A | P40227 | TCPZ_HUMAN | Unmodified | 0 | 0 | 0 | 0 | 0 | 0 | 0 | 0 | 347610 | 0 | 488250 | 2004700 | 394240 | 378320 | 0 | 0 | 143970 | 275410 |
| KRVYLATDI | FUT8; FUT8; FU | Q9BYC5; A0 | FUT8_HUMAN | Unmodified | 0 | 0 | 0 | 0 | 0 | 0 | 0 | 0 | 1516500 | 620740 | 0 | 0 | 0 | 0 | 0 | 50013 | 0 | 0 |
| KRYDLSAL | IGF2R; IGF2R; I | P11717; A0A | MPRI_HUMAN | Unmodified | 0 | 0 | 0 | 0 | 0 | 0 | 0 | 0 | 2006400 | 255120 | 191590 | 174030 | 0 | 0 | 0 | 0 | 0 | 0 |
| KSADTLWC | LDHB; LDHB | P07195; A0A | LDHB_HUMAN | Unmodified | 124710 | 0 | 0 | 0 | 8622900 | 1548300 | 0 | 0 | 0 | 0 | 3640100 | 18105000 | 3934000 | 1714700 | 0 | 0 | 1357400 | 3383400 |
| KSADTLWC | LDHA | P00338 | LDHA_HUMAN | Unmodified | 115430 | 0 | 0 | 0 | 13753000 | 480720 | 2055500 | 1668700 | 1238400 | 1162900 | 1207100 | 11495000 | 2581400 | 3014900 | 210610 | 204470 | 13846000 | 15188000 |
| KSADTLWC | LDHA | P00338 | LDHA_HUMAN | Unmodified | 44016 | 0 | 0 | 0 | 0 | 0 | 0 | 0 | 198270 | 0 | 86661 | 0 | 0 | 0 | 0 | 0 | 8601100 | 1215700 |
| KSAEFLLHI | RPL17; RPL17; | P18621; A0A | RL17_HUMAN | Unmodified | 0 | 0 | 0 | 0 | 868670 | 0 | 706060 | 0 | 122640 | 0 | 0 | 908210 | 793570 | 0 | 0 | 0 | 0 | 342130 |
| KSAEFLLHI | RPL17; RPL17; | P18621; A0A | RL17_HUMAN | Oxidation (M) | 120010 | 95125 | 0 | 0 | 204210 | 0 | 196370 | 119290 | 0 | 36933 | 0 | 0 | 159780 | 0 | 0 | 0 | 596750 | 1535800 |
| KSAEIDSDI | KIF5B | P33176 | KINH_HUMAN | Unmodified | 0 | 0 | 0 | 0 | 4522400 | 0 | 0 | 911520 | 0 | 0 | 0 | 0 | 0 | 0 | 0 | 0 | 212300 | 0 |
| KSAVADKH | EIF3H; EIF3H; E | O15372; E5F | EIF3H_HUMAN | Unmodified | 243310 | 0 | 0 | 674760 | 2034600 | 0 | 583320 | 0 | 0 | 459860 | 0 | 3926700 | 3284800 | 1273200 | 0 | 0 | 893870 | 871900 |
| KSAYCPYSI | CDA | P32320 | CDD_HUMAN | Unmodified | 0 | 0 | 0 | 0 | 1282000 | 0 | 0 | 0 | 0 | 0 | 0 | 0 | 0 | 0 | 0 | 0 | 0 | 0 |
| KSCVEEPEI | HDGF | P51858 | HDGF_HUMAN | Unmodified | 0 | 0 | 0 | 0 | 0 | 0 | 0 | 0 | 0 | 0 | 0 | 422800 | 0 | 0 | 0 | 0 | 0 | 0 |
| KSCVSSAW | NOSTRIN | Q8IVI9 | NOSTN_HUMA | Oxidation (M) | 0 | 0 | 0 | 0 | 0 | 0 | 0 | 0 | 0 | 0 | 0 | 0 | 0 | 0 | 0 | 766300 | 0 | 0 |
| KSDFQVNL | CTNND1; CTNI | O60716; H0 | CTND1_HUMAI | Unmodified | 754590 | 724590 | 1915600 | 3903600 | 2947900 | 757230 | 459510 | 12138 | 0 | 60214 | 1346200 | 8056500 | 2029300 | 3048500 | 6920300 | 5450700 | 573960 | 3268200 |
| KSDGIYIINI | RPSA; RPSA; RI | P08865; A0A | RSSA_HUMAN | Unmodified | 0 | 0 | 0 | 0 | 0 | 0 | 0 | 0 | 0 | 0 | 0 | 1970500 | 0 | 0 | 0 | 0 | 0 | 0 |
| KSDGIYIINI | RPSA; RPSA; RI | P08865; A0A | RSSA_HUMAN | Unmodified | 388150 | 396010 | 0 | 0 | 0 | 0 | 1847600 | 0 | 0 | 0 | 0 | 3148600 | 0 | 0 | 249230 | 284540 | 619200 | 0 |
| KSDGKIDV | DLD; DLD | P09622; E9P | DLDH_HUMAN | Unmodified | 0 | 0 | 0 | 0 | 1350600 | 0 | 0 | 0 | 441560 | 0 | 0 | 0 | 0 | 0 | 0 | 0 | 84633 | 98591 |
| KSDIDEIVL | HSPA5; HSPA5 | P11021; A0A | BIP_HUMAN Er | Unmodified | 365550 | 414270 | 2629000 | 658160 | 0 | 0 | 178070 | 278610 | 1.77E+08 | 3647100 | 9812500 | 8647700 | 0 | 0 | 3036600 | 1132100 | 0 | 0 |
| KSDIDEIVL | HSPA5; HSPA5 | P11021; A0A | BIP_HUMAN Er | Unmodified | 0 | 0 | 1992200 | 3566900 | 2122500 | 120980 | 4394400 | 610720 | 2.08E+08 | 53014000 | 26160000 | 1443700 | 1185700 | 0 | 1772800 | 0 | 185780 | 0 |
| KSDIYVCM | GDI2 | P50395 | GDIB_HUMAN | Unmodified | 663060 | 0 | 0 | 0 | 0 | 5216800 | 145870 | 0 | 6205100 | 1205200 | 2238400 | 4551700 | 0 | 0 | 1153300 | 958930 | 3121500 | 11591000 |
| KSDIYVCM | GDI1 | P31150 | GDIA_HUMAN | Unmodified | 0 | 0 | 0 | 0 | 0 | 0 | 0 | 0 | 1675800 | 0 | 0 | 0 | 0 | 0 | 0 | 0 | 67273 | 0 |
| KSDLHPTLI | TMEM30A; TME | Q9NV96; A0 | CC50A_HUMAI | Unmodified | 0 | 0 | 761600 | 0 | 0 | 0 | 0 | 0 | 3761000 | 661870 | 278870 | 2341700 | 0 | 0 | 687010 | 0 | 46330 | 0 |
| KSDPVVSY | EEF2 | P13639 | EF2_HUMAN EI | Unmodified | 88040 | 0 | 0 | 0 | 0 | 0 | 0 | 0 | 82780 | 0 | 112670 | 3547500 | 0 | 0 | 132130 | 229860 | 0 | 0 |
| KSDPVVSY | EEF2 | P13639 | EF2_HUMAN EI | Unmodified | 0 | 0 | 0 | 681990 | 7249300 | 1013500 | 1252900 | 175110 | 0 | 0 | 1202200 | 4739200 | 1741100 | 1375700 | 61986 | 0 | 0 | 4138700 |
| KSDSNPLT | RBM17 | Q96I25 | SPF45_HUMAN | Unmodified | 0 | 0 | 0 | 0 | 0 | 0 | 0 | 0 | 0 | 0 | 0 | 1111400 | 0 | 0 | 0 | 0 | 0 | 0 |
| KSDVEAIFS | HNRNPCL3; HN | B7ZW38; P0 | HNRC3_HUMA | Unmodified | 0 | 0 | 0 | 0 | 0 | 0 | 3918500 | 1210500 | 0 | 0 | 0 | 0 | 1076800 | 0 | 0 | 0 | 0 | 0 |
| KSDVEDHS | DTYMK | P23919 | KTHY_HUMAN | Unmodified | 0 | 0 | 0 | 0 | 13750000 | 2249300 | 3983300 | 0 | 1404400 | 1157600 | 603450 | 1906300 | 7281500 | 2636000 | 0 | 0 | 4876700 | 5801700 |
| KSEALESIK | EEA1 | Q15075 | EEA1_HUMAN | Unmodified | 0 | 0 | 0 | 0 | 413410 | 42418 | 0 | 0 | 248340 | 120160 | 174170 | 354980 | 0 | 50667 | 0 | 0 | 0 | 48759 |
| KSEEEIDFL | EIF3L; EIF3L; EI | Q9Y262; B0 | EIF3L_HUMAN | Unmodified | 0 | 156640 | 0 | 0 | 0 | 0 | 0 | 0 | 0 | 196130 | 0 | 3407400 | 0 | 0 | 0 | 151930 | 0 | 0 |
| KSEEEIDFL | EIF3L; EIF3L; EI | Q9Y262; B0 | EIF3L_HUMAN | Unmodified | 0 | 0 | 0 | 0 | 0 | 0 | 0 | 0 | 0 | 159280 | 471520 | 1083400 | 0 | 0 | 0 | 0 | 0 | 0 |
| KSEHLSVR | MRPS23 | Q9Y3D9 | RT23_HUMAN | Unmodified | 0 | 0 | 0 | 0 | 0 | 0 | 0 | 0 | 369620 | 0 | 0 | 0 | 0 | 0 | 0 | 0 | 0 | 0 |
| KSEIEYYAM | RPL30; RPL30 | P62888; E5R | RL30_HUMAN | Oxidation (M) | 9204900 | 6791300 | 1859100 | 3199000 | 0 | 0 | 0 | 0 | 497940 | 0 | 0 | 15010000 | 0 | 0 | 14145000 | 7361600 | 0 | 0 |
| KSEIEYYAM | RPL30; RPL30 | P62888; E5R | RL30_HUMAN | Unmodified | 2805300 | 1851900 | 741020 | 1928300 | 0 | 0 | 0 | 0 | 477790 | 0 | 0 | 13845000 | 335130 | 197680 | 1522300 | 563380 | 0 | 0 |
| KSESEVEE | DBN1; DBN1; D | Q16643; D6 | DREB_HUMAN | Unmodified | 0 | 0 | 700840 | 410090 | 0 | 0 | 0 | 0 | 558340 | 0 | 2144300 | 21806000 | 0 | 6022400 | 987490 | 836850 | 0 | 0 |
| KSEYTQPTI | DDX42; DDX42 | Q86XP3; A0 | DDX42_HUMAI | Unmodified | 0 | 0 | 0 | 0 | 0 | 0 | 0 | 0 | 118670 | 0 | 0 | 834930 | 0 | 0 | 0 | 0 | 0 | 0 |
| KSFYGSTLI | FASN; FASN | P49327; A0A | FAS_HUMAN F | Unmodified | 0 | 0 | 0 | 103910 | 0 | 0 | 0 | 0 | 8806600 | 527030 | 0 | 1369400 | 0 | 0 | 0 | 55457 | 0 | 0 |
| KSGALPVA | GPAA1; GPAA1 | O43292; E9F | GPAA1_HUMAI | Unmodified | 0 | 0 | 0 | 0 | 0 | 0 | 0 | 0 | 2352600 | 0 | 0 | 0 | 0 | 0 | 0 | 0 | 0 | 0 |
| KSGNYFFL | ABCE1 | P61221 | ABCE1_HUMAI | Unmodified | 0 | 0 | 0 | 0 | 1440500 | 0 | 0 | 0 | 0 | 0 | 0 | 1242800 | 407020 | 0 | 0 | 107470 | 350300 | 0 |
| KSGVGNIF | PABPC5; PABP | Q96DU9; P1 | PABP5_HUMAI | Unmodified | 11904000 | 3968200 | 3362300 | 4353000 | 0 | 0 | 0 | 0 | 0 | 0 | 0 | 800950 | 136190 | 0 | 2366200 | 3849600 | 0 | 0 |
| KSGVGNVF | PABPC4; PABP | Q13310; Q5 | PABP4_HUMAI | Unmodified | 3622800 | 0 | 2174900 | 1014100 | 0 | 0 | 0 | 0 | 0 | 0 | 0 | 0 | 0 | 0 | 1207500 | 1043800 | 0 | 0 |
| KSHQIEELE | KIF20B | Q96Q89 | KI20B_HUMAN | Unmodified | 0 | 0 | 0 | 0 | 696320 | 153490 | 0 | 0 | 0 | 0 | 0 | 0 | 0 | 0 | 0 | 0 | 159310 | 107340 |
| KSIDTGMG | AARS1; AARS1 | P49588; A0A | SYAC_HUMAN | Oxidation (M) | 0 | 0 | 0 | 0 | 1199200 | 927900 | 367460 | 0 | 0 | 708740 | 0 | 0 | 3043700 | 2072100 | 0 | 0 | 509120 | 585050 |
| KSIPHITSD | CRMP1 | Q14194 | DPYL1_HUMAI | Unmodified | 0 | 0 | 0 | 0 | 0 | 0 | 729650 | 489100 | 242120 | 356530 | 49185 | 441830 | 0 | 0 | 0 | 0 | 0 | 0 |
| KSISHYHE | TM9SF3; TM9SI | Q9HD45; Q5 | TM9S3_HUMAI | Unmodified | 0 | 0 | 0 | 0 | 66933 | 0 | 0 | 0 | 4029700 | 0 | 0 | 0 | 0 | 0 | 0 | 0 | 0 | 0 |
| KSLADEAE | GAS7 | O60861 | GAS7_HUMAN | Unmodified | 0 | 189640 | 0 | 42744 | 0 | 0 | 0 | 0 | 0 | 0 | 313120 | 0 | 0 | 0 | 0 | 0 | 0 | 0 |
| KSLDKDPL | ATP2B1 | P20020 | AT2B1_HUMAI | Unmodified | 0 | 0 | 0 | 0 | 0 | 0 | 0 | 0 | 0 | 0 | 0 | 0 | 1608000 | 0 | 214190 | 0 | 0 | 0 |
| KSLDQDPV | HSPH1; HSPH1 | Q92598; Q5 | HS105_HUMAI | Unmodified | 0 | 0 | 0 | 0 | 429880 | 0 | 839550 | 231020 | 0 | 200600 | 0 | 0 | 977150 | 425720 | 0 | 0 | 726550 | 686390 |
| KSLDSDES | PDAP1 | Q13442 | HAP28_HUMAI | Unmodified | 0 | 0 | 0 | 0 | 385620 | 79763 | 0 | 0 | 203620 | 166450 | 99927 | 0 | 0 | 0 | 0 | 0 | 96450 | 290530 |
| KSLEDQVE | PRKCSH; PRKC | P14314; K7E | GLU2B_HUMAI | Unmodified | 0 | 0 | 0 | 0 | 0 | 0 | 0 | 0 | 4179900 | 690740 | 0 | 0 | 0 | 0 | 232940 | 0 | 0 | 0 |
| KSLEDVVIC | MANSC1; MAN | Q9H8J5; F5I | MANS1_HUMA | Unmodified | 0 | 0 | 0 | 0 | 0 | 0 | 0 | 0 | 645470 | 1175000 | 0 | 0 | 0 | 0 | 0 | 0 | 0 | 0 |
| KSLESINSI | RPL30; RPL30; | P62888; A0A | RL30_HUMAN | Unmodified | 481490 | 528950 | 0 | 0 | 0 | 0 | 56650 | 0 | 0 | 0 | 0 | 791320 | 108080 | 0 | 0 | 188030 | 0 | 0 |
| KSLNQSLA | ATP1B3; ATP1B | P54709; F8V | AT1B3_HUMAI | Unmodified | 0 | 0 | 0 | 0 | 2264300 | 43800 | 2703900 | 1820000 | 0 | 0 | 0 | 3696800 | 0 | 0 | 874950 | 637820 | 0 | 0 |
| KSNFAEAL | P4HB; P4HB; P | P07237; A0A | PDIA1_HUMAN | Unmodified | 0 | 0 | 0 | 0 | 108750 | 0 | 0 | 166940 | 3713900 | 1303000 | 1371100 | 0 | 375140 | 0 | 0 | 27039 | 0 | 0 |
| KSPCPSVV | AGRN; AGRN; A | O00468; A0 | AGRIN_HUMAN | Unmodified | 0 | 0 | 0 | 0 | 0 | 0 | 0 | 0 | 0 | 0 | 0 | 0 | 660580 | 899260 | 0 | 0 | 0 | 0 |
| KSPLIESTA | PPP1R12A; PPP | O14974; F8V | MYPT1_HUMAI | Unmodified | 0 | 0 | 0 | 112290 | 0 | 0 | 0 | 0 | 80869 | 0 | 0 | 1333000 | 0 | 206420 | 0 | 0 | 0 | 282820 |
| KSPLIESTA | PPP1R12A; PPP | O14974; F8V | MYPT1_HUMAI | Oxidation (M) | 0 | 0 | 53497 | 61944 | 242660 | 0 | 0 | 0 | 0 | 0 | 26913 | 302710 | 86178 | 89845 | 84472 | 61723 | 0 | 0 |
| KSPLSWIE | BLVRA | P53004 | BIEA_HUMAN E | Unmodified | 0 | 0 | 0 | 0 | 2451000 | 0 | 0 | 0 | 0 | 0 | 0 | 0 | 0 | 0 | 0 | 0 | 0 | 0 |
| KSPPHLIK | GALNT2 | Q10471 | GALT2_HUMAN | Unmodified | 0 | 0 | 0 | 0 | 0 | 0 | 0 | 0 | 3042300 | 436460 | 0 | 0 | 0 | 0 | 0 | 0 | 36756 | 0 |
| KSPQIFDDI | MYO1D; MYO1I | O94832; J3Q | MYO1D_HUMA | Unmodified | 604030 | 783410 | 0 | 0 | 0 | 0 | 8351.8 | 0 | 196410 | 0 | 0 | 0 | 0 | 0 | 870610 | 664670 | 0 | 0 |
| KSPSDTEG | MPZL1 | O95297 | MPZL1_HUMAN | Unmodified | 289650 | 376320 | 1967900 | 0 | 1258500 | 265320 | 0 | 0 | 0 | 99853 | 0 | 780790 | 0 | 0 | 0 | 0 | 102220 | 558880 |
| KSPSPEFS | FOLH1B; FOLH | Q9HBA9; Q0 | FOH1B_HUMAI | Unmodified | 2185700 | 2012700 | 16334000 | 29083000 | 0 | 0 | 0 | 45462 | 0 | 0 | 0 | 0 | 1856700 | 543560 | 12164000 | 9988900 | 0 | 0 |
| KSPSPEFS | FOLH1B; FOLH | Q9HBA9; Q0 | FOH1B_HUMAI | Oxidation (M) | 1080800 | 773440 | 3526700 | 4923100 | 0 | 0 | 0 | 0 | 0 | 0 | 0 | 0 | 0 | 0 | 3527400 | 3665500 | 0 | 0 |
| KSPVAKDV | VCP; VCP; VCP | P55072; C9J | TERA_HUMAN | Unmodified | 0 | 0 | 0 | 0 | 908070 | 0 | 0 | 0 | 237930 | 249790 | 0 | 0 | 887290 | 209160 | 0 | 0 | 346340 | 0 |
| KSQIFSTAS | HSPA5; HSPA5 | P11021; A0A | BIP_HUMAN Er | Unmodified | 372140 | 500600 | 1847300 | 502890 | 0 | 0 | 19032000 | 5873700 | 1.44E+08 | 1.98E+08 | 10395000 | 5882500 | 1736200 | 2385200 | 2606100 | 745700 | 742630 | 1177100 |
| KSQVFSTA | HSPA9; HSPA9 | P38646; A0A | GRP75_HUMAN | Unmodified | 0 | 0 | 0 | 0 | 1032500 | 0 | 0 | 0 | 12395000 | 2134500 | 0 | 0 | 0 | 0 | 0 | 0 | 235500 | 0 |
| KSSGEIVYC | RPL18A; RPL18 | Q02543; M0 | RL18A_HUMAN | Unmodified | 0 | 0 | 0 | 0 | 0 | 0 | 0 | 58506 | 0 | 0 | 0 | 787810 | 0 | 0 | 0 | 0 | 0 | 0 |
| KSSGGKG | HLA-A; HLA-B; | A0A1W2PRT | A0A1W2PRT9_ | Unmodified | 0 | 0 | 0 | 0 | 1362600 | 0 | 0 | 0 | 0 | 0 | 0 | 0 | 0 | 0 | 0 | 0 | 0 | 516540 |
| KSSPSVKP | CIAPIN1; CIAPI | Q6FI81; H3B | CPIN1_HUMAN | Unmodified | 0 | 0 | 0 | 0 | 0 | 0 | 90661 | 68046 | 79420 | 0 | 125260 | 585900 | 314590 | 0 | 45378 | 0 | 269730 | 279250 |
| KSSSYSSS | F2RL1 | P55085 | PAR2_HUMAN | Unmodified | 0 | 0 | 0 | 592760 | 0 | 277770 | 0 | 0 | 0 | 0 | 0 | 0 | 64991 | 51211 | 0 | 0 | 230570 | 374800 |

| | | | | | | | | | | | | | | | | | | | | | | |
|---|---|---|---|---|---|---|---|---|---|---|---|---|---|---|---|---|---|---|---|---|---|---|
| KSSTVGEIV | ABCC1; ABCC1 | P33527; I3L4 | MRP1_HUMAN | Unmodified | 0 | 0 | 0 | 28459 | 0 | 0 | 0 | 0 | 0 | 0 | 0 | 551260 | 0 | 277210 | 0 | 47780 | 0 | 0 |
| KSTGSPTQ | ZNF185 | H0Y704 | H0Y704_HUMA | Unmodified | 53128 | 0 | 0 | 0 | 288200 | 0 | 48459 | 0 | 0 | 65739 | 0 | 0 | 0 | 0 | 0 | 0 | 745780 | 0 |
| KSVEVNFTI | PDCD10; PDCD | Q9BUL8; C9. | PDC10_HUMAI | Unmodified | 0 | 0 | 0 | 0 | 0 | 0 | 0 | 0 | 0 | 0 | 0 | 0 | 0 | 0 | 0 | 0 | 0 | 0 |
| KSVKVIVVG | MDH1; MDH1; I | P40925; C9J | MDHC_HUMAN | Unmodified | 0 | 0 | 0 | 0 | 278270 | 0 | 4659200 | 2986700 | 0 | 404670 | 0 | 0 | 431760 | 842430 | 0 | 0 | 9814300 | 2042000 |
| KSVQYDDV | ALCAM; ALCAM | Q13740; F5C | CD166_HUMAI | Unmodified | 2579400 | 2141900 | 0 | 803110 | 0 | 0 | 0 | 0 | 0 | 0 | 0 | 0 | 0 | 0 | 0 | 915840 | 0 | 787840 |
| KSVSSENP | SH3D19; SH3D | Q5HYK7; A0 | SH319_HUMAN | Unmodified | 0 | 0 | 0 | 1168000 | 31840000 | 12108000 | 826270 | 520090 | 0 | 10457000 | 1402600 | 0 | 0 | 4324700 | 0 | 0 | 408640 | 1.13E+08 |
| KSVTDSIRD | DNM1L; DNM1I | O00429; F8V | DNM1L_HUMA | Unmodified | 0 | 0 | 0 | 0 | 0 | 0 | 0 | 0 | 0 | 0 | 0 | 870240 | 170360 | 0 | 0 | 0 | 0 | 0 |
| KSVVFTSA( | CA14 | Q9ULX7 | CAH14_HUMAI | Unmodified | 0 | 0 | 0 | 0 | 0 | 0 | 0 | 0 | 0 | 0 | 0 | 1263800 | 0 | 0 | 0 | 0 | 0 | 0 |
| KSWHPVC( | TMPRSS2; TMPI | O15393; C9J | TMPS2_HUMAI | Unmodified | 0 | 0 | 0 | 504970 | 361780 | 0 | 166950 | 0 | 0 | 0 | 1363000 | 0 | 0 | 0 | 0 | 0 | 0 | 212180 |
| KSYGVENR | GPRC5A; GPRC | Q8NFJ5; A0A | RAI3_HUMAN F | Unmodified | 0 | 0 | 0 | 0 | 2259000 | 0 | 0 | 120640 | 575570 | 0 | 0 | 307460 | 0 | 0 | 0 | 0 | 0 | 0 |
| KSYIATQG( | PTPN11 | Q06124 | PTN11_HUMAN | Unmodified | 94909 | 0 | 0 | 54125 | 451660 | 50806 | 0 | 0 | 0 | 0 | 64175 | 0 | 212450 | 149950 | 0 | 39206 | 160650 | 1504300 |
| KSYSLQVP! | CDCP1 | Q9H5V8 | CDCP1_HUMA | Unmodified | 79501 | 87667 | 0 | 1684600 | 1352000 | 70642 | 0 | 0 | 115840 | 284470 | 0 | 443170 | 1988500 | 956910 | 929490 | 485330 | 3610100 | 1067300 |
| KTAGLSDL\ | TSG101; TSG10 | Q99816; J3Q | TS101_HUMAN | Unmodified | 12424000 | 10283000 | 5909100 | 8316200 | 0 | 0 | 245230 | 103410 | 2364300 | 0 | 486730 | 507480 | 0 | 1426200 | 12249000 | 13082000 | 0 | 0 |
| KTANDMIH | MSN | P26038 | MOES_HUMAN | Unmodified | 85799 | 119990 | 0 | 601720 | 9499900 | 2759900 | 146050 | 47647 | 0 | 0 | 0 | 240960 | 1408300 | 49302 | 237300 | 126030 | 1615400 | 4786200 |
| KTANDMIH | MSN | P26038 | MOES_HUMAN | 2 Oxidation (M) | 0 | 0 | 0 | 0 | 0 | 309760 | 0 | 0 | 0 | 0 | 0 | 0 | 0 | 84162 | 0 | 50486 | 0 | 0 |
| KTCRPSTQ | MGAT2 | Q10469 | MGAT2_HUMAI | Unmodified | 0 | 0 | 0 | 0 | 0 | 0 | 0 | 0 | 1129500 | 0 | 0 | 0 | 0 | 0 | 0 | 0 | 0 | 0 |
| KTCTTVAFT | RPL7A; RPL7A | P62424; Q5T | RL7A_HUMAN ( | Unmodified | 1231500 | 515320 | 0 | 308970 | 0 | 0 | 0 | 0 | 0 | 0 | 0 | 8720000 | 0 | 0 | 468790 | 434970 | 0 | 0 |
| KTCTTVAFT | RPL7A; RPL7A | P62424; Q5T | RL7A_HUMAN ( | Unmodified | 1400300 | 645660 | 0 | 0 | 0 | 0 | 0 | 0 | 0 | 0 | 0 | 0 | 0 | 0 | 0 | 0 | 0 | 0 |
| KTDEFCQA | LEMD2 | Q8NC56 | LEMD2_HUMAI | Unmodified | 0 | 0 | 0 | 0 | 0 | 0 | 0 | 0 | 0 | 0 | 0 | 0 | 0 | 0 | 0 | 0 | 0 | 0 |
| KTDELPQL\ | FN1; FN1 | P02751; H0Y | FINC_HUMAN I | Unmodified | 0 | 0 | 865580 | 0 | 0 | 0 | 0 | 0 | 0 | 0 | 0 | 0 | 0 | 0 | 0 | 0 | 0 | 0 |
| KTDFFIGGE | NUDC; NUDC | Q9Y266; A0A | NUDC_HUMAN | Unmodified | 0 | 0 | 0 | 0 | 0 | 0 | 0 | 0 | 434140 | 0 | 131480 | 2062400 | 103420 | 0 | 52732 | 0 | 0 | 0 |
| KTDFFIGGE | NUDC; NUDC | Q9Y266; A0A | NUDC_HUMAN | Oxidation (M) | 82856 | 0 | 0 | 0 | 0 | 0 | 0 | 0 | 102110 | 37119 | 231120 | 328770 | 0 | 0 | 0 | 0 | 0 | 0 |
| KTDGYSGA | VPS4B | O75351 | VPS4B_HUMAN | Unmodified | 375130 | 313350 | 246240 | 0 | 2073500 | 471600 | 46445 | 0 | 2485200 | 0 | 0 | 0 | 0 | 0 | 0 | 0 | 68807 | 174400 |
| KTEAPAAP/ | BASP1 | P80723 | BASP1_HUMAN | Unmodified | 0 | 0 | 0 | 0 | 3233400 | 0 | 0 | 0 | 22710000 | 759680 | 0 | 0 | 0 | 8083.4 | 0 | 0 | 44533 | 56739 |
| KTEEIYGFL | CHST1 | O43916 | CHST1_HUMAI | Unmodified | 0 | 0 | 0 | 0 | 0 | 0 | 82085 | 0 | 9596000 | 743490 | 0 | 0 | 0 | 0 | 37924 | 20120 | 0 | 0 |
| KTEGYSGA | VPS4A; N/A | Q9UN37; I3L | VPS4A_HUMAN | Unmodified | 0 | 0 | 0 | 0 | 0 | 0 | 0 | 0 | 0 | 0 | 0 | 0 | 0 | 0 | 0 | 0 | 0 | 0 |
| KTENWLSE | CRAT; CRAT; CI | P43155; A0A | CACP_HUMAN | Unmodified | 0 | 0 | 0 | 0 | 0 | 0 | 0 | 0 | 479520 | 261350 | 0 | 0 | 0 | 0 | 0 | 0 | 0 | 0 |
| KTEPATGFI | DDB1; DDB1; D | Q16531; A0A | DDB1_HUMAN | Unmodified | 0 | 0 | 0 | 0 | 0 | 0 | 0 | 0 | 722600 | 172410 | 262790 | 3472900 | 0 | 0 | 0 | 0 | 0 | 0 |
| KTESAVSQI | IARS1; IARS1; I/ | P41252; A0A | SYIC_HUMAN I | Unmodified | 0 | 0 | 0 | 0 | 0 | 0 | 0 | 0 | 0 | 0 | 0 | 5027300 | 36770 | 0 | 0 | 0 | 0 | 0 |
| KTETQEKNI | TMSB4X | P62328 | TYB4_HUMAN ` | Unmodified | 0 | 0 | 0 | 0 | 0 | 0 | 0 | 313730 | 0 | 1004900 | 0 | 0 | 0 | 0 | 0 | 0 | 1271700 | 1807900 |
| KTEVLMEN | RUVBL1; RUVBI | Q9Y265; E7E | RUVB1_HUMAI | Unmodified | 0 | 0 | 0 | 0 | 0 | 0 | 0 | 0 | 0 | 0 | 0 | 0 | 0 | 0 | 0 | 0 | 0 | 0 |
| KTEVVMNS | PALM | O75781 | PALM_HUMAN | Unmodified | 0 | 0 | 0 | 0 | 0 | 0 | 0 | 0 | 0 | 0 | 0 | 0 | 0 | 0 | 0 | 0 | 87854 | 462740 |
| KTFSHELSI | PDIA3 | P30101 | PDIA3_HUMAN | Unmodified | 0 | 0 | 0 | 0 | 0 | 0 | 0 | 0 | 3.81E+08 | 16257000 | 10556000 | 3097000 | 0 | 0 | 3979000 | 0 | 0 | 0 |
| KTGCNVLL | CCT4 | P50991 | TCPD_HUMAN | Unmodified | 0 | 0 | 0 | 53975 | 0 | 0 | 350570 | 412120 | 0 | 110360 | 188770 | 0 | 342210 | 604290 | 0 | 39152 | 231460 | 0 |
| KTGNFQVT | SNRNP200; SN | O75643; A0/ | U520_HUMAN | Unmodified | 0 | 0 | 0 | 0 | 102620 | 0 | 0 | 0 | 0 | 139360 | 0 | 3535900 | 0 | 0 | 0 | 48084 | 0 | 0 |
| KTGQLDEL | SMAD3 | P84022 | SMAD3_HUMA | Unmodified | 0 | 0 | 0 | 0 | 0 | 0 | 0 | 0 | 0 | 0 | 0 | 0 | 0 | 0 | 0 | 0 | 572900 | 0 |
| KTGTLQPW | RACGAP1; RAC | Q9H0H5; HC | RGAP1_HUMAI | Unmodified | 0 | 0 | 0 | 0 | 367710 | 0 | 0 | 0 | 0 | 0 | 171630 | 0 | 281510 | 0 | 0 | 0 | 241810 | 122940 |
| KTGVAGED | PRKCD | Q05655 | KPCD_HUMAN | Unmodified | 0 | 0 | 0 | 0 | 0 | 0 | 0 | 0 | 0 | 0 | 0 | 0 | 0 | 0 | 0 | 0 | 130970 | 199110 |
| KTHILILATT | CHST1 | O43916 | CHST1_HUMAI | Unmodified | 0 | 0 | 0 | 0 | 0 | 0 | 0 | 0 | 486660 | 0 | 0 | 0 | 0 | 0 | 0 | 0 | 0 | 0 |
| KTHIQDNH | FLNA; FLNA | P21333; Q6( | FLNA_HUMAN | Unmodified | 933690 | 1505600 | 0 | 1919000 | 0 | 0 | 0 | 0 | 0 | 0 | 0 | 1443000 | 0 | 0 | 886810 | 1179300 | 0 | 0 |
| KTIEWAEEI | UBAP1; UBAP1 | Q9NZ09; A0/ | UBAP1_HUMAI | Unmodified | 1386800 | 1496100 | 0 | 0 | 0 | 0 | 0 | 0 | 0 | 0 | 0 | 0 | 0 | 0 | 0 | 0 | 0 | 0 |
| KTIQFDFQI | ITGA5 | P08648 | ITA5_HUMAN I | Unmodified | 0 | 0 | 0 | 48118 | 1020300 | 0 | 0 | 0 | 0 | 0 | 0 | 0 | 103990 | 0 | 0 | 0 | 393290 | 221790 |
| KTIQGDEEI | NAPA; NAPA; N. | P54920; M0I | SNAA_HUMAN | Unmodified | 659800 | 24643 | 216980 | 586190 | 0 | 181390 | 940850 | 640220 | 553860 | 528330 | 260610 | 1299600 | 1019300 | 1030400 | 384370 | 224090 | 625020 | 1000700 |
| KTKFENLCI | HSP90AA5P; H! | Q58FG0; P0 | HS905_HUMAI | Unmodified | 0 | 0 | 0 | 0 | 0 | 0 | 537420 | 0 | 0 | 694070 | 536200 | 1895700 | 0 | 0 | 0 | 0 | 0 | 2445700 |
| KTKPYIQVD | HSPA5; HSPA5; | P11021; A0A | BIP_HUMAN Er | Unmodified | 0 | 0 | 0 | 757140 | 2389100 | 0 | 28143000 | 15964000 | 1.4E+08 | 84923000 | 6495900 | 1850500 | 3091400 | 3980500 | 1586400 | 289420 | 715350 | 5480800 |
| KTKTENPLI | LONP1; LONP1 | P36776; K7E | LONM_HUMAN | Unmodified | 0 | 0 | 0 | 0 | 0 | 0 | 0 | 0 | 200480 | 0 | 0 | 0 | 0 | 0 | 0 | 0 | 0 | 0 |
| KTLAVLLDN | MGAT5 | Q09328 | MGT5A_HUMAI | Unmodified | 0 | 0 | 0 | 0 | 0 | 0 | 0 | 0 | 742630 | 85007 | 0 | 0 | 0 | 0 | 0 | 0 | 0 | 0 |
| KTLDEILEN | UTRN | P46939 | UTRN_HUMAN | Unmodified | 0 | 41573 | 0 | 967220 | 0 | 0 | 0 | 0 | 3111100 | 722920 | 0 | 0 | 573600 | 1379300 | 475080 | 1019000 | 1021600 | 1240300 |
| KTLEEAIRSI | ANXA11 | P50995 | ANX11_HUMAN | Unmodified | 456040 | 544690 | 0 | 1015900 | 941800 | 362630 | 0 | 38681 | 13333000 | 2321400 | 838020 | 1080600 | 187350 | 81856 | 428990 | 0 | 214370 | 1313400 |
| KTLEEEAKT | MYH9 | P35579 | MYH9_HUMAN | Unmodified | 0 | 0 | 0 | 0 | 1402100 | 0 | 0 | 0 | 0 | 0 | 0 | 0 | 0 | 0 | 0 | 0 | 27832 | 0 |
| KTLFEDSF( | ITCH; ITCH; ITC | Q96J02; A0A | ITCH_HUMAN I | Oxidation (M) | 736970 | 1628500 | 0 | 183670 | 0 | 0 | 0 | 0 | 0 | 0 | 0 | 35209 | 0 | 0 | 345940 | 136870 | 0 | 0 |
| KTLQALEFI | LGALS3BP | Q08380 | LG3BP_HUMAN | Unmodified | 0 | 0 | 0 | 0 | 0 | 794860 | 387040 | 183010 | 21337000 | 11063000 | 15284000 | 5632300 | 5945700 | 1939800 | 0 | 0 | 0 | 11404000 |
| KTLQALEFI | LGALS3BP | Q08380 | LG3BP_HUMAN | Unmodified | 0 | 0 | 0 | 0 | 0 | 0 | 750850 | 0 | 0 | 1923100 | 0 | 0 | 1499500 | 1617400 | 0 | 0 | 0 | 6827900 |
| KTNPFPLLE | WASHC2A; WA! | Q641Q2; Q9 | WAC2A_HUMA | Unmodified | 0 | 52318 | 0 | 407790 | 0 | 0 | 0 | 0 | 0 | 0 | 0 | 293560 | 281050 | 0 | 0 | 0 | 0 | 5216000 |
| KTPESFLGI | EPN1 | Q9Y6I3 | EPN1_HUMAN | Unmodified | 0 | 0 | 0 | 1079800 | 498330 | 0 | 612860 | 0 | 848370 | 0 | 501630 | 0 | 3981700 | 0 | 0 | 0 | 0 | 0 |
| KTPNFPSS\ | MYOF; MYOF | Q9NZM1; HC | MYOF_HUMAN | Unmodified | 0 | 0 | 0 | 0 | 3209400 | 0 | 0 | 0 | 0 | 230110 | 0 | 0 | 0 | 0 | 82114 | 0 | 180480 | 0 |
| KTPNFPSS\ | MYOF; MYOF | Q9NZM1; HC | MYOF_HUMAN | Oxidation (M) | 0 | 0 | 0 | 0 | 7067800 | 1183500 | 0 | 0 | 0 | 0 | 0 | 0 | 0 | 0 | 0 | 0 | 0 | 510240 |
| KTPNNPSC | CACNA2D1; CA | P54289; H0Y | CA2D1_HUMAI | Unmodified | 494200 | 223830 | 0 | 0 | 0 | 0 | 0 | 0 | 651420 | 146930 | 0 | 0 | 0 | 0 | 0 | 0 | 0 | 0 |
| KTPPAPSPI | KIF4A | O95239 | KIF4A_HUMAN | Unmodified | 0 | 0 | 0 | 0 | 0 | 0 | 89480 | 0 | 0 | 0 | 0 | 0 | 829970 | 0 | 0 | 0 | 929410 | 0 |
| KTPQGPPE | CDV3; CDV3; C | Q9UKY7; D6 | CDV3_HUMAN | Unmodified | 0 | 250620 | 0 | 0 | 1223900 | 0 | 0 | 0 | 609680 | 0 | 718420 | 652940 | 2482700 | 0 | 0 | 150750 | 0 | 933370 |
| KTPTEAPAI | HUWE1; HUWE | Q7Z6Z7; A0/ | HUWE1_HUMA | Unmodified | 0 | 0 | 0 | 0 | 0 | 0 | 0 | 0 | 165870 | 136810 | 0 | 115520 | 0 | 0 | 0 | 0 | 0 | 0 |
| KTPTFLGAT | ENTPD8; ENTPI | Q5MY95; H7 | ENTP8_HUMAN | Unmodified | 0 | 0 | 0 | 0 | 0 | 0 | 0 | 0 | 4109300 | 0 | 0 | 0 | 0 | 0 | 0 | 0 | 0 | 0 |
| KTPVKEPN! | RAB10 | P61026 | RAB10_HUMAN | Unmodified | 1969100 | 1484600 | 0 | 0 | 0 | 0 | 0 | 0 | 0 | 0 | 0 | 1930400 | 0 | 0 | 641410 | 0 | 0 | 0 |
| KTQDQISN | LASP1; LASP1 | Q14847; C9J | LASP1_HUMAN | Unmodified | 480210 | 1464600 | 0 | 309910 | 1487500 | 0 | 0 | 0 | 382850 | 244400 | 397350 | 258220 | 0 | 0 | 387120 | 181690 | 0 | 387920 |
| KTQDQISN | LASP1; LASP1 | Q14847; C9J | LASP1_HUMAN | Unmodified | 0 | 416460 | 0 | 0 | 6663100 | 870640 | 1001600 | 485510 | 0 | 766730 | 0 | 0 | 3369300 | 1544000 | 0 | 0 | 2624500 | 2531400 |
| KTQEQLALI | MSN | P26038 | MOES_HUMAN | Unmodified | 0 | 0 | 454940 | 1364100 | 39468000 | 6411600 | 0 | 0 | 0 | 0 | 0 | 0 | 3449400 | 2388600 | 923600 | 953590 | 3691600 | 14231000 |
| KTQEQLALI | MSN | P26038 | MOES_HUMAN | Oxidation (M) | 0 | 94189 | 328690 | 0 | 1483700 | 839040 | 0 | 0 | 0 | 0 | 0 | 0 | 644280 | 476480 | 518490 | 363850 | 367830 | 1080200 |
| KTQEQLALI | MSN | P26038 | MOES_HUMAN | Oxidation (M) | 0 | 0 | 0 | 0 | 22201000 | 2742400 | 0 | 0 | 0 | 0 | 0 | 0 | 8505400 | 5143400 | 0 | 0 | 241080 | 13529000 |
| KTQEQLALI | MSN | P26038 | MOES_HUMAN | Unmodified | 0 | 0 | 0 | 0 | 0 | 0 | 0 | 0 | 0 | 130290 | 0 | 0 | 0 | 123250 | 0 | 0 | 0 | 1261500 |
| KTQEQLALI | MSN | P26038 | MOES_HUMAN | 2 Oxidation (M) | 0 | 0 | 0 | 0 | 5610400 | 0 | 0 | 0 | 0 | 0 | 0 | 0 | 0 | 578790 | 0 | 0 | 0 | 3787900 |
| KTQETLSQ/ | TPD52L2; TPD5 | O43399; A0/ | TPD54_HUMAN | Unmodified | 0 | 0 | 0 | 0 | 280690 | 0 | 123380 | 125000 | 0 | 58586 | 26786 | 0 | 237760 | 100890 | 0 | 0 | 80491 | 137860 |
| KTQHGVLS | STX4 | Q12846 | STX4_HUMAN ! | Unmodified | 953310 | 0 | 0 | 0 | 0 | 0 | 0 | 0 | 2069500 | 705350 | 0 | 673440 | 584060 | 0 | 686170 | 640510 | 550730 | 0 |
| KTQNDVLH | RDX; RDX; RDX | P35241; A0A | RADI_HUMAN I | Unmodified | 78873 | 0 | 2746100 | 3898600 | 20771000 | 4442600 | 461350 | 495970 | 3367400 | 1201900 | 1929800 | 7906200 | 13468000 | 12996000 | 4502100 | 2940600 | 2794500 | 7386300 |
| KTQTVCNF | FABP5; FABP5 | Q01469; I6L! | FABP5_HUMAN | Unmodified | 0 | 0 | 0 | 0 | 0 | 0 | 0 | 0 | 0 | 0 | 1474400 | 0 | 0 | 0 | 0 | 3651700 | 0 | 0 |
| KTRLPLELE | TMEM237; TMEI | Q96Q45; A0/ | TM237_HUMAN | Unmodified | 0 | 0 | 0 | 0 | 604830 | 98020 | 38853 | 0 | 0 | 27870 | 0 | 0 | 0 | 152830 | 0 | 0 | 128290 | 208430 |
| KTSATVGPI | H2AX | P16104 | H2AX_HUMAN | Unmodified | 136740 | 560170 | 0 | 0 | 1049500 | 340920 | 301030 | 1689600 | 0 | 36776 | 0 | 0 | 0 | 0 | 0 | 0 | 0 | 1397500 |
| KTSEDASE\ | PARP4 | Q9UKK3 | PARP4_HUMAN | Unmodified | 0 | 0 | 0 | 0 | 0 | 0 | 0 | 0 | 0 | 0 | 0 | 699140 | 0 | 0 | 0 | 0 | 0 | 0 |
| KTSEVDLAI | PDCD6IP; PDCI | Q8WUM4; F! | PDC6I_HUMAN | Unmodified | 8156200 | 8292900 | 4231800 | 4402300 | 0 | 0 | 131370 | 97675 | 2868900 | 0 | 0 | 2018200 | 4068900 | 1122000 | 4499800 | 3535100 | 122510 | 2238600 |
| KTSEVDLAI | PDCD6IP; PDCI | Q8WUM4; F! | PDC6I_HUMAN | Unmodified | 1808900 | 649240 | 0 | 1017300 | 25066000 | 1212200 | 878700 | 436420 | 731490 | 3185900 | 0 | 207150 | 8189800 | 2951700 | 0 | 303750 | 1502300 | 2959200 |
| KTSSFLDQI | ECE1; ECE1; EC | P42892; B4D | ECE1_HUMAN | Unmodified | 0 | 0 | 0 | 0 | 0 | 0 | 0 | 0 | 2934800 | 534420 | 0 | 0 | 0 | 0 | 0 | 0 | 0 | 0 |
| KTSYAQHQ | RPS3A; RPS3A; | P61247; D6F | RS3A_HUMAN | Unmodified | 169960 | 278910 | 0 | 0 | 0 | 0 | 0 | 0 | 86543 | 0 | 0 | 2316500 | 0 | 0 | 228330 | 268380 | 0 | 0 |
| KTSYLTELII | STK26; STK26; ! | Q9P289; B4I | STK26_HUMAN | Unmodified | 0 | 0 | 0 | 0 | 1222700 | 683980 | 0 | 0 | 0 | 0 | 711590 | 721280 | 1915500 | 1055400 | 0 | 0 | 493040 | 1098900 |
| KTSYLTELII | STK24; STK24; ! | Q9Y6E0; H0 | STK24_HUMAN | Unmodified | 0 | 0 | 0 | 0 | 2897800 | 417030 | 0 | 0 | 0 | 403630 | 0 | 0 | 1168500 | 736100 | 0 | 0 | 502930 | 1791600 |
| KTTARDQD | EPRS1 | P07814 | SYEP_HUMAN | Unmodified | 0 | 0 | 0 | 0 | 1654300 | 464240 | 520110 | 0 | 0 | 331960 | 399090 | 552420 | 0 | 0 | 0 | 0 | 595650 | 1430400 |
| KTTDTASV( | CORO1C; COR( | Q9ULV4; H0 | COR1C_HUMA | Unmodified | 0 | 0 | 0 | 0 | 0 | 0 | 0 | 0 | 0 | 0 | 0 | 93090 | 0 | 0 | 13942 | 0 | 0 | 0 |
| KTTDTASV( | CORO1C; COR( | Q9ULV4; H0 | COR1C_HUMA | Unmodified | 214290 | 118880 | 0 | 114270 | 70726000 | 12442000 | 3517600 | 2776300 | 5025800 | 5230200 | 3801500 | 26184000 | 6794500 | 3682700 | 233930 | 136390 | 12183000 | 8378300 |
| KTTHFVEG( | RPL7; RPL7 | P18124; A8N | RL7_HUMAN 6( | Unmodified | 3132600 | 3677300 | 1396300 | 1228700 | 230040 | 0 | 0 | 88093 | 344160 | 48346 | 1546700 | 18126000 | 449020 | 0 | 2010500 | 1538300 | 0 | 0 |
| KTTPTPSTN | MARK2; MARK2 | Q7KZI7; E7E | MARK2_HUMAI | Unmodified | 0 | 0 | 0 | 196880 | 0 | 0 | 0 | 0 | 66359 | 0 | 127750 | 195730 | 347330 | 0 | 0 | 0 | 0 | 110790 |
| KTTTYTQG\ | JUP; JUP; JUP; J | P14923; C9J | PLAK_HUMAN . | Unmodified | 0 | 0 | 0 | 403120 | 1899100 | 0 | 0 | 0 | 563500 | 0 | 0 | 692870 | 244160 | 0 | 326830 | 444060 | 526490 | 0 |
| KTTTYTQG\ | JUP; JUP; JUP; J | P14923; C9J | PLAK_HUMAN . | Unmodified | 0 | 0 | 0 | 0 | 1855700 | 0 | 0 | 0 | 0 | 0 | 0 | 0 | 2682300 | 1626400 | 0 | 0 | 597730 | 0 |
| KTVAYHEA( | AFG3L2 | Q9Y4W6 | AFG32_HUMAN | Unmodified | 0 | 0 | 0 | 0 | 2683900 | 0 | 2465800 | 310200 | 1244500 | 578830 | 0 | 0 | 0 | 0 | 0 | 0 | 941930 | 0 |
| KTVTAMDV | H4C1 | P62805 | H4_HUMAN Hi: | Oxidation (M) | 6438000 | 3226400 | 0 | 0 | 0 | 0 | 0 | 633240 | 0 | 0 | 0 | 69428000 | 0 | 0 | 0 | 0 | 0 | 0 |
| KTVTAMDV | H4C1 | P62805 | H4_HUMAN Hi: | Unmodified | 3036900 | 1495700 | 0 | 0 | 0 | 0 | 0 | 0 | 0 | 0 | 0 | 32080000 | 0 | 400710 | 0 | 0 | 0 | 0 |
| KTVTAMDV | H4C1 | P62805 | H4_HUMAN Hi: | Oxidation (M) | 30776000 | 14720000 | 0 | 0 | 0 | 0 | 0 | 1055900 | 0 | 0 | 0 | 88948000 | 0 | 0 | 0 | 0 | 0 | 0 |
| KTVTAMDV | H4C1 | P62805 | H4_HUMAN Hi: | Unmodified | 26752000 | 15535000 | 0 | 0 | 0 | 0 | 0 | 0 | 0 | 0 | 0 | 1.39E+08 | 0 | 0 | 0 | 0 | 0 | 0 |
| KTYTLTDYL | DPP4; DPP4; DI | P27487; A0A | DPP4_HUMAN | Unmodified | 0 | 0 | 821420 | 434180 | 0 | 0 | 0 | 0 | 0 | 0 | 0 | 0 | 0 | 0 | 565390 | 524390 | 0 | 0 |
| KVAAIEALN | ST13P4; ST13; ! | Q8IZP2; P50 | ST134_HUMAN | Unmodified | 371940 | 453590 | 0 | 1597600 | 618960 | 0 | 0 | 192380 | 5292200 | 1424100 | 130350 | 12801000 | 4323100 | 1701800 | 598080 | 713490 | 236580 | 422010 |
| KVACIGAW | RPL3L; RPL3; RI | Q92901; P39 | RL3L_HUMAN ( | Unmodified | 0 | 440820 | 0 | 534630 | 0 | 0 | 0 | 0 | 74911 | 0 | 495910 | 499260 | 0 | 131820 | 0 | 0 | 194620 | 0 |
| KVAETAVQI | ETF1; ETF1; ETF | P62495; B7Z | ERF1_HUMAN I | Unmodified | 93046 | 117980 | 0 | 0 | 0 | 0 | 0 | 0 | 353700 | 0 | 85636 | 735450 | 0 | 0 | 0 | 274560 | 0 | 0 |
| KVAGMDVE | YWHAE; YWHAI | P62258; I3L3 | 1433E_HUMAN | Oxidation (M) | 0 | 0 | 0 | 0 | 0 | 0 | 0 | 0 | 506130 | 80577 | 937190 | 2045500 | 0 | 0 | 680360 | 898720 | 0 | 0 |
| KVAGMDVE | YWHAE; YWHAI | P62258; I3L3 | 1433E_HUMAN | Unmodified | 0 | 0 | 0 | 0 | 0 | 0 | 0 | 0 | 0 | 0 | 106860 | 2027800 | 0 | 0 | 0 | 261400 | 0 | 0 |
| KVCGEEGV | EPS8L2; EPS8L: | Q9H6S3; A0 | ES8L2_HUMAN | Unmodified | 0 | 0 | 0 | 0 | 478490 | 0 | 0 | 0 | 0 | 52718 | 0 | 0 | 216310 | 118580 | 0 | 0 | 0 | 0 |
| KVCIESEHS | ATOX1; ATOX1; | O00244; E5F | ATOX1_HUMAN | Unmodified | 0 | 0 | 0 | 630660 | 0 | 0 | 0 | 0 | 401260 | 0 | 0 | 0 | 2246100 | 922260 | 0 | 0 | 366300 | 1416400 |
| KVCIESEHS | ATOX1; ATOX1; | O00244; E5F | ATOX1_HUMAN | Unmodified | 0 | 0 | 0 | 224600 | 0 | 0 | 0 | 0 | 0 | 0 | 0 | 0 | 1072500 | 660220 | 0 | 348360 | 0 | 0 |
| KVDEAIALF | VARS1; VARS1 | P26640; A0A | SYVC_HUMAN | Unmodified | 19067 | 0 | 0 | 338610 | 0 | 0 | 0 | 0 | 0 | 690850 | 337250 | 3868700 | 0 | 0 | 0 | 0 | 0 | 0 |
| KVDIIADAA | HSDL2 | Q6YN16 | HSDL2_HUMAI | Unmodified | 0 | 0 | 0 | 0 | 169250 | 559280 | 0 | 0 | 1975100 | 0 | 0 | 0 | 0 | 0 | 0 | 0 | 11020 | 0 |
| KVDIIADAA | HSDL2 | Q6YN16 | HSDL2_HUMAI | Unmodified | 0 | 0 | 0 | 0 | 0 | 123460 | 0 | 54845 | 16264000 | 0 | 0 | 200740 | 0 | 0 | 0 | 0 | 0 | 0 |
| KVDKNDEA | NCAM1; NCAM | P13591; A0A | NCAM1_HUMA | Unmodified | 0 | 0 | 0 | 562110 | 0 | 0 | 100330 | 166340 | 1605500 | 89884 | 0 | 82491 | 0 | 76365 | 0 | 0 | 0 | 0 |
| KVDLSQPLI | TST | Q16762 | THTR_HUMAN | Unmodified | 0 | 0 | 0 | 0 | 0 | 0 | 0 | 0 | 674050 | 0 | 0 | 0 | 0 | 0 | 0 | 0 | 0 | 0 |

| | | | | | | | | | | | | | | | | | | | | | | |
|---|---|---|---|---|---|---|---|---|---|---|---|---|---|---|---|---|---|---|---|---|---|---|
| KVDNELNP | MYOF | Q9NZM1 | MYOF_HUMAN | Unmodified | 0 | 0 | 0 | 189970 | 332080 | 0 | 0 | 0 | 0 | 0 | 0 | 0 | 0 | 0 | 0 | 0 | 0 | 0 |
| KVDPVSSD | GPRIN1 | Q7Z2K8 | GRIN1_HUMAN | Unmodified | 0 | 0 | 0 | 0 | 0 | 0 | 0 | 0 | 958290 | 0 | 0 | 0 | 0 | 0 | 0 | 0 | 0 | 0 |
| KVDQGLFV | SDCBP2 | Q9H190 | SDCB2_HUMA | Unmodified | 0 | 28257 | 0 | 568850 | 0 | 0 | 0 | 0 | 235800 | 0 | 0 | 0 | 2623900 | 580490 | 411010 | 0 | 0 | 0 |
| KVDTHIHA | AMPD1; AMPD3 | P23109; Q01 | AMPD1_HUMA | Unmodified | 0 | 99708 | 0 | 0 | 0 | 0 | 551300 | 0 | 0 | 0 | 0 | 1200900 | 0 | 0 | 107030 | 79252 | 0 | 0 |
| KVDVDEYD | ARPC5; ARPC5 | O15511; B1A | ARPC5_HUMA | Unmodified | 1151200 | 395010 | 0 | 0 | 0 | 0 | 0 | 0 | 271750 | 0 | 207000 | 3048800 | 36322 | 0 | 325390 | 233390 | 0 | 0 |
| KVDVTSKA | SH3GL1; SH3G | Q99961; M0 | SH3G1_HUMA | Unmodified | 0 | 0 | 0 | 0 | 0 | 0 | 169420 | 101730 | 0 | 47461 | 0 | 0 | 839490 | 432270 | 0 | 0 | 0 | 0 |
| KVDWLTEK | EIF4A3; EIF4A1; | P38919; P60 | IF4A3_HUMAN | Unmodified | 909990 | 0 | 0 | 0 | 0 | 0 | 55345 | 0 | 0 | 148420 | 0 | 4328500 | 0 | 0 | 0 | 0 | 0 | 29387 |
| KVEAQLQE | MYH9 | P35579 | MYH9_HUMAN | Unmodified | 602980 | 696180 | 3597200 | 2662500 | 16110000 | 3219600 | 1316200 | 572110 | 2168900 | 1191100 | 1017800 | 7424200 | 4083200 | 3386000 | 3553700 | 3652200 | 4366600 | 7882200 |
| KVEDLFLTF | SPTAN1; SPTAN | Q13813; A0A | SPTN1_HUMA | Unmodified | 0 | 0 | 0 | 0 | 0 | 0 | 0 | 0 | 0 | 0 | 0 | 1181200 | 0 | 0 | 0 | 0 | 0 | 0 |
| KVEEAEPEE | CBX3; CBX3; CI | Q13185; S4F | CBX3_HUMAN | Unmodified | 341930 | 266350 | 0 | 0 | 0 | 0 | 0 | 0 | 982190 | 480160 | 1296600 | 8098400 | 0 | 0 | 0 | 171090 | 205140 | 0 |
| KVEEEEDES | MCM6 | Q14566 | MCM6_HUMAN | Unmodified | 0 | 0 | 0 | 0 | 0 | 0 | 0 | 37034 | 165920 | 28084 | 44767 | 1559400 | 0 | 13188 | 0 | 0 | 30738 | 37989 |
| KVEELEGEI | BST2 | Q10589 | BST2_HUMAN | Unmodified | 0 | 0 | 0 | 539090 | 0 | 0 | 0 | 0 | 0 | 0 | 0 | 207690 | 2191700 | 1182100 | 6806500 | 8025500 | 0 | 0 |
| KVEELEGEI | BST2 | Q10589 | BST2_HUMAN | Unmodified | 0 | 0 | 0 | 195730 | 0 | 0 | 0 | 0 | 0 | 0 | 0 | 0 | 594790 | 2615600 | 0 | 0 | 0 | 118580 |
| KVEEVVYDI | TSN; TSN | Q15631; E9F | TSN_HUMAN T | Unmodified | 0 | 0 | 38002 | 65217 | 0 | 0 | 0 | 142960 | 2166900 | 0 | 785090 | 11751000 | 0 | 0 | 88144 | 269360 | 34072 | 0 |
| KVEEVVYDI | TSN; TSN | Q15631; E9F | TSN_HUMAN T | Unmodified | 0 | 0 | 0 | 0 | 224610 | 0 | 182270 | 84764 | 0 | 145960 | 555680 | 0 | 548960 | 0 | 0 | 0 | 254350 | 362300 |
| KVEILENEI | CCDC88A; CCD | Q3V6T2; A0A | GRDN_HUMAN | Unmodified | 0 | 0 | 0 | 0 | 529710 | 0 | 165310 | 151620 | 252790 | 109980 | 0 | 312630 | 168020 | 0 | 0 | 0 | 959890 | 411330 |
| KVEITTETIK | PIGK; PIGK; PIG | Q92643; B1A | GPI8_HUMAN | Unmodified | 0 | 0 | 0 | 0 | 0 | 0 | 0 | 0 | 3917900 | 151340 | 192660 | 0 | 0 | 0 | 0 | 0 | 0 | 0 |
| KVELCSFS | RPL24; RPL24; | P83731; C9J | RL24_HUMAN | Acetyl (Protein N | 0 | 0 | 0 | 0 | 0 | 0 | 0 | 0 | 0 | 0 | 0 | 0 | 0 | 0 | 0 | 0 | 0 | 0 |
| KVELFHYQ | PDIA5; PDIA5 | Q14554; C9J | PDIA5_HUMAN | Unmodified | 0 | 0 | 0 | 0 | 0 | 95561 | 0 | 0 | 25680000 | 1428500 | 0 | 0 | 0 | 0 | 0 | 0 | 0 | 0 |
| KVELFHYQ | PDIA5; PDIA5 | Q14554; C9J | PDIA5_HUMAN | Unmodified | 0 | 0 | 0 | 0 | 0 | 0 | 0 | 0 | 0 | 4099700 | 0 | 0 | 0 | 0 | 0 | 0 | 1510000 | 0 |
| KVENEDMN | SEPTIN2; SEPTI | Q15019; H7C | SEPT2_HUMAN | Unmodified | 0 | 0 | 0 | 366890 | 3197000 | 964110 | 0 | 0 | 0 | 0 | 737060 | 1324600 | 1129500 | 0 | 0 | 0 | 235770 | 1126100 |
| KVENTRWD | CACYBP | Q9HB71 | CYBP_HUMAN | Unmodified | 0 | 0 | 0 | 0 | 0 | 638120 | 380620 | 78436 | 1304800 | 944570 | 1987500 | 5829000 | 4677900 | 1266300 | 0 | 0 | 525550 | 423390 |
| KVEQLQQE | TRIM25; TRIM25 | Q14258; A0A | TRI25_HUMAN | Unmodified | 0 | 0 | 0 | 125620 | 857860 | 121290 | 0 | 0 | 0 | 0 | 254700 | 253220 | 0 | 0 | 111970 | 0 | 103060 | 0 |
| KVEQLQQE | TRIM25; TRIM25 | Q14258; A0A | TRI25_HUMAN | Oxidation (M) | 0 | 0 | 0 | 0 | 0 | 0 | 0 | 0 | 0 | 0 | 0 | 283060 | 0 | 0 | 0 | 0 | 0 | 0 |
| KVESLQEEI | VIM; VIM; VIM | P08670; B0Y | VIME_HUMAN | Unmodified | 128890 | 135770 | 0 | 0 | 3919200 | 2550800 | 0 | 90759 | 0 | 0 | 0 | 0 | 0 | 0 | 0 | 0 | 767540 | 1952300 |
| KVESTGAVI | HSDL2 | Q6YN16 | HSDL2_HUMA | Unmodified | 0 | 0 | 0 | 0 | 0 | 0 | 0 | 0 | 9358100 | 1787900 | 0 | 0 | 0 | 0 | 0 | 0 | 0 | 0 |
| KVETDHIVA | AIFM1; AIFM1; A | O95831; E9F | AIFM1_HUMAN | Unmodified | 0 | 0 | 0 | 0 | 0 | 0 | 0 | 0 | 711080 | 0 | 0 | 1017300 | 0 | 0 | 0 | 0 | 0 | 0 |
| KVFASLPQ | WDR1; WDR1 | O75083; D6I | WDR1_HUMAN | Unmodified | 698650 | 692880 | 760260 | 707860 | 1619200 | 357980 | 0 | 31820 | 1329900 | 297850 | 918050 | 6217700 | 1022000 | 502210 | 750620 | 812850 | 0 | 0 |
| KVFASLPQ | WDR1; WDR1 | O75083; D6I | WDR1_HUMAN | Unmodified | 0 | 0 | 0 | 219300 | 28748000 | 4915700 | 4387600 | 3006100 | 0 | 2482600 | 462400 | 0 | 19172000 | 15621000 | 0 | 0 | 6225300 | 10569000 |
| KVFSDVME | RRM1 | P23921 | RIR1_HUMAN | Unmodified | 0 | 0 | 0 | 0 | 0 | 0 | 0 | 0 | 0 | 14311 | 0 | 101810 | 0 | 0 | 0 | 0 | 0 | 0 |
| KVFVGGLS | HNRNPDL; HNI | O14979; A0A | HNRDL_HUMA | Unmodified | 0 | 0 | 0 | 0 | 0 | 0 | 0 | 74423 | 0 | 0 | 426030 | 1034700 | 0 | 0 | 0 | 0 | 0 | 0 |
| KVGAENVA | SQOR; SQOR; S | Q9Y6N5; H3 | SQOR_HUMAN | Unmodified | 0 | 0 | 0 | 0 | 0 | 0 | 0 | 42425 | 2443900 | 696380 | 0 | 0 | 0 | 1302500 | 0 | 0 | 0 | 0 |
| KVGAERNV | HSPA8; HSPA8; | P11142; E9P | HSP7C_HUMA | Unmodified | 0 | 0 | 0 | 0 | 108440 | 26807 | 0 | 0 | 0 | 112850 | 0 | 0 | 0 | 877800 | 0 | 0 | 0 | 116950 |
| KVGIRETPE | FARSB | Q9NSD9 | SYFB_HUMAN | Unmodified | 0 | 0 | 0 | 0 | 371640 | 58758 | 316780 | 173660 | 0 | 210630 | 45174 | 0 | 1053400 | 516840 | 0 | 0 | 192740 | 323050 |
| KVGPGYLG | RET | P07949 | RET_HUMAN P | Unmodified | 0 | 0 | 26015 | 0 | 891430 | 91484 | 0 | 19164 | 6425200 | 67950 | 0 | 0 | 213900 | 0 | 0 | 11971 | 418180 | 382910 |
| KVGYTPDW | CLTCL1; CLTC; | P53675; Q00 | CLH2_HUMAN | Unmodified | 14887000 | 18797000 | 2167000 | 3295100 | 775660 | 300250 | 409130 | 0 | 1025600 | 2230500 | 491870 | 37310000 | 760910 | 155500 | 9860500 | 6717700 | 790140 | 0 |
| KVHVIFNYK | CALR; CALR; CA | P27797; A0A | CALR_HUMAN | Unmodified | 0 | 0 | 0 | 0 | 439590 | 0 | 0 | 903510 | 23081000 | 20338000 | 2406000 | 930380 | 900650 | 0 | 0 | 0 | 135540 | 272620 |
| KVHVIFNYK | CALR; CALR; CA | P27797; A0A | CALR_HUMAN | Unmodified | 0 | 0 | 0 | 0 | 147560 | 0 | 1074400 | 317940 | 292140 | 2103300 | 0 | 0 | 0 | 0 | 0 | 0 | 0 | 140230 |
| KVIDDTNIT | KRT18; KRT18 | P05783; F8V | K1C18_HUMAN | Unmodified | 775240 | 0 | 521500 | 828450 | 0 | 0 | 0 | 0 | 0 | 0 | 0 | 544010 | 606890 | 0 | 668850 | 446470 | 0 | 0 |
| KVIEDRDH | MYOF; MYOF | Q9NZM1; HC | MYOF_HUMAN | Unmodified | 0 | 0 | 0 | 0 | 23406000 | 3522400 | 0 | 0 | 0 | 0 | 0 | 0 | 0 | 0 | 0 | 0 | 0 | 1245200 |
| KVIIDDQLP | CAPN7 | Q9Y6W3 | CAN7_HUMAN | Unmodified | 0 | 0 | 0 | 0 | 0 | 0 | 0 | 0 | 0 | 0 | 0 | 591290 | 0 | 0 | 0 | 0 | 922360 | 0 |
| KVIPHFTAT | PDIA5; PDIA5 | Q14554; H7C | PDIA5_HUMAN | Unmodified | 0 | 0 | 0 | 0 | 0 | 0 | 0 | 0 | 699010 | 0 | 0 | 0 | 0 | 0 | 0 | 0 | 0 | 0 |
| KVIQDCED | ITGA1 | P56199 | ITA1_HUMAN I | Unmodified | 146120 | 0 | 0 | 0 | 253700 | 0 | 0 | 46205 | 0 | 0 | 0 | 0 | 0 | 0 | 0 | 0 | 0 | 347350 |
| KVIQWCTH | SKP1; SKP1; SK | P63208; E7E | SKP1_HUMAN | Unmodified | 183800 | 0 | 0 | 74993 | 665150 | 0 | 0 | 0 | 190650 | 0 | 162730 | 967770 | 0 | 199870 | 0 | 0 | 0 | 277030 |
| KVIQYLAH | MYH10 | P35580 | MYH10_HUMA | Unmodified | 29382 | 104740 | 0 | 260890 | 2169200 | 163800 | 624390 | 0 | 0 | 448930 | 0 | 1212200 | 1366500 | 1934200 | 267260 | 0 | 0 | 743370 |
| KVIQYLAH | MYH14; MYH14 | Q7Z406; M0 | MYH14_HUMA | Unmodified | 54753 | 39156 | 0 | 38670 | 361420 | 0 | 0 | 0 | 279410 | 73205 | 0 | 0 | 295710 | 197840 | 0 | 30041 | 43193 | 112440 |
| KVIQYLAYV | MYH9; MYH9 | P35579; Q5E | MYH9_HUMAN | Unmodified | 0 | 0 | 1011000 | 820660 | 12796000 | 2540900 | 639950 | 0 | 778520 | 585900 | 430580 | 5388300 | 1383700 | 1254800 | 1066400 | 1077400 | 4995900 | 1729400 |
| KVIYPTDSR | SLC44A2; SLC4 | Q8IWA5; K7E | CTL2_HUMAN | Unmodified | 149060 | 384800 | 80039 | 140050 | 0 | 0 | 0 | 0 | 185440 | 0 | 0 | 706150 | 0 | 0 | 113480 | 102960 | 0 | 0 |
| KVIYSQPSA | F11R; F11R | Q9Y624; A0A | JAM1_HUMAN | Unmodified | 1262800 | 1192800 | 958740 | 756450 | 100810 | 0 | 0 | 0 | 1356700 | 264650 | 250570 | 1889300 | 0 | 0 | 947680 | 1307100 | 0 | 0 |
| KVIYSQPSA | F11R; F11R | Q9Y624; A0A | JAM1_HUMAN | Unmodified | 637910 | 0 | 71217 | 35478 | 2796500 | 282500 | 0 | 0 | 544200 | 340610 | 104580 | 0 | 1590700 | 302040 | 0 | 0 | 142380 | 668300 |
| KVKYEETVF | NAMPT; NAMPT | P43490; A0A | NAMPT_HUMA | Unmodified | 0 | 0 | 0 | 0 | 715520 | 0 | 0 | 0 | 0 | 0 | 0 | 0 | 0 | 0 | 0 | 0 | 0 | 316840 |
| KVLDHDVS | NF2; NF2 | P35240; A0A | MERL_HUMAN | Unmodified | 0 | 0 | 0 | 0 | 1054600 | 268190 | 0 | 0 | 0 | 0 | 0 | 0 | 0 | 481330 | 0 | 0 | 333950 | 817940 |
| KVLFEDPLF | CAPN5; CAPN5 | O15484; E7E | CAN5_HUMAN | Unmodified | 0 | 0 | 70366 | 165040 | 121860 | 0 | 0 | 0 | 228730 | 0 | 0 | 143150 | 445770 | 0 | 0 | 159930 | 0 | 0 |
| KVLHMDRN | GDI2; GDI2; GD | P50395; V9C | GDIB_HUMAN | Unmodified | 0 | 0 | 0 | 0 | 1331700 | 0 | 0 | 0 | 0 | 279870 | 0 | 0 | 0 | 0 | 0 | 0 | 0 | 0 |
| KVLINLFVG | MME; MME; MM | P08473; A0A | NEP_HUMAN | Unmodified | 0 | 0 | 0 | 0 | 0 | 0 | 0 | 0 | 642170 | 0 | 0 | 0 | 526780 | 0 | 0 | 418590 | 0 | 0 |
| KVLINLFVG | MME; MME; MM | P08473; A0A | NEP_HUMAN | Unmodified | 0 | 0 | 0 | 6939900 | 0 | 0 | 0 | 0 | 1293400 | 0 | 0 | 0 | 4411700 | 1122600 | 0 | 0 | 0 | 0 |
| KVLVDNPK | ITGB4 | P16144 | ITB4_HUMAN I | Unmodified | 0 | 0 | 0 | 0 | 0 | 0 | 0 | 0 | 0 | 0 | 0 | 0 | 0 | 0 | 0 | 0 | 2467400 | 1288000 |
| KVLVEQTK | SLITRK6 | Q9H5Y7 | SLIK6_HUMAN | Unmodified | 0 | 0 | 0 | 0 | 0 | 0 | 0 | 0 | 0 | 2816500 | 0 | 0 | 0 | 0 | 0 | 0 | 87790 | 0 |
| KVMGVLDI | MYO1A; MYO1A | Q9UBC5; G3 | MYO1A_HUMA | Unmodified | 0 | 0 | 0 | 0 | 675240 | 0 | 1440100 | 0 | 7139300 | 17903000 | 0 | 0 | 0 | 0 | 0 | 0 | 0 | 0 |
| KVMSQEIQ | DYNC1H1; DYN | Q14204; A0A | DYHC1_HUMA | Unmodified | 0 | 163630 | 0 | 0 | 0 | 0 | 0 | 0 | 370240 | 25282 | 84980 | 907590 | 0 | 0 | 0 | 0 | 0 | 20379 |
| KVMVIVTD | ITGA1 | P56199 | ITA1_HUMAN I | Unmodified | 0 | 0 | 0 | 0 | 0 | 0 | 0 | 0 | 0 | 0 | 0 | 0 | 0 | 0 | 0 | 0 | 2347800 | 4150100 |
| KVMVLDFV | TXNRD1; TXNRI | Q16881; A0A | TRXR1_HUMAN | Oxidation (M) | 101400 | 106030 | 128910 | 270650 | 807990 | 201210 | 0 | 48752 | 1920800 | 353800 | 1294300 | 5503300 | 670020 | 0 | 397570 | 371220 | 334930 | 252820 |
| KVNAEGSV | AK1; AK1 | P00568; Q5T | KAD1_HUMAN | Unmodified | 0 | 0 | 0 | 0 | 0 | 0 | 0 | 0 | 0 | 2197500 | 0 | 240730 | 1711700 | 0 | 0 | 0 | 0 | 0 |
| KVNAVFEW | HSD17B4; HSD | P51659; E7E | DHB4_HUMAN | Unmodified | 73343 | 0 | 16164 | 0 | 0 | 0 | 0 | 0 | 1671900 | 208320 | 0 | 0 | 0 | 0 | 0 | 0 | 0 | 0 |
| KVNELRAF | ST13P4; ST13; | Q8IZP2; P50 | ST134_HUMAN | Unmodified | 0 | 0 | 0 | 0 | 0 | 0 | 0 | 0 | 0 | 0 | 0 | 0 | 1431600 | 958050 | 0 | 0 | 0 | 0 |
| KVNESSLN | CNN3; CNN3 | Q15417; E9F | CNN3_HUMAN | Unmodified | 149650 | 0 | 18183 | 0 | 0 | 1582800 | 0 | 0 | 758920 | 1017300 | 465610 | 3215200 | 0 | 0 | 0 | 0 | 3407000 | 0 |
| KVNVTVDY | SND1 | Q7KZF4 | SND1_HUMAN | Unmodified | 758040 | 0 | 0 | 4393700 | 0 | 0 | 0 | 0 | 2942600 | 0 | 0 | 17572000 | 1257600 | 0 | 4928700 | 3101600 | 0 | 0 |
| KVPDKLLD | PSMA6; PSMA6 | P60900; G3 | PSA6_HUMAN | Unmodified | 0 | 0 | 0 | 0 | 1903300 | 0 | 1184700 | 0 | 0 | 842150 | 0 | 0 | 2429800 | 1423700 | 0 | 0 | 1280500 | 998790 |
| KVPEFQFLI | MPI; MPI; MPI; | P34949; H3E | MPI_HUMAN M | Unmodified | 1518200 | 5322600 | 0 | 0 | 1415800 | 0 | 0 | 0 | 0 | 0 | 373380 | 2229700 | 491580 | 292660 | 0 | 0 | 164790 | 0 |
| KVPEPEAIP | ATP2B1; ATP2B | P20020; E7E | AT2B1_HUMAN | Unmodified | 0 | 0 | 0 | 210340 | 252850 | 0 | 0 | 0 | 0 | 0 | 0 | 0 | 1003800 | 0 | 798400 | 0 | 0 | 0 |
| KVPGVTAIE | FXR1; FXR1; FXI | P51114; E7E | FXR1_HUMAN | Unmodified | 147140 | 0 | 0 | 0 | 0 | 0 | 0 | 0 | 0 | 0 | 0 | 0 | 0 | 0 | 0 | 142130 | 0 | 0 |
| KVPILVAQE | PTGES2; PTGES | Q9H7Z7; A0A | PGES2_HUMA | Unmodified | 0 | 0 | 0 | 0 | 0 | 0 | 311710 | 0 | 0 | 0 | 641800 | 0 | 0 | 0 | 0 | 0 | 0 | 0 |
| KVPIVQPSF | DST; DST; DST; | Q03001; A0A | DYST_HUMAN | Unmodified | 0 | 0 | 0 | 0 | 1634100 | 0 | 0 | 32280 | 0 | 30371 | 0 | 0 | 0 | 0 | 0 | 0 | 112720 | 0 |
| KVPLDERIV | NIBAN2 | Q96TA1 | NIBA2_HUMAN | Unmodified | 0 | 0 | 0 | 0 | 1055200 | 0 | 0 | 0 | 0 | 0 | 0 | 0 | 0 | 0 | 0 | 0 | 260480 | 0 |
| KVPSFTFTP | ROBO1; ROBO1 | Q9Y6N7; A0A | ROBO1_HUMA | Unmodified | 0 | 0 | 0 | 0 | 0 | 0 | 0 | 39405 | 1567900 | 0 | 0 | 317620 | 0 | 0 | 0 | 0 | 74354 | 301710 |
| KVPSSLQTI | KIAA1109; KIAA | Q2LD37; A0A | K1109_HUMAN | Unmodified | 0 | 0 | 0 | 0 | 0 | 0 | 0 | 0 | 0 | 0 | 430450 | 2575900 | 0 | 0 | 0 | 0 | 30491 | 0 |
| KVPTVSRP | SEC22B | O75396 | SC22B_HUMA | Unmodified | 55689 | 0 | 0 | 1114900 | 83430 | 0 | 438880 | 0 | 1129100 | 481810 | 417480 | 1870000 | 2542500 | 2156400 | 644170 | 309290 | 0 | 205670 |
| KVPVVQAY | NT5E; NT5E | P21589; H0Y | 5NTD_HUMAN | Unmodified | 0 | 0 | 0 | 0 | 0 | 0 | 0 | 0 | 0 | 0 | 0 | 0 | 0 | 0 | 0 | 0 | 0 | 0 |
| KVQFAKDE | ELFN2; ELFN1 | Q5R3F8; P0 | PPR29_HUMA | Unmodified | 0 | 0 | 0 | 0 | 1926300 | 0 | 0 | 0 | 524980 | 315300 | 0 | 0 | 0 | 0 | 0 | 0 | 0 | 0 |
| KVQGGALE | CCT7 | Q99832 | TCPH_HUMAN | Unmodified | 0 | 0 | 355160 | 0 | 0 | 0 | 0 | 0 | 0 | 0 | 0 | 13476000 | 561580 | 599730 | 0 | 0 | 0 | 0 |
| KVQLEHLE | PLCB4; PLCB4 | Q15147; A0A | PLCB4_HUMA | Unmodified | 0 | 0 | 292950 | 364290 | 167750 | 0 | 0 | 0 | 0 | 0 | 0 | 0 | 0 | 0 | 388860 | 527900 | 96802 | 1839200 |
| KVQLEHLE | PLCB4 | A0A7P0MRI8 | A0A7P0MRI8_H | Unmodified | 0 | 0 | 0 | 0 | 0 | 0 | 0 | 0 | 0 | 0 | 0 | 0 | 0 | 0 | 0 | 0 | 1315800 | 2327000 |
| KVQLVTDR | P4HTM; P4HTM | Q9NXG6; A0 | P4HTM_HUMA | Unmodified | 0 | 0 | 0 | 0 | 430810 | 0 | 0 | 0 | 4770000 | 253350 | 0 | 0 | 0 | 0 | 0 | 0 | 0 | 0 |
| KVQQELSR | GSS; GSS; GSS | P48637; A0A | GSHB_HUMAN | 2 Oxidation (M) | 0 | 1073900 | 0 | 0 | 0 | 0 | 0 | 0 | 0 | 0 | 0 | 2082800 | 0 | 0 | 0 | 0 | 0 | 0 |
| KVQSDGQI | DHX9 | Q08211 | DHX9_HUMAN | Unmodified | 1419200 | 9.39E+08 | 4013200 | 1.84E+09 | 0 | 0 | 1277300 | 119730 | 0 | 177450 | 0 | 660830 | 5985200 | 3723400 | 25011000 | 1.03E+09 | 0 | 0 |
| KVQVAALQ | TJP2; TJP2; TJP2 | Q9UDY2; A0 | ZO2_HUMAN T | Unmodified | 0 | 0 | 0 | 0 | 0 | 754620 | 0 | 0 | 516400 | 0 | 0 | 0 | 0 | 0 | 0 | 62382 | 0 | 0 |
| KVSDVTSS | FAM131B | Q86XD5 | F131B_HUMAN | Unmodified | 0 | 0 | 0 | 0 | 0 | 0 | 0 | 0 | 1452000 | 0 | 0 | 0 | 0 | 0 | 0 | 0 | 0 | 933380 |
| KVSEGVEQ | CNOT3; CNOT3 | O75175; B7Z | CNOT3_HUMA | Unmodified | 58500 | 0 | 0 | 80118 | 1075300 | 455330 | 92519 | 19099 | 396290 | 98832 | 272640 | 729810 | 307720 | 716530 | 0 | 0 | 405220 | 256740 |
| KVSGVGGT | ZDHHC5; ZDHI | Q9C0B5; Q9 | ZDHC5_HUMA | Unmodified | 0 | 0 | 794350 | 893430 | 4863000 | 0 | 0 | 0 | 861890 | 758380 | 360850 | 1382200 | 3547900 | 3558700 | 0 | 1318500 | 5507100 | 5676500 |
| KVSKQEEA | VASP; VASP | P50552; K7E | VASP_HUMAN | Unmodified | 0 | 0 | 0 | 0 | 0 | 0 | 0 | 0 | 13405 | 0 | 147470 | 203020 | 0 | 0 | 0 | 0 | 45601 | 0 |
| KVSLLQNE | TRAF3; TRAF3; | Q13114; A6N | TRAF3_HUMAN | Unmodified | 0 | 0 | 0 | 0 | 10383000 | 2704900 | 861430 | 1587500 | 349010 | 4432700 | 0 | 0 | 0 | 0 | 0 | 0 | 944060 | 1001100 |
| KVSMSSSE | AHNAK2 | Q8IVF2 | AHNK2_HUMA | Unmodified | 0 | 0 | 0 | 0 | 571160 | 0 | 0 | 0 | 0 | 0 | 0 | 0 | 0 | 0 | 0 | 0 | 231520 | 572200 |
| KVSPHSGV | FBLN1 | B1AHL2 | B1AHL2_HUMA | Unmodified | 0 | 0 | 0 | 0 | 0 | 0 | 0 | 52156 | 1145400 | 0 | 0 | 0 | 0 | 0 | 0 | 0 | 0 | 0 |
| KVSQNSQL | ROCK1; ROCK1 | Q13464; A0A | ROCK1_HUMA | Unmodified | 0 | 0 | 0 | 0 | 395540 | 0 | 0 | 231710 | 76655 | 0 | 0 | 0 | 0 | 0 | 0 | 0 | 191590 | 193970 |
| KVSQNSQL | ROCK1; ROCK1 | Q13464; A0A | ROCK1_HUMA | Unmodified | 0 | 0 | 0 | 0 | 0 | 0 | 0 | 174050 | 0 | 0 | 0 | 0 | 0 | 0 | 0 | 0 | 194680 | 0 |
| KVSQPIEGI | CLTC; CLTC | Q00610; A0A | CLH1_HUMAN | Unmodified | 7952200 | 3201600 | 2928800 | 3699700 | 88456 | 0 | 87696 | 0 | 363890 | 0 | 0 | 12038000 | 206970 | 0 | 3401400 | 6505200 | 0 | 0 |
| KVSQQILEK | ACAD9; ACAD9 | Q9H845; H0 | ACAD9_HUMA | Unmodified | 0 | 0 | 0 | 0 | 101250 | 0 | 0 | 0 | 421190 | 36637 | 24582 | 229760 | 0 | 1095900 | 0 | 0 | 0 | 671690 |
| KVSSQRFE | PTPRF; PTPRF | P10586; A2A | PTPRF_HUMAN | Unmodified | 391720 | 658420 | 0 | 197440 | 1055700 | 0 | 0 | 0 | 2228600 | 283650 | 0 | 0 | 0 | 0 | 135810 | 98451 | 397440 | 0 |
| KVSVYLED | MPZL2 | O60487 | MPZL2_HUMA | Unmodified | 0 | 0 | 0 | 0 | 0 | 0 | 0 | 0 | 0 | 0 | 408340 | 0 | 2433600 | 0 | 0 | 0 | 171220 | 0 |
| KVSYLPFTE | SELENON; SELI | Q9NZV5; H9 | SELN_HUMAN | Unmodified | 0 | 0 | 0 | 0 | 0 | 0 | 0 | 0 | 919820 | 0 | 0 | 0 | 0 | 0 | 0 | 0 | 41209 | 0 |
| KVTAMQLY | DSP | P15924 | DESP_HUMAN | Unmodified | 0 | 0 | 0 | 0 | 0 | 0 | 0 | 0 | 0 | 0 | 0 | 0 | 0 | 0 | 0 | 0 | 539270 | 0 |
| KVTEEDFYK | CEL | P19835 | CEL_HUMAN B | Unmodified | 0 | 0 | 0 | 0 | 0 | 0 | 0 | 0 | 2709700 | 91661 | 2053000 | 0 | 0 | 0 | 0 | 0 | 0 | 0 |
| KVTEEDFYK | CEL | P19835 | CEL_HUMAN B | Unmodified | 0 | 0 | 0 | 0 | 0 | 0 | 0 | 0 | 326160 | 1688400 | 52336 | 0 | 0 | 0 | 0 | 0 | 0 | 0 |
| KVTELLRTF | STXBP2; STXBP | Q15833; M0I | STXB2_HUMAN | Unmodified | 0 | 0 | 0 | 0 | 2471700 | 0 | 0 | 0 | 0 | 256510 | 0 | 0 | 896300 | 0 | 0 | 0 | 1435500 | 2345700 |
| KVTGTLDAI | FSCN1 | Q16658 | FSCN1_HUMA | Unmodified | 0 | 0 | 615260 | 0 | 2090900 | 765290 | 0 | 0 | 488600 | 0 | 1093700 | 0 | 0 | 0 | 0 | 0 | 1005300 | 1391200 |
| KVTHAVVT | HSPA5; HSPA5; | P11021; A0A | BIP_HUMAN Er | Unmodified | 264280 | 202050 | 7064600 | 4305700 | 2736400 | 152830 | 1922900 | 700440 | 3.12E+08 | 25353000 | 25128000 | 12108000 | 1573800 | 1152200 | 11506000 | 2824100 | 0 | 1099500 |
| KVTHAVVT | HSPA5; HSPA5; | P11021; A0A | BIP_HUMAN Er | Unmodified | 0 | 0 | 0 | 302170 | 19516000 | 2799300 | 54022000 | 22675000 | 15013000 | 2.02E+08 | 6659200 | 0 | 8658300 | 18295000 | 0 | 0 | 2652500 | 15616000 |
| KVTLSVLSF | PIR | O00625 | PIR_HUMAN Pi | Unmodified | 0 | 0 | 0 | 180030 | 3799800 | 265720 | 304810 | 62504 | 0 | 812330 | 1971400 | 433080 | 1754700 | 2499600 | 0 | 0 | 1594800 | 2212400 |
| KVTQLDLD | PDAP1 | Q13442 | HAP28_HUMA | Unmodified | 0 | 0 | 685110 | 0 | 1017400 | 0 | 0 | 0 | 566080 | 0 | 0 | 0 | 0 | 0 | 0 | 0 | 0 | 709400 |

| | | | | | | | | | | | | | | | | | | | | | | |
|---|---|---|---|---|---|---|---|---|---|---|---|---|---|---|---|---|---|---|---|---|---|---|
| KVTSVVFHI | PRPF19 | Q9UMS4 | PRP19_HUMAN | Unmodified | 0 | 0 | 0 | 0 | 0 | 0 | 0 | 0 | 867440 | 0 | 516220 | 9516900 | 400280 | 0 | 0 | 0 | 1209800 | 0 |
| KVTTTVHNI | OSBP | P22059 | OSBP1_HUMAN | Unmodified | 47735 | 0 | 38274 | 33738 | 33117 | 0 | 0 | 19096 | 0 | 100990 | 119100 | 1629600 | 0 | 0 | 239800 | 219030 | 78980 | 0 |
| KVTVGVIGS | STEAP2; STEAP | Q8NFT2; C9 | STEA2_HUMAN | Unmodified | 1900500 | 1992400 | 7008200 | 8338000 | 781720 | 0 | 0 | 903310 | 0 | 0 | 0 | 814960 | 783250 | 0 | 14246000 | 9486200 | 0 | 0 |
| KVVATTQM | ANPEP | P15144 | AMPN_HUMAN | Unmodified | 0 | 0 | 0 | 0 | 43086 | 0 | 0 | 0 | 0 | 0 | 30463 | 0 | 0 | 0 | 0 | 0 | 2121900 | 1205300 |
| KVVATTQM | ANPEP | P15144 | AMPN_HUMAN | Unmodified | 0 | 0 | 0 | 0 | 0 | 0 | 0 | 0 | 0 | 0 | 0 | 0 | 154520 | 0 | 0 | 0 | 10663000 | 17194000 |
| KVVATTQM | ANPEP | P15144 | AMPN_HUMAN | Oxidation (M) | 0 | 0 | 0 | 0 | 102680 | 19359 | 0 | 0 | 0 | 0 | 0 | 0 | 160030 | 0 | 0 | 0 | 976980 | 3070400 |
| KVVDNTTA | RAB1C; RAB1B; | Q92928; Q9 | RAB1C_HUMAN | Unmodified | 0 | 0 | 0 | 0 | 12246000 | 304830 | 1759500 | 47883 | 0 | 3507200 | 0 | 0 | 8272200 | 2290800 | 0 | 0 | 4276400 | 1469800 |
| KVVDYSQF | NUCKS1 | Q9H1E3 | NUCKS_HUMA | Unmodified | 0 | 0 | 0 | 0 | 328850 | 0 | 160670 | 0 | 507370 | 598990 | 0 | 0 | 0 | 0 | 0 | 0 | 581610 | 0 |
| KVVDYTTA | RAB1A; RAB1A | P62820; E7E | RAB1A_HUMAN | Unmodified | 812450 | 0 | 1094700 | 1450800 | 0 | 0 | 0 | 1153200 | 1687400 | 0 | 1484000 | 8016200 | 3642400 | 0 | 837960 | 887990 | 0 | 0 |
| KVVDYTTA | RAB1A; RAB1A | P62820; E7E | RAB1A_HUMAN | Unmodified | 0 | 385300 | 0 | 0 | 6839900 | 531190 | 5253600 | 0 | 0 | 1723700 | 0 | 0 | 5286700 | 5296000 | 0 | 0 | 4052800 | 4683200 |
| KVVEINHE | SLC39A10; SLC | Q9ULF5; E7E | S39AA_HUMAN | Unmodified | 0 | 0 | 0 | 0 | 1381300 | 0 | 0 | 0 | 0 | 0 | 0 | 0 | 0 | 0 | 0 | 0 | 326810 | 1063300 |
| KVVENGAL | DAG1 | Q14118 | DAG1_HUMAN | Unmodified | 0 | 0 | 309710 | 0 | 0 | 0 | 0 | 0 | 544580 | 80263 | 62306 | 0 | 0 | 0 | 57368 | 36094 | 0 | 0 |
| KVVETEDA | RAB35; RAB35 | Q15286; F5H | RAB35_HUMAN | Unmodified | 926030 | 647020 | 824550 | 449740 | 0 | 0 | 0 | 0 | 0 | 28401 | 358590 | 1482200 | 0 | 0 | 787280 | 981910 | 0 | 0 |
| KVVGCSCV | RPS12 | P25398 | RS12_HUMAN | Unmodified | 0 | 0 | 0 | 154510 | 600850 | 100480 | 125680 | 59993 | 0 | 240740 | 636200 | 1746000 | 1623200 | 409300 | 0 | 0 | 272410 | 871970 |
| KVVKQASE | GAPDH; GAPDH | P04406; E7E | G3P_HUMAN G | Unmodified | 0 | 0 | 0 | 0 | 0 | 0 | 5187100 | 1452700 | 0 | 0 | 0 | 73039 | 0 | 0 | 0 | 0 | 9589200 | 77313 |
| KVVLEAPDI | PTRHD1 | Q6GMV3 | PTRD1_HUMAN | Unmodified | 0 | 0 | 0 | 0 | 1526300 | 0 | 0 | 0 | 0 | 0 | 0 | 0 | 94273 | 0 | 0 | 0 | 613350 | 0 |
| KVVLLTGET | SNRNP200 | O75643 | U520_HUMAN | Unmodified | 0 | 0 | 0 | 0 | 0 | 0 | 0 | 0 | 384440 | 0 | 0 | 2065900 | 0 | 0 | 0 | 0 | 0 | 0 |
| KVVNPLFEI | RPL7A; RPL7A | P62424; Q5 | RL7A_HUMAN | Unmodified | 1657200 | 1644300 | 763720 | 431620 | 276590 | 0 | 0 | 239280 | 149860 | 106130 | 606600 | 2699100 | 844040 | 206200 | 1118300 | 1528200 | 0 | 0 |
| KVVPCLVT | FLNB; FLNB | O75369; E7E | FLNB_HUMAN | Unmodified | 0 | 0 | 0 | 0 | 1316100 | 250220 | 512310 | 0 | 0 | 0 | 0 | 0 | 804820 | 0 | 0 | 0 | 593090 | 263980 |
| KVVPCRHD | CHORDC1; CH | Q9UHD1; E9 | CHRD1_HUMA | Unmodified | 0 | 0 | 0 | 0 | 364350 | 0 | 128430 | 0 | 311570 | 730210 | 0 | 0 | 1564600 | 822580 | 0 | 0 | 1013600 | 964650 |
| KVVQSLEQ | TSN; TSN; TSN; | Q15631; H7 | TSN_HUMAN T | Unmodified | 0 | 0 | 0 | 0 | 0 | 0 | 54341 | 0 | 34072 | 0 | 0 | 1247600 | 0 | 0 | 0 | 0 | 0 | 31259 |
| KVVVCDN | ACTR2; ACTR2; | P61160; A0A | ARP2_HUMAN | Unmodified | 0 | 0 | 0 | 0 | 0 | 0 | 437530 | 197710 | 0 | 363710 | 355250 | 0 | 1505900 | 0 | 0 | 0 | 557040 | 0 |
| KVVVEQLN | MPI; MPI; MPI; | P34949; H3E | MPI_HUMAN M | Unmodified | 0 | 0 | 0 | 58429 | 84528 | 0 | 0 | 0 | 0 | 33330 | 48859 | 478920 | 72214 | 68062 | 289730 | 69407 | 500190 | 0 |
| KVVVYLQK | CTSC; CTSC; C | P53634; A0A | CATC_HUMAN | Unmodified | 0 | 0 | 0 | 0 | 0 | 0 | 0 | 0 | 0 | 0 | 1526600 | 0 | 0 | 0 | 0 | 0 | 0 | 0 |
| KVVYENAY | TXNRD1; TXNR | Q16881; A0A | TRXR1_HUMAN | Unmodified | 193620 | 0 | 0 | 0 | 1917600 | 0 | 0 | 0 | 4720100 | 515340 | 3659300 | 11230000 | 1204800 | 0 | 0 | 748220 | 288480 | 0 |
| KVVYENAY | TXNRD1; TXNR | Q16881; A0A | TRXR1_HUMAN | Unmodified | 0 | 0 | 0 | 0 | 22471000 | 2309700 | 1572300 | 0 | 458490 | 2391600 | 1585400 | 0 | 4790100 | 3192800 | 0 | 0 | 5293000 | 10164000 |
| KVWLDPNE | RPL19; RPL19; | P84098; J3K | RL19_HUMAN | Unmodified | 1916500 | 1923400 | 0 | 0 | 0 | 0 | 0 | 0 | 0 | 0 | 1239800 | 7421400 | 0 | 0 | 0 | 1408100 | 0 | 246060 |
| KVWLDPNE | RPL19; RPL19; | P84098; J3K | RL19_HUMAN | Unmodified | 0 | 0 | 0 | 0 | 0 | 0 | 0 | 0 | 0 | 0 | 0 | 0 | 2211700 | 0 | 0 | 0 | 0 | 0 |
| KVYEEVLSV | ASPH | Q12797 | ASPH_HUMAN | Unmodified | 0 | 0 | 0 | 0 | 0 | 0 | 0 | 0 | 9281200 | 2940700 | 0 | 0 | 0 | 0 | 0 | 0 | 0 | 0 |
| KWAEQYLK | ERP29; ERP29 | P30040; F8V | ERP29_HUMAN | Unmodified | 0 | 0 | 105000 | 0 | 0 | 0 | 105320 | 69048 | 37223000 | 8878700 | 945040 | 0 | 0 | 0 | 0 | 0 | 0 | 0 |
| KWAEQYLK | ERP29; ERP29 | P30040; F8V | ERP29_HUMAN | Unmodified | 0 | 0 | 0 | 0 | 0 | 0 | 18178 | 0 | 0 | 243150 | 0 | 0 | 0 | 0 | 0 | 0 | 0 | 0 |
| KWDEFEDI | GNPAT; GNPAT | O15228; A0A | GNPAT_HUMA | Unmodified | 0 | 0 | 0 | 0 | 0 | 0 | 0 | 0 | 2559300 | 0 | 0 | 0 | 0 | 0 | 0 | 0 | 0 | 0 |
| KWDICAGN | BPNT2 | Q9NX62 | IMPA3_HUMAN | Unmodified | 0 | 0 | 0 | 0 | 0 | 0 | 0 | 328900 | 42667000 | 23127000 | 0 | 2010400 | 0 | 0 | 0 | 895870 | 0 | 0 |
| KWDLSELP | DDX17; DDX17 | Q92841; A0A | DDX17_HUMA | Unmodified | 0 | 0 | 0 | 0 | 219250 | 0 | 0 | 0 | 0 | 142030 | 0 | 1544900 | 0 | 0 | 0 | 0 | 0 | 0 |
| KWDPYKQ | FGB; FGB | P02675; D6F | FIBB_HUMAN F | Unmodified | 0 | 796100 | 0 | 0 | 0 | 0 | 0 | 0 | 0 | 0 | 0 | 0 | 0 | 0 | 0 | 0 | 0 | 0 |
| KWDTCAPE | BPNT1; BPNT1 | O95861; A6 | BPNT1_HUMAN | Unmodified | 103620 | 0 | 21484 | 0 | 0 | 0 | 0 | 0 | 76561 | 0 | 323350 | 660990 | 0 | 0 | 0 | 0 | 0 | 0 |
| KWELICDQ | FMNL2; FMNL1 | Q96PY5; O9 | FMNL2_HUMA | Unmodified | 0 | 0 | 0 | 36580 | 2525700 | 319470 | 0 | 0 | 180920 | 0 | 107490 | 456430 | 0 | 0 | 0 | 0 | 0 | 1048800 |
| KWENPTQ | PRKACB; PRKA | P22694; B1A | KAPCB_HUMA | Unmodified | 0 | 0 | 0 | 1165700 | 0 | 0 | 0 | 0 | 763450 | 0 | 0 | 918420 | 67211 | 0 | 0 | 1422400 | 0 | 0 |
| KWESPAQ | PRKACA; PRKA | P17612; Q1 | KAPCA_HUMA | Unmodified | 0 | 0 | 0 | 0 | 0 | 0 | 0 | 0 | 813750 | 0 | 0 | 2129800 | 0 | 0 | 0 | 0 | 51720 | 0 |
| KWESPAQ | PRKACA; PRKA | P17612; Q1 | KAPCA_HUMA | Unmodified | 0 | 0 | 0 | 0 | 6660600 | 496470 | 4274800 | 289000 | 1570900 | 3751200 | 78614 | 0 | 2903400 | 2270300 | 0 | 0 | 359530 | 1929200 |
| KWGTDEV | ANXA4; ANXA4 | P09525; Q6 | ANXA4_HUMAN | Unmodified | 0 | 0 | 284980 | 0 | 1960900 | 494750 | 259280 | 203860 | 7413800 | 3638700 | 4447800 | 1730200 | 0 | 177910 | 0 | 242550 | 0 | 666330 |
| KWISDWNI | EIF3M; EIF3M; | Q7L2H7; E9 | EIF3M_HUMAN | Unmodified | 0 | 0 | 0 | 0 | 0 | 0 | 231980 | 62952 | 0 | 0 | 0 | 689700 | 0 | 0 | 0 | 0 | 0 | 0 |
| KWLPELVE | ALDH6A1; ALD | Q02252; G3 | MMSA_HUMAN | Unmodified | 0 | 0 | 0 | 0 | 0 | 0 | 0 | 0 | 834960 | 142720 | 0 | 0 | 0 | 0 | 0 | 0 | 0 | 0 |
| KWMEVVVI | TRRAP; TRRAP; | Q9Y4A5; F2 | TRRAP_HUMAN | Oxidation (M) | 0 | 563860 | 0 | 0 | 0 | 0 | 122710 | 2794500 | 0 | 0 | 0 | 0 | 0 | 0 | 0 | 0 | 0 | 0 |
| KWPEVLLS | RELT | Q969Z4 | TR19L_HUMAN | Unmodified | 0 | 0 | 0 | 0 | 0 | 0 | 0 | 0 | 0 | 0 | 227960 | 0 | 0 | 0 | 0 | 0 | 0 | 0 |
| KWPTVDAS | ANTXR1; ANTX | Q9H6X2; P5 | ANTR1_HUMAN | Unmodified | 0 | 0 | 0 | 0 | 2051300 | 773830 | 678580 | 0 | 0 | 0 | 0 | 0 | 0 | 0 | 0 | 0 | 263150 | 1976100 |
| KWQTMIEA | RPS3A; RPS3A; | P61247; D6 | RS3A_HUMAN | Oxidation (M) | 0 | 0 | 202760 | 62970 | 102630 | 0 | 141740 | 0 | 207250 | 0 | 252880 | 1079000 | 0 | 0 | 53575 | 0 | 123660 | 47152 |
| KWQTMIEA | RPS3A; RPS3A; | P61247; D6 | RS3A_HUMAN | Unmodified | 108510 | 0 | 0 | 0 | 412350 | 251820 | 764280 | 35120 | 0 | 0 | 0 | 1695800 | 0 | 0 | 0 | 58753 | 0 | 581330 |
| KWRVENQ | CAP1 | Q01518 | CAP1_HUMAN | Unmodified | 0 | 0 | 0 | 586060 | 39055000 | 5433600 | 2811600 | 38477 | 118270 | 6940200 | 4364200 | 4211400 | 11849000 | 5716800 | 264750 | 0 | 9068000 | 19449000 |
| KWVPAGG | ENAH; ENAH; E | Q8N8S7; A0 | ENAH_HUMAN | Unmodified | 0 | 0 | 80564 | 143100 | 1504900 | 264420 | 0 | 0 | 356410 | 592560 | 265810 | 2104300 | 0 | 0 | 246790 | 167560 | 127360 | 298840 |
| KYADALQE | SQOR; N/A | Q9Y6N5; H3 | SQOR_HUMAN | Unmodified | 0 | 0 | 0 | 0 | 0 | 0 | 0 | 0 | 3674700 | 0 | 0 | 0 | 647060 | 0 | 0 | 0 | 0 | 0 |
| KYAEAVTR | PGK1 | P00558 | PGK1_HUMAN | Unmodified | 0 | 0 | 0 | 513180 | 2222100 | 117730 | 97222 | 0 | 1709900 | 0 | 0 | 19213000 | 0 | 0 | 527780 | 0 | 3156700 | 0 |
| KYAELLEE | TXLNA | P40222 | TXLNA_HUMAN | Unmodified | 0 | 0 | 0 | 0 | 444190 | 0 | 179210 | 96535 | 284960 | 81468 | 0 | 0 | 832870 | 320380 | 0 | 0 | 516990 | 0 |
| KYAPTEAQ | XRCC5 | P13010 | XRCC5_HUMA | Unmodified | 0 | 0 | 0 | 0 | 0 | 0 | 0 | 0 | 0 | 0 | 0 | 0 | 0 | 0 | 0 | 0 | 0 | 0 |
| KYAPTEAQ | XRCC5 | P13010 | XRCC5_HUMA | Oxidation (M) | 0 | 0 | 0 | 0 | 0 | 0 | 0 | 0 | 0 | 0 | 0 | 2522700 | 0 | 0 | 0 | 0 | 0 | 0 |
| KYAVTDDY | MAPKAPK3; MA | Q16644; C9 | MAPK3_HUMA | Unmodified | 0 | 0 | 0 | 0 | 0 | 0 | 0 | 0 | 0 | 180280 | 0 | 676010 | 0 | 0 | 0 | 0 | 413430 | 273340 |
| KYDAFLASI | RPL10A | P62906 | RL10A_HUMAN | Unmodified | 5610100 | 5951700 | 4004900 | 2920300 | 3653300 | 0 | 3485100 | 901030 | 1143400 | 1147600 | 5498200 | 78022000 | 5804900 | 2023000 | 4919900 | 3573300 | 290760 | 1537600 |
| KYDEVLHM | COPA; COPA; C | P53621; A0A | COPA_HUMAN | Unmodified | 0 | 0 | 0 | 0 | 0 | 0 | 0 | 0 | 0 | 0 | 0 | 1577900 | 0 | 0 | 0 | 0 | 0 | 0 |
| KYDFYINV | IGF2R; IGF2R; I | P11717; A0A | MPRI_HUMAN | Unmodified | 0 | 0 | 0 | 0 | 0 | 0 | 0 | 0 | 1362100 | 0 | 0 | 719960 | 0 | 0 | 0 | 0 | 0 | 0 |
| KYDGYTSC | SQOR; SQOR | Q9Y6N5; H3 | SQOR_HUMAN | Unmodified | 0 | 0 | 0 | 0 | 0 | 0 | 0 | 0 | 1522800 | 0 | 0 | 0 | 0 | 0 | 0 | 0 | 0 | 0 |
| KYDLESAS | RAB11FIP5; RA | Q9BXF6; A0 | RFIP5_HUMAN | Unmodified | 0 | 0 | 0 | 0 | 2507000 | 0 | 0 | 0 | 0 | 0 | 0 | 0 | 0 | 0 | 0 | 0 | 0 | 0 |
| KYDVDTLD | COMT; COMT; | P21964; E7E | COMT_HUMAN | Oxidation (M) | 0 | 0 | 468250 | 485700 | 0 | 815220 | 0 | 0 | 0 | 0 | 0 | 0 | 0 | 0 | 691730 | 980870 | 0 | 1555900 |
| KYEAFLQR | MYO1C; MYO1 | O00159; F5 | MYO1C_HUMA | Unmodified | 775100 | 580530 | 293190 | 887190 | 1269000 | 79325 | 0 | 0 | 0 | 30957 | 0 | 832720 | 0 | 0 | 2075400 | 1686300 | 265830 | 0 |
| KYEDICPST | EIF5A; EIF5AL1; | P63241; Q6 | IF5A1_HUMAN | Unmodified | 5093600 | 3545500 | 1830600 | 2950600 | 13206000 | 3453400 | 707330 | 180950 | 8655100 | 1657300 | 6375800 | 28821000 | 8263200 | 2982400 | 3377100 | 2002400 | 2728200 | 8408700 |
| KYEDICPST | EIF5A; EIF5AL1; | P63241; Q6 | IF5A1_HUMAN | Oxidation (M) | 1604100 | 671070 | 1316900 | 699430 | 11914000 | 976520 | 106180 | 103770 | 1432200 | 162770 | 1439200 | 6726400 | 1452600 | 589530 | 726680 | 1216700 | 1082600 | 1559100 |
| KYEDICPST | EIF5A; EIF5AL1; | P63241; Q6 | IF5A1_HUMAN | Unmodified | 0 | 82589 | 0 | 109440 | 333950 | 0 | 27393 | 0 | 0 | 58621 | 556500 | 1181900 | 198550 | 0 | 0 | 0 | 65357 | 129300 |
| KYEDICPST | EIF5A; EIF5AL1; | P63241; Q6 | IF5A1_HUMAN | Oxidation (M) | 0 | 0 | 0 | 0 | 0 | 0 | 0 | 0 | 0 | 0 | 536900 | 628390 | 0 | 0 | 0 | 0 | 0 | 290320 |
| KYEEIDNA | TUFM | P49411 | EFTU_HUMAN | Unmodified | 0 | 0 | 0 | 0 | 0 | 0 | 0 | 0 | 8576800 | 0 | 201100 | 0 | 0 | 0 | 0 | 0 | 0 | 71793 |
| KYEEIDNA | TUFM | P49411 | EFTU_HUMAN | Unmodified | 0 | 0 | 0 | 0 | 0 | 0 | 0 | 39479 | 1233900 | 607050 | 105190 | 0 | 237660 | 0 | 0 | 0 | 0 | 0 |
| KYEETQKE | CEP55; CEP55 | Q53EZ4; H0 | CEP55_HUMAN | Unmodified | 0 | 0 | 0 | 0 | 895430 | 342060 | 0 | 0 | 0 | 1180900 | 0 | 0 | 0 | 0 | 0 | 0 | 0 | 87313 |
| KYEGGREL | PDIA3 | P30101 | PDIA3_HUMAN | Unmodified | 0 | 0 | 325880 | 1040700 | 0 | 0 | 717810 | 0 | 96141000 | 26111000 | 33893000 | 3114400 | 1059300 | 692870 | 1032600 | 0 | 0 | 746040 |
| KYELGRPA | RPS8; RPS8 | P62241; Q5 | RS8_HUMAN 4 | Unmodified | 1591400 | 1263200 | 404470 | 1462100 | 101750 | 112010 | 0 | 684460 | 0 | 20007 | 775650 | 4362700 | 0 | 0 | 447190 | 587170 | 34823 | 87992 |
| KYELRAPV | PEBP1 | P30086 | PEBP1_HUMAN | Unmodified | 0 | 270640 | 0 | 0 | 7128800 | 1088400 | 616610 | 0 | 0 | 5133300 | 6382600 | 0 | 31350000 | 16235000 | 0 | 0 | 9035900 | 12858000 |
| KYEMFAQT | VCP; VCP; VCP; | P55072; C9 | TERA_HUMAN | Unmodified | 668270 | 516790 | 0 | 753510 | 1261900 | 481490 | 0 | 92211 | 5634000 | 0 | 0 | 12038000 | 1493500 | 0 | 459370 | 0 | 0 | 240020 |
| KYEMFAQT | VCP; VCP; VCP; | P55072; C9 | TERA_HUMAN | Oxidation (M) | 782560 | 271230 | 675850 | 894240 | 44542 | 187760 | 0 | 0 | 1841200 | 161550 | 866770 | 10708000 | 405750 | 0 | 627800 | 1193700 | 51232 | 927390 |
| KYEPPVPTF | PSMC1 | P62191 | PRS4_HUMAN | Unmodified | 0 | 0 | 0 | 0 | 1114200 | 243530 | 228000 | 102590 | 317640 | 57894 | 0 | 531020 | 1119400 | 268460 | 0 | 0 | 0 | 362770 |
| KYEQGFITD | IMPDH2; IMPDH | P12268; A0A | IMDH2_HUMA | Unmodified | 0 | 0 | 0 | 0 | 2712600 | 477430 | 2039900 | 1061400 | 0 | 0 | 0 | 0 | 2699900 | 0 | 0 | 0 | 2413900 | 2153100 |
| KYETVIMPV | SUPT16H | Q9Y5B9 | SP16H_HUMAN | Oxidation (M) | 32227 | 0 | 17330 | 0 | 981600 | 72836 | 3308800 | 0 | 451860 | 1179600 | 792390 | 4615100 | 0 | 0 | 0 | 0 | 0 | 0 |
| KYETVIMPV | SUPT16H | Q9Y5B9 | SP16H_HUMAN | Unmodified | 0 | 0 | 0 | 0 | 0 | 0 | 382070 | 0 | 362840 | 104340 | 0 | 153830 | 0 | 0 | 0 | 0 | 0 | 196230 |
| KYFAATQFE | LNPEP | Q9UIQ6 | LCAP_HUMAN | Unmodified | 0 | 0 | 0 | 0 | 0 | 0 | 0 | 0 | 2154900 | 137140 | 0 | 0 | 113880 | 0 | 0 | 0 | 0 | 0 |
| KYFELEPLA | ARL15; ARL15 | Q9NXU5; A0 | ARL15_HUMAN | Unmodified | 0 | 0 | 0 | 0 | 0 | 0 | 0 | 19227 | 925280 | 152590 | 0 | 1542600 | 0 | 0 | 0 | 0 | 0 | 0 |
| KYFPIPVEH | PTPRE; PTPRE; | P23469; S4F | PTPRE_HUMAN | Unmodified | 0 | 0 | 0 | 0 | 2485500 | 762780 | 0 | 0 | 0 | 0 | 0 | 0 | 0 | 0 | 0 | 0 | 0 | 0 |
| KYGETANE | NASP; NASP; N | P49321; E9P | NASP_HUMAN | Unmodified | 0 | 0 | 51142 | 0 | 1212000 | 0 | 1033100 | 0 | 1372300 | 1294900 | 239320 | 1698400 | 1053000 | 307490 | 169500 | 0 | 642960 | 337650 |
| KYGGPPPD | SYNCRIP; SYN | O60506; B7 | HNRPQ_HUMA | Unmodified | 0 | 0 | 0 | 0 | 0 | 0 | 0 | 0 | 0 | 0 | 0 | 6763000 | 0 | 0 | 0 | 0 | 0 | 0 |
| KYGGPPPD | HNRNPR; HNR | O43390; A0 | HNRPR_HUMA | Unmodified | 0 | 0 | 0 | 0 | 0 | 0 | 0 | 0 | 0 | 0 | 0 | 1585000 | 0 | 0 | 0 | 0 | 0 | 0 |
| KYGLFKEE | PTTG1IP; PTTG | P53801; A8 | PTTG_HUMAN | Unmodified | 179290 | 0 | 0 | 0 | 0 | 0 | 0 | 0 | 5335600 | 468900 | 0 | 729000 | 0 | 0 | 810070 | 791450 | 0 | 0 |
| KYGTDLSR | ATP1A1; ATP1A | P05023; Q5 | AT1A1_HUMAN | Unmodified | 382980 | 59442 | 1653400 | 3506400 | 0 | 0 | 901640 | 0 | 2063100 | 442370 | 399520 | 8057000 | 0 | 140460 | 2258300 | 1718300 | 103180 | 0 |
| KYHEVLQF | PDGFD | Q9GZP0 | PDGFD_HUMA | Unmodified | 0 | 0 | 0 | 0 | 1408500 | 0 | 0 | 0 | 0 | 0 | 0 | 0 | 0 | 0 | 0 | 0 | 0 | 0 |
| KYHNVGLS | HNRNPD; HNR | Q14103; H0 | HNRPD_HUMA | Unmodified | 0 | 0 | 0 | 0 | 483470 | 0 | 0 | 320430 | 0 | 0 | 0 | 0 | 574590 | 0 | 0 | 0 | 0 | 0 |
| KYILGNPLT | ALDH1A1 | P00352 | AL1A1_HUMAN | Unmodified | 0 | 0 | 0 | 0 | 0 | 0 | 0 | 0 | 0 | 0 | 7154300 | 93949000 | 0 | 0 | 0 | 0 | 0 | 0 |
| KYILGNPLT | ALDH1A1 | P00352 | AL1A1_HUMAN | Unmodified | 0 | 0 | 0 | 0 | 0 | 0 | 0 | 0 | 3735000 | 2154700 | 12742000 | 4.8E+08 | 0 | 0 | 0 | 0 | 64212 | 0 |
| KYILYDFSV | MGAM2 | Q2M2H8 | MGAL_HUMAN | Unmodified | 0 | 0 | 0 | 0 | 0 | 0 | 0 | 0 | 0 | 0 | 658570 | 1470400 | 0 | 0 | 0 | 0 | 0 | 0 |
| KYLQPDNL | MTTP; MTTP | P55157; E9P | MTP_HUMAN M | Unmodified | 0 | 0 | 417540 | 394980 | 0 | 346420 | 135340 | 556550 | 3087400 | 1229300 | 166360 | 1673900 | 1350600 | 670900 | 753080 | 484670 | 0 | 0 |
| KYNPTWH | DYNLL2; DYNL | Q96FJ2; P63 | DYL2_HUMAN | Unmodified | 4448300 | 1735900 | 693930 | 951790 | 0 | 137750 | 84832 | 32132 | 1762400 | 0 | 286210 | 7608500 | 8039500 | 6894000 | 1018600 | 1089400 | 165660 | 0 |
| KYNPTWH | DYNLL2; DYNL | Q96FJ2; P63 | DYL2_HUMAN | Unmodified | 0 | 0 | 0 | 0 | 0 | 0 | 2907400 | 0 | 0 | 0 | 0 | 0 | 0 | 0 | 0 | 54396 | 758780 | 737120 |
| KYNTYAYV | SFTPA2; SFTPA | Q8IWL1; Q8 | SFPA2_HUMAN | Unmodified | 0 | 0 | 0 | 0 | 0 | 0 | 0 | 0 | 1890600 | 0 | 0 | 0 | 0 | 0 | 84347 | 0 | 0 | 0 |
| KYPDYESK | HYOU1; HYOU | Q9Y4L1; E9 | HYOU1_HUMA | Unmodified | 0 | 0 | 175080 | 0 | 0 | 0 | 198570 | 0 | 8906100 | 1950600 | 0 | 0 | 0 | 0 | 0 | 0 | 0 | 0 |
| KYPEAAAC | STUB1; STUB1 | Q9UNE7; H3 | CHIP_HUMAN | Unmodified | 430210 | 340410 | 0 | 446470 | 150920 | 73224 | 0 | 0 | 207160 | 50881 | 0 | 717660 | 452640 | 0 | 0 | 0 | 104580 | 461370 |
| KYPLLLDV | DPP4; DPP4; D | P27487; A0A | DPP4_HUMAN | Unmodified | 0 | 0 | 179060 | 257240 | 278860 | 0 | 0 | 0 | 0 | 0 | 0 | 0 | 0 | 0 | 1489800 | 952990 | 0 | 0 |
| KYPYVEVN | UGGT1 | Q9NYU2 | UGGG1_HUMA | Unmodified | 0 | 379950 | 0 | 0 | 0 | 0 | 0 | 0 | 3198600 | 896410 | 0 | 0 | 1116400 | 0 | 0 | 0 | 0 | 0 |
| KYPYWPH | ATP5PD | O75947 | ATP5H_HUMAN | Unmodified | 0 | 0 | 0 | 0 | 0 | 0 | 103580 | 0 | 7894500 | 936470 | 204200 | 449310 | 0 | 0 | 0 | 0 | 0 | 0 |
| KYQDWGW | MAN1B1; MAN1 | Q9UKM7; H | MA1B1_HUMA | Unmodified | 0 | 0 | 0 | 0 | 0 | 0 | 0 | 0 | 327270 | 0 | 0 | 0 | 0 | 0 | 0 | 0 | 0 | 0 |
| KYQEQELP | RRAS | P10301 | RRAS_HUMAN | Unmodified | 244310 | 140010 | 321890 | 252890 | 0 | 0 | 164000 | 0 | 269300 | 0 | 0 | 0 | 0 | 358780 | 628220 | 326460 | 0 | 0 |
| KYQILNNE | CYFIP2; CYFIP2 | Q96F07; H7 | CYFP2_HUMAN | Unmodified | 298230 | 0 | 147600 | 639320 | 294360 | 0 | 510690 | 43219 | 2319600 | 764480 | 475210 | 5055900 | 872020 | 511720 | 1032000 | 499800 | 0 | 0 |
| KYQKPIIQS | GUSB | P08236 | BGLR_HUMAN | Oxidation (M) | 0 | 0 | 0 | 0 | 0 | 0 | 0 | 0 | 15638000 | 16764000 | 0 | 0 | 0 | 0 | 0 | 0 | 0 | 0 |
| KYQKPIIQS | GUSB | P08236 | BGLR_HUMAN | Unmodified | 0 | 0 | 0 | 0 | 0 | 0 | 0 | 0 | 19068000 | 1662900 | 0 | 0 | 0 | 0 | 0 | 0 | 0 | 0 |
| KYQLNLPP | NAXE; NAXE | Q8NCW5; A | NNRE_HUMAN | Unmodified | 0 | 0 | 0 | 0 | 759010 | 184560 | 0 | 0 | 250680 | 0 | 0 | 1078400 | 0 | 0 | 301660 | 0 | 0 | 0 |
| KYQLNLPP | NAXE; NAXE | Q8NCW5; A | NNRE_HUMAN | Unmodified | 0 | 0 | 0 | 0 | 1220200 | 0 | 454980 | 0 | 0 | 1395600 | 286160 | 0 | 0 | 881510 | 0 | 0 | 2447600 | 3936200 |
| KYQPNIDV | CD109 | Q6YHK3 | CD109_HUMA | Unmodified | 0 | 366680 | 48898 | 0 | 272760 | 291520 | 209480 | 0 | 1285000 | 0 | 0 | 0 | 0 | 696650 | 0 | 0 | 141810 | 4164800 |

| | | | | | | | | | | | | | | | | | | | | | | |
|---|---|---|---|---|---|---|---|---|---|---|---|---|---|---|---|---|---|---|---|---|---|---|
| KYRPLFDYI | IARS1; IARS1; IA | P41252; A0A | SYIC_HUMAN I | Unmodified | 0 | 0 | 0 | 0 | 0 | 0 | 0 | 0 | 139640 | 0 | 0 | 3325500 | 0 | 0 | 0 | 0 | 0 | 0 |
| KYSHHRNI | TNIK; MINK1; M | Q9UKE5; Q8 | TNIK_HUMAN T | Unmodified | 0 | 0 | 0 | 0 | 0 | 0 | 0 | 0 | 0 | 0 | 0 | 0 | 0 | 0 | 0 | 0 | 641370 | 167720 |
| KYSPSDPA | RAB11FIP1; RAI | Q6WKZ4; E7 | RFIP1_HUMAN | Unmodified | 0 | 0 | 0 | 0 | 7199800 | 192330 | 0 | 0 | 0 | 429300 | 0 | 0 | 0 | 0 | 0 | 0 | 0 | 0 |
| KYSQFINFF | HSP90B1; HSP9 | P14625; A0A | ENPL_HUMAN | Unmodified | 0 | 0 | 0 | 0 | 0 | 0 | 0 | 0 | 35975000 | 4988700 | 3270300 | 9039600 | 0 | 0 | 933850 | 0 | 0 | 0 |
| KYTELPHG | DCTPP1; DCTPI | Q9H773; H3 | DCTP1_HUMAI | Unmodified | 0 | 0 | 0 | 123760 | 0 | 0 | 0 | 0 | 0 | 0 | 1931000 | 2541900 | 0 | 0 | 0 | 0 | 0 | 0 |
| KYTGEDFD | DYNC1H1; DYN | Q14204; A0A | DYHC1_HUMA | Unmodified | 0 | 0 | 0 | 0 | 0 | 0 | 0 | 0 | 0 | 0 | 0 | 0 | 0 | 0 | 0 | 0 | 0 | 0 |
| KYTLPPGVI | HSPB1; HSPB1 | P04792; A0A | HSPB1_HUMAI | Oxidation (M) | 1096900 | 0 | 1098800 | 4536100 | 1123000 | 0 | 0 | 0 | 4724100 | 5915800 | 0 | 0 | 7873700 | 0 | 5217300 | 2571700 | 3460500 | 0 |
| KYTLPPGVI | HSPB1; HSPB1 | P04792; A0A | HSPB1_HUMAI | Unmodified | 0 | 0 | 0 | 0 | 0 | 0 | 0 | 0 | 7636000 | 0 | 0 | 0 | 0 | 0 | 0 | 0 | 1034000 | 1091700 |
| KYVAAQLH | CA14 | Q9ULX7 | CAH14_HUMAI | Unmodified | 0 | 0 | 0 | 0 | 0 | 0 | 0 | 0 | 0 | 0 | 0 | 1060800 | 0 | 0 | 0 | 0 | 0 | 0 |
| KYVATLGVI | RAN; RAN; RAN | P62826; H0Y | RAN_HUMAN G | Unmodified | 1323100 | 994120 | 576990 | 313910 | 0 | 0 | 0 | 0 | 369440 | 640950 | 714380 | 32570000 | 0 | 0 | 3227800 | 2225300 | 321000 | 293450 |
| KYVATLGVI | RAN; RAN; RAN | P62826; H0Y | RAN_HUMAN G | Unmodified | 691630 | 640210 | 589720 | 770020 | 15899000 | 1986600 | 6463300 | 0 | 3984600 | 2948200 | 5983700 | 28831000 | 11628000 | 9996000 | 654780 | 516990 | 4180500 | 6855500 |
| KYVDAETSI | SLC4A7; SLC4A | Q9Y6M7; A0 | S4A7_HUMAN | Unmodified | 0 | 0 | 0 | 0 | 0 | 0 | 0 | 0 | 0 | 0 | 0 | 0 | 0 | 0 | 0 | 0 | 922200 | 2566900 |
| KYVFFLDP( | SLC44A1 | Q8WWI5 | CTL1_HUMAN | Unmodified | 543110 | 446130 | 0 | 654880 | 0 | 0 | 0 | 0 | 319340 | 0 | 0 | 326030 | 0 | 0 | 359370 | 179720 | 0 | 0 |
| KYVFFLDP( | SLC44A1 | Q8WWI5 | CTL1_HUMAN | Unmodified | 0 | 0 | 0 | 771950 | 427020 | 0 | 0 | 0 | 547420 | 335320 | 0 | 603630 | 949930 | 0 | 0 | 0 | 0 | 0 |
| KYVFLDPL | WASHC1; WASI | A8K0Z3; Q6\ | WASH1_HUMA | Unmodified | 0 | 0 | 0 | 0 | 1056600 | 0 | 178530 | 53700 | 0 | 0 | 0 | 0 | 217860 | 325670 | 0 | 0 | 208050 | 410120 |
| KYVLENHP | HP1BP3; HP1BI | Q5SSJ5; B0C | HP1B3_HUMAI | Unmodified | 0 | 0 | 0 | 0 | 83708 | 0 | 569410 | 0 | 0 | 0 | 0 | 0 | 0 | 0 | 0 | 0 | 0 | 0 |
| KYVPTPDR | CEMIP2 | Q9UHN6 | CEIP2_HUMAN | Unmodified | 0 | 0 | 0 | 103910 | 5263800 | 267060 | 154310 | 0 | 1007000 | 168960 | 0 | 0 | 102750 | 0 | 89675 | 0 | 222660 | 0 |
| KYVPTPDR | CEMIP2 | Q9UHN6 | CEIP2_HUMAN | Oxidation (M) | 0 | 0 | 132940 | 149090 | 1032100 | 147540 | 286080 | 0 | 277210 | 250370 | 0 | 0 | 57388 | 0 | 83106 | 77839 | 0 | 0 |
| KYVPYKVP | GALNT10 | Q86SR1 | GLT10_HUMAN | Unmodified | 0 | 0 | 0 | 0 | 0 | 0 | 0 | 0 | 671330 | 0 | 0 | 0 | 0 | 0 | 0 | 0 | 0 | 0 |
| KYWLLPIFS | ZDHHC20; ZDH | Q5W0Z9; B4 | ZDH20_HUMAI | Unmodified | 0 | 0 | 22467 | 100050 | 0 | 0 | 0 | 0 | 0 | 0 | 0 | 0 | 0 | 0 | 130710 | 37811 | 0 | 0 |
| KYYGTKDP | NAMPT; NAMPT | P43490; A0A | NAMPT_HUMAI | Unmodified | 0 | 0 | 0 | 0 | 0 | 0 | 0 | 0 | 0 | 0 | 0 | 1129700 | 0 | 0 | 0 | 0 | 0 | 0 |
| LAAAAAAA( | GPX1; GPX1; GI | P07203; A0A | GPX1_HUMAN | Unmodified | 0 | 0 | 0 | 0 | 0 | 0 | 0 | 0 | 0 | 2016300 | 0 | 0 | 0 | 0 | 0 | 0 | 0 | 0 |
| LAAAILGGV | FBL; FBL; FBL; F | P22087; M0F | FBRL_HUMAN | Unmodified | 535860 | 259000 | 695960 | 293000 | 767640 | 323810 | 3389200 | 631410 | 273640 | 333690 | 411010 | 3779100 | 1631000 | 706480 | 468400 | 338900 | 644680 | 754750 |
| LAACGPPP | IGFBP2 | P18065 | IBP2_HUMAN I | Unmodified | 0 | 0 | 0 | 0 | 0 | 0 | 0 | 33826 | 46234000 | 732120 | 0 | 0 | 0 | 0 | 0 | 0 | 0 | 169590 |
| LAADEDDD | NPM1; NPM1; N | P06748; A0A | NPM_HUMAN N | Unmodified | 0 | 0 | 0 | 0 | 0 | 0 | 0 | 0 | 0 | 0 | 870190 | 0 | 0 | 0 | 0 | 0 | 0 | 0 |
| LAADEDDD | NPM1; NPM1; N | P06748; A0A | NPM_HUMAN N | Unmodified | 0 | 0 | 0 | 0 | 1664000 | 0 | 0 | 0 | 1702800 | 1860500 | 7451500 | 2758500 | 6783600 | 0 | 0 | 0 | 0 | 0 |
| LAADPDPN | VAC14; VAC14 | Q08AM6; H3 | VAC14_HUMAI | Unmodified | 0 | 0 | 0 | 12741 | 0 | 0 | 0 | 0 | 0 | 0 | 0 | 554370 | 0 | 0 | 51288 | 0 | 0 | 0 |
| LAADPSVVI | KYAT3; KYAT3 | Q6YP21; A0A | KAT3_HUMAN I | Unmodified | 0 | 0 | 0 | 0 | 0 | 0 | 0 | 0 | 607740 | 0 | 182530 | 472460 | 0 | 0 | 0 | 0 | 0 | 0 |
| LAAEAGNV | DTNB; DTNB; D | O60941; E7E | DTNB_HUMAN | Unmodified | 0 | 0 | 0 | 813270 | 0 | 0 | 0 | 0 | 1696300 | 516730 | 0 | 388150 | 1068000 | 352530 | 1580200 | 1272800 | 0 | 0 |
| LAAEQELIR | PLEC | Q15149 | PLEC_HUMAN | Unmodified | 0 | 0 | 0 | 70405 | 0 | 0 | 0 | 20285 | 0 | 0 | 62904 | 1059400 | 219550 | 0 | 241930 | 66476 | 0 | 0 |
| LAAFGQLH | ILF3 | Q12906 | ILF3_HUMAN Ir | Unmodified | 496630 | 281880 | 735420 | 391390 | 0 | 0 | 0 | 0 | 0 | 0 | 0 | 1705400 | 164430 | 0 | 0 | 149530 | 0 | 0 |
| LAAGDQLL | AFDN; AFDN; A | P55196; Q5T | AFAD_HUMAN | Unmodified | 325290 | 0 | 219730 | 0 | 0 | 0 | 0 | 0 | 0 | 0 | 457810 | 2033600 | 0 | 0 | 0 | 254540 | 0 | 0 |
| LAAGGPWA | DLL3 | Q9NYJ7 | DLL3_HUMAN I | Unmodified | 0 | 0 | 0 | 0 | 0 | 0 | 0 | 0 | 2545100 | 83488 | 0 | 1154900 | 0 | 0 | 0 | 0 | 0 | 0 |
| LAAGRPFN | PAK4 | O96013 | PAK4_HUMAN | Unmodified | 308180 | 318220 | 0 | 468660 | 0 | 0 | 0 | 0 | 0 | 0 | 0 | 1720800 | 0 | 0 | 0 | 0 | 0 | 0 |
| LAAGSAIVK | PDS5B | Q9NTI5 | PDS5B_HUMAI | Unmodified | 0 | 0 | 0 | 0 | 0 | 0 | 0 | 0 | 0 | 0 | 0 | 318800 | 0 | 0 | 0 | 0 | 0 | 0 |
| LAAIAESGV | PSMB6 | P28072 | PSB6_HUMAN | Unmodified | 66983 | 105850 | 0 | 1571900 | 0 | 0 | 0 | 151340 | 30875000 | 9767500 | 1363200 | 20250000 | 0 | 0 | 1393800 | 992480 | 0 | 0 |
| LAAIAESGV | PSMB6 | P28072 | PSB6_HUMAN | Unmodified | 0 | 0 | 0 | 0 | 2138400 | 0 | 0 | 0 | 763020 | 2540600 | 0 | 0 | 3092200 | 1262200 | 0 | 0 | 985590 | 0 |
| LAAIGEATR | PLEC | Q15149 | PLEC_HUMAN | Unmodified | 0 | 0 | 0 | 0 | 896740 | 190290 | 0 | 0 | 0 | 0 | 0 | 0 | 0 | 0 | 0 | 0 | 1007900 | 1109700 |
| LAALADQW | SPTAN1; SPTAN | Q13813; A0A | SPTN1_HUMAI | Unmodified | 60608 | 0 | 204790 | 46616 | 354400 | 96099 | 89394 | 0 | 277250 | 141770 | 225040 | 2116600 | 114910 | 98467 | 154080 | 164200 | 351930 | 110680 |
| LAALNPESI | CLIC1 | O00299 | CLIC1_HUMAN | Unmodified | 23511000 | 19413000 | 10038000 | 24278000 | 3.12E+08 | 73638000 | 31757000 | 14113000 | 53350000 | 32544000 | 16181000 | 83374000 | 1.15E+08 | 1.42E+08 | 44468000 | 18945000 | 3E+08 | 1.16E+08 |
| LAALPENPI | ATP5PD; ATP5P | O75947; F5H | ATP5H_HUMAI | Unmodified | 0 | 0 | 0 | 0 | 0 | 0 | 0 | 0 | 8581900 | 1703400 | 0 | 0 | 0 | 157750 | 32026 | 70435 | 64539 | 0 |
| LAALPNVYI | MCM5; MCM5 | P33992; B1A | MCM5_HUMAN | Unmodified | 0 | 0 | 0 | 0 | 0 | 0 | 0 | 0 | 462500 | 0 | 0 | 1365900 | 0 | 0 | 0 | 0 | 0 | 0 |
| LAALQGRLI | CKAP4 | Q07065 | CKAP4_HUMAI | Unmodified | 0 | 0 | 0 | 0 | 0 | 0 | 190270 | 88270 | 17965000 | 120410 | 0 | 203330 | 0 | 0 | 0 | 0 | 0 | 0 |
| LAALTAGDI | CPT1A | P50416 | CPT1A_HUMAI | Unmodified | 0 | 0 | 0 | 0 | 0 | 0 | 50486 | 38643 | 580720 | 69867 | 283010 | 1372500 | 52951 | 0 | 0 | 0 | 0 | 0 |
| LAAMPALR | LRRC20; LRRC2 | Q8TCA0; X6I | LRC20_HUMAI | Unmodified | 0 | 0 | 0 | 0 | 749320 | 117160 | 51837 | 0 | 0 | 269690 | 0 | 0 | 114790 | 66082 | 0 | 0 | 231820 | 228630 |
| LAANAFLA( | UGDH | O60701 | UGDH_HUMAI | Unmodified | 0 | 0 | 0 | 0 | 8826400 | 2099500 | 110730 | 0 | 0 | 0 | 0 | 2162600 | 2465600 | 1154000 | 0 | 809520 | 0 | 974750 |
| LAANSSLFI | CIT; CIT; CIT; CI | O14578; A0A | CTRO_HUMAN | Unmodified | 0 | 0 | 0 | 414550 | 1138800 | 409250 | 0 | 0 | 187320 | 0 | 187250 | 498860 | 0 | 0 | 0 | 0 | 0 | 0 |
| LAANSSLFI | CIT; CIT; CIT; CI | O14578; A0A | CTRO_HUMAN | Oxidation (M) | 0 | 0 | 0 | 0 | 0 | 0 | 0 | 0 | 0 | 73122 | 0 | 0 | 0 | 0 | 0 | 0 | 209670 | 101340 |
| LAANSSLFI | CIT; CIT; CIT; CI | O14578; A0A | CTRO_HUMAN | Unmodified | 0 | 0 | 0 | 0 | 3739000 | 322420 | 0 | 0 | 0 | 0 | 0 | 747430 | 292830 | 0 | 494000 | 0 | 481850 | 613500 |
| LAAQENRP | TBCD; TBCD; TI | Q9BTW9; I3L | TBCD_HUMAN | Unmodified | 0 | 0 | 186860 | 0 | 0 | 0 | 431500 | 0 | 0 | 0 | 0 | 6278200 | 122210 | 0 | 0 | 336220 | 0 | 0 |
| LAAQSCAL | DHX9 | Q08211 | DHX9_HUMAN | Unmodified | 0 | 0 | 0 | 0 | 0 | 0 | 0 | 0 | 0 | 0 | 343870 | 1439500 | 244380 | 0 | 0 | 0 | 0 | 0 |
| LAASASTQ( | KIF20A | O95235 | KI20A_HUMAN | Unmodified | 0 | 0 | 682220 | 1423100 | 1644300 | 283430 | 0 | 37891 | 1444100 | 147930 | 478780 | 0 | 0 | 0 | 424340 | 298180 | 400410 | 416850 |
| LAASASTQ( | KIF20A | O95235 | KI20A_HUMAN | Unmodified | 0 | 0 | 0 | 0 | 317630 | 0 | 0 | 0 | 0 | 0 | 0 | 0 | 264830 | 1334800 | 0 | 0 | 538630 | 0 |
| LAASGEGG | DBN1; DBN1 | Q16643; D6I | DREB_HUMAN | Unmodified | 433270 | 434040 | 2107800 | 3617500 | 19063000 | 2351800 | 1671300 | 1365900 | 5729700 | 1730800 | 5317000 | 35794000 | 8117700 | 0 | 5651100 | 3019000 | 6344300 | 6698800 |
| LAASLDGA | LAMA5 | O15230 | LAMA5_HUMAI | Unmodified | 0 | 0 | 1172100 | 0 | 0 | 0 | 0 | 0 | 0 | 0 | 0 | 0 | 0 | 0 | 0 | 0 | 0 | 0 |
| LAATGRED | RABGAP1L; RAE | B7ZAP0; F5H | RBG10_HUMAI | Unmodified | 0 | 0 | 0 | 0 | 0 | 0 | 0 | 0 | 127160 | 0 | 69608 | 401460 | 0 | 0 | 0 | 0 | 0 | 0 |
| LAATNALLN | KPNB1; KPNB1 | Q14974; J3K | IMB1_HUMAN I | Unmodified | 1127500 | 1857300 | 1170800 | 2502000 | 14442000 | 2569300 | 3410100 | 299570 | 4996700 | 3860500 | 2698100 | 13701000 | 11503000 | 4869700 | 3738400 | 1906700 | 7723500 | 7224700 |
| LAATPSPG | XYLB | O75191 | XYLB_HUMAN | Unmodified | 0 | 0 | 0 | 150640 | 0 | 0 | 0 | 0 | 0 | 0 | 0 | 153700 | 3536000 | 272150 | 0 | 0 | 1269900 | 464390 |
| LAAVALINA | IQGAP1; IQGAP | P46940; A0A | IQGA1_HUMAI | Unmodified | 284230 | 841730 | 43962 | 0 | 1712200 | 871810 | 58228 | 59413 | 678600 | 1415400 | 69556 | 3507300 | 81277 | 0 | 99710 | 334290 | 839500 | 0 |
| LAAVDATVI | PDIA6 | Q15084 | PDIA6_HUMAN | Unmodified | 76191 | 30676 | 96273 | 102700 | 259960 | 97808 | 117750 | 1500900 | 1.47E+08 | 29233000 | 19514000 | 4291000 | 137480 | 86995 | 283990 | 1127600 | 191840 | 140490 |
| LAAVDATVI | PDIA6 | Q15084 | PDIA6_HUMAN | Unmodified | 0 | 0 | 0 | 0 | 0 | 337390 | 337770 | 0 | 2670200 | 20855000 | 1268500 | 0 | 545920 | 0 | 0 | 0 | 0 | 497210 |
| LAAVDATVI | PDIA6 | Q15084 | PDIA6_HUMAN | Unmodified | 0 | 0 | 0 | 0 | 6869200 | 783040 | 6262400 | 0 | 1882400 | 1.09E+08 | 1990900 | 0 | 1772800 | 0 | 0 | 0 | 0 | 2815800 |
| LAAVQLLQI | TEX10 | Q9NXF1 | TEX10_HUMAN | Unmodified | 0 | 0 | 0 | 0 | 0 | 0 | 0 | 0 | 0 | 0 | 0 | 75500 | 0 | 0 | 0 | 11353 | 10053 | 0 |
| LAAVSDEN | RIPOR2; RIPOR | Q9Y4F9; F5C | RIPR2_HUMAN | Unmodified | 0 | 0 | 0 | 0 | 0 | 0 | 0 | 0 | 0 | 0 | 0 | 882480 | 0 | 1792200 | 0 | 0 | 0 | 0 |
| LAAVTYNG | SEPTIN7; SEPTI | Q16181; E7E | SEPT7_HUMAN | Unmodified | 0 | 19535 | 231890 | 283230 | 8428500 | 0 | 129710 | 0 | 278680 | 227240 | 89746 | 4151400 | 761870 | 541730 | 457080 | 624950 | 2016600 | 2139100 |
| LAAVTYNG | SEPTIN7; SEPTI | Q16181; E7E | SEPT7_HUMAN | Unmodified | 0 | 0 | 0 | 0 | 0 | 0 | 0 | 0 | 0 | 0 | 0 | 0 | 0 | 0 | 0 | 0 | 0 | 0 |
| LAAVTYNG | SEPTIN7; SEPTI | Q16181; E7E | SEPT7_HUMAN | Unmodified | 0 | 0 | 0 | 0 | 676590 | 68085 | 159990 | 115500 | 0 | 0 | 0 | 0 | 111390 | 135500 | 0 | 0 | 1408100 | 416780 |
| LACGIIAR | CCS; CCS; CCS | O14618; J3K | CCS_HUMAN C | Unmodified | 0 | 197240 | 0 | 0 | 79968 | 20734 | 0 | 0 | 0 | 0 | 254010 | 1033600 | 0 | 0 | 0 | 0 | 0 | 63503 |
| LACGKLTG | OLFM1; OLFM1 | Q99784; F5H | NOE1_HUMAN | Unmodified | 0 | 0 | 0 | 0 | 125320 | 0 | 0 | 0 | 0 | 563390 | 0 | 0 | 0 | 0 | 0 | 0 | 55799 | 70682 |
| LACLAGELI | CD320 | Q9NPF0 | CD320_HUMAI | Unmodified | 387200 | 248530 | 384010 | 837880 | 0 | 0 | 0 | 0 | 3087600 | 211820 | 0 | 0 | 170670 | 0 | 213240 | 219120 | 0 | 0 |
| LADAAENF | CACNA2D2; CA | Q9NY47; C9 | CA2D2_HUMAI | Unmodified | 0 | 0 | 0 | 0 | 549260 | 950070 | 0 | 396830 | 301050 | 463660 | 0 | 0 | 0 | 0 | 0 | 0 | 0 | 0 |
| LADALQELI | LMNA; LMNA; L | P02545; A0A | LMNA_HUMAN | Unmodified | 0 | 0 | 0 | 0 | 1694600 | 0 | 0 | 0 | 0 | 49127 | 0 | 0 | 0 | 0 | 0 | 0 | 0 | 792660 |
| LADCDCD( | EHD4 | Q9H223 | EHD4_HUMAN | Oxidation (M) | 483520 | 177990 | 0 | 0 | 939380 | 0 | 0 | 0 | 0 | 0 | 0 | 0 | 703820 | 0 | 0 | 0 | 524980 | 2811300 |
| LADCDCD( | EHD4 | Q9H223 | EHD4_HUMAN | Unmodified | 0 | 0 | 0 | 0 | 422100 | 0 | 0 | 0 | 0 | 0 | 0 | 0 | 0 | 0 | 0 | 0 | 526730 | 849790 |
| LADDLGFE | RAB3D | O95716 | RAB3D_HUMAI | Unmodified | 0 | 180650 | 267600 | 1340400 | 0 | 0 | 0 | 0 | 0 | 0 | 0 | 0 | 0 | 0 | 1050400 | 405550 | 0 | 0 |
| LADDLSTL( | NUMA1; NUMA | Q14980; A0A | NUMA1_HUMA | Unmodified | 285570 | 0 | 0 | 0 | 0 | 0 | 0 | 0 | 705530 | 1736400 | 0 | 1246600 | 2232100 | 663540 | 0 | 0 | 153910 | 0 |
| LADDVDLE | VCP; VCP; VCP; | P55072; C9J | TERA_HUMAN | Unmodified | 0 | 0 | 0 | 3269600 | 7431600 | 0 | 0 | 0 | 0 | 2207400 | 0 | 0 | 0 | 1763600 | 0 | 0 | 0 | 0 |
| LADEQLSS | ABHD11 | Q8NFV4 | ABHDB_HUMA | Unmodified | 0 | 0 | 0 | 0 | 1545600 | 0 | 0 | 0 | 732440 | 0 | 28070 | 0 | 0 | 65554 | 0 | 0 | 0 | 0 |
| LADFGFAQ | ULK3 | Q6PHR2 | ULK3_HUMAN | Oxidation (M) | 0 | 0 | 0 | 0 | 0 | 0 | 48686 | 95881 | 0 | 0 | 0 | 285020 | 0 | 110160 | 0 | 239330 | 59626 | 0 |
| LADFGLAR | CDK13; CDK12 | Q14004; Q9I | CDK13_HUMAI | Unmodified | 6797600 | 5437300 | 7098200 | 9705300 | 3729900 | 2149000 | 899260 | 718740 | 15352000 | 3572800 | 2871600 | 18278000 | 2591300 | 1465200 | 12233000 | 7987800 | 494710 | 3284400 |
| LADFGTCM | ROCK2; ROCK1 | O75116; Q1 | ROCK2_HUMAI | Unmodified | 0 | 0 | 0 | 0 | 1158600 | 0 | 0 | 0 | 0 | 170600 | 0 | 927980 | 0 | 310230 | 0 | 0 | 893610 | 1288700 |
| LADFGVAG | STK24; STK25; | Q9Y6E0; O0 | STK24_HUMAN | Unmodified | 384790 | 584930 | 847370 | 843320 | 19155000 | 5468000 | 579760 | 677740 | 4639900 | 4193700 | 4758500 | 9748300 | 13413000 | 6871700 | 822600 | 877270 | 3907900 | 9757100 |
| LADFGVAG | STK24; STK25; | Q9Y6E0; O0 | STK24_HUMAN | Unmodified | 0 | 0 | 0 | 0 | 1521900 | 0 | 0 | 0 | 1076200 | 0 | 6827900 | 1651700 | 1651800 | 1827600 | 0 | 0 | 526170 | 0 |
| LADFGVLH | TARS1 | P26639 | SYTC_HUMAN | Unmodified | 263820 | 199290 | 1007000 | 3230700 | 0 | 134180 | 0 | 0 | 5705000 | 217380 | 1585100 | 27417000 | 0 | 0 | 3510300 | 4269100 | 0 | 0 |
| LADFGVSA | STK10; SLK | O94804; Q9I | STK10_HUMAN | Unmodified | 126470 | 116780 | 615000 | 753880 | 1375400 | 408290 | 77275 | 57633 | 923210 | 0 | 2846200 | 5857000 | 3554100 | 2869900 | 460940 | 339130 | 0 | 0 |
| LADGGATN | LGALS3BP; LGA | Q08380; K7E | LG3BP_HUMAI | Unmodified | 0 | 0 | 326970 | 0 | 0 | 0 | 0 | 0 | 43316000 | 416650 | 16542000 | 19690000 | 0 | 0 | 237800 | 357410 | 0 | 0 |
| LADIQIEQL | PFDN1; PFDN1 | O60925; E5F | PFD1_HUMAN | Unmodified | 0 | 0 | 0 | 87540 | 0 | 0 | 0 | 0 | 2820100 | 208710 | 2397500 | 8945100 | 332090 | 0 | 53124 | 388500 | 0 | 0 |
| LADIQIEQL | PFDN1; PFDN1 | O60925; E5F | PFD1_HUMAN | Unmodified | 0 | 0 | 638630 | 0 | 3347800 | 763980 | 0 | 457330 | 0 | 539690 | 0 | 0 | 1699100 | 2236400 | 0 | 0 | 2307200 | 2732200 |
| LADIQIEQL | PFDN1; PFDN1 | O60925; E5F | PFD1_HUMAN | Unmodified | 0 | 0 | 0 | 0 | 2807100 | 0 | 0 | 220400 | 0 | 249680 | 0 | 0 | 1900500 | 1102500 | 0 | 0 | 1922900 | 2999200 |
| LADKYPVVI | PLOD2; PLOD2; | O00469; C9J | PLOD2_HUMAI | Unmodified | 0 | 0 | 0 | 0 | 38570 | 0 | 0 | 0 | 226100 | 296990 | 0 | 0 | 0 | 0 | 0 | 0 | 0 | 0 |
| LADLEQEQ | SARG | Q9BW04 | SARG_HUMAN | Unmodified | 112870 | 0 | 0 | 0 | 271200 | 0 | 0 | 0 | 0 | 0 | 0 | 0 | 1401300 | 1850900 | 384770 | 300030 | 0 | 0 |
| LADLFGWS | ADD1; ADD1; A | P35611; E7E | ADDA_HUMAN | Unmodified | 0 | 0 | 0 | 0 | 434270 | 0 | 0 | 0 | 0 | 0 | 0 | 0 | 29059 | 47694 | 531190 | 0 | 0 | 379220 |
| LADLIER | ALDH2; ALDH1 | P05091; P00 | ALDH2_HUMAI | Unmodified | 0 | 376890 | 142560 | 50132 | 257880 | 0 | 93417 | 0 | 7706700 | 483750 | 5242400 | 74510000 | 0 | 0 | 115670 | 0 | 58921 | 282480 |
| LADLIERDF | ALDH2; ALDH1 | P05091; P00 | ALDH2_HUMAI | Unmodified | 0 | 0 | 0 | 0 | 0 | 0 | 0 | 0 | 134970 | 73867 | 462330 | 54094000 | 0 | 0 | 0 | 0 | 0 | 557290 |
| LADLIERDF | ALDH1A1; ALDI | P00352; Q5S | AL1A1_HUMAN | Oxidation (M) | 0 | 0 | 0 | 0 | 0 | 0 | 0 | 0 | 0 | 0 | 2115700 | 15764000 | 180360 | 0 | 0 | 0 | 0 | 0 |
| LADLIERDF | ALDH1A1; ALDI | P00352; Q5S | AL1A1_HUMAN | 2 Oxidation (M) | 0 | 0 | 0 | 0 | 0 | 0 | 0 | 0 | 0 | 100120 | 2023800 | 13869000 | 476410 | 0 | 0 | 0 | 0 | 0 |
| LADLIERDF | ALDH1A1; ALDI | P00352; Q5S | AL1A1_HUMAN | Unmodified | 0 | 0 | 0 | 0 | 0 | 0 | 0 | 0 | 647310 | 0 | 33073 | 1353900 | 0 | 0 | 0 | 0 | 0 | 0 |
| LADLYGSKI | CDH11; CDH11 | P55287; H3E | CAD11_HUMAI | Unmodified | 0 | 0 | 0 | 0 | 0 | 0 | 0 | 0 | 0 | 0 | 0 | 0 | 0 | 0 | 0 | 0 | 15046000 | 13661000 |
| LADMGIAIF | RTCB | Q9Y3I0 | RTCB_HUMAN | Oxidation (M) | 89287 | 172000 | 228580 | 175790 | 101780 | 0 | 0 | 0 | 98625 | 0 | 248080 | 2643100 | 852480 | 107260 | 298820 | 185050 | 0 | 0 |
| LADMGIAIF | RTCB | Q9Y3I0 | RTCB_HUMAN | Unmodified | 0 | 0 | 95590 | 157470 | 0 | 0 | 0 | 0 | 212490 | 0 | 0 | 1468000 | 390800 | 0 | 121500 | 0 | 0 | 0 |
| LADMYGG( | CDH2; CDH2 | P19022; C9J | CADH2_HUMA | Oxidation (M) | 0 | 0 | 0 | 0 | 0 | 0 | 0 | 0 | 271720 | 0 | 0 | 0 | 0 | 0 | 0 | 0 | 845750 | 791210 |
| LADMYGG( | CDH1; CDH3 | P12830; P22 | CADH1_HUMA | Oxidation (M) | 123940 | 126120 | 393790 | 641630 | 4652100 | 621670 | 0 | 0 | 0 | 52533 | 166580 | 532880 | 1625500 | 1880400 | 893780 | 729020 | 0 | 0 |
| LADNSDW | ADGRG6 | Q86SQ4 | AGRG6_HUMA | Unmodified | 823470 | 576150 | 1244900 | 1509400 | 1614100 | 35406 | 0 | 0 | 343800 | 58084 | 0 | 1028500 | 0 | 0 | 0 | 3034800 | 0 | 0 |
| LADPGDRE | PLSCR3; PLSCF | Q9NRY6; I3L | PLS3_HUMAN I | Unmodified | 11720000 | 7108800 | 0 | 1125200 | 81662 | 0 | 0 | 0 | 1948400 | 681400 | 210950 | 7610700 | 0 | 0 | 4428900 | 4700200 | 0 | 0 |
| LADQFLEE( | GOLM2; GOLM | Q6P4E1; H0 | GOLM2_HUMA | Unmodified | 0 | 0 | 0 | 0 | 0 | 0 | 88030 | 34930 | 3288000 | 351770 | 0 | 0 | 0 | 0 | 0 | 0 | 0 | 0 |
| LADQIFAYY | CYFIP1; CYFIP2 | Q7L576; Q9 | CYFP1_HUMAN | Unmodified | 0 | 0 | 350070 | 0 | 181370 | 0 | 700400 | 125020 | 0 | 124950 | 0 | 2410900 | 0 | 0 | 859650 | 554070 | 0 | 0 |
| LADVDKDG | EHD1; EHD1 | Q9H4M9; A0 | EHD1_HUMAN | Unmodified | 11318000 | 12467000 | 1254100 | 7997000 | 2.03E+08 | 34535000 | 1362400 | 62344 | 8587500 | 6202500 | 24969000 | 41868000 | 43136000 | 26001000 | 21736000 | 6772200 | 85374000 | 2.04E+08 |
| LADYYPINS | SLC4A4 | Q9Y6R1 | S4A4_HUMAN | Unmodified | 0 | 0 | 221640 | 0 | 0 | 0 | 0 | 0 | 0 | 0 | 0 | 0 | 0 | 0 | 445410 | 581180 | 0 | 0 |
| LAEAGESA( | DCHS1 | Q96JQ0 | PCD16_HUMAI | Unmodified | 0 | 0 | 0 | 0 | 3915400 | 1536100 | 10710000 | 27346 | 801620 | 410760 | 205380 | 89250 | 0 | 0 | 0 | 0 | 0 | 1699900 |

| | | | | | | | | | | | | | | | | | | | | | | |
|---|---|---|---|---|---|---|---|---|---|---|---|---|---|---|---|---|---|---|---|---|---|---|
| LAEAPSPAF | SCRIB; SCRIB; S | Q14160; A0A | SCRIB_HUMAN | Unmodified | 0 | 0 | 0 | 0 | 0 | 0 | 0 | 0 | 3585100 | 0 | 0 | 1154700 | 0 | 0 | 981130 | 0 | 2497200 | 0 |
| LAEAQIEEL | DIABLO; DIABL( | Q9NR28; A0, | DBLOH_HUMA | Unmodified | 0 | 0 | 0 | 0 | 0 | 0 | 0 | 0 | 2060200 | 0 | 0 | 0 | 0 | 0 | 0 | 0 | 0 | 0 |
| LAEAQIEEL | DIABLO; DIABL( | Q9NR28; A0, | DBLOH_HUMA | Unmodified | 0 | 0 | 0 | 0 | 0 | 0 | 0 | 45056 | 4435200 | 811750 | 340420 | 1961300 | 0 | 0 | 0 | 113880 | 38238 | 91539 |
| LAEDAPNF | EIF4G2; EIF4G2 | P78344; H0Y | IF4G2_HUMAN | Unmodified | 0 | 131230 | 0 | 0 | 183650 | 0 | 0 | 0 | 0 | 0 | 0 | 1508500 | 0 | 0 | 0 | 255660 | 0 | 0 |
| LAEEAFFR | FURIN; FURIN; | P09958; A0A | FURIN_HUMAN | Unmodified | 0 | 0 | 212990 | 0 | 0 | 0 | 0 | 0 | 1530600 | 58183 | 1803200 | 0 | 65618 | 0 | 152220 | 91736 | 0 | 0 |
| LAEEESCRI | GLG1; GLG1; G | Q92896; H3I | GSLG1_HUMAI | Unmodified | 82001 | 78523 | 0 | 0 | 0 | 939950 | 0 | 0 | 9642400 | 319510 | 1143700 | 3694200 | 0 | 0 | 0 | 0 | 121040 | 0 |
| LAEEESCRI | GLG1; GLG1; G | Q92896; H3I | GSLG1_HUMAI | Unmodified | 0 | 0 | 0 | 0 | 0 | 0 | 0 | 0 | 5416000 | 1298000 | 0 | 0 | 0 | 0 | 0 | 0 | 0 | 0 |
| LAEEFPGQ | SLC38A10; SLC | Q9HBR0; H0 | S38AA_HUMAN | Unmodified | 0 | 0 | 0 | 418510 | 0 | 0 | 94651 | 62901 | 3449500 | 0 | 0 | 0 | 0 | 182250 | 0 | 373770 | 0 | 0 |
| LAEEQQK | UBTF; UBTF; UE | P17480; H0Y | UBF1_HUMAN | Unmodified | 53478 | 398380 | 0 | 0 | 479950 | 95097 | 170420 | 194960 | 1236100 | 162540 | 278510 | 0 | 40462 | 0 | 0 | 0 | 50409 | 0 |
| LAEEVPPPF | CASC3; CASC3 | O15234; J3Q | CASC3_HUMAI | Unmodified | 0 | 0 | 544360 | 0 | 0 | 0 | 0 | 0 | 0 | 0 | 0 | 0 | 0 | 0 | 0 | 1239800 | 0 | 0 |
| LAEEVSGR | IGSF3 | O75054 | IGSF3_HUMAN | Unmodified | 0 | 0 | 245290 | 196820 | 0 | 0 | 0 | 0 | 0 | 0 | 0 | 968700 | 0 | 0 | 0 | 0 | 0 | 0 |
| LAEEVSGR | IGSF3 | O75054 | IGSF3_HUMAN | Unmodified | 0 | 0 | 0 | 1396500 | 0 | 147860 | 0 | 0 | 552470 | 0 | 68448 | 1284000 | 316450 | 68615 | 0 | 0 | 49506 | 0 |
| LAEFIDFLD | FUT11 | Q495W5 | FUT11_HUMAN | Oxidation (M) | 0 | 0 | 0 | 0 | 0 | 0 | 0 | 0 | 2953300 | 408610 | 201960 | 0 | 0 | 0 | 0 | 0 | 0 | 0 |
| LAEFIDFLD | FUT11 | Q495W5 | FUT11_HUMAN | Unmodified | 0 | 0 | 0 | 0 | 0 | 0 | 0 | 0 | 2302300 | 82365 | 0 | 0 | 0 | 0 | 0 | 0 | 0 | 0 |
| LAEFKLENI | VPS13A | Q96RL7 | VP13A_HUMAN | Unmodified | 0 | 0 | 0 | 0 | 520070 | 90310 | 123890 | 234910 | 0 | 116290 | 300290 | 869100 | 164000 | 141170 | 0 | 0 | 517450 | 128920 |
| LAEFQTDS | DPP9; DPP9 | Q86TI2; M0R | DPP9_HUMAN | Unmodified | 0 | 0 | 261700 | 0 | 1749300 | 142620 | 0 | 0 | 279390 | 0 | 195000 | 647980 | 168240 | 0 | 0 | 0 | 557980 | 1076700 |
| LAEFQTDS | DPP9; DPP9 | Q86TI2; M0R | DPP9_HUMAN | Unmodified | 0 | 0 | 0 | 0 | 1135800 | 0 | 0 | 0 | 0 | 0 | 0 | 0 | 380280 | 0 | 0 | 0 | 726490 | 0 |
| LAEIGAPIQ | SF3B2; SF3B2; | Q13435; E9F | SF3B2_HUMAN | Unmodified | 2253100 | 1525800 | 580330 | 0 | 0 | 0 | 0 | 0 | 582720 | 63387 | 1568000 | 8601600 | 0 | 0 | 546550 | 906650 | 0 | 0 |
| LAEIPYDVQ | MGAM2 | Q2M2H8 | MGAL_HUMAN | Oxidation (M) | 0 | 0 | 0 | 0 | 0 | 0 | 0 | 0 | 0 | 0 | 0 | 8112600 | 0 | 0 | 0 | 0 | 0 | 0 |
| LAEISDVWI | SPTBN1; SPTBN | Q01082; A0A | SPTB2_HUMAN | Unmodified | 0 | 21364 | 0 | 0 | 625330 | 56989 | 48024 | 0 | 0 | 165870 | 0 | 0 | 0 | 0 | 0 | 0 | 0 | 0 |
| LAEKEETGI | TGM2 | P21980 | TGM2_HUMAN | Oxidation (M) | 0 | 0 | 0 | 0 | 1212800 | 781230 | 0 | 0 | 0 | 0 | 12222 | 0 | 0 | 0 | 0 | 0 | 0 | 0 |
| LAEKEETGI | TGM2 | P21980 | TGM2_HUMAN | Unmodified | 0 | 0 | 0 | 0 | 2892500 | 630580 | 0 | 0 | 0 | 0 | 0 | 86535 | 0 | 0 | 0 | 0 | 0 | 0 |
| LAEKEETGI | TGM2 | P21980 | TGM2_HUMAN | Unmodified | 0 | 0 | 0 | 0 | 1627100 | 330420 | 0 | 0 | 0 | 0 | 0 | 0 | 0 | 0 | 0 | 0 | 0 | 0 |
| LAEKFPVIS | SNX18 | Q96RF0 | SNX18_HUMAN | Unmodified | 3560300 | 4025700 | 0 | 6807700 | 0 | 0 | 0 | 0 | 0 | 0 | 0 | 0 | 8318100 | 5321600 | 2485800 | 4196800 | 0 | 0 |
| LAEKLTILNI | NCKAP1 | Q9Y2A7 | NCKP1_HUMAI | Unmodified | 0 | 0 | 0 | 0 | 550060 | 0 | 0 | 0 | 0 | 68406 | 0 | 0 | 136280 | 55561 | 0 | 0 | 91354 | 161570 |
| LAELEEFIN | CLTC; CLTC; Cl | Q00610; J3K | CLH1_HUMAN | Unmodified | 18859000 | 18772000 | 8316700 | 5875000 | 0 | 0 | 0 | 65833 | 3458800 | 1874200 | 1019100 | 1.4E+08 | 0 | 0 | 12579000 | 13171000 | 0 | 0 |
| LAELEEFIN | CLTC; CLTC; Cl | Q00610; J3K | CLH1_HUMAN | Unmodified | 10795000 | 7807200 | 2573900 | 6742100 | 0 | 0 | 0 | 0 | 1844400 | 0 | 494410 | 22327000 | 631670 | 0 | 5760300 | 6548100 | 0 | 0 |
| LAELHADLI | LRPAP1; LRPAP | P30533; A0A | AMRP_HUMAN | Unmodified | 0 | 0 | 3218200 | 0 | 860100 | 0 | 0 | 124930 | 1505000 | 233290 | 0 | 0 | 296290 | 1748300 | 0 | 0 | 0 | 1528500 |
| LAELKNLG | HSPA4; HSPA4 | P34932; A0A | HSP74_HUMAI | Unmodified | 0 | 0 | 0 | 0 | 213690 | 0 | 0 | 31305 | 0 | 0 | 0 | 0 | 0 | 0 | 0 | 0 | 225390 | 0 |
| LAELLSGK | CAPN15; CAPN | O75808; A0A | CAN15_HUMAI | Unmodified | 0 | 258880 | 0 | 0 | 0 | 0 | 0 | 0 | 0 | 0 | 0 | 0 | 0 | 0 | 0 | 0 | 0 | 0 |
| LAELLVSVL | RIC8A; RIC8A | Q9NPQ8; E9 | RIC8A_HUMAN | Unmodified | 0 | 0 | 0 | 0 | 0 | 0 | 0 | 0 | 386430 | 78190 | 19307 | 153670 | 0 | 0 | 0 | 0 | 0 | 0 |
| LAELNRNF | UTRN | P46939 | UTRN_HUMAN | Unmodified | 0 | 0 | 0 | 726300 | 1706400 | 240190 | 0 | 58149 | 288390 | 640330 | 0 | 303690 | 526970 | 459310 | 0 | 0 | 873030 | 437010 |
| LAELNRNF | UTRN | P46939 | UTRN_HUMAN | Unmodified | 7495600 | 0 | 0 | 3190100 | 292650 | 99631 | 0 | 0 | 330610 | 0 | 0 | 783250 | 1787700 | 335900 | 0 | 0 | 0 | 115040 |
| LAELRGVPI | DPYSL2; DPYSL | Q16555; A0A | DPYL2_HUMAN | Unmodified | 0 | 0 | 0 | 0 | 1924300 | 372940 | 0 | 0 | 0 | 253140 | 0 | 0 | 0 | 0 | 0 | 0 | 0 | 0 |
| LAENIDAQI | NUP62; NUP62 | P37198; M0( | NUP62_HUMAI | Unmodified | 0 | 0 | 0 | 0 | 455060 | 0 | 105840 | 70914 | 0 | 0 | 0 | 0 | 457170 | 143370 | 0 | 0 | 213080 | 276710 |
| LAENNINLI | ITGB5; ITGB5; I | P18084; H7( | ITB5_HUMAN I | Unmodified | 0 | 0 | 0 | 0 | 418120 | 172740 | 0 | 0 | 257210 | 695230 | 418590 | 2421300 | 0 | 0 | 0 | 0 | 0 | 0 |
| LAENSDWS | ADGRG2 | Q8IZP9 | AGRG2_HUMA | Unmodified | 0 | 27497 | 0 | 0 | 119900 | 0 | 0 | 0 | 435290 | 161470 | 37674 | 0 | 0 | 0 | 0 | 0 | 74340 | 0 |
| LAEPGQSC | MGAT5 | Q09328 | MGT5A_HUMAI | Unmodified | 0 | 0 | 0 | 0 | 0 | 0 | 0 | 0 | 2368600 | 0 | 0 | 526720 | 0 | 0 | 0 | 0 | 0 | 0 |
| LAEPSQLLK | JUP; JUP; JUP; J | P14923; C9J | PLAK_HUMAN | Unmodified | 1034000 | 790990 | 7032600 | 9794600 | 21145000 | 6899400 | 0 | 0 | 6140700 | 2121600 | 6381000 | 17093000 | 13993000 | 6677200 | 11076000 | 9527100 | 3246100 | 5024800 |
| LAEPSQML | CTNNB1; CTNN | P35222; A0A | CTNB1_HUMAI | Unmodified | 764800 | 754170 | 2073000 | 4073200 | 0 | 3034800 | 0 | 0 | 1634800 | 324740 | 1505600 | 6737500 | 11193000 | 6830200 | 3757500 | 3030500 | 3753200 | 7080000 |
| LAEPSQML | CTNNB1; CTNN | P35222; A0A | CTNB1_HUMAI | Oxidation (M) | 0 | 0 | 752770 | 1535800 | 3654300 | 0 | 0 | 0 | 0 | 0 | 369200 | 1196200 | 2757800 | 493610 | 1261600 | 790740 | 0 | 2147600 |
| LAEQAAQL | TRIM41; TRIM4 | Q8WV44; H( | TRI41_HUMAN | Unmodified | 32425000 | 772510 | 114230 | 78473 | 0 | 0 | 0 | 0 | 0 | 0 | 0 | 0 | 0 | 0 | 446460 | 137210 | 0 | 0 |
| LAEQAERYI | YWHAB; YWHA | P31946; P61 | 1433B_HUMAN | Oxidation (M) | 660200 | 636740 | 5904700 | 3756800 | 10956000 | 3722800 | 515140 | 915680 | 3494700 | 537530 | 2076500 | 19917000 | 6829800 | 835650 | 5372900 | 5020100 | 17050000 | 16870000 |
| LAEQAERYI | YWHAB; YWHA | P31946; P61 | 1433B_HUMAN | Unmodified | 1142200 | 1120300 | 2686500 | 4807300 | 62172000 | 6985700 | 5058000 | 1221500 | 19160000 | 5244400 | 7038900 | 40184000 | 13756000 | 10474000 | 8342600 | 3914800 | 26717000 | 72027000 |
| LAEQAERYI | YWHAB; YWHA | P31946; P61 | 1433B_HUMAN | 2 Oxidation (M) | 0 | 0 | 2192400 | 1155100 | 3156600 | 794800 | 835240 | 509860 | 903580 | 330050 | 1303500 | 4443700 | 1970000 | 1217700 | 3272300 | 3267200 | 3079000 | 3611700 |
| LAEQAERYI | YWHAZ; YWHA | P63104; E5R | 1433Z_HUMAN | Oxidation (M) | 660490 | 958550 | 2599100 | 4635600 | 13287000 | 2241100 | 4342200 | 3046700 | 6304300 | 4599400 | 2584900 | 9946500 | 2448600 | 5582200 | 3480600 | 3138000 | 21466000 | 22318000 |
| LAEQAERYI | YWHAZ; YWHA | P63104; E5R | 1433Z_HUMAN | 2 Oxidation (M) | 0 | 138710 | 0 | 753560 | 3133700 | 762770 | 1232300 | 695940 | 2845700 | 743540 | 398280 | 2803800 | 2267500 | 2468200 | 406050 | 0 | 6796400 | 6782200 |
| LAEQAERYI | YWHAZ; YWHA | P63104; E5R | 1433Z_HUMAN | Unmodified | 1191900 | 1342600 | 761480 | 3183700 | 29540000 | 6081600 | 8373100 | 3182300 | 34265000 | 10058000 | 3731500 | 22993000 | 9773000 | 12274000 | 2111200 | 1320900 | 58099000 | 44615000 |
| LAEQAERYI | YWHAH; YWHA | Q04917; F8V | 1433F_HUMAN | Oxidation (M) | 55102 | 26271 | 39655 | 993610 | 5086100 | 631710 | 474730 | 761530 | 1183400 | 312160 | 408420 | 1438000 | 1777500 | 702440 | 281040 | 1158500 | 3612200 | 1547900 |
| LAEQAERYI | YWHAH; YWHA | Q04917; F8V | 1433F_HUMAN | 2 Oxidation (M) | 69950 | 103210 | 121100 | 156600 | 964190 | 184710 | 149410 | 208390 | 132590 | 92285 | 375220 | 706250 | 338370 | 219000 | 140880 | 176390 | 472440 | 1225400 |
| LAEQAERYI | YWHAH; YWHA | Q04917; F8V | 1433F_HUMAN | Unmodified | 651460 | 0 | 687780 | 4541800 | 25143000 | 4328700 | 4549500 | 1164300 | 7361100 | 3185200 | 2470200 | 11692000 | 6334800 | 6248100 | 1925300 | 3232100 | 7491300 | 9165200 |
| LAEQAERYI | YWHAQ; YWHA | P27348; E9P | 1433T_HUMAN | 2 Oxidation (M) | 84306 | 56182 | 391070 | 431690 | 1397000 | 300530 | 1054600 | 168230 | 575830 | 310840 | 446370 | 2685300 | 1098100 | 509820 | 291510 | 980730 | 1723200 | 1943100 |
| LAEQAERYI | YWHAQ; YWHA | P27348; E9P | 1433T_HUMAN | Oxidation (M) | 609130 | 370030 | 295020 | 2968000 | 7392200 | 508970 | 420530 | 995510 | 1967700 | 1112800 | 2266300 | 12545000 | 5599200 | 4347300 | 2417500 | 1190100 | 7067800 | 6735000 |
| LAEQAERYI | YWHAQ; YWHA | P27348; E9P | 1433T_HUMAN | Unmodified | 432820 | 273720 | 0 | 1425700 | 19110000 | 3731500 | 3481500 | 686520 | 9589100 | 2142900 | 765330 | 24041000 | 8279900 | 8824400 | 1483400 | 1116200 | 23142000 | 18812000 |
| LAEQAERYI | YWHAE; YWHA | P62258; I3L | 1433E_HUMAN | Unmodified | 518780 | 632730 | 382740 | 4227200 | 11601000 | 2434600 | 2224500 | 373320 | 5035100 | 3436100 | 5507500 | 16039000 | 12316000 | 10680000 | 2921700 | 1598100 | 5018500 | 3256700 |
| LAEQAERYI | YWHAE; YWHA | P62258; I3L | 1433E_HUMAN | Oxidation (M) | 526430 | 257640 | 1933400 | 2904100 | 5109900 | 1500900 | 695810 | 158460 | 2127600 | 1736500 | 1958400 | 21493000 | 6651600 | 2498800 | 3903200 | 5024700 | 2077000 | 588210 |
| LAEQAERYI | YWHAE; YWHA | P62258; I3L | 1433E_HUMAN | 2 Oxidation (M) | 158050 | 124840 | 530610 | 72741 | 484590 | 0 | 0 | 90185 | 0 | 389560 | 785520 | 2979100 | 1317900 | 455800 | 589960 | 0 | 330730 | 0 |
| LAEQAERYI | YWHAE; YWHA | P62258; I3L | 1433E_HUMAN | Oxidation (M) | 3043600 | 817540 | 5350000 | 5384300 | 7577200 | 6019000 | 3085100 | 1729800 | 3130800 | 4807000 | 7045100 | 50785000 | 13654000 | 9834000 | 9808000 | 8455900 | 7998800 | 2581400 |
| LAEQAERYI | YWHAE; YWHA | P62258; I3L | 1433E_HUMAN | Unmodified | 2004200 | 1820700 | 2708000 | 6066000 | 34025000 | 11052000 | 8749400 | 1214400 | 4301000 | 14125000 | 9274900 | 60227000 | 25671000 | 30199000 | 8695600 | 4561800 | 19462000 | 9465700 |
| LAEQAERYI | YWHAE; YWHA | P62258; I3L | 1433E_HUMAN | 2 Oxidation (M) | 0 | 0 | 0 | 0 | 0 | 0 | 0 | 52275 | 0 | 0 | 398050 | 1085800 | 0 | 0 | 231630 | 132260 | 0 | 0 |
| LAEQAERYI | SFN | P31947 | 1433S_HUMAN | 2 Oxidation (M) | 0 | 0 | 0 | 0 | 3462300 | 1893100 | 0 | 0 | 501150 | 228090 | 0 | 0 | 0 | 0 | 0 | 0 | 1121500 | 712580 |
| LAEQAERYI | SFN | P31947 | 1433S_HUMAN | Oxidation (M) | 755560 | 0 | 0 | 0 | 13449000 | 2614900 | 0 | 0 | 3237200 | 736540 | 0 | 0 | 0 | 0 | 0 | 0 | 4071700 | 566510 |
| LAEQAERYI | SFN | P31947 | 1433S_HUMAN | Unmodified | 115980 | 140480 | 0 | 0 | 16895000 | 1198700 | 0 | 0 | 6765200 | 1289700 | 0 | 0 | 55800 | 0 | 0 | 0 | 5310800 | 4841600 |
| LAEQFPR | BTF3L4; BTF3L | Q96K17; E9F | BT3L4_HUMAN | Unmodified | 332630 | 436580 | 1146700 | 1884800 | 4317000 | 0 | 0 | 0 | 1086900 | 125470 | 83303 | 2403100 | 0 | 0 | 1249100 | 1267500 | 0 | 379120 |
| LAEQIATLC | STXBP1; STXBP | P61764; A0A | STXB1_HUMAN | Unmodified | 110440 | 0 | 0 | 110060 | 0 | 0 | 0 | 39871 | 183800 | 38203 | 61859 | 1663000 | 17000 | 0 | 38818 | 230830 | 0 | 0 |
| LAERESQE | HIP1R | O75146 | HIP1R_HUMAN | Unmodified | 0 | 91032 | 0 | 0 | 0 | 18254 | 0 | 0 | 0 | 0 | 0 | 580310 | 0 | 0 | 0 | 251980 | 0 | 0 |
| LAERPYIIA | PCYT2; PCYT2; | Q99447; I3L | PCY2_HUMAN | Unmodified | 0 | 0 | 0 | 0 | 0 | 0 | 0 | 0 | 170320 | 0 | 0 | 2908400 | 0 | 0 | 0 | 0 | 0 | 0 |
| LAERQLQE | INCENP | Q9NQS7 | INCE_HUMAN I | Unmodified | 0 | 0 | 0 | 0 | 560870 | 803610 | 66207 | 0 | 0 | 0 | 0 | 0 | 0 | 0 | 0 | 0 | 143410 | 156710 |
| LAERTSNLI | IGSF3 | O75054 | IGSF3_HUMAN | Unmodified | 0 | 0 | 0 | 122080 | 367970 | 0 | 0 | 0 | 0 | 0 | 0 | 0 | 345940 | 0 | 0 | 0 | 0 | 0 |
| LAERTSNLI | IGSF3 | O75054 | IGSF3_HUMAN | Unmodified | 0 | 0 | 0 | 0 | 0 | 0 | 0 | 0 | 0 | 0 | 0 | 0 | 758880 | 610890 | 0 | 0 | 0 | 0 |
| LAETMNNII | FNBP1L; FNBP1 | Q5T0N5; A0 | FBP1L_HUMAN | Unmodified | 1104000 | 1255800 | 219290 | 183720 | 57812 | 0 | 0 | 0 | 443620 | 0 | 0 | 380650 | 64980 | 0 | 362660 | 181070 | 0 | 0 |
| LAETVFNF | DYNC1H1; DYN | Q14204; A0A | DYHC1_HUMA | Unmodified | 0 | 0 | 0 | 74968 | 0 | 0 | 0 | 0 | 0 | 0 | 267780 | 2552600 | 0 | 0 | 0 | 0 | 0 | 0 |
| LAETYLHR | DDX6 | P26196 | DDX6_HUMAN | Unmodified | 184510 | 87986 | 0 | 0 | 0 | 0 | 0 | 65703 | 43453 | 56675 | 56588 | 335670 | 0 | 0 | 121970 | 98915 | 0 | 0 |
| LAEVEAALE | PLEC | Q15149 | PLEC_HUMAN | Unmodified | 0 | 0 | 0 | 0 | 0 | 0 | 0 | 0 | 0 | 0 | 0 | 1535200 | 0 | 0 | 0 | 0 | 0 | 0 |
| LAEVGQYE | CNOT1 | A5YKK6 | CNOT1_HUMA | Unmodified | 0 | 0 | 0 | 0 | 0 | 0 | 0 | 0 | 0 | 0 | 0 | 1365000 | 0 | 0 | 0 | 0 | 0 | 0 |
| LAEVKDSLI | SH3GL1; SH3G | Q99961; M0I | SH3G1_HUMAI | Unmodified | 589710 | 0 | 0 | 0 | 0 | 0 | 0 | 0 | 0 | 0 | 0 | 0 | 0 | 0 | 0 | 641060 | 0 | 0 |
| LAEVSQNIE | FNBP1; FNBP1; | Q96RU3; H0 | FNBP1_HUMAI | Unmodified | 375400 | 337870 | 82369 | 0 | 0 | 0 | 0 | 0 | 0 | 0 | 0 | 0 | 0 | 0 | 134010 | 168830 | 0 | 0 |
| LAEVWMDI | GALNT1; GALN | Q10472; Q8I | GALT1_HUMAN | Oxidation (M) | 0 | 0 | 0 | 0 | 0 | 0 | 0 | 0 | 2417200 | 0 | 373220 | 1757800 | 0 | 0 | 0 | 0 | 0 | 0 |
| LAEVWMDI | GALNT1; GALN | Q10472; Q8I | GALT1_HUMAN | Unmodified | 0 | 0 | 0 | 0 | 0 | 0 | 0 | 0 | 1399200 | 0 | 0 | 0 | 0 | 0 | 0 | 0 | 0 | 0 |
| LAEVWMDI | GALNT3; GALN | Q14435; E7E | GALT3_HUMAN | Unmodified | 0 | 0 | 0 | 0 | 0 | 0 | 0 | 0 | 611900 | 0 | 0 | 0 | 0 | 0 | 0 | 0 | 0 | 0 |
| LAFETGPPF | NOTCH1; NOT( | P46531; A0A | NOTC1_HUMA | Unmodified | 203540 | 119770 | 159500 | 195390 | 0 | 0 | 0 | 0 | 476780 | 0 | 142630 | 1765300 | 0 | 0 | 333090 | 286490 | 0 | 0 |
| LAFLNVQA | GPD2 | P43304 | GPDM_HUMAN | Unmodified | 0 | 0 | 0 | 0 | 0 | 0 | 0 | 0 | 10970000 | 2087700 | 0 | 0 | 0 | 0 | 0 | 0 | 0 | 0 |
| LAFLPPEAT | ABHD17A; ABH | Q96GS6; A0, | AB17A_HUMAN | Unmodified | 0 | 0 | 0 | 0 | 0 | 0 | 0 | 0 | 815940 | 0 | 0 | 0 | 0 | 0 | 0 | 168350 | 0 | 0 |
| LAFPSPFGI | SQSTM1; SQST | Q13501; E9F | SQSTM_HUMAI | Unmodified | 1840600 | 1462500 | 3569500 | 3228000 | 18857000 | 3548500 | 4398100 | 41615 | 14445000 | 6717500 | 472500 | 818470 | 13571000 | 4604300 | 1725700 | 2661200 | 4946300 | 9126500 |
| LAFSSTAR | PMPCA | Q10713 | MPPA_HUMAN | Unmodified | 0 | 204360 | 0 | 0 | 0 | 35447 | 44431 | 1223200 | 1605400 | 89709 | 76166 | 111680 | 0 | 0 | 0 | 0 | 183410 | 50515 |
| LAFYPNQD | GAPVD1; GAPV | Q14C86; F8 | GAPD1_HUMAI | Unmodified | 0 | 0 | 0 | 0 | 0 | 0 | 0 | 0 | 496120 | 51170 | 0 | 3549300 | 0 | 0 | 0 | 0 | 0 | 0 |
| LAGANPAV | PRKDC | P78527 | PRKDC_HUMAI | Unmodified | 226530 | 189230 | 933590 | 240620 | 254800 | 0 | 826090 | 163070 | 0 | 0 | 131000 | 2985100 | 288610 | 381810 | 0 | 0 | 0 | 0 |
| LAGAPLPEI | STS; STS; STS; S | P08842; A0A | STS_HUMAN S | Unmodified | 0 | 0 | 0 | 0 | 0 | 0 | 0 | 0 | 1593600 | 355070 | 0 | 0 | 0 | 0 | 0 | 0 | 0 | 0 |
| LAGAPLPEI | STS; STS; STS; S | P08842; A0A | STS_HUMAN S | Unmodified | 0 | 0 | 0 | 0 | 0 | 0 | 0 | 0 | 1121900 | 38674 | 0 | 0 | 0 | 0 | 0 | 0 | 0 | 0 |
| LAGAPLPEI | STS; STS; STS; S | P08842; A0A | STS_HUMAN S | Unmodified | 0 | 0 | 0 | 0 | 0 | 0 | 0 | 0 | 3038800 | 726610 | 0 | 0 | 0 | 0 | 0 | 0 | 0 | 0 |
| LAGAPSED | QSOX1 | O00391 | QSOX1_HUMAI | Unmodified | 0 | 0 | 1019200 | 421110 | 586920 | 1572200 | 0 | 0 | 4808600 | 1024100 | 1149000 | 457930 | 107050 | 165170 | 0 | 0 | 130510 | 372170 |
| LAGATISEE | ITGB6; ITGB6; I | P18564; A0A | ITB6_HUMAN I | Unmodified | 0 | 0 | 0 | 0 | 612750 | 0 | 0 | 0 | 0 | 0 | 0 | 2538000 | 0 | 0 | 0 | 0 | 0 | 0 |
| LAGCTVFIT | HSDL2 | Q6YN16 | HSDL2_HUMAI | Unmodified | 0 | 0 | 0 | 0 | 0 | 0 | 0 | 16479 | 8812100 | 780270 | 0 | 0 | 0 | 0 | 0 | 0 | 0 | 0 |
| LAGDPSAG | PDXP | Q96GD0 | PLPP_HUMAN | Unmodified | 0 | 0 | 0 | 0 | 0 | 0 | 0 | 0 | 0 | 0 | 191100 | 514160 | 0 | 0 | 0 | 87359 | 0 | 0 |
| LAGENMTG | AIFM1; AIFM1; | O95831; E9F | AIFM1_HUMAN | Unmodified | 0 | 0 | 0 | 0 | 0 | 0 | 0 | 0 | 0 | 0 | 0 | 700510 | 0 | 0 | 0 | 0 | 0 | 0 |
| LAGESESN | VCP; VCP; VCP; | P55072; C9J | TERA_HUMAN | Unmodified | 0 | 332900 | 0 | 364340 | 2110600 | 681100 | 0 | 75081 | 1514100 | 0 | 2355200 | 2315500 | 0 | 0 | 601050 | 495560 | 0 | 434880 |
| LAGESESN | VCP; VCP; VCP; | P55072; C9J | TERA_HUMAN | Unmodified | 1393300 | 756730 | 909830 | 3030200 | 22449000 | 4969300 | 547740 | 212290 | 1714900 | 10374000 | 4981300 | 13408000 | 17508000 | 12749000 | 1058800 | 97853 | 306910 | 0 |
| LAGESESN | VCP; VCP; VCP; | P55072; C9J | TERA_HUMAN | Unmodified | 0 | 0 | 0 | 0 | 3030700 | 0 | 1002500 | 696320 | 526290 | 2114500 | 0 | 0 | 4998600 | 2668900 | 0 | 0 | 0 | 338920 |
| LAGFLDLTE | EIF3L; EIF3L; EI | Q9Y262; B0( | EIF3L_HUMAN | Unmodified | 43439 | 80819 | 22130 | 46910 | 0 | 0 | 12216 | 0 | 618230 | 28782 | 494260 | 2871900 | 0 | 0 | 84519 | 79476 | 0 | 0 |
| LAGGYENV | PLOD3 | O60568 | PLOD3_HUMAI | Oxidation (M) | 0 | 0 | 319680 | 0 | 0 | 0 | 0 | 0 | 4012200 | 586220 | 8076300 | 0 | 0 | 0 | 0 | 0 | 0 | 0 |
| LAGGYENV | PLOD3 | O60568 | PLOD3_HUMAI | Unmodified | 0 | 0 | 0 | 0 | 0 | 0 | 0 | 0 | 16082000 | 744770 | 0 | 213680 | 0 | 0 | 0 | 0 | 0 | 0 |
| LAGHLSED | QSOX2 | Q6ZRP7 | QSOX2_HUMAI | Unmodified | 0 | 0 | 0 | 0 | 0 | 0 | 0 | 0 | 13998000 | 298670 | 459110 | 674990 | 0 | 0 | 0 | 0 | 0 | 0 |
| LAGIINFQR | PSMD12 | O00232 | PSD12_HUMAN | Unmodified | 64821 | 0 | 0 | 0 | 0 | 0 | 0 | 0 | 0 | 0 | 0 | 2902900 | 0 | 0 | 256060 | 206470 | 0 | 0 |
| LAGIINFQR | PSMD12 | O00232 | PSD12_HUMAN | Unmodified | 0 | 0 | 0 | 45015 | 0 | 0 | 0 | 0 | 0 | 0 | 0 | 535680 | 0 | 0 | 0 | 0 | 0 | 0 |
| LAGIPELLN | ANKRD13D | Q6ZTN6 | AN13D_HUMAI | Unmodified | 762610 | 551850 | 274620 | 0 | 0 | 0 | 0 | 0 | 0 | 0 | 0 | 535950 | 0 | 0 | 0 | 305820 | 0 | 0 |
| LAGIVIPND | ITGB6; ITGB6; I | P18564; A0A | ITB6_HUMAN I | Unmodified | 0 | 0 | 0 | 0 | 0 | 0 | 0 | 0 | 0 | 0 | 0 | 498260 | 0 | 0 | 0 | 0 | 0 | 0 |
| LAGIVQPNI | ITGB3; N/A; ITG | P05106; H3E | ITB3_HUMAN I | 2 Oxidation (M) | 5887600 | 0 | 0 | 0 | 0 | 0 | 570300 | 275900 | 0 | 0 | 0 | 0 | 1143300 | 966710 | 4011000 | 4018700 | 0 | 0 |
| LAGIVQPNI | ITGB3; N/A; ITG | P05106; H3E | ITB3_HUMAN I | Oxidation (M) | 1088100 | 1458500 | 0 | 7358300 | 667240 | 0 | 0 | 0 | 0 | 0 | 0 | 0 | 0 | 0 | 322830 | 280340 | 0 | 0 |
| LAGKENITE | BROX; BROX; B | Q5VW32; F5 | BROX_HUMAN | Unmodified | 2014300 | 2449800 | 0 | 1165700 | 0 | 0 | 0 | 145150 | 0 | 479750 | 0 | 1221800 | 148700 | 0 | 881660 | 1756900 | 0 | 0 |

| | | | | | | | | | | | | | | | | | | | | | | |
|---|---|---|---|---|---|---|---|---|---|---|---|---|---|---|---|---|---|---|---|---|---|---|
| LAGKENITE | BROX; BROX; B | Q5VW32; F5 | BROX_HUMAN | Unmodified | 876330 | 521680 | 0 | 859840 | 0 | 0 | 0 | 0 | 0 | 128000 | 0 | 0 | 0 | 0 | 0 | 0 | 0 | 0 |
| LAGLALPR | DUSP23 | Q9BVJ7 | DUS23_HUMAI | Unmodified | 0 | 0 | 0 | 0 | 0 | 0 | 0 | 0 | 128010 | 0 | 0 | 1236300 | 0 | 0 | 0 | 30279 | 0 | 0 |
| LAGLEYVLI | RPF2; RPF2 | Q9H7B2; Q5 | RPF2_HUMAN | Unmodified | 0 | 0 | 0 | 0 | 0 | 0 | 0 | 0 | 0 | 0 | 0 | 95211 | 0 | 0 | 0 | 0 | 0 | 0 |
| LAGLFNEQ | HYOU1; HYOU1 | Q9Y4L1; E9F | HYOU1_HUMA | Unmodified | 56069 | 0 | 169400 | 0 | 0 | 0 | 0 | 0 | 9289200 | 1257600 | 457460 | 0 | 365040 | 0 | 349250 | 225660 | 0 | 0 |
| LAGLFNEQ | HYOU1; HYOU1 | Q9Y4L1; E9F | HYOU1_HUMA | Unmodified | 0 | 0 | 0 | 0 | 0 | 0 | 0 | 0 | 5667900 | 1520600 | 313120 | 0 | 0 | 0 | 0 | 0 | 0 | 0 |
| LAGLHLEK | SELENON; N/A; | Q9NZV5; H3 | SELN_HUMAN | Unmodified | 0 | 0 | 0 | 0 | 0 | 0 | 0 | 0 | 504300 | 23903 | 0 | 0 | 0 | 0 | 0 | 0 | 0 | 0 |
| LAGLKPGT | CRLF1; CRLF1 | O75462; M0 | CRLF1_HUMAN | Unmodified | 0 | 0 | 0 | 0 | 0 | 0 | 0 | 0 | 5861100 | 257450 | 0 | 0 | 0 | 0 | 0 | 0 | 0 | 0 |
| LAGLLQHP | SMARCD2; SMA | Q92925; J3K | SMRD2_HUMA | Unmodified | 0 | 0 | 0 | 0 | 0 | 0 | 0 | 0 | 0 | 0 | 0 | 2119700 | 0 | 0 | 0 | 0 | 0 | 3150500 |
| LAGLNQRV | UTRN | P46939 | UTRN_HUMAN | Unmodified | 37099 | 0 | 529450 | 1692500 | 2923400 | 652060 | 65753 | 0 | 1173100 | 2122900 | 0 | 0 | 0 | 509910 | 957740 | 770690 | 2628900 | 1315800 |
| LAGPQLVQ | PSMC3; PSMC3 | P17980; E9P | PRS6A_HUMAN | Unmodified | 0 | 0 | 0 | 0 | 337540 | 108550 | 378550 | 0 | 464440 | 405930 | 0 | 0 | 714790 | 322490 | 0 | 0 | 930640 | 2204200 |
| LAGPQLVQ | PSMC3; PSMC3 | P17980; E9P | PRS6A_HUMAN | Oxidation (M) | 98675 | 80239 | 204570 | 315640 | 2253400 | 358200 | 2341300 | 375400 | 808120 | 1277400 | 555440 | 0 | 1954800 | 1365100 | 176810 | 156860 | 3005700 | 1703900 |
| LAGPSPYL | EEPD1 | Q7L9B9 | EEPD1_HUMAN | Unmodified | 0 | 0 | 0 | 0 | 0 | 0 | 0 | 0 | 0 | 116460 | 0 | 0 | 0 | 0 | 0 | 0 | 2131100 | 4045600 |
| LAGQRFTA | FHL1; FHL1; FH | Q13642; A0A | FHL1_HUMAN | Unmodified | 0 | 0 | 0 | 0 | 596860 | 0 | 1153200 | 0 | 0 | 0 | 0 | 0 | 0 | 0 | 0 | 0 | 344380 | 0 |
| LAGSLLTQ | MOGS; MOGS; | Q13724; A0A | MOGS_HUMAN | Unmodified | 0 | 0 | 0 | 0 | 0 | 63122 | 0 | 0 | 20973000 | 2076200 | 64212 | 0 | 0 | 0 | 0 | 0 | 0 | 0 |
| LAGSLLTQ | MOGS; MOGS; | Q13724; A0A | MOGS_HUMAN | Unmodified | 0 | 0 | 886100 | 0 | 0 | 0 | 0 | 0 | 35863000 | 3030700 | 601100 | 0 | 0 | 0 | 0 | 0 | 570580 | 0 |
| LAGSSGRG | PKP2 | Q99959 | PKP2_HUMAN | Unmodified | 0 | 0 | 0 | 0 | 252300 | 74068 | 0 | 0 | 0 | 0 | 0 | 0 | 282690 | 161760 | 0 | 0 | 1443600 | 1268200 |
| LAGTESPVI | TFRC; TFRC | P02786; G3 | TFR1_HUMAN | Unmodified | 0 | 65055 | 0 | 0 | 78257 | 0 | 0 | 0 | 1642900 | 140650 | 0 | 305470 | 0 | 0 | 0 | 0 | 35001 | 0 |
| LAGTQPLE | UBA1; UBA1 | P22314; Q5J | UBA1_HUMAN | Unmodified | 337800 | 500620 | 985230 | 673130 | 759420 | 301400 | 464760 | 132660 | 6372200 | 2300200 | 4205600 | 29076000 | 930970 | 0 | 1888500 | 0 | 737150 | 845280 |
| LAGVTALS | LYPLA1; LYPLA | O75608; A0A | LYPA1_HUMAN | Unmodified | 763330 | 354340 | 0 | 207100 | 0 | 0 | 0 | 0 | 1612100 | 227630 | 635250 | 4569300 | 0 | 0 | 122470 | 314990 | 0 | 0 |
| LAGWVLSK | CLGN; CLGN | O14967; D6 | CLGN_HUMAN | Unmodified | 0 | 0 | 0 | 0 | 105450 | 0 | 0 | 0 | 15843000 | 696540 | 898580 | 1476100 | 0 | 0 | 0 | 0 | 0 | 0 |
| LAHEVGWI | RPL13A; RPL13 | P40429; A0A | RL13A_HUMAN | Unmodified | 763580 | 905560 | 376580 | 235580 | 57675 | 0 | 0 | 0 | 184050 | 20952 | 367190 | 3921900 | 0 | 20554 | 497040 | 340820 | 0 | 73836 |
| LAHEVGWI | RPL13A; RPL13 | P40429; A0A | RL13A_HUMAN | Unmodified | 1339800 | 1222800 | 0 | 686350 | 214660 | 51125 | 169700 | 25959 | 0 | 172060 | 0 | 0 | 328960 | 108040 | 0 | 0 | 0 | 120330 |
| LAHLGVQV | RPN1; RPN1 | P04843; F8V | RPN1_HUMAN | Unmodified | 0 | 0 | 0 | 0 | 409200 | 2760100 | 0 | 0 | 1.73E+08 | 8131500 | 4758100 | 4262800 | 0 | 92210 | 0 | 0 | 266400 | 0 |
| LAHLGVQV | RPN1; RPN1 | P04843; F8V | RPN1_HUMAN | Unmodified | 0 | 0 | 0 | 856870 | 0 | 0 | 0 | 0 | 2.26E+08 | 26608000 | 11277000 | 0 | 242090 | 421290 | 0 | 0 | 0 | 0 |
| LAHPFSLA | LDLR; LDLR; LD | P01130; J3K | LDLR_HUMAN | Unmodified | 45429 | 84473 | 0 | 0 | 0 | 56968 | 0 | 0 | 1725500 | 39845 | 150250 | 0 | 0 | 0 | 0 | 13702 | 0 | 0 |
| LAHQSWAI | FUT11 | Q495W5 | FUT11_HUMAN | Unmodified | 0 | 0 | 0 | 0 | 0 | 0 | 0 | 0 | 5139000 | 0 | 0 | 0 | 0 | 0 | 0 | 0 | 0 | 0 |
| LAIAERYLV | LONP1; LONP1 | P36776; K7E | LONM_HUMAN | Unmodified | 0 | 0 | 0 | 0 | 0 | 0 | 308580 | 0 | 0 | 1435600 | 0 | 0 | 308600 | 257170 | 0 | 0 | 0 | 290760 |
| LAIEAER | CCDC50 | Q8IVM0 | CCD50_HUMA | Unmodified | 53085 | 2902300 | 1961200 | 2054000 | 345930 | 0 | 0 | 14888 | 218570 | 50218 | 0 | 169020 | 824560 | 0 | 3849500 | 0 | 202720 | 0 |
| LAIEAGFR | AKR1C4; AKR1C | P17516; P42 | AK1C4_HUMAN | Unmodified | 54804 | 220400 | 34564 | 0 | 2812000 | 0 | 0 | 99172 | 0 | 0 | 0 | 609140 | 0 | 0 | 18792 | 925240 | 0 | 0 |
| LAIKLPDD | PFKP; PFKP; PF | Q01813; A0A | PFKAP_HUMAN | Unmodified | 0 | 0 | 0 | 0 | 611550 | 0 | 0 | 0 | 553960 | 170430 | 1411600 | 5776200 | 0 | 0 | 0 | 0 | 500700 | 652190 |
| LAILGIHNE | ACTN1; ACTN1 | P12814; A0A | ACTN1_HUMAI | Unmodified | 351830 | 550580 | 474700 | 206190 | 1170100 | 0 | 0 | 0 | 389150 | 157080 | 493830 | 21292000 | 130680 | 0 | 1416800 | 1455400 | 473600 | 0 |
| LAILQVGNI | MTHFD1; MTHF | P11586; V9C | C1TC_HUMAN | Unmodified | 0 | 0 | 0 | 0 | 1564800 | 178700 | 541080 | 0 | 0 | 967020 | 287300 | 288550 | 2824600 | 335810 | 159200 | 54701 | 1411600 | 1102400 |
| LAINLLAK | EPS8L2; EPS8L | Q9H6S3; A0 | ES8L2_HUMAN | Unmodified | 0 | 0 | 0 | 0 | 0 | 0 | 0 | 0 | 0 | 0 | 0 | 0 | 281910 | 466350 | 0 | 0 | 0 | 0 |
| LAIQGPED | MLF2 | Q15773 | MLF2_HUMAN | Unmodified | 5113500 | 7640600 | 2584000 | 2044100 | 0 | 0 | 0 | 0 | 0 | 0 | 0 | 0 | 252960 | 0 | 1904000 | 1602800 | 0 | 0 |
| LAIQGPED | MLF2 | Q15773 | MLF2_HUMAN | Unmodified | 2351300 | 1860100 | 637330 | 5164800 | 0 | 0 | 0 | 0 | 0 | 0 | 0 | 0 | 2495700 | 739500 | 1169100 | 0 | 0 | 0 |
| LAIRESIESE | VCP; VCP; VCP; | P55072; C9J | TERA_HUMAN | Unmodified | 0 | 0 | 0 | 0 | 947060 | 239560 | 0 | 0 | 3098400 | 511720 | 0 | 0 | 0 | 0 | 0 | 0 | 0 | 72862 |
| LAISREANS | PACSIN2; PACS | Q9UNF0; A0 | PACN2_HUMAI | Unmodified | 0 | 0 | 0 | 360480 | 1380100 | 0 | 0 | 0 | 0 | 208970 | 0 | 0 | 3240100 | 1119500 | 0 | 0 | 1006700 | 0 |
| LAISREANS | PACSIN2; PACS | Q9UNF0; A0 | PACN2_HUMAI | Unmodified | 0 | 0 | 0 | 729460 | 19793000 | 4128600 | 6519200 | 3119300 | 0 | 2506600 | 0 | 0 | 1.51E+08 | 92619000 | 0 | 0 | 4332400 | 9892700 |
| LAIWDTAG | RAB21; RAB18 | Q9UL25; Q9 | RAB21_HUMAN | Unmodified | 471810 | 1007300 | 493310 | 583200 | 1932500 | 0 | 757440 | 540470 | 1187600 | 108680 | 973710 | 10232000 | 0 | 0 | 748640 | 0 | 473820 | 802380 |
| LAIWDTAG | RAB18 | Q9NP72 | RAB18_HUMAN | Unmodified | 0 | 0 | 0 | 0 | 971910 | 0 | 324630 | 51681 | 359040 | 426840 | 738730 | 0 | 1101400 | 387410 | 0 | 0 | 0 | 820810 |
| LAKPGEGL | SHANK2; SHAN | Q9UPX8; A6 | SHAN2_HUMAI | Unmodified | 0 | 0 | 0 | 59628 | 0 | 0 | 0 | 0 | 0 | 23689 | 0 | 0 | 737360 | 905210 | 0 | 0 | 52180 | 92595 |
| LAKPLSSLT | PAK2; PAK2 | Q13177; H7 | PAK2_HUMAN | Oxidation (M) | 0 | 0 | 0 | 0 | 1211100 | 0 | 0 | 0 | 0 | 532990 | 0 | 374520 | 1183500 | 830180 | 0 | 0 | 1982900 | 1069100 |
| LAKVDATE | P4HB; P4HB; P4 | P07237; A0A | PDIA1_HUMAN | Unmodified | 0 | 0 | 0 | 0 | 0 | 0 | 58925 | 110710 | 2.03E+08 | 3139200 | 3268600 | 3957600 | 28415 | 0 | 1416400 | 0 | 0 | 0 |
| LAKVDATE | P4HB; P4HB; P4 | P07237; A0A | PDIA1_HUMAN | Unmodified | 0 | 0 | 0 | 283350 | 0 | 0 | 788750 | 0 | 23668000 | 23278000 | 820710 | 0 | 1577100 | 931080 | 0 | 0 | 0 | 0 |
| LAKVETPLE | NAA15; NAA15 | Q9BXJ9; A0A | NAA15_HUMAN | Unmodified | 0 | 0 | 0 | 0 | 0 | 0 | 0 | 0 | 177960 | 0 | 0 | 1983300 | 0 | 0 | 0 | 0 | 148670 | 0 |
| LAKYNQLLI | ENO1; ENO1 | P06733; A0A | ENOA_HUMAN | Unmodified | 0 | 0 | 0 | 0 | 1308900 | 0 | 328090 | 0 | 0 | 663750 | 0 | 0 | 0 | 0 | 0 | 0 | 8409500 | 1416500 |
| LALASLGYE | PDCD6IP; PDCI | Q8WUM4; A | PDC6I_HUMAN | Unmodified | 46893000 | 45205000 | 16063000 | 16216000 | 13752000 | 3837000 | 2727700 | 1440800 | 14122000 | 14631000 | 2682100 | 11703000 | 14534000 | 10900000 | 28983000 | 27364000 | 3442400 | 7007500 |
| LALDGETL | SCRIB; SCRIB; S | Q14160; A0A | SCRIB_HUMAN | Unmodified | 0 | 0 | 0 | 934470 | 0 | 0 | 0 | 0 | 6043700 | 1925000 | 1586400 | 1077300 | 0 | 0 | 969070 | 0 | 926610 | 1291500 |
| LALDYGIK | RAB8A | P61006 | RAB8A_HUMAN | Unmodified | 620500 | 3739300 | 5280200 | 6009700 | 19099000 | 2357600 | 5940900 | 588180 | 3788500 | 2073900 | 3837700 | 10916000 | 2149200 | 6077400 | 6173300 | 7420700 | 128240 | 8958200 |
| LALDYGIKF | RAB8A | P61006 | RAB8A_HUMAN | Unmodified | 0 | 0 | 0 | 0 | 753750 | 0 | 1461100 | 0 | 753920 | 36549 | 0 | 0 | 341070 | 322500 | 0 | 0 | 520260 | 0 |
| LALDYGIKF | RAB8A | P61006 | RAB8A_HUMAN | Oxidation (M) | 0 | 0 | 439630 | 0 | 0 | 0 | 592300 | 0 | 0 | 0 | 0 | 0 | 0 | 0 | 0 | 0 | 0 | 0 |
| LALEGVEK | MAPRE2 | Q15555 | MARE2_HUMAI | Unmodified | 322290 | 1100400 | 2089700 | 766350 | 512820 | 304910 | 903190 | 1014500 | 1382900 | 263750 | 287910 | 13040000 | 468470 | 4739100 | 1625100 | 8627000 | 308270 | 2072100 |
| LALEGVEKI | MAPRE2 | Q15555 | MARE2_HUMAI | Unmodified | 0 | 0 | 0 | 0 | 0 | 0 | 2032600 | 69523 | 0 | 0 | 0 | 0 | 309790 | 0 | 0 | 0 | 0 | 0 |
| LALELPDVT | RMDN3; RMDN | Q96TC7; H0 | RMD3_HUMAN | Unmodified | 0 | 0 | 0 | 61745 | 0 | 0 | 0 | 0 | 0 | 0 | 182370 | 479860 | 222460 | 33281 | 70931 | 0 | 0 | 0 |
| LALENYITA | APP; APP; APP; | P05067; H7 | A4_HUMAN Am | Unmodified | 0 | 0 | 0 | 0 | 0 | 0 | 0 | 0 | 7500100 | 1690000 | 50523 | 0 | 0 | 0 | 0 | 0 | 0 | 0 |
| LALEPGENI | CHSY1 | Q86X52 | CHSS1_HUMAI | Oxidation (M) | 0 | 0 | 0 | 0 | 0 | 0 | 0 | 0 | 127470 | 0 | 0 | 0 | 0 | 0 | 0 | 0 | 0 | 0 |
| LALEQER | PKP3 | Q9Y446 | PKP3_HUMAN | Unmodified | 3849600 | 3284100 | 2052700 | 2029400 | 0 | 0 | 0 | 20597 | 953180 | 27995 | 272980 | 4401400 | 73187 | 264370 | 2345200 | 2420900 | 25906 | 0 |
| LALFAHREI | VPS35 | Q96QK1 | VPS35_HUMAN | Unmodified | 0 | 0 | 0 | 0 | 0 | 0 | 0 | 0 | 0 | 0 | 0 | 2059400 | 0 | 0 | 0 | 0 | 0 | 0 |
| LALFGEHV | RPAP1 | Q9BWH6 | RPAP1_HUMAN | Unmodified | 0 | 0 | 0 | 0 | 0 | 0 | 0 | 0 | 0 | 0 | 0 | 442670 | 0 | 0 | 0 | 0 | 0 | 0 |
| LALFNSAIV | TMTC3 | Q6ZXV5 | TMTC3_HUMAI | Unmodified | 0 | 0 | 0 | 0 | 0 | 0 | 0 | 0 | 806900 | 0 | 85128 | 948710 | 0 | 0 | 0 | 0 | 0 | 0 |
| LALFNSAIV | TMTC3 | Q6ZXV5 | TMTC3_HUMAI | Oxidation (M) | 0 | 0 | 0 | 0 | 0 | 918350 | 0 | 0 | 1830600 | 300740 | 401680 | 0 | 0 | 0 | 0 | 0 | 123730 | 0 |
| LALGDDSP | GOT1 | P17174 | AATC_HUMAN | Unmodified | 693750 | 494540 | 970560 | 2198600 | 2989400 | 0 | 0 | 0 | 1858200 | 698850 | 5242800 | 26225000 | 3873400 | 1828100 | 2457300 | 1593000 | 0 | 471440 |
| LALGIPLPE | CAD; CAD | P27708; F8V | PYR1_HUMAN | Unmodified | 0 | 0 | 197030 | 110720 | 0 | 0 | 0 | 0 | 0 | 141410 | 228180 | 1559000 | 0 | 0 | 0 | 180830 | 0 | 164390 |
| LALGSFVEI | DCAF7; DCAF7 | P61962; A0A | DCAF7_HUMAI | Unmodified | 0 | 0 | 0 | 0 | 0 | 0 | 0 | 0 | 0 | 0 | 0 | 1340800 | 0 | 0 | 0 | 0 | 0 | 0 |
| LALIGQSLF | ALDH1L2; ALDH | Q3SY69; A0A | AL1L2_HUMAN | Unmodified | 0 | 0 | 0 | 0 | 0 | 0 | 0 | 0 | 3246500 | 1127900 | 0 | 0 | 0 | 0 | 0 | 0 | 0 | 0 |
| LALIQEK | GSTA2 | P09210 | GSTA2_HUMAN | Unmodified | 0 | 0 | 0 | 0 | 0 | 0 | 0 | 0 | 5078400 | 0 | 0 | 0 | 0 | 0 | 0 | 0 | 0 | 0 |
| LALIQLQIS | NIT2; NIT2; NIT | Q9NQR4; F8 | NIT2_HUMAN | Unmodified | 0 | 0 | 0 | 0 | 0 | 0 | 0 | 0 | 0 | 19134 | 54165 | 68283 | 23225 | 0 | 32902 | 0 | 0 | 0 |
| LALIQPSRE | PSPH; PSPH | P78330; C9J | SERB_HUMAN | Unmodified | 185160 | 0 | 65525 | 0 | 0 | 0 | 0 | 25725 | 2667800 | 0 | 839970 | 2690500 | 0 | 0 | 819830 | 674620 | 125590 | 0 |
| LALLEEAR | CROCC; CROC | Q5TZA2; B1A | CROCC_HUMA | Unmodified | 2623400 | 1281000 | 2346500 | 4117900 | 19413000 | 4155000 | 1545200 | 282740 | 19379000 | 3063000 | 3992200 | 28956000 | 6185100 | 3754100 | 3052200 | 3443900 | 1528500 | 3179900 |
| LALLNVAT | WDR26; WDR2 | Q9H7D7; H0 | WDR26_HUMA | Unmodified | 53519 | 0 | 18550 | 48585 | 0 | 0 | 0 | 39264 | 177130 | 0 | 84878 | 944420 | 0 | 0 | 261150 | 43063 | 30004 | 0 |
| LALLQPALI | FKRP; FKRP; FK | Q9H9S5; M0 | FKRP_HUMAN | Unmodified | 0 | 0 | 0 | 0 | 0 | 0 | 0 | 0 | 3125200 | 0 | 0 | 0 | 0 | 0 | 0 | 0 | 0 | 0 |
| LALPADSVI | CUL5 | Q93034 | CUL5_HUMAN | Unmodified | 0 | 0 | 0 | 0 | 918030 | 0 | 0 | 0 | 0 | 0 | 0 | 1393000 | 0 | 0 | 0 | 0 | 0 | 1259200 |
| LALPVNSSI | CHAT | P28329 | CLAT_HUMAN | Unmodified | 0 | 0 | 0 | 0 | 0 | 0 | 0 | 0 | 0 | 0 | 370290 | 5951200 | 0 | 0 | 0 | 0 | 0 | 0 |
| LALQLHPD | DNAJB11; DNA | Q9UBS4; A0 | DJB11_HUMAN | Unmodified | 0 | 0 | 399810 | 0 | 0 | 0 | 187190 | 0 | 23251000 | 5694000 | 2101400 | 2463300 | 0 | 0 | 0 | 209150 | 0 | 0 |
| LALQLHPD | DNAJB11; DNA | Q9UBS4; A0 | DJB11_HUMAN | Unmodified | 0 | 0 | 0 | 0 | 939780 | 0 | 0 | 0 | 2648200 | 0 | 0 | 0 | 0 | 92712 | 0 | 0 | 0 | 0 |
| LALQQDLT | ERLIN2; ERLIN2 | O94905; E5F | ERLN2_HUMAN | Oxidation (M) | 0 | 0 | 0 | 0 | 0 | 0 | 0 | 0 | 7300500 | 242280 | 0 | 0 | 0 | 0 | 0 | 0 | 0 | 17106 |
| LALQQDLT | ERLIN2; ERLIN2 | O94905; E5F | ERLN2_HUMAN | Unmodified | 0 | 0 | 0 | 0 | 0 | 0 | 0 | 0 | 1854000 | 0 | 0 | 0 | 0 | 0 | 0 | 0 | 0 | 0 |
| LALQTIPED | PAPOLA; PAPOI | P51003; Q9 | PAPOA_HUMAI | Unmodified | 0 | 0 | 0 | 0 | 0 | 0 | 0 | 0 | 0 | 0 | 0 | 925140 | 0 | 0 | 377460 | 0 | 0 | 0 |
| LALQWHPI | DNAJC3; DNAJC | Q13217; X6F | DNJC3_HUMAI | Unmodified | 0 | 0 | 0 | 0 | 0 | 0 | 0 | 0 | 13048000 | 2247600 | 863540 | 0 | 0 | 0 | 0 | 0 | 0 | 0 |
| LALQWHPI | DNAJC3; DNAJC | Q13217; X6F | DNJC3_HUMAI | Unmodified | 0 | 0 | 0 | 0 | 0 | 0 | 0 | 0 | 2731800 | 296590 | 0 | 0 | 0 | 0 | 0 | 0 | 0 | 0 |
| LALSEDTEF | FAM171A2 | A8MVW0 | F1712_HUMAN | Unmodified | 0 | 0 | 0 | 0 | 0 | 0 | 0 | 0 | 586670 | 0 | 0 | 164320 | 0 | 0 | 0 | 0 | 0 | 0 |
| LALTEWLQ | OGA | O60502 | OGA_HUMAN F | Unmodified | 0 | 0 | 38103 | 220300 | 0 | 67139 | 0 | 0 | 0 | 166900 | 32116 | 0 | 1097500 | 274360 | 215170 | 94143 | 147800 | 282560 |
| LALTMDLAI | C8orf82; C8orf | Q6P1X6; H0 | CH082_HUMAI | Oxidation (M) | 0 | 0 | 0 | 0 | 0 | 0 | 0 | 0 | 233400 | 0 | 0 | 0 | 0 | 0 | 0 | 0 | 0 | 0 |
| LALVEENGI | PKP3 | Q9Y446 | PKP3_HUMAN | Unmodified | 0 | 0 | 0 | 294660 | 0 | 0 | 0 | 0 | 0 | 0 | 0 | 17261 | 97071 | 0 | 1463600 | 231860 | 0 | 0 |
| LALVISELE | TRIM26; TRIM2 | Q12899; A2A | TRI26_HUMAN | Unmodified | 10265000 | 16005000 | 8491200 | 9496100 | 0 | 0 | 0 | 0 | 0 | 0 | 0 | 0 | 2962000 | 0 | 14855000 | 9324300 | 0 | 0 |
| LALVTGGEI | CCT2; CCT2; C | P78371; F5G | TCPB_HUMAN | Unmodified | 1125800 | 77439 | 97404 | 2819500 | 1377100 | 201930 | 0 | 30296 | 9017800 | 1485500 | 7804700 | 54700000 | 3152300 | 0 | 2998900 | 2493600 | 0 | 419050 |
| LALWEGPS | FARP2; FARP2 | O94887; H7 | FARP2_HUMAN | Unmodified | 0 | 0 | 0 | 0 | 0 | 0 | 0 | 0 | 0 | 0 | 0 | 0 | 0 | 0 | 0 | 0 | 0 | 0 |
| LALYQTAIE | CC2D1A | Q6P1N0 | C2D1A_HUMAI | Unmodified | 1691600 | 0 | 0 | 301370 | 0 | 0 | 0 | 131040 | 0 | 0 | 0 | 886330 | 0 | 0 | 1049500 | 972240 | 0 | 0 |
| LAMQEFMI | ENO1; ENO1; E | P06733; K7E | ENOA_HUMAN | 2 Oxidation (M) | 0 | 0 | 2278700 | 0 | 17954000 | 0 | 337290 | 0 | 13619000 | 6659400 | 6040200 | 48664000 | 18092000 | 0 | 12820000 | 12503000 | 3067100 | 7464500 |
| LAMQEFMI | ENO1; ENO1; E | P06733; K7E | ENOA_HUMAN | Oxidation (M) | 0 | 0 | 0 | 673820 | 6782100 | 806340 | 0 | 0 | 17504000 | 1758800 | 128320 | 4232500 | 2721400 | 650500 | 1625600 | 279020 | 472330 | 3701800 |
| LAMQEFMI | ENO1; ENO1; E | P06733; K7E | ENOA_HUMAN | Unmodified | 0 | 412100 | 0 | 0 | 616170 | 0 | 0 | 0 | 3034600 | 211580 | 0 | 183560 | 0 | 0 | 0 | 0 | 0 | 0 |
| LAMQEFMI | ENO1; ENO1; E | P06733; K7E | ENOA_HUMAN | 3 Oxidation (M) | 0 | 0 | 0 | 0 | 18799000 | 677020 | 0 | 0 | 0 | 0 | 0 | 0 | 69466 | 1225700 | 0 | 0 | 0 | 2926100 |
| LAMTREMN | PACSIN1; PACS | Q9BY11; F6 | PACN1_HUMAI | Unmodified | 0 | 0 | 0 | 0 | 153210 | 0 | 63268 | 0 | 0 | 0 | 0 | 0 | 4998100 | 2252900 | 0 | 0 | 36650 | 199270 |
| LAMTREMN | PACSIN1; PACS | Q9BY11; F6 | PACN1_HUMAI | Oxidation (M) | 0 | 0 | 0 | 0 | 0 | 0 | 0 | 0 | 0 | 0 | 0 | 0 | 1023600 | 132930 | 0 | 0 | 0 | 0 |
| LANDIEATA | H6PD | O95479 | G6PE_HUMAN | Unmodified | 0 | 0 | 0 | 0 | 0 | 107890 | 0 | 54597 | 1659800 | 197330 | 0 | 145460 | 0 | 0 | 0 | 0 | 0 | 0 |
| LANELPDW | TJP2; TJP2; TJP2 | Q9UDY2; A0 | ZO2_HUMAN T | Unmodified | 0 | 0 | 167280 | 0 | 170100 | 0 | 0 | 0 | 6068200 | 224450 | 60448 | 483670 | 0 | 0 | 0 | 0 | 397500 | 0 |
| LANELPDW | TJP2; TJP2; TJP2 | Q9UDY2; A0 | ZO2_HUMAN T | Unmodified | 0 | 0 | 0 | 274900 | 6812100 | 1356200 | 0 | 0 | 4699100 | 3743100 | 0 | 979490 | 0 | 582360 | 0 | 164520 | 306360 | 0 |
| LANELPDW | TJP2; TJP2; TJP2 | Q9UDY2; A0 | ZO2_HUMAN T | Unmodified | 0 | 0 | 0 | 0 | 3014100 | 620120 | 0 | 0 | 0 | 1961000 | 0 | 0 | 371700 | 0 | 0 | 0 | 689810 | 0 |
| LANEQIQK | ANTXR2; ANTXF | P58335; A0A | ANTR2_HUMAN | Unmodified | 181110 | 305630 | 222220 | 818250 | 0 | 0 | 278010 | 29167 | 239720 | 0 | 0 | 0 | 228600 | 368270 | 0 | 316640 | 0 | 1693500 |
| LANEQIQK | ANTXR2; ANTXF | P58335; A0A | ANTR2_HUMAN | Unmodified | 0 | 0 | 0 | 0 | 179150 | 0 | 197820 | 165190 | 200370 | 37921 | 298240 | 292350 | 345670 | 200730 | 0 | 0 | 273530 | 449110 |
| LANFASLSI | NUDT19 | A8MXV4 | NUD19_HUMA | Unmodified | 0 | 0 | 0 | 0 | 0 | 0 | 0 | 0 | 1581800 | 0 | 358170 | 0 | 0 | 0 | 0 | 0 | 0 | 446420 |
| LANIDEEMI | RARS1 | P54136 | SYRC_HUMAN | Unmodified | 0 | 0 | 0 | 0 | 0 | 0 | 225140 | 0 | 1201100 | 0 | 1146200 | 2675400 | 0 | 0 | 0 | 39422 | 0 | 0 |
| LANLAATIC | ANLN | Q9NQW6 | ANLN_HUMAN | Unmodified | 0 | 0 | 0 | 379380 | 6821800 | 1431100 | 0 | 0 | 0 | 0 | 0 | 0 | 0 | 604520 | 0 | 0 | 5858400 | 2187300 |
| LANLNRKY | UACA; UACA; U | Q9BZF9; H0 | UACA_HUMAN | Unmodified | 0 | 0 | 0 | 0 | 778010 | 0 | 0 | 0 | 0 | 0 | 0 | 0 | 0 | 0 | 0 | 0 | 2765000 | 0 |
| LANQAADY | PDCD6IP; PDCI | Q8WUM4; F | PDC6I_HUMAN | Unmodified | 64341000 | 58031000 | 25844000 | 24243000 | 9992600 | 5127600 | 703450 | 708060 | 17463000 | 8144000 | 0 | 14891000 | 19789000 | 7963600 | 50493000 | 40669000 | 1372000 | 5091800 |
| LANQAADY | PDCD6IP; PDCI | Q8WUM4; F | PDC6I_HUMAN | Unmodified | 1029100 | 328850 | 0 | 0 | 3132600 | 0 | 0 | 0 | 0 | 0 | 0 | 0 | 6358400 | 0 | 0 | 0 | 1313800 | 0 |
| LANQAADY | PDCD6IP; PDCI | Q8WUM4; F | PDC6I_HUMAN | Unmodified | 587880 | 0 | 0 | 493680 | 7246900 | 641540 | 0 | 0 | 0 | 3993600 | 0 | 0 | 10756000 | 9377900 | 0 | 0 | 4025000 | 4203400 |
| LANQDEGF | CC2D1A | A0A7P0Z4M | A0A7P0Z4M5_I | Unmodified | 724230 | 0 | 0 | 0 | 752680 | 0 | 0 | 0 | 0 | 0 | 0 | 0 | 0 | 0 | 553730 | 359260 | 0 | 0 |
| LANQDEGF | CC2D1A | Q6P1N0 | C2D1A_HUMAI | Unmodified | 738200 | 599070 | 371850 | 335220 | 814320 | 119570 | 0 | 0 | 392220 | 72558 | 77480 | 0 | 390690 | 104840 | 482750 | 460320 | 72838 | 247570 |

| | | | | | | | | | | | | | | | | | | | | | | |
|---|---|---|---|---|---|---|---|---|---|---|---|---|---|---|---|---|---|---|---|---|---|---|
| LANQDEGF | CC2D1A | Q6P1N0 | C2D1A_HUMAI | Unmodified | 0 | 0 | 0 | 0 | 1001600 | 0 | 0 | 0 | 0 | 0 | 0 | 0 | 466630 | 0 | 0 | 0 | 0 | 0 |
| LANQDSRF | SEZ6; SEZ6; SE | Q53EL9; C9J | SEZ6_HUMAN | Unmodified | 0 | 0 | 0 | 0 | 0 | 0 | 0 | 0 | 1690200 | 0 | 0 | 0 | 0 | 0 | 0 | 0 | 0 | 0 |
| LANRAPEP | CHD4; CHD4; C | Q14839; A0A | CHD4_HUMAN | Unmodified | 0 | 0 | 0 | 0 | 0 | 0 | 72464 | 93879 | 0 | 0 | 0 | 684090 | 221790 | 0 | 0 | 0 | 0 | 0 |
| LANVMMGI | VCL | P18206 | VINC_HUMAN | Unmodified | 0 | 0 | 0 | 0 | 0 | 0 | 118160 | 0 | 386790 | 0 | 0 | 0 | 0 | 0 | 0 | 0 | 416990 | 0 |
| LANVMMGI | VCL | P18206 | VINC_HUMAN | Oxidation (M) | 359920 | 434880 | 0 | 239430 | 622220 | 97932 | 281500 | 26666 | 0 | 910190 | 0 | 5806000 | 153450 | 95195 | 274280 | 241060 | 1452100 | 174820 |
| LANVQLLD | PAFAH1B2; PAF | P68402; J3K | PA1B2_HUMAN | Oxidation (M) | 0 | 0 | 0 | 0 | 1439700 | 0 | 519880 | 0 | 0 | 0 | 0 | 0 | 2083100 | 1192500 | 0 | 0 | 1782700 | 1077500 |
| LANVVFFPI | PDGFD | Q9GZP0 | PDGFD_HUMA | Unmodified | 0 | 0 | 0 | 0 | 1028900 | 498400 | 0 | 0 | 0 | 0 | 632810 | 0 | 0 | 0 | 0 | 0 | 380920 | 1298300 |
| LANYTNLS | SLC12A7; SLC1 | Q9Y666; A0A | S12A7_HUMAN | Unmodified | 0 | 0 | 328690 | 0 | 0 | 0 | 0 | 212550 | 0 | 252540 | 0 | 5407500 | 0 | 0 | 1188700 | 330020 | 29814 | 0 |
| LANYTNLS | SLC12A7; SLC1 | Q9Y666; A0A | S12A7_HUMAN | Unmodified | 720860 | 930770 | 1577200 | 0 | 0 | 0 | 0 | 0 | 6181900 | 351010 | 0 | 7296200 | 0 | 0 | 3193000 | 3410400 | 27929 | 0 |
| LAPAVLLTG | SLC4A7; SLC4A | Q9Y6M7; C9 | S4A7_HUMAN | Unmodified | 0 | 0 | 129960 | 0 | 0 | 0 | 0 | 0 | 211630 | 0 | 447860 | 3613000 | 0 | 0 | 0 | 226150 | 0 | 0 |
| LAPDYDALI | RPL23A; RPL23 | P62750; K7E | RL23A_HUMAN | Unmodified | 8809300 | 5560300 | 3509800 | 3917900 | 0 | 0 | 0 | 105330 | 1297900 | 810560 | 4971900 | 18948000 | 0 | 0 | 4626200 | 2652600 | 516730 | 205910 |
| LAPDYDALI | RPL23A; RPL23 | P62750; K7E | RL23A_HUMAN | Unmodified | 0 | 0 | 0 | 0 | 224650 | 0 | 89411 | 0 | 0 | 81933 | 0 | 0 | 539810 | 134980 | 0 | 0 | 169740 | 0 |
| LAPEFAKRI | PRDX6 | P30041 | PRDX6_HUMAI | Unmodified | 0 | 0 | 0 | 0 | 177080 | 0 | 170640 | 0 | 0 | 0 | 0 | 0 | 265740 | 467210 | 0 | 0 | 1287700 | 960550 |
| LAPEYEAA | PDIA3; PDIA3 | P30101; H7E | PDIA3_HUMAN | Unmodified | 0 | 0 | 0 | 0 | 0 | 0 | 0 | 0 | 7461000 | 561190 | 2853000 | 1272800 | 0 | 0 | 0 | 0 | 0 | 0 |
| LAPEYEAA | PDIA3; PDIA3 | P30101; H7E | PDIA3_HUMAN | Unmodified | 0 | 0 | 0 | 0 | 0 | 312350 | 751120 | 602040 | 2400600 | 4248400 | 0 | 0 | 0 | 0 | 0 | 0 | 1698400 | 0 |
| LAPEYEK | PDIA4; PDIA4 | P13667; A0A | PDIA4_HUMAN | Unmodified | 0 | 0 | 0 | 0 | 0 | 0 | 20806 | 747450 | 6708600 | 2907500 | 1090800 | 47943 | 0 | 0 | 0 | 0 | 39707 | 326790 |
| LAPGFDAE | FKBP9; FKBP9P | O95302; Q7 | FKBP9_HUMAN | Unmodified | 0 | 0 | 0 | 0 | 0 | 0 | 0 | 0 | 0 | 3162800 | 2885000 | 308350 | 0 | 0 | 0 | 0 | 0 | 0 |
| LAPGTIVEV | HEXB; HEXB; H | P07686; H0 | HEXB_HUMAN | Unmodified | 0 | 0 | 0 | 0 | 0 | 0 | 863530 | 0 | 10012000 | 0 | 0 | 0 | 0 | 104040 | 0 | 0 | 0 | 0 |
| LAPITSDPT | PKM; PKM | P14618; B4D | KPYM_HUMAN | Unmodified | 153750 | 0 | 9469100 | 0 | 25883000 | 7350700 | 8188800 | 2238500 | 21910000 | 6643200 | 48806000 | 1.04E+08 | 0 | 0 | 0 | 0 | 5883700 | 10166000 |
| LAPNQTKE | EPB41; EPB41; | P11171; A0A | EPB41_HUMAN | Unmodified | 0 | 0 | 0 | 0 | 0 | 0 | 140000 | 0 | 0 | 0 | 0 | 0 | 0 | 0 | 0 | 0 | 0 | 0 |
| LAPPLVTLL | AP2B1; AP2B1; | P63010; A0A | AP2B1_HUMAN | Unmodified | 131590 | 364060 | 0 | 0 | 0 | 0 | 0 | 0 | 122640 | 120280 | 15917 | 461210 | 0 | 0 | 62603 | 157330 | 0 | 0 |
| LAPSEYR | ERP44 | Q9BS26 | ERP44_HUMAN | Unmodified | 205670 | 0 | 308840 | 176940 | 572550 | 39916 | 60402 | 153060 | 57803000 | 6631100 | 1005100 | 739260 | 993590 | 1111800 | 371810 | 312260 | 12920 | 389580 |
| LAPVVIPAQ | DLG5; DLG5 | Q8TDM6; A0 | DLG5_HUMAN | Unmodified | 0 | 0 | 83534 | 0 | 748840 | 486090 | 0 | 24705 | 388470 | 198330 | 28023 | 885650 | 148680 | 134190 | 0 | 0 | 1070300 | 1475100 |
| LAPWLLR | B3GNT8 | Q7Z7M8 | B3GN8_HUMAI | Unmodified | 0 | 0 | 0 | 0 | 0 | 0 | 0 | 0 | 937210 | 0 | 0 | 0 | 0 | 0 | 0 | 0 | 0 | 0 |
| LAQAAQSS | TLN1 | Q9Y490 | TLN1_HUMAN | Unmodified | 545030 | 625590 | 1223000 | 1405100 | 637680 | 665180 | 212620 | 28733 | 518200 | 1008000 | 0 | 5546000 | 252820 | 232790 | 1883600 | 1190300 | 148080 | 0 |
| LAQAAQSS | TLN1 | Q9Y490 | TLN1_HUMAN | Unmodified | 0 | 0 | 0 | 0 | 4499900 | 2066600 | 2949700 | 0 | 0 | 705000 | 0 | 0 | 0 | 1431100 | 0 | 0 | 1043200 | 723280 |
| LAQAEEQLI | MYH14; MYH14 | Q7Z406; A1L | MYH14_HUMAI | Unmodified | 0 | 0 | 0 | 0 | 0 | 0 | 0 | 0 | 2171700 | 123130 | 0 | 0 | 824250 | 392400 | 0 | 0 | 0 | 104250 |
| LAQAHGFC | SPTBN2 | O15020 | SPTN2_HUMAN | Unmodified | 0 | 0 | 0 | 366710 | 0 | 0 | 0 | 0 | 0 | 0 | 0 | 2487300 | 0 | 0 | 1021400 | 0 | 0 | 0 |
| LAQAHPAG | SKIV2L; SKIV2L | Q15477; H7 | SKIV2_HUMAN | Unmodified | 0 | 0 | 0 | 0 | 0 | 0 | 0 | 0 | 0 | 0 | 0 | 1358200 | 0 | 0 | 0 | 0 | 0 | 0 |
| LAQANGW | ENO1; ENO1 | P06733; A0A | ENOA_HUMAN | Unmodified | 1014000 | 777690 | 1615400 | 2983900 | 14026000 | 2742800 | 300740 | 9826.9 | 69976000 | 5654100 | 3783100 | 1.01E+08 | 5708000 | 2985100 | 3956200 | 2250600 | 3480000 | 10055000 |
| LAQANGW | ENO1; ENO1 | P06733; A0A | ENOA_HUMAN | Oxidation (M) | 905550 | 914510 | 2193300 | 1964100 | 3643400 | 1013400 | 0 | 0 | 9221400 | 1911300 | 2588900 | 36778000 | 2980800 | 843290 | 3556500 | 3863800 | 1041900 | 3179500 |
| LAQAQEVF | PDCD6IP; PDCI | Q8WUM4; F | PDC6I_HUMAN | Unmodified | 172100 | 949480 | 0 | 0 | 0 | 0 | 0 | 0 | 0 | 0 | 0 | 0 | 0 | 0 | 0 | 56976 | 0 | 0 |
| LAQATQER | SMC3 | Q9UQE7 | SMC3_HUMAN | Unmodified | 0 | 0 | 77776 | 42359 | 1269100 | 1162500 | 297860 | 0 | 0 | 696440 | 630710 | 1005900 | 353840 | 0 | 0 | 0 | 80865 | 69809 |
| LAQAWFN | CCDC47 | Q96A33 | CCD47_HUMA | Unmodified | 0 | 0 | 0 | 0 | 0 | 0 | 0 | 0 | 0 | 106160 | 0 | 618380 | 0 | 0 | 0 | 0 | 0 | 0 |
| LAQDAEVE | MGAT1; MGAT1 | P26572; D6F | MGAT1_HUMAI | Unmodified | 0 | 0 | 0 | 0 | 0 | 0 | 0 | 0 | 2631700 | 384410 | 0 | 0 | 0 | 0 | 0 | 0 | 0 | 0 |
| LAQDAEVE | MGAT1; MGAT1 | P26572; D6F | MGAT1_HUMAI | Unmodified | 45131 | 0 | 355470 | 0 | 0 | 0 | 0 | 0 | 17026000 | 226610 | 54714 | 0 | 0 | 0 | 0 | 52537 | 0 | 0 |
| LAQDFLDS | DPP3; DPP3; DI | Q9NY33; E9I | DPP3_HUMAN | Unmodified | 0 | 0 | 0 | 0 | 0 | 0 | 0 | 0 | 1135900 | 0 | 1215400 | 8172400 | 0 | 0 | 0 | 0 | 0 | 0 |
| LAQDPFPL | MVP; MVP; MVP | Q14764; H3I | MVP_HUMAN | Unmodified | 2199000 | 1587200 | 0 | 0 | 0 | 0 | 0 | 0 | 4407200 | 629900 | 1697700 | 23791000 | 31143 | 0 | 79766 | 0 | 0 | 0 |
| LAQEGIYTL | NSUN2 | Q08J23 | NSUN2_HUMA | Unmodified | 0 | 0 | 228530 | 357810 | 0 | 0 | 0 | 0 | 0 | 0 | 101500 | 1491900 | 0 | 0 | 412410 | 0 | 0 | 0 |
| LAQENGW | ENO2; ENO2 | P09104; F5H | ENOG_HUMAN | Unmodified | 65489 | 79296 | 0 | 0 | 0 | 0 | 0 | 0 | 2131600 | 0 | 0 | 719450 | 0 | 0 | 0 | 0 | 0 | 158400 |
| LAQENMDI | CTNNA1; CTNN | P35221; E5R | CTNA1_HUMAI | Unmodified | 0 | 0 | 0 | 108230 | 0 | 362150 | 0 | 0 | 227960 | 0 | 0 | 1038500 | 299060 | 0 | 4825100 | 1652400 | 136030 | 0 |
| LAQENMDI | CTNNA1; CTNN | P35221; E5R | CTNA1_HUMAI | Oxidation (M) | 0 | 0 | 0 | 0 | 0 | 130190 | 898330 | 0 | 0 | 0 | 0 | 0 | 0 | 0 | 2958300 | 954790 | 0 | 354420 |
| LAQENMDI | CTNNA1; CTNN | P35221; E5R | CTNA1_HUMAI | Unmodified | 103220 | 229710 | 0 | 478830 | 9735200 | 3107400 | 0 | 0 | 0 | 662860 | 0 | 335540 | 1958100 | 1026800 | 470460 | 1008700 | 0 | 0 |
| LAQENMDI | CTNNA1; CTNN | P35221; E5R | CTNA1_HUMAI | Oxidation (M) | 317770 | 398890 | 0 | 465480 | 2388900 | 1878200 | 0 | 17490 | 0 | 0 | 0 | 536250 | 682390 | 390770 | 920380 | 1555400 | 0 | 0 |
| LAQERAEIV | USP6NL; USP6I | Q92738; X6F | US6NL_HUMAI | Unmodified | 0 | 0 | 0 | 0 | 0 | 0 | 0 | 0 | 0 | 0 | 159950 | 0 | 385270 | 0 | 243720 | 0 | 238870 | 1666200 |
| LAQFEPSQ | DHX9 | Q08211 | DHX9_HUMAN | Unmodified | 154450 | 131020 | 0 | 0 | 0 | 0 | 0 | 0 | 199270 | 71186 | 229590 | 1474500 | 0 | 0 | 0 | 0 | 0 | 0 |
| LAQGPELA | WASF1 | Q92558 | WASF1_HUMAI | Unmodified | 0 | 0 | 0 | 0 | 1684400 | 254030 | 190700 | 140500 | 1404400 | 475700 | 585290 | 3388800 | 0 | 0 | 0 | 0 | 0 | 413130 |
| LAQILAAR | OTUD6B; OTUD | Q8N6M0; A0 | OTU6B_HUMAI | Unmodified | 2576200 | 1681000 | 712690 | 2732200 | 0 | 0 | 0 | 14463 | 261530 | 139200 | 0 | 1762400 | 0 | 0 | 4471300 | 1432900 | 0 | 0 |
| LAQKNPMY | SFPQ; SFPQ | P23246; H0 | SFPQ_HUMAN | Unmodified | 0 | 0 | 0 | 0 | 0 | 75024 | 429160 | 374420 | 0 | 595840 | 0 | 0 | 1209600 | 460470 | 0 | 0 | 0 | 0 |
| LAQLEEAK | ATP5PB; ATP5P | P24539; Q5 | AT5F1_HUMAN | Unmodified | 0 | 0 | 0 | 0 | 357110 | 0 | 46989 | 0 | 24473000 | 3461600 | 79800 | 290270 | 202730 | 0 | 0 | 0 | 123480 | 97323 |
| LAQLEEAK | ATP5PB; ATP5P | P24539; Q5 | AT5F1_HUMAN | Unmodified | 0 | 0 | 0 | 0 | 0 | 0 | 0 | 0 | 0 | 876990 | 0 | 0 | 790460 | 0 | 111660 | 185640 | 161070 | 0 |
| LAQLQVAY | IKBKG; IKBKG; I | Q9Y6K9; C9 | NEMO_HUMAN | Unmodified | 0 | 0 | 0 | 0 | 3758900 | 764550 | 0 | 0 | 313020 | 0 | 0 | 605520 | 0 | 696150 | 0 | 0 | 0 | 0 |
| LAQMFSDN | TFRC; TFRC | P02786; G3 | TFR1_HUMAN | Oxidation (M) | 0 | 0 | 0 | 0 | 0 | 0 | 0 | 0 | 1720500 | 420240 | 0 | 1504800 | 0 | 0 | 1690600 | 391030 | 0 | 0 |
| LAQMFSDN | TFRC; TFRC | P02786; G3 | TFR1_HUMAN | Unmodified | 0 | 0 | 0 | 0 | 0 | 0 | 0 | 0 | 2859200 | 0 | 120560 | 832150 | 0 | 0 | 0 | 0 | 0 | 0 |
| LAQMFSDN | TFRC; TFRC | P02786; G3 | TFR1_HUMAN | 2 Oxidation (M) | 2038200 | 552950 | 1269300 | 293270 | 0 | 0 | 0 | 0 | 0 | 0 | 0 | 0 | 0 | 0 | 2387500 | 1599300 | 0 | 0 |
| LAQMISDH | UBAP2L; UBAP | Q14157; Q5 | UBP2L_HUMAN | Unmodified | 0 | 0 | 0 | 0 | 0 | 0 | 0 | 0 | 418390 | 0 | 0 | 1432700 | 0 | 0 | 522210 | 0 | 0 | 0 |
| LAQNILSYL | BAG5 | Q9UL15 | BAG5_HUMAN | Unmodified | 0 | 0 | 164740 | 0 | 0 | 0 | 424340 | 0 | 857760 | 97432 | 68281 | 348750 | 0 | 0 | 0 | 0 | 0 | 0 |
| LAQQCLER | VPS9D1; VPS9D | Q9Y2B5; H3I | VP9D1_HUMAN | Unmodified | 0 | 0 | 0 | 0 | 656610 | 114160 | 59756 | 32871 | 0 | 92246 | 0 | 0 | 0 | 0 | 0 | 0 | 45198 | 0 |
| LAQQIANC | MID1; MID1; MI | O15344; A0A | TRI18_HUMAN | Unmodified | 0 | 0 | 0 | 0 | 175030 | 41817 | 0 | 0 | 90426 | 51599 | 0 | 0 | 138530 | 254920 | 0 | 0 | 342460 | 304780 |
| LAQQIKQE | MTHFD2; MTHF | P13995; B8Z | MTDC_HUMAN | Unmodified | 0 | 0 | 0 | 0 | 0 | 0 | 0 | 0 | 1417900 | 45659 | 0 | 0 | 0 | 0 | 0 | 0 | 0 | 0 |
| LAQQISDE | USP9X | Q93008 | USP9X_HUMAI | Unmodified | 0 | 0 | 0 | 0 | 0 | 0 | 0 | 0 | 0 | 57147 | 0 | 1132000 | 0 | 0 | 0 | 0 | 0 | 0 |
| LAQQQAA | TBL1XR1; TBL1 | Q9BZK7; A0 | TBL1R_HUMAN | Unmodified | 0 | 0 | 0 | 0 | 0 | 0 | 0 | 0 | 0 | 0 | 342070 | 1006200 | 0 | 0 | 0 | 0 | 0 | 340200 |
| LAQQQAAL | SF3B2; SF3B2; | Q13435; E9F | SF3B2_HUMAN | Unmodified | 467160 | 0 | 0 | 0 | 0 | 0 | 0 | 80986 | 0 | 0 | 0 | 0 | 0 | 0 | 0 | 0 | 0 | 0 |
| LAQQYYLV | PSMA2; N/A; N | P25787; A0A | PSA2_HUMAN | Unmodified | 0 | 0 | 0 | 0 | 0 | 0 | 0 | 29018 | 1504800 | 1776900 | 527750 | 3308900 | 0 | 0 | 0 | 0 | 194140 | 0 |
| LAQSAEMY | NPDC1; NPDC | Q9NQX5; Q5 | NPDC1_HUMA | Unmodified | 300520 | 606150 | 845920 | 1923400 | 0 | 0 | 0 | 0 | 457170 | 0 | 0 | 0 | 0 | 136520 | 1031200 | 686950 | 0 | 0 |
| LAQSEYQLI | EXOC7; EXOC7 | Q9UPT5; B4I | EXOC7_HUMAI | Unmodified | 0 | 0 | 0 | 0 | 3560600 | 2670200 | 0 | 0 | 1691900 | 617570 | 0 | 2311200 | 558960 | 0 | 0 | 0 | 40287 | 1698300 |
| LAQSEYQLI | EXOC7; EXOC7 | Q9UPT5; B4I | EXOC7_HUMAI | Unmodified | 0 | 0 | 0 | 0 | 1396200 | 0 | 0 | 0 | 0 | 0 | 0 | 0 | 707720 | 0 | 0 | 0 | 370200 | 0 |
| LAQSNGW | ENO3 | P13929 | ENOB_HUMAN | Unmodified | 0 | 0 | 0 | 521220 | 0 | 0 | 0 | 0 | 0 | 0 | 0 | 0 | 0 | 0 | 0 | 855310 | 0 | 0 |
| LAQSVQKL | HYOU1; HYOU | Q9Y4L1; A0A | HYOU1_HUMA | Unmodified | 0 | 0 | 0 | 0 | 0 | 0 | 0 | 0 | 0 | 1157900 | 0 | 0 | 0 | 0 | 0 | 0 | 0 | 0 |
| LAQTFSPSI | THNSL1 | Q8IYQ7 | THNS1_HUMAI | Unmodified | 0 | 0 | 0 | 0 | 0 | 0 | 0 | 0 | 1733000 | 0 | 0 | 0 | 0 | 0 | 0 | 0 | 0 | 0 |
| LAQTLSTQI | ARFIP1 | P53367 | ARFP1_HUMAN | Oxidation (M) | 0 | 0 | 260450 | 0 | 0 | 0 | 0 | 519260 | 0 | 0 | 0 | 979600 | 187510 | 0 | 0 | 0 | 0 | 0 |
| LAQVEPGE | PTPRS; PTPRS | Q13332; G8 | PTPRS_HUMAN | Unmodified | 0 | 0 | 0 | 0 | 1186900 | 160230 | 330630 | 0 | 621070 | 286750 | 0 | 0 | 281720 | 182300 | 0 | 0 | 0 | 0 |
| LAQVEPGE | PTPRS; PTPRS | Q13332; G8 | PTPRS_HUMAN | Oxidation (M) | 0 | 0 | 0 | 619080 | 0 | 262200 | 0 | 0 | 225740 | 0 | 0 | 0 | 0 | 0 | 0 | 0 | 78425 | 0 |
| LAQVSPELI | XPOT | O43592 | XPOT_HUMAN | Unmodified | 0 | 0 | 0 | 0 | 0 | 0 | 0 | 0 | 0 | 0 | 0 | 1760100 | 0 | 0 | 0 | 0 | 0 | 0 |
| LAQVSPELI | XPOT | O43592 | XPOT_HUMAN | Unmodified | 0 | 0 | 0 | 0 | 0 | 0 | 0 | 0 | 0 | 0 | 0 | 1203500 | 0 | 0 | 0 | 0 | 0 | 0 |
| LAQYEPPQ | MYCBP; N/A | Q99417; A0A | MYCBP_HUMA | Unmodified | 275450 | 0 | 0 | 0 | 787470 | 300240 | 0 | 149640 | 46853 | 152070 | 236020 | 1256900 | 660040 | 351580 | 0 | 0 | 88284 | 235000 |
| LAQYEPPQ | MYCBP; N/A | Q99417; A0A | MYCBP_HUMA | Unmodified | 0 | 0 | 0 | 0 | 752690 | 76950 | 454990 | 324630 | 82413 | 162350 | 0 | 0 | 394230 | 423360 | 0 | 0 | 234590 | 285650 |
| LARASGNY | RPL8; RPL8; RP | P62917; E9P | RL8_HUMAN 6 | Unmodified | 0 | 0 | 0 | 0 | 2005000 | 0 | 1244600 | 878400 | 0 | 0 | 0 | 0 | 594090 | 0 | 0 | 0 | 0 | 0 |
| LARASGNY | RPL8; RPL8; RP | P62917; E9P | RL8_HUMAN 6 | Unmodified | 0 | 0 | 0 | 0 | 59883 | 0 | 324680 | 302280 | 0 | 0 | 0 | 0 | 423380 | 0 | 0 | 0 | 99476 | 244530 |
| LARDDQIH | ALOX5 | P09917 | LOX5_HUMAN | Unmodified | 0 | 0 | 0 | 0 | 0 | 0 | 0 | 0 | 933170 | 63163 | 0 | 1158400 | 0 | 0 | 0 | 0 | 0 | 0 |
| LARDEGWI | PCK2; PCK2; P | Q16822; H0 | PCKGM_HUMA | Unmodified | 0 | 0 | 0 | 0 | 0 | 0 | 0 | 0 | 1217900 | 0 | 0 | 0 | 0 | 0 | 0 | 0 | 0 | 0 |
| LAREGENP | PAPSS2 | O95340 | PAPS2_HUMAN | Oxidation (M) | 0 | 0 | 0 | 0 | 0 | 0 | 0 | 0 | 0 | 0 | 0 | 0 | 0 | 0 | 0 | 0 | 472170 | 437080 |
| LAREGENP | PAPSS2 | O95340 | PAPS2_HUMAN | Unmodified | 0 | 0 | 0 | 0 | 417890 | 794540 | 0 | 0 | 0 | 0 | 0 | 0 | 0 | 0 | 0 | 0 | 593170 | 945320 |
| LAREGQKP | PAPSS1 | O43252 | PAPS1_HUMAN | Unmodified | 0 | 0 | 0 | 0 | 0 | 0 | 228070 | 0 | 0 | 0 | 0 | 0 | 0 | 0 | 0 | 0 | 0 | 0 |
| LAREHGIR | RAB13; RAB13 | P51153; A0A | RAB13_HUMAN | Unmodified | 0 | 0 | 0 | 0 | 7059100 | 0 | 423770 | 80542 | 0 | 1679700 | 0 | 0 | 19838000 | 6032100 | 0 | 0 | 1325900 | 11424000 |
| LAREIGLLS | MTHFD1; MTHF | P11586; V9 | C1TC_HUMAN | Unmodified | 0 | 0 | 0 | 0 | 4122400 | 0 | 114900 | 0 | 0 | 0 | 0 | 0 | 0 | 0 | 0 | 0 | 0 | 3077900 |
| LAREIGLLS | MTHFD1; MTHF | P11586; V9 | C1TC_HUMAN | Unmodified | 0 | 0 | 0 | 0 | 0 | 0 | 1737700 | 0 | 0 | 0 | 0 | 0 | 0 | 0 | 0 | 0 | 1860000 | 0 |
| LARGWPE | MIA3 | Q5JRA6 | TGO1_HUMAN | Unmodified | 0 | 0 | 0 | 0 | 0 | 0 | 0 | 49329 | 84017 | 1078800 | 50484 | 0 | 70416 | 122090 | 54414 | 0 | 0 | 0 |
| LARWEYDE | FAM20B | O75063 | XYLK_HUMAN | Unmodified | 0 | 0 | 0 | 0 | 0 | 0 | 0 | 0 | 3376400 | 470550 | 0 | 0 | 0 | 0 | 0 | 0 | 0 | 0 |
| LARWEYDE | FAM20B | O75063 | XYLK_HUMAN | Unmodified | 0 | 0 | 0 | 0 | 0 | 0 | 0 | 0 | 3703000 | 795460 | 0 | 0 | 0 | 0 | 0 | 0 | 23894 | 0 |
| LASAAYPDI | YARS1; YARS1 | P54577; A0A | SYYC_HUMAN | Unmodified | 0 | 0 | 0 | 0 | 202460 | 41068 | 0 | 0 | 107600 | 44508 | 252790 | 1020900 | 416110 | 500990 | 76819 | 117070 | 129330 | 275520 |
| LASAEDSAI | CC2D1B; CC2D | H0Y517; H7 | H0Y517_HUMA | Unmodified | 91330 | 277350 | 91585 | 214040 | 436010 | 0 | 0 | 0 | 620110 | 0 | 0 | 185010 | 0 | 216000 | 83364 | 49075 | 0 | 0 |
| LASAEMQG | BCAM; BCAM | P50895; A0A | BCAM_HUMAN | Unmodified | 472920 | 789780 | 0 | 185630 | 0 | 0 | 0 | 0 | 815500 | 0 | 0 | 0 | 0 | 0 | 0 | 0 | 0 | 0 |
| LASAEMQG | BCAM; BCAM | P50895; A0A | BCAM_HUMAN | 2 Oxidation (M) | 412350 | 380040 | 0 | 0 | 0 | 0 | 0 | 0 | 0 | 0 | 0 | 0 | 0 | 0 | 0 | 0 | 0 | 0 |
| LASAEMQG | BCAM; BCAM | P50895; A0A | BCAM_HUMAN | Oxidation (M) | 0 | 1244000 | 0 | 0 | 358430 | 0 | 0 | 0 | 311660 | 0 | 0 | 0 | 0 | 0 | 0 | 0 | 0 | 0 |
| LASAFTEEC | SRP68 | Q9UHB9 | SRP68_HUMAN | Unmodified | 0 | 0 | 0 | 0 | 0 | 0 | 1185500 | 0 | 0 | 0 | 0 | 4329700 | 0 | 0 | 0 | 0 | 0 | 0 |
| LASALAHN | CARMIL1 | Q5VZK9 | CARL1_HUMAN | Unmodified | 0 | 0 | 224640 | 308390 | 0 | 0 | 0 | 0 | 0 | 0 | 0 | 13105000 | 0 | 0 | 446130 | 359670 | 0 | 0 |
| LASALAHN | CARMIL1 | Q5VZK9 | CARL1_HUMAN | Unmodified | 0 | 0 | 0 | 0 | 3470500 | 0 | 0 | 0 | 0 | 0 | 0 | 0 | 933380 | 0 | 0 | 0 | 1206300 | 0 |
| LASALAPSI | MCM4; MCM4; | P33991; A0A | MCM4_HUMAN | Unmodified | 0 | 0 | 0 | 0 | 0 | 0 | 0 | 89543 | 348800 | 74284 | 0 | 2737200 | 114240 | 0 | 0 | 0 | 0 | 0 |
| LASALDLPL | WASHC5; WASI | Q12768; E7E | WASC5_HUMA | Unmodified | 0 | 0 | 32700 | 0 | 0 | 0 | 0 | 0 | 78633 | 0 | 0 | 1025100 | 0 | 0 | 0 | 0 | 0 | 0 |
| LASAYGAR | LGALS3BP; LGA | Q08380; K7E | LG3BP_HUMAN | Unmodified | 1974600 | 0 | 1072900 | 3205900 | 174160 | 0 | 0 | 0 | 2041300 | 632480 | 504390 | 1510400 | 540230 | 109840 | 1081100 | 1231200 | 860190 | 1152000 |
| LASDLLEW | ACTN4; ACTN1 | O43707; P12 | ACTN4_HUMAI | Unmodified | 290260 | 191050 | 167930 | 0 | 0 | 729470 | 0 | 0 | 128210 | 74659 | 0 | 4612500 | 0 | 0 | 948420 | 0 | 302740 | 0 |
| LASDLLEW | ACTN4; ACTN1 | O43707; P12 | ACTN4_HUMAI | Unmodified | 0 | 0 | 3060800 | 3546700 | 8046200 | 4856600 | 0 | 0 | 8795900 | 4517000 | 6232500 | 50555000 | 0 | 0 | 5237200 | 10594000 | 1741100 | 0 |
| LASEAKPA | TLN1 | Q9Y490 | TLN1_HUMAN | Unmodified | 160530 | 0 | 143870 | 496660 | 1911500 | 0 | 0 | 0 | 1795500 | 208010 | 1066000 | 7357800 | 0 | 0 | 606630 | 379880 | 260900 | 463890 |
| LASEEPPDI | ATXN10; ATXN1 | Q9UBB4; B1 | ATX10_HUMAN | Unmodified | 0 | 0 | 0 | 0 | 0 | 0 | 0 | 0 | 0 | 0 | 0 | 1164700 | 0 | 0 | 0 | 151480 | 0 | 0 |
| LASEQQHS | SEL1L | Q9UBV2 | SE1L1_HUMAN | Unmodified | 0 | 0 | 0 | 0 | 0 | 0 | 0 | 0 | 4988900 | 0 | 0 | 0 | 0 | 0 | 0 | 0 | 0 | 0 |
| LASGCDGS | STK39 | Q9UEW8 | STK39_HUMAN | Unmodified | 0 | 0 | 401380 | 0 | 0 | 0 | 0 | 0 | 218180 | 0 | 0 | 758090 | 345730 | 0 | 385610 | 432310 | 0 | 0 |
| LASGEHIA | ACAD9; ACAD9 | Q9H845; A0 | ACAD9_HUMAI | Unmodified | 0 | 0 | 0 | 0 | 0 | 0 | 0 | 0 | 5863400 | 159800 | 0 | 2063100 | 0 | 0 | 0 | 0 | 0 | 0 |
| LASGETVA | ACADVL | P49748 | ACADV_HUMAI | Unmodified | 0 | 0 | 0 | 0 | 0 | 0 | 0 | 0 | 3294500 | 0 | 0 | 0 | 0 | 0 | 0 | 0 | 0 | 0 |

| | | | | | | | | | | | | | | | | | | | | | | |
|---|---|---|---|---|---|---|---|---|---|---|---|---|---|---|---|---|---|---|---|---|---|---|
| LASGSFDK | TBL1Y; THOC3; | Q9BQ87; Q9 | TBL1Y_HUMAN | Unmodified | 190220 | 0 | 304190 | 756070 | 317320 | 0 | 32689 | 75594 | 2693200 | 93301 | 103730 | 5998400 | 1360800 | 397920 | 531650 | 433070 | 297420 | 293020 |
| LASHEYLK | MFGE8; MFGE8 | Q08431; H0Y | MFGM_HUMAN | Unmodified | 0 | 634260 | 2201100 | 2130000 | 3705800 | 2620000 | 0 | 0 | 2972000 | 0 | 0 | 2001300 | 0 | 173300 | 2090500 | 1564900 | 587940 | 495660 |
| LASHLNLAI | FKBP4 | Q02790 | FKBP4_HUMAN | Unmodified | 0 | 0 | 0 | 0 | 292000 | 162210 | 307130 | 71795 | 110260 | 0 | 511560 | 2801900 | 0 | 680950 | 0 | 0 | 150350 | 0 |
| LASHLNLAI | FKBP4 | Q02790 | FKBP4_HUMAN | Oxidation (M) | 0 | 0 | 194400 | 0 | 0 | 0 | 0 | 13667 | 0 | 0 | 238600 | 3602500 | 545310 | 0 | 0 | 310290 | 0 | 0 |
| LASLEELKR | DSP | P15924 | DESP_HUMAN | Unmodified | 0 | 0 | 0 | 0 | 0 | 0 | 0 | 0 | 0 | 0 | 0 | 0 | 0 | 142270 | 0 | 0 | 953130 | 0 |
| LASLQQEN | CCDC6 | Q16204 | CCDC6_HUMA | Unmodified | 0 | 0 | 0 | 0 | 0 | 0 | 0 | 42035 | 706130 | 34243 | 137620 | 2295100 | 0 | 0 | 0 | 0 | 0 | 0 |
| LASLVSINA | PCDHB4; PCDH | Q9Y5E5; Q9Y | PCDB4_HUMAI | Unmodified | 0 | 0 | 0 | 0 | 0 | 0 | 0 | 0 | 542950 | 261870 | 80306 | 0 | 0 | 0 | 0 | 154550 | 14553 | 0 |
| LASMPAWF | ESPN; ESPN; ES | B1AK53; A0A | ESPN_HUMAN | Unmodified | 32029 | 0 | 0 | 0 | 0 | 0 | 0 | 0 | 32305 | 0 | 0 | 1245000 | 0 | 0 | 0 | 0 | 0 | 0 |
| LASNYISK | MBLAC2 | Q68D91 | MBLC2_HUMAI | Unmodified | 0 | 0 | 165730 | 0 | 0 | 0 | 0 | 0 | 0 | 0 | 70771 | 3106800 | 527370 | 145970 | 337770 | 0 | 0 | 0 |
| LASQANIA( | AP1B1; AP2B1; | Q10567; P63 | AP1B1_HUMAN | Unmodified | 337430 | 289440 | 1063700 | 122380 | 0 | 0 | 0 | 7963.9 | 168590 | 67457 | 745710 | 6249900 | 46140 | 0 | 3183600 | 1541700 | 18726 | 0 |
| LASQGNPF | EHHADH | Q08426 | ECHP_HUMAN | Unmodified | 0 | 0 | 0 | 0 | 52697 | 0 | 0 | 0 | 1187000 | 328810 | 0 | 0 | 0 | 0 | 0 | 0 | 0 | 0 |
| LASRNVES | RPN1; RPN1 | P04843; B7Z | RPN1_HUMAN | Unmodified | 0 | 0 | 0 | 0 | 0 | 0 | 311920 | 0 | 242100 | 3764000 | 30939 | 297080 | 0 | 120220 | 0 | 0 | 0 | 0 |
| LASTLVHL( | CLTC; CLTC; CL | Q00610; J3K | CLH1_HUMAN | Unmodified | 20305000 | 11449000 | 4876100 | 7992400 | 0 | 466910 | 162820 | 36102 | 2951700 | 348880 | 745790 | 54065000 | 138030 | 235790 | 7474000 | 6848300 | 0 | 0 |
| LASTLVHL( | CLTC; CLTC; CL | Q00610; J3K | CLH1_HUMAN | Unmodified | 12706000 | 14105000 | 5733800 | 8790300 | 2893400 | 2546400 | 1047100 | 227910 | 3005400 | 836690 | 1802200 | 68444000 | 2983700 | 2089900 | 12574000 | 14150000 | 669710 | 1315500 |
| LASVLGSE | DPP3; DPP3; DI | Q9NY33; G3 | DPP3_HUMAN | Unmodified | 0 | 0 | 0 | 0 | 170790 | 0 | 0 | 0 | 4664300 | 492090 | 1214300 | 7107400 | 0 | 0 | 0 | 103600 | 1018200 | 222940 |
| LASVPAGG | RPLP2; RPLP2 | P05387; H0Y | RLA2_HUMAN ( | Unmodified | 24277000 | 20760000 | 19833000 | 6354800 | 1653900 | 0 | 355230 | 0 | 0 | 0 | 7995900 | 53282000 | 1572500 | 0 | 51267000 | 47894000 | 0 | 0 |
| LASVPAGG | RPLP2; RPLP2 | P05387; H0Y | RLA2_HUMAN ( | Unmodified | 1783200 | 1851000 | 688830 | 0 | 0 | 0 | 0 | 0 | 0 | 0 | 0 | 0 | 0 | 0 | 1410900 | 1540500 | 0 | 0 |
| LASVPAGG | RPLP2; RPLP2 | P05387; H0Y | RLA2_HUMAN ( | Unmodified | 2621500 | 420390 | 4020400 | 2848900 | 1024600 | 0 | 0 | 0 | 1134100 | 193450 | 12431000 | 27421000 | 586320 | 429230 | 3153900 | 10039000 | 2081200 | 994460 |
| LASVPGSQ | PODXL | O00592 | PODXL_HUMAI | Unmodified | 645750 | 325600 | 928940 | 1748000 | 1113900 | 0 | 427320 | 168430 | 56298000 | 5835900 | 9194200 | 6392200 | 5978400 | 1962900 | 941040 | 1139100 | 424430 | 1375700 |
| LASYLDR | KRT18; KRT18 | P05783; F8V | K1C18_HUMAN | Unmodified | 73254 | 523470 | 73534 | 565530 | 0 | 0 | 0 | 0 | 186630 | 0 | 0 | 9639700 | 182260 | 0 | 134400 | 280490 | 0 | 0 |
| LATCSSDR | SEC13; SEC13; | P55735; A8N | SEC13_HUMAN | Unmodified | 0 | 0 | 0 | 0 | 0 | 0 | 0 | 0 | 0 | 0 | 0 | 0 | 96969 | 0 | 0 | 0 | 0 | 0 |
| LATDKNDP | FTH1; FTH1; FTI | P02794; G3V | FRIH_HUMAN I | Unmodified | 102280 | 0 | 0 | 175920 | 2324100 | 172030 | 1485200 | 433450 | 1.01E+08 | 47583000 | 0 | 2860500 | 5299700 | 6137000 | 213880 | 205240 | 4542000 | 4238200 |
| LATEEPPPF | CDCP1 | Q9H5V8 | CDCP1_HUMA | Unmodified | 0 | 0 | 0 | 0 | 216220 | 0 | 0 | 0 | 0 | 0 | 53884 | 241820 | 0 | 0 | 0 | 0 | 1430800 | 1348700 |
| LATEEPPPF | CDCP1 | Q9H5V8 | CDCP1_HUMA | Unmodified | 0 | 0 | 4254900 | 773870 | 956240 | 0 | 0 | 0 | 0 | 0 | 0 | 0 | 230000 | 134160 | 3397900 | 2480400 | 1063800 | 4174400 |
| LATELPDAE | CUL5 | Q93034 | CUL5_HUMAN | Unmodified | 0 | 0 | 0 | 0 | 0 | 0 | 51436 | 40277 | 0 | 0 | 0 | 0 | 146440 | 0 | 0 | 848180 | 203670 | 0 |
| LATFWYYAI | ATP5MG; N/A; A | O75964; A0A | ATP5L_HUMAN | Unmodified | 0 | 0 | 0 | 0 | 0 | 0 | 0 | 0 | 4563500 | 0 | 0 | 0 | 0 | 0 | 0 | 0 | 0 | 0 |
| LATLEGDD | PSD2 | Q9BQI7 | PSD2_HUMAN | Unmodified | 0 | 0 | 194010 | 0 | 0 | 0 | 0 | 0 | 192940 | 0 | 498930 | 275980 | 0 | 0 | 0 | 0 | 0 | 0 |
| LATLEGDD | PSD2 | Q9BQI7 | PSD2_HUMAN | Unmodified | 0 | 0 | 0 | 0 | 0 | 0 | 0 | 0 | 0 | 0 | 3724100 | 6395800 | 0 | 0 | 0 | 0 | 0 | 0 |
| LATLGALET | ENTPD5; ENTPI | O75356; G3Y | ENTP5_HUMAN | Unmodified | 0 | 0 | 0 | 0 | 0 | 0 | 0 | 0 | 1207600 | 0 | 0 | 0 | 248860 | 0 | 0 | 0 | 0 | 0 |
| LATLLGLQ/ | EIF3A | Q14152 | EIF3A_HUMAN | Unmodified | 0 | 82409 | 855710 | 0 | 0 | 0 | 0 | 0 | 0 | 110020 | 0 | 8547200 | 0 | 0 | 565420 | 808100 | 0 | 0 |
| LATLNEKLT | BRK1 | Q8WUW1 | BRK1_HUMAN | Unmodified | 0 | 0 | 0 | 0 | 1517000 | 0 | 193910 | 66558 | 0 | 883480 | 0 | 0 | 1059000 | 521290 | 0 | 0 | 1147400 | 561330 |
| LATLNIHDY | ANAPC1; ANAP | Q9H1A4; H0 | APC1_HUMAN | Unmodified | 0 | 0 | 0 | 0 | 0 | 0 | 0 | 0 | 0 | 0 | 0 | 1351300 | 0 | 0 | 128010 | 0 | 0 | 0 |
| LATLSDQVI | MYO18A; MYO1 | Q92614; A0A | MY18A_HUMAI | Unmodified | 0 | 0 | 0 | 0 | 0 | 0 | 0 | 0 | 0 | 0 | 0 | 817120 | 0 | 0 | 0 | 0 | 0 | 0 |
| LATMMVQ( | PCYOX1; PCYO | Q9UHG3; F8 | PCYOX_HUMAI | Oxidation (M) | 0 | 0 | 0 | 0 | 0 | 336190 | 0 | 0 | 5846600 | 0 | 0 | 0 | 0 | 0 | 0 | 0 | 0 | 0 |
| LATMMVQ( | PCYOX1; PCYO | Q9UHG3; F8 | PCYOX_HUMAI | Unmodified | 0 | 0 | 0 | 0 | 0 | 0 | 0 | 0 | 2760700 | 0 | 0 | 0 | 0 | 0 | 0 | 0 | 289980 | 0 |
| LATMMVQ( | PCYOX1; PCYO | Q9UHG3; F8 | PCYOX_HUMAI | 2 Oxidation (M) | 0 | 0 | 0 | 0 | 0 | 0 | 0 | 0 | 1089600 | 0 | 0 | 0 | 0 | 0 | 0 | 0 | 0 | 0 |
| LATMMVQ( | PCYOX1; PCYO | Q9UHG3; F8 | PCYOX_HUMAI | Unmodified | 0 | 0 | 0 | 0 | 0 | 0 | 0 | 0 | 2517100 | 279750 | 0 | 0 | 0 | 0 | 0 | 0 | 1372700 | 0 |
| LATNAAVT\ | CCT8; CCT8 | P50990; H7( | TCPQ_HUMAN | Unmodified | 211070 | 212060 | 0 | 192680 | 0 | 0 | 0 | 0 | 488050 | 0 | 444250 | 11558000 | 0 | 0 | 675240 | 593350 | 0 | 0 |
| LATNAAVT\ | CCT8; CCT8 | P50990; H7( | TCPQ_HUMAN | Oxidation (M) | 0 | 0 | 0 | 0 | 814710 | 0 | 0 | 164650 | 937370 | 0 | 974290 | 7064400 | 0 | 0 | 0 | 0 | 0 | 0 |
| LATNTSAPI | MAP4; MAP4; M | P27816; H7( | MAP4_HUMAN | Unmodified | 0 | 0 | 0 | 0 | 0 | 0 | 0 | 0 | 0 | 0 | 96933 | 675350 | 0 | 0 | 0 | 0 | 0 | 111030 |
| LATPFTAAM | VPS37B; VPS37 | Q9H9H4; F5 | VP37B_HUMAN | Oxidation (M) | 5588400 | 0 | 0 | 470380 | 0 | 0 | 0 | 0 | 0 | 0 | 0 | 0 | 0 | 0 | 1933500 | 2021100 | 315340 | 0 |
| LATQLTGP\ | RPL13 | P26373 | RL13_HUMAN ( | Unmodified | 3185000 | 2337800 | 0 | 0 | 0 | 0 | 0 | 0 | 0 | 0 | 2201800 | 10027000 | 0 | 0 | 0 | 0 | 0 | 0 |
| LATQLTGP\ | RPL13 | P26373 | RL13_HUMAN ( | Oxidation (M) | 4531300 | 3599900 | 1836700 | 1126400 | 0 | 0 | 0 | 0 | 442960 | 0 | 436350 | 10153000 | 0 | 0 | 2211900 | 2585900 | 0 | 0 |
| LATQLTGP\ | RPL13 | P26373 | RL13_HUMAN ( | Unmodified | 0 | 0 | 0 | 0 | 1060400 | 187780 | 620320 | 263630 | 0 | 138770 | 0 | 0 | 1271300 | 922980 | 0 | 0 | 139840 | 502930 |
| LATQSNEIT | HSPB1; HSPB1; | P04792; A0A | HSPB1_HUMAI | Unmodified | 3453800 | 3104500 | 8916500 | 15474000 | 1812900 | 1034600 | 1370200 | 642800 | 30513000 | 0 | 0 | 0 | 11512000 | 4564200 | 8970900 | 7962800 | 1736000 | 0 |
| LATQSNEIT | HSPB1; HSPB1; | P04792; A0A | HSPB1_HUMAI | Unmodified | 0 | 0 | 0 | 0 | 6770000 | 1077700 | 5590400 | 0 | 0 | 14871000 | 0 | 0 | 18521000 | 10180000 | 0 | 0 | 20618000 | 7652900 |
| LATVGELQ/ | VCAN; VCAN; V | P13611; Q86 | CSPG2_HUMAI | Unmodified | 0 | 0 | 0 | 0 | 0 | 0 | 0 | 0 | 4074800 | 0 | 0 | 0 | 0 | 0 | 0 | 0 | 0 | 0 |
| LATVHMSR | CLPTM1; CLPTN | O96005; K7E | CLPT1_HUMAN | Unmodified | 0 | 0 | 0 | 0 | 0 | 0 | 0 | 0 | 538790 | 0 | 0 | 0 | 0 | 0 | 0 | 0 | 0 | 0 |
| LATVVGEN | INCENP; INCEN | Q9NQS7; E9 | INCE_HUMAN I | Unmodified | 0 | 0 | 0 | 563870 | 937600 | 0 | 0 | 0 | 0 | 0 | 0 | 0 | 952460 | 0 | 0 | 0 | 173530 | 0 |
| LATWNLQ( | EEPD1 | Q7L9B9 | EEPD1_HUMAN | Unmodified | 0 | 0 | 0 | 209690 | 0 | 0 | 0 | 0 | 0 | 0 | 0 | 0 | 0 | 0 | 234420 | 0 | 1069000 | 0 |
| LAVAWDR\ | TRPM5; TRPM5; | Q9NZQ8; A0 | TRPM5_HUMAI | Unmodified | 0 | 0 | 0 | 0 | 0 | 0 | 0 | 151300 | 0 | 0 | 0 | 2933400 | 0 | 0 | 0 | 0 | 0 | 0 |
| LAVCQHCI | TRAFD1; TRAFD | O14545; F8V | TRAD1_HUMAN | Unmodified | 256120 | 187740 | 0 | 0 | 0 | 0 | 0 | 0 | 0 | 0 | 0 | 0 | 0 | 0 | 0 | 0 | 46035 | 0 |
| LAVDEEEN. | TFRC; TFRC | P02786; F8V | TFR1_HUMAN ` | Unmodified | 4393400 | 4839100 | 22821000 | 21994000 | 13931000 | 0 | 1648200 | 287200 | 2480400 | 518340 | 984010 | 9043400 | 3661500 | 1410200 | 14748000 | 11475000 | 476260 | 5895500 |
| LAVEALSSL | CKB; CKB | P12277; G3\ | KCRB_HUMAN | Unmodified | 660840 | 909530 | 1261500 | 3108900 | 1726200 | 172110 | 0 | 110030 | 2341600 | 735900 | 706190 | 11483000 | 928940 | 0 | 5423600 | 4933100 | 168400 | 78701 |
| LAVEALSSL | CKB; CKB | P12277; G3\ | KCRB_HUMAN | Unmodified | 0 | 0 | 660640 | 17038000 | 1.28E+08 | 10860000 | 3.11E+08 | 711080 | 3834700 | 43017000 | 947320 | 432590 | 2.54E+08 | 50264000 | 0 | 0 | 0 | 498780 |
| LAVEAVLR | CCT2; CCT2; C( | P78371; F5C | TCPB_HUMAN | Unmodified | 4372300 | 2624800 | 1487500 | 752220 | 2055700 | 255310 | 511110 | 333580 | 3160800 | 1728800 | 5176600 | 21297000 | 5355400 | 2050200 | 1706600 | 1482300 | 1313900 | 1069200 |
| LAVEILNK | EIF2AK2; EIF2AI | P19525; A0A | E2AK2_HUMAN | Unmodified | 229750 | 249650 | 0 | 0 | 682120 | 30894 | 0 | 88437 | 140310 | 398380 | 660260 | 2040000 | 456640 | 411200 | 0 | 43270 | 137830 | 349780 |
| LAVGDNNS | IQGAP1; IQGAP | P46940; A0A | IQGA1_HUMAN | Unmodified | 0 | 0 | 0 | 0 | 1841900 | 928260 | 380370 | 71665 | 56462 | 797920 | 0 | 82519 | 0 | 0 | 0 | 0 | 379720 | 0 |
| LAVHPSGV | ISG15; ISG15; I: | P05161; A0A | ISG15_HUMAN | Unmodified | 863110 | 0 | 490800 | 0 | 0 | 0 | 0 | 0 | 0 | 0 | 0 | 1448600 | 0 | 0 | 1197600 | 310020 | 0 | 0 |
| LAVLITNSN | GALK1 | P51570 | GALK1_HUMAI | Unmodified | 295590 | 983080 | 798400 | 467810 | 471670 | 0 | 0 | 120410 | 0 | 348660 | 0 | 1216600 | 0 | 0 | 830470 | 552870 | 178390 | 332650 |
| LAVLPPELA | SCRIB; SCRIB; S | Q14160; A0A | SCRIB_HUMAN | Unmodified | 970040 | 529930 | 528090 | 981060 | 0 | 0 | 0 | 0 | 6680500 | 412290 | 759760 | 6559100 | 0 | 0 | 2791000 | 2386500 | 0 | 0 |
| LAVLPPELA | SCRIB; SCRIB; S | Q14160; A0A | SCRIB_HUMAN | Unmodified | 0 | 0 | 0 | 711090 | 1921200 | 171010 | 0 | 0 | 0 | 421710 | 0 | 0 | 2921400 | 0 | 1349100 | 0 | 3867900 | 3274600 |
| LAVLSSSLT | LAMTOR1; LAM | Q6IAA8; F5G | LTOR1_HUMAN | Unmodified | 0 | 0 | 1681500 | 1101500 | 1884400 | 0 | 220860 | 0 | 2882600 | 845820 | 1787000 | 1886200 | 0 | 1393900 | 936540 | 626950 | 0 | 1171800 |
| LAVNCFVN | EPCAM; EPCAM | P16422; C9J | EPCAM_HUMA | Unmodified | 34708000 | 13413000 | 22720000 | 24332000 | 1795100 | 365290 | 0 | 0 | 47326000 | 4270000 | 8565800 | 36775000 | 10173000 | 721090 | 34524000 | 37438000 | 56711 | 525240 |
| LAVNCFVN | EPCAM; EPCAM | P16422; C9J | EPCAM_HUMA | Unmodified | 364950 | 0 | 0 | 2658400 | 7489600 | 665680 | 0 | 0 | 2849700 | 16099000 | 0 | 0 | 38959000 | 53717000 | 0 | 0 | 7682500 | 4211600 |
| LAVNMVPF | TUBB2A; TUBB( | Q13885; A6N | TBB2A_HUMAN | 2 Oxidation (M) | 0 | 0 | 0 | 0 | 37789 | 0 | 0 | 0 | 0 | 262790 | 0 | 0 | 144480 | 0 | 0 | 0 | 0 | 49612 |
| LAVNNIAGI | BTF3L4; BTF3L4 | Q96K17; E9F | BT3L4_HUMAN | Oxidation (M) | 588270 | 0 | 0 | 0 | 837370 | 0 | 0 | 0 | 0 | 959000 | 0 | 2054400 | 4342800 | 0 | 0 | 0 | 0 | 0 |
| LAVQKYEEI | NAPA; NAPA; N. | P54920; M0F | SNAA_HUMAN | Unmodified | 1017200 | 1063400 | 0 | 332870 | 0 | 0 | 234640 | 0 | 819240 | 0 | 393700 | 1537200 | 142550 | 0 | 481510 | 578310 | 0 | 56154 |
| LAVQKYEEI | NAPA; NAPA; N. | P54920; M0F | SNAA_HUMAN | Unmodified | 647190 | 1003800 | 0 | 0 | 1483500 | 0 | 3837100 | 0 | 0 | 727930 | 0 | 0 | 3150900 | 4326300 | 0 | 311670 | 1715400 | 2417100 |
| LAVQQVEE | GOLIM4; GOLIN | O00461; F8V | GOLI4_HUMAN | Unmodified | 0 | 0 | 0 | 0 | 4777200 | 625460 | 0 | 441760 | 10458000 | 914860 | 591320 | 797370 | 0 | 380450 | 0 | 0 | 0 | 0 |
| LAVQQVEE | GOLIM4; GOLIN | O00461; F8V | GOLI4_HUMAN | Unmodified | 0 | 0 | 0 | 0 | 0 | 0 | 0 | 0 | 71862000 | 424660 | 1556100 | 2439300 | 0 | 0 | 0 | 0 | 0 | 0 |
| LAVVDPLF( | ASNS; ASNS; AS | P08243; C9J | ASNS_HUMAN | Oxidation (M) | 0 | 0 | 0 | 0 | 0 | 0 | 0 | 0 | 0 | 0 | 0 | 12035000 | 0 | 0 | 0 | 0 | 0 | 0 |
| LAVVDPLF( | ASNS; ASNS; AS | P08243; C9J | ASNS_HUMAN | Unmodified | 0 | 0 | 0 | 0 | 0 | 0 | 0 | 0 | 0 | 0 | 0 | 343450 | 0 | 0 | 0 | 0 | 0 | 0 |
| LAVVDPLF( | ASNS; ASNS; AS | P08243; C9J | ASNS_HUMAN | Oxidation (M) | 0 | 0 | 0 | 0 | 0 | 0 | 219600 | 0 | 0 | 207250 | 1216600 | 4854200 | 279390 | 0 | 0 | 0 | 0 | 0 |
| LAWGQPV( | QARS1; QARS1; | P47897; H7( | SYQ_HUMAN G | Unmodified | 0 | 0 | 0 | 0 | 0 | 0 | 0 | 0 | 0 | 0 | 0 | 2009500 | 0 | 0 | 0 | 0 | 0 | 0 |
| LAWVSHDS | ARPC1A; ARPC | Q92747; E9F | ARC1A_HUMAI | Unmodified | 1660600 | 1749400 | 2011400 | 2107300 | 0 | 0 | 410160 | 139020 | 1288800 | 0 | 1520700 | 19132000 | 423440 | 0 | 2399600 | 2627500 | 101890 | 0 |
| LAYARIPAH | MYOF; MYOF | Q9NZM1; H( | MYOF_HUMAN | Unmodified | 0 | 1122700 | 0 | 0 | 0 | 235460 | 0 | 206850 | 313910 | 0 | 0 | 0 | 0 | 0 | 0 | 52342 | 0 | 236090 |
| LAYELYTEA | DNAJC7; DNAJ( | Q99615; A0A | DNJC7_HUMAI | Unmodified | 0 | 0 | 0 | 0 | 2219200 | 128400 | 0 | 0 | 966430 | 901810 | 0 | 0 | 3046500 | 0 | 0 | 0 | 0 | 0 |
| LAYEQLDK | RIPOR2; RIPOR | Q9Y4F9; F5( | RIPR2_HUMAN | Unmodified | 0 | 0 | 0 | 0 | 0 | 0 | 0 | 0 | 0 | 0 | 0 | 379410 | 0 | 0 | 0 | 0 | 0 | 0 |
| LAYEQLDKI | RIPOR2; RIPOR | Q9Y4F9; F5( | RIPR2_HUMAN | Unmodified | 0 | 0 | 0 | 0 | 0 | 0 | 73213 | 0 | 0 | 22276 | 0 | 1136900 | 0 | 0 | 0 | 0 | 0 | 0 |
| LAYFNDIAV | NAA50; NAA50 | Q9GZZ1; E7I | NAA50_HUMAI | Unmodified | 254610 | 231900 | 0 | 0 | 0 | 0 | 0 | 0 | 0 | 0 | 0 | 1598400 | 0 | 0 | 0 | 136220 | 0 | 0 |
| LAYFNDIAV | NAA50; NAA50 | Q9GZZ1; E7I | NAA50_HUMAI | Unmodified | 0 | 0 | 0 | 0 | 0 | 0 | 1174300 | 0 | 0 | 0 | 0 | 0 | 0 | 0 | 0 | 78811 | 1316900 | 0 |
| LAYINPDLA | STIP1 | P31948 | STIP1_HUMAN | Unmodified | 0 | 780370 | 918680 | 1472400 | 0 | 0 | 0 | 0 | 11860000 | 952910 | 8371800 | 52688000 | 1750000 | 843080 | 1903600 | 1718900 | 0 | 802810 |
| LAYKVEQLI | PLXNA2 | O75051 | PLXA2_HUMAN | Unmodified | 0 | 15548 | 0 | 0 | 531740 | 0 | 0 | 0 | 607460 | 312640 | 0 | 521330 | 431750 | 0 | 14330 | 0 | 16067 | 0 |
| LAYSNDGE | EDIL3 | O43854 | EDIL3_HUMAN | Unmodified | 1987500 | 0 | 0 | 0 | 6.51E+08 | 1.61E+08 | 10744000 | 0 | 19327000 | 4186200 | 0 | 0 | 0 | 0 | 0 | 0 | 11519000 | 2.38E+08 |
| LAYSNDGE | EDIL3 | O43854 | EDIL3_HUMAN | Unmodified | 0 | 0 | 0 | 0 | 1273400 | 0 | 212720 | 267450 | 0 | 539300 | 0 | 0 | 0 | 0 | 0 | 0 | 109920 | 1320600 |
| LAYSNDGE | EDIL3 | O43854 | EDIL3_HUMAN | Unmodified | 0 | 0 | 0 | 0 | 43712000 | 8682200 | 1005600 | 0 | 0 | 66278 | 0 | 0 | 0 | 0 | 68533 | 0 | 3209500 | 23118000 |
| LAYTLLNK | DIP2A; DIP2B; [ | Q14689; Q9I | DIP2A_HUMAN | Unmodified | 0 | 0 | 0 | 0 | 1035600 | 3656000 | 0 | 35190 | 0 | 0 | 0 | 3638400 | 0 | 0 | 1419400 | 445660 | 206570 | 3435400 |
| LAYVWNNI | DPP4; DPP4; DI | P27487; A0A | DPP4_HUMAN | Unmodified | 478470 | 0 | 7472200 | 2514200 | 0 | 0 | 0 | 0 | 215740 | 0 | 0 | 0 | 0 | 0 | 5136300 | 3165400 | 191320 | 0 |
| LAYYYQR | SRP54; SRP54; | P61011; G3\ | SRP54_HUMAN | Unmodified | 0 | 0 | 0 | 0 | 0 | 0 | 0 | 0 | 0 | 0 | 69827 | 1151500 | 0 | 0 | 0 | 0 | 0 | 0 |
| LCAAAASIL | DDT; DDTL; DD | P30046; A6N | DOPD_HUMAN | Unmodified | 317180 | 0 | 0 | 0 | 0 | 0 | 0 | 0 | 0 | 0 | 0 | 4010400 | 0 | 0 | 1091800 | 935040 | 0 | 0 |
| LCAEEDQG | H6PD | O95479 | G6PE_HUMAN | Unmodified | 0 | 0 | 0 | 0 | 203310 | 60672 | 0 | 0 | 1675200 | 0 | 0 | 0 | 0 | 0 | 36118 | 0 | 0 | 21228 |
| LCAGASED | MYO6; MYO6; N | Q9UM54; A0 | MYO6_HUMAN | Unmodified | 0 | 0 | 655520 | 1832200 | 336050 | 0 | 0 | 0 | 559640 | 144690 | 145420 | 1123700 | 0 | 0 | 2229800 | 2571000 | 92140 | 225040 |
| LCDEQLSS | DYNC1H1; DYN | Q14204; A0A | DYHC1_HUMA | Unmodified | 0 | 0 | 0 | 0 | 0 | 0 | 0 | 0 | 820940 | 304390 | 0 | 3860100 | 137290 | 0 | 0 | 466410 | 0 | 0 |
| LCDFGSAK | GSK3B; GSK3A; | P49841; P49 | GSK3B_HUMAI | Unmodified | 0 | 0 | 0 | 0 | 181430 | 107140 | 0 | 0 | 1522200 | 149710 | 0 | 0 | 0 | 0 | 0 | 0 | 164240 | 150390 |
| LCDFGVSG | MAP2K2; MAP2 | P36507; Q02 | MP2K2_HUMAI | Unmodified | 0 | 0 | 0 | 0 | 143380 | 0 | 0 | 0 | 286730 | 0 | 0 | 0 | 0 | 0 | 0 | 0 | 41536 | 43048 |
| LCDFNPK | CLTB; CLTA; CL | P09497; P09 | CLCB_HUMAN | Unmodified | 8295800 | 195090 | 1594500 | 2190700 | 5049900 | 0 | 615900 | 338650 | 957500 | 0 | 460220 | 24585000 | 7133800 | 0 | 1726900 | 3043600 | 1372900 | 1815400 |
| LCDSGELV | GSK3B; GSK3B; | P49841; A0A | GSK3B_HUMAI | Unmodified | 117050 | 186560 | 0 | 0 | 339450 | 0 | 386170 | 112580 | 171210 | 0 | 189030 | 765620 | 444170 | 506160 | 199740 | 0 | 314570 | 272740 |
| LCDSYEIRF | HNRNPR; HNRI | O43390; B4I | HNRPR_HUMA | Unmodified | 0 | 0 | 0 | 0 | 0 | 0 | 0 | 0 | 0 | 0 | 829950 | 3881400 | 554610 | 0 | 0 | 0 | 0 | 0 |
| LCEAICPA( | NDUFS8; NDUI | O00217; E9F | NDUS8_HUMA | Unmodified | 0 | 0 | 0 | 0 | 0 | 0 | 0 | 0 | 1358500 | 0 | 0 | 0 | 0 | 0 | 0 | 0 | 0 | 0 |
| LCENGNMI | USP15; USP15; | Q9Y4E8; H0Y | UBP15_HUMAI | Unmodified | 0 | 0 | 0 | 0 | 0 | 0 | 0 | 0 | 0 | 0 | 0 | 128300 | 0 | 0 | 0 | 0 | 0 | 0 |
| LCERETPPI | ABHD12; ABHD | Q8N2K0; A0, | ABD12_HUMAI | Unmodified | 0 | 0 | 0 | 0 | 0 | 0 | 65877 | 0 | 8498100 | 1365400 | 0 | 0 | 0 | 0 | 0 | 0 | 0 | 0 |
| LCEVVDHV | LPIN1 | Q14693 | LPIN1_HUMAN | Unmodified | 0 | 0 | 0 | 0 | 0 | 0 | 0 | 0 | 0 | 0 | 53471 | 408460 | 0 | 10980 | 0 | 0 | 0 | 0 |
| LCFDNSFS | TMED1 | Q13445 | TMED1_HUMAI | Unmodified | 0 | 0 | 0 | 0 | 0 | 0 | 0 | 0 | 8002000 | 988490 | 0 | 0 | 0 | 0 | 0 | 0 | 64734 | 0 |
| LCFKENAE( | CD99 | P14209 | CD99_HUMAN | Unmodified | 0 | 0 | 0 | 0 | 438060 | 0 | 0 | 0 | 0 | 0 | 0 | 0 | 0 | 77142 | 0 | 0 | 461710 | 988830 |
| LCFLDKVEI | TECR; TECR; TE | Q9NZ01; M0 | TECR_HUMAN | Unmodified | 0 | 0 | 0 | 0 | 1154200 | 0 | 1651100 | 191180 | 1304400 | 531690 | 0 | 0 | 1374500 | 865540 | 0 | 0 | 0 | 0 |
| LCFSTAQH | HNRNPL; HNRI | P14866; M0( | HNRPL_HUMAI | Unmodified | 311760 | 152860 | 0 | 152760 | 651350 | 0 | 7180800 | 2588300 | 1049700 | 1589200 | 218640 | 7199400 | 2259400 | 1122600 | 0 | 140240 | 374260 | 582180 |
| LCGDDHF( | SUSD2 | Q9UGT4 | SUSD2_HUMAI | Unmodified | 0 | 0 | 0 | 0 | 0 | 0 | 0 | 0 | 2708800 | 0 | 0 | 0 | 0 | 0 | 0 | 0 | 0 | 0 |
| LCGDLVSC | PACSIN3; PACS | Q9UKS6; E9I | PACN3_HUMAI | Unmodified | 5109300 | 1502200 | 641690 | 0 | 0 | 0 | 0 | 0 | 0 | 0 | 0 | 0 | 0 | 0 | 2517000 | 1672100 | 0 | 0 |

| | | | | | | | | | | | | | | | | | | | | | | |
|---|---|---|---|---|---|---|---|---|---|---|---|---|---|---|---|---|---|---|---|---|---|---|
| LCHLRKGP | SLC9A3R2; SLC | Q15599; H3I | NHRF2_HUMAI | Unmodified | 0 | 0 | 0 | 0 | 1740100 | 146040 | 0 | 0 | 0 | 116510 | 0 | 0 | 1993900 | 1024000 | 0 | 0 | 452230 | 2337700 |
| LCIEVTPQS | ABCE1; ABCE1; | P61221; D6F | ABCE1_HUMAI | Unmodified | 0 | 0 | 0 | 0 | 1086000 | 0 | 0 | 0 | 0 | 0 | 0 | 2435000 | 0 | 0 | 0 | 0 | 0 | 0 |
| LCKTEQDL | STXBP3 | O00186 | STXB3_HUMAN | Unmodified | 168380 | 0 | 0 | 203520 | 729280 | 0 | 0 | 0 | 0 | 208840 | 0 | 386110 | 921120 | 855580 | 0 | 0 | 516150 | 1095200 |
| LCKTEQDL | STXBP3 | O00186 | STXB3_HUMAN | Unmodified | 0 | 0 | 0 | 0 | 284340 | 0 | 485270 | 109130 | 0 | 0 | 0 | 0 | 0 | 489220 | 0 | 0 | 773360 | 272430 |
| LCMHQLIH | ZNF311 | Q5JNZ3 | ZN311_HUMAN | Unmodified | 0 | 0 | 0 | 0 | 19914000 | 5119300 | 10197000 | 5327100 | 0 | 5094500 | 0 | 0 | 15462000 | 10372000 | 0 | 0 | 18064000 | 10720000 |
| LCNDLMN( | PACSIN1 | Q9BY11 | PACN1_HUMAI | Unmodified | 931530 | 238190 | 35453 | 0 | 156350 | 0 | 0 | 0 | 0 | 0 | 0 | 0 | 0 | 0 | 130620 | 66217 | 0 | 0 |
| LCNDLMN( | PACSIN1 | Q9BY11 | PACN1_HUMAI | Oxidation (M) | 240880 | 0 | 0 | 0 | 0 | 0 | 0 | 0 | 0 | 0 | 0 | 0 | 0 | 0 | 0 | 0 | 0 | 0 |
| LCNYLSHH | PAIP1; PAIP1; P | Q9H074; D6 | PAIP1_HUMAN | Unmodified | 2649300 | 1109500 | 721190 | 1263600 | 0 | 0 | 0 | 0 | 0 | 0 | 0 | 1359500 | 0 | 0 | 600900 | 526330 | 0 | 0 |
| LCPFKLVS( | PRKCSH; PRKC | P14314; K7E | GLU2B_HUMAI | Unmodified | 0 | 0 | 0 | 0 | 0 | 0 | 0 | 0 | 0 | 1828000 | 0 | 0 | 0 | 0 | 0 | 0 | 0 | 0 |
| LCPGGQLF | CLIC1 | O00299 | CLIC1_HUMAN | Unmodified | 0 | 0 | 0 | 0 | 51520 | 0 | 0 | 0 | 0 | 0 | 0 | 0 | 71617 | 164980 | 0 | 0 | 75958 | 0 |
| LCPNSTGA | PSMC2; PSMC2 | P35998; C9J | PRS7_HUMAN | Unmodified | 0 | 0 | 0 | 0 | 0 | 0 | 0 | 0 | 740650 | 0 | 418890 | 2573300 | 0 | 0 | 0 | 0 | 0 | 0 |
| LCPQFLQL | STAMBP; STAMI | O95630; C9J | STABP_HUMAN | Unmodified | 775940 | 0 | 0 | 0 | 0 | 0 | 0 | 0 | 1908500 | 165320 | 0 | 429840 | 0 | 0 | 0 | 0 | 90408 | 0 |
| LCQFEEAK( | SPAG1 | Q07617 | SPAG1_HUMAI | Unmodified | 0 | 0 | 0 | 0 | 1248900 | 0 | 0 | 0 | 0 | 0 | 0 | 0 | 0 | 0 | 0 | 0 | 1626100 | 1031600 |
| LCQIPVHG | LTBP1; LTBP1; I | Q14766; E7E | LTBP1_HUMAN | Unmodified | 0 | 0 | 0 | 0 | 0 | 0 | 0 | 17977 | 0 | 0 | 0 | 0 | 0 | 0 | 0 | 0 | 319130 | 1929400 |
| LCREALDVI | MRPS27; MRPS | Q92552; D6I | RT27_HUMAN : | Unmodified | 0 | 0 | 0 | 0 | 0 | 0 | 23658 | 0 | 74762 | 110860 | 0 | 0 | 0 | 0 | 0 | 0 | 0 | 0 |
| LCRGDSS( | MELTF | P08582 | TRFM_HUMAN | Unmodified | 0 | 0 | 0 | 0 | 0 | 0 | 0 | 0 | 1388300 | 0 | 0 | 0 | 0 | 0 | 0 | 0 | 0 | 0 |
| LCRLVRGE | SLC9A3R2 | Q15599 | NHRF2_HUMAI | Unmodified | 0 | 0 | 0 | 0 | 533030 | 0 | 0 | 0 | 0 | 0 | 0 | 0 | 455670 | 802540 | 0 | 0 | 362990 | 904570 |
| LCRTNQIG | IST1; IST1; IST1 | P53990; H3E | IST1_HUMAN I | Unmodified | 0 | 0 | 0 | 0 | 6380900 | 1214500 | 214860 | 0 | 0 | 2252500 | 0 | 0 | 2598400 | 3704800 | 0 | 0 | 0 | 1050900 |
| LCRVEQDL | STXBP1; STXBP | P61764; A0A | STXB1_HUMAN | Oxidation (M) | 130080 | 0 | 0 | 0 | 107930 | 37019 | 74092 | 86365 | 170410 | 75331 | 0 | 470600 | 0 | 0 | 163110 | 0 | 0 | 0 |
| LCSAHGVL | CTPS1; CTPS1; | P17812; B4E | PYRG1_HUMAI | Unmodified | 0 | 0 | 0 | 0 | 0 | 0 | 0 | 28177 | 586630 | 87117 | 0 | 5225200 | 0 | 0 | 0 | 0 | 0 | 0 |
| LCSDLMN( | PACSIN2; PACS | Q9UNF0; B0 | PACN2_HUMAI | Unmodified | 6950700 | 2048300 | 1114000 | 747220 | 0 | 0 | 0 | 0 | 0 | 0 | 0 | 878990 | 0 | 0 | 4141100 | 1702800 | 0 | 0 |
| LCSDLMN( | PACSIN2; PACS | Q9UNF0; B0 | PACN2_HUMAI | Oxidation (M) | 211900 | 0 | 593110 | 0 | 0 | 0 | 0 | 0 | 0 | 0 | 0 | 290610 | 0 | 0 | 3686600 | 2091500 | 0 | 0 |
| LCSGVLGT | MTCH2; MTCH2 | Q9Y6C9; E9I | MTCH2_HUMA | Unmodified | 0 | 0 | 0 | 0 | 0 | 0 | 78514 | 0 | 3390700 | 861500 | 0 | 0 | 0 | 0 | 71555 | 0 | 0 | 0 |
| LCSKRTDL( | CNPY2; CNPY2 | Q9Y2B0; H0 | CNPY2_HUMAI | Unmodified | 0 | 0 | 0 | 0 | 155400 | 0 | 132670 | 0 | 0 | 3214600 | 0 | 123630 | 0 | 0 | 0 | 0 | 98140 | 0 |
| LCSLFYTNI | MATR3; MATR3; | P43243; B3K | MATR3_HUMAI | Unmodified | 0 | 0 | 0 | 0 | 0 | 0 | 0 | 0 | 0 | 0 | 0 | 3721300 | 1434400 | 0 | 0 | 0 | 0 | 0 |
| LCSVEQDL | STXBP2; STXBP | Q15833; M0I | STXB2_HUMAN | Unmodified | 0 | 0 | 0 | 0 | 881080 | 0 | 0 | 0 | 0 | 0 | 0 | 0 | 284100 | 334800 | 0 | 0 | 278400 | 492940 |
| LCTFILK | EIF5; EIF5; EIF5 | P55010; H0Y | IF5_HUMAN Eu | Unmodified | 0 | 0 | 0 | 0 | 1565400 | 0 | 0 | 0 | 0 | 0 | 0 | 1313500 | 733310 | 0 | 0 | 0 | 677610 | 1329700 |
| LCTFILKNP | EIF5; EIF5 | P55010; H0Y | IF5_HUMAN Eu | Unmodified | 0 | 0 | 0 | 0 | 3500700 | 0 | 0 | 0 | 0 | 0 | 0 | 0 | 356980 | 486670 | 0 | 0 | 0 | 0 |
| LCTSATESE | UQCRC1 | P31930 | QCR1_HUMAN | Unmodified | 0 | 0 | 0 | 0 | 0 | 0 | 0 | 0 | 8954300 | 478140 | 0 | 0 | 0 | 0 | 0 | 0 | 0 | 0 |
| LCTSATESE | UQCRC1 | P31930 | QCR1_HUMAN | Unmodified | 0 | 0 | 0 | 0 | 226270 | 0 | 0 | 0 | 2400500 | 2067200 | 0 | 0 | 0 | 0 | 0 | 0 | 124080 | 0 |
| LCTSVTESE | PMPCB; PMPCE | O75439; G3 | MPPB_HUMAN | Unmodified | 0 | 2734600 | 0 | 0 | 0 | 0 | 0 | 0 | 768100 | 0 | 0 | 0 | 0 | 0 | 0 | 0 | 399000 | 0 |
| LCVLNAGV | UTRN | P46939 | UTRN_HUMAN | Unmodified | 0 | 0 | 308130 | 906660 | 0 | 0 | 0 | 0 | 0 | 0 | 0 | 0 | 0 | 0 | 572910 | 442720 | 0 | 0 |
| LCVPAMN\ | AHCYL1; AHCY | O43865; Q9 | SAHH2_HUMAI | Oxidation (M) | 42069 | 145970 | 0 | 529980 | 1216900 | 396200 | 53587 | 0 | 0 | 0 | 0 | 0 | 253160 | 214510 | 236030 | 123600 | 723580 | 375540 |
| LCVPAMN\ | AHCYL1; AHCY | O43865; Q9 | SAHH2_HUMAI | Unmodified | 0 | 0 | 0 | 0 | 1279300 | 343340 | 0 | 0 | 0 | 0 | 0 | 0 | 374680 | 337470 | 1489200 | 0 | 608920 | 605580 |
| LCVQNSP( | CSTF2; CSTF2; | P33240; E7E | CSTF2_HUMAN | Unmodified | 0 | 0 | 0 | 0 | 0 | 0 | 0 | 0 | 0 | 0 | 69154 | 437200 | 0 | 0 | 0 | 0 | 0 | 0 |
| LCWEANPI | RIPK1 | Q13546 | RIPK1_HUMAN | Unmodified | 0 | 0 | 0 | 0 | 0 | 0 | 0 | 114790 | 0 | 392390 | 0 | 0 | 0 | 0 | 0 | 0 | 820400 | 0 |
| LCWFLDEA | PGLS; PGLS | O95336; M0I | 6PGL_HUMAN | Unmodified | 18186 | 0 | 0 | 0 | 407470 | 0 | 0 | 0 | 1488000 | 585570 | 904660 | 1132000 | 0 | 0 | 0 | 0 | 142110 | 576670 |
| LCYVALDFI | POTEE; ACTG1; | Q6S8J3; P63 | POTEE_HUMAN | Oxidation (M) | 5293500 | 2702500 | 1522400 | 10681000 | 37641000 | 11136000 | 6308600 | 3389200 | 3934400 | 4430000 | 2990400 | 7947500 | 16596000 | 11382000 | 9606200 | 2420800 | 22237000 | 16751000 |
| LCYVALDFI | POTEE; ACTG1; | Q6S8J3; P63 | POTEE_HUMAN | Unmodified | 0 | 6085200 | 317180 | 8900000 | 78939000 | 15432000 | 14240000 | 0 | 8710000 | 6754200 | 1732400 | 2855000 | 27368000 | 23396000 | 12258000 | 1213500 | 50835000 | 30739000 |
| LCYVGYNII | ACTR2; ACTR2 | P61160; F5H | ARP2_HUMAN | Unmodified | 490080 | 591060 | 129590 | 554910 | 335450 | 0 | 36491 | 0 | 0 | 0 | 0 | 4484700 | 0 | 0 | 24736 | 1080600 | 0 | 0 |
| LCYYIGATD | MANF | P55145 | MANF_HUMAN | Unmodified | 144360 | 0 | 0 | 0 | 0 | 0 | 0 | 0 | 25329000 | 816960 | 1992100 | 1270300 | 0 | 0 | 0 | 0 | 0 | 0 |
| LDADKYEN | ADI1 | Q9BV57 | MTND_HUMAN | Unmodified | 0 | 216730 | 560970 | 495830 | 333990 | 74092 | 0 | 21219 | 827820 | 81613 | 1154700 | 5741400 | 169120 | 0 | 662090 | 533240 | 0 | 59457 |
| LDAEPRPPI | FARSA; FARSA | Q9Y285; K7E | SYFA_HUMAN I | Unmodified | 0 | 0 | 0 | 0 | 0 | 0 | 0 | 0 | 0 | 0 | 332270 | 2360000 | 0 | 0 | 0 | 0 | 0 | 0 |
| LDAFIEALH | OPA1; OPA1; OI | O60313; E5K | OPA1_HUMAN | Unmodified | 0 | 0 | 0 | 0 | 46256 | 176380 | 0 | 0 | 278280 | 0 | 1001200 | 6638300 | 85906 | 43528 | 63863 | 81301 | 70498 | 0 |
| LDAGRLEG | AHNAK2 | Q8IVF2 | AHNK2_HUMAI | Unmodified | 0 | 0 | 0 | 0 | 743150 | 0 | 0 | 0 | 0 | 0 | 0 | 0 | 0 | 0 | 0 | 0 | 140210 | 0 |
| LDAYKADD | STXBP1; STXBP | P61764; A0A | STXB1_HUMAN | Unmodified | 173770 | 0 | 0 | 945030 | 0 | 2048400 | 0 | 0 | 171420 | 56977 | 0 | 601260 | 0 | 0 | 0 | 119260 | 23471 | 53813 |
| LDAYKADD | STXBP1; STXBP | P61764; A0A | STXB1_HUMAN | Oxidation (M) | 0 | 0 | 0 | 0 | 0 | 0 | 0 | 0 | 24493 | 0 | 0 | 262560 | 0 | 0 | 0 | 0 | 0 | 0 |
| LDCAQLK | CNP; CNP; CNI | P09543; K7E | CN37_HUMAN | Unmodified | 0 | 0 | 0 | 0 | 0 | 0 | 0 | 0 | 189480 | 314200 | 858640 | 340620 | 0 | 0 | 0 | 0 | 0 | 0 |
| LDCPLAME | VPS28; VPS28; | Q9UK41; E9I | VPS28_HUMAN | Oxidation (M) | 666170 | 367230 | 514830 | 172290 | 0 | 0 | 0 | 0 | 0 | 0 | 0 | 230160 | 0 | 0 | 525890 | 468410 | 0 | 0 |
| LDCPLAME | VPS28; VPS28; | Q9UK41; E9I | VPS28_HUMAN | Unmodified | 912120 | 0 | 0 | 0 | 264200 | 0 | 0 | 0 | 0 | 283060 | 0 | 380880 | 0 | 0 | 0 | 480360 | 159680 | 0 |
| LDDCGLTE | RNH1; RNH1; R | P13489; E9P | RINI_HUMAN R | Unmodified | 48141 | 0 | 0 | 0 | 0 | 0 | 0 | 0 | 0 | 0 | 0 | 1826800 | 0 | 0 | 105660 | 0 | 0 | 0 |
| LDDLVSKS | YKT6; YKT6 | O15498; A0/ | YKT6_HUMAN ! | Unmodified | 210560 | 0 | 0 | 0 | 0 | 0 | 0 | 74273 | 495760 | 209200 | 368070 | 1224600 | 116040 | 0 | 0 | 215580 | 189900 | 0 |
| LDDMSSIV( | NXF1; NXF1; N) | Q9UBU9; E9 | NXF1_HUMAN | Unmodified | 253930 | 259810 | 0 | 908530 | 13799000 | 2347000 | 3114100 | 2656000 | 797110 | 1163000 | 613460 | 1144800 | 4289200 | 4097100 | 0 | 353050 | 17909000 | 20320000 |
| LDDPSCPR | EIF2S3; EIF2S3I | P41091; Q2\ | IF2G_HUMAN E | Unmodified | 0 | 0 | 89295 | 0 | 0 | 0 | 0 | 0 | 101080 | 0 | 0 | 661800 | 0 | 0 | 71095 | 74495 | 0 | 0 |
| LDDTVHVV | DDX6 | P26196 | DDX6_HUMAN | Unmodified | 748900 | 306900 | 268480 | 430060 | 0 | 0 | 0 | 0 | 0 | 0 | 241730 | 2123200 | 214450 | 0 | 478260 | 546440 | 0 | 0 |
| LDEAEDDF | DNAJC3; DNAJ( | Q13217; X6F | DNJC3_HUMAI | Unmodified | 0 | 0 | 0 | 0 | 0 | 0 | 0 | 0 | 0 | 110250 | 168480 | 48113 | 0 | 0 | 0 | 0 | 0 | 0 |
| LDEDLAAY( | CNP; CNP; CNI | P09543; K7E | CN37_HUMAN | Unmodified | 0 | 0 | 0 | 0 | 0 | 0 | 0 | 0 | 0 | 0 | 0 | 676400 | 0 | 0 | 0 | 0 | 0 | 0 |
| LDEEAENL | PTPA; PTPA; PTI | Q15257; A6F | PTPA_HUMAN ! | Unmodified | 0 | 0 | 0 | 0 | 2187600 | 0 | 0 | 0 | 0 | 0 | 0 | 0 | 595070 | 0 | 0 | 0 | 1197000 | 0 |
| LDEHDFLG | CPNE2; CPNE2 | Q96FN4; H3 | CPNE2_HUMAI | Unmodified | 0 | 0 | 0 | 0 | 0 | 0 | 0 | 0 | 978450 | 145380 | 0 | 0 | 0 | 0 | 0 | 0 | 0 | 0 |
| LDEIYVAGL | SYNCRIP; SYN( | O60506; A0/ | HNRPQ_HUMA | Unmodified | 0 | 0 | 0 | 0 | 0 | 0 | 0 | 51491 | 0 | 0 | 0 | 1403500 | 0 | 0 | 0 | 0 | 0 | 0 |
| LDELYGTW | RPL4; RPL4 | P36578; H3E | RL4_HUMAN 6( | Unmodified | 0 | 0 | 0 | 0 | 0 | 0 | 0 | 0 | 0 | 0 | 0 | 644790 | 0 | 0 | 0 | 68317 | 0 | 0 |
| LDEREAGII | TPI1; TPI1; TPI1 | P60174; U3k | TPIS_HUMAN T | Unmodified | 1168400 | 106400 | 836460 | 639170 | 866250 | 0 | 0 | 0 | 1671300 | 44511 | 459210 | 12575000 | 0 | 0 | 518300 | 835450 | 143480 | 368780 |
| LDEREAGII | TPI1; TPI1; TPI1 | P60174; U3k | TPIS_HUMAN T | Unmodified | 0 | 0 | 0 | 0 | 1790700 | 0 | 0 | 114070 | 0 | 0 | 0 | 0 | 964570 | 827150 | 0 | 0 | 1485500 | 0 |
| LDFGNSQ( | GGCT; GGCT; ( | O75223; H7I | GGCT_HUMAN | Unmodified | 0 | 0 | 0 | 0 | 0 | 0 | 0 | 0 | 0 | 0 | 0 | 900380 | 0 | 0 | 63362 | 0 | 0 | 0 |
| LDFLPLK | ZFAND2B; ZFAN | Q8WV99; B4 | ZFN2B_HUMAN | Unmodified | 84046 | 114660 | 0 | 0 | 0 | 0 | 0 | 0 | 0 | 0 | 0 | 0 | 0 | 0 | 0 | 38630 | 0 | 0 |
| LDFNLIR | ATIC; ATIC | P31939; C9J | PUR9_HUMAN | Unmodified | 0 | 0 | 538120 | 0 | 0 | 0 | 0 | 0 | 446830 | 0 | 807080 | 6135100 | 0 | 0 | 0 | 490600 | 0 | 0 |
| LDFNPDRY | CYP51A1; CYP5 | Q16850; A0/ | CP51A_HUMAN | Unmodified | 0 | 0 | 0 | 0 | 0 | 2546000 | 0 | 0 | 0 | 0 | 0 | 500600 | 0 | 0 | 0 | 0 | 0 | 0 |
| LDFYQHSV | MYG1; MYG1; M | Q9HB07; F8 | MYG1_HUMAN | Unmodified | 0 | 0 | 0 | 137660 | 0 | 0 | 0 | 0 | 431570 | 0 | 0 | 2336100 | 0 | 0 | 0 | 0 | 0 | 0 |
| LDGAQLEG | AHNAK2 | Q8IVF2 | AHNK2_HUMAI | Unmodified | 0 | 0 | 0 | 0 | 0 | 0 | 196060 | 0 | 0 | 1183000 | 0 | 127630 | 0 | 0 | 0 | 0 | 0 | 304360 |
| LDGARLEG | AHNAK2 | Q8IVF2 | AHNK2_HUMAI | Unmodified | 0 | 0 | 0 | 0 | 0 | 214210 | 0 | 0 | 0 | 0 | 0 | 0 | 0 | 0 | 0 | 0 | 118500 | 2196200 |
| LDGARLEG | AHNAK2 | Q8IVF2 | AHNK2_HUMAI | Unmodified | 0 | 0 | 0 | 0 | 1395300 | 285250 | 0 | 0 | 0 | 0 | 0 | 0 | 0 | 0 | 0 | 0 | 194480 | 1559100 |
| LDGARLEG | AHNAK2 | Q8IVF2 | AHNK2_HUMAI | Unmodified | 0 | 0 | 0 | 0 | 639200 | 0 | 0 | 0 | 0 | 0 | 0 | 0 | 0 | 0 | 121530 | 0 | 97610 | 735930 |
| LDGARLEG | AHNAK2 | Q8IVF2 | AHNK2_HUMAI | Unmodified | 0 | 0 | 0 | 0 | 657990 | 0 | 121130 | 0 | 168970 | 0 | 0 | 0 | 0 | 0 | 0 | 0 | 46475 | 749370 |
| LDGEIIPHII | ADSS2 | P30520 | PURA2_HUMAI | Unmodified | 0 | 0 | 0 | 0 | 0 | 0 | 0 | 0 | 0 | 0 | 0 | 1368400 | 0 | 0 | 0 | 0 | 0 | 0 |
| LDGHGTVF | ATP8A1 | Q9Y2Q0 | AT8A1_HUMAN | Unmodified | 0 | 0 | 0 | 0 | 0 | 0 | 0 | 0 | 0 | 138700 | 0 | 367660 | 0 | 0 | 0 | 0 | 0 | 0 |
| LDGLVETP | NAP1L1; NAP1L | P55209; F8V | NP1L1_HUMAN | Unmodified | 1034000 | 1821300 | 944740 | 0 | 0 | 0 | 0 | 0 | 849240 | 304390 | 942080 | 0 | 1528000 | 0 | 1410900 | 1150000 | 1371500 | 1787300 |
| LDGNQDLI | IDH2 | P48735 | IDHP_HUMAN | Unmodified | 0 | 49021 | 148790 | 0 | 0 | 0 | 0 | 0 | 2231800 | 0 | 0 | 0 | 131830 | 601010 | 0 | 95812 | 0 | 1071100 |
| LDGQTPHI | SMARCA5 | O60264 | SMCA5_HUMA | Unmodified | 0 | 0 | 0 | 0 | 0 | 0 | 0 | 0 | 0 | 0 | 0 | 1228500 | 0 | 0 | 0 | 0 | 0 | 0 |
| LDHHPEWI | PCBD1 | P61457 | PHS_HUMAN P | Unmodified | 80313 | 0 | 0 | 0 | 0 | 0 | 0 | 0 | 902780 | 275150 | 0 | 3186200 | 898070 | 118470 | 0 | 300460 | 266430 | 0 |
| LDHKFDLM | TUBA3C; TUBA: | P0DPH7; Q7 | TBA3C_HUMAN | Oxidation (M) | 2562300 | 1187700 | 3284000 | 2445800 | 4519900 | 1405700 | 2967300 | 1820700 | 10107000 | 2994900 | 5106600 | 81383000 | 3363300 | 1651900 | 2598900 | 4218300 | 1122900 | 2279200 |
| LDHKFDLM | TUBA3C; TUBA: | P0DPH7; Q7 | TBA3C_HUMAN | Unmodified | 990920 | 0 | 275990 | 722390 | 790110 | 187500 | 969640 | 191950 | 9871000 | 1097900 | 1870400 | 30817000 | 1446400 | 662890 | 1027700 | 0 | 127080 | 589460 |
| LDHKFDLM | TUBA3C; TUBA: | P0DPH7; Q7 | TBA3C_HUMAN | Unmodified | 0 | 0 | 0 | 0 | 0 | 0 | 0 | 215920 | 700230 | 0 | 0 | 0 | 0 | 0 | 0 | 0 | 0 | 0 |
| LDHTLQSP | FAM20B; FAM2( | O75063; X6F | XYLK_HUMAN ( | Unmodified | 0 | 0 | 0 | 0 | 0 | 0 | 0 | 0 | 7101400 | 652290 | 0 | 0 | 0 | 0 | 0 | 0 | 129070 | 0 |
| LDHVVTIIK | PRMT1; PRMT1; | Q99873; E9F | ANM1_HUMAN | Unmodified | 0 | 0 | 0 | 0 | 0 | 0 | 0 | 0 | 0 | 0 | 0 | 314390 | 0 | 0 | 0 | 0 | 0 | 0 |
| LDHYAIIK | RPL23A; RPL23. | P62750; K7E | RL23A_HUMAN | Unmodified | 0 | 0 | 270920 | 0 | 0 | 0 | 0 | 0 | 0 | 0 | 0 | 783220 | 0 | 0 | 0 | 0 | 0 | 0 |
| LDIDPETIT | MTHFD1; MTHF | P11586; V9G | C1TC_HUMAN | Unmodified | 0 | 0 | 0 | 0 | 0 | 0 | 0 | 0 | 3034800 | 0 | 0 | 7035700 | 0 | 0 | 648320 | 0 | 0 | 0 |
| LDIDSPPIT | PKM; PKM; PKM | P14618; B4D | KPYM_HUMAN | Unmodified | 354560 | 479440 | 1387200 | 648870 | 0 | 165130 | 0 | 0 | 2582100 | 109730 | 15576000 | 59079000 | 0 | 0 | 1572700 | 1414200 | 120120 | 0 |
| LDIIETNVK | ELP1; ELP1; ELF | O95163; A0/ | ELP1_HUMAN I | Unmodified | 0 | 0 | 0 | 0 | 0 | 0 | 0 | 0 | 0 | 0 | 0 | 0 | 0 | 0 | 494460 | 0 | 0 | 0 |
| LDIKLDSEI | EEF2 | P13639 | EF2_HUMAN EI | Unmodified | 0 | 0 | 1337900 | 0 | 0 | 0 | 871410 | 430500 | 212400 | 0 | 512050 | 5691300 | 0 | 3243800 | 0 | 0 | 0 | 0 |
| LDIKLDSEI | EEF2 | P13639 | EF2_HUMAN EI | Unmodified | 442070 | 0 | 0 | 0 | 0 | 0 | 35510 | 0 | 695580 | 0 | 1661300 | 12458000 | 344160 | 0 | 0 | 444290 | 0 | 0 |
| LDILDTAG( | RRAS; RRAS2; F | P10301; P62 | RRAS_HUMAN | Oxidation (M) | 0 | 327960 | 0 | 0 | 0 | 0 | 0 | 0 | 0 | 0 | 0 | 0 | 0 | 0 | 10598 | 840930 | 0 | 0 |
| LDILHQVAI | MRPL4; MRPL4; | Q9BYD3; K7I | RM04_HUMAN | Oxidation (M) | 0 | 33258 | 0 | 0 | 0 | 0 | 0 | 0 | 221710 | 0 | 0 | 0 | 0 | 0 | 0 | 0 | 0 | 0 |
| LDILHQVAI | MRPL4; MRPL4; | Q9BYD3; K7I | RM04_HUMAN | Unmodified | 0 | 0 | 0 | 0 | 0 | 0 | 0 | 0 | 287020 | 0 | 0 | 0 | 0 | 0 | 0 | 0 | 0 | 0 |
| LDILQGITP | N/A | N/A | N/A | Oxidation (M) | 6078400 | 12774000 | 1120600 | 11316000 | 18255000 | 7010500 | 3776100 | 0 | 77793000 | 6200100 | 3635000 | 0 | 12379000 | 5942200 | 5216600 | 1379900 | 0 | 0 |
| LDIPSAVPL | RHBDF1; RHBD | Q96CC6; F8 | RHDF1_HUMAI | Unmodified | 172580 | 0 | 195990 | 488120 | 0 | 0 | 0 | 0 | 3541600 | 650170 | 0 | 0 | 237830 | 0 | 0 | 473230 | 0 | 842480 |
| LDKESYPVI | ERP29; ERP29 | P30040; F8V | ERP29_HUMAN | Unmodified | 0 | 0 | 0 | 0 | 0 | 0 | 0 | 0 | 610700 | 0 | 0 | 0 | 0 | 0 | 0 | 0 | 0 | 0 |
| LDKESYPVI | ERP29; ERP29 | P30040; F8V | ERP29_HUMAN | Unmodified | 0 | 0 | 0 | 0 | 0 | 0 | 0 | 0 | 2039000 | 735450 | 0 | 0 | 0 | 0 | 0 | 0 | 0 | 0 |
| LDKSQIHD | HSPA8; HSPA8; | P11142; E9P | HSP7C_HUMAI | Unmodified | 0 | 0 | 0 | 0 | 4955800 | 0 | 878810 | 0 | 0 | 593320 | 0 | 0 | 3922000 | 1289600 | 0 | 0 | 973550 | 668130 |
| LDLAGRDL | ACTG1; ACTC1; | P63261; P68 | ACTG_HUMAN | Oxidation (M) | 67559000 | 59597000 | 48729000 | 18050000 | 19197000 | 9450200 | 9110100 | 981290 | 1251400 | 12277000 | 18383000 | 6.17E+08 | 5659500 | 1435800 | 1.2E+08 | 92878000 | 6518400 | 5335300 |
| LDLAGRDL | ACTG1; ACTC1; | P63261; P68 | ACTG_HUMAN | Unmodified | 4872200 | 5960200 | 1291400 | 1703800 | 8165700 | 1392500 | 1865800 | 26112 | 34306000 | 0 | 1328700 | 22598000 | 841980 | 381940 | 5507700 | 3719000 | 1575800 | 1778100 |
| LDLERETIEI | TWF2; TWF2; T\ | Q6IBS0; D6F | TWF2_HUMAN | Unmodified | 0 | 0 | 0 | 0 | 3588400 | 0 | 0 | 0 | 183490 | 0 | 535680 | 1520300 | 0 | 338730 | 0 | 0 | 0 | 0 |
| LDLERETIEI | TWF2; TWF2; T\ | Q6IBS0; D6F | TWF2_HUMAN | Unmodified | 395030 | 0 | 0 | 0 | 532270 | 0 | 0 | 0 | 2567700 | 0 | 3553300 | 18759000 | 0 | 0 | 0 | 0 | 0 | 0 |
| LDLERKVE! | VIM; VIM; VIM | P08670; B0Y | VIME_HUMAN \ | Unmodified | 0 | 55281 | 0 | 0 | 3509300 | 531380 | 0 | 0 | 0 | 0 | 0 | 0 | 0 | 0 | 0 | 0 | 1151300 | 0 |
| LDLHVIPV\ | PCSK1; PCSK1 | P29120; D6F | NEC1_HUMAN | Unmodified | 0 | 0 | 0 | 0 | 0 | 0 | 0 | 0 | 1721000 | 317820 | 0 | 0 | 0 | 0 | 0 | 0 | 0 | 0 |
| LDLISKGEE | SLC29A1 | Q99808 | S29A1_HUMAN | Unmodified | 5613900 | 3706300 | 5069600 | 2646300 | 174120 | 0 | 0 | 0 | 1679500 | 393960 | 2650900 | 33176000 | 0 | 0 | 8052800 | 4668000 | 40155 | 32055 |
| LDLLEANS\ | NOSTRIN; NOS | Q8IVI9; A0A( | NOSTN_HUMA | Unmodified | 0 | 0 | 4092900 | 385480 | 0 | 0 | 0 | 0 | 0 | 0 | 0 | 0 | 0 | 0 | 3970100 | 1675900 | 0 | 0 |
| LDLMDEGT | DNM3; DNM2; I | Q9UQ16; P5 | DYN3_HUMAN | Oxidation (M) | 196760 | 156160 | 494450 | 232000 | 0 | 0 | 0 | 0 | 0 | 0 | 0 | 0 | 0 | 0 | 335210 | 0 | 0 | 0 |
| LDLQEINN\ | SERPINF1 | P36955 | PEDF_HUMAN | Oxidation (M) | 0 | 0 | 0 | 0 | 0 | 0 | 0 | 0 | 454800 | 27055 | 532470 | 361770 | 0 | 0 | 0 | 0 | 0 | 0 |
| LDLQQGQI | MOV10; MOV1( | Q9HCE1; Q5 | MOV10_HUMA | Unmodified | 7058900 | 8140800 | 2226800 | 1306100 | 0 | 0 | 0 | 0 | 0 | 0 | 0 | 189120 | 284500 | 62458 | 4080900 | 4710400 | 0 | 0 |

| | | | | | | | | | | | | | | | | | | | | | | |
|---|---|---|---|---|---|---|---|---|---|---|---|---|---|---|---|---|---|---|---|---|---|---|
| LDLRDDKD | RAC2; RAC1; R | P15153; P63 | RAC2_HUMAN | Unmodified | 7954200 | 8933800 | 14217000 | 6769800 | 2413000 | 589460 | 764620 | 0 | 1265300 | 214160 | 5847200 | 22308000 | 727720 | 510350 | 8750600 | 8707300 | 515520 | 1042800 |
| LDLRDDKD | RAC3; RAC3; R | P60763; J3K | RAC3_HUMAN | Unmodified | 0 | 794600 | 0 | 0 | 0 | 0 | 0 | 0 | 766800 | 0 | 0 | 992520 | 0 | 0 | 1473800 | 1540300 | 712840 | 2311100 |
| LDLSNNNL | FLRT1; FLRT1 | Q9NZU1; A0 | FLRT1_HUMAN | Unmodified | 0 | 0 | 4631500 | 2945000 | 0 | 0 | 0 | 0 | 0 | 0 | 0 | 0 | 0 | 0 | 8059100 | 10343000 | 0 | 0 |
| LDLTEKDYI | GGH; GGH; GG | Q92820; A0 | GGH_HUMAN | Unmodified | 0 | 0 | 0 | 0 | 0 | 0 | 0 | 0 | 411100 | 283080 | 0 | 0 | 0 | 0 | 0 | 0 | 0 | 0 |
| LDNGGYYI | YES1; FYN; FYN | P07947; P06 | YES_HUMAN T | Unmodified | 0 | 0 | 0 | 0 | 0 | 0 | 0 | 0 | 193290 | 0 | 0 | 477910 | 0 | 0 | 0 | 0 | 0 | 0 |
| LDNGGYYI | YES1; YES1 | P07947; J3Q | YES_HUMAN T | Unmodified | 0 | 0 | 0 | 0 | 155410 | 1054700 | 0 | 0 | 0 | 0 | 0 | 210320 | 103930 | 0 | 0 | 0 | 172300 | 0 |
| LDNGQVGI | PACSIN2 | Q9UNF0 | PACN2_HUMA | Unmodified | 2048900 | 1609200 | 82423 | 413810 | 0 | 0 | 29591 | 0 | 2301700 | 633220 | 0 | 0 | 0 | 0 | 1273800 | 1219300 | 0 | 0 |
| LDNIYSSDI | APOB | P04114 | APOB_HUMAN | Unmodified | 0 | 0 | 1227700 | 0 | 0 | 0 | 238000 | 0 | 0 | 104640 | 0 | 218730 | 0 | 0 | 0 | 0 | 0 | 0 |
| LDNLVAILD | TKT; TKT; TKT | P29401; A0 | TKT_HUMAN Tr | Unmodified | 0 | 0 | 0 | 0 | 0 | 0 | 0 | 0 | 0 | 0 | 152880 | 603200 | 0 | 0 | 0 | 0 | 0 | 0 |
| LDNMNVSI | HUWE1; HUWE | Q7Z6Z7; H0 | HUWE1_HUMA | Unmodified | 0 | 0 | 0 | 2925800 | 69542 | 0 | 0 | 0 | 626230 | 0 | 0 | 9926000 | 0 | 0 | 0 | 0 | 4984200 | 47480 |
| LDNVPHTP | NAP1L4; NAP1L | Q99733; C9J | NP1L4_HUMAN | Unmodified | 0 | 2146500 | 1213400 | 819090 | 132530 | 0 | 141880 | 66813 | 2280800 | 282170 | 1855800 | 25498000 | 0 | 200750 | 4335200 | 2198000 | 108830 | 491600 |
| LDPALQDK | GLG1; GLG1 | Q92896; H3I | GSLG1_HUMA | Unmodified | 83371 | 0 | 318880 | 0 | 189090 | 264710 | 0 | 0 | 7005100 | 49753 | 828760 | 1999600 | 0 | 0 | 236120 | 0 | 39937 | 0 |
| LDPGSEET | EEF1G | P26641 | EF1G_HUMAN | Unmodified | 0 | 0 | 304660 | 0 | 0 | 0 | 0 | 0 | 310500 | 0 | 378450 | 7238400 | 0 | 0 | 362190 | 305600 | 0 | 0 |
| LDPHLVLD | MYH9; MYH10 | P35579; P35 | MYH9_HUMAN | Unmodified | 0 | 670910 | 3919600 | 671470 | 0 | 0 | 0 | 42727 | 657360 | 179380 | 1703400 | 22852000 | 0 | 0 | 2753600 | 1886900 | 0 | 0 |
| LDPHNHVI | STIP1; STIP1; S | P31948; F5H | STIP1_HUMAN | Unmodified | 50357 | 0 | 0 | 179000 | 0 | 0 | 0 | 0 | 343840 | 0 | 179410 | 4262300 | 0 | 0 | 155120 | 188690 | 0 | 0 |
| LDPIQPSD | QSOX2 | Q6ZRP7 | QSOX2_HUMA | Unmodified | 0 | 0 | 25822 | 0 | 0 | 0 | 0 | 0 | 18370000 | 2218100 | 59741 | 527740 | 0 | 0 | 0 | 48094 | 0 | 0 |
| LDPSIFESL | RUVBL1; RUVBI | Q9Y265; E7E | RUVB1_HUMA | Unmodified | 996090 | 416890 | 1267500 | 896870 | 1959500 | 261330 | 0 | 87104 | 1470900 | 269660 | 2166800 | 15121000 | 1359700 | 0 | 0 | 0 | 0 | 565970 |
| LDQEVAEV | TSG101; TSG10 | Q99816; F5H | TS101_HUMAN | Unmodified | 2972300 | 2221200 | 4225100 | 0 | 1382600 | 0 | 0 | 0 | 746800 | 0 | 241320 | 2249800 | 351830 | 0 | 7926500 | 4607000 | 0 | 0 |
| LDQEVAEV | TSG101; TSG10 | Q99816; F5H | TS101_HUMAN | Unmodified | 12552000 | 13108000 | 172770 | 6755300 | 0 | 0 | 0 | 0 | 0 | 0 | 734620 | 693950 | 0 | 0 | 10872000 | 10917000 | 0 | 0 |
| LDQEVAEV | TSG101; TSG10 | Q99816; F5H | TS101_HUMAN | Unmodified | 476660 | 349610 | 0 | 418750 | 0 | 0 | 0 | 0 | 0 | 0 | 0 | 0 | 0 | 0 | 563520 | 688470 | 0 | 0 |
| LDQLIYIPLI | VCP; VCP; VCP; | P55072; C9J | TERA_HUMAN | Unmodified | 0 | 0 | 33023 | 42399 | 0 | 0 | 0 | 0 | 182520 | 0 | 202340 | 850780 | 0 | 0 | 0 | 0 | 0 | 0 |
| LDQRVPQI | COL6A3; COL6 | P12111; E7E | CO6A3_HUMA | Unmodified | 53771 | 141270 | 188290 | 237430 | 0 | 0 | 0 | 0 | 0 | 218030 | 0 | 0 | 0 | 51783 | 0 | 67692 | 0 | 0 |
| LDRDSLVH | PTGFRN | Q9P2B2 | FPRP_HUMAN | Unmodified | 31824000 | 25000000 | 75666000 | 54590000 | 216690 | 697570 | 1181300 | 223450 | 517270 | 0 | 0 | 92358000 | 8829700 | 3346700 | 85596000 | 1.02E+08 | 0 | 0 |
| LDRETEELI | EDF1 | O60869 | EDF1_HUMAN | Unmodified | 749100 | 630670 | 2234500 | 419710 | 122700 | 0 | 0 | 0 | 839050 | 0 | 327070 | 1092800 | 435610 | 0 | 2230900 | 1344700 | 705290 | 564440 |
| LDRLTSTAT | P4HA1 | P13674 | P4HA1_HUMA | Unmodified | 0 | 0 | 0 | 0 | 470290 | 0 | 0 | 0 | 27212000 | 3949600 | 838260 | 0 | 0 | 0 | 0 | 0 | 0 | 1444300 |
| LDRQHVQI | CEP55 | Q53EZ4 | CEP55_HUMA | Unmodified | 385360 | 26300 | 93414 | 2382000 | 15444000 | 2525000 | 0 | 0 | 0 | 0 | 311570 | 325750 | 0 | 0 | 1559300 | 0 | 556800 | 1048100 |
| LDRQSSSA | VPS37B; VPS37 | Q9H9H4; F5 | VP37B_HUMA | Unmodified | 0 | 0 | 99797 | 0 | 0 | 0 | 0 | 0 | 0 | 0 | 0 | 0 | 0 | 0 | 0 | 0 | 0 | 0 |
| LDSEQLPLI | CD2AP | Q9Y5K6 | CD2AP_HUMA | Unmodified | 756010 | 819090 | 3460900 | 1049000 | 0 | 0 | 0 | 0 | 0 | 0 | 0 | 0 | 1136000 | 127640 | 6324300 | 1613400 | 0 | 0 |
| LDSGDLLQ | NAPRT; NAPRT; | Q6XQN6; G5 | PNCB_HUMAN | Unmodified | 0 | 0 | 0 | 0 | 0 | 0 | 0 | 0 | 0 | 0 | 0 | 0 | 0 | 0 | 137750 | 0 | 0 | 0 |
| LDSGQLGL | PACSIN1; PACS | Q9BY11; F6 | PACN1_HUMA | Unmodified | 356920 | 381750 | 0 | 33101 | 0 | 0 | 0 | 0 | 0 | 0 | 0 | 0 | 0 | 0 | 119690 | 0 | 0 | 0 |
| LDSLQDIGI | NUCB2; NUCB | P80303; Q2L | NUCB2_HUMA | Oxidation (M) | 0 | 0 | 233020 | 123780 | 0 | 0 | 50596 | 17632 | 6562900 | 570760 | 743680 | 0 | 0 | 0 | 184020 | 58340 | 0 | 0 |
| LDSLQDIGI | NUCB2; NUCB | P80303; Q2L | NUCB2_HUMA | Unmodified | 0 | 0 | 0 | 0 | 0 | 0 | 0 | 0 | 12757000 | 0 | 0 | 0 | 0 | 0 | 0 | 0 | 0 | 0 |
| LDSSETTM | SAE1; SAE1; SA | Q9UBE0; B3 | SAE1_HUMAN | Unmodified | 0 | 0 | 0 | 0 | 0 | 0 | 0 | 0 | 0 | 190050 | 0 | 549340 | 0 | 0 | 0 | 0 | 0 | 0 |
| LDSSGGAV | SH3BP4 | Q9P0V3 | SH3B4_HUMA | Oxidation (M) | 0 | 0 | 0 | 0 | 0 | 0 | 0 | 0 | 931990 | 0 | 404600 | 0 | 0 | 0 | 0 | 0 | 0 | 0 |
| LDSTDFTG | TFRC; TFRC | P02786; G3 | TFR1_HUMAN | Unmodified | 0 | 0 | 1374600 | 381340 | 375360 | 0 | 0 | 0 | 9939000 | 1071500 | 589770 | 1080600 | 0 | 0 | 0 | 539700 | 0 | 0 |
| LDSTRLEGI | AHNAK2 | Q8IVF2 | AHNK2_HUMA | Unmodified | 0 | 0 | 0 | 0 | 1761800 | 0 | 526960 | 0 | 0 | 0 | 0 | 0 | 0 | 0 | 0 | 0 | 0 | 466460 |
| LDTIFFAK | UCHL5; UCHL5 | Q9Y5K5; Q5 | UCHL5_HUMA | Unmodified | 0 | 0 | 0 | 0 | 0 | 0 | 0 | 0 | 0 | 0 | 0 | 0 | 0 | 0 | 0 | 0 | 0 | 0 |
| LDTNSPSIF | NBEAL2; NBEA | Q6ZNJ1; H0 | NBEL2_HUMA | Unmodified | 0 | 0 | 0 | 0 | 0 | 0 | 0 | 0 | 0 | 0 | 174580 | 1239000 | 0 | 0 | 0 | 0 | 25527 | 0 |
| LDTPLYFSY | P3H1 | Q32P28 | P3H1_HUMAN | Unmodified | 0 | 0 | 11195 | 0 | 0 | 0 | 0 | 0 | 1964400 | 0 | 0 | 0 | 0 | 0 | 0 | 0 | 0 | 0 |
| LDTSQWPL | DKC1; DKC1; D | O60832; H7 | DKC1_HUMAN | Unmodified | 301410 | 0 | 0 | 0 | 0 | 38872 | 0 | 0 | 370700 | 0 | 307050 | 1907100 | 0 | 0 | 0 | 0 | 0 | 0 |
| LDTVGSDA | PTGFRN | Q9P2B2 | FPRP_HUMAN | Unmodified | 2986000 | 5072400 | 8566300 | 4948400 | 0 | 0 | 113300 | 0 | 0 | 0 | 109740 | 9706700 | 101250 | 107380 | 9498700 | 8914200 | 0 | 0 |
| LDTVGSDA | PTGFRN | Q9P2B2 | FPRP_HUMAN | Unmodified | 115360 | 71005 | 0 | 4798400 | 303670 | 421980 | 121540 | 21765 | 0 | 0 | 0 | 0 | 91569 | 0 | 824260 | 0 | 148150 | 0 |
| LDVAPTGL | PLA2G3 | Q9NZ20 | PA2G3_HUMA | Unmodified | 982670 | 502530 | 0 | 0 | 0 | 0 | 0 | 0 | 1305300 | 0 | 0 | 0 | 0 | 0 | 0 | 0 | 0 | 0 |
| LDVATDNF | DCUN1D1; DCI | Q96GG9; C9 | DCNL1_HUMA | Unmodified | 153890 | 0 | 0 | 485610 | 0 | 0 | 0 | 0 | 0 | 0 | 0 | 191470 | 0 | 0 | 0 | 66326 | 0 | 0 |
| LDVGNAEV | BCAP31; BCAP | P51572; C9J | BAP31_HUMA | Unmodified | 39946 | 0 | 316350 | 42681 | 652970 | 0 | 299730 | 119750 | 151170 | 0 | 303610 | 425480 | 0 | 163860 | 276160 | 140500 | 152430 | 112470 |
| LDVGNFSV | RPS8 | P62241 | RS8_HUMAN 4 | Unmodified | 382320 | 0 | 66736 | 0 | 0 | 0 | 68904 | 0 | 316890 | 0 | 0 | 1017700 | 0 | 0 | 0 | 0 | 0 | 0 |
| LDVGNFSV | RPS8 | P62241 | RS8_HUMAN 4 | Unmodified | 1978500 | 0 | 0 | 0 | 0 | 0 | 0 | 1150400 | 0 | 0 | 0 | 10485000 | 105060 | 0 | 1153900 | 679620 | 0 | 0 |
| LDVPQNLM | ALCAM | Q13740 | CD166_HUMA | Oxidation (M) | 224250 | 282940 | 0 | 0 | 0 | 0 | 0 | 0 | 0 | 0 | 0 | 0 | 0 | 0 | 138590 | 248070 | 0 | 0 |
| LDVSNVATI | LRBA; LRBA; LR | P50851; A0 | LRBA_HUMAN | Unmodified | 0 | 0 | 0 | 96212 | 0 | 0 | 0 | 0 | 282480 | 0 | 709900 | 1333800 | 0 | 0 | 0 | 156880 | 0 | 0 |
| LDVTSVED | CCT5; CCT5; C | P48643; B7Z | TCPE_HUMAN | Unmodified | 0 | 0 | 0 | 0 | 0 | 0 | 0 | 0 | 0 | 0 | 1066100 | 3579000 | 0 | 0 | 0 | 0 | 0 | 0 |
| LDYDEDAS | ST13; ST13P5; | P50502; Q8 | F10A1_HUMAN | Unmodified | 0 | 0 | 185960 | 0 | 0 | 0 | 0 | 0 | 303920 | 0 | 205300 | 792670 | 0 | 0 | 0 | 0 | 0 | 0 |
| LDYDEDAS | ST13; ST13P5; | P50502; Q8 | F10A1_HUMAN | Oxidation (M) | 0 | 0 | 310700 | 0 | 0 | 0 | 0 | 0 | 0 | 0 | 0 | 0 | 0 | 0 | 0 | 224450 | 0 | 0 |
| LDYFLLSHS | APEX1 | P27695 | APEX1_HUMAN | Unmodified | 0 | 0 | 30540 | 0 | 36777 | 0 | 0 | 0 | 0 | 102170 | 0 | 2371200 | 0 | 0 | 62421 | 0 | 25538 | 0 |
| LDYGQHV | EIF4A3; EIF4A3 | P38919; I3L | IF4A3_HUMAN | Unmodified | 0 | 0 | 0 | 0 | 0 | 0 | 0 | 0 | 0 | 0 | 0 | 1708700 | 0 | 0 | 0 | 194390 | 0 | 0 |
| LDYLDLYLI | AKR1B1; AKR1E | P15121; E9P | ALDR_HUMAN | Unmodified | 0 | 0 | 0 | 0 | 0 | 0 | 0 | 0 | 0 | 0 | 706550 | 1518500 | 0 | 0 | 0 | 0 | 0 | 0 |
| LDYLSSLK | SLC3A2; SLC3A | P08195; A0 | 4F2_HUMAN 4I | Unmodified | 1175100 | 865090 | 1162200 | 1846200 | 0 | 0 | 0 | 0 | 932610 | 335090 | 2865200 | 17954000 | 0 | 0 | 1574600 | 1094600 | 0 | 0 |
| LDYRALAA | SCRIB; SCRIB; | Q14160; A0 | SCRIB_HUMAN | Unmodified | 0 | 0 | 0 | 0 | 0 | 0 | 0 | 0 | 0 | 560290 | 0 | 0 | 641540 | 354830 | 0 | 0 | 1135900 | 2304100 |
| LEAAAQVR | NPEPPSL1; NPI | A6NEC2; P5 | PSAL_HUMAN | Unmodified | 0 | 970390 | 1291100 | 0 | 0 | 17684 | 0 | 0 | 0 | 35484 | 105510 | 2322700 | 291820 | 482530 | 166710 | 131850 | 0 | 0 |
| LEAARPGD | IGSF8 | Q969P0 | IGSF8_HUMAN | Unmodified | 3919600 | 3825600 | 6088300 | 0 | 181650 | 0 | 0 | 0 | 265770 | 60052 | 52875 | 2222600 | 0 | 66207 | 1184100 | 3880000 | 0 | 0 |
| LEAASSAL | USP24 | Q9UPU5 | UBP24_HUMA | Unmodified | 0 | 0 | 0 | 0 | 0 | 0 | 0 | 0 | 0 | 0 | 0 | 834240 | 0 | 0 | 0 | 81587 | 0 | 0 |
| LEAEGVPE | GLRX3 | O76003 | GLRX3_HUMA | Unmodified | 0 | 0 | 0 | 0 | 0 | 0 | 0 | 91443 | 0 | 0 | 0 | 873100 | 0 | 0 | 0 | 0 | 0 | 0 |
| LEAEIATYR | KRT18; KRT18 | P05783; F8V | K1C18_HUMAN | Unmodified | 0 | 493690 | 0 | 172120 | 936200 | 253380 | 0 | 741880 | 0 | 0 | 562870 | 614590 | 366460 | 0 | 0 | 0 | 501590 | 746130 |
| LEAELEK | MYO5C; RRBP1 | Q9NQX4; Q | MYO5C_HUMA | Unmodified | 0 | 0 | 0 | 0 | 0 | 0 | 0 | 0 | 2239000 | 737460 | 0 | 0 | 0 | 0 | 0 | 0 | 0 | 0 |
| LEAEMEAAI | PSPC1 | Q8WXF1 | PSPC1_HUMA | 3 Oxidation (M) | 0 | 0 | 0 | 0 | 0 | 1015600 | 923870 | 0 | 0 | 0 | 2437700 | 0 | 0 | 0 | 0 | 0 | 0 | 0 |
| LEAFNSRYI | UTRN | P46939 | UTRN_HUMAN | Unmodified | 0 | 0 | 1507300 | 150720 | 0 | 0 | 0 | 0 | 0 | 0 | 0 | 0 | 0 | 82864 | 1143800 | 1399700 | 0 | 0 |
| LEAGLNRV | GNG5 | P63218 | GBG5_HUMAN | Unmodified | 0 | 0 | 0 | 0 | 0 | 0 | 305130 | 93609 | 0 | 0 | 0 | 0 | 0 | 0 | 0 | 0 | 1014100 | 0 |
| LEAIEDDS | USP14; USP14 | P54578; A6 | UBP14_HUMA | Unmodified | 0 | 0 | 0 | 0 | 0 | 0 | 0 | 0 | 0 | 0 | 0 | 0 | 0 | 0 | 0 | 0 | 0 | 0 |
| LEAIEDDS | USP14; USP14 | P54578; A6 | UBP14_HUMA | Unmodified | 0 | 0 | 0 | 0 | 0 | 0 | 0 | 0 | 0 | 0 | 143780 | 1434900 | 0 | 0 | 0 | 0 | 0 | 0 |
| LEAIPNSGI | SLC44A1 | Q8WWI5 | CTL1_HUMAN | Oxidation (M) | 0 | 0 | 0 | 0 | 0 | 0 | 0 | 0 | 513350 | 142980 | 0 | 2028900 | 0 | 0 | 0 | 251220 | 0 | 0 |
| LEAIPNSGI | SLC44A1 | Q8WWI5 | CTL1_HUMAN | Unmodified | 0 | 429780 | 708040 | 962530 | 0 | 0 | 0 | 0 | 712350 | 27027 | 0 | 5748200 | 0 | 0 | 614840 | 472940 | 0 | 0 |
| LEAIPNSGI | SLC44A1 | Q8WWI5 | CTL1_HUMAN | Unmodified | 126700 | 0 | 749700 | 0 | 0 | 0 | 0 | 0 | 222570 | 0 | 0 | 1687900 | 0 | 0 | 238450 | 363510 | 0 | 63915 |
| LEAIPNSGI | SLC44A1 | Q8WWI5 | CTL1_HUMAN | Oxidation (M) | 143630 | 67695 | 527500 | 176510 | 0 | 0 | 0 | 0 | 0 | 0 | 0 | 434380 | 0 | 0 | 366200 | 247630 | 0 | 0 |
| LEALAAEFS | SUN2 | Q9UH99 | SUN2_HUMAN | Unmodified | 20772 | 0 | 0 | 0 | 0 | 0 | 0 | 0 | 948370 | 0 | 0 | 0 | 0 | 0 | 0 | 0 | 0 | 0 |
| LEALFPDLI | PRUNE1; PRUN | Q86TP1; A0 | PRUN1_HUMA | Unmodified | 0 | 0 | 0 | 0 | 0 | 0 | 0 | 0 | 0 | 0 | 180470 | 2057900 | 0 | 0 | 0 | 0 | 0 | 0 |
| LEALITQTR | SRP68 | Q9UHB9 | SRP68_HUMA | Unmodified | 0 | 0 | 0 | 0 | 0 | 0 | 46799 | 19464 | 0 | 0 | 118810 | 1539600 | 0 | 0 | 0 | 0 | 0 | 0 |
| LEALREAAI | TMCO3; TMCO | Q6UWJ1; A0 | TMCO3_HUMA | Unmodified | 0 | 0 | 0 | 126950 | 0 | 0 | 0 | 0 | 1332000 | 570990 | 0 | 0 | 0 | 0 | 0 | 0 | 0 | 0 |
| LEALSVKEE | RANBP1; RANB | P43487; F6 | RANG_HUMAN | Unmodified | 0 | 0 | 0 | 0 | 0 | 0 | 0 | 0 | 0 | 0 | 0 | 832310 | 0 | 0 | 0 | 0 | 0 | 0 |
| LEAQETLNI | GOLIM4; GOLI | O00461; F8 | GOLI4_HUMAN | Unmodified | 0 | 0 | 23490 | 0 | 153550 | 0 | 30334 | 35535 | 5938200 | 380920 | 0 | 449330 | 0 | 0 | 0 | 0 | 0 | 0 |
| LEAQLTENI | PFDN6; PFDN6 | O15212; A2 | PFD6_HUMAN | Unmodified | 0 | 0 | 162110 | 0 | 0 | 0 | 0 | 0 | 0 | 0 | 0 | 1461400 | 0 | 0 | 0 | 0 | 0 | 0 |
| LEASELFK | MYO1B; MYO1E | O43795; H7 | MYO1B_HUMA | Unmodified | 207890 | 214700 | 1363900 | 119980 | 2843000 | 609360 | 139080 | 0 | 1105700 | 333920 | 4176800 | 17492000 | 245240 | 0 | 1032200 | 615280 | 514730 | 591180 |
| LEASIER | GNG12 | Q9UBI6 | GBG12_HUMA | Unmodified | 1083800 | 866210 | 833900 | 530170 | 0 | 0 | 0 | 0 | 3038000 | 62548 | 37903 | 0 | 0 | 83680 | 2730600 | 2066400 | 0 | 0 |
| LEATAASS | ANLN; ANLN | Q9NQW6; C | ANLN_HUMAN | Unmodified | 58962 | 30566 | 100800 | 106140 | 197060 | 18079 | 18913 | 0 | 1812100 | 0 | 33663 | 108610 | 0 | 0 | 61269 | 42565 | 34278 | 17743 |
| LEAYQHLF | IQGAP1; IQGAP | P46940; A0 | IQGA1_HUMAN | Unmodified | 0 | 0 | 319950 | 0 | 1353300 | 373350 | 816340 | 0 | 545940 | 0 | 140030 | 330100 | 0 | 0 | 1225600 | 338490 | 488510 | 0 |
| LECAATGH | LRIG1 | Q96JA1 | LRIG1_HUMAN | Unmodified | 0 | 0 | 0 | 0 | 0 | 0 | 0 | 0 | 553250 | 56969 | 0 | 0 | 86136 | 0 | 0 | 0 | 0 | 0 |
| LECGGGP | ALDH1A1 | P00352 | AL1A1_HUMAN | Unmodified | 0 | 0 | 0 | 0 | 0 | 0 | 0 | 0 | 0 | 0 | 0 | 81882000 | 73009 | 0 | 0 | 0 | 0 | 0 |
| LECSEELGI | CLTCL1; CLTC; | P53675; Q0 | CLH2_HUMAN | Unmodified | 372310 | 191270 | 0 | 278770 | 0 | 0 | 0 | 0 | 0 | 0 | 0 | 1516600 | 0 | 0 | 0 | 331010 | 0 | 0 |
| LECVEPNC | RPL36A; RPL36 | P83881; Q9 | RL36A_HUMAN | Unmodified | 462700 | 265990 | 0 | 55750 | 0 | 0 | 0 | 0 | 0 | 0 | 284160 | 3984000 | 0 | 0 | 183900 | 227440 | 0 | 0 |
| LEDAADVY | NAA15; NAA15 | Q9BXJ9; A0 | NAA15_HUMA | Unmodified | 0 | 0 | 0 | 0 | 0 | 133780 | 0 | 0 | 732840 | 0 | 0 | 8053600 | 0 | 0 | 850650 | 0 | 0 | 0 |
| LEDCTPQR | TSPAN6; TSPAN | O43657; A0 | TSN6_HUMAN | Unmodified | 300550 | 0 | 1187100 | 130040 | 0 | 0 | 0 | 0 | 0 | 0 | 0 | 4160700 | 0 | 0 | 1225400 | 571250 | 0 | 0 |
| LEDEIDFLA | NPDC1; NPDC | Q9NQX5; Q | NPDC1_HUMA | Unmodified | 0 | 0 | 0 | 81683 | 0 | 0 | 0 | 0 | 677510 | 185550 | 0 | 0 | 0 | 0 | 0 | 0 | 0 | 0 |
| LEDEIDFLA | NPDC1; NPDC | Q9NQX5; Q | NPDC1_HUMA | Unmodified | 429420 | 695620 | 1487200 | 2127500 | 0 | 0 | 0 | 0 | 9189400 | 0 | 0 | 0 | 0 | 0 | 3268600 | 902120 | 32995 | 0 |
| LEDGTEFD | FKBP2; FKBP2; | P26885; A0 | FKBP2_HUMAN | Unmodified | 0 | 0 | 0 | 0 | 0 | 0 | 0 | 0 | 803980 | 0 | 0 | 0 | 0 | 0 | 0 | 0 | 0 | 0 |
| LEDLATLIQ | MYO1B; MYO1E | O43795; E9 | MYO1B_HUMA | Unmodified | 0 | 0 | 0 | 0 | 0 | 0 | 0 | 0 | 0 | 0 | 0 | 6049600 | 0 | 0 | 0 | 0 | 0 | 0 |
| LEDLSESIV | TMED10 | P49755 | TMEDA_HUMA | Unmodified | 0 | 0 | 0 | 0 | 0 | 0 | 0 | 0 | 1922000 | 268560 | 0 | 0 | 0 | 0 | 0 | 0 | 0 | 0 |
| LEDLSESIV | TMED10 | P49755 | TMEDA_HUMA | Oxidation (M) | 0 | 0 | 0 | 0 | 78607 | 0 | 0 | 0 | 2326800 | 640190 | 0 | 0 | 0 | 0 | 0 | 28202 | 0 | 0 |
| LEDLSESIV | TMED10 | P49755 | TMEDA_HUMA | Unmodified | 0 | 0 | 0 | 0 | 0 | 0 | 53824 | 0 | 257520 | 0 | 0 | 0 | 0 | 222040 | 0 | 0 | 0 | 12418 |
| LEDPHVDII | TMEM43 | Q9BTV4 | TMM43_HUMA | Unmodified | 0 | 0 | 0 | 0 | 0 | 0 | 0 | 0 | 2828000 | 593530 | 125990 | 343480 | 0 | 0 | 0 | 0 | 0 | 0 |
| LEDQNKEN | MYO5C | Q9NQX4 | MYO5C_HUMA | Unmodified | 0 | 0 | 0 | 0 | 471500 | 625440 | 392810 | 0 | 0 | 174530 | 0 | 0 | 1005900 | 0 | 0 | 0 | 443110 | 0 |
| LEDRDNWI | DSG2 | Q14126 | DSG2_HUMAN | Unmodified | 0 | 0 | 261020 | 0 | 0 | 0 | 0 | 23931 | 2337000 | 0 | 0 | 858040 | 0 | 0 | 315200 | 148120 | 0 | 0 |
| LEDSSNQV | UTRN | P46939 | UTRN_HUMAN | Unmodified | 0 | 0 | 0 | 0 | 0 | 0 | 0 | 0 | 0 | 0 | 0 | 96091 | 0 | 0 | 353570 | 270590 | 0 | 0 |
| LEDSTRET | DSP | P15924 | DESP_HUMAN | Unmodified | 263270 | 0 | 106460 | 0 | 0 | 0 | 0 | 70825 | 0 | 0 | 0 | 178220 | 0 | 0 | 64759 | 62131 | 0 | 0 |
| LEDTENWL | HSPA4; HSPA4I | P34932; O9 | HSP74_HUMA | Unmodified | 0 | 0 | 0 | 0 | 0 | 0 | 0 | 0 | 0 | 0 | 0 | 4830700 | 0 | 0 | 0 | 0 | 0 | 0 |
| LEDTVKHH | MYO1C; MYO1 | O00159; F5H | MYO1C_HUMA | Unmodified | 0 | 0 | 0 | 60624 | 1773000 | 226980 | 0 | 0 | 0 | 0 | 0 | 0 | 4662600 | 993330 | 0 | 0 | 368670 | 176970 |
| LEDTYFDRI | HMGCS1 | Q01581 | HMCS1_HUMA | Unmodified | 0 | 0 | 0 | 0 | 0 | 0 | 0 | 0 | 0 | 0 | 236250 | 1499000 | 0 | 0 | 0 | 0 | 0 | 0 |
| LEDVLPLAF | HP1BP3; HP1BI | Q5SSJ5; B0 | HP1B3_HUMA | Unmodified | 671000 | 376700 | 0 | 0 | 0 | 0 | 0 | 98766 | 0 | 0 | 0 | 0 | 0 | 0 | 0 | 0 | 0 | 0 |
| LEDYDKAE | UNC45A; UNC | Q9H3U1; A0 | UN45A_HUMA | Unmodified | 0 | 0 | 86858 | 90338 | 0 | 0 | 0 | 0 | 0 | 0 | 152450 | 583620 | 14966 | 0 | 209940 | 193520 | 18755 | 0 |
| LEEDAEMK | SSB | P05455 | LA_HUMAN Lu | Unmodified | 0 | 0 | 0 | 0 | 0 | 0 | 0 | 0 | 0 | 0 | 180300 | 0 | 0 | 0 | 0 | 0 | 0 | 0 |
| LEEDISSSM | PCBP2; PCBP2; | Q15366; F8 | PCBP2_HUMA | Unmodified | 0 | 519510 | 1134000 | 0 | 0 | 0 | 0 | 0 | 0 | 0 | 0 | 804430 | 0 | 0 | 483600 | 232670 | 0 | 0 |
| LEEDISSSM | PCBP2; PCBP2; | Q15366; F8 | PCBP2_HUMA | Oxidation (M) | 0 | 332420 | 1296800 | 167200 | 0 | 0 | 0 | 0 | 0 | 0 | 0 | 0 | 0 | 0 | 291630 | 532020 | 0 | 0 |

| | | | | | | | | | | | | | | | | | | | | | | |
|---|---|---|---|---|---|---|---|---|---|---|---|---|---|---|---|---|---|---|---|---|---|---|
| LEEENLTF | LUZP1; LUZP1 | Q86V48; E5F | LUZP1_HUMAN | Unmodified | 0 | 0 | 0 | 0 | 0 | 0 | 0 | 0 | 2004600 | 0 | 0 | 0 | 0 | 0 | 0 | 0 | 0 | 0 |
| LEEEGEQF | SCP2; SCP2 | P22307; H0 | SCP2_HUMAN | Unmodified | 0 | 0 | 0 | 0 | 0 | 0 | 0 | 0 | 1050100 | 358700 | 0 | 0 | 0 | 0 | 0 | 0 | 0 | 0 |
| LEEEGEQF | SCP2; SCP2 | P22307; H0 | SCP2_HUMAN | Unmodified | 0 | 0 | 0 | 0 | 0 | 0 | 0 | 0 | 1474100 | 691460 | 0 | 0 | 0 | 0 | 0 | 0 | 0 | 0 |
| LEEESAQLI | RABGAP1 | Q9Y3P9 | RBGP1_HUMAI | Unmodified | 0 | 0 | 0 | 0 | 0 | 0 | 0 | 0 | 0 | 0 | 0 | 0 | 0 | 0 | 0 | 0 | 0 | 0 |
| LEEGLVNN | WDR36; WDR3 | Q8NI36; A0A | WDR36_HUMA | Unmodified | 0 | 0 | 0 | 0 | 0 | 0 | 0 | 0 | 0 | 0 | 0 | 167900 | 0 | 0 | 0 | 0 | 0 | 0 |
| LEEGPPVTI | PDHA1; PDHA1 | P08559; Q5J | ODPA_HUMAN | Unmodified | 0 | 0 | 0 | 0 | 0 | 0 | 0 | 0 | 2929700 | 239380 | 0 | 0 | 0 | 0 | 0 | 0 | 0 | 0 |
| LEEMVTR | TSG101; TSG1 | Q99816; F5I | TS101_HUMAN | Unmodified | 1113700 | 729680 | 1988000 | 460770 | 418720 | 953990 | 79319 | 223950 | 356470 | 128930 | 167960 | 255080 | 0 | 0 | 3011700 | 1980500 | 77652 | 0 |
| LEEMVTR | TSG101; TSG1 | Q99816; F5I | TS101_HUMAN | Oxidation (M) | 0 | 0 | 577750 | 161730 | 0 | 118120 | 107020 | 0 | 0 | 0 | 45240 | 0 | 0 | 0 | 179320 | 1110400 | 0 | 0 |
| LEEMVTRLI | TSG101; TSG1 | Q99816; F5I | TS101_HUMAN | Oxidation (M) | 5244500 | 2799400 | 28910000 | 2482100 | 0 | 0 | 0 | 0 | 654920 | 0 | 0 | 584900 | 0 | 0 | 31388000 | 12022000 | 0 | 0 |
| LEEMVTRLI | TSG101; TSG1 | Q99816; F5I | TS101_HUMAN | Unmodified | 2180900 | 1775300 | 5779600 | 1205600 | 0 | 0 | 0 | 0 | 2098800 | 0 | 169700 | 608950 | 0 | 0 | 10005000 | 2637200 | 0 | 0 |
| LEEMVTRLI | TSG101; TSG1 | Q99816; F5I | TS101_HUMAN | Oxidation (M) | 1286400 | 11269000 | 0 | 3388500 | 0 | 0 | 285720 | 0 | 207060 | 0 | 0 | 0 | 2383200 | 0 | 12423000 | 707310 | 0 | 0 |
| LEEMVTRLI | TSG101; TSG1 | Q99816; F5I | TS101_HUMAN | Unmodified | 49688 | 1160400 | 0 | 1143000 | 0 | 0 | 0 | 0 | 0 | 0 | 0 | 0 | 285200 | 0 | 1592300 | 0 | 0 | 0 |
| LEENLQETI | SLC35F2 | Q8IXU6 | S35F2_HUMAN | Unmodified | 0 | 0 | 743910 | 0 | 0 | 0 | 0 | 0 | 0 | 0 | 0 | 676360 | 0 | 0 | 287380 | 0 | 0 | 0 |
| LEEPSLR | DNM1L | O00429 | DNM1L_HUMA | Unmodified | 0 | 0 | 0 | 0 | 0 | 0 | 0 | 0 | 0 | 0 | 0 | 726130 | 0 | 0 | 0 | 0 | 0 | 0 |
| LEEQLGR | CORO1B; FER1 | Q9BR76; A9 | COR1B_HUMA | Unmodified | 0 | 0 | 0 | 0 | 0 | 0 | 0 | 0 | 0 | 0 | 0 | 1439600 | 159140 | 228390 | 0 | 0 | 0 | 0 |
| LEEREVLKS | MTHFD2; MTHF | P13995; A0A | MTDC_HUMAN | Unmodified | 1765500 | 1272400 | 0 | 518370 | 42817 | 0 | 436380 | 0 | 0 | 105720 | 0 | 0 | 4529500 | 1275700 | 782210 | 1136600 | 0 | 0 |
| LEESTNYN | CCDC88A; CCD | Q3V6T2; A0A | GRDN_HUMAN | Unmodified | 0 | 0 | 0 | 0 | 0 | 0 | 0 | 0 | 100790 | 0 | 0 | 375040 | 0 | 0 | 0 | 0 | 0 | 0 |
| LEESVALR | PPP3CA; PPP3C | Q08209; E9P | PP2BA_HUMAN | Unmodified | 352890 | 367500 | 0 | 37133 | 193800 | 0 | 0 | 0 | 54293 | 0 | 0 | 494020 | 0 | 0 | 0 | 0 | 228980 | 147180 |
| LEETQALLF | DCTN1; DCTN1 | Q14203; A0A | DCTN1_HUMAI | Unmodified | 0 | 0 | 0 | 0 | 0 | 0 | 0 | 0 | 0 | 0 | 0 | 980080 | 0 | 0 | 0 | 0 | 0 | 0 |
| LEETVQAK | CORO1A; COR | P31146; H3E | COR1A_HUMA | Unmodified | 0 | 0 | 64760 | 0 | 0 | 60952 | 67823 | 21842 | 960300 | 253830 | 366300 | 273910 | 0 | 0 | 46708 | 64694 | 819660 | 0 |
| LEEWLGR | MARS1 | P56192 | SYMC_HUMAN | Unmodified | 0 | 0 | 0 | 0 | 0 | 0 | 0 | 0 | 0 | 0 | 0 | 1328000 | 0 | 0 | 0 | 0 | 0 | 0 |
| LEFENLVEE | GPC6 | Q9Y625 | GPC6_HUMAN | Unmodified | 522240 | 1088200 | 251480 | 0 | 0 | 0 | 0 | 0 | 0 | 0 | 0 | 0 | 0 | 0 | 0 | 113430 | 0 | 0 |
| LEGALGAD | MYOF; MYOF | Q9NZM1; H0 | MYOF_HUMAN | Unmodified | 0 | 0 | 0 | 0 | 1562900 | 231930 | 0 | 0 | 217620 | 0 | 0 | 0 | 0 | 0 | 0 | 0 | 0 | 231950 |
| LEGALGAD | MYOF; MYOF | Q9NZM1; H0 | MYOF_HUMAN | Unmodified | 0 | 46995 | 154630 | 0 | 1062800 | 0 | 40891 | 0 | 0 | 0 | 0 | 286790 | 0 | 0 | 0 | 0 | 0 | 0 |
| LEGDLSLAI | AHNAK2 | Q8IVF2 | AHNK2_HUMA | Unmodified | 0 | 50490 | 0 | 0 | 1016200 | 82859 | 175500 | 0 | 0 | 0 | 0 | 693170 | 0 | 0 | 0 | 0 | 122560 | 344730 |
| LEGDNVNF | USP9X | Q93008 | USP9X_HUMAI | Unmodified | 0 | 0 | 0 | 0 | 0 | 0 | 0 | 0 | 0 | 0 | 0 | 1602400 | 0 | 0 | 0 | 121480 | 0 | 0 |
| LEGDSTDL | MYH9 | P35579 | MYH9_HUMAN | Unmodified | 0 | 0 | 0 | 166650 | 0 | 0 | 0 | 0 | 0 | 0 | 0 | 0 | 0 | 0 | 745330 | 0 | 0 | 0 |
| LEGDTLIIPI | HSPG2 | P98160 | PGBM_HUMAN | Unmodified | 0 | 0 | 1907400 | 0 | 0 | 0 | 0 | 0 | 131570 | 0 | 0 | 0 | 0 | 0 | 0 | 891180 | 0 | 0 |
| LEGEACGV | IGFBP2 | P18065 | IBP2_HUMAN I | Unmodified | 0 | 0 | 0 | 0 | 0 | 0 | 0 | 0 | 29863000 | 2069600 | 0 | 0 | 0 | 0 | 0 | 0 | 0 | 0 |
| LEGEACGV | IGFBP2 | P18065 | IBP2_HUMAN I | Unmodified | 93398 | 91070 | 253730 | 0 | 0 | 0 | 0 | 0 | 99491 | 758940 | 0 | 0 | 0 | 0 | 0 | 0 | 0 | 0 |
| LEGELQEAI | ATP6V0A4; ATP | Q9HBG4; A0 | VPP4_HUMAN | Unmodified | 0 | 0 | 0 | 0 | 431900 | 0 | 0 | 0 | 0 | 0 | 0 | 685850 | 0 | 0 | 0 | 0 | 0 | 0 |
| LEGELVPCI | SORL1 | Q92673 | SORL_HUMAN | Unmodified | 0 | 0 | 0 | 0 | 0 | 0 | 0 | 0 | 1076200 | 0 | 0 | 0 | 0 | 0 | 0 | 0 | 0 | 0 |
| LEGFHTQIS | PSME1; PSME1 | Q06323; H0 | PSME1_HUMAI | Unmodified | 0 | 70282 | 0 | 0 | 0 | 0 | 0 | 0 | 0 | 0 | 0 | 316310 | 0 | 0 | 0 | 0 | 0 | 0 |
| LEGFYGER | DOCK7; DOCK | Q96N67; A0 | DOCK7_HUMA | Unmodified | 0 | 0 | 0 | 1597500 | 3186700 | 719220 | 815990 | 0 | 0 | 0 | 0 | 0 | 0 | 10290000 | 119930 | 0 | 0 | 0 |
| LEGHGLPA | EHD2 | Q9NZN4 | EHD2_HUMAN | Unmodified | 67984 | 56036 | 0 | 0 | 3561300 | 0 | 97076 | 0 | 422860 | 0 | 0 | 0 | 0 | 162630 | 0 | 0 | 67888 | 570640 |
| LEGPGEQE | SLC29A1 | Q99808 | S29A1_HUMAN | Unmodified | 1244400 | 685860 | 2858300 | 274730 | 0 | 0 | 0 | 0 | 0 | 0 | 335560 | 15491000 | 0 | 0 | 2816100 | 1792000 | 0 | 0 |
| LEGPGEQE | SLC29A1 | Q99808 | S29A1_HUMAN | Unmodified | 1580800 | 0 | 3783400 | 415850 | 0 | 0 | 352660 | 0 | 0 | 1449600 | 0 | 7999600 | 0 | 0 | 3175700 | 2194900 | 214420 | 0 |
| LEGPGEQE | SLC29A1 | Q99808 | S29A1_HUMAN | Unmodified | 2671200 | 2869000 | 1687600 | 1927000 | 0 | 0 | 0 | 0 | 1087900 | 1154100 | 0 | 16534000 | 0 | 0 | 5432700 | 3187500 | 0 | 0 |
| LEGQEEEE | CHGA; CHGA | P10645; G5E | CMGA_HUMAN | Unmodified | 0 | 0 | 0 | 0 | 0 | 0 | 0 | 0 | 6679700 | 2807700 | 0 | 0 | 0 | 0 | 0 | 0 | 0 | 0 |
| LEGQEEEE | CHGA; CHGA | P10645; G5E | CMGA_HUMAN | Unmodified | 0 | 0 | 0 | 256910 | 0 | 0 | 0 | 91065 | 30526000 | 17961000 | 0 | 0 | 0 | 0 | 0 | 0 | 0 | 0 |
| LEGQEEEE | CHGA; CHGA | P10645; G5E | CMGA_HUMAN | Oxidation (M) | 0 | 0 | 0 | 0 | 0 | 0 | 0 | 24732 | 2769000 | 1416800 | 0 | 0 | 0 | 0 | 0 | 0 | 0 | 0 |
| LEGQGDVF | SUGT1 | Q9Y2Z0 | SGT1_HUMAN | Unmodified | 0 | 0 | 0 | 0 | 0 | 390760 | 0 | 0 | 0 | 61190 | 305750 | 942690 | 0 | 0 | 0 | 0 | 0 | 0 |
| LEGTNVQE | SUCLG2; SUCL | Q96I99; H0 | SUCB2_HUMAI | Unmodified | 0 | 0 | 0 | 0 | 0 | 0 | 0 | 0 | 623910 | 0 | 0 | 0 | 0 | 0 | 0 | 0 | 0 | 0 |
| LEGVLAEV | IQGAP1; IQGAP | P46940; A0A | IQGA1_HUMAN | Unmodified | 1609400 | 1643600 | 6853000 | 306380 | 2174600 | 1047200 | 213660 | 0 | 2637500 | 498290 | 0 | 5366800 | 171820 | 0 | 2470700 | 1186500 | 0 | 0 |
| LEHEYIQNI | MAPRE1 | Q15691 | MARE1_HUMAN | Unmodified | 263930 | 0 | 633610 | 0 | 0 | 0 | 0 | 0 | 0 | 0 | 736740 | 7255700 | 0 | 0 | 564200 | 0 | 0 | 0 |
| LEHLQSAE | STON1; STON1 | Q9Y6Q2; Q5 | STON1_HUMAI | Unmodified | 0 | 0 | 0 | 0 | 205090 | 60275 | 0 | 0 | 0 | 0 | 156300 | 743460 | 0 | 0 | 0 | 0 | 0 | 302470 |
| LEHPVLHV | SUMF2; SUMF2 | Q8NBJ7; C9 | SUMF2_HUMAN | Unmodified | 0 | 0 | 266580 | 0 | 755600 | 51406 | 0 | 0 | 17006000 | 807130 | 0 | 0 | 0 | 0 | 0 | 0 | 0 | 638560 |
| LEIEPEWAY | FKBP3 | Q00688 | FKBP3_HUMAN | Unmodified | 0 | 0 | 0 | 0 | 0 | 0 | 0 | 0 | 0 | 0 | 422750 | 0 | 0 | 0 | 0 | 0 | 0 | 0 |
| LEIERLEK | KIF23; KIF23; KI | Q02241; A0 | KIF23_HUMAN | Unmodified | 0 | 0 | 0 | 569250 | 1107500 | 0 | 0 | 57324 | 0 | 0 | 0 | 329480 | 0 | 3763600 | 351450 | 107990 | 682010 | 0 |
| LEIPYQAEV | TMEM131 | Q92545 | TM131_HUMAN | Unmodified | 0 | 0 | 0 | 0 | 0 | 0 | 0 | 130220 | 0 | 0 | 0 | 0 | 0 | 0 | 0 | 0 | 87170 | 0 |
| LEKEDENY | NOSTRIN; NOS | Q8IVI9; A0A | NOSTN_HUMA | Unmodified | 1064500 | 632700 | 2426900 | 1466600 | 0 | 0 | 0 | 0 | 0 | 0 | 13153 | 0 | 0 | 0 | 5644600 | 2635500 | 0 | 0 |
| LEKPAKYDI | GAPDH; GAPDI | P04406; E7E | G3P_HUMAN G | Unmodified | 498900 | 272160 | 75709 | 321720 | 0 | 0 | 0 | 22101 | 0 | 92555 | 1747400 | 5353200 | 608510 | 0 | 394450 | 626750 | 0 | 258640 |
| LEKTIDDLE | TPM3; TPM3 | J3KN67; A0A | J3KN67_HUMA | Unmodified | 0 | 0 | 0 | 0 | 296470 | 0 | 0 | 0 | 0 | 0 | 0 | 0 | 37239 | 0 | 0 | 0 | 292060 | 0 |
| LELAQYR | ATP5F1A | P25705 | ATPA_HUMAN , | Unmodified | 0 | 0 | 0 | 0 | 0 | 0 | 0 | 0 | 10311000 | 0 | 0 | 0 | 0 | 0 | 0 | 0 | 0 | 0 |
| LELASNHFI | TPBG | Q13641 | TPBG_HUMAN | Unmodified | 0 | 0 | 0 | 0 | 0 | 48791 | 0 | 0 | 150500 | 0 | 0 | 4786800 | 0 | 0 | 0 | 0 | 0 | 0 |
| LELCDER | UQCRH | P07919 | QCR6_HUMAN | Unmodified | 0 | 0 | 0 | 0 | 0 | 0 | 0 | 0 | 4201900 | 0 | 0 | 0 | 0 | 0 | 0 | 0 | 0 | 0 |
| LELCDERV | UQCRH | P07919 | QCR6_HUMAN | Unmodified | 0 | 0 | 0 | 0 | 0 | 0 | 19188 | 0 | 4733500 | 0 | 107870 | 0 | 0 | 0 | 0 | 0 | 0 | 0 |
| LELEITDSK | USO1 | O60763 | USO1_HUMAN | Unmodified | 0 | 0 | 0 | 0 | 0 | 0 | 0 | 0 | 0 | 0 | 0 | 0 | 0 | 0 | 0 | 0 | 0 | 0 |
| LELFPVELE | PPP1R9A; PPP1 | Q9ULJ8; Q9 | NEB1_HUMAN | Unmodified | 0 | 0 | 0 | 0 | 0 | 0 | 0 | 0 | 0 | 0 | 0 | 226520 | 0 | 0 | 0 | 0 | 0 | 0 |
| LELGIGPG | AGRN; AGRN; A | O00468; A0A | AGRIN_HUMAN | Unmodified | 754740 | 0 | 0 | 0 | 0 | 0 | 0 | 0 | 800540 | 0 | 0 | 0 | 0 | 0 | 1057300 | 0 | 0 | 0 |
| LELGNDWI | PARP4 | Q9UKK3 | PARP4_HUMAN | Unmodified | 0 | 0 | 0 | 0 | 0 | 0 | 0 | 0 | 0 | 0 | 0 | 2245200 | 0 | 0 | 0 | 0 | 0 | 0 |
| LELMEKDL | SSRP1 | Q08945 | SSRP1_HUMAN | Oxidation (M) | 0 | 0 | 0 | 1476900 | 2066400 | 170420 | 375150 | 218130 | 1055400 | 545790 | 31154 | 294910 | 0 | 1373200 | 0 | 0 | 0 | 4968000 |
| LELNITTIKE | N/A | N/A | N/A | Unmodified | 0 | 0 | 0 | 0 | 1540400 | 0 | 0 | 0 | 902140 | 0 | 0 | 777400 | 0 | 0 | 0 | 0 | 2018700 | 0 |
| LELQGPR | NCL; NCL; NCL | P19338; C9J | NUCL_HUMAN | Unmodified | 0 | 0 | 0 | 0 | 119960 | 141790 | 222530 | 0 | 290710 | 0 | 340050 | 10353000 | 0 | 0 | 0 | 82780 | 172830 | 0 |
| LELVHHGP | CTSC; CTSC; C | P53634; A0A | CATC_HUMAN | Unmodified | 0 | 0 | 0 | 0 | 0 | 0 | 0 | 0 | 0 | 0 | 760140 | 0 | 0 | 0 | 0 | 0 | 0 | 0 |
| LELVHHGP | CTSC; CTSC; C | P53634; A0A | CATC_HUMAN | Oxidation (M) | 0 | 0 | 0 | 0 | 0 | 0 | 0 | 0 | 0 | 0 | 2496100 | 974790 | 0 | 0 | 0 | 0 | 0 | 0 |
| LELYSNR | TM9SF4; TM9SF | Q92544; A0 | TM9S4_HUMAI | Unmodified | 0 | 0 | 0 | 0 | 0 | 0 | 0 | 0 | 662810 | 0 | 0 | 0 | 0 | 0 | 0 | 0 | 0 | 0 |
| LELYSNRD | TM9SF4; TM9SF | Q92544; A0 | TM9S4_HUMAI | Unmodified | 0 | 0 | 0 | 0 | 0 | 0 | 0 | 0 | 634020 | 0 | 0 | 0 | 0 | 0 | 0 | 0 | 0 | 0 |
| LEMNLGLQ | SPTBN1; SPTBN | Q01082; A0 | SPTB2_HUMAN | Unmodified | 0 | 0 | 0 | 0 | 0 | 0 | 0 | 0 | 0 | 0 | 0 | 572480 | 0 | 0 | 0 | 0 | 0 | 0 |
| LENDQIESI | MARS1; MARS1 | P56192; H0 | SYMC_HUMAN | Unmodified | 0 | 0 | 0 | 0 | 0 | 0 | 0 | 0 | 108170 | 0 | 0 | 5684400 | 0 | 0 | 0 | 0 | 0 | 0 |
| LENEHLNN | XPR1 | Q9UBH6 | XPR1_HUMAN | Unmodified | 0 | 0 | 0 | 0 | 0 | 0 | 0 | 0 | 0 | 0 | 0 | 255080 | 0 | 0 | 0 | 0 | 0 | 0 |
| LENFCRDF | FARP1; FARP1; | Q9Y4F1; C9 | FARP1_HUMAN | Unmodified | 0 | 0 | 0 | 0 | 0 | 0 | 0 | 0 | 468310 | 0 | 0 | 0 | 0 | 0 | 0 | 0 | 0 | 0 |
| LENGEIETI | APMAP; APMAP | Q9HDC9; H0 | APMAP_HUMAI | Unmodified | 0 | 0 | 0 | 411430 | 0 | 0 | 0 | 0 | 29871000 | 1157900 | 681530 | 0 | 0 | 0 | 0 | 32133 | 0 | 0 |
| LENGELEH | PAFAH1B3; PAF | Q15102; M0 | PA1B3_HUMAN | Unmodified | 122850 | 0 | 0 | 24973 | 277470 | 0 | 0 | 33221 | 0 | 0 | 6497900 | 12914000 | 0 | 0 | 131180 | 170550 | 0 | 48323 |
| LENIIVTDVI | VPS13C | Q709C8 | VP13C_HUMAN | Unmodified | 0 | 0 | 0 | 0 | 0 | 0 | 0 | 0 | 0 | 0 | 0 | 10389000 | 0 | 0 | 0 | 0 | 0 | 0 |
| LENLPDTEI | SLK | Q9H2G2 | SLK_HUMAN S | Unmodified | 0 | 0 | 308200 | 0 | 0 | 94970 | 0 | 0 | 0 | 0 | 0 | 1760600 | 0 | 0 | 447110 | 209710 | 0 | 0 |
| LENNDNKF | YTHDF1; YTHD | Q9BYJ9; Q7 | YTHD1_HUMAN | Unmodified | 314760 | 257800 | 190270 | 189370 | 0 | 0 | 0 | 0 | 163260 | 17358 | 0 | 380600 | 0 | 0 | 120860 | 167290 | 0 | 0 |
| LENNENKP | YTHDF2 | Q9Y5A9 | YTHD2_HUMAI | Unmodified | 0 | 449750 | 563400 | 0 | 0 | 0 | 0 | 0 | 0 | 0 | 0 | 0 | 0 | 0 | 413390 | 359860 | 0 | 0 |
| LENSIIPVH | EXOC7; EXOC7 | Q9UPT5; B5 | EXOC7_HUMAI | Unmodified | 0 | 0 | 0 | 0 | 1325400 | 0 | 0 | 0 | 0 | 133640 | 0 | 0 | 0 | 0 | 0 | 0 | 0 | 0 |
| LENTEDIEE | LMBRD1; LMBR | Q9NUN5; A0 | LMBD1_HUMAI | Unmodified | 0 | 0 | 0 | 0 | 0 | 0 | 0 | 0 | 668890 | 491600 | 0 | 110900 | 0 | 0 | 34421 | 0 | 0 | 0 |
| LENTLQDII | L1RE1 | Q9UN81 | LORF1_HUMAN | Unmodified | 243580 | 248070 | 67185 | 29512 | 0 | 0 | 0 | 0 | 0 | 0 | 0 | 0 | 0 | 0 | 0 | 0 | 0 | 0 |
| LENYPIPEP | SORD; SORD; S | Q00796; H0 | DHSO_HUMAN | Unmodified | 0 | 62659 | 1230700 | 1951800 | 403560 | 118530 | 0 | 0 | 0 | 84545 | 3727000 | 10052000 | 239670 | 0 | 2606800 | 189120 | 0 | 151480 |
| LEPDILLR | AMPD2; AMPD2 | Q01433; H0 | AMPD2_HUMAN | Unmodified | 0 | 0 | 39641 | 0 | 0 | 0 | 0 | 26152 | 0 | 0 | 69391 | 285030 | 0 | 56569 | 0 | 0 | 0 | 0 |
| LEPHEGLLI | SLC3A2; SLC3A | P08195; A0 | 4F2_HUMAN 4I | Unmodified | 0 | 0 | 420450 | 0 | 0 | 0 | 0 | 0 | 0 | 0 | 0 | 4125400 | 0 | 0 | 239920 | 223240 | 0 | 0 |
| LEPHEGLLI | SLC3A2; SLC3A | P08195; A0 | 4F2_HUMAN 4I | Unmodified | 9977200 | 5362700 | 20174000 | 5527000 | 9419400 | 2281200 | 0 | 0 | 13554000 | 1803600 | 22974000 | 75533000 | 985710 | 0 | 18466000 | 9556900 | 846210 | 0 |
| LEPKPQPP | SEPTIN9; SEPTI | Q9UHD8; K7 | SEPT9_HUMAN | Unmodified | 0 | 0 | 0 | 2402800 | 0 | 0 | 21662 | 0 | 1303600 | 0 | 3776000 | 3393100 | 0 | 0 | 0 | 602230 | 671520 | 1303900 |
| LEPLGSYR | PTPRJ; PTPRJ; P | Q12913; A0 | PTPRJ_HUMAN | Unmodified | 0 | 0 | 0 | 223710 | 0 | 0 | 0 | 0 | 0 | 0 | 266750 | 1335700 | 0 | 0 | 0 | 0 | 0 | 0 |
| LEPNAQAQ | AP2A1 | O95782 | AP2A1_HUMAN | Unmodified | 0 | 0 | 0 | 0 | 0 | 0 | 0 | 0 | 0 | 0 | 0 | 892790 | 0 | 0 | 0 | 0 | 0 | 0 |
| LEQAAIVK | FKBP5 | Q13451 | FKBP5_HUMAN | Unmodified | 0 | 0 | 373550 | 119470 | 0 | 0 | 54347 | 0 | 66086 | 0 | 125340 | 465010 | 504480 | 0 | 0 | 235500 | 8739400 | 1022800 |
| LEQDILEK | KIF20B | Q96Q89 | KI20B_HUMAN | Unmodified | 205360 | 0 | 0 | 0 | 0 | 647680 | 651080 | 329970 | 0 | 0 | 475100 | 0 | 51100 | 0 | 0 | 0 | 0 | 0 |
| LEQELTEGI | TRIM26; TRIM2 | Q12899; A0 | TRI26_HUMAN | Unmodified | 2251000 | 3435500 | 15635000 | 1832800 | 331860 | 0 | 0 | 0 | 126340 | 0 | 0 | 0 | 362720 | 0 | 10663000 | 5518200 | 0 | 93850 |
| LEQENDDL | RABGAP1L; RAE | B7ZAP0; Q9 | RBG10_HUMAI | Unmodified | 0 | 167750 | 0 | 185240 | 0 | 861170 | 36793 | 0 | 578130 | 0 | 475680 | 2302400 | 183640 | 0 | 245870 | 875110 | 0 | 0 |
| LEQGQAID | CKB; CKB | P12277; H0 | KCRB_HUMAN | Unmodified | 0 | 0 | 0 | 78059 | 0 | 0 | 0 | 0 | 967410 | 0 | 0 | 8785800 | 300720 | 0 | 0 | 0 | 0 | 0 |
| LEQGQAID | CKB; CKB | P12277; H0 | KCRB_HUMAN | Oxidation (M) | 121140 | 0 | 645910 | 315930 | 0 | 128060 | 0 | 0 | 0 | 0 | 0 | 6444500 | 0 | 0 | 1252000 | 1656700 | 0 | 0 |
| LEQNTCSV | WDFY1; WDFY1 | Q8IWB7; C9 | WDFY1_HUMA | Unmodified | 7537500 | 6141500 | 12503000 | 9724200 | 2281700 | 0 | 0 | 0 | 0 | 1092600 | 0 | 8118500 | 0 | 944290 | 22469000 | 14485000 | 0 | 0 |
| LEQPDPGA | TKFC | Q3LXA3 | TKFC_HUMAN | Unmodified | 0 | 0 | 0 | 0 | 0 | 0 | 0 | 0 | 24109 | 0 | 180230 | 2647400 | 0 | 0 | 0 | 215890 | 0 | 0 |
| LEQQLVHV | DNPEP; DNPEP | Q9ULA0; E5I | DNPEP_HUMAI | Unmodified | 0 | 0 | 0 | 0 | 0 | 0 | 0 | 0 | 0 | 0 | 0 | 1559700 | 0 | 0 | 119950 | 79778 | 0 | 0 |
| LEQQVPVN | RPL3; RPL3; RP | P39023; B5 | RL3_HUMAN 6 | Oxidation (M) | 6612300 | 4134300 | 296930 | 837360 | 0 | 0 | 0 | 0 | 566400 | 0 | 5421900 | 9499700 | 0 | 0 | 20980000 | 0 | 0 | 0 |
| LEQSEAQL | COPS5 | Q92905 | CSN5_HUMAN | Unmodified | 0 | 0 | 0 | 0 | 69123 | 87883 | 0 | 0 | 400500 | 0 | 41723 | 2198100 | 0 | 0 | 0 | 0 | 0 | 0 |
| LEQSTIVK | FKBP4 | Q02790 | FKBP4_HUMAN | Unmodified | 501780 | 261800 | 1226900 | 1466000 | 560160 | 367960 | 407850 | 0 | 1270500 | 284450 | 4332200 | 22591000 | 4049200 | 1675500 | 1728800 | 2030200 | 348350 | 644310 |
| LERENAEVI | EIF2S1 | P05198 | IF2A_HUMAN E | Unmodified | 114650 | 0 | 0 | 0 | 0 | 0 | 0 | 0 | 0 | 0 | 0 | 3763400 | 0 | 0 | 0 | 0 | 0 | 0 |
| LESDYEILE | IARS1; IARS1; I | P41252; A0 | SYIC_HUMAN I | Unmodified | 0 | 0 | 134610 | 682490 | 8411500 | 0 | 0 | 0 | 73429 | 0 | 0 | 923040 | 0 | 0 | 0 | 0 | 0 | 0 |
| LESEGSPE | HACD3; HACD | Q9P035; H3 | HACD3_HUMA | Unmodified | 0 | 0 | 0 | 0 | 0 | 0 | 0 | 0 | 403640 | 0 | 0 | 0 | 0 | 0 | 0 | 0 | 0 | 0 |
| LESEHPDQ | SPTBN1; SPTBN | Q01082; A0 | SPTB2_HUMAN | Unmodified | 0 | 0 | 0 | 0 | 0 | 195360 | 0 | 0 | 0 | 0 | 0 | 1225900 | 0 | 0 | 0 | 0 | 0 | 0 |
| LESIISGAAI | CTNNA1; CTNN | P35221; P26 | CTNA1_HUMAI | Oxidation (M) | 0 | 0 | 0 | 0 | 0 | 0 | 0 | 0 | 0 | 0 | 0 | 973870 | 0 | 0 | 236090 | 454700 | 0 | 0 |
| LETETHWV | MPP7 | Q5T2T1 | MPP7_HUMAN | Unmodified | 0 | 0 | 228060 | 38968 | 0 | 93170 | 0 | 0 | 0 | 0 | 0 | 454720 | 0 | 0 | 201920 | 118270 | 0 | 0 |
| LETFIQEHL | CD151; CD151 | P48509; E9P | CD151_HUMAI | Unmodified | 1572000 | 1246900 | 0 | 0 | 0 | 0 | 0 | 0 | 0 | 0 | 0 | 0 | 0 | 0 | 1615500 | 21959 | 167900 | 0 |
| LETGQFLTF | UBA6 | A0AVT1 | UBA6_HUMAN | Unmodified | 0 | 0 | 0 | 0 | 0 | 0 | 0 | 0 | 0 | 0 | 0 | 0 | 0 | 0 | 0 | 0 | 0 | 0 |
| LETHMTPEI | DHX9 | Q08211 | DHX9_HUMAN | Unmodified | 0 | 0 | 0 | 0 | 0 | 0 | 0 | 71452 | 0 | 41902 | 0 | 1361700 | 0 | 0 | 0 | 0 | 0 | 0 |

| | | | | | | | | | | | | | | | | | | | | | | |
|---|---|---|---|---|---|---|---|---|---|---|---|---|---|---|---|---|---|---|---|---|---|---|
| LETKYPVVS | PLOD1; PLOD1; | Q02809; Q5 | PLOD1_HUMAI | Unmodified | 0 | 0 | 0 | 0 | 0 | 0 | 0 | 0 | 2425400 | 0 | 0 | 0 | 0 | 0 | 0 | 0 | 0 | 0 |
| LETLGIGQF | MAT2B | Q9NZL9 | MAT2B_HUMAI | Unmodified | 0 | 0 | 0 | 0 | 0 | 0 | 0 | 0 | 0 | 113050 | 520170 | 587490 | 196480 | 0 | 0 | 0 | 286480 | 587080 |
| LETNEFQQ | IKBIP | Q70UQ0 | IKIP_HUMAN In | Unmodified | 0 | 0 | 0 | 0 | 0 | 333580 | 0 | 0 | 1612600 | 0 | 0 | 0 | 0 | 0 | 0 | 0 | 0 | 0 |
| LETTSNQD | ITGA6 | P23229 | ITA6_HUMAN In | Unmodified | 0 | 0 | 2072000 | 725020 | 1285500 | 249460 | 0 | 0 | 0 | 0 | 128810 | 9288700 | 1016000 | 0 | 4411300 | 4299700 | 632500 | 1727500 |
| LEVAESEFT | PIP5K1A; PIP5K | Q99755; A6F | PI51A_HUMAN | Unmodified | 145170 | 0 | 0 | 0 | 0 | 0 | 0 | 0 | 848650 | 0 | 314070 | 4747200 | 635900 | 0 | 100440 | 0 | 0 | 1211500 |
| LEVANLLLC | ANK3 | Q12955 | ANK3_HUMAN | Unmodified | 0 | 0 | 0 | 0 | 0 | 0 | 0 | 0 | 0 | 0 | 0 | 0 | 0 | 0 | 0 | 0 | 0 | 0 |
| LEVAPISDII | ETFA; ETFA; ETF | P13804; H0Y | ETFA_HUMAN I | Unmodified | 89121 | 57129 | 90994 | 0 | 0 | 44343 | 0 | 47163 | 3451500 | 209740 | 0 | 32129 | 0 | 0 | 108350 | 83961 | 82992 | 0 |
| LEVEANNA | CAPZB; CAPZB; | P47756; A0A | CAPZB_HUMAI | Unmodified | 220470 | 0 | 85225 | 0 | 201380 | 468150 | 0 | 0 | 0 | 0 | 0 | 1013300 | 0 | 0 | 202880 | 166910 | 0 | 454600 |
| LEVLDSTK | UQCRFS1P1; U | P0C7P4; P47 | UCRIL_HUMAN | Unmodified | 0 | 0 | 1142300 | 226550 | 0 | 0 | 70553 | 76471 | 2504000 | 592010 | 248540 | 0 | 157440 | 104920 | 272690 | 0 | 35724 | 0 |
| LEVQATDR | TOP1; TOP1 | P11387; A0A | TOP1_HUMAN | Unmodified | 234670 | 48556 | 0 | 35984 | 0 | 0 | 0 | 14500 | 13244 | 0 | 34378 | 2057000 | 0 | 0 | 39821 | 779410 | 0 | 0 |
| LEVSVDSD | ITGAV | P06756 | ITAV_HUMAN In | Unmodified | 274680 | 0 | 929180 | 713770 | 1133200 | 0 | 0 | 0 | 1232100 | 317460 | 117990 | 2071800 | 172020 | 0 | 757660 | 448590 | 111590 | 0 |
| LEVSVDSD | ITGAV | P06756 | ITAV_HUMAN In | Unmodified | 0 | 0 | 355990 | 0 | 0 | 0 | 0 | 0 | 827200 | 307410 | 0 | 0 | 0 | 0 | 664300 | 518120 | 0 | 0 |
| LEWAIDED | PSD2; PSD2 | Q9BQI7; A0A | PSD2_HUMAN | Unmodified | 0 | 0 | 0 | 0 | 0 | 0 | 0 | 0 | 0 | 0 | 702790 | 767300 | 0 | 0 | 0 | 0 | 0 | 0 |
| LEWLGNCS | ATP1B1; ATP1B | P05026; V9C | AT1B1_HUMAN | Unmodified | 0 | 1343800 | 611550 | 0 | 769690 | 0 | 0 | 0 | 0 | 0 | 0 | 377790 | 0 | 0 | 1727300 | 124370 | 0 | 0 |
| LEYCEALAI | DARS1; DARS1 | P14868; H7E | SYDC_HUMAN | Oxidation (M) | 0 | 0 | 0 | 0 | 0 | 0 | 0 | 0 | 0 | 0 | 0 | 1228300 | 0 | 0 | 68966 | 0 | 0 | 0 |
| LEYHQVIQ | NUCB2; NUCB2 | P80303; Q2L | NUCB2_HUMA | Unmodified | 14080 | 0 | 659420 | 0 | 0 | 0 | 174990 | 107340 | 10611000 | 0 | 0 | 0 | 0 | 0 | 344750 | 78061 | 0 | 0 |
| LEYHQVIQ | NUCB2; NUCB2 | P80303; Q2L | NUCB2_HUMA | Unmodified | 0 | 0 | 0 | 0 | 0 | 0 | 0 | 0 | 3089000 | 0 | 0 | 0 | 0 | 0 | 0 | 0 | 0 | 0 |
| LEYKFCTRE | ITGA6 | P23229 | ITA6_HUMAN In | Unmodified | 0 | 0 | 0 | 0 | 0 | 0 | 150550 | 72900 | 0 | 0 | 0 | 0 | 0 | 0 | 0 | 0 | 31542 | 168750 |
| LFAALLIK | ARFGEF1; ARFG | Q9Y6D6; E5I | BIG1_HUMAN I | Unmodified | 0 | 0 | 0 | 0 | 45203 | 0 | 92466 | 21156 | 52755 | 106900 | 39237 | 0 | 0 | 0 | 0 | 0 | 0 | 64549 |
| LFADAVQE | MCM7 | P33993 | MCM7_HUMAN | Unmodified | 126990 | 0 | 44937 | 0 | 1006100 | 123890 | 551340 | 69166 | 502220 | 350900 | 400940 | 508750 | 0 | 104520 | 0 | 49043 | 160730 | 172850 |
| LFAPQQILC | TK1; TK1 | P04183; K7E | KITH_HUMAN T | Unmodified | 0 | 44048 | 29103 | 0 | 324310 | 88078 | 0 | 0 | 0 | 0 | 0 | 0 | 206880 | 137600 | 0 | 0 | 597970 | 365110 |
| LFAQQCK | NNT; NNT; NNT | Q13423; E9F | NNTM_HUMAN | Unmodified | 0 | 0 | 0 | 0 | 0 | 0 | 42115 | 0 | 2481000 | 613040 | 303120 | 0 | 0 | 0 | 0 | 0 | 0 | 0 |
| LFCIPVHGI | IMPA1; IMPA1 | P29218; H0Y | IMPA1_HUMAN | Unmodified | 0 | 0 | 0 | 0 | 0 | 0 | 0 | 0 | 0 | 0 | 0 | 1935700 | 0 | 0 | 0 | 0 | 335800 | 0 |
| LFCVGFTK | RPS3A; RPS3A; | P61247; D6F | RS3A_HUMAN | Unmodified | 0 | 1071900 | 0 | 0 | 523590 | 0 | 419640 | 390830 | 513160 | 0 | 1311800 | 20693000 | 387590 | 304090 | 982340 | 0 | 0 | 0 |
| LFDANKAE | SPTBN1; SPTBN | Q01082; A0A | SPTB2_HUMAN | Unmodified | 0 | 0 | 0 | 0 | 0 | 0 | 0 | 0 | 0 | 0 | 0 | 1400300 | 0 | 0 | 0 | 234930 | 0 | 0 |
| LFDAPEAPI | MTOR | P42345 | MTOR_HUMAN | Unmodified | 0 | 166830 | 0 | 0 | 0 | 0 | 0 | 0 | 794660 | 195730 | 0 | 663060 | 0 | 0 | 0 | 1458200 | 0 | 0 |
| LFDDDETG | CETN2; CETN1 | P41208; Q12 | CETN2_HUMAI | Unmodified | 0 | 279650 | 0 | 0 | 567680 | 0 | 429810 | 105060 | 0 | 0 | 0 | 0 | 0 | 0 | 0 | 0 | 569910 | 1079100 |
| LFDFPGR | RCC2 | Q9P258 | RCC2_HUMAN | Unmodified | 0 | 0 | 0 | 0 | 0 | 0 | 0 | 0 | 308130 | 0 | 196420 | 8980100 | 0 | 0 | 0 | 0 | 0 | 0 |
| LFDHAVSK | ACSL4; ACSL4; | O60488; D6I | ACSL4_HUMAN | Unmodified | 0 | 0 | 0 | 0 | 5303000 | 171220 | 0 | 0 | 0 | 0 | 624480 | 1871700 | 0 | 0 | 0 | 0 | 1188900 | 3465300 |
| LFDHAVSK | ACSL4; ACSL4; | O60488; D6I | ACSL4_HUMAN | Unmodified | 0 | 0 | 0 | 0 | 0 | 0 | 0 | 0 | 0 | 0 | 0 | 0 | 0 | 0 | 0 | 0 | 267070 | 462380 |
| LFDHPESP | FASN; FASN | P49327; A0A | FAS_HUMAN F | Unmodified | 0 | 0 | 3206800 | 3430000 | 623330 | 0 | 0 | 0 | 12345000 | 377780 | 15994000 | 1.58E+08 | 0 | 0 | 20129000 | 10435000 | 0 | 0 |
| LFDIFSQQ | VPS35 | Q96QK1 | VPS35_HUMAN | Unmodified | 0 | 0 | 48129 | 0 | 86557 | 0 | 63567 | 0 | 129800 | 0 | 65120 | 1717800 | 0 | 0 | 501100 | 94055 | 0 | 30543 |
| LFDLDEK | SAAL1; SAAL1; | Q96ER3; E9F | SAAL1_HUMAN | Unmodified | 0 | 0 | 0 | 0 | 0 | 0 | 0 | 0 | 0 | 0 | 67342 | 2026200 | 0 | 0 | 0 | 0 | 0 | 0 |
| LFDLGETY | SLC4A7; SLC4A | Q9Y6M7; C9 | S4A7_HUMAN | Oxidation (M) | 0 | 0 | 0 | 0 | 0 | 0 | 0 | 0 | 359330 | 0 | 131390 | 343410 | 0 | 0 | 0 | 0 | 66835 | 0 |
| LFDPQLK | SEC16A; SEC16 | O15027; F1T | SC16A_HUMAI | Unmodified | 0 | 0 | 0 | 0 | 0 | 0 | 186020 | 0 | 0 | 25405 | 0 | 1375000 | 0 | 0 | 0 | 0 | 0 | 0 |
| LFDQAFGL | HSPB1; HSPB1; | P04792; A0A | HSPB1_HUMAI | Unmodified | 0 | 0 | 0 | 3577100 | 0 | 0 | 0 | 0 | 6484600 | 0 | 0 | 0 | 0 | 0 | 11232000 | 7840500 | 0 | 0 |
| LFDQANK | TJP2; TJP2; TJP2 | Q9UDY2; A0 | ZO2_HUMAN T | Unmodified | 0 | 0 | 0 | 0 | 3409000 | 1062700 | 0 | 0 | 0 | 0 | 0 | 0 | 0 | 0 | 0 | 0 | 0 | 0 |
| LFDQQLSP | INPP5D | Q92835 | SHIP1_HUMAN | Oxidation (M) | 0 | 0 | 0 | 0 | 0 | 0 | 0 | 0 | 0 | 0 | 0 | 4655300 | 0 | 0 | 0 | 0 | 0 | 0 |
| LFDSDPITV | CLU; CLU | P10909; H0Y | CLUS_HUMAN | Unmodified | 0 | 0 | 0 | 0 | 127490 | 1483500 | 0 | 0 | 8023100 | 0 | 0 | 0 | 0 | 1259700 | 0 | 0 | 0 | 0 |
| LFDSDPITV | CLU; CLU | P10909; H0Y | CLUS_HUMAN | Unmodified | 0 | 0 | 0 | 0 | 11922000 | 9070300 | 0 | 0 | 801550 | 4217600 | 0 | 0 | 0 | 0 | 0 | 0 | 818020 | 15277000 |
| LFDSICNNI | GNAI3; GNAI2; | P08754; P04 | GNAI3_HUMAN | Unmodified | 2815900 | 3468800 | 1406400 | 2559200 | 1910100 | 1389000 | 2292800 | 605840 | 4113700 | 0 | 8358100 | 46153000 | 1074200 | 1666400 | 3944000 | 4109700 | 1423700 | 3604000 |
| LFDSICNNI | GNAI2; GNAI1; | P04899; P63 | GNAI2_HUMAN | Unmodified | 0 | 0 | 0 | 0 | 181140 | 0 | 454320 | 0 | 0 | 0 | 0 | 0 | 151570 | 177850 | 0 | 0 | 739770 | 627330 |
| LFDSTIADE | NUDCD2 | Q8WVJ2 | NUDC2_HUMA | Unmodified | 378750 | 672170 | 735350 | 513570 | 4648600 | 1960300 | 690540 | 90440 | 833520 | 631020 | 1805400 | 3015100 | 2249200 | 3717300 | 793780 | 657820 | 1104300 | 110060 |
| LFDSTTLEH | EIF3I; EIF3I; EIF | Q13347; A0A | EIF3I_HUMAN I | Unmodified | 0 | 0 | 2159400 | 0 | 3897200 | 744500 | 1030000 | 636580 | 897170 | 976850 | 1676800 | 8481200 | 8316000 | 3090900 | 2256800 | 2475600 | 2418700 | 2519400 |
| LFDTLNEDI | HSPA13 | P48723 | HSP13_HUMAI | Unmodified | 0 | 0 | 0 | 0 | 0 | 0 | 0 | 0 | 2394500 | 599020 | 0 | 427370 | 0 | 0 | 0 | 0 | 0 | 0 |
| LFDVGGQF | GNAO1; GNAO1 | P09471; H3E | GNAO_HUMAN | Unmodified | 0 | 0 | 0 | 0 | 0 | 0 | 105830 | 0 | 1318500 | 55907 | 0 | 231050 | 0 | 0 | 0 | 0 | 0 | 0 |
| LFDWANTS | ATAD3A; ATAD3 | Q9NVI7; Q51 | ATD3A_HUMAN | Unmodified | 0 | 0 | 0 | 0 | 0 | 0 | 0 | 0 | 1398400 | 0 | 0 | 0 | 0 | 0 | 0 | 0 | 0 | 0 |
| LFDYFPKPY | CYC1 | P08574 | CY1_HUMAN C | Unmodified | 0 | 0 | 0 | 0 | 0 | 0 | 0 | 0 | 55379000 | 2721600 | 789070 | 429540 | 0 | 0 | 0 | 0 | 32257 | 132880 |
| LFEAEAQDI | EHD4 | Q9H223 | EHD4_HUMAN | Unmodified | 244800 | 259710 | 1461100 | 182880 | 0 | 0 | 0 | 0 | 0 | 0 | 0 | 3180300 | 0 | 0 | 652630 | 414900 | 0 | 0 |
| LFEAEAQDI | EHD4 | Q9H223 | EHD4_HUMAN | Unmodified | 0 | 210850 | 0 | 2910500 | 1399200 | 0 | 0 | 0 | 0 | 0 | 0 | 0 | 0 | 0 | 2846700 | 0 | 0 | 1360200 |
| LFEAEEQDI | EHD1; EHD1; EI | Q9H4M9; C9 | EHD1_HUMAN | Unmodified | 378480 | 91900 | 81251 | 190740 | 0 | 0 | 0 | 13457 | 50400 | 0 | 0 | 4663500 | 65976 | 63574 | 841380 | 489200 | 0 | 962810 |
| LFEAEEQDI | EHD1; EHD1; EI | Q9H4M9; C9 | EHD1_HUMAN | Unmodified | 0 | 0 | 0 | 0 | 0 | 0 | 0 | 0 | 0 | 0 | 0 | 0 | 1472700 | 0 | 0 | 0 | 0 | 1157500 |
| LFEAEEQDI | EHD1; EHD1; EI | Q9H4M9; C9 | EHD1_HUMAN | Unmodified | 0 | 0 | 0 | 0 | 949370 | 0 | 128490 | 0 | 0 | 94779 | 0 | 0 | 1118500 | 0 | 0 | 0 | 1986300 | 749840 |
| LFEAEEQDI | EHD3 | Q9NZN3 | EHD3_HUMAN | Unmodified | 38975 | 87195 | 589610 | 517310 | 0 | 0 | 0 | 145980 | 0 | 294060 | 0 | 509040 | 0 | 0 | 1277300 | 797950 | 0 | 0 |
| LFEAQEKLY | CARS1 | P49589 | SYCC_HUMAN | Unmodified | 0 | 0 | 0 | 0 | 95180 | 0 | 0 | 0 | 360850 | 0 | 0 | 0 | 0 | 0 | 0 | 0 | 207490 | 0 |
| LFEASIETG | LAP3; LAP3 | P28838; H0Y | AMPL_HUMAN | Unmodified | 0 | 0 | 0 | 0 | 0 | 0 | 0 | 0 | 0 | 0 | 0 | 0 | 0 | 0 | 0 | 0 | 0 | 0 |
| LFECDRDQ | LRPPRC | P42704 | LPPRC_HUMAI | Oxidation (M) | 0 | 0 | 0 | 0 | 0 | 0 | 0 | 0 | 1274900 | 852030 | 0 | 0 | 0 | 0 | 0 | 0 | 0 | 0 |
| LFECSNK | VIL1; AVIL; VIL1 | P09327; O75 | VILI_HUMAN Vi | Unmodified | 0 | 0 | 0 | 0 | 0 | 0 | 0 | 0 | 0 | 0 | 0 | 20635000 | 0 | 0 | 0 | 0 | 0 | 0 |
| LFEDGGG( | SLITRK3 | O94933 | SLIK3_HUMAN | Unmodified | 0 | 0 | 47099 | 69249 | 0 | 0 | 0 | 0 | 640650 | 52130 | 0 | 0 | 0 | 0 | 0 | 352610 | 22978 | 0 |
| LFEDTKYTT | TES | Q9UGI8 | TES_HUMAN Te | Unmodified | 0 | 0 | 0 | 834130 | 5063500 | 911850 | 351720 | 136800 | 0 | 1216300 | 209300 | 929560 | 3994700 | 0 | 0 | 0 | 265050 | 7182700 |
| LFEELGK | CAMK2D; HSPA | Q13557; P34 | KCC2D_HUMA | Unmodified | 136690 | 0 | 254660 | 823320 | 3538400 | 691220 | 346080 | 322840 | 1406300 | 1190500 | 2630000 | 12032000 | 4123700 | 1740600 | 480350 | 618450 | 1538300 | 1218900 |
| LFEELGKQI | HSPA4 | P34932 | HSP74_HUMAI | Unmodified | 0 | 0 | 0 | 0 | 239040 | 0 | 0 | 0 | 0 | 0 | 0 | 0 | 154310 | 0 | 0 | 0 | 161350 | 0 |
| LFEELVR | PKM; PKM | H3BTN5; H3 | H3BTN5_HUM | Unmodified | 0 | 0 | 82200 | 0 | 0 | 0 | 0 | 0 | 279320 | 0 | 1164100 | 11004000 | 0 | 0 | 509070 | 0 | 0 | 83634 |
| LFEEPEDPS | PPP2R2C; PPP2 | Q9Y2T4; P63 | 2ABG_HUMAN | Unmodified | 0 | 0 | 0 | 0 | 0 | 0 | 0 | 0 | 0 | 0 | 0 | 2995800 | 0 | 0 | 0 | 764680 | 0 | 0 |
| LFEFAGYD | RARS1 | P54136 | SYRC_HUMAN | Unmodified | 0 | 0 | 0 | 0 | 0 | 0 | 0 | 0 | 0 | 0 | 0 | 5100000 | 0 | 0 | 0 | 0 | 0 | 0 |
| LFEGENEH | IQGAP2; IQGAP | Q13576; E7E | IQGA2_HUMAN | Oxidation (M) | 0 | 0 | 928930 | 586220 | 0 | 0 | 0 | 0 | 0 | 0 | 0 | 0 | 625050 | 0 | 1212000 | 454620 | 0 | 0 |
| LFEGIIK | CASK; CASK; C | O14936; A0A | CSKP_HUMAN | Unmodified | 0 | 0 | 0 | 0 | 16054 | 0 | 259460 | 148040 | 4479600 | 1211700 | 1423700 | 3746000 | 0 | 0 | 0 | 165060 | 225460 | 257070 |
| LFEGNALLI | RPS9; RPS9; RP | P46781; B5N | RS9_HUMAN 4 | Unmodified | 2078600 | 2159600 | 1400000 | 330830 | 35531 | 0 | 76682 | 0 | 116630 | 0 | 941150 | 35207000 | 82927 | 0 | 0 | 1438700 | 0 | 0 |
| LFEGNMHY | NRP2 | O60462 | NRP2_HUMAN | Oxidation (M) | 130390 | 0 | 0 | 0 | 0 | 0 | 0 | 0 | 1088000 | 0 | 932850 | 0 | 0 | 0 | 557960 | 0 | 0 | 0 |
| LFEGNMHY | NRP2 | O60462 | NRP2_HUMAN | Unmodified | 0 | 0 | 0 | 0 | 0 | 0 | 39034 | 0 | 1194200 | 131710 | 0 | 0 | 0 | 0 | 228640 | 258440 | 0 | 0 |
| LFEISDIVIK | FARSB | Q9NSD9 | SYFB_HUMAN | Unmodified | 126150 | 111770 | 142670 | 128800 | 0 | 0 | 212260 | 0 | 411090 | 328600 | 833360 | 9572000 | 0 | 240300 | 0 | 210100 | 0 | 0 |
| LFEISDIVIK | FARSB | Q9NSD9 | SYFB_HUMAN | Unmodified | 0 | 0 | 0 | 0 | 1077800 | 0 | 1853600 | 254290 | 2255700 | 783490 | 120680 | 1440000 | 3525600 | 0 | 0 | 0 | 739020 | 787890 |
| LFEKPLDS( | SCG2 | P13521 | SCG2_HUMAN | Unmodified | 0 | 0 | 0 | 0 | 0 | 0 | 0 | 0 | 1429000 | 697740 | 0 | 0 | 0 | 0 | 0 | 0 | 0 | 0 |
| LFELLEK | SMARCA1; SMA | P28370; O60 | SMCA1_HUMA | Unmodified | 473220 | 428480 | 0 | 0 | 0 | 0 | 0 | 0 | 0 | 0 | 0 | 0 | 0 | 0 | 0 | 0 | 0 | 0 |
| LFELTVERT | LMAN2L | Q9H0V9 | LMA2L_HUMAN | Unmodified | 0 | 105250 | 0 | 144250 | 0 | 0 | 0 | 94171 | 30306000 | 396990 | 0 | 103320 | 0 | 0 | 84531 | 74207 | 0 | 0 |
| LFEMAYK | HSPA9; HSPA9; | P38646; A0A | GRP75_HUMAI | Unmodified | 0 | 0 | 0 | 0 | 0 | 0 | 0 | 0 | 1439000 | 0 | 0 | 0 | 0 | 0 | 0 | 0 | 27086 | 0 |
| LFENQLVG | APMAP; APMAP | Q9HDC9; H( | APMAP_HUMAI | Oxidation (M) | 0 | 0 | 0 | 0 | 0 | 0 | 0 | 0 | 55967000 | 4321800 | 0 | 0 | 0 | 0 | 0 | 0 | 0 | 0 |
| LFENQLVG | APMAP; APMAP | Q9HDC9; H( | APMAP_HUMAI | Unmodified | 0 | 0 | 0 | 0 | 0 | 0 | 0 | 0 | 40117000 | 2279400 | 0 | 0 | 0 | 0 | 0 | 0 | 0 | 0 |
| LFENQLVG | APMAP; APMAP | Q9HDC9; H( | APMAP_HUMAI | Unmodified | 0 | 0 | 0 | 559560 | 0 | 0 | 0 | 0 | 1290100 | 0 | 0 | 0 | 0 | 0 | 0 | 0 | 0 | 0 |
| LFENYIELK | EXOC2 | Q96KP1 | EXOC2_HUMAI | Unmodified | 0 | 0 | 0 | 0 | 0 | 14254 | 0 | 0 | 434970 | 619810 | 210130 | 838340 | 0 | 0 | 0 | 0 | 0 | 0 |
| LFEQNVQR | UBE2M; UBE2M | P61081; M0( | UBC12_HUMAI | Unmodified | 509460 | 1149400 | 925710 | 0 | 534760 | 0 | 239320 | 0 | 2673100 | 933080 | 941690 | 4607000 | 0 | 0 | 1109500 | 882150 | 982100 | 285150 |
| LFERYGEN | SLC39A10; SLC | Q9ULF5; E7E | S39AA_HUMAN | Unmodified | 0 | 13672 | 0 | 0 | 7504900 | 0 | 1327200 | 0 | 0 | 0 | 0 | 0 | 2628100 | 2315800 | 0 | 0 | 4616400 | 6126700 |
| LFEVIETEK | MARK3; MARK2 | P27448; Q7I | MARK3_HUMAI | Unmodified | 0 | 0 | 0 | 0 | 0 | 0 | 368250 | 790380 | 627020 | 456390 | 326360 | 2201000 | 0 | 397020 | 0 | 0 | 0 | 0 |
| LFEYGGFPI | PPP1CA; PPP1C | P62136; P36 | PP1A_HUMAN | Unmodified | 0 | 0 | 0 | 0 | 0 | 0 | 0 | 0 | 0 | 0 | 0 | 359390 | 0 | 0 | 877420 | 0 | 0 | 0 |
| LFEYRNQG | POLR1C; POLR1 | O15160; A0A | RPAC1_HUMAI | Unmodified | 0 | 0 | 0 | 0 | 0 | 0 | 0 | 0 | 0 | 0 | 253910 | 0 | 0 | 0 | 0 | 0 | 0 | 0 |
| LFFDKRDN | EIF3D | O15371 | EIF3D_HUMAN | Unmodified | 0 | 0 | 0 | 0 | 1729100 | 0 | 0 | 0 | 3636900 | 1364900 | 0 | 0 | 2764600 | 0 | 0 | 0 | 327280 | 722460 |
| LFFHCSQY | CSDE1; CSDE1 | O75534; E9F | CSDE1_HUMAI | Unmodified | 0 | 0 | 190540 | 0 | 0 | 0 | 0 | 0 | 0 | 0 | 0 | 659040 | 0 | 0 | 239030 | 413520 | 0 | 0 |
| LFFILLTPVI | VPS13A; VPS13 | Q96RL7; H0Y | VP13A_HUMAN | Unmodified | 0 | 0 | 0 | 0 | 0 | 0 | 0 | 0 | 438960 | 157430 | 319360 | 815640 | 0 | 0 | 0 | 25232 | 0 | 0 |
| LFFLQVK | RDX; EZR; MSN | P35241; P15 | RADI_HUMAN I | Unmodified | 8332400 | 7732100 | 14916000 | 7192700 | 36565000 | 19779000 | 0 | 0 | 62736000 | 15394000 | 0 | 1.02E+08 | 5974900 | 11641000 | 24700000 | 18904000 | 10367000 | 1581500 |
| LFFQPHQA | UBA5; UBA5; UI | Q9GZZ9; E7I | UBA5_HUMAN | Unmodified | 0 | 0 | 0 | 0 | 0 | 0 | 104500 | 0 | 146790 | 12533 | 0 | 2735400 | 0 | 0 | 0 | 0 | 0 | 0 |
| LFFVGSR | SLC7A7; SLC7A | Q9UM01; QC | YLAT1_HUMAN | Unmodified | 4551100 | 5838700 | 8797000 | 0 | 0 | 0 | 0 | 0 | 0 | 247850 | 172010 | 1E+08 | 0 | 0 | 12520000 | 3621700 | 0 | 0 |
| LFFWNSK | ACO1 | P21399 | ACOC_HUMAN | Unmodified | 0 | 0 | 111050 | 110370 | 0 | 0 | 0 | 0 | 0 | 0 | 0 | 4084200 | 179760 | 0 | 0 | 205970 | 0 | 0 |
| LFGAGGGI | CHMP4B | Q9H444 | CHM4B_HUMA | Unmodified | 0 | 0 | 0 | 0 | 1366200 | 0 | 140850 | 0 | 0 | 812410 | 0 | 0 | 0 | 0 | 0 | 0 | 561780 | 485820 |
| LFGAPEPS | LARP1; LARP1; | Q6PKG0; H0 | LARP1_HUMAN | Unmodified | 2480500 | 3070300 | 0 | 4287200 | 0 | 708570 | 326260 | 0 | 0 | 0 | 206070 | 1831300 | 1671300 | 0 | 1076700 | 1221600 | 324830 | 0 |
| LFGAVEHL( | LMF2 | Q9BU23 | LMF2_HUMAN | Unmodified | 0 | 0 | 0 | 0 | 0 | 0 | 769640 | 0 | 523360 | 513210 | 0 | 0 | 0 | 0 | 0 | 0 | 0 | 0 |
| LFGAVEHL( | LMF2 | Q9BU23 | LMF2_HUMAN | Unmodified | 881660 | 2459000 | 0 | 563010 | 0 | 0 | 0 | 0 | 3773100 | 0 | 0 | 0 | 0 | 0 | 0 | 271140 | 0 | 0 |
| LFGFVLR | APPL1 | Q9UKG1 | DP13A_HUMAN | Unmodified | 201400 | 160470 | 0 | 0 | 0 | 0 | 0 | 0 | 233680 | 44973 | 24852 | 750250 | 0 | 0 | 128860 | 0 | 0 | 0 |
| LFGGFNSS | SRC | P12931 | SRC_HUMAN P | Unmodified | 197210 | 170630 | 119420 | 281730 | 0 | 0 | 0 | 0 | 626460 | 78684 | 0 | 0 | 0 | 0 | 257730 | 338050 | 221920 | 272790 |
| LFGGLILDI | SLC4A7; SLC4A | Q9Y6M7; C9 | S4A7_HUMAN | Unmodified | 0 | 26673 | 61949 | 80088 | 63541 | 0 | 0 | 0 | 0 | 214910 | 45130 | 1784100 | 0 | 0 | 112230 | 107770 | 0 | 0 |
| LFGGNFAH | TPP1; TPP1; TPI | O14773; A0A | TPP1_HUMAN | Unmodified | 50047 | 0 | 142060 | 223560 | 0 | 0 | 0 | 0 | 3805000 | 186570 | 0 | 0 | 448340 | 0 | 598150 | 279610 | 151510 | 0 |
| LFGGQIVG | ACOT8; ACOT8; | O14734; E9F | ACOT8_HUMAI | Unmodified | 0 | 0 | 0 | 0 | 0 | 0 | 0 | 0 | 11076000 | 279490 | 51048 | 0 | 0 | 0 | 0 | 0 | 0 | 15792 |
| LFGHSSTS | HUWE1 | Q7Z6Z7 | HUWE1_HUMA | Unmodified | 45090 | 0 | 47352 | 0 | 0 | 0 | 0 | 0 | 541200 | 0 | 0 | 1890200 | 0 | 0 | 86272 | 110540 | 0 | 0 |
| LFGIESSG | EMC1 | Q8N766 | EMC1_HUMAN | Unmodified | 0 | 0 | 0 | 0 | 0 | 0 | 0 | 0 | 0 | 904150 | 285690 | 0 | 0 | 0 | 0 | 0 | 0 | 0 |
| LFGNMEGI | TFRC; TFRC | P02786; G3V | TFR1_HUMAN | Oxidation (M) | 173540 | 0 | 0 | 1548400 | 575490 | 254450 | 0 | 0 | 3280500 | 460240 | 0 | 0 | 396420 | 0 | 0 | 0 | 0 | 512150 |
| LFGNMEGI | TFRC; TFRC | P02786; G3V | TFR1_HUMAN | Unmodified | 0 | 0 | 0 | 0 | 933680 | 0 | 0 | 0 | 4790900 | 341110 | 0 | 0 | 0 | 0 | 0 | 0 | 0 | 450990 |
| LFGPYVWG | RNPEP; RNPEP; | Q9H4A4; H7 | AMPB_HUMAN | Unmodified | 0 | 0 | 0 | 0 | 0 | 0 | 0 | 0 | 0 | 0 | 0 | 3960500 | 0 | 0 | 117340 | 76568 | 0 | 0 |
| LFGQETPE | SNX4 | O95219 | SNX4_HUMAN | Unmodified | 0 | 0 | 0 | 0 | 0 | 2595700 | 0 | 0 | 262000 | 602600 | 0 | 0 | 241170 | 127330 | 0 | 0 | 36119 | 90470 |
| LFGQETPE | SNX4 | O95219 | SNX4_HUMAN | Unmodified | 0 | 0 | 0 | 0 | 2489600 | 0 | 193810 | 116100 | 0 | 149890 | 0 | 0 | 358070 | 0 | 0 | 0 | 0 | 426500 |
| LFGQHLAT | TOR1B; TOR1B | O14657; H0Y | TOR1B_HUMAI | Unmodified | 18770 | 0 | 0 | 0 | 422660 | 0 | 0 | 0 | 1824700 | 1769800 | 0 | 0 | 0 | 0 | 0 | 0 | 0 | 0 |

| | | | | | | | | | | | | | | | | | | | | | | |
|---|---|---|---|---|---|---|---|---|---|---|---|---|---|---|---|---|---|---|---|---|---|---|
| LFGSPLSD' | H6PD | O95479 | G6PE_HUMAN | Unmodified | 0 | 0 | 0 | 0 | 0 | 64183 | 0 | 0 | 4326900 | 0 | 0 | 0 | 0 | 0 | 0 | 0 | 86976 | 0 |
| LFGTSGQK | EGFR; EGFR; E( | P00533; Q5( | EGFR_HUMAN | Unmodified | 0 | 153580 | 0 | 830880 | 13710000 | 3068100 | 24742 | 0 | 252560 | 117290 | 1713100 | 0 | 7983400 | 1867800 | 0 | 803030 | 2070900 | 2846200 |
| LFGYEPTIY | NAXE; NAXE; N | Q8NCW5; A( | NNRE_HUMAN | Unmodified | 42816 | 0 | 0 | 0 | 0 | 0 | 0 | 0 | 2859300 | 1366500 | 1953300 | 4504200 | 0 | 0 | 0 | 0 | 1235700 | 0 |
| LFHRFLQEI | UGP2; UGP2; U | Q16851; A0/ | UGPA_HUMAN | Unmodified | 0 | 0 | 0 | 0 | 1026800 | 0 | 750700 | 0 | 0 | 562800 | 0 | 0 | 341680 | 524760 | 0 | 0 | 838370 | 431920 |
| LFHTAPNVI | METAP1; METAI | P53582; H0\ | MAP11_HUMAI | Unmodified | 0 | 0 | 108790 | 0 | 1254000 | 0 | 0 | 0 | 0 | 0 | 0 | 872880 | 0 | 436370 | 0 | 0 | 0 | 306640 |
| LFHVLDCT' | XXYLT1; XXYLT1 | Q8NBI6; A0/ | XXLT1_HUMAN | Unmodified | 0 | 0 | 0 | 0 | 0 | 0 | 0 | 0 | 2196700 | 0 | 0 | 0 | 0 | 0 | 0 | 0 | 0 | 0 |
| LFIGGLNTE | RBMX; RBMXL1 | P38159; Q9( | RBMX_HUMAN | Unmodified | 0 | 0 | 0 | 0 | 646510 | 0 | 33011 | 295640 | 1866200 | 1577500 | 3228500 | 9533200 | 794750 | 292860 | 0 | 0 | 233750 | 0 |
| LFIGGLNTE | RBMX; RBMX; R | P38159; H3E | RBMX_HUMAN | Unmodified | 0 | 0 | 0 | 0 | 97884 | 0 | 0 | 0 | 0 | 140590 | 0 | 0 | 93138 | 0 | 0 | 0 | 0 | 0 |
| LFIGGLNV( | HNRNPA0 | Q13151 | ROA0_HUMAN | Unmodified | 0 | 0 | 0 | 0 | 0 | 0 | 0 | 0 | 0 | 2239900 | 212010 | 821830 | 0 | 0 | 0 | 0 | 0 | 0 |
| LFIGGLPN\ | U2AF2; U2AF2 | P26368; K7E | U2AF2_HUMAI | Unmodified | 909350 | 3117300 | 746510 | 1719100 | 3292900 | 346290 | 848850 | 22135 | 1102000 | 1827400 | 1571100 | 7458200 | 3489000 | 1613600 | 1126700 | 1402900 | 543150 | 536840 |
| LFIGGLSFE | HNRNPA3; HNI | P51991; A0A | ROA3_HUMAN | Unmodified | 0 | 0 | 0 | 615340 | 4673400 | 0 | 17411000 | 0 | 3828300 | 3056700 | 1936000 | 5667900 | 11501000 | 4507600 | 0 | 0 | 0 | 8631700 |
| LFIGGLSFE | HNRNPA1L2; H | Q32P51; P09 | RA1L2_HUMAI | Unmodified | 45280 | 61146 | 116460 | 34949 | 0 | 31448 | 32474 | 35989 | 747570 | 60876 | 356870 | 1964300 | 46641 | 31003 | 46936 | 39249 | 0 | 0 |
| LFIGGLSFE | HNRNPA2B1; H | P22626; A0A | ROA2_HUMAN | Unmodified | 23316 | 0 | 0 | 0 | 0 | 0 | 13228 | 0 | 937280 | 33627 | 157770 | 2397900 | 31555 | 0 | 0 | 61371 | 0 | 0 |
| LFIGNLPR | RBM4; RBM4B; | Q9BWF3; Q9 | RBM4_HUMAN | Unmodified | 247730 | 303490 | 0 | 219820 | 0 | 0 | 0 | 0 | 0 | 0 | 0 | 743530 | 0 | 0 | 261360 | 295040 | 0 | 0 |
| LFIGTAVDC | PLXNA2 | O75051 | PLXA2_HUMAI | Unmodified | 0 | 0 | 0 | 0 | 0 | 0 | 0 | 0 | 2159900 | 0 | 0 | 0 | 0 | 0 | 0 | 0 | 0 | 0 |
| LFIHESIHD | ALDH7A1; ALDI | P49419; A0A | AL7A1_HUMAI | Unmodified | 0 | 0 | 0 | 0 | 671840 | 75966 | 0 | 0 | 0 | 0 | 0 | 0 | 0 | 0 | 0 | 0 | 97963 | 268200 |
| LFIHNHEQ | PLOD1 | Q02809 | PLOD1_HUMAI | Unmodified | 0 | 0 | 29075 | 0 | 0 | 0 | 0 | 0 | 901630 | 16361 | 48717 | 0 | 20031 | 69134 | 0 | 0 | 0 | 121760 |
| LFIHNKEV\ | PLOD2; PLOD2 | O00469; E7E | PLOD2_HUMAI | Unmodified | 0 | 0 | 0 | 0 | 0 | 0 | 0 | 0 | 0 | 476670 | 0 | 0 | 0 | 0 | 0 | 0 | 0 | 0 |
| LFIIQACR | CASP3 | P42574 | CASP3_HUMAI | Unmodified | 0 | 0 | 0 | 0 | 0 | 0 | 0 | 0 | 120560 | 0 | 0 | 4570400 | 0 | 0 | 0 | 0 | 0 | 0 |
| LFIIRGSPQ | KHSRP; KHSRP | Q92945; M0I | FUBP2_HUMAI | Unmodified | 0 | 0 | 0 | 0 | 1985400 | 0 | 1778400 | 1001500 | 0 | 3123200 | 0 | 0 | 1746400 | 335160 | 0 | 0 | 1141700 | 1977000 |
| LFIPDRDLII | IL1RAPL1; IL1R/ | Q9NZN1; A0 | IRPL1_HUMAN | Unmodified | 0 | 0 | 158660 | 511540 | 0 | 0 | 0 | 0 | 0 | 0 | 0 | 0 | 0 | 0 | 0 | 210500 | 0 | 0 |
| LFIRELPQP | ARHGAP18 | Q8N392 | RHG18_HUMAI | Unmodified | 0 | 0 | 0 | 8897.4 | 534410 | 69161 | 347050 | 0 | 0 | 0 | 0 | 0 | 94896 | 241440 | 0 | 0 | 362780 | 27931 |
| LFISTRLGTI | MYO1C; MYO1( | O00159; F5I | MYO1C_HUMA | Unmodified | 0 | 0 | 0 | 0 | 12138000 | 3417900 | 0 | 0 | 0 | 1017100 | 0 | 498270 | 11174000 | 5219600 | 0 | 0 | 3352900 | 4038900 |
| LFIVESNSS | GCLM | P48507 | GSH0_HUMAN | Unmodified | 0 | 0 | 0 | 0 | 0 | 0 | 0 | 61538 | 0 | 0 | 0 | 590730 | 765050 | 335770 | 0 | 0 | 0 | 0 |
| LFIYNPTTG | ATP1B3; ATP1B | P54709; F8V | AT1B3_HUMAI | Unmodified | 0 | 0 | 3194200 | 0 | 1354900 | 0 | 0 | 0 | 0 | 67533 | 0 | 14676000 | 0 | 0 | 7789800 | 5397900 | 0 | 0 |
| LFKPGQEA' | TXNDC5 | Q8NBS9 | TXND5_HUMAI | Unmodified | 0 | 0 | 0 | 0 | 0 | 0 | 0 | 0 | 5608700 | 2171200 | 47387 | 0 | 0 | 0 | 0 | 0 | 0 | 0 |
| LFKPGQEA' | TXNDC5 | Q8NBS9 | TXND5_HUMAI | Unmodified | 0 | 0 | 0 | 0 | 0 | 0 | 0 | 0 | 108570 | 392570 | 0 | 0 | 0 | 0 | 0 | 0 | 0 | 0 |
| LFKPPQPP/ | EIF3H; EIF3H; E | O15372; B3I | EIF3H_HUMAN | Oxidation (M) | 0 | 0 | 0 | 0 | 0 | 0 | 0 | 0 | 0 | 0 | 0 | 0 | 2629900 | 0 | 0 | 0 | 0 | 0 |
| LFKQDTSS! | ANLN | Q9NQW6 | ANLN_HUMAN | Unmodified | 0 | 0 | 0 | 0 | 3313500 | 0 | 0 | 294250 | 0 | 0 | 0 | 0 | 0 | 0 | 0 | 0 | 871010 | 1386200 |
| LFKTALQEE | IQGAP3; IQGAP | Q86VI3; P46 | IQGA3_HUMAI | Unmodified | 0 | 0 | 0 | 0 | 0 | 0 | 0 | 0 | 0 | 0 | 1157900 | 1321900 | 0 | 0 | 0 | 0 | 0 | 0 |
| LFLAGYELT | VPS26B | Q4G0F5 | VP26B_HUMAI | Oxidation (M) | 0 | 0 | 0 | 0 | 0 | 0 | 0 | 0 | 140620 | 291560 | 0 | 704950 | 0 | 0 | 0 | 0 | 0 | 0 |
| LFLAPLPSS | AGAP3; AGAP3 | Q96P47; H0' | AGAP3_HUMAI | Unmodified | 0 | 0 | 0 | 0 | 0 | 0 | 0 | 0 | 109000 | 0 | 0 | 215690 | 0 | 0 | 0 | 0 | 0 | 0 |
| LFLDDRDLI | SIGIRR; SIGIRR; | Q6IA17; C9JI | SIGIR_HUMAN | Unmodified | 0 | 0 | 1101100 | 3422200 | 0 | 0 | 0 | 0 | 0 | 358600 | 0 | 3441300 | 0 | 0 | 6813000 | 2651600 | 0 | 0 |
| LFLDRDLM' | FECH; FECH; FI | P22830; K7E | HEMH_HUMAI | Oxidation (M) | 0 | 0 | 0 | 0 | 0 | 0 | 0 | 0 | 438680 | 0 | 65771 | 0 | 0 | 0 | 0 | 0 | 0 | 0 |
| LFLDRDLM' | FECH; FECH; FI | P22830; K7E | HEMH_HUMAI | Unmodified | 0 | 0 | 0 | 0 | 0 | 0 | 0 | 0 | 626370 | 0 | 0 | 0 | 0 | 0 | 0 | 0 | 0 | 0 |
| LFLEGEK | SYT4 | Q9H2B2 | SYT4_HUMAN S | Unmodified | 94350 | 0 | 2546100 | 3246000 | 0 | 0 | 355470 | 49065 | 181240 | 85178 | 160620 | 0 | 293450 | 0 | 6423400 | 4479400 | 50389 | 0 |
| LFLEGEKES | SYT4 | Q9H2B2 | SYT4_HUMAN S | Unmodified | 0 | 0 | 0 | 458790 | 0 | 0 | 656690 | 0 | 0 | 0 | 112080 | 366310 | 0 | 0 | 0 | 0 | 0 | 0 |
| LFLEQEDEI | CUL9; CUL9; CI | Q8IWT3; E9F | CUL9_HUMAN | Unmodified | 0 | 0 | 0 | 261550 | 0 | 282320 | 0 | 0 | 484060 | 0 | 0 | 880070 | 0 | 608330 | 162110 | 0 | 0 | 344120 |
| LFLFHTSLP | SEC24C; SEC2 | P53992; G5E | SC24C_HUMAI | Unmodified | 0 | 0 | 0 | 0 | 0 | 0 | 0 | 0 | 0 | 0 | 0 | 0 | 0 | 0 | 0 | 0 | 0 | 2257700 |
| LFLIDFGLA | CSNK1A1L; CSI | Q8N752; P4 | KC1AL_HUMAI | Unmodified | 0 | 0 | 0 | 0 | 0 | 0 | 0 | 0 | 295930 | 235450 | 129590 | 955100 | 0 | 0 | 572920 | 279640 | 0 | 0 |
| LFLIDFGLA | CSNK1A1L; CSI | Q8N752; P4 | KC1AL_HUMAI | Unmodified | 625290 | 0 | 398290 | 3861200 | 0 | 0 | 79561 | 0 | 652930 | 173720 | 0 | 1024000 | 0 | 0 | 980920 | 1063300 | 0 | 0 |
| LFLLQQPL/ | RHBDD2; RHBI | Q6NTF9; Q9 | RHBD2_HUMA | Unmodified | 0 | 0 | 36808 | 0 | 0 | 0 | 0 | 0 | 636960 | 151140 | 0 | 255270 | 0 | 0 | 0 | 49201 | 0 | 0 |
| LFLLTLIHTI | PLXNB3; PLXNE | Q9ULL4; H7I | PLXB3_HUMAI | Unmodified | 0 | 0 | 0 | 0 | 0 | 0 | 0 | 0 | 0 | 0 | 0 | 0 | 0 | 0 | 91729 | 0 | 0 | 0 |
| LFLNETQT( | IARS1 | P41252 | SYIC_HUMAN I | Unmodified | 0 | 0 | 0 | 723080 | 2626000 | 0 | 2939700 | 231270 | 1373000 | 1243000 | 3389800 | 8615600 | 2983200 | 724480 | 913980 | 566870 | 0 | 1816400 |
| LFLPLILK | GXYLT1 | Q4G148 | GXLT1_HUMAI | Unmodified | 0 | 0 | 35218 | 0 | 0 | 0 | 0 | 0 | 1880700 | 700020 | 210870 | 322200 | 0 | 0 | 0 | 0 | 0 | 0 |
| LFLQFTTGT | UBE3A; UBE3A; | Q05086; A0/ | UBE3A_HUMAI | Unmodified | 0 | 129970 | 0 | 0 | 0 | 0 | 0 | 0 | 0 | 0 | 0 | 519940 | 0 | 0 | 0 | 182490 | 0 | 0 |
| LFLQNNLIF | RTN4RL2; RTN | Q86UN3; G3 | R4RL2_HUMAI | Unmodified | 84083 | 0 | 0 | 0 | 0 | 0 | 0 | 74584 | 1810400 | 59847 | 0 | 0 | 0 | 0 | 142350 | 95085 | 0 | 0 |
| LFLSYDYA\ | PCYOX1 | Q9UHG3 | PCYOX_HUMAI | Unmodified | 0 | 0 | 0 | 0 | 0 | 0 | 0 | 0 | 0 | 2724500 | 0 | 0 | 0 | 0 | 0 | 0 | 187410 | 338690 |
| LFLTDFFR | NDST2 | P52849 | NDST2_HUMAI | Unmodified | 0 | 0 | 0 | 0 | 0 | 0 | 0 | 0 | 4386800 | 171200 | 46241 | 247580 | 0 | 0 | 0 | 0 | 0 | 0 |
| LFLVQLQEI | NUDT21; NUDT | O43809; H3I | CPSF5_HUMAI | Unmodified | 12496000 | 1781000 | 1289200 | 1015700 | 0 | 0 | 136030 | 222010 | 0 | 306690 | 378550 | 6218300 | 288280 | 0 | 625850 | 1202000 | 0 | 0 |
| LFLYDLPEC | CDC42BPB; CD | Q9Y5S2; H0' | MRCKB_HUMA | Unmodified | 0 | 0 | 0 | 0 | 0 | 0 | 0 | 0 | 1130700 | 0 | 0 | 847390 | 0 | 0 | 362410 | 324600 | 322160 | 299220 |
| LFMAQALQ | EIF3H; EIF3H; E | O15372; B3I | EIF3H_HUMAN | Oxidation (M) | 339100 | 0 | 0 | 0 | 0 | 0 | 0 | 275040 | 0 | 0 | 0 | 2700400 | 0 | 1993400 | 0 | 0 | 541730 | 0 |
| LFMAQALQ | EIF3H; EIF3H; E | O15372; B3I | EIF3H_HUMAN | Unmodified | 0 | 0 | 0 | 0 | 1915500 | 0 | 0 | 166760 | 0 | 0 | 0 | 1319300 | 0 | 0 | 0 | 0 | 0 | 0 |
| LFMILWLK | CLIC5; CLIC6; ( | Q9NZA1; Q9 | CLIC5_HUMAN | Oxidation (M) | 63493 | 0 | 13207 | 0 | 2507200 | 41588 | 260110 | 0 | 589280 | 458200 | 225620 | 346420 | 960350 | 0 | 532100 | 95740 | 564520 | 80347 |
| LFMVLWLK | CLIC1 | O00299 | CLIC1_HUMAN | Oxidation (M) | 377730 | 312370 | 250370 | 207840 | 4114700 | 854750 | 457500 | 0 | 2987400 | 1066700 | 632510 | 1527500 | 4130400 | 2987900 | 1145700 | 498220 | 1843800 | 1959400 |
| LFNDYGG( | ANPEP | P15144 | AMPN_HUMAN | Unmodified | 0 | 0 | 0 | 0 | 0 | 0 | 0 | 0 | 0 | 0 | 0 | 0 | 0 | 0 | 0 | 0 | 522350 | 338390 |
| LFNESHGII | NUP85; NUP85 | Q9BW27; J3I | NUP85_HUMAI | Unmodified | 0 | 0 | 0 | 0 | 0 | 0 | 83916 | 0 | 0 | 0 | 0 | 244120 | 0 | 0 | 0 | 0 | 0 | 0 |
| LFNHANQE | PKP3 | Q9Y446 | PKP3_HUMAN | Unmodified | 1226800 | 1293000 | 2606400 | 5383900 | 1377500 | 568260 | 0 | 0 | 2730600 | 275540 | 2578600 | 6995700 | 2206600 | 810550 | 5725500 | 4918100 | 765800 | 1180800 |
| LFNHANQE | PKP3 | Q9Y446 | PKP3_HUMAN | Unmodified | 0 | 0 | 0 | 0 | 1219300 | 333090 | 0 | 0 | 0 | 0 | 81313 | 0 | 3598400 | 529880 | 0 | 0 | 925550 | 798870 |
| LFNHLSAV! | CAP1; CAP1; C/ | Q01518; Q5' | CAP1_HUMAN | Oxidation (M) | 0 | 0 | 0 | 0 | 5947600 | 0 | 2309900 | 0 | 0 | 3214800 | 284300 | 917950 | 0 | 0 | 0 | 0 | 5299300 | 0 |
| LFNLREGD | AHNAK2 | Q8IVF2 | AHNK2_HUMAI | Unmodified | 0 | 0 | 0 | 0 | 2073200 | 0 | 0 | 0 | 0 | 0 | 0 | 0 | 0 | 0 | 0 | 0 | 562280 | 3397500 |
| LFNLSKEDI | RPS6; RPS6 | P62753; A2A | RS6_HUMAN 4 | Unmodified | 4583700 | 3389300 | 1192500 | 2891800 | 1248000 | 256090 | 55939 | 90420 | 1984000 | 0 | 726210 | 10037000 | 2723400 | 259080 | 2006200 | 2356200 | 73178 | 1052400 |
| LFNTAVCE! | NAA15 | Q9BXJ9 | NAA15_HUMAI | Unmodified | 76563 | 0 | 223030 | 0 | 0 | 0 | 0 | 0 | 0 | 0 | 0 | 1262000 | 0 | 0 | 0 | 0 | 0 | 0 |
| LFPAPSEK | SCG3 | Q8WXD2 | SCG3_HUMAN | Unmodified | 0 | 0 | 0 | 0 | 0 | 0 | 0 | 0 | 5908700 | 1824800 | 0 | 0 | 0 | 0 | 0 | 0 | 0 | 0 |
| LFPDTPLAI | ILF3 | Q12906 | ILF3_HUMAN Ir | Unmodified | 1651000 | 1241800 | 1278700 | 482190 | 0 | 0 | 0 | 0 | 0 | 0 | 0 | 1657900 | 198970 | 0 | 317170 | 619720 | 0 | 0 |
| LFPDTPLAI | ILF3 | Q12906 | ILF3_HUMAN Ir | Unmodified | 0 | 0 | 0 | 0 | 0 | 0 | 0 | 0 | 0 | 0 | 0 | 0 | 0 | 0 | 0 | 0 | 0 | 0 |
| LFPGFEIET' | ASNS; ASNS | P08243; F8V | ASNS_HUMAN | Unmodified | 0 | 0 | 1306900 | 933630 | 0 | 568470 | 0 | 536000 | 1669900 | 687200 | 4733800 | 21570000 | 1495300 | 0 | 2154100 | 1736400 | 0 | 535570 |
| LFPHEVAD' | TBCD; TBCD | Q9BTW9; J3I | TBCD_HUMAN | Unmodified | 30092 | 29741 | 0 | 0 | 0 | 0 | 0 | 0 | 0 | 0 | 0 | 366330 | 0 | 0 | 37274 | 0 | 0 | 0 |
| LFPLIQAMI | PABPC1; PABP( | P11940; Q9I | PABP1_HUMAI | Oxidation (M) | 6569100 | 12744000 | 1498800 | 0 | 0 | 0 | 0 | 0 | 0 | 0 | 0 | 0 | 0 | 0 | 4222000 | 3768700 | 0 | 0 |
| LFPLIQAMI | PABPC1; PABP( | P11940; Q9I | PABP1_HUMAI | Unmodified | 441560 | 1011600 | 120700 | 0 | 0 | 0 | 0 | 33312 | 0 | 0 | 0 | 0 | 0 | 0 | 255520 | 155770 | 0 | 0 |
| LFPLIQTMI | PABPC4; PABP( | Q13310; B1/ | PABP4_HUMAI | Unmodified | 1520700 | 1332800 | 791380 | 410370 | 0 | 0 | 0 | 0 | 508760 | 0 | 0 | 482240 | 0 | 0 | 600670 | 474280 | 0 | 0 |
| LFPNCVPS | ITGB5; ITGB5; I | P18084; H7( | ITB5_HUMAN I | Unmodified | 0 | 0 | 172400 | 1485100 | 0 | 0 | 0 | 0 | 592780 | 0 | 190820 | 1448200 | 0 | 0 | 0 | 366170 | 0 | 183270 |
| LFPNSLDQ | CALR; CALR; C/ | P27797; A0A | CALR_HUMAN | 2 Oxidation (M) | 0 | 0 | 0 | 0 | 827260 | 0 | 18110000 | 1782900 | 69583000 | 95102000 | 14179000 | 1082500 | 0 | 0 | 0 | 0 | 805590 | 1103900 |
| LFPNSLDQ | CALR; CALR; C/ | P27797; A0A | CALR_HUMAN | Oxidation (M) | 0 | 0 | 0 | 0 | 0 | 0 | 912660 | 0 | 56534000 | 16687000 | 0 | 0 | 0 | 0 | 0 | 0 | 0 | 845120 |
| LFPNSLDQ | CALR; CALR; C/ | P27797; A0A | CALR_HUMAN | Unmodified | 0 | 0 | 0 | 0 | 0 | 0 | 0 | 0 | 15130000 | 732470 | 0 | 0 | 0 | 0 | 0 | 0 | 0 | 0 |
| LFPNSLDQ | CALR; CALR; C/ | P27797; A0A | CALR_HUMAN | 2 Oxidation (M) | 0 | 0 | 0 | 0 | 0 | 0 | 0 | 499260 | 17136000 | 33483000 | 11556000 | 0 | 0 | 0 | 0 | 0 | 0 | 0 |
| LFPNSLDQ | CALR; CALR; C/ | P27797; A0A | CALR_HUMAN | Oxidation (M) | 0 | 0 | 0 | 0 | 0 | 0 | 0 | 0 | 30101000 | 3759800 | 360350 | 0 | 0 | 0 | 0 | 0 | 0 | 0 |
| LFPNSLDQ | CALR; CALR; C/ | P27797; A0A | CALR_HUMAN | Unmodified | 0 | 0 | 0 | 0 | 0 | 0 | 0 | 0 | 6880100 | 1249800 | 0 | 0 | 0 | 0 | 0 | 0 | 0 | 0 |
| LFPPWIKP/ | PRPF8; PRPF8 | Q6P2Q9; I3L | PRP8_HUMAN | Unmodified | 0 | 0 | 0 | 0 | 0 | 0 | 0 | 0 | 0 | 0 | 0 | 563350 | 0 | 0 | 0 | 0 | 0 | 0 |
| LFPYALAFI | NAA15; NAA15 | Q9BXJ9; A0A | NAA15_HUMAI | 2 Oxidation (M) | 195720 | 419460 | 0 | 0 | 0 | 0 | 0 | 11033 | 0 | 0 | 136090 | 431820 | 0 | 0 | 0 | 50471 | 0 | 0 |
| LFQECCPH | HNRNPK; HNRI | P61978; Q5I | HNRPK_HUMA | Unmodified | 176760 | 0 | 0 | 0 | 0 | 0 | 264480 | 168670 | 157920 | 133700 | 150730 | 3793700 | 0 | 0 | 0 | 0 | 0 | 0 |
| LFQECCPH | HNRNPK; HNRI | P61978; Q5I | HNRPK_HUMA | Unmodified | 297650 | 0 | 0 | 0 | 0 | 0 | 64425 | 0 | 0 | 0 | 0 | 6605800 | 0 | 0 | 0 | 0 | 0 | 0 |
| LFQENSVL! | ATP6AP2; ATP6. | O75787; A0/ | RENR_HUMAN | Unmodified | 322320 | 202140 | 0 | 646910 | 51522000 | 5571800 | 0 | 0 | 5074000 | 410850 | 539210 | 330420 | 19847 | 784780 | 0 | 0 | 11741000 | 32826000 |
| LFQEWEAI | TCF25; TCF25; | Q9BQ70; H3 | TCF25_HUMAI | Unmodified | 111730 | 0 | 67980 | 0 | 0 | 0 | 0 | 0 | 0 | 0 | 0 | 0 | 0 | 0 | 677480 | 426820 | 0 | 0 |
| LFQIHGND | AVIL | O75366 | AVIL_HUMAN A | Unmodified | 0 | 0 | 0 | 0 | 0 | 51906 | 0 | 0 | 0 | 0 | 690350 | 5807400 | 0 | 819740 | 0 | 0 | 68127 | 0 |
| LFQLMVEH | LMAN2; LMAN2 | Q12907; D6I | LMAN2_HUMAI | Oxidation (M) | 651850 | 0 | 406100 | 0 | 0 | 0 | 0 | 0 | 33586000 | 4701900 | 2472900 | 1221800 | 0 | 0 | 1338400 | 112960 | 0 | 0 |
| LFQLMVEH | LMAN2; LMAN2 | Q12907; D6I | LMAN2_HUMAI | Unmodified | 0 | 0 | 0 | 0 | 0 | 0 | 0 | 0 | 25856000 | 661850 | 0 | 0 | 0 | 0 | 0 | 0 | 0 | 0 |
| LFQLMVEH | LMAN2; LMAN2 | Q12907; D6I | LMAN2_HUMAI | Oxidation (M) | 0 | 0 | 0 | 0 | 0 | 0 | 0 | 0 | 1850800 | 1809100 | 0 | 0 | 0 | 0 | 0 | 0 | 0 | 0 |
| LFQPSIQLA | GGT1; GGT2; G | P19440; P36 | GGT1_HUMAN | Unmodified | 5375600 | 3976200 | 11065000 | 15188000 | 65193 | 89878 | 43283 | 0 | 12328000 | 4945900 | 743130 | 5400200 | 1185000 | 391910 | 15378000 | 9557000 | 0 | 87157 |
| LFQPSIQLA | GGT1; GGT2; G | P19440; P36 | GGT1_HUMAN | Unmodified | 0 | 0 | 0 | 0 | 0 | 0 | 0 | 0 | 0 | 1587200 | 0 | 0 | 621000 | 0 | 0 | 0 | 0 | 0 |
| LFQTNTELI | ATP11C; ATP11 | Q8NB49; A0. | AT11C_HUMAI | Unmodified | 0 | 0 | 182060 | 0 | 0 | 0 | 0 | 0 | 0 | 0 | 368440 | 2018200 | 0 | 0 | 717980 | 357310 | 117370 | 0 |
| LFQVEYAIE | PSMA5 | P28066 | PSA5_HUMAN | Unmodified | 0 | 0 | 239730 | 0 | 0 | 0 | 0 | 0 | 2292400 | 1164900 | 2385400 | 8239700 | 2700100 | 0 | 831630 | 547580 | 0 | 0 |
| LFQVHTFEI | TMCO3 | Q6UWJ1 | TMCO3_HUMA | Unmodified | 25890 | 0 | 0 | 0 | 0 | 0 | 149120 | 0 | 23291000 | 2782000 | 598160 | 0 | 0 | 0 | 0 | 0 | 0 | 0 |
| LFQVQGTG | VIL1; VIL1; VIL1 | P09327; H7( | VILI_HUMAN Vi | Unmodified | 497250 | 215840 | 5872000 | 3178500 | 0 | 0 | 0 | 0 | 1536300 | 535180 | 2329900 | 11368000 | 0 | 0 | 0 | 3760100 | 0 | 0 |
| LFREEFNAI | PTPRA | P18433 | PTPRA_HUMAI | Unmodified | 0 | 0 | 0 | 0 | 0 | 0 | 0 | 0 | 0 | 0 | 0 | 0 | 0 | 0 | 0 | 0 | 0 | 5563300 |
| LFRFASENI | RANBP1; RANB | P43487; C9J | RANG_HUMAN | Unmodified | 0 | 0 | 0 | 0 | 22490000 | 0 | 1955500 | 325570 | 0 | 1120300 | 0 | 0 | 6000000 | 3937800 | 0 | 0 | 1665300 | 8178700 |
| LFRFASENI | RANBP1; RANB | P43487; C9J | RANG_HUMAN | Unmodified | 0 | 0 | 0 | 0 | 1626600 | 0 | 246120 | 0 | 0 | 0 | 0 | 0 | 0 | 0 | 0 | 0 | 0 | 461350 |
| LFRPVAAD( | ATG5; ATG5; AT | Q9H1Y0; A0, | ATG5_HUMAN | Unmodified | 40524 | 27026 | 0 | 148000 | 0 | 0 | 626650 | 0 | 926490 | 742470 | 349560 | 2669800 | 782230 | 392300 | 179850 | 98898 | 707010 | 0 |
| LFSELGSR | FAM3A; FAM3A; | P98173; D3I | FAM3A_HUMAI | Unmodified | 367090 | 0 | 0 | 0 | 0 | 0 | 0 | 0 | 10209000 | 487640 | 0 | 3320200 | 0 | 0 | 0 | 0 | 0 | 0 |
| LFSELGSRI | FAM3A; FAM3A; | P98173; D3I | FAM3A_HUMAI | Unmodified | 0 | 0 | 0 | 0 | 0 | 0 | 427510 | 0 | 0 | 655460 | 388560 | 0 | 76208 | 0 | 137700 | 98599 | 0 | 0 |
| LFSEVENK | IDE; IDE; IDE; ID | P14735; A0A | IDE_HUMAN In | Unmodified | 0 | 0 | 0 | 0 | 498780 | 0 | 0 | 0 | 232950 | 0 | 216860 | 3556800 | 0 | 82484 | 0 | 0 | 107360 | 48159 |
| LFSHEGSS | MELTF | P08582 | TRFM_HUMAN | Unmodified | 0 | 0 | 0 | 0 | 1306600 | 696810 | 0 | 0 | 6680900 | 252420 | 46458 | 0 | 0 | 0 | 0 | 88044 | 441150 | 0 |
| LFSHEGSS | MELTF | P08582 | TRFM_HUMAN | Oxidation (M) | 0 | 0 | 0 | 0 | 475510 | 0 | 0 | 0 | 2599400 | 0 | 0 | 1035700 | 0 | 0 | 0 | 0 | 0 | 0 |
| LFSLTPAEV | SNX17; SNX17 | Q15036; F8V | SNX17_HUMAI | Unmodified | 96867 | 110320 | 0 | 197730 | 4532500 | 928680 | 920670 | 0 | 0 | 0 | 0 | 0 | 0 | 0 | 0 | 0 | 0 | 0 |
| LFSQDHLV | NUDT9; NUDT9 | Q9BW91; H7 | NUDT9_HUMAI | Unmodified | 0 | 0 | 0 | 159400 | 1111800 | 0 | 0 | 91283 | 1308500 | 795650 | 233840 | 1026500 | 448220 | 224430 | 358780 | 0 | 199730 | 0 |
| LFSQETVMI | APMAP; APMAP | Q9HDC9; H( | APMAP_HUMAI | Unmodified | 0 | 0 | 0 | 0 | 0 | 0 | 0 | 0 | 2584300 | 1443600 | 0 | 245980 | 0 | 0 | 0 | 0 | 0 | 0 |
| LFSQETVMI | APMAP; APMAP | Q9HDC9; H( | APMAP_HUMAI | Oxidation (M) | 0 | 0 | 0 | 0 | 0 | 0 | 0 | 0 | 934750 | 492140 | 0 | 0 | 0 | 0 | 0 | 0 | 0 | 0 |
| LFSQGQD\ | AFDN; AFDN; A | P55196; H0\ | AFAD_HUMAN | Unmodified | 0 | 0 | 148690 | 41894 | 0 | 0 | 0 | 0 | 0 | 0 | 1238000 | 535150 | 0 | 945610 | 0 | 0 | 0 | 72493 |
| LFSQGQD\ | AFDN; AFDN; A | P55196; H0\ | AFAD_HUMAN | Unmodified | 0 | 0 | 0 | 0 | 599550 | 0 | 0 | 152850 | 0 | 120470 | 0 | 0 | 1717300 | 1217200 | 0 | 0 | 0 | 0 |

| | | | | | | | | | | | | | | | | | | | | | | |
|---|---|---|---|---|---|---|---|---|---|---|---|---|---|---|---|---|---|---|---|---|---|---|
| LFSSFLK | EPB41; EPB41; | P11171; A0A | EPB41_HUMAN | Unmodified | 86386 | 0 | 608890 | 0 | 0 | 0 | 436670 | 158010 | 654880 | 485470 | 392060 | 13671000 | 608690 | 465340 | 0 | 0 | 464530 | 1525900 |
| LFSVPDFV( | MVP | Q14764 | MVP_HUMAN N | Unmodified | 0 | 0 | 0 | 0 | 1191400 | 0 | 0 | 0 | 1814500 | 828870 | 0 | 22760000 | 0 | 0 | 0 | 0 | 0 | 760330 |
| LFSWLVNR | MYO1B; MYO1E | O43795; E9F | MYO1B_HUMA | Unmodified | 0 | 0 | 0 | 0 | 0 | 0 | 0 | 0 | 0 | 0 | 145860 | 10497000 | 0 | 0 | 0 | 0 | 0 | 0 |
| LFTAESLIG | ARCN1; ARCN1 | P48444; B0Y | COPD_HUMAN | Unmodified | 0 | 0 | 0 | 0 | 0 | 0 | 0 | 0 | 158620 | 0 | 642430 | 2350600 | 0 | 0 | 0 | 0 | 637380 | 0 |
| LFTAESLIG | ARCN1; ARCN1 | P48444; B0Y | COPD_HUMAN | Unmodified | 0 | 0 | 0 | 0 | 1077300 | 0 | 0 | 136160 | 344910 | 357040 | 0 | 1049000 | 1297600 | 0 | 0 | 0 | 612370 | 1283900 |
| LFTAHNNN | DHX9 | Q08211 | DHX9_HUMAN | Unmodified | 0 | 0 | 0 | 0 | 0 | 0 | 348080 | 19603 | 0 | 0 | 62984 | 520290 | 0 | 0 | 0 | 0 | 0 | 121090 |
| LFTAIGYSE | VPS13C | Q709C8 | VP13C_HUMAN | Unmodified | 0 | 0 | 0 | 0 | 0 | 0 | 0 | 0 | 0 | 0 | 0 | 2577700 | 0 | 0 | 0 | 0 | 0 | 0 |
| LFTDLFDR | CSNK1G2; CSN | P78368; Q9Y | KC1G2_HUMAI | Unmodified | 0 | 33318 | 353800 | 0 | 11817000 | 360450 | 0 | 0 | 451410 | 0 | 0 | 971230 | 33122 | 0 | 815570 | 439590 | 3337500 | 13524000 |
| LFTDLREIG | TAOK1; TAOK1 | Q7L7X3; J3Q | TAOK1_HUMAN | Unmodified | 0 | 0 | 0 | 212230 | 0 | 0 | 0 | 0 | 568760 | 123430 | 102660 | 764430 | 0 | 0 | 0 | 0 | 0 | 0 |
| LFTEEELAR | NENF | Q9UMX5 | NENF_HUMAN | Unmodified | 78926 | 0 | 0 | 219930 | 444680 | 0 | 0 | 779750 | 9683000 | 0 | 0 | 1281400 | 0 | 0 | 184650 | 0 | 0 | 0 |
| LFTEEELAR | NENF | Q9UMX5 | NENF_HUMAN | Unmodified | 0 | 1071600 | 0 | 0 | 0 | 0 | 0 | 0 | 15283000 | 1489900 | 0 | 0 | 0 | 0 | 0 | 0 | 0 | 0 |
| LFTIRGTPQ | FUBP1; FUBP1 | Q96AE4; E9F | FUBP1_HUMAN | Unmodified | 0 | 0 | 0 | 0 | 0 | 0 | 0 | 0 | 0 | 324960 | 175370 | 395770 | 0 | 0 | 0 | 0 | 0 | 0 |
| LFTLTALR | QARS1; QARS1; | P47897; H7C | SYQ_HUMAN G | Unmodified | 0 | 0 | 95426 | 0 | 0 | 0 | 0 | 0 | 0 | 0 | 0 | 5707300 | 0 | 0 | 195480 | 200060 | 0 | 0 |
| LFTPDVIR | ME2; ME2; ME2 | P23368; A0A | MAOM_HUMAN | Unmodified | 17929 | 0 | 0 | 0 | 259800 | 0 | 41947 | 0 | 3255800 | 129140 | 0 | 209850 | 48775 | 209890 | 0 | 11082 | 57013 | 0 |
| LFTQILR | RRAS | P10301 | RRAS_HUMAN | Unmodified | 544940 | 533650 | 1716500 | 575820 | 0 | 0 | 0 | 0 | 0 | 0 | 0 | 740740 | 40219 | 0 | 3222800 | 2335700 | 148690 | 307150 |
| LFTSTGLK | AKAP12 | Q02952 | AKA12_HUMAN | Unmodified | 0 | 0 | 0 | 227270 | 1186000 | 66134 | 144310 | 96074 | 984550 | 271490 | 0 | 244520 | 0 | 0 | 0 | 1091700 | 1939900 | 2125300 |
| LFVDNGAD | TRPM5; TRPM5; | Q9NZQ8; A0 | TRPM5_HUMAI | Unmodified | 0 | 0 | 0 | 0 | 0 | 0 | 0 | 0 | 0 | 0 | 0 | 307730 | 0 | 0 | 0 | 0 | 0 | 0 |
| LFVEESIHD | ALDH1L2; ALDH | Q3SY69; A0A | AL1L2_HUMAN | Unmodified | 0 | 0 | 0 | 0 | 0 | 0 | 0 | 0 | 344020 | 19227 | 0 | 0 | 126300 | 0 | 0 | 0 | 0 | 573390 |
| LFVESYELII | MUC5B | Q9HC84 | MUC5B_HUMA | Unmodified | 0 | 0 | 0 | 0 | 0 | 0 | 19978 | 0 | 0 | 0 | 0 | 0 | 0 | 0 | 0 | 0 | 9507.1 | 495120 |
| LFVGEPVW | ADI1; ADI1; ADI | Q9BV57; H7 | MTND_HUMAN | Unmodified | 0 | 0 | 0 | 0 | 0 | 0 | 0 | 0 | 0 | 0 | 1659800 | 5690100 | 0 | 0 | 0 | 0 | 0 | 0 |
| LFVGEPVW | ADI1; ADI1; ADI | Q9BV57; H7 | MTND_HUMAN | Unmodified | 0 | 0 | 277500 | 0 | 5076400 | 1079100 | 0 | 0 | 0 | 774040 | 575600 | 0 | 3481600 | 1346600 | 0 | 0 | 2819600 | 1409300 |
| LFVGGIKED | HNRNPA2B1; H | P22626; A0A | ROA2_HUMAN | Unmodified | 0 | 0 | 0 | 0 | 0 | 0 | 0 | 0 | 0 | 0 | 2437600 | 8811500 | 0 | 0 | 420890 | 0 | 0 | 0 |
| LFVGGIKED | HNRNPA2B1; H | P22626; A0A | ROA2_HUMAN | Unmodified | 0 | 0 | 0 | 83391 | 0 | 3009100 | 189300 | 0 | 0 | 0 | 0 | 1937300 | 0 | 0 | 0 | 0 | 0 | 0 |
| LFVGGLDW | DAZAP1; DAZAF | Q96EP5; K7E | DAZP1_HUMAN | Unmodified | 396840 | 610220 | 0 | 166700 | 0 | 0 | 0 | 0 | 1339400 | 89253 | 1205400 | 8582300 | 145070 | 0 | 0 | 104820 | 0 | 45874 |
| LFVGGLSFI | CIRBP; CIRBP; ( | Q14011; K7E | CIRBP_HUMAN | Unmodified | 0 | 478210 | 0 | 203430 | 0 | 0 | 0 | 0 | 0 | 0 | 0 | 0 | 432350 | 0 | 1730500 | 0 | 0 | 0 |
| LFVGNLPAI | SFPQ | P23246 | SFPQ_HUMAN | Unmodified | 0 | 115790 | 0 | 0 | 0 | 0 | 0 | 24753 | 501990 | 162770 | 1051100 | 4235800 | 0 | 0 | 0 | 0 | 0 | 0 |
| LFVGNLPAI | SFPQ | P23246 | SFPQ_HUMAN | Unmodified | 0 | 0 | 0 | 0 | 168280 | 0 | 0 | 0 | 3724700 | 0 | 58650 | 11469000 | 0 | 0 | 0 | 0 | 0 | 0 |
| LFVGNLPAI | SFPQ | P23246 | SFPQ_HUMAN | Unmodified | 0 | 0 | 0 | 0 | 0 | 0 | 649690 | 0 | 0 | 977020 | 123780 | 0 | 1115500 | 197980 | 0 | 0 | 0 | 0 |
| LFVGNLPPI | NONO; NONO; | Q15233; C9I | NONO_HUMAN | Oxidation (M) | 0 | 0 | 0 | 0 | 797020 | 0 | 0 | 0 | 32209 | 0 | 90760 | 827310 | 0 | 0 | 0 | 0 | 101520 | 53714 |
| LFVGNLPPI | NONO; NONO; | Q15233; C9I | NONO_HUMAN | Oxidation (M) | 0 | 0 | 0 | 0 | 0 | 0 | 0 | 0 | 0 | 0 | 350270 | 1521900 | 0 | 0 | 0 | 0 | 0 | 0 |
| LFVGNLPPI | NONO; NONO; | Q15233; C9I | NONO_HUMAN | Unmodified | 0 | 0 | 0 | 0 | 763140 | 0 | 1329200 | 0 | 821180 | 1349400 | 78739 | 0 | 740530 | 784750 | 0 | 0 | 0 | 1596000 |
| LFVGNLPPI | NONO; NONO; | Q15233; C9I | NONO_HUMAN | Oxidation (M) | 0 | 0 | 0 | 0 | 50164 | 0 | 573620 | 0 | 0 | 333220 | 0 | 0 | 361980 | 0 | 0 | 0 | 0 | 0 |
| LFVGNLPTI | PSPC1; PSPC1 | Q8WXF1; X6 | PSPC1_HUMAN | Unmodified | 73599 | 0 | 0 | 0 | 0 | 0 | 0 | 0 | 264530 | 0 | 0 | 987360 | 0 | 0 | 0 | 0 | 0 | 0 |
| LFVGNLPTI | PSPC1; PSPC1 | Q8WXF1; X6 | PSPC1_HUMAN | Unmodified | 0 | 0 | 0 | 0 | 0 | 0 | 0 | 0 | 0 | 188180 | 713860 | 4797000 | 23585 | 0 | 0 | 0 | 0 | 0 |
| LFVGTPIDC | PLXNA1 | Q9UIW2 | PLXA1_HUMAN | Unmodified | 0 | 0 | 0 | 0 | 2368700 | 183680 | 0 | 0 | 843860 | 0 | 0 | 0 | 0 | 0 | 0 | 0 | 1562200 | 1620100 |
| LFVLFGAEI | TALDO1; TALDO | P37837; F2Z | TALDO_HUMAN | Unmodified | 0 | 0 | 0 | 0 | 0 | 0 | 0 | 0 | 0 | 0 | 380970 | 531610 | 0 | 0 | 0 | 0 | 0 | 0 |
| LFVLLPEQS | USP47 | Q96K76 | UBP47_HUMAN | Unmodified | 18474 | 0 | 0 | 0 | 29796 | 0 | 0 | 0 | 0 | 50669 | 85224 | 270500 | 61415 | 19926 | 0 | 45985 | 0 | 31555 |
| LFVLTTK | NUDCD1 | Q96RS6 | NUDC1_HUMA | Unmodified | 0 | 32185 | 0 | 84595 | 0 | 0 | 0 | 0 | 401640 | 0 | 449130 | 1389800 | 0 | 0 | 58883 | 144300 | 0 | 0 |
| LFVNAQR | RAB3GAP1 | Q15042 | RB3GP_HUMAI | Unmodified | 0 | 0 | 0 | 0 | 321720 | 0 | 0 | 0 | 0 | 51979 | 101470 | 1630200 | 146040 | 0 | 0 | 0 | 0 | 0 |
| LFVPALDKI | DUSP12; DUSP | Q9UNI6; V9C | DUS12_HUMAI | Unmodified | 0 | 0 | 0 | 0 | 0 | 0 | 0 | 0 | 0 | 0 | 0 | 1718600 | 0 | 0 | 206140 | 89506 | 0 | 0 |
| LFVPNTQS | OCRL; OCRL | Q01968; A0A | OCRL_HUMAN | Unmodified | 176490 | 107580 | 117340 | 133010 | 597450 | 104640 | 225030 | 80618 | 125840 | 163870 | 0 | 0 | 296520 | 249520 | 0 | 0 | 523450 | 606880 |
| LFVQDEK | DPP3; DPP3; DI | Q9NY33; G3 | DPP3_HUMAN | Unmodified | 74584 | 0 | 0 | 0 | 642280 | 0 | 213190 | 0 | 1976900 | 841080 | 818350 | 3780000 | 0 | 0 | 0 | 271690 | 546120 | 1312500 |
| LFVQDLATI | KIF5B | P33176 | KINH_HUMAN | Unmodified | 0 | 0 | 0 | 0 | 1555900 | 0 | 601700 | 0 | 2238000 | 1434100 | 0 | 7223000 | 1340300 | 838170 | 0 | 0 | 45859 | 1073800 |
| LFVQDLATI | KIF5B | P33176 | KINH_HUMAN | Unmodified | 0 | 0 | 0 | 0 | 5204200 | 0 | 3298100 | 1523900 | 407580 | 1521200 | 0 | 0 | 2434300 | 0 | 0 | 0 | 2839400 | 3379300 |
| LFVQDLATI | KIF5B | P33176 | KINH_HUMAN | Unmodified | 0 | 0 | 0 | 0 | 2646900 | 0 | 1079900 | 158450 | 0 | 0 | 0 | 0 | 991540 | 0 | 0 | 0 | 1680900 | 636050 |
| LFVQDLTTI | KIF5C; KIF5C; K | O60282; A0A | KIF5C_HUMAN | Unmodified | 0 | 0 | 0 | 0 | 0 | 0 | 0 | 0 | 0 | 0 | 0 | 0 | 537440 | 0 | 0 | 0 | 0 | 0 |
| LFVSDGVP | MARS1; MARS1 | P56192; F8V | SYMC_HUMAN | Unmodified | 0 | 0 | 0 | 0 | 0 | 0 | 67516 | 0 | 190180 | 0 | 197510 | 2595100 | 68030 | 0 | 146210 | 136370 | 0 | 0 |
| LFVSGACD | GNB1; GNB1 | P62873; B3K | GBB1_HUMAN | Unmodified | 8009600 | 2665900 | 0 | 3682800 | 7423300 | 0 | 2103700 | 365770 | 51591000 | 4001300 | 418020 | 1.43E+08 | 4889700 | 4587700 | 3502100 | 7791400 | 6870200 | 18782000 |
| LFVSGACD | GNB1; GNB1 | P62873; B3K | GBB1_HUMAN | Unmodified | 0 | 0 | 0 | 0 | 582650 | 0 | 0 | 0 | 0 | 0 | 0 | 0 | 0 | 0 | 0 | 0 | 1085300 | 0 |
| LFVTNDAA | CCT8; CCT8 | P50990; H7C | TCPQ_HUMAN | Unmodified | 0 | 352190 | 527990 | 1025600 | 0 | 0 | 0 | 0 | 3726700 | 325060 | 4288500 | 41531000 | 292990 | 0 | 1220500 | 990990 | 73523 | 0 |
| LFVTNDAA | CCT8; CCT8 | P50990; H7C | TCPQ_HUMAN | Unmodified | 1909000 | 1506600 | 0 | 10072000 | 35699000 | 4545700 | 12543000 | 42305 | 12795000 | 14363000 | 7143000 | 20538000 | 71312000 | 38101000 | 7055900 | 2498800 | 24396000 | 19978000 |
| LFVVPADE | PLD3 | Q8IV08 | PLD3_HUMAN | Unmodified | 0 | 0 | 0 | 0 | 0 | 0 | 0 | 0 | 1485200 | 279410 | 529560 | 732450 | 656190 | 0 | 0 | 231710 | 0 | 0 |
| LFVVPADE | PLD3 | Q8IV08 | PLD3_HUMAN | Unmodified | 0 | 0 | 0 | 0 | 537710 | 251310 | 0 | 0 | 1585800 | 795200 | 1414600 | 0 | 0 | 0 | 0 | 0 | 0 | 0 |
| LFVWEILHS | NCBP1; NCBP1 | Q09161; X6F | NCBP1_HUMAI | Unmodified | 0 | 0 | 0 | 0 | 0 | 0 | 0 | 0 | 0 | 0 | 0 | 349600 | 0 | 0 | 0 | 0 | 0 | 0 |
| LFVWEILHS | NCBP1; NCBP1 | Q09161; X6F | NCBP1_HUMAI | Unmodified | 0 | 0 | 0 | 0 | 0 | 0 | 0 | 0 | 0 | 0 | 0 | 781240 | 0 | 0 | 0 | 0 | 0 | 0 |
| LFVYDPNN | PON2; PON2 | Q15165; A0A | PON2_HUMAN | Unmodified | 58578 | 0 | 0 | 0 | 0 | 0 | 0 | 0 | 10375000 | 577530 | 414300 | 70932 | 0 | 0 | 0 | 69772 | 0 | 0 |
| LFYADHPFI | SERPINH1; SER | P50454; E9P | SERPH_HUMAI | Unmodified | 0 | 96726 | 0 | 0 | 0 | 0 | 0 | 0 | 30233000 | 3377700 | 90311 | 0 | 0 | 0 | 0 | 0 | 0 | 0 |
| LFYAPAAG | COL12A1; COL | Q99715; D6I | COCA1_HUMA | Unmodified | 0 | 0 | 0 | 0 | 0 | 0 | 0 | 0 | 96059 | 213760 | 0 | 0 | 0 | 0 | 0 | 0 | 0 | 0 |
| LFYEDGSG | PLXNB2; PLXNE | O15031; A60 | PLXB2_HUMAN | Unmodified | 690370 | 484840 | 1749600 | 1398600 | 1426000 | 684760 | 0 | 0 | 527280 | 0 | 0 | 0 | 812970 | 0 | 2384300 | 1980800 | 146210 | 257940 |
| LFYLALPPT | G6PD; G6PD; G | P11413; E7E | G6PD_HUMAN | Unmodified | 88534 | 0 | 33827 | 167660 | 0 | 104400 | 0 | 0 | 942390 | 0 | 400310 | 540800 | 0 | 0 | 328440 | 169780 | 0 | 0 |
| LFYLDPDA | DOCK9; DOCKS | Q9BZ29; A0A | DOCK9_HUMA | Unmodified | 0 | 203160 | 0 | 0 | 1259400 | 0 | 0 | 25341 | 0 | 0 | 0 | 0 | 643210 | 0 | 0 | 322540 | 0 | 0 |
| LFYRVLTSD | ME1 | P48163 | MAOX_HUMAN | Unmodified | 0 | 0 | 0 | 0 | 324800 | 308170 | 0 | 0 | 0 | 54881 | 0 | 0 | 218900 | 189620 | 0 | 0 | 423450 | 105260 |
| LFYSTFATD | ECHS1 | P30084 | ECHM_HUMAN | Unmodified | 0 | 0 | 52766 | 334970 | 0 | 0 | 0 | 0 | 708260 | 53224 | 0 | 0 | 0 | 0 | 0 | 0 | 0 | 0 |
| LGAAAGFN | TP53I3; TP53I3 | Q53FA7; H7I | QORX_HUMAN | Unmodified | 0 | 0 | 0 | 62264 | 1419400 | 782400 | 0 | 0 | 1126900 | 0 | 192160 | 627690 | 478290 | 0 | 0 | 0 | 248380 | 0 |
| LGAAPEEE | NSFL1C; NSFL1 | Q9UNZ2; F2 | NSF1C_HUMAI | Unmodified | 915660 | 0 | 0 | 0 | 0 | 0 | 0 | 0 | 468860 | 0 | 0 | 1944700 | 910900 | 0 | 0 | 0 | 269790 | 0 |
| LGAASLGA | TLN1 | Q9Y490 | TLN1_HUMAN | Unmodified | 0 | 0 | 0 | 0 | 3915700 | 4725700 | 0 | 814460 | 0 | 665170 | 1041100 | 7758500 | 0 | 0 | 4471500 | 3058300 | 226480 | 0 |
| LGADAVGN | PNP | P00491 | PNPH_HUMAN | Oxidation (M) | 0 | 0 | 0 | 625470 | 1191300 | 0 | 0 | 108410 | 953810 | 573950 | 1600500 | 7003100 | 0 | 253230 | 0 | 469920 | 0 | 693430 |
| LGADAVGN | PNP | P00491 | PNPH_HUMAN | Unmodified | 32337 | 0 | 0 | 0 | 0 | 0 | 36564 | 16453 | 918010 | 210850 | 0 | 980880 | 447210 | 0 | 99145 | 31243 | 118270 | 419920 |
| LGADFHGT | UBAP1 | Q9NZ09 | UBAP1_HUMAI | Unmodified | 315660 | 909160 | 0 | 0 | 0 | 59909 | 0 | 0 | 0 | 0 | 0 | 0 | 0 | 0 | 0 | 0 | 0 | 0 |
| LGAEMESL | PTPRB; PTPRB | P23467; F8V | PTPRB_HUMAN | Unmodified | 266410 | 0 | 0 | 0 | 0 | 0 | 0 | 0 | 0 | 0 | 0 | 0 | 0 | 0 | 0 | 0 | 0 | 0 |
| LGAEPFPLF | TCAF1 | Q9Y4C2 | TCAF1_HUMAN | Unmodified | 398300 | 370950 | 124660 | 84447 | 0 | 0 | 0 | 0 | 0 | 0 | 0 | 500570 | 0 | 0 | 223410 | 311030 | 0 | 0 |
| LGAGGGSI | STUB1; STUB1 | Q9UNE7; H3 | CHIP_HUMAN | Unmodified | 0 | 120130 | 0 | 93458 | 378440 | 68179 | 56103 | 17967 | 0 | 64036 | 67194 | 0 | 627330 | 216080 | 0 | 0 | 0 | 472660 |
| LGAGPDG\ | RPS6KA3; RPS6 | P51812; B7Z | KS6A3_HUMAN | Unmodified | 0 | 0 | 0 | 353840 | 0 | 0 | 0 | 0 | 346490 | 32963 | 97612 | 457090 | 1887200 | 0 | 0 | 0 | 70990 | 0 |
| LGAGQTPG | TUBGCP6; TUB | Q96RT7; H7 | GCP6_HUMAN | Unmodified | 0 | 0 | 0 | 0 | 0 | 0 | 0 | 0 | 0 | 0 | 0 | 426540 | 0 | 0 | 0 | 0 | 0 | 0 |
| LGAHQLDS | PRSS8; PRSS8 | Q16651; H3I | PRSS8_HUMAN | Unmodified | 0 | 0 | 1335800 | 2319200 | 0 | 0 | 0 | 20521 | 6073400 | 1272200 | 941540 | 611410 | 621960 | 491010 | 778680 | 768080 | 0 | 0 |
| LGAHQLDS | PRSS8 | Q16651 | PRSS8_HUMAN | Unmodified | 0 | 0 | 0 | 265920 | 727060 | 0 | 85506 | 0 | 0 | 863460 | 0 | 0 | 1414400 | 961150 | 0 | 0 | 313630 | 481430 |
| LGALSGAA | TMEM256; TME | Q8N2U0; K7 | TM256_HUMAN | Unmodified | 0 | 0 | 0 | 0 | 0 | 0 | 0 | 0 | 15829000 | 4016500 | 0 | 0 | 0 | 0 | 0 | 0 | 22869 | 0 |
| LGANSLLD | SDHA; SDHA; S | P31040; D6F | SDHA_HUMAN | Unmodified | 0 | 0 | 0 | 0 | 0 | 0 | 0 | 0 | 231830 | 177310 | 0 | 0 | 0 | 0 | 0 | 0 | 0 | 0 |
| LGAPALTSI | SHMT2; SHMT2 | P34897; H0Y | GLYM_HUMAN | Unmodified | 4513500 | 5900100 | 3542500 | 3341700 | 3447100 | 90850 | 0 | 25175 | 48968000 | 2164900 | 450500 | 1366500 | 0 | 0 | 3447400 | 3210000 | 1562000 | 4318400 |
| LGAPENSG | FARP1; FARP1; | Q9Y4F1; A0A | FARP1_HUMAN | Unmodified | 253360 | 0 | 0 | 487380 | 0 | 0 | 0 | 0 | 6295400 | 344880 | 2500500 | 5782500 | 0 | 0 | 1562900 | 981550 | 0 | 0 |
| LGAPENSG | FARP1; FARP1; | Q9Y4F1; A0A | FARP1_HUMAN | Unmodified | 0 | 0 | 0 | 414570 | 687710 | 0 | 0 | 0 | 1411500 | 1102700 | 736230 | 0 | 403970 | 410070 | 0 | 0 | 267310 | 0 |
| LGAPLPFR | ARRDC1; ARRD | Q8N5I2; Q51 | ARRD1_HUMAI | Unmodified | 4998300 | 3428400 | 4074600 | 0 | 389690 | 0 | 0 | 0 | 3871500 | 708980 | 530430 | 5084100 | 604780 | 0 | 7410600 | 6188600 | 267690 | 237870 |
| LGAPQTHL | EFHD1; EFHD2; | Q9BUP0; Q9 | EFHD1_HUMAI | Unmodified | 0 | 0 | 0 | 0 | 4159100 | 0 | 0 | 0 | 466430 | 268670 | 0 | 502290 | 0 | 0 | 0 | 0 | 273660 | 669640 |
| LGAQALLG | PYCR1; PYCR2; | P32322; Q96 | P5CR1_HUMAI | Unmodified | 0 | 0 | 0 | 51800 | 0 | 0 | 77927 | 17732 | 2231700 | 639970 | 50943 | 0 | 51157 | 0 | 329410 | 466650 | 0 | 0 |
| LGAQLADL | FN3KRP; FN3KI | Q9HA64; I3L | KT3K_HUMAN I | Unmodified | 0 | 353370 | 381120 | 206320 | 0 | 339550 | 76964 | 24551 | 819200 | 361220 | 241530 | 1607700 | 759330 | 396480 | 722170 | 110170 | 661120 | 1762300 |
| LGAQSVYH | HINT2 | Q9BX68 | HINT2_HUMAN | Unmodified | 0 | 0 | 0 | 0 | 0 | 0 | 0 | 0 | 0 | 0 | 0 | 0 | 0 | 0 | 0 | 0 | 0 | 0 |
| LGAQTPVG | FARP2; FARP2; | O94887; C9J | FARP2_HUMAN | Unmodified | 0 | 0 | 0 | 3423700 | 0 | 0 | 0 | 95123 | 2081300 | 0 | 0 | 0 | 269630 | 0 | 4656300 | 2031200 | 0 | 0 |
| LGASPLHV | ATP6AP1; ATP6 | Q15904; A0A | VAS1_HUMAN | Unmodified | 0 | 0 | 0 | 0 | 239580 | 0 | 111330 | 0 | 6660900 | 349000 | 99481 | 1496000 | 0 | 0 | 0 | 0 | 18025 | 0 |
| LGASPLHV | ATP6AP1; ATP6 | Q15904; A0A | VAS1_HUMAN | Unmodified | 0 | 0 | 0 | 0 | 0 | 0 | 0 | 0 | 1859900 | 0 | 0 | 0 | 0 | 0 | 0 | 0 | 0 | 0 |
| LGATAEKA | VCL | P18206 | VINC_HUMAN | Unmodified | 0 | 0 | 0 | 0 | 0 | 0 | 0 | 0 | 0 | 73154 | 0 | 0 | 0 | 0 | 0 | 0 | 149310 | 0 |
| LGAVDESL | ERO1A; ERO1A; | Q96HE7; G3 | ERO1A_HUMAI | Unmodified | 0 | 78186 | 0 | 0 | 0 | 0 | 0 | 0 | 10133000 | 492200 | 2089300 | 182910 | 0 | 0 | 0 | 0 | 36655 | 0 |
| LGAVFNQV | SF3B3 | Q15393 | SF3B3_HUMAN | Unmodified | 2350600 | 902190 | 198890 | 206420 | 0 | 0 | 0 | 0 | 0 | 0 | 0 | 6275600 | 0 | 0 | 0 | 790160 | 0 | 0 |
| LGAVGGS( | ELP1; ELP1; ELF | O95163; A0A | ELP1_HUMAN I | Unmodified | 0 | 254260 | 0 | 0 | 236510 | 0 | 11780 | 428040 | 172480 | 0 | 0 | 781740 | 0 | 0 | 0 | 52237 | 0 | 0 |
| LGAVVAVT( | ATP1A4 | Q13733 | AT1A4_HUMAN | Unmodified | 0 | 0 | 0 | 624030 | 492720 | 0 | 1589400 | 3399600 | 0 | 0 | 0 | 0 | 0 | 1587900 | 0 | 0 | 38785 | 1020300 |
| LGAVVAVT( | ATP1A4 | Q13733 | AT1A4_HUMAN | Unmodified | 0 | 0 | 0 | 0 | 0 | 0 | 0 | 2541500 | 0 | 0 | 0 | 0 | 0 | 0 | 0 | 0 | 0 | 0 |
| LGAVYDVS | LRPPRC; LRPPF | P42704; B8Z | LPPRC_HUMAN | Unmodified | 0 | 0 | 0 | 125700 | 0 | 0 | 501510 | 0 | 6153300 | 2198500 | 0 | 471870 | 0 | 0 | 0 | 0 | 0 | 0 |
| LGAVYTEG | CEL | P19835 | CEL_HUMAN B | Unmodified | 0 | 0 | 0 | 0 | 0 | 0 | 42051 | 0 | 12720000 | 2002400 | 16667000 | 1349300 | 0 | 0 | 0 | 0 | 0 | 0 |
| LGAVYTEG | CEL | P19835 | CEL_HUMAN B | Unmodified | 0 | 0 | 0 | 0 | 0 | 0 | 0 | 0 | 0 | 0 | 0 | 0 | 0 | 0 | 0 | 0 | 0 | 0 |
| LGAYYVTG | COA7 | Q96BR5 | COA7_HUMAN | Unmodified | 0 | 0 | 0 | 0 | 0 | 0 | 368470 | 164640 | 0 | 2528200 | 0 | 0 | 2324900 | 0 | 0 | 0 | 6305000 | 397840 |
| LGCCVVEK | CDC42SE1 | Q9NRR8 | C42S1_HUMAN | Unmodified | 851230 | 240900 | 0 | 0 | 1330800 | 535330 | 284620 | 185800 | 53036 | 214930 | 0 | 381850 | 2593800 | 2369500 | 0 | 276000 | 2432900 | 0 |
| LGCCVVEK | CDC42SE1 | Q9NRR8 | C42S1_HUMAN | Unmodified | 0 | 0 | 0 | 0 | 494360 | 0 | 61073 | 0 | 0 | 248430 | 0 | 0 | 0 | 66225 | 0 | 0 | 601340 | 427510 |
| LGCEVLGV | PRDX2; PRDX2 | P32119; A6N | PRDX2_HUMAN | Unmodified | 0 | 0 | 0 | 0 | 985170 | 0 | 2400300 | 0 | 1066900 | 0 | 462620 | 2411700 | 0 | 0 | 0 | 0 | 0 | 0 |
| LGCINAEYI | MGAT2 | Q10469 | MGAT2_HUMAI | Unmodified | 0 | 0 | 405670 | 504180 | 0 | 0 | 0 | 0 | 4383500 | 613300 | 0 | 0 | 0 | 0 | 0 | 28017 | 0 | 0 |
| LGCINAEYI | MGAT2 | Q10469 | MGAT2_HUMAI | Unmodified | 0 | 0 | 0 | 0 | 0 | 0 | 0 | 0 | 4713300 | 0 | 281640 | 0 | 0 | 0 | 0 | 0 | 0 | 0 |
| LGCPGFTLI | GCLC; GCLC; G | P48506; A0A | GSH1_HUMAN | Unmodified | 218210 | 0 | 0 | 0 | 5902700 | 0 | 0 | 0 | 0 | 0 | 831000 | 4298800 | 0 | 0 | 436930 | 0 | 0 | 0 |
| LGCRQFVT | PLS3; PLS3; PLS | P13797; H7C | PLST_HUMAN I | Unmodified | 0 | 0 | 0 | 0 | 397010 | 237800 | 0 | 0 | 0 | 45810 | 0 | 0 | 0 | 0 | 0 | 0 | 447510 | 0 |
| LGCSASNF | IGF1R; IGF1R | P08069; C9J | IGF1R_HUMAN | Unmodified | 0 | 0 | 0 | 0 | 827130 | 325330 | 0 | 0 | 225540 | 0 | 0 | 0 | 0 | 1205900 | 0 | 0 | 0 | 0 |
| LGCSLNQN | DAG1 | Q14118 | DAG1_HUMAN | Unmodified | 0 | 0 | 0 | 0 | 0 | 0 | 0 | 0 | 1456300 | 0 | 0 | 0 | 0 | 0 | 185880 | 0 | 0 | 0 |
| LGDAILSVN | SNTB2; SNTB2; | Q13425; A0A | SNTB2_HUMAN | Unmodified | 0 | 0 | 151200 | 0 | 0 | 0 | 0 | 0 | 288790 | 0 | 0 | 0 | 0 | 0 | 116920 | 184340 | 0 | 0 |

| | | | | | | | | | | | | | | | | | | | | | | |
|---|---|---|---|---|---|---|---|---|---|---|---|---|---|---|---|---|---|---|---|---|---|---|
| LGDAVEQG | GARS1; GARS1 | P41250; A0A | GARS_HUMAN | Unmodified | 0 | 0 | 0 | 0 | 0 | 0 | 0 | 0 | 0 | 0 | 0 | 173030 | 0 | 0 | 0 | 0 | 0 | 0 |
| LGDDEVRT | CAPRIN1; CAPF | Q14444; G3V | CAPR1_HUMAN | Unmodified | 0 | 0 | 0 | 0 | 0 | 0 | 0 | 0 | 0 | 76409 | 143760 | 3239500 | 0 | 0 | 0 | 278210 | 0 | 0 |
| LGDHGYDI | ENPP4 | Q9Y6X5 | ENPP4_HUMAN | Unmodified | 0 | 0 | 0 | 0 | 0 | 0 | 0 | 139990 | 25127000 | 628570 | 2009000 | 0 | 0 | 0 | 420800 | 0 | 0 | 0 |
| LGDHGYDI | ENPP4 | Q9Y6X5 | ENPP4_HUMAN | Oxidation (M) | 0 | 0 | 677420 | 217400 | 0 | 0 | 0 | 0 | 1291500 | 0 | 0 | 885760 | 0 | 0 | 201970 | 0 | 0 | 0 |
| LGDLEEAPI | SLC20A1; SLC2 | Q8WUM9; H | S20A1_HUMAN | Unmodified | 0 | 0 | 0 | 0 | 0 | 0 | 0 | 0 | 0 | 0 | 407660 | 2546200 | 0 | 0 | 0 | 425480 | 0 | 0 |
| LGDLEEAPI | SLC20A1; SLC2 | Q8WUM9; H | S20A1_HUMAN | Unmodified | 0 | 0 | 0 | 241640 | 192170 | 0 | 0 | 64686 | 0 | 0 | 289480 | 786420 | 0 | 0 | 190310 | 0 | 0 | 97873 |
| LGDPHFYQ | TMEM138; TMEI | Q9NPI0; J3Q | TM138_HUMAN | Unmodified | 0 | 0 | 89383 | 43702 | 0 | 0 | 0 | 0 | 0 | 0 | 0 | 0 | 0 | 0 | 208140 | 124460 | 0 | 0 |
| LGDRYNPF | DCAF11; DCAF | Q8TEB1; H0Y | DCA11_HUMAI | Unmodified | 0 | 0 | 0 | 0 | 532970 | 0 | 0 | 0 | 0 | 0 | 0 | 0 | 0 | 0 | 0 | 0 | 469150 | 580680 |
| LGDSMANY | NRN1; NRN1 | Q9NPD7; A0 | NRN1_HUMAN | Oxidation (M) | 0 | 0 | 0 | 0 | 0 | 529740 | 632850 | 654470 | 0 | 88662 | 0 | 0 | 0 | 0 | 0 | 0 | 311580 | 0 |
| LGDSMANY | NRN1; NRN1 | Q9NPD7; A0 | NRN1_HUMAN | Unmodified | 0 | 0 | 76191 | 0 | 363950 | 0 | 848340 | 334190 | 0 | 0 | 0 | 0 | 213960 | 0 | 0 | 0 | 0 | 0 |
| LGDSSLSR | EIF3A | Q14152 | EIF3A_HUMAN | Unmodified | 0 | 0 | 0 | 0 | 91133 | 31386 | 0 | 0 | 134010 | 0 | 145350 | 611300 | 0 | 0 | 0 | 0 | 65109 | 84894 |
| LGDTALHA | OSTF1 | Q92882 | OSTF1_HUMAN | Unmodified | 564120 | 356390 | 1743100 | 1947100 | 1379000 | 0 | 195530 | 127510 | 3421300 | 0 | 4380200 | 8335700 | 1613900 | 1567800 | 1507600 | 1711200 | 289330 | 0 |
| LGDTALHA | OSTF1 | Q92882 | OSTF1_HUMAN | Unmodified | 0 | 0 | 0 | 0 | 131270 | 0 | 176030 | 0 | 0 | 79854 | 0 | 0 | 287910 | 1079000 | 0 | 0 | 0 | 0 |
| LGDTYPSIS | RNPEP; RNPEP | Q9H4A4; A6I | AMPB_HUMAN | Unmodified | 202440 | 0 | 0 | 0 | 0 | 0 | 0 | 0 | 0 | 0 | 730440 | 2714300 | 0 | 0 | 0 | 0 | 0 | 0 |
| LGDVGMAI | RNH1; RNH1 | P13489; H0Y | RINI_HUMAN R | Oxidation (M) | 0 | 0 | 0 | 399300 | 0 | 0 | 0 | 0 | 0 | 0 | 0 | 3921800 | 0 | 0 | 729600 | 652660 | 0 | 0 |
| LGDVGMAI | RNH1; RNH1 | P13489; H0Y | RINI_HUMAN R | Unmodified | 0 | 0 | 0 | 0 | 0 | 197700 | 0 | 27651 | 0 | 210240 | 0 | 771590 | 0 | 0 | 0 | 0 | 417380 | 0 |
| LGDVISIQP | VCP; VCP; VCP; | P55072; C9J | TERA_HUMAN | Unmodified | 401210 | 396460 | 0 | 0 | 0 | 0 | 0 | 0 | 2191800 | 399660 | 1047600 | 8112300 | 752610 | 0 | 461940 | 560070 | 0 | 0 |
| LGDVISIQP | VCP; VCP; VCP; | P55072; C9J | TERA_HUMAN | Unmodified | 0 | 0 | 0 | 0 | 6636500 | 908910 | 0 | 184540 | 299140 | 3285000 | 0 | 0 | 8389600 | 6122200 | 0 | 0 | 2634500 | 1185900 |
| LGDVYVND | PGK1; PGK2 | P00558; P07 | PGK1_HUMAN | Unmodified | 5332000 | 5583100 | 16037000 | 18530000 | 9533000 | 2911400 | 1369500 | 0 | 43410000 | 7334200 | 51909000 | 2.5E+08 | 8204800 | 4609800 | 22395000 | 16702000 | 5379100 | 14208000 |
| LGDVYVND | PGK1 | P00558 | PGK1_HUMAN | Oxidation (M) | 142670 | 0 | 175540 | 3137200 | 10858000 | 6385700 | 8974600 | 100530 | 0 | 11082000 | 2231400 | 0 | 23024000 | 19933000 | 0 | 0 | 52233000 | 78468000 |
| LGDVYVND | PGK1 | P00558 | PGK1_HUMAN | Unmodified | 0 | 0 | 0 | 123590 | 54907000 | 5938800 | 10952000 | 0 | 6311700 | 19751000 | 0 | 0 | 47357000 | 4876000 | 0 | 0 | 98755000 | 2.14E+08 |
| LGDYQTQL | UTRN | P46939 | UTRN_HUMAN | Unmodified | 0 | 0 | 0 | 773740 | 1135500 | 0 | 0 | 0 | 839550 | 451060 | 0 | 106930 | 442640 | 447300 | 0 | 0 | 571890 | 488540 |
| LGEAAVLEI | SMPDL3B; SMP | Q92485; Q5 | ASM3B_HUMAI | Unmodified | 340600 | 0 | 5230300 | 11893000 | 0 | 0 | 0 | 0 | 3386600 | 0 | 1881600 | 5992200 | 0 | 0 | 14176000 | 1039900 | 0 | 0 |
| LGEAAVLEI | SMPDL3B; SMP | Q92485; Q5 | ASM3B_HUMAI | Unmodified | 168810 | 0 | 727140 | 2327700 | 771660 | 0 | 0 | 0 | 1181900 | 711430 | 427500 | 29653 | 4424800 | 1471700 | 3601300 | 362130 | 0 | 922330 |
| LGEAEGLM | SLC38A10 | Q9HBR0 | S38AA_HUMAN | Unmodified | 182620 | 0 | 582220 | 0 | 302820 | 0 | 10899 | 0 | 3766700 | 163710 | 295490 | 62515 | 638090 | 0 | 549030 | 383250 | 47341 | 98203 |
| LGEAEGLM | SLC38A10 | Q9HBR0 | S38AA_HUMAN | Unmodified | 0 | 0 | 0 | 0 | 0 | 0 | 0 | 0 | 1615600 | 0 | 0 | 0 | 0 | 0 | 0 | 0 | 0 | 0 |
| LGEEAFFYI | THYN1 | Q9P016 | THYN1_HUMAI | Unmodified | 0 | 0 | 0 | 0 | 0 | 0 | 0 | 0 | 0 | 0 | 51840 | 973550 | 0 | 0 | 0 | 0 | 0 | 0 |
| LGEEDEQG | PACSIN1; PACS | Q9BY11; F6L | PACN1_HUMAI | Unmodified | 1537500 | 1637500 | 874270 | 0 | 0 | 0 | 0 | 0 | 0 | 0 | 0 | 0 | 0 | 0 | 0 | 0 | 0 | 0 |
| LGEGEGSN | SNX5; SNX5; SI | Q9Y5X3; Q5 | SNX5_HUMAN | Oxidation (M) | 0 | 0 | 0 | 0 | 0 | 0 | 17442 | 20343 | 90570 | 31263 | 299610 | 0 | 0 | 0 | 0 | 73450 | 0 | 0 |
| LGEGEGSN | SNX5; SNX5; SI | Q9Y5X3; Q5 | SNX5_HUMAN | Unmodified | 109250 | 0 | 30899 | 89072 | 0 | 0 | 0 | 0 | 0 | 0 | 538060 | 1551900 | 0 | 244040 | 114780 | 97175 | 0 | 0 |
| LGEGEGSN | SNX6; SNX6; SI | Q9UNH7; A0 | SNX6_HUMAN | Unmodified | 98770 | 0 | 47731 | 128150 | 2795900 | 0 | 664540 | 378350 | 502500 | 216760 | 560700 | 1941700 | 652310 | 350530 | 203840 | 145290 | 284340 | 998470 |
| LGEGEGSN | SNX6; SNX6; SI | Q9UNH7; A0 | SNX6_HUMAN | Oxidation (M) | 0 | 0 | 0 | 133450 | 963230 | 178780 | 258730 | 123470 | 154220 | 103380 | 202200 | 859740 | 271790 | 186510 | 83067 | 100060 | 129130 | 201620 |
| LGEGGPDS | FAM234A; FAM | Q9H0X4; H7 | F234A_HUMAN | Unmodified | 0 | 0 | 0 | 0 | 0 | 0 | 0 | 0 | 0 | 40969 | 0 | 1020900 | 0 | 0 | 0 | 0 | 0 | 0 |
| LGEGGPDS | FAM234A; FAM | Q9H0X4; H7 | F234A_HUMAN | Unmodified | 893510 | 0 | 427360 | 361940 | 0 | 0 | 0 | 0 | 702840 | 49278 | 151070 | 4324600 | 55791 | 0 | 633800 | 541900 | 0 | 0 |
| LGEIVTTIPT | ARF4; ARF5; AR | P18085; P84 | ARF4_HUMAN | Unmodified | 0 | 0 | 38979 | 113830 | 0 | 0 | 166880 | 0 | 32978 | 0 | 0 | 107570 | 90556 | 75539 | 1283200 | 37919 | 70533 | 0 |
| LGEMWNN | HMGB1; HMGB | P09429; Q5 | HMGB1_HUMA | Oxidation (M) | 0 | 0 | 0 | 0 | 189890 | 0 | 345520 | 0 | 93861 | 0 | 358300 | 2584500 | 0 | 0 | 0 | 0 | 0 | 0 |
| LGEMWNN | HMGB1; HMGB | P09429; Q5 | HMGB1_HUMA | Unmodified | 0 | 0 | 0 | 88318 | 0 | 0 | 704530 | 17341 | 0 | 0 | 0 | 5786000 | 0 | 0 | 0 | 0 | 0 | 0 |
| LGEMWNN | HMGB1; HMGB | P09429; Q5 | HMGB1_HUMA | Unmodified | 0 | 0 | 0 | 0 | 0 | 0 | 0 | 0 | 0 | 0 | 0 | 2191800 | 0 | 0 | 0 | 0 | 0 | 0 |
| LGEMWNN | HMGB1; HMGB | P09429; Q5 | HMGB1_HUMA | Oxidation (M) | 0 | 0 | 0 | 0 | 0 | 0 | 0 | 0 | 0 | 0 | 0 | 1318000 | 0 | 0 | 0 | 0 | 0 | 0 |
| LGEPDYIPS | GNA13 | Q14344 | GNA13_HUMAI | Unmodified | 0 | 0 | 0 | 0 | 0 | 0 | 0 | 0 | 2787400 | 130390 | 2107800 | 10434000 | 0 | 0 | 0 | 0 | 0 | 0 |
| LGEQVFGT | SLC38A1; SLC3 | Q9H2H9; F8 | S38A1_HUMAN | Unmodified | 0 | 0 | 145270 | 160980 | 0 | 1068000 | 0 | 0 | 504280 | 0 | 0 | 6766600 | 0 | 0 | 231960 | 317990 | 123930 | 487850 |
| LGESLQSA | KIF20A | O95235 | KI20A_HUMAN | Unmodified | 0 | 0 | 366740 | 1860300 | 392540 | 0 | 0 | 0 | 993750 | 80752 | 0 | 0 | 379890 | 67437 | 0 | 0 | 0 | 0 |
| LGESLQSA | KIF20A | O95235 | KI20A_HUMAN | Unmodified | 0 | 0 | 0 | 0 | 1268900 | 0 | 35763 | 0 | 0 | 0 | 0 | 0 | 823680 | 810630 | 0 | 0 | 867820 | 493640 |
| LGESQTLQ | SPTAN1; SPTAN | Q13813; A0 | SPTN1_HUMAN | Unmodified | 0 | 0 | 0 | 0 | 0 | 0 | 0 | 0 | 0 | 0 | 0 | 3691700 | 0 | 0 | 0 | 0 | 0 | 0 |
| LGESVQDL | ATP6V1H; ATP6 | Q9UI12; G3 | VATH_HUMAN | Unmodified | 0 | 0 | 0 | 0 | 0 | 0 | 0 | 39215 | 0 | 0 | 0 | 2583800 | 0 | 0 | 0 | 0 | 0 | 0 |
| LGETYKDH | P4HB; P4HB; P | P07237; A0A | PDIA1_HUMAN | Unmodified | 0 | 0 | 0 | 0 | 3635700 | 0 | 4042800 | 2594000 | 2.42E+08 | 84378000 | 27303000 | 3857000 | 3591200 | 1180400 | 777280 | 1055300 | 0 | 4738500 |
| LGETYKDH | P4HB; P4HB; P | P07237; A0A | PDIA1_HUMAN | Unmodified | 0 | 0 | 0 | 0 | 1301500 | 0 | 1836800 | 0 | 1374100 | 16978000 | 0 | 0 | 494410 | 0 | 0 | 0 | 270320 | 0 |
| LGETYKDH | P4HB; P4HB; P | P07237; A0A | PDIA1_HUMAN | Oxidation (M) | 0 | 0 | 0 | 0 | 0 | 0 | 0 | 0 | 0 | 542830 | 0 | 0 | 0 | 0 | 0 | 0 | 0 | 0 |
| LGEVVNTH | NMT1; NMT1; N | P30419; A0A | NMT1_HUMAN | Unmodified | 0 | 0 | 0 | 0 | 0 | 0 | 0 | 0 | 744470 | 44692 | 0 | 2538700 | 0 | 0 | 0 | 0 | 0 | 0 |
| LGEVVNTH | NMT1; NMT1; N | P30419; A0A | NMT1_HUMAN | Unmodified | 0 | 0 | 0 | 0 | 0 | 195100 | 0 | 0 | 0 | 0 | 0 | 6882700 | 0 | 0 | 0 | 0 | 0 | 0 |
| LGEWNEK | TPD52L2; TPD5 | O43399; A0 | TPD54_HUMAN | Unmodified | 0 | 0 | 0 | 0 | 638050 | 126300 | 106920 | 82994 | 0 | 139170 | 81894 | 0 | 406890 | 0 | 0 | 0 | 183340 | 0 |
| LGEWVGL | RPS12 | P25398 | RS12_HUMAN | Unmodified | 0 | 0 | 0 | 0 | 0 | 0 | 0 | 0 | 0 | 0 | 0 | 4627400 | 1006000 | 0 | 0 | 0 | 0 | 0 |
| LGEYEDVS | TBCB; TBCB; TE | Q99426; K7E | TBCB_HUMAN | Unmodified | 0 | 0 | 527600 | 0 | 0 | 0 | 274470 | 0 | 3344900 | 0 | 319500 | 3994000 | 0 | 0 | 0 | 0 | 0 | 0 |
| LGFAGLVQ | CCT5; CCT5; C | P48643; B7Z | TCPE_HUMAN | Unmodified | 770830 | 308390 | 635700 | 1842900 | 7154600 | 1136000 | 2157900 | 48278 | 4960800 | 4720600 | 3264900 | 17426000 | 17764000 | 6300000 | 1879600 | 1118400 | 3722900 | 5907700 |
| LGFAGLVQ | CCT5; CCT5; C | P48643; B7Z | TCPE_HUMAN | Unmodified | 0 | 0 | 0 | 0 | 4395100 | 0 | 4790600 | 47946 | 0 | 2798900 | 73097 | 255360 | 8025500 | 5405700 | 0 | 0 | 3836800 | 6259100 |
| LGFDKENV | SMARCA5 | O60264 | SMCA5_HUMA | Unmodified | 0 | 136950 | 0 | 0 | 0 | 0 | 0 | 0 | 121670 | 0 | 0 | 0 | 0 | 0 | 0 | 170820 | 0 | 0 |
| LGFEDGSV | UCHL1; UCHL1 | P09936; D6F | UCHL1_HUMAI | Unmodified | 0 | 229270 | 0 | 498960 | 28289000 | 8868700 | 0 | 1437100 | 17186000 | 5085700 | 38710000 | 95519000 | 0 | 0 | 415890 | 0 | 639960 | 0 |
| LGFEDGSV | UCHL1; UCHL1 | P09936; D6F | UCHL1_HUMAI | Unmodified | 0 | 0 | 0 | 0 | 3772100 | 265690 | 990920 | 0 | 0 | 426870 | 92784 | 134810 | 0 | 0 | 0 | 0 | 0 | 0 |
| LGFEEFK | CAPNS2; CAPN | Q96L46; P04 | CPNS2_HUMAI | Unmodified | 216400 | 0 | 557260 | 0 | 0 | 0 | 0 | 0 | 297420 | 0 | 222710 | 1897900 | 0 | 0 | 761330 | 581190 | 0 | 0 |
| LGFEGGQT | MRPL15; MRPL | Q9P015; E5F | RM15_HUMAN | Unmodified | 0 | 0 | 0 | 0 | 0 | 0 | 0 | 0 | 1960900 | 0 | 0 | 125590 | 0 | 0 | 0 | 0 | 0 | 0 |
| LGFGRADG | MFAP4; MFAP4 | P55083; K7E | MFAP4_HUMAI | Oxidation (M) | 0 | 167830 | 0 | 661550 | 0 | 745730 | 884440 | 0 | 0 | 485010 | 0 | 0 | 1836400 | 1041300 | 0 | 0 | 542300 | 4082900 |
| LGFGRADG | MFAP4; MFAP4 | P55083; K7E | MFAP4_HUMAI | Unmodified | 0 | 0 | 0 | 0 | 0 | 0 | 0 | 0 | 0 | 0 | 0 | 0 | 652280 | 276180 | 0 | 0 | 0 | 0 |
| LGFGSFVD | ITGB5; ITGB5; I | P18084; V9G | ITB5_HUMAN I | Unmodified | 878520 | 478490 | 1465700 | 705970 | 0 | 0 | 0 | 0 | 489520 | 0 | 511360 | 7928900 | 0 | 0 | 2417300 | 1460900 | 0 | 0 |
| LGFGSFVEI | ITGB6; ITGB6; I | P18564; A0A | ITB6_HUMAN I | Unmodified | 0 | 0 | 9402500 | 15930000 | 78702000 | 13594000 | 0 | 518950 | 42670000 | 85675 | 4751500 | 19481000 | 6642800 | 2657600 | 16053000 | 12550000 | 12855000 | 69104000 |
| LGFGSFVEI | ITGB6; ITGB6; I | P18564; A0A | ITB6_HUMAN I | Unmodified | 0 | 0 | 0 | 0 | 0 | 0 | 0 | 0 | 0 | 10848 | 0 | 4465600 | 11729 | 0 | 0 | 0 | 0 | 0 |
| LGFHLPLE | PLEC; PLEC | Q15149; H0Y | PLEC_HUMAN | Unmodified | 0 | 65707 | 165580 | 161990 | 310430 | 160470 | 0 | 17268 | 759940 | 90665 | 0 | 0 | 0 | 0 | 145680 | 184200 | 819220 | 449780 |
| LGFHLPLE | PLEC; PLEC | Q15149; H0Y | PLEC_HUMAN | Unmodified | 0 | 0 | 0 | 0 | 451350 | 157020 | 0 | 0 | 0 | 0 | 0 | 0 | 0 | 0 | 0 | 0 | 522670 | 449620 |
| LGFMSAFV | DLST; DLST | P36957; Q8 | ODO2_HUMAN | Oxidation (M) | 24575 | 0 | 81953 | 135330 | 0 | 0 | 600980 | 242890 | 1729200 | 880020 | 0 | 139580 | 93368 | 194260 | 144390 | 51730 | 0 | 0 |
| LGFMSAFV | DLST; DLST | P36957; Q8 | ODO2_HUMAN | Unmodified | 0 | 0 | 0 | 0 | 0 | 0 | 1159000 | 125880 | 5467400 | 1320500 | 0 | 0 | 0 | 67720 | 0 | 0 | 0 | 0 |
| LGFSEVEL | CKB; CKB | P12277; H0Y | KCRB_HUMAN | Oxidation (M) | 0 | 0 | 0 | 36860 | 772130 | 31952 | 0 | 0 | 0 | 0 | 0 | 51662 | 27150 | 0 | 418460 | 0 | 0 | 0 |
| LGFTGSTPI | ALDH1L2; ALDH | Q3SY69; A0 | AL1L2_HUMAN | Unmodified | 0 | 0 | 281570 | 0 | 784020 | 644520 | 0 | 0 | 821990 | 68002 | 0 | 0 | 0 | 0 | 0 | 0 | 0 | 229310 |
| LGFTVTVGI | TPP2; TPP2; TPI | P29144; A0A | TPP2_HUMAN | Unmodified | 0 | 0 | 0 | 0 | 0 | 0 | 0 | 0 | 0 | 0 | 0 | 1558800 | 0 | 0 | 0 | 0 | 0 | 0 |
| LGFVAGVII | ATP6V0A1; ATP | Q93050; K7E | VPP1_HUMAN | Unmodified | 0 | 0 | 446870 | 0 | 0 | 0 | 0 | 0 | 2457300 | 306030 | 420630 | 2181900 | 0 | 0 | 1234800 | 0 | 0 | 0 |
| LGFVAGVII | ATP6V0A1; ATP | Q93050; K7E | VPP1_HUMAN | Unmodified | 0 | 0 | 84057 | 0 | 0 | 0 | 0 | 0 | 1474300 | 0 | 0 | 0 | 0 | 0 | 0 | 0 | 33782 | 0 |
| LGFYGLDE | OGDH; OGDH; | Q02218; A0 | ODO1_HUMAN | Unmodified | 0 | 0 | 0 | 0 | 0 | 0 | 0 | 0 | 1111700 | 237040 | 0 | 0 | 0 | 0 | 0 | 0 | 0 | 0 |
| LGFYNAETI | TRIM41 | Q8WV44 | TRI41_HUMAN | Unmodified | 246490 | 793060 | 41015 | 0 | 0 | 0 | 0 | 0 | 0 | 0 | 0 | 0 | 0 | 0 | 249860 | 0 | 0 | 0 |
| LGGAIAAIN | GLT8D1; GLT8D | Q68CQ7; C9 | GL8D1_HUMAI | Unmodified | 0 | 0 | 0 | 0 | 0 | 0 | 0 | 149880 | 5044000 | 955040 | 0 | 558780 | 0 | 0 | 0 | 0 | 0 | 0 |
| LGGAVLSF | CAD; CAD; CAD | P27708; F8V | PYR1_HUMAN | Unmodified | 0 | 12566 | 0 | 0 | 0 | 0 | 70152 | 0 | 0 | 165320 | 0 | 701260 | 0 | 0 | 24511 | 44176 | 0 | 272270 |
| LGGEAGAA | FAM171A2 | A8MVW0 | F1712_HUMAN | Unmodified | 0 | 0 | 0 | 0 | 287360 | 0 | 0 | 0 | 1486700 | 236990 | 149950 | 0 | 0 | 0 | 0 | 69837 | 395880 | 897820 |
| LGGEVSCL | ETFA; ETFA; ETF | P13804; H0Y | ETFA_HUMAN | Unmodified | 0 | 0 | 0 | 0 | 907860 | 0 | 0 | 0 | 1283300 | 0 | 0 | 0 | 0 | 0 | 0 | 0 | 0 | 0 |
| LGGHGPSI | XRCC5 | P13010 | XRCC5_HUMAI | Unmodified | 98300 | 37335 | 0 | 0 | 0 | 0 | 85021 | 164490 | 0 | 0 | 193690 | 3128000 | 0 | 0 | 0 | 0 | 0 | 0 |
| LGGIGQFL | DHX9 | Q08211 | DHX9_HUMAN | Unmodified | 0 | 0 | 0 | 0 | 0 | 0 | 0 | 0 | 0 | 0 | 0 | 9537400 | 0 | 0 | 0 | 0 | 0 | 0 |
| LGGIPVGV | ACACA; ACACA | Q13085; Q5 | ACACA_HUMAI | Unmodified | 0 | 0 | 0 | 0 | 0 | 0 | 0 | 0 | 0 | 0 | 0 | 1285800 | 0 | 0 | 0 | 0 | 0 | 0 |
| LGGIVLPNI | ITGB1; ITGB1; I | P05556; H7 | ITB1_HUMAN I | 2 Oxidation (M) | 9150500 | 4508100 | 1627100 | 7408500 | 39997000 | 19587000 | 499000 | 0 | 9694600 | 0 | 0 | 1723800 | 0 | 1179500 | 19348000 | 6592400 | 6796600 | 54424000 |
| LGGIVLPNI | ITGB1; ITGB1; I | P05556; H7 | ITB1_HUMAN I | Oxidation (M) | 0 | 0 | 0 | 3109200 | 17801000 | 0 | 0 | 0 | 15183000 | 0 | 0 | 0 | 688180 | 0 | 4260100 | 689490 | 2737900 | 18173000 |
| LGGIVLPNI | ITGB1; ITGB1; I | P05556; H7 | ITB1_HUMAN I | Unmodified | 0 | 0 | 0 | 0 | 7165100 | 0 | 0 | 0 | 8025200 | 0 | 0 | 0 | 0 | 0 | 0 | 0 | 1134900 | 6885400 |
| LGGKLSSE | HSPA5; HSPA5; | P11021; A0A | BIP_HUMAN Er | Unmodified | 0 | 0 | 0 | 0 | 0 | 0 | 0 | 0 | 7483400 | 3257700 | 34520 | 2044000 | 0 | 0 | 124830 | 0 | 70949 | 0 |
| LGGLDVLE | PPM1L | Q5SGD2 | PPM1L_HUMAN | Unmodified | 0 | 4970.7 | 0 | 0 | 0 | 0 | 0 | 0 | 3273700 | 0 | 0 | 170820 | 0 | 0 | 0 | 0 | 0 | 0 |
| LGGLQVLR | SIL1; SIL1 | Q9H173; D6 | SIL1_HUMAN N | Unmodified | 0 | 0 | 407240 | 0 | 0 | 0 | 0 | 0 | 1147700 | 746280 | 0 | 0 | 0 | 0 | 488780 | 0 | 108740 | 0 |
| LGGLTQAP | U2AF2; U2AF2 | P26368; K7E | U2AF2_HUMAN | Unmodified | 3305200 | 2238300 | 0 | 0 | 0 | 0 | 0 | 0 | 0 | 0 | 0 | 4218100 | 0 | 0 | 0 | 268000 | 0 | 0 |
| LGGLVQPH | ITGB5; ITGB5; I | P18084; H7 | ITB5_HUMAN I | Oxidation (M) | 1481600 | 142270 | 1004000 | 1000900 | 0 | 0 | 0 | 0 | 0 | 0 | 0 | 8040100 | 57129 | 419830 | 2150700 | 1318300 | 0 | 0 |
| LGGNVVVS | CFL2 | Q9Y281 | COF2_HUMAN | Unmodified | 0 | 0 | 0 | 0 | 26941000 | 4636300 | 0 | 0 | 0 | 144200 | 0 | 610340 | 1462700 | 1097800 | 0 | 0 | 3698000 | 2198100 |
| LGGPEAGL | FTL | P02792 | FRIL_HUMAN F | Unmodified | 0 | 0 | 0 | 0 | 0 | 0 | 0 | 0 | 16191000 | 393180 | 168030 | 0 | 0 | 0 | 874700 | 493910 | 0 | 0 |
| LGGPEAGL | FTL | P02792 | FRIL_HUMAN F | Unmodified | 0 | 0 | 0 | 15985 | 577240 | 126820 | 1083700 | 0 | 1167200 | 4984400 | 184510 | 0 | 161560 | 0 | 0 | 0 | 123390 | 117730 |
| LGGPNQDI | OS9 | Q13438 | OS9_HUMAN P | Unmodified | 0 | 0 | 0 | 0 | 0 | 0 | 28674 | 36692 | 937810 | 0 | 0 | 0 | 518900 | 0 | 0 | 0 | 0 | 0 |
| LGGPQASL | DECR2; DECR2 | Q9NUI1; Q4 | DECR2_HUMAI | Unmodified | 0 | 0 | 0 | 0 | 0 | 0 | 0 | 0 | 2108100 | 0 | 0 | 0 | 0 | 0 | 0 | 0 | 0 | 0 |
| LGGQDFN | HSPA13 | P48723 | HSP13_HUMAN | Unmodified | 0 | 0 | 0 | 0 | 0 | 0 | 70117 | 0 | 0 | 620710 | 0 | 0 | 0 | 0 | 0 | 0 | 0 | 171270 |
| LGGSAVISI | CFL1; CFL1; CF | P23528; G3 | COF1_HUMAN | Unmodified | 15469000 | 17550000 | 9702200 | 13691000 | 2.62E+08 | 35460000 | 47630000 | 14213000 | 20542000 | 21206000 | 21644000 | 51013000 | 1.18E+08 | 65599000 | 12117000 | 12200000 | 86166000 | 65517000 |
| LGGSLADS | CCT2; CCT2; C | P78371; F5G | TCPB_HUMAN | Unmodified | 94130 | 111030 | 0 | 221840 | 1442000 | 0 | 46772 | 50472 | 0 | 53666 | 290120 | 1036600 | 2236300 | 2368600 | 431970 | 856970 | 863890 | 0 |
| LGGSLADS | CCT2; CCT2; C | P78371; F5G | TCPB_HUMAN | Unmodified | 0 | 0 | 0 | 0 | 690880 | 0 | 567740 | 20276 | 1199000 | 0 | 4545000 | 7753900 | 0 | 0 | 0 | 0 | 788180 | 0 |
| LGGSLADS | CCT2; CCT2; C | P78371; F5G | TCPB_HUMAN | Unmodified | 0 | 0 | 0 | 0 | 744910 | 0 | 2362000 | 37453 | 0 | 369400 | 0 | 0 | 1839700 | 2462100 | 0 | 0 | 2752300 | 1029300 |
| LGGSPFGP | DYNC1H1; DYN | Q14204; A0 | DYHC1_HUMA | Unmodified | 0 | 0 | 0 | 0 | 0 | 0 | 0 | 0 | 0 | 0 | 0 | 1014500 | 0 | 0 | 0 | 0 | 0 | 0 |
| LGGSPFGP | DYNC1H1; DYN | Q14204; A0 | DYHC1_HUMA | Unmodified | 0 | 0 | 0 | 1580600 | 0 | 0 | 0 | 0 | 1422000 | 0 | 780920 | 11489000 | 1337900 | 0 | 1007000 | 749360 | 0 | 0 |
| LGGSPTSL | PRKCSH; PRKC | P14314; K7E | GLU2B_HUMAI | Unmodified | 0 | 0 | 875770 | 0 | 1107100 | 0 | 0 | 0 | 4.45E+08 | 17246000 | 22587000 | 7484800 | 0 | 0 | 1394300 | 1067700 | 0 | 563260 |
| LGGSPTSL | PRKCSH; PRKC | P14314; K7E | GLU2B_HUMAI | Unmodified | 0 | 0 | 0 | 0 | 0 | 17221000 | 0 | 0 | 1.75E+08 | 3828800 | 1922500 | 0 | 0 | 0 | 0 | 0 | 0 | 0 |
| LGGSPTSL | PRKCSH; PRKC | P14314; K7E | GLU2B_HUMAI | Oxidation (M) | 0 | 793110 | 0 | 0 | 292560 | 0 | 0 | 0 | 67604000 | 13792000 | 8447700 | 1339900 | 0 | 0 | 0 | 0 | 0 | 0 |
| LGGSPTSL | PRKCSH; PRKC | P14314; K7E | GLU2B_HUMAI | Oxidation (M) | 0 | 0 | 0 | 0 | 0 | 0 | 0 | 0 | 20338000 | 8466400 | 0 | 0 | 0 | 0 | 0 | 0 | 0 | 0 |
| LGGSPTSL | PRKCSH; PRKC | P14314; K7E | GLU2B_HUMAI | Unmodified | 0 | 0 | 0 | 0 | 0 | 0 | 0 | 0 | 8995600 | 455040 | 0 | 0 | 0 | 0 | 0 | 0 | 0 | 0 |
| LGGSQED | TALDO1; TALD | P37837; F2Z | TALDO_HUMAN | Unmodified | 3327800 | 2410400 | 2775700 | 2400100 | 0 | 0 | 316470 | 99982 | 611370 | 187120 | 2240900 | 18293000 | 0 | 13202 | 3501800 | 2100000 | 0 | 0 |

| | | | | | | | | | | | | | | | | | | | | | | |
|---|---|---|---|---|---|---|---|---|---|---|---|---|---|---|---|---|---|---|---|---|---|---|
| LGGSRALS | SULF2 | Q8IWU5 | SULF2_HUMAN | Unmodified | 0 | 0 | 0 | 0 | 0 | 0 | 0 | 0 | 0 | 0 | 0 | 0 | 72288 | 0 | 0 | 0 | 83044 | 820560 |
| LGGSTSDP | SNTB1; SNTB1 | Q13884; A0A | SNTB1_HUMAN | Unmodified | 0 | 0 | 0 | 0 | 0 | 0 | 0 | 0 | 1360600 | 46779 | 0 | 0 | 0 | 0 | 0 | 0 | 0 | 0 |
| LGGTIDDC | CCT4 | P50991 | TCPD_HUMAN | Unmodified | 0 | 0 | 0 | 0 | 0 | 0 | 0 | 0 | 866860 | 0 | 791360 | 1057100 | 1252200 | 0 | 0 | 0 | 0 | 0 |
| LGGVKYDII | ATOX1; ATOX1; | O00244; E5F | ATOX1_HUMAN | Unmodified | 1162400 | 922340 | 1068100 | 1734300 | 3059900 | 498420 | 89998 | 67871 | 810230 | 274350 | 832650 | 1592000 | 5998000 | 4048600 | 2302100 | 3520900 | 1984100 | 2849400 |
| LGGVKYDII | ATOX1; ATOX1; | O00244; E5F | ATOX1_HUMAN | Unmodified | 0 | 0 | 478020 | 778590 | 505780 | 0 | 0 | 0 | 0 | 82564 | 362680 | 0 | 2529100 | 1511100 | 1232700 | 1414600 | 0 | 808360 |
| LGGYLAEE( | DNAJC13 | O75165 | DJC13_HUMAN | Unmodified | 0 | 0 | 0 | 0 | 0 | 0 | 0 | 0 | 212380 | 0 | 0 | 3255300 | 0 | 0 | 0 | 0 | 0 | 0 |
| LGHAEQKD | RCC2 | Q9P258 | RCC2_HUMAN | Unmodified | 27235 | 0 | 0 | 0 | 0 | 109640 | 15053 | 0 | 334190 | 0 | 272210 | 7871400 | 0 | 0 | 0 | 0 | 122990 | 0 |
| LGHAETDE | ECPAS; ECPAS; | Q5VYK3; R4( | ECM29_HUMAI | Unmodified | 74623 | 95525 | 0 | 100570 | 0 | 0 | 0 | 0 | 0 | 0 | 604830 | 3725300 | 0 | 0 | 302680 | 166930 | 0 | 0 |
| LGHCFAPS | IQGAP3; IQGAP | Q86VI3; F2Z: | IQGA3_HUMAN | Unmodified | 0 | 0 | 0 | 0 | 1304200 | 0 | 0 | 0 | 202250 | 150920 | 0 | 0 | 0 | 0 | 0 | 0 | 569330 | 170440 |
| LGHGLLSG | USP5 | P45974 | UBP5_HUMAN | Unmodified | 7617600 | 4946900 | 1368000 | 1822900 | 742700 | 0 | 0 | 0 | 1279100 | 164490 | 396520 | 5252900 | 3530500 | 769880 | 2021400 | 2249500 | 282120 | 979430 |
| LGHGLLSG | USP13; USP13; | Q92995; A0A | UBP13_HUMAN | Unmodified | 502510 | 206820 | 0 | 219390 | 0 | 0 | 0 | 0 | 0 | 0 | 0 | 0 | 0 | 0 | 151550 | 205680 | 0 | 0 |
| LGHGYHTL | ADA; ADA; ADA | P00813; F5G | ADA_HUMAN A | Unmodified | 55174 | 0 | 0 | 0 | 0 | 0 | 0 | 0 | 0 | 0 | 0 | 1429600 | 0 | 0 | 0 | 0 | 0 | 0 |
| LGHPDTLN | S100A9 | P06702 | S10A9_HUMAN | Unmodified | 0 | 291480 | 0 | 4938700 | 1372600 | 291640 | 0 | 242760 | 462590 | 66155 | 0 | 0 | 310620 | 0 | 0 | 156740 | 137810 | 661210 |
| LGHPEALS. | ZYX; ZYX; ZYX | Q15942; H0Y | ZYX_HUMAN Zy | Unmodified | 0 | 0 | 0 | 0 | 1100700 | 0 | 0 | 0 | 0 | 0 | 0 | 0 | 172250 | 0 | 514750 | 210600 | 485330 | 6427700 |
| LGHPEALS. | ZYX; ZYX; ZYX | Q15942; H0Y | ZYX_HUMAN Zy | Unmodified | 0 | 0 | 0 | 751200 | 0 | 0 | 0 | 0 | 0 | 0 | 0 | 0 | 0 | 0 | 0 | 0 | 0 | 2570400 |
| LGHVVMGI | EIF3E; EIF3E; EI | P60228; B3K | EIF3E_HUMAN | Unmodified | 0 | 0 | 204570 | 154890 | 0 | 0 | 0 | 0 | 1065600 | 0 | 0 | 8890500 | 419770 | 0 | 0 | 227430 | 0 | 214340 |
| LGHVVMGI | EIF3E; EIF3E; EI | P60228; B3K | EIF3E_HUMAN | Oxidation (M) | 1295600 | 1123100 | 958150 | 1927400 | 0 | 0 | 0 | 109580 | 0 | 0 | 565300 | 15611000 | 706070 | 0 | 1495000 | 888740 | 0 | 0 |
| LGIAPQIQD | IQGAP2; IQGAP | Q13576; D6I | IQGA2_HUMAN | Unmodified | 0 | 0 | 311970 | 0 | 0 | 0 | 0 | 0 | 0 | 0 | 0 | 0 | 0 | 0 | 527470 | 0 | 0 | 0 |
| LGIEGLSLH | UGGT1 | Q9NYU2 | UGGG1_HUMA | Unmodified | 0 | 0 | 0 | 0 | 0 | 0 | 0 | 0 | 26125000 | 1228800 | 781940 | 2051000 | 0 | 0 | 0 | 0 | 0 | 0 |
| LGIHEDSQ | HSP90AA1 | P07900 | HS90A_HUMAN | Unmodified | 453990 | 0 | 449710 | 680210 | 0 | 0 | 0 | 0 | 1923600 | 186960 | 1708200 | 16382000 | 0 | 0 | 712400 | 445030 | 0 | 0 |
| LGIHEDSQ | HSP90AA1 | P07900 | HS90A_HUMAN | Unmodified | 918160 | 563140 | 5861600 | 1927100 | 876830 | 1520600 | 0 | 0 | 2980400 | 9938300 | 15884000 | 50042000 | 0 | 0 | 1441600 | 2279100 | 2072100 | 5838700 |
| LGIHEDSQ | HSP90AA1 | P07900 | HS90A_HUMAN | Unmodified | 0 | 0 | 0 | 0 | 8118000 | 1318300 | 1177500 | 570910 | 1705600 | 3106400 | 1040900 | 1390300 | 1450700 | 898290 | 0 | 59614 | 2656000 | 3696500 |
| LGIHEDSTI | HSP90AB1 | P08238 | HS90B_HUMAN | Unmodified | 1379800 | 891210 | 3263700 | 2156500 | 1028500 | 241890 | 72409 | 0 | 2861300 | 594510 | 4233400 | 54293000 | 931650 | 298160 | 3663200 | 2775900 | 287870 | 499740 |
| LGIHEDSTI | HSP90AB1 | P08238 | HS90B_HUMAN | Unmodified | 23712 | 370070 | 0 | 1398000 | 5104300 | 0 | 0 | 75604 | 0 | 615080 | 3363300 | 16378000 | 0 | 0 | 0 | 565860 | 413540 | 4860100 |
| LGIHSALLD | GLRX5 | Q86SX6 | GLRX5_HUMAN | Unmodified | 0 | 0 | 0 | 0 | 0 | 0 | 286970 | 0 | 908870 | 381190 | 0 | 0 | 0 | 0 | 0 | 0 | 29440 | 0 |
| LGIIGGNLY | SLC22A3 | O75751 | S22A3_HUMAN | Unmodified | 0 | 0 | 0 | 0 | 0 | 0 | 0 | 0 | 0 | 0 | 0 | 0 | 0 | 0 | 101210 | 206440 | 0 | 0 |
| LGIIGVVNF | DNM1L; DNM1L | O00429; F8V | DNM1L_HUMA | Unmodified | 0 | 0 | 0 | 0 | 0 | 0 | 0 | 0 | 189620 | 0 | 396770 | 4482300 | 0 | 0 | 115610 | 80409 | 0 | 0 |
| LGIKPESVC | CSRP2; CSRP2 | Q16527; F8V | CSRP2_HUMAN | Unmodified | 0 | 0 | 0 | 0 | 0 | 0 | 0 | 0 | 0 | 0 | 621350 | 1473600 | 0 | 0 | 0 | 0 | 0 | 0 |
| LGIKPSINY | RNASET2; RNAS | O00584; D6I | RNT2_HUMAN | Unmodified | 0 | 0 | 172030 | 0 | 0 | 0 | 0 | 0 | 3437900 | 1083100 | 2288100 | 496190 | 2909900 | 1612400 | 0 | 0 | 0 | 0 |
| LGIKPSINY | RNASET2; RNAS | O00584; D6I | RNT2_HUMAN | Unmodified | 0 | 0 | 0 | 0 | 0 | 0 | 0 | 0 | 1430800 | 1263000 | 0 | 0 | 1185300 | 819330 | 0 | 0 | 0 | 0 |
| LGINKTDPS | MTHFD1L; MTH | Q6UB35; A0. | C1TM_HUMAN | Unmodified | 0 | 0 | 0 | 0 | 0 | 0 | 0 | 0 | 768200 | 68392 | 0 | 0 | 0 | 0 | 0 | 0 | 0 | 0 |
| LGINLLGGI | NEK9; NEK9; NI | Q8TD19; A0A | NEK9_HUMAN | Unmodified | 0 | 0 | 0 | 0 | 0 | 639020 | 0 | 0 | 0 | 0 | 0 | 718150 | 0 | 0 | 0 | 0 | 0 | 0 |
| LGINSLQEI | TPD52; TPD52; | P55327; H0Y | TPD52_HUMAN | Unmodified | 0 | 102590 | 0 | 378220 | 1715900 | 0 | 648900 | 240280 | 6703800 | 1714400 | 0 | 6468600 | 1200700 | 0 | 270100 | 374980 | 504550 | 902440 |
| LGIQTDDK | GSR | P00390 | GSHR_HUMAN | Unmodified | 0 | 0 | 0 | 0 | 1347600 | 0 | 0 | 0 | 0 | 0 | 0 | 3459800 | 0 | 0 | 0 | 0 | 0 | 0 |
| LGIQTDDK( | GSR | P00390 | GSHR_HUMAN | Unmodified | 0 | 180590 | 0 | 0 | 30612000 | 5204300 | 1160600 | 0 | 0 | 1982000 | 1608800 | 3243200 | 14090000 | 8151100 | 0 | 189560 | 624790 | 2640700 |
| LGISSLIR | AGL | P35573 | GDE_HUMAN ( | Unmodified | 0 | 0 | 0 | 0 | 0 | 0 | 0 | 0 | 0 | 0 | 0 | 2452200 | 0 | 0 | 0 | 0 | 0 | 0 |
| LGISTFWSI | KIF14 | Q15058 | KIF14_HUMAN | Unmodified | 0 | 0 | 0 | 0 | 805460 | 0 | 525720 | 0 | 0 | 132010 | 0 | 0 | 62009 | 0 | 0 | 0 | 411030 | 0 |
| LGITHVLN/ | DUSP3; DUSP3 | P51452; K7E | DUS3_HUMAN | Unmodified | 502050 | 0 | 0 | 0 | 0 | 0 | 0 | 0 | 0 | 0 | 0 | 782930 | 428620 | 245150 | 0 | 0 | 0 | 0 |
| LGIVQGIVC | LAMB2 | P55268 | LAMB2_HUMAN | Unmodified | 0 | 0 | 0 | 0 | 0 | 0 | 0 | 0 | 2230800 | 0 | 0 | 0 | 0 | 0 | 0 | 0 | 0 | 0 |
| LGIWGGN\ | VPS13C | Q709C8 | VP13C_HUMAN | Unmodified | 0 | 0 | 0 | 0 | 0 | 0 | 0 | 0 | 0 | 0 | 0 | 544580 | 0 | 0 | 0 | 0 | 0 | 0 |
| LGIYDADG | ASPH; ASPH; A | Q12797; E5F | ASPH_HUMAN | Unmodified | 0 | 0 | 0 | 0 | 0 | 0 | 0 | 0 | 8343200 | 582160 | 2266600 | 0 | 0 | 0 | 0 | 0 | 0 | 0 |
| LGKADETKI | BIN1 | O00499 | BIN1_HUMAN I | Unmodified | 0 | 0 | 0 | 0 | 311670 | 0 | 0 | 0 | 0 | 0 | 0 | 0 | 1788200 | 0 | 200440 | 47285 | 1404000 | 734940 |
| LGKDAVED | DCD | P81605 | DCD_HUMAN I | Unmodified | 123940 | 69677 | 0 | 27925 | 2227300 | 0 | 354860 | 0 | 0 | 0 | 101730 | 0 | 96821 | 0 | 0 | 0 | 0 | 0 |
| LGKDPNTY | DDB1; DDB1; D | Q16531; F5C | DDB1_HUMAN | Oxidation (M) | 0 | 0 | 0 | 0 | 0 | 0 | 0 | 0 | 0 | 0 | 0 | 3870200 | 0 | 0 | 0 | 0 | 0 | 0 |
| LGKPGLGC | MGMT; MGMT | P16455; B4D | MGMT_HUMAN | Unmodified | 0 | 0 | 0 | 0 | 0 | 0 | 0 | 0 | 0 | 0 | 0 | 175610 | 212640 | 0 | 630920 | 307680 | 0 | 0 |
| LGLDEYLDI | FMNL2 | Q96PY5 | FMNL2_HUMAI | Unmodified | 0 | 0 | 0 | 0 | 0 | 0 | 717030 | 101370 | 0 | 0 | 0 | 1511000 | 0 | 0 | 0 | 0 | 0 | 0 |
| LGLEDFESI | STK38 | Q15208 | STK38_HUMAN | Unmodified | 0 | 0 | 0 | 0 | 0 | 0 | 0 | 0 | 0 | 0 | 0 | 1165700 | 0 | 0 | 0 | 0 | 0 | 0 |
| LGLENAEAI | RO60 | P10155 | RO60_HUMAN | Unmodified | 51289000 | 38467000 | 8961400 | 14904000 | 0 | 0 | 0 | 0 | 1159700 | 0 | 735130 | 2750500 | 0 | 0 | 31032000 | 13539000 | 0 | 0 |
| LGLERNPQ | MYO1C; MYO1( | O00159; F5H | MYO1C_HUMA | Unmodified | 0 | 0 | 2039300 | 2038800 | 2091500 | 594880 | 149230 | 19165 | 325610 | 0 | 0 | 236030 | 302730 | 305720 | 0 | 1664200 | 940240 | 322750 |
| LGLFPSNF\ | CD2AP | Q9Y5K6 | CD2AP_HUMAI | Unmodified | 31071000 | 41311000 | 17758000 | 23078000 | 5068400 | 1475900 | 0 | 0 | 1368600 | 745070 | 1237200 | 0 | 16762000 | 9244200 | 55788000 | 32334000 | 0 | 0 |
| LGLGEGAE | RCC1; RCC1 | P18754; C9J | RCC1_HUMAN | Unmodified | 1479600 | 362360 | 36150 | 2942700 | 1053800 | 166020 | 84937 | 0 | 938010 | 256980 | 455600 | 1178700 | 1389300 | 381830 | 192420 | 201610 | 614950 | 710840 |
| LGLGIDEDI | HSP90AA1 | P07900 | HS90A_HUMAI | 2 Oxidation (M) | 0 | 0 | 0 | 0 | 4604900 | 0 | 0 | 1803500 | 0 | 527900 | 0 | 0 | 1274500 | 0 | 0 | 0 | 0 | 5292600 |
| LGLGIDEDI | HSP90AB1 | P08238 | HS90B_HUMAI | Unmodified | 0 | 0 | 0 | 0 | 0 | 0 | 0 | 0 | 0 | 651960 | 3178200 | 0 | 0 | 0 | 0 | 0 | 0 | 0 |
| LGLGLEFQ. | VDAC1 | P21796 | VDAC1_HUMAI | Unmodified | 0 | 0 | 0 | 0 | 0 | 0 | 0 | 0 | 0 | 1341900 | 0 | 0 | 0 | 0 | 0 | 0 | 0 | 0 |
| LGLHQVLQ | DIAPH1; DIAPH | O60610; A0A | DIAP1_HUMAN | Unmodified | 0 | 0 | 0 | 0 | 0 | 0 | 0 | 0 | 0 | 0 | 126100 | 2593600 | 0 | 0 | 258250 | 257420 | 0 | 0 |
| LGLIPLISDI | POLR2C | P19387 | RPB3_HUMAN | Unmodified | 185830 | 0 | 75841 | 0 | 0 | 0 | 0 | 0 | 222280 | 0 | 0 | 824930 | 0 | 0 | 0 | 76238 | 0 | 0 |
| LGLKSLVSH | H1-3; H1-4; H1- | P16402; P10 | H13_HUMAN H | Unmodified | 0 | 0 | 0 | 0 | 0 | 0 | 0 | 3474500 | 0 | 0 | 0 | 0 | 0 | 0 | 0 | 0 | 0 | 0 |
| LGLLDNHS | LONP1; LONP1 | P36776; K7E | LONM_HUMAN | Unmodified | 1912800 | 1953300 | 1140300 | 319500 | 0 | 0 | 2326100 | 285350 | 6435100 | 132960 | 0 | 0 | 0 | 0 | 1546700 | 1664000 | 0 | 0 |
| LGLLEALLK | THOC2; THOC2 | Q8NI27; A0A | THOC2_HUMA | Unmodified | 38422 | 0 | 32393 | 0 | 0 | 0 | 37617 | 0 | 30383 | 19117 | 59939 | 685540 | 0 | 0 | 0 | 161270 | 0 | 0 |
| LGLLGDSV | CEL | P19835 | CEL_HUMAN B | Unmodified | 0 | 0 | 0 | 0 | 0 | 0 | 0 | 0 | 11721000 | 1081500 | 5456300 | 483860 | 0 | 0 | 0 | 0 | 0 | 0 |
| LGLLGDSV | CEL | P19835 | CEL_HUMAN B | Unmodified | 0 | 0 | 0 | 0 | 0 | 0 | 0 | 0 | 127570 | 2854500 | 48916 | 0 | 0 | 0 | 0 | 0 | 0 | 0 |
| LGLLGLAN: | HSD17B4; HSD | P51659; A0A | DHB4_HUMAN | Unmodified | 0 | 0 | 0 | 0 | 0 | 0 | 0 | 0 | 3328100 | 996690 | 35268 | 0 | 156820 | 0 | 0 | 0 | 0 | 0 |
| LGLLGLAN: | HSD17B4; HSD | P51659; A0A | DHB4_HUMAN | Unmodified | 0 | 0 | 0 | 0 | 928850 | 99581 | 2291300 | 34348 | 75777000 | 14645000 | 2700800 | 343670 | 0 | 0 | 0 | 0 | 0 | 285280 |
| LGLMRDDT | UQCRB; UQCRI | P14927; B7Z | QCR7_HUMAN | Oxidation (M) | 0 | 0 | 0 | 0 | 0 | 0 | 0 | 0 | 9241400 | 2152600 | 898910 | 0 | 0 | 0 | 0 | 0 | 0 | 0 |
| LGLMRDDT | UQCRB; UQCRI | P14927; B7Z | QCR7_HUMAN | Unmodified | 111400 | 0 | 43885 | 0 | 0 | 0 | 0 | 0 | 18044000 | 1116200 | 224230 | 0 | 0 | 0 | 0 | 0 | 0 | 0 |
| LGLMRDDT | UQCRB; UQCRI | P14927; B7Z | QCR7_HUMAN | Unmodified | 0 | 0 | 0 | 0 | 0 | 0 | 7224700 | 4921700 | 4539700 | 627960 | 0 | 0 | 0 | 0 | 0 | 0 | 2683800 | 0 |
| LGLMRDDT | UQCRB; UQCRI | P14927; B7Z | QCR7_HUMAN | Oxidation (M) | 0 | 0 | 0 | 0 | 0 | 0 | 0 | 95579 | 2819000 | 934690 | 0 | 0 | 0 | 0 | 0 | 0 | 0 | 0 |
| LGLNDRVL | AP1M2; AP1M2; | Q9Y6Q5; K7I | AP1M2_HUMAI | Unmodified | 62810 | 377050 | 0 | 352000 | 0 | 0 | 0 | 0 | 0 | 0 | 0 | 0 | 0 | 0 | 0 | 0 | 0 | 0 |
| LGLPPLTPE | PUF60; PUF60; | Q9UHX1; E9 | PUF60_HUMAN | Unmodified | 0 | 0 | 0 | 0 | 991460 | 0 | 0 | 108660 | 771360 | 1015000 | 979630 | 1222600 | 707010 | 174930 | 0 | 0 | 628000 | 0 |
| LGLPQDQD | SARG | Q9BW04 | SARG_HUMAN | Unmodified | 1306200 | 383070 | 0 | 2793100 | 549930 | 0 | 0 | 0 | 0 | 0 | 0 | 0 | 22544000 | 19152000 | 0 | 2034000 | 0 | 0 |
| LGLPSRDY | MME; MME; MM | P08473; A0A | NEP_HUMAN N | Unmodified | 0 | 0 | 17103000 | 57170000 | 0 | 0 | 0 | 0 | 0 | 1854700 | 0 | 996440 | 12035000 | 2194400 | 56111000 | 34293000 | 0 | 0 |
| LGLQCLPS | SLC19A1 | P41440 | S19A1_HUMAN | Unmodified | 0 | 0 | 341800 | 464250 | 0 | 0 | 0 | 0 | 0 | 0 | 159020 | 469140 | 832670 | 603410 | 0 | 484230 | 0 | 0 |
| LGLQHLR | ARF5; ARF5 | P84085; C9J | ARF5_HUMAN . | Unmodified | 329310 | 535970 | 765540 | 151150 | 0 | 0 | 0 | 0 | 0 | 0 | 221110 | 2100300 | 0 | 0 | 687830 | 432240 | 0 | 0 |
| LGLQNDLF | NPEPPS; NPEPI | P55786; A0A | PSA_HUMAN P | Unmodified | 0 | 0 | 0 | 0 | 0 | 0 | 0 | 0 | 0 | 0 | 0 | 7296600 | 0 | 0 | 0 | 0 | 0 | 0 |
| LGLQYMEL | LMCD1; LMCD1 | Q9NZU5; H7 | LMCD1_HUMA | Oxidation (M) | 0 | 0 | 0 | 0 | 0 | 0 | 0 | 0 | 0 | 0 | 0 | 2504500 | 0 | 0 | 0 | 0 | 0 | 0 |
| LGLSDMNI | BLMH; BLMH; B | Q13867; K7E | BLMH_HUMAN | Oxidation (M) | 0 | 0 | 0 | 0 | 1968100 | 0 | 1986300 | 0 | 0 | 1202800 | 0 | 0 | 2407900 | 1605500 | 0 | 0 | 97267 | 2151500 |
| LGLSLVVP( | NAPG | Q99747 | SNAG_HUMAN | Unmodified | 0 | 0 | 0 | 0 | 2466700 | 0 | 0 | 0 | 213960 | 361440 | 287260 | 422510 | 345930 | 305680 | 198530 | 0 | 313610 | 1780000 |
| LGLSLVVP( | NAPG | Q99747 | SNAG_HUMAN | Unmodified | 0 | 0 | 0 | 0 | 5954700 | 0 | 0 | 284450 | 1109300 | 803250 | 547690 | 0 | 1734600 | 1829400 | 0 | 0 | 4073300 | 3880800 |
| LGLSTLGEI | TPD52L2; TPD5 | O43399; A0A | TPD54_HUMAN | Unmodified | 0 | 0 | 0 | 0 | 1517800 | 42151 | 0 | 0 | 540650 | 167880 | 339130 | 2034300 | 0 | 0 | 0 | 0 | 416980 | 0 |
| LGLVGVNL | ACLY; ACLY | P53396; K7E | ACLY_HUMAN . | Unmodified | 0 | 0 | 0 | 23900 | 3250700 | 0 | 137750 | 0 | 563490 | 875360 | 0 | 2658100 | 1583500 | 36332 | 0 | 0 | 26518 | 0 |
| LGMLSPEG | FASN; FASN | P49327; A0A | FAS_HUMAN F | Oxidation (M) | 0 | 0 | 302910 | 0 | 0 | 0 | 0 | 0 | 0 | 0 | 111220 | 5165300 | 0 | 0 | 418440 | 400910 | 0 | 0 |
| LGMLSPEG | FASN; FASN | P49327; A0A | FAS_HUMAN F | Unmodified | 0 | 0 | 0 | 677130 | 0 | 185450 | 127120 | 46219 | 823450 | 0 | 0 | 14919000 | 0 | 0 | 270300 | 327060 | 0 | 0 |
| LGMSQIPQ | UTRN | P46939 | UTRN_HUMAN | Unmodified | 0 | 0 | 206260 | 333730 | 570380 | 125460 | 0 | 150680 | 481020 | 268050 | 434150 | 0 | 0 | 263320 | 209020 | 202920 | 508040 | 929260 |
| LGMSQIPQ | UTRN | P46939 | UTRN_HUMAN | Oxidation (M) | 0 | 0 | 133310 | 189440 | 341650 | 0 | 48393 | 0 | 327930 | 0 | 0 | 0 | 86162 | 0 | 168200 | 275460 | 71236 | 483020 |
| LGMSSDLV | PARP12; PARP1 | Q9H0J9; H7( | PAR12_HUMAN | Oxidation (M) | 0 | 288000 | 790210 | 441020 | 0 | 0 | 0 | 0 | 0 | 0 | 0 | 0 | 350640 | 0 | 690390 | 377440 | 0 | 0 |
| LGNDFHTN | RPS17; RPS17; | P08708; H0Y | RS17_HUMAN . | Unmodified | 633820 | 0 | 294060 | 0 | 0 | 0 | 0 | 0 | 0 | 0 | 0 | 4592700 | 0 | 0 | 315490 | 338470 | 0 | 0 |
| LGNFFSPK | IQGAP1; IQGAP | P46940; A0A | IQGA1_HUMAN | Unmodified | 3973600 | 4367800 | 3862600 | 8110000 | 54622000 | 11869000 | 6584000 | 8684300 | 4390300 | 3761600 | 1309000 | 11078000 | 14238000 | 14544000 | 4618300 | 3932800 | 11813000 | 8529900 |
| LGNIEFKPE | MYO1B; MYO1E | O43795; E9F | MYO1B_HUMA | Unmodified | 395240 | 400060 | 236150 | 451020 | 2928800 | 491460 | 404970 | 155250 | 552490 | 0 | 4357300 | 23175000 | 0 | 0 | 1786500 | 394440 | 367650 | 0 |
| LGNIEFKPE | MYO1B; MYO1E | O43795; E9F | MYO1B_HUMA | Unmodified | 0 | 0 | 0 | 0 | 2803000 | 1691400 | 0 | 113910 | 0 | 0 | 0 | 0 | 0 | 0 | 0 | 0 | 1650900 | 883680 |
| LGNKVQEL | STX4; STX4; ST) | Q12846; A0A | STX4_HUMAN S | Unmodified | 110640 | 80044 | 0 | 374380 | 802520 | 185620 | 72665 | 0 | 231170 | 175380 | 75511 | 794660 | 464960 | 0 | 72392 | 0 | 239150 | 286680 |
| LGNPTRSE | RPN1; RPN1 | P04843; B7Z | RPN1_HUMAN | Unmodified | 53502 | 0 | 122440 | 173940 | 0 | 1559700 | 605210 | 472530 | 1.11E+08 | 2402200 | 1040000 | 2947000 | 0 | 1738500 | 151690 | 0 | 0 | 1716300 |
| LGNPTRSE | RPN1; RPN1 | P04843; B7Z | RPN1_HUMAN | Unmodified | 0 | 0 | 527180 | 0 | 2768700 | 1531900 | 1677100 | 0 | 3.51E+08 | 36582000 | 16981000 | 1482100 | 67577000 | 3553700 | 0 | 0 | 0 | 9006700 |
| LGNSADAL | PEX11B | O96011 | PX11B_HUMAN | Unmodified | 0 | 0 | 0 | 0 | 0 | 0 | 0 | 0 | 2095200 | 141700 | 389460 | 0 | 0 | 0 | 0 | 0 | 0 | 137760 |
| LGNSADAL | PEX11B | O96011 | PX11B_HUMAN | Unmodified | 0 | 59056 | 0 | 0 | 0 | 0 | 0 | 0 | 3829300 | 323040 | 0 | 0 | 0 | 0 | 0 | 0 | 0 | 0 |
| LGNSLLLK | CPSF1 | Q10570 | CPSF1_HUMAN | Unmodified | 0 | 9803000 | 0 | 0 | 0 | 0 | 0 | 0 | 0 | 0 | 0 | 244930 | 0 | 0 | 0 | 52506 | 0 | 0 |
| LGNWLAVF | TJP2; TJP2; TJP2 | Q9UDY2; A0 | ZO2_HUMAN T | Unmodified | 63732 | 0 | 0 | 184620 | 482160 | 147130 | 0 | 0 | 4872400 | 63275 | 682910 | 1862600 | 0 | 0 | 217190 | 219070 | 0 | 0 |
| LGNWLAVF | TJP2; TJP2; TJP2 | Q9UDY2; A0 | ZO2_HUMAN T | Unmodified | 0 | 0 | 0 | 0 | 9242100 | 1403200 | 0 | 0 | 0 | 3601400 | 0 | 0 | 852230 | 354740 | 0 | 0 | 0 | 0 |
| LGNYAGAV | SGTA; SGTA; S( | O43765; K7E | SGTA_HUMAN | Unmodified | 0 | 0 | 0 | 0 | 0 | 0 | 0 | 0 | 795660 | 0 | 745370 | 2488900 | 313620 | 0 | 0 | 0 | 0 | 0 |
| LGNYAGAV | SGTA; SGTA; S( | O43765; K7E | SGTA_HUMAN | Unmodified | 0 | 0 | 0 | 0 | 832010 | 0 | 783540 | 0 | 0 | 0 | 0 | 0 | 0 | 0 | 0 | 0 | 1432000 | 0 |
| LGPARHGE | ERBIN | Q96RT1 | ERBIN_HUMAN | Oxidation (M) | 0 | 0 | 0 | 0 | 2984800 | 0 | 0 | 0 | 0 | 0 | 0 | 0 | 0 | 0 | 0 | 0 | 0 | 0 |
| LGPDDPN\ | KLC1; KLC2; KL | Q07866; Q9I | KLC1_HUMAN | Unmodified | 0 | 0 | 0 | 0 | 2612200 | 598790 | 450620 | 0 | 1109300 | 652190 | 408050 | 3118900 | 1194700 | 722570 | 0 | 241590 | 594530 | 0 |
| LGPEAFWF | MAN1C1; MAN: | Q9NR34; A6 | MA1C1_HUMAI | Unmodified | 0 | 0 | 0 | 0 | 0 | 252020 | 0 | 0 | 2416300 | 0 | 0 | 0 | 0 | 0 | 0 | 0 | 0 | 647460 |
| LGPESFKFI | MAN1A2; MAN1 | O60476; H0Y | MA1A2_HUMAI | Unmodified | 0 | 0 | 0 | 0 | 0 | 0 | 0 | 11911 | 5553900 | 0 | 0 | 0 | 0 | 0 | 0 | 0 | 0 | 46613 |
| LGPGGLDD | PKP3; PKP3; PK | Q9Y446; E9F | PKP3_HUMAN | Unmodified | 2560300 | 3769400 | 3144800 | 11679000 | 1111800 | 0 | 0 | 0 | 762030 | 0 | 1183500 | 10644000 | 2867000 | 0 | 12571000 | 9167700 | 636180 | 0 |
| LGPGGLDF | CDC37; CDC37 | Q16543; K7E | CDC37_HUMA | Unmodified | 0 | 0 | 0 | 364980 | 1149600 | 0 | 0 | 0 | 1310600 | 689010 | 1326700 | 546660 | 843110 | 482880 | 0 | 0 | 1279800 | 1419200 |
| LGPLALPEI | PCDH1 | D6RAX3 | D6RAX3_HUM/ | Unmodified | 282620 | 0 | 0 | 0 | 132300 | 258960 | 0 | 0 | 1068300 | 0 | 2903500 | 0 | 50687 | 0 | 508840 | 599980 | 492460 | 210050 |
| LGPLFSLLN | PSMD5; PSMD5 | Q16401; F2Z | PSMD5_HUMA | Unmodified | 0 | 0 | 0 | 0 | 0 | 0 | 0 | 0 | 0 | 0 | 0 | 1044600 | 0 | 0 | 0 | 0 | 0 | 0 |
| LGPLLDILA | MOGS; MOGS; | Q13724; A0A | MOGS_HUMAN | Unmodified | 0 | 0 | 0 | 0 | 0 | 0 | 0 | 0 | 2432600 | 0 | 127950 | 894990 | 0 | 0 | 0 | 0 | 0 | 0 |
| LGPQPLPIH | KDR | P35968 | VGFR2_HUMAN | Unmodified | 0 | 0 | 0 | 0 | 0 | 0 | 0 | 0 | 0 | 0 | 0 | 0 | 0 | 0 | 0 | 0 | 161200 | 875910 |

| | | | | | | | | | | | | | | | | | | | | | | |
|---|---|---|---|---|---|---|---|---|---|---|---|---|---|---|---|---|---|---|---|---|---|---|
| LGPSDYFG | PRKAR1A | P10644 | KAP0_HUMAN | Unmodified | 0 | 168480 | 59041 | 110840 | 0 | 0 | 0 | 0 | 1155100 | 0 | 0 | 0 | 0 | 0 | 743490 | 0 | 0 | 0 |
| LGQALTEV | HDLBP; HDLBP | Q00341; A0A | VIGLN_HUMAN | Unmodified | 326620 | 334750 | 382160 | 297890 | 670540 | 0 | 0 | 0 | 223490 | 0 | 339020 | 1584700 | 810850 | 539380 | 416520 | 410380 | 0 | 446360 |
| LGQEATVG | ACLY; ACLY | P53396; K7E | ACLY_HUMAN | Unmodified | 0 | 0 | 3885000 | 4798900 | 4795600 | 0 | 3375300 | 0 | 1515000 | 0 | 292940 | 20521000 | 3115500 | 570020 | 488590 | 406200 | 0 | 0 |
| LGQEATVG | ACLY; ACLY | P53396; K7E | ACLY_HUMAN | Unmodified | 0 | 0 | 0 | 0 | 1355600 | 0 | 3019800 | 667500 | 0 | 312960 | 0 | 0 | 720850 | 0 | 0 | 0 | 388970 | 0 |
| LGQEQQTL | CGN | Q9P2M7 | CING_HUMAN | Unmodified | 0 | 0 | 0 | 109590 | 0 | 0 | 0 | 0 | 305080 | 0 | 41112 | 1590000 | 0 | 0 | 90226 | 74346 | 0 | 0 |
| LGQGCFGI | YES1; YES1 | P07947; J3Q | YES_HUMAN Ty | Oxidation (M) | 0 | 0 | 0 | 0 | 0 | 0 | 0 | 0 | 0 | 0 | 0 | 0 | 0 | 0 | 0 | 0 | 730420 | 56931 |
| LGQGIGPIH | LOXL2; LOXL2 | Q9Y4K0; H0Y | LOXL2_HUMAN | Unmodified | 0 | 0 | 0 | 0 | 422320 | 1676200 | 0 | 0 | 13984000 | 2703700 | 0 | 0 | 0 | 0 | 0 | 0 | 0 | 1127600 |
| LGQGVLVQ | EXOSC2 | Q13868 | EXOS2_HUMAN | Unmodified | 0 | 0 | 0 | 0 | 0 | 0 | 0 | 0 | 0 | 0 | 0 | 495430 | 0 | 0 | 0 | 0 | 0 | 0 |
| LGQHLYGT | RET; RET; RET; R | P07949; Q9E | RET_HUMAN Pr | Unmodified | 0 | 0 | 0 | 0 | 20300 | 0 | 0 | 0 | 3475900 | 0 | 0 | 0 | 0 | 0 | 0 | 0 | 0 | 0 |
| LGQHVVGI | TSFM | P43897 | EFTS_HUMAN E | Unmodified | 0 | 0 | 0 | 0 | 0 | 0 | 0 | 314970 | 4270400 | 0 | 0 | 0 | 0 | 0 | 0 | 0 | 0 | 0 |
| LGQMIVDY | NCKAP1 | Q9Y2A7 | NCKP1_HUMAN | Oxidation (M) | 361520 | 310540 | 513720 | 656330 | 0 | 0 | 0 | 0 | 428710 | 165340 | 221170 | 2867400 | 0 | 0 | 837860 | 966010 | 0 | 0 |
| LGQMIVDY | NCKAP1 | Q9Y2A7 | NCKP1_HUMAN | Unmodified | 0 | 0 | 0 | 0 | 0 | 0 | 0 | 0 | 1474000 | 0 | 81522 | 1551600 | 0 | 0 | 0 | 0 | 51279 | 0 |
| LGQPVPSW | NDST2 | P52849 | NDST2_HUMAN | Unmodified | 0 | 21908 | 0 | 0 | 0 | 3023900 | 676600 | 0 | 10576000 | 1908400 | 0 | 0 | 0 | 2042400 | 0 | 0 | 292850 | 0 |
| LGQSDPAF | TKT; TKT; TKT | P29401; A0A | TKT_HUMAN Tr | Unmodified | 0 | 0 | 0 | 0 | 0 | 0 | 0 | 0 | 2319800 | 489550 | 1584900 | 8938200 | 0 | 0 | 122090 | 142250 | 197760 | 0 |
| LGQSDPAF | TKT; TKT; TKT | P29401; A0A | TKT_HUMAN Tr | Oxidation (M) | 0 | 75368 | 0 | 0 | 0 | 0 | 0 | 0 | 484820 | 229870 | 0 | 3965900 | 0 | 0 | 143890 | 0 | 0 | 0 |
| LGQSESQG | PSMD9; PSMD9 | O00233; F5H | PSMD9_HUMA | Unmodified | 0 | 49377 | 0 | 0 | 186490 | 41315 | 115240 | 0 | 0 | 0 | 0 | 333140 | 0 | 0 | 243430 | 176320 | 266970 | 517520 |
| LGQSESQG | PSMD9; PSMD9 | O00233; F5H | PSMD9_HUMA | Unmodified | 0 | 0 | 0 | 0 | 672800 | 143490 | 195090 | 0 | 0 | 0 | 0 | 0 | 1229100 | 500660 | 0 | 0 | 276460 | 510270 |
| LGQSVTTIP | ARF6 | P62330 | ARF6_HUMAN | Unmodified | 0 | 0 | 0 | 0 | 1269900 | 0 | 0 | 0 | 943110 | 623790 | 0 | 2446300 | 0 | 0 | 0 | 0 | 0 | 0 |
| LGQTIPLPF | B3GNT8 | Q7Z7M8 | B3GN8_HUMAN | Unmodified | 0 | 0 | 0 | 0 | 0 | 0 | 148260 | 0 | 1277900 | 59970 | 0 | 0 | 0 | 0 | 0 | 0 | 0 | 0 |
| LGQYIRLVE | SLC9A3R1; SLC | O14745; J3Q | NHRF1_HUMAN | Unmodified | 0 | 0 | 0 | 0 | 3116900 | 0 | 0 | 722950 | 0 | 644530 | 0 | 0 | 7531400 | 9399200 | 0 | 0 | 1500900 | 1128100 |
| LGRLDQAV | UNC45A; UNC4 | Q9H3U1; A0 | UN45A_HUMAN | Unmodified | 0 | 0 | 142720 | 480370 | 325290 | 0 | 90677 | 0 | 0 | 137930 | 193350 | 694300 | 2209500 | 708450 | 371330 | 309000 | 0 | 315930 |
| LGRSEIQFL | DSG2 | Q14126 | DSG2_HUMAN | Unmodified | 0 | 0 | 0 | 0 | 0 | 0 | 0 | 0 | 1303200 | 0 | 0 | 0 | 0 | 0 | 0 | 0 | 0 | 0 |
| LGRSPPNV | PLXNB2 | O15031 | PLXB2_HUMAN | Unmodified | 0 | 0 | 318510 | 285570 | 343240 | 249400 | 186380 | 0 | 0 | 0 | 0 | 0 | 0 | 0 | 0 | 0 | 91725 | 0 |
| LGRSVSFV | JAM2; JAM2 | P57087; H0Y | JAM2_HUMAN | Unmodified | 0 | 0 | 0 | 0 | 0 | 0 | 441570 | 0 | 0 | 0 | 0 | 0 | 0 | 0 | 0 | 0 | 0 | 0 |
| LGRTIADH | CTNNA1; CTNN | P35221; G3X | CTNA1_HUMAN | Unmodified | 0 | 0 | 0 | 0 | 1657600 | 0 | 0 | 0 | 543110 | 0 | 0 | 0 | 1102500 | 1741200 | 0 | 0 | 0 | 0 |
| LGRYNEEE | TBCB; TBCB; TE | Q99426; K7E | TBCB_HUMAN | Unmodified | 0 | 0 | 1073200 | 0 | 176650 | 0 | 0 | 0 | 3073000 | 286650 | 308480 | 2250200 | 0 | 0 | 0 | 0 | 102740 | 0 |
| LGSAFQAIS | SNX33 | Q8WV41 | SNX33_HUMAN | Oxidation (M) | 0 | 0 | 1944100 | 9447200 | 0 | 0 | 0 | 0 | 0 | 0 | 0 | 0 | 0 | 0 | 29991000 | 0 | 0 | 0 |
| LGSAGGTD | UNC45A; UNC4 | Q9H3U1; A0 | UN45A_HUMAN | Unmodified | 0 | 0 | 0 | 0 | 0 | 0 | 166770 | 53991 | 0 | 0 | 0 | 722450 | 512700 | 428960 | 144440 | 0 | 136290 | 0 |
| LGSALGGP | POFUT2; POFUT | Q9Y2G5; A0A | OFUT2_HUMAN | Unmodified | 0 | 0 | 0 | 0 | 0 | 0 | 0 | 0 | 8148700 | 1305700 | 0 | 0 | 0 | 0 | 0 | 0 | 0 | 0 |
| LGSAVVTR | ALDH18A1 | P54886 | P5CS_HUMAN | Unmodified | 0 | 0 | 0 | 0 | 0 | 0 | 0 | 0 | 809440 | 0 | 0 | 0 | 0 | 230150 | 0 | 0 | 0 | 0 |
| LGSCKLIEE | CCT2; CCT2; CC | P78371; F5G | TCPB_HUMAN | Unmodified | 0 | 0 | 0 | 0 | 0 | 0 | 0 | 0 | 0 | 0 | 0 | 0 | 218210 | 229230 | 0 | 0 | 132640 | 0 |
| LGSEGEDL | SLC1A4 | P43007 | SATT_HUMAN N | Unmodified | 0 | 0 | 0 | 0 | 0 | 0 | 0 | 0 | 379810 | 0 | 0 | 1186200 | 0 | 0 | 0 | 0 | 0 | 0 |
| LGSEIRDS | SEMA4B | Q9NPR2 | SEM4B_HUMAN | Unmodified | 0 | 0 | 0 | 0 | 2312200 | 483190 | 0 | 0 | 0 | 0 | 0 | 0 | 0 | 0 | 0 | 0 | 0 | 152130 |
| LGSELIQGI | CEMIP2; CEMIP | Q9UHN6; F5 | CEIP2_HUMAN | Unmodified | 684250 | 0 | 1245800 | 942650 | 0 | 422810 | 24638 | 0 | 7609500 | 287530 | 81925 | 484100 | 0 | 96954 | 1048800 | 1423900 | 0 | 66008 |
| LGSGDDIS | PPM1H | Q9ULR3 | PPM1H_HUMA | Unmodified | 0 | 0 | 0 | 0 | 0 | 0 | 0 | 0 | 1002700 | 0 | 0 | 849530 | 0 | 0 | 0 | 0 | 0 | 0 |
| LGSGGHLH | GDF15; GDF15 | Q99988; A0A | GDF15_HUMAN | Unmodified | 0 | 0 | 68285 | 61288 | 92543 | 0 | 0 | 0 | 288440 | 65450 | 0 | 0 | 0 | 0 | 234120 | 163560 | 0 | 0 |
| LGSGNDFE | FOLH1B; FOLH | Q9HBA9; Q0 | FOH1B_HUMAN | Unmodified | 30463000 | 22603000 | 41399000 | 1.08E+08 | 0 | 0 | 0 | 0 | 0 | 0 | 0 | 0 | 0 | 0 | 79378000 | 51640000 | 0 | 0 |
| LGSGNDFE | FOLH1B; FOLH | Q9HBA9; Q0 | FOH1B_HUMAN | Unmodified | 0 | 0 | 651550 | 4390200 | 0 | 0 | 0 | 0 | 0 | 0 | 0 | 0 | 0 | 0 | 0 | 0 | 0 | 0 |
| LGSGPDGA | RPS6KA1; RPS6 | Q15418; E9P | KS6A1_HUMAN | Unmodified | 0 | 0 | 0 | 185700 | 0 | 0 | 0 | 0 | 617910 | 37056 | 0 | 946910 | 39396 | 7737.1 | 310350 | 236450 | 0 | 0 |
| LGSIAIQGA | ACAT1; ACAT1; | P24752; E9P | THIL_HUMAN A | Unmodified | 0 | 0 | 0 | 0 | 0 | 0 | 464100 | 0 | 3417300 | 1386900 | 0 | 0 | 878600 | 0 | 0 | 0 | 0 | 0 |
| LGSLVDEFI | XRCC6; XRCC6 | P12956; B1A | XRCC6_HUMAN | Unmodified | 0 | 0 | 0 | 0 | 0 | 0 | 0 | 0 | 0 | 0 | 0 | 0 | 0 | 0 | 0 | 0 | 0 | 0 |
| LGSLVENN | EIF3CL; EIF3C | B5ME19; Q9 | EIFCL_HUMAN | Unmodified | 220550 | 165550 | 339240 | 224240 | 0 | 0 | 0 | 0 | 0 | 0 | 922090 | 5044600 | 0 | 0 | 180580 | 200670 | 0 | 0 |
| LGSPDEEFI | STX4; STX4; STX | Q12846; H3B | STX4_HUMAN S | Unmodified | 149720 | 789220 | 1467900 | 1211500 | 1315100 | 0 | 122130 | 56188 | 2283000 | 808390 | 1038000 | 5227700 | 1391600 | 813300 | 1397200 | 1076800 | 556500 | 772680 |
| LGSPDEEFI | STX4; STX4; STX | Q12846; A0A | STX4_HUMAN S | Unmodified | 0 | 0 | 0 | 0 | 259150 | 804040 | 44818 | 85721 | 0 | 0 | 0 | 0 | 564060 | 0 | 0 | 0 | 503980 | 0 |
| LGSQIFVK | TJP2; TJP2; TJP2 | Q9UDY2; A0 | ZO2_HUMAN T | Unmodified | 115200 | 81425 | 0 | 0 | 2355500 | 0 | 0 | 0 | 5082000 | 1570100 | 0 | 5447000 | 327630 | 0 | 147480 | 387250 | 0 | 0 |
| LGSQRELA | DIPK2A | Q8NDZ4 | DIK2A_HUMAN | Unmodified | 0 | 0 | 0 | 0 | 0 | 0 | 0 | 0 | 692240 | 0 | 0 | 0 | 0 | 0 | 0 | 0 | 417730 | 0 |
| LGSRAEITE | TMPPE | Q6ZT21 | TMPPE_HUMAN | Unmodified | 0 | 0 | 0 | 0 | 0 | 0 | 148120 | 0 | 1305500 | 248820 | 0 | 71813 | 0 | 0 | 0 | 0 | 0 | 0 |
| LGSRLYGP | SERPINH1; SER | P50454; E9P | SERPH_HUMAN | Unmodified | 0 | 0 | 0 | 0 | 232940 | 0 | 0 | 0 | 0 | 4609600 | 0 | 192820 | 0 | 0 | 0 | 0 | 0 | 93602 |
| LGSRLYGP | SERPINH1; SER | P50454; E9P | SERPH_HUMAN | Unmodified | 0 | 0 | 0 | 0 | 819240 | 0 | 0 | 0 | 0 | 3184500 | 0 | 0 | 0 | 0 | 0 | 0 | 0 | 407490 |
| LGSSGLGS | HUWE1; HUWE | Q7Z6Z7; H0Y | HUWE1_HUMA | Unmodified | 0 | 0 | 42897 | 53318 | 0 | 0 | 0 | 0 | 0 | 0 | 105810 | 1901700 | 0 | 0 | 70400 | 89919 | 0 | 0 |
| LGSTAPQV | NSFL1C | Q9UNZ2 | NSF1C_HUMAN | Unmodified | 940160 | 697520 | 706290 | 603670 | 0 | 0 | 0 | 0 | 1811000 | 0 | 556210 | 3024300 | 1098600 | 0 | 0 | 467540 | 374310 | 0 |
| LGSTEVASI | YTHDF2; YTHDI | Q9Y5A9; S4F | YTHD2_HUMAN | Unmodified | 1463200 | 0 | 0 | 0 | 372650 | 0 | 0 | 88561 | 369140 | 111550 | 187190 | 544150 | 0 | 1205900 | 0 | 0 | 0 | 210440 |
| LGSTVFVAI | HNRNPM; HNR | P52272; M0F | HNRPM_HUMA | Unmodified | 0 | 0 | 0 | 0 | 953810 | 0 | 0 | 0 | 0 | 0 | 0 | 2243600 | 0 | 0 | 0 | 0 | 817830 | 923770 |
| LGSVPVTVI | TCAF1 | Q9Y4C2 | TCAF1_HUMAN | Unmodified | 0 | 420990 | 0 | 0 | 0 | 0 | 0 | 0 | 0 | 0 | 0 | 0 | 0 | 0 | 0 | 0 | 0 | 0 |
| LGSWLAIR | TJP1; TJP1; TJP1 | Q07157; A0A | ZO1_HUMAN T | Unmodified | 0 | 0 | 0 | 670590 | 1101600 | 0 | 0 | 0 | 0 | 0 | 0 | 0 | 809070 | 0 | 0 | 0 | 0 | 0 |
| LGTDEISPF | MYO1C; MYO1C | O00159; F5H | MYO1C_HUMA | Unmodified | 5815900 | 108470 | 225230 | 8525800 | 3211100 | 2388900 | 0 | 0 | 892510 | 504880 | 589630 | 258540 | 3918600 | 999980 | 13173000 | 11961000 | 685830 | 1280400 |
| LGTDEISPF | MYO1C; MYO1C | O00159; F5H | MYO1C_HUMA | Unmodified | 0 | 0 | 0 | 0 | 1944200 | 0 | 0 | 0 | 0 | 0 | 0 | 0 | 934310 | 0 | 0 | 0 | 0 | 763190 |
| LGTDESCF | ANXA7 | P20073 | ANXA7_HUMAN | Oxidation (M) | 385150 | 215070 | 709990 | 0 | 0 | 0 | 112750 | 26487 | 15635000 | 445690 | 0 | 768830 | 0 | 0 | 242900 | 449560 | 0 | 0 |
| LGTDESCF | ANXA7 | P20073 | ANXA7_HUMAN | Unmodified | 0 | 0 | 0 | 0 | 0 | 0 | 0 | 0 | 25760000 | 0 | 0 | 270940 | 494550 | 0 | 0 | 0 | 0 | 0 |
| LGTFEVED | DPP4; DPP4; DI | P27487; A0A | DPP4_HUMAN | Unmodified | 1331000 | 714480 | 11205000 | 13414000 | 0 | 0 | 0 | 0 | 0 | 0 | 0 | 670680 | 0 | 0 | 13603000 | 7641800 | 0 | 0 |
| LGTFEVED | DPP4; DPP4; DI | P27487; A0A | DPP4_HUMAN | Unmodified | 0 | 0 | 2254000 | 11122000 | 0 | 0 | 0 | 69625 | 0 | 0 | 0 | 0 | 460890 | 0 | 2018600 | 0 | 5180100 | 6959000 |
| LGTGENLA | NPTX2 | P47972 | NPTX2_HUMAN | Unmodified | 0 | 0 | 0 | 0 | 3134200 | 0 | 0 | 0 | 1562400 | 0 | 0 | 0 | 0 | 0 | 0 | 0 | 0 | 0 |
| LGTGFNPN | DOCK7; DOCK7 | Q96N67; A0A | DOCK7_HUMA | Unmodified | 0 | 0 | 0 | 0 | 1509400 | 0 | 0 | 118600 | 0 | 204470 | 241710 | 940780 | 0 | 0 | 0 | 0 | 0 | 0 |
| LGTKNEPV | DIP2B | Q9P265 | DIP2B_HUMAN | Unmodified | 0 | 0 | 0 | 0 | 0 | 0 | 0 | 0 | 0 | 0 | 0 | 580900 | 0 | 0 | 151320 | 107790 | 0 | 0 |
| LGTLFFSLE | SYT4 | Q9H2B2 | SYT4_HUMAN S | Unmodified | 0 | 0 | 0 | 223000 | 0 | 0 | 473410 | 0 | 187520 | 84423 | 218140 | 1314000 | 0 | 0 | 681840 | 0 | 0 | 0 |
| LGTLSALDI | CAND1; CAND1 | Q86VP6; A0A | CAND1_HUMA | Unmodified | 0 | 0 | 0 | 0 | 0 | 0 | 0 | 0 | 111950 | 137750 | 67462 | 2802900 | 0 | 0 | 0 | 0 | 0 | 0 |
| LGTLSPAFS | SND1 | Q7KZF4 | SND1_HUMAN | Unmodified | 1244600 | 368510 | 1004500 | 2190400 | 193780 | 0 | 0 | 197420 | 807610 | 0 | 2481800 | 11515000 | 2180000 | 0 | 2264600 | 1968500 | 0 | 189390 |
| LGTPQQIAI | CYFIP1; CYFIP2 | Q7L576; Q96 | CYFP1_HUMAN | Unmodified | 813350 | 589820 | 2343600 | 2105400 | 0 | 0 | 0 | 0 | 2913500 | 743980 | 0 | 8065300 | 0 | 0 | 2835100 | 2443600 | 0 | 0 |
| LGTPTLLAK | CDCP1 | Q9H5V8 | CDCP1_HUMA | Unmodified | 0 | 0 | 0 | 0 | 0 | 0 | 0 | 0 | 0 | 0 | 0 | 0 | 0 | 0 | 0 | 0 | 0 | 739800 |
| LGTPTLLAK | CDCP1 | Q9H5V8 | CDCP1_HUMA | Unmodified | 97170 | 65698 | 39546 | 1141500 | 3983600 | 1647400 | 0 | 0 | 266160 | 739540 | 1433200 | 5098600 | 3864900 | 3137800 | 4542800 | 2104400 | 15741000 | 7808100 |
| LGTPTLLAK | CDCP1 | Q9H5V8 | CDCP1_HUMA | Unmodified | 0 | 0 | 0 | 837690 | 0 | 497560 | 0 | 0 | 0 | 0 | 0 | 3633900 | 0 | 0 | 155570 | 185190 | 1520300 | 0 |
| LGTPVLQAI | PCK2; PCK2; PC | Q16822; H0Y | PCKGM_HUMA | Unmodified | 0 | 0 | 0 | 0 | 0 | 0 | 0 | 0 | 4067700 | 0 | 0 | 0 | 0 | 0 | 0 | 0 | 0 | 0 |
| LGTQQLIPK | ARHGEF16 | Q5VV41 | ARHGG_HUMA | Unmodified | 136770 | 0 | 0 | 112050 | 0 | 0 | 0 | 0 | 658850 | 347370 | 97859 | 446300 | 685440 | 606430 | 0 | 127490 | 0 | 0 |
| LGTVYEK | HSPA4L; HSPA4 | O95757; E9F | HS74L_HUMAN | Unmodified | 68685 | 929090 | 0 | 0 | 79818 | 207340 | 0 | 0 | 95067 | 0 | 0 | 2188500 | 0 | 0 | 2477500 | 0 | 0 | 0 |
| LGVAGQW | UCHL1; UCHL1 | P09936; D6F | UCHL1_HUMAN | Unmodified | 0 | 0 | 0 | 0 | 9058200 | 3424500 | 0 | 0 | 18404000 | 5471500 | 11396000 | 67535000 | 0 | 0 | 0 | 0 | 0 | 0 |
| LGVDISATK | FARSB | Q9NSD9 | SYFB_HUMAN | Unmodified | 69461 | 132410 | 581900 | 55774 | 0 | 0 | 0 | 0 | 1293200 | 674640 | 390600 | 1335900 | 0 | 0 | 7913.5 | 85149 | 0 | 0 |
| LGVDLDAQ | GART | P22102 | PUR2_HUMAN | Unmodified | 0 | 0 | 0 | 0 | 0 | 0 | 0 | 0 | 0 | 0 | 226280 | 1841100 | 0 | 0 | 0 | 0 | 0 | 0 |
| LGVDTIAK | UBXN6; UBXN6 | Q9BZV1; K7E | UBXN6_HUMAN | Unmodified | 330730 | 1033300 | 237230 | 47937 | 238460 | 194950 | 120100 | 154010 | 710600 | 288380 | 107450 | 960850 | 933330 | 676940 | 338740 | 818740 | 0 | 949970 |
| LGVFRAESI | SEC23B; SEC23 | Q15437; A0A | SC23B_HUMAN | Unmodified | 0 | 0 | 0 | 0 | 0 | 0 | 0 | 0 | 0 | 0 | 0 | 639990 | 0 | 0 | 0 | 0 | 0 | 0 |
| LGVHPLSC | LDHA | P00338 | LDHA_HUMAN | Oxidation (M) | 0 | 0 | 0 | 0 | 3156300 | 0 | 596090 | 0 | 1294600 | 0 | 5693200 | 17173000 | 1020500 | 0 | 0 | 1583500 | 0 | 2314900 |
| LGVHPLSC | LDHA | P00338 | LDHA_HUMAN | Unmodified | 0 | 0 | 0 | 0 | 0 | 0 | 0 | 0 | 4778100 | 0 | 0 | 3017800 | 0 | 0 | 0 | 0 | 0 | 3167100 |
| LGVIEDHSI | HSP90B2P; HSI | Q58FF3; P14 | ENPLL_HUMAN | Unmodified | 0 | 0 | 814870 | 0 | 0 | 0 | 0 | 0 | 17872000 | 710310 | 3993200 | 6130600 | 0 | 0 | 0 | 216020 | 0 | 0 |
| LGVIEDHSI | HSP90B1; HSP9 | P14625; A0A | ENPL_HUMAN | Unmodified | 0 | 0 | 0 | 0 | 0 | 0 | 595120 | 0 | 87850000 | 57264000 | 16854000 | 2094000 | 0 | 32549 | 0 | 0 | 0 | 0 |
| LGVIEDHSI | HSP90B1; HSP9 | P14625; A0A | ENPL_HUMAN | Unmodified | 0 | 0 | 0 | 0 | 777690 | 0 | 0 | 0 | 903470 | 79800000 | 1318700 | 0 | 1110800 | 0 | 0 | 0 | 225520 | 538560 |
| LGVIGDKV | IGF2R; IGF2R | P11717; S4R | MPRI_HUMAN | Unmodified | 60031 | 0 | 0 | 319610 | 327880 | 0 | 592940 | 77119 | 19676000 | 13683000 | 1562500 | 479120 | 853440 | 714260 | 0 | 0 | 0 | 158410 |
| LGVLYDLD | RIPOR2; RIPOR | Q9Y4F9; B7Z | RIPR2_HUMAN | Unmodified | 0 | 0 | 0 | 0 | 0 | 0 | 0 | 0 | 0 | 0 | 676220 | 3290200 | 1636200 | 815950 | 0 | 140980 | 0 | 0 |
| LGVLYWKL | ADI1 | Q9BV57 | MTND_HUMAN | Unmodified | 0 | 93591 | 0 | 365070 | 1583600 | 0 | 0 | 0 | 280610 | 264380 | 507360 | 211960 | 1723900 | 407910 | 87336 | 0 | 530740 | 631690 |
| LGVNNNQ | ANKFY1; ANKFY | Q9P2R3; K7E | ANFY1_HUMAN | Unmodified | 0 | 0 | 0 | 0 | 0 | 0 | 68530 | 138640 | 172750 | 0 | 30068 | 930820 | 0 | 0 | 0 | 0 | 0 | 0 |
| LGVQDLFN | SERPINB1 | P30740 | ILEU_HUMAN L | Unmodified | 0 | 0 | 0 | 0 | 323930 | 0 | 289690 | 0 | 1905900 | 312920 | 0 | 351780 | 0 | 0 | 0 | 0 | 0 | 0 |
| LGVQVREE | SLC9A3R1; SLC | O14745; J3Q | NHRF1_HUMAN | Unmodified | 206370 | 153390 | 724290 | 1945900 | 0 | 140600 | 0 | 0 | 1273500 | 0 | 1407900 | 2552600 | 3709900 | 2382600 | 1870700 | 1789600 | 259760 | 163250 |
| LGVQVVITD | TCP1; TCP1; TC | P17987; F5G | TCPA_HUMAN | Unmodified | 0 | 0 | 0 | 0 | 0 | 0 | 0 | 0 | 916400 | 0 | 1051200 | 3022100 | 0 | 0 | 0 | 0 | 0 | 0 |
| LGVRPSQG | FN1 | P02751 | FINC_HUMAN F | Unmodified | 55457 | 217260 | 316910 | 0 | 0 | 0 | 0 | 0 | 0 | 0 | 0 | 0 | 0 | 0 | 81201 | 242230 | 0 | 0 |
| LGVTANDV | MDH1; MDH1; M | P40925; C9J | MDHC_HUMAN | Unmodified | 1628500 | 845670 | 93118 | 149250 | 28366000 | 3740000 | 2931700 | 1972600 | 6879700 | 5930600 | 5329800 | 59285000 | 14878000 | 8724300 | 1503000 | 1556700 | 12187000 | 16566000 |
| LGVTANDV | MDH1; MDH1; M | P40925; C9J | MDHC_HUMAN | Unmodified | 0 | 0 | 0 | 0 | 5638600 | 0 | 6588400 | 148180 | 733640 | 1113200 | 0 | 0 | 1236500 | 3056500 | 0 | 0 | 11872000 | 3108700 |
| LGVTVLSR | CBR3 | O75828 | CBR3_HUMAN | Unmodified | 289700 | 414890 | 1095100 | 1550200 | 11318000 | 4162400 | 1040900 | 787970 | 1294600 | 1196300 | 64130 | 5735800 | 2847000 | 1543300 | 1903300 | 1979000 | 4192500 | 9881000 |
| LGWDPKP | NPEPPS; NPEPI | P55786; A0A | PSA_HUMAN P | Unmodified | 0 | 303960 | 438950 | 0 | 0 | 0 | 307590 | 0 | 591290 | 0 | 426540 | 10868000 | 0 | 0 | 762970 | 499840 | 0 | 0 |
| LGWYHSM | MMP15 | P51511 | MMP15_HUMA | Unmodified | 114280 | 70356 | 202440 | 222940 | 0 | 0 | 0 | 0 | 0 | 0 | 0 | 0 | 0 | 0 | 190080 | 120390 | 0 | 0 |
| LGYAGNTE | ACTR3 | P61158 | ARP3_HUMAN | Unmodified | 0 | 2357700 | 1229300 | 4717700 | 8705700 | 3232800 | 1359600 | 446050 | 3738600 | 1511200 | 1262700 | 19358000 | 6019500 | 8120300 | 1842800 | 2237000 | 3410600 | 6586500 |
| LGYAGNTE | ACTR3 | P61158 | ARP3_HUMAN | Unmodified | 0 | 0 | 0 | 0 | 3413000 | 0 | 0 | 0 | 0 | 0 | 0 | 0 | 0 | 0 | 0 | 0 | 2572500 | 1637100 |
| LGYAVYETI | TOLLIP; TOLLIP | Q9H0E2; E9I | TOLIP_HUMAN | Unmodified | 10895000 | 11488000 | 6922600 | 6145000 | 2773900 | 1280300 | 0 | 0 | 1747000 | 0 | 746520 | 0 | 5757200 | 3333600 | 11516000 | 13741000 | 847900 | 4270400 |
| LGYAVYETI | TOLLIP; TOLLIP | Q9H0E2; E9I | TOLIP_HUMAN | Unmodified | 186170 | 429450 | 0 | 492690 | 0 | 0 | 1481400 | 0 | 0 | 0 | 0 | 0 | 1089000 | 469280 | 483810 | 393410 | 0 | 0 |
| LGYFSVDP | QARS1; QARS1 | P47897; A0A | SYQ_HUMAN G | Unmodified | 0 | 0 | 0 | 0 | 0 | 0 | 0 | 16641 | 433280 | 0 | 271250 | 5181200 | 0 | 0 | 176940 | 0 | 99281 | 94367 |
| LGYILTCPS | CKB; CKMT1A; | P12277; P12 | KCRB_HUMAN | Unmodified | 0 | 0 | 0 | 0 | 0 | 0 | 52999 | 0 | 0 | 0 | 30176 | 2979000 | 0 | 0 | 0 | 0 | 0 | 0 |
| LGYISVVDT | ANK3; ANK2; AN | Q12955; Q01 | ANK3_HUMAN | Unmodified | 0 | 0 | 0 | 681900 | 0 | 0 | 0 | 0 | 0 | 0 | 0 | 0 | 823220 | 738280 | 1384200 | 1233700 | 0 | 0 |
| LGYLLFR | GRK2; GRK2 | P25098; E9P | ARBK1_HUMAN | Unmodified | 0 | 0 | 0 | 0 | 0 | 0 | 0 | 0 | 0 | 0 | 0 | 1236500 | 0 | 0 | 0 | 0 | 0 | 0 |
| LGYPQAPP | KIRREL1; KIRRE | Q96J84; Q5V | KIRR1_HUMAN | Unmodified | 0 | 0 | 0 | 0 | 5964600 | 898720 | 0 | 0 | 0 | 0 | 0 | 0 | 869340 | 0 | 0 | 0 | 0 | 1031500 |
| LGYSEPTV | NCAM2; NCAM | O15394; H9I | NCAM2_HUMA | Oxidation (M) | 0 | 0 | 0 | 0 | 0 | 0 | 0 | 0 | 0 | 0 | 0 | 0 | 0 | 0 | 954670 | 680290 | 0 | 0 |
| LGYSPLHQ | ANK1; ANK1 | P16157; H0Y | ANK1_HUMAN | Unmodified | 0 | 0 | 0 | 0 | 174170 | 0 | 409020 | 0 | 0 | 0 | 121700 | 226940 | 0 | 0 | 0 | 0 | 0 | 0 |
| LGYTHLYS | DDOST; DDOST | P39656; A0A | OST48_HUMAN | Unmodified | 0 | 0 | 0 | 0 | 0 | 0 | 0 | 0 | 1475400 | 0 | 0 | 0 | 0 | 0 | 0 | 0 | 0 | 0 |

| | | | | | | | | | | | | | | | | | | | | | | |
|---|---|---|---|---|---|---|---|---|---|---|---|---|---|---|---|---|---|---|---|---|---|---|
| LGYTPVCR | PAFAH1B3; PAF | Q15102; M0I | PA1B3_HUMAN | Unmodified | 275310 | 0 | 0 | 244100 | 0 | 0 | 0 | 0 | 2353100 | 0 | 681770 | 6029400 | 0 | 0 | 132280 | 146900 | 0 | 0 |
| LHAEFAAEI | DCTPP1 | Q9H773 | DCTP1_HUMAN | Unmodified | 230580 | 91190 | 350420 | 422480 | 0 | 0 | 0 | 0 | 856270 | 0 | 454820 | 6988400 | 0 | 0 | 485480 | 355390 | 0 | 0 |
| LHATHPAG | TMEM132A | Q24JP5 | T132A_HUMAN | Unmodified | 0 | 0 | 0 | 0 | 0 | 0 | 0 | 0 | 146050 | 33846 | 0 | 116640 | 0 | 0 | 0 | 0 | 0 | 0 |
| LHAVNAEE | ECI2; ECI2 | O75521; A0A | ECI2_HUMAN E | Unmodified | 0 | 0 | 0 | 0 | 2131100 | 0 | 0 | 0 | 4037800 | 734790 | 222120 | 0 | 0 | 0 | 0 | 0 | 0 | 0 |
| LHAWFSWI | MOGS; MOGS; | Q13724; A0A | MOGS_HUMAN | Unmodified | 936020 | 0 | 205300 | 2715500 | 0 | 0 | 716660 | 42137 | 10111000 | 0 | 188570 | 0 | 1135700 | 0 | 0 | 0 | 0 | 0 |
| LHCWSLTC | GYG1; GYG1; G | P46976; C9J | GLYG_HUMAN | Unmodified | 42820 | 0 | 126820 | 73678 | 0 | 0 | 283430 | 68207 | 303090 | 0 | 267530 | 1847500 | 351530 | 0 | 151390 | 51982 | 0 | 0 |
| LHDAIVEVV | DNM1L; DNM1L | O00429; H0Y | DNM1L_HUMA | Unmodified | 0 | 0 | 0 | 0 | 0 | 0 | 981680 | 0 | 0 | 0 | 141000 | 468650 | 0 | 0 | 0 | 50220 | 0 | 0 |
| LHDALDAK | FH | P07954 | FUMH_HUMAN | Unmodified | 0 | 0 | 0 | 53738 | 0 | 0 | 0 | 0 | 1246700 | 228990 | 108120 | 517200 | 0 | 72261 | 95936 | 89579 | 0 | 0 |
| LHDETLTYL | UBP1; TFCP2; U | Q9NZI7; Q12 | UBIP1_HUMAN | Unmodified | 66353 | 168920 | 0 | 114790 | 399100 | 0 | 0 | 0 | 530590 | 0 | 688020 | 2892800 | 0 | 0 | 99691 | 75962 | 0 | 0 |
| LHDFGYRG | NAMPT; NAMPT | P43490; A0A | NAMPT_HUMAI | Unmodified | 0 | 0 | 0 | 1046600 | 3210400 | 0 | 0 | 0 | 0 | 0 | 0 | 0 | 0 | 494690 | 0 | 0 | 0 | 0 |
| LHDINAQL | AP1B1; AP1B1 | Q10567; C9J | AP1B1_HUMAN | Unmodified | 196360 | 0 | 554260 | 1851900 | 0 | 0 | 0 | 0 | 2053800 | 1499900 | 2047800 | 6239400 | 1520100 | 1017300 | 1384200 | 316700 | 0 | 1224800 |
| LHDINAQM | AP2B1; AP2B1; | P63010; A0A | AP2B1_HUMAN | Oxidation (M) | 154410 | 0 | 0 | 0 | 0 | 0 | 0 | 0 | 0 | 0 | 0 | 778550 | 0 | 291700 | 125310 | 0 | 0 | 0 |
| LHDNPLTD | NAA15; NAA15 | Q9BXJ9; A0A | NAA15_HUMAN | Oxidation (M) | 0 | 0 | 0 | 0 | 0 | 0 | 0 | 0 | 0 | 0 | 52855 | 607850 | 0 | 0 | 0 | 0 | 0 | 0 |
| LHDNPLTD | NAA15; NAA15 | Q9BXJ9; A0A | NAA15_HUMAN | Unmodified | 0 | 0 | 0 | 0 | 356340 | 52735 | 89522 | 48843 | 68936 | 54401 | 115390 | 1429500 | 0 | 265420 | 28650 | 0 | 48107 | 34820 |
| LHDSVVMV | C20orf27; C20c | Q9GZN8; H7 | CT027_HUMAN | Unmodified | 0 | 0 | 0 | 0 | 272800 | 0 | 0 | 0 | 0 | 0 | 42360 | 138340 | 0 | 0 | 0 | 0 | 265500 | 275630 |
| LHDSVVMV | C20orf27; C20c | Q9GZN8; H7 | CT027_HUMAN | Oxidation (M) | 0 | 0 | 0 | 0 | 400790 | 0 | 118130 | 43101 | 99989 | 50377 | 325830 | 0 | 236250 | 108430 | 0 | 0 | 0 | 360260 |
| LHDVEHVL | PFDN5; PFDN5 | Q99471; A0A | PFD5_HUMAN | Unmodified | 0 | 0 | 0 | 0 | 741430 | 0 | 0 | 0 | 866490 | 140760 | 1115400 | 2949300 | 1939400 | 0 | 0 | 0 | 367240 | 189530 |
| LHDVNSD( | NUCB2; NUCB2 | P80303; Q2L | NUCB2_HUMA | Unmodified | 671630 | 663440 | 1122500 | 5156600 | 1890200 | 477900 | 5754000 | 42890 | 2.02E+08 | 82504000 | 870690 | 549310 | 15856000 | 19408000 | 7656500 | 1216800 | 161430 | 5439200 |
| LHDVNSD( | NUCB2; NUCB2 | P80303; Q2L | NUCB2_HUMA | Unmodified | 0 | 0 | 0 | 0 | 0 | 0 | 671830 | 0 | 0 | 1063900 | 0 | 0 | 0 | 1761100 | 0 | 0 | 0 | 0 |
| LHEATDYP | LAMC1 | P11047 | LAMC1_HUMAI | Unmodified | 1455800 | 0 | 399100 | 774700 | 0 | 0 | 0 | 0 | 5809100 | 85101 | 0 | 471680 | 0 | 0 | 471950 | 222650 | 0 | 0 |
| LHEEEPPQ | EIF2A; EIF2A; EI | Q9BY44; F8V | EIF2A_HUMAN | Unmodified | 0 | 0 | 0 | 0 | 336310 | 0 | 0 | 0 | 0 | 0 | 383110 | 519640 | 231970 | 0 | 0 | 0 | 69739 | 275750 |
| LHEEGIIYR | VARS1; VARS1 | P26640; A0A | SYVC_HUMAN | Unmodified | 0 | 0 | 0 | 0 | 0 | 0 | 0 | 0 | 0 | 0 | 0 | 1810600 | 0 | 0 | 0 | 120580 | 0 | 0 |
| LHELNQKV | SPTAN1; SPTAN | Q13813; A0A | SPTN1_HUMAN | Unmodified | 0 | 0 | 0 | 0 | 0 | 0 | 0 | 0 | 0 | 132600 | 0 | 1479500 | 0 | 0 | 0 | 0 | 182050 | 0 |
| LHELTVMQ | KIF5B | P33176 | KINH_HUMAN | Unmodified | 0 | 0 | 0 | 0 | 0 | 0 | 37741 | 0 | 866700 | 0 | 0 | 1733700 | 0 | 0 | 0 | 0 | 0 | 0 |
| LHENADF( | TNFSF11 | O14788 | TNF11_HUMAN | Unmodified | 0 | 0 | 0 | 0 | 0 | 0 | 0 | 0 | 994180 | 0 | 0 | 0 | 0 | 0 | 0 | 0 | 0 | 0 |
| LHERKCEP | CORO1B | Q9BR76 | COR1B_HUMA | Unmodified | 0 | 0 | 0 | 0 | 0 | 0 | 0 | 0 | 0 | 97492 | 0 | 0 | 319660 | 424660 | 0 | 0 | 175610 | 187390 |
| LHEVGHER | PPFIA1; PPFIA1 | Q13136; E9F | LIPA1_HUMAN | Unmodified | 0 | 0 | 0 | 0 | 0 | 0 | 0 | 0 | 0 | 0 | 0 | 0 | 0 | 201870 | 0 | 0 | 195540 | 510420 |
| LHEWTKPE | SERPINB1 | P30740 | ILEU_HUMAN L | Unmodified | 70770 | 0 | 0 | 0 | 0 | 0 | 0 | 0 | 0 | 362220 | 0 | 0 | 0 | 0 | 0 | 0 | 0 | 0 |
| LHFAFRLY | CHP1; CHP1; C | Q99653; H0Y | CHP1_HUMAN | Unmodified | 0 | 0 | 0 | 0 | 601990 | 0 | 0 | 0 | 0 | 0 | 0 | 0 | 2357900 | 821230 | 0 | 0 | 0 | 349090 |
| LHFFMPGF | TUBB3; N/A | Q13509; A0A | TBB3_HUMAN | Unmodified | 0 | 0 | 0 | 0 | 0 | 0 | 0 | 0 | 3377500 | 343700 | 0 | 544710 | 0 | 0 | 0 | 0 | 0 | 21002 |
| LHFFMPGF | TUBB3; N/A | Q13509; A0A | TBB3_HUMAN | Oxidation (M) | 0 | 0 | 0 | 0 | 0 | 0 | 0 | 0 | 0 | 1053500 | 0 | 3903900 | 0 | 0 | 0 | 0 | 0 | 0 |
| LHFFMPGF | TUBB2A; TUBB | Q13885; A6N | TBB2A_HUMAN | Oxidation (M) | 4131000 | 3251800 | 4968400 | 4490000 | 10346000 | 0 | 0 | 0 | 11306000 | 6649100 | 19233000 | 1.9E+08 | 3364100 | 2296200 | 7206600 | 5358300 | 1174000 | 0 |
| LHFFMPGF | TUBB2A; TUBB | Q13885; A6N | TBB2A_HUMAN | Unmodified | 0 | 0 | 0 | 0 | 348660 | 0 | 0 | 0 | 4659000 | 0 | 637060 | 5059700 | 296160 | 0 | 0 | 0 | 0 | 0 |
| LHFLMTGY | TUBG1; TUBG2 | P23258; Q9N | TBG1_HUMAN | Oxidation (M) | 516290 | 170400 | 62447 | 86706 | 0 | 0 | 0 | 338340 | 0 | 0 | 0 | 1817500 | 0 | 0 | 247820 | 209110 | 0 | 0 |
| LHFLMTGY | TUBG1; TUBG2 | P23258; Q9N | TBG1_HUMAN | Unmodified | 0 | 0 | 133830 | 0 | 0 | 0 | 0 | 0 | 0 | 220230 | 0 | 840950 | 39485 | 0 | 525530 | 0 | 0 | 0 |
| LHGDDNG | NRP2 | O60462 | NRP2_HUMAN | Unmodified | 0 | 0 | 0 | 0 | 0 | 0 | 0 | 0 | 1000900 | 84956 | 0 | 0 | 0 | 0 | 18561 | 0 | 0 | 0 |
| LHGLQAIL | NUP210 | Q8TEM1 | PO210_HUMAN | Unmodified | 0 | 0 | 0 | 0 | 0 | 0 | 0 | 0 | 27967000 | 1612700 | 862240 | 0 | 0 | 0 | 0 | 0 | 558870 | 0 |
| LHGQHLV( | TOR3A; TOR3A | Q9H497; Q5 | TOR3A_HUMAN | Unmodified | 0 | 0 | 0 | 0 | 931370 | 0 | 246640 | 0 | 1226500 | 0 | 0 | 0 | 0 | 0 | 0 | 0 | 72183 | 1908400 |
| LHGRSDES | EIF2B2; EIF2B2 | P49770; H0Y | EI2BB_HUMAN | Unmodified | 75760 | 798050 | 0 | 0 | 0 | 0 | 149690 | 0 | 0 | 0 | 0 | 193680 | 0 | 0 | 447680 | 0 | 82837 | 0 |
| LHGTAQQL | PKN2; PKN2 | Q16513; B1A | PKN2_HUMAN | Unmodified | 0 | 171530 | 223360 | 330710 | 856250 | 257990 | 24368 | 68423 | 825420 | 193520 | 441520 | 1129900 | 560290 | 455480 | 0 | 0 | 568210 | 602670 |
| LHGVRDYA | GLRX5 | Q86SX6 | GLRX5_HUMAN | Unmodified | 0 | 0 | 0 | 0 | 0 | 0 | 0 | 55748 | 3832700 | 0 | 0 | 0 | 0 | 0 | 0 | 0 | 0 | 0 |
| LHGVRDYA | GLRX5 | Q86SX6 | GLRX5_HUMAN | Unmodified | 0 | 0 | 0 | 0 | 0 | 0 | 486740 | 0 | 5967900 | 723030 | 0 | 0 | 0 | 0 | 0 | 0 | 1164100 | 0 |
| LHHVSPAD | HSPG2 | P98160 | PGBM_HUMAN | Unmodified | 195050 | 0 | 1133600 | 270130 | 0 | 146480 | 0 | 0 | 159320 | 0 | 77728 | 117140 | 0 | 0 | 0 | 208350 | 0 | 48205 |
| LHHVSSLA | WDR1 | O75083 | WDR1_HUMAN | Unmodified | 4601500 | 4894400 | 3164800 | 4605300 | 1331300 | 772870 | 0 | 0 | 10170000 | 613970 | 2192200 | 48283000 | 479180 | 0 | 5981700 | 5609500 | 146300 | 0 |
| LHIDHEIET | SULF2 | Q8IWU5 | SULF2_HUMAN | Unmodified | 0 | 0 | 0 | 0 | 0 | 0 | 0 | 0 | 0 | 0 | 0 | 0 | 0 | 0 | 0 | 0 | 122700 | 1349200 |
| LHIEALLHC | IDE; IDE; IDE; ID | P14735; A0A | IDE_HUMAN In | Unmodified | 0 | 0 | 0 | 0 | 0 | 0 | 0 | 0 | 0 | 0 | 0 | 1961700 | 0 | 0 | 92814 | 0 | 0 | 0 |
| LHIIEVGTP | CLTC; CLTC | Q00610; A0A | CLH1_HUMAN | Unmodified | 36658000 | 18836000 | 15175000 | 18574000 | 6617800 | 3342200 | 8619800 | 2966300 | 4258700 | 1503700 | 4212900 | 78170000 | 6224000 | 3217200 | 13270000 | 20715000 | 2307800 | 3751700 |
| LHIIEVGTP | CLTC; CLTC | Q00610; A0A | CLH1_HUMAN | Unmodified | 3513400 | 3993900 | 1688000 | 2155800 | 1747700 | 309070 | 0 | 81652 | 655240 | 143700 | 0 | 9347100 | 908520 | 327770 | 3236800 | 4109100 | 397250 | 465010 |
| LHIPWGDN | GNA13 | Q14344 | GNA13_HUMAI | Unmodified | 590380 | 472900 | 0 | 0 | 7402600 | 1928300 | 2115900 | 538160 | 5174900 | 2377500 | 3641900 | 14594000 | 0 | 0 | 0 | 0 | 2569500 | 5062100 |
| LHISPSNM | HDAC2 | Q92769 | HDAC2_HUMA | Oxidation (M) | 0 | 0 | 0 | 0 | 0 | 0 | 0 | 0 | 0 | 0 | 0 | 2718400 | 0 | 0 | 0 | 0 | 0 | 0 |
| LHISQLQH | DCTN1; DCTN1 | Q14203; A0A | DCTN1_HUMAI | Unmodified | 42590 | 0 | 123130 | 107850 | 0 | 52560 | 0 | 0 | 466070 | 0 | 0 | 3106400 | 397830 | 221170 | 116990 | 122990 | 89969 | 0 |
| LHIVQVVCI | CLIC1 | O00299 | CLIC1_HUMAN | Unmodified | 543220 | 333970 | 248040 | 409160 | 6651300 | 1400400 | 649910 | 429610 | 1303300 | 818600 | 0 | 2663800 | 3774500 | 4836900 | 654830 | 550880 | 4553800 | 0 |
| LHIVQVVCI | CLIC1 | O00299 | CLIC1_HUMAN | Unmodified | 1527200 | 945300 | 615390 | 1397500 | 26544000 | 4515200 | 1919800 | 1003600 | 3783100 | 3033000 | 1103900 | 7008900 | 17071000 | 9548900 | 1780700 | 1292500 | 16113000 | 11472000 |
| LHKPPADS | SLC16A3; SLC1 | O15427; J3Q | MOT4_HUMAN | Unmodified | 264120 | 247360 | 985110 | 347300 | 486800 | 118310 | 0 | 14409 | 635250 | 303590 | 331690 | 5318200 | 0 | 0 | 1383400 | 1263900 | 0 | 230260 |
| LHKPPADS | SLC16A3; SLC1 | O15427; J3Q | MOT4_HUMAN | Unmodified | 0 | 0 | 0 | 0 | 10197000 | 0 | 0 | 0 | 421760 | 0 | 0 | 0 | 0 | 0 | 0 | 0 | 1084100 | 3548000 |
| LHLESPSP | IGSF3 | O75054 | IGSF3_HUMAN | Unmodified | 0 | 0 | 0 | 0 | 0 | 552680 | 0 | 274690 | 0 | 0 | 0 | 716220 | 0 | 150500 | 0 | 0 | 0 | 574370 |
| LHLFASEQI | DNAJC13; DNA | O75165; A0A | DJC13_HUMAN | Unmodified | 0 | 0 | 0 | 0 | 0 | 0 | 0 | 0 | 0 | 0 | 0 | 2851500 | 0 | 0 | 0 | 0 | 0 | 0 |
| LHLGFIEIRI | LUC7L2; LUC7L | Q9Y383; A0A | LC7L2_HUMAN | Unmodified | 0 | 0 | 0 | 0 | 622840 | 0 | 368590 | 0 | 0 | 626950 | 0 | 0 | 999540 | 615480 | 0 | 0 | 85153 | 700000 |
| LHLGHTFS | LARS1; LARS1; | Q9P2J5; A0A | SYLC_HUMAN | Unmodified | 320680 | 7379.6 | 107750 | 0 | 135860 | 0 | 0 | 0 | 0 | 0 | 326200 | 3432600 | 83343 | 0 | 80097 | 0 | 0 | 0 |
| LHLGSTPH | VPS4A; N/A | Q9UN37; I3L | VPS4A_HUMAN | Unmodified | 696750 | 2465900 | 2287900 | 3799200 | 0 | 0 | 0 | 0 | 10211000 | 1410400 | 960510 | 14587000 | 334790 | 0 | 2042300 | 2384700 | 0 | 0 |
| LHLGTTQN | VPS4B | O75351 | VPS4B_HUMAN | Unmodified | 2461600 | 2981500 | 0 | 0 | 0 | 0 | 0 | 0 | 7590800 | 0 | 0 | 16648000 | 0 | 768370 | 1951400 | 3640500 | 0 | 0 |
| LHLGTTQN | VPS4B | O75351 | VPS4B_HUMAN | Unmodified | 3281300 | 2570000 | 1475300 | 2494300 | 0 | 0 | 0 | 0 | 11204000 | 1388900 | 1876700 | 3089900 | 1014100 | 0 | 1256800 | 909980 | 0 | 321870 |
| LHLIAQELP | EXOC5; EXOC5 | O00471; F8V | EXOC5_HUMAI | Unmodified | 0 | 0 | 0 | 0 | 5328500 | 605740 | 0 | 0 | 1181800 | 1094900 | 0 | 0 | 0 | 0 | 0 | 0 | 891060 | 716770 |
| LHLITRNL( | YARS1; YARS1; | P54577; A0A | SYYC_HUMAN | Unmodified | 0 | 0 | 0 | 0 | 467490 | 0 | 157770 | 0 | 0 | 0 | 0 | 0 | 592680 | 766140 | 0 | 0 | 0 | 548140 |
| LHLPADDV | VGF | O15240 | VGF_HUMAN N | Unmodified | 116050 | 977420 | 0 | 0 | 0 | 0 | 0 | 0 | 0 | 1319200 | 0 | 15518 | 0 | 0 | 0 | 0 | 0 | 0 |
| LHLPQQPL | TRIM41 | Q8WV44 | TRI41_HUMAN | Unmodified | 156980 | 192710 | 282430 | 0 | 0 | 0 | 0 | 0 | 0 | 0 | 0 | 0 | 0 | 0 | 88261 | 69589 | 0 | 0 |
| LHLSGIDAI | FASN; FASN | P49327; A0A | FAS_HUMAN F | Unmodified | 1628700 | 1484400 | 2244100 | 4492700 | 0 | 0 | 354390 | 0 | 6087900 | 989970 | 19145000 | 1.29E+08 | 0 | 0 | 13040000 | 4542100 | 0 | 0 |
| LHLSSPDN | MYO6; MYO6; M | Q9UM54; A0 | MYO6_HUMAN | Unmodified | 0 | 0 | 0 | 0 | 0 | 0 | 0 | 0 | 0 | 0 | 0 | 0 | 0 | 0 | 0 | 643970 | 0 | 0 |
| LHLVSPAD | HSPG2 | P98160 | PGBM_HUMAN | Unmodified | 67529 | 155150 | 3066100 | 1981300 | 0 | 0 | 0 | 0 | 782420 | 586960 | 0 | 0 | 0 | 0 | 366310 | 825080 | 0 | 0 |
| LHNAIEGG | DKC1; DKC1; D | O60832; H7I | DKC1_HUMAN | Unmodified | 0 | 0 | 0 | 0 | 0 | 0 | 0 | 0 | 0 | 0 | 0 | 0 | 0 | 0 | 0 | 0 | 0 | 0 |
| LHNFHQLS | PRMT5; PRMT5; | O14744; H0Y | ANM5_HUMAN | Unmodified | 0 | 0 | 0 | 0 | 0 | 0 | 0 | 0 | 0 | 0 | 0 | 744780 | 0 | 0 | 0 | 0 | 0 | 0 |
| LHNLAQFE | TRRAP; TRRAP; | Q9Y4A5; A0A | TRRAP_HUMAN | Unmodified | 55992 | 32059 | 0 | 0 | 80692 | 0 | 107540 | 16096 | 0 | 0 | 0 | 1160200 | 0 | 0 | 0 | 0 | 0 | 116880 |
| LHPDKNPN | DNAJC10; DNA | Q8IXB1; A0A | DJC10_HUMAN | Unmodified | 0 | 0 | 0 | 0 | 0 | 0 | 0 | 0 | 335880 | 112650 | 0 | 0 | 0 | 0 | 0 | 0 | 53225 | 0 |
| LHPEMSNL | UBTF; UBTF; UE | P17480; E9P | UBF1_HUMAN | Unmodified | 0 | 0 | 0 | 0 | 0 | 0 | 224400 | 139480 | 0 | 0 | 0 | 0 | 0 | 0 | 0 | 0 | 0 | 0 |
| LHPGLLEVI | TOR3A; TOR3A | Q9H497; Q5 | TOR3A_HUMAN | Unmodified | 0 | 0 | 0 | 0 | 0 | 0 | 0 | 0 | 3764200 | 226250 | 0 | 0 | 0 | 0 | 0 | 0 | 0 | 0 |
| LHPGLPEN | CARMIL1; CARN | Q5VZK9; A0A | CARL1_HUMAN | Unmodified | 0 | 0 | 0 | 0 | 2369200 | 256890 | 0 | 0 | 0 | 0 | 0 | 0 | 429170 | 536020 | 0 | 0 | 522530 | 1164300 |
| LHPVILASI | EIF3F; EIF3F; EI | O00303; A0A | EIF3F_HUMAN | Unmodified | 378730 | 0 | 0 | 0 | 0 | 0 | 0 | 0 | 0 | 0 | 0 | 6653100 | 0 | 0 | 0 | 386090 | 0 | 0 |
| LHPVILASI | EIF3F; EIF3F; EI | O00303; A0A | EIF3F_HUMAN | Unmodified | 0 | 0 | 0 | 0 | 0 | 0 | 0 | 0 | 0 | 0 | 0 | 3363600 | 0 | 0 | 0 | 0 | 0 | 0 |
| LHQEDND | PTPN1 | P18031 | PTN1_HUMAN | Unmodified | 0 | 0 | 0 | 0 | 333030 | 0 | 0 | 0 | 205160 | 0 | 211820 | 1383400 | 0 | 197850 | 0 | 0 | 0 | 0 |
| LHQGITVSI | GRAMD1A; GRA | Q96CP6; M0 | ASTRA_HUMAN | Unmodified | 0 | 0 | 0 | 0 | 0 | 0 | 0 | 0 | 1773100 | 0 | 0 | 0 | 0 | 0 | 0 | 0 | 0 | 0 |
| LHQHVSTH | TEP1; TEP1; TEF | Q99973; G3Y | TEP1_HUMAN | Unmodified | 456340 | 298910 | 0 | 131790 | 295830 | 0 | 0 | 53038 | 577920 | 0 | 781180 | 5450600 | 0 | 0 | 221940 | 275470 | 0 | 245830 |
| LHQLAMQ( | PCBP2; PCBP2; | Q15366; F8V | PCBP2_HUMAI | Unmodified | 3667500 | 2199200 | 0 | 2825200 | 1777100 | 0 | 952710 | 0 | 0 | 0 | 0 | 236680 | 7057300 | 2128400 | 462350 | 910710 | 593100 | 2401100 |
| LHQLAMQ( | PCBP2; PCBP2; | Q15366; F8V | PCBP2_HUMAI | 2 Oxidation (M) | 13248000 | 6180200 | 9664800 | 9358500 | 2288000 | 0 | 1052800 | 416520 | 870520 | 293850 | 0 | 1870800 | 9992200 | 3035800 | 3200400 | 6074700 | 396990 | 0 |
| LHQLAMQ( | PCBP2; PCBP2; | Q15366; F8V | PCBP2_HUMAI | Oxidation (M) | 13831000 | 13347000 | 1657900 | 12920000 | 4857900 | 0 | 680180 | 0 | 0 | 541810 | 0 | 1453400 | 16259000 | 7041200 | 3677600 | 1653400 | 1242600 | 768220 |
| LHQLLENQ | UACA; UACA; U | Q9BZF9; H0Y | UACA_HUMAN | Unmodified | 0 | 0 | 0 | 0 | 0 | 0 | 0 | 0 | 0 | 0 | 0 | 0 | 0 | 0 | 0 | 0 | 103170 | 0 |
| LHQLSGSD | SPAG9; SPAG9 | O60271; A0A | JIP4_HUMAN C | Unmodified | 0 | 0 | 0 | 0 | 0 | 0 | 0 | 0 | 0 | 32304 | 0 | 0 | 0 | 0 | 0 | 0 | 287960 | 1648600 |
| LHQLSGSD | SPAG9; SPAG9 | O60271; A0A | JIP4_HUMAN C | Unmodified | 0 | 0 | 0 | 0 | 0 | 0 | 0 | 0 | 0 | 0 | 0 | 0 | 0 | 0 | 0 | 0 | 162620 | 0 |
| LHQLSSIDI | LRP8; LRP8; LRI | Q14114; A0A | LRP8_HUMAN | Unmodified | 0 | 0 | 0 | 0 | 0 | 0 | 0 | 0 | 1987100 | 243350 | 0 | 0 | 0 | 0 | 0 | 0 | 0 | 0 |
| LHQMSVAI | HSPG2 | P98160 | PGBM_HUMAN | Unmodified | 0 | 0 | 726750 | 500550 | 0 | 0 | 0 | 0 | 0 | 0 | 0 | 0 | 0 | 0 | 0 | 306370 | 0 | 0 |
| LHQMWLS | PLIN3; PLIN3 | O60664; K7E | PLIN3_HUMAN | Oxidation (M) | 0 | 0 | 0 | 0 | 274910 | 214820 | 0 | 0 | 0 | 0 | 0 | 0 | 85882 | 0 | 0 | 75671 | 570450 | 0 |
| LHQVYFDA | SYP | P08247 | SYPH_HUMAN | Unmodified | 0 | 0 | 0 | 0 | 0 | 0 | 12666 | 0 | 4761500 | 871150 | 0 | 293920 | 0 | 0 | 0 | 0 | 0 | 0 |
| LHQYDGSI | CADM4 | Q8NFZ8 | CADM4_HUMA | Unmodified | 0 | 0 | 0 | 0 | 0 | 0 | 0 | 0 | 1067900 | 0 | 0 | 535830 | 0 | 0 | 1235300 | 0 | 0 | 0 |
| LHREFHHG | ERP44 | Q9BS26 | ERP44_HUMAN | Unmodified | 0 | 0 | 0 | 0 | 0 | 0 | 0 | 84219 | 4778700 | 917540 | 1031200 | 0 | 0 | 0 | 0 | 0 | 0 | 0 |
| LHRTELAT( | PLEC | Q15149 | PLEC_HUMAN | Unmodified | 0 | 0 | 0 | 0 | 42237 | 0 | 0 | 0 | 0 | 0 | 0 | 0 | 0 | 0 | 0 | 0 | 818950 | 864090 |
| LHSELDMV | TOM1L1; TOM1 | O75674; J3K | TM1L1_HUMAN | Unmodified | 1654300 | 1850300 | 1502800 | 753480 | 837750 | 0 | 0 | 0 | 0 | 0 | 165660 | 497960 | 139780 | 0 | 2444800 | 1161200 | 0 | 0 |
| LHSELDMV | TOM1L1; TOM1 | O75674; J3K | TM1L1_HUMAN | Oxidation (M) | 1433900 | 1233300 | 15163000 | 228630 | 0 | 0 | 191420 | 11353 | 118650 | 0 | 0 | 141560 | 0 | 0 | 27429000 | 1633400 | 0 | 0 |
| LHSEPILQI | POLRMT; POLRI | O00411; K7E | RPOM_HUMAN | Unmodified | 599660 | 0 | 574330 | 322230 | 1140800 | 384170 | 0 | 0 | 437630 | 0 | 0 | 1049200 | 55059 | 0 | 0 | 550420 | 0 | 195940 |
| LHSFLGDD | PYGL; PYGL | P06737; E9P | PYGL_HUMAN | Unmodified | 0 | 0 | 0 | 0 | 9837300 | 1599800 | 0 | 0 | 0 | 335240 | 0 | 0 | 0 | 0 | 0 | 0 | 2262500 | 4977500 |
| LHSISSIDV | LDLR; LDLR; LD | P01130; J3K | LDLR_HUMAN | Unmodified | 0 | 0 | 0 | 0 | 368140 | 115740 | 0 | 0 | 2994700 | 0 | 112540 | 0 | 0 | 0 | 0 | 0 | 0 | 0 |
| LHSLGPED | IGSF8 | Q969P0 | IGSF8_HUMAN | Unmodified | 16738000 | 15072000 | 4265600 | 10213000 | 0 | 0 | 0 | 0 | 812960 | 0 | 369800 | 7172900 | 0 | 0 | 13634000 | 10522000 | 0 | 0 |
| LHSLLGDY | EIF3L; EIF3L; EI | Q9Y262; B0( | EIF3L_HUMAN | Unmodified | 0 | 0 | 0 | 0 | 0 | 0 | 0 | 0 | 0 | 0 | 0 | 1474000 | 0 | 0 | 0 | 0 | 0 | 0 |
| LHSLQLLPI | HPS3; HPS3 | Q969F9; G5I | HPS3_HUMAN | Unmodified | 0 | 0 | 40567 | 0 | 0 | 0 | 0 | 0 | 73934 | 0 | 32169 | 4659000 | 0 | 0 | 104510 | 57052 | 0 | 0 |
| LHSLVKPS | LANCL1 | O43813 | LANC1_HUMAI | Unmodified | 216730 | 674850 | 0 | 0 | 0 | 0 | 0 | 0 | 0 | 0 | 0 | 5452100 | 0 | 0 | 340910 | 857750 | 0 | 121220 |
| LHTDDELN | TLN1 | Q9Y490 | TLN1_HUMAN | Unmodified | 423680 | 2137500 | 3350800 | 5716000 | 663690 | 0 | 0 | 0 | 1324600 | 0 | 0 | 2906900 | 0 | 0 | 2268700 | 1286000 | 0 | 0 |
| LHTDDELN | TLN1 | Q9Y490 | TLN1_HUMAN | Unmodified | 0 | 0 | 0 | 0 | 2502600 | 951600 | 0 | 0 | 0 | 0 | 0 | 0 | 0 | 1264900 | 0 | 0 | 424660 | 1498900 |
| LHTDRSAG | TRAF3; TRAF3; | Q13114; A6N | TRAF3_HUMAN | Unmodified | 0 | 0 | 0 | 0 | 10000000 | 496340 | 0 | 0 | 0 | 0 | 0 | 0 | 58443 | 0 | 0 | 0 | 14364 | 0 |
| LHTDRSAG | TRAF3; TRAF3; | Q13114; A6N | TRAF3_HUMAN | Unmodified | 0 | 0 | 0 | 0 | 7365100 | 2453700 | 0 | 0 | 0 | 0 | 0 | 0 | 0 | 0 | 0 | 0 | 0 | 0 |
| LHTLEEEKE | SMC3 | Q9UQE7 | SMC3_HUMAN | Unmodified | 48596 | 0 | 0 | 0 | 0 | 0 | 0 | 0 | 0 | 0 | 0 | 395490 | 0 | 0 | 0 | 0 | 0 | 0 |
| LHTLGDNL | LAMB1; LAMB1; | P07942; A0A | LAMB1_HUMAI | Unmodified | 0 | 0 | 1054100 | 0 | 0 | 0 | 0 | 0 | 441410 | 84662 | 0 | 0 | 0 | 0 | 0 | 0 | 0 | 0 |

| | | | | | | | | | | | | | | | | | | | | | | |
|---|---|---|---|---|---|---|---|---|---|---|---|---|---|---|---|---|---|---|---|---|---|---|
| LHVAILERE | PLEC; PLEC; PL | Q15149; H0Y | PLEC_HUMAN | Unmodified | 0 | 0 | 0 | 0 | 0 | 0 | 0 | 0 | 121130 | 0 | 0 | 187470 | 0 | 0 | 0 | 0 | 678930 | 0 |
| LHVGEESK | PIP4K2B | P78356 | PI42B_HUMAN | Unmodified | 0 | 0 | 0 | 0 | 238620 | 0 | 138460 | 90341 | 0 | 0 | 0 | 0 | 0 | 270450 | 0 | 0 | 285370 | 202160 |
| LHVGYLQP | ATP1B3 | P54709 | AT1B3_HUMAN | Unmodified | 0 | 0 | 54398 | 153240 | 0 | 0 | 0 | 0 | 0 | 0 | 0 | 67378 | 61808 | 0 | 813880 | 78848 | 0 | 0 |
| LHVPEKFEI | SLC4A7; SLC4A | Q9Y6M7; C9 | S4A7_HUMAN | Unmodified | 0 | 0 | 0 | 0 | 0 | 0 | 0 | 0 | 0 | 0 | 0 | 4194100 | 113560 | 0 | 0 | 0 | 51218 | 0 |
| LHYCVSCA | RPS26; RPS26P | P62854; Q5J | RS26_HUMAN | Unmodified | 906180 | 375740 | 468830 | 0 | 0 | 0 | 1227800 | 840420 | 274370 | 180390 | 550070 | 5102200 | 0 | 512950 | 0 | 254250 | 0 | 574780 |
| LHYGDLTD | GMDS | O60547 | GMDS_HUMAN | Unmodified | 0 | 0 | 55607 | 0 | 0 | 0 | 116060 | 31842 | 0 | 0 | 0 | 631400 | 0 | 0 | 0 | 0 | 0 | 0 |
| LHYGLPVV | CTNNB1; CTNN | P35222; A0A | CTNB1_HUMAI | Unmodified | 264460 | 190080 | 828600 | 380380 | 0 | 0 | 0 | 0 | 0 | 0 | 0 | 1715000 | 0 | 0 | 999930 | 1090000 | 58151 | 0 |
| LHYYCEDD | TRIM26; TRIM26 | Q12899; A2A | TRI26_HUMAN | Unmodified | 3612100 | 2413200 | 5066800 | 3445200 | 0 | 0 | 42484 | 27517 | 0 | 47863 | 0 | 102800 | 1336900 | 0 | 4313100 | 3053900 | 33711 | 0 |
| LIAAQSGL | REPS2 | Q8NFH8 | REPS2_HUMAN | Unmodified | 0 | 0 | 0 | 0 | 0 | 0 | 0 | 0 | 0 | 0 | 0 | 0 | 0 | 0 | 0 | 0 | 0 | 0 |
| LIAAQTGTR | UBL5 | Q9BZL1 | UBL5_HUMAN | Unmodified | 239180 | 149840 | 1853600 | 1466800 | 0 | 0 | 0 | 0 | 0 | 0 | 0 | 340510 | 0 | 0 | 1829900 | 2461300 | 0 | 0 |
| LIAAQTGTR | UBL5 | Q9BZL1 | UBL5_HUMAN | Unmodified | 0 | 0 | 0 | 0 | 330760 | 0 | 408120 | 452120 | 37752 | 67224 | 0 | 0 | 1361900 | 725560 | 0 | 0 | 815590 | 764500 |
| LIADLGSTS | FAM3C | Q92520 | FAM3C_HUMAI | Unmodified | 0 | 0 | 0 | 0 | 0 | 0 | 0 | 11594 | 5116200 | 483220 | 0 | 567690 | 0 | 0 | 0 | 0 | 0 | 0 |
| LIADVAPSA | FEN1; FEN1 | P39748; F5H | FEN1_HUMAN | Unmodified | 245550 | 264030 | 363940 | 305130 | 2790500 | 434520 | 3021700 | 2757200 | 692240 | 1424900 | 2190900 | 9442300 | 1164500 | 0 | 249880 | 163380 | 884590 | 693360 |
| LIAEQPPHL | PSPH; PSPH | P78330; C9J | SERB_HUMAN | Unmodified | 111350 | 0 | 707660 | 0 | 0 | 0 | 0 | 0 | 0 | 0 | 434200 | 1962900 | 0 | 0 | 580050 | 568420 | 0 | 0 |
| LIAHAGSLL | NOP58 | Q9Y2X3 | NOP58_HUMAI | Unmodified | 0 | 0 | 0 | 0 | 0 | 0 | 250380 | 0 | 351480 | 0 | 0 | 820740 | 0 | 0 | 0 | 0 | 46162 | 0 |
| LIAHAGSLT | NOP56; NOP56 | O00567; A0A | NOP56_HUMAI | Unmodified | 460690 | 171500 | 0 | 290980 | 0 | 0 | 0 | 0 | 333100 | 0 | 0 | 0 | 112330 | 0 | 165450 | 0 | 0 | 0 |
| LIAHAGSLT | NOP56; NOP56 | O00567; A0A | NOP56_HUMAI | Unmodified | 0 | 0 | 0 | 0 | 0 | 0 | 1199400 | 0 | 0 | 149370 | 0 | 0 | 198970 | 134880 | 0 | 0 | 0 | 273010 |
| LIAINANDP | PPA2; PPA2; PP | Q9H2U2; D6 | IPYR2_HUMAN | Unmodified | 641030 | 0 | 0 | 0 | 0 | 0 | 0 | 0 | 666200 | 0 | 0 | 1347800 | 577150 | 0 | 605040 | 380270 | 0 | 0 |
| LIAINANDP | PPA2; PPA2; PP | Q9H2U2; D6 | IPYR2_HUMAN | Unmodified | 0 | 457810 | 0 | 0 | 744550 | 0 | 0 | 153590 | 2115800 | 61240 | 0 | 2021100 | 189190 | 0 | 0 | 0 | 0 | 0 |
| LIAINANDP | PPA2; PPA2; PP | Q9H2U2; D6 | IPYR2_HUMAN | Unmodified | 0 | 0 | 0 | 0 | 790620 | 0 | 0 | 0 | 867400 | 0 | 0 | 1174700 | 0 | 0 | 0 | 0 | 0 | 0 |
| LIAKVEGKG | EIF5; EIF5; EIF5 | P55010; H0Y | IF5_HUMAN Eu | Unmodified | 0 | 0 | 0 | 0 | 270230 | 0 | 160910 | 0 | 0 | 0 | 0 | 0 | 380040 | 0 | 0 | 0 | 243250 | 263190 |
| LIALLEVLS | FLNC; FLNB; FL | Q14315; O75 | FLNC_HUMAN | Unmodified | 341270 | 594250 | 437600 | 1760000 | 456910 | 106600 | 0 | 0 | 296670 | 87773 | 293490 | 1342400 | 255060 | 234430 | 1173900 | 770880 | 0 | 109340 |
| LIALLEVLS | FLNA; FLNA; FL | P21333; A0A | FLNA_HUMAN | Unmodified | 0 | 0 | 1667200 | 8415300 | 734240 | 0 | 0 | 0 | 0 | 0 | 0 | 7317900 | 0 | 0 | 4824400 | 3536000 | 0 | 0 |
| LIALLEVLS | FLNC; FLNB | Q14315; O75 | FLNC_HUMAN | Unmodified | 0 | 0 | 1818400 | 1224000 | 0 | 0 | 0 | 0 | 309420 | 0 | 215460 | 3702200 | 0 | 0 | 1293100 | 2238300 | 0 | 0 |
| LIALSIDSVE | PRDX6 | P30041 | PRDX6_HUMAN | Unmodified | 3373600 | 3888100 | 2636700 | 18792000 | 58383000 | 5773600 | 3294000 | 105240 | 6933900 | 10741000 | 5739300 | 10251000 | 2.09E+08 | 71619000 | 8793700 | 3323700 | 1.34E+08 | 2.57E+08 |
| LIANNTTVE | ACTL6A | O96019 | ACL6A_HUMAN | Unmodified | 280430 | 155510 | 180620 | 150080 | 0 | 0 | 0 | 0 | 464000 | 259720 | 315350 | 4717500 | 233190 | 203390 | 156620 | 177980 | 110590 | 0 |
| LIAPVAEEE | LDHB; LDHB; LI | P07195; A0A | LDHB_HUMAN | Unmodified | 1610700 | 1420800 | 2318400 | 2778500 | 0 | 1624700 | 0 | 0 | 0 | 0 | 32274000 | 1.36E+08 | 2808000 | 704630 | 3681700 | 2539300 | 960940 | 1386700 |
| LIAQDYK | GLG1; GLG1 | Q92896; H3I | GSLG1_HUMAI | Unmodified | 0 | 0 | 0 | 320990 | 0 | 0 | 0 | 0 | 10942000 | 1600900 | 2234400 | 933710 | 0 | 0 | 0 | 0 | 0 | 0 |
| LIAQDYKVS | GLG1; GLG1 | Q92896; H3I | GSLG1_HUMAI | Unmodified | 0 | 0 | 0 | 0 | 0 | 129690 | 223540 | 168260 | 0 | 2674100 | 0 | 0 | 0 | 141360 | 0 | 0 | 0 | 0 |
| LIASSLVAG | GAPVD1; GAPV | Q14C86; C9 | GAPD1_HUMAI | Unmodified | 80894 | 0 | 0 | 102240 | 0 | 0 | 0 | 0 | 305920 | 0 | 0 | 4357100 | 0 | 0 | 215830 | 126940 | 0 | 0 |
| LIASVAGSV | EXOSC2; EXOS | Q13868; A3K | EXOS2_HUMAN | Unmodified | 0 | 0 | 75782 | 147280 | 0 | 0 | 0 | 0 | 0 | 0 | 1095500 | 596560 | 0 | 0 | 0 | 0 | 0 | 0 |
| LIASVAGSV | EXOSC2; EXOS | Q13868; A3K | EXOS2_HUMAN | Unmodified | 0 | 0 | 0 | 204950 | 516060 | 0 | 0 | 0 | 0 | 0 | 0 | 61927 | 597320 | 0 | 0 | 0 | 0 | 0 |
| LIASYCNVC | LRPPRC; LRPPF | P42704; B8Z | LPPRC_HUMAN | Unmodified | 0 | 0 | 0 | 0 | 0 | 0 | 0 | 18636 | 3636200 | 288960 | 0 | 0 | 0 | 0 | 0 | 0 | 0 | 0 |
| LIATFPDTL | ITGA6 | P23229 | ITA6_HUMAN I | Unmodified | 699120 | 1423000 | 5092000 | 16934000 | 1206200 | 0 | 0 | 0 | 1897900 | 0 | 1038300 | 28646000 | 856850 | 0 | 12583000 | 11136000 | 1479900 | 8034800 |
| LIATFPDTL | ITGA6 | P23229 | ITA6_HUMAN I | Unmodified | 0 | 0 | 631510 | 0 | 21501000 | 0 | 0 | 0 | 0 | 0 | 1533800 | 0 | 19308000 | 5812000 | 5734800 | 1198100 | 11921000 | 61128000 |
| LIATFPDTL | ITGA6 | P23229 | ITA6_HUMAN I | Unmodified | 0 | 0 | 0 | 0 | 77707000 | 12714000 | 15752000 | 82728 | 0 | 648410 | 0 | 0 | 28887000 | 29307000 | 0 | 0 | 1.02E+08 | 5.34E+08 |
| LICCDILDV | YWHAE; YWHAI | P62258; K7E | 1433E_HUMAN | Unmodified | 2701200 | 997990 | 1405200 | 7117800 | 2855600 | 2778100 | 1887500 | 32612 | 7628800 | 6343700 | 6077000 | 20381000 | 6896200 | 545720 | 7689400 | 5841000 | 6825000 | 785820 |
| LICCDILDV | YWHAE | P62258 | 1433E_HUMAN | Unmodified | 37784 | 0 | 0 | 513790 | 1008500 | 0 | 0 | 0 | 273080 | 774750 | 166220 | 46841 | 2208800 | 2235300 | 0 | 0 | 6015700 | 369320 |
| LICDVSRQI | RARS1 | P54136 | SYRC_HUMAN | Unmodified | 199590 | 0 | 0 | 0 | 0 | 0 | 196950 | 86503 | 416640 | 0 | 483630 | 4767400 | 0 | 231160 | 299730 | 287410 | 0 | 0 |
| LICGFMDD | GLG1; GLG1; G | Q92896; H3I | GSLG1_HUMAI | Oxidation (M) | 0 | 0 | 0 | 0 | 0 | 0 | 0 | 0 | 926750 | 264600 | 377790 | 0 | 0 | 0 | 0 | 0 | 0 | 0 |
| LICGFMDD | GLG1; GLG1; G | Q92896; H3I | GSLG1_HUMAI | Unmodified | 0 | 0 | 0 | 0 | 0 | 0 | 0 | 61109 | 2562300 | 148980 | 30802000 | 0 | 0 | 0 | 0 | 0 | 0 | 0 |
| LICSELNGI | CLSTN1; CLSTN | O94985; Q5 | CSTN1_HUMAI | Unmodified | 0 | 0 | 0 | 0 | 0 | 0 | 76491 | 0 | 2161200 | 58540 | 0 | 0 | 598740 | 0 | 0 | 0 | 0 | 0 |
| LIDDEVRILI | AFG3L2 | Q9Y4W6 | AFG32_HUMAN | Unmodified | 0 | 0 | 0 | 0 | 0 | 0 | 0 | 0 | 0 | 467370 | 0 | 0 | 0 | 0 | 0 | 0 | 0 | 0 |
| LIDDMVAQ | IDH1 | O75874 | IDHC_HUMAN | Unmodified | 0 | 0 | 0 | 168540 | 272160 | 0 | 0 | 0 | 1745700 | 0 | 0 | 0 | 0 | 0 | 368130 | 0 | 0 | 0 |
| LIDDMVAQ | IDH1 | O75874 | IDHC_HUMAN | Oxidation (M) | 0 | 0 | 0 | 545130 | 0 | 0 | 0 | 0 | 0 | 0 | 367440 | 2029800 | 0 | 0 | 1611700 | 121650 | 0 | 0 |
| LIDDMVAQ | IDH2; IDH2 | P48735; H0Y | IDHP_HUMAN | Oxidation (M) | 0 | 0 | 0 | 0 | 0 | 0 | 0 | 0 | 7447400 | 218590 | 0 | 0 | 0 | 0 | 0 | 0 | 0 | 0 |
| LIDDMVAQ | IDH2; IDH2 | P48735; H0Y | IDHP_HUMAN | Unmodified | 0 | 0 | 0 | 0 | 0 | 0 | 0 | 0 | 11720000 | 745740 | 37343 | 0 | 0 | 0 | 0 | 0 | 0 | 0 |
| LIDDYDSTK | SCG3 | Q8WXD2 | SCG3_HUMAN | Unmodified | 0 | 0 | 0 | 0 | 0 | 0 | 0 | 52149 | 43536000 | 6961400 | 64756 | 0 | 0 | 0 | 0 | 0 | 0 | 0 |
| LIDDYGVEI | PURA | Q00577 | PURA_HUMAN | Unmodified | 160360 | 0 | 0 | 0 | 0 | 0 | 0 | 0 | 0 | 0 | 0 | 1708900 | 0 | 0 | 0 | 0 | 0 | 0 |
| LIDEMVGK | RP2 | O75695 | XRP2_HUMAN | Unmodified | 375880 | 0 | 314470 | 415270 | 1782700 | 237610 | 250170 | 0 | 2260400 | 491860 | 928010 | 6749900 | 672140 | 533920 | 0 | 0 | 1576500 | 2259200 |
| LIDEVIEDTI | MYOF; MYOF | Q9NZM1; HC | MYOF_HUMAN | Unmodified | 0 | 0 | 0 | 0 | 15830000 | 5737500 | 0 | 0 | 0 | 419400 | 0 | 0 | 0 | 0 | 0 | 0 | 359740 | 0 |
| LIDFLECGK | DDX5; DDX5; D | P17844; A0A | DDX5_HUMAN | Unmodified | 0 | 0 | 0 | 0 | 0 | 0 | 0 | 0 | 0 | 0 | 0 | 2381000 | 0 | 0 | 0 | 0 | 0 | 0 |
| LIDFLESGK | DDX17; DDX17; | Q92841; A0A | DDX17_HUMAI | Unmodified | 0 | 0 | 618780 | 0 | 0 | 0 | 0 | 0 | 621500 | 0 | 1052600 | 8638100 | 0 | 0 | 142250 | 0 | 0 | 0 |
| LIDFLQAK | CIT; CIT; CIT; CI | O14578; A0A | CTRO_HUMAN | Unmodified | 0 | 0 | 0 | 0 | 10660000 | 1126800 | 0 | 0 | 769970 | 0 | 0 | 953910 | 0 | 0 | 0 | 0 | 2171500 | 1008600 |
| LIDFLQAKN | CIT; CIT; CIT; CI | O14578; A0A | CTRO_HUMAN | Oxidation (M) | 0 | 0 | 0 | 0 | 796980 | 0 | 165370 | 109670 | 0 | 0 | 0 | 0 | 0 | 0 | 0 | 0 | 960420 | 0 |
| LIDFLQAKN | CIT; CIT; CIT; CI | O14578; A0A | CTRO_HUMAN | Unmodified | 0 | 0 | 0 | 0 | 1473500 | 0 | 547420 | 0 | 0 | 0 | 290620 | 0 | 0 | 0 | 0 | 0 | 530830 | 0 |
| LIDIFYPGD | ALDH18A1 | P54886 | P5CS_HUMAN | Unmodified | 25765 | 28088 | 19537 | 0 | 36607 | 39398 | 83616 | 63875 | 2413700 | 324890 | 0 | 248060 | 0 | 0 | 0 | 0 | 40151 | 581160 |
| LIDIKDTLP | MVB12B; MVB1 | Q9H7P6; F8 | MB12B_HUMAI | Unmodified | 0 | 0 | 0 | 0 | 0 | 0 | 0 | 0 | 0 | 0 | 0 | 0 | 0 | 0 | 808680 | 0 | 0 | 0 |
| LIDIKDTLP | MVB12B; MVB1 | Q9H7P6; F8 | MB12B_HUMAI | Unmodified | 305380 | 1767500 | 0 | 2573800 | 0 | 0 | 0 | 0 | 364550 | 1049300 | 0 | 0 | 1918400 | 1741500 | 4948300 | 0 | 0 | 0 |
| LIDLHSPSE | RPS20; RPS20; | P60866; A0A | RS20_HUMAN | Unmodified | 1077300 | 1358600 | 546800 | 1344100 | 0 | 0 | 0 | 0 | 720160 | 222200 | 8782100 | 17658000 | 445000 | 0 | 606900 | 407450 | 146610 | 801470 |
| LIDNISSRE | MRPS27; MRPS | Q92552; D6I | RT27_HUMAN | Unmodified | 0 | 0 | 0 | 0 | 0 | 0 | 0 | 0 | 1180600 | 0 | 0 | 0 | 0 | 0 | 0 | 0 | 0 | 0 |
| LIDPQTQV | APPL1 | Q9UKG1 | DP13A_HUMAN | Unmodified | 329190 | 67903 | 0 | 0 | 626990 | 0 | 172510 | 61257 | 472360 | 235100 | 2831200 | 3046200 | 70941 | 0 | 0 | 0 | 1025000 | 0 |
| LIDREIISHD | CYB5R3; CYB5I | P00387; B1A | NB5R3_HUMAI | Unmodified | 0 | 0 | 0 | 0 | 0 | 0 | 0 | 0 | 122360 | 0 | 0 | 484100 | 0 | 0 | 0 | 0 | 0 | 0 |
| LIDREIISHD | CYB5R3; CYB5I | P00387; B1A | NB5R3_HUMAI | Unmodified | 0 | 0 | 139930 | 0 | 0 | 0 | 0 | 0 | 332630 | 0 | 0 | 966480 | 0 | 0 | 0 | 0 | 0 | 0 |
| LIDTVKDPV | CSNK2A2; CSN | P19784; H3E | CSK22_HUMAN | Unmodified | 92979 | 74543 | 0 | 0 | 116520 | 0 | 0 | 16207 | 52177 | 23995 | 92752 | 1826100 | 0 | 0 | 161490 | 69598 | 0 | 13264 |
| LIDVNHYAI | SPTAN1; SPTAN | Q13813; A0A | SPTN1_HUMAN | Unmodified | 0 | 0 | 0 | 0 | 0 | 0 | 0 | 0 | 0 | 181470 | 0 | 2056400 | 0 | 0 | 0 | 0 | 0 | 0 |
| LIDWGLAEI | CSNK2A2; CSN | P19784; H3E | CSK22_HUMAN | Unmodified | 176820 | 501690 | 0 | 187870 | 0 | 0 | 38398 | 0 | 772230 | 0 | 0 | 2049200 | 38896 | 61953 | 0 | 116350 | 0 | 0 |
| LIDWGLAEI | CSNK2A3; CSN | Q8NEV1; P6 | CSK23_HUMAN | Unmodified | 1337800 | 928180 | 206590 | 1790300 | 335590 | 0 | 42210 | 0 | 1103900 | 54017 | 268970 | 4086100 | 0 | 0 | 883890 | 791100 | 0 | 0 |
| LIEALEK | KIF23; LRRFIP2; | Q02241; H7 | KIF23_HUMAN | Unmodified | 377620 | 36760 | 1089800 | 1471800 | 15248000 | 2359400 | 197450 | 48022 | 7325200 | 2564600 | 2294500 | 4181300 | 910500 | 483240 | 1055500 | 346920 | 1359300 | 433220 |
| LIEAVDNMI | EHD4 | Q9H223 | EHD4_HUMAN | Oxidation (M) | 786340 | 604690 | 741630 | 806970 | 1391000 | 305200 | 0 | 0 | 0 | 0 | 0 | 577060 | 785760 | 0 | 1514400 | 1372600 | 451530 | 1390300 |
| LIEAVDNMI | EHD4 | Q9H223 | EHD4_HUMAN | Unmodified | 218920 | 1674800 | 711980 | 2121400 | 9113000 | 995320 | 0 | 0 | 1206300 | 0 | 0 | 1394700 | 3214600 | 1836900 | 2317200 | 1029300 | 2277700 | 9480200 |
| LIEAYVIQC | BROX; BROX; B | Q5VW32; F5 | BROX_HUMAN | Unmodified | 7225200 | 11322000 | 0 | 6960700 | 42029 | 0 | 0 | 0 | 0 | 0 | 276670 | 0 | 0 | 0 | 5292800 | 4634900 | 0 | 0 |
| LIEDFLAR | TRAPPC3; TRAF | O43617; A0A | TPPC3_HUMAN | Unmodified | 162490 | 172040 | 490100 | 581630 | 0 | 0 | 0 | 0 | 500300 | 0 | 214320 | 6061400 | 117100 | 0 | 377220 | 0 | 20097 | 0 |
| LIEDGRGC | RO60; RO60 | P10155; H0Y | RO60_HUMAN | Unmodified | 29416000 | 14983000 | 3954900 | 7038100 | 0 | 0 | 0 | 0 | 600490 | 189530 | 1048400 | 2014600 | 2161000 | 0 | 8597600 | 4083300 | 154880 | 1722700 |
| LIEDNEYTA | YES1; SRC; FYN | P07947; P12 | YES_HUMAN Ty | Unmodified | 589360 | 548390 | 1663900 | 1278300 | 0 | 0 | 0 | 0 | 2894300 | 468900 | 418630 | 3491300 | 87129 | 0 | 2279500 | 1759300 | 0 | 542470 |
| LIEDNEYTA | YES1; SRC; FYN | P07947; P12 | YES_HUMAN Ty | Unmodified | 0 | 0 | 0 | 203120 | 0 | 0 | 0 | 0 | 391700 | 527560 | 0 | 0 | 385570 | 0 | 0 | 0 | 911920 | 0 |
| LIEEAWSPI | NAA25 | Q14CX7 | NAA25_HUMAN | Unmodified | 0 | 91096 | 0 | 0 | 0 | 0 | 0 | 0 | 0 | 0 | 0 | 1380800 | 0 | 0 | 0 | 0 | 0 | 0 |
| LIEECISQI | DIAPH1; DIAPH | O60610; A0A | DIAP1_HUMAN | Unmodified | 0 | 0 | 0 | 511390 | 0 | 0 | 747800 | 0 | 382780 | 426280 | 0 | 3631300 | 650220 | 0 | 727950 | 319340 | 1147100 | 0 |
| LIEELAIAK | RUFY2; RUFY2 | Q8WXA3; HC | RUFY2_HUMAN | Unmodified | 0 | 0 | 0 | 0 | 280820 | 0 | 87845 | 58321 | 0 | 0 | 0 | 0 | 54030 | 0 | 0 | 0 | 61054 | 183330 |
| LIEENEK | STK31; NCAM2; | Q9BXU1; O1 | STK31_HUMAN | Unmodified | 0 | 0 | 2139700 | 33660 | 0 | 0 | 15860 | 0 | 418760 | 77856 | 167080 | 0 | 0 | 111700 | 2591200 | 135710 | 0 | 0 |
| LIEENEKYIL | NCAM2; NCAM | O15394; H9I | NCAM2_HUMA | Unmodified | 0 | 0 | 0 | 171970 | 0 | 0 | 156230 | 54372 | 0 | 0 | 0 | 0 | 0 | 0 | 0 | 0 | 0 | 120820 |
| LIEESSVDE | PRPSAP1; PRPS | Q14558; B4I | KPRA_HUMAN | Unmodified | 0 | 0 | 0 | 0 | 0 | 0 | 0 | 0 | 0 | 0 | 0 | 1322900 | 0 | 0 | 453190 | 0 | 0 | 0 |
| LIEEVHAVV | AP2S1; AP2S1; | P53680; M0 | AP2S1_HUMAN | Unmodified | 0 | 0 | 0 | 0 | 0 | 0 | 0 | 0 | 144660 | 0 | 104840 | 1090100 | 0 | 0 | 0 | 0 | 0 | 0 |
| LIEEVMIGE | CCT2; CCT2; C | P78371; F5G | TCPB_HUMAN | Unmodified | 0 | 0 | 0 | 0 | 0 | 0 | 0 | 0 | 2339800 | 366950 | 1018900 | 4899400 | 1768900 | 528160 | 175030 | 0 | 24848 | 0 |
| LIEEVMIGE | CCT2; CCT2; C | P78371; F5G | TCPB_HUMAN | Oxidation (M) | 0 | 0 | 258230 | 417050 | 0 | 0 | 0 | 81744 | 0 | 0 | 1426500 | 12493000 | 902710 | 525880 | 319360 | 354540 | 0 | 0 |
| LIEFLSK | CAB39; CAB39 | Q9Y376; A0A | CAB39_HUMAN | Unmodified | 308970 | 0 | 439960 | 0 | 435820 | 0 | 0 | 0 | 0 | 315270 | 0 | 2760400 | 0 | 0 | 775280 | 687800 | 0 | 0 |
| LIEFLSKFQ | CAB39; CAB39 | Q9Y376; A0A | CAB39_HUMAN | Unmodified | 0 | 0 | 0 | 0 | 306630 | 0 | 0 | 0 | 0 | 0 | 0 | 0 | 0 | 0 | 0 | 0 | 911080 | 0 |
| LIEFLSSFQ | CAB39L; CAB39 | Q9H9S4; B7 | CB39L_HUMAN | Unmodified | 0 | 0 | 0 | 0 | 0 | 0 | 0 | 0 | 0 | 0 | 856510 | 2522800 | 0 | 0 | 0 | 0 | 0 | 0 |
| LIEGGAHIY | POR; POR | P16435; E7E | NCPR_HUMAN | Unmodified | 0 | 0 | 0 | 0 | 0 | 0 | 676150 | 514550 | 477840 | 46683 | 0 | 675770 | 163260 | 818880 | 0 | 0 | 489170 | 0 |
| LIEGLSHEV | RCC2 | Q9P258 | RCC2_HUMAN | Unmodified | 0 | 0 | 0 | 0 | 0 | 0 | 0 | 0 | 2164200 | 0 | 2373400 | 10596000 | 0 | 0 | 0 | 0 | 0 | 0 |
| LIEHGVNT | CRLF3 | Q8IUI8 | CRLF3_HUMAN | Unmodified | 13907 | 0 | 0 | 0 | 0 | 0 | 0 | 0 | 0 | 0 | 138050 | 622640 | 0 | 0 | 34625 | 52192 | 0 | 0 |
| LIEIASR | LAMC1 | P11047 | LAMC1_HUMAI | Unmodified | 0 | 0 | 0 | 0 | 637450 | 0 | 498450 | 854570 | 667180 | 0 | 0 | 61384 | 181530 | 145630 | 0 | 0 | 65977 | 287140 |
| LIEIENFK | SMC1A; SMC1A | Q14683; V9 | SMC1A_HUMA | Unmodified | 0 | 0 | 0 | 0 | 0 | 0 | 440360 | 490860 | 0 | 147590 | 0 | 1074000 | 96531 | 0 | 0 | 0 | 0 | 606090 |
| LIELQAGK | PRKCSH; PRKC | P14314; K7E | GLU2B_HUMAI | Unmodified | 236850 | 240100 | 0 | 0 | 152950 | 0 | 128390 | 135720 | 17292000 | 6055700 | 2631400 | 0 | 0 | 0 | 0 | 201130 | 268170 | 66299 |
| LIELVDR | MBLAC2 | Q68D91 | MBLC2_HUMAI | Unmodified | 135430 | 0 | 792860 | 143300 | 0 | 0 | 0 | 0 | 0 | 0 | 0 | 3274700 | 347020 | 299680 | 582080 | 960050 | 0 | 0 |
| LIELVDRGL | MBLAC2 | Q68D91 | MBLC2_HUMAI | Unmodified | 221480 | 0 | 546960 | 500630 | 0 | 0 | 0 | 0 | 0 | 0 | 295410 | 2079500 | 0 | 42953 | 667120 | 655560 | 0 | 0 |
| LIENPLLR | SHCBP1 | Q8NEM2 | SHCBP_HUMA | Unmodified | 0 | 0 | 0 | 0 | 2763000 | 860930 | 0 | 0 | 0 | 142700 | 0 | 0 | 0 | 0 | 0 | 0 | 0 | 0 |
| LIENPLLRY | SHCBP1 | Q8NEM2 | SHCBP_HUMA | Unmodified | 0 | 0 | 0 | 0 | 3897600 | 347360 | 234490 | 0 | 0 | 69482 | 0 | 0 | 101380 | 0 | 0 | 0 | 20074 | 203970 |
| LIENVDPEK | NAPG; NAPG | Q99747; J3K | SNAG_HUMAN | Unmodified | 286640 | 584220 | 854140 | 375230 | 3750100 | 764020 | 218330 | 0 | 2535200 | 346390 | 1942700 | 6145400 | 563740 | 1034200 | 1504300 | 1271600 | 520060 | 2999700 |
| LIERDPYEH | PTPN23 | Q9H3S7 | PTN23_HUMAN | Unmodified | 0 | 449160 | 0 | 0 | 0 | 0 | 73017 | 0 | 336540 | 44975 | 0 | 0 | 302870 | 0 | 0 | 0 | 0 | 0 |
| LIESAEELIR | DNAJC3; DNAJ | Q13217; X6F | DNJC3_HUMAN | Unmodified | 0 | 0 | 0 | 0 | 0 | 0 | 374120 | 0 | 5780500 | 0 | 0 | 333030 | 0 | 0 | 0 | 0 | 0 | 0 |
| LIESAEELIR | DNAJC3; DNAJ | Q13217; X6F | DNJC3_HUMAN | Unmodified | 0 | 0 | 0 | 0 | 0 | 0 | 0 | 0 | 9447900 | 71839 | 51309 | 112880 | 117680 | 88487 | 0 | 0 | 0 | 0 |
| LIESSLGSV | PIK3C2B; PIK3C | O00750; F5C | P3C2B_HUMAN | Unmodified | 128100 | 177730 | 123110 | 0 | 0 | 0 | 117140 | 0 | 236760 | 0 | 0 | 0 | 0 | 0 | 349580 | 333590 | 0 | 0 |
| LIETELLQL | LRSAM1; LRSAN | Q6UWE0; AC | LRSM1_HUMAN | Unmodified | 0 | 0 | 0 | 0 | 471340 | 17098 | 0 | 0 | 0 | 43902 | 0 | 0 | 99789 | 0 | 87406 | 0 | 46253 | 68123 |
| LIETVVK | EXOC6; EXOC6 | Q8TAG9; E7I | EXOC6_HUMAI | Unmodified | 0 | 0 | 0 | 47771 | 464130 | 0 | 0 | 20247 | 0 | 0 | 0 | 469730 | 67378 | 69680 | 52621 | 0 | 24081 | 69234 |
| LIEVANLAC | CTNNA1; CTNN | P35221; E5R | CTNA1_HUMAI | Unmodified | 542120 | 1011700 | 668690 | 602670 | 1606800 | 0 | 0 | 0 | 0 | 0 | 619270 | 2923400 | 526370 | 264000 | 7964800 | 6329000 | 0 | 0 |
| LIEVDDERK | RPS6; RPS6 | P62753; A2A | RS6_HUMAN 4 | Unmodified | 0 | 0 | 0 | 0 | 762800 | 0 | 58452 | 258000 | 0 | 0 | 79930 | 7012100 | 1092000 | 0 | 0 | 0 | 0 | 0 |
| LIEVNGQN | SLC9A3R2; SLC | Q15599; H3I | NHRF2_HUMAI | Unmodified | 245690 | 179150 | 605180 | 317190 | 0 | 0 | 0 | 0 | 0 | 0 | 0 | 1771200 | 0 | 0 | 556280 | 562740 | 0 | 0 |
| LIEVQSNSI | ERMP1; ERMP1 | Q7Z2K6; E7E | ERMP1_HUMAI | Unmodified | 215380 | 0 | 0 | 0 | 344640 | 0 | 0 | 0 | 6746400 | 374650 | 0 | 412550 | 0 | 0 | 0 | 0 | 0 | 0 |

| | | | | | | | | | | | | | | | | | | | | | | |
|---|---|---|---|---|---|---|---|---|---|---|---|---|---|---|---|---|---|---|---|---|---|---|
| LIEVTETICK | CNPY3; CNPY3 | Q9BT09; A0A | CNPY3_HUMAI | Unmodified | 0 | 0 | 0 | 0 | 0 | 0 | 0 | 0 | 15102000 | 2436900 | 0 | 0 | 0 | 0 | 293500 | 0 | 0 | 0 |
| LIEVTETICK | CNPY3; CNPY3 | Q9BT09; A0A | CNPY3_HUMAI | Unmodified | 0 | 0 | 0 | 0 | 0 | 0 | 0 | 0 | 716650 | 0 | 0 | 0 | 0 | 0 | 0 | 0 | 0 | 0 |
| LIEWLPLHL | ACAD10; ACAD | Q6JQN1; F8\ | ACD10_HUMAI | Unmodified | 0 | 0 | 0 | 0 | 0 | 0 | 0 | 0 | 594930 | 0 | 0 | 0 | 0 | 0 | 0 | 0 | 0 | 0 |
| LIEWLPQNI | UGT8 | Q16880 | CGT_HUMAN 2 | Unmodified | 0 | 0 | 0 | 0 | 0 | 0 | 0 | 0 | 4392300 | 0 | 195350 | 968420 | 0 | 0 | 0 | 0 | 0 | 0 |
| LIFDNLK | ATP1A1; ATP1A | P05023; P50 | AT1A1_HUMAN | Unmodified | 172510 | 175780 | 2708300 | 462640 | 0 | 0 | 28652 | 11442 | 732970 | 725720 | 0 | 21046000 | 85880 | 0 | 3512300 | 2997100 | 0 | 0 |
| LIFIVDVWH | ASPH | Q12797 | ASPH_HUMAN | Unmodified | 0 | 0 | 0 | 0 | 0 | 0 | 0 | 0 | 463360 | 138350 | 25485 | 0 | 0 | 0 | 0 | 0 | 0 | 0 |
| LIFKPDLTLI | B4GALT4; B4GA | O60513; C9J | B4GT4_HUMAN | Unmodified | 0 | 0 | 0 | 0 | 0 | 0 | 0 | 0 | 15713000 | 3378200 | 0 | 0 | 0 | 0 | 0 | 0 | 0 | 0 |
| LIFLFRQDP | CLSTN1; CLSTN | O94985; Q5 | CSTN1_HUMAI | Unmodified | 0 | 41437 | 2306100 | 285670 | 0 | 149410 | 2852500 | 0 | 4370100 | 588590 | 111510 | 108610 | 81989 | 90792 | 248520 | 31814 | 0 | 0 |
| LIFNENDF\ | CSGALNACT2 | Q8N6G5 | CGAT2_HUMAI | Unmodified | 0 | 3443100 | 0 | 0 | 1989100 | 0 | 0 | 0 | 1246600 | 0 | 0 | 0 | 0 | 0 | 0 | 0 | 1971800 | 0 |
| LIFPERIDPI | TIPRL | O75663 | TIPRL_HUMAN | Unmodified | 1144400 | 0 | 0 | 2666900 | 4784500 | 981250 | 1597700 | 1030200 | 508550 | 5901800 | 2406700 | 14357000 | 7336700 | 2336700 | 626200 | 0 | 2225100 | 3668300 |
| LIFPERIDPI | TIPRL | O75663 | TIPRL_HUMAN | Unmodified | 1919000 | 0 | 0 | 0 | 1848100 | 0 | 812410 | 73990 | 0 | 0 | 0 | 0 | 0 | 0 | 0 | 0 | 1619500 | 0 |
| LIFPITSQNI | SH3BP4 | Q9P0V3 | SH3B4_HUMAI | Unmodified | 26957 | 18664 | 74347 | 571790 | 19160 | 0 | 0 | 0 | 4112200 | 266140 | 221710 | 1002300 | 0 | 0 | 1639400 | 771850 | 0 | 0 |
| LIFPYVELDI | IDH1; IDH1; IDH | O75874; C9J | IDHC_HUMAN | Unmodified | 0 | 0 | 0 | 0 | 0 | 0 | 0 | 0 | 0 | 0 | 0 | 0 | 6773500 | 1683300 | 593320 | 0 | 0 | 0 |
| LIFQMPQN | IQGAP2; IQGAP | Q13576; Q8 | IQGA2_HUMAN | Oxidation (M) | 0 | 0 | 0 | 0 | 1081800 | 0 | 0 | 0 | 0 | 262500 | 0 | 2071100 | 0 | 0 | 0 | 133650 | 0 | 0 |
| LIFQMPQN | IQGAP2; IQGAP | Q13576; Q8 | IQGA2_HUMAN | Unmodified | 87113 | 45167 | 0 | 0 | 2179400 | 931460 | 0 | 102430 | 2385400 | 309520 | 0 | 3333900 | 0 | 0 | 1148000 | 420760 | 129160 | 0 |
| LIFYDLR | KARS1; KARS1 | Q15046; H3I | SYK_HUMAN Ly | Unmodified | 16237 | 24871 | 79486 | 73176 | 0 | 0 | 55113 | 43700 | 0 | 0 | 281610 | 5046200 | 0 | 43925 | 70447 | 37915 | 0 | 0 |
| LIFYDLRGE | KARS1; KARS1 | Q15046; H3I | SYK_HUMAN Ly | Unmodified | 0 | 0 | 1335300 | 490790 | 778400 | 2161500 | 521540 | 418150 | 645110 | 0 | 824530 | 7379200 | 0 | 718660 | 0 | 183610 | 224520 | 424930 |
| LIGAYLPQF | DIP2C; DIP2C | Q9Y2E4; E7E | DIP2C_HUMAN | Unmodified | 0 | 46585 | 0 | 0 | 0 | 0 | 0 | 0 | 0 | 422020 | 0 | 175510 | 992390 | 0 | 249370 | 0 | 0 | 0 |
| LIGDAAKN( | HSPA5; HSPA5; | P11021; A0A | BIP_HUMAN Er | Unmodified | 0 | 0 | 0 | 0 | 0 | 0 | 516760 | 160050 | 0 | 1301100 | 0 | 0 | 0 | 0 | 0 | 0 | 0 | 0 |
| LIGDAAKN( | HSPA8; HSPA8; | P11142; E9P | HSP7C_HUMAI | Oxidation (M) | 0 | 0 | 0 | 0 | 796470 | 0 | 295380 | 181410 | 0 | 0 | 0 | 0 | 1128700 | 493600 | 0 | 0 | 422360 | 0 |
| LIGDAAKN( | HSPA8; HSPA8; | P11142; E9P | HSP7C_HUMAI | Unmodified | 0 | 0 | 0 | 0 | 1992700 | 0 | 0 | 0 | 0 | 0 | 0 | 0 | 1239800 | 1000000 | 0 | 0 | 603390 | 0 |
| LIGEELAQL | ATXN3; ATXN3; | P54252; A0A | ATX3_HUMAN / | Unmodified | 864160 | 836210 | 904190 | 0 | 0 | 0 | 0 | 0 | 0 | 348120 | 0 | 0 | 923030 | 0 | 718490 | 586250 | 0 | 0 |
| LIGEYGLR | RPS9; RPS9; RP | P46781; F2Z | RS9_HUMAN 4 | Unmodified | 6065100 | 6793600 | 4622900 | 2464300 | 0 | 0 | 2012700 | 254760 | 836560 | 166040 | 1419400 | 55105000 | 2235100 | 1139800 | 4234300 | 0 | 0 | 0 |
| LIGGWEDN | PKHD1L1; PKHI | Q86WI1; H0 | PKHL1_HUMAN | Unmodified | 0 | 0 | 0 | 0 | 0 | 0 | 0 | 0 | 760120 | 36487 | 0 | 57533 | 0 | 0 | 0 | 0 | 0 | 0 |
| LIGNLSEK | CKAP5 | Q14008 | CKAP5_HUMAN | Unmodified | 0 | 0 | 0 | 0 | 1078000 | 0 | 14816 | 278650 | 658110 | 1964700 | 220720 | 2084400 | 0 | 0 | 0 | 120390 | 28009 | 2674200 |
| LIGNMALLI | ARPC3; ARPC3; | O15145; F8\ | ARPC3_HUMAN | Unmodified | 228340 | 654430 | 148370 | 0 | 113530 | 0 | 0 | 266540 | 1297500 | 0 | 124710 | 2989500 | 0 | 0 | 262980 | 132940 | 0 | 0 |
| LIGNMALLI | ARPC3; ARPC3; | O15145; F8\ | ARPC3_HUMAN | Oxidation (M) | 306900 | 3542500 | 5336800 | 4641900 | 0 | 0 | 0 | 14737 | 622940 | 0 | 695490 | 16611000 | 0 | 0 | 5385300 | 5722800 | 0 | 0 |
| LIGPLSHSII | POP1 | Q99575 | POP1_HUMAN | Unmodified | 89088 | 106200 | 0 | 0 | 317210 | 69489 | 0 | 0 | 0 | 0 | 0 | 658410 | 0 | 0 | 0 | 0 | 0 | 177240 |
| LIGQLNLK | USP9X; USP9Y | Q93008; O0 | USP9X_HUMAN | Unmodified | 0 | 0 | 0 | 0 | 0 | 0 | 0 | 0 | 86099 | 0 | 120330 | 1593700 | 0 | 0 | 83704 | 0 | 0 | 0 |
| LIGRKFEDA | HSPA2 | P54652 | HSP72_HUMAN | Oxidation (M) | 0 | 0 | 0 | 0 | 0 | 360260 | 108270 | 0 | 0 | 2041100 | 0 | 0 | 157420 | 0 | 0 | 0 | 1024300 | 0 |
| LIGRRFDDA | HSPA8; HSPA8; | P11142; E9P | HSP7C_HUMAI | Unmodified | 0 | 0 | 0 | 0 | 341830 | 0 | 662710 | 0 | 0 | 0 | 0 | 0 | 0 | 0 | 0 | 0 | 505920 | 141630 |
| LIGRTWND | HSPA5; HSPA5; | P11021; A0A | BIP_HUMAN Er | Unmodified | 0 | 0 | 0 | 0 | 0 | 0 | 1611200 | 394520 | 386790 | 6373300 | 0 | 0 | 0 | 0 | 0 | 0 | 0 | 0 |
| LIGTATVALI | MYOF | Q9NZM1 | MYOF_HUMAN | Unmodified | 0 | 0 | 0 | 0 | 38420000 | 4382900 | 0 | 0 | 218140 | 34691 | 0 | 0 | 0 | 0 | 0 | 0 | 919070 | 4136200 |
| LIHEQEQQ | S100A16 | Q96FQ6 | S10AG_HUMAN | Unmodified | 0 | 333030 | 394530 | 0 | 45666000 | 6212200 | 1250600 | 11960 | 2033700 | 0 | 0 | 0 | 0 | 0 | 0 | 0 | 28016000 | 1E+08 |
| LIHPDEDIS | ZNF207; ZNF20 | O43670; X6F | ZN207_HUMAN | Unmodified | 0 | 0 | 0 | 239120 | 0 | 0 | 201660 | 119830 | 0 | 0 | 1807200 | 6231100 | 0 | 0 | 164190 | 0 | 0 | 0 |
| LIHPDEDIS | ZNF207; ZNF20 | O43670; X6F | ZN207_HUMAN | Unmodified | 0 | 0 | 0 | 0 | 0 | 0 | 0 | 162720 | 0 | 0 | 0 | 2392900 | 86313 | 0 | 466760 | 0 | 0 | 0 |
| LIHPDEDIS | ZNF207; ZNF20 | O43670; X6F | ZN207_HUMAN | Unmodified | 0 | 0 | 0 | 0 | 2296200 | 0 | 211580 | 0 | 0 | 557060 | 0 | 0 | 788360 | 312740 | 0 | 0 | 224420 | 0 |
| LIHQTNLIL( | PDCD10; PDCD | Q9BUL8; C9 | PDC10_HUMAI | Unmodified | 0 | 0 | 0 | 0 | 0 | 0 | 0 | 0 | 0 | 0 | 0 | 3011200 | 0 | 0 | 0 | 0 | 0 | 0 |
| LIHYHQHN | BMX | P51813 | BMX_HUMAN ( | Unmodified | 0 | 0 | 0 | 0 | 35762 | 0 | 0 | 0 | 0 | 0 | 1107900 | 7114100 | 0 | 0 | 0 | 0 | 0 | 0 |
| LIHYHQHN | BMX | P51813 | BMX_HUMAN ( | Oxidation (M) | 0 | 0 | 0 | 0 | 0 | 0 | 0 | 0 | 0 | 0 | 471620 | 4593600 | 34420 | 0 | 0 | 0 | 0 | 0 |
| LIIAGTSAY/ | SHMT2; SHMT2 | P34897; H0 | GLYM_HUMAN | Unmodified | 0 | 303830 | 0 | 196850 | 0 | 0 | 227520 | 0 | 32195000 | 9968700 | 0 | 1965100 | 0 | 0 | 0 | 0 | 0 | 0 |
| LIIAGTSCY | SHMT1 | P34896 | GLYC_HUMAN | Unmodified | 0 | 0 | 207390 | 194980 | 0 | 0 | 0 | 0 | 0 | 229640 | 0 | 2915600 | 0 | 0 | 232860 | 249140 | 0 | 0 |
| LIIFDARQN | MTMR1; MTMR1 | Q13613; H0 | MTMR1_HUMA | Unmodified | 0 | 0 | 0 | 0 | 545540 | 0 | 0 | 0 | 0 | 0 | 0 | 0 | 0 | 0 | 0 | 0 | 0 | 0 |
| LIILANGGP | JUP | P14923 | PLAK_HUMAN . | Oxidation (M) | 0 | 0 | 0 | 849710 | 0 | 0 | 0 | 0 | 0 | 0 | 0 | 0 | 0 | 0 | 2904500 | 1527400 | 0 | 0 |
| LIILANGGP | JUP | P14923 | PLAK_HUMAN . | Unmodified | 0 | 0 | 0 | 161840 | 0 | 0 | 0 | 0 | 0 | 0 | 0 | 0 | 0 | 0 | 521500 | 103050 | 0 | 0 |
| LIILASGGP | CTNNB1; CTNN | P35222; A0A | CTNB1_HUMAI | Oxidation (M) | 182000 | 335910 | 211690 | 0 | 0 | 0 | 0 | 0 | 0 | 0 | 0 | 100790 | 0 | 0 | 5961200 | 2363100 | 0 | 0 |
| LIILASGGP | CTNNB1; CTNN | P35222; A0A | CTNB1_HUMAI | Unmodified | 0 | 0 | 0 | 0 | 0 | 0 | 0 | 175210 | 101060 | 0 | 0 | 0 | 0 | 0 | 150980 | 81853 | 0 | 0 |
| LIILEHYSQ/ | GNPDA1; GNPI | P46926; D6F | GNPI1_HUMAN | Unmodified | 0 | 0 | 142900 | 319460 | 482660 | 207100 | 959230 | 0 | 347140 | 317920 | 1497100 | 1728800 | 499160 | 995180 | 409410 | 252410 | 479350 | 454370 |
| LIINSLYK | HSP90B1; HSP | P14625; A0A | ENPL_HUMAN | Unmodified | 1175900 | 484770 | 2409500 | 1386400 | 5021400 | 787470 | 31388000 | 12473000 | 3.9E+08 | 4.9E+08 | 41261000 | 59851000 | 5562200 | 8320700 | 3350700 | 2163300 | 1708100 | 6312900 |
| LIIQWNGPI | VIL1 | P09327 | VILI_HUMAN Vi | Unmodified | 0 | 0 | 0 | 0 | 0 | 0 | 0 | 0 | 0 | 0 | 820400 | 5665700 | 0 | 0 | 0 | 0 | 0 | 0 |
| LIITEETAK | FARSB | Q9NSD9 | SYFB_HUMAN | Unmodified | 0 | 0 | 0 | 0 | 921030 | 0 | 522550 | 342970 | 0 | 281630 | 0 | 7410000 | 1513600 | 0 | 0 | 0 | 0 | 0 |
| LIITQVR | LRPPRC | P42704 | LPPRC_HUMAN | Unmodified | 0 | 0 | 138700 | 54521 | 132970 | 0 | 340140 | 216580 | 2792500 | 0 | 0 | 0 | 25637 | 0 | 119530 | 0 | 133170 | 57005 |
| LIIVEGCQR | ATP1A1; ATP1A | P05023; P50 | AT1A1_HUMAN | Unmodified | 3172800 | 0 | 21059000 | 30019000 | 8154800 | 1879000 | 0 | 757510 | 26556000 | 4369100 | 19786000 | 1.23E+08 | 14170000 | 7670200 | 32041000 | 0 | 1949400 | 0 |
| LIIVEGCQR | ATP1A1; ATP1A | P05023; P50 | AT1A1_HUMAN | Unmodified | 0 | 0 | 0 | 0 | 20314000 | 3671800 | 6597700 | 0 | 0 | 4409000 | 1273900 | 0 | 19050000 | 32210000 | 0 | 0 | 3134300 | 3460400 |
| LIIVEGCQR | ATP1A1; ATP1A | P05023; P50 | AT1A1_HUMAN | Unmodified | 0 | 0 | 0 | 0 | 6831400 | 0 | 973620 | 0 | 0 | 662410 | 386770 | 0 | 5621700 | 1974500 | 0 | 0 | 888000 | 1015600 |
| LIIWDSYTT | GNB2; GNB4; G | P62879; Q9 | GBB2_HUMAN | Unmodified | 25898000 | 24327000 | 0 | 11355000 | 8492900 | 3692400 | 5192500 | 2504200 | 24680000 | 7927900 | 30937000 | 2.95E+08 | 546580 | 9307600 | 47999000 | 24124000 | 5357700 | 6552500 |
| LIIWDSYTT | GNB2; GNB1; G | P62879; P62 | GBB2_HUMAN | Unmodified | 0 | 0 | 0 | 0 | 0 | 0 | 0 | 0 | 307140 | 0 | 0 | 613360 | 0 | 0 | 0 | 0 | 0 | 0 |
| LIKDDFLQ( | ATP6V1A | P38606 | VATA_HUMAN | Unmodified | 0 | 0 | 0 | 0 | 1020000 | 0 | 439850 | 0 | 195390 | 0 | 0 | 4442200 | 0 | 0 | 622560 | 579860 | 310190 | 0 |
| LIKEDQLEY | DOCK9; DOCK | Q9BZ29; A0A | DOCK9_HUMA | Unmodified | 32851 | 0 | 95752 | 0 | 0 | 0 | 0 | 0 | 729110 | 0 | 0 | 432930 | 52011 | 0 | 20049 | 0 | 0 | 0 |
| LIKEPAPDS | ISOC2 | Q96AB3 | ISOC2_HUMAN | Unmodified | 215780 | 0 | 0 | 122380 | 0 | 0 | 839200 | 0 | 6219900 | 2137800 | 0 | 0 | 310980 | 0 | 0 | 33819 | 171420 | 0 |
| LILDVFCGS | G6PD | P11413 | G6PD_HUMAN | Oxidation (M) | 0 | 0 | 0 | 0 | 0 | 0 | 852630 | 396800 | 262980 | 0 | 274010 | 1124400 | 0 | 0 | 0 | 0 | 0 | 176820 |
| LILEQMQK | STIP1; STIP1 | P31948; H0 | STIP1_HUMAN | Unmodified | 0 | 0 | 0 | 0 | 0 | 0 | 0 | 0 | 0 | 0 | 1419900 | 2240400 | 0 | 0 | 0 | 0 | 182620 | 0 |
| LILEQMQK | STIP1; STIP1 | P31948; H0 | STIP1_HUMAN | Oxidation (M) | 0 | 17575 | 0 | 0 | 0 | 0 | 0 | 0 | 0 | 0 | 2971500 | 633110 | 0 | 0 | 0 | 0 | 0 | 0 |
| LILEQMQKI | STIP1 | P31948 | STIP1_HUMAN | Oxidation (M) | 0 | 0 | 0 | 0 | 0 | 0 | 0 | 0 | 0 | 0 | 409390 | 1009400 | 0 | 0 | 0 | 0 | 0 | 0 |
| LILEVHQFS | SPTBN1; SPTBN | Q01082; A0A | SPTB2_HUMAN | Unmodified | 77372 | 0 | 0 | 0 | 0 | 0 | 0 | 0 | 0 | 0 | 0 | 2832800 | 0 | 0 | 0 | 0 | 0 | 0 |
| LILGLMMPI | ANXA6; ANXA6; | P08133; E7E | ANXA6_HUMAN | 2 Oxidation (M) | 0 | 0 | 0 | 984170 | 718130 | 0 | 0 | 128350 | 1589700 | 1332000 | 0 | 0 | 2022600 | 745440 | 1611800 | 281290 | 278050 | 1511700 |
| LILGLMMPI | ANXA6; ANXA6; | P08133; E7E | ANXA6_HUMAN | Oxidation (M) | 0 | 0 | 0 | 591780 | 360640 | 243730 | 514920 | 38130 | 8670500 | 2281800 | 0 | 0 | 0 | 1039600 | 415500 | 0 | 358010 | 2416900 |
| LILGLMMPI | ANXA6; ANXA6; | P08133; E7E | ANXA6_HUMAN | Unmodified | 0 | 0 | 0 | 0 | 551010 | 0 | 0 | 0 | 8470900 | 304510 | 0 | 0 | 0 | 116570 | 0 | 0 | 0 | 1303800 |
| LILGNYK | ACSL4; ACSL4; | O60488; D6I | ACSL4_HUMAN | Unmodified | 0 | 312410 | 7940000 | 15358000 | 0 | 0 | 1519500 | 1371400 | 0 | 0 | 0 | 2750800 | 11305000 | 16380000 | 8816800 | 5113100 | 0 | 0 |
| LILIACGTSY | GFPT1; GFPT1; | Q06210; A0A | GFPT1_HUMAN | Unmodified | 131260 | 107460 | 0 | 391990 | 104710 | 0 | 178020 | 28131 | 0 | 0 | 0 | 546680 | 761600 | 1066400 | 223000 | 223640 | 135770 | 0 |
| LILIESR | RPS13 | P62277 | RS13_HUMAN | Unmodified | 0 | 2919400 | 1940700 | 2500600 | 0 | 0 | 0 | 0 | 0 | 289960 | 0 | 56010000 | 270890 | 0 | 1943600 | 2147100 | 0 | 0 |
| LILISTNGSI | WDR82 | Q6UXN9 | WDR82_HUMA | Unmodified | 0 | 0 | 4349400 | 0 | 0 | 0 | 0 | 462990 | 0 | 0 | 0 | 1584400 | 0 | 0 | 0 | 0 | 0 | 0 |
| LILPELGPV | SCYL2; SCYL2; | Q6P3W7; A0 | SCYL2_HUMAN | Unmodified | 0 | 0 | 0 | 0 | 0 | 0 | 0 | 0 | 0 | 0 | 140330 | 1183800 | 0 | 0 | 0 | 0 | 0 | 0 |
| LILPHVDIQ | IDH2 | P48735 | IDHP_HUMAN | Unmodified | 0 | 0 | 0 | 0 | 0 | 0 | 1252000 | 0 | 24523000 | 5939200 | 1228600 | 336530 | 458570 | 0 | 0 | 0 | 0 | 0 |
| LILPVGPAG | PCMT1; PCMT1 | P22061; F6S | PIMT_HUMAN I | Oxidation (M) | 957620 | 638660 | 1237300 | 0 | 0 | 0 | 50217 | 42011 | 2226800 | 4887700 | 2426400 | 9617300 | 0 | 197960 | 1411900 | 1341600 | 0 | 0 |
| LILPVGPAG | PCMT1; PCMT1 | P22061; F6S | PIMT_HUMAN I | Unmodified | 0 | 34730 | 0 | 0 | 235210 | 0 | 0 | 0 | 2583700 | 113880 | 147580 | 961010 | 103890 | 34138 | 164360 | 0 | 0 | 101880 |
| LILPVGPAG | PCMT1; PCMT1 | P22061; F6S | PIMT_HUMAN I | Oxidation (M) | 902980 | 797100 | 0 | 0 | 0 | 0 | 0 | 0 | 2235700 | 2282100 | 0 | 1599900 | 2163200 | 0 | 0 | 0 | 0 | 0 |
| LILPVGPAG | PCMT1; PCMT1 | P22061; F6S | PIMT_HUMAN I | Unmodified | 0 | 0 | 0 | 0 | 0 | 0 | 0 | 0 | 2478000 | 1098300 | 0 | 0 | 0 | 0 | 0 | 0 | 0 | 0 |
| LILSTDTLEI | UQCC2; UQCC | Q9BRT2; Q5 | UQCC2_HUMA | Unmodified | 0 | 0 | 0 | 0 | 0 | 0 | 0 | 0 | 2700600 | 789080 | 108990 | 35264 | 0 | 0 | 0 | 0 | 0 | 0 |
| LIMEFLPSG | JAK1; JAK1; JAK | P23458; A0A | JAK1_HUMAN T | Unmodified | 0 | 0 | 0 | 0 | 1229800 | 0 | 0 | 0 | 277440 | 0 | 0 | 0 | 0 | 0 | 0 | 0 | 0 | 271540 |
| LIMGIGHR | ACLY | P53396 | ACLY_HUMAN . | Unmodified | 0 | 0 | 0 | 0 | 96288 | 0 | 0 | 0 | 0 | 0 | 0 | 5495200 | 25039 | 0 | 0 | 0 | 17280 | 0 |
| LIMRFDDTI | EPRS1; EPRS1 | P07814; V9C | SYEP_HUMAN | Unmodified | 0 | 0 | 175080 | 228190 | 0 | 0 | 0 | 0 | 868850 | 383750 | 0 | 0 | 654190 | 0 | 0 | 0 | 0 | 0 |
| LIMSERDQ' | DLG5 | Q8TDM6 | DLG5_HUMAN | Unmodified | 0 | 0 | 0 | 0 | 0 | 0 | 0 | 0 | 0 | 0 | 0 | 0 | 240620 | 0 | 0 | 0 | 0 | 0 |
| LIMTQVSK | GPC3 | P51654 | GPC3_HUMAN | Oxidation (M) | 0 | 0 | 59359 | 0 | 0 | 0 | 0 | 0 | 366020 | 111180 | 0 | 0 | 0 | 0 | 0 | 0 | 0 | 0 |
| LIMVQHWI | RNASET2; RNA | O00584; A0A | RNT2_HUMAN | Unmodified | 0 | 0 | 83332 | 0 | 0 | 0 | 0 | 0 | 1565700 | 0 | 0 | 2373800 | 0 | 280000 | 0 | 116430 | 0 | 0 |
| LIMVQHWI | RNASET2; RNA | O00584; A0A | RNT2_HUMAN | Oxidation (M) | 0 | 0 | 0 | 0 | 0 | 0 | 0 | 0 | 0 | 0 | 0 | 0 | 520790 | 442900 | 0 | 0 | 0 | 0 |
| LIMWDPDF | MPP2; MPP2; M | Q14168; A0A | MPP2_HUMAN | Oxidation (M) | 0 | 0 | 183180 | 271780 | 0 | 0 | 0 | 0 | 0 | 441580 | 0 | 0 | 0 | 0 | 386200 | 0 | 0 | 0 |
| LINANMMV | PSMD14 | O00487 | PSDE_HUMAN | Oxidation (M) | 0 | 0 | 0 | 0 | 0 | 0 | 0 | 0 | 0 | 0 | 0 | 2223100 | 0 | 0 | 0 | 0 | 0 | 0 |
| LINAVELK | MARS1 | P56192 | SYMC_HUMAN | Unmodified | 0 | 0 | 181020 | 0 | 0 | 0 | 0 | 0 | 0 | 0 | 0 | 2234500 | 0 | 0 | 0 | 0 | 0 | 0 |
| LINDCHGS | LRBA; LRBA; LR | P50851; A0A | LRBA_HUMAN | Unmodified | 0 | 0 | 35779 | 0 | 0 | 0 | 0 | 0 | 47712 | 28708 | 112350 | 274890 | 136810 | 68174 | 0 | 51367 | 37474 | 42336 |
| LINDGILSQ | DAGLB; DAGLB | Q8NCG7; E7 | DGLB_HUMAN | Unmodified | 0 | 0 | 0 | 0 | 0 | 0 | 0 | 0 | 0 | 252550 | 0 | 270620 | 0 | 0 | 0 | 0 | 0 | 0 |
| LINKLDTEP | PALS1 | Q8N3R9 | PALS1_HUMAN | Unmodified | 99525 | 201230 | 59824 | 1762400 | 6083500 | 1146300 | 0 | 0 | 1623900 | 690910 | 170900 | 797030 | 4123800 | 320760 | 619300 | 307020 | 0 | 4184500 |
| LINLVGESL | BAG6; BAG6; B/ | P46379; F6S | BAG6_HUMAN | Unmodified | 0 | 0 | 0 | 0 | 0 | 0 | 0 | 0 | 0 | 0 | 328440 | 4395400 | 27653 | 0 | 67658 | 0 | 0 | 0 |
| LINNAIKNE | AP2A2; AP2A1; | O94973; O9 | AP2A2_HUMAN | Unmodified | 0 | 0 | 0 | 0 | 0 | 0 | 103910 | 0 | 0 | 0 | 0 | 1454200 | 0 | 0 | 0 | 0 | 182600 | 0 |
| LINRNTTIP | HSPA9; HSPA9; | P38646; A0A | GRP75_HUMAN | Unmodified | 0 | 0 | 0 | 0 | 1458100 | 59005 | 2529700 | 0 | 1860600 | 2095800 | 0 | 0 | 1272700 | 0 | 0 | 0 | 20000 | 0 |
| LINSSDEL( | MTMR12 | Q9C0I1 | MTMRC_HUMA | Unmodified | 0 | 0 | 0 | 0 | 1520200 | 0 | 0 | 0 | 0 | 0 | 0 | 826030 | 1977200 | 0 | 0 | 0 | 496200 | 739220 |
| LIPDSIGK | RPS3A; RPS3A; | P61247; D6F | RS3A_HUMAN | Unmodified | 780710 | 297850 | 0 | 0 | 0 | 0 | 0 | 0 | 0 | 0 | 0 | 0 | 0 | 0 | 0 | 0 | 0 | 0 |
| LIPDSIGKD | RPS3A; RPS3A; | P61247; D6F | RS3A_HUMAN | Unmodified | 4014800 | 5795000 | 0 | 4404200 | 3532600 | 0 | 3429100 | 2936300 | 1669800 | 1544700 | 9635800 | 33960000 | 3004000 | 0 | 2576300 | 1782100 | 467250 | 2581600 |
| LIPEMDQIF | HYOU1; HYOU | Q9Y4L1; A0A | HYOU1_HUMA | Oxidation (M) | 0 | 0 | 0 | 0 | 0 | 0 | 0 | 0 | 0 | 1045000 | 0 | 0 | 0 | 0 | 34950 | 0 | 0 | 0 |
| LIPGATVGL | ITGB6; ITGB6; I | P18564; A0A | ITB6_HUMAN I | Unmodified | 0 | 0 | 0 | 0 | 0 | 0 | 0 | 0 | 0 | 0 | 144590 | 1717200 | 0 | 0 | 0 | 0 | 0 | 0 |
| LIPGSNLDS | GOLM2; GOLM2 | Q6P4E1; A0A | GOLM2_HUMA | Unmodified | 0 | 0 | 0 | 0 | 0 | 0 | 0 | 0 | 2769400 | 131040 | 146310 | 303350 | 0 | 0 | 0 | 0 | 0 | 0 |
| LIPGSNLDS | GOLM2; GOLM2 | Q6P4E1; A0A | GOLM2_HUMA | Unmodified | 0 | 0 | 0 | 0 | 0 | 0 | 0 | 0 | 0 | 1725000 | 0 | 0 | 0 | 486370 | 0 | 0 | 0 | 0 |
| LIPHAVLPP | KIAA1522 | Q9P206 | K1522_HUMAN | Unmodified | 0 | 0 | 203900 | 712670 | 1926200 | 242840 | 0 | 0 | 658390 | 449110 | 219020 | 513820 | 1286000 | 1188400 | 887240 | 553770 | 634340 | 1657200 |
| LIPNATGTC | PKN1; PKN1 | Q16512; K7E | PKN1_HUMAN | Unmodified | 0 | 0 | 0 | 0 | 0 | 0 | 0 | 0 | 666680 | 0 | 0 | 0 | 0 | 0 | 0 | 0 | 0 | 0 |
| LIPPQDILRI | EXOC4 | Q96A65 | EXOC4_HUMAI | Unmodified | 0 | 0 | 378900 | 762510 | 2184500 | 282490 | 0 | 0 | 3078600 | 225760 | 379130 | 0 | 0 | 0 | 0 | 304400 | 144580 | 424320 |
| LIPRNTVVP | HSPA5; HSPA5; | P11021; A0A | BIP_HUMAN Er | Unmodified | 0 | 0 | 0 | 0 | 0 | 0 | 0 | 0 | 0 | 2130000 | 153300 | 0 | 0 | 0 | 0 | 0 | 0 | 0 |
| LIPVSDIND | PPAT | Q06203 | PUR1_HUMAN | Unmodified | 0 | 0 | 0 | 0 | 0 | 0 | 0 | 0 | 0 | 0 | 1127400 | 11662000 | 96764 | 103410 | 0 | 0 | 0 | 0 |
| LIPVSDIND | PPAT | Q06203 | PUR1_HUMAN | Unmodified | 0 | 0 | 0 | 0 | 0 | 0 | 0 | 0 | 0 | 0 | 440500 | 1656100 | 73876 | 0 | 0 | 0 | 231660 | 0 |
| LIQDQQEQ | RAD50; N/A; RA | Q92878; A0A | RAD50_HUMAN | Unmodified | 0 | 0 | 0 | 0 | 0 | 0 | 0 | 132630 | 234130 | 219520 | 295290 | 1423100 | 0 | 0 | 0 | 0 | 21419 | 0 |

| | | | | | | | | | | | | | | | | | | | | | | |
|---|---|---|---|---|---|---|---|---|---|---|---|---|---|---|---|---|---|---|---|---|---|---|
| LIQEEQILR | TTC17; TTC17 | Q96AE7; A0A | TTC17_HUMAN | Unmodified | 587440 | 221060 | 365480 | 1423400 | 192020 | 0 | 0 | 213230 | 1834200 | 97984 | 0 | 0 | 0 | 0 | 294740 | 1711900 | 68218 | 0 |
| LIQELAK | NMRK1; GET3; | Q9NWW6; C | NRK1_HUMAN | Unmodified | 36705 | 15971 | 0 | 0 | 151830 | 0 | 0 | 17838 | 107530 | 33421 | 250450 | 1823500 | 0 | 0 | 182680 | 0 | 58903 | 0 |
| LIQEQHPEI | SPTAN1; SPTAN | Q13813; A0A | SPTN1_HUMAN | Unmodified | 0 | 0 | 0 | 0 | 1210300 | 0 | 0 | 0 | 230510 | 340760 | 0 | 1161800 | 0 | 0 | 0 | 0 | 42553 | 0 |
| LIQGAPTIR | IGFBP2; IGFBP2 | P18065; C9J | IBP2_HUMAN I | Unmodified | 0 | 0 | 732800 | 230220 | 0 | 0 | 0 | 0 | 43729000 | 1737600 | 0 | 0 | 0 | 0 | 326350 | 0 | 0 | 0 |
| LIQGAPTIR | IGFBP2; IGFBP2 | P18065; C9J | IBP2_HUMAN I | Unmodified | 67900 | 0 | 0 | 0 | 0 | 0 | 0 | 0 | 2.48E+08 | 20180000 | 0 | 0 | 0 | 0 | 0 | 0 | 0 | 0 |
| LIQGAPTIR | IGFBP2; IGFBP2 | P18065; C9J | IBP2_HUMAN I | Unmodified | 0 | 0 | 0 | 0 | 0 | 0 | 0 | 0 | 301490 | 4723600 | 0 | 0 | 0 | 0 | 0 | 0 | 0 | 0 |
| LIQHANVQ | STXBP2; STXBP | Q15833; M0I | STXB2_HUMAN | Unmodified | 751010 | 559660 | 802150 | 959100 | 422320 | 88184 | 0 | 0 | 592960 | 115770 | 556620 | 4053300 | 387030 | 0 | 1002800 | 1348400 | 159060 | 460050 |
| LIQNLDANI | S100A16 | Q96FQ6 | S10AG_HUMAN | Unmodified | 0 | 0 | 0 | 0 | 220840 | 15555 | 0 | 0 | 4043100 | 124950 | 0 | 0 | 0 | 0 | 0 | 0 | 0 | 243040 |
| LIQNLDANI | S100A16 | Q96FQ6 | S10AG_HUMAN | Unmodified | 0 | 0 | 0 | 0 | 887030 | 0 | 0 | 0 | 0 | 0 | 0 | 0 | 0 | 79979 | 0 | 0 | 0 | 10313000 |
| LIQNMDAT | MME; MME; MM | P08473; C9I | NEP_HUMAN N | Oxidation (M) | 0 | 0 | 6004000 | 12784000 | 0 | 0 | 0 | 0 | 3513900 | 730910 | 0 | 0 | 9310800 | 3747000 | 13289000 | 9350700 | 0 | 0 |
| LIQNMDAT | MME; MME; MM | P08473; C9I | NEP_HUMAN N | Unmodified | 0 | 0 | 425850 | 3362300 | 0 | 0 | 0 | 0 | 6336100 | 459110 | 0 | 0 | 4001000 | 2326900 | 3074000 | 2039800 | 0 | 0 |
| LIQNMDAT | MME; MME; MM | P08473; C9I | NEP_HUMAN N | Oxidation (M) | 0 | 0 | 0 | 0 | 0 | 0 | 0 | 0 | 0 | 0 | 0 | 0 | 2628100 | 2154500 | 0 | 0 | 0 | 0 |
| LIQNVFEIK | EXOC5; EXOC5 | O00471; F8V | EXOC5_HUMAN | Unmodified | 0 | 0 | 0 | 0 | 0 | 0 | 0 | 0 | 0 | 0 | 0 | 2018900 | 0 | 0 | 0 | 0 | 0 | 0 |
| LIQNVKIEN | STXBP3 | O00186 | STXB3_HUMAN | Oxidation (M) | 198610 | 128700 | 160630 | 0 | 0 | 0 | 0 | 0 | 113220 | 0 | 0 | 1749300 | 0 | 0 | 356890 | 399890 | 157440 | 0 |
| LIQNVKIEN | STXBP3 | O00186 | STXB3_HUMAN | Unmodified | 160580 | 131870 | 0 | 37163 | 572100 | 0 | 0 | 0 | 0 | 180060 | 113100 | 1742500 | 0 | 0 | 194260 | 133320 | 284250 | 0 |
| LIQPDHPQ | HERC4 | Q5GLZ8 | HERC4_HUMAN | Unmodified | 157100 | 154200 | 401670 | 267270 | 1154900 | 212950 | 177230 | 0 | 737860 | 301820 | 0 | 482450 | 342450 | 122110 | 188170 | 179570 | 415420 | 0 |
| LIQQFLTLR | CDS2 | O95674 | CDS2_HUMAN | Unmodified | 0 | 0 | 0 | 0 | 0 | 0 | 0 | 0 | 1933000 | 478210 | 0 | 0 | 0 | 0 | 0 | 0 | 0 | 0 |
| LIQQLEK | SRPRB; SRPRB | Q9Y5M8; H7 | SRPRB_HUMAN | Unmodified | 1457800 | 4834400 | 131840 | 109550 | 0 | 0 | 373840 | 0 | 2071700 | 134040 | 339520 | 349240 | 0 | 763720 | 602310 | 316410 | 0 | 0 |
| LIQSHPESA | SPTAN1; SPTAN | Q13813; A0A | SPTN1_HUMAN | Unmodified | 0 | 250940 | 248960 | 54998 | 0 | 0 | 0 | 0 | 0 | 0 | 2136800 | 8998100 | 0 | 0 | 459610 | 0 | 0 | 0 |
| LIQYCHSK | AKR1B15; AKR1 | C9JRZ8; O60 | AK1BF_HUMAN | Unmodified | 0 | 0 | 0 | 0 | 0 | 0 | 120310 | 207500 | 0 | 0 | 0 | 275010 | 0 | 0 | 0 | 0 | 0 | 0 |
| LIQYCQSK | AKR1B1; AKR1B | P15121; E9P | ALDR_HUMAN | Unmodified | 102550 | 0 | 0 | 457590 | 7062500 | 1708500 | 0 | 58244 | 0 | 0 | 1887700 | 7454900 | 0 | 0 | 105890 | 106790 | 0 | 0 |
| LIRAPDSLK | EPHA2 | P29317 | EPHA2_HUMAN | Unmodified | 24737 | 0 | 0 | 0 | 639430 | 0 | 0 | 0 | 0 | 0 | 0 | 726700 | 0 | 0 | 0 | 0 | 2247900 | 1507700 |
| LIRDEGLAV | ASPH | Q12797 | ASPH_HUMAN | Unmodified | 0 | 0 | 0 | 0 | 0 | 0 | 0 | 0 | 701160 | 141640 | 0 | 0 | 0 | 0 | 0 | 0 | 0 | 0 |
| LIRELKAEIA | KIF14 | Q15058 | KIF14_HUMAN | Unmodified | 0 | 0 | 0 | 0 | 2675200 | 470740 | 0 | 30414 | 0 | 0 | 0 | 0 | 0 | 0 | 0 | 0 | 223940 | 0 |
| LIRETEATFI | GPC2; GPC2 | Q8N158; A0A | GPC2_HUMAN | Unmodified | 0 | 0 | 0 | 0 | 46658 | 0 | 22197 | 0 | 775640 | 283470 | 0 | 0 | 0 | 0 | 0 | 0 | 0 | 0 |
| LIREVFPDT | SMPDL3B; SMP | Q92485; Q5T | ASM3B_HUMAN | Unmodified | 0 | 0 | 5778500 | 8122700 | 6085700 | 1389200 | 0 | 178600 | 4986900 | 1696400 | 1276000 | 3919700 | 14574000 | 7536200 | 5890600 | 3438600 | 13463000 | 985430 |
| LIRGPAETE | TWF1 | Q12792 | TWF1_HUMAN | Unmodified | 660380 | 815000 | 461980 | 457830 | 4451400 | 0 | 186610 | 262440 | 173450 | 161630 | 0 | 1072800 | 3529200 | 5635200 | 438450 | 405920 | 1329600 | 2375900 |
| LIRGVFVNV | EXOC1 | Q9NV70 | EXOC1_HUMAN | Unmodified | 0 | 0 | 0 | 0 | 877980 | 0 | 324070 | 0 | 0 | 132100 | 0 | 0 | 0 | 125930 | 0 | 0 | 211370 | 293130 |
| LIRQEERLP | MME; MME; MM | P08473; A0A | NEP_HUMAN N | Unmodified | 0 | 0 | 8858600 | 34287000 | 0 | 0 | 0 | 23069 | 23742000 | 4389300 | 0 | 0 | 20928000 | 5221100 | 25572000 | 11913000 | 174540 | 0 |
| LIRQEERLP | MME; MME; MM | P08473; A0A | NEP_HUMAN N | Oxidation (M) | 0 | 0 | 26889000 | 62195000 | 0 | 0 | 0 | 0 | 7822800 | 5600600 | 783000 | 0 | 32877000 | 5137600 | 59114000 | 43678000 | 0 | 0 |
| LIRQQMEE | SWAP70; SWAP | Q9UH65; E7 | SWP70_HUMAN | Unmodified | 0 | 0 | 0 | 0 | 824350 | 187210 | 133900 | 0 | 94698 | 0 | 0 | 0 | 0 | 0 | 0 | 0 | 0 | 0 |
| LIRSNLVDN | DDX19B; DDX19 | Q9UMR2; Q9 | DD19B_HUMAN | Unmodified | 0 | 0 | 0 | 0 | 11345000 | 0 | 0 | 0 | 0 | 0 | 0 | 0 | 0 | 3483900 | 0 | 0 | 0 | 9803100 |
| LIRVENFEA | PTPRJ; PTPRJ; P | Q12913; A0A | PTPRJ_HUMAN | Unmodified | 1138900 | 0 | 0 | 0 | 2618700 | 789600 | 0 | 0 | 0 | 0 | 2322900 | 8092400 | 2294100 | 0 | 0 | 456960 | 322660 | 980500 |
| LIRVENFEA | PTPRJ; PTPRJ; P | Q12913; A0A | PTPRJ_HUMAN | Unmodified | 66840 | 0 | 0 | 0 | 2822300 | 532190 | 599910 | 0 | 79051 | 0 | 3047900 | 3724400 | 98501 | 0 | 34708 | 26585 | 0 | 581620 |
| LIRVLQDQI | GOLM1 | Q8NBJ4 | GOLM1_HUMA | Unmodified | 0 | 0 | 0 | 0 | 0 | 0 | 0 | 0 | 0 | 587870 | 0 | 0 | 0 | 0 | 0 | 0 | 0 | 0 |
| LIRVNPEST | EXOC8 | Q8IYI6 | EXOC8_HUMAN | Unmodified | 0 | 0 | 0 | 177880 | 7493000 | 702960 | 280850 | 271420 | 0 | 606160 | 0 | 0 | 1270400 | 926900 | 95701 | 48497 | 1576400 | 2057500 |
| LISDINK | SPTBN2; SPTBN | O15020; Q01 | SPTN2_HUMAN | Unmodified | 705460 | 0 | 690060 | 44284 | 365490 | 211700 | 0 | 645830 | 472950 | 25757 | 71588 | 2221300 | 76366 | 692380 | 939130 | 575020 | 267100 | 381770 |
| LISETAQR | ARHGEF39 | Q8N4T4 | ARG39_HUMAN | Unmodified | 485520 | 565470 | 853970 | 1074000 | 0 | 0 | 122090 | 0 | 0 | 0 | 135760 | 589990 | 0 | 0 | 1123100 | 160450 | 61167 | 226210 |
| LISETAQRV | ARHGEF39 | Q8N4T4 | ARG39_HUMAN | Unmodified | 0 | 0 | 0 | 0 | 1421700 | 258420 | 0 | 0 | 0 | 197810 | 0 | 0 | 119780 | 73935 | 0 | 0 | 0 | 0 |
| LISETTSVC | KDSR; KDSR | Q06136; K7E | KDSR_HUMAN | Unmodified | 0 | 0 | 0 | 0 | 0 | 0 | 38624 | 281370 | 1211200 | 596830 | 227950 | 0 | 0 | 0 | 0 | 0 | 53368 | 148650 |
| LISHRECQ | PLAU; PLAU | P00749; E7E | UROK_HUMAN | Unmodified | 0 | 0 | 0 | 0 | 1562600 | 558550 | 0 | 0 | 0 | 0 | 0 | 0 | 0 | 0 | 0 | 0 | 1121600 | 1725800 |
| LISLICR | PODXL | O00592 | PODXL_HUMAN | Unmodified | 0 | 0 | 0 | 363370 | 1076300 | 0 | 0 | 0 | 25477000 | 2487700 | 6251200 | 1521700 | 2343700 | 1088700 | 0 | 0 | 362790 | 1598400 |
| LISLLNEK | MYOF | Q9NZM1 | MYOF_HUMAN | Unmodified | 0 | 0 | 0 | 0 | 15257000 | 3167700 | 0 | 0 | 425270 | 0 | 0 | 1404200 | 0 | 0 | 0 | 0 | 0 | 3017500 |
| LISLQMETT | MACF1; MACF1 | Q9UPN3; H3 | MACF1_HUMAN | Oxidation (M) | 0 | 0 | 0 | 211230 | 0 | 0 | 0 | 0 | 0 | 0 | 0 | 0 | 0 | 0 | 0 | 0 | 0 | 0 |
| LISLTDENA | HSP90B1; HSP9 | P14625; A0A | ENPL_HUMAN | Unmodified | 0 | 0 | 0 | 0 | 0 | 0 | 167600 | 127870 | 16420000 | 7994600 | 7069900 | 1574700 | 0 | 0 | 0 | 0 | 0 | 0 |
| LISNPTNFN | CDC42BPA; CD | Q5VT25; A0A | MRCKA_HUMA | Oxidation (M) | 0 | 0 | 0 | 424000 | 706640 | 214300 | 0 | 0 | 0 | 0 | 0 | 708420 | 1039000 | 891560 | 0 | 237430 | 0 | 1692300 |
| LISQDIHSN | NMD3; NMD3 | Q96D46; C9 | NMD3_HUMAN | Unmodified | 224450 | 415090 | 138890 | 145940 | 0 | 0 | 0 | 0 | 105800 | 0 | 104450 | 2032300 | 0 | 37003 | 236330 | 189950 | 0 | 0 |
| LISQIVSSIT | TUBA4A; TUBA1 | P68366; Q9E | TBA4A_HUMAN | Unmodified | 0 | 0 | 0 | 46009 | 0 | 0 | 19781 | 0 | 98754 | 73184 | 0 | 0 | 0 | 0 | 172350 | 12860 | 0 | 0 |
| LISSDGHEI | ELOC; ELOC; EL | Q15369; R40 | ELOC_HUMAN | Unmodified | 307650 | 258680 | 1007400 | 744300 | 2953600 | 0 | 0 | 0 | 1579600 | 0 | 0 | 8503900 | 2807200 | 0 | 1048400 | 1276400 | 0 | 1491700 |
| LISSDGHEI | ELOC; ELOC; EL | Q15369; R40 | ELOC_HUMAN | Unmodified | 143550 | 136040 | 0 | 234600 | 311600 | 0 | 548450 | 85940 | 849520 | 0 | 0 | 1843200 | 514940 | 192170 | 70369 | 126430 | 0 | 130350 |
| LISSDGHEI | ELOC; ELOC; EL | Q15369; R40 | ELOC_HUMAN | Unmodified | 0 | 0 | 0 | 180020 | 15073000 | 2649900 | 2747000 | 125480 | 0 | 2498800 | 0 | 0 | 8271700 | 6999900 | 0 | 0 | 8375400 | 7983800 |
| LISSLLR | ELAC2; ELAC2; | Q9BQ52; E7I | RNZ2_HUMAN | Unmodified | 0 | 0 | 0 | 37321 | 142850 | 0 | 0 | 0 | 78790 | 0 | 0 | 568940 | 0 | 0 | 0 | 0 | 250220 | 0 |
| LISSNAAFR | ARHGEF16; ARI | Q5VV41; B0 | ARHGG_HUMA | Unmodified | 0 | 0 | 0 | 0 | 0 | 0 | 0 | 0 | 0 | 0 | 0 | 353270 | 0 | 0 | 0 | 0 | 0 | 0 |
| LISSVDPQF | PBDC1; PBDC1 | Q9BVG4; A6 | PBDC1_HUMAN | Unmodified | 0 | 0 | 182380 | 176000 | 885700 | 412740 | 564250 | 305450 | 388210 | 461370 | 376120 | 2443500 | 447340 | 721500 | 0 | 0 | 0 | 1206500 |
| LISTLIYK | XPO1; XPO1; XF | O14980; A0A | XPO1_HUMAN | Unmodified | 0 | 0 | 0 | 0 | 0 | 0 | 0 | 0 | 0 | 18178 | 0 | 4199500 | 0 | 0 | 0 | 0 | 0 | 0 |
| LISVEDLWI | STIM1; STIM1; S | Q13586; E9F | STIM1_HUMAN | Unmodified | 0 | 0 | 0 | 0 | 0 | 0 | 0 | 0 | 31308000 | 0 | 1514700 | 106650 | 0 | 0 | 0 | 0 | 19893 | 0 |
| LISWYDNE | GAPDH; GAPDH | P04406; E7E | G3P_HUMAN G | Unmodified | 8438700 | 8997300 | 5233400 | 5539700 | 3172100 | 680830 | 324300 | 93253 | 13189000 | 1253000 | 9306400 | 1.4E+08 | 2319200 | 437400 | 19665000 | 14500000 | 984620 | 2128900 |
| LISWYDNE | GAPDH; GAPDH | P04406; E7E | G3P_HUMAN G | 2 Oxidation (M) | 0 | 317190 | 0 | 2005700 | 49787000 | 6768000 | 10966000 | 0 | 1535600 | 3653500 | 0 | 1166600 | 21812000 | 12871000 | 0 | 0 | 11014000 | 49037000 |
| LISWYDNE | GAPDH; GAPDH | P04406; E7E | G3P_HUMAN G | Oxidation (M) | 0 | 0 | 0 | 2255200 | 55068000 | 0 | 14262000 | 0 | 0 | 2823900 | 0 | 0 | 10178000 | 0 | 0 | 0 | 65300000 | 90766000 |
| LISWYDNE | GAPDH; GAPDH | P04406; E7E | G3P_HUMAN G | Unmodified | 0 | 0 | 0 | 0 | 27568000 | 469980 | 9875200 | 0 | 429560 | 493850 | 0 | 0 | 8273300 | 5451400 | 0 | 0 | 18263000 | 33853000 |
| LISYSYMVF | EPHX1 | P07099 | HYEP_HUMAN | Unmodified | 0 | 0 | 0 | 0 | 0 | 0 | 0 | 0 | 6339400 | 0 | 0 | 0 | 822860 | 0 | 378330 | 0 | 0 | 0 |
| LITDALNKN | PRKG2 | Q13237 | KGP2_HUMAN | Unmodified | 0 | 0 | 0 | 0 | 0 | 0 | 224830 | 433870 | 0 | 0 | 0 | 0 | 0 | 0 | 0 | 0 | 0 | 0 |
| LITDLQDQI | KIF5B | P33176 | KINH_HUMAN | Unmodified | 0 | 0 | 0 | 248020 | 2408200 | 409410 | 525870 | 412450 | 0 | 0 | 0 | 0 | 874930 | 873070 | 329180 | 238380 | 506490 | 1023200 |
| LITEDVQGI | RPS3A; RPS3A; | P61247; D6F | RS3A_HUMAN | Unmodified | 3824400 | 2988800 | 3205000 | 3088600 | 0 | 0 | 0 | 1328300 | 1132100 | 333720 | 4280000 | 54247000 | 0 | 0 | 3195800 | 3149000 | 0 | 0 |
| LITEGASK | IDH3A; IDH3A; | P50213; H0Y | IDH3A_HUMAN | Unmodified | 0 | 0 | 0 | 0 | 0 | 0 | 0 | 0 | 0 | 0 | 0 | 0 | 0 | 0 | 0 | 0 | 0 | 0 |
| LITGIIYPQN | CAPN7; CAPN7 | Q9Y6W3; H7 | CAN7_HUMAN | Unmodified | 158980 | 0 | 0 | 0 | 0 | 0 | 0 | 0 | 0 | 0 | 0 | 2256100 | 0 | 0 | 0 | 0 | 0 | 0 |
| LITGIIYPQN | CAPN7; CAPN7 | Q9Y6W3; H7 | CAN7_HUMAN | Unmodified | 0 | 0 | 0 | 0 | 0 | 0 | 0 | 0 | 0 | 0 | 0 | 917930 | 0 | 0 | 0 | 0 | 0 | 0 |
| LITGMDR | FKBP10; FKBP1 | Q96AY3; C9J | FKB10_HUMAN | Unmodified | 0 | 0 | 0 | 0 | 562380 | 8028800 | 689160 | 0 | 0 | 0 | 1200300 | 1282700 | 0 | 0 | 0 | 0 | 0 | 0 |
| LITLEEEMTI | P4HB; P4HB; P4 | P07237; A0A | PDIA1_HUMAN | Oxidation (M) | 0 | 0 | 0 | 0 | 85771 | 0 | 0 | 52568 | 28140000 | 5470000 | 3216200 | 817190 | 0 | 0 | 0 | 0 | 0 | 0 |
| LITLEEEMTI | P4HB; P4HB; P4 | P07237; A0A | PDIA1_HUMAN | Unmodified | 0 | 0 | 0 | 0 | 311380 | 0 | 0 | 56374 | 79950000 | 14377000 | 5386100 | 1753100 | 0 | 201250 | 413090 | 0 | 0 | 159730 |
| LITLEQGK | ALDH6A1 | Q02252 | MMSA_HUMAN | Unmodified | 0 | 0 | 0 | 0 | 0 | 0 | 0 | 0 | 2133100 | 0 | 0 | 0 | 0 | 0 | 0 | 0 | 0 | 0 |
| LITNDTFQF | ERBIN | Q96RT1 | ERBIN_HUMAN | Oxidation (M) | 0 | 0 | 0 | 160980 | 938360 | 218630 | 0 | 0 | 0 | 0 | 0 | 0 | 146280 | 0 | 209140 | 0 | 0 | 0 |
| LITNDTFQF | ERBIN | Q96RT1 | ERBIN_HUMAN | Unmodified | 0 | 0 | 0 | 0 | 726060 | 0 | 0 | 0 | 690570 | 0 | 98004 | 0 | 0 | 352570 | 0 | 0 | 50630 | 109380 |
| LITNDTFQF | ERBIN | Q96RT1 | ERBIN_HUMAN | Unmodified | 989680 | 0 | 0 | 0 | 9679900 | 2795300 | 0 | 0 | 0 | 0 | 0 | 0 | 0 | 0 | 0 | 0 | 491780 | 3051900 |
| LITNDTFQF | ERBIN | Q96RT1 | ERBIN_HUMAN | Oxidation (M) | 0 | 0 | 0 | 0 | 0 | 1150700 | 0 | 0 | 285010 | 0 | 0 | 51858 | 336150 | 0 | 0 | 0 | 0 | 0 |
| LITPAVVSEI | RPS25 | P62851 | RS25_HUMAN | Unmodified | 5481300 | 4058000 | 6030300 | 838150 | 830900 | 254520 | 0 | 0 | 1639500 | 131220 | 5179900 | 50142000 | 585900 | 124550 | 4068600 | 3154900 | 370240 | 1136800 |
| LITPAVVSEI | RPS25 | P62851 | RS25_HUMAN | Unmodified | 0 | 0 | 0 | 691990 | 466110 | 0 | 711400 | 770820 | 0 | 201520 | 484160 | 1849400 | 2022300 | 1620900 | 0 | 0 | 0 | 0 |
| LITQTFSHH | NUDC; NUDC; | Q9Y266; A0A | NUDC_HUMAN | Unmodified | 267830 | 0 | 930010 | 2093700 | 3813300 | 468940 | 520650 | 1177100 | 8745500 | 2842800 | 3237200 | 23063000 | 4931400 | 2685100 | 1663800 | 1004800 | 1751700 | 1853800 |
| LITWSPVCI | GPX1; GPX1 | P07203; A0A | GPX1_HUMAN | Unmodified | 0 | 0 | 0 | 0 | 488500 | 0 | 651610 | 0 | 1540600 | 1099200 | 414710 | 699960 | 0 | 0 | 64104 | 0 | 393060 | 2297700 |
| LIVDAFLQC | UPF2 | Q9HAU5 | RENT2_HUMAN | Unmodified | 0 | 0 | 0 | 0 | 0 | 0 | 0 | 0 | 0 | 19325 | 0 | 218840 | 0 | 0 | 0 | 0 | 0 | 264660 |
| LIVDEAINE | VCP; VCP; VCP; | P55072; A0A | TERA_HUMAN T | Unmodified | 545280 | 0 | 792720 | 2941700 | 0 | 0 | 0 | 0 | 2109300 | 0 | 1782400 | 8132000 | 0 | 0 | 1598800 | 3307600 | 730170 | 0 |
| LIVDGRGKA | CCT7; CCT7 | Q99832; F8V | TCPH_HUMAN | Unmodified | 0 | 0 | 0 | 0 | 1996000 | 0 | 1932800 | 765400 | 0 | 1294800 | 0 | 0 | 4708900 | 4022900 | 30490 | 0 | 3417300 | 1910500 |
| LIVDHNIAD | PRPF8; PRPF8 | Q6P2Q9; I3L | PRP8_HUMAN | Unmodified | 0 | 0 | 0 | 0 | 0 | 0 | 0 | 0 | 0 | 0 | 0 | 2025100 | 0 | 0 | 0 | 0 | 0 | 0 |
| LIVDVIR | IQGAP1; IQGAP | P46940; H0Y | IQGA1_HUMAN | Unmodified | 0 | 0 | 73098 | 0 | 0 | 0 | 0 | 0 | 771510 | 74690 | 0 | 200610 | 0 | 0 | 0 | 0 | 0 | 0 |
| LIVDVIRFQ | IQGAP1; IQGAP | P46940; H0Y | IQGA1_HUMAN | Unmodified | 0 | 0 | 0 | 151090 | 434110 | 0 | 0 | 0 | 0 | 0 | 0 | 0 | 314120 | 0 | 0 | 0 | 0 | 0 |
| LIVENLSSR | SRSF6; SRSF4; | Q13247; Q08 | SRSF6_HUMAN | Unmodified | 1117100 | 547130 | 426790 | 465630 | 0 | 0 | 0 | 0 | 808720 | 0 | 389790 | 3388000 | 621200 | 0 | 376420 | 633090 | 0 | 0 |
| LIVENLSSR | SRSF6; SRSF4; | Q13247; Q08 | SRSF6_HUMAN | Unmodified | 0 | 0 | 0 | 0 | 0 | 0 | 3423700 | 47085 | 0 | 0 | 0 | 0 | 536670 | 335180 | 0 | 0 | 0 | 0 |
| LIVFENPNII | IGSF21 | Q96ID5 | IGS21_HUMAN | Unmodified | 0 | 0 | 0 | 0 | 0 | 0 | 0 | 77512 | 910260 | 0 | 0 | 0 | 0 | 0 | 0 | 0 | 0 | 0 |
| LIVGLMRPF | ANXA6; ANXA6; | P08133; E5R | ANXA6_HUMAN | Oxidation (M) | 93685 | 0 | 854680 | 1178800 | 479320 | 523320 | 217020 | 68584 | 4696700 | 1192200 | 0 | 0 | 622030 | 0 | 1051300 | 455800 | 151730 | 1486900 |
| LIVGLMRPF | ANXA6; ANXA6; | P08133; E5R | ANXA6_HUMAN | Unmodified | 0 | 0 | 0 | 0 | 802340 | 244660 | 80448 | 0 | 16995000 | 1352000 | 0 | 0 | 548490 | 0 | 0 | 0 | 0 | 0 |
| LIVLDRLIEL | COPB1; COPB1 | P53618; E9P | COPB_HUMAN | Unmodified | 0 | 0 | 0 | 0 | 0 | 0 | 0 | 0 | 168800 | 43139 | 88090 | 1923600 | 0 | 0 | 0 | 0 | 0 | 0 |
| LIVLVPPSKI | F11R; F11R | Q9Y624; A0A | JAM1_HUMAN . | Unmodified | 19119000 | 18146000 | 21659000 | 31563000 | 5979200 | 0 | 0 | 0 | 24068000 | 2057200 | 4119100 | 14841000 | 9233100 | 0 | 19631000 | 14354000 | 618600 | 0 |
| LIVPSEFKPT | SLC4A4 | Q9Y6R1 | S4A4_HUMAN | Unmodified | 0 | 0 | 101010 | 0 | 0 | 0 | 0 | 0 | 0 | 0 | 0 | 0 | 0 | 0 | 118640 | 299100 | 0 | 0 |
| LIVPVLLDA | STXBP2; STXBP | Q15833; M0I | STXB2_HUMAN | Unmodified | 416580 | 368540 | 314610 | 564030 | 1005800 | 269900 | 273830 | 0 | 278740 | 219350 | 512490 | 1151200 | 697440 | 414170 | 909610 | 831740 | 448600 | 677670 |
| LIVQPNTRL | DPP3; DPP3; DI | Q9NY33; G3 | DPP3_HUMAN | Unmodified | 0 | 0 | 0 | 122780 | 0 | 0 | 0 | 0 | 0 | 0 | 0 | 0 | 406950 | 0 | 480770 | 0 | 0 | 0 |
| LIVTGHAPF | HMCN2; HMCN | Q8NDA2; A0 | HMCN2_HUMA | Unmodified | 0 | 0 | 0 | 0 | 0 | 0 | 0 | 0 | 356760 | 28230 | 0 | 0 | 0 | 0 | 0 | 0 | 0 | 0 |
| LIYEDYVSIL | RGS19 | P49795 | RGS19_HUMAN | Unmodified | 0 | 0 | 0 | 0 | 0 | 0 | 0 | 0 | 0 | 382730 | 0 | 0 | 0 | 0 | 0 | 0 | 206660 | 130280 |
| LIYEEKLEAS | MYO1B; MYO1B | O43795; H7 | MYO1B_HUMA | Unmodified | 332020 | 0 | 0 | 291020 | 15374000 | 2667500 | 730880 | 58763 | 0 | 1545600 | 1878100 | 7071000 | 770220 | 373480 | 0 | 0 | 3039800 | 4361700 |
| LIYQWVSR | VPS25; VPS25 | Q9BRG1; K7 | VPS25_HUMAN | Unmodified | 7261300 | 5136200 | 3674900 | 285880 | 0 | 0 | 0 | 0 | 0 | 0 | 0 | 846870 | 0 | 0 | 14979000 | 8785300 | 0 | 0 |
| LKADASGF | MST1R; MST1R | Q04912; J3K | RON_HUMAN N | Unmodified | 0 | 10588 | 0 | 0 | 116040 | 0 | 0 | 0 | 0 | 0 | 0 | 0 | 0 | 0 | 0 | 0 | 147360 | 0 |
| LKAEAYQK | FLOT2; FLOT2; I | Q14254; J3Q | FLOT2_HUMAN | Unmodified | 0 | 0 | 0 | 0 | 0 | 0 | 221740 | 88612 | 0 | 106720 | 0 | 0 | 0 | 0 | 0 | 0 | 0 | 0 |
| LKAEEMDT | EHBP1L1; EHBP | Q8N3D4; E9 | EH1L1_HUMAN | Unmodified | 0 | 0 | 0 | 0 | 706830 | 0 | 0 | 0 | 430270 | 116710 | 0 | 0 | 0 | 0 | 0 | 0 | 0 | 0 |
| LKAEGSEIR | P4HB; P4HB; P4 | P07237; A0A | PDIA1_HUMAN | Unmodified | 0 | 0 | 0 | 0 | 0 | 0 | 0 | 0 | 1824100 | 2645100 | 0 | 0 | 357010 | 0 | 0 | 0 | 0 | 0 |
| LKAESLSDI | CIT; CIT; CIT; CI | O14578; A0A | CTRO_HUMAN | Unmodified | 0 | 0 | 0 | 266850 | 3556100 | 383990 | 202280 | 34903 | 0 | 0 | 78581 | 182250 | 413130 | 128490 | 0 | 0 | 508220 | 361770 |
| LKAGDTVIF | ADH5; ADH5 | P11766; D6F | ADHX_HUMAN | Unmodified | 0 | 0 | 0 | 0 | 0 | 0 | 806080 | 441960 | 0 | 253960 | 0 | 261570 | 913970 | 807090 | 0 | 0 | 0 | 2641900 |
| LKAQALAIE | MVP | Q14764 | MVP_HUMAN N | Unmodified | 0 | 0 | 0 | 0 | 3628300 | 1646700 | 0 | 796580 | 0 | 308740 | 0 | 0 | 0 | 0 | 0 | 0 | 0 | 691160 |
| LKAQTYELC | SEPTIN10; SEPT | Q9P0V9; B5I | SEP10_HUMAN | Unmodified | 0 | 0 | 0 | 0 | 1968700 | 0 | 0 | 0 | 0 | 0 | 0 | 0 | 0 | 0 | 0 | 0 | 990260 | 139140 |
| LKCEAVVAI | HSD17B4; HSD | P51659; E7E | DHB4_HUMAN | Unmodified | 49729 | 0 | 0 | 0 | 0 | 0 | 0 | 0 | 7432000 | 785490 | 679460 | 0 | 0 | 0 | 96499 | 0 | 0 | 0 |
| LKDAIEEK | MYO1B; MYO1B | O43795; H7 | MYO1B_HUMA | Unmodified | 0 | 0 | 0 | 324140 | 5652200 | 741420 | 0 | 0 | 0 | 203910 | 0 | 3167900 | 0 | 0 | 0 | 0 | 752360 | 0 |

| | | | | | | | | | | | | | | | | | | | | | | |
|---|---|---|---|---|---|---|---|---|---|---|---|---|---|---|---|---|---|---|---|---|---|---|
| LKDAIEEKII | MYO1B; MYO1E | O43795; H7( | MYO1B_HUMA | Unmodified | 0 | 0 | 0 | 0 | 2645700 | 120650 | 916920 | 0 | 0 | 0 | 0 | 0 | 0 | 0 | 0 | 0 | 2114900 | 624990 |
| LKDDEVAQ | LDHB; LDHB | P07195; A0A | LDHB_HUMAN | Unmodified | 156940 | 109990 | 0 | 279300 | 2483800 | 393690 | 0 | 0 | 0 | 0 | 2207200 | 17284000 | 1361200 | 0 | 291910 | 229580 | 671190 | 945790 |
| LKDDEVAQ | LDHB; LDHB | P07195; A0A | LDHB_HUMAN | Unmodified | 1678200 | 1124400 | 974030 | 2649000 | 24343000 | 3462200 | 0 | 0 | 106270 | 64075 | 24177000 | 1.15E+08 | 6885800 | 3083000 | 1995700 | 1579700 | 5476900 | 6563800 |
| LKDDSIVD\ | SUCO | Q9UBS9 | SUCO_HUMAN | Unmodified | 0 | 0 | 0 | 0 | 0 | 0 | 0 | 0 | 421080 | 68445 | 0 | 0 | 0 | 0 | 0 | 0 | 0 | 0 |
| LKDELASTK | BCAP31 | P51572 | BAP31_HUMAN | Unmodified | 0 | 28494 | 243060 | 0 | 337560 | 0 | 443310 | 136680 | 0 | 0 | 0 | 58443 | 0 | 593760 | 149810 | 120180 | 63462 | 470280 |
| LKDGKLVV | FABP5; FABP5 | Q01469; I6L | FABP5_HUMAN | Unmodified | 0 | 0 | 0 | 0 | 3146300 | 176920 | 0 | 28608 | 0 | 0 | 0 | 0 | 58659 | 0 | 0 | 0 | 3642100 | 9399300 |
| LKDIEHALS | TOR1A; TOR1A | O14656; A0A | TOR1A_HUMAN | Unmodified | 0 | 0 | 0 | 0 | 0 | 0 | 0 | 0 | 3988900 | 175430 | 0 | 0 | 0 | 0 | 0 | 0 | 0 | 0 |
| LKDISTLEP | ANP32B | Q92688 | AN32B_HUMAN | Unmodified | 0 | 0 | 95003 | 0 | 3148300 | 1430200 | 823090 | 422100 | 324580 | 1561700 | 2472900 | 3030200 | 1919800 | 824320 | 0 | 0 | 648410 | 0 |
| LKDQEAKH | FKBP9 | O95302 | FKBP9_HUMAN | Unmodified | 0 | 0 | 0 | 0 | 0 | 0 | 0 | 0 | 0 | 4361500 | 1761300 | 1260000 | 0 | 0 | 0 | 0 | 0 | 0 |
| LKDSEAEL( | SEPTIN7; SEPTI | Q16181; E7E | SEPT7_HUMAN | Unmodified | 0 | 0 | 0 | 216730 | 443210 | 0 | 89709 | 0 | 171510 | 0 | 134800 | 749020 | 0 | 96598 | 85038 | 80616 | 79985 | 127720 |
| LKDTENAIE | OPA1; OPA1; OI | O60313; E5K | OPA1_HUMAN | Oxidation (M) | 0 | 0 | 0 | 0 | 0 | 0 | 0 | 0 | 0 | 0 | 0 | 600050 | 0 | 0 | 0 | 0 | 0 | 0 |
| LKDVLLQVI | MYH9 | P35579 | MYH9_HUMAN | Unmodified | 53053 | 0 | 254950 | 159460 | 0 | 0 | 194130 | 0 | 0 | 210420 | 0 | 6455500 | 0 | 0 | 184200 | 424040 | 0 | 0 |
| LKDWQEG( | ACSL4; ACSL4 | O60488; H0 | ACSL4_HUMAN | Unmodified | 0 | 0 | 0 | 0 | 231190 | 0 | 0 | 56519 | 0 | 0 | 0 | 1615400 | 0 | 0 | 0 | 0 | 124480 | 0 |
| LKDYAFIHF | SYNCRIP; SYN( | O60506; B7Z | HNRPQ_HUMA | Unmodified | 44680 | 0 | 0 | 0 | 0 | 0 | 0 | 0 | 0 | 0 | 0 | 434650 | 0 | 0 | 0 | 71237 | 0 | 0 |
| LKEAELLEP | COPB1; COPB1 | P53618; E9P | COPB_HUMAN | Oxidation (M) | 0 | 0 | 0 | 0 | 0 | 0 | 130900 | 0 | 0 | 0 | 0 | 1065800 | 0 | 0 | 0 | 78454 | 0 | 0 |
| LKEALNSLS | RAI14 | Q9P0K7 | RAI14_HUMAN | Unmodified | 0 | 0 | 0 | 0 | 352980 | 0 | 16264 | 37355 | 668500 | 0 | 0 | 0 | 0 | 0 | 0 | 0 | 276170 | 216300 |
| LKEASRSSI | CPNE3; CPNE3 | O75131; H0 | CPNE3_HUMAI | Unmodified | 0 | 0 | 0 | 0 | 2037700 | 0 | 5256500 | 3602000 | 0 | 2263700 | 0 | 0 | 2664100 | 2998500 | 0 | 0 | 4164900 | 4973800 |
| LKEDKAPQ | GARS1; GARS1 | P41250; F8V | GARS_HUMAN | Unmodified | 0 | 0 | 20821 | 89387 | 164610 | 0 | 0 | 118290 | 795570 | 307720 | 475710 | 2630100 | 118600 | 91280 | 44146 | 270220 | 0 | 94515 |
| LKEEIEVMA | ZMPSTE24; ZMI | O75844; A0A | FACE1_HUMAN | Unmodified | 0 | 0 | 0 | 0 | 0 | 0 | 0 | 0 | 432530 | 0 | 0 | 0 | 0 | 0 | 0 | 0 | 0 | 0 |
| LKEEITQLTI | KIF20B | Q96Q89 | KI20B_HUMAN | Unmodified | 0 | 0 | 0 | 0 | 1299700 | 126270 | 0 | 0 | 0 | 0 | 0 | 0 | 0 | 0 | 0 | 0 | 0 | 0 |
| LKEELIDAIF | ENAH; ENAH | Q8N8S7; A0 | ENAH_HUMAN | Unmodified | 322680 | 372840 | 1366300 | 4225000 | 37633000 | 6704300 | 3922400 | 26174 | 14711000 | 8505900 | 10790000 | 20252000 | 37289000 | 20459000 | 4722700 | 2090600 | 12201000 | 9567600 |
| LKEEYQSLI | UFC1 | Q9Y3C8 | UFC1_HUMAN | Unmodified | 0 | 0 | 0 | 0 | 106780 | 0 | 128280 | 139540 | 0 | 0 | 618500 | 2876000 | 0 | 0 | 0 | 0 | 0 | 0 |
| LKEGDTIIVI | EIF5B; EIF5B | O60841; A0A | IF2P_HUMAN E | Unmodified | 346780 | 287220 | 84651 | 176720 | 0 | 0 | 0 | 0 | 369280 | 125600 | 1015400 | 2695400 | 1406200 | 0 | 134890 | 120810 | 153170 | 148660 |
| LKEGDTMN | PTGR1; PTGR1 | Q14914; F2Z | PTGR1_HUMAN | Unmodified | 0 | 0 | 0 | 0 | 420530 | 0 | 0 | 0 | 216370 | 0 | 0 | 0 | 0 | 0 | 297950 | 279230 | 0 | 0 |
| LKEGLDISH | SLC12A2; SLC1 | P55011; G3 | S12A2_HUMAN | Unmodified | 0 | 63936 | 0 | 0 | 0 | 0 | 0 | 0 | 0 | 0 | 0 | 0 | 0 | 0 | 1396800 | 0 | 0 | 0 |
| LKEINFRGN | LRRC47 | Q8N1G4 | LRC47_HUMAN | Unmodified | 0 | 0 | 0 | 0 | 37375 | 0 | 74271 | 0 | 0 | 50194 | 0 | 0 | 246220 | 179870 | 0 | 0 | 557380 | 163680 |
| LKEIVTNFL | ATP5F1A | P25705 | ATPA_HUMAN | Unmodified | 0 | 0 | 0 | 0 | 0 | 0 | 0 | 0 | 721570 | 942150 | 0 | 0 | 0 | 0 | 0 | 0 | 0 | 0 |
| LKEIYENMA | PSMD2; PSMD2 | Q13200; H7( | PSMD2_HUMA | Unmodified | 0 | 0 | 0 | 0 | 1127600 | 460620 | 0 | 23415 | 515400 | 307460 | 0 | 1327500 | 0 | 0 | 0 | 0 | 717980 | 552170 |
| LKEIYENMA | PSMD2; PSMD2 | Q13200; H7( | PSMD2_HUMA | Oxidation (M) | 0 | 0 | 0 | 0 | 0 | 77033 | 0 | 0 | 0 | 0 | 0 | 276840 | 0 | 87578 | 0 | 0 | 201880 | 57301 |
| LKELDEEHS | SLK | Q9H2G2 | SLK_HUMAN S | Unmodified | 0 | 0 | 0 | 182440 | 1173400 | 0 | 0 | 0 | 1302600 | 192200 | 380750 | 893100 | 1042700 | 165080 | 0 | 105070 | 307210 | 0 |
| LKELESQVS | RRBP1; RRBP1; | Q9P2E9; A0A | RRBP1_HUMAN | Unmodified | 0 | 0 | 0 | 0 | 0 | 0 | 0 | 0 | 0 | 0 | 0 | 0 | 0 | 0 | 0 | 0 | 0 | 0 |
| LKEPWPNS | ITGB4; ITGB4 | P16144; J3Q | ITB4_HUMAN I | Unmodified | 0 | 0 | 0 | 0 | 4831000 | 725360 | 0 | 0 | 0 | 0 | 0 | 0 | 0 | 0 | 0 | 0 | 7182000 | 14109000 |
| LKEQGQAF | RHOG | P84095 | RHOG_HUMAN | Unmodified | 208870 | 310160 | 0 | 0 | 1593800 | 451970 | 441890 | 230610 | 0 | 0 | 650190 | 389940 | 1093300 | 949920 | 0 | 0 | 1036400 | 1935100 |
| LKEQSIFGE | DBN1; DBN1 | Q16643; D6I | DREB_HUMAN | Unmodified | 0 | 0 | 0 | 0 | 0 | 0 | 0 | 0 | 0 | 0 | 461180 | 0 | 0 | 0 | 0 | 0 | 0 | 0 |
| LKESQTQD | UPF1 | Q92900 | RENT1_HUMAN | Unmodified | 435440 | 327780 | 0 | 0 | 3306100 | 797010 | 123110 | 189490 | 0 | 32778 | 1297400 | 5897400 | 0 | 0 | 0 | 529860 | 2284600 | 1718100 |
| LKETAESVL | HSPA4; HSPA4; | P34932; A0A | HSP74_HUMAN | Unmodified | 0 | 0 | 0 | 0 | 116460 | 0 | 0 | 0 | 361540 | 54643 | 485510 | 3735200 | 72214 | 0 | 0 | 60576 | 0 | 0 |
| LKETCVSGI | AKAP12 | Q02952 | AKA12_HUMAN | Unmodified | 86591 | 0 | 0 | 0 | 2194000 | 0 | 0 | 0 | 0 | 0 | 0 | 0 | 0 | 0 | 0 | 0 | 2505300 | 5393100 |
| LKETIDNNS | EIF3E; EIF3E; EI | P60228; B3K | EIF3E_HUMAN | Unmodified | 0 | 0 | 0 | 0 | 0 | 0 | 0 | 0 | 0 | 0 | 145480 | 1567100 | 0 | 150330 | 0 | 45136 | 0 | 0 |
| LKETVYNLF | GOLIM4; GOLIM | O00461; F8V | GOLI4_HUMAN | Unmodified | 0 | 0 | 0 | 0 | 71252 | 0 | 91169 | 0 | 26733000 | 2214700 | 802360 | 0 | 78918 | 63499 | 0 | 0 | 0 | 0 |
| LKEVLEYNA | FDPS; FDPS; FD | P14324; A0A | FPPS_HUMAN | Unmodified | 0 | 171210 | 0 | 87032 | 0 | 0 | 0 | 0 | 0 | 0 | 0 | 4134400 | 0 | 0 | 0 | 361200 | 0 | 0 |
| LKEYDEPY\ | ABCC4 | O15439 | MRP4_HUMAN | Unmodified | 0 | 0 | 0 | 684480 | 0 | 0 | 0 | 0 | 214430 | 156710 | 0 | 1775700 | 0 | 0 | 0 | 0 | 0 | 0 |
| LKFGVYDIE | CPNE3; CPNE3 | O75131; A0A | CPNE3_HUMAI | Unmodified | 0 | 0 | 0 | 1003300 | 1755500 | 0 | 1460800 | 72333 | 11288000 | 3714100 | 221640 | 160590 | 1135700 | 1329900 | 0 | 290610 | 1703000 | 3413300 |
| LKFPSGSVI | TBCE; TBCE; TB | Q15813; A0A | TBCE_HUMAN | Unmodified | 0 | 0 | 0 | 0 | 108620 | 0 | 82956 | 0 | 21800 | 34018 | 25836 | 300410 | 32560 | 0 | 0 | 0 | 0 | 0 |
| LKGDDLQA | HNRNPC; HNR | P07910; G3 | HNRPC_HUMA | Unmodified | 0 | 0 | 0 | 0 | 6227500 | 0 | 606630 | 92949 | 2260000 | 3080800 | 3482000 | 3067600 | 3583700 | 0 | 0 | 0 | 373300 | 1109700 |
| LKGEATVSI | FUS; FUS | P35637; H3E | FUS_HUMAN R | Unmodified | 0 | 0 | 0 | 0 | 897370 | 0 | 0 | 13569 | 2302900 | 1117600 | 1447400 | 2305500 | 994930 | 0 | 0 | 160500 | 189200 | 378280 |
| LKGEMMDI | LDHB; LDHB; LI | P07195; A0A | LDHB_HUMAN | Oxidation (M) | 0 | 303010 | 247220 | 1293100 | 1674500 | 0 | 0 | 0 | 0 | 119890 | 607010 | 45273000 | 1673700 | 0 | 1135200 | 794470 | 0 | 1017700 |
| LKGEMMDI | LDHB; LDHB; LI | P07195; A0A | LDHB_HUMAN | Unmodified | 0 | 0 | 0 | 0 | 2504100 | 0 | 0 | 0 | 0 | 146340 | 0 | 1451100 | 0 | 0 | 0 | 0 | 2525500 | 645010 |
| LKGEMMDI | LDHB; LDHB; LI | P07195; A0A | LDHB_HUMAN | 2 Oxidation (M) | 1628500 | 469890 | 3623500 | 0 | 0 | 0 | 0 | 129630 | 0 | 0 | 3167200 | 40517000 | 0 | 468940 | 1608800 | 528300 | 0 | 0 |
| LKGEMMDI | LDHA; LDHA; LI | P00338; F5G | LDHA_HUMAN | 2 Oxidation (M) | 0 | 0 | 367870 | 396750 | 0 | 0 | 0 | 0 | 0 | 0 | 0 | 26622000 | 0 | 0 | 1030300 | 1407300 | 0 | 0 |
| LKGEMMDI | LDHA; LDHA; LI | P00338; F5G | LDHA_HUMAN | Oxidation (M) | 555310 | 0 | 0 | 0 | 0 | 0 | 0 | 0 | 482480 | 0 | 0 | 31525000 | 0 | 0 | 193750 | 373060 | 0 | 0 |
| LKGEMMDI | LDHA; LDHA; LI | P00338; F5G | LDHA_HUMAN | Unmodified | 0 | 0 | 0 | 0 | 0 | 0 | 0 | 0 | 0 | 0 | 0 | 1360700 | 0 | 0 | 66758 | 39779 | 0 | 0 |
| LKGEMMDI | LDHA; LDHA; LI | P00338; F5G | LDHA_HUMAN | Oxidation (M) | 0 | 0 | 0 | 189350 | 1892500 | 244890 | 0 | 0 | 499430 | 283230 | 833620 | 3727500 | 295750 | 0 | 0 | 98407 | 1260600 | 3066900 |
| LKGEMMDI | LDHA; LDHA; LI | P00338; F5G | LDHA_HUMAN | Unmodified | 511240 | 0 | 0 | 0 | 3942200 | 0 | 0 | 77191 | 611790 | 101190 | 553520 | 0 | 68424 | 88595 | 0 | 0 | 1243000 | 5051500 |
| LKGEMMDI | LDHA; LDHA; LI | P00338; F5G | LDHA_HUMAN | 2 Oxidation (M) | 0 | 0 | 186090 | 0 | 342850 | 0 | 0 | 0 | 130280 | 0 | 678620 | 1924700 | 438280 | 59836 | 142800 | 207030 | 162690 | 904940 |
| LKGPQITGI | AHNAK | Q09666 | AHNK_HUMAN | Unmodified | 181730 | 133500 | 130580 | 222080 | 51653000 | 5902400 | 0 | 0 | 0 | 0 | 0 | 0 | 0 | 1971900 | 229520 | 127690 | 18516000 | 29725000 |
| LKGTEDELI | TPM1; TPM3; TF | P09493; P06 | TPM1_HUMAN | Unmodified | 0 | 0 | 0 | 0 | 1548600 | 212020 | 0 | 0 | 0 | 0 | 0 | 0 | 0 | 0 | 1748400 | 134420 | 0 | 0 |
| LKHYGPGV | PPIB; PPIB; PPII | P23284; A0A | PPIB_HUMAN F | Oxidation (M) | 0 | 0 | 0 | 0 | 313470 | 0 | 0 | 0 | 0 | 722840 | 0 | 0 | 0 | 0 | 0 | 0 | 0 | 0 |
| LKHYGPGV | PPIB; PPIB; PPII | P23284; A0A | PPIB_HUMAN F | Unmodified | 0 | 0 | 0 | 0 | 0 | 0 | 552200 | 0 | 0 | 823430 | 0 | 0 | 106400 | 0 | 0 | 0 | 0 | 0 |
| LKIDFGDS/ | GNAI1 | P63096 | GNAI1_HUMAN | Unmodified | 0 | 0 | 0 | 0 | 0 | 0 | 0 | 0 | 203230 | 0 | 0 | 3187200 | 0 | 0 | 0 | 0 | 0 | 0 |
| LKIDFGEA/ | GNAI3 | P08754 | GNAI3_HUMAN | Unmodified | 0 | 0 | 431320 | 1662300 | 0 | 0 | 0 | 0 | 0 | 0 | 0 | 193470 | 186260 | 0 | 0 | 0 | 0 | 0 |
| LKIPEGLFD | ACTL6A | O96019 | ACL6A_HUMAN | Unmodified | 0 | 0 | 0 | 0 | 0 | 0 | 0 | 0 | 254850 | 191890 | 0 | 1132200 | 0 | 0 | 0 | 0 | 0 | 0 |
| LKITGNMG | SCP2 | P22307 | SCP2_HUMAN | Unmodified | 0 | 0 | 0 | 0 | 0 | 0 | 90506 | 0 | 3208500 | 2441000 | 0 | 0 | 0 | 0 | 0 | 0 | 0 | 0 |
| LKITGNMG | SCP2 | P22307 | SCP2_HUMAN | Oxidation (M) | 0 | 0 | 0 | 140670 | 0 | 0 | 0 | 0 | 2252100 | 2570000 | 0 | 0 | 0 | 0 | 0 | 0 | 0 | 0 |
| LKKDVEFL/ | PIP4K2A; PIP4K | P48426; S4F | PI42A_HUMAN | Unmodified | 0 | 0 | 0 | 0 | 397220 | 0 | 3811200 | 349290 | 0 | 0 | 0 | 0 | 0 | 0 | 0 | 0 | 785330 | 697570 |
| LKKEDIYAV | HSPA4; HSPA4; | P34932; A0A | HSP74_HUMAN | Unmodified | 0 | 0 | 0 | 0 | 0 | 0 | 0 | 0 | 0 | 0 | 267600 | 2489700 | 0 | 0 | 0 | 0 | 0 | 0 |
| LKKLEEEQI | MYH9 | P35579 | MYH9_HUMAN | Unmodified | 0 | 0 | 0 | 0 | 7960700 | 4285800 | 4047400 | 84977 | 0 | 1104800 | 0 | 0 | 4619600 | 5615600 | 0 | 0 | 11611000 | 2966400 |
| LKLDGLDE | LRPAP1; LRPAP | P30533; A0A | AMRP_HUMAN | Unmodified | 0 | 0 | 0 | 0 | 0 | 0 | 0 | 0 | 558540 | 290610 | 0 | 0 | 0 | 0 | 0 | 0 | 0 | 0 |
| LKLDSEQLI | CD2AP | Q9Y5K6 | CD2AP_HUMAI | Unmodified | 6856300 | 8885100 | 8614100 | 6881900 | 1748900 | 463000 | 514490 | 393630 | 569180 | 302830 | 0 | 1160100 | 14959000 | 5289800 | 10098000 | 5939000 | 133820 | 0 |
| LKLEDFFAF | G6PD; G6PD; G | P11413; E7E | G6PD_HUMAN | Unmodified | 0 | 0 | 0 | 1004400 | 0 | 500830 | 0 | 0 | 1053400 | 0 | 0 | 0 | 0 | 0 | 0 | 0 | 0 | 0 |
| LKLEPHEGI | SLC3A2; SLC3A | P08195; A0A | 4F2_HUMAN 4I | Unmodified | 327600 | 258300 | 104930 | 90757 | 0 | 0 | 0 | 0 | 535860 | 166740 | 404430 | 4515300 | 0 | 1973000 | 243910 | 216290 | 0 | 0 |
| LKLEPHEGI | SLC3A2; SLC3A | P08195; A0A | 4F2_HUMAN 4I | Unmodified | 21748000 | 21111000 | 8331900 | 22361000 | 7611900 | 6113100 | 902020 | 0 | 74694000 | 6469700 | 50350000 | 1.15E+08 | 1763300 | 0 | 19994000 | 12008000 | 2111200 | 2887000 |
| LKLEQIDGI | SMC4; SMC4 | Q9NTJ3; E9F | SMC4_HUMAN | Unmodified | 0 | 0 | 0 | 769110 | 882880 | 431140 | 1055300 | 0 | 498540 | 211510 | 266690 | 772110 | 484870 | 0 | 86282 | 99185 | 248970 | 345730 |
| LKLGEIVTTI | ARF4; ARF5; AR | P18085; P84 | ARF4_HUMAN | Unmodified | 0 | 0 | 0 | 0 | 0 | 0 | 0 | 0 | 0 | 0 | 0 | 0 | 0 | 0 | 0 | 0 | 0 | 238550 |
| LKLGQSVT | ARF6 | P62330 | ARF6_HUMAN | Unmodified | 609790 | 0 | 0 | 0 | 2094800 | 0 | 190670 | 0 | 0 | 467800 | 0 | 181310 | 0 | 178760 | 0 | 0 | 61073 | 241840 |
| LKLIENPLLI | SHCBP1 | Q8NEM2 | SHCBP_HUMA | Unmodified | 112520 | 113380 | 0 | 0 | 1403600 | 0 | 0 | 174790 | 0 | 0 | 0 | 0 | 0 | 0 | 0 | 0 | 0 | 0 |
| LKLIPDPKN | PRKCA; PRKCA | P17252; J3K | KPCA_HUMAN | Unmodified | 0 | 0 | 0 | 0 | 1390700 | 252550 | 2929000 | 1145600 | 0 | 0 | 0 | 0 | 182470 | 359420 | 0 | 0 | 345890 | 865460 |
| LKLIPDPKS | PRKCB; PRKCB | P05771; I3L1 | KPCB_HUMAN | Unmodified | 0 | 0 | 0 | 63682 | 789930 | 0 | 3674100 | 2376900 | 0 | 0 | 0 | 0 | 691020 | 1560100 | 0 | 0 | 0 | 0 |
| LKLPSIPLVI | TOMM34 | Q15785 | TOM34_HUMAI | Unmodified | 0 | 0 | 0 | 266330 | 379340 | 0 | 0 | 0 | 0 | 122170 | 754370 | 1185400 | 404790 | 314130 | 0 | 89040 | 332830 | 0 |
| LKLQVNQD | PLCB3 | Q01970 | PLCB3_HUMAN | Unmodified | 0 | 0 | 0 | 0 | 708300 | 0 | 313550 | 273650 | 0 | 0 | 0 | 0 | 89192 | 0 | 0 | 0 | 143300 | 142030 |
| LKLTPEYLQ | ERLIN2; ERLIN2 | O94905; E5F | ERLN2_HUMAN | Unmodified | 0 | 0 | 0 | 0 | 0 | 0 | 476490 | 0 | 15999000 | 2154600 | 0 | 0 | 0 | 467820 | 0 | 0 | 0 | 0 |
| LKLTPEYLQ | ERLIN2; ERLIN2 | O94905; E5F | ERLN2_HUMAN | Oxidation (M) | 0 | 0 | 0 | 0 | 0 | 0 | 898300 | 0 | 7800700 | 6998400 | 0 | 0 | 0 | 0 | 0 | 0 | 0 | 0 |
| LKMEVEQL | GNG11 | P61952 | GBG11_HUMAI | Unmodified | 0 | 0 | 0 | 1371400 | 1534300 | 0 | 1886600 | 991840 | 0 | 0 | 0 | 0 | 2051500 | 2958100 | 0 | 0 | 0 | 2752100 |
| LKMEVEQL | GNG11 | P61952 | GBG11_HUMAI | Oxidation (M) | 0 | 0 | 0 | 658250 | 543420 | 0 | 0 | 608080 | 0 | 0 | 0 | 0 | 526220 | 1236800 | 0 | 218130 | 89614 | 0 |
| LKMPDMH\ | AHNAK | Q09666 | AHNK_HUMAN | Unmodified | 0 | 0 | 334700 | 0 | 721350 | 226480 | 0 | 0 | 0 | 0 | 133030 | 0 | 0 | 0 | 0 | 0 | 0 | 0 |
| LKMPEMNI | AHNAK | Q09666 | AHNK_HUMAN | Unmodified | 0 | 0 | 0 | 0 | 672740 | 262060 | 0 | 0 | 0 | 0 | 0 | 0 | 0 | 0 | 0 | 0 | 0 | 0 |
| LKNDNTEA | DNAJC3; DNAJ( | Q13217; X6F | DNJC3_HUMAN | Unmodified | 250720 | 0 | 0 | 0 | 202210 | 0 | 824370 | 389530 | 8588300 | 3468600 | 560570 | 185240 | 0 | 0 | 40454 | 0 | 0 | 0 |
| LKNEIPNSI | CBS; CBSL | P35520; P0D | CBS_HUMAN C | Unmodified | 0 | 0 | 0 | 0 | 0 | 0 | 0 | 0 | 0 | 49861 | 478790 | 2339900 | 0 | 0 | 90723 | 0 | 0 | 0 |
| LKNELLEIK | SYTL4 | Q96C24 | SYTL4_HUMAN | Unmodified | 0 | 2440600 | 0 | 0 | 0 | 0 | 0 | 0 | 0 | 0 | 0 | 0 | 0 | 0 | 0 | 0 | 0 | 0 |
| LKPDEGGE | ABCE1; ABCE1 | P61221; D6F | ABCE1_HUMAN | Unmodified | 396010 | 660890 | 0 | 739680 | 1153200 | 0 | 0 | 0 | 805750 | 0 | 2424900 | 10461000 | 218360 | 0 | 2678300 | 827140 | 215310 | 807430 |
| LKPDTVPAI | GDF15 | Q99988 | GDF15_HUMAI | Unmodified | 0 | 0 | 0 | 0 | 0 | 0 | 0 | 0 | 1002800 | 0 | 0 | 0 | 0 | 0 | 0 | 0 | 0 | 0 |
| LKPEDITQI( | ITGB1; ITGB1; I | P05556; E7E | ITB1_HUMAN I | Unmodified | 24290000 | 19600000 | 24737000 | 36775000 | 1.26E+09 | 2.64E+08 | 6646600 | 1604900 | 3.06E+08 | 53446000 | 15741000 | 80251000 | 74055000 | 23897000 | 59197000 | 48674000 | 1.67E+08 | 9.09E+08 |
| LKPEDITQI( | ITGB1; ITGB1; I | P05556; E7E | ITB1_HUMAN I | Unmodified | 0 | 0 | 0 | 0 | 9.65E+08 | 75804000 | 6117100 | 0 | 0 | 15207000 | 86224 | 0 | 22453000 | 12590000 | 0 | 0 | 1.58E+08 | 8.05E+08 |
| LKPEGAPEI | INPPL1; INPPL1 | O15357; A0A | SHIP2_HUMAN | Unmodified | 0 | 0 | 0 | 0 | 0 | 0 | 0 | 0 | 0 | 0 | 0 | 968770 | 0 | 0 | 0 | 0 | 0 | 0 |
| LKPEGLHQ | ITPA | Q9BY32 | ITPA_HUMAN I | Unmodified | 233390 | 156210 | 0 | 239750 | 1698000 | 0 | 497620 | 0 | 485660 | 97880 | 1250600 | 5849800 | 1042900 | 335820 | 113150 | 0 | 942850 | 1209500 |
| LKPEIQCVS | MELTF | P08582 | TRFM_HUMAN | Unmodified | 59518 | 41385 | 0 | 0 | 1588300 | 657310 | 0 | 0 | 1784900 | 738730 | 217220 | 0 | 0 | 0 | 0 | 0 | 406490 | 370060 |
| LKPEVVEEL | NCS1 | P62166 | NCS1_HUMAN | Unmodified | 0 | 0 | 0 | 1112100 | 0 | 0 | 152010 | 171030 | 0 | 390800 | 0 | 2142500 | 1870800 | 853410 | 0 | 0 | 0 | 0 |
| LKPFLGSSI | OSBPL8; OSBPI | Q9BZF1; F8V | OSBL8_HUMAN | Unmodified | 0 | 0 | 0 | 0 | 0 | 0 | 0 | 0 | 467170 | 31425 | 0 | 718420 | 0 | 0 | 0 | 0 | 0 | 350820 |
| LKPGKQCS | ADAM10 | O14672 | ADA10_HUMAN | Unmodified | 0 | 0 | 0 | 0 | 498300 | 51924 | 184530 | 83328 | 0 | 0 | 0 | 0 | 1005400 | 1429400 | 0 | 0 | 1143300 | 300040 |
| LKPGYLEA1 | PPA1 | Q15181 | IPYR_HUMAN I | Unmodified | 197550 | 0 | 0 | 0 | 0 | 0 | 0 | 0 | 0 | 0 | 562600 | 2883700 | 0 | 0 | 0 | 0 | 0 | 0 |
| LKPGYLEA1 | PPA1 | Q15181 | IPYR_HUMAN I | Unmodified | 0 | 0 | 0 | 0 | 0 | 0 | 0 | 0 | 0 | 0 | 0 | 1550400 | 501280 | 0 | 0 | 0 | 0 | 0 |
| LKPKPIDV( | ACTR3; ACTR3 | P61158; B4D | ARP3_HUMAN | Unmodified | 732010 | 4265000 | 0 | 4317900 | 341290 | 0 | 224900 | 0 | 2598600 | 98284 | 1744400 | 11396000 | 707640 | 0 | 819350 | 596620 | 88303 | 0 |
| LKPKPIDV( | ACTR3; ACTR3 | P61158; B4D | ARP3_HUMAN | Oxidation (M) | 0 | 0 | 0 | 691630 | 0 | 0 | 0 | 0 | 0 | 0 | 203670 | 1284200 | 0 | 0 | 601740 | 649720 | 0 | 0 |
| LKPLEVELR | TMED10 | P49755 | TMEDA_HUMAI | Unmodified | 0 | 0 | 0 | 0 | 0 | 0 | 0 | 0 | 4976600 | 652090 | 0 | 0 | 0 | 0 | 0 | 0 | 0 | 0 |
| LKPLGEAEI | NLN; NLN; NLN | Q9BYT8; E9F | NEUL_HUMAN | Unmodified | 0 | 0 | 0 | 0 | 1389800 | 0 | 0 | 0 | 1022000 | 0 | 0 | 1751800 | 0 | 0 | 0 | 0 | 0 | 0 |
| LKPLPGETI | TLN1 | Q9Y490 | TLN1_HUMAN | Unmodified | 256100 | 207850 | 101470 | 0 | 2158700 | 645060 | 780300 | 237120 | 320170 | 306620 | 92831 | 1217500 | 0 | 559420 | 404670 | 250240 | 295660 | 0 |
| LKPMYGPG | USP13; USP13; | Q92995; A0A | UBP13_HUMAN | Oxidation (M) | 416720 | 0 | 0 | 0 | 0 | 0 | 0 | 0 | 0 | 0 | 137390 | 0 | 0 | 0 | 0 | 472240 | 141090 | 0 |
| LKPNLGNG | NUDC; NUDC; | Q9Y266; A0A | NUDC_HUMAN | Unmodified | 117740 | 59131 | 416200 | 102010 | 166090 | 0 | 0 | 0 | 1983000 | 0 | 451960 | 11348000 | 0 | 0 | 1911500 | 283290 | 224670 | 0 |
| LKPPTLIHG | RANBP3; RANB | Q9H6Z4; B7 | RANB3_HUMAI | Unmodified | 0 | 0 | 0 | 0 | 0 | 0 | 0 | 0 | 0 | 0 | 418440 | 1337100 | 0 | 0 | 0 | 0 | 0 | 0 |
| LKPQYLEEI | GSTM3; GSTM3 | P21266; A0A | GSTM3_HUMAI | Unmodified | 0 | 111360 | 269100 | 392480 | 110600 | 0 | 319120 | 225350 | 172070 | 0 | 0 | 0 | 964950 | 612820 | 1824900 | 1031000 | 885540 | 140120 |

| | | | | | | | | | | | | | | | | | | | | | | |
|---|---|---|---|---|---|---|---|---|---|---|---|---|---|---|---|---|---|---|---|---|---|---|
| LKPSNAQS | ARL6; ARL6 | Q9H0F7; C9 | ARL6_HUMAN | Unmodified | 302820 | 282390 | 0 | 0 | 731260 | 0 | 0 | 0 | 446280 | 229890 | 0 | 1025000 | 742500 | 763350 | 535290 | 384550 | 513410 | 1390400 |
| LKPTMAFM | HSDL2 | Q6YN16 | HSDL2_HUMAN | Unmodified | 0 | 0 | 0 | 0 | 0 | 0 | 0 | 0 | 1026700 | 0 | 0 | 0 | 0 | 0 | 0 | 0 | 86973 | 0 |
| LKPVDGHC | HABP2 | Q14520 | HABP2_HUMAN | Unmodified | 0 | 0 | 0 | 0 | 0 | 0 | 132150 | 161480 | 0 | 0 | 0 | 0 | 0 | 304950 | 0 | 0 | 0 | 0 |
| LKPVDGHC | HABP2 | Q14520 | HABP2_HUMAN | Unmodified | 0 | 0 | 0 | 0 | 0 | 0 | 398330 | 313060 | 0 | 0 | 0 | 0 | 0 | 0 | 0 | 0 | 0 | 0 |
| LKPVHGLIF | UCHL5; UCHL5 | Q9Y5K5; Q5 | UCHL5_HUMAN | Unmodified | 0 | 0 | 0 | 0 | 0 | 0 | 0 | 0 | 181250 | 0 | 0 | 1766000 | 377600 | 0 | 0 | 0 | 0 | 0 |
| LKPWGLFE | SRPK1; SRPK1 | Q96SB4; H3 | SRPK1_HUMAN | Unmodified | 0 | 0 | 0 | 0 | 0 | 0 | 0 | 0 | 115240 | 0 | 146070 | 1168500 | 0 | 0 | 0 | 49568 | 0 | 0 |
| LKPYFLTD( | ACAT2 | Q9BWD1 | THIC_HUMAN | Oxidation (M) | 0 | 0 | 0 | 0 | 9061200 | 1718200 | 0 | 0 | 2946900 | 2896300 | 1675000 | 10835000 | 12219000 | 0 | 0 | 0 | 0 | 0 |
| LKPYFLTD( | ACAT2 | Q9BWD1 | THIC_HUMAN | Unmodified | 0 | 0 | 0 | 0 | 3090500 | 0 | 0 | 0 | 5088800 | 267520 | 0 | 0 | 0 | 0 | 0 | 0 | 0 | 0 |
| LKPYFLTD( | ACAT2 | Q9BWD1 | THIC_HUMAN | Oxidation (M) | 0 | 0 | 0 | 0 | 0 | 0 | 0 | 0 | 761010 | 531440 | 213810 | 2121200 | 0 | 0 | 0 | 0 | 0 | 0 |
| LKPYVSYL | MRPL41 | Q8IXM3 | RM41_HUMAN | Unmodified | 0 | 0 | 0 | 0 | 0 | 0 | 0 | 0 | 259160 | 0 | 0 | 0 | 0 | 0 | 0 | 0 | 0 | 0 |
| LKQEEETL | EXOC7; EXOC7 | Q9UPT5; B5 | EXOC7_HUMAN | Unmodified | 0 | 0 | 0 | 273590 | 7757700 | 0 | 0 | 0 | 1872500 | 254040 | 538270 | 618080 | 1222600 | 0 | 0 | 0 | 0 | 1335100 |
| LKQEMGGI | CD82; CD82 | P27701; E9 | CD82_HUMAN | Unmodified | 141840 | 82356 | 0 | 0 | 0 | 0 | 0 | 0 | 0 | 215860 | 0 | 137010 | 0 | 0 | 0 | 71275 | 0 | 0 |
| LKQEMGGI | CD82; CD82 | P27701; E9 | CD82_HUMAN | Oxidation (M) | 0 | 136870 | 235640 | 0 | 104180 | 0 | 0 | 0 | 75881 | 134760 | 0 | 480090 | 58041 | 0 | 448240 | 180500 | 95730 | 40938 |
| LKQETPAV | ATP13A3; ATP1 | Q9H7F0; A0 | AT133_HUMAN | Unmodified | 0 | 0 | 0 | 0 | 1267500 | 0 | 0 | 0 | 856220 | 0 | 0 | 0 | 0 | 0 | 0 | 0 | 0 | 0 |
| LKQFVFDL | ERP44 | Q9BS26 | ERP44_HUMAN | Unmodified | 0 | 0 | 0 | 0 | 0 | 0 | 1304600 | 0 | 0 | 520880 | 0 | 0 | 0 | 0 | 82346 | 0 | 0 | 0 |
| LKQIEEQTI | RDX; RDX; RDX | P35241; A0 | RADI_HUMAN | Unmodified | 0 | 0 | 0 | 0 | 428300 | 0 | 0 | 0 | 0 | 105770 | 0 | 0 | 0 | 0 | 0 | 0 | 0 | 0 |
| LKQIEHTLN | PRKACA; PRKA | P17612; P2 | KAPCA_HUMAN | Unmodified | 0 | 0 | 0 | 0 | 712690 | 0 | 1416400 | 430230 | 99989 | 277380 | 0 | 0 | 525850 | 681830 | 0 | 0 | 0 | 0 |
| LKQNLESN | MID1; MID1; MI | O15344; A0 | TRI18_HUMAN | Unmodified | 0 | 0 | 0 | 0 | 766850 | 0 | 94570 | 0 | 0 | 91315 | 0 | 0 | 275150 | 266350 | 0 | 0 | 510730 | 391420 |
| LKQQIAEDI | RAB23 | Q9ULC3 | RAB23_HUMAN | Unmodified | 0 | 928220 | 0 | 1134300 | 2691400 | 748420 | 196230 | 405440 | 1128000 | 1069900 | 0 | 0 | 2388700 | 593930 | 0 | 0 | 740300 | 2111900 |
| LKRDVEFL | PIP4K2B | P78356 | PI42B_HUMAN | Unmodified | 0 | 0 | 0 | 0 | 0 | 0 | 0 | 0 | 0 | 294540 | 0 | 0 | 1373000 | 808180 | 0 | 0 | 509130 | 1303500 |
| LKRDVEFL | PIP4K2C; PIP4 | Q8TBX8; H0 | PI42C_HUMAN | Unmodified | 0 | 0 | 0 | 0 | 1299100 | 0 | 703700 | 0 | 0 | 476270 | 0 | 0 | 1015800 | 1080900 | 0 | 0 | 999470 | 1478000 |
| LKSEDEEI | FKBP10; FKBP1 | Q96AY3; H0 | FKB10_HUMAN | Unmodified | 0 | 0 | 0 | 89732 | 0 | 0 | 0 | 0 | 0 | 54803 | 3181900 | 1580600 | 0 | 65232 | 0 | 292630 | 0 | 0 |
| LKSEDEPTI | TNPO2; TNPO1 | O14787; Q9 | TNPO2_HUMAN | Unmodified | 0 | 0 | 0 | 169070 | 3528300 | 0 | 685330 | 1056800 | 0 | 292010 | 294310 | 0 | 1282200 | 578110 | 0 | 0 | 746710 | 0 |
| LKSEDGVE | AHNAK | Q09666 | AHNK_HUMAN | Unmodified | 0 | 0 | 0 | 0 | 0 | 121840 | 0 | 0 | 134680 | 73773 | 0 | 0 | 0 | 0 | 0 | 0 | 0 | 414980 |
| LKSEDGVE | AHNAK | Q09666 | AHNK_HUMAN | Unmodified | 0 | 0 | 0 | 0 | 1319700 | 0 | 0 | 0 | 0 | 0 | 0 | 0 | 0 | 0 | 0 | 0 | 1272300 | 0 |
| LKSIDNGIF | SDCBP; SDCBF | O00560; G5 | SDCB1_HUMAN | Unmodified | 0 | 241300 | 0 | 198560 | 178880 | 0 | 0 | 0 | 50876 | 731900 | 0 | 0 | 344650 | 163030 | 363660 | 0 | 0 | 0 |
| LKSPQAVL( | COMMD10; CO | Q9Y6G5; D6 | COMDA_HUMA | Unmodified | 0 | 0 | 0 | 0 | 1340100 | 0 | 0 | 0 | 0 | 0 | 0 | 0 | 0 | 0 | 0 | 0 | 332760 | 0 |
| LKSPSQDN | RASAL2; RASAL | Q9UJF2; H0 | NGAP_HUMAN | Unmodified | 0 | 0 | 0 | 0 | 893580 | 0 | 0 | 32448 | 0 | 106520 | 0 | 0 | 0 | 0 | 0 | 0 | 667160 | 398890 |
| LKTDVDVS | AHNAK | Q09666 | AHNK_HUMAN | Unmodified | 0 | 0 | 0 | 0 | 814570 | 0 | 0 | 0 | 0 | 0 | 0 | 0 | 0 | 0 | 0 | 0 | 124690 | 0 |
| LKTESEMH | ADGRE2; ADGR | Q9UHX3; M | AGRE2_HUMAN | Unmodified | 0 | 0 | 0 | 0 | 0 | 0 | 0 | 0 | 54996 | 0 | 0 | 556210 | 0 | 0 | 0 | 0 | 0 | 0 |
| LKTSPNEG | ATP2B1 | P20020 | AT2B1_HUMAN | Unmodified | 0 | 0 | 0 | 3040200 | 1531100 | 0 | 0 | 248370 | 2000700 | 0 | 1813600 | 5177800 | 5720200 | 0 | 1497600 | 292830 | 279630 | 0 |
| LKTSPVEG | ATP2B4 | P23634 | AT2B4_HUMAN | Unmodified | 0 | 0 | 0 | 0 | 1770800 | 0 | 694710 | 319060 | 985790 | 787810 | 741060 | 1694800 | 0 | 0 | 0 | 149700 | 972820 | 2032500 |
| LKVINLSDI | ERBIN; ERBIN | Q96RT1; B4 | ERBIN_HUMAN | Unmodified | 0 | 0 | 0 | 0 | 3650400 | 827220 | 0 | 0 | 0 | 0 | 0 | 0 | 368720 | 370800 | 0 | 0 | 894540 | 420270 |
| LKVLGTAF | HSPH1 | Q92598 | HS105_HUMAN | Unmodified | 0 | 0 | 0 | 0 | 0 | 0 | 908000 | 22965 | 0 | 0 | 0 | 0 | 0 | 0 | 0 | 0 | 279150 | 44596 |
| LKVNFLPE | DYNC1H1; DYN | Q14204; A0 | DYHC1_HUMA | Unmodified | 35333 | 0 | 0 | 0 | 0 | 0 | 381130 | 0 | 0 | 213500 | 0 | 259300 | 0 | 88769 | 0 | 0 | 0 | 0 |
| LKVPEWVD | RPS19; RPS19 | P39019; M0 | RS19_HUMAN | Unmodified | 3080200 | 2945800 | 294400 | 2057100 | 3441700 | 0 | 1585400 | 436100 | 1072400 | 942070 | 1576500 | 14693000 | 6535400 | 2061900 | 983480 | 1548500 | 471870 | 1311800 |
| LKVPPAIN( | RPL7A; RPL7A | P62424; Q5 | RL7A_HUMAN | Unmodified | 6762300 | 3597300 | 448020 | 1371400 | 0 | 0 | 0 | 0 | 917000 | 0 | 4278500 | 12807000 | 327430 | 0 | 1684600 | 2322900 | 235190 | 0 |
| LKVPPAIN( | RPL7A; RPL7A | P62424; Q5 | RL7A_HUMAN | Unmodified | 0 | 2879800 | 0 | 1895900 | 0 | 0 | 30855 | 0 | 0 | 0 | 367380 | 1352300 | 5393300 | 2204200 | 0 | 0 | 0 | 0 |
| LKVSELKEE | HNRNPU; HNR | Q00839; A0 | HNRPU_HUMA | Unmodified | 0 | 0 | 0 | 0 | 0 | 0 | 24798 | 0 | 409760 | 244990 | 0 | 661950 | 198800 | 161930 | 0 | 0 | 0 | 0 |
| LKVTEGGE | GANAB; GANAE | Q14697; F5 | GANAB_HUMA | Unmodified | 0 | 0 | 51561 | 0 | 0 | 0 | 0 | 0 | 8336900 | 438600 | 278900 | 0 | 0 | 0 | 0 | 0 | 0 | 0 |
| LKWENTLE | NOSTRIN; NOS | Q8IVI9; E9 | NOSTN_HUMA | Unmodified | 551140 | 517260 | 0 | 2741700 | 0 | 0 | 0 | 0 | 0 | 0 | 0 | 0 | 0 | 85887 | 2953800 | 589150 | 0 | 0 |
| LKYPLEEFV | ABAT; ABAT; AB | P80404; H3 | GABT_HUMAN | Unmodified | 0 | 0 | 0 | 0 | 0 | 0 | 0 | 0 | 604750 | 0 | 0 | 0 | 0 | 0 | 0 | 0 | 0 | 0 |
| LLAACSQR | RTKN | Q9BST9 | RTKN_HUMAN | Unmodified | 65305 | 0 | 0 | 60663 | 1630800 | 276880 | 26388 | 0 | 0 | 238910 | 0 | 0 | 0 | 0 | 0 | 0 | 757000 | 614450 |
| LLAAEPLSK | COLGALT1 | Q8NBJ5 | GT251_HUMAN | Unmodified | 0 | 0 | 110810 | 0 | 0 | 0 | 359160 | 0 | 15799000 | 5917900 | 405620 | 230560 | 0 | 0 | 0 | 0 | 0 | 0 |
| LLAAERATT | EPPK1; EPPK1 | P58107; A0 | EPIPL_HUMAN | Unmodified | 0 | 0 | 0 | 0 | 0 | 0 | 0 | 0 | 0 | 0 | 0 | 0 | 878310 | 2009000 | 0 | 0 | 0 | 0 |
| LLAAGQLAI | DNAJC3; DNAJ | Q13217; X6 | DNJC3_HUMAN | Unmodified | 848450 | 0 | 0 | 0 | 699740 | 0 | 0 | 0 | 443950 | 3551100 | 0 | 0 | 0 | 0 | 0 | 0 | 0 | 0 |
| LLAALLEDE | TLN1 | Q9Y490 | TLN1_HUMAN | Unmodified | 0 | 2802200 | 6389400 | 6907900 | 14111000 | 2398100 | 17767000 | 346950 | 3361800 | 3942500 | 2662500 | 10789000 | 6413000 | 3501000 | 2834300 | 3114400 | 4859900 | 1442900 |
| LLAAVKED | KTN1 | Q86UP2 | KTN1_HUMAN | Unmodified | 0 | 0 | 0 | 0 | 0 | 0 | 0 | 0 | 0 | 0 | 0 | 555430 | 0 | 0 | 0 | 0 | 0 | 0 |
| LLACIASRP | RPS8; RPS8 | P62241; Q5 | RS8_HUMAN | Unmodified | 769510 | 0 | 191960 | 853030 | 3703700 | 0 | 3087700 | 733370 | 0 | 947060 | 519630 | 861930 | 2745800 | 0 | 0 | 0 | 700360 | 3579800 |
| LLADPTGAI | PRDX5 | P30044 | PRDX5_HUMAN | Unmodified | 2067100 | 286060 | 2669500 | 680330 | 206600 | 274430 | 0 | 0 | 3848100 | 0 | 1278800 | 6135100 | 0 | 248380 | 5903900 | 2946000 | 76497 | 0 |
| LLADQAEA | RPL19; RPL19; | P84098; J3 | RL19_HUMAN | Unmodified | 2407800 | 2532500 | 1569700 | 1790500 | 0 | 0 | 0 | 0 | 0 | 0 | 2527000 | 8901200 | 1425800 | 168880 | 0 | 0 | 0 | 0 |
| LLADSKTA( | FMNL1; FMNL1 | O95466; K7 | FMNL1_HUMAN | Unmodified | 0 | 0 | 0 | 0 | 611280 | 0 | 0 | 0 | 0 | 25876 | 0 | 0 | 0 | 0 | 0 | 0 | 0 | 0 |
| LLADTDDLI | CLGN; CLGN | O14967; D6 | CLGN_HUMAN | Unmodified | 0 | 37587 | 0 | 132780 | 0 | 0 | 234650 | 0 | 12254000 | 2413200 | 991820 | 304820 | 113610 | 98062 | 171130 | 18624 | 0 | 0 |
| LLADVTHL | ZYG11B; ZYG1 | Q9C0D3; A8 | ZY11B_HUMAN | Unmodified | 0 | 0 | 24043 | 0 | 0 | 0 | 0 | 0 | 0 | 0 | 0 | 599210 | 0 | 0 | 29220 | 25319 | 0 | 0 |
| LLAEALNQ | PRKCD | Q05655 | KPCD_HUMAN | Unmodified | 0 | 0 | 0 | 217710 | 917330 | 350700 | 61633 | 0 | 1506300 | 747600 | 2525600 | 3945700 | 1736300 | 0 | 167190 | 225160 | 815310 | 2844600 |
| LLAEALNQ | PRKCD | Q05655 | KPCD_HUMAN | Unmodified | 0 | 0 | 0 | 0 | 970870 | 0 | 61197 | 83420 | 0 | 481580 | 0 | 0 | 1039600 | 479180 | 0 | 0 | 890940 | 2068100 |
| LLAEAPLVL | ADGRE2; ADGR | Q9UHX3; M | AGRE2_HUMAN | Unmodified | 0 | 0 | 0 | 0 | 736390 | 0 | 0 | 0 | 0 | 0 | 3687700 | 6904200 | 0 | 0 | 0 | 0 | 0 | 0 |
| LLAEILR | LRPPRC | P42704 | LPPRC_HUMAN | Unmodified | 0 | 53085 | 90388 | 140470 | 0 | 0 | 0 | 0 | 2559400 | 0 | 0 | 0 | 0 | 0 | 387010 | 0 | 0 | 0 |
| LLAELPASV | FLOT2; FLOT2; | Q14254; J3 | FLOT2_HUMAN | Unmodified | 1360800 | 1904700 | 3815000 | 8691700 | 2304800 | 2879700 | 2682600 | 629940 | 25500000 | 1885100 | 3950400 | 7133200 | 547820 | 291870 | 17710000 | 4681500 | 758600 | 918690 |
| LLAENNEII | MAN2A1 | Q16706 | MA2A1_HUMAN | Unmodified | 0 | 0 | 0 | 0 | 0 | 0 | 0 | 0 | 784040 | 171410 | 0 | 1674800 | 0 | 0 | 0 | 0 | 0 | 0 |
| LLAEPVPGI | UBE2NL; UBE2I | Q5JXB2; P6 | UE2NL_HUMAN | Unmodified | 2301200 | 1456100 | 2533200 | 0 | 0 | 0 | 0 | 0 | 1004500 | 0 | 7415100 | 9600200 | 0 | 0 | 2754100 | 2515600 | 0 | 0 |
| LLAEPVPGI | UBE2NL; UBE2I | Q5JXB2; P6 | UE2NL_HUMAN | Unmodified | 2755400 | 4085200 | 645100 | 593260 | 0 | 0 | 0 | 0 | 1690600 | 0 | 1348300 | 14883000 | 0 | 0 | 2750800 | 2940200 | 0 | 0 |
| LLAETVAPA | ESYT1; ESYT1 | Q9BSJ8; F8 | ESYT1_HUMAN | Unmodified | 324050 | 0 | 347820 | 0 | 0 | 0 | 5340300 | 0 | 0 | 299130 | 0 | 621090 | 0 | 0 | 290470 | 0 | 0 | 0 |
| LLAEVTLK | FARSA; FARSA; | Q9Y285; K7 | SYFA_HUMAN | Unmodified | 0 | 0 | 0 | 0 | 0 | 0 | 547220 | 0 | 0 | 0 | 189750 | 1522300 | 0 | 0 | 0 | 0 | 0 | 0 |
| LLAGEDGT | LIMCH1; LIMCH | Q9UPQ0; H | LIMC1_HUMAN | Unmodified | 0 | 0 | 0 | 128120 | 48161 | 0 | 0 | 0 | 0 | 320070 | 0 | 125920 | 0 | 68845 | 473610 | 332600 | 0 | 0 |
| LLAGTPEP | DLL3 | Q9NYJ7 | DLL3_HUMAN | Unmodified | 0 | 0 | 0 | 0 | 0 | 0 | 112220 | 0 | 2469000 | 216330 | 165990 | 829910 | 0 | 0 | 0 | 0 | 0 | 0 |
| LLAHQKPP | PFKL | P17858 | PFKAL_HUMAN | Unmodified | 0 | 0 | 0 | 0 | 667370 | 0 | 0 | 270800 | 118520 | 0 | 0 | 0 | 399970 | 264030 | 0 | 0 | 584530 | 954910 |
| LLAHVRPP | PFKM; PFKM; P | P08237; A0 | PFKAM_HUMAN | Unmodified | 0 | 0 | 0 | 0 | 102150 | 0 | 0 | 0 | 0 | 0 | 83352 | 678620 | 953070 | 500290 | 0 | 321050 | 284200 | 261730 |
| LLAHVTLEL | MARS1; MARS1 | P56192; H0 | SYMC_HUMAN | Unmodified | 0 | 28089 | 69424 | 0 | 0 | 0 | 0 | 0 | 0 | 0 | 0 | 2550600 | 0 | 0 | 120480 | 42645 | 0 | 0 |
| LLALGDSG | RAB27B; RAB2 | O00194; K7 | RB27B_HUMAN | Unmodified | 797370 | 827650 | 589050 | 1264400 | 1205800 | 0 | 1014500 | 884700 | 0 | 0 | 0 | 0 | 747450 | 916940 | 738310 | 794430 | 182090 | 1091900 |
| LLALGDSG | RAB27B; RAB2 | O00194; K7 | RB27B_HUMAN | Unmodified | 1148900 | 0 | 0 | 221520 | 399070 | 0 | 637830 | 61342 | 0 | 0 | 0 | 0 | 680800 | 528760 | 0 | 0 | 0 | 512610 |
| LLALNPDA | UBAC1 | Q9BSL1 | UBAC1_HUMAN | Unmodified | 230420 | 0 | 72661 | 0 | 0 | 0 | 0 | 0 | 0 | 0 | 113930 | 534820 | 0 | 0 | 0 | 0 | 0 | 0 |
| LLALNPDA | UBAC1 | Q9BSL1 | UBAC1_HUMAN | Unmodified | 0 | 0 | 0 | 0 | 0 | 0 | 0 | 0 | 0 | 0 | 0 | 527420 | 0 | 0 | 0 | 0 | 0 | 0 |
| LLALNSLYS | PRKDC | P78527 | PRKDC_HUMA | Unmodified | 0 | 0 | 0 | 0 | 680910 | 0 | 0 | 0 | 0 | 0 | 0 | 2708700 | 0 | 0 | 0 | 0 | 0 | 0 |
| LLANMVYQ | PSMB5; PSMB5 | P28074; H0 | PSB5_HUMAN | Unmodified | 0 | 0 | 0 | 0 | 0 | 0 | 50818 | 66581 | 0 | 317900 | 0 | 1661300 | 0 | 0 | 0 | 0 | 297110 | 0 |
| LLANQQVF | LIMA1; LIMA1; | Q9UHB6; F8 | LIMA1_HUMAN | Unmodified | 4329800 | 1293600 | 4968500 | 1942300 | 1620200 | 0 | 104640 | 0 | 0 | 0 | 223960 | 2496000 | 1833700 | 0 | 4187300 | 1159100 | 308680 | 0 |
| LLAQDQG( | PAFAH1B3; PA | Q15102; M0 | PA1B3_HUMAN | Unmodified | 0 | 0 | 0 | 0 | 293530 | 0 | 8747.9 | 0 | 1638500 | 942670 | 1901900 | 5122100 | 62686 | 0 | 0 | 0 | 0 | 0 |
| LLAQHATS | TRAF4; TRAF4; | Q9BUZ4; K7 | TRAF4_HUMAN | Unmodified | 491220 | 0 | 0 | 0 | 0 | 0 | 0 | 0 | 0 | 0 | 0 | 0 | 645990 | 250100 | 0 | 739470 | 0 | 0 |
| LLAQQLVS | MCAT | Q8IVS2 | FABD_HUMAN | Unmodified | 0 | 0 | 0 | 0 | 0 | 0 | 0 | 0 | 634540 | 69684 | 0 | 0 | 0 | 0 | 0 | 0 | 0 | 0 |
| LLAQRLYA( | TXNRD1; TXNR | Q16881; A0 | TRXR1_HUMAN | Unmodified | 0 | 0 | 0 | 0 | 3559900 | 0 | 0 | 0 | 0 | 466000 | 0 | 0 | 614540 | 400250 | 0 | 0 | 0 | 1784300 |
| LLAQTTLR | STOML3; STOM | Q8TAV4; P2 | STML3_HUMAN | Unmodified | 3138200 | 4870300 | 3519500 | 7229500 | 0 | 0 | 427830 | 301160 | 0 | 179420 | 0 | 0 | 0 | 0 | 4408400 | 4432800 | 0 | 0 |
| LLASADDF | EML2; EML2 | O95834; K7 | EMAL2_HUMAN | Unmodified | 0 | 264750 | 616420 | 338930 | 0 | 0 | 427520 | 338670 | 0 | 0 | 0 | 1015100 | 558280 | 574020 | 0 | 234050 | 0 | 1342400 |
| LLASCCK | STXBP3 | O00186 | STXB3_HUMAN | Unmodified | 587330 | 16831 | 0 | 387030 | 0 | 0 | 343080 | 337830 | 30194 | 28648 | 117700 | 0 | 0 | 2254200 | 387340 | 379110 | 2613400 | 2494400 |
| LLASCSAD | PAFAH1B1; PA | P43034; A0 | LIS1_HUMAN | Oxidation (M) | 0 | 316840 | 455820 | 0 | 0 | 0 | 0 | 0 | 0 | 0 | 0 | 680000 | 0 | 0 | 342430 | 0 | 0 | 0 |
| LLASINSTV | DDB1; DDB1; D | Q16531; A0 | DDB1_HUMAN | Unmodified | 55491 | 88702 | 0 | 0 | 0 | 0 | 0 | 0 | 429810 | 54727 | 0 | 3570500 | 0 | 0 | 0 | 0 | 0 | 0 |
| LLATEQED | RRBP1; RRBP1; | Q9P2E9; A0 | RRBP1_HUMAN | Unmodified | 0 | 0 | 0 | 0 | 0 | 0 | 0 | 0 | 0 | 0 | 0 | 120970 | 913510 | 501900 | 0 | 0 | 0 | 0 |
| LLAVAATAF | VCL | P18206 | VINC_HUMAN | Unmodified | 584970 | 0 | 0 | 383310 | 204000 | 0 | 0 | 0 | 1125300 | 0 | 0 | 14332000 | 0 | 0 | 939160 | 717980 | 425470 | 173360 |
| LLAVIEEQH | SLC38A10; SLC | Q9HBR0; H | S38AA_HUMAN | Unmodified | 0 | 0 | 0 | 0 | 0 | 0 | 0 | 0 | 1973300 | 0 | 0 | 0 | 0 | 0 | 0 | 0 | 0 | 0 |
| LLAVNNVC | DLG1; DLG1; D | Q12959; H7 | DLG1_HUMAN | Unmodified | 0 | 0 | 0 | 0 | 0 | 1352800 | 0 | 0 | 3237300 | 1220700 | 679390 | 0 | 0 | 0 | 0 | 0 | 581950 | 0 |
| LLAVQELLE | EEPD1 | Q7L9B9 | EEPD1_HUMAN | Unmodified | 125010 | 0 | 1418100 | 1777000 | 1477400 | 136270 | 898230 | 143180 | 3796500 | 1833700 | 338200 | 2497700 | 5659200 | 4744700 | 2548300 | 1417700 | 12663000 | 18534000 |
| LLCGATLIA | KLK11; KLK11; | Q9UBX7; M0 | KLK11_HUMAN | Unmodified | 0 | 0 | 0 | 0 | 0 | 0 | 489800 | 0 | 1178700 | 0 | 0 | 0 | 0 | 0 | 0 | 0 | 0 | 0 |
| LLCGLLAEF | MIF | P14174 | MIF_HUMAN | Unmodified | 17181000 | 5837500 | 9772700 | 12849000 | 10300000 | 4946800 | 1930600 | 565570 | 16949000 | 6038600 | 23945000 | 1.3E+08 | 24675000 | 9123800 | 21449000 | 15699000 | 12525000 | 9933400 |
| LLCGLLAEF | MIF | P14174 | MIF_HUMAN | Unmodified | 0 | 0 | 0 | 5025800 | 23791000 | 8090600 | 4132700 | 42459 | 3084900 | 22205000 | 9701100 | 0 | 69472000 | 36545000 | 263360 | 0 | 54334000 | 25057000 |
| LLCMQRQ | ARHGAP29; AR | Q52LW3; F8 | RHG29_HUMAN | Unmodified | 0 | 0 | 0 | 0 | 526540 | 0 | 0 | 0 | 0 | 0 | 0 | 0 | 0 | 0 | 0 | 0 | 1808300 | 2610900 |
| LLCMQRQ | ARHGAP29; AR | Q52LW3; F8 | RHG29_HUMAN | Oxidation (M) | 0 | 0 | 0 | 0 | 123150 | 37521 | 0 | 9989 | 0 | 0 | 0 | 0 | 0 | 0 | 0 | 0 | 706630 | 670670 |
| LLCNFMR | PSMA1; N/A; PS | P25786; B4 | PSA1_HUMAN | Unmodified | 0 | 0 | 0 | 0 | 0 | 0 | 0 | 0 | 0 | 0 | 0 | 2207500 | 0 | 0 | 0 | 48767 | 0 | 0 |
| LLCQCLGF | FN1; FN1 | P02751; H0 | FINC_HUMAN | Unmodified | 0 | 0 | 10670000 | 9823800 | 0 | 0 | 0 | 0 | 1166200 | 1012800 | 0 | 0 | 2561600 | 619390 | 249300 | 2484300 | 563820 | 984160 |
| LLCSNSAF | NAE1; NAE1 | Q13564; J3 | ULA1_HUMAN | Unmodified | 0 | 0 | 0 | 0 | 0 | 0 | 0 | 0 | 0 | 0 | 0 | 1232400 | 0 | 0 | 0 | 0 | 0 | 0 |
| LLCVMCR | TRIM26; TRIM2 | Q12899; Q5 | TRI26_HUMAN | Unmodified | 277990 | 0 | 0 | 229550 | 790510 | 2818300 | 0 | 228540 | 0 | 0 | 0 | 932700 | 0 | 0 | 0 | 0 | 0 | 1395100 |
| LLDAAYKG | PLXNA1 | Q9UIW2 | PLXA1_HUMAN | Unmodified | 0 | 0 | 0 | 0 | 2747400 | 0 | 200480 | 0 | 0 | 0 | 0 | 0 | 845750 | 2758200 | 0 | 0 | 2394900 | 2582300 |
| LLDAESEDI | KIF4A; KIF4B | O95239; Q2 | KIF4A_HUMAN | Unmodified | 0 | 0 | 0 | 0 | 1110400 | 0 | 596850 | 231340 | 0 | 0 | 0 | 0 | 0 | 27697 | 0 | 0 | 1013700 | 0 |
| LLDAVDTYI | TUFM | P49411 | EFTU_HUMAN | Unmodified | 0 | 0 | 0 | 0 | 0 | 0 | 0 | 0 | 5540900 | 0 | 0 | 0 | 0 | 0 | 0 | 0 | 0 | 0 |
| LLDAVDTYI | TUFM | P49411 | EFTU_HUMAN | Unmodified | 0 | 0 | 0 | 0 | 0 | 0 | 4198900 | 0 | 14808000 | 1080400 | 0 | 501090 | 0 | 0 | 0 | 0 | 73355 | 279810 |
| LLDAVVPFI | TOR2A | Q5JU69 | TOR2A_HUMAN | Unmodified | 0 | 0 | 0 | 0 | 0 | 0 | 0 | 0 | 10317000 | 226080 | 0 | 44533 | 0 | 0 | 0 | 0 | 209080 | 0 |
| LLDDAMAA | GLOD4; GLOD4 | Q9HC38; I3 | GLOD4_HUMA | Unmodified | 0 | 0 | 0 | 0 | 0 | 0 | 0 | 53798 | 0 | 0 | 0 | 1423300 | 969630 | 1134600 | 0 | 0 | 0 | 0 |
| LLDDAMAA | GLOD4; GLOD4 | Q9HC38; I3 | GLOD4_HUMA | Oxidation (M) | 249690 | 0 | 211070 | 235160 | 1554500 | 215720 | 240620 | 279280 | 517420 | 278260 | 1032800 | 5483900 | 2983000 | 929530 | 491750 | 432500 | 0 | 438830 |
| LLDDGTFM | HSH2D; HSH2D | Q96JZ2; K7 | HSH2D_HUMA | Oxidation (M) | 0 | 0 | 0 | 0 | 0 | 0 | 0 | 0 | 0 | 0 | 0 | 0 | 0 | 0 | 1390300 | 1028800 | 0 | 0 |
| LLDDNGNI | SELENOF; SELE | O60613; A0 | SEP15_HUMAN | Unmodified | 0 | 0 | 0 | 0 | 0 | 0 | 0 | 0 | 3119000 | 0 | 1073900 | 0 | 0 | 0 | 0 | 0 | 134310 | 0 |
| LLDDVSFH | AP3M1 | Q9Y2T2 | AP3M1_HUMAN | Unmodified | 460040 | 184170 | 0 | 0 | 0 | 0 | 0 | 0 | 142290 | 0 | 0 | 1417900 | 0 | 0 | 0 | 157440 | 0 | 0 |

| | | | | | | | | | | | | | | | | | | | | | | |
|---|---|---|---|---|---|---|---|---|---|---|---|---|---|---|---|---|---|---|---|---|---|---|
| LLDEAIQAV | VPS35; VPS35; | Q96QK1; A0 | VPS35_HUMAN | Unmodified | 0 | 0 | 1116200 | 627570 | 1417400 | 308390 | 221850 | 0 | 3651700 | 0 | 2264600 | 15108000 | 752300 | 0 | 0 | 578200 | 0 | 574690 |
| LLDEALKEV | EXOC1 | Q9NV70 | EXOC1_HUMAI | Unmodified | 0 | 0 | 0 | 0 | 786120 | 215620 | 0 | 0 | 0 | 386640 | 0 | 0 | 313250 | 108890 | 0 | 0 | 648560 | 286610 |
| LLDEDEIRG | HTATSF1; HTAT | O43719; Q5I | HTSF1_HUMAN | Unmodified | 0 | 0 | 0 | 0 | 0 | 0 | 0 | 0 | 0 | 0 | 0 | 0 | 0 | 0 | 0 | 0 | 0 | 0 |
| LLDEEEATD | PDCD6IP | Q8WUM4 | PDC6I_HUMAN | Unmodified | 9805100 | 8833100 | 677500 | 253520 | 0 | 0 | 0 | 0 | 1170000 | 230460 | 739570 | 3637000 | 114950 | 0 | 1208500 | 1379300 | 0 | 0 |
| LLDEEEATD | PDCD6IP | Q8WUM4 | PDC6I_HUMAN | Unmodified | 20096000 | 26371000 | 7037400 | 3940000 | 336230 | 0 | 0 | 0 | 4064600 | 237960 | 441690 | 6776300 | 550950 | 0 | 9712500 | 10602000 | 0 | 0 |
| LLDEVFFSE | PSMA4; PSMA4 | P25789; H0Y | PSA4_HUMAN | Unmodified | 0 | 0 | 2638500 | 1932700 | 6093200 | 981860 | 1110400 | 549900 | 4750200 | 5116200 | 4448000 | 16797000 | 5308400 | 3454300 | 371980 | 1792600 | 2984100 | 4324300 |
| LLDEYNLVI | NOTCH1; NOTC | P46531; A0A | NOTC1_HUMA | Unmodified | 330220 | 400350 | 0 | 412680 | 0 | 0 | 0 | 0 | 1200700 | 0 | 0 | 3061500 | 0 | 0 | 0 | 0 | 0 | 0 |
| LLDFEFSSC | H6PD | O95479 | G6PE_HUMAN | Unmodified | 0 | 0 | 0 | 0 | 0 | 0 | 0 | 0 | 0 | 0 | 0 | 0 | 0 | 0 | 0 | 0 | 0 | 0 |
| LLDFGSLSI | RPS17; N/A | P08708; H3E | RS17_HUMAN | Oxidation (M) | 3190500 | 5608400 | 366700 | 2567900 | 4555400 | 0 | 0 | 0 | 0 | 359340 | 0 | 1330400 | 4562400 | 2697300 | 2667300 | 0 | 915670 | 2253300 |
| LLDFGSLSI | RPS17; N/A | P08708; H3E | RS17_HUMAN | Oxidation (M) | 0 | 0 | 0 | 0 | 6999300 | 1459600 | 0 | 0 | 0 | 0 | 0 | 0 | 9775100 | 4022900 | 0 | 0 | 874100 | 3462800 |
| LLDFLNVK | DIPK2A | Q8NDZ4 | DIK2A_HUMAN | Unmodified | 0 | 0 | 0 | 0 | 0 | 0 | 0 | 0 | 2074200 | 737780 | 0 | 0 | 0 | 0 | 0 | 57500 | 0 | 0 |
| LLDFSYEQ | NCLN; NCLN | Q969V3; K7E | NCLN_HUMAN | Unmodified | 0 | 0 | 0 | 0 | 0 | 0 | 0 | 0 | 4756400 | 1026900 | 0 | 0 | 0 | 0 | 0 | 0 | 0 | 0 |
| LLDGFFIRP | NEDD4; NEDD4 | P46934; Q96 | NEDD4_HUMA | Unmodified | 1232000 | 3856500 | 973870 | 347850 | 0 | 0 | 325580 | 0 | 44643 | 245370 | 0 | 750090 | 0 | 0 | 4897600 | 3228800 | 136480 | 0 |
| LLDGPSTEI | TCEA1; TCEA1; | P23193; A0A | TCEA1_HUMAN | Unmodified | 0 | 0 | 0 | 0 | 1127100 | 0 | 0 | 0 | 0 | 0 | 0 | 0 | 0 | 0 | 0 | 0 | 0 | 0 |
| LLDGPSTEI | TCEA1; TCEA1; | P23193; A0A | TCEA1_HUMAN | Unmodified | 0 | 0 | 0 | 0 | 2390400 | 218090 | 10473 | 68894 | 0 | 225990 | 0 | 822880 | 227690 | 243280 | 0 | 73686 | 0 | 0 |
| LLDGSAEFI | IGSF21 | Q96ID5 | IGS21_HUMAN | Unmodified | 0 | 0 | 0 | 0 | 0 | 0 | 1572500 | 0 | 3027300 | 0 | 0 | 0 | 0 | 1670200 | 0 | 0 | 0 | 0 |
| LLDIDETEY | ERBB2; ERBB2; | P04626; J3Q | ERBB2_HUMAN | Unmodified | 389920 | 482610 | 4430500 | 1567300 | 0 | 0 | 0 | 0 | 0 | 0 | 0 | 0 | 0 | 0 | 4022900 | 2914800 | 0 | 0 |
| LLDKVELMI | MPC2; MPC2 | O95563; Q5I | MPC2_HUMAN | Unmodified | 0 | 0 | 0 | 0 | 0 | 0 | 0 | 0 | 1316200 | 0 | 0 | 0 | 0 | 0 | 0 | 0 | 0 | 0 |
| LLDLDEAAY | GAS6 | Q14393 | GAS6_HUMAN | Unmodified | 0 | 0 | 0 | 0 | 0 | 0 | 0 | 0 | 13857000 | 0 | 0 | 0 | 0 | 0 | 0 | 0 | 226000 | 0 |
| LLDLDPRP | CAMP; CAMP | P49913; J3K | CAMP_HUMAN | Unmodified | 0 | 0 | 0 | 0 | 0 | 0 | 0 | 0 | 1687100 | 0 | 0 | 0 | 0 | 0 | 0 | 0 | 0 | 0 |
| LLDLLEGLT | UTRN; UTRN; U | P46939; A0A | UTRN_HUMAN | Unmodified | 0 | 0 | 0 | 402300 | 261940 | 0 | 0 | 0 | 572420 | 531660 | 0 | 41736 | 0 | 0 | 794130 | 269180 | 558120 | 0 |
| LLDLSVFR | SLC16A3; SLC1 | O15427; J3Q | MOT4_HUMAN | Unmodified | 211790 | 0 | 2279400 | 3767400 | 1248800 | 9944500 | 3932600 | 0 | 482370 | 228120 | 104270 | 6945900 | 0 | 2653400 | 3667000 | 3425600 | 670260 | 3261800 |
| LLDLSWTE | PARP4 | Q9UKK3 | PARP4_HUMAN | Unmodified | 0 | 0 | 0 | 34311 | 594500 | 0 | 0 | 0 | 0 | 0 | 0 | 438230 | 350950 | 55712 | 0 | 0 | 0 | 1401900 |
| LLDLVQQS | SNU13; SNU13 | P55769; A0A | NH2L1_HUMAI | Unmodified | 0 | 0 | 0 | 0 | 0 | 0 | 0 | 382170 | 0 | 0 | 0 | 536030 | 582640 | 0 | 205700 | 0 | 0 | 334380 |
| LLDMDGIIV | EIF3A | Q14152 | EIF3A_HUMAN | Oxidation (M) | 120960 | 0 | 0 | 252460 | 0 | 0 | 0 | 0 | 0 | 0 | 251480 | 2050500 | 0 | 0 | 0 | 495230 | 0 | 0 |
| LLDMDGIIV | EIF3A | Q14152 | EIF3A_HUMAN | Unmodified | 0 | 0 | 0 | 0 | 0 | 0 | 0 | 0 | 0 | 0 | 0 | 2119300 | 0 | 0 | 0 | 0 | 0 | 0 |
| LLDMLSK | ANKFY1; ANKFY | Q9P2R3; K7E | ANFY1_HUMAN | Unmodified | 0 | 0 | 0 | 0 | 0 | 0 | 0 | 0 | 0 | 0 | 0 | 0 | 0 | 2035800 | 0 | 0 | 0 | 0 |
| LLDNTMEIF | FARP2; FARP2; | O94887; C9J | FARP2_HUMAN | Oxidation (M) | 0 | 0 | 509350 | 0 | 214150 | 0 | 0 | 0 | 0 | 0 | 0 | 0 | 0 | 0 | 1579900 | 0 | 0 | 0 |
| LLDPEDISV | SPTBN1; SPTBN | Q01082; A0A | SPTB2_HUMAN | Unmodified | 0 | 0 | 0 | 0 | 781700 | 1093000 | 313250 | 197650 | 1222500 | 668860 | 0 | 9424200 | 0 | 0 | 0 | 0 | 218230 | 438120 |
| LLDPEDVA | UTRN | P46939 | UTRN_HUMAN | Unmodified | 0 | 0 | 1761700 | 933140 | 0 | 0 | 0 | 0 | 1885300 | 1279000 | 0 | 0 | 0 | 0 | 3478900 | 3182100 | 0 | 0 |
| LLDPEDVN | SPTBN2; SPTBN | O15020; A4C | SPTN2_HUMAN | Unmodified | 0 | 0 | 96879 | 136300 | 36530 | 0 | 0 | 0 | 0 | 0 | 0 | 751500 | 238250 | 0 | 188130 | 169640 | 79068 | 0 |
| LLDQIVEK | FUBP1; FUBP1; | Q96AE4; C9J | FUBP1_HUMAN | Unmodified | 140960 | 0 | 0 | 0 | 0 | 0 | 0 | 0 | 0 | 0 | 1085800 | 248250 | 0 | 0 | 0 | 0 | 0 | 0 |
| LLDRLPSF | COG8; COG8; ( | Q96MW5; H | COG8_HUMAN | Unmodified | 0 | 35622 | 0 | 0 | 0 | 0 | 0 | 0 | 0 | 0 | 0 | 432670 | 0 | 0 | 0 | 0 | 0 | 109570 |
| LLDRTDSRI | PLXNB2; PLXNE | O15031; E2F | PLXB2_HUMAN | Unmodified | 211920 | 79574 | 850160 | 2205800 | 7264100 | 1670700 | 312410 | 0 | 1445800 | 214580 | 0 | 0 | 4343200 | 2409900 | 1489500 | 165700 | 1401200 | 3941900 |
| LLDSEIK | PSMC3; PSMC3 | P17980; E9P | PRS6A_HUMAN | Unmodified | 0 | 0 | 0 | 0 | 0 | 0 | 0 | 0 | 0 | 0 | 0 | 3985700 | 0 | 0 | 0 | 0 | 0 | 0 |
| LLDSITVPV | AP3B1 | O00203 | AP3B1_HUMAN | Unmodified | 0 | 0 | 0 | 0 | 0 | 0 | 0 | 0 | 0 | 0 | 0 | 3583400 | 0 | 0 | 0 | 0 | 0 | 0 |
| LLDSLEPP | TFG; TFG; TFG; | Q92734; A0A | TFG_HUMAN P | Unmodified | 0 | 0 | 0 | 0 | 0 | 0 | 0 | 0 | 794900 | 0 | 0 | 0 | 0 | 0 | 0 | 0 | 0 | 0 |
| LLDSLEPP | TFG; TFG; TFG; | Q92734; A0A | TFG_HUMAN P | Unmodified | 0 | 0 | 0 | 0 | 781460 | 0 | 0 | 0 | 4384800 | 217690 | 0 | 1932600 | 0 | 0 | 0 | 1480400 | 0 | 0 |
| LLDSSTVTI | PSMA6; PSMA6 | P60900; G3V | PSA6_HUMAN | Unmodified | 59219 | 7589500 | 2226700 | 5290900 | 10070000 | 1987600 | 1468600 | 468690 | 7473100 | 4558200 | 4810800 | 34228000 | 6777400 | 4622700 | 3795100 | 2260700 | 2634800 | 6209300 |
| LLDTIGISQ | CAD; CAD | P27708; F8V | PYR1_HUMAN | Unmodified | 0 | 0 | 0 | 0 | 0 | 0 | 0 | 0 | 496210 | 0 | 0 | 1641500 | 0 | 0 | 0 | 0 | 0 | 0 |
| LLDTVDDM | EHD1; EHD1; E | Q9H4M9; C9 | EHD1_HUMAN | Oxidation (M) | 509380 | 575930 | 0 | 1225600 | 1647100 | 443910 | 152220 | 38368 | 395060 | 306170 | 1094500 | 2399100 | 1174700 | 308450 | 828040 | 569740 | 1811800 | 4551100 |
| LLDTVDDM | EHD1; EHD1; E | Q9H4M9; C9 | EHD1_HUMAN | Unmodified | 108800 | 399130 | 36048 | 309430 | 2170000 | 349670 | 0 | 30912 | 1438000 | 0 | 248550 | 586320 | 663230 | 287380 | 283360 | 100640 | 1155000 | 1730900 |
| LLDVDNRV | MT-CO2 | P00403 | COX2_HUMAN | Unmodified | 0 | 0 | 0 | 0 | 0 | 0 | 0 | 0 | 4703200 | 1563300 | 0 | 0 | 0 | 0 | 0 | 0 | 0 | 0 |
| LLDVNNQF | AGRN; AGRN; A | O00468; A0A | AGRIN_HUMAN | Unmodified | 694370 | 99434 | 1584000 | 460370 | 0 | 136060 | 0 | 0 | 709450 | 0 | 0 | 924010 | 0 | 0 | 594930 | 552510 | 0 | 57983 |
| LLDVNNQF | AGRN; AGRN; A | O00468; A0A | AGRIN_HUMAN | Unmodified | 0 | 0 | 920170 | 3761300 | 0 | 0 | 0 | 0 | 1229800 | 0 | 393480 | 0 | 642560 | 0 | 989130 | 461980 | 0 | 0 |
| LLDVTQKY | UACA; UACA; U | Q9BZF9; H0Y | UACA_HUMAN | Unmodified | 0 | 0 | 0 | 0 | 400540 | 0 | 0 | 16571 | 0 | 0 | 0 | 0 | 227280 | 0 | 0 | 0 | 653150 | 321110 |
| LLDVVHPA | CCT7; CCT7 | Q99832; F8V | TCPH_HUMAN | Unmodified | 0 | 0 | 147980 | 120170 | 896310 | 0 | 393370 | 356800 | 778810 | 523290 | 813820 | 3406000 | 1421500 | 0 | 106190 | 115940 | 0 | 475870 |
| LLEAAAQS | PLEC; PLEC | Q15149; H0Y | PLEC_HUMAN | Unmodified | 0 | 0 | 0 | 0 | 0 | 0 | 0 | 0 | 0 | 0 | 0 | 0 | 0 | 0 | 0 | 0 | 190180 | 0 |
| LLEAGNFIC | HMGCL; HMGC | P35914; H0Y | HMGCL_HUMA | Unmodified | 0 | 0 | 0 | 0 | 0 | 0 | 0 | 0 | 740550 | 232990 | 0 | 0 | 0 | 0 | 0 | 0 | 0 | 0 |
| LLEAGNFIC | HMGCL; HMGC | P35914; H0Y | HMGCL_HUMA | Unmodified | 0 | 0 | 0 | 0 | 0 | 0 | 0 | 0 | 6123800 | 2421600 | 0 | 176690 | 0 | 0 | 0 | 0 | 0 | 0 |
| LLEAHEEQI | NAPA; NAPA; N | P54920; M0F | SNAA_HUMAN | Unmodified | 4940800 | 3776100 | 1660800 | 1912300 | 1402000 | 0 | 806660 | 947040 | 4399400 | 806430 | 3133800 | 10594000 | 4836500 | 873430 | 2376000 | 2412500 | 1322800 | 2673600 |
| LLEAKGPV | WDR1 | O75083 | WDR1_HUMAN | Unmodified | 0 | 0 | 0 | 0 | 4892700 | 0 | 0 | 0 | 0 | 532870 | 0 | 0 | 0 | 1456300 | 0 | 0 | 1668700 | 0 |
| LLEALDEMI | EHD2 | Q9NZN4 | EHD2_HUMAN | Oxidation (M) | 0 | 0 | 0 | 0 | 2349300 | 0 | 0 | 0 | 0 | 0 | 0 | 0 | 0 | 0 | 0 | 0 | 142840 | 0 |
| LLEALDEMI | EHD2 | Q9NZN4 | EHD2_HUMAN | Unmodified | 0 | 0 | 0 | 0 | 1315900 | 237260 | 11155 | 0 | 0 | 0 | 0 | 0 | 0 | 0 | 0 | 0 | 0 | 0 |
| LLEALLR | C2orf72 | A6NCS6 | CB072_HUMAN | Unmodified | 114820 | 0 | 0 | 0 | 0 | 0 | 0 | 0 | 0 | 0 | 0 | 0 | 0 | 0 | 0 | 294810 | 0 | 0 |
| LLEAQACT | DSP | P15924 | DESP_HUMAN | Unmodified | 0 | 0 | 136100 | 0 | 0 | 0 | 0 | 0 | 1658700 | 0 | 0 | 0 | 0 | 0 | 0 | 0 | 0 | 0 |
| LLEASTQF | SLC12A2; SLC1 | P55011; G3X | S12A2_HUMAN | Unmodified | 459980 | 219490 | 950030 | 657260 | 1365900 | 0 | 411280 | 197870 | 392650 | 99912 | 0 | 1232800 | 632900 | 394510 | 1241300 | 1015900 | 438710 | 6262500 |
| LLEASTQF | SLC12A2; SLC1 | P55011; G3X | S12A2_HUMAN | Unmodified | 1032300 | 861490 | 161670 | 647950 | 902570 | 0 | 128770 | 92757 | 0 | 29690 | 0 | 1230900 | 813130 | 107130 | 1621700 | 719240 | 4742700 | 4043300 |
| LLEATELK | SPTAN1; SPTAN | Q13813; A0A | SPTN1_HUMAN | Unmodified | 0 | 0 | 0 | 0 | 584420 | 0 | 0 | 0 | 0 | 315480 | 0 | 0 | 0 | 0 | 308120 | 0 | 457210 | 0 |
| LLEDKNGE | CAND2; CAND | O75155; Q8 | CAND2_HUMA | Unmodified | 0 | 0 | 0 | 284300 | 6056300 | 798190 | 2675900 | 1572700 | 631940 | 1765400 | 518590 | 839490 | 7288500 | 4840400 | 0 | 0 | 5693500 | 5450300 |
| LLEDRIAEF | MYH9 | P35579 | MYH9_HUMAN | Unmodified | 0 | 0 | 0 | 1273500 | 19926000 | 4178800 | 1162100 | 49559 | 1840400 | 1438500 | 2707500 | 0 | 7612400 | 5574200 | 0 | 0 | 5125900 | 6223900 |
| LLEDRIAEF | MYH9 | P35579 | MYH9_HUMAN | Unmodified | 0 | 0 | 0 | 0 | 12797000 | 3959200 | 172850 | 0 | 0 | 1368400 | 0 | 0 | 5165000 | 5356600 | 0 | 0 | 6534300 | 4241800 |
| LLEEALLR | PRKDC | P78527 | PRKDC_HUMAI | Unmodified | 444100 | 223330 | 610650 | 0 | 0 | 24595 | 0 | 15391 | 0 | 59517 | 25487 | 4352700 | 0 | 0 | 89902 | 133200 | 21762 | 37609 |
| LLEEAVR | LSR; ARHGEF1 | Q86X29; O1 | LSR_HUMAN Li | Unmodified | 2335500 | 2796200 | 3839800 | 3498600 | 315020 | 43540 | 216490 | 99997 | 17071000 | 1567200 | 1936600 | 6797200 | 0 | 386390 | 6086600 | 4638700 | 315510 | 14729 |
| LLEEAVRK | LSR; LSR | Q86X29; S4F | LSR_HUMAN Li | Unmodified | 0 | 0 | 483230 | 0 | 0 | 0 | 0 | 0 | 0 | 0 | 0 | 0 | 0 | 0 | 732010 | 0 | 0 | 0 |
| LLEEEDSK | CCDC124; CCI | Q96CT7; M0 | CC124_HUMAI | Unmodified | 504820 | 6136200 | 6243500 | 947190 | 0 | 0 | 0 | 0 | 0 | 0 | 287550 | 681760 | 4501200 | 2235800 | 2175200 | 5194500 | 0 | 0 |
| LLEEEEK | RTN2; SPECC1; | O75298; Q5I | RTN2_HUMAN | Unmodified | 0 | 0 | 0 | 0 | 0 | 0 | 0 | 0 | 0 | 0 | 0 | 0 | 0 | 0 | 0 | 0 | 0 | 0 |
| LLEEENAM | ZC3H10 | Q96K80 | ZC3HA_HUMAI | Unmodified | 1877500 | 1220300 | 1045900 | 1979000 | 0 | 0 | 0 | 0 | 0 | 0 | 0 | 0 | 889430 | 0 | 498620 | 747140 | 0 | 0 |
| LLEEENAM | ZC3H10 | Q96K80 | ZC3HA_HUMAI | Oxidation (M) | 0 | 0 | 0 | 0 | 0 | 0 | 0 | 55366 | 0 | 0 | 0 | 0 | 651770 | 65100 | 0 | 137740 | 24420 | 0 |
| LLEEHKSL | UTRN | P46939 | UTRN_HUMAN | Unmodified | 0 | 0 | 0 | 0 | 0 | 0 | 0 | 0 | 0 | 0 | 0 | 0 | 0 | 0 | 0 | 0 | 310360 | 0 |
| LLEELEEG | UBE2V2; UBE2 | Q15819; Q1 | UB2V2_HUMAN | Unmodified | 3088600 | 2918800 | 2387600 | 1068900 | 332950 | 0 | 0 | 0 | 1925100 | 0 | 2840300 | 13753000 | 406880 | 0 | 3547600 | 3207800 | 0 | 224270 |
| LLEENETE | BAIAP2L1 | Q9UHR4 | BI2L1_HUMAN | Unmodified | 0 | 0 | 0 | 0 | 0 | 0 | 0 | 69485 | 643890 | 0 | 0 | 1343500 | 0 | 0 | 2606900 | 1447600 | 0 | 0 |
| LLEENVSA | SMC4; SMC4 | Q9NTJ3; E9F | SMC4_HUMAN | Unmodified | 0 | 0 | 0 | 204010 | 1476000 | 1150600 | 1133800 | 451720 | 1539300 | 1132900 | 1120700 | 1351000 | 1219000 | 0 | 0 | 0 | 845490 | 868370 |
| LLEEQAKL | CIT; CIT; CIT; C | O14578; A0A | CTRO_HUMAN | Unmodified | 0 | 0 | 0 | 0 | 937900 | 0 | 0 | 0 | 0 | 0 | 0 | 0 | 0 | 0 | 0 | 0 | 120960 | 0 |
| LLEEQLQH | KTN1 | Q86UP2 | KTN1_HUMAN | Unmodified | 0 | 0 | 0 | 0 | 0 | 0 | 0 | 0 | 0 | 0 | 0 | 503140 | 0 | 0 | 0 | 0 | 0 | 0 |
| LLEEYGSDI | UTRN | P46939 | UTRN_HUMAN | Unmodified | 0 | 0 | 1987200 | 0 | 0 | 0 | 0 | 0 | 0 | 0 | 0 | 0 | 0 | 0 | 1928300 | 3212700 | 0 | 0 |
| LLEEYGSDI | UTRN | P46939 | UTRN_HUMAN | Unmodified | 0 | 0 | 0 | 1440800 | 537030 | 0 | 0 | 0 | 596460 | 588600 | 0 | 0 | 187490 | 0 | 342050 | 155950 | 270090 | 0 |
| LLEEYGSDI | UTRN | P46939 | UTRN_HUMAN | Unmodified | 0 | 0 | 0 | 689390 | 8734700 | 0 | 237590 | 238440 | 0 | 5183600 | 0 | 0 | 3992100 | 1249300 | 0 | 0 | 6049500 | 0 |
| LLEEYTQLA | NIT1 | Q86X76 | NIT1_HUMAN [ | Unmodified | 0 | 0 | 0 | 63277 | 0 | 0 | 299890 | 0 | 0 | 0 | 217580 | 1820600 | 0 | 0 | 0 | 0 | 0 | 0 |
| LLEFNQGK | GARS1; GARS1 | P41250; A0A | GARS_HUMAN | Unmodified | 0 | 0 | 3298200 | 0 | 0 | 0 | 0 | 0 | 2998500 | 1130800 | 1475500 | 2987400 | 0 | 0 | 0 | 0 | 0 | 0 |
| LLEFQLALK | STIP1; STIP1 | P31948; H0Y | STIP1_HUMAN | Unmodified | 0 | 0 | 0 | 0 | 0 | 0 | 0 | 0 | 0 | 432130 | 932710 | 404730 | 284520 | 0 | 0 | 0 | 0 | 0 |
| LLEGDEK | ERBB4; ERBB4 | Q15303; A0A | ERBB4_HUMAN | Unmodified | 27051 | 0 | 82110 | 0 | 2619700 | 375110 | 0 | 0 | 556600 | 0 | 0 | 541520 | 0 | 94337 | 0 | 0 | 261200 | 1514100 |
| LLEGEEER | LMNB1; LMNB2 | P20700; Q03 | LMNB1_HUMAI | Unmodified | 0 | 0 | 0 | 0 | 0 | 0 | 0 | 0 | 0 | 0 | 0 | 444610 | 0 | 0 | 0 | 0 | 0 | 0 |
| LLEGEEERI | LMNB1; LMNB2 | P20700; Q03 | LMNB1_HUMAI | Unmodified | 0 | 115290 | 0 | 0 | 4607500 | 0 | 197380 | 28123 | 1515100 | 1673000 | 1664200 | 3125900 | 4077200 | 0 | 0 | 0 | 0 | 1264200 |
| LLEGEESRI | VIM; VIM | P08670; B0Y | VIME_HUMAN | Unmodified | 0 | 0 | 0 | 0 | 196150 | 0 | 0 | 0 | 0 | 0 | 0 | 0 | 0 | 0 | 0 | 0 | 289390 | 720570 |
| LLEGEESRI | VIM | P08670 | VIME_HUMAN | Unmodified | 0 | 0 | 0 | 195410 | 0 | 0 | 0 | 0 | 0 | 0 | 0 | 0 | 0 | 0 | 0 | 0 | 1178900 | 2837200 |
| LLEGEESRI | PRPH; PRPH | P41219; H7C | PERI_HUMAN F | Unmodified | 0 | 0 | 0 | 0 | 0 | 0 | 3866400 | 188620 | 0 | 0 | 0 | 0 | 0 | 0 | 0 | 0 | 0 | 0 |
| LLEGEGLPI | RET; RET | P07949; Q9E | RET_HUMAN P | Unmodified | 0 | 0 | 0 | 0 | 0 | 0 | 0 | 0 | 4748900 | 57986 | 0 | 0 | 0 | 0 | 0 | 0 | 0 | 0 |
| LLEGHVPE | AHNAK2 | Q8IVF2 | AHNK2_HUMAI | Unmodified | 0 | 0 | 0 | 0 | 0 | 433010 | 0 | 0 | 0 | 221070 | 0 | 0 | 0 | 0 | 0 | 0 | 131620 | 1246300 |
| LLEGHVPE | AHNAK2 | Q8IVF2 | AHNK2_HUMAI | Unmodified | 0 | 0 | 0 | 0 | 637830 | 216000 | 0 | 0 | 0 | 0 | 0 | 0 | 0 | 0 | 0 | 0 | 116820 | 751230 |
| LLEGIGLEN | DNAJC13; DNA | O75165; H0Y | DJC13_HUMAN | Unmodified | 0 | 0 | 0 | 0 | 44344 | 0 | 0 | 0 | 379750 | 0 | 0 | 2631800 | 0 | 0 | 0 | 0 | 0 | 0 |
| LLEGPVPEE | AHNAK2 | Q8IVF2 | AHNK2_HUMAI | Unmodified | 0 | 0 | 0 | 0 | 3472700 | 620230 | 0 | 0 | 0 | 0 | 0 | 0 | 0 | 0 | 0 | 0 | 657470 | 4469400 |
| LLEHLPQE | ERAL1; ERAL1 | O75616; K7E | ERAL1_HUMAN | Unmodified | 0 | 0 | 0 | 0 | 900660 | 0 | 0 | 228960 | 2903500 | 118360 | 0 | 0 | 0 | 0 | 0 | 0 | 845740 | 456340 |
| LLEIDPYLK | GBE1 | Q04446 | GLGB_HUMAN | Unmodified | 0 | 0 | 0 | 0 | 0 | 0 | 0 | 19932 | 1827700 | 1315500 | 0 | 977890 | 0 | 0 | 0 | 0 | 0 | 0 |
| LLEIPAVSH | ZBTB7A | O95365 | ZBT7A_HUMAN | Unmodified | 0 | 0 | 0 | 0 | 0 | 0 | 0 | 0 | 0 | 0 | 0 | 149920 | 0 | 69397 | 38567 | 0 | 0 | 0 |
| LLEIVER | TWF1 | Q12792 | TWF1_HUMAN | Unmodified | 943330 | 1415700 | 195710 | 143670 | 0 | 0 | 0 | 0 | 0 | 0 | 0 | 2210100 | 103750 | 74254 | 1176600 | 1266600 | 0 | 0 |
| LLEIVSYK | USP7; USP7; U | Q93009; A0A | UBP7_HUMAN | Unmodified | 0 | 0 | 0 | 0 | 0 | 0 | 0 | 0 | 0 | 0 | 0 | 898020 | 0 | 0 | 0 | 0 | 0 | 0 |
| LLELDPEH | P4HA1 | P13674 | P4HA1_HUMAN | Unmodified | 0 | 0 | 0 | 0 | 188550 | 0 | 0 | 22890 | 736890 | 670580 | 407670 | 0 | 0 | 0 | 0 | 0 | 26986 | 0 |
| LLELDSFLK | PSME3; PSME3 | P61289; A0A | PSME3_HUMAI | Unmodified | 113350 | 0 | 112440 | 56684 | 98735 | 0 | 0 | 35081 | 249550 | 0 | 522910 | 960580 | 379010 | 0 | 0 | 0 | 58563 | 0 |
| LLELFPVNF | BZW2; BZW2; B | Q9Y6E2; B5N | BZW2_HUMAN | Unmodified | 0 | 0 | 0 | 0 | 0 | 0 | 0 | 0 | 0 | 0 | 1024400 | 6344100 | 0 | 0 | 0 | 0 | 0 | 0 |
| LLELFTDLS | BROX; BROX; B | Q5VW32; F5 | BROX_HUMAN | 2 Oxidation (M) | 0 | 0 | 0 | 793450 | 0 | 0 | 0 | 0 | 0 | 0 | 0 | 0 | 0 | 0 | 0 | 449310 | 0 | 0 |
| LLELGPKP | COPA; COPA; C | P53621; A0A | COPA_HUMAN | Unmodified | 0 | 0 | 416250 | 201770 | 0 | 0 | 0 | 0 | 424070 | 274710 | 1478700 | 3026300 | 0 | 0 | 341260 | 326550 | 105960 | 247840 |
| LLELGVKP | FAM171A2 | A8MVW0 | F1712_HUMAN | Unmodified | 0 | 0 | 0 | 0 | 458530 | 0 | 0 | 0 | 218100 | 172770 | 0 | 0 | 0 | 0 | 0 | 0 | 137960 | 525270 |
| LLELIGR | BMPR2; BMPR2 | Q13873; A0A | BMPR2_HUMAI | Unmodified | 42062 | 0 | 176610 | 93743 | 0 | 0 | 0 | 0 | 352750 | 60806 | 0 | 261130 | 0 | 0 | 271140 | 124650 | 0 | 0 |
| LLELLPGK | CPSF7; CPSF7; | Q8N684; F5I | CPSF7_HUMAN | Unmodified | 547350 | 0 | 1974700 | 1506600 | 0 | 0 | 0 | 0 | 290210 | 38983 | 292810 | 1631900 | 1809900 | 720680 | 726250 | 1302600 | 0 | 0 |
| LLELLQEAF | PROM2 | Q8N271 | PROM2_HUMA | Unmodified | 1576200 | 1257600 | 0 | 0 | 0 | 0 | 245240 | 127590 | 0 | 0 | 0 | 0 | 0 | 0 | 1658700 | 772490 | 0 | 0 |
| LLELQEVD | ITGB4 | P16144 | ITB4_HUMAN I | Unmodified | 19217 | 16081 | 0 | 11779 | 3003100 | 478180 | 0 | 0 | 0 | 0 | 129840 | 0 | 84336 | 19883 | 73725 | 30835 | 3817900 | 5137000 |
| LLELQEVD | ITGB4 | P16144 | ITB4_HUMAN I | Unmodified | 0 | 0 | 0 | 0 | 11172000 | 1391500 | 229300 | 0 | 417910 | 411230 | 0 | 0 | 2184400 | 1047300 | 0 | 0 | 26857000 | 32591000 |
| LLELQYFISI | GLG1; GLG1 | Q92896; H3I | GSLG1_HUMAI | Unmodified | 0 | 0 | 0 | 0 | 0 | 0 | 0 | 0 | 6240900 | 0 | 699490 | 1178800 | 0 | 0 | 0 | 0 | 0 | 0 |

| | | | | | | | | | | | | | | | | | | | | | | |
|---|---|---|---|---|---|---|---|---|---|---|---|---|---|---|---|---|---|---|---|---|---|---|
| LLEMILNKF | AKR1C2; AKR1C | P52895; B4D | AK1C2_HUMAN | Oxidation (M) | 508500 | 228640 | 0 | 0 | 1234300 | 0 | 0 | 0 | 0 | 0 | 0 | 0 | 43475 | 37997 | 0 | 0 | 0 | 0 |
| LLEMMEEV | MTMR12 | Q9C0I1 | MTMRC_HUMA | Oxidation (M) | 0 | 136840 | 0 | 0 | 0 | 1184600 | 0 | 0 | 0 | 0 | 0 | 0 | 0 | 0 | 0 | 0 | 0 | 0 |
| LLEMNLMH | CLTC; CLTC | Q00610; A0A | CLH1_HUMAN | 3 Oxidation (M) | 0 | 0 | 1476900 | 1270400 | 0 | 0 | 0 | 0 | 0 | 0 | 0 | 3348400 | 0 | 0 | 4333400 | 0 | 0 | 0 |
| LLEPAQVQ | XXYLT1; XXYLT1 | Q8NBI6; A0A | XXLT1_HUMAN | Unmodified | 0 | 0 | 0 | 0 | 0 | 0 | 0 | 0 | 2230200 | 0 | 0 | 0 | 0 | 0 | 0 | 0 | 0 | 0 |
| LLEPAQVQ | XXYLT1; XXYLT1 | Q8NBI6; A0A | XXLT1_HUMAN | Unmodified | 0 | 0 | 0 | 0 | 0 | 0 | 0 | 0 | 3313200 | 0 | 0 | 0 | 0 | 0 | 0 | 0 | 0 | 0 |
| LLEPEAPGI | AFDN; AFDN; A | P55196; H0Y | AFAD_HUMAN | Unmodified | 0 | 0 | 0 | 0 | 0 | 0 | 0 | 0 | 0 | 0 | 480170 | 1717400 | 150490 | 0 | 0 | 0 | 0 | 0 |
| LLEPFEVR | THEM6 | Q8WUY1 | THEM6_HUMAI | Unmodified | 2.9E+08 | 2.7E+08 | 2.02E+08 | 1.96E+08 | 0 | 0 | 0 | 0 | 361430 | 378010 | 0 | 0 | 3923300 | 825490 | 2.88E+08 | 2.28E+08 | 0 | 0 |
| LLEPLVTQV | CTNNA1; CTNN | P35221; E5R | CTNA1_HUMAI | Unmodified | 1244500 | 0 | 2490400 | 0 | 0 | 0 | 0 | 0 | 0 | 0 | 3800900 | 0 | 0 | 0 | 17141000 | 8417900 | 0 | 0 |
| LLEPLVTQV | CTNNA1; CTNN | P35221; E5R | CTNA1_HUMAI | Unmodified | 0 | 2997600 | 0 | 18243000 | 26967000 | 7108200 | 167880 | 0 | 616570 | 1047700 | 1543400 | 1496900 | 17067000 | 6565800 | 9901700 | 8572800 | 0 | 0 |
| LLEPLVTQV | CTNNA1; CTNN | P35221; E5R | CTNA1_HUMAI | Unmodified | 0 | 0 | 0 | 3292400 | 5856700 | 1830700 | 0 | 0 | 0 | 0 | 0 | 0 | 7733400 | 2723200 | 0 | 1289000 | 0 | 0 |
| LLEPLVTQV | CTNNA2; CTNN | P26232; C9J | CTNA2_HUMAI | Unmodified | 0 | 1487400 | 2490400 | 6964500 | 1684500 | 0 | 1056200 | 0 | 1146000 | 0 | 3800900 | 5034800 | 1298400 | 97779 | 17364 | 8417900 | 0 | 0 |
| LLEPSVGSI | WASHC2A; WA | Q641Q2; Q9 | WAC2A_HUMA | Unmodified | 0 | 0 | 0 | 0 | 2065500 | 0 | 78716 | 0 | 0 | 174480 | 0 | 0 | 267180 | 179870 | 105290 | 0 | 183500 | 629810 |
| LLEPVDHG | DDX46; DDX46 | Q7L014; A0A | DDX46_HUMAI | Unmodified | 0 | 0 | 0 | 0 | 0 | 0 | 467310 | 165710 | 0 | 252040 | 218690 | 1585500 | 2005100 | 674370 | 0 | 76208 | 0 | 0 |
| LLEPVLLLG | RPS16; RPS16; | P62249; M0R | RS16_HUMAN | Unmodified | 16265000 | 22048000 | 8217800 | 4260400 | 6941800 | 883580 | 27011000 | 23577000 | 3834600 | 2943500 | 9427600 | 2.35E+08 | 2762900 | 4323000 | 9568300 | 10819000 | 1057700 | 4055900 |
| LLEPVVCM | VPS4A | Q9UN37 | VPS4A_HUMAN | Unmodified | 0 | 0 | 0 | 0 | 59542 | 0 | 0 | 0 | 377900 | 0 | 0 | 0 | 37332 | 34026 | 0 | 0 | 0 | 0 |
| LLEPVVSMS | VPS4B | O75351 | VPS4B_HUMAN | Oxidation (M) | 0 | 0 | 0 | 35697 | 70359 | 0 | 0 | 0 | 397890 | 106890 | 0 | 0 | 41588 | 42506 | 0 | 0 | 0 | 22742 |
| LLEPVVSMS | VPS4B | O75351 | VPS4B_HUMAN | Unmodified | 0 | 0 | 0 | 0 | 162700 | 0 | 0 | 0 | 3715300 | 310530 | 0 | 0 | 60656 | 0 | 0 | 257750 | 0 | 0 |
| LLEQALVIE | CHD4; CHD3; C | Q14839; Q1 | CHD4_HUMAN | Unmodified | 0 | 0 | 0 | 0 | 0 | 0 | 0 | 0 | 0 | 0 | 232540 | 2078900 | 0 | 0 | 0 | 0 | 0 | 0 |
| LLEQEALMI | FMNL2; FMNL2 | Q96PY5; C9I | FMNL2_HUMAI | Unmodified | 0 | 0 | 0 | 0 | 2404700 | 988280 | 0 | 0 | 389310 | 325300 | 0 | 0 | 0 | 0 | 0 | 0 | 497350 | 1628800 |
| LLEQEALMI | FMNL2; FMNL2 | Q96PY5; C9I | FMNL2_HUMAI | Oxidation (M) | 0 | 0 | 0 | 0 | 881560 | 0 | 0 | 38110 | 0 | 0 | 221780 | 335310 | 118640 | 0 | 0 | 0 | 0 | 0 |
| LLEQGLR | FASN; FASN | P49327; A0A | FAS_HUMAN F | Unmodified | 0 | 0 | 4315700 | 0 | 0 | 0 | 0 | 0 | 0 | 0 | 2853100 | 1.14E+08 | 0 | 0 | 5022500 | 4135100 | 0 | 0 |
| LLEQLTDTH | TRIM59; N/A; TF | Q8IWR1; H3 | TRI59_HUMAN | Unmodified | 0 | 0 | 0 | 20801 | 496030 | 59818 | 0 | 0 | 0 | 0 | 0 | 23158 | 37290 | 0 | 0 | 0 | 0 | 74928 |
| LLEQWEK | DCAF1; MESD | Q9Y4B6; Q1 | DCAF1_HUMAI | Unmodified | 0 | 0 | 0 | 0 | 0 | 0 | 0 | 0 | 2565600 | 0 | 417530 | 0 | 0 | 0 | 0 | 0 | 0 | 0 |
| LLEQWEKD | MESD | Q14696 | MESD_HUMAN | Unmodified | 0 | 0 | 0 | 0 | 1183800 | 0 | 1523300 | 0 | 13877000 | 0 | 2381400 | 557860 | 0 | 732150 | 0 | 0 | 0 | 0 |
| LLEQYKEES | HNRNPU; HNR | Q00839; A0A | HNRPU_HUMA | Unmodified | 425360 | 199220 | 121480 | 0 | 284220 | 44739 | 600890 | 643400 | 246420 | 338860 | 468000 | 1685900 | 540970 | 257420 | 48492 | 56120 | 0 | 103560 |
| LLEQYKEES | HNRNPU; HNR | Q00839; A0A | HNRPU_HUMA | Unmodified | 386050 | 0 | 0 | 0 | 1738000 | 0 | 0 | 0 | 588840 | 1268800 | 689970 | 0 | 1522700 | 0 | 0 | 0 | 0 | 587210 |
| LLERLPEVE | NUCB1 | Q02818 | NUCB1_HUMA | Unmodified | 0 | 0 | 0 | 0 | 607200 | 272410 | 2096500 | 79368 | 13687000 | 1930900 | 2344300 | 1579600 | 542570 | 310190 | 0 | 0 | 83956 | 949310 |
| LLESDYFR | ERO1A; ERO1A; | Q96HE7; G3 | ERO1A_HUMAI | Unmodified | 0 | 0 | 0 | 0 | 0 | 0 | 0 | 44946 | 23640000 | 2975500 | 201370 | 1046000 | 0 | 0 | 0 | 0 | 0 | 0 |
| LLESDYFRY | ERO1A; ERO1A; | Q96HE7; G3 | ERO1A_HUMAI | Unmodified | 0 | 0 | 0 | 0 | 1032700 | 0 | 1269100 | 0 | 0 | 25199000 | 0 | 0 | 0 | 224710 | 0 | 0 | 377790 | 1085400 |
| LLESLTPLC | IRAG2; IRAG2; I | Q12912; F5H | IRAG2_HUMAN | Unmodified | 0 | 0 | 0 | 0 | 0 | 0 | 0 | 0 | 0 | 0 | 51787 | 2871700 | 0 | 308000 | 0 | 0 | 0 | 0 |
| LLESSLSSS | ITGA5 | P08648 | ITA5_HUMAN I | Unmodified | 1261100 | 845130 | 789010 | 0 | 878000 | 791070 | 0 | 0 | 0 | 0 | 0 | 0 | 0 | 0 | 2387700 | 2153200 | 889300 | 538420 |
| LLETCSIAL | CTPS1; CTPS1; | P17812; B4E | PYRG1_HUMAI | Unmodified | 0 | 0 | 0 | 0 | 0 | 0 | 0 | 0 | 0 | 0 | 195390 | 1656800 | 0 | 0 | 0 | 0 | 0 | 0 |
| LLETECPQ | S100A8 | P05109 | S10A8_HUMAN | Unmodified | 0 | 0 | 0 | 292530 | 0 | 0 | 0 | 0 | 188140 | 0 | 0 | 0 | 0 | 0 | 0 | 0 | 0 | 458770 |
| LLETELLQE | DTX3L | Q8TDB6 | DTX3L_HUMAN | Unmodified | 0 | 25799 | 0 | 0 | 71414 | 0 | 0 | 0 | 18115 | 0 | 0 | 0 | 227240 | 124060 | 54024 | 0 | 0 | 202440 |
| LLETERQLC | EEA1 | Q15075 | EEA1_HUMAN | Unmodified | 0 | 0 | 0 | 0 | 3314500 | 233370 | 171670 | 31267 | 0 | 0 | 0 | 0 | 487600 | 313050 | 0 | 0 | 124650 | 338520 |
| LLETHIHN | CUL1; CUL1; C | Q13616; A0A | CUL1_HUMAN | Unmodified | 0 | 0 | 0 | 0 | 0 | 441360 | 83015 | 0 | 0 | 134590 | 378590 | 2782600 | 189430 | 0 | 0 | 61795 | 193900 | 123400 |
| LLETIDQLY | ACTN1; ACTN1 | P12814; A0A | ACTN1_HUMAI | Unmodified | 0 | 0 | 0 | 0 | 2235300 | 502900 | 359420 | 0 | 0 | 362950 | 0 | 1687500 | 0 | 0 | 507080 | 262320 | 1660800 | 0 |
| LLETIDQLY | ACTN1; ACTN1 | P12814; A0A | ACTN1_HUMAI | Unmodified | 0 | 0 | 0 | 0 | 0 | 267240 | 0 | 0 | 0 | 0 | 0 | 2975400 | 0 | 0 | 0 | 301140 | 125360 | 0 |
| LLETPSTLH | EXOC2 | Q96KP1 | EXOC2_HUMAI | Unmodified | 0 | 0 | 376180 | 0 | 992570 | 0 | 132630 | 0 | 379420 | 312550 | 0 | 924260 | 169890 | 0 | 194660 | 82524 | 263820 | 406940 |
| LLETTDRPD | KPNB1; KPNB1 | Q14974; J3K | IMB1_HUMAN | Unmodified | 2447800 | 2168700 | 2930400 | 1821800 | 315170 | 0 | 244510 | 0 | 1225500 | 603740 | 0 | 16411000 | 0 | 0 | 4455600 | 2519700 | 221490 | 162980 |
| LLETTEEDK | TRMT10C; TRM | Q7L0Y3; C9J | TM10C_HUMAI | Unmodified | 100370 | 615180 | 83835 | 369240 | 0 | 97621 | 0 | 0 | 678510 | 24836 | 0 | 0 | 0 | 0 | 227480 | 101710 | 0 | 0 |
| LLEVDLK | SRSF6; SRSF6; | Q13247; A0A | SRSF6_HUMAN | Unmodified | 299610 | 308270 | 0 | 0 | 0 | 0 | 0 | 0 | 0 | 178240 | 0 | 605960 | 0 | 0 | 223090 | 0 | 0 | 0 |
| LLEVEHPA | TCP1; TCP1; TC | P17987; F5H | TCPA_HUMAN | Unmodified | 0 | 0 | 136530 | 0 | 652840 | 0 | 326900 | 317230 | 552590 | 446410 | 769990 | 4202600 | 1454700 | 597510 | 158900 | 142470 | 303330 | 572200 |
| LLEVLADLD | RASAL1 | O95294 | RASL1_HUMAN | Unmodified | 0 | 0 | 0 | 0 | 0 | 0 | 179070 | 0 | 785160 | 85686 | 0 | 67629 | 0 | 0 | 0 | 0 | 0 | 0 |
| LLEVLADLD | RASAL1 | O95294 | RASL1_HUMAN | Unmodified | 0 | 0 | 0 | 0 | 6174000 | 0 | 1096600 | 0 | 878160 | 2390500 | 0 | 0 | 7257800 | 0 | 0 | 0 | 598580 | 1194000 |
| LLEVLSGEF | SPTBN1; SPTBN | Q01082; A0A | SPTB2_HUMAN | Unmodified | 0 | 0 | 0 | 0 | 0 | 0 | 0 | 0 | 0 | 0 | 0 | 5664100 | 0 | 0 | 0 | 0 | 0 | 0 |
| LLEVNGVA | SCRIB; SCRIB; | Q14160; H0Y | SCRIB_HUMAN | Unmodified | 0 | 0 | 0 | 292710 | 0 | 0 | 0 | 0 | 1125600 | 0 | 0 | 0 | 0 | 0 | 0 | 271670 | 0 | 0 |
| LLEVPVML | ISOC2; ISOC2; | Q96AB3; K7E | ISOC2_HUMAN | Unmodified | 0 | 0 | 0 | 0 | 0 | 0 | 0 | 0 | 636330 | 0 | 0 | 0 | 0 | 0 | 0 | 0 | 0 | 0 |
| LLEVPVML | ISOC2; ISOC2; | Q96AB3; K7E | ISOC2_HUMAN | Oxidation (M) | 0 | 0 | 0 | 0 | 0 | 0 | 0 | 0 | 0 | 2959400 | 0 | 0 | 0 | 0 | 0 | 0 | 0 | 0 |
| LLEVQPQV | EHBP1; EHBP1 | Q8NDI1; H7I | EHBP1_HUMAI | Unmodified | 0 | 0 | 0 | 0 | 723490 | 0 | 0 | 0 | 223020 | 0 | 0 | 442010 | 264870 | 0 | 0 | 0 | 357550 | 0 |
| LLEYDTVTR | APMAP; APMAP | Q9HDC9; H0 | APMAP_HUMAI | Unmodified | 0 | 0 | 0 | 0 | 257670 | 0 | 0 | 0 | 4750900 | 952220 | 0 | 0 | 134520 | 0 | 0 | 0 | 228580 | 0 |
| LLEYDTVTR | APMAP; APMAP | Q9HDC9; H0 | APMAP_HUMAI | Unmodified | 0 | 0 | 0 | 0 | 0 | 0 | 0 | 0 | 1115400 | 0 | 0 | 0 | 0 | 0 | 0 | 0 | 0 | 0 |
| LLFAIEETE | WWP1; ITCH; IT | Q9H0M0; Q9 | WWP1_HUMAN | Unmodified | 0 | 491200 | 0 | 0 | 467490 | 0 | 0 | 0 | 0 | 0 | 0 | 0 | 219190 | 0 | 0 | 0 | 323970 | 0 |
| LLFEGAGSI | ATP6V0D1; ATP | P61421; R4C | VA0D1_HUMAN | Unmodified | 607340 | 884210 | 1679200 | 1198100 | 665040 | 0 | 215160 | 164400 | 9995200 | 658340 | 4698600 | 5693700 | 259550 | 280150 | 720240 | 467820 | 0 | 0 |
| LLFEGAGSI | ATP6V0D1; ATP | P61421; R4C | VA0D1_HUMAN | Unmodified | 0 | 0 | 0 | 0 | 0 | 0 | 1091300 | 0 | 0 | 1749200 | 0 | 0 | 63504 | 1207800 | 0 | 0 | 0 | 0 |
| LLFEVFDEN | NEDD4L; NEDD | Q96PU5; A0A | NED4L_HUMAN | Unmodified | 695750 | 611510 | 739560 | 518960 | 0 | 0 | 0 | 0 | 366300 | 0 | 0 | 1232500 | 0 | 0 | 1823700 | 836190 | 11124 | 0 |
| LLFGHSTE | ENOPH1; ENOF | Q9UHY7; D6 | ENOPH_HUMA | Unmodified | 0 | 0 | 0 | 0 | 0 | 0 | 0 | 0 | 0 | 903640 | 1182000 | 715720 | 2491600 | 0 | 0 | 279210 | 567240 | 0 |
| LLFPAVDDI | TOP2B; TOP2B | Q02880; E9F | TOP2B_HUMAN | Unmodified | 252330 | 0 | 0 | 0 | 0 | 0 | 0 | 0 | 0 | 0 | 0 | 2054700 | 0 | 0 | 0 | 0 | 0 | 0 |
| LLFPSSHLF | PIK3C2B; PIK3C | O00750; F5C | P3C2B_HUMAN | Unmodified | 998380 | 255860 | 246220 | 536910 | 150860 | 0 | 0 | 0 | 0 | 23184 | 0 | 229190 | 0 | 0 | 2180700 | 2695100 | 0 | 0 |
| LLFQLAQLI | MAU2; MAU2; M | Q9Y6X3; V9C | SCC4_HUMAN | Unmodified | 47887 | 0 | 0 | 0 | 0 | 0 | 0 | 0 | 0 | 0 | 0 | 533190 | 0 | 0 | 0 | 0 | 0 | 0 |
| LLFQPDQN | KIF23; KIF23; KI | Q02241; A0A | KIF23_HUMAN | Unmodified | 0 | 0 | 0 | 205440 | 0 | 506510 | 0 | 0 | 432870 | 0 | 0 | 1135200 | 0 | 0 | 0 | 0 | 0 | 0 |
| LLFQPDQN | KIF23; KIF23; KI | Q02241; A0A | KIF23_HUMAN | Unmodified | 0 | 0 | 0 | 172590 | 2701300 | 532960 | 0 | 0 | 251210 | 354100 | 309790 | 0 | 174090 | 0 | 0 | 0 | 532930 | 691410 |
| LLFQSYNVI | NCOA4; NCOA4 | Q13772; B4D | NCOA4_HUMA | Unmodified | 0 | 0 | 0 | 11553 | 185850 | 24407 | 813310 | 0 | 2809900 | 190640 | 0 | 0 | 243010 | 470390 | 0 | 0 | 105540 | 1761800 |
| LLFRNTNVI | CCDC43; CCDC | Q96MW1; Q | CCD43_HUMA | Unmodified | 270570 | 0 | 107400 | 123880 | 0 | 0 | 0 | 0 | 280830 | 0 | 0 | 241710 | 0 | 0 | 215970 | 0 | 161920 | 335650 |
| LLFSEDQQ | SUN1; SUN1; S | O94901; H0Y | SUN1_HUMAN | Unmodified | 0 | 0 | 0 | 0 | 0 | 0 | 0 | 0 | 24660000 | 1057800 | 0 | 0 | 0 | 0 | 0 | 0 | 0 | 0 |
| LLFSGGHW | NBEAL2; NBEAI | Q6ZNJ1; H0Y | NBEL2_HUMAN | Unmodified | 0 | 0 | 0 | 45222 | 0 | 0 | 0 | 0 | 0 | 0 | 55362 | 849240 | 0 | 0 | 216180 | 115570 | 0 | 0 |
| LLFSHDLVS | UBE2O | Q9C0C9 | UBE2O_HUMAI | Unmodified | 0 | 0 | 440760 | 0 | 0 | 0 | 0 | 0 | 239240 | 0 | 566170 | 4211800 | 102790 | 0 | 313400 | 189340 | 46071 | 0 |
| LLFYATSR | ZC3HAV1; ZC3H | Q7Z2W4; C9 | ZCCHV_HUMA | Unmodified | 1512800 | 840520 | 1347800 | 3555300 | 4662300 | 1744700 | 2529600 | 1326300 | 0 | 507700 | 18533 | 1204800 | 262380 | 0 | 1709900 | 1236000 | 0 | 232620 |
| LLGASELPI | KPNA2; KPNA2; | P52292; A0A | IMA1_HUMAN | Unmodified | 0 | 0 | 0 | 0 | 1564500 | 830940 | 0 | 0 | 4600700 | 1459800 | 1928100 | 6251900 | 0 | 0 | 0 | 0 | 1570800 | 0 |
| LLGELLLDF | CAB39; CAB39 | Q9Y376; A0A | CAB39_HUMAI | Oxidation (M) | 1813000 | 1356700 | 0 | 1731800 | 0 | 0 | 0 | 0 | 0 | 0 | 0 | 421670 | 0 | 0 | 450220 | 244220 | 99093 | 0 |
| LLGELLQDI | TALDO1; TALDC | P37837; F2Z | TALDO_HUMAN | Unmodified | 0 | 0 | 579740 | 0 | 13178000 | 2759000 | 413970 | 272950 | 3109100 | 2644500 | 11701000 | 38351000 | 8090400 | 3460400 | 3832700 | 0 | 3144800 | 3306700 |
| LLGELLQDI | TALDO1; TALDC | P37837; F2Z | TALDO_HUMAN | Unmodified | 0 | 0 | 0 | 0 | 6376800 | 773840 | 0 | 0 | 0 | 1598800 | 0 | 0 | 5895700 | 5274100 | 0 | 0 | 5871400 | 1668300 |
| LLGFGSALI | AP2A1 | O95782 | AP2A1_HUMAN | Unmodified | 0 | 176740 | 0 | 95716 | 0 | 0 | 325330 | 0 | 0 | 0 | 0 | 0 | 421790 | 0 | 0 | 0 | 1175500 | 960260 |
| LLGGQIGLI | GIPC1; GIPC1; | O14908; K7E | GIPC1_HUMAN | Unmodified | 0 | 0 | 0 | 0 | 0 | 0 | 0 | 0 | 0 | 0 | 0 | 0 | 406550 | 394500 | 0 | 0 | 2124300 | 0 |
| LLGGQLGL | GIPC2 | Q8TF65 | GIPC2_HUMAN | Unmodified | 0 | 0 | 0 | 129780 | 8253900 | 287680 | 250340 | 0 | 0 | 396250 | 0 | 0 | 4247700 | 1339500 | 780140 | 0 | 2895300 | 6262500 |
| LLGGVTIAQ | H2AX; H2AC1; I | P16104; Q96 | H2AX_HUMAN | Unmodified | 574320 | 16675000 | 0 | 0 | 876440 | 71434 | 582680 | 0 | 0 | 0 | 0 | 613440 | 0 | 0 | 123790 | 0 | 0 | 626940 |
| LLGGVTIAQ | H2AX; H2AC1; I | P16104; Q96 | H2AX_HUMAN | Unmodified | 2538700 | 7095100 | 0 | 0 | 1021400 | 34536 | 83752 | 0 | 0 | 0 | 0 | 169950 | 72300 | 0 | 0 | 0 | 0 | 166920 |
| LLGHEVED | UPF1 | Q92900 | RENT1_HUMAN | Unmodified | 709440 | 1643200 | 858980 | 905010 | 189370 | 0 | 804880 | 0 | 128710 | 0 | 0 | 724380 | 536750 | 295290 | 719330 | 600320 | 215750 | 238880 |
| LLGIETPLPI | ICAM1; ICAM1 | P05362; K7E | ICAM1_HUMAN | Unmodified | 0 | 0 | 0 | 0 | 3181200 | 368750 | 0 | 0 | 0 | 0 | 0 | 0 | 0 | 0 | 0 | 0 | 201880 | 445040 |
| LLGIETPLPI | ICAM1; ICAM1 | P05362; K7E | ICAM1_HUMAN | Unmodified | 0 | 0 | 1088700 | 1575500 | 2.32E+08 | 26323000 | 0 | 0 | 448110 | 359720 | 0 | 0 | 1181300 | 0 | 1091600 | 1025900 | 17911000 | 27609000 |
| LLGIFENQI | NUDT4; NUDT4 | Q9NZJ9; F8V | NUDT4_HUMAI | Unmodified | 0 | 0 | 0 | 0 | 0 | 0 | 0 | 0 | 0 | 0 | 0 | 0 | 0 | 0 | 386280 | 214140 | 0 | 0 |
| LLGKVTIAQ | H2AC18; H2AC | Q6FI13; Q99 | H2A2A_HUMAN | Unmodified | 307560 | 6538700 | 0 | 0 | 1442500 | 95298 | 2124700 | 0 | 0 | 59401 | 0 | 36051 | 190650 | 0 | 0 | 0 | 0 | 1573500 |
| LLGNTFVAI | BAG6; BAG6; BA | P46379; F6S | BAG6_HUMAN | Unmodified | 0 | 0 | 0 | 0 | 0 | 0 | 0 | 0 | 0 | 0 | 0 | 1866200 | 0 | 0 | 0 | 0 | 0 | 0 |
| LLGPDAAIN | DCTN2; DCTN2 | Q13561; A8K | DCTN2_HUMAI | Unmodified | 0 | 0 | 0 | 583800 | 3004000 | 735510 | 0 | 593110 | 787900 | 0 | 510100 | 117090 | 3270000 | 2356100 | 0 | 0 | 0 | 2298200 |
| LLGPGPAA | NAGLU | P54802 | ANAG_HUMAN | Unmodified | 0 | 0 | 0 | 0 | 0 | 0 | 192830 | 0 | 450840 | 0 | 0 | 0 | 0 | 0 | 0 | 0 | 0 | 0 |
| LLGPGPAA | NAGLU | P54802 | ANAG_HUMAN | Unmodified | 0 | 0 | 0 | 0 | 0 | 0 | 0 | 0 | 0 | 1192800 | 0 | 0 | 0 | 0 | 0 | 0 | 0 | 0 |
| LLGPSAAAI | HUWE1 | Q7Z6Z7 | HUWE1_HUMA | Unmodified | 0 | 0 | 0 | 0 | 0 | 0 | 0 | 0 | 614510 | 467780 | 0 | 326030 | 0 | 0 | 0 | 0 | 245100 | 534950 |
| LLGPSAAAI | HUWE1 | Q7Z6Z7 | HUWE1_HUMA | Unmodified | 0 | 0 | 0 | 0 | 106050 | 0 | 49947 | 0 | 0 | 154120 | 0 | 0 | 348160 | 124930 | 0 | 0 | 213940 | 193940 |
| LLGPSETSS | FARP2 | H7C3M7 | H7C3M7_HUM | Unmodified | 0 | 0 | 451680 | 590910 | 561430 | 0 | 0 | 0 | 379060 | 0 | 0 | 0 | 0 | 0 | 695090 | 687490 | 237310 | 0 |
| LLGPVDKG | VSIR | Q9H7M9 | VISTA_HUMAN | Unmodified | 0 | 0 | 0 | 0 | 2336100 | 0 | 0 | 0 | 287450 | 734760 | 0 | 0 | 0 | 0 | 0 | 0 | 860580 | 1839500 |
| LLGPVLVK | PFDN6; PFDN6 | O15212; A2A | PFD6_HUMAN | Unmodified | 0 | 0 | 0 | 0 | 2031800 | 0 | 0 | 0 | 759340 | 736150 | 0 | 0 | 6634800 | 0 | 0 | 0 | 1213100 | 627000 |
| LLGPVLVK | PFDN6; PFDN6 | O15212; A2A | PFD6_HUMAN | Unmodified | 0 | 0 | 0 | 0 | 6149100 | 708930 | 1470100 | 0 | 0 | 1869700 | 0 | 0 | 7030400 | 2115200 | 0 | 0 | 9421300 | 3735200 |
| LLGQFSEKI | BLVRA | P53004 | BIEA_HUMAN E | Unmodified | 0 | 0 | 0 | 73259 | 474490 | 0 | 732570 | 642440 | 0 | 0 | 0 | 0 | 0 | 405170 | 0 | 0 | 951470 | 336360 |
| LLGQFTLIG | HSPA9; HSPA9; | P38646; H0Y | GRP75_HUMAN | Unmodified | 0 | 0 | 0 | 0 | 317650 | 0 | 172630 | 0 | 8888100 | 1501300 | 395000 | 188460 | 566700 | 0 | 0 | 0 | 0 | 177300 |
| LLGREVPEI | TRMT10C; TRM | Q7L0Y3; C9J | TM10C_HUMAI | Unmodified | 0 | 0 | 0 | 0 | 0 | 0 | 0 | 0 | 468710 | 0 | 0 | 0 | 0 | 0 | 0 | 0 | 0 | 0 |
| LLGRQQPD | IARS2 | Q9NSE4 | SYIM_HUMAN I | Unmodified | 0 | 0 | 0 | 0 | 0 | 0 | 827220 | 224300 | 0 | 1602800 | 0 | 0 | 350430 | 195750 | 0 | 0 | 197390 | 528420 |
| LLGSGYGG | CLASP1; CLASF | Q7Z460; F8V | CLAP1_HUMAN | Unmodified | 0 | 0 | 0 | 0 | 287250 | 0 | 0 | 0 | 0 | 0 | 0 | 223760 | 0 | 0 | 0 | 0 | 236990 | 0 |
| LLGSVDYD | PIK3C2B; PIK3C | O00750; F5C | P3C2B_HUMAN | Unmodified | 0 | 0 | 59814 | 74094 | 0 | 0 | 0 | 0 | 0 | 0 | 0 | 0 | 0 | 0 | 0 | 214420 | 0 | 0 |
| LLGSVQQD | PBXIP1 | Q96AQ6 | PBIP1_HUMAN | Unmodified | 0 | 0 | 0 | 0 | 0 | 0 | 0 | 96743 | 5214000 | 0 | 1466500 | 813070 | 0 | 0 | 0 | 0 | 0 | 0 |
| LLGTIYTAAI | HSDL2 | Q6YN16 | HSDL2_HUMAI | Unmodified | 0 | 0 | 0 | 0 | 428230 | 0 | 1054700 | 0 | 18363000 | 16941000 | 0 | 0 | 292040 | 0 | 0 | 0 | 0 | 246710 |
| LLGVAHLFI | LIPE; LIPE; LIPE | Q05469; M0 | LIPS_HUMAN H | Unmodified | 0 | 0 | 0 | 0 | 0 | 0 | 0 | 0 | 0 | 0 | 0 | 147560 | 0 | 0 | 0 | 0 | 0 | 0 |
| LLGWIQNK | FLNC; FLNB; FL | Q14315; O7 | FLNC_HUMAN | Unmodified | 0 | 281090 | 1857900 | 0 | 0 | 0 | 0 | 0 | 0 | 0 | 0 | 2708000 | 0 | 0 | 1546600 | 798160 | 0 | 0 |
| LLGWIQNK | FLNA; FLNA; FL | P21333; A0A | FLNA_HUMAN | Unmodified | 200970 | 168260 | 0 | 282490 | 0 | 0 | 0 | 0 | 107740 | 0 | 0 | 0 | 0 | 0 | 1279800 | 50729 | 0 | 0 |
| LLHDGIQP | PDE2A; PDE2A; | O00408; E9F | PDE2A_HUMAN | Unmodified | 0 | 0 | 0 | 0 | 0 | 0 | 0 | 0 | 0 | 0 | 79349 | 3973900 | 0 | 0 | 0 | 0 | 0 | 0 |
| LLHDQVGV | COPA; COPA; C | P53621; A0A | COPA_HUMAN | Unmodified | 0 | 0 | 0 | 0 | 0 | 0 | 0 | 0 | 2782700 | 431950 | 681630 | 8790500 | 0 | 0 | 0 | 0 | 0 | 633730 |
| LLHEVQEL | DCTN2; DCTN2 | Q13561; A8K | DCTN2_HUMAI | Unmodified | 0 | 0 | 0 | 402410 | 334430 | 0 | 0 | 0 | 1080100 | 197860 | 1283700 | 0 | 452670 | 0 | 629210 | 0 | 0 | 0 |
| LLHIEELRE | RTRAF | Q9Y224 | RTRAF_HUMAN | Unmodified | 546870 | 191800 | 822100 | 1896400 | 3306200 | 2677400 | 0 | 0 | 822240 | 494340 | 51257000 | 7385000 | 4558000 | 431550 | 639570 | 851480 | 308940 | 3003400 |
| LLHKYPFIL | MCTS1; MCTS2 | Q9ULC4; A0 | MCTS1_HUMAI | Unmodified | 301610 | 277850 | 0 | 500660 | 0 | 0 | 0 | 0 | 407500 | 497740 | 0 | 2163200 | 886190 | 695910 | 266110 | 0 | 452270 | 332930 |
| LLHLPLSLT | LRRC1 | Q9BTT6 | LRRC1_HUMAN | Unmodified | 0 | 0 | 0 | 0 | 0 | 0 | 0 | 0 | 2507600 | 0 | 1950900 | 6520400 | 0 | 0 | 0 | 1453900 | 0 | 0 |

| | | | | | | | | | | | | | | | | | | | | | | |
|---|---|---|---|---|---|---|---|---|---|---|---|---|---|---|---|---|---|---|---|---|---|---|
| LLHPPSHW | CTNNB1; CTNN | P35222; A0A | CTNB1_HUMAI | Unmodified | 470030 | 0 | 1141800 | 738350 | 390000 | 0 | 0 | 0 | 0 | 0 | 0 | 2199900 | 405000 | 0 | 2085000 | 2147100 | 1273800 | 0 |
| LLHPSPDL | CMIP; CMIP; CN | Q8IY22; A0A | CMIP_HUMAN | Unmodified | 0 | 0 | 0 | 0 | 0 | 0 | 0 | 0 | 1292600 | 46082 | 323340 | 1400400 | 0 | 0 | 0 | 0 | 30879 | 0 |
| LLHQLGFS | USP24 | Q9UPU5 | UBP24_HUMAI | Unmodified | 0 | 0 | 0 | 61488 | 0 | 0 | 0 | 0 | 177760 | 0 | 35193 | 312540 | 0 | 0 | 0 | 51840 | 0 | 0 |
| LLHQTGLS | ECHDC1; ECHD | Q9NTX5; J3K | ECHD1_HUMA | Unmodified | 0 | 0 | 0 | 0 | 499110 | 0 | 0 | 0 | 0 | 0 | 328460 | 3344700 | 392420 | 0 | 0 | 0 | 0 | 0 |
| LLHSPPDM | ADGRG2 | Q8IZP9 | AGRG2_HUMA | Unmodified | 0 | 0 | 0 | 0 | 1294300 | 209310 | 0 | 0 | 8038500 | 444960 | 0 | 0 | 0 | 0 | 0 | 0 | 73580 | 0 |
| LLHSPPDM | ADGRG2 | Q8IZP9 | AGRG2_HUMA | Oxidation (M) | 0 | 0 | 0 | 0 | 0 | 0 | 0 | 0 | 733650 | 179470 | 0 | 0 | 0 | 0 | 0 | 0 | 67093 | 0 |
| LLHSPPDM | ADGRG2 | Q8IZP9 | AGRG2_HUMA | Unmodified | 0 | 0 | 0 | 0 | 1697500 | 0 | 0 | 0 | 0 | 799040 | 0 | 0 | 0 | 0 | 0 | 0 | 0 | 0 |
| LLHSPPDM | ADGRG2 | Q8IZP9 | AGRG2_HUMA | Oxidation (M) | 0 | 0 | 0 | 0 | 626170 | 0 | 0 | 0 | 0 | 765630 | 0 | 0 | 0 | 0 | 0 | 0 | 0 | 0 |
| LLHVAVSD | PSMD1; PSMD1 | Q99460; A0A | PSMD1_HUMA | Unmodified | 0 | 0 | 109720 | 0 | 103480 | 0 | 0 | 52359 | 0 | 0 | 115920 | 1491600 | 0 | 0 | 87964 | 0 | 0 | 0 |
| LLHVAVSD | PSMD1; PSMD1 | Q99460; A0A | PSMD1_HUMA | Unmodified | 0 | 0 | 0 | 0 | 0 | 0 | 0 | 115230 | 0 | 0 | 0 | 6906100 | 0 | 0 | 0 | 0 | 0 | 0 |
| LLHWNGH | GLT8D1 | Q68CQ7 | GL8D1_HUMAI | Unmodified | 0 | 0 | 0 | 151660 | 0 | 0 | 72093 | 0 | 6684700 | 357560 | 0 | 0 | 0 | 0 | 122720 | 0 | 0 | 0 |
| LLHYAGEV | MYO1C; MYO1C | O00159; F5H | MYO1C_HUMA | Unmodified | 40795 | 0 | 0 | 2406600 | 67804 | 705830 | 146320 | 0 | 0 | 0 | 0 | 0 | 0 | 0 | 0 | 1441000 | 639500 | 0 |
| LLIDEAILK | SARS1; SARS1 | P49591; Q5T | SYSC_HUMAN | Unmodified | 0 | 0 | 622740 | 996440 | 947980 | 0 | 403210 | 252260 | 3344600 | 789200 | 1539800 | 9993800 | 2411000 | 1271800 | 1401600 | 908430 | 203540 | 85342 |
| LLIDGDGA | THOC7; THOC7 | Q6I9Y2; H7C | THOC7_HUMA | Unmodified | 0 | 0 | 0 | 0 | 0 | 0 | 0 | 0 | 0 | 0 | 65201 | 396710 | 0 | 0 | 0 | 78935 | 0 | 0 |
| LLIEMEQR | CKB; CKB | P12277; H0Y | KCRB_HUMAN | Oxidation (M) | 169450 | 0 | 1046600 | 0 | 0 | 0 | 0 | 0 | 204490 | 85049 | 0 | 10893000 | 0 | 0 | 3305500 | 3303300 | 0 | 0 |
| LLIEMEQR | CKB; CKB | P12277; H0Y | KCRB_HUMAN | Unmodified | 0 | 0 | 0 | 761410 | 0 | 0 | 0 | 0 | 0 | 0 | 0 | 0 | 0 | 0 | 1917400 | 1088300 | 0 | 0 |
| LLIGADGH | COQ6; COQ6; C | Q9Y2Z9; A0A | COQ6_HUMAN | Unmodified | 0 | 0 | 0 | 0 | 0 | 0 | 0 | 0 | 269160 | 65977 | 0 | 0 | 0 | 0 | 0 | 0 | 0 | 0 |
| LLIHQSLAG | HNRNPK; HNRI | P61978; Q5T | HNRPK_HUMA | Unmodified | 4563400 | 4307700 | 2675600 | 2524200 | 4334300 | 897030 | 1480200 | 278100 | 7608100 | 2468000 | 11781000 | 76107000 | 4891100 | 0 | 2673800 | 0 | 0 | 1302700 |
| LLIIDYSENI | PLXNA2 | O75051 | PLXA2_HUMAN | Unmodified | 182160 | 0 | 0 | 0 | 217930 | 809910 | 0 | 0 | 1015300 | 0 | 0 | 629680 | 0 | 0 | 338520 | 0 | 0 | 0 |
| LLIIGDSGV | RAB35; RAB35 | Q15286; F5H | RAB35_HUMAN | Unmodified | 0 | 0 | 0 | 0 | 0 | 0 | 222750 | 0 | 0 | 0 | 0 | 0 | 415540 | 119730 | 0 | 0 | 0 | 0 |
| LLINAHEQ | PLCB4; PLCB4 | Q15147; A0A | PLCB4_HUMAN | Unmodified | 0 | 0 | 34977 | 0 | 0 | 0 | 0 | 0 | 0 | 0 | 0 | 0 | 0 | 0 | 0 | 84434 | 151480 | 909510 |
| LLINAHEQ | PLCB4; PLCB4 | Q15147; A0A | PLCB4_HUMAN | Unmodified | 0 | 0 | 0 | 0 | 0 | 0 | 0 | 0 | 0 | 0 | 0 | 0 | 0 | 0 | 0 | 0 | 1153600 | 982700 |
| LLINESLIK | ABCA12 | Q86UK0 | ABCAC_HUMAI | Unmodified | 0 | 0 | 0 | 0 | 0 | 0 | 0 | 118020 | 42380 | 0 | 0 | 198760 | 0 | 0 | 0 | 0 | 0 | 0 |
| LLINLLK | PSME4; PSME4 | Q14997; F8W | PSME4_HUMAI | Unmodified | 0 | 0 | 0 | 0 | 0 | 0 | 0 | 32449 | 0 | 0 | 0 | 452840 | 0 | 0 | 0 | 0 | 0 | 0 |
| LLIPERDPL | TNS3 | Q68CZ2 | TENS3_HUMAN | Unmodified | 0 | 1455400 | 0 | 0 | 4431100 | 0 | 0 | 0 | 0 | 289030 | 0 | 0 | 0 | 0 | 0 | 0 | 2936000 | 1805900 |
| LLIPGNYK | CPE | P16870 | CBPE_HUMAN | Unmodified | 0 | 0 | 0 | 0 | 0 | 0 | 0 | 0 | 93709000 | 3392200 | 0 | 0 | 0 | 0 | 0 | 26306 | 0 | 0 |
| LLIPGNYKL | CPE | P16870 | CBPE_HUMAN | Unmodified | 0 | 0 | 0 | 0 | 0 | 0 | 0 | 0 | 0 | 719450 | 0 | 0 | 0 | 0 | 0 | 0 | 0 | 0 |
| LLIPGNYKL | CPE | P16870 | CBPE_HUMAN | Unmodified | 0 | 0 | 0 | 0 | 0 | 0 | 0 | 0 | 711940 | 1374500 | 0 | 0 | 0 | 0 | 0 | 0 | 0 | 0 |
| LLIPVAEG | FERMT2; FERM | Q96AC1; A0A | FERM2_HUMAN | Unmodified | 0 | 0 | 0 | 0 | 4463400 | 283640 | 0 | 0 | 0 | 0 | 0 | 0 | 13831 | 0 | 0 | 0 | 2113200 | 3608600 |
| LLIPVAEG | FERMT2; FERM | Q96AC1; A0A | FERM2_HUMAN | Oxidation (M) | 0 | 0 | 0 | 0 | 7703800 | 613320 | 0 | 0 | 0 | 0 | 0 | 0 | 0 | 0 | 0 | 0 | 3268600 | 3302700 |
| LLIQAAIVR | CUL1; CUL1; CI | Q13616; A0A | CUL1_HUMAN | Unmodified | 0 | 0 | 0 | 0 | 0 | 0 | 0 | 0 | 0 | 0 | 96840 | 415950 | 0 | 0 | 0 | 0 | 0 | 0 |
| LLIQAEQTS | CACNA2D1 | P54289 | CA2D1_HUMAI | Unmodified | 626930 | 237950 | 0 | 0 | 0 | 0 | 0 | 0 | 2401700 | 95822 | 0 | 0 | 0 | 0 | 0 | 0 | 0 | 0 |
| LLIQAEQTS | CACNA2D1 | P54289 | CA2D1_HUMAI | Oxidation (M) | 491860 | 276290 | 0 | 0 | 0 | 0 | 0 | 0 | 0 | 0 | 0 | 0 | 0 | 0 | 0 | 0 | 0 | 0 |
| LLISDTQHY | MPP7; MPP7; M | Q5T2T1; U5C | MPP7_HUMAN | Unmodified | 0 | 0 | 0 | 0 | 1315300 | 0 | 0 | 0 | 0 | 0 | 0 | 0 | 469030 | 0 | 0 | 0 | 232370 | 702250 |
| LLISLSQGN | ANXA11 | P50995 | ANX11_HUMAN | Oxidation (M) | 0 | 0 | 0 | 0 | 0 | 0 | 0 | 0 | 8475200 | 0 | 0 | 753270 | 768520 | 0 | 0 | 0 | 0 | 0 |
| LLIVSTTPY | RUVBL2; RUVBI | Q9Y230; M0I | RUVB2_HUMAI | Unmodified | 0 | 0 | 0 | 0 | 0 | 0 | 0 | 0 | 0 | 0 | 283040 | 9839600 | 0 | 0 | 0 | 303070 | 0 | 0 |
| LLIYETEAK | RBM25 | P49756 | RBM25_HUMAI | Unmodified | 0 | 0 | 0 | 0 | 0 | 0 | 0 | 97219 | 0 | 0 | 0 | 769580 | 0 | 0 | 0 | 0 | 0 | 0 |
| LLKFPLGP | ACP2; ACP2; AC | P11117; B7Z | PPAL_HUMAN I | Unmodified | 0 | 0 | 0 | 0 | 0 | 0 | 0 | 0 | 566870 | 0 | 0 | 0 | 0 | 0 | 0 | 0 | 0 | 0 |
| LLKGDLAYI | TMX3 | Q96JJ7 | TMX3_HUMAN | Unmodified | 0 | 0 | 0 | 0 | 114550 | 0 | 0 | 0 | 846010 | 265560 | 0 | 65915 | 0 | 0 | 0 | 0 | 17238 | 0 |
| LLKGDLAYI | TMX3 | Q96JJ7 | TMX3_HUMAN | Unmodified | 0 | 0 | 0 | 0 | 0 | 0 | 0 | 0 | 5865000 | 1192900 | 298800 | 0 | 0 | 0 | 0 | 0 | 0 | 0 |
| LLKGEGPK | CRABP2 | P29373 | RABP2_HUMAN | Unmodified | 0 | 0 | 0 | 0 | 0 | 367980 | 1252200 | 496870 | 0 | 106910 | 0 | 0 | 198070 | 538830 | 0 | 0 | 561730 | 1184800 |
| LLKIPVDTY | VPS35 | Q96QK1 | VPS35_HUMAN | Unmodified | 0 | 0 | 0 | 26416 | 0 | 0 | 67040 | 0 | 0 | 0 | 0 | 286230 | 0 | 0 | 0 | 0 | 0 | 0 |
| LLKPGEEP | PGRMC2; PGRM | O15173; U3I | PGRC2_HUMAI | Unmodified | 0 | 0 | 0 | 0 | 0 | 0 | 0 | 0 | 229030 | 0 | 0 | 2578500 | 0 | 0 | 0 | 0 | 0 | 0 |
| LLKYPDVIQ | CNNM4 | Q6P4Q7 | CNNM4_HUMA | Unmodified | 0 | 0 | 0 | 0 | 0 | 0 | 0 | 0 | 0 | 0 | 0 | 1183100 | 0 | 0 | 0 | 0 | 0 | 0 |
| LLLAALVQD | CALCA; CALCA | P01258; P06 | CALC_HUMAN | Oxidation (M) | 0 | 0 | 0 | 0 | 0 | 0 | 0 | 0 | 650860 | 145240 | 0 | 0 | 0 | 0 | 0 | 0 | 0 | 0 |
| LLLADNPLI | TPBGL | P0DKB5 | TPBGL_HUMAN | Unmodified | 0 | 0 | 0 | 0 | 0 | 0 | 0 | 0 | 796050 | 0 | 0 | 540640 | 0 | 0 | 0 | 0 | 0 | 0 |
| LLLAEALLE | TTC7A; TTC7A; | Q9ULT0; G5I | TTC7A_HUMAN | Unmodified | 0 | 0 | 0 | 0 | 346580 | 0 | 0 | 0 | 0 | 0 | 0 | 0 | 0 | 0 | 0 | 0 | 0 | 0 |
| LLLAGYDD | GNB1; GNB1 | P62873; B3K | GBB1_HUMAN | Unmodified | 0 | 211440 | 0 | 0 | 225340 | 0 | 0 | 0 | 56568 | 146120 | 0 | 0 | 196170 | 249440 | 219020 | 0 | 163800 | 0 |
| LLLAIVGQ | ANXA7 | P20073 | ANXA7_HUMAN | Unmodified | 0 | 0 | 0 | 0 | 34465 | 0 | 67613 | 0 | 2214400 | 334710 | 0 | 0 | 0 | 0 | 0 | 0 | 0 | 13298 |
| LLLAPSSK | MOV10; MOV10 | Q9HCE1; Q5 | MOV10_HUMA | Unmodified | 2038500 | 2374400 | 630750 | 1143200 | 0 | 0 | 0 | 0 | 0 | 0 | 0 | 1876500 | 0 | 0 | 927670 | 1489700 | 0 | 0 |
| LLLASFKNE | STX8; STX8 | Q9UNK0; I3L | STX8_HUMAN | Unmodified | 0 | 0 | 0 | 0 | 0 | 0 | 0 | 0 | 578700 | 76022 | 252810 | 855370 | 0 | 0 | 326810 | 161650 | 0 | 0 |
| LLLASTSEV | UXS1; UXS1; U) | Q8NBZ7; C9 | UXS1_HUMAN | Unmodified | 0 | 0 | 0 | 0 | 0 | 0 | 0 | 0 | 11556000 | 1413000 | 0 | 0 | 0 | 0 | 0 | 0 | 0 | 0 |
| LLLATMESI | ALDH1A1; ALDI | P00352; Q59 | AL1A1_HUMAN | 2 Oxidation (M) | 85478 | 322570 | 0 | 0 | 0 | 0 | 0 | 0 | 0 | 0 | 0 | 12462000 | 0 | 0 | 0 | 0 | 0 | 0 |
| LLLATMESI | ALDH1A1; ALDI | P00352; Q59 | AL1A1_HUMAN | Oxidation (M) | 0 | 0 | 0 | 0 | 0 | 0 | 0 | 0 | 0 | 0 | 476660 | 42013000 | 0 | 120990 | 0 | 0 | 220930 | 0 |
| LLLATMESI | ALDH1A1; ALDI | P00352; Q59 | AL1A1_HUMAN | Unmodified | 0 | 0 | 0 | 0 | 0 | 0 | 0 | 0 | 0 | 0 | 0 | 16566000 | 542290 | 0 | 0 | 0 | 0 | 331650 |
| LLLDGAPLI | HDHD2; HDHD | Q9H0R4; K7 | HDHD2_HUMA | Unmodified | 0 | 0 | 0 | 0 | 0 | 0 | 0 | 0 | 617390 | 291270 | 2175900 | 3839400 | 0 | 0 | 0 | 0 | 0 | 0 |
| LLLDIPLQT | RGPD3; RGPD2 | A6NKT7; P0I | RGPD3_HUMAI | Unmodified | 0 | 0 | 0 | 0 | 0 | 0 | 0 | 0 | 0 | 10634 | 0 | 1081700 | 0 | 0 | 0 | 0 | 0 | 0 |
| LLLDLFR | VIPAS39; VIPAS | Q9H9C1; G3 | SPE39_HUMAN | Unmodified | 0 | 0 | 0 | 0 | 0 | 0 | 0 | 0 | 0 | 0 | 10707 | 536920 | 0 | 0 | 0 | 0 | 0 | 0 |
| LLLDPEFTL | WDR11 | Q9BZH6 | WDR11_HUMA | Unmodified | 0 | 0 | 0 | 0 | 0 | 0 | 0 | 0 | 0 | 0 | 0 | 160310 | 0 | 0 | 0 | 0 | 0 | 0 |
| LLLDPSSPI | LAMTOR1; LAM | Q6IAA8; H0Y | LTOR1_HUMAN | Unmodified | 0 | 250720 | 1355200 | 0 | 2119000 | 0 | 811190 | 425180 | 1742800 | 844790 | 923250 | 5299000 | 2175300 | 1933900 | 850850 | 373470 | 0 | 1228000 |
| LLLDQEQK | DNAJC8 | O75937 | DNJC8_HUMAN | Unmodified | 0 | 0 | 0 | 0 | 0 | 0 | 0 | 0 | 0 | 142620 | 0 | 2580600 | 0 | 0 | 0 | 0 | 0 | 0 |
| LLLDRGGQ | ANK2; ANK2; AI | Q01484; A0A | ANK2_HUMAN | Unmodified | 0 | 0 | 0 | 0 | 0 | 0 | 0 | 0 | 0 | 258630 | 0 | 0 | 0 | 0 | 0 | 0 | 0 | 0 |
| LLLDTFEYQ | CENPV; CENPV | Q7Z7K6; A0A | CENPV_HUMAI | Unmodified | 201690 | 20213 | 0 | 0 | 0 | 0 | 2272900 | 205160 | 0 | 20638 | 73037 | 2109800 | 0 | 0 | 0 | 0 | 0 | 0 |
| LLLEEALQD | APPL2; APPL2 | Q8NEU8; F8 | DP13B_HUMAN | Unmodified | 68195 | 0 | 67349 | 173880 | 127260 | 0 | 0 | 0 | 142880 | 164550 | 0 | 0 | 0 | 287960 | 153840 | 254110 | 0 | 71016 |
| LLLEEDENI | DNAJC13 | O75165 | DJC13_HUMAN | Unmodified | 0 | 0 | 4296800 | 0 | 0 | 0 | 0 | 29566 | 0 | 0 | 0 | 4203500 | 4673300 | 1052300 | 0 | 3984500 | 0 | 0 |
| LLLEFTDTS | GSTM3; GSTM3 | P21266; A0A | GSTM3_HUMAI | Unmodified | 0 | 80444 | 13674 | 54878 | 0 | 0 | 0 | 0 | 0 | 0 | 0 | 0 | 0 | 0 | 62852 | 85948 | 0 | 0 |
| LLLENNAN | ANK1 | P16157 | ANK1_HUMAN | Unmodified | 588490 | 1234300 | 342730 | 447720 | 0 | 0 | 188910 | 125000 | 0 | 0 | 0 | 0 | 0 | 0 | 789900 | 724300 | 0 | 0 |
| LLLFSDGN | COL6A1; COL6A | P12109; A0A | CO6A1_HUMAI | Unmodified | 0 | 0 | 1092800 | 510270 | 627440 | 237650 | 0 | 109680 | 931710 | 45020 | 0 | 320670 | 42991 | 0 | 0 | 385840 | 502350 | 0 |
| LLLGNKCD | RAB13; RAB13 | P51153; A0A | RAB13_HUMAN | Oxidation (M) | 1477200 | 765350 | 312920 | 541930 | 307320 | 0 | 0 | 0 | 480590 | 75206 | 0 | 0 | 247800 | 102400 | 418360 | 403110 | 179390 | 183730 |
| LLLGNKCD | RAB13; RAB13 | P51153; A0A | RAB13_HUMAN | Unmodified | 3029300 | 3875600 | 550080 | 2073300 | 780310 | 0 | 0 | 0 | 1878600 | 220430 | 154330 | 859380 | 649890 | 0 | 1367400 | 875480 | 144650 | 249690 |
| LLLHLEELQ | MOV10; MOV10 | Q9HCE1; Q5 | MOV10_HUMA | Oxidation (M) | 1416400 | 0 | 0 | 0 | 0 | 0 | 0 | 0 | 0 | 0 | 0 | 0 | 0 | 0 | 0 | 2005900 | 0 | 0 |
| LLLIGDSG | RAB1C; RAB10; | Q92928; P61 | RAB1C_HUMAI | Unmodified | 42108000 | 37310000 | 35620000 | 24154000 | 12504000 | 4054100 | 8874300 | 6698300 | 25741000 | 8983800 | 22767000 | 1.2E+08 | 11139000 | 6752200 | 51754000 | 40094000 | 4951500 | 6550800 |
| LLLIGDSG | RAB1C; RAB1A; | Q92928; P62 | RAB1C_HUMAI | Unmodified | 0 | 0 | 0 | 0 | 0 | 0 | 226300 | 0 | 0 | 408080 | 0 | 0 | 455180 | 203270 | 0 | 0 | 324750 | 0 |
| LLLIGDSG | RAB10; RAB8A | P61026; P61 | RAB10_HUMAN | Unmodified | 0 | 0 | 0 | 0 | 0 | 0 | 139330 | 0 | 0 | 0 | 0 | 0 | 0 | 0 | 0 | 0 | 0 | 0 |
| LLLIGNSSV | RAB3D | O95716 | RAB3D_HUMAI | Unmodified | 1449700 | 0 | 5385100 | 6066100 | 212800 | 0 | 0 | 163300 | 0 | 1108500 | 849840 | 4251500 | 0 | 0 | 4387300 | 3942600 | 109630 | 0 |
| LLLINNAGS | SPR | P35270 | SPRE_HUMAN | Unmodified | 0 | 0 | 0 | 0 | 0 | 0 | 0 | 200290 | 651970 | 0 | 0 | 1078300 | 945710 | 631970 | 247790 | 325940 | 0 | 0 |
| LLLIQAFRP | DYNC1H1; DYN | Q14204; A0A | DYHC1_HUMA | Unmodified | 0 | 0 | 0 | 0 | 73903 | 0 | 0 | 0 | 0 | 0 | 0 | 1965400 | 0 | 0 | 0 | 0 | 0 | 0 |
| LLLLAGVDI | PPME1 | Q9Y570 | PPME1_HUMAI | Unmodified | 0 | 0 | 238610 | 0 | 1245300 | 0 | 0 | 66557 | 0 | 625690 | 0 | 4036200 | 3248200 | 1644100 | 0 | 255740 | 971320 | 0 |
| LLLLGAGES | GNAS; GNAI3; ( | Q5JWF2; P08 | GNAS1_HUMAI | Unmodified | 13400000 | 18179000 | 15544000 | 36964000 | 61760000 | 13538000 | 20117000 | 13619000 | 52201000 | 16393000 | 77392000 | 4.26E+08 | 59437000 | 31635000 | 31869000 | 31182000 | 17534000 | 78901000 |
| LLLLGAGES | GNAS; GNAI3; ( | Q5JWF2; P08 | GNAS1_HUMAI | Unmodified | 0 | 0 | 0 | 3416300 | 82221000 | 11228000 | 47523000 | 40218000 | 794450 | 17871000 | 808130 | 877150 | 57914000 | 56530000 | 0 | 0 | 97296000 | 1.2E+08 |
| LLLLGAGES | GNAI3; GNAT1; | P08754; P11 | GNAI3_HUMAN | Oxidation (M) | 0 | 0 | 0 | 0 | 0 | 757500 | 753270 | 344600 | 0 | 0 | 0 | 0 | 0 | 0 | 0 | 0 | 1309300 | 0 |
| LLLLGAGES | GNAI3; GNAT1; | P08754; P11 | GNAI3_HUMAN | Unmodified | 0 | 0 | 0 | 0 | 1098500 | 0 | 1532100 | 256240 | 0 | 0 | 0 | 0 | 792220 | 1116900 | 0 | 0 | 3546000 | 3694900 |
| LLLLGAGES | GNAS; GNAS; G | Q5JWF2; P63 | GNAS1_HUMAI | Unmodified | 0 | 0 | 0 | 0 | 0 | 0 | 0 | 0 | 0 | 0 | 0 | 0 | 224470 | 0 | 0 | 0 | 2843400 | 0 |
| LLLLGTGES | GNAQ; GNA11; | P50148; P29 | GNAQ_HUMAN | Unmodified | 2804400 | 3045300 | 1986800 | 1031300 | 5759800 | 1236900 | 1992300 | 1095100 | 5166800 | 1890600 | 4305300 | 23269000 | 0 | 492860 | 1011500 | 994720 | 1015500 | 1654600 |
| LLLLGTGES | GNAQ; GNA11; | P50148; P29 | GNAQ_HUMAN | Unmodified | 0 | 0 | 0 | 0 | 0 | 1585900 | 7628000 | 3642500 | 561050 | 2699500 | 0 | 0 | 0 | 0 | 0 | 0 | 2245600 | 3997300 |
| LLLLGTSNS | GNAZ | P19086 | GNAZ_HUMAN | Unmodified | 0 | 0 | 0 | 0 | 0 | 0 | 0 | 288340 | 0 | 0 | 0 | 0 | 0 | 0 | 0 | 0 | 0 | 0 |
| LLLNAENPI | SRC | P12931 | SRC_HUMAN P | Unmodified | 811760 | 3.36E+09 | 7062200 | 6300900 | 436780 | 281010 | 19315000 | 4242700 | 369340 | 0 | 0 | 567870 | 0 | 0 | 2.82E+09 | 3.14E+09 | 0 | 0 |
| LLLNLSENI | ARMC10; ARMC | Q8N2F6; A0A | ARM10_HUMAI | Oxidation (M) | 0 | 0 | 157560 | 286650 | 0 | 0 | 0 | 185520 | 0 | 0 | 0 | 171370 | 0 | 0 | 0 | 230320 | 1695100 | 0 |
| LLLNNDNL | CAPZA2; CAPZA | P47755; P52 | CAZA2_HUMAN | Unmodified | 10009000 | 9594900 | 3819600 | 4606700 | 0 | 0 | 0 | 0 | 7491100 | 0 | 3979600 | 54820000 | 0 | 0 | 9524100 | 10033000 | 0 | 0 |
| LLLNNGAK | AKR1B1; AKR1E | P15121; E9P | ALDR_HUMAN | Unmodified | 0 | 0 | 0 | 0 | 463620 | 792020 | 0 | 0 | 0 | 0 | 1773600 | 12820000 | 0 | 0 | 0 | 0 | 0 | 0 |
| LLLNPGNQ | YES1; YES1 | P07947; J3Q | YES_HUMAN Ty | Unmodified | 2899700 | 0 | 849240 | 0 | 315980 | 143720 | 0 | 0 | 2742600 | 373440 | 562010 | 5405900 | 0 | 0 | 0 | 1555700 | 381400 | 615170 |
| LLLNVTPTV | SCYL2; SCYL2; | Q6P3W7; A0 | SCYL2_HUMAN | Oxidation (M) | 0 | 0 | 0 | 0 | 0 | 0 | 0 | 0 | 0 | 0 | 371620 | 2614300 | 0 | 0 | 0 | 0 | 0 | 0 |
| LLLNVTPTV | SCYL2; SCYL2; | Q6P3W7; A0 | SCYL2_HUMAN | Unmodified | 0 | 0 | 0 | 0 | 62326 | 0 | 0 | 0 | 0 | 0 | 0 | 294440 | 0 | 0 | 0 | 0 | 0 | 0 |
| LLLPGELAK | H2BE1; H2BC4 | A0A2R8Y619 | H2BE1_HUMAN | Unmodified | 1.07E+09 | 6.4E+08 | 9437600 | 4588700 | 39760000 | 43516000 | 19283000 | 1.07E+08 | 15740000 | 5678900 | 36279000 | 5.89E+09 | 15115000 | 4671800 | 3720000 | 6172700 | 1967900 | 22945000 |
| LLLPGELAK | H2BE1; H2BC4 | A0A2R8Y619 | H2BE1_HUMAN | Unmodified | 1933800 | 1617800 | 0 | 0 | 598090 | 336320 | 5440900 | 14789000 | 0 | 130770 | 0 | 5756600 | 2195100 | 0 | 0 | 0 | 0 | 6627500 |
| LLLPGWCH | NCDN; NCDN | Q9UBB6; H7 | NCDN_HUMAN | Unmodified | 0 | 106000 | 0 | 0 | 0 | 0 | 0 | 0 | 0 | 0 | 0 | 972590 | 0 | 0 | 0 | 0 | 0 | 0 |
| LLLPTPTVK | CCAR1; CCAR1 | Q8IX12; A0A | CCAR1_HUMAI | Unmodified | 0 | 0 | 0 | 0 | 0 | 0 | 0 | 0 | 0 | 0 | 759050 | 230750 | 0 | 0 | 0 | 0 | 0 | 0 |
| LLLPWLEAI | CLTC; CLTC | Q00610; A0A | CLH1_HUMAN | Unmodified | 57367000 | 35493000 | 22435000 | 14530000 | 2882100 | 1532700 | 725210 | 0 | 10601000 | 1599200 | 2445400 | 2.73E+08 | 3317500 | 1514200 | 21968000 | 38678000 | 684660 | 1632900 |
| LLLPWLEAI | CLTC; CLTC | Q00610; A0A | CLH1_HUMAN | Unmodified | 0 | 0 | 0 | 0 | 4234300 | 2628100 | 4690800 | 0 | 0 | 1110400 | 0 | 0 | 11243000 | 7480300 | 0 | 0 | 5084700 | 8116000 |
| LLLQIPSTIF | EXT1 | Q16394 | EXT1_HUMAN I | Unmodified | 0 | 0 | 1431400 | 0 | 0 | 0 | 0 | 0 | 1884300 | 122730 | 0 | 0 | 0 | 0 | 161740 | 0 | 0 | 0 |
| LLLQQLEAI | NAA25 | Q14CX7 | NAA25_HUMAN | Unmodified | 0 | 0 | 0 | 0 | 0 | 0 | 130490 | 30789 | 0 | 0 | 0 | 1138300 | 0 | 0 | 0 | 0 | 0 | 0 |
| LLLQVQHA | SLC25A6 | P12236 | ADT3_HUMAN | Unmodified | 0 | 0 | 0 | 0 | 80868 | 0 | 73271 | 27130 | 0 | 280310 | 0 | 0 | 0 | 0 | 0 | 0 | 0 | 0 |
| LLLQVQHA | SLC25A4; SLC2 | P12235; V9C | ADT1_HUMAN | Unmodified | 0 | 0 | 0 | 0 | 0 | 0 | 723790 | 0 | 0 | 0 | 0 | 0 | 0 | 0 | 0 | 0 | 0 | 0 |
| LLLQVQHA | SLC25A5 | P05141 | ADT2_HUMAN | Unmodified | 0 | 0 | 0 | 0 | 0 | 0 | 1860200 | 776270 | 0 | 1082700 | 0 | 0 | 0 | 22542 | 0 | 0 | 117420 | 0 |
| LLLSIPHSD | XPO7; XPO7; XF | Q9UIA9; E5F | XPO7_HUMAN | Unmodified | 0 | 0 | 0 | 0 | 0 | 0 | 0 | 0 | 0 | 0 | 0 | 2801800 | 0 | 0 | 0 | 0 | 0 | 0 |
| LLLSSETPI | APMAP; APMAP | Q9HDC9; H( | APMAP_HUMAI | Unmodified | 0 | 0 | 423390 | 0 | 0 | 0 | 0 | 0 | 2.46E+08 | 10581000 | 2898500 | 1771900 | 0 | 0 | 421930 | 0 | 0 | 0 |
| LLLSTLTLL | GNPNAT1; GNF | Q96EK6; G3 | GNA1_HUMAN | Unmodified | 230490 | 0 | 209800 | 170030 | 1052400 | 151460 | 0 | 0 | 111290 | 470320 | 258200 | 385920 | 806560 | 944200 | 590510 | 489230 | 1043000 | 705910 |
| LLLTPWVK | EIF3A | Q14152 | EIF3A_HUMAN | Unmodified | 102210 | 209880 | 275260 | 22991 | 32919 | 0 | 190210 | 10584 | 0 | 132180 | 207860 | 4973100 | 0 | 0 | 282520 | 364980 | 80592 | 0 |
| LLLTTPSK | SLIT2; SLIT2; SL | O94813; H0 | SLIT2_HUMAN | Unmodified | 0 | 0 | 0 | 0 | 0 | 0 | 62484 | 0 | 974780 | 31383 | 0 | 742890 | 0 | 0 | 0 | 0 | 0 | 2150200 |

| | | | | | | | | | | | | | | | | | | | | | | |
|---|---|---|---|---|---|---|---|---|---|---|---|---|---|---|---|---|---|---|---|---|---|---|
| LLLVGAPR | ITGA6 | P23229 | ITA6_HUMAN I | Unmodified | 170590 | 198090 | 242210 | 583270 | 382160 | 237480 | 229260 | 38229 | 0 | 0 | 0 | 1337500 | 1297000 | 438010 | 625370 | 330470 | 256500 | 1294400 |
| LLLVSFDGI | ENPP4; ENPP7 | Q9Y6X5; Q6I | ENPP4_HUMAN | Unmodified | 0 | 0 | 116010 | 0 | 0 | 0 | 0 | 0 | 4177100 | 363350 | 415550 | 952940 | 0 | 0 | 0 | 0 | 0 | 0 |
| LLLVSFDGI | ENPP4 | Q9Y6X5 | ENPP4_HUMAN | Unmodified | 115410 | 0 | 0 | 823150 | 2006100 | 285090 | 1312700 | 0 | 46285000 | 21032000 | 5660100 | 649270 | 1855000 | 851740 | 590110 | 217410 | 130420 | 350450 |
| LLLWAQNE | GID8 | Q9NWU2 | GID8_HUMAN | Unmodified | 0 | 0 | 0 | 0 | 611340 | 0 | 211250 | 0 | 0 | 203940 | 0 | 1638600 | 458260 | 353090 | 0 | 0 | 0 | 339760 |
| LLMAVENA | NEDD4L; NEDD | Q96PU5; K7I | NED4L_HUMAN | Oxidation (M) | 742310 | 880090 | 641880 | 402610 | 0 | 0 | 0 | 0 | 443400 | 330920 | 0 | 535200 | 0 | 0 | 593120 | 565930 | 0 | 0 |
| LLMFPESRI | EXOC8 | Q8IYI6 | EXOC8_HUMAN | Oxidation (M) | 0 | 0 | 155090 | 0 | 892600 | 114420 | 323860 | 88991 | 146810 | 713290 | 0 | 0 | 0 | 233790 | 0 | 0 | 533040 | 237120 |
| LLMFPESRI | EXOC8 | Q8IYI6 | EXOC8_HUMAN | Unmodified | 0 | 0 | 0 | 0 | 356550 | 0 | 0 | 0 | 0 | 0 | 0 | 0 | 0 | 0 | 0 | 0 | 75986 | 0 |
| LLMHGKEV | PCBP1; PCBP3; | Q15365; P57 | PCBP1_HUMAN | Oxidation (M) | 208380 | 182750 | 0 | 0 | 0 | 0 | 55057 | 23566 | 0 | 0 | 0 | 0 | 601150 | 108460 | 0 | 0 | 0 | 0 |
| LLMIAGEEF | CIT; CIT; CIT; CI | O14578; A0A | CTRO_HUMAN | Oxidation (M) | 0 | 0 | 0 | 0 | 1297300 | 0 | 0 | 0 | 0 | 56731 | 0 | 0 | 0 | 0 | 0 | 0 | 506290 | 0 |
| LLMLDDQF | GBA; GBA; GBA | P04062; A0A | GLCM_HUMAN | Unmodified | 0 | 0 | 72572 | 34925 | 0 | 0 | 0 | 0 | 1563000 | 0 | 0 | 0 | 0 | 0 | 0 | 34239 | 30535 | 0 |
| LLMLDDQF | GBA; GBA; GBA | P04062; A0A | GLCM_HUMAN | Oxidation (M) | 0 | 0 | 0 | 0 | 0 | 0 | 0 | 0 | 571820 | 1399900 | 0 | 0 | 0 | 0 | 0 | 0 | 0 | 0 |
| LLMSEDYF | LRPPRC | P42704 | LPPRC_HUMAN | 2 Oxidation (M) | 0 | 0 | 0 | 0 | 730890 | 0 | 0 | 2684100 | 322540 | 1160500 | 0 | 0 | 0 | 0 | 0 | 0 | 881070 | 0 |
| LLMTEVAW | IDE; IDE; IDE; ID | P14735; A0A | IDE_HUMAN In | Oxidation (M) | 0 | 0 | 0 | 0 | 0 | 0 | 0 | 11908 | 0 | 0 | 0 | 1370200 | 0 | 0 | 0 | 0 | 0 | 0 |
| LLMTEVAW | IDE; IDE; IDE; ID | P14735; A0A | IDE_HUMAN In | Unmodified | 0 | 0 | 0 | 0 | 0 | 0 | 0 | 0 | 0 | 0 | 0 | 1086200 | 0 | 0 | 46001 | 22072 | 0 | 0 |
| LLNASAYR | GIT1; GIT1 | Q9Y2X7; A0A | GIT1_HUMAN A | Unmodified | 0 | 0 | 0 | 0 | 0 | 0 | 0 | 0 | 0 | 0 | 0 | 1414500 | 430140 | 134410 | 0 | 0 | 0 | 112070 |
| LLNDEDPV | JUP; JUP; JUP; J | P14923; C9J | PLAK_HUMAN | Unmodified | 2067000 | 1456800 | 15505000 | 25046000 | 77670000 | 13947000 | 167310 | 61713 | 9564000 | 5059700 | 16973000 | 43865000 | 35817000 | 25859000 | 19893000 | 23851000 | 14764000 | 17381000 |
| LLNDEDQV | CTNNB1; CTNN | P35222; A0A | CTNB1_HUMAN | Unmodified | 3274500 | 3140800 | 7568200 | 16483000 | 51734000 | 10423000 | 97937 | 120830 | 2451300 | 1745800 | 5749900 | 16589000 | 23998000 | 4504400 | 13569000 | 13524000 | 23896000 | 20103000 |
| LLNDEDQV | CTNNB1; CTNN | P35222; A0A | CTNB1_HUMAN | Unmodified | 0 | 0 | 0 | 0 | 1335800 | 0 | 0 | 0 | 0 | 0 | 0 | 0 | 798170 | 0 | 0 | 0 | 1402600 | 2272500 |
| LLNENSYV | TFRC; TFRC | P02786; G3V | TFR1_HUMAN | Unmodified | 3416100 | 2494600 | 6097200 | 4008200 | 564100 | 315220 | 0 | 214260 | 29244000 | 2861900 | 1634600 | 8180000 | 332610 | 0 | 3161900 | 3864900 | 93173 | 0 |
| LLNENSYV | TFRC; TFRC | P02786; G3V | TFR1_HUMAN | Unmodified | 0 | 0 | 1435500 | 340200 | 0 | 0 | 0 | 0 | 1036700 | 165510 | 0 | 0 | 0 | 0 | 272000 | 111860 | 0 | 231600 |
| LLNENSYV | TFRC; TFRC | P02786; G3V | TFR1_HUMAN | Unmodified | 0 | 0 | 0 | 2559400 | 0 | 0 | 0 | 0 | 10595000 | 0 | 0 | 0 | 0 | 0 | 0 | 0 | 0 | 0 |
| LLNFPTIVE | GET3 | O43681 | GET3_HUMAN | Unmodified | 0 | 0 | 0 | 0 | 0 | 0 | 0 | 0 | 342690 | 0 | 271090 | 6221900 | 0 | 0 | 370500 | 0 | 0 | 0 |
| LLNGPGDV | TMEM165; TME | Q9HC07; V9 | TM165_HUMAN | Unmodified | 0 | 0 | 0 | 0 | 0 | 0 | 0 | 0 | 0 | 0 | 0 | 5338000 | 0 | 0 | 0 | 0 | 0 | 0 |
| LLNHHPLL | SEZ6; SEZ6; SEZ | Q53EL9; C9J | SEZ6_HUMAN | Unmodified | 0 | 0 | 0 | 0 | 0 | 0 | 0 | 0 | 1004300 | 0 | 0 | 0 | 0 | 0 | 758030 | 0 | 0 | 0 |
| LLNHYQAV | EXTL2; EXTL2; E | Q9UBQ6; C9 | EXTL2_HUMAN | Unmodified | 0 | 0 | 0 | 0 | 0 | 0 | 0 | 0 | 1397900 | 0 | 151780 | 145300 | 0 | 108870 | 0 | 0 | 0 | 0 |
| LLNIYGRD | MYO1D; MYO1D | O94832; J3Q | MYO1D_HUMA | Unmodified | 2576300 | 4329100 | 1883200 | 2229200 | 289470 | 0 | 0 | 0 | 1128600 | 898730 | 580670 | 769860 | 1983100 | 1958700 | 3896600 | 2569300 | 1917900 | 4924000 |
| LLNLYPR | DPYSL5 | Q9BPU6 | DPYL5_HUMAN | Unmodified | 0 | 0 | 0 | 0 | 406240 | 0 | 0 | 0 | 93244 | 0 | 0 | 1579300 | 0 | 0 | 0 | 0 | 80979 | 0 |
| LLNNAFEEI | MCM3; MCM3; | P25205; A0A | MCM3_HUMAN | Unmodified | 0 | 0 | 0 | 0 | 0 | 0 | 0 | 0 | 0 | 0 | 155920 | 429930 | 0 | 0 | 0 | 0 | 0 | 0 |
| LLNPRYNLI | TOM1L1; TOM1 | O75674; J3K | TM1L1_HUMAN | Unmodified | 0 | 0 | 0 | 0 | 385750 | 0 | 206840 | 0 | 0 | 442800 | 0 | 0 | 2402600 | 2099800 | 0 | 0 | 586580 | 621850 |
| LLNPRYNLI | TOM1L1; TOM1 | O75674; J3K | TM1L1_HUMAN | Unmodified | 0 | 0 | 0 | 0 | 0 | 0 | 282330 | 0 | 0 | 0 | 0 | 0 | 1845000 | 2954200 | 0 | 0 | 385730 | 0 |
| LLNQEEGE | PRKCA | P17252 | KPCA_HUMAN | Oxidation (M) | 0 | 0 | 0 | 0 | 3743800 | 0 | 3565000 | 551720 | 0 | 0 | 0 | 0 | 1757700 | 0 | 0 | 0 | 0 | 3586500 |
| LLNQPNQV | JUP | P14923 | PLAK_HUMAN | Unmodified | 450940 | 305280 | 5786500 | 3128800 | 4383300 | 2019700 | 0 | 31437 | 918270 | 1373000 | 0 | 10166000 | 1951500 | 3452900 | 8461300 | 5245300 | 3801500 | 1064700 |
| LLNRNNEV | HEBP2 | Q9Y5Z4 | HEBP2_HUMAN | Unmodified | 314870 | 303350 | 336200 | 243190 | 177000 | 0 | 109180 | 0 | 1605500 | 1062100 | 1716600 | 979260 | 821230 | 632940 | 426900 | 321830 | 306930 | 196340 |
| LLNRNNEV | HEBP2 | Q9Y5Z4 | HEBP2_HUMAN | Unmodified | 0 | 0 | 0 | 0 | 0 | 0 | 0 | 0 | 0 | 489340 | 0 | 0 | 425070 | 223920 | 0 | 0 | 0 | 440580 |
| LLNTFLER | DYNC1H1; DYN | Q14204; A0A | DYHC1_HUMA | Unmodified | 0 | 0 | 0 | 0 | 0 | 0 | 0 | 0 | 1216200 | 0 | 225550 | 0 | 0 | 0 | 0 | 0 | 0 | 0 |
| LLNTHHR | SDF2L1 | Q9HCN8 | SDF2L_HUMAN | Unmodified | 0 | 14269 | 0 | 0 | 201100 | 0 | 0 | 0 | 1302100 | 0 | 0 | 0 | 0 | 0 | 0 | 0 | 65622 | 94299 |
| LLNTVEETT | KIF11; KIF11; KI | P52732; A0A | KIF11_HUMAN | Unmodified | 0 | 0 | 0 | 0 | 440300 | 0 | 0 | 0 | 107950 | 0 | 0 | 0 | 0 | 0 | 0 | 0 | 417480 | 763750 |
| LLNTWTNR | PRKDC | P78527 | PRKDC_HUMAN | Unmodified | 180480 | 97171 | 296990 | 0 | 0 | 423700 | 0 | 0 | 93117 | 43750 | 0 | 913380 | 0 | 0 | 82857 | 124060 | 0 | 522110 |
| LLNVGFQE | B4GALT1 | P15291 | B4GT1_HUMAN | Unmodified | 0 | 0 | 30729 | 0 | 0 | 331570 | 70408 | 23766 | 10535000 | 563530 | 0 | 1317300 | 0 | 49582 | 646260 | 0 | 0 | 0 |
| LLNWAYQ | STEAP1B; STEA | Q6NZ63; Q9 | STEAL_HUMAN | Unmodified | 396330 | 160040 | 5019300 | 137410 | 0 | 0 | 0 | 52140 | 0 | 0 | 0 | 0 | 0 | 0 | 9598400 | 6464300 | 0 | 0 |
| LLPAFHTPS | MAN1A1 | P33908 | MA1A1_HUMAN | Oxidation (M) | 0 | 189500 | 0 | 0 | 0 | 0 | 0 | 0 | 0 | 98384 | 158890 | 337790 | 0 | 0 | 0 | 0 | 0 | 0 |
| LLPAFNTPT | MAN1A2; MAN1 | O60476; H0Y | MA1A2_HUMAN | Oxidation (M) | 0 | 0 | 0 | 0 | 0 | 0 | 0 | 0 | 2132800 | 208870 | 0 | 0 | 0 | 0 | 0 | 0 | 0 | 0 |
| LLPAFNTPT | MAN1A2; MAN1 | O60476; H0Y | MA1A2_HUMAN | Unmodified | 0 | 0 | 0 | 0 | 0 | 0 | 0 | 0 | 343260 | 31230 | 0 | 0 | 0 | 0 | 0 | 0 | 0 | 326800 |
| LLPAITILG | GMPPA | Q96IJ6 | GMPPA_HUMA | Unmodified | 0 | 0 | 0 | 0 | 0 | 0 | 0 | 0 | 346120 | 0 | 0 | 641810 | 0 | 0 | 0 | 0 | 0 | 0 |
| LLPALASVF | ACY1; ABHD14 | Q03154; C9J | ACY1_HUMAN | Unmodified | 0 | 0 | 304730 | 279090 | 0 | 0 | 0 | 0 | 214400 | 0 | 1601900 | 4670300 | 0 | 0 | 370710 | 0 | 0 | 0 |
| LLPDDPYEI | GSTO1; GSTO1 | P78417; Q5T | GSTO1_HUMAN | Unmodified | 0 | 0 | 0 | 668360 | 0 | 0 | 0 | 0 | 0 | 0 | 1520000 | 7472700 | 0 | 0 | 0 | 315450 | 0 | 0 |
| LLPDIYGWI | MME; MME; MM | P08473; A0A | NEP_HUMAN N | Unmodified | 0 | 0 | 7810400 | 38532000 | 0 | 0 | 0 | 0 | 22054000 | 4124900 | 258740 | 230240 | 12496000 | 4723700 | 64366000 | 16931000 | 0 | 0 |
| LLPDIYGWI | MME; MME; MM | P08473; A0A | NEP_HUMAN N | Unmodified | 0 | 0 | 0 | 1472900 | 0 | 0 | 0 | 0 | 0 | 635990 | 0 | 0 | 9412300 | 7178300 | 0 | 0 | 0 | 0 |
| LLPEKDDA | EXTL3 | O43909 | EXTL3_HUMAN | Unmodified | 0 | 0 | 0 | 0 | 0 | 2850600 | 0 | 0 | 529410 | 0 | 0 | 0 | 591350 | 2261700 | 0 | 0 | 0 | 0 |
| LLPERKEVE | DNAJA1 | P31689 | DNJA1_HUMAN | Unmodified | 0 | 0 | 0 | 1647100 | 0 | 0 | 595940 | 174600 | 13566000 | 1843900 | 248650 | 0 | 1937900 | 976100 | 347350 | 0 | 0 | 0 |
| LLPERKEVE | DNAJA1 | P31689 | DNJA1_HUMAN | Oxidation (M) | 0 | 2267000 | 0 | 7813600 | 0 | 0 | 870800 | 0 | 8718400 | 1446700 | 0 | 0 | 9730900 | 1437600 | 0 | 0 | 0 | 0 |
| LLPEYPGVI | DHFR | P00374 | DYR_HUMAN D | Unmodified | 0 | 0 | 0 | 382270 | 2362500 | 0 | 0 | 202330 | 0 | 0 | 0 | 497670 | 873170 | 531990 | 0 | 0 | 0 | 250180 |
| LLPEYRDAI | MYG1; MYG1 | Q9HB07; F8Y | MYG1_HUMAN | Unmodified | 208240 | 363690 | 0 | 0 | 0 | 0 | 219930 | 66208 | 1328900 | 161660 | 0 | 820130 | 0 | 0 | 0 | 116100 | 0 | 0 |
| LLPFLSNPY | TP53BP2 | Q13625 | ASPP2_HUMAN | Unmodified | 0 | 0 | 0 | 0 | 1059100 | 0 | 661870 | 0 | 0 | 0 | 0 | 0 | 0 | 0 | 0 | 0 | 895350 | 855710 |
| LLPFNPQLI | ENDOD1 | O94919 | ENDD1_HUMA | Unmodified | 0 | 0 | 0 | 0 | 426310 | 0 | 0 | 0 | 18832000 | 2801600 | 0 | 0 | 1387900 | 1308100 | 0 | 0 | 0 | 0 |
| LLPFPRAEE | RASAL1 | O95294 | RASL1_HUMAN | Unmodified | 0 | 0 | 0 | 0 | 0 | 0 | 0 | 0 | 1840200 | 145990 | 43335 | 350870 | 0 | 0 | 0 | 0 | 0 | 0 |
| LLPGIEVLW | OGA | O60502 | OGA_HUMAN F | Unmodified | 0 | 0 | 0 | 0 | 0 | 0 | 270710 | 105810 | 218230 | 165640 | 129700 | 397430 | 0 | 460040 | 0 | 0 | 0 | 511750 |
| LLPGLLPAA | IGSF9; IGSF9 | Q9P2J2; A0A | TUTLA_HUMAN | Unmodified | 0 | 0 | 0 | 0 | 0 | 0 | 0 | 0 | 0 | 0 | 0 | 844140 | 0 | 0 | 0 | 0 | 0 | 0 |
| LLPGYYMN | PIP5K1A; PIP5K | Q99755; O6 | PI51A_HUMAN | Oxidation (M) | 277500 | 0 | 255810 | 0 | 0 | 0 | 0 | 0 | 0 | 0 | 0 | 1479700 | 0 | 0 | 433060 | 0 | 0 | 0 |
| LLPIDGANI | OGA | O60502 | OGA_HUMAN F | Unmodified | 0 | 0 | 125680 | 0 | 155650 | 170390 | 0 | 0 | 835700 | 151800 | 119900 | 446330 | 371120 | 0 | 2272200 | 586380 | 191770 | 201970 |
| LLPLEEHYF | EHD1; EHD3; E | Q9H4M9; Q9 | EHD1_HUMAN | Unmodified | 0 | 0 | 250760 | 0 | 0 | 0 | 0 | 0 | 0 | 0 | 0 | 1677300 | 0 | 0 | 0 | 0 | 0 | 0 |
| LLPLEEHYF | EHD1; EHD3; E | Q9H4M9; Q9 | EHD1_HUMAN | Oxidation (M) | 0 | 0 | 0 | 0 | 14430000 | 0 | 0 | 0 | 0 | 0 | 1507700 | 0 | 0 | 0 | 0 | 0 | 0 | 0 |
| LLPLEEHYF | EHD1; EHD3; E | Q9H4M9; Q9 | EHD1_HUMAN | Unmodified | 0 | 0 | 0 | 0 | 15794000 | 0 | 910950 | 0 | 0 | 0 | 0 | 0 | 0 | 0 | 0 | 0 | 0 | 4886200 |
| LLPLGATD | MVB12A; MVB1 | Q96EY5; E9F | MB12A_HUMAN | Unmodified | 51683000 | 58101000 | 7685200 | 29157000 | 430450 | 0 | 0 | 0 | 4617900 | 734390 | 1657400 | 10656000 | 11389000 | 2579100 | 58702000 | 34569000 | 268340 | 630120 |
| LLPLGATD | MVB12A; MVB1 | Q96EY5; E9F | MB12A_HUMAN | Unmodified | 0 | 0 | 368750 | 1310500 | 4781300 | 168200 | 0 | 0 | 0 | 2359100 | 218280 | 739340 | 42771000 | 13378000 | 0 | 0 | 1519500 | 5121400 |
| LLPLLRSSC | RAB12 | A0A3B3ITT1 | A0A3B3ITT1_H | Unmodified | 0 | 0 | 0 | 0 | 2393000 | 0 | 0 | 0 | 0 | 0 | 0 | 0 | 0 | 0 | 0 | 0 | 0 | 0 |
| LLPLNDCR | CFL2 | Q9Y281 | COF2_HUMAN | Unmodified | 1496000 | 665410 | 434660 | 447120 | 4336100 | 1378000 | 0 | 67171 | 283850 | 0 | 0 | 1087800 | 356850 | 131830 | 0 | 0 | 0 | 509600 |
| LLPLNDCR | CFL2 | Q9Y281 | COF2_HUMAN | Unmodified | 0 | 0 | 0 | 0 | 87486000 | 17887000 | 2705600 | 381950 | 374020 | 0 | 0 | 0 | 0 | 2610900 | 0 | 0 | 10548000 | 8806000 |
| LLPLVSDE | PYGB | P11216 | PYGB_HUMAN | Unmodified | 0 | 0 | 146750 | 0 | 1315200 | 0 | 578810 | 0 | 1683800 | 1025500 | 381390 | 1994600 | 2699100 | 2231600 | 443680 | 0 | 2362500 | 11638000 |
| LLPPFKPQ | AKT2; AKT2; AKT | P31751; M0F | AKT2_HUMAN | Unmodified | 0 | 56433 | 1021200 | 606690 | 0 | 0 | 0 | 0 | 0 | 0 | 0 | 0 | 84224 | 0 | 1629800 | 2443600 | 0 | 0 |
| LLPQDPK | GSTZ1; GSTZ1; | O43708; G3V | MAAI_HUMAN | Unmodified | 0 | 0 | 0 | 0 | 0 | 0 | 0 | 0 | 802320 | 0 | 0 | 0 | 0 | 0 | 0 | 0 | 0 | 0 |
| LLPQLTYLD | ANP32A; ANP32 | P39687; H0Y | AN32A_HUMAN | Unmodified | 0 | 0 | 0 | 0 | 0 | 0 | 0 | 98738 | 0 | 0 | 503300 | 5621300 | 0 | 0 | 0 | 0 | 0 | 0 |
| LLPQNPEA | VPS9D1; VPS9D | Q9Y2B5; H3I | VP9D1_HUMAN | Unmodified | 0 | 0 | 0 | 0 | 370470 | 0 | 0 | 0 | 0 | 0 | 0 | 0 | 0 | 0 | 0 | 0 | 0 | 0 |
| LLPQRVLE | MSN | P26038 | MOES_HUMAN | Unmodified | 0 | 0 | 0 | 0 | 677490 | 0 | 0 | 0 | 0 | 0 | 0 | 0 | 0 | 0 | 0 | 0 | 771700 | 159570 |
| LLPRHLEIY | PYGL; PYGL | P06737; E9P | PYGL_HUMAN | Unmodified | 0 | 0 | 0 | 0 | 0 | 0 | 0 | 0 | 0 | 0 | 0 | 726540 | 0 | 0 | 0 | 0 | 443660 | 0 |
| LLPSLIGR | RSL1D1; RSL1D | O76021; J3Q | RL1D1_HUMAN | Unmodified | 0 | 0 | 0 | 0 | 0 | 0 | 0 | 0 | 0 | 0 | 0 | 3768100 | 0 | 0 | 0 | 0 | 0 | 0 |
| LLPTGAGLI | SLC4A2; SLC4A | P04920; A0A | B3A2_HUMAN | Unmodified | 525590 | 429430 | 2774800 | 2184500 | 0 | 0 | 64790 | 0 | 0 | 121440 | 0 | 2867700 | 324160 | 0 | 5114100 | 3104800 | 0 | 0 |
| LLPVDCERI | LEMD2 | Q8NC56 | LEMD2_HUMAN | Unmodified | 0 | 0 | 0 | 0 | 0 | 0 | 0 | 0 | 2531100 | 435980 | 0 | 0 | 0 | 0 | 0 | 0 | 0 | 0 |
| LLPVLDRIY | TP53I3 | Q53FA7 | QORX_HUMAN | Unmodified | 0 | 0 | 0 | 0 | 0 | 0 | 0 | 0 | 0 | 0 | 490820 | 3248800 | 0 | 0 | 0 | 0 | 0 | 0 |
| LLPVLLSTA | IPO4; IPO4; IPO | Q8TEX9; H0Y | IPO4_HUMAN | Unmodified | 56025 | 0 | 0 | 0 | 0 | 0 | 0 | 0 | 348220 | 0 | 0 | 1959800 | 0 | 0 | 187080 | 81861 | 250630 | 0 |
| LLPVLVHTF | MTX1 | Q13505 | MTX1_HUMAN | Unmodified | 34257 | 0 | 161000 | 0 | 0 | 0 | 213930 | 0 | 16261 | 0 | 0 | 1107600 | 0 | 0 | 0 | 12103 | 0 | 0 |
| LLPVYLNC | SEC24C; SEC24 | P53992; G5E | SC24C_HUMAN | Unmodified | 0 | 0 | 0 | 0 | 0 | 0 | 0 | 0 | 894100 | 0 | 0 | 1270700 | 0 | 0 | 0 | 0 | 0 | 0 |
| LLPYIVGVA | COL6A3; COL6 | P12111; C9J | CO6A3_HUMAN | Unmodified | 0 | 0 | 0 | 0 | 0 | 0 | 0 | 0 | 222120 | 36093 | 0 | 0 | 0 | 0 | 0 | 0 | 0 | 0 |
| LLPYLAAVY | ACOX3 | O15254 | ACOX3_HUMAN | Unmodified | 0 | 0 | 0 | 0 | 0 | 0 | 0 | 0 | 512360 | 0 | 0 | 0 | 0 | 0 | 0 | 0 | 0 | 0 |
| LLPYLAEDE | GP1BB | P13224 | GP1BB_HUMAN | Unmodified | 4223100 | 6022600 | 5701600 | 6184200 | 0 | 0 | 0 | 0 | 0 | 0 | 0 | 0 | 0 | 0 | 7042600 | 5447000 | 0 | 0 |
| LLQAAAGA | MMS19; MMS19 | Q96T76; H7C | MMS19_HUMA | Unmodified | 0 | 0 | 0 | 0 | 0 | 0 | 0 | 0 | 74886 | 0 | 163520 | 2100200 | 159330 | 0 | 0 | 0 | 0 | 0 |
| LLQAFLER | CSE1L | P55060 | XPO2_HUMAN | Unmodified | 0 | 0 | 0 | 0 | 0 | 0 | 0 | 0 | 0 | 0 | 0 | 18206000 | 0 | 0 | 0 | 0 | 0 | 0 |
| LLQAGFSEI | DNPEP; DNPEP | Q9ULA0; E5I | DNPEP_HUMAN | Unmodified | 0 | 0 | 0 | 0 | 0 | 0 | 0 | 0 | 381890 | 0 | 0 | 2483700 | 0 | 0 | 0 | 0 | 0 | 0 |
| LLQAGFSEI | DNPEP; DNPEP | Q9ULA0; E5I | DNPEP_HUMAN | Unmodified | 69859 | 0 | 0 | 0 | 0 | 99040 | 105380 | 148200 | 928590 | 0 | 0 | 1932300 | 0 | 431400 | 0 | 0 | 0 | 64124 |
| LLQAQGVE | EIF5B; EIF5B | O60841; A0A | IF2P_HUMAN E | Unmodified | 0 | 0 | 0 | 0 | 0 | 0 | 0 | 0 | 93037 | 0 | 747760 | 1558700 | 225690 | 123960 | 0 | 0 | 0 | 225210 |
| LLQAQGVE | EIF5B; EIF5B | O60841; A0A | IF2P_HUMAN E | Unmodified | 0 | 0 | 0 | 0 | 0 | 0 | 254610 | 0 | 0 | 0 | 0 | 0 | 1694500 | 0 | 0 | 0 | 0 | 0 |
| LLQAQLQA | PBXIP1 | Q96AQ6 | PBIP1_HUMAN | Unmodified | 0 | 0 | 0 | 0 | 548360 | 0 | 0 | 0 | 962450 | 0 | 259210 | 263400 | 0 | 0 | 0 | 0 | 0 | 0 |
| LLQAQLQA | PBXIP1 | Q96AQ6 | PBIP1_HUMAN | Unmodified | 0 | 0 | 0 | 0 | 0 | 0 | 0 | 0 | 8033400 | 361480 | 492180 | 0 | 0 | 0 | 0 | 0 | 0 | 0 |
| LLQASFK | MAPRE2; MAPR | Q15555; K7E | MARE2_HUMAN | Unmodified | 155400 | 714660 | 14035000 | 3549700 | 0 | 5306000 | 2127800 | 2171500 | 15573000 | 1468300 | 259980 | 673650 | 1320100 | 3134500 | 1723400 | 442320 | 219440 | 1121600 |
| LLQCDPSS | HPCA; HPCAL1 | P84074; P37 | HPCA_HUMAN | Unmodified | 0 | 0 | 0 | 0 | 664590 | 0 | 0 | 0 | 0 | 0 | 0 | 0 | 0 | 0 | 0 | 0 | 559070 | 483920 |
| LLQDAIR | ARHGEF2; ARH | Q92974; V9C | ARHG2_HUMA | Unmodified | 0 | 0 | 0 | 0 | 0 | 0 | 0 | 0 | 151180 | 0 | 81555 | 770770 | 0 | 0 | 365670 | 273570 | 0 | 0 |
| LLQDAIREV | ARHGEF2; ARH | Q92974; V9C | ARHG2_HUMA | Unmodified | 0 | 0 | 182980 | 185670 | 231960 | 0 | 62108 | 0 | 1236700 | 246650 | 610260 | 1202100 | 136110 | 78044 | 152420 | 0 | 0 | 0 |
| LLQDANYN | CLPX | O76031 | CLPX_HUMAN | Unmodified | 0 | 0 | 0 | 0 | 1122300 | 3777800 | 1853400 | 1382700 | 0 | 66422 | 0 | 0 | 273320 | 3318800 | 0 | 0 | 512580 | 3724500 |
| LLQDFFNG | HSPA6; HSPA8; | P17066; P11 | HSP76_HUMAN | Unmodified | 41898000 | 54015000 | 64907000 | 96681000 | 2.71E+08 | 37861000 | 53279000 | 13834000 | 89905000 | 70318000 | 23654000 | 1.78E+08 | 3.04E+08 | 1.42E+08 | 46428000 | 45195000 | 39249000 | 66038000 |
| LLQDFFNG | HSPA6; HSPA8; | P17066; P11 | HSP76_HUMAN | Unmodified | 0 | 187740 | 0 | 231750 | 10442000 | 866280 | 14147000 | 2590500 | 0 | 2668100 | 0 | 0 | 15218000 | 15065000 | 0 | 55927 | 12396000 | 6419800 |
| LLQDFFNG | HSPA1A; HSPA | P0DMV8; P0 | HS71A_HUMAN | Unmodified | 6094200 | 7682500 | 3511800 | 2185200 | 0 | 0 | 0 | 0 | 8082100 | 512600 | 267560 | 11289000 | 754400 | 0 | 90028 | 3251700 | 0 | 0 |
| LLQDFFNG | HSPA1A; HSPA | P0DMV8; P0 | HS71A_HUMAN | Unmodified | 17074000 | 21742000 | 10885000 | 19965000 | 1.01E+08 | 15735000 | 30157000 | 4259800 | 68385000 | 64176000 | 5118700 | 14616000 | 1.26E+08 | 52772000 | 8687800 | 7009500 | 37102000 | 35325000 |
| LLQDHPWI | ACLY; ACLY | P53396; K7E | ACLY_HUMAN | Unmodified | 0 | 0 | 0 | 0 | 2360300 | 0 | 1523800 | 56711 | 4782700 | 0 | 1336600 | 9101500 | 512100 | 0 | 0 | 0 | 509260 | 0 |
| LLQDHPWI | ACLY; ACLY | P53396; K7E | ACLY_HUMAN | Unmodified | 0 | 0 | 0 | 0 | 1788700 | 0 | 0 | 0 | 1934300 | 0 | 0 | 2781100 | 0 | 0 | 0 | 0 | 0 | 0 |
| LLQDPLGL | RGS14 | O43566 | RGS14_HUMAN | Unmodified | 0 | 0 | 0 | 0 | 0 | 0 | 0 | 0 | 0 | 0 | 123280 | 93438 | 0 | 0 | 0 | 0 | 0 | 0 |
| LLQDYFNG | HSPA1L; HSPA1 | P34931; Q53 | HS71L_HUMAN | Unmodified | 0 | 0 | 0 | 0 | 0 | 0 | 0 | 0 | 0 | 0 | 0 | 0 | 0 | 0 | 0 | 0 | 0 | 0 |
| LLQDYPITD | TBC1D13 | Q9NVG8 | TBC13_HUMAN | Unmodified | 17932 | 0 | 0 | 33307 | 0 | 0 | 0 | 0 | 0 | 0 | 135990 | 74278 | 21069 | 0 | 56903 | 28113 | 0 | 0 |
| LLQEAAAN | ATP6V0A4; ATP | Q9HBG4; A0 | VPP4_HUMAN | Unmodified | 0 | 0 | 0 | 0 | 0 | 0 | 0 | 0 | 0 | 0 | 0 | 515120 | 0 | 0 | 0 | 0 | 0 | 0 |
| LLQEKELQI | CCDC50 | Q8IVM0 | CCD50_HUMA | Unmodified | 0 | 0 | 0 | 0 | 20336 | 35708 | 0 | 0 | 0 | 33839 | 0 | 0 | 481650 | 183260 | 34317 | 0 | 0 | 43400 |

| | | | | | | | | | | | | | | | | | | | | | | |
|---|---|---|---|---|---|---|---|---|---|---|---|---|---|---|---|---|---|---|---|---|---|---|
| LLQENVPS | PIGG; PIGG; PI | Q5H8A4; E7I | PIGG_HUMAN | Unmodified | 0 | 0 | 0 | 0 | 0 | 0 | 0 | 0 | 1409300 | 0 | 0 | 0 | 0 | 0 | 0 | 0 | 45157 | 0 |
| LLQERELVE | EGFR; EGFR; E | P00533; Q5 | EGFR_HUMAN | Unmodified | 0 | 0 | 0 | 0 | 6600600 | 3446300 | 0 | 0 | 0 | 0 | 0 | 0 | 1672500 | 0 | 0 | 0 | 0 | 0 |
| LLQESYK | KNTC1 | P50748 | KNTC1_HUMA | Unmodified | 0 | 0 | 0 | 0 | 0 | 0 | 0 | 0 | 0 | 0 | 0 | 303580 | 0 | 0 | 0 | 0 | 0 | 0 |
| LLQETELVE | ERBB2; ERBB2; | P04626; J3Q | ERBB2_HUMA | 2 Oxidation (M) | 168650 | 0 | 0 | 334200 | 0 | 0 | 0 | 0 | 0 | 0 | 0 | 0 | 0 | 0 | 843790 | 537970 | 0 | 0 |
| LLQEYCME | PLCG2; PLCG2 | P16885; A0A | PLCG2_HUMA | Oxidation (M) | 0 | 0 | 545490 | 0 | 0 | 510140 | 0 | 0 | 0 | 0 | 241460 | 2061800 | 0 | 0 | 0 | 0 | 0 | 0 |
| LLQEYCME | PLCG2; PLCG2 | P16885; A0A | PLCG2_HUMA | Unmodified | 0 | 0 | 0 | 0 | 0 | 0 | 0 | 0 | 108200 | 123780 | 0 | 2779700 | 0 | 0 | 0 | 0 | 0 | 0 |
| LLQFLAGQ | HUS1 | O60921 | HUS1_HUMAN | Unmodified | 0 | 0 | 0 | 0 | 155480 | 0 | 0 | 36429 | 0 | 0 | 0 | 139700 | 212580 | 0 | 0 | 0 | 0 | 28913 |
| LLQFVTGS | SMURF1 | Q9HCE7 | SMUF1_HUMA | Unmodified | 591580 | 2900100 | 0 | 0 | 0 | 0 | 0 | 0 | 0 | 0 | 0 | 0 | 0 | 0 | 1698700 | 1057900 | 0 | 0 |
| LLQFVTGT | WWP1; ITCH; W | Q9H0M0; Q | WWP1_HUMA | Unmodified | 4621400 | 7287400 | 0 | 0 | 0 | 0 | 0 | 0 | 0 | 0 | 0 | 677600 | 0 | 0 | 0 | 0 | 0 | 0 |
| LLQFVTGT | NEDD4; NEDD4 | P46934; Q9 | NEDD4_HUMA | Unmodified | 1211200 | 0 | 0 | 0 | 0 | 0 | 0 | 0 | 0 | 0 | 0 | 0 | 0 | 0 | 0 | 1486200 | 0 | 0 |
| LLQFYAET | MYO1C; MYO1 | O00159; I3L | MYO1C_HUMA | Unmodified | 350060 | 138100 | 244000 | 472710 | 0 | 0 | 0 | 0 | 0 | 0 | 0 | 176070 | 35870 | 0 | 397680 | 423930 | 0 | 0 |
| LLQFYAET | MYO1C; MYO1 | O00159; I3L | MYO1C_HUMA | Unmodified | 2408000 | 772320 | 1151000 | 2710700 | 480700 | 0 | 0 | 0 | 0 | 0 | 0 | 0 | 331420 | 0 | 3271700 | 2188400 | 212910 | 0 |
| LLQFYPSLE | LAT2; LAT2; LAT | Q9GZY6; C9 | NTAL_HUMAN | Unmodified | 323580 | 352100 | 0 | 0 | 0 | 0 | 0 | 0 | 480510 | 16522 | 0 | 0 | 0 | 0 | 0 | 0 | 37391 | 0 |
| LLQGGGV | GRTP1; GRTP1 | Q5TC63; A0 | GRTP1_HUMA | Unmodified | 0 | 0 | 625760 | 5249000 | 0 | 0 | 0 | 0 | 0 | 0 | 0 | 0 | 0 | 0 | 0 | 0 | 0 | 0 |
| LLQHHITE | PPM1H | Q9ULR3 | PPM1H_HUMA | Unmodified | 0 | 0 | 0 | 0 | 0 | 111820 | 0 | 0 | 137940 | 369520 | 0 | 0 | 59535 | 0 | 0 | 0 | 0 | 0 |
| LLQLGQQF | PKP3; PKP3 | Q9Y446; E9F | PKP3_HUMAN | Unmodified | 0 | 0 | 0 | 4653300 | 1107600 | 0 | 0 | 0 | 123000 | 0 | 2506400 | 4147800 | 0 | 0 | 3178200 | 4399000 | 0 | 876850 |
| LLQLTEK | SPTBN1; SPTBN | Q01082; A0 | SPTB2_HUMA | Unmodified | 0 | 0 | 0 | 0 | 363010 | 0 | 829350 | 0 | 275950 | 0 | 0 | 2424100 | 0 | 0 | 0 | 0 | 0 | 0 |
| LLQLVEDR | DDX5; DDX5; D | P17844; A0A | DDX5_HUMAN | Unmodified | 0 | 51254 | 0 | 169000 | 0 | 0 | 7117900 | 32247 | 0 | 0 | 349700 | 2026400 | 206340 | 0 | 0 | 63217 | 0 | 0 |
| LLQLVEDR | DDX5; DDX5; D | P17844; A0A | DDX5_HUMAN | Unmodified | 144950 | 0 | 0 | 112840 | 915390 | 222480 | 545420 | 454230 | 804460 | 690440 | 232070 | 816020 | 1072500 | 331600 | 73122 | 48354 | 140480 | 321340 |
| LLQMQPAC | AP3B2; AP3B1; | Q13367; O0 | AP3B2_HUMA | Unmodified | 0 | 0 | 0 | 0 | 0 | 0 | 0 | 38158 | 0 | 0 | 0 | 2549600 | 0 | 0 | 0 | 0 | 0 | 0 |
| LLQNIGIDK | SLC39A6 | Q13433 | S39A6_HUMA | Unmodified | 0 | 531980 | 961000 | 987960 | 10275000 | 2199400 | 105770 | 56546 | 1539500 | 318220 | 1026600 | 11726000 | 3779400 | 1438000 | 1001000 | 1200600 | 2832300 | 9769300 |
| LLQNIGIDK | SLC39A6 | Q13433 | S39A6_HUMA | Unmodified | 0 | 0 | 0 | 0 | 1160600 | 0 | 547280 | 481900 | 0 | 0 | 0 | 0 | 440650 | 494300 | 0 | 0 | 1564300 | 1423500 |
| LLQPGDKI | ERBIN; ERBIN | Q96RT1; B4 | ERBIN_HUMA | Unmodified | 0 | 0 | 0 | 0 | 3076000 | 0 | 0 | 0 | 0 | 0 | 0 | 0 | 0 | 0 | 0 | 0 | 340640 | 0 |
| LLQPPAPIM | COPA; COPA; C | P53621; A0A | COPA_HUMAN | Oxidation (M) | 345410 | 80613 | 119570 | 487720 | 3195800 | 501550 | 4582700 | 0 | 1490300 | 1511600 | 1674400 | 1778400 | 5998700 | 2585000 | 392740 | 277500 | 1765200 | 4390700 |
| LLQQEEEIK | RARS1; RARS1; | P54136; E5R | SYRC_HUMAN | Unmodified | 0 | 0 | 0 | 0 | 576540 | 74256 | 0 | 0 | 775590 | 367360 | 1519100 | 8570800 | 1105800 | 442860 | 0 | 0 | 253810 | 520460 |
| LLQQGLAC | VGF; VGF | O15240; Q0 | VGF_HUMAN | Unmodified | 29825 | 53088 | 0 | 0 | 367980 | 0 | 0 | 0 | 1364100 | 0 | 0 | 0 | 0 | 0 | 0 | 0 | 0 | 0 |
| LLQQGLAC | VGF; VGF | O15240; Q0 | VGF_HUMAN | Unmodified | 3913600 | 2705100 | 0 | 0 | 0 | 0 | 0 | 0 | 17221000 | 475020 | 0 | 0 | 0 | 0 | 0 | 0 | 0 | 0 |
| LLQQLFLK | BCCIP | Q9P287 | BCCIP_HUMA | Unmodified | 0 | 0 | 0 | 0 | 603020 | 0 | 161320 | 41955 | 0 | 0 | 1716700 | 6111300 | 497810 | 0 | 0 | 0 | 95880 | 0 |
| LLQQQNLF | INPPL1; INPPL1 | O15357; F5 | SHIP2_HUMA | Unmodified | 0 | 0 | 0 | 0 | 0 | 0 | 0 | 0 | 0 | 0 | 1212200 | 2425200 | 0 | 0 | 0 | 0 | 0 | 0 |
| LLQQVSQI | EIF3A | Q14152 | EIF3A_HUMA | Unmodified | 0 | 0 | 0 | 484460 | 0 | 0 | 0 | 0 | 113120 | 18140 | 0 | 1138800 | 0 | 0 | 185560 | 0 | 0 | 0 |
| LLQSAFSK | PCSK1 | P29120 | NEC1_HUMAN | Unmodified | 0 | 0 | 0 | 0 | 0 | 0 | 0 | 0 | 1726600 | 100630 | 0 | 74381 | 0 | 0 | 35594 | 0 | 0 | 0 |
| LLQSENYV | CAB39L; CAB3 | Q9H9S4; B7 | CB39L_HUMA | Unmodified | 0 | 0 | 0 | 0 | 0 | 0 | 0 | 0 | 0 | 0 | 0 | 1146100 | 0 | 0 | 0 | 0 | 0 | 0 |
| LLQSENYV | CAB39L; CAB3 | Q9H9S4; B7 | CB39L_HUMA | Unmodified | 47393 | 22447 | 107220 | 60984 | 0 | 0 | 0 | 0 | 268150 | 39357 | 0 | 688410 | 0 | 77617 | 251000 | 371920 | 0 | 0 |
| LLQSGNSD | PKP1 | Q13835 | PKP1_HUMAN | Unmodified | 194170 | 0 | 0 | 0 | 0 | 617320 | 0 | 0 | 0 | 0 | 0 | 0 | 0 | 0 | 0 | 0 | 141010 | 0 |
| LLQSIGQA | NAE1 | Q13564 | ULA1_HUMAN | Unmodified | 33286 | 0 | 40329 | 36443 | 0 | 0 | 0 | 175150 | 0 | 113160 | 623780 | 3796900 | 394390 | 0 | 0 | 31744 | 0 | 0 |
| LLQSLGLK | MVP | Q14764 | MVP_HUMAN | Unmodified | 1281800 | 0 | 0 | 0 | 5692400 | 0 | 1421500 | 307820 | 2438600 | 0 | 245730 | 29607000 | 0 | 0 | 0 | 0 | 459310 | 1415600 |
| LLQSLLDTF | FLII | Q13045 | FLII_HUMAN P | Unmodified | 0 | 0 | 0 | 0 | 0 | 0 | 0 | 0 | 0 | 106150 | 0 | 454410 | 0 | 0 | 0 | 0 | 0 | 0 |
| LLQSNPVL | MYO1C; MYO1 | O00159; O4 | MYO1C_HUMA | Unmodified | 3364200 | 0 | 5429100 | 0 | 4786700 | 2142600 | 287810 | 48385 | 719400 | 103280 | 259160 | 1102300 | 2233000 | 2115600 | 15908000 | 11034000 | 932350 | 1272800 |
| LLQSTIR | ABCA12 | Q86UK0 | ABCAC_HUMA | Unmodified | 0 | 0 | 0 | 0 | 0 | 59659 | 114600 | 0 | 2397500 | 0 | 0 | 0 | 0 | 0 | 0 | 0 | 0 | 0 |
| LLQTAATA | STAT3; STAT3; | P40763; A0A | STAT3_HUMA | Unmodified | 127220 | 113350 | 0 | 0 | 0 | 0 | 0 | 0 | 0 | 0 | 0 | 3588600 | 0 | 0 | 878550 | 611670 | 0 | 0 |
| LLQTCFSS | HSPBP1 | Q9NZL4 | HPBP1_HUMA | Oxidation (M) | 0 | 0 | 0 | 0 | 0 | 0 | 122720 | 57584 | 0 | 0 | 0 | 0 | 49666 | 0 | 0 | 0 | 628730 | 411980 |
| LLQTCFSS | HSPBP1 | Q9NZL4 | HPBP1_HUMA | Unmodified | 0 | 0 | 0 | 0 | 0 | 0 | 79064 | 0 | 0 | 0 | 0 | 0 | 0 | 0 | 0 | 0 | 433940 | 200090 |
| LLQTDDEE | CSE1L | P55060 | XPO2_HUMAN | Unmodified | 0 | 0 | 0 | 12068 | 0 | 0 | 0 | 0 | 259860 | 127290 | 0 | 2078300 | 0 | 0 | 0 | 0 | 63535 | 0 |
| LLQTIGKG | CSK | P41240 | CSK_HUMAN T | Oxidation (M) | 0 | 171030 | 0 | 0 | 0 | 0 | 0 | 0 | 0 | 0 | 0 | 1008800 | 0 | 0 | 0 | 0 | 0 | 0 |
| LLQTIGKG | CSK | P41240 | CSK_HUMAN T | Oxidation (M) | 310300 | 0 | 0 | 0 | 0 | 0 | 1179200 | 0 | 0 | 235070 | 6099600 | 2101300 | 942180 | 1738200 | 177890 | 181610 | 775410 | 0 |
| LLQTSNITK | PAK2; PAK3; PA | Q13177; O7 | PAK2_HUMAN | Unmodified | 0 | 0 | 0 | 0 | 0 | 0 | 0 | 0 | 0 | 0 | 692750 | 0 | 0 | 0 | 0 | 0 | 0 | 0 |
| LLQTSNITK | PAK2; PAK3; PA | Q13177; O7 | PAK2_HUMAN | Unmodified | 629160 | 395290 | 53172 | 506660 | 3361700 | 979270 | 133640 | 57202 | 1306400 | 681980 | 819920 | 2324900 | 2608100 | 1046700 | 249270 | 314390 | 705310 | 2218600 |
| LLQTSNITK | PAK2; PAK3; PA | Q13177; O7 | PAK2_HUMAN | Unmodified | 0 | 0 | 0 | 146010 | 592200 | 385350 | 0 | 0 | 0 | 0 | 0 | 175580 | 486310 | 215740 | 0 | 0 | 70415 | 324760 |
| LLQTSNITK | PAK1; PAK1; PA | Q13153; E9F | PAK1_HUMAN | Unmodified | 0 | 0 | 0 | 0 | 0 | 60247 | 0 | 0 | 0 | 124460 | 0 | 0 | 0 | 0 | 0 | 0 | 0 | 121750 |
| LLQTTNNS | GTPBP1; GTPB | O00178; B0 | GTPB1_HUMA | Unmodified | 325320 | 319490 | 212840 | 322370 | 0 | 0 | 222310 | 192840 | 151720 | 101470 | 327710 | 2096400 | 315380 | 257170 | 294010 | 406420 | 59388 | 216470 |
| LLQTTNNS | GTPBP1; GTPB | O00178; B0 | GTPB1_HUMA | Oxidation (M) | 0 | 0 | 0 | 0 | 40097 | 0 | 0 | 0 | 0 | 0 | 0 | 319990 | 0 | 0 | 176120 | 141420 | 0 | 0 |
| LLQVSVDD | UTRN; UTRN | P46939; Q5 | UTRN_HUMAN | Unmodified | 0 | 0 | 948010 | 1312600 | 0 | 3300900 | 73262 | 837690 | 0 | 608610 | 0 | 0 | 0 | 0 | 0 | 3672400 | 0 | 0 |
| LLQVTPAD | HSPG2 | P98160 | PGBM_HUMAN | Unmodified | 0 | 0 | 318120 | 947640 | 0 | 0 | 30899 | 0 | 374650 | 0 | 0 | 0 | 0 | 0 | 0 | 553500 | 0 | 0 |
| LLQYLYK | HSPA13 | P48723 | HSP13_HUMA | Unmodified | 0 | 0 | 294300 | 169160 | 0 | 0 | 0 | 0 | 10198000 | 0 | 0 | 0 | 0 | 0 | 0 | 118540 | 0 | 0 |
| LLRDPADA | CHGB; CHGB | P05060; A0A | SCG1_HUMAN | Unmodified | 0 | 0 | 0 | 0 | 0 | 0 | 0 | 0 | 800070 | 0 | 0 | 0 | 0 | 0 | 0 | 0 | 0 | 0 |
| LLREDKNH | KIF23; KIF23; K | Q02241; A0 | KIF23_HUMAN | Oxidation (M) | 0 | 0 | 0 | 0 | 1125300 | 0 | 0 | 0 | 381910 | 44512 | 0 | 0 | 0 | 0 | 0 | 0 | 346370 | 332530 |
| LLREDKNH | KIF23; KIF23; K | Q02241; A0 | KIF23_HUMAN | Unmodified | 0 | 0 | 0 | 0 | 1474800 | 0 | 0 | 0 | 0 | 0 | 0 | 0 | 0 | 0 | 0 | 0 | 392460 | 0 |
| LLRFQSSH | HSP90B1; HSP | P14625; A0A | ENPL_HUMAN | Unmodified | 0 | 0 | 0 | 0 | 93792 | 0 | 267540 | 0 | 82270000 | 39831000 | 20249000 | 0 | 0 | 0 | 0 | 0 | 0 | 0 |
| LLRFQSSH | HSP90B1; HSP | P14625; A0A | ENPL_HUMAN | Oxidation (M) | 0 | 0 | 0 | 0 | 0 | 0 | 4058300 | 0 | 9425900 | 98860000 | 22116000 | 0 | 603250 | 657700 | 0 | 0 | 99435 | 334340 |
| LLRFQSSH | HSP90B1; HSP | P14625; A0A | ENPL_HUMAN | Unmodified | 0 | 0 | 0 | 0 | 0 | 0 | 31299000 | 0 | 22592000 | 1.06E+08 | 13951000 | 0 | 0 | 219880 | 0 | 0 | 474510 | 0 |
| LLRGHDQS | HLA-B; HLA-B; | A0A1W2PNI | A0A1W2PNR7_ | Unmodified | 0 | 0 | 0 | 0 | 0 | 0 | 0 | 0 | 0 | 0 | 0 | 0 | 0 | 0 | 0 | 0 | 0 | 0 |
| LLRGYDQS | HLA-C; HLA-C; | P10321; Q5 | HLAC_HUMAN | Unmodified | 0 | 0 | 0 | 0 | 0 | 0 | 0 | 0 | 1789400 | 0 | 0 | 0 | 0 | 0 | 0 | 0 | 0 | 0 |
| LLRIYDLNK | STRAP | Q9Y3F4 | STRAP_HUMA | Unmodified | 0 | 0 | 0 | 0 | 8892600 | 8096700 | 1233500 | 422790 | 0 | 867410 | 204780 | 0 | 4024800 | 3572800 | 0 | 0 | 2284700 | 4880600 |
| LLRLQLPNI | FERMT2; FERM | Q96AC1; G3 | FERM2_HUMA | Unmodified | 0 | 0 | 0 | 0 | 911570 | 0 | 0 | 0 | 0 | 0 | 0 | 0 | 0 | 0 | 0 | 0 | 0 | 315280 |
| LLRQSPPL | ZDHHC5 | Q9C0B5 | ZDHC5_HUMA | Unmodified | 0 | 0 | 0 | 0 | 955660 | 0 | 0 | 0 | 0 | 310940 | 0 | 0 | 1606800 | 1293000 | 0 | 0 | 486460 | 1500200 |
| LLRSEESVS | CKAP4 | Q07065 | CKAP4_HUMA | Unmodified | 0 | 0 | 102380 | 0 | 0 | 0 | 0 | 0 | 6714200 | 100800 | 0 | 0 | 0 | 0 | 146150 | 0 | 0 | 0 |
| LLRTNLDE | RTCB | Q9Y3I0 | RTCB_HUMAN | Unmodified | 130740 | 98413 | 294740 | 511070 | 0 | 0 | 0 | 68020 | 0 | 0 | 903430 | 0 | 1353500 | 235770 | 224720 | 161880 | 0 | 0 |
| LLRYESSAL | TRAP1; TRAP1 | Q12931; I3L | TRAP1_HUMA | Unmodified | 0 | 0 | 0 | 0 | 0 | 0 | 0 | 167760 | 6651700 | 1224600 | 0 | 0 | 0 | 0 | 0 | 0 | 0 | 0 |
| LLSAAKILAI | TLN1 | Q9Y490 | TLN1_HUMAN | Unmodified | 0 | 0 | 0 | 0 | 57445 | 0 | 207940 | 78689 | 0 | 0 | 0 | 0 | 0 | 0 | 0 | 0 | 127250 | 0 |
| LLSAEERIS | SEPTIN5; SEPT | Q99719; F8 | SEPT5_HUMA | Unmodified | 0 | 0 | 0 | 0 | 0 | 0 | 824770 | 0 | 0 | 0 | 0 | 0 | 614970 | 0 | 0 | 0 | 1119400 | 0 |
| LLSAERAVT | PLEC; PLEC | Q15149; H0 | PLEC_HUMAN | Unmodified | 0 | 0 | 0 | 0 | 908820 | 0 | 0 | 0 | 0 | 0 | 0 | 0 | 0 | 0 | 0 | 0 | 1764200 | 2323500 |
| LLSAERAVT | PLEC; PLEC | Q15149; H0 | PLEC_HUMAN | Unmodified | 0 | 0 | 0 | 0 | 1953300 | 0 | 0 | 0 | 0 | 0 | 0 | 0 | 0 | 0 | 0 | 0 | 2422700 | 1839500 |
| LLSALCPEE | EIF2B3; EIF2B3 | Q9NR50; H0 | EI2BG_HUMA | Unmodified | 0 | 0 | 0 | 0 | 366900 | 0 | 494710 | 386760 | 225280 | 172330 | 0 | 1933000 | 752740 | 412610 | 0 | 0 | 935660 | 1067000 |
| LLSASMDK | ELP2 | Q6IA86 | ELP2_HUMAN | Unmodified | 0 | 0 | 0 | 0 | 44449 | 0 | 0 | 0 | 56131 | 0 | 0 | 769700 | 0 | 0 | 0 | 0 | 106980 | 0 |
| LLSATGESE | TSPAN8 | P19075 | TSN8_HUMAN | Unmodified | 3.21E+08 | 2.03E+08 | 151460 | 110750 | 0 | 0 | 70616 | 0 | 11720000 | 3015700 | 0 | 0 | 0 | 0 | 731690 | 928800 | 182980 | 9667700 |
| LLSATGESE | TSPAN8 | P19075 | TSN8_HUMAN | Unmodified | 0 | 0 | 0 | 0 | 0 | 0 | 0 | 0 | 537200 | 410180 | 0 | 0 | 0 | 0 | 0 | 0 | 0 | 1361400 |
| LLSAVPEE | PSD2 | Q9BQI7 | PSD2_HUMAN | Unmodified | 0 | 0 | 0 | 0 | 0 | 0 | 0 | 0 | 0 | 0 | 3505900 | 2290800 | 0 | 0 | 0 | 0 | 0 | 0 |
| LLSDATVE | FAM50A; FAM5 | Q14320; B0 | FA50A_HUMA | Unmodified | 216820 | 305140 | 0 | 425440 | 0 | 0 | 0 | 0 | 0 | 0 | 431910 | 750230 | 0 | 0 | 636960 | 408480 | 0 | 0 |
| LLSDATVE | FAM50A | Q14320 | FA50A_HUMA | Unmodified | 0 | 0 | 0 | 0 | 0 | 0 | 0 | 0 | 34714 | 0 | 187770 | 108220 | 0 | 0 | 0 | 0 | 0 | 0 |
| LLSDDRSN | PIGK; PIGK; PIG | Q92643; B1 | GPI8_HUMAN | Unmodified | 0 | 0 | 0 | 0 | 0 | 0 | 0 | 0 | 2760500 | 0 | 0 | 0 | 0 | 0 | 0 | 0 | 0 | 0 |
| LLSDPNYG | TMEM43 | Q9BTV4 | TMM43_HUMA | Unmodified | 0 | 0 | 0 | 0 | 0 | 0 | 89937 | 0 | 13590000 | 3786700 | 0 | 0 | 0 | 0 | 0 | 0 | 0 | 0 |
| LLSDSLPP | TLN1 | Q9Y490 | TLN1_HUMAN | Unmodified | 212550 | 119360 | 126410 | 0 | 0 | 0 | 0 | 0 | 0 | 0 | 174670 | 1275200 | 0 | 0 | 0 | 0 | 0 | 0 |
| LLSDTVAS | MACF1; MACF1 | Q9UPN3; H | MACF1_HUMA | Unmodified | 0 | 0 | 0 | 0 | 0 | 0 | 0 | 0 | 675230 | 543570 | 0 | 544980 | 0 | 0 | 0 | 0 | 0 | 0 |
| LLSEAEVLD | POMGNT1 | Q8WZA1 | PMGT1_HUMA | Unmodified | 0 | 0 | 0 | 0 | 0 | 0 | 0 | 0 | 667080 | 76844 | 0 | 0 | 0 | 0 | 0 | 0 | 124750 | 0 |
| LLSEAQR | ZNF552; ZNF58 | Q9H707; Q9 | ZN552_HUMA | Unmodified | 0 | 0 | 0 | 0 | 0 | 0 | 36581 | 0 | 1871600 | 0 | 0 | 0 | 0 | 0 | 0 | 0 | 0 | 0 |
| LLSEEVFDF | XPO1; XPO1; XI | O14980; A0 | XPO1_HUMAN | Unmodified | 0 | 0 | 0 | 0 | 0 | 0 | 0 | 0 | 0 | 0 | 0 | 2566300 | 0 | 0 | 0 | 0 | 0 | 0 |
| LLSEGADV | CLPB; CLPB; CI | Q9H078; A0 | CLPB_HUMAN | Unmodified | 0 | 0 | 0 | 0 | 0 | 0 | 0 | 0 | 0 | 0 | 204470 | 862180 | 0 | 0 | 0 | 0 | 0 | 0 |
| LLSESNERI | PPFIA3; PPFIA1 | O75145; Q1 | LIPA3_HUMAN | Unmodified | 0 | 0 | 0 | 0 | 975140 | 0 | 0 | 0 | 0 | 0 | 0 | 0 | 355860 | 0 | 0 | 0 | 0 | 0 |
| LLSFGER | PAAT | Q9H8K7 | PAAT_HUMAN | Unmodified | 0 | 965080 | 0 | 0 | 0 | 0 | 0 | 0 | 0 | 0 | 0 | 0 | 0 | 0 | 0 | 0 | 0 | 0 |
| LLSFTSQLK | SLC12A7; SLC1 | Q9Y666; A0 | S12A7_HUMA | Unmodified | 885370 | 0 | 1304800 | 1045500 | 223050 | 0 | 435540 | 0 | 1818300 | 464790 | 752350 | 4526600 | 624650 | 0 | 1935900 | 1941000 | 0 | 0 |
| LLSGGSVLI | OXSR1; OXSR1 | O95747; C9 | OXSR1_HUMA | Unmodified | 0 | 0 | 0 | 0 | 0 | 0 | 0 | 0 | 0 | 0 | 376050 | 633720 | 0 | 0 | 0 | 0 | 0 | 0 |
| LLSGPLDQ | PTPN23 | Q9H3S7 | PTN23_HUMA | Unmodified | 0 | 0 | 0 | 0 | 1900400 | 536780 | 0 | 0 | 0 | 591630 | 0 | 0 | 0 | 0 | 0 | 0 | 0 | 0 |
| LLSGPYFW | HSPG2 | P98160 | PGBM_HUMAN | Unmodified | 0 | 0 | 842450 | 0 | 0 | 0 | 0 | 0 | 0 | 0 | 0 | 0 | 0 | 0 | 0 | 870310 | 0 | 0 |
| LLSGVLFH | MAN1A1 | P33908 | MA1A1_HUMA | Unmodified | 0 | 0 | 0 | 0 | 0 | 0 | 0 | 0 | 0 | 0 | 0 | 1446200 | 0 | 0 | 0 | 0 | 0 | 0 |
| LLSITVR | PRKDC | P78527 | PRKDC_HUMA | Unmodified | 0 | 0 | 0 | 0 | 0 | 0 | 0 | 0 | 1648400 | 536030 | 0 | 0 | 304630 | 0 | 25263 | 45963 | 0 | 0 |
| LLSIVNVCK | EXOC1 | Q9NV70 | EXOC1_HUMA | Unmodified | 0 | 0 | 0 | 0 | 0 | 0 | 0 | 0 | 363200 | 0 | 0 | 612310 | 0 | 0 | 0 | 0 | 0 | 0 |
| LLSKEPNPI | KPNA1; KPNA1; | P52294; C9 | IMA5_HUMAN | Unmodified | 224820 | 706530 | 98251 | 851740 | 0 | 0 | 0 | 149940 | 0 | 152600 | 113800 | 1195300 | 1621900 | 951320 | 192640 | 238320 | 556760 | 1149500 |
| LLSKEPSPI | KPNA6; KPNA6; | O60684; Q5 | IMA7_HUMAN | Unmodified | 878690 | 0 | 0 | 0 | 0 | 0 | 0 | 0 | 850990 | 357740 | 0 | 0 | 0 | 0 | 0 | 0 | 993210 | 0 |
| LLSKTPELN | CANX; CANX; C | P27824; A0 | CALX_HUMAN | Unmodified | 0 | 0 | 0 | 0 | 551410 | 0 | 909580 | 148930 | 52705000 | 20680000 | 2866800 | 0 | 0 | 0 | 0 | 0 | 0 | 0 |
| LLSKTPELN | CANX; CANX; C | P27824; A0 | CALX_HUMAN | Oxidation (M) | 0 | 0 | 0 | 0 | 4187100 | 0 | 5244900 | 0 | 64529000 | 1.07E+08 | 1977600 | 0 | 828360 | 999090 | 0 | 0 | 0 | 0 |
| LLSKTPELN | CANX; CANX; C | P27824; A0 | CALX_HUMAN | Unmodified | 0 | 0 | 0 | 0 | 3150300 | 0 | 2483900 | 0 | 60622000 | 22475000 | 0 | 0 | 0 | 0 | 0 | 0 | 0 | 0 |
| LLSLCLQK | STK39 | Q9UEW8 | STK39_HUMA | Unmodified | 0 | 0 | 0 | 0 | 0 | 0 | 0 | 0 | 0 | 0 | 0 | 0 | 601320 | 0 | 0 | 0 | 0 | 0 |
| LLSLGAQA | GIT2; GIT1; GIT | Q14161; Q9 | GIT2_HUMAN | Unmodified | 0 | 0 | 0 | 136910 | 579250 | 0 | 0 | 0 | 941990 | 453140 | 369280 | 2863200 | 0 | 0 | 290350 | 102590 | 166460 | 0 |
| LLSLISIALP | HUWE1; HUWE | Q7Z6Z7; H0 | HUWE1_HUMA | Unmodified | 0 | 0 | 0 | 0 | 0 | 0 | 0 | 0 | 0 | 0 | 0 | 172060 | 0 | 0 | 0 | 0 | 0 | 0 |
| LLSLLEKDE | SPR | P35270 | SPRE_HUMAN | Unmodified | 0 | 0 | 0 | 0 | 1154900 | 0 | 375770 | 78265 | 560620 | 187010 | 0 | 0 | 1232200 | 580560 | 0 | 0 | 0 | 942520 |
| LLSLLEKDE | SPR | P35270 | SPRE_HUMAN | Unmodified | 0 | 0 | 0 | 0 | 1312700 | 0 | 316590 | 0 | 0 | 321040 | 0 | 0 | 0 | 0 | 0 | 0 | 548800 | 675050 |
| LLSLLPLSC | SLC5A6 | Q9Y289 | SC5A6_HUMA | Unmodified | 0 | 0 | 0 | 0 | 0 | 0 | 0 | 0 | 243260 | 0 | 1086800 | 825650 | 0 | 0 | 0 | 0 | 0 | 0 |
| LLSNDEVTI | TFG; TFG; TFG; | Q92734; A0 | TFG_HUMAN P | Unmodified | 682530 | 613790 | 133280 | 33940 | 1092300 | 718860 | 261960 | 0 | 3287700 | 681680 | 447470 | 9943000 | 654930 | 0 | 979990 | 954310 | 207290 | 485420 |
| LLSNTVMP | CTNNA1; CTNN | P35221; G3 | CTNA1_HUMA | Oxidation (M) | 0 | 0 | 0 | 0 | 3164700 | 1387000 | 0 | 0 | 218320 | 0 | 0 | 2886200 | 2701600 | 2225000 | 205870 | 0 | 289230 | 0 |

| | | | | | | | | | | | | | | | | | | | | | | |
|---|---|---|---|---|---|---|---|---|---|---|---|---|---|---|---|---|---|---|---|---|---|---|
| LLSNTVMP | CTNNA1; CTNN | P35221; G3 | CTNA1_HUMA | Unmodified | 1340400 | 1472200 | 2417100 | 3883800 | 10523000 | 4525200 | 0 | 0 | 0 | 89909 | 2232000 | 14948000 | 7305800 | 5850600 | 6319900 | 3510300 | 0 | 0 |
| LLSPSILKP | POLR1A; POLR | O95602; B9 | RPA1_HUMAN | Unmodified | 0 | 0 | 30637 | 41870 | 165070 | 0 | 384040 | 0 | 0 | 0 | 108340 | 777730 | 0 | 0 | 0 | 0 | 65183 | 189440 |
| LLSPTER | DPP3; DPP3; D | Q9NY33; G3 | DPP3_HUMAN | Unmodified | 0 | 0 | 599760 | 93275 | 365100 | 0 | 0 | 0 | 527120 | 349140 | 0 | 1965400 | 286020 | 0 | 0 | 34097 | 0 | 0 |
| LLSPVVPQ | NUP93; NUP93 | Q8N1F7; H3 | NUP93_HUMA | Unmodified | 0 | 0 | 0 | 0 | 0 | 0 | 0 | 0 | 0 | 0 | 0 | 3920700 | 0 | 0 | 23678 | 151210 | 0 | 0 |
| LLSPYSPQ | DIP2B | Q9P265 | DIP2B_HUMAN | Unmodified | 290800 | 220020 | 269130 | 1941400 | 3675000 | 0 | 275110 | 402740 | 3753600 | 1283500 | 10834000 | 11089000 | 6391600 | 2159500 | 2835000 | 810740 | 1139900 | 16871000 |
| LLSPYSPQ | DIP2B | Q9P265 | DIP2B_HUMAN | Unmodified | 0 | 0 | 0 | 0 | 0 | 0 | 0 | 0 | 0 | 0 | 0 | 0 | 0 | 0 | 0 | 0 | 2199600 | 1433100 |
| LLSQCKFD | MAIP1; MAIP1 | Q8WWC4; H | MAIP1_HUMAN | Unmodified | 0 | 0 | 0 | 0 | 0 | 0 | 0 | 0 | 318870 | 0 | 0 | 0 | 0 | 2359500 | 0 | 0 | 0 | 0 |
| LLSRPQDA | ATAD3A | Q9NVI7 | ATD3A_HUMAN | Unmodified | 0 | 0 | 0 | 0 | 11711000 | 5380900 | 46419000 | 0 | 1757200 | 0 | 0 | 0 | 0 | 0 | 0 | 0 | 2111700 | 11803000 |
| LLSSDRNP | KPNA4; KPNA3; | O00629; O0 | IMA3_HUMAN | Unmodified | 336770 | 687840 | 147920 | 239900 | 834960 | 816640 | 0 | 93181 | 197870 | 315240 | 201290 | 1148700 | 639790 | 957420 | 237230 | 294700 | 1810500 | 705580 |
| LLSSYLQK | DTYMK; DTYMK | P23919; G5 | KTHY_HUMAN | Unmodified | 159020 | 114010 | 0 | 0 | 3720900 | 0 | 1362900 | 1030400 | 862460 | 676040 | 0 | 609230 | 900570 | 0 | 0 | 132150 | 1363400 | 3353500 |
| LLSTDPVA | AP1B1; AP1B1 | Q10567; C9 | AP1B1_HUMAN | Unmodified | 0 | 0 | 889460 | 0 | 0 | 0 | 0 | 0 | 680890 | 220350 | 569050 | 6759500 | 0 | 0 | 0 | 0 | 0 | 0 |
| LLSTDPVT | AP2B1; AP2B1; | P63010; A0 | AP2B1_HUMAN | Unmodified | 564540 | 619960 | 1346100 | 1403800 | 0 | 366060 | 0 | 0 | 2451500 | 924290 | 715090 | 8281300 | 0 | 0 | 1079500 | 1130200 | 0 | 0 |
| LLSTELQAE | CNPY4; CNPY4 | Q8N129; A0 | CNPY4_HUMA | Unmodified | 0 | 0 | 0 | 0 | 0 | 0 | 0 | 0 | 3046500 | 426590 | 137520 | 0 | 0 | 0 | 0 | 0 | 0 | 0 |
| LLSTELQAE | CNPY4; CNPY4 | Q8N129; A0 | CNPY4_HUMA | Unmodified | 0 | 0 | 0 | 0 | 0 | 0 | 0 | 0 | 1648400 | 4300600 | 0 | 0 | 0 | 0 | 0 | 0 | 0 | 0 |
| LLSTVAQA | PDIA5 | Q14554 | PDIA5_HUMAN | Unmodified | 0 | 0 | 0 | 0 | 0 | 0 | 0 | 0 | 22521000 | 1123600 | 0 | 378730 | 0 | 0 | 0 | 0 | 0 | 0 |
| LLSVEGSTI | MYO1C; MYO1 | O00159; F5 | MYO1C_HUMA | Unmodified | 5672300 | 6339000 | 8925500 | 7250400 | 0 | 0 | 0 | 0 | 929920 | 0 | 0 | 3968500 | 0 | 0 | 26675000 | 16140000 | 0 | 0 |
| LLSVEGSTI | MYO1C; MYO1 | O00159; F5 | MYO1C_HUMA | Unmodified | 0 | 0 | 0 | 6285800 | 2082700 | 995860 | 0 | 0 | 569080 | 220190 | 0 | 0 | 0 | 0 | 0 | 0 | 870570 | 520210 |
| LLSVEGSTI | MYO1C; MYO1 | O00159; F5 | MYO1C_HUMA | Unmodified | 884860 | 599940 | 839030 | 3631800 | 7000100 | 4761400 | 0 | 0 | 331820 | 206450 | 146690 | 0 | 1677400 | 1320300 | 2334900 | 1182300 | 2234300 | 998070 |
| LLSVLATVK | FAM20B | O75063 | XYLK_HUMAN | Unmodified | 0 | 0 | 0 | 0 | 0 | 0 | 0 | 0 | 24859000 | 1389500 | 668920 | 422100 | 0 | 0 | 0 | 0 | 0 | 0 |
| LLSVNIR | EPRS1 | P07814 | SYEP_HUMAN | Unmodified | 0 | 0 | 0 | 0 | 0 | 0 | 0 | 0 | 0 | 0 | 0 | 6503000 | 0 | 0 | 0 | 0 | 0 | 0 |
| LLSVVPVPE | C20orf27 | Q9GZN8 | CT027_HUMAN | Unmodified | 1022000 | 0 | 1098100 | 0 | 789350 | 0 | 0 | 0 | 0 | 0 | 0 | 3356900 | 651920 | 0 | 0 | 0 | 0 | 864890 |
| LLSYASYCI | IMMT; IMMT; IM | Q16891; A0 | MIC60_HUMAN | Unmodified | 0 | 0 | 0 | 0 | 0 | 0 | 77220 | 0 | 0 | 137680 | 0 | 0 | 0 | 0 | 0 | 0 | 0 | 0 |
| LLTDDGNK | PSMD2 | Q13200 | PSMD2_HUMA | Unmodified | 0 | 0 | 0 | 0 | 407000 | 137110 | 204230 | 136400 | 606240 | 39778 | 0 | 1733900 | 32130 | 0 | 0 | 0 | 0 | 0 |
| LLTDDGNK | PSMD2 | Q13200 | PSMD2_HUMA | Unmodified | 0 | 0 | 0 | 0 | 1180700 | 305930 | 397430 | 127360 | 109550 | 300790 | 0 | 0 | 1331100 | 1054500 | 0 | 0 | 1644700 | 1288800 |
| LLTDSDLK | PALS2; PALS2 | Q9NZW5; B | PALS2_HUMAN | Unmodified | 0 | 0 | 0 | 0 | 1387400 | 0 | 0 | 0 | 0 | 0 | 0 | 0 | 0 | 0 | 0 | 0 | 0 | 0 |
| LLTEEGQK | SLC25A24 | Q6NUK1 | SCMC1_HUMA | Unmodified | 0 | 0 | 0 | 0 | 0 | 0 | 0 | 0 | 2940400 | 0 | 0 | 0 | 0 | 0 | 0 | 0 | 0 | 0 |
| LLTEEGQK | SLC25A24 | Q6NUK1 | SCMC1_HUMA | Unmodified | 0 | 0 | 0 | 0 | 0 | 0 | 0 | 0 | 1155800 | 242010 | 0 | 0 | 150950 | 0 | 0 | 0 | 0 | 0 |
| LLTETEDW | HSPH1; HSPH1 | Q92598; A0 | HS105_HUMAN | Unmodified | 0 | 0 | 2311200 | 0 | 0 | 0 | 0 | 0 | 1469700 | 351210 | 0 | 1673400 | 321380 | 0 | 0 | 0 | 138170 | 265120 |
| LLTFNPTY | UBE4A | Q14139 | UBE4A_HUMAN | Unmodified | 0 | 0 | 0 | 0 | 0 | 0 | 0 | 0 | 0 | 0 | 0 | 993270 | 0 | 0 | 0 | 0 | 0 | 0 |
| LLTGSSYK | CNRIP1; CNRIP | Q96F85; B8 | CNRP1_HUMA | Unmodified | 0 | 0 | 0 | 0 | 0 | 0 | 243530 | 0 | 0 | 0 | 0 | 0 | 0 | 173980 | 0 | 0 | 410900 | 0 |
| LLTHHKDH | CCT2; CCT2; C | P78371; F5 | TCPB_HUMAN | Unmodified | 0 | 0 | 0 | 0 | 1061700 | 0 | 914480 | 53837 | 0 | 285120 | 0 | 0 | 821320 | 941440 | 0 | 0 | 914600 | 691990 |
| LLTHNLLS | TRMT112; TRM | Q9UI30; F5 | TR112_HUMAN | Unmodified | 0 | 0 | 0 | 0 | 0 | 0 | 0 | 0 | 0 | 0 | 0 | 1554100 | 651230 | 0 | 0 | 0 | 0 | 0 |
| LLTHTFVA | MAP4K5 | Q9Y4K4 | M4K5_HUMAN | Unmodified | 0 | 90458 | 108430 | 140700 | 434350 | 0 | 0 | 0 | 607550 | 0 | 513320 | 4010800 | 0 | 0 | 0 | 0 | 78741 | 156690 |
| LLTLEVPG | MOV10; MOV1 | Q9HCE1; Q | MOV10_HUMA | Unmodified | 28528000 | 12509000 | 6663800 | 0 | 0 | 0 | 0 | 112200 | 0 | 0 | 0 | 755340 | 0 | 0 | 5267800 | 22738000 | 0 | 0 |
| LLTLEVPG | MOV10; MOV1 | Q9HCE1; Q | MOV10_HUMA | Unmodified | 0 | 2462300 | 0 | 0 | 0 | 0 | 0 | 0 | 0 | 0 | 0 | 0 | 0 | 0 | 1983000 | 0 | 0 | 0 |
| LLTLPNGE | DYNC1H1; DYN | Q14204; A0 | DYHC1_HUMA | Unmodified | 0 | 0 | 158390 | 0 | 0 | 0 | 0 | 42001 | 0 | 0 | 0 | 8032800 | 0 | 0 | 380330 | 382370 | 0 | 0 |
| LLTNLGLG | SLC39A10; SLC | Q9ULF5; E7 | S39AA_HUMAN | Unmodified | 0 | 162980 | 0 | 0 | 3578700 | 659220 | 0 | 61541 | 0 | 126340 | 96183 | 3311700 | 863400 | 0 | 0 | 0 | 957320 | 2313700 |
| LLTNLGLG | SLC39A10; SLC | Q9ULF5; E7 | S39AA_HUMAN | Unmodified | 0 | 0 | 0 | 637270 | 2822800 | 0 | 0 | 0 | 515990 | 146110 | 0 | 8603600 | 0 | 0 | 1140900 | 803490 | 625560 | 2049000 |
| LLTNLGLG | SLC39A10; SLC | Q9ULF5; E7 | S39AA_HUMAN | Unmodified | 0 | 0 | 0 | 0 | 33515000 | 2984800 | 925860 | 0 | 0 | 0 | 0 | 0 | 8793100 | 2645800 | 0 | 0 | 3155000 | 16297000 |
| LLTPEAVE | DIPK2A | Q8NDZ4 | DIK2A_HUMAN | Unmodified | 0 | 0 | 0 | 0 | 0 | 0 | 257130 | 0 | 69762000 | 2401400 | 331640 | 775730 | 200110 | 0 | 0 | 0 | 0 | 0 |
| LLTQENR | AGL | P35573 | GDE_HUMAN | Unmodified | 0 | 0 | 0 | 0 | 0 | 0 | 0 | 0 | 0 | 0 | 0 | 944080 | 0 | 0 | 0 | 0 | 0 | 0 |
| LLTQGFTN | SMARCA5 | O60264 | SMCA5_HUMA | Unmodified | 572280 | 278740 | 0 | 0 | 0 | 0 | 0 | 0 | 0 | 0 | 0 | 790390 | 162320 | 184000 | 0 | 0 | 0 | 0 |
| LLTQHENI | SPTBN1; SPTBN | Q01082; A0 | SPTB2_HUMAN | Unmodified | 93754 | 0 | 0 | 0 | 1101500 | 406090 | 0 | 0 | 0 | 363870 | 301040 | 3075200 | 0 | 0 | 0 | 1035800 | 411670 | 0 |
| LLTQYILNL | AP3B1 | O00203 | AP3B1_HUMAN | Unmodified | 0 | 0 | 0 | 0 | 0 | 0 | 1066500 | 16769 | 0 | 412940 | 0 | 2827500 | 582920 | 0 | 0 | 0 | 22949000 | 3003500 |
| LLTSFLPAQ | SLC3A2; SLC3A | P08195; A0 | 4F2_HUMAN 4 | Unmodified | 17438000 | 6978300 | 3066800 | 4528600 | 1719700 | 1267400 | 55410 | 0 | 9655700 | 2342000 | 5777400 | 3.89E+08 | 109790 | 0 | 37759000 | 16793000 | 108710 | 262470 |
| LLTSGYLQ | OTUB1; OTUB1 | Q96FW1; J3 | OTUB1_HUMA | Unmodified | 324460 | 282560 | 0 | 0 | 0 | 0 | 0 | 0 | 285080 | 0 | 688310 | 7816600 | 0 | 0 | 643240 | 652430 | 0 | 0 |
| LLTSSLQK | PPFIBP1; PPFIB | Q86W92; H | LIPB1_HUMAN | Unmodified | 25819 | 0 | 212640 | 244440 | 1527000 | 0 | 0 | 41724 | 0 | 0 | 0 | 94693 | 901000 | 1224600 | 154710 | 151630 | 1374900 | 1638700 |
| LLTSVEVD | USP54; USP54; | Q70EL1; A0 | UBP54_HUMAN | Unmodified | 0 | 0 | 0 | 0 | 0 | 0 | 858560 | 0 | 0 | 0 | 0 | 0 | 1565900 | 0 | 0 | 26418 | 0 | 0 |
| LLTSYGFIQ | CSDE1; CSDE1 | O75534; E9 | CSDE1_HUMA | Unmodified | 0 | 0 | 0 | 148510 | 214220 | 0 | 0 | 0 | 135530 | 0 | 134600 | 818350 | 157140 | 117440 | 116580 | 101640 | 74603 | 0 |
| LLTSYGFIQ | CSDE1; CSDE1 | O75534; E9 | CSDE1_HUMA | Unmodified | 0 | 0 | 0 | 0 | 0 | 0 | 0 | 159450 | 0 | 522960 | 0 | 0 | 2745300 | 0 | 0 | 0 | 333970 | 0 |
| LLTTGVSR | CORO2B | Q9UQ03 | COR2B_HUMA | Unmodified | 0 | 0 | 0 | 0 | 0 | 101380 | 542090 | 0 | 0 | 0 | 54651 | 983990 | 0 | 0 | 0 | 0 | 53575 | 0 |
| LLTTIGKDL | EIF4G1; EIF4G3 | Q04637; O4 | IF4G1_HUMAN | Unmodified | 128690 | 66054 | 0 | 179450 | 1126400 | 0 | 76755 | 15974 | 300000 | 327000 | 98988 | 303490 | 1111800 | 415070 | 0 | 30607 | 0 | 489530 |
| LLTTILHSD | PDS5B | Q9NTI5 | PDS5B_HUMAN | Unmodified | 0 | 0 | 0 | 0 | 0 | 0 | 0 | 0 | 0 | 0 | 0 | 0 | 0 | 0 | 0 | 0 | 0 | 0 |
| LLTVPFEK | PGLS; PGLS | O95336; M0 | 6PGL_HUMAN | Unmodified | 0 | 0 | 0 | 0 | 0 | 721710 | 315370 | 35167 | 0 | 1964200 | 0 | 1231500 | 0 | 0 | 0 | 0 | 1404100 | 0 |
| LLTYNSLYK | MOV10; MOV1 | Q9HCE1; Q | MOV10_HUMA | Unmodified | 4545500 | 4474700 | 2156200 | 1869500 | 0 | 0 | 0 | 0 | 0 | 0 | 0 | 0 | 0 | 0 | 2087700 | 2223300 | 0 | 0 |
| LLTYNSLYK | MOV10; MOV1 | Q9HCE1; Q | MOV10_HUMA | Unmodified | 568980 | 611520 | 170160 | 117530 | 0 | 0 | 0 | 0 | 0 | 0 | 0 | 0 | 0 | 0 | 224690 | 311320 | 0 | 0 |
| LLVATSVA | DDX46; DDX46 | Q7L014; A0 | DDX46_HUMA | Unmodified | 0 | 0 | 0 | 0 | 0 | 0 | 959170 | 91498 | 0 | 0 | 0 | 396340 | 0 | 0 | 332710 | 0 | 0 | 0 |
| LLVATTFTH | IARS1; IARS1; I | P41252; A0 | SYIC_HUMAN | Unmodified | 0 | 0 | 0 | 0 | 0 | 0 | 0 | 0 | 0 | 0 | 167260 | 2458400 | 0 | 0 | 0 | 0 | 0 | 0 |
| LLVCDLGN | ITGA5 | P08648 | ITA5_HUMAN I | Oxidation (M) | 395740 | 137550 | 0 | 273500 | 0 | 0 | 0 | 0 | 0 | 0 | 0 | 0 | 0 | 175220 | 86104 | 455630 | 0 | 0 |
| LLVDVDES | SF3B1 | O75533 | SF3B1_HUMAN | Unmodified | 0 | 0 | 51576 | 0 | 0 | 0 | 17728 | 0 | 0 | 414360 | 3363400 | 40456 | 0 | 0 | 0 | 0 | 0 | 0 |
| LLVDVDES | SF3B1 | O75533 | SF3B1_HUMAN | Unmodified | 0 | 0 | 0 | 0 | 0 | 7692400 | 0 | 0 | 0 | 609940 | 0 | 1960600 | 0 | 0 | 0 | 0 | 2041600 | 0 |
| LLVEDINK | SLC39A10 | Q9ULF5 | S39AA_HUMAN | Unmodified | 287200 | 0 | 720490 | 49050 | 0 | 0 | 86801 | 78238 | 573810 | 278720 | 711530 | 4491200 | 0 | 245300 | 964760 | 586400 | 0 | 0 |
| LLVFATDD | ITGB2; ITGB2; I | P05107; D3 | ITB2_HUMAN I | Unmodified | 262430 | 414530 | 262150 | 1014200 | 138300 | 0 | 79989 | 0 | 496350 | 0 | 139470 | 0 | 202680 | 181840 | 295510 | 326730 | 0 | 98880 |
| LLVFSTDA | ITGB1; ITGB1; I | P05556; H7 | ITB1_HUMAN I | Unmodified | 3015200 | 4633800 | 2281000 | 5049500 | 18089000 | 3669400 | 576380 | 0 | 16392000 | 1555100 | 2301400 | 6167600 | 1736600 | 748030 | 7901000 | 4017800 | 2466900 | 11570000 |
| LLVGNDDV | HUWE1 | Q7Z6Z7 | HUWE1_HUMA | Unmodified | 0 | 0 | 0 | 0 | 0 | 0 | 0 | 0 | 685980 | 0 | 0 | 4749400 | 0 | 0 | 0 | 0 | 0 | 0 |
| LLVGNKCD | RAB1A; RAB1A | P62820; E7 | RAB1A_HUMAN | Unmodified | 2563200 | 1672500 | 73127 | 4138300 | 485300 | 1484500 | 4051500 | 2258300 | 295190 | 2999800 | 4187300 | 16824000 | 6768700 | 6073000 | 1256200 | 1341500 | 2892800 | 0 |
| LLVGNKCD | RAB1A; RAB1A | P62820; E7 | RAB1A_HUMAN | Unmodified | 375070 | 0 | 0 | 392960 | 702270 | 0 | 700000 | 0 | 596320 | 323970 | 484110 | 1288500 | 1508600 | 1328100 | 0 | 166970 | 544400 | 824240 |
| LLVGNKSD | RAB1C; RAB1B; | Q92928; Q9 | RAB1C_HUMA | Unmodified | 3175100 | 2890200 | 827260 | 6651600 | 9938300 | 2397900 | 4411900 | 2434000 | 5382500 | 5248800 | 2934800 | 13452000 | 12021000 | 7947800 | 2922800 | 2553200 | 3442800 | 11030000 |
| LLVGNKSD | RAB1C; RAB1B; | Q92928; Q9 | RAB1C_HUMA | Unmodified | 0 | 721430 | 0 | 1532900 | 2585000 | 296890 | 0 | 22788 | 1117700 | 609570 | 427630 | 2865500 | 1880000 | 0 | 0 | 464790 | 512500 | 1693000 |
| LLVGPTTLA | GGT6; GGT6; G | Q6P531; B4 | GGT6_HUMAN | Unmodified | 0 | 0 | 0 | 0 | 0 | 0 | 0 | 0 | 327800 | 198980 | 0 | 0 | 0 | 0 | 0 | 0 | 0 | 0 |
| LLVGSEDY | SPTAN1; SPTAN | Q13813; A0 | SPTN1_HUMAN | Unmodified | 0 | 0 | 0 | 0 | 167830 | 197750 | 0 | 0 | 365280 | 0 | 0 | 2044900 | 934980 | 0 | 0 | 0 | 0 | 0 |
| LLVGSEDY | SPTAN1; SPTAN | Q13813; A0 | SPTN1_HUMAN | Unmodified | 0 | 0 | 0 | 0 | 715740 | 0 | 0 | 0 | 0 | 0 | 0 | 1834400 | 0 | 0 | 0 | 0 | 0 | 0 |
| LLVGVDEK | ACLY | P53396 | ACLY_HUMAN | Unmodified | 0 | 0 | 0 | 0 | 0 | 0 | 0 | 0 | 0 | 0 | 0 | 19979000 | 0 | 0 | 0 | 0 | 0 | 0 |
| LLVGVDEK | ACLY | P53396 | ACLY_HUMAN | Unmodified | 0 | 0 | 0 | 0 | 0 | 197270 | 364830 | 123740 | 0 | 0 | 0 | 0 | 0 | 0 | 0 | 0 | 0 | 0 |
| LLVGVDEK | ACLY | P53396 | ACLY_HUMAN | Unmodified | 0 | 0 | 0 | 0 | 18811000 | 0 | 7717800 | 254720 | 0 | 0 | 0 | 0 | 3518300 | 0 | 0 | 0 | 1513700 | 0 |
| LLVITVATA | PLOD3; PLOD3 | O60568; C9 | PLOD3_HUMA | Unmodified | 0 | 0 | 0 | 0 | 0 | 0 | 0 | 0 | 281400 | 776510 | 0 | 0 | 0 | 0 | 0 | 0 | 0 | 0 |
| LLVITVATK | PLOD2; PLOD2 | O00469; C9 | PLOD2_HUMA | Unmodified | 0 | 0 | 0 | 0 | 106120 | 0 | 0 | 0 | 7932800 | 4983300 | 403090 | 0 | 0 | 0 | 0 | 0 | 0 | 0 |
| LLVLDATK | PDPK2P; PDPK | Q6A1A2; O1 | PDPK2_HUMA | Unmodified | 103990 | 0 | 1138400 | 375940 | 185210 | 0 | 204780 | 188250 | 0 | 135540 | 75500 | 418130 | 2918000 | 688390 | 1965500 | 1414300 | 117210 | 0 |
| LLVLEAFQ | GSTZ1; GSTZ1; | O43708; G3 | MAAI_HUMAN | Unmodified | 0 | 0 | 0 | 0 | 288220 | 0 | 0 | 0 | 917320 | 150680 | 0 | 0 | 0 | 0 | 0 | 0 | 0 | 0 |
| LLVLYSGE | UNC45A; UNC4 | Q9H3U1; A0 | UN45A_HUMA | Unmodified | 92692 | 0 | 0 | 0 | 0 | 0 | 74375 | 80278 | 252280 | 107560 | 76100 | 1839500 | 322070 | 0 | 480020 | 0 | 82216 | 0 |
| LLVPANEG | TNFRSF10B | O14763 | TR10B_HUMAN | Unmodified | 0 | 0 | 0 | 0 | 0 | 0 | 0 | 0 | 295590 | 0 | 0 | 0 | 0 | 0 | 1570900 | 0 | 0 | 0 |
| LLVPGKIQ | VPS29; VPS29 | Q9UBQ0; F8 | VPS29_HUMAN | Unmodified | 0 | 0 | 0 | 0 | 0 | 0 | 0 | 0 | 0 | 410900 | 0 | 0 | 2809400 | 1621200 | 0 | 0 | 2149100 | 0 |
| LLVPGTYK | CPD; CPD | O75976; J3 | CBPD_HUMAN | Unmodified | 108790 | 90382 | 518630 | 217460 | 0 | 0 | 0 | 0 | 36985000 | 5743100 | 390480 | 397210 | 121010 | 0 | 149910 | 333190 | 0 | 0 |
| LLVPGTYK | CPD; CPD | O75976; J3 | CBPD_HUMAN | Unmodified | 0 | 0 | 0 | 264720 | 0 | 0 | 206960 | 196120 | 1295900 | 3075500 | 0 | 0 | 0 | 0 | 0 | 0 | 0 | 0 |
| LLVPILLPE | MPDU1; MPDU | O75352; J3 | MPU1_HUMAN | Unmodified | 20200 | 0 | 54045 | 0 | 0 | 0 | 0 | 0 | 3959300 | 799380 | 765300 | 562770 | 0 | 0 | 44344 | 39332 | 0 | 0 |
| LLVPLVPD | ESYT1 | Q9BSJ8 | ESYT1_HUMAN | Unmodified | 164940 | 127980 | 22308 | 0 | 37553 | 0 | 0 | 0 | 40314 | 0 | 0 | 172710 | 0 | 0 | 276990 | 0 | 0 | 0 |
| LLVPSPEG | FERMT3 | Q86UX7 | URP2_HUMAN | Oxidation (M) | 439810 | 210840 | 496900 | 2206400 | 0 | 0 | 74835 | 69209 | 0 | 0 | 0 | 0 | 432290 | 132990 | 0 | 472420 | 0 | 0 |
| LLVPTQFV | IGF2BP3 | O00425 | IF2B3_HUMAN | Unmodified | 2100100 | 1563100 | 152560 | 493730 | 0 | 0 | 0 | 451050 | 207620 | 86193 | 458810 | 3435900 | 278360 | 183560 | 255570 | 112110 | 0 | 164030 |
| LLVPTQYV | IGF2BP1 | Q9NZI8 | IF2B1_HUMAN | Unmodified | 611970 | 419470 | 0 | 0 | 0 | 0 | 0 | 0 | 0 | 0 | 0 | 0 | 0 | 0 | 0 | 0 | 0 | 421660 |
| LLVPYLME | HADH; HADH; | Q16836; A0 | HCDH_HUMAN | Oxidation (M) | 0 | 0 | 0 | 0 | 0 | 0 | 0 | 0 | 1359100 | 0 | 0 | 0 | 0 | 0 | 0 | 0 | 0 | 0 |
| LLVPYLME | HADH; HADH; | Q16836; A0 | HCDH_HUMAN | Unmodified | 0 | 0 | 0 | 0 | 0 | 0 | 0 | 0 | 5378900 | 61788 | 0 | 0 | 0 | 0 | 0 | 0 | 0 | 0 |
| LLVQEQQ | RASAL2; RASAL | Q9UJF2; H0 | NGAP_HUMAN | Unmodified | 0 | 0 | 0 | 0 | 521590 | 212610 | 47419 | 0 | 0 | 0 | 0 | 0 | 0 | 192110 | 0 | 0 | 0 | 0 |
| LLVQNYR | CA14; CA14 | Q9ULX7; U3 | CAH14_HUMA | Unmodified | 0 | 0 | 0 | 0 | 0 | 0 | 0 | 0 | 0 | 0 | 0 | 6100400 | 0 | 0 | 0 | 0 | 0 | 0 |
| LLVRGLGE | LNPEP | Q9UIQ6 | LCAP_HUMAN | Unmodified | 0 | 0 | 0 | 0 | 2122800 | 0 | 0 | 0 | 0 | 0 | 0 | 0 | 0 | 0 | 0 | 0 | 0 | 0 |
| LLVRHAYS | BTBD17 | A6NE02 | BTBDH_HUMA | Unmodified | 0 | 0 | 0 | 0 | 0 | 0 | 1073700 | 0 | 0 | 195200 | 0 | 0 | 0 | 0 | 0 | 0 | 0 | 0 |
| LLVSASQD | GNB2; GNB4; G | P62879; Q9 | GBB2_HUMAN | Unmodified | 12872000 | 15567000 | 14284000 | 7443200 | 52592 | 0 | 0 | 0 | 21978000 | 1317100 | 21725000 | 4.79E+08 | 712020 | 57651 | 24722000 | 21428000 | 0 | 187780 |
| LLVSGSSD | NLE1; NLE1; NL | Q9NVX2; A0 | NLE1_HUMAN | Unmodified | 0 | 0 | 0 | 0 | 0 | 0 | 0 | 0 | 0 | 0 | 0 | 1175200 | 0 | 0 | 0 | 0 | 0 | 0 |
| LLVSHGAE | ANKRD28; ANK | O15084; B4 | ANR28_HUMA | Unmodified | 0 | 0 | 0 | 156650 | 64918 | 0 | 47112 | 27879 | 160260 | 85131 | 0 | 1605100 | 90191 | 0 | 162200 | 0 | 78049 | 0 |
| LLVSMCQ | ANXA7 | P20073 | ANXA7_HUMA | Oxidation (M) | 0 | 0 | 0 | 0 | 0 | 0 | 0 | 0 | 540770 | 186390 | 0 | 0 | 0 | 0 | 0 | 0 | 0 | 0 |
| LLVSMCQ | ANXA7 | P20073 | ANXA7_HUMA | Unmodified | 0 | 0 | 0 | 0 | 0 | 0 | 0 | 0 | 2994400 | 0 | 0 | 0 | 274280 | 0 | 0 | 0 | 109830 | 0 |
| LLVSMCQ | ANXA7 | P20073 | ANXA7_HUMA | Oxidation (M) | 1146000 | 0 | 1397600 | 1857000 | 0 | 0 | 0 | 0 | 1.2E+08 | 1058900 | 7179500 | 3248700 | 0 | 0 | 36281 | 2961100 | 0 | 182150 |
| LLVSMCQ | ANXA7 | P20073 | ANXA7_HUMA | Unmodified | 487660 | 0 | 0 | 1143300 | 0 | 0 | 0 | 0 | 2.27E+08 | 1292800 | 0 | 2472500 | 0 | 0 | 1211700 | 397680 | 0 | 0 |
| LLVSMCQ | ANXA7 | P20073 | ANXA7_HUMA | 2 Oxidation (M) | 405400 | 166420 | 434850 | 377350 | 0 | 0 | 0 | 0 | 11706000 | 270220 | 0 | 0 | 0 | 0 | 534680 | 38622 | 0 | 55623 |
| LLVSMCQ | ANXA7 | P20073 | ANXA7_HUMA | Oxidation (M) | 0 | 0 | 0 | 0 | 0 | 0 | 0 | 0 | 4957500 | 0 | 0 | 0 | 0 | 0 | 0 | 0 | 0 | 0 |
| LLVSMCQ | ANXA7 | P20073 | ANXA7_HUMA | Unmodified | 0 | 0 | 0 | 0 | 0 | 0 | 0 | 0 | 30921000 | 0 | 0 | 0 | 0 | 0 | 0 | 0 | 0 | 0 |
| LLVSQGAS | PSMD10; PSMD | O75832; B1 | PSD10_HUMAN | Unmodified | 407020 | 261890 | 391240 | 0 | 870940 | 0 | 353700 | 0 | 1874700 | 0 | 565280 | 0 | 3856200 | 0 | 847410 | 1225200 | 1545600 | 2135000 |
| LLVSQGAS | PSMD10; PSMD | O75832; B1 | PSD10_HUMAN | Unmodified | 0 | 0 | 0 | 0 | 0 | 0 | 1584100 | 0 | 0 | 1448400 | 0 | 0 | 2934200 | 3100400 | 0 | 0 | 5544200 | 2843000 |
| LLVTTSTSL | HIP1R | O75146 | HIP1R_HUMAN | Unmodified | 0 | 0 | 0 | 0 | 0 | 0 | 0 | 0 | 0 | 0 | 0 | 718500 | 0 | 0 | 0 | 0 | 0 | 0 |
| LLVVDPET | SLC9A3R2; SLC | Q15599; H3 | NHRF2_HUMA | Unmodified | 1719700 | 1120100 | 1248600 | 2809700 | 3796100 | 1555700 | 0 | 0 | 1671000 | 462100 | 0 | 8510600 | 4842200 | 1251700 | 3666600 | 3014900 | 1340600 | 0 |

| | | | | | | | | | | | | | | | | | | | | | | |
|---|---|---|---|---|---|---|---|---|---|---|---|---|---|---|---|---|---|---|---|---|---|---|
| LLVVDPETI | SLC9A3R1; SLC | O14745; J3Q | NHRF1_HUMAI | Unmodified | 65519 | 0 | 4589200 | 8871200 | 9745200 | 1400300 | 0 | 1131200 | 2926000 | 1856300 | 6582100 | 23895000 | 29475000 | 21127000 | 9365300 | 7683600 | 2293500 | 1737100 |
| LLVVDPETI | SLC9A3R1; SLC | O14745; J3Q | NHRF1_HUMAI | Unmodified | 0 | 0 | 0 | 0 | 0 | 0 | 0 | 0 | 0 | 85002 | 0 | 0 | 2917100 | 1246900 | 0 | 0 | 294650 | 0 |
| LLVVDQETI | SLC9A3R2; SLC | Q15599; H3I | NHRF2_HUMAI | Unmodified | 1488500 | 851810 | 0 | 0 | 0 | 0 | 0 | 0 | 3662000 | 0 | 0 | 2832400 | 1336600 | 0 | 2101700 | 1406400 | 0 | 1730900 |
| LLVVDRETI | SLC9A3R1; SLC | O14745; J3Q | NHRF1_HUMAI | Unmodified | 0 | 0 | 0 | 0 | 12288000 | 1403200 | 265270 | 0 | 0 | 86490 | 27938 | 0 | 29453000 | 473040 | 0 | 0 | 4600900 | 2310200 |
| LLVVDRETI | SLC9A3R1; SLC | O14745; J3Q | NHRF1_HUMAI | Unmodified | 650080 | 516120 | 1775100 | 4573300 | 23445000 | 4641500 | 9900900 | 1559500 | 5236300 | 2074300 | 1840100 | 11249000 | 47654000 | 60996000 | 3216200 | 4662600 | 14531000 | 7427400 |
| LLVVTDPR | RPSA; RPSA; RF | P08865; A0A | RSSA_HUMAN | Unmodified | 2735100 | 2411300 | 3089500 | 1584300 | 169460 | 0 | 113170 | 149660 | 1155600 | 67771 | 3017500 | 45415000 | 352070 | 0 | 2424000 | 2825500 | 0 | 0 |
| LLVVWNN( | EXT2 | Q93063 | EXT2_HUMAN I | Unmodified | 0 | 0 | 0 | 0 | 0 | 0 | 0 | 0 | 1208600 | 0 | 0 | 0 | 0 | 0 | 0 | 0 | 0 | 0 |
| LLWEQELY | WASL | O00401 | WASL_HUMAN | Unmodified | 614570 | 0 | 95023 | 628120 | 0 | 0 | 0 | 0 | 0 | 0 | 0 | 100880 | 0 | 0 | 3295800 | 63959 | 0 | 0 |
| LLWFVTESI | DIP2C; DIP2C; I | Q9Y2E4; A0/ | DIP2C_HUMAN | Unmodified | 0 | 0 | 382950 | 0 | 0 | 0 | 0 | 0 | 772220 | 402700 | 0 | 0 | 0 | 0 | 1070000 | 774590 | 0 | 0 |
| LLWHPVMI | MYOF; MYOF | Q9NZM1; HC | MYOF_HUMAN | Oxidation (M) | 0 | 0 | 0 | 0 | 1395500 | 0 | 0 | 0 | 0 | 0 | 0 | 0 | 0 | 0 | 0 | 0 | 0 | 102940 |
| LLWHPVMI | MYOF; MYOF | Q9NZM1; HC | MYOF_HUMAN | Unmodified | 0 | 0 | 0 | 0 | 449640 | 1112800 | 0 | 0 | 0 | 0 | 0 | 0 | 0 | 0 | 0 | 0 | 0 | 0 |
| LLWTLESL\ | ALDH16A1 | Q8IZ83 | A16A1_HUMAN | Unmodified | 103660 | 0 | 0 | 0 | 0 | 0 | 0 | 0 | 0 | 0 | 0 | 230610 | 0 | 0 | 0 | 0 | 0 | 0 |
| LLWTTSR | JUP; CTNNB1; ( | P14923; P35 | PLAK_HUMAN . | Unmodified | 1187800 | 879150 | 4306800 | 2994000 | 1238700 | 470430 | 0 | 0 | 7250100 | 2209500 | 0 | 0 | 65405 | 0 | 9656200 | 9701600 | 391600 | 473810 |
| LLWTTSRVI | JUP; CTNNB1; ( | P14923; P35 | PLAK_HUMAN . | Unmodified | 0 | 0 | 0 | 0 | 1380900 | 494810 | 0 | 0 | 0 | 195840 | 0 | 0 | 1024900 | 0 | 0 | 0 | 1391100 | 686140 |
| LLWVIRED( | NRP2 | O60462 | NRP2_HUMAN | Unmodified | 0 | 0 | 0 | 0 | 0 | 0 | 1010300 | 0 | 2053000 | 261690 | 0 | 0 | 0 | 0 | 0 | 0 | 400660 | 0 |
| LLYAIEETE( | WWP2 | O00308 | WWP2_HUMAN | Unmodified | 428380 | 504420 | 0 | 199750 | 0 | 0 | 0 | 0 | 0 | 0 | 0 | 0 | 159330 | 143310 | 392580 | 211490 | 0 | 0 |
| LLYDLADQ | ETFA; ETFA; ETF | P13804; H0Y | ETFA_HUMAN I | Unmodified | 0 | 0 | 0 | 0 | 275770 | 0 | 0 | 0 | 4321600 | 711690 | 0 | 0 | 0 | 0 | 0 | 0 | 0 | 0 |
| LLYEALVD( | EIF3M; EIF3M; E | Q7L2H7; E9I | EIF3M_HUMAN | Unmodified | 0 | 0 | 0 | 0 | 0 | 0 | 0 | 0 | 0 | 0 | 0 | 991390 | 0 | 0 | 0 | 919890 | 0 | 0 |
| LLYGFLIK | NCSTN; NCSTN | Q92542; H0Y | NICA_HUMAN | Unmodified | 325670 | 0 | 1005600 | 0 | 0 | 0 | 0 | 0 | 19195000 | 2220700 | 0 | 19172000 | 0 | 0 | 265010 | 1256600 | 0 | 0 |
| LLYNNVSN | CLTC; CLTC; CI | Q00610; J3K | CLH1_HUMAN | Unmodified | 27235000 | 27429000 | 15846000 | 14128000 | 2358400 | 1782100 | 519240 | 0 | 4612500 | 1268300 | 2648100 | 1.14E+08 | 2778600 | 817310 | 13742000 | 14681000 | 670380 | 1225500 |
| LLYNNVSN | CLTC; CLTC; CI | Q00610; J3K | CLH1_HUMAN | Unmodified | 0 | 0 | 0 | 183600 | 14390000 | 5052400 | 9228500 | 0 | 0 | 1682300 | 0 | 0 | 7000600 | 11767000 | 0 | 0 | 7299000 | 10697000 |
| LLYNRPGT\ | DEK; DEK; DEK; | P35659; B4D | DEK_HUMAN P | Unmodified | 0 | 0 | 0 | 0 | 0 | 0 | 0 | 756580 | 0 | 0 | 0 | 0 | 0 | 0 | 0 | 0 | 0 | 0 |
| LLYPPETGI | CSK; CSK; CSK | P41240; H3E | CSK_HUMAN T | Unmodified | 0 | 0 | 0 | 0 | 0 | 0 | 0 | 0 | 0 | 0 | 515540 | 6673900 | 0 | 0 | 0 | 212050 | 0 | 0 |
| LLYSGVPFI | ST14 | Q9Y5Y6 | ST14_HUMAN ! | Unmodified | 0 | 8038800 | 19249000 | 12935000 | 0 | 215360 | 0 | 15249 | 4433500 | 0 | 923320 | 0 | 4580200 | 0 | 31983000 | 16285000 | 0 | 0 |
| LMAEGAPK | L1CAM; L1CAM | P32004; E7E | L1CAM_HUMAI | Unmodified | 0 | 0 | 0 | 0 | 0 | 0 | 0 | 0 | 741900 | 0 | 0 | 0 | 0 | 0 | 0 | 0 | 0 | 0 |
| LMAFQPAP | PYCR2; PYCR2; | Q96C36; A0/ | P5CR2_HUMAN | Unmodified | 0 | 0 | 0 | 21814 | 208150 | 0 | 0 | 0 | 920210 | 128420 | 0 | 0 | 0 | 0 | 44437 | 0 | 0 | 0 |
| LMDDDAN( | STIM1; STIM1; S | Q13586; G0 | STIM1_HUMAN | Oxidation (M) | 0 | 0 | 0 | 0 | 0 | 0 | 0 | 0 | 1439900 | 0 | 0 | 0 | 0 | 0 | 0 | 0 | 0 | 0 |
| LMDEVAGI\ | ACOT7; ACOT7 | O00154; K7E | BACH_HUMAN | Oxidation (M) | 0 | 0 | 0 | 0 | 0 | 0 | 0 | 118570 | 436990 | 0 | 351060 | 1678800 | 0 | 0 | 0 | 0 | 449150 | 402170 |
| LMDEVAGI\ | ACOT7; ACOT7 | O00154; K7E | BACH_HUMAN | Unmodified | 0 | 0 | 0 | 0 | 0 | 0 | 0 | 0 | 0 | 0 | 0 | 0 | 0 | 0 | 0 | 0 | 583560 | 0 |
| LMDHVGTE | RPN2; RPN2; RI | P04844; F2Z | RPN2_HUMAN | Oxidation (M) | 0 | 0 | 0 | 0 | 0 | 0 | 0 | 0 | 1974400 | 2588000 | 116840 | 0 | 0 | 0 | 0 | 0 | 0 | 0 |
| LMDHVGTE | RPN2; RPN2; RI | P04844; F2Z | RPN2_HUMAN | Unmodified | 0 | 0 | 0 | 0 | 0 | 0 | 0 | 0 | 142320 | 202130 | 0 | 0 | 0 | 0 | 0 | 0 | 0 | 0 |
| LMDHVGTE | RPN2; RPN2; RI | P04844; F2Z | RPN2_HUMAN | Oxidation (M) | 0 | 0 | 0 | 0 | 0 | 0 | 0 | 0 | 7410600 | 0 | 270570 | 0 | 0 | 0 | 0 | 0 | 0 | 0 |
| LMDHVGTE | RPN2; RPN2; RI | P04844; F2Z | RPN2_HUMAN | Unmodified | 0 | 0 | 0 | 0 | 0 | 0 | 0 | 0 | 980660 | 440670 | 0 | 0 | 0 | 0 | 0 | 0 | 371480 | 0 |
| LMDKEQQ\ | EEA1 | Q15075 | EEA1_HUMAN I | Oxidation (M) | 0 | 0 | 0 | 0 | 1256400 | 0 | 235730 | 100640 | 0 | 593930 | 0 | 431620 | 0 | 0 | 0 | 0 | 0 | 0 |
| LMDKEQQ\ | EEA1 | Q15075 | EEA1_HUMAN I | Unmodified | 0 | 0 | 0 | 338890 | 993990 | 0 | 0 | 0 | 0 | 154140 | 0 | 0 | 0 | 0 | 0 | 0 | 0 | 0 |
| LMDLDVEC | PCNA | P12004 | PCNA_HUMAN | Oxidation (M) | 0 | 0 | 0 | 0 | 0 | 0 | 0 | 0 | 0 | 0 | 0 | 1154500 | 0 | 0 | 0 | 0 | 0 | 0 |
| LMDLLADS | USO1 | O60763 | USO1_HUMAN | Unmodified | 0 | 0 | 0 | 0 | 0 | 0 | 0 | 0 | 0 | 0 | 0 | 1750200 | 0 | 0 | 0 | 0 | 0 | 0 |
| LMDLLGEG | RARS1 | P54136 | SYRC_HUMAN | Oxidation (M) | 0 | 0 | 0 | 0 | 0 | 0 | 0 | 0 | 0 | 0 | 0 | 1114600 | 0 | 0 | 0 | 0 | 0 | 0 |
| LMDLLGEG | RARS1 | P54136 | SYRC_HUMAN | Unmodified | 0 | 0 | 0 | 0 | 0 | 0 | 0 | 0 | 0 | 0 | 163050 | 622260 | 0 | 0 | 0 | 0 | 0 | 0 |
| LMDLLSK | MYO3B; CHPF; | Q8WXR4; Q8 | MYO3B_HUMA | Unmodified | 74257 | 83265 | 208380 | 0 | 2272300 | 446590 | 487840 | 0 | 1805800 | 44130 | 0 | 9292100 | 59650 | 1124700 | 0 | 182790 | 0 | 2224700 |
| LMDQLEAL | VTA1; VTA1 | Q9NP79; A0. | VTA1_HUMAN \ | Unmodified | 0 | 0 | 0 | 0 | 0 | 0 | 0 | 0 | 759670 | 786440 | 0 | 484630 | 0 | 0 | 0 | 0 | 0 | 0 |
| LMDQLEAL | VTA1; VTA1 | Q9NP79; A0. | VTA1_HUMAN \ | Oxidation (M) | 289720 | 386470 | 0 | 687800 | 1404800 | 0 | 0 | 120400 | 948820 | 1021500 | 0 | 2342100 | 904850 | 196160 | 649320 | 668280 | 35594 | 0 |
| LMDQLEAL | VTA1; VTA1 | Q9NP79; A0. | VTA1_HUMAN \ | Unmodified | 61431 | 0 | 0 | 82319 | 0 | 0 | 0 | 0 | 0 | 293130 | 158710 | 974760 | 0 | 174150 | 0 | 0 | 0 | 0 |
| LMDQNLK | TPM3; TPM3; TF | J3KN67; A0A | J3KN67_HUMA | Unmodified | 0 | 0 | 1292600 | 1371500 | 3887900 | 866300 | 91531 | 44583 | 4135400 | 0 | 1878400 | 23440000 | 0 | 0 | 2291100 | 1471700 | 596690 | 660570 |
| LMDVGLIAI | STIP1 | P31948 | STIP1_HUMAN | Unmodified | 0 | 0 | 0 | 0 | 0 | 0 | 0 | 0 | 1147600 | 204710 | 304780 | 1235300 | 0 | 0 | 0 | 0 | 0 | 0 |
| LMDVGLIAI | STIP1 | P31948 | STIP1_HUMAN | Oxidation (M) | 0 | 0 | 366130 | 74078 | 0 | 0 | 0 | 0 | 0 | 592680 | 1448600 | 6364300 | 972150 | 119000 | 407560 | 434490 | 0 | 0 |
| LMEEFVK | LMCD1; LMCD1 | Q9NZU5; H7 | LMCD1_HUMA | Unmodified | 45700 | 0 | 0 | 0 | 0 | 0 | 0 | 0 | 0 | 0 | 421220 | 2215300 | 0 | 0 | 0 | 0 | 0 | 0 |
| LMELFPANI | BZW1; BZW1 | Q7L1Q6; C9I | BZW1_HUMAN | Oxidation (M) | 79572 | 113920 | 95147 | 86120 | 0 | 0 | 0 | 0 | 0 | 0 | 0 | 4351700 | 0 | 0 | 68567 | 123410 | 0 | 0 |
| LMELHGEG | RPS3A; RPS3A; | P61247; D6F | RS3A_HUMAN | Oxidation (M) | 606870 | 1716500 | 2155500 | 0 | 88460 | 0 | 0 | 0 | 0 | 115850 | 1141700 | 14195000 | 0 | 0 | 0 | 1680700 | 0 | 0 |
| LMELHGEG | RPS3A; RPS3A; | P61247; D6F | RS3A_HUMAN | Unmodified | 962780 | 664520 | 477990 | 413040 | 105770 | 0 | 0 | 0 | 254600 | 33829 | 1440500 | 10368000 | 0 | 0 | 173870 | 363000 | 0 | 0 |
| LMELRSRD | ATP2B4 | P23634 | AT2B4_HUMAN | Unmodified | 0 | 0 | 0 | 0 | 506210 | 0 | 193100 | 0 | 0 | 0 | 0 | 0 | 0 | 0 | 0 | 0 | 1741900 | 391190 |
| LMELYERLE | ABCF2; ABCF2; | Q9UG63; C9 | ABCF2_HUMAN | 2 Oxidation (M) | 200310 | 0 | 86968 | 0 | 0 | 0 | 0 | 0 | 0 | 0 | 0 | 0 | 0 | 0 | 341030 | 347520 | 0 | 0 |
| LMEPIYLVE | EEF2 | P13639 | EF2_HUMAN EI | Oxidation (M) | 0 | 0 | 0 | 0 | 777080 | 0 | 0 | 0 | 0 | 0 | 0 | 0 | 779830 | 0 | 293880 | 0 | 0 | 380680 |
| LMEPIYLVE | EEF2 | P13639 | EF2_HUMAN EI | Oxidation (M) | 0 | 0 | 0 | 0 | 0 | 0 | 0 | 0 | 0 | 0 | 0 | 0 | 3444200 | 0 | 0 | 0 | 0 | 0 |
| LMETQEED | DARS2; DARS2; | Q6PI48; A0A | SYDM_HUMAN | Unmodified | 0 | 0 | 0 | 104180 | 0 | 0 | 0 | 0 | 2550500 | 0 | 0 | 0 | 0 | 0 | 0 | 0 | 0 | 0 |
| LMEVMNH\ | VIL1 | P09327 | VILI_HUMAN Vi | Unmodified | 63903 | 45515 | 0 | 0 | 0 | 0 | 0 | 0 | 1760600 | 548340 | 0 | 2002600 | 0 | 0 | 0 | 0 | 0 | 0 |
| LMEVMNH\ | VIL1 | P09327 | VILI_HUMAN Vi | Oxidation (M) | 0 | 0 | 0 | 0 | 732060 | 0 | 0 | 0 | 123710 | 0 | 0 | 530100 | 0 | 0 | 0 | 0 | 0 | 0 |
| LMFDRSEV | ANPEP | P15144 | AMPN_HUMAN | 2 Oxidation (M) | 0 | 0 | 0 | 0 | 0 | 0 | 0 | 0 | 0 | 0 | 0 | 0 | 0 | 0 | 0 | 19392 | 6157400 | 1389500 |
| LMFDRSEV | ANPEP | P15144 | AMPN_HUMAN | Oxidation (M) | 0 | 0 | 0 | 0 | 0 | 0 | 0 | 0 | 0 | 0 | 0 | 0 | 0 | 0 | 0 | 0 | 4761500 | 2713100 |
| LMFDRSEV | ANPEP | P15144 | AMPN_HUMAN | Unmodified | 0 | 0 | 1531500 | 0 | 0 | 0 | 1562400 | 0 | 1034100 | 347780 | 1206100 | 0 | 0 | 11001000 | 0 | 0 | 8291900 | 3556400 |
| LMFEQQK | PLCG2; PLCG2 | P16885; A0A | PLCG2_HUMAI | Unmodified | 0 | 0 | 0 | 0 | 0 | 0 | 0 | 0 | 195990 | 0 | 201820 | 3221700 | 0 | 0 | 0 | 0 | 0 | 0 |
| LMGEDEKF | PFDN2 | Q9UHV9 | PFD2_HUMAN | Oxidation (M) | 0 | 0 | 0 | 0 | 0 | 13013 | 0 | 0 | 59012 | 0 | 629820 | 292340 | 45858 | 0 | 13123 | 0 | 0 | 0 |
| LMGEDEKF | PFDN2 | Q9UHV9 | PFD2_HUMAN | Unmodified | 0 | 0 | 0 | 0 | 0 | 0 | 0 | 0 | 155840 | 0 | 281170 | 1513800 | 0 | 0 | 0 | 29929 | 0 | 0 |
| LMHQAALL | TXNRD1; TXNRI | Q16881; A0/ | TRXR1_HUMAN | Oxidation (M) | 131550 | 0 | 54862 | 0 | 0 | 68545 | 0 | 0 | 131980 | 0 | 0 | 2303900 | 0 | 0 | 53709 | 46626 | 0 | 0 |
| LMHSFCAF | TFPI; TFPI; TFPI; | P10646; C9J | TFPI1_HUMAN | Unmodified | 0 | 0 | 0 | 353060 | 818510 | 0 | 0 | 17695 | 0 | 0 | 0 | 0 | 214020 | 270940 | 0 | 0 | 914460 | 1906400 |
| LMIEMDGT | ENO1; ENO1; E | P06733; A0A | ENOA_HUMAN | 2 Oxidation (M) | 153700 | 90791 | 0 | 852100 | 9598600 | 1524300 | 157900 | 85400 | 4419100 | 871120 | 4388700 | 8573700 | 2391300 | 858610 | 189920 | 354520 | 574380 | 1000700 |
| LMIEMDGT | ENO1; ENO1; E | P06733; A0A | ENOA_HUMAN | Oxidation (M) | 780100 | 1052300 | 1107500 | 2894700 | 27626000 | 5930200 | 834880 | 389190 | 21047000 | 5740500 | 19000000 | 43846000 | 10424000 | 4634400 | 4035100 | 2446500 | 3346700 | 16513000 |
| LMIEMDGT | ENO1; ENO1; E | P06733; A0A | ENOA_HUMAN | Unmodified | 423430 | 259520 | 281960 | 1153900 | 38084000 | 3668800 | 706060 | 99228 | 30261000 | 4608800 | 8074300 | 52380000 | 9502100 | 3008200 | 965920 | 669050 | 5034400 | 18866000 |
| LMIEMDGT | ENO1; ENO1; E | P06733; A0A | ENOA_HUMAN | Oxidation (M) | 0 | 0 | 0 | 0 | 2488300 | 136180 | 45941 | 67898 | 0 | 171450 | 0 | 0 | 622750 | 301340 | 0 | 0 | 822310 | 3359300 |
| LMLAQDEE | XPOT | O43592 | XPOT_HUMAN | Unmodified | 0 | 0 | 0 | 0 | 138430 | 0 | 195910 | 0 | 0 | 0 | 0 | 1856300 | 0 | 0 | 0 | 0 | 213200 | 0 |
| LMLENLSN | N/A | N/A | N/A | Oxidation (M) | 0 | 0 | 0 | 0 | 0 | 3581900 | 1640100 | 286070 | 0 | 219990 | 0 | 0 | 0 | 1235500 | 0 | 0 | 352400 | 975430 |
| LMLESQEFI | DHTKD1 | Q96HY7 | DHTK1_HUMAI | Oxidation (M) | 0 | 0 | 0 | 46067 | 0 | 0 | 0 | 0 | 1003400 | 450010 | 0 | 187980 | 0 | 0 | 0 | 0 | 0 | 0 |
| LMLESQEFI | DHTKD1 | Q96HY7 | DHTK1_HUMAI | Unmodified | 0 | 0 | 0 | 0 | 0 | 0 | 0 | 0 | 1618200 | 74812 | 0 | 0 | 0 | 0 | 0 | 0 | 0 | 0 |
| LMLGWEP\ | UXS1 | Q8NBZ7 | UXS1_HUMAN | Oxidation (M) | 0 | 0 | 0 | 8598 | 451500 | 133030 | 476340 | 32259 | 5728500 | 2167100 | 214610 | 111240 | 0 | 0 | 0 | 0 | 161950 | 952120 |
| LMLGWEP\ | UXS1 | Q8NBZ7 | UXS1_HUMAN | Unmodified | 0 | 0 | 0 | 228450 | 215180 | 0 | 0 | 203980 | 1030200 | 30792 | 0 | 0 | 0 | 0 | 322950 | 0 | 0 | 0 |
| LMLWDTA( | RAB23 | Q9ULC3 | RAB23_HUMAN | Oxidation (M) | 321000 | 664820 | 144010 | 569030 | 268000 | 51849 | 0 | 0 | 960750 | 0 | 0 | 0 | 0 | 51948 | 1207900 | 439410 | 44646 | 108060 |
| LMMDGHE | UXS1; UXS1; U) | Q8NBZ7; A0. | UXS1_HUMAN | 2 Oxidation (M) | 0 | 0 | 0 | 0 | 0 | 0 | 0 | 0 | 717820 | 90905 | 27401 | 0 | 0 | 0 | 0 | 0 | 0 | 93183 |
| LMMDGHE | UXS1; UXS1; U) | Q8NBZ7; A0. | UXS1_HUMAN | Oxidation (M) | 0 | 0 | 0 | 0 | 0 | 0 | 0 | 0 | 379650 | 65248 | 0 | 0 | 0 | 0 | 0 | 0 | 0 | 0 |
| LMMDGHE | UXS1; UXS1; U) | Q8NBZ7; A0. | UXS1_HUMAN | 2 Oxidation (M) | 0 | 0 | 0 | 0 | 0 | 0 | 542300 | 0 | 1088400 | 0 | 0 | 0 | 0 | 0 | 0 | 0 | 0 | 0 |
| LMMDGHE | UXS1; UXS1; U) | Q8NBZ7; A0. | UXS1_HUMAN | Oxidation (M) | 0 | 0 | 0 | 178060 | 0 | 0 | 0 | 0 | 1191800 | 0 | 0 | 0 | 0 | 0 | 0 | 0 | 0 | 0 |
| LMMDPLS( | HNRNPR; HNRI | O43390; B4I | HNRPR_HUMA | 2 Oxidation (M) | 0 | 0 | 0 | 0 | 0 | 0 | 0 | 0 | 0 | 0 | 0 | 621180 | 0 | 0 | 0 | 0 | 0 | 0 |
| LMMDPLS( | HNRNPR; HNRI | O43390; B4I | HNRPR_HUMA | Oxidation (M) | 0 | 0 | 0 | 0 | 0 | 0 | 335080 | 0 | 194300 | 0 | 0 | 881200 | 0 | 0 | 0 | 0 | 0 | 0 |
| LMMDPLS( | HNRNPR; HNRI | O43390; B4I | HNRPR_HUMA | Unmodified | 0 | 0 | 0 | 0 | 0 | 0 | 0 | 0 | 0 | 0 | 173770 | 899210 | 0 | 0 | 0 | 0 | 0 | 0 |
| LMMDPLTC | SYNCRIP; SYN( | O60506; B7Z | HNRPQ_HUMA | 2 Oxidation (M) | 576550 | 0 | 0 | 0 | 0 | 0 | 0 | 69887 | 0 | 0 | 0 | 1954000 | 0 | 0 | 0 | 0 | 0 | 0 |
| LMMDPLTC | SYNCRIP; SYN( | O60506; B7Z | HNRPQ_HUMA | Oxidation (M) | 0 | 0 | 0 | 0 | 0 | 0 | 0 | 0 | 0 | 0 | 0 | 3752800 | 0 | 0 | 0 | 0 | 0 | 0 |
| LMMPSWM | TRABD2B | A6NFA1 | TIKI2_HUMAN I | Oxidation (M) | 0 | 0 | 8645800 | 0 | 0 | 0 | 0 | 0 | 0 | 0 | 0 | 0 | 0 | 0 | 1542500 | 0 | 0 | 0 |
| LMNETTAV. | HSPA4; HSPA4I | P34932; O9! | HSP74_HUMAN | Oxidation (M) | 0 | 26429000 | 0 | 0 | 0 | 0 | 121890 | 61474 | 0 | 0 | 312580 | 856900 | 0 | 0 | 0 | 0 | 0 | 0 |
| LMNETTAV. | HSPA4; HSPA4; | P34932; A0A | HSP74_HUMAN | Oxidation (M) | 0 | 0 | 0 | 0 | 0 | 0 | 223530 | 0 | 0 | 0 | 544410 | 3753700 | 0 | 0 | 0 | 0 | 0 | 0 |
| LMNIFLK | C3 | P01024 | CO3_HUMAN ( | Oxidation (M) | 0 | 76270 | 111380 | 243670 | 0 | 374880 | 285320 | 177540 | 294010 | 136660 | 490310 | 0 | 0 | 0 | 218820 | 0 | 0 | 0 |
| LMPGQDPI | FKBP10; FKBP1 | Q96AY3; H0' | FKB10_HUMAN | Unmodified | 0 | 0 | 0 | 0 | 0 | 0 | 0 | 0 | 0 | 0 | 1781900 | 391130 | 0 | 0 | 0 | 0 | 0 | 0 |
| LMPMQQIA | OXCT1; OXCT1 | P55809; E9P | SCOT1_HUMAI | Unmodified | 0 | 0 | 0 | 0 | 0 | 0 | 0 | 0 | 1475400 | 0 | 0 | 104310 | 0 | 0 | 0 | 0 | 0 | 0 |
| LMPSNQQI | SMCHD1; SMCI | A6NHR9; J3I | SMHD1_HUMA | Unmodified | 0 | 0 | 0 | 0 | 0 | 0 | 227670 | 118650 | 0 | 0 | 0 | 0 | 0 | 0 | 0 | 0 | 0 | 0 |
| LMQDDNR( | MAN2A1 | Q16706 | MA2A1_HUMAI | Unmodified | 0 | 0 | 0 | 0 | 0 | 0 | 0 | 0 | 1902000 | 0 | 0 | 0 | 0 | 0 | 0 | 0 | 0 | 0 |
| LMQDDNR( | MAN2A1 | Q16706 | MA2A1_HUMAI | Oxidation (M) | 0 | 0 | 0 | 0 | 0 | 0 | 0 | 0 | 2110000 | 0 | 0 | 0 | 0 | 0 | 0 | 0 | 0 | 0 |
| LMQITSLHS | GMPS | P49915 | GUAA_HUMAN | Oxidation (M) | 296930 | 320570 | 334010 | 370770 | 3015700 | 760790 | 1955800 | 0 | 1570500 | 2375700 | 2517900 | 5541800 | 3154400 | 539510 | 1119400 | 388980 | 1041700 | 1157700 |
| LMQITSLHS | GMPS | P49915 | GUAA_HUMAN | Unmodified | 0 | 0 | 0 | 0 | 75310 | 174000 | 0 | 0 | 231620 | 56402 | 0 | 67737 | 59347 | 0 | 0 | 0 | 0 | 0 |
| LMQLQHEI | CCDC6 | Q16204 | CCDC6_HUMA | Unmodified | 0 | 0 | 0 | 0 | 0 | 0 | 0 | 20555 | 0 | 0 | 0 | 250260 | 0 | 0 | 0 | 0 | 0 | 0 |
| LMQLQHEI | CCDC6 | Q16204 | CCDC6_HUMA | Oxidation (M) | 0 | 0 | 0 | 0 | 0 | 0 | 0 | 0 | 0 | 0 | 0 | 0 | 269910 | 0 | 0 | 0 | 85191 | 427990 |
| LMQLVDHI | DDX17; DDX17; | Q92841; A0/ | DDX17_HUMAI | Unmodified | 0 | 0 | 0 | 0 | 571050 | 98969 | 621930 | 0 | 24558 | 0 | 0 | 1002000 | 0 | 0 | 0 | 0 | 0 | 231340 |
| LMQVVPQE | SLC7A5; SLC7A | Q01650; A0/ | LAT1_HUMAN I | Unmodified | 536080 | 208750 | 0 | 0 | 844160 | 0 | 0 | 0 | 416700 | 93852 | 461590 | 1297000 | 0 | 0 | 366250 | 0 | 159200 | 0 |
| LMQVVPQE | SLC7A5; SLC7A | Q01650; A0/ | LAT1_HUMAN I | Oxidation (M) | 398210 | 1079900 | 0 | 0 | 3198300 | 0 | 0 | 0 | 0 | 0 | 89077 | 2474400 | 0 | 0 | 363410 | 571690 | 0 | 0 |
| LMRFDDPL | LARS1; LARS1; | Q9P2J5; A0A | SYLC_HUMAN | Unmodified | 0 | 0 | 0 | 0 | 891630 | 0 | 121130 | 0 | 244370 | 315370 | 156630 | 0 | 268720 | 141310 | 0 | 0 | 0 | 174810 |
| LMRFDDPL | LARS1; LARS1; | Q9P2J5; A0A | SYLC_HUMAN | Oxidation (M) | 0 | 0 | 79199 | 92529 | 0 | 115150 | 112750 | 35930 | 0 | 167860 | 728850 | 1614500 | 339870 | 149160 | 201240 | 152750 | 0 | 0 |
| LMRNEFTII | STAMBP; STAMI | O95630; C9J | STABP_HUMAN | Oxidation (M) | 0 | 0 | 408440 | 1345400 | 3151800 | 1248100 | 846370 | 0 | 1267500 | 1055600 | 153680 | 624280 | 3308100 | 1577800 | 712420 | 441820 | 834130 | 952020 |
| LMRNEFTII | STAMBP; STAMI | O95630; C9J | STABP_HUMAN | Unmodified | 0 | 0 | 0 | 0 | 2117100 | 0 | 0 | 0 | 884880 | 191390 | 0 | 0 | 0 | 216760 | 0 | 0 | 0 | 412620 |
| LMSALLSGI | ECPAS; ECPAS | Q5VYK3; J3K | ECM29_HUMAI | Unmodified | 0 | 0 | 0 | 34270 | 112270 | 0 | 1196200 | 129230 | 321440 | 135140 | 0 | 180700 | 76425 | 1038000 | 0 | 23863 | 964110 | 0 |
| LMSELVSN | SEPTIN10; SEP1 | Q9P0V9; B5I | SEP10_HUMAN | Oxidation (M) | 0 | 0 | 0 | 0 | 4315000 | 0 | 0 | 0 | 0 | 0 | 0 | 0 | 0 | 0 | 0 | 0 | 1296200 | 0 |
| LMSENTEG | MAP4K5 | Q9Y4K4 | M4K5_HUMAN | Unmodified | 0 | 0 | 0 | 0 | 880220 | 0 | 0 | 0 | 0 | 0 | 0 | 0 | 0 | 0 | 0 | 0 | 0 | 241840 |
| LMSEVQTLI | TAX1BP1; TAX1I | Q86VP1; B8Z | TAXB1_HUMAN | Unmodified | 58396 | 0 | 0 | 0 | 1130900 | 121970 | 2095800 | 0 | 934240 | 413280 | 0 | 0 | 883360 | 218300 | 0 | 0 | 656550 | 1875100 |

| | | | | | | | | | | | | | | | | | | | | | | |
|---|---|---|---|---|---|---|---|---|---|---|---|---|---|---|---|---|---|---|---|---|---|---|
| LMSEVQTLI | TAX1BP1; TAX1 | Q86VP1; B8 | TAXB1_HUMAN | Oxidation (M) | 0 | 0 | 0 | 0 | 491090 | 0 | 1023500 | 129490 | 352010 | 119380 | 0 | 0 | 0 | 214340 | 0 | 0 | 630090 | 0 |
| LMSIVSALF | FRYL; FRYL; FRY | O94915; F22 | FRYL_HUMAN I | Oxidation (M) | 0 | 0 | 0 | 0 | 0 | 0 | 0 | 0 | 0 | 0 | 0 | 1031300 | 0 | 0 | 0 | 0 | 0 | 0 |
| LMSNALST | MTCH1; MTCH1 | Q9NZJ7; H0Y | MTCH1_HUMA | Unmodified | 0 | 0 | 0 | 0 | 0 | 0 | 0 | 0 | 1444000 | 0 | 0 | 0 | 0 | 0 | 0 | 0 | 0 | 0 |
| LMSSGNEE | CPT2; CPT2; CF | P23786; A0A | CPT2_HUMAN | Unmodified | 0 | 0 | 0 | 0 | 0 | 0 | 73436 | 0 | 847500 | 0 | 0 | 0 | 0 | 0 | 0 | 0 | 0 | 0 |
| LMSSGNEE | CPT2; CPT2; CF | P23786; A0A | CPT2_HUMAN | Unmodified | 0 | 0 | 0 | 0 | 0 | 0 | 0 | 67765 | 509590 | 0 | 0 | 0 | 0 | 0 | 0 | 0 | 0 | 0 |
| LMTDTINEF | NARS1 | O43776 | SYNC_HUMAN | Oxidation (M) | 0 | 0 | 0 | 0 | 0 | 0 | 0 | 0 | 408360 | 0 | 0 | 3748800 | 0 | 0 | 0 | 0 | 0 | 0 |
| LMTEILGR | NUDT21; NUDT | O43809; H3 | CPSF5_HUMAN | Unmodified | 1951700 | 1013400 | 90545 | 0 | 0 | 0 | 0 | 0 | 0 | 0 | 0 | 1555500 | 0 | 0 | 0 | 164660 | 0 | 0 |
| LMVCGLLL | N/A | N/A | N/A | Oxidation (M) | 131500 | 0 | 47642 | 687980 | 204380 | 0 | 114730 | 0 | 0 | 122880 | 0 | 415270 | 386180 | 53408 | 374580 | 0 | 574600 | 0 |
| LMVELSK | CCT5; CCT5; C | P48643; B7Z | TCPE_HUMAN | Unmodified | 82513 | 59560 | 0 | 356470 | 2640500 | 0 | 0 | 281300 | 1269700 | 792410 | 1221700 | 8174000 | 7933300 | 0 | 586580 | 0 | 1064700 | 0 |
| LMVELSK | CCT5; CCT5; C | P48643; B7Z | TCPE_HUMAN | Oxidation (M) | 79464 | 98257 | 208010 | 0 | 1177800 | 132860 | 366750 | 245860 | 467350 | 401870 | 561170 | 1633100 | 2459800 | 623000 | 236020 | 232670 | 497530 | 1273900 |
| LMVEPLQA | CUL2; CUL2; C | Q13617; Q5 | CUL2_HUMAN | Oxidation (M) | 0 | 0 | 94120 | 0 | 0 | 0 | 0 | 0 | 192550 | 113120 | 150850 | 2092500 | 0 | 0 | 194410 | 98300 | 0 | 0 |
| LMVFDKTE | HPN; HPN; HPN | P05981; M0 | HEPS_HUMAN | Oxidation (M) | 0 | 59689 | 0 | 658600 | 0 | 0 | 42862 | 0 | 0 | 0 | 0 | 0 | 0 | 0 | 0 | 229110 | 0 | 0 |
| LMVFDKTE | HPN; HPN; HPN | P05981; M0 | HEPS_HUMAN | Unmodified | 60496 | 122280 | 0 | 207390 | 355150 | 0 | 0 | 0 | 0 | 0 | 0 | 0 | 0 | 0 | 100820 | 67268 | 0 | 0 |
| LMVHTVAT | SPTAN1; SPTAN | Q13813; A0 | SPTN1_HUMAN | Unmodified | 0 | 0 | 0 | 0 | 1232200 | 157540 | 0 | 0 | 573560 | 118970 | 0 | 725500 | 337670 | 0 | 0 | 23839 | 292170 | 448330 |
| LMVIFSK | SDF4; SDF4; SD | Q9BRK5; H0 | CAB45_HUMAN | Unmodified | 0 | 0 | 0 | 44449 | 0 | 0 | 0 | 0 | 6876600 | 294450 | 135900 | 178640 | 134040 | 68774 | 44367 | 24159 | 0 | 0 |
| LMVIFSKVD | SDF4; SDF4; SD | Q9BRK5; H0 | CAB45_HUMAN | Unmodified | 0 | 0 | 0 | 0 | 214250 | 0 | 0 | 0 | 3663200 | 934310 | 0 | 0 | 0 | 0 | 0 | 0 | 0 | 0 |
| LMVIFSKVD | SDF4; SDF4; SD | Q9BRK5; H0 | CAB45_HUMAN | Oxidation (M) | 0 | 0 | 0 | 0 | 0 | 0 | 0 | 0 | 0 | 1920900 | 196700 | 0 | 0 | 1218200 | 0 | 0 | 0 | 0 |
| LMVPRFSF | AHNAK2 | Q8IVF2 | AHNK2_HUMA | Oxidation (M) | 0 | 0 | 0 | 0 | 692810 | 109140 | 0 | 0 | 0 | 0 | 0 | 0 | 0 | 0 | 0 | 0 | 0 | 0 |
| LMVPVTVV | PIGS | Q96S52 | PIGS_HUMAN | Unmodified | 0 | 0 | 0 | 0 | 0 | 0 | 0 | 0 | 1395000 | 0 | 0 | 0 | 0 | 0 | 0 | 0 | 0 | 0 |
| LMVSMLDF | SRI; SRI; SRI | P30626; B4 | SORCN_HUMA | 2 Oxidation (M) | 0 | 0 | 0 | 0 | 0 | 0 | 0 | 0 | 1070000 | 0 | 0 | 115730 | 0 | 0 | 0 | 0 | 0 | 0 |
| LMWIPDTK | DCAF7 | P61962 | DCAF7_HUMAN | Oxidation (M) | 0 | 37992 | 0 | 0 | 0 | 0 | 0 | 0 | 0 | 32377 | 0 | 486970 | 0 | 0 | 46145 | 0 | 0 | 0 |
| LMYKESGV | SCARB1; SCAR | Q8WTV0; A0 | SCRB1_HUMAI | Oxidation (M) | 0 | 0 | 0 | 0 | 817200 | 0 | 0 | 0 | 0 | 0 | 0 | 0 | 0 | 0 | 0 | 0 | 0 | 0 |
| LMYKESGV | SCARB1; SCAR | Q8WTV0; A0 | SCRB1_HUMAI | Unmodified | 0 | 0 | 0 | 0 | 377780 | 0 | 0 | 0 | 0 | 0 | 0 | 0 | 0 | 0 | 0 | 0 | 222250 | 0 |
| LMYKESGV | SCARB1; SCAR | Q8WTV0; A0 | SCRB1_HUMAI | Unmodified | 0 | 0 | 0 | 0 | 971090 | 0 | 0 | 0 | 0 | 0 | 0 | 0 | 2908900 | 0 | 0 | 0 | 0 | 0 |
| LMYNSSNF | MYO1D; MYO1 | O94832; J3 | MYO1D_HUMA | Oxidation (M) | 2620600 | 1963800 | 0 | 0 | 0 | 0 | 0 | 85754 | 0 | 0 | 0 | 451850 | 0 | 0 | 225200 | 0 | 38208 | 0 |
| LMYNSSNF | MYO1D; MYO1 | O94832; J3 | MYO1D_HUMA | Unmodified | 1554200 | 685790 | 248620 | 248860 | 0 | 0 | 0 | 0 | 0 | 0 | 0 | 489840 | 0 | 0 | 382110 | 151360 | 0 | 0 |
| LNAAEEML | PPFIBP2; PPFIB | Q8ND30; B8 | LIPB2_HUMAN | Unmodified | 15311 | 0 | 20008 | 69816 | 50886 | 0 | 0 | 0 | 0 | 0 | 23683 | 403390 | 23013 | 0 | 122980 | 39279 | 62932 | 31569 |
| LNAFGNAF | EHD1; EHD3; E | Q9H4M9; Q | EHD1_HUMAN | Unmodified | 0 | 52075 | 3939000 | 4859100 | 0 | 0 | 0 | 0 | 850280 | 0 | 1247000 | 13829000 | 373340 | 0 | 5379800 | 6514800 | 0 | 0 |
| LNAFKADT | STXBP2; STXBP | Q15833; M0 | STXB2_HUMAN | Unmodified | 752790 | 696050 | 234870 | 2276800 | 12862000 | 2539600 | 0 | 0 | 1359100 | 0 | 1511300 | 2892000 | 7744500 | 6321800 | 673680 | 886070 | 7858100 | 10565000 |
| LNAILFK | DIAPH1; DIAPH | O60610; A0 | DIAP1_HUMAN | Unmodified | 69447 | 55956 | 200690 | 215780 | 0 | 72709 | 52512 | 0 | 428330 | 220420 | 172310 | 3105500 | 0 | 0 | 553870 | 261360 | 0 | 0 |
| LNAIYQNN | GSR; GSR; GSR | P00390; H0 | GSHR_HUMAN | Unmodified | 0 | 0 | 0 | 0 | 855430 | 171900 | 0 | 0 | 1204400 | 195130 | 513650 | 4399500 | 1348000 | 0 | 0 | 1822400 | 0 | 0 |
| LNATGRDL | CDK5 | Q00535 | CDK5_HUMAN | Unmodified | 97169 | 65009 | 130420 | 417630 | 2347500 | 702810 | 0 | 122370 | 266990 | 0 | 222350 | 0 | 1567700 | 1677500 | 167440 | 69203 | 2284100 | 2790700 |
| LNATNIELA | PSMA5 | P28066 | PSA5_HUMAN | Oxidation (M) | 0 | 0 | 0 | 0 | 0 | 0 | 0 | 0 | 0 | 0 | 0 | 5004800 | 0 | 0 | 0 | 0 | 0 | 0 |
| LNDALDRL | MACF1; MACF1 | Q9UPN3; H3 | MACF1_HUMAI | Unmodified | 0 | 0 | 193370 | 75358 | 0 | 0 | 0 | 0 | 0 | 245700 | 269780 | 946970 | 0 | 0 | 0 | 0 | 482700 | 0 |
| LNDFASTV | COX5A; COX5A | P20674; H3 | COX5A_HUMAI | Unmodified | 0 | 0 | 0 | 0 | 0 | 0 | 0 | 59533 | 7608900 | 1641900 | 1697600 | 189230 | 191090 | 0 | 0 | 0 | 0 | 0 |
| LNDFASTV | COX5A; COX5A | P20674; H3 | COX5A_HUMAI | Unmodified | 0 | 0 | 0 | 0 | 0 | 0 | 0 | 0 | 0 | 1792600 | 0 | 0 | 0 | 0 | 0 | 0 | 0 | 0 |
| LNDGHFMI | AKR1C2; AKR1 | P52895; Q0 | AK1C2_HUMAI | Oxidation (M) | 386220 | 0 | 0 | 0 | 9452500 | 0 | 0 | 0 | 0 | 0 | 0 | 0 | 0 | 0 | 0 | 0 | 0 | 0 |
| LNDGHFMI | AKR1C4; AKR1 | P17516; P42 | AK1C4_HUMAI | Oxidation (M) | 1151600 | 0 | 0 | 0 | 0 | 0 | 0 | 0 | 0 | 0 | 0 | 0 | 0 | 0 | 0 | 0 | 0 | 0 |
| LNDGSQIT | AIFM1; AIFM1; | O95831; A0 | AIFM1_HUMAN | Unmodified | 0 | 0 | 0 | 0 | 0 | 0 | 415360 | 82645 | 1115200 | 845000 | 316550 | 936890 | 865770 | 531530 | 0 | 0 | 45790 | 172370 |
| LNDMEPSK | VAPA | Q9P0L0 | VAPA_HUMAN | Unmodified | 0 | 0 | 0 | 0 | 252270 | 0 | 126970 | 0 | 166000 | 0 | 91439 | 1032300 | 0 | 49393 | 0 | 0 | 0 | 0 |
| LNDMYGD | SLC16A1; SLC1 | P53985; Q5 | MOT1_HUMAN | Oxidation (M) | 0 | 0 | 0 | 0 | 0 | 0 | 0 | 0 | 0 | 0 | 0 | 964240 | 0 | 0 | 0 | 0 | 0 | 0 |
| LNDRIEMW | GLG1 | Q92896 | GSLG1_HUMAI | Unmodified | 0 | 0 | 0 | 0 | 0 | 0 | 0 | 0 | 5205400 | 749760 | 122150 | 795350 | 0 | 0 | 0 | 0 | 0 | 0 |
| LNDRIEMW | GLG1 | Q92896 | GSLG1_HUMAI | Oxidation (M) | 0 | 0 | 0 | 0 | 0 | 0 | 0 | 0 | 836900 | 233240 | 0 | 0 | 0 | 0 | 0 | 0 | 0 | 0 |
| LNDRVMR | TFRC; TFRC | P02786; G3 | TFR1_HUMAN | Unmodified | 0 | 0 | 0 | 0 | 1195200 | 0 | 261050 | 0 | 1353500 | 420510 | 0 | 0 | 675620 | 0 | 0 | 0 | 0 | 456180 |
| LNDYIFSF | RARS1 | P54136 | SYRC_HUMAN | Unmodified | 77040 | 0 | 163670 | 0 | 0 | 0 | 0 | 0 | 0 | 0 | 380030 | 5255600 | 0 | 0 | 0 | 108950 | 0 | 0 |
| LNEAAAGL | TLN1 | Q9Y490 | TLN1_HUMAN | Unmodified | 4285600 | 0 | 0 | 0 | 0 | 0 | 0 | 0 | 0 | 0 | 0 | 7230400 | 0 | 0 | 2324000 | 2726100 | 0 | 0 |
| LNEAAVTV | CCNDBP1; CCN | O95273; H3 | CCDB1_HUMA | Unmodified | 0 | 0 | 0 | 0 | 0 | 0 | 0 | 0 | 0 | 0 | 0 | 0 | 0 | 0 | 0 | 0 | 0 | 0 |
| LNEAKEEF | PAFAH1B1; PAF | P43034; A0 | LIS1_HUMAN P | Unmodified | 0 | 44189 | 0 | 58005 | 0 | 0 | 0 | 0 | 189720 | 0 | 327690 | 1172600 | 0 | 0 | 114040 | 115670 | 0 | 0 |
| LNEAQPST | EIF6; N/A | P56537; A0 | IF6_HUMAN Eu | Oxidation (M) | 120840 | 157350 | 0 | 0 | 0 | 0 | 0 | 0 | 0 | 0 | 0 | 82348 | 0 | 0 | 0 | 95967 | 0 | 0 |
| LNEAQPST | EIF6 | P56537 | IF6_HUMAN Eu | Unmodified | 606230 | 1195400 | 76404 | 434910 | 2024300 | 300260 | 0 | 0 | 581240 | 349550 | 0 | 0 | 984690 | 876940 | 688010 | 0 | 960720 | 2644600 |
| LNECVDHT | GSTO1; GSTO1 | P78417; Q5 | GSTO1_HUMAI | Unmodified | 0 | 701240 | 0 | 0 | 0 | 0 | 386480 | 0 | 0 | 0 | 0 | 6135400 | 0 | 0 | 1471800 | 238630 | 0 | 0 |
| LNEELLSK | EFCAB14; EFCA | O75071; A0 | EFC14_HUMAN | Unmodified | 0 | 0 | 0 | 38881 | 236640 | 99994 | 0 | 0 | 8074500 | 747090 | 0 | 484160 | 0 | 0 | 0 | 0 | 0 | 0 |
| LNEHFLNT | IDH2; IDH2 | P48735; H0 | IDHP_HUMAN | Unmodified | 0 | 0 | 225740 | 536550 | 508680 | 123270 | 4818800 | 58570 | 1.31E+08 | 13091000 | 2173700 | 526650 | 1367200 | 1559100 | 226410 | 0 | 0 | 53069 |
| LNEIEGTLN | LAMC1 | P11047 | LAMC1_HUMAI | Unmodified | 0 | 0 | 499310 | 0 | 0 | 0 | 0 | 0 | 650740 | 0 | 0 | 0 | 0 | 0 | 264070 | 0 | 0 | 649590 |
| LNEILFPFY | PLCB4; PLCB4 | Q15147; A0 | PLCB4_HUMAI | Unmodified | 0 | 0 | 285960 | 274710 | 0 | 0 | 0 | 0 | 0 | 0 | 0 | 0 | 0 | 0 | 742490 | 852140 | 0 | 0 |
| LNEILFPFY | PLCB4; PLCB4 | Q15147; A0 | PLCB4_HUMAI | Unmodified | 343820 | 0 | 461710 | 947940 | 400910 | 474490 | 0 | 0 | 0 | 0 | 0 | 0 | 0 | 0 | 1485600 | 921380 | 0 | 2254900 |
| LNELLVK | TRAP1; TRAP1; | Q12931; I3L | TRAP1_HUMAN | Unmodified | 0 | 0 | 0 | 0 | 0 | 0 | 0 | 0 | 1822800 | 42022 | 132900 | 526390 | 0 | 0 | 0 | 0 | 113140 | 428530 |
| LNENAISRL | TRIM16; TRIM1 | O95361; J3Q | TRI16_HUMAN | Unmodified | 0 | 0 | 0 | 0 | 1054700 | 0 | 166210 | 139270 | 0 | 0 | 0 | 0 | 0 | 0 | 0 | 0 | 0 | 0 |
| LNENHSGE | ILK; ILK | Q13418; A0 | ILK_HUMAN Int | Unmodified | 0 | 0 | 1381500 | 1539100 | 0 | 0 | 3252500 | 0 | 0 | 0 | 538510 | 0 | 0 | 2467800 | 0 | 0 | 0 | 0 |
| LNEQHQLI | SMAP1 | Q8IYB5 | SMAP1_HUMAI | Unmodified | 268640 | 0 | 0 | 0 | 697140 | 0 | 224220 | 195410 | 1308600 | 562110 | 0 | 2134700 | 987000 | 841860 | 0 | 0 | 0 | 0 |
| LNEQYEHA | CAPRIN1; CAPF | Q14444; G3 | CAPR1_HUMAI | Unmodified | 0 | 0 | 0 | 0 | 0 | 0 | 0 | 0 | 0 | 58958 | 3639500 | 5303900 | 523700 | 0 | 0 | 0 | 754980 | 0 |
| LNERHYGG | PGAM1; PGAM4 | P18669; Q8 | PGAM1_HUMA | Unmodified | 0 | 0 | 0 | 0 | 1656900 | 0 | 0 | 0 | 0 | 333150 | 104160 | 0 | 676360 | 443180 | 0 | 0 | 818700 | 0 |
| LNERIIATS | SEMA3C | Q99985 | SEM3C_HUMAI | Unmodified | 0 | 0 | 0 | 0 | 0 | 318350 | 0 | 0 | 0 | 0 | 0 | 0 | 0 | 0 | 0 | 0 | 0 | 0 |
| LNESLDEN | BAIAP2L1 | Q9UHR4 | BI2L1_HUMAN | Unmodified | 0 | 0 | 0 | 0 | 0 | 0 | 0 | 0 | 0 | 0 | 0 | 0 | 0 | 0 | 0 | 0 | 0 | 0 |
| LNESLDEN | BAIAP2L1 | Q9UHR4 | BI2L1_HUMAN | Unmodified | 0 | 0 | 0 | 0 | 2099200 | 0 | 124370 | 0 | 4749600 | 466800 | 319370 | 0 | 1339400 | 573210 | 0 | 0 | 0 | 0 |
| LNESTFDT | PRKDC | P78527 | PRKDC_HUMAI | Unmodified | 61803 | 0 | 0 | 0 | 6945400 | 0 | 0 | 0 | 0 | 0 | 0 | 1728800 | 0 | 0 | 0 | 65254 | 23119 | 7032200 |
| LNEVIRQV | AHCYL1 | O43865 | SAHH2_HUMA | Unmodified | 0 | 0 | 0 | 0 | 3490500 | 987080 | 465760 | 77149 | 0 | 271680 | 0 | 0 | 848000 | 0 | 0 | 0 | 1590900 | 1609900 |
| LNEVIVNK | SLC12A6; SLC1 | Q9UHW9; B | S12A6_HUMAN | Unmodified | 42279 | 99022 | 0 | 0 | 1632100 | 0 | 0 | 0 | 344300 | 0 | 641240 | 1136800 | 414290 | 0 | 149130 | 134170 | 86585 | 533840 |
| LNEVSSDA | RANBP3; RANB | Q9H6Z4; B7 | RANB3_HUMAI | Unmodified | 0 | 0 | 0 | 0 | 0 | 0 | 0 | 0 | 183600 | 0 | 0 | 543900 | 0 | 0 | 0 | 0 | 0 | 0 |
| LNFAVASR | PDIA3 | P30101 | PDIA3_HUMAN | Unmodified | 541560 | 158330 | 993730 | 378330 | 0 | 0 | 120040 | 272030 | 60298000 | 11302000 | 5647600 | 8226000 | 458200 | 214910 | 2443000 | 1753000 | 0 | 0 |
| LNFAVASR | PDIA3 | P30101 | PDIA3_HUMAN | Unmodified | 0 | 0 | 0 | 0 | 0 | 0 | 200570 | 25056 | 6799400 | 4966000 | 3953600 | 873840 | 0 | 375800 | 0 | 0 | 0 | 221220 |
| LNFELTDA | CPT2; CPT2; CF | P23786; A0 | CPT2_HUMAN | Unmodified | 0 | 0 | 0 | 0 | 0 | 563750 | 0 | 0 | 2110600 | 627610 | 0 | 0 | 0 | 1376900 | 0 | 0 | 0 | 0 |
| LNFGDDIP | STUB1; STUB1; | Q9UNE7; H3 | CHIP_HUMAN | Unmodified | 0 | 1037300 | 1494600 | 0 | 0 | 0 | 0 | 0 | 1585800 | 0 | 390900 | 4392000 | 289450 | 0 | 2320400 | 1582200 | 0 | 0 |
| LNFLESK | AMPD3; AMPD3 | Q01432; E9 | AMPD3_HUMA | Unmodified | 0 | 0 | 0 | 0 | 0 | 0 | 0 | 0 | 0 | 0 | 0 | 0 | 0 | 0 | 0 | 0 | 0 | 0 |
| LNFSHGTH | PKM; PKM; PKM | P14618; H3 | KPYM_HUMAN | Unmodified | 2658300 | 1578100 | 7256900 | 10336000 | 72522000 | 28479000 | 2448600 | 855300 | 15334000 | 5076600 | 24277000 | 1.57E+08 | 13383000 | 5412900 | 10497000 | 7582300 | 8815600 | 25948000 |
| LNFSHGTH | PKM; PKM; PKM | P14618; H3 | KPYM_HUMAN | Unmodified | 0 | 0 | 0 | 0 | 6953200 | 0 | 187900 | 0 | 0 | 0 | 0 | 0 | 2543800 | 180090 | 0 | 0 | 4536300 | 4298700 |
| LNFSTPTST | PFAS; PFAS; PF | O15067; J3Q | PUR4_HUMAN | Unmodified | 0 | 0 | 0 | 0 | 102760 | 0 | 0 | 21981 | 0 | 56297 | 107310 | 1004000 | 0 | 0 | 0 | 0 | 0 | 0 |
| LNGFEVFA | ITGAV | P06756 | ITAV_HUMAN I | Unmodified | 296280 | 37097 | 1380000 | 986950 | 86169 | 49658 | 0 | 0 | 2324600 | 811340 | 437650 | 6174500 | 268340 | 185370 | 3971200 | 1720700 | 0 | 368610 |
| LNGGLGTS | UGP2; UGP2; U | Q16851; A0 | UGPA_HUMAN | Unmodified | 0 | 0 | 0 | 0 | 0 | 0 | 0 | 0 | 551200 | 238480 | 0 | 659150 | 0 | 0 | 0 | 0 | 203040 | 119570 |
| LNGGLGTS | UGP2; UGP2; U | Q16851; A0 | UGPA_HUMAN | Unmodified | 0 | 89628 | 0 | 0 | 695050 | 0 | 274430 | 120340 | 91007 | 0 | 160210 | 0 | 366950 | 249840 | 0 | 0 | 426290 | 376460 |
| LNGHVVVK | MLEC; MLEC; M | Q14165; H0 | MLEC_HUMAN | Unmodified | 0 | 0 | 0 | 0 | 0 | 0 | 0 | 0 | 817190 | 0 | 0 | 286140 | 0 | 0 | 0 | 0 | 0 | 0 |
| LNGLENRV | FLNB; FLNB | O75369; E7 | FLNB_HUMAN | Unmodified | 193500 | 133780 | 1261900 | 648720 | 938620 | 0 | 0 | 0 | 558740 | 77638 | 344320 | 5088100 | 0 | 0 | 1099600 | 945770 | 171510 | 127090 |
| LNGPQDH | CTNND1; CTNN | O60716; E9 | CTND1_HUMAI | Unmodified | 0 | 0 | 0 | 684630 | 855570 | 0 | 0 | 0 | 122050 | 0 | 306350 | 1577700 | 0 | 0 | 666600 | 663140 | 837930 | 1712300 |
| LNGPVTML | GALNT1 | Q10472 | GALT1_HUMAN | Oxidation (M) | 0 | 0 | 0 | 0 | 0 | 0 | 0 | 83885 | 602540 | 125400 | 0 | 0 | 0 | 0 | 0 | 0 | 0 | 0 |
| LNGPVTML | GALNT1 | Q10472 | GALT1_HUMAN | Unmodified | 0 | 0 | 0 | 0 | 0 | 0 | 0 | 0 | 2971400 | 0 | 0 | 1263600 | 0 | 0 | 0 | 0 | 0 | 0 |
| LNGTDPED | MYL12A; MYL12 | P19105; O1 | ML12A_HUMAI | Unmodified | 248950 | 264480 | 717090 | 379600 | 0 | 0 | 0 | 0 | 877460 | 0 | 1088100 | 21587000 | 111000 | 0 | 1112800 | 2268600 | 0 | 0 |
| LNGVVLNK | SLC12A7; SLC1 | Q9Y666; A0 | S12A7_HUMAN | Unmodified | 0 | 199340 | 391730 | 0 | 790970 | 166860 | 1056700 | 2043900 | 1130500 | 107970 | 0 | 4560400 | 2875300 | 2288800 | 854380 | 710740 | 624890 | 2720200 |
| LNGYVDAG | ACE; ACE; ACE; | P12821; J3K | ACE_HUMAN A | Unmodified | 442310 | 353300 | 0 | 0 | 0 | 0 | 0 | 0 | 0 | 0 | 0 | 101710 | 0 | 0 | 0 | 286740 | 0 | 0 |
| LNHANVVF | PTK7; PTK7; PTI | Q13308; H0 | PTK7_HUMAN I | Oxidation (M) | 0 | 0 | 0 | 0 | 0 | 0 | 0 | 0 | 0 | 0 | 0 | 0 | 827770 | 949150 | 0 | 0 | 923060 | 1062100 |
| LNHFFEGV | EXOC1 | Q9NV70 | EXOC1_HUMAI | Unmodified | 0 | 0 | 0 | 0 | 0 | 0 | 0 | 0 | 960580 | 370360 | 0 | 1210800 | 0 | 0 | 0 | 0 | 0 | 0 |
| LNHNAAFN | GFM1; GFM1; G | Q96RP9; C9 | EFGM_HUMAN | Oxidation (M) | 17769 | 27604 | 0 | 71033 | 0 | 0 | 0 | 0 | 698620 | 60135 | 0 | 0 | 0 | 0 | 0 | 0 | 0 | 0 |
| LNHNAAFN | GFM1; GFM1; G | Q96RP9; C9 | EFGM_HUMAN | Unmodified | 0 | 0 | 0 | 0 | 0 | 0 | 0 | 0 | 601580 | 34467 | 0 | 0 | 0 | 0 | 0 | 0 | 0 | 0 |
| LNHQVSEV | RAB17 | Q9H0T7 | RAB17_HUMAN | Unmodified | 412640 | 664500 | 84439 | 614020 | 0 | 0 | 0 | 0 | 674220 | 82088 | 0 | 0 | 0 | 0 | 874920 | 221290 | 0 | 0 |
| LNHVAIAVF | MCEE; MCEE | Q96PE7; H7 | MCEE_HUMAN | Unmodified | 0 | 0 | 0 | 0 | 0 | 0 | 0 | 0 | 0 | 0 | 0 | 0 | 0 | 0 | 1548800 | 0 | 0 | 0 |
| LNHYLEDK | EXOC8 | Q8IYI6 | EXOC8_HUMAI | Unmodified | 0 | 0 | 0 | 0 | 646080 | 0 | 75208 | 0 | 722060 | 0 | 0 | 713670 | 0 | 0 | 0 | 0 | 0 | 0 |
| LNIANMK | NECAP2; NECA | Q9NVZ3; D6 | NECP2_HUMAI | Unmodified | 0 | 52283 | 230060 | 0 | 623570 | 73625 | 59931 | 72190 | 275520 | 176980 | 0 | 181820 | 201670 | 221880 | 58715 | 0 | 160730 | 883250 |
| LNIDSIIQR | PPP1CC; PPP1 | P36873; F8V | PP1G_HUMAN | Unmodified | 0 | 0 | 0 | 0 | 0 | 0 | 0 | 0 | 2410700 | 325090 | 1176100 | 7639800 | 0 | 0 | 0 | 58727 | 0 | 0 |
| LNIDTRNC | ACTR2; ACTR2 | P61160; F5 | ARP2_HUMAN | Unmodified | 0 | 0 | 0 | 0 | 582510 | 248510 | 238220 | 0 | 28878 | 0 | 0 | 0 | 360930 | 392450 | 0 | 0 | 250920 | 452990 |
| LNIGDLQV | PSMD13; PSMD | Q9UNM6; A | PSD13_HUMAN | Unmodified | 0 | 0 | 0 | 0 | 0 | 0 | 0 | 0 | 0 | 0 | 0 | 0 | 0 | 0 | 0 | 0 | 0 | 0 |
| LNIIIAEGA | PFKL | P17858 | PFKAL_HUMAN | Unmodified | 113140 | 263560 | 59473 | 195940 | 66130 | 32002 | 64182 | 0 | 176850 | 41593 | 37218 | 401870 | 26865 | 0 | 657140 | 119830 | 119090 | 129140 |
| LNILDTLSK | PSMD2; PSMD2 | Q13200; H7 | PSMD2_HUMA | Unmodified | 0 | 0 | 0 | 0 | 0 | 0 | 0 | 0 | 244620 | 58532 | 79319 | 5599500 | 0 | 0 | 0 | 0 | 0 | 0 |
| LNIPATNVF | PSPH; PSPH | P78330; C9J | SERB_HUMAN | Unmodified | 0 | 0 | 0 | 0 | 0 | 0 | 0 | 0 | 489340 | 0 | 0 | 4313600 | 0 | 0 | 0 | 0 | 0 | 0 |
| LNIPATNVF | PSPH; PSPH | P78330; C9J | SERB_HUMAN | Unmodified | 0 | 0 | 0 | 249910 | 471940 | 0 | 389440 | 95807 | 180600 | 1923100 | 410330 | 0 | 5408900 | 2615600 | 0 | 0 | 776140 | 233240 |
| LNIPVSQV | ATP1A1; ATP1A | P05023; P13 | AT1A1_HUMAN | Unmodified | 1173900 | 2997900 | 45119000 | 32159000 | 2783100 | 5144300 | 99739 | 1280100 | 32931000 | 5563300 | 24686000 | 2.74E+08 | 3845200 | 1132200 | 76464000 | 61234000 | 2072900 | 579650 |
| LNIPVSQV | ATP1A1; ATP1A | P05023; P13 | AT1A1_HUMAN | Unmodified | 0 | 0 | 0 | 0 | 11202000 | 0 | 731450 | 319370 | 0 | 1694600 | 1743200 | 0 | 14240000 | 6305800 | 0 | 0 | 1340700 | 0 |
| LNIPYENF | PCSK1 | P29120 | NEC1_HUMAN | Unmodified | 0 | 0 | 0 | 0 | 0 | 0 | 0 | 0 | 1047500 | 1575200 | 0 | 0 | 0 | 0 | 0 | 0 | 0 | 0 |
| LNIQPSEA | UGGT1 | Q9NYU2 | UGGG1_HUMA | Unmodified | 0 | 0 | 0 | 0 | 353990 | 0 | 0 | 18383 | 3719400 | 244760 | 33594 | 98204 | 0 | 64634 | 0 | 0 | 0 | 228320 |
| LNISFPATG | RPS6; RPS6 | P62753; A2 | RS6_HUMAN 4 | Unmodified | 3991500 | 0 | 0 | 0 | 0 | 0 | 0 | 0 | 0 | 0 | 1506600 | 15662000 | 0 | 0 | 337920 | 2745000 | 0 | 0 |
| LNITVVQA | TOLLIP; TOLLIP | Q9H0E2; E7 | TOLIP_HUMAN | Unmodified | 14098000 | 16771000 | 7149800 | 0 | 0 | 0 | 286100 | 69871 | 1655500 | 0 | 87679 | 5367500 | 0 | 0 | 19785000 | 17740000 | 0 | 0 |

| | | | | | | | | | | | | | | | | | | | | | | |
|---|---|---|---|---|---|---|---|---|---|---|---|---|---|---|---|---|---|---|---|---|---|---|
| LNIWDVGG | ARL2; ARL2-SN | P36404; V9G | ARL2_HUMAN | Unmodified | 0 | 356450 | 0 | 0 | 0 | 0 | 0 | 0 | 0 | 0 | 0 | 3292500 | 0 | 0 | 0 | 0 | 0 | 0 |
| LNKDQWEI | MSN | P26038 | MOES_HUMAN | Unmodified | 0 | 0 | 0 | 815320 | 1228900 | 852230 | 0 | 0 | 0 | 0 | 0 | 0 | 218690 | 0 | 0 | 0 | 279120 | 0 |
| LNKPFLFD | CANX; CANX; C | P27824; A0A | CALX_HUMAN | Unmodified | 0 | 0 | 39249 | 0 | 0 | 0 | 0 | 0 | 33613000 | 1558600 | 1086900 | 1329100 | 0 | 0 | 0 | 0 | 0 | 0 |
| LNKPFLFD | CANX; CANX; C | P27824; A0A | CALX_HUMAN | Unmodified | 0 | 73837 | 128540 | 437420 | 0 | 0 | 1518500 | 0 | 1088100 | 8912200 | 0 | 0 | 0 | 0 | 842740 | 1393400 | 0 | 0 |
| LNLDSIIGR | PPP1CA; PPP1C | P62136; E9P | PP1A_HUMAN | Unmodified | 0 | 0 | 0 | 0 | 2561000 | 871280 | 0 | 0 | 4622100 | 226690 | 1578700 | 11832000 | 0 | 0 | 1806900 | 345960 | 1229100 | 433680 |
| LNLDSIIGR | PPP1CA; PPP1C | P62136; E9P | PP1A_HUMAN | Unmodified | 0 | 0 | 0 | 34545 | 12954000 | 2904500 | 784180 | 0 | 0 | 3026600 | 267220 | 22306 | 3252500 | 1845000 | 0 | 0 | 1975100 | 3784000 |
| LNLEHIATS | VPS35 | Q96QK1 | VPS35_HUMAN | Unmodified | 0 | 0 | 0 | 0 | 0 | 0 | 0 | 0 | 0 | 0 | 0 | 2251400 | 0 | 0 | 0 | 0 | 0 | 0 |
| LNLFLEK | PNMA2 | Q9UL42 | PNMA2_HUMA | Unmodified | 6330000 | 7568900 | 0 | 0 | 0 | 0 | 0 | 0 | 2694800 | 604670 | 93055 | 159180 | 0 | 0 | 0 | 0 | 0 | 0 |
| LNLGTDSD | PKN1; PKN1 | Q16512; K7E | PKN1_HUMAN | Unmodified | 0 | 0 | 0 | 0 | 0 | 0 | 0 | 0 | 0 | 97055 | 134520 | 522180 | 302920 | 0 | 0 | 0 | 175340 | 296130 |
| LNLGTVGF | NPEPPS; NPEPI | P55786; A0A | PSA_HUMAN P | Unmodified | 0 | 0 | 0 | 0 | 0 | 0 | 0 | 0 | 520090 | 146720 | 305820 | 7674800 | 6810600 | 0 | 0 | 403060 | 0 | 0 |
| LNLLGRCN | ZC3HAV1; ZC3I | Q7Z2W4; C9 | ZCCHV_HUMA | Unmodified | 0 | 0 | 0 | 0 | 2096700 | 378530 | 831660 | 201750 | 0 | 0 | 0 | 0 | 0 | 1743300 | 0 | 0 | 1288300 | 0 |
| LNLLHEFL( | DNMT1; DNMT | P26358; A0A | DNMT1_HUMA | Unmodified | 0 | 0 | 0 | 31620 | 47051 | 0 | 0 | 0 | 169160 | 54493 | 154770 | 282760 | 0 | 0 | 0 | 0 | 0 | 0 |
| LNLNNTVL | EPRS1; EPRS1 | P07814; V9G | SYEP_HUMAN | Unmodified | 0 | 0 | 342260 | 0 | 230500 | 0 | 0 | 79920 | 712300 | 0 | 707920 | 10407000 | 0 | 0 | 206010 | 232510 | 0 | 0 |
| LNLNNTVL | EPRS1; EPRS1 | P07814; V9G | SYEP_HUMAN | Unmodified | 143870 | 0 | 0 | 187920 | 0 | 0 | 0 | 0 | 682900 | 92821 | 208710 | 3847300 | 0 | 0 | 620040 | 255010 | 101300 | 409240 |
| LNLPINIIGL | LAP3 | P28838 | AMPL_HUMAN | Oxidation (M) | 0 | 0 | 0 | 0 | 0 | 0 | 868670 | 0 | 596550 | 626910 | 180970 | 458670 | 2235200 | 1072700 | 0 | 0 | 1076300 | 1118100 |
| LNLPINIIGL | LAP3 | P28838 | AMPL_HUMAN | Unmodified | 0 | 0 | 0 | 0 | 0 | 0 | 0 | 0 | 83908 | 0 | 0 | 0 | 78848 | 0 | 0 | 0 | 0 | 0 |
| LNLPSDMH | CA12; CA12 | O43570; B3I | CAH12_HUMA | Unmodified | 710340 | 570820 | 0 | 0 | 0 | 0 | 0 | 0 | 0 | 63798 | 0 | 836140 | 0 | 0 | 261750 | 0 | 44699 | 0 |
| LNLQTGER | SIL1; SIL1; SIL1 | Q9H173; D6 | SIL1_HUMAN N | Unmodified | 0 | 0 | 0 | 0 | 48933 | 0 | 0 | 0 | 5460200 | 0 | 0 | 0 | 66930 | 0 | 0 | 0 | 34558 | 0 |
| LNLQTSTSI | ITGA2; ITGA2; I | P17301; D6F | ITA2_HUMAN I | Oxidation (M) | 0 | 173850 | 320690 | 214370 | 0 | 0 | 0 | 0 | 0 | 0 | 0 | 0 | 19546 | 0 | 1196400 | 1704900 | 346860 | 0 |
| LNLRGCEV | FERMT1; FERM | Q9BQL6; G3 | FERM1_HUMA | Unmodified | 0 | 0 | 0 | 0 | 0 | 0 | 0 | 0 | 0 | 0 | 0 | 0 | 0 | 0 | 0 | 0 | 308810 | 628930 |
| LNLSQVR | DDX1; DDX1; D | Q92499; A0A | DDX1_HUMAN | Unmodified | 331880 | 0 | 550920 | 990470 | 320430 | 0 | 40888 | 49197 | 561050 | 76388 | 2061200 | 7212700 | 583110 | 206830 | 405970 | 515420 | 46749 | 138520 |
| LNLSSNELI | LRRC40 | Q9H9A6 | LRC40_HUMAN | Unmodified | 0 | 0 | 138730 | 0 | 0 | 0 | 0 | 0 | 0 | 0 | 0 | 1408700 | 0 | 0 | 447740 | 0 | 0 | 0 |
| LNLVQRNV | LDHA; LDHA; LI | P00338; F5G | LDHA_HUMAN | Unmodified | 0 | 0 | 0 | 0 | 255110 | 0 | 0 | 0 | 0 | 83928 | 0 | 0 | 38156 | 41685 | 0 | 0 | 151650 | 76512 |
| LNLWISR | LAD1; LAD1 | O00515; E9F | LAD1_HUMAN | Unmodified | 0 | 0 | 68298 | 0 | 0 | 0 | 0 | 0 | 0 | 0 | 0 | 993960 | 0 | 0 | 0 | 0 | 0 | 0 |
| LNLWISRT( | LAD1; LAD1 | O00515; E9F | LAD1_HUMAN | Unmodified | 0 | 0 | 0 | 0 | 0 | 0 | 0 | 0 | 0 | 0 | 0 | 0 | 476200 | 0 | 0 | 0 | 0 | 0 |
| LNLWRYGI | SND1 | Q7KZF4 | SND1_HUMAN | Unmodified | 0 | 0 | 0 | 0 | 470570 | 0 | 105060 | 0 | 0 | 0 | 0 | 0 | 876710 | 736270 | 0 | 0 | 0 | 407780 |
| LNLYELK | PSMB3; PSMB3 | P49720; A0A | PSB3_HUMAN | Unmodified | 894430 | 0 | 1321600 | 765160 | 0 | 0 | 606980 | 2905100 | 0 | 0 | 0 | 6509200 | 8306000 | 1652300 | 950020 | 717720 | 0 | 0 |
| LNMATLR | DHX9 | Q08211 | DHX9_HUMAN | Unmodified | 25860 | 56611 | 0 | 0 | 0 | 0 | 0 | 0 | 0 | 0 | 0 | 984380 | 0 | 0 | 0 | 0 | 0 | 0 |
| LNMELSEK | ERP29 | P30040 | ERP29_HUMAN | Oxidation (M) | 0 | 0 | 0 | 0 | 0 | 0 | 0 | 64424 | 4627200 | 540760 | 0 | 102300 | 0 | 0 | 0 | 0 | 119170 | 0 |
| LNMELSEK | ERP29 | P30040 | ERP29_HUMAN | Unmodified | 0 | 0 | 816140 | 0 | 0 | 0 | 0 | 63586 | 57184000 | 9268500 | 3051200 | 0 | 0 | 0 | 424420 | 0 | 0 | 0 |
| LNMEQLLQ | GPC3 | P51654 | GPC3_HUMAN | Oxidation (M) | 0 | 0 | 29434 | 903710 | 0 | 140150 | 0 | 0 | 769500 | 1998200 | 0 | 0 | 0 | 0 | 925150 | 0 | 14045 | 0 |
| LNMHMNV | APP; APP | P05067; A0A | A4_HUMAN Am | Oxidation (M) | 0 | 0 | 0 | 0 | 0 | 137150 | 0 | 0 | 611620 | 0 | 0 | 0 | 0 | 0 | 56941 | 0 | 0 | 0 |
| LNMHMNV | APP; APP | P05067; A0A | A4_HUMAN Am | Unmodified | 0 | 0 | 0 | 0 | 0 | 0 | 0 | 0 | 2707200 | 0 | 173290 | 0 | 0 | 0 | 0 | 0 | 0 | 0 |
| LNMHNLVE | DCP1A; DCP1A | Q9NPI6; A0A | DCP1A_HUMA | Unmodified | 0 | 0 | 0 | 0 | 0 | 0 | 0 | 0 | 0 | 0 | 0 | 702030 | 0 | 0 | 0 | 0 | 0 | 0 |
| LNMHVNIQ | APLP2; APLP2; | Q06481; E9F | APLP2_HUMAN | Unmodified | 0 | 0 | 0 | 0 | 2107200 | 440980 | 0 | 0 | 11419000 | 2190600 | 0 | 0 | 1145200 | 0 | 0 | 0 | 0 | 0 |
| LNMILVQIL | XPO1; XPO1; XI | O14980; A0A | XPO1_HUMAN | Oxidation (M) | 0 | 0 | 0 | 0 | 0 | 0 | 0 | 0 | 0 | 0 | 0 | 693320 | 0 | 0 | 0 | 0 | 0 | 0 |
| LNMTPEEA | EIF3E; EIF3E; EI | P60228; B3K | EIF3E_HUMAN | Unmodified | 0 | 0 | 0 | 0 | 0 | 0 | 0 | 0 | 0 | 0 | 0 | 513280 | 130380 | 0 | 0 | 0 | 0 | 0 |
| LNNAPVAG | ST3GAL4; ST3G | Q11206; E9F | SIA4C_HUMAN | Unmodified | 0 | 0 | 0 | 0 | 0 | 0 | 0 | 0 | 1012900 | 69885 | 284410 | 747580 | 0 | 0 | 0 | 0 | 0 | 0 |
| LNNDDNVI | MTHFD2; MTHF | P13995; A0A | MTDC_HUMAN | Unmodified | 0 | 0 | 0 | 0 | 912420 | 0 | 0 | 0 | 5452400 | 25326 | 35056 | 0 | 0 | 0 | 0 | 0 | 0 | 0 |
| LNNDDNVI | MTHFD2; MTHF | P13995; A0A | MTDC_HUMAN | Unmodified | 0 | 0 | 0 | 0 | 975060 | 0 | 0 | 0 | 2071400 | 0 | 0 | 0 | 0 | 0 | 0 | 0 | 20172 | 0 |
| LNNEYLELI | MME; MME; MM | P08473; A0A | NEP_HUMAN N | Unmodified | 0 | 0 | 0 | 526500 | 0 | 0 | 0 | 0 | 0 | 0 | 0 | 0 | 0 | 0 | 3450800 | 0 | 0 | 0 |
| LNNLVLFDI | RPS25 | P62851 | RS25_HUMAN | Unmodified | 6671200 | 4714100 | 3467900 | 3056600 | 0 | 0 | 789820 | 402580 | 0 | 0 | 5292600 | 48497000 | 2721500 | 1105800 | 4135400 | 2863600 | 0 | 0 |
| LNNLVLFDI | RPS25 | P62851 | RS25_HUMAN | Unmodified | 0 | 0 | 0 | 0 | 548450 | 0 | 2706400 | 1584400 | 0 | 0 | 0 | 0 | 2780100 | 1554900 | 0 | 0 | 0 | 590760 |
| LNPEDIKK | ACLY | P53396 | ACLY_HUMAN | Unmodified | 0 | 0 | 132970 | 0 | 220880 | 0 | 0 | 0 | 0 | 44454 | 0 | 2483400 | 0 | 0 | 0 | 0 | 0 | 36791 |
| LNPHWDG | PXDN | Q92626 | PXDN_HUMAN | Unmodified | 0 | 0 | 0 | 0 | 0 | 0 | 0 | 0 | 1105400 | 44227 | 0 | 0 | 0 | 0 | 0 | 0 | 0 | 0 |
| LNPINTTLL | NDRG3; NDRG | Q9UGV2; F8 | NDRG3_HUMA | Unmodified | 202180 | 127060 | 769810 | 1184000 | 2739700 | 0 | 0 | 0 | 2876600 | 0 | 6058400 | 46727000 | 1654100 | 0 | 1527700 | 1350700 | 279080 | 0 |
| LNPNFLVD | SELENBP1 | Q13228 | SBP1_HUMAN | Unmodified | 303570 | 0 | 400770 | 515210 | 0 | 0 | 0 | 0 | 0 | 0 | 0 | 0 | 0 | 0 | 3812100 | 2052400 | 0 | 0 |
| LNPRLAILY | ST13P4; ST13; | Q8IZP2; P50 | ST134_HUMAN | Unmodified | 0 | 0 | 0 | 0 | 0 | 0 | 1785700 | 0 | 0 | 0 | 0 | 0 | 0 | 0 | 0 | 0 | 714190 | 0 |
| LNPTWNHI | MYG1; MYG1 | Q9HB07; F8 | MYG1_HUMAN | Unmodified | 61178 | 0 | 252380 | 156180 | 0 | 0 | 0 | 0 | 915110 | 0 | 908560 | 4014900 | 392760 | 0 | 218330 | 191050 | 0 | 0 |
| LNPTWNHI | MYG1; MYG1 | Q9HB07; F8 | MYG1_HUMAN | Unmodified | 0 | 0 | 0 | 0 | 0 | 0 | 0 | 0 | 484880 | 793380 | 296860 | 773540 | 0 | 0 | 0 | 0 | 0 | 0 |
| LNPVGAKF | GRPEL1 | Q9HAV7 | GRPE1_HUMA | Unmodified | 0 | 0 | 0 | 0 | 3416400 | 362660 | 3387700 | 0 | 9178000 | 1692200 | 0 | 0 | 938900 | 784590 | 0 | 0 | 857110 | 1511500 |
| LNQALLDLI | FTL | P02792 | FRIL_HUMAN F | Unmodified | 0 | 0 | 0 | 0 | 0 | 0 | 0 | 0 | 4402700 | 0 | 0 | 0 | 0 | 0 | 0 | 0 | 0 | 0 |
| LNQDQLD | CAPRIN1; CAPI | Q14444; G3 | CAPR1_HUMA | Unmodified | 217390 | 424120 | 0 | 0 | 687650 | 0 | 174770 | 190970 | 261660 | 0 | 1108600 | 5817700 | 0 | 0 | 275270 | 102400 | 0 | 0 |
| LNQLKPGL | ILF3 | Q12906 | ILF3_HUMAN I | Unmodified | 3313200 | 5935500 | 761910 | 2898900 | 0 | 0 | 325690 | 91418 | 0 | 312310 | 474770 | 2526100 | 3309800 | 877470 | 1118800 | 1521300 | 0 | 0 |
| LNQLWEK | LRPAP1; LRPAP | P30533; D6F | AMRP_HUMAN | Unmodified | 0 | 0 | 0 | 0 | 198520 | 301680 | 0 | 0 | 4911100 | 2469400 | 0 | 0 | 0 | 0 | 0 | 0 | 0 | 0 |
| LNQNLND | BIN1 | O00499 | BIN1_HUMAN I | Unmodified | 0 | 0 | 0 | 0 | 0 | 0 | 0 | 0 | 0 | 0 | 0 | 0 | 0 | 104930 | 0 | 0 | 638990 | 94877 |
| LNQPPEDG | BUB3; BUB3; BI | O43684; J3Q | BUB3_HUMAN | Unmodified | 0 | 0 | 0 | 0 | 0 | 0 | 0 | 0 | 0 | 0 | 0 | 740030 | 0 | 0 | 0 | 0 | 0 | 0 |
| LNQPQPDI | MYO1D; MYO1I | O94832; K7E | MYO1D_HUMA | Unmodified | 10715000 | 11558000 | 4563500 | 3754900 | 764670 | 391910 | 110110 | 0 | 2129800 | 0 | 1068700 | 2131100 | 2021000 | 0 | 6491600 | 5445800 | 789600 | 3745600 |
| LNQQFVHI | MGAT1 | P26572 | MGAT1_HUMA | Unmodified | 0 | 0 | 0 | 0 | 0 | 0 | 0 | 0 | 8557900 | 274360 | 0 | 0 | 0 | 0 | 0 | 0 | 0 | 0 |
| LNQSQREE | PRPF8; PRPF8 | Q6P2Q9; I3L | PRP8_HUMAN | Unmodified | 0 | 0 | 0 | 0 | 0 | 0 | 0 | 0 | 0 | 0 | 0 | 2931800 | 0 | 0 | 0 | 0 | 0 | 0 |
| LNQVARQ( | PCBP1 | Q15365 | PCBP1_HUMA | 2 Oxidation (M) | 0 | 0 | 0 | 0 | 2327500 | 0 | 713060 | 600170 | 0 | 186100 | 0 | 0 | 4790600 | 0 | 0 | 0 | 829970 | 1207900 |
| LNQVARQ( | PCBP1 | Q15365 | PCBP1_HUMA | Oxidation (M) | 0 | 0 | 0 | 0 | 761270 | 0 | 0 | 0 | 0 | 0 | 0 | 0 | 1457600 | 1090200 | 0 | 0 | 263370 | 428710 |
| LNQVCATH | MYO1B; MYO1E | O43795; E9F | MYO1B_HUMA | Unmodified | 0 | 0 | 117430 | 0 | 800320 | 2809300 | 769700 | 429520 | 1660700 | 1488600 | 3433500 | 18005000 | 203480 | 0 | 648040 | 577250 | 1858400 | 1358500 |
| LNQVCATH | MYO1B; MYO1E | O43795; E9F | MYO1B_HUMA | Unmodified | 0 | 177580 | 0 | 0 | 5034900 | 409060 | 845540 | 616720 | 0 | 538090 | 0 | 0 | 777740 | 148410 | 0 | 0 | 2952700 | 690060 |
| LNQVLVDII | ANLN | Q9NQW6 | ANLN_HUMAN | Unmodified | 0 | 0 | 0 | 549950 | 3743500 | 1229800 | 296940 | 0 | 530540 | 0 | 203610 | 353190 | 281400 | 0 | 0 | 0 | 791690 | 0 |
| LNQWCNV | EPRS1 | P07814 | SYEP_HUMAN | Unmodified | 76356 | 0 | 0 | 0 | 0 | 0 | 0 | 0 | 0 | 0 | 0 | 3249200 | 0 | 0 | 0 | 92411 | 0 | 0 |
| LNQYFQK | DHX9 | Q08211 | DHX9_HUMAN | Unmodified | 96115 | 26374 | 274340 | 0 | 0 | 0 | 0 | 0 | 396290 | 144650 | 74662 | 2644200 | 0 | 0 | 0 | 0 | 0 | 0 |
| LNRAGMVI | MFGE8; MFGE8 | Q08431; H0 | MFGM_HUMAN | Oxidation (M) | 0 | 560670 | 0 | 0 | 5308400 | 882030 | 476260 | 0 | 0 | 1050800 | 0 | 0 | 224490 | 0 | 0 | 0 | 0 | 0 |
| LNRLPAAG | RPL23; RPL23; I | P62829; J3K | RL23_HUMAN | 2 Oxidation (M) | 352480 | 287190 | 0 | 192430 | 0 | 0 | 0 | 0 | 0 | 0 | 0 | 2570600 | 0 | 0 | 350430 | 0 | 67292 | 0 |
| LNRLPAAG | RPL23; RPL23; I | P62829; J3K | RL23_HUMAN | Oxidation (M) | 279890 | 481350 | 0 | 299230 | 0 | 0 | 0 | 0 | 0 | 0 | 0 | 3556700 | 0 | 0 | 0 | 312580 | 0 | 0 |
| LNRNHLQI | SLIT2; SLIT2; SL | O94813; X6F | SLIT2_HUMAN | Unmodified | 0 | 119340 | 0 | 0 | 0 | 0 | 0 | 0 | 0 | 0 | 0 | 0 | 0 | 0 | 0 | 0 | 0 | 129510 |
| LNRYNYIE( | CNDP2 | Q96KP4 | CNDP2_HUMA | Unmodified | 0 | 0 | 709580 | 1325700 | 2325900 | 531940 | 787790 | 636220 | 1861900 | 1410900 | 4271900 | 16232000 | 19848000 | 10751000 | 1960900 | 1784500 | 43629 | 1908500 |
| LNRYPASSI | TMEM59; TMEM | Q9BXS4; Q5 | TMM59_HUMA | Unmodified | 0 | 0 | 0 | 0 | 0 | 0 | 232270 | 0 | 4729500 | 641410 | 0 | 0 | 0 | 0 | 0 | 0 | 0 | 0 |
| LNSAIIYDR | RRM1 | P23921 | RIR1_HUMAN F | Unmodified | 0 | 0 | 0 | 0 | 0 | 0 | 0 | 0 | 0 | 0 | 0 | 800450 | 0 | 0 | 0 | 0 | 0 | 0 |
| LNSEGMEE | BPGM; BPGM | P07738; C9J | PMGE_HUMAN | Unmodified | 49745 | 81870 | 0 | 34550 | 0 | 0 | 0 | 0 | 0 | 0 | 53372 | 454870 | 0 | 0 | 0 | 31332 | 0 | 0 |
| LNSELQQC | REPS2; REPS1; | Q8NFH8; Q9 | REPS2_HUMAN | Unmodified | 924030 | 902600 | 0 | 754440 | 0 | 0 | 0 | 0 | 657540 | 0 | 0 | 1159500 | 263360 | 0 | 887620 | 840650 | 0 | 0 |
| LNSEMDITI | MYOF; MYOF | Q9NZM1; H( | MYOF_HUMAN | Unmodified | 0 | 0 | 0 | 0 | 16342000 | 3562200 | 0 | 0 | 0 | 0 | 0 | 0 | 0 | 0 | 0 | 0 | 405700 | 1848800 |
| LNSEMDITI | MYOF; MYOF | Q9NZM1; H( | MYOF_HUMAN | Oxidation (M) | 0 | 0 | 0 | 0 | 2446700 | 439590 | 0 | 0 | 0 | 0 | 0 | 0 | 0 | 0 | 0 | 0 | 0 | 324550 |
| LNSGGRW | NAXE; NAXE; N | Q8NCW5; A | NNRE_HUMAN | Unmodified | 0 | 0 | 0 | 0 | 0 | 0 | 0 | 0 | 429110 | 0 | 0 | 0 | 180960 | 0 | 0 | 0 | 0 | 0 |
| LNSGLGQL | ABLIM1; ABLIM | O14639; J3Q | ABLM1_HUMA | Unmodified | 1.04E+08 | 79814000 | 63437000 | 49497000 | 0 | 0 | 0 | 0 | 0 | 0 | 0 | 0 | 6422800 | 1618500 | 1.54E+08 | 94117000 | 0 | 0 |
| LNSGVDYF | LSM6 | P62312 | LSM6_HUMAN | Unmodified | 0 | 0 | 0 | 0 | 265480 | 40353 | 154770 | 0 | 0 | 117350 | 245100 | 2331900 | 0 | 0 | 0 | 0 | 172520 | 0 |
| LNSHMNAI | G6PD; G6PD; G | P11413; E7E | G6PD_HUMAN | Unmodified | 0 | 0 | 0 | 0 | 117160 | 165500 | 0 | 0 | 1130400 | 0 | 0 | 3777400 | 0 | 0 | 0 | 330970 | 0 | 0 |
| LNSIGNYYI | DHCR24; DHCI | Q15392; A0A | DHC24_HUMA | Unmodified | 72752 | 0 | 98391 | 145330 | 0 | 0 | 0 | 0 | 439480 | 0 | 107060 | 2233800 | 0 | 0 | 431950 | 177850 | 0 | 0 |
| LNSLAIGLF | ALDH18A1 | P54886 | P5CS_HUMAN | Unmodified | 0 | 113520 | 0 | 54396 | 0 | 0 | 0 | 0 | 707360 | 0 | 0 | 2771000 | 0 | 0 | 101510 | 69302 | 0 | 0 |
| LNSLLFPPA | PLCB2 | Q00722 | PLCB2_HUMAN | Unmodified | 0 | 0 | 0 | 0 | 0 | 0 | 0 | 0 | 0 | 0 | 189700 | 4546300 | 0 | 0 | 0 | 0 | 0 | 0 |
| LNSLLFPPA | PLCB2 | Q00722 | PLCB2_HUMAN | Unmodified | 0 | 0 | 0 | 0 | 0 | 0 | 0 | 0 | 0 | 0 | 489590 | 2406400 | 0 | 0 | 0 | 0 | 0 | 0 |
| LNSLLNIPE | C1GALT1C1 | Q96EU7 | C1GLC_HUMA | Unmodified | 0 | 0 | 0 | 0 | 2660300 | 242210 | 324840 | 313680 | 1527200 | 393990 | 347730 | 886260 | 1269000 | 536610 | 0 | 0 | 1787100 | 0 |
| LNSLMSLV | PSMD12 | O00232 | PSD12_HUMAN | Oxidation (M) | 0 | 0 | 0 | 0 | 0 | 0 | 0 | 0 | 0 | 0 | 0 | 1087900 | 0 | 0 | 0 | 363650 | 0 | 0 |
| LNSLMSLV | PSMD12 | O00232 | PSD12_HUMAN | Unmodified | 0 | 0 | 0 | 0 | 0 | 0 | 0 | 0 | 0 | 0 | 0 | 1338600 | 0 | 0 | 0 | 0 | 0 | 0 |
| LNSNDEDI | ASL; ASL; ASL; | P04424; F8V | ARLY_HUMAN | Unmodified | 90346 | 0 | 0 | 195330 | 0 | 0 | 0 | 0 | 220470 | 0 | 0 | 510510 | 0 | 0 | 0 | 177150 | 0 | 0 |
| LNSNTQVV | EIF4A1; EIF4A1 | P60842; J3K | IF4A1_HUMAN | Oxidation (M) | 0 | 0 | 765900 | 0 | 679680 | 0 | 0 | 0 | 0 | 0 | 0 | 3304400 | 1570800 | 0 | 3036900 | 1187200 | 0 | 0 |
| LNSVGNDI | ITGB8 | P26012 | ITB8_HUMAN I | Unmodified | 12439 | 171110 | 0 | 0 | 0 | 0 | 0 | 0 | 242280 | 0 | 0 | 13388 | 0 | 0 | 0 | 0 | 331890 | 0 |
| LNSVQSSE | FASN; FASN | P49327; A0A | FAS_HUMAN F | Unmodified | 8682900 | 1165000 | 16786000 | 38055000 | 0 | 707100 | 372720 | 0 | 20716000 | 0 | 24974000 | 8.1E+08 | 599000 | 201860 | 49152000 | 29034000 | 1237300 | 412080 |
| LNSYIQGK | HEBP2; HEBP2 | Q9Y5Z4; Q5 | HEBP2_HUMA | Unmodified | 0 | 0 | 145200 | 0 | 0 | 0 | 0 | 0 | 0 | 0 | 0 | 200710 | 0 | 0 | 0 | 0 | 0 | 0 |
| LNTAEDAKI | RANGAP1; RAN | P46060; B0( | RAGP1_HUMA | Unmodified | 0 | 0 | 0 | 0 | 0 | 0 | 0 | 0 | 0 | 0 | 0 | 329650 | 0 | 0 | 0 | 0 | 0 | 0 |
| LNTDPMLL | USP7; USP7; U | Q93009; A0A | UBP7_HUMAN | Oxidation (M) | 0 | 469560 | 88694 | 0 | 0 | 0 | 0 | 0 | 0 | 0 | 0 | 595320 | 0 | 0 | 0 | 0 | 0 | 0 |
| LNTEWSEL | P4HA1 | P13674 | P4HA1_HUMA | Unmodified | 0 | 0 | 0 | 0 | 0 | 0 | 10512 | 0 | 165590 | 132630 | 200000 | 0 | 0 | 0 | 0 | 0 | 0 | 0 |
| LNTHVKPLI | AP1M1; AP1M1 | Q9BXS5; K7E | AP1M1_HUMA | Unmodified | 46495 | 0 | 0 | 78896 | 0 | 0 | 0 | 0 | 193890 | 40594 | 0 | 233180 | 0 | 0 | 165600 | 42462 | 0 | 0 |
| LNTIIDSDK | ABCC4 | O15439 | MRP4_HUMAN | Unmodified | 323840 | 284100 | 4108600 | 961350 | 11057000 | 0 | 0 | 0 | 221770 | 92717 | 285960 | 4843700 | 0 | 0 | 3226000 | 2838900 | 0 | 0 |
| LNTIIDSDK | ABCC4 | O15439 | MRP4_HUMAN | Oxidation (M) | 0 | 0 | 0 | 607450 | 0 | 0 | 0 | 0 | 0 | 0 | 0 | 0 | 71368 | 0 | 0 | 0 | 0 | 0 |
| LNTIMDYTI | ABCC3; ABCC1 | O15438; P3 | MRP3_HUMAN | Oxidation (M) | 0 | 0 | 251990 | 0 | 126650 | 0 | 0 | 0 | 0 | 0 | 0 | 608490 | 208550 | 0 | 142160 | 0 | 48863 | 0 |
| LNTIMDYTI | ABCC3; ABCC1 | O15438; P3 | MRP3_HUMAN | Unmodified | 0 | 0 | 0 | 345780 | 0 | 0 | 0 | 0 | 0 | 0 | 0 | 1703600 | 0 | 0 | 0 | 31911 | 0 | 0 |
| LNTLETEEV | LYN; LYN | P07948; E5R | LYN_HUMAN T | Unmodified | 0 | 0 | 0 | 0 | 1376800 | 457020 | 0 | 0 | 0 | 1327900 | 0 | 0 | 0 | 0 | 1940900 | 3134300 | 0 | 0 |
| LNTLLQR | MYO18B; HNRN | Q8IUG5; Q0 | MY18B_HUMA | Unmodified | 1092700 | 785190 | 387670 | 125860 | 0 | 0 | 85196 | 114030 | 1021200 | 204190 | 869640 | 9319500 | 104300 | 0 | 1018900 | 758710 | 31704 | 42865 |
| LNTLLQRAI | HNRNPU; HNR | Q00839; A0A | HNRPU_HUMA | Unmodified | 684190 | 0 | 0 | 0 | 557150 | 0 | 1631400 | 742280 | 0 | 434450 | 239580 | 0 | 754210 | 474410 | 0 | 0 | 55415 | 503800 |
| LNTLMHYQ | PLXNA2 | O75051 | PLXA2_HUMAN | Unmodified | 0 | 0 | 0 | 0 | 0 | 0 | 18017 | 0 | 2136900 | 0 | 0 | 638640 | 0 | 0 | 0 | 53664 | 0 | 0 |
| LNTQEIFDI | DYNC1H1; DYN | Q14204; A0A | DYHC1_HUMA | Unmodified | 0 | 0 | 0 | 0 | 0 | 0 | 0 | 0 | 533910 | 0 | 0 | 1883500 | 0 | 0 | 0 | 0 | 0 | 0 |
| LNTQEIFDI | DYNC1H1; DYN | Q14204; A0A | DYHC1_HUMA | Unmodified | 0 | 0 | 0 | 0 | 0 | 0 | 0 | 0 | 0 | 0 | 50049 | 2183600 | 0 | 0 | 0 | 0 | 0 | 0 |
| LNTQSTYSI | NRP1; NRP1; N | O14786; E9F | NRP1_HUMAN | Unmodified | 0 | 0 | 227810 | 0 | 1223200 | 526800 | 0 | 0 | 0 | 0 | 0 | 0 | 0 | 0 | 164440 | 529690 | 327500 | 2451400 |

| | | | | | | | | | | | | | | | | | | | | | | |
|---|---|---|---|---|---|---|---|---|---|---|---|---|---|---|---|---|---|---|---|---|---|---|
| LNTSELIK | CLCN3; CLCN3 | P51790; H0Y | CLCN3_HUMAI | Unmodified | 0 | 0 | 0 | 0 | 0 | 0 | 0 | 0 | 1490300 | 15497 | 333360 | 2272700 | 0 | 0 | 330000 | 242350 | 0 | 0 |
| LNVAEVTQ | PARVA; PARVA | Q9NVD7; E9 | PARVA_HUMAN | Unmodified | 0 | 0 | 0 | 0 | 403570 | 0 | 0 | 0 | 0 | 0 | 0 | 0 | 0 | 0 | 0 | 0 | 121960 | 272680 |
| LNVAEVTQ | PARVB; PARVB; | Q9HBI1; A0A | PARVB_HUMAN | Unmodified | 580940 | 679860 | 1418000 | 1062100 | 228740 | 0 | 139070 | 11177 | 521200 | 0 | 0 | 0 | 172180 | 164860 | 1436400 | 1101300 | 0 | 75589 |
| LNVCVSK | HNRNPL; HNRN | P14866; Q8V | HNRPL_HUMAI | Unmodified | 104770 | 49199 | 0 | 0 | 427140 | 0 | 695770 | 312900 | 652910 | 384620 | 1253600 | 9106600 | 278190 | 0 | 0 | 49349 | 0 | 277350 |
| LNVDEAFE | RRAS | P10301 | RRAS_HUMAN | Unmodified | 163490 | 374320 | 74185 | 148700 | 66293 | 14934 | 0 | 0 | 0 | 21771 | 0 | 49948 | 0 | 0 | 459420 | 433300 | 34308 | 61093 |
| LNVEDVDS | SCG3 | Q8WXD2 | SCG3_HUMAN | Unmodified | 0 | 0 | 0 | 0 | 0 | 0 | 0 | 0 | 17087000 | 3284900 | 56565 | 300720 | 0 | 0 | 0 | 0 | 0 | 0 |
| LNVEEGLY | GLT8D1 | Q68CQ7 | GL8D1_HUMAI | Unmodified | 0 | 0 | 0 | 0 | 0 | 0 | 0 | 0 | 15092000 | 2040100 | 0 | 0 | 0 | 0 | 0 | 0 | 0 | 0 |
| LNVEERSV | EPHA2 | P29317 | EPHA2_HUMAI | Unmodified | 0 | 0 | 0 | 0 | 580990 | 0 | 0 | 0 | 0 | 0 | 0 | 0 | 0 | 0 | 0 | 0 | 488860 | 1605200 |
| LNVEERSV | EPHA2 | P29317 | EPHA2_HUMAI | Unmodified | 0 | 0 | 0 | 0 | 859590 | 0 | 129960 | 51008 | 0 | 0 | 0 | 0 | 0 | 0 | 0 | 0 | 1181700 | 2012100 |
| LNVFAKPE | BCAM; BCAM; E | P50895; A0A | BCAM_HUMAN | Unmodified | 4990500 | 505190 | 0 | 0 | 0 | 0 | 0 | 0 | 2044500 | 0 | 0 | 0 | 0 | 0 | 0 | 0 | 99099 | 0 |
| LNVFKNDQ | HNRNPL; HNRN | P14866; M0( | HNRPL_HUMAI | Unmodified | 0 | 0 | 0 | 0 | 0 | 0 | 0 | 0 | 342830 | 0 | 0 | 5177800 | 0 | 0 | 0 | 0 | 0 | 0 |
| LNVFKNDQ | HNRNPL; HNRN | P14866; M0( | HNRPL_HUMAI | Unmodified | 0 | 0 | 0 | 0 | 0 | 0 | 0 | 0 | 0 | 0 | 0 | 2311000 | 0 | 0 | 0 | 0 | 0 | 0 |
| LNVHMDSI | SELENOT; SELE | P62341; A0A | SELT_HUMAN T | Unmodified | 0 | 0 | 0 | 0 | 0 | 0 | 0 | 0 | 1019600 | 0 | 0 | 0 | 0 | 0 | 48811 | 38075 | 0 | 0 |
| LNVKLTKLT | AHCY | P23526 | SAHH_HUMAN | Unmodified | 0 | 0 | 0 | 0 | 1378900 | 0 | 259550 | 114920 | 0 | 0 | 0 | 0 | 246880 | 272680 | 0 | 0 | 1382600 | 839120 |
| LNVTEQEK | ENO1; ENO1; E | P06733; A0A | ENOA_HUMAN | Unmodified | 525750 | 429070 | 3880200 | 1790500 | 6442400 | 2993300 | 247720 | 186660 | 10485000 | 1431500 | 2191500 | 25304000 | 5361000 | 1890100 | 4737000 | 2881500 | 1023300 | 7944100 |
| LNVTEQEKI | ENO1; ENO1; E | P06733; A0A | ENOA_HUMAN | Unmodified | 212830 | 0 | 0 | 1054500 | 23126000 | 4300600 | 723220 | 250330 | 4836400 | 3026600 | 1437300 | 3995800 | 9960600 | 2177800 | 239640 | 374180 | 3772400 | 15329000 |
| LNVTEQEKI | ENO1; ENO1; E | P06733; A0A | ENOA_HUMAN | Unmodified | 0 | 0 | 0 | 0 | 18207000 | 1934900 | 0 | 0 | 0 | 2181800 | 0 | 0 | 1013800 | 2769300 | 0 | 0 | 5196300 | 15212000 |
| LNVTEQEKI | ENO1; ENO1; E | P06733; A0A | ENOA_HUMAN | Oxidation (M) | 0 | 0 | 0 | 0 | 9953600 | 1183700 | 128920 | 0 | 0 | 1858700 | 0 | 0 | 2934900 | 1866700 | 0 | 0 | 4864500 | 5443400 |
| LNVTEQEKI | ENO1; ENO1; E | P06733; A0A | ENOA_HUMAN | 2 Oxidation (M) | 0 | 0 | 0 | 0 | 1569600 | 0 | 121240 | 0 | 0 | 690910 | 0 | 0 | 137180 | 361690 | 0 | 0 | 1261700 | 1795400 |
| LNVTVQDQ | ETNK1; ETNK1; | Q9HBU6; H( | EKI1_HUMAN E | Unmodified | 0 | 0 | 0 | 0 | 0 | 0 | 0 | 0 | 0 | 0 | 0 | 0 | 0 | 0 | 0 | 0 | 0 | 0 |
| LNVVYTEDI | RASA3 | Q14644 | RASA3_HUMAI | Unmodified | 304170 | 339310 | 0 | 817670 | 0 | 0 | 0 | 0 | 347730 | 0 | 0 | 77445 | 0 | 0 | 543670 | 426680 | 0 | 778080 |
| LNVWDIGG | ARL3 | P36405 | ARL3_HUMAN | Unmodified | 0 | 0 | 597570 | 1901400 | 1754000 | 0 | 683720 | 353200 | 1473900 | 450390 | 0 | 6143100 | 4895100 | 1943300 | 0 | 0 | 0 | 2417500 |
| LNVWDLSK | RBBP7; RBBP4; | Q16576; Q0 | RBBP7_HUMAI | Unmodified | 0 | 0 | 0 | 0 | 43624 | 13579 | 0 | 0 | 1714800 | 29412 | 525760 | 6876100 | 117150 | 0 | 0 | 0 | 0 | 0 |
| LNWGHIPS | HSPG2 | P98160 | PGBM_HUMAN | Unmodified | 0 | 0 | 3579500 | 817460 | 0 | 0 | 0 | 0 | 363650 | 0 | 0 | 0 | 0 | 0 | 0 | 423150 | 0 | 0 |
| LNWTGTSK | AHSA1; AHSA1; | O95433; H0Y | AHSA1_HUMAI | Unmodified | 31757 | 23217 | 44479 | 0 | 587510 | 194850 | 186600 | 180480 | 818670 | 818520 | 989160 | 2093000 | 1079500 | 256900 | 83409 | 0 | 953150 | 563470 |
| LNYGIPAIV | JUP | P14923 | PLAK_HUMAN | Unmodified | 1613700 | 423800 | 5773100 | 3628900 | 0 | 0 | 0 | 0 | 0 | 0 | 331430 | 21960000 | 0 | 0 | 9126600 | 7401900 | 0 | 0 |
| LNYLDILMF | ITGA6; ITGA6 | P23229; H7E | ITA6_HUMAN I | Oxidation (M) | 0 | 0 | 0 | 523290 | 0 | 0 | 0 | 0 | 0 | 0 | 0 | 6578900 | 0 | 0 | 2252900 | 12879000 | 0 | 0 |
| LNYLDILMF | ITGA6; ITGA6 | P23229; H7E | ITA6_HUMAN I | Unmodified | 0 | 0 | 0 | 0 | 0 | 0 | 0 | 0 | 0 | 0 | 0 | 0 | 0 | 0 | 0 | 0 | 0 | 0 |
| LNYSDHDV | AP2M1; AP2M1; | Q96CW1; E9 | AP2M1_HUMAI | Unmodified | 0 | 0 | 962750 | 0 | 583340 | 411990 | 236900 | 79709 | 0 | 0 | 431700 | 4211200 | 181590 | 928060 | 959650 | 75068 | 126820 | 464910 |
| LPAAGVGD | RPL23; RPL23; | P62829; J3K | RL23_HUMAN ( | 2 Oxidation (M) | 0 | 0 | 0 | 0 | 9034200 | 3883200 | 0 | 0 | 0 | 0 | 0 | 734930 | 0 | 0 | 0 | 464680 | 169660 | 230630 |
| LPAAGVGD | RPL23; RPL23; | P62829; J3K | RL23_HUMAN ( | Oxidation (M) | 271820 | 223740 | 166650 | 0 | 0 | 0 | 0 | 20927 | 0 | 0 | 0 | 1695400 | 0 | 0 | 162540 | 118650 | 0 | 0 |
| LPAELGPQ | COMMD8 | Q9NX08 | COMD8_HUMA | Unmodified | 0 | 0 | 0 | 0 | 0 | 0 | 0 | 0 | 171430 | 199500 | 0 | 1153000 | 358820 | 0 | 0 | 0 | 0 | 0 |
| LPAELQELF | APEX1; APEX1; | P27695; A0A | APEX1_HUMAN | Unmodified | 0 | 0 | 0 | 0 | 0 | 0 | 0 | 0 | 0 | 0 | 0 | 1497000 | 0 | 0 | 0 | 0 | 0 | 0 |
| LPAHTVGD | GRN; GRN | P28799; K7E | GRN_HUMAN F | Unmodified | 153360 | 0 | 0 | 0 | 0 | 0 | 0 | 0 | 799300 | 0 | 0 | 1109200 | 0 | 0 | 0 | 0 | 0 | 0 |
| LPANHPLL | ATP6V1A | P38606 | VATA_HUMAN | Unmodified | 0 | 0 | 0 | 0 | 0 | 0 | 0 | 0 | 0 | 0 | 0 | 1767000 | 0 | 0 | 223080 | 0 | 0 | 0 |
| LPANQYTW | TPP2; TPP2; TPI | P29144; A0A | TPP2_HUMAN | Unmodified | 0 | 0 | 65666 | 0 | 0 | 0 | 0 | 0 | 284180 | 0 | 0 | 4027700 | 0 | 0 | 71163 | 0 | 0 | 0 |
| LPASFDARI | CTSB; CTSB; C1 | P07858; E9P | CATB_HUMAN | Unmodified | 0 | 0 | 0 | 0 | 840910 | 141430 | 153280 | 0 | 0 | 0 | 115950 | 521420 | 0 | 0 | 0 | 0 | 337060 | 0 |
| LPASVFVV | DOP1B | Q9Y3R5 | DOP2_HUMAN | Unmodified | 0 | 0 | 0 | 0 | 0 | 0 | 0 | 0 | 0 | 0 | 0 | 871930 | 0 | 0 | 0 | 99382 | 0 | 0 |
| LPATAAEPE | HMGCS1 | Q01581 | HMCS1_HUMA | Unmodified | 0 | 32586 | 0 | 0 | 0 | 0 | 218320 | 0 | 0 | 0 | 116490 | 1959400 | 0 | 0 | 0 | 0 | 0 | 0 |
| LPATEKPVL | HYOU1; HYOU1 | Q9Y4L1; A0A | HYOU1_HUMA | Unmodified | 0 | 0 | 441510 | 0 | 0 | 0 | 0 | 0 | 4269700 | 0 | 0 | 615440 | 0 | 0 | 246210 | 0 | 0 | 0 |
| LPAVVTADI | ETFB; ETFB | P38117; M0( | ETFB_HUMAN I | Unmodified | 0 | 0 | 0 | 278260 | 314190 | 0 | 0 | 0 | 1924600 | 0 | 590550 | 3429700 | 163780 | 0 | 0 | 302610 | 190260 | 0 |
| LPAYFVVPT | MTMR12 | Q9C0I1 | MTMRC_HUMA | Unmodified | 16893 | 17381 | 61056 | 0 | 0 | 0 | 0 | 0 | 0 | 0 | 0 | 375330 | 0 | 0 | 0 | 0 | 0 | 0 |
| LPDERGPL | IGFBP2; IGFBP2 | P18065; C9J | IBP2_HUMAN I | Unmodified | 0 | 0 | 0 | 0 | 0 | 0 | 0 | 0 | 32390000 | 1353800 | 0 | 0 | 0 | 0 | 0 | 0 | 0 | 0 |
| LPDGFTQL | SCRIB; SCRIB; | Q14160; A0A | SCRIB_HUMAN | Unmodified | 208110 | 0 | 0 | 0 | 0 | 0 | 0 | 0 | 0 | 0 | 0 | 3182700 | 0 | 0 | 332740 | 344960 | 0 | 0 |
| LPDGTSLT | UBXN1; UBXN1 | Q04323; E9F | UBXN1_HUMAI | Unmodified | 0 | 0 | 199380 | 428110 | 0 | 0 | 0 | 0 | 273640 | 0 | 285290 | 4508800 | 0 | 0 | 0 | 623430 | 0 | 0 |
| LPDYKPYPI | CDK5 | Q00535 | CDK5_HUMAN | Oxidation (M) | 0 | 0 | 451800 | 0 | 0 | 0 | 0 | 0 | 0 | 0 | 0 | 2262700 | 0 | 0 | 633360 | 785690 | 0 | 0 |
| LPEAVQLIE | NAPG; NAPG | Q99747; J3K | SNAG_HUMAN | Unmodified | 0 | 0 | 0 | 0 | 903290 | 0 | 0 | 0 | 0 | 0 | 0 | 3146700 | 0 | 0 | 0 | 0 | 210390 | 1489000 |
| LPEDPLLS | FASN; FASN | P49327; A0A | FAS_HUMAN F | Unmodified | 0 | 0 | 188420 | 0 | 0 | 0 | 0 | 0 | 0 | 0 | 943130 | 5831100 | 0 | 0 | 687550 | 314390 | 0 | 0 |
| LPEEIQR | MAN1A1 | P33908 | MA1A1_HUMAI | Unmodified | 1.83E+08 | 1.9E+08 | 0 | 1.15E+08 | 0 | 440460 | 49773 | 21015 | 0 | 67941 | 752210 | 327940 | 0 | 88809 | 2552500 | 77269 | 49649 | 0 |
| LPEEIQRDI | MAN1A1 | P33908 | MA1A1_HUMAI | Unmodified | 0 | 0 | 0 | 0 | 0 | 0 | 0 | 0 | 1196700 | 0 | 162230 | 150810 | 0 | 0 | 0 | 0 | 0 | 0 |
| LPEFADWA | HLA-DMA; HLA- | P28067; A0A | DMA_HUMAN I | Unmodified | 0 | 0 | 19845 | 0 | 0 | 0 | 0 | 0 | 137630 | 0 | 0 | 0 | 0 | 0 | 334380 | 161830 | 0 | 0 |
| LPEFIVQTA | LDOC1 | O95751 | LDOC1_HUMAI | Unmodified | 0 | 137210 | 0 | 0 | 20867 | 0 | 0 | 0 | 0 | 23228 | 0 | 0 | 0 | 0 | 429480 | 0 | 0 | 0 |
| LPEHCIEYV | UBA3; UBA3 | Q8TBC4; F8V | UBA3_HUMAN | Unmodified | 0 | 0 | 0 | 52846 | 0 | 0 | 0 | 0 | 992990 | 41020 | 698400 | 3643000 | 118550 | 0 | 0 | 0 | 0 | 0 |
| LPEHPEGG | LYAR | Q9NX58 | LYAR_HUMAN ( | Unmodified | 0 | 0 | 0 | 0 | 0 | 0 | 0 | 0 | 0 | 0 | 343230 | 524020 | 0 | 0 | 0 | 0 | 0 | 0 |
| LPELAPTAP | VPS37B; VPS37 | Q9H9H4; F5 | VP37B_HUMAN | Unmodified | 2938400 | 2893500 | 1322200 | 731090 | 0 | 0 | 0 | 0 | 0 | 0 | 0 | 897200 | 0 | 0 | 3678100 | 5349900 | 0 | 0 |
| LPEMTAPLF | LMAN2L | Q9H0V9 | LMA2L_HUMAN | Unmodified | 0 | 0 | 0 | 0 | 0 | 0 | 0 | 0 | 0 | 0 | 0 | 0 | 1879600 | 0 | 0 | 0 | 0 | 0 |
| LPENKLEG | DAAM1 | Q9Y4D1 | DAAM1_HUMA | Unmodified | 251410 | 0 | 0 | 0 | 485840 | 420600 | 469650 | 0 | 996630 | 0 | 1219600 | 4740100 | 0 | 0 | 1862000 | 1621100 | 43728 | 487070 |
| LPETNLFET | KARS1 | Q15046 | SYK_HUMAN Ly | Unmodified | 0 | 0 | 370240 | 0 | 0 | 0 | 0 | 0 | 338720 | 0 | 1707600 | 11400000 | 0 | 0 | 241570 | 251410 | 0 | 0 |
| LPEVEVPQI | NUCB1 | Q02818 | NUCB1_HUMA | Unmodified | 0 | 0 | 0 | 0 | 0 | 1400700 | 0 | 0 | 767930 | 0 | 647310 | 0 | 0 | 0 | 0 | 0 | 428950 | 0 |
| LPFAAAQIG | GARS1; GARS1; | P41250; A0A | GARS_HUMAN | Unmodified | 0 | 0 | 0 | 0 | 0 | 0 | 0 | 0 | 0 | 0 | 0 | 3072500 | 0 | 0 | 0 | 0 | 0 | 0 |
| LPFAAAQIG | GARS1; GARS1; | P41250; A0A | GARS_HUMAN | Unmodified | 0 | 0 | 0 | 0 | 0 | 0 | 2073600 | 0 | 401320 | 0 | 0 | 2192000 | 0 | 0 | 0 | 0 | 0 | 0 |
| LPFPIIDDR | PRDX6 | P30041 | PRDX6_HUMAI | Unmodified | 151630 | 0 | 1170300 | 0 | 0 | 0 | 156340 | 0 | 200180 | 0 | 2633400 | 8896500 | 0 | 0 | 596220 | 1294700 | 0 | 0 |
| LPFPIIDDRI | PRDX6 | P30041 | PRDX6_HUMAI | Unmodified | 0 | 0 | 0 | 0 | 0 | 0 | 0 | 41153 | 482570 | 478900 | 0 | 22380000 | 0 | 0 | 0 | 4342300 | 0 | 0 |
| LPFTIPELV | ERN1; ERN1 | O75460; A0A | ERN1_HUMAN | Unmodified | 0 | 0 | 0 | 0 | 0 | 0 | 0 | 0 | 296040 | 64018 | 0 | 0 | 0 | 0 | 0 | 0 | 10557 | 0 |
| LPFTPLSYI | TMCO1; TMCO1 | Q9UM00; J3I | TMCO1_HUMA | Unmodified | 0 | 0 | 0 | 0 | 0 | 67195 | 0 | 0 | 6825100 | 1724100 | 0 | 2032200 | 14061 | 0 | 224940 | 0 | 0 | 0 |
| LPFTVVHEF | PIGS | Q96S52 | PIGS_HUMAN ( | Unmodified | 0 | 0 | 0 | 0 | 0 | 0 | 0 | 0 | 1410700 | 0 | 0 | 0 | 0 | 0 | 0 | 0 | 0 | 0 |
| LPFVINDGI | NPTX1 | Q15818 | NPTX1_HUMAI | Unmodified | 0 | 0 | 0 | 0 | 0 | 0 | 0 | 0 | 20014000 | 407830 | 0 | 285630 | 0 | 0 | 0 | 0 | 0 | 0 |
| LPFVINDGI | NPTX1 | Q15818 | NPTX1_HUMAI | Unmodified | 0 | 0 | 0 | 0 | 0 | 0 | 0 | 0 | 1175400 | 0 | 207400 | 0 | 0 | 78852 | 0 | 0 | 0 | 0 |
| LPFYNQDH | AKT2; AKT2; AKT | P31751; J3Q | AKT2_HUMAN I | Unmodified | 0 | 0 | 291700 | 0 | 0 | 228730 | 37472 | 0 | 0 | 174780 | 0 | 0 | 0 | 0 | 0 | 334120 | 0 | 226040 |
| LPGDKGLV | CANX; CANX; C | P27824; A0A | CALX_HUMAN | Unmodified | 0 | 0 | 0 | 0 | 0 | 0 | 0 | 0 | 0 | 148170 | 0 | 0 | 263400 | 218720 | 0 | 0 | 123740 | 0 |
| LPGGELNP | NUDT21; NUDT | O43809; H3I | CPSF5_HUMAI | Unmodified | 0 | 135430 | 0 | 0 | 0 | 0 | 0 | 0 | 0 | 21969 | 0 | 2815900 | 0 | 0 | 0 | 57728 | 0 | 0 |
| LPGGELNP | NUDT21; NUDT | O43809; H3I | CPSF5_HUMAI | Unmodified | 1600600 | 984200 | 2443600 | 0 | 0 | 0 | 0 | 0 | 0 | 0 | 0 | 5165200 | 0 | 0 | 452780 | 1223200 | 0 | 0 |
| LPGGVQNI | HUWE1 | Q7Z6Z7 | HUWE1_HUMA | Unmodified | 23161 | 0 | 0 | 0 | 0 | 0 | 0 | 12286 | 97972 | 0 | 0 | 876280 | 0 | 0 | 49388 | 50340 | 0 | 0 |
| LPGHAGSII | SNRNP40; SNR | Q96DI7; Q9N | SNR40_HUMAI | Unmodified | 0 | 0 | 0 | 0 | 0 | 0 | 99339 | 0 | 0 | 0 | 0 | 673340 | 0 | 0 | 0 | 0 | 0 | 0 |
| LPGKWPFS | SNX27; SNX27; | Q96L92; A0A | SNX27_HUMAI | Unmodified | 386060 | 247250 | 160590 | 116580 | 0 | 0 | 0 | 0 | 226910 | 0 | 3380100 | 1160200 | 0 | 0 | 458090 | 326840 | 0 | 0 |
| LPGLLGNF | ASAH1; ASAH1; | Q13510; A0A | ASAH1_HUMAI | Oxidation (M) | 0 | 0 | 106700 | 220630 | 0 | 0 | 0 | 0 | 2106300 | 1648300 | 336700 | 134360 | 124230 | 122930 | 195500 | 80660 | 0 | 0 |
| LPGMEYVD | IGFBP5 | P24593 | IBP5_HUMAN I | Unmodified | 0 | 0 | 0 | 0 | 0 | 0 | 0 | 0 | 2168100 | 235610 | 0 | 0 | 0 | 0 | 0 | 0 | 0 | 0 |
| LPGPSGAG | SMPDL3B; SMP | Q92485; Q5T | ASM3B_HUMAI | Unmodified | 0 | 1435300 | 14047000 | 6427300 | 176240 | 128520 | 0 | 0 | 5973300 | 535900 | 1740700 | 10715000 | 2196700 | 0 | 10625000 | 13622000 | 0 | 0 |
| LPHCDPKT | TJP1 | Q07157 | ZO1_HUMAN T | Unmodified | 0 | 0 | 0 | 0 | 0 | 0 | 65150 | 0 | 0 | 104310 | 0 | 0 | 0 | 201380 | 0 | 0 | 304170 | 1135200 |
| LPHSVLLEI | PSAT1 | Q9Y617 | SERC_HUMAN | Unmodified | 220320 | 135410 | 1015900 | 1191000 | 0 | 0 | 0 | 60954 | 0 | 255030 | 2713300 | 14092000 | 919390 | 482170 | 2262000 | 1741000 | 0 | 0 |
| LPICAQCG | ZNF330; ZNF33 | Q9Y3S2; D6I | ZN330_HUMAN | Unmodified | 0 | 0 | 0 | 0 | 54610 | 0 | 0 | 0 | 0 | 34932 | 0 | 225670 | 0 | 0 | 0 | 0 | 0 | 0 |
| LPIDENQLA | MME; MME; MM | P08473; A0A | NEP_HUMAN N | Oxidation (M) | 0 | 0 | 2666600 | 614600 | 0 | 391230 | 0 | 0 | 0 | 0 | 0 | 0 | 0 | 0 | 3435800 | 2168700 | 0 | 0 |
| LPIDENQLA | MME; MME; MM | P08473; A0A | NEP_HUMAN N | Unmodified | 0 | 1347800 | 0 | 0 | 0 | 0 | 0 | 0 | 0 | 0 | 0 | 0 | 0 | 0 | 1359000 | 0 | 0 | 0 |
| LPIGDVATC | CCT7 | Q99832 | TCPH_HUMAN | Oxidation (M) | 0 | 0 | 0 | 0 | 0 | 0 | 0 | 0 | 1339100 | 0 | 1218100 | 7405300 | 33585000 | 20919000 | 758380 | 0 | 0 | 0 |
| LPIPESQVI | PGLS; PGLS; PC | O95336; M0I | 6PGL_HUMAN | Unmodified | 0 | 0 | 0 | 0 | 0 | 0 | 0 | 0 | 784820 | 0 | 348120 | 724930 | 0 | 0 | 0 | 0 | 0 | 0 |
| LPIPRPGEV | ABCE1; ABCE1; | P61221; H0Y | ABCE1_HUMAI | Unmodified | 0 | 0 | 0 | 43958 | 0 | 0 | 0 | 0 | 0 | 0 | 76363 | 810150 | 0 | 0 | 0 | 0 | 0 | 0 |
| LPISSSTSN | TTC7B; TTC7B | Q86TV6; G3V | TTC7B_HUMAI | Unmodified | 0 | 0 | 437990 | 0 | 0 | 0 | 292290 | 91790 | 0 | 0 | 0 | 503520 | 276810 | 0 | 1150600 | 796520 | 0 | 659770 |
| LPKEEDSAI | CD2AP | Q9Y5K6 | CD2AP_HUMAI | Unmodified | 817540 | 698880 | 925850 | 632350 | 0 | 0 | 0 | 0 | 0 | 0 | 0 | 117730 | 0 | 0 | 1485200 | 897400 | 0 | 0 |
| LPLDVLGR | EPS15L1; EPS1 | Q9UBC2; M( | EP15R_HUMAN | Unmodified | 209530 | 366320 | 0 | 0 | 0 | 0 | 0 | 0 | 0 | 0 | 0 | 568350 | 0 | 0 | 0 | 0 | 0 | 0 |
| LPLGLLLHF | SEC24A | O95486 | SC24A_HUMAI | Unmodified | 0 | 0 | 0 | 0 | 0 | 0 | 0 | 0 | 0 | 0 | 0 | 856790 | 0 | 0 | 0 | 0 | 0 | 0 |
| LPLGLLLHF | SEC24B | O95487 | SC24B_HUMAI | Unmodified | 0 | 0 | 0 | 0 | 0 | 0 | 0 | 0 | 0 | 0 | 0 | 429600 | 0 | 0 | 0 | 0 | 0 | 0 |
| LPLMECVQ | PFKL | P17858 | PFKAL_HUMAN | Unmodified | 0 | 0 | 0 | 109160 | 0 | 0 | 0 | 0 | 0 | 0 | 1245300 | 882080 | 0 | 0 | 38647 | 0 | 0 | 0 |
| LPLPYGFS | PPM1G | O15355 | PPM1G_HUMA | Oxidation (M) | 0 | 0 | 0 | 0 | 0 | 0 | 0 | 0 | 0 | 0 | 0 | 0 | 0 | 0 | 0 | 0 | 0 | 0 |
| LPLQDVYK | EEF1A2; EEF1A | Q05639; Q5V | EF1A2_HUMAN | Unmodified | 11449000 | 10459000 | 6345100 | 3829800 | 713180 | 175960 | 0 | 165880 | 199960 | 0 | 1473300 | 12963000 | 0 | 0 | 9737500 | 9793200 | 369330 | 413410 |
| LPLQLDDA | DARS1; DARS1; | P14868; C9J | SYDC_HUMAN | Unmodified | 0 | 0 | 0 | 0 | 0 | 0 | 0 | 0 | 391960 | 147040 | 2172200 | 13390000 | 0 | 0 | 0 | 0 | 0 | 0 |
| LPLSPSLLV | PIK3C2B; PIK3C | O00750; F5C | P3C2B_HUMAI | Unmodified | 0 | 0 | 672400 | 0 | 21813 | 23444 | 0 | 0 | 0 | 0 | 0 | 582610 | 0 | 0 | 58758 | 0 | 0 | 0 |
| LPLVTPHT | PSMC1 | P62191 | PRS4_HUMAN | Unmodified | 0 | 0 | 393540 | 0 | 0 | 0 | 0 | 0 | 0 | 0 | 0 | 2212900 | 0 | 0 | 218690 | 0 | 0 | 0 |
| LPLVVEDF | GATD1; GATD1; | Q8NB37; H0 | GALD1_HUMAI | Unmodified | 0 | 0 | 0 | 48596 | 0 | 0 | 0 | 0 | 481700 | 0 | 224200 | 1307900 | 0 | 0 | 122160 | 146950 | 0 | 0 |
| LPLWQQA | BAIAP2; BAIAP2 | Q9UQB8; I3L | BAIP2_HUMAN | Unmodified | 0 | 0 | 0 | 0 | 0 | 0 | 0 | 0 | 0 | 0 | 0 | 1221800 | 0 | 0 | 1231300 | 0 | 0 | 0 |
| LPNFGFVV | G3BP1; G3BP1; | Q13283; A0A | G3BP1_HUMAI | Unmodified | 0 | 0 | 0 | 0 | 533950 | 109410 | 0 | 415150 | 19492 | 0 | 251920 | 391820 | 0 | 0 | 0 | 0 | 1132400 | 0 |
| LPNFGFVV | G3BP2; G3BP2; | Q9UN86; A0 | G3BP2_HUMAI | Unmodified | 0 | 0 | 0 | 0 | 0 | 0 | 0 | 0 | 86277 | 0 | 0 | 227940 | 0 | 0 | 0 | 0 | 0 | 0 |
| LPNGLVIAS | UQCRC2; UQCI | P22695; H3E | QCR2_HUMAN | Unmodified | 35744 | 20291 | 78095 | 49406 | 33338 | 0 | 32209 | 36523 | 11198000 | 1051100 | 332490 | 158050 | 0 | 0 | 38838 | 32700 | 30681 | 27003 |
| LPNGVFTPI | MAP2K2; MAP2 | P36507; G5E | MP2K2_HUMAI | Unmodified | 0 | 0 | 96187 | 0 | 0 | 0 | 15776 | 0 | 474690 | 0 | 26199 | 130890 | 0 | 0 | 231690 | 124320 | 0 | 0 |
| LPNLGKHE | CKB; CKB; CKB | P12277; H0Y | KCRB_HUMAN | Unmodified | 0 | 0 | 0 | 0 | 359280 | 66459 | 0 | 0 | 0 | 0 | 0 | 1718300 | 0 | 0 | 0 | 0 | 22071 | 288510 |
| LPNLLEHE | SFRP5 | Q5T4F7 | SFRP5_HUMAN | Unmodified | 0 | 0 | 0 | 0 | 0 | 0 | 0 | 0 | 6294500 | 125000 | 0 | 0 | 0 | 0 | 0 | 0 | 0 | 0 |
| LPNLTHLN | ANP32B | Q92688 | AN32B_HUMAI | Unmodified | 0 | 450060 | 417270 | 0 | 0 | 0 | 0 | 0 | 79867 | 0 | 622480 | 3663300 | 0 | 0 | 0 | 0 | 0 | 0 |
| LPNNHIGIS | FLNB; FLNB | O75369; E7E | FLNB_HUMAN | Unmodified | 0 | 0 | 0 | 0 | 0 | 0 | 0 | 38109 | 0 | 53055 | 0 | 335730 | 0 | 0 | 164720 | 130940 | 0 | 0 |
| LPNSVLGK | EHD4; EHD3 | Q9H223; Q9 | EHD4_HUMAN | Unmodified | 736480 | 640750 | 3086300 | 1715900 | 1139600 | 0 | 486990 | 237110 | 223140 | 0 | 0 | 11951000 | 666590 | 703620 | 5095500 | 2418000 | 384950 | 886860 |
| LPNTVLGK | EHD1; EHD1 | Q9H4M9; A0 | EHD1_HUMAN | Unmodified | 156810 | 888890 | 545600 | 0 | 3105100 | 129700 | 0 | 89323 | 0 | 66756 | 268760 | 7244900 | 981600 | 0 | 352410 | 431690 | 279330 | 396610 |

| | | | | | | | | | | | | | | | | | | | | | | |
|---|---|---|---|---|---|---|---|---|---|---|---|---|---|---|---|---|---|---|---|---|---|---|
| LPPEEASPY | SEC24C; SEC24 | P53992; G5E | SC24C_HUMAI | Unmodified | 271750 | 0 | 0 | 0 | 0 | 0 | 0 | 102530 | 0 | 0 | 0 | 2458100 | 0 | 0 | 0 | 0 | 0 | 0 |
| LPPHQPGF | VPS37B | Q9H9H4 | VP37B_HUMAN | Unmodified | 5254800 | 2641300 | 879860 | 362300 | 0 | 0 | 0 | 0 | 0 | 0 | 0 | 628270 | 0 | 0 | 2596500 | 2754900 | 0 | 0 |
| LPPLPPREI | CREG1; CREG1 | O75629; A0A | CREG1_HUMAI | Unmodified | 0 | 0 | 0 | 96038 | 0 | 0 | 0 | 0 | 2456200 | 364140 | 0 | 0 | 316900 | 118370 | 0 | 0 | 0 | 0 |
| LPPNTNDE | SF3B3 | Q15393 | SF3B3_HUMAN | Unmodified | 0 | 0 | 0 | 0 | 0 | 0 | 0 | 0 | 0 | 0 | 96281 | 738770 | 0 | 0 | 0 | 0 | 0 | 0 |
| LPPPKPLP( | ADAM10 | O14672 | ADA10_HUMAN | Unmodified | 6518500 | 3943000 | 11662000 | 2548400 | 0 | 0 | 0 | 0 | 1671600 | 371130 | 0 | 22633000 | 119350 | 0 | 28150000 | 20009000 | 0 | 0 |
| LPPQDFLD | CKAP4 | Q07065 | CKAP4_HUMAN | Unmodified | 88443 | 44372 | 190370 | 119950 | 130240 | 0 | 0 | 0 | 8973100 | 785200 | 2198600 | 76330 | 0 | 0 | 125220 | 140240 | 0 | 0 |
| LPPVLNLQI | VPS50; VPS50 | Q96JG6; C9J | VPS50_HUMAN | Unmodified | 0 | 0 | 0 | 0 | 0 | 0 | 0 | 0 | 0 | 0 | 0 | 2039500 | 0 | 0 | 0 | 0 | 0 | 0 |
| LPQALAPLI | VPS37B; VPS37 | Q9H9H4; F5 | VP37B_HUMAN | Unmodified | 4719800 | 0 | 0 | 1378900 | 0 | 0 | 0 | 0 | 0 | 0 | 558380 | 4605800 | 0 | 0 | 9145100 | 5900000 | 0 | 0 |
| LPQFGISTF | AHNAK | Q09666 | AHNK_HUMAN | Unmodified | 71975 | 60769 | 0 | 0 | 637530 | 263820 | 0 | 18575 | 0 | 0 | 0 | 0 | 0 | 0 | 0 | 0 | 0 | 782610 |
| LPQKEPVG | KIAA1522 | Q9P206 | K1522_HUMAN | Unmodified | 0 | 80540 | 273610 | 839750 | 710470 | 0 | 0 | 0 | 22633 | 0 | 224110 | 2397300 | 0 | 205080 | 0 | 0 | 0 | 1208000 |
| LPQLPITNF | FLNA; FLNA; FL | P21333; A0A | FLNA_HUMAN | Unmodified | 712950 | 0 | 2003400 | 0 | 345560 | 0 | 0 | 0 | 0 | 0 | 0 | 2455200 | 0 | 0 | 1759000 | 1247900 | 0 | 0 |
| LPQLPITNF | FLNA; FLNA; FL | P21333; A0A | FLNA_HUMAN | Unmodified | 42250 | 89287 | 0 | 656020 | 0 | 0 | 0 | 0 | 0 | 0 | 0 | 0 | 206640 | 0 | 117010 | 0 | 0 | 0 |
| LPQPEGCP | PTK7; PTK7 | Q13308; H0Y | PTK7_HUMAN I | Unmodified | 107330 | 83414 | 0 | 127920 | 0 | 0 | 0 | 0 | 316610 | 71507 | 445010 | 41837 | 0 | 0 | 65971 | 57353 | 0 | 0 |
| LPQVAEEIS | FLOT1; FLOT1; I | O75955; A0A | FLOT1_HUMAN | Unmodified | 0 | 0 | 2291400 | 1660100 | 0 | 0 | 337570 | 54681 | 9865700 | 0 | 1920600 | 8293900 | 0 | 0 | 3197900 | 1459400 | 0 | 0 |
| LPQVVTDV | LRRC8A | Q8IWT6 | LRC8A_HUMAN | Unmodified | 107750 | 0 | 0 | 0 | 0 | 0 | 27269 | 0 | 0 | 0 | 0 | 477820 | 0 | 0 | 0 | 0 | 0 | 0 |
| LPSANVYG | MGAM2 | Q2M2H8 | MGAL_HUMAN | Unmodified | 0 | 0 | 0 | 0 | 0 | 0 | 0 | 0 | 0 | 0 | 0 | 2093900 | 0 | 0 | 0 | 97608 | 0 | 0 |
| LPSDVVTA\ | OAT | P04181 | OAT_HUMAN C | Unmodified | 0 | 0 | 0 | 0 | 74086 | 0 | 0 | 0 | 4062000 | 947710 | 0 | 355280 | 0 | 0 | 0 | 0 | 165720 | 93220 |
| LPSEPGMT | AIDA | Q96BJ3 | AIDA_HUMAN / | Oxidation (M) | 0 | 0 | 0 | 1866000 | 0 | 0 | 0 | 0 | 0 | 0 | 0 | 303020 | 0 | 0 | 0 | 2155100 | 0 | 0 |
| LPSEYGVIP | CSGALNACT2 | Q8N6G5 | CGAT2_HUMAI | Oxidation (M) | 0 | 0 | 0 | 0 | 63911 | 29091 | 36326 | 0 | 671640 | 160730 | 0 | 115490 | 0 | 0 | 0 | 0 | 38616 | 0 |
| LPSFWNNI | CPM; CPM | P14384; H0Y | CBPM_HUMAN | Unmodified | 0 | 0 | 0 | 0 | 1288400 | 0 | 0 | 0 | 2383100 | 82012 | 0 | 0 | 0 | 0 | 0 | 0 | 0 | 0 |
| LPSIPLVPV | TOMM34 | Q15785 | TOM34_HUMAI | Unmodified | 0 | 0 | 0 | 0 | 0 | 0 | 0 | 0 | 0 | 0 | 627080 | 2232900 | 0 | 0 | 0 | 0 | 0 | 0 |
| LPSLGNKEI | SLC7A11 | Q9UPY5 | XCT_HUMAN C | Unmodified | 0 | 0 | 0 | 0 | 45183 | 34826 | 0 | 0 | 74466 | 19244 | 0 | 538020 | 0 | 9366 | 33489 | 591410 | 0 | 0 |
| LPSLSGSD | EPHA1 | P21709 | EPHA1_HUMAN | Unmodified | 11823000 | 6997600 | 0 | 0 | 0 | 0 | 0 | 0 | 0 | 235810 | 0 | 0 | 0 | 0 | 741800 | 0 | 0 | 0 |
| LPSLWADN | CPD | O75976 | CBPD_HUMAN | Unmodified | 0 | 0 | 0 | 0 | 0 | 0 | 0 | 0 | 2485800 | 0 | 0 | 0 | 0 | 0 | 0 | 0 | 0 | 513120 |
| LPSPLDITA | EFL1 | Q7Z2Z2 | EFL1_HUMAN I | Unmodified | 0 | 0 | 56196 | 69601 | 0 | 0 | 0 | 0 | 822190 | 0 | 0 | 375400 | 0 | 0 | 50199 | 0 | 0 | 0 |
| LPSQYIYGF | MGAM2 | Q2M2H8 | MGAL_HUMAN | Unmodified | 0 | 0 | 0 | 0 | 0 | 0 | 0 | 0 | 168950 | 0 | 0 | 2345100 | 0 | 0 | 0 | 0 | 0 | 0 |
| LPSQYNFA | NUP210 | Q8TEM1 | PO210_HUMAN | Unmodified | 0 | 0 | 0 | 0 | 0 | 0 | 0 | 0 | 1719100 | 0 | 0 | 0 | 0 | 0 | 0 | 0 | 0 | 0 |
| LPSSVFASE | LTV1 | Q96GA3 | LTV1_HUMAN I | Unmodified | 0 | 0 | 0 | 0 | 0 | 0 | 0 | 0 | 0 | 0 | 0 | 0 | 0 | 0 | 402270 | 0 | 0 | 0 |
| LPSTSGSE( | EPHA2 | P29317 | EPHA2_HUMAN | Unmodified | 90534 | 106460 | 116360 | 0 | 0 | 0 | 168120 | 59496 | 0 | 0 | 0 | 1982900 | 0 | 0 | 0 | 527670 | 0 | 0 |
| LPSVPSTHI | CHMP4A; CHM | Q9BY43; H0 | CHM4A_HUMA | Unmodified | 599680 | 402100 | 213260 | 0 | 0 | 0 | 0 | 0 | 0 | 0 | 0 | 577230 | 0 | 0 | 296210 | 342170 | 0 | 0 |
| LPSVPSTHI | CHMP4A; CHM | Q9BY43; H0 | CHM4A_HUMA | Unmodified | 4681300 | 3567400 | 0 | 403020 | 0 | 0 | 0 | 0 | 0 | 0 | 0 | 3891100 | 0 | 0 | 1729900 | 1420600 | 0 | 0 |
| LPTAADLL( | ATP9A | O75110 | ATP9A_HUMAN | Unmodified | 128950 | 149480 | 120500 | 0 | 0 | 0 | 0 | 0 | 537260 | 481840 | 0 | 0 | 0 | 0 | 415350 | 203850 | 0 | 0 |
| LPTAVFEGI | DTNB; DTNB; D | O60941; E7E | DTNB_HUMAN | Unmodified | 0 | 698760 | 0 | 0 | 0 | 0 | 0 | 0 | 0 | 0 | 0 | 0 | 0 | 0 | 462780 | 223440 | 0 | 43800 |
| LPTEEEWEI | SUMF2; SUMF2 | Q8NBJ7; C9 | SUMF2_HUMAI | Unmodified | 0 | 0 | 0 | 0 | 0 | 0 | 0 | 0 | 7127500 | 0 | 0 | 0 | 0 | 0 | 0 | 0 | 0 | 0 |
| LPTEEEWEI | SUMF2; SUMF2 | Q8NBJ7; C9 | SUMF2_HUMAI | Unmodified | 0 | 0 | 0 | 0 | 57985 | 0 | 0 | 0 | 2175000 | 346910 | 0 | 0 | 0 | 0 | 0 | 0 | 0 | 0 |
| LPTFGAPE( | TRIM25; TRIM25 | Q14258; A0A | TRI25_HUMAN | Unmodified | 0 | 0 | 85085 | 19789 | 89943 | 30089 | 169750 | 0 | 0 | 0 | 57535 | 258220 | 0 | 0 | 77726 | 51157 | 0 | 29841 |
| LPTGINPLT | WDR44; WDR44 | Q5JSH3; H7I | WDR44_HUMA | Oxidation (M) | 89371 | 0 | 48514 | 29921 | 0 | 0 | 0 | 0 | 0 | 0 | 37026 | 162260 | 0 | 0 | 177250 | 76149 | 0 | 2309900 |
| LPTGQISGI | AHNAK | Q09666 | AHNK_HUMAN | Unmodified | 0 | 0 | 0 | 0 | 1140300 | 0 | 0 | 0 | 0 | 0 | 0 | 0 | 0 | 0 | 0 | 0 | 277630 | 857040 |
| LPTGYYFG/ | LMAN2; LMAN2 | Q12907; D6I | LMAN2_HUMAI | Oxidation (M) | 0 | 0 | 0 | 0 | 0 | 278740 | 0 | 0 | 6597500 | 2192700 | 0 | 0 | 0 | 0 | 0 | 0 | 0 | 0 |
| LPTGYYFG/ | LMAN2; LMAN2 | Q12907; D6I | LMAN2_HUMAI | Unmodified | 0 | 0 | 0 | 0 | 0 | 0 | 0 | 0 | 8779900 | 1356900 | 0 | 0 | 0 | 0 | 0 | 0 | 0 | 0 |
| LPTKNEFIP | IDE; IDE; IDE; ID | P14735; A0A | IDE_HUMAN In | Unmodified | 0 | 0 | 0 | 0 | 0 | 0 | 0 | 0 | 0 | 0 | 31849 | 218540 | 0 | 0 | 0 | 0 | 0 | 0 |
| LPTPVLLPY | MMS19; MMS19 | Q96T76; H0Y | MMS19_HUMA | Unmodified | 0 | 0 | 0 | 0 | 0 | 0 | 0 | 0 | 0 | 0 | 0 | 3912800 | 0 | 0 | 0 | 0 | 0 | 0 |
| LPVACPNQ | TRAF4; TRAF4 | Q9BUZ4; F6 | TRAF4_HUMAN | Unmodified | 0 | 0 | 0 | 0 | 0 | 0 | 0 | 0 | 0 | 0 | 0 | 0 | 0 | 0 | 0 | 1497800 | 0 | 0 |
| LPVALDPG | RPN1 | P04843 | RPN1_HUMAN | Unmodified | 61098 | 0 | 293330 | 159000 | 0 | 0 | 109460 | 32389 | 1.22E+08 | 6708200 | 4295400 | 3184800 | 0 | 103490 | 0 | 259590 | 0 | 0 |
| LPVDFSNIF | BABAM2; BABA | Q9NXR7; F8 | BABA2_HUMAN | Unmodified | 0 | 0 | 0 | 0 | 0 | 0 | 0 | 0 | 0 | 0 | 0 | 255150 | 0 | 0 | 0 | 0 | 0 | 0 |
| LPVESIQIVI | VPS25; VPS25; | Q9BRG1; K7 | VPS25_HUMAN | Unmodified | 0 | 0 | 0 | 0 | 0 | 0 | 0 | 0 | 0 | 0 | 0 | 0 | 0 | 0 | 1111600 | 0 | 0 | 0 |
| LPVGGFAD | ITCH; ITCH; ITC | Q96J02; A0A | ITCH_HUMAN I | Unmodified | 558690 | 0 | 0 | 0 | 21347 | 0 | 0 | 0 | 0 | 0 | 0 | 0 | 0 | 0 | 0 | 0 | 0 | 0 |
| LPVGGFAE | WWP2 | O00308 | WWP2_HUMAN | Unmodified | 1581100 | 2922100 | 1008200 | 768390 | 0 | 0 | 43341 | 0 | 0 | 110720 | 0 | 0 | 0 | 0 | 3563100 | 1216300 | 0 | 0 |
| LPVLLLGR | HTRA1; HTRA1; | Q92743; A0A | HTRA1_HUMAN | Unmodified | 14126 | 0 | 86555 | 0 | 0 | 0 | 0 | 0 | 1140100 | 0 | 0 | 0 | 0 | 0 | 0 | 0 | 27916 | 0 |
| LPVLYQVEI | ACOT8; ACOT8 | O14734; E9F | ACOT8_HUMAI | Unmodified | 909450 | 903860 | 190490 | 443800 | 0 | 0 | 0 | 130880 | 5482100 | 155580 | 646770 | 1487900 | 349340 | 0 | 366730 | 271140 | 0 | 0 |
| LPVPASAPI | SLC44A1 | Q8WWI5 | CTL1_HUMAN | Unmodified | 15818000 | 4657400 | 16061000 | 7537700 | 0 | 0 | 0 | 0 | 10208000 | 1451600 | 2413100 | 16014000 | 119240 | 0 | 23997000 | 17445000 | 0 | 0 |
| LPVQLQR | STOM | P27105 | STOM_HUMAN | Unmodified | 3432800 | 2567700 | 5060300 | 3455800 | 3279400 | 1272800 | 801210 | 312390 | 0 | 0 | 0 | 0 | 901370 | 0 | 3916600 | 2712800 | 632800 | 0 |
| LPVSEGVF | MAN2B1; MAN2 | O00754; M0I | MA2B1_HUMAI | Unmodified | 168110 | 5670100 | 267980 | 0 | 0 | 0 | 0 | 0 | 0 | 344220 | 0 | 428580 | 198400 | 0 | 0 | 218450 | 0 | 0 |
| LPVVIGGLI | CLTC; CLTC | Q00610; A0A | CLH1_HUMAN | Unmodified | 327810 | 153960 | 0 | 0 | 0 | 0 | 70947 | 0 | 0 | 0 | 0 | 0 | 0 | 0 | 105760 | 0 | 230480 | 444590 |
| LPYTASSGI | ATP6AP1; ATP6 | Q15904; A0A | VAS1_HUMAN | Unmodified | 0 | 0 | 0 | 0 | 0 | 0 | 0 | 0 | 1236300 | 0 | 0 | 0 | 0 | 0 | 0 | 0 | 127540 | 0 |
| LQAALDDE | HNRNPU; HNR | Q00839; A0A | HNRPU_HUMA | Oxidation (M) | 0 | 0 | 0 | 0 | 0 | 0 | 0 | 0 | 0 | 0 | 4973500 | 11176000 | 0 | 0 | 0 | 0 | 0 | 0 |
| LQAALDDE | HNRNPU; HNR | Q00839; A0A | HNRPU_HUMA | Unmodified | 0 | 0 | 0 | 0 | 0 | 0 | 0 | 0 | 0 | 0 | 2067900 | 4384800 | 0 | 0 | 0 | 0 | 0 | 0 |
| LQAANAED | NUCB1; NUCB1 | Q02818; H7I | NUCB1_HUMA | Unmodified | 0 | 0 | 0 | 0 | 787740 | 694720 | 0 | 90873 | 6279000 | 634080 | 2637700 | 1257600 | 0 | 0 | 0 | 415200 | 44283 | 1966400 |
| LQAAVDGF | BCAP31; BCAP3 | P51572; A0A | BAP31_HUMAN | Unmodified | 38981 | 43089 | 449620 | 341670 | 331890 | 0 | 1597700 | 350910 | 860480 | 225890 | 0 | 0 | 643480 | 992380 | 347970 | 0 | 0 | 695120 |
| LQAAVDGF | BCAP31; BCAP3 | P51572; A0A | BAP31_HUMAN | Oxidation (M) | 0 | 0 | 0 | 0 | 145500 | 0 | 279330 | 0 | 0 | 0 | 0 | 0 | 194100 | 209510 | 187890 | 0 | 0 | 0 |
| LQAAYAGD | SPTBN1; SPTBN | Q01082; A0A | SPTB2_HUMAN | Unmodified | 0 | 0 | 0 | 0 | 0 | 0 | 0 | 0 | 0 | 0 | 0 | 3503500 | 0 | 0 | 0 | 0 | 0 | 0 |
| LQADFKISE | YARS1; YARS1 | P54577; A0A | SYYC_HUMAN | Unmodified | 0 | 0 | 0 | 0 | 372170 | 0 | 67749 | 0 | 0 | 28780 | 0 | 0 | 415440 | 0 | 0 | 0 | 218130 | 0 |
| LQADPKPIS | COBL; COBL; C | O75128; J3K | COBL_HUMAN | Unmodified | 832730 | 1013600 | 302130 | 215310 | 67654 | 21122 | 0 | 0 | 0 | 0 | 0 | 175660 | 139970 | 43751 | 194220 | 293660 | 0 | 0 |
| LQAEAPHI\ | EIF4A2; EIF4A2 | Q14240; E7E | IF4A2_HUMAN | Unmodified | 255120 | 598850 | 543090 | 165950 | 0 | 0 | 0 | 0 | 0 | 0 | 0 | 6383900 | 0 | 0 | 403210 | 390940 | 0 | 0 |
| LQAEAQQL | CDC37; CDC37 | Q16543; K7E | CDC37_HUMA | Unmodified | 0 | 0 | 0 | 0 | 0 | 0 | 0 | 0 | 0 | 0 | 0 | 357330 | 0 | 0 | 0 | 0 | 0 | 0 |
| LQAEEVAQ | PLEC | Q15149 | PLEC_HUMAN | Unmodified | 0 | 75457 | 158480 | 60023 | 468660 | 0 | 0 | 0 | 213660 | 0 | 101530 | 0 | 0 | 0 | 261800 | 0 | 899130 | 795260 |
| LQAENLQL | GIT1; GIT1; GIT | Q9Y2X7; J3Q | GIT1_HUMAN / | Unmodified | 0 | 158340 | 0 | 275570 | 0 | 0 | 0 | 53078 | 0 | 0 | 0 | 1376000 | 0 | 0 | 0 | 0 | 138450 | 0 |
| LQAFGFCE | RBM25; RBM25 | P49756; E9P | RBM25_HUMAI | Unmodified | 0 | 0 | 0 | 0 | 0 | 0 | 0 | 0 | 0 | 0 | 297460 | 1920400 | 116620 | 0 | 251480 | 0 | 0 | 548120 |
| LQAFSAAIE | FKBP4 | Q02790 | FKBP4_HUMAN | Unmodified | 238510 | 357490 | 390340 | 0 | 607260 | 0 | 764160 | 678640 | 0 | 437750 | 2828800 | 11180000 | 10342000 | 4869200 | 982750 | 722800 | 1064500 | 938820 |
| LQAFSAAIE | FKBP4 | Q02790 | FKBP4_HUMAN | Unmodified | 0 | 0 | 0 | 0 | 0 | 0 | 0 | 310840 | 0 | 0 | 0 | 570240 | 0 | 0 | 0 | 0 | 0 | 0 |
| LQAGEETA | NCDN | Q9UBB6 | NCDN_HUMAN | Unmodified | 100480 | 156840 | 303850 | 24326 | 0 | 0 | 0 | 0 | 767180 | 0 | 0 | 4756100 | 0 | 0 | 450520 | 796810 | 0 | 0 |
| LQAGEYVS | ACSL4; ACSL3; | O60488; O9 | ACSL4_HUMAN | Unmodified | 0 | 0 | 0 | 0 | 0 | 147270 | 0 | 0 | 404000 | 0 | 1226900 | 5332200 | 0 | 188280 | 0 | 0 | 324500 | 370320 |
| LQAGEYVS | ACSL4; ACSL3; | O60488; O9 | ACSL4_HUMAN | Unmodified | 0 | 0 | 0 | 0 | 11692000 | 0 | 2326300 | 239430 | 79932 | 500500 | 4180200 | 9326800 | 4628500 | 2782000 | 0 | 0 | 11296000 | 12747000 |
| LQAHLEQL | EPHA1 | P21709 | EPHA1_HUMAN | Unmodified | 262070 | 236570 | 1161400 | 1037600 | 584140 | 0 | 323380 | 0 | 0 | 0 | 0 | 0 | 214970 | 0 | 2751500 | 1532900 | 0 | 605300 |
| LQAISWAR | CTPS2; CTPS2 | Q9NRF8; H0 | PYRG2_HUMAN | Unmodified | 0 | 0 | 0 | 0 | 0 | 0 | 0 | 0 | 0 | 0 | 0 | 880940 | 0 | 0 | 0 | 0 | 0 | 0 |
| LQALAEEPS | ARHGEF16 | Q5VV41 | ARHGG_HUMA | Unmodified | 0 | 0 | 0 | 244060 | 0 | 0 | 0 | 0 | 0 | 0 | 0 | 0 | 0 | 1495400 | 0 | 0 | 0 | 0 |
| LQALANEQ | KTN1 | Q86UP2 | KTN1_HUMAN | Unmodified | 0 | 0 | 0 | 0 | 0 | 0 | 0 | 0 | 538040 | 0 | 856520 | 3289800 | 423830 | 0 | 0 | 0 | 0 | 0 |
| LQALKDTAI | TKT; TKT; TKT; T | P29401; F8V | TKT_HUMAN Tr | Unmodified | 0 | 0 | 0 | 0 | 1944900 | 0 | 0 | 0 | 0 | 0 | 0 | 0 | 620310 | 275900 | 0 | 0 | 684710 | 710670 |
| LQALTGNE | CIAPIN1; CIAPI | Q6FI81; H3B | CPIN1_HUMAN | Unmodified | 0 | 0 | 0 | 0 | 0 | 0 | 0 | 0 | 0 | 0 | 0 | 0 | 0 | 0 | 0 | 0 | 0 | 0 |
| LQALTGNE | CIAPIN1; CIAPI | Q6FI81; H3B | CPIN1_HUMAN | Unmodified | 0 | 0 | 0 | 0 | 94353 | 0 | 0 | 0 | 0 | 0 | 0 | 668440 | 0 | 0 | 1388700 | 92404 | 0 | 0 |
| LQANWNR | SPART | Q8N0X7 | SPART_HUMAN | Unmodified | 0 | 0 | 135690 | 239380 | 0 | 0 | 0 | 0 | 0 | 0 | 0 | 0 | 0 | 0 | 2877500 | 92902 | 0 | 0 |
| LQAQDAGI | IGSF8; IGSF8 | Q969P0; C9 | IGSF8_HUMAN | Unmodified | 31169000 | 20720000 | 2880000 | 9445600 | 0 | 0 | 0 | 0 | 1339700 | 0 | 311510 | 0 | 669230 | 0 | 11728000 | 14845000 | 0 | 0 |
| LQAQDAGI | IGSF8; IGSF8 | Q969P0; C9 | IGSF8_HUMAN | Unmodified | 2059400 | 497360 | 0 | 5559500 | 561230 | 123350 | 0 | 0 | 0 | 55499 | 263890 | 0 | 3622100 | 0 | 261290 | 243560 | 181780 | 494790 |
| LQAQQDA\ | PSMD4; PSMD4 | P55036; A6P | PSMD4_HUMA | Unmodified | 0 | 0 | 0 | 0 | 0 | 0 | 0 | 0 | 0 | 0 | 0 | 3862700 | 0 | 0 | 0 | 0 | 0 | 0 |
| LQAQRNEL | PSMC5; PSMC5 | P62195; J3Q | PRS8_HUMAN | Unmodified | 0 | 0 | 0 | 278510 | 1152100 | 0 | 0 | 0 | 0 | 265670 | 0 | 2151500 | 2607300 | 628960 | 0 | 0 | 571930 | 0 |
| LQAQVTLSI | TNIP1; TNIP1; T | Q15025; A0A | TNIP1_HUMAN | Unmodified | 0 | 0 | 0 | 0 | 306910 | 0 | 0 | 0 | 0 | 0 | 0 | 0 | 0 | 0 | 0 | 0 | 181320 | 481070 |
| LQASGDAL | COLEC12 | Q5KU26 | COL12_HUMAI | Unmodified | 218590 | 0 | 1033100 | 794770 | 0 | 0 | 385590 | 172350 | 0 | 0 | 0 | 0 | 0 | 0 | 506430 | 0 | 0 | 0 |
| LQASISAFR | INF2; INF2; INF | Q27J81; A0A | INF2_HUMAN I | Unmodified | 0 | 0 | 0 | 0 | 0 | 0 | 0 | 462120 | 806410 | 94464 | 3299500 | 857720 | 0 | 0 | 0 | 0 | 0 | 0 |
| LQATEVR | TRMT112; TRM | Q9UI30; F5G | TR112_HUMAN | Unmodified | 0 | 0 | 0 | 0 | 0 | 0 | 0 | 0 | 0 | 0 | 0 | 1358200 | 0 | 0 | 0 | 0 | 0 | 0 |
| LQATLDMQ | PSMD11; PSMD | O00231; J3Q | PSD11_HUMAN | Unmodified | 0 | 0 | 0 | 0 | 791330 | 0 | 238270 | 0 | 320130 | 0 | 380220 | 2894600 | 593530 | 670310 | 132170 | 0 | 0 | 0 |
| LQATLDMQ | PSMD11; PSMD | O00231; J3Q | PSD11_HUMAN | Oxidation (M) | 0 | 0 | 0 | 0 | 0 | 0 | 0 | 26428 | 156070 | 0 | 0 | 812110 | 0 | 0 | 0 | 0 | 644470 | 0 |
| LQATLQGL | IQGAP3; IQGAP | Q86VI3; F2Z | IQGA3_HUMAN | Unmodified | 0 | 0 | 0 | 411940 | 2920000 | 399300 | 283360 | 177610 | 844330 | 531080 | 320100 | 1412300 | 0 | 0 | 0 | 0 | 183570 | 0 |
| LQATMETD | LRBA; LRBA; LR | P50851; A0A | LRBA_HUMAN | Unmodified | 0 | 0 | 0 | 0 | 0 | 0 | 0 | 0 | 0 | 0 | 0 | 558870 | 0 | 0 | 0 | 0 | 0 | 0 |
| LQATVQEL( | CORO1A; COR( | P31146; H3E | COR1A_HUMAI | Unmodified | 0 | 0 | 763720 | 453110 | 0 | 0 | 267520 | 369320 | 230740 | 297910 | 5801300 | 15766000 | 680680 | 334750 | 309760 | 228390 | 462820 | 507790 |
| LQAVEVVIT | CARHSP1 | Q9Y2V2 | CHSP1_HUMAI | Unmodified | 179200 | 128490 | 347530 | 118430 | 0 | 0 | 180760 | 50540 | 2821200 | 310580 | 635170 | 2506600 | 274860 | 0 | 485530 | 0 | 0 | 0 |
| LQAVEVVIT | CARHSP1 | Q9Y2V2 | CHSP1_HUMAI | Unmodified | 373750 | 0 | 0 | 656300 | 6753100 | 0 | 4142900 | 0 | 2888100 | 5448000 | 976560 | 1524000 | 0 | 4257000 | 0 | 0 | 14565000 | 7420600 |
| LQAWMTR | SNX9; SNX9; SI | Q9Y5X1; A0A | SNX9_HUMAN | Unmodified | 0 | 0 | 0 | 0 | 0 | 0 | 0 | 0 | 0 | 55285 | 226020 | 0 | 0 | 0 | 241400 | 0 | 0 | 0 |
| LQAYHTQT | AK2; AK2; AK2; | P54819; A0A | KAD2_HUMAN | Unmodified | 335640 | 127210 | 92705 | 0 | 0 | 0 | 0 | 0 | 1397400 | 0 | 0 | 7580500 | 199780 | 0 | 360390 | 208690 | 78844 | 90732 |
| LQAYHTQT | AK2; AK2; AK2; | P54819; A0A | KAD2_HUMAN | Unmodified | 0 | 0 | 0 | 0 | 0 | 0 | 0 | 0 | 685410 | 183770 | 857690 | 5132600 | 0 | 0 | 0 | 0 | 0 | 0 |
| LQAYTSVSI | NCKAP1 | Q9Y2A7 | NCKP1_HUMAI | Unmodified | 443120 | 0 | 346280 | 0 | 0 | 0 | 25420 | 0 | 0 | 30108 | 0 | 2439400 | 0 | 0 | 1252700 | 780500 | 0 | 0 |
| LQCENVQE | ADAM9; ADAM9 | Q13443; A0A | ADAM9_HUMA | Unmodified | 0 | 0 | 67820 | 0 | 0 | 0 | 0 | 0 | 81531 | 0 | 0 | 0 | 0 | 0 | 411010 | 0 | 0 | 0 |
| LQCENVQE | ADAM9; ADAM9 | Q13443; A0A | ADAM9_HUMA | Unmodified | 2341500 | 967150 | 923050 | 4031300 | 16520000 | 0 | 0 | 0 | 2807900 | 559720 | 144370 | 0 | 4452200 | 1712900 | 6582900 | 1997300 | 612460 | 1236400 |
| LQCGENGI | TANC1 | Q9C0D5 | TANC1_HUMAI | Unmodified | 0 | 0 | 0 | 208370 | 0 | 0 | 0 | 0 | 0 | 0 | 0 | 0 | 297890 | 82929 | 152050 | 144340 | 0 | 0 |
| LQCKPFSW | GALNT1 | Q10472 | GALT1_HUMAN | Unmodified | 0 | 0 | 0 | 0 | 0 | 0 | 0 | 0 | 1808400 | 0 | 0 | 0 | 0 | 0 | 0 | 0 | 0 | 0 |
| LQDAEIAR | S100A6; S100A | P06703; R4C | S10A6_HUMAN | Unmodified | 68718 | 2040200 | 369200 | 0 | 1284200 | 533660 | 0 | 0 | 6317500 | 598090 | 0 | 242030 | 0 | 0 | 83415 | 216470 | 590060 | 1436300 |
| LQDAINILK | LRPPRC | P42704 | LPPRC_HUMAN | Unmodified | 0 | 0 | 0 | 0 | 0 | 0 | 0 | 0 | 1783100 | 0 | 0 | 0 | 0 | 0 | 0 | 0 | 0 | 0 |
| LQDASAEV | BST2 | Q10589 | BST2_HUMAN I | Unmodified | 511410 | 0 | 158600 | 0 | 0 | 0 | 0 | 0 | 637730 | 0 | 199250 | 596630 | 123450 | 0 | 3939700 | 3216900 | 0 | 0 |
| LQDAYYIF( | COPE; COPE | O14579; M0 | COPE_HUMAN | Unmodified | 0 | 0 | 0 | 0 | 0 | 0 | 0 | 0 | 392550 | 0 | 96612 | 0 | 167860 | 0 | 0 | 0 | 0 | 0 |
| LQDFKLDF | GGCT; GGCT | O75223; B8Z | GGCT_HUMAN | Unmodified | 827240 | 493340 | 734670 | 0 | 638180 | 0 | 203470 | 0 | 3232800 | 2890800 | 3110100 | 6208400 | 2439000 | 1343500 | 1530700 | 1251500 | 834230 | 264570 |

| | | | | | | | | | | | | | | | | | | | | | | |
|---|---|---|---|---|---|---|---|---|---|---|---|---|---|---|---|---|---|---|---|---|---|---|
| LQDLELK | BRINP2; HDLBF | Q9C0B6; Q0 | BRNP2_HUMAI | Unmodified | 2021600 | 0 | 1040300 | 1324300 | 891710 | 177050 | 329780 | 598560 | 1227300 | 25825 | 643780 | 2917300 | 1871600 | 0 | 466210 | 0 | 469820 | 447800 |
| LQDLTLR | HYOU1; TEX15; | Q9Y4L1; Q9I | HYOU1_HUMA | Unmodified | 0 | 0 | 0 | 0 | 0 | 0 | 14938 | 0 | 4834300 | 147050 | 530630 | 2134600 | 0 | 0 | 98713 | 198320 | 0 | 0 |
| LQDLTLRDI | HYOU1; HYOU1 | Q9Y4L1; A0A | HYOU1_HUMA | Unmodified | 147050 | 93507 | 1214200 | 437450 | 3255300 | 118600 | 337990 | 513390 | 88931000 | 26471000 | 7558300 | 4805400 | 5183600 | 1492700 | 1371400 | 551740 | 237130 | 4005700 |
| LQDNVRED | PTGES2; PTGES | Q9H7Z7; A0A | PGES2_HUMAI | Unmodified | 0 | 0 | 0 | 0 | 0 | 0 | 0 | 0 | 666640 | 93229 | 0 | 0 | 58365 | 0 | 0 | 0 | 0 | 0 |
| LQDPFSLYI | SDHB | P21912 | SDHB_HUMAN | Unmodified | 0 | 0 | 0 | 0 | 0 | 0 | 0 | 0 | 5637500 | 1019000 | 0 | 0 | 0 | 0 | 0 | 0 | 0 | 180010 |
| LQDREWLT | EIF4G2; EIF4G2 | P78344; H0Y | IF4G2_HUMAN | Unmodified | 250140 | 0 | 0 | 0 | 0 | 0 | 0 | 0 | 0 | 0 | 63308 | 1485200 | 0 | 0 | 0 | 182370 | 0 | 0 |
| LQDTATATI | FUT11 | Q495W5 | FUT11_HUMAN | Unmodified | 0 | 0 | 0 | 0 | 0 | 0 | 0 | 0 | 1667100 | 0 | 0 | 0 | 0 | 0 | 0 | 0 | 0 | 0 |
| LQDTENEA | FMNL2 | Q96PY5 | FMNL2_HUMAI | Unmodified | 0 | 0 | 0 | 0 | 753630 | 108780 | 0 | 20243 | 473530 | 69040 | 179900 | 1539500 | 0 | 0 | 0 | 0 | 0 | 468950 |
| LQDTSSYAI | DSP | P15924 | DESP_HUMAN | Unmodified | 0 | 0 | 0 | 0 | 0 | 0 | 57079 | 0 | 653490 | 0 | 0 | 0 | 0 | 0 | 0 | 0 | 0 | 0 |
| LQDTYNLD | P4HA1 | P13674 | P4HA1_HUMAI | Unmodified | 0 | 0 | 0 | 0 | 0 | 0 | 0 | 0 | 3981600 | 894470 | 900020 | 627340 | 0 | 0 | 0 | 0 | 0 | 0 |
| LQDYEEK | EZR; EZR | P15311; E7E | EZRI_HUMAN E | Unmodified | 832850 | 589470 | 693520 | 1036800 | 3318100 | 320530 | 136160 | 0 | 2409300 | 167580 | 0 | 12055000 | 205180 | 721230 | 1577300 | 1044700 | 319180 | 147210 |
| LQEACKDII | PRKAR2A; PRKA | P13861; P31 | KAP2_HUMAN | Unmodified | 0 | 0 | 0 | 0 | 0 | 0 | 0 | 0 | 0 | 0 | 0 | 0 | 0 | 797090 | 0 | 0 | 0 | 288490 |
| LQEALNLFI | GNAS; GNAS; G | Q5JWF2; P63 | GNAS1_HUMAI | Unmodified | 1625800 | 0 | 4157000 | 0 | 0 | 0 | 0 | 431580 | 0 | 0 | 0 | 29914000 | 0 | 0 | 4167800 | 2027600 | 0 | 0 |
| LQEEAQGL | PROM2 | Q8N271 | PROM2_HUMA | Unmodified | 1735700 | 1396800 | 1006500 | 1193400 | 2514200 | 909280 | 0 | 64618 | 0 | 0 | 0 | 0 | 381580 | 0 | 0 | 1593400 | 511670 | 993510 |
| LQEEELLAT | CCDC50 | Q8IVM0 | CCD50_HUMA | Oxidation (M) | 0 | 0 | 811370 | 0 | 0 | 0 | 0 | 0 | 0 | 0 | 0 | 0 | 0 | 0 | 1511000 | 1001700 | 0 | 0 |
| LQEEGWLE | CD2AP | Q9Y5K6 | CD2AP_HUMAI | Unmodified | 0 | 0 | 158870 | 0 | 0 | 0 | 0 | 0 | 0 | 0 | 0 | 0 | 0 | 0 | 0 | 160950 | 0 | 0 |
| LQEEGWLE | CD2AP | Q9Y5K6 | CD2AP_HUMAI | Unmodified | 920730 | 713070 | 0 | 0 | 0 | 0 | 0 | 0 | 0 | 0 | 0 | 0 | 0 | 0 | 979340 | 0 | 0 | 0 |
| LQEEIIK | SETD2; SETD2; | Q9BYW2; A0 | SETD2_HUMAN | Unmodified | 0 | 0 | 0 | 0 | 0 | 0 | 0 | 0 | 0 | 1807700 | 0 | 0 | 0 | 0 | 0 | 0 | 0 | 0 |
| LQEEITK | USP14; RUFY1; | P54578; H0Y | UBP14_HUMAN | Unmodified | 64418 | 60939 | 0 | 235680 | 0 | 0 | 29697 | 0 | 135980 | 81552 | 0 | 1369300 | 49448 | 0 | 128830 | 93220 | 0 | 0 |
| LQEEITKQS | USP14; USP14 | P54578; A6N | UBP14_HUMAN | Unmodified | 0 | 0 | 0 | 0 | 0 | 15918 | 19450 | 0 | 0 | 0 | 0 | 0 | 188380 | 72692 | 0 | 0 | 82964 | 72157 |
| LQEENLVIT | VIL1; VIL1; VIL1 | P09327; H7C | VILI_HUMAN Vi | Unmodified | 0 | 0 | 0 | 0 | 0 | 0 | 0 | 0 | 0 | 0 | 1499900 | 999380 | 0 | 0 | 0 | 0 | 152650 | 0 |
| LQEESDLEI | EIF3J; EIF3J | O75822; H0Y | EIF3J_HUMAN I | Unmodified | 0 | 0 | 0 | 0 | 364110 | 0 | 0 | 0 | 890810 | 171910 | 1778800 | 3544100 | 0 | 0 | 0 | 0 | 66020 | 0 |
| LQEESDYIT | NAALADL2; NA | Q58DX5; C9 | NADL2_HUMAI | Unmodified | 0 | 0 | 691570 | 913060 | 75599 | 0 | 0 | 0 | 0 | 0 | 313880 | 0 | 0 | 0 | 1439800 | 1379500 | 0 | 0 |
| LQEESEVL( | QTRT2 | Q9H974 | QTRT2_HUMAN | Unmodified | 0 | 0 | 0 | 0 | 0 | 0 | 0 | 0 | 0 | 0 | 0 | 6918500 | 0 | 0 | 0 | 0 | 0 | 0 |
| LQEFFTQQ | KIAA1109; KIAA | Q2LD37; A0A | K1109_HUMAN | Unmodified | 0 | 420910 | 0 | 0 | 746070 | 0 | 0 | 0 | 0 | 0 | 0 | 611440 | 0 | 0 | 85480 | 0 | 0 | 0 |
| LQEGQTLE | LAMB2 | P55268 | LAMB2_HUMAI | Unmodified | 0 | 0 | 0 | 0 | 0 | 0 | 0 | 0 | 683580 | 0 | 0 | 0 | 1901400 | 0 | 0 | 0 | 0 | 0 |
| LQEGRPEF | ARHGEF39 | Q8N4T4 | ARG39_HUMAI | Unmodified | 1263000 | 286340 | 143310 | 2072700 | 0 | 0 | 0 | 0 | 1544200 | 303260 | 249430 | 316500 | 0 | 0 | 2335200 | 524970 | 0 | 0 |
| LQEGYDHS | ESD; ESD | P10768; X6R | ESTD_HUMAN | Unmodified | 0 | 0 | 0 | 1480000 | 16411 | 0 | 0 | 0 | 759590 | 100110 | 185080 | 677880 | 0 | 0 | 7695900 | 0 | 0 | 0 |
| LQEIYNR | ARHGEF2; ARH | Q92974; V9C | ARHG2_HUMA | Unmodified | 0 | 0 | 545170 | 29856 | 0 | 0 | 94989 | 220140 | 164110 | 0 | 0 | 1917900 | 0 | 0 | 0 | 0 | 0 | 0 |
| LQEKEELRI | LMNB1; LMNB2 | P20700; Q03 | LMNB1_HUMAI | Unmodified | 0 | 0 | 0 | 0 | 0 | 49242 | 0 | 0 | 576330 | 0 | 841500 | 1737900 | 0 | 41537 | 0 | 0 | 0 | 0 |
| LQELDAASI | CLTB | P09497 | CLCB_HUMAN | Unmodified | 560580 | 935390 | 0 | 0 | 179400 | 37922 | 0 | 0 | 0 | 278030 | 0 | 273040 | 835650 | 148990 | 0 | 105690 | 0 | 147200 |
| LQELDAASI | CLTB | P09497 | CLCB_HUMAN | Unmodified | 0 | 0 | 0 | 0 | 2851300 | 589890 | 0 | 128150 | 0 | 0 | 0 | 0 | 7907100 | 1974200 | 0 | 0 | 568720 | 806780 |
| LQELEGAVI | MYH10 | P35580 | MYH10_HUMAI | Unmodified | 26672 | 516460 | 920200 | 0 | 4482300 | 805600 | 0 | 0 | 346050 | 0 | 120860 | 6178100 | 0 | 472780 | 0 | 719930 | 277580 | 279970 |
| LQELEGTYI | LAMB2 | P55268 | LAMB2_HUMAI | Unmodified | 237220 | 368400 | 196410 | 317790 | 0 | 51702 | 0 | 0 | 2673100 | 97878 | 0 | 200180 | 0 | 0 | 114610 | 672440 | 0 | 0 |
| LQELIQK | IPO5; IPO5 | O00410; H0Y | IPO5_HUMAN I | Unmodified | 3533200 | 9020700 | 0 | 0 | 34361000 | 6541400 | 23687000 | 57831 | 4518200 | 4824800 | 0 | 6169300 | 0 | 0 | 2449900 | 0 | 0 | 13502000 |
| LQELLLCNI | RABGGTA; RAB | Q92696; H0Y | PGTA_HUMAN | Unmodified | 0 | 0 | 0 | 0 | 0 | 0 | 0 | 0 | 0 | 0 | 0 | 791240 | 0 | 0 | 0 | 0 | 0 | 0 |
| LQELSAEEF | OLA1; OLA1 | Q9NTK5; J3K | OLA1_HUMAN | Unmodified | 0 | 0 | 0 | 0 | 0 | 0 | 0 | 0 | 0 | 0 | 852940 | 12337000 | 0 | 0 | 948540 | 1362200 | 0 | 0 |
| LQELSAEEF | OLA1; OLA1 | Q9NTK5; J3K | OLA1_HUMAN | Unmodified | 1250200 | 1364200 | 643110 | 1336700 | 1131600 | 22857 | 212910 | 150960 | 1274700 | 17348 | 1003600 | 3548500 | 877740 | 654680 | 1172100 | 911310 | 273310 | 449160 |
| LQEMEGTV | MYH9 | P35579 | MYH9_HUMAN | Unmodified | 442100 | 521380 | 1389400 | 1232200 | 14487000 | 2925000 | 2919000 | 604660 | 1090400 | 510620 | 551220 | 4156600 | 11876000 | 3142500 | 2333700 | 978400 | 4960500 | 4268300 |
| LQEMEGTV | MYH9 | P35579 | MYH9_HUMAN | Oxidation (M) | 0 | 0 | 649410 | 0 | 1821800 | 322040 | 336520 | 118860 | 58489 | 253080 | 0 | 1318200 | 0 | 0 | 0 | 1317200 | 811110 | 591630 |
| LQEMEGTV | MYH9 | P35579 | MYH9_HUMAN | Unmodified | 0 | 0 | 0 | 0 | 276520 | 0 | 187180 | 0 | 0 | 0 | 0 | 0 | 0 | 0 | 0 | 0 | 579570 | 365600 |
| LQEMVLK | VPS37B; VPS37 | Q9H9H4; F5 | VP37B_HUMAN | Unmodified | 6899900 | 7349300 | 7002200 | 9224900 | 3627700 | 0 | 0 | 184490 | 2270700 | 1375500 | 0 | 4480900 | 9107900 | 4957200 | 15865000 | 8901200 | 436190 | 5486000 |
| LQEMVLK | VPS37B; VPS37 | Q9H9H4; F5 | VP37B_HUMAN | Oxidation (M) | 0 | 0 | 5706000 | 4609000 | 0 | 0 | 0 | 0 | 0 | 296940 | 533990 | 1626300 | 0 | 0 | 8733800 | 6288500 | 0 | 0 |
| LQENLQQL | STX12; STX12 | Q86Y82; B1A | STX12_HUMAN | Unmodified | 740930 | 784130 | 4574800 | 2276800 | 523360 | 0 | 452660 | 58666 | 3550500 | 0 | 0 | 1682500 | 622950 | 0 | 2258700 | 2403200 | 0 | 213690 |
| LQEPTPSS( | TRIM25; TRIM25 | Q14258; A0A | TRI25_HUMAN | Unmodified | 0 | 0 | 0 | 0 | 0 | 0 | 0 | 0 | 104240 | 0 | 0 | 1446800 | 0 | 0 | 0 | 0 | 0 | 0 |
| LQEQEELEI | VGF | O15240 | VGF_HUMAN N | Unmodified | 243430 | 121880 | 0 | 0 | 0 | 0 | 0 | 0 | 499770 | 159620 | 0 | 0 | 0 | 0 | 0 | 0 | 0 | 0 |
| LQEQLMGA | VPS4A; N/A | Q9UN37; I3L | VPS4A_HUMAN | Unmodified | 0 | 0 | 0 | 0 | 0 | 12926000 | 1629300 | 0 | 0 | 1378700 | 179090 | 140920 | 0 | 2591500 | 0 | 0 | 785790 | 1282200 |
| LQEQVTDL | TPR | P12270 | TPR_HUMAN N | Unmodified | 0 | 0 | 0 | 0 | 462470 | 0 | 0 | 0 | 253460 | 12449 | 0 | 0 | 655260 | 127010 | 0 | 0 | 35938 | 0 |
| LQESGEVP | LIN7A; LIN7A; L | O14910; H0Y | LIN7A_HUMAN | Unmodified | 0 | 0 | 0 | 0 | 0 | 0 | 0 | 0 | 2841500 | 116720 | 229740 | 0 | 0 | 0 | 0 | 0 | 0 | 0 |
| LQESGEVP | LIN7A; LIN7A; L | O14910; H0Y | LIN7A_HUMAN | Unmodified | 0 | 0 | 0 | 0 | 0 | 0 | 160350 | 265500 | 0 | 332110 | 0 | 0 | 0 | 0 | 0 | 0 | 0 | 0 |
| LQESGEVP | LIN7A; LIN7A; L | O14910; H0Y | LIN7A_HUMAN | Unmodified | 0 | 0 | 0 | 0 | 0 | 0 | 1239100 | 694880 | 0 | 469320 | 0 | 0 | 0 | 0 | 0 | 0 | 0 | 0 |
| LQESTQTV( | CDC42BPB | Q9Y5S2 | MRCKB_HUMA | Unmodified | 0 | 0 | 0 | 0 | 444840 | 149850 | 0 | 0 | 267080 | 0 | 0 | 0 | 0 | 0 | 0 | 0 | 236500 | 427210 |
| LQETEMMI | GLG1 | Q92896 | GSLG1_HUMAI | 2 Oxidation (M) | 0 | 0 | 0 | 0 | 0 | 0 | 0 | 0 | 2392100 | 0 | 447020 | 0 | 0 | 0 | 0 | 0 | 0 | 0 |
| LQETEMMI | GLG1 | Q92896 | GSLG1_HUMAI | Oxidation (M) | 0 | 0 | 0 | 0 | 0 | 0 | 0 | 0 | 2398300 | 0 | 0 | 0 | 0 | 0 | 0 | 0 | 0 | 0 |
| LQETLSAAI | PRKDC | P78527 | PRKDC_HUMAI | Unmodified | 0 | 0 | 0 | 0 | 0 | 0 | 0 | 0 | 0 | 0 | 0 | 1387600 | 0 | 0 | 0 | 0 | 0 | 0 |
| LQETTLVAN | MAN2B1; N/A | O00754; M0 | MA2B1_HUMAI | Unmodified | 0 | 0 | 0 | 0 | 0 | 0 | 0 | 0 | 1031800 | 0 | 0 | 0 | 0 | 0 | 100170 | 67427 | 0 | 0 |
| LQEVECEE | SNX1; SNX1 | Q13596; H0Y | SNX1_HUMAN | Unmodified | 0 | 0 | 0 | 0 | 0 | 0 | 0 | 0 | 0 | 0 | 0 | 0 | 0 | 0 | 0 | 0 | 0 | 0 |
| LQEVEVPEI | EML2; EML2; EN | O95834; A0A | EMAL2_HUMAN | Unmodified | 288890 | 0 | 680250 | 0 | 0 | 0 | 0 | 0 | 0 | 21445 | 0 | 256570 | 0 | 0 | 0 | 0 | 0 | 0 |
| LQEVPHEG | PPP2CB; PPP2C | P62714; P67 | PP2AB_HUMAN | Oxidation (M) | 0 | 0 | 0 | 0 | 0 | 0 | 0 | 0 | 0 | 0 | 0 | 1647800 | 0 | 0 | 0 | 298820 | 0 | 0 |
| LQEVSLR | TBCE; CHTF18; | Q15813; Q8 | TBCE_HUMAN | Unmodified | 0 | 109500 | 359410 | 0 | 420810 | 397220 | 129690 | 53384 | 225370 | 44489 | 891370 | 1855000 | 88953 | 0 | 0 | 120070 | 163490 | 354630 |
| LQFAPFSS/ | ATG7; ATG7; AT | O95352; F8V | ATG7_HUMAN | Unmodified | 0 | 0 | 0 | 0 | 317130 | 0 | 46452 | 0 | 0 | 85883 | 0 | 0 | 480290 | 430310 | 0 | 0 | 239830 | 975680 |
| LQFHDVAG | G6PD | P11413 | G6PD_HUMAN | Unmodified | 0 | 0 | 0 | 1553300 | 4967800 | 584840 | 0 | 0 | 0 | 103210 | 661350 | 1267900 | 409980 | 0 | 0 | 0 | 589690 | 1123700 |
| LQFHNVKP | NIPSNAP2; NIP | O75323; F8V | NIPS2_HUMAN | Unmodified | 0 | 0 | 0 | 0 | 317770 | 0 | 0 | 0 | 4101300 | 877720 | 0 | 740220 | 0 | 0 | 0 | 0 | 0 | 0 |
| LQGAHLTS | VCAN; VCAN | P13611; E9P | CSPG2_HUMAI | Unmodified | 0 | 0 | 0 | 0 | 0 | 0 | 0 | 0 | 1760600 | 0 | 0 | 0 | 0 | 0 | 0 | 0 | 0 | 0 |
| LQGDANNI | POLR2H; POLR | P52434; C9J | RPAB3_HUMAN | Unmodified | 0 | 0 | 0 | 0 | 0 | 0 | 0 | 0 | 0 | 571950 | 0 | 1281900 | 0 | 0 | 0 | 0 | 0 | 0 |
| LQGDAVVL | IGSF8; IGSF8 | Q969P0; C9J | IGSF8_HUMAN | Unmodified | 2026400 | 1688500 | 1058900 | 1167100 | 0 | 0 | 0 | 0 | 0 | 0 | 0 | 0 | 339660 | 0 | 1079300 | 1289800 | 0 | 0 |
| LQGEFQLR | DYNC1H1; DYN | Q14204; A0A | DYHC1_HUMA | Unmodified | 0 | 64508 | 351900 | 0 | 0 | 575320 | 0 | 0 | 1252800 | 0 | 0 | 5001800 | 0 | 0 | 317810 | 437750 | 0 | 0 |
| LQGEGAEV | SDSL; SDSL | Q96GA7; F8Y | SDSL_HUMAN | Unmodified | 304690 | 0 | 0 | 0 | 0 | 0 | 0 | 0 | 215330 | 0 | 0 | 657770 | 0 | 0 | 538470 | 292200 | 0 | 97618 |
| LQGEVEKY | PFDN6; PFDN6 | O15212; A2A | PFD6_HUMAN | Unmodified | 0 | 0 | 0 | 0 | 434800 | 0 | 204080 | 140930 | 0 | 271840 | 0 | 0 | 308710 | 276410 | 0 | 0 | 554640 | 398520 |
| LQGEVVAF | LSM12; LSM12 | Q3MHD2; K7 | LSM12_HUMAN | Unmodified | 0 | 1142700 | 0 | 0 | 0 | 0 | 0 | 0 | 0 | 0 | 369620 | 6508100 | 0 | 0 | 0 | 0 | 0 | 0 |
| LQGHLLRQ | GOLIM4; GOLIN | O00461; F8V | GOLI4_HUMAN | Unmodified | 0 | 0 | 0 | 0 | 0 | 0 | 0 | 0 | 971550 | 312360 | 0 | 0 | 0 | 0 | 0 | 0 | 0 | 0 |
| LQGMVTET | KIF23; KIF23; KI | Q02241; A0A | KIF23_HUMAN | Unmodified | 0 | 0 | 0 | 0 | 8740300 | 757050 | 0 | 0 | 2367100 | 525360 | 0 | 1031100 | 0 | 0 | 0 | 0 | 543460 | 849250 |
| LQGMVTET | KIF23; KIF23; KI | Q02241; A0A | KIF23_HUMAN | 2 Oxidation (M) | 0 | 0 | 0 | 0 | 1456400 | 321880 | 0 | 0 | 0 | 191240 | 0 | 126460 | 0 | 0 | 0 | 0 | 333440 | 0 |
| LQGMVTET | KIF23; KIF23; KI | Q02241; A0A | KIF23_HUMAN | Oxidation (M) | 0 | 0 | 0 | 157160 | 4952400 | 827410 | 0 | 0 | 196140 | 256260 | 301700 | 598650 | 429180 | 0 | 414470 | 0 | 0 | 662890 |
| LQGNSEIT( | KRIT1; KRIT1; KI | O00522; C9J | KRIT1_HUMAN | Unmodified | 0 | 0 | 99064 | 306660 | 0 | 0 | 0 | 0 | 342450 | 62664 | 0 | 39966 | 123700 | 228000 | 0 | 0 | 0 | 0 |
| LQGNSEIT( | KRIT1; KRIT1; KI | O00522; C9J | KRIT1_HUMAN | Unmodified | 0 | 0 | 0 | 0 | 942040 | 0 | 307940 | 0 | 0 | 250110 | 0 | 0 | 613770 | 694500 | 0 | 0 | 121000 | 591280 |
| LQGPQTSA | GPD1L | Q8N335 | GPD1L_HUMAI | Unmodified | 0 | 0 | 0 | 0 | 0 | 2201300 | 0 | 1266700 | 885550 | 0 | 0 | 1273500 | 784030 | 2023900 | 0 | 0 | 257170 | 2142200 |
| LQGQLEQ( | ABCF1; ABCF1; | Q8NE71; H0 | ABCF1_HUMAN | Unmodified | 0 | 0 | 0 | 0 | 1101200 | 0 | 0 | 0 | 358810 | 0 | 518060 | 1989900 | 0 | 0 | 0 | 0 | 0 | 0 |
| LQGTLFSTI | CA14; CA14 | Q9ULX7; U3I | CAH14_HUMAI | Unmodified | 0 | 0 | 0 | 0 | 0 | 0 | 0 | 0 | 0 | 0 | 1608000 | 8678400 | 0 | 0 | 0 | 0 | 0 | 0 |
| LQHAAELIK | PDCD6IP | Q8WUM4 | PDC6I_HUMAN | Unmodified | 12892000 | 10391000 | 5242300 | 5472300 | 698960 | 0 | 0 | 0 | 6978500 | 1064200 | 629270 | 3476500 | 1426900 | 0 | 5657200 | 6946000 | 90121 | 313220 |
| LQHPDML\ | EPB41L1; EPB4 | Q9H4G0; H0 | E41L1_HUMAN | Unmodified | 0 | 0 | 0 | 0 | 3222900 | 0 | 0 | 0 | 0 | 0 | 169450 | 319050 | 0 | 0 | 0 | 0 | 303360 | 684970 |
| LQHPLGVT | NHLRC2 | Q8NBF2 | NHLC2_HUMA | Unmodified | 0 | 0 | 0 | 0 | 632740 | 98257 | 369540 | 0 | 0 | 0 | 3326700 | 5921800 | 0 | 0 | 0 | 0 | 231510 | 276380 |
| LQHPLTILP | SPRY4; SPRY4; | Q9C004; D6 | SPY4_HUMAN | Unmodified | 29286 | 0 | 0 | 0 | 0 | 0 | 0 | 0 | 304380 | 0 | 0 | 0 | 0 | 0 | 0 | 0 | 63568 | 483070 |
| LQHPNVV( | ROR2; ROR2 | Q01974; B1A | ROR2_HUMAN | Unmodified | 0 | 0 | 0 | 0 | 0 | 0 | 0 | 0 | 812740 | 170110 | 0 | 693720 | 0 | 0 | 0 | 0 | 0 | 0 |
| LQHQTELE | TAOK3; TAOK3 | Q9H2K8; G3 | TAOK3_HUMAN | Unmodified | 1162500 | 1109900 | 3018200 | 2449500 | 2672900 | 203190 | 236080 | 115580 | 194580 | 86160 | 372000 | 402420 | 1853000 | 508430 | 2048600 | 1449700 | 79535 | 554910 |
| LQHQTELG | TAOK2 | Q9UL54 | TAOK2_HUMAN | Unmodified | 193930 | 100020 | 405410 | 386230 | 0 | 0 | 73099 | 70400 | 1226000 | 84350 | 168180 | 795040 | 0 | 0 | 614250 | 780690 | 0 | 0 |
| LQHQTELTI | TAOK1 | Q7L7X3 | TAOK1_HUMAN | Unmodified | 680350 | 0 | 473830 | 1645900 | 659480 | 237440 | 0 | 52877 | 2545600 | 317980 | 500430 | 3240500 | 190020 | 155860 | 618560 | 874780 | 55666 | 0 |
| LQHVEDG\ | CKAP4 | Q07065 | CKAP4_HUMAN | Unmodified | 0 | 0 | 0 | 0 | 0 | 0 | 0 | 0 | 7476100 | 178710 | 0 | 0 | 0 | 0 | 0 | 0 | 0 | 0 |
| LQHVEDG\ | CKAP4 | Q07065 | CKAP4_HUMAN | Oxidation (M) | 0 | 0 | 0 | 0 | 0 | 0 | 130260 | 0 | 1616300 | 0 | 133250 | 0 | 0 | 0 | 0 | 114380 | 0 | 0 |
| LQIAQLYPI | DLG1; DLG1; D | Q12959; A0A | DLG1_HUMAN | Unmodified | 232690 | 0 | 193470 | 352370 | 1070400 | 0 | 0 | 0 | 738760 | 1050400 | 645190 | 566060 | 0 | 0 | 841200 | 489550 | 0 | 1331600 |
| LQIASDEN' | SPTAN1; SPTAN | Q13813; A0A | SPTN1_HUMAN | Unmodified | 0 | 0 | 0 | 0 | 220400 | 228120 | 35863 | 0 | 0 | 253350 | 103660 | 2171000 | 302430 | 0 | 0 | 0 | 203890 | 0 |
| LQILELR | ERBIN; ERBIN | Q96RT1; B4D | ERBIN_HUMAN | Unmodified | 0 | 0 | 916460 | 179150 | 331910 | 0 | 0 | 0 | 420630 | 17027 | 127850 | 5281800 | 0 | 0 | 90420 | 574000 | 0 | 0 |
| LQILELREN | ERBIN; ERBIN | Q96RT1; B4D | ERBIN_HUMAN | Unmodified | 0 | 0 | 969820 | 1726400 | 26622000 | 6612700 | 1183900 | 462530 | 1078000 | 1060400 | 1784400 | 2652400 | 2678900 | 2675200 | 0 | 0 | 2351800 | 1489700 |
| LQILNLGAI | AP3B1 | O00203 | AP3B1_HUMAN | Unmodified | 0 | 0 | 0 | 0 | 314580 | 0 | 0 | 48836 | 0 | 0 | 154830 | 1941300 | 0 | 0 | 0 | 63921 | 0 | 0 |
| LQILSLR | RSU1 | Q15404 | RSU1_HUMAN | Unmodified | 108390 | 175150 | 138970 | 109620 | 0 | 0 | 0 | 0 | 114330 | 0 | 0 | 291360 | 0 | 0 | 133580 | 204970 | 0 | 0 |
| LQILSLRDN | RSU1 | Q15404 | RSU1_HUMAN | Unmodified | 1522200 | 5006200 | 1856600 | 7535200 | 5548500 | 1144400 | 4482700 | 209890 | 2274800 | 951310 | 618160 | 1437600 | 3623100 | 6895200 | 7370900 | 1853300 | 1809500 | 1806900 |
| LQILSLRDN | RSU1 | Q15404 | RSU1_HUMAN | Unmodified | 0 | 0 | 0 | 0 | 216990 | 0 | 432370 | 0 | 0 | 0 | 0 | 0 | 270060 | 610380 | 0 | 0 | 356230 | 105820 |
| LQIRHEQIS | TPP2; TPP2; TPI | P29144; A0A | TPP2_HUMAN | Unmodified | 0 | 0 | 0 | 0 | 0 | 0 | 0 | 0 | 384300 | 0 | 0 | 2904500 | 0 | 0 | 0 | 0 | 0 | 0 |
| LQISHEAA | DHX9 | Q08211 | DHX9_HUMAN | Unmodified | 0 | 0 | 0 | 0 | 0 | 0 | 284930 | 97786 | 0 | 69094 | 0 | 773590 | 20965 | 0 | 0 | 0 | 0 | 0 |
| LQITPDSN( | NEDD4L; NEDD | Q96PU5; A0A | NED4L_HUMAI | Unmodified | 0 | 1841600 | 10098000 | 0 | 0 | 0 | 0 | 0 | 0 | 0 | 0 | 0 | 0 | 0 | 0 | 1122500 | 0 | 0 |
| LQIVEMPLA | SERPINH1; SER | P50454; E9P | SERPH_HUMAI | Unmodified | 0 | 0 | 0 | 0 | 0 | 0 | 0 | 0 | 28235000 | 3989600 | 0 | 0 | 0 | 0 | 0 | 0 | 0 | 0 |
| LQIVEMPLA | SERPINH1; SER | P50454; E9P | SERPH_HUMAI | Oxidation (M) | 0 | 0 | 0 | 0 | 0 | 0 | 0 | 0 | 1956100 | 614060 | 0 | 0 | 0 | 0 | 0 | 0 | 0 | 0 |
| LQIWDTAG | RAB10 | P61026 | RAB10_HUMAN | Unmodified | 811720 | 994790 | 931020 | 0 | 0 | 0 | 0 | 277890 | 0 | 0 | 2136000 | 826080 | 0 | 0 | 978800 | 604140 | 0 | 0 |
| LQIWDTAG | RAB2A; RAB2B; | P61019; Q8V | RAB2A_HUMAN | Unmodified | 0 | 319540 | 883580 | 1090900 | 0 | 0 | 0 | 0 | 5645700 | 454330 | 915790 | 696430 | 0 | 0 | 1656100 | 2215200 | 0 | 0 |
| LQKADIEEI | NUCB2; NUCB2 | P80303; Q2L | NUCB2_HUMA | Unmodified | 0 | 0 | 0 | 0 | 0 | 0 | 0 | 0 | 4811400 | 1127300 | 0 | 0 | 263100 | 0 | 0 | 0 | 0 | 0 |
| LQKEEEIEF | IARS1; IARS1; IA | P41252; A0A | SYIC_HUMAN I | Unmodified | 0 | 0 | 0 | 0 | 0 | 0 | 0 | 0 | 0 | 0 | 0 | 16753000 | 0 | 0 | 0 | 0 | 0 | 0 |
| LQKLGEGE | SNX6; SNX6; SN | Q9UNH7; A0 | SNX6_HUMAN | Unmodified | 0 | 0 | 0 | 31721 | 1560400 | 0 | 0 | 0 | 0 | 0 | 0 | 0 | 340240 | 64603 | 0 | 0 | 214020 | 0 |
| LQKSEQSV | KLC1; KLC1; KL | Q07866; G3Y | KLC1_HUMAN | Unmodified | 0 | 0 | 0 | 0 | 336250 | 0 | 128200 | 97907 | 0 | 135630 | 0 | 0 | 0 | 0 | 0 | 0 | 245890 | 0 |
| LQLDSPED | PSMD3 | O43242 | PSMD3_HUMA | Unmodified | 0 | 0 | 232870 | 110860 | 0 | 0 | 0 | 43314 | 1200000 | 0 | 0 | 2492700 | 0 | 0 | 0 | 0 | 0 | 0 |

| | | | | | | | | | | | | | | | | | | | | | | |
|---|---|---|---|---|---|---|---|---|---|---|---|---|---|---|---|---|---|---|---|---|---|---|
| LQLEAIHK | GTF3C2; GTF3C | Q8WUA4; H | TF3C2_HUMAN | Unmodified | 40057000 | 38079000 | 18335000 | 11968000 | 0 | 0 | 0 | 15351 | 207240 | 0 | 0 | 707890 | 428090 | 0 | 30817000 | 27867000 | 0 | 0 |
| LQLEETDH | PLEC | Q15149 | PLEC_HUMAN | Unmodified | 0 | 0 | 0 | 0 | 34399 | 0 | 0 | 0 | 191050 | 54844 | 0 | 424170 | 0 | 0 | 0 | 0 | 0 | 21802 |
| LQLEKVTTE | MYH14; MYH9 | Q7Z406; P35 | MYH14_HUMA | Unmodified | 174720 | 0 | 0 | 836000 | 168450 | 47236 | 1725700 | 0 | 0 | 0 | 0 | 0 | 0 | 379250 | 0 | 240550 | 714940 | 452050 |
| LQLELNQE | ROCK1; ROCK1 | Q13464; A0A | ROCK1_HUMA | Unmodified | 0 | 0 | 0 | 724420 | 1392600 | 0 | 0 | 0 | 464400 | 0 | 0 | 0 | 0 | 0 | 439510 | 456500 | 463760 | 1134100 |
| LQLELSK | VARS2; LMNA; | Q5ST30; P02 | SYVM_HUMAN | Unmodified | 0 | 0 | 0 | 0 | 0 | 0 | 0 | 0 | 0 | 0 | 0 | 0 | 1128300 | 0 | 0 | 0 | 0 | 0 |
| LQLEQQVA | PLXNB2; PLXNE | O15031; A6 | PLXB2_HUMAN | Unmodified | 292900 | 123130 | 482120 | 629010 | 1403600 | 297330 | 39618 | 0 | 418760 | 0 | 0 | 89806 | 251280 | 0 | 1164000 | 920200 | 0 | 1086500 |
| LQLEQQVA | PLXNB2; PLXNE | O15031; A6 | PLXB2_HUMAN | Unmodified | 2059600 | 1198100 | 2468000 | 2972200 | 7179300 | 1936700 | 0 | 299440 | 2924600 | 0 | 0 | 1738300 | 3794800 | 1784100 | 5225700 | 5499900 | 2006200 | 6437000 |
| LQLERPVSI | KIAA1522 | Q9P206 | K1522_HUMAN | Unmodified | 0 | 0 | 0 | 4198000 | 0 | 0 | 0 | 0 | 2862900 | 0 | 0 | 2132700 | 1412000 | 0 | 4847000 | 4045000 | 201930 | 0 |
| LQLESIK | N/A | N/A | N/A | Unmodified | 274860 | 69022 | 0 | 0 | 6452500 | 0 | 419590 | 248090 | 387830 | 0 | 236520 | 719770 | 0 | 0 | 0 | 0 | 551940 | 2441900 |
| LQLHESQK | CTTN | Q14247 | SRC8_HUMAN | Unmodified | 0 | 0 | 0 | 0 | 0 | 0 | 0 | 0 | 0 | 0 | 0 | 0 | 0 | 0 | 0 | 0 | 0 | 0 |
| LQLLEDDK | PARP1; PARP1 | P09874; A0A | PARP1_HUMAN | Unmodified | 410660 | 156100 | 0 | 0 | 0 | 0 | 0 | 0 | 619760 | 48538 | 0 | 1092500 | 0 | 0 | 102370 | 36199 | 0 | 0 |
| LQLLEDDK | PARP1; PARP1 | P09874; A0A | PARP1_HUMAN | Unmodified | 0 | 961620 | 0 | 0 | 0 | 0 | 0 | 0 | 542490 | 290880 | 347870 | 1243600 | 0 | 0 | 0 | 0 | 0 | 0 |
| LQLLEEKTD | CCAR1 | Q8IX12 | CCAR1_HUMA | Unmodified | 33237 | 0 | 0 | 0 | 0 | 0 | 0 | 0 | 0 | 0 | 0 | 491510 | 58991 | 16121 | 0 | 19897 | 0 | 0 |
| LQLLEPFD | ACO2; ACO2; A | Q99798; A2A | ACON_HUMAN | Unmodified | 0 | 0 | 0 | 0 | 0 | 0 | 0 | 0 | 4807100 | 186520 | 452950 | 0 | 0 | 0 | 0 | 0 | 0 | 0 |
| LQLLLLNPL | MON2; MON2; | Q7Z3U7; A0 | MON2_HUMAN | Unmodified | 0 | 0 | 0 | 0 | 0 | 0 | 0 | 0 | 395230 | 0 | 133210 | 1333200 | 0 | 0 | 0 | 0 | 0 | 0 |
| LQLLNPEIE | NUP210 | Q8TEM1 | PO210_HUMAN | Oxidation (M) | 0 | 0 | 0 | 0 | 0 | 0 | 0 | 0 | 3069400 | 0 | 0 | 0 | 0 | 0 | 0 | 0 | 0 | 0 |
| LQLLNPEIE | NUP210 | Q8TEM1 | PO210_HUMAN | Unmodified | 0 | 0 | 0 | 0 | 0 | 0 | 0 | 0 | 559820 | 0 | 0 | 0 | 0 | 0 | 0 | 0 | 0 | 0 |
| LQLLSNLF | EIF3M; EIF3M; E | Q7L2H7; E9 | EIF3M_HUMAN | Oxidation (M) | 192560 | 235950 | 320010 | 309280 | 0 | 0 | 0 | 0 | 0 | 0 | 0 | 2266400 | 0 | 0 | 657720 | 621100 | 0 | 0 |
| LQLLSNLF | EIF3M; EIF3M; E | Q7L2H7; E9 | EIF3M_HUMAN | Unmodified | 63689 | 80277 | 0 | 0 | 0 | 0 | 0 | 0 | 191150 | 0 | 49968 | 538860 | 0 | 0 | 125510 | 0 | 0 | 0 |
| LQLLSPQV | FRYL; FRYL; FR | O94915; F22 | FRYL_HUMAN | Unmodified | 0 | 0 | 0 | 0 | 0 | 0 | 0 | 0 | 0 | 0 | 51578 | 417950 | 0 | 0 | 0 | 0 | 0 | 0 |
| LQLNEDEL | GALNT4; POC1 | Q8N4A0; F8 | GALT4_HUMAN | Unmodified | 0 | 0 | 0 | 0 | 0 | 0 | 0 | 0 | 1164200 | 0 | 0 | 0 | 0 | 0 | 0 | 0 | 0 | 0 |
| LQLNIVEMI | DDAH1; DDAH | O94760; B4 | DDAH1_HUMA | Oxidation (M) | 0 | 0 | 1698800 | 0 | 0 | 0 | 0 | 0 | 0 | 0 | 0 | 0 | 0 | 0 | 0 | 0 | 0 | 0 |
| LQLNYLGN | PLOD1; PLOD1 | Q02809; Q5 | PLOD1_HUMA | Unmodified | 0 | 0 | 0 | 0 | 0 | 0 | 0 | 0 | 3212500 | 0 | 0 | 0 | 0 | 0 | 0 | 0 | 0 | 0 |
| LQLNYLGN | PLOD3; PLOD3 | O60568; H7 | PLOD3_HUMA | Unmodified | 0 | 0 | 0 | 0 | 0 | 0 | 0 | 0 | 3037900 | 0 | 0 | 0 | 0 | 0 | 0 | 0 | 0 | 0 |
| LQLPAGLE | FAM171A1 | Q5VUB5 | F1711_HUMAN | Unmodified | 0 | 0 | 0 | 131260 | 0 | 0 | 0 | 0 | 0 | 0 | 0 | 587070 | 0 | 0 | 174370 | 205300 | 205310 | 0 |
| LQLPVWEY | DHX15 | O43143 | DHX15_HUMA | Unmodified | 0 | 0 | 0 | 0 | 0 | 0 | 0 | 0 | 0 | 0 | 416940 | 2713800 | 0 | 0 | 0 | 0 | 0 | 0 |
| LQLSVYTTT | CLPTM1L | Q96KA5 | CLP1L_HUMAN | Unmodified | 143930 | 0 | 0 | 0 | 0 | 0 | 0 | 0 | 5286100 | 863200 | 0 | 0 | 0 | 0 | 0 | 0 | 0 | 0 |
| LQLTTER | CUL5 | Q93034 | CUL5_HUMAN | Unmodified | 0 | 0 | 0 | 0 | 0 | 0 | 38273 | 0 | 0 | 0 | 0 | 810450 | 0 | 0 | 0 | 252650 | 0 | 0 |
| LQLVERLQ | SOGA1; SOGA1 | O94964; X6 | SOGA1_HUMA | Unmodified | 0 | 0 | 0 | 0 | 0 | 0 | 39681 | 18109 | 0 | 221540 | 0 | 0 | 2548000 | 861060 | 0 | 0 | 267520 | 327300 |
| LQLVQLSK | POGLUT3; POG | Q7Z4H8; H0 | PLGT3_HUMAN | Unmodified | 0 | 0 | 0 | 56226 | 0 | 565760 | 33570 | 0 | 12985000 | 520390 | 0 | 0 | 0 | 0 | 0 | 0 | 59299 | 0 |
| LQLVVENS | CCDC22 | O60826 | CCD22_HUMA | Unmodified | 0 | 0 | 0 | 0 | 2254000 | 156960 | 290950 | 0 | 0 | 0 | 0 | 0 | 511990 | 296060 | 0 | 0 | 968710 | 774600 |
| LQLVVLR | TJP2; TJP2; TJP | Q9UDY2; A0 | ZO2_HUMAN T | Unmodified | 0 | 0 | 0 | 0 | 0 | 0 | 0 | 0 | 867030 | 0 | 46704 | 414040 | 0 | 0 | 0 | 0 | 0 | 0 |
| LQLWDIAG | RAB29; RAB38; | O14966; P5 | RAB7L_HUMAN | Unmodified | 0 | 0 | 0 | 0 | 0 | 0 | 0 | 117080 | 0 | 0 | 0 | 1561700 | 0 | 0 | 691740 | 869590 | 0 | 0 |
| LQLWDTAC | RAB6A; RAB39A | P20340; Q1 | RAB6A_HUMAN | Unmodified | 3128500 | 24949000 | 17540000 | 9914500 | 396920 | 280360 | 604690 | 226450 | 17764000 | 2424600 | 13126000 | 79069000 | 348700 | 0 | 33495000 | 30260000 | 196850 | 0 |
| LQLWDTAC | RAB6A; RAB39A | P20340; Q1 | RAB6A_HUMAN | Unmodified | 4141600 | 3120700 | 5983700 | 18524000 | 4302400 | 1812200 | 2618000 | 489350 | 9596600 | 2821200 | 11311000 | 6717500 | 4298600 | 2428500 | 7177500 | 2197000 | 801250 | 2203600 |
| LQMAIEES | EPN1 | Q9Y6I3 | EPN1_HUMAN | Oxidation (M) | 1102200 | 493680 | 0 | 0 | 0 | 0 | 42406 | 20879 | 0 | 0 | 0 | 0 | 0 | 57063 | 0 | 1156700 | 0 | 0 |
| LQMAIEES | EPN1 | Q9Y6I3 | EPN1_HUMAN | Unmodified | 2471200 | 0 | 0 | 0 | 0 | 0 | 0 | 0 | 0 | 198450 | 0 | 0 | 0 | 0 | 0 | 0 | 19686 | 0 |
| LQMEAPHI | EIF4A1; EIF4A1; | P60842; J3K | IF4A1_HUMAN | Unmodified | 1854600 | 0 | 0 | 190060 | 0 | 0 | 0 | 0 | 684140 | 0 | 0 | 3955700 | 0 | 0 | 1040300 | 0 | 39449 | 0 |
| LQMEVNDI | SH3KBP1; SH3I | Q96B97; Q5 | SH3K1_HUMAN | Unmodified | 115240 | 64913 | 257170 | 226510 | 1399000 | 0 | 0 | 0 | 0 | 0 | 0 | 0 | 224800 | 135360 | 0 | 0 | 0 | 0 |
| LQMVTVPP | DOP1B | Q9Y3R5 | DOP2_HUMAN | Oxidation (M) | 0 | 0 | 0 | 0 | 0 | 0 | 0 | 0 | 0 | 0 | 0 | 401200 | 0 | 0 | 0 | 0 | 0 | 0 |
| LQMWVDV | MYOF | Q9NZM1 | MYOF_HUMAN | Oxidation (M) | 0 | 0 | 81629 | 424880 | 1924100 | 0 | 0 | 0 | 0 | 0 | 0 | 176990 | 37256 | 0 | 0 | 0 | 0 | 0 |
| LQNALNEC | NUMA1; NUMA | Q14980; A0 | NUMA1_HUMA | Unmodified | 0 | 0 | 0 | 0 | 0 | 0 | 0 | 0 | 670180 | 152560 | 538640 | 550300 | 461300 | 0 | 497990 | 0 | 684960 | 0 |
| LQNEILK | CKAP4 | Q07065 | CKAP4_HUMAN | Unmodified | 1789100 | 187350 | 893370 | 588180 | 153010 | 0 | 465650 | 221410 | 3542600 | 54232 | 0 | 186800 | 96834 | 0 | 1061900 | 1281600 | 0 | 334320 |
| LQNEILKDI | CKAP4 | Q07065 | CKAP4_HUMAN | Unmodified | 0 | 0 | 0 | 0 | 0 | 0 | 213580 | 0 | 0 | 460810 | 0 | 0 | 0 | 0 | 0 | 0 | 0 | 0 |
| LQNELDNV | MYH10 | P35580 | MYH10_HUMA | Unmodified | 0 | 0 | 570940 | 0 | 0 | 0 | 0 | 0 | 0 | 0 | 0 | 784060 | 0 | 0 | 216100 | 98051 | 0 | 0 |
| LQNELDNV | MYH10 | P35580 | MYH10_HUMA | Unmodified | 0 | 0 | 0 | 945550 | 3894900 | 0 | 0 | 0 | 0 | 0 | 793770 | 3131900 | 246290 | 1772900 | 0 | 987000 | 0 | 0 |
| LQNELDTLI | CLIP1; CLIP1; C | P30622; F5H | CLIP1_HUMAN | Unmodified | 0 | 0 | 0 | 0 | 215970 | 0 | 572340 | 8747.2 | 0 | 0 | 0 | 0 | 0 | 294240 | 0 | 0 | 0 | 0 |
| LQNLDALT | PPP1R7; PPP1R | Q15435; C9 | PP1R7_HUMAN | Oxidation (M) | 0 | 0 | 0 | 0 | 0 | 0 | 56773 | 0 | 0 | 0 | 0 | 3185300 | 0 | 0 | 320800 | 138500 | 0 | 0 |
| LQNLQLQF | SCP2 | P22307 | SCP2_HUMAN | Unmodified | 956150 | 0 | 0 | 0 | 0 | 0 | 0 | 0 | 4833400 | 1361800 | 948640 | 276290 | 0 | 0 | 0 | 0 | 0 | 0 |
| LQNLQLQF | SCP2 | P22307 | SCP2_HUMAN | Unmodified | 0 | 0 | 0 | 0 | 499320 | 0 | 160420 | 70245 | 6118500 | 2441100 | 586440 | 1093700 | 346170 | 275610 | 19698 | 23633 | 0 | 85849 |
| LQNNNVY | AP2B1; AP2B1 | P63010; A0A | AP2B1_HUMAN | Unmodified | 0 | 0 | 0 | 0 | 2399700 | 549080 | 195590 | 80587 | 992060 | 485110 | 643050 | 3663400 | 0 | 987380 | 1413300 | 0 | 1278100 | 2727400 |
| LQNNNVY | AP2B1; AP2B1 | P63010; A0A | AP2B1_HUMAN | Unmodified | 0 | 0 | 0 | 0 | 292970 | 0 | 72502 | 29926 | 968340 | 64986 | 0 | 0 | 0 | 0 | 0 | 0 | 333730 | 560160 |
| LQNPAITG | NEDD4L; NEDD | Q96PU5; A0 | NED4L_HUMAN | Unmodified | 0 | 385400 | 0 | 0 | 0 | 0 | 215510 | 123350 | 0 | 0 | 0 | 879360 | 75888 | 0 | 662550 | 403190 | 0 | 0 |
| LQNQLQG | VPS4B | O75351 | VPS4B_HUMAN | Unmodified | 1081800 | 567480 | 709920 | 881930 | 0 | 0 | 0 | 0 | 0 | 0 | 1226200 | 1368700 | 0 | 0 | 730420 | 686020 | 0 | 0 |
| LQNREDSS | CIT; CIT; CIT; C | O14578; A0 | CTRO_HUMAN | Unmodified | 33002 | 0 | 0 | 0 | 837610 | 363090 | 51669 | 52537 | 24736 | 50073 | 0 | 109470 | 73922 | 0 | 0 | 25703 | 0 | 0 |
| LQNSYQPT | DDX46; DDX46 | Q7L014; A0 | DDX46_HUMA | Unmodified | 0 | 0 | 0 | 0 | 26355 | 0 | 0 | 0 | 21255 | 25094 | 0 | 1006200 | 0 | 0 | 0 | 0 | 0 | 0 |
| LQPAPVIPS | ADAM17 | P78536 | ADA17_HUMAN | Unmodified | 0 | 0 | 0 | 388230 | 0 | 0 | 0 | 0 | 1403300 | 289350 | 0 | 378290 | 0 | 0 | 385430 | 284370 | 0 | 0 |
| LQPASVALI | IQSEC2; IQSEC | Q5JU85; A0 | IQEC2_HUMAN | Unmodified | 0 | 0 | 0 | 0 | 2145600 | 0 | 0 | 0 | 0 | 454870 | 0 | 0 | 0 | 0 | 0 | 0 | 0 | 0 |
| LQPFLQR | BAG6; BAG6; B | P46379; F6S | BAG6_HUMAN | Unmodified | 0 | 0 | 117220 | 118300 | 0 | 0 | 0 | 0 | 87826 | 0 | 0 | 3233400 | 0 | 0 | 118100 | 129610 | 2657000 | 2089500 |
| LQPHEFQG | DLAT; DLAT; DL | P10515; A0A | ODP2_HUMAN | Unmodified | 0 | 0 | 0 | 0 | 0 | 0 | 0 | 0 | 1469600 | 0 | 0 | 0 | 0 | 0 | 0 | 0 | 0 | 0 |
| LQPHPGLE | MLEC | Q14165 | MLEC_HUMAN | Unmodified | 0 | 0 | 0 | 0 | 0 | 0 | 0 | 0 | 22796000 | 660800 | 0 | 605830 | 0 | 0 | 0 | 0 | 0 | 0 |
| LQPSSTTTN | AOC1; AOC1; A | P19801; C9 | AOC1_HUMAN | Unmodified | 0 | 0 | 0 | 0 | 0 | 0 | 0 | 0 | 3107300 | 173680 | 0 | 0 | 239880 | 0 | 0 | 0 | 0 | 0 |
| LQPTWNDI | TXNDC5 | Q8NBS9 | TXND5_HUMA | Unmodified | 0 | 0 | 0 | 0 | 0 | 0 | 0 | 0 | 9673800 | 183090 | 736330 | 0 | 0 | 0 | 0 | 0 | 0 | 0 |
| LQPTWNDI | TXNDC5 | Q8NBS9 | TXND5_HUMA | Oxidation (M) | 0 | 0 | 0 | 0 | 0 | 0 | 0 | 0 | 9729900 | 2101700 | 0 | 0 | 0 | 0 | 0 | 0 | 0 | 0 |
| LQPTWNDI | TXNDC5 | Q8NBS9 | TXND5_HUMA | Unmodified | 0 | 0 | 0 | 0 | 178250 | 2569400 | 0 | 0 | 41749000 | 4678000 | 0 | 141110 | 0 | 2240000 | 0 | 0 | 567510 | 0 |
| LQQAIAMH | VPS37A; VPS37 | Q8NEZ2; H0 | VP37A_HUMAN | Oxidation (M) | 2639300 | 2287500 | 919650 | 0 | 1993400 | 495260 | 0 | 0 | 0 | 0 | 0 | 1322600 | 0 | 0 | 70940 | 57415 | 109790 | 0 |
| LQQAIAMH | VPS37A; VPS37 | Q8NEZ2; H0 | VP37A_HUMAN | Unmodified | 2480200 | 1991200 | 135820 | 328690 | 3815200 | 538550 | 65494 | 0 | 673880 | 164540 | 0 | 0 | 796430 | 419770 | 320660 | 523890 | 124930 | 288530 |
| LQQAKDEI | SNX1; SNX1 | Q13596; H0 | SNX1_HUMAN | Unmodified | 96897 | 0 | 147510 | 86274 | 448860 | 0 | 297180 | 0 | 1511500 | 41784 | 164690 | 0 | 194380 | 296690 | 275810 | 127610 | 0 | 0 |
| LQQAQEMI | TXLNA | P40222 | TXLNA_HUMAN | Unmodified | 0 | 410200 | 0 | 0 | 0 | 0 | 0 | 0 | 379710 | 0 | 0 | 552760 | 0 | 0 | 0 | 0 | 0 | 1294300 |
| LQQAQKG | ICAM1; ICAM1 | P05362; E7E | ICAM1_HUMAN | Unmodified | 0 | 0 | 0 | 0 | 3174500 | 1835700 | 0 | 0 | 0 | 0 | 0 | 0 | 0 | 0 | 0 | 0 | 0 | 1379900 |
| LQQAQKG | ICAM1; ICAM1 | P05362; E7E | ICAM1_HUMAN | Oxidation (M) | 0 | 0 | 0 | 0 | 706350 | 0 | 0 | 0 | 0 | 0 | 0 | 0 | 0 | 0 | 0 | 0 | 67092 | 139600 |
| LQQCKAEL | KIF20A | O95235 | KI20A_HUMAN | Unmodified | 0 | 0 | 0 | 0 | 1445700 | 0 | 134010 | 0 | 0 | 134910 | 0 | 0 | 0 | 0 | 0 | 0 | 1098300 | 0 |
| LQQCKAEL | KIF20A | O95235 | KI20A_HUMAN | Unmodified | 0 | 0 | 0 | 0 | 240590 | 0 | 0 | 0 | 0 | 72421 | 0 | 114960 | 276870 | 0 | 0 | 0 | 1240400 | 0 |
| LQQDSEIM | PIGK; PIGK | Q92643; B1 | GPI8_HUMAN ( | 2 Oxidation (M) | 0 | 0 | 0 | 0 | 0 | 0 | 0 | 0 | 571430 | 0 | 0 | 0 | 0 | 0 | 0 | 0 | 0 | 0 |
| LQQDSEIM | PIGK; PIGK | Q92643; B1 | GPI8_HUMAN ( | Oxidation (M) | 0 | 0 | 0 | 0 | 0 | 0 | 0 | 0 | 2607200 | 0 | 0 | 0 | 0 | 0 | 0 | 0 | 0 | 0 |
| LQQDSEIM | PIGK; PIGK | Q92643; B1 | GPI8_HUMAN ( | Unmodified | 0 | 0 | 0 | 0 | 0 | 0 | 0 | 0 | 5240200 | 0 | 0 | 0 | 0 | 0 | 0 | 0 | 0 | 0 |
| LQQDVLQF | GOLM1 | Q8NBJ4 | GOLM1_HUMA | Unmodified | 0 | 0 | 0 | 0 | 0 | 0 | 0 | 404480 | 42286000 | 37081 | 0 | 0 | 0 | 0 | 0 | 0 | 300600 | 190010 |
| LQQEDGIS | CPE; CPE | P16870; H0 | CBPE_HUMAN | Unmodified | 0 | 0 | 0 | 0 | 0 | 0 | 0 | 0 | 24824000 | 507230 | 0 | 0 | 0 | 3509600 | 0 | 0 | 0 | 0 |
| LQQEILDV | AVIL | O75366 | AVIL_HUMAN A | Unmodified | 0 | 0 | 0 | 0 | 0 | 0 | 0 | 0 | 0 | 0 | 2272100 | 6779200 | 0 | 0 | 0 | 0 | 0 | 0 |
| LQQELDDL | MYH9 | P35579 | MYH9_HUMAN | Unmodified | 0 | 0 | 0 | 0 | 0 | 0 | 0 | 0 | 0 | 0 | 0 | 0 | 0 | 0 | 0 | 0 | 0 | 0 |
| LQQELDDL | MYH9 | P35579 | MYH9_HUMAN | Unmodified | 0 | 0 | 0 | 0 | 1921000 | 97491 | 0 | 0 | 0 | 0 | 0 | 0 | 753290 | 220480 | 0 | 0 | 1983700 | 637960 |
| LQQELDDL | MYH9 | P35579 | MYH9_HUMAN | Unmodified | 0 | 0 | 0 | 0 | 170170 | 35519 | 30314 | 0 | 0 | 0 | 0 | 0 | 0 | 0 | 0 | 0 | 279250 | 575680 |
| LQQELEFLI | PSMC4 | P43686 | PRS6B_HUMAN | Unmodified | 595100 | 0 | 0 | 469420 | 4734700 | 1383100 | 2283200 | 0 | 1126400 | 3378800 | 1198000 | 1994400 | 8864300 | 4699300 | 0 | 0 | 5072200 | 4812000 |
| LQQENLQI | NEK9; NEK9; N | Q8TD19; A0 | NEK9_HUMAN | Unmodified | 0 | 0 | 0 | 0 | 897760 | 0 | 354670 | 116400 | 0 | 266640 | 0 | 427630 | 677070 | 517760 | 0 | 0 | 0 | 0 |
| LQQETAELI | EPS15 | P42566 | EPS15_HUMAN | Unmodified | 484840 | 459760 | 0 | 0 | 0 | 0 | 0 | 0 | 0 | 0 | 0 | 0 | 0 | 0 | 0 | 273380 | 0 | 0 |
| LQQEYAAN | SLC43A1 | O75387 | LAT3_HUMAN | Oxidation (M) | 0 | 0 | 0 | 0 | 207010 | 0 | 0 | 0 | 0 | 0 | 0 | 0 | 0 | 0 | 670910 | 531690 | 363350 | 91921 |
| LQQEYAAN | SLC43A1 | O75387 | LAT3_HUMAN | Unmodified | 0 | 0 | 204000 | 229910 | 0 | 0 | 0 | 0 | 0 | 0 | 0 | 182610 | 0 | 0 | 562760 | 0 | 0 | 0 |
| LQQFFNER | MYO6; MYO6; | Q9UM54; A0 | MYO6_HUMAN | Unmodified | 258390 | 219260 | 1900600 | 1057700 | 0 | 0 | 0 | 66037 | 564980 | 230230 | 317950 | 1772000 | 168700 | 0 | 2701000 | 1548600 | 0 | 0 |
| LQQFFNER | MYO6; MYO6; | Q9UM54; A0 | MYO6_HUMAN | Unmodified | 0 | 0 | 0 | 563100 | 1307600 | 599980 | 240280 | 0 | 0 | 917440 | 91247 | 0 | 1609800 | 0 | 0 | 0 | 376390 | 536640 |
| LQQGIPPS | NUCB2; NUCB | P80303; Q2 | NUCB2_HUMA | Unmodified | 0 | 0 | 0 | 0 | 0 | 0 | 0 | 384330 | 4619000 | 1270000 | 0 | 0 | 510680 | 510320 | 0 | 0 | 0 | 0 |
| LQQHCVAS | ADGRE2; ADGR | Q9UHX3; B4 | AGRE2_HUMA | Unmodified | 0 | 0 | 0 | 0 | 0 | 0 | 0 | 0 | 0 | 0 | 177960 | 1483200 | 0 | 0 | 0 | 0 | 0 | 0 |
| LQQIAAALE | PLXNB2 | O15031 | PLXB2_HUMAN | Unmodified | 0 | 0 | 928970 | 216360 | 0 | 0 | 34128 | 27232 | 261650 | 0 | 196490 | 2372800 | 308740 | 103480 | 970260 | 810850 | 0 | 0 |
| LQQIAAALE | PLXNB2 | O15031 | PLXB2_HUMAN | Unmodified | 415980 | 347840 | 575480 | 2976400 | 271710 | 0 | 109230 | 63730 | 1637400 | 291890 | 589150 | 3383600 | 246080 | 0 | 2037000 | 1300600 | 0 | 0 |
| LQQLFNHT | MYH9; MYH10; | P35579; P35 | MYH9_HUMAN | Oxidation (M) | 0 | 0 | 0 | 0 | 0 | 0 | 0 | 0 | 0 | 53225 | 0 | 5392800 | 0 | 0 | 2366600 | 0 | 0 | 0 |
| LQQLGEAH | NUMA1; NUMA | Q14980; F5 | NUMA1_HUMA | Unmodified | 0 | 77566 | 0 | 0 | 0 | 0 | 0 | 0 | 637660 | 90435 | 536390 | 1632900 | 0 | 0 | 0 | 0 | 0 | 0 |
| LQQLKDET | UPF1 | Q92900 | RENT1_HUMAN | Unmodified | 2123200 | 2475600 | 0 | 2462700 | 0 | 0 | 481030 | 214850 | 559930 | 374530 | 403750 | 1548700 | 2468600 | 0 | 0 | 0 | 163400 | 319030 |
| LQQLKDET | UPF1 | Q92900 | RENT1_HUMAN | Unmodified | 744070 | 0 | 0 | 928300 | 0 | 0 | 0 | 0 | 0 | 0 | 126730 | 0 | 0 | 0 | 0 | 328290 | 0 | 0 |
| LQQLPADF | LRRC59 | Q96AG4 | LRC59_HUMAN | Unmodified | 0 | 0 | 0 | 0 | 1114800 | 152540 | 227410 | 0 | 0 | 176770 | 0 | 0 | 387420 | 534070 | 0 | 0 | 231980 | 1180400 |
| LQQLQEEK | DEF6 | Q9H4E7 | DEFI6_HUMAN | Unmodified | 623440 | 0 | 0 | 0 | 388310 | 0 | 100440 | 0 | 4329100 | 235850 | 0 | 301600 | 0 | 0 | 0 | 0 | 0 | 0 |
| LQQLTAMV | ERBIN | Q96RT1 | ERBIN_HUMAN | Oxidation (M) | 0 | 0 | 0 | 1553200 | 15712000 | 5401900 | 0 | 0 | 954920 | 365900 | 1036800 | 2085200 | 2979700 | 0 | 0 | 79403 | 917480 | 2312900 |
| LQQLTAMV | ERBIN | Q96RT1 | ERBIN_HUMAN | Unmodified | 0 | 0 | 0 | 0 | 2195100 | 0 | 0 | 0 | 0 | 0 | 0 | 0 | 0 | 0 | 0 | 0 | 0 | 0 |
| LQQLTAMV | ERBIN | Q96RT1 | ERBIN_HUMAN | Oxidation (M) | 0 | 0 | 0 | 0 | 26744000 | 3926100 | 1488900 | 141270 | 0 | 0 | 0 | 0 | 0 | 0 | 0 | 0 | 0 | 1709500 |
| LQQNPGKI | TBC1D10A; TB | Q9BXI6; F8 | TB10A_HUMAN | Oxidation (M) | 0 | 0 | 0 | 0 | 0 | 0 | 0 | 0 | 0 | 0 | 0 | 215820 | 0 | 0 | 0 | 182330 | 0 | 0 |
| LQQQMDL | CIT; CIT; CIT; C | O14578; A0 | CTRO_HUMAN | Unmodified | 0 | 0 | 0 | 397270 | 2242000 | 340080 | 133390 | 0 | 262490 | 0 | 0 | 397820 | 167410 | 0 | 0 | 0 | 308700 | 0 |
| LQQSDIWK | HS2ST1 | Q7LGA3 | HS2ST_HUMAN | Unmodified | 0 | 0 | 0 | 0 | 0 | 0 | 0 | 0 | 3132500 | 204990 | 0 | 0 | 0 | 0 | 0 | 0 | 0 | 0 |
| LQQSSHG | GPR180 | Q86V85 | GP180_HUMAN | Unmodified | 0 | 0 | 0 | 0 | 0 | 0 | 26039 | 0 | 2629300 | 473310 | 0 | 46269 | 0 | 0 | 0 | 0 | 0 | 0 |
| LQQTEAELI | VARS1; VARS1 | P26640; A0A | SYVC_HUMAN | Unmodified | 132810 | 119090 | 288930 | 138290 | 100670 | 0 | 0 | 60061 | 806810 | 40829 | 747880 | 5060000 | 0 | 40575 | 418660 | 283930 | 0 | 0 |
| LQQTLNAL | IQGAP2; IQGAP | Q13576; E7E | IQGA2_HUMAN | Unmodified | 0 | 0 | 1114100 | 308080 | 161140 | 0 | 0 | 0 | 0 | 0 | 0 | 0 | 230920 | 189510 | 448660 | 444350 | 0 | 0 |
| LQQTQNQ | VAMP3 | Q15836 | VAMP3_HUMA | Unmodified | 0 | 0 | 0 | 2049500 | 3553300 | 0 | 0 | 0 | 2559200 | 459570 | 0 | 0 | 2098000 | 0 | 0 | 0 | 0 | 4113000 |

| | | | | | | | | | | | | | | | | | | | | | | |
|---|---|---|---|---|---|---|---|---|---|---|---|---|---|---|---|---|---|---|---|---|---|---|
| LQQTYAALI | IQGAP1; IQGAP | P46940; H0Y | IQGA1_HUMAN | Unmodified | 1033200 | 0 | 603310 | 1617500 | 33529000 | 333780 | 1302100 | 529010 | 2772700 | 1696300 | 552460 | 6382000 | 2922800 | 1432100 | 1116400 | 1067900 | 735420 | 1816100 |
| LQQTYAALI | IQGAP1; IQGAP | P46940; H0Y | IQGA1_HUMAN | Unmodified | 0 | 0 | 0 | 0 | 1091800 | 0 | 0 | 0 | 0 | 0 | 0 | 0 | 0 | 0 | 0 | 0 | 0 | 0 |
| LQQVLQME | KPNB1; KPNB1; | Q14974; J3Q | IMB1_HUMAN I | Oxidation (M) | 70927 | 0 | 0 | 0 | 0 | 0 | 0 | 0 | 0 | 0 | 0 | 1020300 | 0 | 0 | 62266 | 90626 | 0 | 0 |
| LQQVLQME | KPNB1; KPNB1; | Q14974; J3Q | IMB1_HUMAN I | Unmodified | 0 | 0 | 0 | 0 | 0 | 0 | 0 | 0 | 0 | 0 | 185490 | 1885500 | 0 | 0 | 326420 | 201620 | 0 | 0 |
| LQQVLTGL | TBCD; TBCD; TI | Q9BTW9; A0 | TBCD_HUMAN | Unmodified | 1779700 | 0 | 1079900 | 989420 | 0 | 0 | 0 | 0 | 160290 | 0 | 0 | 1549700 | 0 | 0 | 2877700 | 1666600 | 0 | 0 |
| LQQYENLV | ARHGEF39 | Q8N4T4 | ARG39_HUMAN | Unmodified | 1267500 | 0 | 446980 | 2133400 | 0 | 0 | 1814600 | 0 | 1763600 | 273850 | 0 | 0 | 0 | 0 | 0 | 545350 | 77866 | 0 |
| LQQYIAPGI | EPHB3 | P54753 | EPHB3_HUMAN | Oxidation (M) | 542780 | 471140 | 0 | 0 | 0 | 0 | 0 | 71342 | 0 | 0 | 0 | 1235900 | 0 | 0 | 0 | 0 | 0 | 395590 |
| LQQYIAPGI | EPHB3 | P54753 | EPHB3_HUMAN | Unmodified | 0 | 0 | 0 | 0 | 0 | 0 | 0 | 0 | 0 | 0 | 384090 | 3047600 | 0 | 0 | 171110 | 0 | 99569 | 1599600 |
| LQRDLEAE | BOLA2; BOLA2E | Q9H3K6; H3 | BOLA2_HUMAN | Unmodified | 1287700 | 764210 | 435460 | 1537700 | 4626300 | 0 | 0 | 155390 | 1630400 | 351200 | 1742500 | 969750 | 2341600 | 518630 | 491720 | 236200 | 199060 | 868310 |
| LQRELDDA | MYH10 | P35580 | MYH10_HUMA | Unmodified | 0 | 0 | 277650 | 250540 | 296890 | 0 | 0 | 0 | 0 | 0 | 0 | 4427100 | 0 | 0 | 450620 | 420210 | 0 | 403270 |
| LQRELDDA | MYH10 | P35580 | MYH10_HUMA | Unmodified | 0 | 0 | 0 | 1091000 | 23226000 | 4203700 | 2162100 | 508200 | 0 | 216810 | 0 | 0 | 5997300 | 5445600 | 0 | 0 | 1454200 | 1636500 |
| LQRELEDA | MYH9 | P35579 | MYH9_HUMAN | Oxidation (M) | 0 | 0 | 0 | 0 | 1396800 | 0 | 0 | 0 | 0 | 0 | 0 | 423900 | 0 | 0 | 0 | 0 | 616980 | 0 |
| LQRELEDA | MYH9 | P35579 | MYH9_HUMAN | Unmodified | 0 | 723080 | 0 | 3184200 | 76968000 | 10407000 | 5070900 | 0 | 2048100 | 0 | 0 | 2223200 | 11546000 | 8800000 | 1663800 | 1219300 | 9715600 | 22384000 |
| LQREPQVC | FURIN; FURIN; | P09958; A0A | FURIN_HUMAN | Unmodified | 0 | 0 | 0 | 0 | 0 | 0 | 0 | 0 | 200480 | 0 | 554820 | 0 | 0 | 0 | 0 | 0 | 0 | 0 |
| LQREVQEA | RASSF8; RASSF | Q8NHQ8; H( | RASF8_HUMAN | Unmodified | 0 | 0 | 0 | 0 | 92083 | 0 | 0 | 0 | 0 | 0 | 0 | 0 | 47785 | 50573 | 0 | 0 | 166390 | 446410 |
| LQRLEESM | VPS37D | Q86XT2 | VP37D_HUMAN | Unmodified | 101980 | 53322 | 152250 | 0 | 0 | 0 | 148620 | 127240 | 0 | 45806 | 0 | 0 | 0 | 0 | 464050 | 132350 | 0 | 0 |
| LQSECHPD | MOB4; HSPE1-I | Q9Y3A3; S4F | PHOCN_HUMA | Unmodified | 0 | 0 | 0 | 0 | 0 | 0 | 0 | 0 | 0 | 0 | 0 | 0 | 0 | 535030 | 0 | 535110 | 0 | 0 |
| LQSEPESIR | CLTA; CLTA | P09496; F8W | CLCA_HUMAN | Unmodified | 1453300 | 829710 | 365450 | 200050 | 0 | 0 | 0 | 0 | 88228 | 0 | 0 | 3972500 | 0 | 0 | 618370 | 693120 | 13610 | 0 |
| LQSEVAELI | IST1; IST1; IST1 | P53990; H3E | IST1_HUMAN IS | Unmodified | 1949100 | 1463700 | 3833500 | 5303500 | 2950400 | 1591000 | 0 | 0 | 24907000 | 4219600 | 7837500 | 50309000 | 1308800 | 340600 | 2372100 | 2511300 | 602030 | 0 |
| LQSEVAELI | IST1; IST1; IST1 | P53990; H3E | IST1_HUMAN IS | Unmodified | 0 | 0 | 0 | 0 | 0 | 0 | 0 | 0 | 0 | 1492900 | 0 | 0 | 2417400 | 464140 | 0 | 0 | 0 | 0 |
| LQSGICHLI | DARS1; DARS1; | P14868; C9J | SYDC_HUMAN | Unmodified | 0 | 0 | 0 | 229740 | 0 | 0 | 0 | 0 | 0 | 0 | 608160 | 6553900 | 0 | 0 | 341250 | 406560 | 0 | 0 |
| LQSITQAAC | N/A | N/A | N/A | Unmodified | 0 | 0 | 0 | 0 | 0 | 0 | 0 | 0 | 0 | 0 | 0 | 0 | 0 | 0 | 4377500 | 0 | 0 | 0 |
| LQSLDGPG | QSOX2 | Q6ZRP7 | QSOX2_HUMA | Unmodified | 0 | 0 | 0 | 0 | 215000 | 2852100 | 0 | 147770 | 832590 | 60462 | 0 | 0 | 0 | 0 | 0 | 0 | 315270 | 1349600 |
| LQSLDTSFI | MVK; MVK; MVK | Q03426; F5H | KIME_HUMAN I | Unmodified | 0 | 0 | 0 | 0 | 0 | 0 | 0 | 0 | 982340 | 0 | 0 | 485590 | 0 | 0 | 0 | 0 | 0 | 0 |
| LQSLFDSPI | SERPINF1 | P36955 | PEDF_HUMAN | Unmodified | 0 | 0 | 0 | 0 | 0 | 0 | 0 | 0 | 2260800 | 139650 | 18396000 | 1369100 | 0 | 0 | 0 | 0 | 0 | 0 |
| LQSLPFALT | SCRIB; SCRIB; S | Q14160; A0A | SCRIB_HUMAN | Unmodified | 0 | 0 | 0 | 0 | 0 | 0 | 0 | 0 | 3520600 | 377160 | 327340 | 5935500 | 0 | 0 | 0 | 0 | 367110 | 0 |
| LQSLYFK | DAAM1 | Q9Y4D1 | DAAM1_HUMA | Unmodified | 0 | 0 | 0 | 0 | 266910 | 0 | 117390 | 56980 | 1015000 | 153870 | 172160 | 2757000 | 307190 | 0 | 281180 | 0 | 30657 | 0 |
| LQSPEFQS | TRNT1; TRNT1; | Q96Q11; A0A | TRNT1_HUMAN | Unmodified | 0 | 0 | 0 | 0 | 409360 | 0 | 479650 | 0 | 220010 | 85247 | 782560 | 1347700 | 0 | 0 | 0 | 0 | 0 | 0 |
| LQSQLLSLI | DNM2; DNM2 | P50570; K7E | DYN2_HUMAN | Unmodified | 1555900 | 646520 | 1235600 | 790500 | 2770500 | 532970 | 249970 | 313810 | 1643200 | 1049700 | 380240 | 2840600 | 3431200 | 1677900 | 945390 | 1158600 | 445960 | 924760 |
| LQSQLLSLI | DNM2; DNM2 | P50570; K7E | DYN2_HUMAN | Unmodified | 0 | 0 | 0 | 0 | 0 | 0 | 0 | 0 | 0 | 271910 | 0 | 0 | 2614800 | 1915100 | 0 | 0 | 0 | 0 |
| LQSRPAAP | VAT1; VAT1 | Q99536; K7E | VAT1_HUMAN S | Unmodified | 0 | 53599 | 0 | 0 | 573640 | 211220 | 0 | 0 | 3080100 | 0 | 395070 | 899760 | 120510 | 0 | 0 | 71071 | 117250 | 142750 |
| LQSSASSIH | UEVLD | Q8IX04 | UEVLD_HUMAN | Unmodified | 288870 | 0 | 454090 | 2426800 | 30527000 | 4378800 | 220180 | 0 | 8529400 | 4548600 | 2538000 | 7250000 | 5763600 | 4527500 | 0 | 0 | 1863400 | 3353700 |
| LQSSQEPE | MAGED2; MAGI | Q9UNF1; Q5 | MAGD2_HUMA | Unmodified | 0 | 0 | 0 | 0 | 0 | 0 | 0 | 0 | 1141300 | 0 | 0 | 0 | 0 | 0 | 0 | 0 | 0 | 0 |
| LQSTFVFEE | HSD17B4; HSD | P51659; E7E | DHB4_HUMAN | Unmodified | 0 | 0 | 0 | 0 | 0 | 0 | 0 | 0 | 2421100 | 0 | 0 | 0 | 0 | 0 | 0 | 0 | 266250 | 0 |
| LQSTFVFEE | HSD17B4; HSD | P51659; E7E | DHB4_HUMAN | Unmodified | 0 | 0 | 0 | 0 | 0 | 0 | 0 | 0 | 4697300 | 488890 | 166430 | 0 | 0 | 0 | 0 | 0 | 0 | 0 |
| LQTASDES | SPTAN1; SPTAN | A0A0D9SGF | A0A0D9SGF6_I | Unmodified | 105590 | 36502 | 0 | 0 | 220960 | 0 | 117000 | 111250 | 2384300 | 629600 | 504470 | 3587600 | 0 | 0 | 0 | 0 | 85276 | 221890 |
| LQTASDES | SPTAN1 | Q13813 | SPTN1_HUMAN | Unmodified | 0 | 0 | 347590 | 114260 | 1138300 | 517510 | 24702 | 0 | 369980 | 164830 | 1204300 | 11977000 | 382920 | 22220 | 375100 | 277670 | 878070 | 0 |
| LQTDIKEN( | PARP4 | Q9UKK3 | PARP4_HUMAN | Unmodified | 2323100 | 0 | 0 | 0 | 0 | 0 | 0 | 0 | 0 | 0 | 0 | 0 | 0 | 0 | 0 | 0 | 0 | 0 |
| LQTEKQELI | USO1 | O60763 | USO1_HUMAN | Unmodified | 0 | 0 | 0 | 0 | 293550 | 15741 | 0 | 94715 | 224530 | 46978 | 384630 | 318600 | 269490 | 51437 | 0 | 0 | 0 | 274730 |
| LQTELLAR | DDO | Q99489 | OXDD_HUMAN | Unmodified | 0 | 0 | 0 | 0 | 268870 | 752520 | 2317100 | 1023800 | 3001700 | 162870 | 0 | 332390 | 207920 | 0 | 0 | 0 | 75061 | 1957600 |
| LQTINLSFN | LRRC40 | Q9H9A6 | LRC40_HUMAN | Unmodified | 0 | 0 | 0 | 0 | 0 | 0 | 0 | 0 | 0 | 0 | 0 | 1762000 | 0 | 0 | 0 | 0 | 0 | 0 |
| LQTLQSEN | GIT2; GIT2; GIT | Q14161; R4( | GIT2_HUMAN A | Unmodified | 0 | 0 | 0 | 0 | 191030 | 0 | 0 | 0 | 0 | 38744 | 0 | 0 | 0 | 0 | 0 | 0 | 121170 | 183560 |
| LQTLVSEQI | KTN1 | Q86UP2 | KTN1_HUMAN | Unmodified | 0 | 0 | 0 | 0 | 0 | 0 | 0 | 0 | 0 | 0 | 0 | 1743600 | 327010 | 0 | 0 | 0 | 0 | 0 |
| LQTMKEELI | LMNA; LMNA; L | P02545; A0A | LMNA_HUMAN | Unmodified | 0 | 0 | 0 | 0 | 0 | 0 | 0 | 0 | 0 | 0 | 0 | 926150 | 0 | 0 | 0 | 0 | 23142 | 0 |
| LQTMKEELI | LMNA; LMNA; L | P02545; A0A | LMNA_HUMAN | Oxidation (M) | 0 | 0 | 0 | 0 | 0 | 0 | 0 | 0 | 0 | 0 | 0 | 763620 | 0 | 0 | 0 | 0 | 0 | 0 |
| LQTNRDGA | NUP210 | Q8TEM1 | PO210_HUMAN | Unmodified | 99790 | 0 | 0 | 0 | 0 | 0 | 0 | 0 | 4515100 | 0 | 0 | 0 | 0 | 0 | 0 | 0 | 0 | 0 |
| LQTQASAT | HDLBP; HDLBP | Q00341; A0A | VIGLN_HUMAN | Unmodified | 311310 | 249670 | 231530 | 326290 | 382200 | 0 | 0 | 0 | 267260 | 0 | 0 | 991050 | 1053000 | 408230 | 225510 | 244890 | 85040 | 239130 |
| LQTQGLGT | ERO1B; ERO1A; | Q86YB8; Q9( | ERO1B_HUMA | Unmodified | 0 | 0 | 0 | 0 | 0 | 0 | 0 | 0 | 2382900 | 3874700 | 876860 | 0 | 0 | 0 | 0 | 0 | 0 | 0 |
| LQTQVFK | RPS9; RPS9; RP | P46781; B5N | RS9_HUMAN 4 | Unmodified | 2236700 | 1830700 | 0 | 0 | 746540 | 147600 | 681350 | 100550 | 475630 | 431280 | 0 | 19500000 | 0 | 0 | 0 | 1803600 | 656190 | 530020 |
| LQTSSVLVS | UBA1; UBA1; UI | P22314; Q5J | UBA1_HUMAN | Unmodified | 0 | 0 | 0 | 0 | 0 | 0 | 0 | 0 | 0 | 0 | 1357700 | 3339400 | 786950 | 0 | 0 | 0 | 0 | 0 |
| LQTVLEAVI | PARVB; PARVB; | Q9HBI1; A0A | PARVB_HUMAN | Unmodified | 0 | 0 | 166030 | 0 | 0 | 0 | 0 | 0 | 0 | 0 | 0 | 704460 | 0 | 0 | 181660 | 236570 | 0 | 0 |
| LQTYKTAVE | CCT3; CCT3 | P49368; B4D | TCPG_HUMAN | Unmodified | 0 | 0 | 0 | 0 | 122500 | 0 | 212480 | 0 | 0 | 0 | 0 | 0 | 383430 | 426240 | 0 | 0 | 873900 | 225130 |
| LQVEELLK | NOL6; NOL6 | Q9H6R4; A0 | NOL6_HUMAN | Unmodified | 0 | 0 | 25337 | 0 | 0 | 0 | 0 | 0 | 0 | 0 | 272560 | 236360 | 0 | 0 | 0 | 11827 | 0 | 0 |
| LQVEEVHQ | DIABLO; DIABL( | Q9NR28; A0 | DBLOH_HUMA | Unmodified | 0 | 0 | 0 | 0 | 0 | 0 | 0 | 0 | 2105700 | 0 | 0 | 0 | 0 | 0 | 0 | 0 | 0 | 0 |
| LQVELDNV | MYH9 | P35579 | MYH9_HUMAN | Unmodified | 0 | 0 | 0 | 0 | 2962400 | 0 | 0 | 0 | 530340 | 0 | 0 | 48146 | 1807500 | 0 | 0 | 0 | 868900 | 1431000 |
| LQVELDNV | MYH9 | P35579 | MYH9_HUMAN | Unmodified | 0 | 0 | 0 | 0 | 0 | 0 | 0 | 0 | 232580 | 0 | 0 | 0 | 368590 | 0 | 0 | 0 | 2364200 | 0 |
| LQVEPAVD | FLNA; FLNA | P21333; Q6( | FLNA_HUMAN | Unmodified | 0 | 1956500 | 0 | 0 | 257080 | 0 | 0 | 36129 | 0 | 0 | 0 | 1512100 | 0 | 0 | 0 | 0 | 0 | 704130 |
| LQVEQAPH | HDDC3; HDDC | Q8N4P3; H0 | MESH1_HUMA | Unmodified | 0 | 0 | 225650 | 0 | 0 | 200190 | 0 | 0 | 0 | 22745 | 275780 | 325540 | 0 | 0 | 37924 | 29242 | 0 | 0 |
| LQVFQALR | H6PD | O95479 | G6PE_HUMAN | Unmodified | 0 | 0 | 0 | 0 | 0 | 0 | 0 | 0 | 1105400 | 0 | 0 | 0 | 0 | 0 | 0 | 0 | 0 | 0 |
| LQVLGTNS | EFR3A; EFR3A; | Q14156; E5F | EFR3A_HUMAN | Unmodified | 0 | 0 | 316420 | 0 | 0 | 0 | 0 | 119290 | 226080 | 0 | 0 | 1426900 | 0 | 0 | 726740 | 402540 | 0 | 0 |
| LQVLQVLD | PATJ; PATJ; PAT | Q8NI35; A0A | INADL_HUMAN | Unmodified | 0 | 0 | 465420 | 0 | 1048800 | 0 | 10809 | 0 | 1034400 | 0 | 0 | 0 | 369530 | 366540 | 327610 | 365680 | 131220 | 962180 |
| LQVMANSF | KARS1; KARS1 | Q15046; H3I | SYK_HUMAN Ly | Unmodified | 23888 | 17141 | 0 | 0 | 543370 | 0 | 0 | 7111.2 | 320100 | 66279 | 635820 | 6862500 | 0 | 0 | 42398 | 37584 | 134620 | 0 |
| LQVNQDGI | PLCB3 | Q01970 | PLCB3_HUMAN | Unmodified | 275310 | 145090 | 0 | 0 | 1529200 | 0 | 286070 | 98729 | 200550 | 42695 | 439590 | 1930900 | 147480 | 427240 | 92322 | 124970 | 260980 | 118800 |
| LQVSQQED | IMPA1; IMPA1; I | P29218; E5R | IMPA1_HUMAN | Unmodified | 115380 | 101810 | 0 | 139200 | 848600 | 0 | 0 | 262770 | 1340200 | 1175600 | 0 | 1127700 | 693950 | 0 | 0 | 128890 | 641430 | 1983700 |
| LQVTMPGII | AHNAK | Q09666 | AHNK_HUMAN | Oxidation (M) | 0 | 0 | 0 | 0 | 395010 | 0 | 0 | 0 | 0 | 0 | 0 | 0 | 0 | 0 | 0 | 0 | 314400 | 870920 |
| LQVTMPGII | AHNAK | Q09666 | AHNK_HUMAN | Unmodified | 0 | 0 | 0 | 0 | 10686000 | 1690300 | 0 | 95287 | 0 | 0 | 0 | 0 | 0 | 89847 | 0 | 0 | 984420 | 8486700 |
| LQVTMPGII | AHNAK | Q09666 | AHNK_HUMAN | Unmodified | 0 | 0 | 0 | 0 | 1247000 | 0 | 0 | 0 | 0 | 0 | 0 | 0 | 0 | 65231 | 0 | 0 | 1244900 | 800210 |
| LQVTNVLS( | RPN2; RPN2; RI | P04844; Q5J | RPN2_HUMAN | Unmodified | 0 | 0 | 0 | 0 | 141350 | 1517800 | 0 | 0 | 55686000 | 12792000 | 5513400 | 1518900 | 0 | 0 | 256990 | 109950 | 0 | 682330 |
| LQVTNVLS( | RPN2; RPN2; RI | P04844; Q5J | RPN2_HUMAN | Unmodified | 0 | 0 | 0 | 0 | 0 | 0 | 0 | 0 | 1.08E+08 | 11727000 | 0 | 0 | 32830 | 0 | 0 | 0 | 0 | 0 |
| LQVVDQPL | FASN; FASN | P49327; A0A | FAS_HUMAN Fa | Unmodified | 0 | 0 | 4268500 | 0 | 0 | 0 | 0 | 0 | 0 | 0 | 4538300 | 83250000 | 0 | 0 | 7361300 | 4474500 | 0 | 0 |
| LQVVGRAG | PHGDH; PHGD | O43175; A0A | SERA_HUMAN | Unmodified | 0 | 0 | 0 | 0 | 0 | 0 | 42220 | 0 | 0 | 0 | 0 | 0 | 1068400 | 621110 | 0 | 0 | 35199 | 129510 |
| LQVVGRAG | PHGDH; PHGD | O43175; A0A | SERA_HUMAN | Unmodified | 0 | 0 | 92644 | 0 | 2135900 | 0 | 1830200 | 336490 | 0 | 1615000 | 0 | 0 | 11972000 | 8648700 | 0 | 0 | 2765300 | 2430700 |
| LQVVLNES | SMCHD1; SMCI | A6NHR9; J3K | SMHD1_HUMA | Unmodified | 0 | 0 | 0 | 0 | 0 | 0 | 1058100 | 280610 | 0 | 0 | 0 | 0 | 0 | 0 | 0 | 0 | 0 | 0 |
| LQVWDTA( | RAB13 | P51153 | RAB13_HUMAN | Unmodified | 5506000 | 5369400 | 2620000 | 1185200 | 0 | 0 | 0 | 0 | 0 | 0 | 0 | 2991700 | 0 | 0 | 4488000 | 2737400 | 0 | 0 |
| LQVWDTA( | RAB13 | P51153 | RAB13_HUMAN | Unmodified | 4032300 | 3756500 | 2883800 | 2315000 | 4075800 | 1517200 | 0 | 89366 | 1447200 | 434030 | 0 | 892400 | 919780 | 658510 | 2606600 | 1494800 | 674140 | 1265200 |
| LQWFCDR( | LRPPRC | P42704 | LPPRC_HUMAN | Unmodified | 0 | 0 | 0 | 0 | 0 | 0 | 0 | 0 | 531940 | 651300 | 0 | 0 | 0 | 0 | 0 | 0 | 0 | 66739 |
| LQWPTPDL | QSOX2 | Q6ZRP7 | QSOX2_HUMA | Unmodified | 0 | 0 | 0 | 0 | 0 | 0 | 0 | 0 | 3952600 | 419650 | 0 | 0 | 0 | 0 | 0 | 0 | 0 | 0 |
| LQYFRDHII | IQGAP1; IQGAP | P46940; A0A | IQGA1_HUMAN | Unmodified | 188360 | 130320 | 177980 | 122920 | 371250 | 0 | 0 | 0 | 348730 | 0 | 0 | 2313300 | 59416 | 0 | 341070 | 317640 | 0 | 0 |
| LQYSQDIPS | DTNB; DTNB | O60941; E7E | DTNB_HUMAN | Unmodified | 26066 | 0 | 0 | 0 | 0 | 0 | 0 | 0 | 0 | 215410 | 0 | 0 | 0 | 107660 | 268720 | 109400 | 0 | 0 |
| LRAANEKY | CLIP2 | Q9UDT6 | CLIP2_HUMAN | Unmodified | 0 | 0 | 0 | 0 | 3483900 | 0 | 777450 | 0 | 0 | 0 | 0 | 0 | 0 | 582040 | 0 | 0 | 0 | 535070 |
| LRADEAKQ | LAMB1; LAMB1; | P07942; A0A | LAMB1_HUMAN | Unmodified | 0 | 0 | 0 | 0 | 0 | 92798 | 0 | 0 | 0 | 183850 | 1995200 | 0 | 191200 | 118800 | 0 | 0 | 267060 | 614580 |
| LRAEIAEIQ | RALBP1 | Q15311 | RBP1_HUMAN | Unmodified | 124200 | 79734 | 356610 | 292740 | 0 | 112980 | 1267100 | 554540 | 221030 | 0 | 0 | 330850 | 0 | 0 | 259650 | 455340 | 732800 | 0 |
| LRAETEQG | PLEC | Q15149 | PLEC_HUMAN | Unmodified | 606160 | 0 | 0 | 0 | 0 | 0 | 277270 | 0 | 108780 | 0 | 35493 | 522640 | 0 | 0 | 134690 | 106350 | 0 | 0 |
| LRAFHNEA | TCP1; TCP1; TC | P17987; F5H | TCPA_HUMAN | Unmodified | 0 | 0 | 33423 | 0 | 646870 | 59195 | 120150 | 172320 | 118760 | 0 | 0 | 0 | 0 | 35826 | 0 | 0 | 188170 | 382280 |
| LRAFHNEA | TCP1; TCP1; TC | P17987; F5H | TCPA_HUMAN | Unmodified | 0 | 0 | 0 | 0 | 1600400 | 0 | 0 | 0 | 0 | 228300 | 196050 | 2505800 | 750040 | 1118400 | 0 | 0 | 450400 | 160360 |
| LRAILQNQ( | APOBEC3B; AP( | Q9UH17; P3 | ABC3B_HUMA | Unmodified | 0 | 0 | 291500 | 496220 | 0 | 0 | 0 | 0 | 0 | 0 | 0 | 0 | 663970 | 0 | 353680 | 160250 | 0 | 0 |
| LRALTDSFF | SRI; SRI; SRI | P30626; B4D | SORCN_HUMA | Unmodified | 0 | 276540 | 298060 | 0 | 531740 | 0 | 116890 | 68332 | 2612300 | 791650 | 898340 | 566040 | 440300 | 468340 | 198550 | 167090 | 0 | 730950 |
| LRAPIICVL( | EIF5B; EIF5B | O60841; A0A | IF2P_HUMAN E | Unmodified | 0 | 0 | 0 | 0 | 0 | 0 | 0 | 0 | 0 | 0 | 0 | 0 | 1290200 | 330900 | 0 | 0 | 0 | 0 |
| LRAVDFAEI | RPL8; RPL8; RP | P62917; E9P | RL8_HUMAN 6( | Unmodified | 0 | 0 | 0 | 182230 | 2313400 | 0 | 1415800 | 0 | 0 | 546720 | 374020 | 0 | 4512400 | 2415300 | 0 | 0 | 205390 | 1277900 |
| LRAWNDVI | VIPAS39; VIPAS | Q9H9C1; G3 | SPE39_HUMAN | Unmodified | 0 | 0 | 0 | 0 | 357820 | 92396 | 272720 | 0 | 178510 | 0 | 46395 | 281420 | 182030 | 70467 | 0 | 40424 | 0 | 0 |
| LRCVFEMP | VAPA | Q9P0L0 | VAPA_HUMAN | Unmodified | 0 | 0 | 329230 | 0 | 1588200 | 0 | 0 | 0 | 0 | 0 | 0 | 0 | 0 | 0 | 0 | 0 | 0 | 1544400 |
| LRDFFGSIS | GLB1L2 | Q8IW92 | GLBL2_HUMAN | Unmodified | 0 | 0 | 0 | 0 | 0 | 0 | 0 | 0 | 2457300 | 568220 | 0 | 159770 | 0 | 0 | 0 | 0 | 0 | 0 |
| LRDLHPDL | UGGT1 | Q9NYU2 | UGGG1_HUMA | Unmodified | 0 | 0 | 0 | 0 | 0 | 0 | 77921 | 0 | 4175100 | 976620 | 0 | 0 | 0 | 0 | 0 | 0 | 0 | 0 |
| LRDLQGAN | UTRN; UTRN | P46939; Q5T | UTRN_HUMAN | Unmodified | 0 | 0 | 0 | 0 | 0 | 0 | 0 | 0 | 439920 | 77004 | 14092 | 0 | 0 | 0 | 175840 | 0 | 0 | 0 |
| LRDLQGAN | UTRN; UTRN | P46939; Q5T | UTRN_HUMAN | Oxidation (M) | 0 | 0 | 214780 | 0 | 47942 | 0 | 0 | 0 | 111010 | 0 | 0 | 0 | 0 | 0 | 0 | 274810 | 83671 | 0 |
| LRDMLSDV | EPS15L1; EPS1 | Q9UBC2; M( | EP15R_HUMAN | Unmodified | 0 | 87889 | 0 | 0 | 437700 | 0 | 149250 | 0 | 64894 | 155090 | 0 | 0 | 0 | 0 | 0 | 0 | 192360 | 385420 |
| LRDNGLLA | OAT | P04181 | OAT_HUMAN C | Unmodified | 0 | 0 | 0 | 0 | 0 | 0 | 0 | 0 | 4973100 | 0 | 0 | 192360 | 0 | 0 | 0 | 0 | 0 | 0 |
| LRDQPSLV | ANKRD28 | O15084 | ANR28_HUMA | Unmodified | 0 | 0 | 417350 | 1159300 | 0 | 0 | 0 | 0 | 0 | 0 | 0 | 1290900 | 0 | 0 | 0 | 964400 | 0 | 0 |
| LRDTADGT | PIK3R1; PIK3R1 | P27986; H0Y | P85A_HUMAN | Unmodified | 0 | 0 | 0 | 100530 | 0 | 0 | 47769 | 0 | 0 | 0 | 0 | 0 | 196200 | 81280 | 0 | 0 | 0 | 0 |
| LRDTEEML | CHMP4B | Q9H444 | CHM4B_HUMA | Unmodified | 12725000 | 8927600 | 5476000 | 7829800 | 5380200 | 210600 | 31780 | 0 | 18469000 | 360430 | 2627000 | 5385400 | 1995400 | 572080 | 0 | 5220100 | 155140 | 946670 |
| LRDVAALN | EMC10; EMC10 | Q5UCC4; M( | EMC10_HUMA | Unmodified | 0 | 0 | 0 | 0 | 0 | 0 | 0 | 0 | 4021300 | 116700 | 633780 | 353970 | 0 | 0 | 0 | 0 | 0 | 0 |
| LRDVFNMD | PLCB2; PLCB2 | Q00722; Q9I | PLCB2_HUMAN | Oxidation (M) | 23291 | 0 | 0 | 0 | 0 | 0 | 0 | 0 | 589640 | 0 | 0 | 943580 | 0 | 0 | 0 | 0 | 0 | 0 |
| LRDYIWNT | CS; CS; CS; CS | O75390; A0A | CISY_HUMAN ( | Unmodified | 0 | 0 | 0 | 0 | 0 | 0 | 0 | 0 | 9339300 | 748740 | 0 | 0 | 0 | 0 | 0 | 0 | 0 | 0 |
| LREAGLAP | UROD; UROD; U | P06132; A0A | DCUP_HUMAN | Oxidation (M) | 0 | 0 | 0 | 0 | 0 | 0 | 0 | 0 | 0 | 0 | 0 | 1949200 | 0 | 0 | 0 | 0 | 0 | 0 |
| LREAGNIN( | KIF23; KIF23; KI | Q02241; A0A | KIF23_HUMAN | Unmodified | 36471 | 34358 | 138000 | 244340 | 1643700 | 330110 | 0 | 78627 | 1846500 | 167120 | 253390 | 504940 | 431720 | 141590 | 95515 | 39107 | 283520 | 597220 |
| LREAGNIN( | KIF23; KIF23; KI | Q02241; A0A | KIF23_HUMAN | Oxidation (M) | 50226 | 73606 | 226820 | 353190 | 477390 | 0 | 0 | 0 | 284720 | 0 | 0 | 601510 | 0 | 0 | 157800 | 92041 | 0 | 0 |
| LREANQQC | SPTAN1; SPTAN | Q13813; A0A | SPTN1_HUMAN | Unmodified | 0 | 0 | 0 | 0 | 90849 | 0 | 0 | 18748 | 116030 | 0 | 0 | 2013200 | 0 | 0 | 0 | 0 | 0 | 21330 |
| LRECEGLV | ARVCF; ARVCF; | O00192; E9F | ARVC_HUMAN | Unmodified | 0 | 0 | 0 | 85657 | 0 | 0 | 0 | 0 | 0 | 0 | 0 | 0 | 0 | 0 | 544100 | 0 | 0 | 0 |
| LRECLPLIIF | RPS4X | P62701 | RS4X_HUMAN | Unmodified | 1473700 | 899720 | 356800 | 907130 | 262900 | 117250 | 783870 | 15047 | 0 | 292650 | 1712300 | 7191800 | 153340 | 0 | 734920 | 527900 | 0 | 298740 |

| | | | | | | | | | | | | | | | | | | | | | | |
|---|---|---|---|---|---|---|---|---|---|---|---|---|---|---|---|---|---|---|---|---|---|---|
| LRECLPLIV | RPS4Y2; RPS4Y | Q8TD47; P2 | RS4Y2_HUMAN | Unmodified | 0 | 0 | 0 | 0 | 0 | 0 | 0 | 0 | 0 | 0 | 84884 | 2148200 | 0 | 0 | 0 | 0 | 0 | 0 |
| LREDENAE | DBN1; DBN1 | Q16643; D6I | DREB_HUMAN | Unmodified | 0 | 0 | 2322900 | 1682600 | 3841000 | 737040 | 203200 | 144630 | 817380 | 291910 | 542990 | 11164000 | 70097 | 0 | 2354800 | 1253100 | 1125700 | 2013600 |
| LREDENAE | DBN1; DBN1 | Q16643; D6I | DREB_HUMAN | Unmodified | 0 | 0 | 0 | 0 | 2095300 | 0 | 0 | 0 | 0 | 235130 | 0 | 0 | 2169000 | 1388400 | 0 | 0 | 3891600 | 324280 |
| LREENDIAF | CEP55; CEP55 | Q53EZ4; H0Y | CEP55_HUMAN | Unmodified | 0 | 0 | 0 | 0 | 0 | 0 | 0 | 0 | 2229200 | 0 | 50152 | 0 | 0 | 0 | 0 | 0 | 0 | 35516 |
| LREGEPGW | B3GAT3; B3GAT | O94766; E9I | B3GA3_HUMAN | Unmodified | 0 | 0 | 0 | 0 | 0 | 0 | 0 | 0 | 2465400 | 432520 | 99864 | 0 | 0 | 0 | 0 | 0 | 0 | 0 |
| LREIELLCQ | MAPRE2 | Q15555 | MARE2_HUMAI | Unmodified | 271360 | 208500 | 224990 | 794090 | 0 | 0 | 0 | 0 | 0 | 0 | 0 | 0 | 0 | 0 | 244160 | 288090 | 0 | 0 |
| LREIFGLYG | SAR1A; SAR1A | Q9NR31; H0 | SAR1A_HUMAN | Unmodified | 0 | 0 | 0 | 0 | 0 | 38177 | 289800 | 52308 | 1642700 | 82487 | 0 | 4070700 | 0 | 247330 | 0 | 0 | 0 | 0 |
| LREIFGLYG | SAR1A; SAR1A | Q9NR31; H0 | SAR1A_HUMAN | Unmodified | 0 | 0 | 0 | 0 | 0 | 0 | 626550 | 0 | 0 | 483510 | 0 | 0 | 299760 | 0 | 0 | 0 | 324010 | 602470 |
| LRELLIIGG | DDX1; DDX1; D | Q92499; A0A | DDX1_HUMAN | Unmodified | 0 | 0 | 0 | 0 | 0 | 0 | 0 | 0 | 0 | 0 | 0 | 1040000 | 0 | 0 | 0 | 0 | 0 | 0 |
| LRELMGEL | TRIM32; TRIM32 | Q13049; Q5 | TRI32_HUMAN | Oxidation (M) | 0 | 0 | 664820 | 97810 | 53155 | 0 | 246170 | 0 | 0 | 0 | 0 | 0 | 0 | 0 | 97409 | 96076 | 0 | 0 |
| LRELPLAAD | PKP3 | Q9Y446 | PKP3_HUMAN | Unmodified | 0 | 0 | 0 | 0 | 0 | 0 | 0 | 0 | 603890 | 517090 | 0 | 1980400 | 0 | 0 | 0 | 0 | 567140 | 0 |
| LRELSELSL | ACP3 | P15309 | PPAP_HUMAN | Unmodified | 0 | 784650 | 0 | 0 | 0 | 0 | 0 | 0 | 5346300 | 3091300 | 0 | 0 | 0 | 0 | 0 | 716130 | 0 | 43403 |
| LREMEHYF | PGM3; PGM3; F | O95394; J3K | AGM1_HUMAN | Oxidation (M) | 0 | 0 | 0 | 0 | 0 | 0 | 0 | 0 | 0 | 0 | 0 | 0 | 1878100 | 414940 | 0 | 0 | 0 | 0 |
| LRENELTYY | CD59; CD59; N | P13987; A0A | CD59_HUMAN | Unmodified | 3987100 | 2243800 | 2719400 | 2043500 | 894400 | 0 | 0 | 0 | 1435900 | 143180 | 0 | 2688800 | 2100700 | 0 | 5361200 | 4374500 | 1140500 | 6550000 |
| LRENELTYY | CD59; CD59; N | P13987; A0A | CD59_HUMAN | Unmodified | 9351500 | 3986600 | 2922700 | 4663300 | 9746100 | 4195900 | 1275300 | 331330 | 5432600 | 1549000 | 154490 | 6392700 | 5869800 | 8978500 | 7115700 | 6695200 | 4862700 | 47637000 |
| LRENVFQE | CTTN | Q14247 | SRC8_HUMAN | Unmodified | 3145000 | 876110 | 0 | 3185500 | 17096000 | 3031900 | 0 | 121120 | 1469600 | 822450 | 0 | 6324600 | 10631000 | 9434200 | 2619800 | 2715300 | 1548900 | 7870300 |
| LRENVFQE | CTTN | Q14247 | SRC8_HUMAN | Unmodified | 0 | 0 | 118840 | 0 | 0 | 0 | 0 | 0 | 0 | 0 | 0 | 0 | 719270 | 0 | 0 | 0 | 621440 | 292250 |
| LREQILDLS | CCDC126; MG/ | Q96EE4; Q9 | CC126_HUMAI | Unmodified | 0 | 0 | 0 | 0 | 0 | 0 | 0 | 0 | 2874700 | 0 | 0 | 0 | 0 | 0 | 0 | 0 | 0 | 0 |
| LREQSEHY | CDC42BPA; CD | Q5VT25; A0A | MRCKA_HUMA | Unmodified | 0 | 0 | 11314 | 0 | 0 | 0 | 49106 | 0 | 518200 | 177890 | 0 | 357060 | 0 | 1393600 | 0 | 0 | 0 | 2157800 |
| LREQTLSPT | SEC31A; SEC31 | O94979; H0Y | SC31A_HUMAN | Unmodified | 0 | 0 | 0 | 126500 | 995450 | 0 | 0 | 0 | 0 | 197320 | 0 | 3153300 | 1372100 | 1084700 | 0 | 63962 | 228910 | 840250 |
| LRESAELFC | NAALADL2 | Q58DX5 | NADL2_HUMAI | Oxidation (M) | 0 | 0 | 475770 | 266440 | 0 | 0 | 0 | 0 | 0 | 0 | 264980 | 0 | 255810 | 0 | 355290 | 221250 | 0 | 0 |
| LRESAELFC | NAALADL2 | Q58DX5 | NADL2_HUMAI | Unmodified | 0 | 0 | 574880 | 538490 | 0 | 0 | 0 | 0 | 0 | 0 | 0 | 0 | 647400 | 0 | 816420 | 499640 | 0 | 0 |
| LRESVVERI | N/A | N/A | N/A | Oxidation (M) | 0 | 0 | 0 | 1002400 | 4106700 | 1433600 | 3774700 | 2699200 | 0 | 0 | 0 | 0 | 0 | 4749000 | 0 | 0 | 0 | 0 |
| LRETAEAD | TPM1; TPM1; TF | H0YKJ4; H0Y | H0YKJ4_HUMA | Unmodified | 43048 | 0 | 236190 | 218780 | 253210 | 0 | 0 | 0 | 311680 | 0 | 0 | 1107500 | 652510 | 0 | 373600 | 424360 | 0 | 0 |
| LRETEEML | CHMP4C | Q96CF2 | CHM4C_HUMA | Unmodified | 882180 | 1425400 | 357620 | 0 | 145300 | 0 | 0 | 0 | 2411700 | 317800 | 35300 | 190830 | 0 | 0 | 566820 | 377370 | 0 | 279440 |
| LRETEEML | CHMP4C | Q96CF2 | CHM4C_HUMA | Oxidation (M) | 646920 | 479580 | 212010 | 84576 | 0 | 0 | 0 | 0 | 823910 | 123040 | 0 | 0 | 0 | 0 | 157000 | 284180 | 0 | 0 |
| LRETLNISG | LAP3; LAP3 | P28838; H0 | AMPL_HUMAN | Unmodified | 0 | 97439 | 0 | 0 | 3664200 | 0 | 826900 | 461120 | 1363100 | 852770 | 0 | 2193500 | 2664200 | 1195500 | 456880 | 233290 | 1317600 | 1626800 |
| LRETQGIEK | CTNND1; CTNN | O60716; A0 | CTND1_HUMAI | Unmodified | 0 | 0 | 0 | 0 | 0 | 0 | 0 | 0 | 0 | 0 | 0 | 0 | 814200 | 256920 | 0 | 0 | 2004500 | 1024900 |
| LREVVLDAI | PLCB3 | Q01970 | PLCB3_HUMAN | Unmodified | 0 | 0 | 0 | 0 | 1610600 | 353580 | 0 | 0 | 0 | 0 | 175560 | 0 | 0 | 0 | 0 | 0 | 526600 | 478230 |
| LREYEAALN | LMNB1; LMNB1 | P20700; A0A | LMNB1_HUMAI | Unmodified | 0 | 0 | 0 | 0 | 2723300 | 1567600 | 411290 | 0 | 1189000 | 726860 | 0 | 5019600 | 1371200 | 0 | 0 | 0 | 573170 | 0 |
| LREYLEGSI | SRGAP1; SRGA | Q7Z6B7; G5I | SRGP1_HUMAN | Unmodified | 112330 | 97333 | 89021 | 159970 | 2839800 | 647300 | 711940 | 336910 | 0 | 1096800 | 0 | 1113200 | 413850 | 489610 | 69679 | 64993 | 460000 | 491930 |
| LRFGIYDID | CPNE1; CPNE1 | Q99829; Q5 | CPNE1_HUMAI | Unmodified | 85130 | 0 | 0 | 0 | 0 | 0 | 0 | 0 | 0 | 0 | 0 | 1020800 | 0 | 0 | 0 | 0 | 0 | 0 |
| LRFLPEFPS | LRRC40 | Q9H9A6 | LRC40_HUMAN | Unmodified | 54123 | 0 | 0 | 0 | 0 | 59964 | 241270 | 0 | 249110 | 176500 | 110220 | 249390 | 197340 | 0 | 50697 | 0 | 226300 | 182410 |
| LRFPAEDEI | CKB; CKB | P12277; G3\ | KCRB_HUMAN | Oxidation (M) | 3967500 | 3055200 | 10715000 | 16721000 | 60670000 | 9119800 | 1.31E+08 | 22081000 | 4580300 | 7319100 | 5538300 | 47812000 | 79594000 | 25441000 | 15899000 | 16830000 | 366040 | 0 |
| LRFPAEDEI | CKB; CKB | P12277; G3\ | KCRB_HUMAN | Unmodified | 0 | 0 | 705930 | 1922500 | 49503000 | 3115400 | 35506000 | 1591900 | 11566000 | 3121600 | 1922300 | 33752000 | 26401000 | 13834000 | 2877200 | 700250 | 0 | 0 |
| LRGDAAAG | NXN; NXN | Q6DKJ4; A0A | NXN_HUMAN N | Unmodified | 0 | 0 | 0 | 0 | 0 | 0 | 0 | 0 | 98782 | 0 | 0 | 1916500 | 0 | 0 | 0 | 37920 | 0 | 0 |
| LRGEDDYN | GOLM1 | Q8NBJ4 | GOLM1_HUMA | Oxidation (M) | 0 | 0 | 0 | 0 | 0 | 0 | 0 | 0 | 2055400 | 0 | 0 | 0 | 0 | 0 | 0 | 0 | 0 | 0 |
| LRGEDDYN | GOLM1 | Q8NBJ4 | GOLM1_HUMA | Unmodified | 0 | 0 | 0 | 0 | 82732 | 0 | 0 | 0 | 5816300 | 368130 | 0 | 0 | 0 | 0 | 0 | 0 | 0 | 0 |
| LRGGAYIGI | LOXL2; LOXL2 | Q9Y4K0; H0Y | LOXL2_HUMAN | Unmodified | 0 | 0 | 0 | 0 | 840200 | 0 | 0 | 0 | 3591300 | 1857300 | 0 | 0 | 0 | 0 | 0 | 0 | 0 | 980570 |
| LRGIEQAV( | IMMT; IMMT; IM | Q16891; B9/ | MIC60_HUMAN | Unmodified | 0 | 0 | 0 | 0 | 0 | 0 | 193530 | 0 | 1286600 | 290090 | 0 | 0 | 0 | 0 | 0 | 0 | 0 | 0 |
| LRGISTKPV | TRIM25; TRIM25 | Q14258; A0A | TRI25_HUMAN | Unmodified | 0 | 0 | 0 | 0 | 23842000 | 1279800 | 0 | 0 | 0 | 0 | 292120 | 1724200 | 4045100 | 518620 | 0 | 253130 | 3046400 | 3941600 |
| LRGMVFGA | SMCHD1; SMCI | A6NHR9; A0 | SMHD1_HUMAN | Unmodified | 0 | 0 | 0 | 0 | 0 | 0 | 0 | 0 | 0 | 0 | 0 | 0 | 1046400 | 0 | 0 | 0 | 0 | 0 |
| LRGPSGG( | NUP88; NUP88 | Q99567; J3K | NUP88_HUMAI | Unmodified | 0 | 0 | 0 | 0 | 0 | 0 | 0 | 0 | 0 | 0 | 0 | 1293800 | 0 | 67323 | 0 | 0 | 0 | 0 |
| LRGQPFYA | LPP; LPP | Q93052; A0A | LPP_HUMAN Li | Unmodified | 0 | 46726 | 0 | 60429 | 1080100 | 0 | 0 | 0 | 0 | 0 | 0 | 0 | 1195700 | 557120 | 0 | 0 | 260080 | 502010 |
| LRGQYQGI | UGGT1 | Q9NYU2 | UGGG1_HUMA | Unmodified | 0 | 0 | 0 | 0 | 0 | 0 | 0 | 0 | 4817600 | 0 | 0 | 0 | 0 | 0 | 0 | 0 | 0 | 0 |
| LRGTADVTI | SLC2A1; SLC2A | P11166; A0A | GTR1_HUMAN | Unmodified | 151520 | 0 | 1611700 | 1268400 | 1867200 | 945470 | 0 | 0 | 2838600 | 677900 | 875740 | 21015000 | 134940 | 52632 | 1388000 | 845010 | 447030 | 219040 |
| LRGWFDAI | RPA1; RPA1 | P27694; I3L | RFA1_HUMAN | Unmodified | 0 | 0 | 0 | 0 | 2627900 | 0 | 522840 | 0 | 0 | 163520 | 0 | 0 | 1176100 | 0 | 0 | 44954 | 2647800 | 0 |
| LRGYDYGF | FASN; FASN | P49327; A0A | FAS_HUMAN F | Unmodified | 0 | 0 | 0 | 0 | 2196900 | 0 | 489750 | 0 | 1411400 | 567430 | 1549100 | 11768000 | 0 | 1575700 | 492910 | 0 | 0 | 1393200 |
| LRHAESVG | LRPAP1 | P30533 | AMRP_HUMAN | Unmodified | 0 | 0 | 0 | 0 | 0 | 0 | 0 | 0 | 2804500 | 438650 | 0 | 0 | 0 | 0 | 0 | 0 | 0 | 0 |
| LRHAESVG | LRPAP1 | P30533 | AMRP_HUMAN | Unmodified | 0 | 0 | 0 | 0 | 0 | 0 | 0 | 0 | 1445400 | 1760500 | 0 | 0 | 0 | 0 | 0 | 0 | 0 | 0 |
| LRHEILLSQ | ALDH18A1 | P54886 | P5CS_HUMAN | Unmodified | 0 | 0 | 390150 | 0 | 0 | 0 | 0 | 0 | 365650 | 0 | 0 | 0 | 0 | 0 | 0 | 0 | 0 | 0 |
| LRHPVLGTI | MAN1C1 | Q9NR34 | MA1C1_HUMAN | Unmodified | 0 | 0 | 0 | 0 | 0 | 0 | 0 | 0 | 539660 | 148990 | 0 | 0 | 0 | 0 | 0 | 0 | 0 | 0 |
| LRHTCEQC | MOGS; MOGS; | Q13724; A0A | MOGS_HUMAN | Unmodified | 0 | 0 | 0 | 0 | 0 | 0 | 0 | 0 | 6032900 | 0 | 0 | 0 | 0 | 0 | 0 | 0 | 0 | 0 |
| LRHTHVQE | EIF5B; EIF5B | O60841; A0A | IF2P_HUMAN E | Unmodified | 0 | 0 | 297830 | 0 | 2437700 | 0 | 0 | 0 | 0 | 0 | 0 | 1165000 | 5628500 | 2298000 | 0 | 860550 | 0 | 2861500 |
| LRHVGSNL | GALNT2 | Q10471 | GALT2_HUMAN | Unmodified | 0 | 0 | 0 | 0 | 0 | 0 | 0 | 0 | 615110 | 102730 | 0 | 0 | 0 | 0 | 0 | 0 | 0 | 0 |
| LRHVGSNL | GALNT2 | Q10471 | GALT2_HUMAN | Unmodified | 0 | 0 | 0 | 0 | 616630 | 15364 | 0 | 0 | 0 | 3217000 | 0 | 0 | 620140 | 0 | 0 | 0 | 0 | 0 |
| LRIAWPPP | LIMA1; LIMA1; L | Q9UHB6; F8 | LIMA1_HUMAN | Unmodified | 0 | 0 | 0 | 679160 | 6260300 | 513590 | 0 | 0 | 0 | 0 | 0 | 0 | 1592400 | 1347500 | 0 | 0 | 378830 | 421300 |
| LRIPQVTPA | HSPG2 | P98160 | PGBM_HUMAN | Unmodified | 0 | 0 | 721550 | 0 | 0 | 0 | 0 | 0 | 0 | 0 | 0 | 0 | 0 | 0 | 0 | 440880 | 0 | 0 |
| LRISPDRVY | MIF | P14174 | MIF_HUMAN M | Oxidation (M) | 0 | 0 | 0 | 0 | 0 | 0 | 0 | 0 | 2114400 | 0 | 0 | 0 | 0 | 0 | 0 | 0 | 4148800 | 6584600 |
| LRITESEEV | LMNA; LMNA; L | P02545; A0A | LMNA_HUMAN | Unmodified | 0 | 0 | 0 | 0 | 5275600 | 550360 | 0 | 0 | 116650 | 339840 | 0 | 0 | 1553000 | 0 | 0 | 0 | 799910 | 1398400 |
| LRIWDTTQ | WDR1; WDR1 | O75083; D6I | WDR1_HUMAN | Unmodified | 296150 | 0 | 228800 | 437200 | 6013300 | 2079300 | 318810 | 127990 | 639430 | 485380 | 870840 | 3488700 | 1664900 | 1376300 | 359080 | 276830 | 393360 | 942730 |
| LRIWDTTQ | WDR1; WDR1 | O75083; D6I | WDR1_HUMAN | Unmodified | 0 | 0 | 0 | 0 | 4231300 | 298370 | 478830 | 0 | 0 | 329270 | 0 | 0 | 2623400 | 1565400 | 0 | 0 | 997600 | 766180 |
| LRIWRDYV | SND1 | Q7KZF4 | SND1_HUMAN | Unmodified | 0 | 0 | 0 | 34754 | 0 | 0 | 108590 | 0 | 0 | 0 | 0 | 0 | 408610 | 0 | 28502 | 0 | 0 | 0 |
| LRKDDPLTI | ACTN1; ACTN1 | P12814; G3\ | ACTN1_HUMAI | Unmodified | 438840 | 322400 | 0 | 2252100 | 39283000 | 8714100 | 6410600 | 1397900 | 1808700 | 1392900 | 4063200 | 19861000 | 11127000 | 9366500 | 1528900 | 947620 | 12721000 | 23502000 |
| LRKDDPVT | ACTN4; ACTN4 | O43707; F5C | ACTN4_HUMAI | Unmodified | 241000 | 276810 | 1083100 | 2071800 | 28487000 | 8685700 | 2291000 | 1100500 | 5733200 | 3621400 | 3782900 | 13716000 | 9151500 | 10941000 | 1669900 | 430790 | 9372100 | 8565300 |
| LRKLELSDI | ANP32E; ANP3 | Q9BTT0; Q5 | AN32E_HUMAN | Unmodified | 0 | 0 | 0 | 0 | 6433800 | 93665 | 162830 | 0 | 0 | 1369700 | 466140 | 0 | 1906700 | 259560 | 0 | 0 | 203890 | 337670 |
| LRLDSIVIQ | SMARCA1; SMA | P28370; O6C | SMCA1_HUMAN | Unmodified | 985730 | 1459300 | 167370 | 0 | 0 | 0 | 105660 | 0 | 0 | 95757 | 0 | 0 | 0 | 0 | 0 | 0 | 0 | 0 |
| LRLECADLI | DARS2; DARS2 | Q6PI48; A0A | SYDM_HUMAN | Unmodified | 0 | 0 | 0 | 0 | 0 | 0 | 0 | 0 | 0 | 0 | 0 | 0 | 0 | 0 | 0 | 0 | 0 | 0 |
| LRLEHEQA | DTNA; DTNB; D | Q9Y4J8; O6C | DTNA_HUMAN | Unmodified | 91546 | 40076 | 375440 | 451080 | 237810 | 0 | 17396 | 0 | 3135100 | 305720 | 287310 | 682590 | 305180 | 143470 | 430740 | 855880 | 71724 | 123990 |
| LRLEVNLQ | MYH9 | P35579 | MYH9_HUMAN | Oxidation (M) | 108510 | 53796 | 0 | 0 | 601890 | 0 | 0 | 0 | 0 | 322560 | 0 | 294680 | 301520 | 0 | 182640 | 0 | 120040 | 155060 |
| LRLEWPYQ | HPSE; HPSE; HI | Q9Y251; D6I | HPSE_HUMAN | Unmodified | 0 | 0 | 0 | 0 | 0 | 0 | 0 | 0 | 513190 | 21074 | 0 | 0 | 0 | 0 | 0 | 0 | 0 | 0 |
| LRLGGPNC | OS9 | Q13438 | OS9_HUMAN P | Unmodified | 0 | 0 | 0 | 0 | 0 | 0 | 0 | 0 | 1264400 | 0 | 0 | 0 | 0 | 0 | 0 | 0 | 0 | 0 |
| LRLGYAVYI | TOLLIP; TOLLIP | Q9H0E2; E9I | TOLIP_HUMAN | Unmodified | 1721400 | 1495800 | 2117300 | 2054200 | 0 | 0 | 0 | 0 | 0 | 0 | 0 | 0 | 3275500 | 289760 | 3216100 | 2225500 | 0 | 1715700 |
| LRLHSLGPI | IGSF8 | Q969P0 | IGSF8_HUMAN | Unmodified | 4481000 | 0 | 285210 | 1297600 | 0 | 0 | 0 | 0 | 0 | 0 | 0 | 591240 | 286020 | 0 | 1206600 | 1045500 | 59732 | 0 |
| LRLNVDEA | RRAS | P10301 | RRAS_HUMAN | Unmodified | 277790 | 0 | 178810 | 652530 | 0 | 108690 | 0 | 0 | 2694000 | 0 | 0 | 0 | 0 | 0 | 1209700 | 301410 | 1109700 | 1501400 |
| LRLQAEEV/ | PLEC | Q15149 | PLEC_HUMAN | Unmodified | 0 | 0 | 42058 | 46427 | 570420 | 229080 | 0 | 0 | 24775 | 146780 | 45657 | 377040 | 182040 | 0 | 0 | 0 | 730150 | 1115400 |
| LRLQEEITK | USP14; USP14 | P54578; A6N | UBP14_HUMAN | Unmodified | 0 | 0 | 0 | 0 | 0 | 0 | 0 | 0 | 0 | 0 | 0 | 1741400 | 0 | 0 | 0 | 0 | 0 | 0 |
| LRLQEEITK | USP14; USP14 | P54578; A6N | UBP14_HUMAN | Unmodified | 0 | 0 | 0 | 0 | 825680 | 0 | 0 | 0 | 0 | 234120 | 0 | 0 | 1639700 | 574370 | 0 | 0 | 375670 | 188940 |
| LRLQMAAD | TAX1BP1; TAX1 | Q86VP1; B8 | TAXB1_HUMAN | Unmodified | 0 | 0 | 0 | 0 | 70273 | 0 | 257620 | 0 | 155340 | 0 | 0 | 0 | 0 | 0 | 0 | 0 | 0 | 0 |
| LRLREDEN | DBN1; DBN1 | Q16643; D6I | DREB_HUMAN | Unmodified | 0 | 0 | 0 | 0 | 2154600 | 150700 | 0 | 0 | 146440 | 211160 | 261360 | 1721500 | 0 | 850050 | 0 | 0 | 0 | 659000 |
| LRLTPQHD | DCP1A | Q9NPI6 | DCP1A_HUMAI | Unmodified | 0 | 0 | 0 | 0 | 0 | 0 | 0 | 0 | 0 | 0 | 0 | 0 | 0 | 156820 | 0 | 0 | 0 | 103570 |
| LRLTVVDTF | SEPTIN2; SEPTI | Q15019; C9I | SEPT2_HUMAN | Unmodified | 0 | 0 | 451640 | 0 | 0 | 0 | 0 | 31452 | 0 | 0 | 154360 | 1852600 | 0 | 0 | 393460 | 293740 | 0 | 0 |
| LRLTVVDTF | SEPTIN2; SEPTI | Q15019; C9I | SEPT2_HUMAN | Unmodified | 0 | 0 | 0 | 0 | 2779700 | 568350 | 0 | 0 | 0 | 553470 | 0 | 0 | 1485500 | 0 | 0 | 0 | 2585500 | 0 |
| LRLVEIAQV | UGP2; UGP2; U | Q16851; E7E | UGPA_HUMAN | Unmodified | 0 | 51555 | 0 | 0 | 4499100 | 0 | 659950 | 490050 | 4723300 | 2666900 | 592860 | 2321900 | 1705600 | 864260 | 208280 | 132330 | 867830 | 1364100 |
| LRMDLIRL( | TAOK3; TAOK3 | Q9H2K8; G3 | TAOK3_HUMAN | Unmodified | 0 | 0 | 0 | 0 | 2106200 | 0 | 736260 | 0 | 0 | 0 | 0 | 0 | 567410 | 580520 | 0 | 0 | 0 | 430610 |
| LRMDLIRL( | TAOK3; TAOK3 | Q9H2K8; G3 | TAOK3_HUMAN | Oxidation (M) | 0 | 0 | 0 | 0 | 5009900 | 0 | 0 | 0 | 0 | 2346800 | 0 | 0 | 2107600 | 4244200 | 0 | 0 | 1661800 | 0 |
| LRNAGNEC | CTNNA1; CTNN | P35221; E5R | CTNA1_HUMAI | Unmodified | 739000 | 0 | 2932000 | 3141400 | 7708400 | 3086600 | 44553 | 0 | 476740 | 600090 | 1016200 | 8363100 | 3957700 | 4882400 | 3643900 | 6084900 | 0 | 329410 |
| LRNEDYHE | SLC36A1; SLC3 | Q7Z2H8; E5I | S36A1_HUMAN | Unmodified | 0 | 0 | 0 | 87874 | 0 | 0 | 0 | 0 | 3455100 | 0 | 0 | 0 | 0 | 0 | 179860 | 0 | 0 | 0 |
| LRNELQTD | ARMT1 | Q9H993 | ARMT1_HUMAI | Unmodified | 0 | 0 | 0 | 0 | 0 | 0 | 0 | 0 | 0 | 0 | 0 | 4827300 | 0 | 0 | 0 | 298700 | 0 | 0 |
| LRNIELICQ | MAPRE1 | Q15691 | MARE1_HUMAI | Unmodified | 636140 | 0 | 220180 | 704140 | 683930 | 0 | 0 | 0 | 0 | 341850 | 1503400 | 7729500 | 1066300 | 0 | 270270 | 302790 | 1026200 | 725490 |
| LRNWDPG( | CEMIP2; CEMIP | Q9UHN6; F5 | CEIP2_HUMAN | Unmodified | 0 | 0 | 0 | 0 | 265620 | 119850 | 40525 | 29909 | 19618000 | 1876500 | 0 | 0 | 0 | 0 | 0 | 0 | 0 | 0 |
| LRPDANSA | FARP1; FARP1; | Q9Y4F1; A0A | FARP1_HUMAN | Unmodified | 0 | 0 | 0 | 0 | 0 | 76537 | 0 | 0 | 615040 | 0 | 0 | 91808 | 0 | 0 | 2958800 | 0 | 0 | 0 |
| LRPDANSA | FARP1; FARP1; | Q9Y4F1; A0A | FARP1_HUMAN | Oxidation (M) | 143030 | 322130 | 69899 | 462090 | 0 | 0 | 0 | 0 | 0 | 0 | 0 | 329320 | 0 | 0 | 3935800 | 1828500 | 0 | 0 |
| LRPEVLQD | HPCAL1; HPCA | P37235; E9P | HPCL1_HUMAI | Unmodified | 0 | 0 | 0 | 0 | 3637100 | 0 | 0 | 0 | 0 | 0 | 0 | 0 | 1.07E+08 | 66564000 | 0 | 0 | 0 | 0 |
| LRPGALGG | PLCB3 | Q01970 | PLCB3_HUMAN | Unmodified | 0 | 0 | 0 | 0 | 1489300 | 271670 | 0 | 0 | 424070 | 0 | 201730 | 2424400 | 0 | 0 | 0 | 0 | 0 | 575140 |
| LRPGSIETY | DUSP23 | Q9BVJ7 | DUS23_HUMAI | Unmodified | 323970 | 313460 | 0 | 133660 | 58264 | 0 | 0 | 0 | 0 | 0 | 0 | 0 | 0 | 0 | 345350 | 0 | 140390 | 0 |
| LRPHFLDV | PLPP1 | O14494 | PLPP1_HUMAN | Unmodified | 0 | 0 | 3697000 | 3044400 | 0 | 0 | 0 | 14671 | 0 | 0 | 0 | 0 | 0 | 0 | 2532300 | 1584300 | 0 | 0 |
| LRPIPITASV | ITGA6 | P23229 | ITA6_HUMAN I | Unmodified | 0 | 0 | 42587 | 289340 | 1949600 | 0 | 0 | 0 | 0 | 0 | 41598 | 994730 | 1204800 | 1042900 | 625530 | 302550 | 2527600 | 12827000 |
| LRPIPITASV | ITGA6 | P23229 | ITA6_HUMAN I | Unmodified | 0 | 0 | 0 | 1011900 | 3725000 | 0 | 0 | 0 | 0 | 0 | 0 | 0 | 3419300 | 1688300 | 0 | 2343000 | 4245500 | 19261000 |
| LRPLYDIPY | SND1 | Q7KZF4 | SND1_HUMAN | Oxidation (M) | 300360 | 605830 | 300360 | 428700 | 0 | 0 | 0 | 0 | 0 | 0 | 0 | 2761700 | 0 | 0 | 511410 | 435430 | 0 | 0 |
| LRPNFLAV( | PLPP2; PLPP2; | O43688; K7I | PLPP2_HUMAN | Unmodified | 0 | 0 | 0 | 0 | 0 | 0 | 0 | 0 | 0 | 0 | 0 | 0 | 0 | 0 | 0 | 0 | 0 | 0 |
| LRPNIIPVLI | PROM1 | O43490 | PROM1_HUMA | Unmodified | 0 | 0 | 0 | 0 | 0 | 0 | 0 | 0 | 0 | 0 | 0 | 15972000 | 0 | 0 | 0 | 0 | 0 | 0 |
| LRPNIPNW | ACVR1B | P36896 | ACV1B_HUMAN | Unmodified | 444580 | 489390 | 0 | 0 | 0 | 0 | 0 | 0 | 0 | 0 | 0 | 0 | 0 | 0 | 856610 | 773530 | 0 | 0 |
| LRPPPARTI | IGFBP2; IGFBP2 | P18065; C9J | IBP2_HUMAN I | Unmodified | 0 | 0 | 0 | 0 | 0 | 0 | 0 | 0 | 34159000 | 11819000 | 0 | 0 | 0 | 0 | 0 | 0 | 0 | 0 |
| LRPQTYDL( | SEPTIN8; SEPTI | Q92599; C9J | SEPT8_HUMAN | Unmodified | 121270 | 0 | 0 | 0 | 1344200 | 0 | 0 | 0 | 657200 | 194240 | 0 | 4723900 | 1332900 | 0 | 0 | 0 | 172720 | 272500 |
| LRPYGAPG | KDF1 | Q8NAX2 | KDF1_HUMAN | Unmodified | 0 | 0 | 0 | 0 | 0 | 0 | 0 | 0 | 0 | 0 | 0 | 0 | 300820 | 459730 | 67316 | 41482 | 0 | 0 |
| LRQAERLFI | APMAP; APMAP | Q9HDC9; H( | APMAP_HUMAI | Unmodified | 0 | 0 | 0 | 0 | 0 | 0 | 0 | 0 | 1988100 | 80421 | 0 | 0 | 0 | 0 | 0 | 0 | 0 | 1286300 |
| LRQAERLFI | APMAP; APMAP | Q9HDC9; H( | APMAP_HUMAI | Oxidation (M) | 0 | 0 | 0 | 0 | 0 | 0 | 0 | 0 | 573970 | 2386000 | 0 | 0 | 0 | 0 | 0 | 0 | 0 | 0 |
| LRQDPLIF | IGSF9 | Q9P2J2 | TUTLA_HUMAN | Unmodified | 0 | 0 | 0 | 0 | 0 | 0 | 0 | 0 | 0 | 76316 | 0 | 900590 | 0 | 0 | 0 | 0 | 0 | 0 |

| | | | | | | | | | | | | | | | | | | | | | | |
|---|---|---|---|---|---|---|---|---|---|---|---|---|---|---|---|---|---|---|---|---|---|---|
| LRQEFAQH | SPTAN1; SPTAN | Q13813; A0A | SPTN1_HUMAN | Unmodified | 0 | 0 | 0 | 0 | 0 | 0 | 0 | 0 | 282060 | 0 | 78870 | 4511500 | 0 | 0 | 0 | 0 | 0 | 0 |
| LRQENMEL | TXLNA | P40222 | TXLNA_HUMAN | Unmodified | 0 | 0 | 0 | 0 | 628310 | 0 | 0 | 0 | 0 | 0 | 0 | 0 | 390000 | 0 | 0 | 0 | 0 | 347850 |
| LRQHHDEY | CPD | O75976 | CBPD_HUMAN | Unmodified | 0 | 0 | 0 | 0 | 0 | 0 | 0 | 0 | 0 | 0 | 0 | 0 | 0 | 0 | 0 | 0 | 0 | 0 |
| LRQIFNGTI | RPL7; RPL7; RP | P18124; C9J | RL7_HUMAN 60 | Unmodified | 361230 | 244520 | 175610 | 0 | 0 | 0 | 0 | 0 | 0 | 0 | 0 | 7312700 | 0 | 0 | 436230 | 0 | 0 | 0 |
| LRQIGPGES | CDCP1 | Q9H5V8 | CDCP1_HUMA | Unmodified | 64097 | 0 | 43980 | 30598 | 0 | 69493 | 0 | 0 | 0 | 88362 | 0 | 2766600 | 0 | 61467 | 836470 | 265270 | 0 | 0 |
| LRQIGPGES | CDCP1 | Q9H5V8 | CDCP1_HUMA | Unmodified | 0 | 0 | 0 | 0 | 634840 | 0 | 0 | 0 | 0 | 0 | 0 | 0 | 0 | 0 | 0 | 0 | 672630 | 757830 |
| LRQLAEEDI | PLEC | Q15149 | PLEC_HUMAN | Unmodified | 0 | 0 | 0 | 0 | 296780 | 0 | 0 | 0 | 0 | 0 | 0 | 0 | 0 | 0 | 0 | 0 | 606000 | 848750 |
| LRQLLEQPI | UTRN; UTRN | P46939; Q5T | UTRN_HUMAN | Unmodified | 0 | 0 | 0 | 0 | 0 | 0 | 230820 | 17043 | 0 | 150850 | 0 | 0 | 0 | 0 | 324400 | 162890 | 0 | 0 |
| LRQLTAELA | PSD2 | Q9BQI7 | PSD2_HUMAN | Unmodified | 0 | 0 | 0 | 0 | 0 | 0 | 0 | 0 | 0 | 0 | 250470 | 0 | 0 | 0 | 0 | 0 | 0 | 0 |
| LRQMEAQL | PPFIA1; PPFIA1 | Q13136; E9F | LIPA1_HUMAN | Unmodified | 0 | 0 | 95572 | 0 | 0 | 0 | 0 | 0 | 291110 | 22475 | 68938 | 0 | 0 | 0 | 101810 | 18358 | 0 | 0 |
| LRQMHLMI | SMG9 | Q9H0W8 | SMG9_HUMAN | Unmodified | 0 | 0 | 0 | 0 | 0 | 0 | 0 | 0 | 0 | 0 | 91852 | 0 | 0 | 0 | 0 | 310680 | 0 | 0 |
| LRQNLDAL | DYNC1H1; DYN | Q14204; A0A | DYHC1_HUMA | Unmodified | 0 | 0 | 0 | 0 | 0 | 0 | 0 | 0 | 0 | 0 | 0 | 908430 | 0 | 0 | 0 | 66039 | 0 | 0 |
| LRQVEELQ | PYM1 | Q9BRP8 | PYM1_HUMAN | Unmodified | 0 | 0 | 81095 | 77682 | 0 | 0 | 0 | 0 | 0 | 0 | 0 | 0 | 0 | 0 | 179350 | 429630 | 0 | 0 |
| LRSELEMVS | TOM1; TOM1 | O60784; B0 | TOM1_HUMAN | Unmodified | 786510 | 728100 | 770620 | 928200 | 162600 | 0 | 0 | 0 | 663690 | 52856 | 0 | 454590 | 243650 | 0 | 889970 | 434680 | 0 | 0 |
| LRSGEPQT | ITGB1; ITGB1; I | P05556; E7E | ITB1_HUMAN I | Unmodified | 1569900 | 1465000 | 1610800 | 3136200 | 1.66E+08 | 35319000 | 1072200 | 492430 | 13693000 | 5302000 | 1363700 | 11044000 | 10078000 | 7289300 | 3241700 | 1898500 | 46279000 | 1.59E+08 |
| LRSGEPQT | ITGB1; ITGB1; I | P05556; E7E | ITB1_HUMAN I | Unmodified | 0 | 0 | 0 | 0 | 762250 | 0 | 0 | 0 | 0 | 0 | 0 | 0 | 0 | 0 | 0 | 0 | 1327600 | 1088900 |
| LRSIPLDEG | DYNC1H1; DYN | Q14204; A0A | DYHC1_HUMA | Unmodified | 0 | 0 | 0 | 74355 | 0 | 147200 | 0 | 0 | 245170 | 0 | 122090 | 2139600 | 221180 | 45744 | 71701 | 62184 | 0 | 0 |
| LRSPVISIDI | HSPG2 | P98160 | PGBM_HUMAN | Unmodified | 0 | 0 | 7693200 | 1978700 | 797530 | 1530600 | 0 | 0 | 864290 | 0 | 0 | 415200 | 0 | 1788200 | 523940 | 1522000 | 0 | 6073300 |
| LRSQALNA | DNAJC3 | Q13217 | DNJC3_HUMAI | Unmodified | 0 | 0 | 0 | 0 | 0 | 0 | 0 | 0 | 7241200 | 2143900 | 0 | 0 | 0 | 0 | 0 | 0 | 0 | 0 |
| LRSRGPAT | GIPC1; GIPC1 | O14908; K7E | GIPC1_HUMAN | Unmodified | 86137 | 0 | 0 | 0 | 2792500 | 451470 | 0 | 0 | 0 | 0 | 0 | 0 | 574380 | 468940 | 0 | 0 | 0 | 2853700 |
| LRSYEDMI( | CTNND1; CTNN | O60716; A0A | CTND1_HUMAI | Oxidation (M) | 269200 | 0 | 0 | 3571600 | 0 | 0 | 0 | 0 | 0 | 0 | 0 | 0 | 0 | 0 | 2602600 | 1778400 | 0 | 0 |
| LRTAGPLES | RRBP1; RRBP1; | Q9P2E9; A0A | RRBP1_HUMAN | Unmodified | 0 | 0 | 0 | 0 | 0 | 0 | 0 | 0 | 0 | 0 | 0 | 0 | 2346300 | 381710 | 0 | 0 | 0 | 0 |
| LRTAGTICL | GSPT1; GSPT2; | P15170; Q8I | ERF3A_HUMAN | Unmodified | 215750 | 0 | 0 | 0 | 565920 | 0 | 0 | 0 | 0 | 49548 | 0 | 955220 | 466080 | 0 | 63936 | 0 | 0 | 0 |
| LRTIVSIGDI | PAK2 | Q13177 | PAK2_HUMAN | Unmodified | 0 | 0 | 0 | 0 | 2318400 | 208320 | 0 | 0 | 156600 | 132070 | 0 | 0 | 735450 | 392520 | 0 | 0 | 399720 | 461940 |
| LRTLHPAPI | PLCD3 | Q8N3E9 | PLCD3_HUMAI | Unmodified | 0 | 0 | 0 | 0 | 669770 | 0 | 452350 | 0 | 0 | 0 | 0 | 0 | 2201900 | 375430 | 0 | 0 | 782890 | 1382900 |
| LRTLWIWE | RNH1; RNH1 | P13489; H0Y | RINI_HUMAN R | Unmodified | 0 | 0 | 0 | 0 | 998510 | 136290 | 0 | 0 | 0 | 169700 | 279810 | 0 | 0 | 0 | 0 | 0 | 0 | 992210 |
| LRTYLPDFF | GFUS; GFUS | Q13630; H0Y | FCL_HUMAN G | Unmodified | 0 | 0 | 0 | 0 | 1722300 | 413780 | 0 | 0 | 0 | 906790 | 0 | 0 | 1727000 | 0 | 0 | 0 | 3481700 | 0 |
| LRVDKAAA | RPL4; RPL4; RP | P36578; H3E | RL4_HUMAN 60 | Unmodified | 0 | 0 | 0 | 0 | 0 | 0 | 0 | 0 | 0 | 0 | 0 | 0 | 594790 | 0 | 0 | 0 | 0 | 0 |
| LRVHLDIQ\ | TMED4 | Q7Z7H5 | TMED4_HUMAI | Unmodified | 0 | 0 | 0 | 0 | 2584100 | 0 | 12884000 | 541900 | 6523100 | 36891000 | 0 | 0 | 0 | 0 | 0 | 0 | 449270 | 1604300 |
| LRVKIEPGV | RNPEP; RNPEP; | Q9H4A4; A6I | AMPB_HUMAN | Unmodified | 0 | 0 | 0 | 0 | 1677200 | 0 | 0 | 268030 | 0 | 0 | 0 | 0 | 0 | 0 | 0 | 0 | 0 | 0 |
| LRVSEWEV | USP13; USP13; | Q92995; A0A | UBP13_HUMAN | Unmodified | 625330 | 609970 | 670120 | 774100 | 232490 | 0 | 0 | 0 | 0 | 0 | 0 | 0 | 826460 | 0 | 370470 | 360580 | 197220 | 174290 |
| LRYEIDTGE | HSPA14; HSPA1 | Q0VDF9; H7 | HSP7E_HUMAN | Unmodified | 0 | 0 | 0 | 0 | 0 | 0 | 0 | 0 | 0 | 0 | 0 | 1031800 | 0 | 0 | 0 | 0 | 0 | 0 |
| LRYPINEEA | PLCG1 | P19174 | PLCG1_HUMAI | Unmodified | 0 | 0 | 0 | 0 | 0 | 24667 | 0 | 0 | 377690 | 0 | 616980 | 1428600 | 519270 | 0 | 0 | 0 | 606740 | 446450 |
| LRYPVTPEL | PLCG2; PLCG2 | P16885; A0A | PLCG2_HUMAI | Unmodified | 0 | 0 | 0 | 0 | 0 | 0 | 1150300 | 0 | 0 | 0 | 0 | 8670700 | 0 | 0 | 0 | 0 | 0 | 0 |
| LRYVANLFI | PPA1; PPA1 | Q15181; Q5 | IPYR_HUMAN I | Unmodified | 390210 | 317860 | 650090 | 2253700 | 21194000 | 2708500 | 1829600 | 188000 | 3671900 | 2054800 | 5519400 | 9028300 | 10856000 | 4154600 | 588890 | 468720 | 9821200 | 7763700 |
| LRYVDQVL | IGF2R | P11717 | MPRI_HUMAN | Unmodified | 0 | 0 | 0 | 115890 | 0 | 0 | 0 | 0 | 5135000 | 667890 | 543810 | 224570 | 39674 | 0 | 87812 | 0 | 0 | 0 |
| LRYVDQVL | IGF2R | P11717 | MPRI_HUMAN | Unmodified | 0 | 0 | 0 | 0 | 0 | 0 | 0 | 0 | 1139200 | 220320 | 0 | 755160 | 1368300 | 0 | 0 | 0 | 0 | 0 |
| LSAAGGGT | OLFM1 | Q99784 | NOE1_HUMAN | Unmodified | 0 | 402220 | 0 | 0 | 125390 | 0 | 0 | 45261 | 1517900 | 51147 | 0 | 152870 | 0 | 0 | 0 | 0 | 213040 | 275680 |
| LSAASGYSI | PMS2P11 | Q13670 | PM2PB_HUMAI | Unmodified | 0 | 0 | 0 | 0 | 919800 | 0 | 0 | 0 | 1744900 | 1284400 | 0 | 0 | 0 | 0 | 0 | 0 | 243000 | 536640 |
| LSAASTWLI | HYOU1; HYOU1 | Q9Y4L1; A0A | HYOU1_HUMA | Oxidation (M) | 0 | 0 | 0 | 0 | 0 | 24816 | 0 | 57504 | 8070200 | 5508900 | 2430300 | 0 | 0 | 0 | 0 | 0 | 0 | 0 |
| LSAASTWLI | HYOU1; HYOU1 | Q9Y4L1; A0A | HYOU1_HUMA | Unmodified | 0 | 0 | 0 | 0 | 931240 | 0 | 0 | 0 | 4509100 | 111880 | 0 | 0 | 0 | 0 | 0 | 0 | 194710 | 0 |
| LSAAVTEAF | VARS1; VARS1 | P26640; A0A | SYVC_HUMAN | Unmodified | 289260 | 0 | 0 | 0 | 0 | 0 | 0 | 0 | 0 | 0 | 921020 | 12725000 | 0 | 0 | 0 | 531140 | 0 | 0 |
| LSADTYEDI | CMIP; CMIP; CN | Q8IY22; A0A | CMIP_HUMAN | Unmodified | 0 | 0 | 0 | 0 | 2288700 | 384810 | 0 | 371030 | 431980 | 124920 | 0 | 4284200 | 0 | 0 | 0 | 0 | 264760 | 695390 |
| LSAEDLVLE | LAMC1 | P11047 | LAMC1_HUMAI | Unmodified | 262500 | 115760 | 117680 | 65446 | 0 | 0 | 0 | 0 | 870630 | 0 | 73854 | 139660 | 0 | 0 | 145680 | 182900 | 0 | 0 |
| LSAEERDQ | PCBD1 | P61457 | PHS_HUMAN P | Unmodified | 0 | 0 | 1064300 | 0 | 656000 | 0 | 0 | 0 | 14178000 | 0 | 2643300 | 12384000 | 0 | 0 | 3844600 | 0 | 0 | 192620 |
| LSAEKVEIA | PSMA4; PSMA4 | P25789; H0Y | PSA4_HUMAN | Unmodified | 0 | 345300 | 0 | 0 | 431500 | 0 | 0 | 0 | 1296000 | 0 | 0 | 3176800 | 53275 | 0 | 249730 | 761050 | 0 | 0 |
| LSAIEIDIPV | CLDN12 | P56749 | CLD12_HUMAN | Unmodified | 3199900 | 2325600 | 1821600 | 4790700 | 2444700 | 0 | 0 | 0 | 5270000 | 2097900 | 0 | 0 | 3969600 | 1766600 | 8000900 | 4530100 | 0 | 0 |
| LSAIFRDFL | PAK1; PAK1; PA | Q13153; E9F | PAK1_HUMAN | Unmodified | 0 | 0 | 0 | 0 | 0 | 0 | 0 | 0 | 318270 | 102700 | 72526 | 320220 | 0 | 0 | 0 | 0 | 0 | 0 |
| LSAIYGGTY | GDI2; GDI2; GD | P50395; Q5 | GDIB_HUMAN | Oxidation (M) | 0 | 0 | 0 | 0 | 0 | 0 | 0 | 0 | 2008500 | 726090 | 327610 | 2340100 | 0 | 0 | 0 | 0 | 0 | 0 |
| LSALHLEVI | PACSIN3; PACS | Q9UKS6; E9I | PACN3_HUMAI | Unmodified | 1921300 | 2015200 | 658660 | 346500 | 75868 | 0 | 0 | 0 | 0 | 0 | 0 | 0 | 0 | 0 | 2695000 | 1328400 | 0 | 0 |
| LSALHLEVI | PACSIN3; PACS | Q9UKS6; E9I | PACN3_HUMAI | Unmodified | 3121700 | 3163100 | 1207200 | 242180 | 132690 | 0 | 218250 | 0 | 0 | 0 | 0 | 0 | 218380 | 239080 | 2391900 | 106580 | 0 | 0 |
| LSALLQETK | USO1 | O60763 | USO1_HUMAN | Unmodified | 0 | 0 | 118800 | 0 | 615100 | 1450200 | 0 | 328180 | 0 | 293290 | 371340 | 5974200 | 425050 | 1340700 | 0 | 0 | 0 | 1238300 |
| LSALRPIVN | DPY19L1; DPY1 | Q2PZI1; A0A | D19L1_HUMAN | Unmodified | 0 | 0 | 0 | 0 | 409210 | 0 | 0 | 0 | 3857800 | 364080 | 0 | 0 | 0 | 0 | 0 | 0 | 0 | 0 |
| LSALYKLES | IARS1; IARS1; IA | P41252; A0A | SYIC_HUMAN I | Unmodified | 0 | 0 | 0 | 0 | 0 | 351250 | 0 | 0 | 0 | 47584 | 0 | 558140 | 0 | 0 | 0 | 0 | 0 | 0 |
| LSANQQNI | TMEM33; TMEM | P57088; H0Y | TMM33_HUMAI | Unmodified | 0 | 0 | 73434 | 0 | 0 | 0 | 0 | 0 | 1918700 | 639470 | 270090 | 155440 | 0 | 0 | 97757 | 79307 | 0 | 0 |
| LSAQDPVV | SLC38A10 | Q9HBR0 | S38AA_HUMAN | Unmodified | 0 | 0 | 0 | 0 | 0 | 0 | 0 | 0 | 737520 | 127560 | 0 | 0 | 0 | 0 | 0 | 0 | 0 | 0 |
| LSASLPDT\ | UBTD1 | Q9HAC8 | UBTD1_HUMAI | Unmodified | 0 | 0 | 0 | 0 | 1452900 | 0 | 0 | 0 | 697370 | 191620 | 0 | 3424100 | 0 | 0 | 0 | 0 | 783700 | 1923300 |
| LSATEPELG | MST1R; MST1R | Q04912; J3K | RON_HUMAN N | Unmodified | 32705 | 0 | 32398 | 0 | 0 | 0 | 0 | 0 | 1004300 | 0 | 0 | 0 | 0 | 0 | 0 | 0 | 0 | 0 |
| LSATEPELG | MST1R; MST1R | Q04912; J3K | RON_HUMAN N | Unmodified | 0 | 0 | 0 | 0 | 1234800 | 395470 | 0 | 0 | 0 | 339080 | 98394 | 0 | 105370 | 223140 | 0 | 0 | 0 | 981170 |
| LSATSVLYY | BMP8B; BMP8A | P34820; Q7Z | BMP8B_HUMAI | Unmodified | 0 | 0 | 0 | 0 | 0 | 0 | 0 | 0 | 826440 | 565410 | 0 | 0 | 0 | 0 | 0 | 0 | 0 | 0 |
| LSAVSLRGI | SNX17 | Q15036 | SNX17_HUMAN | Unmodified | 0 | 0 | 0 | 0 | 2543300 | 0 | 0 | 0 | 0 | 0 | 0 | 0 | 0 | 2225800 | 0 | 536830 | 0 | 0 |
| LSCAPPPPI | SLC23A2; SLC2 | Q9UGH3; H | S23A2_HUMAN | Unmodified | 83697 | 0 | 0 | 0 | 0 | 0 | 0 | 0 | 140980 | 598590 | 0 | 596640 | 0 | 0 | 54101 | 142560 | 145680 | 35976 |
| LSCAYSGF | F11R; F11R | Q9Y624; A0A | JAM1_HUMAN | Unmodified | 5485400 | 3918200 | 3356900 | 2112100 | 425190 | 141310 | 0 | 0 | 2706100 | 0 | 577730 | 3216400 | 969590 | 219420 | 4726300 | 4667300 | 150140 | 0 |
| LSCAYSGF | F11R | Q9Y624 | JAM1_HUMAN | Unmodified | 630900 | 356490 | 1482200 | 878460 | 0 | 0 | 0 | 0 | 0 | 0 | 351100 | 317520 | 0 | 52626 | 547310 | 413390 | 46126 | 0 |
| LSCFAQTV | FSCN1 | Q16658 | FSCN1_HUMAI | Unmodified | 0 | 0 | 0 | 0 | 0 | 0 | 0 | 0 | 0 | 0 | 340000 | 7907800 | 0 | 0 | 0 | 0 | 0 | 0 |
| LSCLGASL | MUC5B | Q9HC84 | MUC5B_HUMA | Unmodified | 231780 | 38052 | 70155 | 332360 | 2867900 | 1877200 | 864700 | 964840 | 0 | 0 | 686530 | 0 | 0 | 0 | 173960 | 287550 | 174960 | 9229700 |
| LSCSLEDLI | PPFIBP2; PPFIB | Q8ND30; H | LIPB2_HUMAN | Unmodified | 0 | 0 | 0 | 0 | 0 | 0 | 0 | 0 | 0 | 0 | 0 | 0 | 0 | 367720 | 0 | 0 | 0 | 0 |
| LSCVPVLIF | ARL3 | P36405 | ARL3_HUMAN | Unmodified | 0 | 0 | 0 | 0 | 0 | 0 | 0 | 0 | 0 | 0 | 0 | 1297700 | 0 | 0 | 0 | 0 | 0 | 0 |
| LSDDNTIGI | SPTAN1; SPTAN | Q13813; A0A | SPTN1_HUMAN | Unmodified | 0 | 0 | 0 | 0 | 76159 | 0 | 0 | 0 | 53635 | 0 | 0 | 1880600 | 0 | 0 | 0 | 0 | 58912 | 425550 |
| LSDFWQQ | UBASH3B | Q8TF42 | UBS3B_HUMAI | Unmodified | 0 | 0 | 0 | 0 | 932670 | 200810 | 58573 | 777650 | 0 | 0 | 0 | 0 | 0 | 0 | 0 | 0 | 650920 | 62296 |
| LSDGFNGA | PSMC6; PSMC6 | P62333; A0A | PRS10_HUMAN | Unmodified | 0 | 0 | 0 | 0 | 0 | 0 | 0 | 0 | 0 | 0 | 0 | 3017400 | 0 | 0 | 0 | 0 | 0 | 0 |
| LSDGGLLL | ACACA | Q13085 | ACACA_HUMAI | Oxidation (M) | 0 | 17329000 | 18359000 | 11861000 | 0 | 0 | 0 | 0 | 0 | 0 | 0 | 0 | 0 | 0 | 32895000 | 20468000 | 0 | 0 |
| LSDGVAVL | HSPD1; HSPD1 | P10809; A0A | CH60_HUMAN | Unmodified | 637400 | 506100 | 316250 | 688520 | 2543100 | 312310 | 5832800 | 1788700 | 28138000 | 10669000 | 2406700 | 3375600 | 6706200 | 6298800 | 494600 | 706490 | 811250 | 2698000 |
| LSDGVAVL | HSPD1; HSPD1 | P10809; A0A | CH60_HUMAN | Unmodified | 0 | 0 | 0 | 0 | 0 | 0 | 631890 | 0 | 0 | 0 | 0 | 0 | 0 | 0 | 0 | 0 | 0 | 0 |
| LSDIPEGK | UQCRFS1P1; U | P0C7P4; P47 | UCRIL_HUMAN | Unmodified | 0 | 0 | 0 | 0 | 0 | 0 | 26290 | 0 | 10696000 | 0 | 238090 | 262340 | 0 | 321720 | 0 | 43363 | 57605 | 0 |
| LSDLDSETI | NUDC | Q9Y266 | NUDC_HUMAN | Unmodified | 0 | 0 | 103650 | 0 | 0 | 0 | 0 | 0 | 2543600 | 0 | 2171900 | 5477700 | 0 | 0 | 842760 | 0 | 71610 | 0 |
| LSDLDSETI | NUDC | Q9Y266 | NUDC_HUMAN | Unmodified | 0 | 0 | 0 | 0 | 325450 | 56222 | 251360 | 62153 | 703770 | 727130 | 0 | 0 | 2308000 | 1021800 | 87590 | 0 | 638330 | 1205400 |
| LSDLDSETI | NUDC | Q9Y266 | NUDC_HUMAN | Oxidation (M) | 0 | 0 | 42301 | 0 | 507500 | 27708 | 112020 | 35664 | 0 | 69954 | 0 | 34774 | 276190 | 715840 | 0 | 0 | 354840 | 88758 |
| LSDLLAPIS | CAP1; CAP1; CA | Q01518; Q5 | CAP1_HUMAN | Unmodified | 0 | 0 | 748340 | 0 | 1039800 | 249340 | 0 | 0 | 3211700 | 368780 | 1562400 | 18477000 | 409470 | 0 | 2040900 | 1035000 | 0 | 508020 |
| LSDLLDWK | GYS1; GYS1 | P13807; M0 | GYS1_HUMAN | Unmodified | 0 | 0 | 0 | 0 | 0 | 0 | 0 | 0 | 0 | 0 | 76816 | 1901200 | 0 | 0 | 0 | 0 | 93197 | 0 |
| LSDMPLK | UGGT1 | Q9NYU2 | UGGG1_HUMA | Unmodified | 0 | 0 | 0 | 0 | 0 | 0 | 0 | 0 | 2497900 | 833520 | 0 | 229200 | 0 | 0 | 0 | 0 | 0 | 0 |
| LSDNTPEH | PALD1 | Q9ULE6 | PALD_HUMAN | Unmodified | 0 | 0 | 0 | 0 | 0 | 0 | 0 | 0 | 0 | 0 | 399330 | 4099800 | 0 | 0 | 0 | 0 | 0 | 0 |
| LSDPANWI | CDH2; CDH2; ( | P19022; C9J | CADH2_HUMA | Unmodified | 0 | 0 | 0 | 0 | 0 | 0 | 0 | 0 | 896390 | 262030 | 653690 | 0 | 0 | 206830 | 0 | 0 | 0 | 0 |
| LSDQFHDII | PDCD6; PDCD6 | O75340; H0Y | PDCD6_HUMA | Unmodified | 17300000 | 10286000 | 4083200 | 2702900 | 0 | 0 | 38778 | 0 | 27100000 | 521340 | 192880 | 14670000 | 154690 | 81478 | 13147000 | 10895000 | 0 | 0 |
| LSDQFHDII | PDCD6; PDCD6 | O75340; H0Y | PDCD6_HUMA | Unmodified | 294550 | 721570 | 196280 | 0 | 0 | 0 | 0 | 95342 | 880380 | 0 | 0 | 343010 | 0 | 0 | 285570 | 351960 | 0 | 0 |
| LSDQSANN | TAX1BP1; TAX1 | Q86VP1; B8 | TAXB1_HUMAN | Unmodified | 0 | 0 | 90014 | 0 | 1026900 | 134900 | 526290 | 57310 | 374430 | 120840 | 49941 | 0 | 376150 | 322160 | 45816 | 0 | 463570 | 1896100 |
| LSDSGRYD | NCAM2; NCAM | O15394; H9I | NCAM2_HUMA | Unmodified | 0 | 280190 | 2760200 | 3819900 | 0 | 0 | 0 | 0 | 0 | 0 | 0 | 123720 | 68343 | 0 | 3752200 | 3296200 | 0 | 0 |
| LSDSIAATS | TRIP11; TRIP11 | Q15643; H0Y | TRIPB_HUMAN | Unmodified | 0 | 0 | 0 | 0 | 0 | 0 | 1170800 | 0 | 0 | 0 | 0 | 0 | 0 | 0 | 0 | 0 | 0 | 0 |
| LSDSLGQR | CAST; CAST; CA | P20810; H0Y | ICAL_HUMAN ( | Unmodified | 42803 | 0 | 0 | 0 | 0 | 44271 | 0 | 0 | 219980 | 0 | 241620 | 1729500 | 62838 | 0 | 103810 | 73400 | 0 | 254470 |
| LSDSQKMF | CXADR | P78310 | CXAR_HUMAN | Oxidation (M) | 0 | 0 | 0 | 3632800 | 1095500 | 199210 | 0 | 0 | 0 | 0 | 0 | 0 | 3224700 | 0 | 0 | 0 | 669940 | 0 |
| LSDSWQPI | CD109 | Q6YHK3 | CD109_HUMAI | Unmodified | 0 | 88315 | 0 | 0 | 0 | 0 | 0 | 152170 | 2691200 | 316810 | 0 | 0 | 0 | 0 | 0 | 0 | 0 | 0 |
| LSDSYSNTI | BAIAP2; BAIAP2 | Q9UQB8; I3I | BAIP2_HUMAN | Unmodified | 403780 | 482090 | 1233400 | 2903800 | 6850500 | 2936500 | 504640 | 785660 | 1529800 | 0 | 794490 | 2510300 | 8397000 | 7164100 | 3506100 | 3133900 | 2557800 | 5021600 |
| LSDVDRDG | EHD2 | Q9NZN4 | EHD2_HUMAN | Oxidation (M) | 0 | 0 | 0 | 0 | 3120900 | 0 | 0 | 0 | 0 | 0 | 0 | 0 | 0 | 0 | 0 | 0 | 0 | 0 |
| LSDYLFTLA | MMAB; MMAB | Q96EY8; S4F | MMAB_HUMAN | Unmodified | 0 | 0 | 0 | 0 | 0 | 0 | 0 | 0 | 3188800 | 0 | 0 | 0 | 0 | 0 | 0 | 0 | 0 | 0 |
| LSECLQK | EXOC5; EXOC5 | O00471; F8V | EXOC5_HUMAI | Unmodified | 0 | 0 | 520750 | 0 | 775560 | 118530 | 251330 | 0 | 67739 | 0 | 0 | 281090 | 0 | 0 | 633900 | 461300 | 41858 | 294050 |
| LSEDSLTK( | STK3; STK3; STI | Q13188; Q8I | STK3_HUMAN | Unmodified | 0 | 167990 | 0 | 0 | 510960 | 0 | 164670 | 0 | 441960 | 256270 | 104570 | 83784 | 451090 | 137130 | 0 | 0 | 0 | 0 |
| LSEDYGVLI | PRDX2; PRDX2 | P32119; A6N | PRDX2_HUMAN | Unmodified | 830810 | 417590 | 135800 | 1413300 | 0 | 0 | 0 | 0 | 4342800 | 239440 | 6086200 | 56167000 | 0 | 0 | 316150 | 188140 | 0 | 187070 |
| LSEDYGVLI | PRDX2 | P32119 | PRDX2_HUMAN | Unmodified | 1726200 | 1958500 | 0 | 4728100 | 805590 | 3158000 | 0 | 0 | 16282000 | 880890 | 9779500 | 48406000 | 0 | 0 | 1317300 | 2252900 | 394430 | 2021100 |
| LSEEFLTAH | TBCE; TBCE; TB | Q15813; A0A | TBCE_HUMAN | Unmodified | 0 | 0 | 0 | 0 | 0 | 0 | 0 | 0 | 0 | 0 | 364800 | 9608300 | 0 | 0 | 52947 | 73145 | 0 | 0 |
| LSEEFVDPI | VANGL2 | Q9ULK5 | VANG2_HUMAI | Unmodified | 0 | 0 | 0 | 0 | 723400 | 0 | 0 | 0 | 0 | 0 | 0 | 2066300 | 0 | 324670 | 0 | 0 | 0 | 0 |
| LSEELSGGI | ACTR3B; ACTR3 | Q9P1U1; P6 | ARP3B_HUMAN | Unmodified | 3246000 | 2192300 | 1999900 | 1330300 | 1156500 | 125120 | 150460 | 102780 | 2173200 | 301210 | 1062900 | 26381000 | 1322600 | 384480 | 2884400 | 4682200 | 143120 | 228500 |
| LSEEMTQR | VPS36; VPS36 | Q86VN1; A0 | VPS36_HUMAN | Unmodified | 1549800 | 3233800 | 0 | 0 | 138820 | 350300 | 35151 | 314540 | 77197 | 0 | 0 | 0 | 0 | 384290 | 1075600 | 776420 | 91720 | 3273200 |
| LSEGVTISY | ITGB1; ITGB1; I | P05556; H7 | ITB1_HUMAN I | Unmodified | 5373100 | 4612800 | 8585500 | 5924800 | 59695000 | 10665000 | 0 | 0 | 27569000 | 3425900 | 4231400 | 13519000 | 4529200 | 1195700 | 13674000 | 13301000 | 10692000 | 31927000 |
| LSEHSSPEI | PPP1R1B; PPP1 | Q9UD71; J3I | PPR1B_HUMAN | Unmodified | 29517 | 0 | 0 | 0 | 0 | 0 | 0 | 0 | 355040 | 0 | 2651000 | 2435200 | 0 | 0 | 90444 | 0 | 82587 | 0 |
| LSEIDVSSE | EFHD2; EFHD2 | Q96C19; H0 | EFHD2_HUMAI | Unmodified | 0 | 112030 | 0 | 149150 | 6648000 | 3038200 | 0 | 0 | 3676800 | 1061200 | 791930 | 1580900 | 115240 | 254150 | 105210 | 122870 | 847310 | 1169900 |
| LSEIDVSSE | EFHD2; EFHD2 | Q96C19; H0 | EFHD2_HUMAI | Unmodified | 0 | 0 | 0 | 0 | 0 | 428330 | 0 | 0 | 0 | 167980 | 0 | 214890 | 136920 | 74144 | 0 | 0 | 504820 | 591810 |
| LSEIFAAR | TTYH2; TTYH2 | Q9BSA4; B4I | TTYH2_HUMAN | Unmodified | 0 | 89920 | 193560 | 153950 | 0 | 0 | 0 | 0 | 1417300 | 748120 | 37573 | 675360 | 0 | 0 | 932620 | 1157800 | 0 | 0 |
| LSEIFAARG | TTYH2; TTYH2 | Q9BSA4; B4I | TTYH2_HUMAN | Unmodified | 0 | 0 | 161230 | 0 | 0 | 0 | 0 | 0 | 0 | 0 | 0 | 0 | 0 | 0 | 447360 | 479670 | 0 | 0 |
| LSEIHHLHI | ZC3HAV1; ZC3I | Q7Z2W4; C9 | ZCCHV_HUMA | Unmodified | 0 | 0 | 0 | 4864100 | 0 | 0 | 0 | 0 | 1280600 | 0 | 217610 | 922230 | 0 | 0 | 3735500 | 1794700 | 0 | 0 |

| | | | | | | | | | | | | | | | | | | | | | | |
|---|---|---|---|---|---|---|---|---|---|---|---|---|---|---|---|---|---|---|---|---|---|---|
| LSEIHHLHI | ZC3HAV1; ZC3I | Q7Z2W4; C9 | ZCCHV_HUMA | Unmodified | 0 | 0 | 0 | 0 | 10980000 | 0 | 0 | 0 | 0 | 0 | 0 | 0 | 0 | 208260 | 0 | 0 | 0 | 0 |
| LSEIPLTLH | BAZ1B | Q9UIG0 | BAZ1B_HUMAN | Unmodified | 1113600 | 1040100 | 58301 | 125870 | 0 | 0 | 0 | 67256 | 0 | 84257 | 0 | 697490 | 0 | 0 | 51263 | 23545 | 0 | 0 |
| LSEIVTLAK | SUCLA2; SUCL | Q9P2R7; A0 | SUCB1_HUMA | Unmodified | 0 | 5289900 | 0 | 2427100 | 110580 | 0 | 131220 | 37841 | 2153400 | 350970 | 0 | 1425800 | 73524 | 68395 | 2734500 | 4122600 | 36107 | 110150 |
| LSEKLDSTD | TFRC; TFRC | P02786; G3 | TFR1_HUMAN | Unmodified | 789030 | 789690 | 0 | 1499600 | 1581400 | 923770 | 580640 | 0 | 11822000 | 1331700 | 0 | 1608700 | 951830 | 334730 | 502510 | 875910 | 234910 | 1442900 |
| LSELDDR | VAMP3; VAMP2 | Q15836; P6 | VAMP3_HUMA | Unmodified | 663830 | 686510 | 1829500 | 633790 | 0 | 0 | 0 | 0 | 0 | 0 | 89441 | 1462500 | 0 | 0 | 1856700 | 0 | 0 | 0 |
| LSELDDRA | VAMP3; VAMP2 | Q15836; P6 | VAMP3_HUMA | Unmodified | 0 | 0 | 11425000 | 4268000 | 1572300 | 0 | 155700 | 136880 | 5457700 | 258570 | 409560 | 14265000 | 1695500 | 0 | 399320 | 188050 | 0 | 2681100 |
| LSELLRYHT | HSP90AB1 | P08238 | HS90B_HUMA | Oxidation (M) | 0 | 0 | 0 | 0 | 3739000 | 1792200 | 0 | 0 | 0 | 3076200 | 2205000 | 0 | 7645600 | 1560900 | 0 | 0 | 1515000 | 1948000 |
| LSELLRYHT | HSP90AB1 | P08238 | HS90B_HUMA | Unmodified | 0 | 0 | 0 | 0 | 6702200 | 3049200 | 3149800 | 0 | 0 | 0 | 0 | 0 | 10549000 | 0 | 0 | 0 | 0 | 0 |
| LSELLRYHT | HSP90AB1 | P08238 | HS90B_HUMA | Oxidation (M) | 0 | 0 | 0 | 0 | 4873700 | 0 | 195370 | 0 | 0 | 818930 | 0 | 0 | 2708600 | 0 | 0 | 0 | 1668400 | 0 |
| LSELLRYHT | HSP90AB1 | P08238 | HS90B_HUMA | Unmodified | 0 | 0 | 0 | 0 | 4417100 | 0 | 158370 | 0 | 0 | 263680 | 0 | 0 | 0 | 1730100 | 0 | 0 | 544730 | 1836400 |
| LSELQETSE | FKBP15 | Q5T1M5 | FKB15_HUMAN | Unmodified | 0 | 0 | 0 | 0 | 1821100 | 0 | 0 | 31049 | 0 | 41497 | 0 | 976330 | 0 | 0 | 0 | 0 | 0 | 242950 |
| LSELQLR | TMED9; HERC1 | Q9BVK6; Q1 | TMED9_HUMA | Unmodified | 0 | 0 | 0 | 202560 | 0 | 0 | 197310 | 82107 | 33538000 | 1493300 | 1425500 | 2753200 | 0 | 0 | 0 | 417920 | 0 | 221810 |
| LSELVQAV | TPP1; TPP1; TP | O14773; A0 | TPP1_HUMAN | Unmodified | 0 | 0 | 0 | 0 | 0 | 0 | 0 | 0 | 1418200 | 0 | 0 | 0 | 0 | 0 | 0 | 0 | 0 | 0 |
| LSEMTEAE | CKMT1A; CKMT | P12532; F8 | KCRU_HUMAN | 2 Oxidation (M) | 0 | 0 | 0 | 0 | 0 | 0 | 4219600 | 0 | 0 | 0 | 0 | 1172300 | 0 | 0 | 0 | 0 | 0 | 0 |
| LSENAGIQ | RAI14 | Q9P0K7 | RAI14_HUMAN | Unmodified | 0 | 0 | 0 | 0 | 280340 | 0 | 0 | 0 | 0 | 0 | 0 | 4938300 | 0 | 0 | 0 | 49566 | 1785200 | 237570 |
| LSENNIQTI | ITGB1; ITGB1; I | P05556; H7 | ITB1_HUMAN I | Unmodified | 1161800 | 4086500 | 0 | 5186500 | 3.08E+08 | 1896000 | 9908400 | 0 | 8390000 | 8740300 | 622700 | 695360 | 12970000 | 14598000 | 13045000 | 1052700 | 67420000 | 2.57E+08 |
| LSENNIQTI | ITGB1; ITGB1; I | P05556; H7 | ITB1_HUMAN I | Unmodified | 0 | 0 | 0 | 0 | 1488400 | 0 | 539470 | 0 | 0 | 0 | 0 | 0 | 0 | 0 | 0 | 0 | 2406100 | 2156600 |
| LSEPAELTD | KLK3; KLK3; KL | P07288; M0 | KLK3_HUMAN | Unmodified | 0 | 0 | 4933300 | 3151700 | 0 | 0 | 0 | 0 | 0 | 0 | 0 | 0 | 0 | 0 | 4084600 | 2375600 | 0 | 0 |
| LSEPVPQTI | PLCG1 | P19174 | PLCG1_HUMA | Unmodified | 43170 | 0 | 101230 | 95863 | 110330 | 74304 | 25812 | 0 | 409790 | 20180 | 0 | 2399100 | 0 | 0 | 151180 | 116830 | 42586 | 62381 |
| LSEVLQAV | USP7; USP7; U | Q93009; A0 | UBP7_HUMAN | Unmodified | 0 | 0 | 0 | 0 | 0 | 0 | 0 | 0 | 0 | 0 | 0 | 3063200 | 0 | 0 | 0 | 51280 | 0 | 0 |
| LSEVQASLI | DNAJB12; DNA | Q9NXW2; V | DJB12_HUMAN | Unmodified | 0 | 0 | 0 | 0 | 0 | 150740 | 0 | 0 | 10575000 | 1069800 | 0 | 0 | 0 | 0 | 0 | 0 | 0 | 0 |
| LSFAVPFR | PRKDC | P78527 | PRKDC_HUMA | Unmodified | 0 | 0 | 0 | 0 | 0 | 0 | 0 | 0 | 832280 | 0 | 0 | 1736300 | 0 | 0 | 0 | 0 | 0 | 0 |
| LSFEAVR | STIM1; STIM1; S | Q13586; G0 | STIM1_HUMAN | Unmodified | 112050 | 96947 | 0 | 0 | 0 | 0 | 0 | 0 | 14011000 | 45011 | 264510 | 360800 | 0 | 0 | 0 | 23829 | 0 | 0 |
| LSFEAVRNI | STIM1; STIM1; S | Q13586; G0 | STIM1_HUMAN | Unmodified | 0 | 0 | 0 | 0 | 0 | 0 | 0 | 0 | 0 | 461830 | 0 | 0 | 0 | 0 | 0 | 0 | 0 | 0 |
| LSFEDFVTI | PEF1 | Q9UBV8 | PEF1_HUMAN | Oxidation (M) | 0 | 573910 | 70987 | 840200 | 0 | 0 | 0 | 0 | 3176000 | 86800 | 48305 | 216020 | 60970 | 0 | 1367600 | 153430 | 0 | 19789 |
| LSFEDFVTI | PEF1 | Q9UBV8 | PEF1_HUMAN | Unmodified | 0 | 0 | 0 | 0 | 0 | 0 | 0 | 0 | 1526900 | 0 | 0 | 0 | 0 | 0 | 0 | 0 | 0 | 0 |
| LSFFGLEK | SLC39A10; SLC | Q9ULF5; E7 | S39AA_HUMAN | Unmodified | 0 | 0 | 0 | 0 | 0 | 0 | 0 | 0 | 0 | 60995 | 0 | 14798000 | 0 | 0 | 947410 | 149200 | 0 | 0 |
| LSFGDAAQ | DIP2B | Q9P265 | DIP2B_HUMAN | Unmodified | 0 | 0 | 0 | 0 | 0 | 0 | 0 | 0 | 0 | 0 | 0 | 3383500 | 0 | 0 | 0 | 845100 | 0 | 0 |
| LSFGDTLQ | ERGIC1 | Q969X5 | ERGI1_HUMAN | Unmodified | 0 | 0 | 0 | 0 | 0 | 0 | 0 | 0 | 0 | 2221800 | 0 | 0 | 1900200 | 0 | 0 | 0 | 0 | 0 |
| LSFGDTLQ | ERGIC1 | Q969X5 | ERGI1_HUMAN | Unmodified | 0 | 0 | 0 | 0 | 0 | 0 | 0 | 0 | 651500 | 3191400 | 0 | 0 | 0 | 0 | 1255900 | 0 | 0 | 0 |
| LSFGLEDEI | ERLIN2; ERLIN2 | O94905; E5 | ERLN2_HUMAN | Unmodified | 0 | 0 | 0 | 0 | 0 | 0 | 520800 | 0 | 13167000 | 0 | 664950 | 304820 | 0 | 0 | 0 | 0 | 0 | 0 |
| LSFGLEDEI | ERLIN2 | O94905 | ERLN2_HUMAN | Unmodified | 0 | 0 | 0 | 0 | 0 | 88197 | 0 | 59393 | 5560300 | 666750 | 0 | 0 | 0 | 0 | 0 | 0 | 0 | 0 |
| LSFISEGEI | L1RE1 | Q9UN81 | LORF1_HUMAN | Unmodified | 2919200 | 2436000 | 731090 | 782880 | 0 | 0 | 0 | 0 | 0 | 0 | 0 | 38783 | 0 | 0 | 685200 | 857050 | 0 | 0 |
| LSFISVGN | MYO6; MYO6; M | Q9UM54; A0 | MYO6_HUMAN | Unmodified | 0 | 0 | 1201800 | 59337 | 0 | 0 | 0 | 0 | 0 | 0 | 0 | 0 | 0 | 0 | 787840 | 1174900 | 0 | 0 |
| LSFMPFFL | DBT; DBT; DBT; | P11182; A0A | ODB2_HUMAN | Unmodified | 0 | 0 | 0 | 0 | 0 | 0 | 0 | 0 | 329970 | 0 | 0 | 0 | 0 | 0 | 0 | 0 | 0 | 0 |
| LSFQHDPE | GANAB; GANAE | Q14697; F5 | GANAB_HUMA | Unmodified | 0 | 0 | 0 | 0 | 0 | 0 | 0 | 0 | 19367000 | 0 | 478800 | 614160 | 0 | 0 | 0 | 0 | 0 | 0 |
| LSFQHDPE | GANAB; GANAE | Q14697; F5 | GANAB_HUMA | Unmodified | 0 | 0 | 339390 | 0 | 0 | 0 | 0 | 0 | 11797000 | 0 | 0 | 0 | 0 | 0 | 0 | 0 | 0 | 0 |
| LSFQHDPE | GANAB; GANAE | Q14697; F5 | GANAB_HUMA | Unmodified | 0 | 0 | 0 | 0 | 0 | 0 | 0 | 0 | 6301700 | 11820000 | 0 | 0 | 0 | 0 | 0 | 0 | 0 | 0 |
| LSGAAPAH | PIMREG; PIMRE | Q9BSJ6; I3L | PIMRE_HUMAN | Unmodified | 0 | 86517 | 0 | 0 | 156900 | 0 | 0 | 86177 | 0 | 108750 | 0 | 0 | 0 | 161080 | 36449 | 0 | 0 | 404500 |
| LSGAHLPD | POMT1; POMT1 | Q9Y6A1; A0 | POMT1_HUMA | Unmodified | 0 | 0 | 0 | 0 | 0 | 0 | 0 | 0 | 4233400 | 0 | 0 | 0 | 0 | 0 | 0 | 0 | 0 | 0 |
| LSGEAFDI | GEMIN5 | Q8TEQ6 | GEMI5_HUMAN | Unmodified | 0 | 0 | 0 | 0 | 0 | 0 | 0 | 0 | 0 | 0 | 275500 | 4241500 | 0 | 0 | 0 | 0 | 0 | 0 |
| LSGEQVDG | FZD6 | O60353 | FZD6_HUMAN | Unmodified | 65939 | 0 | 101300 | 0 | 0 | 0 | 0 | 0 | 830790 | 0 | 0 | 0 | 0 | 0 | 80717 | 100270 | 0 | 0 |
| LSGFSFKK | MARCKS | P29966 | MARCS_HUMA | Unmodified | 0 | 0 | 0 | 2771400 | 0 | 0 | 0 | 2708400 | 7385800 | 7000800 | 0 | 2956100 | 0 | 0 | 0 | 0 | 3294400 | 10829000 |
| LSGGHSLH | FLNC | Q14315 | FLNC_HUMAN | Unmodified | 0 | 0 | 0 | 0 | 1439500 | 0 | 0 | 0 | 0 | 0 | 0 | 0 | 0 | 0 | 0 | 0 | 591720 | 0 |
| LSGIEER | SPTBN1; SPTBN | Q01082; A0 | SPTB2_HUMAN | Unmodified | 0 | 0 | 500260 | 61293 | 181930 | 0 | 720420 | 0 | 147900 | 0 | 0 | 1219600 | 0 | 0 | 0 | 0 | 0 | 474540 |
| LSGKLYLVI | KIF5B; KIF5C; K | P33176; O6 | KINH_HUMAN | Unmodified | 0 | 0 | 0 | 0 | 163610 | 0 | 501350 | 124410 | 0 | 71496 | 0 | 0 | 126320 | 0 | 0 | 0 | 364930 | 39243 |
| LSGLALDH | SLC43A1; SLC4 | O75387; E9 | LAT3_HUMAN | Oxidation (M) | 0 | 0 | 0 | 5099200 | 0 | 0 | 0 | 0 | 0 | 0 | 0 | 0 | 0 | 0 | 0 | 1540200 | 0 | 0 |
| LSGLHGQI | ACP3 | P15309 | PPAP_HUMAN | Unmodified | 676270 | 696940 | 329600 | 170850 | 0 | 0 | 0 | 0 | 7501000 | 2681000 | 83084 | 168930 | 273100 | 283230 | 948650 | 921870 | 0 | 0 |
| LSGLNKET | ORC6; ORC6; C | Q9Y5N6; H3 | ORC6_HUMAN | Unmodified | 0 | 0 | 0 | 0 | 0 | 0 | 0 | 141990 | 0 | 0 | 0 | 0 | 0 | 0 | 0 | 0 | 0 | 0 |
| LSGLSFKR | MARCKSL1 | P49006 | MRP_HUMAN | Unmodified | 0 | 0 | 1009000 | 1404500 | 991320 | 0 | 0 | 53945 | 1903900 | 547780 | 1417900 | 2250800 | 2049800 | 1184100 | 1077900 | 1269000 | 866930 | 4696700 |
| LSGLWSR | TOR1AIP2 | Q8NFQ8 | TOIP2_HUMAN | Unmodified | 0 | 0 | 0 | 0 | 0 | 0 | 0 | 0 | 3607300 | 85209 | 0 | 0 | 0 | 0 | 0 | 0 | 0 | 0 |
| LSGPDDDF | DPP9; DPP9 | Q86TI2; M0 | DPP9_HUMAN | Unmodified | 0 | 0 | 0 | 0 | 0 | 0 | 0 | 0 | 0 | 0 | 0 | 226570 | 257140 | 0 | 0 | 0 | 0 | 44877 |
| LSGPSLKM | AHNAK | Q09666 | AHNK_HUMAN | Unmodified | 0 | 0 | 0 | 0 | 986420 | 147480 | 0 | 0 | 0 | 266750 | 0 | 0 | 0 | 0 | 0 | 0 | 0 | 1065800 |
| LSGSHSQ | HSPG2 | P98160 | PGBM_HUMAN | Unmodified | 0 | 0 | 3271000 | 0 | 0 | 0 | 0 | 0 | 278040 | 167780 | 0 | 0 | 0 | 0 | 0 | 0 | 205540 | 147250 |
| LSGSNPYT | ACTN4; ACTN4 | O43707; F5 | ACTN4_HUMA | Unmodified | 0 | 0 | 307410 | 0 | 750660 | 0 | 0 | 0 | 357160 | 0 | 0 | 7235500 | 0 | 0 | 965330 | 1081000 | 0 | 0 |
| LSGSNPYT | ACTN4; ACTN4 | O43707; F5 | ACTN4_HUMA | Unmodified | 1306600 | 851550 | 3204900 | 2042900 | 33851000 | 11754000 | 2279100 | 816230 | 6122700 | 6958700 | 3286500 | 19550000 | 0 | 0 | 4036300 | 2824000 | 11748000 | 1239700 |
| LSGTGSAG | PGM1; PGM1 | P36871; A0A | PGM1_HUMAN | Unmodified | 0 | 0 | 0 | 0 | 0 | 0 | 0 | 44607 | 695170 | 0 | 0 | 1212300 | 0 | 79567 | 0 | 137560 | 92080 | 0 |
| LSGVEDHV | SLC12A2; SLC1 | P55011; G3 | S12A2_HUMAN | Unmodified | 0 | 0 | 261750 | 1278400 | 158240 | 111120 | 447490 | 0 | 70506 | 0 | 4346700 | 361670 | 0 | 280490 | 940470 | 0 | 0 | 0 |
| LSGVMVPA | ATXN10; ATXN1 | Q9UBB4; A0 | ATX10_HUMAN | Oxidation (M) | 303890 | 0 | 155360 | 431630 | 0 | 0 | 0 | 188280 | 0 | 0 | 429480 | 2204100 | 0 | 0 | 1036300 | 688790 | 0 | 0 |
| LSGWGNV | GIGYF2; GIGYF | Q6Y7W6; I1 | GGYF2_HUMA | Unmodified | 0 | 0 | 0 | 172820 | 569800 | 137320 | 0 | 0 | 0 | 0 | 0 | 397570 | 406740 | 1294100 | 0 | 0 | 0 | 0 |
| LSHANSAV | AP1B1; AP2B1; | Q10567; P6 | AP1B1_HUMAN | Unmodified | 0 | 0 | 0 | 0 | 0 | 0 | 0 | 0 | 0 | 0 | 0 | 0 | 0 | 0 | 0 | 0 | 0 | 0 |
| LSHEGPGS | DCTN1; DCTN1 | Q14203; A0 | DCTN1_HUMA | Unmodified | 0 | 0 | 0 | 0 | 2583600 | 0 | 1422700 | 694440 | 462300 | 436800 | 0 | 1533400 | 1004800 | 0 | 0 | 0 | 970310 | 2048600 |
| LSHETVTIE | SNRPD1; SNRP | P62314; J3Q | SMD1_HUMAN | Unmodified | 1458100 | 1484300 | 506210 | 256000 | 1425700 | 553320 | 3303400 | 2330000 | 640580 | 1775600 | 3735200 | 10860000 | 646940 | 270190 | 362050 | 581450 | 742360 | 556650 |
| LSHLKPSSI | RO60 | P10155 | RO60_HUMAN | Unmodified | 33816000 | 28418000 | 4777100 | 4441600 | 0 | 0 | 0 | 0 | 0 | 0 | 199530 | 2708200 | 253740 | 89624 | 13839000 | 7372300 | 0 | 0 |
| LSIAEVVHC | GAK | O14976 | GAK_HUMAN C | Unmodified | 0 | 55799 | 0 | 0 | 0 | 0 | 0 | 18459 | 0 | 0 | 0 | 42547 | 0 | 0 | 85922 | 0 | 0 | 0 |
| LSIHSLEAC | NOXO1 | Q8NFA2 | NOXO1_HUMA | Unmodified | 0 | 0 | 0 | 0 | 0 | 0 | 0 | 71612 | 0 | 0 | 1487700 | 2405400 | 0 | 207880 | 0 | 0 | 0 | 0 |
| LSIIFPLWK | PIK3C2A | O00443 | P3C2A_HUMAN | Unmodified | 0 | 0 | 0 | 0 | 0 | 0 | 0 | 0 | 0 | 0 | 0 | 539970 | 0 | 0 | 29349 | 0 | 0 | 0 |
| LSIIPQEPVI | ABCC5 | O15440 | MRP5_HUMAN | Unmodified | 0 | 0 | 0 | 0 | 0 | 0 | 0 | 0 | 127870 | 0 | 308940 | 3429300 | 0 | 0 | 0 | 0 | 0 | 0 |
| LSIKHPSVN | PLAA; PLAA | Q9Y263; H0 | PLAP_HUMAN | Unmodified | 0 | 0 | 0 | 0 | 0 | 0 | 0 | 0 | 0 | 43490 | 0 | 546630 | 0 | 0 | 0 | 0 | 0 | 0 |
| LSILYPATT | PRDX6 | P30041 | PRDX6_HUMA | Unmodified | 2103600 | 1276400 | 3717100 | 0 | 0 | 0 | 215180 | 94247 | 1260300 | 314930 | 0 | 6328000 | 1120400 | 0 | 5340600 | 0 | 0 | 0 |
| LSIPGTGVS | UNC5B | Q8IZJ1 | UNC5B_HUMA | Unmodified | 0 | 0 | 0 | 0 | 0 | 0 | 0 | 0 | 519690 | 0 | 0 | 881200 | 0 | 0 | 0 | 0 | 40548 | 0 |
| LSIPTYGLQ | FASN; FASN; FA | P49327; J3K | FAS_HUMAN F | Unmodified | 0 | 0 | 0 | 0 | 0 | 0 | 0 | 0 | 0 | 0 | 0 | 11760000 | 0 | 0 | 0 | 0 | 0 | 0 |
| LSIQCYLSA | HMGCS1 | Q01581 | HMCS1_HUMA | Unmodified | 0 | 0 | 0 | 0 | 0 | 0 | 0 | 0 | 1080500 | 0 | 0 | 2630700 | 0 | 0 | 0 | 0 | 0 | 0 |
| LSITEVTK | MYO1D; MYO1I | O94832; J3C | MYO1D_HUMA | Unmodified | 4437400 | 2232400 | 4175000 | 743000 | 0 | 0 | 0 | 0 | 0 | 0 | 206550 | 1525500 | 0 | 0 | 4301000 | 4141100 | 0 | 0 |
| LSITEVTKRI | MYO1D; MYO1I | O94832; J3C | MYO1D_HUMA | Unmodified | 19401000 | 13731000 | 1031400 | 10981000 | 418620 | 730930 | 0 | 0 | 5762000 | 192350 | 2307100 | 2315600 | 3012800 | 1228400 | 6548400 | 6649700 | 1302600 | 3115400 |
| LSITKQDD | NCAM2; NCAM | O15394; H9 | NCAM2_HUMA | Unmodified | 0 | 0 | 0 | 2051500 | 0 | 0 | 0 | 0 | 0 | 0 | 0 | 0 | 0 | 0 | 1715100 | 796850 | 0 | 0 |
| LSKDPNIVI | PDIA3 | P30101 | PDIA3_HUMAN | Unmodified | 0 | 0 | 0 | 0 | 1168200 | 0 | 440370 | 481320 | 22205000 | 7776400 | 3242000 | 2461500 | 0 | 570080 | 204810 | 0 | 69802 | 850380 |
| LSKDPNIVI | PDIA3 | P30101 | PDIA3_HUMAN | Unmodified | 0 | 0 | 0 | 0 | 0 | 0 | 0 | 0 | 620100 | 4171900 | 0 | 0 | 0 | 0 | 0 | 0 | 0 | 0 |
| LSKDPNIVI | PDIA3 | P30101 | PDIA3_HUMAN | Oxidation (M) | 0 | 0 | 0 | 0 | 0 | 0 | 0 | 0 | 0 | 1256200 | 0 | 0 | 210500 | 0 | 0 | 0 | 0 | 0 |
| LSKEDIER | HSPA8; HSPA8; | P11142; E9P | HSP7C_HUMA | Unmodified | 3353600 | 2894600 | 843630 | 339590 | 0 | 0 | 196240 | 512330 | 1253300 | 115040 | 368100 | 5167100 | 0 | 285490 | 1438400 | 2588100 | 0 | 0 |
| LSKEETVLA | RPN2 | P04844 | RPN2_HUMAN | Unmodified | 0 | 0 | 0 | 0 | 0 | 0 | 0 | 0 | 20345000 | 6953800 | 1080500 | 578450 | 0 | 0 | 0 | 0 | 0 | 0 |
| LSKIYSQST | CAMSAP3 | Q9P1Y5 | CAMP3_HUMA | Unmodified | 0 | 0 | 0 | 0 | 0 | 0 | 0 | 0 | 0 | 0 | 0 | 0 | 1856900 | 0 | 0 | 0 | 0 | 0 |
| LSKLTPGLY | KIAA0319L; KIA | Q8IZA0; B1A | K319L_HUMAN | Unmodified | 0 | 0 | 0 | 0 | 109200 | 0 | 0 | 0 | 0 | 655100 | 0 | 0 | 0 | 0 | 0 | 0 | 0 | 0 |
| LSKNEVLM | EIF2S3; EIF2S3 | P41091; Q2 | IF2G_HUMAN | Unmodified | 0 | 0 | 0 | 0 | 0 | 0 | 0 | 0 | 383180 | 0 | 0 | 212770 | 0 | 0 | 0 | 0 | 0 | 0 |
| LSKNEVLM | EIF2S3; EIF2S3 | P41091; Q2 | IF2G_HUMAN | Oxidation (M) | 240690 | 143270 | 0 | 853770 | 0 | 0 | 0 | 0 | 0 | 46855 | 269590 | 2821900 | 92733 | 0 | 155060 | 266310 | 0 | 66153 |
| LSLAQEDLI | EEA1 | Q15075 | EEA1_HUMAN | Unmodified | 0 | 0 | 0 | 0 | 5386200 | 667470 | 192810 | 55729 | 299900 | 163460 | 0 | 710050 | 787470 | 0 | 241460 | 0 | 0 | 928330 |
| LSLDDLLQI | CAD; CAD; CAD | P27708; F8 | PYR1_HUMAN | Unmodified | 0 | 0 | 0 | 0 | 0 | 0 | 0 | 0 | 0 | 0 | 0 | 1859400 | 0 | 0 | 0 | 0 | 0 | 0 |
| LSLDELHR | ATP1A1; ATP1A | P05023; Q5 | AT1A1_HUMAN | Unmodified | 0 | 0 | 642140 | 689690 | 244860 | 20831 | 0 | 0 | 762980 | 0 | 484320 | 4131000 | 0 | 0 | 1982700 | 1331000 | 0 | 0 |
| LSLDELHRI | ATP1A1; ATP1A | P05023; Q5 | AT1A1_HUMAN | Unmodified | 0 | 0 | 1385200 | 279610 | 0 | 0 | 327840 | 230500 | 1099500 | 0 | 591440 | 3980000 | 0 | 0 | 830660 | 777630 | 0 | 0 |
| LSLDGQNI | PTBP1; PTBP1; | P26599; A0A | PTBP1_HUMAN | Unmodified | 0 | 0 | 0 | 0 | 0 | 0 | 0 | 0 | 0 | 0 | 25848 | 593220 | 0 | 0 | 0 | 0 | 0 | 0 |
| LSLEEFIR | NCALD; HPCA; | P61601; P8 | NCALD_HUMA | Unmodified | 209110 | 138640 | 342460 | 0 | 136510 | 0 | 0 | 0 | 2139600 | 0 | 356780 | 2678800 | 0 | 0 | 406770 | 447530 | 0 | 0 |
| LSLEEFIRG | NCALD; HPCA; | P61601; P8 | NCALD_HUMA | Unmodified | 0 | 0 | 16919 | 0 | 1496000 | 334440 | 48379 | 0 | 563340 | 87739 | 54318 | 0 | 158970 | 41727 | 0 | 0 | 179350 | 758450 |
| LSLEGDHS | ANXA2P2; ANX | A6NMY6; P0 | AXA2L_HUMAN | Unmodified | 1196800 | 851450 | 246170 | 783290 | 4.4E+08 | 75146000 | 5754800 | 3405500 | 9225000 | 8245200 | 91440 | 0 | 1068800 | 1406500 | 385350 | 228860 | 91855000 | 1.54E+08 |
| LSLEGWFL | LPAR2 | Q9HBW0 | LPAR2_HUMAN | Unmodified | 0 | 0 | 0 | 0 | 0 | 0 | 0 | 0 | 0 | 0 | 0 | 0 | 0 | 0 | 0 | 261640 | 0 | 0 |
| LSLEQALPI | FAF1; FAF1; FAI | Q9UNN5; B1 | FAF1_HUMAN | Unmodified | 238980 | 293900 | 0 | 0 | 0 | 0 | 0 | 0 | 0 | 0 | 0 | 930310 | 0 | 0 | 0 | 66861 | 0 | 0 |
| LSLGTYASI | LIMA1; LIMA1; L | Q9UHB6; F8 | LIMA1_HUMAN | Unmodified | 701200 | 852650 | 1275700 | 1131000 | 4856500 | 1028700 | 0 | 0 | 0 | 0 | 0 | 3056500 | 2862400 | 3882900 | 1130300 | 872270 | 1529800 | 991980 |
| LSLHEEEG | ANK3; ANK3; A | Q12955; H0 | ANK3_HUMAN | Unmodified | 0 | 0 | 414540 | 495460 | 100400 | 135610 | 0 | 0 | 3220600 | 0 | 0 | 0 | 124810 | 79973 | 732750 | 607260 | 0 | 0 |
| LSLHEEEG | ANK3; ANK3; A | Q12955; H0 | ANK3_HUMAN | Unmodified | 0 | 0 | 0 | 0 | 0 | 0 | 0 | 0 | 0 | 0 | 0 | 0 | 2590000 | 604360 | 0 | 0 | 0 | 0 |
| LSLHRTPA | NAMPT; NAMPT | P43490; A0A | NAMPT_HUMA | Unmodified | 0 | 0 | 0 | 0 | 3229400 | 0 | 0 | 0 | 0 | 0 | 0 | 0 | 1304600 | 0 | 0 | 0 | 0 | 0 |
| LSLLSFWTI | RIPOR2; RIPOR | Q9Y4F9; F5 | RIPR2_HUMAN | Unmodified | 0 | 0 | 296550 | 0 | 346270 | 81348 | 0 | 0 | 104840 | 93852 | 96782 | 794340 | 0 | 0 | 0 | 10067 | 0 | 0 |
| LSLMPWFH | MATK; CSK; MA | P42679; P41 | MATK_HUMAN | Oxidation (M) | 82590 | 0 | 0 | 231950 | 0 | 0 | 0 | 0 | 0 | 0 | 270310 | 3423900 | 0 | 0 | 0 | 113020 | 0 | 0 |
| LSLNIDPDA | HSP90B1; HSP | P14625; A0A | ENPL_HUMAN | Unmodified | 1086100 | 0 | 1149900 | 0 | 1186000 | 0 | 381080 | 722990 | 5346500 | 843570 | 4094000 | 1201200 | 0 | 0 | 802940 | 177300 | 844000 | 756660 |
| LSLNLDHK | PLOD3; PLOD3 | O60568; H7 | PLOD3_HUMA | Unmodified | 0 | 0 | 0 | 0 | 0 | 0 | 0 | 0 | 1650100 | 0 | 0 | 0 | 0 | 0 | 0 | 0 | 0 | 0 |
| LSLNRQFFI | DYNC1I2; DYN | Q13409; E7 | DC1I2_HUMAN | Unmodified | 0 | 0 | 0 | 0 | 599120 | 0 | 0 | 0 | 0 | 749290 | 0 | 0 | 480200 | 0 | 0 | 0 | 1135800 | 274730 |
| LSLPDNDL | ERBIN; ERBIN | Q96RT1; B4 | ERBIN_HUMAN | Unmodified | 0 | 0 | 0 | 0 | 117950 | 0 | 0 | 0 | 0 | 0 | 0 | 0 | 0 | 0 | 0 | 0 | 0 | 0 |
| LSLPPNVR | DYNC1H1; DYN | Q14204; A0 | DYHC1_HUMA | Unmodified | 0 | 0 | 47426 | 0 | 0 | 0 | 0 | 0 | 196210 | 109550 | 0 | 1290400 | 0 | 0 | 0 | 0 | 0 | 0 |
| LSLPWLLN | PLXNA1 | Q9UIW2 | PLXA1_HUMAN | Unmodified | 0 | 0 | 0 | 123780 | 0 | 0 | 0 | 0 | 0 | 0 | 0 | 416500 | 337610 | 0 | 292420 | 295840 | 0 | 0 |
| LSLRDTTEC | AHNAK2 | Q8IVF2 | AHNK2_HUMA | Unmodified | 0 | 0 | 0 | 0 | 629370 | 124320 | 0 | 0 | 0 | 0 | 0 | 0 | 64703 | 0 | 0 | 0 | 65425 | 529290 |
| LSLRQPGL | RIPOR1; RIPOR | Q6ZS17; A0 | RIPR1_HUMAN | Unmodified | 0 | 0 | 0 | 0 | 0 | 0 | 0 | 0 | 0 | 0 | 0 | 636980 | 0 | 0 | 324110 | 142860 | 0 | 0 |

| | | | | | | | | | | | | | | | | | | | | | | |
|---|---|---|---|---|---|---|---|---|---|---|---|---|---|---|---|---|---|---|---|---|---|---|
| LSLSNAIST | DYNC1H1; DYN | Q14204; A0A | DYHC1_HUMA | Unmodified | 48800 | 134440 | 189460 | 331450 | 214580 | 52076 | 68220 | 0 | 679800 | 553970 | 367600 | 1744100 | 607990 | 0 | 848280 | 291980 | 116440 | 771850 |
| LSLSQLNSI | PROM1 | O43490 | PROM1_HUMA | Unmodified | 0 | 0 | 0 | 0 | 0 | 0 | 0 | 0 | 0 | 0 | 2316100 | 18784000 | 0 | 0 | 0 | 0 | 0 | 0 |
| LSLTQDPV | HSPA4L; HSPA | O95757; E9F | HS74L_HUMAN | Unmodified | 0 | 0 | 0 | 0 | 298350 | 0 | 0 | 0 | 381880 | 188850 | 1800100 | 5650600 | 1634600 | 947530 | 0 | 0 | 137030 | 0 |
| LSLTQDPV | HSPA4L; HSPA | O95757; E9F | HS74L_HUMAN | Unmodified | 0 | 0 | 0 | 0 | 346840 | 54027 | 242280 | 40858 | 0 | 569540 | 0 | 0 | 1523600 | 1239800 | 0 | 0 | 607360 | 190430 |
| LSLVLVPAC | CDCP1 | Q9H5V8 | CDCP1_HUMA | Unmodified | 3578200 | 0 | 0 | 260320 | 0 | 0 | 0 | 0 | 0 | 0 | 0 | 4170700 | 0 | 0 | 702840 | 724250 | 0 | 0 |
| LSMINTMS | SH3GL2 | Q99962 | SH3G2_HUMA | Unmodified | 0 | 0 | 0 | 0 | 0 | 0 | 0 | 0 | 1612900 | 1034400 | 0 | 186540 | 0 | 0 | 0 | 0 | 0 | 0 |
| LSMKEGDF | TES; TES | Q9UGI8; H7 | TES_HUMAN T | Oxidation (M) | 0 | 0 | 0 | 0 | 0 | 0 | 0 | 0 | 0 | 0 | 0 | 0 | 234640 | 0 | 0 | 0 | 580660 | 0 |
| LSMKEGDF | TES; TES | Q9UGI8; H7 | TES_HUMAN T | Unmodified | 0 | 0 | 0 | 0 | 1042300 | 0 | 0 | 0 | 0 | 236760 | 0 | 0 | 0 | 0 | 0 | 0 | 3101100 | 1853000 |
| LSMKPEPLI | MAP1B; MAP1B | P46821; D6F | MAP1B_HUMA | Oxidation (M) | 0 | 0 | 0 | 0 | 0 | 0 | 1055500 | 0 | 0 | 438930 | 0 | 1521100 | 0 | 0 | 0 | 0 | 0 | 0 |
| LSNFKTAA | P4HB; P4HB; P | P07237; A0A | PDIA1_HUMAN | Unmodified | 0 | 0 | 0 | 0 | 0 | 0 | 0 | 0 | 0 | 242880 | 0 | 0 | 0 | 0 | 0 | 0 | 0 | 0 |
| LSNHISSLF | G6PD; G6PD; G | P11413; E7E | G6PD_HUMAN | Unmodified | 0 | 0 | 0 | 0 | 0 | 0 | 0 | 0 | 0 | 0 | 0 | 4091200 | 0 | 0 | 0 | 270390 | 0 | 0 |
| LSNIFVIGK | RPS4X | P62701 | RS4X_HUMAN | Unmodified | 3028100 | 2367200 | 6878900 | 5089700 | 563040 | 220570 | 584540 | 165860 | 4565400 | 696440 | 4899400 | 1.33E+08 | 700950 | 284180 | 5154100 | 4563100 | 88958 | 817150 |
| LSNIFVIGN | RPS4Y1; RPS4Y | P22090; C9J | RS4Y1_HUMAN | Unmodified | 679030 | 1418200 | 0 | 0 | 0 | 0 | 0 | 0 | 0 | 0 | 246400 | 5503300 | 0 | 0 | 0 | 89350 | 0 | 0 |
| LSNKTEEDI | GAS7 | O60861 | GAS7_HUMAN | Unmodified | 0 | 0 | 0 | 0 | 0 | 0 | 0 | 0 | 0 | 0 | 281130 | 1185100 | 0 | 0 | 0 | 0 | 0 | 0 |
| LSNLALVKI | PDCD5; PDCD5 | O14737; K7E | PDCD5_HUMA | Unmodified | 0 | 0 | 369870 | 501630 | 227940 | 0 | 0 | 0 | 358560 | 173500 | 0 | 0 | 559440 | 188710 | 658480 | 0 | 118460 | 255510 |
| LSNLEGLG | CHPF2; CHPF2 | Q9P2E5; G5I | CHPF2_HUMA | Unmodified | 0 | 0 | 30752 | 0 | 139810 | 223910 | 0 | 0 | 705940 | 0 | 0 | 1901100 | 0 | 0 | 0 | 0 | 131860 | 0 |
| LSNLEVK | SH3BP4; FERM | Q9P0V3; Q8 | SH3B4_HUMA | Unmodified | 0 | 0 | 0 | 0 | 776480 | 122230 | 160830 | 13881 | 679230 | 570860 | 156340 | 2368900 | 412690 | 229490 | 117330 | 0 | 338710 | 1194000 |
| LSNRPAFM | ACTN4; ACTN4 | O43707; F5C | ACTN4_HUMA | Unmodified | 471940 | 451030 | 2584000 | 1759800 | 3149900 | 953770 | 0 | 0 | 4168800 | 0 | 1311100 | 12700000 | 846390 | 509780 | 1981700 | 1142300 | 907020 | 340210 |
| LSNRPAFM | ACTN1; ACTN1 | P12814; A0A | ACTN1_HUMA | Unmodified | 0 | 0 | 0 | 0 | 0 | 0 | 68060 | 0 | 0 | 0 | 0 | 10106000 | 0 | 0 | 1129400 | 586600 | 353090 | 0 |
| LSNSDSQD | TRAFD1 | O14545 | TRAD1_HUMAN | Unmodified | 1685600 | 1803000 | 346760 | 77454 | 181570 | 0 | 0 | 0 | 44277 | 0 | 0 | 0 | 0 | 0 | 195390 | 209400 | 252150 | 0 |
| LSNVATSV | SLC32A1 | Q9H598 | VIAAT_HUMAN | Unmodified | 0 | 0 | 0 | 0 | 0 | 0 | 153260 | 28614 | 0 | 55819 | 0 | 0 | 0 | 0 | 0 | 0 | 0 | 0 |
| LSPDAIPGI | FASN; FASN | P49327; A0A | FAS_HUMAN F | Unmodified | 135170 | 0 | 1739300 | 121420 | 1083900 | 9593000 | 33026 | 0 | 196240 | 34815 | 78440000 | 49001000 | 0 | 0 | 2321200 | 2404900 | 0 | 4134900 |
| LSPDAIPGI | FASN; FASN | P49327; A0A | FAS_HUMAN F | Oxidation (M) | 899630 | 0 | 0 | 1370700 | 0 | 0 | 0 | 0 | 211440 | 0 | 343750 | 12055000 | 0 | 0 | 0 | 2626500 | 0 | 0 |
| LSPDASPQ | ABCA2; ABCA2 | Q9BZC7; A0 | ABCA2_HUMA | Unmodified | 0 | 0 | 0 | 0 | 0 | 0 | 0 | 0 | 5516600 | 0 | 0 | 0 | 0 | 0 | 0 | 0 | 0 | 0 |
| LSPEEVQK | PAM16; PAM16 | Q9Y3D7; I3L | TIM16_HUMAN | Unmodified | 0 | 0 | 0 | 0 | 0 | 0 | 0 | 0 | 1176700 | 0 | 0 | 0 | 0 | 0 | 0 | 0 | 0 | 0 |
| LSPEIVELSI | P4HTM | Q9NXG6 | P4HTM_HUMA | Unmodified | 0 | 0 | 0 | 0 | 0 | 0 | 0 | 0 | 12868000 | 0 | 0 | 0 | 0 | 0 | 0 | 0 | 0 | 0 |
| LSPELLNTC | HK2; HK2 | P52789; E9P | HXK2_HUMAN | Unmodified | 457400 | 1089900 | 5088600 | 620790 | 0 | 0 | 39729 | 33336 | 0 | 0 | 105030 | 1714100 | 0 | 0 | 2103300 | 868650 | 0 | 0 |
| LSPEPWTP | ACSL4; ACSL3; | O60488; O9 | ACSL4_HUMAN | Unmodified | 0 | 0 | 0 | 0 | 0 | 5156200 | 0 | 0 | 0 | 0 | 239250 | 1711800 | 0 | 0 | 0 | 0 | 0 | 7835900 |
| LSPEPWTP | ACSL4; ACSL3; | O60488; O9 | ACSL4_HUMAN | Unmodified | 0 | 0 | 0 | 0 | 1975800 | 0 | 0 | 0 | 0 | 0 | 0 | 0 | 0 | 0 | 0 | 0 | 3842200 | 3204900 |
| LSPFMADIF | FLNA; FLNA; FL | P21333; A0A | FLNA_HUMAN | Oxidation (M) | 0 | 0 | 0 | 0 | 786380 | 484800 | 132790 | 0 | 0 | 0 | 0 | 0 | 0 | 271850 | 0 | 399250 | 562530 | 362760 |
| LSPGESAY | PPP1R18; PPP1 | Q6NYC8; A0 | PPR18_HUMAN | Unmodified | 0 | 0 | 0 | 0 | 430660 | 0 | 0 | 0 | 0 | 0 | 0 | 0 | 0 | 0 | 0 | 0 | 0 | 0 |
| LSPGGYFI | RNMT; RNMT | O43148; K7E | MCES_HUMAN | Unmodified | 0 | 0 | 0 | 0 | 0 | 0 | 0 | 0 | 286640 | 0 | 359560 | 1213200 | 0 | 0 | 0 | 0 | 0 | 0 |
| LSPHAFKTI | TGOLN2; TGOL | O43493; A0 | TGON2_HUMA | Unmodified | 0 | 0 | 0 | 0 | 0 | 0 | 0 | 0 | 10500000 | 0 | 0 | 648130 | 0 | 0 | 0 | 0 | 0 | 0 |
| LSPHKVQG | IMUP; N/A | Q9GZP8; K7I | IMUP_HUMAN | Unmodified | 0 | 0 | 0 | 0 | 3124900 | 0 | 0 | 0 | 0 | 55694 | 0 | 0 | 0 | 0 | 0 | 0 | 0 | 485480 |
| LSPIFRDFL | PAK2 | Q13177 | PAK2_HUMAN | Unmodified | 0 | 0 | 0 | 1164700 | 1248800 | 617620 | 188780 | 0 | 2607600 | 0 | 1425900 | 2159100 | 2969000 | 0 | 0 | 0 | 504800 | 421540 |
| LSPIFRDFL | PAK2 | Q13177 | PAK2_HUMAN | Oxidation (M) | 0 | 0 | 0 | 0 | 0 | 0 | 0 | 0 | 0 | 0 | 0 | 0 | 2017300 | 0 | 0 | 0 | 3642300 | 0 |
| LSPISHGN | TCF25 | Q9BQ70 | TCF25_HUMAN | Unmodified | 243220 | 0 | 557590 | 0 | 66663 | 0 | 0 | 0 | 0 | 0 | 0 | 133380 | 0 | 0 | 922520 | 466380 | 0 | 0 |
| LSPKAEEV | TOP1 | P11387 | TOP1_HUMAN | Unmodified | 402730 | 307710 | 0 | 0 | 939810 | 549330 | 695870 | 706730 | 0 | 0 | 0 | 2020300 | 352220 | 223060 | 0 | 0 | 0 | 765570 |
| LSPKHPESI | CLIC4 | Q9Y696 | CLIC4_HUMAN | Unmodified | 53706 | 78404 | 0 | 190800 | 36192000 | 2559500 | 8315300 | 4232100 | 2706600 | 4598900 | 291600 | 0 | 8609300 | 13434000 | 34615 | 0 | 26883000 | 17969000 |
| LSPKHPESI | CLIC4 | Q9Y696 | CLIC4_HUMAN | Oxidation (M) | 0 | 0 | 0 | 0 | 4362000 | 1215000 | 1306800 | 21159 | 0 | 432500 | 80350 | 0 | 1980500 | 2318400 | 0 | 0 | 4759800 | 4124600 |
| LSPKHPESI | CLIC4 | Q9Y696 | CLIC4_HUMAN | Oxidation (M) | 0 | 0 | 0 | 0 | 0 | 0 | 0 | 0 | 0 | 0 | 0 | 0 | 0 | 0 | 0 | 0 | 1685900 | 0 |
| LSPLEACA | GSK3A; GSK3A | P49840; A0A | GSK3A_HUMA | Unmodified | 209140 | 230990 | 0 | 277830 | 0 | 0 | 0 | 0 | 360890 | 34007 | 220690 | 1189600 | 31229 | 0 | 591920 | 250550 | 0 | 0 |
| LSPNEYFSI | PCDHA4; PCDH | Q9UN74; A0 | PCDA4_HUMA | Unmodified | 0 | 0 | 0 | 0 | 0 | 0 | 0 | 0 | 2496500 | 174940 | 0 | 0 | 0 | 0 | 78189 | 0 | 0 | 0 |
| LSPPYSSP | TRIM28; TRIM2 | Q13263; M0 | TIF1B_HUMAN | Unmodified | 578210 | 0 | 0 | 0 | 141620 | 0 | 662600 | 491270 | 2196900 | 1295300 | 42132 | 2956400 | 233410 | 0 | 0 | 661140 | 485090 | 0 |
| LSPQAVNS | SRI; SRI | P30626; C9J | SORCN_HUMA | Unmodified | 3477400 | 3929900 | 0 | 5388500 | 852620 | 618440 | 493970 | 128260 | 33222000 | 1469700 | 8367500 | 11236000 | 913830 | 0 | 4719400 | 2505700 | 336210 | 2363300 |
| LSPQAVNS | SRI; SRI | P30626; C9J | SORCN_HUMA | Unmodified | 1761200 | 1896300 | 933600 | 2199300 | 1411600 | 34755 | 4176600 | 1209800 | 19370000 | 616100 | 1439700 | 3565600 | 78972 | 2794300 | 1655100 | 1369800 | 248480 | 266060 |
| LSPQFPNE | EIF2B2 | P49770 | EI2BB_HUMAN | Unmodified | 0 | 0 | 0 | 0 | 0 | 0 | 0 | 0 | 367690 | 157120 | 706250 | 5971000 | 0 | 0 | 0 | 0 | 22972 | 0 |
| LSPQTLTTI | GCA; GCA; GC | P28676; H7 | GRAN_HUMAN | Unmodified | 0 | 0 | 316610 | 144360 | 0 | 0 | 0 | 0 | 1937700 | 622430 | 638120 | 1711600 | 0 | 0 | 0 | 320680 | 0 | 0 |
| LSPQTLTTI | GCA; GCA; GC | P28676; H7 | GRAN_HUMAN | Unmodified | 309030 | 158410 | 0 | 435230 | 0 | 0 | 0 | 0 | 1762400 | 0 | 183890 | 3214300 | 0 | 0 | 334220 | 196710 | 67893 | 0 |
| LSPSARES | N/A | N/A | N/A | Unmodified | 0 | 0 | 0 | 0 | 0 | 0 | 0 | 0 | 1611600 | 0 | 0 | 0 | 0 | 0 | 0 | 0 | 0 | 0 |
| LSPSCPDA | MUC5B; MUC5 | Q9HC84; P9 | MUC5B_HUMA | Unmodified | 0 | 478520 | 0 | 0 | 0 | 0 | 0 | 0 | 0 | 425790 | 83779 | 0 | 0 | 0 | 0 | 331030 | 0 | 3994000 |
| LSPSTNSIN | PLCB2 | Q00722 | PLCB2_HUMAN | Unmodified | 0 | 0 | 0 | 1093600 | 0 | 0 | 0 | 0 | 0 | 0 | 2464000 | 17178000 | 0 | 2203400 | 0 | 0 | 0 | 0 |
| LSPSTSGPI | MOV10; MOV1 | Q9HCE1; Q5 | MOV10_HUMA | Unmodified | 2471000 | 2871600 | 177950 | 123520 | 0 | 0 | 0 | 0 | 0 | 0 | 619550 | 0 | 0 | 0 | 1401700 | 1966800 | 0 | 0 |
| LSPSTSGPI | MOV10; MOV1 | Q9HCE1; Q5 | MOV10_HUMA | Unmodified | 15846000 | 16022000 | 0 | 10255000 | 0 | 0 | 7821800 | 0 | 0 | 0 | 0 | 0 | 20272000 | 6095900 | 8920400 | 6447600 | 0 | 0 |
| LSPTDNLPI | CNP | P09543 | CN37_HUMAN | Unmodified | 0 | 0 | 0 | 0 | 0 | 1135800 | 360520 | 206050 | 1495000 | 439830 | 6392700 | 24618000 | 0 | 0 | 2442900 | 0 | 1959300 | 0 |
| LSPVTACA | NHLRC2 | Q8NBF2 | NHLC2_HUMA | Unmodified | 0 | 0 | 0 | 0 | 0 | 0 | 0 | 0 | 0 | 0 | 0 | 1209800 | 0 | 0 | 0 | 0 | 0 | 0 |
| LSQADGSF | UTRN | P46939 | UTRN_HUMAN | Unmodified | 0 | 0 | 449750 | 493370 | 1490100 | 306720 | 0 | 0 | 0 | 0 | 0 | 0 | 230910 | 196310 | 0 | 763650 | 0 | 152070 |
| LSQAQGNI | IGSF3 | O75054 | IGSF3_HUMAN | Unmodified | 0 | 126490 | 152490 | 0 | 56550 | 0 | 1630400 | 115720 | 0 | 0 | 0 | 0 | 66785 | 0 | 273300 | 164190 | 0 | 0 |
| LSQDPEPD | SIT1; SIT1 | Q9Y3P8; A0 | SIT1_HUMAN S | Unmodified | 0 | 0 | 0 | 0 | 0 | 0 | 0 | 0 | 1537200 | 0 | 0 | 0 | 0 | 0 | 0 | 0 | 0 | 0 |
| LSQECPIK | BZW2; BZW2; B | Q9Y6E2; B5 | BZW2_HUMAN | Unmodified | 0 | 655260 | 184040 | 0 | 606570 | 453690 | 172160 | 0 | 172300 | 67554 | 162560 | 5134800 | 959030 | 0 | 0 | 0 | 0 | 634310 |
| LSQETEAL | NUCB1 | Q02818 | NUCB1_HUMA | Unmodified | 146660 | 113540 | 153640 | 0 | 0 | 0 | 0 | 0 | 9778400 | 105140 | 2739600 | 326760 | 0 | 0 | 0 | 73733 | 0 | 0 |
| LSQEYESIE | PTPRS; PTPRS | Q13332; G8 | PTPRS_HUMAN | Unmodified | 0 | 0 | 0 | 0 | 2033000 | 1234900 | 923160 | 0 | 2244700 | 529220 | 0 | 2414200 | 0 | 0 | 0 | 0 | 610370 | 1174700 |
| LSQIGVEN | ALDOC; ALDOC | P09972; J3Q | ALDOC_HUMA | Unmodified | 0 | 0 | 0 | 0 | 0 | 0 | 0 | 0 | 0 | 0 | 0 | 0 | 0 | 0 | 0 | 0 | 0 | 0 |
| LSQKAPSLI | SLC43A1; SLC4 | O75387; E9F | LAT3_HUMAN | Oxidation (M) | 0 | 0 | 0 | 2695800 | 0 | 0 | 0 | 0 | 0 | 0 | 0 | 0 | 0 | 0 | 0 | 1069700 | 0 | 0 |
| LSQLEGVN | VPS35 | Q96QK1 | VPS35_HUMAN | Unmodified | 0 | 0 | 0 | 0 | 891460 | 0 | 0 | 0 | 0 | 0 | 0 | 3114900 | 0 | 0 | 0 | 0 | 899480 | 1160400 |
| LSQLEGVN | VPS35 | Q96QK1 | VPS35_HUMAN | Unmodified | 0 | 0 | 168080 | 99938 | 0 | 0 | 0 | 0 | 362380 | 0 | 0 | 1802100 | 250160 | 0 | 264360 | 181630 | 0 | 65218 |
| LSQLTSLLS | INPP5D | Q92835 | SHIP1_HUMAN | Unmodified | 0 | 0 | 0 | 0 | 0 | 0 | 0 | 0 | 0 | 0 | 228290 | 840220 | 0 | 0 | 0 | 0 | 0 | 0 |
| LSQMILDK | PSMD11 | O00231 | PSD11_HUMAN | Unmodified | 0 | 0 | 0 | 0 | 0 | 0 | 0 | 0 | 0 | 0 | 257090 | 2637200 | 0 | 437300 | 0 | 0 | 0 | 190030 |
| LSQNNFAL | VDAC3 | Q9Y277 | VDAC3_HUMA | Unmodified | 0 | 0 | 0 | 0 | 0 | 0 | 0 | 73974 | 46713000 | 3334400 | 511140 | 0 | 0 | 0 | 0 | 0 | 479260 | 604770 |
| LSQPITVQ | CAPN5; CAPN5 | O15484; E7E | CAN5_HUMAN | Unmodified | 0 | 0 | 345940 | 239670 | 321680 | 0 | 0 | 0 | 1899800 | 0 | 0 | 1043900 | 1081500 | 1262000 | 341810 | 554030 | 184880 | 263770 |
| LSQQSKLE | GPC6 | Q9Y625 | GPC6_HUMAN | Unmodified | 5870200 | 11799000 | 3899400 | 8868100 | 0 | 0 | 0 | 0 | 0 | 0 | 0 | 0 | 0 | 0 | 0 | 1162300 | 0 | 0 |
| LSQRPTPEI | PHACTR4 | Q8IZ21 | PHAR4_HUMA | Unmodified | 114810 | 50854 | 76489 | 82795 | 310410 | 152490 | 0 | 0 | 1147200 | 120880 | 178490 | 365930 | 199920 | 91858 | 106250 | 113270 | 63587 | 299310 |
| LSQRPTVAI | PHACTR4 | Q8IZ21 | PHAR4_HUMA | Unmodified | 0 | 0 | 0 | 0 | 596910 | 33886 | 0 | 0 | 0 | 0 | 0 | 0 | 0 | 85038 | 0 | 0 | 0 | 0 |
| LSQSVYHG | WASF3 | Q9UPY6 | WASF3_HUMA | Unmodified | 331530 | 0 | 0 | 186570 | 0 | 0 | 0 | 0 | 0 | 0 | 0 | 0 | 0 | 0 | 232910 | 270200 | 0 | 0 |
| LSQTLSLVF | B3GAT3; B3GA | O94766; E9F | B3GA3_HUMA | Unmodified | 0 | 0 | 0 | 0 | 0 | 0 | 0 | 0 | 974770 | 171040 | 0 | 846240 | 0 | 0 | 29391 | 14752 | 0 | 0 |
| LSQTVAPN | CEP55 | Q53EZ4 | CEP55_HUMAN | Unmodified | 0 | 0 | 0 | 0 | 850020 | 0 | 0 | 0 | 3093300 | 0 | 0 | 535110 | 0 | 0 | 0 | 0 | 0 | 0 |
| LSQTVAPN | CEP55 | Q53EZ4 | CEP55_HUMAN | Oxidation (M) | 0 | 0 | 0 | 1905100 | 0 | 0 | 0 | 0 | 0 | 0 | 0 | 0 | 0 | 0 | 0 | 0 | 0 | 0 |
| LSRADLTE | UQCRC1 | P31930 | QCR1_HUMAN | Unmodified | 0 | 0 | 0 | 0 | 0 | 0 | 0 | 0 | 12855000 | 3299600 | 431140 | 0 | 0 | 0 | 0 | 0 | 0 | 0 |
| LSRADLTE | UQCRC1 | P31930 | QCR1_HUMAN | Unmodified | 0 | 0 | 0 | 0 | 0 | 0 | 239620 | 0 | 0 | 2425000 | 0 | 0 | 0 | 0 | 0 | 0 | 0 | 0 |
| LSRDATFH | IGSF3 | O75054 | IGSF3_HUMAN | Unmodified | 0 | 0 | 0 | 0 | 603980 | 205070 | 0 | 0 | 0 | 0 | 0 | 50728 | 0 | 305840 | 0 | 0 | 0 | 51503 |
| LSREHGAF | MTHFD1; MTHF | P11586; V9G | C1TC_HUMAN | Unmodified | 0 | 0 | 196320 | 0 | 371780 | 306850 | 0 | 0 | 0 | 0 | 0 | 2767700 | 169100 | 72659 | 442150 | 0 | 0 | 0 |
| LSRELDFV | NUCB1; NUCB | Q02818; H7I | NUCB1_HUMA | Unmodified | 0 | 0 | 0 | 0 | 0 | 0 | 0 | 0 | 232920 | 52105 | 0 | 0 | 0 | 0 | 0 | 0 | 127050 | 0 |
| LSRELQVLI | EXOC1 | Q9NV70 | EXOC1_HUMA | Unmodified | 0 | 0 | 0 | 0 | 2305900 | 0 | 4084900 | 0 | 20916000 | 465920 | 0 | 0 | 0 | 0 | 0 | 0 | 492630 | 0 |
| LSRQEAVA | CRK; CRK | P46108; I3L | CRK_HUMAN A | Unmodified | 552300 | 597950 | 0 | 0 | 0 | 0 | 0 | 0 | 0 | 0 | 0 | 563300 | 0 | 0 | 494300 | 514030 | 0 | 0 |
| LSRRDLPA | TRAF4; TRAF4; | Q9BUZ4; A0 | TRAF4_HUMAN | Unmodified | 0 | 0 | 0 | 0 | 0 | 0 | 0 | 15084 | 0 | 0 | 0 | 0 | 411530 | 275960 | 0 | 0 | 35235 | 38276 |
| LSSADDHR | HSPA13 | P48723 | HSP13_HUMAN | Unmodified | 1517900 | 0 | 177040 | 85071 | 0 | 0 | 0 | 0 | 1377900 | 69656 | 69710 | 0 | 0 | 0 | 0 | 0 | 0 | 0 |
| LSSAIAFTYI | LRBA; LRBA; LR | P50851; A0A | LRBA_HUMAN | Unmodified | 0 | 0 | 0 | 0 | 0 | 0 | 0 | 0 | 246890 | 0 | 0 | 1427700 | 78291 | 0 | 0 | 0 | 0 | 714050 |
| LSSAMSAA | MDH1; MDH1; | P40925; B8Z | MDHC_HUMAN | Oxidation (M) | 0 | 0 | 54171 | 0 | 564530 | 0 | 144730 | 103080 | 0 | 213580 | 0 | 0 | 0 | 0 | 41878 | 0 | 222650 | 293230 |
| LSSAMSAA | MDH1; MDH1; | P40925; B8Z | MDHC_HUMAN | Unmodified | 426500 | 462800 | 0 | 935620 | 19929000 | 0 | 3451600 | 2011900 | 4116100 | 2748800 | 1110100 | 24537000 | 7122100 | 7080300 | 619550 | 463000 | 9766800 | 11077000 |
| LSSCDSFT | EPRS1; EPRS1; | P07814; V9G | SYEP_HUMAN | Unmodified | 0 | 0 | 0 | 0 | 0 | 0 | 0 | 0 | 0 | 0 | 0 | 2510700 | 0 | 0 | 0 | 0 | 0 | 0 |
| LSSDCEDC | GLG1 | Q92896 | GSLG1_HUMA | Unmodified | 0 | 0 | 0 | 0 | 0 | 0 | 0 | 0 | 4231300 | 111430 | 423680 | 1229800 | 0 | 0 | 0 | 0 | 0 | 0 |
| LSSDQND | LMNB2 | Q03252 | LMNB2_HUMA | Unmodified | 0 | 0 | 0 | 0 | 607720 | 0 | 0 | 0 | 0 | 133530 | 0 | 0 | 0 | 0 | 0 | 0 | 0 | 0 |
| LSSDVCPT | IGF2R; IGF2R | P11717; A0A | MPRI_HUMAN | Unmodified | 134910 | 0 | 411420 | 184120 | 0 | 0 | 16368 | 0 | 15355000 | 2856200 | 1021200 | 2108700 | 93954 | 145550 | 221180 | 240550 | 85310 | 139900 |
| LSSDVLTLL | LONP1; LONP1 | P36776; K7E | LONM_HUMAN | Unmodified | 0 | 0 | 0 | 0 | 0 | 0 | 385880 | 0 | 5597900 | 1116100 | 0 | 66857 | 0 | 0 | 0 | 0 | 0 | 0 |
| LSSEQEKE | KIF20B | Q96Q89 | KI20B_HUMAN | Unmodified | 0 | 0 | 84948 | 0 | 327860 | 0 | 27063 | 11787 | 347980 | 0 | 0 | 433970 | 0 | 0 | 0 | 0 | 120530 | 0 |
| LSSGFDDI | PKP3; PKP3 | Q9Y446; E9F | PKP3_HUMAN | Unmodified | 1207200 | 2168200 | 3004700 | 6335600 | 319880 | 0 | 0 | 0 | 2569900 | 1321900 | 5567700 | 6495300 | 5343800 | 345870 | 6463600 | 4810600 | 92553 | 0 |
| LSSHPVLSI | SNX5; SNX5; S | Q9Y5X3; Q5 | SNX5_HUMAN | Unmodified | 0 | 0 | 0 | 0 | 0 | 175400 | 0 | 0 | 47010 | 38090 | 56810 | 398870 | 0 | 0 | 0 | 0 | 0 | 0 |
| LSSLIILMP | SERPINH1; SER | P50454; E9P | SERPH_HUMA | Oxidation (M) | 0 | 0 | 0 | 0 | 0 | 0 | 0 | 0 | 1463700 | 0 | 0 | 0 | 0 | 0 | 0 | 0 | 0 | 0 |
| LSSLIILMP | SERPINH1; SER | P50454; E9P | SERPH_HUMA | Unmodified | 0 | 0 | 0 | 0 | 0 | 0 | 0 | 0 | 2719800 | 0 | 0 | 0 | 0 | 0 | 0 | 0 | 0 | 0 |
| LSSLIILMP | SERPINH1; SER | P50454; E9P | SERPH_HUMA | Oxidation (M) | 0 | 0 | 0 | 0 | 0 | 0 | 0 | 0 | 16438000 | 10071000 | 0 | 0 | 0 | 0 | 147680 | 0 | 0 | 0 |
| LSSLIILMP | SERPINH1; SER | P50454; E9P | SERPH_HUMA | Unmodified | 0 | 0 | 0 | 0 | 0 | 0 | 0 | 0 | 17198000 | 4214000 | 0 | 0 | 0 | 0 | 0 | 0 | 0 | 0 |
| LSSLNLDD | MRPS35 | P82673 | RT35_HUMAN | Unmodified | 0 | 0 | 0 | 0 | 0 | 0 | 13408 | 0 | 477650 | 0 | 0 | 73938 | 0 | 0 | 0 | 0 | 0 | 0 |
| LSSLPFQK | NUTF2; NUTF2 | P61970; H3 | NTF2_HUMAN | Unmodified | 231600 | 1279100 | 2250100 | 1770500 | 9756300 | 1506100 | 3552900 | 0 | 3518600 | 0 | 3518500 | 12695000 | 10878000 | 6471500 | 0 | 2623700 | 6295100 | 4835700 |
| LSSLRASTS | RPS6; RPS6 | P62753; A2A | RS6_HUMAN 4 | Unmodified | 0 | 0 | 0 | 0 | 0 | 0 | 820090 | 0 | 0 | 0 | 0 | 0 | 1676400 | 0 | 0 | 0 | 0 | 955760 |
| LSSMLAEL | NOSTRIN; NOS | Q8IVI9; A0A | NOSTN_HUMA | Oxidation (M) | 3266900 | 511630 | 3456000 | 4593100 | 0 | 0 | 0 | 0 | 0 | 0 | 0 | 505510 | 0 | 0 | 10000000 | 4404400 | 0 | 0 |
| LSSMLAEL | NOSTRIN; NOS | Q8IVI9; A0A | NOSTN_HUMA | Unmodified | 1242700 | 715360 | 1007100 | 2064900 | 0 | 0 | 0 | 0 | 0 | 0 | 0 | 0 | 0 | 0 | 3164700 | 815900 | 0 | 0 |
| LSSQISAGI | MTDH; MTDH; | Q86UE4; E5I | LYRIC_HUMAN | Unmodified | 0 | 0 | 0 | 0 | 0 | 0 | 0 | 0 | 0 | 0 | 170340 | 210800 | 0 | 117050 | 109520 | 57786 | 0 | 0 |
| LSSSDRYS | CTPS2 | Q9NRF8 | PYRG2_HUMA | Unmodified | 0 | 0 | 0 | 0 | 0 | 0 | 0 | 0 | 206880 | 0 | 94631 | 711810 | 0 | 0 | 111980 | 0 | 0 | 0 |
| LSSSGIWA | TRIM26 | Q12899 | TRI26_HUMAN | Unmodified | 17921000 | 20200000 | 6063700 | 11813000 | 1692500 | 0 | 0 | 0 | 352130 | 98485 | 0 | 0 | 61899 | 0 | 20563000 | 17031000 | 0 | 0 |

| | | | | | | | | | | | | | | | | | | | | | | |
|---|---|---|---|---|---|---|---|---|---|---|---|---|---|---|---|---|---|---|---|---|---|---|
| LSSSQGTIE | TBK1; TBK1; TBI | Q9UHD2; A0 | TBK1_HUMAN ! | Unmodified | 0 | 0 | 0 | 0 | 0 | 0 | 0 | 0 | 309230 | 0 | 0 | 0 | 0 | 0 | 0 | 0 | 0 | 0 |
| LSSTSSEET | PROM2 | Q8N271 | PROM2_HUMA | Unmodified | 0 | 0 | 0 | 0 | 0 | 0 | 0 | 0 | 0 | 0 | 0 | 1248500 | 0 | 0 | 0 | 0 | 0 | 0 |
| LSSTWEGI( | TALDO1; TALDO | P37837; F2Z | TALDO_HUMAI | Unmodified | 0 | 0 | 0 | 0 | 1746100 | 0 | 351740 | 0 | 0 | 277670 | 3083500 | 14265000 | 0 | 0 | 0 | 0 | 1095900 | 620960 |
| LSSVQQIL( | KIAA2013 | Q8IYS2 | K2013_HUMAN | Unmodified | 0 | 14800 | 0 | 0 | 0 | 0 | 0 | 0 | 1776700 | 1326500 | 0 | 16030 | 0 | 0 | 48078 | 0 | 0 | 0 |
| LSTASGYSI | DTX2; DTX2; DT | Q86UW9; Q7 | DTX2_HUMAN | Unmodified | 0 | 0 | 0 | 0 | 0 | 226070 | 0 | 0 | 0 | 0 | 0 | 0 | 0 | 0 | 0 | 54122 | 0 | 0 |
| LSTDHIPIL\ | GFPT1; GFPT1; | Q06210; A0/ | GFPT1_HUMAN | Unmodified | 0 | 0 | 2299000 | 1608600 | 178110 | 0 | 0 | 22974 | 6728200 | 375030 | 739890 | 4206600 | 2143600 | 964210 | 0 | 0 | 64647 | 143310 |
| LSTDHIPIL\ | GFPT1; GFPT1 | A0A6I8PTT9; | A0A6I8PTT9_HI | Unmodified | 0 | 0 | 0 | 138770 | 1477000 | 630160 | 3251600 | 1607200 | 224240 | 1468000 | 0 | 0 | 10764000 | 4472900 | 317410 | 0 | 928570 | 1772100 |
| LSTDHIPIL\ | GFPT1; GFPT1 | A0A6I8PTT9; | A0A6I8PTT9_HI | Unmodified | 0 | 0 | 0 | 0 | 0 | 0 | 0 | 0 | 0 | 104830 | 0 | 0 | 1428400 | 760890 | 0 | 0 | 0 | 471400 |
| LSTDVCEQ | CARMIL1 | Q5VZK9 | CARL1_HUMAN | Unmodified | 0 | 0 | 0 | 0 | 0 | 0 | 0 | 261290 | 0 | 0 | 0 | 1086800 | 0 | 0 | 0 | 0 | 104210 | 397970 |
| LSTGCTGTI | PRSS23 | O95084 | PRS23_HUMAN | Unmodified | 494620 | 207610 | 0 | 0 | 6512000 | 2168700 | 0 | 0 | 171730 | 70736 | 0 | 227480 | 312020 | 0 | 221970 | 126470 | 0 | 1975700 |
| LSTGVFRP( | CRYZL1; CRYZL | O95825; A6N | QORL1_HUMAI | Unmodified | 208970 | 0 | 202190 | 363520 | 6280100 | 1100200 | 863010 | 134270 | 1634400 | 1175400 | 904710 | 3998700 | 1899600 | 899100 | 460920 | 258690 | 3111300 | 1937600 |
| LSTHELTSL | MPRIP; MPRIP; | Q6WCQ1; J3 | MPRIP_HUMAN | Unmodified | 0 | 0 | 0 | 0 | 0 | 0 | 0 | 0 | 0 | 0 | 50444 | 538080 | 0 | 0 | 20032 | 0 | 0 | 0 |
| LSTKDPLPT | MRPL37; MRPL: | Q9BZE1; H0' | RM37_HUMAN | Unmodified | 0 | 0 | 0 | 0 | 0 | 0 | 0 | 0 | 1199800 | 0 | 0 | 0 | 0 | 0 | 0 | 0 | 0 | 0 |
| LSTKDVEEL | PALM2; PALM2/ | Q8IXS6; B1A | PALM2_HUMAN | Unmodified | 0 | 0 | 404210 | 0 | 415800 | 0 | 0 | 37182 | 212220 | 227930 | 0 | 0 | 0 | 0 | 0 | 0 | 0 | 0 |
| LSTLILHGG | TERF2IP; TERF2 | Q9NYB0; A0. | TE2IP_HUMAN | Unmodified | 0 | 0 | 0 | 0 | 0 | 0 | 0 | 0 | 0 | 0 | 0 | 1068400 | 0 | 0 | 0 | 0 | 0 | 0 |
| LSTLKIENE | SRGAP2; SRGA | O75044; P0I | SRGP2_HUMAI | Unmodified | 0 | 0 | 0 | 0 | 631120 | 0 | 75075 | 65962 | 0 | 91643 | 80391 | 0 | 1777300 | 0 | 0 | 0 | 160030 | 255710 |
| LSTMETGT( | TPP2; TPP2; TPI | P29144; A0A | TPP2_HUMAN ' | Unmodified | 0 | 0 | 0 | 0 | 845830 | 0 | 0 | 54101 | 521160 | 0 | 0 | 2467600 | 0 | 0 | 0 | 0 | 0 | 349100 |
| LSTMETGT( | TPP2; TPP2; TPI | P29144; A0A | TPP2_HUMAN ' | Oxidation (M) | 0 | 0 | 0 | 0 | 0 | 0 | 0 | 327140 | 0 | 0 | 0 | 3987600 | 0 | 0 | 102950 | 92655 | 0 | 0 |
| LSTQVKPLI | AP1M2; AP1M2 | Q9Y6Q5; K7I | AP1M2_HUMAI | Unmodified | 0 | 110430 | 16151 | 190100 | 0 | 0 | 0 | 0 | 38041 | 0 | 60282 | 0 | 20078 | 0 | 141960 | 150870 | 0 | 0 |
| LSTRSDLSI | SNX18 | Q96RF0 | SNX18_HUMAN | Unmodified | 0 | 121500 | 134470 | 993920 | 0 | 0 | 39671 | 0 | 0 | 28077 | 52730 | 0 | 0 | 201010 | 159910 | 78921 | 0 | 0 |
| LSTRSDLSI | SNX18 | Q96RF0 | SNX18_HUMAN | Unmodified | 0 | 0 | 0 | 0 | 0 | 0 | 0 | 0 | 0 | 0 | 0 | 0 | 3449600 | 2341600 | 0 | 0 | 0 | 0 |
| LSTSGNRP | SMCHD1; SMCI | A6NHR9; J3I | SMHD1_HUMA | Unmodified | 112730 | 23623 | 0 | 0 | 340350 | 0 | 398610 | 188650 | 0 | 0 | 0 | 111260 | 162400 | 123800 | 28106 | 0 | 37933 | 140160 |
| LSTSIRDLQ | ERO1B | Q86YB8 | ERO1B_HUMAI | Unmodified | 0 | 0 | 0 | 0 | 0 | 0 | 0 | 0 | 841300 | 358690 | 0 | 0 | 0 | 0 | 0 | 0 | 0 | 0 |
| LSTTECVD/ | PXDN | Q92626 | PXDN_HUMAN | Unmodified | 0 | 0 | 0 | 0 | 0 | 0 | 0 | 0 | 801790 | 0 | 0 | 0 | 0 | 0 | 0 | 0 | 0 | 0 |
| LSVAAQEA/ | PLEC | Q15149 | PLEC_HUMAN | Unmodified | 14907000 | 18887000 | 0 | 0 | 811170 | 25782000 | 140950 | 24246 | 421150 | 0 | 0 | 501780 | 2112500 | 0 | 11093000 | 3031100 | 0 | 1.04E+08 |
| LSVDEDFC | TMEM87B; TME | Q96K49; A0/ | TM87B_HUMAN | Unmodified | 0 | 0 | 0 | 0 | 0 | 0 | 0 | 0 | 2783300 | 0 | 0 | 0 | 0 | 0 | 0 | 0 | 0 | 0 |
| LSVDYGKK | TUBA3C; TUBA: | P0DPH7; Q7 | TBA3C_HUMAN | Unmodified | 0 | 0 | 1651200 | 1202500 | 44858 | 532840 | 155520 | 0 | 1900100 | 671880 | 24850 | 36286000 | 0 | 0 | 1597800 | 1174500 | 0 | 0 |
| LSVEAPPK | IST1; IST1; IST1 | P53990; H3E | IST1_HUMAN I: | Unmodified | 587320 | 663820 | 0 | 2787700 | 12506000 | 3296300 | 0 | 0 | 9215600 | 5221800 | 4823400 | 14537000 | 4029700 | 5160700 | 0 | 714080 | 865250 | 2318200 |
| LSVEAPPKI | IST1; IST1; IST1 | P53990; H3E | IST1_HUMAN I: | Unmodified | 0 | 0 | 0 | 0 | 12371000 | 0 | 2044900 | 0 | 0 | 1572300 | 0 | 0 | 2827300 | 4726900 | 0 | 0 | 1532000 | 2605300 |
| LSVEGFAVI | NPEPPS; NPEPI | P55786; E9P | PSA_HUMAN P | Unmodified | 0 | 0 | 0 | 0 | 0 | 0 | 0 | 0 | 0 | 0 | 0 | 95519000 | 0 | 0 | 0 | 430250 | 0 | 0 |
| LSVEIWDW | PRKCA; PRKCA | P17252; J3KI | KPCA_HUMAN | Unmodified | 44768 | 0 | 0 | 0 | 0 | 0 | 0 | 0 | 0 | 0 | 0 | 493020 | 0 | 0 | 0 | 0 | 0 | 0 |
| LSVGDIENI | SLC20A1 | Q8WUM9 | S20A1_HUMAN | Unmodified | 483240 | 504340 | 213830 | 0 | 0 | 0 | 0 | 0 | 0 | 0 | 270660 | 883250 | 0 | 0 | 253710 | 304310 | 0 | 0 |
| LSVGDIENI | SLC20A1 | Q8WUM9 | S20A1_HUMAN | Unmodified | 0 | 0 | 0 | 0 | 0 | 0 | 0 | 0 | 2744500 | 0 | 3146700 | 6510900 | 0 | 0 | 0 | 0 | 0 | 0 |
| LSVIQNSA\ | MGAT4A | Q9UM21 | MGT4A_HUMAI | Unmodified | 0 | 0 | 0 | 0 | 0 | 0 | 0 | 0 | 860710 | 138750 | 0 | 0 | 0 | 0 | 0 | 0 | 0 | 0 |
| LSVISVEDP | ETFB | P38117 | ETFB_HUMAN I | Unmodified | 0 | 0 | 0 | 0 | 0 | 0 | 0 | 0 | 1644100 | 0 | 0 | 0 | 0 | 0 | 0 | 0 | 0 | 0 |
| LSVLEEEQI | PTPRF; PTPRF | P10586; A2A | PTPRF_HUMAN | Oxidation (M) | 530150 | 682980 | 0 | 0 | 2452900 | 107160 | 0 | 0 | 700020 | 1057200 | 83761 | 0 | 933940 | 0 | 416410 | 0 | 0 | 0 |
| LSVLGAITS | ETF1; ETF1; ETF | P62495; B7Z | ERF1_HUMAN I | Unmodified | 1836800 | 352340 | 825250 | 1205400 | 0 | 83360 | 0 | 0 | 1197300 | 0 | 2188400 | 10488000 | 1166000 | 0 | 1174600 | 1193700 | 0 | 0 |
| LSVLLLEK | TKFC; TKFC; TK | Q3LXA3; I3L: | TKFC_HUMAN ' | Unmodified | 0 | 0 | 0 | 0 | 0 | 0 | 0 | 0 | 0 | 390710 | 0 | 0 | 1147200 | 588970 | 481930 | 0 | 0 | 409320 |
| LSVLQEEK | KANK2 | Q63ZY3 | KANK2_HUMAI | Unmodified | 0 | 0 | 0 | 0 | 840520 | 0 | 0 | 0 | 0 | 0 | 0 | 0 | 0 | 0 | 0 | 0 | 0 | 0 |
| LSVLQQELI | SOGA1; SOGA1 | O94964; X6F | SOGA1_HUMAI | Unmodified | 0 | 0 | 0 | 0 | 0 | 0 | 0 | 0 | 0 | 0 | 0 | 0 | 341190 | 0 | 0 | 47173 | 0 | 0 |
| LSVPSGFS\ | SLC4A2 | P04920 | B3A2_HUMAN . | Unmodified | 568200 | 518390 | 2987900 | 1493300 | 0 | 0 | 0 | 0 | 756560 | 0 | 0 | 2239900 | 0 | 0 | 3749200 | 2026500 | 0 | 0 |
| LSVPTSDEI | ABCF1; ABCF1; | Q8NE71; A0. | ABCF1_HUMAN | Unmodified | 0 | 0 | 0 | 0 | 0 | 0 | 0 | 0 | 0 | 0 | 1111600 | 1268600 | 0 | 0 | 0 | 0 | 0 | 0 |
| LSVPYVPQ' | EFR3A; EFR3A | Q14156; A0/ | EFR3A_HUMAN | Unmodified | 520030 | 0 | 765470 | 1484800 | 0 | 0 | 0 | 0 | 3209600 | 0 | 229330 | 0 | 1398000 | 0 | 1224700 | 1156600 | 0 | 0 |
| LSVQGEVS | MLEC; MLEC; M | Q14165; H0' | MLEC_HUMAN | Unmodified | 140410 | 116700 | 0 | 0 | 0 | 0 | 0 | 0 | 9888300 | 739720 | 0 | 233510 | 0 | 0 | 259810 | 134420 | 0 | 0 |
| LSVQTHSTI | PTPRJ; PTPRJ; P | Q12913; A0/ | PTPRJ_HUMAN | Unmodified | 0 | 0 | 0 | 0 | 0 | 0 | 0 | 0 | 0 | 0 | 752110 | 2383500 | 0 | 0 | 0 | 0 | 0 | 0 |
| LSVQTHSTI | PTPRJ; PTPRJ; P | Q12913; A0/ | PTPRJ_HUMAN | Oxidation (M) | 0 | 0 | 0 | 0 | 636980 | 0 | 0 | 0 | 0 | 0 | 0 | 201010 | 501120 | 0 | 0 | 0 | 0 | 0 |
| LSVQTHSTI | PTPRJ; PTPRJ; P | Q12913; A0/ | PTPRJ_HUMAN | Unmodified | 439910 | 458660 | 0 | 398570 | 7808800 | 0 | 435410 | 0 | 440830 | 0 | 649240 | 7547000 | 1643300 | 0 | 165730 | 384320 | 0 | 2746400 |
| LSVSGRDE | GANAB; GANAE | Q14697; F5H | GANAB_HUMA | Oxidation (M) | 0 | 0 | 0 | 0 | 0 | 0 | 0 | 0 | 9819700 | 2154200 | 0 | 0 | 0 | 0 | 0 | 0 | 0 | 0 |
| LSVSGRDE | GANAB; GANAE | Q14697; F5H | GANAB_HUMA | Unmodified | 0 | 0 | 0 | 0 | 0 | 0 | 0 | 0 | 50933000 | 4333600 | 1358700 | 0 | 0 | 0 | 0 | 0 | 0 | 445400 |
| LSVSRALS/ | PTGFRN | Q9P2B2 | FPRP_HUMAN | Unmodified | 0 | 0 | 0 | 0 | 347500 | 154350 | 0 | 0 | 0 | 0 | 0 | 0 | 789440 | 371350 | 0 | 0 | 0 | 0 |
| LSVTNLEDI | DAGLB; DAGLB | Q8NCG7; E7 | DGLB_HUMAN | Unmodified | 0 | 0 | 0 | 0 | 0 | 0 | 0 | 0 | 0 | 0 | 0 | 0 | 0 | 0 | 0 | 0 | 0 | 0 |
| LSVTSTCGI | LAMB1; LAMB1; | P07942; A0A | LAMB1_HUMAI | Unmodified | 186510 | 43084 | 0 | 116680 | 0 | 0 | 0 | 0 | 0 | 0 | 0 | 0 | 0 | 0 | 0 | 0 | 0 | 649400 |
| LSVTVDPK | HDLBP; HDLBP | Q00341; A0/ | VIGLN_HUMAN | Unmodified | 0 | 0 | 0 | 118460 | 95528 | 0 | 0 | 0 | 0 | 103680 | 1158200 | 2221800 | 0 | 0 | 0 | 0 | 9612.1 | 164500 |
| LSVYAALQI | PLEC; PLEC | Q15149; H0' | PLEC_HUMAN | Unmodified | 0 | 0 | 55356 | 0 | 0 | 283750 | 0 | 0 | 0 | 132640 | 78712 | 0 | 0 | 0 | 237570 | 51863 | 2104500 | 0 |
| LSWATHYV | FOXRED2 | Q8IWF2 | FXRD2_HUMAN | Unmodified | 0 | 0 | 33235 | 0 | 0 | 0 | 0 | 0 | 522240 | 169360 | 0 | 0 | 0 | 0 | 0 | 38837 | 0 | 0 |
| LSWHLPGI | LIFR | P42702 | LIFR_HUMAN L | Unmodified | 0 | 0 | 0 | 0 | 169720 | 0 | 0 | 0 | 646270 | 246770 | 0 | 0 | 0 | 0 | 0 | 0 | 0 | 0 |
| LSWSQLG( | BCAM; BCAM | P50895; A0A | BCAM_HUMAN | Unmodified | 1332000 | 1597700 | 0 | 0 | 0 | 0 | 0 | 0 | 508980 | 0 | 0 | 162460 | 0 | 0 | 84345 | 224150 | 0 | 0 |
| LSYEHAQS | DIS3L2 | Q8IYB7 | DI3L2_HUMAN | Unmodified | 0 | 0 | 0 | 0 | 0 | 0 | 0 | 0 | 0 | 0 | 0 | 705860 | 0 | 0 | 280430 | 0 | 0 | 0 |
| LSYIQPR | TC2N | Q8N9U0 | TAC2N_HUMAI | Unmodified | 0 | 0 | 0 | 336600 | 0 | 0 | 0 | 0 | 0 | 0 | 0 | 0 | 0 | 0 | 0 | 0 | 0 | 0 |
| LSYLDVYLI | AKR1B15; AKR1 | C9JRZ8; O6C | AK1BF_HUMAN | Unmodified | 0 | 0 | 14854 | 0 | 212430 | 0 | 594380 | 0 | 1825300 | 518910 | 55282 | 0 | 0 | 0 | 0 | 0 | 0 | 0 |
| LSYLLMCLI | GLG1; GLG1 | Q92896; H3I | GSLG1_HUMAI | Oxidation (M) | 0 | 0 | 42894 | 0 | 0 | 0 | 0 | 0 | 10560000 | 639620 | 415800 | 1337500 | 0 | 1303700 | 171160 | 0 | 0 | 0 |
| LSYNTASNI | RPL34 | P49207 | RL34_HUMAN ( | Unmodified | 1687900 | 1614500 | 1669000 | 2750700 | 286470 | 0 | 1002500 | 581500 | 437480 | 24389 | 10032000 | 3580900 | 1375700 | 2170600 | 1617900 | 0 | 17991 | 1752700 |
| LSYQNDPP | DSC2; DSC2 | Q02487; A0/ | DSC2_HUMAN | Unmodified | 0 | 0 | 0 | 0 | 0 | 0 | 0 | 0 | 3296200 | 0 | 0 | 0 | 0 | 0 | 0 | 0 | 0 | 0 |
| LSYQNDPP | DSC2; DSC2 | Q02487; A0/ | DSC2_HUMAN | Unmodified | 0 | 0 | 0 | 0 | 0 | 0 | 0 | 0 | 1005600 | 0 | 0 | 0 | 0 | 0 | 0 | 0 | 46859 | 0 |
| LSYYDGMI( | IGF2R; IGF2R | P11717; A0A | MPRI_HUMAN ( | Oxidation (M) | 0 | 0 | 0 | 0 | 0 | 189290 | 442250 | 0 | 3878400 | 595300 | 441530 | 1643200 | 0 | 0 | 272500 | 0 | 0 | 0 |
| LSYYDGMI( | IGF2R; IGF2R | P11717; A0A | MPRI_HUMAN ( | Unmodified | 0 | 0 | 0 | 0 | 0 | 0 | 0 | 0 | 16806000 | 0 | 0 | 0 | 3060700 | 0 | 0 | 0 | 0 | 0 |
| LSYYDGMI( | IGF2R; IGF2R | P11717; A0A | MPRI_HUMAN ( | Unmodified | 0 | 0 | 0 | 0 | 0 | 0 | 0 | 166000 | 1506300 | 0 | 0 | 0 | 0 | 0 | 0 | 0 | 0 | 0 |
| LTAALLEST. | HOMER1; HOM | Q86YM7; Q8 | HOME1_HUMA | Unmodified | 0 | 0 | 0 | 0 | 0 | 0 | 0 | 0 | 0 | 0 | 0 | 0 | 1992000 | 971100 | 0 | 0 | 0 | 0 |
| LTADLSAET | L1RE1 | Q9UN81 | LORF1_HUMAN | Unmodified | 4260200 | 3147800 | 538210 | 765150 | 112620 | 0 | 158410 | 58279 | 0 | 0 | 0 | 0 | 0 | 0 | 664090 | 1241100 | 176140 | 171930 |
| LTADLSAET | L1RE1 | Q9UN81 | LORF1_HUMAN | Unmodified | 1795700 | 1354500 | 287970 | 666970 | 0 | 0 | 0 | 0 | 130760 | 0 | 0 | 0 | 0 | 0 | 302010 | 264930 | 0 | 0 |
| LTAEAIQTP | CDCA8 | Q53HL2 | BOREA_HUMAI | Unmodified | 0 | 0 | 1946200 | 368670 | 0 | 0 | 0 | 0 | 0 | 0 | 0 | 0 | 0 | 0 | 610220 | 185380 | 0 | 0 |
| LTAEEMDE( | IQGAP3 | Q86VI3 | IQGA3_HUMAN | Unmodified | 0 | 0 | 0 | 97672 | 0 | 0 | 0 | 0 | 0 | 0 | 0 | 374120 | 0 | 0 | 0 | 0 | 0 | 0 |
| LTAEEMDEI | IQGAP1; IQGAP | P46940; A0A | IQGA1_HUMAN | Unmodified | 0 | 0 | 0 | 0 | 0 | 0 | 0 | 0 | 0 | 128050 | 0 | 1261500 | 0 | 0 | 77997 | 0 | 0 | 0 |
| LTAELIEQA | SNRPA1; SNRP, | P09661; H0' | RU2A_HUMAN | Unmodified | 574070 | 1128700 | 220850 | 0 | 0 | 0 | 0 | 0 | 297360 | 0 | 0 | 1234400 | 0 | 0 | 463460 | 535430 | 0 | 0 |
| LTAFVNTLI | PSMD4; PIPSL; | P55036; A2A | PSMD4_HUMA | Unmodified | 0 | 137220 | 194420 | 237050 | 0 | 0 | 0 | 0 | 0 | 0 | 106040 | 2434600 | 0 | 0 | 0 | 0 | 763470 | 0 |
| LTAGEACA( | ECI2; ECI2 | O75521; A0/ | ECI2_HUMAN E | Unmodified | 0 | 0 | 0 | 545920 | 0 | 0 | 357120 | 0 | 2063200 | 0 | 0 | 0 | 0 | 0 | 0 | 0 | 0 | 0 |
| LTAGLGSLI | CPD | O75976 | CBPD_HUMAN | Unmodified | 0 | 0 | 0 | 0 | 886500 | 0 | 0 | 0 | 1.38E+08 | 15400000 | 1427200 | 773540 | 1557600 | 0 | 0 | 0 | 194600 | 0 |
| LTAGVPDTI | ITGB4 | P16144 | ITB4_HUMAN I | Unmodified | 0 | 0 | 0 | 0 | 0 | 0 | 0 | 0 | 130140 | 0 | 0 | 0 | 0 | 0 | 294220 | 0 | 779830 | 0 |
| LTAIDILTTC | SNRNP200 | O75643 | U520_HUMAN | Unmodified | 253830 | 303240 | 58676 | 0 | 0 | 0 | 0 | 0 | 0 | 57574 | 157930 | 1395200 | 0 | 0 | 101920 | 0 | 111760 | 0 |
| LTAIELFDIL | UGT8; UGT8 | Q16880; D6I | CGT_HUMAN 2 | Unmodified | 0 | 0 | 0 | 0 | 0 | 0 | 0 | 0 | 242340 | 105580 | 0 | 0 | 0 | 0 | 0 | 0 | 0 | 0 |
| LTALDYHN | RTRAF; RTRAF; | Q9Y224; G3' | RTRAF_HUMAN | Unmodified | 0 | 228290 | 0 | 0 | 0 | 0 | 0 | 0 | 0 | 0 | 0 | 2906100 | 0 | 0 | 0 | 428320 | 0 | 0 |
| LTALEGSR | LLGL2; LLGL2 | Q6P1M3; J3( | L2GL2_HUMAN | Unmodified | 0 | 0 | 0 | 0 | 208270 | 0 | 0 | 0 | 1184300 | 182320 | 115700 | 1679800 | 0 | 0 | 0 | 17175 | 197920 | 66210 |
| LTANETQS/ | MAP4K4 | E7EN19 | E7EN19_HUM/ | Unmodified | 0 | 0 | 0 | 0 | 0 | 0 | 0 | 0 | 0 | 0 | 0 | 909640 | 0 | 0 | 0 | 0 | 0 | 0 |
| LTAQFVAR | SF3A1 | Q15459 | SF3A1_HUMAN | Unmodified | 957700 | 735530 | 595320 | 323700 | 256520 | 147500 | 924550 | 322890 | 462400 | 0 | 132960 | 312740 | 337070 | 478530 | 166280 | 295070 | 0 | 303410 |
| LTASAPGYI | CPE | P16870 | CBPE_HUMAN | Unmodified | 0 | 0 | 0 | 0 | 0 | 0 | 0 | 340560 | 60409000 | 7096800 | 0 | 0 | 0 | 0 | 0 | 0 | 0 | 0 |
| LTASAPGYI | CPE | P16870 | CBPE_HUMAN | Unmodified | 0 | 0 | 0 | 0 | 0 | 0 | 0 | 244140 | 32817000 | 5092500 | 0 | 0 | 0 | 0 | 0 | 0 | 0 | 0 |
| LTAVCMEC | TK1; TK1; TK1; T | P04183; K7E | KITH_HUMAN T | Unmodified | 0 | 0 | 0 | 0 | 440390 | 0 | 0 | 0 | 0 | 0 | 157720 | 0 | 0 | 0 | 0 | 0 | 941430 | 0 |
| LTAYAMTIP | HADHA; HADH/ | P40939; A0A | ECHA_HUMAN | Unmodified | 0 | 0 | 0 | 0 | 0 | 0 | 0 | 0 | 1711100 | 0 | 0 | 0 | 0 | 0 | 0 | 0 | 0 | 0 |
| LTAYAMTIP | HADHA; HADH/ | P40939; A0A | ECHA_HUMAN | Oxidation (M) | 29771000 | 22928000 | 23587000 | 16670000 | 0 | 0 | 783730 | 0 | 0 | 691960 | 0 | 0 | 6254600 | 1408400 | 25422000 | 22923000 | 0 | 0 |
| LTAYTATPE | LARS2; LARS2; | Q15031; A0/ | SYLM_HUMAN | Unmodified | 0 | 0 | 0 | 0 | 0 | 0 | 0 | 0 | 4227800 | 0 | 0 | 0 | 0 | 0 | 0 | 0 | 0 | 0 |
| LTCLQNLK | CD55; CD55; C | P08174; H3E | DAF_HUMAN C | Unmodified | 2443600 | 1670400 | 861110 | 1370100 | 4786400 | 1168000 | 0 | 55221 | 1514200 | 427690 | 2147400 | 15421000 | 526780 | 1008500 | 1142300 | 827340 | 4637700 | 10118000 |
| LTCNDTSA. | FARP1; FARP1; | Q9Y4F1; A0/ | FARP1_HUMAN | Unmodified | 0 | 0 | 0 | 0 | 0 | 0 | 0 | 0 | 0 | 0 | 0 | 0 | 0 | 0 | 682510 | 0 | 0 | 0 |
| LTDEEAQH | MLPH; MLPH; M | Q9BV36; C9. | MELPH_HUMAI | Unmodified | 0 | 0 | 0 | 658450 | 0 | 0 | 0 | 0 | 0 | 0 | 0 | 0 | 0 | 0 | 0 | 44340 | 0 | 0 |
| LTDEEVEM' | PMPCA | Q10713 | MPPA_HUMAN | Unmodified | 0 | 0 | 0 | 0 | 0 | 0 | 0 | 0 | 690810 | 25739 | 0 | 0 | 0 | 0 | 0 | 0 | 0 | 186740 |
| LTDFGFAK | PRKX; MAPKAPI | P51817; Q16 | PRKX_HUMAN | Unmodified | 0 | 0 | 111080 | 147610 | 652760 | 0 | 0 | 103190 | 0 | 91144 | 0 | 1241200 | 361590 | 0 | 196810 | 174400 | 142830 | 910310 |
| LTDFGFCA( | PAK2; PAK3; PA | Q13177; O7! | PAK2_HUMAN | Unmodified | 47634 | 59435 | 0 | 58030 | 0 | 43709 | 0 | 0 | 400030 | 59370 | 110000 | 642280 | 370670 | 63114 | 0 | 90442 | 87816 | 118830 |
| LTDFNFLM' | PRKCB; PRKCA | P05771; P17 | KPCB_HUMAN | Oxidation (M) | 0 | 94214 | 0 | 827940 | 811140 | 0 | 409240 | 0 | 0 | 0 | 0 | 174720 | 0 | 0 | 111580 | 0 | 0 | 0 |
| LTDFNFLM' | PRKCB; PRKCA | P05771; P17 | KPCB_HUMAN | Oxidation (M) | 0 | 0 | 0 | 0 | 651830 | 0 | 1890400 | 0 | 109960 | 169290 | 0 | 27362 | 551830 | 758980 | 0 | 0 | 252680 | 358040 |
| LTDHGLEEI | STAMBP | O95630 | STABP_HUMAN | Unmodified | 589280 | 0 | 649690 | 529480 | 0 | 0 | 0 | 0 | 754690 | 0 | 0 | 954740 | 0 | 0 | 301750 | 314510 | 0 | 0 |
| LTDHGLEEI | STAMBP | O95630 | STABP_HUMAN | Unmodified | 0 | 0 | 0 | 0 | 1026900 | 285080 | 0 | 0 | 0 | 0 | 0 | 0 | 0 | 0 | 0 | 0 | 38330 | 0 |
| LTDIHGNVI | BPNT1; BPNT1 | O95861; A6N | BPNT1_HUMAN | Unmodified | 0 | 0 | 333130 | 0 | 0 | 79965 | 0 | 0 | 0 | 103730 | 254700 | 2709900 | 0 | 195140 | 0 | 0 | 0 | 0 |
| LTDISVTDP | LSM2 | Q9Y333 | LSM2_HUMAN | Unmodified | 0 | 0 | 0 | 0 | 0 | 0 | 0 | 0 | 0 | 0 | 276650 | 1717700 | 0 | 0 | 0 | 0 | 0 | 0 |
| LTDPVPNPI | PLCG2; PLCG2 | P16885; A0A | PLCG2_HUMAI | Unmodified | 0 | 0 | 0 | 0 | 0 | 0 | 0 | 0 | 1083800 | 0 | 0 | 8695200 | 0 | 0 | 0 | 0 | 84316 | 0 |
| LTDQVMQI | NAP1L4; NAP1I | Q99733; C9J | NP1L4_HUMAN | Unmodified | 0 | 0 | 661440 | 0 | 0 | 103230 | 0 | 0 | 789530 | 0 | 637860 | 4553900 | 0 | 0 | 0 | 0 | 89220 | 238810 |
| LTDQVMQI | NAP1L4; NAP1I | Q99733; C9J | NP1L4_HUMAN | Oxidation (M) | 0 | 0 | 313450 | 0 | 0 | 0 | 0 | 0 | 0 | 26668 | 125610 | 488180 | 0 | 0 | 0 | 523340 | 0 | 0 |
| LTDQVMQI | NAP1L4; NAP1I | Q99733; C9J | NP1L4_HUMAN | Oxidation (M) | 0 | 0 | 0 | 0 | 0 | 132190 | 0 | 0 | 0 | 519960 | 0 | 0 | 0 | 0 | 0 | 0 | 0 | 1881100 |
| LTDTLVSK | KTN1 | Q86UP2 | KTN1_HUMAN | Unmodified | 0 | 0 | 30471 | 0 | 0 | 23740 | 45148 | 0 | 211970 | 0 | 207720 | 2795600 | 141260 | 602440 | 0 | 0 | 0 | 128710 |
| LTDVAEEC( | PLCB1; PLCB1; | Q9NQ66; A0 | PLCB1_HUMAN | Unmodified | 0 | 0 | 0 | 0 | 0 | 0 | 17373 | 0 | 415250 | 69213 | 0 | 0 | 326350 | 41938 | 0 | 0 | 0 | 37338 |
| LTDYVAFLE | COPS8; COPS8 | Q99627; H7( | CSN8_HUMAN | Unmodified | 0 | 0 | 0 | 0 | 0 | 0 | 0 | 0 | 0 | 0 | 91416 | 273670 | 0 | 0 | 0 | 0 | 0 | 0 |

| | | | | | | | | | | | | | | | | | | | | | | |
|---|---|---|---|---|---|---|---|---|---|---|---|---|---|---|---|---|---|---|---|---|---|---|
| LTEADYLSS | NUDT16L1; NU | Q9BRJ7; K7E | TIRR_HUMAN T | Unmodified | 221920 | 0 | 0 | 0 | 0 | 0 | 0 | 0 | 0 | 0 | 0 | 1587600 | 0 | 0 | 607920 | 358490 | 0 | 139540 |
| LTECLETILI | AP2A2; AP2A2; | O94973; A0A | AP2A2_HUMAN | Unmodified | 0 | 0 | 0 | 0 | 0 | 0 | 0 | 0 | 0 | 0 | 0 | 1449200 | 0 | 0 | 0 | 0 | 0 | 0 |
| LTEDKADV | TRIM28; TRIM28 | Q13263; M0 | TIF1B_HUMAN | Unmodified | 0 | 0 | 0 | 0 | 0 | 0 | 0 | 0 | 0 | 0 | 0 | 3264800 | 655180 | 0 | 0 | 0 | 0 | 684360 |
| LTEDLEYHE | OSBP; OSBP | P22059; H0Y | OSBP1_HUMAN | Unmodified | 0 | 289730 | 564580 | 339240 | 106130 | 0 | 0 | 16083 | 0 | 0 | 523360 | 8947500 | 464990 | 0 | 698330 | 457940 | 47771 | 0 |
| LTEDPAER | CHDH; CHDH; | Q8NE62; C9 | CHDH_HUMAN | Unmodified | 0 | 0 | 0 | 0 | 0 | 0 | 0 | 0 | 634000 | 0 | 0 | 0 | 0 | 0 | 0 | 0 | 0 | 0 |
| LTEDPEKQ | NAPRT; NAPRT; | Q6XQN6; G5 | PNCB_HUMAN | Unmodified | 0 | 0 | 0 | 113250 | 708210 | 177490 | 68952 | 37885 | 0 | 282310 | 0 | 0 | 1112200 | 346610 | 0 | 0 | 0 | 55453 |
| LTEEESAPI | IGSF3 | O75054 | IGSF3_HUMAN | Unmodified | 211580 | 0 | 0 | 379590 | 0 | 0 | 0 | 872040 | 0 | 0 | 423900 | 530560 | 240730 | 731200 | 0 | 1094500 | 0 | 0 |
| LTEEVPLG | RNF31; RNF31; | Q96EP0; H0 | RNF31_HUMAN | Unmodified | 0 | 1234300 | 0 | 0 | 0 | 0 | 0 | 0 | 0 | 0 | 0 | 0 | 0 | 0 | 0 | 0 | 0 | 0 |
| LTEGCSFR | RPS27L; RPS27 | Q71UM5; P4 | RS27L_HUMAN | Unmodified | 1235200 | 368520 | 0 | 0 | 319220 | 0 | 0 | 109240 | 0 | 44783 | 336650 | 5893600 | 0 | 0 | 0 | 0 | 198040 | 0 |
| LTEGMSGR | ATAD3A; ATAD3 | Q9NVI7; Q5 | ATD3A_HUMAN | Unmodified | 0 | 0 | 0 | 0 | 0 | 0 | 0 | 0 | 2101700 | 0 | 0 | 178750 | 0 | 0 | 0 | 0 | 15128 | 0 |
| LTEIIPK | ITGB2; ITGB2; I | P05107; D3 | ITB2_HUMAN I | Unmodified | 0 | 0 | 0 | 389280 | 639750 | 0 | 1892700 | 1614400 | 0 | 0 | 0 | 0 | 772450 | 1110300 | 0 | 0 | 0 | 327610 |
| LTEKEEEIV | MYOF; MYOF | Q9NZM1; H0 | MYOF_HUMAN | Unmodified | 52422 | 0 | 0 | 0 | 9218000 | 720130 | 0 | 104280 | 253790 | 242580 | 0 | 0 | 0 | 0 | 0 | 0 | 0 | 887620 |
| LTEKQVEQ | ITGB4 | P16144 | ITB4_HUMAN I | Unmodified | 0 | 0 | 0 | 0 | 0 | 0 | 0 | 0 | 0 | 0 | 0 | 0 | 0 | 0 | 0 | 0 | 780490 | 530970 |
| LTELGTVDI | IQGAP1; IQGAP | P46940; H0Y | IQGA1_HUMAN | Unmodified | 0 | 0 | 0 | 0 | 2065500 | 504030 | 0 | 0 | 794990 | 259270 | 444280 | 806320 | 107310 | 0 | 0 | 0 | 88771 | 119170 |
| LTELGTVDI | IQGAP1; IQGAP | P46940; H0Y | IQGA1_HUMAN | Unmodified | 0 | 0 | 0 | 0 | 287490 | 38908 | 0 | 0 | 0 | 46323 | 0 | 0 | 0 | 0 | 0 | 0 | 0 | 0 |
| LTELLVK | SGPL1; SGPL1; | O95470; H7 | SGPL1_HUMAN | Unmodified | 0 | 72917 | 0 | 0 | 989950 | 0 | 0 | 206130 | 1349500 | 355560 | 0 | 900710 | 0 | 0 | 0 | 0 | 354570 | 0 |
| LTELMFEH | ACTL6A; ACTL6 | O96019; H7 | ACL6A_HUMAN | Oxidation (M) | 141420 | 0 | 0 | 102370 | 855300 | 0 | 0 | 0 | 78950 | 1275600 | 128260 | 405080 | 259230 | 0 | 44977 | 10737 | 580390 | 0 |
| LTELQLR | TMED4; LRTM2 | Q7Z7H5; Q8 | TMED4_HUMAN | Unmodified | 38133 | 16879 | 208080 | 42418 | 0 | 0 | 9291.1 | 0 | 11255000 | 756520 | 106450 | 1065400 | 0 | 0 | 132690 | 150250 | 0 | 0 |
| LTEPRKAFI | GIPC1; GIPC1 | O14908; K7E | GIPC1_HUMAN | Unmodified | 0 | 0 | 0 | 0 | 709780 | 0 | 53671 | 0 | 0 | 76458 | 0 | 0 | 330320 | 225970 | 0 | 0 | 1374000 | 714670 |
| LTEPVVPK | ACP2; ACP2; A | P11117; B7Z | PPAL_HUMAN | Unmodified | 0 | 0 | 0 | 0 | 0 | 0 | 0 | 0 | 1341400 | 0 | 0 | 785560 | 0 | 0 | 0 | 0 | 0 | 0 |
| LTEQLAGPI | MTREX | P42285 | MTREX_HUMAN | Unmodified | 0 | 0 | 0 | 0 | 394290 | 0 | 0 | 0 | 0 | 136220 | 128290 | 1399700 | 0 | 213320 | 0 | 0 | 593220 | 0 |
| LTESPCAL | HSP90B1; HSP | P14625; A0A | ENPL_HUMAN | Oxidation (M) | 462720 | 230260 | 596980 | 1878800 | 0 | 0 | 0 | 71053 | 52674000 | 8087800 | 7229600 | 11786000 | 0 | 0 | 0 | 0 | 0 | 0 |
| LTESPCAL | HSP90B1; HSP | P14625; A0A | ENPL_HUMAN | Unmodified | 0 | 0 | 0 | 0 | 0 | 0 | 0 | 0 | 79303000 | 1171600 | 601940 | 493160 | 0 | 0 | 0 | 0 | 0 | 0 |
| LTESPCAL | HSP90B1; HSP | P14625; A0A | ENPL_HUMAN | 2 Oxidation (M) | 0 | 0 | 0 | 0 | 0 | 0 | 0 | 0 | 0 | 2007500 | 477030 | 0 | 0 | 0 | 0 | 0 | 0 | 0 |
| LTESPCAL | HSP90B1; HSP | P14625; A0A | ENPL_HUMAN | Oxidation (M) | 0 | 0 | 0 | 0 | 0 | 0 | 0 | 0 | 0 | 3310300 | 0 | 0 | 0 | 0 | 0 | 0 | 0 | 0 |
| LTESTYR | BAIAP2L1 | Q9UHR4 | BI2L1_HUMAN | Unmodified | 137780 | 0 | 88443 | 0 | 0 | 0 | 0 | 0 | 111270 | 0 | 0 | 22634 | 0 | 0 | 0 | 129070 | 0 | 0 |
| LTEVLTDS | GCN1 | Q92616 | GCN1_HUMAN | Unmodified | 0 | 0 | 0 | 0 | 275680 | 0 | 0 | 80669 | 76009 | 0 | 0 | 470830 | 173990 | 0 | 0 | 0 | 0 | 290790 |
| LTEVSISSD | ATIC; ATIC | P31939; H7 | PUR9_HUMAN | Unmodified | 0 | 0 | 0 | 0 | 0 | 0 | 0 | 0 | 0 | 0 | 0 | 924660 | 0 | 0 | 0 | 0 | 0 | 0 |
| LTEVSISSD | ATIC; ATIC | P31939; H7 | PUR9_HUMAN | Unmodified | 0 | 0 | 0 | 0 | 14119000 | 0 | 0 | 0 | 922860 | 0 | 1018300 | 1875800 | 13050000 | 0 | 0 | 0 | 0 | 0 |
| LTFDEEAY | HTATIP2; HTATI | Q9BUP3; E9 | HTAI2_HUMAN | Unmodified | 0 | 0 | 0 | 0 | 325970 | 0 | 0 | 0 | 4267900 | 177640 | 0 | 122370 | 0 | 0 | 0 | 0 | 0 | 0 |
| LTFDEYRNI | ANXA3; ANXA3 | P12429; D6 | ANXA3_HUMAN | Unmodified | 0 | 0 | 0 | 34473 | 2911100 | 147370 | 0 | 56078 | 0 | 93346 | 0 | 0 | 62365 | 117030 | 0 | 0 | 99607 | 480620 |
| LTFDSSFSF | VDAC1; VDAC1 | P21796; C9J | VDAC1_HUMAN | Unmodified | 0 | 0 | 0 | 0 | 0 | 0 | 0 | 0 | 22432000 | 4003600 | 0 | 515450 | 0 | 0 | 0 | 0 | 0 | 0 |
| LTFDSSFSF | VDAC1; VDAC1 | P21796; C9J | VDAC1_HUMAN | Unmodified | 0 | 0 | 0 | 0 | 0 | 0 | 0 | 584580 | 3.16E+08 | 47349000 | 5247000 | 0 | 0 | 1985200 | 0 | 0 | 0 | 0 |
| LTFDTTFSP | VDAC2; VDAC2 | P45880; Q5J | VDAC2_HUMAN | Unmodified | 0 | 0 | 0 | 0 | 0 | 0 | 0 | 0 | 1.49E+08 | 11187000 | 0 | 0 | 0 | 0 | 0 | 0 | 610040 | 0 |
| LTFDTTFSP | VDAC2; VDAC2 | P45880; Q5J | VDAC2_HUMAN | Unmodified | 0 | 0 | 0 | 0 | 84279 | 0 | 773570 | 310730 | 5.24E+08 | 60699000 | 3097300 | 0 | 0 | 724600 | 0 | 0 | 0 | 786500 |
| LTFDTTFSP | VDAC2; VDAC2 | P45880; Q5J | VDAC2_HUMAN | Unmodified | 0 | 0 | 0 | 0 | 0 | 0 | 63136 | 0 | 0 | 512820 | 0 | 0 | 0 | 0 | 0 | 0 | 0 | 0 |
| LTFEATGPS | ERMP1 | Q7Z2K6 | ERMP1_HUMAN | Unmodified | 0 | 0 | 0 | 0 | 101750 | 0 | 0 | 0 | 1661400 | 0 | 0 | 0 | 0 | 0 | 0 | 0 | 0 | 0 |
| LTFEQVYSI | PROM1 | O43490 | PROM1_HUMA | Unmodified | 0 | 0 | 0 | 0 | 0 | 0 | 0 | 0 | 0 | 0 | 0 | 1452100 | 111640 | 0 | 0 | 0 | 0 | 0 |
| LTFEQVYSI | PROM1 | O43490 | PROM1_HUMA | Unmodified | 1050500 | 0 | 1157800 | 0 | 0 | 0 | 0 | 0 | 0 | 0 | 0 | 10138000 | 102500 | 170610 | 0 | 367780 | 0 | 0 |
| LTFIPGFIG | ERBIN; ERBIN | Q96RT1; B4 | ERBIN_HUMAN | Unmodified | 0 | 252900 | 388270 | 382070 | 1081000 | 181070 | 0 | 0 | 808090 | 0 | 564610 | 9635100 | 0 | 0 | 985360 | 867350 | 0 | 0 |
| LTFLYLAN | RPRD1A; RPRD | Q96P16; Q9 | RPR1A_HUMAN | Unmodified | 0 | 0 | 0 | 0 | 0 | 0 | 0 | 0 | 0 | 0 | 144170 | 454120 | 0 | 0 | 0 | 0 | 0 | 0 |
| LTFQLEPNI | ERBB3; ERBB3 | P21860; B4 | ERBB3_HUMAN | Unmodified | 246170 | 191360 | 0 | 125030 | 0 | 0 | 0 | 0 | 0 | 0 | 284770 | 0 | 53801 | 38446 | 0 | 0 | 0 | 0 |
| LTFREVVSF | NCAM2; NCAM | O15394; H9 | NCAM2_HUMA | Unmodified | 209450 | 33510 | 632030 | 941320 | 0 | 0 | 0 | 0 | 0 | 0 | 0 | 0 | 0 | 0 | 1165500 | 715680 | 0 | 0 |
| LTFREVVSF | NCAM2; NCAM | O15394; H9 | NCAM2_HUMA | Unmodified | 0 | 0 | 0 | 3756300 | 3909800 | 8758900 | 3127800 | 751250 | 0 | 0 | 0 | 0 | 0 | 8300500 | 0 | 0 | 0 | 5330400 |
| LTFSCLGG | PTGES3; PTGES | Q15185; A0 | TEBP_HUMAN | Unmodified | 0 | 0 | 0 | 475560 | 0 | 0 | 0 | 0 | 0 | 0 | 0 | 6333400 | 797010 | 0 | 0 | 797190 | 0 | 0 |
| LTFSVTLK | ITGA2; ITGA2 | P17301; E7E | ITA2_HUMAN I | Unmodified | 114750 | 0 | 0 | 158550 | 0 | 0 | 0 | 16115 | 0 | 0 | 0 | 0 | 0 | 0 | 586240 | 103360 | 24135 | 0 |
| LTFYAVSAF | EFR3A; EFR3A; | Q14156; E5 | EFR3A_HUMAN | Unmodified | 0 | 0 | 0 | 981540 | 0 | 0 | 0 | 19181 | 0 | 0 | 0 | 833880 | 0 | 0 | 0 | 0 | 0 | 0 |
| LTGADGTP | SLC25A11; SLC | Q02978; I3L | M2OM_HUMAN | Unmodified | 0 | 0 | 0 | 0 | 0 | 90909 | 0 | 0 | 1181500 | 0 | 0 | 0 | 0 | 0 | 0 | 0 | 0 | 1226300 |
| LTGDVEEL | CPSF3; CPSF3 | Q9UKF6; G5 | CPSF3_HUMAN | Unmodified | 0 | 0 | 0 | 0 | 0 | 0 | 127190 | 121380 | 0 | 120960 | 0 | 1412300 | 0 | 0 | 0 | 0 | 169190 | 0 |
| LTGFHETSI | GLUL; GLUL | P15104; A0A | GLNA_HUMAN | Unmodified | 54387 | 0 | 0 | 225440 | 278680 | 0 | 0 | 0 | 1067500 | 358500 | 362260 | 2828800 | 322890 | 0 | 0 | 0 | 126190 | 293650 |
| LTGIKHELC | CFL1; CFL1; CF | P23528; G3 | COF1_HUMAN | Unmodified | 4273400 | 4585500 | 3147500 | 4903900 | 27748000 | 8994500 | 2736600 | 1141000 | 9996400 | 1764600 | 5851500 | 47164000 | 9465700 | 5307400 | 3847100 | 5891400 | 5654000 | 12197000 |
| LTGIKHELC | CFL1; CFL1; CF | P23528; G3 | COF1_HUMAN | Unmodified | 3345000 | 2635100 | 0 | 1407700 | 2009900 | 0 | 0 | 0 | 718830 | 0 | 122860 | 6106100 | 1267700 | 210760 | 2006700 | 0 | 0 | 653450 |
| LTGISDPVT | OLFM1; OLFM1 | Q99784; F5 | NOE1_HUMAN | Unmodified | 0 | 0 | 0 | 0 | 0 | 0 | 0 | 0 | 4092400 | 207540 | 0 | 0 | 0 | 0 | 0 | 0 | 0 | 0 |
| LTGKDVNF | RPS7 | P62081 | RS7_HUMAN 4 | Unmodified | 113450 | 141720 | 0 | 162100 | 584370 | 0 | 523910 | 40061 | 0 | 289630 | 0 | 0 | 1230000 | 560610 | 0 | 0 | 310490 | 363230 |
| LTGLACDR | PAN2 | Q504Q3 | PAN2_HUMAN | Unmodified | 283230 | 158430 | 0 | 131160 | 0 | 0 | 0 | 0 | 0 | 0 | 0 | 0 | 0 | 0 | 0 | 0 | 0 | 0 |
| LTGMAFR | GAPDHS; GAPD | O14556; P0 | G3PT_HUMAN | Oxidation (M) | 578010 | 494010 | 717640 | 329150 | 203610 | 0 | 0 | 30523 | 395800 | 11009 | 1077000 | 11170000 | 0 | 0 | 1403900 | 1769900 | 0 | 0 |
| LTGMAFRV | GAPDH; GAPD | P04406; E7E | G3P_HUMAN G | Oxidation (M) | 7889500 | 3461900 | 584750 | 1696000 | 0 | 0 | 574550 | 0 | 1133100 | 1869600 | 1185500 | 34105000 | 0 | 0 | 6365700 | 7501800 | 0 | 0 |
| LTGMAFRV | GAPDH; GAPD | P04406; E7E | G3P_HUMAN G | Unmodified | 573840 | 482470 | 0 | 0 | 0 | 0 | 0 | 0 | 1325100 | 0 | 0 | 2052100 | 0 | 0 | 28861 | 224190 | 0 | 0 |
| LTGMAFRV | GAPDH; GAPD | P04406; E7E | G3P_HUMAN G | Oxidation (M) | 16722000 | 4037200 | 8792400 | 24620000 | 15518000 | 6351300 | 0 | 0 | 12135000 | 0 | 24214000 | 92527000 | 10494000 | 0 | 0 | 17319000 | 4168000 | 16188000 |
| LTGMAFRV | GAPDH; GAPD | P04406; E7E | G3P_HUMAN G | Unmodified | 0 | 0 | 406740 | 1268600 | 4350200 | 0 | 0 | 0 | 32395000 | 0 | 2043400 | 0 | 0 | 0 | 0 | 0 | 0 | 4122900 |
| LTGPIQVIH | FYB1 | O15117 | FYB1_HUMAN | Unmodified | 0 | 0 | 0 | 0 | 0 | 0 | 0 | 0 | 0 | 0 | 0 | 1883900 | 0 | 0 | 0 | 0 | 87663 | 0 |
| LTGSLSGW | ACO2; ACO2; A | Q99798; A2 | ACON_HUMAN | Unmodified | 0 | 0 | 0 | 0 | 0 | 0 | 0 | 0 | 2007400 | 430380 | 0 | 0 | 0 | 0 | 0 | 0 | 0 | 0 |
| LTGSSGFV | ATRN | O75882 | ATRN_HUMAN | Unmodified | 0 | 0 | 0 | 0 | 0 | 0 | 0 | 0 | 2074400 | 291410 | 286580 | 422420 | 0 | 0 | 0 | 0 | 0 | 0 |
| LTGSVLED | PAIP1; PAIP1 | Q9H074; D6 | PAIP1_HUMAN | Unmodified | 301690 | 0 | 648050 | 0 | 0 | 959290 | 0 | 0 | 0 | 0 | 0 | 866530 | 0 | 0 | 269570 | 223520 | 0 | 0 |
| LTGSVLED | PAIP1; PAIP1 | Q9H074; D6 | PAIP1_HUMAN | Unmodified | 1417500 | 1027600 | 0 | 360940 | 0 | 0 | 0 | 0 | 265770 | 269130 | 0 | 1206600 | 960350 | 0 | 0 | 531960 | 0 | 0 |
| LTGTIQNDI | MMUT | P22033 | MUTA_HUMAN | Unmodified | 0 | 0 | 0 | 0 | 0 | 0 | 0 | 0 | 2038800 | 256300 | 102100 | 0 | 0 | 0 | 0 | 0 | 0 | 0 |
| LTGVFAPR | RPS4Y2; RPS4X | Q8TD47; P6 | RS4Y2_HUMAN | Unmodified | 0 | 0 | 0 | 0 | 0 | 0 | 0 | 0 | 0 | 0 | 0 | 5551400 | 0 | 0 | 0 | 0 | 0 | 0 |
| LTGVFAPR | RPS4Y2; RPS4X | Q8TD47; P6 | RS4Y2_HUMAN | Unmodified | 1655500 | 162780 | 1379600 | 1066500 | 0 | 0 | 0 | 0 | 0 | 0 | 0 | 27796000 | 0 | 0 | 1575900 | 0 | 0 | 0 |
| LTGVPAAL | EHHADH | Q08426 | ECHP_HUMAN | Unmodified | 0 | 0 | 0 | 0 | 0 | 0 | 0 | 0 | 4589000 | 638560 | 0 | 78967 | 0 | 0 | 20518 | 13484 | 30093 | 0 |
| LTGVPAAL | EHHADH | Q08426 | ECHP_HUMAN | Unmodified | 0 | 0 | 0 | 0 | 0 | 0 | 0 | 59171 | 6791500 | 746430 | 0 | 0 | 0 | 0 | 0 | 0 | 332750 | 0 |
| LTGVSISQ | FLOT1; FLOT1; | O75955; A0 | FLOT1_HUMAN | Unmodified | 117200 | 259430 | 1030500 | 669600 | 112660 | 0 | 81815 | 0 | 3449700 | 707520 | 1311900 | 11468000 | 0 | 0 | 2044800 | 2057800 | 0 | 0 |
| LTGVSISQ | FLOT1; FLOT1; | O75955; A0 | FLOT1_HUMAN | Unmodified | 0 | 0 | 0 | 676510 | 1956000 | 352110 | 0 | 21180 | 586930 | 836170 | 246140 | 0 | 554200 | 463040 | 0 | 0 | 328430 | 671550 |
| LTHDVELN | TFRC; TFRC; TF | P02786; H7 | TFR1_HUMAN | Unmodified | 694470 | 674640 | 0 | 146150 | 0 | 0 | 42115 | 39054 | 918670 | 1434800 | 0 | 694370 | 0 | 0 | 1306600 | 527010 | 0 | 0 |
| LTHDVELN | TFRC; TFRC; TF | P02786; H7 | TFR1_HUMAN | Unmodified | 0 | 0 | 0 | 452440 | 0 | 0 | 0 | 0 | 0 | 2379700 | 0 | 0 | 0 | 0 | 2139000 | 0 | 0 | 0 |
| LTHLHEGL | PLOD2; PLOD2 | O00469; E7E | PLOD2_HUMAN | Unmodified | 0 | 0 | 0 | 0 | 0 | 0 | 0 | 0 | 6030300 | 0 | 109990 | 0 | 0 | 0 | 0 | 0 | 0 | 0 |
| LTHLLGPD | TMEM132A | Q24JP5 | T132A_HUMAN | Unmodified | 102320 | 0 | 0 | 99692 | 0 | 0 | 0 | 0 | 2577800 | 0 | 0 | 0 | 0 | 0 | 109920 | 49744 | 0 | 0 |
| LTHVDSPLI | ESYT1 | Q9BSJ8 | ESYT1_HUMAN | Unmodified | 297560 | 0 | 191170 | 171040 | 522290 | 231050 | 270220 | 97975 | 1136400 | 336920 | 0 | 0 | 0 | 185230 | 360230 | 253070 | 43606 | 183860 |
| LTHVLTGK | SDF2L1 | Q9HCN8 | SDF2L_HUMAN | Unmodified | 108890 | 0 | 34609 | 171680 | 0 | 0 | 0 | 0 | 2631600 | 517930 | 469150 | 290540 | 35266 | 0 | 0 | 139650 | 0 | 0 |
| LTHYDHVL | MCM6 | Q14566 | MCM6_HUMAN | Unmodified | 102670 | 0 | 0 | 0 | 188190 | 0 | 0 | 0 | 455560 | 78060 | 1323400 | 6787500 | 264200 | 0 | 0 | 0 | 0 | 0 |
| LTHYHEGL | PLOD1 | Q02809 | PLOD1_HUMAN | Unmodified | 781980 | 0 | 0 | 0 | 0 | 0 | 0 | 0 | 1111200 | 0 | 35365 | 327870 | 0 | 0 | 582850 | 559590 | 0 | 0 |
| LTHYHEGL | PLOD3; PLOD3 | O60568; H7 | PLOD3_HUMAN | Unmodified | 0 | 0 | 0 | 0 | 0 | 0 | 0 | 0 | 2941400 | 0 | 0 | 0 | 0 | 0 | 0 | 0 | 0 | 0 |
| LTIAEERDK | RPS4Y2; RPS4X | Q8TD47; P6 | RS4Y2_HUMAN | Unmodified | 0 | 0 | 0 | 0 | 0 | 0 | 0 | 0 | 0 | 0 | 0 | 3480600 | 0 | 0 | 1532800 | 11619 | 0 | 0 |
| LTIAFQAK | CSDE1; CSDE1 | O75534; E9 | CSDE1_HUMAN | Unmodified | 98612 | 152680 | 174830 | 263040 | 0 | 0 | 0 | 0 | 0 | 0 | 471910 | 0 | 419980 | 122330 | 242530 | 211830 | 0 | 0 |
| LTIGEGQQ | FGG; FGG; FGG | P02679; C9J | FIBG_HUMAN | Unmodified | 0 | 707810 | 0 | 0 | 0 | 0 | 0 | 0 | 0 | 0 | 0 | 0 | 0 | 0 | 0 | 0 | 0 | 0 |
| LTIGSNLSI | XRCC5 | P13010 | XRCC5_HUMAN | Unmodified | 0 | 0 | 0 | 0 | 0 | 0 | 0 | 0 | 0 | 0 | 0 | 1702800 | 0 | 0 | 0 | 0 | 0 | 0 |
| LTIHAENPI | MPDZ; MPDZ; M | O75970; B7 | MPDZ_HUMAN | Unmodified | 0 | 0 | 0 | 0 | 1015200 | 0 | 158730 | 0 | 0 | 0 | 0 | 0 | 0 | 0 | 89500 | 0 | 202070 | 338480 |
| LTIHGDLY | SF3B2; SF3B2; | Q13435; H0 | SF3B2_HUMAN | Unmodified | 869720 | 689830 | 315520 | 518150 | 455380 | 78371 | 39247 | 0 | 0 | 127230 | 575470 | 1731300 | 836740 | 244440 | 0 | 361410 | 74516 | 211340 |
| LTILNDR | NCKAP1L; NCK | P55160; Q9 | NCKPL_HUMAN | Unmodified | 194200 | 0 | 222880 | 162500 | 2153300 | 423180 | 117000 | 0 | 1808000 | 434970 | 320330 | 5221200 | 204110 | 0 | 1352200 | 2040400 | 1095000 | 0 |
| LTIQEHLYP | FAM171A1 | Q5VUB5 | F1711_HUMAN | Unmodified | 0 | 0 | 0 | 0 | 0 | 0 | 0 | 0 | 0 | 0 | 1236500 | 2353600 | 0 | 0 | 0 | 0 | 0 | 0 |
| LTISNGPK | USP10; USP10; | Q14694; H3 | UBP10_HUMAN | Unmodified | 0 | 0 | 0 | 0 | 0 | 0 | 0 | 0 | 0 | 0 | 0 | 0 | 0 | 0 | 0 | 0 | 0 | 0 |
| LTIYNANIE | NCAM2; NCAM | O15394; H9 | NCAM2_HUMA | Unmodified | 597950 | 374750 | 1425200 | 3415200 | 0 | 0 | 0 | 0 | 0 | 0 | 0 | 0 | 0 | 0 | 8308500 | 5181600 | 670680 | 0 |
| LTKEEILEN | RCN1 | Q15293 | RCN1_HUMAN | Oxidation (M) | 0 | 0 | 0 | 0 | 0 | 642710 | 0 | 0 | 3945800 | 482320 | 0 | 0 | 0 | 0 | 0 | 0 | 0 | 0 |
| LTKVDDQI | PBDC1; PBDC1 | Q9BVG4; A6 | PBDC1_HUMAN | Unmodified | 0 | 0 | 0 | 0 | 0 | 0 | 0 | 0 | 0 | 0 | 0 | 0 | 0 | 0 | 0 | 0 | 1559900 | 0 |
| LTLAELNQI | AGL | P35573 | GDE_HUMAN ( | Unmodified | 0 | 0 | 0 | 0 | 0 | 0 | 0 | 0 | 19738 | 0 | 0 | 979690 | 0 | 0 | 0 | 0 | 0 | 0 |
| LTLAGLGT | TRPM5; TRPM5; | Q9NZQ8; A0 | TRPM5_HUMAN | Unmodified | 138610 | 151320 | 0 | 513750 | 0 | 0 | 0 | 0 | 4422100 | 0 | 115420 | 1395000 | 0 | 0 | 598990 | 275500 | 0 | 0 |
| LTLAPVYTS | NUP210 | Q8TEM1 | PO210_HUMAN | Oxidation (M) | 0 | 0 | 0 | 0 | 0 | 0 | 0 | 0 | 45131000 | 1837800 | 541180 | 0 | 0 | 0 | 0 | 0 | 0 | 891000 |
| LTLAPVYTS | NUP210 | Q8TEM1 | PO210_HUMAN | Unmodified | 0 | 0 | 0 | 0 | 0 | 0 | 0 | 0 | 23576000 | 0 | 0 | 0 | 0 | 0 | 0 | 0 | 1953100 | 0 |
| LTLDKLDV | PGK1 | P00558 | PGK1_HUMAN | Unmodified | 0 | 154210 | 0 | 0 | 693630 | 0 | 0 | 302610 | 1106100 | 298180 | 1159600 | 4290300 | 876900 | 0 | 248750 | 383880 | 0 | 1992900 |
| LTLDKLDV | PGK1 | P00558 | PGK1_HUMAN | Unmodified | 0 | 0 | 0 | 0 | 159800 | 0 | 0 | 131930 | 0 | 150720 | 0 | 0 | 168520 | 0 | 0 | 0 | 619170 | 593810 |
| LTLDTIFVPI | VDAC3; VDAC3 | Q9Y277; E5 | VDAC3_HUMAN | Unmodified | 0 | 0 | 0 | 0 | 603880 | 0 | 0 | 0 | 14520000 | 970990 | 0 | 0 | 0 | 0 | 0 | 0 | 0 | 0 |
| LTLDTIFVPI | VDAC3; VDAC3 | Q9Y277; E5 | VDAC3_HUMAN | Unmodified | 0 | 0 | 0 | 0 | 0 | 0 | 0 | 0 | 9360600 | 1107900 | 0 | 0 | 0 | 0 | 0 | 0 | 0 | 0 |
| LTLEDQATI | DAG1 | Q14118 | DAG1_HUMAN | Unmodified | 5053100 | 1609400 | 2282100 | 0 | 0 | 0 | 0 | 0 | 115300 | 0 | 0 | 3680700 | 0 | 0 | 1199100 | 1403100 | 0 | 0 |
| LTLEDQATI | DAG1 | Q14118 | DAG1_HUMAN | Unmodified | 6514100 | 4854300 | 1634500 | 1289900 | 0 | 0 | 0 | 0 | 739990 | 202460 | 0 | 7882300 | 379370 | 157480 | 3819000 | 2459800 | 99122 | 164950 |
| LTLFVNGQ | TFG; TFG; TFG; | Q92734; A0 | TFG_HUMAN P | Unmodified | 0 | 71158 | 73333 | 0 | 229080 | 0 | 0 | 0 | 326370 | 0 | 27377 | 786840 | 76172 | 0 | 94356 | 0 | 0 | 78091 |
| LTLHGLQQ | DDX39B; DDX3 | Q13838; O0 | DX39B_HUMAN | Unmodified | 0 | 0 | 738190 | 0 | 0 | 0 | 0 | 0 | 826480 | 387270 | 1934000 | 17459000 | 0 | 0 | 0 | 0 | 113230 | 0 |
| LTLIDPETL | CAND1; CAND | Q86VP6; A0 | CAND1_HUMA | Unmodified | 0 | 0 | 526730 | 0 | 0 | 0 | 0 | 175230 | 659550 | 46262 | 784050 | 8635700 | 0 | 0 | 966960 | 66881 | 0 | 0 |
| LTLIDPETL | CAND1; CAND | Q86VP6; A0 | CAND1_HUMA | Unmodified | 0 | 0 | 0 | 0 | 0 | 61393 | 0 | 0 | 0 | 224960 | 0 | 105960 | 85046 | 94456 | 0 | 0 | 0 | 0 |
| LTLLAQQK | ACSL4 | O60488 | ACSL4_HUMAN | Unmodified | 70383 | 52620000 | 0 | 0 | 0 | 18766000 | 7234600 | 5045900 | 70908000 | 12161000 | 0 | 4169900 | 82026000 | 29584000 | 0 | 0 | 44942000 | 78552000 |
| LTLLQLK | PLBD2 | Q8NHP8 | PLBL2_HUMAN | Unmodified | 0 | 0 | 0 | 0 | 0 | 0 | 0 | 0 | 486660 | 0 | 0 | 0 | 0 | 0 | 0 | 0 | 0 | 0 |

| | | | | | | | | | | | | | | | | | | | | | | |
|---|---|---|---|---|---|---|---|---|---|---|---|---|---|---|---|---|---|---|---|---|---|---|
| LTLLQVASF | KIF4A | O95239 | KIF4A_HUMAN | Unmodified | 0 | 0 | 0 | 0 | 146890 | 0 | 0 | 0 | 0 | 0 | 0 | 0 | 0 | 581190 | 0 | 0 | 516750 | 131140 |
| LTLQEFQE( | NCS1 | P62166 | NCS1_HUMAN | Unmodified | 0 | 0 | 0 | 0 | 0 | 0 | 0 | 0 | 143920 | 0 | 0 | 0 | 0 | 0 | 593030 | 739610 | 172680 | 76640 |
| LTLQPGVT1 | SORCS2; SORC | Q96PQ0; B5 | SORC2_HUMAI | Unmodified | 79157 | 0 | 0 | 0 | 0 | 0 | 157620 | 0 | 2086100 | 1555600 | 0 | 0 | 0 | 0 | 119860 | 0 | 0 | 0 |
| LTLSALIDG | VDAC3 | Q9Y277 | VDAC3_HUMAI | Unmodified | 0 | 0 | 0 | 0 | 0 | 1969100 | 0 | 0 | 427520 | 960550 | 0 | 0 | 0 | 353170 | 0 | 0 | 308970 | 0 |
| LTLSALLDG | VDAC1 | P21796 | VDAC1_HUMAI | Unmodified | 450530 | 0 | 721870 | 393230 | 941090 | 149040 | 2567000 | 269840 | 2.53E+08 | 68137000 | 8302000 | 2276900 | 1084200 | 2025300 | 722130 | 276260 | 102390 | 565830 |
| LTLSALLDG | VDAC1 | P21796 | VDAC1_HUMAI | Unmodified | 0 | 0 | 0 | 0 | 403060 | 0 | 483560 | 0 | 6462900 | 15045000 | 0 | 0 | 0 | 0 | 0 | 0 | 67109 | 0 |
| LTLSALVD( | VDAC2; VDAC2 | P45880; A0A | VDAC2_HUMAI | Unmodified | 0 | 141230 | 274890 | 363690 | 1106500 | 0 | 1691200 | 920020 | 5.43E+08 | 1.39E+08 | 3414100 | 876700 | 597350 | 1037100 | 196340 | 0 | 233850 | 2063000 |
| LTLTFNR | ATP5F1C | P36542 | ATPG_HUMAN | Unmodified | 0 | 0 | 0 | 0 | 0 | 0 | 0 | 0 | 17171000 | 0 | 0 | 0 | 0 | 0 | 0 | 0 | 0 | 0 |
| LTLTFSAQT | DIAPH1; DIAPH | O60610; A0/ | DIAP1_HUMAN | Unmodified | 0 | 0 | 0 | 0 | 648570 | 0 | 156160 | 0 | 312360 | 267360 | 194680 | 172930 | 821110 | 481590 | 0 | 0 | 511570 | 422900 |
| LTLTILR | SCRIB; SCRIB; | Q14160; H0 | SCRIB_HUMAN | Unmodified | 58536 | 71095 | 0 | 272970 | 0 | 0 | 0 | 0 | 9172900 | 639900 | 372410 | 0 | 0 | 0 | 425600 | 66663 | 0 | 585610 |
| LTLTPWVG | ATRN | O75882 | ATRN_HUMAN | Unmodified | 0 | 0 | 182870 | 0 | 230290 | 0 | 0 | 0 | 3551400 | 1210700 | 640110 | 1379000 | 0 | 0 | 326460 | 198410 | 0 | 0 |
| LTLTPWVG | ATRN | O75882 | ATRN_HUMAN | Unmodified | 0 | 0 | 0 | 0 | 0 | 0 | 0 | 0 | 855550 | 178870 | 0 | 0 | 0 | 0 | 0 | 0 | 0 | 0 |
| LTLTSDEST | SERINC1 | Q9NRX5 | SERC1_HUMAI | Unmodified | 0 | 972940 | 0 | 0 | 0 | 0 | 0 | 0 | 0 | 0 | 0 | 1005300 | 0 | 0 | 0 | 0 | 0 | 0 |
| LTLVVVEDI | EPB41L4B | Q9H329 | E41LB_HUMAN | Unmodified | 0 | 0 | 0 | 0 | 0 | 0 | 0 | 0 | 0 | 0 | 0 | 0 | 0 | 0 | 1827600 | 0 | 0 | 0 |
| LTLWYYSEI | ESYT1 | Q9BSJ8 | ESYT1_HUMAN | Unmodified | 0 | 0 | 0 | 0 | 1383300 | 0 | 0 | 0 | 0 | 447260 | 0 | 0 | 0 | 0 | 0 | 0 | 0 | 0 |
| LTLYDIAHT | MDH2 | P40926 | MDHM_HUMAI | Unmodified | 0 | 0 | 364860 | 0 | 950010 | 0 | 0 | 0 | 42804000 | 4500400 | 0 | 1201400 | 0 | 0 | 0 | 700110 | 0 | 0 |
| LTMLNTVSI | SH3GL1; SH3G | Q99961; M0I | SH3G1_HUMAI | Unmodified | 20335000 | 18193000 | 13781000 | 9190900 | 4543000 | 0 | 2602800 | 1129800 | 2681900 | 1461200 | 0 | 0 | 35450000 | 12110000 | 15141000 | 14272000 | 540470 | 1114300 |
| LTMLNTVSI | SH3GL1; SH3G | Q99961; M0I | SH3G1_HUMAI | Oxidation (M) | 17616000 | 17547000 | 14567000 | 14273000 | 3000000 | 390360 | 1223200 | 322350 | 0 | 359860 | 0 | 1019200 | 18144000 | 5289900 | 17569000 | 15844000 | 353780 | 349800 |
| LTNAEGVEI | SRP72 | O76094 | SRP72_HUMAN | Unmodified | 0 | 0 | 0 | 0 | 0 | 0 | 0 | 0 | 0 | 0 | 0 | 0 | 0 | 0 | 0 | 0 | 0 | 0 |
| LTNLSSNSI | TPBG | Q13641 | TPBG_HUMAN | Unmodified | 205700 | 182390 | 0 | 0 | 0 | 0 | 0 | 0 | 0 | 0 | 856500 | 6531900 | 0 | 0 | 0 | 0 | 0 | 0 |
| LTNSMMMI | RPS5; RPS5; RP | P46782; M0I | RS5_HUMAN 4 | Unmodified | 0 | 0 | 0 | 0 | 0 | 0 | 0 | 0 | 241650 | 0 | 0 | 1006100 | 0 | 78993 | 0 | 0 | 0 | 0 |
| LTNSMMMI | RPS5; RPS5; RP | P46782; M0I | RS5_HUMAN 4 | 3 Oxidation (M) | 0 | 0 | 0 | 0 | 0 | 0 | 0 | 0 | 0 | 0 | 0 | 0 | 0 | 0 | 0 | 0 | 0 | 0 |
| LTNTNLLF\ | CACNA2D2; CA | Q9NY47; C9 | CA2D2_HUMAI | Unmodified | 0 | 0 | 0 | 0 | 0 | 0 | 0 | 0 | 5266200 | 0 | 200930 | 0 | 0 | 0 | 0 | 0 | 0 | 0 |
| LTNTNLLF\ | CACNA2D2; CA | Q9NY47; C9 | CA2D2_HUMAI | Unmodified | 0 | 0 | 0 | 0 | 0 | 0 | 0 | 0 | 6141900 | 0 | 61100 | 172700 | 0 | 240190 | 0 | 0 | 0 | 0 |
| LTNVAATS( | P3H1; P3H1 | Q32P28; H7 | P3H1_HUMAN | Unmodified | 0 | 0 | 0 | 0 | 0 | 0 | 0 | 0 | 6888400 | 396770 | 351400 | 0 | 0 | 0 | 0 | 0 | 0 | 0 |
| LTNVAVVR | SBDS; SBDS; SI | Q9Y3A5; A0/ | SBDS_HUMAN | Unmodified | 6331900 | 799870 | 486490 | 930460 | 286610 | 0 | 88122 | 87422 | 1649800 | 1642300 | 282650 | 2417100 | 645780 | 272710 | 985940 | 638910 | 0 | 605090 |
| LTPASSTHI | PRUNE1; PRUN | Q86TP1; A0/ | PRUN1_HUMAI | Unmodified | 0 | 0 | 38267 | 0 | 187150 | 0 | 0 | 0 | 534770 | 0 | 0 | 1289600 | 0 | 0 | 121330 | 0 | 0 | 0 |
| LTPDERSKI | LMAN2; LMAN2 | Q12907; D6I | LMAN2_HUMAI | Unmodified | 0 | 0 | 0 | 0 | 0 | 0 | 0 | 0 | 2476600 | 4871200 | 487280 | 1650800 | 0 | 0 | 0 | 0 | 0 | 0 |
| LTPDMQSK | LMAN2L; LMAN | Q9H0V9; F8\ | LMA2L_HUMAN | Unmodified | 0 | 0 | 0 | 0 | 0 | 0 | 0 | 0 | 5422300 | 699800 | 20502 | 0 | 0 | 0 | 0 | 0 | 0 | 0 |
| LTPEDFAR | VTA1; VTA1; VT/ | Q9NP79; Q5 | VTA1_HUMAN \ | Unmodified | 2125600 | 1662300 | 1814800 | 3218800 | 0 | 0 | 0 | 0 | 12726000 | 633820 | 2147000 | 11644000 | 527320 | 136720 | 2118300 | 1579900 | 0 | 0 |
| LTPEDFAR/ | VTA1; VTA1; VT/ | Q9NP79; Q5 | VTA1_HUMAN \ | Unmodified | 0 | 0 | 0 | 0 | 1090000 | 634810 | 115000 | 0 | 0 | 553720 | 0 | 0 | 1265300 | 824590 | 0 | 0 | 131910 | 1183500 |
| LTPEEEEILI | RPS8; RPS8 | P62241; Q5J | RS8_HUMAN 4 | Unmodified | 6858800 | 4647800 | 8152400 | 441520 | 0 | 523750 | 86107 | 150680 | 2647800 | 246640 | 1551300 | 54047000 | 952840 | 0 | 3219300 | 164130 | 259510 | 9930.6 |
| LTPEEEEILI | RPS8; RPS8 | P62241; Q5J | RS8_HUMAN 4 | Unmodified | 1109600 | 460620 | 2597100 | 469490 | 0 | 0 | 0 | 0 | 0 | 0 | 1090500 | 14337000 | 0 | 0 | 0 | 419640 | 0 | 0 |
| LTPEEIERM | HSPA5; HSPA5; | P11021; A0A | BIP_HUMAN Er | Oxidation (M) | 0 | 0 | 0 | 0 | 0 | 0 | 0 | 0 | 970980 | 250740 | 0 | 0 | 0 | 0 | 0 | 0 | 0 | 0 |
| LTPEEIERM | HSPA5; HSPA5; | P11021; A0A | BIP_HUMAN Er | Unmodified | 22712 | 12193 | 0 | 38226 | 0 | 0 | 0 | 20332 | 10630000 | 0 | 0 | 0 | 0 | 0 | 125510 | 0 | 0 | 0 |
| LTPEEIERM | HSPA5; HSPA5; | P11021; A0A | BIP_HUMAN Er | Unmodified | 0 | 0 | 0 | 0 | 0 | 0 | 0 | 0 | 1361100 | 270920 | 0 | 0 | 0 | 0 | 46475 | 0 | 0 | 0 |
| LTPEYLQLI | ERLIN2; ERLIN2 | O94905; E5I | ERLN2_HUMAN | Oxidation (M) | 0 | 0 | 0 | 0 | 0 | 0 | 0 | 0 | 3294700 | 891220 | 288220 | 0 | 0 | 0 | 0 | 0 | 0 | 0 |
| LTPEYLQLI | ERLIN2; ERLIN2 | O94905; E5I | ERLN2_HUMAN | Unmodified | 0 | 0 | 0 | 0 | 0 | 0 | 0 | 0 | 9639400 | 0 | 0 | 0 | 0 | 0 | 0 | 0 | 0 | 0 |
| LTPFMLGAI | GPI; GPI | P06744; A0A | G6PI_HUMAN ( | 2 Oxidation (M) | 0 | 0 | 0 | 0 | 1205600 | 1477700 | 0 | 0 | 466620 | 1817400 | 2619300 | 2966900 | 943800 | 0 | 1335400 | 1431700 | 730730 | 669810 |
| LTPGEVYNI | NPTX1 | Q15818 | NPTX1_HUMAN | Unmodified | 0 | 0 | 0 | 0 | 2149900 | 948330 | 0 | 86912 | 22708000 | 2191200 | 0 | 0 | 714540 | 0 | 0 | 202430 | 0 | 0 |
| LTPITYPQG | RAC1 | P63000 | RAC1_HUMAN | Oxidation (M) | 806700 | 501800 | 1474300 | 520180 | 892750 | 64345 | 165080 | 230340 | 0 | 0 | 274420 | 1277700 | 231380 | 201070 | 1096300 | 500300 | 0 | 0 |
| LTPLSHEVI | EIF2S3; EIF2S3I | P41091; Q2\ | IF2G_HUMAN E | Unmodified | 0 | 135940 | 621470 | 453010 | 0 | 12330 | 340920 | 487170 | 537660 | 0 | 1193800 | 14925000 | 0 | 0 | 522200 | 441930 | 225080 | 81159 |
| LTPRPTAAE | ESYT1 | Q9BSJ8 | ESYT1_HUMAN | Unmodified | 0 | 0 | 0 | 0 | 1130100 | 0 | 0 | 0 | 1351400 | 473410 | 0 | 557280 | 0 | 0 | 0 | 0 | 0 | 0 |
| LTPSYDLAF | PIPOX | Q9P0Z9 | SOX_HUMAN P | Unmodified | 0 | 0 | 0 | 0 | 0 | 0 | 0 | 0 | 4787800 | 701840 | 0 | 0 | 0 | 0 | 0 | 0 | 0 | 0 |
| LTPTHAASI | SNX33; SNX33 | Q8WV41; H3 | SNX33_HUMAN | Unmodified | 1894700 | 1669700 | 10856000 | 5687300 | 0 | 0 | 99694 | 0 | 0 | 0 | 0 | 0 | 0 | 0 | 20930000 | 13170000 | 0 | 0 |
| LTPTHAASI | SNX33; SNX33 | Q8WV41; H3 | SNX33_HUMAN | Unmodified | 0 | 0 | 3499800 | 1841900 | 0 | 0 | 0 | 0 | 0 | 0 | 0 | 0 | 0 | 0 | 3223000 | 1839800 | 0 | 0 |
| LTPVLLAK | DCTN2; DCTN2 | Q13561; A8I | DCTN2_HUMAI | Unmodified | 97008 | 87279 | 0 | 0 | 0 | 0 | 0 | 148880 | 0 | 0 | 0 | 1107600 | 0 | 1161600 | 361030 | 0 | 0 | 0 |
| LTPVSPESS | SQSTM1; SQSTI | Q13501; E7E | SQSTM_HUMAI | Unmodified | 637120 | 505950 | 805850 | 1155900 | 706240 | 479700 | 682900 | 106500 | 5432100 | 1031800 | 207370 | 317850 | 607060 | 754920 | 426830 | 462280 | 0 | 1994900 |
| LTPYEVDEL | DDX17; DDX17 | Q92841; A0/ | DDX17_HUMAI | Unmodified | 0 | 0 | 193800 | 0 | 0 | 0 | 82143 | 0 | 717500 | 0 | 12047000 | 7681000 | 0 | 0 | 309380 | 0 | 0 | 0 |
| LTQDEIISK | KLC1; KLC1; KL | Q07866; G3 | KLC1_HUMAN | Unmodified | 47365 | 35681 | 139910 | 203970 | 5010300 | 308700 | 257300 | 224180 | 119890 | 0 | 715470 | 5839600 | 822930 | 500320 | 174560 | 420380 | 2465600 | 8468300 |
| LTQDEIISK | KLC1; KLC1; KL | Q07866; G3 | KLC1_HUMAN | Unmodified | 0 | 0 | 0 | 0 | 0 | 0 | 0 | 0 | 0 | 0 | 0 | 0 | 0 | 0 | 0 | 0 | 102890 | 0 |
| LTQDQDVI | PPP2R1A; PPP2 | P30153; B3K | 2AAA_HUMAN | Unmodified | 374050 | 393520 | 0 | 0 | 3943000 | 1062600 | 0 | 0 | 3050500 | 842610 | 3606400 | 12386000 | 3448400 | 5606600 | 1467300 | 0 | 901610 | 3666200 |
| LTQEETNFI | SMC4; SMC4 | Q9NTJ3; E9F | SMC4_HUMAN | Unmodified | 0 | 0 | 0 | 0 | 140020 | 0 | 166020 | 19263 | 245040 | 0 | 182780 | 776950 | 0 | 0 | 0 | 0 | 133070 | 166910 |
| LTQEPESIR | CLTB; CLTB | P09497; H0 | CLCB_HUMAN | Unmodified | 507660 | 212230 | 118310 | 0 | 0 | 0 | 0 | 0 | 0 | 0 | 0 | 1382300 | 22276 | 0 | 302410 | 0 | 0 | 0 |
| LTQGETLR | DHCR24; DHCI | Q15392; A0/ | DHC24_HUMA | Unmodified | 0 | 0 | 0 | 0 | 0 | 0 | 0 | 0 | 0 | 0 | 0 | 724940 | 0 | 0 | 0 | 0 | 0 | 0 |
| LTQGLQEA | CIT; CIT; CIT; CI | O14578; A0/ | CTRO_HUMAN | Unmodified | 67861 | 84192 | 480910 | 770770 | 974630 | 0 | 28387 | 0 | 787990 | 23577 | 640030 | 1873500 | 68164 | 0 | 1127100 | 176810 | 0 | 0 |
| LTQLGTFEI | EGFR; EGFR; E( | P00533; Q5( | EGFR_HUMAN | Unmodified | 753880 | 616000 | 2896300 | 1636800 | 425110 | 256000 | 0 | 0 | 755500 | 0 | 0 | 0 | 0 | 0 | 5483600 | 4748900 | 36521 | 45687 |
| LTQNVDLL( | DOCK1; DOCK1 | Q14185; A0/ | DOCK1_HUMA | Unmodified | 0 | 0 | 126800 | 0 | 0 | 0 | 0 | 0 | 0 | 0 | 0 | 0 | 0 | 0 | 346820 | 0 | 0 | 0 |
| LTQQLEEEI | SHTN1 | A0MZ66 | SHOT1_HUMAI | Unmodified | 0 | 0 | 0 | 0 | 1061200 | 0 | 0 | 59969 | 1619900 | 0 | 639570 | 3160100 | 915860 | 137130 | 344230 | 422680 | 268210 | 136720 |
| LTQQMIK | ROCK2; ROCK2 | O75116; Q1 | ROCK2_HUMAI | Unmodified | 0 | 0 | 0 | 0 | 0 | 0 | 46466 | 0 | 246240 | 68134 | 65219 | 322740 | 238260 | 155790 | 0 | 0 | 167860 | 581700 |
| LTQRLEEM | MATN2; MATN2 | O00339; H0 | MATN2_HUMAI | Unmodified | 0 | 0 | 0 | 0 | 0 | 0 | 0 | 0 | 656420 | 0 | 0 | 0 | 0 | 0 | 0 | 0 | 0 | 0 |
| LTQYIDGQ( | CAMK2G; CAMI | Q13555; Q1 | KCC2G_HUMA | Unmodified | 0 | 0 | 0 | 0 | 82536 | 0 | 0 | 0 | 1828800 | 135990 | 99134 | 561430 | 0 | 0 | 0 | 0 | 194970 | 74574 |
| LTQYMDGS | CAMK2D; CAMI | Q13557; E9F | KCC2D_HUMA | Unmodified | 0 | 0 | 0 | 0 | 0 | 0 | 0 | 0 | 918500 | 0 | 0 | 0 | 0 | 0 | 0 | 0 | 0 | 0 |
| LTRAQFEGI | HSPA9; HSPA9; | P38646; H0 | GRP75_HUMAI | Unmodified | 0 | 0 | 0 | 0 | 0 | 0 | 0 | 0 | 2427500 | 4463700 | 0 | 0 | 0 | 0 | 0 | 0 | 98175 | 0 |
| LTRDETNY( | RACK1; RACK1; | P63244; D6I | RACK1_HUMAI | Unmodified | 1016000 | 0 | 771230 | 0 | 1240600 | 0 | 0 | 194490 | 202130 | 30452 | 564020 | 5914100 | 0 | 0 | 554150 | 739570 | 0 | 15167 |
| LTRDLLNC: | NAMPT; NAMPT | P43490; A0A | NAMPT_HUMAI | Unmodified | 0 | 112830 | 0 | 0 | 1815400 | 0 | 533800 | 0 | 0 | 308460 | 0 | 0 | 212900 | 266210 | 0 | 0 | 0 | 206180 |
| LTRDQWEI | EZR; EZR | P15311; E7E | EZRI_HUMAN E | Unmodified | 536110 | 592670 | 0 | 0 | 0 | 0 | 21311 | 0 | 5597200 | 0 | 3025300 | 22991 | 0 | 0 | 0 | 0 | 0 | 0 |
| LTRDQWEI | EZR; EZR | P15311; E7E | EZRI_HUMAN E | Unmodified | 596950 | 852550 | 1331800 | 2919400 | 5445500 | 2137800 | 57009 | 0 | 7481100 | 610860 | 4028100 | 6231200 | 460690 | 418440 | 1431500 | 1353800 | 0 | 0 |
| LTRIPAEVS | LRRC1 | Q9BTT6 | LRRC1_HUMAN | Unmodified | 0 | 0 | 0 | 299110 | 505850 | 0 | 0 | 0 | 1661800 | 0 | 0 | 3267100 | 0 | 0 | 690720 | 518920 | 0 | 0 |
| LTRIPAEVS | LRRC1 | Q9BTT6 | LRRC1_HUMAN | Unmodified | 0 | 0 | 0 | 0 | 0 | 0 | 0 | 0 | 0 | 0 | 0 | 0 | 675040 | 0 | 0 | 0 | 0 | 0 |
| LTRLSPEIVI | P4HTM | Q9NXG6 | P4HTM_HUMAI | Unmodified | 0 | 0 | 0 | 0 | 0 | 0 | 0 | 0 | 1144500 | 713520 | 0 | 0 | 0 | 0 | 0 | 0 | 0 | 0 |
| LTRNNFAV( | VDAC2; VDAC2 | P45880; A0A | VDAC2_HUMAI | Unmodified | 0 | 0 | 0 | 0 | 0 | 0 | 0 | 0 | 0 | 15095000 | 0 | 0 | 0 | 0 | 0 | 0 | 0 | 0 |
| LTSADALRI | GCN1 | Q92616 | GCN1_HUMAN | Unmodified | 70346 | 0 | 0 | 0 | 0 | 0 | 0 | 0 | 0 | 0 | 0 | 1835800 | 0 | 0 | 84205 | 250130 | 0 | 0 |
| LTSALDELL | PTX3 | P26022 | PTX3_HUMAN I | Unmodified | 0 | 0 | 0 | 0 | 0 | 0 | 0 | 0 | 1751300 | 0 | 0 | 0 | 0 | 0 | 0 | 0 | 0 | 0 |
| LTSAVSSLF | SCFD1; SCFD1; | Q8WVM8; A( | SCFD1_HUMAI | Unmodified | 0 | 0 | 0 | 0 | 1065600 | 0 | 0 | 0 | 767310 | 0 | 312850 | 1099100 | 0 | 0 | 0 | 0 | 0 | 0 |
| LTSAVSSLF | SCFD1; SCFD1; | Q8WVM8; A( | SCFD1_HUMAI | Unmodified | 0 | 0 | 0 | 40668 | 623350 | 135280 | 0 | 0 | 0 | 0 | 0 | 428280 | 90666 | 0 | 0 | 33441 | 0 | 0 |
| LTSDDVKEI | RPS13; RPS13; | P62277; J3K | RS13_HUMAN | Unmodified | 606730 | 1035700 | 0 | 351920 | 935360 | 0 | 91366 | 231080 | 0 | 0 | 102590 | 5446900 | 168030 | 63627 | 259550 | 184430 | 0 | 0 |
| LTSDLTFAY | HEXA; HEXA; N/ | P06865; H3E | HEXA_HUMAN | Unmodified | 1021300 | 0 | 1159800 | 938270 | 0 | 0 | 0 | 0 | 1316500 | 0 | 0 | 0 | 0 | 0 | 936440 | 0 | 32954 | 0 |
| LTSEIQSR | NAA25 | Q14CX7 | NAA25_HUMAI | Unmodified | 0 | 0 | 0 | 0 | 0 | 0 | 0 | 0 | 0 | 0 | 485560 | 1986200 | 0 | 0 | 224590 | 0 | 0 | 0 |
| LTSELITHIC | MX1; MX1; MX1 | P20591; A0A | MX1_HUMAN Ir | Unmodified | 0 | 0 | 0 | 0 | 0 | 0 | 0 | 0 | 0 | 0 | 0 | 1007900 | 0 | 0 | 0 | 0 | 0 | 839020 |
| LTSEPQPQ | RAB14 | P61106 | RAB14_HUMAN | Unmodified | 466970 | 0 | 420150 | 0 | 696540 | 381180 | 36428 | 0 | 1279100 | 246610 | 0 | 3532600 | 0 | 0 | 173700 | 290080 | 0 | 130660 |
| LTSFIGAIAI | CCT2; CCT2; C( | P78371; F5G | TCPB_HUMAN | Unmodified | 0 | 0 | 0 | 0 | 281400 | 0 | 0 | 0 | 298290 | 298490 | 48653 | 911040 | 894740 | 0 | 24415 | 0 | 141690 | 140130 |
| LTSGDSESI | TEP1; TEP1; TEI | Q99973; G3 | TEP1_HUMAN | Unmodified | 0 | 0 | 0 | 0 | 0 | 0 | 0 | 0 | 0 | 0 | 0 | 406160 | 0 | 0 | 0 | 0 | 0 | 0 |
| LTSLAAALD | LETM1 | O95202 | LETM1_HUMAN | Unmodified | 0 | 0 | 0 | 0 | 0 | 0 | 77986 | 0 | 1655100 | 1035700 | 0 | 0 | 181720 | 0 | 0 | 0 | 0 | 0 |
| LTSLAALRA | MYO5C | Q9NQX4 | MYO5C_HUMA | Unmodified | 40439 | 95073 | 0 | 0 | 635620 | 107570 | 0 | 0 | 0 | 0 | 0 | 0 | 0 | 136910 | 0 | 0 | 0 | 80004 |
| LTSLLEETLI | NRBP1; NRBP1 | Q9UHY1; F8 | NRBP_HUMAN | Unmodified | 0 | 0 | 0 | 0 | 0 | 0 | 0 | 0 | 0 | 0 | 0 | 8919700 | 0 | 0 | 0 | 0 | 0 | 0 |
| LTSLNVKYI | PTBP1; PTBP3; | P26599; O9 | PTBP1_HUMAN | Unmodified | 0 | 0 | 0 | 0 | 0 | 0 | 0 | 0 | 0 | 141760 | 0 | 0 | 0 | 0 | 0 | 0 | 0 | 0 |
| LTSLSDRY\ | C4A; C4B; C4B_ | P0C0L4; P0( | CO4A_HUMAN | Unmodified | 155960 | 352300 | 4936700 | 87876000 | 6863500 | 3094300 | 507360 | 0 | 0 | 0 | 0 | 0 | 16089000 | 8015800 | 71372000 | 15463000 | 1271800 | 9433700 |
| LTSLVPFVI | EIF3A | Q14152 | EIF3A_HUMAN | Unmodified | 337670 | 204950 | 0 | 185230 | 0 | 0 | 0 | 0 | 106840 | 0 | 233250 | 3241200 | 0 | 0 | 322750 | 331370 | 0 | 0 |
| LTSMHFYG | RRM1; RRM1 | P23921; E9P | RIR1_HUMAN F | Unmodified | 0 | 0 | 0 | 0 | 214960 | 0 | 0 | 0 | 343430 | 0 | 0 | 2462400 | 0 | 0 | 0 | 0 | 99830 | 143420 |
| LTSNPLASI | ERGIC1 | Q969X5 | ERGI1_HUMAN | Unmodified | 34496 | 0 | 0 | 0 | 0 | 0 | 0 | 0 | 3630600 | 1304300 | 0 | 0 | 0 | 0 | 157350 | 325300 | 0 | 0 |
| LTSPIVSTH | ANKRD13D | Q6ZTN6 | AN13D_HUMAI | Unmodified | 1063000 | 1213300 | 0 | 336560 | 0 | 0 | 111260 | 0 | 0 | 0 | 0 | 0 | 0 | 0 | 498340 | 658330 | 0 | 0 |
| LTSPVINTS | ANKRD13A; AN | Q8IZ07; H0Y | AN13A_HUMAN | Unmodified | 773310 | 975840 | 1129600 | 0 | 2244200 | 0 | 0 | 0 | 0 | 0 | 0 | 0 | 0 | 0 | 1155000 | 492710 | 0 | 0 |
| LTSSLTTK | UQCRC2; UQCI | P22695; H3E | QCR2_HUMAN | Unmodified | 0 | 0 | 0 | 0 | 0 | 0 | 0 | 12107 | 8757800 | 1980000 | 0 | 61624 | 0 | 0 | 0 | 0 | 0 | 0 |
| LTSSRTQLF | TM9SF4; TM9SI | Q92544; F2Z | TM9S4_HUMAN | Unmodified | 0 | 0 | 0 | 0 | 0 | 0 | 0 | 0 | 0 | 1336200 | 0 | 0 | 0 | 0 | 0 | 0 | 0 | 0 |
| LTSTATKDF | P4HA1 | P13674 | P4HA1_HUMAN | Unmodified | 0 | 0 | 0 | 0 | 0 | 0 | 0 | 0 | 815810 | 148820 | 0 | 0 | 0 | 0 | 0 | 0 | 0 | 0 |
| LTSTDTIPK | EPB41; EPB41; | P11171; A0A | EPB41_HUMAN | Unmodified | 0 | 0 | 310990 | 799840 | 0 | 0 | 0 | 0 | 523890 | 0 | 292230 | 1431800 | 0 | 0 | 0 | 2009900 | 0 | 0 |
| LTTDFGNA | TFRC; TFRC; TF | P02786; H7( | TFR1_HUMAN | Unmodified | 2620100 | 1560000 | 4849600 | 4883200 | 99931 | 52898 | 0 | 82389 | 20667000 | 2613600 | 974850 | 980630 | 0 | 0 | 4271800 | 3992300 | 0 | 45113 |
| LTTDFGNA | TFRC; TFRC; TF | P02786; H7( | TFR1_HUMAN | Unmodified | 833420 | 729150 | 0 | 491600 | 0 | 0 | 0 | 0 | 1808400 | 158730 | 0 | 567610 | 0 | 0 | 158600 | 287870 | 0 | 0 |
| LTTDFGNA | TFRC; TFRC; TF | P02786; H7( | TFR1_HUMAN | Unmodified | 218190 | 343850 | 0 | 1011300 | 899500 | 0 | 0 | 0 | 4336400 | 1171700 | 0 | 0 | 179210 | 0 | 0 | 0 | 125100 | 386150 |
| LTTDFGNA | TFRC; TFRC; TF | P02786; H7( | TFR1_HUMAN | Oxidation (M) | 0 | 0 | 0 | 560600 | 117080 | 0 | 74068 | 30150 | 783250 | 420430 | 0 | 732440 | 0 | 0 | 0 | 92253 | 68677 | 250050 |
| LTTLELLEVI | SPTBN1; SPTBI | Q01082; A0/ | SPTB2_HUMAN | Unmodified | 0 | 0 | 0 | 0 | 0 | 0 | 0 | 0 | 0 | 0 | 1435100 | 2926400 | 0 | 0 | 0 | 0 | 0 | 0 |
| LTTPTYGDI | TUBB2A; TUBB4 | Q13885; P0 | TBB2A_HUMAN | Oxidation (M) | 0 | 0 | 2654100 | 5977500 | 0 | 0 | 0 | 0 | 391420 | 0 | 0 | 3970700 | 0 | 0 | 12367000 | 5349800 | 0 | 0 |
| LTTPTYGDI | TUBB2A; TUBB4 | Q13885; P0 | TBB2A_HUMAN | Oxidation (M) | 0 | 0 | 0 | 0 | 0 | 0 | 0 | 0 | 0 | 3129600 | 0 | 0 | 19621000 | 5885800 | 0 | 0 | 4685600 | 7362300 |
| LTTPTYGDI | TUBB2A; TUBB4 | Q13885; P0 | TBB2A_HUMAN | Unmodified | 0 | 0 | 0 | 396380 | 9371500 | 462420 | 17704000 | 0 | 0 | 3271900 | 0 | 0 | 9532800 | 9171000 | 0 | 0 | 0 | 7235500 |
| LTTPVFGK | CYP51A1; CYP5 | Q16850; A0/ | CP51A_HUMAN | Unmodified | 43717 | 0 | 63710 | 36447 | 0 | 0 | 460280 | 208140 | 159030 | 129160 | 160720 | 501360 | 0 | 0 | 0 | 87738 | 0 | 0 |
| LTTVCPTSK | SRC | P12931 | SRC_HUMAN P | Unmodified | 1033600 | 549940 | 296630 | 425610 | 237100 | 151610 | 0 | 470550 | 1751400 | 191290 | 638980 | 1219800 | 0 | 0 | 880760 | 733970 | 626780 | 716100 |
| LTTVCPTVK | YES1; YES1 | P07947; J3Q | YES_HUMAN Ty | Unmodified | 2726500 | 1013300 | 1845800 | 2697500 | 1978900 | 1269500 | 0 | 148100 | 2971400 | 855650 | 776510 | 5956200 | 1592900 | 1987600 | 5205100 | 3531800 | 2327400 | 6532200 |
| LTTVLTTHH | HAGH; HAGH; I | Q16775; H3I | GLO2_HUMAN | Unmodified | 0 | 0 | 0 | 0 | 624510 | 0 | 142430 | 0 | 2252600 | 513970 | 740600 | 869680 | 1712000 | 1213000 | 341380 | 232190 | 226840 | 769280 |

| | | | | | | | | | | | | | | | | | | | | | | |
|---|---|---|---|---|---|---|---|---|---|---|---|---|---|---|---|---|---|---|---|---|---|---|
| LTVADALEF | PRKAR1A; PRKA | P10644; K7E | KAP0_HUMAN | Unmodified | 0 | 0 | 0 | 0 | 406340 | 0 | 0 | 48065 | 3493000 | 0 | 0 | 2633100 | 0 | 0 | 0 | 0 | 0 | 0 |
| LTVAENEAE | EPRS1 | P07814 | SYEP_HUMAN | Unmodified | 0 | 0 | 754230 | 0 | 0 | 0 | 0 | 0 | 963610 | 105960 | 0 | 9885700 | 0 | 0 | 772900 | 315600 | 0 | 216540 |
| LTVDSAIAR | DSP | P15924 | DESP_HUMAN | Unmodified | 0 | 0 | 0 | 0 | 0 | 0 | 0 | 0 | 776710 | 61662 | 0 | 282980 | 0 | 0 | 130470 | 0 | 100680 | 0 |
| LTVEDLEK | MAPRE1 | Q15691 | MARE1_HUMAI | Unmodified | 0 | 0 | 442040 | 484310 | 1198700 | 0 | 0 | 0 | 1286500 | 0 | 3983900 | 19239000 | 495810 | 0 | 0 | 0 | 290060 | 740030 |
| LTVEDLEKE | MAPRE1 | Q15691 | MARE1_HUMAI | Unmodified | 0 | 0 | 0 | 0 | 981340 | 0 | 0 | 0 | 0 | 0 | 0 | 0 | 907380 | 0 | 0 | 0 | 430380 | 0 |
| LTVEDPVTV | PSMA7; PSMA8 | O14818; Q8 | PSA7_HUMAN | Unmodified | 0 | 0 | 0 | 0 | 0 | 0 | 0 | 0 | 2395900 | 51672 | 2450600 | 16047000 | 0 | 0 | 0 | 0 | 0 | 0 |
| LTVENSPK( | DENR; DENR | O43583; F8V | DENR_HUMAN | Unmodified | 493650 | 396040 | 0 | 880280 | 2955000 | 171800 | 206360 | 0 | 964950 | 0 | 622030 | 2943200 | 4220400 | 222670 | 332430 | 440520 | 466550 | 4840400 |
| LTVGPVTLF | PGAP6; PGAP6 | Q9HCN3; K4 | PGAP6_HUMAI | Unmodified | 0 | 0 | 142060 | 0 | 0 | 0 | 0 | 0 | 0 | 0 | 239520 | 1914500 | 0 | 0 | 669130 | 0 | 0 | 0 |
| LTVIDTPGF | SEPTIN9; SEPTI | Q9UHD8; K7 | SEPT9_HUMAN | Oxidation (M) | 0 | 0 | 0 | 0 | 6058200 | 0 | 0 | 0 | 0 | 0 | 0 | 2982900 | 1774400 | 0 | 0 | 0 | 2335500 | 5017400 |
| LTVIDTPGF | SEPTIN9; SEPTI | Q9UHD8; K7 | SEPT9_HUMAN | Unmodified | 0 | 0 | 0 | 0 | 2688400 | 0 | 0 | 0 | 0 | 0 | 0 | 0 | 286610 | 0 | 0 | 0 | 979250 | 1398800 |
| LTVLPDEIC | LRRC57; N/A; L | Q8N9N7; A0 | LRC57_HUMAN | Unmodified | 0 | 0 | 0 | 0 | 0 | 0 | 0 | 0 | 312590 | 0 | 0 | 1293800 | 0 | 0 | 0 | 0 | 0 | 0 |
| LTVLPPELG | RSU1 | Q15404 | RSU1_HUMAN | Unmodified | 622210 | 852450 | 489980 | 3075700 | 144050 | 73664 | 0 | 0 | 645530 | 296410 | 140680 | 791100 | 0 | 0 | 1742900 | 957410 | 59211 | 0 |
| LTVMTDLEI | LMAN2; LMAN2 | Q12907; D6I | LMAN2_HUMAI | Unmodified | 0 | 0 | 0 | 0 | 0 | 0 | 391200 | 221410 | 494610 | 79521 | 0 | 0 | 0 | 0 | 0 | 0 | 0 | 0 |
| LTVMTDLEI | LMAN2; LMAN2 | Q12907; D6I | LMAN2_HUMAI | Oxidation (M) | 0 | 0 | 99460 | 207430 | 0 | 0 | 0 | 0 | 11774000 | 999140 | 310760 | 758060 | 202060 | 0 | 336350 | 145940 | 0 | 0 |
| LTVMTDLEI | LMAN2; LMAN2 | Q12907; D6I | LMAN2_HUMAI | Unmodified | 0 | 0 | 0 | 0 | 0 | 0 | 0 | 0 | 69313000 | 1192000 | 0 | 0 | 0 | 0 | 0 | 0 | 0 | 0 |
| LTVNDFVRI | SLC25A12; SLC | O75746; Q9I | CMC1_HUMAN | Unmodified | 0 | 0 | 0 | 0 | 0 | 0 | 0 | 0 | 2532500 | 326680 | 74719 | 50150 | 0 | 0 | 0 | 0 | 0 | 0 |
| LTVNEAVKI | PLEC; PLEC | Q15149; H0Y | PLEC_HUMAN | Unmodified | 0 | 0 | 0 | 0 | 0 | 0 | 0 | 0 | 0 | 0 | 0 | 0 | 0 | 0 | 0 | 0 | 1889800 | 1567800 |
| LTVPDDIAA | DDAH1; DDAH1 | O94760; B4I | DDAH1_HUMA | Unmodified | 0 | 0 | 0 | 0 | 0 | 0 | 0 | 0 | 272150 | 0 | 33228 | 0 | 1317500 | 0 | 0 | 0 | 307630 | 54887 |
| LTVPFLLLQ | MGLL; MGLL; M | Q99685; H7 | MGLL_HUMAN | Unmodified | 0 | 0 | 0 | 0 | 2808000 | 0 | 0 | 0 | 0 | 0 | 0 | 0 | 0 | 0 | 0 | 0 | 1133900 | 2519400 |
| LTVPGLSEI | ITGB4 | P16144 | ITB4_HUMAN I | Unmodified | 0 | 0 | 0 | 0 | 0 | 0 | 0 | 0 | 0 | 0 | 0 | 0 | 0 | 0 | 0 | 0 | 764230 | 910470 |
| LTVPGLSEI | ITGB4 | P16144 | ITB4_HUMAN I | Unmodified | 0 | 0 | 0 | 0 | 0 | 0 | 0 | 0 | 0 | 0 | 0 | 0 | 0 | 0 | 0 | 0 | 2227400 | 605060 |
| LTVSNVLK | TFRC; TFRC | P02786; G3\ | TFR1_HUMAN | Unmodified | 4194900 | 3895900 | 6137600 | 4801000 | 4766200 | 0 | 1145800 | 372890 | 6306000 | 5240900 | 2240700 | 4404000 | 1061300 | 431540 | 7244300 | 4870300 | 0 | 0 |
| LTVSNVLKE | TFRC; TFRC | P02786; G3\ | TFR1_HUMAN | Unmodified | 0 | 0 | 0 | 0 | 0 | 0 | 0 | 0 | 0 | 377560 | 0 | 0 | 0 | 0 | 0 | 0 | 0 | 0 |
| LTVSWLKD | NFASC; NFASC | O94856; H7I | NFASC_HUMAI | Unmodified | 0 | 0 | 430750 | 856650 | 0 | 0 | 0 | 0 | 836600 | 0 | 0 | 557300 | 0 | 0 | 897090 | 0 | 0 | 0 |
| LTVTAYDC( | CLSTN1; CLSTN | O94985; A0A | CSTN1_HUMAI | Unmodified | 0 | 0 | 178470 | 0 | 0 | 0 | 0 | 0 | 1891200 | 445630 | 0 | 0 | 0 | 0 | 0 | 0 | 0 | 0 |
| LTVTSQNL( | CD74; CD74; C | P04233; H0Y | HG2A_HUMAN | Unmodified | 0 | 0 | 0 | 0 | 0 | 0 | 0 | 0 | 8187700 | 424700 | 0 | 0 | 0 | 0 | 0 | 0 | 0 | 0 |
| LTVVILEAK | SYT1; SYT1 | P21579; J3K | SYT1_HUMAN | Unmodified | 0 | 0 | 0 | 0 | 0 | 0 | 0 | 0 | 1392900 | 0 | 0 | 0 | 0 | 0 | 0 | 0 | 0 | 0 |
| LTVYTTLID\ | ASAH1; ASAH1; | Q13510; A0A | ASAH1_HUMAI | Unmodified | 0 | 0 | 0 | 0 | 0 | 0 | 0 | 0 | 1690700 | 0 | 631320 | 0 | 0 | 0 | 0 | 0 | 0 | 0 |
| LTWHAYPE | CBX5 | P45973 | CBX5_HUMAN | Unmodified | 237400 | 0 | 0 | 0 | 0 | 0 | 0 | 0 | 0 | 28517 | 157280 | 1457300 | 0 | 0 | 0 | 0 | 0 | 0 |
| LTWHSCPE | CBX3 | Q13185 | CBX3_HUMAN | Unmodified | 0 | 0 | 0 | 0 | 0 | 0 | 0 | 35533 | 355830 | 0 | 0 | 3996200 | 0 | 0 | 0 | 0 | 0 | 0 |
| LTYDEAVQ | HAPLN1 | P10915 | HPLN1_HUMAI | Unmodified | 0 | 0 | 0 | 0 | 0 | 0 | 0 | 0 | 1422000 | 0 | 0 | 0 | 0 | 0 | 0 | 0 | 0 | 0 |
| LTYENGLLI | IGF2R; IGF2R | P11717; A0A | MPRI_HUMAN | Unmodified | 0 | 0 | 0 | 0 | 281260 | 0 | 0 | 0 | 25810000 | 69413 | 207530 | 4606200 | 0 | 0 | 0 | 0 | 0 | 0 |
| LTYLSNHD | MAPKAP1; MAP | Q9BPZ7; H0 | SIN1_HUMAN | Unmodified | 0 | 0 | 0 | 132420 | 486240 | 0 | 72252 | 0 | 0 | 0 | 93226 | 296930 | 392860 | 0 | 59088 | 0 | 0 | 0 |
| LTYTAEVSV | LTA4H | P09960 | LKHA4_HUMAI | Unmodified | 0 | 0 | 0 | 0 | 604680 | 0 | 0 | 0 | 840670 | 0 | 1642200 | 6559000 | 0 | 0 | 0 | 0 | 0 | 176790 |
| LVAAPVATA | FKRP; FKRP | Q9H9S5; M0 | FKRP_HUMAN | Unmodified | 0 | 0 | 0 | 0 | 0 | 0 | 0 | 0 | 6139900 | 141870 | 0 | 0 | 0 | 0 | 0 | 0 | 0 | 0 |
| LVAEDLSQ | DIAPH1; DIAPH | O60610; A0A | DIAP1_HUMAN | Unmodified | 0 | 0 | 0 | 0 | 2950000 | 0 | 987560 | 293810 | 763860 | 226610 | 0 | 909370 | 1128400 | 883380 | 0 | 0 | 1857400 | 1204500 |
| LVAEFDFR | GAS6 | Q14393 | GAS6_HUMAN | Unmodified | 0 | 0 | 0 | 0 | 0 | 0 | 0 | 18310 | 772570 | 0 | 0 | 0 | 0 | 0 | 0 | 0 | 0 | 0 |
| LVAEWEG( | PLOD1; PLOD1 | Q02809; Q5 | PLOD1_HUMAI | Unmodified | 0 | 0 | 0 | 0 | 0 | 0 | 0 | 0 | 13127000 | 5779700 | 0 | 0 | 0 | 0 | 0 | 0 | 0 | 1847600 |
| LVAFGTSH( | LLGL2; LLGL2 | Q6P1M3; J3 | L2GL2_HUMAN | Unmodified | 861220 | 786320 | 0 | 887380 | 0 | 0 | 0 | 0 | 0 | 0 | 266130 | 1981300 | 0 | 0 | 1828400 | 1490500 | 0 | 0 |
| LVAGEMG( | HSD17B10; HS | Q99714; Q5I | HCD2_HUMAN | Oxidation (M) | 0 | 0 | 0 | 0 | 0 | 0 | 0 | 0 | 1855100 | 0 | 0 | 0 | 0 | 0 | 22376 | 0 | 0 | 0 |
| LVAGEMG( | HSD17B10; HS | Q99714; Q5I | HCD2_HUMAN | Unmodified | 0 | 0 | 0 | 0 | 0 | 0 | 122230 | 85989 | 8897800 | 174150 | 355820 | 0 | 0 | 123470 | 0 | 0 | 0 | 0 |
| LVAGQQQ\ | PLCB3 | Q01970 | PLCB3_HUMAN | Unmodified | 0 | 370290 | 0 | 0 | 0 | 0 | 0 | 0 | 0 | 0 | 0 | 1209500 | 0 | 0 | 0 | 0 | 0 | 0 |
| LVAIVDPHI | GANAB; GANAE | Q14697; F5H | GANAB_HUMA | Unmodified | 0 | 0 | 0 | 0 | 0 | 0 | 306150 | 161360 | 17682000 | 1319100 | 0 | 229740 | 0 | 0 | 0 | 0 | 0 | 0 |
| LVAIVDPHI | GANAB; GANAE | Q14697; F5H | GANAB_HUMA | Unmodified | 0 | 0 | 0 | 90716 | 0 | 0 | 0 | 0 | 45532000 | 2597800 | 0 | 0 | 0 | 0 | 0 | 0 | 0 | 0 |
| LVAIVDVID | RPL14; RPL14 | P50914; E7E | RL14_HUMAN | Unmodified | 8646900 | 2254500 | 3749600 | 2146600 | 0 | 0 | 0 | 356480 | 979570 | 457060 | 4210000 | 30659000 | 0 | 0 | 4297800 | 2961900 | 0 | 0 |
| LVAIVDVID | RPL14; RPL14 | P50914; E7E | RL14_HUMAN | Unmodified | 90005 | 0 | 0 | 241710 | 122730 | 0 | 204000 | 0 | 0 | 74304 | 309860 | 0 | 113710 | 111110 | 0 | 0 | 0 | 1007000 |
| LVAIVDVID | RPL14; RPL14 | P50914; E7E | RL14_HUMAN | Unmodified | 0 | 0 | 0 | 0 | 915910 | 0 | 2484600 | 0 | 175930 | 493290 | 975020 | 0 | 0 | 1373300 | 0 | 0 | 0 | 1138700 |
| LVALAVIDE | TMX3 | Q96JJ7 | TMX3_HUMAN | Unmodified | 0 | 0 | 0 | 0 | 0 | 0 | 0 | 0 | 841150 | 0 | 108260 | 1059800 | 0 | 0 | 0 | 0 | 0 | 0 |
| LVALHTFGI | FAM111B | Q6SJ93 | F111B_HUMAN | Unmodified | 216160 | 72927 | 0 | 0 | 0 | 0 | 0 | 0 | 0 | 28043 | 0 | 55226 | 0 | 0 | 46398 | 33260 | 0 | 0 |
| LVALLNTLD | PTPA; PTPA; PTI | Q15257; A6F | PTPA_HUMAN | Unmodified | 0 | 0 | 0 | 0 | 0 | 0 | 0 | 0 | 0 | 0 | 0 | 795400 | 0 | 0 | 135610 | 0 | 0 | 0 |
| LVALLNTLD | PTPA; PTPA; PTI | Q15257; A6F | PTPA_HUMAN | Unmodified | 0 | 0 | 0 | 0 | 477390 | 0 | 359430 | 0 | 222710 | 374920 | 280960 | 73088 | 626260 | 454100 | 0 | 0 | 0 | 0 |
| LVANHVAS | SEL1L | Q9UBV2 | SE1L1_HUMAN | Unmodified | 0 | 0 | 0 | 0 | 0 | 0 | 0 | 0 | 11278000 | 483390 | 0 | 380610 | 0 | 0 | 0 | 0 | 0 | 0 |
| LVAPLAPD | PDE2A; PDE2A | O00408; F5H | PDE2A_HUMAN | Oxidation (M) | 0 | 0 | 0 | 0 | 0 | 0 | 0 | 0 | 0 | 0 | 1504100 | 8807200 | 0 | 0 | 0 | 0 | 0 | 0 |
| LVAPLAPD | PDE2A; PDE2A | O00408; F5H | PDE2A_HUMAN | Unmodified | 0 | 0 | 0 | 0 | 0 | 0 | 0 | 0 | 0 | 0 | 65345 | 247520 | 0 | 0 | 0 | 0 | 0 | 0 |
| LVAQLDTE( | IGSF8 | Q969P0 | IGSF8_HUMAN | Unmodified | 28267000 | 33671000 | 11628000 | 16504000 | 0 | 0 | 0 | 0 | 2360200 | 291930 | 1442400 | 10550000 | 0 | 0 | 19125000 | 15873000 | 0 | 0 |
| LVAQLDTE( | IGSF8 | Q969P0 | IGSF8_HUMAN | Oxidation (M) | 0 | 0 | 303120 | 0 | 0 | 0 | 0 | 0 | 689980 | 0 | 289080 | 1832200 | 3981800 | 2570000 | 0 | 0 | 0 | 0 |
| LVAQLYK | CS; CS; CS; CS | O75390; A0A | CISY_HUMAN | Unmodified | 0 | 0 | 0 | 0 | 0 | 0 | 339400 | 65850 | 19165000 | 2051100 | 324350 | 3275400 | 0 | 0 | 162910 | 0 | 57180 | 300150 |
| LVARPEPA | FSCN1 | Q16658 | FSCN1_HUMAI | Unmodified | 0 | 0 | 0 | 0 | 955400 | 578320 | 230370 | 0 | 2972500 | 0 | 10162000 | 20031000 | 0 | 0 | 0 | 36461 | 742260 | 0 |
| LVARPEPA | FSCN1 | Q16658 | FSCN1_HUMAI | Unmodified | 0 | 0 | 0 | 0 | 3175400 | 2012000 | 0 | 100010 | 0 | 349110 | 266930 | 0 | 0 | 0 | 0 | 0 | 1050300 | 1963100 |
| LVASAEQLI | DPP3; DPP3; DI | Q9NY33; G3 | DPP3_HUMAN | Unmodified | 0 | 0 | 125960 | 204560 | 312810 | 87096 | 0 | 0 | 0 | 224060 | 410830 | 8419900 | 632990 | 165220 | 0 | 252310 | 215330 | 300190 |
| LVASAYSIA | BPNT1; BPNT1; | O95861; F8V | BPNT1_HUMAN | Unmodified | 0 | 0 | 0 | 0 | 2352100 | 155600 | 1832400 | 404750 | 422100 | 1655000 | 653880 | 5349600 | 1366800 | 521500 | 353970 | 508280 | 1219900 | 646460 |
| LVATSKITL( | SNX9; SNX9; SI | Q9Y5X1; A0A | SNX9_HUMAN | Unmodified | 0 | 0 | 0 | 0 | 82482 | 0 | 0 | 0 | 0 | 0 | 0 | 0 | 144440 | 42665 | 0 | 0 | 14594 | 0 |
| LVAVAQSG | REPS1; REPS1; | Q96D71; H0 | REPS1_HUMAN | Unmodified | 0 | 0 | 0 | 0 | 0 | 0 | 0 | 109720 | 0 | 0 | 0 | 0 | 0 | 52729 | 338250 | 0 | 0 | 0 |
| LVAYSSDQ | DDC; DDC; DD | P20711; A0A | DDC_HUMAN | Unmodified | 1105000 | 1252700 | 9820300 | 1962900 | 0 | 18927 | 48165 | 43507 | 9430700 | 1580700 | 0 | 0 | 42709000 | 307340 | 422660 | 1138800 | 0 | 0 |
| LVAYSSDQ | DDC; DDC; DD | P20711; A0A | DDC_HUMAN | Unmodified | 0 | 0 | 0 | 0 | 0 | 0 | 0 | 0 | 0 | 7174600 | 0 | 0 | 0 | 0 | 0 | 0 | 135330 | 0 |
| LVCLNTK | PTGS1; PTGS1 | P23219; A0A | PGH1_HUMAN | Unmodified | 0 | 0 | 0 | 0 | 0 | 0 | 0 | 0 | 0 | 0 | 936170 | 526900 | 0 | 0 | 0 | 0 | 0 | 0 |
| LVCSGLLQ | PCYOX1; PCYO | Q9UHG3; F8 | PCYOX_HUMAI | Unmodified | 0 | 0 | 0 | 0 | 0 | 0 | 0 | 0 | 5065600 | 1356400 | 0 | 0 | 0 | 0 | 0 | 0 | 0 | 0 |
| LVCYNNK | F11R | Q9Y624 | JAM1_HUMAN | Unmodified | 0 | 1383200 | 2751500 | 1986700 | 66763 | 128790 | 0 | 0 | 3279800 | 50873 | 43728 | 967360 | 0 | 0 | 2976800 | 1709900 | 16261 | 70132 |
| LVDADGPL | ADAM10 | O14672 | ADA10_HUMAN | Unmodified | 591140 | 88794 | 9196500 | 4880400 | 385890 | 0 | 0 | 0 | 610680 | 446390 | 470090 | 10401000 | 0 | 0 | 12630000 | 12130000 | 0 | 0 |
| LVDAEYTK | POGLUT1 | Q8NBL1 | PGLT1_HUMAN | Unmodified | 0 | 0 | 0 | 0 | 27713 | 145670 | 25459 | 80978 | 3987700 | 999750 | 255560 | 296460 | 0 | 0 | 0 | 0 | 726690 | 807680 |
| LVDAGEEC | ADAM9; ADAM9 | Q13443; A0A | ADAM9_HUMA | Unmodified | 252380 | 0 | 63747 | 1813700 | 0 | 0 | 0 | 0 | 433220 | 0 | 0 | 184830 | 0 | 0 | 121960 | 109530 | 54534 | 0 |
| LVDFGSAA | CIT | O14578 | CTRO_HUMAN | Unmodified | 0 | 0 | 0 | 447000 | 3341200 | 1178700 | 609140 | 402360 | 295300 | 983520 | 347430 | 1318700 | 372070 | 271470 | 0 | 90629 | 2422200 | 1218700 |
| LVDGLATS\ | CLASP1; CLASP | Q7Z460; F8V | CLAP1_HUMAN | Unmodified | 0 | 0 | 0 | 26960 | 0 | 0 | 0 | 0 | 0 | 226210 | 0 | 317970 | 0 | 0 | 0 | 0 | 59146 | 0 |
| LVDLLRSP | PKP1 | Q13835 | PKP1_HUMAN | Unmodified | 0 | 0 | 0 | 0 | 0 | 0 | 0 | 0 | 0 | 0 | 0 | 0 | 0 | 0 | 0 | 0 | 614180 | 0 |
| LVDNIFPEI | EFR3B | Q9Y2G0 | EFR3B_HUMAN | Unmodified | 16175 | 124310 | 0 | 0 | 0 | 0 | 0 | 0 | 0 | 0 | 120860 | 214680 | 128770 | 0 | 0 | 0 | 135290 | 0 |
| LVDNTDPH | EVPL; EVPL | Q92817; K7E | EVPL_HUMAN | Unmodified | 0 | 0 | 0 | 0 | 2242600 | 354640 | 0 | 0 | 893300 | 0 | 0 | 0 | 168040 | 173390 | 0 | 0 | 0 | 126160 |
| LVDPERDM | CAPRIN1; CAPF | Q14444; G3 | CAPR1_HUMAN | Unmodified | 0 | 0 | 0 | 0 | 0 | 0 | 0 | 108600 | 0 | 0 | 606590 | 16348000 | 0 | 0 | 0 | 0 | 0 | 0 |
| LVDPLYSIK | GNPDA1 | P46926 | GNPI1_HUMAN | Unmodified | 0 | 0 | 276870 | 0 | 0 | 520960 | 182950 | 0 | 711780 | 0 | 813850 | 0 | 0 | 1013200 | 0 | 277760 | 0 | 0 |
| LVDPQEMI | FDXR; FDXR; FD | P22570; A0A | ADRO_HUMAN | Unmodified | 0 | 0 | 0 | 0 | 0 | 0 | 906640 | 0 | 0 | 395960 | 0 | 0 | 0 | 0 | 0 | 0 | 46344 | 167580 |
| LVDQNLNK | SMARCA5 | O60264 | SMCA5_HUMA | Unmodified | 982770 | 698770 | 0 | 0 | 743400 | 186690 | 144200 | 0 | 0 | 0 | 356400 | 951600 | 0 | 0 | 0 | 0 | 1425800 | 153200 |
| LVDQSGPF | ADAR; ADAR; AI | P55265; A0A | DSRAD_HUMAI | Unmodified | 608780 | 169660 | 0 | 0 | 0 | 0 | 0 | 464460 | 999230 | 132390 | 240340 | 761360 | 251900 | 57313 | 0 | 526050 | 0 | 0 |
| LVDRDSVV | MYORG | Q6NSJ0 | MYORG_HUMA | Unmodified | 56788 | 71063 | 0 | 0 | 0 | 0 | 0 | 0 | 0 | 0 | 0 | 541790 | 0 | 0 | 171140 | 43645 | 0 | 0 |
| LVDSKGFD | FABP5 | Q01469 | FABP5_HUMAN | Unmodified | 0 | 0 | 0 | 0 | 4773900 | 0 | 0 | 0 | 0 | 0 | 0 | 0 | 0 | 0 | 0 | 0 | 8748700 | 14623000 |
| LVDSKGFD | FABP5 | Q01469 | FABP5_HUMAN | Oxidation (M) | 0 | 0 | 0 | 0 | 2054900 | 0 | 0 | 0 | 0 | 39890 | 0 | 0 | 1024100 | 0 | 0 | 0 | 2898700 | 3688100 |
| LVDVICEK | RAB3D; RAB3A; | O95716; P2 | RAB3D_HUMAI | Unmodified | 144790 | 490680 | 1503000 | 4083900 | 0 | 0 | 0 | 0 | 0 | 0 | 280480 | 2605600 | 0 | 0 | 1250900 | 1052200 | 0 | 0 |
| LVDVLERTI | B4GALNT1 | Q00973 | B4GN1_HUMAI | Unmodified | 0 | 0 | 0 | 0 | 0 | 0 | 0 | 0 | 2559100 | 0 | 0 | 0 | 0 | 0 | 0 | 0 | 0 | 0 |
| LVDVVAVIL | TBCD; TBCD | Q9BTW9; J3I | TBCD_HUMAN | Unmodified | 0 | 0 | 0 | 0 | 0 | 0 | 0 | 0 | 0 | 0 | 0 | 448270 | 0 | 0 | 0 | 0 | 0 | 0 |
| LVDYDSAR | BIN1 | O00499 | BIN1_HUMAN I | Unmodified | 0 | 0 | 0 | 237410 | 987060 | 0 | 0 | 0 | 0 | 0 | 0 | 0 | 2291600 | 556620 | 0 | 0 | 436530 | 932360 |
| LVEAFGGA | NPTXR; NPTXR | O95502; A0A | NPTXR_HUMAN | Unmodified | 0 | 0 | 0 | 0 | 841040 | 0 | 1540100 | 36470 | 2180100 | 242280 | 0 | 0 | 0 | 0 | 1905500 | 0 | 313050 | 662670 |
| LVEAISRDL | DYNC1H1; DYN | Q14204; A0A | DYHC1_HUMA | Unmodified | 21681 | 0 | 36751 | 0 | 0 | 0 | 70679 | 0 | 0 | 19814 | 0 | 425150 | 0 | 0 | 0 | 26552 | 0 | 0 |
| LVEALAAAT | NUDT16L1; NU | Q9BRJ7; K7E | TIRR_HUMAN | Unmodified | 0 | 0 | 231380 | 0 | 0 | 0 | 0 | 0 | 0 | 0 | 0 | 0 | 0 | 0 | 377080 | 0 | 0 | 0 |
| LVEALCAEI | RPS12 | P25398 | RS12_HUMAN | Unmodified | 3169300 | 1108400 | 1919300 | 926420 | 771700 | 516410 | 807490 | 350230 | 2051700 | 0 | 5220100 | 25989000 | 1973200 | 826270 | 2992200 | 1721100 | 0 | 850370 |
| LVEALCAEI | RPS12 | P25398 | RS12_HUMAN | Unmodified | 2097000 | 1856800 | 0 | 0 | 671000 | 0 | 543650 | 427030 | 0 | 164320 | 1757600 | 10400000 | 9076300 | 800180 | 0 | 805750 | 3654200 | 4774600 |
| LVEALCAEI | RPS12 | P25398 | RS12_HUMAN | Unmodified | 542580 | 564600 | 0 | 856520 | 0 | 0 | 1425400 | 209280 | 0 | 262520 | 751580 | 1916900 | 3990700 | 2033900 | 0 | 0 | 891360 | 0 |
| LVEARPMII | KPNB1; KPNB1 | Q14974; J3K | IMB1_HUMAN | Unmodified | 0 | 0 | 0 | 0 | 317570 | 0 | 0 | 0 | 855450 | 101590 | 0 | 1005100 | 0 | 0 | 0 | 0 | 94016 | 97432 |
| LVEARPMII | KPNB1; KPNB1 | Q14974; J3K | IMB1_HUMAN | Oxidation (M) | 0 | 17609 | 104890 | 0 | 0 | 0 | 0 | 0 | 0 | 0 | 184060 | 1508100 | 0 | 0 | 161920 | 401710 | 0 | 0 |
| LVEAVDGK | COLGALT1; CO | Q8NBJ5; M0 | GT251_HUMAN | Unmodified | 0 | 0 | 0 | 0 | 238030 | 152560 | 0 | 0 | 842340 | 31610 | 131670 | 762050 | 0 | 0 | 15658 | 0 | 0 | 0 |
| LVECGSLFI | PLXNB2; PLXNE | O15031; A6 | PLXB2_HUMAN | Unmodified | 0 | 0 | 0 | 0 | 0 | 0 | 0 | 0 | 0 | 0 | 0 | 0 | 637030 | 507470 | 0 | 0 | 0 | 0 |
| LVECLETVL | AP2A1 | O95782 | AP2A1_HUMAN | Unmodified | 0 | 0 | 0 | 0 | 0 | 0 | 0 | 0 | 0 | 0 | 0 | 1201300 | 0 | 0 | 0 | 0 | 0 | 0 |
| LVEDHLAV | OGDHL; OGDH | Q9ULD0; Q0 | OGDHL_HUMA | Unmodified | 0 | 108650 | 0 | 0 | 0 | 0 | 0 | 13497 | 3954400 | 0 | 122520 | 0 | 0 | 0 | 45241 | 0 | 0 | 0 |
| LVEDMENK | CAPZB; CAPZB; | P47756; B1A | CAPZB_HUMAN | Unmodified | 0 | 2320500 | 0 | 1332100 | 214100 | 224440 | 0 | 0 | 2727200 | 155930 | 996820 | 13601000 | 0 | 0 | 1867200 | 1615800 | 0 | 68103 |
| LVEDMENK | CAPZB; CAPZB; | P47756; B1A | CAPZB_HUMAN | Unmodified | 397620 | 286510 | 0 | 0 | 0 | 0 | 0 | 0 | 0 | 0 | 0 | 0 | 0 | 0 | 0 | 0 | 0 | 0 |
| LVEIAQVPK | UGP2; UGP2; U | Q16851; E7E | UGPA_HUMAN | Unmodified | 446690 | 1883200 | 14829000 | 7862300 | 8854300 | 42058000 | 228910 | 172290 | 3812000 | 3651300 | 30617000 | 6908200 | 3339000 | 12901000 | 3819900 | 0 | 2267800 | 18177000 |
| LVEIDNGK | LMNA; LMNA; L | P02545; A0A | LMNA_HUMAN | Unmodified | 0 | 0 | 0 | 0 | 728750 | 103110 | 1060000 | 805220 | 195340 | 80191 | 665580 | 839160 | 0 | 0 | 49535 | 304980 | 779570 | 397330 |
| LVEIIGSR | ECHDC1; ECHI | Q9NTX5; H0 | ECHD1_HUMA | Unmodified | 535660 | 0 | 68219 | 0 | 0 | 0 | 0 | 0 | 167950 | 0 | 898130 | 4733900 | 0 | 0 | 0 | 0 | 0 | 0 |
| LVEIVHPS( | PPP6R3; PPP6F | Q5H9R7; E9I | PP6R3_HUMAN | Unmodified | 0 | 0 | 0 | 0 | 0 | 133890 | 0 | 0 | 0 | 140810 | 177870 | 1003800 | 0 | 0 | 0 | 0 | 0 | 93988 |
| LVEIVHPS( | PPP6R3; PPP6F | Q5H9R7; E9I | PP6R3_HUMAN | Unmodified | 0 | 0 | 0 | 0 | 0 | 0 | 0 | 0 | 0 | 0 | 0 | 4729400 | 0 | 0 | 116890 | 104270 | 0 | 236450 |
| LVELLMHN | KPNA6; KPNA1; | O60684; P5 | IMA7_HUMAN | Oxidation (M) | 227890 | 784120 | 0 | 231750 | 0 | 0 | 143010 | 0 | 0 | 0 | 140500 | 386860 | 0 | 0 | 0 | 249250 | 0 | 0 |
| LVELQNTR\ | TARS2; TARS2; | Q9BW92; F6 | SYTM_HUMAN | Unmodified | 0 | 0 | 0 | 0 | 0 | 26899 | 0 | 0 | 446300 | 0 | 0 | 0 | 0 | 0 | 0 | 0 | 0 | 0 |

| | | | | | | | | | | | | | | | | | | | | | | |
|---|---|---|---|---|---|---|---|---|---|---|---|---|---|---|---|---|---|---|---|---|---|---|
| LVELSGGC | ABCC5 | O15440 | MRP5_HUMAN | Unmodified | 0 | 0 | 0 | 0 | 0 | 0 | 0 | 0 | 0 | 0 | 0 | 0 | 0 | 0 | 0 | 0 | 0 | 0 |
| LVELTQK | LRPPRC; PNPL | P42704; Q9 | LPPRC_HUMAN | Unmodified | 0 | 0 | 0 | 0 | 204350 | 95000 | 0 | 0 | 5607700 | 1056400 | 176710 | 99939 | 0 | 0 | 203460 | 0 | 70570 | 167050 |
| LVEMGPAG | ANK3; ANK3 | Q12955; A0 | ANK3_HUMAN | 2 Oxidation (M) | 0 | 0 | 102630 | 0 | 0 | 0 | 0 | 0 | 0 | 0 | 0 | 124280 | 0 | 0 | 1209700 | 0 | 0 | 0 |
| LVENCVCL | CTNND1; CTN | O60716; A0 | CTND1_HUMA | Unmodified | 721060 | 256930 | 1329100 | 1616700 | 0 | 0 | 0 | 0 | 0 | 0 | 0 | 3793200 | 0 | 1113100 | 2694800 | 2923800 | 880360 | 0 |
| LVEPGSPAI | SLC9A3R1; SLC | O14745; J3C | NHRF1_HUMA | Unmodified | 1456700 | 1303500 | 4010200 | 7736400 | 11679000 | 3148600 | 2901100 | 1559100 | 5416700 | 1524200 | 2374300 | 19828000 | 39162000 | 31414000 | 9527200 | 7494100 | 3404900 | 4204600 |
| LVEQHGSD | IARS2 | Q9NSE4 | SYIM_HUMAN | Unmodified | 0 | 0 | 0 | 0 | 218050 | 0 | 0 | 0 | 2540500 | 0 | 76555 | 0 | 0 | 0 | 0 | 0 | 0 | 0 |
| LVEQLKLE | GNG10 | P50151 | GBG10_HUMA | Unmodified | 0 | 0 | 0 | 0 | 2491800 | 0 | 526140 | 654240 | 0 | 217650 | 0 | 0 | 0 | 0 | 0 | 0 | 0 | 1465000 |
| LVEQLKME | GNG2; GNG2; | P59768; G3 | GBG2_HUMAN | Unmodified | 0 | 0 | 0 | 0 | 162800 | 194020 | 659380 | 306550 | 0 | 0 | 0 | 0 | 0 | 1222300 | 0 | 0 | 2104700 | 6484300 |
| LVEQLKME | GNG2; GNG2; | P59768; G3 | GBG2_HUMAN | Oxidation (M) | 0 | 0 | 0 | 0 | 0 | 0 | 0 | 28956 | 0 | 0 | 0 | 0 | 241010 | 0 | 0 | 0 | 0 | 1772600 |
| LVEQYKLPI | HACL1; HACL1 | Q9UJ83; C9 | HACL1_HUMA | Unmodified | 0 | 0 | 0 | 0 | 32223 | 147220 | 0 | 0 | 256310 | 0 | 0 | 0 | 0 | 0 | 0 | 0 | 0 | 49991 |
| LVEQYTK | SF3A1 | Q15459 | SF3A1_HUMAN | Unmodified | 0 | 105560 | 0 | 0 | 0 | 0 | 0 | 0 | 508550 | 0 | 297080 | 776170 | 0 | 0 | 0 | 137460 | 7958.2 | 0 |
| LVERCQEQ | VPS37C; VPS37 | A5D8V6; F5 | VP37C_HUMAN | Unmodified | 0 | 0 | 0 | 0 | 0 | 0 | 0 | 0 | 0 | 262480 | 0 | 0 | 1038600 | 723760 | 0 | 0 | 59584 | 167260 |
| LVERGAPQ | DIP2B | Q9P265 | DIP2B_HUMAN | Unmodified | 0 | 0 | 0 | 120330 | 696020 | 0 | 48471 | 23345 | 1067500 | 622490 | 639050 | 0 | 402570 | 370810 | 0 | 0 | 207550 | 637920 |
| LVEVDSGR | LMNB1; LMNB1 | P20700; A0A | LMNB1_HUMA | Unmodified | 46148 | 0 | 165230 | 444660 | 0 | 98609 | 0 | 0 | 378010 | 0 | 0 | 0 | 0 | 0 | 146550 | 0 | 0 | 0 |
| LVEVIEEVN | SHTN1 | A0MZ66 | SHOT1_HUMA | Unmodified | 0 | 0 | 0 | 0 | 488740 | 0 | 0 | 0 | 489570 | 0 | 325370 | 1007400 | 0 | 0 | 0 | 0 | 0 | 0 |
| LVEVIKTPM | HADH; HADH; | Q16836; A0 | HCDH_HUMAN | Unmodified | 0 | 0 | 0 | 0 | 0 | 0 | 2249400 | 466690 | 2602600 | 1920300 | 0 | 0 | 0 | 0 | 0 | 0 | 0 | 0 |
| LVEVIKTPM | HADH; HADH; | Q16836; A0 | HCDH_HUMAN | Oxidation (M) | 0 | 0 | 0 | 0 | 0 | 0 | 563120 | 148130 | 0 | 730660 | 0 | 0 | 237660 | 0 | 0 | 0 | 0 | 0 |
| LVEVIKTPM | HADH; HADH; | Q16836; A0 | HCDH_HUMAN | Oxidation (M) | 0 | 0 | 0 | 0 | 0 | 0 | 1450800 | 0 | 0 | 0 | 0 | 0 | 0 | 0 | 0 | 0 | 67320 | 0 |
| LVEVLEGV | CRELD1; CREL | Q96HD1; A0 | CREL1_HUMAN | Unmodified | 0 | 0 | 0 | 0 | 0 | 0 | 0 | 0 | 1949600 | 0 | 145480 | 0 | 0 | 0 | 0 | 0 | 0 | 0 |
| LVEVNGVN | SLC9A3R2 | Q15599 | NHRF2_HUMA | Unmodified | 0 | 0 | 784980 | 276730 | 0 | 0 | 0 | 0 | 0 | 0 | 804550 | 2443800 | 0 | 0 | 410200 | 534950 | 0 | 0 |
| LVEVPNDG | PARD3; PARD3; | Q8TEW0; Q5 | PARD3_HUMA | Unmodified | 0 | 0 | 0 | 0 | 0 | 436820 | 0 | 0 | 4329200 | 0 | 0 | 2354300 | 0 | 0 | 0 | 0 | 513090 | 0 |
| LVEVPNDG | PARD3; PARD3; | Q8TEW0; Q5 | PARD3_HUMA | Unmodified | 0 | 0 | 0 | 0 | 2636400 | 316450 | 293210 | 33645 | 0 | 494940 | 0 | 0 | 0 | 0 | 0 | 0 | 1704000 | 3097500 |
| LVEVPNDG | PARD3; PARD3; | Q8TEW0; Q5 | PARD3_HUMA | Unmodified | 0 | 0 | 0 | 0 | 3407200 | 0 | 0 | 0 | 0 | 0 | 0 | 0 | 0 | 0 | 0 | 0 | 3243500 | 3479200 |
| LVFDEYLK | ANXA6 | P08133 | ANXA6_HUMA | Unmodified | 1389800 | 1713200 | 5580500 | 6635100 | 2203200 | 3406300 | 1769600 | 632640 | 62230000 | 14882000 | 0 | 0 | 1844300 | 1629400 | 6150900 | 1865100 | 1087000 | 3152300 |
| LVFGFLNG | PNP; PNP; PNP | P00491; G3 | PNPH_HUMAN | Unmodified | 204250 | 246650 | 0 | 0 | 0 | 0 | 0 | 0 | 0 | 0 | 0 | 16935000 | 0 | 0 | 396050 | 495720 | 0 | 0 |
| LVFHTQLAI | GIPC1 | O14908 | GIPC1_HUMAN | Unmodified | 0 | 0 | 0 | 0 | 0 | 0 | 0 | 0 | 0 | 0 | 0 | 1210500 | 0 | 0 | 0 | 0 | 0 | 0 |
| LVFHTQLAI | GIPC1 | O14908 | GIPC1_HUMAN | Unmodified | 332250 | 217250 | 905440 | 1705500 | 37961000 | 6558900 | 2454400 | 75233 | 3905300 | 5273800 | 2091100 | 2810600 | 19265000 | 13766000 | 0 | 0 | 26746000 | 19346000 |
| LVFLGEQS | RAB6A; RAB6B; | P20340; Q9 | RAB6A_HUMAN | Unmodified | 0 | 0 | 0 | 0 | 695470 | 0 | 702960 | 557840 | 1278600 | 681550 | 14261000 | 13750000 | 0 | 0 | 1446700 | 833740 | 0 | 0 |
| LVFLGLDN | SAR1B; SAR1A; | Q9Y6B6; Q9 | SAR1B_HUMAN | Unmodified | 369140 | 0 | 520310 | 278680 | 3015300 | 24040000 | 1104000 | 95397 | 1963600 | 2528800 | 478260 | 7942700 | 0 | 0 | 1399100 | 873180 | 2481000 | 0 |
| LVFPQDLL | HM13; HM13; | Q8TCT9; A0 | HM13_HUMAN | Unmodified | 0 | 0 | 0 | 0 | 0 | 0 | 0 | 0 | 4589100 | 485250 | 332500 | 1637000 | 0 | 0 | 0 | 0 | 0 | 0 |
| LVFRDVKD | GNA13 | Q14344 | GNA13_HUMA | Unmodified | 0 | 0 | 0 | 0 | 888460 | 294670 | 0 | 0 | 1204300 | 916370 | 0 | 0 | 0 | 0 | 0 | 0 | 435570 | 674070 |
| LVFSNVNL | DDB1; DDB1; D | Q16531; A0 | DDB1_HUMAN | Unmodified | 0 | 0 | 0 | 0 | 0 | 0 | 0 | 35435 | 0 | 677390 | 281370 | 4526400 | 0 | 0 | 0 | 0 | 0 | 0 |
| LVFVHSAE | CAPN1; CAPN1 | P07384; E9P | CAN1_HUMAN | Unmodified | 462260 | 724990 | 113760 | 125480 | 0 | 0 | 0 | 0 | 0 | 0 | 0 | 1873900 | 0 | 0 | 1270100 | 324710 | 752160 | 0 |
| LVGAIVYY | GPD2; GPD2 | P43304; F5G | GPDM_HUMAN | Unmodified | 0 | 0 | 0 | 0 | 0 | 0 | 0 | 0 | 6221900 | 0 | 0 | 0 | 0 | 0 | 0 | 0 | 0 | 0 |
| LVGDVDFE | MTHFD2; MTHF | P13995; A0A | MTDC_HUMAN | Unmodified | 0 | 0 | 0 | 0 | 0 | 0 | 0 | 366040 | 4985700 | 1759500 | 475910 | 895080 | 0 | 0 | 0 | 0 | 210680 | 188050 |
| LVGFISAIP | NMT1; NMT1; N | P30419; A0A | NMT1_HUMAN | Unmodified | 0 | 0 | 0 | 0 | 0 | 0 | 0 | 0 | 0 | 0 | 0 | 1334500 | 0 | 0 | 0 | 0 | 0 | 0 |
| LVGGKDFE | PDLIM1 | O00151 | PDLI1_HUMAN | Unmodified | 0 | 0 | 0 | 0 | 1434600 | 350790 | 0 | 0 | 243120 | 325120 | 0 | 0 | 0 | 410670 | 0 | 0 | 945180 | 1775000 |
| LVGGPMD | CST3 | P01034 | CYTC_HUMAN | Unmodified | 0 | 0 | 0 | 0 | 0 | 0 | 0 | 0 | 1367200 | 189650 | 0 | 0 | 879010 | 0 | 0 | 0 | 0 | 0 |
| LVGGPMD | CST3 | P01034 | CYTC_HUMAN | Oxidation (M) | 0 | 0 | 974600 | 0 | 0 | 0 | 0 | 0 | 9070900 | 157030 | 0 | 0 | 0 | 0 | 127490 | 89732 | 0 | 0 |
| LVGGPMD | CST3 | P01034 | CYTC_HUMAN | Unmodified | 0 | 0 | 832720 | 394010 | 138720 | 0 | 28692 | 0 | 36594000 | 1109200 | 0 | 0 | 0 | 0 | 168610 | 0 | 0 | 0 |
| LVGGPMD | CST3 | P01034 | CYTC_HUMAN | Oxidation (M) | 0 | 0 | 0 | 0 | 0 | 0 | 0 | 0 | 0 | 8405600 | 0 | 0 | 0 | 106680 | 0 | 0 | 0 | 0 |
| LVGGTTPG | SUCLG1; SUCL | P53597; A0A | SUCA_HUMAN | Unmodified | 0 | 0 | 0 | 0 | 139320 | 0 | 4046100 | 113160 | 6227100 | 5000500 | 0 | 0 | 222800 | 205600 | 0 | 0 | 0 | 1681600 |
| LVGIFENQ | NUDT3; RPS10- | O95989; S4 | NUDT3_HUMA | Unmodified | 0 | 0 | 0 | 0 | 0 | 531840 | 0 | 0 | 348540 | 0 | 338790 | 1557700 | 0 | 0 | 0 | 18976 | 0 | 0 |
| LVGIISSR | IMPDH2; IMPD | P12268; A0A | IMDH2_HUMA | Unmodified | 308410 | 0 | 0 | 406400 | 0 | 0 | 0 | 0 | 1580500 | 0 | 1451200 | 1669100 | 76277 | 266870 | 430310 | 204360 | 0 | 0 |
| LVGILDILD | MYO6; MYO6; | Q9UM54; A0 | MYO6_HUMAN | Unmodified | 1980900 | 2172000 | 5915900 | 29512000 | 3357900 | 878570 | 782440 | 0 | 9630100 | 4906600 | 4325600 | 10405000 | 11085000 | 3985600 | 24472000 | 21610000 | 0 | 0 |
| LVGIVTSRD | IMPDH1; IMPD | P20839; C9J | IMDH1_HUMA | Unmodified | 23783 | 0 | 32208 | 0 | 846250 | 0 | 0 | 0 | 0 | 0 | 0 | 0 | 913160 | 282280 | 0 | 0 | 0 | 405560 |
| LVGMSVLL | VPS36 | Q86VN1 | VPS36_HUMAN | Unmodified | 1054800 | 1493600 | 0 | 835540 | 796330 | 491220 | 1290000 | 3602600 | 537730 | 778990 | 0 | 88612 | 1364500 | 544370 | 1549000 | 983370 | 1467200 | 1170800 |
| LVGMSVLL | VPS36 | Q86VN1 | VPS36_HUMAN | Oxidation (M) | 2204700 | 2040700 | 1885600 | 1266900 | 144060 | 0 | 0 | 0 | 0 | 72300 | 0 | 0 | 1119400 | 483100 | 3014900 | 2737500 | 0 | 110890 |
| LVGNMHG | CPD | O75976 | CBPD_HUMAN | Oxidation (M) | 0 | 0 | 0 | 0 | 0 | 0 | 0 | 0 | 24725000 | 2201000 | 0 | 0 | 0 | 0 | 0 | 0 | 0 | 0 |
| LVGNMHG | CPD | O75976 | CBPD_HUMAN | Unmodified | 179570 | 181940 | 156010 | 0 | 0 | 0 | 108580 | 0 | 10889000 | 965540 | 45780 | 0 | 0 | 0 | 210100 | 236390 | 0 | 0 |
| LVGNMHG | CPD | O75976 | CBPD_HUMAN | Oxidation (M) | 0 | 0 | 0 | 0 | 0 | 0 | 0 | 0 | 0 | 4482700 | 0 | 0 | 0 | 0 | 0 | 0 | 0 | 0 |
| LVGPEEAL | PLOD3; PLOD3 | O60568; H7 | PLOD3_HUMA | Unmodified | 0 | 0 | 0 | 0 | 0 | 0 | 278240 | 0 | 13041000 | 128370 | 356980 | 401100 | 0 | 0 | 0 | 0 | 0 | 0 |
| LVGQGAS | HSD17B10; HS | Q99714; Q5 | HCD2_HUMAN | Unmodified | 90835 | 281610 | 215610 | 0 | 0 | 0 | 0 | 0 | 29878000 | 0 | 0 | 0 | 0 | 0 | 0 | 0 | 0 | 0 |
| LVGQGAS | HSD17B10; HS | Q99714; Q5 | HCD2_HUMAN | Unmodified | 0 | 0 | 0 | 0 | 0 | 0 | 0 | 0 | 705850 | 0 | 0 | 0 | 0 | 0 | 0 | 0 | 0 | 0 |
| LVGQQYLH | RASAL2 | Q9UJF2 | NGAP_HUMAN | Unmodified | 0 | 166300 | 27083 | 378060 | 463160 | 195310 | 77318 | 0 | 74425 | 329470 | 0 | 431670 | 0 | 0 | 0 | 0 | 117420 | 0 |
| LVGQSIIAY | COPA; COPA; C | P53621; A0A | COPA_HUMAN | Unmodified | 0 | 0 | 0 | 501560 | 0 | 0 | 0 | 0 | 0 | 0 | 0 | 3545900 | 0 | 0 | 893330 | 966620 | 0 | 0 |
| LVGQSIIAY | COPA; COPA; C | P53621; A0A | COPA_HUMAN | Unmodified | 0 | 0 | 0 | 0 | 159380 | 0 | 248770 | 0 | 0 | 155230 | 0 | 3538300 | 0 | 0 | 0 | 0 | 0 | 0 |
| LVGRPPLP | DHTKD1 | Q96HY7 | DHTK1_HUMA | Unmodified | 0 | 0 | 0 | 0 | 0 | 0 | 0 | 0 | 2059200 | 0 | 0 | 0 | 0 | 0 | 0 | 0 | 0 | 0 |
| LVGSQEEL | PSMD2; PSMD2 | Q13200; H7 | PSMD2_HUMA | Unmodified | 0 | 0 | 0 | 0 | 0 | 0 | 872350 | 459830 | 0 | 0 | 896670 | 4403100 | 0 | 0 | 0 | 0 | 0 | 0 |
| LVGTPGAE | MAPK14; MAPK | Q16539; B4 | MK14_HUMAN | Unmodified | 0 | 0 | 0 | 0 | 0 | 0 | 0 | 0 | 0 | 0 | 0 | 457280 | 0 | 0 | 0 | 0 | 0 | 84419 |
| LVGVFHTE | NIPSNAP3A | Q9UFN0 | NPS3A_HUMA | Unmodified | 0 | 0 | 0 | 0 | 0 | 0 | 0 | 0 | 772840 | 634360 | 0 | 0 | 0 | 0 | 0 | 0 | 0 | 0 |
| LVGVLVHS | USP9X; USP9Y | Q93008; O0 | USP9X_HUMA | Unmodified | 0 | 0 | 0 | 0 | 0 | 0 | 0 | 0 | 0 | 0 | 0 | 1859300 | 0 | 0 | 0 | 0 | 0 | 0 |
| LVGYLDR | PSAP; PSAP; PS | P07602; Q5 | SAP_HUMAN P | Unmodified | 21247 | 22295 | 0 | 496120 | 0 | 0 | 0 | 28287 | 6618900 | 0 | 292310 | 910300 | 0 | 0 | 0 | 148270 | 0 | 0 |
| LVGYLDRN | PSAP; PSAP; PS | P07602; Q5 | SAP_HUMAN P | Unmodified | 30370 | 0 | 1823000 | 918660 | 4191200 | 1462800 | 2564900 | 3350200 | 74843000 | 23812000 | 2620600 | 690950 | 8386900 | 10166000 | 1544500 | 402970 | 400030 | 1689900 |
| LVHANFGT | TFRC; TFRC | P02786; G3 | TFR1_HUMAN | Unmodified | 102220 | 0 | 0 | 0 | 0 | 0 | 0 | 0 | 906980 | 171780 | 0 | 0 | 0 | 0 | 0 | 0 | 0 | 0 |
| LVHIIGALR | TMEM43 | Q9BTV4 | TMM43_HUMA | Unmodified | 0 | 0 | 0 | 0 | 0 | 0 | 0 | 0 | 4548600 | 287350 | 0 | 0 | 0 | 0 | 0 | 0 | 0 | 0 |
| LVHPGVAE | POLDIP3; POLD | Q9BY77; F6 | PDIP3_HUMAN | Unmodified | 0 | 0 | 0 | 0 | 0 | 0 | 0 | 0 | 0 | 0 | 0 | 1098900 | 0 | 0 | 0 | 0 | 0 | 0 |
| LVHSGPGK | SNTB1 | Q13884 | SNTB1_HUMAN | Unmodified | 0 | 0 | 0 | 0 | 0 | 0 | 0 | 0 | 0 | 861010 | 0 | 0 | 0 | 0 | 0 | 0 | 0 | 0 |
| LVHSGSGC | SNTB2 | Q13425 | SNTB2_HUMAN | Unmodified | 0 | 0 | 0 | 440020 | 0 | 0 | 113150 | 0 | 0 | 0 | 0 | 0 | 0 | 0 | 0 | 0 | 0 | 0 |
| LVHSLFESI | PFAS | O15067 | PUR4_HUMAN | Oxidation (M) | 0 | 0 | 0 | 0 | 0 | 0 | 0 | 0 | 0 | 0 | 581050 | 521230 | 0 | 0 | 0 | 0 | 0 | 0 |
| LVICPDEG | UBE2M; UBE2M | P61081; M0 | UBC12_HUMA | Unmodified | 446400 | 279860 | 425090 | 479640 | 0 | 0 | 258390 | 204080 | 2234400 | 0 | 592820 | 3334800 | 0 | 600500 | 709800 | 635850 | 1346200 | 0 |
| LVIDQIDN | PYGL; PYGL | P06737; E9P | PYGL_HUMAN | Unmodified | 0 | 0 | 0 | 0 | 155450 | 0 | 14401 | 0 | 0 | 0 | 0 | 0 | 0 | 0 | 0 | 0 | 23678 | 113740 |
| LVIEEAER | CCT4 | P50991 | TCPD_HUMAN | Unmodified | 0 | 0 | 480750 | 0 | 0 | 0 | 0 | 10595 | 0 | 0 | 3014700 | 21490000 | 0 | 0 | 720860 | 690480 | 0 | 0 |
| LVIEEQSGF | DLG2; DLG2; D | Q15700; E9 | DLG2_HUMAN | Unmodified | 0 | 248610 | 0 | 0 | 0 | 0 | 0 | 0 | 0 | 0 | 0 | 45839 | 64070000 | 3395600 | 555960 | 86718 | 45767 | 0 |
| LVIFEGGTS | AVIL | O75366 | AVIL_HUMAN A | Unmodified | 0 | 0 | 0 | 0 | 0 | 0 | 221160 | 0 | 0 | 0 | 2920500 | 26818000 | 0 | 0 | 0 | 0 | 0 | 0 |
| LVIFEGGTS | AVIL | O75366 | AVIL_HUMAN A | Unmodified | 0 | 0 | 0 | 0 | 0 | 0 | 0 | 0 | 0 | 0 | 43072 | 5939200 | 0 | 0 | 0 | 0 | 428940 | 580130 |
| LVIIEGDLE | TPM3; TPM3; T | P06753; J3K | TPM3_HUMAN | Unmodified | 0 | 140490 | 0 | 0 | 0 | 0 | 0 | 0 | 0 | 0 | 568780 | 5592000 | 0 | 0 | 1078900 | 926000 | 0 | 0 |
| LVIIEGDLE | TPM3; TPM3; T | J3KN67; A0A | J3KN67_HUMA | Unmodified | 512140 | 0 | 308130 | 1665200 | 8695200 | 698870 | 0 | 0 | 3512400 | 0 | 340960 | 10254000 | 2434100 | 0 | 1680800 | 1054600 | 525350 | 0 |
| LVIIESDLE | TPM1; TPM1; T | P09493; H0 | TPM1_HUMAN | Unmodified | 142780 | 0 | 0 | 0 | 0 | 0 | 0 | 0 | 0 | 0 | 0 | 1067300 | 0 | 0 | 0 | 0 | 0 | 0 |
| LVIITAGAR | LDHAL6A; LDH | Q6ZMR3; P0 | LDH6A_HUMA | Unmodified | 5298500 | 5321600 | 6221800 | 12344000 | 2955800 | 353300 | 132330 | 101300 | 22240000 | 2693200 | 15662000 | 3.78E+08 | 573540 | 288690 | 15258000 | 15117000 | 1937000 | 2207900 |
| LVIITAGAR | LDHA; LDHA; L | P00338; F5G | LDHA_HUMAN | Unmodified | 205200 | 132720 | 620100 | 1824400 | 888360 | 0 | 144290 | 36121 | 1399800 | 462920 | 1899100 | 2653600 | 476660 | 152060 | 257700 | 161740 | 739910 | 1427800 |
| LVIKNQQF | NONO; NONO; | Q15233; A0 | NONO_HUMAN | Unmodified | 0 | 0 | 0 | 0 | 0 | 0 | 406140 | 381430 | 0 | 0 | 0 | 0 | 0 | 0 | 0 | 0 | 0 | 0 |
| LVILANNC | RPL30; RPL30; | P62888; E5R | RL30_HUMAN | Unmodified | 7624200 | 3471600 | 2976800 | 3181700 | 0 | 0 | 0 | 0 | 566230 | 147520 | 1673700 | 28780000 | 0 | 0 | 5141200 | 4431800 | 0 | 0 |
| LVILANNC | RPL30; RPL30 | P62888; E5R | RL30_HUMAN | Unmodified | 1173600 | 506140 | 1845100 | 408970 | 365640 | 0 | 0 | 0 | 296110 | 0 | 383020 | 13132000 | 0 | 597030 | 1331000 | 1002500 | 0 | 0 |
| LVILEGELE | TPM4; TPM4; T | P67936; A0A | TPM4_HUMAN | Unmodified | 613720 | 661550 | 0 | 1089400 | 35489000 | 4451500 | 456350 | 0 | 0 | 0 | 0 | 5872700 | 2589600 | 0 | 0 | 646950 | 1242300 | 0 |
| LVILEGELE | TPM2; TPM2; T | P07951; U3K | TPM2_HUMAN | Unmodified | 0 | 0 | 0 | 0 | 0 | 0 | 0 | 0 | 0 | 0 | 0 | 9082300 | 0 | 0 | 0 | 0 | 0 | 0 |
| LVINGNPIT | GAPDH | P04406 | G3P_HUMAN G | Unmodified | 1961900 | 1359900 | 2041900 | 879470 | 1613200 | 2627600 | 1568100 | 1557900 | 2730500 | 1101300 | 3760300 | 21126000 | 655830 | 883490 | 2060300 | 2589100 | 1212600 | 1434600 |
| LVINGNPIT | GAPDH | P04406 | G3P_HUMAN G | Unmodified | 7181700 | 5777100 | 0 | 1377000 | 22619000 | 3764200 | 0 | 427160 | 11695000 | 2846800 | 28221000 | 1.47E+08 | 10145000 | 4299400 | 1472100 | 1900100 | 4251400 | 26243000 |
| LVINGNPIT | GAPDH | P04406 | G3P_HUMAN G | Unmodified | 0 | 0 | 0 | 0 | 9256100 | 835530 | 742240 | 131830 | 459140 | 885980 | 0 | 0 | 3915100 | 1503700 | 0 | 0 | 1444400 | 3341400 |
| LVINSGNG | CD44; CD44; C | P16070; H0 | CD44_HUMAN | Unmodified | 750230 | 738270 | 0 | 0 | 0 | 0 | 0 | 0 | 0 | 0 | 0 | 0 | 113930 | 0 | 0 | 0 | 0 | 0 |
| LVINSGNG | CD44; CD44; C | P16070; H0 | CD44_HUMAN | Unmodified | 740080 | 825850 | 0 | 0 | 2418500 | 712880 | 0 | 0 | 0 | 0 | 0 | 0 | 0 | 0 | 0 | 0 | 630370 | 2331400 |
| LVINTEEVF | PRKDC | P78527 | PRKDC_HUMA | Unmodified | 0 | 0 | 0 | 0 | 757350 | 0 | 0 | 110270 | 19273 | 0 | 0 | 2321900 | 0 | 0 | 0 | 0 | 0 | 0 |
| LVIPSELGY | FKBP2; FKBP2 | P26885; A0A | FKBP2_HUMAN | Unmodified | 0 | 0 | 0 | 0 | 0 | 217270 | 0 | 0 | 76696000 | 1392400 | 3371500 | 0 | 0 | 0 | 0 | 0 | 0 | 0 |
| LVIPSELGY | FKBP2; FKBP2 | P26885; A0A | FKBP2_HUMAN | Unmodified | 0 | 0 | 0 | 0 | 0 | 0 | 0 | 0 | 2775800 | 499120 | 481480 | 0 | 0 | 0 | 0 | 0 | 0 | 0 |
| LVISASWAI | EXOC4 | Q96A65 | EXOC4_HUMA | Unmodified | 136080 | 0 | 0 | 0 | 0 | 0 | 0 | 0 | 577660 | 0 | 0 | 2660700 | 0 | 0 | 242850 | 145400 | 0 | 0 |
| LVIVGDGA | RHOA; RHOC; | P61586; P08 | RHOA_HUMAN | Unmodified | 4095300 | 3868000 | 2066900 | 5026800 | 13493000 | 2887300 | 2167100 | 0 | 10775000 | 5318100 | 5035100 | 20731000 | 6247200 | 4784900 | 6987400 | 6523700 | 5557100 | 10096000 |
| LVKEVPEE | GRK6; GRK6; G | P43250; D6 | GRK6_HUMAN | Unmodified | 0 | 0 | 0 | 0 | 0 | 0 | 0 | 0 | 0 | 40415 | 0 | 0 | 435470 | 141720 | 0 | 0 | 0 | 169720 |
| LVKGHAYS | CAPN2 | P17655 | CAN2_HUMAN | Unmodified | 0 | 0 | 0 | 0 | 0 | 0 | 0 | 0 | 0 | 0 | 0 | 0 | 0 | 0 | 0 | 0 | 640070 | 0 |
| LVKPGNQ | PA2G4; PA2G4; | Q9UQ80; F8 | PA2G4_HUMA | Unmodified | 5481500 | 5700500 | 2013000 | 5464100 | 3205800 | 752030 | 613930 | 310770 | 7424800 | 1423200 | 6151500 | 30382000 | 4482600 | 0 | 3378900 | 2800000 | 597820 | 793870 |
| LVLCEVFK | GLUL; GLUL | P15104; A0A | GLNA_HUMAN | Unmodified | 0 | 0 | 114430 | 0 | 0 | 0 | 0 | 0 | 425430 | 0 | 0 | 2114600 | 0 | 0 | 0 | 0 | 0 | 0 |
| LVLDQVHT | FAT1; FAT1 | Q14517; A0 | FAT1_HUMAN | Unmodified | 98821 | 99687 | 107230 | 51037 | 0 | 0 | 0 | 0 | 0 | 0 | 0 | 247670 | 0 | 0 | 183210 | 0 | 0 | 0 |
| LVLDSVKL | TBCA; TBCA; T | O75347; E5 | TBCA_HUMAN | Unmodified | 5143100 | 200810 | 2745400 | 4231000 | 4104100 | 0 | 0 | 358380 | 6771100 | 1925400 | 0 | 17156000 | 8288300 | 3454900 | 5355100 | 5167300 | 981240 | 1215800 |
| LVLDWDS | ARHGAP17; AR | Q68EM7; C9 | RHG17_HUMA | Unmodified | 0 | 0 | 0 | 0 | 1225800 | 0 | 0 | 0 | 0 | 192980 | 0 | 0 | 1218300 | 1171800 | 0 | 0 | 0 | 0 |
| LVLEMVHH | BAZ1B | Q9UIG0 | BAZ1B_HUMAN | Unmodified | 0 | 0 | 0 | 0 | 0 | 0 | 569320 | 223510 | 0 | 0 | 0 | 0 | 0 | 0 | 0 | 0 | 0 | 0 |
| LVLEQVVT | IPO5; IPO5 | O00410; H0 | IPO5_HUMAN | Unmodified | 0 | 0 | 64500 | 0 | 0 | 0 | 0 | 0 | 20432 | 114560 | 74265 | 694450 | 0 | 0 | 306810 | 71064 | 0 | 0 |
| LVLEVAQH | ATP5F1B | P06576 | ATPB_HUMAN | Unmodified | 661520 | 0 | 784650 | 0 | 0 | 623460 | 0 | 399430 | 67961000 | 4021900 | 1693400 | 1022200 | 0 | 0 | 527430 | 531180 | 1480100 | 3113300 |
| LVLIGSNH | SEC16A; SEC1 | O15027; F1 | SC16A_HUMAN | Unmodified | 669450 | 0 | 168700 | 0 | 336260 | 0 | 413220 | 41257 | 0 | 0 | 0 | 3082600 | 315620 | 0 | 0 | 227280 | 254370 | 0 |
| LVLINKSG | CTNND1; CTN | O60716; A0 | CTND1_HUMA | Unmodified | 0 | 0 | 0 | 0 | 491530 | 0 | 0 | 0 | 0 | 0 | 0 | 0 | 0 | 945870 | 0 | 0 | 2422900 | 831240 |
| LVLIRHGE | PGAM1 | P18669 | PGAM1_HUMA | Unmodified | 0 | 0 | 0 | 0 | 17998000 | 3985500 | 0 | 0 | 0 | 700250 | 0 | 0 | 15899000 | 5697900 | 0 | 0 | 5506500 | 14817000 |

| | | | | | | | | | | | | | | | | | | | | | | |
|---|---|---|---|---|---|---|---|---|---|---|---|---|---|---|---|---|---|---|---|---|---|---|
| LVLLDLSEG | UEVLD | Q8IX04 | UEVLD_HUMAN | Unmodified | 0 | 0 | 0 | 0 | 0 | 0 | 0 | 0 | 0 | 0 | 0 | 768540 | 0 | 0 | 0 | 0 | 0 | 0 |
| LVLLFSGK | PMVK | Q15126 | PMVK_HUMAN | Unmodified | 0 | 0 | 0 | 0 | 2471100 | 0 | 477040 | 45585 | 985440 | 804480 | 39795 | 1250500 | 2378600 | 351080 | 0 | 0 | 640290 | 1768500 |
| LVLLGESAV | RAB5B; RAB5C; | P61020; P51 | RAB5B_HUMAN | Unmodified | 21617000 | 25205000 | 10551000 | 10872000 | 9264600 | 1707300 | 10325000 | 5474300 | 13392000 | 13798000 | 20708000 | 38233000 | 4815100 | 5187000 | 34305000 | 25905000 | 2314600 | 2152400 |
| LVLLGESAV | RAB5B; RAB5C; | P61020; P51 | RAB5B_HUMAN | Unmodified | 0 | 0 | 0 | 0 | 265190 | 0 | 381250 | 0 | 0 | 275570 | 0 | 0 | 0 | 214360 | 0 | 0 | 31442 | 0 |
| LVLNYKPPS | USP25; USP25 | Q9UHP3; Q9 | UBP25_HUMAN | Unmodified | 0 | 0 | 0 | 0 | 0 | 0 | 0 | 131070 | 1209700 | 96381 | 132720 | 0 | 14696 | 0 | 0 | 0 | 91688 | 0 |
| LVLPAPQIS | CDC5L | Q99459 | CDC5L_HUMAN | Unmodified | 0 | 0 | 0 | 14125000 | 0 | 0 | 0 | 0 | 0 | 0 | 499730 | 232440 | 0 | 0 | 0 | 57607 | 0 | 0 |
| LVLPPPDPI | SKIV2L; SKIV2L; | Q15477; B4E | SKIV2_HUMAN | Unmodified | 0 | 0 | 0 | 0 | 0 | 0 | 0 | 0 | 0 | 0 | 0 | 297190 | 0 | 0 | 0 | 0 | 0 | 0 |
| LVLPSALPA | LARGE2; LARGE | Q8N3Y3; E9I | LARG2_HUMAN | Unmodified | 0 | 0 | 0 | 0 | 0 | 0 | 0 | 0 | 0 | 0 | 0 | 1092200 | 0 | 0 | 0 | 0 | 0 | 0 |
| LVLPSLISSI | SELENBP1; SEL | Q13228; A6F | SBP1_HUMAN | Unmodified | 0 | 0 | 5008900 | 3393000 | 0 | 0 | 0 | 42783 | 1938100 | 717000 | 0 | 120760 | 18104000 | 11040000 | 5395200 | 4228300 | 1282900 | 16097000 |
| LVLPSLLAA | GCN1 | Q92616 | GCN1_HUMAN | Unmodified | 0 | 0 | 0 | 0 | 374420 | 0 | 0 | 0 | 0 | 0 | 0 | 0 | 390850 | 0 | 0 | 19202 | 0 | 226400 |
| LVLSDLHLI | L1CAM | P32004 | L1CAM_HUMAN | Unmodified | 44874 | 71541 | 0 | 0 | 266380 | 0 | 0 | 0 | 7360200 | 438080 | 0 | 279810 | 0 | 0 | 0 | 0 | 0 | 0 |
| LVLSLEPLE | RIF1; RIF1 | Q5UIP0; H7E | RIF1_HUMAN T | Unmodified | 0 | 0 | 0 | 0 | 0 | 0 | 104570 | 0 | 0 | 0 | 0 | 164450 | 0 | 0 | 0 | 0 | 0 | 0 |
| LVLTQEQLI | DCTN1; DCTN1 | Q14203; A0A | DCTN1_HUMAN | Unmodified | 0 | 0 | 0 | 0 | 593100 | 0 | 70226 | 0 | 264220 | 165370 | 0 | 1746100 | 1177800 | 0 | 232270 | 363520 | 48665 | 0 |
| LVLVGDGG | RAN; RAN; RAN | P62826; H0Y | RAN_HUMAN G | Unmodified | 8638300 | 10647000 | 7785400 | 8625200 | 22860000 | 4900300 | 1489400 | 0 | 11864000 | 4088300 | 25888000 | 1.74E+08 | 6464100 | 7239800 | 8613000 | 7988400 | 2574600 | 12468000 |
| LVLVGDGG | RAN; RAN; RAN | P62826; H0Y | RAN_HUMAN G | Unmodified | 0 | 0 | 0 | 0 | 23994000 | 77551 | 8830500 | 4760100 | 0 | 5359400 | 107550 | 68320 | 10716000 | 16691000 | 0 | 0 | 6341000 | 13382000 |
| LVLVGDGG | RAN; RAN; RAN | P62826; H0Y | RAN_HUMAN G | Unmodified | 0 | 0 | 0 | 0 | 2780800 | 0 | 370310 | 287520 | 384760 | 158450 | 0 | 0 | 1385200 | 0 | 0 | 0 | 615720 | 1973200 |
| LVLVLGDLI | VPS29; VPS29 | Q9UBQ0; F8 | VPS29_HUMAN | Unmodified | 0 | 0 | 304670 | 0 | 0 | 0 | 0 | 0 | 0 | 0 | 0 | 1448300 | 0 | 0 | 389690 | 0 | 0 | 0 |
| LVLVNAIYF | SERPINB8; SER | P50452; P30 | SPB8_HUMAN | Unmodified | 0 | 0 | 0 | 0 | 997080 | 0 | 719420 | 0 | 1319700 | 1242300 | 0 | 0 | 745520 | 0 | 0 | 0 | 1083000 | 0 |
| LVLVNAVYI | SERPINB6; SER | P35237; A0A | SPB6_HUMAN | Unmodified | 94340 | 0 | 0 | 418170 | 0 | 0 | 0 | 0 | 380730 | 0 | 454570 | 11710000 | 0 | 0 | 747780 | 337500 | 0 | 0 |
| LVLVSPTSE | GTPBP1 | O00178 | GTPB1_HUMAN | Unmodified | 537210 | 1013000 | 299860 | 466570 | 0 | 0 | 0 | 24865 | 275480 | 0 | 193060 | 1816700 | 0 | 0 | 950360 | 636350 | 0 | 0 |
| LVMAAANR | HADHB; HADHI | P55084; F5G | ECHB_HUMAN | Unmodified | 0 | 0 | 0 | 0 | 0 | 0 | 0 | 0 | 4351800 | 0 | 0 | 0 | 0 | 0 | 0 | 0 | 0 | 0 |
| LVMEDGKN | QARS1; QARS1; | P47897; A0A | SYQ_HUMAN G | Oxidation (M) | 0 | 0 | 0 | 0 | 0 | 0 | 0 | 0 | 324360 | 0 | 64512 | 2332700 | 0 | 0 | 0 | 0 | 0 | 0 |
| LVMEDGKN | QARS1; QARS1; | P47897; A0A | SYQ_HUMAN G | 2 Oxidation (M) | 0 | 0 | 0 | 0 | 0 | 0 | 0 | 0 | 0 | 0 | 0 | 0 | 0 | 0 | 0 | 0 | 0 | 0 |
| LVMEEAPES | RTCB | Q9Y3I0 | RTCB_HUMAN | Oxidation (M) | 43185 | 0 | 157820 | 0 | 0 | 0 | 0 | 0 | 0 | 0 | 0 | 2581300 | 0 | 0 | 0 | 81186 | 0 | 0 |
| LVMEEAPES | RTCB | Q9Y3I0 | RTCB_HUMAN | Unmodified | 0 | 0 | 0 | 0 | 761070 | 0 | 0 | 0 | 0 | 0 | 0 | 0 | 678250 | 0 | 0 | 0 | 0 | 610720 |
| LVMLGAGG | RIT1; RIT1; RIT1 | Q92963; A0A | RIT1_HUMAN G | Unmodified | 0 | 0 | 0 | 0 | 100710 | 0 | 39433 | 45318 | 197720 | 75861 | 320400 | 0 | 0 | 0 | 0 | 0 | 330640 | 950970 |
| LVMSYVAA | MYO1B; MYO1A | O43795; Q9I | MYO1B_HUMA | Oxidation (M) | 0 | 0 | 0 | 0 | 0 | 0 | 0 | 0 | 0 | 0 | 272580 | 1212100 | 0 | 0 | 0 | 0 | 0 | 0 |
| LVMSYVAA | MYO1B; MYO1A | O43795; Q9I | MYO1B_HUMA | Unmodified | 0 | 0 | 0 | 0 | 658500 | 333260 | 0 | 0 | 271880 | 0 | 135750 | 1629800 | 0 | 0 | 0 | 0 | 170480 | 0 |
| LVNACLAEI | BOLA2; BOLA2E | Q9H3K6; A0A | BOLA2_HUMAN | Unmodified | 620290 | 436150 | 0 | 1061300 | 3958700 | 0 | 972360 | 0 | 1314100 | 652940 | 0 | 1318900 | 2568800 | 1150200 | 1414200 | 860300 | 1888900 | 1637200 |
| LVNEAPVYS | STAM2 | O75886 | STAM2_HUMAN | Unmodified | 1004600 | 901850 | 0 | 0 | 1871900 | 0 | 1059900 | 399340 | 1332500 | 836470 | 0 | 454810 | 1159000 | 1068000 | 0 | 0 | 1551500 | 2669900 |
| LVNEERTLE | DNAJB11; N/A | Q9UBS4; H7 | DJB11_HUMAN | Unmodified | 0 | 0 | 0 | 105080 | 0 | 0 | 0 | 0 | 2501500 | 0 | 0 | 0 | 0 | 0 | 0 | 0 | 0 | 0 |
| LVNEERTLE | DNAJB11; N/A | Q9UBS4; H7 | DJB11_HUMAN | Unmodified | 0 | 0 | 0 | 0 | 0 | 0 | 0 | 0 | 1786200 | 736750 | 0 | 0 | 0 | 6929200 | 0 | 0 | 0 | 0 |
| LVNELTEK | KTN1 | Q86UP2 | KTN1_HUMAN | Unmodified | 0 | 0 | 0 | 0 | 94143 | 93347 | 0 | 0 | 0 | 91329 | 371420 | 517190 | 0 | 0 | 0 | 0 | 0 | 0 |
| LVNHFVEEI | HSPA1A; HSPA1 | P0DMV8; P0 | HS71A_HUMAN | Unmodified | 1117500 | 1269800 | 715520 | 467830 | 0 | 253570 | 0 | 0 | 1184700 | 0 | 204440 | 1721700 | 0 | 0 | 1154700 | 832830 | 0 | 0 |
| LVNHFVEEI | HSPA1A; HSPA1 | P0DMV8; P0 | HS71A_HUMAN | Unmodified | 1559300 | 1585600 | 0 | 429140 | 309880 | 0 | 0 | 58165 | 739520 | 41075 | 0 | 887590 | 0 | 0 | 0 | 476510 | 46611 | 0 |
| LVNIMPYEI | PTPRF; PTPRF; I | P10586; H0Y | PTPRF_HUMAN | Unmodified | 0 | 0 | 0 | 0 | 0 | 0 | 0 | 0 | 1939800 | 0 | 0 | 0 | 0 | 0 | 0 | 0 | 0 | 0 |
| LVNIMPYEI | PTPRF; PTPRF; I | P10586; H0Y | PTPRF_HUMAN | Oxidation (M) | 1195300 | 722170 | 907940 | 0 | 0 | 0 | 0 | 0 | 895590 | 0 | 319640 | 1290200 | 0 | 0 | 1498400 | 1618200 | 0 | 0 |
| LVNITIIK | ITGB4 | P16144 | ITB4_HUMAN I | Unmodified | 120530 | 0 | 66093 | 81436 | 4363000 | 649810 | 0 | 0 | 0 | 0 | 105460 | 323190 | 938630 | 406120 | 265620 | 280400 | 4046300 | 10233000 |
| LVNIYDSMI | EXTL2 | Q9UBQ6 | EXTL2_HUMAN | Unmodified | 0 | 0 | 0 | 0 | 0 | 0 | 0 | 0 | 1961700 | 0 | 0 | 0 | 0 | 0 | 0 | 0 | 0 | 0 |
| LVNLQHLD | LRRC59 | Q96AG4 | LRC59_HUMAN | Unmodified | 385340 | 94145 | 464840 | 475990 | 1207500 | 0 | 139550 | 58190 | 1114300 | 276210 | 593560 | 1543600 | 1389700 | 460150 | 359310 | 220460 | 99479 | 777950 |
| LVNLQHLD | LRRC59 | Q96AG4 | LRC59_HUMAN | Unmodified | 0 | 0 | 0 | 0 | 0 | 0 | 0 | 423760 | 0 | 0 | 0 | 0 | 869360 | 590260 | 0 | 106510 | 317840 | 0 |
| LVNMLDAV | CAVIN2 | O95810 | CAVN2_HUMAI | Unmodified | 18877 | 0 | 0 | 0 | 2167900 | 0 | 0 | 30983 | 1007300 | 429190 | 0 | 0 | 56082 | 0 | 0 | 31027 | 0 | 0 |
| LVNMLDAV | CAVIN2 | O95810 | CAVN2_HUMAI | Unmodified | 0 | 0 | 0 | 0 | 992940 | 234580 | 0 | 0 | 0 | 0 | 0 | 0 | 0 | 0 | 0 | 0 | 0 | 0 |
| LVNMLDAV | CAVIN2 | O95810 | CAVN2_HUMAI | Unmodified | 0 | 0 | 0 | 0 | 1880000 | 0 | 0 | 0 | 0 | 0 | 0 | 0 | 0 | 0 | 0 | 0 | 0 | 0 |
| LVNQQLLA | NDUFS1; NDUF | P28331; B4D | NDUS1_HUMA | Unmodified | 0 | 0 | 0 | 0 | 0 | 0 | 0 | 0 | 3392200 | 590270 | 202140 | 91677 | 0 | 0 | 0 | 232970 | 0 | 0 |
| LVNTGLLTL | NLN; NLN; NLN | Q9BYT8; H0Y | NEUL_HUMAN | Unmodified | 0 | 0 | 0 | 0 | 0 | 0 | 0 | 0 | 1100100 | 0 | 0 | 2213300 | 0 | 0 | 0 | 0 | 0 | 0 |
| LVNTINPGA | UPF1 | Q92900 | RENT1_HUMAN | Unmodified | 923500 | 1019600 | 887800 | 269730 | 0 | 0 | 0 | 0 | 0 | 41098 | 0 | 669370 | 160630 | 0 | 382450 | 0 | 0 | 0 |
| LVNTTITPEI | LUZP1 | Q86V48 | LUZP1_HUMAN | Unmodified | 0 | 0 | 0 | 0 | 1582400 | 0 | 0 | 0 | 0 | 0 | 0 | 0 | 549560 | 236400 | 0 | 0 | 0 | 448220 |
| LVNWSCTL | VARS1; VARS1 | P26640; A0A | SYVC_HUMAN | Unmodified | 50278 | 0 | 50037 | 0 | 0 | 0 | 0 | 472330 | 0 | 19498 | 0 | 2253800 | 0 | 0 | 797750 | 0 | 1936100 | 181050 |
| LVPEAVGD | CARS1; CARS1 | P49589; B4D | SYCC_HUMAN | Unmodified | 0 | 0 | 178920 | 0 | 986510 | 167760 | 171570 | 0 | 210720 | 196930 | 1405500 | 4868400 | 831360 | 0 | 0 | 0 | 35868 | 558290 |
| LVPFDHAES | PGLS; PGLS; PG | O95336; M0I | 6PGL_HUMAN | Unmodified | 438150 | 0 | 0 | 0 | 0 | 0 | 0 | 0 | 1735900 | 0 | 0 | 4572800 | 0 | 0 | 0 | 0 | 0 | 0 |
| LVPGHAPG | HYAL2 | Q12891 | HYAL2_HUMAN | Unmodified | 6742300 | 0 | 2034000 | 0 | 0 | 0 | 0 | 0 | 1209800 | 0 | 0 | 2306400 | 0 | 0 | 17432000 | 0 | 0 | 0 |
| LVPGPVFG | LMAN2; LMAN2 | Q12907; D6I | LMAN2_HUMAI | Unmodified | 397810 | 384360 | 2449300 | 1344300 | 0 | 0 | 0 | 0 | 1.55E+08 | 5568700 | 2351400 | 2737100 | 0 | 0 | 1283700 | 670310 | 0 | 0 |
| LVPGPVFG | LMAN2; LMAN2 | Q12907; D6I | LMAN2_HUMAI | Unmodified | 0 | 0 | 0 | 0 | 0 | 0 | 0 | 0 | 620090 | 1146600 | 0 | 0 | 0 | 0 | 0 | 0 | 0 | 0 |
| LVPGRHDI | SNRPB2 | P08579 | RU2B_HUMAN | Unmodified | 1368100 | 2061100 | 0 | 0 | 2316900 | 0 | 0 | 0 | 0 | 0 | 0 | 0 | 360380 | 0 | 1131000 | 309800 | 0 | 239300 |
| LVPGWTKP | IDH2 | P48735 | IDHP_HUMAN | Unmodified | 0 | 0 | 0 | 0 | 0 | 0 | 350010 | 0 | 33261000 | 3209700 | 0 | 464540 | 0 | 0 | 0 | 0 | 0 | 0 |
| LVPLLDTGI | PGD; PGD; PGD | P52209; K7E | 6PGD_HUMAN | Unmodified | 0 | 0 | 0 | 180490 | 0 | 0 | 0 | 0 | 0 | 0 | 535850 | 2061500 | 0 | 0 | 445360 | 292440 | 0 | 0 |
| LVPLLDTGI | PGD; PGD; PGD | P52209; K7E | 6PGD_HUMAN | Unmodified | 0 | 0 | 0 | 0 | 4705400 | 0 | 75434 | 0 | 5625900 | 2265500 | 5249800 | 6555900 | 3272200 | 0 | 0 | 0 | 701350 | 0 |
| LVPLLDTGI | PGD; PGD; PGD | P52209; K7E | 6PGD_HUMAN | Unmodified | 1005700 | 0 | 787970 | 4140200 | 0 | 0 | 1971000 | 0 | 16999000 | 17034000 | 13984000 | 30825000 | 15396000 | 6631500 | 4627900 | 1479800 | 0 | 0 |
| LVPLLLEDG | DYNC1H1; DYN | Q14204; A0A | DYHC1_HUMA | Unmodified | 0 | 0 | 0 | 503580 | 4742000 | 150870 | 645300 | 0 | 322200 | 755700 | 70919 | 211640 | 2611800 | 1964800 | 1081700 | 0 | 756800 | 1510000 |
| LVPLNQES | NUP93; NUP93 | Q8N1F7; H3 | NUP93_HUMAI | Unmodified | 0 | 0 | 0 | 0 | 0 | 0 | 0 | 0 | 0 | 0 | 0 | 1092400 | 0 | 0 | 0 | 137980 | 0 | 0 |
| LVPLVQATI | ANK1; ANK1 | P16157; H0Y | ANK1_HUMAN | Unmodified | 0 | 0 | 0 | 0 | 0 | 0 | 371300 | 157820 | 0 | 0 | 0 | 0 | 0 | 0 | 0 | 0 | 0 | 0 |
| LVPMAAHE | PTPN23 | Q9H3S7 | PTN23_HUMAN | Unmodified | 917180 | 0 | 1240200 | 0 | 2488200 | 0 | 0 | 0 | 1404100 | 974750 | 0 | 0 | 0 | 0 | 0 | 0 | 278530 | 0 |
| LVPMAAHE | PTPN23 | Q9H3S7 | PTN23_HUMAN | Oxidation (M) | 478540 | 744050 | 0 | 0 | 0 | 416620 | 157400 | 73438 | 158120 | 0 | 0 | 0 | 18585 | 0 | 0 | 0 | 0 | 0 |
| LVPNMTPE | CNDP2 | Q96KP4 | CNDP2_HUMA | Oxidation (M) | 0 | 0 | 0 | 0 | 985170 | 0 | 0 | 0 | 0 | 0 | 0 | 4324200 | 8064400 | 0 | 0 | 2151600 | 0 | 0 |
| LVPNMTPE | CNDP2 | Q96KP4 | CNDP2_HUMA | Unmodified | 0 | 0 | 0 | 67815 | 0 | 0 | 0 | 0 | 456800 | 0 | 100950 | 628990 | 311460 | 0 | 311350 | 0 | 0 | 0 |
| LVPNMTPE | CNDP2 | Q96KP4 | CNDP2_HUMA | Oxidation (M) | 0 | 0 | 0 | 0 | 0 | 0 | 0 | 0 | 0 | 298750 | 0 | 3075900 | 2876600 | 1199300 | 203970 | 840030 | 0 | 0 |
| LVPNMTPE | CNDP2 | Q96KP4 | CNDP2_HUMA | Unmodified | 0 | 0 | 0 | 0 | 0 | 0 | 0 | 0 | 0 | 0 | 0 | 691140 | 0 | 0 | 0 | 0 | 0 | 0 |
| LVPPGPAD | N/A | N/A | N/A | Unmodified | 0 | 0 | 362700 | 0 | 33571 | 0 | 0 | 0 | 232730 | 84477 | 0 | 4257700 | 0 | 0 | 1428700 | 325820 | 0 | 0 |
| LVPQQLAH | COX5B | P10606 | COX5B_HUMAI | Unmodified | 0 | 0 | 0 | 0 | 0 | 0 | 0 | 0 | 6605000 | 2671000 | 1397000 | 0 | 0 | 0 | 0 | 0 | 0 | 0 |
| LVPRPGSG | MAP4K4; MAP4 | O95819; C9J | M4K4_HUMAN | Unmodified | 0 | 0 | 0 | 0 | 0 | 0 | 0 | 0 | 0 | 0 | 2427500 | 4492000 | 0 | 0 | 619900 | 0 | 0 | 0 |
| LVPSVNGV | MTHFD1; MTHF | P11586; V9G | C1TC_HUMAN | Unmodified | 0 | 0 | 0 | 0 | 0 | 0 | 0 | 0 | 0 | 0 | 0 | 3856000 | 0 | 0 | 0 | 0 | 0 | 0 |
| LVPTHTQVI | SNX18 | Q96RF0 | SNX18_HUMAN | Unmodified | 0 | 0 | 2315500 | 1998500 | 0 | 0 | 0 | 0 | 0 | 0 | 0 | 0 | 0 | 0 | 2632400 | 4255200 | 0 | 0 |
| LVPVGYGII | EEF1D; EEF1D; | P29692; E9P | EF1D_HUMAN | Unmodified | 0 | 0 | 223820 | 0 | 1259800 | 0 | 0 | 0 | 673960 | 0 | 982500 | 15794000 | 0 | 616560 | 658830 | 610800 | 1730100 | 1047700 |
| LVPVLSAK | TALDO1; TALDO | P37837; F2Z | TALDO_HUMAN | Unmodified | 0 | 0 | 0 | 0 | 8302400 | 0 | 0 | 0 | 2065900 | 2559700 | 0 | 23014000 | 6945300 | 2257400 | 0 | 0 | 1480200 | 2389300 |
| LVQAAQML | VCL | P18206 | VINC_HUMAN | Oxidation (M) | 0 | 0 | 0 | 0 | 0 | 0 | 0 | 0 | 0 | 0 | 0 | 642500 | 0 | 0 | 111580 | 145960 | 0 | 0 |
| LVQAAQML | VCL | P18206 | VINC_HUMAN | Oxidation (M) | 0 | 0 | 280900 | 3027300 | 1316700 | 0 | 0 | 0 | 0 | 0 | 1114000 | 8886400 | 0 | 0 | 1057100 | 0 | 1305800 | 0 |
| LVQAAQML | VCL | P18206 | VINC_HUMAN | Unmodified | 0 | 0 | 0 | 0 | 2497400 | 0 | 0 | 0 | 4660600 | 0 | 0 | 0 | 416570 | 0 | 0 | 0 | 0 | 0 |
| LVQAFQYTI | PRDX2 | P32119 | PRDX2_HUMAN | Unmodified | 14368000 | 5092000 | 4499900 | 12537000 | 0 | 0 | 0 | 0 | 25114000 | 2354500 | 17353000 | 3.53E+08 | 1066700 | 0 | 9184200 | 7727400 | 1084600 | 0 |
| LVQAFQYTI | PRDX4; PRDX4 | Q13162; H7 | PRDX4_HUMAN | Unmodified | 0 | 0 | 0 | 217280 | 0 | 0 | 0 | 0 | 1.39E+08 | 1787600 | 11585000 | 11015000 | 0 | 0 | 559330 | 414990 | 0 | 0 |
| LVQAFQYTI | PRDX4 | Q13162 | PRDX4_HUMAN | Unmodified | 0 | 0 | 0 | 0 | 0 | 0 | 0 | 0 | 4024500 | 0 | 0 | 0 | 0 | 0 | 0 | 0 | 0 | 2852000 |
| LVQAPLDAI | KIF23; KIF23; KI | Q02241; A0A | KIF23_HUMAN | Unmodified | 0 | 0 | 1145600 | 0 | 8219700 | 1837800 | 273850 | 64462 | 2166200 | 1559300 | 769190 | 2375400 | 0 | 0 | 789450 | 368390 | 1285100 | 0 |
| LVQDVANN | HSPD1; HSPD1 | P10809; A0A | CH60_HUMAN | Unmodified | 0 | 0 | 0 | 0 | 0 | 0 | 0 | 190140 | 12792000 | 2424100 | 211170 | 0 | 1223300 | 812100 | 0 | 0 | 0 | 0 |
| LVQDVANN | HSPD1; HSPD1 | P10809; A0A | CH60_HUMAN | Unmodified | 0 | 0 | 0 | 0 | 0 | 0 | 2231800 | 766250 | 0 | 0 | 0 | 0 | 0 | 308710 | 0 | 0 | 0 | 0 |
| LVQDVANN | HSPD1; HSPD1 | P10809; A0A | CH60_HUMAN | Unmodified | 0 | 0 | 0 | 0 | 0 | 1036900 | 6156500 | 0 | 0 | 0 | 0 | 0 | 0 | 0 | 0 | 0 | 626980 | 0 |
| LVQDYGLE | GPD2 | P43304 | GPDM_HUMAN | Unmodified | 0 | 50275 | 0 | 0 | 0 | 0 | 0 | 0 | 23493000 | 5742600 | 0 | 0 | 0 | 0 | 0 | 0 | 0 | 0 |
| LVQGDQDI | CALCOCO2; CA | Q13137; D6I | CACO2_HUMA | Unmodified | 1139400 | 707140 | 149770 | 410990 | 2433100 | 367720 | 613150 | 392020 | 3007400 | 795490 | 151440 | 0 | 1305700 | 547550 | 222050 | 206280 | 788400 | 6381100 |
| LVQGSILK | PCNA | P12004 | PCNA_HUMAN | Unmodified | 0 | 430050 | 549490 | 557140 | 1831400 | 326760 | 718700 | 690800 | 1690300 | 1863300 | 1503100 | 3297900 | 4493400 | 1156900 | 781160 | 467750 | 990390 | 1058800 |
| LVQHDLQV | CCDC50 | Q8IVM0 | CCD50_HUMA | Unmodified | 2302800 | 2560000 | 2788200 | 3152300 | 0 | 0 | 0 | 0 | 0 | 0 | 0 | 0 | 1051300 | 431980 | 3287400 | 3756900 | 0 | 0 |
| LVQIEYALA | PSMA2; N/A; PS | P25787; A0A | PSA2_HUMAN | Unmodified | 0 | 269300 | 0 | 249780 | 2446000 | 0 | 0 | 0 | 293180 | 1027000 | 0 | 0 | 2588800 | 1035200 | 348480 | 0 | 0 | 2458800 |
| LVQILHR | RAP1GDS1 | P52306 | GDS1_HUMAN | Unmodified | 0 | 0 | 0 | 0 | 0 | 0 | 0 | 0 | 0 | 0 | 0 | 384790 | 0 | 0 | 0 | 0 | 0 | 0 |
| LVQLLGGV | PIK3C2A | O00443 | P3C2A_HUMAN | Unmodified | 0 | 0 | 0 | 0 | 0 | 0 | 0 | 0 | 0 | 0 | 0 | 966280 | 0 | 0 | 0 | 0 | 0 | 0 |
| LVQLLVK | JUP | P14923 | PLAK_HUMAN . | Unmodified | 1937600 | 983360 | 7817800 | 4443700 | 2966400 | 2555100 | 0 | 0 | 2386200 | 1400600 | 766710 | 26497000 | 1089800 | 401680 | 11884000 | 18933000 | 1445900 | 86171 |
| LVQLLVR | RNF123; CTNN | Q5XPI4; P35 | RN123_HUMAN | Unmodified | 3502900 | 2909400 | 5937200 | 4756300 | 682930 | 836400 | 261280 | 0 | 2331200 | 373860 | 1007500 | 21175000 | 567210 | 183510 | 10184000 | 11924000 | 1562000 | 330490 |
| LVQLQEK | NUDT21; FKBP | O43809; Q5 | CPSF5_HUMAN | Unmodified | 0 | 0 | 0 | 0 | 1202200 | 125520 | 0 | 0 | 2462400 | 367670 | 104480 | 463990 | 0 | 24174 | 0 | 0 | 0 | 0 |
| LVQNCLW | JUP; CTNNB1; | P14923; P35 | PLAK_HUMAN . | Unmodified | 0 | 0 | 2431500 | 6870000 | 1924500 | 0 | 0 | 0 | 3604900 | 260570 | 2593200 | 8250100 | 1787400 | 801050 | 8387000 | 7227800 | 624990 | 863670 |
| LVQNCLW | JUP | P14923 | PLAK_HUMAN . | Unmodified | 0 | 0 | 0 | 0 | 45349000 | 8563400 | 0 | 0 | 0 | 1551200 | 0 | 0 | 40907000 | 33535000 | 0 | 0 | 4656300 | 21669000 |
| LVQNGTEP | CAD; CAD | P27708; F8V | PYR1_HUMAN | Unmodified | 0 | 0 | 82837 | 0 | 0 | 0 | 0 | 0 | 0 | 0 | 0 | 1629500 | 258290 | 0 | 0 | 47817 | 0 | 347480 |
| LVQSPNSY | RPS27L; RPS27 | Q71UM5; P4 | RS27L_HUMAN | Oxidation (M) | 319930 | 276890 | 0 | 0 | 0 | 0 | 0 | 0 | 0 | 0 | 647190 | 7329800 | 0 | 0 | 0 | 0 | 0 | 0 |
| LVQTAAQQ | HIKESHI; HIKES | Q53FT3; E9F | HIKES_HUMAN | Unmodified | 0 | 180770 | 0 | 0 | 0 | 0 | 0 | 0 | 0 | 0 | 0 | 929470 | 0 | 0 | 0 | 0 | 0 | 0 |
| LVQTAELTK | GMFB; GMFG; ( | P60983; O6( | GMFB_HUMAN | Unmodified | 687960 | 383790 | 1805600 | 1477200 | 3011500 | 708620 | 492640 | 294420 | 1917400 | 847460 | 1039300 | 7045900 | 2874300 | 2854900 | 2220600 | 1318000 | 1585100 | 1801500 |
| LVQTPLTDI | DYNC1H1; DYN | Q14204; A0A | DYHC1_HUMA | Unmodified | 0 | 0 | 0 | 0 | 0 | 0 | 0 | 0 | 0 | 0 | 0 | 3846500 | 0 | 0 | 0 | 0 | 0 | 0 |
| LVQTVRDY | ARHGAP5; ARH | Q13017; A0A | RHG05_HUMAI | Unmodified | 0 | 0 | 0 | 0 | 671930 | 0 | 0 | 0 | 0 | 0 | 0 | 0 | 0 | 0 | 0 | 0 | 61287 | 62668 |
| LVREIAQDI | H3-3A; H3C15; | P84243; Q71 | H33_HUMAN H | Unmodified | 2649900 | 1326600 | 0 | 0 | 1870400 | 324420 | 500430 | 1340900 | 155040 | 858180 | 337120 | 4722900 | 60130 | 0 | 0 | 141740 | 0 | 260340 |
| LVREIAQDI | H3-3A; H3C15; | P84243; Q71 | H33_HUMAN H | Unmodified | 6995000 | 2365300 | 0 | 0 | 2906200 | 646650 | 542130 | 0 | 293990 | 1244100 | 131780 | 21078000 | 147610 | 0 | 0 | 0 | 0 | 0 |
| LVREVAAPI | FLOT2; FLOT2; I | Q14254; J3Q | FLOT2_HUMAN | Unmodified | 0 | 0 | 0 | 0 | 627930 | 131930 | 89651 | 0 | 902210 | 669600 | 249830 | 0 | 238670 | 0 | 0 | 0 | 0 | 0 |
| LVREVAAPI | FLOT2; FLOT2; I | Q14254; J3Q | FLOT2_HUMAN | Unmodified | 238020 | 148100 | 0 | 0 | 582840 | 0 | 148440 | 0 | 0 | 378700 | 0 | 0 | 0 | 109430 | 0 | 0 | 0 | 0 |
| LVRGEQGY | SLC9A3R2 | Q15599 | NHRF2_HUMAI | Unmodified | 511150 | 247330 | 0 | 1183300 | 4567800 | 580870 | 171100 | 0 | 615550 | 374880 | 209970 | 586160 | 3635300 | 1983800 | 619100 | 0 | 809960 | 1970400 |
| LVRPPVQV | ATP5PO; ATP5P | P48047; H7( | ATPO_HUMAN | Unmodified | 90581 | 0 | 0 | 0 | 150670 | 0 | 0 | 0 | 14560000 | 2231500 | 0 | 422330 | 0 | 0 | 0 | 0 | 0 | 0 |

| | | | | | | | | | | | | | | | | | | | | | | |
|---|---|---|---|---|---|---|---|---|---|---|---|---|---|---|---|---|---|---|---|---|---|---|
| LVRPPVQV | ATP5PO; ATP5P | P48047; H7( | ATPO_HUMAN | Unmodified | 0 | 0 | 0 | 0 | 0 | 0 | 1448700 | 0 | 0 | 2979800 | 0 | 0 | 0 | 0 | 0 | 0 | 0 | 0 |
| LVRTGGSA | LAMB2 | P55268 | LAMB2_HUMAN | Unmodified | 0 | 0 | 0 | 0 | 0 | 0 | 0 | 0 | 0 | 564390 | 0 | 0 | 0 | 0 | 0 | 0 | 0 | 0 |
| LVSAVVEY( | VPS45; VPS45; | Q9NRW7; A( | VPS45_HUMAN | Unmodified | 0 | 0 | 0 | 0 | 638310 | 0 | 265790 | 83808 | 0 | 0 | 0 | 0 | 0 | 0 | 0 | 0 | 0 | 159930 |
| LVSDEMVV | AK2; AK2; AK2; | P54819; G3\ | KAD2_HUMAN | Oxidation (M) | 0 | 0 | 0 | 0 | 0 | 0 | 0 | 0 | 0 | 0 | 0 | 0 | 0 | 0 | 0 | 0 | 0 | 0 |
| LVSDGNIN | SPTBN1; SPTBN | Q01082; A0A | SPTB2_HUMAN | Unmodified | 0 | 0 | 0 | 0 | 0 | 0 | 0 | 53256 | 0 | 0 | 0 | 2394700 | 0 | 0 | 0 | 0 | 0 | 0 |
| LVSDGNIN | SPTBN1; SPTBN | Q01082; A0A | SPTB2_HUMAN | Unmodified | 0 | 0 | 0 | 0 | 0 | 0 | 0 | 0 | 56011 | 0 | 0 | 851910 | 0 | 0 | 0 | 0 | 0 | 0 |
| LVSECVHA | DDO | Q99489 | OXDD_HUMAN | Unmodified | 0 | 0 | 0 | 0 | 0 | 0 | 0 | 0 | 941410 | 450050 | 0 | 0 | 0 | 0 | 0 | 0 | 0 | 0 |
| LVSEDPINI | HSPG2 | P98160 | PGBM_HUMAN | Unmodified | 0 | 0 | 2254000 | 0 | 0 | 0 | 0 | 0 | 0 | 0 | 0 | 0 | 0 | 0 | 0 | 0 | 0 | 0 |
| LVSEDPINI | HSPG2 | P98160 | PGBM_HUMAN | Unmodified | 0 | 0 | 283870 | 182380 | 1003500 | 0 | 303470 | 0 | 0 | 0 | 0 | 0 | 0 | 0 | 0 | 0 | 369970 | 204570 |
| LVSEFPYIE | GPD2 | P43304 | GPDM_HUMAN | Unmodified | 0 | 0 | 0 | 0 | 0 | 0 | 0 | 0 | 6175200 | 1674700 | 0 | 0 | 0 | 0 | 0 | 0 | 0 | 0 |
| LVSEFTITK | CEL | P19835 | CEL_HUMAN B | Unmodified | 0 | 0 | 0 | 0 | 0 | 0 | 0 | 0 | 46062000 | 7753100 | 24752000 | 2277400 | 0 | 0 | 0 | 0 | 0 | 0 |
| LVSEYMSK | ADH5 | P11766 | ADHX_HUMAN | Unmodified | 0 | 0 | 0 | 0 | 3225900 | 0 | 215540 | 0 | 0 | 0 | 1331300 | 13127000 | 450640 | 1405500 | 167410 | 0 | 1575900 | 0 |
| LVSEYMSK | ADH5 | P11766 | ADHX_HUMAN | Oxidation (M) | 0 | 0 | 0 | 0 | 0 | 0 | 0 | 0 | 627890 | 91990 | 0 | 1422700 | 0 | 0 | 0 | 0 | 0 | 0 |
| LVSFHDDS | IGF2R; IGF2R | P11717; A0A | MPRI_HUMAN ( | Unmodified | 288700 | 135720 | 281880 | 0 | 2515800 | 232430 | 457430 | 46553 | 1621900 | 1350600 | 186450 | 0 | 4623000 | 3639600 | 0 | 208430 | 301390 | 698300 |
| LVSFPIGIYI | LRRC20; LRRC2 | Q8TCA0; X6I | LRC20_HUMAN | Unmodified | 0 | 0 | 0 | 0 | 1477200 | 483460 | 0 | 212990 | 160470 | 586550 | 0 | 1501400 | 0 | 0 | 0 | 0 | 0 | 0 |
| LVSFTLSDL | HECW2; HECW | Q9P2P5; A0A | HECW2_HUMA | Unmodified | 0 | 0 | 0 | 0 | 0 | 0 | 44077 | 0 | 0 | 101070 | 0 | 0 | 0 | 0 | 0 | 0 | 0 | 0 |
| LVSGMQHI | LMAN1 | P49257 | LMAN1_HUMAI | Oxidation (M) | 231460 | 1693800 | 1511100 | 2919100 | 0 | 0 | 0 | 0 | 1.03E+08 | 2638000 | 5581800 | 4281700 | 0 | 0 | 2451300 | 1204500 | 0 | 0 |
| LVSGMQHI | LMAN1 | P49257 | LMAN1_HUMAI | Unmodified | 0 | 220420 | 944900 | 0 | 299030 | 0 | 0 | 0 | 1.47E+08 | 1242900 | 0 | 419520 | 0 | 0 | 0 | 0 | 0 | 0 |
| LVSGMQHI | LMAN1 | P49257 | LMAN1_HUMAI | Oxidation (M) | 0 | 0 | 0 | 0 | 0 | 0 | 0 | 0 | 4662300 | 2276900 | 0 | 0 | 0 | 0 | 0 | 0 | 0 | 0 |
| LVSGMQHI | LMAN1 | P49257 | LMAN1_HUMAI | Unmodified | 0 | 0 | 0 | 0 | 0 | 0 | 0 | 0 | 5900900 | 0 | 0 | 0 | 0 | 0 | 0 | 0 | 0 | 0 |
| LVSGMQHI | LMAN1 | P49257 | LMAN1_HUMAI | Oxidation (M) | 0 | 0 | 0 | 0 | 0 | 0 | 0 | 0 | 3640000 | 0 | 0 | 0 | 0 | 0 | 0 | 0 | 0 | 0 |
| LVSGMQHI | LMAN1 | P49257 | LMAN1_HUMAI | Unmodified | 0 | 0 | 0 | 0 | 0 | 0 | 0 | 0 | 29124000 | 0 | 0 | 0 | 0 | 0 | 0 | 0 | 0 | 0 |
| LVSGWVKF | IDH1; IDH1 | O75874; C9J | IDHC_HUMAN | Unmodified | 1033600 | 919770 | 1222000 | 2983400 | 2313500 | 1097300 | 198290 | 0 | 7974500 | 1401300 | 2679800 | 15942000 | 14460000 | 2917400 | 5557400 | 2767900 | 638090 | 904810 |
| LVSGYDSY | SLC44A1 | Q8WWI5 | CTL1_HUMAN ( | Unmodified | 1856800 | 260910 | 901670 | 228110 | 0 | 0 | 0 | 0 | 0 | 0 | 0 | 3748200 | 0 | 0 | 2562700 | 2190400 | 135020 | 109200 |
| LVSIGAEEI | ACTN4; ACTN4 | O43707; F5( | ACTN4_HUMAI | Unmodified | 0 | 0 | 388300 | 354520 | 2077000 | 1363000 | 0 | 17326 | 109070 | 759620 | 1625300 | 7249700 | 67691 | 1014800 | 1179700 | 971130 | 1276200 | 128860 |
| LVSLIGSK | PSMA1; PSMA1 | P25786; F5G | PSA1_HUMAN | Unmodified | 0 | 353920 | 1520600 | 735570 | 367660 | 0 | 112000 | 266820 | 4457700 | 2754700 | 2497700 | 19881000 | 1920300 | 1673400 | 1407200 | 1044800 | 1641400 | 0 |
| LVSLIGSKT | PSMA1; PSMA1 | P25786; F5G | PSA1_HUMAN | Unmodified | 0 | 0 | 0 | 0 | 0 | 0 | 151770 | 47731 | 0 | 411720 | 0 | 0 | 318270 | 260880 | 0 | 0 | 365660 | 131190 |
| LVSLSAQN | CTSS; CTSS; CT | P25774; A0A | CATS_HUMAN | Unmodified | 0 | 0 | 0 | 0 | 0 | 0 | 0 | 0 | 1464200 | 535090 | 90935 | 0 | 834300 | 0 | 0 | 0 | 0 | 0 |
| LVSLYFDTK | PSMD11; PSMD | O00231; J3Q | PSD11_HUMAN | Unmodified | 0 | 0 | 0 | 0 | 0 | 0 | 0 | 0 | 0 | 0 | 171600 | 1933400 | 0 | 0 | 0 | 0 | 0 | 0 |
| LVSLYFDTK | PSMD11; PSMD | O00231; J3Q | PSD11_HUMAN | Unmodified | 15407 | 0 | 0 | 0 | 0 | 0 | 0 | 0 | 0 | 0 | 0 | 937540 | 0 | 0 | 0 | 0 | 0 | 0 |
| LVSNHSLH | FLNA; FLNA; FL | P21333; A0A | FLNA_HUMAN | Unmodified | 1347600 | 1911100 | 2456800 | 3152900 | 12494000 | 2100700 | 1528600 | 186400 | 1304400 | 1330000 | 695880 | 12489000 | 5353300 | 3361300 | 3110000 | 2651600 | 7772800 | 4018700 |
| LVSPEPPPI | EPB41L1; EPB4 | Q9H4G0; A0 | E41L1_HUMAN | Unmodified | 102280 | 0 | 216900 | 311610 | 111740 | 0 | 0 | 0 | 0 | 0 | 320510 | 1144200 | 0 | 0 | 441510 | 364980 | 0 | 0 |
| LVSPEQPPI | EPB41L2; EPB4 | O43491; A0A | E41L2_HUMAN | Unmodified | 0 | 0 | 0 | 0 | 205270 | 511950 | 0 | 24570 | 0 | 0 | 0 | 330080 | 0 | 244120 | 0 | 0 | 46848 | 0 |
| LVSPGSAN | FLNB; FLNB; FL | O75369; A0A | FLNB_HUMAN | Unmodified | 0 | 481860 | 0 | 624790 | 0 | 298640 | 0 | 0 | 428070 | 0 | 1179000 | 5545000 | 0 | 0 | 0 | 809230 | 0 | 0 |
| LVSQDNFC | SPTBN1; SPTBN | Q01082; A0A | SPTB2_HUMAN | Unmodified | 0 | 0 | 0 | 0 | 0 | 0 | 0 | 0 | 545970 | 0 | 1259200 | 3237700 | 0 | 0 | 0 | 0 | 0 | 0 |
| LVSSDPEIN | NUDC | Q9Y266 | NUDC_HUMAN | Unmodified | 0 | 0 | 0 | 0 | 52136 | 0 | 0 | 0 | 1651600 | 573550 | 2576800 | 13478000 | 4525000 | 0 | 905000 | 849150 | 962330 | 0 |
| LVSSDPEIN | NUDC | Q9Y266 | NUDC_HUMAN | Unmodified | 0 | 0 | 0 | 0 | 0 | 0 | 0 | 0 | 0 | 0 | 0 | 365700 | 0 | 0 | 0 | 0 | 0 | 0 |
| LVSSPELGI | SPTBN2 | O15020 | SPTN2_HUMAN | Unmodified | 0 | 0 | 809750 | 412950 | 0 | 0 | 0 | 0 | 0 | 0 | 0 | 1507900 | 161450 | 0 | 1347700 | 884190 | 0 | 0 |
| LVSVLQNK | AARS1; AARS1; | P49588; A0A | SYAC_HUMAN | Unmodified | 0 | 0 | 0 | 238060 | 1000600 | 317290 | 0 | 0 | 4251100 | 406810 | 1562900 | 12586000 | 1663100 | 526670 | 746060 | 1192000 | 319600 | 860580 |
| LVSVNFINI | SLIT2; SLIT2; SL | O94813; X6F | SLIT2_HUMAN | Unmodified | 35891 | 76146 | 0 | 126360 | 0 | 0 | 515910 | 1235000 | 0 | 287640 | 1032200 | 0 | 0 | 0 | 0 | 0 | 672200 | 3797900 |
| LVSWEGGI | FKRP | Q9H9S5 | FKRP_HUMAN | Unmodified | 0 | 0 | 0 | 0 | 0 | 55107 | 0 | 0 | 508100 | 0 | 0 | 0 | 0 | 94032 | 0 | 0 | 0 | 0 |
| LVSYYYFLR | HS2ST1 | Q7LGA3 | HS2ST_HUMAN | Unmodified | 0 | 0 | 0 | 0 | 0 | 0 | 0 | 0 | 1961900 | 0 | 0 | 2171200 | 0 | 0 | 0 | 0 | 0 | 0 |
| LVTDLLQRI | IKBIP | Q70UQ0 | IKIP_HUMAN In | Unmodified | 0 | 0 | 0 | 0 | 0 | 0 | 1370700 | 0 | 818280 | 226650 | 61742 | 290040 | 209760 | 735730 | 0 | 0 | 0 | 0 |
| LVTDQNISI | ACP1; ACP1; A( | P24666; F2Z | PPAC_HUMAN | Unmodified | 0 | 0 | 0 | 0 | 0 | 0 | 0 | 0 | 0 | 0 | 0 | 0 | 0 | 0 | 0 | 0 | 0 | 0 |
| LVTEMGTY | COPB1 | P53618 | COPB_HUMAN | Unmodified | 0 | 0 | 0 | 0 | 183600 | 0 | 0 | 0 | 0 | 0 | 0 | 948900 | 0 | 0 | 0 | 0 | 0 | 0 |
| LVTGGGEE | NXN; NXN | Q6DKJ4; A0A | NXN_HUMAN N | Unmodified | 0 | 0 | 0 | 0 | 357480 | 0 | 138360 | 0 | 0 | 73260 | 391140 | 4840600 | 0 | 0 | 134680 | 208180 | 34062 | 0 |
| LVTLEEFLA | NUCB1; NUCB1 | Q02818; H7I | NUCB1_HUMA | Unmodified | 0 | 0 | 0 | 0 | 0 | 0 | 0 | 0 | 0 | 61621 | 510550 | 0 | 0 | 0 | 0 | 0 | 0 | 0 |
| LVTLEEFLA | NUCB1; NUCB1 | Q02818; H7I | NUCB1_HUMA | Unmodified | 0 | 0 | 601830 | 0 | 978980 | 0 | 0 | 0 | 15405000 | 726080 | 2453000 | 3370600 | 0 | 57826 | 0 | 0 | 838680 | 552680 |
| LVTLEEFLK | NUCB2; NUCB2 | P80303; H0Y | NUCB2_HUMA | Unmodified | 0 | 0 | 0 | 799800 | 0 | 0 | 0 | 0 | 50992000 | 4299700 | 1519100 | 1104400 | 0 | 0 | 681150 | 0 | 0 | 0 |
| LVTLESWAI | RSPRY1 | Q96DX4 | RSPRY_HUMAN | Unmodified | 0 | 516190 | 0 | 0 | 0 | 0 | 0 | 0 | 0 | 0 | 0 | 1280800 | 0 | 0 | 291680 | 159790 | 0 | 0 |
| LVTLKDVEI | KIAA1217; KIAA | Q5T5P2; Q5T | SKT_HUMAN Si | Unmodified | 0 | 0 | 0 | 0 | 0 | 0 | 0 | 0 | 152410 | 0 | 0 | 0 | 0 | 0 | 0 | 0 | 0 | 0 |
| LVTLPDITS | SDSL; SDSL; SD | Q96GA7; F8Y | SDSL_HUMAN | Unmodified | 434460 | 0 | 326270 | 0 | 0 | 0 | 0 | 0 | 997650 | 0 | 0 | 0 | 988230 | 0 | 756030 | 541280 | 161790 | 220610 |
| LVTLPVSFA | LRRC59 | Q96AG4 | LRC59_HUMAN | Unmodified | 0 | 0 | 266710 | 0 | 2780200 | 343630 | 989610 | 203390 | 1098800 | 520630 | 460270 | 1633800 | 1481300 | 1448100 | 0 | 0 | 925890 | 2289700 |
| LVTMQIWD | RAB7A; RAB7A; | P51149; A0A | RAB7A_HUMAN | Oxidation (M) | 0 | 0 | 0 | 0 | 1396000 | 0 | 0 | 0 | 0 | 0 | 0 | 4670900 | 0 | 0 | 1609800 | 164180 | 0 | 0 |
| LVTMQIWD | RAB7A; RAB7A; | P51149; A0A | RAB7A_HUMAN | Unmodified | 0 | 0 | 0 | 0 | 0 | 0 | 0 | 0 | 0 | 0 | 0 | 1326600 | 0 | 504670 | 0 | 0 | 0 | 0 |
| LVTPHGES | DLG3; DLG3 | Q92796; Q5J | DLG3_HUMAN | Unmodified | 143460 | 122240 | 609710 | 222180 | 198260 | 0 | 167420 | 126320 | 2505500 | 656130 | 1130800 | 5003100 | 868990 | 428190 | 959750 | 1263200 | 0 | 274310 |
| LVTPHGES | DLG3; DLG3 | Q92796; Q5J | DLG3_HUMAN | Unmodified | 0 | 0 | 0 | 1097500 | 0 | 0 | 0 | 53958 | 1903300 | 211880 | 993370 | 0 | 0 | 0 | 1078000 | 597630 | 0 | 0 |
| LVTQLDRN | PCNX3 | Q9H6A9 | PCX3_HUMAN | Unmodified | 0 | 0 | 0 | 0 | 0 | 0 | 0 | 0 | 0 | 0 | 0 | 1315500 | 0 | 0 | 0 | 0 | 0 | 0 |
| LVTQRDAD | ECI1; ECI1; ECI | P42126; Q9( | ECI1_HUMAN E | Unmodified | 0 | 0 | 0 | 0 | 0 | 0 | 0 | 21320 | 467180 | 1018800 | 0 | 111040 | 0 | 0 | 0 | 0 | 0 | 48104 |
| LVTSGAES( | NEDD1 | Q8NHV4 | NEDD1_HUMA | Unmodified | 0 | 0 | 0 | 0 | 0 | 0 | 0 | 0 | 0 | 0 | 189060 | 305410 | 0 | 0 | 0 | 0 | 0 | 0 |
| LVTSIGDVV | PYGB; PYGB | P11216; H0Y | PYGB_HUMAN | Unmodified | 0 | 0 | 0 | 0 | 0 | 0 | 0 | 0 | 124260 | 0 | 0 | 830250 | 0 | 0 | 0 | 179030 | 0 | 0 |
| LVTSIGDVV | PYGB; PYGB | P11216; H0Y | PYGB_HUMAN | Unmodified | 0 | 0 | 0 | 383480 | 1841000 | 0 | 0 | 92141 | 0 | 1907000 | 0 | 1454000 | 920940 | 864560 | 0 | 0 | 2396700 | 13389000 |
| LVTSPCCIV | HSP90AA1 | P07900 | HS90A_HUMAN | Unmodified | 0 | 0 | 0 | 0 | 0 | 0 | 0 | 0 | 2282000 | 0 | 0 | 0 | 0 | 0 | 0 | 0 | 0 | 0 |
| LVTSPCCIV | HSP90AA1 | P07900 | HS90A_HUMAN | Oxidation (M) | 0 | 80424 | 0 | 0 | 0 | 0 | 0 | 0 | 2019200 | 0 | 0 | 7887200 | 0 | 0 | 0 | 0 | 0 | 0 |
| LVTTALRGE | CCPG1 | Q9ULG6 | CCPG1_HUMA | Unmodified | 0 | 0 | 0 | 0 | 0 | 0 | 0 | 0 | 1746600 | 0 | 0 | 0 | 0 | 0 | 0 | 0 | 0 | 0 |
| LVTTGVLK | H1-0 | P07305 | H10_HUMAN H | Unmodified | 4032800 | 4666300 | 0 | 0 | 0 | 0 | 0 | 248170 | 0 | 0 | 0 | 0 | 0 | 0 | 0 | 0 | 0 | 323900 |
| LVTTVTEIA( | CAP1 | Q01518 | CAP1_HUMAN | Unmodified | 202620 | 153610 | 164800 | 556530 | 9690900 | 1181800 | 6887000 | 2358600 | 2045100 | 1391000 | 1238900 | 3149800 | 1614900 | 2189500 | 295990 | 263170 | 3645500 | 4293900 |
| LVTVWSAP | PPP6C | O00743 | PPP6_HUMAN | Unmodified | 0 | 0 | 0 | 0 | 0 | 0 | 52762 | 0 | 0 | 0 | 0 | 501660 | 0 | 0 | 0 | 0 | 0 | 0 |
| LVTVYCEH( | DHTKD1; DHTK | Q96HY7; C9 | DHTK1_HUMAI | Unmodified | 0 | 455790 | 0 | 0 | 0 | 0 | 0 | 0 | 518480 | 255620 | 0 | 0 | 0 | 0 | 0 | 136430 | 42898 | 0 |
| LVVAVEEAF | VANGL2 | Q9ULK5 | VANG2_HUMAI | Unmodified | 0 | 0 | 0 | 0 | 117640 | 0 | 0 | 0 | 265110 | 343400 | 898390 | 1983200 | 0 | 0 | 0 | 0 | 0 | 0 |
| LVVDLTDIE | LTA4H | P09960 | LKHA4_HUMAN | Unmodified | 0 | 0 | 0 | 0 | 0 | 0 | 0 | 0 | 0 | 102230 | 215140 | 910500 | 0 | 0 | 0 | 0 | 0 | 0 |
| LVVDSHVQ | PRPF8; PRPF8 | Q6P2Q9; I3L | PRP8_HUMAN | Unmodified | 0 | 0 | 30494 | 52366 | 809350 | 196710 | 101570 | 149850 | 0 | 31795 | 0 | 845500 | 441300 | 271180 | 0 | 52061 | 299390 | 294510 |
| LVVECVMN | FABP5; FABP5 | Q01469; I6L | FABP5_HUMAN | Unmodified | 304760 | 0 | 0 | 0 | 910320 | 213520 | 43731 | 0 | 315800 | 0 | 1777000 | 4434300 | 0 | 0 | 0 | 0 | 539910 | 1168600 |
| LVVECVMN | FABP5; FABP5 | Q01469; I6L | FABP5_HUMAN | Unmodified | 0 | 0 | 0 | 0 | 635730 | 0 | 3225000 | 0 | 0 | 0 | 0 | 0 | 0 | 0 | 0 | 0 | 1890300 | 2579800 |
| LVVEEYYEV | AIP; AIP | O00170; E9F | AIP_HUMAN AH | Unmodified | 0 | 0 | 0 | 0 | 1667500 | 0 | 250170 | 0 | 0 | 0 | 0 | 0 | 0 | 0 | 0 | 0 | 0 | 4317000 |
| LVVEWQLC | NPC2; NPC2; N | P61916; J3K | NPC2_HUMAN | Unmodified | 0 | 0 | 0 | 0 | 872560 | 214160 | 0 | 135990 | 887490 | 0 | 0 | 0 | 0 | 0 | 0 | 0 | 370780 | 0 |
| LVVHGLLEI | HERC1 | Q15751 | HERC1_HUMAI | Unmodified | 0 | 0 | 0 | 0 | 0 | 0 | 0 | 0 | 62261 | 0 | 0 | 253130 | 1062800 | 601570 | 0 | 0 | 0 | 0 |
| LVVLATPQ\ | GYG1; GYG1; G | P46976; C9J | GLYG_HUMAN | Oxidation (M) | 0 | 0 | 0 | 0 | 0 | 0 | 0 | 0 | 0 | 0 | 170420 | 1078800 | 0 | 0 | 0 | 0 | 0 | 0 |
| LVVLDYIIR | PI4K2A; N/A | Q9BTU6; E9I | P4K2A_HUMAN | Unmodified | 0 | 0 | 142180 | 156320 | 0 | 0 | 0 | 0 | 564400 | 274720 | 0 | 625600 | 0 | 0 | 361830 | 266370 | 0 | 0 |
| LVVLECYVI | SPRED2; SPREI | Q7Z698; H7( | SPRE2_HUMAN | Unmodified | 44886 | 25378 | 48619 | 38580 | 90478 | 0 | 0 | 0 | 0 | 0 | 0 | 0 | 0 | 65124 | 36171 | 33382 | 155710 | 291500 |
| LVVLGSGG | RAP1B; RAP1A; | P61224; P62 | RAP1B_HUMAN | Unmodified | 38217000 | 70025000 | 44712000 | 1.06E+08 | 88157000 | 21264000 | 59083000 | 44185000 | 16958000 | 8985800 | 52158000 | 2.35E+08 | 66351000 | 70758000 | 62172000 | 63649000 | 18286000 | 41206000 |
| LVVNFESDI | TGM2 | P21980 | TGM2_HUMAN | Unmodified | 0 | 671420 | 0 | 121700 | 7566300 | 1231200 | 616620 | 177500 | 4228500 | 973020 | 814360 | 287370 | 0 | 262970 | 181700 | 0 | 211530 | 413690 |
| LVVNFESDI | TGM2 | P21980 | TGM2_HUMAN | Unmodified | 0 | 0 | 0 | 0 | 11517000 | 955460 | 268410 | 0 | 0 | 0 | 0 | 0 | 0 | 0 | 0 | 0 | 824980 | 744730 |
| LVVPASQC | PCBP3; PCBP2; | P57721; Q15 | PCBP3_HUMAI | Unmodified | 51350000 | 16652000 | 13326000 | 20346000 | 1125600 | 641910 | 602530 | 472440 | 2395200 | 666170 | 2424400 | 16757000 | 6915100 | 1951300 | 10730000 | 13931000 | 335120 | 1189300 |
| LVVPASQC | PCBP2; PCBP2; | Q15366; F8V | PCBP2_HUMAN | Unmodified | 6825900 | 0 | 0 | 3162800 | 859940 | 0 | 2074800 | 1259500 | 0 | 503260 | 96421 | 0 | 16846000 | 4829500 | 0 | 0 | 1737200 | 0 |
| LVVPASQC | PCBP3; PCBP3; | P57721; E9P | PCBP3_HUMAI | Unmodified | 196900 | 762580 | 0 | 1138700 | 0 | 0 | 972310 | 598090 | 1252800 | 121210 | 0 | 0 | 2984700 | 1108400 | 0 | 0 | 96164 | 0 |
| LVVPATQC | PCBP1 | Q15365 | PCBP1_HUMAN | Unmodified | 25390000 | 0 | 0 | 6731400 | 1482900 | 2047600 | 431870 | 227250 | 2295500 | 581600 | 6759300 | 30571000 | 5851300 | 245160 | 0 | 0 | 306800 | 797220 |
| LVVPATQC | PCBP1 | Q15365 | PCBP1_HUMAN | Unmodified | 1999500 | 327820 | 0 | 1104400 | 1630900 | 0 | 754720 | 549500 | 0 | 560890 | 0 | 0 | 7263700 | 3000600 | 0 | 0 | 2215400 | 1175700 |
| LVVTPASS( | BTBD17 | A6NE02 | BTBDH_HUMAI | Unmodified | 0 | 0 | 0 | 0 | 3820800 | 0 | 2799700 | 1773900 | 1300000 | 1371400 | 0 | 0 | 0 | 0 | 0 | 0 | 1617200 | 1704300 |
| LVVVGDGA | RHOB | P62745 | RHOB_HUMAN | Unmodified | 0 | 0 | 0 | 0 | 0 | 0 | 0 | 0 | 0 | 0 | 0 | 0 | 0 | 0 | 0 | 462070 | 0 | 0 |
| LVVVGGG( | RRAS; RRAS2; F | P10301; P62 | RRAS_HUMAN | Unmodified | 3140700 | 4231900 | 3887400 | 0 | 6198000 | 1828700 | 0 | 0 | 812410 | 424830 | 0 | 0 | 2009500 | 0 | 7734500 | 8045600 | 147650 | 6925000 |
| LVVVLCNLI | YARS1; YARS1 | P54577; A0A | SYYC_HUMAN | Unmodified | 100130 | 73908 | 77191 | 73228 | 0 | 0 | 100810 | 0 | 416860 | 0 | 0 | 2453500 | 65028 | 0 | 209320 | 163950 | 65570 | 0 |
| LVWVPSDK | MYH9; MYH9; M | P35579; Q5E | MYH9_HUMAN | Unmodified | 0 | 72455 | 0 | 157990 | 8738500 | 1072600 | 303120 | 188620 | 696810 | 289940 | 1246000 | 7792600 | 0 | 0 | 1159100 | 239720 | 1091500 | 1620300 |
| LVYDGIR | CTNNA1; CTNN | P35221; G3X | CTNA1_HUMAI | Unmodified | 1600500 | 1518200 | 3017300 | 2082300 | 0 | 0 | 0 | 400290 | 0 | 644830 | 296070 | 9753900 | 0 | 0 | 9691400 | 10930000 | 0 | 0 |
| LVYDGIRDI | CTNNA1; CTNN | P35221; G3X | CTNA1_HUMAI | Unmodified | 342400 | 203950 | 679060 | 858990 | 0 | 380630 | 0 | 0 | 0 | 0 | 0 | 2602600 | 87795 | 0 | 1230800 | 1768700 | 0 | 128860 |
| LVYILNRDA | SF3B3 | Q15393 | SF3B3_HUMAN | Unmodified | 106850 | 151640 | 0 | 307730 | 0 | 0 | 0 | 0 | 0 | 0 | 0 | 0 | 107540 | 0 | 0 | 0 | 0 | 0 |
| LVYLVENP( | TFRC; TFRC | P02786; G3\ | TFR1_HUMAN | Unmodified | 3100200 | 2887400 | 4208500 | 5298700 | 526120 | 394480 | 0 | 79915 | 23240000 | 4471600 | 0 | 4822700 | 322150 | 0 | 4799500 | 3775300 | 0 | 0 |
| LWADHGV( | GNAI2 | P04899 | GNAI2_HUMAN | Unmodified | 3692700 | 1758300 | 600100 | 1921600 | 1306400 | 0 | 203000 | 0 | 5750300 | 0 | 4362000 | 47314000 | 0 | 0 | 1592400 | 1675500 | 1453900 | 1340400 |
| LWAGDQIL | PATJ; PATJ; PAT | Q8NI35; A0A | INADL_HUMAN | Unmodified | 0 | 0 | 0 | 90689 | 0 | 0 | 0 | 0 | 0 | 0 | 0 | 0 | 0 | 0 | 62837 | 0 | 0 | 0 |
| LWCAVGVI | SPAG9; SPAG9 | O60271; A0A | JIP4_HUMAN C | Unmodified | 0 | 0 | 0 | 0 | 301570 | 0 | 154800 | 0 | 0 | 104040 | 0 | 457300 | 0 | 0 | 0 | 0 | 431180 | 838490 |
| LWCSTTAD | IGF2R | P11717 | MPRI_HUMAN ( | Unmodified | 0 | 0 | 0 | 0 | 0 | 0 | 0 | 0 | 9645600 | 1382400 | 0 | 0 | 0 | 0 | 0 | 0 | 0 | 0 |
| LWCTYHEK | AKR1B1; AKR1E | P15121; E9P | ALDR_HUMAN | Unmodified | 0 | 0 | 0 | 0 | 293420 | 0 | 0 | 0 | 0 | 0 | 0 | 1045100 | 0 | 0 | 0 | 0 | 0 | 0 |
| LWDHATM | STRAP | Q9Y3F4 | STRAP_HUMAN | Oxidation (M) | 0 | 0 | 0 | 0 | 0 | 0 | 0 | 0 | 0 | 0 | 0 | 523350 | 0 | 0 | 0 | 0 | 0 | 0 |
| LWDIGGQI | ARL8A; ARL8A | Q96BM9; A0 | ARL8A_HUMAN | Unmodified | 82665 | 0 | 0 | 0 | 999300 | 1022500 | 0 | 0 | 348080 | 187780 | 170220 | 2510300 | 0 | 0 | 258310 | 137660 | 784760 | 0 |
| LWDLTTGT | RACK1; RACK1; | P63244; D6F | RACK1_HUMAN | Unmodified | 0 | 0 | 0 | 0 | 0 | 85859 | 0 | 50162 | 978410 | 92329 | 2559300 | 0 | 0 | 0 | 0 | 2293000 | 0 | 0 |
| LWDLTTGT | RACK1; RACK1; | P63244; D6F | RACK1_HUMAN | Unmodified | 917810 | 1144100 | 909330 | 779570 | 0 | 0 | 0 | 186630 | 0 | 0 | 1312700 | 18569000 | 0 | 0 | 703350 | 1092300 | 0 | 0 |
| LWDPRTPC | BUB3; BUB3; BI | O43684; J3Q | BUB3_HUMAN | Unmodified | 0 | 0 | 0 | 0 | 1024300 | 0 | 768000 | 741560 | 0 | 730320 | 0 | 0 | 247330 | 0 | 0 | 0 | 234840 | 667810 |
| LWDSGPEI | RNF31; RNF31; | Q96EP0; H0Y | RNF31_HUMAN | Unmodified | 0 | 172950 | 0 | 294140 | 0 | 0 | 0 | 0 | 600200 | 0 | 0 | 0 | 0 | 892470 | 0 | 0 | 0 | 0 |
| LWDTAGQI | RAB6A; RAB39A | P20340; Q14 | RAB6A_HUMAN | Unmodified | 3151800 | 1970200 | 1631200 | 1211100 | 0 | 0 | 0 | 0 | 0 | 0 | 4168700 | 1520400 | 432150 | 0 | 1427600 | 2104500 | 0 | 0 |

| | | | | | | | | | | | | | | | | | | | | | | |
|---|---|---|---|---|---|---|---|---|---|---|---|---|---|---|---|---|---|---|---|---|---|---|
| LWEAGWK | XRN2 | Q9H0D6 | XRN2_HUMAN | Unmodified | 0 | 0 | 0 | 0 | 0 | 0 | 0 | 0 | 0 | 0 | 44905 | 647160 | 0 | 0 | 0 | 0 | 0 | 0 |
| LWEEQLAA | PRKCSH; PRKC | P14314; K7E | GLU2B_HUMAI | Unmodified | 0 | 0 | 344160 | 176200 | 0 | 0 | 841480 | 499710 | 3.93E+08 | 95382000 | 7400100 | 4520000 | 0 | 328980 | 345660 | 351520 | 0 | 0 |
| LWEISTGR | CSTF1; CSTF1 | Q05048; A0/ | CSTF1_HUMAN | Unmodified | 76511 | 68571 | 100800 | 104460 | 0 | 0 | 0 | 0 | 0 | 0 | 0 | 1129100 | 0 | 0 | 161240 | 69685 | 0 | 0 |
| LWGDRYFI | EEF2 | P13639 | EF2_HUMAN El | Unmodified | 0 | 67943 | 919370 | 1474300 | 1413500 | 0 | 0 | 0 | 598000 | 0 | 2737100 | 15418000 | 585710 | 0 | 1888600 | 1131700 | 162440 | 409030 |
| LWGDSGIC | GNAO1; GNAO | P09471; H3E | GNAO_HUMAN | Unmodified | 0 | 0 | 0 | 0 | 106810 | 0 | 0 | 0 | 2664100 | 0 | 0 | 0 | 0 | 0 | 0 | 0 | 0 | 0 |
| LWGGRFV( | ASL; ASL; ASL | P04424; A0A | ARLY_HUMAN / | Oxidation (M) | 0 | 0 | 0 | 0 | 554170 | 0 | 107020 | 171320 | 0 | 0 | 273800 | 0 | 0 | 0 | 0 | 0 | 0 | 0 |
| LWGVIDVK | CHORDC1; CH | Q9UHD1; E9 | CHRD1_HUMA | Unmodified | 62669 | 0 | 0 | 0 | 1579600 | 0 | 0 | 0 | 1659700 | 1222900 | 2816600 | 7613100 | 4252500 | 0 | 0 | 222200 | 1572900 | 0 |
| LWHLDPD1 | PTPRK; PTPRK; | Q15262; Q5 | PTPRK_HUMAN | Unmodified | 0 | 0 | 0 | 0 | 0 | 0 | 0 | 0 | 2124900 | 14503 | 0 | 0 | 0 | 0 | 0 | 129690 | 0 | 0 |
| LWKEEEDC | SEC13; SEC13 | P55735; A8N | SEC13_HUMAN | Unmodified | 0 | 0 | 0 | 0 | 0 | 0 | 72432 | 0 | 0 | 0 | 122060 | 1265400 | 0 | 0 | 0 | 0 | 0 | 0 |
| LWKKGEGI | DNAJB11; N/A | Q9UBS4; H7 | DJB11_HUMAN | Unmodified | 0 | 0 | 0 | 0 | 0 | 0 | 746520 | 117220 | 0 | 3723700 | 0 | 0 | 0 | 0 | 0 | 0 | 0 | 0 |
| LWLDNTEN | ILK; ILK; ILK | Q13418; A0/ | ILK_HUMAN Int | Unmodified | 1067100 | 0 | 0 | 2226800 | 0 | 0 | 0 | 0 | 0 | 0 | 0 | 0 | 0 | 0 | 1343200 | 959400 | 0 | 0 |
| LWNLNNYI | PRPF8; PRPF8 | Q6P2Q9; I3L | PRP8_HUMAN | Unmodified | 0 | 0 | 0 | 0 | 0 | 0 | 0 | 0 | 0 | 0 | 0 | 1190700 | 0 | 0 | 0 | 0 | 0 | 0 |
| LWNNVGH | TMTC3 | Q6ZXV5 | TMTC3_HUMAI | Unmodified | 0 | 0 | 0 | 0 | 0 | 0 | 0 | 0 | 4498100 | 637750 | 339510 | 0 | 0 | 0 | 0 | 0 | 260300 | 50743 |
| LWNTLGV( | RACK1; RACK1; | P63244; D6F | RACK1_HUMAI | Unmodified | 2015200 | 1208400 | 1010200 | 964850 | 0 | 0 | 1652500 | 860090 | 0 | 438220 | 1193100 | 16620000 | 1013300 | 964740 | 1039600 | 1338300 | 0 | 1000700 |
| LWNTTEVA | STRN | O43815 | STRN_HUMAN | Unmodified | 0 | 0 | 0 | 0 | 0 | 0 | 0 | 0 | 0 | 0 | 1666300 | 585070 | 0 | 0 | 0 | 0 | 0 | 0 |
| LWPSGDR | PAICS; PAICS; F | P22234; D6F | PUR6_HUMAN | Unmodified | 0 | 0 | 120930 | 0 | 0 | 0 | 0 | 0 | 0 | 0 | 0 | 5829300 | 0 | 0 | 0 | 140750 | 0 | 0 |
| LWPSGDRS | PAICS; PAICS; F | P22234; D6F | PUR6_HUMAN | Unmodified | 0 | 0 | 0 | 383310 | 0 | 0 | 483080 | 190420 | 917810 | 0 | 0 | 6930300 | 0 | 0 | 253000 | 0 | 51525 | 210700 |
| LWQEELNE | ABCC5; ABCC5 | O15440; F8V | MRP5_HUMAN | Unmodified | 0 | 0 | 0 | 0 | 0 | 0 | 0 | 0 | 0 | 0 | 0 | 1628200 | 0 | 0 | 0 | 0 | 0 | 0 |
| LWQPDAC' | ANLN | Q9NQW6 | ANLN_HUMAN | Unmodified | 0 | 0 | 0 | 0 | 1398700 | 1997100 | 0 | 30973 | 0 | 0 | 0 | 374620 | 50226 | 0 | 0 | 0 | 880680 | 276910 |
| LWQTTEEIF | MRPL37; MRPL: | Q9BZE1; S4F | RM37_HUMAN | Unmodified | 0 | 0 | 3366900 | 0 | 0 | 0 | 0 | 0 | 0 | 0 | 0 | 0 | 0 | 0 | 0 | 0 | 0 | 0 |
| LWQTVVGI | STRAP; STRAP | Q9Y3F4; H0' | STRAP_HUMAN | Unmodified | 264920 | 150310 | 200310 | 244380 | 0 | 0 | 0 | 11267 | 2388300 | 79416 | 539220 | 6717600 | 0 | 0 | 332140 | 497330 | 0 | 0 |
| LWRDGGV | GNAI3 | P08754 | GNAI3_HUMAN | Unmodified | 115610 | 0 | 0 | 461970 | 0 | 0 | 0 | 0 | 311370 | 111790 | 0 | 244590 | 0 | 126180 | 0 | 0 | 161380 | 73272 |
| LWRDPGIC | GNAT3 | A8MTJ3 | GNAT3_HUMAI | Unmodified | 72688 | 0 | 0 | 0 | 0 | 283330 | 0 | 0 | 0 | 0 | 113880 | 1025100 | 0 | 0 | 0 | 0 | 0 | 0 |
| LWRQDGV | ACOT8; ACOT8; | O14734; H7( | ACOT8_HUMAI | Unmodified | 0 | 0 | 0 | 0 | 76317 | 0 | 0 | 0 | 2352400 | 0 | 0 | 0 | 0 | 0 | 0 | 0 | 60581 | 0 |
| LWRVPAEG | CORO7; CORO | P57737; A0A | CORO7_HUMA | Unmodified | 0 | 0 | 0 | 0 | 0 | 0 | 0 | 0 | 0 | 0 | 363540 | 583100 | 0 | 0 | 0 | 0 | 0 | 0 |
| LWSLDSDE | PRPF4 | O43172 | PRP4_HUMAN | Unmodified | 0 | 0 | 0 | 0 | 0 | 0 | 0 | 0 | 0 | 0 | 0 | 1532100 | 0 | 0 | 0 | 0 | 0 | 0 |
| LWSLGQQI | WDR48 | Q8TAF3 | WDR48_HUMA | Unmodified | 0 | 0 | 0 | 0 | 0 | 0 | 0 | 0 | 0 | 0 | 0 | 1007700 | 0 | 0 | 0 | 0 | 0 | 0 |
| LWSNFWG | PLOD3; PLOD3; | O60568; H7( | PLOD3_HUMAI | Unmodified | 0 | 0 | 0 | 0 | 0 | 0 | 0 | 0 | 246030 | 0 | 0 | 0 | 0 | 0 | 0 | 0 | 0 | 0 |
| LWSNFWG | PLOD2; PLOD2 | O00469; E7E | PLOD2_HUMAI | Unmodified | 0 | 0 | 0 | 0 | 0 | 0 | 0 | 0 | 516570 | 49644 | 0 | 0 | 0 | 0 | 0 | 0 | 0 | 0 |
| LWSPSDEF | SORL1 | Q92673 | SORL_HUMAN | Unmodified | 0 | 0 | 0 | 0 | 0 | 0 | 0 | 0 | 2376700 | 0 | 0 | 0 | 0 | 0 | 0 | 0 | 0 | 0 |
| LWSTFHRP | AKR1C3; AKR1( | P42330; S4F | AK1C3_HUMAN | Unmodified | 0 | 0 | 0 | 0 | 4006200 | 426340 | 0 | 0 | 0 | 0 | 0 | 0 | 0 | 0 | 0 | 0 | 3313800 | 0 |
| LWTLVSEQ | RPL27A; RPL27 | P46776; E9P | RL27A_HUMAN | Unmodified | 3800900 | 2539300 | 1499600 | 730680 | 0 | 0 | 0 | 0 | 546460 | 0 | 1622200 | 27441000 | 0 | 0 | 2431700 | 2219800 | 0 | 0 |
| LWTLVSEQ | RPL27A; RPL27 | P46776; E9P | RL27A_HUMAN | Unmodified | 1709600 | 1294100 | 1489400 | 2225600 | 0 | 0 | 7243100 | 1792100 | 0 | 0 | 0 | 4693900 | 4695300 | 2207200 | 829350 | 480870 | 0 | 0 |
| LWVLETK | WDR43; WDR4: | Q15061; C9I | WDR43_HUMA | Unmodified | 0 | 0 | 0 | 0 | 0 | 0 | 0 | 33780 | 160960 | 0 | 0 | 1254800 | 0 | 0 | 0 | 0 | 0 | 0 |
| LWWADQV | LRP1 | Q07954 | LRP1_HUMAN I | Unmodified | 0 | 0 | 0 | 0 | 0 | 0 | 0 | 0 | 1096500 | 169250 | 0 | 0 | 0 | 0 | 0 | 0 | 0 | 0 |
| LWWLDLK | HPX | P02790 | HEMO_HUMAN | Unmodified | 0 | 0 | 0 | 0 | 0 | 0 | 0 | 0 | 0 | 0 | 0 | 946610 | 0 | 0 | 0 | 0 | 0 | 0 |
| LWYCDLQ( | PREP; PREP | P48147; A0A | PPCE_HUMAN | Unmodified | 0 | 0 | 0 | 0 | 0 | 0 | 0 | 0 | 0 | 2674900 | 0 | 0 | 0 | 0 | 0 | 0 | 0 | 0 |
| LWYGTGR | CHST1 | O43916 | CHST1_HUMAI | Unmodified | 0 | 0 | 0 | 0 | 363130 | 0 | 0 | 0 | 521730 | 193490 | 0 | 0 | 0 | 0 | 0 | 0 | 82372 | 0 |
| LYAGSTVK | TXNRD1; TXNRI | Q16881; A0/ | TRXR1_HUMAN | Unmodified | 0 | 0 | 0 | 167600 | 1916600 | 648190 | 0 | 0 | 4262100 | 383730 | 2007800 | 1757900 | 777060 | 0 | 0 | 735460 | 412290 | 934450 |
| LYAMQTGN | VTA1; VTA1 | Q9NP79; A0, | VTA1_HUMAN ' | Oxidation (M) | 0 | 0 | 0 | 0 | 0 | 0 | 0 | 0 | 1539100 | 0 | 0 | 153440 | 0 | 0 | 0 | 0 | 0 | 0 |
| LYAMQTGN | VTA1; VTA1 | Q9NP79; A0, | VTA1_HUMAN ' | Unmodified | 0 | 183180 | 0 | 0 | 0 | 0 | 0 | 0 | 0 | 0 | 129460 | 2797700 | 0 | 0 | 0 | 0 | 0 | 0 |
| LYANMFER | FKBP4; FKBP4 | Q02790; H0' | FKBP4_HUMAN | Unmodified | 0 | 0 | 91995 | 106310 | 0 | 0 | 0 | 0 | 206060 | 0 | 0 | 1882000 | 0 | 0 | 105200 | 53619 | 0 | 233390 |
| LYAQHALIF | GPC2 | Q8N158 | GPC2_HUMAN | Unmodified | 0 | 0 | 0 | 0 | 0 | 0 | 0 | 0 | 39872 | 0 | 0 | 496800 | 0 | 0 | 17629 | 0 | 31974 | 0 |
| LYASILQQK | AP3D1; AP3D1 | O14617; A0/ | AP3D1_HUMAI | Unmodified | 0 | 0 | 0 | 0 | 0 | 0 | 0 | 0 | 0 | 0 | 81870 | 737690 | 0 | 0 | 0 | 0 | 0 | 0 |
| LYASYEK | RRM1; RRM1 | P23921; E9P | RIR1_HUMAN F | Unmodified | 0 | 0 | 0 | 176830 | 0 | 0 | 0 | 0 | 246240 | 147540 | 524240 | 2935900 | 0 | 0 | 63208 | 0 | 0 | 0 |
| LYAVHQEG | CCT8; CCT8 | P50990; H7( | TCPQ_HUMAN | Unmodified | 0 | 0 | 0 | 0 | 1700000 | 0 | 0 | 0 | 0 | 152220 | 0 | 3152100 | 2855800 | 493520 | 0 | 183790 | 307940 | 0 |
| LYAVHQEG | CCT8; CCT8 | P50990; H7( | TCPQ_HUMAN | Unmodified | 0 | 0 | 0 | 0 | 0 | 0 | 0 | 0 | 0 | 0 | 0 | 0 | 1808700 | 0 | 0 | 0 | 1456000 | 961650 |
| LYCIYVAIG | ATP5F1A; ATP5I | P25705; K7E | ATPA_HUMAN / | Unmodified | 0 | 0 | 0 | 0 | 0 | 0 | 35176 | 0 | 1312300 | 223820 | 0 | 0 | 0 | 0 | 0 | 0 | 0 | 0 |
| LYCPVEFSI | SMS | P52788 | SPSY_HUMAN | Unmodified | 0 | 0 | 0 | 548980 | 0 | 0 | 224950 | 0 | 0 | 0 | 0 | 3018000 | 42643 | 0 | 367450 | 0 | 0 | 0 |
| LYCYYYQVI | POGLUT3; POG | Q7Z4H8; H0 | PLGT3_HUMAN | Unmodified | 0 | 0 | 0 | 0 | 0 | 0 | 0 | 0 | 3474400 | 93355 | 0 | 0 | 0 | 0 | 0 | 0 | 0 | 0 |
| LYDEMPPS | PKP3 | Q9Y446 | PKP3_HUMAN | Oxidation (M) | 0 | 0 | 1464200 | 0 | 0 | 0 | 0 | 0 | 0 | 0 | 307540 | 1372800 | 0 | 0 | 0 | 0 | 0 | 0 |
| LYDEMPPS | PKP3 | Q9Y446 | PKP3_HUMAN | Oxidation (M) | 553280 | 31686 | 264380 | 554030 | 0 | 0 | 0 | 0 | 0 | 0 | 405970 | 0 | 0 | 0 | 0 | 0 | 0 | 0 |
| LYDIDVAK | RPL23A; RPL23 | P62750; K7E | RL23A_HUMAN | Unmodified | 1329600 | 1177000 | 0 | 734660 | 0 | 0 | 293360 | 0 | 0 | 0 | 0 | 9955600 | 867900 | 421650 | 786120 | 631940 | 0 | 0 |
| LYDILGVPF | DNAJA2; DNAJ/ | O60884; I3L: | DNJA2_HUMAN | Unmodified | 0 | 0 | 0 | 0 | 0 | 0 | 0 | 0 | 0 | 0 | 0 | 327340 | 0 | 0 | 432900 | 0 | 0 | 0 |
| LYDLDNDC | NCS1 | P62166 | NCS1_HUMAN | Unmodified | 0 | 0 | 0 | 0 | 0 | 0 | 0 | 0 | 0 | 0 | 0 | 0 | 0 | 0 | 332880 | 0 | 0 | 0 |
| LYDLHGDC | MUC5B | Q9HC84 | MUC5B_HUMA | Unmodified | 0 | 0 | 0 | 0 | 0 | 0 | 0 | 0 | 0 | 0 | 0 | 0 | 0 | 0 | 0 | 0 | 0 | 3520500 |
| LYDLNHNE | SNRNP200; SN | O75643; A0/ | U520_HUMAN | Unmodified | 672390 | 363710 | 361880 | 259860 | 0 | 25197 | 0 | 18825 | 992720 | 0 | 0 | 1445500 | 0 | 0 | 176490 | 265880 | 20871 | 0 |
| LYDNLLEQ | PSMD11 | O00231 | PSD11_HUMAN | Unmodified | 0 | 26003 | 293050 | 0 | 0 | 0 | 0 | 0 | 55638 | 0 | 1440400 | 18002000 | 0 | 0 | 65014 | 590620 | 0 | 0 |
| LYDPGQEY | ARHGAP29; ARI | Q52LW3; F8' | RHG29_HUMAI | Unmodified | 0 | 523340 | 0 | 0 | 1235200 | 0 | 0 | 0 | 0 | 0 | 0 | 0 | 0 | 0 | 0 | 0 | 1213400 | 1835000 |
| LYDRDVAS | POLDIP2; POLD | Q9Y2S7; B4I | PDIP2_HUMAN | Unmodified | 0 | 0 | 0 | 0 | 45590 | 0 | 0 | 0 | 731650 | 0 | 0 | 0 | 0 | 0 | 0 | 0 | 49762 | 0 |
| LYDRSLESI | MOV10; MOV1( | Q9HCE1; Q5 | MOV10_HUMA | Oxidation (M) | 479970 | 253460 | 0 | 518760 | 0 | 0 | 0 | 0 | 0 | 0 | 0 | 0 | 350330 | 0 | 0 | 0 | 0 | 0 |
| LYDRSLESI | MOV10; MOV1( | Q9HCE1; Q5 | MOV10_HUMA | Unmodified | 0 | 205670 | 0 | 458330 | 0 | 0 | 394920 | 138760 | 1892400 | 282850 | 0 | 0 | 613610 | 0 | 166310 | 0 | 653550 | 0 |
| LYDRYSTV( | BMP3 | P12645 | BMP3_HUMAN | Unmodified | 0 | 0 | 0 | 0 | 0 | 0 | 72724 | 0 | 550490 | 0 | 0 | 0 | 0 | 0 | 0 | 0 | 0 | 0 |
| LYDTAMDK | DLG5 | Q8TDM6 | DLG5_HUMAN | Unmodified | 0 | 0 | 0 | 0 | 1761600 | 0 | 0 | 0 | 167470 | 0 | 0 | 0 | 0 | 0 | 0 | 0 | 832570 | 1635300 |
| LYEEVSKCI | DNAH17 | Q9UFH2 | DYH17_HUMAI | Unmodified | 0 | 0 | 0 | 0 | 0 | 0 | 0 | 0 | 1556400 | 1104200 | 0 | 0 | 0 | 0 | 0 | 0 | 532090 | 0 |
| LYEHIVADI | LSS; LSS | P48449; A0A | LSS_HUMAN La | Unmodified | 0 | 0 | 0 | 0 | 302650 | 0 | 336360 | 49320 | 651070 | 0 | 0 | 627500 | 0 | 0 | 0 | 0 | 148050 | 0 |
| LYELIITR | CAPN1 | P07384 | CAN1_HUMAN | Unmodified | 0 | 0 | 0 | 0 | 0 | 0 | 41914 | 255520 | 0 | 0 | 0 | 490630 | 0 | 0 | 0 | 0 | 27018 | 0 |
| LYENFISEF | PSMD13; PSMD | Q9UNM6; E9 | PSD13_HUMAN | Unmodified | 0 | 0 | 0 | 0 | 0 | 0 | 0 | 0 | 0 | 0 | 0 | 2962000 | 0 | 0 | 0 | 0 | 0 | 0 |
| LYEPDQLQ | FAM120A; FAM: | Q9NZB2; A0, | F120A_HUMAN | Unmodified | 0 | 0 | 0 | 0 | 0 | 0 | 0 | 0 | 0 | 0 | 0 | 715260 | 0 | 0 | 0 | 0 | 0 | 0 |
| LYEQILAEN | PPP1R12A; PPP | O14974; A0/ | MYPT1_HUMAN | Unmodified | 0 | 0 | 0 | 0 | 1077700 | 0 | 73322 | 0 | 0 | 0 | 213410 | 0 | 0 | 0 | 0 | 0 | 0 | 0 |
| LYEQLSGK | PEBP1 | P30086 | PEBP1_HUMAN | Unmodified | 0 | 0 | 3077500 | 3210500 | 13326000 | 2236200 | 2790400 | 0 | 9324300 | 11363000 | 7461200 | 26491000 | 23736000 | 14889000 | 5940000 | 5220300 | 14771000 | 15833000 |
| LYERGDAS | HADH; HADH; I | Q16836; A0/ | HCDH_HUMAN | Unmodified | 0 | 0 | 0 | 0 | 97654 | 0 | 0 | 0 | 10295000 | 221780 | 213750 | 106460 | 0 | 155210 | 234450 | 0 | 0 | 0 |
| LYERRGGG | PURB | Q96QR8 | PURB_HUMAN | Unmodified | 0 | 0 | 0 | 0 | 0 | 56539 | 51097 | 0 | 0 | 0 | 0 | 0 | 552020 | 161560 | 0 | 0 | 0 | 0 |
| LYEWISIDK | INPPL1; INPPL1 | O15357; A0/ | SHIP2_HUMAN | Unmodified | 0 | 0 | 0 | 0 | 0 | 0 | 0 | 0 | 0 | 0 | 669280 | 1806100 | 0 | 0 | 0 | 0 | 0 | 0 |
| LYFLQCET( | EIF2S2 | P20042 | IF2B_HUMAN E | Unmodified | 0 | 140070 | 132170 | 0 | 0 | 0 | 0 | 0 | 0 | 0 | 0 | 2703200 | 0 | 0 | 0 | 288610 | 0 | 0 |
| LYFSDATLI | LRP1; LRP1 | Q07954; H0' | LRP1_HUMAN I | Unmodified | 0 | 0 | 0 | 0 | 0 | 0 | 0 | 0 | 2083200 | 0 | 0 | 0 | 0 | 0 | 0 | 0 | 0 | 0 |
| LYFTNLQSI | TMEM63B; TME | Q5T3F8; H0' | CSCL2_HUMAI | Unmodified | 116990 | 0 | 245910 | 130830 | 0 | 0 | 0 | 0 | 338210 | 0 | 263390 | 3664400 | 0 | 0 | 408540 | 227660 | 0 | 0 |
| LYFYSDAYI | CNPY1; CNPY1 | Q3B7I2; A0A | CNPY1_HUMAI | Unmodified | 0 | 0 | 345950 | 0 | 0 | 0 | 0 | 0 | 0 | 41623 | 1201700 | 1086800 | 0 | 0 | 0 | 13567 | 0 | 0 |
| LYGDADYL | CTPS1; CTPS1; | P17812; B4E | PYRG1_HUMAI | Unmodified | 0 | 0 | 0 | 0 | 0 | 0 | 0 | 0 | 1066800 | 0 | 190130 | 2669200 | 0 | 0 | 227820 | 0 | 0 | 0 |
| LYGDDALD | XPO7; XPO7; XI | Q9UIA9; E5F | XPO7_HUMAN | Unmodified | 0 | 0 | 0 | 0 | 0 | 0 | 0 | 0 | 0 | 0 | 0 | 227490 | 0 | 0 | 40213 | 0 | 0 | 0 |
| LYGDVPFIE | CTPS2 | Q9NRF8 | PYRG2_HUMAI | Unmodified | 0 | 0 | 0 | 0 | 0 | 0 | 0 | 0 | 0 | 0 | 0 | 1544400 | 0 | 0 | 0 | 0 | 0 | 0 |
| LYGMSDPN | RET | P07949 | RET_HUMAN Pi | Oxidation (M) | 195000 | 90855 | 0 | 0 | 0 | 0 | 0 | 0 | 400340 | 0 | 0 | 0 | 0 | 0 | 0 | 0 | 0 | 0 |
| LYGPNFIL( | TMPRSS2; TMPI | O15393; C9J | TMPS2_HUMAI | Unmodified | 0 | 0 | 125470 | 1006300 | 0 | 0 | 0 | 0 | 0 | 0 | 0 | 0 | 0 | 0 | 2080900 | 1225300 | 0 | 0 |
| LYGPNFIL( | TMPRSS2; TMPI | O15393; C9J | TMPS2_HUMAI | Unmodified | 0 | 0 | 4440600 | 15463000 | 0 | 0 | 0 | 0 | 0 | 0 | 0 | 0 | 0 | 0 | 3211000 | 4179800 | 0 | 0 |
| LYGPSSVSI | SERPINH1; SER | P50454; E9P | SERPH_HUMAI | Unmodified | 18468 | 17580 | 0 | 0 | 0 | 0 | 0 | 22946 | 16797000 | 964830 | 0 | 0 | 0 | 0 | 28849 | 12632 | 25213 | 34879 |
| LYGPSSVSI | SERPINH1; SER | P50454; E9P | SERPH_HUMAI | Unmodified | 0 | 0 | 0 | 0 | 2853400 | 0 | 252130 | 115610 | 0 | 11568000 | 0 | 0 | 351110 | 0 | 0 | 0 | 0 | 631260 |
| LYGPTNFAI | CPNE1; CPNE1 | Q99829; Q5. | CPNE1_HUMAI | Unmodified | 0 | 1577500 | 1826300 | 3983300 | 231080 | 158910 | 0 | 0 | 8637800 | 795420 | 662160 | 9946100 | 0 | 0 | 3889100 | 4272600 | 576670 | 252540 |
| LYGPTNFSI | CPNE3 | O75131 | CPNE3_HUMAI | Unmodified | 1729700 | 1800000 | 16923000 | 11456000 | 0 | 0 | 779240 | 503630 | 57173000 | 5663400 | 688840 | 13000000 | 0 | 0 | 16357000 | 8969600 | 1191700 | 1929500 |
| LYGSAGPP | HSPA5; HSPA5; | P11021; A0A | BIP_HUMAN Er | Unmodified | 0 | 0 | 949250 | 0 | 0 | 0 | 11027000 | 8364200 | 89348000 | 48197000 | 5603300 | 3855600 | 0 | 0 | 2197800 | 697840 | 0 | 0 |
| LYGSGDQE | LAP3; LAP3 | P28838; H0' | AMPL_HUMAN | Unmodified | 7735.9 | 0 | 0 | 36686 | 4236500 | 488980 | 933010 | 589890 | 724880 | 511570 | 348580 | 3052700 | 330770 | 252220 | 33297 | 23328 | 96308 | 1496500 |
| LYGVYCFR | HAPLN1 | P10915 | HPLN1_HUMAI | Unmodified | 0 | 0 | 0 | 0 | 0 | 0 | 0 | 0 | 10178000 | 667570 | 0 | 0 | 0 | 0 | 0 | 0 | 0 | 0 |
| LYGYLPQP: | MMP15 | P51511 | MMP15_HUMA | Unmodified | 555890 | 647140 | 898240 | 662610 | 0 | 0 | 0 | 0 | 0 | 0 | 0 | 0 | 0 | 0 | 1950300 | 1386400 | 117040 | 0 |
| LYHDIAQQ | EIF3A | Q14152 | EIF3A_HUMAN | Unmodified | 364220 | 343950 | 388670 | 388020 | 0 | 0 | 0 | 0 | 0 | 0 | 762660 | 8748300 | 0 | 0 | 566900 | 663040 | 0 | 0 |
| LYHIQGIEC | MAN2A1 | Q16706 | MA2A1_HUMAI | Unmodified | 0 | 0 | 0 | 0 | 0 | 0 | 0 | 0 | 0 | 601030 | 0 | 0 | 0 | 0 | 0 | 0 | 0 | 0 |
| LYHNEVEIE | SMC1A; SMC1A | Q14683; G8. | SMC1A_HUMA | Unmodified | 0 | 0 | 0 | 0 | 338680 | 0 | 394740 | 0 | 0 | 402690 | 164330 | 525610 | 0 | 184360 | 0 | 0 | 0 | 54477 |
| LYHSCADP | DPP7; DPP7 | Q9UHL4; R4 | DPP2_HUMAN | Unmodified | 83610 | 0 | 31749 | 0 | 0 | 0 | 0 | 0 | 758290 | 0 | 17806 | 0 | 0 | 0 | 94137 | 73600 | 382470 | 138940 |
| LYHVSDSE | VIL1 | P09327 | VILI_HUMAN Vi | Unmodified | 1001200 | 495920 | 0 | 0 | 0 | 0 | 0 | 0 | 3698900 | 332170 | 2360300 | 39914000 | 0 | 0 | 0 | 0 | 0 | 0 |
| LYHVSDSE | VIL1 | P09327 | VILI_HUMAN Vi | Unmodified | 0 | 0 | 0 | 0 | 0 | 0 | 0 | 0 | 0 | 960970 | 0 | 0 | 0 | 0 | 0 | 0 | 0 | 0 |
| LYIEFVK | MLEC; MLEC; M | Q14165; H0' | MLEC_HUMAN | Unmodified | 0 | 0 | 0 | 0 | 0 | 0 | 65374 | 21785 | 6256200 | 1697700 | 632720 | 0 | 0 | 0 | 366390 | 0 | 0 | 0 |
| LYIFQASPA | HSPG2 | P98160 | PGBM_HUMAN | Unmodified | 0 | 0 | 1144900 | 0 | 0 | 0 | 0 | 0 | 0 | 0 | 0 | 0 | 0 | 0 | 255890 | 714100 | 0 | 0 |
| LYIGLAGLA | PSMB3; PSMB3 | P49720; A0A | PSB3_HUMAN | Unmodified | 0 | 0 | 54192 | 0 | 140510 | 0 | 0 | 0 | 45146 | 0 | 113990 | 1312700 | 41293 | 0 | 310330 | 93563 | 0 | 0 |
| LYIGNLSEN | IGF2BP3; IGF2E | O00425; F8V | IF2B3_HUMAN | Unmodified | 648550 | 2352000 | 0 | 317640 | 115440 | 178690 | 0 | 0 | 0 | 107670 | 0 | 183770 | 3375300 | 0 | 0 | 0 | 268110 | 0 |
| LYILQQAR | UTRN | P46939 | UTRN_HUMAN | Unmodified | 49111 | 0 | 1187300 | 501460 | 0 | 0 | 0 | 0 | 1043100 | 178080 | 0 | 0 | 0 | 0 | 1667500 | 1047100 | 43962 | 0 |
| LYILQQARF | UTRN | P46939 | UTRN_HUMAN | Unmodified | 0 | 0 | 0 | 0 | 0 | 515500 | 0 | 0 | 0 | 0 | 0 | 0 | 680530 | 0 | 0 | 0 | 729490 | 738760 |
| LYIYHVTG( | FAM91A1; FAM! | Q658Y4; E7E | F91A1_HUMAN | Unmodified | 0 | 0 | 0 | 0 | 0 | 0 | 246420 | 0 | 516080 | 483720 | 187780 | 3452900 | 0 | 0 | 326830 | 259160 | 0 | 0 |
| LYKDDQLL | ELOB; ELOB | Q15370; B82 | ELOB_HUMAN | Unmodified | 687940 | 356000 | 1341800 | 619320 | 395710 | 1560000 | 211630 | 37498 | 1793900 | 98780 | 1719500 | 7389700 | 406350 | 0 | 1323200 | 1254800 | 272050 | 226720 |
| LYKEELEQ1 | LMNB1; LMNB1 | P20700; A0A | LMNB1_HUMAI | Unmodified | 190350 | 210620 | 92997 | 348750 | 0 | 0 | 0 | 0 | 646920 | 37911 | 397190 | 2182800 | 0 | 0 | 0 | 245850 | 0 | 0 |
| LYKQEFDD | UTRN | P46939 | UTRN_HUMAN | Unmodified | 0 | 0 | 803500 | 485650 | 422350 | 0 | 0 | 0 | 594140 | 64379 | 0 | 228140 | 106560 | 405680 | 1347500 | 1120800 | 118670 | 221160 |
| LYKQEFDD | UTRN | P46939 | UTRN_HUMAN | Unmodified | 0 | 0 | 0 | 0 | 922190 | 0 | 0 | 0 | 0 | 382780 | 181500 | 358610 | 414870 | 359410 | 0 | 0 | 321020 | 0 |

| | | | | | | | | | | | | | | | | | | | | | | |
|---|---|---|---|---|---|---|---|---|---|---|---|---|---|---|---|---|---|---|---|---|---|---|
| LYKQLDDK | KIF5B | P33176 | KINH_HUMAN | Unmodified | 0 | 0 | 0 | 0 | 0 | 84144 | 655860 | 301750 | 0 | 34325 | 0 | 0 | 0 | 0 | 0 | 0 | 413940 | 0 |
| LYKYDYNS | STRAP; STRAP | Q9Y3F4; H0Y | STRAP_HUMAN | Unmodified | 465630 | 481920 | 0 | 1032200 | 643560 | 0 | 60088 | 0 | 775640 | 99337 | 1464900 | 5427700 | 732650 | 278330 | 535900 | 277920 | 0 | 0 |
| LYLADPMK | SRP9; SRP9; SR | P49458; Q65 | SRP09_HUMAN | Oxidation (M) | 1188300 | 1767300 | 1641900 | 2403600 | 1425500 | 133040 | 336320 | 260060 | 287750 | 219520 | 258110 | 2309200 | 466530 | 0 | 4246600 | 4554200 | 255780 | 535930 |
| LYLDGSEK | RANBP2 | P49792 | RBP2_HUMAN | Unmodified | 0 | 0 | 0 | 0 | 0 | 0 | 0 | 0 | 0 | 0 | 0 | 783740 | 0 | 0 | 0 | 0 | 0 | 0 |
| LYLDPGLRI | PLOD3; PLOD3 | O60568; H7 | PLOD3_HUMAN | Unmodified | 0 | 0 | 0 | 0 | 0 | 0 | 0 | 0 | 2461800 | 541790 | 0 | 247330 | 0 | 0 | 0 | 0 | 0 | 0 |
| LYLEQQVQ | SNX9; SNX9; SN | Q9Y5X1; A0 | SNX9_HUMAN | Unmodified | 0 | 0 | 0 | 0 | 0 | 0 | 0 | 0 | 0 | 0 | 0 | 0 | 0 | 0 | 0 | 0 | 0 | 0 |
| LYLEREEY | COPS2; COPS2 | P61201; B4D | CSN2_HUMAN | Unmodified | 0 | 0 | 0 | 0 | 0 | 0 | 0 | 0 | 719730 | 138400 | 336290 | 2865600 | 0 | 0 | 22508 | 0 | 73401 | 0 |
| LYLFQAQE | GPR180 | Q86V85 | GP180_HUMAN | Unmodified | 0 | 0 | 0 | 67353 | 216440 | 0 | 0 | 0 | 7099800 | 2169900 | 201890 | 0 | 0 | 0 | 0 | 0 | 0 | 0 |
| LYLINSPVV | IARS1; IARS1; I | P41252; A0A | SYIC_HUMAN I | Unmodified | 0 | 0 | 0 | 0 | 0 | 0 | 0 | 0 | 743820 | 0 | 0 | 16723000 | 0 | 0 | 0 | 0 | 0 | 0 |
| LYLLTEK | TPMT | P51580 | TPMT_HUMAN | Unmodified | 0 | 0 | 0 | 0 | 1252500 | 267940 | 0 | 0 | 0 | 153600 | 0 | 924340 | 0 | 0 | 0 | 0 | 1211100 | 0 |
| LYLNSNKLI | FLII; FLII; FLII | Q13045; J3K | FLII_HUMAN Pr | Unmodified | 46362 | 44944 | 165720 | 0 | 0 | 0 | 0 | 0 | 177620 | 0 | 102790 | 1589900 | 0 | 0 | 96309 | 130980 | 99813 | 50369 |
| LYLTNSK | AP3B2; AP3B1; | Q13367; O0 | AP3B2_HUMAN | Unmodified | 0 | 0 | 0 | 0 | 0 | 0 | 0 | 0 | 0 | 0 | 0 | 0 | 0 | 0 | 0 | 0 | 0 | 0 |
| LYLTSSTER | ADAM17; ADAM | P78536; A0A | ADA17_HUMAN | Unmodified | 0 | 0 | 0 | 139170 | 0 | 0 | 0 | 0 | 426880 | 314010 | 159690 | 0 | 0 | 0 | 0 | 0 | 0 | 578500 |
| LYLVDLAG | KIF5B; KIF5C; K | P33176; O6 | KINH_HUMAN | Unmodified | 362000 | 0 | 409810 | 0 | 5840200 | 1524600 | 1381200 | 917990 | 2620400 | 2058000 | 3495500 | 6490500 | 0 | 1492900 | 698650 | 491900 | 1786500 | 2324000 |
| LYNLFLK | SUCLG2; SUCL | Q96I99; E9P | SUCB2_HUMAI | Unmodified | 0 | 0 | 0 | 0 | 0 | 0 | 127130 | 0 | 3147300 | 916080 | 0 | 0 | 0 | 0 | 0 | 0 | 0 | 191990 |
| LYNLLLLR | USP15 | Q9Y4E8 | UBP15_HUMAN | Unmodified | 0 | 39136 | 41060 | 0 | 129320 | 0 | 0 | 0 | 250640 | 124860 | 548650 | 3931000 | 213280 | 0 | 0 | 71426 | 73746 | 308950 |
| LYNNHEIR | SYNCRIP; SYNC | O60506; B7 | HNRPQ_HUMA | Unmodified | 0 | 0 | 67538 | 37167 | 0 | 214990 | 138450 | 0 | 840800 | 59583 | 0 | 4265600 | 0 | 0 | 0 | 57710 | 26145 | 183970 |
| LYNPENIYL | TUBG1; TUBG2 | P23258; Q9N | TBG1_HUMAN | Unmodified | 0 | 0 | 0 | 0 | 1135700 | 0 | 0 | 0 | 0 | 0 | 519580 | 2331400 | 0 | 0 | 0 | 0 | 0 | 0 |
| LYNTCAVF | NIT2; NIT2 | Q9NQR4; H7 | NIT2_HUMAN ( | Unmodified | 0 | 0 | 0 | 0 | 0 | 0 | 0 | 0 | 0 | 0 | 0 | 0 | 0 | 0 | 0 | 0 | 0 | 0 |
| LYPAAVDTI | MCTS1 | Q9ULC4 | MCTS1_HUMAI | Unmodified | 0 | 0 | 0 | 0 | 388750 | 0 | 108650 | 0 | 336750 | 133820 | 52952 | 0 | 198400 | 0 | 403860 | 0 | 0 | 465260 |
| LYPEGLAQ | ATP6V0D1; ATP | P61421; R4 | VA0D1_HUMAN | Unmodified | 0 | 0 | 0 | 0 | 1071700 | 0 | 1765100 | 0 | 6508100 | 151490 | 1044300 | 6485600 | 0 | 0 | 1674700 | 923310 | 0 | 0 |
| LYPEGLAQ | ATP6V0D1; ATP | P61421; R4 | VA0D1_HUMAN | Unmodified | 0 | 0 | 0 | 0 | 0 | 0 | 0 | 0 | 3134200 | 578150 | 0 | 0 | 0 | 0 | 0 | 0 | 0 | 0 |
| LYPELSQY | SDCBP; SDCBP | O00560; G5 | SDCB1_HUMAI | Oxidation (M) | 1424600 | 0 | 0 | 0 | 0 | 0 | 84813 | 0 | 4827700 | 0 | 0 | 3393900 | 0 | 0 | 0 | 0 | 0 | 0 |
| LYPGGAEN | H6PD | O95479 | G6PE_HUMAN | Unmodified | 0 | 35498 | 0 | 596880 | 0 | 0 | 0 | 0 | 216800 | 0 | 0 | 0 | 0 | 0 | 0 | 0 | 650990 | 0 |
| LYPNMTAV | PNPT1 | Q8TCS8 | PNPT1_HUMAN | Unmodified | 0 | 0 | 0 | 0 | 0 | 0 | 188000 | 0 | 1397100 | 0 | 0 | 0 | 0 | 0 | 0 | 0 | 0 | 0 |
| LYPPLNTIR | ME1 | P48163 | MAOX_HUMAN | Unmodified | 0 | 0 | 14973 | 0 | 0 | 0 | 0 | 0 | 0 | 0 | 0 | 1345000 | 0 | 0 | 0 | 0 | 0 | 0 |
| LYPPSAEYF | CKMT1A; CKMT | P12532; C9J | KCRU_HUMAN | Unmodified | 0 | 0 | 0 | 0 | 0 | 0 | 0 | 0 | 0 | 0 | 0 | 2376800 | 0 | 0 | 0 | 0 | 0 | 0 |
| LYPQVCR | EDIL3 | O43854 | EDIL3_HUMAN | Unmodified | 427980 | 142040 | 0 | 0 | 5961500 | 3898000 | 0 | 0 | 8277000 | 1066000 | 75489 | 0 | 0 | 0 | 0 | 0 | 341290 | 1506400 |
| LYPTLVIR | ELP3; ELP3 | Q9H9T3; B4 | ELP3_HUMAN I | Unmodified | 0 | 0 | 0 | 0 | 0 | 0 | 0 | 0 | 0 | 0 | 0 | 1600300 | 0 | 0 | 0 | 0 | 0 | 0 |
| LYPTSCHT | MFGE8; MFGE8 | Q08431; H0 | MFGM_HUMAN | Unmodified | 739970 | 149110 | 1008500 | 1194600 | 65837 | 0 | 0 | 0 | 3653200 | 0 | 0 | 1618800 | 372400 | 0 | 1868000 | 1679700 | 55067 | 0 |
| LYPWGVVE | SEPTIN2; SEPTI | Q15019; H7 | SEPT2_HUMAN | Unmodified | 0 | 0 | 0 | 0 | 15256000 | 0 | 0 | 0 | 0 | 0 | 1102300 | 3742400 | 2608000 | 0 | 0 | 0 | 0 | 0 |
| LYQAGEGR | ANXA7 | P20073 | ANXA7_HUMAI | Unmodified | 826040 | 1759800 | 2752400 | 589760 | 0 | 0 | 0 | 0 | 67623000 | 1191300 | 0 | 0 | 39541 | 0 | 767080 | 2615600 | 0 | 0 |
| LYQAIQRA | COG4; COG4; ( | Q9H9E3; A0 | COG4_HUMAN | Unmodified | 52093 | 0 | 0 | 0 | 0 | 0 | 0 | 0 | 0 | 0 | 0 | 198710 | 0 | 0 | 64604 | 57242 | 0 | 0 |
| LYQASPAD | HSPG2 | P98160 | PGBM_HUMAN | Unmodified | 0 | 0 | 575180 | 379570 | 0 | 0 | 0 | 0 | 0 | 0 | 0 | 0 | 0 | 0 | 0 | 422980 | 0 | 0 |
| LYQDQLQK | TOR1B | O14657 | TOR1B_HUMAI | Unmodified | 0 | 0 | 0 | 0 | 0 | 0 | 0 | 0 | 827710 | 0 | 33516 | 0 | 0 | 0 | 0 | 0 | 0 | 0 |
| LYQECEK | HSPH1; HSPA4 | Q92598; O9 | HS105_HUMAI | Unmodified | 0 | 0 | 0 | 0 | 117200 | 0 | 0 | 0 | 159620 | 59864 | 0 | 2175800 | 0 | 0 | 0 | 0 | 88470 | 0 |
| LYQEVEIAS | UPF1 | Q92900 | RENT1_HUMAI | Unmodified | 1690100 | 0 | 937290 | 1923600 | 0 | 0 | 711780 | 0 | 483310 | 306920 | 0 | 646920 | 0 | 0 | 976660 | 1248500 | 0 | 0 |
| LYQGGPG | HSPA2 | P54652 | HSP72_HUMAI | Unmodified | 1046800 | 1068300 | 0 | 0 | 3030500 | 334560 | 0 | 0 | 0 | 0 | 563970 | 1174000 | 0 | 0 | 0 | 0 | 1857300 | 0 |
| LYQGINQL | MSH2; MSH2; N | P43246; A0A | MSH2_HUMAN | Unmodified | 8421.1 | 0 | 0 | 0 | 0 | 0 | 0 | 0 | 0 | 0 | 68362 | 151050 | 0 | 0 | 0 | 0 | 0 | 0 |
| LYQHAVEY | VPS4A; N/A | Q9UN37; I3L | VPS4A_HUMAN | Unmodified | 0 | 0 | 0 | 0 | 342300 | 0 | 0 | 0 | 0 | 0 | 0 | 2533700 | 0 | 0 | 0 | 0 | 0 | 0 |
| LYQHAVEY | VPS4A; N/A | Q9UN37; I3L | VPS4A_HUMAN | Unmodified | 1902400 | 1828100 | 0 | 191200 | 1547200 | 0 | 462060 | 44101 | 0 | 0 | 0 | 4256200 | 474400 | 1276000 | 570970 | 716040 | 0 | 0 |
| LYQHSQQF | LTBP1; LTBP1; l | Q14766; E7E | LTBP1_HUMAN | Unmodified | 0 | 0 | 0 | 0 | 0 | 0 | 0 | 0 | 0 | 0 | 0 | 0 | 0 | 0 | 0 | 0 | 77142 | 422700 |
| LYQPEYQE | HYOU1; HYOU1 | Q9Y4L1; A0A | HYOU1_HUMA | Unmodified | 0 | 0 | 2016900 | 94840 | 0 | 0 | 142880 | 0 | 66457000 | 10891000 | 9185700 | 5224900 | 0 | 151320 | 1384800 | 753560 | 0 | 1037700 |
| LYQSAGGN | HSPA8; HSPA8; | P11142; E9P | HSP7C_HUMAI | Oxidation (M) | 0 | 0 | 0 | 0 | 0 | 177380 | 3339700 | 849430 | 9064800 | 4138400 | 0 | 0 | 6232100 | 5897900 | 0 | 0 | 3530200 | 2864300 |
| LYQSATQA | RABGGTA; RAB | Q92696; H0 | PGTA_HUMAN | Unmodified | 0 | 0 | 0 | 0 | 0 | 0 | 0 | 0 | 0 | 0 | 0 | 1110500 | 242920 | 0 | 0 | 0 | 0 | 0 |
| LYQTDPSG | PSMA7; PSMA8 | O14818; Q8 | PSA7_HUMAN | Unmodified | 507810 | 0 | 0 | 2798900 | 2059800 | 0 | 225420 | 91227 | 2440100 | 733600 | 0 | 15529000 | 2002200 | 471700 | 1334100 | 1872400 | 527750 | 0 |
| LYQVEYAF | PSMA6; PSMA6 | P60900; G3 | PSA6_HUMAN | Unmodified | 0 | 0 | 0 | 0 | 0 | 0 | 286390 | 0 | 3828800 | 1695700 | 0 | 11338000 | 0 | 0 | 0 | 0 | 0 | 0 |
| LYQYPNFA | EIF2A; EIF2A; EI | Q9BY44; C9 | EIF2A_HUMAN | Unmodified | 236780 | 215540 | 607250 | 450830 | 0 | 154830 | 0 | 81476 | 335780 | 0 | 0 | 7915100 | 380730 | 129830 | 446400 | 357100 | 0 | 0 |
| LYRETNEP | XRCC6; XRCC6 | P12956; B1A | XRCC6_HUMAI | Unmodified | 658660 | 127850 | 358310 | 0 | 77022 | 0 | 0 | 0 | 763430 | 497610 | 1147000 | 7690100 | 612510 | 908330 | 201130 | 263150 | 66322 | 636200 |
| LYRGKSDE | MYOF; MYOF | Q9NZM1; H | MYOF_HUMAN | Unmodified | 0 | 0 | 0 | 0 | 9792800 | 2168000 | 0 | 52098 | 0 | 195690 | 0 | 0 | 0 | 0 | 0 | 0 | 0 | 347690 |
| LYRLDLSE | SLIT2; SLIT2; SL | O94813; X6 | SLIT2_HUMAN | Unmodified | 0 | 0 | 0 | 0 | 0 | 0 | 0 | 0 | 405480 | 195690 | 0 | 0 | 0 | 0 | 0 | 0 | 401410 | 1612800 |
| LYRPGSVA | ACLY | P53396 | ACLY_HUMAN | Unmodified | 428470 | 332180 | 817880 | 1225700 | 1964800 | 1553700 | 2555100 | 592120 | 2055100 | 0 | 196970 | 36882000 | 1876200 | 501130 | 1269700 | 1108700 | 406010 | 803070 |
| LYRPPPVL | GCN1 | Q92616 | GCN1_HUMAN | Unmodified | 45042 | 110110 | 277460 | 304580 | 0 | 0 | 0 | 85227 | 0 | 0 | 41352 | 2273100 | 0 | 0 | 220370 | 0 | 0 | 0 |
| LYRSIGEY | PRKDC | P78527 | PRKDC_HUMAI | Unmodified | 0 | 0 | 0 | 0 | 0 | 64490 | 2502900 | 0 | 0 | 0 | 0 | 0 | 169230 | 64777 | 0 | 0 | 47674 | 2127700 |
| LYRTTLSQ | MYO1B; MYO1E | O43795; H7 | MYO1B_HUMA | Unmodified | 0 | 0 | 0 | 0 | 18390000 | 3922000 | 4066900 | 3709100 | 0 | 154070 | 444230 | 0 | 1431100 | 1162300 | 0 | 0 | 7841100 | 5037000 |
| LYSEMEQL | SHCBP1 | Q8NEM2 | SHCBP_HUMA | Oxidation (M) | 0 | 0 | 0 | 0 | 344640 | 0 | 0 | 0 | 154090 | 0 | 0 | 0 | 0 | 0 | 0 | 0 | 0 | 115260 |
| LYSEMEQL | SHCBP1 | Q8NEM2 | SHCBP_HUMA | Unmodified | 100090 | 0 | 76351 | 0 | 2414900 | 0 | 74371 | 0 | 0 | 0 | 0 | 326870 | 0 | 0 | 0 | 0 | 0 | 0 |
| LYSESLAR | GDI1; GDI2; GD | P31150; P50 | GDIA_HUMAN | Unmodified | 3839500 | 3345600 | 6590100 | 8817800 | 5099400 | 2667200 | 1480400 | 401230 | 26276000 | 7352600 | 9336400 | 50196000 | 5597800 | 3820900 | 9023100 | 8235300 | 3001100 | 3307300 |
| LYSESLARY | GDI1; GDI2; GD | P31150; P50 | GDIA_HUMAN | Unmodified | 0 | 0 | 0 | 132540 | 7123500 | 3036700 | 3113300 | 1600300 | 237420 | 6498700 | 1087500 | 0 | 4226500 | 4249800 | 0 | 0 | 4389800 | 3691600 |
| LYSGSPTR | WDR59; WDR5 | Q6PJI9; H3B | WDR59_HUMA | Unmodified | 0 | 162850 | 182160 | 398400 | 0 | 0 | 0 | 0 | 0 | 0 | 521990 | 1426500 | 0 | 0 | 482610 | 412940 | 0 | 0 |
| LYSIAAPAR | ABHD12; ABHD | Q8N2K0; A0 | ABD12_HUMAN | Unmodified | 21782 | 31317 | 30213 | 0 | 0 | 0 | 56878 | 0 | 20985000 | 2628000 | 309580 | 238520 | 0 | 0 | 0 | 0 | 0 | 0 |
| LYSILGTTL | WDR1 | O75083 | WDR1_HUMAN | Unmodified | 0 | 0 | 2649100 | 0 | 0 | 0 | 0 | 0 | 467380 | 0 | 800330 | 6566300 | 0 | 0 | 2296200 | 0 | 0 | 0 |
| LYSILGTTL | WDR1 | O75083 | WDR1_HUMAN | Unmodified | 5539300 | 6546200 | 0 | 8280900 | 0 | 0 | 0 | 0 | 8813000 | 0 | 0 | 30531000 | 0 | 0 | 5756500 | 5364800 | 0 | 0 |
| LYSLFLK | SUCLA2; SUCL | Q9P2R7; A0 | SUCB1_HUMAI | Unmodified | 0 | 0 | 0 | 0 | 0 | 0 | 102700 | 33575 | 2593200 | 521770 | 0 | 0 | 0 | 0 | 0 | 0 | 134410 | 143520 |
| LYSLGNGR | CNP | P09543 | CN37_HUMAN | Unmodified | 0 | 0 | 517700 | 492730 | 0 | 0 | 444700 | 470230 | 956110 | 0 | 2143100 | 9659100 | 0 | 0 | 2007600 | 1346100 | 123710 | 0 |
| LYSLGNGR | CNP | P09543 | CN37_HUMAN | Oxidation (M) | 23790 | 0 | 130480 | 416130 | 1048900 | 157670 | 1018200 | 0 | 1198600 | 804680 | 2192000 | 749120 | 721750 | 344420 | 69554 | 78880 | 129610 | 224170 |
| LYSLGNGR | CNP | P09543 | CN37_HUMAN | Unmodified | 0 | 0 | 0 | 0 | 1563700 | 71090 | 0 | 0 | 409480 | 112790 | 329490 | 53464 | 134330 | 124390 | 0 | 0 | 66945 | 512520 |
| LYSLSVLYK | YKT6; YKT6; YK | O15498; A0 | YKT6_HUMAN | Unmodified | 0 | 0 | 0 | 0 | 0 | 0 | 304980 | 0 | 0 | 471960 | 613270 | 353430 | 0 | 0 | 0 | 0 | 0 | 0 |
| LYSNAYLN | ALDH1A1; ALDH | P00352; Q5 | AL1A1_HUMAN | Unmodified | 0 | 0 | 0 | 0 | 0 | 0 | 0 | 0 | 0 | 0 | 2363600 | 1.03E+08 | 0 | 0 | 0 | 0 | 0 | 0 |
| LYSNRAAC | STIP1 | P31948 | STIP1_HUMAN | Unmodified | 0 | 0 | 0 | 0 | 451190 | 89780 | 0 | 0 | 0 | 0 | 0 | 0 | 0 | 939410 | 0 | 0 | 1026300 | 474670 |
| LYSPSQIG | HSPA9; HSPA9; | P38646; H0 | GRP75_HUMAI | Oxidation (M) | 0 | 0 | 0 | 0 | 1487200 | 0 | 0 | 0 | 4873000 | 2610600 | 58148 | 273210 | 0 | 0 | 0 | 0 | 0 | 0 |
| LYSPSQIG | HSPA9; HSPA9; | P38646; H0 | GRP75_HUMAI | Unmodified | 0 | 0 | 0 | 0 | 0 | 0 | 0 | 0 | 9436500 | 713400 | 0 | 0 | 0 | 29020 | 0 | 0 | 0 | 0 |
| LYSQKAPN | TMEM260 | Q9NX78 | TM260_HUMAN | Unmodified | 0 | 0 | 0 | 0 | 0 | 0 | 0 | 0 | 1820600 | 230540 | 0 | 1835800 | 0 | 0 | 0 | 0 | 0 | 0 |
| LYSVDDRT | CANT1; CANT1 | Q8WVQ1; K7 | CANT1_HUMAI | Unmodified | 0 | 0 | 0 | 274000 | 0 | 0 | 0 | 0 | 40353000 | 6663200 | 824810 | 514690 | 0 | 0 | 130110 | 0 | 874250 | 715010 |
| LYSVSYLLK | FLNA; FLNA; FL | P21333; A0A | FLNA_HUMAN | Unmodified | 133060 | 1956200 | 2131600 | 6636100 | 16993000 | 4566600 | 8989900 | 3342900 | 1448100 | 1299900 | 758240 | 4288400 | 21735000 | 17558000 | 3881200 | 5209400 | 7764500 | 7006900 |
| LYTGKPNL | ERG28 | Q9UKR5 | ERG28_HUMAN | Unmodified | 0 | 0 | 0 | 0 | 0 | 0 | 0 | 0 | 6155600 | 556160 | 0 | 1089100 | 0 | 0 | 0 | 0 | 0 | 0 |
| LYTLQDK | FASN; FASN | P49327; A0A | FAS_HUMAN F | Unmodified | 125610 | 37055 | 0 | 3467200 | 1954800 | 536130 | 1237100 | 250880 | 237020 | 357790 | 2440500 | 35212000 | 1244900 | 0 | 4206200 | 0 | 1052200 | 633000 |
| LYTLQDKA | FASN; FASN | P49327; A0A | FAS_HUMAN F | Unmodified | 0 | 0 | 0 | 0 | 0 | 0 | 184390 | 0 | 0 | 77978 | 0 | 0 | 525790 | 0 | 0 | 0 | 156240 | 0 |
| LYTLVLTDP | PEBP1 | P30086 | PEBP1_HUMAN | Unmodified | 0 | 0 | 0 | 0 | 0 | 0 | 0 | 0 | 485330 | 0 | 0 | 2739900 | 0 | 0 | 0 | 0 | 0 | 0 |
| LYTLVLTDP | PEBP1 | P30086 | PEBP1_HUMAN | Unmodified | 0 | 185860 | 1685600 | 0 | 0 | 0 | 0 | 0 | 0 | 121470 | 0 | 0 | 0 | 0 | 2556700 | 837230 | 0 | 0 |
| LYTLVLTDP | PEBP1 | P30086 | PEBP1_HUMAN | Unmodified | 6550200 | 4425900 | 3730700 | 3561200 | 4418200 | 0 | 0 | 0 | 13164000 | 3232100 | 15558000 | 31660000 | 9254200 | 4161500 | 7520800 | 6151800 | 1984300 | 7533100 |
| LYTLVTYVP | RPL31; RPL31; | P62899; H7 | RL31_HUMAN ( | Unmodified | 1902700 | 1219500 | 614380 | 0 | 665600 | 191090 | 0 | 0 | 0 | 0 | 0 | 12784000 | 0 | 0 | 793990 | 756400 | 163950 | 311410 |
| LYTTMPVA | EIF3L; EIF3L; EI | Q9Y262; B0 | EIF3L_HUMAN | Oxidation (M) | 0 | 0 | 0 | 0 | 0 | 0 | 0 | 64576 | 0 | 0 | 64275 | 1114700 | 0 | 0 | 0 | 0 | 0 | 0 |
| LYTTMPVA | EIF3L; EIF3L; EI | Q9Y262; B0 | EIF3L_HUMAN | Unmodified | 0 | 0 | 0 | 0 | 0 | 0 | 0 | 22841 | 0 | 0 | 0 | 5686100 | 0 | 0 | 0 | 0 | 0 | 0 |
| LYTVEYLSN | BLMH; BLMH; B | Q13867; K7E | BLMH_HUMAN | Unmodified | 0 | 0 | 0 | 0 | 0 | 0 | 0 | 0 | 2581200 | 871240 | 0 | 589370 | 0 | 0 | 0 | 0 | 0 | 0 |
| LYTVTITTR | PTPRB; PTPRB; | P23467; F8V | PTPRB_HUMAN | Unmodified | 284330 | 0 | 0 | 0 | 0 | 0 | 0 | 0 | 0 | 99288 | 157160 | 0 | 0 | 0 | 0 | 0 | 0 | 0 |
| LYTWVEW | LARS1; LARS1; | Q9P2J5; A0A | SYLC_HUMAN | Unmodified | 0 | 0 | 0 | 0 | 0 | 0 | 0 | 0 | 0 | 0 | 0 | 6497400 | 0 | 0 | 0 | 0 | 0 | 0 |
| LYVANAGD | PPM1H | Q9ULR3 | PPM1H_HUMA | Unmodified | 42926 | 0 | 46222 | 0 | 0 | 0 | 0 | 214640 | 182450 | 92237 | 0 | 1685200 | 279930 | 0 | 0 | 0 | 262430 | 0 |
| LYVDQAAG | RET; RET; RET; | P07949; Q9 | RET_HUMAN P | Unmodified | 0 | 0 | 0 | 0 | 0 | 0 | 0 | 0 | 8653100 | 370060 | 130320 | 0 | 0 | 0 | 0 | 0 | 0 | 0 |
| LYVEAAFA | MME; MME; MM | P08473; A0A | NEP_HUMAN N | Unmodified | 0 | 0 | 7910800 | 17253000 | 0 | 0 | 0 | 0 | 7040700 | 0 | 0 | 500250 | 0 | 0 | 24345000 | 16663000 | 174380 | 423790 |
| LYVIGTGEF | NUDCD1; NUD | Q96RS6; A0 | NUDC1_HUMA | Unmodified | 0 | 0 | 0 | 0 | 0 | 0 | 0 | 0 | 0 | 0 | 0 | 3220900 | 0 | 0 | 0 | 0 | 0 | 0 |
| LYVIYKVPQ | SNRPA1; SNRP | P09661; H0 | RU2A_HUMAN | Unmodified | 0 | 0 | 0 | 0 | 0 | 0 | 0 | 0 | 0 | 0 | 0 | 3116200 | 0 | 0 | 0 | 0 | 0 | 0 |
| LYVPHSLW | ALDH4A1 | P30038 | AL4A1_HUMAN | Unmodified | 0 | 0 | 0 | 0 | 0 | 0 | 0 | 0 | 1023300 | 518590 | 0 | 0 | 0 | 0 | 0 | 0 | 0 | 0 |
| LYVRRVFIT | HSP90B1; HSP | P14625; A0A | ENPL_HUMAN | Oxidation (M) | 0 | 0 | 199100 | 0 | 0 | 0 | 1791500 | 0 | 0 | 1537900 | 0 | 0 | 0 | 0 | 0 | 0 | 0 | 0 |
| LYVSNLGI | PPAT; PPAT | Q06203; D6 | PUR1_HUMAN | Unmodified | 0 | 0 | 0 | 0 | 0 | 0 | 0 | 0 | 0 | 0 | 0 | 6706400 | 0 | 0 | 0 | 0 | 0 | 0 |
| LYVSSDLG | SORCS2; SORC | Q96PQ0; B5 | SORC2_HUMAI | Unmodified | 0 | 0 | 0 | 0 | 0 | 0 | 0 | 0 | 5708700 | 1790900 | 0 | 162320 | 0 | 0 | 0 | 0 | 0 | 0 |
| LYYEGLQD | HGS; HGS; HG | O14964; A0 | HGS_HUMAN H | Unmodified | 1256600 | 0 | 0 | 0 | 0 | 0 | 0 | 0 | 1459200 | 1672300 | 747580 | 1255300 | 0 | 0 | 67144 | 54395 | 0 | 0 |
| LYYYWER | APOBEC3B; AP | Q9UH17; Q8 | ABC3B_HUMAI | Unmodified | 0 | 347590 | 0 | 0 | 0 | 0 | 0 | 0 | 0 | 0 | 0 | 0 | 0 | 0 | 867940 | 453110 | 0 | 0 |
| MAAAGGG | HNRNPL | P14866 | HNRPL_HUMAI | Unmodified | 0 | 0 | 0 | 0 | 0 | 0 | 247690 | 1073900 | 0 | 506230 | 0 | 0 | 1134800 | 423880 | 0 | 0 | 0 | 0 |
| MAAERGAF | RRM1; RRM1 | P23921; E9P | RIR1_HUMAN F | Oxidation (M) | 0 | 0 | 0 | 0 | 0 | 0 | 874900 | 0 | 0 | 0 | 0 | 0 | 1556700 | 0 | 0 | 0 | 1076500 | 0 |
| MADCGGL | NDRG1; NDRG | Q92597; E7E | NDRG1_HUMA | Oxidation (M) | 0 | 0 | 0 | 0 | 4357500 | 1253000 | 0 | 0 | 1913400 | 2601100 | 0 | 647060 | 230790 | 161010 | 0 | 284550 | 9126700 | 7379500 |
| MADCGGL | NDRG1; NDRG | Q92597; E7E | NDRG1_HUMA | Unmodified | 0 | 0 | 0 | 0 | 4930500 | 1144400 | 37090 | 0 | 4622900 | 2149100 | 120300 | 1825500 | 746290 | 809860 | 0 | 527850 | 10420000 | 10648000 |
| MADCGGL | NDRG1; NDRG | Q92597; E7E | NDRG1_HUMA | Oxidation (M) | 0 | 0 | 0 | 0 | 912870 | 0 | 0 | 0 | 0 | 1111500 | 0 | 0 | 0 | 0 | 0 | 0 | 11495000 | 2754600 |
| MADCGGL | NDRG1; NDRG | Q92597; E7E | NDRG1_HUMA | Unmodified | 0 | 0 | 0 | 0 | 310990 | 0 | 0 | 0 | 0 | 268180 | 0 | 648730 | 0 | 0 | 0 | 0 | 3328800 | 3238200 |
| MADCGGL | NDRG3; NDRG | Q9UGV2; F8 | NDRG3_HUMA | Oxidation (M) | 0 | 0 | 0 | 0 | 0 | 0 | 0 | 0 | 0 | 0 | 512110 | 3791400 | 0 | 0 | 0 | 70412 | 0 | 0 |
| MADCGGL | NDRG3; NDRG | Q9UGV2; F8 | NDRG3_HUMA | Unmodified | 0 | 0 | 0 | 0 | 0 | 0 | 0 | 0 | 0 | 0 | 465760 | 4777100 | 0 | 0 | 0 | 0 | 0 | 0 |
| MADCGGL | NDRG3; NDRG | Q9UGV2; F8 | NDRG3_HUMA | Oxidation (M) | 65057 | 0 | 0 | 0 | 13322000 | 742140 | 868070 | 0 | 0 | 772590 | 3209800 | 12468000 | 9683500 | 10124000 | 0 | 113240 | 2878700 | 3791500 |
| MADCGGL | NDRG3; NDRG | Q9UGV2; F8 | NDRG3_HUMA | Unmodified | 0 | 0 | 0 | 0 | 2220800 | 0 | 0 | 0 | 0 | 0 | 31640 | 102910 | 1111500 | 0 | 0 | 0 | 0 | 1096000 |
| MADGAPLA | SV2A | Q7L0J3 | SV2A_HUMAN | Unmodified | 0 | 0 | 0 | 0 | 0 | 0 | 0 | 0 | 293180 | 0 | 0 | 3780400 | 0 | 89894 | 0 | 0 | 0 | 0 |

| | | | | | | | | | | | | | | | | | | | | | | |
|---|---|---|---|---|---|---|---|---|---|---|---|---|---|---|---|---|---|---|---|---|---|---|
| MADKAKPA | TPPP | O94811 | TPPP_HUMAN | Acetyl (Protein N | 0 | 0 | 0 | 2797000 | 0 | 0 | 0 | 0 | 0 | 0 | 0 | 0 | 0 | 0 | 0 | 0 | 0 | 0 |
| MADKDGD | CALU | O43852 | CALU_HUMAN | 2 Oxidation (M) | 0 | 0 | 0 | 0 | 0 | 0 | 0 | 0 | 5996500 | 3455700 | 1604500 | 0 | 0 | 0 | 0 | 0 | 0 | 558740 |
| MADKDGD | CALU | O43852 | CALU_HUMAN | Oxidation (M) | 0 | 0 | 0 | 0 | 0 | 0 | 0 | 0 | 11659000 | 0 | 0 | 0 | 0 | 0 | 0 | 0 | 0 | 0 |
| MADLHAVF | DPYSL3; DPYSL | Q14195; H0 | DPYL3_HUMAN | 2 Oxidation (M) | 0 | 0 | 0 | 0 | 0 | 0 | 985180 | 183010 | 0 | 0 | 496190 | 0 | 0 | 0 | 0 | 0 | 0 | 0 |
| MADPQSIQ | CYFIP1; CYFIP1 | Q7L576; A0A | CYFP1_HUMAN | 2 Oxidation (M) | 0 | 0 | 0 | 456830 | 0 | 0 | 0 | 0 | 0 | 0 | 0 | 1720100 | 0 | 0 | 253720 | 955000 | 0 | 0 |
| MADPQSIQ | CYFIP2; CYFIP2 | Q96F07; A0A | CYFP2_HUMAN | 2 Oxidation (M) | 0 | 0 | 0 | 0 | 0 | 0 | 0 | 0 | 0 | 0 | 0 | 1789300 | 0 | 0 | 1531300 | 433400 | 0 | 0 |
| MADQAPFD | FBP1 | P09467 | F16P1_HUMAN | Acetyl (Protein N | 0 | 0 | 0 | 0 | 0 | 0 | 27026000 | 918940 | 0 | 0 | 815490 | 492140 | 0 | 0 | 0 | 0 | 61908 | 0 |
| MADSGGLI | NDRG4; NDRG4 | Q9ULP0; B7Z | NDRG4_HUMA | Oxidation (M) | 0 | 0 | 0 | 0 | 0 | 0 | 651110 | 380770 | 0 | 170210 | 379570 | 1062600 | 0 | 0 | 0 | 0 | 0 | 0 |
| MADSGGQ | NDRG2; NDRG2 | Q9UN36; G3 | NDRG2_HUMA | Unmodified | 0 | 0 | 0 | 0 | 0 | 0 | 0 | 76110 | 0 | 9941.9 | 177270 | 983840 | 0 | 33419 | 0 | 131830 | 0 | 20181 |
| MADSGGQ | NDRG2; NDRG2 | Q9UN36; G3 | NDRG2_HUMA | Oxidation (M) | 91900 | 263200 | 0 | 925920 | 1589200 | 279290 | 367100 | 288650 | 0 | 67232 | 729590 | 1574400 | 6022800 | 1283600 | 602070 | 865190 | 0 | 132910 |
| MADSGGQ | NDRG2; NDRG2 | Q9UN36; G3 | NDRG2_HUMA | Unmodified | 0 | 0 | 0 | 0 | 0 | 0 | 179610 | 0 | 0 | 0 | 296190 | 0 | 2338400 | 678270 | 0 | 0 | 0 | 0 |
| MAEAASGA | ANO8; ANO8 | Q9HCE9; M0 | ANO8_HUMAN | Acetyl (Protein N | 0 | 11396000 | 0 | 0 | 41882000 | 0 | 0 | 1.83E+08 | 0 | 0 | 0 | 0 | 43551000 | 0 | 0 | 0 | 48049000 | 0 |
| MAEEKLTH | STMN1 | P16949 | STMN1_HUMA | Unmodified | 0 | 0 | 0 | 0 | 0 | 0 | 1647200 | 1053500 | 0 | 3004400 | 0 | 0 | 0 | 272440 | 0 | 0 | 0 | 0 |
| MAEGDELV | CDC42BPA; CD | Q5VT25; A0A | MRCKA_HUMA | Unmodified | 0 | 0 | 0 | 299260 | 247610 | 0 | 0 | 0 | 524790 | 181350 | 58188 | 0 | 0 | 0 | 0 | 0 | 359600 | 375540 |
| MAEGGVTS | ATXN3; ATXN3; | P54252; D3V | ATX3_HUMAN | Oxidation (M) | 112670 | 136440 | 207130 | 0 | 0 | 0 | 0 | 0 | 0 | 0 | 0 | 0 | 0 | 0 | 84783 | 130850 | 0 | 0 |
| MAEHVSCH | LPAR2 | Q9HBW0 | LPAR2_HUMAN | Unmodified | 0 | 0 | 0 | 98808 | 0 | 0 | 0 | 0 | 213920 | 0 | 15402 | 127980 | 0 | 0 | 288240 | 0 | 0 | 0 |
| MAEIGEHV | CAD; CAD | P27708; F8V | PYR1_HUMAN | Oxidation (M) | 0 | 0 | 0 | 0 | 0 | 0 | 0 | 0 | 177000 | 0 | 0 | 2898400 | 0 | 0 | 316370 | 251200 | 0 | 0 |
| MAEMGSK | BIN1 | O00499 | BIN1_HUMAN | Acetyl (Protein N | 0 | 769910 | 0 | 1310100 | 0 | 725480 | 0 | 0 | 695610 | 711550 | 4672700 | 0 | 0 | 0 | 372740 | 0 | 0 | 0 |
| MAEMRPPL | ERBIN | Q96RT1 | ERBIN_HUMAN | Oxidation (M) | 0 | 0 | 0 | 51156 | 4331100 | 0 | 0 | 0 | 0 | 0 | 191730 | 0 | 136490 | 0 | 103660 | 0 | 0 | 0 |
| MAEMRPPL | ERBIN | Q96RT1 | ERBIN_HUMAN | 2 Oxidation (M) | 371660 | 188850 | 0 | 0 | 1510700 | 0 | 0 | 0 | 0 | 0 | 0 | 0 | 0 | 0 | 0 | 97471 | 112240 | 0 |
| MAEMRPPL | ERBIN | Q96RT1 | ERBIN_HUMAN | Unmodified | 0 | 0 | 0 | 0 | 1233600 | 0 | 0 | 0 | 0 | 0 | 0 | 0 | 0 | 0 | 0 | 0 | 0 | 0 |
| MAENLGFV | SUCLG2; SUCL | Q96I99; E9P | SUCB2_HUMA | Oxidation (M) | 0 | 0 | 0 | 0 | 0 | 0 | 0 | 0 | 1019800 | 0 | 0 | 0 | 0 | 0 | 263090 | 0 | 0 | 200790 |
| MAENLGFV | SUCLG2; SUCL | Q96I99; E9P | SUCB2_HUMA | Unmodified | 0 | 0 | 0 | 0 | 0 | 0 | 0 | 0 | 2178200 | 0 | 0 | 0 | 0 | 0 | 0 | 0 | 0 | 0 |
| MAESLRIPE | FUT8; FUT8 | Q9BYC5; A0 | FUT8_HUMAN | Oxidation (M) | 0 | 0 | 0 | 0 | 862760 | 0 | 240740 | 0 | 11774000 | 1670700 | 0 | 0 | 0 | 460000 | 0 | 0 | 415640 | 1736100 |
| MAESLRIPE | FUT8; FUT8 | Q9BYC5; A0 | FUT8_HUMAN | Unmodified | 0 | 0 | 0 | 0 | 0 | 0 | 0 | 0 | 13413000 | 596390 | 0 | 93776 | 0 | 0 | 0 | 0 | 0 | 0 |
| MAESLYEIF | TM9SF1; TM9SF | O15321; E9F | TM9S1_HUMAN | Unmodified | 0 | 0 | 0 | 0 | 0 | 0 | 0 | 0 | 806780 | 0 | 0 | 132920 | 0 | 0 | 0 | 0 | 0 | 0 |
| MAESPCSP | SCRIB; SCRIB; | Q14160; A0A | SCRIB_HUMAN | Oxidation (M) | 0 | 0 | 0 | 0 | 0 | 0 | 0 | 0 | 344350 | 0 | 0 | 0 | 0 | 0 | 0 | 0 | 1718800 | 0 |
| MAEVGYDF | PDE2A; PDE2A; | O00408; C9J | PDE2A_HUMAN | Unmodified | 0 | 0 | 8475.4 | 0 | 0 | 0 | 0 | 0 | 0 | 0 | 0 | 1454000 | 0 | 0 | 89338 | 0 | 0 | 0 |
| MAGAMSTI | CHMP2B; CHM | Q9UQN3; A0 | CHM2B_HUMA | Unmodified | 1952500 | 779820 | 0 | 1605000 | 12162000 | 1635900 | 677350 | 722910 | 6101500 | 3308200 | 383850 | 3715900 | 10060000 | 7096700 | 1621800 | 699040 | 2115400 | 6291300 |
| MAGAMSTI | CHMP2B; CHM | Q9UQN3; A0 | CHM2B_HUMA | Oxidation (M) | 0 | 0 | 0 | 0 | 364710 | 0 | 0 | 0 | 0 | 128780 | 0 | 152280 | 266260 | 232770 | 0 | 484530 | 51698 | 119130 |
| MAGATPVF | KYAT3; KYAT3 | Q6YP21; A0A | KAT3_HUMAN | Oxidation (M) | 0 | 0 | 0 | 0 | 0 | 0 | 0 | 0 | 0 | 0 | 0 | 0 | 0 | 0 | 0 | 0 | 0 | 0 |
| MAGDPVAI | PPP2R1A; PPP2 | P30153; B3K | 2AAA_HUMAN | Oxidation (M) | 258010 | 0 | 313850 | 388930 | 0 | 0 | 0 | 63443 | 799330 | 0 | 802030 | 1392200 | 0 | 0 | 506650 | 537110 | 0 | 0 |
| MAGDPVAI | PPP2R1A; PPP2 | P30153; B3K | 2AAA_HUMAN | Oxidation (M) | 0 | 0 | 0 | 327150 | 2375800 | 885950 | 674080 | 691310 | 371970 | 897280 | 533700 | 0 | 4860500 | 2234500 | 0 | 0 | 2563700 | 1981300 |
| MAGDPVAI | PPP2R1A; PPP2 | P30153; B3K | 2AAA_HUMAN | Unmodified | 0 | 0 | 0 | 181470 | 7533000 | 917820 | 1645500 | 462180 | 1595100 | 1362100 | 334330 | 0 | 6685500 | 3596500 | 0 | 134110 | 3683600 | 4855100 |
| MAGEVSAA | CYP7B1 | O75881 | CP7B1_HUMAN | Acetyl (Protein N | 0 | 0 | 219710 | 0 | 0 | 852460 | 0 | 0 | 0 | 0 | 258640 | 0 | 0 | 0 | 0 | 0 | 0 | 0 |
| MAGQAFRI | HSPE1; HSPE1- | P61604; S4F | CH10_HUMAN | Acetyl (Protein N | 0 | 0 | 0 | 0 | 0 | 0 | 0 | 0 | 5396400 | 0 | 0 | 0 | 0 | 0 | 0 | 0 | 0 | 0 |
| MAGRAVLL | RUVBL1; RUVBI | Q9Y265; E7E | RUVB1_HUMA | Oxidation (M) | 0 | 0 | 0 | 0 | 354650 | 0 | 373360 | 316520 | 227940 | 0 | 0 | 0 | 0 | 0 | 0 | 0 | 101540 | 0 |
| MAGRAVLL | RUVBL1; RUVBI | Q9Y265; E7E | RUVB1_HUMA | Unmodified | 0 | 0 | 0 | 0 | 302260 | 0 | 175710 | 29215 | 0 | 30348 | 0 | 0 | 135700 | 105420 | 0 | 0 | 217890 | 195690 |
| MAHGYGEI | CHGB | P05060 | SCG1_HUMAN | Unmodified | 0 | 0 | 0 | 0 | 0 | 0 | 0 | 0 | 696370 | 149870 | 0 | 0 | 0 | 0 | 0 | 0 | 0 | 0 |
| MAHRDQE | CCDC50 | Q8IVM0 | CCD50_HUMA | Oxidation (M) | 2181400 | 3244900 | 4100700 | 3724600 | 0 | 0 | 0 | 0 | 176730 | 0 | 0 | 0 | 0 | 0 | 6249100 | 3454300 | 0 | 0 |
| MAHRDQE | CCDC50 | Q8IVM0 | CCD50_HUMA | Unmodified | 1024800 | 1259100 | 858510 | 1602100 | 0 | 0 | 0 | 0 | 424500 | 0 | 0 | 104740 | 385470 | 0 | 1857200 | 1218500 | 0 | 0 |
| MAHRDQE | CCDC50 | Q8IVM0 | CCD50_HUMA | Unmodified | 866670 | 646980 | 2168100 | 1207800 | 0 | 0 | 0 | 0 | 152270 | 0 | 0 | 0 | 0 | 0 | 1852500 | 891590 | 0 | 0 |
| MAHRDQE | CCDC50 | Q8IVM0 | CCD50_HUMA | Oxidation (M) | 0 | 0 | 3344500 | 1841200 | 0 | 0 | 0 | 0 | 0 | 0 | 0 | 0 | 0 | 0 | 2939400 | 2400500 | 98926 | 0 |
| MAIESAGFI | CAPN1 | P07384 | CAN1_HUMAN | Unmodified | 0 | 0 | 0 | 0 | 0 | 0 | 0 | 0 | 0 | 0 | 0 | 1088200 | 0 | 0 | 0 | 0 | 0 | 0 |
| MAIITTHLQ | BORCS7; BORC | Q96B45; A0A | BORC7_HUMA | Unmodified | 0 | 0 | 0 | 0 | 412530 | 0 | 0 | 0 | 0 | 67942 | 0 | 0 | 218460 | 388490 | 0 | 0 | 0 | 0 |
| MAILQIMK | TRIM28; TRIM28 | Q13263; M0 | TIF1B_HUMAN | Unmodified | 0 | 0 | 0 | 0 | 0 | 0 | 321800 | 46313 | 544370 | 199690 | 0 | 0 | 0 | 0 | 0 | 0 | 0 | 509680 |
| MAILQIMK | TRIM28; TRIM28 | Q13263; M0 | TIF1B_HUMAN | Oxidation (M) | 0 | 0 | 0 | 0 | 28095 | 0 | 171670 | 338460 | 0 | 212300 | 0 | 0 | 0 | 0 | 0 | 0 | 0 | 85406 |
| MAIQQADK | NAA25; NAA25 | Q14CX7; F8 | NAA25_HUMAN | Unmodified | 0 | 0 | 0 | 0 | 0 | 20171 | 30782 | 0 | 600150 | 53887 | 0 | 288710 | 0 | 0 | 0 | 0 | 0 | 415650 |
| MAIQTQQS | TOMM40; TOM | O96008; K7E | TOM40_HUMA | Unmodified | 0 | 0 | 0 | 300940 | 0 | 0 | 92585 | 91946 | 1612000 | 285370 | 0 | 0 | 398190 | 0 | 0 | 0 | 0 | 0 |
| MAITVSLVN | FUT8 | G3XAD2 | G3XAD2_HUM | Acetyl (Protein N | 0 | 0 | 0 | 0 | 0 | 0 | 0 | 0 | 0 | 0 | 0 | 0 | 0 | 0 | 0 | 0 | 0 | 0 |
| MAIVADHL | ANK3; ANK3; A | Q12955; H0 | ANK3_HUMAN | Oxidation (M) | 0 | 0 | 0 | 0 | 0 | 0 | 75858 | 0 | 0 | 0 | 0 | 0 | 0 | 0 | 1248000 | 546120 | 0 | 0 |
| MAKPEEVL | XPO1; XPO1; XF | O14980; A0A | XPO1_HUMAN | Oxidation (M) | 0 | 0 | 0 | 0 | 0 | 0 | 0 | 0 | 0 | 0 | 0 | 6793600 | 0 | 0 | 0 | 329030 | 0 | 0 |
| MAKPEEVL | XPO1; XPO1; XF | O14980; A0A | XPO1_HUMAN | Unmodified | 0 | 0 | 0 | 0 | 0 | 0 | 0 | 0 | 0 | 0 | 0 | 1130100 | 0 | 0 | 0 | 0 | 0 | 0 |
| MAKPLTDQ | PYGL; PYGL | P06737; E9P | PYGL_HUMAN | Acetyl (Protein N | 0 | 1687300 | 15542000 | 54231000 | 21218000 | 2.68E+08 | 12497000 | 31077000 | 4369400 | 6514800 | 1.13E+08 | 2586200 | 8198700 | 1.46E+08 | 15028000 | 1550100 | 1796500 | 2.28E+08 |
| MAKPLTDS | PYGB | P11216 | PYGB_HUMAN | Acetyl (Protein N | 0 | 0 | 0 | 0 | 137820 | 0 | 438300 | 346230 | 0 | 0 | 127440 | 0 | 0 | 0 | 0 | 0 | 0 | 446290 |
| MALDIEIAT | PRPH; VIM; VIM | P41219; P08 | PERI_HUMAN | Oxidation (M) | 0 | 0 | 0 | 0 | 1209700 | 0 | 452210 | 0 | 0 | 0 | 0 | 0 | 0 | 0 | 0 | 0 | 1091200 | 996620 |
| MALEFNRL | TNIP1; TNIP1; T | Q15025; A0A | TNIP1_HUMAN | Unmodified | 0 | 0 | 0 | 0 | 555940 | 0 | 29538 | 0 | 129620 | 150840 | 0 | 0 | 78642 | 0 | 0 | 0 | 318900 | 661600 |
| MALELLTQI | AARS1; AARS1; | P49588; A0A | SYAC_HUMAN | Oxidation (M) | 0 | 0 | 0 | 0 | 0 | 0 | 65341 | 0 | 309080 | 276710 | 111430 | 992460 | 104250 | 0 | 417060 | 60668 | 0 | 0 |
| MALEVGDY | CDH1 | P12830 | CADH1_HUMA | Unmodified | 469350 | 188180 | 278700 | 593370 | 482100 | 201300 | 0 | 16765 | 4054400 | 0 | 0 | 1043000 | 768840 | 873710 | 290090 | 304000 | 0 | 0 |
| MALEVGDY | CDH1 | P12830 | CADH1_HUMA | Oxidation (M) | 730640 | 0 | 0 | 0 | 0 | 0 | 0 | 0 | 728100 | 91342 | 0 | 628020 | 329940 | 304970 | 0 | 430420 | 0 | 0 |
| MALGVAAII | IQGAP3; IQGAP | Q86VI3; F2Z | IQGA3_HUMAN | Unmodified | 0 | 0 | 0 | 0 | 1178900 | 0 | 0 | 0 | 485200 | 0 | 0 | 0 | 0 | 0 | 0 | 0 | 0 | 0 |
| MALHLPWI | CDCP1 | Q9H5V8 | CDCP1_HUMA | Oxidation (M) | 0 | 0 | 643550 | 111020 | 0 | 0 | 0 | 0 | 0 | 0 | 0 | 1668000 | 0 | 0 | 364870 | 423930 | 0 | 0 |
| MALHLPWI | CDCP1 | Q9H5V8 | CDCP1_HUMA | Unmodified | 0 | 0 | 0 | 0 | 54447 | 0 | 0 | 0 | 0 | 0 | 0 | 837390 | 0 | 0 | 0 | 0 | 0 | 64839 |
| MALIGLGV | UQCRC2; UQC | P22695; H3E | QCR2_HUMAN | Oxidation (M) | 0 | 0 | 0 | 0 | 0 | 0 | 0 | 0 | 1833200 | 1368300 | 0 | 0 | 0 | 0 | 0 | 0 | 0 | 0 |
| MALIGLGV | UQCRC2; UQC | P22695; H3E | QCR2_HUMAN | Unmodified | 0 | 0 | 0 | 0 | 0 | 0 | 0 | 0 | 4273500 | 2057400 | 0 | 0 | 0 | 0 | 0 | 0 | 0 | 0 |
| MALKRMGI | UBA5; UBA5; U | Q9GZZ9; E7I | UBA5_HUMAN | Acetyl (Protein N | 0 | 0 | 0 | 0 | 0 | 0 | 0 | 0 | 0 | 0 | 0 | 0 | 0 | 0 | 0 | 0 | 0 | 0 |
| MALNHPYF | CDK1; CDK1 | P06493; A0A | CDK1_HUMAN | Unmodified | 363240 | 365480 | 638520 | 1014800 | 9145400 | 612550 | 1268000 | 106550 | 2437700 | 1182100 | 0 | 0 | 7784300 | 4114000 | 644530 | 485410 | 3835600 | 5971500 |
| MALNHPYF | CDK1; CDK1 | P06493; A0A | CDK1_HUMAN | Oxidation (M) | 257530 | 759800 | 1217900 | 1591700 | 3919500 | 1240400 | 0 | 443490 | 842330 | 903630 | 553030 | 502380 | 5951400 | 2900600 | 1760800 | 1189500 | 1755900 | 3209700 |
| MALPMPTK | DIP2B | Q9P265 | DIP2B_HUMAN | Oxidation (M) | 466130 | 46164 | 0 | 91074 | 313930 | 0 | 0 | 29403 | 0 | 100030 | 236770 | 483910 | 0 | 429130 | 45455 | 207220 | 483810 | 416270 |
| MALSIVDFF | EEPD1 | Q7L9B9 | EEPD1_HUMAN | Oxidation (M) | 0 | 0 | 50126 | 0 | 0 | 0 | 0 | 0 | 0 | 0 | 0 | 47682 | 0 | 0 | 280270 | 83494 | 0 | 0 |
| MALVLEALI | FLOT2; FLOT2; | Q14254; J3Q | FLOT2_HUMAN | Oxidation (M) | 0 | 0 | 2473400 | 0 | 11268000 | 0 | 4674900 | 1869900 | 6286100 | 10549000 | 4390000 | 4118500 | 0 | 2977400 | 3591100 | 1589600 | 0 | 0 |
| MALVLEALI | FLOT2; FLOT2; | Q14254; J3Q | FLOT2_HUMAN | Unmodified | 0 | 0 | 0 | 0 | 6058600 | 136030 | 510030 | 0 | 7690100 | 1491000 | 138600 | 0 | 0 | 0 | 0 | 0 | 287790 | 0 |
| MANASEPC | GPR27 | Q9NS67 | GPR27_HUMA | Unmodified | 358710 | 379190 | 318110 | 640550 | 1356600 | 219070 | 0 | 0 | 0 | 240190 | 0 | 273280 | 161650 | 0 | 1109000 | 741040 | 246010 | 0 |
| MANRDLLS | ANXA7 | P20073 | ANXA7_HUMA | Unmodified | 0 | 0 | 0 | 0 | 0 | 0 | 0 | 0 | 2746000 | 0 | 0 | 0 | 0 | 0 | 0 | 0 | 0 | 0 |
| MANRDLLS | ANXA7 | P20073 | ANXA7_HUMA | Oxidation (M) | 0 | 0 | 0 | 59956 | 1017400 | 822350 | 765660 | 0 | 8683700 | 22483000 | 0 | 0 | 849610 | 1866700 | 0 | 0 | 3680000 | 12912000 |
| MANRDLLS | ANXA7 | P20073 | ANXA7_HUMA | Unmodified | 0 | 0 | 0 | 249470 | 0 | 0 | 0 | 37257 | 2169900 | 2162600 | 0 | 0 | 0 | 0 | 0 | 0 | 0 | 0 |
| MAPEEDAG | PACSIN3; PACS | Q9UKS6; E9I | PACN3_HUMA | Acetyl (Protein N | 0 | 29383000 | 0 | 10470000 | 0 | 0 | 0 | 0 | 272390 | 0 | 0 | 0 | 12523000 | 3756800 | 0 | 0 | 1930700 | 0 |
| MAPRAPWI | HLA-B; HLA-B; | D9J307; A0A | D9J307_HUMA | Unmodified | 0 | 0 | 0 | 0 | 0 | 0 | 0 | 0 | 108600 | 0 | 0 | 0 | 0 | 0 | 0 | 0 | 0 | 8223300 |
| MAPRAPWI | HLA-B; HLA-B; | D9J307; A0A | D9J307_HUMA | Oxidation (M) | 0 | 0 | 0 | 0 | 0 | 0 | 0 | 0 | 0 | 0 | 1945700 | 0 | 0 | 0 | 0 | 0 | 0 | 11488000 |
| MAPSETHG | MYOF; MYOF | Q9NZM1; H0 | MYOF_HUMAN | Unmodified | 0 | 0 | 0 | 0 | 0 | 0 | 0 | 0 | 0 | 0 | 0 | 0 | 0 | 0 | 0 | 0 | 0 | 0 |
| MAPSETHG | MYOF; MYOF | Q9NZM1; H0 | MYOF_HUMAN | Oxidation (M) | 0 | 0 | 0 | 0 | 33350000 | 2434500 | 0 | 0 | 0 | 0 | 0 | 0 | 0 | 0 | 0 | 0 | 0 | 0 |
| MAPSETHG | MYOF; MYOF | Q9NZM1; H0 | MYOF_HUMAN | Unmodified | 0 | 0 | 0 | 0 | 8315300 | 0 | 0 | 0 | 0 | 0 | 0 | 0 | 0 | 0 | 0 | 0 | 0 | 1505300 |
| MAPSETHG | MYOF; MYOF | Q9NZM1; H0 | MYOF_HUMAN | Oxidation (M) | 0 | 0 | 0 | 0 | 2944400 | 591550 | 0 | 0 | 0 | 0 | 0 | 0 | 0 | 0 | 0 | 0 | 0 | 0 |
| MAPTPIPTF | DBN1; DBN1 | Q16643; D6I | DREB_HUMAN | Unmodified | 0 | 0 | 0 | 498210 | 0 | 0 | 0 | 0 | 1033100 | 0 | 0 | 3425700 | 0 | 0 | 0 | 125910 | 0 | 0 |
| MAPTPIPTF | DBN1; DBN1 | Q16643; D6I | DREB_HUMAN | Oxidation (M) | 0 | 0 | 646990 | 1054400 | 0 | 0 | 0 | 0 | 0 | 0 | 1097500 | 9710400 | 0 | 0 | 1020500 | 703020 | 0 | 0 |
| MAPYQGPI | ACTN4; ACTN4 | O43707; F5C | ACTN4_HUMA | Unmodified | 0 | 0 | 0 | 0 | 10102000 | 1777900 | 0 | 42556 | 3239600 | 1041700 | 0 | 0 | 2570000 | 0 | 0 | 0 | 2397800 | 2390300 |
| MAPYQGPI | ACTN4; ACTN4 | O43707; F5C | ACTN4_HUMA | Oxidation (M) | 403360 | 0 | 771320 | 382030 | 8076000 | 0 | 0 | 757190 | 1381600 | 1434000 | 1614700 | 6612900 | 4360600 | 3014700 | 618960 | 630090 | 566360 | 0 |
| MAQEFVNC | GLRX | P35754 | GLRX1_HUMAN | Acetyl (Protein N | 172180 | 73706 | 169970 | 517850 | 461560 | 0 | 0 | 0 | 930670 | 0 | 906490 | 4528400 | 0 | 0 | 195820 | 268590 | 337770 | 487330 |
| MAQELYME | EIF3B | P55884 | EIF3B_HUMAN | Oxidation (M) | 0 | 0 | 0 | 61609 | 0 | 0 | 0 | 0 | 0 | 136630 | 66323 | 473420 | 0 | 0 | 0 | 0 | 0 | 0 |
| MAQELYME | EIF3B | P55884 | EIF3B_HUMAN | Unmodified | 0 | 0 | 0 | 0 | 0 | 303160 | 0 | 0 | 0 | 0 | 0 | 1433600 | 0 | 0 | 0 | 0 | 0 | 0 |
| MAQERLDL | GJC1; GJC1; GJ | P36383; Q5I | CXG1_HUMAN | Unmodified | 0 | 0 | 0 | 0 | 10830000 | 1235000 | 0 | 0 | 0 | 0 | 0 | 0 | 0 | 0 | 0 | 0 | 0 | 0 |
| MAQFDAK | EEF1G | P26641 | EF1G_HUMAN | Oxidation (M) | 0 | 17787 | 154020 | 269240 | 652580 | 286250 | 112290 | 65159 | 118470 | 80586 | 140870 | 1769600 | 367940 | 231000 | 52825 | 69658 | 388610 | 478090 |
| MAQFDAK | EEF1G | P26641 | EF1G_HUMAN | Unmodified | 0 | 118720 | 385810 | 0 | 10264000 | 1049400 | 558160 | 619740 | 2635400 | 979110 | 0 | 23534000 | 2841000 | 0 | 394370 | 390730 | 2146900 | 5667300 |
| MAQFDAKI | EEF1G | P26641 | EF1G_HUMAN | Unmodified | 0 | 0 | 0 | 0 | 0 | 0 | 0 | 0 | 0 | 0 | 0 | 1423500 | 0 | 0 | 0 | 41710 | 0 | 0 |
| MAQTQGTI | HDAC1; HDAC1 | Q13547; Q5 | HDAC1_HUMA | Acetyl (Protein N | 0 | 0 | 0 | 0 | 0 | 0 | 0 | 0 | 0 | 0 | 0 | 0 | 0 | 0 | 0 | 0 | 0 | 0 |
| MAQWGNI | GSTM3 | P21266 | GSTM3_HUMA | Unmodified | 0 | 0 | 0 | 0 | 0 | 0 | 0 | 0 | 0 | 0 | 0 | 0 | 837330 | 0 | 0 | 0 | 147890 | 83482 |
| MARDPQRI | ERBB2; ERBB2; | P04626; J3Q | ERBB2_HUMAN | Oxidation (M) | 0 | 0 | 0 | 1391400 | 602080 | 0 | 0 | 0 | 0 | 0 | 1382600 | 0 | 0 | 0 | 0 | 0 | 175370 | 0 |
| MARVFSVLI | PPP3CA; PPP3 | Q08209; P16 | PP2BA_HUMAN | Oxidation (M) | 0 | 0 | 0 | 0 | 0 | 0 | 488120 | 0 | 0 | 0 | 0 | 0 | 355150 | 570330 | 0 | 0 | 879250 | 989810 |
| MASAPASY | SCFD1; SCFD1 | Q8WVM8; A | SCFD1_HUMA | Unmodified | 0 | 0 | 0 | 0 | 671970 | 0 | 1717300 | 0 | 0 | 53044 | 0 | 96647 | 106340 | 0 | 0 | 0 | 0 | 100750 |
| MASGAAN | FAM3C; FAM3C | Q92520; C9J | FAM3C_HUMA | Unmodified | 373440 | 0 | 218260 | 155160 | 524530 | 176530 | 167820 | 145530 | 38321000 | 6223400 | 2771800 | 2914300 | 494790 | 918910 | 0 | 0 | 0 | 1314800 |
| MASGAAN | FAM3C; FAM3C | Q92520; C9J | FAM3C_HUMA | Oxidation (M) | 0 | 0 | 0 | 0 | 164910 | 0 | 84394 | 0 | 8763600 | 1175300 | 445840 | 434860 | 164840 | 0 | 0 | 0 | 0 | 995250 |
| MASGAAN | FAM3C; FAM3C | Q92520; C9J | FAM3C_HUMA | 2 Oxidation (M) | 0 | 270980 | 0 | 0 | 0 | 0 | 42161 | 0 | 1081000 | 0 | 0 | 0 | 0 | 0 | 0 | 0 | 0 | 0 |
| MASIMEGP | OSBPL9; OSBPI | Q96SU4; G8 | OSBL9_HUMAN | Acetyl (Protein N | 0 | 0 | 582810 | 294460 | 0 | 587930 | 0 | 8883.6 | 0 | 104820 | 2171800 | 50560 | 0 | 69495 | 256870 | 0 | 0 | 0 |
| MASIYSEEC | STAMBP; STAM | O95630; C9J | STABP_HUMAN | Oxidation (M) | 2823900 | 3613100 | 0 | 0 | 0 | 0 | 0 | 0 | 1553600 | 671100 | 550370 | 347240 | 1803000 | 0 | 0 | 0 | 0 | 0 |
| MASNEDEF | CHAT | P28329 | CLAT_HUMAN | Oxidation (M) | 194210 | 0 | 0 | 404330 | 759170 | 0 | 213310 | 0 | 774840 | 0 | 0 | 642500 | 2125100 | 448820 | 0 | 123480 | 195670 | 222500 |
| MASNIFGTI | JPT1; JPT1; JPT1 | Q9UK76; J3K | JUPI1_HUMAN | Oxidation (M) | 0 | 216580 | 615340 | 803350 | 1320400 | 0 | 216710 | 260620 | 0 | 128750 | 385320 | 588840 | 1456400 | 1307500 | 878930 | 643230 | 762760 | 1059600 |
| MASNIFGTI | JPT1; JPT1; JPT1 | Q9UK76; J3K | JUPI1_HUMAN | Unmodified | 0 | 83307 | 0 | 223860 | 1635200 | 143360 | 105230 | 0 | 0 | 0 | 53296 | 164310 | 1227000 | 787910 | 155530 | 99011 | 1040800 | 1770300 |
| MASPATVH | MYOF | H0YD14 | H0YD14_HUM | Unmodified | 0 | 0 | 0 | 0 | 403340 | 0 | 0 | 0 | 0 | 0 | 0 | 0 | 0 | 0 | 0 | 0 | 0 | 0 |
| MASPVAAC | SUCLG2; SUCL | Q96I99; E9P | SUCB2_HUMA | Oxidation (M) | 0 | 0 | 0 | 0 | 1415600 | 475420 | 300100 | 0 | 0 | 0 | 0 | 361380 | 0 | 449490 | 0 | 128650 | 227990 | 195110 |

| | | | | | | | | | | | | | | | | | | | | | | |
|---|---|---|---|---|---|---|---|---|---|---|---|---|---|---|---|---|---|---|---|---|---|---|
| MASRVDAV | CHMP1A; CHM | Q9HD42; F8 | CHM1A_HUMA | Unmodified | 427040 | 0 | 0 | 0 | 2854300 | 264610 | 0 | 0 | 1773900 | 49782 | 17734 | 0 | 0 | 395720 | 0 | 0 | 0 | 1224500 |
| MASRVDAV | CHMP1A; CHM | Q9HD42; A0 | CHM1A_HUMA | Unmodified | 0 | 0 | 0 | 0 | 1930300 | 0 | 0 | 0 | 0 | 244160 | 0 | 0 | 663670 | 2612200 | 0 | 0 | 0 | 244200 |
| MASTFIGN | TUBB6 | Q9BUF5 | TBB6_HUMAN | Oxidation (M) | 20516 | 0 | 0 | 0 | 616030 | 0 | 0 | 0 | 539350 | 67715 | 0 | 25262000 | 321710 | 119430 | 0 | 27264 | 0 | 30329 |
| MASVGLSD | HSPG2 | P98160 | PGBM_HUMAN | Unmodified | 0 | 0 | 0 | 0 | 0 | 0 | 0 | 0 | 638850 | 0 | 0 | 0 | 0 | 0 | 0 | 0 | 0 | 905920 |
| MASVGLSD | HSPG2 | P98160 | PGBM_HUMAN | 2 Oxidation (M) | 0 | 0 | 3814500 | 0 | 0 | 0 | 0 | 0 | 0 | 0 | 0 | 0 | 0 | 0 | 225460 | 896010 | 0 | 0 |
| MASVGLSD | HSPG2 | P98160 | PGBM_HUMAN | Oxidation (M) | 0 | 0 | 1751900 | 573500 | 0 | 0 | 0 | 0 | 0 | 0 | 0 | 0 | 0 | 0 | 0 | 248530 | 0 | 0 |
| MASVRAVP | PABPC1; PABP | P11940; A0A | PABP1_HUMAN | 2 Oxidation (M) | 0 | 0 | 0 | 0 | 0 | 0 | 0 | 0 | 0 | 0 | 0 | 0 | 1760400 | 0 | 0 | 0 | 0 | 0 |
| MATDPENII | SYP; SYP | P08247; H7 | SYPH_HUMAN | Unmodified | 418900 | 393290 | 0 | 0 | 0 | 0 | 140240 | 0 | 26471000 | 2030000 | 0 | 733080 | 0 | 0 | 0 | 0 | 0 | 0 |
| MATDPENII | SYP; SYP | P08247; H7 | SYPH_HUMAN | Oxidation (M) | 1404000 | 437400 | 0 | 0 | 0 | 0 | 0 | 0 | 9085900 | 1341100 | 0 | 0 | 0 | 0 | 0 | 0 | 186970 | 0 |
| MATEVAAD | RPS6; RPS6; RP | P62753; A2A | RS6_HUMAN 4 | Oxidation (M) | 0 | 260240 | 0 | 0 | 0 | 9470400 | 880490 | 478660 | 0 | 377810 | 1051400 | 3993500 | 0 | 0 | 128770 | 265610 | 276740 | 457700 |
| MATFEIDEK | IDE; IDE; IDE; ID | P14735; A0A | IDE_HUMAN In | Oxidation (M) | 0 | 0 | 0 | 0 | 0 | 0 | 0 | 0 | 0 | 0 | 0 | 1296500 | 0 | 0 | 0 | 0 | 0 | 0 |
| MATGIEDSI | CUL4A; CUL4A | Q13619; A0A | CUL4A_HUMAN | Unmodified | 0 | 0 | 0 | 0 | 0 | 0 | 0 | 0 | 0 | 0 | 0 | 938610 | 0 | 0 | 0 | 246110 | 0 | 0 |
| MATNAAAC | TLN1 | Q9Y490 | TLN1_HUMAN | Unmodified | 0 | 0 | 0 | 0 | 1276400 | 233230 | 477480 | 265890 | 0 | 0 | 0 | 303680 | 80850 | 251510 | 0 | 0 | 152660 | 330880 |
| MATNAAAC | TLN1 | Q9Y490 | TLN1_HUMAN | Unmodified | 0 | 0 | 0 | 0 | 1220300 | 0 | 1768800 | 625570 | 0 | 0 | 0 | 1967200 | 862740 | 1814100 | 0 | 0 | 259830 | 391000 |
| MATQADLN | CTNNB1; CTNN | P35222; A0A | CTNB1_HUMAN | 2 Oxidation (M) | 0 | 0 | 0 | 0 | 0 | 0 | 0 | 0 | 0 | 0 | 0 | 23355000 | 0 | 0 | 0 | 0 | 0 | 0 |
| MATQASTL | NNT; NNT; NNT | Q13423; E9F | NNTM_HUMAN | Oxidation (M) | 0 | 0 | 0 | 0 | 0 | 0 | 0 | 0 | 448910 | 0 | 0 | 0 | 0 | 0 | 0 | 0 | 0 | 0 |
| MATQASTL | NNT; NNT; NNT | Q13423; E9F | NNTM_HUMAN | Unmodified | 0 | 0 | 0 | 0 | 0 | 0 | 0 | 0 | 2819600 | 296630 | 0 | 0 | 0 | 0 | 0 | 0 | 0 | 0 |
| MAVDQDW | MRPS35; MRPS | P82673; H0 | RT35_HUMAN | Oxidation (M) | 0 | 0 | 0 | 0 | 0 | 0 | 0 | 0 | 998720 | 0 | 0 | 0 | 0 | 0 | 0 | 0 | 0 | 0 |
| MAVLNEQV | NASP; NASP; N | P49321; E9P | NASP_HUMAN | Unmodified | 76700 | 0 | 79128 | 110110 | 3761200 | 148270 | 120710 | 534730 | 2472600 | 1482800 | 104640 | 2228700 | 1187900 | 540460 | 84952 | 211260 | 364980 | 3367400 |
| MAVLYPQV | SERPINI1; SERP | Q99574; H7 | NEUS_HUMAN | Oxidation (M) | 0 | 0 | 0 | 0 | 0 | 0 | 0 | 0 | 74407 | 293160 | 0 | 0 | 0 | 0 | 0 | 0 | 0 | 0 |
| MAVRNNL | CLTC; CLTC | Q00610; A0 | CLH1_HUMAN | Oxidation (M) | 0 | 0 | 0 | 2204600 | 434100 | 312990 | 0 | 20606 | 0 | 177930 | 0 | 0 | 839610 | 317610 | 0 | 0 | 198860 | 201820 |
| MAVRNNL | CLTC; CLTC | Q00610; A0 | CLH1_HUMAN | Unmodified | 0 | 0 | 0 | 388610 | 1302100 | 1097700 | 197880 | 24725 | 210910 | 218730 | 0 | 0 | 811590 | 254100 | 0 | 0 | 312710 | 612340 |
| MAVTFIGN | TUBB; TUBB; TU | P07437; Q5J | TBB5_HUMAN | Oxidation (M) | 0 | 0 | 0 | 0 | 2295400 | 673660 | 10718000 | 700010 | 354040 | 1120900 | 308930 | 5082600 | 2805600 | 4275900 | 43837 | 761120 | 1428300 | 2238200 |
| MAVTFIGN | TUBB; TUBB; TU | P07437; Q5J | TBB5_HUMAN | Oxidation (M) | 296030 | 445020 | 624630 | 783420 | 1975300 | 487660 | 2939700 | 140740 | 0 | 1369500 | 727880 | 28663000 | 3111300 | 1711600 | 2202200 | 1332900 | 1480300 | 1594600 |
| MAVTFIGN | TUBB; TUBB; TU | P07437; Q5J | TBB5_HUMAN | Unmodified | 0 | 0 | 0 | 0 | 548330 | 0 | 0 | 0 | 289330 | 113950 | 0 | 1098200 | 0 | 0 | 0 | 39577 | 0 | 0 |
| MCDLVSDF | ADSS2 | P30520 | PURA2_HUMAN | Oxidation (M) | 0 | 0 | 0 | 0 | 0 | 0 | 0 | 0 | 0 | 0 | 0 | 1327600 | 0 | 0 | 0 | 0 | 0 | 0 |
| MCDLVSDF | ADSS2 | P30520 | PURA2_HUMAN | Oxidation (M) | 0 | 0 | 0 | 0 | 0 | 0 | 0 | 0 | 0 | 0 | 0 | 2992800 | 0 | 0 | 0 | 0 | 0 | 0 |
| MCFNYEIR | MUC5B | Q9HC84 | MUC5B_HUMA | Unmodified | 0 | 0 | 0 | 0 | 0 | 0 | 0 | 0 | 0 | 0 | 0 | 0 | 0 | 0 | 0 | 0 | 0 | 8974900 |
| MCFNYEIR | MUC5B | Q9HC84 | MUC5B_HUMA | Oxidation (M) | 0 | 0 | 0 | 0 | 0 | 0 | 0 | 0 | 0 | 0 | 0 | 0 | 0 | 0 | 0 | 0 | 0 | 2583800 |
| MCKQDPS | ST13; ST13P5; | P50502; Q8 | F10A1_HUMAN | Unmodified | 0 | 0 | 0 | 0 | 168290 | 0 | 133430 | 0 | 0 | 0 | 0 | 0 | 767190 | 522390 | 0 | 0 | 0 | 304690 |
| MCLFAGFC | HNRNPU; HNR | Q00839; A0 | HNRPU_HUMA | Oxidation (M) | 0 | 0 | 0 | 0 | 0 | 0 | 85929 | 0 | 0 | 0 | 0 | 710800 | 0 | 0 | 0 | 0 | 0 | 0 |
| MCLFAGFC | HNRNPU; HNR | Q00839; A0 | HNRPU_HUMA | Unmodified | 0 | 0 | 0 | 0 | 0 | 0 | 2907400 | 376430 | 0 | 441380 | 229130 | 613960 | 926880 | 780520 | 0 | 0 | 0 | 792950 |
| MCLFAGFC | HNRNPU; HNR | Q00839; A0 | HNRPU_HUMA | Oxidation (M) | 0 | 0 | 0 | 0 | 0 | 0 | 1312700 | 551380 | 0 | 0 | 0 | 246200 | 374860 | 0 | 0 | 0 | 0 | 185040 |
| MCLNYEVF | MUC5AC | P98088 | MUC5A_HUMA | Unmodified | 0 | 0 | 0 | 0 | 0 | 0 | 0 | 0 | 0 | 90899 | 0 | 0 | 0 | 0 | 0 | 0 | 0 | 2337900 |
| MCNFHHQ | EXOC6; EXOC6 | Q8TAG9; E7 | EXOC6_HUMAN | Oxidation (M) | 0 | 0 | 0 | 0 | 413670 | 63331 | 0 | 0 | 0 | 0 | 0 | 0 | 0 | 288570 | 0 | 0 | 74219 | 79697 |
| MCNFHYQ | EXOC6B; EXOC | Q9Y2D4; A0 | EXC6B_HUMAN | Oxidation (M) | 0 | 0 | 0 | 65947 | 955160 | 73512 | 159700 | 34330 | 37688 | 148470 | 0 | 0 | 259860 | 195350 | 38882 | 0 | 397870 | 377700 |
| MCRHPVIS | SNX9; SNX9; SN | Q9Y5X1; A0 | SNX9_HUMAN | Oxidation (M) | 15971000 | 3767400 | 218150 | 800060 | 414330 | 92467 | 0 | 0 | 215670 | 248850 | 372550 | 1995700 | 1787800 | 0 | 2276200 | 1545400 | 0 | 0 |
| MCSKLPK | PSAP; PSAP; PS | P07602; A0A | SAP_HUMAN P | Unmodified | 0 | 0 | 699860 | 0 | 46918 | 0 | 0 | 0 | 6502800 | 5575900 | 147690 | 142300 | 0 | 1605000 | 0 | 0 | 302270 | 0 |
| MCTLIDKF | COPA; COPA; C | P53621; A0A | COPA_HUMAN | Oxidation (M) | 0 | 0 | 0 | 0 | 0 | 0 | 0 | 0 | 0 | 0 | 0 | 1254500 | 0 | 0 | 0 | 0 | 0 | 0 |
| MCVDVNE | FBLN1; FBLN1 | P23142; B1A | FBLN1_HUMAN | Unmodified | 0 | 0 | 0 | 0 | 0 | 0 | 19231 | 0 | 3091200 | 186700 | 0 | 0 | 0 | 0 | 0 | 0 | 0 | 0 |
| MDADDSR | PFKP | Q01813 | PFKAP_HUMAN | Acetyl (Protein N | 0 | 0 | 0 | 0 | 0 | 0 | 0 | 0 | 0 | 0 | 0 | 641340 | 0 | 0 | 0 | 0 | 82710 | 967600 |
| MDADINVT | FNBP1; FNBP1; | Q96RU3; H0 | FNBP1_HUMAN | Unmodified | 0 | 0 | 0 | 0 | 1537000 | 0 | 0 | 0 | 0 | 0 | 0 | 139330 | 382240 | 0 | 0 | 0 | 0 | 0 |
| MDAEQDPI | NUCB1; NUCB | Q02818; C9J | NUCB1_HUMA | Unmodified | 0 | 0 | 0 | 0 | 88312 | 0 | 0 | 0 | 2418400 | 198500 | 787430 | 0 | 0 | 44321 | 0 | 0 | 0 | 0 |
| MDAEQDPI | NUCB1; NUCB | Q02818; C9J | NUCB1_HUMA | Oxidation (M) | 0 | 0 | 0 | 0 | 0 | 0 | 0 | 0 | 1152800 | 0 | 1786000 | 0 | 0 | 0 | 0 | 0 | 0 | 0 |
| MDAGIICD | ITGB4 | P16144 | ITB4_HUMAN I | Unmodified | 0 | 0 | 0 | 0 | 0 | 0 | 0 | 0 | 124260 | 0 | 0 | 0 | 0 | 0 | 0 | 0 | 268190 | 619130 |
| MDAILTEAI | HP1BP3; HP1B | Q5SSJ5; B0 | HP1B3_HUMAN | Oxidation (M) | 0 | 0 | 0 | 0 | 0 | 0 | 490120 | 524650 | 0 | 281350 | 0 | 0 | 0 | 0 | 0 | 0 | 0 | 0 |
| MDAPGALA | RHOF; RHOF | Q9HBH0; F5 | RHOF_HUMAN | Acetyl (Protein N | 0 | 0 | 0 | 0 | 3605600 | 0 | 0 | 0 | 132960 | 0 | 0 | 0 | 0 | 0 | 0 | 0 | 1202400 | 1680600 |
| MDAPGALA | RHOF; RHOF | Q9HBH0; F5 | RHOF_HUMAN | Acetyl (Protein N | 0 | 0 | 0 | 0 | 219750 | 0 | 0 | 0 | 0 | 0 | 0 | 0 | 0 | 0 | 0 | 0 | 650870 | 0 |
| MDAPGALA | RHOF; RHOF | Q9HBH0; F5 | RHOF_HUMAN | Acetyl (Protein N | 0 | 0 | 0 | 0 | 1154400 | 0 | 0 | 0 | 0 | 0 | 0 | 0 | 0 | 0 | 0 | 0 | 1026400 | 697900 |
| MDAPRQV | PSAT1 | Q9Y617 | SERC_HUMAN | Acetyl (Protein N | 0 | 0 | 0 | 0 | 25211000 | 3467800 | 2978300 | 65552 | 0 | 3370800 | 0 | 0 | 33381000 | 20266000 | 0 | 0 | 6587500 | 3852400 |
| MDAPRQV | PSAT1 | Q9Y617 | SERC_HUMAN | Acetyl (Protein N | 0 | 0 | 0 | 0 | 999300 | 0 | 0 | 0 | 0 | 255910 | 0 | 0 | 3482100 | 2372500 | 0 | 0 | 0 | 0 |
| MDAPSSST | BMX | P51813 | BMX_HUMAN C | Unmodified | 0 | 0 | 0 | 0 | 0 | 0 | 0 | 0 | 0 | 0 | 844150 | 1662900 | 0 | 0 | 0 | 0 | 0 | 0 |
| MDASTGD | FERMT2; FERM | Q96AC1; H0 | FERM2_HUMAN | Unmodified | 0 | 80900 | 0 | 0 | 98097 | 0 | 0 | 0 | 80022 | 0 | 0 | 100560 | 20331 | 17349 | 86914 | 147060 | 0 | 0 |
| MDATAND | PDIA4; PDIA4 | P13667; A0A | PDIA4_HUMAN | Unmodified | 0 | 0 | 0 | 0 | 248110 | 3074700 | 0 | 109340 | 1146900 | 0 | 0 | 0 | 0 | 0 | 0 | 0 | 142000 | 0 |
| MDATAND | PDIA4; PDIA4 | P13667; A0A | PDIA4_HUMAN | Unmodified | 0 | 0 | 0 | 0 | 0 | 0 | 0 | 28987 | 11198000 | 560870 | 1441200 | 0 | 0 | 0 | 42241 | 0 | 0 | 0 |
| MDATAND | PDIA4; PDIA4 | P13667; A0A | PDIA4_HUMAN | Oxidation (M) | 409270 | 211990 | 0 | 1169900 | 2302800 | 730870 | 1834700 | 438900 | 91923000 | 25252000 | 19550000 | 2718000 | 3186900 | 1787300 | 2326600 | 594630 | 127270 | 1378600 |
| MDATAND | PDIA4; PDIA4 | P13667; A0A | PDIA4_HUMAN | Unmodified | 0 | 0 | 0 | 0 | 491600 | 0 | 468290 | 0 | 52175000 | 3806800 | 0 | 324370 | 0 | 325060 | 0 | 0 | 0 | 0 |
| MDATAND | PDIA3 | P30101 | PDIA3_HUMAN | Oxidation (M) | 0 | 0 | 295840 | 0 | 0 | 0 | 0 | 0 | 19146000 | 2263400 | 3756400 | 2460100 | 190080 | 0 | 826930 | 0 | 0 | 0 |
| MDATAND | PDIA3 | P30101 | PDIA3_HUMAN | Unmodified | 0 | 0 | 0 | 0 | 0 | 0 | 0 | 0 | 94788000 | 5865300 | 9955100 | 3398400 | 492590 | 0 | 598270 | 0 | 0 | 256090 |
| MDATAND | PDIA3 | P30101 | PDIA3_HUMAN | Oxidation (M) | 0 | 0 | 0 | 0 | 10501000 | 2055800 | 10796000 | 2262400 | 6115200 | 1.12E+08 | 6038500 | 0 | 2473000 | 12288000 | 0 | 0 | 2810200 | 10143000 |
| MDATAND | PDIA3 | P30101 | PDIA3_HUMAN | Unmodified | 0 | 8151700 | 0 | 0 | 1590800 | 0 | 0 | 0 | 2819800 | 24030000 | 0 | 0 | 0 | 0 | 0 | 0 | 0 | 0 |
| MDATAND | PDIA3 | P30101 | PDIA3_HUMAN | Oxidation (M) | 0 | 0 | 0 | 0 | 0 | 5805400 | 0 | 149140 | 489200 | 7308000 | 453250 | 0 | 0 | 0 | 0 | 0 | 0 | 533750 |
| MDATAND | PDIA3 | P30101 | PDIA3_HUMAN | Unmodified | 0 | 0 | 0 | 0 | 0 | 0 | 0 | 0 | 0 | 1440200 | 0 | 0 | 0 | 1200400 | 0 | 0 | 2468700 | 0 |
| MDATSYSS | TMX3; TMX3; TM | Q96JJ7; B4D | TMX3_HUMAN | Oxidation (M) | 0 | 0 | 0 | 0 | 0 | 0 | 0 | 0 | 1011100 | 0 | 0 | 0 | 0 | 0 | 0 | 0 | 0 | 0 |
| MDATSYSS | TMX3; TMX3; TM | Q96JJ7; B4D | TMX3_HUMAN | Unmodified | 0 | 0 | 0 | 0 | 0 | 0 | 0 | 0 | 2616100 | 0 | 0 | 0 | 0 | 0 | 0 | 0 | 0 | 0 |
| MDATSYSS | TMX3; TMX3; TM | Q96JJ7; B4D | TMX3_HUMAN | Unmodified | 0 | 0 | 0 | 0 | 0 | 0 | 0 | 0 | 1305900 | 0 | 0 | 0 | 0 | 0 | 0 | 0 | 0 | 0 |
| MDDASISR | PARP4 | Q9UKK3 | PARP4_HUMAN | Unmodified | 19724 | 0 | 0 | 0 | 0 | 0 | 0 | 0 | 0 | 0 | 0 | 371900 | 0 | 0 | 0 | 8799.9 | 0 | 0 |
| MDDKAFTK | PPP2CB; PPP2C | P62714; E5R | PP2AB_HUMAN | Acetyl (Protein N | 0 | 0 | 0 | 0 | 0 | 0 | 0 | 0 | 0 | 0 | 0 | 0 | 0 | 0 | 0 | 0 | 192670 | 0 |
| MDDPTVN | GSPT1; GSPT1 | P15170; H3E | ERF3A_HUMAN | Oxidation (M) | 181060 | 0 | 0 | 0 | 0 | 0 | 0 | 0 | 0 | 0 | 0 | 1846400 | 0 | 0 | 0 | 311150 | 0 | 0 |
| MDDPTVN | GSPT1; GSPT1 | P15170; H3E | ERF3A_HUMAN | Unmodified | 0 | 0 | 0 | 0 | 0 | 0 | 0 | 0 | 0 | 0 | 282280 | 2023100 | 0 | 0 | 0 | 0 | 0 | 0 |
| MDDREDL | YWHAE; YWHA | P62258; I3L | 1433E_HUMAN | Acetyl (Protein N | 1560600 | 1511200 | 7638800 | 17324000 | 23718000 | 10351000 | 7396900 | 7123800 | 7158300 | 6347500 | 12226000 | 50016000 | 35201000 | 27122000 | 9517100 | 16867000 | 19635000 | 9382200 |
| MDDREDL | YWHAE; YWHA | P62258; I3L | 1433E_HUMAN | Acetyl (Protein N | 1141600 | 1202800 | 3184600 | 8711000 | 30577000 | 7245400 | 6775300 | 2713400 | 15158000 | 10036000 | 10751000 | 36715000 | 21192000 | 6557000 | 6958900 | 4696200 | 23430000 | 10616000 |
| MDDSALQI | SNX18 | Q96RF0 | SNX18_HUMAN | Unmodified | 0 | 0 | 221200 | 0 | 0 | 0 | 0 | 0 | 0 | 0 | 0 | 0 | 0 | 0 | 276630 | 342500 | 0 | 0 |
| MDDSALQI | SNX18 | Q96RF0 | SNX18_HUMAN | Oxidation (M) | 0 | 0 | 682230 | 311450 | 0 | 0 | 0 | 0 | 0 | 0 | 0 | 0 | 0 | 0 | 0 | 1042900 | 0 | 0 |
| MDDSEVES | CTNND1; CTNN | O60716; A0A | CTND1_HUMA | Acetyl (Protein N | 0 | 0 | 0 | 0 | 0 | 0 | 0 | 0 | 0 | 0 | 0 | 0 | 0 | 0 | 0 | 0 | 5486400 | 9854100 |
| MDDTLFQL | CHMP1A; CHM | Q9HD42; F8 | CHM1A_HUMA | Acetyl (Protein N | 2098000 | 2610000 | 705360 | 1319800 | 4678000 | 0 | 1477200 | 1460000 | 1553800 | 1936000 | 762460 | 1322100 | 5175200 | 2631700 | 1125100 | 1006100 | 974640 | 2048400 |
| MDEDVLTT | RAB18; RAB18; | Q9NP72; A0 | RAB18_HUMAN | Acetyl (Protein N | 0 | 36883 | 0 | 0 | 0 | 0 | 0 | 414500 | 0 | 0 | 0 | 1711300 | 0 | 954180 | 0 | 0 | 0 | 0 |
| MDEKVFTK | PPP2CA; N/A | P67775; E5R | PP2AA_HUMAN | Acetyl (Protein N | 0 | 0 | 0 | 0 | 0 | 0 | 288810 | 303300 | 0 | 0 | 0 | 0 | 139630 | 0 | 0 | 0 | 570520 | 0 |
| MDELQDV | NDRG3; NDRG | Q9UGV2; Q5 | NDRG3_HUMA | Acetyl (Protein N | 18136 | 27390 | 0 | 0 | 2776700 | 0 | 0 | 0 | 0 | 0 | 0 | 2111800 | 0 | 0 | 0 | 0 | 0 | 0 |
| MDELQLFR | VCP; VCP; VCP; | P55072; A0A | TERA_HUMAN | Unmodified | 0 | 0 | 0 | 0 | 0 | 0 | 0 | 0 | 300380 | 0 | 0 | 1213100 | 0 | 0 | 0 | 0 | 0 | 0 |
| MDELQLFR | VCP; VCP; VCP; | P55072; A0A | TERA_HUMAN | Oxidation (M) | 0 | 0 | 0 | 314300 | 1075400 | 190420 | 102040 | 67352 | 577990 | 258290 | 395040 | 1649100 | 1218600 | 285150 | 0 | 0 | 0 | 0 |
| MDENQFV | CRMP1; DPYSL | Q14194; Q1 | DPYL1_HUMAN | Unmodified | 0 | 0 | 0 | 0 | 0 | 0 | 0 | 0 | 962650 | 0 | 3181600 | 10016000 | 0 | 0 | 0 | 0 | 0 | 0 |
| MDENQFV | CRMP1; DPYSL | Q14194; Q1 | DPYL1_HUMAN | Oxidation (M) | 0 | 0 | 0 | 0 | 0 | 0 | 0 | 0 | 518410 | 0 | 6009800 | 14785000 | 0 | 0 | 0 | 113250 | 0 | 0 |
| MDENRFV | DPYSL5 | Q9BPU6 | DPYL5_HUMAN | Unmodified | 0 | 0 | 0 | 0 | 0 | 0 | 0 | 0 | 337760 | 0 | 500270 | 2413300 | 0 | 0 | 0 | 0 | 107390 | 0 |
| MDERPIHA | CDH2; CDH2; C | P19022; C9J | CADH2_HUMA | Unmodified | 0 | 0 | 0 | 0 | 445190 | 55119 | 0 | 0 | 0 | 0 | 280300 | 682090 | 0 | 0 | 0 | 0 | 571070 | 1156100 |
| MDERPIHA | CDH2; CDH2 | P19022; C9J | CADH2_HUMA | Oxidation (M) | 0 | 0 | 0 | 0 | 0 | 0 | 0 | 0 | 0 | 0 | 0 | 0 | 0 | 0 | 0 | 0 | 2005400 | 0 |
| MDERPIHA | CDH2; CDH2 | P19022; C9J | CADH2_HUMA | Unmodified | 0 | 0 | 0 | 0 | 0 | 0 | 0 | 0 | 0 | 0 | 0 | 0 | 0 | 0 | 0 | 0 | 1058100 | 2244200 |
| MDEVYKVII | NT5E; NT5E | P21589; H0 | 5NTD_HUMAN | Oxidation (M) | 0 | 0 | 0 | 0 | 0 | 0 | 0 | 0 | 0 | 0 | 0 | 0 | 3321500 | 0 | 0 | 0 | 0 | 0 |
| MDFILSHA | UGGT1 | Q9NYU2 | UGGG1_HUMA | Unmodified | 0 | 0 | 0 | 0 | 0 | 0 | 0 | 0 | 1345500 | 0 | 0 | 0 | 0 | 0 | 0 | 0 | 0 | 0 |
| MDFLLGNF | TOM1; TOM1; T | O60784; F8V | TOM1_HUMAN | Acetyl (Protein N | 43281 | 88180 | 46855 | 22307 | 0 | 0 | 0 | 0 | 38219 | 0 | 0 | 50622 | 0 | 0 | 281740 | 130900 | 23891 | 0 |
| MDFLLGNF | TOM1; TOM1; T | O60784; F8V | TOM1_HUMAN | Acetyl (Protein N | 154100 | 713390 | 0 | 65658 | 0 | 0 | 929320 | 0 | 198870 | 222340 | 0 | 0 | 20603 | 461700 | 0 | 0 | 237750 | 513610 |
| MDFLLGNF | TOM1; TOM1; T | O60784; F8V | TOM1_HUMAN | Acetyl (Protein N | 843740 | 1219200 | 235680 | 1553700 | 1067800 | 286960 | 0 | 0 | 0 | 767710 | 0 | 148880 | 6170300 | 2986700 | 697120 | 478580 | 708640 | 909150 |
| MDFSKLPK | ATP6V1A; ATP6 | P38606; F8V | VATA_HUMAN | Acetyl (Protein N | 0 | 0 | 0 | 0 | 1785100 | 0 | 881670 | 339980 | 0 | 0 | 0 | 0 | 0 | 2198100 | 0 | 0 | 1006600 | 0 |
| MDFTAAR | DNAJC3; DNAJ | Q13217; X6F | DNJC3_HUMAN | Oxidation (M) | 0 | 0 | 0 | 0 | 0 | 0 | 0 | 0 | 493100 | 147200 | 0 | 468630 | 0 | 0 | 0 | 0 | 0 | 0 |
| MDFTAAR | DNAJC3; DNAJ | Q13217; X6F | DNJC3_HUMAN | Unmodified | 0 | 0 | 0 | 0 | 0 | 0 | 34075 | 13541 | 3946300 | 1228800 | 454660 | 0 | 0 | 0 | 57640 | 0 | 0 | 0 |
| MDGPAIIT | CTDSPL; CTDS | O15194; F8V | CTDSL_HUMAN | Acetyl (Protein N | 1229900 | 1004800 | 0 | 0 | 600060 | 0 | 2717200 | 438540 | 0 | 346250 | 291160 | 0 | 3279000 | 0 | 0 | 208660 | 0 | 0 |
| MDGSHSA | NIPAL3; NIPAL3 | Q6P499; A6 | NPAL3_HUMAN | Acetyl (Protein N | 0 | 0 | 0 | 0 | 0 | 0 | 0 | 0 | 620430 | 246630 | 137100 | 0 | 0 | 0 | 0 | 0 | 0 | 0 |
| MDGTEGS | FRMD8; FRMD8 | Q9BZ67; Q8 | FRMD8_HUMA | Acetyl (Protein N | 0 | 0 | 0 | 0 | 313740 | 84100 | 0 | 0 | 0 | 0 | 0 | 0 | 0 | 0 | 0 | 0 | 617070 | 273940 |
| MDHYDSQ | ACTN1; ACTN1 | P12814; A0A | ACTN1_HUMA | Acetyl (Protein N | 0 | 0 | 0 | 0 | 3125600 | 0 | 0 | 0 | 0 | 0 | 0 | 0 | 0 | 0 | 0 | 0 | 1950300 | 0 |
| MDKNELV | YWHAZ; YWHA | P63104; E5R | 1433Z_HUMAN | Acetyl (Protein N | 2811100 | 2680600 | 611550 | 7401400 | 55813000 | 8286200 | 29106000 | 31474000 | 10530000 | 19906000 | 4116100 | 10140000 | 17992000 | 22249000 | 3180900 | 4379200 | 1.3E+08 | 1.09E+08 |
| MDKNELV | YWHAZ; YWHA | P63104; E5R | 1433Z_HUMAN | Acetyl (Protein N | 2659300 | 3292500 | 1324900 | 13142000 | 1.93E+08 | 22592000 | 80740000 | 60737000 | 60726000 | 70024000 | 16349000 | 43036000 | 47348000 | 67631000 | 3925200 | 5811100 | 3.81E+08 | 2.72E+08 |
| MDKNELV | YWHAZ; YWHA | P63104; E5R | 1433Z_HUMAN | Acetyl (Protein N | 0 | 0 | 0 | 26167 | 2831000 | 311710 | 4360100 | 2546100 | 0 | 560320 | 0 | 0 | 1028800 | 1270200 | 0 | 0 | 18058000 | 3090000 |
| MDKNELV | YWHAZ; YWHA | P63104; E5R | 1433Z_HUMAN | Acetyl (Protein N | 0 | 0 | 0 | 0 | 12590000 | 0 | 16911000 | 11803000 | 290630 | 2382000 | 139210 | 126400 | 4256200 | 2679200 | 0 | 0 | 84225000 | 20753000 |
| MDLAAAAE | MCM6 | Q14566 | MCM6_HUMAN | Acetyl (Protein N | 0 | 0 | 0 | 0 | 0 | 0 | 333100 | 368920 | 0 | 0 | 219080 | 704470 | 0 | 0 | 0 | 0 | 277530 | 0 |
| MDLEKPNY | DYNC1H1; DYN | Q14204; A0 | DYHC1_HUMA | 2 Oxidation (M) | 0 | 0 | 0 | 0 | 0 | 0 | 0 | 0 | 0 | 0 | 0 | 1504700 | 0 | 0 | 0 | 0 | 0 | 0 |
| MDLFSALIK | UBE2O; UBE2O | Q9C0C9; K7 | UBE2O_HUMA | Oxidation (M) | 0 | 0 | 0 | 0 | 0 | 0 | 0 | 0 | 0 | 0 | 0 | 778830 | 0 | 0 | 18721 | 16848 | 0 | 0 |
| MDLPVGPC | HSF1; HSF1; HS | Q00613; A0 | HSF1_HUMAN | Acetyl (Protein N | 0 | 0 | 0 | 0 | 0 | 0 | 310730 | 296260 | 0 | 0 | 0 | 0 | 0 | 0 | 0 | 0 | 0 | 1128800 |
| MDLSRHG | FBLN1 | B1AHL2 | B1AHL2_HUMA | Unmodified | 0 | 0 | 0 | 0 | 0 | 0 | 0 | 0 | 0 | 362440 | 0 | 0 | 0 | 0 | 0 | 0 | 0 | 0 |
| MDLTAKEL | MDH1; MDH1; I | P40925; B8Z | MDHC_HUMAN | Unmodified | 0 | 0 | 0 | 0 | 926530 | 0 | 196200 | 84446 | 0 | 254270 | 0 | 0 | 356690 | 497180 | 0 | 0 | 1383000 | 0 |

| | | | | | | | | | | | | | | | | | | | | | | |
|---|---|---|---|---|---|---|---|---|---|---|---|---|---|---|---|---|---|---|---|---|---|---|
| MDLTAKEL | MDH1; MDH1; I | P40925; B8Z | MDHC_HUMAN | Oxidation (M) | 508690 | 0 | 0 | 0 | 2699700 | 0 | 0 | 0 | 0 | 0 | 0 | 0 | 0 | 0 | 0 | 0 | 7956600 | 0 |
| MDLVTYFG | CNP | P09543 | CN37_HUMAN | Unmodified | 0 | 0 | 835000 | 1081600 | 680040 | 0 | 0 | 14752 | 3007300 | 0 | 5120200 | 11955000 | 0 | 0 | 964940 | 79154 | 0 | 2419000 |
| MDLVTYFG | CNP | P09543 | CN37_HUMAN | Oxidation (M) | 91899 | 0 | 2133900 | 1601700 | 0 | 0 | 0 | 0 | 0 | 0 | 3242900 | 18746000 | 0 | 0 | 643820 | 2278300 | 0 | 0 |
| MDLVTYFG | CNP | P09543 | CN37_HUMAN | Unmodified | 0 | 0 | 0 | 865240 | 6785000 | 1110900 | 268260 | 0 | 1836700 | 783010 | 2712300 | 3076900 | 3418000 | 1469900 | 0 | 0 | 1120500 | 5772900 |
| MDNLSDTL | RAP1GDS1; RAI | P52306; D6F | GDS1_HUMAN | Oxidation (M) | 3609600 | 947200 | 0 | 399600 | 0 | 0 | 0 | 0 | 2511800 | 82680 | 0 | 253670 | 266110 | 773370 | 480070 | 761670 | 296680 | 0 |
| MDNLSSEE | SNAP23; SNAP2 | O00161; H3I | SNP23_HUMAN | Acetyl (Protein N | 0 | 0 | 0 | 0 | 404370 | 134820 | 0 | 0 | 0 | 0 | 0 | 0 | 380620 | 0 | 0 | 0 | 230600 | 546690 |
| MDNLSSEE | SNAP23; SNAP2 | O00161; H3I | SNP23_HUMAN | Acetyl (Protein N | 0 | 0 | 0 | 0 | 6798600 | 0 | 0 | 0 | 0 | 0 | 0 | 0 | 4044000 | 1684500 | 0 | 0 | 0 | 0 |
| MDNSGKE/ | NAPA; NAPA; N. | P54920; M0( | SNAA_HUMAN | Acetyl (Protein N | 257170 | 0 | 0 | 0 | 1101400 | 0 | 2446000 | 0 | 654890 | 0 | 0 | 0 | 55964 | 0 | 0 | 0 | 953610 | 0 |
| MDPASSGE | GPRIN1 | Q7Z2K8 | GRIN1_HUMAN | Unmodified | 0 | 0 | 0 | 0 | 0 | 0 | 0 | 0 | 1745800 | 208780 | 0 | 0 | 0 | 0 | 0 | 0 | 437240 | 0 |
| MDPNCSC. | MT2A; MT2A | P02795; H3E | MT2_HUMAN M | Acetyl (Protein N | 0 | 0 | 0 | 0 | 0 | 0 | 0 | 0 | 0 | 0 | 0 | 0 | 0 | 0 | 0 | 0 | 2065400 | 0 |
| MDPNCSC. | MT2A; MT2A | P02795; H3E | MT2_HUMAN M | Acetyl (Protein N | 0 | 0 | 0 | 0 | 0 | 0 | 0 | 0 | 0 | 0 | 0 | 0 | 0 | 0 | 0 | 0 | 7556400 | 0 |
| MDPNCSC. | MT2A; MT2A | P02795; H3E | MT2_HUMAN M | Acetyl (Protein N | 0 | 0 | 0 | 0 | 0 | 0 | 0 | 0 | 0 | 0 | 0 | 0 | 0 | 0 | 0 | 0 | 480420 | 0 |
| MDPNCSC | MT1X; MT1X | P80297; U3K | MT1X_HUMAN | Acetyl (Protein N | 0 | 0 | 0 | 0 | 0 | 0 | 0 | 0 | 0 | 0 | 0 | 0 | 0 | 0 | 0 | 0 | 225530 | 0 |
| MDPVNNIN | LRBA; LRBA; LR | P50851; A0A | LRBA_HUMAN | Unmodified | 0 | 668010 | 0 | 131990 | 0 | 0 | 0 | 0 | 0 | 0 | 0 | 1146900 | 0 | 0 | 0 | 0 | 0 | 0 |
| MDPVPGTD | MVB12A; MVB1 | Q96EY5; E9F | MB12A_HUMAI | Acetyl (Protein N | 3769300 | 4637200 | 697800 | 1628500 | 0 | 0 | 0 | 0 | 0 | 0 | 76834 | 164470 | 0 | 0 | 6347000 | 2915200 | 585340 | 593750 |
| MDQARSAI | TFRC; TFRC | P02786; F8V | TFR1_HUMAN | Acetyl (Protein N | 215750 | 0 | 0 | 0 | 0 | 281330 | 536970 | 0 | 0 | 0 | 0 | 0 | 796080 | 0 | 0 | 0 | 192890 | 662210 |
| MDQLEGQI | NPTXR; NPTXR | O95502; A0A | NPTXR_HUMAN | Oxidation (M) | 58209 | 133980 | 0 | 0 | 0 | 0 | 0 | 0 | 0 | 0 | 0 | 0 | 0 | 0 | 0 | 0 | 0 | 0 |
| MDRGEQG | TPD52 | E5RFR7 | E5RFR7_HUMA | Acetyl (Protein N | 0 | 0 | 0 | 0 | 0 | 0 | 0 | 0 | 1245200 | 195570 | 475310 | 1685000 | 0 | 0 | 0 | 0 | 0 | 0 |
| MDSAGQD | TPD52L2; TPD5 | O43399; A0A | TPD54_HUMAN | Acetyl (Protein N | 0 | 0 | 0 | 0 | 458220 | 0 | 35512 | 58952 | 0 | 53413 | 130330 | 0 | 65816 | 0 | 54781 | 59030 | 173030 | 189740 |
| MDSGTEEY | NINJ1 | Q92982 | NINJ1_HUMAN | Acetyl (Protein N | 0 | 0 | 0 | 0 | 0 | 0 | 0 | 0 | 0 | 0 | 0 | 0 | 0 | 0 | 0 | 2426400 | 0 | 0 |
| MDSGTRP\ | RIT1; RIT1; RIT1 | Q92963; A0A | RIT1_HUMAN C | Acetyl (Protein N | 0 | 0 | 0 | 0 | 0 | 0 | 0 | 0 | 0 | 0 | 555880 | 0 | 0 | 0 | 0 | 0 | 599330 | 0 |
| MDSKYQC\ | AKR1C2; AKR1C | P52895; Q0A | AK1C2_HUMAN | Acetyl (Protein N | 0 | 0 | 14588000 | 0 | 1962300 | 80838 | 0 | 0 | 0 | 0 | 0 | 0 | 0 | 0 | 0 | 0 | 0 | 0 |
| MDSQGRK' | ACTR2; ACTR2; | P61160; A0A | ARP2_HUMAN | Acetyl (Protein N | 0 | 0 | 0 | 0 | 1460500 | 0 | 2135600 | 1751800 | 0 | 0 | 0 | 0 | 2319800 | 2720500 | 0 | 0 | 4331400 | 962590 |
| MDSQGRK' | ACTR2; ACTR2; | P61160; A0A | ARP2_HUMAN | Acetyl (Protein N | 0 | 0 | 0 | 0 | 3950800 | 0 | 4395000 | 1776800 | 271520 | 810530 | 0 | 0 | 1999900 | 3599000 | 0 | 0 | 6591000 | 4319900 |
| MDSTANE\ | P4HB; P4HB; P4 | P07237; A0A | PDIA1_HUMAN | Unmodified | 0 | 0 | 736160 | 0 | 3682400 | 362080 | 3545700 | 0 | 86089000 | 17286000 | 5140600 | 1877000 | 0 | 0 | 1144100 | 140690 | 195740 | 3965900 |
| MDSTANE\ | P4HB; P4HB; P4 | P07237; A0A | PDIA1_HUMAN | Oxidation (M) | 0 | 0 | 0 | 0 | 0 | 0 | 0 | 313750 | 28934000 | 5453300 | 2247200 | 194180 | 311290 | 178520 | 191240 | 0 | 58319 | 40615 |
| MDSTANE\ | P4HB; P4HB; P4 | P07237; A0A | PDIA1_HUMAN | Oxidation (M) | 0 | 126950 | 0 | 0 | 0 | 0 | 2631300 | 2436400 | 0 | 7575000 | 0 | 0 | 0 | 0 | 0 | 0 | 0 | 0 |
| MDSTANE\ | P4HB; P4HB; P4 | P07237; A0A | PDIA1_HUMAN | Unmodified | 0 | 0 | 705940 | 595430 | 1484400 | 0 | 0 | 0 | 495210 | 6166700 | 0 | 0 | 0 | 0 | 0 | 0 | 0 | 0 |
| MDSTEPAY | EEF1A2; EEF1A | Q05639; A0A | EF1A2_HUMAN | Unmodified | 0 | 0 | 1159300 | 0 | 0 | 0 | 0 | 0 | 0 | 0 | 0 | 0 | 0 | 0 | 1658000 | 1395200 | 0 | 0 |
| MDSTEPAY | EEF1A2; EEF1A | Q05639; A0A | EF1A2_HUMAN | Oxidation (M) | 152210 | 0 | 633900 | 129950 | 0 | 0 | 0 | 0 | 0 | 0 | 0 | 0 | 0 | 0 | 444010 | 308810 | 0 | 0 |
| MDSTEPPY | EEF1A1P5; EEF | Q5VTE0; P68 | EF1A3_HUMAN | Unmodified | 0 | 1816000 | 5472300 | 2163700 | 520050 | 0 | 284130 | 0 | 1035600 | 238640 | 0 | 2711600 | 298170 | 0 | 7249800 | 2822500 | 0 | 0 |
| MDSTEPPY | EEF1A1P5; EEF | Q5VTE0; P68 | EF1A3_HUMAN | Oxidation (M) | 817020 | 642270 | 1784000 | 643370 | 0 | 0 | 0 | 0 | 229470 | 0 | 208180 | 727320 | 73118 | 0 | 1516500 | 1106000 | 0 | 0 |
| MDSTLTASI | AARS1; AARS1; | P49588; A0A | SYAC_HUMAN | Acetyl (Protein N | 0 | 0 | 0 | 0 | 327190 | 134270 | 0 | 0 | 0 | 107320 | 0 | 0 | 342360 | 322670 | 0 | 0 | 118910 | 121990 |
| MDSTLTASI | AARS1; AARS1; | P49588; A0A | SYAC_HUMAN | Acetyl (Protein N | 0 | 0 | 0 | 0 | 2188000 | 0 | 0 | 0 | 0 | 0 | 0 | 0 | 0 | 2934700 | 0 | 0 | 0 | 978380 |
| MDSTNAD/ | DNAJC7; DNAJC | Q99615; A0A | DNJC7_HUMAN | Unmodified | 0 | 0 | 0 | 0 | 0 | 0 | 0 | 0 | 568560 | 0 | 0 | 691200 | 0 | 0 | 0 | 0 | 0 | 0 |
| MDSVRSGI | TUBB2A; TUBB4 | Q13885; P04 | TBB2A_HUMAN | Oxidation (M) | 0 | 0 | 0 | 0 | 0 | 0 | 0 | 0 | 0 | 557140 | 0 | 0 | 0 | 0 | 0 | 0 | 471660 | 0 |
| MDTAEDPA | GPRIN1 | Q7Z2K8 | GRIN1_HUMAN | Acetyl (Protein N | 0 | 0 | 0 | 0 | 37077 | 0 | 0 | 36449 | 74575 | 71786 | 0 | 0 | 45126 | 181510 | 0 | 0 | 173970 | 161570 |
| MDTELAES | ABCC4 | O15439 | MRP4_HUMAN | Unmodified | 0 | 0 | 0 | 0 | 0 | 0 | 0 | 0 | 751640 | 198740 | 0 | 0 | 0 | 0 | 0 | 0 | 0 | 0 |
| MDTESTYS | VANGL1 | Q8TAA9 | VANG1_HUMAI | Acetyl (Protein N | 0 | 0 | 0 | 0 | 535460 | 0 | 0 | 0 | 0 | 0 | 0 | 0 | 0 | 0 | 0 | 0 | 201280 | 198120 |
| MDTESTYS | VANGL1 | Q8TAA9 | VANG1_HUMAI | Acetyl (Protein N | 0 | 0 | 0 | 0 | 3369300 | 0 | 494700 | 0 | 347210 | 659000 | 0 | 0 | 0 | 0 | 0 | 0 | 1605800 | 1078900 |
| MDTESTYS | VANGL1 | Q8TAA9 | VANG1_HUMAI | Acetyl (Protein N | 0 | 0 | 0 | 0 | 1660200 | 0 | 0 | 101280 | 0 | 86721 | 0 | 0 | 0 | 886340 | 0 | 0 | 0 | 0 |
| MDTSRVQF | RPS28 | P62857 | RS28_HUMAN | Acetyl (Protein N | 1113800 | 397730 | 298140 | 1703600 | 2019900 | 0 | 0 | 463740 | 110780 | 359130 | 69540 | 910960 | 3044200 | 1422800 | 456650 | 246790 | 1345000 | 1586400 |
| MDTSRVQF | RPS28 | P62857 | RS28_HUMAN | Acetyl (Protein N | 1126900 | 858660 | 0 | 1554900 | 2533300 | 289480 | 994570 | 805830 | 0 | 548540 | 0 | 214510 | 3984100 | 1413300 | 839740 | 0 | 1731400 | 1552600 |
| MDVANLAK | RACGAP1; RAC | Q9H0H5; HC | RGAP1_HUMAI | Unmodified | 0 | 0 | 406510 | 794650 | 349870 | 718300 | 77640 | 0 | 17240000 | 1602000 | 535710 | 2124300 | 0 | 0 | 766020 | 355140 | 738340 | 128820 |
| MDVCETHL | APP; APP; APP; | P05067; H7C | A4_HUMAN Am | Unmodified | 0 | 0 | 0 | 0 | 0 | 0 | 0 | 0 | 2253100 | 102010 | 0 | 0 | 0 | 0 | 0 | 0 | 0 | 0 |
| MEAAAEPG | NUBP2; NUBP2 | Q9Y5Y2; H3I | NUBP2_HUMAI | Acetyl (Protein N | 0 | 0 | 0 | 0 | 1351200 | 29829 | 360550 | 0 | 0 | 270930 | 0 | 0 | 355310 | 341040 | 0 | 0 | 455910 | 643300 |
| MEAAAEPG | NUBP2; NUBP2 | Q9Y5Y2; H3I | NUBP2_HUMAI | Acetyl (Protein N | 0 | 6790000 | 0 | 0 | 0 | 152240 | 3729500 | 0 | 0 | 353530 | 0 | 0 | 2202600 | 1909300 | 0 | 0 | 1946300 | 812570 |
| MEAAHLLP | ATP2A3 | Q93084 | AT2A3_HUMAN | Acetyl (Protein N | 0 | 0 | 0 | 0 | 0 | 0 | 0 | 0 | 0 | 0 | 451470 | 1090400 | 0 | 0 | 0 | 0 | 0 | 0 |
| MEAAPSRF | ERP27 | Q96DN0 | ERP27_HUMAN | Acetyl (Protein N | 0 | 0 | 0 | 0 | 8452400 | 459780 | 4029600 | 0 | 0 | 0 | 0 | 0 | 1420100 | 4673000 | 0 | 0 | 5446100 | 4599800 |
| MEADASVD | TXNDC9; TXND | O14530; G3 | TXND9_HUMAN | Acetyl (Protein N | 0 | 0 | 0 | 0 | 0 | 0 | 0 | 0 | 0 | 1210100 | 0 | 0 | 0 | 0 | 0 | 0 | 0 | 0 |
| MEALIPVIN | DNM1L; DNM1L | O00429; F8V | DNM1L_HUMA | Acetyl (Protein N | 0 | 0 | 0 | 0 | 0 | 0 | 0 | 0 | 0 | 0 | 305720 | 1014500 | 0 | 0 | 0 | 0 | 0 | 0 |
| MEANIDRIK | GNG2; GNG2; ( | P59768; G3\ | GBG2_HUMAN | Oxidation (M) | 0 | 0 | 0 | 0 | 217720 | 0 | 45652 | 0 | 64276 | 42542 | 133770 | 0 | 0 | 56223 | 0 | 0 | 111910 | 505750 |
| MEANIDRIK | GNG2; GNG2; ( | P59768; G3\ | GBG2_HUMAN | Unmodified | 0 | 0 | 0 | 0 | 106890 | 0 | 169220 | 65462 | 0 | 0 | 0 | 0 | 0 | 0 | 0 | 0 | 137630 | 2871900 |
| MEATGVLP | FLII; FLII | Q13045; J3K | FLII_HUMAN Pr | Acetyl (Protein N | 0 | 0 | 0 | 0 | 0 | 0 | 0 | 0 | 0 | 0 | 0 | 0 | 0 | 0 | 0 | 0 | 1468700 | 1001900 |
| MEDATANC | SLC26A6; SLC2 | Q9BXS9; A0A | S26A6_HUMAN | Unmodified | 0 | 0 | 0 | 0 | 97074 | 0 | 0 | 0 | 0 | 0 | 0 | 261290 | 0 | 0 | 0 | 0 | 0 | 0 |
| MEDAVQILI | GNAS | A0A0A0MR1 | A0A0A0MR13_I | Acetyl (Protein N | 0 | 0 | 0 | 398360 | 0 | 0 | 0 | 0 | 0 | 0 | 0 | 2250300 | 0 | 0 | 710280 | 0 | 0 | 0 |
| MEDAVQILI | GNAS | A0A0A0MR1 | A0A0A0MR13_I | Oxidation (M) | 129930 | 0 | 607460 | 1180500 | 0 | 0 | 0 | 0 | 0 | 0 | 0 | 0 | 0 | 0 | 0 | 66900 | 0 | 0 |
| MEDEDEQ( | PACSIN2 | Q9UNF0 | PACN2_HUMAI | Oxidation (M) | 339010 | 207270 | 2493100 | 0 | 0 | 0 | 0 | 0 | 0 | 0 | 0 | 0 | 0 | 0 | 106000 | 615390 | 0 | 0 |
| MEDEDEQ( | PACSIN2 | Q9UNF0 | PACN2_HUMAI | Unmodified | 827450 | 225500 | 1664000 | 0 | 0 | 0 | 0 | 0 | 0 | 0 | 0 | 0 | 0 | 0 | 1479300 | 865700 | 0 | 0 |
| MEDIKESIE | TMED1; TMED1 | Q13445; K7E | TMED1_HUMAI | Unmodified | 0 | 0 | 0 | 158490 | 52328 | 0 | 0 | 0 | 1247000 | 0 | 0 | 152240 | 181730 | 1026300 | 0 | 0 | 0 | 0 |
| MEDIKESIE | TMED1; TMED1 | Q13445; K7E | TMED1_HUMAI | Oxidation (M) | 0 | 222450 | 0 | 0 | 1293200 | 666940 | 0 | 186250 | 535420 | 219450 | 0 | 0 | 0 | 1236000 | 24981 | 144420 | 165460 | 0 |
| MEDLDQSF | RTN4 | Q9NQC3 | RTN4_HUMAN | Acetyl (Protein N | 0 | 0 | 0 | 0 | 1940800 | 0 | 0 | 0 | 0 | 0 | 510220 | 196340 | 0 | 0 | 0 | 0 | 634720 | 868450 |
| MEDYTKIEK | CDK1; CDK1; C | P06493; A0A | CDK1_HUMAN | Acetyl (Protein N | 0 | 0 | 0 | 0 | 0 | 0 | 1147400 | 851410 | 0 | 0 | 0 | 0 | 0 | 0 | 0 | 0 | 734650 | 427140 |
| MEEANIQP | LRPPRC; LRPPF | P42704; B8Z | LPPRC_HUMAN | Oxidation (M) | 0 | 0 | 0 | 0 | 0 | 0 | 0 | 0 | 1401700 | 166420 | 0 | 219150 | 0 | 0 | 0 | 0 | 0 | 0 |
| MEEANIQP | LRPPRC; LRPPF | P42704; B8Z | LPPRC_HUMAN | Unmodified | 0 | 0 | 0 | 0 | 0 | 0 | 0 | 0 | 2205900 | 225710 | 91484 | 0 | 0 | 0 | 0 | 0 | 0 | 0 |
| MEEASEGC | VAMP8; VAMP8 | Q9BV40; B8Z | VAMP8_HUMAI | Acetyl (Protein N | 638670 | 0 | 0 | 1496700 | 885230 | 324780 | 0 | 0 | 2568600 | 408460 | 0 | 2605500 | 649240 | 393870 | 479990 | 0 | 207730 | 486070 |
| MEEASEGC | VAMP8; VAMP8 | Q9BV40; B8Z | VAMP8_HUMAI | Acetyl (Protein N | 0 | 0 | 853720 | 1217200 | 0 | 0 | 0 | 0 | 0 | 0 | 874170 | 0 | 0 | 0 | 0 | 601710 | 0 | 0 |
| MEEASEGC | VAMP8; VAMP8 | Q9BV40; B8Z | VAMP8_HUMAI | Acetyl (Protein N | 0 | 0 | 0 | 0 | 6975500 | 575700 | 0 | 0 | 0 | 1023600 | 347590 | 0 | 3892000 | 2907000 | 0 | 0 | 2695500 | 2369000 |
| MEEASEGC | VAMP8; VAMP8 | Q9BV40; B8Z | VAMP8_HUMAI | Acetyl (Protein N | 508810 | 0 | 0 | 0 | 2778100 | 0 | 0 | 0 | 0 | 1929100 | 0 | 0 | 4198300 | 0 | 0 | 0 | 601290 | 0 |
| MEEDPDD\ | NAA10; NAA10; | P41227; F8V | NAA10_HUMAN | Oxidation (M) | 0 | 0 | 0 | 0 | 0 | 0 | 0 | 0 | 0 | 0 | 403340 | 2684900 | 0 | 0 | 0 | 0 | 0 | 0 |
| MEEDPDD\ | NAA10; NAA10; | P41227; F8V | NAA10_HUMAN | Unmodified | 0 | 0 | 0 | 0 | 0 | 0 | 0 | 0 | 0 | 0 | 0 | 4471700 | 0 | 0 | 0 | 0 | 0 | 0 |
| MEEDPDD\ | NAA10; NAA10; | P41227; F8V | NAA10_HUMAN | Oxidation (M) | 0 | 0 | 0 | 0 | 0 | 0 | 0 | 0 | 68635 | 0 | 0 | 706780 | 0 | 0 | 0 | 0 | 0 | 0 |
| MEEDPDD\ | NAA10; NAA10; | P41227; F8V | NAA10_HUMAN | Unmodified | 0 | 0 | 0 | 0 | 0 | 0 | 0 | 0 | 0 | 0 | 0 | 1717600 | 0 | 0 | 0 | 0 | 0 | 0 |
| MEEIVEGC | JUP; CTNNB1; ( | P14923; P35 | PLAK_HUMAN | Oxidation (M) | 0 | 0 | 540570 | 0 | 0 | 0 | 0 | 0 | 0 | 0 | 0 | 3520800 | 0 | 204230 | 3182300 | 4318800 | 0 | 0 |
| MEEIVEGC | JUP | P14923 | PLAK_HUMAN | 2 Oxidation (M) | 0 | 992650 | 331720 | 0 | 0 | 0 | 0 | 0 | 0 | 0 | 0 | 2722400 | 406760 | 0 | 1899300 | 715210 | 0 | 0 |
| MEEKDYSE | EPB41L1; EPB4 | Q9H4G0; Q4 | E41L1_HUMAN | Acetyl (Protein N | 102040 | 0 | 0 | 0 | 2085400 | 0 | 0 | 167610 | 0 | 0 | 0 | 0 | 0 | 0 | 0 | 61341 | 793010 | 1297300 |
| MEEKDYSE | EPB41L1; EPB4 | Q9H4G0; Q4 | E41L1_HUMAN | Acetyl (Protein N | 0 | 0 | 0 | 0 | 1824800 | 330640 | 0 | 0 | 0 | 0 | 0 | 0 | 0 | 762270 | 0 | 0 | 1725400 | 3233000 |
| MEELHNQI | SFPQ; SFPQ | P23246; H0Y | SFPQ_HUMAN | Unmodified | 0 | 0 | 0 | 0 | 0 | 0 | 10117 | 0 | 236620 | 0 | 517620 | 529140 | 0 | 0 | 0 | 0 | 0 | 0 |
| MEELHNQI | NONO; NONO; | Q15233; A0A | NONO_HUMAN | Unmodified | 0 | 0 | 0 | 0 | 0 | 0 | 0 | 0 | 0 | 0 | 699610 | 0 | 0 | 0 | 0 | 0 | 0 | 22994 |
| MEEQGLAD | ATXN10; ATXN1 | Q9UBB4; B1 | ATX10_HUMAN | Oxidation (M) | 0 | 0 | 144360 | 0 | 0 | 0 | 0 | 0 | 0 | 0 | 0 | 1217800 | 0 | 0 | 352150 | 172900 | 0 | 0 |
| MEESVNQI | RO60; RO60; R( | P10155; D6F | RO60_HUMAN | Acetyl (Protein N | 682280 | 711470 | 160300 | 281990 | 0 | 0 | 34100 | 0 | 0 | 0 | 0 | 0 | 301930 | 110440 | 559420 | 282070 | 0 | 0 |
| MEESVNQI | RO60; RO60; R( | P10155; D6F | RO60_HUMAN | Acetyl (Protein N | 505410 | 1206000 | 94267 | 268770 | 0 | 0 | 0 | 29537 | 0 | 0 | 0 | 173480 | 962850 | 0 | 655630 | 127030 | 0 | 0 |
| MEETIKDPI | CALCOCO2; CA | Q13137; D6I | CACO2_HUMA | Acetyl (Protein N | 0 | 0 | 0 | 0 | 0 | 632360 | 0 | 0 | 0 | 0 | 0 | 0 | 0 | 0 | 0 | 0 | 2025700 | 0 |
| MEETQNVC | VPS37B | Q9H9H4 | VP37B_HUMAN | Unmodified | 8473400 | 6756100 | 14364000 | 7663800 | 850740 | 0 | 67988 | 21136 | 1281000 | 349380 | 1948000 | 5082400 | 1716100 | 1042500 | 22710000 | 11148000 | 62908 | 1054000 |
| MEETQNVC | VPS37B | Q9H9H4 | VP37B_HUMAN | Oxidation (M) | 5241800 | 4143200 | 8599700 | 4693200 | 236900 | 400770 | 158580 | 127410 | 275900 | 97076 | 528080 | 934180 | 725080 | 310040 | 13605000 | 8995700 | 53566 | 293040 |
| MEEVYLPD | PTGR2; N/A | Q8N8N7; G3 | PTGR2_HUMAN | Oxidation (M) | 0 | 319180 | 41803 | 171610 | 0 | 0 | 0 | 0 | 0 | 0 | 0 | 659630 | 145820 | 35778 | 1154300 | 1577300 | 0 | 0 |
| MEEWDVP( | GNG13 | Q9P2W3 | GBG13_HUMAI | Acetyl (Protein N | 0 | 0 | 0 | 0 | 0 | 0 | 0 | 0 | 0 | 0 | 0 | 1356000 | 0 | 0 | 0 | 0 | 0 | 0 |
| MEEWDVP( | GNG13 | Q9P2W3 | GBG13_HUMAI | Acetyl (Protein N | 0 | 0 | 0 | 0 | 0 | 0 | 0 | 0 | 0 | 0 | 174360 | 4746400 | 0 | 0 | 0 | 0 | 0 | 961120 |
| MEFDLGAA | IMUP | Q9GZP8 | IMUP_HUMAN | Acetyl (Protein N | 0 | 0 | 0 | 0 | 5160700 | 0 | 0 | 0 | 0 | 0 | 0 | 0 | 0 | 0 | 0 | 0 | 0 | 1317000 |
| MEFGTAGL | PGM2; PGM2; F | Q96G03; E9I | PGM2_HUMAN | Oxidation (M) | 94392 | 95354 | 0 | 129950 | 0 | 0 | 0 | 0 | 0 | 0 | 0 | 286820 | 0 | 0 | 0 | 92397 | 0 | 0 |
| MEFGTAGL | PGM2; PGM2; F | Q96G03; E9I | PGM2_HUMAN | Unmodified | 0 | 0 | 0 | 500480 | 88465 | 0 | 0 | 0 | 77717 | 0 | 0 | 853150 | 50214 | 0 | 0 | 0 | 177190 | 0 |
| MEFLLGNP | TOM1L2; TOM1 | Q6ZVM7; F5I | TM1L2_HUMAN | Acetyl (Protein N | 199150 | 203400 | 0 | 58453 | 0 | 0 | 95253 | 0 | 25692 | 24130 | 0 | 0 | 161950 | 463490 | 479620 | 0 | 226160 | 280030 |
| MEFPDLGA | ZFAND2B; ZFAI | Q8WV99; B4 | ZFN2B_HUMAN | Acetyl (Protein N | 1891100 | 0 | 0 | 0 | 0 | 0 | 0 | 0 | 0 | 0 | 0 | 0 | 0 | 0 | 0 | 0 | 0 | 1766400 |
| MEGAKPTL | TNPO3; TNPO3 | Q9Y5L0; C9J | TNPO3_HUMAI | Acetyl (Protein N | 36284 | 166820 | 0 | 0 | 0 | 0 | 0 | 0 | 0 | 0 | 0 | 0 | 0 | 0 | 166740 | 0 | 0 | 0 |
| MEGISNFK | PBK; PBK | Q96KB5; E5F | TOPK_HUMAN | Acetyl (Protein N | 0 | 0 | 0 | 0 | 813310 | 0 | 0 | 106970 | 0 | 0 | 0 | 0 | 0 | 0 | 0 | 0 | 1053400 | 0 |
| MEGLDDGI | SNX6; SNX6; SI | Q9UNH7; AC | SNX6_HUMAN | Acetyl (Protein N | 0 | 0 | 0 | 0 | 1917000 | 109300 | 455320 | 206420 | 0 | 0 | 0 | 0 | 0 | 0 | 0 | 0 | 598440 | 1108800 |
| MEGPLSVF | TCP1; TCP1; TC | P17987; F5G | TCPA_HUMAN | Acetyl (Protein N | 0 | 0 | 0 | 0 | 0 | 0 | 0 | 0 | 148160 | 0 | 0 | 218810 | 0 | 0 | 0 | 0 | 0 | 0 |
| MEGPLSVF | TCP1; TCP1; TC | P17987; F5G | TCPA_HUMAN | Acetyl (Protein N | 0 | 0 | 200870 | 0 | 0 | 0 | 0 | 0 | 366420 | 26350 | 0 | 6127700 | 0 | 0 | 0 | 169410 | 0 | 0 |
| MEGPLSVF | TCP1; TCP1; TC | P17987; F5G | TCPA_HUMAN | Acetyl (Protein N | 0 | 965010 | 119450 | 0 | 0 | 0 | 0 | 0 | 736430 | 0 | 0 | 3830800 | 0 | 0 | 0 | 591660 | 0 | 0 |
| MEGPLSVF | TCP1; TCP1; TC | P17987; F5G | TCPA_HUMAN | Acetyl (Protein N | 0 | 0 | 0 | 0 | 0 | 0 | 25663 | 0 | 0 | 0 | 0 | 617480 | 0 | 0 | 0 | 0 | 0 | 0 |
| MEGPLSVF | TCP1; TCP1; TC | P17987; F5G | TCPA_HUMAN | Acetyl (Protein N | 0 | 0 | 0 | 0 | 0 | 0 | 5270600 | 0 | 0 | 0 | 0 | 0 | 0 | 8455700 | 0 | 0 | 5809500 | 8777100 |
| MEGQRWL | UCHL3 | P15374 | UCHL3_HUMAI | Acetyl (Protein N | 0 | 0 | 0 | 0 | 2349100 | 0 | 237390 | 0 | 0 | 810640 | 0 | 0 | 5419500 | 5976500 | 0 | 0 | 4903700 | 1310100 |
| MEGQRWL | UCHL3 | P15374 | UCHL3_HUMAI | Acetyl (Protein N | 0 | 0 | 0 | 0 | 161210 | 0 | 0 | 0 | 0 | 0 | 0 | 0 | 283340 | 0 | 0 | 0 | 236100 | 0 |
| MEGRQGLI | N/A | N/A | N/A | Oxidation (M) | 102030 | 3286000 | 0 | 0 | 17688000 | 8562800 | 34634000 | 63913000 | 0 | 5518100 | 991860 | 2913100 | 19853000 | 13662000 | 0 | 0 | 37938000 | 32240000 |
| MEGRSAAA | PSD3 | Q9NYI0 | PSD3_HUMAN | Acetyl (Protein N | 0 | 1242500 | 1610700 | 3708500 | 1502900 | 877460 | 0 | 0 | 0 | 0 | 0 | 1269800 | 0 | 0 | 0 | 0 | 811480 | 0 |
| MEGVEEKK | RPL7 | P18124 | RL7_HUMAN 60 | Acetyl (Protein N | 212110 | 615990 | 0 | 4788500 | 0 | 0 | 0 | 0 | 2117400 | 1861700 | 0 | 9752700 | 8043800 | 26595000 | 206320 | 248920 | 0 | 0 |
| MEHFDASL | ELOVL5; ELOVL | Q9NYP7; A0 | ELOV5_HUMAN | Acetyl (Protein N | 0 | 0 | 0 | 0 | 0 | 569200 | 0 | 0 | 1234000 | 417660 | 0 | 0 | 0 | 0 | 0 | 0 | 0 | 0 |
| MEIATKDPI | PPA1; PPA1 | Q15181; Q5 | IPYR_HUMAN I | Oxidation (M) | 0 | 0 | 0 | 0 | 0 | 0 | 0 | 0 | 0 | 0 | 1127700 | 5851300 | 0 | 0 | 0 | 0 | 0 | 0 |
| MEIATKDPI | PPA1; PPA1 | Q15181; Q5 | IPYR_HUMAN I | Unmodified | 0 | 0 | 0 | 0 | 0 | 0 | 0 | 0 | 0 | 0 | 0 | 25955000 | 0 | 0 | 3034000 | 662800 | 0 | 0 |

| | | | | | | | | | | | | | | | | | | | | | | |
|---|---|---|---|---|---|---|---|---|---|---|---|---|---|---|---|---|---|---|---|---|---|---|
| MEIATKDPL | PPA1; PPA1 | Q15181; Q5 | IPYR_HUMAN I | Unmodified | 0 | 0 | 0 | 121310 | 957990 | 0 | 0 | 0 | 0 | 0 | 675030 | 1719300 | 816010 | 0 | 0 | 0 | 415410 | 0 |
| MEIGLPDEI | NSF; NSF | P46459; I3L | NSF_HUMAN V | Unmodified | 0 | 180000 | 179460 | 0 | 0 | 0 | 0 | 0 | 451210 | 0 | 0 | 1603700 | 442830 | 0 | 226650 | 173620 | 0 | 0 |
| MEIHKNEA | FNBP1L; FNBP1 | Q5T0N5; A0 | FBP1L_HUMAN | Oxidation (M) | 182980 | 389950 | 0 | 0 | 89626 | 0 | 0 | 0 | 0 | 0 | 0 | 0 | 0 | 0 | 0 | 0 | 0 | 0 |
| MEILLLMVD | RABGGTA; RAB | Q92696; H0 | PGTA_HUMAN | Oxidation (M) | 0 | 0 | 0 | 0 | 0 | 0 | 0 | 0 | 184310 | 1450600 | 0 | 0 | 143440 | 0 | 0 | 0 | 0 | 0 |
| MEIVDHALI | CTNND1; CTNN | O60716; A0 | CTND1_HUMA | Oxidation (M) | 0 | 0 | 0 | 677060 | 0 | 0 | 0 | 0 | 0 | 0 | 0 | 0 | 0 | 0 | 1340800 | 0 | 0 | 0 |
| MEKTELIQK | YWHAQ; YWHA | P27348; E9P | 1433T_HUMAN | Acetyl (Protein N | 0 | 0 | 0 | 0 | 21138000 | 2256600 | 6300700 | 5793100 | 83335 | 2901500 | 0 | 0 | 9615300 | 6205300 | 0 | 0 | 33095000 | 13138000 |
| MEKTELIQK | YWHAQ; YWHA | P27348; E9P | 1433T_HUMAN | Acetyl (Protein N | 0 | 0 | 0 | 0 | 38115000 | 5965500 | 14503000 | 11537000 | 289550 | 7159700 | 0 | 0 | 15277000 | 21470000 | 0 | 0 | 81258000 | 30908000 |
| MEKTELIQK | YWHAQ; YWHA | P27348; E9P | 1433T_HUMAN | Acetyl (Protein N | 0 | 0 | 0 | 0 | 913750 | 0 | 485780 | 612860 | 0 | 0 | 0 | 0 | 0 | 0 | 0 | 0 | 1806100 | 571810 |
| MEKTELIQK | YWHAQ; YWHA | P27348; E9P | 1433T_HUMAN | Acetyl (Protein N | 0 | 0 | 0 | 0 | 2785900 | 0 | 1887200 | 792730 | 136820 | 21348 | 0 | 0 | 2020100 | 0 | 0 | 0 | 11186000 | 4411800 |
| MELEADNA | ING3; ING3 | Q9NXR8; E7 | ING3_HUMAN | Oxidation (M) | 0 | 43892000 | 0 | 0 | 0 | 0 | 0 | 0 | 0 | 9249.2 | 0 | 0 | 0 | 0 | 0 | 0 | 0 | 0 |
| MELEAMSR | TMEM258; TME | P61165; F5G | TM258_HUMAN | Acetyl (Protein N | 0 | 0 | 0 | 0 | 0 | 0 | 0 | 0 | 2044600 | 257530 | 0 | 0 | 0 | 0 | 0 | 0 | 0 | 0 |
| MELEAMSR | TMEM258; TME | P61165; F5G | TM258_HUMAN | Acetyl (Protein N | 0 | 0 | 0 | 0 | 0 | 0 | 0 | 0 | 10530000 | 999470 | 0 | 0 | 0 | 0 | 0 | 0 | 0 | 0 |
| MELEAMSR | TMEM258; TME | P61165; F5G | TM258_HUMAN | Acetyl (Protein N | 0 | 0 | 0 | 0 | 0 | 0 | 0 | 0 | 11122000 | 0 | 0 | 0 | 0 | 0 | 0 | 0 | 0 | 0 |
| MELEAMSR | TMEM258; TME | P61165; F5G | TM258_HUMAN | Oxidation (M) | 0 | 0 | 0 | 0 | 0 | 0 | 0 | 0 | 1025000 | 25630 | 0 | 0 | 0 | 0 | 0 | 0 | 0 | 0 |
| MELEAMSR | TMEM258; TME | P61165; F5G | TM258_HUMAN | Unmodified | 0 | 0 | 0 | 0 | 0 | 0 | 0 | 0 | 5068500 | 151890 | 0 | 0 | 0 | 0 | 0 | 0 | 0 | 0 |
| MELERPGG | FGA | P02671 | FIBA_HUMAN F | Oxidation (M) | 0 | 3525700 | 0 | 0 | 0 | 0 | 0 | 0 | 0 | 0 | 0 | 0 | 0 | 0 | 0 | 0 | 0 | 0 |
| MELITILEK | KPNB1; KPNB1 | Q14974; J3Q | IMB1_HUMAN I | Acetyl (Protein N | 0 | 0 | 458670 | 0 | 5101900 | 464480 | 2005100 | 0 | 1115100 | 2011000 | 953080 | 2155700 | 0 | 0 | 1060600 | 0 | 3393500 | 1437000 |
| MELPQRDV | PLOD3; PLOD3 | O60568; H7 | PLOD3_HUMA | Unmodified | 0 | 0 | 0 | 0 | 0 | 0 | 0 | 0 | 1646500 | 0 | 0 | 0 | 0 | 0 | 0 | 0 | 0 | 0 |
| MELQEIQLI | TPM3; TPM3; TF | P06753; J3K | TPM3_HUMAN | Unmodified | 0 | 0 | 0 | 0 | 4970500 | 1288600 | 0 | 0 | 3633700 | 162690 | 1368000 | 6433300 | 1100500 | 0 | 0 | 0 | 0 | 0 |
| MELQPPEA | SLC3A2; SLC3A | P08195; A0A | 4F2_HUMAN 4I | Acetyl (Protein N | 0 | 0 | 0 | 0 | 546500 | 0 | 0 | 0 | 309110 | 0 | 96725 | 87383 | 0 | 0 | 0 | 0 | 0 | 0 |
| MELQPPEA | SLC3A2; SLC3A | P08195; A0A | 4F2_HUMAN 4I | Acetyl (Protein N | 297960 | 0 | 0 | 0 | 0 | 0 | 0 | 28580 | 1283500 | 0 | 4157200 | 3102200 | 0 | 0 | 777910 | 467840 | 0 | 0 |
| MELSAEYLI | BOLA2; BOLA2E | Q9H3K6; H3 | BOLA2_HUMA | Acetyl (Protein N | 0 | 0 | 0 | 0 | 0 | 0 | 0 | 0 | 0 | 0 | 147830 | 1880400 | 0 | 0 | 0 | 0 | 0 | 0 |
| MELSDANL | CSE1L | P55060 | XPO2_HUMAN | Acetyl (Protein N | 12254 | 0 | 0 | 0 | 0 | 17747 | 179460 | 0 | 59440 | 263560 | 43814 | 138600 | 617210 | 63713 | 38409 | 0 | 76626 | 61236 |
| MELSDANL | CSE1L | P55060 | XPO2_HUMAN | Acetyl (Protein N | 0 | 0 | 47549 | 456120 | 2651400 | 203130 | 1952700 | 62500 | 633290 | 1884000 | 641090 | 2343600 | 1189100 | 661600 | 0 | 0 | 1198300 | 967800 |
| MENAELDV | THBS1 | P07996 | TSP1_HUMAN | Oxidation (M) | 0 | 0 | 166680 | 0 | 0 | 0 | 0 | 0 | 0 | 0 | 0 | 0 | 0 | 0 | 168300 | 61302 | 0 | 0 |
| MENDLPQK | CD58; CD58 | P19256; B1A | LFA3_HUMAN I | Unmodified | 0 | 0 | 0 | 0 | 0 | 0 | 0 | 0 | 0 | 0 | 0 | 0 | 0 | 0 | 0 | 0 | 0 | 0 |
| MENGAVYS | CMTM6 | Q9NX76 | CKLF6_HUMAN | Acetyl (Protein N | 0 | 0 | 0 | 0 | 0 | 0 | 0 | 0 | 1184300 | 0 | 0 | 938860 | 0 | 0 | 0 | 0 | 0 | 0 |
| MENGAVYS | CMTM6 | Q9NX76 | CKLF6_HUMAN | Acetyl (Protein N | 814600 | 667910 | 721800 | 1014100 | 0 | 0 | 0 | 0 | 0 | 0 | 0 | 3470700 | 0 | 0 | 681570 | 807800 | 0 | 0 |
| MENGYTYE | PNP; PNP | P00491; G3 | PNPH_HUMAN | Acetyl (Protein N | 0 | 0 | 0 | 0 | 0 | 0 | 0 | 0 | 0 | 0 | 0 | 0 | 0 | 0 | 0 | 0 | 0 | 0 |
| MENQSENI | TSG101; TSG10 | Q99816; F5H | TS101_HUMAN | Oxidation (M) | 7492800 | 37358000 | 29802000 | 31980000 | 3228800 | 1145500 | 3894400 | 0 | 9961600 | 1653200 | 3644100 | 8553400 | 9218800 | 4290200 | 1.19E+08 | 43962000 | 0 | 2517200 |
| MENQSENI | TSG101; TSG10 | Q99816; F5H | TS101_HUMAN | Unmodified | 0 | 0 | 0 | 0 | 0 | 0 | 0 | 0 | 3460700 | 607600 | 0 | 0 | 0 | 0 | 0 | 0 | 0 | 863760 |
| MENSQLCH | HNRNPA0 | Q13151 | ROA0_HUMAN | Acetyl (Protein N | 0 | 0 | 0 | 0 | 0 | 0 | 559780 | 453060 | 0 | 0 | 0 | 130370 | 0 | 79858 | 0 | 0 | 0 | 0 |
| MENSQLCH | HNRNPA0 | Q13151 | ROA0_HUMAN | Acetyl (Protein N | 0 | 0 | 0 | 0 | 0 | 0 | 0 | 0 | 0 | 0 | 0 | 0 | 0 | 0 | 0 | 0 | 0 | 0 |
| MEPPASKV | SEPTIN9; SEPTI | Q9UHD8; K7 | SEPT9_HUMAN | Oxidation (M) | 0 | 0 | 0 | 0 | 1236500 | 206410 | 0 | 42117 | 0 | 0 | 0 | 112310 | 0 | 0 | 0 | 0 | 1024900 | 341500 |
| MEPPASKV | SEPTIN9; SEPTI | Q9UHD8; K7 | SEPT9_HUMAN | Unmodified | 0 | 0 | 0 | 0 | 2907500 | 655900 | 91460 | 0 | 354890 | 178030 | 0 | 0 | 177870 | 0 | 0 | 0 | 1544300 | 1684800 |
| MEPRAPWI | HLA-A; HLA-A; I | A0A0G2JL56 | A0A0G2JL56_H | Oxidation (M) | 0 | 0 | 0 | 0 | 0 | 0 | 0 | 0 | 0 | 0 | 0 | 0 | 0 | 0 | 0 | 0 | 0 | 1403300 |
| MEPRAPWI | HLA-A; HLA-A; I | A0A1W2PR6 | A0A1W2PR61_ | Unmodified | 0 | 0 | 0 | 0 | 3876200 | 1449100 | 0 | 0 | 0 | 0 | 0 | 0 | 0 | 0 | 157470 | 0 | 0 | 0 |
| MEPRAPWI | HLA-A; HLA-A; I | A0A1W2PR6 | A0A1W2PR61_ | Oxidation (M) | 0 | 0 | 0 | 9151300 | 0 | 1733700 | 0 | 0 | 4479000 | 192210 | 94790 | 0 | 0 | 0 | 0 | 0 | 0 | 0 |
| MEPSSLELI | KYNU; KYNU; K | Q16719; B8 | KYNU_HUMAN | Acetyl (Protein N | 0 | 0 | 0 | 0 | 2497600 | 0 | 0 | 0 | 0 | 0 | 0 | 0 | 0 | 0 | 0 | 0 | 0 | 0 |
| MEQPTSST | SPANXN3; SPAI | Q5MJ09; Q5I | SPXN3_HUMA | Acetyl (Protein N | 0 | 0 | 0 | 0 | 297830 | 0 | 0 | 0 | 0 | 1994100 | 859900 | 0 | 0 | 221010 | 0 | 0 | 0 | 683870 |
| MEQVNELK | STIP1; STIP1; ST | P31948; F5H | STIP1_HUMAN | Acetyl (Protein N | 130060 | 106220 | 0 | 0 | 1673900 | 356620 | 273620 | 151900 | 0 | 707660 | 1697900 | 3891800 | 1738400 | 856680 | 803350 | 195030 | 1065600 | 1500800 |
| MEQVNELK | STIP1; STIP1; ST | P31948; F5H | STIP1_HUMAN | Acetyl (Protein N | 83395 | 0 | 0 | 200450 | 3548000 | 0 | 363980 | 0 | 2159700 | 1355900 | 2364400 | 4915000 | 2733600 | 1555100 | 140440 | 86343 | 2385400 | 2721300 |
| MEQVNELK | STIP1; STIP1; ST | P31948; F5H | STIP1_HUMAN | Acetyl (Protein N | 0 | 0 | 0 | 0 | 0 | 0 | 479300 | 172170 | 0 | 0 | 0 | 0 | 0 | 0 | 0 | 0 | 801740 | 0 |
| MEQVNELK | STIP1; STIP1; ST | P31948; F5H | STIP1_HUMAN | Acetyl (Protein N | 0 | 0 | 0 | 0 | 140480 | 0 | 0 | 0 | 182920 | 0 | 0 | 571630 | 0 | 0 | 0 | 221190 | 2652900 | 0 |
| MERASLIQI | SFN | P31947 | 1433S_HUMAN | Acetyl (Protein N | 0 | 0 | 0 | 0 | 58913000 | 11312000 | 0 | 0 | 0 | 5385900 | 0 | 0 | 0 | 0 | 0 | 0 | 38349000 | 22808000 |
| MERASLIQI | SFN | P31947 | 1433S_HUMAN | Acetyl (Protein N | 52455 | 0 | 0 | 0 | 1501600 | 0 | 0 | 0 | 0 | 0 | 0 | 0 | 0 | 0 | 0 | 0 | 2685600 | 1718100 |
| MERPYTFK | NEDD4L; NEDD | Q96PU5; A0 | NED4L_HUMA | Acetyl (Protein N | 1756100 | 1135100 | 678530 | 838070 | 233580 | 0 | 0 | 21530 | 0 | 0 | 0 | 607170 | 0 | 0 | 1502400 | 851960 | 200810 | 490490 |
| MERPYTFK | NEDD4L; NEDD | Q96PU5; A0 | NED4L_HUMA | Acetyl (Protein N | 0 | 0 | 0 | 641600 | 0 | 0 | 0 | 0 | 0 | 0 | 0 | 0 | 0 | 0 | 0 | 0 | 0 | 0 |
| MESALDQL | TALDO1; TALDO | P37837; E9P | TALDO_HUMA | Unmodified | 0 | 0 | 0 | 0 | 504390 | 0 | 243820 | 0 | 1277800 | 0 | 668360 | 0 | 0 | 0 | 0 | 0 | 0 | 0 |
| MESALDQL | TALDO1; TALDO | P37837; E9P | TALDO_HUMA | Oxidation (M) | 164300 | 220840 | 0 | 0 | 0 | 0 | 870790 | 571420 | 0 | 100840 | 281470 | 1511600 | 0 | 0 | 0 | 0 | 127420 | 0 |
| MESALTAR | MYO1C; MYO1C | O00159; I3L | MYO1C_HUMA | Acetyl (Protein N | 0 | 0 | 108410 | 653570 | 0 | 0 | 0 | 0 | 0 | 0 | 0 | 0 | 0 | 0 | 186290 | 741920 | 0 | 0 |
| MESGAYGA | SYNGR2; SYNG | O43760; K7E | SNG2_HUMAN | Acetyl (Protein N | 0 | 0 | 899940 | 1405600 | 788270 | 0 | 0 | 59498 | 211590 | 87573 | 0 | 376210 | 1038100 | 517690 | 595440 | 435150 | 134390 | 464110 |
| MESGAYGA | SYNGR2; SYNG | O43760; K7E | SNG2_HUMAN | Acetyl (Protein N | 0 | 0 | 300790 | 318060 | 461430 | 0 | 0 | 0 | 0 | 0 | 0 | 0 | 285660 | 0 | 197380 | 207640 | 0 | 536550 |
| MESGFTSK | NNMT | P40261 | NNMT_HUMAN | Acetyl (Protein N | 0 | 0 | 0 | 0 | 15209000 | 2492500 | 0 | 0 | 0 | 0 | 0 | 0 | 0 | 0 | 0 | 0 | 34947 | 665910 |
| MESGFTSK | NNMT | P40261 | NNMT_HUMAN | Acetyl (Protein N | 0 | 0 | 0 | 0 | 7769100 | 0 | 0 | 0 | 0 | 0 | 0 | 0 | 0 | 0 | 0 | 0 | 0 | 0 |
| MESISMMG | STEAP2; STEAP | Q8NFT2; C9 | STEA2_HUMAN | Acetyl (Protein N | 80971 | 0 | 563470 | 320140 | 0 | 0 | 0 | 0 | 0 | 0 | 66404 | 0 | 0 | 190140 | 396240 | 333250 | 0 | 0 |
| MESISMMG | STEAP2; STEAP | Q8NFT2; C9 | STEA2_HUMAN | Acetyl (Protein N | 0 | 0 | 0 | 0 | 0 | 0 | 0 | 0 | 0 | 0 | 0 | 0 | 250570 | 0 | 271220 | 213090 | 0 | 0 |
| MESLRLDG | NIBAN2 | Q96TA1 | NIBA2_HUMAN | Unmodified | 141660 | 189620 | 212550 | 184840 | 0 | 0 | 0 | 0 | 1167800 | 0 | 0 | 952550 | 0 | 0 | 390160 | 325950 | 0 | 0 |
| MESLRLDG | NIBAN2 | Q96TA1 | NIBA2_HUMAN | Unmodified | 100300 | 111650 | 0 | 152060 | 5969900 | 434650 | 0 | 0 | 0 | 1328900 | 0 | 0 | 0 | 55269 | 47596 | 0 | 459080 | 1104600 |
| MESLRLDG | NIBAN2 | Q96TA1 | NIBA2_HUMAN | Oxidation (M) | 345560 | 897670 | 0 | 0 | 15620000 | 2838000 | 132350 | 0 | 828680 | 1750600 | 0 | 101050 | 1347700 | 1264800 | 566730 | 198450 | 1429700 | 1175700 |
| MESYDVIAI | ACTR1A; ACTR1 | P61163; A0A | ACTZ_HUMAN | Acetyl (Protein N | 0 | 0 | 0 | 0 | 3871600 | 0 | 0 | 842520 | 699200 | 654120 | 731890 | 2071400 | 0 | 0 | 0 | 0 | 1892200 | 2243900 |
| MESYHKPD | TKT; TKT; TKT; T | P29401; F8V | TKT_HUMAN Tr | Acetyl (Protein N | 0 | 0 | 0 | 0 | 9190700 | 1738100 | 0 | 0 | 806340 | 1622900 | 0 | 5853600 | 2411100 | 0 | 0 | 0 | 2444700 | 3246300 |
| MESYHKPD | TKT; TKT; TKT; T | P29401; F8V | TKT_HUMAN Tr | Acetyl (Protein N | 0 | 0 | 18454 | 0 | 1881700 | 0 | 0 | 0 | 0 | 0 | 588990 | 1209800 | 0 | 0 | 0 | 24458 | 323380 | 0 |
| METEQPEE | HNRNPK; HNRI | P61978; Q5T | HNRPK_HUMA | Acetyl (Protein N | 0 | 0 | 0 | 0 | 0 | 0 | 0 | 0 | 0 | 0 | 1200300 | 824280 | 0 | 0 | 0 | 0 | 0 | 0 |
| METEQPEE | HNRNPK; HNRI | P61978; Q5T | HNRPK_HUMA | Acetyl (Protein N | 0 | 0 | 0 | 0 | 797700 | 0 | 0 | 147290 | 0 | 0 | 0 | 3803800 | 0 | 0 | 0 | 0 | 0 | 0 |
| METKTESS | EPB41L3; EPB4 | Q9Y2J2; A0A | E41L3_HUMAN | Unmodified | 0 | 0 | 0 | 0 | 0 | 0 | 0 | 0 | 0 | 0 | 0 | 0 | 58685 | 0 | 0 | 0 | 0 | 1534200 |
| METLKDK | VPS37C; VPS37 | A5D8V6; F5H | VP37C_HUMAN | Acetyl (Protein N | 0 | 0 | 0 | 15208 | 994370 | 94298 | 501870 | 172650 | 0 | 33403 | 0 | 136600 | 2623500 | 1495900 | 0 | 0 | 414590 | 957950 |
| METTISEIH | AKAP5 | P24588 | AKAP5_HUMAN | Acetyl (Protein N | 0 | 0 | 184990 | 0 | 0 | 0 | 0 | 0 | 567660 | 0 | 0 | 0 | 3861000 | 2170700 | 0 | 0 | 0 | 2226500 |
| METVSGYE | ANKRD13D | Q6ZTN6 | AN13D_HUMA | Oxidation (M) | 113320 | 90482 | 47656 | 0 | 0 | 0 | 0 | 0 | 0 | 0 | 0 | 0 | 0 | 0 | 61748 | 104770 | 0 | 0 |
| MEVPRLDH | PHYHIPL | Q96FC7 | PHIPL_HUMAN | Acetyl (Protein N | 0 | 0 | 0 | 0 | 0 | 0 | 0 | 323810 | 0 | 0 | 0 | 0 | 0 | 0 | 0 | 0 | 0 | 0 |
| MFDRHSSL | CLTC; CLTC | Q00610; A0 | CLH1_HUMAN | Oxidation (M) | 1819500 | 1370100 | 208190 | 935540 | 274420 | 0 | 0 | 0 | 0 | 0 | 0 | 1769700 | 0 | 0 | 850840 | 1121100 | 0 | 160650 |
| MFDRHSSL | CLTC; CLTC | Q00610; A0 | CLH1_HUMAN | Unmodified | 267240 | 253350 | 0 | 126530 | 0 | 0 | 0 | 0 | 0 | 0 | 0 | 0 | 0 | 0 | 160450 | 0 | 0 | 0 |
| MFDRHSSL | CLTC; CLTC | Q00610; A0 | CLH1_HUMAN | Unmodified | 0 | 0 | 0 | 0 | 1470000 | 737460 | 3047900 | 0 | 0 | 1111200 | 0 | 0 | 797750 | 0 | 0 | 0 | 2180200 | 113270 |
| MFDRHSSL | CLTC; CLTC | Q00610; A0 | CLH1_HUMAN | Oxidation (M) | 0 | 0 | 0 | 0 | 0 | 0 | 1332200 | 252690 | 0 | 207390 | 0 | 0 | 1007600 | 0 | 0 | 0 | 409110 | 327530 |
| MFDSSNYH | MELTF | P08582 | TRFM_HUMAN | Oxidation (M) | 103550 | 90022 | 0 | 0 | 8173400 | 2353500 | 0 | 0 | 2406800 | 1595500 | 444670 | 1253000 | 71258 | 0 | 0 | 0 | 1186800 | 893710 |
| MFDSSNYH | MELTF | P08582 | TRFM_HUMAN | Unmodified | 0 | 0 | 0 | 0 | 5772000 | 977090 | 0 | 0 | 3976800 | 1191100 | 123010 | 275100 | 0 | 0 | 0 | 0 | 557800 | 1089400 |
| MFDVGGQ | GNAS; GNAI3; G | Q5JWF2; P0 | GNAS1_HUMA | Unmodified | 1902700 | 2776500 | 1087700 | 2184900 | 566140 | 0 | 278920 | 186210 | 2316300 | 793180 | 4386400 | 56384000 | 0 | 0 | 2845500 | 1946300 | 192340 | 441650 |
| MFDVGGQ | GNAS; GNAI3; G | Q5JWF2; P0 | GNAS1_HUMA | Oxidation (M) | 2102400 | 3085600 | 1177600 | 793950 | 64851 | 58374 | 55785 | 0 | 1577800 | 321530 | 2121800 | 37936000 | 134710 | 113460 | 3185100 | 2801800 | 147790 | 148870 |
| MFDVGGQ | GNAI3; GNAT1; | P08754; P11 | GNAI3_HUMAN | Unmodified | 71297 | 0 | 0 | 187190 | 0 | 0 | 36794 | 0 | 0 | 0 | 0 | 1689800 | 72878 | 0 | 246620 | 58626 | 0 | 0 |
| MFEARLVQ | PCNA | P12004 | PCNA_HUMAN | Unmodified | 0 | 0 | 0 | 0 | 271600 | 0 | 456760 | 81096 | 0 | 287590 | 0 | 0 | 217280 | 183420 | 0 | 0 | 593810 | 0 |
| MFEDPETTI | EIF3E; EIF3E; EI | P60228; A0A | EIF3E_HUMAN | Oxidation (M) | 0 | 0 | 0 | 0 | 0 | 0 | 0 | 0 | 0 | 152520 | 127180 | 1463100 | 0 | 0 | 0 | 107140 | 0 | 0 |
| MFEELGQR | HSPH1; HSPH1 | Q92598; A0 | HS105_HUMAN | Oxidation (M) | 0 | 0 | 50976 | 186260 | 0 | 0 | 0 | 329780 | 174230 | 0 | 277950 | 1838800 | 0 | 0 | 64791 | 117910 | 0 | 0 |
| MFEELGQR | HSPH1; HSPH1 | Q92598; A0 | HS105_HUMAN | Unmodified | 0 | 0 | 0 | 229130 | 0 | 350350 | 0 | 0 | 712150 | 0 | 647760 | 2645300 | 0 | 0 | 0 | 110490 | 0 | 0 |
| MFEIDYSRI | GLB1; GLB1; GI | P16278; C9J | BGAL_HUMAN | Oxidation (M) | 0 | 0 | 0 | 0 | 83783 | 0 | 0 | 0 | 1888600 | 490820 | 0 | 0 | 0 | 0 | 0 | 0 | 0 | 0 |
| MFEIDYSRI | GLB1; GLB1; GI | P16278; C9J | BGAL_HUMAN | Unmodified | 0 | 0 | 42867 | 0 | 0 | 0 | 0 | 0 | 1407800 | 0 | 0 | 0 | 0 | 0 | 0 | 0 | 0 | 0 |
| MFEIVFEDF | EPRS1 | P07814 | SYEP_HUMAN | Oxidation (M) | 285470 | 122880 | 440600 | 832710 | 3769200 | 442530 | 2136000 | 255590 | 1450100 | 1322100 | 1728500 | 7482900 | 4431100 | 2599300 | 914790 | 456400 | 1512800 | 0 |
| MFESFIESV | PRKAR2A; PRKA | P13861; H7 | KAP2_HUMAN | Oxidation (M) | 74798 | 0 | 0 | 703310 | 3629100 | 402920 | 0 | 0 | 960770 | 699810 | 377060 | 929820 | 6494400 | 5246000 | 1462200 | 1626200 | 521490 | 2273000 |
| MFESFIESV | PRKAR2A; PRKA | P13861; H7 | KAP2_HUMAN | Unmodified | 0 | 0 | 0 | 0 | 0 | 0 | 0 | 0 | 304340 | 125390 | 0 | 0 | 0 | 513880 | 0 | 0 | 0 | 0 |
| MFGIDRDA | TKT; TKT | P29401; A0A | TKT_HUMAN Tr | Oxidation (M) | 0 | 0 | 0 | 0 | 870120 | 0 | 0 | 0 | 554480 | 471680 | 1159000 | 2435100 | 351350 | 0 | 0 | 0 | 199510 | 0 |
| MFGIDRDA | TKT; TKT | P29401; A0A | TKT_HUMAN Tr | Unmodified | 151040 | 83332 | 0 | 169840 | 6150500 | 762970 | 0 | 0 | 5202900 | 1710700 | 490750 | 4498900 | 1459300 | 324100 | 119240 | 56769 | 648280 | 2602900 |
| MFGIPVVA | MTHFD1; MTHF | P11586; V9C | C1TC_HUMAN | Oxidation (M) | 0 | 0 | 78637 | 0 | 0 | 0 | 0 | 0 | 0 | 0 | 257280 | 867800 | 26721 | 0 | 339620 | 132240 | 0 | 0 |
| MFGMVLEK | CSE1L | P55060 | XPO2_HUMAN | Oxidation (M) | 0 | 0 | 0 | 0 | 0 | 0 | 0 | 0 | 1885200 | 0 | 0 | 3234100 | 0 | 0 | 0 | 0 | 0 | 0 |
| MFGMVLEK | CSE1L | P55060 | XPO2_HUMAN | Unmodified | 0 | 0 | 0 | 0 | 0 | 0 | 0 | 0 | 140650 | 0 | 0 | 888660 | 0 | 0 | 0 | 0 | 0 | 0 |
| MFGSSVDL | PRKAR2A; PRKA | P13861; H7 | KAP2_HUMAN | Oxidation (M) | 0 | 0 | 0 | 0 | 0 | 0 | 0 | 142180 | 0 | 0 | 0 | 477040 | 2155500 | 0 | 0 | 0 | 704870 | 611380 |
| MFHGIPATI | VPS28; VPS28; | Q9UK41; E9I | VPS28_HUMAN | Acetyl (Protein N | 1.37E+08 | 1.07E+08 | 98125000 | 1.18E+08 | 50556000 | 12874000 | 6746300 | 1936200 | 20615000 | 28423000 | 10505000 | 2440600 | 1.75E+08 | 59575000 | 2.34E+08 | 1.32E+08 | 8753700 | 41024000 |
| MFHGIPATI | VPS28; VPS28; | Q9UK41; E9I | VPS28_HUMAN | Acetyl (Protein N | 13298000 | 11783000 | 5069500 | 11442000 | 11859000 | 0 | 0 | 0 | 24105000 | 5112200 | 0 | 0 | 4868700 | 16430000 | 11277000 | 6478400 | 1906400 | 10041000 |
| MFHGIPATI | VPS28; VPS28; | Q9UK41; E9I | VPS28_HUMAN | Acetyl (Protein N | 0 | 0 | 0 | 1852500 | 10503000 | 1856300 | 9024500 | 0 | 0 | 6588600 | 0 | 0 | 58043000 | 30868000 | 0 | 0 | 11196000 | 11595000 |
| MFHGIPATI | VPS28; VPS28; | Q9UK41; E9I | VPS28_HUMAN | Acetyl (Protein N | 0 | 0 | 0 | 0 | 0 | 0 | 0 | 0 | 0 | 1503600 | 0 | 0 | 18510000 | 9854800 | 0 | 0 | 4028500 | 6340100 |
| MFHGIQDY | PHYH; PHYH; P | O14832; B1 | PAHX_HUMAN | Unmodified | 0 | 0 | 0 | 0 | 0 | 0 | 0 | 0 | 4477600 | 839060 | 0 | 0 | 0 | 0 | 0 | 0 | 0 | 0 |
| MFHGIQDY | PHYH; PHYH; P | O14832; B1 | PAHX_HUMAN | Oxidation (M) | 0 | 0 | 0 | 0 | 0 | 0 | 0 | 0 | 5526700 | 976580 | 0 | 0 | 0 | 11233 | 0 | 0 | 0 | 0 |
| MFIGGLSW | HNRNPDL; HNI | O14979; A0 | HNRDL_HUMA | Oxidation (M) | 0 | 0 | 0 | 0 | 699940 | 0 | 307620 | 147300 | 0 | 276100 | 424010 | 3151200 | 0 | 0 | 0 | 0 | 0 | 225520 |
| MFIGGLSW | HNRNPDL; HNI | O14979; A0 | HNRDL_HUMA | Unmodified | 0 | 0 | 0 | 0 | 0 | 0 | 0 | 0 | 0 | 0 | 0 | 0 | 0 | 0 | 0 | 0 | 0 | 0 |
| MFIGGLSW | HNRNPD; HNR | Q14103; H0 | HNRPD_HUMA | Oxidation (M) | 0 | 0 | 0 | 0 | 0 | 0 | 0 | 0 | 0 | 0 | 0 | 1751900 | 0 | 0 | 0 | 0 | 0 | 0 |
| MFIGGLSW | HNRNPD; HNR | Q14103; H0 | HNRPD_HUMA | Unmodified | 0 | 0 | 191370 | 0 | 215990 | 0 | 385130 | 0 | 1784400 | 0 | 1031600 | 5295500 | 1044800 | 0 | 0 | 75265 | 0 | 0 |
| MFIGGLSW | HNRNPD; HNR | Q14103; H0 | HNRPD_HUMA | Oxidation (M) | 0 | 0 | 0 | 0 | 0 | 0 | 596020 | 231770 | 647510 | 642370 | 2203100 | 11687000 | 0 | 443340 | 0 | 0 | 0 | 0 |
| MFILQSK | PFDN1; PFDN1 | O60925; E5F | PFD1_HUMAN | Unmodified | 0 | 0 | 0 | 0 | 0 | 0 | 123750 | 29428 | 578350 | 489540 | 312860 | 2258400 | 807890 | 302020 | 0 | 0 | 557930 | 1389600 |
| MFILQSK | PFDN1; PFDN1 | O60925; E5F | PFD1_HUMAN | Oxidation (M) | 0 | 0 | 0 | 157470 | 624670 | 0 | 0 | 38492 | 166150 | 0 | 0 | 1594000 | 387980 | 230720 | 0 | 166320 | 356260 | 356460 |
| MFKPQALL | FERMT2; FERM | Q96AC1; A0 | FERM2_HUMA | Oxidation (M) | 0 | 69626 | 0 | 0 | 4690700 | 0 | 170230 | 193150 | 0 | 0 | 0 | 0 | 161570 | 0 | 0 | 145040 | 1798100 | 6688600 |
| MFKPQALL | FERMT2; FERM | Q96AC1; A0 | FERM2_HUMA | Unmodified | 0 | 0 | 0 | 0 | 2256500 | 0 | 0 | 0 | 0 | 0 | 0 | 0 | 0 | 0 | 0 | 0 | 555930 | 1542700 |

| | | | | | | | | | | | | | | | | | | | | | | |
|---|---|---|---|---|---|---|---|---|---|---|---|---|---|---|---|---|---|---|---|---|---|---|
| MFKPQALL | FERMT2; FERM | Q96AC1; A0 | FERM2_HUMAN | Unmodified | 0 | 0 | 0 | 0 | 619180 | 0 | 0 | 0 | 0 | 0 | 0 | 0 | 0 | 0 | 0 | 0 | 877190 | 0 |
| MFLGDNAI | IQGAP1; IQGAP | P46940; H0 | IQGA1_HUMAN | Oxidation (M) | 0 | 0 | 0 | 0 | 8735400 | 3773400 | 0 | 0 | 0 | 2174300 | 0 | 0 | 0 | 0 | 0 | 0 | 1765500 | 0 |
| MFLLVGAP | ITGAV | P06756 | ITAV_HUMAN I | Unmodified | 0 | 0 | 0 | 208460 | 0 | 0 | 0 | 0 | 1429200 | 0 | 0 | 590480 | 0 | 0 | 214140 | 0 | 0 | 193990 |
| MFLNLLSPI | GTPBP1 | O00178 | GTPB1_HUMAN | Oxidation (M) | 190630 | 0 | 0 | 573860 | 0 | 0 | 0 | 0 | 0 | 0 | 0 | 483250 | 0 | 0 | 366920 | 0 | 0 | 0 |
| MFLQYYLN | NOP10; NOP10 | Q9NPE3; H0 | NOP10_HUMAI | Oxidation (M) | 242180 | 0 | 0 | 0 | 0 | 0 | 0 | 0 | 0 | 0 | 0 | 0 | 1644300 | 1044000 | 0 | 0 | 0 | 0 |
| MFLRGDSV | SNRPD2; SNRP | P62316; K7E | SMD2_HUMAN | Oxidation (M) | 0 | 0 | 0 | 0 | 0 | 0 | 0 | 0 | 0 | 0 | 60810 | 343110 | 0 | 0 | 0 | 0 | 0 | 0 |
| MFLRGDSV | SNRPD2; SNRP | P62316; K7E | SMD2_HUMAN | Oxidation (M) | 27855 | 0 | 0 | 0 | 304540 | 0 | 0 | 0 | 0 | 376530 | 291440 | 0 | 1350700 | 0 | 0 | 0 | 0 | 0 |
| MFLTRSEYI | PSMA5 | P28066 | PSA5_HUMAN | Acetyl (Protein N | 0 | 37850 | 24691 | 0 | 5080300 | 0 | 0 | 0 | 0 | 205950 | 0 | 0 | 2263900 | 0 | 0 | 0 | 319800 | 1222700 |
| MFLTRSEYI | PSMA5 | P28066 | PSA5_HUMAN | Acetyl (Protein N | 0 | 0 | 0 | 0 | 4415100 | 1049300 | 0 | 0 | 0 | 1402500 | 0 | 0 | 5585600 | 2701700 | 0 | 0 | 2437900 | 2546500 |
| MFLVNSFLI | CAPNS1; CAPN | P04632; A0A | CPNS1_HUMAI | Acetyl (Protein N | 0 | 23711 | 14366 | 0 | 1126200 | 0 | 104290 | 0 | 0 | 151180 | 0 | 0 | 0 | 100190 | 46644 | 0 | 1689500 | 402350 |
| MFLYADNE | VTA1; VTA1; VT/ | Q9NP79; Q5 | VTA1_HUMAN | Unmodified | 206110 | 224750 | 135900 | 192130 | 0 | 0 | 0 | 0 | 1123700 | 0 | 0 | 2560500 | 0 | 0 | 169150 | 172930 | 0 | 0 |
| MFLYADNE | VTA1; VTA1; VT/ | Q9NP79; Q5 | VTA1_HUMAN | Oxidation (M) | 0 | 0 | 0 | 0 | 0 | 0 | 0 | 0 | 335210 | 0 | 144750 | 1235900 | 0 | 0 | 0 | 0 | 0 | 0 |
| MFLYADNE | VTA1; VTA1; VT/ | Q9NP79; Q5 | VTA1_HUMAN | Unmodified | 323160 | 489770 | 353220 | 899340 | 0 | 0 | 0 | 0 | 3194000 | 0 | 317860 | 3799100 | 0 | 0 | 1153900 | 320800 | 0 | 0 |
| MFLYADNE | VTA1; VTA1; VT/ | Q9NP79; Q5 | VTA1_HUMAN | Oxidation (M) | 220410 | 385580 | 207350 | 452890 | 39185 | 0 | 0 | 0 | 961540 | 54324 | 141270 | 1610000 | 0 | 0 | 438610 | 226830 | 0 | 0 |
| MFLYADNE | VTA1; VTA1; VT/ | Q9NP79; Q5 | VTA1_HUMAN | Unmodified | 0 | 0 | 0 | 0 | 2360400 | 3475400 | 833560 | 615000 | 1420400 | 4202700 | 556000 | 158750 | 1605400 | 1991900 | 0 | 0 | 907990 | 489560 |
| MFNAENGI | TALDO1 | P37837 | TALDO_HUMAI | Unmodified | 0 | 0 | 0 | 172090 | 1146400 | 323280 | 17808 | 14746 | 97964 | 0 | 1065300 | 5252300 | 897030 | 185820 | 175720 | 9556.2 | 16487 | 209480 |
| MFQDIGVS | CUL1; CUL1; CI | Q13616; A0/ | CUL1_HUMAN | Oxidation (M) | 0 | 0 | 0 | 0 | 0 | 0 | 0 | 0 | 424950 | 0 | 120920 | 599140 | 0 | 0 | 28320 | 0 | 24208 | 0 |
| MFQVPDSE | JPT2; JPT2; JPT2 | Q9H910; B4I | JUPI2_HUMAN | Acetyl (Protein N | 0 | 0 | 0 | 53657 | 268930 | 27265 | 29589 | 0 | 0 | 0 | 0 | 0 | 277320 | 103530 | 0 | 11461 | 123890 | 158960 |
| MFRNQYDI | PSMA1; N/A | P25786; B4D | PSA1_HUMAN | Acetyl (Protein N | 0 | 0 | 0 | 327120 | 2144000 | 1578800 | 4660700 | 2624700 | 0 | 701330 | 193890 | 0 | 1578500 | 2841200 | 0 | 0 | 609030 | 1427900 |
| MFRNQYDI | PSMA1; N/A | P25786; B4D | PSA1_HUMAN | Acetyl (Protein N | 0 | 0 | 0 | 0 | 3071300 | 0 | 561140 | 0 | 0 | 0 | 0 | 0 | 1311300 | 1044000 | 0 | 0 | 802950 | 910300 |
| MFSCVKPY | CAPN5; CAPN5 | O15484; E7E | CAN5_HUMAN | Acetyl (Protein N | 0 | 0 | 0 | 0 | 0 | 0 | 0 | 0 | 0 | 0 | 0 | 0 | 1151100 | 871740 | 0 | 91244 | 227930 | 889330 |
| MFSDEILLS | POGLUT3 | Q7Z4H8 | PLGT3_HUMAN | Oxidation (M) | 0 | 0 | 0 | 9921.7 | 0 | 0 | 0 | 0 | 2312200 | 3100800 | 0 | 0 | 0 | 0 | 0 | 0 | 0 | 0 |
| MFSDEILLS | POGLUT3 | Q7Z4H8 | PLGT3_HUMAN | Unmodified | 0 | 0 | 0 | 0 | 0 | 0 | 0 | 0 | 915870 | 336620 | 0 | 0 | 0 | 0 | 0 | 0 | 0 | 0 |
| MFSGTPVII | GAS6 | Q14393 | GAS6_HUMAN | Unmodified | 0 | 0 | 0 | 0 | 0 | 0 | 0 | 0 | 4933100 | 300310 | 0 | 0 | 0 | 0 | 0 | 0 | 107870 | 0 |
| MFSWVSK | EHD1; EHD1; EI | Q9H4M9; C9 | EHD1_HUMAN | Acetyl (Protein N | 91201 | 110910 | 120970 | 0 | 0 | 395970 | 186400 | 174200 | 0 | 0 | 0 | 1272200 | 721100 | 504060 | 0 | 355910 | 2065900 | 2330000 |
| MFSWVSK | EHD1; EHD1; EI | Q9H4M9; C9 | EHD1_HUMAN | Acetyl (Protein N | 0 | 0 | 0 | 0 | 1559100 | 52839 | 47782 | 0 | 224040 | 20564 | 14571 | 67021 | 185220 | 58442 | 0 | 0 | 1642000 | 1793300 |
| MFTQQQP( | KCTD12 | Q96CX2 | KCD12_HUMAI | Unmodified | 0 | 0 | 0 | 0 | 0 | 0 | 0 | 0 | 736950 | 0 | 0 | 0 | 0 | 0 | 0 | 0 | 0 | 0 |
| MFVDLNPD | GNAQ; GNA11; | P50148; P29 | GNAQ_HUMAN | Unmodified | 0 | 0 | 0 | 0 | 0 | 0 | 0 | 0 | 0 | 0 | 0 | 887740 | 0 | 0 | 0 | 0 | 0 | 0 |
| MFVEEVST( | AARS1; AARS1; | P49588; A0A | SYAC_HUMAN | Oxidation (M) | 0 | 0 | 0 | 0 | 1474700 | 50929 | 987440 | 0 | 543500 | 219770 | 280200 | 4044600 | 2824900 | 0 | 0 | 0 | 1517700 | 0 |
| MFVEEVST( | AARS1; AARS1; | P49588; A0A | SYAC_HUMAN | Unmodified | 0 | 0 | 0 | 0 | 2511700 | 82711 | 0 | 26838 | 1174000 | 562000 | 91000 | 758080 | 1777500 | 1347300 | 0 | 0 | 155910 | 291500 |
| MFVGGLSV | HNRNPAB; HNI | Q99729; D6I | ROAA_HUMAN | Unmodified | 0 | 0 | 0 | 0 | 953990 | 0 | 0 | 0 | 1239200 | 791010 | 908110 | 7792000 | 0 | 0 | 0 | 0 | 0 | 0 |
| MFVGGLSV | HNRNPAB; HNI | Q99729; D6I | ROAA_HUMAN | Oxidation (M) | 146500 | 55105 | 0 | 0 | 0 | 0 | 128980 | 0 | 1570000 | 349400 | 815220 | 4444500 | 561330 | 0 | 0 | 0 | 0 | 0 |
| MFVGQVPI | CELF1; CELF1; | Q92879; G5I | CELF1_HUMAN | Oxidation (M) | 619250 | 0 | 185850 | 144980 | 0 | 0 | 0 | 0 | 0 | 0 | 0 | 0 | 71868 | 0 | 141080 | 140890 | 43382 | 0 |
| MFVGQVPI | CELF1; CELF1; | Q92879; G5I | CELF1_HUMAN | Unmodified | 0 | 0 | 0 | 0 | 0 | 0 | 0 | 0 | 0 | 0 | 0 | 0 | 1085700 | 0 | 0 | 0 | 0 | 0 |
| MFVLDEAD | EIF4A1; EIF4A2; | P60842; Q1 | IF4A1_HUMAN | Unmodified | 28774 | 0 | 0 | 34661 | 24044 | 0 | 0 | 0 | 1272200 | 19042 | 21228 | 107840 | 0 | 0 | 63486 | 14560 | 8727.9 | 0 |
| MFVLDEAD | EIF4A1; EIF4A2; | P60842; Q1 | IF4A1_HUMAN | Oxidation (M) | 168350 | 191530 | 50664 | 237250 | 0 | 1153300 | 0 | 0 | 736510 | 34355 | 77433 | 1381300 | 84042 | 0 | 708650 | 185570 | 29333 | 0 |
| MFVLDEAD | EIF4A1; EIF4A2; | P60842; Q1 | IF4A1_HUMAN | 2 Oxidation (M) | 562060 | 950020 | 170430 | 810000 | 0 | 0 | 37040 | 0 | 88281 | 0 | 153100 | 2656200 | 0 | 31451 | 421970 | 1552400 | 0 | 0 |
| MFVLDEAD | EIF4A1; EIF4A1; | P60842; J3K | IF4A1_HUMAN | 2 Oxidation (M) | 0 | 0 | 0 | 0 | 133340 | 0 | 94309 | 0 | 0 | 47979 | 0 | 0 | 58137 | 183210 | 0 | 0 | 96211 | 78620 |
| MFVSESK | RP2 | O75695 | XRP2_HUMAN | Unmodified | 0 | 0 | 150750 | 688000 | 904930 | 0 | 239060 | 46309 | 2299100 | 593240 | 1466300 | 9158500 | 204260 | 1144500 | 1683300 | 1426400 | 940800 | 2855500 |
| MFVSESK | RP2 | O75695 | XRP2_HUMAN | Oxidation (M) | 0 | 0 | 0 | 0 | 47716 | 38467 | 0 | 0 | 192810 | 40599 | 44987 | 777250 | 0 | 12855 | 171340 | 0 | 124290 | 175620 |
| MFYEEHLH | ADI1; ADI1 | Q9BV57; A0 | MTND_HUMAN | Oxidation (M) | 0 | 0 | 0 | 0 | 0 | 0 | 0 | 0 | 0 | 0 | 0 | 513190 | 0 | 0 | 40072 | 36096 | 0 | 0 |
| MGAIPVMII | CXADR | P78310 | CXAR_HUMAN | 2 Oxidation (M) | 417450 | 495880 | 0 | 0 | 1074900 | 0 | 0 | 410520 | 385750 | 0 | 0 | 0 | 641770 | 0 | 0 | 0 | 0 | 0 |
| MGAIPVMII | CXADR | P78310 | CXAR_HUMAN | Oxidation (M) | 0 | 0 | 0 | 0 | 2336600 | 0 | 0 | 0 | 0 | 0 | 0 | 589750 | 0 | 0 | 0 | 0 | 0 | 0 |
| MGAIPVMII | CXADR | P78310 | CXAR_HUMAN | Unmodified | 0 | 0 | 0 | 0 | 2366400 | 0 | 0 | 0 | 1354200 | 0 | 0 | 0 | 0 | 0 | 0 | 0 | 0 | 0 |
| MGAMAKPI | FABP5; FABP5 | Q01469; I6L | FABP5_HUMAN | Oxidation (M) | 0 | 0 | 0 | 0 | 1182500 | 352350 | 0 | 0 | 0 | 0 | 283240 | 2430300 | 0 | 0 | 0 | 0 | 6578900 | 4248200 |
| MGAMAKPI | FABP5; FABP5 | Q01469; I6L | FABP5_HUMAN | 2 Oxidation (M) | 218600 | 0 | 0 | 0 | 642830 | 116490 | 0 | 0 | 43925 | 0 | 243360 | 2336100 | 0 | 0 | 0 | 0 | 4250200 | 4664900 |
| MGAMAKPI | FABP5; FABP5 | Q01469; I6L | FABP5_HUMAN | Unmodified | 0 | 0 | 0 | 0 | 2149500 | 0 | 0 | 0 | 0 | 0 | 0 | 2389000 | 0 | 0 | 0 | 0 | 9553200 | 7708900 |
| MGANSLEF | HNRNPM; HNR | P52272; M0 | HNRPM_HUMA | Unmodified | 0 | 0 | 397550 | 0 | 0 | 226080 | 0 | 0 | 0 | 112690 | 0 | 288450 | 374910 | 0 | 79318 | 0 | 0 | 0 |
| MGAPESGI | FTH1; FTH1; FTI | P02794; G3 | FRIH_HUMAN | Oxidation (M) | 0 | 0 | 178390 | 0 | 0 | 0 | 0 | 0 | 27943000 | 7489700 | 1180800 | 0 | 0 | 0 | 913000 | 0 | 0 | 0 |
| MGAPESGI | FTH1; FTH1; FTI | P02794; G3 | FRIH_HUMAN | Unmodified | 0 | 0 | 0 | 0 | 81971 | 0 | 0 | 0 | 34621000 | 1453600 | 0 | 0 | 0 | 0 | 0 | 0 | 0 | 0 |
| MGAPESGI | FTH1; FTH1; FTI | P02794; G3 | FRIH_HUMAN | Oxidation (M) | 0 | 0 | 0 | 0 | 0 | 0 | 7102400 | 937340 | 15762000 | 52129000 | 0 | 0 | 1106100 | 0 | 0 | 0 | 3626800 | 2371000 |
| MGAPESGI | FTH1; FTH1; FTI | P02794; G3 | FRIH_HUMAN | Unmodified | 0 | 0 | 0 | 0 | 0 | 0 | 0 | 0 | 0 | 6573800 | 0 | 0 | 0 | 0 | 0 | 0 | 0 | 0 |
| MGATKADF | GSR | P00390 | GSHR_HUMAN | Oxidation (M) | 157370 | 0 | 0 | 677970 | 27933000 | 4396100 | 1779000 | 785250 | 387970 | 2795700 | 296840 | 0 | 9989800 | 7759700 | 0 | 0 | 4105700 | 2258300 |
| MGATKADF | GSR | P00390 | GSHR_HUMAN | Unmodified | 54877 | 0 | 0 | 0 | 22563000 | 1459800 | 617940 | 59264 | 1440000 | 612220 | 173710 | 0 | 2784700 | 1520500 | 0 | 0 | 2286600 | 2474900 |
| MGATPTPF | TJP2; TJP2; TJP2 | Q9UDY2; A0 | ZO2_HUMAN T | Unmodified | 0 | 0 | 0 | 0 | 2072000 | 53810 | 0 | 0 | 3318200 | 1752700 | 0 | 0 | 577920 | 508630 | 0 | 0 | 0 | 0 |
| MGATPTPF | TJP2; TJP2; TJP2 | Q9UDY2; A0 | ZO2_HUMAN T | Oxidation (M) | 0 | 0 | 36351 | 69128 | 1023400 | 192130 | 0 | 0 | 559700 | 915260 | 0 | 199500 | 161190 | 104420 | 29692 | 64619 | 0 | 44877 |
| MGAVFMD/ | HIBADH; HIBAD | P31937; H7E | 3HIDH_HUMAN | Unmodified | 0 | 0 | 0 | 0 | 0 | 0 | 0 | 0 | 176930 | 0 | 0 | 0 | 0 | 0 | 0 | 0 | 326530 | 0 |
| MGDTHTGI | VWA1; VWA1 | Q6PCB0; J3 | VWA1_HUMAN | Oxidation (M) | 0 | 0 | 0 | 0 | 0 | 145520 | 0 | 0 | 0 | 84927 | 0 | 0 | 0 | 0 | 404430 | 133940 | 0 | 0 |
| MGDVLSTH | NIBAN2 | Q96TA1 | NIBA2_HUMAN | Oxidation (M) | 47329 | 18303 | 71963 | 32912 | 0 | 0 | 0 | 0 | 95077 | 0 | 47551 | 602630 | 0 | 0 | 51345 | 72329 | 0 | 0 |
| MGDVYKCI | ITGA2; ITGA2; I | P17301; D6F | ITA2_HUMAN I | Oxidation (M) | 0 | 65492 | 0 | 201110 | 427960 | 0 | 274750 | 195380 | 0 | 0 | 0 | 0 | 0 | 316720 | 0 | 0 | 309260 | 1947900 |
| MGDVYKCI | ITGA2; ITGA2; I | P17301; D6F | ITA2_HUMAN I | Unmodified | 90727 | 0 | 0 | 184100 | 1161000 | 274320 | 0 | 0 | 0 | 0 | 0 | 0 | 256380 | 0 | 0 | 0 | 793230 | 3104800 |
| MGEDMHT | CAPN2 | P17655 | CAN2_HUMAN | 2 Oxidation (M) | 0 | 0 | 0 | 0 | 7808600 | 0 | 0 | 0 | 0 | 622250 | 0 | 0 | 0 | 0 | 0 | 0 | 3406900 | 4024100 |
| MGELASES | BAIAP2; BAIAP2 | Q9UQB8; I3I | BAIP2_HUMAN | Unmodified | 0 | 0 | 955540 | 0 | 1898500 | 364580 | 86171 | 119910 | 0 | 96539 | 50492 | 0 | 2357400 | 974570 | 987180 | 1048300 | 189840 | 359940 |
| MGESDDSI | RPS21; RPS21 | P63220; Q8 | RS21_HUMAN | Oxidation (M) | 861760 | 444020 | 286690 | 345030 | 0 | 0 | 0 | 0 | 0 | 0 | 679920 | 5420700 | 0 | 0 | 0 | 357300 | 58112 | 0 |
| MGESDDSI | RPS21; RPS21 | P63220; Q8 | RS21_HUMAN | Unmodified | 887660 | 604960 | 482930 | 532140 | 0 | 515290 | 0 | 0 | 281470 | 0 | 0 | 6873200 | 0 | 0 | 424170 | 0 | 0 | 0 |
| MGESDDSI | RPS21; RPS21 | P63220; Q8 | RS21_HUMAN | Unmodified | 0 | 0 | 0 | 0 | 0 | 0 | 131050 | 0 | 0 | 148160 | 0 | 0 | 671770 | 243940 | 0 | 0 | 0 | 611170 |
| MGFGNLVI | TAOK3; TAOK3; | Q9H2K8; G3 | TAOK3_HUMAN | Oxidation (M) | 0 | 0 | 0 | 1751200 | 0 | 0 | 0 | 0 | 73107 | 0 | 0 | 0 | 0 | 0 | 0 | 854880 | 0 | 0 |
| MGFNIGVR | TRAPPC3; TRAF | O43617; A6 | TPPC3_HUMAN | Unmodified | 0 | 0 | 405600 | 49995 | 0 | 398970 | 0 | 0 | 224250 | 113340 | 0 | 1085800 | 73687 | 470360 | 0 | 552380 | 0 | 209480 |
| MGFNIGVR | TRAPPC3; TRAF | O43617; A6 | TPPC3_HUMAN | Oxidation (M) | 46912 | 0 | 0 | 39502 | 0 | 0 | 0 | 0 | 666360 | 172940 | 95137 | 382870 | 0 | 0 | 100120 | 126810 | 0 | 0 |
| MGFPEAAS | SCP2; SCP2; S( | P22307; H0 | SCP2_HUMAN | Unmodified | 0 | 0 | 0 | 0 | 0 | 0 | 0 | 0 | 3260700 | 199430 | 211240 | 1945800 | 0 | 0 | 0 | 0 | 0 | 0 |
| MGGDIANI | PA2G4; PA2G4 | Q9UQ80; F8 | PA2G4_HUMAN | Unmodified | 1624800 | 965090 | 0 | 310360 | 0 | 0 | 0 | 0 | 496630 | 24243 | 417500 | 12831000 | 0 | 0 | 1084900 | 0 | 0 | 0 |
| MGGDIANI | PA2G4; PA2G4 | Q9UQ80; F8 | PA2G4_HUMAN | Oxidation (M) | 90886 | 24184 | 0 | 0 | 0 | 0 | 0 | 0 | 0 | 0 | 0 | 970100 | 0 | 0 | 0 | 172990 | 0 | 0 |
| MGGEEAEI | DHX9 | Q08211 | DHX9_HUMAN | Unmodified | 0 | 41085 | 0 | 0 | 0 | 0 | 0 | 0 | 2204000 | 0 | 2543500 | 4578600 | 0 | 0 | 39028 | 0 | 0 | 0 |
| MGGEEAEI | DHX9 | Q08211 | DHX9_HUMAN | Oxidation (M) | 361780 | 0 | 0 | 0 | 0 | 0 | 0 | 0 | 0 | 37116 | 48238 | 1978700 | 0 | 0 | 0 | 157510 | 0 | 0 |
| MGGEEKPI | TARS1; TARS1; | P26639; D6F | SYTC_HUMAN | Unmodified | 0 | 218520 | 545120 | 841080 | 5926100 | 1216900 | 204630 | 0 | 1762300 | 391240 | 1805500 | 9613900 | 2868900 | 3042800 | 615760 | 642180 | 424050 | 956240 |
| MGGNQLFI | GALNT7; GALN | Q86SF2; H0 | GALT7_HUMAN | Unmodified | 0 | 0 | 0 | 0 | 0 | 0 | 0 | 0 | 2996900 | 223970 | 255430 | 543430 | 0 | 0 | 0 | 0 | 0 | 0 |
| MGGNQLFI | GALNT7; GALN | Q86SF2; H0 | GALT7_HUMAN | Oxidation (M) | 0 | 0 | 0 | 0 | 0 | 0 | 0 | 0 | 1022700 | 95347 | 0 | 0 | 0 | 0 | 0 | 0 | 0 | 0 |
| MGGSAPPI | FOLH1B; FOLH | Q9HBA9; Q0 | FOH1B_HUMAI | Oxidation (M) | 1244800 | 672910 | 3289000 | 2819100 | 0 | 0 | 0 | 0 | 0 | 0 | 0 | 0 | 0 | 0 | 2638800 | 3744400 | 0 | 0 |
| MGGSAPPI | FOLH1B; FOLH | Q9HBA9; Q0 | FOH1B_HUMAI | Unmodified | 1587700 | 1018000 | 4313200 | 4626500 | 0 | 0 | 0 | 0 | 0 | 0 | 0 | 0 | 0 | 0 | 3762700 | 6826600 | 0 | 0 |
| MGGSAPPI | FOLH1B; FOLH | Q9HBA9; Q0 | FOH1B_HUMAI | Unmodified | 0 | 0 | 2832600 | 8032000 | 32289 | 0 | 0 | 126180 | 0 | 0 | 0 | 0 | 935630 | 190940 | 3613500 | 0 | 0 | 0 |
| MGGSAPPI | FOLH1B; FOLH | Q9HBA9; Q0 | FOH1B_HUMAI | Oxidation (M) | 323140 | 0 | 1175100 | 2679100 | 0 | 272800 | 0 | 0 | 0 | 0 | 0 | 0 | 0 | 0 | 0 | 629840 | 0 | 0 |
| MGGSSGAI | TKFC; TKFC; TK | Q3LXA3; I3L | TKFC_HUMAN | Oxidation (M) | 0 | 0 | 0 | 445930 | 0 | 0 | 425660 | 0 | 103300 | 0 | 0 | 588320 | 1145000 | 0 | 605860 | 117180 | 701300 | 0 |
| MGHAGAII | SUCLG1 | P53597 | SUCA_HUMAN | Unmodified | 0 | 0 | 0 | 0 | 0 | 0 | 0 | 0 | 522820 | 0 | 0 | 0 | 0 | 0 | 0 | 0 | 0 | 0 |
| MGIEILSFTI | FLOT2; FLOT2; | Q14254; J3Q | FLOT2_HUMAN | Oxidation (M) | 174720 | 154600 | 370590 | 747860 | 0 | 0 | 0 | 0 | 1785600 | 103000 | 353760 | 1008400 | 0 | 0 | 1592700 | 428760 | 0 | 0 |
| MGIEILSFTI | FLOT2; FLOT2; | Q14254; J3Q | FLOT2_HUMAN | Unmodified | 0 | 0 | 0 | 0 | 72408 | 0 | 0 | 0 | 462030 | 0 | 0 | 0 | 0 | 0 | 0 | 0 | 15011 | 0 |
| MGIGGLDK | NSF; NSF | P46459; I3L | NSF_HUMAN V | Oxidation (M) | 37986 | 0 | 0 | 78276 | 0 | 0 | 15367 | 159950 | 80169 | 0 | 0 | 1459300 | 0 | 318880 | 221230 | 48952 | 103120 | 180410 |
| MGIGGLDK | NSF; NSF | P46459; I3L | NSF_HUMAN V | Oxidation (M) | 0 | 0 | 0 | 0 | 0 | 0 | 0 | 0 | 10924 | 0 | 0 | 0 | 2537500 | 229910 | 0 | 0 | 209330 | 88491 |
| MGIRVDQL | CALCOCO2; C/ | Q13137; D6I | CACO2_HUMA | Oxidation (M) | 41790 | 123040 | 52398 | 0 | 505730 | 0 | 1734800 | 272590 | 340160 | 0 | 0 | 0 | 193350 | 381970 | 79541 | 0 | 459220 | 1341400 |
| MGIRVDQL | CALCOCO2; C/ | Q13137; D6I | CACO2_HUMA | Unmodified | 0 | 0 | 0 | 0 | 1641900 | 141650 | 2170000 | 101020 | 1590600 | 503220 | 0 | 0 | 801860 | 439570 | 0 | 0 | 1174000 | 4233800 |
| MGITEYNN | IARS1; IARS1; I/ | P41252; A0A | SYIC_HUMAN I | Unmodified | 0 | 0 | 0 | 0 | 0 | 0 | 0 | 43272 | 0 | 0 | 0 | 665830 | 0 | 0 | 0 | 0 | 0 | 0 |
| MGKIDIDY( | SF3B2; SF3B2 | Q13435; E9F | SF3B2_HUMAN | Unmodified | 0 | 0 | 0 | 113570 | 0 | 0 | 0 | 0 | 0 | 0 | 0 | 308560 | 709740 | 0 | 0 | 0 | 0 | 0 |
| MGLALSSLI | SGTA; SGTA; S( | O43765; A0/ | SGTA_HUMAN | Unmodified | 0 | 0 | 0 | 0 | 787620 | 0 | 0 | 0 | 0 | 0 | 264710 | 1625600 | 725450 | 0 | 0 | 0 | 145410 | 334910 |
| MGLALSSLI | SGTA; SGTA; S( | O43765; A0/ | SGTA_HUMAN | Oxidation (M) | 0 | 0 | 54437 | 315220 | 0 | 0 | 0 | 0 | 0 | 190640 | 219630 | 2801200 | 0 | 0 | 0 | 52712 | 49411 | 0 |
| MGLDGLQ( | ATP2C1; ATP2C | P98194; B4E | AT2C1_HUMAN | Unmodified | 0 | 459800 | 0 | 0 | 241120 | 842620 | 0 | 0 | 1977300 | 0 | 0 | 0 | 0 | 430430 | 0 | 531390 | 137520 | 621090 |
| MGLGHEQ( | TES; TES | Q9UGI8; F8 | TES_HUMAN T | Oxidation (M) | 0 | 0 | 0 | 0 | 353460 | 0 | 64174 | 0 | 0 | 0 | 0 | 0 | 0 | 498990 | 0 | 0 | 583660 | 882920 |
| MGLVDQL\ | HADHA; HADH/ | P40939; A0A | ECHA_HUMAN | Oxidation (M) | 0 | 0 | 0 | 0 | 0 | 0 | 0 | 0 | 2185700 | 0 | 0 | 0 | 441200 | 0 | 0 | 0 | 0 | 0 |
| MGLVDQL\ | HADHA; HADH/ | P40939; A0A | ECHA_HUMAN | Unmodified | 0 | 0 | 0 | 0 | 0 | 0 | 0 | 0 | 0 | 0 | 0 | 0 | 0 | 0 | 0 | 0 | 0 | 0 |
| MGNFPWQ | C1R; C1R; C1R; | P00736; B4D | C1R_HUMAN C | Oxidation (M) | 0 | 0 | 0 | 0 | 0 | 0 | 0 | 0 | 753920 | 222910 | 0 | 0 | 0 | 0 | 0 | 0 | 0 | 0 |
| MGNTPDS/ | SUMF2; SUMF2 | Q8NBJ7; J3Q | SUMF2_HUMAI | Oxidation (M) | 51728 | 69914 | 0 | 0 | 0 | 0 | 0 | 0 | 9978900 | 1115800 | 214170 | 0 | 0 | 0 | 0 | 0 | 0 | 0 |
| MGNTPDS/ | SUMF2; SUMF2 | Q8NBJ7; J3Q | SUMF2_HUMAI | Unmodified | 0 | 0 | 0 | 0 | 0 | 0 | 0 | 0 | 25509000 | 427190 | 0 | 0 | 376980 | 0 | 0 | 0 | 0 | 0 |
| MGNTPDS/ | SUMF2; SUMF2 | Q8NBJ7; J3Q | SUMF2_HUMAI | Oxidation (M) | 0 | 0 | 0 | 0 | 0 | 0 | 0 | 0 | 0 | 4894300 | 110280 | 0 | 0 | 0 | 0 | 0 | 0 | 0 |
| MGNTPDS/ | SUMF2; SUMF2 | Q8NBJ7; J3Q | SUMF2_HUMAI | Unmodified | 0 | 0 | 0 | 0 | 0 | 0 | 0 | 0 | 0 | 3048800 | 0 | 0 | 0 | 0 | 0 | 0 | 0 | 0 |
| MGPGAAS( | RAB1C; RAB1B; | Q92928; Q9I | RAB1C_HUMAI | Unmodified | 4309900 | 3290100 | 3460100 | 4818600 | 33696 | 0 | 296700 | 0 | 3917700 | 1133500 | 4544100 | 26182000 | 0 | 0 | 5148000 | 4095900 | 0 | 0 |
| MGPGAAS( | RAB1C; RAB1B; | Q92928; Q9I | RAB1C_HUMAI | Oxidation (M) | 3702500 | 3609200 | 4207700 | 2998000 | 69460 | 0 | 0 | 44054 | 1381100 | 678670 | 3177100 | 12887000 | 159440 | 77736 | 3750600 | 4572700 | 80117 | 282770 |
| MGPGATA( | RAB1A; RAB1A | P62820; E7E | RAB1A_HUMAN | Oxidation (M) | 0 | 0 | 29580 | 0 | 133830 | 0 | 0 | 0 | 80384 | 88847 | 260620 | 302320 | 177010 | 97839 | 0 | 74613 | 235170 | 109950 |
| MGPGATA( | RAB1A; RAB1A | P62820; E7E | RAB1A_HUMAN | Unmodified | 759370 | 912890 | 538300 | 0 | 318000 | 0 | 0 | 0 | 421090 | 961610 | 1414700 | 9094400 | 234490 | 1368100 | 0 | 1084700 | 0 | 0 |
| MGPSFPSP | NECTIN2 | Q92692 | NECT2_HUMAN | Unmodified | 222110 | 247160 | 0 | 229360 | 0 | 135260 | 0 | 0 | 647610 | 0 | 0 | 0 | 0 | 0 | 0 | 711190 | 0 | 364500 |
| MGPSGGE( | LASP1; LASP1 | Q14847; C9 | LASP1_HUMAN | Unmodified | 430130 | 0 | 201820 | 0 | 1471100 | 0 | 0 | 4486500 | 0 | 63183 | 572190 | 894710 | 0 | 0 | 331720 | 267440 | 0 | 11103000 |
| MGPTELLIE | FGB; FGB | P02675; D6F | FIBB_HUMAN F | 2 Oxidation (M) | 0 | 735970 | 0 | 0 | 0 | 0 | 0 | 0 | 0 | 0 | 0 | 0 | 0 | 0 | 0 | 0 | 0 | 0 |

| | | | | | | | | | | | | | | | | | | | | | | |
|---|---|---|---|---|---|---|---|---|---|---|---|---|---|---|---|---|---|---|---|---|---|---|
| MGQEFVES | WASF2 | Q9Y6W5 | WASF2_HUMAI | Unmodified | 0 | 0 | 0 | 0 | 1264200 | 0 | 151800 | 0 | 700480 | 145960 | 0 | 886390 | 816180 | 703200 | 0 | 0 | 0 | 873600 |
| MGQEFVES | WASF2 | Q9Y6W5 | WASF2_HUMAI | Unmodified | 0 | 0 | 0 | 0 | 0 | 0 | 135200 | 0 | 160350 | 0 | 0 | 0 | 0 | 759210 | 0 | 54504 | 700230 | 115960 |
| MGQLGLGI | RCC2 | Q9P258 | RCC2_HUMAN | 2 Oxidation (M) | 0 | 0 | 0 | 0 | 0 | 0 | 0 | 0 | 0 | 0 | 0 | 1481800 | 0 | 0 | 0 | 0 | 0 | 0 |
| MGQNLNR | ILK; ILK | Q13418; A0A | ILK_HUMAN Int | Unmodified | 0 | 452360 | 573220 | 0 | 1640900 | 53874 | 0 | 0 | 579130 | 0 | 0 | 0 | 0 | 641030 | 844560 | 0 | 0 | 723240 |
| MGQNLNR | ILK; ILK | Q13418; A0A | ILK_HUMAN Int | Unmodified | 0 | 0 | 0 | 0 | 525640 | 0 | 0 | 0 | 0 | 0 | 0 | 0 | 0 | 0 | 0 | 0 | 820620 | 522420 |
| MGQNLNR | ILK; ILK | Q13418; A0A | ILK_HUMAN Int | Oxidation (M) | 0 | 0 | 0 | 0 | 316820 | 0 | 562390 | 30454 | 0 | 0 | 0 | 0 | 453310 | 372820 | 0 | 0 | 91890 | 257040 |
| MGQRPAA( | TNFAIP2 | Q03169 | TNAP2_HUMAN | Oxidation (M) | 0 | 0 | 0 | 0 | 0 | 0 | 0 | 0 | 69875 | 121780 | 0 | 1778800 | 85536 | 0 | 0 | 0 | 0 | 0 |
| MGRYNLSF | SORD; SORD | Q00796; A0A | DHSO_HUMAN | Oxidation (M) | 0 | 0 | 0 | 0 | 0 | 0 | 0 | 0 | 0 | 0 | 0 | 0 | 4686000 | 4904000 | 0 | 0 | 0 | 0 |
| MGTANIQL | BSG; BSG; BSG | P35613; A0A | BASI_HUMAN E | Unmodified | 0 | 0 | 0 | 0 | 0 | 96869 | 346760 | 156040 | 0 | 58616 | 0 | 0 | 214510 | 264740 | 0 | 0 | 664860 | 486300 |
| MGTLVSLEI | PSMG3 | Q9BT73 | PSMG3_HUMA | Oxidation (M) | 0 | 0 | 0 | 0 | 0 | 0 | 0 | 0 | 0 | 336000 | 0 | 1683700 | 0 | 0 | 0 | 0 | 0 | 0 |
| MGTPSCSF | DNAJB12; DNA. | Q9NXW2; V9 | DJB12_HUMAN | Unmodified | 1055800 | 0 | 0 | 0 | 0 | 0 | 0 | 0 | 0 | 1685800 | 0 | 0 | 0 | 0 | 0 | 0 | 0 | 0 |
| MGVGDRC | CLTCL1; CLTCL | A0A087WV7 | A0A087WV74_I | Acetyl (Protein N | 0 | 0 | 0 | 0 | 0 | 553910 | 0 | 0 | 0 | 0 | 0 | 0 | 0 | 0 | 0 | 0 | 0 | 0 |
| MGVGKLEN | MAP1B; MAP1B | P46821; D6F | MAP1B_HUMAI | Oxidation (M) | 0 | 0 | 0 | 0 | 0 | 0 | 333210 | 0 | 0 | 0 | 0 | 0 | 151210 | 0 | 0 | 0 | 0 | 0 |
| MGVPYCIIK | RPL7A; RPL7A | P62424; Q5T | RL7A_HUMAN ( | Oxidation (M) | 1389000 | 0 | 167630 | 0 | 0 | 0 | 1171100 | 837360 | 0 | 151510 | 191080 | 1988000 | 0 | 0 | 250880 | 226620 | 0 | 0 |
| MGYAEEAP | PCMT1; PCMT1 | P22061; F6S | PIMT_HUMAN F | Oxidation (M) | 0 | 0 | 0 | 0 | 0 | 0 | 0 | 0 | 7282000 | 0 | 1818700 | 0 | 0 | 0 | 0 | 742360 | 0 | 0 |
| MGYTPLHV | ANK3 | Q12955 | ANK3_HUMAN | Oxidation (M) | 0 | 0 | 0 | 0 | 0 | 0 | 0 | 0 | 0 | 0 | 186700 | 0 | 0 | 2308600 | 1312700 | 0 | 0 | 0 |
| MGYTPLHV | ANK3 | Q12955 | ANK3_HUMAN | Unmodified | 0 | 0 | 0 | 0 | 0 | 0 | 0 | 0 | 0 | 0 | 206000 | 0 | 1875800 | 1760000 | 0 | 0 | 0 | 0 |
| MHDGTTPL | NOTCH1; NOT( | P46531; A0A | NOTC1_HUMA | Unmodified | 0 | 0 | 0 | 176070 | 0 | 0 | 0 | 0 | 818060 | 0 | 52363 | 217160 | 0 | 0 | 0 | 0 | 0 | 0 |
| MHDGTTPL | NOTCH1; NOT( | P46531; A0A | NOTC1_HUMA | Oxidation (M) | 557570 | 119610 | 487720 | 742100 | 0 | 0 | 209640 | 27382 | 661360 | 27183 | 394070 | 3159500 | 0 | 0 | 0 | 0 | 0 | 0 |
| MHDLNTD( | BUB3; BUB3; BI | O43684; J3Q | BUB3_HUMAN | Oxidation (M) | 761340 | 216200 | 335420 | 0 | 158490 | 0 | 0 | 92199 | 302260 | 0 | 741880 | 16309000 | 0 | 0 | 0 | 0 | 0 | 0 |
| MHDLNTD( | BUB3; BUB3; BI | O43684; J3Q | BUB3_HUMAN | Unmodified | 0 | 0 | 63319 | 0 | 0 | 0 | 0 | 0 | 0 | 0 | 0 | 11940000 | 0 | 0 | 0 | 0 | 0 | 0 |
| MHDYDGN | MCFD2; MCFD2 | Q8NI22; A0A | MCFD2_HUMA | Oxidation (M) | 0 | 0 | 0 | 0 | 0 | 0 | 0 | 0 | 2312000 | 0 | 0 | 0 | 0 | 0 | 0 | 0 | 0 | 0 |
| MHEDINEE | NDUFS1; NDUF | P28331; B4D | NDUS1_HUMA | Oxidation (M) | 0 | 0 | 0 | 0 | 158470 | 0 | 0 | 0 | 678000 | 117730 | 0 | 0 | 0 | 0 | 0 | 0 | 0 | 0 |
| MHEDINEE | NDUFS1; NDUF | P28331; B4D | NDUS1_HUMA | Unmodified | 0 | 0 | 0 | 0 | 0 | 0 | 0 | 0 | 1025200 | 0 | 0 | 0 | 0 | 0 | 0 | 0 | 0 | 0 |
| MHEFSAKD | GPX4; GPX4; GI | P36969; A0A | GPX4_HUMAN | Acetyl (Protein N | 0 | 0 | 91471 | 0 | 0 | 0 | 0 | 0 | 0 | 0 | 0 | 0 | 0 | 0 | 0 | 0 | 0 | 0 |
| MHFESGST | UCHL3; UCHL3 | P15374; Q5T | UCHL3_HUMAI | Unmodified | 0 | 0 | 63423 | 110700 | 0 | 0 | 35720 | 0 | 0 | 0 | 199170 | 1248200 | 525730 | 152050 | 0 | 0 | 354170 | 660540 |
| MHFKDLIG | N/A | N/A | N/A | 2 Oxidation (M) | 0 | 0 | 946180 | 1192200 | 0 | 3991900 | 0 | 171340 | 0 | 0 | 0 | 0 | 0 | 0 | 0 | 0 | 0 | 830950 |
| MHFPRSSL | PTK7; PTK7 | Q13308; C9J | PTK7_HUMAN I | Oxidation (M) | 0 | 0 | 0 | 0 | 4232100 | 0 | 2100100 | 0 | 0 | 1690000 | 0 | 0 | 4489600 | 4163200 | 0 | 0 | 0 | 9129700 |
| MHGGGPT | MTHFD1; MTHF | P11586; V9C | C1TC_HUMAN | Oxidation (M) | 189950 | 0 | 885010 | 644320 | 2673100 | 847810 | 4455000 | 5013000 | 0 | 1440900 | 1078400 | 5807100 | 2149500 | 0 | 519390 | 560470 | 3625500 | 1349600 |
| MHGGGPT | MTHFD1; MTHF | P11586; V9C | C1TC_HUMAN | Unmodified | 0 | 0 | 0 | 0 | 3938400 | 0 | 4341100 | 0 | 1199300 | 670820 | 282240 | 1641900 | 2596500 | 0 | 0 | 0 | 3495100 | 3806200 |
| MHIAQDIN | NDRG3; NDRG3 | Q9UGV2; F8 | NDRG3_HUMA | Oxidation (M) | 0 | 0 | 0 | 0 | 2988000 | 0 | 0 | 0 | 0 | 0 | 0 | 1766800 | 244600 | 0 | 0 | 0 | 0 | 0 |
| MHIAQDIN | NDRG3; NDRG3 | Q9UGV2; F8 | NDRG3_HUMA | Oxidation (M) | 0 | 0 | 0 | 319010 | 0 | 0 | 0 | 0 | 0 | 0 | 0 | 15602000 | 0 | 0 | 1972700 | 462910 | 0 | 0 |
| MHIHSTNE | FOLH1B; FOLH | Q9HBA9; Q0 | FOH1B_HUMAI | Oxidation (M) | 1038000 | 1158900 | 4514100 | 14270000 | 0 | 0 | 0 | 0 | 13747 | 0 | 0 | 0 | 112260 | 0 | 4272800 | 4979400 | 0 | 0 |
| MHIHSTNE | FOLH1B; FOLH | Q9HBA9; Q0 | FOH1B_HUMAI | Unmodified | 0 | 13297 | 0 | 7018500 | 0 | 0 | 0 | 0 | 0 | 0 | 0 | 0 | 0 | 0 | 4467400 | 0 | 0 | 0 |
| MHIHSTNE | FOLH1B; FOLH | Q9HBA9; Q0 | FOH1B_HUMAI | Oxidation (M) | 0 | 0 | 222620 | 1385100 | 0 | 0 | 0 | 0 | 2067200 | 0 | 0 | 0 | 324490 | 128960 | 223450 | 144910 | 94693 | 829150 |
| MHKEEHE\ | IFITM1; IFITM1; | P13164; A0A | IFM1_HUMAN I | Oxidation (M) | 0 | 0 | 0 | 0 | 0 | 0 | 0 | 0 | 0 | 0 | 0 | 0 | 0 | 1018200 | 0 | 0 | 0 | 13184000 |
| MHLAGKTE | PDAP1 | Q13442 | HAP28_HUMAI | Unmodified | 578480 | 328130 | 53159 | 531440 | 993040 | 209950 | 0 | 0 | 87027 | 142120 | 21015 | 161020 | 512900 | 435660 | 255340 | 0 | 465590 | 1133500 |
| MHLGLVIPI | ASPH | Q12797 | ASPH_HUMAN | Unmodified | 0 | 0 | 0 | 0 | 0 | 0 | 0 | 0 | 892530 | 177060 | 0 | 0 | 0 | 0 | 0 | 0 | 0 | 0 |
| MHLPSPTD | EGFR; EGFR; E( | P00533; Q5( | EGFR_HUMAN | Unmodified | 0 | 89561 | 265400 | 695510 | 0 | 0 | 0 | 0 | 0 | 0 | 0 | 0 | 0 | 0 | 672140 | 399770 | 0 | 0 |
| MHLPSPTD | EGFR; EGFR; E( | P00533; Q5( | EGFR_HUMAN | Oxidation (M) | 0 | 0 | 551050 | 152820 | 0 | 0 | 0 | 0 | 0 | 0 | 0 | 0 | 0 | 0 | 158450 | 118820 | 0 | 0 |
| MHNQIPQ\ | DDX1; DDX1; D | Q92499; A0A | DDX1_HUMAN | Unmodified | 68974 | 23157 | 0 | 0 | 0 | 0 | 0 | 0 | 0 | 0 | 209420 | 1414000 | 0 | 0 | 0 | 0 | 0 | 0 |
| MHNQIPQ\ | DDX1; DDX1; D | Q92499; A0A | DDX1_HUMAN | Oxidation (M) | 83792 | 49998 | 132370 | 98278 | 0 | 0 | 28579 | 35880 | 52021 | 0 | 231590 | 992640 | 0 | 0 | 96119 | 86420 | 14736 | 29290 |
| MHRDSCPI | SRSF3; SRSF3 | P84103; A0A | SRSF3_HUMAN | Acetyl (Protein N | 225380 | 0 | 1652400 | 693750 | 220750 | 1769400 | 197900 | 115910 | 0 | 0 | 0 | 0 | 343980 | 385540 | 80162 | 160140 | 35463 | 0 |
| MHRDSCPI | SRSF3; SRSF3 | P84103; A0A | SRSF3_HUMAN | Acetyl (Protein N | 463620 | 163180 | 65068 | 0 | 0 | 0 | 588760 | 334690 | 70456 | 184940 | 39187 | 71894 | 1192300 | 1080300 | 54102 | 33892 | 0 | 0 |
| MHSLLIK | COPA; COPA; C | P53621; A0A | COPA_HUMAN | Unmodified | 0 | 0 | 0 | 0 | 0 | 0 | 0 | 0 | 0 | 0 | 0 | 610550 | 0 | 0 | 0 | 0 | 0 | 0 |
| MHSPQTSA | AARS1; AARS1; | P49588; A0A | SYAC_HUMAN | 2 Oxidation (M) | 82062 | 0 | 0 | 0 | 0 | 0 | 0 | 0 | 0 | 0 | 0 | 1940100 | 0 | 0 | 0 | 331570 | 0 | 0 |
| MHSPQTSA | AARS1; AARS1; | P49588; A0A | SYAC_HUMAN | 2 Oxidation (M) | 0 | 0 | 0 | 0 | 10900000 | 0 | 0 | 0 | 0 | 357300 | 0 | 0 | 0 | 0 | 0 | 0 | 2689500 | 0 |
| MHSVGICG | SORD; SORD; S | Q00796; H0Y | DHSO_HUMAN | Oxidation (M) | 0 | 0 | 977950 | 1360000 | 0 | 0 | 0 | 0 | 0 | 0 | 0 | 3434500 | 0 | 0 | 1375700 | 1198700 | 0 | 0 |
| MHSVGICG | SORD; SORD; S | Q00796; H0Y | DHSO_HUMAN | Oxidation (M) | 0 | 0 | 0 | 0 | 0 | 0 | 0 | 0 | 0 | 0 | 0 | 0 | 3136000 | 1460800 | 0 | 0 | 0 | 0 |
| MHTCFDVY | EXT2; EXT2 | Q93063; E9F | EXT2_HUMAN I | Unmodified | 0 | 0 | 0 | 0 | 0 | 0 | 0 | 0 | 460640 | 0 | 0 | 0 | 0 | 0 | 0 | 0 | 0 | 0 |
| MHTTFEHD | SPTBN1; SPTBN | Q01082; A0A | SPTB2_HUMAN | Oxidation (M) | 67242 | 0 | 0 | 0 | 0 | 0 | 0 | 0 | 0 | 121720 | 0 | 1091600 | 0 | 0 | 0 | 0 | 36864 | 0 |
| MHTTFEHD | SPTBN1; SPTBN | Q01082; A0A | SPTB2_HUMAN | Unmodified | 0 | 0 | 0 | 0 | 0 | 0 | 0 | 0 | 0 | 0 | 0 | 650300 | 359590 | 0 | 0 | 0 | 0 | 0 |
| MIAAVDTD: | BAX; BAX; BAX; | Q07812; I6LI | BAX_HUMAN A | Unmodified | 514150 | 7559100 | 0 | 0 | 0 | 0 | 483850 | 244730 | 0 | 0 | 0 | 794130 | 0 | 0 | 0 | 1569100 | 0 | 300660 |
| MIAAVDTD: | BAX; BAX; BAX; | Q07812; I6LI | BAX_HUMAN A | Oxidation (M) | 75051000 | 290200 | 332710 | 234450 | 0 | 0 | 79881 | 31026 | 0 | 0 | 0 | 716160 | 0 | 0 | 0 | 94825 | 59191 | 0 |
| MIAILTENY( | TARS1 | P26639 | SYTC_HUMAN | Oxidation (M) | 0 | 0 | 0 | 84322 | 0 | 0 | 0 | 0 | 0 | 0 | 0 | 453460 | 0 | 0 | 0 | 0 | 0 | 0 |
| MIAPEGSL\ | PITPNB; PITPNE | P48739; A0A | PIPNB_HUMAN | Oxidation (M) | 0 | 0 | 402990 | 718890 | 1525000 | 569430 | 0 | 0 | 0 | 0 | 694710 | 2599000 | 1857800 | 1352900 | 515820 | 479650 | 837030 | 1380500 |
| MIAPEGSL\ | PITPNB; PITPNE | P48739; A0A | PIPNB_HUMAN | Unmodified | 105960 | 0 | 0 | 0 | 3949500 | 0 | 456830 | 344050 | 764950 | 357430 | 682520 | 1741300 | 1215000 | 811410 | 340540 | 0 | 1225200 | 2479200 |
| MIATDPDD | COL12A1; COL | Q99715; D6I | COCA1_HUMA | Oxidation (M) | 0 | 0 | 82046 | 0 | 0 | 0 | 0 | 0 | 404380 | 141410 | 0 | 0 | 0 | 0 | 156540 | 128680 | 0 | 0 |
| MIATDPDD | COL12A1; COL | Q99715; D6I | COCA1_HUMA | Unmodified | 0 | 0 | 0 | 0 | 0 | 0 | 0 | 0 | 1661400 | 0 | 0 | 0 | 0 | 0 | 0 | 0 | 0 | 0 |
| MICSIQNDI | PCDHB5 | Q9Y5E4 | PCDB5_HUMAI | Oxidation (M) | 0 | 0 | 0 | 0 | 0 | 0 | 27936 | 0 | 369190 | 0 | 0 | 0 | 0 | 0 | 0 | 0 | 0 | 0 |
| MIDERQQE | VCL | P18206 | VINC_HUMAN | Unmodified | 299290 | 497470 | 0 | 1342200 | 1106400 | 374190 | 0 | 0 | 2650600 | 304120 | 0 | 2535800 | 0 | 0 | 702700 | 411820 | 2015900 | 837370 |
| MIDERQQE | VCL | P18206 | VINC_HUMAN | 2 Oxidation (M) | 0 | 0 | 0 | 0 | 1056800 | 0 | 1328400 | 207370 | 0 | 118430 | 0 | 0 | 282080 | 617820 | 0 | 0 | 6304600 | 140710 |
| MIDERQQE | VCL | P18206 | VINC_HUMAN | Oxidation (M) | 0 | 0 | 0 | 0 | 4158500 | 285470 | 2910300 | 0 | 0 | 182920 | 0 | 0 | 949600 | 1589300 | 0 | 0 | 22972000 | 6380700 |
| MIDERQQE | VCL | P18206 | VINC_HUMAN | Unmodified | 0 | 0 | 0 | 0 | 4756600 | 225350 | 2828800 | 0 | 0 | 0 | 0 | 0 | 1156300 | 463720 | 0 | 0 | 12897000 | 8546100 |
| MIDGESGE | CACNA2D1; CA | P54289; H0Y | CA2D1_HUMAI | Unmodified | 819250 | 274720 | 0 | 0 | 0 | 0 | 5250700 | 4168000 | 934360 | 220240 | 0 | 0 | 0 | 0 | 0 | 0 | 0 | 0 |
| MIDGESGE | CACNA2D1; CA | P54289; H0Y | CA2D1_HUMAI | Unmodified | 0 | 0 | 0 | 0 | 0 | 0 | 1138000 | 0 | 0 | 0 | 0 | 0 | 0 | 0 | 0 | 0 | 0 | 0 |
| MIDKNLREI | GNAI2; GNAI2 | P04899; F8W | GNAI2_HUMAN | Unmodified | 0 | 0 | 0 | 0 | 0 | 0 | 3015400 | 650160 | 0 | 0 | 0 | 0 | 0 | 22936 | 0 | 0 | 0 | 0 |
| MIDNVYNP | VLDLR; VLDLR; | P98155; A0A | VLDLR_HUMAN | Oxidation (M) | 0 | 0 | 0 | 0 | 0 | 0 | 23869 | 0 | 312350 | 179970 | 0 | 0 | 0 | 0 | 44517 | 93435 | 39585 | 0 |
| MIDPKACLI | SERPINB5 | P36952 | SPB5_HUMAN | Oxidation (M) | 0 | 0 | 0 | 0 | 0 | 0 | 0 | 0 | 0 | 0 | 0 | 0 | 111270 | 59395 | 0 | 0 | 1564100 | 264560 |
| MIDPKACLI | SERPINB5 | P36952 | SPB5_HUMAN | Unmodified | 0 | 0 | 15350 | 0 | 0 | 0 | 0 | 0 | 0 | 0 | 0 | 0 | 94887 | 0 | 0 | 0 | 1111900 | 514310 |
| MIDRNLREI | GNAI3; GNAI1; | P08754; P63 | GNAI3_HUMAN | Unmodified | 1285300 | 918810 | 2919800 | 3465500 | 11427000 | 3665000 | 0 | 1108700 | 5283300 | 2020900 | 1085700 | 3106200 | 22047000 | 1853800 | 260150 | 309580 | 4188700 | 21330000 |
| MIEAVDNN | ANO6; ANO6; A | Q4KMQ2; A0 | ANO6_HUMAN | Unmodified | 0 | 0 | 0 | 0 | 466220 | 387380 | 36366 | 44094 | 0 | 148410 | 0 | 0 | 0 | 0 | 0 | 0 | 310150 | 426420 |
| MIEAVQNH | TXNRD1; TXNRI | Q16881; A0A | TRXR1_HUMAN | Unmodified | 0 | 0 | 0 | 0 | 0 | 0 | 0 | 0 | 0 | 103420 | 0 | 492500 | 0 | 0 | 0 | 0 | 0 | 0 |
| MIEDAER | CBS; CBSL; CB: | P35520; P0D | CBS_HUMAN C | Unmodified | 176790 | 616960 | 0 | 0 | 0 | 0 | 0 | 0 | 927310 | 73424 | 91397 | 979910 | 0 | 0 | 87504 | 0 | 0 | 0 |
| MIEEAGAII: | ACOT7; ACOT7 | O00154; K7E | BACH_HUMAN | Unmodified | 0 | 0 | 0 | 0 | 0 | 0 | 0 | 0 | 2250500 | 529870 | 345880 | 0 | 0 | 0 | 0 | 0 | 0 | 0 |
| MIEEAGAII: | ACOT7; ACOT7 | O00154; K7E | BACH_HUMAN | Oxidation (M) | 0 | 0 | 0 | 0 | 0 | 0 | 0 | 0 | 689210 | 216540 | 0 | 2461800 | 0 | 0 | 0 | 0 | 0 | 0 |
| MIEEAGAII: | ACOT7; ACOT7 | O00154; K7E | BACH_HUMAN | Unmodified | 0 | 0 | 0 | 0 | 181770 | 0 | 0 | 0 | 0 | 93295 | 0 | 0 | 385730 | 0 | 0 | 0 | 257780 | 668280 |
| MIESLTK | RUVBL2; RUVBI | Q9Y230; M0I | RUVB2_HUMAI | Unmodified | 0 | 0 | 44386 | 0 | 578750 | 0 | 0 | 0 | 0 | 0 | 0 | 746560 | 594050 | 265640 | 71440 | 0 | 78971 | 186400 |
| MIETAQVDI | RAB21 | Q9UL25 | RAB21_HUMAN | Unmodified | 240140 | 219030 | 183840 | 386690 | 0 | 0 | 0 | 0 | 604990 | 91510 | 429340 | 521160 | 0 | 0 | 142280 | 134870 | 0 | 0 |
| MIETAQVDI | RAB21 | Q9UL25 | RAB21_HUMAN | Unmodified | 328720 | 0 | 0 | 0 | 598650 | 0 | 0 | 0 | 0 | 45030 | 0 | 0 | 0 | 0 | 0 | 0 | 46873 | 101210 |
| MIEVVCND | UBL5; UBL5 | Q9BZL1; K7E | UBL5_HUMAN | Unmodified | 502390 | 303140 | 266250 | 1044600 | 704410 | 0 | 0 | 163660 | 123200 | 187120 | 0 | 0 | 1236200 | 1210700 | 408290 | 215510 | 1236800 | 1688300 |
| MIEVVCND | UBL5; UBL5 | Q9BZL1; K7E | UBL5_HUMAN | Oxidation (M) | 215850 | 200110 | 0 | 0 | 0 | 37066 | 0 | 0 | 0 | 94014 | 323010 | 244380 | 381220 | 0 | 0 | 0 | 338800 | 0 |
| MIEYDPERI | DKC1; DKC1; D | O60832; H7I | DKC1_HUMAN | Unmodified | 0 | 0 | 0 | 0 | 0 | 0 | 0 | 0 | 0 | 0 | 0 | 433310 | 0 | 0 | 0 | 0 | 0 | 0 |
| MIGFSETAI | NSF; NSF; NSF | P46459; I3L2 | NSF_HUMAN V | Oxidation (M) | 28053 | 0 | 0 | 0 | 143610 | 0 | 83491 | 58616 | 0 | 0 | 0 | 0 | 1393900 | 417210 | 116860 | 298940 | 79932 | 351010 |
| MIGFSETAI | NSF; NSF; NSF | P46459; I3L2 | NSF_HUMAN V | Unmodified | 0 | 0 | 0 | 0 | 0 | 0 | 288330 | 282660 | 751430 | 0 | 0 | 580670 | 3611700 | 2109100 | 401430 | 185450 | 0 | 0 |
| MIGGLFIYN | AP2M1; AP2M1; | Q96CW1; C9 | AP2M1_HUMAI | Oxidation (M) | 0 | 499680 | 0 | 1045200 | 0 | 0 | 0 | 0 | 0 | 164260 | 214760 | 1187300 | 0 | 0 | 525950 | 1055500 | 0 | 0 |
| MIGGPILPS | ALDH7A1; ALDI | P49419; A0A | AL7A1_HUMAN | Oxidation (M) | 0 | 0 | 0 | 0 | 0 | 0 | 0 | 0 | 0 | 0 | 0 | 2163200 | 0 | 0 | 0 | 0 | 0 | 0 |
| MIGGPILPS | ALDH7A1; ALDI | P49419; A0A | AL7A1_HUMAN | Unmodified | 0 | 0 | 0 | 0 | 0 | 0 | 0 | 0 | 0 | 0 | 0 | 1656300 | 0 | 0 | 0 | 0 | 0 | 0 |
| MIHGNPSG | MVK; MVK; MVK | Q03426; F5H | KIME_HUMAN I | Unmodified | 0 | 0 | 0 | 0 | 0 | 0 | 0 | 0 | 418300 | 0 | 0 | 0 | 2157100 | 0 | 0 | 0 | 0 | 0 |
| MIHILLYYK | PSMD10; PSMD | O75832; B1A | PSD10_HUMAN | Oxidation (M) | 0 | 0 | 0 | 0 | 0 | 0 | 0 | 0 | 0 | 286900 | 0 | 0 | 1010900 | 0 | 126490 | 0 | 687800 | 0 |
| MIIEEAK | CCT5; CCT5; C( | P48643; B7Z | TCPE_HUMAN | Unmodified | 0 | 0 | 315580 | 188070 | 983250 | 0 | 310120 | 260860 | 949590 | 517380 | 1346000 | 7789700 | 1462200 | 711010 | 113270 | 131050 | 331550 | 1309600 |
| MIIEEAK | CCT5; CCT5; C( | P48643; B7Z | TCPE_HUMAN | Oxidation (M) | 0 | 0 | 0 | 0 | 0 | 0 | 623230 | 250500 | 0 | 28264 | 0 | 925740 | 0 | 113230 | 0 | 0 | 0 | 195470 |
| MIIYRDLISI | TPT1; TPT1; TPT | P13693; A0A | TCTP_HUMAN | 2 Oxidation (M) | 459170 | 2325900 | 1144800 | 13473000 | 11718000 | 1311800 | 1711600 | 0 | 348010 | 1476100 | 1533600 | 1485600 | 16321000 | 12438000 | 6358700 | 5790900 | 7343400 | 2079600 |
| MIIYRDLISI | TPT1; TPT1; TPT | P13693; A0A | TCTP_HUMAN | Oxidation (M) | 447860 | 533220 | 0 | 1290300 | 6034300 | 205080 | 448140 | 0 | 862880 | 781140 | 312250 | 140770 | 3963400 | 2438000 | 787000 | 0 | 6632200 | 1884300 |
| MIIYRDLISI | TPT1; TPT1; TPT | P13693; A0A | TCTP_HUMAN | Unmodified | 0 | 51830 | 0 | 96637 | 1667400 | 196130 | 259870 | 0 | 810910 | 890370 | 0 | 380320 | 172500 | 0 | 125050 | 0 | 514100 | 1143500 |
| MIKPQTLAS | PRXL2A | Q9BRX8 | PXL2A_HUMAN | Unmodified | 0 | 0 | 0 | 0 | 0 | 0 | 1727500 | 0 | 10451000 | 1869800 | 114450 | 221330 | 0 | 0 | 0 | 0 | 0 | 0 |
| MIKPQTLAS | PRXL2A | Q9BRX8 | PXL2A_HUMAN | Oxidation (M) | 0 | 0 | 0 | 0 | 0 | 0 | 166460 | 210810 | 0 | 675130 | 0 | 64006 | 0 | 0 | 0 | 0 | 0 | 0 |
| MIKPQTLAS | PRXL2A | Q9BRX8 | PXL2A_HUMAN | Unmodified | 0 | 0 | 0 | 0 | 0 | 0 | 0 | 0 | 1287100 | 331710 | 0 | 59770 | 0 | 0 | 0 | 14810 | 0 | 0 |
| MILELFSK | GSTO1; GSTO1 | P78417; Q5T | GSTO1_HUMAI | Oxidation (M) | 11681 | 0 | 0 | 0 | 0 | 0 | 0 | 0 | 0 | 0 | 0 | 1983200 | 0 | 0 | 57159 | 0 | 0 | 0 |
| MILELFSK | GSTO1; GSTO1 | P78417; Q5T | GSTO1_HUMAI | Unmodified | 0 | 0 | 19034 | 0 | 102720 | 32311 | 0 | 0 | 0 | 0 | 181220 | 1737500 | 21797 | 0 | 0 | 0 | 80016 | 14695 |
| MILGNKCD | RAB8A | P61006 | RAB8A_HUMAN | Unmodified | 0 | 0 | 0 | 433000 | 1017800 | 0 | 161750 | 56398 | 287240 | 36296 | 281650 | 847810 | 0 | 0 | 0 | 0 | 0 | 529030 |
| MILIQDGS( | KHSRP; KHSRP | Q92945; M0I | FUBP2_HUMAN | Unmodified | 0 | 0 | 0 | 0 | 0 | 0 | 0 | 0 | 1916600 | 0 | 948660 | 4312500 | 0 | 0 | 0 | 0 | 0 | 0 |
| MILIQDGS( | KHSRP; KHSRP | Q92945; M0I | FUBP2_HUMAN | Oxidation (M) | 0 | 0 | 0 | 0 | 0 | 0 | 0 | 0 | 557150 | 0 | 856230 | 2741500 | 0 | 0 | 0 | 0 | 0 | 0 |
| MILIQDGS( | KHSRP; KHSRP | Q92945; M0I | FUBP2_HUMAN | Oxidation (M) | 0 | 0 | 0 | 0 | 242470 | 174040 | 0 | 0 | 0 | 0 | 0 | 0 | 0 | 0 | 0 | 0 | 0 | 303190 |
| MILIQDGS( | KHSRP; KHSRP | Q92945; M0I | FUBP2_HUMAN | Unmodified | 0 | 0 | 0 | 0 | 245510 | 0 | 0 | 0 | 0 | 350620 | 0 | 0 | 0 | 0 | 0 | 0 | 222250 | 0 |
| MILIYLLPVI | PCID2 | Q5JVF3 | PCID2_HUMAN | Oxidation (M) | 0 | 0 | 0 | 0 | 0 | 0 | 0 | 0 | 0 | 0 | 0 | 522140 | 0 | 0 | 0 | 0 | 0 | 0 |
| MILLEVNNI | ARPC2; ARPC2; | O15144; G5I | ARPC2_HUMAI | Unmodified | 300720 | 0 | 0 | 0 | 0 | 0 | 369750 | 120320 | 1464700 | 0 | 0 | 7161200 | 0 | 0 | 949020 | 324100 | 0 | 0 |
| MILLEVNNI | ARPC2; ARPC2; | O15144; G5I | ARPC2_HUMAI | Oxidation (M) | 258610 | 0 | 212650 | 0 | 250180 | 0 | 0 | 0 | 0 | 0 | 283870 | 4701600 | 0 | 0 | 439810 | 522480 | 0 | 0 |
| MILTNPEGI | PTPRK; PTPRK; | Q15262; Q5 | PTPRK_HUMAN | Unmodified | 0 | 0 | 0 | 0 | 0 | 0 | 0 | 0 | 951430 | 310930 | 0 | 0 | 342400 | 0 | 0 | 0 | 0 | 1141700 |

| | | | | | | | | | | | | | | | | | | | | | | |
|---|---|---|---|---|---|---|---|---|---|---|---|---|---|---|---|---|---|---|---|---|---|---|
| MIMDDSK | NRP1; NRP1; N | O14786; Q5 | NRP1_HUMAN | Unmodified | 0 | 0 | 0 | 0 | 42747 | 485860 | 0 | 0 | 91389 | 25676 | 0 | 0 | 0 | 0 | 0 | 0 | 188080 | 2463300 |
| MIMMFTDC | CACNA2D2; CA | Q9NY47; C9 | CA2D2_HUMAI | Oxidation (M) | 0 | 0 | 0 | 0 | 0 | 0 | 0 | 0 | 2141700 | 261300 | 0 | 0 | 0 | 0 | 0 | 0 | 0 | 0 |
| MINLMRPG | PTPRG | P23470 | PTPRG_HUMAI | 2 Oxidation (M) | 0 | 0 | 0 | 0 | 3465000 | 0 | 0 | 0 | 0 | 0 | 0 | 0 | 0 | 0 | 0 | 0 | 846250 | 0 |
| MINLSEPD' | PLS1; PLS1 | Q14651; C9J | PLSI_HUMAN F | Unmodified | 0 | 0 | 0 | 0 | 344650 | 0 | 0 | 0 | 650020 | 0 | 157060 | 905120 | 0 | 0 | 0 | 0 | 0 | 0 |
| MINLSEPD' | PLS1; PLS1 | Q14651; C9J | PLSI_HUMAN F | Oxidation (M) | 0 | 0 | 0 | 0 | 0 | 0 | 0 | 0 | 0 | 0 | 283660 | 760550 | 0 | 0 | 0 | 0 | 0 | 0 |
| MINLSEPD' | PLS1; PLS1 | Q14651; C9J | PLSI_HUMAN F | Unmodified | 0 | 0 | 0 | 0 | 3152100 | 0 | 0 | 0 | 0 | 0 | 0 | 0 | 489350 | 0 | 0 | 0 | 141580 | 0 |
| MINLSEPD' | PLS1; PLS1 | Q14651; C9J | PLSI_HUMAN F | Unmodified | 0 | 0 | 0 | 0 | 2131400 | 422350 | 0 | 52751 | 0 | 0 | 0 | 0 | 686480 | 584570 | 0 | 0 | 677660 | 0 |
| MINLSVPD' | LCP1; PLS3; PL: | P13796; P13 | PLSL_HUMAN I | Oxidation (M) | 0 | 0 | 343110 | 224450 | 821280 | 419130 | 0 | 0 | 0 | 0 | 0 | 0 | 0 | 0 | 0 | 525840 | 722150 | 0 |
| MINLSVPD' | LCP1; PLS3; PL: | P13796; P13 | PLSL_HUMAN I | Unmodified | 0 | 0 | 0 | 0 | 0 | 0 | 0 | 0 | 752550 | 0 | 0 | 0 | 0 | 0 | 0 | 0 | 0 | 0 |
| MINLSVPD' | PLS3; PLS3; PL! | P13797; U3k | PLST_HUMAN I | Oxidation (M) | 0 | 0 | 0 | 0 | 3688500 | 2243400 | 0 | 572920 | 0 | 0 | 0 | 0 | 0 | 1196400 | 0 | 0 | 4929000 | 3568100 |
| MINLSVPD' | PLS3; PLS3; PL! | P13797; U3k | PLST_HUMAN I | Unmodified | 0 | 0 | 0 | 0 | 9739900 | 923980 | 255640 | 0 | 0 | 153280 | 0 | 0 | 556050 | 0 | 0 | 0 | 5460900 | 2018400 |
| MINLSVPD' | PLS3; PLS3; PL! | P13797; U3k | PLST_HUMAN I | Unmodified | 0 | 0 | 0 | 0 | 10458000 | 1591400 | 0 | 0 | 87949 | 0 | 0 | 0 | 1062200 | 966530 | 0 | 0 | 11574000 | 4080300 |
| MINLSVPD' | PLS3; PLS3; PL! | P13797; U3k | PLST_HUMAN I | Oxidation (M) | 0 | 0 | 0 | 0 | 2605400 | 952120 | 0 | 435530 | 0 | 312800 | 0 | 0 | 947540 | 752010 | 0 | 0 | 3958600 | 1677600 |
| MINLSVPD' | LCP1; LCP1 | P13796; Q5T | PLSL_HUMAN I | Oxidation (M) | 0 | 0 | 0 | 0 | 0 | 0 | 1010200 | 222220 | 0 | 0 | 0 | 0 | 0 | 0 | 0 | 0 | 1267000 | 0 |
| MINLSVPD' | LCP1; LCP1 | P13796; Q5T | PLSL_HUMAN I | Unmodified | 0 | 0 | 0 | 0 | 0 | 238650 | 0 | 52851 | 0 | 274100 | 0 | 0 | 0 | 185810 | 0 | 0 | 2132000 | 408300 |
| MINTDLSR | RPL4; RPL4 | P36578; H3E | RL4_HUMAN 6( | Unmodified | 3467400 | 5607900 | 1274500 | 1471400 | 134930 | 99826000 | 7652700 | 255570 | 16392000 | 306690 | 59670 | 13695000 | 1787400 | 458950 | 2428900 | 1899900 | 0 | 382700 |
| MINTDLSR | RPL4; RPL4 | P36578; H3E | RL4_HUMAN 6( | Oxidation (M) | 2493800 | 1978500 | 1059400 | 966500 | 230550 | 713340 | 0 | 0 | 0 | 0 | 813240 | 5802700 | 759550 | 48926 | 1320600 | 1518800 | 0 | 365220 |
| MIPCDFLIP | GPI; GPI; GPI; N | P06744; K7E | G6PI_HUMAN ( | Oxidation (M) | 0 | 0 | 823320 | 1867800 | 2738300 | 1939500 | 50257 | 1269400 | 5003700 | 0 | 2580700 | 16910000 | 3833200 | 1543700 | 1686800 | 2641000 | 1842000 | 733620 |
| MIPCDFLIP | GPI; GPI; GPI; N | P06744; K7E | G6PI_HUMAN ( | Unmodified | 0 | 0 | 0 | 292650 | 3692400 | 772430 | 0 | 0 | 2820300 | 1068300 | 0 | 4622100 | 1145400 | 1073500 | 101660 | 0 | 601780 | 1178900 |
| MIPVTEFR | LRRC8C; LRRC | Q8TDW0; E9 | LRC8C_HUMAI | Acetyl (Protein N | 0 | 0 | 0 | 0 | 0 | 0 | 0 | 0 | 71453 | 0 | 0 | 272990 | 0 | 0 | 0 | 0 | 0 | 11418 |
| MIPVTELR | LRRC8A | Q8IWT6 | LRC8A_HUMAI | Acetyl (Protein N | 0 | 0 | 24559 | 35569 | 0 | 14719 | 0 | 0 | 117710 | 0 | 0 | 1418000 | 0 | 0 | 0 | 0 | 0 | 0 |
| MIQDGKGI | CCT4 | P50991 | TCPD_HUMAN | Oxidation (M) | 48983 | 0 | 114540 | 112280 | 2884700 | 247890 | 1680000 | 1242500 | 0 | 1048300 | 1829600 | 8295500 | 7293600 | 2898400 | 109880 | 95667 | 1601800 | 1635200 |
| MIQDGKGI | CCT4 | P50991 | TCPD_HUMAN | Unmodified | 118830 | 101400 | 0 | 302760 | 7700800 | 0 | 1662900 | 350840 | 543020 | 1011600 | 1780800 | 1142700 | 7666300 | 3886200 | 0 | 117300 | 3269800 | 5093600 |
| MIQHAVFK | SRP54; SRP54; | P61011; G3\ | SRP54_HUMAN | Unmodified | 0 | 0 | 0 | 0 | 0 | 0 | 0 | 0 | 0 | 0 | 0 | 312460 | 0 | 0 | 0 | 0 | 0 | 0 |
| MIQNDLDN | N/A | N/A | | Oxidation (M) | 0 | 0 | 0 | 0 | 0 | 0 | 0 | 0 | 0 | 0 | 0 | 0 | 4238500 | 0 | 0 | 0 | 0 | 0 |
| MIREYRQM | YWHAE; YWHA | P62258; K7E | 1433E_HUMAN | Oxidation (M) | 0 | 0 | 125820 | 220060 | 2996700 | 2152200 | 249820 | 0 | 0 | 2028400 | 240880 | 0 | 3896100 | 3381100 | 0 | 0 | 3261000 | 250650 |
| MIREYRQM | YWHAE; YWHA | P62258; K7E | 1433E_HUMAN | Unmodified | 0 | 0 | 0 | 0 | 7980100 | 2662200 | 1519500 | 232070 | 0 | 1086600 | 0 | 0 | 1097300 | 923420 | 60596 | 0 | 7635300 | 920640 |
| MISAEAPVL | NFYC; NFYC; N | Q13952; A0/ | NFYC_HUMAN | Oxidation (M) | 0 | 0 | 0 | 0 | 0 | 0 | 0 | 0 | 0 | 63346 | 0 | 1068700 | 0 | 0 | 0 | 0 | 0 | 0 |
| MISATEGP\ | MEAK7; MEAK7; | Q6P9B6; H3 | MEAK7_HUMAI | Unmodified | 0 | 0 | 0 | 0 | 6182300 | 1183400 | 0 | 0 | 1210800 | 0 | 262330 | 0 | 813470 | 593290 | 0 | 53713 | 1510400 | 2125800 |
| MISATEGP\ | MEAK7; MEAK7; | Q6P9B6; H3 | MEAK7_HUMAI | Oxidation (M) | 82911 | 0 | 95859 | 0 | 1420700 | 425730 | 0 | 0 | 234050 | 94648 | 0 | 133170 | 184970 | 92547 | 119590 | 118550 | 367010 | 389020 |
| MISDAIPEL | MDH2; MDH2 | P40926; G3\ | MDHM_HUMAN | Oxidation (M) | 254330 | 183100 | 671940 | 239490 | 1437700 | 0 | 2678000 | 1144600 | 13786000 | 5720300 | 1751200 | 0 | 1959600 | 5170700 | 321060 | 224360 | 447110 | 706230 |
| MISDAIPEL | MDH2; MDH2 | P40926; G3\ | MDHM_HUMAN | Unmodified | 0 | 0 | 0 | 117480 | 0 | 0 | 0 | 72114 | 14142000 | 2789500 | 0 | 0 | 680290 | 771450 | 0 | 0 | 105890 | 450240 |
| MISDLIAGC | GSTZ1; GSTZ1; | O43708; G3\ | MAAI_HUMAN I | Oxidation (M) | 0 | 0 | 0 | 0 | 0 | 0 | 0 | 0 | 426970 | 723040 | 0 | 0 | 0 | 0 | 0 | 0 | 0 | 0 |
| MISEFTWPI | MYO1D; MYO1I | O94832; J3C | MYO1D_HUMA | Oxidation (M) | 6004700 | 5958400 | 2803200 | 1800100 | 0 | 0 | 0 | 0 | 425250 | 0 | 231060 | 0 | 0 | 0 | 6112000 | 5810600 | 0 | 197450 |
| MISEFTWPI | MYO1D; MYO1I | O94832; J3C | MYO1D_HUMA | Unmodified | 1743100 | 1362200 | 344770 | 465590 | 0 | 43528 | 0 | 0 | 793780 | 75261 | 57173 | 212440 | 96936 | 0 | 1915100 | 475280 | 73661 | 217510 |
| MISEFTWPI | MYO1D; MYO1I | O94832; J3C | MYO1D_HUMA | Unmodified | 1069100 | 1958500 | 0 | 1195800 | 0 | 0 | 121380 | 0 | 0 | 0 | 0 | 0 | 0 | 0 | 0 | 220080 | 309330 | 0 |
| MISGQKLEI | KIF23; KIF23; KI | Q02241; A0/ | KIF23_HUMAN | Unmodified | 0 | 0 | 0 | 0 | 0 | 1382800 | 187040 | 98394 | 320790 | 67744 | 0 | 244060 | 0 | 0 | 0 | 0 | 0 | 0 |
| MISGQKLEI | KIF23; KIF23; KI | Q02241; A0/ | KIF23_HUMAN | Oxidation (M) | 0 | 0 | 0 | 0 | 13921000 | 3605700 | 680960 | 365390 | 859850 | 1746600 | 489010 | 0 | 0 | 104450 | 0 | 0 | 2558200 | 1352900 |
| MISGQKLEI | KIF23; KIF23; KI | Q02241; A0/ | KIF23_HUMAN | Unmodified | 0 | 0 | 0 | 468740 | 54964000 | 6288100 | 1257200 | 0 | 4051000 | 4581100 | 1291900 | 0 | 1150900 | 146030 | 0 | 0 | 8528200 | 4246100 |
| MISLIESGV | RIPK2 | O43353 | RIPK2_HUMAN | Unmodified | 0 | 0 | 0 | 0 | 133430 | 0 | 0 | 0 | 401530 | 4125800 | 0 | 242080 | 0 | 0 | 0 | 0 | 366790 | 200740 |
| MISLTDTQI | GOLT1B; GOLT | Q9Y3E0; F5C | GOT1B_HUMAI | Acetyl (Protein N | 0 | 0 | 0 | 0 | 0 | 0 | 1073700 | 0 | 594350 | 67358 | 0 | 0 | 0 | 0 | 0 | 0 | 40580 | 259190 |
| MITAAYVPL | RBM39 | Q14498 | RBM39_HUMAI | 2 Oxidation (M) | 0 | 0 | 0 | 0 | 0 | 0 | 364740 | 0 | 0 | 1018200 | 0 | 0 | 6239000 | 0 | 110380 | 74466 | 5020100 | 0 |
| MITSAAGII: | PSMD1; PSMD1 | Q99460; C9J | PSMD1_HUMA | Oxidation (M) | 0 | 0 | 45049 | 479490 | 0 | 0 | 0 | 0 | 0 | 0 | 0 | 0 | 957420 | 0 | 0 | 0 | 0 | 0 |
| MITSLSKD( | MCM2; MCM2 | P49736; H0) | MCM2_HUMAN | Unmodified | 0 | 0 | 0 | 0 | 603520 | 0 | 174920 | 65639 | 75292 | 245580 | 0 | 0 | 0 | 0 | 0 | 84434 | 126960 | 0 |
| MIVCVNPK | KIF23; KIF23; KI | Q02241; A0/ | KIF23_HUMAN | Unmodified | 0 | 0 | 0 | 0 | 0 | 0 | 0 | 0 | 558570 | 0 | 0 | 652770 | 0 | 0 | 0 | 0 | 0 | 0 |
| MIVDPVEPI | TRIM28; TRIM2( | Q13263; M0 | TIF1B_HUMAN | Oxidation (M) | 0 | 0 | 0 | 0 | 0 | 0 | 130280 | 159860 | 173110 | 117960 | 0 | 936100 | 0 | 0 | 0 | 0 | 0 | 0 |
| MIVDPVEPI | TRIM28; TRIM2( | Q13263; M0 | TIF1B_HUMAN | Unmodified | 20908 | 0 | 0 | 0 | 0 | 60390 | 171590 | 0 | 0 | 0 | 0 | 712100 | 0 | 0 | 0 | 0 | 0 | 0 |
| MIVDPVEPI | TRIM28; TRIM2( | Q13263; M0 | TIF1B_HUMAN | 2 Oxidation (M) | 0 | 0 | 0 | 0 | 0 | 0 | 784160 | 0 | 0 | 0 | 0 | 0 | 0 | 124230 | 0 | 0 | 0 | 0 |
| MIVTDIYNI | USP15 | Q9Y4E8 | UBP15_HUMAI | Unmodified | 0 | 0 | 0 | 0 | 0 | 0 | 0 | 0 | 0 | 0 | 0 | 1050900 | 0 | 0 | 0 | 0 | 0 | 0 |
| MIVVLGEIG | ACLY | P53396 | ACLY_HUMAN | Oxidation (M) | 0 | 0 | 0 | 0 | 1948200 | 0 | 1725600 | 315860 | 1114200 | 0 | 0 | 4759600 | 0 | 0 | 356100 | 256650 | 0 | 0 |
| MIVVLGEIG | ACLY | P53396 | ACLY_HUMAN | Unmodified | 0 | 0 | 0 | 0 | 0 | 0 | 869100 | 0 | 0 | 0 | 0 | 0 | 0 | 0 | 0 | 0 | 0 | 0 |
| MIYASSK | CFL1; DSTN; CI | P23528; P60 | COF1_HUMAN | Oxidation (M) | 714610 | 84705 | 762570 | 0 | 0 | 0 | 0 | 0 | 0 | 0 | 0 | 873910 | 1942300 | 952090 | 0 | 0 | 0 | 0 |
| MIYASSKD/ | CFL1; DSTN; CI | P23528; P60 | COF1_HUMAN | Oxidation (M) | 61063 | 0 | 0 | 0 | 5189600 | 794390 | 1805200 | 2086800 | 0 | 0 | 0 | 0 | 1581700 | 2007500 | 0 | 0 | 3736600 | 1641800 |
| MIYASSKD/ | CFL1; DSTN; CI | P23528; P60 | COF1_HUMAN | Oxidation (M) | 0 | 0 | 0 | 0 | 36885000 | 3031400 | 10613000 | 4416800 | 146000 | 2078600 | 0 | 0 | 9316700 | 5784500 | 0 | 0 | 21285000 | 9028400 |
| MIYASSKD/ | CFL1; DSTN; CI | P23528; P60 | COF1_HUMAN | Oxidation (M) | 0 | 0 | 0 | 0 | 2875700 | 0 | 1949000 | 3706600 | 669960 | 0 | 0 | 0 | 98466 | 192810 | 0 | 0 | 4165500 | 959160 |
| MIYASSKD/ | CFL1; DSTN; CI | P23528; P60 | COF1_HUMAN | Unmodified | 0 | 0 | 0 | 0 | 5769400 | 0 | 3492400 | 13021000 | 0 | 0 | 0 | 0 | 0 | 6926400 | 0 | 0 | 12159000 | 0 |
| MKAMDVLI | TRIO; TRIO; TRI | O75962; E7E | TRIO_HUMAN 1 | Acetyl (Protein N | 0 | 200310 | 113610 | 0 | 0 | 293610 | 0 | 0 | 3193200 | 642060 | 63979000 | 141410 | 0 | 76306 | 0 | 0 | 0 | 86198 |
| MKDTDSEE | CALM2; CALM3 | P0DP24; P0I | CALM2_HUMAI | Unmodified | 210930 | 98337 | 108300 | 0 | 0 | 0 | 0 | 0 | 0 | 10968 | 0 | 2381700 | 0 | 0 | 1242500 | 506060 | 0 | 0 |
| MKDTDSEE | CALM2; CALM3 | P0DP24; P0I | CALM2_HUMAI | Unmodified | 617440 | 0 | 436410 | 0 | 0 | 0 | 0 | 0 | 219430 | 115760 | 0 | 3921600 | 609000 | 0 | 2427500 | 1133400 | 125490 | 0 |
| MKDTDSEE | CALM2; CALM3 | P0DP24; P0I | CALM2_HUMAI | Oxidation (M) | 274560 | 0 | 1180200 | 0 | 0 | 0 | 0 | 0 | 0 | 0 | 0 | 4966400 | 0 | 0 | 1334700 | 739490 | 0 | 82909 |
| MKEDEPWI | SLC12A2; SLC1 | P55011; G3\ | S12A2_HUMAN | Oxidation (M) | 0 | 131140 | 3949000 | 308380 | 0 | 0 | 0 | 0 | 544080 | 0 | 1615500 | 1779100 | 211740 | 0 | 670000 | 401520 | 0 | 0 |
| MKEEMDR/ | TSG101; TSG1C | Q99816; F5F | TS101_HUMAN | Unmodified | 3049600 | 4561700 | 1176600 | 3393500 | 1430900 | 210830 | 0 | 0 | 1733500 | 305020 | 0 | 1015400 | 1693000 | 499060 | 4296700 | 2030200 | 0 | 1126700 |
| MKEEMDR/ | TSG101; TSG1C | Q99816; F5F | TS101_HUMAN | Oxidation (M) | 7344000 | 6803000 | 1332300 | 6112200 | 0 | 277750 | 135560 | 0 | 1835800 | 206810 | 325090 | 124360 | 1823500 | 677410 | 7093600 | 8871900 | 0 | 318320 |
| MKEEMDR/ | TSG101; TSG1C | Q99816; F5F | TS101_HUMAN | 2 Oxidation (M) | 1229100 | 1715500 | 3507200 | 2925200 | 0 | 0 | 0 | 0 | 69558 | 40420 | 0 | 0 | 224650 | 0 | 2246800 | 2227900 | 0 | 0 |
| MKEGMSN | GNG4 | P50150 | GBG4_HUMAN | Unmodified | 0 | 0 | 0 | 0 | 91556 | 0 | 372610 | 415100 | 0 | 467030 | 0 | 0 | 0 | 0 | 0 | 0 | 0 | 0 |
| MKEGMSN | GNG4 | P50150 | GBG4_HUMAN | Unmodified | 0 | 0 | 0 | 0 | 0 | 0 | 1957400 | 2148800 | 589360 | 603560 | 0 | 0 | 0 | 0 | 0 | 0 | 0 | 0 |
| MKEIAEAYL | HSPA2 | P54652 | HSP72_HUMAN | Oxidation (M) | 0 | 0 | 0 | 0 | 940210 | 0 | 0 | 0 | 0 | 0 | 0 | 0 | 0 | 0 | 0 | 0 | 0 | 0 |
| MKEIAEAYL | HSPA2 | P54652 | HSP72_HUMAN | Unmodified | 0 | 0 | 0 | 0 | 1748700 | 178940 | 0 | 0 | 0 | 0 | 0 | 225740 | 0 | 0 | 0 | 0 | 0 | 0 |
| MKEIAEAYL | HSPA8; HSPA8; | P11142; E9P | HSP7C_HUMAI | Oxidation (M) | 25214000 | 7806900 | 0 | 22886000 | 48124000 | 7566800 | 28652000 | 15105000 | 7908800 | 14538000 | 3073500 | 8283900 | 1.44E+08 | 76653000 | 4323200 | 4395100 | 13922000 | 15639000 |
| MKEIAEAYL | HSPA8; HSPA8; | P11142; E9P | HSP7C_HUMAI | Unmodified | 10577000 | 12882000 | 876370 | 16455000 | 1.35E+08 | 23317000 | 55434000 | 12564000 | 31606000 | 42087000 | 4657600 | 23181000 | 87784000 | 77879000 | 4401500 | 3659800 | 31548000 | 47219000 |
| MKEIAEAYL | HSPA1A; HSPA' | P0DMV8; P0 | HS71A_HUMAN | Oxidation (M) | 0 | 0 | 0 | 0 | 10712000 | 0 | 0 | 0 | 0 | 6543200 | 0 | 0 | 9696000 | 9303700 | 0 | 0 | 4144900 | 4178200 |
| MKETAEAYI | HSPA5; HSPA5; | P11021; A0A | BIP_HUMAN Er | Oxidation (M) | 133450 | 0 | 585570 | 355410 | 2244800 | 0 | 7452300 | 5200200 | 48288000 | 24661000 | 3872600 | 2340600 | 1698400 | 2862000 | 700140 | 379790 | 282840 | 2370300 |
| MKETAEAYI | HSPA5; HSPA5; | P11021; A0A | BIP_HUMAN Er | Unmodified | 243460 | 0 | 1173300 | 615170 | 7658300 | 671980 | 21028000 | 5954000 | 2.4E+08 | 96362000 | 7465200 | 3374000 | 2128700 | 2885600 | 2032700 | 532810 | 975610 | 5281500 |
| MKETAEAYI | HSPA5; HSPA5; | P11021; A0A | BIP_HUMAN Er | Unmodified | 1436400 | 0 | 100090 | 0 | 0 | 0 | 1278100 | 624260 | 10433000 | 5377200 | 664060 | 219820 | 0 | 0 | 0 | 0 | 0 | 506560 |
| MKETAEAYI | HSPA5; HSPA5; | P11021; A0A | BIP_HUMAN Er | Oxidation (M) | 320720 | 0 | 0 | 0 | 1287000 | 0 | 567880 | 344460 | 2210400 | 1898900 | 348820 | 147730 | 0 | 73228 | 79317 | 59119 | 248730 | 0 |
| MKETAENY | HSPA9; HSPA9; | P38646; H0) | GRP75_HUMAI | Unmodified | 0 | 0 | 151950 | 0 | 676690 | 111730 | 511650 | 104960 | 3605300 | 1013400 | 135000 | 142830 | 0 | 264290 | 0 | 13928 | 148170 | 318780 |
| MKETAENY | HSPA9; HSPA9; | P38646; H0) | GRP75_HUMAI | Oxidation (M) | 0 | 0 | 0 | 0 | 355540 | 0 | 626040 | 106320 | 0 | 558900 | 68430 | 71847 | 0 | 185360 | 0 | 0 | 79729 | 141260 |
| MKETIMNQ | BTF3; BTF3; BTI | P20290; A0A | BTF3_HUMAN ' | Unmodified | 0 | 64439 | 0 | 44473 | 0 | 45339 | 0 | 151400 | 86045 | 0 | 329000 | 1063500 | 613540 | 355240 | 26298 | 19268 | 0 | 0 |
| MKFNPFVT | RPL26; RPL26L | P61254; Q9l | RL26_HUMAN ( | Unmodified | 0 | 0 | 0 | 0 | 1162400 | 130670 | 0 | 0 | 0 | 0 | 0 | 0 | 1238100 | 0 | 0 | 0 | 0 | 1200000 |
| MKFVYKEE | GABARAP; GAB | O95166; I3L | GBRAP_HUMAI | Unmodified | 0 | 0 | 0 | 0 | 214410 | 0 | 2543000 | 80988 | 230910 | 1432300 | 0 | 0 | 890740 | 329750 | 0 | 0 | 906870 | 1913800 |
| MKGDYFRY | YWHAB | P31946 | 1433B_HUMAN | Oxidation (M) | 0 | 0 | 0 | 0 | 626800 | 250220 | 450280 | 0 | 0 | 721480 | 0 | 0 | 750610 | 0 | 0 | 0 | 891950 | 996300 |
| MKGDYFRY | YWHAB | P31946 | 1433B_HUMAN | Unmodified | 0 | 0 | 0 | 0 | 684180 | 0 | 238210 | 0 | 97506 | 134500 | 0 | 0 | 181620 | 165970 | 0 | 0 | 404120 | 731470 |
| MKGDYVL( | TAS1R3 | Q7RTX0 | TS1R3_HUMAN | Oxidation (M) | 0 | 23472 | 178140 | 0 | 0 | 0 | 0 | 0 | 0 | 0 | 130410 | 167340 | 0 | 0 | 0 | 0 | 0 | 0 |
| MKGDYYR\ | YWHAZ; YWHA | P63104; B7Z | 1433Z_HUMAN | Unmodified | 0 | 0 | 1176300 | 0 | 335080 | 0 | 94072 | 0 | 0 | 528600 | 923120 | 0 | 0 | 0 | 0 | 0 | 248630 | 281810 |
| MKGDYYR\ | YWHAH; YWHA | Q04917; A2I | 1433F_HUMAN | Oxidation (M) | 0 | 0 | 0 | 0 | 3124100 | 246110 | 2333200 | 0 | 117820 | 469870 | 0 | 0 | 1396400 | 1284400 | 0 | 0 | 2896900 | 1330000 |
| MKGDYYR\ | YWHAH; YWHA | Q04917; A2I | 1433F_HUMAN | Unmodified | 0 | 0 | 0 | 0 | 2407700 | 0 | 703710 | 0 | 0 | 282060 | 0 | 0 | 383950 | 501000 | 0 | 0 | 1342300 | 2659400 |
| MKGDYYR\ | YWHAG | P61981 | 1433G_HUMAI | Oxidation (M) | 0 | 0 | 0 | 100170 | 11256000 | 0 | 12309000 | 0 | 0 | 7942200 | 127270 | 205600 | 14962000 | 26182000 | 0 | 0 | 17730000 | 18268000 |
| MKGDYYR\ | YWHAG | P61981 | 1433G_HUMAI | Unmodified | 0 | 0 | 0 | 0 | 8142100 | 851310 | 1079300 | 0 | 0 | 2222900 | 167310 | 0 | 3032900 | 5888700 | 0 | 0 | 15847000 | 5570000 |
| MKGEAEDI | PLGRKT | Q9HBL7 | PLRKT_HUMAN | Unmodified | 0 | 0 | 0 | 0 | 0 | 0 | 0 | 0 | 645280 | 0 | 0 | 0 | 0 | 0 | 0 | 0 | 0 | 0 |
| MKGEGVEI | PPFIBP1; PPFIB | Q86W92; F5 | LIPB1_HUMAN | Unmodified | 0 | 0 | 0 | 0 | 2241300 | 414210 | 0 | 0 | 0 | 0 | 0 | 347970 | 688820 | 518240 | 0 | 0 | 3044300 | 1067000 |
| MKGEVDVS | AHNAK | Q09666 | AHNK_HUMAN | Unmodified | 0 | 0 | 0 | 0 | 665850 | 0 | 0 | 0 | 0 | 0 | 0 | 0 | 0 | 0 | 0 | 0 | 266580 | 282640 |
| MKGLAFIQI | GLO1 | Q04760 | LGUL_HUMAN | Oxidation (M) | 0 | 0 | 0 | 0 | 895130 | 0 | 0 | 0 | 0 | 0 | 0 | 0 | 927170 | 226960 | 0 | 0 | 0 | 0 |
| MKGPEPV\ | GPC3; GPC3 | P51654; H0) | GPC3_HUMAN | Oxidation (M) | 0 | 0 | 0 | 0 | 0 | 0 | 0 | 0 | 0 | 1920700 | 0 | 86007 | 0 | 0 | 0 | 0 | 0 | 0 |
| MKHYEVEII | TECR; TECR | Q9NZ01; M0 | TECR_HUMAN | Oxidation (M) | 0 | 0 | 0 | 0 | 0 | 0 | 269010 | 0 | 0 | 0 | 0 | 0 | 0 | 0 | 0 | 0 | 0 | 0 |
| MKIEEVKST | RUVBL1 | Q9Y265 | RUVB1_HUMAI | Unmodified | 0 | 0 | 0 | 0 | 0 | 0 | 0 | 1057600 | 0 | 0 | 0 | 0 | 0 | 0 | 0 | 0 | 0 | 0 |
| MKLNISFP/ | RPS6; RPS6 | P62753; A2A | RS6_HUMAN 4 | Unmodified | 0 | 31349 | 0 | 0 | 967680 | 264870 | 997320 | 0 | 167650 | 758750 | 0 | 0 | 1942100 | 802530 | 0 | 0 | 662430 | 1275100 |
| MKLNISFP/ | RPS6; RPS6 | P62753; A2A | RS6_HUMAN 4 | Oxidation (M) | 0 | 0 | 0 | 0 | 1869700 | 0 | 1127300 | 105250 | 0 | 130660 | 0 | 0 | 0 | 852620 | 0 | 0 | 483770 | 0 |
| MKLPQFGI | AHNAK | Q09666 | AHNK_HUMAN | Unmodified | 0 | 0 | 0 | 0 | 4136400 | 0 | 0 | 0 | 0 | 0 | 0 | 0 | 0 | 0 | 0 | 0 | 1185500 | 1471100 |
| MKLPQFGI | AHNAK | Q09666 | AHNK_HUMAN | Oxidation (M) | 0 | 0 | 0 | 0 | 1617700 | 0 | 0 | 0 | 0 | 0 | 59678 | 0 | 0 | 0 | 0 | 0 | 0 | 1053100 |
| MKMGQEF\ | WASF2 | Q9Y6W5 | WASF2_HUMAI | Unmodified | 0 | 0 | 0 | 0 | 0 | 0 | 0 | 0 | 0 | 0 | 0 | 0 | 0 | 0 | 0 | 0 | 303810 | 355300 |
| MKPILLQGI | EIF3I; EIF3I; EIF | Q13347; A0/ | EIF3I_HUMAN I | Oxidation (M) | 0 | 96549 | 0 | 0 | 0 | 0 | 0 | 0 | 0 | 0 | 0 | 659340 | 0 | 0 | 0 | 0 | 16329 | 0 |
| MKPILLQGI | EIF3I; EIF3I; EIF | Q13347; A0/ | EIF3I_HUMAN I | Unmodified | 75839 | 115080 | 204530 | 154370 | 0 | 0 | 0 | 0 | 127980 | 0 | 712730 | 3529500 | 258760 | 0 | 94824 | 106210 | 0 | 246920 |
| MKPLMGVI | PCMT1; PCMT1 | P22061; F6S | PIMT_HUMAN | Unmodified | 0 | 0 | 0 | 0 | 0 | 509810 | 0 | 0 | 1102700 | 0 | 0 | 0 | 0 | 0 | 0 | 0 | 0 | 39249 |
| MKPLVVFVI | CMPK1; CMPK1 | P30085; E9P | KCY_HUMAN U | Unmodified | 0 | 0 | 0 | 0 | 263970 | 0 | 0 | 0 | 360320 | 0 | 0 | 32138 | 309580 | 134810 | 26666 | 15876 | 194950 | 202240 |
| MKPWFEV( | USO1 | O60763 | USO1_HUMAN | Oxidation (M) | 0 | 0 | 0 | 0 | 0 | 0 | 0 | 0 | 0 | 0 | 0 | 825180 | 0 | 0 | 0 | 0 | 0 | 0 |
| MKQELEAE | SNX32; SNX6; S | Q86XE0; Q9I | SNX32_HUMAN | Unmodified | 0 | 0 | 0 | 0 | 2029000 | 0 | 635850 | 0 | 420720 | 100400 | 0 | 330190 | 0 | 0 | 0 | 0 | 0 | 320240 |
| MKQELEAE | SNX32; SNX6; S | Q86XE0; Q9I | SNX32_HUMAN | Oxidation (M) | 0 | 0 | 0 | 0 | 4916900 | 1696400 | 877230 | 181980 | 0 | 0 | 0 | 3858000 | 0 | 696200 | 0 | 0 | 0 | 0 |
| MKQELEAE | SNX5; SNX5; SN | Q9Y5X3; Q5I | SNX5_HUMAN | Oxidation (M) | 0 | 0 | 0 | 0 | 0 | 0 | 0 | 0 | 0 | 247320 | 0 | 1187200 | 256460 | 469460 | 0 | 0 | 0 | 0 |
| MKQEPVKF | RHOA | P61586 | RHOA_HUMAN | Oxidation (M) | 0 | 105050 | 115370 | 0 | 0 | 0 | 47527 | 0 | 265940 | 0 | 432730 | 1040400 | 0 | 0 | 0 | 224710 | 0 | 0 |

| | | | | | | | | | | | | | | | | | | | | | | |
|---|---|---|---|---|---|---|---|---|---|---|---|---|---|---|---|---|---|---|---|---|---|---|
| MKQEPVKF | RHOA | P61586 | RHOA_HUMAN | Unmodified | 0 | 0 | 0 | 0 | 0 | 0 | 0 | 0 | 0 | 0 | 0 | 0 | 0 | 0 | 0 | 0 | 0 | 0 |
| MKQEPVRS | RHOC; RHOC; I | P08134; E9P | RHOC_HUMAN | Unmodified | 158110 | 197070 | 261350 | 271770 | 0 | 0 | 0 | 0 | 0 | 0 | 17871 | 214690 | 51791 | 0 | 465600 | 465720 | 0 | 0 |
| MKQSNNE/ | PDCD6IP | Q8WUM4 | PDC6I_HUMAN | Unmodified | 3546900 | 4659600 | 0 | 5656600 | 28157000 | 605620 | 1967300 | 96335 | 1822400 | 10218000 | 334780 | 280890 | 55916000 | 6034800 | 0 | 231000 | 4078100 | 11016000 |
| MKQSNNE/ | PDCD6IP | Q8WUM4 | PDC6I_HUMAN | Oxidation (M) | 1942200 | 858630 | 0 | 2304800 | 8662500 | 1669500 | 1115200 | 1713500 | 832710 | 3348700 | 175560 | 105530 | 13068000 | 6648000 | 0 | 65016 | 2008300 | 3455900 |
| MKSLDVN( | S100A13 | Q99584 | S10AD_HUMAN | Oxidation (M) | 0 | 0 | 0 | 0 | 197250 | 0 | 48174 | 0 | 0 | 0 | 0 | 0 | 0 | 80384 | 0 | 0 | 520250 | 290340 |
| MKSLDVN( | S100A13 | Q99584 | S10AD_HUMAN | Unmodified | 0 | 0 | 0 | 0 | 1495400 | 0 | 0 | 0 | 0 | 0 | 0 | 0 | 0 | 0 | 0 | 0 | 0 | 1048100 |
| MKSPDEVL | PHLDA2 | Q53GA4 | PHLA2_HUMAN | Acetyl (Protein N | 0 | 144610 | 0 | 0 | 1474000 | 356100 | 685730 | 0 | 0 | 0 | 0 | 0 | 1394500 | 1379600 | 0 | 0 | 810500 | 1342200 |
| MKSPDEVL | PHLDA2 | Q53GA4 | PHLA2_HUMAN | Acetyl (Protein N | 0 | 0 | 0 | 138300 | 2877300 | 212770 | 0 | 73830 | 0 | 0 | 0 | 0 | 4363200 | 79772 | 0 | 0 | 284070 | 3988600 |
| MKSPDEVL | PHLDA2 | Q53GA4 | PHLA2_HUMAN | Unmodified | 0 | 0 | 0 | 0 | 672600 | 0 | 0 | 0 | 0 | 0 | 0 | 0 | 165180 | 0 | 0 | 0 | 0 | 0 |
| MKTILSNQ | RPL9; RPL9 | P32969; D6F | RL9_HUMAN 6 | Oxidation (M) | 0 | 0 | 0 | 0 | 917340 | 0 | 2314700 | 409720 | 0 | 502570 | 0 | 0 | 1632600 | 1086100 | 0 | 0 | 0 | 0 |
| MKVELCSF | RPL24; RPL24; I | P83731; C9J | RL24_HUMAN ( | Unmodified | 1414900 | 935390 | 0 | 375450 | 1034700 | 0 | 2331900 | 811660 | 445720 | 192020 | 0 | 11049000 | 1052800 | 1087000 | 34663 | 278220 | 293460 | 1661700 |
| MKVELCSF | RPL24; RPL24; I | P83731; C9J | RL24_HUMAN ( | Oxidation (M) | 591580 | 407430 | 479990 | 234150 | 0 | 0 | 0 | 676610 | 0 | 0 | 393940 | 9293600 | 688960 | 1516700 | 429490 | 158690 | 172300 | 0 |
| MKVIYPAVE | NT5E | P21589 | 5NTD_HUMAN | Unmodified | 0 | 0 | 0 | 0 | 2275000 | 1143800 | 0 | 0 | 0 | 0 | 0 | 0 | 0 | 0 | 0 | 0 | 338930 | 724840 |
| MKVTNEDF | PSMC1 | P62191 | PRS4_HUMAN | Unmodified | 0 | 179870 | 0 | 0 | 438400 | 0 | 0 | 74451 | 0 | 88605 | 70084 | 0 | 390130 | 425410 | 0 | 0 | 311490 | 262430 |
| MKVVENLC | STAT3; STAT3; S | P40763; A0A | STAT3_HUMAN | Oxidation (M) | 0 | 0 | 0 | 0 | 1018900 | 113250 | 251210 | 0 | 0 | 0 | 0 | 0 | 87517 | 82096 | 0 | 0 | 0 | 0 |
| MKVVENLC | STAT3; STAT3; S | P40763; A0A | STAT3_HUMAN | Unmodified | 0 | 0 | 0 | 0 | 527420 | 0 | 36661 | 0 | 0 | 0 | 0 | 0 | 66411 | 0 | 0 | 0 | 0 | 0 |
| MKVVEVLA | FASN | P49327 | FAS_HUMAN F | Oxidation (M) | 0 | 0 | 0 | 0 | 0 | 0 | 0 | 0 | 0 | 0 | 0 | 1857900 | 0 | 0 | 0 | 87470 | 0 | 0 |
| MKVVEVLA | FASN | P49327 | FAS_HUMAN F | Unmodified | 0 | 45724 | 0 | 0 | 0 | 0 | 0 | 0 | 0 | 0 | 0 | 4777000 | 0 | 0 | 277190 | 0 | 0 | 0 |
| MLAAYLYE | CNDP2 | Q96KP4 | CNDP2_HUMA | Oxidation (M) | 0 | 0 | 0 | 0 | 0 | 0 | 0 | 0 | 0 | 139500 | 265180 | 483890 | 2266700 | 1275200 | 0 | 0 | 312740 | 179640 |
| MLAEDELR | ARF3; ARF1; N/ | P61204; P84 | ARF3_HUMAN . | Unmodified | 381220 | 358260 | 451050 | 0 | 0 | 0 | 0 | 0 | 295580 | 0 | 0 | 10285000 | 0 | 0 | 661610 | 642620 | 0 | 0 |
| MLAEDELR | ARF3; ARF1; N/ | P61204; P84 | ARF3_HUMAN . | Oxidation (M) | 0 | 0 | 0 | 0 | 0 | 0 | 0 | 0 | 0 | 0 | 0 | 2440900 | 0 | 0 | 0 | 0 | 0 | 0 |
| MLAEDELR | ARF3; ARF1; N/ | P61204; P84 | ARF3_HUMAN . | Oxidation (M) | 0 | 160900 | 324240 | 169540 | 0 | 0 | 0 | 0 | 66642 | 138400 | 0 | 133300 | 0 | 0 | 1407600 | 0 | 0 | 0 |
| MLAESDES | OSBP | P22059 | OSBP1_HUMAI | Oxidation (M) | 0 | 0 | 0 | 0 | 0 | 0 | 0 | 0 | 0 | 0 | 0 | 1391600 | 0 | 0 | 0 | 0 | 0 | 0 |
| MLAESDES | OSBP | P22059 | OSBP1_HUMAI | Unmodified | 0 | 0 | 0 | 0 | 0 | 0 | 0 | 0 | 0 | 0 | 0 | 1184800 | 0 | 0 | 0 | 0 | 0 | 0 |
| MLAIYDGFI | PFKP; PFKP; PF | Q01813; A0/ | PFKAP_HUMAN | Oxidation (M) | 0 | 0 | 0 | 0 | 0 | 0 | 0 | 0 | 778680 | 541720 | 1089300 | 4670300 | 0 | 0 | 0 | 0 | 0 | 0 |
| MLALKDPV | IPO11; IPO11 | Q9UI26; E7E | IPO11_HUMAN | Oxidation (M) | 0 | 0 | 0 | 0 | 0 | 31055 | 0 | 0 | 0 | 0 | 0 | 2131800 | 324600 | 0 | 0 | 0 | 0 | 0 |
| MLANSASV | DPYSL5; DPYSL | Q9BPU6; E9I | DPYL5_HUMAN | Acetyl (Protein N | 24995 | 0 | 0 | 0 | 190030 | 0 | 0 | 0 | 0 | 0 | 0 | 1661600 | 0 | 0 | 36522 | 0 | 0 | 0 |
| MLANSASV | DPYSL5; DPYSL | Q9BPU6; E9I | DPYL5_HUMAN | Acetyl (Protein N | 0 | 0 | 0 | 0 | 230750 | 0 | 0 | 0 | 0 | 33536 | 0 | 699570 | 128150 | 0 | 0 | 0 | 0 | 0 |
| MLAPDPDF | SLC16A2 | P36021 | MOT8_HUMAN | Oxidation (M) | 0 | 0 | 0 | 0 | 0 | 0 | 0 | 0 | 0 | 0 | 0 | 297580 | 0 | 0 | 0 | 0 | 0 | 0 |
| MLAPEGAL | PITPNA; PITPN/ | Q00169; I3L | PIPNA_HUMAN | Unmodified | 0 | 0 | 0 | 525550 | 2782100 | 368000 | 700280 | 382320 | 518830 | 174400 | 0 | 0 | 0 | 381790 | 993880 | 0 | 848990 | 2106000 |
| MLAPEGAL | PITPNA; PITPN/ | Q00169; I3L | PIPNA_HUMAN | Oxidation (M) | 0 | 0 | 0 | 0 | 616250 | 0 | 0 | 13056 | 0 | 0 | 0 | 1376700 | 1220800 | 0 | 0 | 0 | 249880 | 626550 |
| MLAQPLKD | SHMT1; SHMT1 | P34896; J3K | GLYC_HUMAN | Oxidation (M) | 293110 | 0 | 0 | 0 | 6660200 | 0 | 60591 | 0 | 0 | 529970 | 0 | 3185400 | 3743500 | 0 | 0 | 477780 | 0 | 0 |
| MLAQPLKD | SHMT1; SHMT1 | P34896; J3K | GLYC_HUMAN | Unmodified | 0 | 0 | 0 | 0 | 1068800 | 0 | 0 | 0 | 0 | 0 | 0 | 0 | 0 | 516920 | 0 | 0 | 0 | 0 |
| MLAQQAVI | RPN2; RPN2 | P04844; Q5J | RPN2_HUMAN | Unmodified | 0 | 0 | 0 | 61045 | 0 | 0 | 0 | 0 | 11165000 | 544370 | 0 | 0 | 0 | 212440 | 0 | 0 | 0 | 0 |
| MLAQQAVI | RPN2; RPN2 | P04844; Q5J | RPN2_HUMAN | Unmodified | 0 | 0 | 0 | 0 | 0 | 0 | 0 | 0 | 625320 | 0 | 0 | 0 | 0 | 0 | 0 | 0 | 0 | 0 |
| MLAVHFDK | TP53I3 | Q53FA7 | QORX_HUMAN | Acetyl (Protein N | 0 | 0 | 0 | 0 | 6003400 | 0 | 0 | 0 | 625060 | 2081300 | 531590 | 890190 | 4363600 | 0 | 0 | 0 | 3163100 | 3961700 |
| MLCSNAE( | KCTD9 | Q7L273 | KCTD9_HUMAI | Unmodified | 0 | 0 | 0 | 0 | 193940 | 0 | 0 | 0 | 0 | 0 | 0 | 0 | 0 | 0 | 0 | 0 | 125700 | 314080 |
| MLDAEDIV | ACTN1; ACTN1 | P12814; G3V | ACTN1_HUMAI | Oxidation (M) | 0 | 0 | 0 | 586060 | 12043000 | 0 | 96130 | 1031300 | 0 | 0 | 3448000 | 7992100 | 1867900 | 1570000 | 672080 | 473590 | 1772700 | 4148500 |
| MLDAEDIV | ACTN1; ACTN1 | P12814; G3V | ACTN1_HUMAI | Unmodified | 0 | 0 | 0 | 380790 | 17323000 | 6099000 | 1551300 | 441260 | 639230 | 195370 | 639210 | 4828400 | 2177900 | 1893400 | 383280 | 239760 | 4553800 | 11285000 |
| MLDAEDIV | ACTN4; ACTN4 | F5GXS2; H7 | F5GXS2_HUM/ | Unmodified | 0 | 0 | 0 | 0 | 0 | 0 | 0 | 0 | 1281700 | 225790 | 0 | 0 | 567490 | 0 | 0 | 0 | 0 | 0 |
| MLDAEDIV | ACTN4; ACTN4 | F5GXS2; H7 | F5GXS2_HUM/ | Oxidation (M) | 0 | 0 | 0 | 0 | 0 | 0 | 0 | 29545 | 352440 | 468350 | 276150 | 0 | 0 | 1264500 | 0 | 0 | 0 | 0 |
| MLDAEDIVI | ACTN4 | O43707 | ACTN4_HUMAI | Oxidation (M) | 91402 | 0 | 2153900 | 3477700 | 16998000 | 4188300 | 1026500 | 1791600 | 604000 | 1777500 | 1723500 | 14429000 | 7889000 | 6143100 | 2029900 | 2195100 | 4427800 | 2128800 |
| MLDAEDIVI | ACTN4 | O43707 | ACTN4_HUMAI | Unmodified | 0 | 0 | 442990 | 824530 | 33311000 | 7041200 | 1530100 | 824910 | 6037500 | 1846800 | 1621800 | 15353000 | 8619400 | 6082200 | 635450 | 372180 | 8696700 | 4338200 |
| MLDDRGN | RARS1 | P54136 | SYRC_HUMAN | Oxidation (M) | 0 | 0 | 0 | 0 | 0 | 0 | 0 | 0 | 0 | 0 | 782170 | 1828900 | 0 | 11697 | 0 | 0 | 0 | 0 |
| MLDGEVTD | FURIN; FURIN | P09958; A0A | FURIN_HUMAN | Unmodified | 0 | 0 | 0 | 0 | 0 | 0 | 0 | 0 | 504540 | 0 | 727100 | 0 | 0 | 0 | 0 | 0 | 0 | 0 |
| MLDHEYTT | TOP1; TOP1MT; | P11387; Q9 | TOP1_HUMAN | Unmodified | 76570 | 0 | 0 | 0 | 0 | 0 | 49101 | 43536 | 0 | 0 | 0 | 1043000 | 0 | 0 | 0 | 0 | 0 | 0 |
| MLDQTLLD | TPM3; TPM3 | J3KN67; A0A | J3KN67_HUMA | 2 Oxidation (M) | 0 | 0 | 245080 | 1187100 | 5211800 | 1567100 | 0 | 361110 | 741520 | 344110 | 0 | 0 | 5040800 | 0 | 763430 | 0 | 1224600 | 0 |
| MLDQTLLD | TPM3; TPM3 | J3KN67; A0A | J3KN67_HUMA | Oxidation (M) | 0 | 0 | 0 | 0 | 771120 | 0 | 0 | 0 | 0 | 0 | 0 | 99896 | 0 | 0 | 0 | 0 | 0 | 0 |
| MLEDRDLF | EIF3A | Q14152 | EIF3A_HUMAN | Oxidation (M) | 0 | 0 | 0 | 0 | 0 | 0 | 0 | 0 | 0 | 0 | 0 | 434900 | 0 | 0 | 0 | 0 | 0 | 0 |
| MLEEIMKYI | FGG; FGG; FGG | P02679; C9J | FIBG_HUMAN | 2 Oxidation (M) | 0 | 5348500 | 0 | 0 | 553340 | 0 | 0 | 0 | 0 | 0 | 0 | 0 | 0 | 0 | 0 | 0 | 281070 | 422330 |
| MLEENTNI | TMOD3; TMOD | Q9NYL9; H0 | TMOD3_HUMA | Unmodified | 917050 | 0 | 0 | 0 | 1664700 | 485810 | 0 | 0 | 0 | 0 | 0 | 492540 | 297190 | 0 | 0 | 366140 | 0 | 0 |
| MLEENTNI | TMOD3; TMOD | Q9NYL9; H0 | TMOD3_HUMA | Oxidation (M) | 0 | 0 | 0 | 0 | 715510 | 0 | 151570 | 111310 | 0 | 0 | 0 | 538070 | 150200 | 0 | 0 | 0 | 0 | 0 |
| MLEGPHYY | SORT1 | Q99523 | SORT_HUMAN | Oxidation (M) | 0 | 0 | 0 | 0 | 0 | 0 | 0 | 0 | 714650 | 3951800 | 243180 | 0 | 0 | 0 | 0 | 0 | 0 | 0 |
| MLEHLSSL | SPCS1; SPCS1 | Q9Y6A9; A0/ | SPCS1_HUMAI | Oxidation (M) | 0 | 121020 | 207750 | 0 | 540270 | 1137500 | 0 | 0 | 2933800 | 0 | 186830 | 423140 | 0 | 31282 | 94671 | 0 | 55129 | 0 |
| MLEHLSSL | SPCS1; SPCS1 | Q9Y6A9; A0/ | SPCS1_HUMAI | Unmodified | 0 | 0 | 0 | 0 | 0 | 0 | 0 | 47780 | 1336400 | 0 | 0 | 0 | 0 | 0 | 0 | 0 | 0 | 0 |
| MLEHLSSL | SPCS1; SPCS1 | Q9Y6A9; A0/ | SPCS1_HUMAI | Oxidation (M) | 0 | 0 | 0 | 0 | 0 | 0 | 0 | 28599 | 1007800 | 465110 | 0 | 0 | 0 | 0 | 0 | 0 | 0 | 1076800 |
| MLEIDPQK | ACAT1; ACAT1; | P24752; A0A | THIL_HUMAN A | Unmodified | 0 | 0 | 0 | 0 | 0 | 0 | 0 | 0 | 3357400 | 0 | 0 | 0 | 1039600 | 0 | 0 | 0 | 0 | 0 |
| MLEIDPQK | ACAT1; ACAT1; | P24752; A0A | THIL_HUMAN A | Oxidation (M) | 0 | 0 | 0 | 0 | 0 | 0 | 0 | 0 | 700550 | 148600 | 0 | 0 | 129620 | 127780 | 0 | 0 | 10494 | 0 |
| MLEMGITG | DPYSL3 | Q14195 | DPYL3_HUMAN | 2 Oxidation (M) | 0 | 0 | 0 | 0 | 0 | 0 | 0 | 0 | 0 | 0 | 3681400 | 0 | 0 | 0 | 0 | 0 | 0 | 0 |
| MLENTDNS | ZC3HAV1L | Q96H79 | ZCCHL_HUMAI | Unmodified | 0 | 0 | 104780 | 675750 | 819660 | 0 | 0 | 0 | 0 | 0 | 0 | 662580 | 0 | 71747 | 268200 | 137250 | 319800 | 233050 |
| MLENTDNS | ZC3HAV1L | Q96H79 | ZCCHL_HUMAI | Oxidation (M) | 113860 | 0 | 106590 | 151660 | 376530 | 56027 | 0 | 16084 | 0 | 0 | 45417 | 138190 | 69282 | 40074 | 296990 | 100740 | 48875 | 68889 |
| MLEPPPSA | KIF20A | O95235 | KI20A_HUMAN | Unmodified | 0 | 0 | 0 | 0 | 916290 | 0 | 0 | 0 | 511690 | 43410 | 0 | 0 | 0 | 0 | 0 | 17376 | 310970 | 0 |
| MLEQLDMF | DDX39B; DDX3 | Q13838; O0 | DX39B_HUMAN | Oxidation (M) | 0 | 24230 | 50675 | 64536 | 0 | 793790 | 0 | 0 | 279410 | 0 | 0 | 554480 | 0 | 0 | 0 | 0 | 0 | 0 |
| MLEQLDMF | DDX39B; DDX3 | Q13838; O0 | DX39B_HUMAN | Unmodified | 0 | 0 | 0 | 0 | 0 | 0 | 0 | 0 | 0 | 0 | 0 | 2507000 | 0 | 0 | 0 | 0 | 0 | 0 |
| MLESSGCK | PHLDA1; PHLD. | Q8WV24; R4 | PHLA1_HUMAN | Unmodified | 0 | 0 | 0 | 0 | 374190 | 0 | 0 | 0 | 0 | 0 | 0 | 371910 | 0 | 0 | 0 | 0 | 2776000 | 0 |
| MLESSGCK | PHLDA1; PHLD. | Q8WV24; R4 | PHLA1_HUMAN | Acetyl (Protein N | 0 | 0 | 0 | 0 | 197580 | 28670 | 0 | 0 | 0 | 0 | 0 | 0 | 0 | 0 | 0 | 0 | 1266700 | 608210 |
| MLESSGCK | PHLDA1; PHLD. | Q8WV24; R4 | PHLA1_HUMAN | Acetyl (Protein N | 0 | 0 | 0 | 0 | 565680 | 0 | 0 | 0 | 0 | 0 | 0 | 0 | 0 | 0 | 0 | 0 | 1270600 | 939530 |
| MLESSGCK | PHLDA1; PHLD. | Q8WV24; R4 | PHLA1_HUMAN | Acetyl (Protein N | 0 | 74175 | 0 | 0 | 0 | 0 | 0 | 0 | 0 | 0 | 0 | 0 | 0 | 0 | 0 | 0 | 2357900 | 282520 |
| MLESYLHA | CARM1; CARM1 | Q86X55; K7E | CARM1_HUMA | Unmodified | 0 | 0 | 0 | 0 | 0 | 0 | 0 | 0 | 0 | 0 | 0 | 595990 | 0 | 0 | 0 | 0 | 0 | 0 |
| MLEYALK | STRN4; STRN; S | Q9NRL3; O4 | STRN4_HUMAI | Unmodified | 0 | 0 | 0 | 0 | 285180 | 40314 | 0 | 0 | 0 | 169120 | 0 | 429100 | 0 | 0 | 0 | 0 | 50815 | 0 |
| MLEYLVTG | SLC43A1 | O75387 | LAT3_HUMAN I | Unmodified | 0 | 0 | 0 | 0 | 0 | 0 | 0 | 0 | 0 | 0 | 0 | 0 | 0 | 0 | 981710 | 274950 | 0 | 0 |
| MLEYLVTG | SLC43A1 | O75387 | LAT3_HUMAN I | Oxidation (M) | 0 | 0 | 356560 | 0 | 0 | 0 | 0 | 0 | 0 | 0 | 0 | 0 | 0 | 0 | 752510 | 853210 | 0 | 0 |
| MLFDYLAD | EIF3E; EIF3E; EI | P60228; A0A | EIF3E_HUMAN | Oxidation (M) | 0 | 0 | 344140 | 0 | 0 | 0 | 0 | 0 | 0 | 0 | 0 | 3626800 | 0 | 0 | 0 | 0 | 0 | 0 |
| MLFSPEEM | P3H1 | Q32P28 | P3H1_HUMAN | Oxidation (M) | 0 | 0 | 0 | 0 | 0 | 0 | 0 | 0 | 0 | 1254000 | 0 | 0 | 0 | 0 | 0 | 0 | 0 | 0 |
| MLFYSFFK | LSM2 | Q9Y333 | LSM2_HUMAN | Oxidation (M) | 225800 | 332020 | 183980 | 0 | 1421700 | 292460 | 915730 | 142920 | 217840 | 1393200 | 754710 | 2859500 | 1324100 | 321370 | 0 | 345860 | 1722800 | 752720 |
| MLGEALSK | PHB2; PHB2 | Q99623; F5 | PHB2_HUMAN | Unmodified | 0 | 0 | 0 | 0 | 254040 | 0 | 685130 | 176080 | 0 | 572180 | 0 | 0 | 219370 | 327310 | 0 | 0 | 144270 | 0 |
| MLGEEEEA | TAOK2 | Q9UL54 | TAOK2_HUMAN | Unmodified | 0 | 0 | 0 | 0 | 0 | 0 | 0 | 0 | 0 | 0 | 0 | 848470 | 0 | 0 | 0 | 0 | 0 | 0 |
| MLGNGRLI | EIF1AY; EIF1AY | O14602; A6 | IF1AY_HUMAN | Oxidation (M) | 193130 | 0 | 66114 | 0 | 120750 | 0 | 167230 | 33213 | 0 | 116200 | 158870 | 650070 | 0 | 0 | 0 | 64447 | 0 | 0 |
| MLGPEGGI | HNRNPF; HNRI | P52597; A0A | HNRPF_HUMAI | Acetyl (Protein N | 0 | 0 | 0 | 0 | 0 | 0 | 0 | 1966000 | 754370 | 929800 | 1998500 | 4857000 | 0 | 0 | 0 | 0 | 882530 | 0 |
| MLGQMTD | VCL | P18206 | VINC_HUMAN | Oxidation (M) | 0 | 0 | 0 | 0 | 0 | 0 | 0 | 0 | 0 | 0 | 0 | 903050 | 0 | 0 | 0 | 0 | 0 | 0 |
| MLGQMTD | VCL | P18206 | VINC_HUMAN | Unmodified | 0 | 0 | 0 | 51465 | 0 | 0 | 0 | 0 | 0 | 163650 | 0 | 0 | 0 | 0 | 0 | 0 | 1714000 | 0 |
| MLGQMTD | VCL | P18206 | VINC_HUMAN | 2 Oxidation (M) | 0 | 0 | 0 | 0 | 263160 | 0 | 0 | 0 | 0 | 0 | 0 | 0 | 0 | 0 | 0 | 0 | 3002600 | 0 |
| MLGQMTD | VCL | P18206 | VINC_HUMAN | Oxidation (M) | 0 | 0 | 0 | 0 | 1219100 | 0 | 154710 | 0 | 0 | 469600 | 0 | 0 | 0 | 0 | 0 | 0 | 10542000 | 0 |
| MLGQMTD | VCL | P18206 | VINC_HUMAN | Unmodified | 0 | 0 | 0 | 0 | 1226100 | 291110 | 1026600 | 0 | 0 | 214440 | 0 | 0 | 571280 | 177730 | 0 | 0 | 7308200 | 716730 |
| MLGSGFKA | IST1; IST1; IST1 | P53990; H3E | IST1_HUMAN I | Acetyl (Protein N | 0 | 0 | 0 | 0 | 0 | 0 | 714160 | 0 | 325670 | 367620 | 655730 | 2735900 | 0 | 0 | 0 | 0 | 0 | 0 |
| MLISILTER | ANXA3; ANXA3; | P12429; D6F | ANXA3_HUMAI | Oxidation (M) | 0 | 0 | 0 | 0 | 1891500 | 0 | 0 | 0 | 0 | 0 | 0 | 0 | 0 | 0 | 0 | 0 | 0 | 0 |
| MLISILTERS | ANXA3; ANXA3; | P12429; D6F | ANXA3_HUMAI | Unmodified | 0 | 99088 | 0 | 0 | 3368100 | 1009900 | 105660 | 0 | 0 | 0 | 0 | 0 | 760280 | 65514 | 0 | 0 | 0 | 1818900 |
| MLITILGTV | LGALS3; LGALS | P17931; G3 | LEG3_HUMAN | Oxidation (M) | 0 | 0 | 0 | 0 | 0 | 0 | 0 | 0 | 0 | 0 | 0 | 856210 | 0 | 0 | 0 | 0 | 0 | 0 |
| MLIYDPAK | CDK1; CDK1 | P06493; A0A | CDK1_HUMAN | Oxidation (M) | 0 | 354600 | 175790 | 265230 | 0 | 0 | 0 | 25389 | 0 | 0 | 59782 | 527090 | 794400 | 344590 | 1317500 | 1791000 | 0 | 0 |
| MLLDGEIG. | AASS; AASS | Q9UDR5; A0 | AASS_HUMAN . | Oxidation (M) | 0 | 0 | 179660 | 0 | 437560 | 783570 | 132330 | 0 | 27077 | 24819 | 0 | 0 | 0 | 443260 | 0 | 0 | 0 | 319770 |
| MLLDIFDEI | CCNY | Q8ND76 | CCNY_HUMAN | Oxidation (M) | 349170 | 385630 | 378100 | 301660 | 0 | 0 | 0 | 0 | 0 | 0 | 569940 | 4146200 | 0 | 0 | 1302400 | 1462900 | 0 | 104990 |
| MLLDPMG( | CCT3; CCT3; C | P49368; Q5 | TCPG_HUMAN | 3 Oxidation (M) | 0 | 1236700 | 0 | 0 | 0 | 0 | 0 | 0 | 0 | 0 | 0 | 1421900 | 0 | 0 | 0 | 0 | 0 | 0 |
| MLLDPMG( | CCT3; CCT3; C | P49368; Q5 | TCPG_HUMAN | 3 Oxidation (M) | 0 | 0 | 0 | 0 | 3224400 | 0 | 3434200 | 0 | 0 | 1716300 | 0 | 0 | 12113000 | 7163300 | 0 | 0 | 3608000 | 3116600 |
| MLLDSEQH | PYCR2; PYCR2; | Q96C36; A0/ | P5CR2_HUMAI | Unmodified | 0 | 0 | 0 | 0 | 0 | 0 | 0 | 0 | 1148900 | 256370 | 0 | 83067 | 0 | 0 | 0 | 0 | 0 | 0 |
| MLLGNKCI | RAB10 | P61026 | RAB10_HUMAN | Unmodified | 128890 | 62982 | 0 | 0 | 0 | 0 | 0 | 0 | 181320 | 0 | 51067 | 586550 | 0 | 0 | 152520 | 121960 | 0 | 0 |
| MLLHSEQH | PYCR1; PYCR1; | P32322; J3Q | P5CR1_HUMAI | Unmodified | 0 | 0 | 0 | 0 | 0 | 0 | 91862 | 0 | 480710 | 124300 | 0 | 0 | 0 | 0 | 0 | 0 | 0 | 0 |
| MLLIENLRE | ITGB4 | P16144 | ITB4_HUMAN I | Oxidation (M) | 0 | 0 | 0 | 0 | 0 | 531420 | 0 | 0 | 0 | 0 | 0 | 0 | 185610 | 0 | 0 | 0 | 9717600 | 9721700 |
| MLLIENLRE | ITGB4 | P16144 | ITB4_HUMAN I | Unmodified | 0 | 0 | 0 | 0 | 1089000 | 379310 | 0 | 0 | 0 | 0 | 0 | 0 | 109830 | 108100 | 0 | 0 | 5862500 | 4522700 |
| MLLLEILHE | ERO1A; ERO1A | Q96HE7; G3 | ERO1A_HUMAI | Oxidation (M) | 0 | 0 | 0 | 0 | 0 | 0 | 0 | 0 | 3127900 | 1640300 | 371350 | 346660 | 0 | 0 | 64851 | 0 | 0 | 0 |
| MLLLEILHE | ERO1A; ERO1A | Q96HE7; G3 | ERO1A_HUMAI | Unmodified | 0 | 0 | 0 | 0 | 0 | 0 | 0 | 0 | 2314100 | 856410 | 210490 | 0 | 0 | 0 | 0 | 0 | 0 | 0 |
| MLLLQPQA | SEMA4B; SEMA | Q9NPR2; H0 | SEM4B_HUMAI | Unmodified | 0 | 0 | 0 | 0 | 0 | 0 | 0 | 0 | 859810 | 0 | 0 | 0 | 0 | 0 | 0 | 0 | 0 | 0 |
| MLLNDFLN | HMGCS1 | Q01581 | HMCS1_HUMA | Unmodified | 40761 | 0 | 0 | 0 | 797900 | 0 | 0 | 0 | 1374900 | 197230 | 0 | 4332700 | 0 | 0 | 0 | 78803 | 0 | 0 |
| MLLNDFLN | HMGCS1 | Q01581 | HMCS1_HUMA | Oxidation (M) | 0 | 0 | 0 | 0 | 890270 | 0 | 0 | 0 | 0 | 0 | 2715000 | 6884700 | 0 | 0 | 0 | 0 | 0 | 0 |
| MLLPSGSL | ROBO1; ROBO2 | Q9Y6N7; Q9 | ROBO1_HUMA | Oxidation (M) | 0 | 0 | 0 | 0 | 0 | 0 | 0 | 0 | 547950 | 59445 | 0 | 0 | 0 | 0 | 0 | 0 | 0 | 0 |
| MLLYTEVTF | GDI1; GDI2; GD | P31150; P50 | GDIA_HUMAN | Oxidation (M) | 0 | 1184700 | 1903300 | 0 | 0 | 0 | 0 | 0 | 1579000 | 0 | 0 | 4759300 | 0 | 0 | 2715500 | 4613000 | 0 | 0 |
| MLLYTEVTF | GDI1; GDI2; GD | P31150; P50 | GDIA_HUMAN | Unmodified | 0 | 0 | 0 | 0 | 0 | 0 | 0 | 0 | 3661300 | 417500 | 897060 | 5074700 | 0 | 0 | 0 | 0 | 0 | 0 |
| MLLYTEVTF | GDI1; GDI2; GD | P31150; P50 | GDIA_HUMAN | Oxidation (M) | 0 | 0 | 0 | 0 | 0 | 0 | 0 | 0 | 0 | 750060 | 0 | 0 | 795420 | 577080 | 0 | 0 | 0 | 0 |
| MLMLDPAE | CASK; CASK; C/ | O14936; A0/ | CSKP_HUMAN | 2 Oxidation (M) | 0 | 0 | 0 | 1037200 | 0 | 0 | 0 | 0 | 0 | 1483000 | 0 | 0 | 2908300 | 4243800 | 375650 | 0 | 822330 | 866300 |
| MLMLILEK | NEDD4L | A0A6Q8PFI7 | A0A6Q8PFI7_H | Acetyl (Protein N | 416230 | 0 | 0 | 0 | 277570 | 0 | 340400 | 447170 | 0 | 0 | 0 | 1730800 | 6110000 | 798710 | 187980 | 175570 | 119310 | 1897200 |

| | | | | | | | | | | | | | | | | | | | | | | |
|---|---|---|---|---|---|---|---|---|---|---|---|---|---|---|---|---|---|---|---|---|---|---|
| MLNAREPK | NSF; NSF | P46459; I3L( | NSF_HUMAN V | Unmodified | 0 | 0 | 0 | 0 | 0 | 0 | 127930 | 50696 | 800030 | 0 | 0 | 0 | 1002100 | 668150 | 0 | 0 | 230930 | 244720 |
| MLNAREPK | NSF; NSF | P46459; I3L( | NSF_HUMAN V | Oxidation (M) | 0 | 0 | 0 | 0 | 0 | 0 | 116280 | 445270 | 0 | 0 | 0 | 0 | 684640 | 342610 | 0 | 0 | 110860 | 0 |
| MLNIHPSLI | GART; GART | P22102; A0A | PUR2_HUMAN | Oxidation (M) | 255050 | 0 | 260070 | 0 | 0 | 0 | 0 | 0 | 0 | 180860 | 0 | 3593500 | 281150 | 0 | 1387700 | 484640 | 0 | 0 |
| MLNSSFED | MLF1; MLF1; MI | P58340; A0A | MLF1_HUMAN | 2 Oxidation (M) | 0 | 0 | 51197 | 0 | 0 | 0 | 0 | 0 | 0 | 0 | 0 | 0 | 334450 | 0 | 0 | 0 | 0 | 0 |
| MLPDKDCI | CFL1; CFL1; CF | P23528; G3\ | COF1_HUMAN | Oxidation (M) | 0 | 0 | 0 | 0 | 67021000 | 12320000 | 13529000 | 9533400 | 0 | 4481400 | 0 | 0 | 28983000 | 24998000 | 0 | 0 | 33504000 | 28651000 |
| MLPDKDCI | CFL1; CFL1; CF | P23528; G3\ | COF1_HUMAN | Unmodified | 0 | 0 | 0 | 0 | 1.69E+08 | 13248000 | 24295000 | 872870 | 0 | 9343200 | 0 | 0 | 31682000 | 42856000 | 0 | 0 | 54315000 | 93438000 |
| MLPLSLLK | LSM4; LSM4; LS | Q9Y4Z0; M0 | LSM4_HUMAN | Oxidation (M) | 87177 | 275620 | 0 | 86665 | 137220 | 0 | 139190 | 131840 | 0 | 343550 | 337370 | 814950 | 641540 | 0 | 0 | 156350 | 0 | 155370 |
| MLPTIIADN | CCT2; CCT2 | P78371; F5G | TCPB_HUMAN | Oxidation (M) | 0 | 0 | 0 | 1415900 | 0 | 0 | 0 | 0 | 238180 | 0 | 278080 | 927060 | 0 | 0 | 0 | 0 | 0 | 0 |
| MLPVDEFLI | COLGALT1 | Q8NBJ5 | GT251_HUMAN | 2 Oxidation (M) | 0 | 0 | 0 | 0 | 0 | 0 | 0 | 0 | 548380 | 131380 | 53942 | 0 | 0 | 0 | 0 | 0 | 0 | 0 |
| MLPVDEFLI | COLGALT1 | Q8NBJ5 | GT251_HUMAN | Oxidation (M) | 0 | 0 | 0 | 0 | 421470 | 0 | 387180 | 0 | 7971300 | 4339400 | 0 | 0 | 0 | 0 | 0 | 216650 | 0 | 0 |
| MLPVDEFLI | COLGALT1 | Q8NBJ5 | GT251_HUMAN | 2 Oxidation (M) | 0 | 0 | 0 | 0 | 0 | 0 | 0 | 0 | 13786000 | 15635000 | 0 | 310670 | 0 | 0 | 0 | 0 | 0 | 0 |
| MLQADPNI | RTCB | Q9Y3I0 | RTCB_HUMAN | Unmodified | 41596 | 256900 | 0 | 0 | 0 | 170990 | 359760 | 287780 | 817590 | 0 | 182980 | 1863300 | 0 | 0 | 366480 | 0 | 0 | 0 |
| MLQDDDD | SLC4A7; SLC4A | Q9Y6M7; A0. | S4A7_HUMAN | Oxidation (M) | 0 | 0 | 39404 | 78685 | 0 | 0 | 0 | 0 | 1282800 | 837370 | 1695400 | 4834500 | 0 | 0 | 508920 | 842470 | 0 | 0 |
| MLQDTEDI | WASF3; WASF1 | Q9UPY6; Q9 | WASF3_HUMA | Unmodified | 0 | 0 | 0 | 0 | 0 | 0 | 0 | 82737 | 0 | 25499 | 0 | 657750 | 0 | 0 | 0 | 0 | 0 | 571840 |
| MLQDTEDI | WASF3; WASF1 | Q9UPY6; Q9 | WASF3_HUMA | Unmodified | 0 | 0 | 32869 | 0 | 563640 | 149470 | 78271 | 162790 | 0 | 101030 | 0 | 0 | 375520 | 201510 | 0 | 203170 | 118630 | 137580 |
| MLQDTKDI | WASF2 | Q9Y6W5 | WASF2_HUMA | Unmodified | 0 | 0 | 0 | 0 | 0 | 0 | 0 | 0 | 77606 | 273830 | 0 | 0 | 0 | 0 | 0 | 0 | 874630 | 0 |
| MLQEDELR | ARF5 | P84085 | ARF5_HUMAN | Unmodified | 789380 | 506240 | 0 | 0 | 0 | 0 | 0 | 0 | 0 | 0 | 0 | 1421300 | 0 | 0 | 0 | 0 | 0 | 0 |
| MLQEDELR | ARF5 | P84085 | ARF5_HUMAN | 3 Oxidation (M) | 38461 | 0 | 0 | 1921900 | 57072 | 0 | 0 | 0 | 0 | 0 | 21011 | 23343 | 0 | 0 | 319450 | 5085100 | 0 | 0 |
| MLQFVLPV | C12orf57; C12c | Q99622; U3I | C10_HUMAN P | Oxidation (M) | 314590 | 368360 | 0 | 649930 | 1995400 | 397170 | 933260 | 0 | 321470 | 494030 | 264250 | 374380 | 3120300 | 2125500 | 882460 | 162900 | 1548600 | 0 |
| MLQHAASI | IQGAP1; IQGAP | P46940; H0 | IQGA1_HUMAN | Unmodified | 0 | 0 | 0 | 0 | 524740 | 919980 | 0 | 0 | 0 | 0 | 0 | 0 | 0 | 0 | 44745 | 0 | 108050 | 3863200 |
| MLQLIQSR | ERGIC2; ERGIC | Q96RQ1; F8 | ERGI2_HUMAN | Unmodified | 0 | 0 | 0 | 0 | 0 | 0 | 0 | 0 | 2128800 | 59977 | 0 | 0 | 0 | 0 | 0 | 0 | 0 | 0 |
| MLQNVQM | DPP4; DPP4; DI | P27487; A0A | DPP4_HUMAN | Oxidation (M) | 0 | 0 | 2513900 | 2921800 | 98050 | 0 | 0 | 0 | 0 | 0 | 49389 | 419900 | 0 | 0 | 2107000 | 1335100 | 850110 | 651610 |
| MLQNVQM | DPP4; DPP4; DI | P27487; A0A | DPP4_HUMAN | Unmodified | 0 | 0 | 3309000 | 5772100 | 182130 | 0 | 0 | 0 | 0 | 136920 | 0 | 0 | 0 | 0 | 3049400 | 1259200 | 1360500 | 1904400 |
| MLQNVQM | DPP4; DPP4; DI | P27487; A0A | DPP4_HUMAN | 2 Oxidation (M) | 0 | 142050 | 850350 | 993770 | 0 | 0 | 0 | 0 | 0 | 417640 | 1435700 | 0 | 0 | 0 | 682290 | 572090 | 105820 | 182800 |
| MLQQELLS | PPFIBP2; PPFIB | Q8ND30; E9 | LIPB2_HUMAN | Acetyl (Protein N | 0 | 0 | 26309 | 1669500 | 0 | 297150 | 213610 | 0 | 0 | 0 | 0 | 0 | 0 | 152760 | 0 | 15937 | 0 | 2920600 |
| MLQQVPEI | IARS1; IARS1; I | P41252; A0A | SYIC_HUMAN I | Acetyl (Protein N | 0 | 0 | 0 | 0 | 268920 | 0 | 426160 | 163390 | 346790 | 344140 | 0 | 0 | 256230 | 0 | 0 | 0 | 0 | 162620 |
| MLQVASQ | WBP2; WBP2; V | Q969T9; K7E | WBP2_HUMAN | Unmodified | 242200 | 234540 | 149940 | 0 | 0 | 0 | 0 | 0 | 0 | 0 | 0 | 0 | 0 | 0 | 432590 | 341940 | 0 | 0 |
| MLRALQA | COLGALT1; CO | Q8NBJ5; M0 | GT251_HUMAN | Oxidation (M) | 0 | 0 | 0 | 0 | 0 | 0 | 193410 | 0 | 0 | 769470 | 0 | 0 | 0 | 0 | 0 | 0 | 0 | 0 |
| MLRLGNAI | MYO5A; MYO5A | Q9Y4I1; G3\ | MYO5A_HUMA | Unmodified | 0 | 0 | 0 | 0 | 488940 | 0 | 0 | 0 | 0 | 0 | 0 | 0 | 0 | 0 | 0 | 0 | 0 | 0 |
| MLRVIVES | MYOF | Q9NZM1 | MYOF_HUMAN | Acetyl (Protein N | 0 | 0 | 0 | 0 | 2863200 | 208860 | 0 | 0 | 0 | 0 | 0 | 0 | 0 | 0 | 0 | 0 | 297000 | 766590 |
| MLRVIVES | MYOF | Q9NZM1 | MYOF_HUMAN | Acetyl (Protein N | 2716300 | 1553800 | 0 | 0 | 1568900 | 283470 | 0 | 0 | 0 | 0 | 0 | 0 | 0 | 0 | 0 | 0 | 0 | 0 |
| MLSASTML | PTPRF; PTPRF; | P10586; H0 | PTPRF_HUMAN | Oxidation (M) | 33186 | 46170 | 0 | 0 | 0 | 0 | 0 | 0 | 122510 | 0 | 0 | 0 | 0 | 0 | 0 | 0 | 0 | 0 |
| MLSASTML | PTPRF; PTPRF; | P10586; H0 | PTPRF_HUMAN | 2 Oxidation (M) | 1033500 | 0 | 0 | 0 | 0 | 0 | 0 | 0 | 0 | 0 | 0 | 0 | 0 | 0 | 0 | 0 | 0 | 0 |
| MLSAVSQC | DYNC1H1; DYN | Q14204; A0 | DYHC1_HUMA | Oxidation (M) | 42937 | 0 | 21691 | 0 | 0 | 0 | 0 | 0 | 0 | 0 | 0 | 701810 | 494380 | 0 | 107370 | 36320 | 0 | 0 |
| MLSAVSQC | DYNC1H1; DYN | Q14204; A0 | DYHC1_HUMA | Unmodified | 0 | 0 | 0 | 0 | 0 | 0 | 0 | 0 | 551160 | 0 | 0 | 203760 | 0 | 0 | 0 | 0 | 0 | 0 |
| MLSAVSQC | DYNC1H1; DYN | Q14204; A0 | DYHC1_HUMA | Unmodified | 0 | 0 | 0 | 0 | 0 | 0 | 0 | 0 | 230370 | 0 | 0 | 844940 | 0 | 0 | 183960 | 129410 | 0 | 0 |
| MLSCAGAI | RPL10; RPL10L; | P27635; Q9 | RL10_HUMAN | Unmodified | 400370 | 0 | 262880 | 101190 | 0 | 0 | 172230 | 0 | 0 | 0 | 183860 | 7277200 | 0 | 0 | 310810 | 198930 | 0 | 0 |
| MLSCAGAI | RPL10; RPL10L; | P27635; Q9 | RL10_HUMAN | Unmodified | 1452600 | 1651100 | 326360 | 899290 | 0 | 0 | 187730 | 151870 | 1973700 | 0 | 733960 | 6733000 | 0 | 135950 | 480250 | 0 | 0 | 0 |
| MLSCAGAI | RPL10; RPL10L; | P27635; Q9 | RL10_HUMAN | 2 Oxidation (M) | 1012900 | 476010 | 213260 | 279670 | 0 | 287310 | 64525 | 76044 | 0 | 50736 | 97805 | 1754500 | 0 | 0 | 245070 | 398910 | 0 | 0 |
| MLSCAGAI | RPL10; RPL10L; | P27635; Q9 | RL10_HUMAN | Oxidation (M) | 1056100 | 335040 | 56527 | 362500 | 0 | 44503 | 0 | 60544 | 0 | 38003 | 476390 | 4764800 | 0 | 0 | 707420 | 240870 | 41321 | 0 |
| MLSGGFG | MLF2; MLF2; MI | Q15773; F5 | MLF2_HUMAN | 2 Oxidation (M) | 23190000 | 25407000 | 6515700 | 10495000 | 0 | 0 | 0 | 0 | 0 | 0 | 0 | 0 | 7037100 | 0 | 0 | 5453100 | 0 | 0 |
| MLSGGFG | MLF2; MLF2; MI | Q15773; F5 | MLF2_HUMAN | Oxidation (M) | 436270 | 912390 | 0 | 183260 | 0 | 0 | 0 | 0 | 0 | 0 | 0 | 0 | 1342400 | 0 | 199520 | 134430 | 0 | 0 |
| MLSLDFLD | SEC11A; SEC11 | P67812; H0 | SC11A_HUMAN | Oxidation (M) | 69026 | 48215 | 50197 | 70193 | 0 | 0 | 0 | 0 | 8450900 | 1153800 | 652330 | 1372600 | 0 | 0 | 68829 | 32913 | 0 | 0 |
| MLSLDFLD | SEC11A; SEC11 | P67812; H0 | SC11A_HUMAN | Unmodified | 0 | 0 | 0 | 0 | 0 | 46855 | 0 | 0 | 3780300 | 207570 | 56322 | 35888 | 0 | 0 | 0 | 0 | 0 | 0 |
| MLSLRVPL | RRM2; RRM2 | P31350; A0A | RIR2_HUMAN | Acetyl (Protein N | 0 | 0 | 0 | 0 | 0 | 0 | 0 | 0 | 0 | 0 | 0 | 0 | 854540 | 764070 | 0 | 0 | 0 | 996960 |
| MLSNKPQI | MYOF | Q9NZM1 | MYOF_HUMAN | Unmodified | 0 | 0 | 0 | 0 | 6351300 | 896350 | 0 | 0 | 0 | 0 | 0 | 0 | 0 | 0 | 0 | 0 | 234160 | 437890 |
| MLSNKPQI | MYOF | Q9NZM1 | MYOF_HUMAN | Unmodified | 0 | 0 | 0 | 0 | 2445400 | 539720 | 0 | 0 | 0 | 0 | 0 | 0 | 0 | 0 | 0 | 0 | 0 | 272740 |
| MLSPEHAP | KIT | P10721 | KIT_HUMAN Ma | Oxidation (M) | 0 | 0 | 0 | 0 | 0 | 622690 | 0 | 0 | 914950 | 0 | 162150 | 702120 | 0 | 0 | 154930 | 0 | 0 | 0 |
| MLTAPFAG | SORCS2; SORC | Q96PQ0; B5 | SORC2_HUMA | Oxidation (M) | 0 | 0 | 0 | 0 | 0 | 0 | 0 | 0 | 1845800 | 1316400 | 0 | 0 | 0 | 0 | 0 | 0 | 0 | 0 |
| MLTAPFAG | SORCS2; SORC | Q96PQ0; B5 | SORC2_HUMA | Unmodified | 0 | 0 | 0 | 0 | 0 | 0 | 0 | 0 | 272710 | 0 | 0 | 0 | 0 | 0 | 0 | 0 | 0 | 0 |
| MLTAQDM | SPTBN1; SPTBN | Q01082; A0 | SPTB2_HUMAN | Unmodified | 0 | 0 | 0 | 0 | 922190 | 220270 | 63345 | 0 | 0 | 132980 | 0 | 0 | 0 | 0 | 0 | 0 | 280990 | 126840 |
| MLTAQDVS | SPTBN2 | O15020 | SPTN2_HUMAN | Unmodified | 0 | 0 | 0 | 0 | 0 | 0 | 0 | 0 | 0 | 0 | 0 | 0 | 289120 | 0 | 0 | 0 | 69836 | 0 |
| MLTERMFN | TALDO1 | P37837 | TALDO_HUMAN | Unmodified | 0 | 0 | 0 | 0 | 1820600 | 0 | 0 | 0 | 0 | 0 | 0 | 0 | 538750 | 232640 | 0 | 0 | 699540 | 0 |
| MLTGPVYS | CAND1; CAND1 | Q86VP6; A0 | CAND1_HUMA | Oxidation (M) | 0 | 0 | 0 | 0 | 0 | 0 | 0 | 0 | 0 | 0 | 0 | 2061400 | 0 | 0 | 0 | 0 | 338710 | 0 |
| MLTPLGPG | EPHA1 | P21709 | EPHA1_HUMAI | Oxidation (M) | 0 | 0 | 574950 | 0 | 0 | 193700 | 0 | 0 | 126590 | 0 | 147660 | 0 | 293440 | 0 | 1451000 | 517980 | 0 | 60918 |
| MLVELSK | CCT4 | P50991 | TCPD_HUMAN | Unmodified | 0 | 0 | 0 | 0 | 0 | 0 | 0 | 0 | 0 | 1412100 | 0 | 6187500 | 0 | 0 | 0 | 139090 | 0 | 2015600 |
| MLVELSK | CCT4 | P50991 | TCPD_HUMAN | Oxidation (M) | 0 | 0 | 0 | 0 | 0 | 0 | 0 | 0 | 0 | 0 | 0 | 2516300 | 0 | 0 | 0 | 0 | 0 | 0 |
| MLVETAQE | PSEN2; PSEN1; | P49810; P49 | PSN2_HUMAN | Oxidation (M) | 0 | 0 | 441700 | 0 | 176790 | 0 | 0 | 0 | 0 | 443020 | 46640 | 2140000 | 0 | 0 | 284410 | 754740 | 76416 | 0 |
| MLVETAQE | PSEN2; PSEN1; | P49810; P49 | PSN2_HUMAN | Unmodified | 0 | 0 | 244280 | 0 | 0 | 0 | 0 | 194450 | 224200 | 100270 | 198450 | 1252000 | 0 | 0 | 295860 | 208660 | 0 | 0 |
| MLVGSQSF | RIPOR2; RIPOR | Q9Y4F9; B7 | RIPR2_HUMAN | Acetyl (Protein N | 0 | 0 | 0 | 0 | 0 | 0 | 0 | 0 | 0 | 0 | 0 | 0 | 258510 | 134230 | 0 | 0 | 0 | 0 |
| MLVIEQCK | CCT5; CCT5; C | P48643; B7 | TCPE_HUMAN | Unmodified | 0 | 0 | 0 | 442950 | 962420 | 0 | 585540 | 325800 | 1028700 | 72717 | 788360 | 6283100 | 1956700 | 1146600 | 0 | 377390 | 1324000 | 740260 |
| MLVIEQCK | CCT5; CCT5; C | P48643; B7 | TCPE_HUMAN | Oxidation (M) | 0 | 0 | 0 | 0 | 353420 | 0 | 0 | 0 | 0 | 244500 | 0 | 2019300 | 743210 | 0 | 0 | 0 | 0 | 383740 |
| MLVNENEV | DAAM1 | Q9Y4D1 | DAAM1_HUMA | Unmodified | 0 | 0 | 0 | 0 | 0 | 0 | 0 | 40001 | 1096200 | 184160 | 636970 | 2130600 | 394320 | 489570 | 0 | 25242 | 0 | 0 |
| MLVNENEV | DAAM1 | Q9Y4D1 | DAAM1_HUMA | Oxidation (M) | 0 | 0 | 0 | 0 | 529060 | 0 | 111840 | 0 | 0 | 133280 | 148890 | 1169400 | 0 | 0 | 0 | 0 | 0 | 549290 |
| MLVSGAGI | CCT6A; CCT6B | P40227; Q9 | TCPZ_HUMAN | Oxidation (M) | 91098 | 103000 | 0 | 245200 | 333510 | 0 | 162960 | 97596 | 60020 | 163420 | 0 | 3151000 | 1153200 | 385550 | 0 | 32602 | 486350 | 141390 |
| MLVSGAGI | CCT6A; CCT6B | P40227; Q9 | TCPZ_HUMAN | Unmodified | 0 | 0 | 0 | 0 | 0 | 0 | 0 | 0 | 0 | 0 | 0 | 9227100 | 0 | 0 | 0 | 0 | 0 | 0 |
| MLVSGAGI | CCT6A; CCT6B | P40227; Q9 | TCPZ_HUMAN | Oxidation (M) | 0 | 0 | 0 | 0 | 1590600 | 0 | 1141000 | 727440 | 0 | 1043000 | 0 | 0 | 2566600 | 2274200 | 0 | 0 | 1211500 | 2176500 |
| MLVSGAGI | CCT6A; CCT6B | P40227; Q9 | TCPZ_HUMAN | Unmodified | 0 | 0 | 0 | 0 | 4902100 | 0 | 2153800 | 484980 | 0 | 1295900 | 0 | 0 | 4931800 | 3002800 | 0 | 0 | 3457000 | 3039100 |
| MLVVGGID | KARS1 | Q15046 | SYK_HUMAN Ly | Unmodified | 0 | 0 | 0 | 0 | 0 | 0 | 0 | 0 | 0 | 0 | 0 | 1605900 | 0 | 0 | 0 | 0 | 0 | 0 |
| MLVVGGID | KARS1 | Q15046 | SYK_HUMAN Ly | Oxidation (M) | 0 | 0 | 652190 | 75993 | 0 | 0 | 0 | 0 | 0 | 0 | 109590 | 6337400 | 0 | 0 | 208370 | 0 | 0 | 0 |
| MLVVGGID | KARS1 | Q15046 | SYK_HUMAN Ly | Unmodified | 282920 | 713490 | 0 | 798330 | 0 | 0 | 0 | 0 | 443240 | 0 | 0 | 1923100 | 0 | 0 | 0 | 410510 | 0 | 0 |
| MLYAATR | TWF2; TWF1; T\ | Q6IBS0; Q12 | TWF2_HUMAN | Unmodified | 631860 | 0 | 282410 | 0 | 558590 | 0 | 0 | 224190 | 294000 | 186160 | 182790 | 2906700 | 0 | 0 | 0 | 0 | 510510 | 0 |
| MLYDDAG\ | SMPDL3B; SMP | Q92485; F8 | ASM3B_HUMAI | 2 Oxidation (M) | 0 | 0 | 890750 | 1402800 | 0 | 0 | 0 | 0 | 677700 | 0 | 0 | 116190 | 0 | 0 | 6809400 | 937050 | 875270 | 0 |
| MLYQNDLI | CNTN2; CNTN2 | Q02246; A0 | CNTN2_HUMA | Oxidation (M) | 0 | 0 | 0 | 0 | 0 | 0 | 262200 | 0 | 0 | 0 | 0 | 0 | 0 | 0 | 0 | 0 | 0 | 0 |
| MMADEAL( | ECH1 | Q13011 | ECH1_HUMAN | 2 Oxidation (M) | 0 | 0 | 0 | 0 | 0 | 0 | 245840 | 0 | 672620 | 0 | 0 | 0 | 0 | 0 | 0 | 0 | 0 | 0 |
| MMADEAL( | ECH1 | Q13011 | ECH1_HUMAN | Oxidation (M) | 0 | 673620 | 0 | 0 | 0 | 0 | 0 | 0 | 4321700 | 0 | 0 | 0 | 0 | 0 | 0 | 0 | 0 | 0 |
| MMADEAL( | ECH1 | Q13011 | ECH1_HUMAN | Unmodified | 0 | 0 | 0 | 0 | 0 | 0 | 45599 | 0 | 4781800 | 0 | 0 | 0 | 236920 | 0 | 0 | 0 | 0 | 0 |
| MMALDRE\ | HYOU1; HYOU1 | Q9Y4L1; A0 | HYOU1_HUMA | 2 Oxidation (M) | 0 | 0 | 0 | 0 | 0 | 0 | 442990 | 0 | 4926000 | 8891600 | 3119100 | 0 | 0 | 0 | 551890 | 254240 | 0 | 0 |
| MMALDRE\ | HYOU1; HYOU1 | Q9Y4L1; A0 | HYOU1_HUMA | Oxidation (M) | 0 | 0 | 0 | 0 | 0 | 0 | 0 | 0 | 17162000 | 13151000 | 0 | 0 | 0 | 0 | 0 | 0 | 0 | 0 |
| MMALDRE\ | HYOU1; HYOU1 | Q9Y4L1; A0 | HYOU1_HUMA | Unmodified | 0 | 0 | 0 | 0 | 31743 | 0 | 0 | 0 | 7427300 | 1567900 | 34578 | 0 | 52770 | 45187 | 0 | 0 | 0 | 0 |
| MMANGILK | AHCY | P23526 | SAHH_HUMAN | Oxidation (M) | 31591 | 0 | 39173 | 73945 | 0 | 0 | 0 | 0 | 0 | 0 | 0 | 2315300 | 82478 | 75414 | 180790 | 107420 | 0 | 0 |
| MMANGILK | AHCY | P23526 | SAHH_HUMAN | 2 Oxidation (M) | 0 | 0 | 0 | 383710 | 2996800 | 544010 | 793360 | 375490 | 248570 | 752320 | 291300 | 1246200 | 1477500 | 709140 | 0 | 0 | 4043900 | 1709000 |
| MMANGILK | AHCY | P23526 | SAHH_HUMAN | Oxidation (M) | 0 | 0 | 0 | 0 | 1080700 | 0 | 438480 | 14462 | 263460 | 737140 | 0 | 160670 | 1465200 | 387640 | 0 | 0 | 2658000 | 2191600 |
| MMANGILK | AHCY | P23526 | SAHH_HUMAN | Unmodified | 0 | 0 | 0 | 0 | 664070 | 0 | 0 | 0 | 151060 | 134660 | 0 | 256220 | 0 | 209100 | 0 | 0 | 485800 | 370410 |
| MMDYLQG | GANAB; GANAE | Q14697; F5 | GANAB_HUMA | 2 Oxidation (M) | 0 | 0 | 226810 | 0 | 0 | 0 | 0 | 0 | 4030000 | 339000 | 260260 | 0 | 0 | 0 | 263770 | 0 | 0 | 176560 |
| MMDYLQG | GANAB; GANAE | Q14697; F5 | GANAB_HUMA | Oxidation (M) | 0 | 0 | 0 | 0 | 0 | 101000 | 0 | 0 | 21776000 | 0 | 184050 | 1114500 | 0 | 0 | 106990 | 35419 | 0 | 0 |
| MMDYLQG | GANAB; GANAE | Q14697; F5 | GANAB_HUMA | Unmodified | 0 | 0 | 0 | 265590 | 0 | 0 | 0 | 0 | 13151000 | 0 | 0 | 0 | 0 | 0 | 339380 | 36371 | 0 | 0 |
| MMEEFVPI | NCKAP1 | Q9Y2A7 | NCKP1_HUMAI | Unmodified | 0 | 0 | 0 | 134400 | 0 | 0 | 0 | 0 | 886120 | 240940 | 0 | 1140000 | 0 | 0 | 230550 | 0 | 0 | 0 |
| MMEEFVPI | NCKAP1 | Q9Y2A7 | NCKP1_HUMAI | Oxidation (M) | 0 | 0 | 131150 | 163510 | 0 | 0 | 0 | 0 | 0 | 0 | 0 | 566960 | 0 | 0 | 228590 | 251680 | 0 | 0 |
| MMEEFVPI | NCKAP1 | Q9Y2A7 | NCKP1_HUMAI | 2 Oxidation (M) | 0 | 0 | 16550 | 0 | 0 | 0 | 0 | 0 | 0 | 0 | 0 | 0 | 0 | 0 | 153570 | 174680 | 0 | 0 |
| MMEIMTR | RPS3A; RPS3A; | P61247; D6 | RS3A_HUMAN | Unmodified | 93271 | 104470 | 66275 | 118990 | 0 | 0 | 65080 | 0 | 340980 | 33196 | 287170 | 3579800 | 151190 | 34289 | 50396 | 17945 | 0 | 0 |
| MMEKLGVI | AIF1L; AIF1L; AI | Q9BQI0; Q5 | AIF1L_HUMAN | 3 Oxidation (M) | 0 | 0 | 0 | 174200 | 0 | 0 | 0 | 0 | 0 | 397320 | 0 | 0 | 583170 | 0 | 0 | 0 | 0 | 0 |
| MMEVAAAI | CNDP2; CNDP2 | Q96KP4; J3C | CNDP2_HUMA | Unmodified | 0 | 0 | 0 | 0 | 407620 | 0 | 0 | 0 | 0 | 0 | 552120 | 520710 | 5258300 | 2345700 | 0 | 0 | 181650 | 318610 |
| MMEVAAAI | CNDP2; CNDP2 | Q96KP4; J3C | CNDP2_HUMA | Oxidation (M) | 0 | 0 | 0 | 186270 | 0 | 0 | 0 | 9037.2 | 157450 | 64880 | 454460 | 2692300 | 1731600 | 945030 | 645350 | 446860 | 63590 | 288040 |
| MMEVAAAI | CNDP2; CNDP2 | Q96KP4; J3C | CNDP2_HUMA | 2 Oxidation (M) | 0 | 0 | 0 | 0 | 0 | 0 | 14789 | 0 | 0 | 0 | 103030 | 272190 | 342780 | 198870 | 0 | 0 | 0 | 132020 |
| MMFEKGEI | LIMA1; LIMA1; L | Q9UHB6; F8 | LIMA1_HUMAN | Unmodified | 0 | 0 | 0 | 0 | 463100 | 0 | 0 | 0 | 0 | 0 | 127480 | 0 | 0 | 312210 | 0 | 0 | 129790 | 59128 |
| MMITSQDV | MT-CO2 | P00403 | COX2_HUMAN | Oxidation (M) | 0 | 0 | 0 | 0 | 0 | 0 | 0 | 0 | 966330 | 0 | 0 | 0 | 0 | 0 | 0 | 0 | 0 | 0 |
| MMITSQDV | MT-CO2 | P00403 | COX2_HUMAN | 2 Oxidation (M) | 0 | 0 | 0 | 32273 | 0 | 371850 | 0 | 0 | 0 | 532860 | 0 | 61489 | 0 | 0 | 0 | 0 | 0 | 0 |
| MMKDLEAI | PGM1; PGM1 | P36871; A0A | PGM1_HUMAN | 3 Oxidation (M) | 0 | 0 | 0 | 0 | 0 | 0 | 0 | 0 | 0 | 0 | 0 | 0 | 0 | 0 | 0 | 0 | 0 | 0 |
| MMLSGGAI | ACSL4; ACSL4 | O60488; H0 | ACSL4_HUMAN | Oxidation (M) | 0 | 0 | 0 | 0 | 0 | 0 | 0 | 0 | 0 | 0 | 502510 | 1990800 | 0 | 0 | 0 | 0 | 0 | 0 |
| MMLSGGAI | ACSL4; ACSL4 | O60488; H0 | ACSL4_HUMAN | 2 Oxidation (M) | 0 | 0 | 240270 | 0 | 0 | 0 | 0 | 0 | 0 | 0 | 0 | 2358000 | 0 | 0 | 0 | 160020 | 0 | 0 |
| MMLSGGAI | ACSL4; ACSL4 | O60488; H0 | ACSL4_HUMAN | Unmodified | 0 | 0 | 0 | 0 | 0 | 0 | 0 | 0 | 0 | 0 | 0 | 3787800 | 0 | 0 | 0 | 0 | 0 | 0 |
| MMMQSGF | SLC25A4; SLC2 | P12235; P12 | ADT1_HUMAN | Oxidation (M) | 0 | 0 | 0 | 0 | 0 | 0 | 0 | 0 | 1167900 | 0 | 0 | 0 | 0 | 0 | 0 | 0 | 0 | 0 |
| MMMQSGF | SLC25A4; SLC2 | P12235; P12 | ADT1_HUMAN | Unmodified | 0 | 0 | 0 | 0 | 0 | 0 | 23429 | 14282 | 6404000 | 747490 | 82706 | 0 | 0 | 0 | 47035 | 0 | 0 | 0 |
| MMNDQLM | FOLH1B; FOLH | Q9HBA9; Q0 | FOH1B_HUMAI | 3 Oxidation (M) | 1118600 | 1109200 | 11929000 | 4679000 | 0 | 0 | 0 | 0 | 0 | 0 | 0 | 0 | 0 | 0 | 6089100 | 6637000 | 0 | 0 |
| MMNDQLM | FOLH1B; FOLH | Q9HBA9; Q0 | FOH1B_HUMAI | Oxidation (M) | 0 | 0 | 1270700 | 0 | 0 | 0 | 0 | 0 | 0 | 0 | 0 | 0 | 0 | 0 | 879400 | 709450 | 0 | 0 |
| MMNGGHY | PARK7; PARK7 | Q99497; K7E | PARK7_HUMAN | Unmodified | 744950 | 276700 | 1171500 | 523110 | 8301900 | 710990 | 70574 | 0 | 3062200 | 709610 | 1301500 | 5912000 | 8410500 | 2966600 | 2911400 | 883160 | 768870 | 4838200 |
| MMNGGHY | PARK7; PARK7 | Q99497; K7E | PARK7_HUMAN | Oxidation (M) | 288730 | 0 | 576970 | 915800 | 2126800 | 173360 | 0 | 0 | 1370800 | 168170 | 636590 | 2437500 | 4765600 | 1448600 | 851000 | 335440 | 249550 | 0 |
| MMPTPVILI | CCT7; CCT7 | Q99832; F8 | TCPH_HUMAN | Acetyl (Protein N | 0 | 0 | 418860 | 0 | 1462800 | 0 | 896560 | 630690 | 106750 | 825890 | 2665800 | 6593900 | 3147100 | 0 | 0 | 414250 | 1132800 | 1836500 |

| | | | | | | | | | | | | | | | | | | | | | | |
|---|---|---|---|---|---|---|---|---|---|---|---|---|---|---|---|---|---|---|---|---|---|---|
| MMQTVRRI | PDHA1; PDHA1 | P08559; Q5J | ODPA_HUMAN | 3 Oxidation (M) | 677400 | 1595700 | 2137600 | 0 | 265360 | 0 | 0 | 0 | 0 | 820450 | 0 | 0 | 684370 | 0 | 1058500 | 725700 | 0 | 472080 |
| MMSETLNF | CPE | P16870 | CBPE_HUMAN | 2 Oxidation (M) | 0 | 0 | 0 | 0 | 0 | 0 | 0 | 84553 | 1084900 | 946340 | 0 | 0 | 0 | 0 | 0 | 0 | 0 | 0 |
| MMSETLNF | CPE | P16870 | CBPE_HUMAN | Oxidation (M) | 0 | 0 | 0 | 0 | 0 | 0 | 0 | 20437 | 1715900 | 1091600 | 0 | 0 | 0 | 0 | 0 | 0 | 0 | 0 |
| MMSLTSSK | HTRA1; HTRA1; | Q92743; A0A | HTRA1_HUMAN | Unmodified | 0 | 0 | 0 | 0 | 480700 | 29416 | 0 | 0 | 749650 | 0 | 0 | 0 | 0 | 0 | 0 | 0 | 371240 | 4326900 |
| MMSLTSSK | HTRA1; HTRA1; | Q92743; A0A | HTRA1_HUMAN | Oxidation (M) | 0 | 0 | 0 | 0 | 0 | 0 | 0 | 0 | 96864 | 43474 | 0 | 0 | 0 | 0 | 0 | 0 | 75667 | 488270 |
| MMTKEELE | MXRA7; MXRA7 | Q6ZR64; K7E | Q6ZR64_HUMA | Unmodified | 0 | 0 | 0 | 0 | 238570 | 75565 | 0 | 0 | 0 | 0 | 0 | 0 | 0 | 0 | 0 | 0 | 190730 | 537980 |
| MMVCQVG | CTNNB1; CTNN | P35222; A0A | CTNB1_HUMA | 2 Oxidation (M) | 0 | 0 | 0 | 364330 | 0 | 0 | 0 | 0 | 0 | 0 | 564650 | 0 | 0 | 0 | 0 | 956120 | 0 | 0 |
| MMVDKDG | CCT5; CCT5; C | P48643; B7Z | TCPE_HUMAN | 3 Oxidation (M) | 0 | 0 | 0 | 0 | 0 | 0 | 0 | 0 | 0 | 0 | 0 | 0 | 1660200 | 0 | 0 | 0 | 1427200 | 622660 |
| MMVGNNL | PPP3R1; PPP3R | P63098; F6U | CANB1_HUMA | Oxidation (M) | 0 | 0 | 0 | 0 | 784530 | 0 | 335420 | 0 | 0 | 0 | 0 | 0 | 823530 | 248940 | 0 | 0 | 570870 | 0 |
| MMVGNNL | PPP3R1; PPP3R | P63098; F6U | CANB1_HUMA | 2 Oxidation (M) | 0 | 0 | 0 | 19195 | 257030 | 0 | 0 | 0 | 0 | 0 | 0 | 0 | 913800 | 734470 | 0 | 0 | 260840 | 0 |
| MMVSSGD | SLC26A6 | G3XAC1 | G3XAC1_HUM | Unmodified | 0 | 0 | 0 | 0 | 1585800 | 0 | 0 | 0 | 0 | 0 | 0 | 514620 | 0 | 0 | 0 | 0 | 0 | 0 |
| MNAEPERK | BLVRA; BLVRA | P53004; C9J | BIEA_HUMAN B | Acetyl (Protein N | 0 | 0 | 0 | 0 | 0 | 0 | 99251 | 0 | 0 | 0 | 0 | 0 | 0 | 0 | 171360 | 63991 | 530680 | 273240 |
| MNAEPERK | BLVRA; BLVRA | P53004; C9J | BIEA_HUMAN B | Acetyl (Protein N | 0 | 0 | 0 | 0 | 0 | 0 | 0 | 0 | 0 | 0 | 0 | 0 | 0 | 0 | 0 | 0 | 282440 | 0 |
| MNAGSDP | ACAT2 | Q9BWD1 | THIC_HUMAN | Acetyl (Protein N | 0 | 0 | 0 | 0 | 0 | 0 | 0 | 0 | 0 | 814510 | 757060 | 2941900 | 0 | 0 | 0 | 0 | 0 | 0 |
| MNAKWDT | ITGB1; ITGB1; I | P05556; H7C | ITB1_HUMAN I | Unmodified | 0 | 0 | 0 | 0 | 4527100 | 1225600 | 0 | 0 | 0 | 0 | 0 | 0 | 0 | 0 | 0 | 0 | 609150 | 4749500 |
| MNAKWDT | ITGB1; ITGB1; I | P05556; H7C | ITB1_HUMAN I | Oxidation (M) | 0 | 0 | 0 | 0 | 2634500 | 0 | 0 | 0 | 0 | 0 | 0 | 0 | 0 | 0 | 0 | 0 | 0 | 1229500 |
| MNDFYIVD | ANO6; ANO6; A | Q4KMQ2; A0 | ANO6_HUMAN | Oxidation (M) | 0 | 0 | 87322 | 0 | 0 | 0 | 0 | 0 | 0 | 0 | 0 | 2249600 | 0 | 0 | 0 | 391790 | 0 | 0 |
| MNDPVESS | ANK3; ANK3; A | Q12955; A0A | ANK3_HUMAN | Oxidation (M) | 0 | 0 | 0 | 374230 | 0 | 0 | 0 | 0 | 0 | 0 | 21345 | 154260 | 188340 | 0 | 1312800 | 663440 | 126660 | 0 |
| MNDSNSA | RAB8B | Q92930 | RAB8B_HUMAN | Unmodified | 202440 | 104320 | 1507500 | 0 | 0 | 0 | 248440 | 50301 | 0 | 0 | 24093 | 0 | 0 | 0 | 1141400 | 219650 | 0 | 0 |
| MNDTVTIR | RPS24; RPS24; | P62847; A0A | RS24_HUMAN | Acetyl (Protein N | 0 | 720620 | 126970 | 170160 | 0 | 0 | 51596 | 0 | 0 | 40696 | 313650 | 18251000 | 0 | 0 | 161970 | 334740 | 116060 | 0 |
| MNDTVTIR | RPS24; RPS24; | P62847; A0A | RS24_HUMAN | Acetyl (Protein N | 0 | 263530 | 0 | 66803 | 262070 | 0 | 0 | 0 | 0 | 0 | 354570 | 3289200 | 475500 | 0 | 0 | 0 | 0 | 0 |
| MNDYKLEE | CNPY1; CNPY1 | Q3B7I2; A0A | CNPY1_HUMA | Unmodified | 0 | 0 | 0 | 0 | 22410 | 0 | 0 | 0 | 0 | 0 | 1605600 | 862580 | 0 | 0 | 0 | 0 | 0 | 0 |
| MNEFLENF | NIPSNAP3A | Q9UFN0 | NPS3A_HUMAN | Oxidation (M) | 0 | 0 | 0 | 0 | 0 | 0 | 0 | 0 | 2477900 | 258110 | 0 | 0 | 0 | 0 | 0 | 0 | 0 | 0 |
| MNEFLENF | NIPSNAP3A | Q9UFN0 | NPS3A_HUMAN | Unmodified | 0 | 0 | 0 | 0 | 0 | 0 | 0 | 0 | 2868000 | 0 | 0 | 0 | 0 | 0 | 0 | 0 | 0 | 0 |
| MNEIMDK | BCCIP | Q9P287 | BCCIP_HUMAN | Unmodified | 0 | 0 | 0 | 0 | 0 | 0 | 0 | 0 | 0 | 0 | 103800 | 515360 | 0 | 0 | 0 | 0 | 0 | 0 |
| MNELNHCI | CASK; CASK; C | O14936; A0A | CSKP_HUMAN | Unmodified | 0 | 0 | 0 | 0 | 1856000 | 320410 | 333050 | 46031 | 2119700 | 1293700 | 0 | 611820 | 1131600 | 780770 | 0 | 394440 | 673620 | 648710 |
| MNELNHCI | CASK; CASK; C | O14936; A0A | CSKP_HUMAN | Oxidation (M) | 0 | 0 | 0 | 0 | 692860 | 292370 | 0 | 22991 | 536980 | 512990 | 0 | 0 | 54861 | 1735400 | 0 | 0 | 334640 | 401600 |
| MNESDAW | SNX2 | O60749 | SNX2_HUMAN | Unmodified | 0 | 0 | 0 | 0 | 768860 | 0 | 178970 | 0 | 0 | 0 | 0 | 0 | 0 | 0 | 0 | 0 | 0 | 397290 |
| MNESLEPS | RAB3D | O95716 | RAB3D_HUMAN | Unmodified | 63440 | 45899 | 535160 | 0 | 0 | 0 | 0 | 0 | 0 | 0 | 0 | 0 | 0 | 0 | 610930 | 216490 | 0 | 0 |
| MNEVISLW | SPTAN1; SPTAN | Q13813; A0A | SPTN1_HUMAN | Oxidation (M) | 0 | 0 | 0 | 0 | 0 | 0 | 0 | 0 | 0 | 0 | 0 | 619070 | 0 | 0 | 0 | 237620 | 0 | 0 |
| MNGDQNS | FDPS; FDPS; FD | P14324; A0A | FPPS_HUMAN | Acetyl (Protein N | 0 | 0 | 0 | 0 | 0 | 0 | 0 | 0 | 0 | 0 | 0 | 884880 | 0 | 0 | 0 | 0 | 0 | 0 |
| MNGDQNS | FDPS; FDPS; FD | P14324; A0A | FPPS_HUMAN | Acetyl (Protein N | 0 | 0 | 0 | 0 | 0 | 0 | 0 | 0 | 0 | 0 | 0 | 2848500 | 0 | 0 | 0 | 0 | 0 | 0 |
| MNGFPNE | B4GALT3 | O60512 | B4GT3_HUMAN | Oxidation (M) | 0 | 0 | 0 | 0 | 0 | 0 | 0 | 0 | 1310200 | 0 | 0 | 0 | 0 | 0 | 0 | 0 | 0 | 0 |
| MNGFPNE | B4GALT3 | O60512 | B4GT3_HUMAN | Unmodified | 0 | 0 | 0 | 0 | 0 | 0 | 0 | 0 | 828160 | 0 | 0 | 0 | 0 | 0 | 0 | 0 | 0 | 0 |
| MNGMLLN | PABPC4; PABP | Q13310; P11 | PABP4_HUMAN | Oxidation (M) | 8391100 | 1638400 | 1192100 | 1597000 | 0 | 0 | 0 | 0 | 0 | 0 | 0 | 0 | 0 | 0 | 1304800 | 1869400 | 0 | 0 |
| MNGMLLN | PABPC4; PABP | Q13310; P11 | PABP4_HUMAN | Unmodified | 2195000 | 0 | 0 | 0 | 0 | 0 | 0 | 0 | 0 | 0 | 0 | 835940 | 0 | 0 | 0 | 0 | 0 | 0 |
| MNGMLLN | PABPC4; PABP | Q13310; P11 | PABP4_HUMAN | 2 Oxidation (M) | 144690 | 0 | 0 | 0 | 0 | 0 | 0 | 0 | 0 | 0 | 0 | 0 | 0 | 0 | 0 | 1274400 | 0 | 0 |
| MNGMLLN | PABPC4; PABP | Q13310; P11 | PABP4_HUMAN | Oxidation (M) | 7212100 | 4801000 | 3224000 | 2129100 | 196040 | 202430 | 85554 | 123170 | 0 | 171270 | 97974 | 529250 | 0 | 231400 | 1276600 | 733470 | 305600 | 0 |
| MNGMLLN | PABPC4; PABP | Q13310; P11 | PABP4_HUMAN | 2 Oxidation (M) | 5639700 | 4824300 | 2370200 | 2230500 | 0 | 0 | 0 | 0 | 0 | 0 | 0 | 435090 | 440970 | 0 | 168280 | 2186700 | 0 | 0 |
| MNGMLLN | PABPC4; PABP | Q13310; P11 | PABP4_HUMAN | Unmodified | 1239400 | 1733300 | 874380 | 0 | 0 | 0 | 0 | 0 | 0 | 0 | 0 | 714020 | 0 | 0 | 0 | 540810 | 62236 | 157250 |
| MNGPEDLI | TXNRD1; TXNR | Q16881; A0A | TRXR1_HUMAN | Acetyl (Protein N | 0 | 0 | 0 | 0 | 2675700 | 0 | 0 | 0 | 279170 | 0 | 465420 | 0 | 879120 | 0 | 0 | 0 | 719390 | 899540 |
| MNGPEDLI | TXNRD1; TXNR | Q16881; A0A | TRXR1_HUMAN | Acetyl (Protein N | 0 | 76583 | 0 | 0 | 4233900 | 614480 | 0 | 0 | 0 | 0 | 0 | 1226500 | 282220 | 161370 | 0 | 0 | 536390 | 1594300 |
| MNGTLDHI | CELF1; CELF1; | Q92879; G5I | CELF1_HUMAN | Acetyl (Protein N | 732370 | 0 | 433480 | 836790 | 0 | 0 | 36151 | 0 | 0 | 0 | 0 | 0 | 495220 | 0 | 290950 | 594670 | 0 | 0 |
| MNHQALVI | BPNT2 | Q9NX62 | IMPA3_HUMAN | Unmodified | 0 | 0 | 0 | 0 | 0 | 0 | 0 | 0 | 17743000 | 2460800 | 0 | 263410 | 0 | 0 | 0 | 0 | 0 | 0 |
| MNHSMSS | DLG2; DLG2; D | Q15700; E9F | DLG2_HUMAN | Acetyl (Protein N | 0 | 0 | 0 | 0 | 0 | 0 | 0 | 0 | 0 | 0 | 0 | 0 | 1554900 | 1266000 | 0 | 0 | 0 | 0 |
| MNHSPLK | PKP1 | Q13835 | PKP1_HUMAN | Acetyl (Protein N | 0 | 0 | 51847 | 0 | 0 | 0 | 0 | 0 | 256060 | 0 | 0 | 0 | 0 | 0 | 0 | 0 | 1296100 | 967010 |
| MNHTVQTI | IFITM3; N/A; IFI | Q01628; C9J | IFM3_HUMAN I | Acetyl (Protein N | 0 | 0 | 2308300 | 762240 | 0 | 645600 | 0 | 0 | 0 | 0 | 0 | 0 | 0 | 901170 | 2024000 | 1857500 | 0 | 0 |
| MNIADTAK | SUCO; SUCO | Q9UBS9; A0 | SUCO_HUMAN | Unmodified | 0 | 0 | 0 | 0 | 0 | 0 | 0 | 0 | 492210 | 0 | 0 | 0 | 0 | 0 | 0 | 0 | 0 | 0 |
| MNIFGGFR | SLC25A24 | Q6NUK1 | SCMC1_HUMA | Unmodified | 0 | 0 | 0 | 0 | 0 | 0 | 0 | 0 | 4706700 | 0 | 0 | 0 | 0 | 0 | 0 | 0 | 0 | 0 |
| MNIIRENK | DNAJC13; DNA | O75165; A0A | DJC13_HUMAN | Acetyl (Protein N | 0 | 0 | 0 | 0 | 0 | 0 | 0 | 0 | 0 | 0 | 0 | 749930 | 0 | 0 | 0 | 0 | 0 | 0 |
| MNIPFRIGI | PSAT1 | Q9Y617 | SERC_HUMAN | Unmodified | 69446 | 0 | 0 | 0 | 7186000 | 755100 | 0 | 0 | 0 | 821590 | 209300 | 0 | 5485500 | 5221100 | 0 | 0 | 1208400 | 1785000 |
| MNIPFRIGI | PSAT1 | Q9Y617 | SERC_HUMAN | Oxidation (M) | 0 | 0 | 0 | 0 | 958420 | 0 | 0 | 59198 | 0 | 0 | 0 | 0 | 1065700 | 792690 | 0 | 0 | 0 | 104230 |
| MNLASEPC | PDLIM1 | O00151 | PDLI1_HUMAN | Unmodified | 0 | 0 | 0 | 0 | 158470 | 0 | 0 | 0 | 0 | 0 | 0 | 0 | 0 | 0 | 0 | 0 | 280280 | 996910 |
| MNLASSFV | PSMD2 | Q13200 | PSMD2_HUMA | Oxidation (M) | 30828 | 62326 | 89995 | 234660 | 0 | 0 | 0 | 0 | 112580 | 0 | 0 | 165850 | 462840 | 106980 | 196220 | 52491 | 123990 | 113760 |
| MNLERVSN | PSENEN; N/A; F | Q9NZ42; K7I | PEN2_HUMAN | Acetyl (Protein N | 0 | 0 | 0 | 0 | 506700 | 66589 | 68130 | 0 | 0 | 0 | 0 | 0 | 0 | 0 | 0 | 0 | 0 | 0 |
| MNLERVSN | PSENEN; N/A; F | Q9NZ42; K7I | PEN2_HUMAN | Acetyl (Protein N | 0 | 0 | 0 | 0 | 664040 | 0 | 0 | 0 | 100790 | 119130 | 0 | 0 | 0 | 0 | 0 | 0 | 117880 | 196960 |
| MNLSAIQD | ARL8B; ARL8A | Q9NVJ2; Q9 | ARL8B_HUMAN | Unmodified | 0 | 0 | 0 | 0 | 0 | 0 | 0 | 0 | 0 | 0 | 0 | 1680100 | 0 | 0 | 0 | 0 | 0 | 0 |
| MNNSGAD | DAZAP1; DAZA | Q96EP5; K7E | DAZP1_HUMAN | Acetyl (Protein N | 0 | 0 | 19829 | 0 | 0 | 0 | 26990 | 0 | 0 | 0 | 0 | 1430500 | 33587 | 0 | 0 | 0 | 0 | 0 |
| MNNSGAD | DAZAP1; DAZA | Q96EP5; K7E | DAZP1_HUMAN | Acetyl (Protein N | 0 | 0 | 0 | 0 | 72515 | 0 | 0 | 0 | 200810 | 106650 | 0 | 1888500 | 0 | 91712 | 0 | 0 | 0 | 0 |
| MNPAAEAE | NAMPT; NAMPT | P43490; A0A | NAMPT_HUMA | Acetyl (Protein N | 0 | 0 | 0 | 0 | 388000 | 67277 | 0 | 0 | 127080 | 0 | 0 | 459790 | 0 | 0 | 0 | 0 | 0 | 186150 |
| MNPEYDYL | RAB1A; RAB1B; | P62820; Q9I | RAB1A_HUMAN | Acetyl (Protein N | 2238600 | 3300700 | 1440300 | 2447200 | 2968300 | 754550 | 2917500 | 1726200 | 0 | 2852800 | 3460500 | 16616000 | 3160200 | 3211200 | 3143300 | 2344100 | 1460900 | 2697000 |
| MNPEYDYL | RAB1A; RAB1B; | P62820; Q9I | RAB1A_HUMAN | Acetyl (Protein N | 0 | 0 | 0 | 0 | 940590 | 0 | 742780 | 0 | 1427100 | 358730 | 175370 | 512540 | 303650 | 0 | 0 | 0 | 0 | 0 |
| MNPIVVVH | ASRGL1; ASRGI | Q7L266; E9F | ASGL1_HUMAN | Acetyl (Protein N | 0 | 0 | 0 | 0 | 120540 | 0 | 61973 | 70225 | 254810 | 665140 | 232340 | 0 | 0 | 0 | 0 | 0 | 137110 | 98745 |
| MNPIVVVH | ASRGL1; ASRGI | Q7L266; E9F | ASGL1_HUMAN | Acetyl (Protein N | 0 | 0 | 0 | 0 | 0 | 0 | 0 | 0 | 851360 | 521560 | 0 | 304610 | 0 | 0 | 0 | 0 | 0 | 108140 |
| MNPLEQYE | APLP1; APLP1; | P51693; B7Z | APLP1_HUMAN | Unmodified | 0 | 0 | 0 | 0 | 0 | 0 | 0 | 0 | 4708700 | 146680 | 0 | 0 | 0 | 0 | 0 | 0 | 0 | 0 |
| MNPNCAR | LASP1; LASP1; | Q14847; K7E | LASP1_HUMAN | Acetyl (Protein N | 0 | 0 | 0 | 0 | 1316900 | 125270 | 298260 | 221510 | 0 | 70681 | 0 | 0 | 372880 | 704720 | 0 | 0 | 1793300 | 854560 |
| MNPNCAR | LASP1; LASP1; | Q14847; K7E | LASP1_HUMAN | Acetyl (Protein N | 0 | 0 | 0 | 0 | 0 | 0 | 329070 | 273490 | 0 | 0 | 0 | 0 | 0 | 154900 | 0 | 0 | 2471600 | 661720 |
| MNPQIRNF | APOBEC3C | Q9NRW3 | ABC3C_HUMA | Acetyl (Protein N | 0 | 0 | 0 | 0 | 0 | 0 | 0 | 0 | 0 | 0 | 0 | 0 | 0 | 0 | 0 | 0 | 450120 | 414170 |
| MNPQIRNF | APOBEC3C | Q9NRW3 | ABC3C_HUMA | Acetyl (Protein N | 0 | 0 | 100960 | 0 | 0 | 0 | 0 | 0 | 0 | 0 | 0 | 0 | 180710 | 0 | 0 | 0 | 0 | 1173100 |
| MNPQSAFF | SCP2 | P22307 | SCP2_HUMAN | Unmodified | 0 | 0 | 0 | 0 | 0 | 0 | 681330 | 147500 | 70950000 | 15509000 | 1025400 | 0 | 396530 | 376650 | 0 | 0 | 129620 | 394380 |
| MNPQSAFF | SCP2 | P22307 | SCP2_HUMAN | Oxidation (M) | 0 | 0 | 0 | 0 | 0 | 0 | 232410 | 0 | 0 | 3876900 | 573070 | 0 | 125660 | 0 | 0 | 0 | 0 | 0 |
| MNPSAPSY | PABPC1; PABP | P11940; A0A | PABP1_HUMAN | Acetyl (Protein N | 5132200 | 7917600 | 0 | 3591600 | 0 | 0 | 489510 | 0 | 0 | 0 | 0 | 0 | 0 | 0 | 2716900 | 0 | 0 | 0 |
| MNPTNTVF | HSPA8; HSPA8; | P11142; E9P | HSP7C_HUMA | Unmodified | 0 | 0 | 0 | 0 | 2874700 | 0 | 580190 | 108140 | 0 | 151270 | 0 | 1476600 | 916520 | 0 | 0 | 0 | 1774800 | 1140400 |
| MNPTNTVF | HSPA8; HSPA8; | P11142; E9P | HSP7C_HUMA | Oxidation (M) | 0 | 0 | 604940 | 0 | 749620 | 0 | 0 | 115600 | 222050 | 0 | 0 | 994850 | 0 | 0 | 792170 | 478760 | 637200 | 1013100 |
| MNPVYSPV | FAM168A | Q92567 | F168A_HUMAN | Acetyl (Protein N | 83542 | 76832 | 56915 | 92118 | 1484200 | 0 | 504080 | 278310 | 426160 | 797830 | 0 | 0 | 433870 | 222500 | 0 | 140050 | 0 | 2941400 |
| MNQAPTV | ST6GALNAC4; | Q9H4F1; A6I | SIA7D_HUMAN | Unmodified | 0 | 0 | 0 | 0 | 0 | 0 | 0 | 0 | 1837400 | 0 | 0 | 0 | 0 | 0 | 0 | 0 | 0 | 0 |
| MNRLFGK | CHMP5 | Q9NZZ3 | CHMP5_HUMA | Acetyl (Protein N | 0 | 0 | 0 | 280480 | 16668000 | 3629300 | 815040 | 402290 | 1745300 | 6065700 | 595550 | 0 | 5580700 | 9858000 | 0 | 0 | 3261000 | 3191700 |
| MNRLFGK | CHMP5 | Q9NZZ3 | CHMP5_HUMA | Acetyl (Protein N | 0 | 0 | 0 | 96634 | 5407600 | 818000 | 281380 | 0 | 69400 | 2085700 | 0 | 0 | 1528700 | 1782400 | 0 | 0 | 606850 | 696160 |
| MNRPAPVE | PTP4A2; PTP4A | Q12974; E9F | TP4A2_HUMAN | Acetyl (Protein N | 0 | 3667600 | 0 | 0 | 0 | 0 | 0 | 0 | 0 | 0 | 0 | 707360 | 12312000 | 0 | 0 | 0 | 0 | 0 |
| MNRTKGDI | VIPAS39; VIPAS | Q9H9C1; G3 | SPE39_HUMAN | Acetyl (Protein N | 0 | 0 | 0 | 0 | 2134000 | 0 | 74914 | 0 | 0 | 0 | 0 | 0 | 0 | 117390 | 0 | 0 | 136420 | 118530 |
| MNSDQDV | USP6NL | Q92738 | US6NL_HUMA | Acetyl (Protein N | 0 | 0 | 0 | 0 | 0 | 0 | 0 | 0 | 126860 | 0 | 0 | 0 | 0 | 0 | 0 | 0 | 0 | 494100 |
| MNSLPAER | NMT1; NMT1; N | P30419; A0A | NMT1_HUMAN | Unmodified | 0 | 0 | 0 | 0 | 1912800 | 0 | 217280 | 0 | 0 | 0 | 0 | 0 | 174510 | 0 | 0 | 0 | 0 | 580360 |
| MNTVLSR | SPCS3 | P61009 | SPCS3_HUMAN | Unmodified | 24885 | 0 | 33918 | 0 | 0 | 0 | 0 | 0 | 3940900 | 281740 | 394870 | 901320 | 0 | 0 | 0 | 0 | 0 | 0 |
| MNVDHEVI | ABRACL; ABRA | Q9P1F3; Q5 | ABRAL_HUMAN | Acetyl (Protein N | 2576100 | 1895400 | 0 | 1317700 | 0 | 0 | 0 | 0 | 2398400 | 415760 | 0 | 0 | 0 | 0 | 775880 | 0 | 0 | 0 |
| MNVDHEVI | ABRACL; ABRA | Q9P1F3; Q5 | ABRAL_HUMAN | Acetyl (Protein N | 490100 | 1830800 | 54065 | 306230 | 2042700 | 882620 | 829150 | 0 | 545740 | 3104000 | 0 | 0 | 1597200 | 985600 | 312000 | 0 | 1532100 | 1140800 |
| MNVDHEVI | ABRACL; ABRA | Q9P1F3; Q5 | ABRAL_HUMAN | Acetyl (Protein N | 0 | 0 | 0 | 0 | 486530 | 0 | 80517 | 0 | 0 | 752050 | 0 | 0 | 210110 | 93058 | 0 | 0 | 184570 | 0 |
| MNVDQAFI | RRAS2; RRAS2 | P62070; E9P | RRAS2_HUMAN | Unmodified | 1007400 | 1785800 | 1418300 | 2281500 | 0 | 0 | 0 | 0 | 1646300 | 207190 | 58844 | 0 | 289700 | 0 | 2733900 | 1480300 | 0 | 0 |
| MNVDQAFI | RRAS2; RRAS2 | P62070; E9P | RRAS2_HUMAN | Oxidation (M) | 1755100 | 1491200 | 0 | 1376800 | 0 | 0 | 0 | 0 | 0 | 0 | 0 | 0 | 0 | 0 | 2749100 | 2406100 | 0 | 0 |
| MNVFDTEL | ATP2A2; ATP2A | P16615; H7C | AT2A2_HUMAN | Oxidation (M) | 0 | 0 | 0 | 0 | 354650 | 0 | 805790 | 732480 | 0 | 92289 | 0 | 515130 | 0 | 0 | 0 | 0 | 0 | 154640 |
| MNVFDTEL | ATP2A2; ATP2A | P16615; H7C | AT2A2_HUMAN | Unmodified | 0 | 0 | 0 | 0 | 770540 | 0 | 1746500 | 380740 | 592060 | 0 | 0 | 975290 | 278000 | 0 | 0 | 0 | 0 | 192870 |
| MNVLADAL | RPS15A; N/A; R | P62244; H3E | RS15A_HUMAN | Unmodified | 0 | 601790 | 0 | 795220 | 101670 | 0 | 68973 | 13582 | 698310 | 72932 | 670040 | 13514000 | 0 | 0 | 715100 | 0 | 0 | 0 |
| MNVLADAL | RPS15A; N/A; R | P62244; H3E | RS15A_HUMAN | Oxidation (M) | 651550 | 607460 | 456560 | 224380 | 38098 | 119020 | 102190 | 26679 | 0 | 34601 | 485620 | 11090000 | 61449 | 0 | 387160 | 635710 | 0 | 0 |
| MNVSPDVI | PSMC3; PSMC3 | P17980; E9P | PRS6A_HUMAN | Unmodified | 0 | 0 | 0 | 0 | 0 | 0 | 0 | 0 | 0 | 0 | 0 | 0 | 0 | 0 | 0 | 0 | 548640 | 651190 |
| MNYDSYLA | PROM1 | O43490 | PROM1_HUMA | Unmodified | 0 | 0 | 0 | 0 | 0 | 0 | 0 | 0 | 191500 | 0 | 440960 | 2041400 | 0 | 0 | 0 | 0 | 0 | 0 |
| MNYDSYLA | PROM1 | O43490 | PROM1_HUMA | Oxidation (M) | 0 | 0 | 0 | 0 | 0 | 0 | 0 | 0 | 192910 | 98655 | 0 | 849310 | 0 | 0 | 0 | 0 | 0 | 0 |
| MPCAELVR | IGFBP2 | P18065 | IBP2_HUMAN I | Unmodified | 0 | 0 | 0 | 0 | 0 | 0 | 0 | 0 | 5224200 | 132590 | 0 | 0 | 0 | 0 | 0 | 0 | 0 | 0 |
| MPDSGPLF | VGF; VGF | O15240; Q0 | VGF_HUMAN N | Unmodified | 0 | 0 | 0 | 0 | 0 | 154590 | 150780 | 83022 | 4435100 | 435770 | 0 | 0 | 0 | 0 | 161960 | 0 | 159290 | 89078 |
| MPDSGPLF | VGF; VGF | O15240; Q0 | VGF_HUMAN N | Unmodified | 0 | 1154300 | 0 | 297920 | 0 | 0 | 143350 | 130400 | 2882300 | 4658100 | 0 | 2191900 | 0 | 351500 | 0 | 0 | 0 | 0 |
| MPDSGPLF | VGF; VGF | O15240; Q0 | VGF_HUMAN N | Oxidation (M) | 94299 | 0 | 0 | 0 | 0 | 0 | 0 | 67342 | 2598600 | 1587600 | 0 | 0 | 0 | 0 | 0 | 0 | 0 | 0 |
| MPDSTLPG | PTGFRN | Q9P2B2 | FPRP_HUMAN | Unmodified | 5116600 | 3207400 | 2410600 | 12003000 | 192600 | 0 | 0 | 0 | 138370 | 0 | 0 | 16765000 | 1139100 | 493400 | 17950000 | 17215000 | 0 | 0 |
| MPDSTLPG | PTGFRN | Q9P2B2 | FPRP_HUMAN | Oxidation (M) | 650680 | 429080 | 2693800 | 1442700 | 0 | 0 | 0 | 0 | 0 | 0 | 0 | 1242500 | 176150 | 0 | 1642300 | 2714000 | 0 | 0 |
| MPDYLGAI | PSMC2; PSMC2 | P35998; C9J | PRS7_HUMAN | Unmodified | 0 | 0 | 0 | 0 | 0 | 2150100 | 0 | 0 | 0 | 0 | 0 | 0 | 0 | 0 | 0 | 0 | 0 | 0 |
| MPEDESTP | NCALD; HPCA; | P61601; P84 | NCALD_HUMA | Unmodified | 59643 | 87344 | 134440 | 108130 | 93694 | 0 | 0 | 0 | 384610 | 60484 | 289610 | 1184900 | 0 | 0 | 108780 | 140730 | 0 | 108660 |
| MPEDGLSI | CALD1; CALD1; | Q05682; C9J | CALD1_HUMAN | Unmodified | 0 | 0 | 89442 | 0 | 229470 | 0 | 0 | 0 | 0 | 0 | 388120 | 2818100 | 2627700 | 0 | 140140 | 0 | 86910 | 0 |
| MPEEMDKI | STEAP3; STEAP | Q658P3; B8Z | STEA3_HUMAN | Oxidation (M) | 1424600 | 11687000 | 0 | 0 | 0 | 0 | 0 | 0 | 0 | 0 | 0 | 0 | 1031200 | 720460 | 0 | 439120 | 0 | 0 |
| MPEKEPPP | LAPTM4A | Q15012 | LAP4A_HUMAN | Unmodified | 0 | 0 | 0 | 0 | 0 | 0 | 0 | 0 | 790090 | 116460 | 0 | 0 | 0 | 0 | 0 | 0 | 0 | 0 |
| MPEMHFK | AHNAK | Q09666 | AHNK_HUMAN | Unmodified | 0 | 63284 | 0 | 0 | 794660 | 0 | 0 | 0 | 0 | 29407 | 0 | 0 | 0 | 0 | 0 | 0 | 201690 | 273300 |
| MPEMNIK | AHNAK | Q09666 | AHNK_HUMAN | Unmodified | 0 | 0 | 0 | 0 | 2188500 | 0 | 0 | 0 | 0 | 0 | 0 | 0 | 0 | 0 | 0 | 0 | 633830 | 1462600 |
| MPEQEYEF | ANTXR1 | Q9H6X2 | ANTR1_HUMAN | Unmodified | 0 | 0 | 0 | 0 | 1079000 | 0 | 0 | 0 | 1048400 | 0 | 0 | 0 | 0 | 0 | 0 | 0 | 0 | 0 |

| | | | | | | | | | | | | | | | | | | | | | | |
|---|---|---|---|---|---|---|---|---|---|---|---|---|---|---|---|---|---|---|---|---|---|---|
| MPETGPVL | PTPRG | P23470 | PTPRG_HUMAN | Unmodified | 53609 | 0 | 0 | 103940 | 65633 | 0 | 139140 | 0 | 223100 | 142440 | 0 | 1274300 | 0 | 0 | 357550 | 163260 | 234330 | 0 |
| MPILGLGT | AKR1B1; AKR1E | P15121; E9P | ALDR_HUMAN | Oxidation (M) | 0 | 0 | 0 | 0 | 11788000 | 1695400 | 0 | 19305 | 0 | 0 | 722920 | 429140 | 0 | 0 | 0 | 0 | 0 | 0 |
| MPILGLGT | AKR1B1; AKR1E | P15121; E9P | ALDR_HUMAN | Unmodified | 0 | 0 | 0 | 0 | 17369000 | 1379000 | 0 | 0 | 0 | 0 | 280410 | 106930 | 0 | 0 | 0 | 0 | 0 | 0 |
| MPIVGLGT | AKR1B10 | O60218 | AK1BA_HUMAN | Oxidation (M) | 0 | 0 | 0 | 0 | 0 | 0 | 1053000 | 102290 | 0 | 0 | 0 | 0 | 334200 | 0 | 0 | 0 | 0 | 0 |
| MPIVGLGT | AKR1B10 | O60218 | AK1BA_HUMAN | Unmodified | 0 | 0 | 0 | 0 | 0 | 0 | 319030 | 0 | 0 | 0 | 0 | 0 | 0 | 0 | 0 | 0 | 73530 | 0 |
| MPLIGLGT | AKR1A1; AKR1A | P14550; V9G | AK1A1_HUMAN | Unmodified | 0 | 0 | 0 | 0 | 1680200 | 332900 | 0 | 0 | 2041900 | 0 | 1053300 | 6939300 | 0 | 0 | 538010 | 0 | 0 | 194360 |
| MPLIGLGT | AKR1A1; AKR1A | P14550; V9G | AK1A1_HUMAN | Unmodified | 0 | 0 | 0 | 105390 | 1702300 | 0 | 287210 | 285920 | 1002500 | 831080 | 0 | 0 | 1459100 | 0 | 0 | 0 | 679370 | 1340800 |
| MPLIGLGT | AKR1A1; AKR1A | P14550; V9G | AK1A1_HUMAN | Oxidation (M) | 0 | 0 | 0 | 0 | 1086100 | 0 | 0 | 0 | 0 | 449500 | 0 | 374430 | 723180 | 449850 | 0 | 0 | 0 | 0 |
| MPMIWLDL | PTPN23; PTPN2 | Q9H3S7; R4 | PTN23_HUMAN | Oxidation (M) | 42702 | 71849 | 0 | 115030 | 0 | 0 | 0 | 0 | 281650 | 0 | 171720 | 0 | 0 | 0 | 0 | 74366 | 0 | 0 |
| MPNVPNT | SCAMP1; SCAM | O15126; A0 | SCAM1_HUMA | Unmodified | 0 | 0 | 759590 | 0 | 1309700 | 0 | 0 | 0 | 0 | 0 | 0 | 0 | 381360 | 0 | 471670 | 308200 | 150280 | 0 |
| MPNVPNT | SCAMP1; SCAM | O15126; A0 | SCAM1_HUMA | Oxidation (M) | 466650 | 304370 | 1260000 | 1589700 | 1554300 | 0 | 196370 | 0 | 0 | 0 | 329200 | 1653500 | 0 | 0 | 1019800 | 1151900 | 0 | 0 |
| MPNVPNT | SCAMP1; SCAM | O15126; A0 | SCAM1_HUMA | 2 Oxidation (M) | 0 | 0 | 1315300 | 0 | 0 | 0 | 0 | 0 | 0 | 0 | 0 | 0 | 0 | 0 | 0 | 0 | 0 | 0 |
| MPPAPSGF | BAIAP2L1 | Q9UHR4 | BI2L1_HUMAN | Unmodified | 0 | 0 | 108270 | 0 | 0 | 0 | 0 | 0 | 0 | 0 | 0 | 0 | 0 | 464580 | 120610 | 0 | 0 | 0 |
| MPPVLEER | MRPS22; MRPS | P82650; A0A | RT22_HUMAN | Oxidation (M) | 0 | 0 | 0 | 0 | 0 | 0 | 0 | 0 | 491220 | 59297 | 0 | 0 | 0 | 0 | 0 | 0 | 0 | 0 |
| MPPVLEER | MRPS22; MRPS | P82650; A0A | RT22_HUMAN | Unmodified | 0 | 0 | 0 | 0 | 16129 | 0 | 773410 | 0 | 1234100 | 0 | 0 | 0 | 0 | 587050 | 0 | 0 | 0 | 0 |
| MPPYDEQT | PRKCSH; PRKC | P14314; K7E | GLU2B_HUMA | Unmodified | 0 | 0 | 0 | 0 | 0 | 0 | 0 | 0 | 3627100 | 94957 | 476100 | 0 | 0 | 0 | 397480 | 0 | 0 | 1624400 |
| MPPYDEQT | PRKCSH; PRKC | P14314; K7E | GLU2B_HUMA | Oxidation (M) | 0 | 0 | 0 | 0 | 0 | 0 | 0 | 0 | 1331100 | 0 | 0 | 0 | 0 | 0 | 0 | 0 | 0 | 0 |
| MPPYTVVY | GSTP1; GSTP1; | P09211; A8 | GSTP1_HUMAN | Oxidation (M) | 0 | 0 | 1087200 | 0 | 0 | 0 | 0 | 0 | 0 | 0 | 0 | 1642200 | 0 | 0 | 0 | 0 | 0 | 0 |
| MPQSMPE | LONP2; LONP2 | Q86WA8; H3 | LONP2_HUMA | Oxidation (M) | 0 | 0 | 0 | 0 | 0 | 0 | 0 | 0 | 0 | 572550 | 0 | 0 | 0 | 0 | 0 | 0 | 0 | 0 |
| MPSFGVSA | AHNAK2 | Q8IVF2 | AHNK2_HUMA | Unmodified | 0 | 0 | 0 | 0 | 4902700 | 512480 | 0 | 0 | 0 | 439530 | 0 | 109650 | 0 | 0 | 0 | 0 | 572380 | 3307400 |
| MPSFGVSA | AHNAK2 | Q8IVF2 | AHNK2_HUMA | Oxidation (M) | 0 | 0 | 0 | 0 | 124750 | 0 | 223440 | 76899 | 0 | 0 | 0 | 0 | 0 | 0 | 0 | 0 | 0 | 218760 |
| MPSGEFAR | PCNA | P12004 | PCNA_HUMAN | Unmodified | 0 | 0 | 0 | 0 | 2016100 | 384550 | 53630 | 15422 | 2751500 | 1135700 | 1506200 | 5490600 | 3926100 | 38474 | 0 | 0 | 306560 | 991710 |
| MPSGEFAR | PCNA | P12004 | PCNA_HUMAN | Oxidation (M) | 0 | 0 | 0 | 51521 | 30410 | 43580 | 23522 | 0 | 525710 | 10904 | 17742 | 525320 | 0 | 38702 | 65553 | 0 | 218520 | 70991 |
| MPSHQGA | SEC31A; SEC31 | O94979; H7 | SC31A_HUMAN | Unmodified | 0 | 0 | 0 | 0 | 0 | 0 | 0 | 0 | 759260 | 0 | 248050 | 2203200 | 0 | 0 | 0 | 0 | 0 | 93759 |
| MPSLPSYK | TKT; TKT; TKT | P29401; A0A | TKT_HUMAN Tr | Unmodified | 83694 | 49283 | 0 | 77425 | 769510 | 179890 | 0 | 0 | 2470800 | 295300 | 3200400 | 7118900 | 97950 | 0 | 0 | 180470 | 88014 | 202740 |
| MPSLPSYK | TKT; TKT; TKT | P29401; A0A | TKT_HUMAN Tr | Oxidation (M) | 0 | 0 | 0 | 0 | 174690 | 0 | 0 | 0 | 144110 | 27759 | 377570 | 2121100 | 0 | 0 | 39676 | 90361 | 0 | 0 |
| MPSLPSYK | TKT; TKT; TKT | P29401; A0A | TKT_HUMAN Tr | Unmodified | 0 | 123850 | 0 | 0 | 2938900 | 1231600 | 0 | 0 | 1479400 | 675950 | 439380 | 2207500 | 466430 | 0 | 0 | 0 | 0 | 776280 |
| MPSLSPID | MYO1B; MYO1E | O43795; E9F | MYO1B_HUMA | Unmodified | 0 | 0 | 0 | 0 | 0 | 0 | 0 | 0 | 34914 | 0 | 127480 | 6297200 | 0 | 0 | 0 | 0 | 0 | 0 |
| MPYQEIK | DIAPH1; DIAPH | O60610; A0 | DIAP1_HUMAN | Unmodified | 0 | 0 | 0 | 0 | 0 | 0 | 0 | 0 | 0 | 0 | 0 | 1362900 | 0 | 0 | 0 | 82869 | 0 | 0 |
| MQASIEK | NPM1; NPM1; N | P06748; A0A | NPM_HUMAN N | Unmodified | 0 | 0 | 199260 | 0 | 346530 | 0 | 43764 | 0 | 0 | 0 | 0 | 2559900 | 2435200 | 0 | 0 | 0 | 23186 | 0 |
| MQAVQEA | PLEC | Q15149 | PLEC_HUMAN | Oxidation (M) | 0 | 0 | 0 | 0 | 0 | 0 | 0 | 0 | 242080 | 0 | 0 | 0 | 88644 | 0 | 0 | 0 | 1328900 | 609870 |
| MQAVQEA | PLEC | Q15149 | PLEC_HUMAN | Unmodified | 0 | 0 | 0 | 0 | 0 | 78934 | 0 | 0 | 0 | 0 | 0 | 0 | 0 | 0 | 0 | 0 | 653280 | 442230 |
| MQAYSNP | RFFL; RFFL; RFF | Q8WZ73; C9 | RFFL_HUMAN | Oxidation (M) | 0 | 0 | 0 | 65166 | 357140 | 0 | 0 | 0 | 0 | 0 | 0 | 0 | 132810 | 93239 | 0 | 0 | 86361 | 337360 |
| MQCFSVTE | GAS6 | Q14393 | GAS6_HUMAN | Unmodified | 0 | 0 | 0 | 0 | 0 | 0 | 0 | 0 | 2198800 | 0 | 0 | 0 | 0 | 0 | 0 | 0 | 0 | 0 |
| MQCFSVTE | GAS6 | Q14393 | GAS6_HUMAN | Oxidation (M) | 0 | 0 | 0 | 0 | 0 | 0 | 0 | 0 | 727480 | 0 | 0 | 0 | 0 | 0 | 0 | 0 | 0 | 0 |
| MQDQLQA | EHD3 | Q9NZN3 | EHD3_HUMAN | Oxidation (M) | 0 | 0 | 0 | 0 | 0 | 0 | 0 | 353660 | 0 | 0 | 0 | 0 | 0 | 0 | 193390 | 0 | 0 | 0 |
| MQEDLVTL | CACNA2D1; CA | P54289; E7E | CA2D1_HUMA | Oxidation (M) | 568850 | 0 | 0 | 0 | 0 | 0 | 0 | 158350 | 856290 | 0 | 0 | 0 | 0 | 0 | 0 | 0 | 0 | 0 |
| MQEDLVTL | CACNA2D1; CA | P54289; E7E | CA2D1_HUMA | Unmodified | 532890 | 529160 | 144890 | 0 | 0 | 0 | 0 | 0 | 0 | 0 | 0 | 0 | 0 | 0 | 670070 | 63768 | 0 | 0 |
| MQEEEARK | SCRIB; SCRIB; | Q14160; A0 | SCRIB_HUMAN | Oxidation (M) | 0 | 298490 | 0 | 0 | 225560 | 0 | 0 | 0 | 0 | 171180 | 0 | 0 | 124400 | 88447 | 0 | 0 | 499850 | 263470 |
| MQELALSE | DNAJC13; DNA | O75165; A0 | DJC13_HUMAN | Oxidation (M) | 0 | 0 | 0 | 0 | 0 | 0 | 0 | 0 | 0 | 211110 | 0 | 864500 | 0 | 0 | 0 | 0 | 226350 | 0 |
| MQELLMAH | EHD2 | Q9NZN4 | EHD2_HUMAN | Oxidation (M) | 0 | 0 | 0 | 41526 | 589130 | 189960 | 0 | 0 | 0 | 0 | 0 | 0 | 0 | 0 | 0 | 0 | 0 | 0 |
| MQELLQTC | EHD1; EHD1; E | Q9H4M9; C9 | EHD1_HUMAN | Oxidation (M) | 762680 | 661610 | 970360 | 1030100 | 8590700 | 1651400 | 575960 | 491030 | 0 | 368870 | 3225000 | 4053100 | 3236600 | 1208500 | 1259100 | 1399100 | 3076500 | 7847800 |
| MQELLQTC | EHD1; EHD1; E | Q9H4M9; C9 | EHD1_HUMAN | Unmodified | 446060 | 289040 | 0 | 0 | 11443000 | 381950 | 0 | 0 | 229340 | 0 | 0 | 5286800 | 13332000 | 0 | 0 | 451540 | 6933900 | 12194000 |
| MQELLQTC | EHD1; EHD1; E | Q9H4M9; C9 | EHD1_HUMAN | Oxidation (M) | 0 | 0 | 0 | 0 | 1575500 | 0 | 637000 | 0 | 0 | 0 | 0 | 0 | 933240 | 1159100 | 0 | 0 | 6034600 | 2347800 |
| MQELLQTC | EHD1; EHD1; E | Q9H4M9; C9 | EHD1_HUMAN | Unmodified | 0 | 0 | 0 | 0 | 0 | 0 | 0 | 0 | 0 | 0 | 0 | 0 | 0 | 0 | 0 | 0 | 3043700 | 2631200 |
| MQELQEAC | TJP2; TJP2; TJP2 | Q9UDY2; A0 | ZO2_HUMAN T | Unmodified | 0 | 0 | 0 | 0 | 0 | 0 | 0 | 0 | 0 | 0 | 0 | 0 | 0 | 0 | 0 | 0 | 0 | 0 |
| MQELQEAC | TJP2; TJP2; TJP2 | Q9UDY2; A0 | ZO2_HUMAN T | Oxidation (M) | 0 | 0 | 0 | 0 | 202000 | 0 | 0 | 0 | 552080 | 75531 | 0 | 0 | 0 | 0 | 0 | 0 | 0 | 0 |
| MQELQEAC | TJP2; TJP2; TJP2 | Q9UDY2; A0 | ZO2_HUMAN T | Unmodified | 0 | 0 | 0 | 0 | 413690 | 0 | 0 | 0 | 1490900 | 172280 | 0 | 0 | 0 | 0 | 0 | 0 | 0 | 0 |
| MQELQEAC | TJP2; TJP2; TJP2 | Q9UDY2; A0 | ZO2_HUMAN T | Oxidation (M) | 0 | 0 | 0 | 0 | 2680700 | 0 | 0 | 0 | 0 | 0 | 0 | 0 | 0 | 0 | 0 | 0 | 0 | 0 |
| MQEMIAR | SEPTIN2; SEPTI | Q15019; H7 | SEPT2_HUMAN | Unmodified | 0 | 0 | 0 | 12017 | 111590 | 76249 | 0 | 0 | 222670 | 0 | 78812 | 547960 | 72223 | 0 | 0 | 0 | 0 | 44190 |
| MQENLILD | N/A | N/A | N/A | Unmodified | 0 | 0 | 0 | 0 | 2099600 | 0 | 0 | 0 | 0 | 108600 | 0 | 0 | 0 | 0 | 0 | 0 | 864920 | 4930300 |
| MQEQKNE | ERGIC3; ERGIC | Q9Y282; A0 | ERGI3_HUMAN | Oxidation (M) | 0 | 0 | 0 | 0 | 0 | 0 | 0 | 0 | 2557300 | 0 | 410060 | 0 | 0 | 0 | 0 | 0 | 0 | 0 |
| MQEQKNE | ERGIC3; ERGIC | Q9Y282; A0 | ERGI3_HUMAN | Unmodified | 0 | 0 | 0 | 0 | 935520 | 0 | 0 | 0 | 3675600 | 514320 | 920000 | 0 | 0 | 0 | 0 | 0 | 0 | 0 |
| MQEQQAD | DCTN1; DCTN1 | Q14203; Q6 | DCTN1_HUMA | Unmodified | 0 | 0 | 0 | 0 | 0 | 0 | 0 | 0 | 0 | 73564 | 0 | 724380 | 0 | 0 | 0 | 0 | 111210 | 0 |
| MQEVVANI | DDB1; DDB1; D | Q16531; A0 | DDB1_HUMAN | Unmodified | 0 | 0 | 0 | 0 | 792720 | 0 | 0 | 0 | 617430 | 0 | 0 | 1304600 | 0 | 0 | 0 | 0 | 0 | 0 |
| MQEVVANI | DDB1; DDB1; D | Q16531; A0 | DDB1_HUMAN | 2 Oxidation (M) | 0 | 0 | 0 | 0 | 0 | 0 | 0 | 0 | 0 | 0 | 0 | 953250 | 0 | 0 | 0 | 0 | 0 | 0 |
| MQEVVANI | DDB1; DDB1; D | Q16531; A0 | DDB1_HUMAN | Oxidation (M) | 0 | 0 | 107390 | 0 | 237890 | 0 | 0 | 0 | 447390 | 0 | 258640 | 1339200 | 0 | 0 | 0 | 0 | 0 | 0 |
| MQEVVANI | DDB1; DDB1; D | Q16531; A0 | DDB1_HUMAN | 2 Oxidation (M) | 0 | 0 | 0 | 73411 | 603420 | 112780 | 0 | 0 | 0 | 252070 | 0 | 0 | 419500 | 137540 | 0 | 0 | 331600 | 458100 |
| MQEVYNFN | LTA4H | P09960 | LKHA4_HUMAN | Oxidation (M) | 0 | 0 | 0 | 0 | 0 | 0 | 132840 | 41248 | 0 | 0 | 0 | 1484900 | 60370 | 149330 | 0 | 0 | 0 | 130470 |
| MQGQSPP | CKAP5 | Q14008 | CKAP5_HUMAN | Unmodified | 0 | 0 | 0 | 0 | 0 | 0 | 0 | 0 | 0 | 29657 | 100990 | 799950 | 0 | 0 | 0 | 0 | 0 | 0 |
| MQGQSPP | CKAP5 | Q14008 | CKAP5_HUMAN | Unmodified | 162410 | 0 | 0 | 0 | 0 | 0 | 0 | 0 | 84104 | 285880 | 0 | 0 | 369160 | 0 | 0 | 0 | 0 | 0 |
| MQGSLEAH | SUPT16H | Q9Y5B9 | SP16H_HUMAN | Unmodified | 0 | 0 | 0 | 0 | 68528 | 0 | 69941 | 8499.1 | 0 | 0 | 0 | 774120 | 24698 | 0 | 0 | 0 | 0 | 0 |
| MQHLIAR | PKM; PKM; PKM | P14618; B4D | KPYM_HUMAN | Unmodified | 102120 | 113180 | 927360 | 290570 | 0 | 59284 | 0 | 0 | 2021600 | 88795 | 1138400 | 32350000 | 66220 | 0 | 544080 | 0 | 0 | 0 |
| MQHLIAR | PKM; PKM; PKM | P14618; B4D | KPYM_HUMAN | Oxidation (M) | 77421 | 0 | 115050 | 139000 | 0 | 0 | 0 | 0 | 987310 | 0 | 738310 | 16418000 | 0 | 0 | 286260 | 335510 | 86320 | 0 |
| MQHLIARE | PKM; PKM | P14618; B4D | KPYM_HUMAN | Oxidation (M) | 0 | 0 | 0 | 278670 | 8457900 | 0 | 573080 | 0 | 0 | 1102000 | 1119500 | 0 | 870460 | 340480 | 0 | 0 | 690520 | 1278300 |
| MQHLNPD | FERMT2; FERM | Q96AC1; H0 | FERM2_HUMA | Oxidation (M) | 0 | 0 | 0 | 415360 | 5608800 | 0 | 0 | 0 | 0 | 0 | 0 | 0 | 0 | 0 | 0 | 0 | 0 | 2309400 |
| MQHNLEQ | SPTAN1; SPTAN | Q13813; A0 | SPTN1_HUMAN | Unmodified | 0 | 0 | 0 | 0 | 237390 | 0 | 0 | 0 | 0 | 0 | 0 | 325540 | 0 | 0 | 0 | 0 | 0 | 0 |
| MQKEIITLA | ACTBL2 | Q562R1 | ACTBL_HUMAN | Oxidation (M) | 0 | 0 | 0 | 0 | 2585900 | 11694000 | 3970900 | 522430 | 0 | 0 | 0 | 0 | 1122500 | 5990100 | 0 | 0 | 1124700 | 8951600 |
| MQKEITALA | ACTG1; ACTC1; | P63261; P68 | ACTG_HUMAN | Unmodified | 0 | 0 | 0 | 0 | 2292500 | 0 | 0 | 0 | 0 | 512880 | 0 | 0 | 1116800 | 0 | 0 | 0 | 2080200 | 1137900 |
| MQKYEKLE | CDK5 | Q00535 | CDK5_HUMAN | Acetyl (Protein N | 0 | 0 | 0 | 0 | 0 | 0 | 693660 | 344810 | 0 | 0 | 0 | 0 | 0 | 231870 | 0 | 0 | 1168000 | 846230 |
| MQLKPMEI | UCHL1; UCHL1 | P09936; D6F | UCHL1_HUMA | 2 Oxidation (M) | 0 | 0 | 103490 | 0 | 19155000 | 8804500 | 4871800 | 3457700 | 1420900 | 5707600 | 5245900 | 16168000 | 0 | 0 | 0 | 0 | 0 | 0 |
| MQLKPMEI | UCHL1; UCHL1 | P09936; D6F | UCHL1_HUMA | Oxidation (M) | 0 | 0 | 0 | 0 | 43960000 | 9277400 | 3863100 | 395200 | 9626200 | 5317900 | 3912100 | 13691000 | 0 | 0 | 0 | 0 | 0 | 0 |
| MQLKPMEI | UCHL1; UCHL1 | P09936; D6F | UCHL1_HUMA | Unmodified | 64758 | 0 | 0 | 0 | 17418000 | 0 | 0 | 0 | 8175700 | 1015700 | 0 | 547480 | 0 | 0 | 0 | 0 | 0 | 0 |
| MQLKPMEI | UCHL1; UCHL1 | P09936; D6F | UCHL1_HUMA | 3 Oxidation (M) | 0 | 0 | 0 | 0 | 4889900 | 196030 | 67345 | 2140000 | 0 | 487260 | 1204800 | 7750400 | 0 | 0 | 0 | 0 | 0 | 0 |
| MQLLEIITT | HADHA; HADH | P40939; A0A | ECHA_HUMAN | Unmodified | 0 | 0 | 0 | 0 | 0 | 0 | 0 | 0 | 2334100 | 0 | 0 | 0 | 0 | 0 | 0 | 0 | 0 | 0 |
| MQLLEIITT | HADHA; HADH | P40939; A0A | ECHA_HUMAN | Oxidation (M) | 0 | 0 | 0 | 0 | 0 | 0 | 0 | 0 | 699910 | 0 | 0 | 0 | 0 | 0 | 0 | 0 | 0 | 0 |
| MQLNSEGI | PLCB2; PLCB2 | Q00722; Q9 | PLCB2_HUMAN | Oxidation (M) | 0 | 0 | 0 | 0 | 0 | 0 | 0 | 0 | 0 | 51955 | 0 | 717710 | 0 | 0 | 0 | 0 | 0 | 0 |
| MQLSGEG | SLC25A11; SLC | Q02978; I3L | M2OM_HUMAN | Unmodified | 0 | 0 | 0 | 0 | 94312 | 0 | 0 | 0 | 4088000 | 248240 | 12076 | 0 | 0 | 0 | 0 | 0 | 11628 | 203510 |
| MQLTHQLD | TPRKB | Q9Y3C4 | TPRKB_HUMAN | Oxidation (M) | 0 | 0 | 0 | 0 | 0 | 0 | 222580 | 0 | 0 | 0 | 0 | 0 | 172830 | 104830 | 0 | 0 | 0 | 0 |
| MQMAVSLF | OSMR | Q99650 | OSMR_HUMAN | Unmodified | 0 | 0 | 0 | 0 | 1765400 | 171320 | 0 | 0 | 0 | 0 | 0 | 0 | 0 | 0 | 0 | 0 | 0 | 0 |
| MQNDAGE | RPS21; RPS21; | P63220; Q9 | RS21_HUMAN | Acetyl (Protein N | 3030400 | 5568200 | 0 | 2415100 | 1810700 | 0 | 2222300 | 686670 | 0 | 693750 | 1510500 | 13574000 | 3941400 | 2174500 | 0 | 0 | 1896700 | 2289500 |
| MQNDVKLI | SLC25A10; SLC | Q9UBX3; F6 | DIC_HUMAN M | Unmodified | 0 | 0 | 0 | 0 | 0 | 0 | 0 | 0 | 470590 | 89932 | 0 | 0 | 0 | 0 | 0 | 0 | 0 | 0 |
| MQNHGYE | APLP2 | Q06481 | APLP2_HUMAN | Unmodified | 0 | 0 | 0 | 0 | 740980 | 0 | 0 | 0 | 260820 | 0 | 0 | 0 | 0 | 0 | 0 | 0 | 0 | 0 |
| MQNSEQV | N/A | N/A | N/A | Oxidation (M) | 0 | 0 | 0 | 0 | 0 | 0 | 0 | 0 | 0 | 1932100 | 0 | 950550 | 0 | 0 | 0 | 0 | 0 | 0 |
| MQNYKYDI | PLPP3 | O14495 | PLPP3_HUMAN | Acetyl (Protein N | 0 | 0 | 0 | 0 | 3293600 | 293420 | 78863 | 52311 | 0 | 0 | 0 | 0 | 0 | 268000 | 0 | 0 | 0 | 1610100 |
| MQNYKYDI | PLPP3 | O14495 | PLPP3_HUMAN | Acetyl (Protein N | 0 | 0 | 0 | 112300 | 9819900 | 0 | 147940 | 41073 | 97481 | 205330 | 0 | 0 | 865080 | 188550 | 0 | 60541 | 930460 | 3309400 |
| MQPGPVF | LMAN2L | Q9H0V9 | LMA2L_HUMAN | Unmodified | 0 | 0 | 0 | 0 | 0 | 0 | 0 | 0 | 3061000 | 0 | 0 | 0 | 0 | 0 | 0 | 0 | 0 | 0 |
| MQQHEQA | STUB1; STUB1 | Q9UNE7; H3 | CHIP_HUMAN | Unmodified | 0 | 0 | 0 | 0 | 2713300 | 0 | 1548200 | 0 | 0 | 0 | 0 | 0 | 1586700 | 1692300 | 0 | 0 | 1921700 | 3463600 |
| MQQHFEF | ASNS; ASNS; A | P08243; C9J | ASNS_HUMAN | Oxidation (M) | 0 | 0 | 0 | 0 | 0 | 0 | 0 | 8719.7 | 0 | 0 | 0 | 1512100 | 0 | 0 | 0 | 388170 | 0 | 0 |
| MQQHFEF | ASNS; ASNS; A | P08243; C9J | ASNS_HUMAN | Unmodified | 0 | 0 | 0 | 0 | 0 | 0 | 0 | 0 | 0 | 0 | 0 | 2845500 | 0 | 0 | 465680 | 360370 | 0 | 0 |
| MQQNGYE | APP; APP; APP | P05067; E9P | A4_HUMAN Am | Unmodified | 0 | 0 | 0 | 0 | 2874300 | 0 | 0 | 0 | 0 | 0 | 0 | 0 | 114810 | 0 | 0 | 0 | 0 | 542200 |
| MQQNGYE | APP; APP; APP | P05067; E9P | A4_HUMAN Am | Oxidation (M) | 0 | 0 | 0 | 0 | 362310 | 68401 | 0 | 0 | 0 | 0 | 0 | 0 | 0 | 0 | 0 | 0 | 0 | 0 |
| MQQNIQEL | MYH9 | P35579 | MYH9_HUMAN | Oxidation (M) | 0 | 0 | 0 | 0 | 0 | 0 | 0 | 0 | 0 | 0 | 0 | 0 | 0 | 0 | 1178400 | 452680 | 0 | 0 |
| MQQNIQEL | MYH9 | P35579 | MYH9_HUMAN | Oxidation (M) | 139530 | 0 | 1122400 | 1144400 | 1102300 | 0 | 0 | 0 | 0 | 427630 | 0 | 4266600 | 532200 | 0 | 1089200 | 1343500 | 0 | 0 |
| MQQNIQEL | MYH9 | P35579 | MYH9_HUMAN | Oxidation (M) | 0 | 0 | 0 | 1018900 | 12093000 | 3374300 | 1189900 | 0 | 0 | 2710200 | 0 | 0 | 6480700 | 5914200 | 0 | 0 | 2216100 | 6176800 |
| MQQNIQEL | MYH9 | P35579 | MYH9_HUMAN | Unmodified | 0 | 0 | 0 | 0 | 7540900 | 0 | 654360 | 0 | 0 | 599300 | 0 | 0 | 0 | 0 | 0 | 0 | 1679500 | 1589100 |
| MQQSGHP | EHHADH | Q08426 | ECHP_HUMAN | Unmodified | 0 | 0 | 0 | 0 | 0 | 0 | 37861 | 0 | 609870 | 103350 | 0 | 0 | 0 | 0 | 0 | 0 | 0 | 0 |
| MQQSVYV | KTN1 | Q86UP2 | KTN1_HUMAN | Unmodified | 0 | 0 | 0 | 0 | 0 | 0 | 0 | 0 | 0 | 0 | 0 | 1349400 | 0 | 0 | 0 | 0 | 0 | 0 |
| MQQYDLQ | NCLN | Q969V3 | NCLN_HUMAN | Oxidation (M) | 0 | 0 | 0 | 0 | 0 | 0 | 0 | 0 | 1527200 | 75361 | 121580 | 207590 | 0 | 0 | 0 | 0 | 0 | 0 |
| MQREEQCI | FKBP5 | Q13451 | FKBP5_HUMAN | Oxidation (M) | 0 | 436190 | 0 | 0 | 0 | 1146800 | 348550 | 19002 | 0 | 0 | 0 | 226470 | 263000 | 0 | 0 | 0 | 0 | 223070 |
| MQREEQCI | FKBP5 | Q13451 | FKBP5_HUMAN | Unmodified | 0 | 0 | 0 | 0 | 0 | 0 | 0 | 0 | 0 | 0 | 0 | 0 | 135670 | 0 | 0 | 0 | 0 | 0 |
| MQRPSLPD | WIPF2 | Q8TF74 | WIPF2_HUMAN | Oxidation (M) | 519560 | 0 | 423190 | 0 | 0 | 0 | 0 | 0 | 0 | 0 | 0 | 0 | 0 | 0 | 870360 | 0 | 0 | 0 |
| MQSLQPV | SUSD2 | Q9UGT4 | SUSD2_HUMA | Oxidation (M) | 0 | 0 | 113490 | 0 | 0 | 0 | 186620 | 0 | 516960 | 247440 | 0 | 0 | 0 | 0 | 0 | 0 | 0 | 0 |
| MQSNKTFN | NONO; NONO; | Q15233; C9 | NONO_HUMAN | Acetyl (Protein N | 0 | 0 | 0 | 0 | 0 | 0 | 0 | 181300 | 0 | 0 | 0 | 0 | 0 | 0 | 0 | 0 | 67549 | 0 |
| MQTFSPAG | LAMA5 | O15230 | LAMA5_HUMAN | Oxidation (M) | 67375 | 0 | 50280 | 0 | 0 | 43648 | 0 | 0 | 0 | 40601 | 0 | 0 | 65758 | 0 | 0 | 0 | 0 | 242400 |
| MQTFSPAG | LAMA5 | O15230 | LAMA5_HUMAN | Unmodified | 0 | 0 | 0 | 0 | 0 | 0 | 0 | 0 | 0 | 0 | 0 | 0 | 0 | 0 | 0 | 0 | 0 | 660530 |
| MQTNQTP | ATP6V0A1; ATP | Q93050; B7 | VPP1_HUMAN | Unmodified | 29890 | 0 | 72823 | 31852 | 0 | 0 | 0 | 0 | 1155300 | 296130 | 89954 | 439590 | 0 | 0 | 0 | 54986 | 0 | 0 |
| MQTQQVN | ROCK2; ROCK2 | O75116; Q1 | ROCK2_HUMA | Unmodified | 0 | 0 | 124710 | 0 | 1378100 | 0 | 95524 | 53025 | 933670 | 112360 | 69368 | 623140 | 565070 | 203990 | 135130 | 114500 | 679830 | 2037700 |
| MQTQQVN | ROCK2; ROCK2 | O75116; Q1 | ROCK2_HUMA | Unmodified | 0 | 0 | 0 | 0 | 0 | 0 | 79672 | 72682 | 0 | 591030 | 0 | 0 | 0 | 33762 | 0 | 0 | 189530 | 0 |

| | | | | | | | | | | | | | | | | | | | | | | |
|---|---|---|---|---|---|---|---|---|---|---|---|---|---|---|---|---|---|---|---|---|---|---|
| MQVDQEEI | HSPA4; HSPA4 | P34932; A0A | HSP74_HUMAN | Unmodified | 0 | 0 | 0 | 0 | 0 | 0 | 0 | 0 | 0 | 0 | 942110 | 1069300 | 0 | 0 | 0 | 0 | 0 | 0 |
| MQVDQEEI | HSPA4; HSPA4 | P34932; A0A | HSP74_HUMAN | Unmodified | 0 | 0 | 0 | 0 | 1622100 | 0 | 0 | 0 | 3793600 | 0 | 1815100 | 12532000 | 674680 | 241820 | 0 | 0 | 0 | 451780 |
| MQVDQEEI | HSPA4; HSPA4 | P34932; A0A | HSP74_HUMAN | Oxidation (M) | 0 | 0 | 0 | 0 | 611690 | 0 | 0 | 0 | 450220 | 0 | 1445600 | 5581200 | 333350 | 0 | 274350 | 344540 | 0 | 292880 |
| MQVDQEEI | HSPA4; HSPA4 | P34932; A0A | HSP74_HUMAN | 2 Oxidation (M) | 0 | 0 | 0 | 0 | 0 | 0 | 0 | 0 | 0 | 0 | 0 | 1122000 | 0 | 0 | 0 | 0 | 0 | 0 |
| MQYAPNTC | CAPG; CAPG | P40121; H7C | CAPG_HUMAN | Oxidation (M) | 0 | 0 | 0 | 0 | 0 | 0 | 0 | 0 | 0 | 0 | 0 | 0 | 0 | 0 | 0 | 0 | 959910 | 0 |
| MRAEDGEN | TBCA; TBCA; TB | O75347; E5F | TBCA_HUMAN | Unmodified | 0 | 0 | 360770 | 0 | 0 | 0 | 0 | 0 | 0 | 0 | 0 | 2656300 | 777960 | 271220 | 0 | 0 | 0 | 0 |
| MRAEDGEN | TBCA; TBCA; TB | O75347; E5F | TBCA_HUMAN | Unmodified | 226230 | 153160 | 288140 | 794030 | 830860 | 0 | 13550 | 0 | 1046400 | 296770 | 304170 | 4768400 | 1633500 | 0 | 632940 | 412260 | 165410 | 510320 |
| MRCELIRL( | TAOK1 | Q7L7X3 | TAOK1_HUMAN | Oxidation (M) | 0 | 0 | 0 | 0 | 278540 | 0 | 307430 | 0 | 0 | 446450 | 0 | 0 | 0 | 187060 | 0 | 0 | 152440 | 0 |
| MRCELIRL( | TAOK1 | Q7L7X3 | TAOK1_HUMAN | Unmodified | 0 | 0 | 0 | 0 | 1202400 | 0 | 276870 | 0 | 0 | 0 | 0 | 0 | 0 | 0 | 0 | 0 | 0 | 0 |
| MRDCYCT\ | RASA3 | Q14644 | RASA3_HUMAN | Unmodified | 0 | 0 | 0 | 0 | 434840 | 0 | 606800 | 0 | 0 | 0 | 0 | 0 | 249020 | 290160 | 375950 | 0 | 1271800 | 332750 |
| MRDFINKQ | SCG3 | Q8WXD2 | SCG3_HUMAN | Oxidation (M) | 0 | 0 | 0 | 0 | 0 | 0 | 0 | 0 | 0 | 1443000 | 0 | 218600 | 0 | 0 | 0 | 0 | 0 | 0 |
| MRDFINKQ | SCG3 | Q8WXD2 | SCG3_HUMAN | Unmodified | 0 | 0 | 0 | 0 | 1905900 | 0 | 189470 | 0 | 0 | 4108300 | 0 | 0 | 0 | 0 | 0 | 0 | 0 | 0 |
| MRDPLTDC | NOSTRIN; NOS | Q8IVI9; F8W | NOSTN_HUMA | Unmodified | 3198800 | 0 | 1778500 | 1694200 | 0 | 0 | 0 | 0 | 0 | 0 | 0 | 0 | 1985000 | 1690700 | 3337500 | 2427700 | 0 | 0 |
| MRDPLTDC | NOSTRIN; NOS | Q8IVI9; F8W | NOSTN_HUMA | Oxidation (M) | 418120 | 1007900 | 1586900 | 1601500 | 0 | 0 | 0 | 0 | 0 | 0 | 0 | 299320 | 773470 | 0 | 3330200 | 634030 | 0 | 0 |
| MRECHGL\ | PKP3 | Q9Y446 | PKP3_HUMAN | Oxidation (M) | 0 | 0 | 0 | 697390 | 758740 | 0 | 0 | 0 | 0 | 0 | 0 | 0 | 0 | 1407400 | 489670 | 690320 | 1499200 | 0 |
| MREDDRV/ | MCM5; MCM5 | P33992; B1A | MCM5_HUMAN | Oxidation (M) | 0 | 0 | 0 | 20055 | 0 | 0 | 0 | 0 | 0 | 0 | 0 | 344020 | 0 | 0 | 0 | 0 | 0 | 0 |
| MREDYDS\ | RBM8A | Q9Y5S9 | RBM8A_HUMA | Oxidation (M) | 2460500 | 1682500 | 2880400 | 819800 | 0 | 0 | 0 | 0 | 482070 | 74269 | 235620 | 1251400 | 0 | 0 | 67829 | 0 | 0 | 0 |
| MREDYDS\ | RBM8A | Q9Y5S9 | RBM8A_HUMA | Unmodified | 1363900 | 1072300 | 523530 | 695000 | 145310 | 57777 | 0 | 0 | 2181300 | 0 | 1528700 | 1735800 | 446310 | 63532 | 443470 | 445340 | 38481 | 0 |
| MREEVITLII | IQGAP1; IQGAP | P46940; A0A | IQGA1_HUMAN | Unmodified | 0 | 0 | 0 | 0 | 1642500 | 0 | 0 | 26759 | 1251400 | 190830 | 0 | 717950 | 0 | 0 | 0 | 90019 | 0 | 0 |
| MREFGIADI | ACOX1; ACOX1 | Q15067; I3L | ACOX1_HUMA | 2 Oxidation (M) | 0 | 0 | 0 | 0 | 0 | 0 | 0 | 0 | 1896700 | 638940 | 213780 | 0 | 0 | 0 | 0 | 0 | 0 | 0 |
| MREFGIADI | ACOX1; ACOX1 | Q15067; I3L | ACOX1_HUMA | Oxidation (M) | 0 | 0 | 0 | 0 | 0 | 0 | 0 | 0 | 633150 | 40578 | 0 | 0 | 0 | 0 | 0 | 0 | 0 | 0 |
| MREHLELF | CLTC; CLTC; CI | Q00610; J3K | CLH1_HUMAN | Oxidation (M) | 0 | 0 | 0 | 0 | 0 | 0 | 0 | 0 | 0 | 0 | 749810 | 1346900 | 0 | 0 | 0 | 0 | 0 | 0 |
| MREHVMN | NUCB2; NUCB2 | P80303; H0Y | NUCB2_HUMA | Oxidation (M) | 0 | 0 | 0 | 0 | 0 | 0 | 0 | 0 | 7282000 | 142920 | 0 | 0 | 0 | 0 | 0 | 0 | 0 | 0 |
| MREHVMN | NUCB2; NUCB2 | P80303; H0Y | NUCB2_HUMA | Unmodified | 0 | 0 | 109060 | 0 | 0 | 0 | 0 | 0 | 7520300 | 89528 | 44895 | 0 | 0 | 0 | 222220 | 0 | 0 | 0 |
| MREHVMN | NUCB2; NUCB2 | P80303; H0Y | NUCB2_HUMA | 2 Oxidation (M) | 0 | 0 | 0 | 0 | 322300 | 0 | 0 | 0 | 1215900 | 0 | 0 | 0 | 0 | 0 | 0 | 0 | 0 | 0 |
| MREHVMN | NUCB2; NUCB2 | P80303; H0Y | NUCB2_HUMA | Oxidation (M) | 0 | 0 | 0 | 0 | 0 | 0 | 0 | 0 | 6880200 | 0 | 0 | 0 | 0 | 0 | 0 | 0 | 0 | 0 |
| MREHVMN | NUCB2; NUCB2 | P80303; H0Y | NUCB2_HUMA | Unmodified | 0 | 0 | 0 | 0 | 0 | 0 | 0 | 0 | 13536000 | 0 | 0 | 129390 | 0 | 0 | 0 | 0 | 44434 | 0 |
| MREIVHIQ/ | TUBB6; TUBB6; | Q9BUF5; K7I | TBB6_HUMAN | Oxidation (M) | 0 | 114560 | 0 | 0 | 660650 | 0 | 0 | 77683 | 0 | 28064 | 0 | 0 | 0 | 0 | 0 | 0 | 0 | 0 |
| MREIVHIQ/ | TUBB6; TUBB6; | Q9BUF5; K7I | TBB6_HUMAN | Unmodified | 0 | 0 | 0 | 0 | 2052400 | 254860 | 169180 | 0 | 0 | 30346 | 0 | 0 | 0 | 0 | 0 | 0 | 0 | 103050 |
| MREIVHLQ | TUBB4A; TUBB4 | P04350; P68 | TBB4A_HUMAN | Unmodified | 1793700 | 1160600 | 2675100 | 4278300 | 99170000 | 15303000 | 75105000 | 15136000 | 38167000 | 29552000 | 17196000 | 92721000 | 35108000 | 29720000 | 4092600 | 2521000 | 26875000 | 64843000 |
| MRETAFEEI | GPRC5B | Q9NZH0 | GPC5B_HUMA | Oxidation (M) | 0 | 0 | 1259600 | 1189100 | 101450 | 0 | 0 | 0 | 0 | 243000 | 581520 | 433100 | 0 | 0 | 0 | 0 | 0 | 0 |
| MREVCEQN | KIF4A | O95239 | KIF4A_HUMAN | Oxidation (M) | 0 | 0 | 0 | 0 | 183120 | 0 | 110000 | 0 | 113410 | 0 | 0 | 0 | 862140 | 0 | 0 | 0 | 381700 | 779160 |
| MREYKLVV | RAP1B; RAP1A; | P61224; P62 | RAP1B_HUMAN | Unmodified | 0 | 0 | 0 | 0 | 7393200 | 543230 | 14601000 | 721810 | 0 | 832790 | 0 | 0 | 2902900 | 5016400 | 0 | 0 | 3858400 | 4854900 |
| MREYKLVV | RAP1B; RAP1A; | P61224; P62 | RAP1B_HUMAN | Oxidation (M) | 0 | 0 | 0 | 18322 | 7056600 | 1019100 | 16230000 | 5705300 | 0 | 668180 | 0 | 0 | 9214100 | 15224000 | 0 | 0 | 5557100 | 4964200 |
| MREYKVVV | RAP2C; RAP2A; | Q9Y3L5; P10 | RAP2C_HUMA | Oxidation (M) | 0 | 0 | 0 | 0 | 0 | 185690 | 962620 | 879010 | 0 | 934930 | 0 | 0 | 1126700 | 1822300 | 0 | 0 | 2544000 | 1324300 |
| MREYKVVV | RAP2C; RAP2A; | Q9Y3L5; P10 | RAP2C_HUMA | Unmodified | 0 | 0 | 0 | 0 | 5297200 | 0 | 1518800 | 239550 | 0 | 661400 | 0 | 0 | 165280 | 1959900 | 0 | 0 | 4277500 | 3261300 |
| MRFVDHVF | RPN1; RPN1 | P04843; B7Z | RPN1_HUMAN | Oxidation (M) | 0 | 0 | 0 | 0 | 0 | 0 | 185680 | 0 | 12922000 | 16677000 | 863840 | 0 | 0 | 86704 | 0 | 0 | 108040 | 0 |
| MRFVDHVF | RPN1; RPN1 | P04843; B7Z | RPN1_HUMAN | Unmodified | 0 | 0 | 0 | 0 | 0 | 0 | 2803600 | 0 | 11449000 | 2413300 | 0 | 621120 | 0 | 0 | 0 | 0 | 228810 | 301590 |
| MRGEAEAF | FLOT1; FLOT1; I | O75955; A0/ | FLOT1_HUMAN | Unmodified | 0 | 0 | 1087500 | 1415600 | 9641800 | 1438000 | 107970 | 0 | 6937400 | 1131100 | 869770 | 1437400 | 1508800 | 723710 | 2380200 | 428840 | 388210 | 1239000 |
| MRGEAEAF | FLOT1; FLOT1; I | O75955; A0/ | FLOT1_HUMAN | Oxidation (M) | 0 | 0 | 0 | 687280 | 1468900 | 484530 | 0 | 0 | 1346400 | 512360 | 0 | 1285100 | 0 | 0 | 3075800 | 2079100 | 61749 | 169810 |
| MRGETTLW | VARS1; VARS1 | P26640; A0A | SYVC_HUMAN | Oxidation (M) | 0 | 0 | 0 | 0 | 125180 | 0 | 679750 | 0 | 0 | 0 | 0 | 1308500 | 0 | 0 | 0 | 0 | 0 | 0 |
| MRGHQLLS | NSF; NSF | P46459; I3L( | NSF_HUMAN V | Oxidation (M) | 0 | 0 | 0 | 0 | 0 | 0 | 0 | 0 | 0 | 0 | 0 | 0 | 322850 | 175340 | 0 | 0 | 0 | 0 |
| MRGHQLLS | NSF; NSF | P46459; I3L( | NSF_HUMAN V | Oxidation (M) | 0 | 0 | 0 | 0 | 0 | 0 | 0 | 0 | 479530 | 0 | 0 | 0 | 2608500 | 1580400 | 0 | 0 | 241980 | 0 |
| MRGVESQ( | YARS1; YARS1 | P54577; A0A | SYYC_HUMAN | 2 Oxidation (M) | 0 | 0 | 0 | 0 | 0 | 0 | 0 | 0 | 0 | 0 | 0 | 0 | 3888800 | 0 | 0 | 0 | 1319700 | 3225200 |
| MRIESLSS( | LMNB1; LMNB1 | P20700; A0A | LMNB1_HUMA | Unmodified | 0 | 0 | 0 | 0 | 1276600 | 0 | 690390 | 0 | 1207900 | 0 | 0 | 333220 | 736250 | 0 | 0 | 0 | 0 | 0 |
| MRIESLSS( | LMNB1; LMNB1 | P20700; A0A | LMNB1_HUMA | Oxidation (M) | 0 | 0 | 0 | 0 | 0 | 0 | 0 | 67047 | 0 | 259990 | 471870 | 538210 | 0 | 0 | 0 | 0 | 0 | 0 |
| MRITLRNLE | AGRN; AGRN | O00468; A0/ | AGRIN_HUMAN | Acetyl (Protein N | 0 | 0 | 0 | 0 | 0 | 0 | 0 | 0 | 141850 | 0 | 0 | 0 | 0 | 0 | 0 | 0 | 0 | 0 |
| MRLESLSY( | LMNB2 | Q03252 | LMNB2_HUMA | Unmodified | 47143 | 0 | 0 | 0 | 2170900 | 477150 | 0 | 0 | 917120 | 581590 | 0 | 339690 | 352940 | 0 | 0 | 0 | 45221 | 0 |
| MRLESLSY( | LMNB2 | Q03252 | LMNB2_HUMA | Oxidation (M) | 85577 | 0 | 0 | 0 | 1120000 | 0 | 0 | 77137 | 0 | 400490 | 0 | 0 | 0 | 0 | 0 | 0 | 0 | 0 |
| MRNIEVVEI | KIDINS220; KID | Q9ULH0; E9 | KDIS_HUMAN | Oxidation (M) | 0 | 0 | 0 | 0 | 2024500 | 0 | 0 | 0 | 0 | 369230 | 0 | 27922 | 557730 | 452930 | 0 | 0 | 0 | 0 |
| MRPQQAP\ | GOLGA7 | Q7Z5G4 | GOGA7_HUMA | Acetyl (Protein N | 113010 | 0 | 299530 | 378030 | 1860500 | 0 | 0 | 0 | 264280 | 145990 | 232290 | 1311500 | 1254600 | 0 | 299620 | 289490 | 243910 | 562670 |
| MRPQQAP\ | GOLGA7 | Q7Z5G4 | GOGA7_HUMA | Acetyl (Protein N | 160850 | 0 | 500990 | 799300 | 5874200 | 395690 | 564230 | 0 | 679620 | 312530 | 644290 | 3268100 | 2096700 | 1188800 | 649880 | 482090 | 975660 | 1675700 |
| MRPQTLKG | EPS8L2; EPS8L2 | Q9H6S3; A0, | ES8L2_HUMAN | Unmodified | 0 | 0 | 0 | 0 | 0 | 0 | 0 | 0 | 0 | 0 | 0 | 0 | 0 | 201000 | 0 | 0 | 0 | 135450 |
| MRQGEEVF | GCN1 | Q92616 | GCN1_HUMAN | Unmodified | 0 | 0 | 0 | 0 | 0 | 0 | 164230 | 14840 | 0 | 0 | 0 | 543390 | 182840 | 0 | 0 | 0 | 119410 | 321330 |
| MRQYPGS\ | SYTL4 | Q96C24 | SYTL4_HUMAN | Unmodified | 0 | 0 | 0 | 0 | 489680 | 0 | 105710 | 0 | 0 | 0 | 0 | 0 | 58630 | 0 | 0 | 0 | 0 | 0 |
| MRSSENFD | CRABP1 | P29762 | RABP1_HUMAN | Oxidation (M) | 0 | 0 | 0 | 0 | 380530 | 0 | 0 | 0 | 187660 | 691140 | 0 | 0 | 0 | 0 | 0 | 0 | 970790 | 2559600 |
| MRSSENFD | CRABP1 | P29762 | RABP1_HUMAN | Unmodified | 0 | 0 | 0 | 0 | 0 | 0 | 0 | 0 | 0 | 0 | 0 | 0 | 0 | 0 | 0 | 0 | 249440 | 1171500 |
| MRVTGVIT( | EDIL3 | O43854 | EDIL3_HUMAN | Unmodified | 78104 | 0 | 0 | 0 | 2.03E+08 | 36011000 | 15431000 | 4176700 | 7724700 | 8143100 | 0 | 0 | 0 | 81540 | 0 | 0 | 5190100 | 31382000 |
| MRVTGVIT( | EDIL3 | O43854 | EDIL3_HUMAN | Oxidation (M) | 0 | 0 | 0 | 0 | 15957000 | 6187000 | 1147000 | 979870 | 1188600 | 736340 | 0 | 0 | 0 | 0 | 0 | 0 | 1007300 | 5507200 |
| MRYVASYL | RPLP2 | P05387 | RLA2_HUMAN | Oxidation (M) | 2022700 | 0 | 533920 | 10076000 | 3115800 | 0 | 1534800 | 0 | 0 | 1269100 | 800620 | 458330 | 24313000 | 6558000 | 1372400 | 1429800 | 1197200 | 3112500 |
| MSAAQALK | PLEC; PLEC | Q15149; H0Y | PLEC_HUMAN | Unmodified | 551670 | 494370 | 0 | 0 | 0 | 0 | 0 | 0 | 0 | 0 | 0 | 0 | 0 | 0 | 389550 | 0 | 464460 | 0 |
| MSAEINEII | VCL | P18206 | VINC_HUMAN | Oxidation (M) | 0 | 0 | 0 | 0 | 415000 | 131720 | 0 | 0 | 0 | 152090 | 0 | 1702400 | 0 | 0 | 457460 | 346770 | 398820 | 350190 |
| MSAEINEII | VCL | P18206 | VINC_HUMAN | Unmodified | 0 | 0 | 0 | 0 | 820620 | 0 | 0 | 0 | 1479200 | 389160 | 247190 | 2548800 | 34027 | 0 | 0 | 0 | 1606400 | 334840 |
| MSASAVYV | AP1M1; AP1M1 | Q9BXS5; K7I | AP1M1_HUMA | Acetyl (Protein N | 0 | 0 | 0 | 0 | 0 | 0 | 0 | 0 | 0 | 0 | 0 | 0 | 899640 | 0 | 0 | 0 | 0 | 0 |
| MSASDPNS | WARS1; WARS1 | P23381; H0Y | SYWC_HUMAN | Unmodified | 0 | 0 | 0 | 0 | 0 | 0 | 289430 | 0 | 959110 | 265520 | 161690 | 0 | 1040200 | 312990 | 0 | 71191 | 605680 | 1577200 |
| MSASILLSK | UNC45A; UNC4 | Q9H3U1; A0 | UN45A_HUMA | Unmodified | 0 | 0 | 0 | 0 | 0 | 0 | 0 | 215390 | 0 | 0 | 0 | 611050 | 136560 | 73748 | 149800 | 65740 | 0 | 0 |
| MSASQLEA | CTNNA1; CTNN | P35221; E5R | CTNA1_HUMA | Oxidation (M) | 0 | 0 | 0 | 0 | 0 | 0 | 0 | 0 | 0 | 0 | 0 | 0 | 0 | 0 | 1190100 | 344200 | 0 | 0 |
| MSATFIGNS | TUBB2A; TUBB2 | Q13885; Q9I | TBB2A_HUMAN | Oxidation (M) | 412460 | 785410 | 521860 | 915700 | 2371700 | 559080 | 14352000 | 1139400 | 761690 | 1475500 | 1005100 | 14597000 | 2370900 | 1838800 | 0 | 668860 | 0 | 1638700 |
| MSATFIGNS | TUBB2A; TUBB2 | Q13885; Q9I | TBB2A_HUMAN | Unmodified | 0 | 0 | 0 | 0 | 0 | 0 | 0 | 0 | 0 | 0 | 0 | 0 | 0 | 0 | 0 | 0 | 0 | 631140 |
| MSATFIGNS | TUBB2A; TUBB2 | Q13885; Q9I | TBB2A_HUMAN | Oxidation (M) | 0 | 300450 | 100570 | 0 | 377360 | 282600 | 1805900 | 62957 | 827290 | 462800 | 785540 | 2398500 | 989640 | 0 | 0 | 510770 | 1127600 | 59154 |
| MSATFIGNS | TUBB2A; TUBB2 | Q13885; Q9I | TBB2A_HUMAN | Unmodified | 0 | 1747600 | 0 | 0 | 0 | 0 | 0 | 0 | 3505300 | 393180 | 0 | 6504100 | 0 | 0 | 801190 | 0 | 0 | 0 |
| MSCQQFIA | PSD2; PSD2 | Q9BQI7; A0A | PSD2_HUMAN | Oxidation (M) | 0 | 0 | 0 | 0 | 0 | 0 | 0 | 0 | 0 | 0 | 779780 | 0 | 0 | 0 | 0 | 0 | 0 | 0 |
| MSDGKKEE | ERP29; ERP29 | P30040; F8V | ERP29_HUMAN | Unmodified | 24484 | 0 | 0 | 0 | 25795 | 23368 | 0 | 180270 | 36106000 | 1343100 | 140050 | 319880 | 0 | 0 | 143520 | 54215 | 26031 | 0 |
| MSDGKKEE | ERP29; ERP29 | P30040; F8V | ERP29_HUMAN | Oxidation (M) | 0 | 0 | 0 | 0 | 0 | 0 | 0 | 0 | 0 | 0 | 0 | 0 | 0 | 0 | 0 | 0 | 0 | 0 |
| MSEDKPQI | CTNNB1; CTNN | P35222; A0A | CTNB1_HUMA | Unmodified | 29798 | 0 | 22461 | 1118100 | 130400 | 116230 | 187070 | 176530 | 26804 | 0 | 16090 | 2101100 | 337120 | 425230 | 1704500 | 1564000 | 150260 | 644410 |
| MSEEDEQ( | PACSIN3 | Q9UKS6 | PACN3_HUMA | Unmodified | 7442100 | 3954200 | 971310 | 2431000 | 0 | 0 | 0 | 0 | 293390 | 0 | 0 | 0 | 1050400 | 408080 | 3722300 | 1896200 | 72620 | 0 |
| MSEEDEQ( | PACSIN3 | Q9UKS6 | PACN3_HUMA | Oxidation (M) | 5086800 | 4260200 | 1159200 | 1226400 | 0 | 0 | 0 | 0 | 0 | 0 | 0 | 0 | 455470 | 0 | 3373300 | 2565100 | 0 | 0 |
| MSERQLAL | CORO1C; COR( | Q9ULV4; B4I | COR1C_HUMA | Unmodified | 0 | 0 | 0 | 0 | 23330000 | 1716700 | 547220 | 283350 | 0 | 1172300 | 0 | 311420 | 1518100 | 962540 | 0 | 0 | 2029200 | 1995900 |
| MSERQVAL | CORO1A; COR( | P31146; H3E | COR1A_HUMA | Unmodified | 0 | 0 | 0 | 0 | 0 | 0 | 203920 | 59237 | 0 | 218730 | 0 | 0 | 215480 | 1282000 | 0 | 0 | 0 | 0 |
| MSESLDTA | RAB3A; RAB3A | P20336; S4F | RAB3A_HUMAN | Oxidation (M) | 215860 | 0 | 337140 | 401580 | 0 | 0 | 0 | 0 | 0 | 0 | 0 | 0 | 0 | 0 | 271130 | 206530 | 0 | 0 |
| MSESLDTA | RAB3A; RAB3A | P20336; S4F | RAB3A_HUMAN | Unmodified | 0 | 0 | 0 | 0 | 0 | 0 | 0 | 0 | 0 | 156410 | 755530 | 536030 | 0 | 0 | 438660 | 467530 | 0 | 184770 |
| MSEWMKK | GEMIN6; GEMI | Q8WXD5; B8 | GEMI6_HUMAN | Acetyl (Protein N | 0 | 0 | 0 | 0 | 0 | 0 | 0 | 0 | 0 | 0 | 0 | 0 | 0 | 0 | 0 | 0 | 0 | 0 |
| MSFETQK | CCPG1 | Q9ULG6 | CCPG1_HUMA | Unmodified | 0 | 0 | 0 | 0 | 0 | 0 | 0 | 0 | 422920 | 43086 | 0 | 112870 | 0 | 0 | 0 | 0 | 0 | 0 |
| MSFPKLGA | N/A | N/A | N/A | Unmodified | 0 | 0 | 0 | 0 | 11916000 | 0 | 0 | 0 | 0 | 0 | 0 | 0 | 0 | 1984600 | 0 | 0 | 1366800 | 5268100 |
| MSGGWEL | C1QBP; C1QBP | Q07021; I3L | C1QBP_HUMA | Unmodified | 1713000 | 4850900 | 0 | 0 | 0 | 0 | 0 | 0 | 1184500 | 0 | 0 | 0 | 0 | 0 | 1134000 | 632330 | 0 | 0 |
| MSGNGREI | ALDH1A1 | P00352 | AL1A1_HUMAN | Oxidation (M) | 0 | 0 | 0 | 0 | 0 | 0 | 0 | 0 | 0 | 0 | 994610 | 422480 | 0 | 0 | 0 | 0 | 0 | 0 |
| MSHLEDIL( | SUN1; SUN1; S | O94901; H0Y | SUN1_HUMAN | Unmodified | 0 | 0 | 0 | 0 | 665200 | 0 | 0 | 0 | 882640 | 0 | 0 | 0 | 0 | 0 | 0 | 0 | 0 | 0 |
| MSHLPPDF | VPS28; VPS28 | Q9UK41; E9I | VPS28_HUMAN | Oxidation (M) | 2443200 | 1649800 | 1112900 | 0 | 0 | 0 | 0 | 0 | 108590 | 0 | 0 | 177030 | 0 | 0 | 1910100 | 2256300 | 0 | 0 |
| MSHLPPDF | VPS28; VPS28 | Q9UK41; E9I | VPS28_HUMAN | Unmodified | 0 | 4503000 | 2046500 | 323310 | 0 | 0 | 0 | 0 | 93951 | 0 | 0 | 560220 | 0 | 0 | 2714200 | 3659000 | 0 | 0 |
| MSIIPQEPV | ABCC4 | O15439 | MRP4_HUMAN | 2 Oxidation (M) | 0 | 0 | 1187200 | 503150 | 0 | 0 | 0 | 0 | 0 | 0 | 0 | 0 | 0 | 29043 | 0 | 0 | 0 | 0 |
| MSIKVTLLN | ARL13B | A0A7P0T9C1 | A0A7P0T9C1_H | Acetyl (Protein N | 0 | 0 | 0 | 0 | 0 | 0 | 0 | 288450 | 0 | 0 | 0 | 0 | 306440 | 0 | 0 | 27699 | 0 | 0 |
| MSINAEEV | ATP1A1 | P05023 | AT1A1_HUMAN | Oxidation (M) | 0 | 0 | 0 | 0 | 910380 | 0 | 0 | 0 | 0 | 656430 | 1578800 | 3520800 | 262360 | 0 | 0 | 0 | 0 | 0 |
| MSINAEEV | ATP1A1 | P05023 | AT1A1_HUMAN | Unmodified | 0 | 0 | 0 | 0 | 0 | 0 | 0 | 0 | 0 | 0 | 0 | 0 | 0 | 0 | 1181100 | 0 | 0 | 0 |
| MSINAEEV | ATP1A1 | P05023 | AT1A1_HUMAN | Oxidation (M) | 0 | 0 | 0 | 4765300 | 30050000 | 833520 | 657550 | 0 | 0 | 1066000 | 768740 | 0 | 23971000 | 7698800 | 0 | 0 | 3302300 | 5779600 |
| MSINAEEV | ATP1A1 | P05023 | AT1A1_HUMAN | Unmodified | 0 | 0 | 0 | 621340 | 13949000 | 454380 | 0 | 0 | 0 | 383890 | 0 | 0 | 7232300 | 1791600 | 0 | 0 | 0 | 0 |
| MSKYCEDL | PLCB4; PLCB4 | Q15147; A0/ | PLCB4_HUMAN | Oxidation (M) | 0 | 0 | 0 | 0 | 154560 | 81709 | 0 | 0 | 0 | 0 | 0 | 0 | 0 | 0 | 0 | 0 | 322940 | 0 |
| MSLDPADL | NENF | Q9UMX5 | NENF_HUMAN | Unmodified | 52439 | 91649 | 0 | 0 | 0 | 0 | 0 | 51235 | 29790000 | 7223900 | 214600 | 0 | 0 | 0 | 0 | 0 | 0 | 0 |
| MSLDPADL | NENF | Q9UMX5 | NENF_HUMAN | Oxidation (M) | 0 | 0 | 0 | 0 | 0 | 0 | 0 | 0 | 10862000 | 8542000 | 688070 | 0 | 174220 | 0 | 0 | 0 | 0 | 0 |
| MSLDPADL | NENF | Q9UMX5 | NENF_HUMAN | Oxidation (M) | 0 | 0 | 0 | 0 | 0 | 0 | 0 | 0 | 0 | 2224100 | 0 | 0 | 0 | 0 | 0 | 0 | 0 | 0 |
| MSLFYAEA | CYRIA; CYRIB; ( | Q9H0Q0; Q9 | CYRIA_HUMAN | 2 Oxidation (M) | 0 | 0 | 0 | 0 | 0 | 0 | 0 | 0 | 0 | 0 | 0 | 1842900 | 0 | 0 | 0 | 0 | 0 | 0 |
| MSLKLQAS | RALY; RALY; RA | Q9UKM9; Q5 | RALY_HUMAN | Acetyl (Protein N | 0 | 0 | 0 | 0 | 0 | 0 | 4674000 | 0 | 0 | 0 | 0 | 0 | 0 | 0 | 683040 | 0 | 1171900 | 158730 |
| MSLLQLVEI | MYO1C; MYO1( | O00159; F5H | MYO1C_HUMA | Oxidation (M) | 31288 | 9004.7 | 0 | 52308 | 62617 | 0 | 0 | 0 | 0 | 0 | 0 | 0 | 0 | 0 | 1427000 | 0 | 35119 | 0 |
| MSLLQLVEI | MYO1C; MYO1( | O00159; F5H | MYO1C_HUMA | Unmodified | 0 | 602320 | 0 | 770060 | 830790 | 258530 | 0 | 0 | 0 | 125340 | 0 | 0 | 138030 | 0 | 4694400 | 0 | 254200 | 0 |
| MSLLQLVEI | MYO1C; MYO1( | O00159; F5H | MYO1C_HUMA | Oxidation (M) | 97112 | 960840 | 0 | 2899400 | 821450 | 216630 | 0 | 0 | 0 | 0 | 0 | 0 | 150450 | 0 | 4664600 | 217080 | 0 | 0 |
| MSLNGQR( | IGFBP2; IGFBP2 | P18065; C9J | IBP2_HUMAN | Oxidation (M) | 0 | 0 | 0 | 0 | 0 | 0 | 0 | 0 | 1065400 | 488250 | 0 | 0 | 0 | 0 | 0 | 0 | 0 | 0 |
| MSLNGQR( | IGFBP2; IGFBP2 | P18065; C9J | IBP2_HUMAN | Unmodified | 0 | 0 | 37451 | 49944 | 0 | 210500 | 0 | 0 | 8357000 | 2364800 | 0 | 0 | 0 | 111660 | 0 | 0 | 0 | 0 |
| MSLPDVDL | AHNAK | Q09666 | AHNK_HUMAN | Oxidation (M) | 0 | 0 | 0 | 0 | 2403000 | 424850 | 0 | 0 | 0 | 0 | 0 | 0 | 0 | 0 | 0 | 0 | 1598600 | 1515500 |
| MSLPDVDL | AHNAK | Q09666 | AHNK_HUMAN | Unmodified | 0 | 0 | 0 | 0 | 10177000 | 0 | 0 | 0 | 0 | 0 | 0 | 0 | 0 | 0 | 0 | 0 | 2306200 | 7490300 |
| MSLSSMEV | AHNAK2 | Q8IVF2 | AHNK2_HUMA | Oxidation (M) | 0 | 0 | 0 | 0 | 733530 | 0 | 0 | 0 | 0 | 0 | 0 | 0 | 0 | 0 | 0 | 125080 | 0 | 509520 |

| | | | | | | | | | | | | | | | | | | | | | | |
|---|---|---|---|---|---|---|---|---|---|---|---|---|---|---|---|---|---|---|---|---|---|---|
| MSLSSMEV | AHNAK2 | Q8IVF2 | AHNK2_HUMAI | Unmodified | 0 | 0 | 0 | 0 | 0 | 0 | 0 | 0 | 0 | 0 | 0 | 0 | 0 | 0 | 0 | 0 | 0 | 0 |
| MSMKEVDE | TUBB6; TUBB3; | Q9BUF5; Q1 | TBB6_HUMAN | Unmodified | 0 | 0 | 0 | 0 | 337060 | 0 | 159140 | 0 | 202890 | 0 | 0 | 0 | 0 | 0 | 0 | 0 | 0 | 270310 |
| MSMKEVDE | TUBB2A; TUBB; | Q13885; P07 | TBB2A_HUMAN | Unmodified | 0 | 0 | 0 | 0 | 4816900 | 0 | 2465700 | 426510 | 3810500 | 1534200 | 0 | 0 | 1900200 | 892980 | 0 | 0 | 1013600 | 4058600 |
| MSMKEVDE | TUBB2A; TUBB; | Q13885; P07 | TBB2A_HUMAN | Oxidation (M) | 0 | 128020 | 0 | 171940 | 2969700 | 990280 | 1895900 | 976630 | 880850 | 2578000 | 400010 | 964530 | 2777100 | 1673100 | 0 | 0 | 1083800 | 2001300 |
| MSMKEVDE | TUBB2A; TUBB; | Q13885; P07 | TBB2A_HUMAN | 2 Oxidation (M) | 0 | 0 | 0 | 37403 | 2922700 | 295780 | 3169900 | 1422100 | 374540 | 2907300 | 291260 | 957940 | 1400000 | 1036900 | 0 | 0 | 491070 | 1021900 |
| MSMKEVDE | TUBB2A; TUBB; | Q13885; P07 | TBB2A_HUMAN | 3 Oxidation (M) | 0 | 0 | 0 | 0 | 0 | 0 | 1890200 | 0 | 0 | 0 | 0 | 0 | 0 | 0 | 0 | 28477000 | 0 | 0 |
| MSMLRLQK | RPL19; RPL19; | P84098; A0A | RL19_HUMAN | Acetyl (Protein N | 0 | 0 | 0 | 0 | 0 | 0 | 61700 | 52249 | 51407 | 0 | 0 | 0 | 0 | 0 | 0 | 0 | 315470 | 277250 |
| MSNNGLDI | PAK1; PAK1; PA | Q13153; E9F | PAK1_HUMAN | Acetyl (Protein N | 0 | 0 | 1239000 | 655470 | 0 | 199860 | 0 | 37992 | 0 | 0 | 0 | 0 | 0 | 91863 | 0 | 0 | 0 | 0 |
| MSNYDTDL | AARS1; AARS1; | P49588; A0A | SYAC_HUMAN | Oxidation (M) | 0 | 0 | 0 | 823990 | 2603000 | 0 | 525860 | 0 | 1470100 | 934040 | 227540 | 2721500 | 3797900 | 2750800 | 0 | 0 | 745820 | 1064200 |
| MSPDEGQI | TMEM165; TMEI | Q9HC07; V9 | TM165_HUMAI | Unmodified | 0 | 0 | 0 | 0 | 0 | 0 | 0 | 0 | 2213700 | 310000 | 0 | 0 | 0 | 0 | 0 | 0 | 0 | 0 |
| MSPIERLEI | UTRN | P46939 | UTRN_HUMAN | Unmodified | 0 | 0 | 0 | 0 | 0 | 37046 | 0 | 0 | 259000 | 197640 | 0 | 0 | 0 | 0 | 0 | 0 | 291790 | 0 |
| MSPNNYK | BMX | P51813 | BMX_HUMAN C | Unmodified | 0 | 0 | 0 | 44063 | 0 | 0 | 0 | 0 | 199350 | 0 | 0 | 2181200 | 0 | 0 | 58492 | 146570 | 53335 | 0 |
| MSQLERNI | S100A9 | P06702 | S10A9_HUMAN | Oxidation (M) | 0 | 0 | 0 | 144140 | 2727200 | 81484 | 0 | 0 | 0 | 0 | 0 | 0 | 249940 | 0 | 0 | 0 | 0 | 0 |
| MSQSIIVSG | MYO6; MYO6; M | Q9UM54; A0 | MYO6_HUMAN | Unmodified | 547320 | 183100 | 804640 | 5094000 | 834440 | 0 | 197940 | 0 | 1844900 | 418980 | 0 | 3259200 | 1760600 | 1110000 | 5336900 | 3211000 | 33670 | 0 |
| MSQSIIVSG | MYO6; MYO6; M | Q9UM54; A0 | MYO6_HUMAN | Oxidation (M) | 0 | 0 | 0 | 1004900 | 0 | 0 | 0 | 0 | 191320 | 0 | 0 | 692580 | 432850 | 0 | 1245800 | 1470200 | 0 | 0 |
| MSQVAPSL | NOP56; NOP56 | O00567; A0A | NOP56_HUMAI | Oxidation (M) | 0 | 0 | 0 | 0 | 60124 | 0 | 87010 | 0 | 0 | 90753 | 0 | 687970 | 0 | 0 | 0 | 0 | 171480 | 0 |
| MSQVMREV | APP; APP; APP; | P05067; H7C | A4_HUMAN Am | Unmodified | 0 | 0 | 0 | 0 | 0 | 0 | 0 | 0 | 1344500 | 120780 | 0 | 0 | 0 | 0 | 0 | 0 | 0 | 0 |
| MSQVMREV | APP; APP; APP; | P05067; H7C | A4_HUMAN Am | Unmodified | 0 | 0 | 0 | 0 | 441580 | 1680900 | 0 | 0 | 2182100 | 901410 | 0 | 0 | 61847 | 0 | 0 | 0 | 0 | 0 |
| MSQVMREV | APP; APP; APP; | P05067; H7C | A4_HUMAN Am | Oxidation (M) | 0 | 0 | 0 | 213330 | 195590 | 258460 | 0 | 0 | 2012200 | 1348400 | 0 | 0 | 0 | 0 | 0 | 0 | 0 | 0 |
| MSSDDGIR | SNTB1 | Q13884 | SNTB1_HUMAN | Unmodified | 0 | 0 | 0 | 0 | 0 | 0 | 0 | 0 | 1051600 | 0 | 0 | 0 | 0 | 0 | 0 | 207660 | 0 | 0 |
| MSSPTDAS | SUSD2 | Q9UGT4 | SUSD2_HUMAI | Oxidation (M) | 860610 | 0 | 0 | 0 | 863080 | 250090 | 0 | 0 | 2948500 | 288170 | 0 | 0 | 0 | 0 | 0 | 0 | 0 | 0 |
| MSSPTDAS | SUSD2 | Q9UGT4 | SUSD2_HUMAI | Unmodified | 669460 | 0 | 0 | 0 | 1790100 | 660090 | 0 | 0 | 7238800 | 617880 | 0 | 0 | 0 | 0 | 0 | 0 | 0 | 0 |
| MSSPTDAS | SUSD2 | Q9UGT4 | SUSD2_HUMAI | Oxidation (M) | 0 | 0 | 0 | 0 | 4826000 | 0 | 0 | 0 | 231240 | 1774600 | 0 | 0 | 0 | 0 | 0 | 0 | 0 | 0 |
| MSSPTDAS | SUSD2 | Q9UGT4 | SUSD2_HUMAI | Unmodified | 0 | 0 | 0 | 0 | 14992000 | 1864900 | 0 | 0 | 0 | 2639500 | 0 | 0 | 0 | 0 | 0 | 0 | 0 | 285160 |
| MSSTFIGNS | TUBB3; N/A | Q13509; A0A | TBB3_HUMAN | Oxidation (M) | 0 | 0 | 71843 | 0 | 723520 | 916100 | 1195500 | 220510 | 35901 | 299040 | 57542 | 1261000 | 0 | 0 | 76194 | 0 | 0 | 56044 |
| MSSTFIGNS | TUBB3; N/A | Q13509; A0A | TBB3_HUMAN | Unmodified | 0 | 0 | 0 | 0 | 0 | 559080 | 0 | 0 | 0 | 0 | 0 | 0 | 0 | 0 | 0 | 0 | 0 | 0 |
| MSSTFIGNS | TUBB3; N/A | Q13509; A0A | TBB3_HUMAN | Oxidation (M) | 0 | 0 | 0 | 0 | 0 | 0 | 0 | 30932 | 340920 | 0 | 260810 | 1893200 | 0 | 0 | 0 | 0 | 0 | 0 |
| MSSTIFSTG | PAK2 | Q13177 | PAK2_HUMAN | Oxidation (M) | 224850 | 0 | 0 | 0 | 0 | 0 | 0 | 0 | 0 | 0 | 1942000 | 3344800 | 674100 | 0 | 0 | 0 | 0 | 0 |
| MSSYAFFV | HMGB2; HMGB | P26583; B2F | HMGB2_HUMA | Unmodified | 0 | 0 | 0 | 104490 | 0 | 0 | 0 | 0 | 0 | 338230 | 0 | 1049400 | 547870 | 0 | 0 | 0 | 323980 | 0 |
| MSSYAFFV | HMGB2; HMGB | P26583; B2F | HMGB2_HUMA | Oxidation (M) | 0 | 0 | 0 | 0 | 151090 | 0 | 152050 | 124180 | 0 | 234600 | 0 | 2959800 | 345070 | 194700 | 0 | 0 | 0 | 0 |
| MSSYAFFV | HMGB2; HMGB | P26583; B2F | HMGB2_HUMA | Unmodified | 0 | 0 | 0 | 0 | 0 | 0 | 114730 | 0 | 278920 | 0 | 0 | 938570 | 339410 | 177250 | 0 | 0 | 0 | 0 |
| MSTEEIIQR | PSMC3; PSMC3 | P17980; E9P | PRS6A_HUMAN | Unmodified | 0 | 0 | 0 | 0 | 0 | 0 | 0 | 0 | 1105400 | 195970 | 0 | 2997400 | 660300 | 0 | 0 | 0 | 0 | 0 |
| MSTEEIIQR | PSMC3; PSMC3 | P17980; E9P | PRS6A_HUMAN | Oxidation (M) | 0 | 0 | 0 | 0 | 0 | 0 | 0 | 0 | 218150 | 52877 | 155780 | 708950 | 94181 | 0 | 0 | 28101 | 72695 | 89053 |
| MSTEEIIQR | PSMC3; PSMC3 | P17980; E9P | PRS6A_HUMAN | Unmodified | 2483200 | 1080400 | 0 | 0 | 923960 | 642170 | 0 | 0 | 359630 | 388770 | 0 | 5050900 | 1127500 | 0 | 0 | 0 | 303560 | 980680 |
| MSTGPDIH | AGPS; AGPS; A( | O00116; A0A | ADAS_HUMAN | 2 Oxidation (M) | 0 | 0 | 0 | 0 | 0 | 0 | 0 | 0 | 3864800 | 3567500 | 0 | 0 | 677710 | 0 | 0 | 0 | 0 | 0 |
| MSTMTPK | TMED2; TMED2 | Q15363; E7E | TMED2_HUMAI | 2 Oxidation (M) | 0 | 0 | 0 | 0 | 0 | 0 | 0 | 0 | 1092800 | 123960 | 0 | 0 | 0 | 0 | 0 | 0 | 0 | 0 |
| MSVFQTQL | SEC24A | O95486 | SC24A_HUMAN | Oxidation (M) | 0 | 0 | 0 | 0 | 0 | 0 | 0 | 0 | 0 | 0 | 0 | 1267600 | 0 | 0 | 0 | 0 | 0 | 0 |
| MSVIEEGD | PYGB | P11216 | PYGB_HUMAN | Unmodified | 0 | 0 | 0 | 0 | 0 | 0 | 0 | 0 | 392330 | 300620 | 373810 | 159840 | 0 | 0 | 0 | 34808 | 317740 | 322730 |
| MSVIWDK | DPYSL3; DPYSL | Q14195; Q1( | DPYL3_HUMAN | Oxidation (M) | 0 | 0 | 59366 | 0 | 574900 | 321470 | 9166.2 | 0 | 279050 | 0 | 1506500 | 7315500 | 0 | 0 | 0 | 0 | 0 | 156890 |
| MSVIWDK | DPYSL3; DPYSL | Q14195; Q1( | DPYL3_HUMAN | Unmodified | 147890 | 88977 | 50471 | 42568 | 0 | 0 | 0 | 163450 | 1855100 | 169010 | 10188000 | 56354000 | 63340 | 0 | 114480 | 0 | 0 | 0 |
| MSVKEGAC | INF2; INF2; INF | Q27J81; A0A | INF2_HUMAN I | Acetyl (Protein N | 0 | 0 | 0 | 0 | 0 | 0 | 0 | 60250 | 0 | 0 | 0 | 877230 | 0 | 0 | 0 | 0 | 0 | 0 |
| MSVQPTVS | NPM1; NPM1; N | P06748; A0A | NPM_HUMAN N | Oxidation (M) | 775650 | 1376700 | 577400 | 2052500 | 2280600 | 1197000 | 601370 | 0 | 2881400 | 4062700 | 5465600 | 2888500 | 3427800 | 576550 | 0 | 903190 | 828800 | 1641500 |
| MSVQPTVS | NPM1; NPM1; N | P06748; A0A | NPM_HUMAN N | Unmodified | 0 | 0 | 0 | 0 | 0 | 0 | 0 | 0 | 1210600 | 815420 | 382440 | 189670 | 392550 | 0 | 0 | 0 | 0 | 0 |
| MSVQPTVS | NPM1; NPM1; N | P06748; A0A | NPM_HUMAN N | Oxidation (M) | 0 | 0 | 0 | 0 | 6750300 | 0 | 4279300 | 0 | 0 | 4299000 | 0 | 0 | 12982000 | 6885700 | 0 | 0 | 2386500 | 3529400 |
| MSVQPTVS | NPM1; NPM1; N | P06748; A0A | NPM_HUMAN N | Unmodified | 0 | 0 | 0 | 0 | 762030 | 0 | 894970 | 0 | 0 | 645800 | 0 | 0 | 1200400 | 560380 | 0 | 0 | 708200 | 653660 |
| MTADGRLF | SLC25A11; SLC | Q02978; I3L | M2OM_HUMAN | Unmodified | 0 | 0 | 0 | 0 | 0 | 20790 | 0 | 0 | 2450000 | 0 | 0 | 0 | 0 | 0 | 120770 | 0 | 0 | 0 |
| MTAEEMAS | TCEA1; TCEA1; | P23193; A0A | TCEA1_HUMAN | Unmodified | 0 | 0 | 0 | 0 | 0 | 0 | 0 | 0 | 933050 | 209820 | 0 | 0 | 0 | 0 | 0 | 0 | 0 | 0 |
| MTAIIFSDY | GLG1; GLG1; G | Q92896; H3I | GSLG1_HUMAI | Oxidation (M) | 0 | 0 | 108030 | 0 | 0 | 0 | 0 | 0 | 4794800 | 560520 | 1192900 | 1676200 | 333970 | 0 | 0 | 219940 | 0 | 0 |
| MTAIIFSDY | GLG1; GLG1; G | Q92896; H3I | GSLG1_HUMAI | Unmodified | 0 | 0 | 0 | 0 | 0 | 0 | 0 | 0 | 17524000 | 0 | 0 | 533960 | 0 | 0 | 0 | 0 | 0 | 0 |
| MTAMDNAS | ATP5F1C | P36542 | ATPG_HUMAN | Oxidation (M) | 0 | 0 | 0 | 0 | 0 | 0 | 0 | 0 | 348760 | 0 | 0 | 0 | 0 | 0 | 0 | 0 | 0 | 0 |
| MTAMDNAS | ATP5F1C | P36542 | ATPG_HUMAN | Unmodified | 0 | 0 | 0 | 0 | 0 | 0 | 0 | 0 | 2217100 | 0 | 0 | 0 | 0 | 0 | 0 | 0 | 24539 | 264050 |
| MTANRKVE | TRIM25; TRIM25 | Q14258; A0A | TRI25_HUMAN | Unmodified | 0 | 0 | 0 | 0 | 3343300 | 0 | 0 | 0 | 0 | 68213 | 0 | 0 | 0 | 0 | 0 | 0 | 133420 | 1144800 |
| MTANSVLE | RO60; RO60 | P10155; G5E | RO60_HUMAN | Unmodified | 2039300 | 466550 | 0 | 0 | 0 | 0 | 0 | 0 | 0 | 0 | 0 | 0 | 0 | 0 | 0 | 0 | 175380 | 51789 |
| MTANSVLE | RO60; RO60 | P10155; G5E | RO60_HUMAN | Oxidation (M) | 13420000 | 4421900 | 3355500 | 3050200 | 0 | 0 | 0 | 0 | 0 | 0 | 0 | 0 | 0 | 0 | 6490400 | 4051500 | 0 | 0 |
| MTATGSGE | COPS6; COPS6 | Q7L5N1; H7 | CSN6_HUMAN | Oxidation (M) | 0 | 0 | 0 | 0 | 0 | 0 | 0 | 0 | 0 | 120200 | 700100 | 2938400 | 258380 | 0 | 0 | 0 | 0 | 0 |
| MTDLLEEG | STXBP3 | O00186 | STXB3_HUMAN | Oxidation (M) | 0 | 0 | 0 | 0 | 365610 | 0 | 0 | 0 | 0 | 0 | 0 | 0 | 0 | 0 | 0 | 0 | 124040 | 136650 |
| MTDQEAIQ | NPM1; NPM1; N | P06748; A0A | NPM_HUMAN N | Oxidation (M) | 0 | 0 | 0 | 0 | 0 | 0 | 46402 | 120590 | 2417200 | 1526800 | 4094100 | 6361500 | 3365600 | 0 | 0 | 0 | 0 | 0 |
| MTDQEAIQ | NPM1; NPM1; N | P06748; A0A | NPM_HUMAN N | Unmodified | 0 | 0 | 0 | 0 | 0 | 0 | 0 | 0 | 2690300 | 1750900 | 1363700 | 2802400 | 0 | 0 | 0 | 0 | 0 | 0 |
| MTDSKYFT | AP2B1; AP2B1; | P63010; K7E | AP2B1_HUMAN | Acetyl (Protein N | 0 | 132630 | 0 | 0 | 0 | 0 | 0 | 0 | 951420 | 0 | 2002400 | 3587300 | 134540 | 0 | 0 | 0 | 0 | 0 |
| MTEEEVEM | MYL6; MYL6; M' | G8JLA2; F8V | G8JLA2_HUMA | Oxidation (M) | 0 | 0 | 0 | 0 | 395380 | 0 | 0 | 0 | 0 | 0 | 0 | 0 | 291800 | 0 | 0 | 0 | 0 | 0 |
| MTEELEALF | CDC42BPA; CD | Q5VT25; Q9' | MRCKA_HUMA | Oxidation (M) | 0 | 0 | 0 | 0 | 0 | 0 | 0 | 0 | 0 | 0 | 0 | 1371300 | 0 | 0 | 350220 | 0 | 0 | 0 |
| MTEIQTPEN | SMC4; SMC4 | Q9NTJ3; E9F | SMC4_HUMAN | Oxidation (M) | 0 | 0 | 0 | 0 | 154390 | 0 | 283170 | 12343 | 0 | 0 | 0 | 0 | 0 | 0 | 0 | 0 | 119990 | 0 |
| MTELFQSL | UTRN; UTRN | P46939; Q5T | UTRN_HUMAN | Oxidation (M) | 0 | 0 | 0 | 0 | 0 | 0 | 0 | 0 | 0 | 0 | 0 | 0 | 0 | 0 | 0 | 0 | 0 | 0 |
| MTEQHFPH | PFAS | O15067 | PUR4_HUMAN | 2 Oxidation (M) | 0 | 0 | 0 | 0 | 0 | 0 | 0 | 0 | 0 | 0 | 0 | 801450 | 0 | 0 | 0 | 0 | 0 | 0 |
| MTEWETAA | RPS5; RPS5; RP | P46782; M0F | RS5_HUMAN 4 | Acetyl (Protein N | 0 | 0 | 0 | 0 | 0 | 0 | 0 | 0 | 0 | 0 | 0 | 0 | 0 | 0 | 0 | 0 | 0 | 0 |
| MTFTSNK | ACOT7L; ACOT | Q6ZUV0; O0 | BACHL_HUMAI | Unmodified | 0 | 0 | 0 | 0 | 0 | 0 | 88913 | 0 | 2206800 | 1100600 | 161580 | 0 | 1097200 | 0 | 20202 | 0 | 542160 | 1021600 |
| MTGESECL | SLK | Q9H2G2 | SLK_HUMAN S | Unmodified | 55943 | 0 | 0 | 52685 | 43002 | 0 | 0 | 0 | 0 | 0 | 115950 | 822790 | 186040 | 324140 | 279600 | 24358 | 22990 | 69341 |
| MTGESECL | SLK | Q9H2G2 | SLK_HUMAN S | Unmodified | 0 | 0 | 0 | 0 | 0 | 0 | 0 | 0 | 0 | 0 | 0 | 0 | 579100 | 913170 | 0 | 0 | 0 | 0 |
| MTGHQALI | NLE1; NLE1; NL | Q9NVX2; A0 | NLE1_HUMAN | Oxidation (M) | 0 | 0 | 0 | 0 | 768580 | 0 | 0 | 0 | 0 | 0 | 0 | 585140 | 0 | 0 | 113830 | 0 | 607530 | 0 |
| MTGLVDEA | VCL | P18206 | VINC_HUMAN | Oxidation (M) | 0 | 0 | 0 | 0 | 0 | 0 | 368900 | 0 | 0 | 255160 | 0 | 0 | 0 | 0 | 0 | 0 | 2614500 | 0 |
| MTGLVDEA | VCL | P18206 | VINC_HUMAN | Unmodified | 0 | 238370 | 0 | 0 | 1075100 | 430100 | 483690 | 0 | 191010 | 579790 | 0 | 1035500 | 146650 | 0 | 0 | 0 | 3669700 | 0 |
| MTGLVDEA | VCL | P18206 | VINC_HUMAN | Oxidation (M) | 0 | 0 | 0 | 0 | 0 | 0 | 0 | 0 | 0 | 0 | 0 | 0 | 0 | 0 | 0 | 0 | 2321100 | 86529 |
| MTHNLLLN | HDAC2; HDAC | Q92769; Q1 | HDAC2_HUMA | Oxidation (M) | 313240 | 0 | 112300 | 0 | 0 | 0 | 0 | 0 | 0 | 194740 | 152390 | 4489000 | 238600 | 0 | 0 | 95618 | 0 | 0 |
| MTHNLLLN | HDAC2; HDAC | Q92769; Q1 | HDAC2_HUMA | Oxidation (M) | 0 | 0 | 0 | 0 | 0 | 0 | 0 | 0 | 0 | 0 | 281860 | 776900 | 0 | 0 | 0 | 0 | 0 | 0 |
| MTKNPDEII | ITGA2 | P17301 | ITA2_HUMAN I | Unmodified | 0 | 0 | 0 | 199720 | 686280 | 0 | 0 | 0 | 0 | 0 | 0 | 0 | 0 | 0 | 0 | 0 | 233210 | 0 |
| MTLDRSEY | LDLR; LDLR; LD | P01130; J3K | LDLR_HUMAN | Unmodified | 0 | 0 | 0 | 0 | 0 | 454280 | 6463600 | 1402900 | 959530 | 0 | 0 | 0 | 0 | 843190 | 0 | 0 | 2246600 | 363920 |
| MTLDTLSIY | EIF3B | P55884 | EIF3B_HUMAN | 2 Oxidation (M) | 0 | 0 | 46414 | 0 | 0 | 0 | 0 | 0 | 0 | 190840 | 564060 | 7279400 | 0 | 0 | 0 | 0 | 0 | 0 |
| MTMDKSEL | YWHAB; YWHA | P31946; Q4\ | 1433B_HUMAN | Acetyl (Protein N | 0 | 0 | 0 | 0 | 0 | 0 | 0 | 0 | 0 | 0 | 0 | 0 | 0 | 0 | 0 | 0 | 0 | 0 |
| MTNGFSGA | VCP; VCP; VCP; | P55072; C9J | TERA_HUMAN | Unmodified | 0 | 0 | 0 | 0 | 647040 | 0 | 0 | 0 | 1602800 | 0 | 0 | 1954000 | 0 | 0 | 0 | 0 | 0 | 0 |
| MTNGFSGA | VCP; VCP; VCP; | P55072; C9J | TERA_HUMAN | Oxidation (M) | 428250 | 155370 | 608270 | 504080 | 0 | 0 | 0 | 0 | 0 | 0 | 0 | 4603800 | 860550 | 0 | 254980 | 350920 | 0 | 0 |
| MTNGFSGA | VCP; VCP; VCP; | P55072; C9J | TERA_HUMAN | Oxidation (M) | 0 | 0 | 0 | 0 | 1585000 | 0 | 0 | 614580 | 0 | 0 | 0 | 0 | 3709900 | 0 | 0 | 0 | 1197500 | 0 |
| MTNGFSGA | VCP; VCP; VCP; | P55072; C9J | TERA_HUMAN | Unmodified | 0 | 0 | 0 | 0 | 2038100 | 464640 | 0 | 0 | 0 | 749910 | 0 | 0 | 2134100 | 0 | 0 | 0 | 978380 | 332140 |
| MTNHCVFS | BZW2; BZW2; B | Q9Y6E2; Q7 | BZW2_HUMAN | Unmodified | 0 | 0 | 149640 | 0 | 0 | 0 | 0 | 0 | 0 | 0 | 0 | 1567300 | 0 | 0 | 0 | 0 | 0 | 0 |
| MTNPAIQN | CYRIB; CYRIB; ( | Q9NUQ9; A0 | CYRIB_HUMAN | Oxidation (M) | 38474 | 0 | 0 | 0 | 0 | 0 | 0 | 0 | 0 | 315720 | 0 | 1885800 | 0 | 0 | 0 | 0 | 0 | 0 |
| MTNPAIQN | CYRIB; CYRIB; ( | Q9NUQ9; A0 | CYRIB_HUMAN | Unmodified | 0 | 0 | 0 | 0 | 420310 | 0 | 0 | 0 | 742900 | 306980 | 0 | 1262800 | 0 | 0 | 0 | 0 | 0 | 488480 |
| MTNSSSSP | PAN3; PAN3 | Q58A45; A0A | PAN3_HUMAN | Oxidation (M) | 2452800 | 0 | 0 | 2059900 | 0 | 0 | 0 | 0 | 0 | 0 | 0 | 0 | 0 | 0 | 0 | 0 | 0 | 0 |
| MTNYDVEH | ANXA6; ANXA6; | P08133; E7E | ANXA6_HUMAN | Oxidation (M) | 0 | 0 | 449380 | 383600 | 0 | 0 | 0 | 0 | 706290 | 363990 | 0 | 0 | 0 | 0 | 370210 | 168290 | 0 | 206230 |
| MTNYDVEH | ANXA6; ANXA6; | P08133; E7E | ANXA6_HUMAN | Unmodified | 219660 | 0 | 971140 | 446600 | 872580 | 0 | 901380 | 289080 | 2982000 | 757740 | 0 | 0 | 172500 | 176710 | 600620 | 528930 | 0 | 133750 |
| MTNYDVEH | ANXA6; ANXA6; | P08133; E7E | ANXA6_HUMAN | Unmodified | 206110 | 265280 | 1526600 | 1845300 | 7416900 | 2116000 | 3147700 | 1184800 | 11325000 | 7827200 | 0 | 0 | 3641200 | 3633800 | 1553400 | 634260 | 1508100 | 6965200 |
| MTNYDVEH | ANXA6; ANXA6; | P08133; E7E | ANXA6_HUMAN | Oxidation (M) | 0 | 0 | 839660 | 1242700 | 1651900 | 768340 | 918670 | 671150 | 1914800 | 1378100 | 0 | 23077 | 2011300 | 1126000 | 1997100 | 911430 | 0 | 2194400 |
| MTPEQMAK | MYO6; MYO6; M | Q9UM54; A0 | MYO6_HUMAN | Unmodified | 0 | 0 | 0 | 0 | 0 | 0 | 0 | 13037 | 0 | 154930 | 0 | 0 | 894620 | 1371500 | 0 | 0 | 0 | 0 |
| MTPSYEIRA | DHX9 | Q08211 | DHX9_HUMAN | Unmodified | 0 | 0 | 0 | 0 | 0 | 0 | 552170 | 244390 | 0 | 118760 | 0 | 0 | 0 | 256330 | 0 | 0 | 0 | 0 |
| MTQNPNY | SNRNP200 | O75643 | U520_HUMAN | Oxidation (M) | 0 | 0 | 227510 | 114090 | 0 | 0 | 0 | 0 | 0 | 0 | 0 | 1322800 | 0 | 0 | 0 | 176920 | 0 | 0 |
| MTRDTGTA | COBLL1; COBL | Q53SF7; A0A | COBL1_HUMAI | Oxidation (M) | 0 | 0 | 0 | 0 | 0 | 0 | 0 | 0 | 0 | 0 | 0 | 0 | 4713500 | 2643200 | 0 | 0 | 0 | 0 |
| MTSFVKDA | DIAPH1; DIAPH | O60610; A0A | DIAP1_HUMAN | Unmodified | 0 | 0 | 0 | 0 | 190800 | 0 | 121750 | 40217 | 0 | 0 | 0 | 0 | 0 | 0 | 0 | 0 | 283340 | 379550 |
| MTSGDVLS | BPNT2; BPNT2 | Q9NX62; H0 | IMPA3_HUMAN | Unmodified | 0 | 0 | 0 | 0 | 0 | 0 | 0 | 0 | 1531100 | 178650 | 0 | 0 | 0 | 0 | 0 | 0 | 0 | 0 |
| MTSGDVLS | BPNT2; BPNT2 | Q9NX62; H0 | IMPA3_HUMAN | Unmodified | 0 | 0 | 289880 | 0 | 250300 | 0 | 0 | 60413 | 84505000 | 4596200 | 946810 | 2621700 | 0 | 0 | 239010 | 272460 | 0 | 0 |
| MTSGDVLS | BPNT2; BPNT2 | Q9NX62; H0 | IMPA3_HUMAN | Oxidation (M) | 0 | 0 | 0 | 0 | 0 | 0 | 0 | 23602 | 20315000 | 2280700 | 0 | 1395400 | 0 | 104780 | 0 | 0 | 0 | 0 |
| MTSGSWTE | PTPRK; PTPRK; | Q15262; Q5 | PTPRK_HUMAN | Oxidation (M) | 143490 | 119110 | 0 | 0 | 0 | 0 | 0 | 0 | 475820 | 0 | 0 | 0 | 0 | 0 | 0 | 0 | 0 | 0 |
| MTSGSWTE | PTPRK; PTPRK; | Q15262; Q5 | PTPRK_HUMAN | Unmodified | 0 | 0 | 0 | 0 | 0 | 0 | 0 | 0 | 1547900 | 0 | 0 | 0 | 0 | 0 | 0 | 0 | 0 | 0 |
| MTTETASEI | GCN1 | Q92616 | GCN1_HUMAN | Unmodified | 0 | 0 | 0 | 0 | 0 | 0 | 0 | 0 | 0 | 0 | 0 | 716270 | 0 | 0 | 0 | 0 | 0 | 0 |
| MTTETGPD | EPB41L1; EPB4 | Q9H4G0; Q4 | E41L1_HUMAN | Unmodified | 0 | 0 | 0 | 0 | 0 | 0 | 0 | 0 | 3120400 | 988420 | 0 | 0 | 0 | 0 | 0 | 0 | 0 | 0 |
| MTTNPIYEH | CHN2; CHN2; ( | P52757; A0A | CHIO_HUMAN | Oxidation (M) | 0 | 1325200 | 0 | 0 | 0 | 0 | 0 | 0 | 0 | 0 | 137690 | 1270000 | 0 | 0 | 0 | 0 | 0 | 0 |
| MTTSGALFI | ADRM1; ADRM1 | Q16186; A0A | ADRM1_HUMA | Oxidation (M) | 0 | 0 | 279610 | 52286 | 0 | 0 | 0 | 0 | 0 | 0 | 0 | 0 | 0 | 0 | 163660 | 26438 | 0 | 0 |
| MTVAHMW | ATP1A1 | P05023 | AT1A1_HUMAN | 2 Oxidation (M) | 551060 | 808480 | 6004900 | 13787000 | 2155300 | 851150 | 0 | 0 | 2214800 | 766610 | 10503000 | 39949000 | 8094500 | 720870 | 14783000 | 13196000 | 0 | 291580 |
| MTVAHMW | ATP1A1 | P05023 | AT1A1_HUMAN | Oxidation (M) | 642690 | 0 | 0 | 7505200 | 9864800 | 1194600 | 0 | 0 | 16836000 | 1993600 | 4209000 | 37924000 | 7052700 | 2990700 | 8281800 | 4752500 | 0 | 0 |
| MTVAHMW | ATP1A1 | P05023 | AT1A1_HUMAN | Unmodified | 0 | 0 | 0 | 0 | 9927000 | 72721 | 0 | 0 | 14157000 | 411030 | 854150 | 4879300 | 0 | 1282700 | 0 | 0 | 0 | 0 |
| MTVCLETEI | BLVRA | P53004 | BIEA_HUMAN E | Unmodified | 0 | 0 | 0 | 0 | 536130 | 212040 | 196740 | 87214 | 0 | 0 | 0 | 0 | 481200 | 338820 | 0 | 69417 | 285270 | 436710 |
| MTVCSPDC | TACSTD2 | P09758 | TACD2_HUMAI | Unmodified | 0 | 0 | 0 | 291420 | 1773400 | 631300 | 0 | 0 | 105140 | 0 | 70031 | 0 | 0 | 0 | 0 | 0 | 37778 | 185830 |

| | | | | | | | | | | | | | | | | | | | | | | |
|---|---|---|---|---|---|---|---|---|---|---|---|---|---|---|---|---|---|---|---|---|---|---|
| MTVDESGC | WDR1 | O75083 | WDR1_HUMAN | Oxidation (M) | 0 | 0 | 0 | 0 | 0 | 0 | 0 | 0 | 0 | 0 | 0 | 844880 | 0 | 0 | 121070 | 95154 | 0 | 0 |
| MTVHEGQI | IGSF8; IGSF8 | Q969P0; C9J | IGSF8_HUMAN | Oxidation (M) | 4590200 | 2884700 | 4409400 | 4760800 | 118790 | 0 | 0 | 0 | 0 | 0 | 0 | 0 | 0 | 0 | 3629400 | 3617100 | 0 | 0 |
| MTVHEGQI | IGSF8; IGSF8 | Q969P0; C9J | IGSF8_HUMAN | Unmodified | 4831400 | 2679500 | 756960 | 2101300 | 0 | 0 | 0 | 102810 | 0 | 0 | 0 | 0 | 57169 | 0 | 1350100 | 2001500 | 0 | 0 |
| MTVPKEMP | ATP8B2 | P98198 | AT8B2_HUMAN | Acetyl (Protein N | 180280 | 0 | 0 | 0 | 717510 | 0 | 115890 | 0 | 0 | 0 | 0 | 709590 | 0 | 0 | 448480 | 628550 | 0 | 0 |
| MTVTDQVN | APOD; APOD | P05090; C9J | APOD_HUMAN | Oxidation (M) | 0 | 0 | 0 | 0 | 0 | 4288400 | 0 | 0 | 648020 | 170680 | 0 | 0 | 0 | 0 | 0 | 130410 | 0 | 0 |
| MTVTDQVN | APOD; APOD | P05090; C9J | APOD_HUMAN | Unmodified | 0 | 0 | 0 | 0 | 0 | 0 | 0 | 0 | 911700 | 201630 | 0 | 0 | 0 | 0 | 0 | 0 | 0 | 54791 |
| MTVTDQVN | APOD | P05090 | APOD_HUMAN | Unmodified | 0 | 0 | 0 | 0 | 0 | 0 | 0 | 81283 | 212690 | 452100 | 0 | 0 | 0 | 0 | 0 | 0 | 0 | 0 |
| MTVVQAYI | ATP2B4 | P23634 | AT2B4_HUMAN | Oxidation (M) | 0 | 73331 | 0 | 0 | 304510 | 0 | 144020 | 40979 | 79899 | 0 | 467080 | 1024500 | 0 | 0 | 0 | 0 | 0 | 100150 |
| MTVVQAYI | ATP2B1; ATP2B | P20020; E7E | AT2B1_HUMAN | Unmodified | 164120 | 0 | 0 | 661170 | 4851900 | 0 | 399720 | 0 | 2158300 | 1371800 | 987680 | 1397900 | 2912800 | 900940 | 2468800 | 0 | 947720 | 1293300 |
| MTVVQAYI | ATP2B1; ATP2B | P20020; E7E | AT2B1_HUMAN | Oxidation (M) | 0 | 109150 | 454900 | 471390 | 2074900 | 668930 | 3052800 | 83577 | 0 | 2000800 | 0 | 483170 | 277730 | 0 | 1074700 | 664590 | 970780 | 1109100 |
| MVAAATNN | TLN1 | Q9Y490 | TLN1_HUMAN | Oxidation (M) | 494950 | 0 | 0 | 1303600 | 2044500 | 511460 | 378690 | 311260 | 0 | 0 | 0 | 1722400 | 1304200 | 1302000 | 0 | 566150 | 680720 | 358890 |
| MVAAATNN | TLN1 | Q9Y490 | TLN1_HUMAN | Unmodified | 432520 | 0 | 0 | 1306600 | 6513800 | 0 | 1093400 | 379820 | 1402700 | 579920 | 280380 | 1819900 | 2491700 | 4142100 | 0 | 456180 | 1221100 | 2852000 |
| MVADLVEN | HDLBP; HDLBP | Q00341; A0A | VIGLN_HUMAN | Oxidation (M) | 0 | 0 | 0 | 0 | 1122300 | 222560 | 167470 | 0 | 0 | 0 | 0 | 0 | 0 | 694330 | 0 | 0 | 344990 | 278260 |
| MVADPPRD | SLC1A5 | Q15758 | AAAT_HUMAN | Acetyl (Protein N | 0 | 0 | 262130 | 281890 | 1520500 | 380070 | 94745 | 29954 | 490930 | 325660 | 1002800 | 4560400 | 1211000 | 573480 | 471120 | 692010 | 271690 | 1809800 |
| MVADPPRD | SLC1A5 | Q15758 | AAAT_HUMAN | Acetyl (Protein N | 88912 | 81339 | 0 | 0 | 1098500 | 0 | 0 | 0 | 378080 | 221460 | 0 | 3049400 | 1590000 | 1058900 | 281400 | 255690 | 291520 | 251280 |
| MVADTISR | VIL1; VIL1; VIL1 | P09327; H7C | VILI_HUMAN Vi | Unmodified | 72458 | 0 | 0 | 0 | 79712 | 0 | 0 | 12069 | 624410 | 0 | 662870 | 2234200 | 0 | 0 | 0 | 0 | 0 | 0 |
| MVADTISRI | VIL1; VIL1; VIL1 | P09327; H7C | VILI_HUMAN Vi | Unmodified | 432750 | 0 | 0 | 0 | 0 | 0 | 0 | 0 | 2849500 | 1481300 | 1480500 | 937660 | 0 | 0 | 46535 | 0 | 0 | 0 |
| MVADTISRI | VIL1; VIL1; VIL1 | P09327; H7C | VILI_HUMAN Vi | Oxidation (M) | 0 | 0 | 0 | 0 | 0 | 377950 | 0 | 0 | 0 | 347790 | 12265 | 0 | 0 | 0 | 0 | 0 | 0 | 0 |
| MVALSTSR | QTRT1 | Q9BXR0 | TGT_HUMAN Q | Unmodified | 0 | 0 | 0 | 0 | 0 | 0 | 0 | 0 | 0 | 0 | 0 | 0 | 0 | 0 | 0 | 0 | 0 | 0 |
| MVCEQKLL | CRABP2 | P29373 | RABP2_HUMAN | Unmodified | 0 | 0 | 0 | 0 | 0 | 0 | 199350 | 0 | 688200 | 0 | 0 | 0 | 0 | 0 | 0 | 0 | 375890 | 0 |
| MVDDGNG | AVIL | O75366 | AVIL_HUMAN A | Unmodified | 0 | 0 | 0 | 0 | 0 | 0 | 0 | 0 | 0 | 0 | 347180 | 21474000 | 0 | 0 | 0 | 0 | 0 | 0 |
| MVDDGNG | AVIL | O75366 | AVIL_HUMAN A | Oxidation (M) | 0 | 0 | 0 | 0 | 0 | 0 | 0 | 0 | 0 | 0 | 201740 | 8381100 | 0 | 0 | 0 | 0 | 0 | 0 |
| MVDDGSG | VIL1; VIL1 | P09327; B4D | VILI_HUMAN Vi | Oxidation (M) | 0 | 0 | 0 | 0 | 0 | 0 | 0 | 0 | 0 | 0 | 0 | 1962000 | 0 | 0 | 0 | 0 | 0 | 0 |
| MVDDGSG | VIL1; VIL1 | P09327; B4D | VILI_HUMAN Vi | Unmodified | 0 | 0 | 0 | 0 | 0 | 0 | 0 | 0 | 0 | 0 | 0 | 1171700 | 0 | 0 | 0 | 0 | 0 | 0 |
| MVDDGSG | VIL1; VIL1 | P09327; B4D | VILI_HUMAN Vi | Oxidation (M) | 0 | 0 | 0 | 0 | 0 | 0 | 0 | 0 | 352180 | 377530 | 0 | 2412100 | 0 | 0 | 0 | 0 | 0 | 0 |
| MVDELKRA | ITGB4 | P16144 | ITB4_HUMAN I | Unmodified | 0 | 0 | 0 | 0 | 494760 | 0 | 0 | 0 | 0 | 0 | 0 | 0 | 0 | 0 | 0 | 0 | 2906700 | 0 |
| MVDEWAQ | N/A | N/A | N/A | Unmodified | 0 | 0 | 0 | 0 | 0 | 0 | 0 | 0 | 0 | 0 | 0 | 0 | 0 | 0 | 0 | 0 | 0 | 923980 |
| MVDNRGFI | SERPINH1; SER | P50454; E9P | SERPH_HUMAI | Unmodified | 0 | 0 | 0 | 0 | 202890 | 0 | 72219 | 20655 | 3325000 | 4786600 | 0 | 0 | 0 | 143130 | 0 | 0 | 0 | 0 |
| MVDNRGFI | SERPINH1; SER | P50454; E9P | SERPH_HUMAI | Oxidation (M) | 0 | 0 | 0 | 20411 | 310560 | 0 | 170580 | 63688 | 743460 | 2417900 | 0 | 0 | 67001 | 400870 | 0 | 0 | 0 | 170940 |
| MVDREQLV | YWHAG | P61981 | 1433G_HUMAN | Acetyl (Protein N | 0 | 0 | 0 | 0 | 212970 | 0 | 0 | 0 | 0 | 0 | 0 | 0 | 140320 | 194450 | 0 | 0 | 431140 | 356170 |
| MVDREQLV | YWHAG | P61981 | 1433G_HUMAN | Unmodified | 0 | 0 | 408250 | 826660 | 2751300 | 888700 | 304000 | 0 | 304470 | 403520 | 577080 | 113760 | 1969300 | 1792800 | 1398900 | 1431900 | 4504500 | 6044200 |
| MVDVGGQ | GNAQ; GNAZ; ( | P50148; P19 | GNAQ_HUMAN | Unmodified | 1083800 | 1509100 | 0 | 0 | 284560 | 133590 | 87521 | 69836 | 0 | 0 | 495510 | 27283000 | 0 | 0 | 329020 | 354500 | 128940 | 413810 |
| MVDVGGQ | GNAQ; GNAZ; ( | P50148; P19 | GNAQ_HUMAN | Unmodified | 0 | 0 | 0 | 0 | 0 | 40931 | 0 | 0 | 1160300 | 0 | 40300 | 24384 | 0 | 0 | 0 | 0 | 236030 | 316580 |
| MVEDCEHI | GLG1; GLG1 | Q92896; H3I | GSLG1_HUMAI | Unmodified | 0 | 0 | 0 | 0 | 0 | 0 | 0 | 0 | 213810 | 0 | 0 | 0 | 0 | 0 | 0 | 0 | 0 | 0 |
| MVEPENAV | SCRIB; SCRIB; ! | Q14160; H0' | SCRIB_HUMAN | Oxidation (M) | 0 | 0 | 0 | 0 | 0 | 0 | 0 | 0 | 2537700 | 0 | 1104600 | 1198800 | 279870 | 0 | 0 | 0 | 972240 | 0 |
| MVEPENAV | SCRIB; SCRIB; ! | Q14160; H0' | SCRIB_HUMAN | Unmodified | 0 | 0 | 0 | 0 | 0 | 0 | 0 | 0 | 6371100 | 0 | 0 | 320950 | 0 | 0 | 206190 | 0 | 0 | 0 |
| MVEPENAV | SCRIB; SCRIB; ! | Q14160; H0' | SCRIB_HUMAN | Unmodified | 0 | 0 | 0 | 0 | 1150600 | 0 | 0 | 0 | 0 | 0 | 0 | 0 | 997270 | 0 | 0 | 0 | 1084800 | 0 |
| MVEPENAV | SCRIB; SCRIB; ! | Q14160; H0' | SCRIB_HUMAN | Oxidation (M) | 0 | 0 | 0 | 0 | 0 | 0 | 0 | 0 | 0 | 1473400 | 0 | 0 | 796450 | 0 | 0 | 0 | 1962700 | 2375200 |
| MVEPQYQE | PIR | O00625 | PIR_HUMAN Pi | Oxidation (M) | 0 | 0 | 0 | 0 | 0 | 0 | 0 | 0 | 0 | 49894 | 482930 | 1586300 | 0 | 0 | 0 | 0 | 0 | 0 |
| MVEPQYQE | PIR | O00625 | PIR_HUMAN Pi | Unmodified | 0 | 0 | 0 | 80101 | 0 | 0 | 0 | 0 | 181390 | 0 | 921600 | 2993400 | 0 | 0 | 0 | 0 | 0 | 0 |
| MVEPQYQE | PIR | O00625 | PIR_HUMAN Pi | Unmodified | 0 | 0 | 0 | 0 | 0 | 0 | 0 | 0 | 0 | 0 | 864010 | 2071900 | 200510 | 0 | 0 | 0 | 0 | 0 |
| MVFFVQNE | KIAA0319L; KIA | Q8IZA0; C9J! | K319L_HUMAN | Oxidation (M) | 111580 | 0 | 196140 | 300930 | 0 | 0 | 0 | 0 | 1249600 | 304700 | 40299 | 0 | 0 | 201870 | 363100 | 199790 | 0 | 0 |
| MVFFVQNE | KIAA0319L; KIA | Q8IZA0; C9J! | K319L_HUMAN | Unmodified | 0 | 0 | 0 | 0 | 0 | 667420 | 0 | 0 | 5546500 | 0 | 0 | 0 | 0 | 0 | 0 | 0 | 0 | 0 |
| MVFINNIAL | LRPPRC | P42704 | LPPRC_HUMAI | Unmodified | 0 | 0 | 0 | 0 | 0 | 0 | 0 | 0 | 2284400 | 416490 | 0 | 0 | 0 | 0 | 0 | 0 | 0 | 0 |
| MVGCTGK\ | PCMT1; PCMT1 | P22061; F6S | PIMT_HUMAN I | Unmodified | 0 | 0 | 0 | 0 | 404330 | 0 | 0 | 0 | 0 | 361790 | 0 | 0 | 0 | 190980 | 0 | 0 | 172350 | 247750 |
| MVGFDMSI | SCP2; SCP2; S( | P22307; H0' | SCP2_HUMAN | Unmodified | 0 | 0 | 0 | 0 | 0 | 0 | 0 | 0 | 1954400 | 1046400 | 0 | 0 | 0 | 0 | 0 | 0 | 0 | 0 |
| MVGFIIGR | FUBP1; FUBP3; | Q96AE4; Q9( | FUBP1_HUMAI | Unmodified | 0 | 0 | 0 | 0 | 0 | 0 | 0 | 0 | 143750 | 0 | 12528 | 431580 | 0 | 0 | 0 | 0 | 0 | 0 |
| MVGGIAQII | TLN1 | Q9Y490 | TLN1_HUMAN | 2 Oxidation (M) | 914580 | 1605100 | 1755700 | 1919100 | 2613600 | 821720 | 1051500 | 55180 | 529710 | 221170 | 228350 | 2376700 | 2076500 | 1706800 | 1316700 | 2148600 | 360520 | 1327500 |
| MVGGIAQII | TLN1 | Q9Y490 | TLN1_HUMAN | Oxidation (M) | 0 | 0 | 101070 | 0 | 1496800 | 0 | 163180 | 0 | 519220 | 0 | 0 | 0 | 275000 | 225370 | 0 | 0 | 115540 | 422710 |
| MVGGIAQII | TLN1 | Q9Y490 | TLN1_HUMAN | Unmodified | 0 | 0 | 0 | 0 | 68199 | 0 | 0 | 0 | 145960 | 0 | 0 | 465670 | 0 | 0 | 0 | 0 | 0 | 50477 |
| MVGGVLVE | PFDN2 | Q9UHV9 | PFD2_HUMAN | Unmodified | 0 | 0 | 0 | 0 | 0 | 0 | 0 | 0 | 1993600 | 0 | 2239000 | 11695000 | 0 | 0 | 377670 | 0 | 0 | 0 |
| MVGGVLVE | PFDN2 | Q9UHV9 | PFD2_HUMAN | Oxidation (M) | 0 | 0 | 0 | 244440 | 173400 | 0 | 0 | 0 | 199120 | 103950 | 529540 | 2519600 | 323840 | 66493 | 0 | 185960 | 54324 | 77118 |
| MVGGVLVE | PFDN2 | Q9UHV9 | PFD2_HUMAN | Unmodified | 0 | 0 | 0 | 0 | 1418000 | 0 | 173120 | 119670 | 0 | 515720 | 0 | 0 | 1523000 | 792250 | 0 | 0 | 803020 | 1379600 |
| MVGGVLVE | PFDN2 | Q9UHV9 | PFD2_HUMAN | Oxidation (M) | 0 | 0 | 0 | 0 | 409060 | 0 | 0 | 0 | 0 | 342020 | 0 | 0 | 410000 | 0 | 0 | 0 | 246450 | 0 |
| MVGMGQK | ACTG1; ACTC1; | P63261; P68 | ACTG_HUMAN | Oxidation (M) | 0 | 0 | 0 | 0 | 0 | 0 | 22914 | 0 | 0 | 0 | 0 | 0 | 0 | 0 | 0 | 0 | 155330 | 0 |
| MVGMGQK | ACTG1; ACTC1; | P63261; P68 | ACTG_HUMAN | Unmodified | 0 | 0 | 0 | 0 | 241230 | 0 | 0 | 0 | 0 | 0 | 0 | 0 | 0 | 0 | 0 | 0 | 383870 | 0 |
| MVHGNITP | SCYL2; SCYL2; | Q6P3W7; A0 | SCYL2_HUMAN | Unmodified | 0 | 0 | 0 | 0 | 277300 | 0 | 0 | 16385 | 907180 | 175990 | 0 | 944940 | 56269 | 0 | 0 | 0 | 0 | 0 |
| MVILLCNLI | AIMP1; AIMP1 | Q12904; A0A | AIMP1_HUMAN | Unmodified | 7399500 | 809530 | 0 | 436680 | 620600 | 66368 | 0 | 21841 | 217260 | 225570 | 243040 | 723030 | 1074500 | 1299800 | 0 | 0 | 304990 | 197490 |
| MVILLCNLI | AIMP1; AIMP1 | Q12904; A0A | AIMP1_HUMAN | Oxidation (M) | 0 | 0 | 71412 | 0 | 286240 | 119580 | 63483 | 40253 | 0 | 290080 | 380120 | 1157700 | 854910 | 366030 | 99161 | 79194 | 0 | 116140 |
| MVINHLEK | CCT8; CCT8 | P50990; H7C | TCPQ_HUMAN | Unmodified | 0 | 0 | 0 | 0 | 0 | 0 | 64749 | 0 | 166830 | 132630 | 0 | 1040100 | 0 | 227260 | 0 | 89661 | 0 | 0 |
| MVIPGGID\ | DPYSL3; DPYSL | Q14195; D6I | DPYL3_HUMAN | Oxidation (M) | 0 | 0 | 25142 | 0 | 1740900 | 177270 | 646480 | 38032 | 0 | 0 | 4611600 | 22897000 | 535450 | 116520 | 66673 | 72818 | 0 | 297110 |
| MVIPGGID\ | DPYSL3; DPYSL | Q14195; D6I | DPYL3_HUMAN | Unmodified | 0 | 0 | 0 | 0 | 1888800 | 0 | 417680 | 0 | 0 | 0 | 713430 | 2991000 | 0 | 0 | 0 | 0 | 212220 | 54633 |
| MVIPGGID\ | DPYSL3; DPYSL | Q14195; D6I | DPYL3_HUMAN | 2 Oxidation (M) | 0 | 0 | 0 | 0 | 297070 | 0 | 266380 | 0 | 0 | 0 | 5245200 | 19214000 | 0 | 0 | 0 | 0 | 307770 | 304630 |
| MVIPGGID\ | DPYSL2; DPYSL | Q16555; E5F | DPYL2_HUMAN | Oxidation (M) | 0 | 0 | 0 | 0 | 0 | 0 | 0 | 0 | 0 | 0 | 0 | 1942600 | 0 | 0 | 0 | 0 | 0 | 0 |
| MVIPGGID\ | DPYSL2; DPYSL | Q16555; E5F | DPYL2_HUMAN | Unmodified | 213000 | 380380 | 0 | 0 | 0 | 0 | 0 | 0 | 0 | 0 | 0 | 5588300 | 0 | 0 | 235980 | 281300 | 0 | 0 |
| MVIPGGID\ | DPYSL2; DPYSL | Q16555; E5F | DPYL2_HUMAN | 3 Oxidation (M) | 0 | 0 | 0 | 0 | 2518100 | 0 | 0 | 0 | 0 | 0 | 0 | 0 | 0 | 0 | 0 | 0 | 0 | 0 |
| MVKLAKAG | NCL; NCL; NCL | P19338; A0A | NUCL_HUMAN | Oxidation (M) | 0 | 0 | 0 | 0 | 0 | 0 | 0 | 0 | 0 | 0 | 0 | 0 | 0 | 0 | 0 | 0 | 0 | 209780 |
| MVLAAAGG | UQCRC1 | P31930 | QCR1_HUMAN | Unmodified | 0 | 0 | 0 | 0 | 116120 | 0 | 0 | 0 | 17495000 | 3876100 | 0 | 0 | 0 | 0 | 0 | 0 | 0 | 0 |
| MVLAAAGG | UQCRC1 | P31930 | QCR1_HUMAN | Oxidation (M) | 0 | 0 | 0 | 0 | 0 | 0 | 0 | 52515 | 16129000 | 12571000 | 1128700 | 652120 | 0 | 0 | 0 | 0 | 0 | 0 |
| MVLSGCAII | SND1 | Q7KZF4 | SND1_HUMAN | Oxidation (M) | 0 | 0 | 0 | 0 | 0 | 0 | 0 | 119400 | 0 | 0 | 0 | 834000 | 0 | 0 | 0 | 0 | 0 | 0 |
| MVLVGDVI | PRPS2; PRPS1; | P11908; P60 | PRPS2_HUMAN | Oxidation (M) | 0 | 17421 | 424570 | 431710 | 3007100 | 302840 | 0 | 0 | 265560 | 0 | 43924 | 1718700 | 650590 | 0 | 0 | 0 | 66512 | 511740 |
| MVMAEQSI | TCF4 | A0A0E3D6N | A0A0E3D6N2_I | Acetyl (Protein N | 0 | 0 | 0 | 0 | 0 | 0 | 0 | 0 | 0 | 0 | 0 | 0 | 0 | 0 | 0 | 0 | 0 | 2967200 |
| MVMETIEK | SF3B1 | O75533 | SF3B1_HUMAN | Unmodified | 0 | 0 | 0 | 0 | 0 | 0 | 32551 | 0 | 194720 | 15706 | 106330 | 2174400 | 0 | 0 | 0 | 0 | 0 | 0 |
| MVMIQDGI | FUBP1; FUBP1 | Q96AE4; E9F | FUBP1_HUMAI | Unmodified | 0 | 0 | 0 | 0 | 0 | 0 | 0 | 0 | 0 | 0 | 0 | 1438600 | 189380 | 0 | 0 | 0 | 0 | 0 |
| MVMNNGL | PLCB4; PLCB4 | Q15147; A0A | PLCB4_HUMAN | 2 Oxidation (M) | 0 | 0 | 0 | 0 | 0 | 0 | 0 | 0 | 0 | 431960 | 333720 | 430500 | 0 | 0 | 0 | 623780 | 0 | 0 |
| MVMNNGL | PLCB4; PLCB4 | Q15147; A0A | PLCB4_HUMAN | 2 Oxidation (M) | 0 | 0 | 0 | 0 | 0 | 0 | 0 | 0 | 0 | 0 | 0 | 0 | 0 | 0 | 0 | 0 | 0 | 0 |
| MVNDAEKI | HSPA5; HSPA5; | P11021; A0A | BIP_HUMAN Er | Unmodified | 0 | 309250 | 0 | 0 | 0 | 0 | 0 | 0 | 15073000 | 2177700 | 373110 | 689760 | 0 | 0 | 181440 | 0 | 0 | 0 |
| MVNDAEKI | HSPA5; HSPA5; | P11021; A0A | BIP_HUMAN Er | Oxidation (M) | 0 | 0 | 0 | 0 | 0 | 0 | 0 | 0 | 3448700 | 352490 | 37000 | 528340 | 0 | 0 | 0 | 0 | 0 | 0 |
| MVNDAEKI | HSPA5; HSPA5; | P11021; A0A | BIP_HUMAN Er | Unmodified | 0 | 0 | 0 | 0 | 0 | 0 | 0 | 0 | 4381700 | 552370 | 0 | 0 | 0 | 0 | 0 | 0 | 0 | 0 |
| MVNHFIAE | HSPA8; HSPA8; | P11142; E9P | HSP7C_HUMAI | Oxidation (M) | 19217000 | 19822000 | 20442000 | 20551000 | 5214100 | 1643900 | 588550 | 152250 | 16812000 | 1948500 | 6782700 | 41158000 | 6330800 | 2530800 | 16534000 | 15081000 | 0 | 2234000 |
| MVNHFIAE | HSPA8; HSPA8; | P11142; E9P | HSP7C_HUMAI | Unmodified | 12615000 | 1619500 | 5868600 | 14167000 | 7755500 | 0 | 0 | 0 | 58395000 | 2115600 | 1230700 | 63225000 | 5554600 | 1788100 | 10300000 | 5613900 | 237080 | 376100 |
| MVNHFIAE | HSPA8; HSPA8; | P11142; E9P | HSP7C_HUMAI | Oxidation (M) | 1154300 | 936920 | 0 | 1140900 | 0 | 85812 | 0 | 0 | 1189700 | 113580 | 0 | 1959900 | 0 | 0 | 949020 | 0 | 0 | 0 |
| MVNHFIAE | HSPA8; HSPA8; | P11142; E9P | HSP7C_HUMAI | Unmodified | 4351900 | 3671900 | 904750 | 1740600 | 0 | 0 | 0 | 0 | 3622900 | 121360 | 0 | 10378000 | 409690 | 0 | 1967600 | 909470 | 0 | 0 |
| MVNYIQMK | TWF2; TWF2; T\ | Q6IBS0; D6F | TWF2_HUMAN | Oxidation (M) | 152360 | 17072 | 0 | 0 | 0 | 0 | 0 | 0 | 2895400 | 187260 | 374760 | 1377400 | 0 | 0 | 0 | 0 | 0 | 0 |
| MVNYIQMK | TWF2; TWF2; T\ | Q6IBS0; D6F | TWF2_HUMAN | Unmodified | 0 | 0 | 0 | 0 | 0 | 503860 | 0 | 0 | 254590 | 0 | 0 | 1967800 | 0 | 0 | 0 | 0 | 40884 | 649380 |
| MVPEDIKP( | MOV10; MOV1( | Q9HCE1; Q5 | MOV10_HUMA | Oxidation (M) | 855240 | 0 | 128570 | 249860 | 0 | 0 | 0 | 0 | 0 | 0 | 0 | 0 | 0 | 0 | 0 | 283800 | 0 | 0 |
| MVPEDIKP( | MOV10; MOV1( | Q9HCE1; Q5 | MOV10_HUMA | Unmodified | 735510 | 340550 | 0 | 296720 | 0 | 0 | 0 | 0 | 0 | 0 | 0 | 0 | 0 | 0 | 0 | 298050 | 0 | 0 |
| MVPKPEEIN | CLPTM1L; CLPT | Q96KA5; G5I | CLP1L_HUMAN | Oxidation (M) | 0 | 0 | 0 | 0 | 0 | 0 | 0 | 0 | 1263500 | 0 | 0 | 0 | 0 | 0 | 0 | 0 | 0 | 0 |
| MVPKPEEIN | CLPTM1L; CLPT | Q96KA5; G5I | CLP1L_HUMAN | Unmodified | 0 | 0 | 0 | 0 | 0 | 0 | 0 | 0 | 1263100 | 0 | 0 | 0 | 0 | 0 | 0 | 0 | 0 | 0 |
| MVPLLNK | JUP; JUP; JUP | P14923; C9J | PLAK_HUMAN . | Oxidation (M) | 378720 | 0 | 2098300 | 638700 | 0 | 138510 | 0 | 0 | 897470 | 76086 | 514400 | 2508400 | 1443600 | 0 | 2060900 | 2741100 | 207830 | 284630 |
| MVPPVQVS | ATP5ME | P56385 | ATP5I_HUMAN | Oxidation (M) | 0 | 0 | 0 | 0 | 0 | 0 | 0 | 0 | 1739700 | 539320 | 0 | 0 | 0 | 0 | 0 | 0 | 0 | 0 |
| MVPPVQVS | ATP5ME | P56385 | ATP5I_HUMAN | Unmodified | 0 | 0 | 0 | 0 | 0 | 0 | 0 | 0 | 3946200 | 723730 | 0 | 0 | 0 | 0 | 0 | 0 | 0 | 0 |
| MVPSSPAV | SLC19A1; SLC1 | P41440; C9J | S19A1_HUMAN | Acetyl (Protein N | 0 | 0 | 0 | 0 | 1542300 | 0 | 0 | 0 | 760570 | 107460 | 339740 | 0 | 0 | 0 | 0 | 0 | 0 | 462520 |
| MVPSSPAV | SLC19A1; SLC1 | P41440; C9J | S19A1_HUMAN | Acetyl (Protein N | 0 | 0 | 177080 | 614940 | 0 | 0 | 0 | 0 | 0 | 0 | 0 | 0 | 0 | 0 | 223290 | 410980 | 0 | 0 |
| MVPVSVQ( | PDCD6IP | Q8WUM4 | PDC6I_HUMAN | Oxidation (M) | 17964000 | 13414000 | 6518000 | 7686000 | 3583500 | 1918000 | 0 | 206020 | 2900500 | 243800 | 1715900 | 4885000 | 6465100 | 2172100 | 15003000 | 10336000 | 0 | 2062100 |
| MVPVSVQ( | PDCD6IP | Q8WUM4 | PDC6I_HUMAN | Unmodified | 7025900 | 5388600 | 1918000 | 4222000 | 9297400 | 1112700 | 0 | 0 | 9064000 | 2596200 | 824900 | 2936500 | 7656500 | 2461300 | 7603900 | 2753300 | 0 | 3806500 |
| MVPVSVQ( | PDCD6IP | Q8WUM4 | PDC6I_HUMAN | Unmodified | 6386400 | 9242500 | 4885400 | 10725000 | 7044800 | 1219500 | 0 | 0 | 17511000 | 2965000 | 888950 | 2653600 | 8573900 | 9226000 | 3785100 | 1353600 | 645970 | 3329000 |
| MVPVSVQ( | PDCD6IP | Q8WUM4 | PDC6I_HUMAN | Oxidation (M) | 9521700 | 8975200 | 6787400 | 6538000 | 2890700 | 0 | 239370 | 0 | 3898600 | 1486800 | 0 | 2554900 | 3277200 | 2548000 | 6383400 | 3870300 | 264040 | 786230 |
| MVPVSVQ( | PDCD6IP | Q8WUM4 | PDC6I_HUMAN | Oxidation (M) | 0 | 0 | 0 | 0 | 0 | 0 | 0 | 0 | 0 | 3395700 | 0 | 0 | 14589000 | 5557700 | 0 | 0 | 0 | 2723800 |
| MVPVSVQ( | PDCD6IP | Q8WUM4 | PDC6I_HUMAN | Unmodified | 0 | 0 | 0 | 0 | 0 | 0 | 0 | 0 | 0 | 5168700 | 0 | 0 | 16052000 | 0 | 0 | 0 | 0 | 0 |
| MVPYRDSK | KIF23; KIF23; KI | Q02241; A0A | KIF23_HUMAN | Unmodified | 0 | 0 | 0 | 0 | 21851000 | 0 | 0 | 0 | 0 | 1471100 | 0 | 0 | 0 | 0 | 0 | 0 | 2211300 | 0 |
| MVQDEAD( | SNX33; SNX33 | Q8WV41; H3 | SNX33_HUMAN | Unmodified | 0 | 532260 | 2612600 | 1649700 | 0 | 0 | 0 | 0 | 0 | 0 | 231960 | 0 | 0 | 0 | 7040800 | 2835100 | 0 | 0 |
| MVQDEAD( | SNX33; SNX33 | Q8WV41; H3 | SNX33_HUMAN | Oxidation (M) | 0 | 0 | 1011400 | 0 | 0 | 0 | 0 | 0 | 225070 | 0 | 0 | 212620 | 0 | 0 | 2516800 | 1264800 | 0 | 0 |
| MVQDEAD( | SNX33; SNX33 | Q8WV41; H3 | SNX33_HUMAN | Unmodified | 0 | 0 | 600470 | 767750 | 0 | 0 | 0 | 0 | 0 | 0 | 0 | 0 | 0 | 0 | 0 | 0 | 0 | 0 |
| MVQEAEKY | HSPA8; HSPA8; | P11142; E9P | HSP7C_HUMAI | Unmodified | 0 | 0 | 0 | 0 | 0 | 0 | 0 | 0 | 0 | 1035500 | 0 | 0 | 0 | 0 | 91675 | 0 | 228140 | 0 |
| MVQEAEKY | HSPA1A; HSPA | P0DMV8; P0 | HS71A_HUMAI | Unmodified | 201040 | 0 | 0 | 0 | 0 | 0 | 0 | 0 | 0 | 0 | 0 | 0 | 0 | 0 | 0 | 54080 | 0 | 0 |
| MVQFEEN( | CCT3; CCT3; C( | P49368; Q5! | TCPG_HUMAN | Unmodified | 0 | 0 | 0 | 0 | 0 | 83615 | 0 | 0 | 0 | 0 | 941420 | 13521000 | 467270 | 0 | 0 | 0 | 128660 | 278550 |
| MVQFEEN( | CCT3; CCT3; C( | P49368; Q5! | TCPG_HUMAN | Oxidation (M) | 0 | 0 | 0 | 0 | 0 | 0 | 0 | 0 | 0 | 0 | 0 | 1104400 | 142700 | 0 | 0 | 0 | 0 | 0 |
| MVQQLGK/ | ABCC4 | O15439 | MRP4_HUMAN | Unmodified | 0 | 0 | 0 | 0 | 2063400 | 0 | 853290 | 0 | 0 | 0 | 0 | 0 | 829020 | 610230 | 0 | 0 | 368120 | 0 |

| | | | | | | | | | | | | | | | | | | | | | | |
|---|---|---|---|---|---|---|---|---|---|---|---|---|---|---|---|---|---|---|---|---|---|---|
| MVRKPVVS | SLC7A11 | Q9UPY5 | XCT_HUMAN C | Acetyl (Protein N | 0 | 0 | 0 | 191030 | 1458000 | 159580 | 0 | 0 | 196680 | 146160 | 0 | 0 | 1243500 | 1038900 | 0 | 0 | 154610 | 344090 |
| MVSADAYK | CNP; CNP; CNI | P09543; K7E | CN37_HUMAN | Unmodified | 889690 | 1114600 | 0 | 2723200 | 14485000 | 2445500 | 716510 | 685130 | 4653900 | 1193500 | 6996500 | 27786000 | 0 | 3732600 | 2328200 | 0 | 2811400 | 10527000 |
| MVSADAYK | CNP; CNP; CNI | P09543; K7E | CN37_HUMAN | Unmodified | 0 | 0 | 178470 | 0 | 3208100 | 127150 | 332980 | 175330 | 0 | 689700 | 31761 | 0 | 2635600 | 2307300 | 0 | 0 | 1708500 | 5259200 |
| MVSADAYK | CNP; CNP; CNI | P09543; K7E | CN37_HUMAN | Oxidation (M) | 0 | 0 | 0 | 0 | 970580 | 121170 | 80989 | 0 | 0 | 395230 | 0 | 0 | 1630400 | 0 | 0 | 0 | 684990 | 891420 |
| MVSADAYK | CNP; CNP; CNI | P09543; K7E | CN37_HUMAN | Unmodified | 0 | 0 | 0 | 0 | 4686100 | 348300 | 875100 | 445510 | 0 | 623550 | 0 | 0 | 1340900 | 3296900 | 0 | 0 | 3020500 | 3329400 |
| MVSADAYK | CNP; CNP; CNI | P09543; K7E | CN37_HUMAN | Oxidation (M) | 0 | 0 | 0 | 196860 | 1919000 | 118220 | 621340 | 663780 | 0 | 0 | 0 | 0 | 1592900 | 4108200 | 0 | 0 | 2099900 | 0 |
| MVSDINNA | ACTN1; ACTN1 | P12814; A0A | ACTN1_HUMAI | Oxidation (M) | 0 | 0 | 0 | 0 | 0 | 0 | 0 | 0 | 0 | 0 | 0 | 1378800 | 0 | 0 | 150300 | 77037 | 208730 | 0 |
| MVSDINNG | ACTN4; ACTN4 | O43707; F5C | ACTN4_HUMAI | Unmodified | 312180 | 694000 | 361440 | 154000 | 12236000 | 3429300 | 122750 | 0 | 5288000 | 2445500 | 2183300 | 9845700 | 495320 | 281050 | 969610 | 210490 | 1663800 | 0 |
| MVSDINNG | ACTN4; ACTN4 | O43707; F5C | ACTN4_HUMAI | Oxidation (M) | 478160 | 334280 | 769630 | 696870 | 4015500 | 3516500 | 0 | 516880 | 1712400 | 1476300 | 1179700 | 10842000 | 771240 | 0 | 1596200 | 2069100 | 849670 | 181610 |
| MVSEIYLTR | PLXNA1; PLXNA | Q9UIW2; O7 | PLXA1_HUMAN | Oxidation (M) | 0 | 0 | 0 | 0 | 0 | 0 | 0 | 0 | 0 | 0 | 0 | 0 | 0 | 0 | 0 | 0 | 0 | 0 |
| MVSHFYGN | ERLEC1 | Q96DZ1 | ERLEC_HUMAN | Unmodified | 0 | 0 | 0 | 0 | 0 | 0 | 0 | 0 | 2654800 | 0 | 0 | 0 | 0 | 0 | 0 | 0 | 0 | 0 |
| MVSVLPIFF | NHLRC2 | Q8NBF2 | NHLC2_HUMA | Oxidation (M) | 0 | 0 | 26646 | 0 | 0 | 0 | 0 | 0 | 0 | 0 | 52066 | 642750 | 0 | 0 | 0 | 522910 | 0 | 0 |
| MVSVLPIFF | NHLRC2 | Q8NBF2 | NHLC2_HUMA | Oxidation (M) | 0 | 0 | 0 | 0 | 2114000 | 277030 | 3167700 | 0 | 246980 | 790850 | 313660 | 0 | 5214700 | 2166100 | 0 | 55012 | 2253400 | 2501600 |
| MVTGDNIN | ATP2B2; ATP2B | Q01814; Q1( | AT2B2_HUMAN | Oxidation (M) | 243450 | 253460 | 0 | 1168700 | 1424300 | 267750 | 231460 | 305130 | 347790 | 237350 | 548460 | 2663200 | 2700100 | 1302300 | 1442200 | 1010200 | 787450 | 735320 |
| MVTGDNIN | ATP2B2; ATP2B | Q01814; Q1( | AT2B2_HUMAN | Unmodified | 0 | 0 | 0 | 0 | 10974000 | 1624300 | 2686200 | 1036400 | 4556200 | 2453600 | 6281500 | 24647000 | 52717000 | 14181000 | 0 | 7081300 | 3538500 | 5399400 |
| MVTGDNIN | ATP2B3 | Q16720 | AT2B3_HUMAN | Oxidation (M) | 0 | 0 | 75696 | 0 | 253020 | 0 | 323810 | 207050 | 0 | 0 | 0 | 0 | 0 | 0 | 0 | 0 | 523210 | 0 |
| MVTGDNIN | ATP2B3 | Q16720 | AT2B3_HUMAN | Unmodified | 0 | 0 | 0 | 0 | 0 | 0 | 1131000 | 231260 | 0 | 0 | 0 | 0 | 0 | 0 | 0 | 0 | 132530 | 0 |
| MVTGDNIN | ATP2B2; ATP2B | Q01814; A0/ | AT2B2_HUMAN | Unmodified | 0 | 0 | 0 | 0 | 0 | 0 | 643350 | 0 | 0 | 0 | 0 | 0 | 0 | 0 | 0 | 0 | 101060 | 0 |
| MVTGDNIN | ATP2B4; ATP2B | P23634; P20 | AT2B4_HUMAN | Oxidation (M) | 0 | 0 | 0 | 0 | 4300800 | 135360 | 2116500 | 986110 | 0 | 294410 | 0 | 0 | 4963500 | 6202500 | 0 | 0 | 1514900 | 2904500 |
| MVTGDNIN | ATP2B4; ATP2B | P23634; P20 | AT2B4_HUMAN | Unmodified | 0 | 0 | 0 | 0 | 29559000 | 1926100 | 8867500 | 1553800 | 0 | 781870 | 0 | 0 | 19523000 | 33680000 | 0 | 0 | 7081400 | 11403000 |
| MVVAQQLA | TYK2; TYK2; TYI | P29597; E9P | TYK2_HUMAN I | Oxidation (M) | 0 | 0 | 0 | 0 | 164960 | 240800 | 0 | 0 | 453290 | 0 | 143500 | 0 | 0 | 0 | 0 | 0 | 228600 | 206300 |
| MVVDQGD | PAK6 | Q9NQU5 | PAK6_HUMAN | Oxidation (M) | 0 | 0 | 0 | 522160 | 590730 | 177670 | 0 | 0 | 116630 | 0 | 0 | 0 | 0 | 0 | 0 | 0 | 181860 | 197620 |
| MVVESAYE | LDHB; LDHB; N | P07195; A0A | LDHB_HUMAN | Unmodified | 0 | 0 | 0 | 282690 | 0 | 0 | 0 | 0 | 0 | 0 | 6113800 | 13889000 | 298640 | 0 | 0 | 0 | 0 | 1141600 |
| MVVESAYE | LDHB; LDHB; N | P07195; A0A | LDHB_HUMAN | Oxidation (M) | 858520 | 583480 | 1072400 | 1364200 | 2038700 | 0 | 0 | 59214 | 0 | 0 | 6618800 | 45836000 | 1019800 | 391660 | 1776800 | 1852100 | 0 | 0 |
| MVVESAYE | LDHB; LDHB; N | P07195; A0A | LDHB_HUMAN | Unmodified | 0 | 0 | 0 | 0 | 913810 | 136330 | 293530 | 0 | 0 | 0 | 0 | 0 | 0 | 0 | 0 | 0 | 840990 | 0 |
| MVVGGAQ | IGSF8; IGSF8 | Q969P0; C9J | IGSF8_HUMAN | Oxidation (M) | 18673000 | 5377800 | 1888400 | 11288000 | 0 | 0 | 0 | 0 | 0 | 0 | 0 | 0 | 19403000 | 8302600 | 9507500 | 4911300 | 0 | 0 |
| MVVGGAQ | IGSF8; IGSF8 | Q969P0; C9J | IGSF8_HUMAN | Oxidation (M) | 0 | 0 | 0 | 3026300 | 0 | 0 | 0 | 0 | 0 | 0 | 0 | 0 | 7125400 | 0 | 0 | 0 | 0 | 0 |
| MVVLSLPR | DYNC1H1; DYN | Q14204; A0/ | DYHC1_HUMA | Oxidation (M) | 0 | 0 | 0 | 0 | 0 | 0 | 0 | 0 | 0 | 0 | 0 | 1321700 | 0 | 0 | 0 | 0 | 0 | 0 |
| MVVMAPR | HLA-G; HLA-G; | P17693; Q5I | HLAG_HUMAN | Acetyl (Protein N | 0 | 0 | 0 | 0 | 0 | 0 | 0 | 0 | 0 | 0 | 0 | 0 | 0 | 0 | 0 | 0 | 0 | 0 |
| MVVNEGSI | HINT1 | P49773 | HINT1_HUMAN | Oxidation (M) | 2514500 | 2700100 | 2395300 | 6277400 | 9932500 | 0 | 0 | 0 | 8052200 | 4161500 | 6315300 | 33982000 | 15682000 | 5457400 | 5430000 | 5628900 | 1763300 | 2918600 |
| MVVNEGSI | HINT1 | P49773 | HINT1_HUMAN | Unmodified | 968420 | 696440 | 1270000 | 2143000 | 12516000 | 0 | 1319500 | 0 | 20368000 | 6025000 | 428040 | 17374000 | 20720000 | 5523000 | 1596800 | 693220 | 2906800 | 4194400 |
| MVVPGLDC | FASN; FASN | P49327; A0A | FAS_HUMAN F | Unmodified | 0 | 0 | 0 | 3662500 | 345710 | 509950 | 319010 | 0 | 679610 | 0 | 529600 | 8466600 | 513720 | 220890 | 0 | 0 | 88828 | 488930 |
| MVVPGLDC | FASN; FASN | P49327; A0A | FAS_HUMAN F | Oxidation (M) | 1105800 | 332670 | 3208900 | 6183400 | 1446100 | 0 | 370900 | 91688 | 1193800 | 420070 | 4656400 | 23073000 | 3116800 | 0 | 7995600 | 4799200 | 805560 | 0 |
| MVVPLACL | SEC23B; SEC2 | Q15437; A0/ | SC23B_HUMAI | Oxidation (M) | 0 | 0 | 0 | 0 | 0 | 0 | 0 | 0 | 399400 | 308820 | 0 | 1183500 | 336150 | 0 | 0 | 0 | 483060 | 77835 |
| MVVPLACL | SEC23B; SEC2 | Q15437; A0/ | SC23B_HUMAI | Unmodified | 19431 | 0 | 0 | 0 | 0 | 0 | 0 | 0 | 313290 | 0 | 0 | 125060 | 0 | 0 | 0 | 0 | 191040 | 144140 |
| MVVPVAAL | SEC23A; SEC2 | Q15436; F5I | SC23A_HUMAI | Oxidation (M) | 110910 | 0 | 73652 | 73097 | 406480 | 334520 | 109480 | 0 | 129290 | 0 | 119960 | 2159100 | 194160 | 0 | 125810 | 80545 | 278640 | 825160 |
| MVVPVAAL | SEC23A; SEC2 | Q15436; F5I | SC23A_HUMAI | Unmodified | 0 | 333870 | 0 | 0 | 0 | 88133 | 0 | 0 | 0 | 184380 | 33249 | 112880 | 0 | 0 | 0 | 0 | 92777 | 0 |
| MVVSFNR | IQGAP2; IQGAP | Q13576; P4( | IQGA2_HUMAN | Unmodified | 250100 | 336620 | 0 | 0 | 0 | 0 | 0 | 0 | 1236000 | 0 | 0 | 4113200 | 0 | 0 | 871950 | 507560 | 0 | 0 |
| MVVVGNG | RAB23 | Q9ULC3 | RAB23_HUMAN | Oxidation (M) | 0 | 0 | 0 | 0 | 596520 | 116070 | 0 | 0 | 67264 | 43993 | 0 | 0 | 0 | 0 | 0 | 0 | 219570 | 141780 |
| MVVVGNG | RAB23 | Q9ULC3 | RAB23_HUMAN | Unmodified | 0 | 0 | 357340 | 370460 | 100810 | 433670 | 61263 | 433550 | 0 | 401150 | 0 | 0 | 0 | 0 | 504050 | 296360 | 928450 | 1602900 |
| MVVYQGG | VIL1; VIL1; VIL1 | P09327; H7( | VILI_HUMAN Vi | Unmodified | 0 | 0 | 0 | 0 | 0 | 40151 | 0 | 0 | 0 | 65426 | 628820 | 6994100 | 0 | 0 | 0 | 0 | 0 | 0 |
| MVVYQGG | VIL1; VIL1; VIL1 | P09327; H7( | VILI_HUMAN Vi | Oxidation (M) | 47256 | 0 | 0 | 0 | 0 | 0 | 0 | 0 | 0 | 0 | 0 | 1726900 | 0 | 0 | 0 | 0 | 0 | 0 |
| MVVYQGG | VIL1; VIL1; VIL1 | P09327; H7( | VILI_HUMAN Vi | Unmodified | 0 | 0 | 0 | 0 | 0 | 0 | 0 | 0 | 0 | 190400 | 282180 | 0 | 0 | 0 | 0 | 0 | 0 | 0 |
| MVWDSQA | CHID1; CHID1; | Q9BWS9; H( | CHID1_HUMAN | Unmodified | 0 | 0 | 0 | 0 | 0 | 0 | 0 | 0 | 5807100 | 437850 | 0 | 0 | 0 | 0 | 0 | 0 | 0 | 0 |
| MVWDSQA | CHID1; CHID1; | Q9BWS9; H( | CHID1_HUMAN | Oxidation (M) | 0 | 0 | 0 | 0 | 0 | 0 | 0 | 0 | 2346200 | 657110 | 97148 | 0 | 0 | 0 | 0 | 0 | 0 | 0 |
| MVYDMEK | NCSTN; NCSTN | Q92542; H0 | NICA_HUMAN | Oxidation (M) | 0 | 0 | 42763 | 0 | 0 | 0 | 0 | 0 | 1248200 | 280490 | 0 | 0 | 0 | 0 | 0 | 64916 | 0 | 0 |
| MVYYLDPS | NAA15; NAA15 | Q9BXJ9; A0A | NAA15_HUMAI | Unmodified | 0 | 0 | 0 | 0 | 0 | 0 | 0 | 0 | 0 | 0 | 0 | 1966500 | 0 | 0 | 0 | 0 | 0 | 0 |
| MWAQAEKI | PPP1R16A; PPP | Q96I34; A0A | PP16A_HUMAI | Unmodified | 0 | 0 | 0 | 0 | 0 | 0 | 0 | 0 | 0 | 0 | 49631 | 0 | 0 | 0 | 0 | 0 | 412020 | 0 |
| MWDLSSN | RAE1; RAE1; RA | P78406; E9P | RAE1L_HUMAN | Oxidation (M) | 755470 | 0 | 0 | 293860 | 554530 | 0 | 48255 | 332500 | 0 | 377500 | 344400 | 1876300 | 0 | 0 | 0 | 318520 | 1377800 | 0 |
| MWDPHNI | SNRNP70; SNR | P08621; M0( | RU17_HUMAN | Oxidation (M) | 122770 | 0 | 0 | 0 | 0 | 0 | 0 | 0 | 223540 | 146890 | 376200 | 1162000 | 0 | 0 | 0 | 0 | 0 | 0 |
| MWDPHNI | SNRNP70; SNR | P08621; M0( | RU17_HUMAN | Unmodified | 205410 | 0 | 64210 | 72802 | 847230 | 0 | 0 | 0 | 708190 | 433780 | 0 | 1139400 | 258020 | 165130 | 0 | 0 | 198370 | 792530 |
| MWITNGGI | ACADM; ACADI | P11310; Q5I | ACADM_HUMA | Unmodified | 0 | 0 | 0 | 0 | 99976 | 0 | 90196 | 0 | 698790 | 0 | 0 | 0 | 0 | 0 | 0 | 29999 | 99538 | 0 |
| MWLKPEEV | TBC1D8B; TBC | Q0IIM8; O95 | TBC8B_HUMAI | Acetyl (Protein N | 0 | 0 | 0 | 0 | 2577600 | 0 | 0 | 0 | 0 | 818310 | 0 | 0 | 0 | 0 | 0 | 0 | 0 | 613930 |
| MWNLLHE | FOLH1; FOLH1; | Q04609; E9I | FOLH1_HUMAI | Acetyl (Protein N | 0 | 0 | 0 | 920950 | 0 | 0 | 0 | 0 | 0 | 0 | 0 | 0 | 0 | 0 | 5025300 | 3537000 | 0 | 0 |
| MWNLLHE | FOLH1; FOLH1; | Q04609; E9I | FOLH1_HUMAI | Acetyl (Protein N | 277980 | 0 | 330740 | 612250 | 0 | 0 | 0 | 0 | 0 | 0 | 0 | 0 | 0 | 0 | 1367300 | 373120 | 0 | 0 |
| MWSIENIAI | NAMPT; NAMPT | P43490; A0A | NAMPT_HUMAI | Oxidation (M) | 0 | 0 | 0 | 0 | 2411200 | 374350 | 0 | 0 | 92837 | 197510 | 162410 | 511740 | 0 | 0 | 446580 | 127810 | 0 | 0 |
| MWSIENIAI | NAMPT; NAMPT | P43490; A0A | NAMPT_HUMAI | Unmodified | 0 | 0 | 0 | 0 | 298170 | 0 | 0 | 0 | 0 | 0 | 50274 | 70196 | 0 | 0 | 0 | 0 | 0 | 0 |
| MWVQLLIP | PSME3; PSME3 | P61289; K7E | PSME3_HUMAI | Oxidation (M) | 0 | 0 | 0 | 0 | 0 | 0 | 0 | 0 | 0 | 0 | 564390 | 1189000 | 0 | 0 | 0 | 0 | 0 | 0 |
| MWVTGVV | MFGE8; MFGE8 | Q08431; H0 | MFGM_HUMAN | Unmodified | 0 | 152660 | 0 | 186790 | 4215900 | 1495200 | 0 | 0 | 3996600 | 0 | 0 | 1190300 | 79792 | 0 | 319980 | 158140 | 0 | 0 |
| MWVTGVV | MFGE8; MFGE8 | Q08431; H0 | MFGM_HUMAN | Oxidation (M) | 0 | 524180 | 0 | 501010 | 1452100 | 577960 | 0 | 0 | 2557700 | 0 | 0 | 0 | 0 | 0 | 1060900 | 2375000 | 194040 | 0 |
| MWVTGVV | MFGE8; MFGE8 | Q08431; H0 | MFGM_HUMAN | Oxidation (M) | 0 | 0 | 0 | 0 | 10815000 | 2549500 | 467810 | 0 | 158860 | 1334300 | 0 | 0 | 1512400 | 1162500 | 0 | 0 | 2361900 | 862300 |
| MWVTGVV | MFGE8; MFGE8 | Q08431; H0 | MFGM_HUMAN | Unmodified | 0 | 0 | 0 | 0 | 16351000 | 866830 | 0 | 0 | 0 | 550000 | 0 | 0 | 171010 | 0 | 0 | 0 | 1872900 | 0 |
| MYAPAWV | ILK; ILK | Q13418; A0/ | ILK_HUMAN Int | Oxidation (M) | 0 | 517930 | 0 | 0 | 0 | 0 | 0 | 0 | 0 | 0 | 0 | 0 | 0 | 0 | 0 | 504870 | 0 | 0 |
| MYCIVIVILI | ITGB1 | C9JPK5 | C9JPK5_HUMA | Unmodified | 433260 | 0 | 1874800 | 2727700 | 0 | 3403300 | 7207700 | 1730000 | 0 | 188040 | 0 | 138420 | 1450700 | 2929800 | 869400 | 0 | 0 | 0 |
| MYDDAIEAI | MAN1A2; MAN1 | O60476; H0 | MA1A2_HUMAI | Oxidation (M) | 0 | 0 | 0 | 0 | 0 | 0 | 0 | 0 | 1973700 | 0 | 0 | 0 | 0 | 0 | 0 | 0 | 0 | 0 |
| MYDDAIEAI | MAN1A2; MAN1 | O60476; H0 | MA1A2_HUMAI | Unmodified | 0 | 0 | 0 | 0 | 0 | 0 | 0 | 0 | 1431700 | 0 | 0 | 0 | 0 | 0 | 0 | 0 | 0 | 0 |
| MYDLNSNI | MLST8; MLST8; | Q9BVC4; H3 | LST8_HUMAN T | Oxidation (M) | 0 | 0 | 0 | 55605 | 0 | 0 | 0 | 0 | 0 | 0 | 0 | 313260 | 0 | 0 | 0 | 178860 | 0 | 0 |
| MYEEFLSK | PRKAR1B; PRK/ | P31321; P10 | KAP1_HUMAN | Oxidation (M) | 0 | 0 | 0 | 127610 | 912240 | 72615 | 599910 | 97007 | 698240 | 485670 | 0 | 240540 | 198170 | 76712 | 52067 | 116000 | 581370 | 679430 |
| MYEEFLSK | PRKAR1B; PRK/ | P31321; P10 | KAP1_HUMAN | Unmodified | 0 | 0 | 0 | 0 | 1090600 | 78128 | 377000 | 51186 | 3122700 | 205980 | 0 | 301720 | 0 | 0 | 0 | 0 | 223940 | 355350 |
| MYELSCYLI | MYOF; MYOF | Q9NZM1; H( | MYOF_HUMAN | Oxidation (M) | 0 | 0 | 0 | 0 | 1428300 | 0 | 0 | 0 | 0 | 0 | 0 | 0 | 0 | 0 | 0 | 0 | 0 | 0 |
| MYELSCYLI | MYOF; MYOF | Q9NZM1; H( | MYOF_HUMAN | Unmodified | 0 | 0 | 0 | 0 | 620650 | 222000 | 0 | 0 | 0 | 0 | 0 | 0 | 0 | 0 | 0 | 0 | 0 | 187110 |
| MYEQQRDI | CHMP5 | Q9NZZ3 | CHMP5_HUMA | 2 Oxidation (M) | 1211700 | 929160 | 296950 | 2996400 | 1612400 | 1154800 | 0 | 0 | 0 | 362570 | 0 | 2947200 | 353340 | 0 | 199510 | 1042800 | 0 | 0 |
| MYEQQRDI | CHMP5 | Q9NZZ3 | CHMP5_HUMA | Oxidation (M) | 0 | 0 | 0 | 750370 | 3705500 | 0 | 0 | 0 | 5804500 | 542930 | 0 | 855820 | 0 | 0 | 0 | 0 | 0 | 0 |
| MYEQQRDI | CHMP5 | Q9NZZ3 | CHMP5_HUMA | Unmodified | 0 | 0 | 0 | 0 | 2687900 | 0 | 0 | 0 | 5668500 | 0 | 0 | 0 | 0 | 0 | 0 | 0 | 0 | 0 |
| MYEQQRDI | CHMP5 | Q9NZZ3 | CHMP5_HUMA | Oxidation (M) | 0 | 0 | 0 | 0 | 17670000 | 3659900 | 441640 | 0 | 0 | 4179100 | 0 | 0 | 915100 | 3293600 | 0 | 0 | 0 | 1377900 |
| MYEQQRDI | CHMP5 | Q9NZZ3 | CHMP5_HUMA | 2 Oxidation (M) | 0 | 0 | 0 | 0 | 3105400 | 0 | 0 | 0 | 0 | 816910 | 0 | 0 | 2134300 | 1956000 | 0 | 0 | 0 | 751140 |
| MYEQQRDI | CHMP5 | Q9NZZ3 | CHMP5_HUMA | Unmodified | 0 | 0 | 0 | 0 | 5368600 | 0 | 0 | 0 | 0 | 0 | 0 | 0 | 0 | 0 | 0 | 0 | 0 | 666260 |
| MYFDAVQA | MAN1A1 | P33908 | MA1A1_HUMAI | Oxidation (M) | 0 | 0 | 0 | 0 | 147320 | 0 | 0 | 0 | 0 | 334270 | 775700 | 0 | 0 | 0 | 0 | 0 | 234080 | 0 |
| MYFPDVEF | AHNAK | Q09666 | AHNK_HUMAN | Oxidation (M) | 0 | 0 | 0 | 0 | 2614200 | 0 | 0 | 0 | 0 | 0 | 0 | 0 | 0 | 0 | 0 | 0 | 1369500 | 2879400 |
| MYFPDVEF | AHNAK | Q09666 | AHNK_HUMAN | Unmodified | 0 | 0 | 0 | 0 | 0 | 0 | 0 | 0 | 0 | 0 | 0 | 0 | 0 | 0 | 0 | 44349 | 952360 | 7885800 |
| MYGCDLGI | HLA-C; HLA-C; | Q5QT33; A0/ | Q5QT33_HUM/ | Oxidation (M) | 0 | 0 | 0 | 0 | 0 | 0 | 0 | 0 | 2422300 | 0 | 0 | 0 | 0 | 0 | 0 | 0 | 0 | 0 |
| MYGCDLGI | HLA-C; HLA-C; | Q5QT33; A0/ | Q5QT33_HUM/ | Unmodified | 0 | 0 | 0 | 0 | 681970 | 0 | 0 | 0 | 3927600 | 227580 | 0 | 0 | 0 | 0 | 0 | 0 | 0 | 0 |
| MYGCDVG | HLA-B; HLA-A; I | P01889; A0A | HLAB_HUMAN | Oxidation (M) | 182570 | 0 | 0 | 0 | 0 | 0 | 0 | 0 | 103900 | 0 | 634580 | 9759000 | 0 | 0 | 0 | 0 | 0 | 0 |
| MYGCDVG | HLA-B; HLA-A; I | P01889; A0A | HLAB_HUMAN | Unmodified | 0 | 0 | 0 | 0 | 0 | 0 | 0 | 0 | 682070 | 0 | 1083800 | 11111000 | 0 | 0 | 0 | 0 | 0 | 0 |
| MYGCDVG | HLA-A; HLA-A; I | Q53Z42; A0/ | Q53Z42_HUM/ | Oxidation (M) | 0 | 0 | 0 | 0 | 0 | 0 | 0 | 0 | 0 | 0 | 0 | 4382500 | 0 | 0 | 0 | 0 | 0 | 0 |
| MYGCDVG | HLA-A; HLA-A; I | Q53Z42; A0/ | Q53Z42_HUM/ | Unmodified | 0 | 0 | 0 | 0 | 0 | 0 | 0 | 0 | 0 | 0 | 0 | 9199100 | 0 | 0 | 0 | 0 | 0 | 0 |
| MYGISLCQ | ANXA1 | P04083 | ANXA1_HUMAI | Oxidation (M) | 662390 | 0 | 0 | 0 | 0 | 0 | 0 | 0 | 0 | 0 | 1177000 | 8554000 | 0 | 0 | 0 | 275320 | 0 | 1275900 |
| MYGISLCQ | ANXA1 | P04083 | ANXA1_HUMAI | Oxidation (M) | 0 | 0 | 0 | 0 | 30834000 | 6640200 | 0 | 0 | 636860 | 39984 | 38382000 | 2.41E+08 | 0 | 0 | 0 | 270600 | 3170600 | 7370200 |
| MYGISLCQ | ANXA1 | P04083 | ANXA1_HUMAI | Unmodified | 0 | 0 | 0 | 0 | 8026100 | 584390 | 0 | 0 | 0 | 0 | 5557700 | 25219000 | 0 | 0 | 0 | 0 | 759850 | 1489100 |
| MYKEEGLK | SLC25A3; SLC2 | Q00325; F8V | MPCP_HUMAN | Unmodified | 0 | 0 | 85197 | 0 | 0 | 0 | 388540 | 0 | 8823000 | 1291000 | 0 | 0 | 0 | 0 | 81070 | 0 | 0 | 0 |
| MYKTTPDV | RPS24; RPS24; | P62847; A0A | RS24_HUMAN | Oxidation (M) | 0 | 0 | 0 | 0 | 0 | 0 | 877780 | 0 | 0 | 0 | 0 | 0 | 0 | 0 | 0 | 0 | 0 | 0 |
| MYLETDPS | SCIN | Q9Y6U3 | ADSV_HUMAN | Unmodified | 50948 | 0 | 0 | 0 | 0 | 0 | 0 | 0 | 579730 | 1002000 | 84130 | 220580 | 0 | 0 | 0 | 0 | 0 | 0 |
| MYLSNNPI | TFAP2A; TFAP2/ | P05549; A0A | AP2A_HUMAN | Unmodified | 1709100 | 2938300 | 0 | 0 | 0 | 0 | 0 | 86068 | 0 | 0 | 0 | 0 | 0 | 0 | 198910 | 0 | 0 | 0 |
| MYLTPSEK | CYFIP1; CYFIP2 | Q7L576; Q9( | CYFP1_HUMAN | Unmodified | 0 | 0 | 196100 | 0 | 0 | 0 | 215560 | 115300 | 0 | 69042 | 153950 | 1583000 | 0 | 0 | 0 | 304470 | 0 | 88110 |
| MYQTQVSI | PTGFRN | Q9P2B2 | FPRP_HUMAN | Oxidation (M) | 4989200 | 6335400 | 9705800 | 6027000 | 0 | 0 | 0 | 0 | 0 | 0 | 0 | 6669700 | 167910 | 0 | 9882700 | 8912800 | 0 | 0 |
| MYQTQVSI | PTGFRN | Q9P2B2 | FPRP_HUMAN | Unmodified | 2586700 | 2137800 | 5930400 | 3946600 | 0 | 0 | 0 | 0 | 0 | 0 | 0 | 12596000 | 0 | 0 | 6879500 | 3937800 | 0 | 0 |
| MYRDTFYD | APOBEC3B; AP( | Q9UH17; Q9 | ABC3B_HUMAI | Oxidation (M) | 480500 | 1153800 | 418190 | 971640 | 0 | 0 | 0 | 0 | 0 | 0 | 0 | 0 | 0 | 0 | 1644200 | 639890 | 0 | 0 |
| MYRDTFYD | APOBEC3B; AP( | Q9UH17; Q9 | ABC3B_HUMAI | Unmodified | 0 | 154610 | 0 | 0 | 0 | 0 | 0 | 0 | 0 | 0 | 0 | 0 | 0 | 0 | 255040 | 0 | 0 | 0 |
| MYSIEPADI | UBA6; UBA6 | A0AVT1; H0 | UBA6_HUMAN | Unmodified | 0 | 0 | 0 | 0 | 642330 | 0 | 207550 | 72824 | 437300 | 156180 | 217910 | 2142300 | 384210 | 171420 | 0 | 0 | 353900 | 994780 |
| MYSNVYYH | CHAMP1; CHAI | Q96JM3; A0/ | CHAP1_HUMAI | Unmodified | 0 | 0 | 0 | 0 | 0 | 0 | 218930 | 78860 | 0 | 0 | 71419 | 0 | 0 | 0 | 0 | 0 | 0 | 0 |
| MYSTDDG\ | EFTUD2 | Q15029 | U5S1_HUMAN | Oxidation (M) | 0 | 0 | 0 | 0 | 1311800 | 0 | 446590 | 0 | 0 | 0 | 0 | 0 | 849910 | 308110 | 0 | 0 | 0 | 0 |
| MYSYVTEEI | ESD; ESD; ESD; | P10768; X6R | ESTD_HUMAN | Oxidation (M) | 0 | 0 | 0 | 93226 | 0 | 0 | 0 | 0 | 93588 | 0 | 315580 | 1264400 | 0 | 0 | 0 | 0 | 0 | 0 |
| MYTAIPQS( | CAPG; CAPG; C | P40121; E7E | CAPG_HUMAN | Acetyl (Protein N | 1147800 | 0 | 0 | 0 | 5237000 | 0 | 0 | 0 | 0 | 0 | 0 | 0 | 0 | 0 | 0 | 6368200 | 0 | 0 |
| MYVDPSEII | PLCG2; PLCG2 | P16885; A0A | PLCG2_HUMAI | 2 Oxidation (M) | 0 | 0 | 0 | 0 | 0 | 0 | 0 | 0 | 0 | 0 | 0 | 975510 | 0 | 0 | 0 | 0 | 0 | 0 |
| MYVDPSEII | PLCG2; PLCG2 | P16885; A0A | PLCG2_HUMAI | Oxidation (M) | 0 | 0 | 0 | 0 | 0 | 0 | 0 | 0 | 0 | 0 | 460340 | 1492400 | 0 | 0 | 0 | 0 | 0 | 0 |
| MYVQTVLD | CUL1; CUL1; CI | Q13616; A0/ | CUL1_HUMAN | Unmodified | 0 | 0 | 0 | 0 | 534900 | 0 | 0 | 0 | 0 | 0 | 117810 | 407290 | 102150 | 0 | 0 | 0 | 140540 | 147790 |
| NAADSSVP | SPINT2; SPINT2 | O43291; K7E | SPIT2_HUMAN | Unmodified | 223350 | 163090 | 389880 | 369150 | 0 | 0 | 0 | 0 | 1.08E+08 | 5128400 | 242550 | 289880 | 0 | 0 | 152020 | 115460 | 0 | 0 |
| NAAEIVQY( | SEMA3C | Q99985 | SEM3C_HUMAI | Unmodified | 0 | 0 | 0 | 1267200 | 0 | 0 | 0 | 0 | 306360 | 435140 | 3450600 | 0 | 0 | 0 | 1244100 | 0 | 0 | 0 |
| NAAENMLE | STAU1; STAU1 | O95793; Q5J | STAU1_HUMAN | Unmodified | 0 | 0 | 0 | 0 | 0 | 0 | 0 | 0 | 0 | 71729 | 0 | 271880 | 0 | 0 | 0 | 0 | 0 | 0 |

| | | | | | | | | | | | | | | | | | | | | | | |
|---|---|---|---|---|---|---|---|---|---|---|---|---|---|---|---|---|---|---|---|---|---|---|
| NAAGALDL | TCEA1; TCEA1; | P23193; E5R | TCEA1_HUMAN | Unmodified | 1167700 | 0 | 312490 | 475530 | 1689300 | 297190 | 335150 | 322720 | 1263100 | 1282700 | 575640 | 1034200 | 332790 | 0 | 496600 | 606920 | 206800 | 50111 |
| NAAHALPT | COLGALT1 | Q8NBJ5 | GT251_HUMAN | Unmodified | 30397 | 0 | 379950 | 112230 | 0 | 0 | 34074 | 0 | 35341000 | 4732100 | 537320 | 0 | 0 | 0 | 194340 | 83245 | 0 | 0 |
| NAALAPTS\ | COBL; COBL; C | O75128; J3K | COBL_HUMAN | Unmodified | 359960 | 0 | 0 | 0 | 0 | 0 | 0 | 24904 | 0 | 0 | 129440 | 0 | 0 | 0 | 93185 | 97164 | 0 | 0 |
| NAALAPTS\ | COBL; COBL; C | O75128; J3K | COBL_HUMAN | Unmodified | 0 | 0 | 0 | 0 | 121190 | 0 | 0 | 0 | 0 | 0 | 0 | 0 | 231500 | 185300 | 0 | 0 | 19469 | 0 |
| NAALDVEP | STRN4; STRN4; | Q9NRL3; M0 | STRN4_HUMAN | Unmodified | 0 | 0 | 0 | 0 | 0 | 0 | 0 | 0 | 0 | 0 | 0 | 701520 | 0 | 0 | 0 | 0 | 0 | 0 |
| NAALERVEI | FMNL2 | Q96PY5 | FMNL2_HUMAI | Unmodified | 0 | 0 | 0 | 0 | 1036700 | 266580 | 0 | 0 | 1190200 | 272590 | 878740 | 3379100 | 0 | 0 | 0 | 0 | 137710 | 1390600 |
| NAALFALG( | IPO4; IPO4 | Q8TEX9; H0\ | IPO4_HUMAN I | Unmodified | 0 | 0 | 0 | 0 | 2252100 | 0 | 348620 | 0 | 0 | 0 | 0 | 675530 | 0 | 0 | 0 | 0 | 0 | 0 |
| NAALNTIVT | CKAP5 | Q14008 | CKAP5_HUMAI | Unmodified | 0 | 0 | 0 | 0 | 0 | 0 | 22655 | 0 | 1023200 | 0 | 0 | 3303000 | 0 | 0 | 339200 | 0 | 0 | 0 |
| NAAQFLLS' | PPP2R2A | P63151 | 2ABA_HUMAN | Unmodified | 311310 | 153240 | 338850 | 524840 | 0 | 0 | 320110 | 309010 | 877760 | 418100 | 632110 | 3431000 | 1415600 | 870720 | 571170 | 613530 | 1292700 | 588790 |
| NAAQFLLS' | PPP2R2A | P63151 | 2ABA_HUMAN | Unmodified | 0 | 0 | 257340 | 0 | 0 | 0 | 0 | 233950 | 0 | 0 | 0 | 0 | 0 | 0 | 0 | 0 | 466300 | 0 |
| NAAQVLVG | COPS4; COPS4 | Q9BT78; D6I | CSN4_HUMAN | Unmodified | 0 | 0 | 0 | 0 | 0 | 0 | 0 | 0 | 0 | 0 | 0 | 1225200 | 0 | 0 | 0 | 0 | 0 | 0 |
| NAAYYSPH | EIF2A; EIF2A; EI | Q9BY44; F8\ | EIF2A_HUMAN | Unmodified | 0 | 0 | 248080 | 0 | 0 | 0 | 0 | 0 | 0 | 0 | 0 | 3630400 | 0 | 0 | 0 | 0 | 0 | 0 |
| NACDHLS( | SF3B6 | Q9Y3B4 | SF3B6_HUMAI | Unmodified | 456620 | 0 | 0 | 229530 | 0 | 0 | 0 | 0 | 0 | 0 | 0 | 1118500 | 0 | 0 | 0 | 0 | 0 | 0 |
| NADGKIISL | EPRS1; EPRS1 | P07814; V9C | SYEP_HUMAN | Unmodified | 0 | 0 | 0 | 0 | 0 | 0 | 0 | 0 | 0 | 0 | 0 | 0 | 1475700 | 412710 | 0 | 0 | 0 | 0 |
| NADGTICYI | AKR1A1; AKR1A | P14550; Q5T | AK1A1_HUMAI | Unmodified | 0 | 0 | 0 | 0 | 0 | 0 | 0 | 0 | 543670 | 300290 | 171450 | 1728700 | 0 | 0 | 799190 | 499540 | 153490 | 125130 |
| NADGTICYI | AKR1A1; AKR1A | P14550; Q5T | AK1A1_HUMAI | Unmodified | 0 | 0 | 0 | 0 | 633070 | 0 | 0 | 0 | 349650 | 765010 | 0 | 0 | 1717100 | 503540 | 0 | 0 | 309140 | 267610 |
| NADGYKLD | EIF3B; EIF3B | P55884; C9J | EIF3B_HUMAN | Unmodified | 0 | 0 | 138330 | 0 | 118360 | 0 | 0 | 16411 | 0 | 0 | 0 | 955860 | 0 | 0 | 0 | 97297 | 0 | 327380 |
| NADHSMN | ILF3; ILF3 | Q12906; K7E | ILF3_HUMAN Ir | Unmodified | 323940 | 275130 | 0 | 250610 | 0 | 0 | 76625 | 0 | 0 | 0 | 0 | 0 | 0 | 0 | 0 | 0 | 0 | 0 |
| NADMSEEN | DYNLL1; DYNLI | P63167; F8V | DYL1_HUMAN | Oxidation (M) | 484970 | 0 | 0 | 362770 | 672000 | 0 | 393290 | 0 | 0 | 579390 | 0 | 1409500 | 696600 | 1013300 | 0 | 167000 | 995500 | 2096000 |
| NADMSEEN | DYNLL1; DYNLI | P63167; F8V | DYL1_HUMAN | Unmodified | 0 | 0 | 0 | 0 | 1264900 | 0 | 315040 | 0 | 1545100 | 294160 | 0 | 0 | 0 | 0 | 0 | 0 | 1234900 | 2572100 |
| NADMSEEN | DYNLL1; DYNLI | P63167; F8V | DYL1_HUMAN | 2 Oxidation (M) | 0 | 0 | 0 | 401480 | 0 | 0 | 0 | 150530 | 0 | 0 | 276760 | 1434300 | 0 | 0 | 0 | 0 | 600810 | 451200 |
| NADNDWF | UFC1 | Q9Y3C8 | UFC1_HUMAN | Unmodified | 0 | 0 | 0 | 0 | 238980 | 104930 | 0 | 0 | 0 | 0 | 0 | 1404900 | 0 | 0 | 0 | 0 | 27619 | 78021 |
| NADPILISLI | CORO1C; COR( | Q9ULV4; H0 | COR1C_HUMA | Unmodified | 0 | 167290 | 338360 | 436340 | 59664000 | 9365600 | 1850000 | 127620 | 6231900 | 3083800 | 7934100 | 41852000 | 8463500 | 3730500 | 355190 | 293470 | 6003900 | 6490500 |
| NADVELQC | AP2A2; AP2A1; | O94973; O9 | AP2A2_HUMAN | Unmodified | 411220 | 444020 | 486020 | 686490 | 0 | 0 | 0 | 0 | 1445900 | 160130 | 0 | 1133100 | 0 | 0 | 784950 | 637690 | 0 | 79697 |
| NAEAVLQS | FLII | Q13045 | FLII_HUMAN Pr | Unmodified | 0 | 0 | 0 | 0 | 0 | 0 | 0 | 0 | 0 | 0 | 0 | 695650 | 0 | 0 | 0 | 0 | 0 | 0 |
| NAEDMEVF | BTN2A2; BTN2/ | Q8WVV5; Q | BT2A2_HUMAN | Unmodified | 0 | 0 | 0 | 0 | 0 | 0 | 0 | 0 | 570140 | 0 | 0 | 779810 | 0 | 0 | 0 | 0 | 0 | 0 |
| NAEEFTK | UBE2L5; UBE2L | A0A1B0GUS | UB2L5_HUMAN | Unmodified | 181420 | 149220 | 585610 | 289760 | 0 | 397460 | 0 | 511510 | 271530 | 238970 | 0 | 2822500 | 916850 | 590090 | 480280 | 376300 | 1180300 | 671390 |
| NAEFDRHE | PALS2; PALS2 | Q9NZW5; B | PALS2_HUMAN | Unmodified | 0 | 0 | 532030 | 183590 | 0 | 0 | 0 | 0 | 410770 | 0 | 1881200 | 2234600 | 0 | 0 | 1086900 | 551970 | 0 | 0 |
| NAEFDRHE | MPP2; MPP2; M | Q14168; A0/ | MPP2_HUMAN | Unmodified | 0 | 0 | 417100 | 0 | 415890 | 0 | 0 | 0 | 399150 | 0 | 0 | 0 | 0 | 0 | 1153000 | 772910 | 0 | 0 |
| NAEIGNKD | STT3A | P46977 | STT3A_HUMAN | Unmodified | 0 | 0 | 0 | 0 | 0 | 308730 | 0 | 0 | 0 | 953680 | 0 | 0 | 0 | 0 | 0 | 0 | 0 | 0 |
| NAEIGNKD | STT3B | Q8TCJ2 | STT3B_HUMAN | Unmodified | 0 | 0 | 0 | 0 | 276390 | 0 | 191140 | 100810 | 1860600 | 21220 | 0 | 0 | 12645 | 25757 | 0 | 0 | 222500 | 0 |
| NAELALFEL | TTC13 | Q8NBP0 | TTC13_HUMAN | Unmodified | 0 | 0 | 0 | 0 | 0 | 0 | 0 | 0 | 1276100 | 0 | 0 | 0 | 0 | 0 | 0 | 0 | 0 | 0 |
| NAENAIEAI | ATP2A1; ATP2A | O14983; P16 | AT2A1_HUMAN | Unmodified | 0 | 0 | 0 | 0 | 0 | 0 | 0 | 0 | 0 | 0 | 0 | 1289500 | 0 | 0 | 140010 | 0 | 0 | 0 |
| NAEPLINLD | RTRAF; RTRAF; | Q9Y224; G3\ | RTRAF_HUMAN | Unmodified | 295850 | 0 | 269290 | 1170600 | 330440 | 0 | 0 | 0 | 341460 | 0 | 1190100 | 3793400 | 634140 | 0 | 507070 | 446460 | 0 | 1168300 |
| NAEQAATQ | KTN1 | Q86UP2 | KTN1_HUMAN | Unmodified | 81611 | 145210 | 0 | 0 | 0 | 0 | 0 | 0 | 0 | 127280 | 1521400 | 1449600 | 298980 | 0 | 0 | 0 | 0 | 0 |
| NAEQNGP\ | HSPA4 | P34932 | HSP74_HUMAI | Unmodified | 0 | 116010 | 0 | 0 | 1848700 | 0 | 0 | 0 | 0 | 0 | 1312400 | 5695900 | 901470 | 312060 | 823320 | 249950 | 332660 | 479830 |
| NAEQNGP\ | HSPA4 | P34932 | HSP74_HUMAI | Unmodified | 0 | 0 | 0 | 0 | 1728200 | 0 | 0 | 0 | 0 | 0 | 0 | 0 | 0 | 0 | 0 | 0 | 1166900 | 0 |
| NAESAIEAL | ATP2A3 | Q93084 | AT2A3_HUMAN | Unmodified | 0 | 0 | 0 | 0 | 0 | 0 | 0 | 0 | 0 | 0 | 0 | 0 | 0 | 0 | 0 | 0 | 0 | 0 |
| NAESNAEL | RPL17; RPL17; I | P18621; A0A | RL17_HUMAN ( | Unmodified | 0 | 0 | 0 | 0 | 618760 | 0 | 0 | 387910 | 32586 | 19187 | 0 | 6628700 | 0 | 0 | 0 | 0 | 0 | 76400 |
| NAESNAEL | RPL17; RPL17; I | P18621; A0A | RL17_HUMAN ( | Unmodified | 0 | 0 | 0 | 0 | 5850500 | 0 | 7318500 | 0 | 0 | 231340 | 0 | 0 | 3411500 | 4524700 | 0 | 0 | 1304200 | 294100 |
| NAFERGED | SRGAP2; SRGA | O75044; Q5\ | SRGP2_HUMAI | Oxidation (M) | 0 | 0 | 0 | 0 | 0 | 0 | 0 | 0 | 0 | 0 | 0 | 1268400 | 0 | 0 | 0 | 0 | 0 | 0 |
| NAFYIGSY( | COPE; COPE; C | O14579; M0I | COPE_HUMAN | Unmodified | 0 | 0 | 0 | 0 | 1070900 | 0 | 0 | 0 | 0 | 678110 | 0 | 0 | 603800 | 0 | 0 | 0 | 842930 | 1262200 |
| NAGFTPQE | APEX1; APEX1; , | P27695; G3\ | APEX1_HUMAN | Unmodified | 19599 | 0 | 0 | 0 | 0 | 0 | 35571 | 0 | 1386000 | 0 | 982110 | 5328900 | 0 | 0 | 0 | 83714 | 0 | 0 |
| NAGGGGLI | IRGQ | Q8WZA9 | IRGQ_HUMAN | Unmodified | 37597 | 0 | 0 | 0 | 1666000 | 0 | 0 | 0 | 0 | 243760 | 0 | 1110700 | 0 | 0 | 0 | 0 | 161190 | 161560 |
| NAGKLTTTI | GPC3; GPC3 | P51654; H0\ | GPC3_HUMAN | Unmodified | 0 | 0 | 0 | 0 | 0 | 0 | 0 | 0 | 633220 | 173820 | 0 | 0 | 0 | 0 | 0 | 0 | 0 | 0 |
| NAGLAFIEL | LRBA; LRBA; LR | P50851; A0A | LRBA_HUMAN | Unmodified | 0 | 0 | 0 | 0 | 0 | 0 | 0 | 0 | 0 | 0 | 49058 | 4351400 | 0 | 0 | 0 | 151850 | 0 | 0 |
| NAGNCLSI | PHGDH; PHGD | O43175; A0/ | SERA_HUMAN | Unmodified | 717050 | 0 | 1120400 | 3256200 | 3477700 | 542180 | 10891000 | 785830 | 4591500 | 4990000 | 2173500 | 11043000 | 27344000 | 20976000 | 3400700 | 3091100 | 4743900 | 4298700 |
| NAGNCLSI | PHGDH; PHGD | O43175; A0/ | SERA_HUMAN | Unmodified | 0 | 0 | 0 | 0 | 0 | 0 | 885180 | 0 | 0 | 0 | 0 | 0 | 561310 | 928990 | 0 | 0 | 841380 | 0 |
| NAGNEQDI | CTNNA1; CTNN | P35221; E5R | CTNA1_HUMAI | Unmodified | 1428200 | 1211000 | 4870900 | 7109700 | 673160 | 713060 | 100590 | 0 | 485470 | 87799 | 977960 | 15469000 | 0 | 718270 | 9944100 | 10044000 | 0 | 0 |
| NAGNPVVI | PATJ; PATJ | Q8NI35; A0A | INADL_HUMAN | Unmodified | 16177 | 0 | 67244 | 158190 | 843490 | 0 | 0 | 0 | 204450 | 62901 | 0 | 42088 | 156400 | 174900 | 104750 | 88543 | 0 | 306640 |
| NAGQAVTI | DLG1; DLG1; D | Q12959; A0/ | DLG1_HUMAN | Unmodified | 0 | 0 | 0 | 0 | 0 | 0 | 0 | 0 | 0 | 0 | 0 | 0 | 0 | 0 | 0 | 180400 | 0 | 0 |
| NAGQAVTI | DLG1; DLG1; D | Q12959; A0/ | DLG1_HUMAN | Unmodified | 517830 | 0 | 747340 | 0 | 4027600 | 958880 | 0 | 0 | 2223900 | 2629700 | 0 | 771660 | 0 | 0 | 1744100 | 1385200 | 2764700 | 0 |
| NAGVEGSL | HSPD1; HSPD1 | P10809; A0A | CH60_HUMAN | Unmodified | 209150 | 0 | 534060 | 721200 | 760020 | 0 | 11506000 | 2921100 | 42915000 | 16547000 | 3449500 | 2994200 | 5657300 | 8755900 | 127120 | 327290 | 1244000 | 2111800 |
| NAIANASTL | SNRPA1; SNRP. | P09661; H0\ | RU2A_HUMAN | Unmodified | 0 | 0 | 0 | 0 | 592930 | 401060 | 154110 | 96272 | 0 | 113260 | 411250 | 0 | 0 | 0 | 75862 | 54073 | 0 | 0 |
| NAIEALGSI | FAM3B; FAM3B | P58499; A8N | FAM3B_HUMAI | Unmodified | 10020000 | 0 | 0 | 0 | 0 | 0 | 0 | 0 | 300850 | 509550 | 1066200 | 0 | 0 | 0 | 0 | 0 | 0 | 0 |
| NAIKEGDN | ALCAM; ALCAM | Q13740; F5( | CD166_HUMAI | Unmodified | 0 | 0 | 0 | 0 | 1240700 | 0 | 20673 | 18433 | 42653 | 36475 | 0 | 0 | 243620 | 0 | 0 | 0 | 0 | 551750 |
| NAIKVPIVIN | PEAK1; PEAK1; | Q9H792; H0 | PEAK1_HUMAN | Unmodified | 0 | 0 | 0 | 0 | 1145300 | 0 | 0 | 0 | 0 | 0 | 0 | 0 | 0 | 0 | 0 | 0 | 437690 | 0 |
| NAILEELPR | DOP1B | Q9Y3R5 | DOP2_HUMAN | Unmodified | 0 | 391950 | 0 | 0 | 0 | 0 | 0 | 62502 | 669080 | 0 | 215460 | 0 | 0 | 0 | 0 | 0 | 0 | 0 |
| NAINIEELF( | RAB31 | Q13636 | RAB31_HUMAN | Unmodified | 120300 | 0 | 0 | 0 | 2315600 | 910410 | 0 | 0 | 0 | 0 | 77790 | 0 | 0 | 0 | 0 | 0 | 0 | 0 |
| NAININELF | RAB22A | Q9UL26 | RB22A_HUMAN | Unmodified | 0 | 0 | 0 | 0 | 2801400 | 0 | 0 | 0 | 0 | 0 | 38734 | 0 | 516050 | 172120 | 34426 | 0 | 0 | 1069900 |
| NAINTEMYI | ECI2; ECI2; ECI | O75521; F1L | ECI2_HUMAN E | Oxidation (M) | 0 | 0 | 0 | 0 | 0 | 0 | 0 | 0 | 4946500 | 251930 | 0 | 0 | 0 | 0 | 0 | 0 | 0 | 0 |
| NAINTEMYI | ECI2; ECI2; ECI | O75521; F1L | ECI2_HUMAN E | Unmodified | 0 | 0 | 0 | 0 | 538790 | 107600 | 0 | 215140 | 13516000 | 554440 | 0 | 0 | 0 | 0 | 0 | 0 | 516180 | 337380 |
| NAINTEMYI | ECI2; ECI2; ECI | O75521; F1L | ECI2_HUMAN E | Unmodified | 0 | 0 | 0 | 0 | 0 | 0 | 79137 | 0 | 0 | 329870 | 0 | 0 | 39493 | 0 | 0 | 0 | 60259 | 204200 |
| NAKISSLLE | RPS8; RPS8 | P62241; Q5J | RS8_HUMAN 4 | Unmodified | 0 | 0 | 0 | 0 | 0 | 0 | 1280800 | 293910 | 0 | 0 | 0 | 0 | 1090900 | 269720 | 0 | 0 | 0 | 0 |
| NAKKEDSD | NCL; NCL; NCL | P19338; C9J | NUCL_HUMAN | Unmodified | 0 | 0 | 0 | 0 | 450180 | 0 | 0 | 0 | 673260 | 0 | 0 | 0 | 1115100 | 143490 | 0 | 0 | 0 | 0 |
| NAKPVSILF | STK3; STK3; STI | Q13188; A0/ | STK3_HUMAN ! | Oxidation (M) | 0 | 0 | 0 | 0 | 0 | 0 | 0 | 0 | 0 | 0 | 0 | 0 | 103400 | 65275 | 0 | 0 | 0 | 0 |
| NALAACVR | GNPAT | O15228 | GNPAT_HUMAI | Unmodified | 0 | 0 | 0 | 0 | 0 | 0 | 0 | 0 | 1648400 | 263160 | 108650 | 277950 | 80425 | 0 | 0 | 0 | 74675 | 79893 |
| NALANPLY( | UQCRC2; UQCI | P22695; H3E | QCR2_HUMAN | Unmodified | 0 | 0 | 0 | 0 | 0 | 0 | 0 | 0 | 771290 | 0 | 0 | 0 | 0 | 0 | 0 | 0 | 0 | 0 |
| NALDLLLPI | ADSL; ADSL; AD | P30566; A0A | PUR8_HUMAN | Unmodified | 0 | 62547 | 0 | 0 | 847130 | 1065100 | 558090 | 403480 | 0 | 0 | 606790 | 5522200 | 0 | 0 | 57515 | 0 | 0 | 513630 |
| NALDPMS\ | PRCP; PRCP; PI | P42785; B7Z | PCP_HUMAN L | Unmodified | 0 | 0 | 0 | 0 | 0 | 0 | 0 | 0 | 2545100 | 639920 | 171310 | 0 | 0 | 0 | 0 | 0 | 0 | 0 |
| NALESYAFI | HSPA1L; HSPA: | P34931; P0D | HS71L_HUMAN | Oxidation (M) | 2329100 | 2459200 | 1130700 | 778820 | 2931600 | 2076600 | 2001600 | 1300700 | 1400300 | 3314000 | 479560 | 2890800 | 2923400 | 2217500 | 1057300 | 1152600 | 2549300 | 1591400 |
| NALESYAFI | HSPA1L; HSPA: | P34931; P0D | HS71L_HUMAN | Unmodified | 0 | 2359400 | 0 | 0 | 8152200 | 1883700 | 2605100 | 523650 | 5081400 | 4875900 | 0 | 1814100 | 7052700 | 0 | 995820 | 0 | 4726700 | 3040100 |
| NALESYAFI | HSPA1A; HSPA: | P0DMV8; P0 | HS71A_HUMAI | Unmodified | 0 | 0 | 0 | 238960 | 0 | 0 | 899370 | 0 | 1303700 | 0 | 0 | 0 | 0 | 0 | 0 | 0 | 1509900 | 0 |
| NALESYAFI | HSPA1A; HSPA: | P0DMV8; P0 | HS71A_HUMAI | Oxidation (M) | 404720 | 0 | 0 | 0 | 449670 | 0 | 765830 | 91074 | 0 | 65516 | 0 | 0 | 98956 | 519100 | 0 | 90230 | 675160 | 0 |
| NALESYTYI | HSPA2 | P54652 | HSP72_HUMAI | Unmodified | 0 | 0 | 0 | 0 | 1110000 | 0 | 0 | 0 | 0 | 0 | 330830 | 1155300 | 0 | 0 | 0 | 0 | 0 | 0 |
| NALFPEVFS | MISP | Q8IVT2 | MISP_HUMAN I | Unmodified | 0 | 0 | 0 | 0 | 0 | 0 | 0 | 0 | 0 | 0 | 0 | 383210 | 0 | 0 | 0 | 0 | 0 | 0 |
| NALGAIHH | NRCAM; NRCAI | Q92823; A0/ | NRCAM_HUMA | Unmodified | 0 | 0 | 0 | 0 | 0 | 0 | 0 | 0 | 843260 | 0 | 0 | 0 | 0 | 0 | 0 | 0 | 0 | 0 |
| NALGPGLS | LRRC47 | Q8N1G4 | LRC47_HUMAN | Unmodified | 552550 | 0 | 286950 | 351930 | 0 | 0 | 0 | 0 | 1274100 | 286750 | 478630 | 2681500 | 643290 | 0 | 470400 | 599440 | 0 | 0 |
| NALKQETE' | RAB7A; RAB7A; | P51149; C9J | RAB7A_HUMAN | Unmodified | 798000 | 169640 | 1053300 | 1220300 | 0 | 0 | 0 | 0 | 2089600 | 0 | 3032200 | 4294400 | 0 | 0 | 0 | 1073500 | 0 | 0 |
| NALKQETE' | RAB7A; RAB7A; | P51149; C9J | RAB7A_HUMAN | Unmodified | 0 | 0 | 0 | 0 | 0 | 0 | 0 | 0 | 6965600 | 0 | 3446900 | 13854000 | 1964300 | 374090 | 0 | 0 | 0 | 0 |
| NALLSLAK | ANXA1; ANXA1 | P04083; Q5T | ANXA1_HUMAI | Unmodified | 177820 | 122310 | 0 | 0 | 1041200 | 417530 | 0 | 0 | 738860 | 119240 | 50781000 | 1.98E+09 | 0 | 0 | 87586 | 605070 | 58232 | 0 |
| NALLSLAK( | ANXA1 | P04083 | ANXA1_HUMAI | Unmodified | 0 | 0 | 0 | 0 | 714580 | 0 | 0 | 0 | 0 | 0 | 12620000 | 84006000 | 0 | 795600 | 0 | 0 | 0 | 1205600 |
| NALLSQNP | MPP1; MPP1 | Q00013; C9J | EM55_HUMAN | Unmodified | 0 | 0 | 0 | 0 | 1112200 | 211650 | 0 | 0 | 263490 | 94997 | 0 | 227210 | 0 | 0 | 0 | 0 | 90137 | 0 |
| NALNVLQR | EXOC2 | Q96KP1 | EXOC2_HUMAI | Unmodified | 0 | 53425 | 0 | 0 | 0 | 0 | 0 | 0 | 0 | 0 | 0 | 959600 | 0 | 0 | 0 | 0 | 0 | 0 |
| NALQELQC | SAMD4B; SAMD | Q5PRF9; M0 | SMAG2_HUMA | Unmodified | 0 | 13977 | 24073 | 25906 | 0 | 0 | 0 | 0 | 51460 | 17723 | 107300 | 181560 | 0 | 0 | 29873 | 28277 | 0 | 0 |
| NALQQENI | SLIRP; SLIRP; S | Q9GZT3; G3' | SLIRP_HUMAN | Unmodified | 0 | 0 | 0 | 0 | 145670 | 0 | 0 | 0 | 3776100 | 0 | 362390 | 563980 | 0 | 0 | 0 | 0 | 0 | 0 |
| NALSSLWG | EIF3E; EIF3E; EI | P60228; B3K | EIF3E_HUMAN | Unmodified | 82772 | 177450 | 89050 | 0 | 0 | 0 | 0 | 0 | 0 | 0 | 0 | 6669900 | 0 | 0 | 216880 | 203750 | 0 | 0 |
| NALVSHLD | UQCRC1 | P31930 | QCR1_HUMAN | Unmodified | 0 | 0 | 0 | 0 | 267700 | 0 | 118720 | 0 | 40018000 | 2471700 | 0 | 0 | 0 | 0 | 0 | 0 | 0 | 316480 |
| NALWHTGI | THBS2; THBS1; | P35442; P07 | TSP2_HUMAN ' | Unmodified | 0 | 0 | 338550 | 197330 | 0 | 0 | 0 | 0 | 0 | 0 | 0 | 0 | 0 | 0 | 211180 | 0 | 0 | 0 |
| NAMDGYR | ATP11C; ATP11 | Q8NB49; A0. | AT11C_HUMAN | Unmodified | 0 | 12398 | 78609 | 185240 | 0 | 0 | 0 | 0 | 0 | 0 | 114530 | 753260 | 0 | 0 | 299190 | 86650 | 0 | 0 |
| NAMNELQI | EVI5; EVI5 | O60447; A0/ | EVI5_HUMAN E | Unmodified | 0 | 0 | 0 | 0 | 0 | 129800 | 0 | 0 | 0 | 84515 | 317880 | 1555700 | 0 | 0 | 0 | 0 | 38617 | 0 |
| NANAEPAV | CD99; CD99 | P14209; A0A | CD99_HUMAN | Unmodified | 3576000 | 5300200 | 13020000 | 10288000 | 867530 | 439030 | 0 | 58063 | 0 | 0 | 0 | 50307 | 0 | 33812 | 18722000 | 14118000 | 1048400 | 675560 |
| NANAEPAV | CD99; CD99 | P14209; A0A | CD99_HUMAN | Unmodified | 0 | 0 | 0 | 0 | 4120900 | 1379100 | 0 | 322320 | 0 | 0 | 0 | 0 | 1688700 | 814580 | 0 | 0 | 5573900 | 7807100 |
| NANAVMEY | DNAJC7; DNAJ( | Q99615; A0/ | DNJC7_HUMAI | Unmodified | 0 | 0 | 232770 | 320570 | 1537200 | 0 | 444170 | 73200 | 899270 | 473390 | 153990 | 0 | 1879600 | 0 | 0 | 0 | 0 | 0 |
| NANEMFR | DYNC1H1; DYN | Q14204; A0/ | DYHC1_HUMA | Unmodified | 0 | 0 | 66997 | 64498 | 99534 | 17090 | 24413 | 0 | 306760 | 94233 | 210820 | 1880200 | 126680 | 0 | 100510 | 72785 | 14301 | 0 |
| NANPEWN | MYOF | Q9NZM1 | MYOF_HUMAN | Unmodified | 0 | 0 | 0 | 0 | 2053000 | 364720 | 0 | 0 | 0 | 0 | 0 | 0 | 0 | 0 | 0 | 0 | 29138 | 0 |
| NANPEWN | MYOF | Q9NZM1 | MYOF_HUMAN | Unmodified | 0 | 0 | 0 | 0 | 13520000 | 458470 | 0 | 0 | 0 | 0 | 0 | 0 | 0 | 0 | 0 | 0 | 495640 | 890580 |
| NANVQDA/ | EPB41L5 | Q9HCM4 | E41L5_HUMAN | Unmodified | 584810 | 0 | 0 | 408450 | 1336200 | 0 | 0 | 3564400 | 25056000 | 88950000 | 4263400 | 0 | 0 | 398490 | 241890 | 0 | 67540 | 2971700 |
| NAPAIIFIDE | VCP; VCP; VCP; | P55072; C9J | TERA_HUMAN ' | Unmodified | 126190 | 0 | 37034 | 128400 | 162580 | 0 | 25825 | 0 | 0 | 154800 | 0 | 0 | 205660 | 0 | 110060 | 82902 | 0 | 33202 |
| NAPAIIFIDE | VCP; VCP; VCP; | P55072; C9J | TERA_HUMAN ' | Unmodified | 2948600 | 2671000 | 361730 | 2464800 | 1297400 | 428350 | 40271 | 0 | 10077000 | 1016600 | 2628600 | 37756000 | 3439800 | 225950 | 2563900 | 2717500 | 0 | 187360 |
| NAPHPDDI | APBB2; APBB2; | Q92870; H0\ | APBB2_HUMAN | Unmodified | 0 | 0 | 0 | 0 | 254570 | 45742 | 0 | 0 | 0 | 0 | 18775 | 0 | 0 | 0 | 0 | 0 | 61895 | 0 |
| NAPPEPVP | VGF | O15240 | VGF_HUMAN N | Unmodified | 0 | 0 | 0 | 0 | 0 | 0 | 103980 | 153150 | 14209000 | 2385500 | 0 | 0 | 0 | 0 | 0 | 0 | 0 | 0 |
| NAQAIEDM | PSMD1; PSMD1 | Q99460; A0/ | PSMD1_HUMA | Unmodified | 0 | 0 | 0 | 0 | 0 | 0 | 0 | 121600 | 1145800 | 0 | 874080 | 1240100 | 0 | 0 | 0 | 0 | 0 | 0 |
| NAQAQQEI | DNAJC7; DNAJ( | Q99615; A0/ | DNJC7_HUMAI | Unmodified | 280470 | 0 | 306110 | 658200 | 1139000 | 254950 | 250560 | 135550 | 761750 | 204820 | 448400 | 1551400 | 2127300 | 910840 | 319770 | 325110 | 0 | 300110 |
| NAQAQQEI | DNAJC7; DNAJ( | Q99615; A0/ | DNJC7_HUMAI | Unmodified | 0 | 0 | 0 | 0 | 610210 | 0 | 767590 | 0 | 0 | 0 | 0 | 0 | 0 | 0 | 0 | 0 | 136470 | 0 |
| NAQGKDQ | ABHD12; ABHD | Q8N2K0; A0. | ABD12_HUMAI | Oxidation (M) | 0 | 0 | 0 | 0 | 0 | 0 | 0 | 0 | 8617900 | 2900200 | 0 | 0 | 0 | 0 | 0 | 23690 | 217030 | 0 |
| NAQGKDQ | ABHD12; ABHD | Q8N2K0; A0. | ABD12_HUMAI | Unmodified | 0 | 0 | 0 | 0 | 0 | 0 | 0 | 0 | 1865500 | 109180 | 0 | 0 | 0 | 0 | 0 | 0 | 0 | 0 |
| NAQLELK | DNAJC7; DNAJ( | Q99615; A0/ | DNJC7_HUMAI | Unmodified | 0 | 0 | 0 | 0 | 0 | 0 | 0 | 0 | 0 | 0 | 0 | 0 | 977410 | 401500 | 0 | 0 | 0 | 0 |
| NAQLNIELI | NAMPT; NAMPT | P43490; A0A | NAMPT_HUMAI | Unmodified | 0 | 0 | 0 | 0 | 22546000 | 5021500 | 1953900 | 219670 | 0 | 1355500 | 229820 | 3222900 | 7812800 | 6943300 | 0 | 290330 | 1460600 | 4364700 |

| | | | | | | | | | | | | | | | | | | | | | | |
|---|---|---|---|---|---|---|---|---|---|---|---|---|---|---|---|---|---|---|---|---|---|---|
| NAQMAQSI | ILVBL; ILVBL; IL' | A1L0T0; E9P | HACL2_HUMAI | Oxidation (M) | 0 | 0 | 11464 | 0 | 0 | 0 | 0 | 0 | 104360 | 0 | 58639 | 218420 | 0 | 0 | 27960 | 0 | 0 | 0 |
| NAQVRGG' | AARS1; AARS1; | P49588; A0A | SYAC_HUMAN | Unmodified | 0 | 0 | 0 | 0 | 4066800 | 794210 | 1181300 | 0 | 0 | 1677600 | 0 | 0 | 3063800 | 3139900 | 0 | 0 | 1485200 | 1177100 |
| NAREITQD' | SIRPA | P78324 | SHPS1_HUMAI | Unmodified | 112420 | 0 | 0 | 0 | 4657300 | 0 | 0 | 0 | 0 | 0 | 0 | 0 | 1413800 | 0 | 78819 | 0 | 3611200 | 8806900 |
| NASCYFDII | FSCN1; FSCN1 | Q16658; C9J | FSCN1_HUMAI | Unmodified | 0 | 170410 | 0 | 0 | 1762300 | 345420 | 0 | 0 | 189400 | 180030 | 822410 | 506690 | 0 | 0 | 0 | 0 | 612730 | 952360 |
| NASCYFDII | FSCN1; FSCN1 | Q16658; C9J | FSCN1_HUMAI | Unmodified | 0 | 0 | 0 | 0 | 2347600 | 1101100 | 0 | 0 | 193960 | 636150 | 0 | 0 | 0 | 0 | 0 | 0 | 1006300 | 2306700 |
| NASDGALN | NOTCH1; NOT( | P46531; A0A | NOTC1_HUMA | Oxidation (M) | 0 | 0 | 0 | 0 | 0 | 0 | 0 | 0 | 0 | 0 | 486350 | 1629000 | 0 | 0 | 84485 | 0 | 0 | 0 |
| NASEMIDK | ATP5F1C | P36542 | ATPG_HUMAN | Unmodified | 436130 | 0 | 204940 | 216620 | 4441200 | 76348 | 0 | 52694 | 3766900 | 0 | 15562 | 0 | 936260 | 336760 | 92448 | 106600 | 795880 | 616800 |
| NASEMIDK | ATP5F1C | P36542 | ATPG_HUMAN | Unmodified | 0 | 0 | 0 | 0 | 0 | 0 | 0 | 0 | 4123200 | 0 | 0 | 0 | 0 | 0 | 0 | 0 | 0 | 0 |
| NASEMIDK | ATP5F1C | P36542 | ATPG_HUMAN | Oxidation (M) | 0 | 0 | 0 | 0 | 0 | 0 | 0 | 0 | 1009200 | 248890 | 0 | 0 | 0 | 0 | 0 | 0 | 0 | 0 |
| NASEMIDK | ATP5F1C | P36542 | ATPG_HUMAN | Oxidation (M) | 0 | 0 | 0 | 0 | 0 | 0 | 0 | 0 | 0 | 823750 | 0 | 0 | 0 | 0 | 0 | 0 | 0 | 0 |
| NASEMIDK | ATP5F1C | P36542 | ATPG_HUMAN | Unmodified | 0 | 0 | 0 | 0 | 0 | 0 | 0 | 0 | 420990 | 907850 | 0 | 0 | 0 | 0 | 0 | 0 | 0 | 0 |
| NASTFEDV | CTTN | Q14247 | SRC8_HUMAN | Unmodified | 2611800 | 1849200 | 1801900 | 2070900 | 13854000 | 2809500 | 514550 | 454610 | 640790 | 574250 | 160380 | 1618200 | 8117600 | 5972200 | 2667700 | 2044400 | 5914100 | 6562000 |
| NATFTWAR | ABCC1; ABCC1 | P33527; I3L4 | MRP1_HUMAN | Unmodified | 0 | 0 | 415080 | 291040 | 0 | 0 | 0 | 0 | 0 | 0 | 232760 | 3812200 | 0 | 0 | 0 | 529740 | 0 | 0 |
| NATLAWDS | ABCC5 | O15440 | MRP5_HUMAN | Unmodified | 0 | 0 | 0 | 0 | 0 | 0 | 0 | 0 | 0 | 0 | 1368800 | 3411900 | 0 | 0 | 0 | 0 | 0 | 0 |
| NATLSLNSI | AVIL | O75366 | AVIL_HUMAN A | Unmodified | 203290 | 0 | 0 | 0 | 2793700 | 0 | 0 | 0 | 0 | 0 | 2902500 | 16004000 | 0 | 0 | 0 | 0 | 0 | 829020 |
| NATNVEQA | RAB1C; RAB1B; | Q92928; Q9I | RAB1C_HUMAI | Oxidation (M) | 1180900 | 1075100 | 515700 | 3866600 | 3975600 | 322020 | 1232700 | 27074 | 4143100 | 1861800 | 786780 | 2171100 | 3908500 | 2462400 | 521530 | 1062800 | 417830 | 3256000 |
| NATNVEQA | RAB1C; RAB1B; | Q92928; Q9I | RAB1C_HUMAI | Unmodified | 0 | 0 | 0 | 0 | 623280 | 0 | 245960 | 0 | 1158000 | 0 | 0 | 0 | 0 | 0 | 0 | 0 | 0 | 345370 |
| NATNVEQA | RAB1C; RAB1B; | Q92928; Q9I | RAB1C_HUMAI | 2 Oxidation (M) | 125870 | 145730 | 0 | 428780 | 538120 | 0 | 140040 | 52620 | 0 | 235940 | 0 | 1789600 | 671750 | 0 | 409160 | 651010 | 80627 | 0 |
| NATNVEQA | RAB1C; RAB1B; | Q92928; Q9I | RAB1C_HUMAI | Unmodified | 0 | 0 | 0 | 0 | 0 | 0 | 0 | 0 | 1616500 | 0 | 209600 | 0 | 0 | 0 | 0 | 0 | 0 | 0 |
| NATNVEQA | RAB1C; RAB1B; | Q92928; Q9I | RAB1C_HUMAI | Oxidation (M) | 0 | 1236200 | 549610 | 1141400 | 1871000 | 0 | 0 | 0 | 2013200 | 552750 | 0 | 4763300 | 2352100 | 1918100 | 2123300 | 881010 | 0 | 1928900 |
| NATNVEQA | RAB1B; RAB1B | Q9H0U4; E9 | RAB1B_HUMAI | Oxidation (M) | 0 | 0 | 0 | 0 | 575300 | 0 | 341600 | 0 | 0 | 339590 | 0 | 0 | 2617200 | 0 | 0 | 0 | 502780 | 1034500 |
| NATNVEQS | RAB1A; RAB1A | P62820; E7E | RAB1A_HUMAI | Oxidation (M) | 1050000 | 0 | 0 | 0 | 5037500 | 1203200 | 1997700 | 35585 | 619320 | 1948900 | 767460 | 3055600 | 3608200 | 0 | 0 | 104700 | 393530 | 1247000 |
| NATNVEQS | RAB1A; RAB1A | P62820; E7E | RAB1A_HUMAI | 2 Oxidation (M) | 659790 | 771350 | 0 | 768890 | 1131200 | 0 | 344840 | 222160 | 71668 | 0 | 1093900 | 1882100 | 505590 | 0 | 0 | 212920 | 0 | 249570 |
| NATNVEQS | RAB1A; RAB1A | P62820; E7E | RAB1A_HUMAI | Unmodified | 12365000 | 486210 | 2740400 | 0 | 1784200 | 0 | 964090 | 33355 | 1705900 | 575800 | 418840 | 0 | 954530 | 0 | 0 | 0 | 0 | 1020600 |
| NATNVEQS | RAB1A; RAB1A | P62820; E7E | RAB1A_HUMAI | Oxidation (M) | 0 | 0 | 0 | 1786500 | 2881600 | 0 | 458680 | 0 | 0 | 0 | 514870 | 3815900 | 939410 | 2196500 | 0 | 733760 | 0 | 0 |
| NATNVEQS | RAB1A; RAB1A | P62820; E7E | RAB1A_HUMAI | Oxidation (M) | 0 | 0 | 0 | 0 | 378070 | 0 | 0 | 29781 | 0 | 0 | 0 | 0 | 740450 | 781690 | 0 | 0 | 561500 | 0 |
| NATSLQHL | NAA16 | Q6N069 | NAA16_HUMAI | Unmodified | 0 | 1464700 | 1523600 | 4651600 | 0 | 0 | 0 | 0 | 0 | 0 | 0 | 0 | 650370 | 1357300 | 2600500 | 1562700 | 0 | 0 |
| NATTDALTS | ANK3; ANK3; AI | Q12955; H0' | ANK3_HUMAN | Unmodified | 0 | 0 | 984990 | 0 | 0 | 0 | 0 | 52989 | 529030 | 0 | 0 | 2178200 | 0 | 0 | 0 | 1296500 | 0 | 0 |
| NATVHPGL | PRPSAP1; PRPS | Q14558; C9J | KPRA_HUMAN | Oxidation (M) | 0 | 0 | 0 | 0 | 149360 | 0 | 55764 | 27684 | 188120 | 55507 | 0 | 0 | 120380 | 74801 | 0 | 0 | 275060 | 260900 |
| NATVHPGL | PRPSAP1; PRPS | Q14558; C9J | KPRA_HUMAN | Unmodified | 0 | 0 | 0 | 0 | 0 | 0 | 0 | 0 | 0 | 0 | 0 | 0 | 0 | 0 | 0 | 0 | 0 | 196820 |
| NATVHPGL | PRPSAP1; PRPS | Q14558; C9J | KPRA_HUMAN | 2 Oxidation (M) | 0 | 0 | 0 | 0 | 181130 | 0 | 260690 | 128060 | 0 | 0 | 0 | 0 | 161410 | 0 | 0 | 0 | 198850 | 251960 |
| NAVEEYVYI | HSPA4L; HSPA4 | O95757; E9F | HS74L_HUMAI | Unmodified | 0 | 0 | 0 | 0 | 0 | 0 | 0 | 0 | 0 | 0 | 0 | 1260300 | 0 | 0 | 0 | 0 | 0 | 0 |
| NAVEEYVYI | HSPA4L; HSPA4 | O95757; E9F | HS74L_HUMAI | Unmodified | 0 | 0 | 57162 | 0 | 0 | 0 | 0 | 0 | 193070 | 0 | 505720 | 4276100 | 103280 | 0 | 0 | 0 | 0 | 0 |
| NAVEEYVYI | HSPA4L; HSPA4 | O95757; E9F | HS74L_HUMAI | Unmodified | 0 | 0 | 0 | 0 | 1264700 | 206150 | 363610 | 0 | 424250 | 882010 | 1537400 | 91175 | 5319100 | 2026400 | 0 | 87948 | 1116400 | 718300 |
| NAVEEYVYI | HSPH1; HSPH1 | Q92598; A0A | HS105_HUMAI | Unmodified | 0 | 0 | 0 | 0 | 0 | 0 | 0 | 0 | 0 | 0 | 0 | 2206600 | 0 | 0 | 0 | 0 | 0 | 0 |
| NAVEEYVYI | HSPH1; HSPH1 | Q92598; A0A | HS105_HUMAI | Unmodified | 0 | 0 | 0 | 1776400 | 1884100 | 0 | 2250000 | 304610 | 3120400 | 1948600 | 2068200 | 5585500 | 3333400 | 2280400 | 0 | 0 | 3886200 | 2748300 |
| NAVEEYVYI | HSPA4; HSPA4 | P34932; A0A | HSP74_HUMAI | Oxidation (M) | 0 | 0 | 225610 | 0 | 0 | 0 | 0 | 0 | 0 | 0 | 0 | 1634700 | 0 | 0 | 0 | 317020 | 0 | 0 |
| NAVEEYVYI | HSPA4; HSPA4 | P34932; A0A | HSP74_HUMAI | Unmodified | 0 | 0 | 0 | 0 | 0 | 0 | 0 | 0 | 408710 | 0 | 0 | 3231800 | 0 | 0 | 0 | 0 | 0 | 0 |
| NAVEEYVYI | HSPA4; HSPA4 | P34932; A0A | HSP74_HUMAI | Unmodified | 0 | 0 | 188540 | 0 | 2137500 | 461270 | 0 | 0 | 446280 | 0 | 929520 | 1788500 | 3164200 | 0 | 0 | 0 | 0 | 1393100 |
| NAVEEYVYI | HSPA4; HSPA4 | P34932; A0A | HSP74_HUMAI | Unmodified | 0 | 0 | 0 | 0 | 2235800 | 459050 | 333640 | 0 | 0 | 525730 | 0 | 0 | 2436500 | 1888900 | 0 | 0 | 640040 | 1603000 |
| NAVHVNLF | MFGE8; MFGE8 | Q08431; H0' | MFGM_HUMAN | Unmodified | 7701600 | 0 | 20307000 | 34474000 | 63689000 | 34008000 | 937550 | 0 | 32978000 | 3225500 | 2572900 | 28388000 | 5143400 | 0 | 30685000 | 17696000 | 2336500 | 0 |
| NAVITVPAY | HSPA1L; HSPA1 | P34931; P0D | HS71L_HUMAI | Unmodified | 0 | 0 | 0 | 0 | 0 | 0 | 0 | 0 | 8686500 | 472890 | 0 | 0 | 0 | 0 | 0 | 0 | 0 | 0 |
| NAVITVPAY | HSPA1L; HSPA1 | P34931; P0D | HS71L_HUMAI | Unmodified | 0 | 0 | 0 | 0 | 3703000 | 0 | 0 | 0 | 0 | 2564700 | 0 | 0 | 0 | 0 | 0 | 0 | 0 | 0 |
| NAVIVALSS | UBE2K | P61086 | UBE2K_HUMAI | Unmodified | 347130 | 0 | 0 | 0 | 0 | 0 | 0 | 0 | 423770 | 0 | 805610 | 2733500 | 0 | 0 | 0 | 397150 | 0 | 0 |
| NAVLNTEA | NCLN | Q969V3 | NCLN_HUMAN | Unmodified | 742660 | 930760 | 1664800 | 590520 | 0 | 0 | 236270 | 142380 | 14078000 | 700280 | 539030 | 743240 | 0 | 0 | 621800 | 696100 | 51905 | 0 |
| NAVQALIDI | TMEM68; TMEM | Q96MH6; Q4 | TMM68_HUMAI | Unmodified | 0 | 103500 | 0 | 0 | 0 | 0 | 0 | 0 | 1009700 | 41543 | 0 | 344160 | 0 | 0 | 0 | 0 | 0 | 0 |
| NAVTQEFG | ACO2; ACO2; A | Q99798; A2A | ACON_HUMAN | Unmodified | 0 | 0 | 0 | 0 | 0 | 0 | 0 | 0 | 5471300 | 221720 | 0 | 0 | 0 | 44680 | 0 | 0 | 0 | 0 |
| NAVTSMSII | FAM126A; FAM1 | Q9BYI3; A0A | HYCCI_HUMAI | Unmodified | 0 | 0 | 139220 | 135910 | 0 | 0 | 0 | 0 | 0 | 0 | 0 | 0 | 0 | 0 | 204750 | 229220 | 0 | 0 |
| NAVVQGLE | LRP1 | Q07954 | LRP1_HUMAN | Unmodified | 0 | 0 | 1063500 | 174320 | 1314100 | 1566500 | 491380 | 0 | 4520700 | 1319300 | 0 | 0 | 0 | 358810 | 0 | 0 | 0 | 1828200 |
| NAWNIIQA | TSPAN9; TSPAN | O75954; B5I | TSN9_HUMAN | Oxidation (M) | 0 | 0 | 850930 | 844420 | 311510 | 832960 | 0 | 0 | 0 | 0 | 0 | 0 | 78379 | 0 | 984480 | 481030 | 0 | 0 |
| NAWQDAH | MBP; MBP; MBF | P02686; J3K | MBP_HUMAN I | Unmodified | 43545 | 488630 | 212260 | 1336900 | 4647900 | 7674900 | 872320 | 16447 | 0 | 0 | 200430 | 1008300 | 5894500 | 3165700 | 1276300 | 1123300 | 582020 | 3813800 |
| NAYAVLYD | PSME1; PSME1 | Q06323; H0' | PSME1_HUMAI | Unmodified | 75119 | 0 | 41846 | 92757 | 0 | 100070 | 0 | 0 | 467740 | 244910 | 286240 | 701400 | 510600 | 125070 | 0 | 52576 | 159100 | 374490 |
| NAYQLTEK | CEP55 | Q53EZ4 | CEP55_HUMAI | Unmodified | 602130 | 1389500 | 3283300 | 3298400 | 324770 | 3484400 | 55176 | 105710 | 7539700 | 1977300 | 656590 | 3817300 | 454910 | 338800 | 3831500 | 990070 | 274890 | 428500 |
| NAYTPQEIN | GALNS; GALNS | P34059; H3E | GALNS_HUMAI | Unmodified | 0 | 0 | 0 | 0 | 0 | 0 | 0 | 0 | 2423200 | 586550 | 0 | 0 | 44133 | 2455400 | 0 | 0 | 0 | 0 |
| NAYVWSQI | ARPC1A; ARPC | Q92747; E9F | ARC1A_HUMAI | Unmodified | 0 | 27151 | 0 | 0 | 0 | 0 | 0 | 0 | 0 | 0 | 0 | 2186200 | 0 | 0 | 0 | 67987 | 0 | 0 |
| NAYWVTLK | ARPC1B; ARPC | O15143; C9J | ARC1B_HUMAI | Unmodified | 1120300 | 527430 | 458650 | 100270 | 0 | 0 | 0 | 0 | 0 | 116050 | 0 | 4497800 | 0 | 0 | 1268200 | 464260 | 0 | 0 |
| NCDGVPAI | CTNND1; CTNN | O60716; A0A | CTND1_HUMAI | Unmodified | 0 | 0 | 1502600 | 3248600 | 7566400 | 2228800 | 0 | 0 | 0 | 0 | 1539800 | 3942200 | 4920400 | 5017200 | 3002000 | 3888600 | 5937600 | 413490 |
| NCFASVFE | ARPC2 | O15144 | ARPC2_HUMAI | Unmodified | 384180 | 152830 | 298270 | 557720 | 180760 | 0 | 323040 | 164150 | 0 | 0 | 486960 | 4248300 | 0 | 0 | 238050 | 242110 | 212910 | 102090 |
| NCFRMTD( | NPM1; NPM1; N | P06748; A0A | NPM_HUMAN I | Oxidation (M) | 0 | 0 | 0 | 0 | 0 | 0 | 860300 | 0 | 0 | 1076800 | 0 | 0 | 2856200 | 1966800 | 0 | 0 | 1708800 | 1185100 |
| NCGCLGA: | RARS1; RARS1; | P54136; E5R | SYRC_HUMAN | Unmodified | 0 | 0 | 0 | 0 | 0 | 0 | 0 | 0 | 0 | 0 | 0 | 1572100 | 0 | 0 | 0 | 0 | 0 | 0 |
| NCGHRFH | TOM1L2; TOM1 | Q6ZVM7; K7 | TM1L2_HUMAI | Unmodified | 0 | 0 | 0 | 0 | 0 | 0 | 0 | 0 | 0 | 0 | 0 | 0 | 1260700 | 1262700 | 0 | 0 | 0 | 0 |
| NCGQMSE | TLN1 | Q9Y490 | TLN1_HUMAN | Unmodified | 0 | 0 | 0 | 2051800 | 2023300 | 0 | 3713500 | 2211400 | 0 | 0 | 0 | 1396300 | 3178300 | 2920800 | 0 | 0 | 227500 | 2276900 |
| NCGQMSE | TLN1 | Q9Y490 | TLN1_HUMAN | Oxidation (M) | 150340 | 0 | 0 | 0 | 851950 | 0 | 898140 | 727920 | 0 | 0 | 0 | 0 | 0 | 0 | 158720 | 340670 | 0 | 453960 |
| NCGQMSE | TLN1 | Q9Y490 | TLN1_HUMAN | Unmodified | 0 | 0 | 0 | 0 | 0 | 0 | 750710 | 517480 | 0 | 0 | 0 | 203510 | 0 | 105710 | 0 | 0 | 49829 | 0 |
| NCGQTVHI | HGS; HGS; HGS | O14964; A0A | HGS_HUMAN I | Unmodified | 0 | 586490 | 0 | 0 | 299210 | 381460 | 953770 | 0 | 741600 | 640170 | 0 | 0 | 0 | 0 | 0 | 0 | 245950 | 0 |
| NCHCENG | ADAM9; ADAMS | Q13443; A0A | ADAM9_HUMA | Unmodified | 511630 | 0 | 171840 | 325150 | 804470 | 0 | 0 | 0 | 286040 | 0 | 0 | 78613 | 298010 | 229470 | 357160 | 364290 | 0 | 0 |
| NCHLNENI | IDI2; IDI1 | Q9BXS1; Q1 | IDI2_HUMAN Is | Unmodified | 0 | 0 | 0 | 0 | 439330 | 0 | 0 | 0 | 95179 | 0 | 0 | 1526300 | 0 | 0 | 0 | 0 | 22432 | 73545 |
| NCIAQTSA | MIX23; MIX23; I | Q4VC31; C9 | MIX23_HUMAN | Unmodified | 0 | 0 | 0 | 0 | 0 | 0 | 0 | 58896 | 0 | 250500 | 757720 | 1519000 | 449580 | 252570 | 0 | 0 | 27197 | 0 |
| NCIDITGVF | LMAN2; LMAN2 | Q12907; D6I | LMAN2_HUMAI | Unmodified | 77572 | 0 | 115340 | 0 | 0 | 59062 | 0 | 132960 | 5934900 | 690550 | 0 | 113410 | 0 | 0 | 104870 | 54717 | 0 | 0 |
| NCIIVSPDA | PRPS2; PRPS1L | P11908; P21 | PRPS2_HUMAI | Unmodified | 0 | 0 | 0 | 0 | 244460 | 0 | 0 | 206580 | 281270 | 429880 | 0 | 912030 | 708560 | 0 | 0 | 0 | 126690 | 0 |
| NCIVLIDST | RPS8; RPS8 | P62241; Q5J | RS8_HUMAN 4 | Unmodified | 1316800 | 403000 | 582790 | 142760 | 0 | 0 | 579230 | 0 | 0 | 0 | 0 | 12540000 | 0 | 0 | 331980 | 367130 | 0 | 0 |
| NCIVLIDST | RPS8; RPS8 | P62241; Q5J | RS8_HUMAN 4 | Unmodified | 2570500 | 604730 | 0 | 1261400 | 0 | 0 | 2861500 | 0 | 0 | 0 | 178560 | 1851700 | 1580800 | 656180 | 1187500 | 0 | 0 | 0 |
| NCLALADD | MACROH2A1; I | O75367; B4I | H2AY_HUMAN | Unmodified | 3605100 | 1182600 | 0 | 0 | 0 | 0 | 0 | 1615800 | 0 | 0 | 151530 | 361000 | 177180 | 0 | 0 | 0 | 0 | 0 |
| NCLALADD | MACROH2A1; I | O75367; B4I | H2AY_HUMAN | Unmodified | 0 | 0 | 0 | 0 | 0 | 0 | 87931 | 1606600 | 0 | 0 | 0 | 0 | 0 | 0 | 0 | 0 | 0 | 0 |
| NCLLETAK | ROCK2; ROCK2 | O75116; E9F | ROCK2_HUMAI | Unmodified | 0 | 0 | 0 | 0 | 0 | 0 | 0 | 0 | 0 | 0 | 0 | 0 | 0 | 0 | 0 | 0 | 0 | 436670 |
| NCLNPQFS | CPNE3; CPNE3 | O75131; A0A | CPNE3_HUMAI | Unmodified | 0 | 0 | 0 | 0 | 2401500 | 369700 | 0 | 1212500 | 9137700 | 7932100 | 0 | 0 | 2143600 | 2145200 | 0 | 0 | 3972100 | 8186200 |
| NCLTNFHC | RPS3A; RPS3A; | P61247; D6F | RS3A_HUMAN | Unmodified | 1713200 | 0 | 0 | 0 | 707110 | 0 | 1002300 | 409900 | 465180 | 140670 | 690160 | 9075400 | 524970 | 607400 | 635380 | 0 | 107940 | 629870 |
| NCLTNFHC | RPS3A; RPS3A; | P61247; D6F | RS3A_HUMAN | Oxidation (M) | 269740 | 204130 | 202310 | 185090 | 199690 | 0 | 243050 | 11746 | 0 | 74102 | 226530 | 3079300 | 154620 | 171330 | 208880 | 170390 | 0 | 208640 |
| NCLVDRDL | BMX | P51813 | BMX_HUMAN ( | Unmodified | 0 | 0 | 0 | 0 | 0 | 0 | 0 | 0 | 0 | 0 | 155690 | 1104400 | 0 | 0 | 0 | 0 | 0 | 0 |
| NCLVSAQF | PTK7; PTK7; PTI | Q13308; H0' | PTK7_HUMAN I | Unmodified | 0 | 0 | 0 | 0 | 525030 | 0 | 0 | 0 | 3170200 | 404680 | 901330 | 3166500 | 631090 | 55307 | 1443400 | 875330 | 0 | 152340 |
| NCLVSAQF | PTK7; PTK7; PTI | Q13308; H0' | PTK7_HUMAN I | Unmodified | 0 | 0 | 0 | 0 | 1149000 | 0 | 83925 | 0 | 0 | 500530 | 0 | 0 | 0 | 0 | 0 | 0 | 579060 | 810610 |
| NCNDFQYI | YWHAH; YWHA | Q04917; A2I | 1433F_HUMAI | Unmodified | 161040 | 0 | 0 | 459340 | 2474900 | 834610 | 512550 | 316280 | 764560 | 343650 | 597010 | 2681100 | 1053900 | 377800 | 313760 | 299530 | 1396900 | 1107500 |
| NCPHIVVG | DDX39B; DDX3 | Q13838; F6I | DX39B_HUMAI | Unmodified | 0 | 0 | 0 | 0 | 0 | 0 | 94968 | 0 | 372760 | 218200 | 775710 | 6687700 | 0 | 0 | 0 | 0 | 0 | 0 |
| NCPHVVVC | DDX39A; DDX3 | O00148; A0A | DX39A_HUMAI | Unmodified | 83250 | 0 | 0 | 0 | 0 | 32031 | 161650 | 101530 | 0 | 0 | 100350 | 667850 | 0 | 0 | 0 | 0 | 0 | 0 |
| NCPISAKLI | MACF1; MACF1 | Q9UPN3; H3 | MACF1_HUMAI | Unmodified | 0 | 0 | 0 | 0 | 0 | 0 | 0 | 0 | 0 | 0 | 0 | 0 | 89404 | 278880 | 0 | 0 | 0 | 0 |
| NCPLIDNIC | ACSL4 | O60488 | ACSL4_HUMAI | Unmodified | 0 | 0 | 0 | 33472 | 2438500 | 476970 | 0 | 0 | 0 | 0 | 857130 | 4433500 | 0 | 0 | 11622 | 0 | 3182400 | 2369200 |
| NCQFETAC | ADAM17 | P78536 | ADA17_HUMAI | Unmodified | 74893000 | 68795000 | 71468000 | 0 | 877280 | 377390 | 0 | 0 | 298750 | 135260 | 38837 | 317450 | 4783800 | 1676500 | 46008000 | 60073000 | 621240 | 1240700 |
| NCQFETAC | ADAM17 | P78536 | ADA17_HUMAI | Unmodified | 0 | 304070 | 0 | 0 | 138590 | 0 | 0 | 0 | 0 | 251770 | 0 | 0 | 110270 | 0 | 0 | 38839 | 900500 | 58002 |
| NCQTLVNL | NOTCH2 | Q04721 | NOTC2_HUMA | Unmodified | 0 | 0 | 0 | 0 | 0 | 0 | 0 | 0 | 0 | 0 | 0 | 0 | 941380 | 1832100 | 106100 | 40220 | 116370 | 0 |
| NCSAACP( | ITGB2; ITGB2; I | P05107; D3I | ITB2_HUMAN I | Unmodified | 0 | 1050300 | 0 | 816830 | 0 | 0 | 114880 | 0 | 0 | 0 | 0 | 0 | 0 | 236130 | 0 | 442530 | 0 | 210430 |
| NCSETQYE | YWHAG | P61981 | 1433G_HUMAI | Unmodified | 0 | 0 | 0 | 0 | 0 | 278520 | 0 | 0 | 0 | 0 | 0 | 1172300 | 0 | 0 | 0 | 0 | 734120 | 1117400 |
| NCSETQYE | YWHAG | P61981 | 1433G_HUMAI | Unmodified | 0 | 0 | 0 | 0 | 0 | 0 | 1085000 | 935390 | 0 | 0 | 0 | 0 | 1550300 | 1382100 | 0 | 0 | 6798500 | 1512900 |
| NCSPHMN | ENPP4 | Q9Y6X5 | ENPP4_HUMAI | Unmodified | 0 | 0 | 0 | 0 | 0 | 0 | 0 | 0 | 3340100 | 0 | 699900 | 951350 | 0 | 0 | 0 | 0 | 184980 | 0 |
| NCSSFLIK | RPL28; RPL28; I | P46779; H0' | RL28_HUMAN I | Unmodified | 520600 | 0 | 0 | 0 | 0 | 0 | 118980 | 0 | 0 | 0 | 0 | 2906700 | 0 | 0 | 0 | 0 | 0 | 0 |
| NCSSPEFS | CPNE1; CPNE1 | Q99829; Q5. | CPNE1_HUMAI | Unmodified | 0 | 0 | 79272 | 215140 | 321730 | 0 | 0 | 0 | 571310 | 117760 | 72761 | 871200 | 0 | 42292 | 105380 | 80338 | 192300 | 557390 |
| NCTYTQVC | TCEA1; TCEA2; | P23193; Q1! | TCEA1_HUMAI | Unmodified | 0 | 0 | 0 | 0 | 0 | 0 | 120930 | 0 | 0 | 153030 | 0 | 165860 | 0 | 37631 | 0 | 0 | 0 | 0 |
| NCWETEAS | TYK2; TYK2; TYK | P29597; E9P | TYK2_HUMAN I | Unmodified | 0 | 0 | 0 | 0 | 853420 | 0 | 40176 | 0 | 0 | 0 | 0 | 0 | 33696 | 37878 | 0 | 0 | 729340 | 1077300 |
| NDAPLHEII | GPC4 | O75487 | GPC4_HUMAN | Unmodified | 976030 | 603860 | 0 | 0 | 0 | 0 | 0 | 0 | 1707100 | 0 | 0 | 230120 | 467910 | 0 | 41146 | 300500 | 0 | 0 |
| NDAPVVQE | ROR1; ROR2; R | Q01973; Q0: | ROR1_HUMAN | Unmodified | 0 | 0 | 0 | 0 | 879120 | 672500 | 0 | 0 | 56152 | 165810 | 0 | 305020 | 0 | 0 | 0 | 0 | 287070 | 0 |
| NDASHLLV | ITGB3; N/A; ITG | P05106; H3E | ITB3_HUMAN I | Unmodified | 5823100 | 9045500 | 8018800 | 12250000 | 399540 | 145510 | 235540 | 115930 | 0 | 0 | 0 | 0 | 601270 | 281440 | 9643800 | 11046000 | 32143 | 0 |
| NDDDQKIE | GAP43 | P17677 | NEUM_HUMAN | Unmodified | 0 | 0 | 0 | 0 | 0 | 0 | 700670 | 430500 | 124060 | 152080 | 0 | 0 | 0 | 0 | 0 | 0 | 0 | 0 |
| NDDGDTV( | STX2 | P32856 | STX2_HUMAN ! | Unmodified | 0 | 0 | 119850 | 75527 | 0 | 0 | 0 | 0 | 0 | 0 | 392900 | 2836000 | 0 | 0 | 0 | 286700 | 0 | 0 |
| NDDRLLQ/ | RAI14; RAI14; R | Q9P0K7; D6I | RAI14_HUMAN | Unmodified | 0 | 0 | 96103 | 171050 | 331240 | 159950 | 0 | 0 | 0 | 0 | 0 | 595800 | 191200 | 0 | 0 | 0 | 127790 | 178020 |
| NDDTPNR( | SCG2 | P13521 | SCG2_HUMAN | Oxidation (M) | 0 | 0 | 0 | 0 | 0 | 0 | 0 | 0 | 1548500 | 290440 | 0 | 0 | 0 | 0 | 0 | 0 | 0 | 0 |
| NDDTPNR( | SCG2 | P13521 | SCG2_HUMAN | Unmodified | 0 | 0 | 0 | 0 | 0 | 0 | 0 | 45817 | 1587400 | 0 | 0 | 0 | 0 | 0 | 0 | 0 | 0 | 0 |
| NDEELNK | H2AX; H2AC6; I | P16104; Q9 | H2AX_HUMAN | Unmodified | 92012 | 95482 | 53315 | 53341 | 44700 | 0 | 21202 | 36977 | 50750 | 46954 | 67344 | 518500 | 106130 | 43122 | 62905 | 0 | 0 | 0 |
| NDEELNKL | H2AC18; H2AC | Q6FI13; Q99 | H2A2A_HUMAI | Unmodified | 2083000 | 3344100 | 0 | 0 | 68230 | 0 | 161860 | 217700 | 0 | 350590 | 0 | 863020 | 0 | 0 | 0 | 0 | 38585 | 0 |
| NDFEPVKQ | MYO9B; MYO9E | Q13459; M0 | MYO9B_HUMAI | Unmodified | 0 | 0 | 0 | 0 | 342340 | 47916 | 0 | 0 | 238850 | 427750 | 0 | 135630 | 1518100 | 5871900 | 0 | 0 | 235590 | 0 |
| NDFFGHYI | JAG2 | Q9Y219 | JAG2_HUMAN I | Unmodified | 0 | 0 | 0 | 137810 | 0 | 0 | 0 | 13725 | 557980 | 37060 | 0 | 219990 | 0 | 0 | 109280 | 119200 | 0 | 0 |

| | | | | | | | | | | | | | | | | | | | | | | |
|---|---|---|---|---|---|---|---|---|---|---|---|---|---|---|---|---|---|---|---|---|---|---|
| NDFTEEEA | SKP1; SKP1 | P63208; E5R | SKP1_HUMAN | Unmodified | 0 | 0 | 0 | 0 | 0 | 0 | 0 | 0 | 0 | 0 | 504050 | 367910 | 0 | 0 | 0 | 0 | 0 | 0 |
| NDFTEEEA | SKP1; SKP1 | P63208; E5R | SKP1_HUMAN | Unmodified | 0 | 0 | 1373800 | 0 | 444870 | 0 | 0 | 0 | 421100 | 0 | 231540 | 3372500 | 0 | 0 | 0 | 0 | 0 | 0 |
| NDGEESTD | SLC17A5 | Q9NRA2 | S17A5_HUMAN | Unmodified | 0 | 0 | 0 | 0 | 0 | 0 | 0 | 0 | 650290 | 0 | 0 | 0 | 0 | 0 | 0 | 0 | 0 | 0 |
| NDGFSSYL | GPR107; GPR10 | Q5VW38; U3 | GP107_HUMAN | Unmodified | 24384 | 0 | 0 | 0 | 0 | 0 | 0 | 0 | 3619600 | 0 | 0 | 0 | 0 | 0 | 0 | 0 | 0 | 0 |
| NDGTFIGYI | AKT1 | P31749 | AKT1_HUMAN | Unmodified | 0 | 0 | 936080 | 61887 | 205730 | 0 | 0 | 115260 | 289950 | 0 | 1299000 | 3338700 | 1876500 | 1273100 | 1663700 | 0 | 0 | 197040 |
| NDGVLLLQ | USO1 | O60763 | USO1_HUMAN | Unmodified | 0 | 0 | 0 | 0 | 0 | 0 | 0 | 0 | 0 | 0 | 0 | 628280 | 0 | 0 | 0 | 0 | 0 | 0 |
| NDHQEDF | RNPEP; RNPEP | Q9H4A4; A6 | AMPB_HUMAN | Unmodified | 0 | 0 | 193090 | 0 | 0 | 0 | 0 | 0 | 0 | 0 | 262780 | 1089100 | 86801 | 0 | 0 | 0 | 0 | 0 |
| NDIASHPP | SND1; SND1 | Q7KZF4; H7 | SND1_HUMAN | Unmodified | 184510 | 327530 | 701500 | 429750 | 86593 | 0 | 0 | 0 | 0 | 0 | 775130 | 10710000 | 0 | 0 | 1164900 | 1245500 | 0 | 0 |
| NDIPEIPES | SCRIB; SCRIB; | Q14160; A0 | SCRIB_HUMAN | Unmodified | 0 | 0 | 0 | 0 | 0 | 0 | 0 | 0 | 931330 | 0 | 0 | 234460 | 0 | 0 | 0 | 0 | 0 | 0 |
| NDIVNMLV | RAB18; RAB18; | Q9NP72; H0 | RAB18_HUMAN | Oxidation (M) | 0 | 0 | 121320 | 0 | 0 | 0 | 0 | 0 | 0 | 0 | 0 | 3566300 | 0 | 0 | 80333 | 75842 | 0 | 0 |
| NDLAVVDV | NCL; NCL; NCL | P19338; C9J | NUCL_HUMAN | Unmodified | 1644600 | 110520 | 0 | 37794000 | 0 | 0 | 779310 | 0 | 2084400 | 386390 | 3084300 | 13552000 | 0 | 0 | 0 | 0 | 78749 | 0 |
| NDLEEAFIH | ADPGK; ADPGK | Q9BRR6; H3 | ADPGK_HUMA | Unmodified | 0 | 0 | 0 | 0 | 0 | 0 | 0 | 0 | 1006700 | 0 | 0 | 49544 | 0 | 0 | 0 | 0 | 0 | 0 |
| NDLGVGYL | ASPH | Q12797 | ASPH_HUMAN | Unmodified | 0 | 0 | 0 | 126510 | 0 | 0 | 0 | 0 | 2532200 | 0 | 67041 | 149980 | 0 | 0 | 0 | 0 | 0 | 0 |
| NDLGVGYL | ASPH | Q12797 | ASPH_HUMAN | Unmodified | 0 | 1824800 | 0 | 0 | 0 | 0 | 0 | 0 | 2076700 | 260660 | 0 | 0 | 0 | 0 | 0 | 0 | 0 | 0 |
| NDLIVFLAD | ITFG1 | Q8TB96 | TIP_HUMAN T- | Unmodified | 0 | 0 | 0 | 0 | 0 | 0 | 0 | 0 | 12394000 | 1342100 | 0 | 0 | 0 | 0 | 0 | 0 | 0 | 0 |
| NDLSPASS | LBR; LBR | Q14739; A0 | LBR_HUMAN D | Unmodified | 0 | 0 | 0 | 0 | 0 | 0 | 0 | 0 | 35623 | 48861 | 0 | 358380 | 0 | 0 | 17232 | 0 | 0 | 0 |
| NDLSPTTVI | BLVRB; BLVRB; | P30043; M0 | BLVRB_HUMAN | Unmodified | 0 | 0 | 0 | 0 | 0 | 0 | 0 | 0 | 0 | 0 | 0 | 0 | 0 | 0 | 596660 | 420290 | 0 | 0 |
| NDLTREEIV | CASP3; CASP3; | P42574; A8 | CASP3_HUMAN | Unmodified | 0 | 0 | 0 | 0 | 0 | 0 | 0 | 0 | 0 | 95457 | 0 | 785360 | 0 | 0 | 0 | 0 | 0 | 0 |
| NDNTEAFY | DNAJC3; DNAJ | Q13217; X6 | DNJC3_HUMAN | Unmodified | 215620 | 0 | 0 | 0 | 0 | 0 | 0 | 0 | 488960 | 342710 | 0 | 0 | 0 | 0 | 0 | 0 | 0 | 0 |
| NDPHLCDI | FTH1; FTH1; FT | P02794; G3 | FRIH_HUMAN | Unmodified | 0 | 0 | 0 | 0 | 0 | 0 | 0 | 0 | 3818500 | 336520 | 0 | 0 | 0 | 0 | 0 | 0 | 0 | 0 |
| NDPSLPEP | NARS1; NARS1 | O43776; K7 | SYNC_HUMAN | Unmodified | 0 | 0 | 0 | 0 | 0 | 0 | 0 | 0 | 0 | 0 | 0 | 5549000 | 0 | 0 | 0 | 0 | 0 | 0 |
| NDQANYSI | ADA; ADA | P00813; F5G | ADA_HUMAN A | Unmodified | 0 | 0 | 0 | 100080 | 0 | 0 | 0 | 0 | 0 | 0 | 0 | 620100 | 51102 | 0 | 124150 | 41449 | 0 | 0 |
| NDQCYDD | CORO1C; COR | Q9ULV4; F8 | COR1C_HUMA | Unmodified | 0 | 0 | 0 | 0 | 0 | 0 | 0 | 0 | 0 | 0 | 0 | 1358800 | 0 | 0 | 0 | 0 | 0 | 0 |
| NDQDTWD | HNRNPL; HNRN | P14866; M0 | HNRPL_HUMA | Unmodified | 0 | 0 | 0 | 0 | 0 | 0 | 0 | 0 | 0 | 0 | 0 | 1908400 | 0 | 0 | 0 | 0 | 0 | 0 |
| NDQDTWD | HNRNPL; HNRN | P14866; M0 | HNRPL_HUMA | Unmodified | 0 | 0 | 0 | 0 | 0 | 0 | 0 | 0 | 0 | 0 | 0 | 1993800 | 0 | 0 | 0 | 0 | 0 | 0 |
| NDQFSYFK | N/A | N/A | N/A | Unmodified | 785070 | 11964000 | 0 | 0 | 0 | 0 | 0 | 0 | 0 | 0 | 0 | 0 | 0 | 0 | 0 | 0 | 166980 | 0 |
| NDQWHWI | ROCK2; ROCK2 | O75116; D6 | ROCK2_HUMA | Unmodified | 0 | 0 | 0 | 0 | 0 | 0 | 0 | 164820 | 3429500 | 614480 | 0 | 282490 | 0 | 0 | 0 | 0 | 0 | 0 |
| NDRDVLGI | CTH | P32929 | CGL_HUMAN C | Unmodified | 0 | 0 | 0 | 290080 | 0 | 0 | 0 | 0 | 0 | 0 | 0 | 874050 | 0 | 0 | 386290 | 0 | 0 | 0 |
| NDSIYEASS | MVB12A; MVB1 | Q96EY5; M0 | MB12A_HUMA | Unmodified | 0 | 0 | 0 | 0 | 0 | 0 | 0 | 0 | 0 | 0 | 0 | 0 | 0 | 0 | 964980 | 0 | 0 | 0 |
| NDSLAGVV | YKT6; YKT6; YK | O15498; A0 | YKT6_HUMAN | Unmodified | 0 | 0 | 0 | 0 | 0 | 0 | 0 | 0 | 0 | 0 | 0 | 1477300 | 0 | 153450 | 0 | 0 | 0 | 0 |
| NDTKEDVF | YBX1; YBX2; YB | P67809; Q9 | YBOX1_HUMAN | Unmodified | 805630 | 288430 | 300950 | 611530 | 0 | 0 | 181150 | 94663 | 0 | 0 | 912570 | 4832800 | 0 | 0 | 0 | 365790 | 0 | 0 |
| NDTKEDVF | YBX1; YBX3; YB | P67809; P16 | YBOX1_HUMAN | Unmodified | 363770 | 177450 | 222170 | 151010 | 0 | 0 | 0 | 0 | 0 | 0 | 283680 | 2914700 | 0 | 0 | 158750 | 306900 | 0 | 0 |
| NDVAPTLM | CDH1 | P12830 | CADH1_HUMA | Unmodified | 0 | 0 | 0 | 393360 | 0 | 0 | 0 | 0 | 0 | 0 | 0 | 1267300 | 0 | 0 | 0 | 0 | 0 | 0 |
| NDVAWNF | GPX1; GPX1 | P07203; A0A | GPX1_HUMAN | Unmodified | 0 | 0 | 0 | 0 | 0 | 0 | 0 | 0 | 1222100 | 0 | 0 | 214420 | 0 | 0 | 0 | 93979 | 0 | 0 |
| NDVGPYEC | CEACAM5 | P06731 | CEAM5_HUMA | Unmodified | 0 | 0 | 0 | 0 | 0 | 0 | 0 | 0 | 15547000 | 490310 | 0 | 0 | 0 | 0 | 0 | 0 | 0 | 0 |
| NDVLAHQ | MACF1; MACF1 | Q9UPN3; H3 | MACF1_HUMA | Unmodified | 0 | 0 | 0 | 956530 | 0 | 0 | 0 | 0 | 135640 | 57401 | 0 | 369320 | 0 | 151870 | 0 | 0 | 0 | 0 |
| NDVPLPTD | JAM3 | Q9BX67 | JAM3_HUMAN | Unmodified | 0 | 0 | 0 | 0 | 0 | 0 | 323140 | 0 | 2704600 | 213470 | 113430 | 3946800 | 0 | 0 | 0 | 14775 | 0 | 0 |
| NDVPLPTD | JAM3 | Q9BX67 | JAM3_HUMAN | Unmodified | 2429500 | 637750 | 1300300 | 2360300 | 251250 | 25447 | 15781 | 0 | 3778100 | 347450 | 0 | 0 | 142540 | 0 | 1138900 | 1478400 | 213270 | 492030 |
| NDVVGTTY | MYOF | Q9NZM1 | MYOF_HUMAN | Unmodified | 0 | 0 | 144130 | 0 | 10229000 | 2095300 | 0 | 0 | 0 | 0 | 0 | 0 | 0 | 0 | 92258 | 0 | 301190 | 1181900 |
| NDYAVESY | NIBAN1 | Q9BZQ8 | NIBA1_HUMAN | Unmodified | 0 | 0 | 167670 | 0 | 0 | 0 | 0 | 0 | 1521900 | 132910 | 0 | 0 | 0 | 0 | 124750 | 105770 | 0 | 0 |
| NDYEARPQ | DIAPH1; DIAPH | O60610; A0 | DIAP1_HUMAN | Unmodified | 0 | 0 | 191870 | 0 | 0 | 0 | 0 | 0 | 0 | 0 | 0 | 1267200 | 0 | 0 | 0 | 0 | 0 | 0 |
| NDYVAAER | NAPG | Q99747 | SNAG_HUMAN | Unmodified | 0 | 0 | 0 | 0 | 0 | 0 | 0 | 0 | 0 | 0 | 0 | 1185900 | 0 | 0 | 0 | 0 | 0 | 0 |
| NDYVHALV | PRMT1; PRMT1 | Q99873; E9 | ANM1_HUMAN | Unmodified | 0 | 0 | 2586800 | 0 | 0 | 0 | 0 | 0 | 0 | 0 | 16821000 | 0 | 0 | 0 | 0 | 0 | 0 | 370790 |
| NEAIQAAH | UCHL1; UCHL1 | P09936; D6 | UCHL1_HUMA | Unmodified | 0 | 0 | 0 | 0 | 274470 | 0 | 102150 | 0 | 2592300 | 299630 | 3067800 | 17931000 | 0 | 0 | 0 | 0 | 0 | 0 |
| NEAIQAAH | UCHL1; UCHL1 | P09936; D6 | UCHL1_HUMA | Unmodified | 0 | 0 | 0 | 0 | 5648300 | 1970200 | 0 | 26854 | 14878000 | 1243500 | 6405500 | 46914000 | 0 | 0 | 0 | 0 | 0 | 0 |
| NEALAPPLI | MGAT2 | Q10469 | MGAT2_HUMA | Unmodified | 0 | 0 | 0 | 0 | 0 | 0 | 0 | 0 | 1270500 | 0 | 0 | 313330 | 0 | 0 | 0 | 0 | 0 | 0 |
| NEALAPPLI | MGAT2 | Q10469 | MGAT2_HUMA | Unmodified | 0 | 0 | 0 | 0 | 0 | 0 | 757140 | 0 | 1995900 | 0 | 0 | 0 | 0 | 0 | 0 | 0 | 0 | 0 |
| NEANADVL | RELL1; RELL1 | Q8IUW5; D6 | RELL1_HUMAN | Unmodified | 423920 | 586110 | 624870 | 620330 | 4927300 | 0 | 689260 | 392890 | 2045900 | 623150 | 0 | 111290 | 2715500 | 1670500 | 424270 | 431370 | 133680 | 515290 |
| NECSLHIH | SORT1 | Q99523 | SORT_HUMAN | Unmodified | 0 | 0 | 0 | 0 | 0 | 0 | 0 | 0 | 872160 | 237090 | 0 | 0 | 0 | 0 | 0 | 0 | 0 | 0 |
| NEDDCVVF | ITGB3; N/A | P05106; H3 | ITB3_HUMAN I | Unmodified | 44595 | 52621 | 653380 | 81353 | 0 | 0 | 0 | 0 | 72319 | 0 | 0 | 0 | 0 | 0 | 379920 | 0 | 0 | 0 |
| NEDDCVVF | ITGB3; N/A | P05106; H3 | ITB3_HUMAN I | Unmodified | 0 | 0 | 754600 | 0 | 0 | 0 | 292790 | 24845 | 0 | 0 | 0 | 0 | 0 | 0 | 0 | 0 | 0 | 0 |
| NEDFAAHF | AKAP12 | Q02952 | AKA12_HUMAN | Unmodified | 0 | 0 | 0 | 0 | 185740 | 59994 | 0 | 0 | 0 | 0 | 0 | 0 | 0 | 119190 | 0 | 0 | 0 | 308090 |
| NEDIPNVA | ATP1B3 | P54709 | AT1B3_HUMAN | Unmodified | 0 | 374790 | 993910 | 999800 | 0 | 0 | 64069 | 23950 | 7097800 | 0 | 1424100 | 11507000 | 1743000 | 173990 | 2245800 | 931230 | 135900 | 0 |
| NEDIPNVA | ATP1B3 | P54709 | AT1B3_HUMAN | Oxidation (M) | 203410 | 0 | 4766500 | 1817100 | 0 | 0 | 0 | 0 | 0 | 0 | 1578700 | 14679000 | 0 | 0 | 4714400 | 4051100 | 0 | 0 |
| NEDQTGRI | MUC5B | Q9HC84 | MUC5B_HUMA | Unmodified | 0 | 0 | 0 | 0 | 0 | 0 | 0 | 0 | 0 | 0 | 0 | 0 | 0 | 0 | 0 | 0 | 0 | 939260 |
| NEDVQLR | PPP2R1A; PPP2 | P30153; P30 | 2AAA_HUMAN | Unmodified | 0 | 47242 | 256990 | 140500 | 0 | 0 | 0 | 1080500 | 154960 | 0 | 142980 | 2420400 | 0 | 0 | 0 | 261470 | 0 | 156200 |
| NEEDAAEL | UBA1 | P22314 | UBA1_HUMAN | Unmodified | 729740 | 545840 | 990050 | 0 | 280660 | 19382 | 0 | 0 | 3428000 | 0 | 3610800 | 13019000 | 0 | 0 | 2597300 | 1129600 | 0 | 10651 |
| NEEKPVQM | SERPINB6; SER | P35237; A0A | SPB6_HUMAN | Oxidation (M) | 0 | 99275 | 0 | 0 | 0 | 1442800 | 0 | 0 | 0 | 0 | 0 | 385230 | 0 | 0 | 0 | 0 | 0 | 0 |
| NEEKPVQM | SERPINB6; SER | P35237; A0A | SPB6_HUMAN | Unmodified | 0 | 0 | 0 | 0 | 0 | 0 | 0 | 0 | 0 | 0 | 0 | 2877700 | 0 | 0 | 0 | 0 | 0 | 134900 |
| NEEVLAVL | ZMPSTE24 | O75844 | FACE1_HUMAN | Unmodified | 23131 | 0 | 78350 | 11745 | 0 | 0 | 0 | 0 | 1192000 | 46269 | 98308 | 959220 | 0 | 0 | 62596 | 40595 | 0 | 0 |
| NEEVTNTE | N/A | N/A | N/A | Unmodified | 9180300 | 20984000 | 287110 | 15190000 | 24421000 | 264320 | 1840900 | 536330 | 76310 | 10959000 | 295680 | 24790000 | 22015000 | 4956400 | 2212700 | 1307700 | 3963800 | 5494000 |
| NEEYGLR | TJP1; TJP2; TJP1 | Q07157; Q9 | ZO1_HUMAN T | Unmodified | 0 | 31999 | 269090 | 0 | 0 | 0 | 0 | 0 | 333710 | 125740 | 0 | 968380 | 405440 | 0 | 21871 | 0 | 168160 | 0 |
| NEFIFCHD | ZC3H10; ZC3H | Q96K80; F8 | ZC3HA_HUMA | Unmodified | 1396300 | 343230 | 1115600 | 547410 | 0 | 0 | 0 | 0 | 0 | 0 | 0 | 0 | 0 | 0 | 397580 | 586790 | 0 | 0 |
| NEFIPTNFE | IDE; IDE; IDE; ID | P14735; A0A | IDE_HUMAN In | Unmodified | 0 | 0 | 38458 | 0 | 0 | 0 | 0 | 0 | 234530 | 0 | 156030 | 834490 | 61112 | 0 | 216210 | 87530 | 0 | 0 |
| NEFNLESK | RAB11A; RAB11 | P62491; Q1 | RB11A_HUMAN | Unmodified | 449660 | 176780 | 1038200 | 0 | 6956500 | 892660 | 894130 | 835870 | 1119200 | 107100 | 280500 | 3692100 | 0 | 0 | 48150 | 64260 | 3817100 | 1949700 |
| NEFQGELE | GOLM1; GOLM | Q8NBJ4; F8 | GOLM1_HUMA | Unmodified | 0 | 0 | 0 | 0 | 0 | 98727 | 0 | 0 | 3747600 | 314760 | 0 | 0 | 0 | 0 | 0 | 0 | 67243 | 0 |
| NEFSQTVG | SLC44A4; SLC4 | Q53GD3; A0 | CTL4_HUMAN | Unmodified | 0 | 0 | 7522100 | 0 | 0 | 0 | 0 | 182230 | 0 | 0 | 0 | 0 | 0 | 0 | 4939900 | 3733100 | 0 | 0 |
| NEGCQVY | ERGIC3; ERGIC | Q9Y282; A0 | ERGI3_HUMAN | Unmodified | 0 | 0 | 0 | 0 | 0 | 0 | 0 | 0 | 3290600 | 0 | 0 | 0 | 0 | 0 | 0 | 0 | 0 | 0 |
| NEGGTWS | SELENBP1; SEL | Q13228; A6 | SBP1_HUMAN | Unmodified | 0 | 0 | 590320 | 470320 | 0 | 0 | 0 | 0 | 0 | 0 | 0 | 0 | 715710 | 0 | 977350 | 632660 | 0 | 0 |
| NEGNIFPN | SUPT16H | Q9Y5B9 | SP16H_HUMAN | Unmodified | 0 | 0 | 0 | 0 | 0 | 0 | 0 | 0 | 0 | 0 | 0 | 1761800 | 0 | 0 | 0 | 0 | 0 | 0 |
| NEGNKPLS | PLCB4; PLCB4 | Q15147; A0 | PLCB4_HUMAN | Unmodified | 90840 | 0 | 0 | 70214 | 0 | 0 | 0 | 0 | 0 | 0 | 0 | 0 | 0 | 0 | 164000 | 244970 | 0 | 0 |
| NEGQDATN | NPTN; NPTN; N | Q9Y639; Q9 | NPTN_HUMAN | Unmodified | 0 | 0 | 0 | 0 | 671650 | 137210 | 0 | 0 | 991850 | 52093 | 453800 | 5860900 | 650810 | 405370 | 1515800 | 0 | 122070 | 965940 |
| NEGQDATN | NPTN; NPTN; N | Q9Y639; Q9 | NPTN_HUMAN | Oxidation (M) | 0 | 0 | 0 | 0 | 538570 | 0 | 0 | 0 | 331960 | 0 | 0 | 1994800 | 0 | 267950 | 706890 | 595670 | 0 | 341720 |
| NEGSESAP | YBX1; YBX1; YB | P67809; C9J | YBOX1_HUMAN | Unmodified | 391560 | 811920 | 1408200 | 0 | 0 | 0 | 0 | 0 | 0 | 272250 | 0 | 12821000 | 0 | 0 | 0 | 0 | 20462 | 0 |
| NEGSLQPA | MGAT4A | Q9UM21 | MGT4A_HUMA | Unmodified | 0 | 0 | 0 | 0 | 0 | 0 | 0 | 0 | 997110 | 0 | 942820 | 2763100 | 0 | 0 | 0 | 0 | 0 | 0 |
| NEGTATYA | JUP | P14923 | PLAK_HUMAN | Unmodified | 0 | 0 | 468950 | 700050 | 0 | 0 | 0 | 0 | 0 | 0 | 0 | 5431000 | 0 | 0 | 3621900 | 4665000 | 0 | 0 |
| NEGVATYA | CTNNB1; CTNN | P35222; A0A | CTNB1_HUMA | Unmodified | 124690 | 174310 | 215980 | 0 | 0 | 0 | 0 | 0 | 53029 | 0 | 0 | 4367500 | 0 | 0 | 3873000 | 1975200 | 0 | 0 |
| NEGVGPFS | CTNNB1 | A0A2R8Y6G | A0A2R8Y6G0_ | Unmodified | 0 | 0 | 0 | 0 | 0 | 0 | 0 | 0 | 0 | 0 | 0 | 13658000 | 0 | 0 | 0 | 0 | 0 | 0 |
| NEGVNWL | CHMP1A; CHM | Q9HD42; F8 | CHM1A_HUMA | Unmodified | 0 | 0 | 0 | 0 | 1567700 | 529040 | 0 | 0 | 1359500 | 884480 | 0 | 0 | 1862400 | 0 | 0 | 0 | 0 | 0 |
| NEHNSILQ | KLC1; KLC1; KL | Q07866; G3 | KLC1_HUMAN | Unmodified | 0 | 0 | 206280 | 249360 | 539640 | 0 | 0 | 0 | 1739100 | 246490 | 784590 | 2640700 | 237650 | 0 | 394910 | 354690 | 108700 | 509500 |
| NEIASVAYF | EIF3D | O15371 | EIF3D_HUMAN | Unmodified | 0 | 0 | 0 | 0 | 0 | 0 | 0 | 0 | 0 | 0 | 0 | 751890 | 0 | 0 | 0 | 0 | 0 | 0 |
| NEILTSGNI | VPS13C | Q709C8 | VP13C_HUMAN | Unmodified | 0 | 0 | 0 | 0 | 0 | 0 | 0 | 0 | 86348 | 429920 | 71061 | 371280 | 0 | 0 | 0 | 0 | 0 | 0 |
| NEINIDTLA | EMC1; EMC1 | Q8N766; A0 | EMC1_HUMAN | Unmodified | 0 | 0 | 0 | 0 | 0 | 0 | 0 | 0 | 3071500 | 0 | 0 | 0 | 0 | 0 | 0 | 0 | 0 | 0 |
| NEIPEEALY | ATP2B1; ATP2B | P20020; E7E | AT2B1_HUMAN | Unmodified | 114370 | 0 | 269340 | 0 | 0 | 0 | 0 | 0 | 0 | 0 | 0 | 1257200 | 0 | 0 | 470240 | 231960 | 0 | 0 |
| NEIPNSHIL | CBS; CBSL | P35520; P0D | CBS_HUMAN C | Unmodified | 0 | 0 | 0 | 0 | 0 | 0 | 0 | 0 | 0 | 0 | 0 | 1423800 | 0 | 0 | 0 | 0 | 0 | 0 |
| NEIVFPAGI | ECE1; ECE1; EC | P42892; B4D | ECE1_HUMAN | Unmodified | 484700 | 222230 | 82415 | 137310 | 0 | 92119 | 0 | 0 | 4531300 | 209280 | 0 | 134430 | 0 | 0 | 0 | 76838 | 0 | 0 |
| NEKDFGLT | N/A | N/A | N/A | Unmodified | 172240 | 0 | 0 | 345730 | 27450000 | 1216400 | 90614000 | 13280000 | 0 | 15811000 | 0 | 0 | 25950000 | 22607000 | 0 | 36484 | 24273000 | 12326000 |
| NELESYAYS | HSPA5; HSPA5; | P11021; A0A | BIP_HUMAN Er | Unmodified | 0 | 0 | 456980 | 0 | 0 | 0 | 0 | 0 | 22071000 | 1542600 | 582090 | 1719500 | 0 | 0 | 0 | 213460 | 0 | 0 |
| NELHNLLD | ARL8B; ARL8A | Q9NVJ2; Q9 | ARL8B_HUMAN | Unmodified | 0 | 0 | 298450 | 0 | 105240 | 63944 | 0 | 0 | 446150 | 663040 | 0 | 2307900 | 0 | 0 | 0 | 0 | 443770 | 0 |
| NELLESEM | GOLGB1 | Q14789 | GOGB1_HUMA | Unmodified | 53179 | 78336 | 0 | 0 | 0 | 0 | 0 | 0 | 0 | 0 | 484900 | 1620200 | 0 | 0 | 0 | 0 | 0 | 0 |
| NELLMAQR | KIF14 | Q15058 | KIF14_HUMAN | Oxidation (M) | 0 | 0 | 0 | 0 | 328080 | 103380 | 0 | 0 | 0 | 0 | 0 | 0 | 0 | 0 | 0 | 0 | 0 | 0 |
| NELQTDKP | ARMT1 | Q9H993 | ARMT1_HUMAN | Unmodified | 0 | 0 | 0 | 0 | 0 | 0 | 0 | 0 | 0 | 0 | 0 | 885710 | 0 | 0 | 0 | 0 | 0 | 0 |
| NELSGALT | TARS1 | P26639 | SYTC_HUMAN | Unmodified | 0 | 0 | 0 | 0 | 0 | 0 | 0 | 0 | 0 | 0 | 371540 | 3635000 | 0 | 0 | 0 | 0 | 0 | 0 |
| NELVIRVQF | G6PD | P11413 | G6PD_HUMAN | Unmodified | 0 | 0 | 0 | 0 | 208880 | 81928 | 0 | 0 | 0 | 0 | 0 | 0 | 0 | 0 | 0 | 73620 | 111730 | 0 |
| NENARQQI | MYH9 | P35579 | MYH9_HUMAN | Unmodified | 0 | 0 | 0 | 15424 | 9761000 | 230750 | 1208000 | 288290 | 0 | 47086 | 0 | 0 | 569790 | 2667400 | 0 | 0 | 2713400 | 3824500 |
| NENEEILEF | LIMA1; LIMA1; L | Q9UHB6; F8 | LIMA1_HUMAN | Unmodified | 246880 | 281370 | 1206800 | 225070 | 924300 | 0 | 0 | 0 | 0 | 273860 | 0 | 1339500 | 0 | 0 | 2480400 | 1494800 | 0 | 0 |
| NENGIDAE | TMEM237; TME | Q96Q45; A0 | TM237_HUMAN | Unmodified | 0 | 0 | 0 | 0 | 752130 | 0 | 0 | 0 | 0 | 0 | 460560 | 1214200 | 0 | 0 | 0 | 0 | 364400 | 0 |
| NENIYQIYS | DPY19L4 | Q7Z388 | D19L4_HUMAN | Unmodified | 0 | 0 | 0 | 0 | 998030 | 0 | 2115200 | 274050 | 833300 | 194490 | 28105 | 0 | 0 | 0 | 0 | 0 | 0 | 0 |
| NENKPYLF | SLC44A2 | Q8IWA5 | CTL2_HUMAN | Unmodified | 19017000 | 13520000 | 22776000 | 6208200 | 445400 | 0 | 262800 | 0 | 10875000 | 1131100 | 0 | 11635000 | 0 | 0 | 32259000 | 24372000 | 192350 | 0 |
| NENLDCNS | TMEM87B; TME | Q96K49; A0 | TM87B_HUMAN | Unmodified | 0 | 0 | 0 | 0 | 0 | 0 | 0 | 0 | 982310 | 0 | 116810 | 0 | 0 | 0 | 0 | 0 | 0 | 0 |
| NENPYPAV | SBF1 | O95248 | MTMR5_HUMA | Unmodified | 0 | 0 | 0 | 0 | 0 | 0 | 0 | 0 | 0 | 0 | 0 | 256280 | 0 | 0 | 0 | 0 | 3037600 | 20911000 |
| NENQEVIG | PDE2A; PDE2A | O00408; E9 | PDE2A_HUMAN | Unmodified | 0 | 0 | 0 | 0 | 0 | 0 | 0 | 0 | 0 | 0 | 0 | 1539200 | 0 | 0 | 0 | 0 | 0 | 0 |
| NENSNDP | DNAJC10; DNA | Q8IXB1; A0A | DJC10_HUMAN | Unmodified | 0 | 53119 | 0 | 0 | 0 | 0 | 0 | 0 | 442050 | 0 | 0 | 0 | 0 | 0 | 0 | 0 | 0 | 0 |
| NENTFLDL | UGP2; UGP2; U | Q16851; E7E | UGPA_HUMAN | Unmodified | 0 | 0 | 0 | 29748 | 0 | 0 | 0 | 0 | 215920 | 0 | 40066 | 282400 | 0 | 0 | 0 | 0 | 0 | 0 |
| NEPCGARI | ITGA1 | P56199 | ITA1_HUMAN I | Unmodified | 0 | 0 | 0 | 0 | 129550 | 0 | 0 | 94796 | 0 | 62465 | 0 | 0 | 0 | 0 | 0 | 0 | 483310 | 1831300 |
| NEPQNATG | RAB5C | P51148 | RAB5C_HUMA | Unmodified | 1810800 | 0 | 4399600 | 0 | 0 | 0 | 0 | 9708.7 | 5251700 | 0 | 0 | 5004300 | 0 | 0 | 3288100 | 3064100 | 0 | 0 |
| NEPQNPG | RAB5A | P20339 | RAB5A_HUMA | Unmodified | 118640 | 13677 | 76347 | 0 | 0 | 0 | 0 | 0 | 376600 | 0 | 0 | 279470 | 14533 | 0 | 16301 | 89742 | 0 | 0 |
| NEQDAYAI | ACAT1; ACAT1; | P24752; A0A | THIL_HUMAN A | Unmodified | 0 | 0 | 0 | 0 | 0 | 0 | 0 | 0 | 853430 | 0 | 0 | 0 | 0 | 0 | 0 | 0 | 93319 | 0 |
| NEQINDEM | SCG2 | P13521 | SCG2_HUMAN | Unmodified | 0 | 0 | 223720 | 0 | 0 | 122410 | 0 | 0 | 190830 | 0 | 0 | 289950 | 0 | 0 | 0 | 0 | 0 | 0 |

| | | | | | | | | | | | | | | | | | | | | | | |
|---|---|---|---|---|---|---|---|---|---|---|---|---|---|---|---|---|---|---|---|---|---|---|
| NERPARPP | CCDC50 | Q8IVM0 | CCD50_HUMA | Oxidation (M) | 5568100 | 4205300 | 11922000 | 10225000 | 0 | 0 | 0 | 0 | 0 | 0 | 0 | 0 | 0 | 0 | 12699000 | 15424000 | 0 | 0 |
| NERPARPP | CCDC50 | Q8IVM0 | CCD50_HUMA | Unmodified | 0 | 5275900 | 6079000 | 5814600 | 0 | 0 | 0 | 0 | 0 | 0 | 0 | 0 | 0 | 0 | 11829000 | 1847900 | 0 | 0 |
| NESGTWIQ | ZC3HAV1; ZC3I | Q7Z2W4; C9 | ZCCHV_HUMA | Unmodified | 499670 | 466420 | 636170 | 977610 | 502340 | 0 | 0 | 51387 | 410440 | 108360 | 132470 | 581800 | 137240 | 0 | 649690 | 625690 | 0 | 0 |
| NETDEDGV | EIF3A | Q14152 | EIF3A_HUMAN | Unmodified | 228880 | 203990 | 989170 | 436420 | 0 | 0 | 0 | 0 | 0 | 758960 | 0 | 0 | 0 | 0 | 0 | 317860 | 0 | 0 |
| NETGGGE( | PITPNB; PITPNI | P48739; A0A | PIPNB_HUMAN | Unmodified | 0 | 0 | 0 | 0 | 0 | 0 | 0 | 0 | 471390 | 0 | 0 | 2530700 | 0 | 0 | 0 | 0 | 0 | 0 |
| NETLGGTC | DLD; DLD; DLD | P09622; A0A | DLDH_HUMAN | Unmodified | 0 | 0 | 0 | 0 | 0 | 0 | 83442 | 0 | 963180 | 103120 | 0 | 56430 | 0 | 31149 | 46066 | 0 | 0 | 78785 |
| NETSATHQ | LMBRD2 | Q68DH5 | LMBD2_HUMAI | Unmodified | 121000 | 0 | 801310 | 0 | 0 | 0 | 0 | 0 | 0 | 37762 | 493880 | 855980 | 0 | 0 | 0 | 363520 | 0 | 0 |
| NEVDMQVI | QARS1; QARS1; | P47897; A0A | SYQ_HUMAN G | Oxidation (M) | 0 | 0 | 0 | 0 | 0 | 0 | 0 | 0 | 0 | 0 | 0 | 508340 | 0 | 0 | 0 | 0 | 0 | 0 |
| NEVIGQLVI | SYT4 | Q9H2B2 | SYT4_HUMAN S | Unmodified | 0 | 0 | 0 | 0 | 0 | 0 | 3650100 | 0 | 0 | 0 | 0 | 0 | 0 | 0 | 15404000 | 10807000 | 0 | 0 |
| NEVLEDHV | COG3 | Q96JB2 | COG3_HUMAN | Unmodified | 0 | 0 | 0 | 497360 | 1693100 | 779050 | 5883000 | 0 | 0 | 0 | 0 | 0 | 0 | 3218100 | 0 | 0 | 0 | 2657500 |
| NEVLMVNI( | EIF2S3; EIF2S3I | P41091; Q2\ | IF2G_HUMAN E | Oxidation (M) | 0 | 0 | 68955 | 0 | 0 | 0 | 0 | 16192 | 0 | 0 | 83614 | 576000 | 0 | 0 | 0 | 64050 | 49100 | 0 |
| NEVSFVIHI | SEPHS2 | Q99611 | SPS2_HUMAN | Unmodified | 35154 | 0 | 0 | 0 | 0 | 0 | 0 | 0 | 0 | 0 | 112500 | 893270 | 0 | 0 | 110210 | 0 | 0 | 13284 |
| NEVSFVIHI | SEPHS1 | P49903 | SPS1_HUMAN | Unmodified | 0 | 0 | 95161 | 83251 | 101150 | 0 | 59483 | 0 | 594910 | 77322 | 337190 | 2625900 | 0 | 106930 | 148560 | 108280 | 0 | 19521 |
| NEVTDSGI | GLT8D1; GLT8D | Q68CQ7; C9 | GL8D1_HUMAI | Unmodified | 0 | 0 | 0 | 0 | 0 | 0 | 0 | 0 | 1742600 | 0 | 0 | 109960 | 75836 | 0 | 0 | 0 | 0 | 0 |
| NFAADHFN | DARS2; DARS2; | Q6PI48; A0A | SYDM_HUMAN | Unmodified | 0 | 0 | 0 | 0 | 32691 | 64030 | 250470 | 0 | 1996100 | 0 | 0 | 0 | 0 | 0 | 0 | 0 | 127150 | 61416 |
| NFAALEVLI | UAP1L1 | Q3KQV9 | UAP1L_HUMAN | Unmodified | 155960 | 339990 | 0 | 0 | 0 | 0 | 1605400 | 0 | 0 | 65657 | 0 | 162410 | 0 | 723260 | 358010 | 148970 | 608210 | 0 |
| NFAELAR | PPIL1 | Q9Y3C6 | PPIL1_HUMAN | Unmodified | 890850 | 0 | 622700 | 494730 | 507710 | 124300 | 96848 | 68252 | 231740 | 0 | 351080 | 1337800 | 637340 | 358690 | 0 | 699170 | 194250 | 236970 |
| NFAEPGSE | AGL | P35573 | GDE_HUMAN ( | Unmodified | 0 | 0 | 0 | 0 | 0 | 0 | 0 | 0 | 0 | 0 | 0 | 1230500 | 131960 | 0 | 0 | 0 | 0 | 0 |
| NFAFLEFR | U2AF2; U2AF2 | P26368; K7E | U2AF2_HUMAN | Unmodified | 1511600 | 2567600 | 0 | 0 | 392180 | 0 | 0 | 0 | 0 | 0 | 0 | 8699500 | 0 | 0 | 697730 | 914160 | 0 | 0 |
| NFALLGVG | UBA6; UBA6 | A0AVT1; H0Y | UBA6_HUMAN | Unmodified | 0 | 0 | 0 | 0 | 401860 | 149820 | 330820 | 0 | 0 | 171790 | 0 | 588510 | 685860 | 187870 | 69116 | 0 | 739820 | 607570 |
| NFALLGVG | UBA6; UBA6 | A0AVT1; H0Y | UBA6_HUMAN | Unmodified | 0 | 248980 | 162500 | 0 | 0 | 200520 | 311160 | 0 | 226460 | 0 | 0 | 0 | 0 | 0 | 186460 | 0 | 671900 | 577180 |
| NFCDANNI | MANEAL; MANE | Q5VSG8; E9I | MANEL_HUMAI | Oxidation (M) | 0 | 0 | 0 | 0 | 0 | 0 | 0 | 0 | 2256300 | 0 | 0 | 0 | 0 | 0 | 0 | 0 | 0 | 0 |
| NFCSAVQY | GLG1 | Q92896 | GSLG1_HUMAI | Unmodified | 0 | 0 | 0 | 0 | 0 | 0 | 57394 | 0 | 2053200 | 3056400 | 0 | 0 | 0 | 0 | 0 | 0 | 0 | 427140 |
| NFDEILR | PRDX6 | P30041 | PRDX6_HUMAN | Unmodified | 0 | 415400 | 756370 | 0 | 0 | 0 | 0 | 0 | 987750 | 0 | 1371900 | 20921000 | 0 | 0 | 0 | 2126900 | 0 | 0 |
| NFDEKLVEI | HSPH1; HSPH1 | Q92598; A0A | HS105_HUMAN | Unmodified | 0 | 0 | 0 | 0 | 0 | 0 | 269380 | 0 | 0 | 0 | 0 | 0 | 0 | 0 | 0 | 0 | 161460 | 45681 |
| NFDIPKEM | CLIC4 | Q9Y696 | CLIC4_HUMAN | Unmodified | 0 | 0 | 0 | 0 | 459280 | 0 | 0 | 0 | 175550 | 0 | 0 | 0 | 0 | 0 | 0 | 0 | 571780 | 551350 |
| NFDLRPGV | MAT1A; MAT2A | Q00266; P31 | METK1_HUMAN | Unmodified | 0 | 187360 | 0 | 324310 | 0 | 0 | 0 | 0 | 0 | 0 | 516100 | 3198800 | 0 | 0 | 0 | 412370 | 0 | 0 |
| NFDQEGLV | RNF114; RNF11 | Q9Y508; A0A | RN114_HUMAN | Unmodified | 1375200 | 0 | 0 | 947680 | 936200 | 206360 | 385170 | 206200 | 215470 | 0 | 175400 | 558150 | 4209900 | 1859200 | 1067100 | 0 | 789970 | 1232400 |
| NFDREIGH | TMEM43 | Q9BTV4 | TMM43_HUMAI | Oxidation (M) | 0 | 0 | 0 | 0 | 5304000 | 0 | 0 | 0 | 10919000 | 276160 | 0 | 0 | 0 | 0 | 0 | 0 | 0 | 0 |
| NFDREIGH | TMEM43 | Q9BTV4 | TMM43_HUMAI | Unmodified | 0 | 0 | 0 | 0 | 0 | 0 | 0 | 0 | 4556400 | 0 | 0 | 0 | 0 | 0 | 0 | 0 | 0 | 0 |
| NFDREIGH | TMEM43 | Q9BTV4 | TMM43_HUMAI | 2 Oxidation (M) | 0 | 0 | 0 | 0 | 0 | 0 | 0 | 0 | 921240 | 230720 | 0 | 0 | 0 | 0 | 0 | 0 | 0 | 0 |
| NFDTGLQE | OCLN; OCLN | Q16625; A0A | OCLN_HUMAN | Unmodified | 1047100 | 935210 | 1258900 | 2399700 | 999950 | 335660 | 0 | 0 | 1322500 | 567270 | 0 | 557880 | 1569300 | 619960 | 2409900 | 1693000 | 0 | 0 |
| NFEATLGW | LARS1; LARS1; | Q9P2J5; A0A | SYLC_HUMAN | Unmodified | 48274 | 25005 | 0 | 0 | 0 | 0 | 0 | 0 | 1136300 | 0 | 0 | 5027900 | 0 | 0 | 444900 | 179790 | 0 | 0 |
| NFEDVAFD | P4HB; P4HB; P | P07237; A0A | PDIA1_HUMAN | Unmodified | 120080 | 0 | 0 | 1020900 | 3283000 | 606020 | 3856700 | 1965100 | 78676000 | 42001000 | 9586400 | 2800800 | 3235500 | 3273700 | 818930 | 512980 | 400950 | 1712400 |
| NFEDVAFD | P4HB; P4HB; P | P07237; A0A | PDIA1_HUMAN | Unmodified | 0 | 0 | 461570 | 0 | 2809500 | 594700 | 7240800 | 3261900 | 86295000 | 44517000 | 7919300 | 2526600 | 4766300 | 4749100 | 0 | 0 | 696570 | 2304900 |
| NFEHLIPD/ | COPB1; COPB1 | P53618; E9P | COPB_HUMAN | Unmodified | 0 | 0 | 0 | 0 | 0 | 0 | 0 | 0 | 0 | 0 | 0 | 349810 | 0 | 0 | 147080 | 0 | 0 | 0 |
| NFELMEAE | ERLIN1; ERLIN1 | O75477; B0( | ERLN1_HUMAN | Oxidation (M) | 0 | 350200 | 0 | 0 | 0 | 0 | 0 | 0 | 880210 | 112590 | 0 | 128220 | 0 | 0 | 0 | 0 | 0 | 0 |
| NFELMEAE | ERLIN1; ERLIN1 | O75477; B0( | ERLN1_HUMAN | Unmodified | 0 | 0 | 0 | 0 | 0 | 0 | 0 | 0 | 5575900 | 748460 | 0 | 0 | 0 | 0 | 190960 | 0 | 0 | 0 |
| NFEQWQS | WWP1 | Q9H0M0 | WWP1_HUMAN | Unmodified | 478620 | 0 | 684480 | 626260 | 0 | 0 | 0 | 0 | 0 | 0 | 0 | 0 | 0 | 0 | 832490 | 663450 | 0 | 0 |
| NFESLSEAI | RPN2; RPN2; RI | P04844; F2Z | RPN2_HUMAN | Unmodified | 0 | 0 | 0 | 0 | 0 | 0 | 0 | 0 | 8387300 | 3479800 | 301050 | 553030 | 0 | 0 | 0 | 0 | 0 | 0 |
| NFESSSEW | DOP1B; DOP1B | Q9Y3R5; F8\ | DOP2_HUMAN | Unmodified | 0 | 0 | 10656 | 41965 | 0 | 150880 | 0 | 0 | 180310 | 41530 | 37099 | 73183 | 0 | 0 | 110730 | 0 | 0 | 0 |
| NFETLRIDV | PBDC1; PBDC1 | Q9BVG4; A6 | PBDC1_HUMAI | Unmodified | 0 | 143000 | 0 | 0 | 0 | 183640 | 0 | 0 | 0 | 912530 | 0 | 3495800 | 760610 | 0 | 0 | 0 | 1243800 | 0 |
| NFEWVAEF | PAICS; PAICS; I | P22234; D6F | PUR6_HUMAN | Unmodified | 80609 | 100720 | 25337 | 119940 | 742460 | 0 | 365520 | 0 | 18851 | 554210 | 0 | 0 | 2837700 | 936460 | 251530 | 26159 | 1096600 | 1372300 |
| NFFLTNR | CAPN2 | P17655 | CAN2_HUMAN | Unmodified | 0 | 0 | 0 | 0 | 4121500 | 0 | 0 | 0 | 1854600 | 0 | 0 | 422320 | 0 | 0 | 0 | 0 | 1177400 | 1278900 |
| NFFYIISPG | GALNT1 | Q10472 | GALT1_HUMAN | Unmodified | 0 | 0 | 0 | 0 | 0 | 0 | 0 | 0 | 1844300 | 0 | 0 | 1484200 | 0 | 0 | 0 | 0 | 0 | 0 |
| NFFYIISPG | GALNT1 | Q10472 | GALT1_HUMAN | Unmodified | 0 | 0 | 0 | 0 | 0 | 0 | 0 | 0 | 13114000 | 615770 | 1802200 | 1598700 | 0 | 0 | 0 | 0 | 0 | 0 |
| NFGDQPDI | SF3B3 | Q15393 | SF3B3_HUMAN | Unmodified | 1191500 | 1080500 | 217860 | 0 | 0 | 0 | 0 | 0 | 0 | 0 | 105780 | 4873300 | 0 | 0 | 0 | 161010 | 0 | 0 |
| NFGEDMDI | PABPC1; PABP( | P11940; Q9I | PABP1_HUMAN | Oxidation (M) | 57841000 | 2694300 | 20819000 | 14621000 | 1481800 | 252320 | 491910 | 10952 | 638750 | 443480 | 435830 | 3720600 | 3584500 | 834380 | 575990 | 18028000 | 333390 | 0 |
| NFGEDMDI | PABPC1; PABP( | P11940; Q9I | PABP1_HUMAN | Unmodified | 30175000 | 21638000 | 11527000 | 11625000 | 870090 | 55212 | 583010 | 301070 | 1053100 | 957980 | 2410500 | 2428400 | 2354600 | 1370600 | 7674400 | 7139200 | 310190 | 1048800 |
| NFGEDMDI | PABPC1; PABP( | P11940; Q9I | PABP1_HUMAN | Oxidation (M) | 35050000 | 18658000 | 26943 | 20207000 | 1119600 | 0 | 619650 | 0 | 0 | 499190 | 0 | 0 | 6581500 | 0 | 0 | 0 | 870400 | 0 |
| NFGEDMDI | PABPC1; PABP( | P11940; Q9I | PABP1_HUMAN | Unmodified | 0 | 1529800 | 0 | 3714600 | 0 | 0 | 316800 | 0 | 0 | 204500 | 0 | 0 | 2054500 | 738850 | 0 | 0 | 0 | 0 |
| NFGEEVDE | PABPC4; PABP( | Q13310; B1/ | PABP4_HUMAN | Unmodified | 15356000 | 16496000 | 10116000 | 4296800 | 309100 | 0 | 434240 | 124740 | 254510 | 221800 | 317320 | 1212700 | 778520 | 0 | 6332700 | 5515600 | 0 | 0 |
| NFGEEVDE | PABPC4; PABP( | Q13310; B1/ | PABP4_HUMAN | Unmodified | 10042000 | 10072000 | 0 | 6956700 | 0 | 0 | 2012000 | 0 | 0 | 345530 | 0 | 0 | 1796200 | 0 | 0 | 0 | 0 | 0 |
| NFGEEVDE | PABPC4; PABP( | Q13310; B1/ | PABP4_HUMAN | Unmodified | 0 | 0 | 0 | 41495 | 0 | 0 | 0 | 0 | 0 | 135530 | 0 | 0 | 325240 | 0 | 0 | 0 | 0 | 0 |
| NFGFAFLAI | ATP8B1; ATP8B | O43520; A0/ | AT8B1_HUMAN | Unmodified | 0 | 0 | 0 | 81413 | 0 | 0 | 0 | 0 | 0 | 0 | 0 | 2557100 | 0 | 0 | 343480 | 217260 | 0 | 0 |
| NFGGGNT/ | CRELD2; CRELI | Q6UXH1; A6 | CREL2_HUMAN | Unmodified | 0 | 0 | 0 | 0 | 0 | 0 | 790570 | 516570 | 1722700 | 1291000 | 0 | 0 | 0 | 0 | 0 | 0 | 0 | 0 |
| NFGGKWEI | SORT1; SORT1 | Q99523; A0/ | SORT_HUMAN | Unmodified | 0 | 0 | 0 | 0 | 0 | 0 | 35106 | 394620 | 5868700 | 1385100 | 881200 | 1582900 | 0 | 0 | 0 | 0 | 0 | 0 |
| NFGGLLGF | XYLT2 | Q9H1B5 | XYLT2_HUMAN | Unmodified | 0 | 0 | 0 | 0 | 0 | 0 | 0 | 0 | 776400 | 197010 | 0 | 84266 | 0 | 0 | 0 | 0 | 0 | 0 |
| NFGGTVSII | KIF14 | Q15058 | KIF14_HUMAN | Unmodified | 0 | 0 | 0 | 0 | 4775000 | 0 | 964380 | 262380 | 297210 | 0 | 0 | 0 | 581920 | 1017300 | 0 | 0 | 1752900 | 0 |
| NFGIGQDI( | RPL7A; RPL7A | P62424; Q5T | RL7A_HUMAN ( | Unmodified | 6575800 | 5513300 | 2780000 | 2213200 | 676180 | 329260 | 0 | 336760 | 597840 | 0 | 5169700 | 21117000 | 2525600 | 438530 | 4381000 | 3548300 | 0 | 0 |
| NFGIGQDI( | RPL7A; RPL7A | P62424; Q5T | RL7A_HUMAN ( | Unmodified | 0 | 0 | 0 | 0 | 0 | 0 | 0 | 0 | 0 | 0 | 0 | 4696300 | 0 | 0 | 0 | 0 | 0 | 0 |
| NFGIGQDI( | RPL7A; RPL7A | P62424; Q5T | RL7A_HUMAN ( | Unmodified | 0 | 0 | 0 | 232660 | 2445600 | 542070 | 1592600 | 225650 | 0 | 70566 | 0 | 0 | 12667000 | 5070200 | 0 | 0 | 0 | 1337300 |
| NFGIWLR | RPL18A; RPL18 | Q02543; M0I | RL18A_HUMAN | Unmodified | 9640100 | 655150 | 0 | 1778300 | 0 | 0 | 0 | 0 | 1639300 | 140150 | 2192200 | 38938000 | 0 | 0 | 463340 | 432440 | 0 | 0 |
| NFGLYNER | GOT1 | P17174 | AATC_HUMAN | Unmodified | 0 | 0 | 0 | 0 | 0 | 0 | 0 | 0 | 1014000 | 319290 | 341840 | 4655600 | 0 | 0 | 0 | 0 | 0 | 0 |
| NFGSVQFV | MFGE8 | Q08431 | MFGM_HUMAN | Unmodified | 4191000 | 3595200 | 3984000 | 17211000 | 26993000 | 8152900 | 0 | 0 | 11845000 | 799760 | 771100 | 5458500 | 1597400 | 0 | 13739000 | 9462700 | 772250 | 2312400 |
| NFGSYVTH | DYNLL2; DYNLI | Q96FJ2; P63 | DYL2_HUMAN | Unmodified | 101060 | 205160 | 1376700 | 625160 | 559490 | 0 | 55961 | 642450 | 0 | 0 | 0 | 2671900 | 639920 | 747560 | 0 | 0 | 301080 | 0 |
| NFGTDPEV | CPD | O75976 | CBPD_HUMAN | Unmodified | 0 | 0 | 0 | 535930 | 0 | 0 | 0 | 0 | 39312000 | 4173700 | 0 | 0 | 0 | 0 | 0 | 0 | 0 | 0 |
| NFGTDPEV | CPD | O75976 | CBPD_HUMAN | 2 Oxidation (M) | 0 | 0 | 0 | 0 | 0 | 0 | 0 | 0 | 0 | 10607000 | 0 | 100780 | 0 | 233050 | 0 | 0 | 0 | 104870 |
| NFHIFYQLI | MYO1C; MYO1( | O00159; F5I | MYO1C_HUMA | Unmodified | 579080 | 838860 | 393940 | 1693600 | 0 | 0 | 0 | 0 | 118860 | 0 | 0 | 0 | 92052 | 0 | 3052900 | 424030 | 0 | 0 |
| NFHIFYQLI | MYO1C; MYO1( | O00159; F5I | MYO1C_HUMA | Unmodified | 3884600 | 3069000 | 0 | 5089300 | 0 | 0 | 0 | 0 | 2239300 | 0 | 272980 | 0 | 219560 | 310360 | 0 | 0 | 0 | 0 |
| NFHVFYQL | MYO1B; MYO1E | O43795; E9F | MYO1B_HUMA | Unmodified | 34821 | 635600 | 0 | 0 | 0 | 199610 | 131330 | 0 | 0 | 0 | 0 | 0 | 0 | 0 | 424240 | 0 | 0 | 0 |
| NFHYPPDG | CTNND1; CTNI | O60716; A0/ | CTND1_HUMAI | Unmodified | 238560 | 0 | 934660 | 2427400 | 0 | 0 | 0 | 0 | 438370 | 0 | 267430 | 683690 | 130400 | 0 | 2693300 | 5058900 | 0 | 70916 |
| NFHYPPDG | CTNND1; CTNI | O60716; A0/ | CTND1_HUMAI | Unmodified | 74450 | 0 | 0 | 998580 | 3096900 | 751590 | 0 | 0 | 0 | 0 | 0 | 0 | 1776600 | 2286500 | 301200 | 61709 | 898520 | 7458900 |
| NFIAQGPYI | ATP6V1B2 | P21281 | VATB2_HUMAN | Unmodified | 0 | 0 | 0 | 0 | 153490 | 0 | 0 | 0 | 520440 | 74176 | 390690 | 2917600 | 0 | 0 | 0 | 0 | 0 | 0 |
| NFIELMEK | CACNA2D2; CA | Q9NY47; C9 | CA2D2_HUMAI | Unmodified | 0 | 0 | 0 | 0 | 0 | 0 | 0 | 25216 | 5512500 | 1487800 | 189710 | 0 | 0 | 0 | 0 | 0 | 0 | 0 |
| NFIELMEK\ | CACNA2D2; CA | Q9NY47; C9 | CA2D2_HUMAI | Unmodified | 0 | 0 | 0 | 38115 | 0 | 0 | 99758 | 7773.9 | 0 | 309280 | 0 | 0 | 31876 | 0 | 0 | 0 | 0 | 0 |
| NFILDQTN\ | HNRNPU; HNR | Q00839; A0/ | HNRPU_HUMA | Unmodified | 0 | 0 | 0 | 0 | 0 | 0 | 1166600 | 873590 | 0 | 0 | 566730 | 4540600 | 1029000 | 0 | 0 | 0 | 0 | 1510200 |
| NFILDQTN\ | HNRNPU; HNR | Q00839; A0/ | HNRPU_HUMA | Unmodified | 0 | 0 | 0 | 0 | 0 | 0 | 661830 | 0 | 0 | 0 | 0 | 2341700 | 1879500 | 0 | 0 | 0 | 0 | 0 |
| NFILDQTN\ | HNRNPU; HNR | Q00839; A0/ | HNRPU_HUMA | Unmodified | 0 | 0 | 0 | 0 | 0 | 217810 | 9003700 | 8315400 | 0 | 512170 | 0 | 627100 | 493320 | 1776300 | 0 | 0 | 0 | 1103200 |
| NFINNPLA( | MYH9; MYH9; I | P35579; Q5E | MYH9_HUMAN | Unmodified | 0 | 0 | 438630 | 606700 | 37670000 | 2752600 | 8296600 | 2418000 | 996810 | 2459300 | 2186300 | 11013000 | 9020400 | 10079000 | 1396700 | 723030 | 21664000 | 15759000 |
| NFINNPLA( | MYH9; MYH9; I | P35579; Q5E | MYH9_HUMAN | Unmodified | 0 | 0 | 0 | 0 | 13567000 | 1995000 | 2439100 | 427980 | 0 | 513710 | 437510 | 4043700 | 3081000 | 58873 | 1156700 | 805020 | 5529500 | 4063200 |
| NFIRNETHI | MET | P08581 | MET_HUMAN H | Unmodified | 941490 | 273450 | 0 | 3454400 | 187700 | 61239 | 200730 | 0 | 556970 | 54446 | 56036 | 306610 | 0 | 0 | 0 | 0 | 97605 | 524990 |
| NFITAEELR | ACTN4; ACTN4; | O43707; F5( | ACTN4_HUMAI | Unmodified | 932470 | 511340 | 1395200 | 1949800 | 8120400 | 2431900 | 0 | 0 | 10215000 | 1316300 | 2777500 | 29924000 | 4582900 | 0 | 2944700 | 2399500 | 946070 | 807970 |
| NFITAEELR | ACTN4; ACTN4; | O43707; F5( | ACTN4_HUMAI | Unmodified | 0 | 0 | 0 | 0 | 19936000 | 2796800 | 0 | 0 | 0 | 1838500 | 0 | 0 | 4841100 | 3067900 | 0 | 0 | 0 | 0 |
| NFITRENLE | MRPS26 | Q9BYN8 | RT26_HUMAN | Unmodified | 0 | 0 | 0 | 0 | 0 | 0 | 0 | 1318900 | 551750 | 0 | 0 | 0 | 0 | 0 | 0 | 0 | 41066 | 0 |
| NFIVWLED | RTRAF; RTRAF; | Q9Y224; G3\ | RTRAF_HUMAN | Unmodified | 0 | 204120 | 0 | 0 | 0 | 0 | 0 | 0 | 0 | 0 | 0 | 1100300 | 0 | 0 | 0 | 0 | 0 | 0 |
| NFLDLISSS | RASA3 | Q14644 | RASA3_HUMAN | Unmodified | 314220 | 469930 | 267170 | 0 | 0 | 0 | 0 | 0 | 0 | 0 | 0 | 0 | 0 | 0 | 568550 | 395030 | 0 | 0 |
| NFLEKFVQ | ITGA2; ITGA2; I | P17301; E9P | ITA2_HUMAN I | Unmodified | 0 | 0 | 0 | 0 | 965290 | 90662 | 407580 | 0 | 0 | 0 | 0 | 0 | 207880 | 162570 | 0 | 0 | 1807000 | 912130 |
| NFLELTELK | ATP6V0A1; ATP | Q93050; K7E | VPP1_HUMAN | Unmodified | 0 | 0 | 268810 | 0 | 0 | 0 | 0 | 0 | 1134300 | 2151900 | 2601000 | 232010 | 0 | 0 | 0 | 0 | 0 | 0 |
| NFLINYYNI | SNX5; SNX5; SI | Q9Y5X3; Q5( | SNX5_HUMAN | Unmodified | 0 | 0 | 0 | 0 | 0 | 0 | 0 | 0 | 0 | 0 | 0 | 1161300 | 0 | 0 | 0 | 0 | 0 | 0 |
| NFLINYYNI | SNX5; SNX5; SI | Q9Y5X3; Q5( | SNX5_HUMAN | Unmodified | 0 | 0 | 0 | 0 | 0 | 0 | 87698 | 0 | 0 | 533770 | 0 | 0 | 977650 | 392600 | 0 | 0 | 0 | 0 |
| NFLNALTSI | TRIO; TRIO; TRI | O75962; E7E | TRIO_HUMAN T | Unmodified | 0 | 0 | 32920 | 0 | 0 | 0 | 91216 | 0 | 143120 | 0 | 35182 | 221940 | 0 | 85202 | 106330 | 82816 | 0 | 0 |
| NFLTISEGA | N/A | N/A | N/A | Unmodified | 0 | 0 | 0 | 0 | 0 | 0 | 0 | 0 | 2406100 | 0 | 0 | 0 | 0 | 0 | 0 | 0 | 0 | 0 |
| NFLYAWC( | DHX9 | Q08211 | DHX9_HUMAN | Unmodified | 0 | 0 | 0 | 0 | 0 | 0 | 0 | 0 | 0 | 0 | 0 | 1093800 | 453680 | 264250 | 0 | 0 | 0 | 0 |
| NFMLHYEF | PNPT1 | Q8TCS8 | PNPT1_HUMAN | Oxidation (M) | 151120 | 0 | 0 | 0 | 0 | 0 | 0 | 0 | 545760 | 0 | 0 | 0 | 0 | 0 | 0 | 0 | 0 | 0 |
| NFMLLIEQI | ANKRD26; ANK | Q9UPS8; E7I | ANR26_HUMAI | Unmodified | 0 | 0 | 0 | 0 | 1293500 | 0 | 0 | 0 | 0 | 139070 | 0 | 0 | 226690 | 0 | 0 | 0 | 975020 | 1112500 |
| NFNIHGTN | GNS; GNS; GNS | P15586; H7( | GNS_HUMAN N | Unmodified | 0 | 0 | 117330 | 46880 | 370260 | 0 | 0 | 28372 | 1449500 | 531700 | 126120 | 254880 | 392580 | 226310 | 0 | 0 | 0 | 0 |
| NFNKDGDI | MME; MME; MM | P08473; A0A | NEP_HUMAN N | Unmodified | 0 | 0 | 0 | 6507100 | 0 | 0 | 0 | 0 | 1477500 | 0 | 0 | 0 | 0 | 0 | 3469600 | 3860800 | 0 | 0 |
| NFNLPMCI | OXCT1; OXCT1 | P55809; E9P | SCOT1_HUMAI | Unmodified | 0 | 0 | 0 | 0 | 0 | 0 | 0 | 0 | 1245000 | 491740 | 0 | 0 | 0 | 0 | 0 | 0 | 0 | 0 |
| NFPDLDRI\ | CPE; CPE; CPE; | P16870; C9J | CBPE_HUMAN | Unmodified | 41786 | 20548 | 71050 | 30593 | 0 | 0 | 0 | 0 | 90715000 | 2030400 | 0 | 0 | 0 | 0 | 72232 | 22124 | 0 | 0 |
| NFPNAIEH | UBA1 | P22314 | UBA1_HUMAN | Unmodified | 0 | 0 | 0 | 0 | 0 | 0 | 0 | 0 | 0 | 0 | 0 | 547560 | 0 | 0 | 0 | 0 | 0 | 0 |
| NFPNEFAK | DENR; DENR | O43583; F8\ | DENR_HUMAN | Unmodified | 0 | 0 | 0 | 0 | 0 | 431130 | 0 | 0 | 415790 | 70299 | 0 | 0 | 0 | 0 | 0 | 0 | 580740 | 936170 |
| NFPQTALV | EMC1 | Q8N766 | EMC1_HUMAN | Unmodified | 0 | 0 | 0 | 0 | 21930000 | 748920 | 1460200 | 1304400 | 11925000 | 1992800 | 0 | 0 | 13540000 | 2390100 | 0 | 0 | 12556000 | 19945000 |
| NFPTITALE | WDR11 | Q9BZH6 | WDR11_HUMA | Unmodified | 79447 | 0 | 0 | 0 | 0 | 0 | 0 | 0 | 0 | 0 | 0 | 1348700 | 0 | 0 | 0 | 0 | 0 | 313490 |
| NFQCVDAS | ADAM9; ADAMS | Q13443; A0/ | ADAM9_HUMA | Unmodified | 1069900 | 442590 | 696760 | 880560 | 854210 | 0 | 0 | 0 | 518730 | 0 | 0 | 0 | 918940 | 0 | 897010 | 1648200 | 0 | 0 |
| NFQDFDC( | NDRG3; NDRG; | Q9UGV2; Q5 | NDRG3_HUMA | Unmodified | 0 | 0 | 0 | 0 | 0 | 0 | 0 | 0 | 241960 | 0 | 2579000 | 19374000 | 0 | 0 | 554730 | 0 | 0 | 0 |
| NFQEEQIN | DNAJC10; DNA | Q8IXB1; A0A | DJC10_HUMAN | Unmodified | 0 | 0 | 0 | 0 | 0 | 0 | 0 | 0 | 3949000 | 235770 | 779310 | 0 | 0 | 0 | 0 | 0 | 0 | 0 |
| NFQEEQIN | DNAJC10; DNA | Q8IXB1; A0A | DJC10_HUMAN | Unmodified | 0 | 0 | 0 | 0 | 0 | 0 | 0 | 0 | 0 | 748980 | 0 | 0 | 0 | 0 | 0 | 0 | 0 | 0 |
| NFQIEVQK | JAK1; JAK1; JAK | P23458; A0A | JAK1_HUMAN T | Unmodified | 0 | 0 | 0 | 0 | 963450 | 0 | 0 | 0 | 0 | 0 | 0 | 0 | 0 | 0 | 0 | 0 | 196460 | 0 |

| | | | | | | | | | | | | | | | | | | | | | | |
|---|---|---|---|---|---|---|---|---|---|---|---|---|---|---|---|---|---|---|---|---|---|---|
| NFQMASIT: | MYOF; MYOF | Q9NZM1; H( | MYOF_HUMAN | Oxidation (M) | 0 | 0 | 0 | 0 | 5652500 | 0 | 6099200 | 0 | 0 | 0 | 0 | 0 | 0 | 0 | 0 | 0 | 0 | 0 |
| NFQNIFPP: | PTBP1; PTBP1; | P26599; A0A | PTBP1_HUMAN | Unmodified | 0 | 0 | 0 | 0 | 0 | 0 | 0 | 0 | 2880500 | 2262100 | 4013100 | 3007200 | 0 | 0 | 0 | 0 | 0 | 0 |
| NFRLLEELE | UBE2V2; UBE2\ | Q15819; Q1: | UB2V2_HUMAN | Unmodified | 97022 | 80186 | 744080 | 687620 | 2410500 | 608000 | 237610 | 0 | 947140 | 3020200 | 1857300 | 0 | 4861300 | 1642200 | 93188 | 0 | 2257400 | 8770700 |
| NFRNIQVD | UBE2L3 | P68036 | UB2L3_HUMAN | Unmodified | 0 | 0 | 0 | 0 | 1373100 | 0 | 0 | 0 | 0 | 549270 | 0 | 0 | 1196000 | 3373700 | 0 | 0 | 0 | 0 |
| NFRNPLAK | ENO1; ENO1 | P06733; A0A | ENOA_HUMAN | Unmodified | 523110 | 932300 | 4046200 | 3399600 | 22249000 | 5357100 | 257300 | 346540 | 11701000 | 7668500 | 10126000 | 41373000 | 56597000 | 21590000 | 3347800 | 4527100 | 5880600 | 13026000 |
| NFSAHLLD | TUBGCP2 | Q9BSJ2 | GCP2_HUMAN | Unmodified | 0 | 0 | 0 | 0 | 0 | 0 | 0 | 0 | 0 | 0 | 81390 | 557870 | 0 | 0 | 0 | 0 | 0 | 0 |
| NFSAIINPP | DLAT; DLAT; DL | P10515; A0A | ODP2_HUMAN | Unmodified | 0 | 0 | 0 | 0 | 0 | 0 | 0 | 0 | 667720 | 584380 | 0 | 0 | 0 | 0 | 0 | 0 | 0 | 0 |
| NFSDNQL( | TAGLN2; TAGLI | P37802; X6R | TAGL2_HUMAN | Unmodified | 9794000 | 5609900 | 2040700 | 1807200 | 3491400 | 1427100 | 117220 | 100000 | 6671600 | 584550 | 4505400 | 15618000 | 922510 | 227160 | 2495600 | 2026600 | 540690 | 2291500 |
| NFSGAELE | NSF; NSF | P46459; I3L( | NSF_HUMAN V | Unmodified | 1025700 | 0 | 0 | 0 | 828540 | 0 | 32682 | 0 | 1199800 | 0 | 418830 | 2658000 | 0 | 2584300 | 189170 | 0 | 389390 | 1039600 |
| NFSGAELE | NSF; NSF | P46459; I3L( | NSF_HUMAN V | Unmodified | 0 | 0 | 0 | 0 | 0 | 0 | 418250 | 0 | 0 | 0 | 0 | 0 | 1250700 | 1483900 | 0 | 0 | 1086200 | 1631700 |
| NFTGRND/ | ACO2; ACO2; A | Q99798; A2/ | ACON_HUMAN | Unmodified | 0 | 0 | 0 | 0 | 0 | 0 | 0 | 0 | 1552200 | 0 | 238520 | 0 | 0 | 0 | 0 | 0 | 0 | 0 |
| NFTLDMI | RO60; RO60 | P10155; G5E | RO60_HUMAN | Oxidation (M) | 4007400 | 2958900 | 1249400 | 910940 | 0 | 0 | 0 | 0 | 0 | 0 | 0 | 613920 | 0 | 0 | 1556300 | 1293300 | 0 | 0 |
| NFTMDTES | SMARCA5 | O60264 | SMCA5_HUMA | Oxidation (M) | 416010 | 0 | 0 | 0 | 0 | 0 | 0 | 0 | 0 | 0 | 0 | 0 | 0 | 0 | 0 | 0 | 0 | 0 |
| NFVDSPIIV | GBA; GBA; GBA | P04062; A0A | GLCM_HUMAN | Unmodified | 0 | 0 | 24537 | 0 | 0 | 0 | 0 | 0 | 3779500 | 0 | 0 | 305930 | 0 | 0 | 0 | 164620 | 0 | 0 |
| NFVDTVRD | SLC12A7; SLC1 | Q9Y666; A0/ | S12A7_HUMAN | Unmodified | 814410 | 664710 | 713400 | 702790 | 1921900 | 0 | 0 | 0 | 4011700 | 0 | 806330 | 5773300 | 653780 | 0 | 4435700 | 3530600 | 0 | 0 |
| NFVSFRVQ | HYAL2 | Q12891 | HYAL2_HUMAN | Unmodified | 0 | 0 | 82268 | 434430 | 0 | 0 | 0 | 0 | 0 | 0 | 0 | 0 | 0 | 0 | 0 | 78693 | 0 | 0 |
| NFVSQIVAI | TOR1B; TOR1B | O14657; H0' | TOR1B_HUMAI | Unmodified | 0 | 0 | 0 | 0 | 739740 | 0 | 0 | 0 | 4653900 | 2983700 | 0 | 0 | 0 | 0 | 0 | 0 | 0 | 0 |
| NFVVVMVT | RAD23B; RAD2: | P54727; Q5\ | RD23B_HUMAI | Unmodified | 996910 | 695370 | 0 | 365480 | 1112800 | 0 | 524890 | 0 | 1061700 | 827370 | 0 | 382730 | 1370400 | 0 | 389410 | 246760 | 300070 | 0 |
| NFVVVMVT | RAD23B; RAD2: | P54727; Q5\ | RD23B_HUMAI | Oxidation (M) | 454630 | 726290 | 0 | 0 | 0 | 0 | 0 | 0 | 0 | 366390 | 338040 | 375450 | 0 | 0 | 497260 | 822840 | 475540 | 0 |
| NFYELSPHI | MYO1B; MYO1E | O43795; C9J | MYO1B_HUMA | Unmodified | 438500 | 502020 | 207700 | 475220 | 0 | 314910 | 0 | 0 | 2126100 | 0 | 3002800 | 19978000 | 68849 | 58706 | 1263900 | 378960 | 272260 | 0 |
| NFYESDDD | AP3B1 | O00203 | AP3B1_HUMAN | Unmodified | 0 | 0 | 0 | 0 | 0 | 0 | 0 | 0 | 0 | 0 | 0 | 878580 | 0 | 0 | 0 | 0 | 0 | 0 |
| NFYNEHEE | DDX42; DDX42 | Q86XP3; A0/ | DDX42_HUMAI | Unmodified | 0 | 0 | 0 | 0 | 0 | 0 | 0 | 0 | 0 | 0 | 897440 | 1177000 | 595160 | 0 | 0 | 0 | 0 | 0 |
| NFYQEHPD | DDX5; DDX5; D | P17844; A0A | DDX5_HUMAN | Unmodified | 0 | 0 | 0 | 0 | 0 | 0 | 63829 | 0 | 157140 | 0 | 0 | 1873400 | 0 | 0 | 0 | 0 | 0 | 0 |
| NFYVEVPEI | DDX46; DDX46 | Q7L014; A0/ | DDX46_HUMAI | Unmodified | 0 | 0 | 0 | 0 | 0 | 0 | 1136700 | 0 | 0 | 0 | 0 | 1127300 | 0 | 0 | 0 | 0 | 0 | 0 |
| NFYYAAVP: | GALNT2 | Q10471 | GALT2_HUMAN | Unmodified | 0 | 166180 | 0 | 0 | 1276400 | 618080 | 211900 | 0 | 56917000 | 4769000 | 1808900 | 1325700 | 0 | 0 | 0 | 0 | 492760 | 0 |
| NGAFVNA/ | ANKFY1 | Q9P2R3 | ANFY1_HUMAN | Unmodified | 0 | 0 | 0 | 447340 | 2832600 | 0 | 510950 | 0 | 0 | 0 | 26673 | 106870 | 0 | 0 | 157910 | 0 | 0 | 0 |
| NGAPIIMSI | SCARB2; SCARI | Q14108; A0/ | SCRB2_HUMAI | Unmodified | 0 | 17638 | 86548 | 138890 | 0 | 0 | 0 | 0 | 6837100 | 547750 | 0 | 0 | 0 | 0 | 364370 | 0 | 0 | 0 |
| NGAVIMAV | PRXL2A | Q9BRX8 | PXL2A_HUMAN | Oxidation (M) | 0 | 0 | 0 | 0 | 29817 | 0 | 0 | 0 | 2134100 | 69559 | 0 | 0 | 0 | 0 | 0 | 0 | 9172.5 | 0 |
| NGAVIMAV | PRXL2A | Q9BRX8 | PXL2A_HUMAN | Unmodified | 0 | 0 | 0 | 0 | 0 | 0 | 0 | 0 | 4927500 | 0 | 53471 | 123790 | 0 | 0 | 0 | 0 | 0 | 0 |
| NGAWEIVI | ATP8A1; ATP8A | Q9Y2Q0; H0 | AT8A1_HUMAN | Unmodified | 0 | 0 | 0 | 0 | 0 | 0 | 0 | 0 | 410870 | 120400 | 0 | 0 | 0 | 0 | 37711 | 0 | 0 | 0 |
| NGCGGEIE | ITGB5 | P18084 | ITB5_HUMAN I | Unmodified | 686670 | 381790 | 571820 | 657760 | 502720 | 0 | 0 | 0 | 830300 | 0 | 0 | 2562700 | 1602700 | 452510 | 916710 | 639840 | 612980 | 316580 |
| NGDLDEVI | MTPN | P58546 | MTPN_HUMAN | Unmodified | 0 | 0 | 0 | 0 | 0 | 0 | 0 | 0 | 0 | 0 | 0 | 1575500 | 0 | 0 | 0 | 0 | 0 | 426660 |
| NGDVIELK' | PTPN11; PTPN1 | Q06124; A0/ | PTN11_HUMAI | Unmodified | 0 | 0 | 0 | 0 | 0 | 0 | 0 | 0 | 0 | 0 | 0 | 219710 | 0 | 0 | 0 | 0 | 0 | 0 |
| NGDYKET( | KIF23; KIF23; KI | Q02241; A0/ | KIF23_HUMAN | Unmodified | 46780 | 37795 | 0 | 229960 | 1040800 | 169600 | 0 | 0 | 414490 | 70046 | 102340 | 234530 | 27537 | 0 | 0 | 44812 | 156090 | 0 |
| NGECFRAI | GARS1; GARS1 | P41250; F8V | GARS_HUMAN | Unmodified | 0 | 0 | 0 | 0 | 0 | 0 | 0 | 0 | 0 | 0 | 0 | 0 | 249400 | 150260 | 0 | 0 | 0 | 0 |
| NGEILQISV | TMX4 | Q9H1E5 | TMX4_HUMAN | Unmodified | 0 | 0 | 0 | 0 | 0 | 0 | 0 | 0 | 2681200 | 0 | 0 | 0 | 0 | 0 | 0 | 0 | 0 | 0 |
| NGELENIKI | PAFAH1B2 | P68402 | PA1B2_HUMAI | Unmodified | 0 | 0 | 0 | 0 | 0 | 0 | 0 | 0 | 0 | 0 | 158330 | 1027500 | 0 | 0 | 0 | 0 | 30653 | 0 |
| NGETELCN | AHSA1; AHSA1 | O95433; H0' | AHSA1_HUMAI | Unmodified | 0 | 0 | 0 | 0 | 0 | 0 | 0 | 0 | 0 | 0 | 0 | 780150 | 0 | 0 | 0 | 0 | 0 | 0 |
| NGETVPIDI | KIF5B | P33176 | KINH_HUMAN | Unmodified | 0 | 0 | 0 | 0 | 0 | 0 | 0 | 0 | 0 | 0 | 188920 | 1774900 | 0 | 0 | 0 | 0 | 0 | 0 |
| NGEVQNL/ | CAND2; CAND: | O75155; Q8( | CAND2_HUMA | Unmodified | 0 | 0 | 0 | 0 | 0 | 0 | 0 | 0 | 468470 | 0 | 439510 | 2622800 | 0 | 0 | 0 | 0 | 0 | 0 |
| NGEVVHTF | SLC16A3; SLC1 | O15427; J3Q | MOT4_HUMAN | Unmodified | 0 | 0 | 401380 | 372660 | 13050000 | 1962200 | 0 | 0 | 74499 | 0 | 257470 | 2146000 | 0 | 0 | 724260 | 441440 | 3993300 | 6823000 |
| NGFLLDGF | AK2; AK2; AK2; | P54819; G3\ | KAD2_HUMAN | Unmodified | 0 | 0 | 0 | 0 | 0 | 0 | 0 | 0 | 1765900 | 0 | 942420 | 4815000 | 0 | 0 | 0 | 0 | 0 | 0 |
| NGFPKQTA | MACROH2A1; I | O75367; B4I | H2AY_HUMAN | Unmodified | 1190900 | 0 | 5757600 | 2990800 | 0 | 7215200 | 5895600 | 2796400 | 0 | 646390 | 65313 | 6753900 | 2022300 | 5748600 | 1904800 | 790850 | 1083500 | 11228000 |
| NGGLGHM | PRDX3 | P30048 | PRDX3_HUMAI | Oxidation (M) | 0 | 0 | 0 | 0 | 0 | 0 | 44020 | 0 | 534160 | 179350 | 0 | 0 | 0 | 0 | 0 | 0 | 0 | 0 |
| NGGLGHM | PRDX3 | P30048 | PRDX3_HUMAI | Unmodified | 0 | 0 | 0 | 0 | 0 | 0 | 0 | 0 | 1727900 | 0 | 0 | 0 | 0 | 0 | 0 | 0 | 0 | 0 |
| NGGLITLEE | SNF8; SNF8; SI | Q96H20; D6 | SNF8_HUMAN | Unmodified | 2128800 | 2003900 | 1097600 | 700080 | 0 | 0 | 0 | 0 | 208850 | 0 | 50114 | 2236200 | 201420 | 0 | 2905900 | 2141600 | 0 | 0 |
| NGGQETIE | DSC2; DSC2 | Q02487; A0/ | DSC2_HUMAN | Unmodified | 0 | 0 | 0 | 0 | 1284600 | 0 | 848690 | 622250 | 269240 | 425980 | 143950 | 257210 | 466610 | 983600 | 0 | 0 | 1763600 | 1568900 |
| NGGQILIAI | SEC13; SEC13; | P55735; A8I | SEC13_HUMAI | Unmodified | 0 | 0 | 0 | 0 | 0 | 0 | 0 | 0 | 0 | 0 | 119700 | 2591700 | 0 | 0 | 0 | 100340 | 0 | 0 |
| NGGTQSVI | MOV10; MOV1( | Q9HCE1; Q5 | MOV10_HUMA | Unmodified | 1933400 | 750320 | 479070 | 0 | 0 | 0 | 0 | 0 | 0 | 0 | 0 | 0 | 0 | 0 | 762370 | 1112200 | 0 | 0 |
| NGHLAVV/ | MYO1C; MYO1( | O00159; F5I | MYO1C_HUMA | Unmodified | 0 | 0 | 0 | 429980 | 874440 | 0 | 0 | 0 | 0 | 0 | 0 | 0 | 414840 | 413790 | 461670 | 339270 | 220220 | 838990 |
| NGHVGISF | FLNA; FLNA | P21333; Q6( | FLNA_HUMAN | Unmodified | 0 | 0 | 0 | 0 | 0 | 0 | 750340 | 0 | 0 | 0 | 0 | 0 | 894860 | 895920 | 0 | 0 | 589880 | 184520 |
| NGIHDLDI | PSMB2; PSMB2 | P49721; A0A | PSB2_HUMAN | Unmodified | 0 | 0 | 0 | 0 | 285250 | 0 | 0 | 0 | 1102700 | 290590 | 516310 | 4512700 | 0 | 0 | 157230 | 0 | 0 | 113980 |
| NGILASPQ: | SLC4A7 | C9JRP1 | C9JRP1_HUMA | Unmodified | 0 | 0 | 0 | 0 | 0 | 0 | 71712 | 0 | 0 | 0 | 0 | 989900 | 0 | 0 | 0 | 0 | 0 | 0 |
| NGKITSIVK | SDCBP; SDCBP | O00560; G5I | SDCB1_HUMAI | Unmodified | 607970 | 551860 | 44652 | 2589900 | 2932600 | 0 | 295690 | 140560 | 6524100 | 6940300 | 0 | 0 | 2359200 | 1790900 | 0 | 0 | 143870 | 343250 |
| NGKITSIVK | SDCBP; SDCBP | O00560; G5I | SDCB1_HUMAI | Unmodified | 0 | 0 | 0 | 0 | 293070 | 0 | 0 | 0 | 1222900 | 1057700 | 0 | 0 | 404580 | 291990 | 0 | 0 | 0 | 0 |
| NGLDEYLE | RIPOR2; RIPOR | Q9Y4F9; B7Z | RIPR2_HUMAN | Unmodified | 0 | 0 | 0 | 0 | 0 | 0 | 0 | 0 | 569730 | 173450 | 649880 | 3544400 | 226950 | 132530 | 0 | 0 | 0 | 0 |
| NGLDEYLE | RIPOR2; RIPOR | Q9Y4F9; B7Z | RIPR2_HUMAN | Unmodified | 0 | 0 | 0 | 0 | 0 | 0 | 0 | 0 | 0 | 0 | 0 | 0 | 1548400 | 1270800 | 0 | 0 | 0 | 0 |
| NGLLFTYG' | KIF23; KIF23; KI | Q02241; A0/ | KIF23_HUMAN | Unmodified | 0 | 0 | 0 | 899390 | 8259200 | 0 | 0 | 0 | 2895000 | 18394 | 1109100 | 1578500 | 0 | 0 | 0 | 0 | 0 | 0 |
| NGLLFTYG' | KIF23; KIF23; KI | Q02241; A0/ | KIF23_HUMAN | Oxidation (M) | 0 | 0 | 0 | 0 | 2272000 | 0 | 0 | 0 | 0 | 419970 | 0 | 0 | 0 | 0 | 0 | 0 | 529150 | 0 |
| NGLLTEHN | SDCBP; SDCBP | O00560; G5I | SDCB1_HUMAI | Unmodified | 2605500 | 18075000 | 14682000 | 2407400 | 0 | 0 | 0 | 0 | 52617000 | 3149400 | 1612700 | 34410000 | 0 | 0 | 2141600 | 31529000 | 0 | 0 |
| NGLPDHTI | GAPVD1; GAPV | Q14C86; F8\ | GAPD1_HUMAI | Unmodified | 0 | 35292 | 0 | 0 | 0 | 0 | 0 | 0 | 128280 | 0 | 0 | 645100 | 0 | 0 | 224340 | 1097900 | 0 | 0 |
| NGLSLAALI | H1-5 | P16401 | H15_HUMAN I | Unmodified | 0 | 2149700 | 0 | 0 | 0 | 0 | 0 | 0 | 0 | 0 | 0 | 0 | 0 | 0 | 0 | 0 | 0 | 0 |
| NGLVINND | ATP7A | Q04656 | ATP7A_HUMAN | Unmodified | 0 | 0 | 0 | 0 | 0 | 0 | 0 | 0 | 0 | 0 | 0 | 0 | 0 | 0 | 185920 | 100560 | 0 | 0 |
| NGLVNSEV | TMEM106B; TM | Q9NUM4; C! | T106B_HUMAN | Unmodified | 0 | 0 | 0 | 0 | 0 | 0 | 0 | 0 | 0 | 0 | 0 | 340970 | 33320 | 0 | 0 | 0 | 0 | 0 |
| NGMAVAQI | CTH | P32929 | CGL_HUMAN ( | Oxidation (M) | 0 | 0 | 0 | 81777 | 0 | 0 | 0 | 0 | 0 | 0 | 22966 | 0 | 392490 | 154660 | 49661 | 30032 | 0 | 0 |
| NGMGEAAI | FAM83F | Q8NEG4 | FA83F_HUMAN | Unmodified | 0 | 521420 | 357900 | 1049400 | 0 | 0 | 0 | 0 | 0 | 0 | 0 | 0 | 0 | 0 | 385440 | 747600 | 0 | 0 |
| NGMVLKPI | RPL13; RPL13; I | P26373; H3E | RL13_HUMAN ( | Unmodified | 0 | 0 | 0 | 161310 | 0 | 0 | 0 | 0 | 0 | 0 | 0 | 752730 | 0 | 0 | 0 | 206910 | 0 | 0 |
| NGNGGPG | CTNND1; CTNI | O60716; A0/ | CTND1_HUMAI | Unmodified | 0 | 0 | 0 | 0 | 0 | 0 | 0 | 0 | 0 | 0 | 0 | 0 | 439020 | 0 | 0 | 0 | 1054500 | 2910300 |
| NGNLPEF( | TLN1 | Q9Y490 | TLN1_HUMAN | Unmodified | 0 | 0 | 0 | 0 | 5504300 | 752600 | 0 | 104260 | 0 | 0 | 247270 | 2589300 | 1121700 | 484850 | 0 | 0 | 1506600 | 0 |
| NGPALQE/ | DBNL; DBNL; D | Q9UJU6; F8\ | DBNL_HUMAN | Unmodified | 828080 | 355710 | 672600 | 295730 | 296810 | 0 | 463630 | 447640 | 262680 | 0 | 0 | 3956500 | 322200 | 0 | 0 | 0 | 881380 | 181470 |
| NGPALQE/ | DBNL; DBNL; D | Q9UJU6; F8\ | DBNL_HUMAN | Unmodified | 0 | 0 | 0 | 0 | 0 | 0 | 0 | 0 | 0 | 0 | 0 | 0 | 1834700 | 0 | 0 | 0 | 0 | 0 |
| NGPFKPNY | AHCYL1; AHCY | O43865; Q9( | SAHH2_HUMAI | Unmodified | 0 | 0 | 777200 | 0 | 12001000 | 2199700 | 839270 | 943250 | 356320 | 259180 | 0 | 1734100 | 3094300 | 2786100 | 982850 | 872220 | 4901100 | 4636300 |
| NGPGTVD: | INPP5D; INPP5I | Q92835; H0' | SHIP1_HUMAN | Unmodified | 0 | 0 | 0 | 0 | 0 | 0 | 0 | 0 | 0 | 0 | 0 | 373200 | 0 | 0 | 0 | 0 | 0 | 0 |
| NGPLEVAG | MACROH2A1; I | O75367; B4I | H2AY_HUMAN | Unmodified | 4774100 | 3535300 | 0 | 0 | 0 | 0 | 1298300 | 4655100 | 0 | 0 | 0 | 0 | 0 | 0 | 0 | 0 | 0 | 0 |
| NGPLNES( | TAOK3 | Q9H2K8 | TAOK3_HUMAI | Unmodified | 0 | 0 | 0 | 0 | 0 | 0 | 0 | 0 | 0 | 0 | 0 | 126990 | 0 | 0 | 0 | 409150 | 0 | 0 |
| NGQDLGV/ | HNRNPU; HNR | Q00839; A0/ | HNRPU_HUMA | Unmodified | 660320 | 0 | 0 | 0 | 0 | 0 | 0 | 0 | 0 | 0 | 0 | 5644700 | 0 | 0 | 0 | 0 | 0 | 0 |
| NGQDLGV/ | HNRNPU; HNR | Q00839; A0/ | HNRPU_HUMA | Unmodified | 90886 | 73256 | 0 | 0 | 0 | 0 | 2845600 | 1741700 | 0 | 347050 | 0 | 2696000 | 200120 | 276760 | 0 | 93061 | 0 | 161390 |
| NGQGFILV | RAP2C; RAP2A; | Q9Y3L5; P1( | RAP2C_HUMAI | Oxidation (M) | 458740 | 648090 | 414430 | 449040 | 392430 | 0 | 99518 | 0 | 193000 | 0 | 0 | 1313300 | 0 | 0 | 3874900 | 792530 | 0 | 0 |
| NGQHVAS: | FLNA; FLNA; FL | P21333; H0' | FLNA_HUMAN | Unmodified | 0 | 921830 | 0 | 0 | 0 | 0 | 0 | 180690 | 0 | 0 | 0 | 0 | 0 | 0 | 0 | 0 | 0 | 0 |
| NGQNAAV' | TOR1AIP1; TOR | Q5JTV8; A0A | TOIP1_HUMAN | Unmodified | 0 | 0 | 0 | 0 | 0 | 0 | 0 | 0 | 2968300 | 0 | 165270 | 0 | 0 | 0 | 0 | 0 | 1755600 | 0 |
| NGQRLEVF | BCAM; BCAM; E | P50895; A0A | BCAM_HUMAN | Unmodified | 0 | 0 | 0 | 0 | 0 | 0 | 0 | 0 | 2110400 | 0 | 0 | 0 | 0 | 0 | 0 | 0 | 0 | 0 |
| NGQTREH/ | EEF1A2; EEF1A: | Q05639; Q5\ | EF1A2_HUMAN | Unmodified | 3082900 | 1871800 | 4012900 | 11875000 | 32219000 | 5723500 | 6998800 | 459290 | 1043900 | 3173000 | 3673400 | 1053100 | 45117000 | 19056000 | 1656700 | 817590 | 14648000 | 24243000 |
| NGQVIGIG. | ATIC; ATIC | P31939; H7( | PUR9_HUMAN | Unmodified | 0 | 0 | 0 | 0 | 0 | 0 | 0 | 0 | 0 | 0 | 0 | 0 | 0 | 464450 | 0 | 0 | 0 | 0 |
| NGRIFFDD | GCA; GCA; GC/ | P28676; H7( | GRAN_HUMAN | Unmodified | 0 | 0 | 0 | 0 | 161320 | 46166 | 171410 | 98300 | 0 | 230030 | 219110 | 87186 | 134670 | 319670 | 662480 | 0 | 127760 | 166700 |
| NGRLVYLV | TFRC; TFRC | P02786; G3\ | TFR1_HUMAN | Unmodified | 0 | 0 | 0 | 1256300 | 1846600 | 1482700 | 1063500 | 0 | 4837900 | 1605900 | 431260 | 0 | 681760 | 828020 | 0 | 0 | 147460 | 1475800 |
| NGRVEIIAN | HSPA5; HSPA5; | P11021; A0A | BIP_HUMAN Er | Unmodified | 0 | 353050 | 2301300 | 954950 | 0 | 0 | 235090 | 149130 | 85426000 | 1658800 | 4784100 | 5731000 | 0 | 0 | 3117900 | 1090400 | 0 | 0 |
| NGRVEIIAN | HSPA5; HSPA5; | P11021; A0A | BIP_HUMAN Er | Unmodified | 0 | 0 | 0 | 0 | 0 | 0 | 0 | 0 | 38210000 | 0 | 1169200 | 0 | 0 | 0 | 0 | 0 | 0 | 0 |
| NGRYSISR' | CD44; CD44; C | P16070; Q8( | CD44_HUMAN | Unmodified | 0 | 0 | 0 | 0 | 0 | 0 | 0 | 0 | 0 | 0 | 0 | 0 | 0 | 0 | 0 | 0 | 2297500 | 6920100 |
| NGSCFLRT | ATP9A | O75110 | ATP9A_HUMAN | Unmodified | 0 | 0 | 0 | 75212 | 0 | 641770 | 0 | 0 | 370610 | 0 | 0 | 0 | 0 | 0 | 0 | 0 | 0 | 0 |
| NGSCGVS' | FLNB; FLNB | O75369; E7E | FLNB_HUMAN | Unmodified | 0 | 0 | 0 | 0 | 0 | 0 | 0 | 0 | 0 | 0 | 0 | 675530 | 0 | 0 | 0 | 0 | 0 | 0 |
| NGSEADID | UBA1; UBA1; UI | P22314; Q5J | UBA1_HUMAN | Unmodified | 0 | 0 | 0 | 0 | 0 | 0 | 0 | 0 | 0 | 0 | 0 | 1097800 | 0 | 0 | 0 | 0 | 0 | 0 |
| NGSFANLF | SMS; SMS | P52788; H7( | SPSY_HUMAN | Unmodified | 0 | 0 | 0 | 0 | 508680 | 0 | 0 | 0 | 0 | 0 | 364850 | 4004800 | 794980 | 234730 | 114750 | 0 | 0 | 158090 |
| NGSLDSP( | NUDC; NUDC; | Q9Y266; A0/ | NUDC_HUMAN | Unmodified | 0 | 0 | 0 | 0 | 0 | 0 | 0 | 0 | 0 | 0 | 123740 | 2196100 | 0 | 0 | 182800 | 0 | 0 | 0 |
| NGSLICTA: | CORO1C; COR( | Q9ULV4; F8\ | COR1C_HUMA | Unmodified | 0 | 0 | 0 | 0 | 0 | 0 | 0 | 0 | 0 | 0 | 0 | 7644500 | 0 | 0 | 0 | 0 | 0 | 0 |
| NGTAGLIP: | CASK; CASK; C/ | O14936; A0/ | CSKP_HUMAN | Unmodified | 0 | 0 | 206090 | 0 | 0 | 32553 | 0 | 0 | 634220 | 0 | 48214 | 146300 | 0 | 0 | 1874300 | 1785200 | 242640 | 95917 |
| NGTQVHG' | SNRPD1 | P62314 | SMD1_HUMAN | Oxidation (M) | 282890 | 0 | 0 | 0 | 201920 | 0 | 190650 | 0 | 0 | 183430 | 0 | 1217900 | 0 | 0 | 0 | 91859 | 0 | 0 |
| NGTQVHG' | SNRPD1 | P62314 | SMD1_HUMAN | Unmodified | 0 | 0 | 0 | 0 | 0 | 0 | 0 | 0 | 0 | 0 | 0 | 3613600 | 0 | 0 | 0 | 0 | 0 | 0 |
| NGVELSATI | NPTN; NPTN | Q9Y639; Q9I | NPTN_HUMAN | Unmodified | 0 | 0 | 0 | 28210 | 273390 | 69120 | 0 | 0 | 0 | 56185 | 0 | 461570 | 0 | 214740 | 41715 | 0 | 214170 | 90598 |
| NGVELSATI | NPTN; NPTN | Q9Y639; Q9I | NPTN_HUMAN | Unmodified | 83175 | 142670 | 362740 | 489900 | 260320 | 319630 | 0 | 0 | 936650 | 0 | 0 | 3300300 | 1084700 | 1449000 | 617920 | 356780 | 143480 | 821010 |
| NGVKITLP\ | PGK1 | P00558 | PGK1_HUMAN | Unmodified | 0 | 0 | 0 | 0 | 7067500 | 0 | 3302400 | 99368 | 0 | 2819100 | 313400 | 0 | 13269000 | 9559200 | 0 | 0 | 28298000 | 19679000 |
| NGVNEAKI | FAS | P25445 | TNR6_HUMAN | Unmodified | 0 | 90393 | 0 | 76109 | 499070 | 133600 | 0 | 0 | 0 | 0 | 0 | 0 | 247600 | 0 | 108380 | 79082 | 389240 | 65755 |
| NGVSGLVT | EPHA2 | P29317 | EPHA2_HUMAI | Unmodified | 0 | 216310 | 0 | 0 | 0 | 0 | 0 | 0 | 0 | 0 | 0 | 0 | 0 | 0 | 0 | 0 | 675800 | 1155500 |
| NGYDFYPK | TPP2; TPP2; TPI | P29144; A0A | TPP2_HUMAN | Unmodified | 0 | 0 | 0 | 0 | 0 | 0 | 0 | 0 | 132560 | 0 | 0 | 1204400 | 0 | 0 | 0 | 0 | 0 | 0 |
| NGYDYGQ | SRSF9; SRSF9 | Q13242; S4F | SRSF9_HUMAN | Unmodified | 490640 | 240080 | 0 | 0 | 0 | 0 | 0 | 0 | 0 | 0 | 0 | 0 | 0 | 0 | 252080 | 0 | 0 | 0 |
| NGYELSPT/ | PSMB2 | P49721 | PSB2_HUMAN | Unmodified | 0 | 0 | 0 | 0 | 0 | 0 | 0 | 0 | 1016600 | 0 | 0 | 2999200 | 0 | 0 | 0 | 0 | 0 | 0 |
| NGYGFINR | YBX1; YBX2; YB | P67809; Q9Y | YBOX1_HUMAI | Unmodified | 0 | 0 | 0 | 0 | 0 | 0 | 0 | 0 | 0 | 0 | 0 | 3550700 | 0 | 0 | 0 | 0 | 0 | 0 |
| NGYGFINR | YBX1; YBX2; YB | P67809; Q9Y | YBOX1_HUMAI | Unmodified | 1642100 | 1359800 | 576140 | 3379500 | 538090 | 0 | 0 | 0 | 253800 | 0 | 0 | 8145100 | 0 | 0 | 1080600 | 1503500 | 0 | 0 |
| NGYGTPM1 | F11R; F11R | Q9Y624; A0/ | JAM1_HUMAN | Unmodified | 689290 | 381490 | 503030 | 717750 | 0 | 0 | 0 | 0 | 448630 | 0 | 0 | 614690 | 0 | 0 | 826220 | 409060 | 0 | 0 |
| NGYGTPM1 | F11R; F11R | Q9Y624; A0/ | JAM1_HUMAN | Oxidation (M) | 505920 | 0 | 380320 | 132840 | 0 | 0 | 0 | 0 | 151790 | 0 | 0 | 291560 | 123240 | 0 | 411190 | 341470 | 0 | 0 |

| | | | | | | | | | | | | | | | | | | | | | | |
|---|---|---|---|---|---|---|---|---|---|---|---|---|---|---|---|---|---|---|---|---|---|---|
| NGYTPLHI/ | ANK3; ANK2; AI | Q12955; Q0: | ANK3_HUMAN | Unmodified | 0 | 0 | 72636 | 88498 | 0 | 0 | 0 | 0 | 0 | 0 | 0 | 0 | 232000 | 344620 | 124780 | 0 | 0 | 0 |
| NHAAPFSK | HSPA4; HSPH1 | P34932; Q92 | HSP74_HUMAI | Unmodified | 1048300 | 0 | 480810 | 438700 | 1993000 | 40517 | 0 | 698280 | 824000 | 67995000 | 518100 | 12445000 | 922280 | 1280100 | 0 | 76487 | 564570 | 783890 |
| NHAVVCQ( | TES | Q9UGI8 | TES_HUMAN Te | Unmodified | 0 | 0 | 0 | 0 | 0 | 0 | 0 | 62870 | 0 | 27579 | 318660 | 1129300 | 0 | 116860 | 0 | 224580 | 33970 | 59683 |
| NHDSSQPT | MVB12B | Q9H7P6 | MB12B_HUMAI | Unmodified | 6341400 | 5557900 | 8190400 | 7932700 | 292520 | 0 | 0 | 0 | 6685300 | 1350400 | 729840 | 1595600 | 2541000 | 413830 | 10770000 | 11564000 | 0 | 0 |
| NHEAQIQD | MYH10 | P35580 | MYH10_HUMAI | Unmodified | 0 | 0 | 260370 | 0 | 401140 | 0 | 0 | 0 | 92944 | 0 | 0 | 0 | 0 | 0 | 297000 | 491240 | 0 | 0 |
| NHEFIATFF | PRKCD; PRKCD | Q05655; C9I | KPCD_HUMAN | Unmodified | 0 | 0 | 0 | 0 | 0 | 0 | 0 | 0 | 827230 | 0 | 198600 | 378800 | 339630 | 0 | 0 | 66607 | 853120 | 1844700 |
| NHELSGLN | ZC3HAV1; ZC3I | Q7Z2W4; C9 | ZCCHV_HUMA | Oxidation (M) | 0 | 0 | 0 | 0 | 2418500 | 0 | 0 | 0 | 0 | 0 | 0 | 0 | 0 | 0 | 0 | 0 | 0 | 0 |
| NHFTVALN | SULT1E1; N/A | P49888; H0Y | ST1E1_HUMAN | Unmodified | 0 | 0 | 718970 | 0 | 0 | 0 | 0 | 0 | 0 | 0 | 0 | 0 | 0 | 0 | 0 | 0 | 0 | 0 |
| NHFWELIE | MKRN1 | Q9UHC7 | MKRN1_HUMA | Unmodified | 2182300 | 1353700 | 665350 | 0 | 0 | 0 | 0 | 0 | 0 | 0 | 0 | 0 | 0 | 0 | 897660 | 884450 | 0 | 0 |
| NHGILSGR | MGAT5 | Q09328 | MGT5A_HUMAI | Unmodified | 0 | 0 | 0 | 0 | 0 | 0 | 59538 | 0 | 570750 | 1729000 | 188690 | 0 | 0 | 47582 | 0 | 0 | 0 | 0 |
| NHIFRLTQ( | CIT; CIT; CIT; CI | O14578; A0/ | CTRO_HUMAN | Unmodified | 0 | 0 | 0 | 0 | 2917000 | 0 | 0 | 0 | 0 | 0 | 0 | 0 | 0 | 154410 | 0 | 0 | 826620 | 0 |
| NHILPPMD | CNN2; CNN2; ( | Q99439; H3I | CNN2_HUMAN | Oxidation (M) | 0 | 0 | 0 | 0 | 4072000 | 0 | 76617 | 0 | 0 | 0 | 0 | 0 | 338100 | 0 | 0 | 0 | 371600 | 411580 |
| NHILPPMD | CNN2; CNN2; ( | Q99439; H3I | CNN2_HUMAN | Unmodified | 0 | 0 | 0 | 0 | 5152300 | 0 | 0 | 0 | 513080 | 0 | 0 | 0 | 97465 | 0 | 0 | 77155 | 308320 | 480360 |
| NHILPPMD | CNN2; CNN2; ( | Q99439; H3I | CNN2_HUMAN | 2 Oxidation (M) | 0 | 0 | 0 | 0 | 544640 | 154830 | 0 | 0 | 0 | 0 | 0 | 0 | 0 | 183320 | 0 | 0 | 103910 | 111940 |
| NHIPITEQ( | GEMIN6 | Q8WXD5 | GEMI6_HUMAI | Unmodified | 67171 | 0 | 0 | 0 | 0 | 62419 | 36333 | 0 | 0 | 204010 | 0 | 706570 | 235500 | 0 | 55714 | 0 | 0 | 0 |
| NHIQLVK | DIABLO; DIABL( | Q9NR28; A0. | DBLOH_HUMA | Unmodified | 0 | 0 | 0 | 0 | 0 | 0 | 72233 | 11739 | 457100 | 0 | 0 | 158110 | 0 | 0 | 0 | 0 | 0 | 0 |
| NHLLADG( | ARHGAP10 | A1A4S6 | RHG10_HUMAI | Unmodified | 0 | 0 | 204520 | 340370 | 0 | 0 | 0 | 0 | 0 | 0 | 0 | 0 | 0 | 0 | 455220 | 303180 | 0 | 0 |
| NHLLHVFD | ANXA4; ANXA4 | P09525; Q6F | ANXA4_HUMAI | Unmodified | 0 | 0 | 0 | 0 | 0 | 0 | 0 | 0 | 3031300 | 0 | 540720 | 1848900 | 0 | 0 | 0 | 0 | 0 | 0 |
| NHLLHVFD | ANXA4; ANXA4 | P09525; Q6F | ANXA4_HUMAI | Unmodified | 476160 | 0 | 0 | 0 | 0 | 0 | 0 | 0 | 2507000 | 428660 | 833050 | 748770 | 0 | 0 | 0 | 0 | 24806 | 0 |
| NHLPQDS( | PTPN23 | Q9H3S7 | PTN23_HUMAI | Unmodified | 0 | 0 | 0 | 0 | 8129700 | 1315200 | 0 | 0 | 1593900 | 839960 | 0 | 0 | 0 | 0 | 0 | 0 | 0 | 3308900 |
| NHNALLAS | ATRN | O75882 | ATRN_HUMAN | Unmodified | 0 | 175270 | 0 | 112350 | 0 | 235650 | 83111 | 0 | 1115100 | 204450 | 0 | 447820 | 0 | 0 | 239890 | 0 | 0 | 0 |
| NHNGPGFI | MGAM2 | Q2M2H8 | MGAL_HUMAN | Unmodified | 373440 | 0 | 697230 | 1565800 | 0 | 0 | 26883 | 0 | 0 | 0 | 406560 | 1400300 | 0 | 0 | 2270600 | 1813300 | 76876 | 0 |
| NHNLELSK | NDST2 | P52849 | NDST2_HUMAI | Unmodified | 0 | 0 | 0 | 0 | 0 | 0 | 0 | 0 | 460960 | 0 | 0 | 469470 | 0 | 0 | 0 | 0 | 0 | 0 |
| NHNMYVA( | KIF23; KIF23; KI | Q02241; A0/ | KIF23_HUMAN | Unmodified | 0 | 0 | 0 | 0 | 4157100 | 0 | 0 | 0 | 1209200 | 296590 | 0 | 725480 | 0 | 0 | 0 | 0 | 141790 | 0 |
| NHNMYVA( | KIF23; KIF23; KI | Q02241; A0/ | KIF23_HUMAN | Oxidation (M) | 0 | 0 | 55596 | 0 | 987300 | 0 | 0 | 0 | 300260 | 0 | 181050 | 384550 | 0 | 0 | 0 | 0 | 0 | 0 |
| NHNSLLSL | GAA | P10253 | LYAG_HUMAN | Unmodified | 0 | 0 | 0 | 0 | 0 | 0 | 710480 | 0 | 1194500 | 0 | 0 | 0 | 0 | 0 | 0 | 0 | 0 | 0 |
| NHNVAVYS | PPM1L | Q5SGD2 | PPM1L_HUMAI | Unmodified | 0 | 0 | 0 | 0 | 0 | 0 | 0 | 0 | 2165700 | 116750 | 0 | 0 | 0 | 0 | 0 | 0 | 0 | 0 |
| NHPAPFSK | HSPA4L; HSPA4 | O95757; E9F | HS74L_HUMAI | Unmodified | 0 | 0 | 0 | 0 | 0 | 0 | 0 | 0 | 0 | 0 | 0 | 0 | 0 | 0 | 0 | 0 | 0 | 0 |
| NHPAVVM\ | GUSB | P08236 | BGLR_HUMAN | Oxidation (M) | 0 | 0 | 0 | 0 | 5629000 | 7071900 | 1402600 | 0 | 10302000 | 8925500 | 2116700 | 6334600 | 5994100 | 3300100 | 2796900 | 0 | 0 | 0 |
| NHPAVVM\ | GUSB | P08236 | BGLR_HUMAN | Unmodified | 0 | 0 | 0 | 0 | 0 | 0 | 0 | 0 | 15439000 | 3281300 | 0 | 0 | 0 | 0 | 0 | 0 | 0 | 0 |
| NHPDLAGI | FURIN; FURIN; | P09958; A0A | FURIN_HUMAN | Unmodified | 0 | 0 | 0 | 0 | 0 | 0 | 0 | 0 | 2341500 | 0 | 5122300 | 756020 | 0 | 0 | 0 | 0 | 0 | 0 |
| NHPMTPEA | HSD17B4; HSD | P51659; A0A | DHB4_HUMAN | Unmodified | 0 | 0 | 0 | 0 | 0 | 0 | 134950 | 52176 | 3864700 | 772690 | 0 | 0 | 0 | 79384 | 0 | 0 | 0 | 0 |
| NHQEEDLT | CNPY3 | Q9BT09 | CNPY3_HUMAI | Unmodified | 132520 | 0 | 0 | 0 | 0 | 0 | 0 | 0 | 32268000 | 0 | 0 | 0 | 0 | 0 | 0 | 0 | 0 | 0 |
| NHQLEVTP | TAOK3; TAOK3 | Q9H2K8; G3 | TAOK3_HUMAI | Unmodified | 0 | 0 | 0 | 82738 | 276680 | 164970 | 0 | 196500 | 0 | 0 | 0 | 0 | 162280 | 422620 | 0 | 0 | 81490 | 286150 |
| NHQLEVTP | TAOK3; TAOK3 | Q9H2K8; G3 | TAOK3_HUMAI | Unmodified | 0 | 0 | 0 | 0 | 0 | 13433 | 251520 | 117840 | 0 | 0 | 0 | 0 | 0 | 168240 | 0 | 0 | 103610 | 0 |
| NHQSAAEY | DPYSL3 | Q14195 | DPYL3_HUMAI | Oxidation (M) | 0 | 0 | 158980 | 0 | 0 | 0 | 0 | 0 | 0 | 0 | 3626900 | 31684000 | 0 | 0 | 0 | 0 | 0 | 0 |
| NHQSAAEY | DPYSL3 | Q14195 | DPYL3_HUMAI | Unmodified | 0 | 415970 | 510400 | 648740 | 0 | 0 | 0 | 0 | 0 | 0 | 2959800 | 16597000 | 0 | 0 | 413720 | 324430 | 0 | 31850 |
| NHQSAAEY | DPYSL3 | Q14195 | DPYL3_HUMAI | Oxidation (M) | 0 | 0 | 0 | 0 | 1044700 | 0 | 2457000 | 0 | 0 | 0 | 0 | 0 | 0 | 244210 | 0 | 0 | 0 | 0 |
| NHSIILSAP | STX2; STX2 | P32856; J3KI | STX2_HUMAN S | Unmodified | 0 | 0 | 0 | 0 | 0 | 0 | 0 | 0 | 0 | 0 | 0 | 953990 | 0 | 0 | 0 | 0 | 0 | 0 |
| NHSQRPAI | TNIK; TNIK | Q9UKE5; C9 | TNIK_HUMAN T | Unmodified | 93234 | 0 | 99677 | 220200 | 1035400 | 693000 | 0 | 0 | 461250 | 340470 | 0 | 1609600 | 229140 | 256850 | 0 | 0 | 274640 | 0 |
| NHSTVTVE | TRIM29 | Q14134 | TRI29_HUMAN | Unmodified | 0 | 0 | 0 | 0 | 0 | 0 | 0 | 0 | 0 | 0 | 353630 | 900650 | 0 | 0 | 0 | 0 | 158630 | 0 |
| NHTLALTE | RCC2 | Q9P258 | RCC2_HUMAN | Unmodified | 0 | 0 | 0 | 0 | 0 | 0 | 0 | 0 | 1587500 | 75472 | 0 | 16107000 | 0 | 0 | 0 | 0 | 0 | 0 |
| NHTLQEQ\ | EEA1 | Q15075 | EEA1_HUMAN I | Unmodified | 0 | 0 | 0 | 0 | 1548000 | 0 | 248760 | 0 | 0 | 0 | 0 | 0 | 621480 | 509880 | 0 | 0 | 211160 | 0 |
| NHVQPYIP | NIBAN2 | Q96TA1 | NIBA2_HUMAN | Oxidation (M) | 374940 | 226690 | 0 | 127370 | 142540 | 0 | 0 | 0 | 333200 | 0 | 0 | 235680 | 0 | 0 | 1912500 | 150420 | 0 | 0 |
| NHYMLYK | ITGB5; ITGB5; I | P18084; H7( | ITB5_HUMAN I | Unmodified | 0 | 0 | 0 | 0 | 140690 | 0 | 0 | 0 | 44259 | 40999 | 52381 | 779160 | 36844 | 23782 | 205340 | 74329 | 0 | 81090 |
| NIAASGKF | PYGL | P06737 | PYGL_HUMAN | Unmodified | 0 | 0 | 0 | 0 | 0 | 0 | 0 | 0 | 0 | 0 | 0 | 0 | 0 | 0 | 0 | 0 | 0 | 5198100 |
| NIAFFSTN( | ATP1A1 | P05023 | AT1A1_HUMAN | Unmodified | 0 | 0 | 13594000 | 14752000 | 357020 | 230250 | 90935 | 298870 | 8137700 | 1605100 | 6498900 | 49511000 | 6905200 | 603270 | 21550000 | 16860000 | 3020300 | 202820 |
| NIAIEFLTLE | SMC4; SMC4; S | Q9NTJ3; E9F | SMC4_HUMAN | Unmodified | 0 | 0 | 0 | 0 | 600310 | 104140 | 874870 | 0 | 0 | 261970 | 0 | 0 | 1224000 | 0 | 0 | 0 | 163830 | 559140 |
| NIALRNDSI | CRLF3 | Q8IUI8 | CRLF3_HUMAN | Unmodified | 0 | 0 | 0 | 0 | 0 | 0 | 0 | 157980 | 0 | 102100 | 0 | 663880 | 0 | 0 | 0 | 0 | 0 | 0 |
| NIAQLQPD | SLITRK6 | Q9H5Y7 | SLIK6_HUMAN | Unmodified | 0 | 0 | 0 | 0 | 0 | 0 | 0 | 0 | 9083300 | 4860100 | 0 | 0 | 0 | 0 | 0 | 0 | 49515 | 1045700 |
| NIAQLQPD | SLITRK6 | Q9H5Y7 | SLIK6_HUMAN | Oxidation (M) | 0 | 6244300 | 0 | 0 | 0 | 0 | 0 | 0 | 2243500 | 1865200 | 0 | 0 | 0 | 0 | 0 | 0 | 243900 | 347310 |
| NIATYYGAF | TNIK; MINK1; M | Q9UKE5; Q8 | TNIK_HUMAN T | Unmodified | 0 | 0 | 514430 | 603630 | 720090 | 0 | 0 | 0 | 2413600 | 748510 | 1162700 | 3190500 | 795150 | 0 | 683450 | 601310 | 991360 | 0 |
| NIATYYGAF | TNIK; MINK1; M | Q9UKE5; Q8 | TNIK_HUMAN T | Unmodified | 0 | 0 | 0 | 0 | 227140 | 0 | 0 | 0 | 612210 | 476550 | 0 | 1348500 | 0 | 0 | 0 | 55892 | 0 | 0 |
| NIAYEHILS | CA14 | Q9ULX7 | CAH14_HUMAI | Unmodified | 0 | 0 | 0 | 0 | 0 | 0 | 0 | 0 | 0 | 51183 | 474490 | 10916000 | 0 | 0 | 0 | 0 | 0 | 0 |
| NICKVVEV( | PKM; PKM; PKM | P14618; B4D | KPYM_HUMAN | Unmodified | 0 | 200860 | 0 | 0 | 586870 | 0 | 0 | 888940 | 0 | 27286 | 0 | 0 | 169270 | 199660 | 0 | 0 | 4171800 | 598120 |
| NICQFLLE\ | EIF1B | O60739 | EIF1B_HUMAN | Unmodified | 0 | 0 | 0 | 293650 | 0 | 0 | 0 | 0 | 0 | 518430 | 0 | 0 | 783050 | 0 | 0 | 0 | 783250 | 612500 |
| NICQFLVEI | EIF1; EIF1 | P41567; K7E | EIF1_HUMAN E | Unmodified | 764180 | 884800 | 0 | 1534400 | 0 | 66329 | 0 | 0 | 463880 | 0 | 150500 | 0 | 14211000 | 5340600 | 1225100 | 226680 | 2373200 | 1877900 |
| NICQFLVEI | EIF1; EIF1 | P41567; K7E | EIF1_HUMAN E | Unmodified | 0 | 0 | 0 | 0 | 300510 | 0 | 357470 | 0 | 0 | 0 | 0 | 0 | 393250 | 556000 | 0 | 0 | 1045100 | 162920 |
| NICQQVNII | EIF3A | Q14152 | EIF3A_HUMAN | Unmodified | 338840 | 0 | 1287500 | 517070 | 1899400 | 0 | 0 | 0 | 87221 | 33807 | 0 | 2471600 | 0 | 0 | 448780 | 334140 | 0 | 0 |
| NIDDGTSD | RPL27; RPL27 | P61353; K7E | RL27_HUMAN ( | Unmodified | 9645100 | 18899000 | 5466300 | 10860000 | 2230500 | 543660 | 1076700 | 114160 | 1522300 | 0 | 0 | 36134000 | 4600100 | 866720 | 6279900 | 4451100 | 0 | 1492300 |
| NIDEHANE | RAB10 | P61026 | RAB10_HUMAN | Unmodified | 14914000 | 12646000 | 17981000 | 18747000 | 401630 | 175510 | 0 | 480820 | 9869000 | 3227400 | 5017600 | 57199000 | 325590 | 222570 | 17128000 | 20789000 | 573680 | 264000 |
| NIDINDVTF | PSMC5; PSMC5 | P62195; J3Q | PRS8_HUMAN | Unmodified | 0 | 0 | 280300 | 0 | 0 | 0 | 0 | 0 | 0 | 0 | 0 | 2104000 | 0 | 0 | 0 | 0 | 0 | 0 |
| NIDNPALAI | WDR1; WDR1 | O75083; D6I | WDR1_HUMAN | Unmodified | 1800900 | 1245300 | 0 | 1063900 | 11489000 | 2143600 | 480900 | 187800 | 4246200 | 908610 | 1770400 | 14953000 | 6849500 | 1925800 | 1887100 | 1120300 | 1699600 | 3113700 |
| NIDQAVTA | GGT1; GGT2; G | P19440; P36 | GGT1_HUMAN | Unmodified | 3161700 | 2033300 | 1132900 | 2824300 | 0 | 0 | 0 | 0 | 4769900 | 0 | 887290 | 2340000 | 0 | 0 | 5116200 | 5471100 | 0 | 0 |
| NIDVFNVEI | GOLM1 | Q8NBJ4 | GOLM1_HUMA | Unmodified | 0 | 0 | 0 | 0 | 0 | 0 | 66615 | 0 | 2625900 | 0 | 0 | 0 | 0 | 0 | 0 | 0 | 0 | 0 |
| NIDVFNVEI | GOLM1 | Q8NBJ4 | GOLM1_HUMA | Unmodified | 0 | 0 | 0 | 0 | 370830 | 0 | 0 | 0 | 2289100 | 262250 | 0 | 0 | 0 | 0 | 0 | 0 | 0 | 85695 |
| NIEDVIAQ( | RPLP2; RPLP2 | P05387; H0Y | RLA2_HUMAN ( | Unmodified | 49573000 | 40272000 | 33895000 | 49844000 | 12822000 | 2185000 | 151910 | 604720 | 8749300 | 3069500 | 25182000 | 1.4E+08 | 24832000 | 14064000 | 70181000 | 42526000 | 1951900 | 5269900 |
| NIEEHASAI | RAB8A | P61006 | RAB8A_HUMAN | Unmodified | 1133900 | 791790 | 4539400 | 5032900 | 382680 | 0 | 54920 | 0 | 686970 | 141870 | 1791300 | 2361400 | 0 | 0 | 905730 | 951160 | 85222 | 0 |
| NIEEHASSI | RAB8B; RAB8B | Q92930; H0Y | RAB8B_HUMAN | Unmodified | 0 | 0 | 932770 | 927720 | 0 | 0 | 0 | 0 | 62417 | 0 | 306790 | 0 | 0 | 0 | 1236000 | 313300 | 0 | 0 |
| NIEIDSPYE | RPN1; RPN1 | P04843; B7Z | RPN1_HUMAN | Unmodified | 0 | 0 | 0 | 0 | 0 | 0 | 0 | 0 | 12564000 | 3432500 | 0 | 1914300 | 0 | 143970 | 0 | 758690 | 0 | 0 |
| NIEIDSPYE | RPN1; RPN1 | P04843; B7Z | RPN1_HUMAN | Unmodified | 0 | 0 | 0 | 0 | 0 | 0 | 0 | 0 | 8545200 | 0 | 0 | 0 | 0 | 0 | 0 | 0 | 0 | 0 |
| NIELAVMR | PSMA7; PSMA7 | O14818; H0Y | PSA7_HUMAN | Unmodified | 0 | 0 | 0 | 88246 | 569960 | 0 | 0 | 0 | 626990 | 136910 | 341190 | 1929200 | 251460 | 60980 | 44098 | 0 | 39482 | 115800 |
| NIELICQEN | MAPRE1 | Q15691 | MARE1_HUMAI | Unmodified | 877380 | 0 | 0 | 0 | 0 | 0 | 0 | 0 | 951770 | 0 | 386600 | 3677300 | 0 | 0 | 0 | 164420 | 286210 | 0 |
| NIEMTQED | SNRNP200; SN | O75643; A0/ | U520_HUMAN | Unmodified | 855280 | 0 | 0 | 0 | 0 | 0 | 0 | 0 | 674910 | 0 | 0 | 875260 | 0 | 324860 | 0 | 0 | 0 | 0 |
| NIEPTHAPI | MACF1; MACF1 | Q9UPN3; H3 | MACF1_HUMAI | Unmodified | 0 | 0 | 0 | 0 | 420710 | 65437 | 0 | 0 | 570180 | 531000 | 0 | 449520 | 418210 | 355990 | 97128 | 117350 | 605660 | 773810 |
| NIESLSDLF | SLITRK6 | Q9H5Y7 | SLIK6_HUMAN | Unmodified | 897860 | 486980 | 0 | 0 | 0 | 0 | 0 | 0 | 554430 | 85308 | 0 | 0 | 0 | 0 | 0 | 0 | 0 | 0 |
| NIESLSDLF | SLITRK6 | Q9H5Y7 | SLIK6_HUMAN | Unmodified | 417570 | 613120 | 0 | 0 | 0 | 0 | 0 | 117750 | 0 | 0 | 0 | 0 | 0 | 0 | 0 | 0 | 0 | 0 |
| NIETIINTFI | S100A9 | P06702 | S10A9_HUMAN | Unmodified | 0 | 0 | 98650 | 6625600 | 955080 | 0 | 0 | 0 | 354520 | 0 | 0 | 0 | 0 | 0 | 0 | 0 | 0 | 0 |
| NIEVVELLL | KIDINS220; KID | Q9ULH0; E9 | KDIS_HUMAN I | Unmodified | 0 | 0 | 149570 | 345340 | 441380 | 0 | 0 | 0 | 0 | 0 | 274100 | 1221700 | 213690 | 0 | 323160 | 230750 | 0 | 0 |
| NIEYPVDLE | USP10; USP10 | Q14694; H3I | UBP10_HUMAI | Unmodified | 0 | 0 | 0 | 0 | 0 | 0 | 0 | 0 | 0 | 0 | 0 | 2588000 | 0 | 0 | 0 | 0 | 0 | 0 |
| NIFDQRAIE | GSS; GSS; GSS | P48637; A0A | GSHB_HUMAN | Unmodified | 0 | 0 | 0 | 169400 | 839850 | 0 | 0 | 0 | 0 | 148150 | 0 | 0 | 0 | 0 | 0 | 0 | 154280 | 218240 |
| NIFIECTGT | FARSB | Q9NSD9 | SYFB_HUMAN | Unmodified | 0 | 0 | 0 | 0 | 0 | 0 | 0 | 0 | 0 | 0 | 0 | 2476600 | 0 | 0 | 0 | 0 | 0 | 0 |
| NIFPIWALC | LSS; LSS | P48449; A0A | LSS_HUMAN La | Unmodified | 0 | 0 | 123390 | 0 | 0 | 0 | 85016 | 0 | 2160100 | 238600 | 230340 | 1201900 | 45317 | 0 | 121620 | 0 | 0 | 86749 |
| NIFPSNLVS | SLC1A5; SLC1A | Q15758; M0 | AAAT_HUMAN I | Unmodified | 5220000 | 3984900 | 5972300 | 526660 | 0 | 0 | 0 | 0 | 8436100 | 6251100 | 1665900 | 2.9E+08 | 0 | 0 | 11606000 | 12366000 | 0 | 0 |
| NIFPYHEVI | AGL | P35573 | GDE_HUMAN ( | Unmodified | 0 | 0 | 0 | 0 | 0 | 0 | 49134 | 0 | 0 | 479360 | 0 | 720400 | 46411 | 24296 | 0 | 0 | 0 | 0 |
| NIFSVPEIVI | EXOC3; EXOC3 | O60645; D6I | EXOC3_HUMAI | Unmodified | 0 | 31980 | 0 | 91513 | 2357200 | 0 | 283810 | 0 | 0 | 518000 | 0 | 0 | 913330 | 908600 | 0 | 0 | 283700 | 0 |
| NIGAKLVQI | HSPD1; HSPD1 | P10809; A0A | CH60_HUMAN | Unmodified | 0 | 0 | 0 | 0 | 0 | 0 | 2013600 | 0 | 0 | 0 | 0 | 0 | 718710 | 1042300 | 0 | 0 | 0 | 0 |
| NIGAKLVQI | HSPD1; HSPD1 | P10809; A0A | CH60_HUMAN | Unmodified | 0 | 0 | 0 | 0 | 0 | 0 | 3469700 | 0 | 0 | 0 | 0 | 0 | 0 | 0 | 0 | 0 | 0 | 0 |
| NIGDINQD | ITGA6 | P23229 | ITA6_HUMAN I | Unmodified | 0 | 0 | 521270 | 4197300 | 44478000 | 7739900 | 0 | 0 | 1952100 | 1024500 | 1972800 | 3830100 | 21223000 | 12439000 | 6043100 | 2286500 | 68898000 | 1.82E+08 |
| NIGDLLSS | IKBIP | Q70UQ0 | IKIP_HUMAN I | Unmodified | 0 | 0 | 0 | 0 | 706000 | 0 | 452750 | 0 | 2026800 | 1073100 | 0 | 0 | 0 | 0 | 0 | 0 | 0 | 0 |
| NIGENEGG | GBE1; GBE1 | Q04446; E9F | GLGB_HUMAN | Unmodified | 0 | 168830 | 0 | 0 | 87423 | 0 | 0 | 17327 | 752590 | 221880 | 0 | 0 | 0 | 0 | 0 | 0 | 257570 | 746320 |
| NIGENEGG | GBE1; GBE1 | Q04446; E9F | GLGB_HUMAN | Unmodified | 0 | 0 | 0 | 44747 | 108140 | 0 | 0 | 76691 | 188110 | 370150 | 0 | 0 | 0 | 31270 | 0 | 0 | 125060 | 295960 |
| NIGENEGG | GBE1; GBE1 | Q04446; E9F | GLGB_HUMAN | Unmodified | 0 | 0 | 0 | 0 | 0 | 0 | 0 | 0 | 0 | 1096700 | 0 | 0 | 0 | 0 | 0 | 0 | 676540 | 0 |
| NIGGQIQE | UNC45A; UNC4 | Q9H3U1; A0 | UN45A_HUMAI | Unmodified | 31195 | 33509 | 0 | 226680 | 0 | 0 | 0 | 0 | 193490 | 0 | 237760 | 1111000 | 447390 | 133660 | 257290 | 112260 | 0 | 126230 |
| NIGVDNPA | CCT2; CCT2; C( | P78371; F5G | TCPB_HUMAN | Unmodified | 784210 | 470780 | 1203500 | 0 | 7389400 | 868720 | 4725200 | 2848400 | 3709000 | 3941800 | 3720800 | 17131000 | 16012000 | 6694300 | 811100 | 1166100 | 4524800 | 7061200 |
| NIHESCMS | G6PD; G6PD | P11413; E9P | G6PD_HUMAN | Unmodified | 0 | 0 | 0 | 0 | 275270 | 193280 | 0 | 0 | 2175400 | 0 | 149000 | 1327300 | 0 | 0 | 145740 | 0 | 107520 | 93494 |
| NIHSINPT( | STARD3NL; STA | O95772; C9J | STR3N_HUMAI | Unmodified | 0 | 0 | 0 | 0 | 0 | 0 | 53601 | 0 | 683520 | 0 | 0 | 0 | 0 | 0 | 0 | 0 | 0 | 0 |
| NIHSSDWI | RTRAF; RTRAF; | Q9Y224; G3 | RTRAF_HUMAN | Unmodified | 0 | 0 | 35942 | 0 | 0 | 0 | 0 | 0 | 0 | 0 | 0 | 258160 | 0 | 0 | 0 | 50322 | 0 | 0 |
| NIIGSSPVA | EPS15; EPS15 | P42566; B1A | EPS15_HUMAN | Unmodified | 1031600 | 824270 | 342160 | 0 | 0 | 349340 | 0 | 126440 | 0 | 0 | 0 | 0 | 2722100 | 0 | 0 | 627060 | 0 | 983660 |
| NIIGYFEQK | ERP44 | Q9BS26 | ERP44_HUMAN | Unmodified | 267140 | 0 | 0 | 0 | 0 | 0 | 0 | 0 | 20498000 | 2258700 | 0 | 0 | 171250 | 356180 | 0 | 0 | 0 | 0 |
| NIIGYFEQK | ERP44 | Q9BS26 | ERP44_HUMAN | Unmodified | 61628 | 620810 | 0 | 0 | 81126 | 0 | 0 | 0 | 2394900 | 0 | 103930 | 111950 | 59960 | 0 | 150710 | 0 | 0 | 0 |
| NIIGYFEQK | ERP44 | Q9BS26 | ERP44_HUMAN | Unmodified | 0 | 0 | 0 | 0 | 0 | 0 | 0 | 0 | 3589900 | 1232100 | 0 | 0 | 0 | 0 | 0 | 0 | 0 | 0 |
| NIIHETQM/ | TTC17; TTC17 | Q96AE7; A0/ | TTC17_HUMAN | Unmodified | 0 | 0 | 0 | 0 | 0 | 0 | 0 | 0 | 530890 | 0 | 0 | 0 | 0 | 0 | 0 | 0 | 0 | 0 |
| NIIHGSDS\ | NME1 | P15531 | NDKA_HUMAN | Unmodified | 891750 | 332740 | 1602600 | 3328800 | 5681400 | 2175000 | 229270 | 67540 | 4317100 | 606760 | 19232000 | 61671000 | 6974200 | 1580600 | 1587100 | 2735800 | 1052200 | 3490800 |
| NIIHGSDS\ | NME2P1; NME2 | O60361; P22 | NDK8_HUMAN | Unmodified | 3502600 | 254110 | 2065300 | 4273200 | 7068700 | 1835900 | 270010 | 207790 | 1924400 | 684010 | 5843600 | 25133000 | 19816000 | 0 | 2708800 | 3394900 | 2526500 | 5243000 |
| NIIHGSDS\ | NME2P1; NME2 | O60361; P22 | NDK8_HUMAN | Unmodified | 0 | 0 | 0 | 0 | 3070300 | 0 | 559060 | 393290 | 200160 | 314670 | 0 | 0 | 2359000 | 1244700 | 0 | 0 | 3106700 | 3227300 |
| NIILEEGK | CFL1; CFL1; CF | P23528; G3\ | COF1_HUMAN | Unmodified | 1509900 | 5321500 | 14909000 | 5177400 | 18118000 | 6922800 | 802610 | 281530 | 8838600 | 1097200 | 12930000 | 66146000 | 7067100 | 2655200 | 11130000 | 11023000 | 1213500 | 5562600 |
| NIILEEGKE | CFL1; CFL1; CF | P23528; G3\ | COF1_HUMAN | Unmodified | 0 | 13503000 | 206030 | 12327000 | 2.22E+08 | 5923500 | 13375000 | 0 | 1693400 | 12034000 | 0 | 568470 | 1.07E+08 | 62317000 | 5555000 | 376650 | 45010000 | 1.05E+08 |
| NIILGGVK | UBA1; UBA1; UI | P22314; Q5J | UBA1_HUMAN | Unmodified | 282950 | 351020 | 475700 | 567670 | 0 | 0 | 4875600 | 2268400 | 2051600 | 2328100 | 1163700 | 7150700 | 3396300 | 2202700 | 647600 | 615130 | 0 | 0 |

| | | | | | | | | | | | | | | | | | | | | | | |
|---|---|---|---|---|---|---|---|---|---|---|---|---|---|---|---|---|---|---|---|---|---|---|
| NIIQQAIDA | DNAJC9 | Q8WXX5 | DNJC9_HUMAI | Unmodified | 301430 | 0 | 0 | 0 | 0 | 0 | 0 | 0 | 0 | 1591600 | 0 | 0 | 0 | 0 | 0 | 0 | 0 | 0 |
| NIISPPWDI | CDC16; CDC16 | Q13042; Q5 | CDC16_HUMA | Unmodified | 56681 | 0 | 0 | 0 | 0 | 256770 | 54959 | 0 | 99910 | 0 | 0 | 219510 | 78147 | 0 | 0 | 30876 | 538370 | 0 |
| NIITNLSK | TOP1 | P11387 | TOP1_HUMAN | Unmodified | 478100 | 0 | 0 | 0 | 0 | 0 | 0 | 0 | 0 | 0 | 0 | 17217000 | 0 | 0 | 432740 | 0 | 66225 | 0 |
| NIIVGFAR | PCCB; PCCB; P | P05166; C9J | PCCB_HUMAN | Unmodified | 0 | 0 | 0 | 0 | 0 | 0 | 0 | 0 | 1335800 | 187440 | 0 | 0 | 22877 | 19425 | 0 | 0 | 0 | 0 |
| NIKLGIHED | HSP90AA1 | P07900 | HS90A_HUMAI | Unmodified | 0 | 0 | 0 | 0 | 289350 | 130170 | 348490 | 0 | 0 | 135540 | 0 | 0 | 257550 | 322780 | 0 | 0 | 1181500 | 924020 |
| NILAFRDQI | PRKDC | P78527 | PRKDC_HUMAI | Unmodified | 0 | 0 | 0 | 0 | 0 | 0 | 0 | 0 | 0 | 939250 | 0 | 2929600 | 0 | 0 | 0 | 0 | 0 | 0 |
| NILASNPR | ASF1B; ASF1A; | Q9NVP2; Q9 | ASF1B_HUMAN | Unmodified | 36944 | 0 | 0 | 109790 | 0 | 0 | 0 | 0 | 84684 | 97382 | 269810 | 1490100 | 0 | 0 | 0 | 26176 | 0 | 0 |
| NILDDFR | STT3B | Q8TCJ2 | STT3B_HUMAN | Unmodified | 0 | 0 | 168900 | 225040 | 0 | 102550 | 0 | 273240 | 2270900 | 0 | 0 | 795000 | 0 | 0 | 535940 | 456530 | 0 | 0 |
| NILDFPQH | THOP1; THOP1 | P52888; K7E | THOP1_HUMAI | Unmodified | 280950 | 0 | 190010 | 246950 | 0 | 0 | 0 | 0 | 0 | 0 | 1312400 | 3423900 | 0 | 0 | 502280 | 318700 | 0 | 0 |
| NILDRQDP | AZGP1 | P25311 | ZA2G_HUMAN | Unmodified | 0 | 0 | 3929200 | 1741100 | 936000 | 0 | 0 | 0 | 34309 | 0 | 17451000 | 1834200 | 0 | 0 | 2925000 | 503100 | 232700 | 0 |
| NILDRQDP | AZGP1 | P25311 | ZA2G_HUMAN | Unmodified | 0 | 0 | 0 | 0 | 0 | 0 | 0 | 0 | 0 | 0 | 17425000 | 794730 | 0 | 0 | 798880 | 0 | 0 | 0 |
| NILDSKPTA | CORO1C; COR | Q9ULV4; H0 | COR1C_HUMA | Unmodified | 0 | 0 | 0 | 0 | 2744700 | 0 | 0 | 111900 | 140240 | 73969 | 289290 | 1512000 | 0 | 0 | 0 | 0 | 245620 | 0 |
| NILDSKPTA | CORO1C; COR | Q9ULV4; H0 | COR1C_HUMA | Unmodified | 0 | 0 | 0 | 0 | 4639200 | 0 | 0 | 0 | 0 | 0 | 0 | 3204600 | 0 | 0 | 0 | 0 | 0 | 0 |
| NILDVARQ | HMBS; HMBS; | P08397; A0A | HEM3_HUMAN | Unmodified | 0 | 0 | 0 | 0 | 796540 | 382700 | 164620 | 46182 | 0 | 220700 | 59066 | 0 | 1047300 | 431710 | 0 | 0 | 720560 | 333620 |
| NILEESLCE | PRKDC | P78527 | PRKDC_HUMAI | Unmodified | 56127 | 19898 | 22106 | 0 | 0 | 0 | 45565 | 0 | 0 | 0 | 23202 | 97386 | 0 | 0 | 19357 | 14452 | 0 | 0 |
| NILETLLQ | MRPS35 | P82673 | RT35_HUMAN | Unmodified | 0 | 16835 | 0 | 0 | 0 | 0 | 0 | 0 | 166000 | 0 | 0 | 0 | 0 | 0 | 0 | 0 | 0 | 0 |
| NILEYLEW | CDC42BPA; CD | Q5VT25; A0A | MRCKA_HUMA | Unmodified | 0 | 0 | 0 | 0 | 0 | 674460 | 447820 | 0 | 23615 | 251760 | 0 | 0 | 588520 | 0 | 0 | 0 | 357680 | 257210 |
| NILFVITKP | NACA; NACA; N | E9PAV3; Q1 | NACAM_HUMA | Unmodified | 0 | 0 | 893250 | 0 | 0 | 0 | 0 | 0 | 0 | 0 | 687320 | 7189400 | 0 | 0 | 900140 | 431040 | 0 | 0 |
| NILGGTVFI | IDH1; IDH2; IDH | O75874; P48 | IDHC_HUMAN | Unmodified | 1508600 | 943580 | 1429100 | 2002600 | 0 | 0 | 0 | 0 | 3598900 | 149840 | 2360100 | 14535000 | 1553500 | 0 | 4112100 | 4063400 | 0 | 0 |
| NILGGTVFI | IDH1; IDH1 | O75874; C9J | IDHC_HUMAN | Unmodified | 0 | 0 | 0 | 390070 | 2342800 | 1655100 | 879970 | 235830 | 299340 | 1301200 | 618220 | 0 | 16915000 | 6693100 | 0 | 0 | 983870 | 1184100 |
| NILGGTVFI | IDH2 | P48735 | IDHP_HUMAN | Unmodified | 0 | 0 | 64403 | 208600 | 0 | 0 | 393760 | 0 | 40528000 | 2757300 | 693590 | 676300 | 230540 | 133080 | 0 | 0 | 273260 | 162710 |
| NILHQGQE | DCPS; DCPS | Q96C86; A0 | DCPS_HUMAN | Unmodified | 0 | 0 | 0 | 0 | 0 | 0 | 0 | 0 | 146080 | 0 | 422110 | 6916900 | 0 | 0 | 0 | 0 | 0 | 0 |
| NILHSFVQI | EXOC5; EXOC5 | O00471; F8 | EXOC5_HUMAI | Unmodified | 0 | 0 | 0 | 196650 | 0 | 0 | 0 | 0 | 0 | 0 | 0 | 640450 | 69550 | 0 | 0 | 0 | 0 | 0 |
| NILIVLTK | THOC2; THOC2 | Q8NI27; H0 | THOC2_HUMA | Unmodified | 0 | 0 | 0 | 0 | 0 | 0 | 0 | 7270.8 | 142640 | 14923 | 330190 | 1518800 | 0 | 0 | 0 | 0 | 0 | 0 |
| NILLSIGELI | MTTP; MTTP | P55157; E9P | MTP_HUMAN | Unmodified | 0 | 0 | 0 | 0 | 0 | 0 | 0 | 0 | 0 | 0 | 0 | 2210200 | 0 | 0 | 0 | 0 | 0 | 0 |
| NILLTHGR | KIT | P10721 | KIT_HUMAN M | Unmodified | 0 | 0 | 0 | 0 | 0 | 0 | 0 | 0 | 788650 | 49884 | 267790 | 931410 | 0 | 0 | 0 | 0 | 0 | 0 |
| NILLTNEQL | SFXN1; SFXN1; | Q9H9B4; D6 | SFXN1_HUMAI | Unmodified | 0 | 0 | 0 | 236280 | 0 | 0 | 0 | 17532 | 5729900 | 191120 | 456120 | 0 | 0 | 0 | 0 | 0 | 0 | 0 |
| NILLTNEQL | SFXN1; SFXN1; | Q9H9B4; D6 | SFXN1_HUMAI | Unmodified | 0 | 0 | 0 | 0 | 0 | 0 | 0 | 0 | 5382200 | 488140 | 0 | 728380 | 0 | 0 | 0 | 0 | 0 | 0 |
| NILMYGPP | ATAD3A | Q9NVI7 | ATD3A_HUMAN | Oxidation (M) | 0 | 0 | 0 | 0 | 0 | 0 | 0 | 0 | 733770 | 0 | 0 | 0 | 0 | 0 | 0 | 0 | 0 | 0 |
| NILMYGPP | ATAD3A | Q9NVI7 | ATD3A_HUMAN | Unmodified | 0 | 0 | 0 | 0 | 0 | 0 | 0 | 0 | 1261200 | 0 | 0 | 0 | 0 | 0 | 0 | 0 | 0 | 0 |
| NILPMTPV | SEC24B | O95487 | SC24B_HUMAI | Oxidation (M) | 0 | 0 | 0 | 0 | 0 | 0 | 0 | 0 | 29983 | 0 | 0 | 351830 | 0 | 0 | 0 | 0 | 0 | 0 |
| NILPRGAAI | PARD3; PARD3; | Q8TEW0; Q5 | PARD3_HUMAI | Unmodified | 0 | 0 | 0 | 0 | 0 | 0 | 0 | 0 | 0 | 0 | 0 | 0 | 0 | 0 | 0 | 0 | 294860 | 0 |
| NILQLHDL | PREP; PREP | P48147; A0A | PPCE_HUMAN | Unmodified | 49116 | 54818 | 47951 | 218440 | 171320 | 131300 | 141960 | 0 | 336750 | 373610 | 822000 | 2156500 | 212030 | 0 | 284330 | 214180 | 277130 | 229390 |
| NILRNALVS | UQCRC1 | P31930 | QCR1_HUMAN | Unmodified | 0 | 0 | 0 | 0 | 0 | 0 | 0 | 0 | 0 | 1792500 | 0 | 0 | 0 | 0 | 0 | 0 | 0 | 0 |
| NILSSADY | STX4 | Q12846 | STX4_HUMAN | Unmodified | 2843600 | 1886300 | 3061400 | 1605000 | 12522000 | 3344200 | 2054500 | 0 | 5493200 | 1762800 | 4108400 | 18494000 | 4605200 | 2911900 | 5192000 | 4279500 | 2158800 | 7817400 |
| NILTEIYR | RAB11B | Q15907 | RB11B_HUMAN | Unmodified | 2705800 | 4542100 | 1745400 | 4207300 | 0 | 0 | 0 | 62747 | 5418300 | 0 | 1247600 | 19505000 | 0 | 0 | 0 | 2092500 | 0 | 0 |
| NILTEIYRI | RAB11B | Q15907 | RB11B_HUMAN | Unmodified | 0 | 0 | 0 | 261990 | 673660 | 231070 | 160290 | 0 | 98994 | 267310 | 77661 | 0 | 267980 | 156290 | 0 | 0 | 89629 | 0 |
| NILTSNNID | APOD; APOD | P05090; C9J | APOD_HUMAN | Unmodified | 0 | 6.59E+09 | 6.95E+09 | 1.01E+10 | 0 | 0 | 3.39E+08 | 320300 | 1.24E+08 | 11934000 | 0 | 0 | 5.4E+09 | 2.66E+09 | 44550 | 81293 | 139890 | 371480 |
| NILTSNNID | APOD; APOD | P05090; C9J | APOD_HUMAN | Unmodified | 749180 | 572870 | 0 | 324060 | 2297100 | 1414200 | 1114100 | 316690 | 13533000 | 2005400 | 0 | 131860 | 1201900 | 1216400 | 0 | 120600 | 1592400 | 2265600 |
| NILTSNNID | APOD; APOD | P05090; C9J | APOD_HUMAN | Unmodified | 0 | 0 | 0 | 0 | 0 | 0 | 0 | 0 | 0 | 390300 | 0 | 0 | 0 | 0 | 0 | 0 | 0 | 0 |
| NILTYNK | AIDA | Q96BJ3 | AIDA_HUMAN | Unmodified | 467380 | 0 | 1018200 | 4139200 | 547600 | 0 | 797820 | 652870 | 1351700 | 507120 | 0 | 0 | 8685900 | 7720600 | 5012100 | 4615000 | 0 | 0 |
| NILTYNKEF | AIDA | Q96BJ3 | AIDA_HUMAN | Unmodified | 0 | 0 | 0 | 0 | 0 | 0 | 0 | 0 | 0 | 0 | 0 | 0 | 446620 | 0 | 0 | 0 | 627470 | 908070 |
| NILTYNKEF | AIDA | Q96BJ3 | AIDA_HUMAN | Unmodified | 0 | 0 | 0 | 0 | 0 | 0 | 0 | 0 | 0 | 0 | 0 | 0 | 5101700 | 0 | 0 | 0 | 2617000 | 4337000 |
| NILVFGED | DYNC1LI2; DYN | O43237; H3 | DC1L2_HUMAI | Unmodified | 0 | 0 | 0 | 0 | 0 | 0 | 0 | 0 | 0 | 0 | 286110 | 1687000 | 0 | 0 | 0 | 79221 | 0 | 0 |
| NILVNSNL | EPHB3; EPHB1; | P54753; P54 | EPHB3_HUMAI | Unmodified | 725310 | 385380 | 0 | 0 | 0 | 0 | 0 | 0 | 488040 | 184720 | 380750 | 1526500 | 0 | 0 | 621480 | 307860 | 593220 | 0 |
| NIMDFVQK | TSPAN15; TSPA | O95858; H7 | TSN15_HUMAN | Unmodified | 32961 | 13621 | 1385800 | 0 | 49680 | 0 | 0 | 0 | 780830 | 153540 | 0 | 730460 | 0 | 0 | 1723300 | 273240 | 0 | 0 |
| NIMTQNVE | VAMP8; VAMP8 | Q9BV40; B8 | VAMP8_HUMAI | Unmodified | 0 | 0 | 389770 | 587920 | 568630 | 209510 | 0 | 26216 | 6638000 | 293110 | 4333300 | 7764600 | 0 | 0 | 438050 | 394940 | 0 | 320640 |
| NIMTQNVE | VAMP8; VAMP8 | Q9BV40; B8 | VAMP8_HUMAI | Oxidation (M) | 0 | 0 | 320460 | 329930 | 262870 | 0 | 0 | 0 | 570460 | 142320 | 610290 | 1757800 | 0 | 0 | 0 | 219870 | 0 | 0 |
| NINADEAA | HYOU1; HYOU | Q9Y4L1; E9 | HYOU1_HUMA | Oxidation (M) | 0 | 0 | 0 | 0 | 0 | 0 | 0 | 108180 | 8633900 | 7879600 | 985060 | 0 | 671220 | 0 | 0 | 0 | 0 | 0 |
| NINADEAA | HYOU1; HYOU | Q9Y4L1; E9 | HYOU1_HUMA | Unmodified | 0 | 0 | 0 | 0 | 0 | 0 | 0 | 0 | 39940000 | 9913900 | 0 | 0 | 1408900 | 3491000 | 0 | 0 | 113540 | 1501200 |
| NINDGLSS | SOGA1; SOGA1 | O94964; H0 | SOGA1_HUMAI | Unmodified | 0 | 0 | 0 | 125310 | 0 | 0 | 0 | 0 | 0 | 0 | 0 | 0 | 362800 | 0 | 62371 | 0 | 0 | 0 |
| NINIKEDMI | ELAPOR2; ELAP | A8MWY0; H7 | ELAP2_HUMAN | Oxidation (M) | 0 | 0 | 0 | 0 | 0 | 644890 | 0 | 0 | 0 | 1913600 | 0 | 0 | 0 | 0 | 0 | 0 | 0 | 0 |
| NINLIVQK | AP1B1; AP2B1; | Q10567; P6 | AP1B1_HUMAN | Unmodified | 843060 | 863220 | 1386600 | 0 | 544810 | 0 | 0 | 0 | 0 | 221270 | 446750 | 10727000 | 271130 | 210980 | 2277800 | 2014500 | 31109 | 250240 |
| NINSINFDI | LDLR; LDLR | P01130; J3K | LDLR_HUMAN | Unmodified | 0 | 0 | 0 | 0 | 2587900 | 661890 | 1796000 | 189110 | 0 | 750780 | 0 | 0 | 0 | 0 | 0 | 0 | 0 | 504620 |
| NIPEGSHQ | MOB1B; MOB1 | Q7L9L4; Q9 | MOB1B_HUMA | Unmodified | 0 | 0 | 0 | 0 | 184130 | 121660 | 0 | 275010 | 194660 | 427980 | 0 | 0 | 928530 | 1022100 | 0 | 0 | 1173300 | 4364300 |
| NIPGITLLN | RPL4; RPL4 | P36578; H3 | RL4_HUMAN 6 | Unmodified | 11972000 | 6567500 | 5426100 | 1581600 | 1451000 | 0 | 3522300 | 0 | 0 | 662520 | 2700600 | 1.4E+08 | 0 | 0 | 6611300 | 4966300 | 338830 | 1184900 |
| NIPGPHM | SMARCA1; SMA | P28370; O6 | SMCA1_HUMA | Unmodified | 0 | 0 | 0 | 0 | 0 | 0 | 0 | 0 | 0 | 0 | 0 | 969470 | 221370 | 0 | 0 | 0 | 0 | 0 |
| NIPIIVLPLK | IGSF3 | O75054 | IGSF3_HUMAN | Unmodified | 6462100 | 0 | 2001000 | 1330600 | 6547500 | 1914000 | 3828100 | 483630 | 1766200 | 484930 | 375390 | 4770900 | 229360 | 0 | 624540 | 3971100 | 41686 | 0 |
| NIPNGLQE | AFDN; AFDN; A | P55196; H0 | AFAD_HUMAN | Unmodified | 30092 | 0 | 0 | 0 | 0 | 0 | 673500 | 0 | 212910 | 17277 | 207440 | 441650 | 0 | 0 | 0 | 20761 | 0 | 846700 |
| NIPPYFVAL | XRCC6; XRCC6 | P12956; B1 | XRCC6_HUMAI | Unmodified | 0 | 0 | 0 | 0 | 0 | 0 | 0 | 0 | 1063700 | 0 | 1014100 | 4278500 | 759270 | 0 | 239290 | 0 | 0 | 0 |
| NIQDTSDL | FAM120A; FAM | Q9NZB2; A0 | F120A_HUMAN | Unmodified | 189870 | 0 | 0 | 0 | 0 | 0 | 0 | 0 | 4083700 | 672060 | 763570 | 771910 | 0 | 0 | 0 | 0 | 0 | 0 |
| NIQEPTETC | RGS13 | O14921 | RGS13_HUMAI | Unmodified | 0 | 0 | 0 | 0 | 0 | 0 | 0 | 0 | 0 | 0 | 524780 | 971160 | 0 | 0 | 0 | 0 | 0 | 0 |
| NIQFEAFH | CAB39; CAB39 | Q9Y376; Q9 | CAB39_HUMAN | Unmodified | 0 | 255620 | 354180 | 0 | 0 | 93885 | 0 | 0 | 0 | 0 | 0 | 4083600 | 0 | 0 | 1161900 | 1085000 | 123110 | 203960 |
| NIQGMKHI | FARP1; FARP1 | Q9Y4F1; C9 | FARP1_HUMAN | Unmodified | 0 | 0 | 0 | 0 | 8412300 | 2066600 | 1304700 | 0 | 0 | 693450 | 0 | 0 | 4819800 | 3612300 | 0 | 0 | 2449000 | 4262200 |
| NIQKGDYD | EXOC2 | Q96KP1 | EXOC2_HUMAI | Unmodified | 0 | 0 | 0 | 0 | 760450 | 0 | 0 | 254510 | 0 | 194870 | 0 | 1051300 | 0 | 0 | 165500 | 0 | 0 | 0 |
| NIQTSESHI | PTPRJ; PTPRJ; | Q12913; A0 | PTPRJ_HUMAN | Unmodified | 157350 | 0 | 0 | 0 | 0 | 0 | 0 | 0 | 0 | 0 | 480090 | 7434200 | 0 | 0 | 152950 | 291150 | 0 | 0 |
| NIQVSHQE | PDCD6IP | Q8WUM4 | PDC6I_HUMAN | Unmodified | 70179000 | 89199000 | 48805000 | 42052000 | 84838000 | 16158000 | 10563000 | 5147200 | 36039000 | 37052000 | 6306900 | 18166000 | 86145000 | 29480000 | 44651000 | 50187000 | 14041000 | 33707000 |
| NIRDDEGF | MYO6; MYO6; | Q9UM54; A0 | MYO6_HUMAN | Unmodified | 0 | 0 | 2496100 | 0 | 0 | 0 | 136540 | 67684 | 1667900 | 0 | 0 | 4659300 | 0 | 0 | 5934800 | 4390900 | 0 | 0 |
| NIRDDEGF | MYO6; MYO6; | Q9UM54; A0 | MYO6_HUMAN | Unmodified | 0 | 0 | 0 | 0 | 378600 | 0 | 0 | 0 | 0 | 287410 | 0 | 0 | 921210 | 457410 | 0 | 0 | 0 | 0 |
| NIRELVLA | MANBA; MANB | O00462; A0 | MANBA_HUMA | Unmodified | 0 | 0 | 0 | 0 | 0 | 0 | 0 | 0 | 175750 | 456550 | 0 | 0 | 198550 | 225390 | 0 | 0 | 0 | 0 |
| NIREQQILT | LACTB2 | Q53H82 | LACB2_HUMAN | Unmodified | 0 | 0 | 0 | 0 | 0 | 0 | 0 | 0 | 1166700 | 0 | 0 | 156880 | 0 | 0 | 0 | 0 | 0 | 0 |
| NIREQQILT | LACTB2 | Q53H82 | LACB2_HUMAN | Unmodified | 0 | 0 | 0 | 0 | 0 | 0 | 0 | 0 | 2285800 | 434470 | 0 | 0 | 0 | 0 | 0 | 0 | 0 | 0 |
| NIRTVVTGI | TUFM | P49411 | EFTU_HUMAN | Oxidation (M) | 0 | 81320 | 0 | 0 | 0 | 0 | 1206800 | 0 | 0 | 680440 | 0 | 0 | 0 | 0 | 0 | 0 | 0 | 0 |
| NIRVNCLAI | DHRS4L1; DHR | P0CG22; Q9 | DR4L1_HUMAN | Unmodified | 0 | 0 | 0 | 0 | 0 | 0 | 336630 | 48673 | 1446000 | 1824900 | 0 | 0 | 0 | 0 | 0 | 0 | 0 | 0 |
| NISAGSLG | SLC4A2; SLC4A | P04920; A0 | B3A2_HUMAN | Oxidation (M) | 588080 | 0 | 1491100 | 747000 | 0 | 0 | 0 | 0 | 0 | 0 | 0 | 1017700 | 0 | 0 | 2038400 | 0 | 0 | 0 |
| NISEAFED | VPS36; VPS36 | Q86VN1; A0 | VPS36_HUMAN | Unmodified | 7818000 | 6069600 | 7782100 | 3679200 | 0 | 0 | 33633 | 94154 | 255880 | 241240 | 1068000 | 931810 | 1207200 | 0 | 12003000 | 7827800 | 0 | 270790 |
| NISFGRDQ | CTNND1; CTN | O60716; E9 | CTND1_HUMAN | Unmodified | 375390 | 1715900 | 5976600 | 10335000 | 3463500 | 756060 | 0 | 0 | 0 | 0 | 5292100 | 2792500 | 38201000 | 26804000 | 9297100 | 12034000 | 32985000 | 53891000 |
| NISFGRDQ | CTNND1; CTN | O60716; E9 | CTND1_HUMAI | Unmodified | 0 | 0 | 0 | 0 | 10701000 | 0 | 327760 | 148660 | 0 | 0 | 0 | 0 | 5511600 | 12199000 | 0 | 0 | 23992000 | 14058000 |
| NISFTVWD | ARF3; ARF1; A | P61204; P84 | ARF3_HUMAN | Unmodified | 1164700 | 2190900 | 3281600 | 625940 | 0 | 184430 | 382010 | 54337 | 1906700 | 782150 | 2224500 | 15238000 | 0 | 0 | 1226600 | 1782800 | 0 | 0 |
| NISGSVTL | PALS2; PALS2; | Q9NZW5; B8 | PALS2_HUMAN | Unmodified | 22968 | 0 | 37989 | 42793 | 1116500 | 394220 | 0 | 9872.5 | 665160 | 188100 | 539300 | 526830 | 222480 | 0 | 0 | 181690 | 979500 | 953230 |
| NISHYEEQ | PRKAR2A; PRK | P13861; H7 | KAP2_HUMAN | Unmodified | 0 | 0 | 0 | 0 | 0 | 0 | 0 | 0 | 0 | 0 | 0 | 0 | 0 | 2253300 | 0 | 0 | 473430 | 0 |
| NISQDMTQ | ATXN3; ATXN3 | P54252; A0A | ATX3_HUMAN | Unmodified | 605130 | 639650 | 0 | 740750 | 0 | 0 | 776510 | 0 | 259370 | 0 | 0 | 0 | 500320 | 0 | 510780 | 331260 | 0 | 0 |
| NISSAQIV | IST1 | P53990 | IST1_HUMAN | Unmodified | 1299300 | 890770 | 0 | 2475000 | 2752300 | 830110 | 0 | 0 | 12011000 | 998920 | 7902300 | 47160000 | 877450 | 0 | 2029800 | 1784000 | 113410 | 472410 |
| NISSEHSM | BMPR2 | Q13873 | BMPR2_HUMAN | Unmodified | 0 | 0 | 0 | 0 | 896160 | 0 | 514350 | 0 | 0 | 0 | 0 | 0 | 0 | 0 | 0 | 0 | 116930 | 461000 |
| NITEYCRD | RECQL | P46063 | RECQ1_HUMAN | Unmodified | 0 | 0 | 0 | 0 | 0 | 0 | 0 | 0 | 0 | 0 | 0 | 0 | 0 | 0 | 0 | 0 | 0 | 0 |
| NITLDDAS | CSDE1; CSDE1 | O75534; E9 | CSDE1_HUMAI | Unmodified | 192390 | 134090 | 343400 | 383620 | 682260 | 0 | 143650 | 135400 | 0 | 150950 | 713830 | 1406300 | 1673000 | 980160 | 287400 | 473440 | 243770 | 0 |
| NITLNFGP | NDUFS2; NDUF | O75306; A0 | NDUS2_HUMA | Unmodified | 0 | 0 | 0 | 0 | 0 | 0 | 0 | 0 | 4979200 | 839950 | 0 | 106040 | 0 | 0 | 0 | 0 | 0 | 345340 |
| NITNELLRI | PPL; PPL | O60437; K7 | PEPL_HUMAN | Unmodified | 0 | 0 | 0 | 0 | 1874100 | 295090 | 0 | 0 | 0 | 0 | 0 | 0 | 446900 | 274020 | 0 | 0 | 0 | 209510 |
| NITRPFED | RPF2; RPF2 | Q9H7B2; Q5 | RPF2_HUMAN | Unmodified | 127710 | 76986 | 0 | 0 | 0 | 0 | 539380 | 0 | 0 | 0 | 289020 | 0 | 0 | 337030 | 62106 | 75109 | 0 | 0 |
| NITVLFLHS | ERBIN; ERBIN | Q96RT1; B4 | ERBIN_HUMAN | Oxidation (M) | 0 | 0 | 0 | 297550 | 16110000 | 3512600 | 0 | 0 | 551060 | 0 | 0 | 0 | 1104300 | 0 | 0 | 0 | 810140 | 1568600 |
| NITVLFLHS | ERBIN; ERBIN | Q96RT1; B4 | ERBIN_HUMAN | 2 Oxidation (M) | 134650 | 0 | 0 | 140670 | 22731000 | 4437900 | 0 | 0 | 379950 | 623840 | 1586000 | 1922100 | 2912100 | 1867900 | 506090 | 536270 | 108580 | 1570000 |
| NITVLFLHS | ERBIN; ERBIN | Q96RT1; B4 | ERBIN_HUMAN | Unmodified | 0 | 0 | 0 | 121340 | 2697600 | 0 | 0 | 0 | 730820 | 0 | 0 | 0 | 0 | 0 | 0 | 0 | 227300 | 1225500 |
| NITYLPAG | PHB; PHB | P35232; C9 | PHB_HUMAN | Unmodified | 0 | 0 | 0 | 0 | 0 | 0 | 660960 | 0 | 1375600 | 1077200 | 184160 | 0 | 644190 | 433440 | 0 | 0 | 0 | 0 |
| NIVHLPVF | KIDINS220; KID | Q9ULH0; E9 | KDIS_HUMAN | Unmodified | 0 | 0 | 779920 | 0 | 214770 | 0 | 0 | 0 | 512090 | 298330 | 0 | 1649100 | 0 | 0 | 0 | 536130 | 0 | 0 |
| NIVHNYSE | EPN1; EPN3; E | Q9Y6I3; Q9 | EPN1_HUMAN | Unmodified | 9166200 | 9886000 | 5351400 | 4648800 | 93363 | 0 | 94894 | 178570 | 1894900 | 363430 | 0 | 0 | 2202700 | 1303900 | 4423300 | 5229500 | 1128000 | 1553900 |
| NIVHNYSE | EPN1; EPN3; E | Q9Y6I3; Q9 | EPN1_HUMAN | Unmodified | 0 | 0 | 0 | 249820 | 0 | 0 | 0 | 0 | 0 | 131340 | 0 | 0 | 468250 | 0 | 0 | 0 | 406980 | 459950 |
| NIVLSGGS | ACTR3; ACTR3 | P61158; B4 | ARP3_HUMAN | Unmodified | 0 | 316200 | 0 | 0 | 136390 | 0 | 0 | 0 | 1263000 | 255260 | 0 | 4160100 | 0 | 0 | 0 | 0 | 60642 | 0 |
| NIVLSGGS | ACTR3; ACTR3 | P61158; B4 | ARP3_HUMAN | Oxidation (M) | 295640 | 345990 | 0 | 0 | 0 | 0 | 0 | 0 | 164130 | 0 | 110540 | 2548500 | 0 | 0 | 656880 | 0 | 0 | 0 |
| NIVLSGGS | ACTR3; ACTR3 | P61158; B4 | ARP3_HUMAN | Unmodified | 178480 | 0 | 0 | 571480 | 4046600 | 0 | 699020 | 0 | 1481600 | 895750 | 0 | 523130 | 2682200 | 1666900 | 0 | 0 | 951020 | 1457000 |
| NIVNNYSE | EPN2; EPN2; E | O95208; J3K | EPN2_HUMAN | Unmodified | 0 | 0 | 0 | 0 | 0 | 0 | 0 | 0 | 0 | 0 | 0 | 0 | 0 | 0 | 0 | 0 | 0 | 0 |
| NIVWIAEC | ATP6V0D1; ATP | P61421; R4 | VA0D1_HUMAN | Unmodified | 0 | 0 | 0 | 0 | 0 | 9523 | 0 | 0 | 1065400 | 224300 | 403490 | 1824400 | 0 | 0 | 0 | 0 | 0 | 0 |
| NIYGNIED | AGPS; AGPS; A | O00116; A0 | ADAS_HUMAN | Unmodified | 0 | 0 | 0 | 0 | 0 | 0 | 3511400 | 0 | 21331000 | 7656100 | 1357800 | 0 | 0 | 0 | 0 | 0 | 0 | 0 |
| NIYKEIENY | NAPG | Q99747 | SNAG_HUMAN | Unmodified | 0 | 0 | 0 | 0 | 2513100 | 0 | 0 | 294680 | 0 | 0 | 0 | 0 | 1817400 | 0 | 0 | 0 | 2582700 | 2044300 |
| NIYLNSGL | ADAM17 | P78536 | ADA17_HUMAN | Unmodified | 0 | 0 | 0 | 0 | 2113500 | 0 | 0 | 92891 | 1296200 | 0 | 0 | 0 | 713420 | 632650 | 0 | 0 | 1410400 | 2182600 |
| NIYSEELRE | LMNA; LMNA; L | P02545; A0A | LMNA_HUMAN | Unmodified | 0 | 0 | 0 | 0 | 7108200 | 0 | 13343 | 140820 | 321000 | 358750 | 0 | 373180 | 1866700 | 371250 | 0 | 0 | 2321000 | 2957300 |
| NIYVLQEL | XRCC6; XRCC6 | P12956; B1A | XRCC6_HUMAI | Unmodified | 153780 | 86247 | 169280 | 0 | 1586900 | 280030 | 0 | 405340 | 1172400 | 466830 | 2367600 | 15404000 | 0 | 92965 | 0 | 0 | 134120 | 297540 |
| NIYVLQEL | XRCC6; XRCC6 | P12956; B1A | XRCC6_HUMAI | Unmodified | 0 | 0 | 0 | 0 | 0 | 0 | 0 | 79331 | 308080 | 0 | 747260 | 2283600 | 0 | 0 | 0 | 0 | 0 | 0 |
| NKAHEAER | LAMC1 | P11047 | LAMC1_HUMAN | Unmodified | 0 | 0 | 0 | 0 | 446080 | 0 | 0 | 0 | 0 | 392760 | 0 | 0 | 0 | 139230 | 0 | 0 | 0 | 748370 |
| NKAPSPDV | ABCC5 | O15440 | MRP5_HUMAN | Oxidation (M) | 0 | 0 | 0 | 0 | 0 | 0 | 0 | 0 | 0 | 0 | 0 | 3373200 | 0 | 0 | 0 | 0 | 0 | 0 |

| | | | | | | | | | | | | | | | | | | | | | | |
|---|---|---|---|---|---|---|---|---|---|---|---|---|---|---|---|---|---|---|---|---|---|---|
| NKASPQSE | ANLN; ANLN | Q9NQW6; H | ANLN_HUMAN | Unmodified | 0 | 0 | 0 | 0 | 1711500 | 381850 | 116700 | 15001 | 0 | 0 | 0 | 724040 | 0 | 0 | 0 | 0 | 542750 | 277440 |
| NKASVDEL | RAP2B | P61225 | RAP2B_HUMAN | Unmodified | 0 | 0 | 0 | 2771500 | 0 | 0 | 0 | 0 | 465750 | 0 | 0 | 1942800 | 0 | 0 | 0 | 0 | 0 | 0 |
| NKDENAET | DNAJC1; DNAJ | Q96KC8; Q5 | DNJC1_HUMAN | Unmodified | 0 | 0 | 0 | 0 | 0 | 0 | 0 | 0 | 535850 | 0 | 0 | 0 | 0 | 0 | 0 | 0 | 0 | 0 |
| NKDGVLVD | CHMP5 | Q9NZZ3 | CHMP5_HUMA | Unmodified | 3469200 | 7942200 | 1384400 | 13723000 | 74714000 | 13683000 | 4791700 | 1437600 | 20794000 | 28062000 | 9012600 | 8523500 | 36831000 | 23254000 | 2747000 | 2866000 | 6007900 | 11988000 |
| NKDLPESA | ATIC; ATIC | P31939; H7 | PUR9_HUMAN | Unmodified | 0 | 0 | 0 | 0 | 715230 | 0 | 0 | 0 | 0 | 0 | 0 | 0 | 0 | 840290 | 0 | 0 | 0 | 0 |
| NKDQGTYE | MYL6; MYL6; M | P60660; G3 | MYL6_HUMAN | Unmodified | 617480 | 877240 | 113890 | 828280 | 1446200 | 1122000 | 0 | 0 | 1735700 | 226970 | 0 | 7640100 | 530780 | 0 | 440050 | 750950 | 222400 | 0 |
| NKDQGTYE | MYL6; MYL6; M | P60660; G3 | MYL6_HUMAN | Unmodified | 0 | 0 | 0 | 4747800 | 79798000 | 16043000 | 2364400 | 231210 | 6417100 | 3100600 | 5053900 | 0 | 12047000 | 6415800 | 1065100 | 1434300 | 9637100 | 11524000 |
| NKDSGSD | RAB11FIP1; RA | Q6WKZ4; E7 | RFIP1_HUMAN | Unmodified | 0 | 0 | 0 | 0 | 2599200 | 0 | 0 | 0 | 0 | 0 | 0 | 0 | 0 | 0 | 0 | 0 | 0 | 0 |
| NKDSGSD | RAB11FIP1; RA | Q6WKZ4; E7 | RFIP1_HUMAN | Unmodified | 0 | 0 | 0 | 0 | 2011300 | 0 | 0 | 0 | 0 | 0 | 0 | 0 | 0 | 0 | 0 | 0 | 0 | 0 |
| NKDTLFQD | MYO1D; MYO1 | O94832; J3Q | MYO1D_HUMA | Unmodified | 2402200 | 1436700 | 731740 | 689910 | 0 | 179670 | 0 | 0 | 0 | 507280 | 0 | 0 | 0 | 0 | 2005500 | 1140700 | 518110 | 569140 |
| NKDTLFQD | MYO1D; MYO1 | O94832; J3Q | MYO1D_HUMA | Unmodified | 2257900 | 1211900 | 209760 | 693400 | 0 | 0 | 31218 | 0 | 171260 | 0 | 0 | 588610 | 0 | 0 | 880360 | 1045500 | 0 | 0 |
| NKDTVFEE | MYO5A; MYO5A | Q9Y4I1; G3V | MYO5A_HUMA | Unmodified | 0 | 0 | 0 | 0 | 472810 | 0 | 0 | 0 | 0 | 0 | 0 | 0 | 0 | 0 | 0 | 0 | 0 | 0 |
| NKDVLQAE | SNX6; SNX6 | Q9UNH7; A0 | SNX6_HUMAN | Unmodified | 0 | 0 | 0 | 0 | 480690 | 0 | 166250 | 123850 | 0 | 0 | 0 | 240810 | 1457000 | 1248600 | 0 | 0 | 922980 | 0 |
| NKEDEDIAI | STAM2 | O75886 | STAM2_HUMAN | Unmodified | 0 | 0 | 0 | 0 | 417780 | 45497 | 0 | 0 | 0 | 92266 | 0 | 0 | 153640 | 121640 | 0 | 0 | 154080 | 1464700 |
| NKEDQYDI | HSPA4 | P34932 | HSP74_HUMAN | Unmodified | 0 | 0 | 476650 | 190240 | 1857900 | 203510 | 119530 | 30168 | 596290 | 266390 | 710990 | 6349100 | 998460 | 243960 | 222050 | 416000 | 652560 | 656710 |
| NKEDQYDI | HSPA4 | P34932 | HSP74_HUMAN | Unmodified | 44453 | 0 | 0 | 3231200 | 5800000 | 1525700 | 1627300 | 363730 | 1795500 | 2001800 | 2229400 | 7027200 | 5238600 | 9929300 | 0 | 0 | 3366400 | 1500100 |
| NKEDQYDI | HSPA4 | P34932 | HSP74_HUMAN | Oxidation (M) | 0 | 0 | 0 | 153590 | 644610 | 0 | 132870 | 0 | 0 | 0 | 0 | 413610 | 0 | 0 | 0 | 0 | 429650 | 0 |
| NKEEAAEY | RPS6; RPS6 | P62753; A2A | RS6_HUMAN 4 | Unmodified | 326060 | 294550 | 260400 | 0 | 186480 | 0 | 59528 | 151350 | 118390 | 0 | 45027 | 981460 | 637700 | 0 | 218410 | 118380 | 18843 | 137730 |
| NKEEAAEY | RPS6; RPS6 | P62753; A2A | RS6_HUMAN 4 | Unmodified | 0 | 0 | 0 | 0 | 0 | 0 | 0 | 96886 | 0 | 0 | 0 | 0 | 1279300 | 169160 | 0 | 0 | 0 | 0 |
| NKEIFLREL | HSP90B1; HSP | P14625; P08 | ENPL_HUMAN | Unmodified | 0 | 0 | 0 | 0 | 7897300 | 1586000 | 27747000 | 31062 | 52334000 | 3.87E+08 | 9260700 | 0 | 7492300 | 9558600 | 0 | 0 | 407190 | 4310800 |
| NKELEQGE | FERMT1; FERM | Q9BQL6; G3 | FERM1_HUMAN | Unmodified | 0 | 0 | 0 | 0 | 0 | 61595 | 0 | 0 | 0 | 0 | 0 | 0 | 159290 | 0 | 0 | 0 | 401260 | 1657500 |
| NKELIGKH | CTNND1; CTNN | O60716; A0 | CTND1_HUMA | Unmodified | 0 | 0 | 0 | 0 | 0 | 0 | 0 | 0 | 0 | 0 | 0 | 0 | 416780 | 176090 | 0 | 0 | 957490 | 491360 |
| NKEPPAPA | TMEM165; TME | Q9HC07; D6 | TM165_HUMAN | Unmodified | 0 | 0 | 0 | 0 | 0 | 0 | 0 | 0 | 5378600 | 3602000 | 349230 | 0 | 0 | 0 | 0 | 0 | 0 | 0 |
| NKEPPAPA | TMEM165 | Q9HC07 | TM165_HUMAN | Unmodified | 0 | 0 | 0 | 0 | 0 | 0 | 0 | 0 | 2.43E+08 | 9090300 | 7812700 | 25665000 | 0 | 0 | 0 | 0 | 0 | 0 |
| NKEQLSDM | IQGAP1; IQGAP | P46940; A0A | IQGA1_HUMAN | Unmodified | 0 | 0 | 0 | 0 | 7616000 | 404380 | 300200 | 281970 | 523940 | 0 | 0 | 0 | 376590 | 114980 | 0 | 0 | 57048 | 273800 |
| NKEQLSDM | IQGAP1; IQGAP | P46940; A0A | IQGA1_HUMAN | Oxidation (M) | 27872 | 0 | 0 | 0 | 6896500 | 389760 | 61024 | 0 | 0 | 43092 | 0 | 197320 | 0 | 0 | 0 | 0 | 0 | 0 |
| NKEQLSDM | IQGAP1; IQGAP | P46940; A0A | IQGA1_HUMAN | 2 Oxidation (M) | 0 | 0 | 0 | 0 | 376410 | 74095 | 16851 | 19730 | 0 | 0 | 0 | 0 | 0 | 0 | 0 | 0 | 0 | 0 |
| NKESKDPA | STMN1 | P16949 | STMN1_HUMA | Unmodified | 92198 | 0 | 0 | 62861 | 3624000 | 804040 | 3143700 | 3083000 | 5888300 | 4471300 | 289990 | 2735400 | 314660 | 133450 | 123300 | 15041 | 1928800 | 4539300 |
| NKESVNKA | SMC2 | O95347 | SMC2_HUMAN | Unmodified | 0 | 0 | 0 | 0 | 312770 | 0 | 337090 | 202060 | 0 | 13817 | 0 | 0 | 0 | 0 | 0 | 0 | 0 | 0 |
| NKETLIIEP | PPP1R12A; PPP | O14974; F8 | MYPT1_HUMAN | Unmodified | 0 | 0 | 0 | 0 | 1224300 | 233490 | 71743 | 67371 | 159020 | 0 | 0 | 0 | 417680 | 505110 | 0 | 0 | 540380 | 682420 |
| NKETNTVL | REPS1; REPS1; | Q96D71; H0 | REPS1_HUMAN | Unmodified | 0 | 0 | 0 | 0 | 0 | 0 | 0 | 0 | 0 | 0 | 0 | 0 | 246410 | 107050 | 0 | 0 | 267760 | 217780 |
| NKFEEAER | PRKCSH; PRKC | P14314; K7E | GLU2B_HUMA | Unmodified | 0 | 0 | 0 | 0 | 0 | 0 | 0 | 0 | 4708100 | 3191500 | 26481 | 0 | 0 | 0 | 0 | 0 | 0 | 0 |
| NKFLIDGF | CMPK1; CMPK1 | P30085; Q5 | KCY_HUMAN U | Unmodified | 0 | 0 | 0 | 171060 | 3143200 | 0 | 2006400 | 51681 | 0 | 1202200 | 0 | 0 | 5014000 | 3400800 | 0 | 0 | 9975200 | 3216400 |
| NKFNQVES | GALNT2 | Q10471 | GALT2_HUMAN | Unmodified | 0 | 0 | 0 | 0 | 0 | 0 | 0 | 0 | 1218500 | 631260 | 0 | 0 | 0 | 0 | 0 | 0 | 0 | 0 |
| NKFVEFDM | LIMS2; LIMS1; L | Q7Z4I7; P48 | LIMS2_HUMAN | Unmodified | 0 | 0 | 0 | 0 | 425150 | 0 | 955240 | 0 | 0 | 160720 | 0 | 0 | 0 | 0 | 0 | 0 | 0 | 213940 |
| NKFVEFDM | LIMS2; LIMS1; L | Q7Z4I7; P48 | LIMS2_HUMAN | Oxidation (M) | 0 | 0 | 0 | 0 | 0 | 0 | 179350 | 199900 | 0 | 0 | 0 | 0 | 0 | 0 | 0 | 0 | 0 | 0 |
| NKFVLSGN | CPD | O75976 | CBPD_HUMAN | Unmodified | 0 | 0 | 0 | 0 | 509270 | 0 | 0 | 0 | 14090000 | 5351100 | 0 | 0 | 0 | 0 | 0 | 0 | 0 | 0 |
| NKFVLSGN | CPD | O75976 | CBPD_HUMAN | Unmodified | 0 | 0 | 0 | 0 | 0 | 0 | 0 | 0 | 360690 | 3342700 | 0 | 0 | 0 | 0 | 0 | 0 | 0 | 0 |
| NKGDSHL | MTDH; MTDH | Q86UE4; E5 | LYRIC_HUMAN | Unmodified | 291650 | 1010300 | 0 | 0 | 74931 | 0 | 0 | 0 | 0 | 0 | 0 | 0 | 209190 | 0 | 288110 | 205160 | 46397 | 85506 |
| NKGNECF | STIP1 | P31948 | STIP1_HUMAN | Oxidation (M) | 0 | 0 | 0 | 0 | 726090 | 0 | 0 | 30411 | 0 | 471480 | 0 | 0 | 0 | 777280 | 0 | 0 | 1027300 | 673610 |
| NKGNECF | STIP1 | P31948 | STIP1_HUMAN | Unmodified | 87221 | 672570 | 184850 | 890470 | 4940100 | 1785500 | 2042900 | 671250 | 506190 | 1059600 | 425180 | 0 | 5855000 | 4516600 | 368180 | 155800 | 4473100 | 16013000 |
| NKITITNDQ | HSPA5; HSPA5; | P11021; A0A | BIP_HUMAN Er | Unmodified | 303060 | 179500 | 0 | 0 | 0 | 0 | 232210 | 22874 | 9471500 | 1364900 | 326860 | 1269900 | 0 | 14158 | 0 | 123760 | 18654 | 0 |
| NKITITNDQ | HSPA5; HSPA5; | P11021; A0A | BIP_HUMAN Er | Unmodified | 235900 | 0 | 0 | 221600 | 0 | 0 | 0 | 0 | 35625000 | 7095400 | 2260000 | 1764000 | 0 | 0 | 0 | 0 | 0 | 0 |
| NKITLQDV | IQGAP1; IQGAP | P46940; A0A | IQGA1_HUMAN | Unmodified | 551000 | 295120 | 0 | 974940 | 19479000 | 4199100 | 1545100 | 783530 | 1392600 | 1643400 | 129380 | 1314400 | 2994900 | 2092200 | 234590 | 127340 | 626020 | 904040 |
| NKITQSNA | GSTM3; GSTM3 | P21266; A0A | GSTM3_HUMA | Unmodified | 0 | 0 | 0 | 0 | 364300 | 63989 | 54359 | 48631 | 303830 | 152920 | 0 | 0 | 1318200 | 473960 | 30585 | 75882 | 191970 | 385150 |
| NKITVVSEK | RPL22L1; RPL2 | Q6P5R6; H0 | RL22L_HUMAN | Unmodified | 305880 | 291240 | 17063 | 0 | 0 | 0 | 334350 | 40197 | 0 | 179800 | 16995 | 416880 | 0 | 18485 | 372890 | 314760 | 0 | 0 |
| NKIWYEHR | IDH2; IDH2 | P48735; H0 | IDHP_HUMAN | Unmodified | 0 | 0 | 0 | 0 | 724430 | 37902 | 0 | 0 | 0 | 401000 | 0 | 0 | 0 | 0 | 0 | 0 | 0 | 0 |
| NKLDFLRP | PSMB7 | Q99436 | PSB7_HUMAN | Unmodified | 0 | 0 | 0 | 432810 | 3443500 | 256170 | 237020 | 166750 | 312340 | 1838500 | 1059900 | 4715700 | 7429800 | 771130 | 0 | 201880 | 594670 | 572560 |
| NKLDHYAI | RPL23A; RPL23 | P62750; K7E | RL23A_HUMAN | Unmodified | 677790 | 429810 | 487320 | 202800 | 0 | 0 | 0 | 0 | 0 | 0 | 0 | 3763400 | 0 | 0 | 595840 | 0 | 0 | 0 |
| NKLDHYAI | RPL23A; RPL23 | P62750; K7E | RL23A_HUMAN | Oxidation (M) | 973190 | 1270600 | 0 | 0 | 0 | 0 | 0 | 0 | 0 | 0 | 0 | 1692300 | 393520 | 0 | 224070 | 0 | 0 | 0 |
| NKLESLEK | BAIAP2L1 | Q9UHR4 | BI2L1_HUMAN | Unmodified | 24534 | 0 | 182730 | 0 | 4648400 | 0 | 9487.5 | 79855 | 0 | 0 | 21017 | 90061 | 1082400 | 970640 | 0 | 0 | 42549 | 350260 |
| NKLESLEK | BAIAP2L1 | Q9UHR4 | BI2L1_HUMAN | Unmodified | 0 | 0 | 0 | 0 | 1363300 | 0 | 0 | 0 | 0 | 0 | 0 | 0 | 0 | 145130 | 0 | 0 | 539010 | 444680 |
| NKLPGLITS | EXOC7; EXOC7 | Q9UPT5; B4 | EXOC7_HUMA | Unmodified | 0 | 0 | 0 | 0 | 631110 | 0 | 0 | 0 | 301750 | 0 | 0 | 0 | 0 | 0 | 0 | 0 | 251430 | 0 |
| NKLQQLPA | LRRC59 | Q96AG4 | LRC59_HUMAN | Unmodified | 0 | 0 | 0 | 0 | 486940 | 117850 | 73015 | 0 | 0 | 88383 | 0 | 0 | 514880 | 365240 | 0 | 0 | 0 | 363460 |
| NKLQQLPA | LRRC59 | Q96AG4 | LRC59_HUMAN | Unmodified | 0 | 0 | 0 | 0 | 3630900 | 149180 | 2094900 | 0 | 0 | 497350 | 0 | 0 | 816600 | 3663500 | 0 | 0 | 979370 | 2893300 |
| NKLVQTAE | GMFB; GMFB | P60983; G3 | GMFB_HUMAN | Unmodified | 0 | 0 | 0 | 0 | 898260 | 0 | 725900 | 0 | 0 | 489500 | 0 | 49677 | 0 | 0 | 0 | 0 | 1431800 | 893310 |
| NKMASNIF | JPT1; JPT1 | Q9UK76; J3 | JUPI1_HUMAN | Oxidation (M) | 0 | 0 | 0 | 0 | 0 | 0 | 0 | 0 | 0 | 0 | 0 | 0 | 0 | 0 | 0 | 0 | 742010 | 0 |
| NKMASNIF | JPT1; JPT1 | Q9UK76; J3 | JUPI1_HUMAN | Unmodified | 0 | 0 | 0 | 0 | 0 | 0 | 0 | 0 | 0 | 0 | 0 | 0 | 0 | 0 | 0 | 0 | 390400 | 0 |
| NKMSAYNL | ARHGAP19; AR | Q14CB8; R4 | RHG19_HUMA | 2 Oxidation (M) | 0 | 0 | 0 | 87081 | 0 | 0 | 0 | 134690 | 64807 | 0 | 0 | 62131 | 0 | 0 | 207870 | 86872 | 0 | 137350 |
| NKNECSLH | SORT1 | Q99523 | SORT_HUMAN | Unmodified | 0 | 0 | 0 | 0 | 0 | 0 | 0 | 0 | 6934700 | 2073600 | 1659400 | 1988400 | 0 | 172590 | 0 | 0 | 0 | 116320 |
| NKNPAPPI | KPNA2; KPNA2; | P52292; A0A | IMA1_HUMAN I | Unmodified | 605460 | 0 | 268390 | 875710 | 269740 | 0 | 0 | 0 | 1713200 | 350640 | 0 | 1308100 | 96670 | 0 | 640430 | 706560 | 257910 | 129210 |
| NKPEWFFK | GSTO1; GSTO1 | P78417; Q5 | GSTO1_HUMA | Unmodified | 0 | 0 | 0 | 0 | 2270800 | 0 | 130630 | 0 | 0 | 0 | 507060 | 4410400 | 651510 | 312710 | 0 | 0 | 487240 | 467320 |
| NKPEWREE | ING3; ING3; IN | Q9NXR8; A0 | ING3_HUMAN | Oxidation (M) | 0 | 0 | 0 | 0 | 0 | 2786000 | 4052000 | 791620 | 0 | 927580 | 0 | 0 | 0 | 0 | 0 | 0 | 4494400 | 0 |
| NKPGPNIE | EIF5B; EIF5B | O60841; A0 | IF2P_HUMAN E | Unmodified | 132970 | 0 | 102890 | 0 | 577120 | 0 | 99830 | 0 | 275980 | 154320 | 798940 | 1784100 | 1209400 | 305130 | 0 | 158470 | 244430 | 885950 |
| NKPGPNIE | EIF5B; EIF5B | O60841; A0 | IF2P_HUMAN E | Unmodified | 0 | 0 | 0 | 0 | 1177400 | 0 | 0 | 0 | 0 | 264110 | 0 | 0 | 1219600 | 841460 | 0 | 0 | 643200 | 848370 |
| NKPIVNIDF | KIF4A | O95239 | KIF4A_HUMAN | Unmodified | 115370 | 0 | 580390 | 440640 | 1585100 | 334810 | 271220 | 0 | 0 | 173280 | 0 | 0 | 3075500 | 1066800 | 0 | 0 | 2320400 | 1164300 |
| NKPLEQSV | FMNL1 | O95466 | FMNL1_HUMA | Unmodified | 0 | 0 | 0 | 0 | 2197500 | 1308500 | 79564 | 84509 | 0 | 734830 | 0 | 0 | 0 | 342150 | 0 | 0 | 0 | 0 |
| NKPLFFADI | ANXA6; ANXA6; | P08133; E7E | ANXA6_HUMAN | Unmodified | 0 | 0 | 0 | 0 | 629790 | 0 | 4792900 | 0 | 228160 | 1159700 | 0 | 0 | 779950 | 1201500 | 0 | 0 | 2881600 | 1899500 |
| NKPLISYVG | EMB; EMB | Q6PCB8; D6 | EMB_HUMAN E | Unmodified | 0 | 0 | 0 | 0 | 1100700 | 0 | 0 | 0 | 214300 | 0 | 0 | 0 | 0 | 0 | 0 | 0 | 0 | 0 |
| NKPYISWP | CDC42BPB | Q9Y5S2 | MRCKB_HUMA | Unmodified | 0 | 0 | 101720 | 253300 | 0 | 90750 | 0 | 0 | 636170 | 28516 | 120270 | 793070 | 0 | 0 | 380280 | 244630 | 136450 | 529660 |
| NKQGLETD | MYH10 | P35580 | MYH10_HUMA | Unmodified | 0 | 303620 | 0 | 0 | 1736000 | 0 | 663190 | 303930 | 0 | 32549 | 0 | 0 | 0 | 0 | 0 | 0 | 678810 | 245150 |
| NKQKPITP | CDC42; CDC42 | P60953; A0A | CDC42_HUMA | Unmodified | 0 | 0 | 0 | 0 | 11076000 | 0 | 12307000 | 5958400 | 0 | 0 | 0 | 0 | 13581000 | 23283000 | 0 | 0 | 18286000 | 3887500 |
| NKQTYSTE | RPL28; RPL28; | P46779; H0 | RL28_HUMAN | Unmodified | 1356200 | 1455300 | 0 | 634780 | 72836 | 10354 | 54238 | 175950 | 191130 | 0 | 0 | 2791200 | 0 | 0 | 0 | 0 | 24721 | 0 |
| NKTEDLEA | VAMP8; VAMP8 | Q9BV40; C9 | VAMP8_HUMA | Unmodified | 0 | 0 | 0 | 0 | 0 | 0 | 0 | 0 | 2926100 | 419520 | 4179100 | 4374600 | 0 | 0 | 0 | 0 | 0 | 0 |
| NKTEYEEA | MSH2; MSH2; | P43246; A0A | MSH2_HUMAN | Unmodified | 0 | 0 | 0 | 0 | 130970 | 143800 | 523300 | 258420 | 0 | 482620 | 0 | 0 | 0 | 0 | 0 | 0 | 58915 | 468940 |
| NKTGAAPI | RPL27A; RPL27 | P46776; E9P | RL27A_HUMAN | Unmodified | 1484600 | 1923500 | 0 | 840460 | 363910 | 0 | 205850 | 266700 | 0 | 276910 | 0 | 591490 | 1098400 | 274370 | 0 | 0 | 0 | 793180 |
| NKTGAAPI | RPL27A; RPL27 | P46776; E9P | RL27A_HUMAN | Unmodified | 0 | 0 | 0 | 0 | 2627900 | 0 | 4425200 | 815680 | 0 | 0 | 0 | 0 | 3051700 | 2515400 | 0 | 0 | 517100 | 322220 |
| NKTLEYKIE | KIF23; KIF23; K | Q02241; A0 | KIF23_HUMAN | Unmodified | 0 | 0 | 0 | 0 | 1487300 | 0 | 0 | 0 | 0 | 24825 | 0 | 0 | 0 | 0 | 0 | 0 | 806290 | 0 |
| NKTSTTSS | PPP1R2; PPP1R | P41236; E7E | IPP2_HUMAN F | Unmodified | 0 | 0 | 0 | 0 | 0 | 0 | 0 | 0 | 0 | 0 | 0 | 0 | 307820 | 0 | 0 | 0 | 315010 | 0 |
| NKVCLLDV | MPP7; MPP7 | Q5T2T1; U5 | MPP7_HUMAN | Unmodified | 0 | 0 | 0 | 0 | 529130 | 0 | 0 | 0 | 0 | 0 | 0 | 0 | 168480 | 449080 | 0 | 0 | 252330 | 0 |
| NKVDENM | APLP2 | Q06481 | APLP2_HUMAN | Unmodified | 0 | 0 | 0 | 0 | 547850 | 0 | 0 | 0 | 4087000 | 0 | 0 | 0 | 245980 | 139430 | 0 | 0 | 0 | 368180 |
| NKVDENM | APLP2 | Q06481 | APLP2_HUMAN | Oxidation (M) | 66653 | 0 | 207750 | 0 | 261950 | 144920 | 0 | 0 | 1389700 | 254320 | 0 | 0 | 226920 | 0 | 0 | 71604 | 0 | 110810 |
| NKVEDSGI | MAN2A1 | Q16706 | MA2A1_HUMA | Unmodified | 0 | 0 | 0 | 0 | 0 | 0 | 0 | 0 | 3908200 | 1218000 | 0 | 1598300 | 0 | 0 | 0 | 0 | 0 | 693150 |
| NKVFQEAL | UNC45A; UNC4 | Q9H3U1; A0 | UN45A_HUMA | Unmodified | 0 | 0 | 0 | 0 | 231920 | 0 | 0 | 0 | 0 | 0 | 0 | 0 | 334770 | 0 | 0 | 0 | 239340 | 0 |
| NKVPWNR | MAPK1 | P28482 | MK01_HUMAN | Unmodified | 0 | 0 | 0 | 0 | 0 | 0 | 0 | 0 | 310220 | 224860 | 0 | 0 | 710400 | 0 | 0 | 0 | 464100 | 0 |
| NKVQDLEN | GULP1; GULP1 | Q9UBP9; H0 | GULP1_HUMA | Unmodified | 0 | 0 | 0 | 702980 | 0 | 0 | 0 | 0 | 0 | 0 | 0 | 0 | 517220 | 0 | 0 | 0 | 0 | 0 |
| NKVSMAPD | UAP1 | Q16222 | UAP1_HUMAN | Unmodified | 0 | 0 | 47690 | 28685 | 305480 | 50247 | 0 | 0 | 0 | 102560 | 0 | 0 | 213070 | 0 | 0 | 0 | 0 | 71508 |
| NKYGDAFI | LSM6 | P62312 | LSM6_HUMAN | Unmodified | 0 | 0 | 0 | 0 | 5430700 | 0 | 1548200 | 0 | 0 | 737140 | 315110 | 0 | 4825900 | 1906200 | 57161 | 0 | 3041900 | 1048800 |
| NKYQELINI | IQGAP1; IQGAP | P46940; H0 | IQGA1_HUMAN | Unmodified | 0 | 0 | 0 | 645170 | 2935800 | 1175000 | 0 | 0 | 4372900 | 192770 | 0 | 2662100 | 657770 | 0 | 0 | 2868900 | 0 | 39753 |
| NKYQELINI | IQGAP1; IQGAP | P46940; H0 | IQGA1_HUMAN | Unmodified | 0 | 0 | 0 | 0 | 2056800 | 1012400 | 0 | 0 | 869290 | 0 | 0 | 0 | 0 | 0 | 0 | 0 | 0 | 0 |
| NLAAFPASI | MINK1 | Q8N4C8 | MINK1_HUMAN | Unmodified | 0 | 0 | 1195300 | 2120800 | 1444800 | 0 | 0 | 0 | 2576500 | 0 | 926850 | 1379400 | 1756200 | 0 | 1994100 | 1470600 | 0 | 0 |
| NLADCLR | PSMB2 | P49721 | PSB2_HUMAN | Unmodified | 0 | 0 | 0 | 0 | 0 | 0 | 10972 | 0 | 0 | 0 | 429980 | 2536500 | 0 | 0 | 0 | 0 | 0 | 0 |
| NLAEDRSG | GPHN; GPHN; | Q9NQX3; G3 | GEPH_HUMAN | Unmodified | 0 | 0 | 0 | 0 | 280740 | 0 | 805510 | 229280 | 297430 | 86119 | 688580 | 334150 | 431960 | 897680 | 0 | 0 | 37565 | 860930 |
| NLAFHTLS | ARMH3 | Q5T2E6 | ARMD3_HUMA | Unmodified | 44229 | 42426 | 28789 | 39966 | 82226 | 25175 | 0 | 55124 | 0 | 91487 | 60497 | 845390 | 0 | 0 | 56671 | 60040 | 0 | 71647 |
| NLAFYWIL | SCPEP1; SCPE | Q9HB40; I3L | RISC_HUMAN I | Unmodified | 0 | 0 | 0 | 0 | 53789 | 0 | 64332 | 0 | 792030 | 1450800 | 23458 | 19118 | 86141 | 89863 | 0 | 0 | 0 | 206320 |
| NLAIPVINK | RAP1GDS1; RA | P52306; H0 | GDS1_HUMAN | Unmodified | 26462 | 0 | 57440 | 0 | 0 | 0 | 0 | 0 | 290020 | 0 | 208760 | 1554100 | 0 | 0 | 41763 | 51276 | 0 | 0 |
| NLALCPAN | CTNNB1; CTNN | P35222; A0A | CTNB1_HUMA | Unmodified | 0 | 0 | 0 | 0 | 0 | 0 | 53808 | 0 | 0 | 0 | 0 | 768820 | 111990 | 0 | 0 | 0 | 0 | 0 |
| NLALCPAN | CTNNB1; CTNN | P35222; A0A | CTNB1_HUMA | Unmodified | 1839100 | 156460 | 0 | 4445600 | 0 | 0 | 0 | 114860 | 1053100 | 159470 | 0 | 10591000 | 0 | 0 | 8509500 | 10237000 | 907440 | 0 |
| NLALDINE | CNOT1 | A5YKK6 | CNOT1_HUMA | Unmodified | 0 | 0 | 80409 | 0 | 150050 | 101010 | 116820 | 0 | 0 | 484050 | 124880 | 1075100 | 0 | 42389 | 195530 | 236710 | 49552 | 20240 |
| NLALSGFR | UBA3; UBA3 | Q8TBC4; F8 | UBA3_HUMAN | Unmodified | 0 | 0 | 0 | 0 | 0 | 0 | 0 | 0 | 203420 | 0 | 487500 | 1403800 | 42143 | 0 | 0 | 73915 | 0 | 0 |
| NLALSRESI | LSR; LSR | Q86X29; S4 | LSR_HUMAN Li | Unmodified | 0 | 191320 | 0 | 0 | 14861000 | 2942200 | 0 | 0 | 0 | 23363000 | 0 | 0 | 5723100 | 5396700 | 0 | 0 | 1386700 | 1622500 |
| NLAMEATY | EIF3D | O15371 | EIF3D_HUMAN | Oxidation (M) | 0 | 0 | 0 | 706500 | 0 | 0 | 0 | 0 | 0 | 0 | 0 | 5520600 | 0 | 0 | 0 | 0 | 0 | 0 |
| NLANTVTE | SYNCRIP; SYN | O60506; B7 | HNRPQ_HUMA | Unmodified | 260790 | 0 | 393540 | 275160 | 0 | 0 | 0 | 0 | 489610 | 147840 | 6450500 | 13375000 | 1964700 | 0 | 320250 | 461500 | 42215 | 0 |
| NLAQLNDI | CCDC134; CCD | Q9H6E4; B0 | CC134_HUMA | Unmodified | 0 | 0 | 33405 | 0 | 0 | 0 | 0 | 43667 | 5056000 | 2052500 | 534730 | 419520 | 0 | 0 | 0 | 0 | 0 | 0 |
| NLATAYDN | PDCD6IP | Q8WUM4 | PDC6I_HUMAN | Unmodified | 0 | 2431200 | 78081 | 1334000 | 2476200 | 203420 | 172210 | 0 | 40975 | 0 | 16009 | 0 | 78104 | 2238900 | 1414300 | 0 | 334920 | 1258000 |
| NLATAYDN | PDCD6IP | Q8WUM4 | PDC6I_HUMAN | Unmodified | 0 | 0 | 0 | 77482 | 9296000 | 833710 | 284340 | 0 | 0 | 5009400 | 0 | 0 | 9577000 | 5811000 | 0 | 0 | 2561500 | 4476800 |
| NLATSLAE | TLN2; TLN2 | Q9Y4G6; A0 | TLN2_HUMAN | Unmodified | 0 | 0 | 0 | 0 | 0 | 0 | 0 | 24096 | 0 | 0 | 0 | 0 | 0 | 67547 | 0 | 0 | 132470 | 467220 |
| NLATTVTEE | HNRNPR; HNR | O43390; B4 | HNRPR_HUMA | Unmodified | 0 | 0 | 144960 | 0 | 253200 | 0 | 887260 | 43783 | 2366100 | 1361900 | 3119600 | 13820000 | 0 | 157480 | 0 | 560150 | 0 | 193440 |
| NLAYDTLP | PLOD1 | Q02809 | PLOD1_HUMA | Unmodified | 0 | 0 | 0 | 0 | 0 | 0 | 0 | 0 | 4750300 | 487230 | 0 | 200480 | 0 | 0 | 0 | 0 | 0 | 21298 |

| | | | | | | | | | | | | | | | | | | | | | |
|---|---|---|---|---|---|---|---|---|---|---|---|---|---|---|---|---|---|---|---|---|---|
| NLCSDDTF | PPP2R1A; PPP2 | P30153; C9J | 2AAA_HUMAN | Unmodified | 0 | 0 | 0 | 0 | 0 | 0 | 0 | 0 | 0 | 0 | 0 | 0 | 0 | 0 | 0 | 0 | 0 | 0 |
| NLCYANTII | EGFR; EGFR; E( | P00533; Q5( | EGFR_HUMAN | Unmodified | 0 | 0 | 0 | 0 | 6138000 | 0 | 0 | 0 | 0 | 0 | 0 | 0 | 0 | 0 | 0 | 0 | 1596200 | 1228700 |
| NLCYANTII | EGFR; EGFR; E( | P00533; Q5( | EGFR_HUMAN | Unmodified | 0 | 0 | 0 | 0 | 6733200 | 687570 | 91082 | 64653 | 0 | 0 | 0 | 0 | 0 | 1008500 | 0 | 0 | 0 | 68848 |
| NLDDGIDD | PABPC1; PABP( | P11940; Q9I | PABP1_HUMAN | Unmodified | 7905100 | 9955800 | 2520700 | 1244600 | 0 | 0 | 0 | 0 | 237220 | 56746 | 555260 | 1434400 | 0 | 0 | 2324500 | 3413200 | 0 | 0 |
| NLDDGIDD | PABPC1; PABP( | P11940; Q9I | PABP1_HUMAN | Unmodified | 35654000 | 30431000 | 6396300 | 6864100 | 0 | 0 | 0 | 0 | 222850 | 0 | 985090 | 5696500 | 382860 | 0 | 5219700 | 8729100 | 0 | 0 |
| NLDDLTLL | MIX23; MIX23; I | Q4VC31; C9 | MIX23_HUMAN | Unmodified | 0 | 0 | 0 | 0 | 221580 | 0 | 0 | 0 | 0 | 0 | 1303600 | 3967700 | 0 | 0 | 0 | 0 | 0 | 0 |
| NLDDTIDD | PABPC4L; PABI | P0CB38; Q1 | PAB4L_HUMAN | Unmodified | 14738000 | 8623400 | 7637600 | 5955600 | 292920 | 66231 | 317140 | 194720 | 0 | 123270 | 229020 | 775060 | 1097300 | 1206600 | 3569500 | 3178300 | 98745 | 0 |
| NLDDTIDD | PABPC4L; PABI | P0CB38; Q1 | PAB4L_HUMAN | Unmodified | 3606900 | 2850500 | 0 | 0 | 3233200 | 1467300 | 572710 | 1348900 | 126130 | 1111600 | 0 | 0 | 0 | 0 | 0 | 0 | 3123300 | 368020 |
| NLDDTIDD | PABPC4; PABP( | Q13310; B1/ | PABP4_HUMAN | Unmodified | 1314600 | 0 | 0 | 1844300 | 93569 | 0 | 1992200 | 0 | 0 | 0 | 0 | 84350 | 1719300 | 0 | 0 | 0 | 213920 | 0 |
| NLDENGLI | CDK1; CDK1 | P06493; A0/ | CDK1_HUMAN | Unmodified | 0 | 0 | 0 | 0 | 6058900 | 0 | 479390 | 432460 | 1025400 | 1026800 | 367620 | 0 | 2721300 | 1287600 | 0 | 0 | 0 | 1728300 |
| NLDEQLSA | CNOT1; CNOT1 | A5YKK6; H3I | CNOT1_HUMA | Unmodified | 0 | 0 | 0 | 0 | 0 | 0 | 0 | 0 | 0 | 0 | 0 | 886200 | 0 | 0 | 0 | 0 | 0 | 0 |
| NLDFIVYKF | CHAT; CHAT | P28329; A0/ | CLAT_HUMAN | Unmodified | 0 | 0 | 0 | 0 | 0 | 0 | 0 | 0 | 0 | 0 | 361440 | 2293200 | 0 | 0 | 0 | 0 | 0 | 0 |
| NLDFQDVL | RTCB | Q9Y3I0 | RTCB_HUMAN | Unmodified | 0 | 0 | 474580 | 622020 | 385680 | 0 | 358350 | 0 | 364190 | 0 | 1128700 | 5501200 | 0 | 0 | 0 | 0 | 0 | 0 |
| NLDFSVIV/ | DSG2 | Q14126 | DSG2_HUMAN | Unmodified | 0 | 0 | 0 | 0 | 952610 | 0 | 0 | 0 | 27715000 | 10463000 | 0 | 0 | 0 | 0 | 0 | 0 | 0 | 0 |
| NLDGISHA | GGH; GGH | Q92820; A0/ | GGH_HUMAN ( | Unmodified | 0 | 0 | 366250 | 11058 | 787050 | 0 | 0 | 36976 | 45653000 | 21612000 | 3185500 | 1495100 | 1096000 | 0 | 349760 | 96064 | 0 | 0 |
| NLDIDPKL( | LARP1 | Q6PKG0 | LARP1_HUMAN | Unmodified | 198850 | 124360 | 96652 | 643900 | 0 | 0 | 0 | 0 | 0 | 0 | 0 | 399320 | 0 | 175370 | 184240 | 0 | 0 | 0 |
| NLDIERPTY | TUBA3C; TUBA | P0DPH7; Q7 | TBA3C_HUMAN | Unmodified | 10801000 | 8220600 | 18295000 | 12436000 | 3376800 | 3340600 | 3032400 | 1910600 | 71995000 | 11662000 | 27217000 | 5.98E+08 | 5674000 | 2500800 | 22309000 | 20221000 | 2772100 | 2476400 |
| NLDKSIDN | PABPC4; PABP( | Q13310; P1 | PABP4_HUMAN | Unmodified | 0 | 0 | 0 | 0 | 0 | 0 | 0 | 0 | 0 | 0 | 0 | 0 | 1159600 | 891100 | 0 | 0 | 133470 | 89362 |
| NLDLAAPT/ | PLCD3; PLCD3; | Q8N3E9; F5( | PLCD3_HUMAN | Unmodified | 0 | 0 | 0 | 0 | 835220 | 0 | 0 | 0 | 0 | 136550 | 0 | 0 | 0 | 0 | 0 | 0 | 1127800 | 0 |
| NLDNIVLQ( | EXOC6B; EXOC | Q9Y2D4; A0/ | EXC6B_HUMAN | Unmodified | 4024100 | 4983000 | 0 | 0 | 994720 | 0 | 0 | 0 | 419360 | 0 | 0 | 0 | 855180 | 0 | 0 | 0 | 218450 | 141070 |
| NLDPDVFN | RPAP3 | Q9H6T3 | RPAP3_HUMAN | Unmodified | 0 | 0 | 0 | 0 | 0 | 0 | 0 | 0 | 0 | 0 | 0 | 1687700 | 0 | 0 | 0 | 0 | 0 | 0 |
| NLDPFNEH | ABCC4 | O15439 | MRP4_HUMAN | Unmodified | 0 | 0 | 0 | 0 | 0 | 0 | 0 | 0 | 0 | 0 | 0 | 0 | 0 | 0 | 1553600 | 0 | 0 | 0 |
| NLDPLVYLI | TUBGCP2; TUB | Q9BSJ2; F2Z | GCP2_HUMAN | Unmodified | 0 | 0 | 0 | 0 | 165240 | 55833 | 0 | 0 | 0 | 37711 | 152580 | 932660 | 0 | 0 | 141850 | 0 | 0 | 89450 |
| NLDSDIEG | DNAH17; DNAH | Q9UFH2; K7 | DYH17_HUMAI | Unmodified | 0 | 0 | 0 | 0 | 0 | 0 | 0 | 0 | 1918900 | 496840 | 0 | 0 | 0 | 0 | 0 | 0 | 0 | 0 |
| NLDSTTVAI | CSRP2; CSRP2; | Q16527; F5I | CSRP2_HUMAI | Unmodified | 497130 | 0 | 0 | 0 | 277710 | 0 | 666850 | 535120 | 153070 | 169530 | 492890 | 5218100 | 0 | 0 | 0 | 0 | 2418700 | 0 |
| NLDSTTVA' | CSRP1; CSRP1; | P21291; E9P | CSRP1_HUMAI | Unmodified | 1032200 | 892950 | 1551700 | 1132400 | 9301200 | 1459500 | 2251600 | 3287000 | 4400000 | 4834800 | 647470 | 2370400 | 6480300 | 5361900 | 2465400 | 1378000 | 5698200 | 2492100 |
| NLDTVKGS | PLXNB2 | O15031 | PLXB2_HUMAN | Unmodified | 0 | 0 | 0 | 0 | 6195300 | 913960 | 1719700 | 629070 | 0 | 206160 | 0 | 0 | 4817600 | 3035100 | 0 | 0 | 5391600 | 8294400 |
| NLDYVATS | GOT1 | P17174 | AATC_HUMAN | Unmodified | 262510 | 466450 | 1634000 | 1743000 | 3050900 | 943400 | 1032500 | 0 | 3010200 | 1212100 | 7715500 | 26737000 | 3513600 | 2086200 | 0 | 4657700 | 347980 | 711500 |
| NLDYVATS | GOT1 | P17174 | AATC_HUMAN | Unmodified | 0 | 0 | 0 | 0 | 7142700 | 2348400 | 2297000 | 0 | 660720 | 4892500 | 1758600 | 1077200 | 9621800 | 10705000 | 1851800 | 0 | 3765100 | 2743500 |
| NLEAQHKE | SEPTIN7; SEPTI | Q16181; E7I | SEPT7_HUMAI | Unmodified | 0 | 0 | 0 | 0 | 0 | 0 | 623760 | 0 | 0 | 67909 | 0 | 0 | 774490 | 1143000 | 0 | 0 | 286340 | 0 |
| NLEAVETL( | ATP1A1; ATP1A | P05023; P50 | AT1A1_HUMAN | Unmodified | 2673700 | 735210 | 16378000 | 7037000 | 1778200 | 0 | 1260500 | 89348 | 6980700 | 834640 | 7510900 | 38061000 | 1471200 | 1304500 | 16834000 | 10562000 | 0 | 0 |
| NLEAVETL( | ATP1A1; ATP1A | P05023; P50 | AT1A1_HUMAN | Unmodified | 0 | 0 | 0 | 10486000 | 5164200 | 1153400 | 548050 | 474640 | 4324600 | 2594400 | 4368900 | 5675200 | 7511900 | 2889500 | 1784000 | 997070 | 3095600 | 1229700 |
| NLEECITR | L1RE1 | Q9UN81 | LORF1_HUMAN | Unmodified | 2751000 | 1263700 | 0 | 522610 | 0 | 0 | 0 | 0 | 0 | 0 | 0 | 0 | 0 | 0 | 586100 | 649690 | 0 | 0 |
| NLEECITR | L1RE1 | Q9UN81 | LORF1_HUMAN | Unmodified | 0 | 0 | 0 | 0 | 0 | 0 | 414660 | 0 | 0 | 544910 | 0 | 0 | 2419100 | 1525000 | 0 | 0 | 0 | 0 |
| NLEEDDYL | STEAP1 | Q9UHE8 | STEA1_HUMAN | Unmodified | 490300 | 305890 | 8422000 | 4876100 | 0 | 464680 | 0 | 0 | 0 | 0 | 0 | 0 | 1393800 | 603670 | 8411000 | 5873300 | 0 | 520960 |
| NLEEDDYL | STEAP1 | Q9UHE8 | STEA1_HUMAN | Oxidation (M) | 0 | 0 | 0 | 2457400 | 0 | 0 | 0 | 0 | 0 | 0 | 63045 | 0 | 1840200 | 1248800 | 921850 | 1185400 | 0 | 0 |
| NLEEDDYL | STEAP1 | Q9UHE8 | STEA1_HUMAN | Unmodified | 0 | 0 | 0 | 2941700 | 0 | 0 | 0 | 0 | 0 | 0 | 0 | 0 | 3911200 | 2307600 | 0 | 0 | 0 | 0 |
| NLEELEEK | G3BP2; G3BP2; | Q9UN86; A0 | G3BP2_HUMAI | Unmodified | 0 | 0 | 0 | 0 | 0 | 0 | 0 | 0 | 0 | 0 | 950490 | 4390700 | 455420 | 0 | 0 | 0 | 0 | 0 |
| NLEEPKHQ | ITGA7; ITGA7 | Q13683; J3K | ITA7_HUMAN I | Unmodified | 0 | 0 | 0 | 0 | 0 | 0 | 2046400 | 1252300 | 0 | 476530 | 0 | 0 | 0 | 0 | 0 | 0 | 0 | 0 |
| NLEFQGPL | VPS37C; VPS37 | A5D8V6; F5I | VP37C_HUMAN | Unmodified | 31062000 | 27302000 | 21024000 | 17525000 | 0 | 0 | 0 | 168360 | 14287000 | 0 | 0 | 0 | 242220 | 0 | 16587000 | 23962000 | 0 | 0 |
| NLEGNELQ | FUT8; FUT8 | Q9BYC5; A0 | FUT8_HUMAN | Unmodified | 0 | 0 | 0 | 0 | 0 | 0 | 0 | 0 | 6805800 | 109030 | 0 | 0 | 0 | 0 | 0 | 0 | 0 | 0 |
| NLEGNELQ | FUT8; FUT8 | Q9BYC5; A0 | FUT8_HUMAN | Unmodified | 0 | 0 | 0 | 0 | 0 | 0 | 0 | 0 | 6746900 | 0 | 0 | 0 | 0 | 0 | 0 | 0 | 0 | 0 |
| NLEGYVGF | SEPTIN7; SEPTI | Q16181; E7I | SEPT7_HUMAI | Unmodified | 102630 | 0 | 0 | 311070 | 384300 | 0 | 0 | 0 | 236710 | 0 | 30853 | 849460 | 0 | 0 | 0 | 1017900 | 345650 | 0 |
| NLEGYVGF | SEPTIN7; SEPTI | Q16181; E7I | SEPT7_HUMAI | Unmodified | 248130 | 0 | 0 | 1255200 | 31569000 | 3541200 | 6665000 | 1043800 | 1825100 | 2119800 | 1305100 | 4161700 | 7461400 | 6758700 | 1325800 | 651210 | 16826000 | 10040000 |
| NLENGALQ | STK24; STK24; | Q9Y6E0; H0 | STK24_HUMAN | Unmodified | 0 | 0 | 206950 | 518870 | 3571100 | 754100 | 82535 | 61284 | 0 | 0 | 0 | 0 | 722280 | 1047800 | 336660 | 347450 | 983060 | 3041000 |
| NLENTQN( | UACA; UACA; U | Q9BZF9; H0 | UACA_HUMAN | Unmodified | 0 | 0 | 0 | 0 | 1115800 | 136450 | 0 | 0 | 459080 | 69440 | 61250 | 234810 | 0 | 0 | 0 | 0 | 0 | 0 |
| NLEPEWAA | PDIA6 | Q15084 | PDIA6_HUMAN | Unmodified | 0 | 0 | 0 | 0 | 1357400 | 250080 | 680330 | 267470 | 1.29E+08 | 19820000 | 12740000 | 1341000 | 548720 | 555590 | 0 | 0 | 362590 | 948220 |
| NLEPEWAA | PDIA6 | Q15084 | PDIA6_HUMAN | Unmodified | 0 | 0 | 0 | 0 | 2272800 | 622040 | 2530400 | 1315100 | 0 | 39743000 | 1492800 | 0 | 1123400 | 1566500 | 0 | 0 | 810720 | 1353300 |
| NLEPKYKEI | PDIA3 | P30101 | PDIA3_HUMAN | Unmodified | 0 | 0 | 0 | 0 | 100980 | 0 | 1276000 | 1486900 | 93813 | 1772800 | 0 | 0 | 0 | 0 | 0 | 0 | 236090 | 2575600 |
| NLEQLGGT | STXBP2; STXBP | Q15833; M0 | STXB2_HUMAN | Unmodified | 495880 | 493310 | 870340 | 1141400 | 376830 | 0 | 0 | 0 | 1775700 | 198170 | 0 | 3106200 | 415590 | 0 | 705590 | 477250 | 0 | 0 |
| NLEQYNKL | SMC3 | Q9UQE7 | SMC3_HUMAN | Unmodified | 96166 | 103900 | 0 | 0 | 1333500 | 254360 | 605950 | 49089 | 272160 | 246470 | 169280 | 2424900 | 0 | 0 | 0 | 0 | 0 | 0 |
| NLESTVSQ | ROCK1; ROCK1 | Q13464; A0/ | ROCK1_HUMAI | Unmodified | 0 | 0 | 0 | 0 | 0 | 0 | 0 | 0 | 0 | 0 | 0 | 682320 | 0 | 0 | 0 | 0 | 0 | 0 |
| NLETGKFIL | ADAMTS2; ADA | O95450; A0/ | ATS2_HUMAN / | Unmodified | 0 | 0 | 0 | 0 | 0 | 0 | 947950 | 455140 | 0 | 0 | 0 | 0 | 0 | 0 | 0 | 0 | 0 | 0 |
| NLETPLCK | AK2; AK2; AK2; | P54819; G3 | KAD2_HUMAN | Unmodified | 0 | 0 | 63135 | 0 | 0 | 0 | 265150 | 0 | 0 | 0 | 997770 | 3584600 | 1139900 | 453510 | 0 | 0 | 0 | 0 |
| NLEVLNFFI | RSU1 | Q15404 | RSU1_HUMAN | Unmodified | 0 | 36888 | 0 | 0 | 86477 | 0 | 503210 | 0 | 0 | 0 | 0 | 0 | 0 | 1018700 | 37885 | 0 | 0 | 0 |
| NLFEDQNT | CSE1L | P55060 | XPO2_HUMAN | Unmodified | 0 | 0 | 0 | 0 | 0 | 0 | 390750 | 356990 | 0 | 0 | 0 | 471500 | 0 | 0 | 0 | 0 | 0 | 0 |
| NLFELAR | VPS4A; N/A | Q9UN37; I3L | VPS4A_HUMAN | Unmodified | 743150 | 0 | 668360 | 983540 | 1957300 | 0 | 0 | 0 | 0 | 0 | 888390 | 3613800 | 3243100 | 0 | 0 | 0 | 0 | 0 |
| NLFEQLVR | RACGAP1; RAC | Q9H0H5; F8 | RGAP1_HUMAI | Unmodified | 247060 | 0 | 593580 | 1939500 | 7542400 | 1589600 | 0 | 0 | 4753400 | 666900 | 2752600 | 4877200 | 535620 | 0 | 1069200 | 733550 | 490910 | 0 |
| NLFETPILA | MFGE8 | Q08431 | MFGM_HUMAN | Unmodified | 12854000 | 6863200 | 12612000 | 26072000 | 58593000 | 13908000 | 2324700 | 634690 | 17478000 | 8302900 | 1996800 | 19880000 | 12309000 | 4895100 | 22317000 | 15527000 | 17794000 | 9835000 |
| NLFETPILA | MFGE8 | Q08431 | MFGM_HUMAN | Unmodified | 0 | 0 | 0 | 0 | 7359700 | 643370 | 1404500 | 0 | 0 | 610420 | 0 | 0 | 522410 | 1545400 | 0 | 0 | 2953300 | 897940 |
| NLFEVQEN | CACNA2D2; CA | Q9NY47; C9 | CA2D2_HUMAI | Unmodified | 0 | 0 | 0 | 0 | 394360 | 0 | 246570 | 177570 | 1731300 | 0 | 226830 | 748170 | 222370 | 0 | 0 | 0 | 0 | 0 |
| NLFEVQEN | CACNA2D2; CA | Q9NY47; C9 | CA2D2_HUMAI | Unmodified | 0 | 0 | 0 | 0 | 0 | 0 | 0 | 0 | 1843700 | 0 | 0 | 0 | 0 | 0 | 0 | 0 | 0 | 0 |
| NLFLTGNQ | TPBG | Q13641 | TPBG_HUMAN | Unmodified | 773520 | 1219300 | 0 | 0 | 3464000 | 1997200 | 0 | 0 | 7938800 | 1734600 | 2106200 | 24328000 | 0 | 0 | 0 | 0 | 0 | 258960 |
| NLFNVVDC | EIF3B | P55884 | EIF3B_HUMAN | Unmodified | 0 | 0 | 0 | 0 | 0 | 0 | 0 | 0 | 0 | 0 | 0 | 5505300 | 0 | 0 | 0 | 0 | 0 | 0 |
| NLFPSNLV | SLC1A4 | P43007 | SATT_HUMAN I | Unmodified | 1227100 | 952290 | 431330 | 76077 | 0 | 0 | 0 | 0 | 17343000 | 12544000 | 0 | 18217000 | 0 | 0 | 2207200 | 1944700 | 0 | 0 |
| NLFQLAR | VPS4B | O75351 | VPS4B_HUMAN | Unmodified | 0 | 0 | 973660 | 2188300 | 580600 | 0 | 0 | 37751 | 5209400 | 2103500 | 867590 | 5325400 | 1210000 | 224070 | 1138500 | 1618400 | 0 | 177610 |
| NLFRPSLP | H6PD | O95479 | G6PE_HUMAN | Unmodified | 0 | 0 | 0 | 0 | 0 | 0 | 0 | 0 | 558370 | 0 | 0 | 0 | 0 | 0 | 0 | 0 | 0 | 0 |
| NLFWTSYD | LRP1 | Q07954 | LRP1_HUMAN | Unmodified | 0 | 0 | 0 | 0 | 0 | 620360 | 0 | 0 | 1445200 | 108970 | 0 | 0 | 0 | 0 | 0 | 0 | 0 | 0 |
| NLGESEIR | PSMD6; PSMD6 | Q15008; C9J | PSMD6_HUMA | Unmodified | 63541 | 39669 | 0 | 0 | 0 | 0 | 0 | 0 | 841410 | 0 | 0 | 1231500 | 0 | 0 | 37313 | 0 | 0 | 0 |
| NLGFSPED | PAPSS1 | O43252 | PAPS1_HUMAN | Unmodified | 0 | 1198500 | 0 | 870780 | 83176 | 0 | 183850 | 76670 | 0 | 0 | 0 | 1819200 | 572470 | 0 | 452520 | 279970 | 48377 | 89689 |
| NLGIGKVS | MDH2; MDH2 | P40926; Q3I | MDHM_HUMAN | Unmodified | 0 | 0 | 0 | 0 | 924550 | 119520 | 6331000 | 1397600 | 0 | 6910200 | 0 | 0 | 1603200 | 1592900 | 0 | 0 | 1132500 | 863840 |
| NLGIPIITVL | CKAP5 | Q14008 | CKAP5_HUMAN | Unmodified | 0 | 57264 | 70590 | 71821 | 627850 | 0 | 0 | 273610 | 114700 | 10837 | 719960 | 2595200 | 1175200 | 0 | 0 | 198800 | 164360 | 431450 |
| NLGLEELG | DLD; DLD | P09622; E9P | DLDH_HUMAN | Unmodified | 0 | 0 | 0 | 0 | 0 | 0 | 0 | 0 | 1305300 | 0 | 0 | 0 | 0 | 0 | 0 | 0 | 0 | 0 |
| NLGLFQH | MAN2A1 | Q16706 | MA2A1_HUMAI | Unmodified | 0 | 0 | 0 | 0 | 0 | 0 | 0 | 0 | 0 | 0 | 0 | 1482200 | 0 | 0 | 0 | 0 | 0 | 0 |
| NLGMTDAF | SERPINB6; SER | P35237; A0A | SPB6_HUMAN | Oxidation (M) | 0 | 0 | 0 | 0 | 294890 | 0 | 845760 | 175990 | 0 | 0 | 0 | 3279800 | 0 | 290610 | 367460 | 245690 | 0 | 0 |
| NLGMTDAF | SERPINB6; SER | P35237; A0A | SPB6_HUMAN | Unmodified | 0 | 0 | 0 | 0 | 0 | 0 | 0 | 0 | 362920 | 0 | 0 | 1636200 | 0 | 0 | 0 | 0 | 0 | 0 |
| NLGMTDAF | SERPINB6; SER | P35237; A0A | SPB6_HUMAN | Oxidation (M) | 0 | 0 | 0 | 1945000 | 7719800 | 0 | 7392100 | 0 | 0 | 3300500 | 0 | 17442000 | 6223900 | 1565600 | 1128500 | 0 | 6813300 | 0 |
| NLGPGMTI | VCL | P18206 | VINC_HUMAN | Unmodified | 0 | 274660 | 411380 | 0 | 5702200 | 3351000 | 2811200 | 1824300 | 1459000 | 1572700 | 663280 | 0 | 0 | 0 | 666250 | 0 | 21396000 | 5552200 |
| NLGQKEN( | ADAM10; ADAM | O14672; H0 | ADA10_HUMAI | Unmodified | 0 | 0 | 0 | 0 | 1279900 | 0 | 78190 | 0 | 0 | 126220 | 0 | 117440 | 2224800 | 946660 | 0 | 0 | 0 | 351570 |
| NLGQKEN( | ADAM10; ADAM | O14672; H0 | ADA10_HUMAI | Oxidation (M) | 0 | 0 | 0 | 0 | 222120 | 82171 | 0 | 17454 | 0 | 0 | 0 | 0 | 646000 | 450050 | 0 | 0 | 0 | 0 |
| NLGQNLW | RCC2 | Q9P258 | RCC2_HUMAN | Unmodified | 60220 | 35975 | 0 | 26903 | 0 | 0 | 0 | 0 | 674370 | 26514 | 848890 | 9715000 | 0 | 0 | 66866 | 0 | 0 | 0 |
| NLGSINTEI | SEC22B; SEC22 | O75396; A0/ | SC22B_HUMAN | Unmodified | 0 | 0 | 0 | 0 | 0 | 0 | 0 | 0 | 1007300 | 0 | 492610 | 1874600 | 0 | 0 | 0 | 0 | 0 | 0 |
| NLGTALAEI | TLN1 | Q9Y490 | TLN1_HUMAN | Unmodified | 810600 | 1047400 | 1639900 | 2772400 | 4137200 | 145410 | 2005700 | 699350 | 2757000 | 341440 | 1281600 | 2947900 | 2601800 | 228350 | 1453900 | 1959000 | 466810 | 2779100 |
| NLGTALAEI | TLN1 | Q9Y490 | TLN1_HUMAN | Unmodified | 0 | 0 | 0 | 0 | 18551000 | 0 | 6884900 | 1026800 | 0 | 0 | 0 | 0 | 9545100 | 8093300 | 0 | 0 | 7294600 | 11386000 |
| NLGTIAKS( | HSP90B1; HSP! | P14625; A0A | ENPL_HUMAN | Unmodified | 0 | 0 | 0 | 0 | 0 | 0 | 759320 | 29297 | 0 | 661620 | 0 | 0 | 2698800 | 0 | 0 | 0 | 0 | 0 |
| NLGVIQEL\ | ANKRD16 | Q6P6B7 | ANR16_HUMAI | Unmodified | 0 | 0 | 0 | 26220 | 502910 | 0 | 75341 | 0 | 23683 | 0 | 45862 | 183090 | 597910 | 185850 | 0 | 0 | 0 | 0 |
| NLGVQQS( | CKS1B; CKS1B | P61024; Q9E | CKS1_HUMAN | Oxidation (M) | 1174800 | 2040300 | 0 | 0 | 0 | 0 | 0 | 0 | 0 | 0 | 0 | 0 | 0 | 0 | 976990 | 0 | 0 | 0 |
| NLHAFSVE | COLGALT1 | Q8NBJ5 | GT251_HUMAN | Unmodified | 0 | 0 | 0 | 0 | 0 | 0 | 0 | 0 | 1707900 | 0 | 0 | 0 | 0 | 0 | 0 | 0 | 0 | 0 |
| NLHALILVN | DCN; DCN; DC | P07585; F8V | PGS2_HUMAN | Unmodified | 0 | 0 | 0 | 0 | 0 | 0 | 0 | 172180 | 0 | 0 | 0 | 0 | 0 | 0 | 0 | 0 | 0 | 0 |
| NLHDALSC | ELP3; ELP3 | Q9H9T3; B4I | ELP3_HUMAN I | Unmodified | 0 | 0 | 0 | 0 | 0 | 0 | 0 | 0 | 0 | 0 | 0 | 899480 | 0 | 0 | 32924 | 47907 | 0 | 0 |
| NLHGDGIA | LMAN2; LMAN2 | Q12907; D6I | LMAN2_HUMAI | Unmodified | 229570 | 328810 | 928960 | 576310 | 0 | 0 | 247000 | 0 | 2.04E+08 | 13278000 | 2593900 | 4664500 | 107140 | 0 | 833470 | 1051500 | 0 | 0 |
| NLHGDGIA | LMAN2; LMAN2 | Q12907; D6I | LMAN2_HUMAI | Unmodified | 0 | 0 | 0 | 0 | 0 | 0 | 0 | 0 | 38234000 | 3146200 | 379460 | 186870 | 0 | 0 | 0 | 0 | 0 | 0 |
| NLHGDGIA | LMAN2; LMAN2 | Q12907; D6I | LMAN2_HUMAI | Unmodified | 0 | 0 | 0 | 0 | 538670 | 0 | 1679800 | 0 | 7259600 | 28839000 | 0 | 0 | 579940 | 597380 | 0 | 0 | 205190 | 0 |
| NLHGDGL/ | LMAN2L; LMAN | Q9H0V9; F8\ | LMA2L_HUMAN | Unmodified | 0 | 0 | 52436 | 0 | 0 | 0 | 71071 | 0 | 9265700 | 1565400 | 0 | 0 | 48219 | 40816 | 0 | 0 | 0 | 0 |
| NLHQSGFS | DPYSL3; DPYSL | Q14195; H0 | DPYL3_HUMAN | Unmodified | 49578 | 171240 | 125200 | 118230 | 0 | 0 | 103140 | 0 | 276530 | 64523 | 737590 | 4181100 | 0 | 0 | 173020 | 167190 | 0 | 0 |
| NLHRFTHII | GALNT7; GALN | Q86SF2; H0 | GALT7_HUMAN | Unmodified | 0 | 0 | 0 | 0 | 0 | 0 | 84902 | 47097 | 207920 | 283100 | 0 | 0 | 0 | 74856 | 0 | 0 | 0 | 0 |
| NLHSHYHE | POMT2 | Q9UKY4 | POMT2_HUMAI | Unmodified | 0 | 0 | 0 | 0 | 0 | 0 | 0 | 0 | 725250 | 0 | 0 | 0 | 0 | 0 | 0 | 0 | 0 | 0 |
| NLHTHHFF | SDF2L1 | Q9HCN8 | SDF2L_HUMAN | Unmodified | 0 | 0 | 0 | 0 | 0 | 0 | 0 | 0 | 34348000 | 0 | 0 | 0 | 0 | 0 | 0 | 0 | 0 | 0 |
| NLHVVFTM | DYNC1H1; DYN | Q14204; A0/ | DYHC1_HUMA | Oxidation (M) | 0 | 0 | 0 | 250910 | 147630 | 0 | 0 | 0 | 224420 | 84862 | 300450 | 1192700 | 130050 | 60652 | 308130 | 230470 | 0 | 0 |
| NLICAFLTD | ROCK2; ROCK1 | O75116; Q1 | ROCK2_HUMAI | Unmodified | 0 | 0 | 0 | 893880 | 998290 | 1137900 | 0 | 0 | 1198100 | 352200 | 55366 | 2725800 | 0 | 0 | 0 | 0 | 752610 | 1081000 |
| NLIDAGVD | IMPDH2; IMPDI | P12268; A0A | IMDH2_HUMAN | Unmodified | 334440 | 395180 | 0 | 420060 | 0 | 0 | 0 | 0 | 830630 | 0 | 403020 | 2659200 | 0 | 0 | 740410 | 498710 | 0 | 0 |
| NLIDEDGN | IPO5; IPO5; IPO | O00410; E7I | IPO5_HUMAN I | Unmodified | 0 | 0 | 133120 | 0 | 142640 | 0 | 0 | 0 | 180590 | 54941 | 164440 | 2082500 | 0 | 0 | 211470 | 107070 | 0 | 0 |
| NLIDYFIPFI | TOR1B | O14657 | TOR1B_HUMAN | Unmodified | 0 | 0 | 0 | 0 | 0 | 0 | 0 | 0 | 0 | 290720 | 0 | 0 | 0 | 0 | 0 | 0 | 0 | 0 |
| NLIIYWKPL | LIFR | P42702 | LIFR_HUMAN L | Unmodified | 0 | 0 | 0 | 0 | 249930 | 0 | 0 | 0 | 636780 | 199470 | 0 | 0 | 0 | 0 | 0 | 0 | 134520 | 0 |
| NLILVVR | CLTC; CLTC | Q00610; A0/ | CLH1_HUMAN | Unmodified | 4182700 | 166850 | 2031600 | 15179 | 0 | 0 | 0 | 0 | 0 | 0 | 0 | 16118000 | 0 | 0 | 3372600 | 5270500 | 0 | 0 |
| NLILYLLTD | PAN3; PAN3 | Q58A45; A0/ | PAN3_HUMAN | Unmodified | 351990 | 303390 | 63601 | 87379 | 0 | 0 | 0 | 0 | 0 | 0 | 0 | 0 | 0 | 0 | 90981 | 77982 | 0 | 0 |
| NLIPFDQM | ATP5PD | O75947 | ATP5H_HUMAI | Oxidation (M) | 0 | 0 | 45073 | 0 | 0 | 141030 | 0 | 0 | 560260 | 208900 | 0 | 0 | 0 | 0 | 0 | 0 | 0 | 0 |
| NLIPFDQM | ATP5PD | O75947 | ATP5H_HUMAI | Unmodified | 0 | 0 | 0 | 0 | 0 | 0 | 0 | 0 | 420030 | 0 | 0 | 0 | 0 | 0 | 0 | 13988 | 0 | 0 |
| NLIPFDQM | ATP5PD | O75947 | ATP5H_HUMAI | Oxidation (M) | 0 | 0 | 0 | 0 | 0 | 0 | 0 | 0 | 3459700 | 6422400 | 0 | 0 | 0 | 0 | 0 | 0 | 0 | 0 |

| | | | | | | | | | | | | | | | | | | | | | | |
|---|---|---|---|---|---|---|---|---|---|---|---|---|---|---|---|---|---|---|---|---|---|---|
| NLIPFDQM | ATP5PD | O75947 | ATP5H_HUMAN | Unmodified | 0 | 0 | 0 | 0 | 0 | 0 | 0 | 0 | 1272100 | 329200 | 0 | 0 | 0 | 0 | 0 | 0 | 0 | 0 |
| NLIPMDPN | PRKCA; PRKCG | P17252; P05 | KPCA_HUMAN | Oxidation (M) | 229720 | 0 | 0 | 0 | 0 | 0 | 0 | 740210 | 0 | 0 | 0 | 0 | 0 | 0 | 0 | 0 | 0 | 0 |
| NLISPDLG\ | MAP1B; MAP1B | P46821; D6F | MAP1B_HUMAI | Unmodified | 69464 | 0 | 0 | 0 | 0 | 0 | 0 | 0 | 104000 | 0 | 0 | 234390 | 177210 | 249760 | 57175 | 19061 | 0 | 0 |
| NLISPDLG\ | MAP1B; MAP1B | P46821; D6F | MAP1B_HUMAI | Unmodified | 0 | 0 | 0 | 0 | 273820 | 0 | 4266300 | 0 | 683720 | 600100 | 450820 | 540560 | 711240 | 2151100 | 0 | 0 | 2015100 | 519980 |
| NLIYDNAD | PKP3 | Q9Y446 | PKP3_HUMAN | Unmodified | 707860 | 729830 | 2442900 | 1906600 | 0 | 68026 | 544500 | 0 | 570480 | 74428 | 1015800 | 0 | 0 | 451140 | 2315000 | 2440100 | 0 | 0 |
| NLKEDGIS/ | GNAO1; GNAO | P09471; A0A | GNAO_HUMAN | Unmodified | 0 | 0 | 0 | 0 | 0 | 0 | 582850 | 466550 | 688580 | 490640 | 0 | 51900 | 0 | 0 | 0 | 0 | 0 | 0 |
| NLKEDGIS/ | GNAO1; GNAO | P09471; A0A | GNAO_HUMAN | Unmodified | 0 | 0 | 0 | 162690 | 3981900 | 0 | 1651100 | 1307800 | 493990 | 688520 | 156020 | 0 | 968150 | 469470 | 0 | 0 | 361500 | 0 |
| NLKEPLLTF | RACGAP1; RAC | Q9H0H5; H0 | RGAP1_HUMAI | Unmodified | 0 | 0 | 0 | 0 | 137650 | 0 | 0 | 0 | 577720 | 0 | 0 | 1276600 | 0 | 0 | 0 | 0 | 0 | 0 |
| NLKEQGVT | STAM; STAM | Q92783; A6N | STAM1_HUMAI | Unmodified | 0 | 0 | 0 | 0 | 2632000 | 0 | 967040 | 1280000 | 0 | 677820 | 0 | 0 | 468740 | 0 | 0 | 0 | 0 | 1751400 |
| NLKPIKPM( | PGAM1 | P18669 | PGAM1_HUMA | Unmodified | 0 | 0 | 0 | 0 | 550640 | 0 | 0 | 0 | 70225 | 0 | 1019600 | 492010 | 0 | 0 | 0 | 0 | 73482 | 56231 |
| NLKVPSGQ | NAA50; NAA50; | Q9GZZ1; A0/ | NAA50_HUMAN | Unmodified | 139810 | 0 | 0 | 0 | 2092700 | 367280 | 691750 | 649540 | 160770 | 453520 | 86892 | 148600 | 2325600 | 974060 | 0 | 0 | 2510700 | 1892100 |
| NLKVTAVPT | TXNDC17; TXNI | Q9BRA2; I3L | TXD17_HUMAN | Unmodified | 0 | 0 | 0 | 0 | 457720 | 0 | 0 | 417590 | 0 | 0 | 0 | 0 | 0 | 0 | 0 | 0 | 1861900 | 0 |
| NLLAEFNL( | ENTPD4; ENTPI | Q9Y227; Q8I | ENTP4_HUMAN | Unmodified | 0 | 0 | 0 | 0 | 0 | 0 | 0 | 0 | 1180500 | 0 | 0 | 0 | 0 | 0 | 0 | 0 | 0 | 0 |
| NLLALDKD | MYOF; MYOF | Q9NZM1; H( | MYOF_HUMAN | Unmodified | 0 | 0 | 0 | 0 | 1509300 | 1244300 | 0 | 0 | 0 | 28411 | 0 | 0 | 0 | 0 | 0 | 0 | 148660 | 208480 |
| NLLCGFYG | APOBEC3B; AP( | Q9UH17; P3 | ABC3B_HUMAI | Unmodified | 555760 | 294360 | 310380 | 192580 | 0 | 0 | 0 | 0 | 0 | 0 | 0 | 0 | 0 | 0 | 970070 | 617780 | 0 | 0 |
| NLLDEIASF | PPL; PPL; PPL | O60437; K7E | PEPL_HUMAN | Unmodified | 0 | 0 | 0 | 0 | 6246200 | 674070 | 0 | 0 | 0 | 0 | 0 | 0 | 1323700 | 768590 | 0 | 0 | 386810 | 453740 |
| NLLEISGPE | CHMP4B | Q9H444 | CHM4B_HUMA | Unmodified | 1368100 | 1223900 | 1460900 | 989520 | 118930 | 78667 | 0 | 0 | 3625600 | 309860 | 323260 | 255400 | 82288 | 29417 | 3820400 | 1147500 | 0 | 32581 |
| NLLEISGPE | CHMP4B | Q9H444 | CHM4B_HUMA | Unmodified | 46520000 | 37206000 | 7700300 | 28487000 | 40100000 | 7886300 | 157280 | 0 | 73253000 | 7786200 | 13168000 | 8732400 | 15345000 | 3666000 | 33792000 | 25991000 | 231520 | 7197300 |
| NLLESYHV | LRPPRC | P42704 | LPPRC_HUMAN | Unmodified | 0 | 0 | 0 | 0 | 0 | 0 | 0 | 0 | 2501900 | 0 | 0 | 0 | 0 | 0 | 0 | 0 | 0 | 0 |
| NLLGALSLI | PLCH1; PLCH1 | Q4KWH8; A( | PLCH1_HUMAN | Unmodified | 0 | 0 | 0 | 0 | 2344000 | 0 | 0 | 0 | 0 | 0 | 0 | 0 | 0 | 0 | 0 | 0 | 5171200 | 0 |
| NLLGLIEAK | GRP; GRP | P07492; H0\ | GRP_HUMAN ( | Unmodified | 0 | 0 | 0 | 0 | 0 | 0 | 0 | 0 | 2091000 | 580680 | 0 | 0 | 0 | 0 | 0 | 0 | 0 | 0 |
| NLLHSLQS | GCN1 | Q92616 | GCN1_HUMAN | Unmodified | 202500 | 0 | 0 | 0 | 55319 | 0 | 218340 | 43320 | 0 | 0 | 0 | 0 | 515200 | 0 | 374880 | 386170 | 124350 | 830790 |
| NLLHVTDT( | HSP90B1; HSP9 | P14625; A0A | ENPL_HUMAN | Unmodified | 0 | 0 | 0 | 0 | 0 | 0 | 212060 | 36391 | 5914000 | 2367700 | 7300100 | 504540 | 0 | 0 | 0 | 0 | 0 | 0 |
| NLLHVTDT( | HSP90B1; HSP9 | P14625; A0A | ENPL_HUMAN | Oxidation (M) | 58516 | 0 | 0 | 0 | 0 | 0 | 200550 | 126740 | 3646500 | 849770 | 11962000 | 869810 | 223230 | 229970 | 20407 | 39072 | 0 | 0 |
| NLLHVTDT( | HSP90B1; HSP9 | P14625; A0A | ENPL_HUMAN | Unmodified | 516660 | 81646 | 0 | 640490 | 2618800 | 1429900 | 23191000 | 1264500 | 1.03E+09 | 2.68E+08 | 40669000 | 31641000 | 5721100 | 9704500 | 898790 | 586560 | 1127500 | 5620700 |
| NLLHVTDT( | HSP90B1; HSP9 | P14625; A0A | ENPL_HUMAN | Oxidation (M) | 2284800 | 0 | 2777100 | 2153700 | 1247200 | 1087200 | 23978000 | 3709500 | 2.84E+08 | 2.42E+08 | 59297000 | 55957000 | 4909400 | 6008400 | 3623400 | 3095600 | 1489100 | 2018600 |
| NLLHVTDT( | HSP90B1; HSP9 | P14625; A0A | ENPL_HUMAN | Unmodified | 0 | 0 | 0 | 0 | 0 | 0 | 2877900 | 0 | 0 | 3934400 | 0 | 0 | 0 | 0 | 0 | 0 | 212090 | 0 |
| NLLHVTDT( | HSP90B1; HSP9 | P14625; A0A | ENPL_HUMAN | Oxidation (M) | 0 | 0 | 0 | 0 | 0 | 0 | 12425000 | 0 | 0 | 10276000 | 0 | 0 | 0 | 0 | 0 | 0 | 0 | 0 |
| NLLIFENLI | PRKDC | P78527 | PRKDC_HUMAI | Unmodified | 0 | 20079 | 0 | 0 | 0 | 0 | 45085 | 0 | 33322 | 96474 | 0 | 219700 | 0 | 0 | 0 | 0 | 0 | 0 |
| NLLLAEVIN | ABAT; ABAT; AB | P80404; H3I | GABT_HUMAN | Unmodified | 0 | 0 | 0 | 0 | 0 | 0 | 0 | 0 | 0 | 225760 | 0 | 0 | 0 | 0 | 0 | 0 | 0 | 0 |
| NLLLEPSLE | VPS37A; VPS37 | Q8NEZ2; H0 | VP37A_HUMAN | Unmodified | 6022800 | 7449500 | 1199600 | 1012800 | 3155000 | 646490 | 0 | 0 | 0 | 166900 | 0 | 0 | 994780 | 495080 | 1538100 | 1681200 | 0 | 229450 |
| NLLLEPSLE | VPS37A; VPS37 | Q8NEZ2; H0 | VP37A_HUMAN | Unmodified | 444070 | 346040 | 0 | 317800 | 0 | 0 | 0 | 0 | 0 | 0 | 0 | 0 | 0 | 0 | 0 | 100540 | 0 | 0 |
| NLLLLANS1 | ROCK2; ROCK2 | O75116; Q1 | ROCK2_HUMAI | Unmodified | 0 | 129410 | 0 | 0 | 328550 | 0 | 0 | 0 | 0 | 0 | 0 | 362180 | 690180 | 0 | 0 | 0 | 899950 | 297160 |
| NLLLLLTG( | FAM83F | Q8NEG4 | FA83F_HUMAN | Unmodified | 0 | 0 | 0 | 0 | 0 | 0 | 0 | 0 | 0 | 0 | 0 | 0 | 0 | 0 | 176360 | 0 | 0 | 0 |
| NLLPRTPT( | KIF20A | O95235 | KI20A_HUMAN | Unmodified | 0 | 0 | 0 | 0 | 474230 | 0 | 0 | 0 | 0 | 0 | 0 | 0 | 1094600 | 211600 | 0 | 0 | 659010 | 73177 |
| NLLQQSWI | DLG5; DLG5 | Q8TDM6; R4 | DLG5_HUMAN | Oxidation (M) | 423790 | 31150 | 0 | 0 | 0 | 58417 | 0 | 0 | 3502900 | 255860 | 0 | 0 | 0 | 0 | 0 | 0 | 0 | 0 |
| NLLSEHEQ | GOLIM4 | O00461 | GOLI4_HUMAN | Unmodified | 0 | 0 | 0 | 0 | 1096400 | 0 | 518820 | 0 | 24869000 | 5282100 | 136900 | 0 | 0 | 0 | 0 | 0 | 0 | 0 |
| NLLSLGHN | CSTF1 | Q05048 | CSTF1_HUMAN | Unmodified | 0 | 0 | 121560 | 116680 | 0 | 0 | 24668 | 55671 | 0 | 0 | 0 | 1345400 | 0 | 0 | 169660 | 143510 | 0 | 0 |
| NLLSVAYK | SFN; YWHAB; Y | P31947; P31 | 1433S_HUMAN | Unmodified | 7323200 | 9491500 | 15881000 | 12773000 | 4738700 | 1802400 | 781680 | 0 | 40751000 | 2733000 | 11594000 | 2.52E+08 | 4343000 | 1672200 | 30673000 | 40208000 | 2957400 | 2298200 |
| NLLSVAYKI | YWHAB; YWHA | P31946; P63 | 1433B_HUMAN | Unmodified | 0 | 0 | 0 | 0 | 0 | 0 | 0 | 0 | 0 | 0 | 0 | 0 | 0 | 0 | 0 | 0 | 973220 | 321450 |
| NLLTAAAD | ABCD1 | P33897 | ABCD1_HUMAI | Unmodified | 0 | 0 | 0 | 0 | 0 | 0 | 0 | 0 | 1536200 | 0 | 0 | 0 | 0 | 0 | 0 | 0 | 0 | 0 |
| NLLTGETE/ | CLPTM1; CLPTN | O96005; K7E | CLPT1_HUMAN | Oxidation (M) | 0 | 0 | 0 | 0 | 0 | 0 | 0 | 0 | 1247700 | 0 | 0 | 0 | 0 | 0 | 0 | 0 | 0 | 0 |
| NLLTGETE/ | CLPTM1; CLPTN | O96005; K7E | CLPT1_HUMAN | Unmodified | 0 | 0 | 0 | 0 | 0 | 140540 | 0 | 0 | 9800300 | 0 | 0 | 431770 | 0 | 0 | 0 | 0 | 0 | 0 |
| NLLTSMPH | TMEM260 | Q9NX78 | TM260_HUMAN | Unmodified | 0 | 0 | 0 | 0 | 0 | 0 | 0 | 0 | 1460400 | 158460 | 0 | 0 | 0 | 0 | 0 | 0 | 0 | 0 |
| NLLVAFR | MMS19; MMS19 | Q96T76; Q5T | MMS19_HUMA | Unmodified | 0 | 0 | 0 | 0 | 0 | 0 | 0 | 0 | 0 | 0 | 0 | 712890 | 0 | 0 | 0 | 0 | 0 | 0 |
| NLLVNTLY1 | SORL1; SORL1; | Q92673; E9F | SORL_HUMAN | Unmodified | 0 | 0 | 0 | 0 | 0 | 0 | 0 | 0 | 2110800 | 316780 | 0 | 0 | 0 | 0 | 0 | 0 | 0 | 0 |
| NLLYIYPQS | DOCK7; DOCK; | Q96N67; A0, | DOCK7_HUMA | Unmodified | 90356 | 0 | 0 | 0 | 0 | 0 | 44178 | 0 | 264460 | 0 | 419430 | 1190000 | 0 | 0 | 206550 | 175040 | 0 | 0 |
| NLMASQP( | PKP1 | Q13835 | PKP1_HUMAN | Unmodified | 0 | 0 | 0 | 0 | 0 | 0 | 0 | 0 | 0 | 0 | 0 | 0 | 0 | 0 | 0 | 0 | 0 | 0 |
| NLMFLVLR | NARS1; NARS1 | O43776; K7E | SYNC_HUMAN | Unmodified | 0 | 0 | 0 | 0 | 0 | 0 | 0 | 0 | 0 | 0 | 13041 | 215880 | 0 | 0 | 0 | 0 | 0 | 0 |
| NLMFLVLR | NARS1; NARS1 | O43776; K7E | SYNC_HUMAN | Oxidation (M) | 0 | 0 | 69148 | 0 | 0 | 0 | 0 | 0 | 0 | 0 | 225740 | 285740 | 0 | 0 | 624770 | 355400 | 0 | 0 |
| NLMNAVV( | CTNNA1; CTNN | P35221; Q9( | CTNA1_HUMAI | Oxidation (M) | 0 | 0 | 812130 | 572190 | 1223700 | 247390 | 0 | 0 | 189160 | 105960 | 693680 | 2902900 | 4292000 | 4254500 | 5455600 | 5479100 | 0 | 0 |
| NLMNAVV( | CTNNA1; CTNN | P35221; Q9( | CTNA1_HUMAI | Unmodified | 0 | 0 | 1293100 | 4029000 | 11489000 | 1980200 | 0 | 0 | 0 | 0 | 839130 | 4209800 | 8690600 | 12866000 | 2909100 | 1666300 | 166590 | 0 |
| NLMQFHT\ | DHX30; DHX30 | Q7L2E3; H7I | DHX30_HUMAI | Unmodified | 91560 | 0 | 0 | 41215 | 0 | 0 | 92690 | 30933 | 1110300 | 0 | 0 | 717940 | 0 | 0 | 0 | 0 | 0 | 0 |
| NLMSQLG1 | STX12; STX12 | Q86Y82; B1/ | STX12_HUMAN | Unmodified | 0 | 0 | 591670 | 539490 | 0 | 0 | 0 | 0 | 1415500 | 0 | 0 | 0 | 0 | 0 | 283660 | 237110 | 0 | 0 |
| NLMSQLG1 | STX12; STX12 | Q86Y82; B1/ | STX12_HUMAN | Unmodified | 0 | 0 | 0 | 0 | 225160 | 0 | 0 | 0 | 0 | 98685 | 0 | 0 | 110700 | 96681 | 0 | 0 | 242140 | 119680 |
| NLMTSEIHI | ZW10 | O43264 | ZW10_HUMAN | Oxidation (M) | 0 | 0 | 0 | 0 | 0 | 0 | 0 | 0 | 0 | 0 | 0 | 215960 | 0 | 0 | 0 | 0 | 0 | 36407 |
| NLNEDDTF | NLN; NLN | Q9BYT8; E9F | NEUL_HUMAN | Unmodified | 267250 | 0 | 0 | 0 | 0 | 0 | 0 | 0 | 1289500 | 0 | 321020 | 2317400 | 0 | 0 | 0 | 0 | 0 | 0 |
| NLNIQLK | PIGT; PIGT; PIG | Q969N2; A0, | PIGT_HUMAN ( | Unmodified | 0 | 0 | 0 | 314770 | 361830 | 0 | 0 | 0 | 11120000 | 764730 | 0 | 0 | 0 | 0 | 799110 | 806950 | 153380 | 0 |
| NLNLNDES | XPO7; XPO7 | Q9UIA9; E7E | XPO7_HUMAN | Oxidation (M) | 0 | 0 | 0 | 0 | 0 | 23013 | 0 | 0 | 0 | 0 | 64580 | 1438200 | 0 | 0 | 107280 | 142880 | 0 | 0 |
| NLNREAQ( | GLIPR2; GLIPR2 | Q9H4G4; Q! | GAPR1_HUMAI | Unmodified | 0 | 0 | 433800 | 1596100 | 5582000 | 1624200 | 0 | 0 | 693770 | 748240 | 0 | 0 | 5695100 | 2462000 | 0 | 0 | 3126300 | 4570800 |
| NLNREAQ( | GLIPR2; GLIPR2 | Q9H4G4; Q! | GAPR1_HUMAI | Unmodified | 0 | 0 | 0 | 0 | 12200000 | 0 | 0 | 0 | 0 | 424570 | 0 | 0 | 7479700 | 6411000 | 0 | 0 | 4994000 | 5718300 |
| NLNSVSVP | MET; MET | P08581; H7( | MET_HUMAN I | Unmodified | 0 | 0 | 0 | 0 | 267800 | 210600 | 0 | 0 | 1197600 | 149890 | 0 | 403870 | 0 | 0 | 0 | 0 | 69434 | 560490 |
| NLNVILAK | LRPAP1; LRPAP | P30533; A0A | AMRP_HUMAN | Unmodified | 0 | 0 | 0 | 0 | 0 | 0 | 0 | 0 | 674440 | 0 | 0 | 0 | 0 | 0 | 0 | 0 | 0 | 0 |
| NLPFDFTW | HNRNPM; HNR | P52272; M0I | HNRPM_HUMA | Unmodified | 370710 | 0 | 356030 | 0 | 1841200 | 267700 | 1736500 | 195970 | 1747800 | 834200 | 698810 | 2547900 | 2302000 | 980440 | 0 | 0 | 238970 | 1930200 |
| NLPFLIEDY | TTC13 | Q8NBP0 | TTC13_HUMAN | Unmodified | 0 | 0 | 0 | 316370 | 0 | 0 | 0 | 0 | 6423600 | 0 | 0 | 0 | 4866900 | 0 | 514850 | 132060 | 0 | 0 |
| NLPFSVEN | COX7C; COX7C | P15954; D6F | COX7C_HUMA | Unmodified | 0 | 0 | 0 | 0 | 447520 | 0 | 4301100 | 98024 | 2096100 | 12791000 | 75012 | 0 | 390330 | 643150 | 0 | 0 | 139460 | 216240 |
| NLPFTPILA | SLC44A1 | Q8WWI5 | CTL1_HUMAN | Unmodified | 2878300 | 1316700 | 1761000 | 174190 | 0 | 0 | 0 | 0 | 283020 | 288060 | 0 | 4179400 | 0 | 0 | 1797700 | 3313700 | 0 | 0 |
| NLPGLVQE | SND1; SND1 | Q7KZF4; H7( | SND1_HUMAN | Unmodified | 934860 | 0 | 376530 | 2374000 | 0 | 0 | 1097500 | 0 | 906510 | 660690 | 0 | 0 | 2187500 | 0 | 2044900 | 1356500 | 0 | 1502900 |
| NLPIYSEEI | MYH9; MYH9 | P35579; Q5E | MYH9_HUMAN | Oxidation (M) | 0 | 0 | 207220 | 410470 | 4349100 | 464330 | 0 | 0 | 0 | 999120 | 0 | 2642200 | 556110 | 1392600 | 2005500 | 1107400 | 2162400 | 0 |
| NLPIYSEEI | MYH9; MYH9 | P35579; Q5E | MYH9_HUMAN | Unmodified | 0 | 0 | 0 | 0 | 1532700 | 0 | 0 | 0 | 0 | 173040 | 0 | 0 | 0 | 0 | 0 | 0 | 431090 | 0 |
| NLPIYSENI | MYH10; MYH10 | P35580; E7E | MYH10_HUMA | Oxidation (M) | 0 | 0 | 0 | 0 | 0 | 0 | 0 | 0 | 0 | 0 | 0 | 961990 | 0 | 0 | 76273 | 0 | 0 | 0 |
| NLPLADQG | DSP | P15924 | DESP_HUMAN | Unmodified | 0 | 0 | 0 | 0 | 0 | 0 | 0 | 0 | 1042700 | 1070600 | 0 | 400860 | 171260 | 0 | 0 | 0 | 667970 | 0 |
| NLPLGWD( | SF3A3 | Q12874 | SF3A3_HUMAN | Unmodified | 1312900 | 791620 | 0 | 872210 | 1223400 | 0 | 813630 | 0 | 470750 | 601020 | 304620 | 395760 | 1966300 | 1236600 | 296020 | 258910 | 498580 | 0 |
| NLPLPPPI | HNRNPK; HNRI | P61978; Q5T | HNRPK_HUMA | Unmodified | 477240 | 0 | 0 | 0 | 0 | 0 | 0 | 0 | 0 | 0 | 1233200 | 3815400 | 0 | 0 | 0 | 357900 | 0 | 0 |
| NLPSLGDN | SLC7A14 | Q8TBB6 | S7A14_HUMAN | Unmodified | 0 | 192840 | 0 | 0 | 0 | 98276 | 0 | 0 | 363660 | 0 | 0 | 0 | 38375 | 0 | 0 | 794140 | 871580 | 0 |
| NLPSSLEYI | TLR9; N/A | Q9NR96; H0 | TLR9_HUMAN T | Unmodified | 0 | 0 | 0 | 0 | 0 | 0 | 0 | 0 | 0 | 0 | 169050 | 223580 | 0 | 0 | 0 | 0 | 0 | 0 |
| NLPSYPGP | RASA3 | Q14644 | RASA3_HUMAN | Unmodified | 0 | 0 | 0 | 0 | 998140 | 0 | 0 | 0 | 0 | 0 | 0 | 0 | 0 | 0 | 0 | 0 | 2008400 | 3359900 |
| NLPVPVYLI | SPAG9; SPAG9 | O60271; A0/ | JIP4_HUMAN C | Unmodified | 0 | 0 | 0 | 0 | 0 | 0 | 0 | 0 | 0 | 0 | 1028200 | 2560800 | 0 | 0 | 0 | 0 | 0 | 0 |
| NLPVPVYLI | SPAG9; SPAG9 | O60271; A0/ | JIP4_HUMAN C | Oxidation (M) | 0 | 0 | 0 | 0 | 1050600 | 0 | 0 | 273270 | 0 | 0 | 0 | 0 | 840880 | 198070 | 0 | 0 | 1224200 | 1440500 |
| NLPWLFTF | EPB41L2; EPB4 | O43491; A0/ | E41L2_HUMAN | Unmodified | 0 | 0 | 36975 | 131450 | 0 | 243570 | 0 | 0 | 0 | 0 | 0 | 0 | 122330 | 177930 | 0 | 202410 | 0 | 0 |
| NLPYKVTQ | NCL; NCL; NCL | P19338; C9J | NUCL_HUMAN | Unmodified | 0 | 0 | 0 | 0 | 0 | 0 | 138750 | 60906 | 0 | 781120 | 0 | 708250 | 1377200 | 0 | 0 | 0 | 312350 | 0 |
| NLQDAMQ' | CCT3; CCT3 | P49368; B4E | TCPG_HUMAN | Oxidation (M) | 0 | 0 | 0 | 0 | 0 | 0 | 0 | 192850 | 0 | 0 | 0 | 1379600 | 0 | 0 | 0 | 0 | 0 | 0 |
| NLQDAMQ' | CCT3; CCT3 | P49368; B4E | TCPG_HUMAN | Unmodified | 0 | 0 | 0 | 0 | 0 | 0 | 0 | 0 | 0 | 0 | 0 | 4408500 | 65519 | 0 | 0 | 0 | 0 | 0 |
| NLQDLVR | AP3D1 | O14617 | AP3D1_HUMAI | Unmodified | 61798 | 131190 | 166720 | 0 | 0 | 0 | 0 | 0 | 304160 | 151410 | 314480 | 3801400 | 180810 | 0 | 0 | 0 | 0 | 196910 |
| NLQEAEEW | PRPH; VIM; VIM | P41219; P08 | PERI_HUMAN F | Unmodified | 0 | 0 | 0 | 0 | 2517500 | 448070 | 1113000 | 1211400 | 0 | 0 | 0 | 0 | 0 | 0 | 0 | 0 | 3214100 | 5193300 |
| NLQEAEEW | PRPH; VIM; VIM | P41219; P08 | PERI_HUMAN F | Unmodified | 0 | 0 | 0 | 0 | 434590 | 80838 | 301250 | 236350 | 0 | 0 | 0 | 0 | 0 | 0 | 0 | 0 | 681100 | 0 |
| NLQEILHG | EGFR; EGFR | P00533; C9J | EGFR_HUMAN | Unmodified | 268840 | 259850 | 316830 | 2606700 | 1092400 | 127770 | 0 | 0 | 913720 | 0 | 0 | 0 | 0 | 0 | 3093800 | 2189100 | 0 | 0 |
| NLQEVLGE | YARS1; YARS1; | P54577; A0A | SYYC_HUMAN | Unmodified | 0 | 0 | 0 | 0 | 0 | 0 | 0 | 0 | 983100 | 207490 | 905090 | 2072900 | 0 | 0 | 0 | 0 | 0 | 0 |
| NLQFEDGK | ALDH1L2; ALDI | Q3SY69; A0A | AL1L2_HUMAN | Unmodified | 0 | 0 | 74743 | 153950 | 0 | 0 | 0 | 0 | 401200 | 25351 | 0 | 0 | 0 | 82352 | 34008 | 0 | 0 | 271030 |
| NLQIPPEDI | SCG2 | P13521 | SCG2_HUMAN | Unmodified | 0 | 0 | 0 | 0 | 0 | 0 | 0 | 0 | 310710 | 46081 | 0 | 0 | 0 | 0 | 0 | 0 | 0 | 0 |
| NLQLDYVD | AKR1C2; AKR1( | P52895; Q0/ | AK1C2_HUMAN | Unmodified | 0 | 0 | 0 | 0 | 2051200 | 0 | 0 | 0 | 0 | 0 | 0 | 0 | 0 | 0 | 0 | 0 | 0 | 0 |
| NLQLDYVD | AKR1C2; AKR1( | P52895; Q0/ | AK1C2_HUMAN | Unmodified | 0 | 0 | 0 | 0 | 3969600 | 0 | 0 | 0 | 0 | 0 | 0 | 0 | 0 | 0 | 0 | 0 | 0 | 0 |
| NLQLFMEN | TOP1; TOP1 | P11387; A0A | TOP1_HUMAN | Oxidation (M) | 0 | 926820 | 0 | 0 | 722380 | 0 | 1517700 | 82862 | 0 | 0 | 0 | 990230 | 0 | 0 | 0 | 0 | 0 | 441230 |
| NLQLSLPR | SORL1; SORL1; | Q92673; E9F | SORL_HUMAN | Unmodified | 0 | 0 | 0 | 0 | 0 | 0 | 0 | 0 | 1312600 | 0 | 0 | 0 | 0 | 0 | 0 | 0 | 0 | 0 |
| NLQNLLILT | CLTCL1; CLTC; | P53675; Q0( | CLH2_HUMAN | Unmodified | 12075000 | 10817000 | 3330400 | 542990 | 0 | 0 | 0 | 0 | 482450 | 0 | 504630 | 37593000 | 0 | 0 | 8849000 | 11297000 | 0 | 0 |
| NLQNLLILT | CLTCL1; CLTC; | P53675; Q0( | CLH2_HUMAN | Unmodified | 13527000 | 11219000 | 156130 | 646860 | 0 | 0 | 0 | 0 | 213850 | 0 | 84171 | 40143000 | 0 | 0 | 3522400 | 6700300 | 0 | 0 |
| NLQNSLAK | MYO5C | Q9NQX4 | MYO5C_HUMA | Unmodified | 2194900 | 226860 | 0 | 7227100 | 1016200 | 0 | 1945200 | 870180 | 1092900 | 0 | 0 | 170630 | 0 | 0 | 3762800 | 4138500 | 186160 | 58112 |
| NLQNVDM | CD44; CD44; C | P16070; H0\ | CD44_HUMAN | Unmodified | 0 | 0 | 0 | 0 | 3088200 | 0 | 0 | 0 | 0 | 0 | 0 | 0 | 0 | 0 | 0 | 0 | 0 | 0 |
| NLQNVDM | CD44; CD44; C | P16070; H0\ | CD44_HUMAN | Oxidation (M) | 889470 | 693120 | 0 | 0 | 327750 | 144670 | 0 | 0 | 268810 | 420240 | 0 | 169720 | 22642 | 9003.6 | 404510 | 0 | 79485 | 307610 |
| NLQNVDM | CD44; CD44; C | P16070; H0\ | CD44_HUMAN | Unmodified | 0 | 0 | 0 | 0 | 1639000 | 0 | 0 | 62615 | 0 | 0 | 0 | 0 | 287970 | 69649 | 0 | 0 | 425760 | 1108900 |
| NLQPASEY | FN1 | P02751 | FINC_HUMAN I | Unmodified | 0 | 0 | 29002000 | 10106000 | 0 | 0 | 0 | 0 | 6969900 | 616380 | 2934000 | 2735700 | 0 | 0 | 950020 | 7392600 | 0 | 0 |
| NLQQELET | KIF23; KIF23; KI | Q02241; A0A | KIF23_HUMAN | Unmodified | 156100 | 0 | 0 | 2103600 | 2359300 | 0 | 0 | 0 | 2404700 | 79079 | 1162100 | 2152900 | 0 | 0 | 0 | 220390 | 0 | 0 |
| NLQQQYL( | GOLGB1 | Q14789 | GOGB1_HUMA | Unmodified | 0 | 0 | 0 | 0 | 0 | 0 | 0 | 0 | 0 | 0 | 0 | 0 | 0 | 125240 | 0 | 0 | 0 | 0 |
| NLQSEVEG | VAMP8; VAMP8 | Q9BV40; B8/ | VAMP8_HUMAI | Unmodified | 1743600 | 2144100 | 1396100 | 2198200 | 1226200 | 287330 | 0 | 113740 | 6580900 | 1285300 | 4451600 | 11357000 | 1538600 | 446910 | 1283800 | 1560000 | 0 | 507490 |
| NLQSQSSV | KIAA0319L; KIA | Q8IZA0; C9J | K319L_HUMAN | Unmodified | 0 | 0 | 0 | 0 | 0 | 0 | 0 | 0 | 1359100 | 0 | 0 | 0 | 0 | 0 | 0 | 0 | 0 | 0 |
| NLQSQSSV | KIAA0319L; KIA | Q8IZA0; C9J | K319L_HUMAN | Unmodified | 0 | 0 | 0 | 0 | 0 | 0 | 0 | 0 | 4772400 | 568880 | 0 | 0 | 0 | 0 | 0 | 0 | 0 | 0 |
| NLQSVVLS | MSH2; MSH2; N | P43246; A0A | MSH2_HUMAN | Unmodified | 0 | 0 | 0 | 0 | 21127 | 0 | 0 | 0 | 0 | 0 | 0 | 943150 | 0 | 0 | 0 | 0 | 0 | 0 |
| NLQTKNPN | MYO1B; MYO1E | O43795; H0\ | MYO1B_HUMAN | Unmodified | 28812 | 0 | 0 | 0 | 5494000 | 3022800 | 565720 | 475830 | 331140 | 485240 | 0 | 0 | 312570 | 0 | 0 | 0 | 1463100 | 2104300 |
| NLQVEYEQ | UTRN; UTRN | P46939; Q5T | UTRN_HUMAN | Unmodified | 0 | 0 | 267810 | 0 | 0 | 0 | 0 | 0 | 497280 | 0 | 79199 | 0 | 0 | 0 | 0 | 395240 | 0 | 0 |

| | | | | | | | | | | | | | | | | | | | | | | |
|---|---|---|---|---|---|---|---|---|---|---|---|---|---|---|---|---|---|---|---|---|---|---|
| NLQVKCAC | NAP1L1; NAP1L | P55209; F8V | NP1L1_HUMAN | Unmodified | 0 | 0 | 0 | 0 | 0 | 0 | 113380 | 96877 | 0 | 0 | 0 | 0 | 0 | 0 | 203780 | 0 | 364920 | 0 |
| NLQYYDISA | RAN; RAN; RAN | P62826; J3K | RAN_HUMAN G | Unmodified | 2282300 | 3844400 | 12142000 | 0 | 31667000 | 8204700 | 20490000 | 11930000 | 7581300 | 6371900 | 16485000 | 1.05E+08 | 16633000 | 18355000 | 10744000 | 3752700 | 7567300 | 20401000 |
| NLRDIDEVS | CCT8; CCT8 | P50990; H7C | TCPQ_HUMAN | Unmodified | 225520 | 0 | 315570 | 1273700 | 3683700 | 885010 | 637760 | 0 | 1998700 | 1908000 | 2493600 | 6304100 | 6472800 | 2295400 | 779580 | 460190 | 1594600 | 2392300 |
| NLRDIDEVS | CCT8; CCT8 | P50990; H7C | TCPQ_HUMAN | Oxidation (M) | 0 | 0 | 0 | 0 | 5461700 | 0 | 8733000 | 0 | 0 | 197510 | 0 | 0 | 3071900 | 4298900 | 0 | 0 | 5391000 | 2644800 |
| NLRDIDEVS | CCT8; CCT8 | P50990; H7C | TCPQ_HUMAN | Unmodified | 0 | 0 | 0 | 0 | 1239600 | 109450 | 565180 | 0 | 0 | 166120 | 0 | 0 | 950130 | 739170 | 0 | 0 | 1041200 | 2934800 |
| NLREDGEK | GNAI2; GNAI1; | P04899; P63 | GNAI2_HUMAN | Unmodified | 49446 | 0 | 0 | 32619 | 690560 | 0 | 0 | 0 | 0 | 249680 | 0 | 21650 | 180190 | 114370 | 90795 | 42350 | 741000 | 1642200 |
| NLRERFGIE | PIP4K2A | P48426 | PI42A_HUMAN | Unmodified | 0 | 0 | 0 | 0 | 645950 | 0 | 0 | 0 | 0 | 0 | 0 | 0 | 365710 | 0 | 0 | 0 | 0 | 0 |
| NLRNLAED | LLGL1 | Q15334 | L2GL1_HUMAN | Unmodified | 0 | 0 | 0 | 0 | 0 | 0 | 0 | 0 | 0 | 0 | 0 | 0 | 0 | 0 | 0 | 0 | 978950 | 1435400 |
| NLRPDDVF | MGAT3 | Q09327 | MGAT3_HUMA | Unmodified | 0 | 0 | 0 | 0 | 0 | 0 | 0 | 0 | 1407000 | 0 | 0 | 0 | 0 | 0 | 0 | 0 | 0 | 0 |
| NLRVGSHI | ATP6V1A; ATP6 | P38606; C9J | VATA_HUMAN | Unmodified | 0 | 0 | 0 | 0 | 649400 | 0 | 1347300 | 0 | 0 | 594110 | 0 | 0 | 2312200 | 1423300 | 0 | 0 | 875770 | 0 |
| NLRVNAGD | ALDH6A1; ALD | Q02252; G3 | MMSA_HUMAN | Unmodified | 0 | 0 | 0 | 0 | 0 | 0 | 342130 | 93441 | 1940400 | 722480 | 0 | 0 | 217350 | 250600 | 0 | 0 | 0 | 0 |
| NLSADQW | NCKAP1 | Q9Y2A7 | NCKP1_HUMA | Unmodified | 0 | 0 | 0 | 413170 | 0 | 0 | 0 | 0 | 0 | 0 | 0 | 1941200 | 0 | 0 | 0 | 527370 | 0 | 0 |
| NLSALENY | NRP1; NRP1; N | O14786; E9F | NRP1_HUMAN | Unmodified | 0 | 0 | 0 | 454820 | 0 | 0 | 0 | 0 | 1301700 | 0 | 0 | 0 | 401380 | 0 | 0 | 819080 | 0 | 0 |
| NLSDVATK | JUP | P14923 | PLAK_HUMAN | Unmodified | 0 | 0 | 0 | 0 | 0 | 0 | 95680 | 77431 | 0 | 0 | 0 | 1262400 | 0 | 0 | 0 | 0 | 0 | 0 |
| NLSDVATK | JUP | P14923 | PLAK_HUMAN | Unmodified | 1395400 | 1790000 | 2664600 | 20911000 | 13201000 | 5508900 | 0 | 0 | 9001800 | 1466500 | 2999400 | 17501000 | 7298900 | 3502300 | 9988900 | 15761000 | 1537500 | 2319900 |
| NLSEIENVI | DLG1; DLG1; D | Q12959; A0A | DLG1_HUMAN | Unmodified | 0 | 0 | 1223900 | 2590700 | 9746400 | 714060 | 0 | 0 | 6656700 | 5427700 | 3274900 | 1784400 | 5529800 | 7256900 | 1987900 | 2125300 | 3869000 | 16856000 |
| NLSGQPNF | TM9SF3 | Q9HD45 | TM9S3_HUMAN | Unmodified | 0 | 0 | 0 | 0 | 0 | 0 | 0 | 0 | 243970 | 104890 | 0 | 908780 | 0 | 0 | 0 | 0 | 0 | 0 |
| NLSLQNEI | PKP2 | Q99959 | PKP2_HUMAN | Unmodified | 0 | 737200 | 191520 | 0 | 0 | 0 | 159920 | 0 | 294910 | 0 | 0 | 2790100 | 0 | 0 | 230890 | 0 | 0 | 429830 |
| NLSMPDLE | SNTB2; SNTB2; | Q13425; H7 | SNTB2_HUMAN | Unmodified | 0 | 0 | 0 | 385640 | 0 | 0 | 0 | 0 | 0 | 0 | 0 | 0 | 0 | 0 | 0 | 0 | 0 | 0 |
| NLSPGAVE | COPA; COPA; C | P53621; A0A | COPA_HUMAN | Unmodified | 0 | 0 | 194010 | 0 | 0 | 0 | 0 | 0 | 392890 | 0 | 327080 | 4460900 | 0 | 0 | 0 | 180130 | 0 | 0 |
| NLSPVVSN | PSPC1; PSPC1 | Q8WXF1; X6 | PSPC1_HUMAN | Unmodified | 0 | 0 | 0 | 0 | 2327500 | 197940 | 918890 | 0 | 257770 | 663850 | 325490 | 595870 | 931020 | 386200 | 0 | 0 | 645080 | 1024200 |
| NLSRIHPVS | LDHB; LDHB; N | P07195; A0A | LDHB_HUMAN | Unmodified | 0 | 0 | 0 | 0 | 2667000 | 0 | 0 | 0 | 0 | 0 | 0 | 0 | 0 | 0 | 0 | 0 | 1792300 | 592520 |
| NLSRIHPVS | LDHB; LDHB; N | P07195; A0A | LDHB_HUMAN | Oxidation (M) | 0 | 0 | 0 | 0 | 0 | 943360 | 157710 | 110390 | 0 | 0 | 0 | 0 | 0 | 0 | 0 | 0 | 713050 | 250030 |
| NLSSASQA | PKP3 | Q9Y446 | PKP3_HUMAN | Unmodified | 0 | 12361 | 0 | 2201900 | 0 | 0 | 0 | 0 | 0 | 0 | 0 | 0 | 0 | 0 | 2662500 | 0 | 0 | 0 |
| NLSSASQA | PKP3 | Q9Y446 | PKP3_HUMAN | Unmodified | 0 | 0 | 0 | 0 | 285170 | 0 | 53645 | 0 | 16353 | 0 | 45645 | 0 | 678300 | 0 | 0 | 0 | 357250 | 0 |
| NLSSNEAIS | PRKDC | P78527 | PRKDC_HUMA | Unmodified | 0 | 0 | 0 | 0 | 0 | 0 | 0 | 0 | 0 | 0 | 0 | 1078000 | 0 | 0 | 0 | 0 | 0 | 0 |
| NLSSPFIFH | SLC7A2; SLC7A | P52569; A0A | CTR2_HUMAN | Unmodified | 0 | 113130 | 0 | 428610 | 0 | 40141 | 0 | 0 | 0 | 309510 | 0 | 1236500 | 447310 | 0 | 225010 | 134370 | 50745 | 0 |
| NLSSTTDD | LAD1; LAD1; LA | O00515; A0A | LAD1_HUMAN | Unmodified | 0 | 0 | 0 | 0 | 0 | 226140 | 0 | 0 | 0 | 0 | 309120 | 633610 | 0 | 0 | 0 | 0 | 0 | 0 |
| NLSYGRDT | ARVCF; ARVCF; | O00192; E9F | ARVC_HUMAN | Unmodified | 0 | 117750 | 0 | 794550 | 0 | 0 | 0 | 0 | 273890 | 0 | 101130 | 51276 | 0 | 223570 | 1090100 | 1220700 | 0 | 0 |
| NLSYQVHF | CTNND1; CTNN | O60716; A0A | CTND1_HUMA | Unmodified | 248280 | 236570 | 561470 | 142140 | 0 | 0 | 0 | 0 | 30622 | 0 | 220370 | 1297000 | 0 | 0 | 1368900 | 1380300 | 0 | 0 |
| NLSYQVHF | CTNND1; CTNN | O60716; A0A | CTND1_HUMA | Unmodified | 58977 | 64611 | 192570 | 501310 | 300030 | 111840 | 0 | 0 | 0 | 0 | 301960 | 165700 | 0 | 0 | 296650 | 306270 | 173020 | 160340 |
| NLSYQVHF | CTNND1; CTNN | O60716; A0A | CTND1_HUMA | Unmodified | 0 | 0 | 0 | 570600 | 9846800 | 2908300 | 0 | 0 | 0 | 0 | 0 | 0 | 6290000 | 10092000 | 0 | 0 | 6897100 | 18231000 |
| NLTALGLN | ATIC; ATIC | P31939; F8V | PUR9_HUMAN | Unmodified | 0 | 0 | 0 | 1877000 | 0 | 0 | 0 | 129230 | 0 | 744030 | 770920 | 2270000 | 2182400 | 768380 | 0 | 0 | 667240 | 877410 |
| NLTDIKDR | N/A | N/A | N/A | Unmodified | 0 | 0 | 0 | 0 | 2606900 | 528260 | 762020 | 48940 | 0 | 57497 | 0 | 388570 | 565770 | 283830 | 0 | 0 | 1784600 | 7204000 |
| NLTEDNSC | ATXN10; ATXN1 | Q9UBB4; B1 | ATX10_HUMAN | Unmodified | 0 | 0 | 816420 | 0 | 0 | 0 | 0 | 0 | 117440 | 0 | 265790 | 1550400 | 0 | 54149 | 674350 | 870870 | 0 | 84810 |
| NLTIKTESTI | FABP5P3; FABP | A8MUU1; Q0 | FB5L3_HUMAN | Unmodified | 0 | 305650 | 0 | 0 | 17372000 | 0 | 389120 | 279930 | 141770 | 342570 | 1123500 | 961910 | 0 | 0 | 0 | 0 | 39462000 | 35302000 |
| NLTLQLEQ | ROCK1; ROCK1 | Q13464; A0A | ROCK1_HUMA | Unmodified | 0 | 60974 | 174800 | 518440 | 1616900 | 152840 | 0 | 13473 | 693100 | 97197 | 0 | 501790 | 178950 | 69613 | 0 | 209610 | 120780 | 972060 |
| NLTLQLEQ | ROCK1; ROCK1 | Q13464; A0A | ROCK1_HUMA | Unmodified | 0 | 0 | 0 | 0 | 704830 | 0 | 0 | 0 | 0 | 0 | 0 | 0 | 0 | 0 | 0 | 0 | 592110 | 1748600 |
| NLTNPNTV | RAB14; RAB14 | P61106; X6R | RAB14_HUMAN | Unmodified | 0 | 0 | 1578800 | 0 | 0 | 0 | 0 | 0 | 1457500 | 310720 | 2302600 | 6551000 | 0 | 0 | 1417200 | 783670 | 0 | 0 |
| NLTNPNTV | RAB14; RAB14 | P61106; X6R | RAB14_HUMAN | Unmodified | 0 | 0 | 0 | 0 | 0 | 0 | 0 | 0 | 0 | 0 | 569640 | 4238500 | 0 | 0 | 0 | 0 | 0 | 0 |
| NLTNPNTV | RAB14; RAB14 | P61106; X6R | RAB14_HUMAN | Unmodified | 0 | 0 | 0 | 7923900 | 0 | 0 | 0 | 0 | 11461000 | 1677600 | 3783300 | 8658000 | 0 | 0 | 0 | 0 | 0 | 0 |
| NLTPGEQI | FARP2 | H7C3M7 | H7C3M7_HUM | Unmodified | 145120 | 0 | 459480 | 941390 | 2944300 | 0 | 0 | 249480 | 459720 | 417970 | 0 | 0 | 2246100 | 2309300 | 667730 | 378130 | 1322900 | 1054300 |
| NLTQDEM | EIF3A | Q14152 | EIF3A_HUMAN | Unmodified | 0 | 0 | 147840 | 181180 | 0 | 0 | 0 | 0 | 0 | 0 | 254130 | 2411700 | 0 | 0 | 179720 | 153290 | 0 | 0 |
| NLTQDEM | EIF3A | Q14152 | EIF3A_HUMAN | Oxidation (M) | 273990 | 0 | 0 | 0 | 0 | 0 | 0 | 0 | 0 | 0 | 0 | 970950 | 0 | 0 | 0 | 0 | 0 | 0 |
| NLTQYSWL | AK3 | Q9UIJ7 | KAD3_HUMAN | Unmodified | 0 | 0 | 0 | 0 | 0 | 0 | 0 | 0 | 1071300 | 398810 | 0 | 0 | 0 | 0 | 0 | 0 | 0 | 0 |
| NLTSSSLNI | SLC4A4 | Q9Y6R1 | S4A4_HUMAN | Unmodified | 0 | 0 | 359700 | 156100 | 0 | 0 | 0 | 0 | 0 | 0 | 0 | 0 | 0 | 0 | 421100 | 1209400 | 0 | 0 |
| NLTSSSLNI | SLC4A4 | Q9Y6R1 | S4A4_HUMAN | Unmodified | 0 | 0 | 0 | 0 | 1147400 | 277930 | 0 | 0 | 0 | 0 | 0 | 0 | 0 | 0 | 0 | 0 | 61912 | 0 |
| NLTVILSDA | XRN2 | Q9H0D6 | XRN2_HUMAN | Unmodified | 0 | 0 | 0 | 0 | 45919 | 0 | 0 | 0 | 172140 | 0 | 1023200 | 3779600 | 0 | 0 | 0 | 0 | 0 | 0 |
| NLVAFVSQ | PSMG3 | Q9BT73 | PSMG3_HUMA | Unmodified | 0 | 0 | 0 | 20614 | 0 | 0 | 0 | 0 | 0 | 0 | 0 | 1098100 | 0 | 0 | 0 | 0 | 0 | 0 |
| NLVCTDLF | DDX6 | P26196 | DDX6_HUMAN | Unmodified | 0 | 6600900 | 0 | 0 | 0 | 0 | 0 | 0 | 0 | 282000 | 0 | 1567200 | 0 | 0 | 0 | 472560 | 0 | 0 |
| NLVDPFVE | DYSF; MYOF | O75923; Q9 | DYSF_HUMAN | Unmodified | 0 | 0 | 0 | 0 | 11751000 | 1377600 | 0 | 0 | 417030 | 0 | 0 | 0 | 0 | 0 | 0 | 0 | 0 | 1039900 |
| NLVDPFVE | MYOF | Q9NZM1 | MYOF_HUMAN | Unmodified | 0 | 0 | 0 | 0 | 34414000 | 3487800 | 0 | 0 | 0 | 0 | 0 | 0 | 0 | 0 | 0 | 0 | 457700 | 3828900 |
| NLVDSYVA | DNM2 | P50570 | DYN2_HUMAN | Unmodified | 0 | 0 | 0 | 0 | 0 | 0 | 0 | 0 | 0 | 0 | 0 | 0 | 0 | 0 | 0 | 1601700 | 0 | 0 |
| NLVELAELE | SNX6; SNX6 | Q9UNH7; A0 | SNX6_HUMAN | Unmodified | 0 | 0 | 0 | 0 | 0 | 533370 | 780190 | 0 | 0 | 295530 | 159840 | 1414700 | 2191900 | 816460 | 0 | 0 | 805500 | 1906200 |
| NLVEQHIQ | RPN1; RPN1 | P04843; B7Z | RPN1_HUMAN | Unmodified | 0 | 0 | 0 | 0 | 0 | 0 | 531640 | 0 | 44834000 | 5550900 | 1014700 | 786370 | 0 | 0 | 0 | 0 | 194390 | 0 |
| NLVFEDND | PKP2 | Q99959 | PKP2_HUMAN | Unmodified | 0 | 0 | 0 | 0 | 0 | 0 | 0 | 0 | 0 | 0 | 0 | 2096800 | 0 | 0 | 0 | 0 | 0 | 0 |
| NLVFSDGY | RPS6KA1; RPS6 | Q15418; E9F | KS6A1_HUMAN | Unmodified | 0 | 0 | 0 | 0 | 418360 | 0 | 0 | 452110 | 879940 | 204290 | 134980 | 1284700 | 704530 | 0 | 374630 | 0 | 476430 | 0 |
| NLVGSGSE | UXS1 | Q8NBZ7 | UXS1_HUMAN | Unmodified | 0 | 0 | 0 | 0 | 0 | 0 | 60990 | 0 | 6273900 | 641900 | 0 | 0 | 0 | 0 | 0 | 0 | 1636800 | 0 |
| NLVLHSAR | HSPG2 | P98160 | PGBM_HUMAN | Unmodified | 0 | 0 | 3979800 | 1861200 | 0 | 0 | 286260 | 0 | 829270 | 0 | 0 | 0 | 936160 | 726650 | 302050 | 1155100 | 0 | 1415500 |
| NLVLLVR | MRPL46 | Q9H2W6 | RM46_HUMAN | Unmodified | 0 | 0 | 0 | 0 | 0 | 0 | 0 | 0 | 624180 | 0 | 0 | 0 | 0 | 0 | 0 | 0 | 0 | 0 |
| NLVNDDD | UBE3A; UBE3A; | Q05086; A0A | UBE3A_HUMA | Unmodified | 0 | 0 | 0 | 889530 | 0 | 0 | 0 | 148000 | 274780 | 0 | 285650 | 1177300 | 0 | 0 | 0 | 0 | 110210 | 0 |
| NLVPDADI | PHYH; PHYH; P | O14832; B1A | PAHX_HUMAN | Unmodified | 0 | 0 | 0 | 0 | 0 | 0 | 0 | 0 | 14817000 | 0 | 0 | 0 | 0 | 0 | 0 | 0 | 0 | 0 |
| NLVPDADI | PHYH; PHYH; P | O14832; B1A | PAHX_HUMAN | Unmodified | 0 | 0 | 0 | 0 | 0 | 0 | 498750 | 0 | 1645500 | 11169000 | 0 | 0 | 77774 | 0 | 0 | 0 | 0 | 0 |
| NLVPEEAD | CHAC2 | Q8WUX2 | CHAC2_HUMA | Unmodified | 0 | 0 | 0 | 0 | 0 | 0 | 98204 | 0 | 166190 | 118660 | 0 | 830560 | 0 | 0 | 0 | 0 | 50688 | 0 |
| NLVPGESV | FBL; FBL; FBL; F | P22087; M0 | FBRL_HUMAN | Unmodified | 0 | 129540 | 105500 | 0 | 0 | 0 | 1105200 | 547450 | 0 | 0 | 0 | 525140 | 351550 | 373580 | 0 | 0 | 103300 | 0 |
| NLVPLLLAF | ATL2; ATL2; ATL | Q8NHH9; B5 | ATLA2_HUMAN | Unmodified | 0 | 0 | 26379 | 0 | 0 | 0 | 0 | 0 | 0 | 0 | 329820 | 484530 | 40952 | 0 | 0 | 20248 | 0 | 0 |
| NLVPMDPN | PRKCB | P05771 | KPCB_HUMAN | Oxidation (M) | 0 | 0 | 0 | 0 | 1020900 | 0 | 565300 | 307180 | 0 | 242030 | 0 | 0 | 905810 | 0 | 0 | 301240 | 0 | 0 |
| NLVTEDVM | GPI; GPI; GPI; G | P06744; K7E | G6PI_HUMAN ( | Unmodified | 389800 | 429480 | 653820 | 1358000 | 3791900 | 626630 | 0 | 70913 | 8405500 | 0 | 7967000 | 38628000 | 0 | 0 | 1173900 | 976140 | 537730 | 0 |
| NLVTEDVM | GPI; GPI; GPI; G | P06744; K7E | G6PI_HUMAN ( | Oxidation (M) | 0 | 0 | 1080900 | 1161900 | 1720800 | 0 | 0 | 0 | 306680 | 1714200 | 4946000 | 21646000 | 718150 | 0 | 796790 | 955990 | 0 | 0 |
| NLVTEDVM | GPI; GPI; GPI; G | P06744; K7E | G6PI_HUMAN ( | Oxidation (M) | 0 | 0 | 0 | 0 | 3641100 | 24526 | 696980 | 0 | 0 | 1772700 | 0 | 0 | 1454000 | 1227300 | 0 | 0 | 551170 | 54691 |
| NLVTEDVM | GPI; GPI; GPI; G | P06744; K7E | G6PI_HUMAN ( | 2 Oxidation (M) | 0 | 0 | 0 | 0 | 715380 | 102680 | 119590 | 12070 | 0 | 741330 | 0 | 0 | 401280 | 0 | 0 | 0 | 0 | 0 |
| NLVTGDHI | ENSA; ENSA; EN | O43768; A0A | ENSA_HUMAN | Unmodified | 0 | 0 | 0 | 0 | 0 | 0 | 0 | 0 | 0 | 0 | 419250 | 0 | 0 | 0 | 0 | 0 | 0 | 0 |
| NLVTGDHI | ENSA; ENSA; EN | O43768; A0A | ENSA_HUMAN | Unmodified | 1708500 | 1348300 | 721710 | 1613600 | 9719500 | 1395000 | 1782700 | 1514400 | 3263700 | 1691700 | 1034000 | 1880500 | 8339700 | 8074700 | 921510 | 1512900 | 6211600 | 13382000 |
| NLYAHIQK | PTPRF; PTPRF; | P10586; H0 | PTPRF_HUMAN | Unmodified | 258800 | 216730 | 1000600 | 553280 | 50783 | 0 | 288340 | 0 | 0 | 0 | 0 | 0 | 0 | 0 | 1011000 | 885390 | 0 | 0 |
| NLYDLYTR | EXOC5; EXOC5 | O00471; F8 | EXOC5_HUMA | Unmodified | 0 | 0 | 0 | 0 | 416050 | 152360 | 0 | 0 | 0 | 321100 | 0 | 0 | 88770 | 0 | 0 | 0 | 119790 | 318190 |
| NLYGTSIDS | PALS1 | Q8N3R9 | PALS1_HUMAN | Unmodified | 0 | 0 | 0 | 0 | 3641100 | 1822400 | 0 | 0 | 0 | 345740 | 0 | 0 | 1095700 | 1303600 | 0 | 0 | 689120 | 2595200 |
| NLYHNLCT | MCM6 | Q14566 | MCM6_HUMAN | Unmodified | 0 | 0 | 0 | 0 | 0 | 172340 | 0 | 0 | 413080 | 0 | 381680 | 1474500 | 0 | 0 | 0 | 0 | 414100 | 0 |
| NLYIISVKG | RPL23; RPL23; | P62829; J3K | RL23_HUMAN ( | Unmodified | 0 | 0 | 0 | 0 | 0 | 0 | 1236600 | 454830 | 0 | 0 | 0 | 0 | 129910 | 132050 | 0 | 0 | 0 | 0 |
| NLYPSSSP | SUGT1 | Q9Y2Z0 | SGT1_HUMAN | Unmodified | 0 | 0 | 7356600 | 0 | 725730 | 0 | 0 | 98384 | 645220 | 325840 | 157880 | 601470 | 0 | 0 | 0 | 10648000 | 0 | 449780 |
| NLYPSSSP | SUGT1 | Q9Y2Z0 | SGT1_HUMAN | Unmodified | 0 | 0 | 0 | 0 | 1574300 | 0 | 84085 | 188780 | 0 | 145890 | 0 | 0 | 0 | 0 | 0 | 0 | 466580 | 0 |
| NLYRVGFE | MIB1 | Q86YT6 | MIB1_HUMAN | Unmodified | 0 | 0 | 0 | 0 | 696380 | 59088 | 373920 | 0 | 0 | 0 | 0 | 0 | 120710 | 112230 | 0 | 0 | 160850 | 346430 |
| NLYSDDIP | EIF3E; EIF3E; EI | P60228; A0A | EIF3E_HUMAN | Unmodified | 0 | 134980 | 0 | 0 | 0 | 0 | 60626 | 48969 | 0 | 0 | 0 | 3011600 | 0 | 0 | 468300 | 0 | 0 | 0 |
| NLYSDDIP | EIF3E; EIF3E; EI | P60228; A0A | EIF3E_HUMAN | Unmodified | 341840 | 513760 | 571770 | 0 | 0 | 200090 | 0 | 211160 | 1069400 | 285590 | 0 | 6434600 | 2206900 | 617060 | 1252000 | 0 | 271240 | 604770 |
| NLYVTFPIP | IDE; IDE; IDE; ID | P14735; A0A | IDE_HUMAN In | Unmodified | 0 | 0 | 0 | 0 | 0 | 0 | 0 | 0 | 0 | 0 | 0 | 544900 | 0 | 0 | 0 | 0 | 0 | 0 |
| NMACVQR | TAGLN2; TAGLN | P37802; X6R | TAGL2_HUMAN | Unmodified | 2980400 | 1113000 | 267620 | 1560000 | 10819000 | 6668400 | 1026700 | 1225600 | 817220 | 1488000 | 123500 | 3804700 | 3119900 | 1560000 | 0 | 158600 | 5411400 | 6614100 |
| NMACVQR | TAGLN2; TAGLN | P37802; X6R | TAGL2_HUMAN | Oxidation (M) | 0 | 0 | 0 | 0 | 586850 | 345940 | 0 | 125760 | 157830 | 0 | 0 | 1042700 | 313910 | 150730 | 0 | 0 | 50027 | 460140 |
| NMADAMR | ACLY | P53396 | ACLY_HUMAN | Unmodified | 0 | 0 | 0 | 0 | 53153 | 20044 | 67156 | 0 | 79820 | 36392 | 45522 | 1000000 | 0 | 0 | 0 | 0 | 17401 | 0 |
| NMAEQIIQ | TKT; TKT; TKT | P29401; A0A | TKT_HUMAN Tr | Oxidation (M) | 0 | 45493 | 0 | 54370 | 3210700 | 274490 | 876470 | 0 | 138010 | 1234400 | 34307 | 323780 | 1549700 | 272800 | 0 | 0 | 1223300 | 1520300 |
| NMAEQIIQ | TKT; TKT; TKT | P29401; A0A | TKT_HUMAN Tr | Unmodified | 0 | 0 | 0 | 0 | 71647 | 0 | 0 | 0 | 0 | 0 | 0 | 0 | 0 | 0 | 0 | 0 | 0 | 21481 |
| NMAEQIIQ | TKT; TKT; TKT | P29401; A0A | TKT_HUMAN Tr | Oxidation (M) | 0 | 0 | 0 | 0 | 554430 | 0 | 146640 | 0 | 0 | 0 | 0 | 0 | 184550 | 0 | 0 | 0 | 742660 | 0 |
| NMAGYSTF | NOSTRIN; NOS | Q8IVI9; A0A | NOSTN_HUMA | Unmodified | 1780100 | 1185200 | 2586500 | 4336900 | 0 | 0 | 0 | 0 | 0 | 0 | 0 | 0 | 672660 | 292780 | 5696100 | 2255700 | 0 | 0 |
| NMAILDPP | CCAR1; CCAR1 | Q8IX12; F5H | CCAR1_HUMA | Oxidation (M) | 0 | 0 | 0 | 0 | 0 | 0 | 275950 | 238460 | 0 | 0 | 0 | 2959900 | 0 | 76879 | 0 | 0 | 0 | 0 |
| NMAIQQEL | STAMBP; STAM | O95630; C9J | STABP_HUMAN | Oxidation (M) | 203290 | 162840 | 75171 | 0 | 0 | 0 | 78203 | 0 | 0 | 0 | 0 | 0 | 0 | 0 | 0 | 91334 | 0 | 0 |
| NMAPSQQ | ABI2; ABI2; ABI2 | Q9NYB9; H7 | ABI2_HUMAN A | Unmodified | 69862 | 34378 | 0 | 0 | 0 | 0 | 0 | 0 | 0 | 0 | 63389 | 379340 | 0 | 0 | 0 | 0 | 0 | 0 |
| NMAQFSVL | MTAP; MTAP | Q13126; B4 | MTAP_HUMAN | Oxidation (M) | 0 | 0 | 0 | 0 | 0 | 0 | 0 | 204390 | 168430 | 0 | 0 | 999420 | 535930 | 0 | 0 | 0 | 1074100 | 0 |
| NMAQFSVL | MTAP; MTAP | Q13126; B4 | MTAP_HUMAN | Unmodified | 0 | 0 | 0 | 0 | 1156500 | 0 | 0 | 0 | 0 | 0 | 0 | 0 | 0 | 0 | 0 | 0 | 486640 | 0 |
| NMDPLNDI | MYH9 | P35579 | MYH9_HUMAN | Oxidation (M) | 0 | 738010 | 5638700 | 1770700 | 2047500 | 0 | 161870 | 71909 | 262720 | 612610 | 3074200 | 17926000 | 222330 | 0 | 4773300 | 2894600 | 0 | 0 |
| NMDPLNDI | MYH9 | P35579 | MYH9_HUMAN | Oxidation (M) | 0 | 0 | 0 | 345410 | 0 | 0 | 0 | 0 | 0 | 0 | 0 | 0 | 421740 | 0 | 0 | 0 | 0 | 0 |
| NMDPLNDI | MYH10 | P35580 | MYH10_HUMA | Oxidation (M) | 0 | 0 | 0 | 83874 | 0 | 0 | 0 | 0 | 0 | 0 | 0 | 157770 | 0 | 0 | 596340 | 0 | 0 | 0 |
| NMDSMPEI | UPF1 | Q92900 | RENT1_HUMAN | Unmodified | 0 | 0 | 0 | 0 | 0 | 0 | 0 | 0 | 0 | 0 | 0 | 0 | 0 | 0 | 0 | 0 | 0 | 0 |
| NMDSMPEI | UPF1 | Q92900 | RENT1_HUMAN | 2 Oxidation (M) | 252520 | 208030 | 0 | 49683 | 0 | 0 | 0 | 0 | 0 | 0 | 0 | 0 | 0 | 0 | 42179 | 104470 | 0 | 0 |
| NMERLNLF | DOCK10; DOCK | Q96BY6; A0A | DOC10_HUMA | Oxidation (M) | 0 | 0 | 0 | 0 | 2152000 | 0 | 31395 | 0 | 0 | 0 | 0 | 0 | 0 | 0 | 0 | 0 | 0 | 584440 |
| NMFLVGEC | PSIP1 | O75475 | PSIP1_HUMAN | Oxidation (M) | 239140 | 0 | 0 | 0 | 0 | 0 | 642140 | 45496 | 0 | 210580 | 45764 | 234440 | 182370 | 0 | 0 | 0 | 0 | 0 |
| NMFTINR | CDH1; CDH1; C | P12830; H3E | CADH1_HUMA | Unmodified | 86678 | 133490 | 0 | 120400 | 0 | 914710 | 0 | 0 | 865370 | 0 | 0 | 0 | 0 | 0 | 173210 | 72902 | 0 | 0 |
| NMFTNQDI | FKBP9; FKBP9P | O95302; Q7 | FKBP9_HUMAN | Unmodified | 0 | 0 | 0 | 0 | 0 | 0 | 0 | 10493 | 5921300 | 1716400 | 214900 | 0 | 0 | 0 | 1210100 | 0 | 0 | 0 |
| NMFTNQDI | FKBP9; FKBP9P | O95302; Q7 | FKBP9_HUMAN | Oxidation (M) | 0 | 0 | 0 | 0 | 0 | 0 | 0 | 0 | 737720 | 219490 | 177720 | 0 | 0 | 0 | 0 | 0 | 0 | 0 |
| NMGGPYG | HNRNPA2B1; H | P22626; A0A | ROA2_HUMAN | Oxidation (M) | 538690 | 0 | 0 | 0 | 0 | 0 | 76129 | 432070 | 471750 | 314790 | 278610 | 297180 | 242420 | 0 | 0 | 0 | 0 | 0 |
| NMGGPYG | HNRNPA2B1; H | P22626; A0A | ROA2_HUMAN | Unmodified | 137300 | 0 | 0 | 0 | 575750 | 0 | 170860 | 233040 | 671800 | 574440 | 0 | 0 | 0 | 0 | 0 | 0 | 0 | 0 |
| NMGMDFC | PLOD2; PLOD2 | O00469; E7E | PLOD2_HUMA | Unmodified | 0 | 0 | 0 | 0 | 0 | 0 | 0 | 0 | 621190 | 32680 | 187720 | 117810 | 0 | 0 | 0 | 0 | 0 | 0 |

| | | | | | | | | | | | | | | | | | | | | | | |
|---|---|---|---|---|---|---|---|---|---|---|---|---|---|---|---|---|---|---|---|---|---|---|
| NMGVIAER | ANO6; ANO6; A | Q4KMQ2; A0 | ANO6_HUMAN | Unmodified | 40362 | 0 | 0 | 786970 | 21266 | 2201100 | 83686 | 0 | 252640 | 304780 | 117980 | 919910 | 109380 | 0 | 175550 | 124340 | 0 | 0 |
| NMGYVQQ | BUB3; BUB3 | O43684; J3Q | BUB3_HUMAN | Unmodified | 0 | 0 | 0 | 51941 | 0 | 0 | 0 | 0 | 0 | 0 | 0 | 1883300 | 0 | 0 | 0 | 0 | 0 | 0 |
| NMHWYSP | CLDND1; CLDN | Q9NY35; D6 | CLDN1_HUMA | Oxidation (M) | 1103200 | 971180 | 518190 | 397540 | 0 | 0 | 0 | 0 | 0 | 0 | 0 | 130450 | 276930 | 0 | 234020 | 359900 | 0 | 0 |
| NMIHETNE | BLOC1S5; BLO | Q8TDH9; G5 | BL1S5_HUMAN | Unmodified | 0 | 0 | 0 | 0 | 0 | 42770 | 0 | 0 | 90340 | 54052 | 0 | 389380 | 0 | 0 | 0 | 0 | 0 | 27618 |
| NMIKEVDE | EFHD2; EFHD2 | Q96C19; H0 | EFHD2_HUMA | Unmodified | 0 | 0 | 0 | 0 | 1067900 | 0 | 0 | 0 | 0 | 0 | 0 | 0 | 0 | 0 | 0 | 0 | 0 | 0 |
| NMINIISRL | RARS1; RARS1; | P54136; E5R | SYRC_HUMAN | Oxidation (M) | 0 | 0 | 0 | 0 | 20400 | 0 | 0 | 0 | 0 | 1540700 | 0 | 0 | 4480700 | 0 | 0 | 0 | 805970 | 1427900 |
| NMINIISRL | RARS1; RARS1; | P54136; E5R | SYRC_HUMAN | Unmodified | 0 | 0 | 0 | 0 | 552720 | 48023 | 258860 | 0 | 0 | 156330 | 0 | 0 | 0 | 378300 | 0 | 0 | 118160 | 515760 |
| NMIREQDL | USO1 | O60763 | USO1_HUMAN | Oxidation (M) | 0 | 0 | 0 | 0 | 0 | 0 | 0 | 0 | 0 | 565720 | 0 | 1152800 | 1459000 | 0 | 0 | 0 | 0 | 0 |
| NMITGTSQ | EEF1A2; EEF1A | Q05639; Q5 | EF1A2_HUMAN | Oxidation (M) | 0 | 158550 | 0 | 0 | 98144 | 0 | 23314 | 0 | 0 | 0 | 0 | 0 | 224770 | 103960 | 202650 | 0 | 0 | 0 |
| NMKEIETNI | UTRN | P46939 | UTRN_HUMAN | Unmodified | 0 | 0 | 0 | 0 | 519480 | 0 | 0 | 0 | 0 | 0 | 0 | 0 | 0 | 0 | 0 | 0 | 510250 | 0 |
| NMLFSGTN | ATP2A1; ATP2A | O14983; P16 | AT2A1_HUMAN | Oxidation (M) | 0 | 0 | 0 | 0 | 655210 | 0 | 514560 | 278290 | 0 | 0 | 0 | 0 | 0 | 0 | 0 | 0 | 317100 | 0 |
| NMLIRTHM | SEPTIN7; SEPTI | Q16181; E7E | SEPT7_HUMAN | Unmodified | 146440 | 0 | 148730 | 334310 | 1005400 | 0 | 0 | 0 | 234480 | 107490 | 197930 | 0 | 135630 | 114650 | 376750 | 111080 | 90125 | 0 |
| NMMAACD | TUBB2A; TUBB | Q13885; A6N | TBB2A_HUMAN | Unmodified | 675040 | 364140 | 0 | 846280 | 2080700 | 0 | 1991900 | 660590 | 11102000 | 1236900 | 554770 | 45016000 | 1247200 | 0 | 1076900 | 125290 | 582360 | 2772600 |
| NMMAACD | TUBB2A; TUBB | Q13885; A6N | TBB2A_HUMAN | Oxidation (M) | 443860 | 260900 | 635090 | 1092900 | 685660 | 0 | 707930 | 188800 | 2981600 | 495190 | 1618700 | 22281000 | 2110800 | 0 | 1049000 | 787640 | 160290 | 1706800 |
| NMMAACD | TUBB2A; TUBB | Q13885; A6N | TBB2A_HUMAN | 2 Oxidation (M) | 51442 | 26368 | 98237 | 0 | 60701 | 0 | 0 | 41697 | 156720 | 49157 | 122630 | 1309000 | 72788 | 0 | 0 | 0 | 25955 | 161630 |
| NMMAACD | TUBB2A; TUBB4 | Q13885; P04 | TBB2A_HUMAN | Unmodified | 0 | 0 | 0 | 0 | 3096500 | 756710 | 2840900 | 1265000 | 114770 | 1857400 | 465710 | 252430 | 2203800 | 772940 | 95864 | 59442 | 709630 | 1796700 |
| NMPLQHLI | DSP | P15924 | DESP_HUMAN | Oxidation (M) | 0 | 0 | 66339 | 39058 | 0 | 0 | 0 | 0 | 234900 | 172580 | 0 | 184970 | 0 | 0 | 0 | 0 | 0 | 27601 |
| NMPNKQPI | SLC30A1 | Q9Y6M5 | ZNT1_HUMAN | Unmodified | 0 | 96310 | 0 | 0 | 2199400 | 0 | 0 | 141810 | 417570 | 0 | 0 | 997450 | 921210 | 0 | 171180 | 179110 | 721090 | 1908000 |
| NMQNVEH | PSMB1 | P20618 | PSB1_HUMAN | Unmodified | 210270 | 184650 | 379530 | 296470 | 2126600 | 265650 | 0 | 39768 | 4738300 | 880550 | 806160 | 7044400 | 734360 | 358980 | 244140 | 189770 | 0 | 957450 |
| NMQNVEH | PSMB1 | P20618 | PSB1_HUMAN | Oxidation (M) | 0 | 0 | 255320 | 0 | 0 | 0 | 0 | 0 | 651040 | 0 | 0 | 5605200 | 0 | 0 | 212570 | 314530 | 0 | 0 |
| NMQNVEH | PSMB1 | P20618 | PSB1_HUMAN | Oxidation (M) | 0 | 0 | 0 | 0 | 965800 | 531770 | 194430 | 39771 | 0 | 471550 | 295980 | 0 | 2750200 | 0 | 0 | 0 | 420460 | 572670 |
| NMQNVEH | PSMB1 | P20618 | PSB1_HUMAN | Unmodified | 0 | 0 | 0 | 0 | 1673800 | 190700 | 0 | 0 | 0 | 436210 | 0 | 0 | 1021600 | 0 | 0 | 0 | 0 | 1016300 |
| NMSVHLSF | RPS11; RPS11; | P62280; M0F | RS11_HUMAN | Oxidation (M) | 45461 | 300750 | 408510 | 493310 | 0 | 0 | 0 | 0 | 0 | 0 | 272510 | 6713300 | 0 | 0 | 0 | 415200 | 0 | 0 |
| NMSVHLSF | RPS11; RPS11; | P62280; M0F | RS11_HUMAN | Unmodified | 756430 | 0 | 334090 | 445380 | 0 | 272130 | 0 | 17876 | 0 | 0 | 598690 | 16546000 | 0 | 0 | 960110 | 0 | 0 | 0 |
| NMSVIAHV | EEF2 | P13639 | EF2_HUMAN EI | Unmodified | 1691800 | 2419600 | 3106600 | 3649400 | 5555500 | 1850400 | 383200 | 206090 | 3805800 | 805200 | 9181300 | 55216000 | 1175300 | 1207300 | 2376700 | 2910600 | 1160400 | 3318600 |
| NMSVIAHV | EEF2 | P13639 | EF2_HUMAN EI | Oxidation (M) | 1532000 | 1312900 | 1860000 | 2427600 | 1547800 | 428680 | 159290 | 108800 | 889000 | 0 | 3398600 | 19845000 | 0 | 189770 | 1865400 | 2426700 | 0 | 546170 |
| NMSYQGF | LARS1; LARS1; | Q9P2J5; A0A | SYLC_HUMAN | Oxidation (M) | 0 | 0 | 0 | 0 | 252110 | 0 | 0 | 0 | 72772 | 0 | 0 | 1094100 | 0 | 0 | 0 | 0 | 57258 | 0 |
| NMSYQGF | LARS1; LARS1; | Q9P2J5; A0A | SYLC_HUMAN | Unmodified | 0 | 0 | 0 | 0 | 406620 | 0 | 127040 | 47841 | 235130 | 168480 | 0 | 3511900 | 287650 | 0 | 0 | 0 | 80411 | 225150 |
| NMVHPNV | SQSTM1; SQSTI | Q13501; E9F | SQSTM_HUMA | Oxidation (M) | 0 | 0 | 349460 | 0 | 0 | 0 | 0 | 0 | 583260 | 0 | 0 | 0 | 0 | 0 | 195690 | 0 | 0 | 0 |
| NMVHPNV | SQSTM1; SQSTI | Q13501; E9F | SQSTM_HUMA | Unmodified | 114710 | 92256 | 0 | 217820 | 0 | 0 | 0 | 0 | 1295400 | 0 | 0 | 0 | 0 | 0 | 0 | 0 | 0 | 0 |
| NMVHPNV | SQSTM1; SQSTI | Q13501; E9F | SQSTM_HUMA | Oxidation (M) | 0 | 0 | 0 | 0 | 0 | 0 | 0 | 0 | 0 | 0 | 0 | 0 | 713660 | 0 | 0 | 0 | 398040 | 659710 |
| NMVHPNV | SQSTM1; SQSTI | Q13501; E9F | SQSTM_HUMA | Unmodified | 0 | 0 | 0 | 0 | 0 | 0 | 0 | 0 | 0 | 0 | 0 | 0 | 0 | 0 | 0 | 0 | 939330 | 1547300 |
| NMVLFTR | THYN1 | Q9P016 | THYN1_HUMA | Unmodified | 0 | 0 | 0 | 0 | 0 | 88966 | 0 | 0 | 0 | 0 | 178340 | 686410 | 130990 | 0 | 0 | 0 | 0 | 0 |
| NMVPQQA | ATP1A1; ATP1A | P05023; P50 | AT1A1_HUMAN | Oxidation (M) | 0 | 0 | 7777600 | 3141600 | 0 | 0 | 0 | 0 | 1884500 | 835060 | 3373000 | 30911000 | 402710 | 112340 | 7726500 | 8230100 | 0 | 0 |
| NMVPQQA | ATP1A1; ATP1A | P05023; P50 | AT1A1_HUMAN | Unmodified | 0 | 504160 | 2440600 | 2607400 | 851850 | 0 | 0 | 0 | 7217000 | 1090100 | 1529800 | 28937000 | 524390 | 142520 | 5319800 | 3319400 | 0 | 0 |
| NMVPQQA | ATP1A2; ATP1A | P50993; P13 | AT1A2_HUMAN | Unmodified | 0 | 0 | 0 | 0 | 656460 | 0 | 0 | 0 | 990490 | 0 | 0 | 0 | 0 | 0 | 0 | 0 | 349460 | 0 |
| NMVPQQA | ATP1A1 | P05023 | AT1A1_HUMAN | Unmodified | 0 | 0 | 497320 | 1185900 | 12120000 | 1134500 | 1301900 | 0 | 2248400 | 1075600 | 3268900 | 0 | 6026300 | 2979400 | 659930 | 239620 | 775260 | 1276400 |
| NMVPQQA | ATP1A1 | P05023 | AT1A1_HUMAN | Oxidation (M) | 79656 | 0 | 1076500 | 1405300 | 3020700 | 0 | 427040 | 107290 | 534700 | 454170 | 3856700 | 317580 | 2735000 | 2259400 | 667060 | 438450 | 315280 | 615790 |
| NMYELHK | EIF3F; EIF3F; EI | O00303; A0/ | EIF3F_HUMAN | Unmodified | 29279 | 0 | 0 | 0 | 0 | 0 | 0 | 0 | 0 | 0 | 0 | 891390 | 0 | 0 | 0 | 0 | 0 | 0 |
| NNAFPCQ | ANLN; ANLN | Q9NQW6; H | ANLN_HUMAN | Unmodified | 0 | 0 | 0 | 0 | 6851000 | 1136000 | 419460 | 149420 | 0 | 0 | 191870 | 0 | 0 | 0 | 0 | 0 | 1987800 | 0 |
| NNAFPCQ | ANLN; ANLN | Q9NQW6; H | ANLN_HUMAN | Unmodified | 0 | 0 | 0 | 0 | 0 | 0 | 0 | 0 | 0 | 0 | 0 | 0 | 0 | 0 | 0 | 0 | 0 | 0 |
| NNAIISLRK | SLC39A10 | Q9ULF5 | S39AA_HUMAN | Unmodified | 0 | 0 | 0 | 0 | 391260 | 0 | 0 | 0 | 0 | 0 | 0 | 0 | 122670 | 0 | 0 | 0 | 0 | 0 |
| NNALNQV | SPCS3 | P61009 | SPCS3_HUMA | Unmodified | 0 | 0 | 0 | 0 | 0 | 0 | 0 | 0 | 9582400 | 3028900 | 0 | 206550 | 317890 | 0 | 0 | 0 | 0 | 0 |
| NNASTDYD | RPL3; RPL3; RP | P39023; B5N | RL3_HUMAN 6 | Unmodified | 7552500 | 4942400 | 3267500 | 2150700 | 1214700 | 0 | 1370200 | 852090 | 256890 | 1861600 | 5210000 | 30730000 | 599920 | 595030 | 3518600 | 4709900 | 204990 | 0 |
| NNAYLAQS | DARS1 | P14868 | SYDC_HUMAN | Unmodified | 0 | 0 | 600580 | 0 | 685690 | 0 | 351350 | 294370 | 1411600 | 405700 | 599340 | 5166500 | 686250 | 494670 | 177700 | 0 | 428920 | 780720 |
| NNCPFSAD | PCM1; PCM1; P | Q15154; A0/ | PCM1_HUMAN | Unmodified | 0 | 0 | 0 | 0 | 195600 | 35972 | 381300 | 0 | 0 | 0 | 0 | 0 | 409680 | 474170 | 0 | 0 | 0 | 204030 |
| NNDALHM | MYO6; MYO6; M | Q9UM54; A0 | MYO6_HUMAN | Oxidation (M) | 0 | 0 | 0 | 0 | 0 | 0 | 0 | 0 | 0 | 0 | 0 | 0 | 0 | 0 | 387600 | 410770 | 0 | 0 |
| NNDGYIDY | MCFD2 | Q8NI22 | MCFD2_HUMA | Unmodified | 0 | 0 | 0 | 0 | 0 | 0 | 0 | 0 | 4575300 | 0 | 0 | 362880 | 0 | 0 | 0 | 235600 | 0 | 0 |
| NNDLLFR | MYO1C; MYO1 | O00159; Q9I | MYO1C_HUMA | Unmodified | 3480100 | 5738500 | 6173800 | 8006400 | 3698600 | 1640000 | 207750 | 32515 | 427750 | 100960 | 0 | 969310 | 444970 | 188190 | 17066000 | 9358500 | 491070 | 282960 |
| NNDLLFRN | MYO1C; MYO1 | O00159; F5I | MYO1C_HUMA | Unmodified | 0 | 133910 | 0 | 0 | 2115000 | 1714600 | 116060 | 25554 | 0 | 0 | 0 | 0 | 259900 | 287190 | 0 | 0 | 645600 | 231240 |
| NNDLLYR | MYO1H; MYO1E | Q8N1T3; O4 | MYO1H_HUMA | Unmodified | 0 | 0 | 0 | 0 | 0 | 0 | 0 | 0 | 0 | 0 | 0 | 6053300 | 0 | 0 | 0 | 0 | 0 | 0 |
| NNDLLYRD | MYO1B; MYO1E | O43795; E9F | MYO1B_HUMA | Oxidation (M) | 0 | 0 | 0 | 0 | 2312100 | 739080 | 0 | 0 | 356460 | 262460 | 1839900 | 819200 | 0 | 0 | 100490 | 57766 | 292180 | 0 |
| NNDLLYRD | MYO1B; MYO1E | O43795; E9F | MYO1B_HUMA | Unmodified | 0 | 0 | 0 | 0 | 1553000 | 1627800 | 451810 | 0 | 426640 | 184500 | 0 | 324040 | 0 | 0 | 180230 | 0 | 469840 | 173090 |
| NNDVWQN | GOLIM4; GOLIM | O00461; F8V | GOLI4_HUMAN | Unmodified | 0 | 0 | 0 | 0 | 0 | 0 | 0 | 0 | 957360 | 0 | 0 | 0 | 0 | 0 | 0 | 0 | 0 | 0 |
| NNEDISIIPI | SSR4; SSR4 | P51571; A6N | SSRD_HUMAN | Unmodified | 0 | 0 | 182110 | 0 | 0 | 0 | 0 | 0 | 14678000 | 1087900 | 587490 | 656860 | 0 | 0 | 0 | 0 | 65420 | 0 |
| NNEEYLALI | QSOX1 | O00391 | QSOX1_HUMA | Unmodified | 0 | 0 | 34970 | 0 | 0 | 0 | 0 | 0 | 556800 | 0 | 0 | 0 | 0 | 0 | 66796 | 0 | 0 | 0 |
| NNEFIVIHN | GFPT1; GFPT1; | Q06210; A0/ | GFPT1_HUMA | Unmodified | 371300 | 0 | 959150 | 314400 | 148010 | 84126 | 0 | 0 | 991670 | 107980 | 395960 | 2731100 | 285330 | 153640 | 1268900 | 0 | 77720 | 106310 |
| NNEVSFSC | PTPRJ | Q12913 | PTPRJ_HUMAN | Unmodified | 655350 | 1451100 | 0 | 202190 | 841090 | 0 | 0 | 0 | 0 | 0 | 605220 | 9276200 | 219920 | 231430 | 0 | 567730 | 83750 | 663070 |
| NNEVSLAA | MON2; MON2 | Q7Z3U7; A0/ | MON2_HUMAN | Unmodified | 0 | 326930 | 0 | 0 | 0 | 0 | 158160 | 19726 | 526170 | 0 | 0 | 871330 | 0 | 0 | 0 | 0 | 0 | 0 |
| NNEVWLIC | HEBP2 | Q9Y5Z4 | HEBP2_HUMA | Unmodified | 245890 | 0 | 0 | 0 | 0 | 0 | 0 | 0 | 600150 | 0 | 0 | 986700 | 0 | 0 | 438390 | 352370 | 0 | 0 |
| NNFAVGYF | VDAC2; VDAC2 | P45880; Q5J | VDAC2_HUMA | Unmodified | 116910 | 130460 | 183530 | 478280 | 0 | 0 | 0 | 0 | 3.69E+08 | 17867000 | 2947700 | 1067300 | 121360 | 0 | 123450 | 95965 | 25007 | 0 |
| NNFAVGYF | VDAC2; VDAC2 | P45880; A0A | VDAC2_HUMA | Unmodified | 0 | 0 | 0 | 0 | 0 | 0 | 0 | 0 | 0 | 15711000 | 0 | 0 | 0 | 0 | 0 | 0 | 0 | 753860 |
| NNFEADAY | ITFG1; ITFG1; I | Q8TB96; F5C | TIP_HUMAN T- | Unmodified | 0 | 0 | 0 | 0 | 0 | 0 | 0 | 0 | 4690900 | 1971300 | 0 | 235260 | 0 | 0 | 0 | 0 | 0 | 0 |
| NNFEEEQC | SPINT1; SPINT1 | O43278; H3I | SPIT1_HUMAN | Unmodified | 760270 | 347830 | 97360 | 212250 | 0 | 0 | 0 | 0 | 862290 | 0 | 0 | 0 | 0 | 0 | 0 | 194590 | 0 | 0 |
| NNFEGEVT | P4HB; P4HB; P4 | P07237; A0A | PDIA1_HUMAN | Unmodified | 0 | 0 | 0 | 288500 | 134330 | 0 | 425730 | 35199 | 1442400 | 337830 | 260130 | 613160 | 0 | 0 | 0 | 0 | 0 | 0 |
| NNFEGEVT | P4HB; P4HB; P4 | P07237; A0A | PDIA1_HUMAN | Unmodified | 0 | 0 | 0 | 0 | 0 | 0 | 0 | 0 | 5488800 | 1828500 | 372420 | 0 | 0 | 0 | 0 | 0 | 0 | 0 |
| NNFGAEPF | HSPH1; HSPH1 | Q92598; A0/ | HS105_HUMA | Unmodified | 0 | 0 | 0 | 0 | 0 | 0 | 0 | 0 | 0 | 0 | 0 | 508800 | 0 | 0 | 0 | 0 | 0 | 0 |
| NNFTDFTN | HINT3 | Q9NQE9 | HINT3_HUMAN | Unmodified | 0 | 246540 | 161790 | 172520 | 0 | 0 | 0 | 0 | 0 | 0 | 241610 | 6444400 | 0 | 0 | 0 | 260560 | 0 | 0 |
| NNGAGYFI | UQCRC1 | P31930 | QCR1_HUMAN | Unmodified | 0 | 0 | 0 | 0 | 0 | 0 | 0 | 0 | 22379000 | 1873600 | 0 | 659530 | 0 | 0 | 0 | 0 | 0 | 0 |
| NNGAGYFI | UQCRC1 | P31930 | QCR1_HUMAN | Unmodified | 0 | 0 | 0 | 0 | 0 | 0 | 198340 | 0 | 0 | 1304300 | 0 | 0 | 0 | 0 | 0 | 0 | 0 | 0 |
| NNGCLVVI | PHYH; PHYH; P | O14832; B1/ | PAHX_HUMAN | Unmodified | 0 | 0 | 0 | 0 | 0 | 132440 | 0 | 0 | 1590400 | 0 | 0 | 275240 | 0 | 0 | 0 | 0 | 0 | 0 |
| NNGKIELIK | EIF3B | P55884 | EIF3B_HUMAN | Unmodified | 0 | 0 | 0 | 0 | 0 | 0 | 0 | 0 | 109520 | 0 | 0 | 1639800 | 0 | 0 | 0 | 0 | 0 | 0 |
| NNGPSSFS | ITGAV | P06756 | ITAV_HUMAN I | Unmodified | 348540 | 0 | 0 | 1323900 | 0 | 0 | 0 | 0 | 281180 | 150980 | 0 | 1686300 | 0 | 0 | 0 | 532800 | 115220 | 0 |
| NNGSLDRI | SLC44A4; SLC4 | Q53GD3; A0 | CTL4_HUMAN | Oxidation (M) | 0 | 0 | 0 | 0 | 0 | 0 | 0 | 0 | 0 | 0 | 84153 | 0 | 0 | 0 | 421140 | 246180 | 0 | 0 |
| NNGSLDRI | SLC44A4; SLC4 | Q53GD3; A0 | CTL4_HUMAN | Unmodified | 0 | 0 | 151570 | 0 | 178190 | 0 | 0 | 50713 | 0 | 0 | 0 | 0 | 0 | 0 | 0 | 337540 | 0 | 0 |
| NNGVLFEN | AP2A2; AP2A2 | O94973; A0/ | AP2A2_HUMAN | Unmodified | 0 | 105730 | 0 | 74059 | 188270 | 56482 | 13150 | 0 | 110250 | 189600 | 92115 | 225570 | 26018 | 164130 | 299660 | 95450 | 81612 | 63884 |
| NNGVLFEN | AP2A1 | O95782 | AP2A1_HUMAN | Unmodified | 199100 | 416400 | 78109 | 251530 | 211930 | 0 | 0 | 0 | 310920 | 94230 | 319790 | 390310 | 0 | 115350 | 287530 | 132160 | 427370 | 343020 |
| NNGYAISTI | BCKDHA; N/A; I | P12694; F5H | ODBA_HUMAN | Unmodified | 0 | 0 | 0 | 0 | 0 | 0 | 0 | 0 | 424200 | 0 | 0 | 0 | 0 | 0 | 0 | 0 | 0 | 0 |
| NNHHFKSI | GGH | Q92820 | GGH_HUMAN | Unmodified | 0 | 0 | 0 | 0 | 0 | 0 | 0 | 0 | 0 | 9232300 | 0 | 0 | 0 | 0 | 0 | 0 | 0 | 0 |
| NNIFEESYF | NEDD4L; NEDD | Q96PU5; A0/ | NED4L_HUMA | Unmodified | 275050 | 436120 | 0 | 159230 | 0 | 0 | 0 | 0 | 0 | 0 | 0 | 0 | 0 | 0 | 450510 | 290110 | 0 | 0 |
| NNIFEESYF | NEDD4L; NEDD | Q96PU5; A0/ | NED4L_HUMA | Unmodified | 1218000 | 1371600 | 1105200 | 0 | 0 | 0 | 0 | 0 | 0 | 0 | 0 | 0 | 0 | 0 | 2467900 | 1758700 | 0 | 0 |
| NNIIQTWR | GARS1; GARS1 | P41250; F8V | GARS_HUMAN | Unmodified | 0 | 0 | 1751500 | 824740 | 0 | 0 | 0 | 0 | 3841500 | 0 | 3448300 | 15348000 | 0 | 0 | 624550 | 896530 | 0 | 1176300 |
| NNIPYFETS | RAB7A; RAB7A; | P51149; A0A | RAB7A_HUMA | Unmodified | 5722300 | 4056900 | 4687300 | 7021900 | 31743000 | 4321800 | 784050 | 4936400 | 11546000 | 7370800 | 8849100 | 38967000 | 17822000 | 10569000 | 5471000 | 4763400 | 4884800 | 14566000 |
| NNIPYFETS | RAB7A; RAB7A; | P51149; A0A | RAB7A_HUMA | Unmodified | 0 | 0 | 0 | 0 | 1427600 | 0 | 0 | 0 | 0 | 397420 | 0 | 0 | 0 | 0 | 0 | 0 | 2336900 | 1651300 |
| NNISSGHV | STAU1; STAU1 | O95793; Q5. | STAU1_HUMA | Unmodified | 363600 | 271840 | 395830 | 419880 | 0 | 0 | 0 | 0 | 0 | 0 | 0 | 1366800 | 0 | 0 | 0 | 347900 | 0 | 0 |
| NNITLVQD | FAT3; FAT3; FA | Q8TDW7; E9 | FAT3_HUMAN | Unmodified | 0 | 0 | 0 | 776120 | 0 | 0 | 0 | 0 | 0 | 0 | 0 | 1662100 | 0 | 0 | 0 | 0 | 0 | 0 |
| NNKGPVK | PDIA4; PDIA4 | P13667; A0A | PDIA4_HUMAN | Unmodified | 0 | 0 | 0 | 0 | 0 | 0 | 216190 | 357990 | 0 | 199890 | 0 | 0 | 0 | 0 | 0 | 0 | 0 | 0 |
| NNLAGAEE | CLTC; CLTC | Q00610; A0/ | CLH1_HUMAN | Unmodified | 11188000 | 372430 | 6622200 | 7789500 | 0 | 0 | 0 | 0 | 3345200 | 579860 | 1551600 | 23479000 | 0 | 0 | 6433700 | 6429900 | 475300 | 0 |
| NNLAGAEE | CLTC; CLTC | Q00610; A0/ | CLH1_HUMAN | Unmodified | 13456000 | 11849000 | 5832900 | 3014200 | 1705700 | 544180 | 268870 | 0 | 0 | 0 | 625790 | 40990000 | 1270900 | 0 | 9878600 | 9949100 | 222800 | 551610 |
| NNLAGAEE | CLTC; CLTC | Q00610; A0/ | CLH1_HUMAN | Unmodified | 0 | 0 | 0 | 0 | 0 | 0 | 911250 | 0 | 0 | 0 | 0 | 0 | 2350600 | 962210 | 0 | 0 | 1589300 | 2060900 |
| NNLASCYL | KLC1; KLC2; KL | Q07866; Q9I | KLC1_HUMAN | Unmodified | 0 | 0 | 253210 | 207080 | 784310 | 61430 | 516160 | 338410 | 0 | 0 | 0 | 1415500 | 459480 | 326830 | 238410 | 0 | 980760 | 823230 |
| NNLCPSGS | CD81; CD81; C | P60033; A6N | CD81_HUMAN | Unmodified | 69968000 | 10736000 | 63012000 | 9191500 | 238770 | 18102 | 0 | 0 | 1759400 | 534890 | 1110700 | 14246000 | 624210 | 397620 | 93675000 | 73318000 | 0 | 125340 |
| NNLCPSGS | CD81; CD81; C | P60033; A6N | CD81_HUMAN | Unmodified | 25658000 | 8264400 | 0 | 6077600 | 0 | 0 | 0 | 0 | 0 | 0 | 0 | 0 | 0 | 0 | 22637000 | 14488000 | 0 | 0 |
| NNLIQQK | MBNL1; MBNL3 | Q9NR56; Q9 | MBNL1_HUMA | Unmodified | 1152700 | 0 | 0 | 624120 | 0 | 0 | 0 | 0 | 0 | 0 | 0 | 0 | 0 | 0 | 0 | 0 | 0 | 0 |
| NNLLNEDL | PACSIN1; PACS | Q9BY11; F6I | PACN1_HUMA | Unmodified | 12847000 | 11034000 | 1879100 | 3038100 | 0 | 0 | 0 | 0 | 0 | 0 | 0 | 1936900 | 1969700 | 725680 | 4335000 | 2494500 | 0 | 0 |
| NNLNPTWI | CPNE1; CPNE1 | Q99829; Q5. | CPNE1_HUMA | Unmodified | 0 | 0 | 0 | 742710 | 2111200 | 762920 | 0 | 0 | 1994800 | 405400 | 0 | 2668000 | 1196700 | 835170 | 1017100 | 842810 | 851850 | 2807000 |
| NNLNPVW | CPNE3; CPNE3 | O75131; H0 | CPNE3_HUMA | Unmodified | 1238100 | 775890 | 6189200 | 4994800 | 11441000 | 0 | 3148400 | 1025400 | 52527000 | 11893000 | 937540 | 4363300 | 5687900 | 3783100 | 5660100 | 4080400 | 3384700 | 14755000 |
| NNLNPVW | CPNE3; CPNE3 | O75131; H0 | CPNE3_HUMA | Oxidation (M) | 0 | 1712600 | 0 | 2331400 | 0 | 0 | 2103100 | 0 | 0 | 1520500 | 0 | 0 | 0 | 1639500 | 0 | 0 | 1088300 | 0 |
| NNLSFIETS | RAB11B; RAB11 | Q15907; M0I | RB11B_HUMA | Unmodified | 1006400 | 0 | 422320 | 2039800 | 1070300 | 0 | 0 | 0 | 1429000 | 595490 | 656320 | 544020 | 0 | 0 | 1130300 | 683640 | 0 | 0 |
| NNLVIFHR | PAM; PAM | P19021; H7E | AMD_HUMAN | Unmodified | 0 | 0 | 0 | 0 | 0 | 53082 | 0 | 0 | 2394400 | 0 | 1901700 | 1141200 | 0 | 0 | 0 | 0 | 0 | 0 |
| NNNAVDS | ATP2B1; ATP2B | P20020; E7E | AT2B1_HUMAN | Unmodified | 0 | 98486 | 0 | 114950 | 850330 | 0 | 0 | 0 | 0 | 0 | 0 | 0 | 1295300 | 1311000 | 0 | 0 | 446820 | 1217000 |
| NNNAVDS | ATP2B1; ATP2B | P20020; E7E | AT2B1_HUMAN | Oxidation (M) | 69242 | 0 | 571760 | 567480 | 972500 | 42161 | 0 | 0 | 97617 | 315630 | 196440 | 255280 | 2878100 | 5529200 | 835920 | 552230 | 604290 | 2327900 |
| NNNIDME | COBLL1; COBL | Q53SF7; A0/ | COBL1_HUMA | Unmodified | 1155200 | 1633700 | 1100000 | 1193600 | 292900 | 0 | 0 | 0 | 0 | 0 | 0 | 139210 | 1033700 | 385920 | 0 | 0 | 0 | 0 |
| NNNLAVTS | ATP6V1C1 | P21283 | VATC1_HUMAN | Unmodified | 18620 | 75047 | 20500 | 155760 | 0 | 0 | 239500 | 31931 | 55001 | 0 | 0 | 362400 | 14204 | 40992 | 95228 | 46996 | 55181 | 134480 |
| NNNLSKD | FAT3; FAT3 | Q8TDW7; E9 | FAT3_HUMAN | Unmodified | 0 | 0 | 0 | 0 | 779490 | 18837 | 0 | 0 | 48244 | 92204 | 0 | 0 | 0 | 0 | 0 | 0 | 45553 | 191690 |
| NNNPISNS | TENM4; TENM4 | Q6N022; H0 | TEN4_HUMAN | Unmodified | 0 | 0 | 0 | 0 | 0 | 0 | 0 | 0 | 0 | 0 | 0 | 0 | 0 | 0 | 0 | 0 | 0 | 0 |
| NNNPPKEI | RIPOR2; RIPOR | Q9Y4F9; B7Z | RIPR2_HUMAN | Unmodified | 58273 | 0 | 40095 | 0 | 0 | 0 | 0 | 15931 | 336020 | 60567 | 0 | 578270 | 522080 | 0 | 0 | 0 | 0 | 64371 |
| NNNQQLA | NCKAP1 | Q9Y2A7 | NCKP1_HUMA | Unmodified | 301940 | 22954 | 493190 | 344090 | 0 | 0 | 0 | 0 | 299410 | 44300 | 330200 | 3656800 | 0 | 0 | 980470 | 1054800 | 0 | 0 |
| NNNSNQN | USO1 | O60763 | USO1_HUMAN | Unmodified | 0 | 0 | 0 | 0 | 0 | 0 | 0 | 0 | 0 | 0 | 0 | 1114400 | 0 | 0 | 0 | 0 | 0 | 0 |

| | | | | | | | | | | | | | | | | | | | | | | |
|---|---|---|---|---|---|---|---|---|---|---|---|---|---|---|---|---|---|---|---|---|---|---|
| NNPAIVIIG | LMAN1 | P49257 | LMAN1_HUMAI | Unmodified | 0 | 0 | 0 | 0 | 0 | 0 | 0 | 0 | 1.27E+08 | 3768500 | 1640000 | 2242400 | 0 | 0 | 1417500 | 376150 | 0 | 0 |
| NNPAIVIIG | LMAN1 | P49257 | LMAN1_HUMAI | Unmodified | 0 | 0 | 0 | 0 | 0 | 0 | 0 | 0 | 3395400 | 0 | 278480 | 0 | 0 | 0 | 0 | 0 | 0 | 0 |
| NNPHYDPS | THYN1 | Q9P016 | THYN1_HUMAI | Unmodified | 0 | 0 | 0 | 0 | 0 | 0 | 0 | 0 | 0 | 0 | 102940 | 1052500 | 0 | 0 | 0 | 0 | 0 | 0 |
| NNPIYQFH | CAPN7 | Q9Y6W3 | CAN7_HUMAN | Unmodified | 352360 | 0 | 0 | 181080 | 0 | 0 | 0 | 49261 | 2419100 | 1961500 | 0 | 7626900 | 0 | 0 | 0 | 0 | 0 | 0 |
| NNPPTIVHI | TOM1; TOM1; T | O60784; V9( | TOM1_HUMAN | Unmodified | 0 | 202860 | 279310 | 670590 | 0 | 0 | 0 | 0 | 0 | 0 | 0 | 0 | 0 | 0 | 360890 | 262230 | 26063 | 91566 |
| NNPPTIVQI | TOM1L2; TOM1 | Q6ZVM7; K7 | TM1L2_HUMAN | Unmodified | 2662600 | 2360600 | 2008600 | 1504300 | 0 | 0 | 204440 | 0 | 150340 | 87360 | 0 | 275660 | 985140 | 548840 | 2394300 | 2636700 | 220250 | 466860 |
| NNPPTLEG | STK24; STK24; | Q9Y6E0; H0 | STK24_HUMAN | Unmodified | 0 | 0 | 2226500 | 137840 | 903370 | 335870 | 0 | 60133 | 707150 | 0 | 234010 | 1016800 | 258280 | 116000 | 231960 | 280760 | 0 | 637700 |
| NNPPTLEG | STK24; STK24; | Q9Y6E0; H0 | STK24_HUMAN | Unmodified | 0 | 0 | 0 | 0 | 5489500 | 2134100 | 0 | 1639700 | 1063700 | 1191400 | 664640 | 0 | 4423400 | 3064000 | 0 | 0 | 0 | 3976600 |
| NNPPTLVG | STK26; STK26; | Q9P289; B4 | STK26_HUMAN | Unmodified | 0 | 0 | 0 | 0 | 6284000 | 3399800 | 698380 | 372680 | 0 | 0 | 0 | 4997700 | 7448600 | 7329500 | 1218600 | 0 | 5198800 | 6038500 |
| NNPSKPLH | WDR1 | O75083 | WDR1_HUMAN | Unmodified | 240480 | 159390 | 60983 | 0 | 287560 | 0 | 0 | 0 | 153520 | 68797 | 255160 | 1701100 | 0 | 0 | 149610 | 385560 | 497240 | 0 |
| NNPSSDYF | MBTPS1 | Q14703 | MBTP1_HUMAI | Unmodified | 0 | 0 | 0 | 0 | 0 | 0 | 0 | 0 | 432020 | 0 | 63044 | 0 | 0 | 0 | 0 | 0 | 0 | 0 |
| NNPVMSL( | HS2ST1; HS2S1 | Q7LGA3; K7 | HS2ST_HUMAN | Oxidation (M) | 0 | 0 | 0 | 0 | 0 | 0 | 0 | 0 | 3668200 | 421250 | 79937 | 0 | 0 | 0 | 0 | 0 | 0 | 0 |
| NNPVMSL( | HS2ST1; HS2S1 | Q7LGA3; K7 | HS2ST_HUMAN | Unmodified | 0 | 0 | 0 | 0 | 0 | 0 | 0 | 0 | 14395000 | 853690 | 142170 | 570080 | 0 | 0 | 0 | 0 | 0 | 0 |
| NNPVMSL( | HS2ST1; HS2S1 | Q7LGA3; K7 | HS2ST_HUMAN | Oxidation (M) | 0 | 0 | 0 | 0 | 0 | 0 | 0 | 0 | 0 | 896420 | 0 | 0 | 0 | 0 | 0 | 0 | 0 | 0 |
| NNPVMSL( | HS2ST1; HS2S1 | Q7LGA3; K7 | HS2ST_HUMAN | Unmodified | 0 | 0 | 0 | 0 | 0 | 0 | 0 | 0 | 0 | 395050 | 0 | 0 | 0 | 0 | 211010 | 0 | 0 | 0 |
| NNPWVEK | GNG13 | Q9P2W3 | GBG13_HUMAI | Unmodified | 0 | 0 | 825350 | 93644 | 1410200 | 0 | 0 | 0 | 0 | 60478 | 4373200 | 28309000 | 0 | 64449 | 0 | 0 | 59108 | 0 |
| NNQFQALL | PTBP1; PTBP1; | P26599; A0A | PTBP1_HUMAN | Unmodified | 0 | 0 | 0 | 0 | 0 | 0 | 0 | 0 | 0 | 0 | 653210 | 6543800 | 0 | 1726300 | 0 | 0 | 0 | 0 |
| NNQIEQFS | TTC39C; TTC39 | Q8N584; G3 | TT39C_HUMAN | Unmodified | 0 | 285410 | 0 | 439490 | 98474 | 0 | 0 | 0 | 1968300 | 0 | 0 | 3478100 | 0 | 0 | 0 | 0 | 52617 | 221600 |
| NNQITNNC | PGK1 | P00558 | PGK1_HUMAN | Unmodified | 0 | 0 | 0 | 0 | 27439 | 0 | 0 | 0 | 59755 | 0 | 80079 | 150920 | 42820 | 0 | 0 | 19451 | 60855 | 97166 |
| NNQITNNC | PGK1 | P00558 | PGK1_HUMAN | Unmodified | 0 | 0 | 0 | 0 | 70120000 | 10988000 | 0 | 11205000 | 0 | 0 | 0 | 0 | 32736000 | 13359000 | 0 | 0 | 1.72E+08 | 61284000 |
| NNQLAFDL | MICAL3; MICAL | Q7RTP6; A0/ | MICA3_HUMAN | Unmodified | 0 | 0 | 0 | 201830 | 3856300 | 2261400 | 0 | 180060 | 148780 | 0 | 0 | 0 | 0 | 0 | 0 | 68667 | 0 | 0 |
| NNQVLGIG | RPIA | P49247 | RPIA_HUMAN F | Unmodified | 0 | 90326 | 0 | 0 | 0 | 0 | 0 | 0 | 780180 | 0 | 1046400 | 8427100 | 0 | 0 | 150710 | 0 | 0 | 0 |
| NNRFSTPE | HNRNPL; HNRI | P14866; M0 | HNRPL_HUMAI | Unmodified | 0 | 0 | 0 | 0 | 0 | 0 | 0 | 0 | 39299 | 0 | 410590 | 2181400 | 0 | 0 | 0 | 0 | 0 | 0 |
| NNRPSEGF | CLTC; CLTC | Q00610; A0/ | CLH1_HUMAN | Unmodified | 2952800 | 1370300 | 1849900 | 1550600 | 215490 | 212510 | 101510 | 41752 | 446640 | 290470 | 141600 | 14377000 | 111200 | 0 | 2104000 | 2959400 | 347140 | 468400 |
| NNRQPYA\ | RPL4; RPL4; RP | P36578; H3 | RL4_HUMAN 6 | Unmodified | 10971000 | 12137000 | 8502100 | 6519900 | 3313200 | 0 | 0 | 1410600 | 0 | 1001300 | 3276700 | 42074000 | 16055000 | 2542000 | 5714400 | 5146400 | 645090 | 1522400 |
| NNSELLNN | MARS1 | P56192 | SYMC_HUMAN | Unmodified | 0 | 0 | 70829 | 17138 | 0 | 0 | 0 | 0 | 0 | 0 | 0 | 3519100 | 16052 | 0 | 75701 | 148020 | 0 | 0 |
| NNSFVNEII | MSH2 | P43246 | MSH2_HUMAN | Unmodified | 0 | 0 | 0 | 0 | 277770 | 128140 | 830240 | 0 | 0 | 230430 | 0 | 110230 | 0 | 0 | 0 | 0 | 0 | 0 |
| NNSFVNEII | MSH2 | P43246 | MSH2_HUMAN | Unmodified | 0 | 0 | 0 | 0 | 0 | 0 | 1214900 | 0 | 0 | 0 | 0 | 0 | 0 | 0 | 0 | 0 | 164120 | 0 |
| NNSIPDKQ | MFGE8 | Q08431 | MFGM_HUMAN | Unmodified | 0 | 0 | 0 | 0 | 1229900 | 240680 | 129990 | 0 | 0 | 0 | 0 | 0 | 0 | 0 | 0 | 0 | 26732 | 298550 |
| NNTALQSV | ARHGEF2; ARH | Q92974; V9( | ARHG2_HUMA | Unmodified | 0 | 0 | 0 | 0 | 0 | 0 | 248880 | 371330 | 0 | 191850 | 0 | 1939500 | 1178600 | 0 | 0 | 0 | 0 | 0 |
| NNTQVLIN | SNRPD2; SNRP | P62316; K7E | SMD2_HUMAN | Unmodified | 859520 | 487980 | 0 | 0 | 0 | 0 | 0 | 0 | 0 | 0 | 1270000 | 4285900 | 0 | 0 | 189200 | 271530 | 0 | 0 |
| NNTQVLIN | SNRPD2; SNRP | P62316; K7E | SMD2_HUMAN | Unmodified | 0 | 0 | 0 | 0 | 3429300 | 0 | 1475100 | 1004900 | 0 | 813910 | 0 | 0 | 2076700 | 2370500 | 0 | 0 | 561940 | 80844 |
| NNTQVLIN | SNRPD2; SNRP | P62316; K7E | SMD2_HUMAN | Unmodified | 0 | 2357400 | 0 | 0 | 1031000 | 116870 | 401100 | 547700 | 0 | 103580 | 0 | 0 | 1584400 | 2699600 | 0 | 0 | 0 | 0 |
| NNTVGLIQ | ECHS1 | P30084 | ECHM_HUMAN | Unmodified | 10962000 | 8273500 | 0 | 0 | 0 | 0 | 1329200 | 150910 | 970040 | 245870 | 0 | 0 | 199860 | 0 | 0 | 0 | 0 | 0 |
| NNTVNTMI | PYGB | P11216 | PYGB_HUMAN | Unmodified | 0 | 0 | 0 | 0 | 104070 | 0 | 0 | 0 | 0 | 0 | 0 | 0 | 131520 | 61602 | 0 | 0 | 144140 | 579170 |
| NNTVTPGC | RPL35A; RPL35 | P18077; C9 | RL35A_HUMAN | Unmodified | 1816500 | 1084800 | 1061200 | 632090 | 348740 | 68272 | 508000 | 316060 | 267500 | 0 | 0 | 12283000 | 73434 | 70514 | 1425300 | 341690 | 18919 | 292830 |
| NNVAPDEL | LSS; LSS | P48449; A0/ | LSS_HUMAN L | Unmodified | 22061 | 0 | 0 | 0 | 0 | 0 | 0 | 0 | 0 | 0 | 0 | 794370 | 0 | 187590 | 0 | 51364 | 0 | 0 |
| NNVEQDLI | EPS15; EPS15 | P42566; B1/ | EPS15_HUMAN | Unmodified | 0 | 0 | 32232 | 0 | 286690 | 0 | 376650 | 0 | 0 | 23810 | 43328 | 0 | 0 | 0 | 0 | 0 | 103210 | 0 |
| NNVEQDLI | EPS15; EPS15 | P42566; B1/ | EPS15_HUMAN | Unmodified | 0 | 61940 | 0 | 26468 | 258870 | 0 | 0 | 0 | 0 | 33663 | 0 | 0 | 918560 | 636390 | 0 | 0 | 0 | 436860 |
| NNVGRGIN | FAM3C; FAM3C | Q92520; C9J | FAM3C_HUMAI | Unmodified | 0 | 0 | 286490 | 0 | 306170 | 73592 | 93628 | 118500 | 0 | 2884200 | 0 | 0 | 323680 | 393290 | 0 | 0 | 32965 | 1127200 |
| NNVGRGIN | FAM3C; FAM3C | Q92520; C9J | FAM3C_HUMAI | Unmodified | 0 | 0 | 0 | 0 | 0 | 0 | 0 | 117950 | 0 | 1313400 | 0 | 0 | 0 | 0 | 0 | 0 | 0 | 102220 |
| NNVKWELI | MRPL9 | Q9BYD2 | RM09_HUMAN | Unmodified | 0 | 0 | 0 | 0 | 0 | 0 | 0 | 0 | 1220000 | 329420 | 0 | 0 | 0 | 0 | 0 | 0 | 0 | 0 |
| NNVLIVCVI | NUP50 | Q9UKX7 | NUP50_HUMAI | Unmodified | 0 | 30006 | 0 | 0 | 0 | 0 | 50364 | 24560 | 0 | 31671 | 0 | 176580 | 0 | 0 | 0 | 32494 | 0 | 0 |
| NNVLVLYS | PDIA5 | Q14554 | PDIA5_HUMAN | Unmodified | 0 | 0 | 0 | 0 | 0 | 0 | 224070 | 56254 | 8598700 | 2579500 | 0 | 0 | 0 | 31016 | 81556 | 0 | 0 | 0 |
| NNVRRQA1 | SSBP1 | Q04837 | SSBP_HUMAN | Unmodified | 0 | 0 | 0 | 0 | 0 | 0 | 0 | 0 | 0 | 2211900 | 0 | 0 | 0 | 0 | 0 | 0 | 0 | 0 |
| NNVSLLQS | SNX5 | Q9Y5X3 | SNX5_HUMAN | Unmodified | 0 | 0 | 0 | 0 | 0 | 0 | 0 | 0 | 0 | 0 | 0 | 0 | 0 | 0 | 0 | 0 | 0 | 0 |
| NNVSPVSR | MAP4K4; MAP4 | H7C3Z6; A0/ | H7C3Z6_HUM/ | Unmodified | 0 | 0 | 53760 | 0 | 1716500 | 342900 | 0 | 0 | 0 | 0 | 147320 | 127460 | 0 | 0 | 0 | 0 | 1018400 | 997270 |
| NNVVGRL( | HSPG2 | P98160 | PGBM_HUMAN | Unmodified | 0 | 0 | 0 | 0 | 0 | 0 | 0 | 0 | 0 | 0 | 0 | 0 | 416370 | 0 | 0 | 0 | 569170 | 4102900 |
| NNWNIEDI | ATP7A | Q04656 | ATP7A_HUMAN | Unmodified | 0 | 0 | 0 | 424390 | 0 | 0 | 0 | 0 | 0 | 0 | 0 | 532360 | 0 | 291820 | 0 | 1294200 | 0 | 0 |
| NNYALNTT | PKP4; PKP4 | Q99569; A0/ | PKP4_HUMAN | Unmodified | 0 | 0 | 0 | 530160 | 0 | 0 | 272800 | 0 | 905390 | 0 | 0 | 0 | 0 | 0 | 0 | 253370 | 400530 | 613330 |
| NNYIATIYK | PLPP1 | O14494 | PLPP1_HUMAN | Unmodified | 0 | 0 | 0 | 0 | 0 | 0 | 0 | 0 | 0 | 0 | 0 | 0 | 0 | 0 | 4785900 | 1484700 | 0 | 0 |
| NNYLREEE | SPINT1; SPINT1 | O43278; H3 | SPIT1_HUMAN | Unmodified | 0 | 0 | 0 | 658090 | 0 | 0 | 0 | 20685 | 1867800 | 0 | 235240 | 1468500 | 0 | 0 | 0 | 0 | 183930 | 166820 |
| NNYSEDLE | PLCG1; PLCG1; | P19174; V9C | PLCG1_HUMAI | Unmodified | 0 | 0 | 0 | 0 | 0 | 0 | 0 | 0 | 0 | 0 | 0 | 439850 | 0 | 0 | 0 | 51218 | 0 | 0 |
| NNYYGTSII | MPP7; MPP7 | Q5T2T1; U5( | MPP7_HUMAN | Unmodified | 0 | 0 | 148740 | 326150 | 1041800 | 0 | 0 | 0 | 647920 | 0 | 0 | 0 | 0 | 0 | 139360 | 0 | 0 | 221650 |
| NNYYGTSII | MPP7; MPP7 | Q5T2T1; U5( | MPP7_HUMAN | Unmodified | 91363 | 0 | 0 | 54774 | 3900400 | 390670 | 0 | 0 | 0 | 442320 | 0 | 202930 | 902730 | 1074600 | 0 | 0 | 496810 | 977950 |
| NPACSGAN | IGF2R; IGF2R | P11717; S4F | MPRI_HUMAN ( | Unmodified | 0 | 0 | 0 | 0 | 0 | 0 | 0 | 0 | 1043000 | 569950 | 0 | 583630 | 0 | 0 | 0 | 0 | 0 | 0 |
| NPAFEGVN | ARHGAP29 | Q52LW3 | RHG29_HUMAI | Unmodified | 0 | 0 | 0 | 0 | 0 | 0 | 0 | 0 | 0 | 0 | 0 | 0 | 1672600 | 101670 | 0 | 0 | 54723 | 179850 |
| NPAPPIDA\ | KPNA2; KPNA2; | P52292; A0/ | IMA1_HUMAN I | Unmodified | 204760 | 198620 | 163930 | 300650 | 734900 | 355400 | 53409 | 0 | 552170 | 418010 | 317710 | 560690 | 345270 | 0 | 830310 | 334780 | 338440 | 0 |
| NPAVLSAA | SEC31A; SEC31 | O94979; H7 | SC31A_HUMAI | Unmodified | 0 | 0 | 0 | 0 | 0 | 0 | 0 | 0 | 0 | 0 | 0 | 2801800 | 0 | 0 | 124650 | 66802 | 0 | 0 |
| NPAVTQLV | CPD | O75976 | CBPD_HUMAN | Unmodified | 0 | 0 | 435780 | 0 | 0 | 0 | 0 | 0 | 6636300 | 705330 | 504320 | 481640 | 0 | 0 | 0 | 0 | 0 | 0 |
| NPAVTQLV | CPD | O75976 | CBPD_HUMAN | Unmodified | 0 | 0 | 0 | 0 | 498910 | 457190 | 0 | 0 | 9751300 | 2152900 | 125040 | 58040 | 0 | 0 | 0 | 0 | 0 | 62022 |
| NPCHNGG | MFGE8; MFGE8 | Q08431; H0 | MFGM_HUMAN | Unmodified | 569290 | 198840 | 430110 | 1347100 | 728090 | 163300 | 101650 | 0 | 2842700 | 262770 | 538430 | 6688000 | 283440 | 0 | 1247500 | 1158000 | 0 | 0 |
| NPCHNGG | MFGE8; MFGE8 | Q08431; H0 | MFGM_HUMAN | Unmodified | 0 | 0 | 0 | 0 | 3910700 | 0 | 0 | 0 | 0 | 0 | 0 | 0 | 0 | 0 | 0 | 0 | 838670 | 0 |
| NPDDITNE | HSP90AA1 | P07900 | HS90A_HUMAI | Unmodified | 581110 | 927020 | 0 | 178370 | 0 | 0 | 0 | 0 | 1877200 | 626630 | 5184100 | 23969000 | 0 | 0 | 1186600 | 1007500 | 50970 | 0 |
| NPDDITQE | HSP90AB1 | P08238 | HS90B_HUMAI | Unmodified | 0 | 283510 | 601370 | 707800 | 0 | 0 | 0 | 0 | 3214300 | 671120 | 4419100 | 29321000 | 0 | 0 | 2776600 | 2783000 | 0 | 0 |
| NPDDVFR | DNAJB6; DNAJE | O75190; A0/ | DNJB6_HUMAN | Unmodified | 1183300 | 1271700 | 0 | 0 | 0 | 0 | 0 | 0 | 0 | 0 | 0 | 151260 | 0 | 0 | 1464400 | 939970 | 0 | 0 |
| NPDEIDETI | ITGA2 | P17301 | ITA2_HUMAN I | Unmodified | 836770 | 586940 | 1215900 | 668740 | 1024000 | 0 | 0 | 0 | 0 | 0 | 0 | 0 | 0 | 0 | 1341000 | 1054900 | 275300 | 0 |
| NPDELAEA | GIPC1; GIPC1 | O14908; K7E | GIPC1_HUMAN | Unmodified | 0 | 0 | 0 | 0 | 0 | 0 | 0 | 0 | 0 | 0 | 0 | 0 | 0 | 0 | 0 | 1793400 | 0 | 0 |
| NPDMVAGI | EIF2S2 | P20042 | IF2B_HUMAN E | Unmodified | 178250 | 138600 | 0 | 0 | 1542100 | 268860 | 0 | 0 | 667920 | 667660 | 157710 | 3619400 | 0 | 0 | 501330 | 0 | 0 | 974520 |
| NPDRVLIG | UGDH | O60701 | UGDH_HUMAN | Unmodified | 307740 | 0 | 436710 | 179710 | 593360 | 349660 | 0 | 0 | 294530 | 0 | 119090 | 1309800 | 234570 | 0 | 507660 | 377930 | 0 | 0 |
| NPDSQYGI | CMPK1; CMPK1 | P30085; Q5T | KCY_HUMAN U | Unmodified | 0 | 0 | 0 | 351670 | 482500 | 0 | 0 | 0 | 178180 | 0 | 0 | 1366100 | 51196 | 0 | 570950 | 0 | 0 | 565200 |
| NPDTQWIT | RPL15; RPL15; | P61313; E7E | RL15_HUMAN ( | Unmodified | 340440 | 990570 | 744480 | 438270 | 834300 | 0 | 401270 | 216900 | 88678 | 0 | 216470 | 6566100 | 0 | 0 | 515940 | 485760 | 0 | 187800 |
| NPEDLSAE | PPME1 | Q9Y570 | PPME1_HUMAI | Unmodified | 0 | 0 | 0 | 0 | 600080 | 0 | 0 | 0 | 150570 | 0 | 172400 | 666310 | 0 | 0 | 0 | 0 | 0 | 0 |
| NPELSTQLI | TEX10 | Q9NXF1 | TEX10_HUMAN | Unmodified | 72759 | 48913 | 36836 | 0 | 0 | 0 | 0 | 0 | 0 | 0 | 0 | 407490 | 0 | 0 | 0 | 27423 | 0 | 0 |
| NPEPELLVI | GANAB; GANAE | Q14697; F5H | GANAB_HUMA | Unmodified | 0 | 0 | 337580 | 0 | 0 | 0 | 0 | 0 | 5103900 | 546300 | 908640 | 1417200 | 0 | 0 | 0 | 458590 | 0 | 1872100 |
| NPEQEPIPI | ARHGAP1; ARH | Q07960; H0 | RHG01_HUMAI | Unmodified | 0 | 0 | 143650 | 0 | 0 | 0 | 0 | 0 | 0 | 0 | 0 | 821630 | 0 | 0 | 0 | 0 | 0 | 0 |
| NPEQVDLY | MARS1 | P56192 | SYMC_HUMAN | Oxidation (M) | 0 | 0 | 0 | 0 | 706980 | 0 | 0 | 0 | 0 | 0 | 0 | 2281500 | 0 | 0 | 0 | 0 | 0 | 0 |
| NPEQVDLY | MARS1 | P56192 | SYMC_HUMAN | Unmodified | 0 | 0 | 1324800 | 0 | 97147 | 0 | 0 | 0 | 0 | 0 | 0 | 1154300 | 0 | 1589300 | 25082 | 0 | 0 | 0 |
| NPEVGLKP | EPRS1; EPRS1 | P07814; V9C | SYEP_HUMAN | Unmodified | 265960 | 0 | 327120 | 0 | 787290 | 6038500 | 0 | 0 | 1361900 | 0 | 1274000 | 10811000 | 1467300 | 0 | 564100 | 309770 | 0 | 0 |
| NPFGNAGI | ACADVL; ACAD | P49748; G3\ | ACADV_HUMAI | Unmodified | 0 | 0 | 0 | 0 | 0 | 0 | 0 | 0 | 8281700 | 1459700 | 0 | 0 | 0 | 0 | 0 | 0 | 0 | 0 |
| NPGFEIIHC | FARSB | Q9NSD9 | SYFB_HUMAN | Unmodified | 0 | 0 | 0 | 0 | 0 | 0 | 0 | 0 | 0 | 0 | 0 | 658710 | 0 | 0 | 0 | 0 | 0 | 0 |
| NPGKPDG\ | JAM3 | Q9BX67 | JAM3_HUMAN | Unmodified | 0 | 0 | 0 | 0 | 0 | 0 | 0 | 10367 | 12277000 | 546810 | 1456600 | 1096200 | 0 | 0 | 0 | 0 | 630700 | 167700 |
| NPGSVAEN | MX1; MX1; MX1 | P20591; A0/ | MX1_HUMAN I | Unmodified | 0 | 0 | 0 | 0 | 0 | 0 | 0 | 0 | 0 | 0 | 0 | 0 | 0 | 0 | 0 | 0 | 0 | 653920 |
| NPGVFLIH | MAN1A2 | O60476 | MA1A2_HUMAI | Unmodified | 0 | 89033 | 0 | 681870 | 0 | 0 | 0 | 0 | 2783300 | 306720 | 0 | 0 | 0 | 76771 | 0 | 0 | 52644 | 0 |
| NPGVGNG | MAPRE1 | Q15691 | MARE1_HUMAI | Oxidation (M) | 2018300 | 1106800 | 1147400 | 0 | 0 | 0 | 0 | 0 | 1084500 | 0 | 0 | 5855200 | 0 | 0 | 1168900 | 1377400 | 0 | 0 |
| NPGYPQSE | SH3GL1; SH3G | Q99961; M0 | SH3G1_HUMAI | Oxidation (M) | 9170700 | 7297300 | 7524300 | 6108100 | 0 | 0 | 0 | 0 | 0 | 0 | 0 | 0 | 656600 | 0 | 12760000 | 9244500 | 0 | 0 |
| NPGYPQSE | SH3GL1; SH3G | Q99961; M0 | SH3G1_HUMAI | Unmodified | 1251600 | 1397600 | 518600 | 964420 | 0 | 0 | 0 | 16867 | 0 | 0 | 0 | 0 | 736230 | 0 | 882090 | 1242900 | 0 | 0 |
| NPGYPQSE | SH3GL1; SH3G | Q99961; M0 | SH3G1_HUMAI | Unmodified | 346790 | 142510 | 0 | 1401300 | 1582900 | 0 | 0 | 0 | 0 | 0 | 0 | 0 | 9637600 | 0 | 197180 | 0 | 107400 | 785210 |
| NPGYPQSE | SH3GL1; SH3G | Q99961; M0 | SH3G1_HUMAI | Oxidation (M) | 0 | 0 | 0 | 268520 | 0 | 0 | 0 | 0 | 0 | 82472 | 0 | 0 | 4118500 | 1169800 | 0 | 0 | 0 | 0 |
| NPHEGYE\ | C1QL1 | O75973 | C1QRF_HUMAI | Unmodified | 0 | 0 | 0 | 0 | 0 | 0 | 0 | 0 | 9088400 | 0 | 0 | 0 | 0 | 0 | 0 | 0 | 0 | 0 |
| NPHPSSAF | BLVRA; BLVRA | P53004; C9J | BIEA_HUMAN E | Unmodified | 278070 | 146020 | 46805 | 22300 | 0 | 0 | 0 | 0 | 208080 | 0 | 0 | 230420 | 0 | 0 | 569300 | 310600 | 0 | 0 |
| NPIHNIPST | NHS; NHS; NHS | Q6T4R5; A0/ | NHS_HUMAN N | Unmodified | 0 | 0 | 0 | 0 | 882690 | 237000 | 0 | 0 | 0 | 153970 | 0 | 0 | 0 | 0 | 0 | 0 | 0 | 796510 |
| NPILWNVA | RPN2; RPN2; RI | P04844; H0 | RPN2_HUMAN | Unmodified | 0 | 0 | 0 | 0 | 0 | 0 | 0 | 0 | 567320 | 0 | 0 | 0 | 0 | 0 | 0 | 0 | 0 | 0 |
| NPIMSQCF | MYO1C; MYO1( | O00159; F5H | MYO1C_HUMA | Unmodified | 431950 | 388020 | 0 | 1032500 | 0 | 0 | 0 | 0 | 0 | 0 | 0 | 541720 | 0 | 0 | 3824000 | 1475600 | 46903 | 0 |
| NPIMSQCF | MYO1C; MYO1( | O00159; F5H | MYO1C_HUMA | Oxidation (M) | 518340 | 28124 | 0 | 0 | 115740 | 0 | 0 | 0 | 0 | 0 | 0 | 0 | 0 | 0 | 1257000 | 894250 | 0 | 0 |
| NPKPATVFS | PIGK; PIGK | Q92643; A6I | GPI8_HUMAN ( | Unmodified | 0 | 0 | 0 | 0 | 0 | 0 | 0 | 0 | 3834100 | 297530 | 0 | 0 | 0 | 0 | 0 | 0 | 88375 | 0 |
| NPLGSTHP | LRBA; LRBA; LR | P50851; A0/ | LRBA_HUMAN | Unmodified | 0 | 0 | 0 | 0 | 0 | 0 | 0 | 0 | 0 | 0 | 0 | 2328700 | 0 | 0 | 0 | 0 | 0 | 0 |
| NPLIINSIID | DIMT1; DIMT1; | Q9UNQ2; B4 | DIM1_HUMAN | Unmodified | 0 | 0 | 34198 | 20424 | 0 | 136910 | 0 | 11379 | 63229 | 0 | 464680 | 1195700 | 0 | 0 | 0 | 0 | 0 | 45734 |
| NPLMYLTG | ITGA2 | P17301 | ITA2_HUMAN I | Oxidation (M) | 0 | 0 | 243830 | 0 | 0 | 0 | 0 | 0 | 0 | 0 | 0 | 0 | 0 | 0 | 270370 | 106960 | 0 | 0 |
| NPLPSKETI | TMSB4X | P62328 | TYB4_HUMAN | Unmodified | 0 | 4135000 | 0 | 7177800 | 1.25E+08 | 22519000 | 16291000 | 15621000 | 1.43E+08 | 83588000 | 3164100 | 1753300 | 35519000 | 26572000 | 0 | 0 | 47898000 | 4.04E+08 |
| NPLPSKETI | TMSB4X | P62328 | TYB4_HUMAN | Unmodified | 0 | 0 | 0 | 0 | 36927000 | 3909400 | 27371000 | 33370000 | 0 | 27565000 | 0 | 0 | 5797200 | 11746000 | 0 | 0 | 86050000 | 1.16E+08 |
| NPLVAVYY | STUB1 | Q9UNE7 | CHIP_HUMAN | Unmodified | 593280 | 0 | 0 | 0 | 0 | 175400 | 0 | 0 | 0 | 0 | 0 | 770290 | 0 | 0 | 483390 | 276900 | 0 | 0 |
| NPNAVLTL | IQGAP2; IQGAP | Q13576; D6I | IQGA2_HUMAN | Unmodified | 0 | 0 | 277360 | 0 | 303270 | 0 | 0 | 0 | 0 | 0 | 483360 | 9938100 | 0 | 0 | 428500 | 276860 | 0 | 0 |
| NPNGPYP | HNRNPL; HNRI | P14866; M0 | HNRPL_HUMAI | Unmodified | 228480 | 215650 | 187600 | 121860 | 1021200 | 143520 | 2104000 | 2368200 | 723900 | 516830 | 3809100 | 17375000 | 2560600 | 637600 | 202350 | 167070 | 104270 | 795440 |
| NPNLPPET | APOD; APOD | P05090; C9J | APOD_HUMAN | Unmodified | 0 | 0 | 0 | 0 | 0 | 564840 | 0 | 0 | 2836600 | 197660 | 0 | 419190 | 0 | 0 | 0 | 0 | 0 | 0 |
| NPNTSEPC | ATP1A1 | P05023 | AT1A1_HUMAN | Oxidation (M) | 1203800 | 429560 | 11661000 | 6561800 | 4238500 | 961630 | 1339200 | 296030 | 4907900 | 1390800 | 9404200 | 57070000 | 5034500 | 2926100 | 19450000 | 19471000 | 1280500 | 2112300 |
| NPNTSEPC | ATP1A1 | P05023 | AT1A1_HUMAN | Unmodified | 1072000 | 891870 | 20609000 | 15617000 | 16864000 | 2003000 | 3433000 | 180640 | 29322000 | 2693900 | 28154000 | 1.87E+08 | 13143000 | 7476400 | 24708000 | 15182000 | 1343400 | 2347200 |
| NPPAAYIQI | FMNL1 | O95466 | FMNL1_HUMAN | Unmodified | 0 | 0 | 0 | 903690 | 3952400 | 790180 | 0 | 0 | 0 | 0 | 0 | 0 | 0 | 0 | 0 | 0 | 292130 | 0 |
| NPPDIVVQ | MPZL1; MPZL1 | O95297; Q9 | MPZL1_HUMAN | Unmodified | 2849100 | 1369700 | 5715900 | 1421500 | 0 | 0 | 79989 | 0 | 3998600 | 54720 | 560210 | 39164000 | 173490 | 0 | 3963700 | 3509700 | 168680 | 0 |

| | | | | | | | | | | | | | | | | | | | | | | |
|---|---|---|---|---|---|---|---|---|---|---|---|---|---|---|---|---|---|---|---|---|---|---|
| NPPDVDG | MPZL2 | O60487 | MPZL2_HUMAN | Unmodified | 0 | 0 | 1693900 | 0 | 0 | 0 | 0 | 0 | 0 | 0 | 0 | 2355300 | 0 | 0 | 1645600 | 749250 | 0 | 0 |
| NPPGFAFV | SRSF3; SRSF3 | P84103; A0A | SRSF3_HUMAN | Unmodified | 7467900 | 17139000 | 1374100 | 2435400 | 0 | 139360 | 0 | 0 | 961420 | 0 | 2157900 | 6906500 | 0 | 0 | 6679500 | 3388500 | 0 | 0 |
| NPPGFAFV | SRSF7; SRSF7; | Q16629; A0A | SRSF7_HUMAN | Unmodified | 4327300 | 5852600 | 592490 | 0 | 0 | 0 | 0 | 0 | 0 | 0 | 0 | 1891200 | 0 | 0 | 3163300 | 0 | 0 | 0 |
| NPPGGDN | TMEM30A; TME | Q9NV96; A0 | CC50A_HUMAI | Unmodified | 0 | 0 | 0 | 0 | 0 | 0 | 0 | 0 | 365760 | 167340 | 153930 | 0 | 0 | 0 | 0 | 0 | 0 | 0 |
| NPPNQSSI | U2SURP; U2SU | O15042; E7E | SR140_HUMAN | Unmodified | 0 | 0 | 0 | 0 | 0 | 0 | 0 | 0 | 0 | 0 | 0 | 0 | 0 | 0 | 0 | 0 | 0 | 0 |
| NPPPPINF | DIMT1; DIMT1; | Q9UNQ2; B4 | DIM1_HUMAN | Unmodified | 0 | 0 | 0 | 0 | 0 | 0 | 0 | 0 | 0 | 0 | 51405 | 899940 | 0 | 0 | 0 | 0 | 0 | 0 |
| NPQAHIEG | GMDS | O60547 | GMDS_HUMAN | Unmodified | 0 | 0 | 0 | 114940 | 0 | 137180 | 0 | 0 | 355770 | 0 | 110440 | 2249000 | 76434 | 0 | 138780 | 0 | 0 | 0 |
| NPQAVLDV | PAK2; PAK3; PA | Q13177; O7 | PAK2_HUMAN | Unmodified | 96294 | 92732 | 0 | 213490 | 2417000 | 213730 | 66401 | 51433 | 402840 | 146170 | 920530 | 11098000 | 521710 | 626890 | 131770 | 0 | 826300 | 939640 |
| NPQMGDP | FNBP1L; FNBP1 | Q5T0N5; A0 | FBP1L_HUMAN | Oxidation (M) | 353420 | 63615 | 0 | 0 | 0 | 0 | 0 | 0 | 0 | 0 | 0 | 83550 | 0 | 0 | 0 | 0 | 0 | 0 |
| NPQMGDP | FNBP1L; FNBP1 | Q5T0N5; A0 | FBP1L_HUMAN | Unmodified | 309940 | 394000 | 0 | 0 | 0 | 0 | 0 | 0 | 0 | 0 | 0 | 0 | 0 | 0 | 0 | 0 | 0 | 0 |
| NPQPAQA | GALNT16 | Q8N428 | GLT16_HUMAN | Unmodified | 0 | 0 | 0 | 0 | 483570 | 0 | 0 | 0 | 19336000 | 2247700 | 0 | 0 | 0 | 0 | 0 | 0 | 103630 | 0 |
| NPQQHLN | PABPC1; PABP | P11940; A0A | PABP1_HUMAN | 2 Oxidation (M) | 8413000 | 0 | 2170600 | 0 | 0 | 0 | 0 | 0 | 0 | 0 | 0 | 0 | 0 | 0 | 1544700 | 4112500 | 0 | 0 |
| NPQQHLN | PABPC1; PABP | P11940; A0A | PABP1_HUMAN | Oxidation (M) | 4386900 | 1694700 | 0 | 0 | 0 | 0 | 0 | 0 | 0 | 0 | 0 | 0 | 0 | 0 | 0 | 635100 | 0 | 0 |
| NPQREEIIG | LACTB2 | Q53H82 | LACB2_HUMAN | Unmodified | 359910 | 0 | 0 | 0 | 0 | 0 | 0 | 0 | 8351200 | 0 | 0 | 0 | 0 | 0 | 0 | 0 | 0 | 0 |
| NPQSYLYL | MYO1C; MYO1 | O00159; F5H | MYO1C_HUMA | Unmodified | 95860 | 0 | 0 | 0 | 0 | 0 | 0 | 0 | 0 | 0 | 0 | 2203200 | 0 | 0 | 15447000 | 6033600 | 0 | 0 |
| NPSAAFFC | GGCT; GGCT; | O75223; B8 | GGCT_HUMAN | Unmodified | 0 | 0 | 0 | 0 | 0 | 0 | 388990 | 0 | 0 | 0 | 0 | 1214600 | 0 | 0 | 0 | 262270 | 0 | 70503 |
| NPSAMAVE | TMEM43 | Q9BTV4 | TMM43_HUMAI | Oxidation (M) | 0 | 0 | 0 | 0 | 0 | 0 | 0 | 0 | 1628900 | 0 | 0 | 0 | 0 | 0 | 0 | 0 | 0 | 0 |
| NPSAMAVE | TMEM43 | Q9BTV4 | TMM43_HUMAI | Unmodified | 0 | 0 | 0 | 0 | 0 | 0 | 0 | 0 | 366920 | 0 | 0 | 0 | 0 | 0 | 0 | 0 | 0 | 0 |
| NPSASTFLI | HYAL2 | Q12891 | HYAL2_HUMAN | Unmodified | 595410 | 1077400 | 0 | 410890 | 0 | 0 | 0 | 0 | 233230 | 430050 | 0 | 1471500 | 0 | 0 | 0 | 119320 | 0 | 0 |
| NPSGSYVS | ADAM15 | Q13444 | ADA15_HUMAN | Unmodified | 664760 | 240900 | 260560 | 0 | 0 | 0 | 0 | 0 | 0 | 0 | 0 | 1479400 | 0 | 0 | 621010 | 503380 | 0 | 0 |
| NPSLPSKLI | EPS15L1; EPS1 | Q9UBC2; M | EP15R_HUMAN | Unmodified | 1925300 | 1567100 | 310950 | 784130 | 0 | 0 | 0 | 0 | 1577900 | 309070 | 890170 | 1663400 | 0 | 0 | 0 | 1016600 | 0 | 0 |
| NPSTNFQE | SRM; SRM | P19623; K7E | SPEE_HUMAN | Oxidation (M) | 0 | 0 | 0 | 0 | 0 | 0 | 0 | 0 | 821760 | 344160 | 0 | 1742400 | 0 | 0 | 0 | 468270 | 0 | 0 |
| NPSTSLGP | HSD17B11; HS | Q8NBQ5; D6 | DHB11_HUMAI | Unmodified | 0 | 0 | 0 | 0 | 94705 | 0 | 0 | 0 | 495640 | 0 | 0 | 135310 | 0 | 0 | 0 | 0 | 94336 | 0 |
| NPSTVEAFI | PFAS | O15067 | PUR4_HUMAN | Unmodified | 0 | 10824 | 0 | 0 | 0 | 0 | 0 | 0 | 244390 | 0 | 0 | 3689300 | 0 | 0 | 167140 | 0 | 152790 | 464700 |
| NPSVNWK | CSE1L | P55060 | XPO2_HUMAN | Unmodified | 68357 | 97809 | 213880 | 0 | 258010 | 0 | 139940 | 55615 | 376890 | 214560 | 508810 | 2990200 | 0 | 0 | 208090 | 28228 | 97940 | 249060 |
| NPSVSLVEI | CHST1 | O43916 | CHST1_HUMAI | Unmodified | 0 | 0 | 0 | 0 | 0 | 0 | 0 | 0 | 803790 | 0 | 0 | 70431 | 0 | 0 | 0 | 0 | 0 | 0 |
| NPTFLDYVI | SOAT1 | P35610 | SOAT1_HUMAN | Unmodified | 120240 | 0 | 0 | 0 | 0 | 0 | 0 | 0 | 6039800 | 142040 | 315340 | 0 | 0 | 0 | 0 | 0 | 0 | 0 |
| NPTNTVFD | HSPA8; HSPA8; | P11142; E9P | HSP7C_HUMAI | Unmodified | 0 | 0 | 0 | 0 | 0 | 0 | 0 | 0 | 0 | 0 | 0 | 0 | 0 | 0 | 0 | 0 | 905210 | 0 |
| NPVLQQD | CD276; CD276 | Q5ZPR3; A0 | CD276_HUMAI | Unmodified | 0 | 42287 | 0 | 0 | 0 | 0 | 0 | 0 | 0 | 0 | 0 | 0 | 0 | 0 | 113390 | 432860 | 0 | 0 |
| NPVTIFSLA | SSBP1; SSBP1; | Q04837; C9 | SSBP_HUMAN | Unmodified | 0 | 0 | 0 | 0 | 0 | 0 | 0 | 0 | 643770 | 0 | 0 | 0 | 0 | 0 | 0 | 0 | 0 | 0 |
| NPVWYQAI | IPO7 | O95373 | IPO7_HUMAN I | Unmodified | 0 | 0 | 444160 | 0 | 0 | 0 | 0 | 0 | 0 | 0 | 0 | 3613500 | 0 | 0 | 0 | 0 | 0 | 0 |
| NPWTGVS | PCYT2; PCYT2; | Q99447; I3L | PCY2_HUMAN | Unmodified | 0 | 0 | 25715 | 0 | 0 | 0 | 0 | 0 | 0 | 0 | 0 | 370010 | 0 | 0 | 0 | 41554 | 0 | 0 |
| NQAAIQGR | FAM120A; FAM | Q9NZB2; A0 | F120A_HUMAN | Unmodified | 853930 | 688510 | 0 | 978680 | 538160 | 0 | 0 | 0 | 0 | 0 | 0 | 443860 | 1473800 | 378720 | 0 | 579030 | 121160 | 750290 |
| NQAEAQK | NECTIN2; NEC | Q92692; K7E | NECT2_HUMAI | Unmodified | 0 | 0 | 0 | 0 | 0 | 0 | 0 | 0 | 0 | 0 | 0 | 0 | 0 | 0 | 0 | 0 | 883010 | 987610 |
| NQAIEELEK | STK26; STK26; | Q9P289; B4 | STK26_HUMAN | Unmodified | 0 | 0 | 0 | 0 | 0 | 0 | 0 | 0 | 0 | 0 | 387920 | 2933500 | 0 | 0 | 0 | 325590 | 0 | 0 |
| NQALFPAC | DDX1; DDX1; D | Q92499; A0 | DDX1_HUMAN | Unmodified | 228520 | 0 | 353650 | 202160 | 732800 | 0 | 0 | 235080 | 806040 | 0 | 725130 | 3088100 | 845040 | 292670 | 0 | 298670 | 319500 | 396760 |
| NQALNTDI | SPTAN1; SPTAI | Q13813; A0 | SPTN1_HUMAN | Unmodified | 0 | 0 | 315510 | 25934 | 0 | 0 | 0 | 0 | 1345800 | 0 | 1071300 | 3933200 | 0 | 113520 | 244310 | 178120 | 0 | 0 |
| NQALNTDI | SPTAN1; SPTAI | Q13813; A0 | SPTN1_HUMAN | Unmodified | 0 | 0 | 0 | 0 | 0 | 0 | 197330 | 28533 | 0 | 0 | 0 | 1059600 | 0 | 0 | 0 | 51801 | 0 | 0 |
| NQANPSR | ADAM9; ADAM | Q13443; A0 | ADAM9_HUMA | Unmodified | 0 | 0 | 0 | 0 | 0 | 0 | 0 | 0 | 0 | 0 | 0 | 0 | 0 | 0 | 539260 | 0 | 0 | 0 |
| NQAVCHD | QSOX2 | Q6ZRP7 | QSOX2_HUMAI | Unmodified | 0 | 0 | 0 | 0 | 0 | 0 | 0 | 0 | 37937000 | 2418200 | 988430 | 1224700 | 0 | 0 | 0 | 0 | 0 | 0 |
| NQAVNFLF | CHMP1B | Q7LBR1 | CHM1B_HUMA | Unmodified | 625990 | 3287300 | 0 | 13907000 | 51832000 | 17101000 | 1777400 | 683780 | 37315000 | 46464000 | 21812000 | 59073000 | 20675000 | 14060000 | 1535000 | 858080 | 713390 | 7786800 |
| NQCTQVV | DSP | P15924 | DESP_HUMAN | Unmodified | 0 | 0 | 0 | 0 | 0 | 0 | 0 | 0 | 0 | 676600 | 0 | 182640 | 763860 | 0 | 0 | 57306 | 780890 | 0 |
| NQDATVYV | SF3B4 | Q15427 | SF3B4_HUMAN | Unmodified | 536320 | 459630 | 0 | 0 | 0 | 0 | 0 | 106130 | 0 | 0 | 0 | 0 | 0 | 0 | 0 | 0 | 240330 | 0 |
| NQDEESQ | ABCF1; ABCF1; | Q8NE71; H0 | ABCF1_HUMAN | Unmodified | 0 | 0 | 0 | 0 | 328690 | 0 | 0 | 0 | 0 | 0 | 382500 | 1539200 | 0 | 0 | 149420 | 0 | 0 | 215030 |
| NQDGKITV | FKBP10; FKBP1 | Q96AY3; H0 | FKB10_HUMAN | Unmodified | 0 | 0 | 42310 | 0 | 0 | 0 | 0 | 0 | 0 | 0 | 926080 | 0 | 0 | 0 | 0 | 0 | 0 | 0 |
| NQDLALSN | UBQLN2 | Q9UHD9 | UBQL2_HUMAI | Unmodified | 1212800 | 0 | 719650 | 0 | 0 | 0 | 0 | 0 | 492040 | 0 | 0 | 405780 | 0 | 0 | 0 | 0 | 386880 | 0 |
| NQDLAPN | ILF2; ILF2; ILF2; | Q12905; B4 | ILF2_HUMAN I | Unmodified | 1518600 | 14428 | 272380 | 750710 | 0 | 0 | 0 | 403290 | 1222000 | 0 | 660020 | 8912800 | 0 | 0 | 0 | 0 | 0 | 0 |
| NQDLCQQ | PDIA5; PDIA5 | Q14554; H7 | PDIA5_HUMAN | Unmodified | 0 | 55596 | 0 | 0 | 0 | 0 | 0 | 0 | 2982500 | 274190 | 0 | 136980 | 0 | 0 | 0 | 0 | 0 | 0 |
| NQDNLQG | CMPK1; CMPK1 | P30085; Q5 | KCY_HUMAN U | Unmodified | 188490 | 0 | 872030 | 1407400 | 263470 | 2302400 | 0 | 60756 | 1096100 | 0 | 115170 | 1176200 | 467060 | 213360 | 476790 | 0 | 133720 | 624250 |
| NQDQQGP | MUC5AC | P98088 | MUC5A_HUMA | Unmodified | 628820 | 0 | 0 | 0 | 191170 | 0 | 0 | 14512 | 89005 | 0 | 0 | 0 | 0 | 0 | 0 | 0 | 0 | 314630 |
| NQEASLGN | SPATS2L; SPAT | Q9NUQ6; B | SPS2L_HUMAN | Unmodified | 29103 | 0 | 0 | 0 | 0 | 0 | 0 | 0 | 0 | 0 | 0 | 271580 | 0 | 0 | 0 | 0 | 0 | 0 |
| NQEQLAAE | RDX; RDX; RDX | P35241; A0A | RADI_HUMAN I | Unmodified | 5242400 | 2037000 | 3698800 | 13548000 | 62070000 | 6664600 | 661120 | 1357600 | 11165000 | 5921600 | 4341600 | 39519000 | 46369000 | 52488000 | 8659100 | 2552500 | 5401800 | 14246000 |
| NQEQLLTL | HEBP2 | Q9Y5Z4 | HEBP2_HUMAI | Unmodified | 0 | 0 | 0 | 789360 | 0 | 0 | 0 | 0 | 2421600 | 815210 | 2308400 | 2827800 | 0 | 0 | 0 | 2184900 | 0 | 0 |
| NQEQLLTL | HEBP2 | Q9Y5Z4 | HEBP2_HUMAI | Unmodified | 648020 | 848170 | 0 | 957980 | 1430700 | 161920 | 474420 | 0 | 3665600 | 8717300 | 746030 | 2217200 | 5580500 | 3707100 | 781710 | 448110 | 1664100 | 765280 |
| NQEQMKP | PSMC1 | P62191 | PRS4_HUMAN | Unmodified | 0 | 0 | 22368 | 0 | 0 | 0 | 0 | 0 | 0 | 0 | 93629 | 1359500 | 0 | 0 | 35270 | 39765 | 0 | 0 |
| NQFPAGS | SMPDL3B; SMP | Q92485; Q5 | ASM3B_HUMAI | Unmodified | 0 | 39048 | 331620 | 92830 | 630400 | 0 | 0 | 0 | 96775 | 107510 | 0 | 0 | 0 | 0 | 794260 | 137790 | 0 | 0 |
| NQFSGPTS | PLCB2 | Q00722 | PLCB2_HUMAN | Unmodified | 0 | 0 | 0 | 0 | 0 | 0 | 0 | 0 | 45726 | 0 | 302370 | 2611300 | 0 | 0 | 0 | 0 | 0 | 0 |
| NQGATIHL | CYFIP1; CYFIP1 | Q7L576; A0 | CYFP1_HUMAN | Unmodified | 360850 | 0 | 463680 | 451200 | 0 | 0 | 0 | 0 | 437330 | 0 | 0 | 1159600 | 0 | 80057 | 584250 | 613310 | 0 | 0 |
| NQGEEQR | DNASE2 | O00115 | DNS2A_HUMAI | Unmodified | 0 | 1512200 | 0 | 0 | 0 | 225150 | 0 | 0 | 4088600 | 199970 | 185820 | 0 | 0 | 0 | 0 | 402600 | 0 | 882270 |
| NQGGGLS | EPRS1 | P07814 | SYEP_HUMAN | Unmodified | 0 | 0 | 0 | 0 | 0 | 0 | 0 | 0 | 305370 | 0 | 1193100 | 2297900 | 0 | 119130 | 262380 | 221170 | 0 | 0 |
| NQGGGLS | EPRS1 | P07814 | SYEP_HUMAN | Unmodified | 0 | 0 | 0 | 0 | 133130 | 48406 | 0 | 15942 | 0 | 61481 | 0 | 0 | 329460 | 153410 | 0 | 0 | 68473 | 175010 |
| NQGGYDR | RBM3 | P98179 | RBM3_HUMAN | Unmodified | 5392300 | 13614000 | 1740600 | 2238100 | 0 | 0 | 0 | 0 | 0 | 0 | 0 | 0 | 569310 | 0 | 5007100 | 7027100 | 0 | 0 |
| NQGIEEALI | VCL | P18206 | VINC_HUMAN | Unmodified | 485950 | 1428300 | 1109700 | 983190 | 6873000 | 2555700 | 2111400 | 1126700 | 1412400 | 1279200 | 173530 | 7593800 | 3034700 | 2852300 | 1320000 | 1120800 | 10477000 | 5498600 |
| NQGIEEALI | VCL | P18206 | VINC_HUMAN | Unmodified | 0 | 0 | 0 | 0 | 0 | 0 | 122450 | 0 | 0 | 0 | 0 | 0 | 0 | 0 | 0 | 0 | 570670 | 12255 |
| NQGIEEALI | VCL | P18206 | VINC_HUMAN | Unmodified | 0 | 0 | 0 | 0 | 3582100 | 0 | 3486000 | 1363500 | 0 | 0 | 0 | 0 | 1849300 | 0 | 0 | 0 | 14966000 | 2862900 |
| NQGNLYDI | STT3B | Q8TCJ2 | STT3B_HUMAN | Unmodified | 0 | 0 | 0 | 0 | 0 | 3244300 | 283680 | 0 | 45321000 | 22420000 | 31435000 | 1069500 | 0 | 171640 | 0 | 0 | 0 | 0 |
| NQGSEPK | ITGA6 | P23229 | ITA6_HUMAN I | Unmodified | 0 | 0 | 0 | 91995 | 7201000 | 1731000 | 0 | 0 | 85441 | 115780 | 0 | 0 | 5205500 | 2047100 | 0 | 0 | 6988000 | 33562000 |
| NQGSEPK | ITGA6 | P23229 | ITA6_HUMAN I | Unmodified | 0 | 0 | 0 | 0 | 131680 | 0 | 0 | 0 | 0 | 0 | 0 | 0 | 137490 | 0 | 0 | 0 | 617760 | 1011700 |
| NQGYYDY | OGDH; OGDH; | Q02218; E9 | ODO1_HUMAN | Unmodified | 0 | 0 | 0 | 0 | 0 | 0 | 33686 | 0 | 1164400 | 187990 | 0 | 109500 | 0 | 0 | 0 | 0 | 57137 | 74233 |
| NQHFDGF | CHID1 | Q9BWS9 | CHID1_HUMAN | Unmodified | 0 | 0 | 0 | 0 | 0 | 0 | 0 | 0 | 0 | 346530 | 0 | 0 | 0 | 0 | 136300 | 0 | 0 | 0 |
| NQIIACNG | RRM1; RRM1 | P23921; E9P | RIR1_HUMAN I | Unmodified | 0 | 0 | 0 | 0 | 0 | 0 | 2062900 | 0 | 0 | 0 | 0 | 2697500 | 0 | 0 | 0 | 0 | 0 | 855560 |
| NQIILKLSN | UBE2O; UBE2O | Q9C0C9; K7 | UBE2O_HUMAI | Unmodified | 0 | 286260 | 1192600 | 5655900 | 7631000 | 22558000 | 3667500 | 881930 | 0 | 0 | 0 | 0 | 2975100 | 2609400 | 1413000 | 0 | 2038600 | 18358000 |
| NQILIIDEA | ABCC4 | O15439 | MRP4_HUMAN | Unmodified | 0 | 0 | 0 | 0 | 2196400 | 0 | 0 | 0 | 0 | 0 | 0 | 0 | 1527100 | 0 | 0 | 222030 | 0 | 0 |
| NQITQLESI | CEP55 | Q53EZ4 | CEP55_HUMAN | Unmodified | 178170 | 196380 | 0 | 421110 | 2645800 | 0 | 0 | 0 | 0 | 1115100 | 0 | 349360 | 0 | 0 | 417840 | 212690 | 0 | 0 |
| NQIVFDNR | ENAH | Q8N8S7 | ENAH_HUMAN | Unmodified | 103560 | 168380 | 0 | 0 | 0 | 0 | 0 | 0 | 0 | 0 | 0 | 3565000 | 0 | 0 | 0 | 0 | 0 | 0 |
| NQIVFDNR | ENAH | Q8N8S7 | ENAH_HUMAN | Unmodified | 0 | 0 | 0 | 0 | 0 | 0 | 0 | 0 | 0 | 0 | 0 | 0 | 2904000 | 0 | 0 | 0 | 0 | 0 |
| NQKDDVA | SLC3A2; SLC3A | P08195; A0A | 4F2_HUMAN 4I | Unmodified | 3108500 | 1563200 | 5773600 | 1922400 | 3121800 | 237860 | 140580 | 0 | 6598700 | 856710 | 11209000 | 45141000 | 0 | 0 | 8043900 | 6045000 | 0 | 92906 |
| NQKDDVA | SLC3A2; SLC3A | P08195; A0A | 4F2_HUMAN 4I | Unmodified | 24837000 | 624460 | 816800 | 17674000 | 1.3E+08 | 29237000 | 7840000 | 0 | 46697000 | 25929000 | 2394700 | 22483000 | 17454000 | 6216600 | 37365000 | 9229300 | 5105700 | 22351000 |
| NQKDPGVI | S100A11 | P31949 | S10AB_HUMAN | Oxidation (M) | 0 | 0 | 0 | 0 | 5395700 | 0 | 0 | 437700 | 0 | 378180 | 0 | 0 | 1628000 | 0 | 0 | 0 | 1059800 | 3477100 |
| NQKDPGVI | S100A11 | P31949 | S10AB_HUMAN | Unmodified | 0 | 0 | 0 | 0 | 34181000 | 4420800 | 2562100 | 879410 | 198410 | 2490100 | 887750 | 0 | 3832900 | 9334700 | 0 | 0 | 7114300 | 16907000 |
| NQLAVLRV | CHMP3; RNF10 | Q9Y3E7; A0 | CHMP3_HUMA | Unmodified | 0 | 0 | 0 | 44776 | 2559800 | 504120 | 1481500 | 652820 | 320050 | 2847500 | 220430 | 0 | 1281100 | 3173300 | 0 | 0 | 957910 | 596550 |
| NQLDQEVE | PFDN5; PFDN5 | Q99471; A0 | PFD5_HUMAN | Unmodified | 0 | 0 | 0 | 0 | 5080600 | 435710 | 1064700 | 0 | 569570 | 4305200 | 0 | 511260 | 6683700 | 2492400 | 0 | 0 | 3344700 | 4339800 |
| NQLIEQFP | MCTS1 | Q9ULC4 | MCTS1_HUMAI | Oxidation (M) | 0 | 0 | 0 | 0 | 0 | 0 | 0 | 0 | 0 | 0 | 0 | 0 | 995610 | 0 | 0 | 0 | 0 | 0 |
| NQLLGPDE | PTPN6 | P29350 | PTN6_HUMAN | Unmodified | 0 | 0 | 0 | 0 | 0 | 203100 | 0 | 27333 | 316400 | 81813 | 96493 | 622610 | 309370 | 342110 | 0 | 112080 | 0 | 0 |
| NQLLLALL | PREX1; PREX1 | Q8TCU6; H0 | PREX1_HUMAN | Unmodified | 46434 | 0 | 0 | 0 | 0 | 0 | 0 | 0 | 0 | 0 | 35642 | 199010 | 0 | 0 | 0 | 0 | 0 | 0 |
| NQLLPELQ | IPO11; IPO11 | Q9UI26; F8V | IPO11_HUMAN | Unmodified | 0 | 1962100 | 0 | 0 | 0 | 0 | 0 | 0 | 0 | 0 | 0 | 4290700 | 0 | 0 | 0 | 0 | 0 | 0 |
| NQLLSCSA | STRN3 | Q13033 | STRN3_HUMAI | Unmodified | 0 | 0 | 0 | 0 | 0 | 0 | 0 | 0 | 0 | 0 | 0 | 1404300 | 0 | 0 | 0 | 0 | 0 | 0 |
| NQLLTLK | TBCE; TBCE; TB | Q15813; A0 | TBCE_HUMAN | Unmodified | 0 | 0 | 0 | 43681 | 1396800 | 0 | 508770 | 286700 | 196800 | 339250 | 583180 | 1622300 | 573360 | 0 | 110870 | 0 | 3349800 | 1822200 |
| NQLQGAM | WWP2 | O00308 | WWP2_HUMAN | Unmodified | 4101500 | 5418600 | 2192200 | 2648200 | 0 | 0 | 0 | 0 | 498770 | 0 | 0 | 845390 | 0 | 0 | 5404400 | 2986200 | 138530 | 0 |
| NQLQGAM | WWP2 | O00308 | WWP2_HUMAN | Oxidation (M) | 980510 | 884890 | 470200 | 321080 | 0 | 0 | 0 | 0 | 0 | 0 | 0 | 66958 | 0 | 0 | 671720 | 393990 | 0 | 0 |
| NQLQGAM | WWP1 | Q9H0M0 | WWP1_HUMAN | Unmodified | 0 | 171960 | 0 | 0 | 0 | 115840 | 0 | 0 | 0 | 28961 | 0 | 0 | 0 | 0 | 806330 | 0 | 62092 | 0 |
| NQLQGAM | WWP1 | Q9H0M0 | WWP1_HUMAN | Oxidation (M) | 0 | 0 | 161440 | 125450 | 0 | 0 | 0 | 0 | 0 | 0 | 0 | 0 | 0 | 0 | 193830 | 226450 | 0 | 0 |
| NQLTAMSS | EIF3A | Q14152 | EIF3A_HUMAN | Unmodified | 0 | 289850 | 0 | 0 | 0 | 0 | 0 | 0 | 0 | 0 | 0 | 5679800 | 0 | 0 | 0 | 0 | 0 | 0 |
| NQLTAMSS | EIF3A | Q14152 | EIF3A_HUMAN | Oxidation (M) | 0 | 0 | 0 | 0 | 0 | 0 | 0 | 0 | 0 | 0 | 0 | 1245400 | 0 | 0 | 0 | 0 | 0 | 0 |
| NQLTSLPS | FLII; FLII; FLII | Q13045; J3K | FLII_HUMAN P | Unmodified | 0 | 0 | 0 | 0 | 0 | 0 | 0 | 0 | 0 | 0 | 0 | 0 | 0 | 0 | 0 | 0 | 0 | 0 |
| NQLTSNPE | HSPA5; HSPA5; | P11021; A0A | BIP_HUMAN Er | Unmodified | 771460 | 282740 | 3459600 | 1706200 | 7898200 | 1339200 | 17381000 | 8261300 | 1.51E+08 | 69472000 | 13900000 | 6350300 | 4521200 | 4417300 | 4051500 | 1382100 | 798770 | 6274700 |
| NQLTSNPE | HSPA5; HSPA5; | P11021; A0A | BIP_HUMAN Er | Unmodified | 0 | 0 | 0 | 214750 | 292200 | 0 | 4468900 | 3141800 | 15729000 | 7596500 | 3605800 | 419030 | 0 | 587370 | 0 | 0 | 0 | 0 |
| NQMLIEEE | UCHL5; UCHL5 | Q9Y5K5; H0 | UCHL5_HUMAI | Oxidation (M) | 0 | 0 | 77609 | 0 | 0 | 0 | 0 | 0 | 0 | 0 | 141760 | 280180 | 0 | 0 | 0 | 0 | 0 | 0 |
| NQMLISED | PTK7; PTK7 | Q13308; H7 | PTK7_HUMAN | Unmodified | 0 | 0 | 136740 | 0 | 0 | 0 | 0 | 0 | 688940 | 0 | 0 | 0 | 0 | 0 | 0 | 0 | 0 | 0 |
| NQNDPLP | CSDE1; CSDE1 | O75534; E9 | CSDE1_HUMAI | Unmodified | 194000 | 155870 | 302210 | 494710 | 0 | 0 | 0 | 0 | 0 | 0 | 0 | 1665600 | 146470 | 61632 | 225690 | 674890 | 0 | 0 |
| NQNQELPE | AVIL | O75366 | AVIL_HUMAN A | Unmodified | 0 | 0 | 0 | 0 | 0 | 0 | 0 | 8376 | 0 | 0 | 357690 | 1379800 | 0 | 0 | 51087 | 0 | 0 | 0 |
| NQNQELPE | AVIL | O75366 | AVIL_HUMAN A | Unmodified | 0 | 0 | 0 | 0 | 139560 | 0 | 0 | 0 | 510190 | 0 | 15931000 | 45377000 | 0 | 0 | 0 | 0 | 136780 | 88244 |
| NQNQELPE | AVIL | O75366 | AVIL_HUMAN A | Unmodified | 0 | 0 | 0 | 0 | 0 | 0 | 0 | 0 | 0 | 0 | 2578900 | 6565800 | 0 | 0 | 0 | 0 | 0 | 0 |
| NQNSWGT | LMNB1 | P20700 | LMNB1_HUMAI | Unmodified | 0 | 0 | 0 | 0 | 0 | 0 | 0 | 0 | 0 | 0 | 0 | 406760 | 0 | 0 | 0 | 0 | 0 | 0 |
| NQPIYIQYS | PTBP2; PTBP2; | Q9UKA9; A0 | PTBP2_HUMAN | Unmodified | 0 | 0 | 0 | 0 | 0 | 0 | 0 | 0 | 0 | 0 | 0 | 692010 | 0 | 0 | 0 | 0 | 0 | 0 |
| NQQDGQL | GMPPA; GMPP | Q96IJ6; A0A | GMPPA_HUMA | Unmodified | 0 | 0 | 0 | 0 | 1660900 | 1371900 | 85008 | 104170 | 0 | 171100 | 0 | 2486000 | 0 | 0 | 0 | 70860 | 3733000 | 0 |
| NQQGIKAE | SPTBN2 | O15020 | SPTN2_HUMAN | Unmodified | 0 | 0 | 0 | 245440 | 0 | 0 | 0 | 0 | 100100 | 0 | 12381 | 0 | 178970 | 0 | 279050 | 421660 | 0 | 0 |
| NQQIFLR | GRB2; GRB2 | P62993; J3Q | GRB2_HUMAN | Unmodified | 4601200 | 4870800 | 2271300 | 0 | 385640 | 2062800 | 894860 | 288760 | 322310 | 136450 | 0 | 5447900 | 0 | 920250 | 5154800 | 4017300 | 338330 | 3298000 |
| NQQITHAN | HSPH1; HSPH1 | Q92598; R4 | HS105_HUMAN | Unmodified | 0 | 0 | 0 | 0 | 429080 | 0 | 188060 | 122870 | 101240 | 0 | 87650 | 628800 | 303720 | 270430 | 0 | 0 | 441620 | 0 |

| | | | | | | | | | | | | | | | | | | | | | | |
|---|---|---|---|---|---|---|---|---|---|---|---|---|---|---|---|---|---|---|---|---|---|---|
| NQQSDFYI | STEAP2; STEAP | Q8NFT2; B5I | STEA2_HUMAN | Unmodified | 0 | 301190 | 0 | 188390 | 85573 | 0 | 0 | 0 | 0 | 0 | 0 | 0 | 0 | 0 | 2106300 | 1429900 | 68010 | 38453 |
| NQQWLVYI | CEP55; CEP55 | Q53EZ4; H0 | CEP55_HUMAN | Unmodified | 0 | 0 | 0 | 591100 | 682490 | 0 | 0 | 0 | 2161700 | 197810 | 0 | 0 | 0 | 0 | 575560 | 0 | 0 | 0 |
| NQQWLVYI | CEP55; CEP55 | Q53EZ4; H0 | CEP55_HUMAN | Unmodified | 1201200 | 977240 | 0 | 8755700 | 61709000 | 17160000 | 3825900 | 397500 | 12075000 | 21295000 | 1983600 | 1096900 | 4498200 | 5615400 | 2325300 | 568740 | 4381100 | 6080800 |
| NQRARQSI | EPHA2 | P29317 | EPHA2_HUMAN | Unmodified | 0 | 0 | 0 | 0 | 156520 | 0 | 0 | 0 | 0 | 0 | 0 | 0 | 469520 | 0 | 0 | 0 | 478490 | 626380 |
| NQRVPADI | ATP9A | O75110 | ATP9A_HUMAN | Unmodified | 0 | 0 | 0 | 0 | 0 | 0 | 62416 | 19181 | 789660 | 0 | 0 | 0 | 0 | 0 | 0 | 0 | 0 | 0 |
| NQSETLNV | CKAP5 | Q14008 | CKAP5_HUMAN | Unmodified | 1153300 | 2321300 | 29092 | 1412900 | 0 | 0 | 0 | 0 | 0 | 0 | 98226 | 985810 | 0 | 0 | 1191000 | 1072800 | 0 | 0 |
| NQSFCPTV | RPL27A; RPL27 | P46776; E9P | RL27A_HUMAN | Unmodified | 0 | 1702800 | 0 | 856860 | 0 | 0 | 817310 | 483700 | 0 | 0 | 968390 | 4554000 | 757080 | 0 | 0 | 0 | 0 | 0 |
| NQSIAIELS | PLCG2; PLCG2 | P16885; A0 | PLCG2_HUMAN | Unmodified | 0 | 0 | 0 | 0 | 0 | 0 | 0 | 0 | 1171600 | 0 | 617280 | 1864600 | 0 | 0 | 0 | 0 | 0 | 0 |
| NQSLVEEN | KIF4A; KIF4B | O95239; Q2 | KIF4A_HUMAN | Unmodified | 0 | 0 | 0 | 0 | 491770 | 100760 | 0 | 70385 | 156040 | 258030 | 0 | 0 | 0 | 684530 | 0 | 0 | 487950 | 487920 |
| NQSPEPSS | RFTN1; RFTN1; | Q14699; G3 | RFTN1_HUMAN | Unmodified | 2273400 | 1959000 | 1290200 | 857710 | 0 | 0 | 79807 | 0 | 0 | 0 | 127640 | 1159100 | 580300 | 298070 | 1180100 | 1239600 | 0 | 0 |
| NQSPVDQ | SDC1 | P18827 | SDC1_HUMAN | Unmodified | 3084100 | 3094300 | 1710300 | 882800 | 582650 | 0 | 0 | 0 | 2318100 | 0 | 0 | 0 | 0 | 0 | 561170 | 1123200 | 170650 | 346450 |
| NQSSEDILI | TP53BP2 | Q13625 | ASPP2_HUMAN | Unmodified | 0 | 0 | 0 | 0 | 980750 | 0 | 0 | 0 | 0 | 0 | 0 | 0 | 0 | 671810 | 0 | 0 | 740120 | 1639700 |
| NQTAEKEE | HSPA8; HSPA8; | P11142; E9P | HSP7C_HUMAN | Unmodified | 75362000 | 1.69E+08 | 78757000 | 1.11E+08 | 2.11E+08 | 1.05E+08 | 44293000 | 2263200 | 1.05E+08 | 78445000 | 61957000 | 2.69E+08 | 30692000 | 29069000 | 48868000 | 37699000 | 0 | 63850000 |
| NQTAEKEE | HSPA8; HSPA8; | P11142; E9P | HSP7C_HUMAN | Unmodified | 794490 | 1225100 | 0 | 1805000 | 79277000 | 5038300 | 14890000 | 5809500 | 3953200 | 14789000 | 946340 | 1125900 | 31109000 | 24999000 | 466420 | 236520 | 14178000 | 16949000 |
| NQTIDFLNI | TSPAN15; TSPA | O95858; H7 | TSN15_HUMAN | Unmodified | 17020 | 0 | 0 | 0 | 2013700 | 0 | 0 | 0 | 0 | 146820 | 0 | 1106200 | 0 | 1142900 | 0 | 0 | 0 | 765110 |
| NQTIDFLNI | TSPAN15; TSPA | O95858; H7 | TSN15_HUMAN | Unmodified | 187760 | 290030 | 0 | 0 | 0 | 0 | 501420 | 0 | 0 | 84424 | 0 | 0 | 109790 | 0 | 929840 | 574110 | 81046 | 0 |
| NQTPAPSA | DIP2B | Q9P265 | DIP2B_HUMAN | Unmodified | 0 | 0 | 45373 | 200180 | 0 | 0 | 0 | 0 | 209480 | 0 | 50767 | 1152600 | 0 | 415370 | 0 | 0 | 202520 | 617620 |
| NQTVVVEY | PELI2 | Q9HAT8 | PELI2_HUMAN | Unmodified | 0 | 0 | 0 | 0 | 0 | 0 | 0 | 0 | 1570700 | 72437 | 0 | 331920 | 0 | 0 | 0 | 0 | 0 | 0 |
| NQVALNP | HSPA1A; HSPA | P0DMV8; P0 | HS71A_HUMAN | Unmodified | 6988600 | 8512800 | 4545400 | 6289300 | 26912000 | 5429600 | 6351900 | 3100100 | 14293000 | 10394000 | 1141200 | 5413600 | 30979000 | 12237000 | 3115800 | 3314300 | 6685300 | 24546000 |
| NQVALNP | HSPA1A; HSPA | P0DMV8; P0 | HS71A_HUMAN | Unmodified | 4646900 | 1327100 | 1681500 | 3538200 | 2181700 | 1753700 | 0 | 0 | 5299500 | 1211900 | 319090 | 2336600 | 3482800 | 0 | 1278900 | 1498000 | 1410600 | 2170000 |
| NQVALNP | HSPA1A; HSPA | P0DMV8; P0 | HS71A_HUMAN | Unmodified | 426040 | 0 | 0 | 0 | 0 | 0 | 1507300 | 0 | 0 | 113210 | 0 | 0 | 0 | 358700 | 338380 | 318940 | 1574400 | 0 |
| NQVAMNP | HSPA1L; HSPA | P34931; Q5 | HS71L_HUMAN | Oxidation (M) | 0 | 0 | 0 | 0 | 0 | 0 | 325390 | 0 | 398740 | 236830 | 0 | 0 | 0 | 0 | 0 | 0 | 0 | 0 |
| NQVAMNP | HSPA2 | P54652 | HSP72_HUMAN | Oxidation (M) | 656350 | 1014500 | 0 | 0 | 1297200 | 0 | 0 | 0 | 0 | 0 | 0 | 0 | 0 | 0 | 0 | 0 | 0 | 213450 |
| NQVAMNP | HSPA8; HSPA8; | P11142; E9P | HSP7C_HUMAN | Unmodified | 1260200 | 1730100 | 1197900 | 3944000 | 23536000 | 2722600 | 5866200 | 727970 | 13857000 | 5056100 | 0 | 7325000 | 13935000 | 9345800 | 2225800 | 843620 | 3059900 | 8920100 |
| NQVAMNP | HSPA8; HSPA8; | P11142; E9P | HSP7C_HUMAN | Oxidation (M) | 8668500 | 10210000 | 12256000 | 14253000 | 28837000 | 5370400 | 8819700 | 4150200 | 9241100 | 8179500 | 3566500 | 19710000 | 32044000 | 18816000 | 10786000 | 7979800 | 5468500 | 8729000 |
| NQVAMNP | HSPA8; HSPA8; | P11142; E9P | HSP7C_HUMAN | Oxidation (M) | 2282900 | 1087400 | 968040 | 1370400 | 695140 | 192150 | 725060 | 548190 | 1127300 | 442900 | 554530 | 3826200 | 760920 | 352440 | 998130 | 0 | 674280 | 678180 |
| NQVAMNP | HSPA8; HSPA8; | P11142; E9P | HSP7C_HUMAN | Unmodified | 2109000 | 1185900 | 1532300 | 2553700 | 2294900 | 980830 | 2079100 | 366480 | 4264800 | 1590300 | 1593100 | 6074300 | 2920800 | 1462900 | 1651300 | 1494200 | 2769800 | 1205600 |
| NQVAMNP | HSPA8 | E9PI65 | E9PI65_HUMAN | Oxidation (M) | 0 | 0 | 0 | 66556 | 170330 | 0 | 0 | 0 | 340890 | 0 | 0 | 0 | 0 | 0 | 0 | 0 | 0 | 0 |
| NQVDVEIK | RACGAP1; RAC | Q9H0H5; F8 | RGAP1_HUMAN | Unmodified | 0 | 0 | 0 | 1615600 | 4059700 | 1173400 | 0 | 0 | 3287200 | 784080 | 1447800 | 1769600 | 392330 | 0 | 0 | 0 | 503380 | 312800 |
| NQVEDLLA | NCSTN; NCSTN | Q92542; H0 | NICA_HUMAN | Unmodified | 362710 | 0 | 234530 | 309010 | 0 | 0 | 0 | 0 | 18323000 | 2531400 | 378900 | 2272200 | 0 | 0 | 871230 | 624480 | 0 | 0 |
| NQVGDEN | TNPO1; TNPO1 | Q92973; A0 | TNPO1_HUMAN | Unmodified | 0 | 0 | 64152 | 0 | 0 | 0 | 0 | 0 | 0 | 0 | 0 | 252530 | 0 | 0 | 56049 | 96708 | 0 | 0 |
| NQVLTLEE | TPMT | P51580 | TPMT_HUMAN | Unmodified | 0 | 0 | 0 | 180010 | 2487800 | 0 | 0 | 0 | 604890 | 0 | 0 | 635310 | 311390 | 0 | 0 | 133070 | 0 | 283320 |
| NQVLVFVH | SNRNP200; SN | O75643; A0 | U520_HUMAN | Unmodified | 0 | 0 | 0 | 0 | 0 | 0 | 0 | 0 | 0 | 54239 | 0 | 1422700 | 0 | 0 | 0 | 0 | 0 | 0 |
| NQVSQLLP | SAMHD1; SAM | Q9Y3Z3; A0 | SAMH1_HUMA | Unmodified | 0 | 0 | 0 | 0 | 353700 | 0 | 974090 | 327920 | 0 | 0 | 0 | 331740 | 0 | 0 | 0 | 0 | 129660 | 0 |
| NQVTATKA | DLD; DLD | P09622; E9P | DLDH_HUMAN | Unmodified | 0 | 0 | 0 | 38923 | 217470 | 0 | 327300 | 0 | 2734300 | 1148100 | 0 | 0 | 617670 | 261890 | 0 | 0 | 48387 | 205050 |
| NQVTQLK | MTHFD1; MTHF | P11586; V9 | C1TC_HUMAN | Unmodified | 118670 | 0 | 326080 | 468950 | 0 | 2210100 | 4124600 | 0 | 2090100 | 869310 | 122930 | 3422600 | 0 | 229860 | 349020 | 0 | 0 | 4629200 |
| NQVTQLKE | MTHFD1; MTHF | P11586; V9 | C1TC_HUMAN | Unmodified | 0 | 0 | 0 | 0 | 0 | 0 | 1330000 | 1046200 | 0 | 0 | 0 | 0 | 0 | 94899 | 0 | 0 | 4331900 | 0 |
| NQVVSSTN | CSNK1G3 | H0YBN6 | H0YBN6_HUM | Unmodified | 122410 | 0 | 0 | 0 | 0 | 0 | 0 | 0 | 0 | 0 | 0 | 627410 | 0 | 0 | 0 | 174720 | 0 | 0 |
| NQWDGTQ | UTRN | P46939 | UTRN_HUMAN | Unmodified | 0 | 0 | 1123500 | 1417800 | 0 | 0 | 0 | 0 | 1750200 | 98935 | 0 | 526000 | 0 | 0 | 2094900 | 2300900 | 34495 | 0 |
| NQWIDNVI | VCL | P18206 | VINC_HUMAN | Unmodified | 0 | 0 | 0 | 0 | 1441700 | 990350 | 0 | 0 | 213000 | 397340 | 143510 | 1880500 | 152690 | 0 | 0 | 0 | 275810 | 0 |
| NQWQLSA | CNP; CNP | P09543; K7 | CN37_HUMAN | Unmodified | 726630 | 764840 | 1368200 | 1547500 | 5088400 | 825760 | 975260 | 687160 | 429080 | 709060 | 7186600 | 17684000 | 2580800 | 2312700 | 2386800 | 1373400 | 1621000 | 4556300 |
| NQWQLSA | CNP; CNP | P09543; K7 | CN37_HUMAN | Unmodified | 0 | 0 | 2458400 | 2632400 | 11548000 | 1982300 | 1802900 | 1290100 | 57380 | 1211400 | 24048000 | 29016000 | 4718400 | 4175200 | 4477500 | 5405800 | 3248600 | 15003000 |
| NQYDNDV | PSMA1; N/A | P25786; B4 | PSA1_HUMAN | Unmodified | 79159 | 0 | 287820 | 285290 | 59906 | 0 | 0 | 488430 | 1657400 | 67646 | 945480 | 6887400 | 104370 | 0 | 364570 | 260330 | 0 | 0 |
| NQYVPYPI | MYO1E | Q12965 | MYO1E_HUMA | Unmodified | 0 | 0 | 0 | 130840 | 0 | 0 | 0 | 0 | 170260 | 0 | 0 | 0 | 0 | 0 | 0 | 105990 | 0 | 0 |
| NQYVSNGI | SPON2; SPON2 | Q9BUD6; D | SPON2_HUMA | Unmodified | 0 | 0 | 0 | 283860 | 0 | 0 | 0 | 0 | 0 | 0 | 0 | 0 | 0 | 0 | 0 | 2591500 | 0 | 0 |
| NRAEQLAS | TJP1; TJP1; TJP | Q07157; A0 | ZO1_HUMAN | Unmodified | 0 | 0 | 0 | 1007700 | 8382100 | 0 | 0 | 293980 | 0 | 0 | 0 | 0 | 3247100 | 3207700 | 634810 | 1311200 | 1110500 | 744480 |
| NRAEQWN | RALA; RALA | P11233; H7 | RALA_HUMAN | Unmodified | 9732900 | 10854000 | 7864400 | 12195000 | 15081000 | 3878900 | 5057000 | 2919800 | 46603000 | 14223000 | 39080000 | 95794000 | 30721000 | 21962000 | 12434000 | 15100000 | 3369700 | 20484000 |
| NRAEQWN | RALA; RALA | P11233; H7 | RALA_HUMAN | Unmodified | 0 | 0 | 0 | 0 | 0 | 0 | 205390 | 0 | 0 | 812770 | 0 | 0 | 564180 | 918570 | 0 | 0 | 0 | 0 |
| NRAGLGEE | PTPRF; PTPRF; | P10586; H0 | PTPRF_HUMAN | Unmodified | 0 | 0 | 0 | 0 | 0 | 0 | 76310 | 0 | 329070 | 517340 | 0 | 269980 | 186710 | 0 | 353360 | 0 | 0 | 0 |
| NRAPTGYD | MVB12B; MVB1 | Q9H7P6; F8 | MB12B_HUMAN | Unmodified | 3794200 | 2805200 | 2015000 | 4161900 | 787530 | 0 | 230800 | 0 | 6181400 | 1135200 | 0 | 490160 | 1311900 | 0 | 5742400 | 5175300 | 0 | 0 |
| NRDAPEG | ITGB3; N/A; ITG | P05106; H3 | ITB3_HUMAN | Oxidation (M) | 0 | 0 | 0 | 107970 | 1044300 | 0 | 921670 | 302130 | 0 | 0 | 0 | 0 | 0 | 0 | 0 | 0 | 0 | 465660 |
| NRDAPEG | ITGB3; N/A; ITG | P05106; H3 | ITB3_HUMAN | Unmodified | 0 | 0 | 0 | 1601500 | 1997000 | 0 | 0 | 0 | 0 | 0 | 0 | 0 | 1165100 | 764350 | 0 | 0 | 0 | 0 |
| NRDAPEG | ITGB5; ITGB5; | P18084; H7 | ITB5_HUMAN | Unmodified | 1733400 | 0 | 0 | 0 | 1712200 | 0 | 0 | 0 | 1028700 | 0 | 0 | 2136800 | 0 | 1647400 | 1559900 | 0 | 0 | 949810 |
| NRDHDTFI | LMAN2; LMAN2 | Q12907; D6 | LMAN2_HUMAN | Unmodified | 349020 | 513850 | 77734 | 390570 | 0 | 0 | 0 | 19194 | 18698000 | 25633 | 0 | 0 | 0 | 0 | 377200 | 0 | 312550 | 0 |
| NRDNDPN | POLR2C | P19387 | RPB3_HUMAN | Unmodified | 0 | 0 | 0 | 818950 | 0 | 405880 | 0 | 36172 | 845480 | 0 | 62529 | 583600 | 0 | 0 | 0 | 0 | 0 | 0 |
| NRDTFRDI | MYO10; MYO10 | Q9HD67; D6 | MYO10_HUMA | Unmodified | 0 | 0 | 160940 | 253720 | 245130 | 0 | 0 | 0 | 370480 | 0 | 0 | 0 | 0 | 0 | 520990 | 142360 | 0 | 0 |
| NRDYDDID | TTYH2; TTYH2 | Q9BSA4; B4 | TTYH2_HUMAN | Unmodified | 84872 | 121150 | 870510 | 1924400 | 0 | 0 | 0 | 0 | 11097000 | 2906600 | 0 | 0 | 0 | 0 | 1646100 | 1199900 | 0 | 0 |
| NREEDPSL | CACNA2D1; CA | P54289; E7 | CA2D1_HUMAI | Unmodified | 1089600 | 897080 | 0 | 0 | 0 | 0 | 889600 | 0 | 1646800 | 1098500 | 0 | 0 | 0 | 0 | 0 | 0 | 0 | 0 |
| NREELGFR | PARP1; PARP1; | P09874; A0 | PARP1_HUMAN | Unmodified | 2617300 | 1936400 | 635260 | 0 | 2867700 | 190480 | 2817200 | 4352600 | 3027200 | 1580600 | 3593400 | 6911500 | 2273700 | 1009300 | 530270 | 174450 | 2031600 | 3790300 |
| NREILDESI | PDCD6IP | Q8WUM4 | PDC6I_HUMAN | Unmodified | 7948600 | 12365000 | 3448100 | 2026700 | 0 | 0 | 0 | 0 | 2192000 | 180490 | 806290 | 3331400 | 0 | 0 | 5819600 | 5761200 | 0 | 0 |
| NREILDESI | PDCD6IP | Q8WUM4 | PDC6I_HUMAN | Unmodified | 25416000 | 18624000 | 360920 | 5117800 | 415880 | 492130 | 0 | 0 | 5834200 | 379500 | 0 | 806710 | 0 | 0 | 16299000 | 4569400 | 0 | 0 |
| NREILQEIQ | DTNA; DTNB; D | Q9Y4J8; O6 | DTNA_HUMAN | Unmodified | 138620 | 71480 | 0 | 580310 | 245400 | 0 | 0 | 0 | 1718500 | 582560 | 0 | 215720 | 367170 | 0 | 2647700 | 911060 | 0 | 116760 |
| NRELAILLG | PRDX6 | P30041 | PRDX6_HUMAI | Oxidation (M) | 34710 | 43623 | 21067 | 49834 | 0 | 0 | 0 | 0 | 90184 | 0 | 0 | 617680 | 0 | 0 | 120880 | 103990 | 0 | 0 |
| NRELEEEL | SPAG9; SPAG9 | O60271; H0 | JIP4_HUMAN C | Unmodified | 0 | 0 | 0 | 0 | 425520 | 67982 | 0 | 0 | 122560 | 21397 | 87798 | 828770 | 0 | 0 | 0 | 58029 | 36442 | 694500 |
| NRENEHIFI | DST | A0A0U1RQJ | A0A0U1RQJ2_ | Unmodified | 0 | 0 | 0 | 180300 | 0 | 0 | 0 | 0 | 1852700 | 0 | 0 | 0 | 228480 | 0 | 228560 | 0 | 0 | 0 |
| NREPVQLE | SNRPD1; SNRP | P62314; J3 | SMD1_HUMAN | Unmodified | 2589700 | 2213100 | 572390 | 804400 | 0 | 0 | 760400 | 0 | 1519800 | 202110 | 1245600 | 16049000 | 92646 | 0 | 1046400 | 1155000 | 0 | 0 |
| NREPVQLE | SNRPD1; SNRP | P62314; J3 | SMD1_HUMAN | Unmodified | 584440 | 0 | 179920 | 834130 | 0 | 533520 | 0 | 0 | 0 | 882230 | 2633500 | 0 | 1534600 | 0 | 0 | 200510 | 418000 | 0 |
| NRETWYLS | GSPT1; GSPT2; | P15170; Q8 | ERF3A_HUMAN | Unmodified | 0 | 0 | 0 | 0 | 829170 | 0 | 0 | 0 | 1182300 | 0 | 880570 | 1636700 | 1945600 | 0 | 0 | 0 | 428920 | 810410 |
| NRFSLVPH | NIBAN2 | Q96TA1 | NIBA2_HUMAN | Unmodified | 581130 | 740730 | 665070 | 2239000 | 3923000 | 2377200 | 0 | 0 | 5916800 | 1485800 | 59176 | 585580 | 205820 | 347010 | 1160400 | 927230 | 1410800 | 436220 |
| NRGFCFLE | HNRNPR; SYN | O43390; O6 | HNRPR_HUMA | Unmodified | 0 | 0 | 0 | 0 | 0 | 0 | 0 | 0 | 0 | 602170 | 1384200 | 0 | 0 | 0 | 0 | 0 | 0 | 628200 |
| NRGQGTYI | MYL6B | P14649 | MYL6B_HUMAN | Unmodified | 0 | 0 | 0 | 0 | 1066400 | 0 | 106770 | 0 | 0 | 0 | 0 | 0 | 0 | 0 | 0 | 0 | 0 | 0 |
| NRGSGFGI | YTHDF2 | Q9Y5A9 | YTHD2_HUMAN | Unmodified | 0 | 0 | 441900 | 0 | 0 | 0 | 0 | 0 | 0 | 0 | 0 | 0 | 921640 | 0 | 0 | 0 | 0 | 0 |
| NRHPNFLV | ATP6V1F | Q16864 | VATF_HUMAN | Unmodified | 0 | 0 | 0 | 0 | 442080 | 0 | 200360 | 0 | 153610 | 73814 | 252410 | 1338600 | 786150 | 544150 | 167000 | 0 | 136680 | 2894100 |
| NRHQDVM | PTPN23 | Q9H3S7 | PTN23_HUMAN | Unmodified | 317810 | 511680 | 0 | 0 | 935810 | 0 | 0 | 0 | 0 | 0 | 0 | 0 | 0 | 0 | 0 | 0 | 0 | 0 |
| NRIIQFNP | GNPDA1; GNP | P46926; D6 | GNPI1_HUMAN | Unmodified | 93421 | 0 | 61188 | 91843 | 290900 | 0 | 57653 | 101600 | 0 | 106850 | 0 | 0 | 408550 | 336680 | 0 | 0 | 347990 | 246820 |
| NRINKAVSI | ABCF1; ABCF1; | Q8NE71; A0 | ABCF1_HUMAN | Unmodified | 0 | 0 | 0 | 0 | 0 | 0 | 85396 | 78018 | 0 | 0 | 0 | 0 | 0 | 0 | 0 | 0 | 139660 | 0 |
| NRITLEDVI | IQGAP2; IQGAP | Q13576; D6 | IQGA2_HUMAN | Unmodified | 72279 | 0 | 808270 | 801550 | 4828200 | 0 | 461970 | 0 | 167280 | 460180 | 0 | 106210 | 971470 | 0 | 670030 | 431390 | 0 | 0 |
| NRKEGGGI | MARCKSL1 | P49006 | MRP_HUMAN | Unmodified | 0 | 0 | 0 | 1286800 | 2329200 | 681450 | 839970 | 1880900 | 5246700 | 8431700 | 828100 | 181080 | 25623000 | 11753000 | 0 | 66979 | 24994000 | 68330000 |
| NRKLIDDYI | SCG3 | Q8WXD2 | SCG3_HUMAN | Unmodified | 0 | 0 | 0 | 0 | 0 | 138700 | 25279 | 92115 | 0 | 1417600 | 0 | 0 | 241390 | 0 | 0 | 0 | 237410 | 33625 |
| NRLAGELQ | CENPF; CENPF | P49454; A0 | CENPF_HUMAI | Unmodified | 0 | 0 | 0 | 0 | 202480 | 0 | 0 | 0 | 93036 | 81953 | 42733 | 0 | 0 | 0 | 0 | 0 | 0 | 36427 |
| NRLCPSLIC | DUS3L | Q96G46 | DUS3L_HUMAI | Unmodified | 0 | 0 | 0 | 0 | 0 | 0 | 392010 | 214300 | 0 | 0 | 0 | 0 | 56509 | 0 | 0 | 0 | 0 | 0 |
| NRLFQENS | ATP6AP2; ATP6 | O75787; A0 | RENR_HUMAN | Unmodified | 0 | 0 | 0 | 0 | 0 | 0 | 0 | 0 | 3023600 | 0 | 0 | 0 | 0 | 0 | 0 | 0 | 0 | 0 |
| NRLLSKYD | CNN2; CNN2; | Q99439; H3 | CNN2_HUMAN | Unmodified | 0 | 0 | 0 | 0 | 1880200 | 0 | 0 | 0 | 0 | 0 | 0 | 0 | 0 | 0 | 0 | 0 | 0 | 0 |
| NRLQQELD | MYH10 | P35580 | MYH10_HUMAI | Unmodified | 0 | 0 | 0 | 0 | 1058700 | 114180 | 0 | 0 | 273300 | 0 | 0 | 4197500 | 420310 | 0 | 0 | 0 | 0 | 0 |
| NRLQQELD | MYH10 | P35580 | MYH10_HUMAI | Unmodified | 0 | 0 | 0 | 394010 | 20671000 | 2969100 | 4370700 | 40301 | 0 | 280810 | 0 | 0 | 9043700 | 13418000 | 0 | 0 | 1941900 | 3265500 |
| NRLVNIMP | PTPRF; PTPRF; | P10586; H0 | PTPRF_HUMAN | Unmodified | 0 | 0 | 0 | 0 | 1109100 | 0 | 0 | 0 | 1532200 | 182230 | 0 | 0 | 0 | 0 | 0 | 0 | 0 | 0 |
| NRLVNIMP | PTPRF; PTPRF; | P10586; H0 | PTPRF_HUMAN | Oxidation (M) | 221780 | 130120 | 178760 | 323970 | 169800 | 238990 | 0 | 0 | 398700 | 547180 | 0 | 0 | 336160 | 0 | 239510 | 161130 | 0 | 0 |
| NRLVQHDI | CCDC50 | Q8IVM0 | CCD50_HUMA | Unmodified | 1367200 | 1045300 | 1740600 | 4365900 | 389530 | 63973 | 122710 | 27689 | 249190 | 444060 | 0 | 0 | 11178000 | 2345900 | 2393400 | 1513800 | 118790 | 886680 |
| NRMDGTY | FLNB; FLNB | O75369; E7 | FLNB_HUMAN | Unmodified | 0 | 0 | 0 | 309050 | 8717800 | 1487700 | 345360 | 88878 | 329080 | 142490 | 90215 | 1826300 | 1275300 | 703700 | 125430 | 109380 | 4675800 | 3314800 |
| NRMDGTY | FLNB; FLNB | O75369; E7 | FLNB_HUMAN | Oxidation (M) | 0 | 8269 | 175680 | 0 | 3229300 | 763570 | 0 | 136520 | 0 | 0 | 123460 | 760100 | 585450 | 379850 | 136690 | 113460 | 2700000 | 1725300 |
| NRNAAFG | YTHDF1 | Q9BYJ9 | YTHD1_HUMAN | Unmodified | 0 | 0 | 0 | 0 | 0 | 0 | 0 | 0 | 1080200 | 0 | 0 | 0 | 791050 | 0 | 0 | 0 | 0 | 401230 |
| NRNEQES | DBNL; DBNL; D | Q9UJU6; B4 | DBNL_HUMAN | Unmodified | 0 | 324520 | 623010 | 1414800 | 0 | 89042 | 0 | 0 | 257740 | 0 | 0 | 1678700 | 228460 | 105100 | 433880 | 0 | 31998 | 77961 |
| NRNEQES | DBNL; DBNL; D | Q9UJU6; B4 | DBNL_HUMAN | Unmodified | 0 | 0 | 0 | 0 | 276470 | 0 | 0 | 0 | 0 | 0 | 0 | 0 | 1299300 | 396970 | 0 | 0 | 0 | 0 |
| NRNFYELS | MYO1B; MYO1 | O43795; C9 | MYO1B_HUMA | Unmodified | 0 | 0 | 0 | 0 | 456370 | 110340 | 0 | 0 | 1768800 | 0 | 2475100 | 8928800 | 0 | 0 | 340180 | 0 | 494880 | 0 |
| NRNHLLH | ANXA4; ANXA4 | P09525; Q6 | ANXA4_HUMAN | Unmodified | 411700 | 146780 | 0 | 385680 | 4964000 | 1424900 | 284220 | 0 | 15743000 | 7673500 | 19467000 | 6636800 | 2825700 | 1376000 | 159990 | 1506700 | 0 | 1564300 |
| NRNHLLH | ANXA4; ANXA4 | P09525; Q6 | ANXA4_HUMAN | Unmodified | 0 | 0 | 0 | 1191300 | 386280 | 0 | 0 | 0 | 3617500 | 0 | 2828200 | 0 | 0 | 0 | 0 | 0 | 0 | 0 |
| NRNNPSFI | MPPE1; MPPE1 | Q53F39; A0 | MPPE1_HUMAI | Oxidation (M) | 0 | 0 | 0 | 0 | 0 | 0 | 0 | 0 | 267510 | 0 | 0 | 0 | 0 | 0 | 0 | 0 | 0 | 0 |
| NRPEDYQ | PDIA6 | Q15084 | PDIA6_HUMAN | Unmodified | 0 | 0 | 0 | 0 | 0 | 0 | 0 | 0 | 5406000 | 246410 | 91904 | 393320 | 0 | 0 | 0 | 0 | 0 | 0 |
| NRPEDYQ | PDIA6 | Q15084 | PDIA6_HUMAN | Unmodified | 319390 | 0 | 165910 | 444120 | 2069000 | 355930 | 575030 | 0 | 3.22E+08 | 46647000 | 28580000 | 1870200 | 1746000 | 594140 | 1021400 | 332400 | 684550 | 0 |
| NRPNRLIVI | VCP; VCP; VCP; | P55072; A0 | TERA_HUMAN | Unmodified | 0 | 0 | 0 | 3676500 | 29237000 | 3343500 | 2449300 | 0 | 9513400 | 5801800 | 6307000 | 2539600 | 28895000 | 7023200 | 0 | 0 | 2581500 | 3546500 |
| NRPPFGQ | TFG; TFG | Q92734; A0 | TFG_HUMAN P | Unmodified | 0 | 0 | 0 | 0 | 1543000 | 255260 | 0 | 60708 | 607450 | 310510 | 0 | 0 | 662170 | 393920 | 0 | 0 | 278030 | 706110 |
| NRPPGAA | CLUH; CLUH; C | O75153; I3 | CLU_HUMAN C | Oxidation (M) | 0 | 0 | 0 | 0 | 0 | 0 | 0 | 0 | 0 | 0 | 0 | 0 | 0 | 0 | 0 | 0 | 0 | 0 |
| NRPPLPAG | TMPO; TMPO; T | P42167; P4 | LAP2B_HUMAN | Unmodified | 92834 | 431680 | 0 | 0 | 101110 | 0 | 376610 | 229060 | 899590 | 0 | 520180 | 1205100 | 141640 | 770030 | 35749 | 44761 | 47735 | 0 |
| NRPSSGSL | QDPR | P09417 | DHPR_HUMAN | Unmodified | 0 | 0 | 0 | 0 | 8915000 | 0 | 3833300 | 0 | 0 | 148650 | 0 | 0 | 10627000 | 248730 | 0 | 0 | 3440800 | 3363500 |
| NRPTSISW | PEBP1 | P30086 | PEBP1_HUMAN | Unmodified | 7170200 | 4789600 | 4044500 | 5544600 | 6113900 | 1688000 | 276830 | 261660 | 11785000 | 5024100 | 11287000 | 31893000 | 15594000 | 8254900 | 6767700 | 6470400 | 5734900 | 8323200 |
| NRQALLGE | STX6 | O43752 | STX6_HUMAN | Unmodified | 0 | 0 | 0 | 0 | 0 | 0 | 551310 | 0 | 1283200 | 0 | 78740 | 203970 | 0 | 0 | 0 | 0 | 0 | 0 |
| NRQGDEN | SLC12A7; SLC1 | Q9Y666; A0 | S12A7_HUMAN | Oxidation (M) | 0 | 0 | 0 | 0 | 0 | 0 | 0 | 0 | 0 | 0 | 0 | 0 | 0 | 0 | 595960 | 0 | 0 | 0 |
| NRQSDENI | ILRUN; ILRUN | Q9H6K1; Q5 | ILRUN_HUMAN | Unmodified | 349810 | 336350 | 0 | 115640 | 0 | 0 | 74638 | 0 | 0 | 0 | 0 | 0 | 394410 | 0 | 0 | 0 | 0 | 103270 |
| NRQVNELV | PAFAH1B3; PAF | Q15102; M0 | PA1B3_HUMAN | Unmodified | 0 | 0 | 0 | 0 | 0 | 0 | 0 | 50488 | 0 | 0 | 0 | 884240 | 0 | 0 | 0 | 0 | 0 | 0 |

| | | | | | | | | | | | | | | | | | | | | | | |
|---|---|---|---|---|---|---|---|---|---|---|---|---|---|---|---|---|---|---|---|---|---|---|
| NRSAEEGE | TNKS1BP1 | Q9C0C2 | TB182_HUMAN | Unmodified | 0 | 12737 | 0 | 0 | 0 | 0 | 0 | 0 | 0 | 0 | 0 | 0 | 581800 | 1296100 | 0 | 0 | 0 | 0 |
| NRSGFILS\ | MYO1D; MYO1D | O94832; K7E | MYO1D_HUMA | Unmodified | 0 | 0 | 0 | 0 | 4654300 | 737170 | 0 | 145240 | 1015100 | 1556400 | 620400 | 0 | 0 | 7872800 | 0 | 0 | 0 | 24559000 |
| NRSNTPIL\ | GPI; GPI; GPI; G | P06744; K7E | G6PI_HUMAN ( | Unmodified | 0 | 0 | 0 | 0 | 2595800 | 615980 | 275450 | 0 | 4007800 | 1327100 | 1159800 | 7123200 | 2108300 | 0 | 0 | 0 | 0 | 1075700 |
| NRSNTPIL\ | GPI; GPI; GPI; G | P06744; K7E | G6PI_HUMAN ( | Oxidation (M) | 0 | 294110 | 0 | 1314100 | 561000 | 0 | 0 | 0 | 1189000 | 501250 | 0 | 6778400 | 514920 | 0 | 652490 | 133070 | 0 | 0 |
| NRTFSVWY | FLNA; FLNA; FL | P21333; A0A | FLNA_HUMAN | Unmodified | 197110 | 426310 | 0 | 874090 | 1102200 | 450530 | 548360 | 0 | 0 | 0 | 0 | 0 | 0 | 0 | 450460 | 211150 | 210600 | 0 |
| NRTFSVWY | FLNA; FLNA; FL | P21333; A0A | FLNA_HUMAN | Unmodified | 0 | 1090300 | 0 | 5077700 | 3135600 | 1610500 | 6212800 | 0 | 0 | 549920 | 0 | 0 | 0 | 1965300 | 0 | 0 | 0 | 626180 |
| NRTPYVTTI | TUBGCP2; TUB | Q9BSJ2; F2Z | GCP2_HUMAN | Unmodified | 0 | 0 | 0 | 0 | 1547300 | 45140 | 558190 | 158170 | 78858 | 0 | 60934 | 1483600 | 0 | 0 | 0 | 145410 | 168390 | 590320 |
| NRVGPQVF | EPS8L2; EPS8L | Q9H6S3; H0 | ES8L2_HUMAN | Unmodified | 0 | 0 | 411280 | 694070 | 2005700 | 667120 | 0 | 0 | 775750 | 253700 | 0 | 166800 | 2640300 | 753060 | 1607300 | 1080300 | 314520 | 544870 |
| NRVIGSGC | LDHAL6A; LDH | Q6ZMR3; P0 | LDH6A_HUMAI | Unmodified | 0 | 951810 | 773640 | 2497100 | 2477100 | 729480 | 330990 | 164300 | 1894300 | 1362900 | 5725200 | 33672000 | 1393800 | 655040 | 2288500 | 2318700 | 2231700 | 2885900 |
| NRVLQIIPY | PTPRA | P18433 | PTPRA_HUMAN | Unmodified | 0 | 0 | 0 | 0 | 765030 | 0 | 0 | 0 | 405070 | 0 | 0 | 0 | 555550 | 0 | 0 | 0 | 472520 | 1490800 |
| NRVLQIIPY | PTPRA | P18433 | PTPRA_HUMAN | Unmodified | 0 | 0 | 0 | 0 | 2306800 | 0 | 0 | 0 | 0 | 32928 | 0 | 0 | 0 | 0 | 0 | 0 | 441670 | 1914400 |
| NRVPDSC( | CD63; CD63; C | P08962; F8V | CD63_HUMAN | Unmodified | 0 | 0 | 0 | 305490 | 0 | 0 | 0 | 0 | 1735700 | 636500 | 0 | 216060 | 0 | 0 | 0 | 0 | 0 | 0 |
| NRVQNMA | SSR4 | P51571 | SSRD_HUMAN | Unmodified | 0 | 0 | 0 | 0 | 0 | 0 | 0 | 0 | 14288000 | 4027700 | 1714100 | 0 | 0 | 0 | 0 | 0 | 0 | 0 |
| NRVQNMA | SSR4 | P51571 | SSRD_HUMAN | Oxidation (M) | 0 | 0 | 0 | 0 | 0 | 0 | 0 | 0 | 1987100 | 432110 | 494350 | 45176 | 104050 | 0 | 0 | 0 | 0 | 0 |
| NRYANVIA | PTPRD; PTPRF; | P23468; P10 | PTPRD_HUMAI | Unmodified | 322870 | 236270 | 234400 | 417840 | 134120 | 90176 | 15058 | 0 | 1543300 | 101440 | 190980 | 426880 | 192450 | 72884 | 557700 | 391420 | 193960 | 117770 |
| NRYGNIIAY | PTPRM; PTPRK; | P28827; Q15 | PTPRM_HUMAI | Unmodified | 0 | 0 | 0 | 0 | 0 | 0 | 0 | 0 | 0 | 120050 | 0 | 308780 | 0 | 0 | 0 | 149200 | 0 | 0 |
| NRYINILAY | PTPRG | P23470 | PTPRG_HUMAI | Unmodified | 0 | 0 | 0 | 0 | 5766900 | 1099300 | 0 | 0 | 0 | 0 | 0 | 0 | 493500 | 618920 | 0 | 0 | 0 | 0 |
| NRYNNVLF | PTPRJ; PTPRJ; P | Q12913; A0A | PTPRJ_HUMAN | Unmodified | 23138 | 0 | 0 | 48971 | 0 | 0 | 0 | 0 | 0 | 0 | 989180 | 5149700 | 0 | 0 | 0 | 0 | 106000 | 0 |
| NSAAEIPVT | ARFIP1 | P53367 | ARFP1_HUMAN | Unmodified | 0 | 0 | 0 | 0 | 0 | 0 | 0 | 0 | 0 | 0 | 0 | 1288500 | 0 | 0 | 0 | 0 | 0 | 291150 |
| NSAEQTVT | ARHGEF7; ARH | Q14155; A0A | ARHG7_HUMA | Unmodified | 0 | 0 | 0 | 0 | 7253000 | 0 | 1165000 | 385590 | 0 | 690540 | 0 | 0 | 1141300 | 826930 | 0 | 0 | 3831300 | 5916300 |
| NSAIPSELN | OPTN | Q96CV9 | OPTN_HUMAN | Unmodified | 0 | 0 | 0 | 0 | 0 | 0 | 408300 | 129330 | 0 | 0 | 0 | 0 | 0 | 0 | 0 | 0 | 0 | 0 |
| NSAKVMVT | TLN1 | Q9Y490 | TLN1_HUMAN | Unmodified | 0 | 0 | 0 | 0 | 0 | 0 | 281210 | 0 | 0 | 0 | 0 | 0 | 58276 | 177120 | 0 | 0 | 115460 | 0 |
| NSAQGNV | RBM39; RBM39 | Q14498; H0 | RBM39_HUMAI | Unmodified | 1160700 | 0 | 0 | 0 | 434970 | 0 | 95455 | 210210 | 1223300 | 29759 | 74702 | 1081500 | 165220 | 0 | 0 | 874670 | 125050 | 204570 |
| NSATSADE | MARK2; MARK2 | Q7KZI7; E7E | MARK2_HUMAI | Unmodified | 0 | 0 | 0 | 0 | 0 | 226090 | 0 | 74895 | 0 | 383740 | 0 | 0 | 2800500 | 853440 | 0 | 0 | 375240 | 0 |
| NSCIVCNN | AHCYL1 | O43865 | SAHH2_HUMAI | Oxidation (M) | 98352 | 0 | 69717 | 0 | 0 | 0 | 0 | 0 | 1316100 | 0 | 0 | 929100 | 0 | 0 | 361930 | 208330 | 0 | 0 |
| NSCNNFIY | SPINT2; SPINT2 | O43291; K7E | SPIT2_HUMAN | Unmodified | 0 | 0 | 0 | 0 | 0 | 0 | 0 | 0 | 7350900 | 411370 | 0 | 0 | 0 | 0 | 0 | 0 | 0 | 0 |
| NSCNNFIY | SPINT2; SPINT2 | O43291; K7E | SPIT2_HUMAN | Unmodified | 0 | 0 | 0 | 0 | 0 | 0 | 0 | 0 | 5874600 | 0 | 0 | 0 | 0 | 0 | 0 | 0 | 0 | 0 |
| NSDGSYRI | ATP2B1; ATP2B | P20020; E7E | AT2B1_HUMAN | Unmodified | 0 | 0 | 68797 | 0 | 2020600 | 0 | 1347400 | 383310 | 0 | 812230 | 0 | 0 | 4708500 | 5626500 | 0 | 0 | 918050 | 711580 |
| NSDLLLLVI | GOLM2 | Q6P4E1 | GOLM2_HUMA | Unmodified | 0 | 0 | 0 | 0 | 93705 | 0 | 0 | 0 | 1433100 | 137230 | 206600 | 137980 | 0 | 0 | 0 | 0 | 48864 | 0 |
| NSDLYGAK | SHCBP1 | Q8NEM2 | SHCBP_HUMA | Unmodified | 0 | 1112900 | 0 | 0 | 7038000 | 14481 | 43988 | 0 | 0 | 20416 | 0 | 526060 | 0 | 0 | 109710 | 0 | 121240 | 235060 |
| NSDPALGL | PCSK1N | Q9UHG2 | PCS1N_HUMAI | Unmodified | 0 | 0 | 0 | 0 | 0 | 0 | 0 | 0 | 0 | 6174100 | 0 | 0 | 0 | 0 | 0 | 0 | 0 | 0 |
| NSDPNDPI | SRPK1; SRPK1 | Q96SB4; H3 | SRPK1_HUMAN | Oxidation (M) | 0 | 0 | 0 | 0 | 0 | 0 | 0 | 0 | 0 | 0 | 0 | 832230 | 0 | 0 | 0 | 0 | 0 | 0 |
| NSDPWAAS | EPN2; EPN2; EF | O95208; F6F | EPN2_HUMAN | Unmodified | 0 | 203770 | 0 | 0 | 250690 | 0 | 0 | 0 | 110060 | 45853 | 0 | 166080 | 1312200 | 777850 | 352730 | 364160 | 62332 | 647820 |
| NSDRTPLV | NSF; NSF | P46459; I3L( | NSF_HUMAN V | Unmodified | 0 | 0 | 940550 | 3460600 | 866580 | 0 | 3540300 | 0 | 830240 | 896670 | 1109200 | 2484900 | 21609000 | 9902200 | 2327700 | 0 | 4003300 | 10830000 |
| NSDSILEAI | NDUFV2; NDUF | P19404; E7E | NDUV2_HUMA | Unmodified | 0 | 0 | 0 | 0 | 3196200 | 0 | 466170 | 514740 | 464750 | 0 | 0 | 0 | 0 | 0 | 0 | 0 | 0 | 0 |
| NSDVLQSF | COLGALT1 | Q8NBJ5 | GT251_HUMAN | Unmodified | 0 | 0 | 0 | 0 | 0 | 0 | 0 | 189610 | 8186000 | 3858400 | 0 | 0 | 0 | 0 | 0 | 0 | 0 | 2904900 |
| NSEELRSRI | PSMF1; PSMF1; | Q92530; Q5( | PSMF1_HUMAI | Unmodified | 0 | 0 | 0 | 0 | 0 | 0 | 0 | 0 | 0 | 0 | 0 | 0 | 0 | 0 | 0 | 0 | 253210 | 0 |
| NSELMDPK | GLG1 | Q92896 | GSLG1_HUMAI | Unmodified | 0 | 0 | 0 | 0 | 626820 | 0 | 131530 | 0 | 12771000 | 4207500 | 2432800 | 2229300 | 0 | 0 | 0 | 0 | 0 | 0 |
| NSEMENEI | APPL2 | Q8NEU8 | DP13B_HUMAI | Unmodified | 0 | 0 | 0 | 0 | 832380 | 0 | 0 | 0 | 0 | 92193 | 0 | 0 | 553460 | 0 | 0 | 0 | 313440 | 1842600 |
| NSEPAGLE | EPRS1; EPRS1 | P07814; V9C | SYEP_HUMAN | Unmodified | 69630 | 0 | 0 | 0 | 0 | 0 | 0 | 0 | 0 | 0 | 708050 | 1768300 | 89870 | 0 | 78403 | 87911 | 0 | 0 |
| NSEPEEVIF | YARS1; YARS1 | P54577; A0A | SYYC_HUMAN | Unmodified | 0 | 0 | 585370 | 779780 | 0 | 0 | 0 | 0 | 0 | 263890 | 1505300 | 9224000 | 3981300 | 1237200 | 1014500 | 842780 | 0 | 0 |
| NSEPEEVIF | YARS1; YARS1 | P54577; A0A | SYYC_HUMAN | Unmodified | 0 | 0 | 0 | 0 | 6692800 | 927650 | 2171400 | 208420 | 228350 | 1890900 | 0 | 0 | 16304000 | 9540100 | 0 | 0 | 2127300 | 3809200 |
| NSEPQDEC | VGF; VGF | O15240; Q0! | VGF_HUMAN N | Unmodified | 0 | 645960 | 0 | 0 | 0 | 0 | 0 | 0 | 8925300 | 3251500 | 75346 | 0 | 0 | 0 | 0 | 0 | 0 | 141010 |
| NSETFPTIL | DIAPH1; DIAPH | O60610; A0A | DIAP1_HUMAN | Unmodified | 0 | 0 | 0 | 0 | 1176400 | 0 | 0 | 0 | 0 | 0 | 0 | 1578700 | 1966300 | 2496800 | 0 | 0 | 710240 | 2096900 |
| NSETFPTIL | DIAPH1; DIAPH | O60610; A0A | DIAP1_HUMAN | Unmodified | 0 | 33146 | 0 | 0 | 468870 | 37480 | 304550 | 0 | 0 | 0 | 0 | 0 | 0 | 0 | 0 | 0 | 1062700 | 826020 |
| NSFEDPCS | MUC5B | Q9HC84 | MUC5B_HUMA | Unmodified | 0 | 0 | 0 | 0 | 0 | 0 | 0 | 0 | 0 | 0 | 0 | 0 | 0 | 0 | 0 | 0 | 0 | 1119800 |
| NSFNNPAY | INPPL1; INPPL1 | O15357; A0A | SHIP2_HUMAN | Unmodified | 0 | 0 | 0 | 0 | 0 | 0 | 0 | 0 | 0 | 0 | 854450 | 1809400 | 0 | 0 | 0 | 0 | 0 | 2350800 |
| NSFREQLE | MYH9 | P35579 | MYH9_HUMAN | Unmodified | 241600 | 185810 | 2183400 | 1695800 | 4448400 | 1554800 | 0 | 0 | 3403000 | 0 | 1316300 | 9141000 | 1379400 | 792750 | 3529700 | 2124600 | 644670 | 1076000 |
| NSFREQLE | MYH9 | P35579 | MYH9_HUMAN | Unmodified | 105560 | 0 | 0 | 1810800 | 42373000 | 14483000 | 134680 | 96918 | 1580300 | 1673600 | 429930 | 891200 | 5879800 | 5715900 | 0 | 0 | 9606600 | 12858000 |
| NSFTVDCS | FLNC | Q14315 | FLNC_HUMAN | Unmodified | 0 | 0 | 0 | 0 | 4792900 | 325900 | 0 | 0 | 0 | 0 | 0 | 31905 | 0 | 0 | 0 | 0 | 3386500 | 693840 |
| NSFYVALEI | RABIF | P47224 | MSS4_HUMAN | Unmodified | 0 | 0 | 0 | 526630 | 1627900 | 0 | 329180 | 0 | 214920 | 211560 | 437930 | 0 | 798390 | 0 | 0 | 0 | 630110 | 1092300 |
| NSGFATCV | GAS6 | Q14393 | GAS6_HUMAN | Unmodified | 0 | 0 | 0 | 0 | 0 | 0 | 0 | 0 | 1072300 | 0 | 0 | 0 | 0 | 0 | 0 | 0 | 0 | 0 |
| NSGIIILR | CPNE2; CPNE2 | Q96FN4; H3 | CPNE2_HUMAI | Unmodified | 0 | 66759 | 61653 | 0 | 0 | 0 | 0 | 25399 | 1201800 | 256800 | 0 | 85090 | 0 | 0 | 99187 | 147760 | 0 | 0 |
| NSGTQSD( | LLGL2; LLGL2 | Q6P1M3; J3( | L2GL2_HUMAN | Oxidation (M) | 92060 | 0 | 0 | 147800 | 266110 | 0 | 0 | 0 | 241510 | 220010 | 60435 | 0 | 216150 | 0 | 0 | 0 | 92165 | 251310 |
| NSGTQSD( | LLGL2; LLGL2 | Q6P1M3; J3( | L2GL2_HUMAN | Unmodified | 62248 | 390140 | 0 | 136730 | 0 | 0 | 0 | 0 | 1083200 | 419920 | 158660 | 278650 | 535670 | 0 | 0 | 0 | 701100 | 0 |
| NSHHTWL | P4HTM; P4HTM | Q9NXG6; V9 | P4HTM_HUMAI | Unmodified | 0 | 0 | 0 | 0 | 0 | 0 | 0 | 0 | 1801200 | 265840 | 0 | 0 | 0 | 0 | 0 | 0 | 0 | 0 |
| NSHLINVLI | EIF3H; EIF3H; E | O15372; E5F | EIF3H_HUMAN | Oxidation (M) | 145470 | 87133 | 80869 | 0 | 893640 | 0 | 195750 | 0 | 0 | 149970 | 380180 | 1172200 | 604580 | 371400 | 490420 | 0 | 119230 | 311060 |
| NSHSLTTEI | PARP12; PARP1 | Q9H0J9; G5I | PAR12_HUMAN | Unmodified | 0 | 554200 | 256560 | 890620 | 0 | 0 | 0 | 0 | 0 | 0 | 0 | 0 | 0 | 0 | 1265300 | 910010 | 0 | 0 |
| NSHSSIAEI | VPS37A; VPS37 | Q8NEZ2; E5I | VP37A_HUMAN | Unmodified | 2481800 | 2341600 | 201160 | 0 | 0 | 0 | 0 | 0 | 0 | 0 | 0 | 0 | 0 | 0 | 704690 | 844640 | 0 | 66590 |
| NSIASCADI | MARK3; MARK3 | P27448; A0A | MARK3_HUMAI | Unmodified | 0 | 0 | 65277 | 191380 | 0 | 0 | 0 | 0 | 0 | 0 | 0 | 0 | 0 | 0 | 0 | 128580 | 0 | 0 |
| NSIASCADI | MARK3; MARK3 | P27448; A0A | MARK3_HUMAI | Unmodified | 0 | 0 | 0 | 0 | 717110 | 0 | 44701 | 0 | 0 | 0 | 0 | 0 | 0 | 0 | 0 | 0 | 0 | 0 |
| NSIAYLDEA | COL2A1 | P02458 | CO2A1_HUMAI | Unmodified | 0 | 0 | 0 | 0 | 0 | 0 | 0 | 422580 | 0 | 98041 | 5626300 | 216200 | 0 | 0 | 0 | 0 | 0 | 0 |
| NSIAYLDEA | COL2A1 | P02458 | CO2A1_HUMAI | Unmodified | 0 | 0 | 255550 | 175620 | 0 | 0 | 435670 | 248770 | 315350 | 529780 | 16061000 | 539230 | 0 | 1186600 | 117140 | 0 | 0 | 869120 |
| NSILAQVLD | PDCD5; PDCD5 | O14737; K7E | PDCD5_HUMA | Unmodified | 1380900 | 0 | 985150 | 1759900 | 0 | 0 | 0 | 0 | 4693500 | 0 | 2131000 | 2514100 | 0 | 0 | 3450600 | 2490100 | 0 | 0 |
| NSILFLNSL | DNAJC10; DNA | Q8IXB1; A0A | DJC10_HUMAN | Unmodified | 0 | 0 | 0 | 0 | 0 | 2896900 | 0 | 0 | 7863300 | 1609100 | 474170 | 422320 | 0 | 0 | 0 | 0 | 0 | 33168 |
| NSIVNSQP | XPO7; XPO7 | Q9UIA9; E7E | XPO7_HUMAN | Unmodified | 0 | 0 | 0 | 0 | 0 | 0 | 0 | 0 | 0 | 0 | 0 | 1134000 | 0 | 0 | 0 | 0 | 0 | 0 |
| NSIYSGLEA | HMGCS1 | Q01581 | HMCS1_HUMA | Unmodified | 0 | 0 | 0 | 0 | 1175400 | 414700 | 347180 | 0 | 0 | 289260 | 0 | 0 | 0 | 0 | 0 | 0 | 0 | 0 |
| NSKHEELN | EPRS1; EPRS1 | P07814; V9C | SYEP_HUMAN | Unmodified | 0 | 0 | 0 | 0 | 0 | 0 | 0 | 0 | 0 | 0 | 0 | 1371300 | 0 | 0 | 31011 | 0 | 192510 | 484120 |
| NSKNQGIE | VCL | P18206 | VINC_HUMAN | Unmodified | 0 | 0 | 0 | 0 | 815350 | 64942 | 2199800 | 914960 | 0 | 239570 | 0 | 0 | 0 | 0 | 0 | 0 | 5871000 | 323720 |
| NSKPNMVI | PTPRK; PTPRK; | Q15262; Q5 | PTPRK_HUMAN | Unmodified | 0 | 0 | 0 | 0 | 0 | 0 | 0 | 0 | 1039100 | 106470 | 196440 | 1312700 | 0 | 0 | 0 | 0 | 0 | 0 |
| NSKRDYTG | MPZL1 | O95297 | MPZL1_HUMAI | Unmodified | 0 | 0 | 0 | 0 | 2334300 | 234990 | 68948 | 57143 | 0 | 73158 | 0 | 0 | 0 | 0 | 0 | 0 | 782760 | 1623600 |
| NSLDCEIVS | CAP1 | Q01518 | CAP1_HUMAN | Unmodified | 757140 | 198480 | 0 | 721810 | 11250000 | 2157400 | 5950600 | 3235000 | 1888200 | 1641400 | 1574300 | 7327400 | 5623800 | 6474400 | 893590 | 584920 | 4633500 | 4226000 |
| NSLESISSII | SCRIB; SCRIB; | Q14160; A0A | SCRIB_HUMAN | Unmodified | 0 | 200540 | 1786000 | 1126900 | 497820 | 752540 | 643390 | 0 | 12349000 | 151670 | 1879800 | 4069300 | 0 | 0 | 1894300 | 1595200 | 350930 | 413610 |
| NSLESISSII | SCRIB; SCRIB; | Q14160; A0A | SCRIB_HUMAN | Unmodified | 0 | 0 | 0 | 281670 | 627870 | 0 | 0 | 0 | 968400 | 526850 | 93501 | 0 | 1537700 | 0 | 0 | 0 | 1447100 | 1539100 |
| NSLESSLR( | IFIT2; IFIT2; IFIT | P09913; A0A | IFIT2_HUMAN I | Unmodified | 0 | 0 | 0 | 0 | 209140 | 0 | 0 | 0 | 0 | 0 | 0 | 0 | 0 | 0 | 0 | 0 | 51648 | 234330 |
| NSLESYAFI | HSPA8; HSPA8; | P11142; E9P | HSP7C_HUMAI | Oxidation (M) | 5421900 | 9030800 | 8346000 | 4063200 | 7450700 | 4389100 | 3339300 | 703110 | 3159100 | 6516500 | 2602800 | 17478000 | 4923400 | 3351700 | 6210400 | 5380200 | 3612500 | 1929600 |
| NSLESYAFI | HSPA8; HSPA8; | P11142; E9P | HSP7C_HUMAI | Unmodified | 4982700 | 9511400 | 7383100 | 6239600 | 23557000 | 8439700 | 4406100 | 420310 | 20874000 | 15736000 | 3820900 | 16378000 | 8536600 | 6512000 | 5882100 | 3661900 | 6229600 | 4566400 |
| NSLESYAFI | HSPA8; HSPA8; | P11142; E9P | HSP7C_HUMAI | Unmodified | 0 | 0 | 0 | 0 | 1839900 | 244650 | 1565100 | 0 | 0 | 1580900 | 0 | 0 | 1325000 | 1116000 | 0 | 0 | 2429800 | 0 |
| NSLESYAFI | HSPA8; HSPA8; | P11142; E9P | HSP7C_HUMAI | Oxidation (M) | 0 | 0 | 0 | 0 | 0 | 0 | 0 | 0 | 0 | 324680 | 0 | 0 | 0 | 0 | 0 | 0 | 467530 | 0 |
| NSLETVGTI | SYT7; SYT7; SY | O43581; F5( | SYT7_HUMAN | Unmodified | 0 | 0 | 0 | 0 | 0 | 0 | 0 | 0 | 172970 | 0 | 0 | 0 | 0 | 0 | 261850 | 95934 | 0 | 0 |
| NSLFGSVE | HEBP1; HEBP1 | Q9NRV9; F5( | HEBP1_HUMAI | Unmodified | 20870 | 90692 | 58412 | 146010 | 194840 | 0 | 569060 | 0 | 0 | 0 | 0 | 35292 | 318260 | 0 | 194340 | 16178 | 61288 | 0 |
| NSLGEGQE | RIPOR2; RIPOR | Q9Y4F9; B7Z | RIPR2_HUMAN | Unmodified | 0 | 0 | 0 | 0 | 0 | 37708 | 0 | 89761 | 0 | 187320 | 0 | 851770 | 0 | 290150 | 0 | 0 | 0 | 0 |
| NSLGGFAS | CD109 | Q6YHK3 | CD109_HUMAI | Unmodified | 0 | 0 | 0 | 0 | 0 | 0 | 0 | 0 | 3283500 | 1103400 | 0 | 0 | 0 | 0 | 0 | 0 | 0 | 0 |
| NSLHLDLE | SYT4 | Q9H2B2 | SYT4_HUMAN | Unmodified | 0 | 139900 | 0 | 0 | 0 | 0 | 0 | 0 | 0 | 0 | 0 | 0 | 0 | 0 | 2432900 | 1530000 | 0 | 0 |
| NSLISSLEE | NUMA1; NUMA | Q14980; H0 | NUMA1_HUMA | Unmodified | 0 | 0 | 0 | 26727 | 18196 | 0 | 0 | 0 | 0 | 0 | 0 | 0 | 497130 | 0 | 34322 | 0 | 0 | 0 |
| NSLISYLEQ | CPE | P16870 | CBPE_HUMAN | Unmodified | 0 | 0 | 0 | 0 | 0 | 0 | 0 | 13455 | 1.77E+08 | 5181700 | 0 | 0 | 0 | 0 | 0 | 0 | 0 | 0 |
| NSLISYLEQ | CPE | P16870 | CBPE_HUMAN | Unmodified | 0 | 0 | 0 | 64106 | 0 | 0 | 0 | 0 | 7331300 | 22185000 | 0 | 0 | 0 | 0 | 0 | 0 | 0 | 0 |
| NSLKEANH | ATP2B1; ATP2B | P20020; F8V | AT2B1_HUMAN | Unmodified | 0 | 0 | 0 | 0 | 0 | 0 | 0 | 0 | 0 | 0 | 0 | 0 | 1264900 | 852540 | 0 | 0 | 210650 | 0 |
| NSLLDDLH | ZYG11B | Q9C0D3 | ZY11B_HUMAN | Unmodified | 0 | 0 | 25266 | 0 | 0 | 0 | 0 | 0 | 0 | 0 | 0 | 402240 | 0 | 0 | 18033 | 32678 | 0 | 0 |
| NSLQEQQE | MYH10 | P35580 | MYH10_HUMAI | Unmodified | 0 | 0 | 1149800 | 0 | 3029900 | 0 | 0 | 0 | 0 | 0 | 0 | 7127200 | 0 | 722650 | 0 | 0 | 0 | 0 |
| NSLQEQQE | MYH10 | P35580 | MYH10_HUMAI | Unmodified | 0 | 0 | 0 | 40947 | 3963000 | 625630 | 314910 | 156520 | 0 | 0 | 0 | 0 | 1676100 | 1708300 | 0 | 0 | 0 | 0 |
| NSLRVEGD | EPB41L2; EPB4 | O43491; Q6 | E41L2_HUMAN | Unmodified | 0 | 0 | 0 | 0 | 2847500 | 790910 | 0 | 0 | 0 | 0 | 0 | 0 | 0 | 0 | 0 | 0 | 788630 | 1647700 |
| NSLSDPNII | ARHGAP35 | Q9NRY4 | RHG35_HUMAI | Unmodified | 154910 | 262960 | 138030 | 0 | 778140 | 202880 | 0 | 0 | 239120 | 0 | 300590 | 820680 | 564380 | 0 | 751560 | 673640 | 177830 | 217350 |
| NSLTFPDD | ROCK1; ROCK1 | Q13464; A0A | ROCK1_HUMAI | Unmodified | 0 | 0 | 0 | 147070 | 1005200 | 259760 | 0 | 82166 | 714300 | 361790 | 130700 | 799680 | 364410 | 314780 | 136510 | 144090 | 613950 | 658260 |
| NSLTFPDD | ROCK1; ROCK1 | Q13464; A0A | ROCK1_HUMAI | Unmodified | 0 | 0 | 0 | 0 | 408060 | 0 | 272170 | 191230 | 0 | 90267 | 0 | 0 | 187590 | 221080 | 0 | 0 | 423860 | 615480 |
| NSLTLEAV | PGM2; PGM2; F | Q96G03; E9I | PGM2_HUMAN | Unmodified | 0 | 110440 | 0 | 0 | 1277700 | 121500 | 0 | 114290 | 85967 | 121030 | 436120 | 858430 | 810900 | 406190 | 138580 | 0 | 172040 | 606600 |
| NSLTSKDP | KPNB1; KPNB1 | Q14974; J3C | IMB1_HUMAN | Unmodified | 0 | 0 | 5938200 | 0 | 937060 | 0 | 283140 | 0 | 507350 | 234890 | 332150 | 2287100 | 692240 | 262490 | 0 | 0 | 274930 | 579640 |
| NSLTSKDP | KPNB1; KPNB1 | Q14974; J3C | IMB1_HUMAN | Unmodified | 0 | 0 | 0 | 0 | 2377900 | 0 | 1075400 | 746240 | 0 | 2034200 | 0 | 0 | 4742200 | 0 | 0 | 0 | 2752600 | 0 |
| NSLWHTGI | THBS4; THBS4 | P35443; E7E | TSP4_HUMAN | Unmodified | 56785 | 137280 | 1412000 | 367440 | 0 | 0 | 0 | 0 | 0 | 0 | 0 | 0 | 0 | 0 | 302900 | 184320 | 0 | 0 |
| NSMIALVDI | MYO1D; MYO1D | O94832; J3C | MYO1D_HUMA | Unmodified | 0 | 3138700 | 0 | 0 | 0 | 0 | 0 | 0 | 0 | 0 | 0 | 0 | 0 | 0 | 3309300 | 0 | 0 | 0 |
| NSMIALVDI | MYO1D; MYO1D | O94832; J3C | MYO1D_HUMA | Oxidation (M) | 1701600 | 2139200 | 556280 | 1219100 | 0 | 0 | 0 | 0 | 0 | 0 | 0 | 0 | 0 | 0 | 1730800 | 648080 | 0 | 0 |
| NSMPASSF | LUC7L2; LUC7L | Q9Y383; Q9I | LC7L2_HUMAN | Unmodified | 0 | 0 | 0 | 0 | 0 | 0 | 201400 | 0 | 344670 | 2053400 | 2118600 | 653920 | 0 | 0 | 0 | 0 | 0 | 0 |
| NSMSETVR | LMAN1 | P49257 | LMAN1_HUMAI | Oxidation (M) | 0 | 0 | 0 | 0 | 0 | 0 | 0 | 72578 | 6543200 | 191090 | 0 | 0 | 0 | 0 | 0 | 0 | 0 | 0 |
| NSMSETVR | LMAN1 | P49257 | LMAN1_HUMAI | Unmodified | 0 | 0 | 12033 | 0 | 0 | 0 | 0 | 0 | 53497000 | 3019000 | 0 | 62210 | 0 | 0 | 0 | 0 | 0 | 0 |
| NSNAYGIG | FAM131B | Q86XD5 | F131B_HUMAN | Unmodified | 0 | 0 | 0 | 0 | 0 | 0 | 0 | 0 | 452860 | 0 | 1115700 | 0 | 0 | 0 | 322420 | 278670 | 0 | 523400 |
| NSNFATVG | MKRN1; MKRN | Q9UHC7; C9 | MKRN1_HUMA | Unmodified | 3696900 | 3528300 | 0 | 1397700 | 0 | 0 | 0 | 0 | 0 | 0 | 0 | 0 | 0 | 0 | 0 | 0 | 0 | 0 |
| NSNILEDLE | ARCN1; ARCN1 | P48444; Q6F | COPD_HUMAN | Unmodified | 0 | 0 | 62711 | 0 | 0 | 0 | 0 | 0 | 0 | 0 | 624260 | 6483400 | 213790 | 0 | 32820 | 21680 | 0 | 0 |
| NSNILEDLE | ARCN1; ARCN1 | P48444; Q6F | COPD_HUMAN | Unmodified | 0 | 0 | 0 | 0 | 0 | 0 | 0 | 0 | 0 | 490840 | 0 | 0 | 0 | 0 | 0 | 0 | 0 | 0 |
| NSNLDRHI | MAP1B; MAP1B | P46821; D6F | MAP1B_HUMAI | Unmodified | 0 | 0 | 0 | 0 | 0 | 0 | 0 | 0 | 0 | 0 | 0 | 2196700 | 336080 | 0 | 0 | 75616 | 0 | 0 |

| | | | | | | | | | | | | | | | | | | | | | | |
|---|---|---|---|---|---|---|---|---|---|---|---|---|---|---|---|---|---|---|---|---|---|---|
| NSNLVGAA | LMNA; LMNA; L | P02545; A0A | LMNA_HUMAN | Unmodified | 455410 | 0 | 0 | 0 | 0 | 0 | 0 | 66591 | 0 | 0 | 0 | 0 | 209240 | 0 | 0 | 0 | 0 | 0 |
| NSNPALNE | CLIC1 | O00299 | CLIC1_HUMAN | Unmodified | 4022800 | 4102000 | 3179000 | 3891500 | 1.2E+08 | 22879000 | 4582300 | 4352600 | 12097000 | 7362700 | 4640900 | 29255000 | 38757000 | 60886000 | 5653500 | 3815500 | 61924000 | 34953000 |
| NSNPALNE | CLIC1 | O00299 | CLIC1_HUMAN | Unmodified | 0 | 0 | 0 | 0 | 5817700 | 1004200 | 1677300 | 1576700 | 0 | 0 | 0 | 0 | 5665100 | 3235900 | 0 | 0 | 16260000 | 3137700 |
| NSNVDSSY | ZC3HAV1; ZC3H | Q7Z2W4; C9 | ZCCHV_HUMA | Unmodified | 0 | 0 | 0 | 1534900 | 0 | 0 | 0 | 0 | 687040 | 0 | 0 | 0 | 478100 | 423330 | 0 | 1409800 | 0 | 0 |
| NSPLNKAE | ACOX1; ACOX1 | Q15067; K7E | ACOX1_HUMAI | Unmodified | 0 | 0 | 0 | 0 | 0 | 0 | 82938 | 32529 | 642790 | 694750 | 0 | 0 | 0 | 0 | 0 | 0 | 109860 | 67656 |
| NSPLSEWL | NCOA4; NCOA4 | Q13772; B4I | NCOA4_HUMA | Unmodified | 0 | 0 | 0 | 0 | 52263 | 0 | 0 | 0 | 1258100 | 332340 | 0 | 64426 | 1226100 | 681020 | 0 | 0 | 0 | 1711600 |
| NSPLVSRL | MDH2 | P40926 | MDHM_HUMAI | Unmodified | 0 | 0 | 0 | 1405400 | 18185000 | 2266600 | 53403000 | 0 | 0 | 67925000 | 0 | 0 | 8431300 | 30646000 | 0 | 0 | 3746800 | 10865000 |
| NSPPTLEG | STK25; STK25; S | O00506; C9J | STK25_HUMAN | Unmodified | 148310 | 35046 | 35855 | 115450 | 1351300 | 154470 | 40253 | 0 | 92884 | 0 | 112930 | 539390 | 706400 | 204560 | 354190 | 90917 | 160040 | 786360 |
| NSPQCPES | LRBA; LRBA; LR | P50851; A0A | LRBA_HUMAN | Unmodified | 0 | 0 | 0 | 0 | 0 | 0 | 0 | 0 | 0 | 0 | 0 | 785830 | 0 | 0 | 0 | 0 | 0 | 0 |
| NSPQCPES | LRBA; LRBA; LR | P50851; A0A | LRBA_HUMAN | Unmodified | 0 | 0 | 0 | 0 | 820970 | 165830 | 0 | 0 | 28610 | 0 | 0 | 660730 | 0 | 44712 | 0 | 0 | 300520 | 360890 |
| NSQDDYDI | SRP72; SRP72; | O76094; D6I | SRP72_HUMAN | Unmodified | 0 | 0 | 0 | 0 | 0 | 0 | 0 | 0 | 0 | 0 | 0 | 246650 | 0 | 0 | 0 | 0 | 0 | 0 |
| NSQDITPVI | NEO1; NEO1 | Q92859; Q5 | NEO1_HUMAN | Unmodified | 0 | 502310 | 177450 | 561690 | 109130 | 0 | 0 | 0 | 0 | 0 | 0 | 0 | 0 | 0 | 0 | 268360 | 0 | 0 |
| NSQGEEVA | LMNB1 | P20700 | LMNB1_HUMAI | Unmodified | 0 | 0 | 0 | 0 | 0 | 0 | 81313 | 89138 | 0 | 937660 | 0 | 0 | 0 | 0 | 0 | 0 | 0 | 0 |
| NSQKIDRLI | GLRX3 | O76003 | GLRX3_HUMAN | Unmodified | 233220 | 0 | 0 | 0 | 3126000 | 0 | 1359600 | 1187100 | 0 | 458660 | 378700 | 1697700 | 1823100 | 1941600 | 170900 | 0 | 1854000 | 0 |
| NSQLEQEN | TSC22D1; TSC2 | Q15714; A0A | T22D1_HUMAN | Unmodified | 0 | 0 | 0 | 0 | 0 | 90174 | 0 | 0 | 0 | 357970 | 0 | 534120 | 1047800 | 539770 | 0 | 0 | 363940 | 593260 |
| NSQPWQV | KLK12; KLK12 | Q9UKR0; E9I | KLK12_HUMAN | Unmodified | 0 | 0 | 3634600 | 0 | 0 | 0 | 0 | 0 | 15715000 | 0 | 0 | 0 | 0 | 0 | 0 | 0 | 0 | 0 |
| NSQSFFSG | NAPA; NAPA; N. | P54920; M0I | SNAA_HUMAN | Unmodified | 41569 | 0 | 49892 | 0 | 0 | 0 | 0 | 0 | 39217 | 0 | 0 | 1205600 | 0 | 0 | 88682 | 52669 | 0 | 0 |
| NSQVFRNF | GPRC5C; GPRC | Q9NQ84; J9J | GPC5C_HUMA | Unmodified | 0 | 0 | 0 | 10473000 | 555420 | 88232 | 0 | 0 | 198200 | 97033 | 0 | 0 | 7881700 | 2689400 | 0 | 4018300 | 1304300 | 3308200 |
| NSQVPKDE | GALNT6 | Q8NCL4 | GALT6_HUMAN | Unmodified | 0 | 0 | 0 | 0 | 0 | 0 | 132250 | 0 | 1469000 | 0 | 0 | 873000 | 0 | 0 | 0 | 0 | 0 | 0 |
| NSRAEGQT | PKP4; PKP4 | Q99569; A0A | PKP4_HUMAN | Unmodified | 0 | 0 | 0 | 195530 | 0 | 0 | 0 | 0 | 125600 | 60571 | 38576 | 0 | 0 | 0 | 0 | 0 | 0 | 0 |
| NSRPEANE | CLIC4 | Q9Y696 | CLIC4_HUMAN | Unmodified | 1251800 | 1475800 | 1582000 | 2629300 | 5368900 | 0 | 1652900 | 744550 | 9440200 | 1544300 | 7323500 | 18025000 | 2683800 | 4744300 | 3215400 | 3110400 | 2071600 | 2427700 |
| NSRPEANE | CLIC4 | Q9Y696 | CLIC4_HUMAN | Unmodified | 0 | 0 | 0 | 633020 | 90437000 | 23787000 | 25403000 | 22513000 | 1546900 | 12852000 | 619890 | 0 | 18559000 | 44360000 | 0 | 0 | 57704000 | 48015000 |
| NSSHAGAF | AARS1; AARS1; | P49588; A0A | SYAC_HUMAN | Unmodified | 301420 | 237150 | 631990 | 691910 | 3168900 | 0 | 383460 | 99679 | 4422300 | 730650 | 1811300 | 12125000 | 1765100 | 779700 | 835030 | 839510 | 0 | 580770 |
| NSSPSDSK | USP54; USP54; | Q70EL1; A0A | UBP54_HUMAI | Unmodified | 0 | 0 | 121960 | 209320 | 0 | 0 | 0 | 0 | 0 | 0 | 0 | 0 | 1646700 | 1343400 | 133190 | 92724 | 0 | 0 |
| NSSQVGM | BMX | P51813 | BMX_HUMAN ( | Oxidation (M) | 0 | 0 | 0 | 0 | 0 | 0 | 0 | 0 | 0 | 0 | 340880 | 5559500 | 0 | 957510 | 0 | 0 | 0 | 0 |
| NSSTEPKG | ANLN | Q9NQW6 | ANLN_HUMAN | Unmodified | 0 | 0 | 0 | 0 | 0 | 0 | 0 | 0 | 711020 | 0 | 0 | 0 | 0 | 0 | 0 | 0 | 1726200 | 0 |
| NSSTYWEC | TARS1 | P26639 | SYTC_HUMAN | Unmodified | 0 | 590310 | 247930 | 0 | 0 | 0 | 0 | 29164 | 176870 | 0 | 306690 | 2346500 | 0 | 0 | 304770 | 423110 | 0 | 0 |
| NSSTYWEC | TARS1 | P26639 | SYTC_HUMAN | Unmodified | 66876 | 0 | 310530 | 163450 | 3323900 | 1408400 | 0 | 0 | 1110700 | 666010 | 662580 | 3043800 | 266460 | 144620 | 0 | 272370 | 0 | 0 |
| NSSTYWEC | TARS1 | P26639 | SYTC_HUMAN | Oxidation (M) | 0 | 0 | 0 | 0 | 291790 | 0 | 0 | 0 | 0 | 0 | 237930 | 1660700 | 0 | 0 | 0 | 609880 | 0 | 0 |
| NSSVPLAEI | UACA; UACA; U | Q9BZF9; H0 | UACA_HUMAN | Unmodified | 0 | 0 | 0 | 0 | 463260 | 0 | 0 | 0 | 0 | 0 | 0 | 0 | 0 | 684870 | 0 | 0 | 608620 | 1050000 |
| NSSVVPSE | PTPRG | P23470 | PTPRG_HUMAI | Unmodified | 151040 | 432490 | 0 | 0 | 0 | 0 | 0 | 0 | 132250 | 0 | 97406 | 290150 | 0 | 0 | 36870 | 26657 | 0 | 0 |
| NSSYFVEW | TUBB2A; TUBB | Q13885; P0 | TBB2A_HUMAN | Unmodified | 33141 | 0 | 0 | 4565400 | 15661000 | 4878400 | 27061000 | 6155000 | 17370000 | 9562200 | 8928300 | 1.04E+08 | 16817000 | 17718000 | 7455600 | 2857900 | 7265500 | 13295000 |
| NSSYVHGC | G3BP1; G3BP1; | Q13283; A0 | G3BP1_HUMAI | Unmodified | 0 | 0 | 0 | 0 | 784250 | 0 | 0 | 0 | 314810 | 0 | 604580 | 12057000 | 2824300 | 907460 | 0 | 0 | 1768500 | 0 |
| NSSYVHGC | G3BP2; G3BP2; | Q9UN86; D6 | G3BP2_HUMAI | Unmodified | 0 | 0 | 0 | 0 | 352370 | 0 | 0 | 0 | 108470 | 87366 | 233290 | 6352300 | 1131700 | 0 | 0 | 0 | 0 | 0 |
| NSTGGTSV | SNTA1 | Q13424 | SNTA1_HUMAI | Unmodified | 116080 | 0 | 137190 | 69785 | 131410 | 0 | 0 | 0 | 0 | 0 | 0 | 0 | 104210 | 0 | 391640 | 230640 | 48624 | 0 |
| NSTGSQDC | PPFIA1; PPFIA1 | Q13136; H0 | LIPA1_HUMAN | Unmodified | 0 | 0 | 238370 | 216490 | 109980 | 0 | 0 | 0 | 355160 | 0 | 0 | 380260 | 0 | 0 | 409090 | 399550 | 0 | 0 |
| NSTIVFPLP | STOM | P27105 | STOM_HUMAN | Oxidation (M) | 0 | 66663 | 0 | 0 | 4485000 | 0 | 0 | 0 | 0 | 0 | 0 | 0 | 0 | 0 | 0 | 0 | 89028 | 0 |
| NSTIVFPLP | STOM | P27105 | STOM_HUMAN | Oxidation (M) | 0 | 0 | 0 | 0 | 622490 | 0 | 0 | 0 | 0 | 0 | 0 | 0 | 0 | 0 | 0 | 0 | 0 | 0 |
| NSTIVFPLP | STOM | P27105 | STOM_HUMAN | Unmodified | 0 | 0 | 0 | 0 | 266460 | 0 | 0 | 0 | 0 | 0 | 0 | 0 | 0 | 0 | 0 | 0 | 0 | 0 |
| NSTLSDSG | SH3BP4; SH3BI | Q9P0V3; C9. | SH3B4_HUMAI | Unmodified | 0 | 0 | 0 | 0 | 0 | 0 | 0 | 0 | 485420 | 0 | 0 | 0 | 0 | 0 | 0 | 0 | 0 | 0 |
| NSTPPSLAI | ZC2HC1A; ZC2I | Q96GY0; H0 | ZC21A_HUMAN | Unmodified | 0 | 0 | 0 | 0 | 280860 | 0 | 36406 | 0 | 324720 | 239090 | 0 | 0 | 451190 | 235000 | 0 | 0 | 142810 | 396830 |
| NSTRTDYE | MVB12B | Q9H7P6 | MB12B_HUMAI | 2 Oxidation (M) | 0 | 496700 | 417310 | 1080100 | 0 | 0 | 0 | 0 | 0 | 0 | 0 | 0 | 0 | 0 | 1648000 | 469620 | 0 | 0 |
| NSTSSIEPR | MAP4K4; MAP4 | C9J840; A0A | C9J840_HUMA | Unmodified | 0 | 0 | 63984 | 0 | 0 | 0 | 0 | 0 | 0 | 5952 | 103280 | 928550 | 298930 | 0 | 62904 | 0 | 56525 | 0 |
| NSTTDQVY | SNX27; SNX27; | Q96L92; A0A | SNX27_HUMAI | Unmodified | 59258 | 103900 | 0 | 0 | 0 | 0 | 0 | 0 | 0 | 0 | 0 | 0 | 0 | 0 | 75992 | 83167 | 0 | 0 |
| NSTVTLVPE | TOM1L1; TOM1 | O75674; J3K | TM1L1_HUMAN | Unmodified | 7022800 | 6528500 | 5138500 | 3102700 | 618800 | 552890 | 0 | 0 | 1845700 | 482500 | 1136700 | 3576800 | 924990 | 525910 | 9512700 | 8026800 | 270500 | 487850 |
| NSTVTLVPE | TOM1L1; TOM1 | O75674; J3K | TM1L1_HUMAN | Oxidation (M) | 5216600 | 2558000 | 0 | 6689000 | 4518700 | 659790 | 1417300 | 51294 | 636760 | 2661700 | 0 | 447690 | 6330500 | 3279300 | 527960 | 1119500 | 2842300 | 3657300 |
| NSTVTLVPE | TOM1L1; TOM1 | O75674; J3K | TM1L1_HUMAN | Unmodified | 280000 | 340280 | 0 | 468190 | 1067100 | 0 | 0 | 0 | 1268700 | 467150 | 0 | 0 | 1502900 | 820510 | 0 | 0 | 394140 | 933000 |
| NSTWSGES | NCL; NCL; NCL | P19338; C9J | NUCL_HUMAN | Unmodified | 0 | 0 | 0 | 0 | 55729 | 2278700 | 777240 | 715690 | 1438000 | 9292800 | 4154900 | 0 | 2866700 | 363680 | 0 | 0 | 0 | 0 |
| NSVCIIPLR | RRP9 | O43818 | U3IP2_HUMAN | Unmodified | 0 | 0 | 0 | 0 | 0 | 0 | 0 | 0 | 0 | 0 | 0 | 243830 | 0 | 0 | 0 | 0 | 0 | 0 |
| NSVEVALN | YARS1; YARS1 | P54577; A0A | SYYC_HUMAN | Unmodified | 1136500 | 958450 | 222390 | 714450 | 2561400 | 186060 | 658080 | 619200 | 3314000 | 1296500 | 1105200 | 4667200 | 5472400 | 4518700 | 1113300 | 1553700 | 605310 | 2021000 |
| NSVEVALN | YARS1; YARS1 | P54577; A0A | SYYC_HUMAN | Unmodified | 0 | 0 | 0 | 0 | 243820 | 0 | 466770 | 0 | 0 | 0 | 0 | 0 | 487770 | 496660 | 0 | 0 | 471470 | 297800 |
| NSVFQQGI | ATP1A1; ATP1A | P05023; P50 | AT1A1_HUMAN | Unmodified | 0 | 0 | 2251900 | 410240 | 0 | 0 | 0 | 69259 | 705830 | 0 | 204570 | 9075300 | 0 | 0 | 1739700 | 4261800 | 38305 | 0 |
| NSVFQQGI | ATP1A1; ATP1A | P05023; P50 | AT1A1_HUMAN | Oxidation (M) | 72525 | 69678 | 1274400 | 189270 | 0 | 0 | 0 | 0 | 0 | 0 | 0 | 4789000 | 0 | 0 | 1738700 | 1557800 | 0 | 0 |
| NSVGRWKI | FN1 | P02751 | FINC_HUMAN I | Unmodified | 0 | 0 | 0 | 0 | 0 | 0 | 0 | 0 | 0 | 378920 | 0 | 0 | 0 | 0 | 0 | 0 | 486740 | 0 |
| NSVLQSGL | AMPD3 | Q01432 | AMPD3_HUMA | Unmodified | 0 | 0 | 0 | 0 | 0 | 0 | 39842 | 41804 | 0 | 0 | 0 | 514600 | 0 | 0 | 0 | 0 | 0 | 0 |
| NSVNHVIH | MME; MME; MM | P08473; A0A | NEP_HUMAN N | Unmodified | 0 | 59785 | 4316000 | 3766000 | 0 | 0 | 0 | 0 | 0 | 0 | 0 | 0 | 0 | 0 | 5690500 | 6373700 | 0 | 0 |
| NSVNPVWI | DYSF | O75923 | DYSF_HUMAN | Unmodified | 0 | 0 | 0 | 0 | 363100 | 35737 | 0 | 0 | 0 | 33778 | 0 | 0 | 0 | 0 | 0 | 0 | 0 | 83886 |
| NSVSLVVVI | ADAMTS1; ADA | Q9UHI8; E5F | ATS1_HUMAN / | Unmodified | 0 | 0 | 556870 | 0 | 0 | 0 | 0 | 0 | 0 | 0 | 0 | 0 | 0 | 0 | 201600 | 264430 | 0 | 0 |
| NSVSNFLH | ACACA | Q13085 | ACACA_HUMAI | Unmodified | 0 | 0 | 248680 | 0 | 0 | 0 | 0 | 0 | 0 | 0 | 0 | 1024000 | 0 | 0 | 0 | 0 | 0 | 0 |
| NSVSQISVI | ARPC1B | O15143 | ARC1B_HUMAI | Unmodified | 1455400 | 835850 | 671520 | 879090 | 12966000 | 3221600 | 2500500 | 2088900 | 1142000 | 819350 | 966690 | 9340600 | 4683800 | 3346700 | 1058500 | 857690 | 5174000 | 6845900 |
| NSVSQISVI | ARPC1B | O15143 | ARC1B_HUMAI | Unmodified | 0 | 0 | 0 | 0 | 0 | 0 | 0 | 0 | 0 | 0 | 0 | 0 | 0 | 0 | 0 | 0 | 584330 | 0 |
| NSVTPDMN | RPL5; RPL5 | P46777; A0A | RL5_HUMAN 6 | Oxidation (M) | 193560 | 25341 | 0 | 0 | 0 | 0 | 0 | 0 | 0 | 0 | 0 | 2492800 | 0 | 0 | 0 | 0 | 0 | 0 |
| NSVVEASE | YWHAH; YWHA | Q04917; A2I | 1433F_HUMAN | Unmodified | 369820 | 336060 | 0 | 660860 | 0 | 0 | 0 | 0 | 1062000 | 167690 | 1024400 | 4681900 | 0 | 469650 | 1204000 | 985920 | 544420 | 701240 |
| NSVVEASE | YWHAH; YWHA | Q04917; A2I | 1433F_HUMAN | Unmodified | 0 | 0 | 0 | 0 | 0 | 394670 | 240980 | 0 | 410950 | 1242200 | 0 | 0 | 0 | 0 | 0 | 0 | 571060 | 895250 |
| NSWGHNF | CTSS; CTSS; CT | P25774; A0A | CATS_HUMAN | Unmodified | 0 | 0 | 0 | 0 | 0 | 162930 | 109630 | 0 | 3823800 | 784810 | 0 | 0 | 0 | 0 | 0 | 0 | 0 | 240180 |
| NSWGTGW | CTSC; CTSC; C | P53634; A0A | CATC_HUMAN | Unmodified | 0 | 0 | 0 | 0 | 0 | 0 | 0 | 0 | 1493900 | 330150 | 9819900 | 6289900 | 0 | 0 | 0 | 0 | 0 | 0 |
| NSWQITPR | CEMIP2; CEMIP | Q9UHN6; H | CEIP2_HUMAN | Unmodified | 0 | 0 | 169010 | 0 | 679750 | 0 | 0 | 0 | 4167900 | 0 | 0 | 0 | 0 | 0 | 715100 | 624940 | 0 | 0 |
| NSYATTENI | BAIAP2; BAIAP2 | Q9UQB8; I3I | BAIP2_HUMAN | Unmodified | 0 | 187220 | 0 | 0 | 272870 | 0 | 18414 | 100640 | 0 | 23408 | 0 | 59122 | 0 | 282480 | 0 | 0 | 222720 | 0 |
| NSYLEVLLK | PDIA6 | Q15084 | PDIA6_HUMAN | Unmodified | 0 | 70406 | 315210 | 43399 | 0 | 96421 | 165340 | 46850 | 10677000 | 4785700 | 3218000 | 1863700 | 0 | 26201 | 489980 | 130050 | 241220 | 0 |
| NSYMNPEK | MME; MME; MM | P08473; A0A | NEP_HUMAN N | Oxidation (M) | 100060 | 0 | 1691700 | 4173500 | 0 | 0 | 0 | 0 | 284060 | 0 | 0 | 0 | 3351600 | 976970 | 3788000 | 2403500 | 0 | 44259 |
| NSYMNPEK | MME; MME; MM | P08473; A0A | NEP_HUMAN N | Unmodified | 0 | 0 | 4219200 | 5399300 | 66678 | 0 | 0 | 0 | 2792300 | 1334100 | 15886 | 0 | 8341600 | 2204500 | 8184200 | 5142800 | 0 | 0 |
| NSYMNPEK | MME; MME; MM | P08473; A0A | NEP_HUMAN N | Unmodified | 0 | 0 | 0 | 0 | 291860 | 0 | 0 | 0 | 2127200 | 0 | 0 | 65351 | 0 | 0 | 4644300 | 0 | 0 | 82971 |
| NSYRSEEA | SPINT2; SPINT2 | O43291; K7E | SPIT2_HUMAN | Unmodified | 0 | 0 | 205520 | 686830 | 617970 | 298970 | 85942 | 29888 | 25189000 | 2683700 | 0 | 0 | 288220 | 339670 | 0 | 0 | 0 | 0 |
| NSYRSEEA | SPINT2; SPINT2 | O43291; K7E | SPIT2_HUMAN | Oxidation (M) | 55082 | 0 | 0 | 194040 | 301260 | 146150 | 16651 | 0 | 6603900 | 580730 | 0 | 0 | 0 | 0 | 0 | 0 | 43824 | 0 |
| NSYVAGQY | G6PD; G6PD; G | P11413; E7E | G6PD_HUMAN | Unmodified | 0 | 0 | 122260 | 0 | 0 | 0 | 0 | 0 | 1662400 | 0 | 403890 | 3976800 | 0 | 0 | 262000 | 0 | 0 | 0 |
| NSYVRDDA | TRAF2 | Q12933 | TRAF2_HUMAN | Unmodified | 0 | 0 | 0 | 0 | 4615600 | 890110 | 0 | 0 | 0 | 0 | 0 | 0 | 0 | 0 | 0 | 0 | 0 | 0 |
| NTAELQPES | PPM1G | O15355 | PPM1G_HUMA | Unmodified | 0 | 0 | 0 | 0 | 438510 | 0 | 0 | 0 | 63078 | 0 | 154190 | 853980 | 0 | 0 | 0 | 0 | 0 | 0 |
| NTAEWLLS | PNP; PNP; PNP | P00491; G3 | PNPH_HUMAN | Unmodified | 139870 | 293220 | 0 | 0 | 0 | 0 | 0 | 0 | 0 | 0 | 0 | 13833000 | 357890 | 0 | 560560 | 524390 | 0 | 0 |
| NTAQRQEII | ANXA4; ANXA4; | P09525; B4E | ANXA4_HUMAI | Unmodified | 0 | 0 | 0 | 0 | 0 | 948490 | 210030 | 229780 | 0 | 5707500 | 0 | 0 | 735680 | 0 | 0 | 0 | 0 | 0 |
| NTASLKYEI | APOB | P04114 | APOB_HUMAN | Unmodified | 158850 | 469290 | 0 | 649570 | 0 | 0 | 4426100 | 0 | 0 | 0 | 0 | 0 | 0 | 0 | 0 | 0 | 40813 | 0 |
| NTCQLYIQ | ADAM10 | O14672 | ADA10_HUMAI | Unmodified | 6532400 | 6611700 | 5011900 | 10078000 | 11962000 | 2159000 | 4059600 | 176710 | 2400700 | 1851400 | 929700 | 6481000 | 15369000 | 13791000 | 11606000 | 8579000 | 3616000 | 4893400 |
| NTCTSVYTI | PRKDC; PRKDC | P78527; F5G | PRKDC_HUMAI | Unmodified | 0 | 33886 | 0 | 0 | 0 | 53260 | 0 | 0 | 0 | 197900 | 1608600 | 683350 | 0 | 0 | 23178 | 18733 | 534710 | 0 |
| NTDEMVEL | HNRNPK; HNRI | P61978; Q5 | HNRPK_HUMA | Unmodified | 611410 | 551830 | 0 | 409760 | 439180 | 132140 | 0 | 0 | 1632600 | 309190 | 3034600 | 8074100 | 700850 | 140340 | 298850 | 178520 | 0 | 0 |
| NTDEMVEL | HNRNPK; HNRI | P61978; Q5 | HNRPK_HUMA | Oxidation (M) | 695850 | 0 | 0 | 0 | 0 | 0 | 0 | 30487 | 443610 | 0 | 421480 | 2523400 | 542000 | 0 | 0 | 0 | 0 | 0 |
| NTDEPMGI | CASK; CASK; C | O14936; A0 | CSKP_HUMAN | Unmodified | 0 | 0 | 139430 | 0 | 1643900 | 0 | 0 | 145310 | 3145500 | 1442000 | 0 | 465150 | 957140 | 1304700 | 566360 | 0 | 821750 | 689310 |
| NTDEPMGI | CASK; CASK; C | O14936; A0 | CSKP_HUMAN | Oxidation (M) | 0 | 0 | 654520 | 0 | 396270 | 418270 | 184710 | 137730 | 749090 | 349070 | 0 | 210030 | 567820 | 619250 | 0 | 0 | 0 | 0 |
| NTDFCHGI | ATP8B1; ATP8B | O43520; A0 | AT8B1_HUMAN | Unmodified | 0 | 0 | 0 | 0 | 0 | 35825 | 0 | 0 | 515570 | 0 | 0 | 529600 | 0 | 97760 | 0 | 134250 | 0 | 0 |
| NTDFFIGKI | MGAT5 | Q09328 | MGT5A_HUMAI | Unmodified | 0 | 61593 | 0 | 0 | 408100 | 0 | 0 | 0 | 10202000 | 0 | 1097200 | 0 | 0 | 0 | 0 | 53281 | 0 | 0 |
| NTDFVPPQ | KIF23; KIF23; KI | A0A2U3TZL8 | A0A2U3TZL8_H | Unmodified | 0 | 0 | 0 | 0 | 718090 | 265360 | 47111 | 45797 | 205630 | 46898 | 216510 | 463350 | 0 | 0 | 0 | 0 | 122680 | 0 |
| NTDQASMI | MYH9; MYH11; | P35579; P35 | MYH9_HUMAN | Unmodified | 119620 | 0 | 421870 | 532360 | 476710 | 135600 | 0 | 0 | 356430 | 0 | 869490 | 4301900 | 107460 | 93114 | 395990 | 345350 | 178350 | 234920 |
| NTDQASMI | MYH9; MYH11; | P35579; P35 | MYH9_HUMAN | Oxidation (M) | 0 | 0 | 0 | 20005 | 0 | 0 | 0 | 0 | 0 | 0 | 82234 | 131700 | 0 | 0 | 34419 | 14211 | 0 | 0 |
| NTDQASMI | MYH10 | P35580 | MYH10_HUMAI | Unmodified | 0 | 0 | 0 | 0 | 0 | 0 | 0 | 0 | 0 | 0 | 0 | 2127300 | 0 | 0 | 0 | 0 | 0 | 0 |
| NTDSPEFK | CC2D1A; CC2D | Q6P1N0; A0 | C2D1A_HUMAI | Unmodified | 873140 | 628430 | 0 | 674090 | 453520 | 203750 | 0 | 0 | 780110 | 304090 | 0 | 670730 | 250910 | 189970 | 373190 | 507490 | 95380 | 29524 |
| NTEAIGNE | PI4KA | P42356 | PI4KA_HUMAN | Unmodified | 0 | 0 | 0 | 0 | 0 | 0 | 0 | 35978 | 759110 | 0 | 0 | 422790 | 0 | 0 | 0 | 0 | 0 | 0 |
| NTEAVDLIT | MRPL27; MRPL | Q9P0M9; H7 | RM27_HUMAN | Unmodified | 67921 | 0 | 0 | 0 | 974120 | 0 | 271470 | 17474 | 2188700 | 591850 | 0 | 0 | 0 | 254330 | 444970 | 0 | 0 | 0 |
| NTEDLSRP | GALNT4; POC1 | Q8N4A0; F8 | GALT4_HUMAN | Unmodified | 0 | 0 | 0 | 0 | 0 | 0 | 0 | 0 | 519900 | 0 | 0 | 0 | 106540 | 0 | 0 | 0 | 0 | 1301600 |
| NTEDLTEE | GMFB; GMFB | P60983; G3 | GMFB_HUMAN | Unmodified | 1243600 | 784530 | 0 | 0 | 0 | 0 | 0 | 0 | 0 | 0 | 0 | 0 | 0 | 0 | 1321500 | 2065500 | 0 | 0 |
| NTEIENTKL | TAX1BP1; TAX1 | Q86VP1; B8 | TAXB1_HUMAN | Unmodified | 0 | 0 | 0 | 0 | 161650 | 761540 | 3616800 | 230300 | 0 | 0 | 0 | 1271500 | 0 | 0 | 0 | 0 | 533460 | 536100 |
| NTEIENTKL | TAX1BP1; TAX1 | Q86VP1; B8 | TAXB1_HUMAN | Oxidation (M) | 0 | 0 | 0 | 0 | 0 | 0 | 783970 | 250590 | 0 | 0 | 0 | 0 | 0 | 0 | 0 | 0 | 0 | 0 |
| NTEIGFLQI | DARS2; DARS2; | Q6PI48; A0A | SYDM_HUMAN | Unmodified | 0 | 0 | 0 | 0 | 73641 | 0 | 0 | 24144 | 4270000 | 231390 | 0 | 0 | 0 | 0 | 0 | 0 | 0 | 0 |
| NTEILAALK | MON2; MON2; | Q7Z3U7; A0 | MON2_HUMAN | Unmodified | 0 | 0 | 0 | 0 | 0 | 0 | 0 | 0 | 0 | 102490 | 250410 | 1648700 | 0 | 0 | 0 | 0 | 0 | 0 |
| NTFAEVTGI | FN1 | P02751 | FINC_HUMAN I | Unmodified | 0 | 0 | 2875200 | 4132700 | 0 | 0 | 0 | 0 | 909970 | 1959000 | 169170 | 0 | 0 | 453280 | 109980 | 2714400 | 0 | 0 |
| NTFEVIPVN | DIP2B | Q9P265 | DIP2B_HUMAN | Unmodified | 355680 | 0 | 329160 | 0 | 0 | 0 | 0 | 0 | 51690 | 0 | 0 | 3123100 | 0 | 0 | 1079600 | 1154700 | 0 | 0 |
| NTFLWTEF | PIP4P1; PIP4P1 | Q86T03; H0 | PP4P1_HUMAN | Unmodified | 0 | 0 | 0 | 119810 | 17532 | 0 | 0 | 0 | 5279500 | 1190100 | 0 | 0 | 0 | 0 | 0 | 52392 | 0 | 0 |
| NTFSSGSD | ECHDC1; ECHI | Q9NTX5; J3K | ECHD1_HUMA | Unmodified | 147660 | 153080 | 132490 | 216220 | 625260 | 0 | 0 | 0 | 84651 | 0 | 346330 | 8124800 | 185520 | 1006800 | 0 | 0 | 0 | 0 |
| NTFTESAG | COL12A1; COL | Q99715; D6I | COCA1_HUMA | Unmodified | 0 | 303610 | 103930 | 94564 | 0 | 0 | 0 | 0 | 485940 | 48360 | 60105 | 38374 | 0 | 0 | 110510 | 344950 | 0 | 0 |
| NTGIICTIGI | PKM; PKM; PKM | P14618; B4I | KPYM_HUMAN | Unmodified | 2193600 | 1464100 | 13018000 | 9194900 | 37687000 | 12378000 | 1750900 | 759890 | 9537000 | 3982000 | 19387000 | 1.02E+08 | 15847000 | 3630400 | 6336500 | 4913200 | 11044000 | 14770000 |
| NTGIICTIGI | PKM; PKM; PKM | P14618; H3E | KPYM_HUMAN | Unmodified | 0 | 0 | 0 | 0 | 34014000 | 11742000 | 3285600 | 3141900 | 438290 | 1394100 | 0 | 0 | 6700200 | 0 | 0 | 0 | 17503000 | 20360000 |
| NTGIICTIGI | PKM; PKM; PKM | P14618; H3E | KPYM_HUMAN | Unmodified | 0 | 0 | 0 | 0 | 0 | 0 | 0 | 0 | 0 | 0 | 0 | 0 | 0 | 0 | 138980 | 0 | 3254600 | 823150 |

| | | | | | | | | | | | | | | | | | | | | | | |
|---|---|---|---|---|---|---|---|---|---|---|---|---|---|---|---|---|---|---|---|---|---|---|
| NTGLITVQ( | PCDH1 | Q08174 | PCDH1_HUMA | Unmodified | 556900 | 883090 | 0 | 192600 | 0 | 0 | 0 | 0 | 479850 | 0 | 0 | 201630 | 0 | 0 | 407150 | 189520 | 0 | 0 |
| NTGNVKW: | NBR1; NBR1 | Q14596; B7Z | NBR1_HUMAN | Unmodified | 0 | 0 | 0 | 0 | 0 | 0 | 1023500 | 0 | 0 | 0 | 0 | 0 | 0 | 0 | 0 | 0 | 0 | 0 |
| NTGNVKW: | NBR1; NBR1 | Q14596; B7Z | NBR1_HUMAN | Unmodified | 0 | 0 | 0 | 0 | 0 | 0 | 285190 | 61472 | 23617 | 0 | 0 | 0 | 0 | 882040 | 0 | 0 | 0 | 0 |
| NTGTEAPD | SELENBP1; SEL | Q13228; C9J | SBP1_HUMAN | Unmodified | 0 | 0 | 0 | 0 | 0 | 0 | 0 | 0 | 0 | 0 | 0 | 0 | 0 | 0 | 0 | 0 | 0 | 0 |
| NTHATTHN | PARP1; PARP1 | P09874; A0A | PARP1_HUMAN | Unmodified | 506950 | 351310 | 58397 | 136860 | 0 | 0 | 0 | 0 | 888070 | 278980 | 497840 | 1850800 | 176310 | 66295 | 0 | 0 | 0 | 159580 |
| NTHEAEVLI | SYTL2; SYTL2; S | A0A0U1RQP | A0A0U1RQP0_ | Unmodified | 0 | 1235300 | 170140 | 5860800 | 25604000 | 74357000 | 68870000 | 40923000 | 0 | 192140 | 228870 | 0 | 15234000 | 43254000 | 184740 | 95279 | 10456000 | 84381000 |
| NTHTLLLA( | COPG1; COPG: | Q9Y678; H0Y | COPG1_HUMA | Unmodified | 0 | 0 | 0 | 0 | 0 | 0 | 0 | 0 | 0 | 0 | 0 | 787920 | 0 | 0 | 0 | 0 | 0 | 0 |
| NTIEETGNI | LAMC1 | P11047 | LAMC1_HUMAI | Unmodified | 0 | 0 | 256780 | 0 | 0 | 0 | 0 | 0 | 1056100 | 0 | 153090 | 0 | 0 | 0 | 0 | 0 | 0 | 0 |
| NTILFLR | FAM91A1; FAM9 | Q658Y4; E7E | F91A1_HUMAN | Unmodified | 0 | 44910 | 0 | 0 | 0 | 0 | 0 | 0 | 0 | 0 | 0 | 889730 | 0 | 0 | 204520 | 188350 | 0 | 0 |
| NTILQLPLR | N/A | N/A | N/A | Unmodified | 0 | 0 | 0 | 0 | 0 | 0 | 2276400 | 0 | 0 | 0 | 0 | 0 | 0 | 1711100 | 0 | 0 | 0 | 0 |
| NTIQWLEN | KIF5B | P33176 | KINH_HUMAN | Unmodified | 83026 | 91379 | 23846 | 0 | 111680 | 0 | 0 | 0 | 608420 | 0 | 309130 | 770080 | 73735 | 0 | 160850 | 116640 | 22014 | 0 |
| NTISLLVAG | DYNC1H1; DYN | Q14204; A0A | DYHC1_HUMA | Unmodified | 0 | 0 | 0 | 0 | 0 | 0 | 0 | 0 | 0 | 0 | 157310 | 3568500 | 0 | 0 | 0 | 0 | 0 | 0 |
| NTIVTSYNF | ACO2; ACO2; A | Q99798; A2A | ACON_HUMAN | Unmodified | 0 | 0 | 0 | 0 | 0 | 0 | 0 | 0 | 2579300 | 229270 | 0 | 0 | 0 | 0 | 0 | 0 | 0 | 0 |
| NTKDSAYN | ITGA1 | P56199 | ITA1_HUMAN In | Unmodified | 0 | 0 | 0 | 0 | 447710 | 0 | 0 | 0 | 0 | 0 | 0 | 0 | 143020 | 0 | 0 | 0 | 1058900 | 0 |
| NTKEPPLSI | ILF3; ILF3; ILF3; | Q12906; K7E | ILF3_HUMAN In | Oxidation (M) | 3506700 | 1684400 | 0 | 3520600 | 109810 | 0 | 0 | 0 | 0 | 0 | 0 | 2098400 | 0 | 286740 | 0 | 0 | 0 | 0 |
| NTKLDDLFI | LRPPRC | P42704 | LPPRC_HUMAN | Unmodified | 0 | 0 | 0 | 0 | 0 | 0 | 0 | 0 | 1424800 | 510650 | 0 | 0 | 0 | 0 | 0 | 0 | 0 | 0 |
| NTKLDDLFI | LRPPRC | P42704 | LPPRC_HUMAN | Unmodified | 0 | 0 | 0 | 0 | 0 | 0 | 113460 | 0 | 470020 | 0 | 0 | 0 | 0 | 0 | 0 | 0 | 205210 | 0 |
| NTKLEAIPN | SLC44A1 | Q8WWI5 | CTL1_HUMAN ( | Unmodified | 441140 | 106930 | 0 | 1119700 | 630340 | 0 | 0 | 0 | 918870 | 486700 | 235390 | 1556400 | 634450 | 0 | 479720 | 0 | 0 | 0 |
| NTKLEAIPN | SLC44A1 | Q8WWI5 | CTL1_HUMAN ( | Unmodified | 0 | 151480 | 743280 | 563840 | 1955300 | 232840 | 0 | 0 | 4710700 | 0 | 755370 | 1009600 | 106770 | 41386 | 830280 | 566800 | 0 | 323740 |
| NTLFNLSN | PICALM; SNAPS | Q13492; O6( | PICAL_HUMAN | Unmodified | 1492900 | 1088600 | 217820 | 139280 | 0 | 0 | 0 | 0 | 186350 | 100580 | 0 | 1118500 | 312580 | 0 | 810680 | 718230 | 0 | 0 |
| NTLIQFEDF | ME2; ME2; ME2 | P23368; A0A | MAOM_HUMAN | Unmodified | 0 | 0 | 0 | 0 | 0 | 0 | 23270 | 0 | 3382700 | 0 | 0 | 0 | 0 | 0 | 0 | 0 | 0 | 0 |
| NTLLIAGLQ | DDOST; DDOST | P39656; A0A | OST48_HUMAN | Unmodified | 0 | 0 | 0 | 0 | 0 | 0 | 0 | 0 | 71999000 | 7657500 | 2107300 | 1546200 | 0 | 0 | 0 | 0 | 0 | 0 |
| NTLLQNWI | PEAK1 | Q9H792 | PEAK1_HUMAN | Unmodified | 0 | 0 | 0 | 0 | 0 | 0 | 0 | 0 | 0 | 0 | 0 | 312390 | 0 | 0 | 0 | 0 | 0 | 0 |
| NTLNPVW( | CPNE8; CPNE8 | Q86YQ8; E7I | CPNE8_HUMAI | Unmodified | 0 | 0 | 0 | 0 | 1569400 | 518640 | 0 | 0 | 0 | 0 | 0 | 0 | 0 | 0 | 0 | 0 | 0 | 0 |
| NTLPSKETI | TMSB15B; TMSI | P0CG35; P0( | TB15B_HUMAN | Unmodified | 0 | 0 | 0 | 0 | 110950 | 0 | 597570 | 715530 | 2143500 | 2946900 | 46834 | 0 | 1550400 | 486920 | 0 | 0 | 120110 | 0 |
| NTLPSKETI | TMSB15B; TMSI | P0CG35; P0( | TB15B_HUMAN | Unmodified | 0 | 0 | 0 | 0 | 0 | 0 | 0 | 771890 | 0 | 330690 | 0 | 0 | 0 | 0 | 0 | 0 | 0 | 0 |
| NTLPTKETII | TMSB10 | P63313 | TYB10_HUMAN | Unmodified | 0 | 0 | 0 | 0 | 370270 | 0 | 0 | 0 | 0 | 1371100 | 140310 | 0 | 0 | 0 | 0 | 0 | 1170900 | 973830 |
| NTLQLHR | RPL15; RPL15 | P61313; E7E | RL15_HUMAN ( | Unmodified | 569230 | 638170 | 0 | 0 | 0 | 0 | 1649800 | 0 | 1334200 | 87275 | 29251 | 1048900 | 0 | 0 | 39661 | 0 | 0 | 41393 |
| NTLTQTTEN | DSP | P15924 | DESP_HUMAN | Unmodified | 0 | 0 | 0 | 0 | 0 | 0 | 0 | 0 | 393010 | 55594 | 0 | 206060 | 0 | 0 | 0 | 0 | 0 | 0 |
| NTMETIDW | CD74 | P04233 | HG2A_HUMAN | Unmodified | 0 | 0 | 0 | 0 | 0 | 0 | 0 | 0 | 1999600 | 77866 | 0 | 0 | 0 | 0 | 0 | 0 | 0 | 0 |
| NTMGDLSF | NPTX1 | Q15818 | NPTX1_HUMAN | Unmodified | 0 | 0 | 0 | 0 | 0 | 0 | 0 | 0 | 38127000 | 961340 | 0 | 0 | 0 | 0 | 0 | 0 | 0 | 0 |
| NTNAGAPF | OGDH; OGDH; | Q02218; E9F | ODO1_HUMAN | Unmodified | 0 | 0 | 0 | 0 | 0 | 0 | 0 | 0 | 1407700 | 147370 | 0 | 0 | 0 | 0 | 0 | 0 | 0 | 0 |
| NTNDANS( | GDI1; GDI2; GD | P31150; P50 | GDIA_HUMAN | Unmodified | 1363500 | 564300 | 1291100 | 2211600 | 742910 | 0 | 0 | 82152 | 3787700 | 724750 | 2541800 | 0 | 1182600 | 432870 | 2343600 | 1676300 | 277770 | 855350 |
| NTNDANS( | GDI1; GDI2; GD | P31150; P50 | GDIA_HUMAN | Unmodified | 502890 | 244070 | 1007200 | 970090 | 800950 | 0 | 0 | 0 | 2678400 | 418330 | 1231800 | 7111100 | 574840 | 0 | 1177000 | 1731600 | 251520 | 0 |
| NTNGSQFF | PPID | Q08752 | PPID_HUMAN I | Unmodified | 0 | 0 | 0 | 114070 | 0 | 0 | 0 | 0 | 0 | 0 | 0 | 4225000 | 0 | 0 | 0 | 44999 | 0 | 0 |
| NTNPNFVR | MYH9; MYH10 | P35579; P35 | MYH9_HUMAN | Unmodified | 0 | 0 | 475870 | 0 | 1069800 | 0 | 252450 | 141670 | 505900 | 3004600 | 1038200 | 10693000 | 0 | 99810 | 0 | 341540 | 231580 | 432070 |
| NTNTINNG | DDX6; DDX6 | P26196; Q8I | DDX6_HUMAN | Unmodified | 0 | 516910 | 287540 | 483890 | 892150 | 0 | 0 | 85225 | 0 | 0 | 0 | 0 | 1163200 | 401750 | 0 | 0 | 0 | 539340 |
| NTNTPNN\ | SNX9; SNX9; SN | Q9Y5X1; A0A | SNX9_HUMAN | Unmodified | 6784800 | 0 | 0 | 1691200 | 0 | 0 | 0 | 0 | 0 | 0 | 0 | 0 | 0 | 0 | 0 | 0 | 0 | 0 |
| NTPAFFAEI | ANXA11; ANXA1 | P50995; H0Y | ANX11_HUMAN | Unmodified | 1318900 | 876640 | 709910 | 2399300 | 46432000 | 5496200 | 24833000 | 10738000 | 57611000 | 42209000 | 5274600 | 2393800 | 36615000 | 38020000 | 1079200 | 650800 | 19928000 | 70977000 |
| NTPASASLI | TMEM237; TMEI | Q96Q45; F2Z | TM237_HUMAN | Unmodified | 0 | 0 | 0 | 0 | 15981000 | 1316900 | 585980 | 699650 | 342520 | 372570 | 0 | 0 | 0 | 2514700 | 0 | 0 | 1703600 | 4924400 |
| NTPCSENK | DHX36; DHX36 | Q9H2U1; E7 | DHX36_HUMAI | Unmodified | 151960 | 102860 | 1024600 | 288010 | 208960 | 31341 | 25213 | 28646 | 0 | 0 | 98730 | 305490 | 188480 | 0 | 174850 | 165040 | 194320 | 193030 |
| NTPSPFIET | GCLC; GCLC; G | P48506; A0A | GSH1_HUMAN | Unmodified | 60912 | 0 | 34243 | 0 | 0 | 0 | 0 | 0 | 79981 | 0 | 123860 | 1722600 | 0 | 0 | 0 | 0 | 0 | 0 |
| NTPVYVSSI | GOT1 | P17174 | AATC_HUMAN | Unmodified | 0 | 0 | 2782000 | 3152600 | 1360900 | 649000 | 190460 | 0 | 7789900 | 2604000 | 0 | 49025000 | 0 | 1698200 | 4663500 | 3041800 | 606300 | 439280 |
| NTQAAEAL' | DST; DST; DST; | Q03001; A0A | DYST_HUMAN | Unmodified | 0 | 0 | 0 | 0 | 817480 | 0 | 0 | 0 | 0 | 0 | 0 | 0 | 0 | 0 | 0 | 0 | 0 | 0 |
| NTQIQVLPI | SCIN | Q9Y6U3 | ADSV_HUMAN | Unmodified | 0 | 0 | 0 | 0 | 0 | 0 | 0 | 0 | 2405300 | 1821400 | 0 | 959910 | 0 | 0 | 67817 | 24508 | 0 | 0 |
| NTSDFVYLI | DLG1; DLG1; D | Q12959; H7( | DLG1_HUMAN | Unmodified | 231470 | 0 | 408090 | 0 | 4084400 | 0 | 0 | 0 | 1816800 | 357740 | 1142400 | 0 | 434940 | 0 | 0 | 0 | 1415700 | 629900 |
| NTSDVISA/ | CTNNA1; CTNN | P35221; E5R | CTNA1_HUMAI | Unmodified | 120660 | 108880 | 582250 | 741940 | 1470300 | 459220 | 0 | 0 | 68016 | 136080 | 439270 | 0 | 1663800 | 1215300 | 810560 | 674760 | 98688 | 0 |
| NTSLNPQE | RBCK1; RBCK1; | Q9BYM8; G3 | HOIL1_HUMAN | Unmodified | 276000 | 250020 | 0 | 0 | 125850 | 0 | 0 | 0 | 0 | 0 | 0 | 204830 | 0 | 0 | 73377 | 75537 | 0 | 0 |
| NTSVVDSE | NEXN; NEXN | Q0ZGT2; H7 | NEXN_HUMAN | Unmodified | 0 | 0 | 0 | 0 | 1586300 | 293560 | 139380 | 0 | 0 | 0 | 388500 | 0 | 0 | 0 | 0 | 0 | 1367300 | 0 |
| NTTGVTEE/ | SPTAN1; SPTAN | Q13813; A0A | SPTN1_HUMAN | Unmodified | 64569 | 0 | 440090 | 0 | 124660 | 0 | 0 | 0 | 0 | 184830 | 1519200 | 3203600 | 0 | 0 | 241360 | 153960 | 0 | 0 |
| NTTIPTKQT | HSPA8; HSPA8; | P11142; E9P | HSP7C_HUMAI | Oxidation (M) | 0 | 0 | 0 | 0 | 0 | 0 | 0 | 0 | 0 | 1766500 | 0 | 0 | 0 | 0 | 0 | 0 | 0 | 0 |
| NTTPMYR | GAK; GAK | O14976; D6I | GAK_HUMAN C | Unmodified | 0 | 0 | 0 | 0 | 0 | 0 | 0 | 0 | 0 | 0 | 0 | 309690 | 0 | 0 | 0 | 0 | 0 | 0 |
| NTTQNTGY | MYO1E | Q12965 | MYO1E_HUMAI | Unmodified | 0 | 0 | 0 | 0 | 0 | 0 | 0 | 0 | 0 | 0 | 0 | 0 | 0 | 0 | 0 | 0 | 0 | 0 |
| NTTVSGAG | DSG2 | Q14126 | DSG2_HUMAN | Unmodified | 0 | 0 | 0 | 0 | 0 | 0 | 0 | 0 | 3720100 | 0 | 5981600 | 5258000 | 0 | 0 | 0 | 0 | 0 | 0 |
| NTTVSGAG | DSG2 | Q14126 | DSG2_HUMAN | Unmodified | 0 | 812140 | 0 | 1137700 | 39997000 | 6413400 | 0 | 0 | 0 | 1936100 | 2134000 | 0 | 6367800 | 5650800 | 0 | 0 | 3454300 | 5950600 |
| NTTYDLIAN | USP39; USP39; | Q53GS9; A0 | SNUT2_HUMAI | Unmodified | 0 | 0 | 0 | 0 | 0 | 695370 | 0 | 0 | 250980 | 0 | 160270 | 1070300 | 0 | 0 | 0 | 28910 | 0 | 0 |
| NTVALDAD | VLDLR; VLDLR; | P98155; A0A | VLDLR_HUMAN | Unmodified | 0 | 0 | 0 | 0 | 0 | 1472500 | 208620 | 0 | 1096000 | 123800 | 0 | 0 | 0 | 0 | 0 | 0 | 0 | 0 |
| NTVANSPQ | ABCE1 | P61221 | ABCE1_HUMAI | Oxidation (M) | 0 | 0 | 0 | 0 | 0 | 0 | 118960 | 0 | 645330 | 0 | 0 | 1032200 | 0 | 0 | 0 | 0 | 0 | 0 |
| NTVANSPQ | ABCE1 | P61221 | ABCE1_HUMAI | Unmodified | 0 | 0 | 0 | 0 | 0 | 0 | 28824 | 0 | 0 | 0 | 0 | 1241400 | 108890 | 84790 | 0 | 0 | 0 | 0 |
| NTVCSLSS | PLAA; PLAA; PL | Q9Y263; H0Y | PLAP_HUMAN I | Unmodified | 0 | 0 | 0 | 0 | 121730 | 0 | 0 | 0 | 0 | 0 | 0 | 716250 | 0 | 0 | 0 | 109580 | 0 | 0 |
| NTVEGYSD | TYRP1; TYRP1 | P17643; E7E | TYRP1_HUMAN | Unmodified | 0 | 0 | 0 | 0 | 0 | 0 | 0 | 0 | 3766600 | 0 | 0 | 0 | 0 | 0 | 0 | 0 | 0 | 0 |
| NTVEIFKFE | PON2; PON2; P | Q15165; G3 | PON2_HUMAN | Unmodified | 0 | 0 | 0 | 0 | 0 | 0 | 0 | 0 | 17830000 | 2202800 | 280410 | 0 | 0 | 0 | 0 | 0 | 0 | 0 |
| NTVELLTDI | CGN | Q9P2M7 | CING_HUMAN | Unmodified | 0 | 0 | 23644 | 40358 | 0 | 0 | 0 | 0 | 929250 | 76722 | 0 | 1881300 | 0 | 3757100 | 0 | 0 | 0 | 0 |
| NTVELLVEI | FLNB; FLNB | O75369; E7E | FLNB_HUMAN | Unmodified | 61360 | 80246 | 0 | 298110 | 3449200 | 872100 | 0 | 0 | 0 | 165980 | 0 | 1131900 | 386930 | 138420 | 186880 | 129820 | 436520 | 568020 |
| NTVHDQEI | MCFD2; MCFD2 | Q8NI22; A0A | MCFD2_HUMA | 2 Oxidation (M) | 0 | 0 | 0 | 0 | 0 | 0 | 0 | 0 | 1433200 | 285770 | 0 | 0 | 0 | 0 | 0 | 0 | 0 | 0 |
| NTVHDQEI | MCFD2; MCFD2 | Q8NI22; A0A | MCFD2_HUMA | Oxidation (M) | 0 | 0 | 0 | 0 | 0 | 0 | 0 | 0 | 7880900 | 375810 | 0 | 0 | 0 | 0 | 0 | 0 | 0 | 0 |
| NTVHDQEI | MCFD2; MCFD2 | Q8NI22; A0A | MCFD2_HUMA | Unmodified | 0 | 0 | 0 | 0 | 0 | 0 | 0 | 0 | 5319700 | 0 | 0 | 0 | 0 | 0 | 0 | 0 | 0 | 0 |
| NTVLATWQ | EIF2A; EIF2A; EI | Q9BY44; H7( | EIF2A_HUMAN | Unmodified | 0 | 0 | 68614 | 0 | 0 | 0 | 0 | 0 | 122940 | 110930 | 0 | 3294500 | 156060 | 36003 | 339660 | 129730 | 253220 | 0 |
| NTVQDPAY | OPRK1; OPRK1 | P41145; A0A | OPRK_HUMAN | Unmodified | 0 | 0 | 0 | 0 | 0 | 0 | 0 | 0 | 2016000 | 427300 | 0 | 0 | 604260 | 0 | 0 | 854450 | 0 | 0 |
| NTVQDPAY | OPRK1; OPRK1 | P41145; A0A | OPRK_HUMAN | Unmodified | 0 | 0 | 0 | 0 | 0 | 0 | 0 | 0 | 1603300 | 0 | 0 | 0 | 0 | 0 | 0 | 0 | 0 | 0 |
| NTVSQSIS( | EIF2A; EIF2A | Q9BY44; F8V | EIF2A_HUMAN | Unmodified | 154790 | 252720 | 0 | 349020 | 0 | 0 | 338520 | 76919 | 604190 | 140150 | 489790 | 1744600 | 466140 | 0 | 0 | 241310 | 0 | 0 |
| NTVTDDILC | SLITRK6 | Q9H5Y7 | SLIK6_HUMAN | Unmodified | 2598100 | 1423600 | 0 | 0 | 0 | 0 | 0 | 0 | 940320 | 480270 | 0 | 0 | 0 | 0 | 0 | 0 | 0 | 0 |
| NTVTLLEEA | ASAP1; ASAP1; | Q9ULH1; E5 | ASAP1_HUMAN | Unmodified | 0 | 0 | 0 | 0 | 0 | 0 | 0 | 0 | 537720 | 0 | 0 | 0 | 0 | 0 | 0 | 0 | 270090 | 271590 |
| NTVVATGG | SDHA; SDHA; N | P31040; D6F | SDHA_HUMAN | Unmodified | 0 | 0 | 0 | 0 | 0 | 0 | 0 | 0 | 4373700 | 0 | 0 | 191780 | 0 | 0 | 0 | 0 | 0 | 0 |
| NTVVLFVP( | STOML2 | Q9UJZ1 | STML2_HUMAN | Unmodified | 0 | 0 | 0 | 0 | 0 | 0 | 0 | 0 | 5339100 | 274300 | 121580 | 169050 | 0 | 0 | 0 | 0 | 0 | 0 |
| NTVVPTKKS | HSPA5; HSPA5; | P11021; A0A | BIP_HUMAN Er | Unmodified | 0 | 0 | 0 | 0 | 0 | 0 | 1012700 | 0 | 614320 | 1225000 | 0 | 0 | 0 | 0 | 0 | 0 | 0 | 0 |
| NTWDCGL | SLC25A1; SLC2 | P53007; B4D | TXTP_HUMAN T | Unmodified | 0 | 0 | 0 | 0 | 0 | 0 | 0 | 0 | 1427000 | 0 | 0 | 0 | 0 | 0 | 57752 | 0 | 0 | 0 |
| NVADYYPE | ATP6V0D1; ATP | P61421; R4C | VA0D1_HUMAN | Unmodified | 0 | 0 | 0 | 1057900 | 391290 | 178430 | 327050 | 0 | 2949400 | 644400 | 582930 | 2574000 | 309430 | 0 | 0 | 0 | 0 | 298540 |
| NVAGEVKT | PXDN; PXDN | Q92626; H7( | PXDN_HUMAN | Unmodified | 0 | 0 | 0 | 0 | 0 | 0 | 0 | 0 | 0 | 65665 | 0 | 0 | 43724 | 0 | 0 | 0 | 109100 | 506780 |
| NVALLSQL' | ELAVL1 | Q15717 | ELAV1_HUMAN | Unmodified | 0 | 0 | 0 | 192280 | 0 | 0 | 0 | 0 | 0 | 0 | 0 | 0 | 0 | 0 | 0 | 186920 | 0 | 0 |
| NVAQYGLE | MICAL1; MICAL | Q8TDZ2; H0Y | MICA1_HUMAN | Unmodified | 0 | 98655 | 223710 | 386170 | 79114 | 0 | 0 | 0 | 755310 | 62551 | 0 | 0 | 758260 | 0 | 213850 | 121530 | 0 | 0 |
| NVASETLLS | RIPOR2; RIPOR | Q9Y4F9; F5C | RIPR2_HUMAN | Unmodified | 0 | 0 | 0 | 0 | 0 | 0 | 0 | 0 | 0 | 25316 | 76987 | 1157100 | 0 | 0 | 0 | 0 | 0 | 0 |
| NVASPSRI( | OPA1; OPA1; OI | O60313; E5K | OPA1_HUMAN | Unmodified | 0 | 0 | 0 | 0 | 280210 | 0 | 292520 | 0 | 2509700 | 1013100 | 0 | 0 | 651850 | 298380 | 0 | 0 | 0 | 162120 |
| NVAVIPPG | IREB2 | P48200 | IREB2_HUMAN | Oxidation (M) | 59483 | 0 | 0 | 0 | 0 | 0 | 0 | 0 | 0 | 0 | 0 | 2858900 | 0 | 0 | 0 | 0 | 0 | 0 |
| NVAYDTLPI | PLOD3; PLOD3 | O60568; H7( | PLOD3_HUMAI | Unmodified | 0 | 151180 | 0 | 0 | 0 | 0 | 0 | 0 | 4590000 | 234930 | 0 | 0 | 127870 | 102190 | 0 | 0 | 0 | 0 |
| NVDCVLLA | MTAP; N/A; MTA | Q13126; F2Z | MTAP_HUMAN | Unmodified | 132490 | 43506 | 0 | 183740 | 622360 | 311990 | 0 | 0 | 0 | 0 | 381030 | 1052300 | 892400 | 396240 | 138640 | 205580 | 471310 | 538310 |
| NVDEAINFI | ALDH3A2; ALDI | P51648; A0A | AL3A2_HUMAN | Unmodified | 0 | 0 | 0 | 70921 | 0 | 0 | 0 | 0 | 3634200 | 0 | 132080 | 0 | 0 | 0 | 0 | 0 | 89754 | 101400 |
| NVDEAINFI | ALDH3A2; ALDI | P51648; A0A | AL3A2_HUMAN | Unmodified | 215040 | 0 | 0 | 0 | 0 | 0 | 0 | 0 | 11750000 | 4190600 | 0 | 0 | 0 | 0 | 0 | 0 | 0 | 1404600 |
| NVDFNYAT | ATXN2L; ATXN2 | Q8WWM7; H | ATX2L_HUMAN | Unmodified | 349090 | 0 | 0 | 324700 | 883010 | 75711 | 0 | 0 | 272560 | 0 | 738420 | 1683900 | 414330 | 0 | 1483500 | 0 | 0 | 0 |
| NVDGVNY/ | CTSZ; CTSZ; CT | Q9UBR2; A0 | CATZ_HUMAN | Unmodified | 0 | 0 | 0 | 0 | 0 | 0 | 0 | 0 | 1837500 | 0 | 0 | 0 | 0 | 0 | 0 | 0 | 0 | 0 |
| NVDILKDPI | XPO1; XPO1; XF | O14980; A0A | XPO1_HUMAN | Unmodified | 391090 | 170520 | 511610 | 159290 | 1589000 | 123840 | 411320 | 144260 | 1395900 | 554870 | 1901800 | 12365000 | 448790 | 4996700 | 317840 | 293780 | 914990 | 201780 |
| NVDLASPE | PALS1; PALS1 | Q8N3R9; G3 | PALS1_HUMAN | Unmodified | 0 | 0 | 0 | 0 | 502090 | 185800 | 0 | 0 | 500370 | 0 | 449530 | 0 | 0 | 0 | 0 | 0 | 104270 | 134300 |
| NVDLLSDM | NAP1L1; NAP1L | P55209; F8V | NP1L1_HUMAN | Oxidation (M) | 8750500 | 7347900 | 6855800 | 17086000 | 8291100 | 4502300 | 5099500 | 1331000 | 3418600 | 5462700 | 7044400 | 30595000 | 10316000 | 5502400 | 1188700 | 5767300 | 5633100 | 5682500 |
| NVDLLSDM | NAP1L1; NAP1L | P55209; F8V | NP1L1_HUMAN | Unmodified | 8013500 | 9246500 | 5115800 | 13432000 | 37400000 | 3315500 | 13354000 | 1010600 | 28665000 | 15555000 | 7462800 | 55600000 | 12577000 | 11607000 | 8022400 | 3916300 | 15428000 | 14134000 |
| NVDMLSEL | NAP1L4; NAP1L | Q99733; C9J | NP1L4_HUMAN | Unmodified | 0 | 0 | 0 | 0 | 0 | 0 | 0 | 0 | 116550 | 0 | 0 | 0 | 0 | 0 | 0 | 0 | 0 | 0 |
| NVDMLSEL | NAP1L4; NAP1L | Q99733; C9J | NP1L4_HUMAN | Oxidation (M) | 0 | 0 | 0 | 0 | 0 | 0 | 0 | 0 | 0 | 0 | 0 | 657220 | 0 | 0 | 0 | 0 | 0 | 0 |
| NVDQAIKN | MACF1; MACF1 | Q9UPN3; H3 | MACF1_HUMAI | Unmodified | 0 | 0 | 0 | 0 | 0 | 0 | 0 | 0 | 0 | 0 | 0 | 0 | 0 | 0 | 0 | 0 | 264990 | 0 |
| NVDSFPQN | SLC12A7; SLC1 | Q9Y666; A0A | S12A7_HUMAN | Unmodified | 0 | 0 | 0 | 0 | 0 | 0 | 0 | 0 | 0 | 0 | 0 | 1132000 | 0 | 0 | 0 | 444670 | 0 | 0 |
| NVDTNQDI | NUCB1; NUCB | Q02818; H7I | NUCB1_HUMA | Unmodified | 159740 | 0 | 0 | 0 | 3578200 | 2075300 | 3823000 | 54574 | 13961000 | 10312000 | 3970600 | 0 | 0 | 211720 | 0 | 0 | 285500 | 6227400 |
| NVDVEFFK | MAP1B | P46821 | MAP1B_HUMAI | Unmodified | 0 | 0 | 0 | 0 | 577380 | 0 | 1672800 | 1325600 | 782740 | 359150 | 147380 | 1535300 | 1046100 | 0 | 0 | 0 | 233790 | 327300 |
| NVDVGDF\ | PIK3C2B; PIK3C | O00750; F5C | P3C2B_HUMAN | Unmodified | 0 | 0 | 0 | 0 | 0 | 0 | 0 | 0 | 0 | 0 | 0 | 0 | 0 | 0 | 0 | 170310 | 0 | 0 |
| NVDWGEIF | GALNT7; GALN | Q86SF2; H0Y | GALT7_HUMAN | Unmodified | 0 | 0 | 127520 | 0 | 0 | 0 | 0 | 0 | 4795100 | 107370 | 0 | 608270 | 0 | 0 | 126910 | 261990 | 0 | 0 |
| NVEAMNFA | DLST | P36957 | ODO2_HUMAN | Oxidation (M) | 0 | 0 | 0 | 0 | 0 | 0 | 0 | 0 | 2193100 | 864350 | 0 | 0 | 0 | 0 | 0 | 0 | 0 | 0 |
| NVEAMNFA | DLST | P36957 | ODO2_HUMAN | Unmodified | 0 | 0 | 0 | 0 | 0 | 0 | 0 | 0 | 2284500 | 0 | 0 | 0 | 0 | 0 | 0 | 0 | 0 | 0 |
| NVECARVY | CHMP1A; CHM | Q9HD42; F8 | CHM1A_HUMA | Unmodified | 0 | 0 | 0 | 0 | 5122900 | 779390 | 1382200 | 385810 | 0 | 1065300 | 0 | 0 | 4176700 | 2159000 | 0 | 0 | 652930 | 2161200 |
| NVEDDYIAI | DNAJB12; DNA | Q9NXW2; V9 | DJB12_HUMAN | Unmodified | 0 | 0 | 60356 | 0 | 0 | 0 | 0 | 0 | 2208400 | 282480 | 0 | 0 | 0 | 0 | 0 | 184220 | 0 | 0 |
| NVEDDYIAI | DNAJB12; DNA | Q9NXW2; V9 | DJB12_HUMAN | Unmodified | 0 | 0 | 0 | 0 | 0 | 0 | 0 | 0 | 3817400 | 0 | 0 | 0 | 0 | 0 | 0 | 0 | 0 | 0 |
| NVEDFTGP | SPCS3 | P61009 | SPCS3_HUMAI | Unmodified | 55910 | 63391 | 0 | 57086 | 0 | 0 | 0 | 0 | 23727000 | 1114800 | 1413800 | 1818800 | 0 | 0 | 80790 | 0 | 123630 | 0 |
| NVEDFTGP | SPCS3 | P61009 | SPCS3_HUMAI | Unmodified | 0 | 0 | 0 | 0 | 485140 | 0 | 0 | 0 | 1196200 | 0 | 0 | 0 | 0 | 0 | 0 | 0 | 0 | 0 |

| | | | | | | | | | | | | | | | | | | | | | | |
|---|---|---|---|---|---|---|---|---|---|---|---|---|---|---|---|---|---|---|---|---|---|---|
| NVEDLSGC | ABCE1; ABCE1 | P61221; D6F | ABCE1_HUMAN | Unmodified | 0 | 0 | 0 | 153970 | 0 | 0 | 0 | 0 | 0 | 0 | 573070 | 3052400 | 0 | 0 | 0 | 0 | 0 | 0 |
| NVEGLDLL | CHD7 | Q9P2D1 | CHD7_HUMAN | Unmodified | 0 | 0 | 0 | 0 | 0 | 193320 | 144610 | 15876 | 0 | 253110 | 0 | 0 | 0 | 0 | 0 | 0 | 59695 | 1258300 |
| NVEHIIDSL | NADK2 | Q4G0N4 | NAKD2_HUMAI | Unmodified | 0 | 0 | 0 | 0 | 0 | 0 | 0 | 0 | 1023900 | 225830 | 0 | 0 | 0 | 0 | 0 | 0 | 0 | 0 |
| NVEPFTSVI | MRPL1; MRPL1 | Q9BYD6; H0 | RM01_HUMAN | Unmodified | 0 | 0 | 0 | 0 | 71657 | 0 | 115400 | 0 | 2515000 | 710320 | 37375 | 0 | 28351 | 0 | 0 | 0 | 0 | 0 |
| NVEQSSDL | BORCS7; BORC | Q96B45; A0A | BORC7_HUMA | Unmodified | 75790 | 0 | 0 | 0 | 1285400 | 2534700 | 0 | 22702 | 0 | 0 | 0 | 1765500 | 338620 | 267750 | 0 | 0 | 343490 | 0 |
| NVESTRTFL | MRPL53 | Q96EL3 | RM53_HUMAN | Unmodified | 0 | 0 | 0 | 0 | 0 | 0 | 47048 | 0 | 419760 | 596840 | 0 | 0 | 0 | 0 | 0 | 0 | 78606 | 354900 |
| NVESYTK | RPN1; RPN1 | P04843; B7Z | RPN1_HUMAN | Unmodified | 0 | 0 | 0 | 0 | 0 | 0 | 0 | 0 | 10323000 | 0 | 0 | 0 | 0 | 0 | 0 | 0 | 0 | 0 |
| NVETNQCL | GALNT1 | Q10472 | GALT1_HUMAN | Unmodified | 0 | 0 | 0 | 0 | 0 | 0 | 0 | 0 | 634310 | 0 | 0 | 329630 | 0 | 0 | 0 | 0 | 0 | 0 |
| NVETNQCL | GALNT1 | Q10472 | GALT1_HUMAN | Unmodified | 0 | 0 | 0 | 0 | 0 | 0 | 0 | 0 | 471740 | 0 | 0 | 494040 | 0 | 0 | 0 | 0 | 0 | 0 |
| NVEVIELAK | AP3B1 | O00203 | AP3B1_HUMAN | Unmodified | 30804 | 0 | 0 | 0 | 0 | 0 | 0 | 0 | 0 | 0 | 187370 | 952910 | 0 | 0 | 0 | 0 | 0 | 0 |
| NVFDEAILA | CDC42 | P60953 | CDC42_HUMA | Unmodified | 6907600 | 9521200 | 2500100 | 4764800 | 5152100 | 1132000 | 3576900 | 333090 | 3681100 | 2450000 | 4696000 | 7648000 | 4451700 | 2668300 | 11029000 | 3855600 | 1808200 | 3397100 |
| NVFDEAILA | CDC42 | P60953 | CDC42_HUMA | Unmodified | 3172700 | 3600500 | 2294500 | 5583200 | 21353000 | 287270 | 7650700 | 0 | 10395000 | 13466000 | 6074600 | 8232200 | 8491300 | 8597900 | 4799400 | 2463500 | 7312900 | 11574000 |
| NVFDEAILA | CDC42 | P60953 | CDC42_HUMA | Unmodified | 0 | 0 | 0 | 0 | 1337000 | 0 | 755120 | 82769 | 0 | 240730 | 0 | 0 | 2607800 | 3402000 | 0 | 0 | 765700 | 1481500 |
| NVFEAVPV | DIP2A; DIP2A | Q14689; A0A | DIP2A_HUMAN | Unmodified | 0 | 0 | 0 | 0 | 0 | 0 | 0 | 0 | 1211800 | 836140 | 154120 | 2449400 | 0 | 0 | 0 | 1895000 | 0 | 0 |
| NVFIAQNV | UBR4 | Q5T4S7 | UBR4_HUMAN | Unmodified | 21145 | 0 | 0 | 0 | 1642700 | 0 | 0 | 44126 | 0 | 204690 | 0 | 2387900 | 0 | 0 | 0 | 21416 | 0 | 0 |
| NVFIIGATN | VCP; VCP; VCP; | P55072; C9J | TERA_HUMAN | Unmodified | 0 | 0 | 0 | 0 | 0 | 0 | 0 | 0 | 793760 | 0 | 859960 | 4435900 | 0 | 0 | 0 | 0 | 0 | 0 |
| NVFLLGFIP | UPF1 | Q92900 | RENT1_HUMAN | Unmodified | 2061400 | 0 | 2245700 | 871950 | 0 | 0 | 0 | 0 | 611120 | 94231 | 613610 | 1478000 | 164690 | 0 | 2058800 | 1274700 | 0 | 0 |
| NVFNALIR | SLC25A11; SLC | Q02978; I3L | M2OM_HUMAN | Unmodified | 0 | 0 | 989370 | 0 | 0 | 0 | 0 | 0 | 4951700 | 1018500 | 58714 | 0 | 0 | 0 | 0 | 0 | 0 | 0 |
| NVFPDNW | STK3; STK3 | Q13188; A0A | STK3_HUMAN | Unmodified | 0 | 0 | 0 | 0 | 3937600 | 235770 | 121330 | 0 | 525380 | 1292500 | 31194 | 0 | 588400 | 355980 | 0 | 0 | 553150 | 1651400 |
| NVFPNLHV | GALNT4; POC1 | Q8N4A0; F8 | GALT4_HUMAN | Unmodified | 0 | 0 | 0 | 198640 | 0 | 0 | 0 | 0 | 1907000 | 204020 | 0 | 0 | 0 | 0 | 0 | 0 | 258940 | 0 |
| NVFQLEQA | NAGLU | P54802 | ANAG_HUMAN | Unmodified | 0 | 0 | 164710 | 13246 | 0 | 751810 | 0 | 0 | 1189000 | 815310 | 80969 | 0 | 0 | 0 | 11070 | 26689 | 0 | 199350 |
| NVFSGIYR | ATP2B4 | P23634 | AT2B4_HUMAN | Unmodified | 0 | 0 | 0 | 0 | 0 | 0 | 0 | 0 | 0 | 0 | 0 | 1385100 | 172960 | 0 | 0 | 96426 | 0 | 0 |
| NVFYELED | SRCIN1; SRCIN | Q9C0H9; A0 | SRCN1_HUMAI | Unmodified | 0 | 0 | 0 | 0 | 0 | 0 | 0 | 0 | 68027 | 0 | 0 | 1193700 | 0 | 0 | 0 | 0 | 0 | 0 |
| NVGCLQEA | AARS1; AARS1; | P49588; A0A | SYAC_HUMAN | Unmodified | 0 | 0 | 0 | 0 | 0 | 0 | 0 | 0 | 0 | 0 | 0 | 0 | 91075 | 55851 | 0 | 0 | 0 | 0 |
| NVGCLQEA | AARS1; AARS1; | P49588; A0A | SYAC_HUMAN | Unmodified | 0 | 0 | 0 | 0 | 272670 | 0 | 53063 | 0 | 0 | 95897 | 0 | 0 | 565760 | 0 | 0 | 0 | 267130 | 93678 |
| NVGEEEDE | CLEC16A | Q2KHT3 | CL16A_HUMAN | Unmodified | 0 | 0 | 0 | 0 | 90326 | 0 | 0 | 0 | 0 | 0 | 78939 | 208310 | 0 | 0 | 0 | 0 | 0 | 134970 |
| NVGGIPALI | PKP4; PKP4 | Q99569; A0A | PKP4_HUMAN | Unmodified | 0 | 0 | 259270 | 97478 | 0 | 0 | 0 | 0 | 2537500 | 425970 | 0 | 0 | 0 | 0 | 0 | 0 | 31932 | 0 |
| NVGKVLLV | VAT1; VAT1 | Q99536; K7E | VAT1_HUMAN | Unmodified | 478240 | 573520 | 0 | 0 | 671000 | 91446 | 153930 | 159720 | 258960 | 44792 | 0 | 0 | 0 | 0 | 0 | 0 | 869020 | 162620 |
| NVGLDIEAI | CCT8; CCT8 | P50990; H7C | TCPQ_HUMAN | Unmodified | 0 | 0 | 0 | 0 | 0 | 0 | 0 | 0 | 383540 | 385160 | 1177400 | 1160800 | 0 | 0 | 0 | 0 | 0 | 0 |
| NVGQFSGI | DEK; DEK; DEK; | P35659; B4D | DEK_HUMAN P | Unmodified | 0 | 0 | 0 | 0 | 0 | 0 | 0 | 428830 | 0 | 0 | 0 | 675570 | 0 | 0 | 0 | 0 | 0 | 0 |
| NVGTGLCA | GALNT10 | Q86SR1 | GLT10_HUMAN | Unmodified | 0 | 0 | 0 | 0 | 0 | 0 | 0 | 0 | 1265500 | 0 | 0 | 0 | 0 | 0 | 0 | 0 | 0 | 0 |
| NVHDNHF | CORO2B | Q9UQ03 | COR2B_HUMA | Unmodified | 0 | 0 | 0 | 0 | 0 | 0 | 0 | 0 | 0 | 0 | 0 | 719490 | 0 | 0 | 0 | 0 | 0 | 0 |
| NVHFASVV | NUDT15 | Q9NV35 | NUD15_HUMA | Unmodified | 0 | 0 | 0 | 0 | 873510 | 0 | 243510 | 37529 | 240310 | 124820 | 0 | 1069500 | 0 | 0 | 115860 | 0 | 343800 | 456550 |
| NVHGINFV | CTSC; CTSC; C | P53634; A0A | CATC_HUMAN | Unmodified | 0 | 0 | 241660 | 0 | 0 | 0 | 0 | 0 | 2299900 | 0 | 6807000 | 24350000 | 0 | 0 | 0 | 0 | 0 | 0 |
| NVHLAPGV | DYNC1H1; DYN | Q14204; A0A | DYHC1_HUMA | Oxidation (M) | 0 | 0 | 423020 | 0 | 0 | 0 | 440420 | 0 | 0 | 0 | 0 | 3051200 | 0 | 0 | 142900 | 71873 | 0 | 0 |
| NVIAALNSI | EMC1; EMC1 | Q8N766; Q5 | EMC1_HUMAN | Unmodified | 0 | 0 | 0 | 0 | 401910 | 0 | 0 | 0 | 22186000 | 3078600 | 1585200 | 306210 | 0 | 0 | 0 | 0 | 0 | 217900 |
| NVIAALNSI | EMC1 | Q8N766 | EMC1_HUMAN | Unmodified | 0 | 0 | 0 | 0 | 0 | 0 | 0 | 0 | 0 | 846560 | 0 | 0 | 0 | 0 | 0 | 0 | 0 | 0 |
| NVIAPLMTI | PLOD1 | Q02809 | PLOD1_HUMAI | Unmodified | 0 | 0 | 30348 | 0 | 0 | 0 | 0 | 0 | 4966000 | 678660 | 78549 | 0 | 0 | 0 | 0 | 0 | 0 | 0 |
| NVIATETSY | DSG2 | Q14126 | DSG2_HUMAN | Oxidation (M) | 0 | 0 | 0 | 2348500 | 0 | 0 | 0 | 0 | 0 | 0 | 0 | 0 | 0 | 0 | 2064800 | 0 | 0 | 0 |
| NVICDKCE | DNAJA1 | P31689 | DNJA1_HUMAN | Unmodified | 321000 | 0 | 0 | 0 | 0 | 0 | 0 | 0 | 0 | 0 | 0 | 0 | 0 | 0 | 0 | 0 | 0 | 0 |
| NVICDKCE | DNAJA1 | P31689 | DNJA1_HUMAN | Unmodified | 0 | 0 | 0 | 0 | 271330 | 0 | 0 | 148450 | 43888 | 346490 | 112850 | 0 | 668340 | 611630 | 0 | 0 | 147190 | 391260 |
| NVIDGDLC | SF3B3 | Q15393 | SF3B3_HUMAN | Oxidation (M) | 256270 | 162480 | 0 | 182190 | 0 | 0 | 0 | 0 | 0 | 63085 | 0 | 1294700 | 185310 | 0 | 0 | 77417 | 0 | 0 |
| NVIDPPIYA | EDIL3 | O43854 | EDIL3_HUMAN | Unmodified | 874850 | 677960 | 0 | 462110 | 5.45E+08 | 1.2E+08 | 10137000 | 7216000 | 47972000 | 25842000 | 520960 | 201470 | 0 | 0 | 0 | 85925 | 15337000 | 2.04E+08 |
| NVIDPPIYA | EDIL3 | O43854 | EDIL3_HUMAN | Unmodified | 0 | 0 | 0 | 0 | 11299000 | 790680 | 2206900 | 608650 | 0 | 0 | 0 | 0 | 0 | 0 | 0 | 0 | 1725700 | 4631600 |
| NVIETCKEA | NSDHL; NSDHL | Q15738; C9J | NSDHL_HUMA | Unmodified | 0 | 0 | 0 | 0 | 0 | 0 | 289340 | 128600 | 0 | 50685 | 0 | 0 | 0 | 0 | 30496 | 0 | 0 | 0 |
| NVIFEDEEK | GGA1; GGA1; G | Q9UJY5; B0C | GGA1_HUMAN | Unmodified | 0 | 0 | 0 | 101570 | 281260 | 0 | 0 | 78693 | 0 | 259480 | 264920 | 686480 | 0 | 110080 | 0 | 0 | 210380 | 262000 |
| NVIFEISPTI | IQGAP1; IQGAP | P46940; H0Y | IQGA1_HUMAN | Unmodified | 189950 | 0 | 72384 | 239500 | 23067 | 62684 | 299880 | 38846 | 0 | 183830 | 32787 | 247130 | 0 | 0 | 470530 | 182990 | 0 | 15465 |
| NVIFQPVAI | PFKP; PFKP; PF | Q01813; A0A | PFKAP_HUMAN | Unmodified | 0 | 0 | 0 | 0 | 1329300 | 615760 | 0 | 0 | 648460 | 0 | 241150 | 1690300 | 0 | 0 | 0 | 0 | 435610 | 1008400 |
| NVIGLQMG | TAGLN2; TAGLN | P37802; X6R | TAGL2_HUMAN | Unmodified | 0 | 2355000 | 527770 | 1050100 | 1768500 | 403170 | 0 | 0 | 0 | 313070 | 1660700 | 3108100 | 566610 | 234880 | 1602000 | 0 | 372630 | 1395400 |
| NVIGLQMG | TAGLN2; TAGLN | P37802; X6R | TAGL2_HUMAN | Oxidation (M) | 1663300 | 1446300 | 0 | 757990 | 742650 | 0 | 0 | 0 | 1019900 | 0 | 1113500 | 2023600 | 244670 | 0 | 764060 | 549130 | 149130 | 651290 |
| NVIGLQMG | TAGLN2 | P37802 | TAGL2_HUMAN | Oxidation (M) | 0 | 0 | 0 | 0 | 721550 | 0 | 0 | 0 | 0 | 0 | 0 | 0 | 87519 | 0 | 0 | 0 | 0 | 257900 |
| NVIIWGNH | MDH1; MDH1; I | P40925; C9J | MDHC_HUMAN | Unmodified | 2064700 | 984390 | 1930900 | 3087800 | 54742000 | 9418900 | 13140000 | 4712900 | 14388000 | 9020900 | 20941000 | 98085000 | 19784000 | 16470000 | 2284000 | 2830700 | 18899000 | 34168000 |
| NVINTFTQ1 | FDXR; FDXR; FD | P22570; A0A | ADRO_HUMAN | Unmodified | 0 | 0 | 0 | 0 | 545110 | 0 | 199170 | 0 | 1118300 | 349060 | 87828 | 0 | 0 | 0 | 198190 | 24300 | 0 | 0 |
| NVIPALELV | PSAP; PSAP; PS | P07602; A0A | SAP_HUMAN P | Unmodified | 0 | 251060 | 302370 | 390620 | 1141400 | 3728600 | 2624800 | 0 | 91674000 | 10025000 | 1434500 | 1109600 | 999110 | 1543600 | 414620 | 266870 | 0 | 4047300 |
| NVIPALELV | PSAP; PSAP; PS | P07602; A0A | SAP_HUMAN P | Unmodified | 0 | 0 | 0 | 0 | 0 | 0 | 0 | 109340 | 5450100 | 1621500 | 0 | 244220 | 0 | 0 | 0 | 0 | 0 | 0 |
| NVIPALELV | PSAP; PSAP; PS | P07602; A0A | SAP_HUMAN P | Unmodified | 0 | 0 | 0 | 0 | 0 | 0 | 0 | 0 | 0 | 1098300 | 0 | 0 | 0 | 0 | 0 | 0 | 0 | 0 |
| NVIQSVLQ | CYFIP1; CYFIP1 | Q7L576; A0A | CYFP1_HUMAN | Unmodified | 372060 | 35203 | 0 | 219880 | 0 | 0 | 0 | 200730 | 346150 | 8975.3 | 0 | 2389000 | 0 | 0 | 630650 | 681170 | 7302.1 | 0 |
| NVIQSVLQ | CYFIP1; CYFIP1 | Q7L576; A0A | CYFP1_HUMAN | Unmodified | 67540 | 0 | 104870 | 136210 | 0 | 0 | 46761 | 0 | 135900 | 24387 | 38871 | 610820 | 29906 | 0 | 305200 | 234740 | 0 | 0 |
| NVISLTEDV | ITGB4; ITGB4 | P16144; J3Q | ITB4_HUMAN I | Unmodified | 0 | 1284800 | 0 | 0 | 4913900 | 4037800 | 0 | 0 | 0 | 0 | 0 | 0 | 961960 | 0 | 515240 | 0 | 5653500 | 11436000 |
| NVISLTEDV | ITGB4; ITGB4 | P16144; J3Q | ITB4_HUMAN I | Unmodified | 0 | 0 | 0 | 0 | 990800 | 217070 | 0 | 0 | 0 | 0 | 0 | 0 | 0 | 0 | 0 | 0 | 1439500 | 2052400 |
| NVISQTEH | EXTL3 | O43909 | EXTL3_HUMAN | Unmodified | 0 | 0 | 156270 | 0 | 127640 | 25940 | 0 | 0 | 508400 | 134000 | 161150 | 47954 | 0 | 0 | 38511 | 0 | 0 | 0 |
| NVKFNVWI | ARF6 | P62330 | ARF6_HUMAN | Unmodified | 0 | 0 | 0 | 232590 | 2254500 | 169070 | 0 | 165620 | 950210 | 723130 | 0 | 0 | 0 | 231560 | 31759 | 0 | 333670 | 548030 |
| NVKLIALSI | PRDX6 | P30041 | PRDX6_HUMAN | Unmodified | 0 | 0 | 0 | 0 | 0 | 0 | 0 | 0 | 0 | 0 | 0 | 0 | 989780 | 585390 | 0 | 0 | 1806600 | 0 |
| NVKLPDAY | G6PD | P11413 | G6PD_HUMAN | Unmodified | 0 | 0 | 0 | 0 | 2104800 | 0 | 0 | 0 | 797950 | 0 | 111770 | 676900 | 0 | 0 | 82956 | 0 | 0 | 0 |
| NVLAAITR | SLC7A6 | Q92536 | YLAT2_HUMAN | Unmodified | 175900 | 467910 | 36814 | 19591 | 0 | 0 | 133180 | 37646 | 0 | 21677 | 0 | 1546500 | 0 | 0 | 105510 | 114010 | 216740 | 0 |
| NVLCSACS | DNAJA2; DNAJ | O60884; A0A | DNJA2_HUMAN | Unmodified | 1138100 | 480980 | 588770 | 1084800 | 5708900 | 1940700 | 389480 | 324270 | 1255000 | 1180300 | 151720 | 1242800 | 4133900 | 2248400 | 748600 | 577140 | 902630 | 1333200 |
| NVLCSACS | DNAJA2; DNAJ | O60884; A0A | DNJA2_HUMAN | Unmodified | 0 | 0 | 0 | 0 | 1398700 | 202210 | 1060800 | 685030 | 0 | 518340 | 0 | 0 | 1824700 | 2062900 | 0 | 0 | 1153300 | 404130 |
| NVLDSEDE | CHID1 | Q9BWS9 | CHID1_HUMAN | Unmodified | 0 | 0 | 0 | 0 | 0 | 0 | 0 | 0 | 1618800 | 0 | 0 | 0 | 0 | 0 | 0 | 0 | 0 | 0 |
| NVLELSNV | PTPRF | P10586 | PTPRF_HUMAN | Unmodified | 1426400 | 998040 | 0 | 0 | 0 | 0 | 0 | 0 | 3070400 | 0 | 146120 | 0 | 0 | 0 | 455690 | 0 | 0 | 0 |
| NVLGHMQ | PFKP; PFKP; PF | Q01813; A0A | PFKAP_HUMAN | Unmodified | 0 | 314760 | 0 | 0 | 13426000 | 0 | 0 | 711940 | 1233700 | 0 | 1179600 | 8362200 | 0 | 0 | 0 | 0 | 3153100 | 6723200 |
| NVLGHMQ | PFKP; PFKP; PF | Q01813; A0A | PFKAP_HUMAN | Oxidation (M) | 0 | 247430 | 0 | 0 | 6005500 | 1256500 | 422340 | 0 | 421020 | 0 | 1124100 | 11960000 | 0 | 730250 | 0 | 0 | 3392600 | 1961800 |
| NVLGHMQ | PFKM; PFKM; PI | P08237; A0A | PFKAM_HUMAI | Unmodified | 0 | 0 | 0 | 0 | 380110 | 0 | 0 | 0 | 0 | 0 | 0 | 3781300 | 147420 | 0 | 0 | 0 | 455630 | 1036800 |
| NVLGHMQ | PFKM; PFKM; PI | P08237; A0A | PFKAM_HUMAI | Oxidation (M) | 175680 | 0 | 228260 | 839900 | 0 | 466810 | 464560 | 418820 | 0 | 0 | 0 | 1818000 | 1481600 | 0 | 731890 | 586550 | 248410 | 0 |
| NVLIEVNP | EMG1; EMG1; E | Q92979; A0A | NEP1_HUMAN | Unmodified | 0 | 0 | 0 | 0 | 0 | 0 | 0 | 0 | 0 | 0 | 0 | 1977200 | 0 | 0 | 0 | 0 | 0 | 0 |
| NVLIQESET | PPFIA2; PPFIA2 | O75334; F8V | LIPA2_HUMAN | Unmodified | 0 | 0 | 0 | 678480 | 0 | 0 | 0 | 0 | 0 | 0 | 0 | 0 | 299330 | 0 | 769820 | 0 | 0 | 0 |
| NVLISVLQA | CYFIP2; CYFIP2 | Q96F07; H7C | CYFP2_HUMAN | Unmodified | 0 | 0 | 0 | 0 | 0 | 0 | 0 | 0 | 113510 | 0 | 0 | 347180 | 0 | 0 | 0 | 0 | 0 | 0 |
| NVLITDFFG | PIGK; PIGK; PIG | Q92643; B1A | GPI8_HUMAN ( | Unmodified | 0 | 0 | 0 | 0 | 0 | 0 | 0 | 0 | 33682000 | 3080800 | 704470 | 0 | 0 | 0 | 0 | 0 | 0 | 0 |
| NVLITDFFG | PIGK; PIGK; PIG | Q92643; B1A | GPI8_HUMAN ( | Unmodified | 0 | 0 | 0 | 0 | 0 | 0 | 0 | 0 | 1085400 | 0 | 40941 | 0 | 0 | 0 | 0 | 0 | 0 | 0 |
| NVLLDNDF | TYK2; TYK2 | P29597; E9P | TYK2_HUMAN I | Unmodified | 0 | 0 | 0 | 0 | 0 | 0 | 0 | 0 | 0 | 0 | 0 | 0 | 0 | 0 | 0 | 0 | 0 | 314090 |
| NVLLDPQL | CCT3; CCT3 | P49368; B4D | TCPG_HUMAN | Oxidation (M) | 0 | 0 | 0 | 0 | 0 | 187570 | 0 | 0 | 1327800 | 1946200 | 2036600 | 11176000 | 0 | 0 | 0 | 0 | 0 | 0 |
| NVLLEQSG | CARMIL1; CARN | Q5VZK9; A0A | CARL1_HUMAN | Unmodified | 11035 | 0 | 0 | 0 | 1144100 | 0 | 0 | 0 | 0 | 0 | 0 | 840710 | 0 | 0 | 0 | 0 | 377870 | 0 |
| NVLLFLQD | ATP6AP1; ATP6 | Q15904; A0A | VAS1_HUMAN | Unmodified | 0 | 0 | 0 | 0 | 0 | 0 | 0 | 0 | 3433600 | 0 | 1709500 | 0 | 0 | 0 | 0 | 0 | 0 | 0 |
| NVLLHQAG | EIPR1; EIPR1 | Q53HC9; D3 | EIPR1_HUMAN | Unmodified | 0 | 0 | 0 | 0 | 3771600 | 197190 | 0 | 0 | 0 | 0 | 0 | 0 | 772820 | 0 | 0 | 0 | 0 | 679590 |
| NVLLLGED | DYNC1LI1 | Q9Y6G9 | DC1L1_HUMAN | Unmodified | 0 | 0 | 0 | 0 | 0 | 0 | 0 | 42933 | 699850 | 132460 | 1127700 | 0 | 948160 | 0 | 123580 | 128540 | 0 | 269620 |
| NVLLLGED | DYNC1LI1 | Q9Y6G9 | DC1L1_HUMAN | Unmodified | 0 | 29534 | 0 | 0 | 0 | 0 | 662890 | 65440 | 0 | 458900 | 0 | 0 | 0 | 753910 | 0 | 0 | 304740 | 1615700 |
| NVLLVTQH | SYK | P43405 | KSYK_HUMAN | Unmodified | 76516 | 0 | 0 | 0 | 0 | 362720 | 0 | 0 | 483020 | 130590 | 376080 | 1231700 | 0 | 0 | 0 | 0 | 0 | 0 |
| NVLRWDLS | NLN | Q9BYT8 | NEUL_HUMAN | Unmodified | 0 | 0 | 0 | 204270 | 1560000 | 307960 | 0 | 0 | 633430 | 410150 | 382460 | 429070 | 538710 | 0 | 0 | 0 | 0 | 0 |
| NVLSLTNK | ITGB1; ITGB1; I | P05556; H7C | ITB1_HUMAN I | Unmodified | 1858200 | 1691000 | 5563600 | 551220 | 3795100 | 1025600 | 0 | 0 | 2986000 | 0 | 630730 | 3160600 | 0 | 0 | 2687000 | 2130200 | 243290 | 3784100 |
| NVLSLTNK | ITGB1; ITGB1; I | P05556; H7C | ITB1_HUMAN I | Unmodified | 15430000 | 15975000 | 54738 | 29232000 | 1.73E+09 | 2.1E+08 | 14953000 | 501660 | 1.04E+08 | 36771000 | 8995900 | 34586000 | 133520 | 28616000 | 36570000 | 41247000 | 4.22E+08 | 1.45E+09 |
| NVLSLTNK | ITGB1; ITGB1; I | P05556; H7C | ITB1_HUMAN I | Unmodified | 0 | 0 | 0 | 0 | 18898000 | 2828800 | 378150 | 0 | 0 | 554080 | 0 | 0 | 0 | 824140 | 0 | 0 | 17385000 | 34048000 |
| NVLTESAR | GATD3B; GATD | A0A0B4J2D5 | GAL3B_HUMAN | Unmodified | 436940 | 614760 | 494990 | 39812 | 502630 | 0 | 0 | 12298 | 1905600 | 223040 | 0 | 729770 | 0 | 0 | 431830 | 431920 | 44014 | 0 |
| NVLVESEH | JAK1; JAK1; JAK | P23458; A0A | JAK1_HUMAN | Unmodified | 0 | 0 | 0 | 0 | 3113400 | 1026800 | 0 | 24622 | 786410 | 0 | 137380 | 433210 | 0 | 0 | 0 | 0 | 21488 | 0 |
| NVLVSEDN | CSK | P41240 | CSK_HUMAN T | Unmodified | 0 | 0 | 378560 | 305600 | 0 | 234080 | 0 | 0 | 176240 | 0 | 1700000 | 4502400 | 242760 | 154350 | 359330 | 321340 | 0 | 0 |
| NVMFVFFQ | NCSTN; NCSTN | Q92542; H0Y | NICA_HUMAN | Oxidation (M) | 0 | 0 | 0 | 0 | 0 | 0 | 0 | 0 | 30700 | 144160 | 0 | 0 | 161010 | 0 | 0 | 0 | 0 | 0 |
| NVMLLPVG | CNDP2 | Q96KP4 | CNDP2_HUMA | Oxidation (M) | 227600 | 267750 | 901170 | 1188300 | 1052500 | 0 | 0 | 0 | 540470 | 397860 | 3792800 | 6621800 | 5174500 | 4369400 | 1332400 | 1712000 | 455590 | 1229300 |
| NVMLLPVG | CNDP2 | Q96KP4 | CNDP2_HUMA | Unmodified | 0 | 0 | 207710 | 369760 | 0 | 0 | 192000 | 0 | 1314800 | 564190 | 2142500 | 7068700 | 5302000 | 2684700 | 442170 | 371550 | 250680 | 733720 |
| NVMNATD | STX2 | P32856 | STX2_HUMAN | Unmodified | 0 | 0 | 0 | 0 | 0 | 0 | 0 | 115120 | 0 | 0 | 0 | 1646200 | 0 | 0 | 0 | 0 | 0 | 0 |
| NVMNATD | STX2 | P32856 | STX2_HUMAN | Unmodified | 0 | 0 | 95131 | 0 | 0 | 0 | 0 | 0 | 245730 | 0 | 0 | 1468800 | 0 | 0 | 0 | 43797 | 0 | 0 |
| NVMNATD | STX2 | P32856 | STX2_HUMAN | Unmodified | 0 | 0 | 0 | 0 | 0 | 0 | 0 | 0 | 304560 | 58835 | 91463 | 3358800 | 0 | 0 | 0 | 0 | 0 | 0 |
| NVMSAFGL | TFG; TFG; TFG; | Q92734; A0A | TFG_HUMAN P | Unmodified | 0 | 0 | 0 | 0 | 0 | 0 | 0 | 0 | 0 | 0 | 0 | 0 | 0 | 0 | 0 | 0 | 0 | 0 |
| NVMSAFGL | TFG; TFG; TFG; | Q92734; A0A | TFG_HUMAN P | Oxidation (M) | 259120 | 0 | 138120 | 0 | 0 | 0 | 0 | 0 | 0 | 0 | 0 | 1510600 | 0 | 0 | 607750 | 649490 | 48580 | 0 |
| NVMSKVQI | ACSL4 | O60488 | ACSL4_HUMAN | Unmodified | 0 | 58322 | 0 | 0 | 1150200 | 0 | 332720 | 35276 | 0 | 86639 | 0 | 0 | 0 | 0 | 0 | 143930 | 850340 | 161740 |
| NVNAGGH | VDAC1 | P21796 | VDAC1_HUMAI | Unmodified | 0 | 0 | 0 | 0 | 0 | 0 | 1023500 | 0 | 0 | 8562000 | 0 | 0 | 231170 | 0 | 0 | 0 | 0 | 119030 |
| NVNELNQ | RASAL1 | O95294 | RASL1_HUMAN | Unmodified | 0 | 0 | 0 | 0 | 8539.2 | 0 | 0 | 0 | 156290 | 48002 | 0 | 0 | 0 | 0 | 0 | 0 | 0 | 0 |
| NVNELNQ | RASAL1 | O95294 | RASL1_HUMAN | Unmodified | 0 | 892720 | 0 | 0 | 0 | 0 | 0 | 0 | 1665900 | 0 | 121310 | 1026600 | 0 | 0 | 124620 | 76828 | 0 | 0 |
| NVNIFKFII | LDHA; LDHA; LI | P00338; F5G | LDHA_HUMAN | Unmodified | 0 | 0 | 0 | 0 | 1021800 | 0 | 155400 | 0 | 182660 | 261800 | 53173 | 1014300 | 335290 | 0 | 0 | 0 | 1655500 | 1082900 |
| NVNIQNFH | ACTN1; ACTN1; | P12814; G3V | ACTN1_HUMAI | Unmodified | 904810 | 0 | 997550 | 1041300 | 5712700 | 1350100 | 467790 | 0 | 862450 | 255470 | 1673800 | 17469000 | 165980 | 791020 | 2035100 | 1959600 | 1381500 | 3720300 |
| NVNIQNFH | ACTN1; ACTN1; | P12814; G3V | ACTN1_HUMAI | Unmodified | 0 | 0 | 0 | 0 | 0 | 0 | 433960 | 0 | 0 | 34339 | 0 | 0 | 463380 | 317570 | 0 | 0 | 649610 | 1003300 |
| NVNNEVHI | EIF2A; EIF2A | Q9BY44; F8V | EIF2A_HUMAN | Unmodified | 418380 | 253680 | 0 | 667490 | 0 | 0 | 0 | 0 | 794090 | 0 | 938590 | 5830900 | 0 | 0 | 822790 | 453710 | 0 | 0 |

| | | | | | | | | | | | | | | | | | | | | | | |
|---|---|---|---|---|---|---|---|---|---|---|---|---|---|---|---|---|---|---|---|---|---|---|
| NVNQCLGI | VLDLR; VLDLR; | P98155; A0A | VLDLR_HUMAN | Unmodified | 0 | 0 | 0 | 0 | 0 | 0 | 0 | 0 | 737790 | 99882 | 0 | 0 | 0 | 0 | 0 | 0 | 0 | 0 |
| NVNQEVVD | HTATIP2; HTATI | Q9BUP3; E9 | HTAI2_HUMAN | Unmodified | 0 | 0 | 0 | 0 | 0 | 0 | 85416 | 0 | 1217200 | 0 | 0 | 147200 | 0 | 0 | 0 | 0 | 0 | 0 |
| NVNQSLLE | FTH1; FTH1; FTI | P02794; G3V | FRIH_HUMAN F | Unmodified | 0 | 0 | 0 | 0 | 3158900 | 930950 | 5805300 | 3234900 | 1.64E+08 | 64272000 | 2387400 | 1928400 | 5858600 | 6467900 | 71903 | 0 | 3609700 | 3720200 |
| NVNTGELA | MAP4K3; MAP4 | Q8IVH8; V9C | M4K3_HUMAN | Unmodified | 0 | 0 | 0 | 0 | 0 | 0 | 0 | 0 | 0 | 0 | 0 | 233940 | 0 | 0 | 0 | 0 | 0 | 0 |
| NVNVFFPH | UEVLD; UEVLD | Q8IX04; A8M | UEVLD_HUMAN | Unmodified | 0 | 0 | 0 | 0 | 0 | 0 | 0 | 0 | 487480 | 0 | 154240 | 2045600 | 0 | 0 | 0 | 0 | 0 | 0 |
| NVNVQNFI | ACTN4; ACTN4; | O43707; F5C | ACTN4_HUMAN | Unmodified | 701350 | 542550 | 1415000 | 1680800 | 8304400 | 4831400 | 226870 | 0 | 6269100 | 732050 | 1383400 | 24500000 | 2132700 | 1662100 | 3159600 | 2219700 | 1461400 | 985130 |
| NVNVQNFI | ACTN4; ACTN4; | O43707; F5C | ACTN4_HUMAN | Unmodified | 0 | 0 | 0 | 660320 | 11184000 | 3598200 | 0 | 0 | 500360 | 2954600 | 140760 | 0 | 8552800 | 5412300 | 0 | 0 | 2802700 | 1580400 |
| NVPAAREP | XPO1; XPO1; XF | O14980; A0A | XPO1_HUMAN | Unmodified | 0 | 0 | 0 | 0 | 0 | 0 | 0 | 28382 | 0 | 0 | 0 | 1816800 | 0 | 0 | 0 | 0 | 0 | 0 |
| NVPEIAVYF | LAPTM4A | Q15012 | LAP4A_HUMAN | Oxidation (M) | 0 | 0 | 0 | 0 | 0 | 0 | 0 | 0 | 2428100 | 0 | 0 | 0 | 0 | 0 | 0 | 0 | 0 | 0 |
| NVPILYTAS | CTNNA1; CTNN | P35221; E5R | CTNA1_HUMAN | Unmodified | 5777300 | 374860 | 8320900 | 30493000 | 36554000 | 25053000 | 0 | 0 | 2267800 | 1256700 | 5820200 | 31427000 | 37817000 | 34519000 | 20869000 | 19348000 | 356760 | 2267100 |
| NVPILYTAS | CTNNA1; CTNN | P35221; E5R | CTNA1_HUMAN | Unmodified | 0 | 0 | 0 | 0 | 0 | 1934300 | 0 | 0 | 0 | 0 | 0 | 0 | 9106400 | 10458000 | 0 | 0 | 535080 | 0 |
| NVPLPEFPI | IDE; IDE; IDE; ID | P14735; A0A | IDE_HUMAN In | Unmodified | 0 | 184570 | 0 | 0 | 986600 | 451110 | 0 | 0 | 0 | 0 | 939080 | 2577700 | 0 | 0 | 0 | 0 | 0 | 270270 |
| NVPNRYTF | RNF114; RNF11 | Q9Y508; A0A | RN114_HUMAN | Unmodified | 261600 | 0 | 0 | 323060 | 0 | 152540 | 72555 | 36848 | 0 | 62615 | 0 | 0 | 1394600 | 528630 | 0 | 165020 | 532690 | 865780 |
| NVPNWHR | RAN; RAN; RAN | P62826; J3K | RAN_HUMAN G | Unmodified | 0 | 260020 | 0 | 0 | 0 | 0 | 0 | 8643.5 | 283030 | 0 | 768880 | 5198800 | 0 | 87062 | 429200 | 195830 | 0 | 0 |
| NVPPGLDE | SCAMP1; SCAM | O15126; A0A | SCAM1_HUMA | Unmodified | 0 | 0 | 2742300 | 1865000 | 0 | 0 | 0 | 142110 | 0 | 0 | 0 | 0 | 0 | 0 | 0 | 2062200 | 0 | 0 |
| NVPPPSIEF | TMEM51 | Q9NW97 | TMM51_HUMA | Unmodified | 0 | 0 | 0 | 0 | 3224200 | 0 | 0 | 0 | 3426400 | 0 | 0 | 0 | 0 | 0 | 115020 | 104360 | 0 | 246020 |
| NVPPPSIEF | TMEM51 | Q9NW97 | TMM51_HUMA | Unmodified | 0 | 0 | 0 | 0 | 4001800 | 0 | 0 | 0 | 0 | 1218300 | 0 | 0 | 891970 | 0 | 0 | 0 | 479180 | 2501200 |
| NVPYGNIQ | GALNT2 | Q10471 | GALT2_HUMAN | Unmodified | 0 | 0 | 0 | 0 | 0 | 425270 | 0 | 0 | 36072000 | 2063500 | 546060 | 177290 | 0 | 0 | 0 | 200720 | 0 | 0 |
| NVPYGNIQ | GALNT2 | Q10471 | GALT2_HUMAN | Unmodified | 0 | 0 | 0 | 0 | 538440 | 0 | 0 | 0 | 0 | 4286000 | 0 | 0 | 382880 | 73054 | 0 | 0 | 0 | 0 |
| NVQAEEM\ | ATP5F1A; ATP5I | P25705; K7E | ATPA_HUMAN / | Oxidation (M) | 0 | 0 | 0 | 0 | 0 | 0 | 0 | 0 | 1555300 | 176920 | 0 | 0 | 0 | 0 | 0 | 0 | 0 | 0 |
| NVQAEEM\ | ATP5F1A; ATP5I | P25705; K7E | ATPA_HUMAN / | Unmodified | 0 | 0 | 0 | 0 | 0 | 0 | 0 | 0 | 1937700 | 133440 | 0 | 19222 | 0 | 0 | 0 | 0 | 137880 | 27046 |
| NVQFVFDA | GNAI3; GNAI2; | P08754; P04 | GNAI3_HUMAN | Unmodified | 0 | 96920 | 0 | 0 | 994840 | 0 | 94858 | 0 | 0 | 0 | 0 | 0 | 320050 | 479220 | 0 | 0 | 1099300 | 3377400 |
| NVQFVFDA | GNAI3; GNAI2; | P08754; P04 | GNAI3_HUMAN | Unmodified | 0 | 0 | 0 | 0 | 794700 | 0 | 878130 | 0 | 0 | 101330 | 0 | 0 | 778010 | 45888 | 0 | 0 | 2428900 | 2456800 |
| NVQGIIEILI | LRPPRC; LRPPF | P42704; B8Z | LPPRC_HUMAN | Unmodified | 0 | 0 | 0 | 0 | 0 | 101550 | 292880 | 53623 | 9634000 | 1190300 | 496370 | 309130 | 0 | 85183 | 0 | 0 | 0 | 0 |
| NVQLLSQF | MRPS18C | Q9Y3D5 | RT18C_HUMAN | Unmodified | 0 | 0 | 0 | 0 | 0 | 0 | 12384 | 0 | 345700 | 89022 | 0 | 0 | 0 | 0 | 0 | 0 | 0 | 0 |
| NVQLQENE | PPP1CC; PPP1C | P36873; F8V | PP1G_HUMAN | Unmodified | 41311 | 0 | 773980 | 733230 | 402020 | 0 | 0 | 0 | 2478600 | 0 | 702160 | 3379700 | 0 | 0 | 1079100 | 597700 | 0 | 0 |
| NVQLQENE | PPP1CC; PPP1C | P36873; F8V | PP1G_HUMAN | Unmodified | 0 | 0 | 0 | 0 | 0 | 0 | 0 | 712530 | 799110 | 33619 | 0 | 0 | 0 | 0 | 0 | 0 | 285630 | 0 |
| NVQLTENE | PPP1CA; PPP1C | P62136; E9P | PP1A_HUMAN | Unmodified | 551660 | 366840 | 0 | 929390 | 0 | 0 | 0 | 0 | 2997500 | 0 | 0 | 10417000 | 0 | 0 | 1031300 | 962990 | 0 | 0 |
| NVQLTENE | PPP1CA; PPP1C | P62136; E9P | PP1A_HUMAN | Unmodified | 0 | 0 | 0 | 166710 | 11178000 | 5029100 | 2662400 | 1039000 | 600580 | 3141500 | 926720 | 0 | 1796700 | 4304800 | 0 | 0 | 2903000 | 4707300 |
| NVQPSDQ( | PTGFRN | Q9P2B2 | FPRP_HUMAN | Unmodified | 8575800 | 6424100 | 25569000 | 26047000 | 48177000 | 2227600 | 7990100 | 1301300 | 0 | 0 | 0 | 10217000 | 79258000 | 76270000 | 25978000 | 37324000 | 766580 | 4297000 |
| NVQPSDQ( | PTGFRN | Q9P2B2 | FPRP_HUMAN | Unmodified | 0 | 0 | 0 | 0 | 0 | 0 | 1711100 | 366600 | 0 | 0 | 0 | 0 | 4575000 | 4785400 | 0 | 0 | 0 | 0 |
| NVQVFLIS( | PSPH; PSPH | P78330; C9J | SERB_HUMAN | Unmodified | 0 | 0 | 0 | 0 | 0 | 0 | 0 | 0 | 0 | 0 | 0 | 779760 | 0 | 0 | 0 | 0 | 0 | 0 |
| NVQVFLIS( | PSPH; PSPH | P78330; C9J | SERB_HUMAN | Unmodified | 0 | 0 | 0 | 0 | 0 | 0 | 0 | 0 | 0 | 0 | 0 | 0 | 563120 | 220410 | 0 | 0 | 0 | 0 |
| NVQVVELP | FUT8; FUT8; FU | Q9BYC5; A0 | FUT8_HUMAN / | Unmodified | 0 | 0 | 0 | 0 | 0 | 0 | 0 | 0 | 1796000 | 0 | 0 | 0 | 0 | 0 | 0 | 0 | 0 | 0 |
| NVRTATESI | PKM; PKM; PKM | P14618; H3E | KPYM_HUMAN | Unmodified | 0 | 0 | 0 | 0 | 14878000 | 1298700 | 889530 | 0 | 0 | 852730 | 0 | 0 | 1720600 | 0 | 0 | 0 | 2433500 | 6831400 |
| NVSHNPLL | PLA2G4A | P47712 | PA24A_HUMAN | Unmodified | 0 | 0 | 483100 | 287040 | 213520 | 1401400 | 236860 | 176750 | 0 | 0 | 546610 | 2869600 | 0 | 311990 | 280360 | 0 | 54146 | 910100 |
| NVSIGIVGK | PSMA1; PSMA1 | P25786; F5G | PSA1_HUMAN | Unmodified | 2869300 | 0 | 893060 | 683170 | 1610200 | 0 | 3437300 | 127130 | 3635600 | 990970 | 0 | 14884000 | 825330 | 853840 | 802300 | 467200 | 665160 | 492960 |
| NVSINTVTY | TES; TES | Q9UGI8; F8V | TES_HUMAN Te | Unmodified | 214090 | 0 | 175570 | 1626200 | 5584000 | 918440 | 539180 | 145350 | 1325000 | 1050600 | 0 | 4211400 | 4807200 | 4136900 | 603710 | 480070 | 2737800 | 6961300 |
| NVSLGNVL | DPP3; DPP3; DI | Q9NY33; G3 | DPP3_HUMAN | Unmodified | 0 | 0 | 220100 | 516580 | 0 | 0 | 0 | 0 | 1632500 | 926190 | 3628100 | 4791400 | 586540 | 0 | 0 | 449770 | 0 | 0 |
| NVSQHLED | UTRN | P46939 | UTRN_HUMAN | Unmodified | 0 | 0 | 0 | 653440 | 0 | 42161 | 0 | 0 | 220270 | 0 | 0 | 0 | 0 | 0 | 247980 | 315070 | 298610 | 0 |
| NVSQHLED | UTRN | P46939 | UTRN_HUMAN | Unmodified | 0 | 0 | 2069100 | 825960 | 314670 | 0 | 0 | 0 | 376160 | 84124 | 0 | 179660 | 0 | 0 | 3988300 | 1146500 | 0 | 202590 |
| NVSQHLED | UTRN | P46939 | UTRN_HUMAN | Unmodified | 0 | 0 | 0 | 0 | 1002100 | 0 | 0 | 0 | 238860 | 157870 | 0 | 0 | 6703700 | 405580 | 0 | 0 | 507430 | 663340 |
| NVTAIQGP | IDH3A; IDH3A; | P50213; H0Y | IDH3A_HUMAN | Unmodified | 142080 | 2197800 | 0 | 0 | 0 | 0 | 5599900 | 813370 | 760690 | 38934 | 0 | 0 | 2528300 | 0 | 0 | 91616 | 0 | 0 |
| NVTDVVNT | RTCB | Q9Y3I0 | RTCB_HUMAN | Unmodified | 393640 | 133220 | 340020 | 668610 | 157320 | 325670 | 204400 | 96207 | 965890 | 702620 | 410400 | 3520000 | 724280 | 632940 | 938910 | 211930 | 272770 | 1779600 |
| NVTDVVNT | RTCB | Q9Y3I0 | RTCB_HUMAN | Unmodified | 0 | 64092 | 133630 | 709740 | 465950 | 0 | 205590 | 62866 | 0 | 0 | 328410 | 3448800 | 787590 | 875740 | 300980 | 288660 | 329980 | 310770 |
| NVTELNEPI | YWHAG | P61981 | 1433G_HUMAN | Unmodified | 177000 | 2103900 | 2689800 | 2776700 | 0 | 0 | 0 | 0 | 791100 | 0 | 4368700 | 28207000 | 89919 | 129900 | 6301400 | 5213600 | 112610 | 0 |
| NVTELNEPI | YWHAG | P61981 | 1433G_HUMAN | Unmodified | 7222500 | 8253100 | 17960000 | 48529000 | 1.09E+08 | 31653000 | 27746000 | 1172000 | 26476000 | 23315000 | 34308000 | 95490000 | 86647000 | 1.18E+08 | 74832000 | 38274000 | 92228000 | 86719000 |
| NVTELNEPI | YWHAG | P61981 | 1433G_HUMAN | Unmodified | 0 | 0 | 0 | 0 | 2364600 | 0 | 1779400 | 0 | 0 | 477780 | 0 | 0 | 4636400 | 2825700 | 0 | 0 | 8554500 | 2259100 |
| NVTGHYISI | PPA2; PPA2; PP | Q9H2U2; A0 | IPYR2_HUMAN | Unmodified | 0 | 0 | 0 | 0 | 445170 | 0 | 430590 | 30896 | 3121700 | 619750 | 651930 | 2030900 | 898510 | 233970 | 0 | 0 | 0 | 0 |
| NVTGILWN | PKP3 | Q9Y446 | PKP3_HUMAN | Unmodified | 0 | 8402300 | 2119800 | 2204800 | 32490 | 0 | 0 | 25582 | 0 | 0 | 909000 | 1712000 | 1728500 | 1142600 | 6496500 | 903550 | 751850 | 153900 |
| NVTICVPDI | SERINC5; SERII | Q86VE9; A0A | SERC5_HUMAN | Unmodified | 0 | 0 | 0 | 0 | 0 | 0 | 0 | 0 | 0 | 0 | 0 | 0 | 0 | 0 | 185620 | 0 | 0 | 0 |
| NVTLNPDP | IDI1 | Q13907 | IDI1_HUMAN Is | Unmodified | 0 | 0 | 0 | 0 | 551390 | 140180 | 395740 | 276080 | 691080 | 314470 | 1072300 | 2618700 | 0 | 0 | 0 | 0 | 0 | 0 |
| NVTLPAVFI | RPL4; RPL4; RP | P36578; H3E | RL4_HUMAN 60 | Unmodified | 5938600 | 6821000 | 0 | 0 | 0 | 0 | 0 | 0 | 0 | 137100 | 0 | 0 | 0 | 0 | 0 | 1947500 | 124170 | 0 |
| NVTNTPIDI | SIPA1L1; SIPA1 | O43166; F5C | SI1L1_HUMAN | Unmodified | 0 | 0 | 111210 | 253470 | 1554100 | 310280 | 0 | 0 | 251990 | 0 | 87925 | 383400 | 373110 | 425270 | 223130 | 176260 | 263710 | 748270 |
| NVTSWFTS | INPPL1; INPPL1 | O15357; A0A | SHIP2_HUMAN | Unmodified | 34404 | 0 | 0 | 0 | 0 | 0 | 0 | 0 | 0 | 156940 | 462400 | 2528500 | 0 | 0 | 0 | 0 | 0 | 244200 |
| NVVALDTE | LDLR; LDLR; LD | P01130; J3K | LDLR_HUMAN | Unmodified | 0 | 0 | 0 | 0 | 0 | 0 | 0 | 0 | 2398500 | 0 | 0 | 311650 | 0 | 0 | 0 | 0 | 0 | 0 |
| NVVDGPYA | MGAT5 | Q09328 | MGT5A_HUMA | Unmodified | 0 | 0 | 0 | 0 | 0 | 0 | 0 | 0 | 2311000 | 0 | 0 | 0 | 0 | 0 | 0 | 0 | 0 | 0 |
| NVVHQLSV | DNAJA1 | P31689 | DNJA1_HUMAN | Unmodified | 2160300 | 7184800 | 0 | 13310000 | 25270000 | 3091500 | 4157000 | 0 | 13049000 | 15453000 | 1107600 | 1647700 | 46364000 | 20838000 | 8287700 | 1499300 | 1484600 | 5994700 |
| NVVHQLSV | DNAJA1 | P31689 | DNJA1_HUMAN | Unmodified | 0 | 0 | 0 | 0 | 1707000 | 0 | 719730 | 0 | 0 | 554610 | 0 | 0 | 1484900 | 3306200 | 0 | 0 | 910930 | 0 |
| NVVLPTETE | MAP4; MAP4 | P27816; E7E | MAP4_HUMAN | Unmodified | 156840 | 0 | 35161 | 0 | 0 | 0 | 0 | 0 | 238910 | 0 | 514970 | 1200500 | 276610 | 0 | 0 | 0 | 0 | 405790 |
| NVVLPTETE | MAP4; MAP4 | P27816; E7E | MAP4_HUMAN | Unmodified | 0 | 0 | 0 | 0 | 758510 | 0 | 0 | 101820 | 0 | 0 | 0 | 0 | 0 | 0 | 0 | 0 | 581470 | 1043000 |
| NVVLQTLE( | FLOT2; FLOT2; | Q14254; J3Q | FLOT2_HUMAN | Unmodified | 1451700 | 809070 | 3958000 | 1855400 | 683750 | 0 | 248540 | 0 | 9732100 | 1447100 | 1629400 | 17928000 | 0 | 376260 | 8175200 | 5374900 | 0 | 0 |
| NVVRMLVN | DAAM1 | Q9Y4D1 | DAAM1_HUMA | Unmodified | 0 | 0 | 0 | 0 | 3677100 | 0 | 411590 | 0 | 0 | 1395400 | 0 | 0 | 0 | 0 | 0 | 0 | 4477400 | 0 |
| NVVSGGFF | FBXO3; FBXO3; | Q9UK99; G3 | FBX3_HUMAN | Unmodified | 18782 | 0 | 0 | 0 | 146430 | 0 | 0 | 10230 | 0 | 0 | 0 | 858440 | 0 | 0 | 0 | 0 | 0 | 0 |
| NVVTIFSAP | PPP2CB; PPP2C | P62714; P67 | PP2AB_HUMAN | Unmodified | 0 | 0 | 0 | 0 | 0 | 0 | 0 | 0 | 0 | 91915 | 0 | 3825900 | 0 | 846080 | 0 | 0 | 0 | 0 |
| NVVVVDGV | HADHB; HADHI | P55084; C9J | ECHB_HUMAN | Unmodified | 0 | 0 | 0 | 0 | 0 | 0 | 0 | 0 | 1908800 | 0 | 0 | 0 | 0 | 0 | 0 | 0 | 0 | 0 |
| NVVVVDGV | HADHB; HADHI | P55084; C9J | ECHB_HUMAN | Unmodified | 0 | 0 | 93890 | 0 | 0 | 0 | 0 | 0 | 2173500 | 0 | 0 | 0 | 0 | 0 | 0 | 0 | 402020 | 34810 |
| NVWSFLK | SEC31A; SEC31 | O94979; H7I | SC31A_HUMAN | Unmodified | 0 | 0 | 0 | 0 | 673460 | 0 | 407950 | 0 | 495770 | 504730 | 0 | 2903300 | 359140 | 389660 | 0 | 0 | 302060 | 965460 |
| NVYFAQYG | DIPK2A | Q8NDZ4 | DIK2A_HUMAN | Unmodified | 0 | 0 | 0 | 0 | 0 | 0 | 0 | 0 | 2733200 | 0 | 0 | 0 | 0 | 0 | 0 | 0 | 0 | 0 |
| NVYFAQYG | DIPK2A | Q8NDZ4 | DIK2A_HUMAN | Unmodified | 0 | 0 | 0 | 0 | 0 | 0 | 0 | 0 | 692510 | 592140 | 0 | 0 | 0 | 0 | 0 | 0 | 0 | 0 |
| NVYKFELD | RPN2 | P04844 | RPN2_HUMAN | Unmodified | 0 | 0 | 0 | 0 | 0 | 0 | 0 | 0 | 53920000 | 1806900 | 0 | 1608800 | 0 | 0 | 0 | 0 | 0 | 0 |
| NWADVQS | SEC23B; SEC23 | Q15437; A0A | SC23B_HUMAN | Unmodified | 0 | 0 | 0 | 0 | 0 | 0 | 0 | 0 | 0 | 0 | 0 | 780030 | 0 | 0 | 0 | 0 | 0 | 0 |
| NWDPGQD | CEMIP2; CEMIP | Q9UHN6; F5 | CEIP2_HUMAN | Unmodified | 611250 | 0 | 0 | 0 | 1852500 | 0 | 0 | 0 | 2886800 | 337750 | 0 | 261940 | 0 | 0 | 941430 | 0 | 0 | 0 |
| NWEQEGV | MUC5B | Q9HC84 | MUC5B_HUMA | Unmodified | 0 | 0 | 0 | 0 | 0 | 0 | 46967 | 0 | 0 | 0 | 0 | 0 | 0 | 0 | 0 | 0 | 0 | 833970 |
| NWFGPSP | ITCH; ITCH; ITC | Q96J02; A0A | ITCH_HUMAN I | Unmodified | 0 | 0 | 0 | 0 | 4114300 | 0 | 406020 | 0 | 0 | 174980 | 0 | 70745 | 992410 | 0 | 83405 | 0 | 1228400 | 2048200 |
| NWKPARP | IGF2R; IGF2R | P11717; A0A | MPRI_HUMAN | Unmodified | 0 | 0 | 0 | 0 | 116970 | 0 | 330020 | 0 | 20430000 | 4963800 | 1802200 | 1560600 | 411190 | 0 | 0 | 0 | 0 | 0 |
| NWLSADTK | ANLN | Q9NQW6 | ANLN_HUMAN | Oxidation (M) | 0 | 0 | 0 | 204440 | 4108100 | 945000 | 26230 | 0 | 1699000 | 202000 | 0 | 0 | 0 | 0 | 0 | 0 | 1758900 | 269200 |
| NWLSADTK | ANLN | Q9NQW6 | ANLN_HUMAN | Unmodified | 0 | 0 | 0 | 0 | 7078800 | 0 | 0 | 0 | 0 | 0 | 0 | 0 | 0 | 0 | 0 | 22252 | 1159600 | 1073800 |
| NWMLVEK | ADK; ADK; ADK; | P55263; A0A | ADK_HUMAN A | Oxidation (M) | 0 | 100890 | 0 | 0 | 1496400 | 0 | 0 | 357630 | 0 | 170090 | 577780 | 5778200 | 343110 | 0 | 0 | 253270 | 825900 | 0 |
| NWNPPLL( | CUEDC1; CUED | Q9NWM3; J3 | CUED1_HUMA | Unmodified | 0 | 0 | 0 | 36347 | 267360 | 0 | 0 | 0 | 54195 | 78824 | 0 | 0 | 0 | 0 | 67783 | 41505 | 0 | 0 |
| NWPSDEM | ARHGAP1; ARH | Q07960; E9F | RHG01_HUMA | Unmodified | 0 | 0 | 0 | 0 | 473930 | 0 | 0 | 0 | 0 | 0 | 0 | 0 | 85654 | 0 | 0 | 0 | 97821 | 0 |
| NWPSRPYL | MYO1B; MYO1E | O43795; E9F | MYO1B_HUMA | Unmodified | 0 | 0 | 0 | 0 | 4859400 | 2002100 | 0 | 130890 | 261760 | 171690 | 2461700 | 19599000 | 0 | 0 | 320570 | 179650 | 1081200 | 1135900 |
| NWQYQETI | PSMD6 | Q15008 | PSMD6_HUMA | Unmodified | 0 | 0 | 0 | 0 | 0 | 0 | 0 | 0 | 0 | 0 | 0 | 3046800 | 0 | 0 | 470880 | 0 | 0 | 0 |
| NWSYLGVI | STXBP3 | O00186 | STXB3_HUMAN | Unmodified | 211790 | 0 | 0 | 0 | 0 | 0 | 0 | 0 | 0 | 0 | 0 | 1735800 | 0 | 0 | 632900 | 567790 | 0 | 0 |
| NWSYLGVI | STXBP3 | O00186 | STXB3_HUMAN | Unmodified | 763570 | 782990 | 0 | 783060 | 0 | 0 | 0 | 0 | 737890 | 0 | 0 | 6668600 | 0 | 0 | 1625900 | 1263700 | 0 | 0 |
| NWTAGTAS | SERINC5; SERII | Q86VE9; A0A | SERC5_HUMAN | Unmodified | 0 | 0 | 0 | 0 | 0 | 0 | 0 | 0 | 0 | 0 | 0 | 0 | 0 | 30565 | 1100000 | 596590 | 0 | 0 |
| NWTPELQK | CAPN7; CAPN7 | Q9Y6W3; H7 | CAN7_HUMAN | Unmodified | 84335 | 0 | 0 | 0 | 238050 | 0 | 0 | 0 | 689410 | 994000 | 287830 | 2282300 | 0 | 0 | 0 | 0 | 0 | 0 |
| NWVVVGG | MRPL24; MRPL2 | Q96A35; X6F | RM24_HUMAN | Unmodified | 1854000 | 0 | 0 | 0 | 0 | 0 | 0 | 288570 | 1389700 | 0 | 0 | 13066000 | 0 | 0 | 0 | 0 | 0 | 0 |
| NWYLPAPE | ERMP1; ERMP1 | Q7Z2K6; E7E | ERMP1_HUMA | Unmodified | 0 | 0 | 0 | 0 | 0 | 0 | 0 | 0 | 1575300 | 0 | 0 | 0 | 0 | 0 | 0 | 0 | 269580 | 0 |
| NYAGAYRE | TSTD1; TSTD1 | Q8NFU3; R4 | TSTD1_HUMAN | Unmodified | 476270 | 375890 | 0 | 0 | 0 | 0 | 0 | 0 | 478770 | 122480 | 0 | 0 | 0 | 0 | 286670 | 177100 | 66662 | 0 |
| NYAHALDC | SLC25A10; SLC | Q9UBX3; F6I | DIC_HUMAN M | Unmodified | 0 | 0 | 0 | 0 | 0 | 0 | 0 | 0 | 3330900 | 0 | 126410 | 0 | 0 | 0 | 0 | 0 | 0 | 0 |
| NYAIACER | TMEM260 | Q9NX78 | TM260_HUMAN | Unmodified | 0 | 63645 | 0 | 0 | 0 | 0 | 0 | 0 | 830860 | 145640 | 0 | 0 | 0 | 0 | 0 | 0 | 0 | 0 |
| NYCGLPGE | FGB; FGB | P02675; D6F | FIBB_HUMAN F | Unmodified | 0 | 372640 | 0 | 0 | 0 | 0 | 0 | 0 | 0 | 0 | 0 | 0 | 0 | 0 | 0 | 0 | 0 | 0 |
| NYCNIQVT | ARCN1; ARCN1 | P48444; B0Y | COPD_HUMAN | Unmodified | 0 | 0 | 0 | 0 | 0 | 0 | 0 | 0 | 0 | 0 | 0 | 1146500 | 314380 | 168780 | 0 | 0 | 0 | 0 |
| NYDFAQVL | PAN2 | Q504Q3 | PAN2_HUMAN | Unmodified | 247930 | 95628 | 98713 | 108300 | 0 | 0 | 0 | 0 | 506180 | 0 | 0 | 0 | 0 | 0 | 220860 | 91206 | 0 | 0 |
| NYDIGAALI | SQSTM1 | Q13501 | SQSTM_HUMA | Unmodified | 0 | 236020 | 0 | 322350 | 9643700 | 1788300 | 0 | 43241 | 2002400 | 4095700 | 0 | 0 | 2708000 | 2961100 | 0 | 0 | 5957200 | 3196700 |
| NYDTTIHGI | MYO1D; MYO1E | O94832; J3Q | MYO1D_HUMA | Unmodified | 1179400 | 764470 | 320700 | 0 | 387100 | 0 | 0 | 0 | 104180 | 128040 | 0 | 29116 | 0 | 0 | 746210 | 0 | 0 | 0 |
| NYDYYASR | BPNT1 | O95861 | BPNT1_HUMAN | Unmodified | 0 | 74068 | 0 | 0 | 0 | 0 | 0 | 0 | 1430000 | 0 | 1151100 | 2853500 | 4330300 | 0 | 0 | 0 | 0 | 2095400 |
| NYEASMSH | INPP5A; INPP5A | Q14642; H0Y | I5P1_HUMAN I | Unmodified | 112760 | 175370 | 671750 | 669060 | 451670 | 79104 | 54832 | 0 | 306210 | 53683 | 0 | 11776 | 344500 | 0 | 640070 | 699430 | 0 | 0 |
| NYEEALR | VPS4A; N/A | Q9UN37; I3L | VPS4A_HUMAN | Unmodified | 727780 | 636200 | 643400 | 550500 | 0 | 0 | 0 | 0 | 0 | 0 | 0 | 0 | 30401 | 0 | 697170 | 651520 | 0 | 0 |
| NYEEALRLY | VPS4A; N/A | Q9UN37; I3L | VPS4A_HUMAN | Unmodified | 0 | 0 | 0 | 0 | 0 | 0 | 0 | 0 | 0 | 55608 | 0 | 0 | 193590 | 420700 | 0 | 0 | 0 | 0 |
| NYEEIAK | TMED10 | P49755 | TMEDA_HUMA | Unmodified | 0 | 0 | 0 | 0 | 0 | 22716 | 82240 | 0 | 44627000 | 0 | 805410 | 1065700 | 0 | 0 | 0 | 0 | 0 | 0 |
| NYEEIAKVE | TMED10 | P49755 | TMEDA_HUMA | Unmodified | 430880 | 0 | 0 | 0 | 4228300 | 723130 | 3548700 | 1526200 | 2.83E+08 | 63869000 | 5439900 | 4131100 | 0 | 1317900 | 734390 | 0 | 30668 | 7528500 |
| NYEFPHLQ | ENPP4 | Q9Y6X5 | ENPP4_HUMAN | Unmodified | 0 | 0 | 0 | 902240 | 1185200 | 0 | 750230 | 0 | 25466000 | 6999800 | 10687000 | 5004500 | 1056600 | 0 | 454810 | 623470 | 0 | 0 |
| NYEFPHLQ | ENPP4 | Q9Y6X5 | ENPP4_HUMAN | Unmodified | 0 | 0 | 0 | 0 | 817610 | 0 | 0 | 0 | 0 | 4533200 | 0 | 0 | 0 | 0 | 0 | 0 | 0 | 0 |
| NYEIKNEAI | ARPC3; ARPC3; | O15145; F8V | ARPC3_HUMAN | Unmodified | 47769 | 61379 | 32123 | 0 | 0 | 0 | 0 | 0 | 0 | 0 | 0 | 0 | 39097 | 122220 | 0 | 0 | 0 | 0 |
| NYELMESE | ERLIN2; ERLIN2 | O94905; E5F | ERLN2_HUMAN | Oxidation (M) | 0 | 0 | 0 | 0 | 0 | 0 | 0 | 0 | 2600900 | 381910 | 0 | 0 | 0 | 0 | 0 | 0 | 0 | 0 |
| NYELMESE | ERLIN2; ERLIN2 | O94905; E5F | ERLN2_HUMAN | Unmodified | 0 | 0 | 0 | 0 | 0 | 0 | 0 | 0 | 10573000 | 978710 | 0 | 174870 | 0 | 0 | 0 | 0 | 0 | 0 |
| NYEQWQL( | ITCH; ITCH; ITC | Q96J02; A0A | ITCH_HUMAN I | Unmodified | 2174300 | 2892000 | 1559500 | 758080 | 0 | 0 | 0 | 0 | 252180 | 0 | 0 | 262220 | 0 | 0 | 1704800 | 1644000 | 0 | 0 |

| | | | | | | | | | | | | | | | | | | | | | | |
|---|---|---|---|---|---|---|---|---|---|---|---|---|---|---|---|---|---|---|---|---|---|---|
| NYEQWQS | WWP2 | O00308 | WWP2_HUMAN | Unmodified | 939410 | 728530 | 679350 | 436540 | 0 | 0 | 0 | 0 | 0 | 0 | 0 | 0 | 0 | 0 | 669100 | 619370 | 0 | 0 |
| NYEYLIR | TCF25; TCF25; | Q9BQ70; H3 | TCF25_HUMAN | Unmodified | 101790 | 83416 | 0 | 0 | 0 | 0 | 0 | 63173 | 0 | 0 | 0 | 0 | 0 | 0 | 377410 | 166700 | 0 | 0 |
| NYFDGEGK | KIF23; KIF23; KI | Q02241; A0A | KIF23_HUMAN | Unmodified | 111110 | 162980 | 609860 | 1043200 | 34515000 | 0 | 1123100 | 206770 | 2224000 | 3539900 | 1273400 | 1404000 | 1281400 | 1390600 | 828240 | 302570 | 3526700 | 384400 |
| NYFENTDIL | ARL3 | P36405 | ARL3_HUMAN | Unmodified | 0 | 0 | 0 | 0 | 0 | 0 | 0 | 0 | 0 | 0 | 0 | 0 | 1735100 | 0 | 0 | 0 | 0 | 0 |
| NYGENPEA | PTPN23; PTPN2 | Q9H3S7; R4 | PTN23_HUMAN | Unmodified | 0 | 0 | 0 | 0 | 1576900 | 362480 | 0 | 0 | 590600 | 283040 | 0 | 0 | 0 | 0 | 0 | 0 | 0 | 0 |
| NYGFQGG | CEMIP2 | Q9UHN6 | CEIP2_HUMAN | Unmodified | 0 | 0 | 0 | 0 | 773920 | 269450 | 0 | 0 | 3529100 | 165120 | 0 | 161760 | 0 | 0 | 0 | 0 | 0 | 0 |
| NYGFVHIEI | RBM4; RBM4B; | Q9BWF3; Q9 | RBM4_HUMAN | Unmodified | 0 | 0 | 0 | 0 | 0 | 0 | 0 | 0 | 0 | 50901 | 115300 | 670470 | 0 | 0 | 0 | 0 | 0 | 0 |
| NYGGVYV | OCC1; C12orf7 | Q8TAD7; I7H | OCC1_HUMAN | Oxidation (M) | 728780 | 2749600 | 0 | 0 | 727960 | 0 | 0 | 0 | 0 | 0 | 0 | 0 | 0 | 0 | 0 | 0 | 1000700 | 4597300 |
| NYGGVYV | OCC1; C12orf7 | Q8TAD7; I7H | OCC1_HUMAN | Unmodified | 0 | 0 | 0 | 0 | 0 | 0 | 0 | 45563 | 0 | 0 | 68420 | 243640 | 0 | 0 | 0 | 0 | 665660 | 1713200 |
| NYGILADA | API5; API5; API5 | Q9BZZ5; E9F | API5_HUMAN A | Unmodified | 0 | 0 | 0 | 0 | 0 | 0 | 0 | 0 | 0 | 0 | 0 | 5802700 | 0 | 0 | 0 | 0 | 0 | 0 |
| NYGRYYIA | GGH; GGH; GG | Q92820; A0A | GGH_HUMAN ( | Unmodified | 0 | 0 | 0 | 0 | 0 | 0 | 0 | 0 | 2965800 | 11527000 | 0 | 0 | 185230 | 0 | 0 | 0 | 0 | 0 |
| NYGSQPPS | BMX | P51813 | BMX_HUMAN ( | Unmodified | 0 | 0 | 0 | 0 | 0 | 0 | 0 | 0 | 0 | 0 | 430680 | 989690 | 0 | 0 | 0 | 0 | 0 | 0 |
| NYGSYSTQ | SCAMP3 | O14828 | SCAM3_HUMA | Unmodified | 0 | 0 | 0 | 0 | 1464000 | 0 | 53974 | 23308 | 0 | 0 | 0 | 972340 | 0 | 0 | 0 | 0 | 0 | 782830 |
| NYGVIIPYP | CYFIP2; CYFIP2 | Q96F07; H7 | CYFP2_HUMAN | Unmodified | 160020 | 126000 | 207070 | 529440 | 0 | 0 | 443020 | 0 | 1392000 | 307320 | 249170 | 4507500 | 0 | 373710 | 1083500 | 639840 | 0 | 0 |
| NYGWKVEI | TXNRD1; TXNRI | Q16881; A0A | TRXR1_HUMAN | Unmodified | 0 | 0 | 0 | 0 | 0 | 0 | 0 | 0 | 0 | 0 | 0 | 1074600 | 0 | 0 | 0 | 0 | 0 | 0 |
| NYHATTEFI | PLD1 | Q13393 | PLD1_HUMAN | Unmodified | 0 | 0 | 0 | 0 | 0 | 0 | 0 | 0 | 193650 | 0 | 152310 | 232320 | 0 | 0 | 0 | 0 | 0 | 0 |
| NYIEMKPH | GRB2; GRB2; G | P62993; J3Q | GRB2_HUMAN | Oxidation (M) | 3378900 | 3239200 | 2226400 | 530030 | 0 | 0 | 0 | 0 | 0 | 0 | 0 | 1843000 | 0 | 0 | 4927500 | 3023400 | 282160 | 0 |
| NYIEMKPH | GRB2; GRB2; G | P62993; J3Q | GRB2_HUMAN | Unmodified | 446440 | 609790 | 0 | 0 | 0 | 0 | 0 | 0 | 0 | 0 | 0 | 446900 | 0 | 0 | 1260900 | 0 | 0 | 0 |
| NYIEMKPH | GRB2; GRB2; G | P62993; J3Q | GRB2_HUMAN | Oxidation (M) | 920880 | 917960 | 0 | 0 | 0 | 0 | 0 | 0 | 0 | 191810 | 0 | 403080 | 0 | 0 | 0 | 336100 | 0 | 0 |
| NYIQGINLV | MRPL12; N/A | P52815; B4D | RM12_HUMAN | Unmodified | 0 | 0 | 0 | 0 | 0 | 0 | 387310 | 130000 | 1461000 | 795910 | 0 | 0 | 0 | 0 | 0 | 0 | 0 | 0 |
| NYIRLEELL | BAG5 | Q9UL15 | BAG5_HUMAN | Unmodified | 0 | 575010 | 0 | 1157900 | 0 | 652790 | 221340 | 0 | 3525600 | 1937900 | 352480 | 902200 | 594770 | 0 | 648220 | 536640 | 0 | 268390 |
| NYISQYSEI | PSAP | P07602 | SAP_HUMAN P | 2 Oxidation (M) | 0 | 0 | 0 | 0 | 0 | 0 | 0 | 0 | 775090 | 967310 | 0 | 0 | 0 | 0 | 0 | 0 | 0 | 0 |
| NYITMDELI | ACTN1; ACTN1; | P12814; H0Y | ACTN1_HUMAI | Unmodified | 0 | 0 | 0 | 0 | 1821400 | 948200 | 0 | 0 | 644980 | 0 | 886400 | 15130000 | 0 | 0 | 523160 | 334780 | 302960 | 582450 |
| NYITMDELI | ACTN1; ACTN1; | P12814; H0Y | ACTN1_HUMAI | Oxidation (M) | 0 | 0 | 0 | 0 | 1555300 | 1272800 | 0 | 0 | 0 | 0 | 0 | 0 | 512160 | 539730 | 0 | 0 | 699560 | 1825100 |
| NYITMDELI | ACTN1; ACTN1; | P12814; H0Y | ACTN1_HUMAI | Unmodified | 0 | 0 | 0 | 0 | 4053000 | 182680 | 0 | 0 | 0 | 0 | 0 | 0 | 280490 | 268030 | 0 | 0 | 664940 | 2135000 |
| NYIVEDGD | OLA1; OLA1 | Q9NTK5; J3K | OLA1_HUMAN | Unmodified | 0 | 18132 | 102800 | 0 | 2526100 | 584650 | 0 | 0 | 1708200 | 547670 | 2793800 | 7371300 | 3582700 | 175130 | 0 | 128190 | 2022600 | 1574800 |
| NYKPVVQT | NCEH1; NCEH1 | Q6PIU2; A0A | NCEH1_HUMA | Unmodified | 0 | 0 | 0 | 0 | 1165900 | 251580 | 64574 | 22338 | 2537900 | 721450 | 186400 | 0 | 0 | 0 | 0 | 0 | 0 | 211430 |
| NYKSEEEFI | KARS1; KARS1; | Q15046; H3I | SYK_HUMAN L | Unmodified | 0 | 0 | 0 | 0 | 0 | 0 | 0 | 0 | 694580 | 0 | 357290 | 14104000 | 823330 | 0 | 0 | 144060 | 0 | 81281 |
| NYLAPAWC | BTBD17 | A6NE02 | BTBDH_HUMAI | Unmodified | 0 | 292950 | 0 | 1530800 | 0 | 0 | 0 | 0 | 1333800 | 0 | 0 | 0 | 2573300 | 4291200 | 8411400 | 6061200 | 0 | 0 |
| NYLHYSLYI | PSMD3 | O43242 | PSMD3_HUMA | Unmodified | 183690 | 186330 | 530080 | 399730 | 715780 | 0 | 0 | 0 | 861820 | 0 | 0 | 9677900 | 0 | 0 | 793990 | 432380 | 0 | 0 |
| NYLPAINGI | SAR1A; SAR1A | Q9NR31; Q5 | SAR1A_HUMAN | Unmodified | 0 | 0 | 0 | 0 | 0 | 0 | 0 | 0 | 228710 | 0 | 0 | 137000 | 0 | 0 | 73533 | 0 | 0 | 0 |
| NYLSCDVE | GPD2 | P43304 | GPDM_HUMAN | Unmodified | 0 | 0 | 0 | 0 | 0 | 0 | 0 | 0 | 1741000 | 0 | 0 | 0 | 0 | 0 | 0 | 0 | 0 | 0 |
| NYLSPTFK | UGGT2; UGGT1 | Q9NYU1; Q9 | UGGG2_HUMA | Unmodified | 0 | 0 | 0 | 0 | 0 | 0 | 294320 | 34999 | 1540900 | 2824000 | 0 | 0 | 0 | 0 | 0 | 105340 | 0 | 0 |
| NYLSQPRL | ATP6V1B2; ATP | P21281; E5R | VATB2_HUMAN | Unmodified | 0 | 0 | 0 | 0 | 0 | 0 | 0 | 0 | 0 | 0 | 0 | 0 | 130670 | 1172600 | 0 | 0 | 177970 | 0 |
| NYMQRPS | MAP4K4; MAP4 | O95819; A0A | M4K4_HUMAN | Unmodified | 145830 | 0 | 556120 | 685680 | 2249200 | 0 | 310450 | 135110 | 1541600 | 847640 | 828920 | 2452800 | 1122500 | 704160 | 798090 | 0 | 3009000 | 7389000 |
| NYMSNPS | DYNC1H1; DYN | Q14204; A0A | DYHC1_HUMA | Oxidation (M) | 0 | 0 | 0 | 0 | 0 | 0 | 0 | 0 | 0 | 0 | 0 | 2206300 | 0 | 0 | 0 | 0 | 0 | 0 |
| NYNFQQP | GLIPR2 | Q9H4G4 | GAPR1_HUMAI | Oxidation (M) | 0 | 0 | 0 | 4314100 | 9443600 | 1682500 | 3850300 | 413940 | 0 | 0 | 0 | 0 | 0 | 0 | 4984100 | 3532100 | 1259600 | 5865300 |
| NYNFQQP | GLIPR2 | Q9H4G4 | GAPR1_HUMAI | Unmodified | 0 | 1590900 | 0 | 0 | 0 | 0 | 1925700 | 0 | 0 | 0 | 0 | 0 | 2340200 | 2527000 | 0 | 0 | 726610 | 61727 |
| NYNFQQP | GLIPR2 | Q9H4G4 | GAPR1_HUMAI | Oxidation (M) | 0 | 0 | 0 | 0 | 697100 | 0 | 2562500 | 0 | 0 | 0 | 0 | 0 | 0 | 0 | 0 | 0 | 0 | 0 |
| NYNFQQP | GLIPR2 | Q9H4G4 | GAPR1_HUMAI | Unmodified | 0 | 0 | 0 | 0 | 5751800 | 0 | 989560 | 0 | 0 | 0 | 0 | 0 | 0 | 0 | 0 | 0 | 0 | 0 |
| NYNLEQTL | WDR62; WDR6 | O43379; A0A | WDR62_HUMA | Unmodified | 0 | 0 | 0 | 0 | 0 | 0 | 165930 | 0 | 0 | 703550 | 0 | 0 | 0 | 0 | 0 | 0 | 0 | 0 |
| NYPATFWV | CAPN1 | P07384 | CAN1_HUMAN | Unmodified | 0 | 0 | 0 | 0 | 0 | 0 | 0 | 0 | 1066000 | 0 | 984350 | 4830700 | 0 | 0 | 0 | 0 | 0 | 0 |
| NYPPWQH | LARS1; LARS1; | Q9P2J5; A0A | SYLC_HUMAN | Unmodified | 29178 | 0 | 101180 | 94318 | 0 | 0 | 1443000 | 0 | 0 | 0 | 193600 | 3840700 | 57451 | 0 | 116850 | 141480 | 0 | 0 |
| NYPPWQH | LARS1; LARS1; | Q9P2J5; A0A | SYLC_HUMAN | Unmodified | 0 | 0 | 0 | 0 | 0 | 0 | 637540 | 98754 | 0 | 533570 | 485760 | 11610000 | 0 | 181300 | 0 | 0 | 156410 | 463700 |
| NYPSSLCA | MELTF | P08582 | TRFM_HUMAN | Unmodified | 0 | 0 | 0 | 0 | 0 | 0 | 0 | 0 | 293260 | 0 | 0 | 0 | 0 | 0 | 0 | 0 | 0 | 0 |
| NYPSSLCA | MELTF | P08582 | TRFM_HUMAN | Unmodified | 0 | 0 | 0 | 0 | 3238200 | 844420 | 0 | 0 | 0 | 1016900 | 0 | 360040 | 0 | 0 | 0 | 0 | 1438800 | 877450 |
| NYQFDFLR | SF3A1 | Q15459 | SF3A1_HUMAN | Unmodified | 534430 | 602150 | 0 | 463630 | 0 | 0 | 0 | 0 | 0 | 0 | 0 | 769530 | 1281200 | 0 | 412520 | 350510 | 98459 | 0 |
| NYQNGDV | C1orf21; C1orf2 | Q9H246; A0 | CA021_HUMAN | Unmodified | 0 | 0 | 0 | 0 | 0 | 0 | 0 | 0 | 0 | 0 | 0 | 0 | 4674600 | 2558600 | 0 | 0 | 0 | 0 |
| NYQQLFDY | MAN2A1 | Q16706 | MA2A1_HUMAI | Unmodified | 0 | 0 | 0 | 0 | 0 | 0 | 0 | 0 | 763800 | 0 | 0 | 385050 | 0 | 0 | 0 | 0 | 0 | 0 |
| NYQQNYQ | YBX1; YBX1; YB | P67809; C9J | YBOX1_HUMAN | Unmodified | 2117900 | 229690 | 3003500 | 1269200 | 0 | 0 | 0 | 0 | 0 | 0 | 0 | 5243800 | 0 | 0 | 1298800 | 1593900 | 0 | 0 |
| NYQQNYQ | YBX1; YBX1; YB | P67809; C9J | YBOX1_HUMAN | Unmodified | 1740900 | 1586700 | 0 | 3644200 | 0 | 0 | 0 | 30980 | 0 | 0 | 411470 | 2083700 | 0 | 0 | 0 | 339180 | 0 | 0 |
| NYSREQHC | DFFA; DFFA | O00273; K7E | DFFA_HUMAN | Unmodified | 0 | 0 | 0 | 0 | 0 | 0 | 0 | 0 | 0 | 0 | 0 | 0 | 845770 | 0 | 0 | 0 | 542200 | 0 |
| NYTDEAIET | PSMA7; PSMA7 | O14818; H0Y | PSA7_HUMAN | Unmodified | 0 | 0 | 1310200 | 1334500 | 4029000 | 0 | 0 | 378600 | 7774100 | 2962400 | 6784600 | 13438000 | 5129100 | 1742000 | 1482700 | 0 | 1262900 | 3085300 |
| NYTEEEDR | SMARCA1; SMA | P28370; O60 | SMCA1_HUMA | Unmodified | 160900 | 306090 | 0 | 0 | 72333 | 0 | 0 | 0 | 0 | 0 | 0 | 0 | 0 | 0 | 0 | 0 | 0 | 41944 |
| NYTGDRLN | TKFC; TKFC; TK | Q3LXA3; H0Y | TKFC_HUMAN | Unmodified | 0 | 0 | 0 | 0 | 0 | 0 | 27078 | 0 | 0 | 32263 | 107050 | 2266900 | 193900 | 84776 | 0 | 383500 | 0 | 0 |
| NYVEELNR | RUFY2; RUFY2 | Q8WXA3; H0 | RUFY2_HUMAN | Unmodified | 0 | 0 | 0 | 0 | 419870 | 83643 | 74322 | 0 | 0 | 534020 | 0 | 0 | 0 | 0 | 0 | 0 | 67337 | 325140 |
| NYVLNYLA | XPO7; XPO7; XF | Q9UIA9; H0Y | XPO7_HUMAN | Unmodified | 0 | 0 | 0 | 0 | 0 | 0 | 0 | 0 | 0 | 0 | 97471 | 1684200 | 0 | 0 | 0 | 62611 | 0 | 0 |
| NYVRDDAV | TRAF4; TRAF4; T | Q9BUZ4; A0 | TRAF4_HUMAN | Unmodified | 0 | 0 | 368050 | 1346500 | 0 | 0 | 0 | 0 | 341730 | 0 | 217600 | 891120 | 605270 | 173760 | 760540 | 632980 | 0 | 0 |
| NYVTPVNR | GRB2 | P62993 | GRB2_HUMAN | Unmodified | 5251100 | 3664500 | 2808300 | 2259500 | 354020 | 0 | 0 | 65047 | 2907700 | 0 | 1575600 | 7472800 | 752540 | 0 | 5491400 | 4014800 | 101510 | 0 |
| NYVTPVNR | GRB2 | P62993 | GRB2_HUMAN | Unmodified | 0 | 0 | 0 | 1049100 | 2169900 | 1488900 | 596510 | 628070 | 249560 | 797210 | 0 | 0 | 6710000 | 0 | 0 | 0 | 1479900 | 3997000 |
| NYVVTDHG | EGFR; EGFR; EC | P00533; Q50 | EGFR_HUMAN | Unmodified | 609990 | 338400 | 1427500 | 1993000 | 7162200 | 1538900 | 0 | 0 | 246840 | 0 | 0 | 0 | 1278400 | 794860 | 1449300 | 1764500 | 904140 | 1865700 |
| NYVVTDHG | EGFR; EGFR; EC | P00533; Q50 | EGFR_HUMAN | Unmodified | 0 | 0 | 0 | 0 | 5479000 | 0 | 0 | 0 | 0 | 0 | 0 | 0 | 0 | 0 | 0 | 0 | 669060 | 1835500 |
| NYVVTDHG | EGFR; EGFR; EC | P00533; Q50 | EGFR_HUMAN | Oxidation (M) | 0 | 0 | 0 | 0 | 964700 | 0 | 0 | 0 | 0 | 0 | 0 | 0 | 0 | 0 | 0 | 0 | 187100 | 212540 |
| NYWLLIGH | ERN1; ERN1 | O75460; A0A | ERN1_HUMAN | Unmodified | 0 | 0 | 0 | 0 | 0 | 0 | 0 | 0 | 2173300 | 0 | 0 | 0 | 0 | 0 | 0 | 0 | 0 | 0 |
| NYYEQWG | HNRNPA2B1; H | P22626; A0A | ROA2_HUMAN | Unmodified | 0 | 0 | 0 | 0 | 0 | 0 | 0 | 0 | 493000 | 60966 | 330980 | 6011900 | 0 | 0 | 0 | 49296 | 0 | 0 |
| PAAAAAAT | PABPC1; PABP | P11940; A0A | PABP1_HUMAN | Unmodified | 167480 | 1823200 | 0 | 0 | 0 | 0 | 0 | 0 | 0 | 0 | 0 | 0 | 0 | 0 | 104090 | 155190 | 0 | 0 |
| PAALAAAP | CUL4A; CUL4A | Q13619; A0A | CUL4A_HUMAN | Unmodified | 0 | 0 | 67646 | 0 | 0 | 0 | 0 | 0 | 201580 | 78073 | 0 | 313820 | 0 | 0 | 0 | 31123 | 0 | 0 |
| PAANFDAIF | ANXA7 | P20073 | ANXA7_HUMAN | Unmodified | 1689800 | 463870 | 1537000 | 31375 | 0 | 0 | 0 | 0 | 2744300 | 424470 | 0 | 1693900 | 209700 | 0 | 3201700 | 2258700 | 0 | 0 |
| PAANFDAIF | ANXA7 | P20073 | ANXA7_HUMAN | Unmodified | 65289 | 57811 | 410150 | 0 | 655830 | 0 | 340470 | 0 | 28184000 | 5478400 | 0 | 0 | 0 | 0 | 581480 | 322400 | 1712000 | 0 |
| PAAVVAPIT | CACYBP | Q9HB71 | CYBP_HUMAN | Unmodified | 0 | 0 | 0 | 0 | 0 | 0 | 0 | 0 | 0 | 0 | 0 | 5109600 | 0 | 0 | 0 | 35762 | 220190 | 68855 |
| PAAVVLQTI | GANAB; GANAE | Q14697; F5H | GANAB_HUMA | Unmodified | 0 | 0 | 0 | 0 | 0 | 0 | 0 | 0 | 3280400 | 687430 | 0 | 0 | 0 | 0 | 0 | 0 | 0 | 0 |
| PADVPLAK | CTPS1 | A0A3B3ITF6 | A0A3B3ITF6_H | Unmodified | 3214300 | 2448500 | 4555300 | 1377500 | 0 | 0 | 15041 | 0 | 0 | 0 | 0 | 0 | 2368700 | 1064200 | 2668200 | 2245300 | 0 | 0 |
| PAEKPAETF | RAD23B; RAD2 | P54727; H0Y | RD23B_HUMAN | Unmodified | 0 | 0 | 0 | 0 | 0 | 0 | 0 | 0 | 0 | 0 | 0 | 0 | 0 | 0 | 0 | 0 | 0 | 0 |
| PAEQTPLTA | ALDH1A1; ALDH | P00352; Q5S | AL1A1_HUMAN | Unmodified | 0 | 0 | 0 | 0 | 0 | 0 | 0 | 0 | 103420 | 75224 | 401940 | 46627000 | 0 | 0 | 0 | 0 | 0 | 0 |
| PAEYSSMV | ST14 | Q9Y5Y6 | ST14_HUMAN S | Oxidation (M) | 0 | 0 | 1716400 | 1217900 | 0 | 0 | 0 | 0 | 0 | 0 | 0 | 0 | 0 | 0 | 2468400 | 1895100 | 0 | 0 |
| PAFFAEK | ANXA1 | P04083 | ANXA1_HUMAN | Unmodified | 0 | 0 | 0 | 0 | 934250 | 0 | 0 | 0 | 0 | 0 | 6120800 | 84764000 | 0 | 0 | 0 | 0 | 0 | 0 |
| PAFGTPLEE | ARHGAP17 | Q68EM7 | RHG17_HUMAI | Unmodified | 325640 | 0 | 268100 | 0 | 0 | 0 | 0 | 0 | 0 | 0 | 0 | 0 | 0 | 0 | 268250 | 528240 | 0 | 0 |
| PAFNQNSA | SHANK2; SHAN | Q9UPX8; A6I | SHAN2_HUMAI | Unmodified | 0 | 217700 | 18682000 | 0 | 0 | 0 | 0 | 0 | 0 | 0 | 0 | 0 | 0 | 0 | 519400 | 0 | 0 | 0 |
| PAGDGTFC | HLA-E; HLA-A; H | P13747; P04 | HLAE_HUMAN | Unmodified | 0 | 0 | 301050 | 0 | 157300 | 29211 | 45695 | 0 | 1552600 | 158370 | 51629 | 78498 | 0 | 0 | 0 | 0 | 259850 | 0 |
| PAHVVVGD | BLVRB; BLVRB; | P30043; M0 | BLVRB_HUMAN | Unmodified | 130380 | 0 | 168160 | 0 | 0 | 0 | 0 | 0 | 0 | 0 | 0 | 0 | 0 | 0 | 293820 | 225150 | 98547 | 0 |
| PAKYDDIK | GAPDH; GAPDH | P04406; E7E | G3P_HUMAN G | Unmodified | 20765 | 0 | 0 | 0 | 1140700 | 0 | 0 | 0 | 549880 | 20529 | 240290 | 4095300 | 285310 | 0 | 0 | 0 | 0 | 984340 |
| PALELLEPI | GDI2 | P50395 | GDIB_HUMAN | Unmodified | 193000 | 0 | 0 | 0 | 0 | 0 | 0 | 0 | 0 | 0 | 0 | 1722000 | 0 | 0 | 0 | 0 | 0 | 0 |
| PALHIEDLF | GNG11 | P61952 | GBG11_HUMAI | Unmodified | 89981 | 162110 | 82984 | 269390 | 0 | 175980 | 1006100 | 601950 | 0 | 252950 | 0 | 0 | 426180 | 521530 | 0 | 108850 | 304590 | 1322500 |
| PALHIEDLF | GNG11 | P61952 | GBG11_HUMAI | Unmodified | 0 | 0 | 0 | 0 | 0 | 0 | 2463100 | 1230400 | 0 | 0 | 0 | 0 | 561180 | 944500 | 0 | 0 | 0 | 887470 |
| PALHIEDLF | GNG11 | P61952 | GBG11_HUMAI | Unmodified | 0 | 0 | 0 | 0 | 0 | 0 | 694850 | 484230 | 0 | 0 | 0 | 0 | 0 | 0 | 0 | 0 | 100250 | 0 |
| PALLHSTFF | WARS1; WARS1 | P23381; H0Y | SYWC_HUMAN | Unmodified | 447280 | 456690 | 0 | 0 | 2225200 | 418950 | 0 | 0 | 7240200 | 481090 | 0 | 6622200 | 985290 | 653340 | 0 | 1514100 | 330140 | 1494400 |
| PALPSYWA | MGAM; MGAM | O43451; E7E | MGA_HUMAN N | Oxidation (M) | 0 | 0 | 0 | 2382800 | 0 | 3180800 | 0 | 0 | 0 | 0 | 0 | 0 | 0 | 0 | 0 | 0 | 0 | 0 |
| PALVVLTEK | SNTB1; SNTB1 | Q13884; A0A | SNTB1_HUMAN | Unmodified | 0 | 0 | 0 | 0 | 0 | 0 | 1236400 | 812760 | 686770 | 64903 | 0 | 0 | 83110 | 44251 | 0 | 0 | 0 | 39826 |
| PANDFNPD | ANXA6; ANXA6 | P08133; E5R | ANXA6_HUMAN | Unmodified | 0 | 0 | 0 | 0 | 159220 | 41319 | 1145700 | 268260 | 7586300 | 1963600 | 0 | 0 | 0 | 0 | 0 | 0 | 404310 | 114580 |
| PANKQEDE | ARPC3 | O15145 | ARPC3_HUMAI | Unmodified | 2285900 | 1892000 | 0 | 1245100 | 0 | 0 | 0 | 0 | 1162900 | 0 | 718770 | 1452000 | 0 | 19171 | 6028300 | 2175400 | 13265 | 0 |
| PANKQEDE | ARPC3 | O15145 | ARPC3_HUMAI | Oxidation (M) | 0 | 129020 | 377620 | 301590 | 0 | 0 | 0 | 0 | 0 | 0 | 138230 | 1753700 | 0 | 0 | 424400 | 595620 | 0 | 0 |
| PANSVFTTI | ZC3HAV1; ZC3H | Q7Z2W4; C9 | ZCCHV_HUMA | Unmodified | 685170 | 799450 | 0 | 857190 | 3035600 | 0 | 0 | 0 | 0 | 140570 | 455940 | 44639 | 1399700 | 0 | 154470 | 84113 | 295600 | 792260 |
| PAPLTAEIQ | CTSC; CTSC; CT | P53634; A0A | CATC_HUMAN | Unmodified | 0 | 0 | 0 | 0 | 0 | 0 | 0 | 0 | 0 | 0 | 992950 | 885540 | 0 | 0 | 0 | 0 | 0 | 0 |
| PAPPQQEC | SEL1L | Q9UBV2 | SE1L1_HUMAN | Unmodified | 0 | 0 | 0 | 0 | 0 | 0 | 0 | 0 | 619850 | 42212 | 0 | 0 | 0 | 0 | 0 | 0 | 0 | 0 |
| PAPYWEGT | PRDX4; PRDX4; | Q13162; A6N | PRDX4_HUMAN | Unmodified | 0 | 0 | 0 | 40234 | 118130 | 72217 | 0 | 0 | 1774800 | 0 | 0 | 275440 | 0 | 0 | 0 | 0 | 68464 | 0 |
| PAQGVVTT | FN1; FN1 | P02751; H0Y | FINC_HUMAN I | Unmodified | 0 | 0 | 4170000 | 0 | 0 | 0 | 0 | 0 | 0 | 0 | 0 | 0 | 0 | 0 | 0 | 806620 | 51518 | 0 |
| PAQGVVTT | FN1; FN1 | P02751; H0Y | FINC_HUMAN I | Unmodified | 0 | 0 | 1913600 | 0 | 264590 | 0 | 0 | 0 | 0 | 0 | 0 | 0 | 0 | 0 | 200740 | 750910 | 0 | 0 |
| PAQPQEHF | SEC23B; SEC23 | Q15437; A0A | SC23B_HUMAN | Unmodified | 0 | 0 | 0 | 0 | 108450 | 34040 | 0 | 0 | 233410 | 101300 | 248520 | 3670000 | 0 | 0 | 0 | 0 | 0 | 45107 |
| PASGCEAE | IGF2R; IGF2R | P11717; A0A | MPRI_HUMAN ( | Unmodified | 0 | 0 | 0 | 0 | 0 | 0 | 0 | 0 | 2279600 | 1281400 | 0 | 205960 | 0 | 0 | 0 | 0 | 0 | 0 |
| PASTNSDD | G6PD; G6PD; G | P11413; E7E | G6PD_HUMAN | Unmodified | 0 | 0 | 0 | 0 | 0 | 0 | 58536 | 0 | 0 | 0 | 199220 | 127170 | 26864 | 0 | 0 | 0 | 0 | 0 |
| PATEKDLAE | FLNA; FLNA; FL | P21333; A0A | FLNA_HUMAN | Unmodified | 0 | 0 | 0 | 0 | 172320 | 0 | 257420 | 0 | 0 | 0 | 0 | 0 | 88705 | 222600 | 0 | 0 | 228820 | 0 |
| PATEKDLAE | FLNA; FLNA; FL | P21333; A0A | FLNA_HUMAN | Unmodified | 0 | 0 | 0 | 0 | 1524000 | 0 | 2277200 | 2191300 | 0 | 285520 | 0 | 0 | 412690 | 1475300 | 0 | 0 | 1584200 | 370960 |
| PATFTVNTK | FLNA; FLNA; FL | P21333; H0Y | FLNA_HUMAN | Unmodified | 0 | 0 | 0 | 0 | 0 | 0 | 0 | 0 | 0 | 0 | 0 | 0 | 0 | 0 | 0 | 0 | 0 | 0 |
| PAVASVPKI | USP8; USP8; US | P40818; H0Y | UBP8_HUMAN | Unmodified | 0 | 0 | 0 | 0 | 696560 | 0 | 98407 | 132930 | 0 | 15659 | 0 | 0 | 0 | 0 | 0 | 0 | 92072 | 0 |
| PAVLGFEG | OSGEP; OSGEP | Q9NPF4; G3 | OSGEP_HUMAI | Unmodified | 0 | 0 | 0 | 0 | 0 | 0 | 0 | 0 | 0 | 0 | 235660 | 0 | 0 | 0 | 0 | 0 | 0 | 0 |
| PAVSLPPK | NAA15; NAA15 | Q9BXJ9; A0A | NAA15_HUMAN | Unmodified | 2274400 | 0 | 2137100 | 2052400 | 0 | 0 | 0 | 0 | 0 | 0 | 0 | 0 | 329010 | 136970 | 1051800 | 1411000 | 0 | 0 |
| PAYFQRPEI | EIF3A | Q14152 | EIF3A_HUMAN | Unmodified | 818240 | 387610 | 1516900 | 0 | 2361900 | 0 | 0 | 779230 | 0 | 0 | 0 | 6164500 | 2284800 | 0 | 1383200 | 476800 | 626890 | 423130 |
| PAYHSSLM | ARPC3; ARPC3 | O15145; C9J | ARPC3_HUMAI | Oxidation (M) | 610590 | 457550 | 662390 | 415620 | 817980 | 0 | 140490 | 0 | 0 | 0 | 15957 | 3063200 | 871930 | 471330 | 594540 | 796800 | 255690 | 523200 |
| PAYHSSLM | ARPC3; ARPC3 | O15145; C9J | ARPC3_HUMAI | Unmodified | 1064700 | 965070 | 1050000 | 0 | 2728400 | 650810 | 423180 | 357760 | 1843600 | 0 | 0 | 10970000 | 1974900 | 868470 | 1030400 | 1028500 | 555930 | 1264000 |

| | | | | | | | | | | | | | | | | | | | | | |
|---|---|---|---|---|---|---|---|---|---|---|---|---|---|---|---|---|---|---|---|---|---|
| PAYSPASW | PKP3; PKP3; PK | Q9Y446; E9F | PKP3_HUMAN | Unmodified | 0 | 0 | 1401200 | 0 | 0 | 416320 | 0 | 0 | 0 | 0 | 0 | 792800 | 0 | 0 | 2241500 | 345420 | 303040 | 0 |
| PCSEETPAI | DUT | H0YNJ9 | H0YNJ9_HUMA | Unmodified | 0 | 0 | 0 | 0 | 877920 | 152910 | 94370 | 96501 | 114790 | 0 | 162510 | 1019000 | 531010 | 209700 | 0 | 0 | 209100 | 116470 |
| PCSEETPAI | DUT | H0YNJ9 | H0YNJ9_HUMA | Unmodified | 0 | 0 | 0 | 0 | 0 | 0 | 0 | 0 | 0 | 0 | 1005900 | 0 | 0 | 0 | 0 | 0 | 103970 | 0 |
| PDGKYVVV | MTHFD1; MTHF | P11586; V9C | C1TC_HUMAN | Unmodified | 0 | 0 | 0 | 0 | 0 | 0 | 0 | 16135 | 0 | 0 | 0 | 0 | 0 | 0 | 0 | 0 | 633540 | 0 |
| PDNTYEVK | CALR; CALR; C/ | P27797; A0A | CALR_HUMAN | Unmodified | 0 | 0 | 0 | 0 | 0 | 0 | 0 | 0 | 0 | 1544500 | 0 | 0 | 0 | 0 | 0 | 0 | 0 | 0 |
| PDPAKSAP | H2BC9; H2BC1 | Q93079; P2 | H2B1H_HUMAI | Unmodified | 0 | 0 | 95730 | 142690 | 0 | 0 | 0 | 1603300 | 0 | 0 | 0 | 1247300 | 0 | 0 | 0 | 0 | 0 | 0 |
| PDPAKSAP | H2BC9; H2BC1 | Q93079; P2 | H2B1H_HUMAI | Unmodified | 0 | 0 | 0 | 0 | 0 | 0 | 60545 | 965490 | 0 | 0 | 0 | 82614 | 0 | 0 | 0 | 0 | 0 | 0 |
| PDPGSGG( | ESYT1 | Q9BSJ8 | ESYT1_HUMAN | Unmodified | 0 | 0 | 0 | 0 | 0 | 0 | 0 | 0 | 239040 | 0 | 0 | 0 | 0 | 0 | 109410 | 0 | 0 | 0 |
| PDPSKSAP | H2BU1 | Q8N257 | H2B3B_HUMAI | Unmodified | 0 | 0 | 0 | 0 | 0 | 0 | 0 | 135360 | 0 | 31404 | 0 | 0 | 0 | 0 | 0 | 0 | 14912 | 0 |
| PDQDIKFTI | EFNB2 | P52799 | EFNB2_HUMAN | Unmodified | 0 | 0 | 41281 | 198270 | 0 | 378640 | 63775 | 0 | 0 | 47099 | 667750 | 0 | 0 | 455200 | 0 | 0 | 0 | 292740 |
| PDQISVSEI | ASAP2 | O43150 | ASAP2_HUMAN | Unmodified | 0 | 0 | 0 | 0 | 2876000 | 569410 | 0 | 0 | 0 | 0 | 336750 | 0 | 2241900 | 979630 | 0 | 0 | 1748200 | 3794100 |
| PDSNFAER | HACL1; HACL1 | Q9UJ83; B3I | HACL1_HUMAN | Unmodified | 0 | 0 | 0 | 0 | 253600 | 0 | 81310 | 82146 | 4910000 | 1192200 | 58977 | 0 | 21089 | 55859 | 0 | 0 | 0 | 33900 |
| PDTIEVQQI | RDX; EZR; MSN | P35241; P15 | RADI_HUMAN I | Unmodified | 0 | 0 | 0 | 0 | 2112500 | 845680 | 170720 | 0 | 200280 | 0 | 0 | 0 | 0 | 0 | 0 | 0 | 1367900 | 0 |
| PDTIEVQQI | RDX; EZR; MSN | P35241; P15 | RADI_HUMAN I | Oxidation (M) | 0 | 0 | 0 | 0 | 645750 | 0 | 0 | 0 | 0 | 80381 | 115460 | 0 | 0 | 0 | 0 | 0 | 403770 | 0 |
| PDTIEVQQI | RDX; EZR; MSN | P35241; P15 | RADI_HUMAN I | Unmodified | 0 | 0 | 0 | 0 | 192260 | 72447 | 110670 | 52607 | 0 | 0 | 0 | 95601 | 0 | 0 | 0 | 0 | 1334000 | 0 |
| PEANEALEI | CLIC4 | Q9Y696 | CLIC4_HUMAN | Unmodified | 0 | 0 | 0 | 0 | 0 | 0 | 0 | 0 | 0 | 0 | 0 | 0 | 0 | 0 | 0 | 0 | 1058800 | 0 |
| PEDENNVL | EXOC5; EXOC5 | O00471; F8V | EXOC5_HUMAI | Unmodified | 27952 | 0 | 30232 | 20949 | 0 | 0 | 0 | 0 | 272230 | 29697 | 173560 | 390800 | 28953 | 0 | 39981 | 21036 | 0 | 0 |
| PEDITQIQP | ITGB1; ITGB1; I | P05556; E7E | ITB1_HUMAN I | Unmodified | 0 | 105290 | 0 | 0 | 686470 | 562200 | 0 | 0 | 0 | 154070 | 0 | 0 | 0 | 0 | 0 | 0 | 1235900 | 0 |
| PEDITQIQP | ITGB1; ITGB1; I | P05556; E7E | ITB1_HUMAN I | Unmodified | 0 | 0 | 0 | 0 | 0 | 227160 | 0 | 0 | 0 | 0 | 0 | 0 | 0 | 0 | 0 | 0 | 1141300 | 0 |
| PEDITQIQP | ITGB1; ITGB1; I | P05556; E7E | ITB1_HUMAN I | Unmodified | 0 | 0 | 0 | 0 | 306460 | 0 | 475030 | 0 | 0 | 0 | 0 | 0 | 0 | 0 | 0 | 0 | 4880100 | 0 |
| PEDWDEEI | CALR; CALR; C/ | P27797; A0A | CALR_HUMAN | Oxidation (M) | 0 | 0 | 0 | 0 | 0 | 0 | 0 | 0 | 0 | 780230 | 0 | 0 | 0 | 0 | 0 | 0 | 0 | 0 |
| PEEMDKPL | STEAP3; STEAP | Q658P3; B8 | STEA3_HUMAN | Oxidation (M) | 0 | 0 | 1134000 | 178890 | 353550 | 0 | 0 | 0 | 0 | 312240 | 344860 | 0 | 0 | 0 | 643940 | 705420 | 0 | 0 |
| PEFLEDPS\ | TMPO; TMPO; T | P42167; P42 | LAP2B_HUMAN | Unmodified | 0 | 0 | 0 | 0 | 0 | 0 | 0 | 222470 | 1453400 | 1094000 | 0 | 2155200 | 0 | 0 | 0 | 0 | 0 | 0 |
| PEGALQNN | EPCAM; EPCAM | P16422; C9J | EPCAM_HUMA | Unmodified | 0 | 0 | 465930 | 0 | 0 | 0 | 0 | 0 | 0 | 0 | 0 | 0 | 0 | 0 | 654970 | 297000 | 0 | 0 |
| PEINTNHLI | IDI1 | Q13907 | IDI1_HUMAN Is | Unmodified | 0 | 0 | 0 | 0 | 150490 | 32115 | 45693 | 245610 | 78668 | 42043 | 96814 | 424360 | 0 | 0 | 0 | 0 | 0 | 33363 |
| PEIVDTCSL | LTA4H; LTA4H | P09960; B4E | LKHA4_HUMAN | Unmodified | 0 | 0 | 0 | 0 | 518440 | 0 | 0 | 0 | 538710 | 459750 | 744530 | 4692600 | 0 | 0 | 0 | 29272 | 166940 | 0 |
| PEIVDTCSL | LTA4H; LTA4H | P09960; B4E | LKHA4_HUMAN | Unmodified | 0 | 0 | 0 | 0 | 2070400 | 0 | 0 | 179910 | 556400 | 0 | 0 | 0 | 0 | 0 | 0 | 0 | 2948300 | 0 |
| PENPATDK | PATJ; PATJ; PAT | Q8NI35; A0A | INADL_HUMAN | Unmodified | 0 | 0 | 0 | 0 | 9021400 | 1143400 | 0 | 0 | 0 | 0 | 0 | 0 | 3629500 | 3731100 | 0 | 0 | 6107200 | 5064300 |
| PENVAPRS | CCT4 | P50991 | TCPD_HUMAN | Unmodified | 0 | 0 | 0 | 0 | 0 | 0 | 0 | 0 | 0 | 0 | 0 | 0 | 31811 | 0 | 0 | 0 | 3200800 | 0 |
| PEPAKSAP | H2BC4; H2BC1 | P62807; P06 | H2B1C_HUMAI | Unmodified | 0 | 0 | 0 | 0 | 0 | 0 | 0 | 5617500 | 0 | 0 | 0 | 0 | 41175 | 0 | 0 | 0 | 0 | 0 |
| PETLMEVV | N/A | N/A | N/A | Oxidation (M) | 0 | 20450 | 0 | 0 | 0 | 0 | 0 | 23418 | 2133000 | 1596100 | 0 | 0 | 0 | 0 | 0 | 0 | 0 | 0 |
| PEVPEDVD | LIMA1; LIMA1; L | Q9UHB6; F8 | LIMA1_HUMAN | Unmodified | 0 | 0 | 0 | 0 | 0 | 0 | 0 | 0 | 1164300 | 74046 | 0 | 0 | 0 | 0 | 0 | 0 | 0 | 0 |
| PFAIDIFPV | SUMF2; SUMF2 | Q8NBJ7; J3C | SUMF2_HUMAI | Unmodified | 0 | 0 | 43205 | 0 | 0 | 0 | 0 | 0 | 1172600 | 0 | 0 | 0 | 0 | 0 | 0 | 0 | 0 | 0 |
| PFATPTGYI | CEL | P19835 | CEL_HUMAN B | Unmodified | 0 | 0 | 0 | 0 | 0 | 40219 | 0 | 0 | 1336600 | 31534 | 465400 | 279890 | 0 | 0 | 0 | 0 | 0 | 0 |
| PFEDASFAI | PIN1; PIN1 | Q13526; K7E | PIN1_HUMAN I | Unmodified | 1380000 | 1060100 | 4024400 | 463310 | 0 | 0 | 0 | 55891 | 0 | 0 | 0 | 298140 | 0 | 0 | 1679300 | 1484500 | 0 | 0 |
| PFEVPFLKF | NUDT5; NUDT5 | Q9UKK9; A6I | NUDT5_HUMAI | Unmodified | 51027 | 0 | 0 | 0 | 592570 | 57688 | 0 | 0 | 535520 | 0 | 389220 | 3628900 | 0 | 0 | 165860 | 0 | 35937 | 67123 |
| PFFAAGNT | PTGFRN | Q9P2B2 | FPRP_HUMAN | Oxidation (M) | 428070 | 110940 | 8128900 | 526090 | 0 | 0 | 0 | 0 | 0 | 0 | 0 | 2861700 | 80752 | 0 | 4318500 | 3383400 | 0 | 0 |
| PFFAAGNT | PTGFRN | Q9P2B2 | FPRP_HUMAN | Unmodified | 0 | 0 | 177930 | 61168 | 0 | 0 | 0 | 0 | 3323000 | 1449600 | 1859000 | 147550 | 0 | 0 | 604660 | 307780 | 0 | 0 |
| PFGSEEQL | SNRNP200; SN | O75643; A0A | U520_HUMAN | Unmodified | 0 | 0 | 0 | 0 | 0 | 0 | 0 | 0 | 0 | 0 | 0 | 0 | 0 | 0 | 0 | 0 | 0 | 0 |
| PFHFQISNI | PLCG1; PLCG1 | P19174; V9C | PLCG1_HUMAN | Unmodified | 0 | 5044800 | 220660 | 1795300 | 0 | 0 | 0 | 0 | 0 | 0 | 173420 | 546610 | 941020 | 0 | 3600600 | 370270 | 0 | 0 |
| PFLAILGGA | PGK1; PGK2 | P00558; P07 | PGK1_HUMAN | Unmodified | 40756 | 90127 | 0 | 0 | 328280 | 0 | 292730 | 97180 | 63046 | 927580 | 20122 | 1131800 | 75238 | 0 | 0 | 70073 | 3572500 | 0 |
| PFLELDTNI | DDT; DDTL; DD | P30046; A6N | DOPD_HUMAN | Unmodified | 1678600 | 3883800 | 4212900 | 4187700 | 8285200 | 3427200 | 3095800 | 305030 | 0 | 3826100 | 6434500 | 14885000 | 12317000 | 10060000 | 12976000 | 7814900 | 4456100 | 2821200 |
| PFLELDTNI | DDT; DDTL; DD | P30046; A6N | DOPD_HUMAN | Unmodified | 0 | 0 | 0 | 0 | 0 | 0 | 0 | 0 | 834350 | 0 | 262270 | 11935000 | 0 | 0 | 12936000 | 252970 | 0 | 0 |
| PFLFDTK | CANX; CANX; C | P27824; A0A | CALX_HUMAN | Unmodified | 0 | 0 | 23726 | 0 | 588640 | 0 | 0 | 93082 | 817940 | 140380 | 21978 | 0 | 0 | 0 | 0 | 0 | 0 | 0 |
| PFLLPVEAV | TUFM | P49411 | EFTU_HUMAN | Unmodified | 0 | 0 | 0 | 0 | 0 | 0 | 0 | 0 | 2481500 | 0 | 45390 | 194920 | 0 | 0 | 0 | 44455 | 0 | 0 |
| PFLLPVEAV | TUFM | P49411 | EFTU_HUMAN | Unmodified | 0 | 0 | 0 | 0 | 0 | 0 | 0 | 0 | 5708400 | 250260 | 162640 | 0 | 0 | 0 | 0 | 0 | 0 | 0 |
| PFLNGLTG | SNRPF; SNRPF | P62306; A0A | RUXF_HUMAN | Oxidation (M) | 0 | 301250 | 1151800 | 0 | 0 | 0 | 88562 | 0 | 0 | 0 | 413640 | 1321400 | 0 | 0 | 1215700 | 839030 | 0 | 0 |
| PFLWLAR | RAN; RAN; RAN | P62826; J3K | RAN_HUMAN C | Unmodified | 2265500 | 1943500 | 3548800 | 1257900 | 1128000 | 297820 | 117580 | 38885 | 6511300 | 306210 | 3759700 | 87723000 | 562230 | 254320 | 6670700 | 4069800 | 0 | 1086400 |
| PFRLDLLEI | GANAB; GANAE | Q14697; F5F | GANAB_HUMA | Unmodified | 0 | 0 | 0 | 0 | 0 | 0 | 0 | 0 | 16174000 | 676340 | 118260 | 871150 | 0 | 0 | 0 | 0 | 0 | 0 |
| PFSAPKPQ | CAP1 | Q01518 | CAP1_HUMAN | Unmodified | 0 | 0 | 0 | 0 | 307540 | 0 | 134500 | 0 | 262050 | 0 | 0 | 1236400 | 0 | 0 | 419200 | 0 | 68395 | 0 |
| PFSEYTAW | BPNT2 | Q9NX62 | IMPA3_HUMAN | Oxidation (M) | 0 | 0 | 0 | 0 | 0 | 655610 | 0 | 0 | 1216600 | 0 | 0 | 0 | 0 | 0 | 0 | 0 | 0 | 0 |
| PFSNSHNA | CKB; CKB; CKB | P12277; G3V | KCRB_HUMAN | Unmodified | 252610 | 216440 | 1062600 | 1613700 | 7415900 | 1553000 | 8628700 | 9533000 | 1293500 | 807990 | 443850 | 4717400 | 6977300 | 3159700 | 1636300 | 1275200 | 78437 | 0 |
| PFTQLVK | CDC42BPB | Q9Y5S2 | MRCKB_HUMA | Unmodified | 0 | 0 | 0 | 23015 | 0 | 0 | 0 | 0 | 0 | 0 | 2492300 | 179030 | 0 | 0 | 352620 | 281940 | 0 | 0 |
| PFVPNVHA | GSPT2 | Q8IYD1 | ERF3B_HUMAN | Unmodified | 1328300 | 745360 | 1454400 | 871570 | 46954 | 30739 | 0 | 0 | 681610 | 0 | 628330 | 3751100 | 0 | 70633 | 2274100 | 1496900 | 0 | 138120 |
| PFYYQHPE | LAMB1; LAMB1 | P07942; A0A | LAMB1_HUMAI | Unmodified | 0 | 0 | 294110 | 0 | 0 | 0 | 97016 | 0 | 0 | 0 | 58611 | 0 | 41731 | 0 | 0 | 0 | 0 | 0 |
| PGALGGA | PLCB3 | Q01970 | PLCB3_HUMAN | Unmodified | 220390 | 0 | 0 | 0 | 0 | 0 | 0 | 0 | 87055 | 0 | 0 | 0 | 0 | 0 | 0 | 244330 | 34471 | 11913 |
| PGDGEFVE | NUDT5; NUDT5 | Q9UKK9; A6I | NUDT5_HUMA | Unmodified | 3644900 | 0 | 89045 | 0 | 0 | 0 | 0 | 158820 | 0 | 0 | 191030 | 885430 | 0 | 0 | 0 | 0 | 0 | 0 |
| PGDLESAP | IGF2R; IGF2R; I | P11717; A0A | MPRI_HUMAN | Unmodified | 0 | 0 | 0 | 0 | 0 | 0 | 0 | 0 | 895200 | 497570 | 0 | 0 | 0 | 0 | 0 | 0 | 112050 | 0 |
| PGETEEPRI | ZFP91; ZFP91-C | Q96JP5; A0A | ZFP91_HUMAN | Unmodified | 0 | 0 | 0 | 131050 | 85087 | 0 | 0 | 0 | 45939 | 0 | 90008 | 0 | 981690 | 245850 | 80591 | 61013 | 28683 | 104320 |
| PGGGPVS\ | IGSF8 | Q969P0 | IGSF8_HUMAN | Unmodified | 435250 | 371070 | 154670 | 0 | 0 | 0 | 0 | 0 | 515740 | 0 | 0 | 0 | 0 | 0 | 262810 | 202800 | 0 | 0 |
| PGGLLLGD | PRDX6 | P30041 | PRDX6_HUMAN | Unmodified | 4049600 | 5290500 | 2501000 | 10430000 | 21124000 | 3433400 | 1090200 | 804520 | 13183000 | 6419300 | 8431400 | 22511000 | 91930000 | 17128000 | 8387400 | 10818000 | 27057000 | 58677000 |
| PGGLLLGD | PRDX6 | P30041 | PRDX6_HUMAN | Unmodified | 0 | 0 | 0 | 0 | 4849600 | 499320 | 0 | 0 | 0 | 1278100 | 0 | 0 | 1284800 | 0 | 0 | 0 | 7477200 | 17802000 |
| PGGVLILG( | NPTX2 | P47972 | NPTX2_HUMAN | Unmodified | 0 | 0 | 0 | 0 | 344680 | 0 | 0 | 0 | 0 | 65319 | 0 | 0 | 0 | 0 | 0 | 0 | 0 | 59480 |
| PGIDKLPIE | APPL1; APPL1 | Q9UKG1; C9 | DP13A_HUMAN | Unmodified | 691930 | 2447700 | 452470 | 685350 | 7058100 | 996020 | 151080 | 79971 | 7187500 | 870730 | 2043900 | 6144400 | 3605200 | 605960 | 450370 | 471200 | 1811500 | 3260000 |
| PGIVELPTL | NDUFA8 | P51970 | NDUA8_HUMA | Unmodified | 1260400 | 0 | 0 | 0 | 703920 | 0 | 1115600 | 0 | 1542800 | 542540 | 558620 | 2674800 | 820680 | 0 | 0 | 0 | 0 | 0 |
| PGIVELPTL | NDUFA8 | P51970 | NDUA8_HUMA | Unmodified | 0 | 0 | 0 | 0 | 0 | 0 | 772120 | 0 | 0 | 100820 | 0 | 0 | 151730 | 387600 | 0 | 0 | 50766 | 0 |
| PGMVVTFA | EEF1A1P5; EEF | Q5VTE0; P68 | EF1A3_HUMAN | Oxidation (M) | 0 | 5183000 | 974340 | 0 | 594990 | 0 | 332630 | 490640 | 802510 | 0 | 308950 | 902070 | 0 | 0 | 1711400 | 641630 | 1760500 | 455640 |
| PGNQNTQ' | PA2G4; PA2G4; | Q9UQ80; F8 | PA2G4_HUMAN | Unmodified | 38878 | 39973 | 0 | 0 | 0 | 0 | 0 | 0 | 0 | 0 | 0 | 663610 | 0 | 0 | 0 | 0 | 0 | 0 |
| PGPALWPL | HEXB | P07686 | HEXB_HUMAN | Unmodified | 0 | 0 | 717120 | 0 | 0 | 0 | 0 | 0 | 7937200 | 0 | 429050 | 0 | 546970 | 0 | 872380 | 0 | 304630 | 0 |
| PGPGAGP( | PDCD6; PDCD6 | O75340; Q8 | PDCD6_HUMA | Unmodified | 0 | 0 | 0 | 0 | 0 | 0 | 0 | 0 | 632140 | 0 | 0 | 0 | 0 | 0 | 0 | 0 | 0 | 0 |
| PGPTPSGT | SEC61B | P60468 | SC61B_HUMAI | Unmodified | 0 | 0 | 0 | 0 | 0 | 0 | 0 | 0 | 137420 | 0 | 364730 | 459910 | 0 | 0 | 0 | 0 | 0 | 0 |
| PGPTPSGT | SEC61B | P60468 | SC61B_HUMAI | Unmodified | 0 | 0 | 0 | 0 | 947250 | 105050 | 0 | 0 | 0 | 0 | 0 | 0 | 2064400 | 838150 | 0 | 0 | 0 | 931170 |
| PGQFIHTN | PGD | P52209 | 6PGD_HUMAN | Unmodified | 176190 | 0 | 0 | 1170300 | 1904800 | 428470 | 0 | 0 | 0 | 0 | 0 | 11519000 | 274020 | 0 | 1413700 | 707700 | 0 | 0 |
| PGRLDQLI | VCP; VCP; VCP; | P55072; C9J | TERA_HUMAN | Unmodified | 46719 | 849020 | 0 | 0 | 304560 | 193390 | 136780 | 0 | 145030 | 667530 | 15877 | 1456400 | 84066 | 0 | 0 | 50229 | 547210 | 0 |
| PGSAAKGS | EIF2B2; EIF2B2 | P49770; G3\ | EI2BB_HUMAN | Unmodified | 0 | 0 | 0 | 0 | 12081 | 945880 | 296680 | 0 | 0 | 0 | 0 | 0 | 514750 | 0 | 0 | 0 | 650340 | 375900 |
| PGSLPLNA | HMGCS1; HMG | Q01581; D6I | HMCS1_HUMA | Unmodified | 0 | 0 | 0 | 0 | 884140 | 0 | 0 | 175380 | 1022900 | 418740 | 0 | 4967800 | 0 | 0 | 0 | 0 | 0 | 0 |
| PGTEQYLL | SNX9; SNX9; SN | Q9Y5X1; A0A | SNX9_HUMAN | Unmodified | 0 | 1651800 | 1152600 | 0 | 0 | 0 | 0 | 0 | 0 | 543920 | 147590 | 0 | 0 | 0 | 612450 | 0 | 0 | 0 |
| PGVPMEIVI | HYOU1; HYOU1 | Q9Y4L1; J3Q | HYOU1_HUMA | Unmodified | 167520 | 0 | 0 | 0 | 0 | 0 | 0 | 0 | 1068600 | 0 | 0 | 146390 | 0 | 0 | 0 | 0 | 0 | 0 |
| PGVTVKDV | RPS19; RPS19; | P39019; M0( | RS19_HUMAN | Unmodified | 0 | 0 | 0 | 0 | 463260 | 0 | 298160 | 239860 | 0 | 363990 | 0 | 0 | 1753200 | 420510 | 0 | 0 | 199770 | 389710 |
| PGVTVKDV | RPS19; RPS19; | P39019; M0( | RS19_HUMAN | Unmodified | 0 | 0 | 0 | 0 | 1556800 | 0 | 3119900 | 0 | 0 | 438590 | 0 | 0 | 4167900 | 1871600 | 0 | 0 | 3856400 | 1635100 |
| PGYHDIITII | ADAMTS1; ADA | Q9UHI8; E5F | ATS1_HUMAN | Unmodified | 0 | 0 | 169940 | 0 | 0 | 0 | 0 | 0 | 0 | 0 | 0 | 0 | 0 | 0 | 0 | 0 | 0 | 0 |
| PGYLEATVI | PPA1 | Q15181 | IPYR_HUMAN I | Unmodified | 0 | 0 | 0 | 0 | 0 | 0 | 0 | 0 | 0 | 0 | 40848 | 1052200 | 0 | 0 | 0 | 0 | 0 | 0 |
| PHLMSQEL | P4HB; P4HB; P4 | P07237; A0A | PDIA1_HUMAN | Unmodified | 0 | 0 | 0 | 0 | 0 | 0 | 0 | 0 | 818720 | 1072200 | 0 | 0 | 0 | 0 | 0 | 0 | 0 | 0 |
| PHSYPALS | ALDOC; ALDOC | P09972; J3Q | ALDOC_HUMA | Unmodified | 0 | 0 | 0 | 0 | 0 | 0 | 124250 | 104760 | 0 | 22658 | 91507 | 1110500 | 0 | 572640 | 0 | 0 | 449190 | 498050 |
| PHSYPALS | ALDOC; ALDOC | P09972; J3Q | ALDOC_HUMA | Unmodified | 0 | 0 | 0 | 0 | 0 | 0 | 0 | 0 | 0 | 0 | 0 | 0 | 176030 | 0 | 0 | 0 | 284940 | 400730 |
| PIDKKAPDI | RDX; EZR; MSN | P35241; P15 | RADI_HUMAN I | Unmodified | 0 | 0 | 0 | 0 | 0 | 0 | 0 | 0 | 0 | 0 | 0 | 0 | 0 | 0 | 0 | 0 | 1052400 | 0 |
| PIDVQVITH | ACTR3; ACTR3 | P61158; B4D | ARP3_HUMAN | Unmodified | 0 | 0 | 0 | 0 | 0 | 0 | 0 | 0 | 0 | 0 | 0 | 889020 | 0 | 0 | 0 | 0 | 0 | 0 |
| PIEASIRGEI | ANXA6 | P08133 | ANXA6_HUMAN | Unmodified | 0 | 0 | 0 | 0 | 0 | 0 | 0 | 0 | 0 | 290180 | 0 | 0 | 0 | 0 | 0 | 0 | 0 | 0 |
| PIGALNPK | NBEA; LRBA; LR | Q8NFP9; P5( | NBEA_HUMAN | Unmodified | 0 | 0 | 0 | 0 | 8716400 | 0 | 2561400 | 2784700 | 0 | 0 | 208240 | 693750 | 0 | 7141000 | 3293200 | 0 | 798360 | 9463100 |
| PIHQGPDA | COPB1 | P53618 | COPB_HUMAN | Unmodified | 0 | 0 | 77287 | 0 | 137150 | 0 | 10980 | 0 | 0 | 0 | 0 | 418410 | 0 | 0 | 0 | 0 | 51799 | 0 |
| PIIHFGSDY | PRNP; PRNP; P | P04156; A2A | PRIO_HUMAN I | Unmodified | 189320 | 243200 | 92442 | 0 | 0 | 371930 | 0 | 0 | 0 | 0 | 0 | 0 | 0 | 0 | 488010 | 719880 | 0 | 569240 |
| PIPPPPPP | WIPF2; WIPF2; | Q8TF74; J3Q | WIPF2_HUMAN | Unmodified | 1166300 | 676030 | 1944200 | 3460600 | 0 | 0 | 0 | 0 | 0 | 0 | 0 | 0 | 1258400 | 0 | 3284400 | 1096300 | 0 | 0 |
| PISTLDNRT | DNAJA1 | P31689 | DNJA1_HUMAN | Unmodified | 0 | 0 | 0 | 0 | 779020 | 0 | 0 | 0 | 0 | 1106000 | 0 | 0 | 246210 | 0 | 0 | 0 | 0 | 0 |
| PIWQRPSK | HSP90B1; HSP | P14625; A0A | ENPL_HUMAN | Unmodified | 0 | 0 | 123530 | 0 | 0 | 0 | 187100 | 0 | 31498000 | 4147800 | 3331700 | 4255200 | 0 | 0 | 0 | 149990 | 0 | 0 |
| PIWTRNPD | HSP90AA1 | P07900 | HS90A_HUMAN | Unmodified | 96739 | 312080 | 772390 | 0 | 2500100 | 916370 | 609330 | 0 | 3391600 | 1010400 | 1543200 | 8272600 | 229250 | 0 | 805570 | 188550 | 1641200 | 296200 |
| PIWTRNPD | HSP90AB1 | P08238 | HS90B_HUMAN | Unmodified | 0 | 0 | 0 | 0 | 2621200 | 0 | 0 | 0 | 0 | 601820 | 979670 | 205360 | 1004000 | 0 | 2510900 | 1315700 | 0 | 0 |
| PKEADESLI | TLN1 | Q9Y490 | TLN1_HUMAN | Unmodified | 0 | 0 | 0 | 0 | 0 | 0 | 260970 | 0 | 0 | 0 | 0 | 0 | 0 | 0 | 0 | 0 | 0 | 0 |
| PKFADIVSII | EPHA2 | P29317 | EPHA2_HUMAN | Unmodified | 0 | 0 | 0 | 0 | 0 | 0 | 0 | 0 | 0 | 0 | 0 | 0 | 0 | 0 | 0 | 0 | 1066200 | 0 |
| PKHEFSVD | ATOX1 | O00244 | ATOX1_HUMAN | Unmodified | 731500 | 0 | 0 | 443200 | 464400 | 0 | 0 | 0 | 0 | 0 | 0 | 0 | 2141900 | 362270 | 603070 | 790340 | 637640 | 0 |
| PKHEFSVD | ATOX1 | O00244 | ATOX1_HUMAN | Unmodified | 0 | 0 | 0 | 0 | 0 | 0 | 0 | 0 | 0 | 0 | 0 | 0 | 2049300 | 1229800 | 0 | 0 | 1526000 | 1082200 |
| PKLPEDPLI | FASN; FASN | P49327; A0A | FAS_HUMAN F | Unmodified | 0 | 0 | 0 | 0 | 112440 | 0 | 0 | 0 | 0 | 0 | 185970 | 461580 | 0 | 0 | 0 | 0 | 142100 | 0 |
| PKMVMTVF | PLS3; PLS3 | P13797; A0A | PLST_HUMAN I | 4 Oxidation (M) | 6978100 | 0 | 1.44E+08 | 85518000 | 0 | 19634000 | 0 | 0 | 0 | 0 | 0 | 0 | 0 | 0 | 41284000 | 8922900 | 0 | 0 |
| PKSMENIM | DLG1; DLG1; D | Q12959; A0A | DLG1_HUMAN | 3 Oxidation (M) | 0 | 0 | 0 | 272740 | 0 | 0 | 0 | 0 | 0 | 0 | 0 | 0 | 0 | 0 | 0 | 0 | 0 | 0 |
| PKVQVEYK | HSPA8; HSPA2; | P11142; P54 | HSP7C_HUMAI | Unmodified | 0 | 274590 | 0 | 0 | 0 | 0 | 804580 | 0 | 0 | 1085700 | 0 | 0 | 42587 | 0 | 0 | 0 | 383700 | 0 |
| PLAAVTCTA | MVD; MVD; MVI | P53602; H3E | MVD1_HUMAN | Unmodified | 0 | 0 | 0 | 0 | 0 | 0 | 0 | 0 | 0 | 0 | 0 | 613830 | 0 | 0 | 0 | 0 | 0 | 0 |
| PLAEGPQV | CRIP2; CRIP2 | P52943; H0' | CRIP2_HUMAN | Unmodified | 0 | 636720 | 0 | 0 | 0 | 0 | 0 | 0 | 5359100 | 0 | 0 | 0 | 0 | 0 | 223240 | 127190 | 0 | 0 |
| PLAGTDDS | SLC39A10 | Q9ULF5 | S39AA_HUMAN | Unmodified | 0 | 0 | 419630 | 0 | 0 | 0 | 860920 | 0 | 0 | 0 | 0 | 1411300 | 0 | 571990 | 0 | 0 | 0 | 0 |
| PLAQLADP' | RPS6KA3 | P51812 | KS6A3_HUMAN | Unmodified | 0 | 0 | 0 | 0 | 0 | 0 | 0 | 0 | 0 | 0 | 0 | 0 | 0 | 0 | 0 | 0 | 0 | 1212500 |
| PLDETLK | ANXA1; ANXA1; | P04083; Q5I | ANXA1_HUMAN | Unmodified | 0 | 0 | 182430 | 0 | 0 | 0 | 16636 | 0 | 0 | 0 | 623970 | 1082000 | 0 | 65367 | 25755 | 0 | 0 | 62824 |

| | | | | | | | | | | | | | | | | | | | | | | |
|---|---|---|---|---|---|---|---|---|---|---|---|---|---|---|---|---|---|---|---|---|---|---|
| PLDLAQGA | OSBPL1A | Q9BXW6 | OSBL1_HUMAN | Oxidation (M) | 0 | 0 | 0 | 6949100 | 0 | 0 | 0 | 0 | 0 | 0 | 0 | 0 | 0 | 2422900 | 0 | 5137900 | 0 | 0 |
| PLDRNDGT | POGLUT3 | Q7Z4H8 | PLGT3_HUMAN | Unmodified | 0 | 0 | 0 | 0 | 211360 | 0 | 0 | 0 | 1381700 | 0 | 0 | 0 | 0 | 0 | 0 | 0 | 0 | 0 |
| PLDSQSIY( | SCG2 | P13521 | SCG2_HUMAN | Unmodified | 0 | 0 | 0 | 0 | 0 | 0 | 53183 | 0 | 1059500 | 91652 | 0 | 21552 | 0 | 103550 | 0 | 0 | 11552 | 0 |
| PLELELCP( | NANS | Q9NR45 | SIAS_HUMAN S | Unmodified | 151320 | 0 | 0 | 0 | 0 | 0 | 0 | 0 | 0 | 0 | 0 | 2929200 | 82104 | 0 | 303860 | 0 | 0 | 0 |
| PLENLEEE( | PSMD6; PSMD6 | Q15008; C9J | PSMD6_HUMA | Unmodified | 99811 | 0 | 569160 | 161170 | 0 | 0 | 0 | 0 | 540380 | 0 | 810670 | 5427700 | 203900 | 0 | 231830 | 302700 | 100340 | 0 |
| PLENLEEE( | PSMD6; PSMD6 | Q15008; C9J | PSMD6_HUMA | Unmodified | 235190 | 299070 | 0 | 0 | 0 | 0 | 0 | 0 | 1976800 | 0 | 2091900 | 6575500 | 0 | 0 | 0 | 272000 | 507080 | 0 |
| PLENLEEE( | PSMD6; PSMD6 | Q15008; C9J | PSMD6_HUMA | Unmodified | 0 | 0 | 0 | 0 | 0 | 0 | 847840 | 0 | 0 | 926020 | 0 | 0 | 3475400 | 2420900 | 0 | 0 | 469360 | 1659400 |
| PLEVELR | TMED10 | P49755 | TMEDA_HUMAI | Unmodified | 0 | 0 | 0 | 182940 | 0 | 0 | 0 | 0 | 1793700 | 171550 | 194140 | 1586800 | 0 | 0 | 0 | 0 | 0 | 0 |
| PLFATNPFI | STAM; STAM | Q92783; C9J | STAM1_HUMAI | Unmodified | 4476700 | 1744800 | 0 | 908400 | 6534300 | 930490 | 1986300 | 767460 | 9239500 | 2158500 | 0 | 585920 | 4328600 | 908230 | 614070 | 1124300 | 628230 | 1546900 |
| PLFTANPFI | STAM2 | O75886 | STAM2_HUMAI | Unmodified | 1065000 | 0 | 0 | 0 | 781970 | 0 | 477050 | 161070 | 786100 | 435740 | 0 | 0 | 0 | 0 | 556020 | 0 | 293340 | 1542200 |
| PLHELIMQI | PAFAH1B2; PAF | P68402; J3K | PA1B2_HUMAN | Oxidation (M) | 869050 | 283340 | 1321900 | 0 | 27577 | 31611 | 0 | 0 | 51344 | 0 | 129280 | 1955000 | 30318 | 0 | 1959900 | 273200 | 0 | 0 |
| PLHEVPSG | SH3KBP1; SH3I | Q96B97; Q5 | SH3K1_HUMAI | Unmodified | 0 | 0 | 695060 | 0 | 0 | 0 | 0 | 0 | 0 | 0 | 0 | 0 | 0 | 0 | 37020 | 0 | 0 | 0 |
| PLHPLLVV/ | ELAC2; ELAC2; | Q9BQ52; E7 | RNZ2_HUMAN | Unmodified | 0 | 0 | 0 | 0 | 0 | 0 | 456800 | 0 | 105530 | 0 | 0 | 524910 | 0 | 0 | 0 | 0 | 0 | 0 |
| PLILQSLGP | CD2AP | Q9Y5K6 | CD2AP_HUMAI | Unmodified | 3569400 | 2604500 | 20216000 | 2447400 | 0 | 0 | 0 | 0 | 0 | 0 | 111320 | 735060 | 0 | 0 | 15085000 | 6912700 | 0 | 0 |
| PLIPLQK | ERBIN | Q96RT1 | ERBIN_HUMAN | Unmodified | 0 | 0 | 445370 | 0 | 0 | 0 | 0 | 0 | 242080 | 0 | 511770 | 2320600 | 0 | 0 | 533840 | 237050 | 0 | 0 |
| PLIPLQKET | ERBIN | Q96RT1 | ERBIN_HUMAN | Unmodified | 0 | 0 | 0 | 151700 | 4056700 | 900360 | 0 | 0 | 0 | 0 | 0 | 0 | 0 | 0 | 0 | 0 | 0 | 782330 |
| PLLDHSYD | FTSJ1; FTSJ1 | Q9UET6; B7 | TRM7_HUMAN | Unmodified | 0 | 0 | 0 | 0 | 0 | 0 | 0 | 0 | 0 | 0 | 0 | 997330 | 0 | 0 | 0 | 0 | 0 | 0 |
| PLLESGTL( | UBA1 | P22314 | UBA1_HUMAN | Unmodified | 225540 | 231880 | 0 | 156900 | 0 | 0 | 0 | 0 | 566170 | 0 | 0 | 3188300 | 0 | 0 | 0 | 230400 | 51097 | 0 |
| PLLMVHGV | EPHX1 | P07099 | HYEP_HUMAN | Oxidation (M) | 0 | 0 | 290140 | 169070 | 0 | 0 | 0 | 0 | 343250 | 0 | 0 | 0 | 0 | 0 | 1180700 | 251770 | 0 | 0 |
| PLLMVHGV | EPHX1 | P07099 | HYEP_HUMAN | Unmodified | 0 | 0 | 0 | 0 | 0 | 0 | 0 | 0 | 126370 | 0 | 0 | 0 | 0 | 0 | 0 | 0 | 0 | 0 |
| PLLTGLMW | MOGS; MOGS; | Q13724; A0/ | MOGS_HUMAN | Unmodified | 0 | 0 | 0 | 0 | 0 | 0 | 0 | 0 | 0 | 0 | 0 | 0 | 0 | 0 | 0 | 0 | 0 | 0 |
| PLNIAYSHI | CYFIP2; CYFIP2 | Q96F07; H7( | CYFP2_HUMAN | Unmodified | 0 | 0 | 36300 | 0 | 0 | 0 | 0 | 0 | 0 | 0 | 0 | 1902300 | 0 | 0 | 63204 | 213650 | 0 | 0 |
| PLNNPWH | TMPRSS2; TMPI | O15393; C9J | TMPS2_HUMAI | Unmodified | 22071 | 17338 | 126650 | 245940 | 0 | 0 | 0 | 0 | 0 | 0 | 0 | 0 | 48873 | 19807 | 848210 | 240980 | 0 | 0 |
| PLNPVPGC | IGF2R; IGF2R | P11717; A0A | MPRI_HUMAN | Unmodified | 0 | 0 | 0 | 0 | 0 | 0 | 0 | 0 | 386530 | 324010 | 0 | 0 | 0 | 0 | 0 | 0 | 0 | 0 |
| PLPPSERLE | DDX3X; DDX3Y | O00571; O15 | DDX3X_HUMAI | Unmodified | 0 | 0 | 749960 | 0 | 0 | 0 | 0 | 0 | 0 | 0 | 0 | 0 | 0 | 0 | 1331800 | 0 | 0 | 0 |
| PLPVPPELA | BAIAP2; BAIAP2 | Q9UQB8; I3L | BAIP2_HUMAN | Unmodified | 0 | 0 | 1207100 | 497460 | 0 | 0 | 0 | 0 | 659730 | 0 | 57882 | 836710 | 0 | 0 | 2577100 | 2422200 | 0 | 0 |
| PLQQLEVP | PRSS8; PRSS8 | Q16651; H3I | PRSS8_HUMAN | Unmodified | 0 | 0 | 0 | 0 | 0 | 0 | 0 | 0 | 780680 | 0 | 0 | 0 | 0 | 0 | 0 | 0 | 0 | 0 |
| PLRAEGTN | PDCD6IP | Q8WUM4 | PDC6I_HUMAN | Unmodified | 105980 | 0 | 846320 | 0 | 3830600 | 0 | 97465 | 0 | 0 | 1375300 | 0 | 0 | 1436800 | 768090 | 0 | 0 | 789830 | 3731500 |
| PLRLPLQD | EEF1A2; EEF1A | Q05639; Q5 | EF1A2_HUMAN | Unmodified | 6873000 | 5994600 | 17027000 | 2257800 | 658990 | 0 | 0 | 0 | 1464600 | 0 | 356420 | 6133300 | 447290 | 0 | 13617000 | 9302200 | 0 | 0 |
| PLSDFILAL | BCS1L; BCS1L; | Q9Y276; C9J | BCS1_HUMAN | Unmodified | 0 | 0 | 0 | 0 | 0 | 0 | 0 | 0 | 211750 | 43999 | 0 | 0 | 0 | 0 | 0 | 0 | 0 | 0 |
| PLSPLHSL | NOL9 | Q5SY16 | NOL9_HUMAN | Unmodified | 0 | 25989 | 59646 | 0 | 0 | 0 | 0 | 0 | 0 | 0 | 0 | 394520 | 0 | 0 | 0 | 0 | 0 | 0 |
| PLSSILGR | SHC1; SHC1; S | P29353; Q5I | SHC1_HUMAN | Unmodified | 0 | 37709 | 0 | 0 | 0 | 0 | 89760 | 0 | 511720 | 0 | 0 | 681430 | 377650 | 0 | 0 | 24399 | 0 | 0 |
| PLSSLTPLI | PAK2; PAK2 | Q13177; H7( | PAK2_HUMAN | Oxidation (M) | 0 | 0 | 0 | 0 | 0 | 0 | 0 | 0 | 0 | 0 | 0 | 8155200 | 0 | 0 | 0 | 0 | 0 | 0 |
| PLTAATLFK | MYO1D; MYO1I | O94832; J3Q | MYO1D_HUMA | Unmodified | 0 | 455170 | 0 | 0 | 0 | 0 | 0 | 46806 | 0 | 0 | 0 | 641180 | 0 | 0 | 0 | 38128 | 41980 | 0 |
| PLTVWRDF | BTN2A1 | Q7KYR7 | BT2A1_HUMAN | Unmodified | 0 | 0 | 652650 | 628490 | 0 | 0 | 0 | 0 | 722250 | 0 | 0 | 2333200 | 0 | 0 | 876770 | 605710 | 175880 | 0 |
| PLYEGLGS | ACTR10; ACTR1 | Q9NZ32; F6 | ARP10_HUMAN | Unmodified | 0 | 55233 | 0 | 0 | 0 | 0 | 0 | 0 | 0 | 0 | 0 | 1846400 | 794240 | 363260 | 0 | 98500 | 0 | 0 |
| PLYFADRL | ANXA2; ANXA2; | P07355; H0 | ANXA2_HUMAN | Unmodified | 0 | 0 | 0 | 0 | 783230 | 0 | 0 | 0 | 0 | 0 | 0 | 0 | 0 | 0 | 0 | 0 | 167640 | 171960 |
| PLYHHFTT/ | GNA13 | Q14344 | GNA13_HUMAI | Unmodified | 136510 | 164900 | 171310 | 393660 | 435900 | 213770 | 0 | 0 | 0 | 0 | 820640 | 6066000 | 0 | 0 | 0 | 149720 | 0 | 631790 |
| PLYSVTVK | USP14; USP14 | P54578; A6N | UBP14_HUMAN | Unmodified | 0 | 0 | 0 | 0 | 0 | 57262 | 0 | 15200 | 438280 | 0 | 450570 | 6642900 | 0 | 101010 | 0 | 0 | 178860 | 163620 |
| PLYVALAQI | PABPC4; PABP( | Q13310; P11 | PABP4_HUMAN | Unmodified | 6709200 | 5433500 | 26734000 | 0 | 0 | 0 | 0 | 0 | 0 | 0 | 0 | 0 | 0 | 0 | 5257300 | 3188300 | 0 | 0 |
| PMCIPPSY/ | VDAC2; VDAC2 | P45880; Q5J | VDAC2_HUMAI | Unmodified | 0 | 0 | 0 | 0 | 1434200 | 14565000 | 377520 | 440870 | 0 | 675000 | 870070 | 270910 | 0 | 0 | 0 | 0 | 0 | 99252 |
| PMFIVNTN | MIF | P14174 | MIF_HUMAN M | Oxidation (M) | 10611000 | 8126500 | 11542000 | 14271000 | 9171200 | 3947300 | 844590 | 776930 | 8398600 | 4039500 | 20043000 | 69605000 | 17677000 | 5415100 | 22981000 | 26319000 | 6778500 | 6801500 |
| PMFIVNTN | MIF | P14174 | MIF_HUMAN M | Unmodified | 1108200 | 1085300 | 1302700 | 2575000 | 4791100 | 625530 | 341160 | 0 | 15248000 | 1969700 | 4022200 | 16406000 | 10899000 | 2547800 | 2134900 | 1054900 | 238220 | 3442900 |
| PMMIITEYM | EPHA2 | P29317 | EPHA2_HUMAI | 3 Oxidation (M) | 0 | 0 | 147540 | 0 | 0 | 456150 | 0 | 45811 | 0 | 0 | 0 | 0 | 0 | 0 | 0 | 0 | 0 | 0 |
| PMQFLGDI | PGAM1; PGAM2 | P18669; P15 | PGAM1_HUMA | Oxidation (M) | 0 | 0 | 0 | 0 | 0 | 0 | 0 | 0 | 0 | 0 | 0 | 977720 | 0 | 0 | 0 | 0 | 0 | 0 |
| PMQFLGDI | PGAM1; PGAM2 | P18669; P15 | PGAM1_HUMA | Unmodified | 0 | 0 | 291240 | 0 | 0 | 0 | 0 | 0 | 0 | 0 | 0 | 1762200 | 0 | 0 | 0 | 55281 | 0 | 0 |
| PMVYLVNL | OLA1; OLA1; OL | Q9NTK5; J3K | OLA1_HUMAN | Oxidation (M) | 0 | 0 | 494770 | 0 | 0 | 0 | 0 | 0 | 0 | 0 | 260340 | 3252200 | 0 | 0 | 212680 | 174690 | 0 | 0 |
| PNAAIRDC | ST13P4; ST13; S | Q8IZP2; P50 | ST134_HUMAN | Unmodified | 240670 | 0 | 319320 | 361740 | 0 | 0 | 0 | 0 | 0 | 0 | 557240 | 0 | 1043200 | 0 | 226840 | 0 | 0 | 0 |
| PNFAGTWI | CRABP1 | P29762 | RABP1_HUMAN | Unmodified | 0 | 0 | 0 | 0 | 0 | 0 | 0 | 0 | 771160 | 853190 | 0 | 0 | 0 | 0 | 0 | 0 | 159790 | 3326300 |
| PNFSGNW | CRABP2; CRAB | P29373; Q5 | RABP2_HUMAN | Unmodified | 0 | 0 | 0 | 0 | 5772800 | 552750 | 1717200 | 842710 | 2739400 | 2676300 | 0 | 0 | 0 | 0 | 0 | 0 | 20771000 | 7482800 |
| PNFSGNW | CRABP2; CRAB | P29373; Q5 | RABP2_HUMAN | Unmodified | 0 | 0 | 0 | 0 | 0 | 0 | 97553 | 0 | 0 | 0 | 0 | 0 | 0 | 0 | 0 | 0 | 228820 | 0 |
| PNFSGNW | CRABP2; CRAB | P29373; Q5 | RABP2_HUMAN | Unmodified | 0 | 0 | 0 | 0 | 0 | 509540 | 2291900 | 0 | 0 | 0 | 0 | 0 | 0 | 0 | 0 | 0 | 15432000 | 0 |
| PNGGNILV | ITCH; ITCH; ITC | Q96J02; A0A | ITCH_HUMAN | Unmodified | 0 | 179380 | 0 | 0 | 0 | 0 | 0 | 0 | 0 | 0 | 0 | 0 | 0 | 0 | 0 | 0 | 0 | 0 |
| PNIAIVEMK | KIAA0586; KIAA | Q9BVV6; A0/ | TALD3_HUMAN | Oxidation (M) | 0 | 3503800 | 0 | 0 | 0 | 0 | 0 | 0 | 0 | 0 | 0 | 0 | 0 | 0 | 0 | 4715200 | 0 | 0 |
| PNIVLFSGS | PRPS2; PRPS2; | P11908; D3Y | PRPS2_HUMAN | Unmodified | 657260 | 89066 | 0 | 1341600 | 0 | 0 | 0 | 0 | 3494700 | 577760 | 1905400 | 8133500 | 1733200 | 0 | 0 | 814090 | 271640 | 0 |
| PNIVLFSGS | PRPS2; PRPS2; | P11908; D3Y | PRPS2_HUMAN | Unmodified | 0 | 0 | 0 | 0 | 11140000 | 0 | 7748700 | 0 | 0 | 2047200 | 0 | 0 | 29676000 | 0 | 0 | 0 | 0 | 6589700 |
| PNNLSLVV | SORD; SORD; S | Q00796; H0Y | DHSO_HUMAN | Unmodified | 0 | 0 | 534970 | 102660 | 0 | 0 | 100910 | 0 | 0 | 0 | 0 | 1030600 | 0 | 0 | 666260 | 370240 | 0 | 0 |
| PNPSSTSS | SKAP2 | O75563 | SKAP2_HUMAN | Unmodified | 149740 | 0 | 0 | 0 | 0 | 0 | 0 | 0 | 0 | 0 | 0 | 543780 | 78248 | 0 | 0 | 0 | 0 | 0 |
| PPPQGDVT | IRGQ; IRGQ | Q8WZA9; M( | IRGQ_HUMAN | Unmodified | 0 | 0 | 65695 | 129530 | 1310700 | 169000 | 1097400 | 736060 | 563040 | 304280 | 889420 | 2935500 | 1251700 | 0 | 132330 | 0 | 1148600 | 1127900 |
| PPVATEGE | DHX29; DHX29 | Q7Z478; A0/ | DHX29_HUMAN | Unmodified | 0 | 0 | 0 | 0 | 166810 | 0 | 0 | 0 | 0 | 0 | 205420 | 0 | 0 | 0 | 0 | 0 | 0 | 1259300 |
| PQEAAVAP | MARCKS | P29966 | MARCS_HUMA | Unmodified | 0 | 0 | 0 | 0 | 137950 | 0 | 0 | 0 | 8453600 | 1879000 | 89936 | 0 | 0 | 0 | 0 | 0 | 37152 | 0 |
| PQPQFGRI | PCBP3; PCBP3 | P57721; J3Q | PCBP3_HUMAI | Acetyl (Protein N | 0 | 0 | 0 | 0 | 0 | 0 | 0 | 0 | 0 | 0 | 0 | 0 | 0 | 0 | 0 | 0 | 0 | 0 |
| PQTTTEAK/ | ATP13A3; ATP1 | Q9H7F0; A0/ | AT133_HUMAN | Unmodified | 0 | 0 | 131640 | 0 | 0 | 0 | 0 | 0 | 0 | 0 | 0 | 0 | 0 | 0 | 0 | 0 | 0 | 0 |
| PQYQTWEI | SRP9; SRP9; SR | P49458; Q6 | SRP09_HUMAN | Unmodified | 6810600 | 6343300 | 5939400 | 2167500 | 166230 | 0 | 0 | 0 | 687220 | 0 | 1725900 | 10766000 | 364570 | 467740 | 20648000 | 19058000 | 10520 | 0 |
| PQYQTWEI | SRP9; SRP9; SR | P49458; Q6 | SRP09_HUMAN | Unmodified | 1078500 | 407150 | 1399300 | 5714100 | 4920800 | 836430 | 4969000 | 1277500 | 219290 | 1810900 | 0 | 0 | 5007800 | 2153700 | 2030100 | 943010 | 2949500 | 1806600 |
| PQYQTWEI | SRP9; SRP9; SR | P49458; Q6 | SRP09_HUMAN | Oxidation (M) | 0 | 0 | 0 | 143220 | 1152000 | 0 | 1309100 | 0 | 0 | 0 | 0 | 0 | 649260 | 326560 | 0 | 0 | 591590 | 441590 |
| PRAEDSGE | NPTN; NPTN | Q9Y639; Q9I | NPTN_HUMAN | Unmodified | 0 | 0 | 0 | 0 | 0 | 0 | 0 | 0 | 0 | 0 | 0 | 394320 | 0 | 0 | 0 | 57520 | 0 | 0 |
| PRDQAVFP | MVP | Q14764 | MVP_HUMAN | Unmodified | 0 | 0 | 0 | 0 | 0 | 0 | 0 | 0 | 0 | 0 | 0 | 1042000 | 0 | 0 | 0 | 0 | 0 | 0 |
| PREIMDAA | DARS1; DARS1 | P14868; H7( | SYDC_HUMAN | Unmodified | 0 | 0 | 142370 | 0 | 0 | 0 | 0 | 0 | 141430 | 0 | 0 | 727910 | 0 | 0 | 203160 | 0 | 0 | 0 |
| PREPCGG( | HAPLN1; HAPL | P10915; D6 | HPLN1_HUMAN | Unmodified | 0 | 0 | 0 | 0 | 0 | 0 | 0 | 0 | 344280 | 0 | 0 | 0 | 0 | 0 | 0 | 0 | 0 | 0 |
| PREPFDLG | SH3GL1 | Q99961 | SH3G1_HUMAI | Unmodified | 13330000 | 7283700 | 9669800 | 7841500 | 0 | 0 | 0 | 0 | 0 | 0 | 194110 | 0 | 2141200 | 0 | 25749000 | 14025000 | 0 | 0 |
| PRHQEGEI | RPL6 | Q02878 | RL6_HUMAN 6 | Unmodified | 0 | 0 | 0 | 0 | 304920 | 1453300 | 0 | 0 | 132560 | 55162 | 0 | 759440 | 0 | 0 | 304230 | 0 | 0 | 0 |
| PRLGQEAT | ACLY; ACLY | P53396; K7E | ACLY_HUMAN | Unmodified | 0 | 0 | 0 | 0 | 0 | 0 | 0 | 15333 | 0 | 0 | 0 | 326650 | 0 | 0 | 0 | 0 | 0 | 0 |
| PRLLEPLD | DOCK10; DOCK | Q96BY6; A0/ | DOC10_HUMA | Unmodified | 0 | 0 | 0 | 0 | 1037000 | 0 | 0 | 0 | 0 | 0 | 0 | 0 | 0 | 0 | 0 | 0 | 0 | 0 |
| PSADLTNS | MARK2; MARK2 | Q7KZI7; E7E | MARK2_HUMAN | Unmodified | 0 | 0 | 0 | 0 | 42802 | 0 | 0 | 0 | 388090 | 0 | 107690 | 284460 | 0 | 0 | 212950 | 0 | 59651 | 0 |
| PSASLSQLI | CGN | Q9P2M7 | CING_HUMAN | Unmodified | 0 | 0 | 0 | 0 | 0 | 0 | 0 | 0 | 0 | 0 | 109750 | 1297000 | 0 | 0 | 0 | 0 | 0 | 0 |
| PSDTIENV | UBA52; RPS27A | P62987; P62 | RL40_HUMAN | Unmodified | 3883600 | 4775100 | 1666900 | 1653600 | 178800 | 0 | 0 | 0 | 0 | 541470 | 0 | 0 | 0 | 0 | 4179500 | 2914800 | 96680 | 0 |
| PSDVMLVF | ATXN2L; ATXN2 | Q8WWM7; F | ATX2L_HUMAN | Unmodified | 0 | 0 | 0 | 0 | 0 | 0 | 0 | 0 | 217000 | 0 | 0 | 0 | 0 | 0 | 0 | 0 | 0 | 0 |
| PSEGPLQT | CLTC; CLTC | Q00610; A0/ | CLH1_HUMAN | Unmodified | 107670 | 230660 | 0 | 0 | 0 | 0 | 50837 | 0 | 0 | 76767 | 0 | 627830 | 0 | 0 | 50843 | 72775 | 0 | 0 |
| PSENLGQV | TM9SF2 | Q99805 | TM9S2_HUMAI | Unmodified | 33957 | 11835 | 0 | 0 | 0 | 0 | 0 | 0 | 0 | 0 | 0 | 470720 | 0 | 0 | 13088 | 24581 | 0 | 0 |
| PSENLGQV | TM9SF2 | Q99805 | TM9S2_HUMAI | Unmodified | 0 | 0 | 0 | 277910 | 0 | 0 | 711500 | 32629 | 5669700 | 0 | 0 | 0 | 0 | 0 | 0 | 0 | 0 | 0 |
| PSETPQAE | CBS; CBSL; CB | P35520; P0D | CBS_HUMAN C | Unmodified | 0 | 0 | 0 | 158860 | 0 | 0 | 0 | 0 | 861290 | 0 | 1354000 | 0 | 0 | 0 | 0 | 322570 | 236690 | 0 |
| PSFDEALQ | SLC22A3 | O75751 | S22A3_HUMAN | Unmodified | 0 | 0 | 403250 | 0 | 574320 | 0 | 0 | 0 | 0 | 0 | 10521000 | 0 | 0 | 0 | 0 | 2935200 | 550130 | 470770 |
| PSGLGSAL | N/A | N/A | N/A | Unmodified | 4801500 | 0 | 0 | 4539300 | 0 | 0 | 0 | 0 | 0 | 0 | 0 | 0 | 548310 | 0 | 0 | 0 | 0 | 0 |
| PSIQALIHG | PSMD14 | O00487 | PSDE_HUMAN | Unmodified | 0 | 0 | 0 | 0 | 0 | 0 | 0 | 0 | 0 | 0 | 0 | 224230 | 0 | 0 | 0 | 0 | 0 | 0 |
| PSKEVEEDI | HSP90B1; HSP9 | P14625; A0A | ENPL_HUMAN | Unmodified | 0 | 1355800 | 54201 | 0 | 0 | 0 | 0 | 0 | 226430 | 1299000 | 0 | 0 | 0 | 0 | 0 | 0 | 0 | 0 |
| PSKEVEEDI | HSP90B1; HSP9 | P14625; A0A | ENPL_HUMAN | Unmodified | 0 | 0 | 0 | 0 | 0 | 0 | 83626 | 54341 | 410640 | 1579600 | 0 | 237460 | 0 | 0 | 0 | 0 | 46141 | 0 |
| PSKGPLQS | RPS16; RPS16; | P62249; M0 | RS16_HUMAN | Unmodified | 0 | 0 | 0 | 0 | 913960 | 191750 | 0 | 1056900 | 0 | 830920 | 0 | 0 | 5279800 | 1706100 | 0 | 0 | 824660 | 1119700 |
| PSLWDRFS | DDIT4 | Q9NX09 | DDIT4_HUMAN | Unmodified | 0 | 554970 | 1109900 | 0 | 1673300 | 0 | 0 | 450470 | 0 | 0 | 0 | 4191500 | 0 | 0 | 160200 | 134630 | 413390 | 284750 |
| PSMTLEPN | CD276; CD276 | Q5ZPR3; H0 | CD276_HUMAI | Unmodified | 475830 | 319950 | 3685900 | 356300 | 0 | 0 | 22623 | 0 | 304910 | 0 | 0 | 720200 | 92494 | 0 | 2522000 | 1663900 | 0 | 0 |
| PSMTLEPN | CD276; CD276 | Q5ZPR3; H0 | CD276_HUMAI | Oxidation (M) | 0 | 0 | 945680 | 86198 | 0 | 0 | 79787 | 0 | 0 | 0 | 0 | 1071000 | 0 | 105460 | 592260 | 735840 | 0 | 0 |
| PSNLLLDQ | EPHA3; EPHA3 | P29320; C9J | EPHA3_HUMAI | Unmodified | 244810 | 210950 | 0 | 0 | 0 | 0 | 0 | 39845 | 0 | 0 | 0 | 0 | 0 | 0 | 0 | 0 | 0 | 0 |
| PSPELKQQ | TLN1 | Q9Y490 | TLN1_HUMAN | Unmodified | 0 | 0 | 0 | 0 | 712390 | 153620 | 1420700 | 594950 | 0 | 32770 | 0 | 0 | 234470 | 957580 | 0 | 0 | 0 | 3457300 |
| PSPLLVGR | G3BP1; G3BP2; | Q13283; Q9I | G3BP1_HUMAN | Unmodified | 0 | 0 | 164190 | 185680 | 679170 | 883130 | 0 | 0 | 863990 | 75594 | 1875900 | 13586000 | 0 | 0 | 174110 | 120820 | 73970 | 0 |
| PSPLTLAP | COPB1 | P53618 | COPB_HUMAN | Unmodified | 0 | 0 | 696100 | 0 | 0 | 0 | 0 | 0 | 0 | 0 | 119860 | 3252500 | 0 | 0 | 0 | 0 | 152110 | 99912 |
| PSQMEHAI | S100A10 | P60903 | S10AA_HUMAN | 2 Oxidation (M) | 0 | 0 | 0 | 246630 | 12506000 | 0 | 0 | 0 | 0 | 0 | 0 | 0 | 146270 | 0 | 0 | 0 | 2957500 | 1341000 |
| PSQMEHAI | S100A10 | P60903 | S10AA_HUMAN | Oxidation (M) | 0 | 0 | 0 | 0 | 25790000 | 4693300 | 0 | 0 | 348010 | 0 | 0 | 0 | 0 | 0 | 0 | 0 | 4766500 | 3615600 |
| PSQMEHAI | S100A10 | P60903 | S10AA_HUMAN | Unmodified | 0 | 104050 | 0 | 0 | 21921000 | 716750 | 0 | 0 | 713710 | 0 | 0 | 0 | 0 | 0 | 0 | 0 | 1400100 | 4357000 |
| PSQMEHAI | S100A10 | P60903 | S10AA_HUMAN | 4 Oxidation (M) | 0 | 0 | 0 | 0 | 0 | 0 | 0 | 498890 | 0 | 0 | 7766900 | 0 | 0 | 0 | 0 | 0 | 0 | 0 |
| PSQVLQPL | ITGA5 | P08648 | ITA5_HUMAN | Unmodified | 0 | 0 | 148220 | 0 | 0 | 0 | 0 | 0 | 0 | 0 | 0 | 0 | 0 | 0 | 2345000 | 639560 | 0 | 0 |
| PSSALPVT/ | GGA3; GGA3; G | Q9NZ52; J3( | GGA3_HUMAN | Unmodified | 0 | 215690 | 7119500 | 591490 | 0 | 0 | 0 | 94137 | 0 | 0 | 0 | 0 | 0 | 0 | 3333000 | 889430 | 0 | 0 |
| PSSINYMV | SDCBP; SDCBP | O00560; G5I | SDCB1_HUMAI | Oxidation (M) | 5907600 | 4606800 | 9298700 | 1325000 | 0 | 0 | 0 | 0 | 3009900 | 0 | 0 | 9701200 | 0 | 659670 | 21990000 | 11703000 | 0 | 0 |
| PSSINYMV | SDCBP; SDCBP | O00560; G5I | SDCB1_HUMAI | Unmodified | 0 | 0 | 0 | 0 | 0 | 0 | 0 | 0 | 0 | 0 | 0 | 0 | 0 | 0 | 2629600 | 0 | 0 | 0 |
| PSSINYMV | SDCBP; SDCBP | O00560; G5I | SDCB1_HUMAI | Unmodified | 691640 | 0 | 0 | 0 | 0 | 0 | 0 | 0 | 2009800 | 0 | 0 | 831200 | 0 | 0 | 3616500 | 0 | 208610 | 0 |
| PSSINYMV | SDCBP; SDCBP | O00560; G5I | SDCB1_HUMAI | Unmodified | 848710 | 0 | 1316600 | 0 | 0 | 0 | 0 | 0 | 0 | 0 | 0 | 0 | 0 | 0 | 2506400 | 2156600 | 0 | 0 |
| PSSINYMV | SDCBP; SDCBP | O00560; G5I | SDCB1_HUMAI | Oxidation (M) | 0 | 0 | 0 | 0 | 544210 | 0 | 0 | 0 | 0 | 1470600 | 0 | 0 | 0 | 0 | 0 | 0 | 0 | 0 |
| PSSSPVIFA | PCBP2 | F8VRH0 | F8VRH0_HUM | Unmodified | 9265800 | 0 | 0 | 4069900 | 0 | 0 | 0 | 0 | 0 | 0 | 0 | 259040 | 0 | 0 | 0 | 4627300 | 0 | 0 |
| PSSSPVIFA | PCBP2; PCBP2 | Q15366; H3I | PCBP2_HUMAN | Unmodified | 959390 | 368630 | 1024400 | 94361 | 0 | 0 | 0 | 0 | 0 | 0 | 0 | 0 | 758710 | 0 | 512830 | 616850 | 174350 | 0 |
| PSSTLNVP | PACSIN2; PACS | Q9UNF0; A0 | PACN2_HUMAI | Unmodified | 0 | 0 | 1989900 | 0 | 0 | 0 | 0 | 0 | 0 | 0 | 0 | 0 | 0 | 0 | 902230 | 286080 | 0 | 0 |

| | | | | | | | | | | | | | | | | | | | | | | |
|---|---|---|---|---|---|---|---|---|---|---|---|---|---|---|---|---|---|---|---|---|---|---|
| PSSVPHGA | ISOC1; ISOC1 | Q96CN7; D6 | ISOC1_HUMAN | Unmodified | 0 | 0 | 0 | 0 | 0 | 0 | 0 | 0 | 0 | 0 | 0 | 1230200 | 0 | 0 | 57895 | 47669 | 0 | 0 |
| PSTDLLMLI | SGPL1 | O95470 | SGPL1_HUMAN | Oxidation (M) | 0 | 0 | 0 | 0 | 0 | 0 | 88587 | 38131 | 32342000 | 4763700 | 160000 | 0 | 0 | 0 | 1230500 | 0 | 0 | 0 |
| PSTDLLMLI | SGPL1 | O95470 | SGPL1_HUMAN | Unmodified | 0 | 0 | 0 | 0 | 0 | 0 | 0 | 0 | 27848000 | 1314700 | 27175 | 0 | 0 | 0 | 0 | 0 | 0 | 0 |
| PSTFAYPAF | PSMD1; PSMD1 | Q99460; A0A | PSMD1_HUMA | Unmodified | 0 | 0 | 0 | 0 | 0 | 0 | 0 | 0 | 0 | 0 | 0 | 595170 | 0 | 0 | 0 | 0 | 0 | 0 |
| PSTPLVSMS | N/A | N/A | N/A | Oxidation (M) | 0 | 0 | 2024600 | 29266000 | 0 | 0 | 3309100 | 199500 | 0 | 0 | 0 | 0 | 0 | 0 | 0 | 2284800 | 0 | 0 |
| PSVLLTLR | PEPD; PEPD; PE | P12955; A0A | PEPD_HUMAN | Unmodified | 0 | 0 | 0 | 0 | 283940 | 0 | 0 | 0 | 155290 | 176950 | 0 | 1753800 | 0 | 0 | 70279 | 0 | 190560 | 141680 |
| PSVYLSTPS | CD2AP | Q9Y5K6 | CD2AP_HUMA | Unmodified | 15166000 | 9764900 | 11031000 | 12898000 | 2725400 | 857580 | 0 | 0 | 923820 | 0 | 0 | 0 | 5077900 | 1601400 | 16930000 | 11792000 | 0 | 0 |
| PTDKPLRLI | EEF1A2; EEF1A | Q05639; Q5 | EF1A2_HUMAN | Unmodified | 1603800 | 7632000 | 0 | 0 | 798370 | 0 | 0 | 0 | 0 | 0 | 0 | 976800 | 46189 | 0 | 1075400 | 676650 | 1079200 | 0 |
| PTGDFDSK | EIF3G; EIF3G; E | O75821; K7E | EIF3G_HUMAN | Unmodified | 0 | 0 | 0 | 0 | 435170 | 0 | 0 | 0 | 299340 | 0 | 812240 | 1900300 | 0 | 0 | 0 | 0 | 0 | 548690 |
| PTGDFDSK | EIF3G; EIF3G; E | O75821; K7E | EIF3G_HUMAN | Unmodified | 0 | 0 | 0 | 1314600 | 0 | 0 | 641370 | 0 | 0 | 0 | 0 | 0 | 3873800 | 2983700 | 0 | 0 | 2651500 | 2279600 |
| PTTQQSPQ | VPS35; VPS35; | Q96QK1; A0 | VPS35_HUMAN | Unmodified | 0 | 0 | 0 | 0 | 316720 | 0 | 0 | 0 | 175670 | 0 | 0 | 835310 | 92538 | 0 | 0 | 0 | 14719 | 109380 |
| PTVEELYR | API5; API5; API5 | Q9BZZ5; E9F | API5_HUMAN A | Unmodified | 0 | 132960 | 0 | 0 | 0 | 0 | 0 | 0 | 0 | 0 | 1212900 | 2682600 | 0 | 0 | 0 | 0 | 0 | 587080 |
| PTVFSEVTD | ALDH1A3; ALDH | P47895; H0Y | AL1A3_HUMAN | Oxidation (M) | 0 | 0 | 0 | 0 | 0 | 0 | 0 | 0 | 0 | 0 | 0 | 2165400 | 0 | 0 | 0 | 0 | 0 | 0 |
| PVAVALDTI | PKM; PKM; PKM | P14618; H3E | KPYM_HUMAN | Unmodified | 0 | 0 | 0 | 0 | 1443400 | 0 | 246600 | 0 | 0 | 255440 | 0 | 0 | 45923 | 0 | 0 | 0 | 1989100 | 0 |
| PVAVGPYG | ROMO1 | P60602 | ROMO1_HUMA | Unmodified | 0 | 0 | 0 | 0 | 0 | 0 | 0 | 0 | 993820 | 0 | 0 | 0 | 0 | 1022100 | 0 | 0 | 0 | 0 |
| PVDLSKWS | PEBP1 | P30086 | PEBP1_HUMAN | Unmodified | 0 | 0 | 0 | 0 | 4054600 | 0 | 787140 | 0 | 0 | 3448300 | 0 | 0 | 2947000 | 4231000 | 0 | 0 | 45783000 | 11614000 |
| PVELSDRE | CCT4 | P50991 | TCPD_HUMAN | Unmodified | 0 | 0 | 0 | 0 | 0 | 0 | 0 | 2512400 | 0 | 0 | 0 | 0 | 0 | 0 | 0 | 0 | 0 | 0 |
| PVFHTRTIE | VCL; VCL | P18206; A0A | VINC_HUMAN | Oxidation (M) | 0 | 0 | 0 | 0 | 587910 | 0 | 0 | 0 | 0 | 0 | 0 | 0 | 0 | 0 | 0 | 0 | 2410700 | 1356300 |
| PVFTEYQAS | RHOBTB3; RHC | O94955; H0Y | RHBT3_HUMAN | Unmodified | 286400 | 0 | 0 | 0 | 0 | 0 | 0 | 0 | 1710800 | 706400 | 0 | 50770 | 0 | 0 | 0 | 0 | 0 | 0 |
| PVGGLLPLI | MAN1A1 | P33908 | MA1A1_HUMAN | Unmodified | 0 | 0 | 0 | 0 | 0 | 0 | 0 | 0 | 0 | 0 | 47364 | 135520 | 0 | 0 | 0 | 0 | 0 | 0 |
| PVGSLAGIC | BANF1 | O75531 | BAF_HUMAN B | Unmodified | 0 | 0 | 1042600 | 0 | 1102700 | 0 | 0 | 0 | 599770 | 169580 | 0 | 9299000 | 0 | 0 | 0 | 0 | 0 | 0 |
| PVHHGVN | RPL15; RPL15; | P61313; E7E | RL15_HUMAN | Unmodified | 0 | 0 | 0 | 0 | 140040 | 0 | 0 | 0 | 0 | 0 | 91680 | 179720 | 0 | 0 | 0 | 0 | 179790 | 0 |
| PVHLTAFLC | RPL3L; RPL3; R | Q92901; P39 | RL3L_HUMAN | Unmodified | 0 | 0 | 0 | 0 | 0 | 0 | 0 | 0 | 0 | 0 | 0 | 1189600 | 0 | 0 | 0 | 0 | 0 | 0 |
| PVIDNPNYI | CANX; CANX; C | P27824; A0A | CALX_HUMAN | Unmodified | 0 | 0 | 0 | 0 | 0 | 0 | 0 | 0 | 994360 | 682620 | 33224 | 0 | 0 | 0 | 0 | 0 | 0 | 0 |
| PVINIEDLTI | GNGT1; GNGT1 | P63211; C9J | GBG1_HUMAN | Unmodified | 0 | 0 | 0 | 0 | 0 | 0 | 0 | 0 | 915060 | 286260 | 0 | 0 | 0 | 0 | 0 | 0 | 0 | 0 |
| PVINIQK | ITGA6 | P23229 | ITA6_HUMAN I | Unmodified | 0 | 0 | 0 | 0 | 209650 | 441710 | 0 | 0 | 0 | 31187 | 0 | 0 | 0 | 0 | 0 | 0 | 1187600 | 707960 |
| PVINIVHK | ITGA3; ITGA3 | P26006; K7E | ITA3_HUMAN I | Unmodified | 0 | 0 | 0 | 0 | 3007500 | 831230 | 0 | 0 | 117660 | 0 | 0 | 145530 | 431350 | 236840 | 0 | 0 | 1073800 | 761790 |
| PVKPVEDE | PCDH1 | Q08174 | PCDH1_HUMA | Unmodified | 0 | 0 | 585500 | 84312 | 77505 | 0 | 0 | 0 | 278910 | 0 | 639600 | 3880200 | 0 | 201550 | 631230 | 459490 | 0 | 2432900 |
| PVLLQVAE | SLC25A13 | Q9UJS0 | CMC2_HUMAN | Unmodified | 0 | 56452000 | 0 | 1435600 | 0 | 62821 | 0 | 0 | 0 | 0 | 0 | 0 | 580700 | 0 | 8733800 | 5041100 | 0 | 0 |
| PVLQKYAEI | OGDH; OGDH; | Q02218; E9F | ODO1_HUMAN | Unmodified | 0 | 0 | 0 | 0 | 34998000 | 1.57E+08 | 16754000 | 0 | 0 | 2300900 | 0 | 0 | 0 | 80781000 | 0 | 0 | 7001000 | 75730000 |
| PVNLGLWI | RAC2; RAC3; R | P15153; P60 | RAC2_HUMAN | Unmodified | 1327500 | 2027300 | 499700 | 0 | 0 | 0 | 0 | 0 | 0 | 0 | 0 | 1140400 | 0 | 0 | 3685400 | 3098300 | 0 | 0 |
| PVQTVENK | CPD; CPD | O75976; J3Q | CBPD_HUMAN | Unmodified | 0 | 0 | 0 | 0 | 0 | 0 | 0 | 0 | 1031900 | 230010 | 0 | 0 | 0 | 0 | 0 | 0 | 0 | 0 |
| PVTALEYTF | ATP1B3; ATP1B | P54709; C9J | AT1B3_HUMAN | Unmodified | 0 | 0 | 7051700 | 0 | 0 | 0 | 0 | 168970 | 53896 | 0 | 102870 | 13024000 | 79598 | 62990 | 6006900 | 3299200 | 0 | 0 |
| PVTALEYTF | ATP1B3; ATP1B | P54709; C9J | AT1B3_HUMAN | Unmodified | 0 | 0 | 480980 | 0 | 0 | 0 | 0 | 0 | 0 | 0 | 0 | 254760 | 0 | 0 | 1249300 | 777130 | 0 | 0 |
| PVTEKDLAI | FLNB | O75369 | FLNB_HUMAN | Unmodified | 0 | 0 | 0 | 0 | 0 | 259640 | 289190 | 0 | 1623500 | 0 | 0 | 2164500 | 0 | 0 | 6104700 | 0 | 1693900 | 0 |
| PVTHRKSD | PROSER2; PRO | Q86WR7; X6 | PRSR2_HUMAN | Unmodified | 0 | 0 | 0 | 0 | 419300 | 70355 | 0 | 0 | 0 | 0 | 0 | 0 | 0 | 0 | 0 | 0 | 0 | 0 |
| PVTLLASLF | TRNT1 | Q96Q11 | TRNT1_HUMAN | Unmodified | 0 | 0 | 0 | 0 | 0 | 0 | 0 | 0 | 44385 | 10717 | 17472 | 410550 | 0 | 0 | 16221 | 9178.4 | 0 | 0 |
| PVWGGGN | CSRP2; CSRP2; | Q16527; F8V | CSRP2_HUMA | Unmodified | 337480 | 333500 | 0 | 0 | 729420 | 69104 | 288170 | 285350 | 114220 | 190240 | 250180 | 3674100 | 0 | 1086600 | 0 | 0 | 447310 | 1006500 |
| PWLSNESI | SMPDL3B; SMP | Q92485; Q5 | ASM3B_HUMA | Unmodified | 0 | 0 | 234090 | 0 | 0 | 0 | 0 | 0 | 0 | 0 | 0 | 0 | 0 | 13241 | 0 | 73847 | 0 | 0 |
| PYANQPTV | POLR2C; POLR | P19387; H3E | RPB3_HUMAN | Unmodified | 56800000 | 51166000 | 15259000 | 13598000 | 0 | 0 | 1392400 | 398000 | 0 | 0 | 0 | 558620 | 13279000 | 761820 | 12838000 | 19503000 | 0 | 0 |
| PYDYNGPF | PDIA4; PDIA4 | P13667; A0A | PDIA4_HUMAN | Unmodified | 0 | 0 | 0 | 0 | 0 | 0 | 30477 | 0 | 363530 | 236060 | 0 | 0 | 0 | 0 | 0 | 0 | 0 | 0 |
| PYFLEAYRF | VCP; VCP; VCP; | P55072; C9J | TERA_HUMAN | Unmodified | 361090 | 432530 | 508650 | 613740 | 0 | 0 | 0 | 17830 | 2283600 | 0 | 804390 | 14867000 | 1962700 | 0 | 777940 | 569040 | 0 | 0 |
| PYISNTLPS | COBLL1; COBL | Q53SF7; A0 | COBL1_HUMA | Unmodified | 239870 | 0 | 866640 | 0 | 0 | 0 | 0 | 0 | 0 | 0 | 0 | 0 | 0 | 0 | 0 | 0 | 84978 | 0 |
| PYISNTLPS | COBLL1; COBL | Q53SF7; A0 | COBL1_HUMA | Unmodified | 253590 | 0 | 283230 | 0 | 0 | 0 | 0 | 0 | 0 | 0 | 0 | 0 | 59312 | 0 | 225380 | 0 | 0 | 0 |
| PYLTPNDR | ANPEP | P15144 | AMPN_HUMAN | Unmodified | 0 | 0 | 0 | 0 | 0 | 0 | 0 | 0 | 0 | 0 | 0 | 295970 | 0 | 0 | 0 | 0 | 285950 | 0 |
| PYQGVGSS | LMAN2; LMAN2 | Q12907; D6I | LMAN2_HUMA | 2 Oxidation (M) | 0 | 0 | 0 | 0 | 0 | 0 | 0 | 0 | 4164100 | 0 | 0 | 0 | 0 | 0 | 0 | 0 | 0 | 1382800 |
| PYQGVGSS | LMAN2; LMAN2 | Q12907; D6I | LMAN2_HUMA | Oxidation (M) | 0 | 0 | 0 | 0 | 0 | 0 | 0 | 0 | 4714900 | 0 | 0 | 0 | 0 | 0 | 0 | 0 | 0 | 0 |
| PYQGVGSS | LMAN2; LMAN2 | Q12907; D6I | LMAN2_HUMA | Unmodified | 0 | 0 | 0 | 0 | 0 | 0 | 0 | 0 | 660000 | 0 | 0 | 0 | 0 | 0 | 0 | 0 | 0 | 0 |
| PYQGWMM | FN1 | P02751 | FINC_HUMAN | 2 Oxidation (M) | 0 | 0 | 684700 | 409400 | 0 | 0 | 0 | 0 | 0 | 0 | 0 | 0 | 0 | 0 | 0 | 408540 | 0 | 0 |
| PYQYPALTI | ALDOA; ALDOA | P04075; J3K | ALDOA_HUMA | Unmodified | 0 | 0 | 0 | 0 | 0 | 0 | 0 | 65660 | 0 | 0 | 716350 | 6760400 | 0 | 0 | 0 | 339240 | 0 | 0 |
| PYQYPALTI | ALDOA; ALDOA | P04075; J3K | ALDOA_HUMA | Unmodified | 2191300 | 1426500 | 2253900 | 4246200 | 1.45E+08 | 19965000 | 9305300 | 7365800 | 19498000 | 11238000 | 26719000 | 1.07E+08 | 14068000 | 8818800 | 3820600 | 4050300 | 40242000 | 35266000 |
| PYQYPALTI | ALDOA; ALDOA | P04075; J3K | ALDOA_HUMA | Unmodified | 0 | 0 | 0 | 0 | 9928700 | 1335100 | 1285700 | 0 | 0 | 1293400 | 0 | 0 | 1631300 | 1212600 | 0 | 0 | 6099700 | 2480000 |
| PYSAHDPL | PAN3; PAN3 | Q58A45; A0 | PAN3_HUMAN | Unmodified | 160470 | 0 | 37727 | 0 | 0 | 0 | 0 | 0 | 0 | 0 | 0 | 0 | 0 | 0 | 0 | 0 | 0 | 0 |
| PYSIPPCEI | CTSB; CTSB; C1 | P07858; E9P | CATB_HUMAN | Unmodified | 0 | 0 | 0 | 0 | 0 | 0 | 0 | 0 | 0 | 0 | 20742000 | 0 | 0 | 0 | 0 | 0 | 0 | 0 |
| PYTLEEQK | ATP1B3; ATP1B | P54709; C9J | AT1B3_HUMAN | Unmodified | 0 | 0 | 2992100 | 0 | 0 | 0 | 0 | 0 | 73990 | 0 | 114680 | 1824100 | 0 | 0 | 2344300 | 0 | 0 | 0 |
| PYTLMSMV | PSMB3; PSMB3 | P49720; A0A | PSB3_HUMAN | Oxidation (M) | 12974000 | 10372000 | 0 | 0 | 0 | 0 | 0 | 0 | 845770 | 0 | 0 | 0 | 2113200 | 0 | 852440 | 3591900 | 0 | 0 |
| PYYEIGER | CD46 | P15529 | MCP_HUMAN | Unmodified | 0 | 879100 | 0 | 0 | 0 | 0 | 0 | 0 | 145960 | 0 | 63657 | 0 | 143580 | 96002 | 551390 | 588260 | 15987 | 0 |
| PYYEIGERV | CD46 | P15529 | MCP_HUMAN | Unmodified | 9328500 | 5300600 | 7908400 | 2171100 | 1393700 | 248840 | 0 | 20936 | 4286500 | 149580 | 1212800 | 21559000 | 842240 | 259970 | 6899300 | 3769900 | 114450 | 1045300 |
| QAAASATQ | TLN1 | Q9Y490 | TLN1_HUMAN | Unmodified | 0 | 0 | 0 | 749910 | 19032000 | 5050800 | 954180 | 912230 | 2179200 | 1168000 | 986500 | 7879500 | 5431000 | 2486200 | 683840 | 545000 | 6539900 | 7494800 |
| QAADLTQE | B3GLCT | Q6Y288 | B3GLT_HUMAN | Unmodified | 0 | 0 | 0 | 0 | 0 | 0 | 0 | 0 | 2024300 | 1246300 | 0 | 0 | 0 | 0 | 0 | 0 | 0 | 0 |
| QAADMILLI | ATP1A1; ATP1A | P05023; P50 | AT1A1_HUMAN | Oxidation (M) | 0 | 0 | 0 | 0 | 0 | 0 | 0 | 0 | 0 | 0 | 0 | 0 | 0 | 0 | 502540 | 0 | 0 | 0 |
| QAADMILLI | ATP1A1; ATP1A | P05023; P50 | AT1A1_HUMAN | Oxidation (M) | 0 | 0 | 0 | 0 | 0 | 0 | 0 | 0 | 0 | 0 | 0 | 0 | 194930 | 0 | 774560 | 0 | 0 | 0 |
| QAADQMKI | RDX; RDX; RDX | P35241; A0A | RADI_HUMAN | Oxidation (M) | 0 | 0 | 0 | 0 | 2554500 | 783160 | 0 | 0 | 0 | 0 | 0 | 0 | 0 | 3282300 | 0 | 0 | 1520200 | 0 |
| QAADQMKI | RDX; RDX; RDX | P35241; A0A | RADI_HUMAN | Unmodified | 0 | 0 | 0 | 0 | 4037400 | 0 | 0 | 0 | 0 | 0 | 0 | 0 | 0 | 2731500 | 0 | 0 | 0 | 1108900 |
| QAAENEKK | SLC39A10; SLC | Q9ULF5; E7E | S39AA_HUMAN | Unmodified | 0 | 0 | 0 | 0 | 0 | 0 | 0 | 0 | 0 | 0 | 0 | 0 | 0 | 0 | 0 | 0 | 159520 | 0 |
| QAALAGNE | GOLM1 | Q8NBJ4 | GOLM1_HUMA | Unmodified | 0 | 0 | 0 | 0 | 0 | 0 | 0 | 0 | 884010 | 0 | 0 | 0 | 58588 | 0 | 0 | 0 | 0 | 0 |
| QAALAGNE | GOLM1 | Q8NBJ4 | GOLM1_HUMA | Unmodified | 0 | 0 | 0 | 0 | 0 | 0 | 0 | 0 | 12199000 | 0 | 0 | 0 | 0 | 0 | 0 | 0 | 0 | 0 |
| QAALEEEQ | PICALM; PICALI | Q13492; H0Y | PICAL_HUMAN | Unmodified | 202060 | 231410 | 0 | 63624 | 0 | 0 | 0 | 18816 | 239060 | 0 | 221560 | 577090 | 0 | 0 | 0 | 262320 | 0 | 0 |
| QAALEEEQ | PICALM; PICALI | Q13492; H0Y | PICAL_HUMAN | Unmodified | 0 | 0 | 0 | 0 | 0 | 0 | 0 | 20132 | 827100 | 473390 | 112010 | 0 | 3934100 | 1517400 | 0 | 0 | 0 | 0 |
| QAAPCVLF | VCP; VCP; VCP; | P55072; C9J | TERA_HUMAN | Unmodified | 220660 | 150630 | 46266 | 109560 | 0 | 0 | 0 | 0 | 411050 | 160960 | 129880 | 803230 | 142320 | 0 | 181700 | 100820 | 164010 | 0 |
| QAASGLVG | RUVBL1 | Q9Y265 | RUVB1_HUMA | Unmodified | 0 | 614800 | 0 | 0 | 0 | 0 | 0 | 0 | 0 | 0 | 278570 | 8664100 | 0 | 0 | 0 | 0 | 0 | 0 |
| QAASSLQC | HSPA9; HSPA9; | P38646; A0A | GRP75_HUMA | Unmodified | 601390 | 10814000 | 128990 | 555570 | 0 | 0 | 0 | 30052 | 4795500 | 137490 | 0 | 0 | 0 | 0 | 9658300 | 828470 | 0 | 0 |
| QAAVDHIN | IQGAP2; IQGAP | Q13576; D6I | IQGA2_HUMAN | Unmodified | 0 | 0 | 8544500 | 0 | 0 | 0 | 0 | 0 | 0 | 0 | 0 | 0 | 0 | 0 | 3527000 | 3279500 | 0 | 0 |
| QACGFEYT | CUL1; CUL1; C | Q13616; A0 | CUL1_HUMAN | Unmodified | 0 | 0 | 0 | 0 | 121990 | 0 | 0 | 0 | 0 | 0 | 0 | 375520 | 0 | 0 | 0 | 0 | 68146 | 109660 |
| QACTPMFF | SHMT2; SHMT2 | P34897; H0Y | GLYM_HUMAN | Unmodified | 0 | 0 | 0 | 0 | 0 | 0 | 0 | 0 | 2553200 | 221070 | 47291 | 222830 | 0 | 0 | 0 | 0 | 0 | 0 |
| QACVLMIK | MYH9 | P35579 | MYH9_HUMAN | Unmodified | 0 | 0 | 0 | 0 | 303640 | 55737 | 419850 | 0 | 129960 | 212570 | 0 | 482020 | 0 | 0 | 0 | 0 | 149090 | 105100 |
| QACVLMIK | MYH9 | P35579 | MYH9_HUMAN | Oxidation (M) | 0 | 0 | 0 | 236520 | 0 | 0 | 411810 | 250430 | 0 | 499480 | 0 | 1738200 | 0 | 0 | 272740 | 226620 | 0 | 0 |
| QADFEAHN | GMPS | P49915 | GUAA_HUMAN | Unmodified | 0 | 0 | 0 | 0 | 0 | 0 | 0 | 0 | 532110 | 0 | 0 | 4618300 | 0 | 0 | 0 | 0 | 0 | 0 |
| QADFVQTP | USP10; USP10 | Q14694; H3I | UBP10_HUMA | Unmodified | 0 | 0 | 0 | 0 | 0 | 0 | 0 | 0 | 0 | 0 | 264540 | 4503200 | 0 | 0 | 0 | 0 | 0 | 0 |
| QADGAASA | FARP1; FARP1; | Q9Y4F1; A0 | FARP1_HUMAN | Unmodified | 0 | 0 | 614600 | 350910 | 551620 | 0 | 0 | 0 | 2562600 | 664750 | 1024100 | 1861900 | 347360 | 0 | 775170 | 485340 | 0 | 1165800 |
| QADGAASA | FARP1; FARP1; | Q9Y4F1; A0 | FARP1_HUMAN | Unmodified | 0 | 0 | 0 | 0 | 0 | 0 | 0 | 0 | 468220 | 170060 | 1016400 | 0 | 0 | 0 | 0 | 0 | 0 | 0 |
| QADLYISEC | CCT6A; CCT6B | P40227; Q92 | TCPZ_HUMAN | Unmodified | 0 | 0 | 708030 | 971970 | 2107700 | 835970 | 0 | 0 | 3096600 | 0 | 1300900 | 16229000 | 8762900 | 2406400 | 698420 | 599730 | 1213800 | 2671400 |
| QADLYISEC | CCT6A | P40227 | TCPZ_HUMAN | Unmodified | 0 | 0 | 0 | 0 | 2758200 | 0 | 0 | 0 | 0 | 640590 | 0 | 0 | 4427000 | 647580 | 0 | 0 | 1598000 | 649990 |
| QADLYISEC | CCT6A | P40227 | TCPZ_HUMAN | Unmodified | 0 | 0 | 0 | 0 | 0 | 0 | 1501200 | 0 | 0 | 0 | 0 | 0 | 0 | 1544200 | 0 | 0 | 1724000 | 0 |
| QADNPHVA | HYOU1; HYOU1 | Q9Y4L1; J3Q | HYOU1_HUMA | Unmodified | 191740 | 0 | 124530 | 0 | 517550 | 0 | 51021 | 0 | 17055000 | 1481900 | 479350 | 2157500 | 0 | 0 | 664090 | 350330 | 859500 | 0 |
| QADNPHVA | HYOU1; HYOU1 | Q9Y4L1; J3Q | HYOU1_HUMA | Unmodified | 0 | 0 | 0 | 36771 | 0 | 0 | 0 | 0 | 2621000 | 387310 | 207730 | 0 | 0 | 0 | 0 | 0 | 0 | 0 |
| QADSTSIGI | GPRIN1 | Q7Z2K8 | GRIN1_HUMAN | Unmodified | 0 | 0 | 0 | 0 | 0 | 0 | 0 | 0 | 2108300 | 0 | 0 | 0 | 0 | 0 | 0 | 0 | 0 | 0 |
| QADVNLVN | PHGDH; PHGD | O43175; A0 | SERA_HUMAN | Unmodified | 335070 | 143520 | 3135400 | 3149500 | 2603200 | 1030100 | 5622700 | 3429700 | 4193100 | 2463400 | 1437700 | 15761000 | 18289000 | 18138000 | 2901700 | 3425500 | 2010800 | 3982200 |
| QADVNLVN | PHGDH; PHGD | O43175; A0 | SERA_HUMAN | Unmodified | 0 | 0 | 0 | 0 | 0 | 0 | 427300 | 192270 | 0 | 0 | 0 | 0 | 0 | 0 | 0 | 0 | 97921 | 0 |
| QADYVPSD | GNAS; GNAS; G | Q5JWF2; P6 | GNAS1_HUMA | Unmodified | 2206100 | 979260 | 2998600 | 3426100 | 611440 | 778710 | 0 | 0 | 1711800 | 0 | 2676400 | 12718000 | 566060 | 317880 | 11125000 | 6606300 | 537530 | 557720 |
| QAEALRPF | GPC1; GPC1; G | P35052; H7 | GPC1_HUMAN | Unmodified | 0 | 0 | 834560 | 872740 | 0 | 0 | 0 | 0 | 126990 | 0 | 0 | 0 | 0 | 0 | 1042200 | 897210 | 0 | 0 |
| QAEETYEN | IPO5; IPO5; IPO | O00410; E7E | IPO5_HUMAN I | Unmodified | 0 | 0 | 0 | 0 | 0 | 0 | 0 | 0 | 501060 | 251850 | 903240 | 2843500 | 0 | 0 | 155350 | 94634 | 0 | 0 |
| QAEIENPLE | ATP6V0A1; ATP | Q93050; B7 | VPP1_HUMAN | Unmodified | 504910 | 0 | 0 | 0 | 0 | 0 | 0 | 87383 | 13637000 | 1292200 | 0 | 3595800 | 0 | 0 | 0 | 478380 | 0 | 0 |
| QAEILQESI | TBCA; TBCA; TB | O75347; E5F | TBCA_HUMAN | Unmodified | 1328200 | 864230 | 1114400 | 1174600 | 2068100 | 403130 | 112520 | 108550 | 2777700 | 990430 | 2468500 | 6931800 | 4232400 | 0 | 1956500 | 1908200 | 479980 | 677430 |
| QAELAIER | PI4K2A | Q9BTU6 | P4K2A_HUMAN | Unmodified | 0 | 0 | 0 | 0 | 0 | 0 | 0 | 0 | 396780 | 0 | 0 | 0 | 0 | 0 | 0 | 0 | 0 | 0 |
| QAELEEGR | GOLIM4; GOLIN | O00461; F8V | GOLI4_HUMAN | Unmodified | 0 | 0 | 0 | 0 | 0 | 0 | 0 | 38485 | 16393000 | 360120 | 422030 | 1583000 | 0 | 0 | 0 | 0 | 45579 | 0 |
| QAELEEIYE | ADGRE5 | P48960 | AGRE5_HUMA | Unmodified | 0 | 0 | 0 | 0 | 7665300 | 1123900 | 0 | 0 | 99130 | 101290 | 0 | 0 | 0 | 0 | 0 | 0 | 1443700 | 10056000 |
| QAELEEIYE | ADGRE5 | P48960 | AGRE5_HUMA | Unmodified | 0 | 0 | 0 | 0 | 4661200 | 0 | 0 | 0 | 0 | 0 | 0 | 0 | 0 | 0 | 0 | 0 | 0 | 0 |
| QAELLLPG | TPP1; TPP1; TP | O14773; E7E | TPP1_HUMAN | Unmodified | 0 | 0 | 0 | 340760 | 0 | 1804100 | 0 | 0 | 1485700 | 0 | 0 | 0 | 0 | 0 | 0 | 0 | 0 | 0 |
| QAELTVQV | PVR; PVR; PVR | P15151; A0A | PVR_HUMAN P | Unmodified | 3448400 | 4386400 | 2627900 | 2869700 | 1065400 | 0 | 216130 | 0 | 758350 | 180920 | 2615400 | 6629800 | 167460 | 243290 | 3243300 | 1844300 | 394380 | 1335600 |
| QAERLFEN | APMAP; APMAP | Q9HDC9; H | APMAP_HUMA | Unmodified | 0 | 0 | 0 | 0 | 0 | 0 | 0 | 0 | 3758300 | 0 | 0 | 0 | 0 | 0 | 0 | 0 | 0 | 0 |
| QAETLYK | KLC1; KLC1; KL | Q07866; G3 | KLC1_HUMAN | Unmodified | 0 | 59386 | 491230 | 0 | 790620 | 169210 | 13678 | 75187 | 0 | 110840 | 67252 | 623760 | 0 | 64239 | 51936 | 0 | 253800 | 602050 |
| QAEVANQE | HMGN1; HMGN | P05114; A6N | HMGN1_HUMA | Unmodified | 0 | 0 | 0 | 0 | 0 | 0 | 0 | 0 | 0 | 0 | 0 | 1268200 | 404620 | 0 | 0 | 0 | 0 | 0 |
| QAFATSPA | RIPOR2; RIPOR | Q9Y4F9; B7 | RIPR2_HUMAN | Unmodified | 287770 | 460320 | 0 | 29526 | 0 | 0 | 1481400 | 0 | 717470 | 358390 | 267880 | 2742800 | 2410700 | 1722100 | 18611 | 139330 | 0 | 0 |
| QAFEADIA | PALS1 | Q8N3R9 | PALS1_HUMAN | Unmodified | 0 | 0 | 146650 | 0 | 0 | 0 | 143320 | 34806 | 553780 | 103100 | 0 | 512300 | 0 | 0 | 0 | 0 | 0 | 0 |
| QAFLQVFV | NCAM2; NCAM | O15394; H9I | NCAM2_HUMA | Unmodified | 0 | 0 | 373720 | 162570 | 0 | 0 | 0 | 82730 | 0 | 0 | 0 | 0 | 0 | 0 | 1022000 | 190530 | 0 | 0 |
| QAFPNTNR | EEF1G | P26641 | EF1G_HUMAN | Unmodified | 204030 | 57137 | 371390 | 0 | 0 | 0 | 0 | 0 | 0 | 0 | 0 | 22619000 | 43146 | 0 | 525300 | 603390 | 0 | 0 |
| QAFQGAVC | TNFSF11 | O14788 | TNF11_HUMAN | Unmodified | 0 | 0 | 0 | 0 | 49097 | 0 | 0 | 14090 | 322230 | 18346 | 0 | 95534 | 0 | 0 | 0 | 0 | 0 | 0 |
| QAFQIGSP | ALDH1A1; ALDH | P00352; Q5 | AL1A1_HUMAN | Unmodified | 0 | 0 | 0 | 0 | 0 | 87694 | 0 | 0 | 2637300 | 299460 | 12333000 | 5.02E+08 | 0 | 0 | 0 | 0 | 0 | 275970 |

| | | | | | | | | | | | | | | | | | | | | | | |
|---|---|---|---|---|---|---|---|---|---|---|---|---|---|---|---|---|---|---|---|---|---|---|
| QAFTDVAT | TKT; TKT; TKT | P29401; F8V | TKT_HUMAN Tr | Oxidation (M) | 0 | 0 | 0 | 0 | 0 | 0 | 0 | 0 | 0 | 827320 | 813580 | 0 | 0 | 0 | 0 | 0 | 1226200 | 0 |
| QAGAEALS | GCN1 | Q92616 | GCN1_HUMAN | Unmodified | 0 | 160610 | 0 | 0 | 0 | 0 | 0 | 0 | 2752000 | 144270 | 0 | 1035800 | 0 | 0 | 0 | 0 | 0 | 0 |
| QAGDAMRI | N/A | N/A | N/A | Oxidation (M) | 0 | 0 | 0 | 0 | 1181400 | 123670 | 0 | 0 | 0 | 209990 | 0 | 0 | 0 | 0 | 0 | 0 | 728350 | 6600200 |
| QAGDAMRI | N/A | N/A | N/A | Unmodified | 13553000 | 41409000 | 0 | 0 | 0 | 0 | 0 | 0 | 0 | 0 | 0 | 0 | 205100 | 0 | 2450900 | 0 | 0 | 0 |
| QAGDSSSI | SPAG1 | Q07617 | SPAG1_HUMAI | Unmodified | 0 | 0 | 0 | 0 | 2234400 | 189610 | 50337 | 42217 | 0 | 0 | 0 | 0 | 440210 | 653450 | 0 | 0 | 1216400 | 2043100 |
| QAGGNPW | PDPK1; PDPK1 | O15530; E9F | PDPK1_HUMAI | Unmodified | 191480 | 166180 | 1304100 | 2226900 | 740050 | 0 | 0 | 19608 | 604620 | 1452100 | 0 | 4903000 | 3771900 | 0 | 2420400 | 3061500 | 0 | 677180 |
| QAGLEAAA | TRIM25; TRIM2 | Q14258; A0A | TRI25_HUMAN | Unmodified | 0 | 0 | 0 | 343410 | 5270800 | 793430 | 6417.6 | 126550 | 0 | 0 | 0 | 119380 | 0 | 0 | 241450 | 165460 | 231630 | 530970 |
| QAGPASVP | PDIA3 | P30101 | PDIA3_HUMAN | Unmodified | 0 | 0 | 0 | 291780 | 1350400 | 0 | 1455900 | 1121200 | 571880 | 14821000 | 9545900 | 6525500 | 1209600 | 1273800 | 733910 | 854300 | 242060 | 1259900 |
| QAGPASVP | PDIA3 | P30101 | PDIA3_HUMAN | Unmodified | 369840 | 0 | 0 | 0 | 551770 | 0 | 1345700 | 315490 | 85323000 | 32017000 | 13484000 | 20842000 | 0 | 0 | 0 | 0 | 0 | 0 |
| QAGVFEPT | TCP1; TCP1; TC | P17987; F5F | TCPA_HUMAN | Unmodified | 626190 | 0 | 550830 | 0 | 1516000 | 0 | 77595 | 51643 | 2328200 | 841830 | 2822700 | 16327000 | 0 | 866290 | 0 | 677710 | 627520 | 1488900 |
| QAGVNHVI | XPR1 | Q9UBH6 | XPR1_HUMAN | Unmodified | 0 | 0 | 0 | 0 | 0 | 0 | 0 | 0 | 0 | 0 | 0 | 665280 | 0 | 0 | 71232 | 90587 | 0 | 0 |
| QAHILLDC | ITGAV | P06756 | ITAV_HUMAN I | Unmodified | 0 | 136480 | 0 | 0 | 0 | 0 | 0 | 69181 | 176220 | 241180 | 0 | 887810 | 0 | 0 | 0 | 0 | 0 | 254110 |
| QAHLTNQY | PABPC1; PABP | P11940; A0A | PABP1_HUMAI | Unmodified | 92832000 | 66515000 | 24815000 | 38929000 | 376030 | 149100 | 71847 | 0 | 948320 | 0 | 0 | 2806000 | 5208100 | 703490 | 13453000 | 19779000 | 124950 | 58949 |
| QAHLTNQY | PABPC1; PABP | P11940; A0A | PABP1_HUMAI | Oxidation (M) | 74439000 | 36728000 | 3507600 | 29592000 | 0 | 0 | 0 | 0 | 152130 | 0 | 0 | 550680 | 3032300 | 744550 | 15283000 | 17892000 | 40233 | 198660 |
| QAHTMDP | FASN; FASN; FA | P49327; A0A | FAS_HUMAN F | Unmodified | 179750 | 0 | 0 | 94127 | 0 | 0 | 0 | 0 | 0 | 17721 | 91147 | 3576800 | 0 | 0 | 0 | 282240 | 0 | 0 |
| QAIELSPDI | IFIT3; IFIT3; IFIT | O14879; A0A | IFIT3_HUMAN I | Unmodified | 0 | 0 | 0 | 0 | 533420 | 0 | 0 | 0 | 0 | 0 | 0 | 0 | 0 | 0 | 0 | 0 | 0 | 745010 |
| QAIGIALETI | PSMD1; PSMD1 | Q99460; C9J | PSMD1_HUMA | Unmodified | 82400 | 0 | 0 | 196460 | 0 | 0 | 0 | 0 | 0 | 0 | 0 | 2985300 | 288090 | 0 | 225610 | 216990 | 0 | 189140 |
| QAIGLMGY | GCA; GCA; GCA | P28676; H7C | GRAN_HUMAN | Oxidation (M) | 0 | 0 | 0 | 0 | 0 | 0 | 0 | 0 | 426780 | 401660 | 0 | 0 | 59314 | 0 | 0 | 0 | 0 | 0 |
| QAILAAQR | EDF1 | O60869 | EDF1_HUMAN | Unmodified | 1488800 | 1132800 | 1173800 | 1879300 | 216980 | 104170 | 368960 | 80768 | 499780 | 131830 | 340100 | 4291100 | 228460 | 660850 | 2131900 | 3372200 | 127500 | 143720 |
| QAILAAQRI | EDF1 | O60869 | EDF1_HUMAN | Unmodified | 0 | 0 | 0 | 0 | 0 | 57130 | 0 | 0 | 0 | 0 | 0 | 0 | 346160 | 0 | 0 | 0 | 133240 | 323930 |
| QAIPLDENI | MVP | Q14764 | MVP_HUMAN N | Unmodified | 1732300 | 1371100 | 0 | 0 | 0 | 0 | 0 | 0 | 1320000 | 0 | 0 | 13954000 | 0 | 0 | 0 | 0 | 0 | 0 |
| QAIQASIK | GALNT7; GALN | Q86SF2; E9F | GALT7_HUMAI | Unmodified | 3943500 | 3234900 | 2679300 | 1922200 | 43037 | 0 | 214580 | 91103 | 4836200 | 1982900 | 421280 | 1505300 | 1561600 | 0 | 1156500 | 1285400 | 209770 | 863600 |
| QAIQEELNI | CDC42BPA; CD | Q5VT25; A0A | MRCKA_HUMA | Unmodified | 112540 | 0 | 0 | 0 | 1482600 | 1011900 | 0 | 119810 | 2243100 | 1104400 | 100860 | 613800 | 1085200 | 0 | 108310 | 0 | 0 | 1068100 |
| QAIQEELNI | CDC42BPA; CD | Q5VT25; A0A | MRCKA_HUMA | Unmodified | 0 | 0 | 0 | 0 | 0 | 0 | 0 | 0 | 0 | 0 | 0 | 0 | 0 | 0 | 0 | 0 | 0 | 0 |
| QAITQVVV | SFXN1 | Q9H9B4 | SFXN1_HUMAI | Unmodified | 0 | 0 | 0 | 0 | 0 | 0 | 0 | 40579 | 2687700 | 0 | 0 | 0 | 0 | 0 | 0 | 0 | 0 | 0 |
| QAIVDKVP | COPG1 | Q9Y678 | COPG1_HUMA | Unmodified | 0 | 0 | 0 | 0 | 0 | 0 | 0 | 0 | 0 | 0 | 0 | 2670500 | 0 | 0 | 0 | 0 | 0 | 0 |
| QAKEEAQA | ATP6V1G1; ATP | O75348; A0A | VATG1_HUMAI | Unmodified | 0 | 0 | 0 | 95316 | 211100 | 0 | 0 | 0 | 0 | 0 | 62139 | 1467500 | 111430 | 0 | 0 | 150950 | 0 | 0 |
| QAKEIEDA | AHCYL1 | O43865 | SAHH2_HUMAI | Unmodified | 0 | 0 | 0 | 0 | 264840 | 0 | 32943 | 0 | 0 | 0 | 0 | 0 | 68389 | 0 | 0 | 634770 | 1080900 | 219690 |
| QALCPGST | ARHGEF2; ARH | Q92974; V9C | ARHG2_HUMA | Unmodified | 0 | 0 | 17763 | 0 | 0 | 0 | 0 | 0 | 113120 | 23037 | 0 | 582700 | 0 | 0 | 23108 | 0 | 0 | 0 |
| QALEASNR | DSP | P15924 | DESP_HUMAN | Unmodified | 0 | 0 | 0 | 0 | 354460 | 0 | 0 | 0 | 315340 | 0 | 0 | 0 | 118880 | 0 | 0 | 0 | 0 | 0 |
| QALEDLEK | KLC1; KLC2; KL | Q07866; Q9I | KLC1_HUMAN | Unmodified | 0 | 0 | 0 | 0 | 1295500 | 0 | 0 | 245290 | 882830 | 373560 | 338290 | 1384700 | 583180 | 274520 | 56243 | 53246 | 513040 | 635090 |
| QALEKFEE | SH3GL1 | Q99961 | SH3G1_HUMAI | Unmodified | 4363600 | 4597500 | 0 | 1829400 | 0 | 0 | 0 | 0 | 0 | 0 | 0 | 0 | 0 | 0 | 2293100 | 2415700 | 0 | 109440 |
| QALESHPL | PLCB4; PLCB4 | Q15147; A0A | PLCB4_HUMAI | Unmodified | 0 | 0 | 306390 | 0 | 0 | 0 | 0 | 0 | 0 | 0 | 0 | 0 | 1171800 | 0 | 0 | 0 | 0 | 4727900 |
| QALETVND | DYNC1H1; DYN | Q14204; A0A | DYHC1_HUMA | Oxidation (M) | 0 | 0 | 75163 | 0 | 0 | 0 | 0 | 0 | 0 | 0 | 0 | 741790 | 0 | 0 | 0 | 104690 | 0 | 0 |
| QALFQQEI | P4HTM; P4HTM | Q9NXG6; V9 | P4HTM_HUMAI | Unmodified | 0 | 0 | 0 | 0 | 0 | 0 | 0 | 0 | 2036400 | 0 | 0 | 0 | 0 | 0 | 0 | 0 | 0 | 0 |
| QALIASAR | MBTPS1; MBTP | Q14703; A0A | MBTP1_HUMAI | Unmodified | 998770 | 0 | 0 | 907890 | 0 | 3223700 | 0 | 0 | 892560 | 0 | 0 | 0 | 0 | 0 | 0 | 0 | 0 | 0 |
| QALLKQGC | ERP29; ERP29 | P30040; F8V | ERP29_HUMAI | Unmodified | 0 | 0 | 0 | 0 | 0 | 0 | 0 | 0 | 0 | 1752400 | 0 | 0 | 0 | 0 | 0 | 0 | 0 | 0 |
| QALLKQGC | ERP29; ERP29 | P30040; F8V | ERP29_HUMAI | Unmodified | 0 | 0 | 0 | 0 | 0 | 0 | 112900 | 68160 | 315610 | 5630600 | 0 | 0 | 0 | 0 | 0 | 0 | 0 | 0 |
| QALLQYK | FKBP4 | Q02790 | FKBP4_HUMAI | Unmodified | 356900 | 0 | 0 | 0 | 0 | 0 | 0 | 644240 | 0 | 0 | 1720300 | 11189000 | 538300 | 1871600 | 0 | 106120 | 0 | 0 |
| QALLQYKK | FKBP4 | Q02790 | FKBP4_HUMAI | Unmodified | 0 | 0 | 0 | 0 | 0 | 0 | 177230 | 0 | 0 | 0 | 0 | 1219900 | 3373400 | 571730 | 0 | 0 | 0 | 0 |
| QALMNILK | RXYLT1; RXYLT | Q9Y2B1; G3 | RXLT1_HUMAN | Unmodified | 0 | 0 | 0 | 0 | 0 | 0 | 0 | 0 | 270530 | 76415 | 0 | 0 | 0 | 0 | 0 | 0 | 0 | 0 |
| QALNEHFC | APLP1; APLP1; | P51693; B7Z | APLP1_HUMAI | Unmodified | 0 | 0 | 0 | 0 | 0 | 0 | 0 | 0 | 848160 | 0 | 0 | 0 | 0 | 0 | 0 | 0 | 0 | 0 |
| QALNEHFC | APLP1; APLP1; | P51693; B7Z | APLP1_HUMAI | Unmodified | 0 | 0 | 0 | 0 | 0 | 721580 | 0 | 0 | 2150100 | 0 | 0 | 0 | 0 | 0 | 0 | 0 | 0 | 0 |
| QALNEIESI | STX2 | P32856 | STX2_HUMAN | Unmodified | 0 | 245440 | 204030 | 417400 | 0 | 0 | 192070 | 0 | 583250 | 163620 | 135390 | 4008500 | 0 | 0 | 330240 | 344270 | 0 | 258260 |
| QALNEISAI | STX4 | Q12846 | STX4_HUMAN | Unmodified | 0 | 0 | 0 | 688140 | 0 | 0 | 0 | 0 | 1077100 | 0 | 0 | 10216000 | 0 | 0 | 0 | 0 | 0 | 0 |
| QALNEISAI | STX4 | Q12846 | STX4_HUMAN | Unmodified | 0 | 0 | 0 | 0 | 446230 | 0 | 0 | 0 | 0 | 97085 | 0 | 0 | 454730 | 0 | 0 | 0 | 0 | 0 |
| QALNTDYL | ENDOD1 | O94919 | ENDD1_HUMA | Unmodified | 0 | 0 | 844950 | 101720 | 0 | 0 | 0 | 0 | 5891600 | 216240 | 0 | 0 | 0 | 0 | 381520 | 402890 | 0 | 0 |
| QALPQVR | CPNE1; CPNE1 | Q99829; Q5 | CPNE1_HUMAI | Unmodified | 77894 | 302350 | 372100 | 0 | 0 | 0 | 232790 | 63710 | 1055300 | 43914 | 40351 | 4652900 | 0 | 0 | 0 | 350540 | 190840 | 0 |
| QALQDLLS | CTNNA1; CTNN | P35221; G3 | CTNA1_HUMAI | Unmodified | 297350 | 441150 | 0 | 1986900 | 1356400 | 118800 | 0 | 31340 | 238770 | 0 | 0 | 2234200 | 943200 | 0 | 2232600 | 1493400 | 0 | 0 |
| QALQDLLS | CTNNA1; CTNN | P35221; G3 | CTNA1_HUMAI | Oxidation (M) | 0 | 0 | 0 | 0 | 0 | 0 | 0 | 66996 | 890750 | 0 | 531510 | 1573100 | 0 | 0 | 902680 | 4609000 | 0 | 0 |
| QALQEIVILI | SPAST; SPAST; | Q9UBP0; A0 | SPAST_HUMAN | Unmodified | 0 | 0 | 0 | 0 | 1843000 | 394220 | 90048 | 0 | 461140 | 385710 | 295930 | 370570 | 0 | 0 | 434960 | 0 | 0 | 0 |
| QALQQLQ | CGN | Q9P2M7 | CING_HUMAN | Unmodified | 0 | 0 | 0 | 0 | 0 | 0 | 0 | 0 | 0 | 0 | 0 | 0 | 0 | 0 | 0 | 0 | 0 | 0 |
| QALSEIEGI | STX3; STX3; ST | Q13277; A0A | STX3_HUMAN | Unmodified | 1117400 | 111880 | 527400 | 0 | 113010 | 0 | 0 | 0 | 4815900 | 1003400 | 125970 | 595270 | 0 | 222020 | 1037400 | 781990 | 0 | 198000 |
| QALSEIETR | STX1A; STX1A; | Q16623; A8I | STX1A_HUMAI | Unmodified | 0 | 0 | 0 | 0 | 0 | 0 | 0 | 0 | 1446500 | 1603000 | 1701100 | 137330 | 0 | 0 | 0 | 0 | 0 | 0 |
| QALSGFGY | PDCD6; PDCD | O75340; A0A | PDCD6_HUMA | Unmodified | 10184000 | 8057200 | 6282400 | 2692100 | 263870 | 105100 | 0 | 0 | 20018000 | 1073900 | 63774 | 7096000 | 99241 | 0 | 9953000 | 8103100 | 0 | 313850 |
| QALSGFGY | PDCD6; PDCD | O75340; A0A | PDCD6_HUMA | Unmodified | 465900 | 0 | 0 | 1389800 | 0 | 0 | 0 | 0 | 0 | 2088000 | 58298 | 0 | 557440 | 347850 | 0 | 0 | 255610 | 785090 |
| QALTEFNT | LACTB2 | Q53H82 | LACB2_HUMAI | Unmodified | 498640 | 0 | 145220 | 570930 | 2725100 | 168410 | 0 | 0 | 26330000 | 720720 | 252610 | 639240 | 0 | 0 | 406000 | 272260 | 168530 | 0 |
| QALTEFNT | LACTB2 | Q53H82 | LACB2_HUMAI | Unmodified | 0 | 0 | 0 | 0 | 1382400 | 0 | 0 | 0 | 0 | 2326600 | 0 | 0 | 0 | 0 | 0 | 0 | 785760 | 2241700 |
| QALVAIFTH | DYNC1H1; DYN | Q14204; A0A | DYHC1_HUMA | Unmodified | 0 | 0 | 0 | 0 | 0 | 0 | 0 | 0 | 0 | 0 | 0 | 1289600 | 0 | 0 | 0 | 0 | 0 | 0 |
| QAMLENAS | ABCF1; ABCF1; | Q8NE71; H0 | ABCF1_HUMAI | Unmodified | 133340 | 325550 | 5541700 | 3779400 | 5292700 | 15653000 | 812150 | 324300 | 0 | 37152 | 0 | 0 | 0 | 2633600 | 0 | 0 | 317360 | 1272400 |
| QAMTLEFK | SHMT1 | P34896 | GLYC_HUMAN | Oxidation (M) | 0 | 0 | 0 | 0 | 0 | 0 | 0 | 0 | 0 | 0 | 0 | 415300 | 0 | 0 | 0 | 77078 | 0 | 0 |
| QANEEYQII | TUSC3; TUSC3; | Q13454; D6I | TUSC3_HUMAI | Unmodified | 0 | 0 | 0 | 0 | 0 | 0 | 0 | 0 | 4562600 | 200250 | 80047 | 380080 | 0 | 0 | 0 | 0 | 0 | 0 |
| QANTGLFN | THOP1; THOP1 | P52888; K7E | THOP1_HUMAI | Unmodified | 0 | 0 | 0 | 0 | 0 | 0 | 0 | 0 | 0 | 135320 | 0 | 1609400 | 0 | 0 | 0 | 0 | 0 | 0 |
| QANVQIQE | L1RE1 | Q9UN81 | LORF1_HUMAI | Unmodified | 3876500 | 4090400 | 560830 | 180470 | 0 | 0 | 0 | 0 | 96130 | 0 | 0 | 0 | 0 | 0 | 501540 | 890360 | 0 | 0 |
| QANVQIQE | L1RE1 | Q9UN81 | LORF1_HUMAI | Unmodified | 1294100 | 806900 | 0 | 958910 | 0 | 0 | 0 | 0 | 0 | 0 | 0 | 0 | 161930 | 0 | 351490 | 0 | 60199 | 0 |
| QAPEENQF | PAFAH2; PAFAI | Q99487; Q5 | PAFA2_HUMAI | Unmodified | 0 | 0 | 0 | 0 | 0 | 0 | 0 | 0 | 1005700 | 0 | 0 | 0 | 0 | 0 | 0 | 0 | 0 | 0 |
| QAPHHAAI | UNC119B | A6NIH7 | U119B_HUMAI | Unmodified | 1966500 | 0 | 96116 | 0 | 0 | 0 | 0 | 0 | 0 | 0 | 0 | 0 | 1441900 | 978530 | 384070 | 0 | 0 | 0 |
| QAPQTIHLI | ACO1 | P21399 | ACOC_HUMAN | Unmodified | 67440 | 0 | 0 | 74918 | 0 | 0 | 0 | 0 | 0 | 0 | 472910 | 1950400 | 0 | 0 | 193300 | 0 | 0 | 0 |
| QAPRKSPS | MPZL1 | O95297 | MPZL1_HUMAI | Unmodified | 50427 | 0 | 0 | 0 | 16990000 | 1339800 | 5258100 | 696320 | 0 | 1065500 | 0 | 0 | 4772500 | 5222500 | 0 | 0 | 4271700 | 16063000 |
| QAQDLARS | KRAS | P01116 | RASK_HUMAN | Unmodified | 0 | 0 | 0 | 0 | 3420000 | 0 | 547700 | 240340 | 0 | 352930 | 0 | 0 | 2331400 | 2585500 | 0 | 0 | 1542700 | 1808900 |
| QAQEYEAL | KRT18; KRT18 | P05783; F8V | K1C18_HUMAI | Unmodified | 0 | 0 | 402400 | 0 | 0 | 0 | 0 | 0 | 3624600 | 222380 | 4725600 | 24002000 | 0 | 0 | 0 | 0 | 0 | 0 |
| QAQIEVVP | RPL12 | P30050 | RL12_HUMAN | Unmodified | 4516600 | 14838000 | 3014200 | 3936700 | 1053800 | 0 | 0 | 411110 | 849200 | 1150500 | 6704800 | 1.02E+08 | 4671500 | 1122200 | 13510000 | 7003900 | 0 | 650980 |
| QAQILASE | STOML2; STOM | Q9UJZ1; A0A | STML2_HUMAI | Unmodified | 0 | 0 | 0 | 0 | 0 | 0 | 0 | 0 | 1101900 | 203170 | 0 | 0 | 0 | 0 | 0 | 0 | 0 | 0 |
| QAQILASE | STOML2; STOM | Q9UJZ1; A0A | STML2_HUMAI | Unmodified | 0 | 0 | 0 | 0 | 0 | 0 | 2917500 | 0 | 4249400 | 824400 | 0 | 0 | 0 | 0 | 0 | 0 | 0 | 0 |
| QAQLAQTL | NUMA1; NUMA | Q14980; F5I | NUMA1_HUMA | Unmodified | 0 | 0 | 0 | 0 | 0 | 0 | 0 | 0 | 0 | 363840 | 0 | 0 | 1369500 | 359660 | 0 | 0 | 0 | 0 |
| QAQNPDQ | NECAP2; NECA | Q9NVZ3; D6 | NECP2_HUMAI | Unmodified | 309770 | 172720 | 0 | 0 | 685600 | 0 | 0 | 0 | 343890 | 63497 | 0 | 0 | 0 | 0 | 0 | 378220 | 0 | 915830 |
| QAQNPDQ | NECAP2; NECA | Q9NVZ3; D6 | NECP2_HUMAI | Unmodified | 0 | 0 | 0 | 0 | 2079900 | 0 | 250320 | 0 | 0 | 0 | 0 | 0 | 0 | 0 | 0 | 0 | 1121600 | 2085300 |
| QAQQERDI | MYH9 | P35579 | MYH9_HUMAN | Unmodified | 0 | 401720 | 1128600 | 1098000 | 0 | 0 | 0 | 0 | 154530 | 0 | 332690 | 4226400 | 0 | 0 | 294780 | 260890 | 0 | 0 |
| QAQVPLAA | SEC24C; SEC2 | P53992; G5E | SC24C_HUMAI | Unmodified | 0 | 0 | 0 | 0 | 0 | 0 | 0 | 1024700 | 0 | 0 | 40101 | 360840 | 0 | 0 | 0 | 0 | 0 | 0 |
| QAQYLGM | AHCY | P23526 | SAHH_HUMAN | Oxidation (M) | 0 | 0 | 824500 | 1249500 | 1672400 | 0 | 424940 | 0 | 0 | 759780 | 1958200 | 5761700 | 1047200 | 636770 | 0 | 442580 | 5042900 | 1790800 |
| QAQYLGM | AHCY | P23526 | SAHH_HUMAN | Unmodified | 0 | 0 | 0 | 0 | 4118100 | 438800 | 0 | 0 | 844180 | 455550 | 226500 | 2978000 | 1074400 | 462100 | 0 | 0 | 3826700 | 2414000 |
| QARANSFV | PDPK2P; PDPK | Q6A1A2; O1 | PDPK2_HUMAI | Unmodified | 0 | 0 | 0 | 0 | 0 | 0 | 91280 | 151340 | 0 | 0 | 0 | 0 | 681670 | 643960 | 0 | 0 | 0 | 0 |
| QASALIDRI | EPB41L1; EPB4 | Q9H4G0; A0 | E41L1_HUMAI | Unmodified | 460530 | 380130 | 1254300 | 1439400 | 1240500 | 678130 | 0 | 0 | 1123100 | 0 | 2474600 | 4482800 | 847270 | 221100 | 2644400 | 2574700 | 0 | 0 |
| QASALIDRI | EPB41; EPB41; | P11171; A0A | EPB41_HUMAI | Unmodified | 1092400 | 0 | 2851900 | 2436100 | 0 | 0 | 223610 | 0 | 1492500 | 243770 | 1164600 | 4483100 | 966340 | 0 | 2513600 | 0 | 0 | 339210 |
| QASALIDRI | EPB41; EPB41; | P11171; A0A | EPB41_HUMAI | Unmodified | 0 | 0 | 0 | 0 | 0 | 0 | 0 | 0 | 0 | 0 | 0 | 0 | 0 | 2637800 | 0 | 0 | 0 | 0 |
| QASALSFP | ADAM15 | Q13444 | ADA15_HUMAI | Unmodified | 0 | 0 | 346610 | 691050 | 0 | 90117 | 0 | 0 | 0 | 0 | 98779 | 0 | 0 | 0 | 0 | 0 | 0 | 0 |
| QASCSGDI | PGM3; PGM3; F | O95394; J3K | AGM1_HUMAN | Unmodified | 0 | 0 | 0 | 0 | 158410 | 0 | 0 | 0 | 0 | 0 | 0 | 0 | 551930 | 864040 | 0 | 0 | 471030 | 440270 |
| QASDGPS | N/A | N/A | N/A | Unmodified | 3186500 | 0 | 0 | 0 | 0 | 0 | 0 | 0 | 0 | 0 | 0 | 0 | 0 | 0 | 0 | 639020 | 0 | 0 |
| QASDYGIN | TRMT10A | Q8TBZ6 | TM10A_HUMAI | Unmodified | 0 | 0 | 0 | 0 | 0 | 0 | 534130 | 0 | 0 | 0 | 0 | 236770 | 0 | 0 | 0 | 0 | 0 | 0 |
| QASEGPLK | GAPDH; GAPDI | P04406; E7E | G3P_HUMAN G | Unmodified | 18582000 | 0 | 6766300 | 16898000 | 69554000 | 9654000 | 17056000 | 11292000 | 6127800 | 4603900 | 1614300 | 21713000 | 14891000 | 7606900 | 1445600 | 9165700 | 44671000 | 55038000 |
| QASEWVR | PARP4 | Q9UKK3 | PARP4_HUMAN | Unmodified | 0 | 0 | 0 | 0 | 0 | 0 | 0 | 0 | 107560 | 0 | 0 | 431590 | 0 | 0 | 0 | 0 | 0 | 0 |
| QASIQHIQI | ATP5PB; ATP5P | P24539; Q5 | AT5F1_HUMAN | Unmodified | 0 | 0 | 0 | 0 | 0 | 0 | 0 | 0 | 28473000 | 5967900 | 0 | 0 | 0 | 0 | 0 | 0 | 0 | 0 |
| QASLADCL | XPOT | O43592 | XPOT_HUMAN | Unmodified | 68843 | 0 | 0 | 0 | 0 | 0 | 0 | 36823 | 0 | 0 | 0 | 645530 | 99045 | 0 | 167030 | 0 | 0 | 0 |
| QASMEDIP | LXN | Q9BS40 | LXN_HUMAN L | Unmodified | 56310 | 0 | 0 | 0 | 1163000 | 638640 | 0 | 423840 | 0 | 0 | 420980 | 156780 | 94936 | 68246 | 0 | 0 | 605270 | 1050900 |
| QASNGVLI | ANPEP | P15144 | AMPN_HUMAN | Unmodified | 317390 | 53769 | 0 | 0 | 79093 | 0 | 34837 | 0 | 287040 | 0 | 77611 | 235440 | 0 | 0 | 168380 | 108900 | 910880 | 503710 |
| QASPNIVIA | RAB5C; N/A; RA | P51148; K7E | RAB5C_HUMAI | Unmodified | 2279300 | 2158400 | 9295900 | 2980400 | 0 | 0 | 0 | 0 | 3573300 | 392420 | 5608100 | 8896700 | 0 | 0 | 8717100 | 4182400 | 0 | 0 |
| QASPNIVIA | RAB5C; N/A; RA | P51148; K7E | RAB5C_HUMAI | Unmodified | 8171800 | 16671000 | 1454600 | 17414000 | 2855000 | 444850 | 460720 | 342130 | 14577000 | 2671000 | 2324300 | 21113000 | 13984000 | 1900300 | 28204000 | 5968200 | 0 | 0 |
| QASPNIVIA | RAB5C; N/A; RA | P51148; K7E | RAB5C_HUMAI | Unmodified | 0 | 0 | 0 | 4555500 | 0 | 0 | 0 | 106670 | 5479200 | 0 | 0 | 15316000 | 1624800 | 0 | 2750400 | 1799100 | 0 | 0 |
| QASPNIVIA | RAB5A | P20339 | RAB5A_HUMAI | Unmodified | 537000 | 458830 | 1082300 | 354740 | 0 | 0 | 0 | 0 | 970560 | 0 | 831380 | 6237000 | 0 | 0 | 746800 | 752940 | 0 | 0 |
| QASPNIVIA | RAB5A | P20339 | RAB5A_HUMAI | Unmodified | 2767100 | 5102600 | 0 | 1995100 | 1081900 | 0 | 0 | 0 | 5130600 | 988300 | 896410 | 6032000 | 0 | 0 | 2198000 | 1651400 | 0 | 634380 |
| QASPNIVIA | RAB5A | P20339 | RAB5A_HUMAI | Unmodified | 918260 | 682900 | 0 | 1135900 | 0 | 0 | 0 | 95687 | 1384200 | 256370 | 846690 | 1266800 | 0 | 0 | 0 | 288550 | 301550 | 0 |
| QASPSIVIA | RAB5B | P61020 | RAB5B_HUMAI | Unmodified | 2210700 | 1377500 | 2228300 | 883180 | 0 | 0 | 0 | 0 | 1588700 | 224760 | 0 | 1857600 | 0 | 0 | 0 | 1508900 | 0 | 0 |
| QASPSIVIA | RAB5B | P61020 | RAB5B_HUMAI | Unmodified | 5033600 | 4206200 | 0 | 3872200 | 2628000 | 0 | 123810 | 0 | 7080600 | 2106400 | 494730 | 3242600 | 0 | 720330 | 2408100 | 2293800 | 0 | 0 |
| QASPSIVIA | RAB5B | P61020 | RAB5B_HUMAI | Unmodified | 2234600 | 2396800 | 0 | 1898500 | 0 | 0 | 0 | 0 | 2123400 | 326280 | 0 | 1110200 | 0 | 0 | 863030 | 1234800 | 0 | 0 |
| QASQGMV | RUVBL2; RUVBI | Q9Y230; X6F | RUVB2_HUMAI | Oxidation (M) | 0 | 0 | 0 | 0 | 0 | 0 | 0 | 0 | 0 | 0 | 0 | 482890 | 0 | 0 | 0 | 0 | 0 | 0 |
| QASQGMV | RUVBL2; RUVBI | Q9Y230; X6F | RUVB2_HUMAI | Unmodified | 0 | 0 | 0 | 0 | 121330 | 0 | 0 | 0 | 317970 | 0 | 348870 | 3369700 | 335390 | 0 | 0 | 58337 | 0 | 0 |
| QASSSSSR | PKP3; PKP3; PK | Q9Y446; E9F | PKP3_HUMAN | Unmodified | 0 | 0 | 0 | 0 | 0 | 0 | 0 | 0 | 0 | 0 | 0 | 0 | 336060 | 241220 | 0 | 166190 | 0 | 0 |
| QASTLIDRI | EPB41L2; EPB4 | O43491; A0A | E41L2_HUMAN | Unmodified | 0 | 0 | 599270 | 472630 | 1112400 | 1175200 | 0 | 34624 | 0 | 52345 | 0 | 0 | 0 | 0 | 949460 | 600520 | 0 | 658800 |
| QASTLIDRI | EPB41L2; EPB4 | O43491; A0A | E41L2_HUMAN | Unmodified | 0 | 0 | 0 | 0 | 25935000 | 0 | 149160 | 0 | 0 | 0 | 0 | 0 | 635460 | 0 | 0 | 0 | 1685400 | 10456000 |

| | | | | | | | | | | | | | | | | | | | | | | |
|---|---|---|---|---|---|---|---|---|---|---|---|---|---|---|---|---|---|---|---|---|---|---|
| QASVADYE | ALDH7A1; ALDI | P49419; A0A | AL7A1_HUMAN | Unmodified | 0 | 0 | 0 | 0 | 0 | 0 | 0 | 0 | 0 | 0 | 104070 | 805420 | 0 | 0 | 0 | 0 | 0 | 0 |
| QASVADYE | ALDH7A1; ALDI | P49419; A0A | AL7A1_HUMAN | Unmodified | 0 | 0 | 0 | 0 | 0 | 0 | 0 | 0 | 0 | 0 | 153150 | 1819500 | 0 | 0 | 0 | 0 | 0 | 0 |
| QATEGPCN | ECI2; ECI2; ECI | O75521; F8W | ECI2_HUMAN E | Unmodified | 0 | 0 | 0 | 0 | 0 | 23893 | 184100 | 0 | 11300000 | 822230 | 0 | 0 | 0 | 0 | 0 | 0 | 0 | 0 |
| QATEGPCN | ECI2; ECI2; ECI | O75521; F8W | ECI2_HUMAN E | Oxidation (M) | 0 | 0 | 0 | 0 | 0 | 0 | 141840 | 54816 | 1546000 | 2565200 | 0 | 0 | 0 | 0 | 0 | 0 | 0 | 0 |
| QATELALLC | ARHGEF2; ARH | Q92974; V9C | ARHG2_HUMA | Unmodified | 0 | 0 | 0 | 0 | 0 | 0 | 0 | 0 | 505490 | 274480 | 0 | 1285300 | 0 | 0 | 0 | 0 | 201870 | 0 |
| QATEPAPAI | POMGNT2 | Q8NAT1 | PMGT2_HUMAI | Unmodified | 0 | 116210 | 0 | 0 | 0 | 0 | 38130 | 24039 | 1235800 | 146200 | 0 | 0 | 0 | 0 | 0 | 0 | 0 | 33339 |
| QATGIEDG | CUL4B; CUL4B | Q13620; A0A | CUL4B_HUMAI | Unmodified | 0 | 0 | 0 | 0 | 0 | 0 | 0 | 0 | 0 | 0 | 0 | 195500 | 0 | 199640 | 85478 | 0 | 86133 | 0 |
| QATGRFEE | SNX33; SNX33 | Q8WV41; H3 | SNX33_HUMAN | Unmodified | 1530500 | 1335800 | 9208300 | 11614000 | 0 | 0 | 0 | 0 | 195800 | 0 | 0 | 166550 | 331470 | 0 | 13789000 | 9038000 | 0 | 0 |
| QATGRFEE | SNX18 | Q96RF0 | SNX18_HUMAN | Unmodified | 0 | 0 | 0 | 0 | 0 | 0 | 0 | 0 | 0 | 0 | 0 | 0 | 0 | 0 | 0 | 2918500 | 0 | 0 |
| QATHDQA\ | SNTB2; SNTB2; | Q13425; A0A | SNTB2_HUMAN E | Unmodified | 0 | 0 | 194060 | 0 | 495280 | 41015 | 0 | 0 | 112730 | 0 | 0 | 0 | 0 | 0 | 200020 | 212790 | 61844 | 0 |
| QATINIGTI( | EIF2S3; EIF2S3I | P41091; Q2\ | IF2G_HUMAN E | Unmodified | 0 | 0 | 0 | 270930 | 551890 | 0 | 0 | 0 | 1346500 | 301660 | 0 | 5189800 | 432940 | 0 | 0 | 785840 | 0 | 0 |
| QATKDAGT | HSPA8; HSPA8; | P11142; E9P | HSP7C_HUMAI | Unmodified | 58549 | 108310 | 0 | 69226 | 513440 | 0 | 634690 | 338070 | 0 | 816960 | 0 | 0 | 980230 | 623420 | 75190 | 33662 | 579110 | 187250 |
| QATKDAGT | HSPA5; HSPA5; | P11021; A0A | BIP_HUMAN Er | Unmodified | 0 | 0 | 0 | 0 | 0 | 0 | 0 | 0 | 690750 | 596320 | 0 | 0 | 0 | 0 | 0 | 0 | 0 | 0 |
| QATLIPDEV | LRRC40 | Q9H9A6 | LRC40_HUMAN | Unmodified | 0 | 0 | 0 | 0 | 0 | 0 | 0 | 0 | 0 | 0 | 0 | 815540 | 0 | 0 | 0 | 0 | 0 | 0 |
| QATLTISSA | KIT | P10721 | KIT_HUMAN Ma | Unmodified | 0 | 0 | 0 | 0 | 0 | 0 | 0 | 0 | 602730 | 0 | 48401 | 883930 | 0 | 0 | 0 | 0 | 0 | 0 |
| QATLTQTLI | ITGA5 | P08648 | ITA5_HUMAN I | Unmodified | 0 | 0 | 0 | 0 | 0 | 0 | 0 | 0 | 0 | 0 | 0 | 0 | 0 | 0 | 622140 | 1022100 | 0 | 0 |
| QATTIIADN | SSBP1 | Q04837 | SSBP_HUMAN | Unmodified | 0 | 0 | 0 | 0 | 128170 | 0 | 0 | 0 | 4086100 | 1589900 | 221240 | 684500 | 65189 | 0 | 0 | 0 | 0 | 0 |
| QATVGDIN | DBI; DBI; DBI | P07108; B8Z | ACBP_HUMAN | Unmodified | 1086100 | 1948600 | 0 | 1007100 | 712320 | 351390 | 0 | 108060 | 6762900 | 1022800 | 1010600 | 3736100 | 266850 | 0 | 5510700 | 2102000 | 0 | 300860 |
| QATVGDIN | DBI; DBI; DBI | P07108; B8Z | ACBP_HUMAN | Oxidation (M) | 4464400 | 2761100 | 9667200 | 997630 | 422140 | 288280 | 1101600 | 1489700 | 1191600 | 556140 | 2594200 | 4625200 | 430850 | 647610 | 12148000 | 10771000 | 1316200 | 2783100 |
| QATYGYYL | RCN1 | Q15293 | RCN1_HUMAN | Unmodified | 0 | 0 | 0 | 0 | 0 | 0 | 195010 | 0 | 6162400 | 2527500 | 1398300 | 0 | 0 | 0 | 0 | 0 | 0 | 296750 |
| QATYGYYL | RCN1 | Q15293 | RCN1_HUMAN | Unmodified | 0 | 0 | 244690 | 0 | 1509900 | 0 | 429260 | 0 | 7262500 | 2052000 | 0 | 0 | 595040 | 0 | 433150 | 0 | 0 | 677560 |
| QAVAQILPI | H6PD | O95479 | G6PE_HUMAN | Unmodified | 0 | 0 | 0 | 0 | 0 | 0 | 0 | 0 | 2982400 | 1102000 | 0 | 0 | 0 | 0 | 0 | 0 | 0 | 0 |
| QAVELQR | WDR26; WDR2( | Q9H7D7; A0 | WDR26_HUMA | Unmodified | 0 | 694840 | 523790 | 602140 | 0 | 203020 | 127430 | 77192 | 599560 | 324260 | 195200 | 938070 | 589250 | 0 | 716360 | 69483 | 0 | 0 |
| QAVEQQIQ | KIF5B | P33176 | KINH_HUMAN | Unmodified | 0 | 0 | 135620 | 0 | 0 | 0 | 0 | 0 | 0 | 0 | 273170 | 2745300 | 0 | 0 | 175250 | 136140 | 0 | 0 |
| QAVKETCA | GFUS | Q13630 | FCL_HUMAN G | Unmodified | 0 | 0 | 0 | 0 | 684980 | 0 | 0 | 0 | 0 | 0 | 0 | 0 | 0 | 295980 | 0 | 0 | 2277100 | 1054700 |
| QAVLGAGL | ACAT1; ACAT1; | P24752; H0\ | THIL_HUMAN A | Unmodified | 0 | 0 | 0 | 0 | 0 | 412560 | 0 | 0 | 2670000 | 333820 | 0 | 0 | 0 | 0 | 0 | 56258 | 0 | 0 |
| QAVMEMM | PFDN5; PFDN5 | Q99471; A0A | PFD5_HUMAN | Unmodified | 0 | 0 | 0 | 0 | 1083100 | 0 | 0 | 0 | 209920 | 89207 | 0 | 1092300 | 476900 | 0 | 0 | 0 | 918880 | 418350 |
| QAVPISVPF | SNX27; SNX27 | Q96L92; A0A | SNX27_HUMAN | Unmodified | 278210 | 252190 | 187900 | 264790 | 64891000 | 291810 | 0 | 0 | 289870 | 186110 | 0 | 594650 | 0 | 0 | 225580 | 0 | 10162000 | 15213000 |
| QAVPISVPF | SNX27; SNX27 | Q96L92; A0A | SNX27_HUMAN | Unmodified | 0 | 0 | 0 | 0 | 1306300 | 298780 | 226320 | 58323 | 0 | 0 | 0 | 238340 | 151320 | 188210 | 0 | 0 | 197790 | 386090 |
| QAVQILDEI | SH3GL1 | Q99961 | SH3G1_HUMAI | Unmodified | 42222000 | 33931000 | 38852000 | 18643000 | 0 | 0 | 0 | 0 | 3705900 | 0 | 0 | 1193600 | 3651800 | 0 | 37723000 | 46819000 | 0 | 0 |
| QAVQILDEI | SH3GL1 | Q99961 | SH3G1_HUMAI | Unmodified | 932680 | 0 | 0 | 0 | 0 | 0 | 0 | 0 | 0 | 0 | 0 | 0 | 2820300 | 0 | 0 | 0 | 0 | 0 |
| QAVQILQQ | SH3GL2 | Q99962 | SH3G2_HUMAI | Unmodified | 136880 | 0 | 0 | 0 | 0 | 0 | 0 | 0 | 1496500 | 43661 | 0 | 0 | 0 | 0 | 0 | 29300 | 0 | 0 |
| QAVQILQQ | SH3GL2 | Q99962 | SH3G2_HUMAI | Unmodified | 0 | 0 | 0 | 0 | 0 | 0 | 0 | 0 | 0 | 695890 | 0 | 0 | 0 | 0 | 0 | 0 | 0 | 0 |
| QAVSELDE | INCENP | Q9NQS7 | INCE_HUMAN I | Unmodified | 0 | 0 | 232090 | 786390 | 1702000 | 0 | 0 | 0 | 0 | 0 | 0 | 0 | 1563900 | 479680 | 0 | 0 | 612870 | 0 |
| QAVSELDE | INCENP | Q9NQS7 | INCE_HUMAN I | Unmodified | 0 | 0 | 0 | 0 | 0 | 0 | 0 | 0 | 0 | 0 | 0 | 0 | 0 | 0 | 0 | 0 | 962720 | 0 |
| QAVSMFLG | ALDH7A1; ALDI | P49419; A0A | AL7A1_HUMAN | Oxidation (M) | 0 | 0 | 0 | 0 | 0 | 0 | 0 | 0 | 0 | 0 | 0 | 1212200 | 0 | 0 | 0 | 0 | 0 | 0 |
| QAVSMFLG | ALDH7A1; ALDI | P49419; A0A | AL7A1_HUMAN | Oxidation (M) | 0 | 0 | 0 | 0 | 1153100 | 365380 | 70488 | 0 | 0 | 0 | 0 | 0 | 0 | 0 | 0 | 0 | 1351000 | 2226300 |
| QAVSMFLG | ALDH7A1; ALDI | P49419; A0A | AL7A1_HUMAN | Unmodified | 0 | 0 | 0 | 0 | 2004900 | 0 | 0 | 0 | 0 | 0 | 0 | 0 | 0 | 0 | 0 | 0 | 1376200 | 1906400 |
| QAVTNPNN | HSPA9; HSPA9; | P38646; H0\ | GRP75_HUMAI | Unmodified | 0 | 0 | 0 | 0 | 0 | 0 | 0 | 0 | 247920 | 210010 | 0 | 0 | 0 | 0 | 0 | 0 | 0 | 0 |
| QAWFIENE | ANXA1; ANXA1; | P04083; Q5T | ANXA1_HUMAN | Unmodified | 0 | 0 | 0 | 0 | 34250000 | 4234100 | 0 | 0 | 0 | 36320 | 26842000 | 2.15E+08 | 0 | 0 | 0 | 0 | 4827200 | 7758800 |
| QAWQKAD | RPL14; RPL14 | P50914; E7E | RL14_HUMAN ( | Unmodified | 0 | 0 | 0 | 0 | 0 | 68375 | 49819 | 60324 | 0 | 0 | 0 | 0 | 472850 | 123660 | 0 | 0 | 0 | 112360 |
| QAWVITGG | WDR6; WDR6; \ | Q9NNW5; E9 | WDR6_HUMAN | Unmodified | 286520 | 475910 | 25251 | 0 | 0 | 0 | 0 | 0 | 0 | 0 | 0 | 247130 | 0 | 0 | 100470 | 42376 | 0 | 0 |
| QAYEPCLE | MYO1B; MYO1E | O43795; E9I | MYO1B_HUMA | Unmodified | 151070 | 87273 | 0 | 0 | 0 | 0 | 0 | 0 | 384350 | 0 | 0 | 4745000 | 0 | 0 | 106760 | 102970 | 0 | 0 |
| QAYTQFGC | PSMA4; PSMA4 | P25789; H0\ | PSA4_HUMAN | Unmodified | 0 | 44774 | 238350 | 163300 | 447280 | 0 | 0 | 0 | 1183000 | 581860 | 673840 | 6177300 | 104820 | 172810 | 551250 | 352280 | 0 | 0 |
| QAYTQFGC | PSMA4; PSMA4 | P25789; H0\ | PSA4_HUMAN | Unmodified | 0 | 0 | 0 | 0 | 2580500 | 450640 | 384060 | 0 | 817090 | 1457000 | 383140 | 471960 | 2491500 | 82971 | 0 | 0 | 1653900 | 1178700 |
| QAYVDKLE | HSPH1; HSPH1 | Q92598; A0A | HS105_HUMAN | Unmodified | 0 | 0 | 0 | 359520 | 358080 | 0 | 380640 | 0 | 793570 | 237400 | 419410 | 1610200 | 692480 | 0 | 0 | 0 | 417740 | 1252100 |
| QAYVDKLE | HSPH1; HSPH1 | Q92598; A0A | HS105_HUMAN | Oxidation (M) | 22400 | 0 | 0 | 78150 | 124830 | 0 | 0 | 0 | 0 | 0 | 258820 | 901350 | 0 | 0 | 180400 | 0 | 107100 | 197130 |
| QCCGTLPC | SLC30A1 | Q9Y6M5 | ZNT1_HUMAN | Unmodified | 110920 | 0 | 0 | 0 | 0 | 0 | 0 | 0 | 0 | 0 | 54018 | 285920 | 0 | 0 | 143640 | 120290 | 0 | 0 |
| QCCGTLPC | SLC30A1 | Q9Y6M5 | ZNT1_HUMAN | Unmodified | 0 | 0 | 0 | 158690 | 1127200 | 150790 | 168010 | 41971 | 0 | 94139 | 0 | 333350 | 0 | 0 | 138990 | 139940 | 222730 | 503570 |
| QCEGITSPI | PPME1 | Q9Y570 | PPME1_HUMAN | Unmodified | 0 | 0 | 0 | 0 | 318800 | 60678 | 56890 | 26996 | 282390 | 81260 | 154930 | 761470 | 593080 | 276280 | 103580 | 115200 | 222260 | 676280 |
| QCFHEEVE | TMED3; TMED3 | Q9Y3Q3; G3 | TMED3_HUMAI | Unmodified | 0 | 0 | 0 | 0 | 0 | 0 | 0 | 89888 | 7183500 | 950560 | 391110 | 169900 | 0 | 0 | 0 | 0 | 0 | 0 |
| QCFHGQV | ADGRG6 | Q86SQ4 | AGRG6_HUMA | Unmodified | 29291 | 0 | 98639 | 33967 | 0 | 94042 | 0 | 0 | 0 | 0 | 0 | 0 | 0 | 0 | 231000 | 347910 | 42699 | 0 |
| QCFYEDIA | TMED7; TMED7 | Q9Y3B3; A0A | TMED7_HUMAI | Unmodified | 0 | 0 | 0 | 0 | 0 | 0 | 326570 | 293950 | 11235000 | 2193400 | 0 | 568010 | 680860 | 0 | 0 | 0 | 0 | 0 |
| QCHQTQN | N/A | N/A | N/A | Unmodified | 10588000 | 13609000 | 2639400 | 0 | 0 | 0 | 0 | 0 | 0 | 0 | 0 | 0 | 10073000 | 0 | 5149700 | 0 | 0 | 0 |
| QCPIMDPA | PCK2; PCK2; P( | Q16822; H0\ | PCKGM_HUMA | Oxidation (M) | 0 | 0 | 0 | 0 | 0 | 0 | 0 | 0 | 1370300 | 0 | 0 | 0 | 0 | 0 | 0 | 0 | 1779300 | 0 |
| QCQCTSV( | EPCAM; EPCAM | P16422; C9J | EPCAM_HUMA | Unmodified | 17524000 | 5269200 | 6557900 | 0 | 0 | 0 | 0 | 0 | 17579000 | 1532100 | 2031000 | 20265000 | 3817500 | 1932800 | 13499000 | 15171000 | 0 | 0 |
| QCQYKDTL | PDCD6IP; PDCI | Q8WUM4; F! | PDC6I_HUMAN | Unmodified | 721520 | 0 | 0 | 845810 | 1108600 | 205880 | 110290 | 59848 | 187880 | 533710 | 26835 | 0 | 1510900 | 0 | 0 | 0 | 16454 | 483400 |
| QCQYKDTL | PDCD6IP; PDCI | Q8WUM4; F! | PDC6I_HUMAN | Unmodified | 0 | 0 | 0 | 0 | 1158700 | 0 | 0 | 0 | 0 | 0 | 0 | 0 | 0 | 1871100 | 0 | 0 | 458400 | 798860 |
| QCSILHGP | MUC5B | Q9HC84 | MUC5B_HUMA | Unmodified | 0 | 0 | 0 | 0 | 0 | 0 | 0 | 0 | 0 | 0 | 0 | 0 | 0 | 0 | 0 | 0 | 0 | 3335000 |
| QCSPSQGI | ADAM10 | O14672 | ADA10_HUMAN | Unmodified | 491660 | 0 | 371710 | 1117600 | 773260 | 222150 | 0 | 0 | 0 | 157180 | 0 | 1449100 | 1838400 | 1711000 | 951000 | 796720 | 674680 | 760800 |
| QCSWSSLI | CTHRC1; CTHR | Q96CG8; E7 | CTHR1_HUMAI | Unmodified | 0 | 0 | 0 | 0 | 0 | 0 | 0 | 0 | 1483500 | 331760 | 0 | 0 | 0 | 0 | 0 | 0 | 0 | 0 |
| QCTDQFGI | FAM20B | O75063 | XYLK_HUMAN ( | Oxidation (M) | 159000 | 56931 | 0 | 0 | 0 | 0 | 0 | 0 | 2461000 | 650380 | 128670 | 0 | 0 | 0 | 0 | 0 | 0 | 0 |
| QCTEVHLK | AGO2 | Q9UKV8 | AGO2_HUMAN | Unmodified | 0 | 0 | 0 | 0 | 61138 | 0 | 0 | 0 | 0 | 0 | 0 | 498620 | 0 | 0 | 475860 | 54641 | 0 | 0 |
| QCVENADI | PSMD1; PSMD1 | Q99460; C9J | PSMD1_HUMA | Unmodified | 0 | 0 | 0 | 0 | 0 | 0 | 0 | 0 | 0 | 0 | 0 | 732710 | 0 | 0 | 0 | 0 | 0 | 0 |
| QCVIDDR | RRAS2; RRAS2 | P62070; E9P | RRAS2_HUMAI | Unmodified | 1329100 | 792990 | 1028600 | 1167800 | 0 | 97497 | 0 | 0 | 789480 | 55262 | 0 | 0 | 236790 | 104110 | 1534700 | 1238000 | 0 | 0 |
| QDAFLLSK | THOP1; THOP1 | P52888; K7E | THOP1_HUMAI | Unmodified | 0 | 0 | 297950 | 22563 | 16790 | 359730 | 0 | 0 | 69070 | 90954 | 166710 | 728750 | 0 | 0 | 0 | 0 | 90003 | 0 |
| QDAQDLYE | ANXA4; ANXA4 | P09525; Q6I | ANXA4_HUMAI | Unmodified | 531910 | 534810 | 660550 | 0 | 0 | 0 | 0 | 0 | 2582500 | 501310 | 639460 | 1540200 | 0 | 0 | 425900 | 305030 | 0 | 0 |
| QDAQDLYE | ANXA4; ANXA4 | P09525; Q6I | ANXA4_HUMAI | Unmodified | 0 | 0 | 0 | 0 | 0 | 0 | 0 | 0 | 533190 | 0 | 0 | 673430 | 0 | 0 | 0 | 0 | 0 | 0 |
| QDAQILYK | ANXA3; ANXA3; | P12429; D6I | ANXA3_HUMAI | Unmodified | 0 | 0 | 114000 | 59056 | 1729900 | 0 | 0 | 0 | 204240 | 49841 | 140620 | 728230 | 0 | 0 | 0 | 121620 | 455770 | 457550 |
| QDAQSLH( | DDX21 | Q9NR30 | DDX21_HUMAN | Unmodified | 0 | 0 | 0 | 0 | 6506200 | 0 | 0 | 0 | 0 | 1221500 | 0 | 0 | 752100 | 2225600 | 0 | 0 | 12434000 | 14672000 |
| QDDELNLE | NOSTRIN; NOS | Q8IVI9; A0A( | NOSTN_HUMA | Unmodified | 1119000 | 858860 | 4703500 | 494370 | 0 | 0 | 0 | 0 | 0 | 0 | 0 | 0 | 0 | 0 | 3959500 | 1947500 | 0 | 0 |
| QDDELNLE | NOSTRIN; NOS | Q8IVI9; A0A( | NOSTN_HUMA | Unmodified | 1456600 | 2249800 | 0 | 0 | 0 | 0 | 0 | 0 | 0 | 0 | 0 | 0 | 0 | 0 | 2367100 | 1324800 | 0 | 0 |
| QDDGGAPI | NCAM2; NCAM: | O15394; H9I | NCAM2_HUMA | Unmodified | 0 | 0 | 3473300 | 1813000 | 0 | 0 | 0 | 0 | 0 | 0 | 0 | 0 | 0 | 0 | 9911700 | 1032000 | 0 | 0 |
| QDDNGVR | TLL2 | Q9Y6L7 | TLL2_HUMAN T | Unmodified | 0 | 0 | 0 | 0 | 0 | 0 | 0 | 0 | 284810 | 0 | 0 | 0 | 0 | 0 | 0 | 0 | 0 | 470390 |
| QDEEKPLH | IGFBP5; IGFBP! | P24593; C9J | IBP5_HUMAN I | Unmodified | 0 | 0 | 0 | 0 | 80032 | 0 | 0 | 0 | 5789600 | 102510 | 0 | 0 | 0 | 0 | 0 | 0 | 0 | 0 |
| QDFCHGQ | MGAT5 | Q09328 | MGT5A_HUMAI | Oxidation (M) | 0 | 0 | 0 | 0 | 0 | 0 | 0 | 0 | 6540100 | 2850000 | 701400 | 909490 | 0 | 0 | 0 | 0 | 0 | 0 |
| QDFCHGQ | MGAT5 | Q09328 | MGT5A_HUMAI | Unmodified | 0 | 0 | 0 | 0 | 0 | 212990 | 0 | 0 | 870860 | 45608 | 0 | 0 | 0 | 0 | 0 | 0 | 0 | 0 |
| QDFVQHFS | FDPS; FDPS; FD | P14324; A0A | FPPS_HUMAN | Unmodified | 1873400 | 0 | 0 | 0 | 0 | 0 | 0 | 0 | 2644800 | 256820 | 2451700 | 19957000 | 458360 | 0 | 767950 | 0 | 0 | 0 |
| QDGESYKN | JAM3 | Q9BX67 | JAM3_HUMAN | Unmodified | 0 | 0 | 0 | 0 | 0 | 0 | 0 | 0 | 797420 | 1507400 | 0 | 2600100 | 0 | 0 | 0 | 0 | 0 | 0 |
| QDGHLWC | FN1 | P02751 | FINC_HUMAN | Unmodified | 0 | 381780 | 2173100 | 0 | 0 | 0 | 0 | 0 | 0 | 0 | 0 | 0 | 0 | 0 | 0 | 1550300 | 0 | 0 |
| QDGVLAVT | ACOT8; ACOT8 | O14734; H7( | ACOT8_HUMAI | Unmodified | 0 | 0 | 0 | 0 | 0 | 0 | 0 | 0 | 864040 | 0 | 0 | 0 | 0 | 0 | 0 | 0 | 0 | 0 |
| QDIAFAYQ | ANXA2; ANXA2; | P07355; H0\ | ANXA2_HUMAI | Unmodified | 190410 | 998370 | 0 | 228850 | 413480 | 959230 | 0 | 0 | 7663100 | 208080 | 0 | 0 | 0 | 0 | 0 | 164600 | 181170 | 0 |
| QDIAFAYQ | ANXA2; ANXA2; | P07355; H0\ | ANXA2_HUMAI | Unmodified | 224160 | 0 | 0 | 0 | 940020 | 901660 | 0 | 0 | 3117600 | 173810 | 0 | 0 | 0 | 0 | 164780 | 0 | 307870 | 0 |
| QDIPAGLY | PIGX; PIGX; PIG | Q8TBF5; C9J | PIGX_HUMAN I | Unmodified | 0 | 0 | 0 | 0 | 0 | 0 | 0 | 0 | 520890 | 196070 | 0 | 0 | 0 | 0 | 0 | 0 | 0 | 0 |
| QDIPAGLY | PIGX; PIGX; PIG | Q8TBF5; C9J | PIGX_HUMAN I | Unmodified | 0 | 0 | 0 | 662300 | 0 | 0 | 0 | 0 | 3615300 | 1573900 | 0 | 0 | 0 | 0 | 0 | 0 | 0 | 0 |
| QDLAQLMI | COPS4; COPS4 | Q9BT78; D6I | CSN4_HUMAN | Unmodified | 0 | 34647 | 0 | 0 | 77840 | 0 | 0 | 0 | 0 | 0 | 0 | 900140 | 0 | 0 | 0 | 0 | 0 | 0 |
| QDLLAYLQ | CTNNA1; CTNN | P35221; P26 | CTNA1_HUMAI | Unmodified | 373060 | 465080 | 2996200 | 2650700 | 1423200 | 505720 | 421710 | 0 | 166880 | 0 | 1793000 | 1733500 | 1468200 | 397100 | 8173000 | 6212700 | 0 | 0 |
| QDLLAYLQ | CTNNA1; CTNN | P35221; P26 | CTNA1_HUMAI | Unmodified | 0 | 0 | 0 | 0 | 0 | 0 | 0 | 0 | 0 | 0 | 0 | 0 | 504080 | 1444500 | 0 | 0 | 0 | 0 |
| QDLLTAAP | ARL3 | P36405 | ARL3_HUMAN | Unmodified | 0 | 0 | 74156 | 0 | 0 | 0 | 0 | 0 | 311830 | 170340 | 0 | 2552100 | 0 | 0 | 1452700 | 1215800 | 0 | 0 |
| QDLLTAAP | ARL3 | P36405 | ARL3_HUMAN | Unmodified | 0 | 0 | 376580 | 0 | 0 | 0 | 177190 | 0 | 0 | 0 | 0 | 1651000 | 0 | 0 | 0 | 871850 | 195570 | 0 |
| QDLPALEEI | HSPA4; HSPA4 | P34932; A0A | HSP74_HUMAI | Unmodified | 0 | 0 | 139120 | 0 | 0 | 0 | 0 | 0 | 0 | 0 | 437070 | 5235900 | 0 | 0 | 107520 | 0 | 0 | 0 |
| QDLPDAMI | ARF6 | P62330 | ARF6_HUMAN | Unmodified | 0 | 0 | 6334800 | 0 | 902020 | 0 | 0 | 0 | 2147200 | 715150 | 419570 | 1776200 | 0 | 0 | 4791400 | 3003600 | 0 | 336390 |
| QDLPDAMI | ARF6 | P62330 | ARF6_HUMAN | Oxidation (M) | 109340 | 381430 | 1724100 | 35671 | 197320 | 0 | 0 | 0 | 285440 | 0 | 55511 | 643330 | 0 | 0 | 966170 | 831340 | 0 | 0 |
| QDLPNAMI | ARF3; ARF1; N/ | P61204; P84 | ARF3_HUMAN | Oxidation (M) | 0 | 188010 | 2215300 | 0 | 0 | 0 | 0 | 0 | 0 | 0 | 1282000 | 2303300 | 0 | 0 | 789360 | 787740 | 0 | 0 |
| QDLPNAMI | ARF3; ARF1; N/ | P61204; P84 | ARF3_HUMAN | Unmodified | 200440 | 317270 | 2178900 | 0 | 0 | 0 | 0 | 0 | 781040 | 0 | 386070 | 3489000 | 0 | 0 | 691230 | 453540 | 0 | 0 |
| QDLPNAMI | ARF3; ARF1; N/ | P61204; P84 | ARF3_HUMAN | Oxidation (M) | 0 | 947130 | 0 | 0 | 0 | 0 | 0 | 0 | 0 | 0 | 0 | 0 | 0 | 0 | 0 | 320220 | 0 | 0 |
| QDLTTALG | ARHGEF2; ARH | Q92974; V9C | ARHG2_HUMA | Unmodified | 0 | 0 | 0 | 0 | 0 | 0 | 0 | 0 | 0 | 0 | 0 | 262090 | 0 | 0 | 0 | 0 | 0 | 0 |
| QDLTTLDV | EIF2S3; EIF2S3I | P41091; Q2\ | IF2G_HUMAN E | Unmodified | 0 | 0 | 0 | 0 | 0 | 0 | 0 | 0 | 0 | 0 | 0 | 3488500 | 142430 | 0 | 0 | 0 | 0 | 0 |
| QDLTTLDV | EIF2S3; EIF2S3I | P41091; Q2\ | IF2G_HUMAN E | Unmodified | 339630 | 332220 | 56228 | 840400 | 0 | 0 | 0 | 0 | 1278200 | 0 | 638770 | 3725900 | 161730 | 0 | 265770 | 0 | 0 | 0 |
| QDPQVHSI | RHBDF1 | Q96CC6 | RHDF1_HUMAI | Unmodified | 158770 | 600940 | 0 | 106340 | 0 | 0 | 0 | 0 | 594940 | 157270 | 0 | 0 | 0 | 0 | 224520 | 283460 | 0 | 0 |
| QDPSVLHT | ST13; ST13P5; | P50502; Q8I | F10A1_HUMAN | Unmodified | 0 | 0 | 0 | 0 | 0 | 0 | 0 | 0 | 680290 | 0 | 109850 | 1091100 | 0 | 0 | 0 | 0 | 0 | 0 |
| QDPSVLHT | ST13; ST13P5; | P50502; Q8I | F10A1_HUMAN | Oxidation (M) | 104340 | 86878 | 103350 | 0 | 0 | 0 | 0 | 0 | 0 | 0 | 0 | 663420 | 85770 | 0 | 154950 | 156210 | 0 | 0 |
| QDPTLLWC | CACNA2D2; CA | Q9NY47; C9 | CA2D2_HUMAI | Unmodified | 36744 | 0 | 0 | 0 | 0 | 0 | 0 | 0 | 246200 | 21699 | 0 | 90894 | 21380 | 0 | 0 | 0 | 0 | 0 |
| QDQVCIAR | GSPT1; GSPT2; | P15170; Q8I | ERF3A_HUMAI | Unmodified | 0 | 112510 | 158590 | 0 | 564180 | 0 | 0 | 0 | 292030 | 36408 | 0 | 2152500 | 0 | 0 | 0 | 0 | 164120 | 0 |
| QDRDLEEP | IL1RAP; IL1RAP | Q9NPH3; C9 | IL1AP_HUMAN | Unmodified | 0 | 0 | 0 | 0 | 0 | 0 | 0 | 0 | 0 | 0 | 0 | 0 | 0 | 0 | 628400 | 125630 | 0 | 0 |
| QDRFAGLH | HADH; HADH; I | Q16836; A0A | HCDH_HUMAN | Unmodified | 0 | 0 | 0 | 1089000 | 0 | 0 | 0 | 0 | 569620 | 0 | 0 | 2251500 | 0 | 289060 | 0 | 0 | 0 | 0 |
| QDRITRYQ | PABPC1; PABP( | P11940; A0A | PABP1_HUMAN | Unmodified | 0 | 0 | 0 | 954820 | 0 | 0 | 341110 | 0 | 0 | 0 | 0 | 0 | 1523000 | 574550 | 0 | 0 | 0 | 110440 |
| QDRPFFLY | ARSA; ARSA | P15289; A0A | ARSA_HUMAN | Unmodified | 0 | 0 | 0 | 0 | 0 | 0 | 0 | 0 | 2067800 | 0 | 0 | 0 | 0 | 449500 | 0 | 0 | 0 | 0 |
| QDTFALAS | ACAA1; ACAA1; | P09110; C9J | THIK_HUMAN 3 | Unmodified | 0 | 0 | 0 | 0 | 140560 | 0 | 0 | 90523 | 5333300 | 991730 | 0 | 0 | 148130 | 0 | 0 | 0 | 0 | 0 |

| | | | | | | | | | | | | | | | | | | | | | | |
|---|---|---|---|---|---|---|---|---|---|---|---|---|---|---|---|---|---|---|---|---|---|---|
| QDTFSGTG | USP15 | Q9Y4E8 | UBP15_HUMAN | Unmodified | 59718 | 0 | 0 | 0 | 0 | 0 | 0 | 0 | 0 | 0 | 0 | 1163000 | 0 | 0 | 0 | 129510 | 0 | 0 |
| QDTSSSTTI | ANLN | Q9NQW6 | ANLN_HUMAN | Unmodified | 0 | 0 | 360960 | 1611900 | 1298600 | 0 | 0 | 0 | 0 | 0 | 0 | 295780 | 0 | 0 | 197250 | 147950 | 289990 | 295000 |
| QDTSSSTTI | ANLN | Q9NQW6 | ANLN_HUMAN | Unmodified | 0 | 0 | 0 | 0 | 0 | 0 | 0 | 0 | 0 | 0 | 0 | 0 | 0 | 0 | 0 | 0 | 515130 | 0 |
| QDVIITALD | UBA6; UBA6 | A0AVT1; H0Y | UBA6_HUMAN | Unmodified | 0 | 101250 | 0 | 35568 | 0 | 0 | 0 | 0 | 0 | 0 | 230030 | 1117800 | 0 | 0 | 52526 | 31725 | 0 | 0 |
| QDVIITALD | UBA6; UBA6 | A0AVT1; H0Y | UBA6_HUMAN | Unmodified | 0 | 0 | 0 | 0 | 0 | 104850 | 0 | 212440 | 1283500 | 0 | 0 | 961020 | 0 | 0 | 0 | 0 | 0 | 0 |
| QDVLNWLI | PPP6R3; PPP6R | Q5H9R7; E9I | PP6R3_HUMAN | Unmodified | 0 | 0 | 0 | 0 | 0 | 0 | 0 | 0 | 0 | 0 | 0 | 1679900 | 0 | 0 | 0 | 0 | 0 | 0 |
| QDVYGPQI | CTNND1; CTNN | O60716; A0A | CTND1_HUMAI | Unmodified | 304110 | 256090 | 1016100 | 610020 | 0 | 0 | 0 | 0 | 0 | 0 | 287960 | 1253300 | 0 | 0 | 0 | 1964300 | 0 | 0 |
| QDYKEGEN | PSMA4; PSMA4 | P25789; H0Y | PSA4_HUMAN | Unmodified | 0 | 0 | 0 | 0 | 114430 | 0 | 0 | 0 | 157620 | 0 | 0 | 423000 | 85366 | 58824 | 0 | 0 | 0 | 0 |
| QDYQAVRN | ATP2B4 | P23634 | AT2B4_HUMAN | Unmodified | 0 | 20600 | 1011600 | 904840 | 316620 | 62918 | 284230 | 30209 | 455600 | 0 | 528540 | 3807100 | 0 | 0 | 696690 | 930900 | 0 | 0 |
| QEAEEAER | N/A | N/A | N/A | Unmodified | 0 | 0 | 0 | 0 | 14340000 | 2514100 | 551080 | 334150 | 0 | 0 | 0 | 0 | 0 | 0 | 0 | 0 | 8515100 | 6476100 |
| QEAGGLGI | SNTB2; SNTB2 | Q13425; A0A | SNTB2_HUMAN | Unmodified | 0 | 0 | 2447300 | 242370 | 44117 | 0 | 0 | 43107 | 934600 | 0 | 90402 | 388300 | 25316 | 0 | 2905100 | 1825500 | 0 | 0 |
| QEAGISEG | DENR; DENR | O43583; F8V | DENR_HUMAN | Unmodified | 0 | 0 | 157860 | 0 | 0 | 0 | 0 | 0 | 186770 | 0 | 540000 | 2342400 | 0 | 0 | 0 | 157890 | 0 | 0 |
| QEALEWLIF | XPNPEP1; XPNI | Q9NQW7; Q | XPP1_HUMAN . | Unmodified | 0 | 0 | 47150 | 0 | 0 | 0 | 0 | 0 | 0 | 0 | 0 | 2457500 | 0 | 0 | 90945 | 54004 | 0 | 0 |
| QEATFCSQ | ANLN; ANLN; A | Q9NQW6; H | ANLN_HUMAN | Unmodified | 0 | 0 | 0 | 0 | 534130 | 0 | 0 | 0 | 21600 | 0 | 0 | 0 | 0 | 0 | 0 | 0 | 264590 | 232660 |
| QEATVNYLI | TTC17 | Q96AE7 | TTC17_HUMAN | Unmodified | 0 | 0 | 0 | 0 | 0 | 0 | 0 | 0 | 1536700 | 160380 | 0 | 0 | 0 | 0 | 0 | 0 | 0 | 0 |
| QEDGSLCL | SRP19; SRP19; | P09132; A0A | SRP19_HUMAN | Unmodified | 0 | 19927 | 110450 | 5817700 | 22985000 | 3867100 | 6904800 | 3691700 | 0 | 0 | 0 | 1347900 | 30477000 | 21496000 | 492250 | 236060 | 1576600 | 2414400 |
| QEDHAEAA | KIT | P10721 | KIT_HUMAN Ma | Unmodified | 0 | 0 | 0 | 0 | 0 | 0 | 244130 | 82044 | 9033000 | 76883 | 2221400 | 0 | 0 | 0 | 0 | 0 | 0 | 0 |
| QEDQLQD | GOLM2; GOLM2 | Q6P4E1; H0Y | GOLM2_HUMA | Unmodified | 0 | 0 | 0 | 0 | 0 | 105610 | 0 | 0 | 1110100 | 150000 | 0 | 0 | 147660 | 219500 | 0 | 0 | 0 | 17401 |
| QEEATATEL | MKRN1; MKRN4 | Q9UHC7; Q1 | MKRN1_HUMA | Unmodified | 389740 | 0 | 546390 | 0 | 0 | 0 | 0 | 0 | 0 | 0 | 0 | 0 | 0 | 0 | 0 | 0 | 0 | 0 |
| QEEELQAK | MYH10 | P35580 | MYH10_HUMAI | Unmodified | 40086 | 0 | 0 | 0 | 0 | 0 | 0 | 0 | 0 | 0 | 164040 | 6020200 | 0 | 0 | 202410 | 103270 | 0 | 132090 |
| QEEEMMAI | MYH9 | P35579 | MYH9_HUMAN | 2 Oxidation (M) | 158420 | 62657 | 46634 | 0 | 0 | 0 | 0 | 0 | 0 | 83793 | 47281 | 736670 | 0 | 0 | 215340 | 145210 | 0 | 0 |
| QEEEMMAI | MYH9 | P35579 | MYH9_HUMAN | Unmodified | 0 | 0 | 0 | 0 | 0 | 0 | 0 | 0 | 0 | 0 | 0 | 2390200 | 0 | 0 | 0 | 0 | 0 | 0 |
| QEEEMMAI | MYH9 | P35579 | MYH9_HUMAN | Oxidation (M) | 0 | 30294 | 161730 | 0 | 0 | 0 | 0 | 0 | 0 | 0 | 224430 | 1249100 | 0 | 0 | 216660 | 412920 | 0 | 0 |
| QEEENPAE | ENSA; ENSA; EN | O43768; A0A | ENSA_HUMAN | Unmodified | 0 | 0 | 0 | 0 | 0 | 0 | 0 | 0 | 0 | 0 | 96431 | 0 | 0 | 0 | 0 | 0 | 0 | 0 |
| QEERLPIDE | MME; MME; MM | P08473; A0A | NEP_HUMAN N | Oxidation (M) | 0 | 0 | 5407500 | 6601000 | 0 | 0 | 0 | 0 | 474170 | 266320 | 0 | 0 | 782310 | 0 | 14213000 | 13420000 | 0 | 0 |
| QEERLPIDE | MME; MME; MM | P08473; A0A | NEP_HUMAN N | Unmodified | 0 | 128460 | 747920 | 1625000 | 286730 | 0 | 0 | 0 | 2187800 | 208950 | 0 | 258360 | 273780 | 0 | 4063600 | 1378000 | 284890 | 4061700 |
| QEESLMPS | EHD1; EHD1; EI | Q9H4M9; C9 | EHD1_HUMAN | Unmodified | 0 | 0 | 0 | 0 | 0 | 0 | 0 | 0 | 0 | 0 | 0 | 4708000 | 0 | 0 | 0 | 258380 | 0 | 0 |
| QEETDHKN | OGA | O60502 | OGA_HUMAN F | Unmodified | 0 | 0 | 0 | 160690 | 0 | 0 | 0 | 0 | 0 | 0 | 0 | 0 | 75378 | 0 | 118300 | 135790 | 0 | 0 |
| QEGGDND | ADSL; ADSL; AD | P30566; A0A | PUR8_HUMAN | Unmodified | 0 | 0 | 0 | 0 | 0 | 0 | 0 | 29356 | 0 | 0 | 373820 | 3443600 | 0 | 0 | 157420 | 0 | 0 | 0 |
| QEGIIFIGPI | MCCC1; MCCC | Q96RQ3; E9I | MCCA_HUMAN | Oxidation (M) | 0 | 0 | 0 | 0 | 0 | 0 | 67343 | 0 | 669330 | 178460 | 0 | 0 | 0 | 0 | 0 | 0 | 0 | 0 |
| QEGLESVLI | JUP | P14923 | PLAK_HUMAN . | Unmodified | 0 | 0 | 4893600 | 823670 | 0 | 0 | 0 | 0 | 0 | 0 | 1426400 | 8097900 | 65199 | 0 | 2366900 | 2480700 | 0 | 0 |
| QEGRKDSL | S100A13 | Q99584 | S10AD_HUMAN | Unmodified | 0 | 0 | 0 | 0 | 827930 | 700230 | 0 | 0 | 0 | 0 | 0 | 0 | 0 | 0 | 0 | 0 | 304640 | 1138700 |
| QEILAALEK | PSAP; PSAP; PS | P07602; Q5E | SAP_HUMAN P | Unmodified | 0 | 0 | 0 | 0 | 197110 | 0 | 0 | 0 | 7218300 | 0 | 0 | 1791400 | 0 | 0 | 0 | 0 | 0 | 0 |
| QEIQHLFR | RPN2; RPN2 | P04844; H0Y | RPN2_HUMAN | Unmodified | 0 | 121360 | 199550 | 303210 | 0 | 0 | 0 | 0 | 599810 | 0 | 0 | 0 | 0 | 0 | 96844 | 0 | 0 | 0 |
| QEIQHLFRI | RPN2; RPN2 | P04844; H0Y | RPN2_HUMAN | Unmodified | 0 | 0 | 0 | 0 | 0 | 0 | 0 | 0 | 3642500 | 87620 | 0 | 0 | 0 | 0 | 0 | 0 | 0 | 0 |
| QEISSLKDE | MPRIP; MPRIP; | Q6WCQ1; H | MPRIP_HUMAN | Unmodified | 0 | 0 | 0 | 0 | 0 | 0 | 0 | 22423 | 0 | 32431 | 0 | 425180 | 0 | 0 | 0 | 0 | 0 | 0 |
| QEIVSLFNA | ERO1A | Q96HE7 | ERO1A_HUMAN | Unmodified | 0 | 0 | 0 | 0 | 0 | 0 | 0 | 0 | 481630 | 0 | 0 | 0 | 0 | 0 | 0 | 0 | 0 | 0 |
| QEKPAEKP | RAD23B; RAD23 | P54727; H0Y | RD23B_HUMAN | Unmodified | 208160 | 179710 | 324680 | 108580 | 0 | 0 | 0 | 0 | 148740 | 0 | 0 | 107610 | 0 | 0 | 322100 | 118600 | 0 | 0 |
| QELGGLGI | SNTB1; SNTB1 | Q13884; A0A | SNTB1_HUMAN | Unmodified | 0 | 0 | 0 | 0 | 0 | 0 | 0 | 0 | 2488000 | 0 | 0 | 0 | 0 | 0 | 0 | 0 | 0 | 0 |
| QELGGLGI | SNTB1; SNTB1 | Q13884; A0A | SNTB1_HUMAN | Unmodified | 0 | 0 | 0 | 0 | 0 | 0 | 0 | 0 | 562390 | 350870 | 0 | 0 | 0 | 0 | 0 | 0 | 0 | 208530 |
| QELILSNSE | COPA; COPA; C | P53621; A0A | COPA_HUMAN | Unmodified | 0 | 0 | 0 | 0 | 244730 | 0 | 0 | 0 | 548150 | 0 | 291710 | 3262200 | 0 | 0 | 0 | 0 | 0 | 0 |
| QELLEALTK | XRCC6; XRCC6 | P12956; B1A | XRCC6_HUMAI | Unmodified | 225580 | 505220 | 0 | 0 | 0 | 0 | 0 | 0 | 0 | 0 | 231920 | 0 | 0 | 0 | 494460 | 0 | 0 | 0 |
| QELLQHSL | PLEKHG3; PLEK | A1L390; A0A | PKHG3_HUMAI | Unmodified | 0 | 0 | 0 | 0 | 1076700 | 164640 | 0 | 0 | 0 | 0 | 0 | 0 | 0 | 0 | 0 | 0 | 124330 | 317330 |
| QELQEMSE | CSGALNACT2 | Q8N6G5 | CGAT2_HUMAI | Unmodified | 0 | 0 | 0 | 0 | 124010 | 0 | 0 | 0 | 320650 | 0 | 290400 | 0 | 0 | 0 | 0 | 0 | 46685 | 108830 |
| QELQNLRD | STX4; STX4; ST | Q12846; A0A | STX4_HUMAN S | Unmodified | 0 | 0 | 0 | 0 | 2127700 | 0 | 0 | 53347 | 476740 | 0 | 0 | 924080 | 507140 | 0 | 0 | 0 | 34557 | 0 |
| QELSFAAR | FASN; FASN | P49327; A0A | FAS_HUMAN Fa | Unmodified | 0 | 0 | 0 | 109340 | 0 | 0 | 0 | 0 | 0 | 157950 | 0 | 4987200 | 259190 | 0 | 212920 | 251620 | 394050 | 0 |
| QELSHALY | PRPF19; PRPF1 | Q9UMS4; F5 | PRP19_HUMAN | Unmodified | 71750 | 0 | 0 | 0 | 0 | 128730 | 0 | 0 | 90414 | 0 | 0 | 2676800 | 162150 | 21919 | 0 | 0 | 0 | 0 |
| QEMQEVQS | HNRNPA2B1; H | P22626; A0A | ROA2_HUMAN | Unmodified | 395870 | 134530 | 0 | 0 | 0 | 0 | 0 | 0 | 0 | 0 | 541110 | 2006900 | 73294 | 0 | 94331 | 0 | 0 | 0 |
| QEMTHMVI | PTPRK; PTPRK; | Q15262; H0Y | PTPRK_HUMAN | Unmodified | 0 | 0 | 0 | 0 | 0 | 0 | 0 | 0 | 0 | 63152 | 56100 | 582510 | 0 | 0 | 0 | 0 | 0 | 0 |
| QENGASVI | EEF1D; EEF1D; | P29692; E9P | EF1D_HUMAN | Unmodified | 362790 | 535830 | 820280 | 0 | 0 | 0 | 0 | 0 | 0 | 0 | 0 | 5450200 | 271600 | 0 | 351350 | 1131100 | 0 | 0 |
| QEPLLIGST | FASN; FASN | P49327; A0A | FAS_HUMAN Fa | Unmodified | 0 | 0 | 0 | 0 | 0 | 0 | 0 | 0 | 0 | 0 | 0 | 20737000 | 0 | 0 | 0 | 0 | 0 | 0 |
| QEPVKPEE | RHOA; RHOA | P61586; A0A | RHOA_HUMAN | Unmodified | 0 | 920330 | 1731700 | 0 | 350260 | 0 | 0 | 0 | 5391500 | 103290 | 1111500 | 10167000 | 0 | 62823 | 920940 | 1188600 | 0 | 0 |
| QEQEEYQF | MYO1D; MYO1D | O94832; J3Q | MYO1D_HUMA | Unmodified | 0 | 0 | 0 | 181240 | 0 | 0 | 126220 | 47502 | 0 | 0 | 0 | 180450 | 0 | 0 | 578940 | 201980 | 0 | 0 |
| QEQETSYTI | EPHA3; EPHA3 | P29320; C9J | EPHA3_HUMAN | Unmodified | 1839900 | 242220 | 0 | 0 | 0 | 0 | 0 | 0 | 0 | 0 | 0 | 3230900 | 0 | 0 | 0 | 0 | 0 | 0 |
| QEQRPLPN | C1orf198; C1or | Q9H425; E5I | CA198_HUMAN | Unmodified | 0 | 0 | 0 | 0 | 548160 | 0 | 0 | 0 | 0 | 34054 | 224090 | 650760 | 0 | 0 | 191710 | 0 | 19695 | 289730 |
| QEREPVGT | ITGAV | P06756 | ITAV_HUMAN In | Unmodified | 495980 | 0 | 741270 | 0 | 0 | 0 | 0 | 0 | 511820 | 0 | 403230 | 1984500 | 0 | 0 | 652390 | 539390 | 0 | 0 |
| QERNFDDA | CNN2; CNN2; C | Q99439; H3I | CNN2_HUMAN | Unmodified | 0 | 0 | 0 | 0 | 3091800 | 632150 | 0 | 0 | 0 | 0 | 0 | 0 | 190630 | 168320 | 0 | 0 | 0 | 0 |
| QESNQIGP | ANK3; ANK3; AN | Q12955; A0A | ANK3_HUMAN | Unmodified | 0 | 0 | 170130 | 0 | 0 | 0 | 0 | 0 | 0 | 0 | 0 | 0 | 369040 | 683830 | 3024700 | 599370 | 0 | 0 |
| QESTSVLL( | DSG2 | Q14126 | DSG2_HUMAN | Unmodified | 0 | 0 | 0 | 0 | 0 | 0 | 0 | 0 | 869040 | 0 | 0 | 303510 | 0 | 0 | 0 | 0 | 0 | 0 |
| QETEVELYI | RAB7A; RAB7A; | P51149; C9J | RAB7A_HUMAN | Unmodified | 0 | 0 | 851360 | 0 | 0 | 0 | 0 | 0 | 0 | 0 | 0 | 1479600 | 0 | 0 | 361170 | 270360 | 0 | 0 |
| QETEVELYI | RAB7A; RAB7A; | P51149; C9J | RAB7A_HUMAN | Unmodified | 21663 | 0 | 1070600 | 0 | 0 | 0 | 0 | 0 | 0 | 0 | 1393900 | 5978900 | 0 | 0 | 4316900 | 1350700 | 0 | 0 |
| QETEVELYI | RAB7A; RAB7A; | P51149; C9J | RAB7A_HUMAN | Unmodified | 0 | 586240 | 0 | 0 | 0 | 0 | 0 | 0 | 880320 | 300010 | 0 | 1251800 | 0 | 708120 | 0 | 918850 | 0 | 0 |
| QETQLLED | MAN1B1; MAN1 | Q9UKM7; H0 | MA1B1_HUMAI | Unmodified | 0 | 1007100 | 0 | 0 | 0 | 0 | 0 | 0 | 216880 | 164140 | 0 | 0 | 0 | 0 | 0 | 0 | 0 | 0 |
| QEVISTSSK | RACK1; RACK1; | P63244; D6F | RACK1_HUMAI | Unmodified | 0 | 0 | 329220 | 0 | 0 | 0 | 0 | 0 | 0 | 0 | 0 | 1238500 | 0 | 0 | 148480 | 0 | 0 | 0 |
| QEVPIIIVFR | POLR2B; POLR2 | P30876; C9J | RPB2_HUMAN | Unmodified | 0 | 0 | 0 | 0 | 0 | 0 | 0 | 0 | 0 | 0 | 0 | 426660 | 0 | 0 | 0 | 0 | 0 | 0 |
| QEVPLATLE | SERPINI1 | Q99574 | NEUS_HUMAN | Unmodified | 0 | 0 | 0 | 0 | 0 | 0 | 0 | 0 | 1425300 | 0 | 0 | 0 | 0 | 0 | 0 | 0 | 0 | 0 |
| QEVQNLIK | DPYSL3 | Q14195 | DPYL3_HUMAN | Unmodified | 0 | 0 | 0 | 0 | 546750 | 0 | 0 | 0 | 170280 | 68770 | 4727900 | 35153000 | 471670 | 199090 | 0 | 0 | 190680 | 0 |
| QEWENNR | CPD | O75976 | CBPD_HUMAN | Unmodified | 0 | 0 | 0 | 26832 | 0 | 0 | 0 | 0 | 3805100 | 369560 | 0 | 0 | 0 | 0 | 33855 | 29993 | 0 | 0 |
| QEWFWGS | ECT2; ECT2 | Q9H8V3; H7 | ECT2_HUMAN | Oxidation (M) | 0 | 0 | 268860 | 1686900 | 0 | 0 | 0 | 0 | 9837700 | 0 | 0 | 0 | 0 | 2569600 | 0 | 0 | 0 | 0 |
| QEYDESGP | POTEJ; POTEE; I | P0CG39; Q6 | POTEJ_HUMAN | Unmodified | 20835000 | 14911000 | 13487000 | 2764700 | 1436900 | 135190 | 608390 | 254980 | 1172700 | 0 | 16569000 | 1.36E+08 | 207040 | 0 | 49184000 | 33129000 | 237010 | 371260 |
| QEYDESGP | POTEJ; POTEE; I | P0CG39; Q6 | POTEJ_HUMAN | Unmodified | 10922000 | 13457000 | 29743000 | 15697000 | 50674000 | 14044000 | 3855500 | 2874100 | 10955000 | 2096500 | 12761000 | 97424000 | 14421000 | 6395500 | 23254000 | 30707000 | 5954400 | 10406000 |
| QFAAATIQT | AP3B1 | O00203 | AP3B1_HUMAN | Unmodified | 0 | 0 | 0 | 0 | 0 | 0 | 0 | 0 | 0 | 0 | 0 | 2285300 | 0 | 0 | 0 | 0 | 0 | 0 |
| QFAAVSITT | EPHA3; EPHA3 | P29320; C9J | EPHA3_HUMAN | Unmodified | 1131100 | 708910 | 0 | 0 | 0 | 0 | 0 | 0 | 0 | 0 | 0 | 0 | 0 | 0 | 0 | 0 | 0 | 0 |
| QFAEENGL | RAB14; RAB14 | P61106; X6R | RAB14_HUMAN | Unmodified | 0 | 0 | 0 | 0 | 584550 | 8183.3 | 0 | 0 | 902050 | 235140 | 1267200 | 3975200 | 0 | 0 | 0 | 147870 | 0 | 485330 |
| QFAEMYVA | EEF2 | P13639 | EF2_HUMAN El | Oxidation (M) | 0 | 0 | 0 | 0 | 541500 | 135260 | 390040 | 209710 | 0 | 301660 | 514220 | 7218300 | 0 | 0 | 1683000 | 0 | 267630 | 0 |
| QFAEMYVA | EEF2 | P13639 | EF2_HUMAN El | Unmodified | 200810 | 0 | 255740 | 160610 | 256340 | 0 | 950100 | 226580 | 0 | 1055200 | 1991200 | 6182100 | 20029 | 0 | 570500 | 193750 | 888850 | 294780 |
| QFAPEYEKI | PDIA4; PDIA4; F | P13667; C9J | PDIA4_HUMAN | Unmodified | 0 | 0 | 0 | 0 | 49902 | 0 | 729780 | 0 | 0 | 222890 | 0 | 0 | 0 | 0 | 0 | 0 | 0 | 57006 |
| QFAQLFGK | TRAF5 | O00463 | TRAF5_HUMAN | Unmodified | 152930 | 0 | 0 | 127150 | 0 | 0 | 0 | 0 | 0 | 0 | 0 | 779800 | 0 | 0 | 139420 | 155560 | 0 | 0 |
| QFASQANV | ACTN4; ACTN4; | O43707; F5C | ACTN4_HUMAI | Unmodified | 246840 | 276410 | 570590 | 0 | 997580 | 701300 | 0 | 0 | 700190 | 345470 | 0 | 7389400 | 0 | 42541 | 831850 | 728880 | 0 | 97503 |
| QFAYQNSV | EPRS1 | P07814 | SYEP_HUMAN | Unmodified | 0 | 0 | 184560 | 85551 | 0 | 0 | 0 | 29003 | 255080 | 259730 | 0 | 5503600 | 137010 | 48600 | 0 | 0 | 0 | 0 |
| QFCLQNA( | GNG5 | P63218 | GBG5_HUMAN | Unmodified | 2602900 | 0 | 7970700 | 0 | 13419000 | 1697900 | 3703400 | 613580 | 16903000 | 10990000 | 7640400 | 60680000 | 9845000 | 7646300 | 3551100 | 2131000 | 5693300 | 17926000 |
| QFCVPGFK | SLC44A2; SLC4 | Q8IWA5; K7I | CTL2_HUMAN ( | Unmodified | 0 | 0 | 1851400 | 2681100 | 548030 | 81933 | 0 | 0 | 1971300 | 987250 | 702450 | 6197900 | 765460 | 375550 | 4424100 | 3334700 | 0 | 1444100 |
| QFDGAAV( | AVIL | O75366 | AVIL_HUMAN A | Unmodified | 71921 | 258150 | 0 | 0 | 0 | 83250 | 0 | 0 | 0 | 0 | 2272700 | 38374000 | 0 | 0 | 0 | 151760 | 124750 | 0 |
| QFDHLNHI | NUCB2; NUCB2 | P80303; Q2L | NUCB2_HUMA | Oxidation (M) | 0 | 0 | 0 | 885270 | 0 | 0 | 0 | 0 | 34934000 | 13499000 | 259070 | 0 | 0 | 997100 | 15263000 | 0 | 0 | 0 |
| QFDHLNHI | NUCB2; NUCB2 | P80303; Q2L | NUCB2_HUMA | Unmodified | 0 | 0 | 0 | 0 | 0 | 0 | 0 | 0 | 23964000 | 2423500 | 0 | 0 | 0 | 0 | 0 | 0 | 0 | 0 |
| QFDIQLLTH | GCLM | P48507 | GSH0_HUMAN | Unmodified | 216880 | 0 | 473800 | 249700 | 1418800 | 0 | 910860 | 0 | 756140 | 210630 | 0 | 0 | 1432800 | 660420 | 553070 | 319170 | 352940 | 634110 |
| QFEDELHP | PPIL1 | Q9Y3C6 | PPIL1_HUMAN | Unmodified | 2143100 | 1681500 | 1118100 | 1033500 | 648500 | 0 | 0 | 0 | 609910 | 0 | 484820 | 3227200 | 1123500 | 0 | 1769800 | 1311400 | 68922 | 260300 |
| QFEEIWER | MYO6; MYO6; M | Q9UM54; A0 | MYO6_HUMAN | Unmodified | 0 | 0 | 0 | 3599900 | 0 | 0 | 0 | 0 | 0 | 0 | 0 | 1459100 | 0 | 0 | 1842000 | 1233300 | 10625 | 0 |
| QFEELTLGI | BIRC5; BIRC5; I | O15392; H3I | BIRC5_HUMAN | Unmodified | 0 | 0 | 0 | 0 | 2551600 | 0 | 0 | 0 | 0 | 191060 | 0 | 0 | 0 | 0 | 0 | 0 | 0 | 0 |
| QFEGLADK | ERLIN2; ERLIN2 | O94905; E5F | ERLN2_HUMAN | Unmodified | 0 | 0 | 0 | 0 | 0 | 0 | 0 | 0 | 7253900 | 378570 | 523800 | 147240 | 0 | 0 | 0 | 0 | 74539 | 0 |
| QFEGLADK | ERLIN2; ERLIN2 | O94905; E5F | ERLN2_HUMAN | Unmodified | 0 | 0 | 0 | 0 | 0 | 0 | 0 | 0 | 3217200 | 1686700 | 0 | 0 | 0 | 0 | 0 | 0 | 0 | 0 |
| QFEGLADK | ERLIN2 | O94905 | ERLN2_HUMAN | Unmodified | 0 | 0 | 0 | 0 | 0 | 0 | 0 | 0 | 1820700 | 5083400 | 0 | 0 | 0 | 0 | 0 | 0 | 0 | 0 |
| QFEHLDPQ | NUCB1; NUCB1 | Q02818; C9J | NUCB1_HUMA | Unmodified | 115900 | 193530 | 466580 | 161700 | 165500 | 0 | 0 | 0 | 7651600 | 0 | 722550 | 1074500 | 0 | 0 | 160790 | 242160 | 0 | 0 |
| QFELALQL | VPS39 | Q96JC1 | VPS39_HUMAN | Unmodified | 0 | 0 | 440950 | 0 | 0 | 1019300 | 1144000 | 267840 | 0 | 0 | 0 | 1592400 | 0 | 0 | 0 | 0 | 0 | 5879000 |
| QFGAQAN | ACTN1; ACTN1; | P12814; H0Y | ACTN1_HUMAI | Unmodified | 0 | 0 | 0 | 0 | 633470 | 0 | 409990 | 0 | 0 | 0 | 0 | 5179800 | 0 | 0 | 0 | 0 | 945390 | 570330 |
| QFGFIVLTT | RPS15A | P62244 | RS15A_HUMAN | Oxidation (M) | 0 | 0 | 0 | 0 | 0 | 0 | 0 | 0 | 0 | 0 | 0 | 3851100 | 0 | 0 | 0 | 0 | 0 | 0 |
| QFGIVTIGE | GPM6A; GPM6A | P51674; D6F | GPM6A_HUMA | Unmodified | 4542100 | 2745100 | 432960 | 0 | 0 | 0 | 0 | 0 | 0 | 0 | 0 | 0 | 0 | 0 | 980900 | 318090 | 0 | 0 |
| QFGIVTIGE | GPM6A; GPM6A | P51674; D6F | GPM6A_HUMA | Unmodified | 1802300 | 1114000 | 0 | 0 | 708550 | 359850 | 117760 | 230830 | 0 | 0 | 0 | 387740 | 0 | 0 | 0 | 0 | 136130 | 0 |
| QFGLQEK | LMF2 | Q9BU23 | LMF2_HUMAN | Unmodified | 0 | 0 | 0 | 0 | 179410 | 0 | 0 | 0 | 4702400 | 269210 | 169410 | 0 | 0 | 0 | 0 | 0 | 64031 | 0 |
| QFHFTGWI | PTPRK; PTPRK; | Q15262; Q5 | PTPRK_HUMAN | Unmodified | 353890 | 623560 | 0 | 0 | 0 | 0 | 0 | 0 | 141280 | 0 | 29117 | 180050 | 0 | 0 | 0 | 0 | 178190 | 0 |
| QFHFTSWF | PTPRJ; PTPRJ; P | Q12913; A0A | PTPRJ_HUMAN | Unmodified | 339510 | 403310 | 0 | 0 | 0 | 0 | 0 | 0 | 0 | 393260 | 0 | 857500 | 0 | 0 | 301230 | 128590 | 0 | 0 |
| QFHNEVLK | GLIPR2; GLIPR2 | Q9H4G4; Q5 | GAPR1_HUMAI | Unmodified | 0 | 0 | 0 | 0 | 209240 | 0 | 0 | 0 | 0 | 0 | 115930 | 0 | 0 | 629710 | 0 | 0 | 191380 | 805890 |
| QFHNEVLK | GLIPR2; GLIPR2 | Q9H4G4; Q5 | GAPR1_HUMAI | Unmodified | 0 | 154890 | 0 | 0 | 524530 | 0 | 0 | 0 | 0 | 0 | 0 | 0 | 0 | 1034200 | 0 | 0 | 410520 | 0 |
| QFHSVAAA | NBEAL2; NBEAL | Q6ZNJ1; H0Y | NBEL2_HUMAN | Unmodified | 0 | 0 | 0 | 0 | 0 | 0 | 0 | 0 | 0 | 0 | 0 | 784030 | 0 | 0 | 104040 | 133300 | 0 | 0 |
| QFIAAQGS | EPRS1; EPRS1 | P07814; V9C | SYEP_HUMAN | Unmodified | 0 | 0 | 106280 | 0 | 380640 | 85592 | 0 | 171520 | 2215700 | 311330 | 764780 | 6383100 | 167570 | 0 | 0 | 65532 | 141520 | 0 |
| QFIDSNPN | AARS1; AARS1; | P49588; A0A | SYAC_HUMAN | Oxidation (M) | 0 | 0 | 0 | 0 | 0 | 0 | 0 | 0 | 0 | 4677600 | 355050 | 1139600 | 0 | 0 | 0 | 0 | 0 | 0 |
| QFIQGPPE | ECHDC1 | Q9NTX5 | ECHD1_HUMA | Unmodified | 0 | 0 | 447400 | 0 | 351570 | 98876 | 0 | 0 | 158020 | 0 | 1815600 | 2659800 | 0 | 0 | 0 | 0 | 0 | 0 |
| QFLEVWEK | APEH; APEH; AI | P13798; C9J | ACPH_HUMAN | Unmodified | 5469200 | 5677400 | 3545500 | 3452300 | 2838800 | 2401200 | 0 | 70071 | 484390 | 879690 | 862150 | 8850500 | 2512200 | 2089800 | 4574700 | 5201800 | 1255800 | 2996000 |
| QFLFRPWD | UBA1 | P22314 | UBA1_HUMAN | Unmodified | 0 | 914120 | 687490 | 0 | 1048300 | 204970 | 0 | 0 | 3041000 | 1247000 | 861100 | 6482700 | 2034500 | 0 | 1113600 | 1081900 | 0 | 0 |

| | | | | | | | | | | | | | | | | | | | | | | |
|---|---|---|---|---|---|---|---|---|---|---|---|---|---|---|---|---|---|---|---|---|---|---|
| QFLPFLQR | SND1; SND1 | Q7KZF4; H7( | SND1_HUMAN | Unmodified | 712750 | 2207800 | 1911800 | 2278100 | 611040 | 442510 | 0 | 0 | 962490 | 0 | 1234800 | 11667000 | 2090800 | 0 | 3280300 | 3374400 | 191370 | 1051700 |
| QFLQAAEA | P4HB; P4HB; P4 | P07237; A0A | PDIA1_HUMAN | Unmodified | 0 | 0 | 0 | 0 | 0 | 0 | 0 | 0 | 326300 | 1190300 | 0 | 0 | 0 | 0 | 281530 | 0 | 0 | 0 |
| QFLRQNTG | TVP23B; TVP23 | Q9NYZ1; J3( | TV23B_HUMAN | Unmodified | 0 | 0 | 0 | 0 | 372780 | 0 | 142440 | 0 | 0 | 0 | 0 | 0 | 501250 | 395230 | 0 | 0 | 0 | 440210 |
| QFLSETEK | UCHL1; UCHL1 | P09936; D6F | UCHL1_HUMAN | Unmodified | 0 | 0 | 1591600 | 610530 | 56337000 | 8344500 | 2846800 | 2550500 | 0 | 12896000 | 1553600 | 35279000 | 0 | 783510 | 0 | 491960 | 0 | 0 |
| QFLSETEKN | UCHL1; UCHL1 | P09936; D6F | UCHL1_HUMAI | Unmodified | 0 | 0 | 0 | 0 | 834810 | 0 | 0 | 104870 | 175700 | 136670 | 0 | 0 | 0 | 0 | 0 | 0 | 0 | 0 |
| QFLSETEKN | UCHL1; UCHL1 | P09936; D6F | UCHL1_HUMAI | Unmodified | 0 | 0 | 0 | 0 | 15049000 | 2360500 | 2498800 | 387960 | 165490 | 2313700 | 0 | 0 | 0 | 0 | 0 | 0 | 0 | 0 |
| QFLSETEKN | UCHL1; UCHL1 | P09936; D6F | UCHL1_HUMAI | Oxidation (M) | 0 | 0 | 0 | 0 | 2604000 | 504810 | 336510 | 275690 | 0 | 399550 | 0 | 0 | 0 | 0 | 0 | 0 | 0 | 0 |
| QFLSQFEM | CPSF6; CPSF6 | Q16630; F8V | CPSF6_HUMAN | Unmodified | 517750 | 321350 | 0 | 0 | 0 | 0 | 0 | 0 | 0 | 0 | 0 | 0 | 0 | 340150 | 0 | 198290 | 0 | 0 |
| QFMETNFK | MGAM2 | Q2M2H8 | MGAL_HUMAN | Unmodified | 0 | 0 | 0 | 0 | 0 | 0 | 0 | 0 | 0 | 0 | 0 | 582890 | 0 | 0 | 0 | 226190 | 0 | 0 |
| QFMHQGN | CC2D1B; CC2D | Q5T0F9; H0Y | C2D1B_HUMAI | Unmodified | 225800 | 0 | 0 | 0 | 0 | 0 | 157910 | 0 | 0 | 0 | 0 | 0 | 0 | 0 | 0 | 0 | 442270 | 0 |
| QFQEAIIVF | TSPAN8 | P19075 | TSN8_HUMAN | Unmodified | 5992200 | 10751000 | 0 | 0 | 0 | 0 | 0 | 28779 | 515110 | 309420 | 0 | 0 | 0 | 0 | 0 | 0 | 0 | 1484800 |
| QFQFGHAI | AGPS; AGPS; A( | O00116; A0A | ADAS_HUMAN | Unmodified | 0 | 0 | 0 | 342660 | 0 | 0 | 0 | 0 | 0 | 171230 | 0 | 0 | 285810 | 0 | 102430 | 0 | 0 | 0 |
| QFQFGHAI | AGPS; AGPS; A( | O00116; A0A | ADAS_HUMAN | Unmodified | 0 | 0 | 0 | 0 | 620540 | 0 | 0 | 0 | 305460 | 2822500 | 0 | 0 | 0 | 0 | 0 | 0 | 0 | 0 |
| QFQFMAW | PTPRF; PTPRF; I | P10586; H0Y | PTPRF_HUMAN | Oxidation (M) | 51774 | 57328 | 0 | 42939 | 0 | 0 | 0 | 0 | 0 | 0 | 0 | 0 | 0 | 0 | 1015000 | 47100 | 0 | 0 |
| QFQFTDWI | PTPRD; PTPRF; | P23468; P10 | PTPRD_HUMAN | Unmodified | 1520200 | 0 | 778810 | 999380 | 1030300 | 351600 | 0 | 35237 | 2019400 | 463080 | 2350900 | 2851700 | 0 | 149630 | 3845700 | 3064800 | 0 | 0 |
| QFQGHTD( | TLE4; TLE1 | Q04727; Q0 | TLE4_HUMAN T | Unmodified | 0 | 0 | 0 | 0 | 0 | 0 | 0 | 0 | 0 | 0 | 0 | 346930 | 0 | 0 | 0 | 0 | 0 | 0 |
| QFQLMENS | AKAP2; PALM2/ | Q9Y2D5; C9 | AKAP2_HUMAN | Unmodified | 0 | 0 | 0 | 0 | 0 | 0 | 0 | 0 | 0 | 0 | 0 | 0 | 0 | 0 | 0 | 0 | 513120 | 0 |
| QFRPAVYA | NUDT14 | O95848 | NUD14_HUMA | Unmodified | 0 | 0 | 0 | 0 | 0 | 14855 | 76006 | 0 | 334220 | 0 | 0 | 1380500 | 0 | 55344 | 215020 | 0 | 0 | 35016 |
| QFRTEMED | MYH9 | P35579 | MYH9_HUMAN | Unmodified | 0 | 0 | 0 | 0 | 9643600 | 2044000 | 0 | 0 | 1729900 | 0 | 0 | 1779700 | 1744600 | 2712800 | 185900 | 0 | 1664900 | 6983900 |
| QFRTEMED | MYH9 | P35579 | MYH9_HUMAN | Oxidation (M) | 0 | 0 | 0 | 0 | 1041800 | 272630 | 312710 | 47091 | 0 | 605830 | 0 | 0 | 0 | 565600 | 0 | 0 | 241140 | 452880 |
| QFSDASQL | PARD3; PARD3; | Q8TEW0; Q5 | PARD3_HUMAI | Unmodified | 0 | 0 | 0 | 0 | 0 | 0 | 0 | 0 | 0 | 0 | 0 | 0 | 0 | 0 | 0 | 0 | 2151900 | 0 |
| QFSGPDPL | BMPR2 | Q13873 | BMPR2_HUMAI | Unmodified | 38020 | 27092 | 39067 | 92332 | 1437600 | 269060 | 1052800 | 0 | 304090 | 292030 | 26029 | 21977 | 323420 | 225160 | 61602 | 67911 | 301100 | 835560 |
| QFSLENVQ | N4BP1 | O75113 | N4BP1_HUMAI | Unmodified | 0 | 47574 | 285030 | 0 | 2344400 | 0 | 0 | 0 | 0 | 0 | 76835 | 336560 | 131020 | 0 | 0 | 0 | 0 | 0 |
| QFSQILK | GBE1; GBE1 | Q04446; E9F | GLGB_HUMAN | Unmodified | 0 | 0 | 0 | 0 | 356200 | 0 | 0 | 0 | 768890 | 556570 | 0 | 350790 | 0 | 101670 | 0 | 0 | 1086900 | 911090 |
| QFSQYIK | RPL5; RPL5 | P46777; A0A | RL5_HUMAN 6( | Unmodified | 7327000 | 5569500 | 1672100 | 1801300 | 4346000 | 836550 | 2234300 | 2011900 | 780070 | 750160 | 1707400 | 21924000 | 0 | 1663100 | 1925700 | 1404500 | 994800 | 2151300 |
| QFSSASSQ | EIF2B3; EIF2B3 | Q9NR50; H0 | EI2BG_HUMAN | Unmodified | 0 | 0 | 0 | 0 | 116610 | 22615 | 48216 | 48047 | 0 | 0 | 98496 | 1988100 | 0 | 244140 | 0 | 0 | 89897 | 272090 |
| QFTPCQLL | HADHA; HADH/ | P40939; A0A | ECHA_HUMAN | Unmodified | 0 | 0 | 0 | 0 | 0 | 0 | 0 | 0 | 1516700 | 0 | 0 | 0 | 0 | 0 | 0 | 0 | 0 | 0 |
| QFTSSTSYI | FGA | P02671 | FIBA_HUMAN F | Unmodified | 0 | 7024000 | 0 | 0 | 0 | 0 | 0 | 0 | 0 | 0 | 0 | 0 | 0 | 1372200 | 0 | 0 | 0 | 0 |
| QFVDGPP( | RACGAP1; RAC | Q9H0H5; F8 | RGAP1_HUMAI | Unmodified | 0 | 0 | 0 | 612500 | 10592000 | 1620600 | 91808 | 0 | 0 | 311490 | 439250 | 0 | 0 | 0 | 256560 | 118020 | 2212400 | 0 |
| QFVDGPP( | RACGAP1; RAC | Q9H0H5; F8 | RGAP1_HUMAI | Unmodified | 0 | 0 | 0 | 1041400 | 15628000 | 2864400 | 263460 | 157890 | 0 | 708870 | 0 | 1744400 | 1568800 | 844030 | 0 | 0 | 1448800 | 1264900 |
| QFVEQHTP | SFTPB; SFTPB; | P07988; U3K | PSPB_HUMAN | Unmodified | 0 | 0 | 0 | 0 | 2048500 | 0 | 0 | 0 | 5261300 | 2306500 | 0 | 0 | 0 | 0 | 0 | 0 | 0 | 0 |
| QFVFDLHS | ERP44 | Q9BS26 | ERP44_HUMAI | Unmodified | 504440 | 484460 | 78687 | 0 | 0 | 0 | 0 | 0 | 1154700 | 217230 | 0 | 2553300 | 0 | 0 | 463810 | 253320 | 0 | 0 |
| QFVPAVEK | GFM1; GFM1 | Q96RP9; C9 | EFGM_HUMAN | Unmodified | 0 | 0 | 0 | 0 | 0 | 0 | 0 | 187210 | 810740 | 263680 | 918900 | 0 | 0 | 0 | 0 | 0 | 0 | 0 |
| QFVPGDEF | CNP | P09543 | CN37_HUMAN | Unmodified | 0 | 0 | 0 | 0 | 2669400 | 0 | 0 | 0 | 2386600 | 16685 | 6477400 | 14329000 | 377770 | 0 | 5238400 | 0 | 229730 | 94441 |
| QFVPIKVE( | TXNRD1; TXNRI | Q16881; A0/ | TRXR1_HUMAN | Unmodified | 272750 | 338820 | 299470 | 1418100 | 990060 | 445470 | 43422 | 21212 | 8359700 | 725540 | 1847400 | 21772000 | 477670 | 161030 | 1192200 | 1125400 | 0 | 388940 |
| QFVPIKVE( | TXNRD1; TXNRI | Q16881; A0/ | TRXR1_HUMAN | Unmodified | 0 | 450790 | 0 | 0 | 4434000 | 0 | 0 | 0 | 0 | 3940200 | 3109300 | 0 | 0 | 527620 | 0 | 0 | 1086800 | 3050200 |
| QFVSQELV | ABHD17B | Q5VST6 | AB17B_HUMAN | Unmodified | 0 | 0 | 0 | 0 | 354230 | 0 | 514040 | 95083 | 289700 | 112060 | 0 | 0 | 0 | 0 | 2277200 | 0 | 118030 | 0 |
| QFVTATDV | LCP1 | P13796 | PLSL_HUMAN I | Unmodified | 0 | 0 | 0 | 0 | 0 | 0 | 0 | 0 | 554570 | 0 | 0 | 396360 | 0 | 0 | 1116900 | 0 | 77658 | 0 |
| QFVTGIIDS | MATN2; MATN2 | O00339; H0Y | MATN2_HUMAI | Unmodified | 0 | 0 | 0 | 0 | 0 | 0 | 0 | 0 | 510970 | 0 | 0 | 0 | 0 | 0 | 0 | 0 | 0 | 236050 |
| QFVTPADV | PLS1; PLS3; PLS | Q14651; P13 | PLS1_HUMAN F | Unmodified | 0 | 0 | 0 | 0 | 987970 | 0 | 0 | 0 | 0 | 0 | 0 | 0 | 0 | 0 | 0 | 0 | 0 | 0 |
| QFVVFEGN | RPN1 | P04843 | RPN1_HUMAN | Unmodified | 157340 | 72049 | 412670 | 535450 | 1501700 | 144500 | 2235600 | 71263 | 4.44E+08 | 1E+08 | 7646700 | 7931200 | 934550 | 1016000 | 509030 | 362310 | 99696 | 237860 |
| QFYDQALC | CD81; CD81; C | P60033; E9P | CD81_HUMAN | Unmodified | 3.42E+08 | 1.55E+08 | 3.08E+08 | 2776900 | 11446000 | 3902100 | 1250700 | 16430 | 21774000 | 6304100 | 18597000 | 2.46E+08 | 17293000 | 6888000 | 7.6E+08 | 4.74E+08 | 1339700 | 4489000 |
| QFYDQALC | CD81; CD81; C | P60033; E9P | CD81_HUMAN | Unmodified | 0 | 118090 | 0 | 0 | 449930 | 0 | 0 | 0 | 0 | 0 | 0 | 1184100 | 658180 | 412810 | 153560 | 202160 | 0 | 0 |
| QFYINVERI | EIF4A1; EIF4A2; | P60842; Q14 | IF4A1_HUMAN | Unmodified | 0 | 0 | 0 | 0 | 0 | 0 | 541730 | 0 | 441540 | 97423 | 0 | 4037800 | 0 | 155350 | 1137500 | 2085000 | 0 | 359400 |
| QFYPDLIRE | SELENBP1 | Q13228 | SBP1_HUMAN | Oxidation (M) | 0 | 0 | 0 | 0 | 0 | 0 | 0 | 0 | 0 | 0 | 0 | 0 | 2977900 | 716500 | 0 | 0 | 501360 | 5496200 |
| QGAIGIPY | CRYZ; CRYZ; CF | Q08257; A6I | QOR_HUMAN ( | Unmodified | 0 | 0 | 0 | 0 | 0 | 0 | 0 | 0 | 1433700 | 0 | 0 | 1234200 | 0 | 0 | 0 | 0 | 0 | 0 |
| QGAFITLVA | KDSR; KDSR; KI | Q06136; K7E | KDSR_HUMAN | Unmodified | 0 | 0 | 0 | 0 | 0 | 0 | 0 | 170730 | 1057900 | 122410 | 0 | 0 | 0 | 0 | 122370 | 0 | 0 | 0 |
| QGAFLVNT | CTBP1; CTBP1; | Q13363; H0Y | CTBP1_HUMAN | Unmodified | 0 | 0 | 0 | 0 | 0 | 0 | 0 | 0 | 0 | 0 | 0 | 276760 | 1209400 | 0 | 0 | 190540 | 0 | 0 |
| QGAIVAVT( | ATP1A1; ATP1A | P05023; P50 | AT1A1_HUMAN | Unmodified | 1805100 | 1132500 | 15077000 | 13056000 | 4535900 | 943830 | 236380 | 34587 | 9746000 | 2202700 | 13428000 | 88625000 | 6078300 | 2568500 | 19579000 | 20817000 | 826260 | 921580 |
| QGAIVAVT( | ATP1A1; ATP1A | P05023; P50 | AT1A1_HUMAN | Unmodified | 509980 | 0 | 5037800 | 6126000 | 1075800 | 218360 | 150860 | 0 | 2392400 | 1510400 | 3478600 | 16643000 | 1682900 | 574620 | 5659200 | 5493500 | 622830 | 198620 |
| QGAKINEIF | PCBP2; PCBP2; | Q15366; F8V | PCBP2_HUMAI | Unmodified | 0 | 0 | 0 | 0 | 0 | 0 | 475290 | 0 | 0 | 125230 | 0 | 0 | 1620400 | 867800 | 0 | 0 | 320830 | 0 |
| QGANINEII | PCBP1 | Q15365 | PCBP1_HUMAI | Unmodified | 10142000 | 12473000 | 2827300 | 5292200 | 0 | 0 | 0 | 0 | 0 | 0 | 0 | 0 | 534140 | 0 | 3872000 | 3113700 | 0 | 0 |
| QGANINEII | PCBP1 | Q15365 | PCBP1_HUMAI | Unmodified | 431320 | 448300 | 0 | 1286900 | 2857700 | 1192000 | 704480 | 363950 | 755770 | 2637500 | 878750 | 957110 | 12300000 | 2580100 | 0 | 0 | 702650 | 2643100 |
| QGANINEII | PCBP1 | Q15365 | PCBP1_HUMAI | Oxidation (M) | 200370 | 0 | 65211 | 396250 | 518770 | 219650 | 0 | 0 | 0 | 0 | 143460 | 71147 | 1426400 | 369430 | 0 | 0 | 0 | 225720 |
| QGAPGQG | SUN2 | Q9UH99 | SUN2_HUMAN | Unmodified | 0 | 0 | 0 | 2127600 | 1026300 | 0 | 0 | 0 | 865880 | 0 | 0 | 0 | 0 | 0 | 0 | 0 | 0 | 0 |
| QGAPGQH | CTH | P32929 | CGL_HUMAN ( | Unmodified | 26216 | 0 | 68631 | 0 | 0 | 21183 | 0 | 0 | 0 | 0 | 52393 | 551110 | 225940 | 91333 | 147420 | 145780 | 37025 | 0 |
| QGCDCEC | PHPT1 | Q9NRX4 | PHP14_HUMAI | Unmodified | 0 | 0 | 0 | 0 | 0 | 0 | 0 | 0 | 0 | 0 | 0 | 407450 | 0 | 0 | 0 | 0 | 0 | 0 |
| QGCDCEC | PHPT1 | Q9NRX4 | PHP14_HUMAI | Unmodified | 0 | 0 | 0 | 173610 | 686680 | 184360 | 0 | 0 | 0 | 265080 | 0 | 0 | 828020 | 772340 | 0 | 0 | 483700 | 655040 |
| QGCPFQP\ | APOBEC3B; AP( | Q9UH17; B0 | ABC3B_HUMAI | Unmodified | 3337800 | 3875000 | 1717400 | 1811900 | 0 | 0 | 0 | 0 | 0 | 0 | 0 | 0 | 0 | 0 | 5399500 | 3340600 | 0 | 0 |
| QGDITEDE | VPS36; VPS36 | Q86VN1; A0 | VPS36_HUMAN | Unmodified | 6709500 | 0 | 5449800 | 0 | 0 | 0 | 0 | 17654 | 311380 | 0 | 152550 | 0 | 603880 | 259800 | 0 | 0 | 0 | 237220 |
| QGDLVTC( | IREB2 | P48200 | IREB2_HUMAN | Unmodified | 0 | 0 | 0 | 0 | 0 | 0 | 0 | 0 | 0 | 0 | 204310 | 618290 | 0 | 0 | 0 | 0 | 0 | 0 |
| QGDMVLEI | AFG3L2 | Q9Y4W6 | AFG32_HUMAN | Oxidation (M) | 0 | 0 | 0 | 0 | 0 | 0 | 0 | 0 | 472680 | 0 | 0 | 0 | 0 | 0 | 0 | 0 | 0 | 0 |
| QGDNCDS | LARS1; LARS1; | Q9P2J5; A0A | SYLC_HUMAN | Unmodified | 0 | 0 | 0 | 0 | 0 | 0 | 0 | 0 | 0 | 0 | 0 | 898010 | 0 | 0 | 0 | 0 | 0 | 0 |
| QGDTGDW | STRAP | Q9Y3F4 | STRAP_HUMAI | Unmodified | 649650 | 608850 | 456460 | 1905700 | 0 | 0 | 0 | 0 | 1216900 | 156270 | 2158900 | 10860000 | 1669600 | 335790 | 666020 | 781840 | 0 | 0 |
| QGDTILVS( | NUP210 | Q8TEM1 | PO210_HUMAI | Unmodified | 0 | 0 | 176060 | 0 | 0 | 0 | 87225 | 0 | 20514000 | 0 | 0 | 0 | 0 | 0 | 875020 | 819460 | 0 | 0 |
| QGDTILVS( | NUP210 | Q8TEM1 | PO210_HUMAI | Oxidation (M) | 0 | 52639 | 0 | 0 | 0 | 0 | 0 | 0 | 0 | 771370 | 0 | 0 | 0 | 0 | 0 | 0 | 0 | 0 |
| QGEDAEV\ | NCAM2; NCAM2 | O15394; H9I | NCAM2_HUMA | Unmodified | 0 | 0 | 743290 | 0 | 0 | 1134700 | 239160 | 31603 | 86236 | 88140 | 29500 | 911920 | 0 | 0 | 1060600 | 954350 | 73242 | 0 |
| QGEDNST/ | MTDH; MTDH | Q86UE4; E5I | LYRIC_HUMAN | Unmodified | 0 | 0 | 0 | 0 | 0 | 0 | 0 | 0 | 0 | 0 | 47891 | 203760 | 0 | 0 | 0 | 0 | 0 | 0 |
| QGELENYV | BAIAP2; BAIAP2 | Q9UQB8; I3I | BAIP2_HUMAN | Unmodified | 0 | 0 | 0 | 0 | 0 | 9053200 | 0 | 0 | 853920 | 0 | 0 | 2116600 | 0 | 0 | 1297000 | 1073100 | 0 | 0 |
| QGEQYMN | INPP5A; INPP5/ | Q14642; B1/ | I5P1_HUMAN I | Unmodified | 47918 | 45425 | 153580 | 0 | 0 | 0 | 0 | 0 | 275280 | 0 | 0 | 494530 | 0 | 0 | 236280 | 194740 | 0 | 0 |
| QGESEVSR | CKAP4 | Q07065 | CKAP4_HUMAI | Unmodified | 0 | 0 | 0 | 0 | 2124400 | 8511900 | 1310500 | 330260 | 1793700 | 846510 | 1831900 | 0 | 0 | 4540700 | 0 | 0 | 891490 | 10823000 |
| QGESFAVH | WDR20; WDR2( | Q8TBZ3; A0/ | WDR20_HUMA | Unmodified | 0 | 77737 | 0 | 157560 | 527210 | 241130 | 0 | 0 | 144630 | 174240 | 0 | 324050 | 381890 | 381270 | 0 | 0 | 619880 | 547000 |
| QGEVIVSIE | NASP; NASP | P49321; E9P | NASP_HUMAN | Unmodified | 0 | 0 | 0 | 33657 | 0 | 121560 | 142860 | 62133 | 466220 | 0 | 584490 | 1131600 | 0 | 0 | 0 | 0 | 0 | 0 |
| QGEVSKAA | MACROH2A1 | O75367 | H2AY_HUMAN | Unmodified | 0 | 0 | 0 | 0 | 0 | 0 | 0 | 2020700 | 0 | 0 | 0 | 0 | 0 | 0 | 0 | 0 | 0 | 0 |
| QGFDFLCN | PRMT5; PRMT5; | O14744; G3\ | ANM5_HUMAN | Oxidation (M) | 0 | 0 | 0 | 0 | 0 | 0 | 0 | 0 | 0 | 110670 | 0 | 1679400 | 829520 | 0 | 0 | 0 | 0 | 0 |
| QGFEFNIM | SEPTIN9; SEPTI | Q9UHD8; K7 | SEPT9_HUMAN | Oxidation (M) | 38430 | 0 | 0 | 0 | 227600 | 0 | 0 | 0 | 0 | 0 | 0 | 0 | 0 | 0 | 0 | 0 | 0 | 132960 |
| QGFGELLQ | APEX1; APEX1; | P27695; G3\ | APEX1_HUMAN | Unmodified | 0 | 0 | 0 | 0 | 0 | 0 | 0 | 0 | 173810 | 0 | 0 | 1960600 | 0 | 0 | 0 | 51540 | 0 | 0 |
| QGFGNLPI | MTHFD1L; MTH | Q6UB35; P1 | C1TM_HUMAN | Oxidation (M) | 0 | 193460 | 0 | 0 | 623760 | 417620 | 1049900 | 1025800 | 3987500 | 343230 | 0 | 1963300 | 789140 | 547390 | 244870 | 0 | 1972900 | 1564500 |
| QGFGNLPI | MTHFD1L; MTH | Q6UB35; P1 | C1TM_HUMAN | Unmodified | 0 | 0 | 0 | 0 | 0 | 0 | 1546200 | 269050 | 1173000 | 157150 | 0 | 862300 | 0 | 599470 | 0 | 0 | 3072600 | 2266800 |
| QGFGNVAT | FGB; FGB | P02675; D6F | FIBB_HUMAN F | Unmodified | 0 | 2225200 | 0 | 0 | 0 | 0 | 0 | 0 | 0 | 0 | 0 | 0 | 0 | 185760 | 0 | 0 | 0 | 0 |
| QGFNVVVE | NNT; NNT; NNT | Q13423; A0/ | NNTM_HUMAN | Unmodified | 0 | 0 | 0 | 0 | 0 | 0 | 260180 | 0 | 2210300 | 175990 | 0 | 0 | 0 | 0 | 0 | 0 | 0 | 0 |
| QGFNVVVE | NNT; NNT; NNT | Q13423; A0/ | NNTM_HUMAN | Unmodified | 0 | 0 | 0 | 0 | 0 | 3752500 | 0 | 0 | 1269700 | 3546900 | 0 | 0 | 0 | 1540000 | 0 | 0 | 669030 | 0 |
| QGFPIIFHC | MOV10; MOV1( | Q9HCE1; QE | MOV10_HUMA | Oxidation (M) | 1194100 | 848590 | 476030 | 0 | 0 | 0 | 0 | 0 | 0 | 0 | 0 | 0 | 0 | 0 | 481370 | 773640 | 0 | 0 |
| QGFPIIFHC | MOV10; MOV1( | Q9HCE1; QE | MOV10_HUMA | Oxidation (M) | 5428600 | 11827000 | 0 | 3636500 | 0 | 0 | 0 | 0 | 0 | 0 | 0 | 0 | 0 | 0 | 5157800 | 0 | 0 | 0 |
| QGFPNRV\ | MYH9 | P35579 | MYH9_HUMAN | Unmodified | 1101600 | 1250900 | 1991900 | 4210900 | 787830 | 358740 | 223380 | 0 | 2383700 | 1984100 | 1507800 | 13950000 | 50018 | 45174 | 2305900 | 2154000 | 441400 | 406060 |
| QGFQCQV( | PRKCB; PRKCA | P05771; P17 | KPCB_HUMAN | Unmodified | 0 | 0 | 0 | 0 | 0 | 0 | 1090600 | 530230 | 0 | 0 | 0 | 0 | 356190 | 1731600 | 0 | 0 | 183540 | 0 |
| QGFYVALR | EPS15L1; EPS1 | Q9UBC2; M( | EP15R_HUMAN | Unmodified | 0 | 0 | 0 | 0 | 0 | 0 | 0 | 0 | 0 | 0 | 0 | 1893500 | 0 | 0 | 0 | 0 | 0 | 0 |
| QGGASQSI | SLC2A1 | P11166 | GTR1_HUMAN | Unmodified | 2636500 | 2705000 | 10528000 | 7929000 | 2938000 | 0 | 0 | 63066 | 10329000 | 1992000 | 2556100 | 24795000 | 0 | 0 | 9772300 | 10053000 | 1517300 | 899860 |
| QGGFLGLS | MUC1; MUC1; I | P15941; B1A | MUC1_HUMAN | Unmodified | 0 | 0 | 0 | 0 | 22515000 | 2912600 | 0 | 0 | 38595000 | 4773900 | 0 | 457400 | 0 | 0 | 0 | 0 | 0 | 0 |
| QGGGGGC | HNRNPM; HNR | P52272; M0I | HNRPM_HUMA | Unmodified | 766170 | 0 | 201340 | 0 | 142610 | 50327 | 0 | 105490 | 340600 | 274610 | 300440 | 388190 | 347950 | 0 | 111570 | 0 | 0 | 166040 |
| QGGLGPIR | PRDX4; PRDX4; | Q13162; A6I | PRDX4_HUMAN | Unmodified | 0 | 0 | 0 | 0 | 0 | 0 | 0 | 0 | 94906000 | 7660300 | 8731300 | 6709500 | 0 | 0 | 385970 | 0 | 0 | 0 |
| QGGPTYYI | ACTL6A; ACTL6 | O96019; H7C | ACL6A_HUMAN | Unmodified | 0 | 0 | 0 | 0 | 0 | 0 | 0 | 0 | 330720 | 0 | 0 | 1313700 | 0 | 0 | 0 | 0 | 0 | 0 |
| QGHNSVFI | HPX; HPX | P02790; Q9I | HEMO_HUMAN | Unmodified | 0 | 0 | 0 | 0 | 0 | 0 | 0 | 0 | 0 | 0 | 0 | 1151100 | 0 | 0 | 0 | 0 | 0 | 0 |
| QGIETPED( | SF1; SF1 | Q15637; A0/ | SF01_HUMAN : | Unmodified | 245730 | 251400 | 328170 | 473680 | 725760 | 305580 | 66769 | 0 | 419160 | 365710 | 223360 | 459930 | 953230 | 242780 | 167590 | 209370 | 184720 | 248020 |
| QGIFPASY\ | SORBS3; SORB | O60504; H0Y | VINEX_HUMAN | Unmodified | 0 | 3208700 | 0 | 0 | 0 | 0 | 0 | 0 | 0 | 0 | 0 | 0 | 0 | 0 | 0 | 0 | 0 | 0 |
| QGIREVILC | SDCBP; SDCBF | O00560; G5I | SDCB1_HUMAI | Unmodified | 639440 | 452880 | 679810 | 2481700 | 3039800 | 1702300 | 166320 | 81256 | 4281600 | 3228300 | 482950 | 0 | 2284900 | 1963700 | 670020 | 403800 | 147270 | 870290 |
| QGIREVILC | SDCBP; SDCBF | O00560; G5I | SDCB1_HUMAI | Unmodified | 252680 | 104020 | 0 | 1250600 | 5244700 | 2174800 | 250290 | 72018 | 4082800 | 2337600 | 0 | 0 | 5326000 | 4825300 | 0 | 36343 | 226050 | 1138000 |
| QGIVPPGL | SFXN1; SFXN1; | Q9H9B4; D6 | SFXN1_HUMAN | Unmodified | 0 | 0 | 0 | 0 | 218020 | 0 | 0 | 0 | 3466700 | 250150 | 0 | 0 | 0 | 0 | 0 | 0 | 0 | 0 |
| QGKIPDEEI | SH3GL2; SH3G | Q99962; Q9 | SH3G2_HUMAI | Unmodified | 476470 | 663540 | 0 | 1159700 | 0 | 0 | 0 | 0 | 156910 | 0 | 0 | 0 | 0 | 0 | 466290 | 460730 | 136200 | 0 |
| QGKLDEAE | DNAJC3; DNAJ( | Q13217; X6F | DNJC3_HUMAI | Unmodified | 0 | 0 | 0 | 0 | 0 | 0 | 0 | 0 | 551470 | 112370 | 33254 | 0 | 0 | 0 | 0 | 0 | 0 | 0 |
| QGKLDEAE | DNAJC3; DNAJ( | Q13217; X6F | DNJC3_HUMAI | Unmodified | 0 | 0 | 0 | 0 | 0 | 0 | 0 | 29847 | 1117300 | 432710 | 0 | 0 | 0 | 0 | 0 | 0 | 0 | 0 |
| QGKPQSAT | MIA3; MIA3 | Q5JRA6; A0A | TGO1_HUMAN | Unmodified | 0 | 0 | 0 | 0 | 0 | 0 | 0 | 0 | 4022600 | 0 | 0 | 0 | 4101700 | 0 | 0 | 0 | 0 | 0 |
| QGKVNAW | EDIL3 | O43854 | EDIL3_HUMAN | Unmodified | 0 | 0 | 0 | 0 | 35096000 | 8157800 | 46912 | 0 | 0 | 0 | 0 | 0 | 0 | 0 | 0 | 0 | 1465000 | 10704000 |
| QGKVNAW | EDIL3 | O43854 | EDIL3_HUMAN | Unmodified | 0 | 0 | 0 | 0 | 1.06E+08 | 0 | 2103300 | 0 | 0 | 0 | 0 | 0 | 0 | 0 | 0 | 0 | 2210300 | 54306000 |
| QGLEQFIN | SNX12; SNX3; S | Q9UMY4; O( | SNX12_HUMAN | Unmodified | 153920 | 217380 | 462810 | 275430 | 0 | 0 | 291080 | 0 | 0 | 45476 | 0 | 2038600 | 537330 | 0 | 725960 | 390480 | 0 | 0 |
| QGLLPLTF/ | ACO2; ACO2; A | Q99798; A2/ | ACON_HUMAN | Unmodified | 0 | 0 | 0 | 0 | 0 | 0 | 0 | 106260 | 2134800 | 0 | 0 | 0 | 0 | 0 | 0 | 0 | 0 | 0 |
| QGLPFTIF( | LLGL2; LLGL2; I | Q6P1M3; J3( | L2GL2_HUMAN | Oxidation (M) | 0 | 0 | 0 | 0 | 0 | 0 | 0 | 0 | 0 | 0 | 0 | 944990 | 0 | 0 | 0 | 0 | 0 | 0 |
| QGLQTTQ/ | GAPVD1; GAPV | Q14C86; F8\ | GAPD1_HUMAN | Unmodified | 43035 | 21520 | 0 | 0 | 0 | 0 | 279640 | 0 | 0 | 0 | 0 | 1044000 | 0 | 0 | 71389 | 0 | 53315 | 2330100 |
| QGLSSSIFT | ALDH7A1; ALDI | P49419; A0A | AL7A1_HUMAN | Unmodified | 0 | 0 | 0 | 0 | 3147200 | 269450 | 75964 | 82542 | 259520 | 0 | 0 | 1511300 | 0 | 43937 | 0 | 0 | 968740 | 3303300 |

| | | | | | | | | | | | | | | | | | | | | | | |
|---|---|---|---|---|---|---|---|---|---|---|---|---|---|---|---|---|---|---|---|---|---|---|
| QGLTVSFIF | CDCP1 | Q9H5V8 | CDCP1_HUMA | Unmodified | 0 | 0 | 1513000 | 310550 | 0 | 0 | 0 | 0 | 303920 | 0 | 340620 | 3876300 | 0 | 0 | 2327000 | 1488000 | 0 | 0 |
| QGLTVSFIF | CDCP1 | Q9H5V8 | CDCP1_HUMA | Unmodified | 0 | 0 | 0 | 379050 | 0 | 0 | 0 | 0 | 0 | 0 | 0 | 0 | 0 | 0 | 278980 | 0 | 540620 | 0 |
| QGLYELSA | TXNDC5 | Q8NBS9 | TXND5_HUMAN | Unmodified | 0 | 0 | 0 | 0 | 845210 | 302710 | 563270 | 0 | 48623000 | 14200000 | 0 | 0 | 478520 | 1299600 | 0 | 839530 | 723380 | 0 |
| QGLYRHPV | ACLY | P53396 | ACLY_HUMAN | 2 Oxidation (M) | 0 | 0 | 0 | 0 | 0 | 0 | 1456500 | 0 | 0 | 0 | 0 | 0 | 0 | 0 | 0 | 0 | 0 | 0 |
| QGMDLSAI | FSCN1 | Q16658 | FSCN1_HUMAI | Unmodified | 0 | 0 | 0 | 0 | 1042600 | 0 | 0 | 0 | 0 | 0 | 0 | 937700 | 0 | 0 | 0 | 0 | 0 | 0 |
| QGMDLSAI | FSCN1 | Q16658 | FSCN1_HUMAI | Oxidation (M) | 0 | 0 | 0 | 0 | 0 | 0 | 0 | 0 | 0 | 0 | 1007200 | 3256900 | 0 | 0 | 0 | 0 | 0 | 0 |
| QGNAVTLG | LSR; LSR; LSR; I | Q86X29; M0 | LSR_HUMAN Li | Unmodified | 1638100 | 2661100 | 4466800 | 5583300 | 888860 | 0 | 186340 | 0 | 11992000 | 4192800 | 1469700 | 10464000 | 844160 | 0 | 7331800 | 8155100 | 0 | 0 |
| QGNAVTLG | LSR; LSR; LSR; I | Q86X29; M0 | LSR_HUMAN Li | Unmodified | 738110 | 636620 | 2785300 | 1288500 | 0 | 0 | 0 | 59902 | 7601100 | 2424400 | 1299500 | 5708300 | 67945 | 0 | 1921200 | 2362100 | 0 | 0 |
| QGNFFASF | RACGAP1 | Q9H0H5 | RGAP1_HUMAI | Unmodified | 0 | 28727 | 0 | 0 | 2449800 | 0 | 0 | 0 | 625120 | 192540 | 0 | 0 | 0 | 0 | 0 | 0 | 0 | 0 |
| QGNGPVL\ | UPF1 | Q92900 | RENT1_HUMAN | Unmodified | 1336700 | 834090 | 379090 | 692120 | 0 | 0 | 0 | 0 | 0 | 0 | 0 | 703320 | 0 | 0 | 519000 | 0 | 0 | 0 |
| QGNLSSQ\ | PRKDC | P78527 | PRKDC_HUMAI | Unmodified | 0 | 0 | 0 | 0 | 531800 | 0 | 0 | 0 | 61230 | 20407 | 0 | 1419700 | 0 | 0 | 0 | 0 | 0 | 0 |
| QGNMTAAI | ARPC5; ARPC5 | O15511; B1/ | ARPC5_HUMAI | Unmodified | 0 | 0 | 0 | 0 | 0 | 0 | 0 | 0 | 380420 | 0 | 0 | 4092600 | 0 | 0 | 0 | 128410 | 0 | 0 |
| QGNMTAAI | ARPC5; ARPC5 | O15511; B1/ | ARPC5_HUMAI | Oxidation (M) | 0 | 0 | 0 | 166800 | 0 | 0 | 0 | 0 | 0 | 0 | 0 | 3998000 | 0 | 0 | 341570 | 220580 | 485310 | 0 |
| QGNPVAPI | GALNT2 | Q10471 | GALT2_HUMAN | Unmodified | 0 | 0 | 90100 | 0 | 0 | 0 | 135670 | 0 | 23250000 | 1474900 | 603160 | 0 | 0 | 0 | 11880 | 0 | 0 | 0 |
| QGNPVAPI | GALNT2 | Q10471 | GALT2_HUMAN | Unmodified | 0 | 0 | 0 | 0 | 0 | 0 | 0 | 0 | 1350100 | 415590 | 0 | 0 | 0 | 0 | 0 | 0 | 0 | 0 |
| QGNPVAPI | GALNT2 | Q10471 | GALT2_HUMAN | 2 Oxidation (M) | 0 | 0 | 0 | 0 | 0 | 0 | 0 | 0 | 1473000 | 856620 | 27267 | 0 | 0 | 0 | 0 | 0 | 0 | 0 |
| QGNPVAPI | GALNT2 | Q10471 | GALT2_HUMAN | Oxidation (M) | 0 | 0 | 0 | 0 | 0 | 0 | 0 | 0 | 2638600 | 214720 | 0 | 0 | 0 | 0 | 0 | 0 | 0 | 0 |
| QGQDNLS | ERP29; ERP29 | P30040; F8V | ERP29_HUMAN | Unmodified | 87191 | 0 | 143570 | 820540 | 0 | 0 | 135760 | 62555 | 72969000 | 3297000 | 341810 | 975910 | 958900 | 652720 | 97480 | 202780 | 0 | 0 |
| QGQDNLS | ERP29; ERP29 | P30040; F8V | ERP29_HUMAN | Unmodified | 1149700 | 0 | 0 | 18093 | 0 | 0 | 0 | 684560 | 29983000 | 24808000 | 751470 | 0 | 0 | 0 | 0 | 61370 | 0 | 0 |
| QGQDNLS | ERP29; ERP29 | P30040; F8V | ERP29_HUMAN | Unmodified | 0 | 0 | 0 | 0 | 0 | 0 | 0 | 0 | 11609000 | 87486 | 0 | 0 | 0 | 0 | 0 | 0 | 0 | 0 |
| QGQELMAI | H6PD | O95479 | G6PE_HUMAN | Oxidation (M) | 0 | 0 | 0 | 0 | 1909000 | 0 | 2687000 | 1079300 | 0 | 761370 | 215190 | 1437800 | 0 | 0 | 0 | 0 | 186260 | 0 |
| QGQETAVA | MAPRE1 | Q15691 | MARE1_HUMAI | Unmodified | 514140 | 574480 | 563350 | 637860 | 0 | 0 | 0 | 0 | 1153000 | 0 | 1743900 | 6121400 | 0 | 0 | 642030 | 655220 | 0 | 0 |
| QGQGQLV | DDB1; DDB1; D | Q16531; A0/ | DDB1_HUMAN | Unmodified | 0 | 0 | 0 | 0 | 0 | 0 | 0 | 0 | 0 | 0 | 0 | 2206300 | 0 | 0 | 0 | 0 | 0 | 0 |
| QGQHWLE | SEC11A; SEC11 | P67812; H0Y | SC11A_HUMAN | Unmodified | 0 | 0 | 0 | 0 | 0 | 0 | 0 | 0 | 757680 | 43124 | 0 | 68890 | 0 | 0 | 0 | 0 | 0 | 0 |
| QGQIINPSI | BCKDHA; N/A; I | P12694; F5H | ODBA_HUMAN | Unmodified | 0 | 0 | 0 | 0 | 0 | 0 | 0 | 0 | 2293600 | 0 | 0 | 0 | 0 | 89694 | 0 | 0 | 0 | 0 |
| QGQMDAV | CHMP2A; CHM | O43633; M0I | CHM2A_HUMA | Oxidation (M) | 0 | 0 | 0 | 0 | 0 | 0 | 0 | 0 | 351980 | 46527 | 0 | 936040 | 0 | 0 | 0 | 137070 | 0 | 0 |
| QGQMDAV | CHMP2A; CHM | O43633; M0I | CHM2A_HUMA | Unmodified | 1132500 | 1426900 | 1227000 | 1548400 | 0 | 318710 | 51801 | 0 | 6613800 | 378140 | 51909 | 8242600 | 0 | 0 | 816800 | 862570 | 22425 | 44113 |
| QGQMDAV | CHMP2A; CHM | O43633; M0I | CHM2A_HUMA | Unmodified | 0 | 0 | 0 | 0 | 3544100 | 661950 | 310210 | 78298 | 0 | 1663000 | 267700 | 0 | 0 | 783550 | 0 | 0 | 209870 | 0 |
| QGQMDAV | CHMP2A; CHM | O43633; M0I | CHM2A_HUMA | Oxidation (M) | 0 | 28122 | 0 | 0 | 952470 | 58278 | 0 | 0 | 0 | 352290 | 80216 | 0 | 54155 | 0 | 0 | 0 | 0 | 0 |
| QGQYSPM | ATP5F1A | P25705 | ATPA_HUMAN | Oxidation (M) | 0 | 0 | 0 | 0 | 0 | 0 | 0 | 0 | 1289400 | 0 | 0 | 0 | 0 | 0 | 0 | 0 | 0 | 0 |
| QGRPVVIC | GFPT1; GFPT1 | Q06210; A0/ | GFPT1_HUMAN | Unmodified | 0 | 0 | 0 | 528240 | 377000 | 0 | 0 | 0 | 1518700 | 293800 | 708270 | 2239200 | 1765000 | 2053300 | 754020 | 841960 | 120840 | 723640 |
| QGRQEALE | XPNPEP1; XPNI | Q9NQW7; Q | XPP1_HUMAN | Unmodified | 0 | 0 | 161040 | 244800 | 0 | 0 | 0 | 0 | 0 | 0 | 166040 | 1846100 | 129060 | 431930 | 0 | 223030 | 0 | 143490 |
| QGRQEALE | XPNPEP1; XPNI | Q9NQW7; Q | XPP1_HUMAN | Unmodified | 0 | 0 | 0 | 0 | 0 | 253710 | 480470 | 0 | 433720 | 191950 | 158260 | 0 | 1335000 | 814690 | 0 | 0 | 1231700 | 1133100 |
| QGSATVGL | PSMA1; PSMA1 | P25786; F5H | PSA1_HUMAN | Unmodified | 140170 | 0 | 0 | 332090 | 2234400 | 6705.9 | 305640 | 199910 | 148390 | 681480 | 162390 | 6191900 | 1329300 | 77616 | 0 | 409070 | 996900 | 485870 |
| QGSGSSQI | TRIM28 | Q13263 | TIF1B_HUMAN | Oxidation (M) | 0 | 0 | 0 | 0 | 0 | 0 | 0 | 431500 | 0 | 0 | 0 | 0 | 0 | 0 | 0 | 0 | 0 | 345890 |
| QGSGVILR | CRK | P46108 | CRK_HUMAN A | Unmodified | 485590 | 309790 | 561900 | 300160 | 170500 | 0 | 0 | 0 | 1143200 | 0 | 0 | 1112000 | 0 | 0 | 574150 | 700470 | 411990 | 0 |
| QGSKDPAE | GANAB; GANAE | Q14697; F5H | GANAB_HUMA | Unmodified | 0 | 0 | 78646 | 0 | 0 | 0 | 0 | 0 | 1711700 | 315470 | 672880 | 0 | 0 | 0 | 0 | 0 | 203210 | 0 |
| QGSLAMEE | SPN | P16150 | LEUK_HUMAN | Unmodified | 0 | 0 | 0 | 0 | 0 | 0 | 0 | 0 | 0 | 0 | 0 | 1340500 | 0 | 119010 | 0 | 26285 | 684900 | 0 |
| QGSPVAAC | IGF2BP1 | Q9NZI8 | IF2B1_HUMAN | Unmodified | 174880 | 0 | 0 | 0 | 0 | 0 | 0 | 0 | 0 | 0 | 0 | 0 | 0 | 0 | 0 | 0 | 0 | 0 |
| QGSVVSSR | FRMD8 | Q9BZ67 | FRMD8_HUMAI | Unmodified | 0 | 0 | 0 | 0 | 334800 | 0 | 0 | 0 | 0 | 0 | 0 | 0 | 0 | 0 | 0 | 0 | 0 | 469820 |
| QGTEIDGR | NCL; NCL; NCL | P19338; C9J | NUCL_HUMAN | Unmodified | 0 | 0 | 0 | 0 | 543080 | 0 | 224760 | 0 | 523670 | 1666500 | 0 | 0 | 227750 | 385630 | 0 | 0 | 1064000 | 172760 |
| QGTHITVV | PDHB; PDHB | P11177; C9J | ODPB_HUMAN | Unmodified | 0 | 0 | 0 | 0 | 68313 | 0 | 0 | 0 | 1542500 | 0 | 0 | 0 | 0 | 0 | 0 | 0 | 0 | 0 |
| QGTLSEFV | RTN4IP1; RTN4 | Q8WWV3; G | RT4I1_HUMAN | Unmodified | 0 | 0 | 0 | 0 | 0 | 0 | 0 | 0 | 1650100 | 0 | 0 | 0 | 0 | 0 | 0 | 0 | 0 | 0 |
| QGTPMCVI | EIF5B; EIF5B | O60841; A0/ | IF2P_HUMAN E | Unmodified | 0 | 0 | 167900 | 0 | 175930 | 0 | 0 | 0 | 0 | 0 | 0 | 1013200 | 292590 | 0 | 0 | 87998 | 0 | 0 |
| QGVDQKL\ | PLIN3; PLIN3 | O60664; K7E | PLIN3_HUMAN | Unmodified | 0 | 0 | 0 | 0 | 417450 | 237190 | 0 | 0 | 0 | 0 | 0 | 0 | 184050 | 79067 | 20311 | 0 | 441260 | 168570 |
| QGVLTHGF | RPS6; RPS6 | P62753; A2A | RS6_HUMAN 4 | Unmodified | 0 | 0 | 0 | 0 | 0 | 0 | 0 | 690570 | 0 | 34955 | 0 | 7450700 | 0 | 0 | 301580 | 0 | 0 | 0 |
| QGVPAFTV | FPGS; FPGS; FF | Q05932; Q5 | FOLC_HUMAN | Unmodified | 0 | 0 | 0 | 32994 | 0 | 0 | 0 | 0 | 197030 | 0 | 0 | 0 | 0 | 0 | 231600 | 71652 | 0 | 0 |
| QGVQVQV | FASN; FASN | P49327; A0A | FAS_HUMAN F | Unmodified | 0 | 0 | 0 | 0 | 547480 | 0 | 0 | 58298 | 0 | 0 | 2214200 | 9895100 | 0 | 0 | 782440 | 0 | 0 | 0 |
| QGVTQQQI | PSME4 | Q14997 | PSME4_HUMAI | Unmodified | 0 | 0 | 0 | 0 | 34944 | 0 | 37419 | 0 | 56653 | 0 | 0 | 474940 | 0 | 0 | 774660 | 0 | 0 | 23639 |
| QGVYVTSP | VPS37A; VPS37 | Q8NEZ2; E5I | VP37A_HUMAN | Oxidation (M) | 2453000 | 3549400 | 309170 | 0 | 0 | 0 | 0 | 0 | 0 | 0 | 0 | 0 | 83986 | 121130 | 363170 | 696240 | 0 | 0 |
| QGWTVLHI | ANKRD13A | Q8IZ07 | AN13A_HUMAN | Oxidation (M) | 738570 | 0 | 0 | 0 | 0 | 0 | 0 | 0 | 0 | 0 | 0 | 0 | 186960 | 0 | 733370 | 0 | 0 | 0 |
| QGWVSSSI | BCAM; BCAM | P50895; A0A | BCAM_HUMAN | Unmodified | 3519300 | 3396500 | 0 | 0 | 213660 | 1792900 | 42733 | 74366 | 546100 | 0 | 753560 | 0 | 0 | 0 | 0 | 0 | 0 | 342460 |
| QGYDVSIP | EXT2 | Q93063 | EXT2_HUMAN I | Unmodified | 0 | 0 | 0 | 0 | 0 | 0 | 0 | 0 | 1561800 | 0 | 0 | 0 | 0 | 0 | 0 | 0 | 0 | 0 |
| QGYMSGG | C1GALT1 | Q9NS00 | C1GLT_HUMAN | Unmodified | 0 | 0 | 0 | 0 | 0 | 0 | 0 | 0 | 3479500 | 0 | 0 | 0 | 0 | 0 | 0 | 0 | 0 | 0 |
| QGYVIYR | RPL15; RPL15; I | P61313; E7E | RL15_HUMAN | Unmodified | 5101900 | 0 | 3279700 | 2099400 | 0 | 0 | 69518 | 62995 | 675610 | 727060 | 1111100 | 30644000 | 671930 | 129910 | 3441900 | 3406900 | 0 | 140930 |
| QGYVLSSII | BUB3; BUB3 | O43684; J3Q | BUB3_HUMAN | Unmodified | 0 | 0 | 0 | 0 | 0 | 0 | 138830 | 225420 | 433160 | 0 | 387030 | 7700400 | 0 | 0 | 0 | 153290 | 0 | 0 |
| QGYWAQS | TBC1D24; TBC | Q9ULP9; A0/ | TBC24_HUMAN | Unmodified | 52671 | 0 | 0 | 0 | 0 | 0 | 0 | 0 | 425330 | 0 | 0 | 0 | 0 | 0 | 0 | 0 | 0 | 17601 |
| QHDAEVD/ | COL2A1 | P02458 | CO2A1_HUMAI | Unmodified | 0 | 0 | 0 | 0 | 0 | 0 | 0 | 0 | 0 | 0 | 704040 | 96074 | 0 | 0 | 0 | 0 | 0 | 0 |
| QHDQLEA( | NUCB2; NUCB | P80303; Q2L | NUCB2_HUMA | Unmodified | 0 | 0 | 0 | 0 | 0 | 0 | 0 | 0 | 65030000 | 943020 | 0 | 342250 | 0 | 0 | 504100 | 117300 | 0 | 0 |
| QHDQLEA( | NUCB2; NUCB | P80303; Q2L | NUCB2_HUMA | Oxidation (M) | 0 | 0 | 0 | 34947 | 0 | 0 | 0 | 0 | 1765400 | 0 | 0 | 0 | 0 | 0 | 0 | 0 | 0 | 0 |
| QHDQLEA( | NUCB2; NUCB | P80303; Q2L | NUCB2_HUMA | Unmodified | 0 | 0 | 0 | 0 | 0 | 0 | 0 | 0 | 6830100 | 0 | 0 | 0 | 0 | 0 | 0 | 0 | 0 | 443100 |
| QHEADADI | OXCT1; OXCT1 | P55809; E9P | SCOT1_HUMAI | Unmodified | 0 | 0 | 0 | 0 | 206410 | 56072 | 0 | 0 | 738350 | 0 | 0 | 0 | 0 | 0 | 0 | 0 | 167010 | 249830 |
| QHELSESC | VPS37A; VPS37 | Q8NEZ2; H0 | VP37A_HUMAN | Unmodified | 2164100 | 872490 | 0 | 0 | 0 | 114910 | 0 | 0 | 0 | 0 | 0 | 0 | 0 | 0 | 0 | 0 | 0 | 0 |
| QHESMQEI | TAOK1 | Q7L7X3 | TAOK1_HUMAN | Unmodified | 0 | 0 | 0 | 0 | 0 | 0 | 0 | 0 | 0 | 0 | 0 | 0 | 0 | 0 | 0 | 0 | 0 | 0 |
| QHFISFDTI | SRI; SRI | P30626; C9J | SORCN_HUMA | Unmodified | 0 | 0 | 118020 | 0 | 0 | 0 | 0 | 0 | 686380 | 0 | 0 | 1431000 | 0 | 0 | 0 | 0 | 0 | 0 |
| QHFISFDTI | SRI; SRI | P30626; C9J | SORCN_HUMA | Unmodified | 2004400 | 1761600 | 2167600 | 1041400 | 638780 | 0 | 165250 | 0 | 38392000 | 2375800 | 7000000 | 12599000 | 0 | 0 | 2428300 | 2615200 | 486050 | 339150 |
| QHFNDVL | GOLM2; GOLM | Q6P4E1; A0/ | GOLM2_HUMA | Unmodified | 0 | 0 | 0 | 252470 | 0 | 0 | 0 | 0 | 1475400 | 161020 | 0 | 0 | 0 | 0 | 0 | 0 | 0 | 0 |
| QHFPATPL | ACLY | P53396 | ACLY_HUMAN | Unmodified | 0 | 0 | 0 | 0 | 0 | 0 | 0 | 0 | 536240 | 0 | 279030 | 3299300 | 0 | 0 | 366820 | 0 | 53327 | 0 |
| QHGDVVS/ | SLC9A3R1 | O14745 | NHRF1_HUMAI | Unmodified | 106150 | 401600 | 112910 | 159180 | 0 | 0 | 0 | 0 | 196760 | 0 | 58761 | 573930 | 131430 | 82026 | 356420 | 344880 | 0 | 0 |
| QHGDVVS/ | SLC9A3R1 | O14745 | NHRF1_HUMAI | Unmodified | 0 | 0 | 180940 | 369680 | 3814900 | 653410 | 266810 | 233580 | 0 | 415030 | 380350 | 0 | 9039500 | 3946300 | 96930 | 0 | 1273100 | 1323400 |
| QHGPPNIV | RAB22A | Q9UL26 | RB22A_HUMAN | Unmodified | 212760 | 0 | 0 | 0 | 0 | 0 | 0 | 0 | 0 | 0 | 0 | 372150 | 0 | 0 | 512750 | 236580 | 0 | 0 |
| QHIEMSTT | DPP4; DPP4; DI | P27487; A0A | DPP4_HUMAN | Oxidation (M) | 0 | 0 | 1277500 | 331540 | 0 | 0 | 0 | 0 | 0 | 0 | 0 | 0 | 0 | 0 | 768930 | 915660 | 0 | 0 |
| QHIEMSTT | DPP4; DPP4; DI | P27487; A0A | DPP4_HUMAN | Unmodified | 0 | 151820 | 2119600 | 1544700 | 0 | 0 | 0 | 0 | 0 | 0 | 0 | 0 | 0 | 0 | 4309700 | 2091200 | 49117 | 0 |
| QHIVNDMI | NDRG1; NDRG | Q92597; A0/ | NDRG1_HUMA | Unmodified | 0 | 0 | 0 | 0 | 637720 | 0 | 0 | 0 | 835850 | 0 | 0 | 0 | 0 | 0 | 0 | 0 | 427640 | 0 |
| QHIVNDMI | NDRG1; NDRG | Q92597; A0/ | NDRG1_HUMA | Unmodified | 0 | 0 | 54671 | 663520 | 0 | 196280 | 0 | 0 | 9038500 | 192190 | 0 | 298990 | 0 | 0 | 96413 | 96463 | 307010 | 0 |
| QHLEITGG | QARS1; QARS1 | P47897; A0A | SYQ_HUMAN G | Unmodified | 0 | 0 | 0 | 0 | 169110 | 107560 | 0 | 0 | 0 | 154540 | 202910 | 2312700 | 0 | 0 | 0 | 0 | 0 | 295330 |
| QHLENDP( | TXNL1; TXNL1; | O43396; K7E | TXNL1_HUMAN | Unmodified | 0 | 0 | 653060 | 328530 | 0 | 0 | 0 | 0 | 616700 | 0 | 786260 | 1935700 | 0 | 181510 | 624610 | 455570 | 276830 | 0 |
| QHLEQEIY | KIF14 | Q15058 | KIF14_HUMAN | Unmodified | 0 | 0 | 0 | 0 | 546500 | 80833 | 0 | 0 | 0 | 0 | 0 | 110420 | 0 | 0 | 0 | 0 | 297620 | 0 |
| QHLFYGQ | IPO9 | Q96P70 | IPO9_HUMAN I | Unmodified | 0 | 0 | 0 | 0 | 0 | 0 | 0 | 0 | 103020 | 112640 | 0 | 1087200 | 0 | 0 | 0 | 0 | 0 | 0 |
| QHLQDYEI | PPM1L | Q5SGD2 | PPM1L_HUMAN | Unmodified | 12091 | 0 | 0 | 0 | 0 | 0 | 8657.1 | 0 | 2494300 | 0 | 0 | 0 | 0 | 0 | 0 | 0 | 0 | 0 |
| QHLRLVFD | ANXA6 | P08133 | ANXA6_HUMAN | Unmodified | 0 | 0 | 0 | 0 | 331790 | 0 | 1075100 | 0 | 0 | 2811200 | 0 | 0 | 725370 | 0 | 0 | 0 | 0 | 329920 |
| QHMDITGE | RNPEP; RNPEP | Q9H4A4; A6I | AMPB_HUMAN | Unmodified | 0 | 63130 | 0 | 158800 | 0 | 62450 | 28986 | 15919 | 457500 | 0 | 407780 | 3286900 | 0 | 0 | 27658 | 136730 | 0 | 0 |
| QHMDITGE | RNPEP; RNPEP | Q9H4A4; A6I | AMPB_HUMAN | Oxidation (M) | 0 | 0 | 1174000 | 0 | 0 | 0 | 117620 | 0 | 0 | 0 | 109250 | 743590 | 0 | 0 | 0 | 801390 | 0 | 0 |
| QHMFNEK | ADGRG2 | Q8IZP9 | AGRG2_HUMA | Oxidation (M) | 5375900 | 3882300 | 4277800 | 3211000 | 3421800 | 381890 | 549580 | 0 | 1188400 | 9618300 | 0 | 4133600 | 280300 | 0 | 5249700 | 5189300 | 87299 | 0 |
| QHNPPSDI | ADAMTS1; ADA | Q9UHI8; E5F | ATS1_HUMAN | Unmodified | 0 | 0 | 626240 | 0 | 0 | 0 | 0 | 0 | 0 | 0 | 0 | 0 | 389110 | 0 | 0 | 0 | 0 | 0 |
| QHPVPPPA | ZYX; ZYX | Q15942; H0Y | ZYX_HUMAN Z | Unmodified | 0 | 0 | 0 | 0 | 629600 | 0 | 0 | 0 | 0 | 0 | 0 | 0 | 0 | 0 | 0 | 0 | 652680 | 5889200 |
| QHPYTFPG | GALNT2 | Q10471 | GALT2_HUMAN | Unmodified | 0 | 0 | 159890 | 211050 | 0 | 133700 | 0 | 0 | 23768000 | 4027800 | 0 | 1456000 | 0 | 0 | 170390 | 121500 | 0 | 0 |
| QHQLYVG\ | DDAH1 | O94760 | DDAH1_HUMA | Unmodified | 94293 | 113590 | 0 | 0 | 0 | 69050 | 0 | 0 | 1533000 | 0 | 241780 | 1755100 | 0 | 0 | 0 | 1127300 | 87421 | 0 |
| QHRGPQLI | PODXL2 | Q9NZ53 | PDXL2_HUMAN | Unmodified | 0 | 0 | 298810 | 350750 | 0 | 0 | 0 | 0 | 1080400 | 0 | 0 | 0 | 0 | 0 | 0 | 0 | 0 | 0 |
| QHRLDQDI | TRAF2 | Q12933 | TRAF2_HUMAN | Unmodified | 0 | 0 | 0 | 0 | 1968600 | 1286400 | 0 | 0 | 405900 | 0 | 0 | 1093700 | 0 | 0 | 0 | 0 | 0 | 0 |
| QHSVVPS( | HNRNPLL; HNF | Q8WVV9; C9 | HNRLL_HUMAI | Unmodified | 33382 | 0 | 0 | 0 | 0 | 0 | 0 | 28594 | 0 | 0 | 52178 | 446430 | 0 | 0 | 0 | 0 | 0 | 0 |
| QHTEQEAS | MYO1D; MYO1I | O94832; J3Q | MYO1D_HUMA | Unmodified | 921790 | 170520 | 534400 | 0 | 0 | 0 | 0 | 0 | 32468 | 0 | 0 | 230090 | 0 | 0 | 114940 | 315270 | 0 | 0 |
| QHTEQEAS | MYO1D; MYO1I | O94832; J3Q | MYO1D_HUMA | Unmodified | 2351400 | 3161000 | 1516600 | 806390 | 0 | 45545 | 0 | 0 | 29326 | 0 | 0 | 0 | 611480 | 0 | 1924200 | 1573000 | 116100 | 118750 |
| QHTFVETE | ARCN1; ARCN1 | P48444; Q6F | COPD_HUMAN | Unmodified | 0 | 0 | 0 | 0 | 0 | 0 | 0 | 0 | 0 | 54836 | 0 | 384410 | 0 | 0 | 0 | 0 | 0 | 0 |
| QHTIMPSK | MTAP; N/A; MT/ | Q13126; F2Z | MTAP_HUMAN | Unmodified | 48918 | 0 | 0 | 0 | 0 | 0 | 0 | 0 | 0 | 0 | 36077 | 460440 | 0 | 0 | 0 | 0 | 0 | 0 |
| QHTVTTTI | PRNP; PRNP; PI | P04156; A2A | PRIO_HUMAN I | Unmodified | 0 | 0 | 334680 | 438560 | 0 | 0 | 0 | 0 | 0 | 0 | 0 | 0 | 469120 | 0 | 616560 | 671870 | 0 | 0 |
| QHVTEAFQ | PHGDH; PHGD | O43175; A0/ | SERA_HUMAN | Unmodified | 0 | 0 | 662460 | 203930 | 0 | 0 | 301400 | 30529 | 1095000 | 153730 | 165340 | 5513300 | 429480 | 200880 | 563020 | 309840 | 78104 | 51471 |
| QHYESGA/ | PCYOX1L; PCY( | Q8NBM8; D6 | PCYXL_HUMAN | Unmodified | 0 | 82128 | 0 | 0 | 0 | 0 | 0 | 0 | 1909700 | 0 | 0 | 0 | 0 | 0 | 0 | 0 | 0 | 0 |
| QHYGRDN | QSOX2 | Q6ZRP7 | QSOX2_HUMAI | Unmodified | 0 | 0 | 0 | 0 | 0 | 0 | 0 | 0 | 6684000 | 0 | 0 | 0 | 0 | 0 | 0 | 0 | 0 | 0 |
| QHYIDLKD | NME2P1; NME2 | O60361; P22 | NDK8_HUMAN | Unmodified | 0 | 0 | 0 | 0 | 45528000 | 4175800 | 8999900 | 0 | 368290 | 2182300 | 0 | 0 | 21119000 | 1530200 | 0 | 0 | 45093000 | 21011000 |
| QIAASSQD | ALDH18A1 | P54886 | P5CS_HUMAN | Unmodified | 0 | 0 | 0 | 0 | 20301 | 0 | 36361 | 23966 | 899580 | 170060 | 0 | 0 | 0 | 0 | 0 | 0 | 0 | 0 |
| QIANSQDG | RO60; RO60; R( | P10155; D6F | RO60_HUMAN | Oxidation (M) | 9039700 | 0 | 2328100 | 1578900 | 162020 | 0 | 172550 | 0 | 0 | 0 | 0 | 676040 | 0 | 0 | 3074900 | 3519000 | 0 | 0 |
| QIANSQDG | RO60; RO60; R( | P10155; D6F | RO60_HUMAN | Oxidation (M) | 9492200 | 236110 | 0 | 4053500 | 0 | 0 | 0 | 0 | 0 | 0 | 0 | 0 | 0 | 0 | 1166800 | 0 | 0 | 0 |
| QIANSQDG | RO60; RO60; R( | P10155; D6F | RO60_HUMAN | Unmodified | 0 | 1954700 | 0 | 1314700 | 0 | 0 | 0 | 0 | 0 | 0 | 0 | 0 | 0 | 0 | 0 | 0 | 0 | 0 |
| QIAQLKQEI | CSGALNACT2 | Q8N6G5 | CGAT2_HUMAI | Unmodified | 240280 | 0 | 246090 | 0 | 0 | 0 | 0 | 0 | 1169500 | 205330 | 0 | 0 | 0 | 0 | 435870 | 310010 | 0 | 0 |
| QIASEKEEII | RALBP1 | Q15311 | RBP1_HUMAN | Unmodified | 0 | 0 | 69662 | 103570 | 0 | 0 | 0 | 0 | 102230 | 0 | 54532 | 275720 | 0 | 0 | 130050 | 193040 | 0 | 0 |
| QIASLTGLV | KIAA1217 | Q5T5P2 | SKT_HUMAN S | Unmodified | 0 | 0 | 0 | 0 | 0 | 0 | 0 | 0 | 219910 | 103140 | 0 | 0 | 0 | 0 | 0 | 0 | 0 | 0 |
| QIASLTGLV | KIAA1217 | Q5T5P2 | SKT_HUMAN S | Unmodified | 0 | 0 | 0 | 0 | 0 | 0 | 0 | 0 | 0 | 351160 | 0 | 0 | 188790 | 1029700 | 0 | 0 | 66254 | 0 |
| QIATLHAQ\ | MYH9 | P35579 | MYH9_HUMAN | Oxidation (M) | 204570 | 391290 | 0 | 0 | 477070 | 105340 | 102810 | 0 | 0 | 0 | 0 | 164230 | 140420 | 0 | 0 | 90367 | 265860 | 753650 |

| | | | | | | | | | | | | | | | | | | | | | | |
|---|---|---|---|---|---|---|---|---|---|---|---|---|---|---|---|---|---|---|---|---|---|---|
| QIATLHAQ\ | MYH9 | P35579 | MYH9_HUMAN | Unmodified | 0 | 0 | 0 | 0 | 4076000 | 599590 | 137840 | 0 | 0 | 114960 | 0 | 1889600 | 906340 | 411860 | 0 | 0 | 1966100 | 1824700 |
| QIATLHAQ\ | MYH9 | P35579 | MYH9_HUMAN | Unmodified | 0 | 0 | 0 | 0 | 4042900 | 0 | 0 | 0 | 0 | 0 | 0 | 694490 | 1123400 | 1051900 | 199760 | 147400 | 1928900 | 1501300 |
| QIATSGPPI | SVIP | Q8NHG7 | SVIP_HUMAN S | Unmodified | 2074400 | 1888200 | 794550 | 503210 | 0 | 0 | 0 | 108830 | 2210800 | 229740 | 454480 | 1228500 | 250020 | 0 | 998510 | 934770 | 0 | 280520 |
| QIATSGPPI | SVIP | Q8NHG7 | SVIP_HUMAN S | Unmodified | 311830 | 309540 | 0 | 374870 | 867180 | 0 | 621880 | 585160 | 717830 | 2682900 | 231710 | 0 | 3138200 | 2156700 | 0 | 0 | 2315800 | 4251500 |
| QIAVEAQEI | FLOT2; FLOT2; I | Q14254; J3Q | FLOT2_HUMAN | Unmodified | 1662400 | 0 | 1886900 | 1980000 | 13002000 | 0 | 2360900 | 391690 | 7012600 | 0 | 0 | 7301400 | 3592500 | 3134600 | 3758900 | 1695000 | 0 | 3327700 |
| QIAVEAQEI | FLOT2; FLOT2; I | Q14254; J3Q | FLOT2_HUMAN | Unmodified | 0 | 0 | 0 | 0 | 610500 | 0 | 0 | 0 | 0 | 383880 | 0 | 0 | 0 | 181400 | 0 | 0 | 76406 | 0 |
| QICSQLSEI | RABGAP1 | Q9Y3P9 | RBGP1_HUMAI | Unmodified | 0 | 0 | 75127 | 0 | 820600 | 0 | 0 | 0 | 0 | 96719 | 0 | 729030 | 62692 | 0 | 0 | 115820 | 45575 | 0 |
| QICVVMLE! | PCBP3; PCBP3; | P57721; E9P | PCBP3_HUMAI | Oxidation (M) | 7138000 | 1565200 | 673080 | 3710900 | 0 | 0 | 0 | 30561 | 0 | 101610 | 0 | 0 | 0 | 0 | 0 | 1529100 | 285130 | 0 |
| QIDATFVRI | PPP1R7; PPP1R | Q15435; H7( | PP1R7_HUMAN | Unmodified | 0 | 0 | 839460 | 660420 | 1538700 | 244430 | 0 | 106620 | 292470 | 400450 | 703860 | 3734600 | 642840 | 364820 | 330740 | 511280 | 349220 | 216280 |
| QIDLSTVDI | MANF | P55145 | MANF_HUMAN | Unmodified | 0 | 0 | 0 | 0 | 955010 | 0 | 1161000 | 861910 | 41421000 | 17005000 | 6678300 | 7327500 | 1737500 | 798030 | 0 | 0 | 492030 | 1128500 |
| QIDLSTVDI | MANF | P55145 | MANF_HUMAN | Unmodified | 260940 | 0 | 0 | 0 | 237460 | 0 | 705220 | 726400 | 22939000 | 14231000 | 3696400 | 1418900 | 945670 | 707130 | 253180 | 428440 | 140820 | 941690 |
| QIDNPDYK | CALR; CALR; C/ | P27797; A0A | CALR_HUMAN | Unmodified | 0 | 159740 | 0 | 0 | 0 | 0 | 0 | 115000 | 29350000 | 811880 | 1382400 | 1846900 | 0 | 0 | 339610 | 363760 | 0 | 0 |
| QIDQFLVV/ | MTA1; MTA3; M | Q13330; Q9I | MTA1_HUMAN | Unmodified | 473990 | 0 | 751410 | 0 | 0 | 0 | 0 | 0 | 0 | 0 | 0 | 1098700 | 0 | 0 | 0 | 0 | 0 | 0 |
| QIDSGSNL | PCYT2; PCYT2; | Q99447; I3L | PCY2_HUMAN | Unmodified | 0 | 0 | 0 | 0 | 0 | 0 | 0 | 0 | 0 | 0 | 0 | 846220 | 0 | 0 | 0 | 0 | 0 | 0 |
| QIEELKGQI | UCHL1; UCHL1 | P09936; D6F | UCHL1_HUMAI | Unmodified | 0 | 0 | 0 | 0 | 52883000 | 8140800 | 4486200 | 2360900 | 5872100 | 9659700 | 2751300 | 13570000 | 0 | 0 | 0 | 0 | 0 | 0 |
| QIEEQTIK | RDX; RDX; RDX | P35241; A0A | RADI_HUMAN I | Unmodified | 242420 | 182420 | 434670 | 773500 | 4721700 | 792540 | 61480 | 0 | 1214900 | 157370 | 263070 | 1884100 | 1527500 | 644610 | 0 | 669100 | 294840 | 609460 |
| QIEEQTIKA | RDX; RDX; RDX | P35241; A0A | RADI_HUMAN I | Unmodified | 0 | 0 | 0 | 0 | 2260800 | 692760 | 0 | 0 | 0 | 0 | 0 | 0 | 3593900 | 2628600 | 0 | 0 | 351000 | 0 |
| QIEINTISAS | GSS; GSS; GSS | P48637; A0A | GSHB_HUMAN | Unmodified | 0 | 0 | 0 | 0 | 0 | 0 | 0 | 0 | 1088800 | 17597 | 322490 | 3676600 | 0 | 0 | 461520 | 410260 | 0 | 0 |
| QIEKLEQEL | CENPF | P49454 | CENPF_HUMAI | Unmodified | 0 | 0 | 0 | 0 | 0 | 0 | 0 | 0 | 87553 | 0 | 0 | 0 | 0 | 0 | 0 | 0 | 401360 | 83858 |
| QIFEDHVF( | CALCOCO2 | Q13137 | CACO2_HUMA | Unmodified | 0 | 0 | 0 | 0 | 0 | 0 | 1770300 | 0 | 0 | 199200 | 0 | 0 | 230730 | 0 | 0 | 0 | 1349100 | 1912800 |
| QIFIGPPDII | AFG3L2 | Q9Y4W6 | AFG32_HUMAN | Unmodified | 0 | 0 | 0 | 0 | 0 | 0 | 0 | 0 | 1389900 | 0 | 0 | 0 | 0 | 0 | 0 | 0 | 0 | 0 |
| QIFILLFQR | CSE1L | P55060 | XPO2_HUMAN | Unmodified | 0 | 0 | 0 | 0 | 0 | 0 | 0 | 0 | 0 | 0 | 0 | 769130 | 0 | 0 | 0 | 0 | 0 | 0 |
| QIFLGGVDI | SLC25A6; SLC2 | P12236; P05 | ADT3_HUMAN | Unmodified | 0 | 0 | 0 | 0 | 0 | 0 | 0 | 0 | 2520500 | 0 | 0 | 0 | 0 | 0 | 0 | 0 | 0 | 0 |
| QIFNGTFVI | RPL7; RPL7; RP | P18124; C9J | RL7_HUMAN 6( | Unmodified | 3116400 | 3127700 | 1390800 | 281570 | 0 | 0 | 0 | 0 | 0 | 0 | 0 | 40226000 | 0 | 0 | 0 | 798870 | 0 | 0 |
| QIGDALPV! | ALCAM; ALCAM | Q13740; F5( | CD166_HUMAI | Unmodified | 1207900 | 880110 | 0 | 0 | 0 | 0 | 0 | 0 | 723570 | 0 | 0 | 574900 | 0 | 0 | 0 | 883940 | 0 | 0 |
| QIGDNLIVI | DPYSL3; DPYSL | Q14195; D6I | DPYL3_HUMAN | Unmodified | 0 | 0 | 0 | 0 | 1921400 | 0 | 766410 | 671070 | 0 | 0 | 29718000 | 58784000 | 0 | 0 | 0 | 0 | 0 | 318790 |
| QIGENLIVP | CRMP1; DPYSL | Q14194; Q1( | DPYL1_HUMAN | Unmodified | 0 | 0 | 0 | 0 | 11380000 | 2876000 | 2650700 | 1741200 | 3938500 | 2790900 | 0 | 7116900 | 1419800 | 813340 | 0 | 0 | 924880 | 2540600 |
| QIGIDPPR( | PSMC4 | P43686 | PRS6B_HUMAN | Oxidation (M) | 53083 | 0 | 0 | 446040 | 0 | 684320 | 1179900 | 510730 | 1666500 | 0 | 0 | 748160 | 1342900 | 766710 | 0 | 96708 | 968930 | 768210 |
| QIGIDPPR( | PSMC4 | P43686 | PRS6B_HUMAN | Unmodified | 0 | 0 | 0 | 0 | 367300 | 0 | 0 | 0 | 171130 | 0 | 0 | 0 | 0 | 475290 | 0 | 0 | 297700 | 397900 |
| QIGNVAALI | RTCB | Q9Y3I0 | RTCB_HUMAN | Unmodified | 0 | 0 | 0 | 0 | 1370500 | 0 | 540860 | 0 | 667020 | 364300 | 1584100 | 10158000 | 3955700 | 1304100 | 0 | 0 | 715230 | 1686000 |
| QIGPGESC | CDCP1 | Q9H5V8 | CDCP1_HUMA | Unmodified | 76760 | 0 | 1072300 | 629100 | 0 | 0 | 68070 | 106740 | 0 | 88771 | 343210 | 10009000 | 0 | 0 | 2138200 | 1421800 | 138970 | 0 |
| QIGPGESC | CDCP1 | Q9H5V8 | CDCP1_HUMA | Unmodified | 361910 | 0 | 846700 | 766960 | 635510 | 0 | 0 | 0 | 0 | 0 | 405450 | 3335800 | 301010 | 0 | 1448400 | 1064800 | 384430 | 1637100 |
| QIHAVRYLI | RHOG | P84095 | RHOG_HUMAN | Unmodified | 0 | 0 | 0 | 0 | 877960 | 24150 | 194830 | 0 | 0 | 0 | 0 | 427750 | 2999100 | 0 | 0 | 0 | 0 | 0 |
| QIHKFEELI | EIF3L; EIF3L; EII | Q9Y262; B0( | EIF3L_HUMAN | Unmodified | 182700 | 242870 | 178680 | 0 | 0 | 0 | 42730 | 0 | 0 | 0 | 0 | 1575200 | 0 | 78385 | 0 | 181420 | 0 | 0 |
| QIIDQDK | TJP1; TJP1; TJP1 | Q07157; A0/ | ZO1_HUMAN T | Unmodified | 0 | 144390 | 0 | 0 | 550110 | 0 | 0 | 0 | 0 | 0 | 0 | 1828200 | 401240 | 0 | 0 | 68978 | 0 | 381020 |
| QIIEDNPLC | PLCG2; PLCG2 | P16885; A0A | PLCG2_HUMAI | Unmodified | 0 | 0 | 0 | 0 | 0 | 0 | 0 | 0 | 0 | 0 | 1179100 | 6527300 | 0 | 0 | 0 | 0 | 0 | 0 |
| QIIEDQSGI | DLG3; DLG3 | Q92796; Q5. | DLG3_HUMAN | Unmodified | 137750 | 93178 | 6436100 | 754460 | 381440 | 0 | 0 | 76577 | 1807200 | 931880 | 0 | 6628500 | 1381400 | 437850 | 2917200 | 1412600 | 2483000 | 605450 |
| QIIEEQSGS | DLG1; DLG1; D | Q12959; A0/ | DLG1_HUMAN | Unmodified | 0 | 0 | 0 | 0 | 0 | 0 | 0 | 0 | 0 | 0 | 0 | 0 | 0 | 0 | 0 | 0 | 1008500 | 472100 |
| QIIEQDKH/ | TJP2; TJP2; TJP2 | Q9UDY2; A0 | ZO2_HUMAN T | Unmodified | 0 | 0 | 0 | 0 | 1022100 | 287480 | 0 | 0 | 0 | 826430 | 0 | 0 | 0 | 258520 | 0 | 0 | 184750 | 0 |
| QIIGYVIGT( | CEACAM5; CEA | P06731; A0A | CEAM5_HUMAI | Unmodified | 0 | 0 | 0 | 0 | 0 | 0 | 0 | 0 | 2077200 | 0 | 0 | 0 | 670740 | 0 | 0 | 0 | 0 | 0 |
| QIILEKEETE | XRCC6; XRCC6 | P12956; B1A | XRCC6_HUMAI | Unmodified | 516190 | 178410 | 0 | 75971 | 653630 | 228740 | 0 | 282600 | 606720 | 395030 | 2571600 | 14448000 | 336850 | 0 | 140810 | 0 | 0 | 0 |
| QIILEKEETE | XRCC6; XRCC6 | P12956; B1A | XRCC6_HUMAI | Unmodified | 367840 | 0 | 0 | 62669 | 61161 | 0 | 0 | 0 | 0 | 0 | 320430 | 3180900 | 0 | 0 | 0 | 96957 | 0 | 0 |
| QIIQQNPSI | RAD23B | P54727 | RD23B_HUMAI | Unmodified | 1128500 | 1008800 | 272820 | 112590 | 0 | 0 | 0 | 0 | 205010 | 0 | 0 | 528650 | 0 | 0 | 1296600 | 1014300 | 0 | 0 |
| QIITDKDDL | VPS37A; VPS37 | Q8NEZ2; H0 | VP37A_HUMAN | Unmodified | 4580000 | 6350400 | 830030 | 1471400 | 2536900 | 897160 | 148520 | 64598 | 456440 | 0 | 255400 | 1118300 | 877820 | 593070 | 823260 | 774000 | 0 | 188920 |
| QIIVDPLSF | CTNNA1; CTNN | P35221; H0Y | CTNA1_HUMAI | Unmodified | 2898400 | 2905400 | 6115600 | 7539100 | 0 | 1346400 | 0 | 0 | 1469900 | 0 | 5538400 | 23674000 | 0 | 0 | 22469000 | 13092000 | 0 | 0 |
| QIKLDPSP( | EIF4G2; EIF4G2 | P78344; H0Y | IF4G2_HUMAN | Unmodified | 0 | 0 | 0 | 0 | 0 | 0 | 0 | 0 | 0 | 0 | 0 | 375420 | 112450 | 0 | 0 | 0 | 0 | 0 |
| QILDEAGK\ | VCL | P18206 | VINC_HUMAN ' | Unmodified | 492490 | 742680 | 60224 | 0 | 126280 | 0 | 0 | 0 | 216440 | 0 | 0 | 4137200 | 0 | 0 | 428120 | 1063000 | 447710 | 0 |
| QILESIESIH | TTBK2; TTBK2; 1 | Q6IQ55; A0A | TTBK2_HUMAN | Unmodified | 75827 | 0 | 0 | 0 | 0 | 0 | 0 | 0 | 0 | 0 | 0 | 0 | 0 | 0 | 0 | 0 | 159550 | 0 |
| QILGVIDK | CAPRIN1; CAPF | Q14444; E9F | CAPR1_HUMAI | Unmodified | 209530 | 564760 | 0 | 0 | 800660 | 217950 | 73128 | 146190 | 139460 | 224940 | 0 | 5628900 | 34623 | 0 | 0 | 156680 | 180010 | 461170 |
| QILLYSATFI | DDX6 | P26196 | DDX6_HUMAN | Unmodified | 0 | 0 | 0 | 0 | 0 | 0 | 0 | 0 | 0 | 0 | 0 | 3721200 | 0 | 0 | 0 | 0 | 0 | 0 |
| QILQNRKSI | MYO6; MYO6; N | Q9UM54; A0 | MYO6_HUMAN | Unmodified | 0 | 0 | 0 | 0 | 0 | 0 | 0 | 0 | 0 | 0 | 0 | 0 | 0 | 136960 | 0 | 0 | 0 | 0 |
| QILSADDK | STRAP | Q9Y3F4 | STRAP_HUMAN | Unmodified | 0 | 85785 | 135600 | 411000 | 713930 | 0 | 308110 | 0 | 1573200 | 69035 | 14164 | 5355800 | 559830 | 883120 | 326410 | 0 | 154190 | 1341800 |
| QILSADLR | NPEPPS; NPEPI | P55786; A0A | PSA_HUMAN P | Unmodified | 0 | 0 | 0 | 0 | 0 | 0 | 0 | 0 | 1620400 | 0 | 0 | 7253500 | 0 | 0 | 0 | 0 | 0 | 0 |
| QILSADLRS | NPEPPS; NPEPI | P55786; A0A | PSA_HUMAN P | Unmodified | 0 | 0 | 0 | 0 | 2510800 | 1177200 | 0 | 0 | 260650 | 918790 | 1126500 | 0 | 2510100 | 713980 | 18216 | 0 | 1546000 | 1979900 |
| QILSLNYDI | WASHC5; WASI | Q12768; E7E | WASC5_HUMA | Unmodified | 0 | 0 | 0 | 0 | 0 | 0 | 0 | 0 | 0 | 0 | 0 | 1348100 | 0 | 0 | 0 | 0 | 0 | 0 |
| QILTKGEV( | SBDS; SBDS | Q9Y3A5; A0/ | SBDS_HUMAN | Unmodified | 625590 | 546020 | 0 | 328070 | 420850 | 0 | 77615 | 27056 | 194590 | 29609 | 358940 | 595080 | 197400 | 72405 | 437900 | 375930 | 0 | 96876 |
| QILVGDIGI | CFL2 | Q9Y281 | COF2_HUMAN | Unmodified | 2734900 | 1884400 | 201510 | 1936900 | 1.32E+08 | 23599000 | 3543700 | 442730 | 986850 | 717380 | 354050 | 1296100 | 5349500 | 5376100 | 2840800 | 1415300 | 19424000 | 10230000 |
| QIMKEIQEI | SEPTIN7; SEPTI | Q16181; E7E | SEPT7_HUMAN | Unmodified | 0 | 0 | 0 | 0 | 1913200 | 0 | 201940 | 108200 | 0 | 0 | 0 | 0 | 328470 | 318800 | 0 | 0 | 1207300 | 0 |
| QIMWFWQ | ITCH; ITCH; ITC | Q96J02; A0A | ITCH_HUMAN I | Oxidation (M) | 572740 | 1372800 | 38891 | 0 | 0 | 0 | 0 | 0 | 0 | 0 | 0 | 0 | 0 | 0 | 1009000 | 165000 | 0 | 0 |
| QINDIQLSI | EIF3I; EIF3I; EIF | Q13347; A0/ | EIF3I_HUMAN I | Unmodified | 402270 | 44533 | 0 | 380000 | 0 | 810840 | 796680 | 264360 | 6363600 | 663860 | 9615600 | 1.24E+08 | 0 | 0 | 339320 | 0 | 359200 | 293460 |
| QINKLSDQ | TAX1BP1; TAX1 | Q86VP1; B8 | TAXB1_HUMAN | Unmodified | 0 | 0 | 0 | 0 | 260240 | 0 | 2110200 | 111420 | 0 | 57818 | 0 | 0 | 0 | 199610 | 0 | 0 | 1348800 | 363360 |
| QINLSNIR | SND1 | Q7KZF4 | SND1_HUMAN | Unmodified | 392960 | 407410 | 1038000 | 848980 | 0 | 93496 | 0 | 12832 | 1586700 | 296740 | 497400 | 5209000 | 620490 | 130570 | 1142400 | 883550 | 0 | 0 |
| QINQEYLIS | TPRKB | Q9Y3C4 | TPRKB_HUMAN | Unmodified | 0 | 0 | 0 | 0 | 0 | 0 | 0 | 0 | 0 | 0 | 0 | 1964500 | 0 | 0 | 0 | 0 | 0 | 0 |
| QINVARLD( | LRP1 | Q07954 | LRP1_HUMAN I | Unmodified | 0 | 0 | 0 | 0 | 0 | 0 | 0 | 0 | 788150 | 0 | 0 | 0 | 0 | 0 | 0 | 0 | 0 | 0 |
| QINWTVLY | RPL24; RPL24; I | P83731; C9J | RL24_HUMAN ( | Unmodified | 7853300 | 8484100 | 2926400 | 2055700 | 1158300 | 220760 | 447870 | 107580 | 979480 | 317240 | 1668500 | 12720000 | 1608000 | 498960 | 3896000 | 3638800 | 3067600 | 776030 |
| QINWTVLY | RPL24; RPL24; I | P83731; C9J | RL24_HUMAN ( | Unmodified | 205070 | 343660 | 0 | 0 | 0 | 0 | 0 | 0 | 0 | 0 | 0 | 597750 | 115230 | 0 | 110120 | 0 | 0 | 0 |
| QIPAAQEQ | ITGA2; ITGA2 | P17301; E7E | ITA2_HUMAN I | Unmodified | 2543300 | 2469100 | 2439200 | 1340600 | 0 | 0 | 0 | 0 | 0 | 0 | 0 | 0 | 0 | 0 | 567120 | 852120 | 364280 | 0 |
| QIPAAQEQ | ITGA2; ITGA2 | P17301; E7E | ITA2_HUMAN I | Unmodified | 0 | 0 | 0 | 759180 | 0 | 0 | 0 | 0 | 0 | 0 | 0 | 505050 | 0 | 0 | 854040 | 0 | 173200 | 161660 |
| QIPAITCIQ! | IQGAP1; IQGAP | P46940; A0A | IQGA1_HUMAN | Unmodified | 0 | 0 | 0 | 0 | 2346600 | 463780 | 0 | 211110 | 0 | 103170 | 0 | 0 | 0 | 0 | 0 | 0 | 0 | 1527000 |
| QIPLQSLDI | EMC1 | Q8N766 | EMC1_HUMAN | Unmodified | 0 | 0 | 0 | 0 | 0 | 0 | 0 | 0 | 4380900 | 744970 | 0 | 0 | 0 | 0 | 0 | 0 | 0 | 0 |
| QIPQATASI | PHGDH; PHGD | O43175; A0/ | SERA_HUMAN | Unmodified | 0 | 0 | 998270 | 2714000 | 1278200 | 0 | 0 | 0 | 2106600 | 470340 | 986130 | 4822400 | 6860300 | 2766400 | 2303600 | 2097700 | 0 | 2108800 |
| QIPQATASI | PHGDH; PHGD | O43175; A0/ | SERA_HUMAN | Unmodified | 0 | 0 | 0 | 0 | 0 | 0 | 0 | 0 | 0 | 0 | 0 | 0 | 2798800 | 701000 | 0 | 0 | 0 | 0 |
| QIPVLQTNI | EEF1E1; EEF1E | O43324; D6I | MCA3_HUMAN | Unmodified | 0 | 0 | 0 | 0 | 0 | 0 | 0 | 0 | 204780 | 0 | 0 | 604790 | 0 | 0 | 0 | 0 | 0 | 0 |
| QIPYTMMK | SLC25A3; SLC2 | Q00325; F8V | MPCP_HUMAN | Oxidation (M) | 0 | 0 | 0 | 0 | 0 | 0 | 0 | 0 | 3438100 | 701050 | 0 | 0 | 0 | 0 | 0 | 0 | 0 | 0 |
| QIQELVEAI | PSMC3; PSMC3 | P17980; E9P | PRS6A_HUMAN | Oxidation (M) | 0 | 0 | 1214400 | 0 | 0 | 0 | 0 | 0 | 517210 | 0 | 465850 | 1754100 | 0 | 134500 | 0 | 0 | 30448 | 0 |
| QIQFADDM | AHCYL1 | O43865 | SAHH2_HUMAI | Unmodified | 0 | 0 | 0 | 0 | 5056500 | 1250100 | 0 | 0 | 0 | 0 | 0 | 1072200 | 573900 | 0 | 0 | 49354 | 1075100 | 3272600 |
| QIQFADDM | AHCYL1 | O43865 | SAHH2_HUMAI | Oxidation (M) | 0 | 0 | 0 | 0 | 2007500 | 0 | 0 | 0 | 0 | 0 | 0 | 0 | 595380 | 381070 | 0 | 0 | 685820 | 0 |
| QIQQLESN | ROCK2; ROCK2 | O75116; E9F | ROCK2_HUMAI | Unmodified | 0 | 0 | 0 | 0 | 1101000 | 424300 | 0 | 0 | 372900 | 0 | 0 | 785330 | 0 | 0 | 0 | 0 | 288550 | 1086800 |
| QIQQLESN | ROCK2; ROCK2 | O75116; E9F | ROCK2_HUMAI | Unmodified | 0 | 0 | 0 | 0 | 625760 | 0 | 0 | 0 | 0 | 0 | 0 | 0 | 1261800 | 0 | 0 | 0 | 1475600 | 1201700 |
| QIQQYMK | HSPA4 | P34932 | HSP74_HUMAN | Unmodified | 0 | 0 | 0 | 0 | 2616000 | 481880 | 352160 | 0 | 625390 | 615350 | 1633700 | 1680700 | 1930800 | 780640 | 0 | 0 | 685920 | 929120 |
| QIQSLCFQ | COPS4; COPS4 | Q9BT78; D6I | CSN4_HUMAN | Unmodified | 0 | 0 | 0 | 0 | 0 | 0 | 0 | 30495 | 0 | 0 | 0 | 0 | 0 | 0 | 728530 | 0 | 1455400 | 0 |
| QIQSLENG | PAN2 | Q504Q3 | PAN2_HUMAN | Unmodified | 188490 | 350710 | 0 | 0 | 533530 | 0 | 0 | 0 | 0 | 0 | 0 | 0 | 20483 | 0 | 40667 | 0 | 117880 | 73266 |
| QIQSLKEEI | MYO5C | Q9NQX4 | MYO5C_HUMA | Unmodified | 0 | 0 | 0 | 0 | 805510 | 75789 | 0 | 0 | 405950 | 233540 | 0 | 0 | 92548 | 0 | 0 | 0 | 37463 | 0 |
| QIQSPFPH | SHISA2 | Q6UWI4 | SHSA2_HUMAN | Unmodified | 0 | 0 | 31610 | 66524 | 0 | 0 | 46355 | 164730 | 2248300 | 542500 | 0 | 0 | 33025 | 0 | 0 | 98079 | 42259 | 171200 |
| QIQSQWK | FOLH1; FOLH1 | Q04609; E9F | FOLH1_HUMAN | Unmodified | 3465500 | 3089100 | 20904000 | 19988000 | 0 | 0 | 0 | 0 | 0 | 0 | 0 | 0 | 2340600 | 587580 | 3735600 | 14340000 | 0 | 0 |
| QIQTEAAQI | RMDN1; RMDN | Q96DB5; H0 | RMD1_HUMAN | Unmodified | 0 | 0 | 0 | 0 | 0 | 0 | 0 | 0 | 423950 | 0 | 0 | 0 | 0 | 0 | 0 | 0 | 0 | 0 |
| QIQVNDED | RAB23 | Q9ULC3 | RAB23_HUMAN | Unmodified | 1550200 | 5270100 | 895820 | 1338200 | 83303 | 0 | 0 | 0 | 1246100 | 0 | 120080 | 1637500 | 0 | 0 | 1752700 | 1425600 | 0 | 0 |
| QIQWFWQ | WWP2 | O00308 | WWP2_HUMAN | Unmodified | 1703200 | 1449100 | 377330 | 815450 | 0 | 0 | 0 | 0 | 0 | 0 | 0 | 0 | 0 | 0 | 2654900 | 1079000 | 0 | 0 |
| QIRSSPWN | EPB41L1; EPB4 | Q9H4G0; Q4 | E41L1_HUMAN | Unmodified | 0 | 0 | 0 | 0 | 52311 | 0 | 0 | 0 | 0 | 0 | 0 | 0 | 0 | 0 | 0 | 0 | 110690 | 0 |
| QISDGEREI | BAG2 | O95816 | BAG2_HUMAN | Unmodified | 0 | 0 | 132350 | 0 | 0 | 0 | 0 | 0 | 1127800 | 0 | 0 | 0 | 0 | 0 | 144700 | 185250 | 0 | 0 |
| QISDVERD | DIAPH1; DIAPH | O60610; A0/ | DIAP1_HUMAN | Unmodified | 83204 | 74161 | 202600 | 284910 | 388660 | 79141 | 621430 | 0 | 475300 | 144920 | 372420 | 2698900 | 824450 | 0 | 725030 | 248360 | 484260 | 573700 |
| QISESTNHI | KIT | P10721 | KIT_HUMAN M | Unmodified | 0 | 0 | 0 | 0 | 0 | 0 | 0 | 0 | 370930 | 42867 | 0 | 1686800 | 0 | 0 | 0 | 0 | 0 | 0 |
| QISLLTSAV | EFCAB14; EFC/ | O75071; A0/ | EFC14_HUMAN | Unmodified | 0 | 0 | 0 | 0 | 0 | 0 | 0 | 0 | 10859000 | 3404600 | 0 | 0 | 0 | 0 | 0 | 0 | 0 | 0 |
| QISPELSAL | PLCD3 | Q8N3E9 | PLCD3_HUMAN | Unmodified | 0 | 0 | 0 | 0 | 994310 | 0 | 0 | 0 | 0 | 0 | 0 | 0 | 159710 | 0 | 0 | 0 | 2058900 | 2742700 |
| QISQAYEVI | DNAJA1 | P31689 | DNJA1_HUMAN | Unmodified | 3054400 | 3087500 | 3093800 | 658550 | 0 | 0 | 0 | 0 | 404980 | 637750 | 0 | 2290300 | 0 | 0 | 4112800 | 3102400 | 0 | 0 |
| QISQDVKLI | AMPD2; AMPD2 | Q01433; A0/ | AMPD2_HUMA | Unmodified | 0 | 0 | 0 | 0 | 0 | 0 | 0 | 0 | 0 | 0 | 0 | 796810 | 190860 | 0 | 0 | 0 | 0 | 0 |
| QISSLRDE\ | KIF5B | P33176 | KINH_HUMAN | Unmodified | 0 | 0 | 0 | 0 | 2872400 | 0 | 29844 | 0 | 645900 | 482170 | 1067000 | 2579600 | 860250 | 1137000 | 0 | 0 | 2026500 | 2103400 |
| QISVRGLA( | PYGB | P11216 | PYGB_HUMAN | Unmodified | 0 | 0 | 0 | 0 | 0 | 0 | 0 | 0 | 0 | 0 | 0 | 0 | 0 | 0 | 0 | 0 | 0 | 1348000 |
| QISYSPGV( | CDC42BPA; CD | Q5VT25; A0A | MRCKA_HUMA | Unmodified | 0 | 0 | 0 | 0 | 0 | 0 | 287700 | 0 | 0 | 0 | 0 | 0 | 0 | 0 | 0 | 0 | 2200100 | 0 |
| QITAQQINI | UPF1 | Q92900 | RENT1_HUMAN | Unmodified | 2345500 | 5410100 | 2569700 | 3570900 | 144070 | 68733 | 0 | 0 | 1095100 | 468900 | 406960 | 3153300 | 414520 | 0 | 2671800 | 4852700 | 0 | 91476 |
| QITASSSYK | MFGE8 | Q08431 | MFGM_HUMAN | Unmodified | 978030 | 1038300 | 4081400 | 3986900 | 66194000 | 25013000 | 345430 | 147730 | 3857800 | 489290 | 151790 | 4381000 | 2480100 | 0 | 5235800 | 4419700 | 3234400 | 8885900 |
| QITIEEGEQ | RAB6B; RAB6B | Q9NRW1; J3 | RAB6B_HUMAN | Unmodified | 0 | 0 | 0 | 0 | 0 | 0 | 0 | 0 | 0 | 0 | 572950 | 0 | 0 | 0 | 0 | 0 | 0 | 0 |
| QITIEEGEQ | RAB6B; RAB6B | Q9NRW1; J3 | RAB6B_HUMAN | Unmodified | 0 | 0 | 0 | 0 | 358740 | 0 | 0 | 0 | 446320 | 65629 | 432680 | 409230 | 0 | 0 | 0 | 0 | 0 | 0 |
| QITLNDLP\ | PRDX4; PRDX4 | Q13162; H7( | PRDX4_HUMAN | Unmodified | 0 | 0 | 0 | 261910 | 0 | 0 | 0 | 0 | 1.31E+08 | 5273600 | 12581000 | 9585900 | 23013 | 0 | 0 | 0 | 0 | 0 |
| QITQNTDYI | GLG1 | Q92896 | GSLG1_HUMAI | Unmodified | 0 | 0 | 0 | 0 | 0 | 0 | 0 | 0 | 309280 | 122060 | 0 | 231410 | 0 | 0 | 0 | 0 | 53013 | 118400 |
| QITSYGET( | CCT8 | P50990 | TCPQ_HUMAN | Unmodified | 0 | 0 | 0 | 0 | 0 | 0 | 0 | 347250 | 0 | 0 | 0 | 5067100 | 3755800 | 0 | 0 | 0 | 0 | 0 |
| QITSYGET( | CCT8 | P50990 | TCPQ_HUMAN | Unmodified | 0 | 0 | 0 | 0 | 4004700 | 634460 | 998220 | 583550 | 567660 | 1653600 | 1137600 | 10545000 | 4658800 | 3276300 | 0 | 0 | 2880600 | 2879500 |
| QIVGYAIGT | CEACAM1; CEA | P13688; M0F | CEAM1_HUMAI | Unmodified | 0 | 0 | 0 | 0 | 0 | 0 | 0 | 0 | 0 | 31267 | 0 | 3124000 | 0 | 0 | 0 | 0 | 0 | 0 |
| QIVHFQQE | KIF20B | Q96Q89 | KI20B_HUMAN | Unmodified | 0 | 0 | 0 | 0 | 979850 | 0 | 0 | 0 | 0 | 0 | 0 | 0 | 755620 | 0 | 0 | 0 | 761630 | 0 |

| | | | | | | | | | | | | | | | | | | | | | | |
|---|---|---|---|---|---|---|---|---|---|---|---|---|---|---|---|---|---|---|---|---|---|---|
| QIVLTGILE( | VPS35 | Q96QK1 | VPS35_HUMAN | Unmodified | 0 | 0 | 0 | 0 | 0 | 0 | 0 | 0 | 10494 | 0 | 0 | 171520 | 0 | 0 | 0 | 0 | 0 | 0 |
| QIVQLIEK | KIT | P10721 | KIT_HUMAN Ma | Unmodified | 0 | 156830 | 0 | 70571 | 185690 | 0 | 24885 | 23252 | 0 | 457560 | 0 | 3085700 | 155820 | 44891 | 0 | 0 | 124430 | 539410 |
| QIVSAVQY( | MARK3; MARK2 | P27448; Q7K | MARK3_HUMAI | Unmodified | 80913 | 0 | 0 | 0 | 0 | 0 | 0 | 0 | 330350 | 0 | 0 | 934730 | 0 | 0 | 1141800 | 656520 | 0 | 0 |
| QIVSEMLR | AP2A1 | O95782 | AP2A1_HUMAN | Unmodified | 479760 | 255110 | 1019100 | 2701900 | 0 | 0 | 0 | 133930 | 0 | 0 | 780500 | 1288400 | 0 | 0 | 2145000 | 2522600 | 48271 | 0 |
| QIVTTDLRC | FURIN; FURIN; | P09958; A0A | FURIN_HUMAN | Unmodified | 0 | 0 | 0 | 0 | 0 | 0 | 0 | 28376 | 0 | 166040 | 0 | 119210 | 0 | 0 | 0 | 0 | 184780 | 122410 |
| QIVWNGP\ | PGK1 | P00558 | PGK1_HUMAN | Unmodified | 0 | 89468 | 73500 | 163990 | 311020 | 50777 | 88362 | 0 | 383250 | 566170 | 171970 | 307680 | 687650 | 157270 | 423400 | 44161 | 642350 | 1860100 |
| QIVWNGP\ | PGK1 | P00558 | PGK1_HUMAN | Unmodified | 0 | 0 | 0 | 186260 | 63187000 | 6750600 | 39081000 | 73443 | 0 | 17212000 | 279550 | 0 | 64432000 | 54385000 | 0 | 0 | 3.06E+08 | 2.29E+08 |
| QIWQNLGI | AARS1; AARS1; | P49588; A0A | SYAC_HUMAN | Unmodified | 0 | 0 | 0 | 0 | 0 | 0 | 0 | 116250 | 1568400 | 588690 | 1038100 | 5750200 | 2897300 | 0 | 0 | 0 | 0 | 0 |
| QIWTLEQP | FSCN1 | Q16658 | FSCN1_HUMAI | Unmodified | 0 | 0 | 0 | 1174000 | 841640 | 0 | 0 | 0 | 790650 | 0 | 1454200 | 3287400 | 0 | 2291500 | 0 | 0 | 0 | 0 |
| QIWTLEQP | FSCN1 | Q16658 | FSCN1_HUMAI | Unmodified | 0 | 0 | 0 | 0 | 2740500 | 0 | 0 | 0 | 0 | 0 | 0 | 0 | 0 | 0 | 0 | 0 | 1610300 | 0 |
| QIYIYNEK | NUP93; NUP93 | Q8N1F7; H3 | NUP93_HUMAI | Unmodified | 0 | 0 | 104630 | 0 | 184280 | 450960 | 51681 | 0 | 0 | 0 | 0 | 1036200 | 90599 | 0 | 0 | 0 | 0 | 208500 |
| QIYPPINVL | ATP6V1B2; ATP | P21281; P15 | VATB2_HUMAN | Unmodified | 0 | 0 | 131710 | 0 | 0 | 0 | 0 | 0 | 617600 | 0 | 268980 | 1873900 | 0 | 0 | 0 | 0 | 0 | 0 |
| QIYQTYGF\ | HSPA13 | P48723 | HSP13_HUMAI | Unmodified | 0 | 0 | 0 | 0 | 0 | 0 | 0 | 0 | 4153600 | 375660 | 0 | 0 | 0 | 0 | 0 | 0 | 0 | 0 |
| QIYSVDQTI | PLCG2; PLCG2 | P16885; A0A | PLCG2_HUMAI | Unmodified | 0 | 0 | 0 | 0 | 0 | 297460 | 0 | 0 | 0 | 0 | 155450 | 2468800 | 0 | 0 | 0 | 0 | 0 | 198750 |
| QIYVGRAQ | PABPC1; PABP( | P11940; Q9I | PABP1_HUMAN | Unmodified | 55664 | 0 | 0 | 2024600 | 0 | 0 | 181030 | 0 | 0 | 0 | 0 | 0 | 1121000 | 561580 | 0 | 0 | 240240 | 118010 |
| QIYYSDK | CKS1B; CKS2; ( | P61024; P33 | CKS1_HUMAN | Unmodified | 484620 | 819750 | 970710 | 0 | 313400 | 0 | 0 | 19095 | 182780 | 44822 | 59733 | 810190 | 626320 | 0 | 1441300 | 1181800 | 30513 | 136940 |
| QIYYSDKYI | CKS1B; CKS1B | P61024; Q5T | CKS1_HUMAN | Unmodified | 1672700 | 1094700 | 331590 | 312750 | 0 | 0 | 0 | 0 | 1004700 | 0 | 0 | 474720 | 0 | 0 | 360460 | 269250 | 0 | 0 |
| QKEEESQG | MYO6; MYO6; M | Q9UM54; A0 | MYO6_HUMAN | Unmodified | 0 | 561260 | 2392500 | 3166900 | 0 | 0 | 0 | 0 | 0 | 0 | 167230 | 874280 | 0 | 0 | 3207400 | 3311800 | 0 | 0 |
| QKEKEDAC | AKAP12 | Q02952 | AKA12_HUMAN | Unmodified | 0 | 0 | 0 | 0 | 143990 | 69639 | 0 | 0 | 0 | 0 | 0 | 0 | 0 | 0 | 0 | 0 | 143400 | 246000 |
| QKEVFLPS1 | TMED4; TMED4 | Q7Z7H5; F8\ | TMED4_HUMAI | Unmodified | 666510 | 0 | 0 | 0 | 0 | 0 | 0 | 0 | 16604000 | 1564800 | 0 | 0 | 0 | 0 | 2008800 | 0 | 0 | 0 |
| QKFDNLYC | AHCYL1; AHCY | O43865; Q9( | SAHH2_HUMAI | Unmodified | 0 | 0 | 0 | 0 | 745390 | 362070 | 132210 | 0 | 0 | 0 | 0 | 0 | 267220 | 301080 | 0 | 0 | 1909600 | 399270 |
| QKGADFLV | PKM; PKM; PKM | P14618; B4D | KPYM_HUMAN | Unmodified | 0 | 0 | 0 | 0 | 12160000 | 219960 | 0 | 48773 | 0 | 0 | 0 | 527850 | 2923300 | 0 | 0 | 0 | 5774800 | 5563200 |
| QKGADFLV | PKM; PKM; PKM | P14618; B4D | KPYM_HUMAN | Unmodified | 0 | 0 | 0 | 0 | 16144000 | 3518200 | 2064100 | 640610 | 780910 | 89171 | 247270 | 923340 | 4326500 | 2854700 | 0 | 0 | 9371100 | 7801600 |
| QKGDVVLC | MYO1C; MYO1( | O00159; F5H | MYO1C_HUMA | Unmodified | 5159700 | 4261300 | 7515200 | 13768000 | 34137000 | 8566200 | 1069100 | 179490 | 578180 | 464620 | 0 | 1484900 | 10855000 | 11114000 | 17923000 | 10495000 | 8890900 | 7572800 |
| QKGHFFVE | PDLIM1 | O00151 | PDLI1_HUMAN | Unmodified | 40144 | 0 | 0 | 0 | 1671600 | 0 | 802090 | 205300 | 0 | 0 | 0 | 0 | 257990 | 192740 | 0 | 0 | 4067600 | 1855500 |
| QKIEDHLLS | TNFRSF10B | O14763 | TR10B_HUMAN | Unmodified | 0 | 0 | 0 | 0 | 76194 | 0 | 0 | 0 | 195550 | 0 | 0 | 165410 | 0 | 0 | 1310600 | 1217900 | 215380 | 842960 |
| QKIEVGLV\ | NSF; NSF | P46459; I3L( | NSF_HUMAN V | Unmodified | 0 | 0 | 0 | 0 | 68131 | 0 | 0 | 0 | 0 | 0 | 82761 | 0 | 530020 | 0 | 0 | 0 | 41953 | 0 |
| QKIHPTSVI | TCP1; TCP1; TC | P17987; F5H | TCPA_HUMAN | Unmodified | 0 | 0 | 0 | 0 | 630620 | 0 | 703100 | 0 | 0 | 0 | 0 | 0 | 1260300 | 961220 | 0 | 0 | 0 | 746170 |
| QKISFLENI | KIF5B; KIF5C; K | P33176; O6( | KINH_HUMAN | Unmodified | 0 | 0 | 0 | 140810 | 4336800 | 417200 | 1186700 | 0 | 354700 | 899620 | 0 | 0 | 1755000 | 1301300 | 0 | 0 | 1487000 | 1355300 |
| QKKESEAV | MSN | P26038 | MOES_HUMAN | Unmodified | 0 | 0 | 0 | 0 | 3377400 | 354870 | 0 | 0 | 0 | 0 | 0 | 0 | 0 | 0 | 0 | 0 | 495670 | 1329500 |
| QKKFDQLL | MYH9; MYH10 | P35579; P35 | MYH9_HUMAN | Unmodified | 0 | 0 | 0 | 0 | 2351800 | 0 | 864980 | 0 | 0 | 223570 | 0 | 0 | 901560 | 1307700 | 0 | 0 | 1523900 | 997500 |
| QKLDILDQI | SPTAN1; SPTAN | Q13813; A0/ | SPTN1_HUMAN | Unmodified | 0 | 0 | 192530 | 0 | 246880 | 0 | 0 | 0 | 0 | 0 | 378630 | 2138100 | 0 | 0 | 386600 | 399950 | 0 | 0 |
| QKLDPKIA\ | TPI1 | P60174 | TPIS_HUMAN T | Unmodified | 0 | 0 | 0 | 0 | 160200 | 0 | 0 | 0 | 0 | 0 | 0 | 0 | 0 | 0 | 0 | 0 | 1513100 | 0 |
| QKLEEDAE | SSB | P05455 | LA_HUMAN Lu | Unmodified | 0 | 0 | 0 | 0 | 89561 | 69633 | 0 | 0 | 0 | 0 | 485710 | 2712200 | 0 | 0 | 0 | 0 | 0 | 0 |
| QKLEELK | NGRN; WWC2 | Q9NPE2; Q6 | NGRN_HUMAN | Unmodified | 221070 | 0 | 0 | 0 | 0 | 0 | 0 | 0 | 0 | 0 | 0 | 4525200 | 0 | 0 | 219100 | 0 | 0 | 0 |
| QKLEFLKK | TM9SF2 | Q99805 | TM9S2_HUMAN | Unmodified | 0 | 0 | 0 | 0 | 0 | 0 | 0 | 0 | 5948600 | 365040 | 150900 | 600080 | 0 | 0 | 0 | 0 | 0 | 92289 |
| QKLPDGSE | CNDP2; CNDP2 | Q96KP4; J3C | CNDP2_HUMA | Unmodified | 0 | 0 | 0 | 522350 | 0 | 0 | 21544 | 319670 | 1028300 | 505150 | 2121400 | 9532100 | 3916900 | 3053700 | 1069000 | 1236100 | 0 | 0 |
| QKPATETSI | DSG2 | Q14126 | DSG2_HUMAN | Unmodified | 0 | 0 | 0 | 0 | 0 | 0 | 0 | 0 | 657000 | 0 | 0 | 0 | 0 | 0 | 0 | 0 | 0 | 0 |
| QKPITPETA | CDC42; CDC42 | P60953; A0A | CDC42_HUMA | Unmodified | 17581000 | 16637000 | 17127000 | 12983000 | 64545000 | 13271000 | 6094500 | 4424800 | 9234000 | 4929600 | 7199800 | 52735000 | 29073000 | 23345000 | 23972000 | 16461000 | 11051000 | 52035000 |
| QKPITPETA | CDC42; CDC42 | P60953; A0A | CDC42_HUMA | Unmodified | 0 | 0 | 0 | 0 | 0 | 0 | 393590 | 0 | 0 | 0 | 0 | 0 | 740450 | 0 | 0 | 0 | 1117600 | 771410 |
| QKPSNTED | RECQL | P46063 | RECQ1_HUMAI | Unmodified | 65366 | 0 | 0 | 0 | 0 | 0 | 0 | 0 | 2425600 | 0 | 0 | 810070 | 0 | 0 | 108960 | 0 | 0 | 0 |
| QKPVQTFTI | TMOD3 | Q9NYL9 | TMOD3_HUMA | Unmodified | 262080 | 0 | 300280 | 0 | 3390000 | 631430 | 189140 | 141200 | 347700 | 160700 | 150560 | 1246400 | 1762900 | 958900 | 455340 | 480160 | 585820 | 0 |
| QKTACGAF | ITGA6 | P23229 | ITA6_HUMAN I | Unmodified | 0 | 0 | 0 | 0 | 0 | 0 | 0 | 0 | 0 | 0 | 0 | 0 | 0 | 0 | 0 | 0 | 3107800 | 1236100 |
| QKTEFVVD | PLCG1; PLCG1 | P19174; V9G | PLCG1_HUMAI | Unmodified | 0 | 0 | 0 | 0 | 0 | 0 | 0 | 0 | 0 | 0 | 0 | 1110800 | 0 | 0 | 0 | 0 | 0 | 0 |
| QKTESTLN | NPTX2 | P47972 | NPTX2_HUMAN | Unmodified | 0 | 0 | 0 | 0 | 0 | 0 | 0 | 0 | 959110 | 47893 | 0 | 0 | 0 | 0 | 0 | 0 | 0 | 0 |
| QKVCMEET | ITGA6 | P23229 | ITA6_HUMAN I | Oxidation (M) | 0 | 0 | 0 | 0 | 757350 | 0 | 0 | 0 | 0 | 249100 | 0 | 0 | 0 | 324890 | 67161 | 0 | 3613600 | 5406600 |
| QKVCMEET | ITGA6 | P23229 | ITA6_HUMAN I | Unmodified | 0 | 0 | 0 | 0 | 2622500 | 0 | 0 | 0 | 0 | 0 | 0 | 0 | 821710 | 421390 | 0 | 0 | 5381100 | 12982000 |
| QKVDLQSL | DPY30 | Q9C005 | DPY30_HUMAN | Unmodified | 0 | 0 | 0 | 40278 | 0 | 0 | 0 | 0 | 433290 | 112090 | 209000 | 1168700 | 188510 | 0 | 0 | 0 | 0 | 0 |
| QKVDSLLE | HNRNPCL3; HN | B7ZW38; P0 | HNRC3_HUMA | Unmodified | 1266200 | 426530 | 317980 | 351200 | 4310600 | 513370 | 0 | 854980 | 3671600 | 10270000 | 3461000 | 7174800 | 2471700 | 583190 | 163270 | 174740 | 192890 | 1270100 |
| QKVDSLLE | HNRNPCL3; HN | B7ZW38; P0 | HNRC3_HUMA | Unmodified | 0 | 0 | 0 | 0 | 2533100 | 0 | 0 | 0 | 149220 | 1016700 | 0 | 0 | 1492100 | 669250 | 0 | 0 | 0 | 0 |
| QKVEGTEP | NCL; NCL; NCL | P19338; C9J | NUCL_HUMAN | Unmodified | 68115 | 0 | 39339 | 0 | 0 | 2658400 | 0 | 0 | 2076000 | 2425900 | 2066600 | 6711300 | 0 | 0 | 0 | 0 | 0 | 0 |
| QKVIGAGE | EPHA2 | P29317 | EPHA2_HUMAN | Unmodified | 0 | 0 | 0 | 0 | 0 | 0 | 0 | 0 | 0 | 0 | 0 | 0 | 0 | 0 | 0 | 0 | 1391300 | 1362800 |
| QKVIGAGE | EPHA2 | P29317 | EPHA2_HUMAN | Unmodified | 0 | 0 | 0 | 0 | 0 | 0 | 0 | 0 | 0 | 0 | 0 | 0 | 0 | 0 | 0 | 0 | 2319500 | 4987600 |
| QKVPDVLF | C2CD5 | Q86YS7 | C2CD5_HUMA | Unmodified | 160730 | 0 | 0 | 0 | 443600 | 0 | 0 | 0 | 2056000 | 184570 | 0 | 445730 | 0 | 0 | 0 | 0 | 0 | 0 |
| QKWEAEP\ | AZGP1; AZGP1 | P25311; C9J | ZA2G_HUMAN | Unmodified | 0 | 0 | 0 | 0 | 0 | 0 | 0 | 0 | 0 | 0 | 573800 | 0 | 0 | 0 | 0 | 0 | 232140 | 166600 |
| QKWEQIEG | GALNT2 | Q10471 | GALT2_HUMAN | Unmodified | 0 | 0 | 0 | 0 | 197400 | 49609 | 446780 | 21408 | 27729000 | 1958700 | 822510 | 1513200 | 0 | 465550 | 0 | 0 | 430210 | 29773 |
| QKYTDHLC | EXOC1 | Q9NV70 | EXOC1_HUMAI | Unmodified | 0 | 0 | 0 | 0 | 1802100 | 0 | 0 | 0 | 0 | 322460 | 0 | 380970 | 0 | 0 | 0 | 0 | 0 | 0 |
| QLAAENRL | MYH9 | P35579 | MYH9_HUMAN | Oxidation (M) | 0 | 0 | 0 | 0 | 5829200 | 0 | 0 | 0 | 0 | 0 | 0 | 0 | 0 | 0 | 0 | 0 | 2196800 | 1795700 |
| QLAAGWGI | MAN2B1 | O00754 | MA2B1_HUMAI | Unmodified | 0 | 0 | 0 | 0 | 0 | 0 | 0 | 0 | 445190 | 393530 | 0 | 37899 | 0 | 0 | 0 | 0 | 0 | 894410 |
| QLAALGQT | ARHGEF2; ARH | Q92974; V9( | ARHG2_HUMA | Unmodified | 0 | 0 | 0 | 0 | 0 | 0 | 0 | 0 | 0 | 0 | 0 | 672680 | 0 | 0 | 0 | 0 | 0 | 0 |
| QLAALKQQ | DCTN2; DCTN2 | Q13561; A8I | DCTN2_HUMAI | Unmodified | 0 | 0 | 0 | 0 | 0 | 0 | 746580 | 60470 | 0 | 0 | 0 | 0 | 70502 | 144810 | 0 | 0 | 560070 | 0 |
| QLAASLPPI | VWDE; VWDE; \ | Q8N2E2; E5I | VWDE_HUMAN | Unmodified | 0 | 0 | 0 | 0 | 0 | 0 | 0 | 0 | 477960 | 0 | 0 | 0 | 0 | 0 | 0 | 0 | 0 | 0 |
| QLADAVAE | TAX1BP1; TAX1 | Q86VP1; B8Z | TAXB1_HUMAN | Unmodified | 943230 | 0 | 198170 | 262870 | 726440 | 0 | 3265100 | 331770 | 1157500 | 989430 | 0 | 1945000 | 896260 | 301930 | 44583 | 91616 | 868320 | 2778100 |
| QLADETLLI | LMNB1; LMNB1 | P20700; A0A | LMNB1_HUMAI | Unmodified | 0 | 0 | 233980 | 0 | 0 | 0 | 0 | 0 | 0 | 0 | 1244200 | 2380100 | 0 | 0 | 0 | 0 | 0 | 0 |
| QLADHLGF | RAB3A; RAB3A | P20336; S4F | RAB3A_HUMAN | Unmodified | 307010 | 0 | 487900 | 1459200 | 0 | 0 | 0 | 0 | 0 | 0 | 0 | 3121300 | 0 | 0 | 613520 | 526730 | 0 | 0 |
| QLAEDFLR | DDX17; DDX17 | Q92841; A0/ | DDX17_HUMAI | Unmodified | 0 | 0 | 108270 | 0 | 0 | 0 | 0 | 0 | 0 | 0 | 46627 | 4704900 | 0 | 0 | 107600 | 110060 | 0 | 0 |
| QLAEEDLA | PLEC | Q15149 | PLEC_HUMAN | Unmodified | 0 | 0 | 0 | 0 | 0 | 0 | 0 | 0 | 0 | 0 | 0 | 0 | 205620 | 0 | 0 | 0 | 0 | 513770 |
| QLAEFVPLI | MRPS30 | Q9NP92 | RT30_HUMAN : | Unmodified | 0 | 0 | 0 | 0 | 0 | 0 | 0 | 0 | 137450 | 107980 | 0 | 0 | 0 | 0 | 0 | 0 | 0 | 0 |
| QLAGILQVI | VPS36; VPS36 | Q86VN1; A0 | VPS36_HUMAN | Unmodified | 5040000 | 5400000 | 2613900 | 518460 | 0 | 0 | 0 | 0 | 195170 | 0 | 0 | 809400 | 0 | 0 | 9440000 | 5768800 | 0 | 0 |
| QLALWNPI | CORO1C; COR( | Q9ULV4; B4I | COR1C_HUMA | Unmodified | 0 | 0 | 651160 | 458270 | 12081000 | 2748700 | 0 | 0 | 0 | 0 | 3580900 | 16616000 | 666570 | 0 | 300180 | 0 | 1085000 | 1521100 |
| QLANSLSG | SPTBN2; SPTBN | O15020; A4( | SPTN2_HUMAN | Unmodified | 0 | 0 | 92893 | 0 | 0 | 0 | 0 | 0 | 0 | 0 | 0 | 652880 | 0 | 721290 | 267590 | 143580 | 0 | 0 |
| QLAPIWDK | P4HB; P4HB; P4 | P07237; A0A | PDIA1_HUMAN | Unmodified | 257590 | 545570 | 790190 | 0 | 2720600 | 2773600 | 0 | 0 | 1.34E+08 | 24774000 | 10457000 | 3490300 | 2200100 | 1968700 | 903270 | 1388300 | 528590 | 2359400 |
| QLAPIWDK | P4HB; P4HB; P4 | P07237; A0A | PDIA1_HUMAN | Unmodified | 0 | 0 | 0 | 0 | 0 | 0 | 179410 | 0 | 436670 | 1816400 | 0 | 0 | 0 | 14465 | 0 | 0 | 0 | 0 |
| QLAPIWDK | P4HB; P4HB; P4 | P07237; A0A | PDIA1_HUMAN | Unmodified | 0 | 0 | 0 | 0 | 5748100 | 0 | 8184100 | 0 | 4050100 | 34163000 | 0 | 0 | 1554600 | 0 | 0 | 0 | 0 | 2414100 |
| QLAQIDGTI | CHMP4B | Q9H444 | CHM4B_HUMA | Unmodified | 3373000 | 8972700 | 1104200 | 2120900 | 0 | 0 | 428290 | 41266 | 3472400 | 0 | 0 | 1944700 | 277010 | 0 | 2491300 | 1967700 | 0 | 586620 |
| QLAQIDGTI | CHMP4B | Q9H444 | CHM4B_HUMA | Unmodified | 0 | 5759400 | 0 | 9762500 | 68140000 | 7643000 | 6594000 | 0 | 7093100 | 12671000 | 647970 | 0 | 38660000 | 21043000 | 10230000 | 0 | 0 | 24813000 |
| QLASALSYI | JAK1; JAK1; JAK | P23458; A0A | JAK1_HUMAN 1 | Unmodified | 0 | 0 | 0 | 0 | 2514700 | 552240 | 1707600 | 0 | 214410 | 28170 | 0 | 0 | 140340 | 0 | 0 | 0 | 1822600 | 2423400 |
| QLASEDISI | ARL3 | P36405 | ARL3_HUMAN / | Unmodified | 1031900 | 861190 | 1870000 | 2389400 | 6515700 | 562220 | 11472000 | 8102100 | 3677900 | 2544000 | 2237800 | 18777000 | 18298000 | 12396000 | 3592200 | 3131300 | 6987800 | 9793500 |
| QLASGLLL\ | RPL6 | Q02878 | RL6_HUMAN 6( | Unmodified | 0 | 0 | 58145 | 0 | 0 | 0 | 0 | 0 | 0 | 0 | 0 | 3510600 | 0 | 0 | 0 | 214120 | 0 | 0 |
| QLCDNAGI | CCT7 | Q99832 | TCPH_HUMAN | Unmodified | 0 | 0 | 0 | 0 | 0 | 0 | 0 | 0 | 0 | 0 | 235810 | 1346200 | 0 | 0 | 0 | 0 | 0 | 0 |
| QLCPYAAV | MKRN1; MKRN1 | Q9UHC7; C9 | MKRN1_HUMA | Unmodified | 2924300 | 1000800 | 388770 | 750980 | 0 | 0 | 0 | 0 | 0 | 0 | 585900 | 0 | 0 | 0 | 485470 | 430620 | 0 | 0 |
| QLDAGEEA | P4HA2 | O15460 | P4HA2_HUMAN | Unmodified | 0 | 0 | 0 | 0 | 390970 | 0 | 0 | 0 | 0 | 449690 | 0 | 0 | 0 | 0 | 0 | 0 | 0 | 0 |
| QLDALLEAI | LAMB2 | P55268 | LAMB2_HUMAN | Unmodified | 97397 | 0 | 0 | 0 | 0 | 90930 | 0 | 0 | 2566300 | 209540 | 0 | 69076 | 271710 | 671450 | 0 | 0 | 0 | 115580 |
| QLDDKDEE | KIF5B | P33176 | KINH_HUMAN | Unmodified | 0 | 227070 | 540760 | 616800 | 7740400 | 1165400 | 1140300 | 580170 | 2303700 | 1380200 | 1200000 | 7979600 | 2653500 | 1543800 | 627000 | 437780 | 1571100 | 2894400 |
| QLDDLKVE | RPL35; RPL35 | P42766; F2Z | RL35_HUMAN ( | Unmodified | 3279300 | 3534900 | 430200 | 1944700 | 0 | 539580 | 0 | 74788 | 1002300 | 535620 | 3239900 | 19004000 | 1086300 | 0 | 1375800 | 1370100 | 0 | 0 |
| QLDEGEIS1 | P4HA1 | P13674 | P4HA1_HUMAN | Unmodified | 0 | 0 | 0 | 0 | 0 | 0 | 0 | 0 | 1459100 | 0 | 0 | 0 | 0 | 0 | 0 | 0 | 0 | 0 |
| QLDELEK | SMAD3; UTRN | P84022; P46 | SMAD3_HUMA | Unmodified | 0 | 0 | 219860 | 347890 | 0 | 72853 | 0 | 0 | 343690 | 220670 | 0 | 141260 | 293160 | 168060 | 402160 | 489080 | 0 | 870520 |
| QLDFNSSK | COPA; COPA; C | P53621; A0A | COPA_HUMAN | Unmodified | 0 | 0 | 288020 | 0 | 775540 | 259100 | 568640 | 1442200 | 183120 | 0 | 0 | 3759600 | 209510 | 492050 | 48638 | 0 | 1080100 | 772340 |
| QLDLELK | ANKFY1 | Q9P2R3 | ANFY1_HUMAN | Unmodified | 0 | 0 | 0 | 0 | 2260400 | 0 | 0 | 208300 | 0 | 0 | 0 | 124530 | 454510 | 0 | 0 | 0 | 0 | 0 |
| QLDLTKNE | TNFRSF10A; TN | O00220; F8U | TR10A_HUMAN | Unmodified | 148960 | 164910 | 0 | 205680 | 0 | 0 | 0 | 0 | 1095400 | 0 | 0 | 1014100 | 0 | 0 | 423120 | 421360 | 0 | 0 |
| QLDMILDE( | LMAN1 | P49257 | LMAN1_HUMAI | Oxidation (M) | 0 | 0 | 0 | 0 | 0 | 0 | 0 | 84078 | 2355400 | 0 | 0 | 0 | 202720 | 0 | 0 | 0 | 0 | 0 |
| QLDMILDE( | LMAN1 | P49257 | LMAN1_HUMAI | Unmodified | 0 | 0 | 0 | 0 | 0 | 0 | 0 | 0 | 4797300 | 177590 | 0 | 0 | 0 | 0 | 0 | 0 | 0 | 0 |
| QLDMILDE( | LMAN1 | P49257 | LMAN1_HUMAI | Unmodified | 0 | 0 | 0 | 0 | 0 | 0 | 0 | 0 | 60275000 | 2849300 | 2624400 | 810950 | 0 | 0 | 0 | 0 | 0 | 0 |
| QLDPGGG( | ITGA3 | P26006 | ITA3_HUMAN I | Unmodified | 0 | 0 | 0 | 0 | 4795200 | 0 | 0 | 0 | 0 | 434760 | 0 | 0 | 619790 | 826730 | 0 | 0 | 1924600 | 6652000 |
| QLDPGGG( | ITGA3 | P26006 | ITA3_HUMAN I | Unmodified | 0 | 0 | 0 | 0 | 5361700 | 1343200 | 0 | 0 | 156010 | 0 | 0 | 0 | 0 | 0 | 0 | 0 | 4848100 | 21013000 |
| QLDQCSAI | UTRN | P46939 | UTRN_HUMAN | Unmodified | 0 | 0 | 0 | 465060 | 0 | 0 | 0 | 0 | 0 | 0 | 0 | 0 | 0 | 0 | 470510 | 0 | 0 | 0 |
| QLDQHLDI | LAMB2 | P55268 | LAMB2_HUMAN | Unmodified | 122220 | 0 | 243230 | 0 | 0 | 0 | 0 | 0 | 601360 | 0 | 0 | 0 | 0 | 0 | 0 | 95272 | 0 | 0 |
| QLEADILD\ | STX12 | Q86Y82 | STX12_HUMAN | Unmodified | 0 | 143490 | 457140 | 511220 | 27247 | 0 | 0 | 0 | 1130000 | 0 | 0 | 64654 | 0 | 0 | 1174000 | 615050 | 0 | 0 |
| QLEADIMD | STX7 | O15400 | STX7_HUMAN ! | Oxidation (M) | 28631 | 0 | 30957 | 62719 | 0 | 0 | 0 | 0 | 0 | 0 | 0 | 42783 | 0 | 0 | 0 | 57970 | 0 | 0 |
| QLEAIDQLI | ACTN4; ACTN4 | O43707; F5( | ACTN4_HUMAI | Unmodified | 496670 | 245990 | 776690 | 533770 | 4758000 | 5808700 | 948710 | 97069 | 2341300 | 2440300 | 861430 | 6205400 | 654160 | 0 | 1098600 | 825620 | 3406200 | 319780 |
| QLEAIDQLI | ACTN4; ACTN4 | O43707; F5( | ACTN4_HUMAI | Unmodified | 267460 | 0 | 0 | 0 | 0 | 0 | 0 | 0 | 438060 | 0 | 0 | 5279800 | 0 | 0 | 114700 | 675570 | 0 | 0 |
| QLEDELVSI | TPM1; TPM1; TF | P09493; A0A | TPM1_HUMAN | Unmodified | 0 | 0 | 0 | 0 | 2149000 | 0 | 0 | 141090 | 0 | 0 | 0 | 0 | 0 | 0 | 0 | 0 | 343760 | 0 |
| QLEDILVLA | MACF1; MACF1 | Q9UPN3; H3 | MACF1_HUMAI | Unmodified | 0 | 0 | 0 | 0 | 0 | 350870 | 0 | 0 | 415220 | 494140 | 122260 | 289110 | 88320 | 0 | 0 | 49690 | 555470 | 52395 |
| QLEEAEEE/ | MYH9 | P35579 | MYH9_HUMAN | Unmodified | 0 | 0 | 0 | 0 | 0 | 0 | 0 | 0 | 0 | 0 | 213460 | 748250 | 0 | 0 | 0 | 0 | 0 | 0 |
| QLEEAEEE/ | MYH9 | P35579 | MYH9_HUMAN | Unmodified | 0 | 0 | 0 | 0 | 0 | 0 | 95802 | 0 | 0 | 387650 | 0 | 683140 | 0 | 0 | 0 | 0 | 573500 | 0 |
| QLEEQLRII | TPM1; TPM1; TF | H0YL42; B7Z | H0YL42_HUMA | Unmodified | 0 | 0 | 0 | 44332 | 572010 | 0 | 0 | 0 | 0 | 0 | 0 | 0 | 241420 | 98538 | 0 | 0 | 0 | 0 |
| QLEEQNAF | DCTN1; DCTN1 | Q14203; Q6/ | DCTN1_HUMAI | Unmodified | 0 | 0 | 0 | 0 | 0 | 109570 | 303980 | 159480 | 0 | 338670 | 0 | 0 | 1828100 | 1037200 | 0 | 0 | 0 | 950640 |
| QLEESVSEI | KIF4A | O95239 | KIF4A_HUMAN | Unmodified | 0 | 0 | 205420 | 0 | 251730 | 0 | 0 | 78019 | 0 | 0 | 0 | 0 | 929050 | 0 | 0 | 0 | 1200700 | 0 |
| QLEIAHEK | LRPAP1 | P30533 | AMRP_HUMAN | Unmodified | 0 | 0 | 0 | 0 | 0 | 0 | 0 | 0 | 1503900 | 1037400 | 0 | 0 | 0 | 0 | 50506 | 0 | 0 | 0 |

| | | | | | | | | | | | | | | | | | | | | | | |
|---|---|---|---|---|---|---|---|---|---|---|---|---|---|---|---|---|---|---|---|---|---|---|
| QLEILQLAQ | CC2D1B; CC2D | Q5T0F9; H0Y | C2D1B_HUMAI | Unmodified | 331280 | 373540 | 0 | 0 | 0 | 0 | 0 | 0 | 1098800 | 478770 | 0 | 0 | 886560 | 0 | 455450 | 245060 | 0 | 0 |
| QLEITGR | STX1B; STX2; ST | P61266; P32 | STX1B_HUMAN | Unmodified | 4350300 | 2753800 | 2831300 | 1472500 | 0 | 0 | 0 | 1111900 | 1183300 | 250910 | 268820 | 2801400 | 1922700 | 7036800 | 110790 | 796860 | 0 | 0 |
| QLEIVGEK | POMT1; POMT1 | Q9Y6A1; A0A | POMT1_HUMAI | Unmodified | 0 | 0 | 0 | 0 | 0 | 0 | 0 | 0 | 1318000 | 0 | 0 | 0 | 0 | 0 | 0 | 0 | 0 | 0 |
| QLELEIK | CHMP2B; COR( | Q9UQN3; Q9 | CHM2B_HUMA | Unmodified | 1320400 | 1144600 | 399960 | 706350 | 3495200 | 275220 | 294230 | 398250 | 2074800 | 63686 | 172260 | 2204500 | 2999000 | 1489600 | 0 | 864170 | 294710 | 1381500 |
| QLELEIKK | CHMP2B; CHM | Q9UQN3; C9 | CHM2B_HUMA | Unmodified | 0 | 0 | 0 | 0 | 1265400 | 0 | 0 | 103270 | 316630 | 309030 | 104560 | 0 | 409540 | 273400 | 0 | 0 | 0 | 376190 |
| QLELENLTI | PTPN1; PTPN1 | P18031; B4D | PTN1_HUMAN | Unmodified | 0 | 378900 | 0 | 0 | 0 | 0 | 0 | 0 | 0 | 0 | 0 | 937820 | 0 | 0 | 0 | 0 | 0 | 0 |
| QLELRQEEI | NONO; NONO; | Q15233; A0A | NONO_HUMAN | Unmodified | 0 | 0 | 0 | 0 | 0 | 0 | 0 | 0 | 625760 | 0 | 0 | 892980 | 0 | 0 | 0 | 0 | 0 | 0 |
| QLEMILNKI | AKR1C4; AKR1( | P17516; P42 | AK1C4_HUMAN | Unmodified | 0 | 0 | 0 | 0 | 4235600 | 0 | 0 | 0 | 0 | 0 | 0 | 0 | 0 | 0 | 0 | 0 | 214230 | 0 |
| QLEMILNKI | AKR1C4; AKR1( | P17516; P42 | AK1C4_HUMAN | Oxidation (M) | 305920 | 93769 | 0 | 0 | 3491200 | 299370 | 0 | 0 | 0 | 0 | 0 | 0 | 25265 | 0 | 0 | 0 | 131400 | 177000 |
| QLEMQNK | KIF23; KIF23; KI | Q02241; H7I | KIF23_HUMAN | Unmodified | 50444 | 0 | 0 | 0 | 6964400 | 987940 | 0 | 194040 | 796890 | 333550 | 23835 | 0 | 0 | 230760 | 0 | 0 | 3351300 | 0 |
| QLENELEG | CDC42BPA; CD | Q5VT25; A0A | MRCKA_HUMA | Unmodified | 0 | 0 | 0 | 0 | 576800 | 0 | 97185 | 90577 | 540440 | 0 | 99795 | 511140 | 893910 | 653660 | 0 | 183080 | 648200 | 1712100 |
| QLENLEQC | ATP6V1H; ATP6 | Q9UI12; G3V | VATH_HUMAN | Unmodified | 0 | 0 | 0 | 0 | 0 | 0 | 0 | 0 | 267430 | 0 | 0 | 6046300 | 0 | 0 | 88824 | 77554 | 0 | 0 |
| QLENSLNE | TARS1 | P26639 | SYTC_HUMAN | Unmodified | 1472000 | 0 | 0 | 0 | 0 | 0 | 0 | 0 | 0 | 0 | 0 | 1852700 | 0 | 0 | 0 | 0 | 0 | 0 |
| QLENSLNE | TARS1 | P26639 | SYTC_HUMAN | Unmodified | 0 | 0 | 0 | 0 | 0 | 0 | 0 | 0 | 0 | 325770 | 0 | 3734900 | 0 | 0 | 0 | 0 | 0 | 0 |
| QLEPRTETI | RACGAP1; RAC | Q9H0H5; HC | RGAP1_HUMAI | Unmodified | 0 | 0 | 0 | 0 | 567670 | 0 | 0 | 0 | 0 | 0 | 0 | 0 | 0 | 0 | 0 | 0 | 0 | 0 |
| QLEPRTETI | RACGAP1; RAC | Q9H0H5; HC | RGAP1_HUMAI | Unmodified | 0 | 0 | 0 | 0 | 5654500 | 0 | 0 | 0 | 0 | 0 | 0 | 0 | 0 | 0 | 0 | 0 | 0 | 0 |
| QLEPVYNS | PDIA4; PDIA4 | P13667; A0A | PDIA4_HUMAN | Unmodified | 0 | 0 | 0 | 0 | 1317100 | 0 | 8897300 | 6675900 | 30142000 | 13788000 | 5330300 | 1627000 | 0 | 0 | 0 | 0 | 0 | 0 |
| QLEPVYNS | PDIA4; PDIA4 | P13667; A0A | PDIA4_HUMAN | Unmodified | 0 | 0 | 0 | 0 | 61289 | 0 | 1610400 | 1392500 | 5623100 | 2496400 | 813840 | 277860 | 0 | 0 | 0 | 0 | 230860 | 200360 |
| QLEQVIAK | FGA; FGA | P02671; A0A | FIBA_HUMAN F | Unmodified | 0 | 1924900 | 0 | 0 | 0 | 0 | 0 | 0 | 0 | 0 | 0 | 0 | 0 | 0 | 0 | 0 | 0 | 0 |
| QLERAQLE | RDX; RDX; RDX | P35241; A0A | RADI_HUMAN I | Unmodified | 8617700 | 0 | 0 | 0 | 23859000 | 7525200 | 2898000 | 934670 | 374700 | 2505700 | 692850 | 0 | 8791800 | 14616000 | 0 | 73136 | 5137700 | 0 |
| QLERAQLE | RDX; RDX; RDX | P35241; A0A | RADI_HUMAN I | Unmodified | 0 | 0 | 0 | 0 | 5231600 | 0 | 29809 | 0 | 0 | 0 | 0 | 315250 | 915950 | 0 | 0 | 0 | 0 | 0 |
| QLERQQLE | EZR; EZR | P15311; E7E | EZRI_HUMAN E | Unmodified | 0 | 0 | 0 | 0 | 8211300 | 1864300 | 435330 | 2674200 | 2759300 | 9958200 | 683030 | 0 | 0 | 3656900 | 0 | 0 | 849290 | 1429000 |
| QLERQQLE | EZR; EZR | P15311; E7E | EZRI_HUMAN E | Unmodified | 0 | 0 | 0 | 0 | 3484400 | 0 | 0 | 0 | 0 | 0 | 0 | 0 | 0 | 401170 | 0 | 0 | 0 | 0 |
| QLESHLSL( | PEX11B | O96011 | PX11B_HUMAN | Unmodified | 0 | 0 | 0 | 0 | 0 | 0 | 0 | 0 | 2562500 | 24056 | 0 | 0 | 0 | 0 | 0 | 0 | 0 | 0 |
| QLETKQETI | ANLN | Q9NQW6 | ANLN_HUMAN | Unmodified | 0 | 0 | 0 | 151190 | 1398100 | 0 | 0 | 0 | 0 | 0 | 0 | 0 | 0 | 0 | 0 | 0 | 1355200 | 644810 |
| QLETVLDD | TMOD3 | Q9NYL9 | TMOD3_HUMA | Unmodified | 0 | 0 | 0 | 0 | 4155000 | 562850 | 0 | 0 | 0 | 0 | 0 | 0 | 1229800 | 1402200 | 0 | 0 | 0 | 0 |
| QLEVEPEEI | LONP1; LONP1 | P36776; K7E | LONM_HUMAN | Unmodified | 0 | 0 | 0 | 0 | 0 | 0 | 0 | 0 | 2013200 | 825880 | 0 | 0 | 0 | 195530 | 0 | 0 | 0 | 0 |
| QLEVYTSG | EIF3L; EIF3L; EI | Q9Y262; B0( | EIF3L_HUMAN | Unmodified | 0 | 0 | 304850 | 0 | 0 | 370540 | 293840 | 0 | 0 | 0 | 0 | 2628000 | 0 | 714610 | 110940 | 0 | 0 | 0 |
| QLEYNARN | PKP1 | Q13835 | PKP1_HUMAN | Unmodified | 0 | 0 | 0 | 89599 | 0 | 0 | 0 | 0 | 0 | 0 | 0 | 0 | 0 | 0 | 0 | 0 | 552290 | 0 |
| QLEYQQLE | EPB41L3; EPB4 | Q9Y2J2; A0A | E41L3_HUMAN | Unmodified | 0 | 0 | 0 | 0 | 0 | 0 | 0 | 0 | 0 | 0 | 0 | 0 | 0 | 66378 | 0 | 0 | 0 | 517990 |
| QLFDQVVK | RDX; EZR; MSN | P35241; P15 | RADI_HUMAN I | Unmodified | 2340900 | 2172100 | 3021900 | 0 | 13981000 | 4190700 | 0 | 910340 | 16010000 | 10752000 | 3538100 | 30661000 | 2792900 | 4859700 | 5677700 | 5954000 | 4132400 | 2926400 |
| QLFDVVIV( | NT5DC2; NT5D | Q9H857; H7 | NT5D2_HUMAI | Unmodified | 0 | 0 | 0 | 0 | 0 | 0 | 25906 | 0 | 1812900 | 0 | 0 | 331710 | 0 | 0 | 0 | 0 | 0 | 0 |
| QLFEIDSVI | BAG5 | Q9UL15 | BAG5_HUMAN | Unmodified | 0 | 0 | 0 | 693360 | 1352000 | 0 | 0 | 0 | 2749700 | 1110800 | 0 | 0 | 1389800 | 0 | 393210 | 0 | 0 | 0 |
| QLFEIDSVI | BAG5 | Q9UL15 | BAG5_HUMAN | Unmodified | 0 | 506440 | 0 | 697280 | 2342600 | 275680 | 251320 | 0 | 3397100 | 987560 | 0 | 0 | 0 | 513820 | 0 | 0 | 0 | 0 |
| QLFFLNFA( | MME; MME; MM | P08473; A0A | NEP_HUMAN N | Unmodified | 0 | 0 | 0 | 125630 | 0 | 0 | 0 | 0 | 0 | 0 | 0 | 0 | 121130 | 45519 | 339240 | 0 | 0 | 864610 |
| QLFHPEQL | TUBA3C; TUBA | P0DPH7; Q7 | TBA3C_HUMAN | Unmodified | 0 | 0 | 6900600 | 1145400 | 0 | 303790 | 0 | 217560 | 4472500 | 888120 | 2249300 | 41738000 | 0 | 0 | 646900 | 716940 | 0 | 0 |
| QLFHPEQL | TUBA3C; TUBA | P0DPH7; Q7 | TBA3C_HUMAN | Unmodified | 18399000 | 15718000 | 21071000 | 20589000 | 1931800 | 462660 | 0 | 218800 | 44438000 | 2836900 | 14012000 | 4.82E+08 | 1149100 | 0 | 30584000 | 21502000 | 1123500 | 0 |
| QLFLQTYAI | COPA; COPA; C | P53621; A0A | COPA_HUMAN | Unmodified | 0 | 251410 | 0 | 0 | 1049400 | 0 | 719130 | 89868 | 387710 | 0 | 0 | 2979700 | 458440 | 368230 | 157450 | 107790 | 475510 | 0 |
| QLFNCQSL | ERBIN; ERBIN | Q96RT1; B4I | ERBIN_HUMAN | Unmodified | 0 | 0 | 251560 | 0 | 8615000 | 1462700 | 345100 | 284430 | 436200 | 0 | 307200 | 1169100 | 1246300 | 1236700 | 0 | 274830 | 854820 | 1759400 |
| QLGDNEAI | VTA1; VTA1; VTA | Q9NP79; Q5 | VTA1_HUMAN | Unmodified | 1963000 | 1225800 | 0 | 1644900 | 2710500 | 0 | 0 | 0 | 6157400 | 7585000 | 885110 | 2627500 | 0 | 0 | 872760 | 906020 | 0 | 2213200 |
| QLGGSVEL | CNDP2; CNDP2 | Q96KP4; J3C | CNDP2_HUMA | Unmodified | 0 | 0 | 703430 | 1801500 | 918850 | 0 | 0 | 346760 | 0 | 0 | 4243500 | 17465000 | 14899000 | 8819300 | 2737400 | 2275000 | 415780 | 609210 |
| QLGGSVEL | CNDP2; CNDP2 | Q96KP4; J3C | CNDP2_HUMA | Unmodified | 0 | 0 | 0 | 0 | 0 | 0 | 0 | 0 | 0 | 0 | 0 | 0 | 3397300 | 2870500 | 0 | 0 | 110770 | 0 |
| QLGLLLHD | UTRN; UTRN; U | P46939; H0Y | UTRN_HUMAN | Unmodified | 75478 | 74593 | 0 | 1576200 | 0 | 0 | 0 | 0 | 1271100 | 90187 | 0 | 187930 | 0 | 0 | 2313800 | 0 | 67629 | 0 |
| QLGLQAFL | SNX16; SNX16; | P57768; E5R | SNX16_HUMAN | Unmodified | 0 | 0 | 0 | 0 | 52607 | 0 | 0 | 0 | 150310 | 0 | 0 | 68561 | 0 | 0 | 0 | 0 | 0 | 0 |
| QLGPQVQ( | USO1 | O60763 | USO1_HUMAN | Oxidation (M) | 0 | 0 | 0 | 0 | 0 | 0 | 0 | 0 | 0 | 0 | 0 | 2756900 | 459390 | 0 | 0 | 0 | 0 | 0 |
| QLGPVLRS | MOV10; MOV1( | Q9HCE1; Q5 | MOV10_HUMA | Unmodified | 1533900 | 1480000 | 539890 | 2763700 | 0 | 0 | 0 | 0 | 0 | 0 | 0 | 0 | 206160 | 0 | 574580 | 401430 | 0 | 0 |
| QLGQDLLN | HSPA4L; HSPA4 | O95757; E9F | HS74L_HUMAN | Unmodified | 0 | 0 | 0 | 0 | 0 | 0 | 0 | 14877 | 0 | 47075 | 222380 | 683040 | 0 | 0 | 0 | 0 | 0 | 0 |
| QLGRIMLK( | SNRPE; SNRPE | P62304; A6N | RUXE_HUMAN | Oxidation (M) | 0 | 0 | 0 | 0 | 945070 | 0 | 0 | 0 | 0 | 289740 | 0 | 0 | 1076700 | 0 | 0 | 0 | 1627300 | 0 |
| QLHAQEGI | UBTD1 | Q9HAC8 | UBTD1_HUMAI | Unmodified | 404260 | 238810 | 333770 | 396060 | 1138000 | 0 | 0 | 0 | 796590 | 197580 | 0 | 2319700 | 257700 | 0 | 561340 | 497970 | 0 | 234420 |
| QLHDDYFY | RCN2; RCN2 | Q14257; A8N | RCN2_HUMAN | Unmodified | 24949 | 0 | 104130 | 0 | 0 | 0 | 0 | 0 | 8576600 | 1345700 | 1401100 | 923260 | 0 | 0 | 147020 | 102170 | 0 | 0 |
| QLHEFAITE | CEP55 | Q53EZ4 | CEP55_HUMAN | Unmodified | 0 | 0 | 0 | 7434000 | 52846000 | 0 | 1184400 | 0 | 0 | 10664000 | 0 | 2881700 | 7742500 | 0 | 0 | 0 | 2477800 | 458930 |
| QLHPAGAT | SEZ6; SEZ6; SE | Q53EL9; C9J | SEZ6_HUMAN | Unmodified | 0 | 0 | 0 | 0 | 0 | 0 | 0 | 0 | 2970600 | 1028500 | 0 | 0 | 0 | 0 | 0 | 0 | 0 | 0 |
| QLHQSCQ | COPS2; COPS2 | P61201; B4D | CSN2_HUMAN | Unmodified | 0 | 0 | 0 | 124900 | 0 | 0 | 0 | 0 | 0 | 0 | 0 | 754890 | 0 | 0 | 0 | 0 | 0 | 0 |
| QLIAELENK | DTNA; DTNB; D | Q9Y4J8; O6C | DTNA_HUMAN | Unmodified | 157520 | 0 | 242650 | 264690 | 640270 | 160790 | 0 | 27264 | 1364600 | 289750 | 0 | 399590 | 391840 | 0 | 572490 | 1020100 | 0 | 0 |
| QLIALENK | TAOK3; TAOK3; | Q9H2K8; G3 | TAOK3_HUMAN | Unmodified | 241410 | 132280 | 0 | 101400 | 1636100 | 0 | 72145 | 457170 | 635920 | 0 | 0 | 783300 | 1081400 | 518970 | 102910 | 0 | 965740 | 539190 |
| QLICDPSYI | GDI1 | P31150 | GDIA_HUMAN | Unmodified | 0 | 0 | 0 | 0 | 0 | 0 | 0 | 0 | 573810 | 433260 | 0 | 0 | 680820 | 1121400 | 0 | 0 | 828580 | 0 |
| QLICDPSY\ | GDI2; GDI2 | P50395; Q59 | GDIB_HUMAN | Unmodified | 0 | 0 | 0 | 0 | 4599700 | 1851500 | 0 | 1497700 | 0 | 0 | 0 | 7120300 | 3808400 | 3223000 | 0 | 1065600 | 3296300 | 4859600 |
| QLICDPSY\ | GDI2; GDI2 | P50395; Q59 | GDIB_HUMAN | Unmodified | 133020 | 0 | 163200 | 283470 | 394890 | 719930 | 253070 | 210360 | 285340 | 0 | 0 | 0 | 184290 | 0 | 0 | 204200 | 747600 | 205540 |
| QLIEFQHRI | HOOK1 | Q9UJC3 | HOOK1_HUMA | Unmodified | 0 | 0 | 0 | 217220 | 0 | 0 | 0 | 0 | 616720 | 0 | 0 | 0 | 309890 | 0 | 0 | 0 | 0 | 0 |
| QLIFTTEDN | TSNAX; TSNAX- | Q99598; C4I | TSNAX_HUMAN | Unmodified | 250060 | 0 | 507920 | 0 | 0 | 0 | 0 | 153510 | 5255400 | 467580 | 653560 | 3489300 | 0 | 0 | 136860 | 0 | 0 | 0 |
| QLIFTTEDN | TSNAX | Q99598 | TSNAX_HUMAN | Unmodified | 0 | 0 | 0 | 0 | 113630 | 30034 | 0 | 0 | 0 | 0 | 0 | 0 | 112820 | 0 | 0 | 0 | 214730 | 55897 |
| QLILKLPVIF | EHD2 | Q9NZN4 | EHD2_HUMAN | Unmodified | 0 | 0 | 0 | 0 | 2943500 | 886370 | 220900 | 0 | 0 | 0 | 0 | 0 | 120740 | 109360 | 0 | 0 | 0 | 0 |
| QLILVGDH( | UPF1 | Q92900 | RENT1_HUMAN | Oxidation (M) | 286580 | 221990 | 0 | 0 | 0 | 0 | 102820 | 49936 | 0 | 0 | 0 | 0 | 0 | 0 | 0 | 0 | 0 | 0 |
| QLIQGGVA | SCP2; SCP2; S( | P22307; H0Y | SCP2_HUMAN | Unmodified | 0 | 0 | 0 | 0 | 393750 | 0 | 1118600 | 0 | 1019400 | 1737200 | 0 | 0 | 602230 | 81839 | 0 | 0 | 0 | 14052 |
| QLIQPIAEK | NXN; NXN; NXN | Q6DKJ4; I3L4 | NXN_HUMAN N | Unmodified | 0 | 0 | 0 | 0 | 858080 | 199050 | 371010 | 0 | 172160 | 0 | 0 | 1145800 | 469850 | 223310 | 0 | 0 | 0 | 120970 |
| QLIVANAGI | PPM1G | O15355 | PPM1G_HUMA | Unmodified | 0 | 0 | 0 | 0 | 0 | 0 | 0 | 116590 | 300110 | 85158 | 331060 | 5070800 | 0 | 0 | 0 | 0 | 0 | 0 |
| QLIVGVNK | EEF1A2; EEF1A | Q05639; Q5 | EF1A2_HUMAN | Unmodified | 1.84E+08 | 1.32E+08 | 1.65E+08 | 64716000 | 30060000 | 5544000 | 8419100 | 4652300 | 30686000 | 4888200 | 18114000 | 2.21E+08 | 17737000 | 9456300 | 2.44E+08 | 1.96E+08 | 7847100 | 15538000 |
| QLIVGVNKI | EEF1A2; EEF1A | Q05639; A0A | EF1A2_HUMAN | Unmodified | 917860 | 1168400 | 263610 | 2222700 | 2668000 | 1001500 | 0 | 0 | 910290 | 661740 | 0 | 0 | 1463300 | 1257800 | 1901700 | 822950 | 201840 | 463000 |
| QLIVGVNKI | EEF1A2; EEF1A | Q05639; A0A | EF1A2_HUMAN | Oxidation (M) | 779490 | 0 | 0 | 1124700 | 0 | 0 | 0 | 0 | 0 | 0 | 0 | 0 | 86085 | 0 | 959940 | 908890 | 0 | 0 |
| QLIVGVNKI | EEF1A1P5; EEF | Q5VTE0; P68 | EF1A3_HUMAN | Oxidation (M) | 4122800 | 5593600 | 0 | 5303600 | 6078100 | 0 | 93264 | 0 | 0 | 0 | 0 | 4120700 | 3463300 | 0 | 2270500 | 2597700 | 807400 | 286470 |
| QLIVGVNKI | EEF1A1P5; EEF | Q5VTE0; P68 | EF1A3_HUMAN | Unmodified | 3655000 | 6701700 | 381570 | 6072200 | 14488000 | 1779800 | 1067700 | 484630 | 3742400 | 3347800 | 1273300 | 769070 | 8740900 | 3671500 | 2455700 | 1119200 | 988900 | 1751300 |
| QLKLDPSIF | RUVBL1; RUVBI | Q9Y265; E7E | RUVB1_HUMAI | Unmodified | 0 | 29498 | 0 | 0 | 0 | 0 | 622560 | 104210 | 0 | 121090 | 0 | 0 | 2817600 | 947980 | 0 | 0 | 0 | 0 |
| QLLADIIQF | IQGAP3; IQGAP | Q86VI3; F2Z | IQGA3_HUMAN | Unmodified | 476680 | 69012 | 658070 | 786020 | 3340600 | 696610 | 385240 | 0 | 0 | 593270 | 289320 | 875030 | 282510 | 0 | 0 | 659970 | 640050 | 284540 |
| QLLADIIQF | IQGAP3; IQGAP | Q86VI3; F2Z | IQGA3_HUMAN | Unmodified | 0 | 0 | 0 | 0 | 515320 | 87554 | 0 | 0 | 0 | 0 | 0 | 0 | 0 | 0 | 0 | 0 | 412870 | 0 |
| QLLAGGIA( | SLC25A24; SLC | Q6NUK1; A0 | SCMC1_HUMA | Unmodified | 0 | 0 | 0 | 0 | 0 | 0 | 0 | 0 | 5200600 | 498050 | 0 | 0 | 0 | 0 | 0 | 0 | 0 | 0 |
| QLLALDAVI | BAG5 | Q9UL15 | BAG5_HUMAN | Unmodified | 0 | 0 | 0 | 0 | 0 | 0 | 0 | 0 | 717990 | 970530 | 0 | 975520 | 0 | 0 | 0 | 0 | 0 | 0 |
| QLLAPGNS | LYN | P07948 | LYN_HUMAN T | Unmodified | 0 | 0 | 0 | 0 | 0 | 0 | 0 | 0 | 1466000 | 0 | 0 | 729600 | 0 | 0 | 0 | 0 | 0 | 0 |
| QLLDQVEC | TMED4 | Q7Z7H5 | TMED4_HUMAI | Unmodified | 0 | 0 | 315090 | 235090 | 0 | 0 | 0 | 0 | 38576000 | 1263100 | 1463900 | 887460 | 0 | 0 | 426940 | 319010 | 0 | 0 |
| QLLDQVEC | TMED4 | Q7Z7H5 | TMED4_HUMAI | Unmodified | 0 | 0 | 0 | 0 | 0 | 0 | 0 | 144660 | 24368000 | 1015700 | 0 | 0 | 0 | 0 | 0 | 0 | 0 | 0 |
| QLLEAENEI | FRYL; FRYL; FRY | O94915; F2Z | FRYL_HUMAN I | Unmodified | 0 | 0 | 0 | 0 | 0 | 0 | 0 | 0 | 0 | 0 | 0 | 0 | 0 | 0 | 0 | 0 | 0 | 0 |
| QLLEQPESI | UTRN; UTRN | P46939; Q5T | UTRN_HUMAN | Unmodified | 0 | 0 | 0 | 0 | 0 | 0 | 0 | 0 | 0 | 0 | 0 | 0 | 0 | 0 | 385310 | 339210 | 0 | 0 |
| QLLETLK | CASP8AP2; CAS | Q9UKL3; A0A | C8AP2_HUMAN | Unmodified | 8320400 | 2520900 | 2659700 | 2020700 | 2888800 | 0 | 697510 | 535930 | 463260 | 367810 | 0 | 1711200 | 1326200 | 209140 | 4909800 | 9684200 | 1515000 | 3379600 |
| QLLFPAEEI | ARHGEF1; ARH | Q92888; M0 | ARHG1_HUMA | Unmodified | 0 | 0 | 0 | 0 | 0 | 0 | 0 | 0 | 0 | 0 | 0 | 696400 | 0 | 0 | 0 | 0 | 0 | 0 |
| QLLGLQPIS | PARP4 | Q9UKK3 | PARP4_HUMAN | Unmodified | 0 | 0 | 0 | 0 | 0 | 0 | 0 | 0 | 326810 | 0 | 0 | 6767600 | 0 | 0 | 0 | 0 | 0 | 0 |
| QLLHDALA | ALDH3B1; ALDI | P43353; A0A | AL3B1_HUMAN | Unmodified | 0 | 0 | 0 | 0 | 2502200 | 555310 | 326520 | 302020 | 0 | 109730 | 0 | 0 | 0 | 0 | 0 | 0 | 0 | 0 |
| QLLHNFPP | UBA1; UBA1 | P22314; Q5J | UBA1_HUMAN | Unmodified | 279270 | 0 | 296120 | 0 | 0 | 68426 | 238580 | 0 | 1533500 | 0 | 896710 | 8471700 | 0 | 44237 | 1486700 | 0 | 217290 | 258100 |
| QLLIDPEDI | CDH4; CDH2; C | P55283; P19 | CADH4_HUMA | Unmodified | 0 | 0 | 0 | 0 | 0 | 0 | 0 | 0 | 0 | 0 | 0 | 2471900 | 0 | 0 | 207970 | 0 | 0 | 0 |
| QLLLDILQF | PHKB | Q93100 | KPBB_HUMAN | Unmodified | 0 | 0 | 0 | 0 | 0 | 0 | 0 | 0 | 0 | 0 | 0 | 173920 | 0 | 0 | 0 | 24857 | 0 | 0 |
| QLLLQLNQ | HIC2 | Q96JB3 | HIC2_HUMAN I | Unmodified | 1.05E+10 | 1.06E+10 | 5.89E+09 | 3.48E+09 | 0 | 0 | 1453800 | 425730 | 0 | 0 | 0 | 0 | 71845000 | 17615000 | 8.58E+09 | 7.55E+09 | 0 | 0 |
| QLLLTEGPI | PDPK1; PDPK1 | O15530; E9F | PDPK1_HUMAN | Unmodified | 250910 | 367020 | 2807000 | 2309000 | 532140 | 0 | 390410 | 98298 | 492270 | 0 | 0 | 626000 | 3339500 | 3397900 | 4407100 | 2558200 | 1477600 | 2166300 |
| QLLLTPNA\ | MYO1C; MYO1( | O00159; F5H | MYO1C_HUMA | Unmodified | 2708900 | 2192600 | 4365800 | 3783900 | 6829500 | 1813900 | 0 | 0 | 558630 | 0 | 0 | 1182700 | 0 | 1618900 | 8744100 | 8192800 | 1718000 | 2467100 |
| QLLLTPNA\ | MYO1C; MYO1( | O00159; F5H | MYO1C_HUMA | Unmodified | 0 | 0 | 0 | 0 | 16278000 | 2486000 | 0 | 0 | 141560 | 0 | 0 | 0 | 5557300 | 8501700 | 0 | 0 | 8980400 | 4811400 |
| QLLMLQSL | AP2S1; AP2S1; | P53680; M0F | AP2S1_HUMAN | Oxidation (M) | 0 | 0 | 298060 | 0 | 0 | 0 | 0 | 68370 | 67803 | 0 | 0 | 3597600 | 0 | 641290 | 0 | 571490 | 200460 | 989960 |
| QLLMVGGI | DARS2; DARS2; | Q6PI48; A0A | SYDM_HUMAN | Oxidation (M) | 0 | 0 | 0 | 0 | 0 | 0 | 0 | 0 | 2308900 | 1359500 | 0 | 0 | 0 | 0 | 0 | 0 | 0 | 0 |
| QLLPGVPFI | DCDC1; DCDC | M0R2J8; H7( | DCDC1_HUMA | Unmodified | 21541000 | 17841000 | 9212800 | 28202000 | 0 | 0 | 327500 | 0 | 0 | 0 | 0 | 0 | 5604600 | 1210500 | 7375800 | 6879700 | 0 | 0 |
| QLLPMLLQ | MOV10; MOV1( | Q9HCE1; Q5 | MOV10_HUMA | Oxidation (M) | 863540 | 1090500 | 256290 | 0 | 0 | 0 | 0 | 0 | 0 | 0 | 0 | 0 | 0 | 0 | 1249900 | 820210 | 0 | 0 |
| QLLQAALR | CC2D1A; CC2D | Q6P1N0; A0 | C2D1A_HUMAI | Unmodified | 894570 | 589110 | 717670 | 764840 | 574530 | 0 | 0 | 0 | 508230 | 220820 | 0 | 456900 | 282480 | 0 | 502000 | 508510 | 70578 | 0 |
| QLLQANPII | MYH14; MYH9; | Q7Z406; P35 | MYH14_HUMAI | Unmodified | 0 | 0 | 188360 | 72905 | 0 | 0 | 0 | 0 | 1327500 | 641110 | 564570 | 2955500 | 0 | 0 | 0 | 462620 | 0 | 0 |
| QLLQANPII | MYH10 | P35580 | MYH10_HUMAI | Unmodified | 0 | 0 | 456490 | 0 | 292490 | 0 | 0 | 0 | 0 | 0 | 208330 | 5845900 | 0 | 0 | 0 | 573670 | 0 | 0 |
| QLLQDEK | TAX1BP1; TAX1 | Q86VP1; B8 | TAXB1_HUMAN | Unmodified | 0 | 0 | 84229 | 488070 | 812670 | 4613300 | 1193400 | 168620 | 106330 | 208610 | 0 | 0 | 0 | 106700 | 0 | 0 | 538140 | 1116900 |
| QLLQTVNV | AKAP12 | Q02952 | AKA12_HUMAN | Unmodified | 0 | 0 | 0 | 0 | 312580 | 0 | 13350 | 0 | 0 | 0 | 0 | 0 | 0 | 0 | 0 | 0 | 1190200 | 1933100 |
| QLLSDLIEK | PLXNA1 | Q9UIW2 | PLXA1_HUMAN | Unmodified | 0 | 0 | 0 | 0 | 0 | 0 | 0 | 0 | 0 | 0 | 0 | 963850 | 0 | 0 | 0 | 0 | 0 | 0 |
| QLLSELDEI | SH3KBP1; SH3I | Q96B97; Q5 | SH3K1_HUMAN | Unmodified | 120620 | 0 | 263120 | 295160 | 0 | 0 | 0 | 0 | 0 | 0 | 0 | 0 | 646210 | 0 | 0 | 315110 | 0 | 0 |
| QLLSLLSA( | ADAMTS13; AD | Q76LX8; E7E | ATS13_HUMAN | Unmodified | 0 | 51789 | 386450 | 75810 | 0 | 0 | 0 | 0 | 0 | 0 | 0 | 0 | 0 | 0 | 0 | 52042 | 0 | 0 |
| QLLTDFCTI | COPS4; COPS4 | Q9BT78; D6I | CSN4_HUMAN | Unmodified | 32746 | 0 | 26149 | 0 | 0 | 0 | 0 | 0 | 0 | 0 | 0 | 1627300 | 0 | 0 | 0 | 109170 | 0 | 0 |
| QLLTLSSEI | EZR; EZR | P15311; E7E | EZRI_HUMAN E | Unmodified | 0 | 0 | 0 | 0 | 0 | 0 | 0 | 0 | 696100 | 0 | 419240 | 1847700 | 0 | 0 | 0 | 0 | 1365200 | 0 |
| QLLTLSSEI | EZR; EZR | P15311; E7E | EZRI_HUMAN E | Unmodified | 0 | 2103200 | 1638200 | 1894600 | 18498000 | 4518300 | 0 | 0 | 23651000 | 1208800 | 5450100 | 75066000 | 6682900 | 0 | 3298700 | 3632400 | 492280 | 2280100 |
| QLLTLSSEI | EZR; EZR | P15311; E7E | EZRI_HUMAN E | Unmodified | 0 | 660240 | 0 | 467690 | 613170 | 804270 | 0 | 0 | 4445500 | 0 | 1159000 | 2673300 | 0 | 0 | 0 | 0 | 0 | 0 |
| QLLTPEHVI | AGRN; AGRN; A | O00468; A0A | AGRIN_HUMAN | Unmodified | 638010 | 523470 | 1264400 | 853010 | 0 | 0 | 0 | 0 | 1305800 | 72331 | 0 | 0 | 260460 | 0 | 538080 | 499590 | 0 | 0 |

| | | | | | | | | | | | | | | | | | | | | | | |
|---|---|---|---|---|---|---|---|---|---|---|---|---|---|---|---|---|---|---|---|---|---|---|
| QLLVEAIDV | ITGB4 | P16144 | ITB4_HUMAN I | Unmodified | 0 | 0 | 0 | 0 | 0 | 0 | 0 | 0 | 0 | 0 | 0 | 0 | 0 | 0 | 0 | 0 | 1175200 | 0 |
| QLLVEAIDV | ITGB4 | P16144 | ITB4_HUMAN I | Unmodified | 0 | 0 | 0 | 0 | 2872200 | 648470 | 0 | 0 | 0 | 0 | 0 | 0 | 343420 | 0 | 0 | 0 | 3383600 | 6700700 |
| QLLYDAIK | UGGT1 | Q9NYU2 | UGGG1_HUMA | Unmodified | 0 | 0 | 0 | 0 | 288930 | 0 | 0 | 5389.7 | 10447000 | 1939000 | 0 | 775200 | 16536 | 0 | 0 | 45167 | 0 | 0 |
| QLLYLEK | PTP4A1; PTP4A | Q93096; Q1 | TP4A1_HUMAN | Unmodified | 854160 | 1312600 | 1154100 | 2131600 | 0 | 0 | 0 | 0 | 2560400 | 646630 | 0 | 5928900 | 1188400 | 1178000 | 1840800 | 1224000 | 0 | 0 |
| QLMETPAN | LAP3; LAP3 | P28838; H0 | AMPL_HUMAN | Oxidation (M) | 0 | 0 | 0 | 0 | 2390200 | 78154 | 86062 | 0 | 818120 | 189670 | 64224 | 1516300 | 1125300 | 321930 | 0 | 79719 | 90399 | 1203200 |
| QLMETPAN | LAP3; LAP3 | P28838; H0 | AMPL_HUMAN | 2 Oxidation (M) | 0 | 0 | 0 | 0 | 0 | 0 | 0 | 0 | 0 | 102810 | 718300 | 0 | 1016200 | 0 | 146050 | 106150 | 0 | 0 |
| QLMHQLPI | LMCD1; LMCD1 | Q9NZU5; H7 | LMCD1_HUMA | Unmodified | 0 | 0 | 0 | 0 | 229220 | 0 | 0 | 0 | 700820 | 0 | 0 | 1716600 | 0 | 0 | 0 | 0 | 0 | 0 |
| QLMHQLPI | LMCD1; LMCD1 | Q9NZU5; H7 | LMCD1_HUMA | Oxidation (M) | 0 | 0 | 0 | 0 | 0 | 0 | 0 | 0 | 0 | 0 | 0 | 0 | 0 | 0 | 0 | 0 | 0 | 0 |
| QLMHSGH | SPTBN1; SPTBN | Q01082; A0 | SPTB2_HUMAN | Unmodified | 0 | 0 | 0 | 0 | 932130 | 0 | 1207500 | 636180 | 16428 | 0 | 633710 | 586830 | 0 | 0 | 0 | 0 | 160390 | 0 |
| QLNENQVI | PPP2CB; PPP2 | P62714; E5 | PP2AB_HUMAN | Unmodified | 0 | 0 | 0 | 0 | 1867500 | 345700 | 340810 | 268670 | 0 | 0 | 0 | 0 | 1219300 | 1354100 | 0 | 0 | 0 | 1164700 |
| QLNFIPIDL | STEAP2; STEAP | Q8NFT2; B5 | STEA2_HUMAN | Unmodified | 3074600 | 2165100 | 10536000 | 10165000 | 22326 | 13460 | 0 | 0 | 0 | 0 | 0 | 62391 | 0 | 0 | 48493000 | 27433000 | 0 | 11622 |
| QLNFIPIDL | STEAP2; STEAP | Q8NFT2; B5 | STEA2_HUMAN | Unmodified | 0 | 0 | 0 | 245830 | 0 | 0 | 0 | 0 | 0 | 0 | 0 | 0 | 0 | 0 | 480920 | 0 | 0 | 0 |
| QLNLWSTC | CMIP; CMIP; CN | Q8IY22; A0 | CMIP_HUMAN | Unmodified | 0 | 162970 | 0 | 0 | 0 | 0 | 0 | 0 | 418980 | 0 | 38710 | 667190 | 0 | 0 | 0 | 0 | 0 | 0 |
| QLNPESIRI | DHX30; DHX30 | Q7L2E3; H7 | DHX30_HUMAI | Unmodified | 166880 | 132810 | 0 | 0 | 0 | 0 | 0 | 0 | 349520 | 0 | 0 | 1407300 | 0 | 0 | 0 | 0 | 0 | 0 |
| QLNQVNFI | RAB11FIP1; RA | Q6WKZ4; E7 | RFIP1_HUMAN | Unmodified | 0 | 0 | 0 | 0 | 979560 | 0 | 0 | 0 | 0 | 0 | 0 | 0 | 383570 | 137710 | 0 | 0 | 0 | 0 |
| QLNRQLDN | LMAN1 | P49257 | LMAN1_HUMA | Unmodified | 0 | 0 | 0 | 0 | 0 | 0 | 0 | 0 | 3575200 | 6444200 | 80995 | 0 | 384970 | 0 | 0 | 0 | 0 | 0 |
| QLNSESLS | SERPINB5 | P36952 | SPB5_HUMAN | Oxidation (M) | 0 | 0 | 0 | 0 | 0 | 0 | 0 | 0 | 4836000 | 1657500 | 0 | 0 | 1015100 | 0 | 0 | 0 | 1081400 | 778920 |
| QLNSESLS | SERPINB5 | P36952 | SPB5_HUMAN | Unmodified | 0 | 0 | 0 | 0 | 0 | 0 | 0 | 0 | 0 | 0 | 0 | 0 | 0 | 0 | 0 | 0 | 1206100 | 3249100 |
| QLPADSPG | DOP1B | Q9Y3R5 | DOP2_HUMAN | Unmodified | 0 | 0 | 0 | 0 | 0 | 0 | 0 | 0 | 1715900 | 102300 | 192840 | 1855700 | 0 | 0 | 0 | 0 | 0 | 0 |
| QLPDAQLL | PTGS1; PTGS1 | P23219; A0 | PGH1_HUMAN | Unmodified | 0 | 0 | 0 | 0 | 0 | 0 | 0 | 0 | 0 | 0 | 0 | 2460400 | 0 | 0 | 0 | 0 | 0 | 0 |
| QLPDAQLL | PTGS1; PTGS1 | P23219; A0 | PGH1_HUMAN | Unmodified | 0 | 0 | 0 | 0 | 0 | 0 | 0 | 0 | 0 | 0 | 1064800 | 450650 | 0 | 0 | 0 | 0 | 0 | 0 |
| QLPFRGDD | SNX3 | O60493 | SNX3_HUMAN | Unmodified | 259710 | 313590 | 0 | 377630 | 0 | 0 | 0 | 0 | 2569700 | 0 | 202870 | 339590 | 0 | 0 | 564490 | 696620 | 0 | 0 |
| QLPFRGDD | SNX3 | O60493 | SNX3_HUMAN | Unmodified | 0 | 284060 | 518580 | 559040 | 433540 | 0 | 0 | 0 | 2554500 | 0 | 142600 | 0 | 153930 | 0 | 1262000 | 539660 | 49499 | 0 |
| QLPIDSDD | GRIK3 | Q13003 | GRIK3_HUMAN | Unmodified | 0 | 299770 | 746350 | 0 | 0 | 0 | 199180 | 0 | 368000 | 0 | 0 | 415950 | 103110 | 0 | 373140 | 0 | 0 | 0 |
| QLPIFCQLI | INF2; INF2; INF | Q27J81; A0 | INF2_HUMAN I | Unmodified | 0 | 0 | 0 | 0 | 0 | 0 | 0 | 0 | 0 | 0 | 0 | 123340 | 0 | 0 | 0 | 0 | 0 | 0 |
| QLPLKFSN | MYO1E | Q12965 | MYO1E_HUMA | Unmodified | 0 | 0 | 0 | 0 | 0 | 0 | 0 | 0 | 0 | 0 | 0 | 0 | 0 | 0 | 0 | 0 | 246840 | 0 |
| QLPLVKPYI | CLTCL1; CLTC | P53675; Q0 | CLH2_HUMAN | Unmodified | 29111000 | 16324000 | 8724500 | 16059000 | 14256000 | 4413800 | 1983600 | 1343900 | 4970500 | 1611900 | 1126300 | 1.12E+08 | 15339000 | 6061000 | 11643000 | 15174000 | 1999400 | 958720 |
| QLPLVKPYI | CLTC; CLTC; CI | Q00610; J3K | CLH1_HUMAN | Unmodified | 70606 | 0 | 0 | 0 | 2898000 | 0 | 2915900 | 238100 | 0 | 153530 | 0 | 0 | 1248300 | 2114300 | 0 | 0 | 1368300 | 2795200 |
| QLPNGDW | LSS; LSS | P48449; A0 | LSS_HUMAN La | Unmodified | 0 | 0 | 0 | 0 | 0 | 0 | 0 | 0 | 0 | 0 | 0 | 452350 | 0 | 0 | 0 | 0 | 0 | 0 |
| QLPPVDAE | PROM1 | O43490 | PROM1_HUMA | Unmodified | 0 | 0 | 0 | 0 | 898140 | 0 | 0 | 0 | 0 | 0 | 519060 | 9429300 | 0 | 0 | 0 | 0 | 0 | 0 |
| QLPQNVLD | MYO1C; MYO1 | O00159; F5 | MYO1C_HUMA | Unmodified | 448220 | 7055400 | 2414100 | 0 | 0 | 0 | 0 | 0 | 0 | 0 | 0 | 1459700 | 0 | 0 | 7248800 | 10580000 | 0 | 0 |
| QLPSAGPD | ENSA; ENSA; EN | O43768; A0 | ENSA_HUMAN | Unmodified | 313790 | 194440 | 0 | 684240 | 0 | 0 | 314690 | 188390 | 916360 | 473260 | 223410 | 689490 | 2385100 | 939030 | 481790 | 512640 | 0 | 0 |
| QLPSAGPD | ENSA; ENSA; EN | O43768; A0 | ENSA_HUMAN | Unmodified | 0 | 0 | 0 | 0 | 464240 | 0 | 0 | 487880 | 0 | 0 | 0 | 0 | 591560 | 645670 | 0 | 0 | 1335600 | 0 |
| QLPTPVGA | CSNK1G3; CSN | Q9Y6M4; H0 | KC1G3_HUMAI | Unmodified | 0 | 0 | 0 | 0 | 0 | 0 | 0 | 0 | 0 | 0 | 291720 | 1261400 | 1211800 | 3456700 | 0 | 0 | 0 | 0 |
| QLPTSEAV | SUN1; SUN1; S | O94901; H0 | SUN1_HUMAN | Unmodified | 0 | 0 | 0 | 0 | 0 | 0 | 0 | 0 | 18332000 | 0 | 0 | 0 | 0 | 0 | 0 | 0 | 0 | 0 |
| QLPYYYLSI | ALOX5 | P09917 | LOX5_HUMAN | Unmodified | 0 | 0 | 0 | 0 | 0 | 0 | 0 | 0 | 0 | 0 | 130220 | 341190 | 0 | 0 | 0 | 0 | 0 | 0 |
| QLQAETEPI | EIF3E; EIF3E; EI | P60228; A0 | EIF3E_HUMAN | Unmodified | 467970 | 315140 | 719210 | 852120 | 1249100 | 0 | 576950 | 160900 | 543190 | 291890 | 1102600 | 5310200 | 1453600 | 0 | 556740 | 594290 | 252790 | 0 |
| QLQAIEFNI | SRP72 | O76094 | SRP72_HUMAN | Unmodified | 108540 | 82974 | 356120 | 0 | 0 | 0 | 352410 | 333860 | 0 | 0 | 0 | 2490900 | 535520 | 0 | 141630 | 139750 | 224030 | 346970 |
| QLQALSSE | RDX; RDX; RDX | P35241; A0 | RADI_HUMAN I | Unmodified | 1345800 | 992570 | 1732000 | 4429000 | 13885000 | 5519100 | 1009800 | 479690 | 3869100 | 1521800 | 1317600 | 12317000 | 17689000 | 18727000 | 4143300 | 2809600 | 2458100 | 4059100 |
| QLQALSSE | RDX; RDX; RDX | P35241; A0 | RADI_HUMAN I | Unmodified | 0 | 0 | 742550 | 1978300 | 6957800 | 1828100 | 0 | 250580 | 1526500 | 598970 | 300610 | 8238200 | 5717100 | 2826200 | 2477600 | 899510 | 862190 | 2105600 |
| QLQAQIDN | GPRASP2 | Q96D09 | GASP2_HUMAI | Unmodified | 389840 | 454690 | 167790 | 154040 | 183470 | 0 | 0 | 0 | 0 | 0 | 0 | 0 | 46390 | 0 | 453010 | 305860 | 0 | 0 |
| QLQDELEN | PSMD2; PSMD2 | Q13200; C9 | PSMD2_HUMAN | Oxidation (M) | 0 | 0 | 22719 | 0 | 0 | 0 | 0 | 0 | 0 | 0 | 0 | 237460 | 0 | 0 | 0 | 0 | 0 | 0 |
| QLQDIATL | IPO7 | O95373 | IPO7_HUMAN I | Unmodified | 0 | 0 | 0 | 0 | 241950 | 84322 | 0 | 0 | 0 | 0 | 0 | 2389900 | 0 | 0 | 0 | 0 | 89744 | 330600 |
| QLQEAHRI | UTRN; UTRN | P46939; Q5 | UTRN_HUMAN | Unmodified | 0 | 0 | 1137900 | 0 | 0 | 0 | 0 | 0 | 1653500 | 0 | 0 | 0 | 0 | 0 | 1121200 | 1127400 | 57586 | 0 |
| QLQEEDLK | CFAP45; CCDC | Q9UL16; Q8 | CFA45_HUMAN | Unmodified | 4735100 | 5471300 | 5163600 | 8137800 | 0 | 0 | 0 | 504270 | 382840 | 260090 | 0 | 228020 | 3178000 | 2464000 | 8854800 | 8517900 | 267970 | 1561700 |
| QLQEEDLK | CCDC50 | Q8IVM0 | CCD50_HUMA | Unmodified | 0 | 912670 | 0 | 369460 | 465300 | 0 | 1259800 | 208350 | 206260 | 1100600 | 0 | 0 | 22130000 | 10527000 | 0 | 533100 | 901860 | 2166400 |
| QLQEETPP | NDUFB9; NDUI | Q9Y6M9; A0 | NDUB9_HUMA | Unmodified | 0 | 0 | 0 | 0 | 0 | 0 | 0 | 0 | 3082000 | 0 | 0 | 0 | 0 | 0 | 0 | 0 | 0 | 0 |
| QLQFYTEA | CC2D1A; CC2D | Q6P1N0; A0 | C2D1A_HUMAI | Unmodified | 434630 | 500680 | 0 | 0 | 0 | 0 | 262500 | 0 | 0 | 0 | 0 | 0 | 0 | 0 | 478490 | 443690 | 93409 | 0 |
| QLQGFPFY | SNRPB2 | P08579 | RU2B_HUMAN | Oxidation (M) | 362430 | 153180 | 0 | 0 | 0 | 0 | 0 | 0 | 0 | 0 | 0 | 0 | 112250 | 0 | 0 | 124720 | 0 | 0 |
| QLQGIQIQI | RABEP1 | Q15276 | RABE1_HUMAN | Unmodified | 0 | 0 | 0 | 0 | 1419600 | 361840 | 0 | 0 | 229260 | 0 | 0 | 365490 | 513180 | 0 | 0 | 0 | 0 | 0 |
| QLQGPEKE | PLIN3; PLIN3 | O60664; K7 | PLIN3_HUMAN | Unmodified | 108510 | 0 | 0 | 112500 | 1616100 | 634240 | 0 | 0 | 128860 | 135240 | 65571 | 648750 | 357160 | 118400 | 171380 | 118010 | 984100 | 742380 |
| QLQIRFQK | UGGT1; UGGT1 | Q9NYU2; H7 | UGGG1_HUMA | Unmodified | 0 | 0 | 0 | 0 | 0 | 0 | 0 | 0 | 0 | 808650 | 0 | 0 | 0 | 0 | 0 | 0 | 0 | 0 |
| QLQISERN | SOGA1; SOGA1 | O94964; X6 | SOGA1_HUMAN | Unmodified | 0 | 0 | 0 | 0 | 0 | 0 | 0 | 0 | 0 | 0 | 0 | 0 | 391820 | 293330 | 0 | 0 | 0 | 0 |
| QLQLEAEE | LRRC47 | Q8N1G4 | LRC47_HUMAN | Unmodified | 0 | 0 | 0 | 0 | 0 | 0 | 0 | 27524 | 567350 | 184180 | 251820 | 1488000 | 356220 | 0 | 169900 | 0 | 0 | 109280 |
| QLQLELK | CORO2B | Q9UQ03 | COR2B_HUMA | Unmodified | 0 | 0 | 0 | 0 | 0 | 0 | 0 | 61564 | 127980 | 0 | 266610 | 1590900 | 0 | 0 | 0 | 0 | 20721 | 0 |
| QLQLVTYS | B4GALNT1; B4 | Q00973; H0 | B4GN1_HUMAI | Unmodified | 0 | 0 | 0 | 0 | 0 | 0 | 0 | 0 | 1606900 | 0 | 0 | 0 | 0 | 0 | 0 | 405680 | 0 | 0 |
| QLQNISLA | DYNC1H1; DYN | Q14204; A0 | DYHC1_HUMA | Unmodified | 0 | 0 | 0 | 0 | 0 | 0 | 0 | 0 | 0 | 0 | 0 | 3534600 | 0 | 0 | 81189 | 0 | 0 | 0 |
| QLQNQLAE | RAI14 | Q9P0K7 | RAI14_HUMAN | Unmodified | 0 | 0 | 0 | 0 | 0 | 0 | 0 | 0 | 0 | 0 | 0 | 141800 | 0 | 0 | 0 | 0 | 0 | 0 |
| QLQQAQA | FEN1; FEN1 | P39748; F5 | FEN1_HUMAN | Unmodified | 0 | 0 | 0 | 0 | 0 | 0 | 0 | 0 | 0 | 0 | 187230 | 921150 | 0 | 0 | 0 | 0 | 0 | 0 |
| QLQQAQA | FEN1; FEN1 | P39748; F5 | FEN1_HUMAN | Unmodified | 0 | 0 | 0 | 0 | 0 | 0 | 630370 | 0 | 0 | 842500 | 0 | 0 | 1150600 | 512120 | 0 | 0 | 779460 | 0 |
| QLQQAVTC | CTNNA1; CTNN | P35221; H0 | CTNA1_HUMAI | Unmodified | 0 | 0 | 0 | 6741300 | 6145600 | 7785600 | 0 | 0 | 0 | 0 | 0 | 1458500 | 5053500 | 6719300 | 7727700 | 0 | 0 | 0 |
| QLQQERPC | LRP11; LRP11 | Q86VZ4; Q5 | LRP11_HUMAN | Unmodified | 0 | 0 | 0 | 0 | 0 | 0 | 0 | 0 | 5928300 | 180770 | 0 | 0 | 259650 | 0 | 0 | 0 | 0 | 0 |
| QLQQREI | EEA1 | Q15075 | EEA1_HUMAN | Unmodified | 0 | 0 | 0 | 0 | 1523100 | 0 | 0 | 0 | 0 | 0 | 0 | 0 | 0 | 0 | 0 | 0 | 212620 | 361530 |
| QLQRQLSS | ABCC1; ABCC1 | P33327; I3L | MRP1_HUMAN | Unmodified | 0 | 0 | 0 | 0 | 223410 | 0 | 0 | 0 | 0 | 0 | 0 | 0 | 0 | 0 | 0 | 0 | 0 | 0 |
| QLQSEQPC | ATP6V1H; ATP6 | Q9UI12; G3 | VATH_HUMAN | Unmodified | 0 | 0 | 0 | 0 | 0 | 14358 | 0 | 105620 | 84102 | 63260 | 0 | 304800 | 0 | 0 | 41058 | 56811 | 0 | 0 |
| QLRFEDVV | TIAL1 | Q01085 | TIAR_HUMAN N | Unmodified | 273420 | 296120 | 523060 | 309090 | 891660 | 109130 | 424220 | 0 | 340100 | 257830 | 231160 | 1647000 | 1764900 | 783970 | 615920 | 761050 | 442680 | 353530 |
| QLSAFGEY | NDUFS3; NDUI | O75489; E9 | NDUS3_HUMA | Unmodified | 0 | 0 | 0 | 0 | 0 | 0 | 0 | 0 | 264440 | 0 | 0 | 0 | 0 | 0 | 0 | 0 | 0 | 0 |
| QLSALSSD | ABCC3; ABCC3 | O15438; H0 | MRP3_HUMAN | Unmodified | 0 | 0 | 0 | 0 | 977250 | 124800 | 0 | 0 | 0 | 0 | 0 | 0 | 0 | 0 | 0 | 0 | 0 | 318590 |
| QLSDAISII | CSE1L | P55060 | XPO2_HUMAN | Unmodified | 0 | 122080 | 128030 | 95452 | 0 | 0 | 0 | 0 | 0 | 224080 | 0 | 2939300 | 0 | 0 | 0 | 98473 | 0 | 0 |
| QLSESQVK | PPP2CA; N/A | P67775; E5 | PP2AA_HUMAN | Unmodified | 138430 | 0 | 173480 | 168950 | 2531400 | 354790 | 595510 | 718910 | 193820 | 176110 | 0 | 1032700 | 1402800 | 786570 | 232910 | 145990 | 1059500 | 2329100 |
| QLSESQVK | PPP2CA; N/A | P67775; E5 | PP2AA_HUMAN | Unmodified | 0 | 0 | 0 | 0 | 73220 | 0 | 0 | 178370 | 0 | 99313 | 0 | 0 | 104520 | 0 | 0 | 0 | 1190100 | 69491 |
| QLSEVFETI | CDCA3; CDCA3 | Q99618; F8 | CDCA3_HUMA | Unmodified | 0 | 0 | 1126400 | 0 | 1868200 | 0 | 0 | 72671 | 0 | 191280 | 0 | 0 | 1221200 | 452500 | 0 | 0 | 295150 | 0 |
| QLSKYESQ | SLC39A6; SLC3 | Q13433; K7 | S39A6_HUMAN | Unmodified | 88875 | 0 | 102300 | 275580 | 0 | 38531 | 0 | 0 | 204900 | 0 | 206640 | 1019300 | 0 | 0 | 389070 | 437520 | 0 | 91381 |
| QLSLGQGS | FRMD8 | Q9BZ67 | FRMD8_HUMA | Unmodified | 241730 | 0 | 323470 | 0 | 740760 | 0 | 0 | 0 | 465340 | 39624 | 82046 | 765600 | 0 | 0 | 366050 | 281770 | 455930 | 823730 |
| QLSLLDSR | DST; DST; DST | Q03001; A0 | DYST_HUMAN | Unmodified | 0 | 0 | 0 | 0 | 627910 | 0 | 0 | 0 | 574360 | 362070 | 0 | 0 | 0 | 0 | 0 | 0 | 200860 | 0 |
| QLSNPQQI | AP2A2; AP2A2 | O94973; A0 | AP2A2_HUMAN | Unmodified | 0 | 0 | 169980 | 172530 | 2975300 | 0 | 270070 | 54066 | 995160 | 302100 | 0 | 2177900 | 2175000 | 811660 | 0 | 195140 | 369240 | 0 |
| QLSQATAA | KARS1; KARS1 | Q15046; H3 | SYK_HUMAN L | Unmodified | 0 | 0 | 0 | 0 | 0 | 0 | 0 | 0 | 0 | 0 | 0 | 1311300 | 0 | 0 | 0 | 0 | 0 | 0 |
| QLSQLGQ | SCFD1; SCFD1 | Q8WVM8; A | SCFD1_HUMAI | Unmodified | 24242 | 48013 | 318410 | 1003400 | 1479800 | 218250 | 302660 | 31470 | 147190 | 100090 | 682930 | 708730 | 498060 | 187660 | 493370 | 494390 | 57104 | 133730 |
| QLSQSLLP | PPP2R1A; PPP2 | P30153; P3 | 2AAA_HUMAN | Unmodified | 0 | 48204 | 0 | 0 | 0 | 0 | 12287 | 109730 | 0 | 0 | 21298 | 434800 | 0 | 0 | 468280 | 0 | 0 | 0 |
| QLSQSLLP | PPP2R1A; PPP2 | P30153; P3 | 2AAA_HUMAN | Unmodified | 0 | 0 | 0 | 0 | 0 | 0 | 0 | 0 | 0 | 0 | 0 | 0 | 0 | 0 | 361280 | 0 | 0 | 0 |
| QLSSEELE | IARS1; IARS1; I | P41252; A0 | SYIC_HUMAN | Unmodified | 0 | 0 | 0 | 0 | 1066500 | 246930 | 480030 | 0 | 782930 | 345510 | 1514200 | 5013400 | 0 | 0 | 0 | 0 | 0 | 406040 |
| QLSSGVSE | HSPB1; HSPB1 | P04792; A0 | HSPB1_HUMAN | Unmodified | 2480800 | 327160 | 8877100 | 4998300 | 0 | 0 | 0 | 0 | 4100300 | 0 | 0 | 0 | 115810 | 0 | 7247800 | 5102200 | 25466 | 0 |
| QLSSSGRP | SQOR; SQOR; S | Q9Y6N5; H3 | SQOR_HUMAN | Unmodified | 3442400 | 0 | 0 | 0 | 0 | 0 | 0 | 0 | 7155500 | 1153000 | 0 | 0 | 0 | 0 | 0 | 0 | 0 | 0 |
| QLSSSSSY | ABCC1; ABCC1 | P33327; I3L | MRP1_HUMAN | Unmodified | 0 | 0 | 0 | 0 | 0 | 0 | 0 | 0 | 0 | 0 | 0 | 0 | 0 | 0 | 146820 | 188600 | 28473 | 0 |
| QLSSSVTG | IQGAP1; IQGAP | P46940; A0 | IQGA1_HUMAN | Unmodified | 139980 | 301550 | 0 | 0 | 0 | 179100 | 0 | 0 | 119340 | 0 | 0 | 1032400 | 0 | 0 | 32074 | 0 | 0 | 0 |
| QLSVPEFIS | SDF4; SDF4; SD | Q9BRK5; H0 | CAB45_HUMAN | Unmodified | 0 | 0 | 0 | 0 | 0 | 0 | 0 | 0 | 2487600 | 0 | 0 | 0 | 0 | 0 | 0 | 0 | 0 | 0 |
| QLTCTEEM | SLC9A3R2; SLC | Q15599; H3 | NHRF2_HUMAN | Unmodified | 0 | 0 | 0 | 0 | 0 | 0 | 126210 | 0 | 0 | 0 | 0 | 352960 | 0 | 0 | 293970 | 232830 | 0 | 0 |
| QLTDNLK | CD109 | Q6YHK3 | CD109_HUMAN | Unmodified | 304020 | 172930 | 239370 | 0 | 89811 | 260140 | 0 | 0 | 3496600 | 438040 | 200270 | 0 | 42894 | 0 | 658400 | 300280 | 421150 | 565830 |
| QLTEEKFIV | PSMC6; PSMC6 | P62333; H0 | PRS10_HUMAN | Unmodified | 0 | 0 | 0 | 0 | 0 | 0 | 0 | 359490 | 0 | 0 | 0 | 0 | 0 | 0 | 0 | 0 | 1665800 | 400020 |
| QLTLEQLH | NUDT16L1; NU | Q9BRJ7; K7 | TIRR_HUMAN T | Unmodified | 0 | 139730 | 0 | 61471 | 0 | 0 | 0 | 0 | 0 | 0 | 0 | 1576800 | 0 | 0 | 0 | 408260 | 0 | 0 |
| QLTQEMM | KTN1 | Q86UP2 | KTN1_HUMAN | Unmodified | 2804900 | 0 | 0 | 0 | 0 | 0 | 0 | 0 | 0 | 0 | 0 | 745020 | 0 | 0 | 0 | 0 | 0 | 0 |
| QLTQIDGTI | CHMP4C | Q96CF2 | CHM4C_HUMA | Unmodified | 1618500 | 3266600 | 0 | 0 | 0 | 0 | 0 | 0 | 2391600 | 0 | 25120 | 0 | 0 | 0 | 1736100 | 1740800 | 0 | 0 |
| QLTQIDGTI | CHMP4C | Q96CF2 | CHM4C_HUMA | Unmodified | 3840500 | 0 | 0 | 1776800 | 1575700 | 0 | 0 | 0 | 6190200 | 1553300 | 0 | 0 | 1337700 | 0 | 0 | 0 | 0 | 0 |
| QLTQIDGTI | CHMP4C | Q96CF2 | CHM4C_HUMA | Oxidation (M) | 0 | 0 | 0 | 0 | 0 | 0 | 0 | 0 | 0 | 1016300 | 0 | 0 | 1692000 | 1532000 | 0 | 0 | 0 | 630470 |
| QLTQNFLS | GRK6; GRK6; G | P43250; D6 | GRK6_HUMAN | Unmodified | 933340 | 263100 | 959710 | 1773300 | 0 | 0 | 0 | 0 | 2055100 | 151230 | 779460 | 3501500 | 118990 | 0 | 1839700 | 2028100 | 0 | 0 |
| QLTTYLTDF | UTRN | P46939 | UTRN_HUMAN | Unmodified | 0 | 0 | 0 | 4668300 | 0 | 0 | 0 | 0 | 0 | 285430 | 0 | 0 | 0 | 0 | 3404800 | 3770700 | 0 | 0 |
| QLTVTQQS | HIP1R | O75146 | HIP1R_HUMAN | Unmodified | 0 | 0 | 0 | 0 | 298790 | 0 | 0 | 0 | 0 | 63464 | 0 | 0 | 1166800 | 790960 | 0 | 0 | 0 | 0 |
| QLTYLDVSI | ERBIN; ERBIN | Q96RT1; B4 | ERBIN_HUMAN | Unmodified | 127090 | 55217 | 0 | 420630 | 10420000 | 4667800 | 0 | 0 | 960270 | 0 | 0 | 4253200 | 569380 | 0 | 602190 | 448390 | 694500 | 274340 |
| QLVAIYEVL | DNAJC1 | Q96KC8 | DNJC1_HUMAI | Unmodified | 0 | 0 | 0 | 0 | 0 | 0 | 0 | 0 | 3693700 | 214070 | 0 | 139850 | 0 | 0 | 0 | 0 | 0 | 0 |
| QLVANLRG | ARRDC3 | Q96B67 | ARRD3_HUMAI | Unmodified | 0 | 0 | 0 | 0 | 1159000 | 0 | 0 | 0 | 0 | 0 | 0 | 0 | 0 | 0 | 0 | 0 | 0 | 0 |
| QLVAREQE | RRBP1; RRBP1 | Q9P2E9; A0 | RRBP1_HUMAI | Unmodified | 0 | 0 | 0 | 0 | 0 | 0 | 0 | 0 | 0 | 0 | 0 | 0 | 823900 | 0 | 0 | 0 | 0 | 0 |
| QLVDEFQA | CRAT; CRAT; CI | P43155; A0 | CACP_HUMAN | Unmodified | 0 | 0 | 0 | 0 | 0 | 0 | 0 | 0 | 1422500 | 537050 | 0 | 0 | 0 | 0 | 0 | 0 | 0 | 0 |
| QLVDIPEG | QSOX1; QSOX1 | O00391; H0 | QSOX1_HUMAI | Unmodified | 0 | 0 | 0 | 0 | 0 | 0 | 0 | 0 | 787550 | 146030 | 0 | 0 | 0 | 0 | 0 | 0 | 0 | 0 |
| QLVDTTVEI | SF3B1 | O75533 | SF3B1_HUMAN | Unmodified | 170560 | 147780 | 0 | 0 | 0 | 0 | 0 | 0 | 0 | 0 | 0 | 4174200 | 0 | 0 | 0 | 0 | 0 | 0 |
| QLVEASERI | CDC42BPB | Q9Y5S2 | MRCKB_HUMA | Unmodified | 0 | 0 | 0 | 0 | 375470 | 0 | 0 | 0 | 0 | 0 | 0 | 0 | 0 | 0 | 0 | 0 | 72949 | 377490 |
| QLVEIAARC | CACNA2D1; CA | P54289; E7 | CA2D1_HUMAN | Unmodified | 0 | 151260 | 0 | 0 | 0 | 0 | 868820 | 546750 | 0 | 675340 | 0 | 0 | 346510 | 0 | 0 | 0 | 0 | 0 |
| QLVEKGPQ | PACSIN2; PACS | Q9UNF0; B0 | PACN2_HUMAI | Unmodified | 2181500 | 3634900 | 0 | 0 | 0 | 0 | 0 | 0 | 0 | 413890 | 0 | 52341 | 23679000 | 8690500 | 0 | 0 | 396450 | 640100 |
| QLVEQVEQ | TMED9 | Q9BVK6 | TMED9_HUMA | Unmodified | 0 | 0 | 513840 | 313290 | 0 | 0 | 0 | 30094 | 92856000 | 5954500 | 3052000 | 3025200 | 0 | 0 | 533400 | 467260 | 0 | 264400 |
| QLVEQVEQ | TMED9 | Q9BVK6 | TMED9_HUMA | Unmodified | 0 | 0 | 0 | 0 | 0 | 0 | 0 | 0 | 12334000 | 2994800 | 0 | 0 | 0 | 0 | 0 | 0 | 0 | 130460 |

| | | | | | | | | | | | | | | | | | | | | | | |
|---|---|---|---|---|---|---|---|---|---|---|---|---|---|---|---|---|---|---|---|---|---|---|
| QLVFHIGH | H6PD | O95479 | G6PE_HUMAN | Unmodified | 0 | 0 | 0 | 0 | 0 | 0 | 0 | 0 | 5837900 | 524140 | 0 | 0 | 0 | 0 | 0 | 0 | 0 | 0 |
| QLVHELDE | PSME1; PSME1 | Q06323; H0Y | PSME1_HUMAI | Unmodified | 0 | 0 | 0 | 0 | 0 | 0 | 0 | 15937 | 0 | 0 | 298370 | 5808200 | 0 | 0 | 0 | 0 | 0 | 0 |
| QLVSEGNII | SPTBN2 | O15020 | SPTN2_HUMAN | Unmodified | 0 | 0 | 371130 | 117970 | 0 | 0 | 0 | 0 | 0 | 0 | 0 | 0 | 25587 | 0 | 290360 | 0 | 0 | 0 |
| QLVTNACL | PRMT1; PRMT1; | Q99873; E9F | ANM1_HUMAN | Unmodified | 0 | 0 | 0 | 0 | 0 | 0 | 0 | 0 | 0 | 0 | 355330 | 10305000 | 0 | 0 | 0 | 0 | 0 | 0 |
| QLWDTANT | GYS1 | P13807 | GYS1_HUMAN | Unmodified | 0 | 0 | 0 | 0 | 0 | 0 | 0 | 0 | 0 | 0 | 0 | 3665600 | 151070 | 0 | 0 | 0 | 0 | 0 |
| QLWGLLIEI | SPTBN1; SPTBN | Q01082; A0/ | SPTB2_HUMAN | Unmodified | 0 | 0 | 0 | 0 | 0 | 0 | 0 | 0 | 0 | 0 | 0 | 1652300 | 0 | 0 | 0 | 0 | 0 | 0 |
| QLYEFDIK | GNS; GNS; GNS | P15586; H7( | GNS_HUMAN N | Unmodified | 0 | 0 | 0 | 0 | 0 | 0 | 0 | 0 | 1259300 | 0 | 0 | 0 | 0 | 0 | 0 | 0 | 0 | 0 |
| QLYEMVLT | TPP2; TPP2; TPI | P29144; A0A | TPP2_HUMAN | Oxidation (M) | 0 | 0 | 0 | 0 | 0 | 0 | 57682 | 31523 | 0 | 0 | 22881 | 1080000 | 121380 | 143430 | 0 | 0 | 0 | 264940 |
| QLYETTDTI | XPO7; XPO7; XF | Q9UIA9; H0Y | XPO7_HUMAN | Unmodified | 0 | 0 | 0 | 0 | 0 | 0 | 0 | 0 | 0 | 0 | 0 | 824830 | 0 | 0 | 0 | 0 | 0 | 0 |
| QLYETTDTI | XPO7; XPO7; XF | Q9UIA9; H0Y | XPO7_HUMAN | Unmodified | 38572 | 0 | 0 | 0 | 1667300 | 0 | 330550 | 71265 | 0 | 231550 | 214100 | 0 | 402040 | 0 | 0 | 0 | 514250 | 492400 |
| QLYLERVLI | ACTR2; ACTR2 | P61160; F5H | ARP2_HUMAN | Unmodified | 0 | 0 | 0 | 53377 | 1564900 | 455070 | 539310 | 0 | 0 | 332750 | 0 | 0 | 738530 | 658000 | 0 | 0 | 731270 | 454950 |
| QLYLPMLFI | XPO1; XPO1; XF | O14980; A0/ | XPO1_HUMAN | Oxidation (M) | 0 | 0 | 0 | 0 | 0 | 0 | 0 | 0 | 0 | 0 | 0 | 2972400 | 0 | 0 | 0 | 0 | 0 | 0 |
| QLYQTLTD | CSNK2A3; CSN | Q8NEV1; P6 | CSK23_HUMAN | Unmodified | 0 | 0 | 0 | 0 | 0 | 0 | 0 | 0 | 0 | 0 | 0 | 0 | 0 | 0 | 0 | 0 | 0 | 0 |
| QLYRPEPLI | DCHS1 | Q96JQ0 | PCD16_HUMAI | Unmodified | 62457 | 0 | 0 | 0 | 352870 | 0 | 0 | 0 | 559210 | 0 | 0 | 0 | 0 | 0 | 0 | 0 | 0 | 0 |
| QLYVLGHE | UBA1; UBA1; UI | P22314; Q5J | UBA1_HUMAN | Unmodified | 0 | 0 | 0 | 0 | 0 | 0 | 0 | 0 | 522700 | 0 | 0 | 1366300 | 0 | 0 | 0 | 0 | 0 | 0 |
| QMADTGKI | HNRNPU; HNR | Q00839; A0/ | HNRPU_HUMA | Oxidation (M) | 0 | 0 | 0 | 0 | 0 | 0 | 3082700 | 1122400 | 0 | 547050 | 0 | 0 | 1115300 | 1386000 | 0 | 0 | 0 | 0 |
| QMCMIMM | CTNNA1; CTNN | P35221; P26 | CTNA1_HUMAI | 3 Oxidation (M) | 0 | 0 | 0 | 0 | 380240 | 0 | 0 | 0 | 0 | 0 | 0 | 0 | 414930 | 599960 | 0 | 0 | 0 | 0 |
| QMDTNND | HPCA; HPCAL1 | P84074; P37 | HPCA_HUMAN | Oxidation (M) | 0 | 0 | 110350 | 153960 | 0 | 0 | 0 | 0 | 430940 | 0 | 242800 | 1036700 | 0 | 0 | 0 | 0 | 0 | 0 |
| QMEGICVK | CRELD1; CRELI | Q96HD1; A0 | CREL1_HUMAN | Unmodified | 0 | 0 | 0 | 0 | 0 | 0 | 0 | 0 | 692710 | 0 | 300840 | 0 | 0 | 0 | 0 | 0 | 0 | 0 |
| QMEKDETV | CAPZB; CAPZB; | P47756; B1A | CAPZB_HUMAN | Unmodified | 464350 | 322150 | 0 | 224140 | 0 | 191930 | 0 | 0 | 862130 | 0 | 297230 | 5292500 | 0 | 164970 | 341220 | 203390 | 0 | 0 |
| QMEKDETV | CAPZB; CAPZB; | P47756; B1A | CAPZB_HUMAN | Oxidation (M) | 787270 | 375390 | 0 | 0 | 0 | 0 | 0 | 0 | 0 | 0 | 0 | 2251600 | 0 | 0 | 382460 | 377210 | 0 | 0 |
| QMELLENK | HTATIP2; HTATI | Q9BUP3; E9I | HTAI2_HUMAN | Unmodified | 0 | 0 | 0 | 0 | 648510 | 0 | 0 | 10082 | 2710000 | 582870 | 39479 | 0 | 0 | 0 | 0 | 0 | 0 | 0 |
| QMENGML | ABCC1; ABCC1 | P33527; I3L4 | MRP1_HUMAN | Unmodified | 0 | 0 | 0 | 0 | 664890 | 0 | 0 | 0 | 0 | 0 | 0 | 0 | 0 | 0 | 0 | 0 | 0 | 0 |
| QMEQISQF | TAGLN2; TAGLN | P37802; X6R | TAGL2_HUMAN | Oxidation (M) | 4891700 | 2373100 | 1207100 | 1023900 | 0 | 190730 | 0 | 0 | 3300300 | 0 | 891720 | 10512000 | 0 | 0 | 2310400 | 1660200 | 0 | 0 |
| QMEQISQF | TAGLN2; TAGLN | P37802; X6R | TAGL2_HUMAN | Unmodified | 119910 | 91140 | 0 | 219100 | 458420 | 0 | 0 | 0 | 2045200 | 0 | 139980 | 4356500 | 169890 | 0 | 0 | 0 | 0 | 0 |
| QMERAMLE | MSN | P26038 | MOES_HUMAN | Oxidation (M) | 0 | 140280 | 0 | 0 | 11171000 | 3272900 | 0 | 0 | 0 | 0 | 0 | 0 | 968970 | 1341800 | 0 | 18384 | 2429200 | 2384300 |
| QMERAMLE | MSN | P26038 | MOES_HUMAN | Unmodified | 0 | 0 | 0 | 33350 | 21651000 | 3909300 | 0 | 0 | 0 | 0 | 9632.7 | 87180 | 0 | 210410 | 0 | 0 | 6493900 | 9634400 |
| QMERAMLE | MSN | P26038 | MOES_HUMAN | Unmodified | 0 | 0 | 0 | 53635 | 5184600 | 1230000 | 296400 | 286260 | 0 | 0 | 0 | 0 | 0 | 166230 | 0 | 0 | 233620 | 1895800 |
| QMESELEA | CDC42BPB | Q9Y5S2 | MRCKB_HUMA | Unmodified | 0 | 0 | 0 | 49948 | 577670 | 0 | 0 | 0 | 221790 | 43295 | 0 | 0 | 0 | 0 | 0 | 0 | 158070 | 488180 |
| QMETGLSP | MAN1B1; MAN1 | Q9UKM7; H( | MA1B1_HUMAI | Oxidation (M) | 0 | 0 | 0 | 0 | 0 | 0 | 1335300 | 0 | 6044800 | 2170800 | 0 | 625060 | 0 | 0 | 97929 | 0 | 0 | 0 |
| QMETGLSP | MAN1B1; MAN1 | Q9UKM7; H( | MA1B1_HUMAI | Unmodified | 0 | 0 | 0 | 0 | 0 | 0 | 6711100 | 0 | 8263700 | 0 | 0 | 0 | 0 | 0 | 0 | 0 | 0 | 0 |
| QMEVAQAI | DCTN1; DCTN1 | Q14203; Q6/ | DCTN1_HUMAI | Unmodified | 0 | 0 | 0 | 0 | 0 | 0 | 0 | 0 | 0 | 0 | 0 | 0 | 0 | 0 | 0 | 0 | 0 | 0 |
| QMFAQMY | ANXA7 | P20073 | ANXA7_HUMAN | Oxidation (M) | 773320 | 237910 | 210840 | 0 | 2329500 | 0 | 596310 | 390840 | 12968000 | 2230000 | 2568400 | 16134000 | 0 | 779700 | 740650 | 65731 | 915090 | 602220 |
| QMFAQMY | ANXA7 | P20073 | ANXA7_HUMAN | 2 Oxidation (M) | 0 | 0 | 45207 | 0 | 0 | 0 | 0 | 0 | 1399300 | 272740 | 0 | 0 | 0 | 0 | 0 | 0 | 59884 | 252350 |
| QMFAQMY | ANXA7 | P20073 | ANXA7_HUMAN | Unmodified | 997990 | 83231 | 0 | 0 | 179230 | 0 | 0 | 0 | 12410000 | 1246700 | 0 | 0 | 0 | 0 | 90710 | 0 | 894440 | 453680 |
| QMFLATWI | AP1B1 | Q10567 | AP1B1_HUMAN | Oxidation (M) | 0 | 0 | 0 | 63110 | 0 | 0 | 0 | 0 | 0 | 0 | 0 | 425480 | 0 | 0 | 0 | 0 | 0 | 0 |
| QMFLATWI | AP1B1 | Q10567 | AP1B1_HUMAN | Unmodified | 0 | 0 | 0 | 0 | 233910 | 0 | 16904 | 0 | 62388 | 0 | 0 | 145680 | 0 | 14685 | 0 | 11591 | 0 | 0 |
| QMGFPLIY | NPEPPS; NPEPI | P55786; A0A | PSA_HUMAN P | Oxidation (M) | 0 | 0 | 0 | 0 | 6549800 | 0 | 0 | 0 | 0 | 845220 | 0 | 0 | 3804200 | 2951700 | 0 | 0 | 6094600 | 8778000 |
| QMGSGDA | TPP2; TPP2; TPI | P29144; A0A | TPP2_HUMAN | Unmodified | 0 | 0 | 0 | 0 | 0 | 0 | 0 | 0 | 0 | 0 | 0 | 959230 | 0 | 0 | 0 | 0 | 0 | 0 |
| QMGSYSCI | EMB; EMB | Q6PCB8; D6 | EMB_HUMAN E | Unmodified | 0 | 0 | 0 | 0 | 1128700 | 274810 | 0 | 0 | 1095500 | 0 | 0 | 0 | 0 | 0 | 0 | 60839 | 0 | 0 |
| QMIAVADE | SDF4; SDF4 | Q9BRK5; A0/ | CAB45_HUMAN | Oxidation (M) | 0 | 0 | 0 | 0 | 0 | 0 | 0 | 0 | 11741000 | 1163400 | 1499200 | 0 | 600600 | 451180 | 0 | 0 | 0 | 0 |
| QMIAVADE | SDF4; SDF4 | Q9BRK5; A0/ | CAB45_HUMAN | Unmodified | 0 | 0 | 0 | 0 | 0 | 0 | 0 | 0 | 29371000 | 1700300 | 298500 | 0 | 0 | 0 | 0 | 0 | 0 | 146420 |
| QMIAVADE | SDF4; SDF4 | Q9BRK5; A0/ | CAB45_HUMAN | Oxidation (M) | 0 | 0 | 0 | 0 | 0 | 0 | 598140 | 0 | 433080 | 7003300 | 0 | 0 | 1464600 | 1352300 | 0 | 0 | 0 | 1363900 |
| QMIGYNLA | SUCLG2; SUCL | Q96I99; E9P | SUCB2_HUMAI | Unmodified | 0 | 0 | 0 | 0 | 0 | 0 | 360580 | 138280 | 869600 | 0 | 0 | 0 | 0 | 0 | 0 | 0 | 0 | 0 |
| QMKQEINT | ROCK1; ROCK1 | Q13464; A0/ | ROCK1_HUMAI | Unmodified | 0 | 0 | 0 | 0 | 898300 | 114990 | 161760 | 20443 | 0 | 175830 | 0 | 0 | 411600 | 0 | 0 | 0 | 652880 | 681550 |
| QMLFDLES | VPS28 | Q9UK41 | VPS28_HUMAN | Oxidation (M) | 4369300 | 0 | 0 | 0 | 0 | 0 | 0 | 0 | 0 | 0 | 0 | 0 | 0 | 0 | 11386000 | 3880600 | 0 | 0 |
| QMLGERLF | PABPC1; PABP( | P11940; Q9I | PABP1_HUMAN | 2 Oxidation (M) | 511520 | 954600 | 560240 | 0 | 0 | 0 | 0 | 0 | 0 | 0 | 0 | 0 | 382610 | 0 | 259390 | 220620 | 0 | 0 |
| QMLPIVLKI | PPP2R1B; PPP2 | P30154; H0Y | 2AAB_HUMAN | Oxidation (M) | 0 | 0 | 0 | 0 | 3225300 | 544510 | 0 | 0 | 364220 | 0 | 0 | 119130 | 287790 | 0 | 0 | 0 | 948990 | 395840 |
| QMLPLNTN | XPO1; XPO1; XF | O14980; A0/ | XPO1_HUMAN | Oxidation (M) | 0 | 0 | 0 | 0 | 0 | 0 | 0 | 0 | 0 | 0 | 0 | 1611500 | 0 | 0 | 0 | 0 | 0 | 0 |
| QMLRNGLI | N/A | N/A | N/A | Oxidation (M) | 0 | 0 | 0 | 0 | 0 | 0 | 0 | 2725300 | 0 | 0 | 0 | 0 | 0 | 0 | 0 | 0 | 0 | 0 |
| QMMGGFK | PACSIN2; PACS | Q9UNF0; B0 | PACN2_HUMAI | Unmodified | 398690 | 2355000 | 0 | 1036700 | 0 | 0 | 0 | 0 | 0 | 0 | 30692 | 0 | 936560 | 0 | 0 | 0 | 0 | 0 |
| QMMGGFK | PACSIN2; PACS | Q9UNF0; B0 | PACN2_HUMAI | Oxidation (M) | 0 | 0 | 0 | 0 | 0 | 0 | 0 | 0 | 0 | 0 | 0 | 0 | 0 | 0 | 0 | 0 | 0 | 0 |
| QMMIDEFN | KIF23; KIF23; KI | Q02241; A0/ | KIF23_HUMAN | Oxidation (M) | 0 | 0 | 1093000 | 424940 | 1697500 | 301310 | 0 | 0 | 347950 | 83164 | 177830 | 1214100 | 0 | 0 | 0 | 0 | 71344 | 0 |
| QMNIPGGI | TES | Q9UGI8 | TES_HUMAN Te | Oxidation (M) | 0 | 0 | 0 | 0 | 717970 | 195050 | 0 | 0 | 0 | 0 | 0 | 450140 | 1359600 | 380660 | 128570 | 87907 | 319410 | 1301900 |
| QMNIPGGI | TES | Q9UGI8 | TES_HUMAN Te | Unmodified | 0 | 0 | 0 | 0 | 1454700 | 236120 | 0 | 0 | 384470 | 155220 | 0 | 0 | 1126700 | 537780 | 0 | 0 | 500810 | 2418200 |
| QMPEPEQI | DIAPH1; DIAPH | O60610; A0/ | DIAP1_HUMAN | Unmodified | 0 | 0 | 0 | 0 | 707370 | 0 | 0 | 0 | 0 | 190100 | 533540 | 0 | 763470 | 719930 | 0 | 105520 | 395720 | 434530 |
| QMQEAEMI | SEL1L | Q9UBV2 | SE1L1_HUMAN | 4 Oxidation (M) | 0 | 0 | 1479800 | 0 | 0 | 1932500 | 0 | 0 | 0 | 0 | 0 | 0 | 0 | 0 | 339910 | 0 | 0 | 1337300 |
| QMQVLHP | CCT4 | P50991 | TCPD_HUMAN | Unmodified | 0 | 0 | 0 | 0 | 328260 | 0 | 68141 | 17815 | 277390 | 167130 | 0 | 1160100 | 1214100 | 273670 | 0 | 0 | 280410 | 0 |
| QMQVSLTS | F2RL1 | P55085 | PAR2_HUMAN | Unmodified | 0 | 0 | 0 | 0 | 0 | 0 | 0 | 0 | 0 | 0 | 0 | 0 | 104630 | 0 | 0 | 0 | 111160 | 401470 |
| QMSCLME | GLG1 | Q92896 | GSLG1_HUMAI | Unmodified | 0 | 0 | 0 | 0 | 0 | 0 | 0 | 0 | 2728600 | 0 | 0 | 0 | 0 | 0 | 0 | 0 | 0 | 0 |
| QMSGAQIK | PCBP1; PCBP3; | Q15365; P57 | PCBP1_HUMAN | Oxidation (M) | 712870 | 680510 | 985880 | 698240 | 38500 | 9547.5 | 71584 | 0 | 240520 | 0 | 9188.8 | 328550 | 171450 | 56138 | 633770 | 1091100 | 26841 | 0 |
| QMSVKEDI | DYNC1H1; DYN | Q14204; A0/ | DYHC1_HUMA | Oxidation (M) | 0 | 0 | 0 | 0 | 0 | 0 | 0 | 0 | 872350 | 821530 | 174410 | 5511100 | 2399700 | 0 | 0 | 0 | 0 | 1839100 |
| QMTEAIGP | MVP | Q14764 | MVP_HUMAN N | Unmodified | 0 | 0 | 0 | 0 | 0 | 0 | 0 | 0 | 0 | 0 | 0 | 2361400 | 0 | 0 | 0 | 0 | 0 | 0 |
| QMTEAIGP | MVP | Q14764 | MVP_HUMAN N | 2 Oxidation (M) | 0 | 0 | 0 | 0 | 0 | 0 | 0 | 0 | 0 | 0 | 0 | 8854800 | 0 | 0 | 0 | 0 | 0 | 0 |
| QMTEAIGP | MVP | Q14764 | MVP_HUMAN N | Oxidation (M) | 0 | 0 | 0 | 0 | 0 | 0 | 0 | 0 | 992920 | 0 | 0 | 1898500 | 0 | 0 | 0 | 0 | 0 | 0 |
| QMTENLER | PLCB2; PLCB2 | Q00722; H0Y | PLCB2_HUMAN | Unmodified | 0 | 0 | 0 | 0 | 0 | 0 | 0 | 0 | 0 | 119470 | 70776 | 222560 | 0 | 0 | 0 | 0 | 0 | 0 |
| QMVEHGG | DYNC1H1; DYN | Q14204; A0/ | DYHC1_HUMA | Oxidation (M) | 0 | 0 | 0 | 92746 | 0 | 0 | 0 | 0 | 0 | 0 | 0 | 541640 | 0 | 0 | 0 | 0 | 0 | 0 |
| QMVEHGG | DYNC1H1; DYN | Q14204; A0/ | DYHC1_HUMA | Unmodified | 0 | 0 | 0 | 0 | 0 | 0 | 0 | 0 | 0 | 0 | 0 | 1385400 | 0 | 0 | 0 | 0 | 0 | 0 |
| QMVEILHK | GLRX3 | O76003 | GLRX3_HUMAN | Unmodified | 0 | 0 | 0 | 0 | 736090 | 0 | 0 | 0 | 0 | 0 | 0 | 523850 | 161050 | 87963 | 0 | 0 | 170790 | 0 |
| QMVETELK | YWHAE; YWHAI | P62258; K7E | 1433E_HUMAN | Unmodified | 703570 | 0 | 2905000 | 3084000 | 0 | 60957 | 0 | 0 | 3344500 | 1040700 | 3150300 | 9252900 | 0 | 0 | 5256600 | 1124900 | 291970 | 0 |
| QMVETELK | YWHAE; YWHAI | P62258; K7E | 1433E_HUMAN | Oxidation (M) | 732510 | 692350 | 3177300 | 1329400 | 0 | 0 | 0 | 0 | 0 | 0 | 0 | 7727400 | 0 | 0 | 4024100 | 3642500 | 0 | 0 |
| QMVIDVLH | RPS24; RPS24; | P62847; A0A | RS24_HUMAN | Oxidation (M) | 2568400 | 1230500 | 2075100 | 1157500 | 285040 | 189500 | 511070 | 441590 | 0 | 0 | 1603700 | 8065900 | 746290 | 651560 | 858710 | 2145500 | 79503 | 321620 |
| QMVIDVLH | RPS24; RPS24; | P62847; A0A | RS24_HUMAN | Unmodified | 555200 | 0 | 0 | 317270 | 768360 | 0 | 0 | 0 | 803870 | 123740 | 816820 | 9255000 | 579200 | 421950 | 0 | 143260 | 76207 | 489790 |
| QMVQVSD | PLXNB2 | O15031 | PLXB2_HUMAN | Oxidation (M) | 152570 | 186570 | 752650 | 365710 | 0 | 0 | 0 | 0 | 624110 | 267750 | 0 | 4566800 | 0 | 0 | 3367700 | 1771100 | 0 | 0 |
| QMVQVSD | PLXNB2 | O15031 | PLXB2_HUMAN | 2 Oxidation (M) | 255510 | 82400 | 408100 | 314560 | 0 | 0 | 83139 | 0 | 67994 | 0 | 0 | 1184700 | 0 | 0 | 1182400 | 1280500 | 0 | 0 |
| QMVQVSD | PLXNB2 | O15031 | PLXB2_HUMAN | Unmodified | 0 | 0 | 0 | 111220 | 0 | 0 | 0 | 0 | 0 | 0 | 0 | 805310 | 0 | 0 | 330500 | 125070 | 0 | 0 |
| QMYPELQI | APLP1; APLP1; | P51693; B7Z | APLP1_HUMAN | Unmodified | 0 | 0 | 0 | 0 | 0 | 0 | 0 | 0 | 1633400 | 0 | 0 | 0 | 0 | 0 | 0 | 0 | 0 | 0 |
| QNAHFSTI | LLGL2; LLGL2 | Q6P1M3; J3( | L2GL2_HUMAN | Oxidation (M) | 0 | 0 | 0 | 0 | 0 | 0 | 0 | 0 | 0 | 0 | 0 | 1010700 | 0 | 0 | 1568300 | 0 | 0 | 0 |
| QNALLFAEI | VGF | O15240 | VGF_HUMAN N | Unmodified | 647430 | 855660 | 0 | 0 | 0 | 0 | 0 | 0 | 0 | 0 | 0 | 0 | 0 | 0 | 0 | 0 | 0 | 0 |
| QNALLFAEI | VGF | O15240 | VGF_HUMAN N | Unmodified | 0 | 95871 | 0 | 0 | 0 | 0 | 0 | 0 | 4210000 | 0 | 0 | 0 | 0 | 0 | 0 | 0 | 0 | 0 |
| QNAQCLH | DDX50 | Q9BQ39 | DDX50_HUMAI | Unmodified | 0 | 0 | 0 | 0 | 0 | 0 | 200440 | 0 | 0 | 0 | 0 | 0 | 0 | 0 | 0 | 0 | 0 | 0 |
| QNASFLYE | EXOC8 | Q8IYI6 | EXOC8_HUMAI | Unmodified | 0 | 0 | 0 | 0 | 0 | 0 | 0 | 0 | 0 | 0 | 0 | 684060 | 0 | 0 | 0 | 0 | 0 | 0 |
| QNAYIATQ | PTPRF; PTPRF; I | P10586; H0Y | PTPRF_HUMAN | Unmodified | 585900 | 279180 | 0 | 60762 | 123810 | 81614 | 0 | 0 | 298790 | 100300 | 0 | 0 | 20705 | 0 | 514190 | 76240 | 0 | 0 |
| QNAYIATQ | PTPRF; PTPRF; I | P10586; H0Y | PTPRF_HUMAN | Oxidation (M) | 557640 | 939330 | 0 | 0 | 349950 | 0 | 0 | 0 | 0 | 158000 | 0 | 0 | 0 | 0 | 1654700 | 1890100 | 0 | 0 |
| QNCSQHE | IL6ST | P40189 | IL6RB_HUMAN | Unmodified | 0 | 0 | 0 | 0 | 460470 | 0 | 0 | 0 | 0 | 0 | 0 | 0 | 0 | 179460 | 0 | 0 | 151320 | 243100 |
| QNCVTELA | P3H1 | Q32P28 | P3H1_HUMAN | Unmodified | 0 | 0 | 0 | 0 | 101240 | 0 | 0 | 0 | 1357700 | 61542 | 0 | 0 | 0 | 0 | 0 | 0 | 0 | 0 |
| QNDEEREL | GOLM2 | H0YM51 | H0YM51_HUM | Oxidation (M) | 0 | 0 | 0 | 0 | 0 | 0 | 0 | 0 | 360510 | 85222 | 0 | 0 | 0 | 0 | 0 | 0 | 0 | 0 |
| QNDQVSF | KTN1; KTN1; KT | Q86UP2; H0 | KTN1_HUMAN | Unmodified | 0 | 0 | 0 | 0 | 0 | 0 | 0 | 0 | 0 | 0 | 0 | 0 | 0 | 0 | 0 | 0 | 0 | 0 |
| QNDVFGE | GDI1 | P31150 | GDIA_HUMAN | Unmodified | 81932 | 0 | 0 | 0 | 0 | 0 | 154620 | 194690 | 316870 | 0 | 0 | 439470 | 348210 | 0 | 100070 | 136270 | 408030 | 0 |
| QNEFSVVR | SNX6; SNX6; SI | Q9UNH7; G | SNX6_HUMAN | Unmodified | 746670 | 478100 | 195170 | 0 | 0 | 0 | 0 | 0 | 223190 | 0 | 111090 | 2143200 | 0 | 0 | 456550 | 179160 | 0 | 0 |
| QNEKEEQ | EEA1 | Q15075 | EEA1_HUMAN I | Unmodified | 0 | 0 | 0 | 149250 | 1553800 | 0 | 0 | 0 | 186100 | 0 | 0 | 0 | 0 | 0 | 157970 | 70164 | 64696 | 0 |
| QNEQLLK | PLCB4 | A0A7P0MRI8 | A0A7P0MRI8_H | Unmodified | 0 | 0 | 0 | 0 | 158870 | 24539 | 0 | 11930 | 0 | 24842 | 17866 | 237690 | 240740 | 0 | 0 | 343300 | 514200 | 960830 |
| QNETTAMK | CALCOCO2; C/ | Q13137; D6I | CACO2_HUMA | Unmodified | 0 | 0 | 0 | 0 | 0 | 0 | 322980 | 0 | 0 | 0 | 0 | 0 | 158330 | 0 | 0 | 0 | 155340 | 0 |
| QNFALGK | APPL2; APPL2; | Q8NEU8; F8 | DP13B_HUMAN | Unmodified | 0 | 59817 | 0 | 4308100 | 0 | 0 | 0 | 0 | 0 | 0 | 0 | 0 | 3235800 | 5167900 | 0 | 0 | 970730 | 0 |
| QNFIDPLQ | SH3GL1; SH3G | Q99961; M0 | SH3G1_HUMAI | Unmodified | 16718000 | 5804400 | 7574600 | 6945300 | 430470 | 88530 | 0 | 0 | 0 | 0 | 0 | 295990 | 8374900 | 2651900 | 11554000 | 7135800 | 47400 | 0 |
| QNFIDPLQ | SH3GL1; SH3G | Q99961; M0 | SH3G1_HUMAI | Unmodified | 0 | 0 | 0 | 0 | 0 | 0 | 0 | 0 | 0 | 0 | 0 | 130180 | 879630 | 0 | 0 | 0 | 0 | 0 |
| QNFIDPLQ | SH3GL2 | Q99962 | SH3G2_HUMAI | Unmodified | 907000 | 1123200 | 0 | 0 | 0 | 0 | 0 | 0 | 3839600 | 609310 | 0 | 0 | 0 | 0 | 0 | 0 | 0 | 0 |
| QNFLLFDM | GOLPH3 | Q9H4A6 | GOLP3_HUMAI | Unmodified | 344570 | 943380 | 0 | 108840 | 610260 | 0 | 0 | 0 | 0 | 0 | 46793 | 360860 | 520860 | 511280 | 203080 | 0 | 0 | 446950 |
| QNFLLFDM | GOLPH3 | Q9H4A6 | GOLP3_HUMAI | Oxidation (M) | 0 | 1836500 | 172670 | 1631600 | 0 | 0 | 0 | 0 | 0 | 0 | 0 | 1112500 | 674970 | 0 | 0 | 369720 | 0 | 0 |
| QNFLLFDM | GOLPH3L; GOL | Q9H4A5; Q5 | GLP3L_HUMAN | Oxidation (M) | 64380 | 89352 | 353800 | 277240 | 297890 | 0 | 53831 | 0 | 0 | 172490 | 78949 | 327420 | 0 | 0 | 689010 | 0 | 286940 | 471260 |
| QNFLLFDM | GOLPH3L; GOL | Q9H4A5; Q5 | GLP3L_HUMAN | Unmodified | 27216 | 0 | 30073 | 0 | 603470 | 0 | 0 | 0 | 175130 | 101710 | 0 | 256590 | 397320 | 247280 | 80439 | 37316 | 453440 | 1125200 |
| QNGSFLDF | EPB41 | A0A2R8Y783 | A0A2R8Y783_H | Unmodified | 0 | 0 | 0 | 0 | 0 | 0 | 0 | 0 | 139080 | 136500 | 27075 | 0 | 348720 | 342050 | 0 | 31257 | 0 | 0 |
| QNHESVQ | DHX15 | O43143 | DHX15_HUMAI | Unmodified | 0 | 0 | 0 | 0 | 0 | 0 | 0 | 0 | 0 | 0 | 0 | 879770 | 0 | 0 | 0 | 0 | 0 | 0 |
| QNHLQADI | CRIM1 | Q9NZV1 | CRIM1_HUMAN | Unmodified | 0 | 0 | 0 | 0 | 719840 | 0 | 147340 | 44804 | 0 | 0 | 0 | 0 | 0 | 0 | 0 | 0 | 30868 | 111380 |
| QNKGIEELI | RAB21 | Q9UL25 | RAB21_HUMAN | Unmodified | 0 | 0 | 0 | 0 | 588610 | 94151 | 0 | 22135 | 0 | 401560 | 287350 | 170150 | 925100 | 638030 | 0 | 0 | 530590 | 610140 |
| QNKPENW | DIPK2A; DIPK2/ | Q8NDZ4; E7 | DIK2A_HUMAN | Unmodified | 0 | 0 | 0 | 0 | 0 | 0 | 0 | 0 | 2408300 | 119220 | 0 | 165280 | 0 | 0 | 0 | 0 | 0 | 0 |
| QNLDALLN | DYNC1H1; DYN | Q14204; A0/ | DYHC1_HUMA | Unmodified | 0 | 0 | 146660 | 0 | 0 | 0 | 0 | 0 | 197120 | 0 | 0 | 2717100 | 0 | 0 | 0 | 0 | 0 | 0 |
| QNLFQEAE | PSME2; PSME2 | Q9UL46; A0/ | PSME2_HUMAI | Unmodified | 32826 | 0 | 0 | 0 | 0 | 0 | 0 | 0 | 113400 | 73076 | 17375 | 163230 | 0 | 0 | 0 | 0 | 13677 | 0 |
| QNLFTTWS | DYNC1H1; DYN | Q14204; A0/ | DYHC1_HUMA | Unmodified | 0 | 0 | 0 | 0 | 8419700 | 64494000 | 0 | 0 | 306250 | 0 | 101500 | 1942300 | 0 | 0 | 228900 | 0 | 0 | 20102000 |

| | | | | | | | | | | | | | | | | | | | | | | |
|---|---|---|---|---|---|---|---|---|---|---|---|---|---|---|---|---|---|---|---|---|---|---|
| QNLLSVGD | SMARCA5 | O60264 | SMCA5_HUMA | Unmodified | 681920 | 0 | 0 | 0 | 0 | 1153100 | 738460 | 552770 | 0 | 0 | 0 | 0 | 0 | 352050 | 0 | 0 | 0 | 280330 |
| QNLPLSSG | MPST; MPST | P25325; B1A | THTM_HUMAN | Unmodified | 0 | 0 | 0 | 0 | 0 | 0 | 0 | 0 | 1387700 | 0 | 0 | 2060600 | 0 | 0 | 0 | 0 | 0 | 0 |
| QNLSKEELI | PHGDH; PHGD | O43175; A0A | SERA_HUMAN | Unmodified | 0 | 0 | 0 | 0 | 0 | 0 | 0 | 0 | 0 | 0 | 0 | 0 | 3708000 | 2438600 | 0 | 0 | 0 | 0 |
| QNLSQFEA | CPSF7; CPSF7; | Q8N684; F5I | CPSF7_HUMAN | Unmodified | 1111900 | 491340 | 224810 | 298910 | 0 | 0 | 0 | 0 | 340370 | 0 | 244740 | 262580 | 0 | 0 | 573350 | 167520 | 0 | 0 |
| QNLSQFEA | CPSF7; CPSF7; | Q8N684; F5I | CPSF7_HUMAN | Unmodified | 3350400 | 4880500 | 5402800 | 6229100 | 733660 | 0 | 0 | 0 | 0 | 0 | 0 | 1275300 | 1680100 | 443260 | 6121700 | 3658700 | 0 | 476250 |
| QNNRCPL( | RNF7 | Q9UBF6 | RBX2_HUMAN | Unmodified | 0 | 0 | 0 | 0 | 0 | 0 | 0 | 0 | 0 | 0 | 0 | 0 | 0 | 0 | 0 | 0 | 1349500 | 0 |
| QNPQIDHA | ERBIN | Q96RT1 | ERBIN_HUMAN | Unmodified | 0 | 0 | 0 | 0 | 3470500 | 0 | 0 | 0 | 0 | 0 | 0 | 0 | 365480 | 0 | 0 | 0 | 420980 | 1260300 |
| QNPQNIGI | ZNF638 | Q14966 | ZN638_HUMAN | Unmodified | 0 | 0 | 0 | 0 | 0 | 0 | 0 | 0 | 0 | 0 | 1890100 | 0 | 0 | 0 | 0 | 0 | 0 | 0 |
| QNPSTREA | MCAM | P43121 | MUC18_HUMAN | Unmodified | 0 | 0 | 0 | 0 | 1061600 | 398800 | 0 | 0 | 0 | 0 | 0 | 0 | 0 | 0 | 0 | 0 | 1392600 | 6060700 |
| QNQTTSAV | RANBP2 | P49792 | RBP2_HUMAN | Unmodified | 131910 | 48077 | 0 | 51649 | 0 | 95300 | 0 | 0 | 67848 | 0 | 135870 | 1251100 | 0 | 365000 | 0 | 0 | 0 | 0 |
| QNQTTSAV | RANBP2 | P49792 | RBP2_HUMAN | Unmodified | 0 | 0 | 0 | 0 | 0 | 0 | 0 | 0 | 0 | 0 | 0 | 0 | 698020 | 0 | 0 | 0 | 0 | 0 |
| QNRFEIVYI | NDUFS3 | O75489 | NDUS3_HUMA | Unmodified | 0 | 0 | 0 | 0 | 0 | 0 | 214380 | 0 | 131170 | 0 | 0 | 0 | 0 | 0 | 0 | 0 | 0 | 0 |
| QNRYDGQ | UBA1 | P22314 | UBA1_HUMAN | Unmodified | 0 | 0 | 158200 | 0 | 1187900 | 151090 | 115590 | 37278 | 107770 | 468570 | 0 | 347730 | 1332500 | 303780 | 0 | 122980 | 0 | 824990 |
| QNSDMQA | CALCOCO2; CA | Q13137; D6I | CACO2_HUMA | Unmodified | 0 | 0 | 0 | 0 | 0 | 0 | 240450 | 0 | 431040 | 0 | 0 | 0 | 0 | 0 | 0 | 0 | 0 | 716920 |
| QNSDMQA | CALCOCO2; CA | Q13137; D6I | CACO2_HUMA | Unmodified | 0 | 0 | 0 | 0 | 1715400 | 0 | 6953600 | 0 | 0 | 424400 | 0 | 0 | 1037200 | 653120 | 0 | 0 | 2846700 | 5507700 |
| QNSDMQA | CALCOCO2; CA | Q13137; D6I | CACO2_HUMA | Oxidation (M) | 0 | 0 | 0 | 0 | 0 | 0 | 1638400 | 0 | 0 | 206150 | 0 | 0 | 703340 | 208530 | 0 | 0 | 531200 | 1789200 |
| QNSDPTSE | TNIK | Q9UKE5 | TNIK_HUMAN | Unmodified | 133010 | 0 | 0 | 106650 | 0 | 0 | 0 | 83875 | 0 | 0 | 0 | 697810 | 0 | 0 | 0 | 0 | 0 | 0 |
| QNSQLPAC | ENAH; ENAH; E | Q8N8S7; A0 | ENAH_HUMAN | Unmodified | 0 | 0 | 1049600 | 905470 | 802730 | 0 | 0 | 0 | 0 | 0 | 0 | 20666000 | 817420 | 582360 | 2389600 | 1163600 | 602440 | 0 |
| QNSVQEQI | ANLN | Q9NQW6 | ANLN_HUMAN | Unmodified | 0 | 0 | 0 | 0 | 1485700 | 665150 | 0 | 79750 | 0 | 0 | 0 | 0 | 0 | 0 | 0 | 0 | 1134800 | 737520 |
| QNSVVLQK | COL12A1; COL | Q99715; D6I | COCA1_HUMA | Unmodified | 0 | 0 | 0 | 0 | 624680 | 0 | 385410 | 33452 | 1397900 | 229650 | 67598 | 1192500 | 59271 | 305370 | 0 | 213520 | 0 | 120440 |
| QNTEESTI( | SLC39A10 | Q9ULF5 | S39AA_HUMAN | Unmodified | 101520 | 0 | 583910 | 171730 | 0 | 0 | 0 | 0 | 0 | 0 | 0 | 2089300 | 32752 | 0 | 509660 | 414990 | 0 | 0 |
| QNTEESTI( | SLC39A10 | Q9ULF5 | S39AA_HUMAN | Unmodified | 0 | 65270 | 218680 | 0 | 0 | 0 | 0 | 0 | 184760 | 39173 | 0 | 1978400 | 0 | 0 | 0 | 0 | 53450 | 0 |
| QNTFLVLR | DARS2; DARS2; | Q6PI48; A0A | SYDM_HUMAN | Unmodified | 0 | 0 | 0 | 0 | 0 | 0 | 0 | 0 | 838690 | 0 | 0 | 0 | 0 | 0 | 0 | 0 | 0 | 0 |
| QNTTLGATI | PI4KA | P42356 | PI4KA_HUMAN | Unmodified | 0 | 0 | 211150 | 0 | 0 | 0 | 0 | 0 | 0 | 0 | 0 | 514280 | 0 | 0 | 341570 | 188130 | 1510900 | 1159000 |
| QNVAYEYL | IQGAP1; IQGAP | P46940; A0A | IQGA1_HUMAN | Unmodified | 0 | 0 | 0 | 0 | 1182700 | 0 | 0 | 0 | 1626000 | 0 | 3451800 | 2995400 | 0 | 0 | 0 | 0 | 0 | 0 |
| QNYHQDS | FTH1; FTH1; FT | P02794; E9P | FRIH_HUMAN | Unmodified | 0 | 0 | 0 | 0 | 0 | 0 | 0 | 0 | 7271400 | 0 | 0 | 0 | 0 | 0 | 0 | 0 | 43974 | 0 |
| QNYVDLVS | ECI2; ECI2; ECI | O75521; F8V | ECI2_HUMAN | Unmodified | 0 | 0 | 0 | 0 | 12209000 | 426100 | 788370 | 0 | 4207300 | 0 | 0 | 0 | 0 | 0 | 0 | 0 | 16114000 | 576810 |
| QPAAFCCC | N/A | N/A | N/A | Oxidation (M) | 0 | 0 | 0 | 0 | 13309000 | 1464200 | 3548100 | 256230 | 0 | 3135100 | 0 | 0 | 14627000 | 8370700 | 0 | 0 | 0 | 13354000 |
| QPAENVN( | UBA1; UBA1 | P22314; Q5J | UBA1_HUMAN | Unmodified | 0 | 282250 | 0 | 167670 | 0 | 31324 | 0 | 163180 | 888890 | 0 | 736500 | 6764300 | 0 | 0 | 444840 | 347940 | 0 | 0 |
| QPAGPGS( | SUMF2; SUMF2 | Q8NBJ7; C9J | SUMF2_HUMA | Unmodified | 47711 | 0 | 46055 | 0 | 0 | 0 | 0 | 0 | 13076000 | 109450 | 97016 | 359220 | 0 | 0 | 81674 | 59634 | 0 | 0 |
| QPAIISQLD | DHX9 | Q08211 | DHX9_HUMAN | Unmodified | 0 | 0 | 416040 | 0 | 0 | 0 | 0 | 0 | 0 | 0 | 396070 | 3584800 | 0 | 0 | 0 | 0 | 0 | 0 |
| QPALSAAC | APEH; APEH; AP | P13798; C9J | ACPH_HUMAN | Unmodified | 0 | 0 | 0 | 0 | 0 | 0 | 0 | 0 | 0 | 0 | 0 | 3324400 | 0 | 0 | 455140 | 208630 | 0 | 0 |
| QPAPKPEP | SIRPA | P78324 | SHPS1_HUMAN | Unmodified | 0 | 0 | 0 | 0 | 692660 | 231180 | 0 | 0 | 0 | 0 | 0 | 814980 | 257670 | 0 | 0 | 0 | 0 | 1126100 |
| QPDFISFGI | HSPG2; HSPG2 | P98160; A0A | PGBM_HUMAN | Unmodified | 0 | 0 | 340290 | 80912 | 0 | 209080 | 182570 | 0 | 0 | 230920 | 0 | 0 | 42239 | 49949 | 292460 | 207870 | 1647600 | 0 |
| QPDLEHAL | GFM2 | Q969S9 | RRF2M_HUMAN | Unmodified | 0 | 0 | 0 | 0 | 492930 | 0 | 0 | 0 | 0 | 147140 | 0 | 0 | 0 | 0 | 0 | 0 | 165330 | 846430 |
| QPDSGISS | RPN1; RPN1 | P04843; B7Z | RPN1_HUMAN | Unmodified | 0 | 1389200 | 0 | 305810 | 0 | 0 | 184770 | 0 | 22577000 | 3690000 | 852470 | 640390 | 0 | 0 | 0 | 0 | 0 | 0 |
| QPELPEVIA | CTNND1; CTNN | O60716; E9F | CTND1_HUMA | Oxidation (M) | 0 | 0 | 12634 | 0 | 0 | 0 | 0 | 0 | 0 | 0 | 0 | 0 | 0 | 0 | 147150 | 73334 | 0 | 0 |
| QPGGITNQ | FUT11 | Q495W5 | FUT11_HUMAN | Unmodified | 0 | 0 | 0 | 0 | 0 | 0 | 0 | 0 | 299430 | 0 | 0 | 0 | 0 | 0 | 0 | 0 | 0 | 0 |
| QPGNETAL | VBP1 | P61758 | PFD3_HUMAN | Unmodified | 0 | 0 | 0 | 0 | 0 | 0 | 0 | 0 | 1363400 | 0 | 527050 | 5649200 | 345720 | 0 | 383330 | 269870 | 0 | 0 |
| QPGPVTN( | SNAP23; SNAP: | O00161; H3I | SNP23_HUMAN | Unmodified | 0 | 0 | 853750 | 653560 | 988030 | 0 | 0 | 0 | 0 | 0 | 343000 | 3624700 | 0 | 409720 | 1634800 | 2006700 | 0 | 558620 |
| QPGRFDEV | MYO1C; MYO1( | O00159; F5I | MYO1C_HUMA | Unmodified | 1835400 | 3223500 | 2929800 | 0 | 0 | 0 | 0 | 0 | 786580 | 0 | 0 | 0 | 374380 | 0 | 13146000 | 10542000 | 0 | 0 |
| QPHYSAFG | EPHB4 | P54760 | EPHB4_HUMAN | Unmodified | 282290 | 215030 | 0 | 0 | 0 | 0 | 0 | 0 | 63992 | 493990 | 0 | 0 | 0 | 0 | 318400 | 139360 | 0 | 0 |
| QPILDAIEA | STK4; STK4 | Q13043; F5I | STK4_HUMAN | Unmodified | 79757 | 0 | 65234 | 65943 | 0 | 0 | 0 | 0 | 225660 | 0 | 65342 | 1930900 | 0 | 0 | 0 | 149760 | 0 | 0 |
| QPIPFFVET | SORCS2; SORC | Q96PQ0; B5 | SORC2_HUMA | Unmodified | 44801 | 5798300 | 0 | 4056700 | 125200 | 0 | 0 | 0 | 415750 | 0 | 0 | 0 | 0 | 0 | 0 | 0 | 176180 | 0 |
| QPLALNVA | ILF2; ILF2 | Q12905; B4I | ILF2_HUMAN | Unmodified | 0 | 0 | 144440 | 0 | 447390 | 0 | 0 | 0 | 0 | 0 | 0 | 2643200 | 0 | 0 | 296330 | 0 | 0 | 0 |
| QPLQKPSH | LARS1; LARS1; | Q9P2J5; A0A | SYLC_HUMAN | Unmodified | 0 | 0 | 0 | 0 | 0 | 0 | 0 | 0 | 0 | 0 | 0 | 776290 | 3786900 | 139720 | 0 | 0 | 0 | 0 |
| QPLSEASN | ANLN; ANLN | Q9NQW6; C | ANLN_HUMAN | Unmodified | 82637 | 65046 | 195960 | 189350 | 393560 | 114400 | 0 | 0 | 150590 | 0 | 0 | 348030 | 0 | 0 | 114240 | 127590 | 105840 | 168890 |
| QPLWNHLI | UPF1 | Q92900 | RENT1_HUMAN | Unmodified | 1003100 | 1072300 | 736030 | 0 | 0 | 0 | 0 | 0 | 0 | 0 | 0 | 0 | 0 | 0 | 0 | 0 | 0 | 0 |
| QPMNAAS( | SLC3A2; SLC3A | P08195; A0A | 4F2_HUMAN | Unmodified | 0 | 0 | 0 | 0 | 1461200 | 0 | 0 | 0 | 0 | 0 | 0 | 736470 | 0 | 0 | 0 | 0 | 163570 | 453090 |
| QPNEGSSS | MAGEA4; MAGE | P43358; C9J | MAGA4_HUMA | Unmodified | 347180 | 434910 | 0 | 0 | 0 | 0 | 0 | 0 | 0 | 0 | 0 | 0 | 0 | 0 | 0 | 46992 | 0 | 0 |
| QPPESLKLI | NDUFS3 | O75489 | NDUS3_HUMA | Unmodified | 0 | 0 | 0 | 0 | 0 | 0 | 0 | 0 | 318140 | 0 | 0 | 0 | 0 | 0 | 855510 | 0 | 0 | 0 |
| QPPPQTVP | SEC16A; SEC16 | O15027; F1T | SC16A_HUMAN | Acetyl (Protein N | 0 | 0 | 0 | 0 | 779050 | 0 | 4470200 | 1163200 | 0 | 0 | 0 | 0 | 0 | 3166100 | 0 | 0 | 1881700 | 2373400 |
| QPPTAAGR | DYNC1LI1; DYN | Q9Y6G9; E9I | DC1L1_HUMAN | Unmodified | 0 | 0 | 0 | 0 | 0 | 0 | 0 | 0 | 0 | 0 | 0 | 0 | 0 | 0 | 0 | 156730 | 0 | 0 |
| QPQAIITW | HSPG2 | P98160 | PGBM_HUMAN | Unmodified | 0 | 0 | 1178300 | 0 | 0 | 0 | 0 | 0 | 223910 | 0 | 0 | 0 | 0 | 0 | 0 | 404480 | 0 | 0 |
| QPQQFPSF | ADAM9; ADAM9 | Q13443; A0A | ADAM9_HUMA | Unmodified | 978370 | 948650 | 1734500 | 1455100 | 3610100 | 618790 | 0 | 0 | 312320 | 0 | 0 | 0 | 1023100 | 230040 | 2751500 | 2802200 | 170000 | 468900 |
| QPSQGPTF | MTHFD1; MTHF | P11586; V9C | C1TC_HUMAN | Unmodified | 126180 | 83682 | 267230 | 94728 | 0 | 73243000 | 32128 | 0 | 1072100 | 0 | 82621000 | 5206200 | 0 | 0 | 357690 | 487470 | 760250 | 0 |
| QPSSQDAI | PTK7; PTK7; PTI | Q13308; Q8I | PTK7_HUMAN | Unmodified | 0 | 0 | 0 | 0 | 373610 | 47938 | 0 | 0 | 613620 | 0 | 29291 | 0 | 0 | 0 | 95837 | 0 | 0 | 153550 |
| QPTIFQNK | RPS11; RPS11; | P62280; M0I | RS11_HUMAN | Unmodified | 1339000 | 1640200 | 1413100 | 950140 | 778720 | 195190 | 37955 | 224940 | 484290 | 0 | 447530 | 8704300 | 863060 | 200920 | 1518600 | 1358600 | 0 | 475590 |
| QPVLSQTE | PSMB4 | P28070 | PSB4_HUMAN | Unmodified | 0 | 0 | 0 | 0 | 304920 | 0 | 0 | 0 | 2079200 | 0 | 78408 | 5204700 | 55150 | 0 | 0 | 0 | 142750 | 201770 |
| QPVLSQTE | PSMB4 | P28070 | PSB4_HUMAN | Unmodified | 0 | 0 | 0 | 0 | 461480 | 0 | 0 | 0 | 1866400 | 937720 | 0 | 374850 | 252760 | 0 | 0 | 0 | 0 | 0 |
| QPYAVSEL | RPL4; RPL4; RP | P36578; H3I | RL4_HUMAN | Unmodified | 760030 | 1838400 | 0 | 0 | 0 | 0 | 0 | 0 | 0 | 0 | 135030 | 2120900 | 0 | 0 | 578440 | 387110 | 0 | 0 |
| QQAAQGD | UTRN | P46939 | UTRN_HUMAN | Unmodified | 0 | 0 | 355400 | 512360 | 0 | 0 | 0 | 0 | 0 | 17068 | 0 | 154950 | 0 | 0 | 473820 | 673370 | 0 | 0 |
| QQAASKEE | RABGAP1L; RAI | B7ZAP0; F5I | RBG10_HUMAI | Unmodified | 0 | 0 | 0 | 0 | 0 | 0 | 17638 | 0 | 0 | 98371 | 76454 | 452690 | 162590 | 0 | 0 | 0 | 0 | 0 |
| QQAEIKDT | CCDC88A; CCI | Q3V6T2; A0A | GRDN_HUMAN | Unmodified | 0 | 0 | 0 | 0 | 0 | 0 | 0 | 490800 | 0 | 0 | 0 | 956150 | 0 | 0 | 0 | 0 | 0 | 0 |
| QQANTIFW | EIF3B | P55884 | EIF3B_HUMAN | Unmodified | 0 | 0 | 0 | 290600 | 0 | 0 | 76290 | 0 | 0 | 0 | 0 | 103430 | 0 | 0 | 0 | 0 | 0 | 0 |
| QQASSWLI | SFRP5 | Q5T4F7 | SFRP5_HUMAN | Unmodified | 0 | 0 | 0 | 0 | 0 | 0 | 0 | 0 | 6976600 | 1230000 | 0 | 0 | 0 | 0 | 0 | 0 | 0 | 0 |
| QQASSWVI | SFRP1 | Q8N474 | SFRP1_HUMAN | Unmodified | 0 | 0 | 0 | 0 | 0 | 0 | 0 | 0 | 1278900 | 1201500 | 0 | 0 | 0 | 0 | 0 | 0 | 0 | 0 |
| QQCFSKDI | TRIM28; TRIM28 | Q13263; M0I | TIF1B_HUMAN | Oxidation (M) | 0 | 0 | 0 | 0 | 0 | 0 | 0 | 0 | 0 | 126290 | 0 | 0 | 91565 | 0 | 0 | 0 | 556490 | 0 |
| QQCFSKDI | TRIM28; TRIM28 | Q13263; M0I | TIF1B_HUMAN | Unmodified | 0 | 0 | 0 | 0 | 94319 | 0 | 152550 | 0 | 0 | 0 | 0 | 0 | 48052 | 46727 | 0 | 0 | 262030 | 283780 |
| QQDHFFQ/ | PLEK | P08567 | PLEK_HUMAN | Unmodified | 1393000 | 2455100 | 1315600 | 7952100 | 1246900 | 0 | 294290 | 0 | 0 | 0 | 0 | 0 | 2070100 | 766580 | 2278100 | 2459400 | 0 | 874340 |
| QQDQRQQ | TP53BP2; TP53I | Q13625; H7I | ASPP2_HUMAN | Unmodified | 0 | 0 | 0 | 0 | 346680 | 0 | 0 | 52060 | 0 | 62159 | 0 | 0 | 0 | 0 | 0 | 0 | 0 | 0 |
| QQDVLGFL | SH3BGRL | O75368 | SH3L1_HUMAN | Unmodified | 0 | 0 | 0 | 0 | 0 | 0 | 0 | 0 | 0 | 0 | 213000 | 841700 | 0 | 0 | 0 | 0 | 0 | 0 |
| QQEAPDLF | CRELD1; CRELI | Q96HD1; A0 | CREL1_HUMAN | Unmodified | 0 | 0 | 0 | 0 | 0 | 0 | 0 | 0 | 944910 | 170110 | 0 | 0 | 0 | 0 | 0 | 0 | 0 | 0 |
| QQEGESRL | LDHA; LDHB; LI | P00338; P07 | LDHA_HUMAN | Unmodified | 0 | 31086 | 1610500 | 0 | 264450 | 470680 | 0 | 0 | 0 | 69964 | 52849 | 0 | 0 | 0 | 0 | 1650300 | 420050 | 1181000 |
| QQEIILWR | VPS26A | O75436 | VP26A_HUMAN | Unmodified | 29661 | 0 | 0 | 0 | 0 | 0 | 0 | 0 | 98276 | 0 | 0 | 408860 | 0 | 0 | 103490 | 74405 | 0 | 0 |
| QQEQQVPI | FASN; FASN | P49327; A0A | FAS_HUMAN | Unmodified | 0 | 0 | 1872000 | 0 | 0 | 0 | 0 | 0 | 342470 | 0 | 1053500 | 3761400 | 0 | 0 | 899660 | 160500 | 0 | 0 |
| QQFVEYDK | RCN2; RCN2 | Q14257; H0Y | RCN2_HUMAN | Unmodified | 0 | 0 | 0 | 0 | 203800 | 0 | 0 | 0 | 13820000 | 1581100 | 851300 | 507170 | 0 | 0 | 46498 | 57105 | 0 | 0 |
| QQGELYM\ | PLCG2; PLCG2 | P16885; A0A | PLCG2_HUMAI | Oxidation (M) | 0 | 0 | 0 | 0 | 0 | 0 | 0 | 0 | 0 | 459730 | 0 | 2353300 | 0 | 0 | 0 | 0 | 0 | 0 |
| QQGVLALR | MARS1; MARS1 | P56192; H0Y | SYMC_HUMAN | Unmodified | 133290 | 0 | 422690 | 0 | 0 | 0 | 0 | 0 | 769170 | 87568 | 0 | 6788600 | 461010 | 152440 | 296930 | 0 | 0 | 343230 |
| QQHHQPM | UNC13D; UNC | Q70J99; K7E | UN13D_HUMA | Unmodified | 0 | 0 | 0 | 0 | 485160 | 0 | 1338700 | 0 | 723180 | 1488600 | 0 | 0 | 3269800 | 0 | 0 | 0 | 394160 | 0 |
| QQHIDNIT( | TRIM14 | Q14142 | TRI14_HUMAN | Unmodified | 0 | 0 | 292010 | 320110 | 670850 | 0 | 0 | 0 | 123120 | 0 | 0 | 0 | 251540 | 54475 | 208710 | 136680 | 0 | 0 |
| QQHVIETLI | CCT5; CCT5; C( | P48643; B7Z | TCPE_HUMAN | Unmodified | 50925 | 0 | 0 | 105120 | 2073700 | 0 | 255900 | 0 | 348410 | 30026 | 2481300 | 13545000 | 4132800 | 1135100 | 0 | 215490 | 343630 | 1965300 |
| QQHVIETLI | CCT5; CCT5; C( | P48643; B7Z | TCPE_HUMAN | Unmodified | 109560 | 86106 | 287640 | 179500 | 1044700 | 0 | 0 | 0 | 497040 | 271420 | 32306 | 5977200 | 1357700 | 427050 | 148430 | 204890 | 189390 | 1838900 |
| QQHVIETLI | CCT5; CCT5; C( | P48643; B7Z | TCPE_HUMAN | Unmodified | 0 | 0 | 0 | 0 | 507750 | 0 | 179140 | 0 | 0 | 293810 | 0 | 0 | 710570 | 367040 | 0 | 0 | 465420 | 582220 |
| QQHVIETLI | CCT5; CCT5; C( | P48643; B7Z | TCPE_HUMAN | Oxidation (M) | 0 | 0 | 0 | 0 | 890340 | 0 | 0 | 0 | 0 | 201170 | 0 | 0 | 252150 | 0 | 0 | 0 | 252710 | 0 |
| QQIAEDPEI | RAB23 | Q9ULC3 | RAB23_HUMAN | Unmodified | 0 | 0 | 126140 | 107350 | 0 | 0 | 65741 | 62964 | 80837 | 0 | 27077 | 133600 | 0 | 0 | 117270 | 309920 | 0 | 0 |
| QQIATEKQI | DIAPH1; DIAPH | O60610; A0A | DIAP1_HUMAN | Unmodified | 0 | 0 | 0 | 0 | 3353800 | 0 | 0 | 0 | 0 | 0 | 0 | 0 | 0 | 0 | 0 | 0 | 2182300 | 0 |
| QQIEEQRV | FLOT1; FLOT1; I | O75955; A0A | FLOT1_HUMAN | Unmodified | 402960 | 167250 | 2253100 | 5849700 | 2991000 | 1308600 | 0 | 81843 | 11819000 | 1536400 | 2013100 | 7989300 | 1215000 | 812210 | 5509100 | 3432500 | 592240 | 1642500 |
| QQIGSSAM | ADSL; ADSL; AD | P30566; A0A | PUR8_HUMAN | Oxidation (M) | 0 | 0 | 0 | 0 | 0 | 0 | 0 | 0 | 0 | 0 | 0 | 366170 | 0 | 0 | 0 | 44263 | 0 | 0 |
| QQIGSSAM | ADSL; ADSL; AD | P30566; A0A | PUR8_HUMAN | Unmodified | 0 | 0 | 0 | 67996 | 0 | 0 | 0 | 0 | 0 | 0 | 142170 | 3723500 | 0 | 0 | 0 | 0 | 0 | 0 |
| QQILFYQR | SNX33; SNX33 | Q8WV41; H3 | SNX33_HUMAN | Unmodified | 907710 | 895010 | 6250400 | 0 | 0 | 0 | 0 | 0 | 0 | 0 | 0 | 0 | 0 | 0 | 13706000 | 7132900 | 0 | 0 |
| QQILHSEEI | DYNC1I2; DYN( | Q13409; E7E | DC1I2_HUMAN | Unmodified | 0 | 0 | 169910 | 0 | 0 | 0 | 0 | 0 | 664770 | 0 | 539110 | 4999600 | 0 | 0 | 0 | 330980 | 0 | 0 |
| QQILLSFK | ANXA11 | P50995 | ANX11_HUMAN | Unmodified | 912370 | 656210 | 244420 | 1038800 | 1188900 | 223360 | 462560 | 347980 | 20300000 | 1488800 | 929750 | 2418700 | 697610 | 943570 | 521860 | 606460 | 1096800 | 2803500 |
| QQIQSIQQ | SNU13; SNU13 | P55769; A0A | NH2L1_HUMAI | Unmodified | 0 | 0 | 0 | 0 | 178370 | 0 | 225130 | 74784 | 0 | 0 | 251150 | 1537400 | 210800 | 0 | 0 | 442470 | 0 | 0 |
| QQIQSIQQ | SNU13; SNU13 | P55769; A0A | NH2L1_HUMAI | Unmodified | 0 | 0 | 0 | 0 | 701370 | 0 | 1625600 | 0 | 10703 | 706660 | 0 | 0 | 1613000 | 1147900 | 0 | 0 | 650850 | 1253800 |
| QQISLATQI | CCT5; CCT5; C( | P48643; B7Z | TCPE_HUMAN | Unmodified | 0 | 0 | 0 | 0 | 1484900 | 0 | 0 | 0 | 602640 | 0 | 0 | 3601300 | 1057800 | 503560 | 0 | 0 | 0 | 824750 |
| QQKPLEGE | AKAP5 | P24588 | AKAP5_HUMAN | Unmodified | 0 | 0 | 0 | 0 | 0 | 0 | 0 | 0 | 0 | 0 | 0 | 0 | 94726 | 1396000 | 0 | 0 | 0 | 726120 |
| QQKSDPIQ | ERP44 | Q9BS26 | ERP44_HUMAN | Unmodified | 0 | 0 | 0 | 0 | 0 | 0 | 0 | 0 | 1138800 | 556870 | 0 | 0 | 0 | 0 | 0 | 0 | 0 | 0 |
| QQLAAYQE | HOMER1; HOM | Q86YM7; Q8 | HOME1_HUMA | Unmodified | 0 | 0 | 0 | 0 | 0 | 0 | 0 | 0 | 0 | 0 | 0 | 0 | 694010 | 0 | 57357 | 120070 | 0 | 0 |
| QQLALYTEI | TXLNA | P40222 | TXLNA_HUMAN | Unmodified | 0 | 0 | 0 | 0 | 0 | 0 | 136040 | 80001 | 0 | 0 | 0 | 0 | 2011000 | 0 | 0 | 0 | 1990700 | 2696800 |
| QQLDYGIY | B4GALT4; B4G/ | O60513; C9J | B4GT4_HUMAN | Unmodified | 0 | 0 | 0 | 0 | 0 | 0 | 0 | 0 | 1306200 | 0 | 0 | 0 | 0 | 0 | 0 | 420240 | 0 | 0 |
| QQLDYGIY | B4GALT4; B4G/ | O60513; C9J | B4GT4_HUMAN | Unmodified | 0 | 0 | 0 | 0 | 0 | 0 | 0 | 0 | 884260 | 319730 | 0 | 0 | 0 | 0 | 0 | 0 | 0 | 0 |
| QQLDYGIY | B4GALT1 | P15291 | B4GT1_HUMAN | Unmodified | 0 | 0 | 0 | 0 | 0 | 0 | 0 | 0 | 495800 | 1202700 | 0 | 0 | 0 | 0 | 0 | 0 | 0 | 0 |
| QQLEELAR | GSS; GSS; GSS | P48637; A0A | GSHB_HUMAN | Unmodified | 0 | 0 | 0 | 0 | 0 | 48616 | 0 | 0 | 598420 | 0 | 137660 | 2985800 | 0 | 0 | 610690 | 485140 | 0 | 0 |
| QQLELLEEI | SHTN1 | A0MZ66 | SHOT1_HUMAI | Unmodified | 148770 | 0 | 2926000 | 0 | 0 | 0 | 0 | 309900 | 1046300 | 0 | 0 | 1059300 | 0 | 0 | 0 | 0 | 0 | 0 |
| QQLLIGAY/ | CCT7 | Q99832 | TCPH_HUMAN | Unmodified | 0 | 462780 | 0 | 766210 | 1444800 | 581780 | 988230 | 0 | 2747000 | 1260200 | 2983300 | 15468000 | 2164800 | 0 | 0 | 1117900 | 884250 | 1380800 |
| QQLLQEQ( | MAP4K4; MAP4 | O95819; C9J | M4K4_HUMAN | Unmodified | 177210 | 0 | 174350 | 482810 | 0 | 0 | 0 | 0 | 1689100 | 54420 | 1082100 | 3831500 | 0 | 0 | 0 | 316380 | 134820 | 0 |
| QQLNIHGL | ME1 | P48163 | MAOX_HUMAN | Unmodified | 0 | 0 | 0 | 0 | 0 | 0 | 0 | 0 | 0 | 0 | 0 | 2831900 | 0 | 0 | 642760 | 1047100 | 0 | 0 |
| QQLQALSE | GOLM1 | Q8NBJ4 | GOLM1_HUMA | Unmodified | 0 | 571620 | 681690 | 4027300 | 1884800 | 473390 | 0 | 0 | 9909000 | 116760 | 0 | 0 | 179550 | 0 | 4725500 | 2800600 | 0 | 515250 |

| | | | | | | | | | | | | | | | | | | | | | | |
|---|---|---|---|---|---|---|---|---|---|---|---|---|---|---|---|---|---|---|---|---|---|---|
| QQLQQVP( | TTLL12 | Q14166 | TTL12_HUMAN | Unmodified | 0 | 0 | 1428800 | 430360 | 0 | 0 | 0 | 0 | 0 | 0 | 741910 | 7360300 | 0 | 0 | 1076200 | 966670 | 0 | 0 |
| QQLSAATS | CEP55 | Q53EZ4 | CEP55_HUMAN | Unmodified | 0 | 0 | 0 | 0 | 17929000 | 0 | 335860 | 0 | 0 | 3945300 | 0 | 0 | 670640 | 1058900 | 0 | 0 | 0 | 135820 |
| QQLSAATS | CEP55 | Q53EZ4 | CEP55_HUMAN | Unmodified | 0 | 0 | 0 | 0 | 941030 | 0 | 0 | 0 | 0 | 0 | 0 | 0 | 0 | 164860 | 0 | 0 | 62159 | 0 |
| QQLTNTEV | OPA1; OPA1; OI | O60313; E5K | OPA1_HUMAN | Unmodified | 0 | 0 | 0 | 0 | 0 | 0 | 0 | 0 | 0 | 0 | 0 | 2893600 | 0 | 0 | 0 | 0 | 0 | 0 |
| QQLYVSSN | SEMA3C | Q99985 | SEM3C_HUMAI | Unmodified | 0 | 0 | 1010600 | 0 | 0 | 0 | 0 | 0 | 0 | 0 | 0 | 0 | 0 | 0 | 516770 | 0 | 0 | 0 |
| QQMENYPI | CD63; CD63; C | P08962; F8V | CD63_HUMAN | Unmodified | 10378000 | 223650 | 458440 | 3422600 | 0 | 177480 | 0 | 24567 | 43984000 | 11545000 | 194060 | 9101500 | 0 | 0 | 17689000 | 9031900 | 0 | 0 |
| QQMILQIG' | DSC2; DSC2 | Q02487; A0/ | DSC2_HUMAN | Oxidation (M) | 0 | 0 | 0 | 68239 | 0 | 0 | 0 | 0 | 1253500 | 1379000 | 0 | 0 | 0 | 0 | 1166900 | 0 | 53271 | 0 |
| QQNAQGG | PZP | P20742 | PZP_HUMAN P | Unmodified | 0 | 0 | 0 | 0 | 0 | 0 | 0 | 0 | 0 | 0 | 0 | 0 | 0 | 0 | 0 | 0 | 0 | 0 |
| QQNELAEL | NUMA1; NUMA | Q14980; A0/ | NUMA1_HUMA | Unmodified | 28474 | 25454 | 0 | 0 | 0 | 0 | 0 | 0 | 251740 | 39396 | 0 | 548540 | 0 | 0 | 0 | 0 | 0 | 0 |
| QQNVDQA' | DHRS4L2; DHR | Q6PKH6; Q9 | DR4L2_HUMAN | Unmodified | 0 | 0 | 0 | 0 | 0 | 0 | 0 | 0 | 1116200 | 0 | 0 | 0 | 0 | 0 | 0 | 0 | 0 | 0 |
| QQPDTEAV | CLNS1A; CLNS | P54105; E9P | ICLN_HUMAN I | Unmodified | 0 | 0 | 0 | 0 | 0 | 0 | 0 | 0 | 0 | 0 | 406890 | 2530000 | 0 | 0 | 0 | 0 | 0 | 0 |
| QQPDTELE | IARS2 | Q9NSE4 | SYIM_HUMAN I | Unmodified | 0 | 0 | 0 | 0 | 0 | 0 | 0 | 0 | 1429400 | 97490 | 0 | 0 | 0 | 0 | 0 | 0 | 0 | 0 |
| QQPLFVSG | COPA; COPA; C | P53621; A0A | COPA_HUMAN | Unmodified | 0 | 0 | 0 | 0 | 0 | 0 | 0 | 0 | 0 | 101390 | 0 | 3858800 | 0 | 0 | 0 | 0 | 0 | 0 |
| QQPLTELA/ | CORO7; CORO | P57737; I3L1 | CORO7_HUMA | Unmodified | 26816 | 0 | 0 | 0 | 0 | 0 | 0 | 0 | 0 | 0 | 0 | 6365500 | 0 | 0 | 0 | 42287 | 0 | 0 |
| QQPMALA\ | GAA | P10253 | LYAG_HUMAN | Unmodified | 0 | 0 | 0 | 0 | 0 | 0 | 0 | 0 | 573560 | 0 | 0 | 0 | 0 | 0 | 0 | 0 | 0 | 0 |
| QQPMALA\ | GAA | P10253 | LYAG_HUMAN | Oxidation (M) | 21051 | 0 | 0 | 0 | 0 | 0 | 0 | 0 | 833200 | 0 | 0 | 0 | 0 | 0 | 0 | 0 | 0 | 0 |
| QQPNAGKI | ITGB4 | P16144 | ITB4_HUMAN I | Unmodified | 0 | 0 | 0 | 0 | 0 | 0 | 0 | 0 | 0 | 0 | 0 | 0 | 0 | 0 | 0 | 0 | 281930 | 1194900 |
| QQPNPGN | TTLL12 | Q14166 | TTL12_HUMAN | Unmodified | 0 | 0 | 0 | 0 | 0 | 0 | 0 | 0 | 0 | 0 | 103570 | 1349200 | 128260 | 0 | 234180 | 202900 | 0 | 0 |
| QQPPAVQI | KDM1A; KDM1A | O60341; R4( | KDM1A_HUMA | Unmodified | 14809 | 0 | 26543 | 0 | 0 | 0 | 0 | 0 | 0 | 0 | 152240 | 1445700 | 0 | 0 | 0 | 0 | 0 | 0 |
| QQPPLGPS | TBC1D10B | Q4KMP7 | TB10B_HUMAN | Unmodified | 0 | 0 | 0 | 0 | 78677 | 0 | 0 | 0 | 0 | 227700 | 142760 | 608340 | 0 | 0 | 0 | 0 | 0 | 48431 |
| QQPTQFIN | SEPTIN2; SEPTI | Q15019; C9I | SEPT2_HUMAN | Unmodified | 0 | 0 | 0 | 0 | 22110000 | 13668000 | 0 | 0 | 1826100 | 1376700 | 0 | 10802000 | 9774500 | 6445000 | 0 | 0 | 9204700 | 7717300 |
| QQQDQVD | NONO; NONO; | Q15233; A0/ | NONO_HUMAN | Unmodified | 0 | 0 | 1687400 | 2567900 | 320650 | 2154500 | 2136700 | 2274600 | 0 | 1466300 | 0 | 0 | 1470500 | 79870 | 4932200 | 3274100 | 452950 | 1689900 |
| QQQDYFH: | USP8; USP8; U: | P40818; H0Y | UBP8_HUMAN | Unmodified | 0 | 0 | 0 | 0 | 261140 | 0 | 0 | 0 | 0 | 0 | 0 | 175830 | 0 | 0 | 0 | 0 | 261850 | 0 |
| QQQELDG: | SLK; SLK | Q9H2G2; A0 | SLK_HUMAN S' | Unmodified | 0 | 497330 | 0 | 0 | 0 | 0 | 0 | 0 | 499320 | 0 | 0 | 994220 | 0 | 656990 | 0 | 0 | 0 | 0 |
| QQQELDG: | SLK; SLK | Q9H2G2; A0 | SLK_HUMAN S' | Unmodified | 3803200 | 103010 | 0 | 253490 | 413860 | 196950 | 219910 | 0 | 536410 | 263820 | 485690 | 1852600 | 953230 | 458120 | 248300 | 272070 | 0 | 0 |
| QQQFENLI | SNX2 | O60749 | SNX2_HUMAN | Unmodified | 0 | 113850 | 324840 | 0 | 3165800 | 0 | 570720 | 271910 | 416570 | 190010 | 700160 | 2462500 | 2080200 | 1231100 | 308420 | 647550 | 0 | 1494800 |
| QQQFNVQ. | DARS1; DARS1; | P14868; H7( | SYDC_HUMAN | Unmodified | 0 | 0 | 0 | 0 | 0 | 0 | 0 | 0 | 0 | 52059 | 1045900 | 4960900 | 0 | 0 | 0 | 505770 | 0 | 0 |
| QQQLEEMI | AFDN; AFDN; A | P55196; H0Y | AFAD_HUMAN | Unmodified | 0 | 0 | 0 | 0 | 0 | 0 | 0 | 0 | 0 | 0 | 0 | 384370 | 0 | 0 | 0 | 0 | 0 | 0 |
| QQQQEGN | RBCK1; RBCK1 | Q9BYM8; G3 | HOIL1_HUMAN | Unmodified | 707720 | 601290 | 0 | 0 | 0 | 0 | 0 | 0 | 0 | 0 | 0 | 0 | 0 | 0 | 0 | 0 | 0 | 0 |
| QQQQLQQ | MINK1; MINK1; | Q8N4C8; A0 | MINK1_HUMAN | Unmodified | 0 | 0 | 0 | 0 | 1520100 | 0 | 0 | 0 | 2804000 | 613170 | 0 | 855450 | 2364400 | 3945200 | 0 | 831470 | 348960 | 0 |
| QQQQQLLI | MINK1; MINK1 | Q8N4C8; A0 | MINK1_HUMAN | Unmodified | 0 | 0 | 0 | 391340 | 450230 | 0 | 0 | 0 | 692060 | 477350 | 195900 | 655530 | 662380 | 838430 | 0 | 0 | 307660 | 640520 |
| QQQQQQN | RAB35; RAB35 | Q15286; F5H | RAB35_HUMAN | Unmodified | 0 | 3385700 | 0 | 21096000 | 4188700 | 0 | 90312 | 638110 | 819250 | 403410 | 531970 | 574100 | 13645000 | 3268700 | 0 | 2007200 | 1214800 | 4470800 |
| QQQQQQC | NUCB1 | Q02818 | NUCB1_HUMA | Unmodified | 0 | 29878 | 0 | 0 | 434210 | 86063 | 192100 | 474410 | 2428000 | 787500 | 378500 | 132350 | 0 | 169270 | 0 | 0 | 0 | 675210 |
| QQQTVFIH | DHX16; DHX16 | O60231; Q5! | DHX16_HUMAI | Unmodified | 0 | 0 | 0 | 0 | 0 | 0 | 0 | 0 | 0 | 0 | 0 | 536880 | 0 | 0 | 0 | 0 | 0 | 0 |
| QQREDITQ | ILF3 | Q12906 | ILF3_HUMAN Ir | Unmodified | 7067100 | 4084500 | 0 | 459220 | 0 | 0 | 115500 | 0 | 0 | 0 | 0 | 2946300 | 0 | 2578400 | 1363300 | 996580 | 0 | 0 |
| QQSEEAFP | GLCE; GLCE | O94923; H0Y | GLCE_HUMAN | Unmodified | 0 | 0 | 0 | 0 | 0 | 0 | 0 | 0 | 404900 | 37182 | 0 | 58237 | 263360 | 0 | 0 | 0 | 0 | 0 |
| QQSEISAA\ | PRRC2C; PRRC | Q9Y520; A0/ | PRC2C_HUMAI | Unmodified | 357220 | 141920 | 0 | 0 | 0 | 0 | 0 | 0 | 0 | 0 | 286120 | 887360 | 0 | 0 | 0 | 27798 | 133450 | 0 |
| QQSELQSC | LASP1; LASP1 | Q14847; C9J | LASP1_HUMAN | Unmodified | 3622800 | 2944700 | 1601000 | 957960 | 0 | 0 | 0 | 0 | 914220 | 0 | 0 | 1242700 | 504230 | 271440 | 2482700 | 1527500 | 0 | 0 |
| QQSELQSC | LASP1; LASP1 | Q14847; C9J | LASP1_HUMAN | Unmodified | 0 | 0 | 0 | 0 | 15304000 | 2463600 | 280510 | 1543500 | 0 | 919320 | 0 | 0 | 7184600 | 8533000 | 0 | 0 | 2960200 | 5735700 |
| QQSFFLAE' | MAN1C1; MAN | Q9NR34; P3 | MA1C1_HUMAI | Unmodified | 0 | 0 | 0 | 0 | 0 | 0 | 0 | 0 | 593750 | 0 | 0 | 0 | 0 | 0 | 0 | 0 | 0 | 0 |
| QQSHFAMI | PCBP1 | Q15365 | PCBP1_HUMAI | 2 Oxidation (M) | 1874100 | 1828500 | 0 | 0 | 111060 | 0 | 0 | 0 | 0 | 0 | 0 | 0 | 992180 | 0 | 965100 | 450400 | 0 | 0 |
| QQSHFAMI | PCBP1 | Q15365 | PCBP1_HUMAI | Oxidation (M) | 2290000 | 2230200 | 0 | 131930 | 0 | 0 | 0 | 0 | 0 | 0 | 0 | 699340 | 0 | 0 | 147730 | 183150 | 0 | 0 |
| QQSHFAMI | PCBP1 | Q15365 | PCBP1_HUMAI | Unmodified | 0 | 0 | 0 | 0 | 0 | 0 | 0 | 0 | 0 | 0 | 167700 | 0 | 1273100 | 0 | 362880 | 0 | 0 | 0 |
| QQSIAGSA | AIMP1; AIMP1 | Q12904; A0/ | AIMP1_HUMAN | Unmodified | 0 | 0 | 0 | 0 | 0 | 0 | 0 | 0 | 0 | 0 | 0 | 1410400 | 0 | 0 | 0 | 0 | 0 | 0 |
| QQSTNDYI | CMTM4; CMTM | Q8IZR5; J3Q | CKLF4_HUMAN | Unmodified | 0 | 0 | 0 | 0 | 0 | 0 | 0 | 0 | 0 | 0 | 0 | 1866800 | 0 | 0 | 0 | 196990 | 0 | 0 |
| QQTANKVE | RABGAP1 | Q9Y3P9 | RBGP1_HUMAI | Unmodified | 0 | 0 | 0 | 0 | 2503300 | 0 | 0 | 0 | 0 | 0 | 0 | 0 | 0 | 107190 | 0 | 54237 | 0 | 0 |
| QQTDMAVI | HDAC2; HDAC2 | Q92769; H3I | HDAC2_HUMA | Unmodified | 0 | 0 | 0 | 0 | 0 | 0 | 0 | 0 | 0 | 0 | 0 | 2509800 | 0 | 0 | 0 | 0 | 0 | 0 |
| QQTDNPK( | UTRN | P46939 | UTRN_HUMAN | Oxidation (M) | 0 | 0 | 98690 | 0 | 0 | 0 | 0 | 0 | 63154 | 0 | 0 | 0 | 0 | 0 | 0 | 133460 | 218200 | 98712 |
| QQTDNPK( | UTRN | P46939 | UTRN_HUMAN | Unmodified | 0 | 0 | 0 | 0 | 0 | 0 | 72685 | 0 | 0 | 173830 | 0 | 0 | 0 | 506070 | 0 | 0 | 656630 | 0 |
| QQTGKLSI | SHPRH; SHPRH | Q149N8; A0 | SHPRH_HUMA | Unmodified | 0 | 0 | 0 | 0 | 3989600 | 785490 | 5923400 | 3235500 | 4.41E+08 | 83738000 | 337150 | 0 | 6935300 | 0 | 0 | 0 | 985750 | 7497100 |
| QQTLEAEE | DBN1; DBN1 | Q16643; D6I | DREB_HUMAN | Unmodified | 66264 | 0 | 1202500 | 1536000 | 1645500 | 0 | 0 | 0 | 0 | 0 | 608140 | 8516100 | 0 | 0 | 1735300 | 1434200 | 0 | 0 |
| QQTNSKG: | AHCYL1 | O43865 | SAHH2_HUMAI | Unmodified | 0 | 0 | 0 | 0 | 1599000 | 742120 | 444840 | 285030 | 0 | 102410 | 0 | 0 | 2093100 | 1294100 | 0 | 0 | 1903800 | 1417800 |
| QQTNTLLN | UBE3C | Q15386 | UBE3C_HUMAI | Unmodified | 0 | 0 | 0 | 0 | 0 | 0 | 0 | 0 | 0 | 0 | 0 | 92148 | 0 | 0 | 16595 | 19003 | 0 | 0 |
| QQTPPGLC | REXO2; REXO2; | Q9Y3B8; F5( | ORN_HUMAN ( | Unmodified | 0 | 0 | 0 | 200290 | 0 | 0 | 0 | 0 | 359850 | 0 | 0 | 1137600 | 0 | 93491 | 0 | 159640 | 0 | 0 |
| QQTSLFIAF | CKAP5 | Q14008 | CKAP5_HUMAN | Unmodified | 0 | 0 | 64003 | 0 | 231720 | 0 | 0 | 16734 | 1757100 | 235250 | 124160 | 2126100 | 257130 | 63718 | 0 | 0 | 690020 | 51727 |
| QQVADLTA | DAAM1 | Q9Y4D1 | DAAM1_HUMA | Unmodified | 0 | 0 | 427310 | 479830 | 0 | 0 | 0 | 0 | 0 | 0 | 0 | 0 | 1602200 | 0 | 400260 | 659200 | 0 | 0 |
| QQVADLTA | DAAM1 | Q9Y4D1 | DAAM1_HUMA | Unmodified | 0 | 326670 | 0 | 719670 | 964600 | 0 | 891250 | 0 | 1463200 | 674720 | 779000 | 4055600 | 1530300 | 0 | 0 | 347960 | 0 | 674470 |
| QQVAPTDI | SPTAN1; SPTAN | Q13813; A0/ | SPTN1_HUMAN | Unmodified | 0 | 0 | 0 | 0 | 0 | 0 | 0 | 0 | 0 | 0 | 0 | 1532500 | 0 | 0 | 0 | 0 | 0 | 0 |
| QQVEGTAC | ADIRF | Q15847 | ADIRF_HUMAN | Unmodified | 396670 | 805740 | 0 | 0 | 0 | 0 | 0 | 0 | 0 | 0 | 0 | 0 | 0 | 0 | 0 | 0 | 1082500 | 10975000 |
| QQVIELAR | STEAP2; STEAP | Q8NFT2; B5I | STEA2_HUMAN | Unmodified | 3044300 | 2242800 | 22218000 | 18754000 | 0 | 0 | 0 | 0 | 291950 | 0 | 0 | 190610 | 465910 | 143320 | 40798000 | 29924000 | 0 | 0 |
| QQVPSEDS | KIF20A; KIF20A | O95235; D6I | KI20A_HUMAN | Unmodified | 0 | 0 | 235070 | 194260 | 0 | 0 | 0 | 0 | 0 | 0 | 0 | 0 | 0 | 0 | 266650 | 82813 | 0 | 0 |
| QQVPSGES | PARP1; PARP1; | P09874; A0A | PARP1_HUMAN | Unmodified | 0 | 0 | 0 | 0 | 0 | 0 | 0 | 0 | 496770 | 600970 | 368250 | 1275300 | 0 | 0 | 0 | 0 | 0 | 0 |
| QQWQQLY | EIF3M; EIF3M | Q7L2H7; H0' | EIF3M_HUMAN | Unmodified | 0 | 0 | 0 | 0 | 0 | 60146 | 0 | 0 | 0 | 0 | 82542 | 655170 | 0 | 0 | 11261 | 0 | 0 | 0 |
| QQYEHSV( | SEC14L2; N/A; | O76054; H7( | S14L2_HUMAN | Unmodified | 0 | 0 | 0 | 0 | 0 | 0 | 21259 | 14325 | 99995 | 20265 | 137880 | 472540 | 0 | 0 | 0 | 0 | 0 | 0 |
| QQYQNGLI | CNNM4 | Q6P4Q7 | CNNM4_HUMA | Unmodified | 0 | 0 | 0 | 0 | 0 | 0 | 0 | 0 | 0 | 0 | 0 | 0 | 0 | 0 | 0 | 0 | 0 | 0 |
| QQYTNQL/ | STX7 | O15400 | STX7_HUMAN S | Unmodified | 0 | 364570 | 0 | 0 | 3172800 | 532800 | 0 | 71388 | 760710 | 296720 | 0 | 747230 | 7173600 | 0 | 0 | 724030 | 0 | 1398700 |
| QQYTNQL/ | STX7 | O15400 | STX7_HUMAN S | Unmodified | 0 | 0 | 0 | 0 | 1329700 | 218960 | 445500 | 146570 | 0 | 0 | 0 | 0 | 0 | 0 | 0 | 0 | 128410 | 294030 |
| QQYWTEVI | XPO4 | Q9C0E2 | XPO4_HUMAN | Unmodified | 0 | 0 | 0 | 0 | 0 | 0 | 0 | 0 | 0 | 0 | 0 | 512530 | 0 | 0 | 0 | 0 | 0 | 0 |
| QRAPGEDI | AKT2; AKT2; AK | P31751; A8N | AKT2_HUMAN I | Oxidation (M) | 38793 | 0 | 105720 | 166290 | 0 | 0 | 0 | 0 | 0 | 0 | 0 | 55814 | 0 | 0 | 148140 | 162230 | 0 | 0 |
| QRAQDGD | PTGFRN | Q9P2B2 | FPRP_HUMAN | Unmodified | 1602600 | 1502100 | 5458400 | 5715600 | 1917100 | 0 | 41603 | 0 | 253950 | 42206 | 523880 | 8858400 | 4476100 | 2140700 | 7122600 | 4584900 | 0 | 0 |
| QRAQDGD | PTGFRN | Q9P2B2 | FPRP_HUMAN | Unmodified | 0 | 0 | 0 | 0 | 0 | 0 | 0 | 0 | 0 | 0 | 0 | 0 | 403860 | 0 | 0 | 35668 | 0 | 0 |
| QRCPPGV\ | FASN; FASN | P49327; A0A | FAS_HUMAN F | Unmodified | 0 | 94176 | 86183 | 0 | 0 | 0 | 0 | 0 | 0 | 0 | 136320 | 2704100 | 0 | 0 | 143750 | 109460 | 0 | 0 |
| QRDSEIMQ | SERF2 | P84101 | SERF2_HUMAN | Oxidation (M) | 0 | 267320 | 0 | 358680 | 0 | 0 | 15566 | 0 | 0 | 0 | 0 | 0 | 0 | 0 | 0 | 0 | 34633 | 130780 |
| QREELGQC | CKAP4 | Q07065 | CKAP4_HUMAN | Unmodified | 43428 | 32991 | 150260 | 88178 | 44011 | 0 | 0 | 0 | 8485000 | 0 | 0 | 0 | 47910 | 30493 | 115840 | 100220 | 0 | 39710 |
| QREEYQPA | TMED9 | Q9BVK6 | TMED9_HUMAI | Unmodified | 0 | 2553500 | 0 | 0 | 322780 | 0 | 0 | 187160 | 1063800 | 0 | 0 | 0 | 0 | 0 | 0 | 0 | 0 | 0 |
| QREEYQPA | TMED9 | Q9BVK6 | TMED9_HUMAI | Oxidation (M) | 120740 | 0 | 0 | 0 | 776580 | 0 | 0 | 0 | 46627000 | 2229600 | 3886300 | 2128400 | 194050 | 0 | 3010300 | 539280 | 0 | 0 |
| QREEYQPA | TMED9 | Q9BVK6 | TMED9_HUMAI | Unmodified | 0 | 0 | 0 | 161470 | 0 | 0 | 0 | 0 | 74751000 | 2243200 | 745530 | 647990 | 0 | 0 | 0 | 0 | 0 | 0 |
| QREMEEEE | OS9; OS9 | Q13438; B4E | OS9_HUMAN P | Oxidation (M) | 0 | 0 | 0 | 0 | 0 | 0 | 0 | 0 | 1315500 | 0 | 0 | 0 | 0 | 0 | 0 | 0 | 0 | 0 |
| QREMEEEE | OS9; OS9 | Q13438; B4E | OS9_HUMAN P | Unmodified | 0 | 0 | 0 | 0 | 0 | 0 | 0 | 0 | 1139600 | 0 | 0 | 0 | 0 | 0 | 0 | 0 | 0 | 0 |
| QREQLDKI( | GOLM1; GOLM | Q8NBJ4; F8V | GOLM1_HUMA | Unmodified | 0 | 0 | 0 | 0 | 1189500 | 485020 | 0 | 0 | 1002500 | 5134000 | 0 | 0 | 0 | 0 | 0 | 0 | 0 | 518580 |
| QRFDQPLE | DAP3; DAP3; D/ | P51398; V9G | RT29_HUMAN : | Unmodified | 0 | 0 | 0 | 0 | 0 | 0 | 0 | 0 | 958820 | 0 | 357880 | 0 | 0 | 0 | 0 | 0 | 0 | 0 |
| QRFPLGGI | APPL1; APPL1 | Q9UKG1; C9 | DP13A_HUMAN | Unmodified | 0 | 0 | 0 | 0 | 1942100 | 311850 | 0 | 0 | 1410300 | 837010 | 0 | 540280 | 858670 | 929430 | 0 | 0 | 0 | 0 |
| QRFPLGGI | APPL1; APPL1 | Q9UKG1; C9 | DP13A_HUMAN | Oxidation (M) | 0 | 0 | 0 | 0 | 0 | 795460 | 0 | 89497 | 381260 | 648710 | 0 | 612130 | 828610 | 0 | 0 | 0 | 226010 | 0 |
| QRGDQLVI | TMEM43; N/A | Q9BTV4; V9( | TMM43_HUMAI | Unmodified | 0 | 0 | 0 | 0 | 0 | 0 | 0 | 0 | 20470000 | 2369800 | 1437100 | 599000 | 0 | 0 | 0 | 0 | 102480 | 0 |
| QRGLLQQI | MGAT1; MGAT1 | P26572; D6F | MGAT1_HUMAI | Unmodified | 0 | 0 | 0 | 0 | 0 | 0 | 0 | 0 | 352310 | 26337 | 0 | 0 | 0 | 0 | 0 | 0 | 0 | 0 |
| QRGPEAAS | TRIM32 | Q13049 | TRI32_HUMAN | Unmodified | 0 | 0 | 0 | 0 | 0 | 0 | 445200 | 0 | 0 | 0 | 0 | 0 | 0 | 0 | 0 | 0 | 0 | 0 |
| QRHFDTYI | PKP2 | Q99959 | PKP2_HUMAN | Unmodified | 0 | 0 | 15643 | 0 | 0 | 0 | 0 | 0 | 0 | 0 | 0 | 128000 | 23386 | 0 | 28274 | 0 | 36393 | 0 |
| QRIDEFEAI | EZR; EZR | P15311; E7E | EZRI_HUMAN E | Unmodified | 0 | 0 | 0 | 0 | 40830000 | 0 | 0 | 0 | 7656700 | 6038400 | 1697900 | 9841900 | 8684500 | 9470200 | 1130100 | 1685500 | 12275000 | 7106600 |
| QRIDEFEAI | RDX; RDX; RDX | P35241; A0A | RADI_HUMAN I | Oxidation (M) | 279540 | 264530 | 0 | 515780 | 5871100 | 977720 | 1666800 | 1326000 | 399860 | 401030 | 163100 | 0 | 4971000 | 5301400 | 436740 | 0 | 2528900 | 1386500 |
| QRIDEFEAI | RDX; RDX; RDX | P35241; A0A | RADI_HUMAN I | Unmodified | 0 | 210390 | 0 | 0 | 15624000 | 1740100 | 0 | 0 | 1190900 | 824610 | 0 | 1807600 | 4679300 | 0 | 396660 | 0 | 4295400 | 2427300 |
| QRIDEFESI | ERMN; MSN; EF | Q8TAM6; P2( | ERMIN_HUMAN | Oxidation (M) | 0 | 0 | 0 | 1193300 | 17760000 | 5239800 | 1566600 | 1445500 | 0 | 0 | 0 | 0 | 3461900 | 5127000 | 0 | 0 | 26503000 | 12447000 |
| QRIDEFESI | ERMN; MSN; EF | Q8TAM6; P2( | ERMIN_HUMAN | Unmodified | 199580 | 208370 | 351020 | 813490 | 50118000 | 7027300 | 3271300 | 1875700 | 0 | 0 | 0 | 0 | 7047100 | 7912700 | 361300 | 268640 | 51315000 | 22490000 |
| QRIDYANL | MYO1C; MYO1( | O00159; F5H | MYO1C_HUMA | Unmodified | 0 | 0 | 0 | 330350 | 0 | 0 | 0 | 0 | 0 | 0 | 0 | 0 | 0 | 0 | 1489000 | 0 | 0 | 0 |
| QRIDYANL | MYO1C; MYO1( | O00159; F5H | MYO1C_HUMA | Unmodified | 0 | 0 | 0 | 0 | 39450000 | 0 | 0 | 0 | 0 | 0 | 0 | 0 | 0 | 0 | 0 | 0 | 3386700 | 12183000 |
| QRITADLLS | SEPTIN9; SEPTI | Q9UHD8; A( | SEPT9_HUMAN | Unmodified | 0 | 0 | 0 | 0 | 0 | 496060 | 275150 | 0 | 910100 | 448360 | 0 | 4858000 | 874940 | 75516 | 0 | 0 | 1171600 | 1557800 |
| QRLEALGT | SPTBN2 | O15020 | SPTN2_HUMAN | Unmodified | 0 | 0 | 423590 | 0 | 0 | 70285 | 0 | 0 | 0 | 0 | 0 | 776200 | 0 | 0 | 234890 | 240310 | 0 | 0 |
| QRLEDLATI | MYO1B; MYO1E | O43795; E9F | MYO1B_HUMA | Unmodified | 82031 | 324740 | 0 | 0 | 181160 | 0 | 0 | 0 | 396540 | 90708 | 695370 | 3465800 | 0 | 0 | 299810 | 181530 | 132050 | 0 |
| QRLLDSSC | PANX1 | Q96RD7 | PANX1_HUMAN | Unmodified | 0 | 200280 | 1580200 | 0 | 299540 | 490240 | 0 | 0 | 0 | 129830 | 1202700 | 311090 | 0 | 0 | 0 | 234140 | 329850 | 0 |
| QRLNPTSN | TJP2; TJP2; TJP2 | Q9UDY2; A0 | ZO2_HUMAN T | Unmodified | 0 | 0 | 0 | 0 | 258900 | 127980 | 0 | 0 | 214620 | 78236 | 0 | 53240 | 28969 | 20751 | 0 | 0 | 22247 | 19731 |
| QRLPGPPD | PDXP | Q96GD0 | PLPP_HUMAN ( | Unmodified | 0 | 0 | 0 | 0 | 0 | 0 | 0 | 0 | 0 | 0 | 0 | 2169300 | 0 | 0 | 0 | 0 | 0 | 0 |
| QRMESALD | TALDO1; TALD( | P37837; E9P | TALDO_HUMAN | Unmodified | 100880 | 0 | 0 | 0 | 492230 | 184660 | 0 | 0 | 0 | 277800 | 1123600 | 2727700 | 319120 | 0 | 0 | 0 | 158350 | 0 |
| QRNSLGGI | CD109 | Q6YHK3 | CD109_HUMAI | Unmodified | 0 | 0 | 0 | 0 | 0 | 0 | 0 | 0 | 533120 | 591900 | 0 | 0 | 0 | 0 | 0 | 0 | 262160 | 0 |
| QRPADAEI | EEF1AKNMT | Q8N6R0 | EFNMT_HUMAI | Unmodified | 0 | 0 | 0 | 88847 | 213260 | 0 | 86855 | 0 | 0 | 46277 | 155160 | 512960 | 149360 | 153090 | 116890 | 103840 | 77845 | 351610 |
| QRPCQSYI | TSPAN8 | P19075 | TSN8_HUMAN | Unmodified | 13891000 | 2213000 | 78413 | 0 | 0 | 219940 | 18952 | 0 | 53349 | 31231 | 0 | 0 | 0 | 0 | 0 | 0 | 0 | 112650 |
| QRPYSVAV | BAIAP2; BAIAP2 | Q9UQB8; I3I | BAIP2_HUMAN | Unmodified | 79771 | 0 | 0 | 665840 | 1185700 | 192300 | 0 | 0 | 1039800 | 587560 | 0 | 1197000 | 791620 | 387090 | 4927200 | 4039300 | 0 | 306730 |
| QRQEVCQ: | ANXA6; ANXA6; | P08133; E5R | ANXA6_HUMAN | Unmodified | 0 | 412590 | 526720 | 0 | 0 | 0 | 187320 | 88544 | 7360900 | 360900 | 0 | 0 | 0 | 201760 | 92108 | 520660 | 0 | 874990 |
| QRQLAEAH | PLEC | Q15149 | PLEC_HUMAN | Unmodified | 0 | 0 | 0 | 0 | 0 | 0 | 0 | 0 | 0 | 0 | 0 | 0 | 0 | 0 | 0 | 0 | 0 | 0 |
| QRQQILLSI | ANXA11 | P50995 | ANX11_HUMAN | Unmodified | 1067900 | 1730200 | 616420 | 920310 | 14855000 | 2549300 | 4619900 | 1987900 | 67895000 | 21444000 | 2668400 | 3778400 | 11630000 | 14262000 | 762650 | 816890 | 5428300 | 15694000 |
| QRQWIQIS | NIBAN2 | Q96TA1 | NIBA2_HUMAN | Unmodified | 0 | 0 | 0 | 0 | 2245300 | 260430 | 0 | 0 | 0 | 0 | 0 | 0 | 9168300 | 1510900 | 0 | 0 | 0 | 0 |

| | | | | | | | | | | | | | | | | | | | | | | |
|---|---|---|---|---|---|---|---|---|---|---|---|---|---|---|---|---|---|---|---|---|---|---|
| QRRFSDQA | MARK2; MARK2 | E7ETY4; A0A | E7ETY4_HUMA | Unmodified | 0 | 0 | 0 | 0 | 408870 | 0 | 0 | 0 | 0 | 0 | 0 | 0 | 216250 | 0 | 0 | 433060 | 285390 | 0 |
| QRTNFHQF | VPS26A | O75436 | VP26A_HUMAN | Unmodified | 0 | 0 | 0 | 0 | 3157700 | 841340 | 0 | 0 | 999530 | 302840 | 0 | 0 | 1587100 | 0 | 0 | 0 | 833490 | 1220200 |
| QRTNPSPT | NUMB; NUMB | P49757; G3\ | NUMB_HUMAN | Unmodified | 0 | 126170 | 0 | 0 | 3605200 | 754920 | 0 | 62901 | 0 | 144990 | 0 | 0 | 514650 | 281890 | 0 | 0 | 549910 | 2136800 |
| QRTSEVQD | EPS15; EPS15 | P42566; B1A | EPS15_HUMAN | Unmodified | 120480 | 162980 | 0 | 464950 | 752960 | 137120 | 161290 | 49358 | 0 | 367470 | 144990 | 0 | 1345300 | 582120 | 0 | 0 | 193360 | 526330 |
| QRVAEVLN | KLC1; KLC1; KL | Q07866; G3\ | KLC1_HUMAN | Unmodified | 0 | 0 | 0 | 0 | 1171700 | 0 | 220620 | 115400 | 599460 | 107570 | 0 | 0 | 0 | 0 | 0 | 0 | 437400 | 1084900 |
| QRVDQILE | ITGA7; ITGA7 | Q13683; J3K | ITA7_HUMAN I | Unmodified | 0 | 0 | 0 | 0 | 0 | 0 | 0 | 0 | 666310 | 0 | 153750 | 0 | 0 | 0 | 0 | 124710 | 0 | 0 |
| QRVEDAFY | KRAS | P01116 | RASK_HUMAN | Unmodified | 376110 | 0 | 85830 | 132250 | 0 | 0 | 0 | 0 | 430890 | 0 | 80357 | 1565300 | 0 | 31373 | 293710 | 193160 | 0 | 0 |
| QRVEHQCI | PIPOX; PIPOX; F | Q9P0Z9; K7E | SOX_HUMAN P | Unmodified | 0 | 0 | 0 | 0 | 0 | 0 | 0 | 0 | 1482700 | 91254 | 0 | 0 | 0 | 0 | 0 | 0 | 0 | 0 |
| QRVEQFAR | SHMT2 | P34897 | GLYM_HUMAN | Unmodified | 0 | 0 | 0 | 0 | 0 | 0 | 0 | 0 | 2010300 | 382880 | 0 | 0 | 0 | 0 | 0 | 0 | 0 | 0 |
| QRWDEAFI | APEX1; APEX1; | P27695; H7C | APEX1_HUMAN | Unmodified | 0 | 0 | 0 | 600290 | 0 | 0 | 0 | 0 | 378810 | 59441 | 289740 | 1277900 | 0 | 0 | 0 | 0 | 31582 | 0 |
| QRWENIAT | KIF4A | O95239 | KIF4A_HUMAN | Unmodified | 0 | 0 | 0 | 0 | 0 | 0 | 0 | 0 | 0 | 0 | 0 | 0 | 0 | 0 | 0 | 0 | 1648000 | 0 |
| QRWFKFDI | USP8 | P40818 | UBP8_HUMAN | Unmodified | 0 | 0 | 0 | 0 | 428610 | 89033 | 0 | 0 | 0 | 0 | 0 | 0 | 0 | 0 | 0 | 0 | 0 | 0 |
| QRYASNAV | DNPEP; DNPEP | Q9ULA0; E5I | DNPEP_HUMAI | Unmodified | 0 | 0 | 0 | 0 | 230580 | 0 | 0 | 0 | 0 | 623510 | 0 | 0 | 906440 | 0 | 0 | 0 | 591170 | 615540 |
| QRYEDRIF | ENTPD4; ENTPI | Q9Y227; Q8I | ENTP4_HUMAN | Unmodified | 0 | 0 | 0 | 257170 | 369650 | 0 | 0 | 0 | 536120 | 159760 | 0 | 336460 | 0 | 0 | 0 | 0 | 0 | 474730 |
| QRYEILTPN | MYH9 | P35579 | MYH9_HUMAN | Unmodified | 288580 | 344290 | 928550 | 882720 | 0 | 0 | 0 | 0 | 0 | 0 | 0 | 4463400 | 0 | 0 | 1874100 | 650570 | 0 | 0 |
| QRYPSQPR | NAGLU | P54802 | ANAG_HUMAN | Unmodified | 0 | 0 | 0 | 0 | 0 | 0 | 0 | 0 | 1073200 | 1978400 | 0 | 0 | 0 | 0 | 0 | 0 | 180000 | 0 |
| QRYSYQYT | ERGIC1 | Q969X5 | ERGI1_HUMAN | Unmodified | 0 | 0 | 129270 | 0 | 0 | 0 | 0 | 0 | 8947000 | 737970 | 117190 | 109080 | 0 | 0 | 219560 | 153590 | 0 | 0 |
| QRYVLGDT | UBA6 | A0AVT1 | UBA6_HUMAN | Unmodified | 0 | 0 | 0 | 0 | 115770 | 0 | 0 | 0 | 0 | 51301 | 0 | 0 | 194570 | 0 | 0 | 0 | 135600 | 236390 |
| QSAALCLLI | AP2A2; AP2A1; | O94973; O9! | AP2A2_HUMAN | Unmodified | 133890 | 88387 | 0 | 108320 | 0 | 0 | 0 | 0 | 247800 | 0 | 0 | 4084000 | 0 | 0 | 246390 | 0 | 0 | 0 |
| QSAEDILLK | LAMB1; LAMB1; | P07942; A0A | LAMB1_HUMAI | Unmodified | 228370 | 122210 | 868660 | 108470 | 0 | 0 | 1026000 | 939720 | 651560 | 0 | 289690 | 1110600 | 390830 | 324340 | 80025 | 84066 | 185180 | 474140 |
| QSAERNSN | LMNA; LMNA; L | P02545; A0A | LMNA_HUMAN | Unmodified | 0 | 0 | 0 | 0 | 439850 | 0 | 0 | 0 | 0 | 0 | 0 | 0 | 279690 | 0 | 0 | 0 | 192880 | 270200 |
| QSAMAICQ | MCOLN3; MCO | Q8TDD5; B1 | MCLN3_HUMA | Unmodified | 0 | 0 | 0 | 0 | 0 | 0 | 0 | 0 | 1015900 | 157170 | 0 | 0 | 0 | 0 | 0 | 0 | 0 | 0 |
| QSAQLTAL | SYAP1 | Q96A49 | SYAP1_HUMAN | Unmodified | 0 | 0 | 0 | 0 | 0 | 0 | 0 | 0 | 0 | 0 | 0 | 925800 | 0 | 0 | 469440 | 285690 | 205940 | 0 |
| QSAYDSKA | STK24; STK24; | Q9Y6E0; H0 | STK24_HUMAN | Oxidation (M) | 0 | 0 | 0 | 0 | 4125300 | 707720 | 315050 | 0 | 0 | 0 | 0 | 0 | 4527800 | 0 | 0 | 0 | 567520 | 2154100 |
| QSCITEQT | CACNA2D1 | P54289 | CA2D1_HUMAI | Unmodified | 466620 | 393640 | 0 | 0 | 0 | 0 | 1002100 | 426780 | 1438500 | 0 | 0 | 0 | 0 | 0 | 0 | 0 | 0 | 0 |
| QSEDGSEF | HTATSF1 | O43719 | HTSF1_HUMAN | Unmodified | 0 | 0 | 0 | 0 | 0 | 0 | 0 | 0 | 0 | 0 | 0 | 297430 | 0 | 0 | 0 | 0 | 0 | 0 |
| QSEDLGSC | USO1 | O60763 | USO1_HUMAN | Unmodified | 0 | 0 | 59236 | 0 | 0 | 0 | 0 | 0 | 0 | 0 | 0 | 1045700 | 0 | 0 | 0 | 0 | 0 | 0 |
| QSEENNNI | PEG10; PEG10; | Q86TG7; A0 | PEG10_HUMAI | Unmodified | 0 | 125760 | 1340000 | 138430 | 0 | 157380 | 0 | 0 | 0 | 0 | 0 | 0 | 0 | 0 | 0 | 0 | 0 | 0 |
| QSEPFFK | G6PD; G6PD; G | P11413; E7E | G6PD_HUMAN | Unmodified | 0 | 0 | 0 | 281970 | 794580 | 305970 | 0 | 0 | 374090 | 0 | 141540 | 1104900 | 0 | 0 | 158840 | 191520 | 0 | 227920 |
| QSEPFFKA | G6PD; G6PD; G | P11413; E7E | G6PD_HUMAN | Unmodified | 0 | 0 | 0 | 0 | 858610 | 231210 | 0 | 0 | 0 | 104000 | 0 | 0 | 0 | 472240 | 0 | 0 | 0 | 0 |
| QSEQQRET | PFDN6 | O15212 | PFD6_HUMAN | Unmodified | 0 | 0 | 0 | 203630 | 0 | 0 | 0 | 0 | 1426600 | 0 | 0 | 6563300 | 0 | 0 | 220510 | 109930 | 400310 | 0 |
| QSFASLALI | ANK3; ANK3; AI | Q12955; H0 | ANK3_HUMAN | Unmodified | 0 | 0 | 412510 | 227810 | 75074 | 1435800 | 0 | 0 | 641260 | 416010 | 5890800 | 174650 | 390930 | 226140 | 545400 | 601940 | 0 | 0 |
| QSFMFYGA | TMPRSS2; TMPI | O15393; C9J | TMPS2_HUMAI | Oxidation (M) | 0 | 0 | 0 | 0 | 0 | 0 | 0 | 0 | 0 | 0 | 0 | 0 | 0 | 0 | 421480 | 402570 | 0 | 0 |
| QSFTMVAD | LASP1; LASP1; I | Q14847; C9J | LASP1_HUMAN | Oxidation (M) | 169840 | 0 | 0 | 0 | 8261600 | 1012800 | 0 | 2162000 | 0 | 1198800 | 0 | 0 | 6985600 | 5394000 | 0 | 0 | 3941300 | 1749800 |
| QSFTMVAD | LASP1; LASP1; I | Q14847; C9J | LASP1_HUMAN | Unmodified | 0 | 0 | 0 | 244910 | 13180000 | 3183300 | 0 | 0 | 0 | 0 | 0 | 0 | 5484500 | 2944000 | 0 | 0 | 4540000 | 9900900 |
| QSFVLKEG | ARHGDIA; ARH | P52565; J3K | GDIR1_HUMAN | Unmodified | 0 | 0 | 0 | 0 | 828250 | 0 | 2600400 | 276680 | 0 | 0 | 0 | 0 | 0 | 0 | 0 | 0 | 1636900 | 0 |
| QSGESIDII | DDB1; DDB1; D | Q16531; A0 | DDB1_HUMAN | Unmodified | 0 | 0 | 0 | 0 | 0 | 0 | 0 | 199410 | 0 | 0 | 0 | 1020600 | 0 | 0 | 0 | 0 | 0 | 0 |
| QSGGTTAL | MRPL44 | Q9H9J2 | RM44_HUMAN | Unmodified | 0 | 0 | 0 | 0 | 639130 | 0 | 0 | 0 | 192800 | 112680 | 0 | 0 | 0 | 0 | 0 | 0 | 0 | 95873 |
| QSGYGGQ | RPL36A; RPL36 | P83881; Q96 | RL36A_HUMAN | Unmodified | 0 | 0 | 0 | 282210 | 0 | 0 | 0 | 0 | 0 | 0 | 0 | 528790 | 0 | 599070 | 82984 | 0 | 0 | 0 |
| QSHTLPFP | FAM120A; FAM | Q9NZB2; A0 | F120A_HUMAN | Unmodified | 1078100 | 126030 | 120890 | 439280 | 37773 | 0 | 0 | 0 | 328430 | 23824 | 147700 | 501140 | 88302 | 0 | 0 | 506540 | 0 | 18159 |
| QSIAIDDC | AP2M1; AP2M1 | Q96CW1; C9 | AP2M1_HUMAI | Unmodified | 239550 | 1657000 | 0 | 293630 | 0 | 85987 | 0 | 0 | 662900 | 114180 | 0 | 2653800 | 104050 | 0 | 207210 | 0 | 131830 | 248000 |
| QSIEVFAG | N/A | N/A | N/A | Unmodified | 0 | 0 | 0 | 0 | 0 | 0 | 2491000 | 0 | 0 | 407800 | 0 | 0 | 0 | 129200 | 0 | 0 | 3872000 | 233670 |
| QSIINPDW | NSF; NSF | P46459; I3L | NSF_HUMAN V | Unmodified | 375980 | 0 | 0 | 0 | 0 | 0 | 0 | 0 | 1609700 | 0 | 0 | 1404300 | 2122900 | 0 | 0 | 0 | 423220 | 0 |
| QSIIVAASS | FARP1; FARP1; | Q9Y4F1; C9J | FARP1_HUMAN | Unmodified | 71580 | 77530 | 0 | 0 | 378540 | 0 | 0 | 0 | 821880 | 149500 | 336890 | 1521500 | 0 | 0 | 0 | 144050 | 19712 | 28756 |
| QSIIVAASS | FARP1; FARP1; | Q9Y4F1; C9J | FARP1_HUMAN | Unmodified | 0 | 0 | 0 | 0 | 0 | 0 | 0 | 203330 | 536600 | 0 | 301610 | 440040 | 0 | 0 | 0 | 0 | 0 | 0 |
| QSINTIESLI | EXOC3; EXOC3 | O60645; D6I | EXOC3_HUMAI | Unmodified | 47879 | 0 | 4490100 | 1375700 | 0 | 0 | 0 | 0 | 504480 | 0 | 329480 | 0 | 0 | 0 | 211550 | 424070 | 0 | 0 |
| QSIWTSTIS | QDPR; QDPR; C | P09417; H0 | DHPR_HUMAN | Unmodified | 0 | 0 | 0 | 0 | 1297200 | 343890 | 0 | 0 | 632920 | 0 | 471330 | 0 | 0 | 680240 | 0 | 0 | 503010 | 479480 |
| QSLADTTKI | HACL1; HACL1 | Q9UJ83; B3K | HACL1_HUMAI | Oxidation (M) | 0 | 0 | 0 | 0 | 0 | 0 | 0 | 0 | 314830 | 0 | 0 | 0 | 0 | 0 | 0 | 0 | 0 | 0 |
| QSLADTTKI | HACL1; HACL1 | Q9UJ83; B3K | HACL1_HUMAI | Unmodified | 0 | 0 | 0 | 0 | 0 | 0 | 0 | 0 | 1432100 | 0 | 0 | 0 | 0 | 0 | 0 | 0 | 0 | 0 |
| QSLAQSHL | CIT; CIT; CIT; CI | O14578; A0 | CTRO_HUMAN | Unmodified | 0 | 0 | 143690 | 0 | 13679000 | 3722400 | 1334000 | 0 | 4854000 | 555320 | 212030 | 2152800 | 476920 | 699260 | 977560 | 0 | 3369700 | 1648400 |
| QSLEDMLE | B3GALT6 | Q96L58 | B3GT6_HUMAN | Unmodified | 0 | 0 | 0 | 0 | 970690 | 0 | 0 | 0 | 3508300 | 237390 | 1147500 | 1828000 | 722920 | 0 | 22845 | 0 | 465550 | 0 |
| QSLFFYPS | ARSA; ARSA | P15289; A0A | ARSA_HUMAN | Unmodified | 0 | 0 | 0 | 0 | 0 | 0 | 0 | 135750 | 1355600 | 0 | 0 | 0 | 0 | 986730 | 0 | 0 | 772930 | 715280 |
| QSLFFYPS | ARSA; ARSA | P15289; A0A | ARSA_HUMAN | Unmodified | 0 | 0 | 0 | 0 | 0 | 0 | 0 | 0 | 1431200 | 646280 | 0 | 0 | 0 | 0 | 0 | 0 | 0 | 0 |
| QSLFFYPS | ARSA; ARSA | P15289; A0A | ARSA_HUMAN | Unmodified | 0 | 0 | 0 | 0 | 0 | 0 | 281060 | 0 | 0 | 1471300 | 0 | 0 | 0 | 0 | 0 | 0 | 0 | 0 |
| QSLGELIG | TPI1 | P60174 | TPIS_HUMAN T | Unmodified | 608590 | 141640 | 5705400 | 2356000 | 1629700 | 236940 | 0 | 0 | 3501700 | 281970 | 1638900 | 24285000 | 501910 | 537790 | 1431600 | 15385000 | 0 | 1737700 |
| QSLGELIG | TPI1 | P60174 | TPIS_HUMAN T | Unmodified | 0 | 0 | 0 | 0 | 1065200 | 0 | 0 | 0 | 0 | 0 | 0 | 0 | 1722100 | 0 | 0 | 0 | 0 | 565850 |
| QSLGELIG | TPI1 | P60174 | TPIS_HUMAN T | Unmodified | 0 | 0 | 0 | 0 | 489920 | 0 | 0 | 0 | 0 | 0 | 0 | 0 | 0 | 485240 | 0 | 0 | 0 | 481310 |
| QSLLTAIR | COBLL1; COBLI | Q53SF7; A0 | COBL1_HUMAI | Unmodified | 0 | 0 | 0 | 0 | 0 | 0 | 0 | 0 | 0 | 0 | 0 | 0 | 0 | 0 | 0 | 0 | 0 | 0 |
| QSLLWGDI | IGSF9 | Q9P2J2 | TUTLA_HUMAN | Unmodified | 0 | 0 | 0 | 0 | 0 | 0 | 0 | 0 | 0 | 0 | 0 | 0 | 0 | 0 | 0 | 0 | 0 | 0 |
| QSLPPGLA | RPL38 | P63173 | RL38_HUMAN ( | Unmodified | 2095300 | 2479600 | 773220 | 601500 | 579510 | 0 | 473950 | 640940 | 402070 | 164440 | 510630 | 8073300 | 1339500 | 231190 | 1209500 | 0 | 0 | 0 |
| QSLQVIES | CLPP; CLPP | Q16740; M0I | CLPP_HUMAN | Unmodified | 0 | 0 | 0 | 0 | 0 | 0 | 0 | 0 | 551740 | 0 | 0 | 0 | 0 | 0 | 0 | 0 | 0 | 0 |
| QSLTMDPV | HSPA4 | P34932 | HSP74_HUMAI | Unmodified | 0 | 51309 | 83309 | 219090 | 1396100 | 0 | 0 | 0 | 738270 | 613170 | 597690 | 3747100 | 1711600 | 1299200 | 223660 | 218530 | 1445300 | 1023900 |
| QSLTMDPV | HSPA4 | P34932 | HSP74_HUMAI | Oxidation (M) | 0 | 0 | 0 | 0 | 942720 | 0 | 0 | 92588 | 415870 | 112300 | 0 | 1237500 | 474220 | 0 | 0 | 0 | 882500 | 336750 |
| QSLVLLR | FAM234B; FAM | A2RU67; Q6 | F234B_HUMAN | Unmodified | 0 | 0 | 0 | 0 | 0 | 0 | 0 | 0 | 1898400 | 0 | 0 | 0 | 0 | 0 | 0 | 0 | 0 | 0 |
| QSNAFKAL | KIF23; KIF23; KI | Q02241; A0 | KIF23_HUMAN | Unmodified | 0 | 0 | 0 | 0 | 901230 | 112700 | 242460 | 0 | 0 | 0 | 0 | 44315 | 65318 | 69236 | 0 | 0 | 592050 | 105350 |
| QSNNEANI | PDCD6IP | Q8WUM4 | PDC6I_HUMAN | Unmodified | 1.03E+08 | 1.14E+08 | 65126000 | 37514000 | 37125000 | 8447900 | 0 | 1297800 | 23414000 | 10615000 | 4601800 | 38204000 | 30790000 | 21004000 | 57208000 | 54282000 | 4500700 | 16125000 |
| QSNPLLIH | PREP; PREP | P48147; A0A | PPCE_HUMAN | Unmodified | 0 | 0 | 0 | 0 | 0 | 0 | 0 | 0 | 0 | 0 | 298930 | 2592000 | 0 | 0 | 0 | 0 | 0 | 0 |
| QSNVEVPF | IREB2; IREB2; IF | P48200; H0 | IREB2_HUMAN | Unmodified | 0 | 0 | 0 | 0 | 0 | 0 | 0 | 0 | 0 | 0 | 0 | 0 | 0 | 0 | 0 | 0 | 0 | 0 |
| QSPANGH | CASK | A0A2U3TZM | A0A2U3TZM1_ | Unmodified | 0 | 0 | 0 | 0 | 0 | 0 | 0 | 0 | 0 | 0 | 0 | 0 | 0 | 0 | 337810 | 107340 | 33309 | 0 |
| QSPASPPP | SCRIB; SCRIB; | Q14160; A0 | SCRIB_HUMAN | Unmodified | 0 | 0 | 195910 | 180940 | 0 | 0 | 0 | 0 | 1193300 | 0 | 0 | 527760 | 0 | 0 | 142650 | 162920 | 0 | 0 |
| QSPASPPP | SCRIB; SCRIB; | Q14160; A0 | SCRIB_HUMAN | Unmodified | 0 | 0 | 0 | 617470 | 1056400 | 0 | 404500 | 324690 | 836560 | 892090 | 0 | 0 | 302660 | 406560 | 69573 | 0 | 753290 | 1635300 |
| QSPASPPP | SCRIB; SCRIB; | Q14160; A0 | SCRIB_HUMAN | Unmodified | 0 | 0 | 0 | 0 | 0 | 0 | 0 | 0 | 0 | 160280 | 0 | 0 | 0 | 0 | 0 | 0 | 0 | 1036100 |
| QSPEDVYF | EPHA2 | P29317 | EPHA2_HUMAI | Unmodified | 0 | 0 | 0 | 0 | 959890 | 641450 | 0 | 0 | 0 | 0 | 253340 | 2196500 | 0 | 0 | 0 | 0 | 912450 | 3718100 |
| QSPEDVYF | EPHA2 | P29317 | EPHA2_HUMAI | Unmodified | 74130 | 92294 | 0 | 0 | 4840100 | 1257600 | 0 | 0 | 0 | 0 | 0 | 0 | 309550 | 0 | 0 | 0 | 8310900 | 4629700 |
| QSPLYSR | XXYLT1; XXYLT1 | Q8NBI6; A0 | XXLT1_HUMAN | Unmodified | 0 | 0 | 0 | 0 | 0 | 0 | 0 | 0 | 1250200 | 0 | 0 | 0 | 0 | 0 | 0 | 0 | 0 | 0 |
| QSPPESPIL | PTPRJ; PTPRJ; P | Q12913; A0 | PTPRJ_HUMAN | Unmodified | 0 | 314130 | 0 | 0 | 0 | 0 | 0 | 0 | 0 | 59958 | 799860 | 0 | 0 | 0 | 0 | 727970 | 0 | 0 |
| QSPPLPGR | ZDHHC5 | Q9C0B5 | ZDHC5_HUMA | Unmodified | 765210 | 429870 | 1404400 | 1594400 | 1038400 | 0 | 0 | 0 | 1044900 | 0 | 1161300 | 3168200 | 1028800 | 978590 | 2414400 | 2845900 | 795700 | 1372500 |
| QSPPLPGR | ZDHHC5 | Q9C0B5 | ZDHC5_HUMA | Unmodified | 0 | 0 | 0 | 0 | 1572200 | 0 | 0 | 0 | 0 | 0 | 0 | 0 | 1433600 | 1954600 | 0 | 0 | 0 | 4020300 |
| QSPTLQRN | USP14; USP14 | P54578; A6N | UBP14_HUMAI | Unmodified | 0 | 0 | 0 | 0 | 1424700 | 0 | 674510 | 572010 | 0 | 0 | 0 | 0 | 1066500 | 896650 | 0 | 0 | 982470 | 918530 |
| QSPVPILH | MPP7; MPP7 | Q5T2T1; S4F | MPP7_HUMAN | Unmodified | 0 | 0 | 0 | 0 | 1530800 | 0 | 0 | 0 | 0 | 0 | 0 | 0 | 0 | 0 | 0 | 0 | 0 | 573180 |
| QSQGILNA | HAUS3; HAUS3 | Q68CZ6; A0 | HAUS3_HUMAI | Oxidation (M) | 0 | 0 | 0 | 0 | 0 | 0 | 0 | 0 | 0 | 1045600 | 0 | 0 | 0 | 0 | 0 | 0 | 0 | 0 |
| QSRPYRLA | WDR1; WDR1 | O75083; D6I | WDR1_HUMAN | Unmodified | 0 | 0 | 0 | 0 | 6123000 | 1978000 | 630280 | 0 | 0 | 777760 | 0 | 0 | 3360100 | 3036400 | 0 | 0 | 1914700 | 2719500 |
| QSSFALLGI | TNPO2; TNPO1 | O14787; Q9 | TNPO2_HUMAI | Unmodified | 644120 | 0 | 0 | 0 | 0 | 0 | 0 | 0 | 0 | 0 | 365330 | 4351900 | 0 | 0 | 0 | 0 | 0 | 0 |
| QSSGASSS | DDX3X; DDX3X | O00571; A0 | DDX3X_HUMAI | Unmodified | 774230 | 578080 | 0 | 671350 | 0 | 0 | 0 | 0 | 0 | 0 | 0 | 164850 | 0 | 0 | 1249100 | 1139500 | 0 | 0 |
| QSSGPGAS | EEF1D; EEF1D | E9PL71; E9P | E9PL71_HUMA | Unmodified | 0 | 0 | 0 | 236110 | 218910 | 0 | 0 | 0 | 1500200 | 120520 | 1196500 | 4785800 | 156630 | 0 | 147880 | 187750 | 189730 | 129070 |
| QSSGPSSS | CLIP2 | Q9UDT6 | CLIP2_HUMAN | Unmodified | 0 | 0 | 0 | 0 | 0 | 0 | 0 | 0 | 0 | 0 | 0 | 0 | 0 | 0 | 0 | 0 | 0 | 329550 |
| QSSLEPVAI | CORO7; CORO | P57737; I3L | CORO7_HUMA | Unmodified | 0 | 0 | 0 | 0 | 0 | 0 | 0 | 0 | 0 | 0 | 0 | 1811200 | 0 | 0 | 0 | 0 | 0 | 0 |
| QSSMTVMI | VPS25; VPS25; | Q9BRG1; K7 | VPS25_HUMAN | 2 Oxidation (M) | 1577700 | 2107500 | 984510 | 677300 | 0 | 0 | 0 | 0 | 0 | 0 | 0 | 0 | 0 | 0 | 2672500 | 2545300 | 374300 | 0 |
| QSSMTVMI | VPS25; VPS25; | Q9BRG1; K7 | VPS25_HUMAN | Oxidation (M) | 821760 | 622950 | 40312 | 438090 | 0 | 0 | 0 | 0 | 0 | 0 | 0 | 1310400 | 0 | 0 | 1356700 | 249410 | 0 | 0 |
| QSSSASLE | VPS37B; VPS37 | Q9H9H4; F5 | VP37B_HUMAN | Unmodified | 648220 | 0 | 605550 | 258260 | 0 | 0 | 0 | 0 | 0 | 0 | 0 | 0 | 0 | 0 | 6991100 | 855210 | 0 | 0 |
| QSSSTNYT | LIMA1; LIMA1; L | Q9UHB6; F8 | LIMA1_HUMAN | Unmodified | 700090 | 0 | 0 | 521440 | 1348200 | 0 | 0 | 0 | 0 | 0 | 0 | 487180 | 619700 | 833750 | 0 | 0 | 239450 | 0 |
| QSTDEEVT | TCEA1; TCEA1; | P23193; E5R | TCEA1_HUMAN | Unmodified | 0 | 0 | 0 | 0 | 490600 | 0 | 0 | 0 | 0 | 486080 | 85632 | 0 | 432500 | 0 | 0 | 0 | 0 | 0 |
| QSTLNFDS | SLIT2; SLIT2; SL | O94813; H0 | SLIT2_HUMAN | Oxidation (M) | 138800 | 210900 | 0 | 0 | 0 | 0 | 440260 | 0 | 0 | 0 | 330600 | 2395500 | 0 | 0 | 0 | 0 | 742010 | 1511800 |
| QSTQVMA | TFG; TFG; TFG; | Q92734; A0 | TFG_HUMAN P | Oxidation (M) | 0 | 0 | 0 | 0 | 2019900 | 0 | 0 | 0 | 0 | 0 | 0 | 0 | 0 | 0 | 0 | 0 | 0 | 452020 |
| QSTSSQEA | COBL; COBL; C | O75128; J3K | COBL_HUMAN | Unmodified | 0 | 0 | 0 | 0 | 0 | 0 | 0 | 0 | 0 | 0 | 0 | 0 | 0 | 0 | 0 | 0 | 0 | 0 |
| QSTWEKPD | PRPF40A; PRPF | O75400; H0 | PR40A_HUMAN | Unmodified | 0 | 0 | 0 | 0 | 0 | 0 | 0 | 0 | 0 | 0 | 513430 | 1702700 | 0 | 0 | 0 | 0 | 0 | 0 |
| QSVANGVE | AP1M2; AP1M2 | Q9Y6Q5; K7I | AP1M2_HUMAI | Unmodified | 0 | 150770 | 0 | 0 | 0 | 0 | 0 | 0 | 0 | 0 | 0 | 932370 | 0 | 0 | 0 | 0 | 0 | 0 |
| QSVDHFAK | BZW2; BZW2; B | Q9Y6E2; B5I | BZW2_HUMAN | Unmodified | 0 | 0 | 121480 | 0 | 696650 | 312920 | 527350 | 263710 | 687680 | 526780 | 0 | 861410 | 263540 | 0 | 0 | 42549 | 0 | 34501 |
| QSVDKVTS | CALD1; CALD1 | Q05682; E7E | CALD1_HUMAI | Unmodified | 0 | 0 | 0 | 0 | 264260 | 483070 | 0 | 0 | 0 | 0 | 0 | 0 | 995270 | 0 | 0 | 0 | 695170 | 307160 |
| QSVELCK | CLTCL1; CLTC; | P53675; Q0 | CLH2_HUMAN | Unmodified | 1079600 | 0 | 506060 | 325460 | 482480 | 0 | 0 | 20323 | 0 | 151820 | 46704 | 0 | 304130 | 0 | 425030 | 593820 | 0 | 0 |
| QSVELCKK | CLTCL1; CLTC; | P53675; Q0 | CLH2_HUMAN | Unmodified | 0 | 296730 | 0 | 0 | 326210 | 0 | 0 | 33829 | 0 | 0 | 237700 | 3559400 | 0 | 0 | 0 | 0 | 100890 | 0 |
| QSVENDIH | KRT18; KRT18 | P05783; F8V | K1C18_HUMAN | Unmodified | 0 | 0 | 0 | 0 | 0 | 0 | 0 | 0 | 732020 | 44355 | 0 | 2784800 | 0 | 0 | 0 | 0 | 0 | 0 |
| QSVENDIH | KRT18; KRT18 | P05783; F8V | K1C18_HUMAN | Unmodified | 263710 | 0 | 373400 | 298910 | 358960 | 163780 | 0 | 33719 | 4158900 | 380400 | 3368600 | 19601000 | 396550 | 88291 | 0 | 328090 | 0 | 0 |
| QSVSRNRI | ITGB3; N/A; ITG | P05106; H3I | ITB3_HUMAN I | Oxidation (M) | 0 | 0 | 0 | 0 | 0 | 0 | 780740 | 0 | 0 | 0 | 0 | 0 | 0 | 0 | 0 | 0 | 0 | 0 |
| QSVSRNRI | ITGB3; N/A; ITG | P05106; H3I | ITB3_HUMAN I | Unmodified | 0 | 0 | 1463300 | 0 | 0 | 0 | 3711800 | 208300 | 0 | 0 | 0 | 0 | 1137200 | 5094500 | 0 | 0 | 0 | 0 |
| QSVSYVNI | N/A | N/A | N/A | Unmodified | 0 | 0 | 0 | 0 | 0 | 0 | 0 | 0 | 0 | 69577000 | 0 | 0 | 0 | 0 | 0 | 0 | 0 | 0 |
| QSVTYACN | CD55; CD55; C | P08174; B1A | DAF_HUMAN C | Unmodified | 1743600 | 977680 | 182840 | 412950 | 0 | 3720300 | 0 | 0 | 685040 | 0 | 834850 | 4099300 | 0 | 0 | 498980 | 556240 | 0 | 1553700 |
| QSYKGSPI | LTA4H | P09960 | LKHA4_HUMAI | Oxidation (M) | 0 | 0 | 0 | 0 | 0 | 0 | 0 | 0 | 0 | 0 | 463250 | 566800 | 0 | 0 | 0 | 0 | 0 | 0 |
| QTAAQLILK | MACROH2A1; I | O75367; B4I | H2AY_HUMAN | Unmodified | 6782100 | 2569000 | 0 | 0 | 0 | 0 | 0 | 2407100 | 0 | 0 | 0 | 0 | 64207 | 90584 | 0 | 22480 | 0 | 0 |

| | | | | | | | | | | | | | | | | | | | | | | |
|---|---|---|---|---|---|---|---|---|---|---|---|---|---|---|---|---|---|---|---|---|---|---|
| QTAAQVTLI | MACROH2A2; N | Q9P0M6; A0. | H2AW_HUMAN | Unmodified | 0 | 0 | 0 | 0 | 0 | 0 | 0 | 0 | 0 | 0 | 0 | 1056900 | 0 | 0 | 0 | 0 | 0 | 273340 |
| QTAEVALAI | BICD2 | Q8TD16 | BICD2_HUMAN | Unmodified | 93666 | 219970 | 0 | 121670 | 0 | 0 | 0 | 0 | 0 | 0 | 0 | 0 | 2049600 | 72734 | 32639 | 0 | 0 | 0 |
| QTALQQEA | NPTXR; NPTXR | O95502; A0A | NPTXR_HUMAN | Unmodified | 0 | 157680 | 0 | 215210 | 236450 | 0 | 0 | 0 | 2557800 | 58000 | 231480 | 0 | 0 | 346510 | 173550 | 0 | 0 | 0 |
| QTAQQIVSI | FAM120A | Q9NZB2 | F120A_HUMAN | Unmodified | 119600 | 123750 | 189130 | 411740 | 0 | 0 | 0 | 0 | 225270 | 0 | 69205 | 1050600 | 0 | 0 | 140990 | 0 | 27393 | 0 |
| QTATQLLK | RPL7A; RPL7A | P62424; Q5T | RL7A_HUMAN ( | Unmodified | 4039400 | 3634100 | 1044400 | 648820 | 0 | 0 | 0 | 0 | 640640 | 158270 | 835250 | 16053000 | 112800 | 59928 | 1842800 | 2114200 | 106440 | 0 |
| QTEIEGKR( | PALM2; PALM2/ | Q8IXS6; C9J. | PALM2_HUMAN | Unmodified | 0 | 0 | 0 | 0 | 305060 | 0 | 0 | 0 | 0 | 185460 | 0 | 0 | 0 | 0 | 0 | 0 | 0 | 97787 |
| QTELFAHFI | SMARCA5 | O60264 | SMCA5_HUMA | Unmodified | 1041900 | 595500 | 97364 | 0 | 0 | 51052 | 276220 | 226850 | 0 | 35904 | 181770 | 958320 | 0 | 130900 | 1034700 | 0 | 0 | 152810 |
| QTENLQRL | EXOC7; EXOC7 | Q9UPT5; B5I | EXOC7_HUMAI | Unmodified | 0 | 0 | 0 | 0 | 3163200 | 0 | 140100 | 78066 | 0 | 0 | 0 | 0 | 0 | 0 | 0 | 0 | 543960 | 939730 |
| QTEQTYW( | THBS4; THBS3; | P35443; P49 | TSP4_HUMAN ' | Unmodified | 0 | 0 | 264950 | 0 | 0 | 0 | 0 | 0 | 117790 | 0 | 0 | 0 | 0 | 0 | 0 | 0 | 0 | 0 |
| QTESLESLL | CKAP4 | Q07065 | CKAP4_HUMAI | Unmodified | 0 | 0 | 0 | 0 | 0 | 0 | 0 | 0 | 6907800 | 67778 | 0 | 0 | 0 | 0 | 0 | 0 | 0 | 0 |
| QTESTSFLE | PSMA6; PSMA6 | P60900; G3\ | PSA6_HUMAN | Unmodified | 0 | 155180 | 649460 | 3961900 | 2658300 | 0 | 0 | 175310 | 5451700 | 908000 | 0 | 11663000 | 1536500 | 801360 | 419950 | 587680 | 945730 | 2051200 |
| QTESTSFLE | PSMA6; PSMA6 | P60900; G3\ | PSA6_HUMAN | Unmodified | 0 | 0 | 0 | 0 | 1441700 | 0 | 0 | 0 | 0 | 113740 | 386120 | 0 | 0 | 0 | 0 | 0 | 231980 | 676780 |
| QTEVLLQPI | SH3GLB1; SH3( | Q9Y371; A0/ | SHLB1_HUMAN | Unmodified | 0 | 0 | 0 | 0 | 741780 | 0 | 0 | 89672 | 0 | 791340 | 0 | 0 | 3522700 | 0 | 0 | 0 | 563400 | 0 |
| QTFEAAILT( | EXOSC4; EXOS( | Q9NPD3; E9 | EXOS4_HUMAN | Unmodified | 0 | 0 | 0 | 0 | 0 | 0 | 0 | 0 | 0 | 0 | 0 | 2378100 | 0 | 0 | 0 | 0 | 0 | 0 |
| QTFGYGAR | AHSA1; AHSA1; | O95433; H0\ | AHSA1_HUMAI | Unmodified | 79632 | 0 | 117450 | 0 | 0 | 0 | 128720 | 5068.9 | 69278 | 269150 | 406880 | 3783700 | 0 | 0 | 314080 | 443630 | 0 | 0 |
| QTFIDPLQL | SH3GL3 | Q99963 | SH3G3_HUMAI | Unmodified | 0 | 0 | 60851 | 71732 | 0 | 0 | 0 | 0 | 121710 | 0 | 0 | 0 | 383670 | 462990 | 234390 | 116170 | 0 | 0 |
| QTFIGHESI | GNB2; GNB2; G | P62879; E7E | GBB2_HUMAN | Unmodified | 3057000 | 866980 | 0 | 1283800 | 0 | 0 | 0 | 0 | 0 | 0 | 2240200 | 3868700 | 0 | 0 | 1570600 | 2304100 | 0 | 0 |
| QTFTGHES | GNB1; GNB1 | P62873; B3K | GBB1_HUMAN | Unmodified | 0 | 0 | 0 | 0 | 0 | 0 | 0 | 0 | 10820000 | 0 | 2175500 | 12582000 | 0 | 0 | 552640 | 0 | 0 | 0 |
| QTGEGVGI | LARS1; LARS1; | Q9P2J5; A0A | SYLC_HUMAN | Unmodified | 0 | 0 | 0 | 0 | 0 | 0 | 0 | 0 | 0 | 0 | 99800 | 1393000 | 0 | 0 | 0 | 0 | 0 | 0 |
| QTGGLGISI | SCRIB; SCRIB; ! | Q14160; H0\ | SCRIB_HUMAN | Unmodified | 0 | 0 | 225120 | 175080 | 505420 | 188580 | 0 | 559120 | 4869900 | 419730 | 245710 | 1128300 | 0 | 0 | 276700 | 189830 | 815920 | 266870 |
| QTGGLGISI | SCRIB; SCRIB; ! | Q14160; H0\ | SCRIB_HUMAN | Unmodified | 0 | 0 | 0 | 0 | 0 | 0 | 0 | 0 | 2994800 | 95874 | 0 | 0 | 0 | 0 | 0 | 0 | 0 | 0 |
| QTGIVLNRI | SF3B2; SF3B2; ! | Q13435; E9I | SF3B2_HUMAN | Unmodified | 0 | 0 | 0 | 313800 | 0 | 0 | 0 | 0 | 116530 | 0 | 0 | 1554600 | 0 | 0 | 0 | 0 | 0 | 0 |
| QTGLLLLGI | PIPOX; PIPOX; F | Q9P0Z9; K7E | SOX_HUMAN P | Unmodified | 0 | 0 | 0 | 0 | 0 | 0 | 0 | 0 | 873950 | 0 | 0 | 0 | 0 | 0 | 0 | 0 | 0 | 0 |
| QTIDNSQG | YWHAQ | P27348 | 1433T_HUMAN | Unmodified | 1802800 | 0 | 6512800 | 4881800 | 4022700 | 3789600 | 289880 | 251410 | 11064000 | 947830 | 11844000 | 70807000 | 2055800 | 1360900 | 12240000 | 7301400 | 1477100 | 2133800 |
| QTIERLEQE | SLK | Q9H2G2 | SLK_HUMAN S' | Unmodified | 61778 | 37516 | 53193 | 69650 | 0 | 0 | 0 | 0 | 978750 | 0 | 239040 | 1941100 | 67188 | 18697 | 126710 | 140600 | 157280 | 0 |
| QTIETEEVN | ARHGAP4; ARH | P98171; E9P | RHG04_HUMAI | Unmodified | 0 | 0 | 0 | 0 | 206040 | 0 | 0 | 0 | 0 | 0 | 14445 | 224780 | 0 | 0 | 0 | 33166 | 0 | 0 |
| QTIGNSCG | UCHL1; UCHL1 | P09936; D6F | UCHL1_HUMAI | Unmodified | 0 | 0 | 0 | 0 | 11769000 | 3715600 | 1005000 | 444200 | 11031000 | 1983400 | 10459000 | 44033000 | 0 | 0 | 0 | 0 | 0 | 0 |
| QTIGNSCG | UCHL1; UCHL1 | P09936; D6F | UCHL1_HUMAI | Unmodified | 0 | 0 | 0 | 0 | 2.03E+08 | 22811000 | 33131000 | 266540 | 10875000 | 32559000 | 5548100 | 23528000 | 0 | 0 | 0 | 0 | 0 | 0 |
| QTIGNSCG | UCHL1; UCHL1 | P09936; D6F | UCHL1_HUMAI | Unmodified | 0 | 0 | 0 | 0 | 3365000 | 0 | 0 | 0 | 0 | 297440 | 0 | 0 | 0 | 0 | 0 | 0 | 0 | 0 |
| QTIMETLE( | EXOC3 | O60645 | EXOC3_HUMAI | Unmodified | 0 | 0 | 0 | 0 | 0 | 500590 | 0 | 0 | 367200 | 0 | 0 | 0 | 0 | 0 | 882600 | 0 | 0 | 0 |
| QTISNACG | UCHL3; UCHL3 | P15374; Q5T | UCHL3_HUMAI | Unmodified | 0 | 0 | 0 | 0 | 0 | 0 | 2042100 | 0 | 0 | 0 | 803260 | 2968100 | 930960 | 681220 | 600170 | 1277500 | 910890 | 601540 |
| QTITKWPG | ABCF2 | Q9UG63 | ABCF2_HUMAN | Unmodified | 0 | 0 | 0 | 0 | 0 | 0 | 0 | 0 | 0 | 0 | 0 | 0 | 0 | 0 | 0 | 0 | 370900 | 0 |
| QTKPEFDQ | EXOC7; EXOC7 | Q9UPT5; B4I | EXOC7_HUMAI | Unmodified | 115500 | 71098 | 153970 | 0 | 3852000 | 1240900 | 508860 | 160740 | 994120 | 720010 | 0 | 1178600 | 647740 | 358930 | 267380 | 218440 | 0 | 607920 |
| QTKVEGELI | MYH10 | P35580 | MYH10_HUMAI | Unmodified | 0 | 0 | 0 | 0 | 0 | 58496 | 0 | 0 | 0 | 0 | 54173 | 519430 | 0 | 0 | 0 | 0 | 43835 | 0 |
| QTLDALNY | SLK | Q9H2G2 | SLK_HUMAN S' | Unmodified | 136750 | 0 | 277910 | 0 | 342470 | 0 | 1592100 | 1558100 | 1446900 | 172040 | 1069600 | 5581900 | 596990 | 545420 | 537840 | 594640 | 0 | 0 |
| QTLENERG | MYH9 | P35579 | MYH9_HUMAN | Unmodified | 414410 | 375280 | 2865900 | 1191000 | 4269200 | 917620 | 1330500 | 365060 | 769620 | 557740 | 1430800 | 4239700 | 1771300 | 1291500 | 2311100 | 2018900 | 1821500 | 1865300 |
| QTLGVTQK | SLC25A24 | Q6NUK1 | SCMC1_HUMA | Unmodified | 0 | 0 | 6401.7 | 0 | 12567000 | 0 | 2872600 | 552300 | 3377800 | 657360 | 0 | 0 | 0 | 0 | 166160 | 0 | 2385200 | 2379900 |
| QTLIQHFQ | APLP2 | Q06481 | APLP2_HUMAN | Unmodified | 0 | 0 | 0 | 0 | 0 | 0 | 0 | 0 | 2087600 | 155910 | 0 | 389000 | 0 | 0 | 0 | 0 | 0 | 104160 |
| QTLMWSAI | DDX5; DDX17; I | P17844; Q92 | DDX5_HUMAN | Oxidation (M) | 0 | 0 | 0 | 0 | 0 | 0 | 0 | 0 | 0 | 0 | 0 | 3058100 | 0 | 0 | 0 | 106980 | 0 | 0 |
| QTLNAIKLE | EIF5B; EIF5B | O60841; A0/ | IF2P_HUMAN E | Unmodified | 0 | 0 | 0 | 0 | 0 | 0 | 0 | 0 | 0 | 0 | 0 | 0 | 0 | 0 | 0 | 0 | 0 | 0 |
| QTLQSEQP | PSMC2; PSMC2 | P35998; C9J | PRS7_HUMAN | Unmodified | 0 | 0 | 693350 | 172050 | 235230 | 0 | 0 | 0 | 622080 | 0 | 323950 | 5577900 | 0 | 0 | 180840 | 0 | 0 | 166470 |
| QTLSTPGTI | NTPCR; NTPCR | Q9BSD7; Q5 | NTPCR_HUMAI | Unmodified | 186150 | 0 | 265810 | 0 | 0 | 0 | 0 | 0 | 618150 | 0 | 0 | 1403700 | 0 | 0 | 0 | 602420 | 0 | 0 |
| QTMEALK | CIT; CIT; CIT; CI | O14578; A0/ | CTRO_HUMAN | Unmodified | 0 | 0 | 0 | 0 | 1929100 | 0 | 0 | 0 | 17214000 | 880950 | 0 | 474920 | 0 | 0 | 0 | 0 | 166250 | 0 |
| QTMQVDEI | ACAA2; ACAA2; | P42765; A0A | THIM_HUMAN : | Unmodified | 0 | 0 | 0 | 0 | 0 | 0 | 0 | 0 | 2232900 | 0 | 0 | 0 | 0 | 0 | 0 | 0 | 0 | 0 |
| QTMQVDEI | ACAA2; ACAA2; | P42765; A0A | THIM_HUMAN : | Oxidation (M) | 0 | 0 | 0 | 0 | 0 | 0 | 0 | 370400 | 2310200 | 158630 | 0 | 0 | 0 | 0 | 0 | 0 | 0 | 0 |
| QTMTALGII | ITGA1 | P56199 | ITA1_HUMAN I | Unmodified | 270200 | 482280 | 0 | 0 | 0 | 0 | 0 | 0 | 561710 | 0 | 0 | 0 | 0 | 0 | 0 | 0 | 529500 | 1204100 |
| QTMTALGII | ITGA1 | P56199 | ITA1_HUMAN I | Oxidation (M) | 489670 | 364420 | 0 | 232430 | 0 | 0 | 0 | 0 | 0 | 0 | 0 | 615250 | 0 | 0 | 0 | 0 | 0 | 0 |
| QTNPSAME | VCP; VCP; VCP; | P55072; C9J | TERA_HUMAN ' | Oxidation (M) | 0 | 0 | 0 | 0 | 0 | 0 | 0 | 0 | 0 | 0 | 1018400 | 3854400 | 0 | 0 | 0 | 0 | 0 | 0 |
| QTPADGEA | SLC12A2; SLC1 | P55011; G3) | S12A2_HUMAN | Unmodified | 0 | 0 | 0 | 0 | 1474600 | 223860 | 0 | 0 | 0 | 0 | 0 | 0 | 0 | 0 | 0 | 0 | 0 | 15235000 |
| QTPSDFLK | LSM6 | P62312 | LSM6_HUMAN | Unmodified | 0 | 0 | 0 | 227630 | 976910 | 0 | 75752 | 60891 | 299860 | 341130 | 0 | 1625100 | 0 | 171880 | 0 | 0 | 0 | 180000 |
| QTPSFWIL/ | NAE1 | Q13564 | ULA1_HUMAN | Unmodified | 0 | 0 | 0 | 206320 | 1333200 | 194220 | 215170 | 87897 | 0 | 0 | 1250500 | 1150700 | 2345600 | 0 | 0 | 0 | 351860 | 708200 |
| QTQTFTTYS | HSPA8; HSPA8; | P11142; E9P | HSP7C_HUMAI | Unmodified | 4498400 | 12373000 | 1730700 | 0 | 0 | 0 | 0 | 0 | 5121200 | 3232700 | 699870 | 7311300 | 0 | 0 | 5582500 | 3947200 | 0 | 0 |
| QTQTFTTYS | HSPA8; HSPA8; | P11142; E9P | HSP7C_HUMAI | Oxidation (M) | 4859000 | 5063200 | 4205900 | 3465000 | 8766400 | 8084100 | 4571700 | 0 | 4523700 | 14583000 | 5153200 | 3841200 | 5819400 | 3503400 | 7358500 | 1200200 | 5339300 | 0 |
| QTQTFTTYS | HSPA8; HSPA8; | P11142; E9P | HSP7C_HUMAI | Unmodified | 0 | 0 | 0 | 0 | 4871300 | 1869200 | 0 | 0 | 0 | 4759600 | 0 | 0 | 0 | 0 | 0 | 0 | 1551900 | 0 |
| QTRTASDM | PSMD12 | O00232 | PSD12_HUMAN | Unmodified | 0 | 0 | 0 | 0 | 145720 | 0 | 203640 | 0 | 0 | 0 | 0 | 0 | 0 | 0 | 0 | 0 | 166820 | 0 |
| QTSAYNISI | PLXNA1 | Q9UIW2 | PLXA1_HUMAN | Unmodified | 0 | 0 | 44124 | 77306 | 1168300 | 0 | 0 | 0 | 57513 | 0 | 0 | 416920 | 728490 | 977260 | 0 | 113830 | 1305500 | 2161100 |
| QTSDGNW | CPNE3; CPNE3 | O75131; A0/ | CPNE3_HUMAI | Unmodified | 165780 | 0 | 0 | 48869 | 0 | 0 | 0 | 0 | 3842200 | 104090 | 0 | 0 | 83200 | 0 | 468650 | 0 | 0 | 0 |
| QTSDGNW | CPNE3; CPNE3 | O75131; A0/ | CPNE3_HUMAI | Oxidation (M) | 0 | 0 | 0 | 679260 | 0 | 0 | 0 | 0 | 1402800 | 272600 | 0 | 361930 | 0 | 0 | 0 | 637860 | 0 | 0 |
| QTSDGNW | CPNE3; CPNE3 | O75131; A0/ | CPNE3_HUMAI | Unmodified | 0 | 0 | 0 | 0 | 2896000 | 0 | 1122700 | 0 | 6968900 | 2572300 | 0 | 309930 | 761530 | 2250100 | 0 | 189210 | 945650 | 5773600 |
| QTSDGNW | CPNE3; CPNE3 | O75131; A0/ | CPNE3_HUMAI | Oxidation (M) | 143180 | 0 | 1445400 | 856380 | 1192500 | 0 | 0 | 0 | 2368800 | 2333600 | 0 | 0 | 1448000 | 1523000 | 953470 | 808920 | 0 | 1434400 |
| QTSGGPVI | ATP5PF; ATP5PI | P18859; A8N | ATP5J_HUMAN | Unmodified | 0 | 0 | 0 | 0 | 0 | 0 | 0 | 0 | 2143700 | 253500 | 0 | 0 | 0 | 0 | 0 | 0 | 0 | 0 |
| QTSSYNIP/ | PLXNA2 | O75051 | PLXA2_HUMAN | Unmodified | 0 | 0 | 0 | 0 | 0 | 0 | 0 | 0 | 1204700 | 383670 | 0 | 459260 | 0 | 0 | 0 | 0 | 0 | 0 |
| QTTSNLGH | PSMD14 | O00487 | PSDE_HUMAN | Unmodified | 0 | 0 | 0 | 0 | 0 | 0 | 62227 | 0 | 0 | 0 | 0 | 5337100 | 0 | 0 | 0 | 322100 | 0 | 0 |
| QTTTVVAGI | PROM1 | O43490 | PROM1_HUMA | Unmodified | 190520 | 0 | 0 | 0 | 1304400 | 0 | 1144800 | 1386000 | 1087000 | 154200 | 424670 | 6287300 | 0 | 0 | 0 | 0 | 63125 | 364080 |
| QTTVSNSQ | YWHAB | P31946 | 1433B_HUMAN | Unmodified | 0 | 0 | 0 | 178420 | 0 | 0 | 0 | 0 | 0 | 0 | 1656500 | 1339000 | 0 | 0 | 0 | 464630 | 0 | 0 |
| QTTVSNSQ | YWHAB | P31946 | 1433B_HUMAN | Unmodified | 2752900 | 2497800 | 6742800 | 5649800 | 7185000 | 1821600 | 611220 | 517490 | 9740200 | 269400 | 6741300 | 43657000 | 3519000 | 2018900 | 18265000 | 11625000 | 3874800 | 3908000 |
| QTVADQVL | EIF6 | P56537 | IF6_HUMAN Eu | Unmodified | 1504000 | 2169100 | 349270 | 0 | 0 | 0 | 0 | 0 | 0 | 0 | 0 | 3731500 | 1544100 | 0 | 3553300 | 600690 | 786510 | 0 |
| QTVAIAELT | PRKG2 | Q13237 | KGP2_HUMAN | Unmodified | 0 | 0 | 0 | 0 | 0 | 0 | 1285700 | 767270 | 0 | 0 | 0 | 0 | 0 | 0 | 0 | 0 | 0 | 0 |
| QTVAVGVII | EEF1A2; EEF1A | Q05639; Q5\ | EF1A2_HUMAN | Unmodified | 46806000 | 0 | 61364000 | 58825000 | 1.31E+08 | 0 | 0 | 0 | 0 | 0 | 39254000 | 1.85E+08 | 0 | 0 | 80765000 | 85142000 | 0 | 0 |
| QTVAVGVII | EEF1A1P5; EEF | Q5VTE0; P68 | EF1A3_HUMAN | Unmodified | 0 | 0 | 0 | 0 | 928100 | 0 | 1095900 | 797140 | 0 | 269480 | 0 | 0 | 601620 | 613740 | 0 | 0 | 4482700 | 887650 |
| QTVAVGVII | EEF1A1P5; EEF | Q5VTE0; P68 | EF1A3_HUMAN | Unmodified | 0 | 0 | 0 | 0 | 0 | 0 | 0 | 202510 | 0 | 0 | 0 | 0 | 0 | 0 | 0 | 0 | 1057300 | 383960 |
| QTVNRGP( | PURA | Q00577 | PURA_HUMAN | Unmodified | 672300 | 720660 | 0 | 0 | 0 | 0 | 0 | 0 | 0 | 0 | 0 | 11256000 | 0 | 0 | 0 | 514120 | 0 | 0 |
| QTVNRGP( | PURA | Q00577 | PURA_HUMAN | Unmodified | 0 | 0 | 0 | 0 | 0 | 0 | 0 | 0 | 0 | 0 | 0 | 1136900 | 0 | 0 | 0 | 0 | 0 | 0 |
| QTYEQANK | DLG3; DLG3 | Q92796; Q5. | DLG3_HUMAN | Unmodified | 0 | 0 | 0 | 0 | 0 | 18815 | 0 | 0 | 64830 | 133460 | 0 | 447810 | 0 | 0 | 0 | 0 | 293220 | 81235 |
| QTYFLPVIG | PSMC3; PSMC3 | P17980; E9P | PRS6A_HUMAN | Unmodified | 3319200 | 0 | 370450 | 509610 | 1411800 | 449210 | 233100 | 0 | 4377600 | 709660 | 705990 | 4762200 | 2277000 | 1087400 | 918640 | 104380 | 882390 | 1424000 |
| QTYSTEPNI | RPL28; RPL28; I | P46779; H0\ | RL28_HUMAN ( | Unmodified | 4487700 | 5712500 | 3531200 | 2173600 | 1859300 | 0 | 935510 | 635110 | 1444200 | 184710 | 1487800 | 16082000 | 0 | 0 | 5362500 | 1544000 | 111760 | 709480 |
| QVAASTAQ | TLN2; TLN1; TLI | Q9Y4G6; Q9 | TLN2_HUMAN ' | Unmodified | 0 | 128330 | 170010 | 923630 | 3496500 | 694620 | 2331500 | 1410600 | 323000 | 467990 | 0 | 655520 | 1760100 | 2456200 | 245750 | 168970 | 740930 | 2167600 |
| QVAEAYEVI | DNAJB6; DNAJE | O75190; Q8\ | DNJB6_HUMAN | Unmodified | 2770400 | 5288000 | 3888600 | 0 | 0 | 0 | 0 | 0 | 1088700 | 0 | 0 | 0 | 5916500 | 0 | 2677000 | 1791800 | 0 | 0 |
| QVAEQFLN | UQCRC2; UQCI | P22695; H3E | QCR2_HUMAN | Unmodified | 0 | 0 | 0 | 0 | 0 | 0 | 0 | 0 | 3613600 | 0 | 0 | 0 | 0 | 0 | 0 | 0 | 0 | 0 |
| QVAEQFLN | UQCRC2; UQCI | P22695; H3E | QCR2_HUMAN | Oxidation (M) | 0 | 0 | 0 | 0 | 0 | 0 | 0 | 0 | 0 | 504960 | 0 | 0 | 0 | 0 | 0 | 0 | 0 | 0 |
| QVAEQFLN | UQCRC2; UQCI | P22695; H3E | QCR2_HUMAN | Unmodified | 0 | 0 | 0 | 0 | 0 | 0 | 66628 | 0 | 0 | 275420 | 0 | 0 | 0 | 0 | 0 | 0 | 0 | 0 |
| QVAEQGGI | STK10 | O94804 | STK10_HUMAN | Unmodified | 0 | 0 | 0 | 0 | 1855900 | 278670 | 0 | 0 | 0 | 0 | 0 | 0 | 511650 | 229540 | 0 | 0 | 733620 | 1588600 |
| QVAGTMKI | ATP5F1A | P25705 | ATPA_HUMAN / | Oxidation (M) | 0 | 0 | 0 | 0 | 0 | 0 | 0 | 0 | 1059300 | 209120 | 0 | 0 | 0 | 0 | 0 | 0 | 0 | 0 |
| QVAGTMKI | ATP5F1A | P25705 | ATPA_HUMAN / | Unmodified | 0 | 46196 | 0 | 0 | 0 | 565330 | 0 | 0 | 7916400 | 277740 | 41958 | 0 | 0 | 0 | 0 | 0 | 0 | 0 |
| QVALWDTI | CORO1A; COR( | P31146; H3E | COR1A_HUMAI | Unmodified | 122160 | 215720 | 327740 | 0 | 0 | 0 | 0 | 0 | 0 | 0 | 2227200 | 10023000 | 139330 | 0 | 659920 | 375030 | 0 | 0 |
| QVAQQEA( | PHB2; PHB2; PI | Q99623; F5( | PHB2_HUMAN | Unmodified | 0 | 0 | 0 | 0 | 270170 | 0 | 745190 | 327410 | 938410 | 2415900 | 0 | 0 | 217190 | 0 | 0 | 0 | 0 | 0 |
| QVASFGSA | PARP4 | Q9UKK3 | PARP4_HUMAN | Unmodified | 0 | 0 | 0 | 0 | 385330 | 0 | 0 | 0 | 0 | 0 | 0 | 462130 | 445650 | 0 | 0 | 0 | 0 | 227790 |
| QVATALQN | VCL | P18206 | VINC_HUMAN ' | Unmodified | 436740 | 0 | 449690 | 0 | 693870 | 584870 | 745290 | 223460 | 0 | 555800 | 0 | 1731600 | 113640 | 0 | 993270 | 913270 | 6863800 | 0 |
| QVATAVQV | ITGA5; ITGA5 | P08648; H0\ | ITA5_HUMAN I | Unmodified | 691060 | 228500 | 266050 | 71679 | 0 | 0 | 0 | 0 | 0 | 0 | 0 | 0 | 0 | 316590 | 1923100 | 1852700 | 0 | 0 |
| QVCPLDNI | RBX1 | P62877 | RBX1_HUMAN | Unmodified | 0 | 0 | 0 | 294400 | 0 | 0 | 0 | 0 | 0 | 0 | 0 | 2669300 | 0 | 0 | 159600 | 122040 | 228350 | 0 |
| QVCQESQI | MGAT5 | Q09328 | MGT5A_HUMAI | Unmodified | 0 | 2464900 | 0 | 0 | 15276000 | 0 | 0 | 0 | 2022000 | 0 | 0 | 0 | 0 | 0 | 0 | 0 | 0 | 3303200 |
| QVCQLPGI | MTCH2; MTCH2 | Q9Y6C9; E9I | MTCH2_HUMA | Unmodified | 0 | 0 | 0 | 0 | 1971200 | 0 | 0 | 0 | 1856200 | 0 | 0 | 0 | 0 | 0 | 0 | 0 | 0 | 0 |
| QVCQLPGI | MTCH2; MTCH2 | Q9Y6C9; E9I | MTCH2_HUMA | Unmodified | 0 | 0 | 0 | 0 | 0 | 0 | 0 | 0 | 3831400 | 1033900 | 60314 | 0 | 0 | 0 | 0 | 0 | 89095 | 0 |
| QVCTDVIR | COL21A1; COL: | Q96P44; F5( | COLA1_HUMAI | Unmodified | 83621 | 0 | 0 | 0 | 0 | 0 | 0 | 0 | 3871000 | 0 | 0 | 0 | 0 | 0 | 0 | 0 | 0 | 0 |
| QVDAESWI | TARS1; TARS1; ' | P26639; D6F | SYTC_HUMAN ' | Unmodified | 0 | 0 | 580510 | 0 | 296410 | 140200 | 0 | 0 | 581720 | 52509 | 836080 | 5205300 | 202240 | 139440 | 609280 | 431970 | 244570 | 182100 |
| QVDGDNSI | TFRC; TFRC | P02786; F8V | TFR1_HUMAN ' | Unmodified | 2218300 | 2484700 | 9467300 | 9920600 | 2161300 | 209800 | 628780 | 115070 | 1572200 | 0 | 0 | 0 | 284190 | 286000 | 6171200 | 7310700 | 0 | 223050 |
| QVDGDNSI | TFRC; TFRC | P02786; F8V | TFR1_HUMAN ' | Oxidation (M) | 0 | 3751700 | 1750400 | 2454700 | 131080 | 1237800 | 0 | 0 | 88134 | 69178 | 0 | 274770 | 297430 | 59993 | 1152600 | 1326100 | 2189200 | 42044000 |
| QVDGDNSI | TFRC; TFRC | P02786; F8V | TFR1_HUMAN ' | Unmodified | 0 | 0 | 0 | 984010 | 4155900 | 0 | 0 | 0 | 0 | 0 | 0 | 0 | 1605600 | 0 | 0 | 0 | 307090 | 3181300 |
| QVDLENVV | PLOD2; PLOD2 | O00469; E7E | PLOD2_HUMAI | Unmodified | 0 | 0 | 0 | 0 | 0 | 0 | 0 | 0 | 700490 | 0 | 0 | 0 | 0 | 0 | 0 | 0 | 0 | 0 |
| QVDLLAYIA | UEVLD | Q8IX04 | UEVLD_HUMAI | Unmodified | 0 | 0 | 0 | 0 | 0 | 29607 | 0 | 0 | 1957100 | 0 | 0 | 2877400 | 0 | 0 | 0 | 0 | 0 | 0 |
| QVDPAYTG | EPS15L1; EPS1 | Q9UBC2; K7 | EP15R_HUMAN | Unmodified | 0 | 377060 | 107820 | 0 | 381010 | 12427 | 0 | 40468 | 0 | 0 | 0 | 1468900 | 0 | 0 | 0 | 210790 | 0 | 0 |
| QVDQLTNI | VIM; VIM | P08670; B0Y | VIME_HUMAN ' | Unmodified | 0 | 0 | 0 | 0 | 1015500 | 75553 | 0 | 0 | 1191200 | 0 | 24913 | 0 | 0 | 0 | 0 | 0 | 0 | 640940 |
| QVDRIDWI | YKT6; YKT6; YK | O15498; A0/ | YKT6_HUMAN ! | Unmodified | 1690200 | 2415900 | 752210 | 593890 | 0 | 0 | 0 | 0 | 2989100 | 0 | 0 | 3530700 | 0 | 0 | 2693400 | 866340 | 0 | 0 |
| QVDVTSFA | AGRN; AGRN; A | O00468; A0/ | AGRIN_HUMAN | Unmodified | 1262800 | 303300 | 1246600 | 670030 | 0 | 0 | 0 | 0 | 0 | 223560 | 0 | 0 | 0 | 306790 | 765060 | 1207900 | 0 | 317330 |
| QVDVVITC | AHCYL1 | O43865 | SAHH2_HUMAI | Unmodified | 0 | 0 | 0 | 579810 | 0 | 0 | 0 | 0 | 0 | 0 | 0 | 0 | 0 | 0 | 1615500 | 1291500 | 123920 | 0 |
| QVEDDIQ( | PSMC2; PSMC2 | P35998; C9J | PRS7_HUMAN | Unmodified | 0 | 0 | 0 | 0 | 956980 | 0 | 0 | 0 | 0 | 0 | 147700 | 0 | 0 | 0 | 0 | 0 | 555700 | 1548300 |
| QVEDEKNS | MYH9 | P35579 | MYH9_HUMAN | Unmodified | 0 | 0 | 0 | 0 | 2167800 | 0 | 0 | 0 | 0 | 0 | 0 | 0 | 377880 | 0 | 0 | 0 | 223940 | 986630 |
| QVEEALHQ | PSMD9; PSMD9 | O00233; F5I | PSMD9_HUMA | Unmodified | 0 | 0 | 0 | 0 | 403020 | 63641 | 0 | 0 | 0 | 0 | 0 | 0 | 907100 | 205220 | 0 | 0 | 21042 | 298140 |
| QVEEELQE | PAIP2B | Q9ULR5 | PAI2B_HUMAN | Unmodified | 1312800 | 717060 | 179550 | 0 | 0 | 0 | 0 | 0 | 0 | 45142 | 0 | 0 | 0 | 0 | 443920 | 344450 | 0 | 0 |

| | | | | | | | | | | | | | | | | | | | | | | |
|---|---|---|---|---|---|---|---|---|---|---|---|---|---|---|---|---|---|---|---|---|---|---|
| QVEELYHS | SPTAN1; SPTAN | Q13813; A0A | SPTN1_HUMAN | Unmodified | 0 | 0 | 0 | 0 | 0 | 0 | 0 | 0 | 948090 | 549360 | 291920 | 1561000 | 0 | 0 | 0 | 0 | 1420400 | 467370 |
| QVEGKNPV | SSBP1; SSBP1; | Q04837; C9J | SSBP_HUMAN | Oxidation (M) | 0 | 0 | 0 | 0 | 0 | 0 | 0 | 0 | 4275700 | 0 | 336730 | 2114200 | 0 | 0 | 0 | 0 | 0 | 0 |
| QVEGMEDV | AZGP1; AZGP1 | P25311; C9J | ZA2G_HUMAN | Oxidation (M) | 0 | 0 | 161620 | 370200 | 0 | 0 | 0 | 0 | 0 | 0 | 3948800 | 305090 | 0 | 0 | 433600 | 226450 | 0 | 0 |
| QVEGMEDV | AZGP1; AZGP1 | P25311; C9J | ZA2G_HUMAN | Unmodified | 0 | 0 | 345020 | 461670 | 0 | 0 | 0 | 0 | 0 | 0 | 5480600 | 627690 | 0 | 0 | 802880 | 0 | 0 | 0 |
| QVEIAQR | ASS1; ASS1; AS | P00966; Q5T | ASSY_HUMAN | Unmodified | 16966 | 0 | 128360 | 0 | 0 | 0 | 40129 | 23614 | 352750 | 26020 | 16645 | 1302400 | 0 | 0 | 1821800 | 75936 | 0 | 0 |
| QVEIAQRE | ASS1; ASS1; AS | P00966; Q5T | ASSY_HUMAN | Unmodified | 201990 | 0 | 0 | 0 | 1189500 | 0 | 2966300 | 1048900 | 432900 | 332590 | 0 | 0 | 0 | 53557 | 293720 | 199620 | 387170 | 842860 |
| QVEILNR | EPHX1; EPHX1 | P07099; B1A | HYEP_HUMAN | Unmodified | 0 | 0 | 296900 | 114700 | 109370 | 0 | 103100 | 0 | 814070 | 32874 | 308110 | 995840 | 0 | 0 | 496000 | 192890 | 114730 | 0 |
| QVELALWD | RHOA; RHOC; F | P61586; P08 | RHOA_HUMAN | Unmodified | 2501400 | 0 | 453160 | 0 | 0 | 0 | 0 | 0 | 0 | 0 | 0 | 0 | 0 | 0 | 4145200 | 1353500 | 0 | 0 |
| QVENAGAI | SEPTIN9; SEPTI | Q9UHD8; K7 | SEPT9_HUMAN | Unmodified | 0 | 0 | 0 | 0 | 2549600 | 0 | 0 | 50280 | 0 | 0 | 0 | 0 | 464470 | 615260 | 0 | 0 | 2192900 | 0 |
| QVEPLDPP | YARS1; YARS1 | P54577; A0A | SYYC_HUMAN | Unmodified | 211590 | 0 | 231850 | 497870 | 0 | 0 | 0 | 0 | 1532100 | 192150 | 714090 | 5695200 | 512910 | 0 | 722650 | 572480 | 0 | 0 |
| QVEQRAFF | ITGB4 | P16144 | ITB4_HUMAN I | Unmodified | 0 | 0 | 0 | 0 | 1763200 | 0 | 0 | 26852 | 0 | 0 | 0 | 0 | 0 | 0 | 0 | 0 | 0 | 4140300 |
| QVEVHQEF | SLC38A10; SLC | Q9HBR0; H0 | S38AA_HUMAN | Unmodified | 0 | 0 | 0 | 0 | 0 | 0 | 0 | 0 | 4689800 | 104100 | 0 | 0 | 9103.2 | 0 | 0 | 0 | 0 | 0 |
| QVEVINFGI | RANGAP1; RAN | P46060; H0Y | RAGP1_HUMAI | Unmodified | 0 | 0 | 0 | 0 | 0 | 0 | 0 | 0 | 0 | 0 | 0 | 1319500 | 0 | 0 | 0 | 0 | 0 | 0 |
| QVEVTVHK | PLEC; PLEC | Q15149; H0Y | PLEC_HUMAN | Unmodified | 0 | 0 | 0 | 0 | 590150 | 211370 | 0 | 0 | 0 | 0 | 0 | 223750 | 0 | 0 | 0 | 0 | 1095100 | 1231700 |
| QVEYLLIHC | DPP4; DPP4; DI | P27487; A0A | DPP4_HUMAN | Unmodified | 1170200 | 0 | 3821600 | 7348600 | 0 | 0 | 0 | 0 | 0 | 0 | 0 | 0 | 88024 | 0 | 9307900 | 4172000 | 2434400 | 1181400 |
| QVEYLVNE | GOT1 | P17174 | AATC_HUMAN | Unmodified | 0 | 0 | 0 | 0 | 927330 | 0 | 904360 | 333740 | 711050 | 648270 | 1272300 | 12964000 | 1549600 | 0 | 0 | 758160 | 576020 | 0 |
| QVFEGQNI | LMAN1 | P49257 | LMAN1_HUMAI | Unmodified | 0 | 0 | 0 | 0 | 0 | 0 | 191620 | 0 | 1.1E+08 | 5595600 | 0 | 0 | 0 | 0 | 610480 | 0 | 0 | 0 |
| QVFEGQNI | LMAN1 | P49257 | LMAN1_HUMAI | Unmodified | 0 | 0 | 0 | 0 | 160060 | 0 | 0 | 0 | 7060800 | 2759300 | 1142100 | 0 | 0 | 0 | 0 | 0 | 0 | 0 |
| QVFERLVD | RAB3D | O95716 | RAB3D_HUMAI | Unmodified | 42429 | 0 | 0 | 530470 | 284890 | 212630 | 48809 | 0 | 0 | 121050 | 0 | 0 | 530030 | 257190 | 0 | 0 | 27503 | 404960 |
| QVFGHNVI | ATP2B4 | P23634 | AT2B4_HUMAN | Unmodified | 0 | 0 | 0 | 0 | 0 | 0 | 0 | 0 | 0 | 0 | 0 | 1020700 | 0 | 0 | 64888 | 53778 | 0 | 0 |
| QVFGSLES | PNMA2 | Q9UL42 | PNMA2_HUMA | Unmodified | 5909500 | 7654100 | 0 | 152840 | 797310 | 0 | 0 | 0 | 0 | 0 | 0 | 3323200 | 0 | 0 | 0 | 0 | 0 | 339120 |
| QVFGTHTT | KIF23; KIF23; KI | Q02241; A0A | KIF23_HUMAN | Unmodified | 0 | 0 | 0 | 0 | 0 | 785350 | 0 | 66935 | 861590 | 633850 | 289120 | 676810 | 0 | 0 | 0 | 0 | 806830 | 244320 |
| QVFGTHTT | KIF23; KIF23; KI | Q02241; A0A | KIF23_HUMAN | Unmodified | 0 | 0 | 0 | 0 | 0 | 0 | 0 | 0 | 0 | 0 | 0 | 0 | 0 | 0 | 0 | 0 | 1638600 | 0 |
| QVGNKTEC | ATP2B4 | P23634 | AT2B4_HUMAN | Unmodified | 0 | 0 | 20306 | 0 | 0 | 0 | 0 | 0 | 0 | 0 | 253850 | 1797500 | 0 | 0 | 60984 | 46413 | 0 | 0 |
| QVGSGVTT | IGHM | P01871 | IGHM_HUMAN | Unmodified | 0 | 182360 | 0 | 0 | 0 | 0 | 0 | 0 | 0 | 0 | 0 | 0 | 0 | 0 | 0 | 0 | 0 | 0 |
| QVGYEDQ | PLOD3 | O60568 | PLOD3_HUMAI | Unmodified | 14750 | 0 | 0 | 0 | 0 | 0 | 0 | 0 | 7740100 | 1867900 | 0 | 0 | 0 | 390620 | 0 | 0 | 0 | 0 |
| QVHLVSPL | CLPTM1L | Q96KA5 | CLP1L_HUMAN | Oxidation (M) | 71470 | 0 | 0 | 0 | 0 | 0 | 0 | 0 | 4551400 | 0 | 0 | 0 | 0 | 7546500 | 0 | 0 | 0 | 0 |
| QVHLVSPL | CLPTM1L | Q96KA5 | CLP1L_HUMAN | Unmodified | 0 | 0 | 0 | 0 | 0 | 0 | 0 | 0 | 2878700 | 0 | 0 | 0 | 0 | 0 | 0 | 0 | 0 | 0 |
| QVHPDTGI | H2BC4; H2BC1 | P62807; P06 | H2B1C_HUMAI | Unmodified | 1.74E+08 | 97668000 | 0 | 207400 | 9351300 | 634130 | 24148000 | 89025000 | 1084500 | 0 | 5818100 | 1.46E+09 | 536080 | 562570 | 0 | 262420 | 111220 | 2150300 |
| QVIAGVGA | RASA3 | Q14644 | RASA3_HUMAN | Unmodified | 0 | 0 | 0 | 0 | 0 | 0 | 0 | 0 | 0 | 128840 | 0 | 0 | 0 | 0 | 0 | 0 | 977910 | 3203300 |
| QVIDVLETD | NUCB2; NUCB2 | P80303; E9P | NUCB2_HUMA | Unmodified | 0 | 0 | 1693600 | 1901300 | 623720 | 0 | 347200 | 264600 | 85389000 | 11142000 | 1386300 | 2639800 | 624020 | 788930 | 3411600 | 1439300 | 0 | 0 |
| QVIDVLETD | NUCB2; NUCB2 | P80303; Q2L | NUCB2_HUMA | Unmodified | 0 | 0 | 0 | 410080 | 716600 | 0 | 517230 | 212280 | 3394700 | 9191400 | 0 | 0 | 323120 | 127510 | 0 | 0 | 0 | 0 |
| QVIEYARQI | PSMD8; PSMD8 | P48556; R4C | PSMD8_HUMA | Oxidation (M) | 0 | 0 | 0 | 0 | 234290 | 25194 | 636890 | 0 | 0 | 69268 | 0 | 0 | 437240 | 287890 | 0 | 0 | 156690 | 600260 |
| QVIQGLEAI | KLC1; KLC1; KL | Q07866; G3V | KLC1_HUMAN | Unmodified | 0 | 0 | 0 | 0 | 2710200 | 0 | 0 | 0 | 0 | 482940 | 0 | 0 | 582720 | 0 | 0 | 0 | 520480 | 2964900 |
| QVIQQNPA | RAD23A; RAD23 | P54725; K7E | RD23A_HUMAI | Unmodified | 0 | 350820 | 0 | 0 | 0 | 0 | 0 | 0 | 0 | 0 | 0 | 0 | 0 | 0 | 153390 | 0 | 0 | 0 |
| QVITLLNEL | ACAA1; ACAA1; | P09110; C9J | THIK_HUMAN 3 | Unmodified | 0 | 2699500 | 3706000 | 68958000 | 936320 | 9189600 | 1653400 | 31950 | 14312000 | 5538500 | 721120 | 46168 | 162650 | 0 | 343450 | 0 | 696150 | 2714400 |
| QVITLLNEL | ACAA1; ACAA1; | P09110; C9J | THIK_HUMAN 3 | Unmodified | 483270 | 1841200 | 0 | 0 | 0 | 0 | 0 | 0 | 842690 | 0 | 0 | 0 | 0 | 0 | 0 | 231740 | 0 | 0 |
| QVKDNEN\ | FGB | P02675 | FIBB_HUMAN F | Unmodified | 0 | 1707400 | 0 | 0 | 0 | 0 | 0 | 0 | 0 | 0 | 0 | 0 | 0 | 977690 | 0 | 0 | 0 | 0 |
| QVKPYVEE | CCT7; CCT7 | Q99832; F8V | TCPH_HUMAN | Unmodified | 372180 | 0 | 862390 | 752330 | 2037300 | 540490 | 0 | 89744 | 2154400 | 1483700 | 5507600 | 32927000 | 5071600 | 2255800 | 1447600 | 0 | 718500 | 1346000 |
| QVLALQSQ | MYH10 | P35580 | MYH10_HUMAI | Unmodified | 172160 | 0 | 77219 | 347170 | 1844000 | 156450 | 0 | 0 | 0 | 269960 | 0 | 1284400 | 1451900 | 527330 | 157660 | 95682 | 0 | 0 |
| QVLDLEDL | SNRNP200; SN | O75643; A0A | U520_HUMAN | Unmodified | 0 | 0 | 26065000 | 1290900 | 0 | 0 | 0 | 422610 | 667400 | 0 | 0 | 1306400 | 933940 | 0 | 996150 | 0 | 0 | 0 |
| QVLEDSPA | PATJ; PATJ; PAT | Q8NI35; A0A | INADL_HUMAN | Unmodified | 0 | 0 | 0 | 0 | 636030 | 103110 | 0 | 0 | 0 | 87500 | 0 | 0 | 0 | 390890 | 0 | 0 | 220250 | 537660 |
| QVLFPPSV | PPP1R16A | Q96I34 | PP16A_HUMAN | Unmodified | 0 | 0 | 0 | 0 | 0 | 0 | 0 | 0 | 121540 | 0 | 0 | 31668 | 0 | 0 | 0 | 0 | 0 | 0 |
| QVLFSADD | ALDOC; ALDOC | P09972; J3Q | ALDOC_HUMA | Unmodified | 0 | 0 | 0 | 781660 | 0 | 0 | 96354 | 121220 | 912030 | 324340 | 0 | 1209500 | 140910 | 0 | 0 | 0 | 131120 | 0 |
| QVLFSADD | ALDOC; ALDOC | P09972; J3Q | ALDOC_HUMA | Unmodified | 320000 | 0 | 89093 | 0 | 0 | 0 | 0 | 0 | 173220 | 0 | 157780 | 1266300 | 0 | 0 | 0 | 50449 | 656400 | 172680 |
| QVLGQMVI | PSMB4 | P28070 | PSB4_HUMAN | Oxidation (M) | 0 | 0 | 0 | 907790 | 0 | 617750 | 0 | 0 | 2323200 | 0 | 0 | 2831900 | 0 | 0 | 0 | 281670 | 0 | 0 |
| QVLIRPCSI | RPS15A; N/A; R | P62244; H3E | RS15A_HUMAN | Unmodified | 105600 | 0 | 0 | 0 | 0 | 0 | 0 | 0 | 0 | 0 | 44401 | 2732000 | 0 | 0 | 0 | 0 | 0 | 0 |
| QVLIYLAR | CPD | O75976 | CBPD_HUMAN | Unmodified | 0 | 0 | 0 | 0 | 0 | 0 | 0 | 0 | 38637000 | 2907200 | 0 | 0 | 0 | 0 | 0 | 0 | 0 | 0 |
| QVLLAQAE | FLOT2; FLOT2; | Q14254; J3Q | FLOT2_HUMAN | Unmodified | 0 | 0 | 3051200 | 2222700 | 8730500 | 1254800 | 1105300 | 0 | 7835200 | 2763400 | 1597000 | 6581700 | 2798200 | 1719100 | 4423900 | 4140400 | 0 | 874250 |
| QVLLAQAE | FLOT2; FLOT2; | Q14254; J3Q | FLOT2_HUMAN | Unmodified | 0 | 0 | 0 | 0 | 654970 | 0 | 224690 | 0 | 0 | 380810 | 0 | 0 | 312350 | 285970 | 0 | 0 | 0 | 86671 |
| QVLLGDQI | PSMB6 | P28072 | PSB6_HUMAN | Unmodified | 131770 | 235090 | 649070 | 504930 | 0 | 0 | 0 | 0 | 2314000 | 358910 | 1034600 | 9598900 | 551320 | 0 | 488930 | 408430 | 0 | 0 |
| QVLLQNER | IRAG2; IRAG2; | Q12912; F5H | IRAG2_HUMAN | Unmodified | 0 | 0 | 0 | 0 | 0 | 0 | 0 | 0 | 0 | 0 | 62458 | 2250300 | 0 | 0 | 0 | 0 | 0 | 0 |
| QVLLSEPEI | APEH; APEH; AI | P13798; C9J | ACPH_HUMAN | Unmodified | 88800 | 0 | 0 | 0 | 0 | 0 | 0 | 0 | 0 | 0 | 814570 | 5537700 | 0 | 0 | 376320 | 539320 | 0 | 0 |
| QVLNQDLT | LYPLAL1 | Q5VWZ2 | LYPL1_HUMAN | Unmodified | 0 | 0 | 195430 | 386110 | 877530 | 198750 | 0 | 0 | 418770 | 0 | 0 | 0 | 1844300 | 934250 | 551200 | 503720 | 366970 | 534170 |
| QVLPDPEV | PBXIP1; PBXIP1 | Q96AQ6; Q5 | PBIP1_HUMAN | Unmodified | 0 | 0 | 0 | 0 | 0 | 0 | 0 | 0 | 1464400 | 0 | 420450 | 612280 | 0 | 0 | 0 | 0 | 0 | 0 |
| QVLQHPW | RPS6KA1; RPS6 | Q15418; Q5 | KS6A1_HUMAN | Unmodified | 0 | 0 | 0 | 0 | 476250 | 0 | 190670 | 103880 | 583500 | 273920 | 0 | 371760 | 792920 | 451770 | 173900 | 132120 | 456690 | 802160 |
| QVLVAPGN | GART; GART; G | P22102; C9J | PUR2_HUMAN | Unmodified | 0 | 0 | 0 | 0 | 756770 | 386700 | 534170 | 184700 | 187990 | 0 | 0 | 1289600 | 1467800 | 618040 | 0 | 71690 | 927330 | 521860 |
| QVMMFSA | DDX39B; DDX3 | Q13838; F6T | DX39B_HUMAN | Oxidation (M) | 0 | 0 | 0 | 0 | 142560 | 0 | 0 | 90947 | 443710 | 0 | 0 | 1174500 | 0 | 0 | 0 | 0 | 0 | 0 |
| QVMVVPVC | TARS1 | P26639 | SYTC_HUMAN | Oxidation (M) | 0 | 0 | 0 | 0 | 299890 | 0 | 0 | 13501 | 0 | 0 | 184120 | 4400200 | 77301 | 0 | 327230 | 429950 | 0 | 0 |
| QVMYQQS | EPDR1 | Q9UM22 | EPDR1_HUMAN | Unmodified | 0 | 0 | 0 | 0 | 0 | 51265 | 0 | 52115 | 429500 | 0 | 41343 | 529090 | 0 | 0 | 0 | 0 | 0 | 0 |
| QVNEMKN | LMAN1 | P49257 | LMAN1_HUMAI | Unmodified | 0 | 0 | 0 | 0 | 0 | 0 | 0 | 0 | 0 | 428210 | 0 | 0 | 21471 | 0 | 0 | 0 | 0 | 29483 |
| QVNILMK | EXOC1 | Q9NV70 | EXOC1_HUMAI | Unmodified | 0 | 0 | 0 | 0 | 406930 | 0 | 105300 | 47059 | 0 | 0 | 0 | 413850 | 107240 | 123780 | 0 | 0 | 132370 | 230450 |
| QVPGGGG | SRP68; SRP68; | Q9UHB9; K7 | SRP68_HUMAN | Unmodified | 0 | 0 | 0 | 0 | 0 | 0 | 0 | 0 | 0 | 0 | 0 | 443430 | 0 | 0 | 14830 | 32339 | 0 | 0 |
| QVPICGDC | STEAP3; STEAP | Q658P3; B8Z | STEA3_HUMAN | Unmodified | 0 | 0 | 0 | 0 | 0 | 0 | 0 | 0 | 0 | 0 | 0 | 509350 | 0 | 0 | 283000 | 0 | 0 | 0 |
| QVPNESFF | NAP1L4; NAP1L | Q99733; H0Y | NP1L4_HUMAN | Unmodified | 0 | 265830 | 144690 | 371830 | 1043000 | 159710 | 205360 | 0 | 758500 | 500320 | 256440 | 1712600 | 1476200 | 665650 | 827340 | 338470 | 597700 | 609350 |
| QVPQKLEQ | EXOC4 | Q96A65 | EXOC4_HUMAI | Unmodified | 0 | 0 | 0 | 0 | 848210 | 129000 | 827910 | 0 | 0 | 0 | 0 | 475710 | 0 | 97529 | 0 | 0 | 0 | 0 |
| QVPVEEAR | RALB; RALB; RA | P11234; C9J | RALB_HUMAN | Unmodified | 664980 | 0 | 1109900 | 1289800 | 0 | 0 | 0 | 0 | 0 | 0 | 944320 | 7116100 | 0 | 0 | 0 | 0 | 0 | 24424 |
| QVPVEEAR | RALB; RALB | P11234; C9J | RALB_HUMAN | Unmodified | 0 | 0 | 0 | 0 | 2294700 | 527720 | 0 | 0 | 0 | 354860 | 0 | 0 | 1058000 | 0 | 0 | 0 | 441630 | 0 |
| QVPVEPGP | SLC19A1; SLC1 | P41440; C9J | S19A1_HUMAN | Unmodified | 0 | 244860 | 557510 | 0 | 0 | 0 | 0 | 0 | 0 | 0 | 0 | 1549100 | 0 | 0 | 971880 | 746180 | 0 | 0 |
| QVPVGPDC | ARVCF; ARVCF; | O00192; E9F | ARVC_HUMAN | Unmodified | 0 | 0 | 0 | 0 | 0 | 0 | 0 | 0 | 51379 | 0 | 0 | 0 | 0 | 0 | 184410 | 0 | 0 | 0 |
| QVPVGPDC | ARVCF; ARVCF; | O00192; E9F | ARVC_HUMAN | Unmodified | 0 | 0 | 113250 | 1126800 | 0 | 0 | 0 | 0 | 0 | 0 | 0 | 0 | 1054000 | 575510 | 3566000 | 232940 | 0 | 0 |
| QVQDLERE | UPF2 | Q9HAU5 | RENT2_HUMAN | Unmodified | 0 | 0 | 0 | 0 | 0 | 0 | 0 | 0 | 0 | 0 | 0 | 564470 | 0 | 0 | 0 | 0 | 0 | 0 |
| QVQELELK | NOS1AP; N/A; N | O75052; H7C | CAPON_HUMA | Unmodified | 123920 | 0 | 0 | 0 | 0 | 0 | 53886 | 18695 | 0 | 0 | 0 | 402670 | 395970 | 687490 | 134730 | 0 | 0 | 0 |
| QVQFHQG | MYO1E | Q12965 | MYO1E_HUMAI | Unmodified | 0 | 0 | 0 | 0 | 0 | 0 | 0 | 0 | 0 | 0 | 0 | 678140 | 0 | 0 | 0 | 0 | 0 | 0 |
| QVQFSEEH | HSPG2 | P98160 | PGBM_HUMAN | Unmodified | 0 | 0 | 1163900 | 0 | 0 | 0 | 0 | 0 | 0 | 0 | 0 | 0 | 0 | 0 | 144960 | 415020 | 0 | 0 |
| QVQGSEIS | VPS28; VPS28; | Q9UK41; E9I | VPS28_HUMAN | Unmodified | 4408200 | 1402600 | 1182400 | 2091800 | 0 | 0 | 0 | 0 | 0 | 0 | 0 | 0 | 0 | 0 | 3946400 | 2465200 | 0 | 0 |
| QVQGSEIS | VPS28; VPS28; | Q9UK41; E9I | VPS28_HUMAN | Unmodified | 44352000 | 23965000 | 13821000 | 22710000 | 1182500 | 0 | 0 | 0 | 4839400 | 350720 | 882270 | 5438200 | 6402000 | 1225800 | 32403000 | 31579000 | 275630 | 753850 |
| QVQHILAS | SEC31A; SEC31 | O94979; D6I | SC31A_HUMAN | Unmodified | 0 | 0 | 0 | 0 | 0 | 0 | 0 | 0 | 0 | 0 | 219510 | 1422000 | 185910 | 0 | 132180 | 0 | 0 | 0 |
| QVQIDGLV | EIF3H; EIF3H; E | O15372; E5F | EIF3H_HUMAN | Unmodified | 254230 | 0 | 0 | 0 | 0 | 0 | 170170 | 0 | 0 | 156420 | 0 | 953490 | 0 | 0 | 329770 | 0 | 0 | 0 |
| QVQLDAQ | TOLLIP; TOLLIP | Q9H0E2; F2Z | TOLIP_HUMAN | Unmodified | 4626600 | 4973700 | 1336600 | 2086700 | 0 | 0 | 0 | 0 | 1487500 | 0 | 0 | 1474500 | 0 | 0 | 4805900 | 7159800 | 0 | 0 |
| QVQLEDAV | RAB20 | Q9NX57 | RAB20_HUMAN | Unmodified | 0 | 0 | 0 | 0 | 0 | 0 | 0 | 0 | 1738900 | 479690 | 0 | 0 | 365580 | 0 | 629800 | 533260 | 0 | 0 |
| QVQPEGPY | FASN; FASN | P49327; A0A | FAS_HUMAN F | Unmodified | 1309600 | 56056 | 754200 | 0 | 0 | 0 | 0 | 0 | 155990 | 0 | 955080 | 37954000 | 0 | 0 | 1916400 | 1773400 | 204020 | 0 |
| QVQQNSLI | ITSN1; ITSN1; I | Q15811; F8V | ITSN1_HUMAN | Unmodified | 0 | 0 | 0 | 0 | 595040 | 0 | 0 | 28419 | 0 | 57046 | 0 | 0 | 523300 | 327070 | 0 | 0 | 99727 | 107600 |
| QVQSLLELI | PLCB3 | Q01970 | PLCB3_HUMAN | Unmodified | 0 | 0 | 0 | 0 | 0 | 0 | 0 | 0 | 0 | 0 | 0 | 834090 | 0 | 0 | 0 | 0 | 0 | 0 |
| QVQVALET | PLEC | Q15149 | PLEC_HUMAN | Unmodified | 0 | 0 | 0 | 0 | 0 | 0 | 0 | 0 | 0 | 0 | 0 | 0 | 0 | 0 | 0 | 0 | 721180 | 0 |
| QVRELEAEI | MYH10 | P35580 | MYH10_HUMAI | Unmodified | 0 | 0 | 0 | 0 | 1777300 | 236390 | 0 | 0 | 0 | 0 | 0 | 0 | 0 | 230200 | 0 | 0 | 0 | 0 |
| QVREMEAE | MYH9 | P35579 | MYH9_HUMAN | Unmodified | 0 | 0 | 180550 | 491250 | 2022300 | 0 | 287140 | 54226 | 132520 | 489730 | 163910 | 0 | 1513700 | 683230 | 0 | 0 | 337320 | 586490 |
| QVRGVPWI | EPB41; EPB41; | P11171; A0A | EPB41_HUMAN | Unmodified | 0 | 0 | 0 | 0 | 0 | 38803 | 3141900 | 0 | 0 | 504040 | 0 | 0 | 0 | 0 | 0 | 0 | 407910 | 0 |
| QVRPPSCP | LDOC1 | O95751 | LDOC1_HUMAI | Unmodified | 1273100 | 765260 | 0 | 0 | 0 | 0 | 0 | 0 | 0 | 0 | 0 | 408520 | 0 | 0 | 2958000 | 1483500 | 0 | 0 |
| QVRPVTEKI | ATP6V1A | P38606 | VATA_HUMAN | Unmodified | 0 | 593560 | 0 | 212450 | 0 | 0 | 0 | 0 | 1477600 | 0 | 709430 | 3251700 | 0 | 0 | 0 | 166490 | 0 | 0 |
| QVSDDLTE | PHB; PHB; PHB | P35232; C9J | PHB_HUMAN F | Unmodified | 0 | 0 | 0 | 0 | 0 | 0 | 0 | 0 | 1708600 | 129960 | 0 | 0 | 0 | 0 | 0 | 0 | 0 | 0 |
| QVSDLISVL | DDX5; DDX5; D | P17844; A0A | DDX5_HUMAN | Unmodified | 101740 | 164360 | 0 | 43965 | 1719900 | 236720 | 0 | 14042 | 318810 | 121550 | 292480 | 3343400 | 0 | 0 | 51385 | 52803 | 0 | 0 |
| QVSIEEGEI | RAB6A; RAB6D | P20340; Q53 | RAB6A_HUMAN | Unmodified | 241980 | 0 | 0 | 250790 | 78851 | 0 | 14811 | 164840 | 503940 | 0 | 0 | 2207000 | 593140 | 551590 | 277070 | 265250 | 161630 | 428600 |
| QVSITASTR | CLUH; CLUH; C | O75153; I3L | CLU_HUMAN C | Unmodified | 0 | 0 | 0 | 0 | 0 | 0 | 0 | 0 | 0 | 0 | 0 | 1083300 | 0 | 0 | 0 | 0 | 0 | 0 |
| QVSMVQTL | RIPOR2; RIPOR | Q9Y4F9; F5C | RIPR2_HUMAN | Unmodified | 0 | 0 | 0 | 0 | 0 | 0 | 0 | 0 | 0 | 0 | 0 | 0 | 0 | 0 | 0 | 0 | 0 | 0 |
| QVSNLLATI | ZC3H10 | Q96K80 | ZC3HA_HUMAI | Unmodified | 2929000 | 0 | 0 | 2292500 | 0 | 0 | 0 | 0 | 0 | 0 | 0 | 0 | 0 | 0 | 0 | 0 | 0 | 0 |
| QVSNLLATI | ZC3H10 | Q96K80 | ZC3HA_HUMAI | Unmodified | 0 | 0 | 0 | 1434100 | 0 | 0 | 523680 | 23977 | 0 | 0 | 0 | 0 | 5440300 | 0 | 0 | 0 | 0 | 0 |
| QVSSECQC | GLG1; GLG1 | Q92896; H3I | GSLG1_HUMAI | Unmodified | 0 | 0 | 0 | 0 | 0 | 0 | 0 | 0 | 7370000 | 722290 | 332700 | 1467700 | 0 | 0 | 0 | 69958 | 0 | 0 |
| QVSSVNEE | IL6ST | P40189 | IL6RB_HUMAN | Unmodified | 63338 | 0 | 283520 | 191540 | 2820600 | 414800 | 266270 | 0 | 883670 | 0 | 0 | 450120 | 0 | 0 | 0 | 0 | 163270 | 1210500 |
| QVTCVAW | PWP1 | Q13610 | PWP1_HUMAN | Unmodified | 0 | 0 | 0 | 0 | 0 | 0 | 0 | 108080 | 0 | 0 | 0 | 0 | 6981.6 | 0 | 0 | 0 | 0 | 0 |
| QVTDLEAEI | TNIP1; TNIP1; T | Q15025; A0A | TNIP1_HUMAN | Unmodified | 0 | 0 | 634590 | 11428000 | 290550 | 50119 | 0 | 0 | 0 | 977270 | 0 | 0 | 166980 | 0 | 0 | 40793 | 0 | 412770 |
| QVTESSHL | CDC42BPA; CD | Q5VT25; A0A | MRCKA_HUMA | Unmodified | 0 | 0 | 0 | 0 | 0 | 0 | 0 | 0 | 0 | 675960 | 0 | 0 | 3494100 | 2634100 | 0 | 2058600 | 0 | 5931800 |
| QVTGRFEE | SNX9; SNX9; SN | Q9Y5X1; A0A | SNX9_HUMAN | Unmodified | 10422000 | 9471300 | 0 | 0 | 0 | 0 | 372680 | 0 | 627090 | 244480 | 740850 | 10497000 | 0 | 0 | 0 | 0 | 0 | 0 |
| QVTHPNYN | KLK14; KLK14 | Q9P0G3; A0 | KLK14_HUMAN | Unmodified | 0 | 0 | 0 | 0 | 0 | 0 | 0 | 0 | 0 | 0 | 0 | 0 | 0 | 0 | 0 | 0 | 0 | 0 |
| QVTPDGES | DLG1; DLG1; D | Q12959; A0A | DLG1_HUMAN | Unmodified | 130910 | 138150 | 0 | 866040 | 735290 | 0 | 0 | 0 | 3879100 | 547830 | 656930 | 2970100 | 468030 | 125360 | 1178700 | 812900 | 96275 | 364580 |
| QVTQEEGC | RRAS2; RRAS2 | P62070; E9P | RRAS2_HUMAN | Unmodified | 2102900 | 3913800 | 2918400 | 3942100 | 0 | 421280 | 0 | 0 | 1483000 | 0 | 0 | 0 | 0 | 0 | 6151600 | 5397700 | 0 | 36783 |
| QVTSNSLS | LRPAP1; LRPAP | P30533; A0A | AMRP_HUMAN | Unmodified | 0 | 0 | 0 | 0 | 0 | 0 | 0 | 0 | 20201000 | 3145800 | 0 | 643760 | 0 | 0 | 0 | 0 | 0 | 0 |
| QVTSVLQQ | NOSTRIN; NOS | Q8IVI9; F8W | NOSTN_HUMA | Unmodified | 3573200 | 2592800 | 5708800 | 5658800 | 0 | 0 | 0 | 0 | 0 | 149650 | 0 | 0 | 1446100 | 0 | 13732000 | 0 | 0 | 0 |
| QVTSVLQQ | NOSTRIN | Q8IVI9 | NOSTN_HUMA | Unmodified | 0 | 0 | 133280 | 0 | 0 | 0 | 0 | 0 | 0 | 0 | 0 | 0 | 5340000 | 4347500 | 0 | 0 | 0 | 0 |

| | | | | | | | | | | | | | | | | | | | | | | |
|---|---|---|---|---|---|---|---|---|---|---|---|---|---|---|---|---|---|---|---|---|---|---|
| QVTVLELFF | SLC2A14; SLC2 | Q8TDB8; P1 | GTR14_HUMAN | Unmodified | 3291600 | 5599900 | 1988900 | 1624900 | 306150 | 0 | 0 | 14180 | 0 | 0 | 0 | 0 | 0 | 0 | 5745300 | 4155100 | 0 | 0 |
| QVVAVTGD | ATP2B2; ATP2B | Q01814; Q1( | AT2B2_HUMAN | Unmodified | 0 | 0 | 6708500 | 9500200 | 0 | 0 | 0 | 0 | 5907200 | 390820 | 4811300 | 44605000 | 2239600 | 1072400 | 12071000 | 7869600 | 718000 | 0 |
| QVVAVTGD | ATP2B2; ATP2B | Q01814; Q1( | AT2B2_HUMAN | Unmodified | 0 | 0 | 0 | 0 | 0 | 0 | 0 | 0 | 0 | 0 | 1107700 | 2209400 | 0 | 0 | 0 | 0 | 0 | 0 |
| QVVCDLGN | ITGAV | P06756 | ITAV_HUMAN I | Unmodified | 0 | 0 | 0 | 0 | 0 | 0 | 0 | 0 | 456360 | 0 | 0 | 0 | 0 | 0 | 0 | 0 | 0 | 0 |
| QVVEQDEE | PSEN1; PSEN1; | P49768; G3 | PSN1_HUMAN | Unmodified | 0 | 0 | 0 | 0 | 0 | 0 | 69484 | 48391 | 0 | 1160200 | 0 | 4493900 | 0 | 0 | 384930 | 633140 | 0 | 0 |
| QVVESAYE | LDHA | P00338 | LDHA_HUMAN | Unmodified | 1021200 | 671960 | 1734900 | 2309600 | 9000100 | 1705900 | 0 | 80342 | 2671800 | 2063000 | 15469000 | 1.16E+08 | 2905000 | 626910 | 3932200 | 5389500 | 3023700 | 9069700 |
| QVVFTSDP | CEMIP2; CEMIP | Q9UHN6; B4 | CEIP2_HUMAN | Unmodified | 0 | 0 | 0 | 0 | 0 | 0 | 31313 | 35217 | 1136500 | 0 | 0 | 0 | 0 | 0 | 0 | 0 | 0 | 0 |
| QVVHLNLA | STXBP3 | O00186 | STXB3_HUMAN | Oxidation (M) | 434290 | 195220 | 497550 | 411850 | 1100500 | 403560 | 509300 | 0 | 632370 | 341310 | 227900 | 1846600 | 2759300 | 1940500 | 461090 | 624490 | 1219600 | 1747500 |
| QVVHLNLA | STXBP3 | O00186 | STXB3_HUMAN | Unmodified | 230630 | 217860 | 1164300 | 957480 | 3459100 | 0 | 2311300 | 665360 | 1281100 | 0 | 0 | 1960400 | 2686100 | 5359200 | 465570 | 207420 | 1998900 | 9498100 |
| QVVILSQD | UCK2; UCK2 | Q9BZX2; A0 | UCK2_HUMAN | Unmodified | 0 | 0 | 0 | 0 | 0 | 0 | 0 | 0 | 0 | 0 | 0 | 0 | 1561700 | 1431200 | 0 | 0 | 0 | 0 |
| QVVLYSIR | FAM91A1; FAM | Q658Y4; E7E | F91A1_HUMAN | Unmodified | 120010 | 84033 | 0 | 0 | 0 | 0 | 140090 | 0 | 276490 | 0 | 21846 | 980420 | 0 | 0 | 0 | 167690 | 0 | 0 |
| QVVNFGPC | PSAT1 | Q9Y617 | SERC_HUMAN | Unmodified | 0 | 0 | 524070 | 0 | 0 | 0 | 0 | 0 | 292630 | 0 | 2090700 | 10488000 | 135700 | 0 | 2540400 | 2449000 | 0 | 0 |
| QVVNFGPC | PSAT1 | Q9Y617 | SERC_HUMAN | Unmodified | 0 | 43765 | 0 | 853050 | 1086800 | 0 | 0 | 0 | 198040 | 553800 | 1370700 | 2158100 | 772520 | 1463500 | 65438 | 340930 | 54577 | 0 |
| QVVNIPSFI | RPS9 | P46781 | RS9_HUMAN 4 | Unmodified | 1264400 | 1535100 | 975880 | 568840 | 0 | 0 | 0 | 50654 | 796650 | 0 | 874700 | 30963000 | 0 | 0 | 1618300 | 1145500 | 0 | 0 |
| QVVNIPSFI | RPS9 | P46781 | RS9_HUMAN 4 | Unmodified | 0 | 0 | 0 | 0 | 1433100 | 320450 | 3744000 | 0 | 1291500 | 2421500 | 2488700 | 3899400 | 0 | 0 | 0 | 0 | 106320 | 1022400 |
| QVVNQVW | HUWE1 | Q7Z6Z7 | HUWE1_HUMA | Unmodified | 0 | 0 | 0 | 0 | 0 | 0 | 0 | 0 | 0 | 0 | 0 | 237950 | 0 | 0 | 0 | 0 | 0 | 0 |
| QVVNVYAD | BTN3A2; BTN3A | P78410; O0( | BT3A2_HUMAN | Unmodified | 0 | 0 | 0 | 0 | 149150 | 0 | 155010 | 294090 | 44673 | 0 | 0 | 1013100 | 0 | 0 | 0 | 90219 | 350180 | 0 |
| QVVPVSSH | NUP210 | Q8TEM1 | PO210_HUMAN | Unmodified | 217720 | 0 | 0 | 0 | 0 | 0 | 0 | 0 | 2114500 | 165990 | 0 | 157020 | 0 | 0 | 0 | 0 | 19423 | 0 |
| QVVTVVFEI | MON2; MON2; | Q7Z3U7; A0 | MON2_HUMAN | Unmodified | 0 | 0 | 0 | 0 | 0 | 0 | 0 | 0 | 0 | 0 | 0 | 342010 | 0 | 0 | 0 | 0 | 0 | 0 |
| QVVVCGY( | AHCYL1; AHCY | O43865; Q9 | SAHH2_HUMA | Unmodified | 0 | 0 | 236880 | 405160 | 964650 | 305530 | 0 | 44003 | 0 | 0 | 0 | 182680 | 261010 | 321780 | 754340 | 577670 | 377400 | 712890 |
| QVVVEGQE | VIL1; VIL1; VIL1 | P09327; H7( | VILI_HUMAN Vi | Oxidation (M) | 0 | 0 | 0 | 0 | 0 | 0 | 0 | 0 | 535570 | 0 | 370490 | 2617600 | 0 | 0 | 0 | 0 | 0 | 0 |
| QVYDAHTK | CAV1; CAV1; C/ | Q03135; C9J | CAV1_HUMAN | Unmodified | 0 | 0 | 0 | 0 | 1124100 | 497540 | 96653 | 0 | 0 | 0 | 0 | 0 | 0 | 0 | 0 | 0 | 245740 | 206230 |
| QVYDGFSF | TSNAX | Q99598 | TSNAX_HUMAN | Unmodified | 0 | 0 | 0 | 0 | 0 | 0 | 0 | 0 | 0 | 0 | 0 | 727600 | 0 | 0 | 0 | 0 | 0 | 0 |
| QVYDGFSF | TSNAX | Q99598 | TSNAX_HUMAN | Unmodified | 0 | 0 | 0 | 0 | 0 | 0 | 0 | 0 | 0 | 0 | 0 | 0 | 0 | 0 | 0 | 0 | 0 | 0 |
| QVYDIAAK | AGPS; AGPS; A( | O00116; A0 | ADAS_HUMAN | Unmodified | 0 | 0 | 0 | 0 | 0 | 0 | 0 | 0 | 6140900 | 0 | 0 | 0 | 0 | 0 | 0 | 0 | 0 | 0 |
| QVYGLNFC | ENAH; ENAH; E | Q8N8S7; A0 | ENAH_HUMAN | Unmodified | 0 | 0 | 0 | 0 | 0 | 0 | 0 | 0 | 0 | 0 | 0 | 0 | 0 | 0 | 0 | 0 | 0 | 0 |
| QVYICSNN | STEAP2; STEAP | Q8NFT2; C9 | STEA2_HUMAN | Unmodified | 1010700 | 866400 | 14089000 | 8728600 | 0 | 0 | 0 | 0 | 0 | 0 | 0 | 0 | 0 | 0 | 22597000 | 12913000 | 0 | 0 |
| QVYLSRPP | DDR1; DDR1; D | Q08345; A0 | DDR1_HUMAN | Unmodified | 133850 | 173950 | 0 | 46870 | 0 | 0 | 0 | 0 | 746510 | 0 | 0 | 0 | 0 | 0 | 0 | 0 | 0 | 0 |
| QVYMSLPC | GLUL; GLUL | P15104; A0A | GLNA_HUMAN | Unmodified | 0 | 0 | 0 | 0 | 0 | 0 | 0 | 0 | 0 | 0 | 0 | 0 | 0 | 0 | 0 | 0 | 276050 | 0 |
| QWCNCAF | RAP1A | P62834 | RAP1A_HUMAN | Unmodified | 1105500 | 878900 | 0 | 2973300 | 159110 | 0 | 0 | 0 | 392510 | 147570 | 0 | 3926800 | 1976200 | 943740 | 2442300 | 1708100 | 428790 | 868200 |
| QWDQVEQ | GNPTG | Q9UJJ9 | GNPTG_HUMA | Unmodified | 0 | 0 | 0 | 0 | 0 | 0 | 0 | 0 | 3090200 | 0 | 0 | 0 | 0 | 0 | 0 | 0 | 0 | 0 |
| QWFINITDI | IARS2 | Q9NSE4 | SYIM_HUMAN I | Unmodified | 0 | 0 | 0 | 0 | 0 | 0 | 0 | 0 | 617660 | 386810 | 0 | 0 | 0 | 0 | 0 | 0 | 0 | 0 |
| QWGHNEL | DHTKD1; DHTK | Q96HY7; H7 | DHTK1_HUMAI | Unmodified | 0 | 0 | 0 | 0 | 0 | 0 | 0 | 0 | 437470 | 0 | 0 | 0 | 0 | 0 | 0 | 0 | 0 | 0 |
| QWGWTQ( | RPL17; RPL17; | P18621; A0A | RL17_HUMAN ( | Unmodified | 0 | 670900 | 0 | 0 | 341000 | 0 | 0 | 0 | 0 | 0 | 0 | 2701300 | 0 | 0 | 0 | 0 | 0 | 29308 |
| QWKPALV\ | SNTB1; SNTB1 | Q13884; A0 | SNTB1_HUMAN | Unmodified | 0 | 0 | 0 | 0 | 0 | 0 | 0 | 0 | 1495500 | 664080 | 0 | 0 | 0 | 0 | 0 | 0 | 116850 | 0 |
| QWLLQTH\ | FERMT3; FERM | Q86UX7; F5I | URP2_HUMAN | Unmodified | 206600 | 427370 | 81733 | 152890 | 0 | 0 | 0 | 0 | 0 | 0 | 0 | 0 | 0 | 0 | 466850 | 450910 | 0 | 0 |
| QWLQEIDF | RAB1C; RAB1A; | Q92928; P62 | RAB1C_HUMAI | Unmodified | 691180 | 0 | 510930 | 237780 | 0 | 0 | 0 | 0 | 0 | 0 | 0 | 4420200 | 0 | 0 | 614420 | 278030 | 0 | 0 |
| QWLQEIDF | RAB1A; RAB1B; | P62820; Q9I | RAB1A_HUMAN | Unmodified | 7143300 | 9449700 | 5094600 | 15605000 | 8055600 | 825280 | 5385600 | 75765 | 17894000 | 6351600 | 11930000 | 70130000 | 6557000 | 3799600 | 10997000 | 11004000 | 2400500 | 5030400 |
| QWLQEVK | GOT2 | P00505 | AATM_HUMAN | Unmodified | 0 | 799330 | 0 | 0 | 0 | 0 | 0 | 0 | 1893300 | 0 | 0 | 0 | 0 | 0 | 0 | 0 | 0 | 0 |
| QWMENPN | ANPEP | P15144 | AMPN_HUMAN | Oxidation (M) | 0 | 0 | 0 | 0 | 0 | 0 | 41512 | 0 | 0 | 0 | 0 | 0 | 299350 | 0 | 0 | 0 | 3054600 | 7132700 |
| QWMENPN | ANPEP | P15144 | AMPN_HUMAN | Unmodified | 0 | 0 | 0 | 0 | 0 | 0 | 0 | 0 | 0 | 0 | 0 | 0 | 0 | 0 | 0 | 0 | 4491500 | 11994000 |
| QWNNCAF | RAP1B; N/A; RA | P61224; A6N | RAP1B_HUMAN | Unmodified | 11124000 | 9663700 | 4456300 | 14752000 | 2572400 | 1148800 | 1714000 | 1051900 | 1819200 | 434440 | 5328600 | 17569000 | 4469900 | 4502300 | 9537500 | 8523600 | 1254000 | 3713000 |
| QWRVDLP/ | GALNT2 | Q10471 | GALT2_HUMAN | Unmodified | 0 | 0 | 0 | 0 | 0 | 0 | 21692 | 0 | 366420 | 1141800 | 0 | 0 | 0 | 0 | 0 | 0 | 0 | 0 |
| QWTFNYIL | RABL6; RABL6; | Q3YEC7; F2 | RABL6_HUMAN | Unmodified | 0 | 0 | 0 | 0 | 258840 | 0 | 0 | 0 | 170980 | 0 | 0 | 1140200 | 0 | 0 | 0 | 0 | 314560 | 0 |
| QWVDTDD | RPA2; RPA2 | P15927; Q5 | RFA2_HUMAN | Unmodified | 0 | 0 | 0 | 0 | 0 | 0 | 0 | 0 | 0 | 242620 | 0 | 7789800 | 0 | 0 | 0 | 0 | 0 | 0 |
| QWVPVLQI | ITFG1; ITFG1 | Q8TB96; F5C | TIP_HUMAN T-( | Unmodified | 0 | 0 | 417610 | 0 | 0 | 0 | 0 | 0 | 12093000 | 163990 | 21208 | 0 | 0 | 0 | 674960 | 31366 | 0 | 0 |
| QYAGFSTV | MMUT | P22033 | MUTA_HUMAN | Unmodified | 0 | 0 | 0 | 0 | 0 | 0 | 0 | 0 | 1140000 | 0 | 0 | 0 | 0 | 0 | 0 | 0 | 0 | 0 |
| QYAGLDHE | SIGMAR1; SIGN | Q99720; B4I | SGMR1_HUMA | Unmodified | 0 | 0 | 0 | 0 | 0 | 0 | 0 | 0 | 3475000 | 584410 | 0 | 837460 | 0 | 0 | 0 | 0 | 0 | 0 |
| QYASLTGT | GANAB; GANAE | Q14697; F5H | GANAB_HUMA | Unmodified | 0 | 0 | 0 | 0 | 0 | 0 | 0 | 0 | 35540000 | 0 | 0 | 3826200 | 0 | 0 | 0 | 0 | 0 | 0 |
| QYDADLEC | TAGLN2; TAGLI | P37802; X6R | TAGL2_HUMAN | Unmodified | 83946 | 412580 | 0 | 20208 | 3762000 | 0 | 635950 | 0 | 67707 | 569920 | 0 | 43333 | 867940 | 958830 | 215110 | 0 | 1590000 | 2344500 |
| QYDAGRD( | EFHD2; EFHD2 | Q96C19; H0 | EFHD2_HUMAI | Unmodified | 0 | 0 | 0 | 0 | 15243000 | 11925000 | 135660 | 0 | 4050000 | 1319000 | 168460 | 583580 | 0 | 0 | 0 | 0 | 895800 | 4007000 |
| QYDAGRD( | EFHD2; EFHD2 | Q96C19; H0 | EFHD2_HUMAI | Oxidation (M) | 0 | 0 | 0 | 0 | 10855000 | 5267500 | 115580 | 0 | 1104100 | 757430 | 347980 | 1806100 | 0 | 216300 | 0 | 0 | 480650 | 750610 |
| QYDGIFYEI | NIPSNAP3A | Q9UFN0 | NPS3A_HUMAI | Unmodified | 0 | 0 | 0 | 0 | 0 | 0 | 0 | 0 | 2805800 | 0 | 0 | 0 | 0 | 0 | 0 | 0 | 0 | 0 |
| QYDQEIEN | SLK | Q9H2G2 | SLK_HUMAN S | Unmodified | 0 | 0 | 0 | 0 | 0 | 0 | 0 | 0 | 0 | 0 | 0 | 715570 | 162910 | 0 | 111280 | 228950 | 0 | 0 |
| QYDQEIEN | SLK | Q9H2G2 | SLK_HUMAN S | Unmodified | 0 | 0 | 0 | 0 | 1093900 | 244600 | 1602800 | 2736300 | 0 | 0 | 0 | 0 | 259660 | 0 | 0 | 0 | 453750 | 0 |
| QYDSFTFT/ | TMED7; TMED7 | Q9Y3B3; A0 | TMED7_HUMAI | Unmodified | 0 | 0 | 167960 | 596270 | 0 | 0 | 390840 | 0 | 20334000 | 1943600 | 580430 | 1015500 | 0 | 122150 | 1529600 | 565520 | 0 | 209150 |
| QYDSFTYR | TMED3; TMED3 | Q9Y3Q3; G3 | TMED3_HUMAI | Unmodified | 0 | 0 | 0 | 122930 | 0 | 0 | 113760 | 11552 | 6805300 | 1725900 | 301010 | 0 | 98407 | 75761 | 108110 | 146140 | 0 | 0 |
| QYDYDSST | NIBAN1; NIBAN | Q9BZQ8; H0 | NIBA1_HUMAN | Unmodified | 0 | 0 | 265270 | 841030 | 0 | 0 | 0 | 34950 | 2855500 | 369480 | 0 | 0 | 0 | 0 | 365340 | 0 | 0 | 909020 |
| QYDYDSST | NIBAN1; NIBAN | Q9BZQ8; H0 | NIBA1_HUMAN | Unmodified | 0 | 0 | 177310 | 287420 | 481270 | 386390 | 359020 | 189330 | 2486400 | 1157900 | 0 | 0 | 0 | 0 | 0 | 85445 | 0 | 192020 |
| QYDYDSST | NIBAN1; NIBAN | Q9BZQ8; H0 | NIBA1_HUMAN | Unmodified | 0 | 375140 | 0 | 61354 | 802830 | 821440 | 1159700 | 438520 | 178480 | 1260200 | 0 | 0 | 128270 | 0 | 0 | 0 | 154280 | 0 |
| QYEMGEC | U2AF1; U2AF1L | Q01081; P0I | U2AF1_HUMA | Unmodified | 469270 | 228980 | 0 | 200850 | 0 | 0 | 0 | 0 | 86649 | 0 | 51876 | 368640 | 0 | 0 | 0 | 164560 | 0 | 0 |
| QYEQNSLT | ARHGAP29 | Q52LW3 | RHG29_HUMAI | Unmodified | 0 | 0 | 0 | 0 | 116270 | 0 | 121790 | 49831 | 0 | 0 | 0 | 71791 | 0 | 0 | 0 | 0 | 347680 | 714620 |
| QYEQQTY( | EIF3L; EIF3L; EII | Q9Y262; B0I | EIF3L_HUMAN | Unmodified | 0 | 0 | 0 | 0 | 0 | 0 | 0 | 0 | 0 | 0 | 0 | 3043300 | 0 | 0 | 0 | 215460 | 0 | 0 |
| QYFGSYSA | IGSF3 | O75054 | IGSF3_HUMAN | Unmodified | 1087000 | 549290 | 0 | 370930 | 1045100 | 358820 | 138730 | 0 | 0 | 0 | 0 | 0 | 756120 | 319140 | 747000 | 703930 | 0 | 0 |
| QYFPNDEE | P4HA1 | P13674 | P4HA1_HUMAN | Unmodified | 0 | 0 | 0 | 0 | 0 | 0 | 0 | 0 | 4261200 | 614910 | 461670 | 268020 | 0 | 0 | 68092 | 0 | 0 | 199060 |
| QYGNEVFL | CCT8; CCT8 | P50990; H7C | TCPQ_HUMAN | Unmodified | 355520 | 274260 | 541130 | 217360 | 4507100 | 1104300 | 0 | 1532700 | 1369300 | 1396800 | 5382800 | 23827000 | 5761800 | 3713400 | 1123000 | 890600 | 3730900 | 4134800 |
| QYHLQQLF | SLC39A6 | Q13433 | S39A6_HUMAN | Unmodified | 0 | 0 | 0 | 0 | 0 | 0 | 0 | 0 | 0 | 0 | 0 | 1176800 | 0 | 0 | 94467 | 130420 | 0 | 0 |
| QYIISEELIS | NUDT5; NUDT5 | Q9UKK9; A6I | NUDT5_HUMA | Unmodified | 0 | 0 | 0 | 373980 | 1266200 | 0 | 0 | 17486 | 1567100 | 0 | 1605700 | 7408900 | 23941 | 0 | 0 | 0 | 0 | 0 |
| QYIISEELIS | NUDT5; NUDT5 | Q9UKK9; A6I | NUDT5_HUMA | Unmodified | 0 | 0 | 0 | 0 | 4925400 | 531550 | 1779400 | 0 | 0 | 662740 | 0 | 0 | 1134200 | 0 | 0 | 0 | 1207800 | 0 |
| QYINAIKDY | PLEC; PLEC | Q15149; H0 | PLEC_HUMAN | Unmodified | 0 | 0 | 0 | 0 | 522010 | 153860 | 864340 | 0 | 0 | 0 | 0 | 0 | 0 | 0 | 0 | 0 | 2557500 | 465250 |
| QYIVAEFEC | TRIM26; TRIM2( | Q12899; A0 | TRI26_HUMAN | Unmodified | 13316000 | 11162000 | 0 | 0 | 0 | 0 | 0 | 0 | 0 | 0 | 0 | 0 | 0 | 0 | 17748000 | 10053000 | 0 | 0 |
| QYKIPDWF | RPS18; RPS18 | P62269; J3J | RS18_HUMAN | Unmodified | 268950 | 351390 | 0 | 205280 | 0 | 0 | 0 | 0 | 0 | 78119 | 0 | 137660 | 100280 | 34472 | 0 | 0 | 0 | 0 |
| QYKPVVYS | GNAO1; GNAO | P09471; A0A | GNAO_HUMAN | Unmodified | 0 | 0 | 0 | 0 | 0 | 0 | 0 | 0 | 1538000 | 625450 | 0 | 0 | 0 | 0 | 0 | 0 | 0 | 0 |
| QYLLLHSL | CAND1; CAND | Q86VP6; A0 | CAND1_HUMA | Unmodified | 0 | 0 | 0 | 0 | 0 | 0 | 0 | 0 | 0 | 0 | 0 | 0 | 0 | 0 | 0 | 0 | 0 | 0 |
| QYLPGNEC | CIAO1 | O76071 | CIAO1_HUMAN | Unmodified | 331320 | 1712300 | 0 | 0 | 0 | 0 | 0 | 0 | 0 | 0 | 0 | 830900 | 681130 | 53734 | 0 | 0 | 0 | 0 |
| QYLPNVKP | EMC1 | Q8N766 | EMC1_HUMAN | Unmodified | 0 | 0 | 0 | 0 | 0 | 0 | 0 | 0 | 1704500 | 382610 | 0 | 0 | 0 | 0 | 0 | 0 | 0 | 0 |
| QYLVFHDC | APEH; APEH; AI | P13798; C9J | ACPH_HUMAN | Unmodified | 7109000 | 17624000 | 0 | 5358200 | 4545500 | 0 | 0 | 0 | 0 | 573120 | 0 | 3708300 | 0 | 0 | 0 | 328540 | 0 | 0 |
| QYMEGFNI | CDC37; CDC37 | Q16543; K7E | CDC37_HUMA | Oxidation (M) | 0 | 0 | 0 | 0 | 0 | 0 | 0 | 0 | 0 | 0 | 0 | 0 | 0 | 0 | 0 | 0 | 0 | 0 |
| QYMFPETL | MGAT2 | Q10469 | MGAT2_HUMAI | Oxidation (M) | 0 | 0 | 0 | 0 | 0 | 0 | 0 | 0 | 5177100 | 736260 | 0 | 0 | 0 | 0 | 0 | 0 | 0 | 0 |
| QYMFPETL | MGAT2 | Q10469 | MGAT2_HUMAI | Unmodified | 0 | 0 | 0 | 0 | 0 | 0 | 0 | 0 | 5758500 | 0 | 0 | 822260 | 0 | 0 | 0 | 0 | 0 | 934440 |
| QYNVDYKL | COPS4; COPS4 | Q9BT78; D6I | CSN4_HUMAN | Unmodified | 0 | 0 | 0 | 0 | 331240 | 0 | 0 | 0 | 0 | 0 | 279810 | 1961300 | 585670 | 0 | 0 | 0 | 0 | 0 |
| QYNVGPS\ | FN1 | P02751 | FINC_HUMAN I | Unmodified | 0 | 0 | 20896000 | 1923700 | 0 | 1152100 | 0 | 0 | 2619500 | 0 | 0 | 0 | 0 | 0 | 2931500 | 0 | 0 | 254770 |
| QYNVGPS\ | FN1 | P02751 | FINC_HUMAN I | Unmodified | 131270 | 0 | 591160 | 5286400 | 262030 | 0 | 0 | 0 | 2608800 | 479290 | 602540 | 378740 | 110450 | 0 | 163150 | 3350500 | 0 | 0 |
| QYNVGPS\ | FN1 | P02751 | FINC_HUMAN I | Unmodified | 0 | 0 | 0 | 329910 | 0 | 0 | 0 | 0 | 0 | 2419600 | 158440 | 0 | 474710 | 182700 | 0 | 0 | 0 | 193070 |
| QYPEFPWT | TTLL12 | Q14166 | TTL12_HUMAN | Unmodified | 0 | 0 | 0 | 0 | 0 | 0 | 0 | 0 | 0 | 0 | 25153 | 147050 | 0 | 0 | 10148 | 39511 | 0 | 0 |
| QYPFQIVY | BMX | P51813 | BMX_HUMAN ( | Unmodified | 0 | 0 | 2177300 | 0 | 0 | 0 | 0 | 0 | 0 | 0 | 448340 | 5508600 | 0 | 0 | 0 | 0 | 0 | 0 |
| QYPISLVLA | DDX3X; DDX3Y | O00571; O1 | DDX3X_HUMAI | Unmodified | 0 | 0 | 0 | 0 | 0 | 0 | 0 | 0 | 0 | 0 | 0 | 2567000 | 0 | 0 | 4076700 | 2846100 | 0 | 0 |
| QYPLFRPP/ | SPON2; SPON2 | Q9BUD6; DE | SPON2_HUMA | Oxidation (M) | 0 | 0 | 437850 | 7548100 | 0 | 0 | 0 | 0 | 0 | 0 | 0 | 0 | 0 | 0 | 4924300 | 2511200 | 0 | 0 |
| QYPLFRPP/ | SPON2; SPON2 | Q9BUD6; DE | SPON2_HUMA | Oxidation (M) | 0 | 0 | 1313500 | 7120900 | 0 | 0 | 0 | 0 | 0 | 0 | 0 | 0 | 0 | 9614900 | 0 | 0 | 0 | 0 |
| QYPLVTPN | COPB2; COPB2 | P35606; H0 | COPB2_HUMAN | Unmodified | 0 | 0 | 0 | 0 | 362150 | 0 | 0 | 0 | 0 | 0 | 0 | 1227800 | 0 | 0 | 0 | 0 | 0 | 0 |
| QYPPSTQV | VIL1; VIL1 | P09327; B4E | VILI_HUMAN Vi | Unmodified | 0 | 0 | 0 | 0 | 0 | 0 | 0 | 0 | 0 | 0 | 620740 | 3083500 | 0 | 0 | 0 | 0 | 0 | 0 |
| QYQATGFL | MOGS; MOGS; | Q13724; A0 | MOGS_HUMAN | Unmodified | 0 | 0 | 0 | 0 | 0 | 0 | 0 | 0 | 5823700 | 406130 | 574220 | 0 | 0 | 0 | 0 | 0 | 0 | 0 |
| QYQGSGG | TMEM222; TME | Q9H0R3; Q5 | TM222_HUMAN | Unmodified | 0 | 0 | 0 | 0 | 0 | 0 | 0 | 0 | 208230 | 0 | 0 | 233130 | 0 | 0 | 361610 | 198200 | 0 | 0 |
| QYQIDAAI\ | CUL4B; CUL4A; | Q13620; Q1 | CUL4B_HUMAI | Unmodified | 673140 | 1031800 | 78960 | 50385 | 741250 | 13983000 | 0 | 0 | 1117100 | 0 | 964710 | 10583000 | 0 | 0 | 175500 | 206230 | 0 | 0 |
| QYSLQNW | GSS; GSS; GSS | P48637; A0A | GSHB_HUMAN | Unmodified | 0 | 0 | 134360 | 357290 | 0 | 0 | 0 | 0 | 0 | 0 | 0 | 2124800 | 0 | 0 | 0 | 482990 | 0 | 0 |
| QYSVAFCN | NIBAN1 | Q9BZQ8 | NIBA1_HUMAN | Unmodified | 0 | 0 | 0 | 0 | 0 | 0 | 0 | 56663 | 1223100 | 64259 | 0 | 0 | 0 | 0 | 94705 | 0 | 0 | 0 |
| QYSVAFCN | NIBAN1 | Q9BZQ8 | NIBA1_HUMAN | Unmodified | 0 | 0 | 0 | 0 | 0 | 0 | 0 | 0 | 1780300 | 177080 | 0 | 0 | 0 | 0 | 0 | 40122 | 0 | 0 |
| QYSWDQR | RP2 | O75695 | XRP2_HUMAN | Unmodified | 104220 | 155490 | 227860 | 405860 | 0 | 0 | 0 | 0 | 681540 | 30069 | 158300 | 1083700 | 0 | 0 | 417090 | 391750 | 0 | 0 |
| QYTGINAIS | ACTR3; ACTR3 | P61158; B4E | ARP3_HUMAN | Unmodified | 174930 | 0 | 0 | 0 | 514810 | 0 | 59631 | 26246 | 110480 | 195500 | 581230 | 3355800 | 0 | 0 | 0 | 0 | 0 | 65038 |
| QYTGINAIS | ACTR3; ACTR3 | P61158; B4E | ARP3_HUMAN | Unmodified | 2689100 | 2644000 | 768930 | 2286700 | 6491800 | 1341400 | 1387100 | 668600 | 1054900 | 510980 | 1215500 | 10014000 | 4918200 | 3833600 | 1625800 | 1941500 | 2564700 | 3676500 |
| QYTSIHHG | APP; APP; APP | P05067; E9P | A4_HUMAN Am | Unmodified | 0 | 0 | 0 | 0 | 0 | 0 | 0 | 0 | 812920 | 0 | 0 | 0 | 0 | 0 | 0 | 0 | 0 | 0 |
| QYTSIHHG | APP; APP; APP | P05067; E9P | A4_HUMAN Am | Unmodified | 0 | 0 | 0 | 0 | 3996000 | 0 | 333570 | 0 | 0 | 1260300 | 0 | 0 | 204590 | 174980 | 0 | 0 | 158700 | 2823200 |
| QYTSPEEIC | PDAP1; PDAP1 | Q13442; F8V | HAP28_HUMAI | Unmodified | 1130800 | 954490 | 578260 | 1501700 | 2397700 | 0 | 4761800 | 147840 | 0 | 226280 | 361350 | 767780 | 1334300 | 302450 | 914380 | 995990 | 386210 | 4014100 |
| QYWCTFKI | FERMT2; FERM | Q96AC1; A0 | FERM2_HUMAN | Unmodified | 0 | 0 | 0 | 0 | 610430 | 0 | 0 | 0 | 0 | 0 | 0 | 0 | 0 | 0 | 0 | 0 | 0 | 663280 |
| QYWFIFKD | FERMT1; FERM | Q9BQL6; G3 | FERM1_HUMAN | Unmodified | 40030 | 0 | 0 | 0 | 0 | 0 | 0 | 0 | 0 | 0 | 0 | 0 | 0 | 21636 | 0 | 0 | 129920 | 0 |
| QYYTLLNQ | G3BP1; G3BP1; | Q13283; A0 | G3BP1_HUMAN | Oxidation (M) | 0 | 0 | 0 | 0 | 0 | 0 | 0 | 0 | 0 | 0 | 0 | 2135000 | 0 | 0 | 0 | 0 | 0 | 0 |
| QYYTLLNQ | G3BP1; G3BP1; | Q13283; A0 | G3BP1_HUMAN | Unmodified | 0 | 0 | 0 | 0 | 0 | 0 | 23664 | 0 | 0 | 265040 | 0 | 1715800 | 0 | 0 | 0 | 0 | 0 | 0 |
| RAAAEVNQ | FH | P07954 | FUMH_HUMAN | Unmodified | 63729 | 60597 | 289240 | 115240 | 0 | 0 | 566600 | 266350 | 4086800 | 991870 | 275280 | 1333400 | 1148900 | 762400 | 135850 | 166100 | 634380 | 0 |
| RAAAEVNQ | FH | P07954 | FUMH_HUMAN | Oxidation (M) | 0 | 0 | 0 | 0 | 0 | 0 | 0 | 266220 | 0 | 249750 | 0 | 0 | 544150 | 306760 | 0 | 0 | 821860 | 0 |

| | | | | | | | | | | | | | | | | | | | | | | |
|---|---|---|---|---|---|---|---|---|---|---|---|---|---|---|---|---|---|---|---|---|---|---|
| RAAAEVNQ | FH | P07954 | FUMH_HUMAN | Unmodified | 0 | 0 | 0 | 0 | 91657 | 0 | 1315700 | 88987 | 0 | 933240 | 0 | 0 | 0 | 0 | 0 | 0 | 1253300 | 0 |
| RAAASKLG | PPP2R1A; PPP2 | P30153; P30 | 2AAA_HUMAN | Unmodified | 0 | 0 | 0 | 0 | 1490400 | 239180 | 287820 | 144840 | 0 | 206160 | 0 | 0 | 1854800 | 987310 | 0 | 0 | 1300100 | 1221400 |
| RAAASKLQ | ARHGAP5; ARH | Q13017; G3 | RHG05_HUMA | Unmodified | 0 | 0 | 0 | 0 | 719590 | 0 | 0 | 370150 | 0 | 0 | 0 | 0 | 0 | 88794 | 0 | 0 | 201380 | 116600 |
| RAAATQPD | SND1 | Q7KZF4 | SND1_HUMAN | Unmodified | 317030 | 522640 | 1190200 | 0 | 0 | 0 | 0 | 0 | 0 | 0 | 0 | 4255900 | 0 | 342780 | 609080 | 570470 | 0 | 0 |
| RAAEDDED | PTMA; PTMA; P | P06454; H7 | PTMA_HUMAN | Unmodified | 35485 | 30385 | 312560 | 0 | 573000 | 0 | 0 | 0 | 396050 | 108930 | 0 | 467880 | 30985 | 0 | 138740 | 0 | 88581 | 0 |
| RAAEDDED | PTMA; PTMA; P | P06454; H7 | PTMA_HUMAN | Unmodified | 0 | 0 | 371920 | 0 | 843460 | 102110 | 0 | 0 | 2591300 | 216250 | 0 | 506680 | 0 | 0 | 0 | 117900 | 82417 | 215570 |
| RAAEEEDE | PTMS | P20962 | PTMS_HUMAN | Unmodified | 63038 | 1150300 | 886460 | 768970 | 172940 | 0 | 373730 | 861350 | 159350 | 75843 | 255930 | 3013800 | 605860 | 71721 | 0 | 0 | 0 | 226790 |
| RAAEEEDE | PTMS | P20962 | PTMS_HUMAN | Unmodified | 201890 | 0 | 0 | 0 | 0 | 0 | 0 | 0 | 0 | 0 | 0 | 1952800 | 0 | 0 | 0 | 0 | 0 | 0 |
| RAAELGAR | GSR; GSR; GSR | P00390; H0 | GSHR_HUMAN | Unmodified | 0 | 0 | 0 | 0 | 2422700 | 0 | 0 | 0 | 0 | 0 | 0 | 0 | 1144700 | 457960 | 0 | 0 | 0 | 168900 |
| RAAEVWMI | GALNT4; GALN | Q8N4A0; Q1 | GALT4_HUMAN | Unmodified | 0 | 2000700 | 1456000 | 2236700 | 3177000 | 18959000 | 2115000 | 0 | 2828600 | 1013200 | 10050000 | 0 | 698540 | 2988800 | 869950 | 0 | 495300 | 8260300 |
| RAAEVWMI | GALNT4; GALN | Q8N4A0; Q1 | GALT4_HUMAN | Oxidation (M) | 0 | 0 | 0 | 0 | 0 | 0 | 0 | 0 | 2731900 | 336600 | 0 | 0 | 0 | 0 | 0 | 0 | 0 | 0 |
| RAAEVWMI | GALNT2 | Q10471 | GALT2_HUMAN | Oxidation (M) | 0 | 0 | 0 | 0 | 0 | 0 | 0 | 0 | 0 | 574810 | 0 | 0 | 0 | 0 | 0 | 0 | 0 | 0 |
| RAAGGGS | DDX1; DDX1; D | Q92499; A0 | DDX1_HUMAN | Unmodified | 0 | 0 | 0 | 0 | 1264900 | 0 | 0 | 0 | 0 | 0 | 0 | 0 | 860720 | 1179500 | 0 | 0 | 714470 | 709150 |
| RAAGVVLE | RUVBL2; RUVB | Q9Y230; M0 | RUVB2_HUMA | Unmodified | 0 | 0 | 0 | 0 | 0 | 0 | 0 | 0 | 0 | 110700 | 0 | 0 | 744220 | 145370 | 0 | 0 | 0 | 0 |
| RAALEGFL | APLP1; APLP1; | P51693; B7Z | APLP1_HUMA | Unmodified | 0 | 0 | 0 | 0 | 0 | 0 | 0 | 0 | 993760 | 0 | 0 | 0 | 0 | 0 | 0 | 0 | 0 | 0 |
| RAASAATA | AGPS; AGPS; A | O00116; A0 | ADAS_HUMAN | Unmodified | 0 | 0 | 0 | 0 | 0 | 0 | 1249100 | 684110 | 8981900 | 2882400 | 673540 | 0 | 0 | 0 | 0 | 0 | 0 | 0 |
| RAASAATA | AGPS; AGPS; A | O00116; A0 | ADAS_HUMAN | Unmodified | 0 | 0 | 0 | 0 | 0 | 0 | 38186 | 0 | 504320 | 0 | 0 | 0 | 0 | 0 | 0 | 0 | 0 | 0 |
| RAAVLLEQI | SF3B2; SF3B2; | Q13435; E9 | SF3B2_HUMA | Unmodified | 1256700 | 850090 | 0 | 0 | 404500 | 0 | 0 | 0 | 0 | 110860 | 0 | 4009900 | 1029800 | 0 | 0 | 0 | 69493 | 0 |
| RADLNQGI | EFHD2 | Q96C19 | EFHD2_HUMA | Unmodified | 0 | 0 | 0 | 71911 | 998690 | 0 | 26149 | 0 | 367890 | 81398 | 418180 | 418540 | 0 | 0 | 0 | 16893 | 0 | 0 |
| RADLNQGI | EFHD2 | Q96C19 | EFHD2_HUMA | Unmodified | 0 | 0 | 0 | 67904 | 4142300 | 2044400 | 0 | 0 | 0 | 60201 | 0 | 207120 | 0 | 0 | 0 | 0 | 0 | 1162200 |
| RADQLADE | SNAP25; SNAP | P60880; A0 | SNP25_HUMA | Unmodified | 0 | 286780 | 0 | 0 | 0 | 0 | 0 | 0 | 3599000 | 182410 | 0 | 0 | 87872 | 0 | 0 | 0 | 0 | 0 |
| RADVQHLE | CEP55; CEP55 | Q53EZ4; H0 | CEP55_HUMA | Unmodified | 46790 | 0 | 0 | 0 | 157970 | 0 | 0 | 0 | 375470 | 72229 | 38893 | 153710 | 0 | 0 | 0 | 0 | 0 | 0 |
| RAEDGSVI | ANXA2; ANXA2; | P07355; H0 | ANXA2_HUMA | Unmodified | 997310 | 651870 | 297580 | 577210 | 4521300 | 0 | 526530 | 182210 | 8206000 | 0 | 0 | 0 | 0 | 0 | 412590 | 0 | 663940 | 0 |
| RAEDGSVI | ANXA2; ANXA2; | P07355; H0 | ANXA2_HUMA | Unmodified | 0 | 0 | 0 | 0 | 20679000 | 4426800 | 0 | 0 | 11216000 | 771970 | 0 | 0 | 0 | 0 | 0 | 0 | 0 | 15227000 |
| RAEDYGEH | LLGL2; LLGL2 | Q6P1M3; J3 | L2GL2_HUMA | Unmodified | 0 | 0 | 0 | 1330800 | 0 | 0 | 0 | 0 | 1369600 | 1094500 | 325990 | 0 | 0 | 0 | 0 | 0 | 0 | 0 |
| RAEDYPIDI | ITGB1; ITGB1; | P05556; E7E | ITB1_HUMAN | 2 Oxidation (M) | 0 | 0 | 0 | 0 | 4191200 | 0 | 0 | 0 | 0 | 0 | 0 | 0 | 0 | 0 | 0 | 0 | 0 | 3120300 |
| RAEDYPIDI | ITGB1; ITGB1; | P05556; E7E | ITB1_HUMAN | Unmodified | 0 | 0 | 0 | 0 | 448730 | 0 | 0 | 0 | 0 | 0 | 0 | 0 | 0 | 0 | 0 | 0 | 0 | 935390 |
| RAEDYPIDI | ITGB1; ITGB1; | P05556; E7E | ITB1_HUMAN | Oxidation (M) | 0 | 0 | 0 | 0 | 1827800 | 0 | 0 | 0 | 0 | 0 | 0 | 0 | 0 | 0 | 0 | 0 | 0 | 1035300 |
| RAEEFLTAS | PPP2R5D; PPP2 | Q14738; H0 | 2A5D_HUMAN | Unmodified | 0 | 0 | 0 | 0 | 640610 | 0 | 0 | 0 | 0 | 0 | 0 | 2197700 | 0 | 0 | 745470 | 0 | 0 | 0 |
| RAEELKTQ | PPP5C; PPP5C | P53041; H0 | PPP5_HUMAN | Unmodified | 0 | 0 | 0 | 0 | 0 | 0 | 461750 | 198190 | 0 | 0 | 0 | 38707 | 0 | 0 | 0 | 0 | 0 | 0 |
| RAEGEAEG | ARHGAP39 | Q9C0H5 | RHG39_HUMA | Unmodified | 0 | 0 | 0 | 0 | 0 | 0 | 0 | 0 | 546590 | 0 | 0 | 0 | 0 | 0 | 0 | 0 | 497780 | 267790 |
| RAEGPPRL | MLF2 | Q15773 | MLF2_HUMAN | Unmodified | 0 | 0 | 0 | 0 | 0 | 0 | 0 | 0 | 0 | 0 | 0 | 122980 | 1887000 | 262460 | 0 | 0 | 0 | 0 |
| RAEISNAID | ATIC; ATIC | P31939; H7 | PUR9_HUMAN | Unmodified | 0 | 0 | 0 | 0 | 2282600 | 0 | 0 | 0 | 514990 | 704960 | 839830 | 0 | 0 | 3244100 | 0 | 0 | 0 | 0 |
| RAEISNAID | ATIC; ATIC | P31939; H7 | PUR9_HUMAN | Unmodified | 0 | 0 | 0 | 0 | 2065700 | 0 | 0 | 0 | 0 | 0 | 0 | 0 | 3450900 | 0 | 0 | 0 | 535100 | 0 |
| RAEPNVLLI | NT5E; NT5E | P21589; Q9 | 5NTD_HUMAN | Unmodified | 0 | 0 | 0 | 0 | 4000400 | 0 | 0 | 0 | 0 | 0 | 0 | 0 | 0 | 0 | 0 | 0 | 254610 | 2598600 |
| RAESMLQC | PLS3; PLS3; PL | P13797; H7 | PLST_HUMAN | Unmodified | 0 | 0 | 0 | 83412 | 197540 | 179640 | 0 | 62812 | 184430 | 46868 | 97728 | 0 | 0 | 0 | 0 | 0 | 82825 | 0 |
| RAEVLGHK | SEPTIN9; SEPT | Q9UHD8; K7 | SEPT9_HUMA | Unmodified | 0 | 0 | 0 | 0 | 1171000 | 0 | 0 | 92556 | 0 | 212870 | 0 | 0 | 3093900 | 73304 | 0 | 0 | 468590 | 877790 |
| RAEVNGLA | ALDOC; ALDOC | P09972; A8 | ALDOC_HUMA | Unmodified | 234490 | 0 | 0 | 349350 | 0 | 0 | 0 | 0 | 0 | 0 | 0 | 5608000 | 1232600 | 649500 | 0 | 0 | 3055400 | 1525400 |
| RAFLIEEQK | RPL34 | P49207 | RL34_HUMAN | Unmodified | 2641300 | 2300100 | 3912300 | 1487000 | 3321100 | 186860 | 177260 | 275790 | 145030 | 0 | 399820 | 14729000 | 893200 | 147460 | 2820200 | 2182500 | 0 | 392910 |
| RAFLIEEQK | RPL34 | P49207 | RL34_HUMAN | Unmodified | 0 | 0 | 0 | 0 | 1404900 | 0 | 1718700 | 0 | 0 | 0 | 0 | 0 | 2860200 | 0 | 0 | 0 | 0 | 529200 |
| RAFVHWY | TUBA3C; TUBA | P0DPH7; Q7 | TBA3C_HUMA | Oxidation (M) | 0 | 0 | 780410 | 0 | 644850 | 0 | 4381900 | 0 | 1270400 | 4797000 | 3474400 | 10096000 | 0 | 0 | 0 | 1251900 | 0 | 0 |
| RAFVHWY | TUBA3C; TUBA | P0DPH7; Q7 | TBA3C_HUMA | Unmodified | 0 | 0 | 0 | 0 | 173220 | 0 | 1893100 | 0 | 1362900 | 2133000 | 225190 | 1510700 | 0 | 0 | 0 | 117520 | 0 | 0 |
| RAFVHWY | TUBA1A; TUBA | Q71U36; P6 | TBA1A_HUMA | Oxidation (M) | 0 | 0 | 0 | 1315700 | 14787000 | 7516100 | 24222000 | 0 | 8966200 | 11705000 | 0 | 3347000 | 25770000 | 12725000 | 0 | 0 | 3186100 | 19199000 |
| RAFVHWY | TUBA1A; TUBA | Q71U36; P6 | TBA1A_HUMA | 2 Oxidation (M) | 0 | 0 | 0 | 2066400 | 8327700 | 574240 | 15287000 | 0 | 2770100 | 8998200 | 366910 | 2795100 | 6640700 | 0 | 754320 | 599840 | 2331500 | 5215300 |
| RAFVHWY | TUBA1A; TUBA | Q71U36; P6 | TBA1A_HUMA | Unmodified | 0 | 0 | 0 | 0 | 5497000 | 0 | 2922300 | 0 | 2583600 | 1457900 | 0 | 0 | 3439700 | 3838800 | 0 | 0 | 349320 | 3582900 |
| RAGELTEDI | RPS18 | P62269 | RS18_HUMAN | Unmodified | 862840 | 864530 | 726130 | 641400 | 0 | 0 | 0 | 0 | 299400 | 0 | 1117400 | 28960000 | 0 | 0 | 1040200 | 1272200 | 0 | 0 |
| RAGELTEDI | RPS18 | P62269 | RS18_HUMAN | Oxidation (M) | 0 | 0 | 0 | 0 | 0 | 0 | 0 | 28456 | 2021100 | 0 | 0 | 1011900 | 0 | 0 | 0 | 0 | 0 | 0 |
| RAGELTEDI | RPS18 | P62269 | RS18_HUMAN | Unmodified | 0 | 0 | 0 | 1192800 | 0 | 0 | 0 | 0 | 0 | 0 | 0 | 0 | 931420 | 0 | 1277500 | 0 | 0 | 0 |
| RAGGPEGI | TTLL12 | Q14166 | TTL12_HUMA | Unmodified | 0 | 0 | 0 | 0 | 0 | 0 | 0 | 0 | 0 | 0 | 0 | 1796000 | 102010 | 0 | 763950 | 120450 | 0 | 0 |
| RAGKEVLLI | SNTB1; SNTB2; | Q13884; Q1 | SNTB1_HUMA | Unmodified | 0 | 0 | 0 | 0 | 2546800 | 0 | 0 | 0 | 0 | 1973000 | 0 | 0 | 627370 | 622360 | 0 | 0 | 2025100 | 2679400 |
| RAGLGSGI | ACADVL; ACAD | P49748; G3 | ACADV_HUMA | Unmodified | 0 | 0 | 0 | 0 | 0 | 0 | 0 | 0 | 3750500 | 182940 | 0 | 0 | 0 | 0 | 0 | 0 | 0 | 0 |
| RAGLLTEQ | VPS33B; VPS33 | Q9H267; F5 | VP33B_HUMA | Unmodified | 91609 | 0 | 0 | 0 | 384520 | 0 | 0 | 0 | 0 | 0 | 0 | 580370 | 207350 | 0 | 100810 | 101380 | 0 | 0 |
| RAGLMLQE | PLS1; PLS1 | Q14651; C9 | PLSI_HUMAN | Unmodified | 49824 | 0 | 0 | 0 | 0 | 0 | 328060 | 205460 | 0 | 0 | 0 | 948610 | 0 | 0 | 0 | 45688 | 0 | 0 |
| RAGVTCIVI | LONP1; LONP1 | P36776; K7E | LONM_HUMAN | Unmodified | 0 | 0 | 0 | 0 | 0 | 0 | 0 | 0 | 547690 | 106180 | 0 | 0 | 0 | 0 | 0 | 0 | 0 | 0 |
| RAHLTVGQ | PKP2 | Q99959 | PKP2_HUMAN | Unmodified | 0 | 0 | 0 | 0 | 0 | 0 | 0 | 0 | 0 | 0 | 0 | 626640 | 0 | 0 | 0 | 0 | 82773 | 0 |
| RAHQLPGI | CCM2; CCM2; C | Q9BSQ5; A0 | CCM2_HUMAN | Unmodified | 0 | 0 | 0 | 351630 | 0 | 0 | 0 | 0 | 1028800 | 367040 | 0 | 0 | 1431700 | 2546500 | 0 | 201570 | 2616200 | 0 |
| RAIELATTLI | NAA15; NAA15 | Q9BXJ9; A0 | NAA15_HUMA | Unmodified | 0 | 0 | 0 | 0 | 111360 | 0 | 0 | 101230 | 0 | 0 | 0 | 1620700 | 0 | 0 | 0 | 0 | 0 | 39548 |
| RAIKNDSV | CCT7 | Q99832 | TCPH_HUMAN | Unmodified | 0 | 0 | 0 | 0 | 967830 | 0 | 111500 | 0 | 0 | 241890 | 168010 | 0 | 1111900 | 603870 | 0 | 0 | 482880 | 272560 |
| RAILTMDE | DAAM1 | Q9Y4D1 | DAAM1_HUMA | Unmodified | 0 | 0 | 0 | 0 | 0 | 0 | 0 | 0 | 0 | 0 | 0 | 2052900 | 0 | 0 | 325760 | 127420 | 0 | 0 |
| RAISQSGV | CEL | P19835 | CEL_HUMAN B | Unmodified | 0 | 0 | 0 | 0 | 0 | 0 | 0 | 0 | 12818000 | 2646400 | 7171300 | 0 | 0 | 0 | 0 | 0 | 0 | 0 |
| RAIVHDNK | FLNB; FLNB | O75369; E7 | FLNB_HUMAN | Unmodified | 0 | 0 | 0 | 0 | 1201100 | 537610 | 0 | 0 | 0 | 0 | 0 | 405770 | 187350 | 0 | 0 | 0 | 205330 | 0 |
| RAKEAAEQ | RPL13; RPL13 | P26373; J3K | RL13_HUMAN | Unmodified | 0 | 0 | 0 | 0 | 1395600 | 0 | 115660 | 394060 | 0 | 0 | 8501.3 | 40165 | 0 | 0 | 0 | 0 | 0 | 0 |
| RAKPEGAL | EPCAM; EPCAM | P16422; C9 | EPCAM_HUMA | Unmodified | 37283000 | 11962000 | 30577000 | 54301000 | 25269000 | 3778300 | 0 | 0 | 36376000 | 41037000 | 14984000 | 29429000 | 1.05E+08 | 88633000 | 60240000 | 71022000 | 3764800 | 5245800 |
| RALANSLA | ALDOA; ALDOA | P04075; J3K | ALDOA_HUMA | Unmodified | 0 | 0 | 0 | 0 | 18894000 | 2756200 | 1999900 | 0 | 0 | 2813700 | 387800 | 0 | 5105500 | 4093200 | 0 | 0 | 20858000 | 9308900 |
| RALASIDSK | VCL | P18206 | VINC_HUMAN | Unmodified | 0 | 0 | 0 | 5738800 | 0 | 1298800 | 57060 | 0 | 15099 | 0 | 0 | 132050 | 166900 | 0 | 116900 | 0 | 460200 | 0 |
| RALASIDSK | VCL | P18206 | VINC_HUMAN | Unmodified | 0 | 0 | 0 | 0 | 1842400 | 453950 | 0 | 406610 | 0 | 35192 | 0 | 139230 | 0 | 0 | 0 | 0 | 5218500 | 255680 |
| RALASNTSI | KIF4A; KIF4B | O95239; Q2 | KIF4A_HUMA | Unmodified | 0 | 0 | 0 | 0 | 0 | 0 | 0 | 0 | 0 | 0 | 0 | 0 | 198250 | 0 | 0 | 0 | 746930 | 0 |
| RALDFAVG | CST3 | P01034 | CYTC_HUMAN | Unmodified | 0 | 0 | 0 | 0 | 0 | 0 | 0 | 0 | 1358000 | 0 | 0 | 0 | 0 | 0 | 0 | 0 | 0 | 0 |
| RALDKLDG | SRSF6; SRSF6; | Q13247; A0 | SRSF6_HUMA | Unmodified | 0 | 0 | 0 | 0 | 102530 | 0 | 0 | 0 | 0 | 0 | 0 | 54296 | 228410 | 56960 | 0 | 0 | 24168 | 0 |
| RALDLNLK | UQCRB; UQCR | P14927; B7Z | QCR7_HUMAN | Unmodified | 3462800 | 0 | 0 | 0 | 168510 | 0 | 676200 | 557570 | 7298000 | 2738100 | 0 | 0 | 70861 | 59797 | 0 | 0 | 0 | 0 |
| RALELEQEI | MSN | P26038 | MOES_HUMAN | Unmodified | 0 | 610180 | 0 | 0 | 747190 | 0 | 1397100 | 0 | 0 | 0 | 0 | 0 | 0 | 0 | 0 | 0 | 354020 | 1006500 |
| RALEQQVE | MYH9 | P35579 | MYH9_HUMAN | Unmodified | 223300 | 175380 | 750160 | 552350 | 1339200 | 451340 | 1510700 | 382080 | 1109900 | 255830 | 336700 | 9423500 | 934180 | 1303800 | 1376600 | 1148200 | 118260 | 722200 |
| RALEQQVE | MYH9 | P35579 | MYH9_HUMAN | Oxidation (M) | 37042 | 73221 | 254150 | 0 | 0 | 173090 | 21682 | 0 | 0 | 105950 | 0 | 1002100 | 82498 | 46165 | 477220 | 510210 | 0 | 209250 |
| RALEQQVE | MYH10 | P35580 | MYH10_HUMA | Unmodified | 0 | 0 | 264160 | 0 | 0 | 0 | 0 | 0 | 0 | 0 | 0 | 1044700 | 0 | 0 | 0 | 0 | 0 | 0 |
| RALIAGGG | CCT4 | P50991 | TCPD_HUMAN | Unmodified | 0 | 0 | 0 | 0 | 0 | 0 | 0 | 0 | 0 | 0 | 156480 | 3262100 | 235970 | 0 | 0 | 300880 | 0 | 0 |
| RALIESYQN | ABI1; ABI1; ABI | Q8IZP0; A0 | ABI1_HUMAN | Unmodified | 473000 | 465260 | 1056500 | 1656100 | 5193300 | 433300 | 0 | 0 | 3227300 | 761820 | 718920 | 6823900 | 3625300 | 1481200 | 2000000 | 2513700 | 363230 | 276560 |
| RALIESYQN | ABI1; ABI1; ABI | Q8IZP0; A0 | ABI1_HUMAN | Unmodified | 0 | 0 | 0 | 0 | 0 | 0 | 0 | 0 | 0 | 0 | 0 | 0 | 0 | 0 | 0 | 0 | 863350 | 1845000 |
| RALIQCAKI | VCL | P18206 | VINC_HUMAN | Unmodified | 0 | 0 | 0 | 0 | 98822 | 0 | 344640 | 0 | 0 | 0 | 0 | 0 | 176840 | 287750 | 0 | 0 | 1012000 | 125170 |
| RALNFLVN | RELCH; RELCH | Q9P260; A0 | RELCH_HUMA | Unmodified | 0 | 0 | 0 | 0 | 2296500 | 0 | 12774 | 0 | 0 | 113320 | 0 | 0 | 1299000 | 705790 | 0 | 0 | 0 | 0 |
| RALSELTQ | KIF20B | Q96Q89 | KI20B_HUMA | Unmodified | 0 | 0 | 0 | 0 | 0 | 0 | 0 | 0 | 0 | 0 | 0 | 0 | 0 | 0 | 0 | 47127 | 840590 | 0 |
| RALSSQHQ | HSPA5; HSPA5 | P11021; A0 | BIP_HUMAN Er | Unmodified | 0 | 0 | 0 | 279970 | 560480 | 0 | 3334300 | 0 | 17124000 | 48940000 | 10116000 | 0 | 0 | 0 | 0 | 0 | 0 | 0 |
| RALTHIDH | MARS1; MARS1 | P56192; F5 | SYMC_HUMAN | Unmodified | 0 | 0 | 56900 | 0 | 0 | 0 | 0 | 0 | 0 | 0 | 0 | 770090 | 320230 | 740690 | 11842 | 21493 | 0 | 0 |
| RALVAIGT | FARSB | Q9NSD9 | SYFB_HUMAN | Unmodified | 0 | 0 | 0 | 0 | 1406800 | 0 | 301230 | 0 | 281580 | 234920 | 385480 | 2702700 | 0 | 339540 | 0 | 0 | 0 | 366070 |
| RANNTFYG | ALDH1A1 | P00352 | AL1A1_HUMA | Unmodified | 0 | 0 | 0 | 0 | 0 | 0 | 0 | 0 | 2522500 | 1808300 | 20566000 | 4.89E+08 | 0 | 0 | 0 | 0 | 0 | 0 |
| RANPQVG | XRCC5 | P13010 | XRCC5_HUMA | Unmodified | 0 | 0 | 0 | 0 | 504610 | 0 | 432760 | 224750 | 276050 | 0 | 0 | 2964100 | 0 | 0 | 0 | 0 | 0 | 0 |
| RANSTDYG | ALDH1A3; ALD | P47895; H0 | AL1A3_HUMA | Unmodified | 0 | 0 | 0 | 0 | 0 | 0 | 0 | 0 | 0 | 0 | 117580 | 0 | 0 | 0 | 0 | 0 | 7981500 | 5494800 |
| RAPFDLFEI | HSP90AB1; HSI | P08238; Q5 | HS90B_HUMA | Unmodified | 731540 | 823460 | 2427600 | 1967800 | 6553500 | 721530 | 0 | 35470 | 3752800 | 1633100 | 6811100 | 43627000 | 3970600 | 1691100 | 3389000 | 2280100 | 430590 | 1836000 |
| RAPFDLFEI | HSP90AB1; HSI | P08238; Q5 | HS90B_HUMA | Unmodified | 156230 | 0 | 227280 | 473180 | 1784300 | 459620 | 214980 | 0 | 922720 | 849880 | 566930 | 8497800 | 811610 | 286330 | 1871200 | 473390 | 0 | 664210 |
| RAPFDLFEI | HSP90AA1 | P07900 | HS90A_HUMA | Unmodified | 400040 | 349140 | 1212000 | 862630 | 0 | 0 | 0 | 0 | 3380500 | 396830 | 1091300 | 29280000 | 101690 | 0 | 1464100 | 730330 | 0 | 0 |
| RAPFDLFEI | HSP90AA1 | P07900 | HS90A_HUMA | Unmodified | 610530 | 425860 | 721800 | 944120 | 2005400 | 1972200 | 684460 | 0 | 2001900 | 1940900 | 525620 | 9507800 | 976600 | 659870 | 985390 | 933720 | 524310 | 1382600 |
| RAPPPPGS | COBL; COBL; C | O75128; J3K | COBL_HUMAN | Unmodified | 179110 | 189200 | 22435 | 0 | 0 | 0 | 0 | 0 | 0 | 0 | 0 | 0 | 0 | 0 | 85365 | 85299 | 0 | 1212300 |
| RAPSSAQY | CORO7; CORO | P57737; I3L | CORO7_HUMA | Unmodified | 0 | 0 | 0 | 88384 | 0 | 0 | 0 | 33912 | 0 | 0 | 3735100 | 0 | 0 | 0 | 0 | 0 | 39962 | 0 |
| RAPSTSPS | MCRIP2 | Q9BUT9 | MCRI2_HUMA | Unmodified | 901150 | 0 | 0 | 0 | 0 | 0 | 0 | 0 | 0 | 0 | 0 | 0 | 0 | 0 | 0 | 0 | 0 | 0 |
| RAPVIWDN | OGA; OGA | O60502; H7 | OGA_HUMAN | Unmodified | 0 | 0 | 0 | 0 | 0 | 0 | 0 | 0 | 0 | 0 | 0 | 0 | 0 | 0 | 0 | 0 | 0 | 0 |
| RAQADLAY | FLOT1; FLOT1; | O75955; A0 | FLOT1_HUMA | Unmodified | 0 | 377450 | 2860800 | 3088200 | 10244000 | 1713000 | 1224300 | 284840 | 16612000 | 3919900 | 1599300 | 10259000 | 3013000 | 316700 | 4480600 | 2638100 | 1017400 | 2428700 |
| RAQAWCY | RAB7A; RAB7A; | P51149; A0 | RAB7A_HUMA | Unmodified | 0 | 0 | 0 | 0 | 0 | 0 | 0 | 0 | 0 | 0 | 0 | 0 | 0 | 0 | 0 | 0 | 0 | 989070 |
| RAQCYMD | DNAJC7; DNAJ | Q99615; A0 | DNJC7_HUMA | Unmodified | 0 | 0 | 0 | 0 | 0 | 0 | 0 | 0 | 0 | 0 | 0 | 845040 | 1937400 | 0 | 0 | 0 | 0 | 0 |
| RAQEEAEA | VGF | O15240 | VGF_HUMAN | Unmodified | 72317 | 79026 | 0 | 0 | 0 | 0 | 0 | 0 | 1782700 | 0 | 0 | 0 | 0 | 0 | 0 | 0 | 0 | 0 |
| RAQEILSQI | GLRX | P35754 | GLRX1_HUMA | Unmodified | 0 | 0 | 0 | 0 | 0 | 0 | 0 | 0 | 0 | 0 | 0 | 5856800 | 797150 | 0 | 0 | 0 | 375150 | 0 |
| RAQGEKPL | AHCYL1; AHCY | O43865; Q9 | SAHH2_HUMA | Unmodified | 0 | 0 | 0 | 452740 | 3200600 | 2112600 | 925690 | 391840 | 6400.1 | 0 | 0 | 1208900 | 2871500 | 873680 | 2092300 | 381240 | 5150500 | 2209200 |
| RAQLHVQC | SPNS1; SPNS1 | Q9H2V7; H3 | SPNS1_HUMA | Unmodified | 85905 | 0 | 0 | 7235600 | 2912800 | 40207000 | 1904600 | 32471 | 0 | 0 | 0 | 0 | 1259300 | 4109800 | 0 | 0 | 1907800 | 2399500 |
| RAQMLREA | SCRIB; SCRIB; | Q14160; A0 | SCRIB_HUMA | Unmodified | 0 | 0 | 0 | 0 | 124640 | 19176 | 0 | 0 | 52555 | 173410 | 117310 | 0 | 0 | 51834 | 0 | 0 | 83808 | 337810 |
| RAQQQLEN | TRIP10; TRIP10 | Q15642; W4 | CIP4_HUMAN | Unmodified | 0 | 50663 | 0 | 0 | 334010 | 228040 | 0 | 96222 | 51554 | 0 | 0 | 0 | 0 | 0 | 0 | 0 | 0 | 0 |
| RAQSEAEK | MSN | P26038 | MOES_HUMAN | Unmodified | 0 | 0 | 0 | 0 | 0 | 507040 | 75350 | 78382 | 0 | 0 | 0 | 0 | 279630 | 181400 | 0 | 0 | 0 | 1078100 |
| RARPSPGH | RPAP1; RPAP1; | Q9BWH6; H | RPAP1_HUMA | Unmodified | 132270 | 4103100 | 587280 | 736560 | 0 | 0 | 0 | 0 | 0 | 0 | 0 | 0 | 0 | 0 | 2135400 | 3082600 | 0 | 0 |
| RARQPLSE | ANLN; ANLN | Q9NQW6; C | ANLN_HUMAN | Unmodified | 0 | 0 | 0 | 314580 | 22935000 | 3806100 | 975900 | 373330 | 231430 | 154710 | 0 | 0 | 1234400 | 947480 | 0 | 0 | 8355500 | 8047700 |
| RASAAAAA | SPAG1 | Q07617 | SPAG1_HUMA | Unmodified | 354600 | 289830 | 309130 | 262570 | 4362600 | 1042400 | 93185 | 0 | 554660 | 185740 | 0 | 0 | 1644900 | 900170 | 141070 | 270860 | 1302200 | 3192300 |
| RASALIDRF | EPB41L3; EPB4 | Q9Y2J2; A0 | E41L3_HUMA | Unmodified | 0 | 0 | 0 | 0 | 0 | 0 | 0 | 0 | 0 | 0 | 0 | 0 | 0 | 0 | 0 | 0 | 160210 | 951810 |
| RASEFPGP | FAM171A1 | Q5VUB5 | F1711_HUMA | Unmodified | 0 | 0 | 0 | 0 | 1445000 | 0 | 0 | 0 | 0 | 0 | 0 | 835510 | 329840 | 0 | 0 | 0 | 164660 | 0 |

| | | | | | | | | | | | | | | | | | | | | | | |
|---|---|---|---|---|---|---|---|---|---|---|---|---|---|---|---|---|---|---|---|---|---|---|
| RASGQAFE | STMN1; STMN1 | P16949; A2A | STMN1_HUMAI | Unmodified | 877120 | 751540 | 547780 | 1278100 | 7451100 | 1715800 | 3639700 | 1536700 | 31045000 | 30209000 | 1786200 | 17880000 | 6142400 | 2061800 | 901480 | 500220 | 3038400 | 5397800 |
| RASGQAFE | STMN1; STMN1 | P16949; A2A | STMN1_HUMAI | Unmodified | 0 | 0 | 0 | 0 | 1599100 | 0 | 4895800 | 932730 | 0 | 2163600 | 0 | 0 | 606910 | 0 | 0 | 0 | 1024500 | 1063900 |
| RASGQSFE | STMN3; STMN3 | Q9NZ72; A0/ | STMN3_HUMAI | Unmodified | 0 | 0 | 170090 | 0 | 0 | 0 | 1210200 | 439830 | 1290700 | 267000 | 0 | 0 | 0 | 0 | 940430 | 0 | 0 | 0 |
| RASGQSFE | STMN3; STMN3 | Q9NZ72; A0/ | STMN3_HUMAI | Oxidation (M) | 0 | 0 | 0 | 0 | 0 | 0 | 873090 | 0 | 0 | 0 | 0 | 0 | 0 | 0 | 2844500 | 0 | 0 | 0 |
| RASHTAPQ | ARPC2; ARPC2 | O15144; H7( | ARPC2_HUMAI | Unmodified | 0 | 0 | 0 | 0 | 524320 | 0 | 0 | 0 | 0 | 0 | 0 | 8301700 | 0 | 0 | 0 | 0 | 0 | 0 |
| RASSDLSIA | SLK | Q9H2G2 | SLK_HUMAN S | Unmodified | 0 | 0 | 0 | 0 | 0 | 0 | 0 | 0 | 0 | 0 | 0 | 1052400 | 0 | 0 | 0 | 0 | 0 | 0 |
| RATASEQPI | LAD1; LAD1 | O00515; E9F | LAD1_HUMAN | Unmodified | 93550 | 0 | 153480 | 182580 | 177900 | 0 | 0 | 0 | 0 | 0 | 829760 | 1966000 | 1957100 | 1024700 | 180240 | 155910 | 42468 | 0 |
| RATPIEVLC | CSNK1G2; CSN | P78368; A0A | KC1G2_HUMAI | Unmodified | 0 | 0 | 0 | 0 | 0 | 0 | 0 | 0 | 226850 | 0 | 45658 | 28882 | 0 | 0 | 687790 | 0 | 0 | 0 |
| RATPIEVLC | CSNK1G3 | Q9Y6M4 | KC1G3_HUMAI | Unmodified | 0 | 794060 | 0 | 20703 | 180490 | 0 | 0 | 0 | 0 | 0 | 0 | 0 | 0 | 0 | 45742 | 0 | 0 | 0 |
| RATTEDRN | ARPC1A | Q92747 | ARC1A_HUMAI | Unmodified | 0 | 166890 | 2459300 | 0 | 0 | 0 | 0 | 0 | 0 | 0 | 0 | 1123700 | 0 | 0 | 1285400 | 958940 | 0 | 172450 |
| RATVAPED | ACAT2 | Q9BWD1 | THIC_HUMAN / | Unmodified | 0 | 0 | 0 | 0 | 0 | 0 | 0 | 0 | 2175700 | 0 | 0 | 6255000 | 0 | 0 | 0 | 0 | 0 | 0 |
| RATVLESE( | STOML2; STOM | Q9UJZ1; A0A | STML2_HUMAN | Unmodified | 0 | 0 | 0 | 0 | 0 | 0 | 0 | 0 | 1289000 | 0 | 0 | 0 | 0 | 0 | 0 | 0 | 0 | 0 |
| RATVNTFG | SF3B1; SF3B1 | O75533; H7( | SF3B1_HUMAN | Unmodified | 35589 | 0 | 0 | 0 | 0 | 104420 | 0 | 24231 | 0 | 0 | 0 | 805790 | 0 | 0 | 0 | 0 | 0 | 0 |
| RATVVESSI | YWHAG | P61981 | 1433G_HUMAN | Unmodified | 0 | 0 | 152380 | 768770 | 5582800 | 948250 | 1343600 | 1468800 | 1083200 | 592810 | 11974 | 3168800 | 3219100 | 3801200 | 0 | 74318 | 6207500 | 3233800 |
| RATVVESSI | YWHAG | P61981 | 1433G_HUMAN | Unmodified | 0 | 0 | 0 | 0 | 1032000 | 0 | 1031100 | 0 | 0 | 229540 | 0 | 0 | 896940 | 2664600 | 0 | 0 | 3000900 | 961140 |
| RAVAGDAS | ATP1A1 | P05023 | AT1A1_HUMAN | Unmodified | 5036900 | 3125500 | 42208000 | 36305000 | 97574000 | 1170900 | 15962000 | 1290500 | 37195000 | 14465000 | 30533000 | 1.86E+08 | 94765000 | 66198000 | 49761000 | 66837000 | 10946000 | 11435000 |
| RAVAGDAS | ATP1A1 | P05023 | AT1A1_HUMAN | Unmodified | 0 | 0 | 0 | 0 | 560670 | 0 | 0 | 0 | 0 | 0 | 0 | 0 | 309960 | 2061900 | 0 | 0 | 944760 | 0 |
| RAVAGLLA! | FXN; N/A; N/A; I | Q16595; A0/ | FRDA_HUMAN | Unmodified | 1063300 | 1787500 | 0 | 3077500 | 0 | 0 | 0 | 0 | 0 | 0 | 0 | 0 | 0 | 0 | 747000 | 505860 | 0 | 0 |
| RAVDFQEA | RAB5A | P20339 | RAB5A_HUMAN | Unmodified | 0 | 0 | 386970 | 0 | 4541600 | 0 | 0 | 0 | 3689800 | 0 | 0 | 0 | 0 | 0 | 0 | 0 | 0 | 0 |
| RAVDFQEA | RAB5A | P20339 | RAB5A_HUMAN | Oxidation (M) | 0 | 1168700 | 0 | 0 | 2042800 | 0 | 0 | 0 | 1793700 | 1030300 | 0 | 0 | 2179400 | 713640 | 0 | 825910 | 0 | 1496200 |
| RAVDGFTF | NCAM2; NCAM | O15394; H9I | NCAM2_HUMA | Unmodified | 0 | 0 | 1831300 | 3490500 | 0 | 0 | 0 | 0 | 264840 | 0 | 0 | 0 | 0 | 0 | 2572400 | 2641100 | 0 | 0 |
| RAVEFQEA( | RAB5C; RAB5C | P51148; K7E | RAB5C_HUMAI | Oxidation (M) | 4783200 | 0 | 0 | 0 | 3891400 | 0 | 1710600 | 0 | 3095200 | 3036300 | 0 | 3587200 | 8120400 | 4798800 | 1222600 | 4808600 | 0 | 0 |
| RAVEFQEA( | RAB5C; RAB5C | P51148; K7E | RAB5C_HUMAI | Unmodified | 0 | 0 | 0 | 0 | 0 | 0 | 2762100 | 0 | 5666400 | 1646600 | 0 | 0 | 2826200 | 0 | 0 | 0 | 0 | 0 |
| RAVGGDH/ | SNX11; SNX11 | Q9Y5W9; B4 | SNX11_HUMAN | Unmodified | 0 | 0 | 0 | 0 | 115160 | 0 | 0 | 0 | 0 | 0 | 0 | 0 | 130030 | 0 | 0 | 0 | 418100 | 793290 |
| RAVGIWHC | RPL37A; RPL37. | P61513; M0I | RL37A_HUMAN | Unmodified | 71547 | 58700 | 0 | 0 | 0 | 0 | 0 | 0 | 0 | 0 | 0 | 400900 | 0 | 0 | 456370 | 0 | 0 | 0 |
| RAVGPSST | CYFIP1; CYFIP1 | Q7L576; A0/ | CYFP1_HUMAN | Unmodified | 0 | 0 | 0 | 0 | 0 | 0 | 0 | 0 | 653330 | 0 | 0 | 1327100 | 0 | 0 | 297020 | 156630 | 0 | 0 |
| RAVLAHVD | IGSF8 | Q969P0 | IGSF8_HUMAN | Unmodified | 0 | 0 | 0 | 0 | 0 | 0 | 0 | 0 | 0 | 0 | 0 | 0 | 5174800 | 0 | 0 | 0 | 0 | 0 |
| RAVVMISC | M6PR; M6PR | P20645; H0 | MPRD_HUMAN | Oxidation (M) | 0 | 0 | 0 | 96133 | 0 | 0 | 0 | 0 | 1278500 | 0 | 0 | 0 | 0 | 0 | 0 | 0 | 0 | 0 |
| RAVVMISC | M6PR; M6PR | P20645; H0 | MPRD_HUMAN | Unmodified | 0 | 0 | 0 | 0 | 0 | 0 | 0 | 0 | 3200400 | 0 | 174000 | 0 | 0 | 0 | 0 | 0 | 0 | 0 |
| RAYIPNFES | LRP2; LRP2 | P98164; A0A | LRP2_HUMAN I | Unmodified | 0 | 0 | 0 | 0 | 0 | 0 | 0 | 0 | 2845600 | 597980 | 0 | 0 | 0 | 0 | 0 | 0 | 0 | 0 |
| RAYQCVVL | RARS1 | P54136 | SYRC_HUMAN | Unmodified | 0 | 0 | 0 | 0 | 0 | 0 | 0 | 384320 | 0 | 0 | 0 | 1081300 | 262570 | 0 | 0 | 0 | 0 | 0 |
| RCADELTPI | CSGALNACT2 | Q8N6G5 | CGAT2_HUMAI | Unmodified | 0 | 0 | 0 | 0 | 0 | 0 | 0 | 0 | 1091500 | 115200 | 0 | 0 | 0 | 0 | 0 | 0 | 0 | 0 |
| RCANYVN( | MME; MME; MM | P08473; A0A | NEP_HUMAN N | Unmodified | 0 | 0 | 0 | 1439900 | 0 | 0 | 0 | 0 | 636220 | 0 | 0 | 0 | 0 | 0 | 1814300 | 951420 | 14958 | 0 |
| RCANYVN( | MME; MME; MM | P08473; A0A | NEP_HUMAN N | Oxidation (M) | 0 | 0 | 88868 | 216980 | 0 | 0 | 0 | 0 | 0 | 0 | 0 | 0 | 0 | 0 | 214940 | 263650 | 0 | 0 |
| RCEAFGWI | TKT; TKT; TKT | P29401; A0A | TKT_HUMAN Tr | Unmodified | 0 | 0 | 0 | 0 | 2404300 | 651150 | 316940 | 0 | 867980 | 4195300 | 0 | 1779100 | 0 | 0 | 0 | 0 | 2029900 | 1506700 |
| RCEPIAMT | CORO1A; COR( | P31146; H3E | COR1A_HUMAI | Unmodified | 0 | 0 | 0 | 0 | 234740 | 0 | 0 | 0 | 0 | 45981 | 0 | 2302200 | 0 | 0 | 0 | 0 | 0 | 0 |
| RCFIEEIPD | TMED4; TMED4 | Q7Z7H5; F8 | TMED4_HUMAI | Oxidation (M) | 0 | 0 | 0 | 0 | 0 | 0 | 0 | 0 | 3010500 | 187500 | 0 | 0 | 0 | 0 | 0 | 0 | 1303600 | 0 |
| RCFIEEIPD | TMED4; TMED4 | Q7Z7H5; F8 | TMED4_HUMAI | Unmodified | 0 | 0 | 0 | 0 | 0 | 0 | 0 | 0 | 5275200 | 0 | 15884 | 0 | 0 | 0 | 0 | 0 | 0 | 0 |
| RCFIEEIPD | TMED4; TMED4 | Q7Z7H5; F8 | TMED4_HUMAI | 2 Oxidation (M) | 0 | 0 | 0 | 0 | 0 | 0 | 0 | 0 | 0 | 8500700 | 0 | 0 | 0 | 0 | 0 | 0 | 0 | 0 |
| RCFIEEIPD | TMED4; TMED4 | Q7Z7H5; F8 | TMED4_HUMAI | Oxidation (M) | 0 | 0 | 0 | 0 | 0 | 0 | 0 | 0 | 1539000 | 1608200 | 0 | 0 | 0 | 0 | 0 | 0 | 0 | 0 |
| RCFPAIHA | SLC44A2; SLC4 | Q8IWA5; K7I | CTL2_HUMAN | Unmodified | 0 | 0 | 392350 | 131830 | 0 | 0 | 0 | 0 | 0 | 0 | 0 | 0 | 107250 | 496900 | 685820 | 181790 | 0 | 0 |
| RCGNLSTC | CALCA | P01258 | CALC_HUMAN | Unmodified | 0 | 0 | 0 | 657680 | 146710 | 0 | 0 | 0 | 1516900 | 0 | 0 | 0 | 591490 | 0 | 0 | 0 | 0 | 144640 |
| RCLYASVL | EEF2 | P13639 | EF2_HUMAN El | Unmodified | 0 | 0 | 0 | 435840 | 2533800 | 596130 | 0 | 148290 | 241500 | 453070 | 790960 | 5616800 | 787410 | 610480 | 435970 | 254300 | 723330 | 672880 |
| RCQEAQN( | SUGT1 | Q9Y2Z0 | SGT1_HUMAN | Unmodified | 0 | 0 | 0 | 0 | 0 | 0 | 0 | 0 | 0 | 0 | 50103 | 922040 | 0 | 0 | 0 | 0 | 0 | 0 |
| RCSDNTEV | ANLN; ANLN | Q9NQW6; C | ANLN_HUMAN | Unmodified | 0 | 0 | 0 | 0 | 1109900 | 0 | 0 | 0 | 0 | 0 | 0 | 0 | 0 | 0 | 0 | 0 | 341540 | 0 |
| RDAESIHQ | NPEPPS; NPEPI | P55786; E9P | PSA_HUMAN P | Unmodified | 0 | 0 | 925640 | 1029300 | 3337200 | 1565400 | 536470 | 276240 | 736190 | 557120 | 1187400 | 2852600 | 1789500 | 1381700 | 0 | 847320 | 1460200 | 3779500 |
| RDAYVSRL | GSR; GSR; GSR | P00390; H0 | GSHR_HUMAN | Unmodified | 0 | 0 | 0 | 0 | 1453500 | 0 | 0 | 0 | 0 | 0 | 0 | 0 | 365500 | 246210 | 0 | 18342 | 0 | 0 |
| RDDDGALI | SOD3 | P08294 | SODE_HUMAN | Unmodified | 0 | 0 | 0 | 0 | 0 | 0 | 0 | 0 | 6595300 | 0 | 0 | 0 | 0 | 0 | 0 | 0 | 0 | 0 |
| RDDGTGQI | PLEC; PLEC | Q15149; H0 | PLEC_HUMAN | Unmodified | 0 | 0 | 0 | 0 | 0 | 0 | 0 | 0 | 0 | 124180 | 25575 | 0 | 333230 | 0 | 0 | 0 | 1948500 | 0 |
| RDEAGHFL | PCK2; PCK2 | Q16822; H0 | PCKGM_HUMA | Unmodified | 0 | 0 | 0 | 0 | 0 | 0 | 0 | 0 | 805340 | 0 | 0 | 0 | 0 | 0 | 0 | 0 | 0 | 0 |
| RDFAPPG( | TMEM109 | Q9BVC6 | TM109_HUMAN | Unmodified | 0 | 20069 | 0 | 0 | 0 | 0 | 0 | 0 | 3811600 | 859560 | 187270 | 0 | 0 | 0 | 0 | 0 | 34659 | 0 |
| RDHALLEE | ACTN4; ACTN4 | O43707; F5( | ACTN4_HUMAI | Unmodified | 526940 | 401660 | 1994200 | 479290 | 993680 | 1142400 | 0 | 0 | 2351800 | 307350 | 916450 | 4878100 | 0 | 0 | 2875300 | 2782100 | 7694.9 | 0 |
| RDIDNLVQ | LMAN1 | P49257 | LMAN1_HUMAI | Unmodified | 0 | 0 | 157410 | 0 | 0 | 0 | 82963 | 68380 | 6082300 | 223560 | 0 | 140740 | 0 | 0 | 114850 | 39415 | 0 | 0 |
| RDIQENDE | PSMD11; PSMD | O00231; J3Q | PSD11_HUMAN | Unmodified | 0 | 0 | 0 | 0 | 208550 | 0 | 0 | 20957 | 0 | 0 | 222150 | 2001200 | 0 | 0 | 0 | 0 | 0 | 0 |
| RDLAKDITS | ANXA1; ANXA1 | P04083; Q5I | ANXA1_HUMAN | Unmodified | 0 | 0 | 0 | 0 | 0 | 0 | 0 | 0 | 624160 | 0 | 0 | 665170 | 0 | 0 | 0 | 0 | 0 | 0 |
| RDLEALQS | PROM2 | Q8N271 | PROM2_HUMA | Unmodified | 2147700 | 1480700 | 744670 | 1137500 | 0 | 0 | 21010 | 0 | 0 | 0 | 0 | 1077400 | 52142 | 0 | 1736000 | 1263900 | 80810 | 329000 |
| RDLEIERPII | NDRG3; NDRG | Q9UGV2; F8 | NDRG3_HUMA | Unmodified | 0 | 0 | 0 | 0 | 0 | 0 | 0 | 0 | 0 | 0 | 0 | 2095300 | 0 | 0 | 0 | 0 | 0 | 0 |
| RDLMACA( | DDX3X; DDX3Y | O00571; O1! | DDX3X_HUMAI | Unmodified | 328750 | 0 | 232700 | 0 | 0 | 0 | 46777 | 29210 | 36907 | 0 | 0 | 232430 | 0 | 85506 | 341290 | 215460 | 0 | 0 |
| RDLVLSGD | PLXNB2; PLXNI | O15031; E2F | PLXB2_HUMAN | Oxidation (M) | 0 | 0 | 0 | 0 | 0 | 0 | 0 | 0 | 0 | 0 | 0 | 264510 | 0 | 0 | 2743600 | 0 | 0 | 0 |
| RDMELLSN | EVA1B | Q9NVM1 | EVA1B_HUMAN | Oxidation (M) | 0 | 0 | 605150 | 0 | 0 | 0 | 0 | 0 | 0 | 0 | 0 | 0 | 0 | 0 | 394840 | 217630 | 0 | 0 |
| RDNMNDN | CHGB | P05060 | SCG1_HUMAN | Unmodified | 326050 | 0 | 3000800 | 0 | 0 | 0 | 0 | 0 | 0 | 0 | 0 | 0 | 0 | 0 | 706270 | 567680 | 0 | 0 |
| RDNTNEIY | ITGB1; ITGB1; I | P05556; H7( | ITB1_HUMAN I | Unmodified | 0 | 137440 | 2153100 | 0 | 2972200 | 452490 | 77848 | 0 | 1520100 | 133890 | 0 | 1895200 | 0 | 0 | 610450 | 1154600 | 126850 | 4379300 |
| RDPGVITY | VARS1; VARS1; | P26640; A0A | SYVC_HUMAN | Unmodified | 0 | 0 | 0 | 0 | 0 | 0 | 0 | 0 | 0 | 0 | 0 | 1084700 | 0 | 0 | 0 | 0 | 0 | 0 |
| RDPGVITY | VARS1; VARS1; | P26640; A0A | SYVC_HUMAN | Unmodified | 0 | 0 | 0 | 0 | 1124400 | 0 | 622680 | 0 | 0 | 0 | 0 | 1868100 | 640500 | 764940 | 0 | 0 | 0 | 179630 |
| RDPHLACV | CLTCL1; CLTC; | P53675; Q0( | CLH2_HUMAN | Unmodified | 4340100 | 1813600 | 1138400 | 943640 | 0 | 0 | 79318 | 0 | 336460 | 67649 | 0 | 14733000 | 0 | 37664 | 1212100 | 2041900 | 0 | 0 |
| RDPHLACV | CLTC; CLTC | Q00610; A0/ | CLH1_HUMAN | Unmodified | 77456000 | 0 | 0 | 1780100 | 1373300 | 0 | 0 | 0 | 0 | 0 | 0 | 1555800 | 3788200 | 0 | 0 | 0 | 0 | 1258000 |
| RDPSPVSG | GNPTG; GNPTG | Q9UJJ9; H0 | GNPTG_HUMA | Unmodified | 0 | 0 | 0 | 262410 | 0 | 0 | 0 | 0 | 1138600 | 0 | 0 | 0 | 0 | 0 | 369160 | 245890 | 0 | 0 |
| RDPSSVPN | COG3 | Q96JB2 | COG3_HUMAN | Unmodified | 0 | 0 | 0 | 0 | 0 | 0 | 0 | 0 | 0 | 0 | 0 | 273440 | 408880 | 100760 | 0 | 0 | 0 | 72782 |
| RDQALTEE | ACTN1; ACTN1 | P12814; A0A | ACTN1_HUMAI | Unmodified | 0 | 0 | 96902 | 0 | 0 | 23009 | 0 | 0 | 189590 | 0 | 35056 | 7573300 | 0 | 27743 | 518820 | 476670 | 0 | 0 |
| RDQALTEE | ACTN1; ACTN1 | P12814; A0A | ACTN1_HUMAI | Unmodified | 0 | 0 | 0 | 0 | 0 | 0 | 0 | 0 | 0 | 0 | 168720 | 1026800 | 0 | 0 | 0 | 0 | 0 | 0 |
| RDQAVMIS | MYO1C; MYO1( | O00159; I3L! | MYO1C_HUMA | Oxidation (M) | 0 | 0 | 104370 | 0 | 0 | 0 | 0 | 0 | 0 | 0 | 0 | 0 | 0 | 13828 | 0 | 0 | 0 | 0 |
| RDQAVMIS | MYO1C; MYO1( | O00159; I3L! | MYO1C_HUMA | Unmodified | 0 | 0 | 0 | 0 | 0 | 0 | 0 | 0 | 0 | 0 | 0 | 0 | 0 | 0 | 0 | 922000 | 0 | 0 |
| RDQDNMQ | KIF5B | P33176 | KINH_HUMAN | Unmodified | 0 | 0 | 0 | 0 | 0 | 0 | 0 | 0 | 0 | 0 | 0 | 260770 | 0 | 0 | 0 | 0 | 0 | 0 |
| RDREEILLC | TOR2A | Q5JU69 | TOR2A_HUMAN | Unmodified | 0 | 0 | 0 | 0 | 0 | 0 | 0 | 0 | 820510 | 0 | 0 | 0 | 0 | 0 | 0 | 0 | 0 | 0 |
| RDSHLTLLI | MRPL17; MRPL | Q9NRX2; E9I | RM17_HUMAN | Unmodified | 0 | 0 | 0 | 0 | 2031900 | 0 | 6145000 | 0 | 578190 | 0 | 178560 | 0 | 0 | 0 | 0 | 0 | 0 | 876060 |
| RDSSHAFT | MELTF | P08582 | TRFM_HUMAN | Unmodified | 0 | 0 | 0 | 0 | 656570 | 0 | 0 | 42760 | 713100 | 217000 | 0 | 0 | 0 | 0 | 0 | 0 | 151100 | 0 |
| RDTFNHLT | RAB2A; RAB2A; | P61019; H7( | RAB2A_HUMAN | Unmodified | 194180 | 24970 | 97303 | 0 | 0 | 0 | 0 | 0 | 457000 | 163980 | 0 | 678420 | 0 | 0 | 144850 | 134780 | 0 | 0 |
| RDTGAALL | QSOX1; QSOX1 | O00391; H0 | QSOX1_HUMAI | Unmodified | 0 | 0 | 0 | 0 | 0 | 0 | 0 | 0 | 423390 | 81818 | 0 | 0 | 0 | 0 | 0 | 0 | 0 | 0 |
| RDTINLLD( | GOLM1 | Q8NBJ4 | GOLM1_HUMA | Unmodified | 0 | 0 | 0 | 0 | 0 | 0 | 0 | 0 | 0 | 0 | 0 | 0 | 0 | 0 | 0 | 0 | 0 | 0 |
| RDVAGDAS | ATP1A3; N/A; A | P13637; A0A | AT1A3_HUMAN | Unmodified | 279940 | 0 | 0 | 0 | 135160 | 0 | 1402300 | 0 | 526310 | 485710 | 0 | 81634 | 0 | 0 | 0 | 0 | 0 | 0 |
| RDVQEIFRI | DDX39A; DDX3 | O00148; K7E | DX39A_HUMAN | Unmodified | 0 | 0 | 0 | 0 | 0 | 0 | 353790 | 0 | 0 | 0 | 0 | 0 | 0 | 0 | 0 | 0 | 0 | 0 |
| READGSET | SND1 | Q7KZF4 | SND1_HUMAN | Unmodified | 0 | 0 | 0 | 0 | 0 | 0 | 0 | 78604 | 0 | 0 | 0 | 985970 | 0 | 0 | 188720 | 0 | 0 | 0 |
| REAPVDVL | TMEM109 | Q9BVC6 | TM109_HUMAN | Unmodified | 0 | 0 | 0 | 0 | 0 | 0 | 0 | 0 | 4318300 | 0 | 0 | 0 | 0 | 0 | 0 | 0 | 0 | 0 |
| REDAGTYW | CEACAM1 | P13688 | CEAM1_HUMAI | Unmodified | 0 | 0 | 0 | 0 | 0 | 0 | 0 | 0 | 0 | 0 | 312800 | 1322700 | 0 | 0 | 0 | 0 | 0 | 0 |
| REDLVVAP | IMPDH2; IMPDI | P12268; A0A | IMDH2_HUMAN | Unmodified | 0 | 0 | 922970 | 0 | 0 | 0 | 0 | 0 | 483440 | 0 | 858810 | 1405800 | 108880 | 0 | 0 | 451380 | 51553 | 230000 |
| REEEEFNT( | SNRPD2; SNRP | P62316; K7E | SMD2_HUMAN | Unmodified | 0 | 0 | 0 | 0 | 0 | 0 | 0 | 0 | 0 | 0 | 1008600 | 1076600 | 0 | 0 | 0 | 0 | 0 | 0 |
| REEGAPGD | MAN1A1 | P33908 | MA1A1_HUMAI | Unmodified | 0 | 0 | 0 | 0 | 0 | 0 | 0 | 0 | 438650 | 0 | 354330 | 0 | 0 | 0 | 0 | 0 | 0 | 0 |
| REELITNWI | SPTAN1; SPTAN | Q13813; A0/ | SPTN1_HUMAN | Unmodified | 0 | 0 | 0 | 0 | 0 | 0 | 0 | 0 | 0 | 0 | 0 | 2049800 | 0 | 0 | 0 | 0 | 0 | 0 |
| REESEAVE/ | PTPN23 | Q9H3S7 | PTN23_HUMAN | Unmodified | 0 | 1109000 | 0 | 0 | 271730 | 0 | 0 | 0 | 1156100 | 0 | 0 | 748270 | 0 | 0 | 0 | 208560 | 58492 | 0 |
| REGPDAHI | ITGA7; ITGA7 | Q13683; J3K | ITA7_HUMAN I | Unmodified | 51812 | 0 | 0 | 0 | 0 | 0 | 486240 | 108980 | 2213900 | 937020 | 0 | 0 | 0 | 0 | 0 | 0 | 0 | 0 |
| REGSFPQG | MAN2A1 | Q16706 | MA2A1_HUMAI | Unmodified | 0 | 0 | 0 | 0 | 41719 | 647200 | 0 | 0 | 1029400 | 113130 | 121670 | 0 | 0 | 0 | 1057700 | 0 | 0 | 0 |
| REGTGTEM | FKBP4 | Q02790 | FKBP4_HUMAN | Oxidation (M) | 0 | 0 | 0 | 0 | 0 | 0 | 0 | 0 | 0 | 0 | 0 | 316110 | 0 | 0 | 0 | 0 | 0 | 0 |
| REHALTSG | ELOC; ELOC; EI | Q15369; R4( | ELOC_HUMAN | Unmodified | 0 | 0 | 0 | 0 | 1856900 | 0 | 0 | 31676 | 0 | 5734.7 | 989430 | 9624400 | 65230 | 393210 | 473760 | 0 | 226410 | 0 |
| REHINLGC | VDAC1; VDAC1 | P21796; C9J | VDAC1_HUMAI | Unmodified | 0 | 0 | 0 | 0 | 0 | 0 | 0 | 0 | 1070000 | 0 | 0 | 0 | 0 | 0 | 0 | 0 | 0 | 0 |
| REHSLIKPY | LMAN2; LMAN2 | Q12907; D6I | LMAN2_HUMAI | 2 Oxidation (M) | 0 | 0 | 0 | 0 | 0 | 0 | 0 | 0 | 25554000 | 442260 | 0 | 0 | 0 | 0 | 0 | 0 | 0 | 0 |
| REHSLIKPY | LMAN2; LMAN2 | Q12907; D6I | LMAN2_HUMAI | Oxidation (M) | 0 | 0 | 0 | 0 | 0 | 0 | 0 | 0 | 36297000 | 0 | 0 | 0 | 0 | 0 | 0 | 0 | 0 | 0 |
| REHSLIKPY | LMAN2; LMAN2 | Q12907; D6I | LMAN2_HUMAI | Unmodified | 0 | 0 | 0 | 0 | 0 | 0 | 0 | 0 | 18058000 | 0 | 0 | 0 | 0 | 0 | 0 | 0 | 0 | 0 |
| REINNSHI( | PLCB2; PLCB2 | Q00722; H0 | PLCB2_HUMAN | Unmodified | 0 | 0 | 0 | 0 | 0 | 0 | 0 | 0 | 0 | 0 | 0 | 1568400 | 0 | 0 | 0 | 0 | 0 | 0 |
| RELHGQNI | CPSF6; CPSF6 | Q16630; F8V | CPSF6_HUMAN | Unmodified | 1075100 | 395750 | 975120 | 814440 | 0 | 54950000 | 0 | 0 | 0 | 15441 | 123970 | 703760 | 1338300 | 0 | 331910 | 0 | 0 | 208890 |
| RELQELVQ | VCP; VCP; VCP; | P55072; C9J | TERA_HUMAN | Unmodified | 0 | 0 | 202510 | 0 | 668870 | 0 | 0 | 0 | 200720 | 0 | 537910 | 752830 | 0 | 0 | 72505 | 0 | 228690 | 0 |
| RELQQAVL | NUCB1 | Q02818 | NUCB1_HUMA | Unmodified | 0 | 0 | 0 | 0 | 339020 | 0 | 0 | 0 | 986310 | 138380 | 0 | 217450 | 0 | 0 | 0 | 198080 | 0 | 0 |
| RELVFKED( | EIF1AY; EIF1AX; | O14602; P47 | IF1AY_HUMAN | Unmodified | 435510 | 168680 | 536130 | 94452 | 0 | 0 | 0 | 0 | 0 | 0 | 0 | 1624400 | 276550 | 0 | 696640 | 677900 | 0 | 0 |
| REMDQTM/ | CMPK1; CMPK1 | P30085; Q5I | KCY_HUMAN U | Oxidation (M) | 0 | 0 | 38597 | 0 | 89065 | 0 | 0 | 0 | 79705 | 25832 | 0 | 49791 | 150430 | 0 | 72011 | 61389 | 0 | 0 |
| REMDQTM/ | CMPK1; CMPK1 | P30085; Q5I | KCY_HUMAN U | Unmodified | 0 | 0 | 0 | 0 | 0 | 0 | 0 | 0 | 0 | 0 | 123040 | 1619600 | 1429600 | 0 | 1085400 | 425170 | 425690 | 0 |
| RENLSDED | PLS1 | Q14651 | PLSI_HUMAN F | Unmodified | 0 | 0 | 0 | 0 | 710540 | 0 | 0 | 0 | 0 | 0 | 0 | 1236300 | 0 | 0 | 0 | 0 | 403170 | 91287 |
| REPAVGEP. | ICAM1; ICAM1 | P05362; K7E | ICAM1_HUMAN | Unmodified | 0 | 0 | 1125200 | 593960 | 0 | 0 | 0 | 168080 | 0 | 0 | 0 | 0 | 0 | 0 | 1110400 | 669490 | 0 | 0 |
| REPDSNVI | LPL; LPL; LPL | P06858; E5R | LIPL_HUMAN L | Unmodified | 0 | 0 | 0 | 0 | 0 | 0 | 0 | 0 | 1628000 | 0 | 0 | 0 | 0 | 0 | 0 | 0 | 0 | 653570 |
| REPIPSLEA | STXBP2; STXBP | Q15833; M0 | STXB2_HUMAN | Unmodified | 0 | 0 | 0 | 0 | 0 | 0 | 24430 | 0 | 199290 | 0 | 0 | 663820 | 0 | 0 | 0 | 0 | 0 | 0 |
| REQEVNILI | MYH9 | P35579 | MYH9_HUMAN | Unmodified | 0 | 0 | 14524 | 0 | 593150 | 0 | 15376 | 0 | 169710 | 29293 | 0 | 603630 | 0 | 103430 | 0 | 296550 | 186440 | 104950 |

| | | | | | | | | | | | | | | | | | | | | | | |
|---|---|---|---|---|---|---|---|---|---|---|---|---|---|---|---|---|---|---|---|---|---|---|
| REQQLAEIE | NOMO1; NOMC | Q15155; P69 | NOMO1_HUMA | Unmodified | 0 | 0 | 0 | 0 | 0 | 0 | 0 | 0 | 2242300 | 46423 | 0 | 124580 | 50751 | 0 | 0 | 0 | 63721 | 103950 |
| RESRDHTT | SRGAP2; SRGA | O75044; P0D | SRGP2_HUMAI | Unmodified | 0 | 0 | 0 | 0 | 478530 | 108460 | 0 | 0 | 0 | 0 | 0 | 0 | 616700 | 0 | 0 | 0 | 0 | 0 |
| RETEFKDD: | CD2AP | Q9Y5K6 | CD2AP_HUMAI | Unmodified | 0 | 0 | 0 | 0 | 0 | 0 | 0 | 0 | 0 | 0 | 0 | 0 | 0 | 0 | 495960 | 0 | 0 | 0 |
| RETNSDSV | MGRN1; MGRN | O60291; K7E | MGRN1_HUMA | Unmodified | 78296 | 0 | 30804 | 77485 | 45742 | 0 | 0 | 0 | 462800 | 0 | 0 | 0 | 0 | 0 | 95328 | 56508 | 0 | 0 |
| REVDDLGP | DHX15 | O43143 | DHX15_HUMAI | Unmodified | 35040 | 34047 | 0 | 0 | 0 | 84108 | 0 | 0 | 278020 | 0 | 176210 | 3865100 | 0 | 0 | 74790 | 0 | 0 | 0 |
| REYTMHDI | FMNL2; FMNL2 | Q96PY5; C9I | FMNL2_HUMAI | Unmodified | 0 | 0 | 0 | 0 | 0 | 0 | 0 | 0 | 0 | 0 | 0 | 428240 | 0 | 0 | 0 | 0 | 42413 | 95223 |
| RFASGGCI | SEC13; SEC13 | P55735; A8N | SEC13_HUMAN | Unmodified | 0 | 0 | 0 | 217900 | 1396300 | 296520 | 365950 | 276810 | 244560 | 0 | 220520 | 2157200 | 1434800 | 852730 | 0 | 49237 | 933300 | 1472700 |
| RFCPEADS | GLG1 | Q92896 | GSLG1_HUMAI | Unmodified | 1.96E+08 | 2.21E+08 | 0 | 0 | 75853 | 0 | 0 | 203330 | 5100300 | 2528300 | 685080 | 1415200 | 0 | 3539600 | 1766800 | 0 | 0 | 0 |
| RFDAMPFT | PA2G4 | Q9UQ80 | PA2G4_HUMAN | Oxidation (M) | 480210 | 516640 | 0 | 0 | 0 | 0 | 0 | 0 | 1120400 | 0 | 149130 | 2288200 | 0 | 0 | 0 | 258250 | 0 | 0 |
| RFDAMPFT | PA2G4 | Q9UQ80 | PA2G4_HUMAN | Unmodified | 0 | 0 | 0 | 0 | 0 | 0 | 0 | 0 | 0 | 0 | 0 | 557150 | 0 | 0 | 0 | 0 | 0 | 0 |
| RFDAQTGA | PSMC4 | P43686 | PRS6B_HUMAN | Unmodified | 0 | 0 | 375710 | 0 | 0 | 0 | 0 | 0 | 628270 | 0 | 197870 | 2247500 | 0 | 0 | 0 | 152820 | 0 | 19490 |
| RFDDAVVQ | HSPA8; HSPA8; | P11142; E9P | HSP7C_HUMAI | Unmodified | 26597000 | 23887000 | 21082000 | 30392000 | 1.19E+08 | 18686000 | 7290900 | 1749800 | 76208000 | 27977000 | 12302000 | 1.03E+08 | 1.19E+08 | 37560000 | 25666000 | 15952000 | 5954600 | 38250000 |
| RFDDAVVQ | HSPA8; HSPA8; | P11142; E9P | HSP7C_HUMAI | Oxidation (M) | 22360000 | 15155000 | 23055000 | 20062000 | 40718000 | 8945300 | 2512500 | 1268600 | 25620000 | 7054800 | 4952100 | 39454000 | 43612000 | 18707000 | 17061000 | 15431000 | 2715700 | 13470000 |
| RFDDPGLN | XRCC6; XRCC6 | P12956; B1A | XRCC6_HUMAI | 2 Oxidation (M) | 0 | 0 | 0 | 0 | 0 | 0 | 0 | 0 | 0 | 0 | 0 | 2301700 | 0 | 0 | 0 | 0 | 0 | 0 |
| RFDEAALSI | NAPG | Q99747 | SNAG_HUMAN | Unmodified | 0 | 0 | 0 | 0 | 3722900 | 0 | 747170 | 289690 | 1466000 | 522960 | 0 | 2685400 | 1314800 | 1972000 | 0 | 0 | 1757900 | 0 |
| RFDEATQLI | PFKL | P17858 | PFKAL_HUMAN | Unmodified | 0 | 0 | 0 | 0 | 696750 | 312570 | 315330 | 0 | 0 | 0 | 223360 | 0 | 564690 | 272640 | 0 | 0 | 0 | 0 |
| RFDEILEAS | PKM; PKM; PKM | P14618; B4D | KPYM_HUMAN | Oxidation (M) | 1630400 | 971260 | 814400 | 1169600 | 143480 | 0 | 0 | 0 | 1994900 | 0 | 5275900 | 52399000 | 0 | 0 | 2551600 | 4126400 | 0 | 0 |
| RFDEILEAS | PKM; PKM; PKM | P14618; B4D | KPYM_HUMAN | Unmodified | 0 | 0 | 0 | 0 | 0 | 0 | 0 | 0 | 3577600 | 386840 | 1233500 | 6207200 | 0 | 0 | 0 | 0 | 0 | 0 |
| RFDLGLED | MAN1A2 | O60476 | MA1A2_HUMAI | Unmodified | 0 | 0 | 0 | 0 | 0 | 0 | 0 | 0 | 11052000 | 681050 | 198830 | 592760 | 2233300 | 0 | 0 | 0 | 0 | 0 |
| RFDPIPAQ | NRP2 | O60462 | NRP2_HUMAN | Unmodified | 0 | 0 | 0 | 0 | 0 | 0 | 0 | 0 | 873620 | 0 | 0 | 0 | 0 | 0 | 0 | 0 | 0 | 0 |
| RFDSELSQ | MYO18A; MYO1 | Q92614; A0A | MY18A_HUMAI | Unmodified | 0 | 0 | 0 | 0 | 935280 | 95055 | 0 | 0 | 0 | 122490 | 0 | 491380 | 0 | 0 | 0 | 0 | 0 | 0 |
| RFDVSGYP | PDIA4; PDIA4 | P13667; A0A | PDIA4_HUMAN | Unmodified | 0 | 0 | 0 | 0 | 1565500 | 196450 | 6421200 | 4426300 | 39581000 | 25644000 | 7727700 | 2891100 | 2044200 | 1821700 | 455680 | 0 | 357190 | 1512900 |
| RFEEALQTI | MYO1D; MYO1D | O94832; J3Q | MYO1D_HUMA | Unmodified | 4457200 | 8425700 | 1603400 | 340070 | 0 | 0 | 0 | 0 | 0 | 0 | 221850 | 3255700 | 0 | 0 | 5154600 | 3760300 | 0 | 0 |
| RFEELTNLI | EIF3CL; EIF3C; | B5ME19; Q9 | EIFCL_HUMAN | Unmodified | 172550 | 280150 | 525550 | 331350 | 345340 | 0 | 0 | 0 | 224650 | 0 | 484370 | 3551500 | 0 | 0 | 458770 | 342100 | 0 | 0 |
| RFEETGQE | ARL3 | P36405 | ARL3_HUMAN , | Unmodified | 0 | 0 | 1179400 | 2929900 | 0 | 0 | 0 | 0 | 0 | 0 | 0 | 8740600 | 0 | 0 | 2569800 | 2171000 | 0 | 0 |
| RFELVHIDT | DAAM1 | Q9Y4D1 | DAAM1_HUMA | Unmodified | 791820 | 544160 | 0 | 0 | 0 | 0 | 291760 | 71719 | 604340 | 0 | 0 | 2464100 | 252000 | 77939 | 438100 | 0 | 0 | 0 |
| RFEQEMMS | SLK | Q9H2G2 | SLK_HUMAN S | Unmodified | 0 | 0 | 0 | 0 | 310500 | 0 | 0 | 215650 | 243370 | 0 | 0 | 334450 | 0 | 0 | 0 | 0 | 0 | 0 |
| RFFFDVGS | PURA | Q00577 | PURA_HUMAN | Unmodified | 253320 | 0 | 0 | 0 | 0 | 0 | 0 | 0 | 112490 | 0 | 0 | 606450 | 0 | 0 | 0 | 101050 | 0 | 0 |
| RFGFPEGS | RPS3; RPS3; RP | P23396; H0Y | RS3_HUMAN 4 | Unmodified | 2334800 | 1134700 | 2661600 | 1566000 | 1340200 | 188200 | 4225800 | 231280 | 0 | 967420 | 2721100 | 36048000 | 1010000 | 701590 | 2797000 | 4485400 | 382370 | 1547100 |
| RFGILSNCI | MKRN1 | Q9UHC7 | MKRN1_HUMA | Unmodified | 2818600 | 1188800 | 148630 | 265710 | 0 | 420030 | 0 | 0 | 0 | 0 | 0 | 0 | 383490 | 196840 | 204460 | 261030 | 0 | 0 |
| RFGMKPYT | PTGS1; PTGS1; | P23219; A0A | PGH1_HUMAN | Oxidation (M) | 0 | 0 | 0 | 0 | 0 | 0 | 0 | 0 | 0 | 0 | 1548700 | 642920 | 108630 | 0 | 0 | 0 | 0 | 0 |
| RFGMKPYT | PTGS1; PTGS1; | P23219; A0A | PGH1_HUMAN | Unmodified | 0 | 0 | 0 | 0 | 0 | 0 | 0 | 0 | 0 | 0 | 353210 | 196070 | 0 | 0 | 0 | 0 | 0 | 0 |
| RFGPYYTEI | PSMB3; PSMB3 | P49720; A0A | PSB3_HUMAN | Unmodified | 314470 | 0 | 908140 | 324430 | 0 | 0 | 25541 | 0 | 1890100 | 1837000 | 1446800 | 4795000 | 811630 | 0 | 392670 | 343440 | 645630 | 0 |
| RFGTVLTEI | DNPH1; DNPH: | O43598; H0Y | DNPH1_HUMA | Unmodified | 132170 | 77055 | 247890 | 57427 | 0 | 0 | 195080 | 0 | 0 | 0 | 360630 | 2320900 | 0 | 0 | 417540 | 312120 | 0 | 0 |
| RFGVPVIAI | IMPDH2; IMPDI | P12268; A0A | IMDH2_HUMAI | Unmodified | 214360 | 230260 | 286380 | 0 | 2931900 | 0 | 0 | 299450 | 3100600 | 949950 | 1585900 | 2305500 | 1166000 | 0 | 492870 | 393690 | 2518900 | 1813000 |
| RFHGRAFS | HSPA4; HSPA4; | P34932; A0A | HSP74_HUMAI | Unmodified | 0 | 0 | 0 | 0 | 1884900 | 0 | 0 | 0 | 0 | 222700 | 0 | 0 | 1420300 | 1764700 | 0 | 0 | 1045800 | 632760 |
| RFHVQLSN | ITGB4 | P16144 | ITB4_HUMAN I | Unmodified | 0 | 0 | 0 | 0 | 3322300 | 1016800 | 0 | 0 | 0 | 0 | 0 | 115730 | 0 | 159570 | 0 | 0 | 6815600 | 7812600 |
| RFHWEQD | ABCF2 | Q9UG63 | ABCF2_HUMAN | Unmodified | 118190 | 0 | 0 | 0 | 1020100 | 0 | 0 | 0 | 0 | 0 | 0 | 336990 | 0 | 210960 | 0 | 69113 | 0 | 0 |
| RFKDPGLV | SARS1; SARS1 | P49591; Q5T | SYSC_HUMAN | Unmodified | 0 | 298020 | 0 | 342790 | 1492100 | 165470 | 532890 | 110020 | 1447100 | 701270 | 0 | 866290 | 6296000 | 3211800 | 178100 | 277970 | 595930 | 1482000 |
| RFKEANNF | RPL7; RPL7 | P18124; A8M | RL7_HUMAN 6 | Unmodified | 0 | 0 | 0 | 0 | 0 | 0 | 275280 | 0 | 659610 | 0 | 0 | 212130 | 0 | 0 | 0 | 0 | 0 | 0 |
| RFLDFIGHI | PIP4K2A; PIP4K | P48426; H7E | PI42A_HUMAN | Unmodified | 0 | 0 | 0 | 0 | 471950 | 36866 | 947870 | 0 | 0 | 63017 | 26692 | 52975 | 328150 | 295500 | 0 | 0 | 622100 | 489850 |
| RFMQTFVL | G3BP1; G3BP1; | Q13283; A0A | G3BP1_HUMAI | Oxidation (M) | 0 | 0 | 0 | 0 | 0 | 0 | 54741 | 0 | 0 | 0 | 287200 | 1489500 | 0 | 0 | 0 | 0 | 0 | 0 |
| RFPDTVST | RAB20 | Q9NX57 | RAB20_HUMAN | Unmodified | 0 | 0 | 0 | 0 | 0 | 0 | 0 | 0 | 557920 | 372210 | 0 | 532480 | 123970 | 0 | 0 | 0 | 0 | 127520 |
| RFPGYDSE | RPL5; RPL5 | P46777; A0A | RL5_HUMAN 6 | Unmodified | 923930 | 598780 | 235310 | 722730 | 861000 | 0 | 75808 | 51903 | 0 | 147090 | 286640 | 3175600 | 632270 | 261150 | 618700 | 440180 | 100740 | 390000 |
| RFPVAPLIP | RBM25; RBM25 | P49756; H0Y | RBM25_HUMAI | Unmodified | 0 | 0 | 0 | 0 | 0 | 0 | 0 | 0 | 0 | 0 | 0 | 663710 | 0 | 0 | 0 | 0 | 0 | 0 |
| RFQLLEGPI | GDI1; GDI1 | P31150; G5E | GDIA_HUMAN | Oxidation (M) | 2076100 | 1221600 | 0 | 0 | 1990200 | 0 | 0 | 0 | 1253300 | 0 | 0 | 1995500 | 1525500 | 0 | 1159300 | 1355100 | 221130 | 46959 |
| RFQLLEGPI | GDI1; GDI1 | P31150; G5E | GDIA_HUMAN | Unmodified | 0 | 0 | 0 | 0 | 78328 | 0 | 0 | 0 | 2065700 | 278450 | 387740 | 1105800 | 225840 | 0 | 0 | 0 | 0 | 172910 |
| RFQPVHDL | RAB2A; RAB2B; | P61019; Q8V | RAB2A_HUMAN | Unmodified | 0 | 0 | 0 | 0 | 0 | 0 | 223200 | 0 | 298200 | 1394500 | 0 | 150540 | 0 | 0 | 0 | 0 | 0 | 0 |
| RFQSVPAQ | CLTC; CLTC | Q00610; A0A | CLH1_HUMAN | Unmodified | 0 | 0 | 185190 | 0 | 0 | 0 | 0 | 0 | 38528 | 0 | 0 | 206650 | 0 | 0 | 565940 | 127650 | 0 | 0 |
| RFSALLSD( | NIBAN1 | Q9BZQ8 | NIBA1_HUMAN | Unmodified | 0 | 0 | 0 | 0 | 0 | 0 | 0 | 0 | 1089000 | 259330 | 0 | 0 | 0 | 0 | 0 | 0 | 0 | 0 |
| RFSDATSK | TAX1BP1; TAX1 | Q86VP1; B8Z | TAXB1_HUMAN | Unmodified | 0 | 0 | 0 | 0 | 0 | 0 | 2203200 | 42966 | 0 | 0 | 0 | 0 | 0 | 0 | 0 | 0 | 437220 | 0 |
| RFSDQAGF | MARK2; MARK2 | E7ETY4; A0A | E7ETY4_HUMA | Unmodified | 0 | 0 | 0 | 143790 | 0 | 0 | 0 | 27633 | 0 | 0 | 0 | 0 | 682520 | 398480 | 0 | 62893 | 0 | 0 |
| RFSEGVLQ | TNKS1BP1 | Q9C0C2 | TB182_HUMAN | Unmodified | 0 | 0 | 0 | 228210 | 0 | 0 | 0 | 0 | 0 | 0 | 0 | 219270 | 429260 | 386910 | 116530 | 102820 | 0 | 0 |
| RFSFSGNT | GANAB; GANAE | Q14697; F5H | GANAB_HUMA | Unmodified | 0 | 0 | 0 | 0 | 0 | 0 | 0 | 0 | 1.59E+08 | 16963000 | 1410900 | 0 | 0 | 0 | 0 | 0 | 0 | 0 |
| RFSMVVQD | PRDX5 | P30044 | PRDX5_HUMAI | Oxidation (M) | 301140 | 269890 | 606530 | 1058500 | 3633900 | 0 | 1650000 | 1071300 | 2906300 | 2293000 | 737550 | 0 | 7162800 | 3403800 | 1211200 | 1263300 | 6718700 | 4498600 |
| RFSMVVQD | PRDX5 | P30044 | PRDX5_HUMAI | Unmodified | 167120 | 55025 | 149330 | 390290 | 18660000 | 536470 | 6711100 | 375630 | 23247000 | 7642200 | 666350 | 2069700 | 11644000 | 6679600 | 755390 | 227830 | 12979000 | 16364000 |
| RFSTEYELC | ANPEP | P15144 | AMPN_HUMAN | Unmodified | 0 | 0 | 0 | 0 | 0 | 0 | 0 | 0 | 0 | 0 | 0 | 0 | 0 | 0 | 0 | 0 | 1261000 | 0 |
| RFTAQGLP | UPF1 | Q92900 | RENT1_HUMAN | Unmodified | 3403400 | 3767700 | 3179200 | 3300300 | 1013500 | 0 | 2694400 | 907100 | 1041500 | 421980 | 0 | 3806100 | 4868500 | 2533200 | 2123900 | 2606800 | 1167900 | 825460 |
| RFTDYCDL | SMOC1 | Q9H4F8 | SMOC1_HUMA | Unmodified | 0 | 0 | 0 | 0 | 0 | 0 | 0 | 0 | 341600 | 315940 | 0 | 0 | 0 | 0 | 0 | 0 | 0 | 0 |
| RFVDAFKTI | C1GALT1 | Q9NS00 | C1GLT_HUMAN | Unmodified | 0 | 0 | 0 | 0 | 0 | 0 | 0 | 0 | 1609500 | 349840 | 0 | 0 | 0 | 91328 | 0 | 0 | 0 | 0 |
| RFVEVGRV | RPL14; RPL14 | P50914; E7E | RL14_HUMAN ( | Unmodified | 0 | 0 | 0 | 0 | 558660 | 0 | 304390 | 0 | 0 | 129220 | 0 | 0 | 0 | 138150 | 0 | 0 | 51057 | 149910 |
| RFYPESSYI | OCLN; OCLN; C | Q16625; A0A | OCLN_HUMAN | Unmodified | 0 | 0 | 606970 | 816580 | 1423500 | 0 | 0 | 0 | 542400 | 150280 | 0 | 0 | 1370200 | 608870 | 1032900 | 711830 | 0 | 305420 |
| RGADFDSV | AIDA | Q96BJ3 | AIDA_HUMAN / | Unmodified | 0 | 0 | 0 | 0 | 128640 | 0 | 0 | 0 | 0 | 0 | 0 | 0 | 629060 | 1235300 | 0 | 0 | 142770 | 0 |
| RGAEILPR | MYO6; MYO6; M | Q9UM54; A0 | MYO6_HUMAN | Unmodified | 0 | 0 | 753590 | 588820 | 0 | 0 | 0 | 0 | 113600 | 0 | 0 | 176190 | 129440 | 173870 | 1321400 | 986660 | 0 | 0 |
| RGAGMPG | LMAN1 | P49257 | LMAN1_HUMAI | Oxidation (M) | 0 | 0 | 463370 | 426550 | 0 | 0 | 208230 | 0 | 2.78E+08 | 9859600 | 2746500 | 4553100 | 741590 | 559120 | 1040000 | 131440 | 18203 | 722240 |
| RGAGMPG | LMAN1 | P49257 | LMAN1_HUMAI | Unmodified | 0 | 0 | 0 | 78888 | 0 | 0 | 50470 | 0 | 13132000 | 0 | 390630 | 0 | 0 | 0 | 105510 | 0 | 0 | 0 |
| RGAGMPG | LMAN1 | P49257 | LMAN1_HUMAI | Oxidation (M) | 0 | 0 | 0 | 0 | 0 | 0 | 0 | 0 | 2762800 | 0 | 0 | 0 | 0 | 0 | 0 | 0 | 0 | 0 |
| RGAISAEV | PRKAR1A; PRKA | P10644; K7E | KAP0_HUMAN | Unmodified | 0 | 0 | 0 | 0 | 365380 | 0 | 0 | 0 | 718990 | 523760 | 0 | 0 | 0 | 0 | 0 | 0 | 549290 | 459220 |
| RGALIVLEG | DTYMK; DTYMK | P23919; G5E | KTHY_HUMAN | Unmodified | 0 | 0 | 0 | 0 | 692360 | 0 | 164350 | 0 | 0 | 149620 | 0 | 0 | 586240 | 289100 | 0 | 0 | 301370 | 639860 |
| RGAPAAAT | H1-10 | Q92522 | H1X_HUMAN H | Unmodified | 0 | 0 | 0 | 560590 | 26837000 | 3780300 | 837810 | 0 | 2869000 | 0 | 973800 | 2589200 | 0 | 5331900 | 0 | 0 | 8121200 | 18896000 |
| RGDFFYHS | TMEM43 | Q9BTV4 | TMM43_HUMAI | Unmodified | 0 | 0 | 0 | 0 | 0 | 0 | 0 | 0 | 10360000 | 0 | 0 | 0 | 0 | 0 | 0 | 0 | 0 | 0 |
| RGDFIHVM | GRB2 | P62993 | GRB2_HUMAN | Unmodified | 223850 | 232250 | 208920 | 159990 | 0 | 0 | 0 | 0 | 898440 | 0 | 0 | 0 | 0 | 0 | 0 | 178390 | 0 | 0 |
| RGDFIHVM | GRB2 | P62993 | GRB2_HUMAN | Oxidation (M) | 2547700 | 2246300 | 843370 | 2203100 | 0 | 0 | 0 | 0 | 0 | 0 | 0 | 1822200 | 111070 | 0 | 4998300 | 1643500 | 0 | 0 |
| RGDIIGVQ( | KARS1; KARS1; | Q15046; H3I | SYK_HUMAN L | Unmodified | 0 | 0 | 0 | 19143 | 0 | 0 | 0 | 0 | 0 | 0 | 384700 | 4017100 | 0 | 0 | 98013 | 79580 | 0 | 0 |
| RGDINLLM | MCM5; MCM5 | P33992; B1A | MCM5_HUMAN | Oxidation (M) | 0 | 0 | 0 | 81479 | 0 | 0 | 288330 | 149260 | 0 | 0 | 257630 | 0 | 69290 | 0 | 0 | 0 | 0 | 0 |
| RGDLAIHL | FURIN; FURIN | P09958; A0A | FURIN_HUMAN | Unmodified | 0 | 0 | 0 | 0 | 0 | 0 | 0 | 0 | 0 | 0 | 412360 | 0 | 0 | 0 | 0 | 0 | 0 | 0 |
| RGDQLLSV | LIN7A; LIN7C; L | O14910; Q9I | LIN7A_HUMAN | Unmodified | 632170 | 715780 | 2942000 | 806780 | 0 | 564800 | 0 | 0 | 2522000 | 1404500 | 1067100 | 2861600 | 0 | 0 | 4778100 | 4029100 | 922280 | 0 |
| RGDQLLSV | LIN7A; LIN7C; L | O14910; Q9I | LIN7A_HUMAN | Unmodified | 0 | 0 | 0 | 0 | 220020 | 0 | 0 | 0 | 0 | 476580 | 0 | 0 | 0 | 0 | 0 | 0 | 872800 | 0 |
| RGDTVATL | GART | P22102 | PUR2_HUMAN | Unmodified | 51201 | 0 | 0 | 0 | 0 | 19126 | 0 | 0 | 0 | 22264 | 0 | 801470 | 0 | 0 | 0 | 182500 | 9619.7 | 0 |
| RGDTVATL | GART | P22102 | PUR2_HUMAN | Unmodified | 0 | 0 | 0 | 0 | 410990 | 0 | 203260 | 227660 | 246690 | 138510 | 254140 | 449740 | 477770 | 215590 | 0 | 0 | 540950 | 0 |
| RGDTYELQ | MAGT1; MAGT1 | Q9H0U3; A0 | MAGT1_HUMAI | Unmodified | 0 | 0 | 0 | 0 | 0 | 0 | 0 | 0 | 12493000 | 1194500 | 204230 | 400330 | 0 | 0 | 18660 | 0 | 0 | 0 |
| RGDVIDYS | KIDINS220 | Q9ULH0 | KDIS_HUMAN I | Unmodified | 0 | 0 | 0 | 752610 | 6358900 | 0 | 1119800 | 533700 | 780650 | 0 | 0 | 0 | 3002500 | 849690 | 0 | 0 | 0 | 0 |
| RGDVIYILS | SKAP2 | O75563 | SKAP2_HUMAN | Unmodified | 0 | 0 | 0 | 0 | 0 | 0 | 28561 | 0 | 0 | 0 | 155270 | 314720 | 0 | 60582 | 0 | 0 | 0 | 0 |
| RGDVLAAV | GPD2 | P43304 | GPDM_HUMAN | Unmodified | 0 | 44405 | 0 | 0 | 0 | 0 | 0 | 0 | 16024000 | 1234600 | 0 | 0 | 0 | 2127300 | 0 | 0 | 0 | 0 |
| RGEAHLAV | FKBP4 | Q02790 | FKBP4_HUMAN | Unmodified | 913270 | 594400 | 1309000 | 1280500 | 0 | 0 | 0 | 0 | 921670 | 97090 | 3519200 | 33934000 | 1722500 | 0 | 1957400 | 1789200 | 0 | 72543 |
| RGEAHLAV | FKBP4 | Q02790 | FKBP4_HUMAN | Unmodified | 509050 | 0 | 2032300 | 4716600 | 4541700 | 613650 | 2552300 | 0 | 1022200 | 1476500 | 5090900 | 3145400 | 34879000 | 15352000 | 930480 | 0 | 3425400 | 2147000 |
| RGEDMMH | DNAJA2; DNAJA | O60884; A0A | DNJA2_HUMAN | Unmodified | 0 | 0 | 0 | 0 | 1298100 | 51368 | 0 | 0 | 715460 | 160600 | 0 | 204220 | 0 | 0 | 0 | 0 | 0 | 0 |
| RGEENTDY | PTPRA | P18433 | PTPRA_HUMAN | Unmodified | 48652 | 39087 | 718860 | 0 | 0 | 0 | 0 | 24739 | 0 | 0 | 0 | 2021800 | 0 | 0 | 757000 | 546840 | 0 | 0 |
| RGEQADHI | AARSD1; PTGES | Q9BTE6; B3K | AASD1_HUMAN | Unmodified | 0 | 0 | 0 | 0 | 0 | 0 | 0 | 430050 | 0 | 329730 | 413550 | 3878400 | 0 | 0 | 0 | 0 | 0 | 0 |
| RGEVGIYQ | ITGB4 | P16144 | ITB4_HUMAN I | Unmodified | 704400 | 312840 | 348700 | 757610 | 1170500 | 0 | 0 | 0 | 0 | 163970 | 0 | 0 | 0 | 0 | 1147200 | 301720 | 1740500 | 2318200 |
| RGFAFVTFI | HNRNPA1L2; H | Q32P51; P09 | RA1L2_HUMAN | Unmodified | 0 | 0 | 96571 | 89917 | 1540200 | 193710 | 201220 | 63965 | 3636000 | 1765500 | 2918500 | 15332000 | 1199200 | 340420 | 249170 | 0 | 190180 | 251250 |
| RGFAFVTFI | HNRNPA1L2; H | Q32P51; P09 | RA1L2_HUMAN | Unmodified | 0 | 0 | 0 | 0 | 4881200 | 0 | 1990400 | 0 | 0 | 5650200 | 590210 | 639220 | 3344900 | 532320 | 0 | 0 | 0 | 1072300 |
| RGFAFVTFI | HNRNPA3; HNI | P51991; A0A | ROA3_HUMAN | Unmodified | 0 | 0 | 0 | 0 | 338570 | 0 | 300760 | 75938 | 416980 | 227420 | 1113700 | 3216000 | 747990 | 136220 | 0 | 0 | 28643 | 427390 |
| RGFAFVTFI | HNRNPA3; HNI | P51991; A0A | ROA3_HUMAN | Unmodified | 0 | 0 | 0 | 0 | 0 | 0 | 2101600 | 49201 | 0 | 1247100 | 0 | 0 | 1476600 | 785680 | 0 | 0 | 776650 | 994040 |
| RGFEEQW | GUSB; GUSB; G | P08236; F8V | BGLR_HUMAN | Unmodified | 0 | 0 | 0 | 0 | 0 | 0 | 0 | 0 | 1198700 | 23379 | 0 | 0 | 0 | 0 | 0 | 0 | 0 | 0 |
| RGFEFTLM | SEPTIN7; SEPTI | Q16181; E7E | SEPT7_HUMAN | Oxidation (M) | 0 | 0 | 0 | 0 | 1595900 | 0 | 0 | 0 | 0 | 242310 | 0 | 40339 | 335080 | 0 | 0 | 0 | 0 | 0 |
| RGFEVVYM | HSP90AB1 | P08238 | HS90B_HUMAI | Oxidation (M) | 0 | 0 | 0 | 0 | 1061800 | 0 | 0 | 0 | 816290 | 0 | 0 | 6194300 | 0 | 0 | 0 | 1261100 | 0 | 0 |
| RGFEVVYM | HSP90AB1 | P08238 | HS90B_HUMAI | Unmodified | 0 | 0 | 0 | 0 | 1650300 | 0 | 0 | 0 | 0 | 0 | 0 | 1465300 | 2218900 | 0 | 0 | 0 | 0 | 0 |
| RGFFICDQ | EPRS1; EPRS1 | P07814; V9G | SYEP_HUMAN | Unmodified | 0 | 0 | 0 | 0 | 0 | 0 | 0 | 0 | 0 | 0 | 0 | 268900 | 410930 | 2253200 | 0 | 0 | 0 | 0 |
| RGFGFVTFI | HNRNPA2B1; H | P22626; A0A | ROA2_HUMAN | Unmodified | 0 | 0 | 0 | 0 | 1236500 | 0 | 0 | 104380 | 9755700 | 873280 | 2661400 | 13013000 | 2139100 | 0 | 0 | 0 | 0 | 0 |
| RGFGFVTFI | HNRNPA2B1; H | P22626; A0A | ROA2_HUMAN | Unmodified | 0 | 0 | 0 | 0 | 5625100 | 447790 | 1978900 | 0 | 0 | 5847900 | 206410 | 0 | 8467800 | 3280800 | 0 | 0 | 641770 | 1625000 |
| RGFGFVYF | HNRNPA0 | Q13151 | ROA0_HUMAN | Unmodified | 0 | 99407 | 0 | 0 | 201350 | 0 | 219800 | 0 | 0 | 224330 | 0 | 0 | 363690 | 0 | 0 | 0 | 0 | 263170 |
| RGFQETLY | ELAPOR2; ELAP | A8MWY0; H7 | ELAP2_HUMAN | Unmodified | 0 | 0 | 0 | 0 | 959610 | 0 | 0 | 0 | 5372500 | 0 | 138800 | 0 | 0 | 0 | 0 | 0 | 0 | 0 |
| RGFVLQDT | MARS1; MARS1 | P56192; H0Y | SYMC_HUMAN | Unmodified | 0 | 0 | 0 | 0 | 0 | 0 | 170930 | 0 | 702060 | 0 | 0 | 3616300 | 0 | 0 | 252120 | 211650 | 0 | 0 |
| RGGAGAG | GLG1; GLG1; G | Q92896; H3I | GSLG1_HUMAI | Unmodified | 0 | 0 | 0 | 0 | 0 | 0 | 92618 | 0 | 3858100 | 271920 | 0 | 0 | 0 | 0 | 0 | 0 | 0 | 0 |
| RGGDAGM | GOLM2; GOLM2 | Q6P4E1; A0A | GOLM2_HUMA | Oxidation (M) | 0 | 0 | 0 | 0 | 0 | 0 | 0 | 0 | 917110 | 0 | 0 | 425780 | 0 | 0 | 0 | 0 | 0 | 0 |
| RGGEERVC | VGF | O15240 | VGF_HUMAN N | Unmodified | 0 | 0 | 0 | 0 | 0 | 0 | 0 | 0 | 909710 | 0 | 0 | 0 | 0 | 0 | 0 | 0 | 0 | 0 |
| RGGGSVVI | DHRS4L2; DHR | Q6PKH6; Q9 | DR4L2_HUMAN | Unmodified | 0 | 0 | 0 | 0 | 0 | 0 | 3323800 | 0 | 19967000 | 10834000 | 0 | 0 | 0 | 0 | 0 | 0 | 240810 | 194860 |
| RGGNRFEF | HNRNPM; HNR | P52272; M0F | HNRPM_HUMA | Unmodified | 0 | 0 | 0 | 58751 | 876740 | 0 | 705370 | 341820 | 0 | 272710 | 0 | 294380 | 1048900 | 0 | 0 | 0 | 376630 | 597090 |

| | | | | | | | | | | | | | | | | | | | | | | |
|---|---|---|---|---|---|---|---|---|---|---|---|---|---|---|---|---|---|---|---|---|---|---|
| RGGQALRI | PGAM1 | P18669 | PGAM1_HUMA | Unmodified | 0 | 0 | 0 | 0 | 1177400 | 0 | 0 | 0 | 0 | 0 | 0 | 0 | 0 | 0 | 0 | 0 | 1431100 | 0 |
| RGGSGSHI | SERBP1 | Q8NC51 | PAIRB_HUMAN | Unmodified | 208360 | 0 | 0 | 266160 | 0 | 0 | 0 | 0 | 0 | 0 | 0 | 861160 | 0 | 0 | 743400 | 443760 | 0 | 0 |
| RGHFDYQS | UBE2Z | Q9H832 | UBE2Z_HUMAN | Unmodified | 0 | 0 | 0 | 0 | 0 | 0 | 0 | 0 | 0 | 0 | 27460 | 0 | 167270 | 0 | 0 | 0 | 177380 | 430380 |
| RGHVFEES | EEF2 | P13639 | EF2_HUMAN El | Oxidation (M) | 983060 | 2056000 | 2784100 | 1580500 | 5464500 | 2412100 | 1074200 | 448050 | 744490 | 1076400 | 5241700 | 23580000 | 2111200 | 1672500 | 923100 | 1816200 | 1564300 | 1999100 |
| RGHVFEES | EEF2 | P13639 | EF2_HUMAN El | Unmodified | 0 | 505320 | 412870 | 865410 | 13731000 | 2276500 | 1406900 | 0 | 4474600 | 1269900 | 2309700 | 13861000 | 2624500 | 1664100 | 195770 | 278690 | 3429400 | 4469600 |
| RGIAEAVGI | FOLH1 | Q04609 | FOLH1_HUMAN | Unmodified | 873490 | 1169100 | 518330 | 19907000 | 0 | 0 | 0 | 0 | 0 | 0 | 0 | 0 | 414420 | 0 | 6595300 | 8624300 | 0 | 0 |
| RGIEKPPFE | SF3B2; SF3B2 | Q13435; E9F | SF3B2_HUMAN | Unmodified | 0 | 455940 | 0 | 0 | 0 | 0 | 0 | 0 | 0 | 0 | 0 | 807820 | 462320 | 0 | 0 | 441950 | 0 | 0 |
| RGIENYYDI | TSPAN15; TSPA | O95858; H7( | TSN15_HUMAN | Unmodified | 0 | 0 | 0 | 0 | 0 | 0 | 0 | 0 | 0 | 0 | 0 | 1715900 | 0 | 0 | 2586600 | 1992800 | 0 | 0 |
| RGIGGIFFD | CPOX | P36551 | HEM6_HUMAN | Unmodified | 0 | 0 | 0 | 0 | 0 | 0 | 0 | 346140 | 0 | 0 | 0 | 0 | 0 | 0 | 0 | 0 | 0 | 0 |
| RGIHSAIDA | AK2; AK2; AK2; / | P54819; A0A | KAD2_HUMAN | Unmodified | 0 | 0 | 0 | 0 | 0 | 0 | 340280 | 0 | 0 | 332110 | 0 | 0 | 565450 | 386210 | 0 | 0 | 0 | 0 |
| RGILTLKYP | ACTC1; ACTA1; | P68032; P68 | ACTC_HUMAN | Oxidation (M) | 39529000 | 1.04E+08 | 0 | 1.62E+08 | 4.9E+09 | 9.01E+08 | 4.18E+08 | 862480 | 68578000 | 2.6E+08 | 43056000 | 25724000 | 1.44E+09 | 8.72E+08 | 6639900 | 7873700 | 7.43E+08 | 9.77E+08 |
| RGILTLKYP | ACTC1; ACTA1; | P68032; P68 | ACTC_HUMAN | Unmodified | 2556100 | 5916900 | 0 | 15009000 | 1.69E+09 | 1.8E+08 | 94693000 | 0 | 69784000 | 67782000 | 1567800 | 0 | 4E+08 | 2.26E+08 | 0 | 96141 | 1.73E+08 | 3.9E+08 |
| RGILTLKYP | ACTG1; ACTB; / | P63261; P60 | ACTG_HUMAN | Oxidation (M) | 0 | 0 | 0 | 217410 | 17506000 | 5305400 | 0 | 0 | 0 | 692960 | 0 | 0 | 0 | 772240 | 0 | 0 | 5396000 | 2527400 |
| RGILTLKYP | ACTG1; ACTB; / | P63261; P60 | ACTG_HUMAN | Unmodified | 0 | 0 | 0 | 0 | 13204000 | 0 | 0 | 0 | 0 | 58409 | 0 | 0 | 0 | 844230 | 850440 | 0 | 1258200 | 175240 |
| RGIPHSLS( | KIDINS220 | Q9ULH0 | KDIS_HUMAN I | Unmodified | 0 | 0 | 122870 | 334710 | 992720 | 0 | 0 | 0 | 173170 | 0 | 0 | 0 | 140510 | 0 | 0 | 112710 | 0 | 0 |
| RGKGDTFV | SHCBP1 | Q8NEM2 | SHCBP_HUMA | Unmodified | 0 | 0 | 0 | 0 | 588730 | 0 | 0 | 0 | 0 | 0 | 0 | 67037 | 0 | 0 | 0 | 0 | 44186 | 0 |
| RGKIEEAVI | ASPH; ASPH; AS | Q12797; E5F | ASPH_HUMAN | Unmodified | 0 | 0 | 0 | 0 | 364800 | 0 | 0 | 0 | 1218500 | 509220 | 138720 | 100410 | 0 | 0 | 0 | 0 | 45949 | 0 |
| RGLDVNKC | CORO1C; COR( | Q9ULV4; H0 | COR1C_HUMA | Unmodified | 0 | 0 | 0 | 157390 | 7757500 | 1482000 | 0 | 0 | 476600 | 513890 | 677170 | 2045600 | 1495300 | 551640 | 0 | 0 | 763510 | 1147600 |
| RGLDVSSC | CORO2A | Q92828 | COR2A_HUMAN | Unmodified | 43790 | 0 | 47551 | 67779 | 0 | 0 | 0 | 0 | 0 | 0 | 0 | 153170 | 0 | 0 | 161780 | 143650 | 0 | 0 |
| RGLFDEEM | PLEC; PLEC | Q15149; H0Y | PLEC_HUMAN | Oxidation (M) | 0 | 0 | 0 | 0 | 0 | 0 | 0 | 0 | 0 | 0 | 0 | 0 | 0 | 0 | 0 | 0 | 571120 | 1016000 |
| RGLFDEEM | PLEC; PLEC | Q15149; H0Y | PLEC_HUMAN | Unmodified | 0 | 0 | 0 | 0 | 447420 | 53087 | 0 | 0 | 0 | 0 | 0 | 0 | 0 | 0 | 0 | 0 | 607370 | 656420 |
| RGLNVDQI | STAT1; STAT1; S | P42224; A0A | STAT1_HUMAN | Unmodified | 0 | 0 | 0 | 0 | 0 | 0 | 0 | 0 | 0 | 0 | 0 | 0 | 0 | 0 | 0 | 0 | 0 | 288990 |
| RGLPVGSP | UTRN; UTRN | P46939; Q5T | UTRN_HUMAN | Unmodified | 0 | 0 | 0 | 731090 | 0 | 0 | 0 | 0 | 731790 | 0 | 0 | 0 | 0 | 0 | 1012100 | 1104600 | 0 | 1801800 |
| RGLSGKYQ | TMEM87A | Q8NBN3 | TM87A_HUMAN | Unmodified | 0 | 0 | 0 | 0 | 1256000 | 155650 | 44478 | 0 | 104050 | 686000 | 0 | 0 | 740920 | 258390 | 0 | 0 | 127310 | 47390 |
| RGMFPDNI | CD2AP | Q9Y5K6 | CD2AP_HUMAN | Oxidation (M) | 593860 | 520640 | 595960 | 109320 | 0 | 0 | 0 | 0 | 0 | 0 | 0 | 0 | 0 | 0 | 844160 | 628680 | 0 | 0 |
| RGMFPDNI | CD2AP | Q9Y5K6 | CD2AP_HUMAN | Unmodified | 435870 | 696150 | 0 | 0 | 0 | 0 | 0 | 0 | 0 | 0 | 0 | 0 | 0 | 0 | 620300 | 243170 | 0 | 0 |
| RGNIFLTSY | PLXNB2 | O15031 | PLXB2_HUMAN | Unmodified | 368460 | 141900 | 589700 | 1143000 | 415390 | 249950 | 0 | 0 | 2663500 | 28646 | 0 | 252510 | 1085500 | 93416 | 806950 | 1247300 | 124430 | 229400 |
| RGNILIPGII | CNDP2 | Q96KP4 | CNDP2_HUMA | Unmodified | 0 | 0 | 892330 | 0 | 0 | 0 | 0 | 0 | 0 | 0 | 2341800 | 6526500 | 0 | 0 | 2218100 | 3226300 | 0 | 0 |
| RGPCIIYNE | RPL4; RPL4 | P36578; H3E | RL4_HUMAN 6( | Unmodified | 3528000 | 2350700 | 805240 | 991600 | 0 | 0 | 0 | 235250 | 0 | 0 | 1052400 | 28797000 | 0 | 0 | 347050 | 919860 | 0 | 0 |
| RGPDAVAA | ADRA2A | P08913 | ADA2A_HUMAN | Unmodified | 0 | 0 | 0 | 0 | 0 | 0 | 21026 | 0 | 0 | 218580 | 0 | 0 | 0 | 0 | 323180 | 956070 | 104610 | 0 |
| RGPDEDGI | TTYH3; TTYH3 | Q9C0H2; H7 | TTYH3_HUMAN | Unmodified | 0 | 0 | 0 | 0 | 0 | 0 | 0 | 0 | 3354400 | 558650 | 0 | 2984600 | 69671 | 0 | 0 | 665880 | 0 | 0 |
| RGPFELEAI | HSPH1; HSPH1 | Q92598; A0/ | HS105_HUMAN | Unmodified | 0 | 0 | 620260 | 2316800 | 0 | 0 | 0 | 109670 | 3431100 | 958210 | 1947300 | 4271500 | 2798100 | 521730 | 0 | 1390200 | 1676900 | 0 |
| RGPGATGI | ADRA2A | P08913 | ADA2A_HUMAN | Unmodified | 173710 | 72686 | 1290200 | 0 | 0 | 0 | 0 | 60754 | 1927400 | 444230 | 0 | 0 | 0 | 0 | 732660 | 3110300 | 0 | 0 |
| RGPGATGI | ADRA2A | P08913 | ADA2A_HUMAN | Unmodified | 0 | 0 | 0 | 0 | 0 | 0 | 89816 | 0 | 891930 | 161530 | 0 | 0 | 245570 | 273490 | 0 | 0 | 0 | 0 |
| RGPQLVCT | SNRNP40 | Q96DI7 | SNR40_HUMAN | Unmodified | 0 | 26127 | 0 | 0 | 0 | 0 | 0 | 0 | 0 | 0 | 0 | 776950 | 0 | 0 | 0 | 0 | 0 | 0 |
| RGQADESI | N/A | N/A | N/A | Unmodified | 0 | 0 | 0 | 0 | 0 | 0 | 0 | 0 | 0 | 0 | 0 | 0 | 0 | 0 | 0 | 1.39E+08 | 0 | 0 |
| RGQSPQSS | CXCL16; CXCL1 | Q9H2A7; A0/ | CXL16_HUMAN | Unmodified | 350530 | 0 | 0 | 0 | 0 | 0 | 0 | 0 | 0 | 0 | 0 | 0 | 0 | 0 | 838720 | 466730 | 0 | 0 |
| RGQTCVVI | FKBP1A; FKBP1 | P62942; F6T | FKB1A_HUMAN | Oxidation (M) | 2494000 | 2017000 | 1122700 | 701860 | 901500 | 1614400 | 180580 | 70343 | 977500 | 267350 | 495220 | 702340 | 1990400 | 1284900 | 1735300 | 938410 | 1039100 | 2373500 |
| RGQTCVVI | FKBP1A; FKBP1 | P62942; F6T | FKB1A_HUMAN | Unmodified | 1985000 | 4122400 | 2920300 | 2609800 | 8005900 | 1519100 | 1275500 | 96782 | 4518200 | 1253500 | 2246200 | 2012900 | 11764000 | 7251400 | 2818300 | 2590400 | 5138500 | 9574000 |
| RGQTCVVI | FKBP1A; FKBP1 | P62942; F6T | FKB1A_HUMAN | Unmodified | 1108200 | 581520 | 228150 | 284790 | 369500 | 0 | 90387 | 0 | 0 | 101340 | 0 | 903570 | 789680 | 325680 | 767580 | 649860 | 635520 | 862030 |
| RGQTCVVI | FKBP1A; FKBP1 | P62942; F6T | FKB1A_HUMAN | Oxidation (M) | 1060400 | 606750 | 246570 | 246800 | 201740 | 32923 | 0 | 0 | 59580 | 0 | 146090 | 321890 | 266030 | 83326 | 451060 | 472290 | 42334 | 0 |
| RGSIQVDG | HSPG2 | P98160 | PGBM_HUMAN | Unmodified | 0 | 0 | 1121600 | 958150 | 0 | 392180 | 0 | 0 | 0 | 0 | 0 | 0 | 1539600 | 962050 | 275490 | 657180 | 0 | 3636800 |
| RGSLLWN( | EMC10; EMC10 | Q5UCC4; M( | EMC10_HUMAN | Unmodified | 0 | 0 | 0 | 0 | 257380 | 0 | 501950 | 263750 | 58685000 | 11096000 | 1475100 | 0 | 0 | 0 | 0 | 0 | 0 | 158780 |
| RGSNTTSH | UQCRC2; UQCI | P22695; H3E | QCR2_HUMAN | Unmodified | 0 | 0 | 0 | 0 | 77487 | 13904 | 455540 | 226320 | 13004000 | 6294700 | 272670 | 0 | 165000 | 80780 | 0 | 0 | 47756 | 206660 |
| RGTDECAII | CTSC; CTSC; C | P53634; A0A | CATC_HUMAN | Unmodified | 0 | 0 | 0 | 0 | 0 | 0 | 0 | 0 | 0 | 0 | 4494900 | 0 | 2231400 | 0 | 0 | 0 | 0 | 0 |
| RGTGGVD1 | CKMT1A | P12532 | KCRU_HUMAN | Unmodified | 0 | 0 | 0 | 0 | 0 | 152110 | 0 | 55364 | 0 | 0 | 0 | 1534400 | 94222 | 0 | 0 | 0 | 0 | 70567 |
| RGTGGVD1 | CKMT1A | P12532 | KCRU_HUMAN | Unmodified | 0 | 0 | 1505400 | 0 | 0 | 0 | 137690 | 0 | 364040 | 0 | 0 | 3335400 | 0 | 0 | 0 | 0 | 174810 | 0 |
| RGTGGVD1 | CKB; CKB | P12277; H0Y | KCRB_HUMAN | Unmodified | 0 | 0 | 0 | 0 | 0 | 0 | 0 | 0 | 186260 | 0 | 0 | 974940 | 0 | 0 | 685170 | 489080 | 899040 | 0 |
| RGTHQLYS | RRP9 | O43818 | U3IP2_HUMAN | Unmodified | 0 | 0 | 0 | 0 | 0 | 0 | 0 | 0 | 0 | 0 | 996050 | 109000 | 0 | 0 | 0 | 0 | 87817 | 0 |
| RGTILDCN | ITGA1 | P56199 | ITA1_HUMAN I | Unmodified | 568310 | 0 | 0 | 0 | 271260 | 25051 | 0 | 0 | 38819 | 28242 | 0 | 0 | 0 | 0 | 215890 | 0 | 698570 | 4598200 |
| RGVDQTC1 | DOCK10; DOCI | Q96BY6; A0/ | DOC10_HUMA | Unmodified | 0 | 0 | 118970 | 128940 | 291000 | 0 | 0 | 0 | 706260 | 0 | 323360 | 2043900 | 18405 | 0 | 200530 | 0 | 219100 | 119390 |
| RGVEGAYS | GNPTAB | Q3T906 | GNPTA_HUMAN | Unmodified | 0 | 0 | 0 | 0 | 0 | 470190 | 0 | 0 | 1464800 | 0 | 0 | 0 | 1431700 | 0 | 0 | 0 | 145690 | 0 |
| RGVFNVQI | EIF2S1 | P05198 | IF2A_HUMAN E | Unmodified | 0 | 0 | 0 | 0 | 258160 | 0 | 0 | 0 | 90069 | 0 | 0 | 556700 | 0 | 0 | 0 | 0 | 0 | 188840 |
| RGVLLEDP | PHACTR4; PHA | Q8IZ21; A0A | PHAR4_HUMAN | Unmodified | 0 | 0 | 0 | 0 | 129930 | 0 | 0 | 0 | 1898900 | 288990 | 0 | 0 | 511650 | 0 | 0 | 0 | 401910 | 1822800 |
| RGVMLAVD | HSPD1; HSPD1 | P10809; A0A | CH60_HUMAN | Oxidation (M) | 0 | 0 | 0 | 0 | 0 | 0 | 0 | 0 | 1404900 | 674830 | 0 | 0 | 0 | 0 | 0 | 0 | 0 | 0 |
| RGVSCQF( | PPP5C; PPP5C; | P53041; A8N | PPP5_HUMAN | Unmodified | 92373 | 0 | 70325 | 107580 | 742910 | 169080 | 597530 | 390130 | 638180 | 452560 | 886300 | 3694500 | 656010 | 332200 | 129640 | 117460 | 372900 | 519650 |
| RGWDENV | NIPSNAP1 | Q9BPW8 | NIPS1_HUMAN | Unmodified | 0 | 0 | 0 | 0 | 0 | 0 | 0 | 35586 | 1370400 | 290390 | 0 | 1963900 | 0 | 0 | 0 | 0 | 0 | 0 |
| RGYFNEEL | DSP | P15924 | DESP_HUMAN | Unmodified | 0 | 0 | 691180 | 0 | 0 | 0 | 0 | 0 | 1009200 | 280060 | 0 | 0 | 0 | 0 | 0 | 0 | 0 | 0 |
| RHAEQERI | MYH10 | P35580 | MYH10_HUMAN | Unmodified | 0 | 0 | 574330 | 272830 | 0 | 0 | 0 | 0 | 144710 | 45248 | 0 | 2584200 | 551770 | 120000 | 437000 | 351450 | 0 | 0 |
| RHCTLRME | EDIL3 | O43854 | EDIL3_HUMAN | Unmodified | 0 | 0 | 0 | 0 | 1355300 | 0 | 0 | 0 | 0 | 0 | 0 | 0 | 0 | 0 | 0 | 0 | 0 | 0 |
| RHDAETNI | KIF1A; KIF1A; KI | Q12756; A0/ | KIF1A_HUMAN | Unmodified | 0 | 0 | 0 | 0 | 0 | 0 | 0 | 0 | 0 | 0 | 17977 | 528820 | 0 | 0 | 0 | 0 | 0 | 0 |
| RHDLYDIYI | ZC3H10 | Q96K80 | ZC3HA_HUMAN | Unmodified | 1447700 | 1193100 | 0 | 0 | 0 | 0 | 0 | 0 | 0 | 0 | 176600 | 0 | 0 | 0 | 243050 | 0 | 0 | 0 |
| RHEEGHMI | FN1 | P02751 | FINC_HUMAN I | Oxidation (M) | 0 | 0 | 622830 | 752870 | 0 | 0 | 0 | 0 | 0 | 71605 | 0 | 0 | 0 | 0 | 0 | 145510 | 0 | 0 |
| RHEEGHMI | FN1 | P02751 | FINC_HUMAN I | Unmodified | 0 | 0 | 1132700 | 438880 | 0 | 0 | 0 | 0 | 0 | 0 | 0 | 0 | 0 | 0 | 0 | 0 | 0 | 0 |
| RHEILQWV | LRRC59 | Q96AG4 | LRC59_HUMAN | Unmodified | 0 | 0 | 0 | 0 | 0 | 0 | 0 | 0 | 41105000 | 2318100 | 1948000 | 3922000 | 0 | 0 | 0 | 77079 | 322020 | 0 |
| RHEVEQQI | TAOK3; TAOK3 | Q9H2K8; G3 | TAOK3_HUMAN | Unmodified | 462970 | 727680 | 864160 | 2831400 | 7999600 | 110320 | 2633900 | 610970 | 0 | 0 | 16334 | 721570 | 4082200 | 3593600 | 1127900 | 1274100 | 585510 | 1764000 |
| RHGYTSSL | SEMA4C | Q9C0C4 | SEM4C_HUMAN | Unmodified | 0 | 0 | 0 | 0 | 0 | 0 | 0 | 0 | 4973900 | 143070 | 0 | 0 | 0 | 0 | 0 | 0 | 0 | 0 |
| RHIEIFTDLS | XRCC5; XRCC5 | P13010; C9J | XRCC5_HUMAN | Unmodified | 0 | 0 | 0 | 0 | 1927100 | 0 | 990960 | 0 | 0 | 851490 | 0 | 0 | 430870 | 0 | 0 | 0 | 384170 | 0 |
| RHLNLAEN | PACSIN1; PACS | Q9BY11; F6U | PACN1_HUMAN | Unmodified | 3382000 | 2325900 | 752510 | 370290 | 0 | 0 | 0 | 0 | 0 | 0 | 0 | 539860 | 0 | 0 | 1570200 | 880240 | 0 | 0 |
| RHLQVNVT | MYO1D; MYO1D | O94832; K7E | MYO1D_HUMA | Unmodified | 278750 | 0 | 0 | 0 | 1086600 | 0 | 0 | 0 | 0 | 613350 | 270120 | 569720 | 0 | 142780 | 253330 | 0 | 1619400 | 3101400 |
| RHLQVNVT | MYO1D; MYO1D | O94832; K7E | MYO1D_HUMA | Unmodified | 1291600 | 0 | 0 | 202070 | 0 | 376250 | 0 | 0 | 0 | 0 | 0 | 0 | 974230 | 1150000 | 0 | 106010 | 685230 | 1728900 |
| RHLTGEFEI | RAN; RAN; RAN | P62826; H0Y | RAN_HUMAN G | Unmodified | 0 | 0 | 0 | 0 | 0 | 0 | 0 | 0 | 0 | 185360 | 850750 | 1747100 | 0 | 1640900 | 0 | 0 | 0 | 0 |
| RHLTGEFEI | RAN; RAN; RAN | P62826; H0Y | RAN_HUMAN G | Unmodified | 0 | 0 | 0 | 0 | 2345900 | 0 | 8202.4 | 0 | 0 | 0 | 0 | 5000100 | 351450 | 0 | 123390 | 0 | 539500 | 1162300 |
| RHMEASLQ | IGFBP5 | P24593 | IBP5_HUMAN I | Unmodified | 0 | 0 | 0 | 0 | 0 | 0 | 0 | 0 | 753190 | 385230 | 0 | 0 | 0 | 0 | 0 | 0 | 0 | 0 |
| RHPGFEAT | ST14 | Q9Y5Y6 | ST14_HUMAN S | Unmodified | 0 | 826320 | 1316500 | 2953600 | 0 | 0 | 0 | 0 | 1025300 | 55137 | 0 | 0 | 0 | 0 | 3631000 | 2855800 | 0 | 0 |
| RHQEAQEA | APLP1; APLP1; / | P51693; B7Z | APLP1_HUMAN | Oxidation (M) | 0 | 0 | 0 | 0 | 0 | 0 | 0 | 0 | 981570 | 0 | 0 | 0 | 0 | 0 | 0 | 0 | 0 | 0 |
| RHQGTFTD | SMARCC1 | Q92922 | SMRC1_HUMA | Unmodified | 0 | 0 | 0 | 0 | 0 | 0 | 0 | 0 | 98288 | 197020 | 60289 | 65009 | 0 | 0 | 0 | 0 | 0 | 0 |
| RHVFGESD | TPI1; TPI1; TPI1 | P60174; U3K | TPIS_HUMAN T | Unmodified | 0 | 0 | 70854 | 404280 | 1814100 | 0 | 68032 | 0 | 462840 | 259420 | 1807900 | 3379700 | 2147700 | 490240 | 276800 | 0 | 0 | 0 |
| RHVPYSVS | CNPY4; CNPY4 | Q8N129; A0/ | CNPY4_HUMAN | Unmodified | 0 | 0 | 0 | 0 | 0 | 418390 | 0 | 0 | 908630 | 0 | 0 | 0 | 0 | 0 | 0 | 0 | 0 | 0 |
| RHWGGNV | RPL7A; RPL7A | P62424; Q5T | RL7A_HUMAN ( | Unmodified | 0 | 0 | 0 | 1861900 | 0 | 0 | 806470 | 1246300 | 112650 | 0 | 64950 | 2818500 | 118800 | 0 | 0 | 0 | 196540 | 0 |
| RHWPGAPI | SULF2 | Q8IWU5 | SULF2_HUMAN | Unmodified | 0 | 0 | 0 | 0 | 0 | 0 | 0 | 0 | 0 | 0 | 0 | 0 | 0 | 0 | 0 | 0 | 0 | 4758800 |
| RHYNGEAY | DNAJA1 | P31689 | DNJA1_HUMAN | Unmodified | 0 | 104610 | 134950 | 109340 | 23747 | 0 | 0 | 0 | 149310 | 0 | 0 | 81421 | 0 | 0 | 216320 | 141630 | 0 | 0 |
| RIALTDNAL | RPL7; RPL7 | P18124; A8N | RL7_HUMAN 6( | Unmodified | 1511400 | 1346300 | 714040 | 494270 | 0 | 0 | 390630 | 114800 | 296470 | 138820 | 441790 | 14060000 | 590630 | 0 | 812130 | 905220 | 0 | 0 |
| RIANLQTDI | FLNA; FLNA; FL | P21333; A0A | FLNA_HUMAN | Unmodified | 507430 | 1759700 | 1233100 | 3113000 | 1333400 | 206420 | 241710 | 43673 | 376530 | 57066 | 180390 | 1696800 | 526910 | 442030 | 1555900 | 1951300 | 96723 | 119380 |
| RIAPCYQD' | ITGA4; ITGA4 | P13612; E7E | ITA4_HUMAN I | Unmodified | 107740 | 0 | 0 | 427830 | 313260 | 0 | 535060 | 457170 | 0 | 0 | 0 | 0 | 507440 | 453080 | 0 | 0 | 0 | 0 |
| RIARELAED | ATP5ME | P56385 | ATP5I_HUMAN | Unmodified | 0 | 0 | 0 | 0 | 120590 | 0 | 574730 | 42361 | 161110 | 551980 | 0 | 0 | 206880 | 252740 | 0 | 0 | 0 | 0 |
| RIATGSFLK | PAICS; PAICS; F | P22234; D6F | PUR6_HUMAN | Unmodified | 24629 | 29776 | 59560 | 123200 | 1471600 | 85694 | 160390 | 58150 | 0 | 241340 | 0 | 3411500 | 898200 | 827290 | 54292 | 90828 | 1594900 | 563390 |
| RIAVVGEGI | WDR1; WDR1 | O75083; D6I | WDR1_HUMAN | Unmodified | 283440 | 1249800 | 1442100 | 2345800 | 20004000 | 2466800 | 499220 | 1334600 | 2490000 | 2252900 | 1630700 | 9456800 | 0 | 8294700 | 1354100 | 1655900 | 720560 | 5706600 |
| RIAYSLLGL | EPHA2 | P29317 | EPHA2_HUMAN | Unmodified | 0 | 0 | 0 | 0 | 1806200 | 626350 | 0 | 0 | 0 | 0 | 0 | 0 | 0 | 0 | 0 | 0 | 14763000 | 13911000 |
| RICNAVSPI | MTHFD2; MTHF | P13995; A0A | MTDC_HUMAN | Unmodified | 0 | 0 | 0 | 0 | 0 | 0 | 0 | 0 | 1715500 | 0 | 0 | 0 | 0 | 0 | 0 | 0 | 0 | 334290 |
| RICSYCNN | LMO7; LMO7; LI | Q8WWI1; J3I | LMO7_HUMAN | Unmodified | 0 | 0 | 0 | 0 | 734470 | 0 | 0 | 0 | 125620 | 162000 | 0 | 0 | 0 | 0 | 0 | 0 | 409670 | 407990 |
| RIDDGHRL | PACSIN1 | Q9BY11 | PACN1_HUMAN | Unmodified | 3398200 | 1416400 | 0 | 476810 | 0 | 0 | 0 | 0 | 0 | 0 | 0 | 0 | 708450 | 177530 | 0 | 348150 | 0 | 0 |
| RIDDGHRL | PACSIN1 | Q9BY11 | PACN1_HUMAN | Oxidation (M) | 888090 | 0 | 0 | 0 | 0 | 0 | 0 | 0 | 0 | 0 | 0 | 0 | 0 | 0 | 0 | 0 | 0 | 0 |
| RIDDGHRL | PACSIN2; PACS | Q9UNF0; B0 | PACN2_HUMAN | Unmodified | 7951300 | 2849200 | 19626 | 4884800 | 343800 | 0 | 0 | 0 | 0 | 66234 | 0 | 465870 | 7107100 | 2358800 | 4008900 | 1286000 | 116430 | 168310 |
| RIDDGHRL | PACSIN2; PACS | Q9UNF0; B0 | PACN2_HUMAN | Oxidation (M) | 2160700 | 1716700 | 1145500 | 4946900 | 0 | 0 | 0 | 0 | 0 | 0 | 0 | 0 | 3198500 | 220380 | 2767600 | 1013700 | 0 | 0 |
| RIDITLSSVI | LGALS3BP; LGA | Q08380; K7E | LG3BP_HUMAN | Unmodified | 0 | 0 | 0 | 0 | 0 | 0 | 0 | 0 | 834150 | 470740 | 1133200 | 732890 | 226150 | 290550 | 0 | 0 | 0 | 596610 |
| RIDTNLNK | SDHB; SDHB; S | P21912; A0A | SDHB_HUMAN | Unmodified | 762290 | 2676500 | 1821200 | 848960 | 3550100 | 2263300 | 1873600 | 0 | 1864800 | 197580 | 267340 | 3138300 | 1469000 | 9415800 | 1328300 | 613120 | 4129900 | 0 |
| RIEDNLPA( | TRAPPC3; TRAF | O43617; A0/ | TPPC3_HUMAN | Unmodified | 0 | 172870 | 0 | 1798400 | 0 | 0 | 0 | 0 | 599760 | 109560 | 0 | 1488700 | 318650 | 179230 | 411050 | 0 | 391200 | 0 |
| RIEESAIDE' | PRPSAP2; PRPS | O60256; I3L( | KPRB_HUMAN | Unmodified | 0 | 0 | 0 | 357010 | 0 | 0 | 0 | 0 | 0 | 0 | 430590 | 13896000 | 0 | 162590 | 740770 | 641490 | 419940 | 0 |
| RIEGSGDQ | DNM1; DNM1; I | Q05193; A0/ | DYN1_HUMAN | Unmodified | 0 | 0 | 0 | 0 | 105290 | 0 | 0 | 0 | 890870 | 206580 | 0 | 0 | 0 | 0 | 0 | 0 | 35554 | 0 |
| RIEGSGDQ | DNM2; DNM2 | P50570; K7E | DYN2_HUMAN | Unmodified | 980110 | 1454900 | 545570 | 430270 | 1157400 | 0 | 0 | 0 | 872990 | 98298 | 0 | 593780 | 378770 | 0 | 479590 | 457120 | 23596 | 0 |
| RIEPADAH' | NAA50; NAA50; | Q9GZZ1; A0/ | NAA50_HUMAN | Unmodified | 133410 | 58554 | 0 | 0 | 1826200 | 0 | 354820 | 303960 | 819240 | 657340 | 589380 | 3549900 | 1121900 | 480040 | 169230 | 161880 | 662290 | 0 |
| RIEVNVELF | KPNA2; KPNA2; | P52292; J3Q | IMA1_HUMAN I | Unmodified | 0 | 0 | 221290 | 309600 | 638210 | 100890 | 84080 | 0 | 193890 | 296840 | 0 | 0 | 848040 | 698350 | 0 | 0 | 501640 | 471320 |
| RIFDANTKF | UGDH | O60701 | UGDH_HUMAN | Unmodified | 0 | 0 | 0 | 0 | 9496100 | 0 | 0 | 0 | 0 | 0 | 0 | 0 | 0 | 0 | 0 | 0 | 0 | 0 |
| RIFHTVTTI | EIF3D; EIF3D; E | O15371; B0( | EIF3D_HUMAN | Unmodified | 258610 | 273090 | 466980 | 405770 | 2287200 | 437030 | 494620 | 0 | 730460 | 709380 | 805310 | 4672700 | 1201500 | 610390 | 435800 | 382030 | 367230 | 850800 |
| RIGDELDSI | BAX; BAX | Q07812; K4J | BAX_HUMAN A | Unmodified | 0 | 265580 | 0 | 71566 | 1840900 | 0 | 30153 | 0 | 739550 | 0 | 339890 | 1612000 | 0 | 0 | 0 | 0 | 0 | 131650 |
| RIGIFGQDI | ATP2A2; ATP2A | P16615; H7( | AT2A2_HUMAN | Unmodified | 164800 | 0 | 89023 | 566700 | 0 | 0 | 549420 | 215200 | 0 | 343120 | 240460 | 596340 | 0 | 246500 | 189930 | 275560 | 0 | 0 |
| RIGNLQTD | FLNB | O75369 | FLNB_HUMAN | Unmodified | 0 | 0 | 0 | 0 | 0 | 0 | 0 | 0 | 0 | 0 | 0 | 836760 | 0 | 0 | 464030 | 418470 | 0 | 0 |
| RIGSPEYIK | EDIL3 | O43854 | EDIL3_HUMAN | Unmodified | 0 | 0 | 0 | 0 | 51571000 | 20046000 | 4646100 | 1822700 | 5602500 | 2168000 | 0 | 0 | 0 | 0 | 0 | 0 | 2054400 | 12909000 |

| | | | | | | | | | | | | | | | | | | | | | |
|---|---|---|---|---|---|---|---|---|---|---|---|---|---|---|---|---|---|---|---|---|---|
| RIGTADREI | SH3GLB1; SH3 | Q9Y371; A0A | SHLB1_HUMAN | Unmodified | 0 | 0 | 0 | 0 | 1029600 | 0 | 0 | 0 | 1246900 | 28298 | 0 | 232470 | 0 | 0 | 0 | 996980 | 0 | 0 |
| RIGVFSYGS | HMGCS1 | Q01581 | HMCS1_HUMA | Unmodified | 0 | 0 | 0 | 0 | 1712600 | 0 | 1544700 | 0 | 881900 | 1444100 | 634680 | 993860 | 0 | 0 | 0 | 0 | 0 | 0 |
| RIHQESELI | STUB1; STUB1; | Q9UNE7; H3 | CHIP_HUMAN | Unmodified | 0 | 0 | 0 | 2270100 | 875590 | 0 | 0 | 35151 | 206990 | 0 | 2119100 | 8092700 | 0 | 0 | 0 | 0 | 408930 | 688100 |
| RIIITGDED\ | EPB41L1; EPB4 | Q9H4G0; H0 | E41L1_HUMAN | Unmodified | 0 | 0 | 0 | 0 | 2886600 | 234860 | 0 | 0 | 0 | 0 | 0 | 165140 | 1067200 | 684190 | 332570 | 0 | 1152900 | 2034800 |
| RIILEAEKMI | RHEB; RHEB | Q15382; C9J | RHEB_HUMAN | Unmodified | 0 | 188980 | 0 | 0 | 2629800 | 0 | 0 | 0 | 130390 | 270620 | 0 | 0 | 1137300 | 0 | 0 | 0 | 243490 | 517960 |
| RIITIPPFLA | FKBP9 | O95302 | FKBP9_HUMAN | Unmodified | 0 | 0 | 0 | 0 | 0 | 0 | 0 | 0 | 1031400 | 0 | 154420 | 64518 | 0 | 0 | 0 | 0 | 0 | 0 |
| RILALCMGI | RDX; MSN; RDX | P35241; P26 | RADI_HUMAN I | Oxidation (M) | 0 | 0 | 0 | 0 | 2940900 | 0 | 0 | 0 | 0 | 0 | 0 | 0 | 244890 | 0 | 0 | 0 | 1114700 | 0 |
| RILALCMGI | RDX; MSN; RDX | P35241; P26 | RADI_HUMAN I | Unmodified | 0 | 0 | 0 | 0 | 5168500 | 0 | 249210 | 0 | 295260 | 153560 | 0 | 0 | 0 | 683590 | 0 | 0 | 0 | 682650 |
| RILDAAGAI | GRHPR; GRHPI | Q9UBQ7; U3 | GRHPR_HUMA | Unmodified | 0 | 0 | 0 | 84942 | 0 | 0 | 0 | 0 | 0 | 0 | 188000 | 1435300 | 297010 | 159300 | 176530 | 283190 | 0 | 0 |
| RILDEIEEHI | SEPTIN2; SEPTI | Q15019; H7( | SEPT2_HUMAN | Unmodified | 122150 | 115140 | 270200 | 446490 | 4226600 | 1055000 | 0 | 0 | 701430 | 250820 | 971120 | 3121200 | 663700 | 778370 | 191040 | 291300 | 872680 | 1196400 |
| RILEDQEEI | PGLS; PGLS | O95336; M0I | 6PGL_HUMAN | Unmodified | 389340 | 250800 | 0 | 0 | 1452100 | 0 | 1006800 | 318370 | 8428400 | 3308500 | 0 | 3932700 | 1010700 | 0 | 0 | 245430 | 1368600 | 5579600 |
| RILELDQFK | XRCC6; XRCC6 | P12956; B1A | XRCC6_HUMAI | Unmodified | 0 | 0 | 0 | 0 | 865770 | 0 | 0 | 351410 | 0 | 519620 | 0 | 0 | 278640 | 0 | 0 | 0 | 0 | 313460 |
| RILELSGSS | PRKDC | P78527 | PRKDC_HUMAI | Unmodified | 0 | 0 | 0 | 0 | 0 | 0 | 0 | 0 | 0 | 0 | 0 | 466880 | 0 | 0 | 0 | 0 | 0 | 0 |
| RILGLAIES( | SNAP23; SNAP: | O00161; H3I | SNP23_HUMAN | Unmodified | 0 | 0 | 0 | 0 | 1745400 | 0 | 0 | 0 | 498700 | 0 | 686600 | 3391400 | 1806700 | 436390 | 0 | 0 | 225190 | 2635300 |
| RILGLLDAY | EEF1G | P26641 | EF1G_HUMAN | Unmodified | 0 | 0 | 0 | 0 | 0 | 0 | 0 | 0 | 0 | 0 | 0 | 7020900 | 0 | 0 | 0 | 0 | 74141 | 0 |
| RILIDTGEP, | LACTB2 | Q53H82 | LACB2_HUMAN | Unmodified | 0 | 0 | 127910 | 916080 | 2320000 | 1115100 | 0 | 191970 | 9633400 | 3936900 | 0 | 1262900 | 1366000 | 987490 | 867600 | 0 | 680500 | 787300 |
| RILISLATGI | ANXA6; ANXA6; | P08133; E7E | ANXA6_HUMAN | Unmodified | 0 | 0 | 140330 | 0 | 0 | 0 | 0 | 0 | 600500 | 135320 | 0 | 0 | 0 | 0 | 332310 | 0 | 0 | 0 |
| RILITGGAG | UXS1; UXS1; U) | Q8NBZ7; C9 | UXS1_HUMAN | Unmodified | 0 | 0 | 0 | 0 | 0 | 0 | 0 | 118640 | 3103100 | 161390 | 0 | 0 | 0 | 0 | 0 | 0 | 0 | 0 |
| RILLLEQFFI | PIPOX | Q9P0Z9 | SOX_HUMAN P | Unmodified | 0 | 0 | 0 | 0 | 0 | 0 | 0 | 0 | 487280 | 99944 | 0 | 0 | 0 | 0 | 0 | 0 | 0 | 0 |
| RILNVPQEI | ACACA | Q13085 | ACACA_HUMAI | Unmodified | 0 | 0 | 197630 | 282670 | 0 | 0 | 0 | 0 | 0 | 0 | 0 | 0 | 444360 | 358920 | 180290 | 0 | 0 | 189740 |
| RILQLCMG | EZR; EZR | P15311; E7E | EZRI_HUMAN E | Oxidation (M) | 0 | 0 | 0 | 0 | 11034000 | 918820 | 113370 | 0 | 2990000 | 4605900 | 0 | 1732100 | 982290 | 381890 | 0 | 0 | 357410 | 0 |
| RILQLCMG | EZR; EZR | P15311; E7E | EZRI_HUMAN E | Unmodified | 0 | 0 | 0 | 0 | 4420900 | 0 | 187510 | 0 | 1367300 | 0 | 0 | 0 | 296790 | 252890 | 0 | 0 | 511410 | 0 |
| RILTQPHFI | CASK; CASK; C/ | O14936; A0A | CSKP_HUMAN | Unmodified | 0 | 0 | 0 | 0 | 0 | 0 | 0 | 0 | 942370 | 0 | 0 | 0 | 0 | 0 | 354580 | 0 | 0 | 0 |
| RILTQPHFI | CASK; CASK; C/ | O14936; A0A | CSKP_HUMAN | Oxidation (M) | 0 | 0 | 637950 | 654810 | 212130 | 0 | 0 | 0 | 520280 | 0 | 0 | 0 | 563660 | 0 | 1714800 | 338730 | 0 | 0 |
| RIMENGQE | DNAJB2; DNAJE | P25686; C9J | DNJB2_HUMAN | Unmodified | 487260 | 775800 | 0 | 0 | 0 | 0 | 0 | 0 | 0 | 0 | 0 | 0 | 0 | 0 | 0 | 454030 | 0 | 0 |
| RIMLFTNEI | XRCC6; XRCC6 | P12956; B1A | XRCC6_HUMAI | Oxidation (M) | 71118 | 67553 | 0 | 0 | 0 | 0 | 71399 | 170690 | 0 | 155960 | 258980 | 1140200 | 333540 | 0 | 0 | 95077 | 0 | 0 |
| RIMLFTNEI | XRCC6; XRCC6 | P12956; B1A | XRCC6_HUMAI | Unmodified | 127710 | 101310 | 0 | 0 | 212930 | 0 | 0 | 225330 | 279030 | 0 | 751750 | 4348900 | 61360 | 0 | 0 | 0 | 434560 | 257580 |
| RINAEIEK | GNA11 | P29992 | GNA11_HUMAI | Unmodified | 149730 | 0 | 243070 | 0 | 3623500 | 0 | 0 | 169380 | 1060700 | 272850 | 261890 | 3831900 | 0 | 0 | 0 | 170570 | 508370 | 51408 |
| RINLAEDVI | NCLN; NCLN; N | Q969V3; K7E | NCLN_HUMAN | Unmodified | 0 | 0 | 0 | 0 | 0 | 0 | 0 | 0 | 1676200 | 141870 | 0 | 0 | 0 | 0 | 0 | 0 | 0 | 0 |
| RINLLSHD\ | MEST; MEST; MI | Q5EB52; C9. | MEST_HUMAN | Unmodified | 0 | 0 | 0 | 0 | 0 | 0 | 0 | 0 | 238140 | 0 | 0 | 0 | 0 | 0 | 0 | 0 | 0 | 0 |
| RINMVEELI | KARS1 | Q15046 | SYK_HUMAN Ly | Unmodified | 0 | 0 | 0 | 0 | 0 | 0 | 0 | 0 | 0 | 0 | 0 | 3437100 | 0 | 0 | 0 | 0 | 0 | 0 |
| RIPADTFAA | IQGAP1; IQGAP | P46940; A0A | IQGA1_HUMAN | Unmodified | 0 | 17735 | 0 | 30101 | 0 | 0 | 0 | 0 | 193630 | 0 | 0 | 864690 | 0 | 150650 | 291680 | 114330 | 0 | 0 |
| RIPEFSSFV | DARS2; DARS2; | Q6PI48; A0A | SYDM_HUMAN | Unmodified | 0 | 0 | 0 | 0 | 0 | 0 | 0 | 0 | 839780 | 57133 | 0 | 0 | 0 | 0 | 0 | 0 | 0 | 0 |
| RIPLAEWES | UQCRC1 | P31930 | QCR1_HUMAN | Unmodified | 0 | 0 | 0 | 0 | 0 | 0 | 0 | 0 | 6268200 | 499940 | 0 | 0 | 0 | 0 | 0 | 0 | 0 | 0 |
| RIPQSTLSE | ATP6V1B2 | P21281 | VATB2_HUMAN | Unmodified | 0 | 0 | 0 | 0 | 0 | 0 | 0 | 0 | 0 | 0 | 382660 | 3725000 | 0 | 0 | 0 | 0 | 0 | 0 |
| RIPSGAFEI | PXDN; PXDN; P | Q92626; H7( | PXDN_HUMAN | Unmodified | 0 | 0 | 0 | 0 | 0 | 0 | 0 | 0 | 932000 | 216420 | 0 | 0 | 0 | 0 | 0 | 0 | 0 | 0 |
| RIPSTDANI | RAB22A | Q9UL26 | RB22A_HUMAN | Unmodified | 459330 | 540870 | 303770 | 305710 | 0 | 0 | 0 | 0 | 502030 | 92919 | 158510 | 430640 | 57777 | 0 | 806790 | 441860 | 0 | 0 |
| RIPWTAASS | SLIRP; SLIRP; S | Q9GZT3; H0 | SLIRP_HUMAN | Unmodified | 0 | 0 | 0 | 0 | 0 | 0 | 105450 | 50136 | 1938800 | 710850 | 0 | 0 | 0 | 0 | 0 | 0 | 0 | 0 |
| RIPYAPSGE | UGDH | O60701 | UGDH_HUMAN | Unmodified | 234900 | 0 | 109890 | 100750 | 5581900 | 0 | 0 | 0 | 0 | 0 | 0 | 0 | 0 | 0 | 0 | 136310 | 0 | 0 |
| RIQAEVEA( | MYO6; MYO6; N | Q9UM54; A0 | MYO6_HUMAN | Unmodified | 23147 | 0 | 0 | 0 | 158650 | 0 | 0 | 0 | 444610 | 0 | 0 | 484910 | 0 | 0 | 4439500 | 820270 | 0 | 0 |
| RIQDLETEN | GULP1 | Q9UBP9 | GULP1_HUMAI | Unmodified | 0 | 0 | 0 | 347550 | 925130 | 0 | 0 | 0 | 0 | 0 | 0 | 0 | 958830 | 1221000 | 0 | 0 | 1164200 | 755350 |
| RIQDLETEN | GULP1 | Q9UBP9 | GULP1_HUMAI | Oxidation (M) | 129180 | 112340 | 1781700 | 531440 | 359740 | 0 | 0 | 0 | 0 | 0 | 78108 | 0 | 628390 | 0 | 0 | 222870 | 0 | 258930 |
| RIQEIIEQLI | HSPD1; HSPD1 | P10809; A0A | CH60_HUMAN | Unmodified | 0 | 0 | 0 | 0 | 0 | 0 | 0 | 0 | 28611000 | 16535000 | 0 | 0 | 6963800 | 6597100 | 0 | 0 | 606760 | 2248200 |
| RIQEIIEQLI | HSPD1; HSPD1 | P10809; A0A | CH60_HUMAN | Unmodified | 0 | 0 | 0 | 0 | 941160 | 0 | 553920 | 0 | 0 | 2404900 | 0 | 0 | 0 | 396490 | 0 | 0 | 554430 | 0 |
| RIQTYLQST | CMPK1; CMPK1 | P30085; A0A | KCY_HUMAN U | Oxidation (M) | 0 | 0 | 0 | 0 | 11660000 | 558230 | 1492100 | 0 | 0 | 887080 | 0 | 0 | 3470200 | 1523400 | 843590 | 0 | 1751200 | 2134900 |
| RIQTYLQST | CMPK1; CMPK1 | P30085; A0A | KCY_HUMAN U | Unmodified | 0 | 0 | 0 | 0 | 3564500 | 0 | 360760 | 0 | 2417300 | 985210 | 0 | 0 | 6736800 | 1781700 | 0 | 0 | 3707400 | 4742500 |
| RIQYQLVDI | SH3BGRL3; SH | Q9H299; Q5 | SH3L3_HUMAN | Oxidation (M) | 5812700 | 5352400 | 0 | 2547000 | 3884600 | 0 | 0 | 0 | 1788900 | 0 | 0 | 0 | 1409400 | 818070 | 3400100 | 2145800 | 3159100 | 8818700 |
| RIQYQLVDI | SH3BGRL3; SH | Q9H299; Q5 | SH3L3_HUMAN | Unmodified | 1183000 | 1040000 | 0 | 0 | 4518700 | 407550 | 281180 | 0 | 10862000 | 395280 | 343320 | 798710 | 627110 | 336520 | 180800 | 188600 | 3539100 | 16259000 |
| RISATSIFFE | SHMT2; SHMT2 | P34897; G3\ | GLYM_HUMAN | Oxidation (M) | 0 | 0 | 0 | 0 | 0 | 0 | 0 | 0 | 3883100 | 1067100 | 0 | 0 | 293230 | 2284300 | 0 | 0 | 0 | 0 |
| RISATSIFFE | SHMT2; SHMT2 | P34897; G3\ | GLYM_HUMAN | Unmodified | 0 | 0 | 0 | 0 | 0 | 0 | 0 | 0 | 15489000 | 868670 | 0 | 0 | 0 | 0 | 0 | 0 | 0 | 0 |
| RISATSIFFE | SHMT2; SHMT2 | P34897; G3\ | GLYM_HUMAN | Oxidation (M) | 0 | 0 | 0 | 0 | 0 | 0 | 0 | 0 | 0 | 828750 | 0 | 0 | 0 | 0 | 0 | 0 | 0 | 0 |
| RISATSIFFE | SHMT2; SHMT2 | P34897; G3\ | GLYM_HUMAN | Unmodified | 0 | 0 | 0 | 0 | 0 | 0 | 0 | 0 | 0 | 1203000 | 0 | 0 | 263250 | 0 | 0 | 0 | 0 | 0 |
| RISEIEDAAI | ERLIN1; ERLIN1 | O75477; B0( | ERLN1_HUMAN | Unmodified | 0 | 0 | 0 | 0 | 0 | 0 | 0 | 0 | 11546000 | 504510 | 1025900 | 195060 | 0 | 0 | 0 | 0 | 0 | 0 |
| RISEIEDAAI | ERLIN1; ERLIN1 | O75477; B0( | ERLN1_HUMAN | Unmodified | 43729 | 33938 | 0 | 49900 | 965280 | 132360 | 881530 | 461080 | 2565500 | 2440800 | 0 | 0 | 0 | 0 | 0 | 0 | 51354 | 120370 |
| RISEMEEEL | PPP1R12A | O14974 | MYPT1_HUMAN | Unmodified | 0 | 0 | 0 | 0 | 430910 | 0 | 0 | 0 | 98596 | 704250 | 3249600 | 166500 | 239400 | 130230 | 0 | 0 | 78365 | 182140 |
| RISEQFTAM | TUBB2A; TUBB4 | Q13885; P04 | TBB2A_HUMAN | Unmodified | 0 | 0 | 0 | 0 | 4470100 | 2527700 | 4829200 | 159880 | 1468100 | 2348400 | 659590 | 31932000 | 1650900 | 1031400 | 0 | 0 | 774280 | 0 |
| RISEQFTAM | TUBB2A; TUBB4 | Q13885; P04 | TBB2A_HUMAN | Oxidation (M) | 804240 | 275830 | 1537800 | 647710 | 1431100 | 463300 | 3614100 | 767960 | 126110 | 2234300 | 268910 | 7670600 | 1772300 | 1503300 | 877480 | 834590 | 304180 | 901310 |
| RISGLIYEET | H4C1 | P62805 | H4_HUMAN Hi | Unmodified | 1.54E+08 | 51820000 | 2745300 | 0 | 2887400 | 524110 | 857240 | 1414600 | 5064300 | 1617400 | 3322100 | 1.27E+09 | 2234500 | 1648000 | 352970 | 584850 | 556610 | 2851200 |
| RISGLIYEET | H4C1 | P62805 | H4_HUMAN Hi | Unmodified | 1.01E+08 | 50616000 | 1494400 | 771110 | 19734000 | 3428000 | 1.35E+08 | 14969000 | 10747000 | 51019000 | 3302900 | 2.03E+08 | 3298200 | 6818200 | 0 | 223630 | 2150300 | 6278100 |
| RISPVEESE | SDC4 | P31431 | SDC4_HUMAN | Unmodified | 1976200 | 2200600 | 8696400 | 4478000 | 718080 | 679460 | 0 | 0 | 2074100 | 0 | 0 | 0 | 0 | 0 | 3383600 | 5776600 | 103700 | 0 |
| RISQTYQQ( | ANXA4; ANXA4; | P09525; B4E | ANXA4_HUMAN | Unmodified | 586570 | 281840 | 297970 | 577280 | 291640 | 5070100 | 0 | 0 | 27419000 | 1721600 | 22865000 | 10571000 | 0 | 0 | 559020 | 357540 | 0 | 164280 |
| RISTLTIEEG | PPP1R8; PPP1R | Q12972; A0A | PP1R8_HUMAN | Unmodified | 0 | 0 | 0 | 0 | 458200 | 0 | 0 | 0 | 0 | 281740 | 0 | 267460 | 482920 | 0 | 0 | 0 | 132470 | 609870 |
| RISVFIQTDI | CD109 | Q6YHK3 | CD109_HUMAI | Unmodified | 213160 | 0 | 0 | 0 | 0 | 0 | 0 | 0 | 5589600 | 1519100 | 0 | 0 | 0 | 0 | 0 | 0 | 339920 | 0 |
| RITNDAREI | SNAP23; SNAP: | O00161; H3I | SNP23_HUMAN | Oxidation (M) | 0 | 1902400 | 2311200 | 3747300 | 25170000 | 2966200 | 0 | 0 | 430840 | 0 | 1277200 | 1884300 | 16356000 | 9888400 | 5257700 | 0 | 6075300 | 15415000 |
| RITNDAREI | SNAP23; SNAP: | O00161; H3I | SNP23_HUMAN | Unmodified | 383030 | 953760 | 132750 | 0 | 19805000 | 1995500 | 0 | 0 | 0 | 0 | 0 | 0 | 5705900 | 2347300 | 1436100 | 0 | 10568000 | 11246000 |
| RITSDFRIG | ITGB1; ITGB1; I | P05556; H7( | ITB1_HUMAN I | Unmodified | 0 | 0 | 0 | 0 | 1.2E+08 | 11456000 | 6320700 | 0 | 503760 | 4878700 | 129980 | 0 | 11021000 | 8514100 | 0 | 0 | 33312000 | 86624000 |
| RITSPLMEP | PSMA5 | P28066 | PSA5_HUMAN | Unmodified | 0 | 0 | 0 | 0 | 1048500 | 0 | 0 | 0 | 2454200 | 1377400 | 266060 | 1146200 | 854500 | 421910 | 0 | 0 | 2383300 | 3619600 |
| RITSPLMEP | PSMA5 | P28066 | PSA5_HUMAN | Oxidation (M) | 419540 | 662960 | 662050 | 712900 | 3419800 | 671960 | 882540 | 908110 | 1110200 | 2009900 | 1557200 | 6493700 | 4850600 | 2348200 | 994760 | 900060 | 2569100 | 2762500 |
| RIVAVTGAE | AARS1; AARS1; | P49588; A0A | SYAC_HUMAN | Unmodified | 88379 | 0 | 178730 | 0 | 3254900 | 516060 | 0 | 0 | 1449300 | 512360 | 646090 | 3937500 | 2391500 | 2505700 | 0 | 265720 | 0 | 917850 |
| RIVDDWAN | CANX; CANX; C | P27824; A0A | CALX_HUMAN | Unmodified | 0 | 0 | 0 | 0 | 0 | 569930 | 0 | 0 | 18432000 | 1140900 | 427770 | 241490 | 0 | 0 | 0 | 0 | 0 | 0 |
| RIVDDWAN | CANX; CANX; C | P27824; A0A | CALX_HUMAN | Unmodified | 0 | 0 | 0 | 0 | 0 | 0 | 0 | 0 | 3811600 | 522880 | 0 | 143790 | 0 | 0 | 0 | 0 | 0 | 0 |
| RIVDGKVVS | KRT18 | P05783 | K1C18_HUMAN | Unmodified | 0 | 0 | 0 | 0 | 0 | 0 | 0 | 0 | 0 | 18069 | 0 | 0 | 957200 | 536920 | 0 | 0 | 533850 | 62970 |
| RIVEGILK | ELOB; ELOB; EL | Q15370; A0A | ELOB_HUMAN | Unmodified | 0 | 240120 | 0 | 0 | 3017900 | 268530 | 0 | 0 | 927530 | 60068 | 0 | 0 | 0 | 0 | 0 | 0 | 656620 | 1204900 |
| RIVENGQE | DNAJB6; DNAJE | O75190; Q8I | DNJB6_HUMAN | Unmodified | 4046500 | 4152200 | 7067400 | 2068000 | 933050 | 0 | 78680 | 0 | 0 | 72661 | 0 | 800170 | 1556400 | 1128800 | 3237300 | 2135100 | 300750 | 0 |
| RIVITGDAD | EPB41L3; EPB4 | Q9Y2J2; A0A | E41L3_HUMAN | Unmodified | 0 | 0 | 0 | 0 | 0 | 0 | 0 | 0 | 0 | 0 | 0 | 0 | 0 | 364530 | 0 | 0 | 3759600 | 16396000 |
| RIVITGDAD | EPB41L3; EPB4 | Q9Y2J2; A0A | E41L3_HUMAN | Unmodified | 0 | 0 | 0 | 0 | 0 | 0 | 0 | 0 | 0 | 0 | 0 | 0 | 0 | 0 | 0 | 0 | 1633800 | 1915100 |
| RIVITGDGI | EPB41L2; EPB4 | O43491; Q6 | E41L2_HUMAN | Unmodified | 0 | 0 | 0 | 0 | 3844200 | 0 | 0 | 0 | 0 | 0 | 0 | 0 | 0 | 261860 | 0 | 322930 | 1236900 | 456260 |
| RIVITGDGI | EPB41L2; EPB4 | O43491; Q6 | E41L2_HUMAN | Unmodified | 0 | 0 | 0 | 300850 | 3262500 | 0 | 0 | 0 | 0 | 0 | 0 | 0 | 503040 | 0 | 0 | 0 | 527490 | 0 |
| RIVLVDDSI | PPAT | Q06203 | PUR1_HUMAN | Unmodified | 0 | 0 | 0 | 0 | 0 | 0 | 20166 | 10313 | 0 | 0 | 0 | 1011400 | 0 | 0 | 0 | 0 | 0 | 0 |
| RIVLVDDSI | PPAT | Q06203 | PUR1_HUMAN | Unmodified | 0 | 0 | 0 | 0 | 0 | 0 | 0 | 0 | 0 | 221480 | 0 | 0 | 662740 | 0 | 0 | 0 | 0 | 0 |
| RIVNHAVVI | KIF13A; KIF13A | Q9H1H9; A0 | KI13A_HUMAN | Unmodified | 0 | 0 | 0 | 111060 | 622780 | 0 | 792140 | 288580 | 334380 | 168690 | 158440 | 253580 | 1.59E+08 | 0 | 0 | 0 | 254610 | 663620 |
| RIVPGQFL/ | SF3B3 | Q15393 | SF3B3_HUMAN | Unmodified | 0 | 0 | 0 | 235310 | 0 | 0 | 0 | 0 | 0 | 0 | 0 | 705740 | 0 | 0 | 0 | 0 | 0 | 0 |
| RIVQNSNG | UGDH; UGDH; | O60701; E7E | UGDH_HUMAN | Unmodified | 0 | 0 | 0 | 0 | 3023800 | 0 | 0 | 0 | 0 | 0 | 0 | 0 | 212030 | 180150 | 0 | 0 | 133560 | 0 |
| RIVSQLLTL | VCP; VCP; VCP; | P55072; C9J | TERA_HUMAN | Unmodified | 146740 | 0 | 0 | 0 | 1472700 | 170900 | 0 | 0 | 1707100 | 969630 | 30277 | 625480 | 2483900 | 122380 | 28965 | 98721 | 175670 | 318030 |
| RIYIPLPEE/ | VPS4A; N/A | Q9UN37; I3L | VPS4A_HUMAN | Unmodified | 563520 | 0 | 637210 | 768130 | 1616000 | 0 | 0 | 0 | 1632900 | 778810 | 2378600 | 2642300 | 0 | 0 | 0 | 0 | 0 | 0 |
| RIYQFTAAS | SKAP2 | O75563 | SKAP2_HUMAN | Unmodified | 0 | 0 | 0 | 0 | 0 | 0 | 0 | 0 | 0 | 0 | 0 | 530540 | 0 | 0 | 0 | 0 | 0 | 0 |
| RKADLNTIT | UTRN | P46939 | UTRN_HUMAN | Unmodified | 0 | 0 | 153490 | 1088100 | 0 | 0 | 0 | 0 | 3117100 | 426420 | 0 | 0 | 251690 | 0 | 2372100 | 292250 | 0 | 0 |
| RKASGPPV | H1-3; H1-4; H1 | P16402; P10 | H13_HUMAN H | Unmodified | 1414400 | 1272300 | 0 | 0 | 1325100 | 161010 | 2124100 | 64408000 | 0 | 82497 | 0 | 0 | 78868 | 0 | 0 | 0 | 0 | 454750 |
| RKATGPPV | H1-5 | P16401 | H15_HUMAN H | Unmodified | 3278700 | 1597000 | 0 | 0 | 633720 | 0 | 357900 | 10967000 | 0 | 18010 | 0 | 0 | 0 | 0 | 0 | 0 | 0 | 0 |
| RKDPEGTP | HDDC3; HDDC | Q8N4P3; H0 | MESH1_HUMA | Unmodified | 0 | 0 | 0 | 0 | 446820 | 279320 | 0 | 0 | 0 | 194040 | 52877 | 695730 | 0 | 237560 | 0 | 0 | 204260 | 0 |
| RKDPELWC | CLTC; CLTC | Q00610; A0A | CLH1_HUMAN | Unmodified | 4517300 | 1550900 | 1128800 | 0 | 8337800 | 512990 | 0 | 0 | 216740 | 0 | 0 | 26910000 | 0 | 0 | 0 | 2427200 | 0 | 1209000 |
| RKDPLTHT | SPAST; SPAST; | Q9UBP0; A0 | SPAST_HUMAN | Unmodified | 122190 | 100850 | 330880 | 253280 | 1608400 | 0 | 64139 | 0 | 0 | 47482 | 0 | 0 | 215350 | 101820 | 0 | 103230 | 0 | 117130 |
| RKEDEVEE\ | EZR; EZR | P15311; E7E | EZRI_HUMAN E | Unmodified | 0 | 1013800 | 24884 | 1006400 | 2160200 | 590350 | 0 | 22879 | 7674000 | 1433400 | 3229400 | 15954000 | 911780 | 0 | 1836800 | 2362500 | 0 | 0 |
| RKEDPALP | CIT | O14578 | CTRO_HUMAN | Unmodified | 0 | 0 | 0 | 222230 | 8617100 | 293870 | 0 | 0 | 0 | 0 | 0 | 0 | 180150 | 0 | 44873 | 33357 | 622610 | 209010 |
| RKEEIEAQII | PSMD9; PSMD9 | O00233; J3K | PSMD9_HUMA | Unmodified | 0 | 0 | 0 | 0 | 0 | 0 | 0 | 0 | 0 | 0 | 297810 | 5747500 | 0 | 0 | 530380 | 415900 | 0 | 0 |
| RKELEQVC | HSPA1A; HSPA | P0DMV8; P0 | HS71A_HUMAN | Unmodified | 244710 | 0 | 0 | 0 | 3088900 | 0 | 0 | 0 | 962620 | 730610 | 0 | 0 | 3639100 | 1023600 | 0 | 0 | 864520 | 1156400 |
| RKEPSSLG | C1orf226; N/A | A1L170; F8W | CA226_HUMAN | Unmodified | 0 | 0 | 0 | 250300 | 648670 | 0 | 0 | 0 | 282840 | 153920 | 0 | 0 | 1273700 | 534570 | 0 | 0 | 0 | 0 |
| RKEQEHMI | ATP5PB; ATP5P | P24539; Q5( | AT5F1_HUMAN | Unmodified | 0 | 0 | 0 | 0 | 0 | 0 | 0 | 0 | 1378500 | 403950 | 0 | 0 | 0 | 0 | 0 | 0 | 0 | 0 |
| RKEVTDEIV | WARS1 | P23381 | SYWC_HUMAN | Unmodified | 91784 | 0 | 0 | 0 | 1027500 | 0 | 16614 | 0 | 87791 | 261800 | 0 | 177730 | 0 | 0 | 0 | 0 | 177230 | 484040 |
| RKGDEVD( | PARP1; PARP1; | P09874; A0A | PARP1_HUMAN | Unmodified | 1127700 | 579420 | 234440 | 0 | 401310 | 48140 | 0 | 26051 | 0 | 82054 | 30740 | 0 | 0 | 138190 | 0 | 0 | 0 | 179870 |
| RKGEGYSN | FKBP5 | Q13451 | FKBP5_HUMAN | Unmodified | 137590 | 0 | 0 | 0 | 905290 | 0 | 0 | 0 | 0 | 0 | 0 | 1286900 | 1791700 | 541600 | 0 | 0 | 0 | 0 |
| RKGFSEGL | HDGF | P51858 | HDGF_HUMAN | Unmodified | 0 | 0 | 0 | 0 | 4144900 | 278620 | 0 | 0 | 312000 | 1081100 | 465840 | 1365100 | 1621100 | 0 | 0 | 0 | 0 | 404640 |
| RKGTDVN\ | ANXA1 | P04083 | ANXA1_HUMAI | Unmodified | 0 | 0 | 0 | 0 | 345310 | 0 | 0 | 0 | 0 | 0 | 6463400 | 50986000 | 0 | 0 | 0 | 0 | 0 | 0 |
| RKGTEVQV | RUVBL2 | Q9Y230 | RUVB2_HUMAI | Unmodified | 0 | 0 | 132410 | 477440 | 476960 | 164180 | 0 | 56045 | 152740 | 0 | 405330 | 3360800 | 623300 | 0 | 179580 | 203460 | 87730 | 0 |
| RKGVEFPM | TOM1L2 | Q6ZVM7 | TM1L2_HUMAN | Oxidation (M) | 871690 | 1068300 | 0 | 1115700 | 0 | 89259 | 0 | 0 | 0 | 0 | 0 | 0 | 358660 | 0 | 553720 | 522150 | 0 | 0 |
| RKIPLAQSE | CNPY1; CNPY1 | A0A1B0GVE | A0A1B0GVE0_I | Unmodified | 0 | 0 | 0 | 0 | 0 | 0 | 0 | 0 | 0 | 0 | 1286400 | 344970 | 0 | 0 | 0 | 0 | 0 | 0 |
| RKIPVFHN( | SLC4A7; SLC4A | Q9Y6M7; C9 | S4A7_HUMAN | Unmodified | 0 | 0 | 199700 | 418180 | 909680 | 0 | 0 | 0 | 146130 | 0 | 299170 | 4118400 | 424360 | 0 | 218750 | 361340 | 202370 | 0 |
| RKIQVLQQ | TPM3; TPM3; TF | D6R904; A0A | D6R904_HUMA | Unmodified | 38449 | 0 | 0 | 0 | 74414 | 0 | 0 | 0 | 108610 | 0 | 0 | 249310 | 0 | 0 | 0 | 37609 | 0 | 0 |
| RKLEGDSTI | MYH9 | P35579 | MYH9_HUMAN | Unmodified | 0 | 0 | 0 | 0 | 956560 | 243570 | 0 | 0 | 325170 | 1066400 | 397530 | 0 | 250470 | 0 | 0 | 0 | 219110 | 0 |

| | | | | | | | | | | | | | | | | | | | | | | |
|---|---|---|---|---|---|---|---|---|---|---|---|---|---|---|---|---|---|---|---|---|---|---|
| RKLGINSL( | TPD52; TPD52; | P55327; H0\ | TPD52_HUMAN | Unmodified | 0 | 0 | 147060 | 136740 | 1252300 | 506320 | 217720 | 92077 | 2204200 | 1593400 | 0 | 3799500 | 738910 | 833040 | 182200 | 292990 | 501290 | 612890 |
| RKLLISDTQ | MPP7; MPP7; M | Q5T2T1; U5( | MPP7_HUMAN | Unmodified | 0 | 0 | 0 | 0 | 1123900 | 0 | 0 | 0 | 0 | 0 | 0 | 0 | 703390 | 397760 | 0 | 0 | 135740 | 0 |
| RKLVIPSEL | FKBP2; FKBP2 | P26885; A0A | FKBP2_HUMAN | Unmodified | 0 | 0 | 0 | 0 | 52319 | 0 | 0 | 0 | 23396000 | 1510000 | 1063700 | 0 | 0 | 0 | 0 | 0 | 0 | 0 |
| RKNPAAYE | UQCRQ | O14949 | QCR8_HUMAN | Unmodified | 0 | 66474 | 32848 | 0 | 0 | 0 | 0 | 0 | 2730100 | 962950 | 59715 | 0 | 0 | 0 | 10812 | 34206 | 0 | 0 |
| RKPADLHN | CLIC5; CLIC5; ( | Q9NZA1; A0, | CLIC5_HUMAN | Unmodified | 0 | 0 | 0 | 0 | 1018700 | 0 | 0 | 0 | 0 | 0 | 0 | 1723000 | 0 | 0 | 0 | 0 | 0 | 0 |
| RKPADLQN | CLIC4 | Q9Y696 | CLIC4_HUMAN | Unmodified | 1072200 | 379550 | 1299200 | 1696800 | 7733600 | 0 | 0 | 0 | 7100500 | 572430 | 733360 | 15184000 | 2050800 | 2548700 | 3027600 | 3331000 | 1466400 | 2589800 |
| RKPDIVTPN | GYS1 | P13807 | GYS1_HUMAN | Unmodified | 25660 | 33973 | 0 | 0 | 71143 | 0 | 44558 | 125450 | 0 | 0 | 0 | 1466900 | 97358 | 65808 | 0 | 72793 | 0 | 98449 |
| RKPDTIEV( | RDX; EZR; MSN | P35241; P15 | RADI_HUMAN I | Unmodified | 1407300 | 1370400 | 1671500 | 3166600 | 41761000 | 8413000 | 1191100 | 607460 | 13193000 | 6949300 | 2206300 | 20886000 | 7098900 | 7517100 | 2969200 | 2727800 | 6173200 | 7250400 |
| RKPDTIEV( | RDX; EZR; MSN | P35241; P15 | RADI_HUMAN I | Oxidation (M) | 0 | 1263900 | 1791300 | 3074800 | 8667200 | 5262700 | 807610 | 107140 | 2517000 | 132860 | 1916500 | 10747000 | 2359500 | 3801800 | 1341500 | 3422700 | 3617300 | 4110400 |
| RKPDYEPV | ARFGAP3; ARF( | Q9NP61; H0 | ARFG3_HUMAN | Unmodified | 0 | 0 | 0 | 135300 | 229990 | 0 | 0 | 0 | 0 | 0 | 0 | 0 | 194150 | 197340 | 156780 | 135780 | 0 | 85966 |
| RKPEDVLD | BSG; BSG; BSG | P35613; A0A | BASI_HUMAN E | Unmodified | 554300 | 267800 | 8997100 | 1395000 | 380770 | 0 | 246720 | 117460 | 1826100 | 526390 | 916580 | 14577000 | 0 | 0 | 8578400 | 8176600 | 156130 | 0 |
| RKQNDVF( | GDI1 | P31150 | GDIA_HUMAN | Unmodified | 343100 | 235920 | 0 | 356660 | 2228000 | 398690 | 3307900 | 2030100 | 3056900 | 2227300 | 128730 | 1511300 | 6876300 | 2452500 | 288070 | 355980 | 7475300 | 6499600 |
| RKQPQEAA | PCYOX1L; PCY( | Q8NBM8; E7 | PCYXL_HUMAN | Unmodified | 0 | 0 | 0 | 0 | 0 | 0 | 0 | 0 | 461120 | 0 | 132700 | 0 | 0 | 0 | 0 | 0 | 0 | 0 |
| RKQYDTYG | DNAJB11; DNA | Q9UBS4; A0, | DJB11_HUMAN | Unmodified | 0 | 128780 | 0 | 0 | 0 | 0 | 0 | 0 | 969080 | 0 | 274340 | 0 | 0 | 0 | 0 | 0 | 0 | 0 |
| RKTDFFIG( | NUDC; NUDC | Q9Y266; A0/ | NUDC_HUMAN | Unmodified | 0 | 0 | 0 | 30345 | 747750 | 0 | 0 | 0 | 2837500 | 1175700 | 310700 | 3493700 | 379340 | 123600 | 0 | 0 | 0 | 471960 |
| RKTGTLQP | RACGAP1; RAC | Q9H0H5; H( | RGAP1_HUMAI | Unmodified | 0 | 0 | 0 | 0 | 398290 | 143920 | 0 | 0 | 95029 | 0 | 0 | 0 | 118390 | 0 | 0 | 0 | 0 | 0 |
| RKTSDFNTI | CHORDC1; CH | Q9UHD1; E9 | CHRD1_HUMA | Unmodified | 0 | 270390 | 0 | 0 | 0 | 1769500 | 53502 | 0 | 0 | 0 | 0 | 0 | 175180 | 0 | 0 | 0 | 152220 | 76238 |
| RKTVTAMD | H4C1 | P62805 | H4_HUMAN Hi | Unmodified | 1372700 | 940980 | 0 | 0 | 5070600 | 1250200 | 16857000 | 6410900 | 0 | 1280200 | 0 | 2565000 | 0 | 653940 | 0 | 0 | 903760 | 3268500 |
| RKTVTAMD | H4C1 | P62805 | H4_HUMAN Hi | Oxidation (M) | 1386400 | 0 | 0 | 0 | 680790 | 0 | 5253700 | 8557300 | 0 | 275600 | 0 | 2048300 | 0 | 143910 | 0 | 0 | 0 | 0 |
| RKVGAENV | SQOR; SQOR; S | Q9Y6N5; H3 | SQOR_HUMAN | Unmodified | 0 | 0 | 0 | 0 | 0 | 0 | 0 | 0 | 2162000 | 0 | 0 | 0 | 0 | 0 | 0 | 0 | 0 | 0 |
| RKWEAVH/ | HLA-A; HLA-A; I | A0A0G2JL56 | A0A0G2JL56_H | Unmodified | 0 | 0 | 234230 | 0 | 0 | 0 | 26317 | 0 | 452370 | 2840000 | 0 | 5409800 | 1072200 | 387300 | 0 | 249240 | 1925500 | 17414000 |
| RLAACVNL | CUTA; CUTA; C | O60888; C9I | CUTA_HUMAN | Unmodified | 0 | 0 | 0 | 0 | 711410 | 0 | 0 | 0 | 206660 | 517010 | 41841 | 54591 | 925040 | 452760 | 114250 | 0 | 758230 | 1209100 |
| RLAAEATEV | SRPK1; SRPK1 | Q96SB4; H3 | SRPK1_HUMAN | Unmodified | 0 | 0 | 454080 | 0 | 227520 | 0 | 0 | 0 | 257750 | 140640 | 152090 | 490860 | 0 | 0 | 52419 | 0 | 251360 | 0 |
| RLADAAEN | CACNA2D2; CA | Q9NY47; C9 | CA2D2_HUMAI | Unmodified | 0 | 0 | 0 | 0 | 0 | 0 | 845060 | 431530 | 4153500 | 2269100 | 128600 | 618150 | 0 | 0 | 0 | 0 | 0 | 0 |
| RLADKYPV | PLOD2; PLOD2; | O00469; C9J | PLOD2_HUMAI | Unmodified | 0 | 0 | 0 | 0 | 40010 | 0 | 0 | 0 | 2242500 | 489240 | 0 | 0 | 0 | 0 | 0 | 0 | 0 | 0 |
| RLAENSAS | ERP29 | P30040 | ERP29_HUMAN | Unmodified | 0 | 0 | 0 | 0 | 0 | 0 | 0 | 0 | 7256800 | 1433800 | 0 | 0 | 0 | 0 | 0 | 0 | 0 | 0 |
| RLAEVKDSI | SH3GL1; SH3G | Q99961; M0I | SH3G1_HUMAI | Unmodified | 3747400 | 4526800 | 0 | 693110 | 0 | 0 | 0 | 0 | 0 | 185660 | 0 | 0 | 2325300 | 0 | 104440 | 587480 | 0 | 307310 |
| RLAGGDW | PPP2R1A; PPP2 | P30153; C9J | 2AAA_HUMAN | Unmodified | 109150 | 0 | 212670 | 377710 | 236780 | 0 | 0 | 0 | 189740 | 299340 | 423360 | 4589200 | 1807100 | 2775700 | 350450 | 338390 | 579690 | 513060 |
| RLAHPFSL | LDLR; LDLR; LD | P01130; J3K | LDLR_HUMAN | Unmodified | 131230 | 0 | 0 | 0 | 752720 | 289040 | 0 | 0 | 3507400 | 307050 | 191440 | 0 | 0 | 0 | 0 | 125470 | 0 | 280090 |
| RLAIKLPDD | PFKP; PFKP; PF | Q01813; A0/ | PFKAP_HUMAN | Unmodified | 0 | 0 | 0 | 0 | 3999600 | 900390 | 161920 | 45954 | 380820 | 311220 | 553450 | 2502800 | 335010 | 213640 | 0 | 130720 | 1259200 | 1460200 |
| RLAPEYEA/ | PDIA3; PDIA3 | P30101; H7E | PDIA3_HUMAN | Unmodified | 0 | 56451 | 579200 | 753170 | 77448 | 268670 | 391550 | 320710 | 2E+08 | 9641200 | 18349000 | 12466000 | 77593 | 128960 | 2066800 | 1794100 | 13425 | 290190 |
| RLAPEYEA/ | PDIA3; PDIA3 | P30101; H7E | PDIA3_HUMAN | Unmodified | 0 | 0 | 0 | 477920 | 12183000 | 1106900 | 6589200 | 5341000 | 20226000 | 55512000 | 10802000 | 293890 | 6763400 | 5550200 | 0 | 0 | 1111600 | 5607700 |
| RLAPITSDP | PKM; PKM | P14618; B4D | KPYM_HUMAN | Unmodified | 0 | 0 | 771510 | 1369800 | 7291500 | 0 | 878580 | 183970 | 4845100 | 713550 | 4214400 | 16454000 | 1161200 | 0 | 2463200 | 0 | 1995500 | 5157600 |
| RLAQDGAH | DHRS4; DHRS4 | Q9BTZ2; F5( | DHRS4_HUMAI | Unmodified | 0 | 0 | 0 | 0 | 0 | 0 | 0 | 0 | 1678300 | 482590 | 0 | 0 | 0 | 0 | 0 | 0 | 19241 | 0 |
| RLAQDGAH | DHRS4; DHRS4 | Q9BTZ2; F5( | DHRS4_HUMAI | Unmodified | 0 | 0 | 0 | 0 | 0 | 0 | 112030 | 0 | 6777700 | 1929100 | 0 | 0 | 0 | 0 | 0 | 0 | 0 | 0 |
| RLATEFELR | CNOT1 | A5YKK6 | CNOT1_HUMA | Unmodified | 0 | 0 | 0 | 0 | 623700 | 81577 | 185760 | 0 | 0 | 102960 | 87748 | 589460 | 219070 | 239100 | 578040 | 0 | 0 | 0 |
| RLATYGITV | SNRNP200; SN | O75643; A0/ | U520_HUMAN | Unmodified | 0 | 0 | 0 | 0 | 0 | 0 | 0 | 0 | 0 | 0 | 0 | 772910 | 244570 | 0 | 0 | 0 | 0 | 0 |
| RLAYARIPA | MYOF; MYOF | Q9NZM1; H( | MYOF_HUMAN | Unmodified | 0 | 0 | 0 | 0 | 6339800 | 2087400 | 0 | 0 | 0 | 0 | 0 | 0 | 0 | 0 | 0 | 0 | 164190 | 185010 |
| RLCQENQ\ | KLC1; KLC1; KL | Q07866; G3\ | KLC1_HUMAN | Unmodified | 0 | 0 | 0 | 0 | 2603300 | 0 | 679090 | 0 | 86599 | 0 | 0 | 1578800 | 1492700 | 523770 | 0 | 0 | 0 | 3057500 |
| RLDEDLAA | CNP; CNP; CNI | P09543; K7E | CN37_HUMAN | Unmodified | 0 | 352940 | 1125700 | 1156700 | 2524100 | 1267100 | 0 | 93544 | 2147900 | 406750 | 6012900 | 30992000 | 1842800 | 0 | 1207200 | 1524600 | 364010 | 3018700 |
| RLDEEEEDI | EIF3C | Q99613 | EIF3C_HUMAN | Unmodified | 0 | 0 | 0 | 242640 | 0 | 0 | 0 | 0 | 184640 | 0 | 0 | 1858800 | 0 | 74981 | 192830 | 0 | 0 | 0 |
| RLDEEEEDI | EIF3C | Q99613 | EIF3C_HUMAN | Unmodified | 295740 | 0 | 246910 | 704380 | 0 | 0 | 0 | 0 | 0 | 0 | 201790 | 2722400 | 0 | 0 | 0 | 384210 | 0 | 0 |
| RLDEELED/ | PSMD6; PSMD6 | Q15008; C9J | PSMD6_HUMA | Unmodified | 0 | 0 | 0 | 0 | 159780 | 0 | 0 | 0 | 430390 | 0 | 3433500 | 16538000 | 0 | 0 | 552830 | 244790 | 0 | 0 |
| RLDFFEKPI | CSNK1G1; CSN | Q9HCP0; U3 | KC1G1_HUMAI | Unmodified | 195750 | 0 | 312190 | 284190 | 0 | 0 | 0 | 105740 | 1269100 | 0 | 971310 | 2281800 | 0 | 0 | 815490 | 252350 | 0 | 0 |
| RLDIDEKPL | AFDN; AFDN; A | P55196; Q5T | AFAD_HUMAN | Unmodified | 0 | 23211 | 20124 | 0 | 72219 | 0 | 222320 | 0 | 0 | 0 | 162310 | 191390 | 0 | 0 | 102660 | 36876 | 0 | 0 |
| RLDINTNTY | SIL1; SIL1; SIL1 | Q9H173; D6 | SIL1_HUMAN N | Unmodified | 0 | 0 | 796530 | 341430 | 0 | 0 | 0 | 0 | 3203100 | 1932000 | 0 | 0 | 0 | 0 | 265960 | 158460 | 0 | 0 |
| RLDKFNEV | MYO6; MYO6; N | Q9UM54; A0 | MYO6_HUMAN | Unmodified | 56873 | 0 | 0 | 262030 | 0 | 0 | 37484 | 0 | 0 | 0 | 0 | 0 | 176070 | 246930 | 0 | 0 | 0 | 0 |
| RLDLDAAK | SH3GLB1; SH3( | Q9Y371; A0/ | SHLB1_HUMAN | Unmodified | 0 | 0 | 0 | 0 | 0 | 0 | 0 | 0 | 0 | 0 | 0 | 0 | 418050 | 0 | 0 | 0 | 0 | 0 |
| RLDLMMN\ | HSDL2 | Q6YN16 | HSDL2_HUMAI | Unmodified | 0 | 0 | 0 | 0 | 0 | 0 | 0 | 0 | 1653300 | 156850 | 0 | 0 | 0 | 0 | 28870 | 0 | 0 | 0 |
| RLDLMMN\ | HSDL2 | Q6YN16 | HSDL2_HUMAI | Oxidation (M) | 0 | 0 | 0 | 0 | 0 | 0 | 0 | 0 | 2463700 | 0 | 0 | 0 | 0 | 0 | 0 | 0 | 0 | 260770 |
| RLDPNKYP | FEN1; FEN1 | P39748; I3L3 | FEN1_HUMAN | Unmodified | 0 | 0 | 0 | 0 | 1093500 | 302220 | 350580 | 0 | 241800 | 247410 | 184220 | 3723000 | 241600 | 0 | 0 | 0 | 0 | 0 |
| RLDRLEET\ | CORO1A; COR( | P31146; H3E | COR1A_HUMAI | Unmodified | 178710 | 0 | 39999 | 0 | 2556800 | 420010 | 4238500 | 2246500 | 1699200 | 1581500 | 6593300 | 23694000 | 0 | 1581500 | 0 | 0 | 0 | 0 |
| RLDRSVSY | PPP1R16A | Q96I34 | PP16A_HUMAN | Unmodified | 0 | 0 | 0 | 0 | 254320 | 0 | 0 | 0 | 0 | 0 | 0 | 0 | 0 | 1109900 | 0 | 0 | 1312400 | 1048100 |
| RLDVSCNS | TLR9; N/A | Q9NR96; H0 | TLR9_HUMAN T | Unmodified | 0 | 0 | 0 | 0 | 0 | 0 | 0 | 0 | 0 | 0 | 243650 | 133750 | 0 | 0 | 0 | 0 | 0 | 0 |
| RLDVTVQS | FAM98A; FAM9 | Q8NCA5; Q5 | FA98A_HUMAN | Unmodified | 0 | 0 | 0 | 0 | 0 | 0 | 0 | 0 | 0 | 0 | 0 | 4247300 | 0 | 0 | 0 | 0 | 0 | 0 |
| RLDVTVQS | FAM98A; FAM9 | Q8NCA5; Q5 | FA98A_HUMAN | Unmodified | 0 | 0 | 0 | 347590 | 2820600 | 0 | 1303400 | 0 | 0 | 0 | 0 | 226770 | 2290200 | 0 | 0 | 0 | 353910 | 417400 |
| RLDYITAEI | PFDN6 | O15212 | PFD6_HUMAN | Unmodified | 101520 | 101930 | 207860 | 0 | 2580400 | 0 | 234350 | 143640 | 2557700 | 606280 | 1418800 | 6844000 | 3207600 | 1621500 | 262470 | 297890 | 1115200 | 2551500 |
| RLEAAYLDI | TBCA; TBCA; TB | O75347; E5F | TBCA_HUMAN | Unmodified | 0 | 639470 | 578490 | 604440 | 423810 | 891600 | 0 | 0 | 8100100 | 238040 | 1845900 | 22930000 | 0 | 0 | 885450 | 6513700 | 0 | 133860 |
| RLEAELAAI | SPTAN1; SPTAN | Q13813; A0/ | SPTN1_HUMAN | Unmodified | 0 | 0 | 0 | 0 | 825080 | 0 | 0 | 0 | 0 | 207250 | 0 | 2944800 | 0 | 58902 | 0 | 0 | 347600 | 0 |
| RLEAGDHF | SPARC; SPARC | P09486; F5G | SPRC_HUMAN | Unmodified | 0 | 0 | 150970 | 153880 | 0 | 1940600 | 744020 | 0 | 0 | 0 | 0 | 0 | 0 | 528650 | 0 | 0 | 180440 | 1367700 |
| RLEALAAEF | SUN2 | Q9UH99 | SUN2_HUMAN | Unmodified | 0 | 0 | 0 | 0 | 0 | 0 | 488310 | 10742 | 20772000 | 980650 | 0 | 0 | 0 | 0 | 0 | 0 | 0 | 0 |
| RLEALGLVS | IQGAP3; IQGAP | Q86VI3; F2Z | IQGA3_HUMAN | Unmodified | 0 | 0 | 0 | 0 | 1481300 | 0 | 0 | 0 | 0 | 0 | 0 | 0 | 0 | 0 | 0 | 0 | 0 | 0 |
| RLEASDGG | FARSA; FARSA; | Q9Y285; K7E | SYFA_HUMAN I | Oxidation (M) | 0 | 0 | 0 | 0 | 0 | 0 | 54267 | 0 | 0 | 0 | 0 | 4816800 | 1734700 | 0 | 0 | 0 | 0 | 0 |
| RLEDLSESI | TMED10 | P49755 | TMEDA_HUMAI | Oxidation (M) | 0 | 0 | 0 | 2066600 | 0 | 0 | 0 | 0 | 88546000 | 13891000 | 5036200 | 2024400 | 0 | 0 | 0 | 0 | 0 | 0 |
| RLEDLSESI | TMED10 | P49755 | TMEDA_HUMAI | Unmodified | 0 | 0 | 0 | 0 | 0 | 0 | 0 | 0 | 1.46E+08 | 9785000 | 591100 | 0 | 0 | 0 | 354880 | 0 | 0 | 0 |
| RLEDLSESI | TMED10 | P49755 | TMEDA_HUMAI | Oxidation (M) | 0 | 0 | 0 | 0 | 0 | 0 | 0 | 0 | 1944900 | 251190 | 0 | 0 | 0 | 0 | 0 | 0 | 0 | 0 |
| RLEDQLAG | SUN2 | Q9UH99 | SUN2_HUMAN | Unmodified | 0 | 0 | 0 | 0 | 0 | 50985 | 0 | 0 | 1447900 | 614990 | 0 | 0 | 0 | 0 | 0 | 0 | 0 | 0 |
| RLEDSSQQ | FOXRED2 | Q8IWF2 | FXRD2_HUMAN | Unmodified | 0 | 0 | 0 | 0 | 0 | 0 | 0 | 0 | 5951600 | 0 | 0 | 0 | 0 | 0 | 0 | 0 | 0 | 0 |
| RLEEDDMY | ABCC4; ABCC4 | O15439; A0/ | MRP4_HUMAN | Oxidation (M) | 0 | 0 | 754360 | 163300 | 0 | 0 | 0 | 0 | 0 | 73864 | 0 | 0 | 0 | 0 | 966180 | 1349000 | 0 | 0 |
| RLEEEADRI | CUL4B; CUL4B; | Q13620; A0/ | CUL4B_HUMAI | Unmodified | 25962 | 42840 | 0 | 162010 | 459060 | 0 | 62887 | 0 | 0 | 0 | 0 | 573140 | 384870 | 113100 | 96967 | 77433 | 94366 | 0 |
| RLEELDLGI | LRRC1 | Q9BTT6 | LRRC1_HUMAN | Unmodified | 0 | 0 | 0 | 0 | 0 | 0 | 0 | 0 | 0 | 0 | 0 | 0 | 197200 | 0 | 55446 | 0 | 0 | 0 |
| RLEENDDD | CFAP298; CFAF | P57076; C9J | CF298_HUMAN | Unmodified | 0 | 0 | 0 | 164570 | 453280 | 0 | 0 | 0 | 0 | 0 | 0 | 389460 | 487930 | 0 | 215180 | 180910 | 0 | 0 |
| RLEEWLGR | MARS1 | P56192 | SYMC_HUMAN | Unmodified | 0 | 0 | 0 | 0 | 0 | 0 | 0 | 0 | 0 | 134910 | 0 | 0 | 272770 | 0 | 0 | 0 | 0 | 270010 |
| RLEFVAAR | PVR; PVR; PVR | P15151; A0A | PVR_HUMAN P | Unmodified | 737330 | 567240 | 1248400 | 313880 | 751930 | 239570 | 0 | 0 | 1266300 | 37993 | 927720 | 5594100 | 94449 | 0 | 995530 | 526870 | 0 | 0 |
| RLEGEDSA | PCK2; PCK2 | Q16822; H0\ | PCKGM_HUMA | Unmodified | 0 | 0 | 0 | 0 | 0 | 0 | 0 | 0 | 5391700 | 0 | 0 | 0 | 0 | 0 | 0 | 0 | 0 | 0 |
| RLEGQEEE | CHGA; CHGA | P10645; G5E | CMGA_HUMAN | Unmodified | 0 | 0 | 0 | 0 | 0 | 0 | 0 | 0 | 355420 | 297910 | 0 | 0 | 0 | 0 | 0 | 0 | 0 | 0 |
| RLEGQEEE | CHGA; CHGA | P10645; G5E | CMGA_HUMAN | Oxidation (M) | 0 | 0 | 0 | 0 | 0 | 0 | 0 | 0 | 2654600 | 2905600 | 0 | 0 | 0 | 0 | 0 | 0 | 0 | 0 |
| RLEGQEEE | CHGA; CHGA | P10645; G5E | CMGA_HUMAN | Unmodified | 0 | 0 | 0 | 0 | 0 | 70758 | 0 | 0 | 27714000 | 14947000 | 0 | 0 | 0 | 0 | 0 | 0 | 0 | 0 |
| RLEHVIFPG | DOCK1; DOCK: | Q14185; A0/ | DOCK1_HUMA | Unmodified | 0 | 0 | 233370 | 161860 | 0 | 0 | 0 | 0 | 0 | 0 | 0 | 0 | 0 | 0 | 646210 | 241030 | 0 | 0 |
| RLELASNH | TPBG | Q13641 | TPBG_HUMAN | Unmodified | 0 | 0 | 0 | 0 | 0 | 0 | 0 | 0 | 1697600 | 0 | 0 | 50426000 | 0 | 0 | 0 | 0 | 0 | 0 |
| RLEQEKLEI | CDC42BPA; CD | Q5VT25; Q9\ | MRCKA_HUMA | Unmodified | 0 | 0 | 0 | 0 | 0 | 0 | 216860 | 52237 | 0 | 0 | 0 | 0 | 0 | 0 | 0 | 0 | 46989 | 198400 |
| RLEQEVDG | CACNA2D2; CA | Q9NY47; C9 | CA2D2_HUMAI | Unmodified | 0 | 117080 | 0 | 0 | 0 | 0 | 122760 | 0 | 6481000 | 1135700 | 219560 | 0 | 0 | 0 | 0 | 0 | 0 | 0 |
| RLEQEVDG | CACNA2D2; CA | Q9NY47; C9 | CA2D2_HUMAI | Oxidation (M) | 55764 | 0 | 21783 | 0 | 0 | 0 | 0 | 0 | 405300 | 194880 | 50167 | 144410 | 0 | 0 | 56579 | 0 | 0 | 0 |
| RLERLDHL | ACTN4; ACTN1; | O43707; P12 | ACTN4_HUMAI | Unmodified | 266610 | 204650 | 0 | 194000 | 1482200 | 0 | 0 | 0 | 1187900 | 312140 | 350350 | 2786800 | 0 | 0 | 563760 | 557260 | 746190 | 0 |
| RLESIATTL | ANKFY1 | Q9P2R3 | ANFY1_HUMAN | Oxidation (M) | 0 | 0 | 0 | 674780 | 0 | 0 | 0 | 0 | 0 | 0 | 95019 | 183390 | 0 | 0 | 320320 | 446250 | 0 | 0 |
| RLETKYPV | PLOD1; PLOD1; | Q02809; Q5. | PLOD1_HUMAI | Unmodified | 152240 | 482190 | 0 | 0 | 764240 | 0 | 0 | 0 | 758110 | 58235 | 0 | 0 | 107100 | 0 | 0 | 33808 | 0 | 112020 |
| RLETPEPG( | PIMREG; PIMRE | Q9BSJ6; I3L4 | PIMRE_HUMAN | Unmodified | 0 | 0 | 0 | 0 | 0 | 0 | 0 | 34545 | 302970 | 302050 | 0 | 0 | 102670 | 0 | 0 | 0 | 0 | 0 |
| RLETYYKAT | AK1; AK1 | P00568; Q5T | KAD1_HUMAN | Unmodified | 0 | 0 | 0 | 0 | 1446500 | 0 | 203240 | 206000 | 0 | 0 | 0 | 0 | 1170400 | 0 | 0 | 0 | 0 | 0 |
| RLEVLDST | UQCRFS1P1; U | P0C7P4; P47 | UCRIL_HUMAN | Unmodified | 0 | 0 | 0 | 0 | 0 | 0 | 580610 | 92626 | 2612400 | 664800 | 0 | 0 | 217670 | 379400 | 0 | 0 | 0 | 0 |
| RLEVTWQT | CNTFR; CNTFR | P26992; Q5T | CNTFR_HUMAI | Unmodified | 2220800 | 2694600 | 0 | 0 | 619070 | 0 | 0 | 0 | 2896400 | 627630 | 0 | 0 | 0 | 2823300 | 0 | 0 | 0 | 0 |
| RLEYESFQ( | GOLM2; GOLM2 | Q6P4E1; H0\ | GOLM2_HUMA | Unmodified | 0 | 0 | 0 | 0 | 532000 | 0 | 0 | 0 | 5561900 | 1266100 | 122800 | 0 | 0 | 0 | 0 | 0 | 0 | 1475700 |
| RLEYESFQ( | GOLM2; GOLM2 | Q6P4E1; H0\ | GOLM2_HUMA | Oxidation (M) | 8635.2 | 0 | 0 | 0 | 405020 | 0 | 0 | 0 | 2243500 | 518000 | 0 | 0 | 414700 | 0 | 0 | 0 | 79201 | 107740 |
| RLFEAEAQI | EHD4 | Q9H223 | EHD4_HUMAN | Unmodified | 296060 | 0 | 226960 | 140290 | 0 | 0 | 0 | 0 | 0 | 0 | 0 | 291400 | 0 | 0 | 663580 | 483200 | 0 | 0 |
| RLFEGNALI | RPS9; RPS9; RP | P46781; B5N | RS9_HUMAN 4 | Unmodified | 227220 | 244640 | 0 | 0 | 0 | 0 | 0 | 0 | 0 | 0 | 0 | 6219600 | 0 | 0 | 347150 | 186970 | 0 | 0 |
| RLFIHESIH | ALDH7A1; ALDI | P49419; A0A | AL7A1_HUMAN | Unmodified | 0 | 0 | 0 | 0 | 0 | 0 | 0 | 0 | 0 | 0 | 147410 | 1888200 | 0 | 0 | 0 | 0 | 488650 | 0 |
| RLGAGQF( | LYN | P07948 | LYN_HUMAN T | Oxidation (M) | 1027700 | 692980 | 363840 | 978690 | 3867500 | 1172900 | 1383200 | 23785 | 2643600 | 1660000 | 0 | 0 | 1011800 | 971870 | 1042400 | 1461900 | 217170 | 202650 |
| RLGAGQF( | LYN | P07948 | LYN_HUMAN T | Unmodified | 0 | 0 | 0 | 1445900 | 8861600 | 13831000 | 4541900 | 870010 | 1037700 | 0 | 0 | 0 | 1047500 | 9097900 | 0 | 0 | 1154900 | 13991000 |
| RLGALSGA | TMEM256; TME | Q8N2U0; K7 | TM256_HUMAN | Unmodified | 0 | 0 | 0 | 0 | 0 | 0 | 0 | 0 | 1867700 | 484920 | 0 | 0 | 0 | 0 | 0 | 0 | 0 | 0 |
| RLGCEEME | PDPK2P; PDPK | Q6A1A2; O1 | PDPK2_HUMAN | Unmodified | 0 | 0 | 45382 | 85083 | 0 | 0 | 0 | 0 | 0 | 0 | 0 | 0 | 287830 | 276180 | 94045 | 59863 | 0 | 0 |
| RLGCEEME | PDPK2P; PDPK | Q6A1A2; O1 | PDPK2_HUMAN | Oxidation (M) | 0 | 0 | 0 | 118180 | 0 | 0 | 0 | 0 | 0 | 0 | 0 | 0 | 235290 | 221660 | 311720 | 0 | 0 | 0 |
| RLGCFDHA | UBE2O; UBE2O | Q9C0C9; K7 | UBE2O_HUMAI | Unmodified | 0 | 39385 | 0 | 0 | 0 | 0 | 0 | 0 | 0 | 0 | 0 | 1198900 | 0 | 0 | 0 | 0 | 0 | 28501 |
| RLGIVWQD | ZW10 | O43264 | ZW10_HUMAN | Unmodified | 0 | 0 | 0 | 0 | 104340 | 0 | 147960 | 0 | 202560 | 0 | 0 | 0 | 0 | 0 | 12444 | 0 | 118520 | 72793 |
| RLGLERNP | MYO1C; MYO1( | O00159; F5H | MYO1C_HUMA | Unmodified | 0 | 0 | 0 | 1913500 | 2071400 | 0 | 979060 | 0 | 0 | 0 | 0 | 0 | 0 | 0 | 0 | 0 | 461660 | 0 |
| RLGLPGDE | PRKDC | P78527 | PRKDC_HUMAI | Unmodified | 0 | 0 | 184250 | 0 | 0 | 0 | 0 | 0 | 0 | 0 | 0 | 0 | 0 | 0 | 0 | 0 | 0 | 0 |
| RLGPGGLD | CDC37; CDC37 | Q16543; K7E | CDC37_HUMA | Unmodified | 0 | 0 | 0 | 2073200 | 14041000 | 2270100 | 1445300 | 0 | 4838900 | 1782500 | 1159900 | 6880200 | 5062500 | 4074200 | 7204100 | 1916200 | 5156500 | 16564000 |
| RLGSGQF( | FRK | P42685 | FRK_HUMAN T | Unmodified | 0 | 0 | 0 | 0 | 0 | 0 | 0 | 0 | 585070 | 1869500 | 0 | 0 | 0 | 0 | 0 | 42384 | 0 | 0 |
| RLGSQREL | DIPK2A | Q8NDZ4 | DIK2A_HUMAN | Unmodified | 0 | 0 | 0 | 0 | 0 | 0 | 87332 | 28644 | 235370 | 2195000 | 0 | 0 | 38416 | 0 | 0 | 0 | 0 | 0 |
| RLGTEKEVI | TK1 | P04183 | KITH_HUMAN T | Unmodified | 0 | 0 | 0 | 0 | 729690 | 0 | 0 | 25725 | 0 | 103670 | 0 | 228860 | 325770 | 0 | 0 | 0 | 193990 | 0 |
| RLGTFEVEI | DPP4; DPP4; DI | P27487; A0A | DPP4_HUMAN | Unmodified | 136900 | 0 | 2045200 | 2238800 | 816690 | 0 | 0 | 0 | 0 | 0 | 0 | 0 | 0 | 0 | 3140000 | 1830100 | 259600 | 0 |

| | | | | | | | | | | | | | | | | | | | | | | |
|---|---|---|---|---|---|---|---|---|---|---|---|---|---|---|---|---|---|---|---|---|---|---|
| RLGTFEVEI | DPP4; DPP4; DI | P27487; A0A | DPP4_HUMAN | Unmodified | 0 | 0 | 97205 | 3840500 | 0 | 0 | 0 | 0 | 0 | 0 | 0 | 0 | 0 | 0 | 0 | 0 | 0 | 0 |
| RLGYISVVD | ANK3; ANK2; AI | Q12955; Q0 | ANK3_HUMAN | Unmodified | 0 | 0 | 414200 | 842570 | 0 | 0 | 0 | 0 | 0 | 197610 | 0 | 169180 | 1541900 | 837640 | 2176700 | 1342900 | 0 | 0 |
| RLHDEKEE | DIAPH1; DIAPH | O60610; A0A | DIAP1_HUMAN | Unmodified | 0 | 33300 | 0 | 0 | 0 | 0 | 0 | 0 | 49829 | 37224 | 19825 | 1143700 | 0 | 0 | 92049 | 32357 | 23115 | 0 |
| RLHGLDEE | MCM4; MCM4; | P33991; A0A | MCM4_HUMAN | Unmodified | 99035 | 53624 | 116820 | 118010 | 1456300 | 0 | 0 | 77587 | 538560 | 390650 | 710380 | 1810100 | 492990 | 172960 | 0 | 0 | 293410 | 233990 |
| RLHTLEEVI | GGA3; GGA3; G | Q9NZ52; J3K | GGA3_HUMAN | Unmodified | 0 | 0 | 0 | 353730 | 63498 | 0 | 0 | 0 | 160760 | 0 | 62341 | 183210 | 0 | 0 | 0 | 0 | 238420 | 364720 |
| RLHTVGEP | IGSF3 | O75054 | IGSF3_HUMAN | Unmodified | 1053800 | 1080700 | 1129700 | 398980 | 645530 | 0 | 24163 | 61694 | 269980 | 0 | 0 | 1764100 | 845350 | 221370 | 1334000 | 964350 | 116300 | 282840 |
| RLIADLGST | FAM3C | Q92520 | FAM3C_HUMAI | Unmodified | 0 | 0 | 0 | 0 | 0 | 25725 | 0 | 0 | 2502900 | 0 | 0 | 0 | 0 | 249310 | 0 | 0 | 0 | 0 |
| RLIADLGST | FAM3C | Q92520 | FAM3C_HUMAI | Unmodified | 0 | 0 | 0 | 0 | 0 | 512330 | 0 | 0 | 305440 | 2307500 | 0 | 0 | 1552000 | 0 | 0 | 0 | 0 | 1499100 |
| RLIDLHSPS | RPS20; RPS20; | P60866; A0A | RS20_HUMAN | Unmodified | 3508700 | 2832800 | 1980500 | 2063900 | 4712300 | 1374300 | 3409700 | 0 | 1752000 | 2007500 | 2067700 | 19458000 | 6934700 | 0 | 2057400 | 2388300 | 394910 | 5915600 |
| RLIDLHTN\ | SCFD1; SCFD1 | Q8WVM8; A( | SCFD1_HUMAI | Unmodified | 0 | 0 | 0 | 0 | 752610 | 0 | 0 | 0 | 0 | 0 | 0 | 287850 | 0 | 0 | 0 | 0 | 0 | 0 |
| RLIGRTWN | HSPA5; HSPA5; | P11021; A0A | BIP_HUMAN Er | Unmodified | 0 | 0 | 0 | 0 | 7428500 | 0 | 30099000 | 7456800 | 0 | 1.16E+08 | 536710 | 0 | 0 | 11391000 | 0 | 0 | 947600 | 6352900 |
| RLIPDGCG | RPL10; RPL10; | P27635; X1V | RL10_HUMAN | Unmodified | 820010 | 545560 | 271960 | 243760 | 0 | 0 | 0 | 0 | 0 | 0 | 0 | 4585700 | 0 | 219130 | 0 | 255140 | 0 | 0 |
| RLIPNATGT | PKN1; PKN1 | Q16512; K7E | PKN1_HUMAN | Unmodified | 0 | 0 | 0 | 0 | 0 | 0 | 0 | 0 | 543960 | 0 | 0 | 519470 | 0 | 0 | 0 | 94862 | 0 | 0 |
| RLIVDVIR | IQGAP1; IQGAP | P46940; H0Y | IQGA1_HUMAN | Unmodified | 334780 | 311540 | 223490 | 251300 | 869710 | 120730 | 0 | 0 | 846310 | 94258 | 72443 | 2400000 | 82364 | 0 | 544600 | 490050 | 0 | 0 |
| RLIVDVIRF | IQGAP1; IQGAP | P46940; H0Y | IQGA1_HUMAN | Unmodified | 0 | 0 | 0 | 382640 | 3892500 | 0 | 0 | 0 | 0 | 204250 | 0 | 0 | 254120 | 0 | 0 | 0 | 0 | 0 |
| RLKEGDTM | PTGFR1; PTGFR1 | Q14914; F22 | PTGR1_HUMAI | Unmodified | 0 | 0 | 0 | 71108 | 4022900 | 0 | 0 | 126380 | 0 | 194370 | 0 | 0 | 0 | 144700 | 0 | 0 | 155670 | 0 |
| RLKEGDTM | PTGR1; PTGR1 | Q14914; F22 | PTGR1_HUMAI | Oxidation (M) | 0 | 0 | 46266 | 0 | 794480 | 0 | 0 | 0 | 0 | 0 | 0 | 90405 | 0 | 0 | 52114 | 66393 | 114730 | 235710 |
| RLKEQGQA | RHOG | P84095 | RHOG_HUMAN | Unmodified | 0 | 0 | 0 | 0 | 637600 | 0 | 148440 | 103050 | 0 | 0 | 0 | 0 | 416300 | 317450 | 0 | 0 | 0 | 1548100 |
| RLKNELLEI | SYTL4 | Q96C24 | SYTL4_HUMAN | Unmodified | 0 | 0 | 0 | 0 | 589520 | 0 | 29259 | 8894.4 | 0 | 196940 | 0 | 0 | 122460 | 0 | 0 | 0 | 82401 | 0 |
| RLKPGYLE/ | PPA1 | Q15181 | IPYR_HUMAN | Unmodified | 0 | 0 | 0 | 696790 | 1680700 | 96208 | 0 | 0 | 181550 | 164020 | 1394600 | 8272800 | 2348000 | 299730 | 711760 | 356750 | 475140 | 330440 |
| RLLDGLAQ | PLCB3 | Q01970 | PLCB3_HUMAN | Unmodified | 0 | 0 | 0 | 0 | 967460 | 221900 | 0 | 0 | 166820 | 149480 | 0 | 1446200 | 0 | 0 | 0 | 0 | 320030 | 278290 |
| RLLDQQEK | SLC38A10; SLC | Q9HBR0; H0 | S38AA_HUMAN | Unmodified | 0 | 0 | 0 | 0 | 0 | 0 | 0 | 0 | 3671500 | 754260 | 0 | 0 | 0 | 0 | 0 | 0 | 0 | 0 |
| RLLDTAGH | INHBE; INHBE | P58166; F8V | INHBE_HUMAN | Unmodified | 266310 | 0 | 0 | 1276900 | 0 | 0 | 0 | 0 | 1436400 | 482060 | 0 | 0 | 0 | 0 | 0 | 584670 | 0 | 0 |
| RLLEFELAC | ROCK1; ROCK1 | Q13464; A0A | ROCK1_HUMAI | Unmodified | 0 | 50062 | 0 | 0 | 2663100 | 895480 | 0 | 0 | 1741200 | 976030 | 0 | 0 | 1111000 | 825520 | 0 | 0 | 2271000 | 1909400 |
| RLLEFNQG | GARS1; GARS1 | P41250; A0A | GARS_HUMAN | Unmodified | 0 | 0 | 146780 | 0 | 0 | 0 | 0 | 0 | 0 | 0 | 0 | 756330 | 0 | 0 | 0 | 37018 | 0 | 0 |
| RLLEGDGC | GART; GART | P22102; F8V | PUR2_HUMAN | Oxidation (M) | 0 | 0 | 0 | 0 | 0 | 0 | 0 | 0 | 0 | 0 | 0 | 1002600 | 0 | 0 | 0 | 0 | 857570 | 0 |
| RLLGSVQQ | PBXIP1 | Q96AQ6 | PBIP1_HUMAN | Unmodified | 0 | 0 | 0 | 0 | 0 | 0 | 0 | 184490 | 2743000 | 132180 | 856300 | 0 | 0 | 0 | 0 | 0 | 108760 | 0 |
| RLLGYVATI | CSDE1; CSDE1 | O75534; E9F | CSDE1_HUMAI | Unmodified | 33532 | 0 | 0 | 0 | 334790 | 0 | 0 | 23630 | 87328 | 0 | 121720 | 725690 | 0 | 0 | 0 | 84020 | 0 | 257600 |
| RLLHVAVSI | PSMD1; PSMD1 | Q99460; A0A | PSMD1_HUMA | Unmodified | 0 | 0 | 0 | 0 | 0 | 0 | 0 | 0 | 0 | 0 | 0 | 2134700 | 0 | 230210 | 0 | 0 | 0 | 0 |
| RLLLAIVGQ | ANXA7 | P20073 | ANXA7_HUMAI | Unmodified | 0 | 0 | 0 | 0 | 176950 | 0 | 56964 | 35951 | 14652000 | 1457200 | 0 | 655600 | 0 | 62036 | 0 | 0 | 0 | 0 |
| RLLLFSDGI | COL6A1; COL6 | P12109; A0A | CO6A1_HUMAI | Unmodified | 0 | 0 | 903360 | 529140 | 0 | 520290 | 937200 | 405760 | 716810 | 1067200 | 29597 | 419540 | 147790 | 0 | 190220 | 130020 | 165560 | 297450 |
| RLLQERELV | EGFR; EGFR; E | P00533; Q5 | EGFR_HUMAN | Unmodified | 0 | 0 | 0 | 0 | 5974900 | 4119000 | 0 | 0 | 0 | 0 | 0 | 0 | 1644900 | 419510 | 0 | 0 | 0 | 480760 |
| RLLQFYAET | MYO1C; MYO1 | O00159; I3L | MYO1C_HUMA | Unmodified | 991780 | 197640 | 179040 | 417340 | 0 | 0 | 0 | 0 | 0 | 0 | 0 | 0 | 0 | 246040 | 2877900 | 1788900 | 0 | 0 |
| RLLSDDRS | PIGK; PIGK; PIG | Q92643; B1 | GPI8_HUMAN | Unmodified | 0 | 0 | 0 | 0 | 0 | 0 | 0 | 0 | 2621500 | 414540 | 0 | 0 | 0 | 0 | 0 | 0 | 0 | 0 |
| RLLSDSLPI | TLN1 | Q9Y490 | TLN1_HUMAN | Unmodified | 78330 | 1356000 | 308990 | 190940 | 0 | 0 | 0 | 0 | 0 | 0 | 0 | 1005100 | 0 | 0 | 939640 | 290540 | 0 | 0 |
| RLLSRPQD | ATAD3A | Q9NVI7 | ATD3A_HUMAN | Unmodified | 0 | 0 | 0 | 0 | 0 | 0 | 0 | 0 | 4677600 | 443570 | 0 | 0 | 0 | 0 | 0 | 0 | 0 | 0 |
| RLLVPILLPI | MPDU1; MPDU | O75352; J3Q | MPU1_HUMAN | Unmodified | 0 | 0 | 0 | 0 | 0 | 0 | 0 | 0 | 3638000 | 2546500 | 465170 | 0 | 0 | 0 | 0 | 0 | 0 | 0 |
| RLLVQEQQ | RASAL2; RASAL | Q9UJF2; H0 | NGAP_HUMAN | Unmodified | 0 | 0 | 0 | 0 | 664890 | 0 | 0 | 0 | 0 | 0 | 0 | 0 | 0 | 0 | 0 | 0 | 0 | 79730 |
| RLMSMEMI | PTGFRN | Q9P2B2 | FPRP_HUMAN | 3 Oxidation (M) | 0 | 32005 | 103480 | 188030 | 0 | 0 | 0 | 0 | 0 | 0 | 20939 | 0 | 0 | 113620 | 126100 | 91060 | 0 | 31342 |
| RLMYNSSN | MYO1D; MYO1 | O94832; J3Q | MYO1D_HUMA | Unmodified | 677290 | 783900 | 0 | 242910 | 34928 | 0 | 0 | 0 | 212750 | 0 | 0 | 0 | 0 | 0 | 756590 | 629900 | 75923 | 622850 |
| RLNDFAST | COX5A; COX5A | P20674; H3E | COX5A_HUMAN | Unmodified | 0 | 0 | 0 | 704730 | 1573600 | 41050 | 713720 | 421150 | 3809300 | 673380 | 0 | 0 | 864380 | 1965300 | 309410 | 0 | 0 | 2628300 |
| RLNDRIEM | GLG1 | Q92896 | GSLG1_HUMAI | Unmodified | 19599 | 0 | 0 | 0 | 0 | 0 | 0 | 0 | 7519300 | 0 | 0 | 0 | 0 | 0 | 0 | 0 | 0 | 0 |
| RLNEAAVT | CCNDBP1; CC | O95273; H3I | CCDB1_HUMA | Unmodified | 0 | 0 | 0 | 0 | 0 | 0 | 0 | 0 | 0 | 0 | 0 | 171800 | 0 | 0 | 996220 | 1029500 | 0 | 0 |
| RLNHYVLY | PSMD3 | O43242 | PSMD3_HUMA | Unmodified | 0 | 0 | 0 | 0 | 0 | 0 | 0 | 0 | 0 | 139000 | 0 | 529040 | 0 | 0 | 0 | 0 | 0 | 0 |
| RLNIIIVAEC | PFKP; PFKP; PF | Q01813; A0A | PFKAP_HUMAN | Unmodified | 0 | 0 | 0 | 0 | 2689800 | 475270 | 136820 | 0 | 0 | 234140 | 337010 | 0 | 0 | 0 | 0 | 0 | 1003400 | 1920600 |
| RLNQVIFP | NAA50; NAA50 | Q9GZZ1; A0 | NAA50_HUMAN | Unmodified | 0 | 0 | 0 | 0 | 0 | 0 | 0 | 484700 | 0 | 0 | 0 | 5118200 | 0 | 0 | 0 | 0 | 0 | 0 |
| RLNTEWSE | P4HA1 | P13674 | P4HA1_HUMAI | Unmodified | 0 | 0 | 0 | 0 | 0 | 0 | 0 | 0 | 10513000 | 5129000 | 1043000 | 49932 | 0 | 13833 | 0 | 0 | 0 | 2895200 |
| RLNTLAHY | PLXNA1 | Q9UIW2 | PLXA1_HUMAI | Unmodified | 0 | 34959 | 365760 | 0 | 4077200 | 237820 | 0 | 0 | 1224800 | 0 | 314010 | 2541300 | 2031500 | 483350 | 643590 | 103790 | 261650 | 777190 |
| RLNTLMHY | PLXNA2 | O75051 | PLXA2_HUMAN | Unmodified | 0 | 0 | 0 | 117340 | 0 | 0 | 0 | 0 | 1078700 | 0 | 0 | 0 | 0 | 0 | 0 | 0 | 0 | 0 |
| RLNYQTPG | PTPRF; PTPRF | P10586; H0 | PTPRF_HUMAN | Unmodified | 0 | 0 | 0 | 0 | 0 | 0 | 0 | 0 | 2369900 | 0 | 0 | 0 | 0 | 0 | 0 | 0 | 0 | 0 |
| RLNYQTPG | PTPRF; PTPRF | P10586; H0 | PTPRF_HUMAN | Oxidation (M) | 549740 | 430140 | 503850 | 309670 | 4572600 | 0 | 0 | 0 | 370160 | 1166400 | 0 | 599330 | 0 | 0 | 1036900 | 858940 | 0 | 0 |
| RLPAFTLSI | NCLN; NCLN; | Q969V3; K7E | NCLN_HUMAN | Unmodified | 0 | 0 | 1544100 | 0 | 0 | 0 | 66859 | 0 | 1587600 | 252010 | 90957 | 0 | 0 | 0 | 0 | 0 | 0 | 0 |
| RLPAFTLSI | NCLN; NCLN; | Q969V3; K7E | NCLN_HUMAN | Unmodified | 0 | 444000 | 0 | 0 | 0 | 0 | 0 | 0 | 6032800 | 83262 | 0 | 0 | 0 | 0 | 0 | 0 | 0 | 0 |
| RLPDGIGQ | SCRIB; SCRIB; | Q14160; A0 | SCRIB_HUMAN | Unmodified | 0 | 0 | 0 | 0 | 227970 | 0 | 0 | 0 | 1895300 | 964610 | 1089600 | 3288400 | 169020 | 0 | 0 | 0 | 681550 | 48645 |
| RLPENLYN | UQCRB; UQCR | P14927; B72 | QCR7_HUMAN | Unmodified | 0 | 0 | 0 | 0 | 0 | 0 | 0 | 0 | 3405200 | 172470 | 0 | 920790 | 0 | 0 | 0 | 24663 | 0 | 0 |
| RLPLNFLS | NAA15; NAA15 | Q9BXJ9; A0 | NAA15_HUMAI | Unmodified | 57419 | 0 | 0 | 0 | 0 | 0 | 0 | 0 | 0 | 0 | 0 | 2242400 | 0 | 0 | 0 | 0 | 0 | 0 |
| RLPPQQIEI | CAPZB; CAPZB; | P47756; A0A | CAPZB_HUMAN | Unmodified | 226090 | 245200 | 164690 | 63951 | 0 | 143110 | 34982 | 0 | 268360 | 136100 | 0 | 934750 | 0 | 0 | 0 | 162110 | 128000 | 259740 |
| RLPSSPASI | EPB41L1; EPB4 | Q9H4G0; H7 | E41L1_HUMAN | Unmodified | 0 | 0 | 0 | 0 | 138890 | 63577 | 29954 | 0 | 0 | 10589 | 675470 | 517070 | 0 | 0 | 0 | 0 | 29814 | 400850 |
| RLPSSPASI | EPB41L1; EPB4 | Q9H4G0; H7 | E41L1_HUMAN | Unmodified | 0 | 0 | 0 | 0 | 602500 | 49433 | 0 | 0 | 0 | 0 | 0 | 0 | 317430 | 199100 | 0 | 0 | 141590 | 322150 |
| RLPTEEEW | SUMF2; SUMF2 | Q8NBJ7; C9 | SUMF2_HUMAI | Unmodified | 0 | 0 | 0 | 0 | 0 | 0 | 0 | 0 | 62955000 | 5581700 | 1007500 | 0 | 0 | 0 | 0 | 0 | 0 | 0 |
| RLPTEEEW | SUMF2; SUMF2 | Q8NBJ7; C9 | SUMF2_HUMAI | Unmodified | 0 | 0 | 0 | 0 | 325810 | 0 | 406240 | 0 | 6220100 | 8252600 | 0 | 0 | 0 | 0 | 0 | 0 | 0 | 0 |
| RLPWATGY | PPM1H | Q9ULR3 | PPM1H_HUMA | Unmodified | 0 | 0 | 0 | 0 | 0 | 0 | 0 | 0 | 1596300 | 0 | 462320 | 3130800 | 0 | 0 | 0 | 0 | 0 | 0 |
| RLQEALER | CALD1; CALD1 | Q05682; C9 | CALD1_HUMAN | Unmodified | 1249600 | 0 | 610400 | 689590 | 1382600 | 378470 | 222100 | 0 | 0 | 458810 | 77991 | 0 | 502830 | 999660 | 0 | 0 | 0 | 613690 |
| RLQEDPNY | BAG6; BAG6; B | P46379; H0 | BAG6_HUMAN | Unmodified | 0 | 0 | 0 | 59038 | 0 | 0 | 0 | 0 | 444270 | 94456 | 249090 | 847220 | 110130 | 0 | 57785 | 58257 | 0 | 24426 |
| RLQEENLV | VIL1; VIL1; VIL1 | P09327; H7 | VILI_HUMAN Vi | Unmodified | 655970 | 1371300 | 0 | 0 | 0 | 0 | 246240 | 0 | 3271300 | 1653600 | 771910 | 15645000 | 0 | 0 | 0 | 0 | 0 | 0 |
| RLQELDAA | CLTB | P09497 | CLCB_HUMAN | Unmodified | 4319600 | 2268700 | 3330400 | 2229000 | 0 | 85526 | 122390 | 123870 | 251540 | 0 | 0 | 2649700 | 2196900 | 279110 | 1871100 | 2497900 | 187080 | 342930 |
| RLQELDAA | CLTB | P09497 | CLCB_HUMAN | Unmodified | 826650 | 222390 | 0 | 163230 | 0 | 0 | 0 | 0 | 0 | 0 | 0 | 0 | 0 | 0 | 0 | 0 | 0 | 0 |
| RLQELEGQ | KIF4A | O95239 | KIF4A_HUMAN | Unmodified | 0 | 0 | 112700 | 139680 | 695850 | 245990 | 0 | 0 | 162560 | 133930 | 0 | 130230 | 808980 | 0 | 0 | 0 | 232840 | 387750 |
| RLQEQEEL | VGF | O15240 | VGF_HUMAN | Unmodified | 29863 | 52606 | 0 | 0 | 30430 | 0 | 0 | 0 | 551350 | 128820 | 0 | 0 | 0 | 0 | 0 | 0 | 0 | 0 |
| RLQGQLEC | ABCF1; ABCF1 | Q8NE71; H0 | ABCF1_HUMAI | Unmodified | 0 | 0 | 0 | 0 | 0 | 0 | 0 | 0 | 0 | 0 | 0 | 1654500 | 0 | 0 | 0 | 0 | 0 | 0 |
| RLQIAQLYF | DLG1; DLG1; D | Q12959; A0 | DLG1_HUMAN | Unmodified | 0 | 0 | 0 | 0 | 0 | 0 | 0 | 0 | 0 | 2057300 | 0 | 0 | 0 | 809590 | 0 | 0 | 1378900 | 515170 |
| RLQITPDSI | NEDD4L; NEDD | Q96PU5; A0 | NED4L_HUMAN | Unmodified | 0 | 0 | 0 | 4687400 | 599740 | 0 | 0 | 0 | 0 | 0 | 1067900 | 2810900 | 1263300 | 165360 | 0 | 0 | 0 | 0 |
| RLQLPVWE | DHX15 | O43143 | DHX15_HUMAI | Unmodified | 0 | 0 | 0 | 0 | 0 | 0 | 0 | 0 | 899610 | 152870 | 328660 | 2023100 | 0 | 0 | 0 | 0 | 0 | 0 |
| RLQMQMD | PLXNA2; PLXN | O75051; Q9 | PLXA2_HUMAN | Oxidation (M) | 0 | 0 | 0 | 0 | 0 | 0 | 0 | 0 | 575100 | 0 | 0 | 180650 | 0 | 0 | 0 | 0 | 0 | 0 |
| RLQMQMD | PLXNA2; PLXN | O75051; Q9 | PLXA2_HUMAN | Unmodified | 67757 | 0 | 0 | 0 | 0 | 0 | 0 | 0 | 1050600 | 0 | 0 | 469820 | 0 | 0 | 0 | 29941 | 128320 | 0 |
| RLQQAQLY | DLG3; DLG3 | Q92796; Q5 | DLG3_HUMAN | Unmodified | 0 | 0 | 0 | 0 | 415390 | 0 | 0 | 0 | 826180 | 1270900 | 0 | 1255000 | 1747700 | 731660 | 184380 | 267620 | 2201800 | 0 |
| RLQQEDGI | CPE; CPE | P16870; H0 | CBPE_HUMAN | Unmodified | 0 | 0 | 0 | 0 | 0 | 0 | 0 | 0 | 30043000 | 1168000 | 0 | 0 | 0 | 0 | 0 | 0 | 0 | 0 |
| RLQQEILD | AVIL | O75366 | AVIL_HUMAN | Unmodified | 0 | 20536 | 0 | 0 | 0 | 0 | 0 | 0 | 0 | 0 | 330310 | 48637000 | 0 | 0 | 0 | 0 | 0 | 0 |
| RLQQTQN | VAMP3 | Q15836 | VAMP3_HUMAI | Unmodified | 0 | 0 | 0 | 0 | 0 | 0 | 70038 | 0 | 983530 | 0 | 0 | 0 | 0 | 0 | 0 | 0 | 0 | 0 |
| RLQQTQN | VAMP3 | Q15836 | VAMP3_HUMAI | Unmodified | 71970 | 0 | 296810 | 676100 | 2732000 | 0 | 0 | 0 | 649700 | 1059700 | 0 | 0 | 973580 | 1364000 | 271340 | 0 | 0 | 1023900 |
| RLQYSQDI | DTNB; DTNB | O60941; E7E | DTNB_HUMAN | Unmodified | 0 | 0 | 0 | 1588800 | 492660 | 0 | 0 | 0 | 5212300 | 2122100 | 0 | 0 | 0 | 0 | 1207600 | 824360 | 161490 | 327560 |
| RLSAPLPS | RNF31; RNF31 | Q96EP0; H0 | RNF31_HUMAN | Unmodified | 197220 | 136310 | 0 | 0 | 659520 | 0 | 0 | 0 | 0 | 12519 | 0 | 0 | 0 | 0 | 0 | 0 | 0 | 0 |
| RLSEDYGV | PRDX2; PRDX2 | P32119; A0 | PRDX2_HUMAI | Unmodified | 167450 | 107950 | 788330 | 288940 | 0 | 0 | 0 | 0 | 1952100 | 113770 | 1515400 | 15273000 | 0 | 0 | 542520 | 594800 | 0 | 0 |
| RLSEDYGV | PRDX2 | P32119 | PRDX2_HUMAI | Unmodified | 1220000 | 1409900 | 0 | 2436200 | 0 | 0 | 189780 | 539190 | 16019000 | 1394700 | 4771400 | 33951000 | 463390 | 0 | 542250 | 1121600 | 267850 | 0 |
| RLSELLRY | HSP90AB1 | P08238 | HS90B_HUMAI | Oxidation (M) | 0 | 0 | 0 | 0 | 1332400 | 593570 | 0 | 0 | 0 | 377980 | 0 | 0 | 1242500 | 368440 | 0 | 0 | 172920 | 280410 |
| RLSENEIIC | CALCOCO2; C | Q13137; D6 | CACO2_HUMA | Unmodified | 493080 | 452920 | 70741 | 142150 | 0 | 0 | 0 | 0 | 1885200 | 444640 | 0 | 0 | 0 | 0 | 0 | 0 | 0 | 1314600 |
| RLSENEIIC | CALCOCO2; C | Q13137; D6 | CACO2_HUMA | Unmodified | 0 | 0 | 0 | 0 | 269790 | 0 | 1809000 | 56888 | 0 | 0 | 0 | 0 | 595810 | 0 | 0 | 0 | 1036200 | 2822200 |
| RLSQIGVE | ALDOC; ALDOC | P09972; J3Q | ALDOC_HUMA | Unmodified | 116480 | 0 | 0 | 0 | 0 | 83518 | 54453 | 0 | 0 | 0 | 691190 | 4950700 | 1046100 | 266340 | 360440 | 474960 | 741860 | 629480 |
| RLSQRPTP | PHACTR4 | Q8IZ21 | PHAR4_HUMAN | Unmodified | 0 | 0 | 0 | 0 | 0 | 0 | 0 | 0 | 568830 | 57493 | 0 | 0 | 0 | 0 | 0 | 0 | 117240 | 187490 |
| RLSVELTSS | CTNNB1; CTNN | P35222; A0 | CTNB1_HUMAN | Unmodified | 0 | 0 | 0 | 391920 | 0 | 0 | 0 | 0 | 0 | 0 | 0 | 0 | 0 | 0 | 553670 | 0 | 0 | 293280 |
| RLSYNTASI | RPL34 | P49207 | RL34_HUMAN | Unmodified | 2307900 | 0 | 0 | 1489300 | 0 | 0 | 0 | 0 | 400080 | 0 | 0 | 0 | 1929900 | 0 | 1536800 | 0 | 0 | 0 |
| RLTGFHET | GLUL; GLUL | P15104; A0 | GLNA_HUMAN | Unmodified | 0 | 0 | 0 | 0 | 0 | 58309 | 0 | 0 | 1471500 | 199130 | 0 | 7090400 | 0 | 0 | 0 | 0 | 0 | 218880 |
| RLTQNADC | TUBG1; TUBG2 | P23258; Q9 | TBG1_HUMAN | Unmodified | 0 | 0 | 0 | 0 | 0 | 0 | 0 | 0 | 0 | 0 | 0 | 1280900 | 370090 | 0 | 0 | 0 | 164680 | 0 |
| RLTSPVINT | ANKRD13A; AN | Q8IZ07; H0 | AN13A_HUMAN | Unmodified | 0 | 602580 | 704480 | 875990 | 1474900 | 0 | 0 | 0 | 791320 | 0 | 0 | 0 | 126020 | 0 | 0 | 0 | 0 | 0 |
| RLTVMSLQ | FLNB; FLNB | O75369; E7E | FLNB_HUMAN | Unmodified | 0 | 0 | 0 | 0 | 1161100 | 339970 | 0 | 0 | 0 | 56190 | 0 | 804650 | 0 | 0 | 0 | 0 | 138630 | 0 |
| RLTVNEAVI | PLEC; PLEC | Q15149; H0 | PLEC_HUMAN | Unmodified | 0 | 0 | 0 | 0 | 0 | 0 | 0 | 0 | 0 | 0 | 0 | 0 | 0 | 0 | 0 | 0 | 946010 | 569750 |
| RLTVSSLQI | FLNA; FLNA; FL | P21333; A0 | FLNA_HUMAN | Unmodified | 744770 | 297440 | 1129500 | 232010 | 3107500 | 389280 | 78078 | 416640 | 380880 | 1166200 | 0 | 2722600 | 827730 | 0 | 1586400 | 1630100 | 2348800 | 0 |
| RLTVTSLQE | FLNC | Q14315 | FLNC_HUMAN | Unmodified | 0 | 0 | 0 | 0 | 658060 | 232890 | 0 | 0 | 0 | 0 | 0 | 0 | 0 | 0 | 0 | 0 | 0 | 0 |
| RLVACFQG | ARHGAP17; AR | Q68EM7; C9 | RHG17_HUMAI | Unmodified | 185010 | 0 | 0 | 119920 | 0 | 0 | 0 | 0 | 203920 | 0 | 35122 | 654570 | 441280 | 0 | 98913 | 68212 | 13716 | 0 |
| RLVECGSL | PLXNB2; PLXN | O15031; A6 | PLXB2_HUMAN | Unmodified | 285460 | 70596 | 170490 | 571040 | 252140 | 259710 | 93703 | 74502 | 0 | 0 | 0 | 0 | 430950 | 0 | 587970 | 674010 | 1148000 | 1712300 |
| RLVHQNSA | CUL1; CUL1; C | Q13616; A0 | CUL1_HUMAN | Unmodified | 0 | 0 | 0 | 0 | 1445900 | 0 | 118010 | 138870 | 57033 | 77819 | 114000 | 2085800 | 0 | 0 | 0 | 0 | 377700 | 1025500 |
| RLVNITIIK | ITGB4 | P16144 | ITB4_HUMAN | Unmodified | 0 | 0 | 0 | 0 | 903450 | 0 | 0 | 0 | 0 | 0 | 0 | 0 | 126990 | 126070 | 0 | 0 | 474460 | 589620 |
| RLVPGGGA | CCT8 | P50990 | TCPQ_HUMAN | Unmodified | 1032700 | 0 | 2119800 | 1729100 | 14926000 | 2028500 | 5369700 | 3230300 | 995670 | 566340 | 5896800 | 35943000 | 22221000 | 10482000 | 2328300 | 2316100 | 6700700 | 11437000 |
| RLVQENQV | KLC2; KLC2; KL | Q9H0B6; C9 | KLC2_HUMAN | Unmodified | 0 | 0 | 0 | 0 | 0 | 0 | 0 | 0 | 0 | 0 | 0 | 0 | 879310 | 711310 | 0 | 61794 | 0 | 0 |
| RLVQSPNS | RPS27L; RPS27 | Q71UM5; P4 | RS27L_HUMAN | Unmodified | 378370 | 564320 | 1458100 | 279930 | 2348300 | 0 | 1318700 | 208320 | 0 | 227320 | 0 | 2342500 | 1436800 | 1107500 | 403920 | 119150 | 0 | 1512100 |
| RLVQSPNS | RPS27L; RPS27 | Q71UM5; P4 | RS27L_HUMAN | Oxidation (M) | 1328000 | 924090 | 940080 | 865340 | 1770500 | 0 | 1397000 | 1058200 | 303430 | 807340 | 722310 | 5445800 | 2234800 | 954860 | 1029400 | 1252200 | 0 | 0 |
| RLVTTGVLI | H1-0 | P07305 | H10_HUMAN | Unmodified | 0 | 0 | 0 | 0 | 0 | 0 | 0 | 219220 | 0 | 0 | 0 | 0 | 0 | 0 | 0 | 0 | 0 | 0 |
| RLVVFPTDI | L1CAM; L1CAM | P32004; E9F | L1CAM_HUMAN | Unmodified | 0 | 0 | 0 | 0 | 0 | 0 | 0 | 0 | 1059600 | 0 | 0 | 0 | 0 | 0 | 0 | 0 | 0 | 0 |
| RLWADHG | GNAI2 | P04899 | GNAI2_HUMAN | Unmodified | 2590600 | 2607800 | 517110 | 2020400 | 629030 | 469160 | 0 | 14143 | 5977000 | 1122600 | 7215100 | 36477000 | 813230 | 915890 | 2633800 | 1035000 | 1823300 | 2613000 |

| | | | | | | | | | | | | | | | | | | | | | | |
|---|---|---|---|---|---|---|---|---|---|---|---|---|---|---|---|---|---|---|---|---|---|---|
| RLYHLEGF( | PSD2 | Q9BQI7 | PSD2_HUMAN | Unmodified | 0 | 0 | 0 | 0 | 0 | 0 | 0 | 0 | 0 | 0 | 863420 | 240630 | 0 | 0 | 0 | 0 | 0 | 0 |
| RLYWDDLK | TFRC; TFRC | P02786; G3\ | TFR1_HUMAN T | Unmodified | 0 | 329980 | 707700 | 558620 | 1733300 | 261990 | 128560 | 0 | 5822400 | 3475300 | 436430 | 0 | 797420 | 0 | 549690 | 398760 | 0 | 426430 |
| RLYYEGLQ | HGS; HGS; HGS | O14964; A0A | HGS_HUMAN H | Unmodified | 587830 | 1392400 | 103880 | 0 | 332390 | 94407 | 0 | 38370 | 2183600 | 169380 | 237520 | 1109700 | 205890 | 669490 | 446510 | 752690 | 0 | 0 |
| RMAESLRIF | FUT8; FUT8 | Q9BYC5; A0A | FUT8_HUMAN A | Oxidation (M) | 0 | 0 | 0 | 0 | 0 | 0 | 0 | 0 | 1393000 | 84793 | 0 | 0 | 0 | 0 | 0 | 0 | 0 | 0 |
| RMAESLRIF | FUT8; FUT8 | Q9BYC5; A0A | FUT8_HUMAN A | Unmodified | 0 | 0 | 0 | 0 | 0 | 0 | 0 | 0 | 463540 | 0 | 0 | 0 | 0 | 0 | 0 | 0 | 0 | 0 |
| RMAGTAFD | GDI1 | P31150 | GDIA_HUMAN | Oxidation (M) | 0 | 0 | 0 | 188930 | 0 | 0 | 197660 | 141160 | 1276800 | 855320 | 0 | 816900 | 1886300 | 1533400 | 104260 | 0 | 914860 | 1842300 |
| RMAGTAFD | GDI1 | P31150 | GDIA_HUMAN | Unmodified | 0 | 0 | 0 | 0 | 678080 | 0 | 271660 | 0 | 1429100 | 601320 | 0 | 0 | 728030 | 549090 | 0 | 0 | 1147000 | 1861600 |
| RMAGTAFD | GDI1 | P31150 | GDIA_HUMAN | 2 Oxidation (M) | 0 | 0 | 0 | 0 | 0 | 0 | 0 | 155570 | 275700 | 149780 | 0 | 0 | 317730 | 0 | 0 | 0 | 200510 | 249640 |
| RMATEVAA | RPS6; RPS6 | P62753; A2A | RS6_HUMAN 4 | Oxidation (M) | 1855300 | 1437800 | 2979600 | 2391500 | 385630 | 833610 | 444740 | 276110 | 286330 | 0 | 1095900 | 15046000 | 939260 | 373070 | 2664300 | 1618900 | 0 | 0 |
| RMATEVAA | RPS6; RPS6 | P62753; A2A | RS6_HUMAN 4 | Unmodified | 0 | 0 | 48542 | 0 | 0 | 0 | 0 | 0 | 0 | 0 | 0 | 1995900 | 0 | 0 | 0 | 0 | 0 | 0 |
| RMDAPASA | AKR7A2; AKR7A | O43488; H3I | ARK72_HUMAN | Unmodified | 0 | 0 | 0 | 0 | 91793 | 0 | 0 | 0 | 793720 | 62486 | 0 | 428740 | 0 | 0 | 0 | 0 | 40740 | 0 |
| RMDERPIH | CDH2; CDH2; C | P19022; C9J | CADH2_HUMA | Unmodified | 0 | 0 | 0 | 0 | 0 | 0 | 0 | 0 | 799220 | 0 | 0 | 0 | 0 | 0 | 0 | 0 | 230810 | 1131400 |
| RMEELHNC | NONO; NONO; | Q15233; A0A | NONO_HUMAN | Oxidation (M) | 0 | 23803 | 0 | 0 | 0 | 0 | 0 | 0 | 105750 | 0 | 99509 | 295510 | 0 | 0 | 33738 | 0 | 0 | 0 |
| RMEPPASK | SEPTIN9; SEPTI | Q9UHD8; K7 | SEPT9_HUMAN | Unmodified | 0 | 0 | 0 | 0 | 3560500 | 114010 | 0 | 0 | 0 | 0 | 340300 | 0 | 0 | 0 | 0 | 0 | 1291300 | 2496500 |
| RMEPPASK | SEPTIN9; SEPTI | Q9UHD8; K7 | SEPT9_HUMAN | Oxidation (M) | 0 | 0 | 0 | 0 | 297020 | 87491 | 0 | 0 | 0 | 0 | 0 | 86198 | 0 | 0 | 47743 | 45924 | 269010 | 265570 |
| RMESSINH | ABI1; ABI2; ABI2 | Q8IZP0; Q9N | ABI1_HUMAN A | Oxidation (M) | 292660 | 79066 | 251320 | 337640 | 592450 | 174760 | 51099 | 50758 | 416860 | 16180 | 651880 | 1709600 | 828350 | 283100 | 395220 | 414320 | 257550 | 228050 |
| RMESSINH | ABI1; ABI2; ABI2 | Q8IZP0; Q9N | ABI1_HUMAN A | Unmodified | 212120 | 190770 | 249770 | 617200 | 2380900 | 359480 | 151570 | 0 | 2391200 | 447310 | 1185800 | 6306200 | 2520000 | 940580 | 774910 | 710790 | 526010 | 756730 |
| RMESSINH | ABI1; ABI2; ABI2 | Q8IZP0; Q9N | ABI1_HUMAN A | Unmodified | 0 | 0 | 0 | 0 | 1494700 | 0 | 0 | 0 | 0 | 206780 | 0 | 0 | 833380 | 0 | 0 | 0 | 663320 | 330350 |
| RMGESDDS | RPS21; RPS21 | P63220; Q8V | RS21_HUMAN | Unmodified | 568570 | 493850 | 0 | 563860 | 0 | 0 | 0 | 0 | 0 | 0 | 217210 | 3353000 | 80856 | 0 | 0 | 0 | 0 | 0 |
| RMGESDDS | RPS21; RPS21 | P63220; Q8V | RS21_HUMAN | Oxidation (M) | 0 | 0 | 0 | 0 | 0 | 0 | 0 | 0 | 0 | 0 | 0 | 1423600 | 0 | 0 | 0 | 0 | 0 | 0 |
| RMGESDDS | RPS21; RPS21 | P63220; Q8V | RS21_HUMAN | Oxidation (M) | 0 | 0 | 0 | 138730 | 375720 | 95784 | 95415 | 0 | 528590 | 88254 | 104680 | 291280 | 1123800 | 205350 | 134100 | 0 | 0 | 0 |
| RMGPGAAS | RAB1B; RAB1B | Q9H0U4; E9 | RAB1B_HUMAN | Unmodified | 0 | 0 | 630010 | 391590 | 0 | 0 | 0 | 0 | 291650 | 0 | 175890 | 1972700 | 0 | 0 | 885200 | 386290 | 0 | 0 |
| RMGPGATA | RAB1A; RAB1A | P62820; E7E | RAB1A_HUMAN | Unmodified | 0 | 0 | 0 | 0 | 1201000 | 0 | 0 | 94753 | 909240 | 0 | 465270 | 1377300 | 0 | 0 | 0 | 0 | 33172 | 984140 |
| RMIETAQVI | RAB21 | Q9UL25 | RAB21_HUMAN | Oxidation (M) | 228360 | 354760 | 278110 | 417720 | 0 | 0 | 498310 | 0 | 440330 | 64426 | 220900 | 700200 | 172630 | 0 | 276360 | 384950 | 0 | 0 |
| RMIETAQVI | RAB21 | Q9UL25 | RAB21_HUMAN | Unmodified | 0 | 0 | 0 | 0 | 0 | 0 | 0 | 0 | 113550 | 0 | 39834 | 151240 | 0 | 0 | 0 | 0 | 0 | 0 |
| RMIETAQVI | RAB21 | Q9UL25 | RAB21_HUMAN | Unmodified | 185990 | 140820 | 284020 | 681960 | 2920500 | 437150 | 203970 | 77961 | 595340 | 880690 | 222780 | 0 | 999440 | 487950 | 237220 | 135360 | 0 | 1006700 |
| RMMEVAAA | CNDP2; CNDP2 | Q96KP4; J3C | CNDP2_HUMA | Oxidation (M) | 0 | 0 | 0 | 0 | 0 | 0 | 0 | 0 | 0 | 0 | 81317 | 239650 | 407570 | 0 | 0 | 0 | 0 | 0 |
| RMMEVAAA | CNDP2; CNDP2 | Q96KP4; J3C | CNDP2_HUMA | Unmodified | 0 | 0 | 0 | 0 | 218660 | 0 | 0 | 0 | 0 | 0 | 218110 | 747200 | 0 | 701470 | 0 | 0 | 0 | 0 |
| RMNYSDAI | NARS1 | O43776 | SYNC_HUMAN | Oxidation (M) | 0 | 34683 | 0 | 0 | 0 | 0 | 0 | 0 | 162990 | 0 | 0 | 1500600 | 0 | 0 | 574560 | 223020 | 0 | 0 |
| RMQDQLQ | EHD3 | Q9NZN3 | EHD3_HUMAN | Oxidation (M) | 249920 | 79149 | 539240 | 1206500 | 2138400 | 379380 | 3093300 | 4173700 | 0 | 60287 | 0 | 0 | 2307700 | 3275600 | 772870 | 941510 | 371220 | 2124000 |
| RMQDQLQ | EHD3 | Q9NZN3 | EHD3_HUMAN | Unmodified | 282310 | 7365600 | 5467300 | 10141000 | 3393500 | 351600 | 3504100 | 937240 | 0 | 89570 | 0 | 0 | 3618800 | 4743900 | 7748300 | 4596900 | 566570 | 3034500 |
| RMQDQLQ | EHD3 | Q9NZN3 | EHD3_HUMAN | Oxidation (M) | 21091 | 0 | 0 | 0 | 0 | 0 | 4771000 | 3854000 | 0 | 0 | 0 | 0 | 249050 | 856610 | 0 | 0 | 0 | 0 |
| RMQDQLQ | EHD3 | Q9NZN3 | EHD3_HUMAN | Unmodified | 0 | 0 | 0 | 0 | 223950 | 0 | 1043700 | 0 | 0 | 0 | 0 | 0 | 216990 | 745670 | 0 | 0 | 0 | 0 |
| RMQEIEEM | SRPK1; SRPK1; | Q96SB4; D6I | SRPK1_HUMAN | Unmodified | 0 | 74829 | 0 | 0 | 0 | 0 | 0 | 0 | 298060 | 0 | 0 | 476780 | 0 | 0 | 0 | 0 | 0 | 0 |
| RMQSLQP\ | SUSD2 | Q9UGT4 | SUSD2_HUMAI | Oxidation (M) | 389550 | 0 | 0 | 0 | 951750 | 0 | 0 | 0 | 679450 | 0 | 0 | 0 | 0 | 0 | 0 | 0 | 0 | 0 |
| RMTGSEFD | GDI2 | P50395 | GDIB_HUMAN | 2 Oxidation (M) | 0 | 0 | 0 | 690570 | 1461300 | 2040000 | 0 | 0 | 0 | 500680 | 0 | 2619800 | 1360100 | 771140 | 839570 | 923130 | 333930 | 1625300 |
| RMTGSEFD | GDI2 | P50395 | GDIB_HUMAN | Oxidation (M) | 250950 | 0 | 405530 | 836790 | 8046900 | 1051600 | 906640 | 303300 | 2712600 | 1540600 | 853880 | 9230100 | 3798400 | 1313100 | 613010 | 454820 | 1635500 | 3041800 |
| RMTGSEFD | GDI2 | P50395 | GDIB_HUMAN | Unmodified | 0 | 0 | 0 | 0 | 3950300 | 0 | 0 | 0 | 3194600 | 561530 | 0 | 1671500 | 909120 | 450510 | 0 | 0 | 559550 | 2376000 |
| RMYEQQRI | CHMP5 | Q9NZZ3 | CHMP5_HUMA | Oxidation (M) | 0 | 0 | 0 | 0 | 8126300 | 0 | 0 | 0 | 1444400 | 997170 | 0 | 0 | 0 | 0 | 0 | 0 | 0 | 1016900 |
| RMYEQQRI | CHMP5 | Q9NZZ3 | CHMP5_HUMA | Unmodified | 0 | 0 | 0 | 0 | 2532400 | 0 | 0 | 0 | 671370 | 0 | 0 | 0 | 0 | 0 | 0 | 0 | 0 | 0 |
| RMYEQQRI | CHMP5 | Q9NZZ3 | CHMP5_HUMA | 2 Oxidation (M) | 0 | 0 | 0 | 0 | 602160 | 0 | 0 | 0 | 0 | 801130 | 1976300 | 283780 | 1214600 | 528220 | 0 | 0 | 0 | 0 |
| RNCFASVF | ARPC2 | O15144 | ARPC2_HUMAI | Unmodified | 0 | 0 | 0 | 0 | 743990 | 290590 | 309570 | 188290 | 213180 | 236800 | 249880 | 1842900 | 642650 | 483550 | 0 | 0 | 1142300 | 314780 |
| RNDFQLIG | EIF5A; EIF5A; EI | P63241; I3L3 | IF5A1_HUMAN | Unmodified | 0 | 781820 | 0 | 0 | 0 | 0 | 0 | 0 | 0 | 0 | 0 | 0 | 0 | 0 | 896180 | 0 | 0 | 0 |
| RNDSIIVDII | USP15 | Q9Y4E8 | UBP15_HUMAI | Unmodified | 0 | 0 | 0 | 0 | 0 | 0 | 0 | 0 | 0 | 0 | 50091 | 881560 | 355820 | 157810 | 0 | 32219 | 0 | 0 |
| RNDSIYEAS | MVB12A; MVB1 | Q96EY5; M0 | MB12A_HUMAI | Unmodified | 0 | 0 | 0 | 0 | 0 | 433710 | 0 | 0 | 1670500 | 0 | 0 | 0 | 0 | 0 | 0 | 1743600 | 77997 | 0 |
| RNDSIYEAS | MVB12A; MVB1 | Q96EY5; M0 | MB12A_HUMAI | Oxidation (M) | 2465800 | 0 | 0 | 3185900 | 0 | 0 | 1683800 | 0 | 0 | 14237000 | 0 | 0 | 0 | 0 | 5714500 | 1626100 | 0 | 0 |
| RNDSIYEAS | MVB12A; MVB1 | Q96EY5; M0 | MB12A_HUMAI | Oxidation (M) | 1061500 | 0 | 0 | 8186000 | 0 | 0 | 0 | 0 | 0 | 0 | 0 | 0 | 17181000 | 4314100 | 1785000 | 0 | 0 | 0 |
| RNELVIRV( | G6PD | P11413 | G6PD_HUMAN | Unmodified | 0 | 0 | 0 | 0 | 86027000 | 8920400 | 2452300 | 1022200 | 0 | 552790 | 276160 | 0 | 8210500 | 7038200 | 0 | 0 | 12326000 | 24986000 |
| RNENIYQIY | DPY19L4 | Q7Z388 | D19L4_HUMAN | Unmodified | 0 | 0 | 0 | 0 | 81636 | 0 | 40684 | 0 | 942000 | 107270 | 0 | 0 | 0 | 0 | 0 | 0 | 117560 | 0 |
| RNENQLIIF | SF3B3 | Q15393 | SF3B3_HUMAN | Unmodified | 0 | 0 | 0 | 0 | 0 | 0 | 0 | 0 | 0 | 0 | 0 | 2121000 | 0 | 0 | 0 | 0 | 0 | 0 |
| RNEQSLQE | L1RE1 | Q9UN81 | LORF1_HUMAN | Unmodified | 9288300 | 5211900 | 2776100 | 1997200 | 0 | 0 | 0 | 0 | 1126800 | 838820 | 0 | 534920 | 36383 | 900930 | 0 | 2333300 | 0 | 0 |
| RNFDTGLC | OCLN; OCLN | Q16625; A0A | OCLN_HUMAN | Unmodified | 0 | 0 | 965130 | 1519700 | 3186400 | 0 | 0 | 0 | 42785 | 0 | 80770 | 0 | 2372600 | 4715000 | 3994000 | 2383400 | 0 | 303700 |
| RNFILDQC | HNRNPUL2; HN | Q1KMD3; H3 | HNRL2_HUMAI | Unmodified | 0 | 0 | 0 | 0 | 0 | 0 | 0 | 0 | 0 | 0 | 0 | 798550 | 871180 | 0 | 0 | 1908000 | 0 | 0 |
| RNFILDQTI | HNRNPU; HNR | Q00839; A0A | HNRPU_HUMA | Unmodified | 238000 | 109610 | 220370 | 0 | 2234800 | 0 | 2939200 | 1371400 | 0 | 992430 | 804370 | 3372700 | 2107300 | 751890 | 0 | 141740 | 0 | 0 |
| RNFLELTEL | ATP6V0A1; ATP | Q93050; K7E | VPP1_HUMAN | Unmodified | 0 | 0 | 0 | 0 | 0 | 0 | 286140 | 0 | 6329400 | 1111900 | 1032000 | 1678800 | 0 | 0 | 0 | 0 | 0 | 0 |
| RNFMESTP | EPB41L2; EPB4 | O43491; Q6 | E41L2_HUMAN | Unmodified | 0 | 0 | 0 | 0 | 4699800 | 788020 | 0 | 0 | 0 | 0 | 3057700 | 0 | 0 | 0 | 0 | 0 | 1105800 | 5270200 |
| RNFMESTP | EPB41L2; EPB4 | O43491; Q6 | E41L2_HUMAN | Oxidation (M) | 0 | 0 | 0 | 0 | 1590800 | 107920 | 0 | 0 | 0 | 0 | 0 | 0 | 0 | 0 | 0 | 0 | 635950 | 1290700 |
| RNFQEEQII | DNAJC10; DNA | Q8IXB1; A0A | DJC10_HUMAN | Unmodified | 0 | 0 | 0 | 1529400 | 0 | 0 | 0 | 0 | 13219000 | 625090 | 370260 | 0 | 0 | 23432 | 395730 | 605740 | 0 | 0 |
| RNFQEEQII | DNAJC10; DNA | Q8IXB1; A0A | DJC10_HUMAN | Unmodified | 0 | 82865 | 0 | 0 | 0 | 0 | 0 | 0 | 410320 | 3448300 | 0 | 0 | 9655.5 | 0 | 0 | 0 | 0 | 139280 |
| RNGLHLPE | NLN; NLN | Q9BYT8; E9F | NEUL_HUMAN | Unmodified | 0 | 0 | 0 | 0 | 159660 | 0 | 0 | 0 | 152340 | 0 | 0 | 231090 | 0 | 0 | 0 | 0 | 0 | 0 |
| RNHFWELI | MKRN1 | Q9UHC7 | MKRN1_HUMA | Unmodified | 1377800 | 584300 | 0 | 0 | 0 | 0 | 0 | 0 | 0 | 0 | 0 | 0 | 0 | 0 | 0 | 138950 | 0 | 0 |
| RNIGVVVV( | SPTLC2 | O15270 | SPTC2_HUMAN | Unmodified | 0 | 0 | 0 | 0 | 135200 | 0 | 65253 | 15278 | 0 | 174180 | 72587 | 112490 | 0 | 52141 | 0 | 0 | 0 | 52475 |
| RNIIGYFEQ | ERP44 | Q9BS26 | ERP44_HUMAN | Unmodified | 80752 | 62253 | 0 | 0 | 0 | 0 | 252590 | 63402 | 50725000 | 9134800 | 1547700 | 574400 | 0 | 0 | 0 | 122530 | 0 | 803410 |
| RNIPPYFVA | XRCC6; XRCC6 | P12956; B1A | XRCC6_HUMAI | Unmodified | 0 | 0 | 0 | 0 | 644730 | 58937 | 467840 | 0 | 0 | 0 | 693280 | 8537400 | 0 | 578650 | 0 | 0 | 229140 | 0 |
| RNIQQYNS | PRKAR1A | P10644 | KAP0_HUMAN | Unmodified | 0 | 0 | 0 | 0 | 0 | 0 | 143180 | 0 | 202210 | 0 | 21503 | 0 | 0 | 0 | 0 | 0 | 989180 | 945240 |
| RNISHYEE( | PRKAR2A; PRKA | P13861; H7C | KAP2_HUMAN | Unmodified | 25842 | 0 | 106140 | 193830 | 0 | 0 | 142220 | 63884 | 398820 | 215100 | 130550 | 973060 | 2727900 | 1588800 | 474710 | 250870 | 155750 | 0 |
| RNLDIERPI | TUBA3C; TUBA3 | P0DPH7; Q7 | TBA3C_HUMAN | Unmodified | 338900 | 0 | 807200 | 294380 | 170260 | 0 | 469900 | 78240 | 3651800 | 988020 | 535440 | 28291000 | 0 | 0 | 1095200 | 276830 | 70190 | 0 |
| RNLEEDDY | STEAP1 | Q9UHE8 | STEA1_HUMAN | Unmodified | 0 | 0 | 12261000 | 4903600 | 0 | 0 | 0 | 0 | 0 | 0 | 0 | 169490 | 510700 | 1607000 | 1990200 | 11758000 | 0 | 0 |
| RNLEEDDY | STEAP1 | Q9UHE8 | STEA1_HUMAN | Oxidation (M) | 0 | 0 | 0 | 3179700 | 0 | 0 | 0 | 0 | 0 | 0 | 0 | 0 | 3369300 | 1635400 | 0 | 0 | 0 | 0 |
| RNLEEDDY | STEAP1 | Q9UHE8 | STEA1_HUMAN | Unmodified | 0 | 0 | 0 | 0 | 0 | 0 | 0 | 22055 | 0 | 0 | 0 | 0 | 5751500 | 4862600 | 0 | 0 | 0 | 0 |
| RNLGSINTI | SEC22B; SEC22 | O75396; A0A | SC22B_HUMAN | Unmodified | 128580 | 0 | 383260 | 419480 | 0 | 0 | 0 | 0 | 340800 | 98959 | 0 | 1095700 | 268490 | 102780 | 185220 | 58967 | 334200 | 0 |
| RNLNPIFNI | SYT7 | O43581 | SYT7_HUMAN S | Unmodified | 0 | 0 | 0 | 0 | 0 | 0 | 0 | 0 | 564600 | 707590 | 0 | 0 | 0 | 0 | 0 | 0 | 0 | 0 |
| RNLQQELE | KIF23; KIF23; KI | Q02241; A0A | KIF23_HUMAN | Unmodified | 209900 | 0 | 1808100 | 109980 | 2400000 | 0 | 0 | 0 | 1306000 | 223420 | 369770 | 1961200 | 0 | 0 | 264450 | 126200 | 206090 | 0 |
| RNMQTPDI | SGCE; SGCE; S | O43556; A0A | SGCE_HUMAN | Unmodified | 0 | 0 | 0 | 0 | 922090 | 0 | 0 | 0 | 0 | 403550 | 0 | 0 | 0 | 0 | 0 | 0 | 0 | 0 |
| RNPDTQW | RPL15; RPL15; I | P61313; E7E | RL15_HUMAN ( | Unmodified | 5251700 | 5462700 | 3125300 | 3383400 | 0 | 0 | 0 | 46303 | 2513200 | 0 | 1305800 | 30429000 | 0 | 0 | 3009600 | 2778900 | 13298 | 200510 |
| RNQDLAPN | ILF2; ILF2; ILF2; | Q12905; B4I | ILF2_HUMAN Ir | Unmodified | 9245800 | 10382000 | 3802500 | 7351800 | 1219900 | 292260 | 155800 | 258980 | 5078800 | 1961400 | 4130900 | 21988000 | 4831200 | 1564300 | 2382200 | 3068300 | 0 | 636930 |
| RNQELLQS | USO1 | O60763 | USO1_HUMAN | Unmodified | 0 | 0 | 0 | 0 | 146750 | 74080 | 682370 | 79738 | 506680 | 31139 | 168690 | 5103900 | 125470 | 0 | 0 | 61238 | 0 | 152460 |
| RNQSFCPT | RPL27A; RPL27A | P46776; E9P | RL27A_HUMAN | Unmodified | 2846600 | 6263900 | 5333300 | 12754000 | 975680 | 0 | 1612000 | 661490 | 639220 | 165090 | 433640 | 11346000 | 4963200 | 9398100 | 3203700 | 4226000 | 200980 | 894080 |
| RNSDLLLL\ | GOLM2 | Q6P4E1 | GOLM2_HUMA | Unmodified | 0 | 0 | 0 | 0 | 0 | 0 | 0 | 0 | 1127400 | 334250 | 0 | 147270 | 24697 | 0 | 0 | 0 | 0 | 0 |
| RNSDLLLL\ | GOLM2 | Q6P4E1 | GOLM2_HUMA | Unmodified | 0 | 0 | 0 | 0 | 458250 | 0 | 0 | 0 | 8514600 | 1209600 | 0 | 2038200 | 0 | 0 | 0 | 0 | 0 | 0 |
| RNSHTDN\ | ADGRG6 | Q86SQ4 | AGRG6_HUMA | Unmodified | 0 | 0 | 935870 | 2185600 | 28821000 | 6597500 | 0 | 0 | 0 | 0 | 0 | 37371 | 0 | 0 | 3117800 | 5201900 | 0 | 0 |
| RNTFVGTP | STK26; STK26; S | Q9P289; B4E | STK26_HUMAN | Oxidation (M) | 0 | 0 | 0 | 0 | 0 | 0 | 0 | 0 | 0 | 0 | 0 | 3437300 | 0 | 0 | 0 | 0 | 0 | 0 |
| RNTTIPTK | HSPA8; HSPA2; | P11142; P54 | HSP7C_HUMAI | Unmodified | 153470 | 0 | 81744 | 0 | 4853500 | 1329600 | 1187600 | 0 | 153540 | 0 | 0 | 0 | 1435800 | 995550 | 64779 | 45293 | 3750200 | 278200 |
| RNVEGQDI | AP1B1; AP2B1; | Q10567; P63 | AP1B1_HUMAN | Unmodified | 0 | 0 | 0 | 341840 | 0 | 0 | 0 | 0 | 0 | 1080600 | 0 | 2504600 | 1549100 | 1056900 | 0 | 297500 | 1438700 | 1619700 |
| RNVEGQDI | AP1B1; AP2B1; | Q10567; P63 | AP1B1_HUMAN | Oxidation (M) | 0 | 0 | 0 | 170950 | 329460 | 79851 | 0 | 0 | 153170 | 109440 | 0 | 0 | 286500 | 0 | 0 | 210880 | 123010 | 547310 |
| RNVIEAVYI | PPM1A | P35813 | PPM1A_HUMAI | Unmodified | 0 | 0 | 0 | 0 | 0 | 0 | 0 | 0 | 0 | 0 | 0 | 0 | 0 | 0 | 0 | 0 | 0 | 0 |
| RNVIFQPV | PFKP; PFKP; PF | Q01813; A0A | PFKAP_HUMAN | Unmodified | 0 | 0 | 0 | 0 | 7438700 | 2240300 | 0 | 116060 | 943320 | 440870 | 1932300 | 874030 | 592000 | 0 | 0 | 467530 | 1327700 | 3423000 |
| RNVIPETSS | MME; MME; MM | P08473; C9I | NEP_HUMAN N | Unmodified | 0 | 0 | 6575300 | 5014700 | 0 | 0 | 0 | 0 | 2290400 | 120400 | 54809 | 260140 | 1097900 | 269420 | 7414300 | 4830600 | 0 | 0 |
| RNVKLIALS | PRDX6 | P30041 | PRDX6_HUMAI | Unmodified | 0 | 0 | 0 | 0 | 1476600 | 0 | 580850 | 0 | 0 | 21792 | 0 | 0 | 2031600 | 3410400 | 0 | 0 | 10125000 | 6404100 |
| RNVMILTNI | TES; TES | Q9UGI8; F8V | TES_HUMAN Te | Oxidation (M) | 0 | 0 | 0 | 63910 | 178970 | 0 | 0 | 0 | 0 | 0 | 0 | 140210 | 299920 | 205340 | 0 | 0 | 205080 | 256120 |
| RNVMILTNI | TES; TES | Q9UGI8; F8V | TES_HUMAN Te | Unmodified | 0 | 0 | 0 | 0 | 514000 | 0 | 0 | 0 | 124700 | 0 | 0 | 0 | 541430 | 297160 | 0 | 0 | 841470 | 685040 |
| RNVTVQPD | COPB1 | P53618 | COPB_HUMAN | Oxidation (M) | 0 | 0 | 0 | 0 | 0 | 0 | 0 | 0 | 0 | 0 | 74440 | 2424300 | 0 | 39151 | 145950 | 232710 | 0 | 0 |
| RNVTVQPD | COPB1 | P53618 | COPB_HUMAN | Unmodified | 0 | 0 | 0 | 0 | 155470 | 0 | 0 | 0 | 581410 | 45704 | 0 | 2364400 | 0 | 0 | 0 | 0 | 0 | 265000 |
| RNVVAGEV | VIL1; VIL1 | P09327; C9J | VILI_HUMAN Vi | Oxidation (M) | 230780 | 0 | 0 | 0 | 108140 | 0 | 0 | 0 | 161020 | 0 | 0 | 453350 | 0 | 0 | 0 | 0 | 0 | 0 |
| RNVVAGEV | VIL1; VIL1 | P09327; C9J | VILI_HUMAN Vi | Unmodified | 0 | 0 | 0 | 0 | 109270 | 0 | 0 | 0 | 111630 | 0 | 0 | 1354500 | 191830 | 0 | 0 | 0 | 55656 | 0 |
| RNYETVPC | ADGRG6 | Q86SQ4 | AGRG6_HUMA | Unmodified | 0 | 0 | 161890 | 244090 | 461490 | 257680 | 0 | 0 | 0 | 22415 | 146480 | 267460 | 0 | 0 | 132880 | 215820 | 0 | 62865 |
| RPAEAEAH | FLOT2; FLOT2; I | Q14254; J3C | FLOT2_HUMAN | Unmodified | 646230 | 880370 | 689010 | 8037200 | 4127700 | 0 | 0 | 0 | 17066000 | 1776000 | 1714400 | 7020100 | 0 | 0 | 3627900 | 1914900 | 0 | 0 |
| RPAEDMEE | HNRNPK; HNRI | P61978; Q5T | HNRPK_HUMA | Unmodified | 81989 | 0 | 0 | 0 | 0 | 0 | 0 | 0 | 0 | 0 | 292170 | 1343600 | 0 | 0 | 0 | 0 | 0 | 0 |
| RPAEDMEE | HNRNPK; HNRI | P61978; Q5T | HNRPK_HUMA | Unmodified | 0 | 0 | 259550 | 0 | 434500 | 0 | 0 | 0 | 0 | 0 | 742160 | 461900 | 0 | 0 | 0 | 162580 | 0 | 0 |
| RPAILTYHD | NDRG2; NDRG2 | Q9UN36; G3 | NDRG2_HUMA | Unmodified | 433130 | 1135800 | 588340 | 2211400 | 2107400 | 162200 | 1486300 | 303570 | 0 | 93244 | 790450 | 7324500 | 891720 | 1162500 | 1055200 | 1503100 | 107340 | 582480 |
| RPAMLDNE | USP7; USP7; US | Q93009; A0A | UBP7_HUMAN | Unmodified | 0 | 0 | 0 | 0 | 143140 | 0 | 0 | 0 | 824330 | 47666 | 0 | 417830 | 0 | 0 | 17274 | 17107 | 0 | 0 |
| RPAYGDVT | SEZ6; SEZ6; SE | Q53EL9; H0Y | SEZ6_HUMAN S | Unmodified | 0 | 0 | 0 | 0 | 0 | 0 | 0 | 23537 | 11214000 | 102930 | 0 | 0 | 0 | 0 | 0 | 0 | 42783 | 0 |
| RPDADLKT | CANX; CANX; C | P27824; A0A | CALX_HUMAN | Unmodified | 0 | 0 | 0 | 0 | 0 | 0 | 0 | 0 | 0 | 300630 | 0 | 0 | 0 | 0 | 0 | 0 | 0 | 0 |
| RPDDTAFQ | SUN1; SUN1; S | O94901; H0Y | SUN1_HUMAN | Unmodified | 0 | 0 | 0 | 0 | 0 | 0 | 0 | 0 | 5841300 | 57498 | 93734 | 0 | 0 | 0 | 0 | 0 | 0 | 0 |
| RPDEGWE | SUN2; SUN2; S | Q9UH99; HC | SUN2_HUMAN | Unmodified | 0 | 0 | 0 | 0 | 0 | 0 | 51074 | 0 | 739020 | 11207 | 0 | 0 | 0 | 0 | 0 | 0 | 0 | 0 |
| RPDEGWE | SUN2 | Q9UH99 | SUN2_HUMAN | Unmodified | 0 | 0 | 0 | 0 | 0 | 0 | 0 | 0 | 1248500 | 0 | 0 | 0 | 0 | 0 | 0 | 0 | 0 | 0 |
| RPDQQLQ( | SERBP1 | Q8NC51 | PAIRB_HUMAN | Unmodified | 0 | 0 | 0 | 0 | 251260 | 0 | 0 | 0 | 0 | 0 | 0 | 1831600 | 0 | 0 | 0 | 0 | 0 | 118210 |
| RPEAAQLLI | CARS1; CARS1 | P49589; B4D | SYCC_HUMAN | Unmodified | 0 | 0 | 0 | 0 | 0 | 0 | 0 | 0 | 404310 | 0 | 0 | 3886400 | 485280 | 0 | 733220 | 0 | 0 | 0 |
| RPEDQELE | CHGA; CHGA | P10645; G5E | CMGA_HUMAN | Unmodified | 0 | 0 | 0 | 0 | 0 | 0 | 0 | 0 | 2142600 | 1715100 | 0 | 371260 | 13581 | 0 | 0 | 0 | 0 | 0 |

| | | | | | | | | | | | | | | | | | | | | | | |
|---|---|---|---|---|---|---|---|---|---|---|---|---|---|---|---|---|---|---|---|---|---|---|
| RPEEVCGP | HSPG2 | P98160 | PGBM_HUMAN | Unmodified | 0 | 0 | 491030 | 0 | 0 | 0 | 0 | 0 | 253500 | 0 | 0 | 0 | 0 | 0 | 0 | 230170 | 0 | 139730 |
| RPEILKQEII | AP2B1; AP2B1; | P63010; A0A | AP2B1_HUMAN | Unmodified | 0 | 0 | 0 | 0 | 0 | 0 | 0 | 0 | 135140 | 64872 | 87815 | 1944400 | 0 | 0 | 264080 | 306970 | 32826 | 0 |
| RPEIPTEQS | EMC1 | Q8N766 | EMC1_HUMAN | Unmodified | 0 | 0 | 0 | 0 | 0 | 0 | 0 | 0 | 38793000 | 1730200 | 405090 | 498040 | 0 | 0 | 0 | 0 | 0 | 0 |
| RPEKPVPPI | CD2AP | Q9Y5K6 | CD2AP_HUMAI | Unmodified | 6122200 | 3674900 | 4975000 | 4555900 | 615140 | 0 | 0 | 0 | 0 | 0 | 408740 | 998310 | 2664000 | 433060 | 7576300 | 4220100 | 0 | 0 |
| RPELLTHST | GTF2I; GTF2I; G | P78347; A0A | GTF2I_HUMAN | Unmodified | 0 | 0 | 0 | 0 | 0 | 0 | 0 | 0 | 135390 | 0 | 90916 | 477010 | 138440 | 0 | 0 | 0 | 0 | 0 |
| RPELPDLAI | B4GALNT1; B4( | Q00973; F8V | B4GN1_HUMAI | Unmodified | 0 | 0 | 0 | 0 | 0 | 0 | 0 | 0 | 2371300 | 0 | 0 | 0 | 0 | 0 | 0 | 0 | 0 | 0 |
| RPEQAVEA | LLGL1; LLGL1 | Q15334; A0A | L2GL1_HUMAN | Unmodified | 101710 | 0 | 0 | 0 | 299410 | 0 | 0 | 358930 | 63156 | 0 | 0 | 434690 | 0 | 0 | 356350 | 458240 | 256610 | 0 |
| RPESALLG( | VGF; VGF | O15240; Q0! | VGF_HUMAN N | Unmodified | 270280 | 259820 | 0 | 0 | 0 | 0 | 28028 | 0 | 4997500 | 676620 | 0 | 0 | 0 | 0 | 0 | 0 | 120050 | 90608 |
| RPGCFLCR | PRXL2A | Q9BRX8 | PXL2A_HUMAN | Unmodified | 0 | 0 | 0 | 0 | 0 | 0 | 0 | 0 | 14575000 | 0 | 0 | 0 | 0 | 0 | 0 | 0 | 0 | 0 |
| RPGGEPSP | FN1; FN1 | P02751; H0Y | FINC_HUMAN I | Unmodified | 0 | 0 | 13508000 | 4278900 | 0 | 0 | 0 | 0 | 0 | 0 | 0 | 0 | 0 | 0 | 98972 | 5304600 | 0 | 0 |
| RPGGQHG | MOGS; MOGS; | Q13724; A0A | MOGS_HUMAN | Unmodified | 0 | 58512 | 0 | 0 | 0 | 0 | 40865 | 20599 | 1941500 | 0 | 85530 | 0 | 0 | 0 | 0 | 0 | 0 | 0 |
| RPGHSLEA | HIP1R | O75146 | HIP1R_HUMAN | Unmodified | 0 | 0 | 77482 | 0 | 0 | 0 | 0 | 0 | 0 | 0 | 98218 | 0 | 0 | 558450 | 0 | 143560 | 0 | 0 |
| RPHDYQPL | U2AF2; U2AF2 | P26368; K7E | U2AF2_HUMAN | Oxidation (M) | 1764900 | 520110 | 0 | 0 | 0 | 0 | 0 | 0 | 0 | 0 | 0 | 3769700 | 0 | 0 | 0 | 0 | 0 | 0 |
| RPHETGGY | FN1 | P02751 | FINC_HUMAN I | Oxidation (M) | 0 | 0 | 230280 | 0 | 0 | 0 | 0 | 25575 | 0 | 0 | 110210 | 0 | 71373 | 0 | 0 | 0 | 0 | 0 |
| RPHFPQFS | AKT1; AKT1 | P31749; G3V | AKT1_HUMAN I | Unmodified | 0 | 0 | 1222500 | 0 | 0 | 0 | 0 | 0 | 0 | 0 | 0 | 0 | 0 | 0 | 920800 | 1742800 | 99755 | 0 |
| RPHKEEEEI | LSR; LSR | Q86X29; S4F | LSR_HUMAN Li | Unmodified | 127690 | 0 | 0 | 222240 | 0 | 0 | 0 | 0 | 2940400 | 0 | 0 | 0 | 0 | 0 | 285070 | 317580 | 0 | 0 |
| RPHLSVILV | MTHFD2; MTHF | P13995; B8Z | MTDC_HUMAN | Unmodified | 0 | 0 | 0 | 0 | 0 | 0 | 0 | 0 | 872100 | 0 | 0 | 0 | 0 | 0 | 0 | 0 | 0 | 0 |
| RPHPQHS( | MAP4K4; MAP4 | O95819; C9J | M4K4_HUMAN | Unmodified | 0 | 0 | 35997 | 110510 | 0 | 0 | 0 | 0 | 99003 | 0 | 91479 | 105540 | 0 | 0 | 54196 | 26963 | 0 | 0 |
| RPHVDEFL | CTDSP1; CTDSI | Q9GZU7; O1 | CTDS1_HUMAI | Unmodified | 1116800 | 601380 | 540640 | 287790 | 0 | 0 | 50058 | 55919 | 0 | 33823 | 145350 | 772770 | 0 | 0 | 743360 | 763370 | 0 | 0 |
| RPIFITFEG( | SIGIRR; SIGIRR; | Q6IA17; C9J | SIGIR_HUMAN | Unmodified | 230040 | 230880 | 343580 | 0 | 0 | 0 | 0 | 0 | 0 | 0 | 0 | 1226900 | 0 | 0 | 609260 | 530680 | 0 | 0 |
| RPILQSLLL | EMC1 | Q8N766 | EMC1_HUMAN | Unmodified | 0 | 0 | 0 | 0 | 0 | 0 | 0 | 0 | 3543500 | 271690 | 27434 | 0 | 0 | 0 | 0 | 0 | 0 | 0 |
| RPISADSAI | CLTC; CLTC; CI | Q00610; J3K | CLH1_HUMAN | Oxidation (M) | 1959300 | 225760 | 1530100 | 0 | 0 | 0 | 0 | 59934 | 0 | 0 | 0 | 1510700 | 0 | 0 | 156050 | 817980 | 0 | 0 |
| RPISADSAI | CLTC; CLTC; CI | Q00610; J3K | CLH1_HUMAN | Unmodified | 468950 | 232990 | 474710 | 0 | 0 | 0 | 0 | 0 | 44246 | 0 | 0 | 1782200 | 184180 | 0 | 207420 | 315280 | 0 | 0 |
| RPKFADIVS | EPHA2 | P29317 | EPHA2_HUMAI | Unmodified | 0 | 0 | 0 | 0 | 0 | 0 | 0 | 0 | 0 | 67052 | 0 | 0 | 0 | 0 | 0 | 0 | 254900 | 92660 |
| RPLDDGV( | FUBP3 | Q96I24 | FUBP3_HUMAI | Unmodified | 0 | 0 | 0 | 0 | 0 | 0 | 0 | 0 | 0 | 0 | 361330 | 1009900 | 0 | 0 | 0 | 0 | 0 | 0 |
| RPLEQQQF | LASP1; LASP1; I | Q14847; C9J | LASP1_HUMAN | Unmodified | 252970 | 0 | 579880 | 0 | 0 | 0 | 0 | 0 | 0 | 0 | 0 | 0 | 0 | 0 | 0 | 0 | 0 | 0 |
| RPLHSAQA | RNPEP; RNPEP | Q9H4A4; A6I | AMPB_HUMAN | Unmodified | 0 | 0 | 0 | 0 | 0 | 0 | 0 | 0 | 412220 | 0 | 218060 | 6220000 | 0 | 0 | 0 | 0 | 0 | 0 |
| RPLILQLVH | DNM1L; DNM1I | O00429; F8V | DNM1L_HUMA | Unmodified | 0 | 0 | 0 | 0 | 0 | 0 | 0 | 0 | 318730 | 0 | 274150 | 8514900 | 0 | 0 | 87627 | 151720 | 0 | 0 |
| RPLNPLAS( | ERO1A | Q96HE7 | ERO1A_HUMAI | Unmodified | 0 | 0 | 0 | 0 | 0 | 0 | 0 | 0 | 418420 | 0 | 0 | 0 | 0 | 0 | 0 | 0 | 0 | 0 |
| RPLRPQVV | APMAP | Q9HDC9 | APMAP_HUMAI | Unmodified | 0 | 0 | 0 | 151060 | 0 | 0 | 0 | 0 | 2625900 | 112090 | 0 | 0 | 0 | 0 | 116460 | 137610 | 0 | 0 |
| RPLSIQDSI | SEMA4B | Q9NPR2 | SEM4B_HUMAI | Unmodified | 0 | 0 | 0 | 0 | 3160100 | 0 | 0 | 0 | 0 | 0 | 0 | 0 | 0 | 0 | 0 | 0 | 0 | 0 |
| RPLSSSHE | PHACTR4; PHA( | Q8IZ21; A0A | PHAR4_HUMAI | Unmodified | 0 | 0 | 93006 | 0 | 0 | 0 | 0 | 0 | 295070 | 0 | 0 | 0 | 0 | 0 | 81562 | 0 | 0 | 0 |
| RPLTAATLF | MYO1D; MYO1I | O94832; J3Q | MYO1D_HUMA | Unmodified | 602880 | 552590 | 973390 | 164160 | 0 | 0 | 0 | 0 | 0 | 0 | 0 | 1254700 | 0 | 0 | 1140200 | 530610 | 0 | 0 |
| RPLWESGF | GUSB; GUSB; C | P08236; F8V | BGLR_HUMAN | Oxidation (M) | 0 | 0 | 0 | 0 | 0 | 0 | 0 | 0 | 1442500 | 0 | 0 | 0 | 0 | 0 | 0 | 0 | 0 | 0 |
| RPMEEDGI | ILF3 | Q12906 | ILF3_HUMAN Ir | Unmodified | 0 | 0 | 0 | 0 | 0 | 0 | 0 | 0 | 0 | 0 | 0 | 417950 | 0 | 0 | 0 | 0 | 0 | 0 |
| RPNQSQPL | NCSTN; NCSTN | Q92542; H0Y | NICA_HUMAN | Unmodified | 0 | 0 | 0 | 0 | 0 | 0 | 0 | 0 | 205880 | 0 | 0 | 0 | 0 | 0 | 0 | 0 | 47141 | 25303 |
| RPPENEAP | PIGT; PIGT; PIG | Q969N2; A0A | PIGT_HUMAN ( | Unmodified | 0 | 0 | 0 | 0 | 0 | 0 | 0 | 0 | 768610 | 156880 | 0 | 0 | 0 | 0 | 0 | 0 | 0 | 0 |
| RPPEVFEQ | UBE2O; UBE2O | Q9C0C9; K7 | UBE2O_HUMAI | Unmodified | 0 | 0 | 0 | 0 | 0 | 0 | 0 | 0 | 0 | 0 | 0 | 1953000 | 0 | 0 | 139550 | 0 | 0 | 0 |
| RPPGREAY | ATP5F1A | P25705 | ATPA_HUMAN / | Unmodified | 0 | 0 | 0 | 0 | 0 | 0 | 0 | 0 | 1957000 | 511020 | 0 | 0 | 0 | 0 | 0 | 0 | 0 | 0 |
| RPPGVLHC | CNP | P09543 | CN37_HUMAN | Unmodified | 0 | 0 | 0 | 0 | 0 | 0 | 0 | 0 | 0 | 186360 | 9605000 | 24394000 | 1031700 | 0 | 1803600 | 0 | 395220 | 621890 |
| RPPPAPAQ | B3GAT3; B3GAT | O94766; E9F | B3GA3_HUMAI | Unmodified | 0 | 0 | 0 | 0 | 0 | 0 | 0 | 0 | 8875400 | 762980 | 0 | 1465000 | 0 | 0 | 0 | 0 | 0 | 0 |
| RPPQPIQQ | ADAM10 | O14672 | ADA10_HUMAI | Unmodified | 478870 | 233300 | 679060 | 1107900 | 0 | 0 | 35921 | 232660 | 498210 | 75639 | 592930 | 937410 | 202830 | 0 | 2151100 | 1211800 | 0 | 0 |
| RPPSAFFLF | HMGB1; HMGB | P09429; Q5T | HMGB1_HUMA | Unmodified | 0 | 0 | 0 | 0 | 0 | 0 | 454460 | 0 | 0 | 0 | 992700 | 4897800 | 0 | 0 | 0 | 0 | 0 | 0 |
| RPPSPIKFD | UBA6; UBA6 | A0AVT1; H0Y | UBA6_HUMAN | Unmodified | 0 | 0 | 0 | 0 | 0 | 0 | 0 | 0 | 0 | 0 | 0 | 7195300 | 0 | 0 | 0 | 0 | 0 | 731570 |
| RPQYSNPP | YBX1; YBX1; YB | P67809; C9J | YBOX1_HUMAI | Oxidation (M) | 1960300 | 1310500 | 2423600 | 2302100 | 0 | 0 | 0 | 0 | 0 | 0 | 1070800 | 3432200 | 0 | 0 | 709040 | 1663900 | 0 | 0 |
| RPQYSNPP | YBX1; YBX1; YB | P67809; C9J | YBOX1_HUMAI | Unmodified | 3554700 | 2798200 | 3448200 | 4511700 | 0 | 0 | 0 | 0 | 417310 | 0 | 2212600 | 8436300 | 0 | 0 | 2008100 | 3008900 | 0 | 0 |
| RPSENLGQ | TM9SF2 | Q99805 | TM9S2_HUMAI | Unmodified | 581140 | 347550 | 337300 | 0 | 0 | 0 | 0 | 0 | 13045000 | 5596000 | 251490 | 13569000 | 0 | 0 | 378030 | 777900 | 0 | 0 |
| RPSENLGQ | TM9SF2 | Q99805 | TM9S2_HUMAI | Unmodified | 0 | 0 | 0 | 0 | 0 | 0 | 0 | 0 | 28724000 | 330430 | 1157500 | 5503500 | 0 | 623980 | 0 | 0 | 92425 | 72198 |
| RPSPDEPM | CHORDC1; CH( | Q9UHD1; E9 | CHRD1_HUMA | Oxidation (M) | 0 | 0 | 0 | 0 | 0 | 0 | 0 | 0 | 0 | 0 | 250490 | 1049200 | 0 | 0 | 0 | 0 | 0 | 0 |
| RPSPDEPM | CHORDC1; CH( | Q9UHD1; E9 | CHRD1_HUMA | Unmodified | 0 | 0 | 0 | 0 | 138950 | 0 | 0 | 0 | 0 | 0 | 0 | 1503400 | 0 | 0 | 0 | 181010 | 126860 | 152380 |
| RPTELLSNI | CAPN1; CAPN1 | P07384; E9P | CAN1_HUMAN | Unmodified | 362830 | 0 | 0 | 0 | 0 | 0 | 0 | 0 | 2091100 | 0 | 108320 | 3087400 | 0 | 0 | 0 | 43205 | 0 | 0 |
| RPTFQQLL! | BMX | P51813 | BMX_HUMAN ( | Unmodified | 0 | 0 | 0 | 0 | 0 | 0 | 0 | 0 | 0 | 0 | 1450700 | 8501400 | 0 | 0 | 0 | 0 | 0 | 0 |
| RPTPNDDT | HMGCS1 | Q01581 | HMCS1_HUMA | Unmodified | 0 | 0 | 0 | 0 | 0 | 0 | 0 | 0 | 1387300 | 0 | 0 | 0 | 0 | 358940 | 0 | 0 | 0 | 926640 |
| RPTPQDSP | FASN; FASN | P49327; A0A | FAS_HUMAN Fi | Unmodified | 1718100 | 1105700 | 5451600 | 1465900 | 0 | 0 | 0 | 0 | 976580 | 0 | 3822900 | 1.34E+08 | 0 | 0 | 7797000 | 5687100 | 0 | 0 |
| RPVGGFLL | QTRT2 | Q9H974 | QTRT2_HUMAI | Unmodified | 598090 | 389650 | 87771 | 0 | 0 | 0 | 0 | 0 | 676550 | 0 | 239110 | 1315800 | 0 | 0 | 77664 | 0 | 0 | 48976 |
| RPVPPEVA( | CC2D1A; CC2D | Q6P1N0; A0 | C2D1A_HUMAI | Unmodified | 0 | 0 | 0 | 107200 | 373670 | 0 | 0 | 0 | 733850 | 529560 | 0 | 0 | 0 | 0 | 347820 | 0 | 0 | 272190 |
| RPVPPEVA( | CC2D1A; CC2D | Q6P1N0; A0 | C2D1A_HUMAI | Oxidation (M) | 190160 | 0 | 0 | 256340 | 0 | 0 | 0 | 0 | 107450 | 0 | 0 | 0 | 0 | 0 | 285260 | 121970 | 0 | 0 |
| RPVVEQAL | PLXNB2 | O15031 | PLXB2_HUMAN | Unmodified | 531000 | 448860 | 821000 | 655520 | 0 | 0 | 277960 | 0 | 1012200 | 186980 | 121370 | 2069700 | 0 | 0 | 4128200 | 2551700 | 0 | 0 |
| RPYFTVDE/ | NECTIN1 | Q15223 | NECT1_HUMAI | Unmodified | 798240 | 580620 | 658150 | 0 | 0 | 0 | 0 | 0 | 307040 | 0 | 846990 | 6429900 | 0 | 0 | 811300 | 626090 | 0 | 0 |
| RQAHLCVL | RPS12 | P25398 | RS12_HUMAN | Oxidation (M) | 734220 | 0 | 0 | 375170 | 0 | 0 | 148110 | 0 | 0 | 0 | 0 | 4059200 | 911750 | 0 | 0 | 0 | 117210 | 0 |
| RQAHLCVL | RPS12 | P25398 | RS12_HUMAN | Unmodified | 0 | 0 | 0 | 0 | 0 | 0 | 0 | 0 | 0 | 0 | 0 | 1790300 | 0 | 0 | 0 | 0 | 189310 | 0 |
| RQAQQERI | MYH9 | P35579 | MYH9_HUMAN | Unmodified | 1143600 | 1470400 | 4423900 | 3235300 | 9284000 | 5121900 | 542550 | 75977 | 4894100 | 2367300 | 2462000 | 28351000 | 2594500 | 1239600 | 8989100 | 5733700 | 1059200 | 2232000 |
| RQAQQMV | FN1 | P02751 | FINC_HUMAN I | Oxidation (M) | 0 | 0 | 2324200 | 0 | 0 | 0 | 0 | 0 | 0 | 0 | 0 | 0 | 0 | 31548 | 0 | 0 | 0 | 0 |
| RQAQQMV | FN1 | P02751 | FINC_HUMAN I | Unmodified | 0 | 0 | 1652300 | 106640 | 930710 | 0 | 0 | 0 | 0 | 0 | 0 | 0 | 0 | 0 | 0 | 0 | 0 | 80093 |
| RQAQQMV | FN1 | P02751 | FINC_HUMAN I | Oxidation (M) | 0 | 0 | 2091900 | 596490 | 0 | 0 | 0 | 0 | 0 | 0 | 0 | 0 | 0 | 0 | 0 | 294210 | 0 | 780560 |
| RQAQQMV | FN1 | P02751 | FINC_HUMAN I | Unmodified | 0 | 0 | 2830500 | 0 | 0 | 0 | 0 | 0 | 0 | 0 | 0 | 0 | 0 | 1773700 | 0 | 0 | 0 | 3461000 |
| RQAQSPA( | PPFIA1; PPFIA1 | Q13136; E9F | LIPA1_HUMAN | Unmodified | 0 | 0 | 0 | 521100 | 5635700 | 791900 | 0 | 0 | 1507300 | 967110 | 0 | 843550 | 1900600 | 0 | 621000 | 476040 | 2839100 | 2748200 |
| RQAQSPS( | PPFIA2; PPFIA4 | O75334; O7! | LIPA2_HUMAN | Unmodified | 0 | 0 | 0 | 0 | 0 | 0 | 0 | 0 | 0 | 0 | 0 | 0 | 2599500 | 0 | 0 | 328000 | 41827 | 0 |
| RQATTIIADI | SSBP1 | Q04837 | SSBP_HUMAN | Unmodified | 0 | 0 | 0 | 0 | 0 | 0 | 0 | 0 | 8552500 | 2236900 | 0 | 1240800 | 0 | 0 | 0 | 0 | 0 | 0 |
| RQAVDVSP | RPS5; RPS5; RP | P46782; M0I | RS5_HUMAN 4 | Unmodified | 223660 | 93737 | 0 | 0 | 0 | 0 | 0 | 0 | 106230 | 0 | 109830 | 1718700 | 86096 | 0 | 0 | 70213 | 0 | 46781 |
| RQAVTNPN | HSPA9; HSPA9; | P38646; H0Y | GRP75_HUMAI | Unmodified | 76172 | 41954 | 82589 | 112010 | 2964600 | 421700 | 1955100 | 32172 | 12534000 | 2872700 | 394840 | 631790 | 1035400 | 613760 | 63214 | 31390 | 468810 | 1245800 |
| RQDIKSDG | CAPN2 | P17655 | CAN2_HUMAN | Unmodified | 0 | 0 | 0 | 0 | 77672 | 0 | 54022 | 0 | 0 | 0 | 0 | 0 | 0 | 0 | 509970 | 0 | 146100 | 316360 |
| RQDPGDN | CDV3; CDV3; C | Q9UKY7; D6 | CDV3_HUMAN | Oxidation (M) | 0 | 0 | 89728 | 90759 | 317200 | 0 | 0 | 0 | 85609 | 0 | 102150 | 368400 | 448430 | 93951 | 329310 | 125580 | 0 | 85106 |
| RQDPTLLW | CACNA2D2; CA | Q9NY47; C9 | CA2D2_HUMAI | Unmodified | 0 | 0 | 0 | 0 | 53104 | 0 | 0 | 0 | 744980 | 0 | 0 | 0 | 0 | 0 | 0 | 0 | 0 | 54333 |
| RQDSEDH! | SPINT2; SPINT2 | O43291; K7E | SPIT2_HUMAN | Unmodified | 0 | 0 | 0 | 0 | 0 | 0 | 0 | 0 | 33640000 | 1488500 | 0 | 0 | 0 | 0 | 0 | 0 | 0 | 0 |
| RQDSEDH! | SPINT2; SPINT2 | O43291; K7E | SPIT2_HUMAN | Oxidation (M) | 0 | 0 | 0 | 0 | 0 | 0 | 0 | 0 | 10594000 | 0 | 0 | 0 | 0 | 0 | 0 | 0 | 0 | 0 |
| RQEEAALL! | ACE; ACE; ACE | P12821; A0A | ACE_HUMAN A | Unmodified | 2819700 | 0 | 0 | 0 | 0 | 0 | 0 | 24838 | 3405300 | 0 | 0 | 0 | 0 | 0 | 0 | 0 | 0 | 0 |
| RQEEEEVV | YTHDF1 | Q9BYJ9 | YTHD1_HUMAI | Unmodified | 0 | 0 | 130770 | 106190 | 0 | 0 | 66791 | 55659 | 0 | 0 | 30990 | 180280 | 425300 | 0 | 0 | 128460 | 0 | 0 |
| RQEELNN( | PLCG2; PLCG2 | P16885; A0A | PLCG2_HUMAI | Unmodified | 0 | 0 | 0 | 0 | 0 | 0 | 0 | 0 | 0 | 0 | 603410 | 6994700 | 0 | 0 | 0 | 0 | 0 | 0 |
| RQELDAFL/ | FAM120A; FAM: | Q9NZB2; A0 | F120A_HUMAN | Unmodified | 150090 | 0 | 134700 | 0 | 0 | 0 | 0 | 0 | 0 | 0 | 0 | 2877600 | 0 | 0 | 0 | 0 | 0 | 0 |
| RQEMDSQ' | HIP1 | O00291 | HIP1_HUMAN I | Oxidation (M) | 0 | 0 | 0 | 0 | 0 | 0 | 78853 | 0 | 0 | 0 | 0 | 0 | 700680 | 0 | 0 | 0 | 0 | 0 |
| RQENEQLL | KRTCAP3; KRT( | Q53RY4; A0 | KCP3_HUMAN | Unmodified | 2557000 | 1892300 | 4187200 | 3029200 | 0 | 0 | 0 | 0 | 380360 | 0 | 1476500 | 5384200 | 241420 | 0 | 5260500 | 4828100 | 0 | 0 |
| RQENEQLL | KRTCAP3; KRT( | Q53RY4; A0 | KCP3_HUMAN | Unmodified | 0 | 0 | 45564 | 575090 | 746320 | 0 | 0 | 0 | 0 | 0 | 370540 | 0 | 432840 | 348140 | 97518 | 0 | 0 | 0 |
| RQFASQAN | ACTN4; ACTN4; | O43707; F5C | ACTN4_HUMAI | Unmodified | 0 | 0 | 267780 | 0 | 239520 | 491340 | 0 | 0 | 896760 | 155050 | 0 | 13929000 | 24258 | 0 | 741500 | 977930 | 77421 | 0 |
| RQFVQSAK | TLN1 | Q9Y490 | TLN1_HUMAN | Unmodified | 0 | 0 | 0 | 0 | 517670 | 0 | 4413300 | 209460 | 0 | 0 | 0 | 0 | 249750 | 2713600 | 0 | 0 | 2344900 | 0 |
| RQGEVTVL | RELL1 | Q8IUW5 | RELL1_HUMAN | Unmodified | 0 | 1843600 | 242990 | 459420 | 922030 | 363440 | 25155 | 0 | 1253900 | 0 | 87455 | 0 | 639120 | 172250 | 0 | 663630 | 0 | 0 |
| RQGGLGPI | PRDX4; PRDX4; | Q13162; A6N | PRDX4_HUMAI | Unmodified | 0 | 0 | 0 | 0 | 0 | 0 | 0 | 0 | 5754000 | 0 | 100380 | 0 | 0 | 0 | 0 | 0 | 0 | 0 |
| RQGVQVQ' | FASN; FASN | P49327; A0A | FAS_HUMAN Fi | Unmodified | 0 | 65706 | 151550 | 729150 | 0 | 0 | 0 | 0 | 0 | 0 | 1092100 | 13639000 | 0 | 0 | 912250 | 706510 | 0 | 0 |
| RQHSSQD' | NSFL1C; NSFL1 | Q9UNZ2; F2 | NSF1C_HUMAI | Unmodified | 137480 | 145850 | 96848 | 0 | 0 | 0 | 0 | 0 | 211500 | 0 | 39729 | 625020 | 0 | 0 | 0 | 123370 | 0 | 150100 |
| RQIAQLKQI | CSGALNACT2 | Q8N6G5 | CGAT2_HUMAI | Unmodified | 0 | 118660 | 740860 | 737130 | 0 | 0 | 0 | 0 | 1140700 | 0 | 0 | 0 | 0 | 0 | 256090 | 118830 | 0 | 0 |
| RQIIVEEAK | CFL2; CFL2 | Q9Y281; F8V | COF2_HUMAN | Unmodified | 150780 | 0 | 0 | 84198 | 1367200 | 0 | 0 | 0 | 0 | 0 | 0 | 309530 | 89818 | 0 | 257490 | 163840 | 29747 | 0 |
| RQITIEEGI | RAB6B; RAB6B | Q9NRW1; J3 | RAB6B_HUMAN | Unmodified | 150980 | 0 | 89072 | 134050 | 126250 | 0 | 0 | 0 | 493020 | 79112 | 35025 | 1611900 | 159510 | 37790 | 50359 | 159570 | 0 | 0 |
| RQITQNTD' | GLG1 | Q92896 | GSLG1_HUMAI | Unmodified | 0 | 0 | 0 | 0 | 0 | 0 | 0 | 0 | 0 | 0 | 0 | 0 | 0 | 0 | 0 | 0 | 0 | 0 |
| RQIYVAAFT | FOLH1 | Q04609 | FOLH1_HUMAI | Unmodified | 0 | 0 | 0 | 1898300 | 0 | 0 | 0 | 0 | 0 | 0 | 0 | 0 | 0 | 0 | 1984900 | 0 | 0 | 0 |
| RQLDPGG( | ITGA3 | P26006 | ITA3_HUMAN Ir | Unmodified | 0 | 0 | 0 | 0 | 0 | 0 | 0 | 0 | 0 | 0 | 0 | 0 | 0 | 0 | 0 | 0 | 150530 | 897520 |
| RQLEDGD( | KHSRP; KHSRP | Q92945; M0I | FUBP2_HUMAI | Unmodified | 0 | 0 | 0 | 0 | 0 | 0 | 0 | 0 | 129030 | 134130 | 36053 | 116600 | 0 | 0 | 0 | 0 | 0 | 0 |
| RQLEEAEEI | MYH9 | P35579 | MYH9_HUMAN | Unmodified | 186380 | 193690 | 185070 | 211230 | 0 | 0 | 0 | 0 | 269940 | 234040 | 0 | 3057200 | 158510 | 0 | 443530 | 335640 | 69677 | 0 |
| RQLEEAEEI | MYH9 | P35579 | MYH9_HUMAN | Unmodified | 1529200 | 1496600 | 1525900 | 5309300 | 14576000 | 3857900 | 1596500 | 100940 | 3711900 | 5407900 | 651780 | 14360000 | 6208100 | 4971700 | 7714900 | 6385700 | 2863400 | 10966000 |
| RQLEEAEEI | MYH10 | P35580 | MYH10_HUMAI | Unmodified | 0 | 0 | 37127 | 34551 | 0 | 0 | 0 | 0 | 0 | 0 | 0 | 1403300 | 23442 | 0 | 60948 | 80324 | 0 | 0 |
| RQLEEAEEI | MYH10 | P35580 | MYH10_HUMAI | Unmodified | 0 | 0 | 681270 | 644570 | 1386900 | 1208400 | 37848 | 0 | 0 | 86766 | 52584 | 7526800 | 1676400 | 0 | 1211000 | 1323900 | 0 | 0 |
| RQLESGGF | EXOC8 | Q8IYI6 | EXOC8_HUMAI | Unmodified | 0 | 0 | 0 | 0 | 348690 | 43408 | 0 | 0 | 21105 | 0 | 43112 | 167130 | 0 | 0 | 0 | 0 | 0 | 0 |
| RQLHAQE( | UBTD1 | Q9HAC8 | UBTD1_HUMAI | Unmodified | 564260 | 0 | 0 | 0 | 1557600 | 0 | 0 | 0 | 450640 | 167070 | 187150 | 1427500 | 906600 | 253840 | 234750 | 188280 | 1394400 | 3575400 |
| RQLHAQE( | UBTD1 | Q9HAC8 | UBTD1_HUMAI | Unmodified | 0 | 0 | 0 | 0 | 1444900 | 0 | 0 | 0 | 0 | 0 | 0 | 0 | 0 | 814870 | 0 | 0 | 1785000 | 0 |
| RQLQERFTI | ROCK2; ROCK2 | O75116; E9F | ROCK2_HUMAI | Unmodified | 0 | 0 | 0 | 0 | 2078900 | 0 | 93207 | 0 | 0 | 144730 | 0 | 0 | 0 | 0 | 0 | 0 | 735350 | 1847000 |
| RQLSFSED | MICAL3; MICAL | Q7RTP6; J3Q | MICA3_HUMAN | Unmodified | 130730 | 420330 | 2683400 | 1574900 | 1570300 | 3364700 | 0 | 404100 | 0 | 78878 | 0 | 0 | 0 | 0 | 697820 | 155040 | 0 | 0 |
| RQMGSGD | TPP2; TPP2; TPI | P29144; A0A | TPP2_HUMAN | Unmodified | 0 | 0 | 0 | 0 | 0 | 0 | 0 | 0 | 0 | 0 | 0 | 437460 | 0 | 0 | 0 | 0 | 0 | 0 |
| RQNIGVATI | SMC4; SMC4 | Q9NTJ3; E9F | SMC4_HUMAN | Unmodified | 0 | 0 | 0 | 0 | 0 | 0 | 1272700 | 93805 | 0 | 0 | 272560 | 536400 | 0 | 0 | 0 | 0 | 272760 | 0 |
| RQQEQQVI | FASN; FASN | P49327; A0A | FAS_HUMAN Fi | Unmodified | 497430 | 261050 | 1816800 | 2377500 | 1256300 | 1361500 | 1620100 | 513360 | 1257000 | 483690 | 2892800 | 56227000 | 1873700 | 448500 | 2623300 | 2030200 | 432160 | 836340 |
| RQQETAAA | VGF; VGF | O15240; Q0! | VGF_HUMAN N | Unmodified | 722340 | 768530 | 0 | 0 | 0 | 0 | 0 | 0 | 6042400 | 320410 | 0 | 0 | 0 | 0 | 0 | 0 | 0 | 0 |

| | | | | | | | | | | | | | | | | | | | | | | |
|---|---|---|---|---|---|---|---|---|---|---|---|---|---|---|---|---|---|---|---|---|---|---|
| RQQETAAA | VGF; VGF | O15240; Q0! | VGF_HUMAN N | Unmodified | 351690 | 0 | 0 | 0 | 0 | 0 | 0 | 0 | 4953800 | 1629100 | 0 | 0 | 0 | 0 | 0 | 0 | 0 | 0 |
| RQQEVETE | SNRNP70; SNR | P08621; M0( | RU17_HUMAN | Unmodified | 492040 | 39586 | 0 | 0 | 22359 | 0 | 89485 | 326970 | 0 | 0 | 153550 | 0 | 0 | 0 | 343210 | 393670 | 0 | 0 |
| RQQLLQEC | MAP4K4; MAP4 | O95819; C9J | M4K4_HUMAN | Unmodified | 24711 | 30420 | 0 | 0 | 0 | 0 | 0 | 89895 | 518930 | 0 | 0 | 1247600 | 0 | 0 | 0 | 139870 | 0 | 0 |
| RQQQLTAN | MYH9; MYH11 | P35579; P35 | MYH9_HUMAN | Unmodified | 0 | 0 | 0 | 0 | 634570 | 0 | 257980 | 0 | 0 | 0 | 0 | 844730 | 0 | 0 | 0 | 0 | 0 | 0 |
| RQSEDPSC | ANO6; ANO6; A | Q4KMQ2; A0 | ANO6_HUMAN | Unmodified | 0 | 0 | 0 | 0 | 0 | 0 | 0 | 0 | 50538 | 0 | 0 | 116800 | 0 | 0 | 0 | 0 | 0 | 0 |
| RQSGQTDI | IQGAP1; IQGAP | P46940; A0A | IQGA1_HUMAN | Unmodified | 0 | 346750 | 0 | 0 | 1102100 | 0 | 0 | 0 | 0 | 594230 | 0 | 0 | 0 | 0 | 0 | 0 | 425980 | 0 |
| RQSHLLLQ | SHTN1 | A0MZ66 | SHOT1_HUMAI | Unmodified | 91202 | 0 | 0 | 0 | 1300300 | 258010 | 238820 | 0 | 1100100 | 181810 | 206840 | 1853200 | 1199300 | 1101100 | 0 | 0 | 402330 | 719450 |
| RQSQQLEA | RAI14 | Q9P0K7 | RAI14_HUMAN | Unmodified | 0 | 0 | 0 | 0 | 1397800 | 308460 | 0 | 0 | 0 | 147360 | 0 | 963820 | 0 | 0 | 0 | 0 | 631850 | 816090 |
| RQTLDQLS | UTRN | P46939 | UTRN_HUMAN | Unmodified | 0 | 0 | 0 | 1373100 | 1219700 | 0 | 0 | 0 | 519640 | 479320 | 0 | 0 | 0 | 0 | 0 | 0 | 763780 | 0 |
| RQTSGGP\ | ATP5PF; ATP5PI | P18859; A8M | ATP5J_HUMAN | Unmodified | 0 | 0 | 0 | 0 | 0 | 0 | 696530 | 0 | 12482000 | 2405100 | 0 | 0 | 0 | 304200 | 0 | 0 | 0 | 0 |
| RQTVLDKY | VPS37A; VPS37 | Q8NEZ2; H0 | VP37A_HUMAN | Unmodified | 589110 | 1167700 | 0 | 0 | 1895300 | 0 | 0 | 0 | 0 | 0 | 0 | 0 | 0 | 0 | 0 | 0 | 0 | 0 |
| RQTVLDKY | VPS37A; VPS37 | Q8NEZ2; H0 | VP37A_HUMAN | Oxidation (M) | 1522300 | 2753400 | 0 | 0 | 584950 | 0 | 0 | 0 | 0 | 0 | 0 | 158520 | 241500 | 0 | 342470 | 0 | 0 | 0 |
| RQVEKEETI | GOLM1 | Q8NBJ4 | GOLM1_HUMA | Unmodified | 0 | 0 | 0 | 0 | 0 | 0 | 0 | 0 | 3811900 | 56411 | 0 | 0 | 0 | 0 | 0 | 0 | 0 | 0 |
| RQVLLSAAI | CCT2 | P78371 | TCPB_HUMAN | Unmodified | 0 | 0 | 0 | 0 | 0 | 0 | 0 | 188570 | 0 | 0 | 0 | 1297100 | 0 | 0 | 0 | 0 | 262230 | 0 |
| RQVLLSAAI | CCT2 | P78371 | TCPB_HUMAN | Unmodified | 0 | 0 | 0 | 0 | 0 | 0 | 0 | 0 | 0 | 0 | 0 | 0 | 2219200 | 542640 | 0 | 0 | 577920 | 408610 |
| RQVLLYEP( | CUL5 | Q93034 | CUL5_HUMAN | Unmodified | 0 | 0 | 0 | 0 | 0 | 0 | 0 | 0 | 0 | 0 | 0 | 3640200 | 101230 | 0 | 0 | 0 | 0 | 0 |
| RQYGTISH( | APLP2 | Q06481 | APLP2_HUMAN | Unmodified | 0 | 0 | 0 | 0 | 905520 | 0 | 0 | 0 | 2358500 | 91906 | 1035600 | 0 | 0 | 0 | 0 | 0 | 0 | 0 |
| RQYGTISH( | APLP2 | Q06481 | APLP2_HUMAN | Oxidation (M) | 0 | 0 | 0 | 0 | 3473500 | 460820 | 780600 | 0 | 0 | 0 | 0 | 0 | 444040 | 587230 | 0 | 0 | 0 | 1806800 |
| RRAADPSS | DST; DST; DST; | Q03001; A0A | DYST_HUMAN | Unmodified | 0 | 0 | 0 | 0 | 826110 | 0 | 0 | 0 | 0 | 263000 | 0 | 0 | 317870 | 157680 | 0 | 0 | 146860 | 198860 |
| RRALAAPA | SLC7A5 | Q01650 | LAT1_HUMAN I | Unmodified | 0 | 0 | 195080 | 0 | 5623700 | 0 | 0 | 0 | 186300 | 0 | 0 | 0 | 0 | 0 | 0 | 0 | 0 | 0 |
| RRANQQET | SRGAP2; SRGA | O75044; Q5\ | SRGP2_HUMAI | Unmodified | 0 | 0 | 0 | 0 | 2536400 | 0 | 503630 | 320760 | 270010 | 177010 | 0 | 1214000 | 0 | 938140 | 308740 | 0 | 851690 | 1185900 |
| RRDEELIYH | ECE1; ECE1; EC | P42892; B4E | ECE1_HUMAN | Unmodified | 157020 | 0 | 517770 | 0 | 1800800 | 0 | 1061700 | 0 | 10218000 | 1199800 | 113740 | 0 | 0 | 0 | 251600 | 224490 | 18537 | 79214 |
| RREAVCIVL | VCP; VCP; VCP; | P55072; A0A | TERA_HUMAN | Unmodified | 0 | 0 | 0 | 0 | 638920 | 571580 | 154310 | 32810 | 0 | 402710 | 171450 | 0 | 2610000 | 2813300 | 0 | 0 | 476280 | 604380 |
| RREEGAPG | MAN1A1 | P33908 | MA1A1_HUMAI | Unmodified | 0 | 0 | 0 | 0 | 0 | 0 | 0 | 0 | 9546000 | 1389700 | 3622900 | 9324000 | 0 | 0 | 0 | 0 | 0 | 0 |
| RREFITGD\ | NAP1L4; NAP1L | Q99733; C9J | NP1L4_HUMAN | Unmodified | 0 | 0 | 0 | 0 | 0 | 0 | 0 | 0 | 0 | 0 | 0 | 10535000 | 0 | 0 | 433010 | 0 | 0 | 0 |
| RREIENMIL | ACOX1; ACOX1 | Q15067; I3L | ACOX1_HUMAN | Unmodified | 0 | 0 | 0 | 0 | 0 | 0 | 0 | 0 | 3140500 | 0 | 0 | 0 | 0 | 0 | 0 | 0 | 0 | 0 |
| RREPIPSLE | STXBP2; STXBP | Q15833; M0 | STXB2_HUMAN | Unmodified | 1503500 | 1209700 | 1387700 | 4055400 | 2744800 | 1156700 | 962370 | 0 | 1977200 | 198360 | 1137800 | 5860800 | 2906100 | 2426700 | 2191300 | 1459700 | 592310 | 1648600 |
| RREPLPSLE | STXBP1; STXBP | P61764; A0A | STXB1_HUMAN | Unmodified | 0 | 1097400 | 0 | 0 | 2241700 | 393180 | 1965800 | 57678 | 4022300 | 1562900 | 280850 | 5089500 | 2030900 | 646730 | 392020 | 0 | 668740 | 1029600 |
| RREPVGQD | NOTCH2 | Q04721 | NOTC2_HUMA | Unmodified | 0 | 0 | 0 | 0 | 1226400 | 0 | 0 | 0 | 0 | 0 | 0 | 0 | 1309000 | 0 | 0 | 0 | 0 | 2927500 |
| RRGDALIEI | GRSF1; GRSF1; | Q12849; H0 | GRSF1_HUMAI | Unmodified | 0 | 0 | 0 | 0 | 32809 | 0 | 0 | 0 | 454030 | 41340 | 0 | 0 | 67298 | 0 | 0 | 68657 | 0 | 0 |
| RRITPLSLQ | EPB41L2; EPB4 | O43491; E9I | E41L2_HUMAN | Unmodified | 0 | 0 | 0 | 0 | 12712000 | 1546700 | 0 | 0 | 0 | 0 | 0 | 0 | 0 | 0 | 0 | 0 | 7799900 | 9869700 |
| RRLEIVFVS | NXN; NXN | Q6DKJ4; A0A | NXN_HUMAN N | Unmodified | 0 | 0 | 0 | 0 | 0 | 0 | 0 | 0 | 0 | 0 | 0 | 768620 | 0 | 0 | 0 | 0 | 0 | 0 |
| RRLLEEEIIA | SEPTIN10; SEPT | Q9P0V9; B5I | SEP10_HUMAN | Unmodified | 0 | 0 | 0 | 0 | 6518700 | 2295100 | 165680 | 0 | 0 | 0 | 0 | 0 | 440670 | 376810 | 0 | 0 | 697630 | 1613900 |
| RRPGASPT | SLC4A2; SLC4A | P04920; C9J | B3A2_HUMAN | Unmodified | 772390 | 607750 | 2018000 | 5115600 | 1297800 | 0 | 0 | 0 | 1247300 | 0 | 358660 | 1514300 | 1100500 | 793480 | 4512400 | 4489600 | 0 | 1065000 |
| RRPSDENT | PPFIBP1; PPFIB | Q86W92; F5 | LIPB1_HUMAN | Unmodified | 0 | 0 | 99150 | 114920 | 0 | 0 | 0 | 0 | 153790 | 0 | 0 | 276710 | 0 | 0 | 95130 | 73457 | 0 | 0 |
| RRQEELNN | PLCG2; PLCG2 | P16885; A0A | PLCG2_HUMAI | Unmodified | 0 | 0 | 0 | 0 | 0 | 0 | 0 | 0 | 0 | 0 | 2938000 | 18097000 | 0 | 0 | 0 | 0 | 0 | 0 |
| RRQLLLTEC | PDPK1; PDPK1 | O15530; B9I | PDPK1_HUMAN | Unmodified | 0 | 0 | 0 | 715210 | 0 | 0 | 490730 | 0 | 0 | 0 | 0 | 0 | 1672800 | 2408200 | 273870 | 318220 | 0 | 0 |
| RRQMQEAI | SEL1L | Q9UBV2 | SE1L1_HUMAN | Oxidation (M) | 0 | 0 | 0 | 0 | 0 | 0 | 0 | 0 | 563430 | 0 | 0 | 0 | 0 | 0 | 0 | 0 | 0 | 0 |
| RRSSTVAP/ | KIF23; KIF23; KI | Q02241; A0A | KIF23_HUMAN | Unmodified | 0 | 0 | 0 | 0 | 3408300 | 359760 | 0 | 0 | 0 | 0 | 0 | 0 | 0 | 0 | 0 | 0 | 0 | 0 |
| RRTTDFSD | CHORDC1; CH | Q9UHD1; E9 | CHRD1_HUMA | Unmodified | 0 | 2301000 | 0 | 95016 | 1561800 | 302990 | 694840 | 65615 | 0 | 2573600 | 0 | 0 | 8297100 | 4767600 | 0 | 0 | 10648000 | 3368600 |
| RRVNEFGE | CLPTM1L; CLPT | Q96KA5; G5I | CLP1L_HUMAN | Unmodified | 0 | 0 | 0 | 0 | 0 | 0 | 0 | 0 | 0 | 0 | 4636700 | 0 | 0 | 0 | 0 | 0 | 0 | 0 |
| RRVNSLPE | ITGA6 | P23229 | ITA6_HUMAN I | Unmodified | 0 | 0 | 356480 | 683990 | 6745400 | 2371200 | 0 | 0 | 0 | 0 | 0 | 0 | 4084600 | 2007200 | 811340 | 1748300 | 5629300 | 42204000 |
| RSALQSINI | SERPINH1; SER | P50454; E9P | SERPH_HUMAI | Unmodified | 0 | 0 | 0 | 0 | 0 | 0 | 0 | 0 | 612440 | 0 | 0 | 0 | 0 | 0 | 0 | 0 | 0 | 0 |
| RSALQSINI | SERPINH1; SER | P50454; E9P | SERPH_HUMAI | Unmodified | 0 | 0 | 0 | 0 | 536990 | 0 | 0 | 0 | 74859000 | 3578200 | 0 | 0 | 0 | 0 | 0 | 0 | 0 | 0 |
| RSAPGGGS | NPM1; NPM1; N | P06748; A0A | NPM_HUMAN N | Unmodified | 0 | 0 | 0 | 0 | 160710 | 215110 | 179770 | 60665 | 0 | 100130 | 86777 | 0 | 0 | 0 | 0 | 0 | 35494 | 795870 |
| RSAQFFNY | PLOD1; PLOD1 | Q02809; Q5. | PLOD1_HUMAN | Unmodified | 101420 | 0 | 0 | 0 | 499150 | 33558 | 63580 | 0 | 1332200 | 692560 | 0 | 0 | 0 | 0 | 0 | 0 | 0 | 2200700 |
| RSAQFFNY | PLOD1; PLOD1 | Q02809; Q5. | PLOD1_HUMAN | Unmodified | 0 | 0 | 0 | 0 | 0 | 0 | 0 | 0 | 0 | 491740 | 0 | 0 | 0 | 0 | 0 | 0 | 0 | 0 |
| RSCHAGF( | MELTF | P08582 | TRFM_HUMAN | Unmodified | 0 | 0 | 0 | 0 | 3956800 | 509880 | 211810 | 0 | 1672600 | 0 | 0 | 219740 | 0 | 0 | 0 | 0 | 53591 | 1417100 |
| RSDELSFC | PLCG2; PLCG2 | P16885; A0A | PLCG2_HUMAI | Unmodified | 0 | 0 | 0 | 0 | 0 | 0 | 0 | 0 | 0 | 0 | 187170 | 838040 | 0 | 0 | 0 | 0 | 0 | 0 |
| RSDGLLQL | PHLDA1; PHLD. | Q8WV24; R4 | PHLA1_HUMAN | Unmodified | 0 | 0 | 0 | 0 | 7105200 | 921150 | 0 | 0 | 0 | 0 | 0 | 0 | 0 | 0 | 0 | 0 | 14127000 | 9999100 |
| RSDGLLQL | PHLDA1; PHLD. | Q8WV24; R4 | PHLA1_HUMAN | Unmodified | 0 | 0 | 0 | 0 | 2790800 | 812200 | 0 | 0 | 0 | 0 | 0 | 0 | 0 | 0 | 0 | 0 | 3923800 | 3065500 |
| RSDNVVTS | PTGFRN | Q9P2B2 | FPRP_HUMAN | Unmodified | 0 | 462280 | 68159 | 772060 | 0 | 0 | 0 | 0 | 0 | 0 | 0 | 0 | 222640 | 0 | 2206700 | 0 | 0 | 0 |
| RSDSASSE | PRKCD | Q05655 | KPCD_HUMAN | Unmodified | 0 | 0 | 547940 | 0 | 0 | 0 | 130920 | 0 | 759260 | 326290 | 505950 | 3860100 | 907340 | 0 | 744960 | 0 | 1234100 | 2696400 |
| RSDSLFQL' | PHLDA2 | Q53GA4 | PHLA2_HUMAI | Unmodified | 0 | 0 | 0 | 0 | 792960 | 0 | 0 | 0 | 0 | 0 | 0 | 0 | 738120 | 0 | 794200 | 302010 | 0 | 641050 |
| RSDSLFQL' | PHLDA2 | Q53GA4 | PHLA2_HUMAI | Unmodified | 0 | 0 | 0 | 0 | 957170 | 349630 | 247950 | 0 | 0 | 0 | 0 | 126820 | 610870 | 533120 | 0 | 0 | 0 | 1474200 |
| RSDSVILN\ | CEACAM5; CEA | P06731; A0A | CEAM5_HUMAI | Unmodified | 0 | 0 | 0 | 0 | 0 | 0 | 0 | 0 | 1.16E+08 | 1288200 | 0 | 0 | 0 | 0 | 0 | 0 | 0 | 0 |
| RSDYDREA | RCN2; RCN2 | Q14257; H0' | RCN2_HUMAN | Unmodified | 0 | 0 | 0 | 0 | 182000 | 0 | 0 | 0 | 36467000 | 8363700 | 912450 | 110540 | 120390 | 96529 | 0 | 0 | 0 | 40438 |
| RSEACPCC | SCRIB; SCRIB; ! | Q14160; A0A | SCRIB_HUMAN | Unmodified | 0 | 0 | 0 | 0 | 404550 | 0 | 0 | 0 | 2754800 | 350860 | 0 | 805970 | 0 | 0 | 149070 | 119460 | 1225900 | 1337400 |
| RSEAEEAIT | ELAVL1 | Q15717 | ELAV1_HUMAN | Unmodified | 0 | 0 | 284420 | 0 | 0 | 0 | 0 | 0 | 0 | 0 | 0 | 2374600 | 0 | 0 | 0 | 195650 | 0 | 0 |
| RSECGALE | CFC1; CFC1B | P0CG37; P0( | CFC1_HUMAN | Unmodified | 1089900 | 1397900 | 0 | 0 | 0 | 0 | 0 | 0 | 244470 | 0 | 0 | 0 | 0 | 0 | 0 | 0 | 0 | 0 |
| RSEDEPPA | PAWR; PAWR | Q96IZ0; H0Y | PAWR_HUMAN | Unmodified | 501850 | 749340 | 0 | 320980 | 0 | 120000 | 0 | 0 | 574290 | 0 | 0 | 333050 | 0 | 0 | 0 | 0 | 117990 | 0 |
| RSEELLSQ\ | CEP55; CEP55 | Q53EZ4; H0 | CEP55_HUMAN | Unmodified | 0 | 0 | 0 | 0 | 623720 | 0 | 0 | 0 | 0 | 242860 | 0 | 0 | 0 | 0 | 0 | 0 | 0 | 0 |
| RSEGAVVN | IGSF3 | O75054 | IGSF3_HUMAN | Unmodified | 2091700 | 2805500 | 822720 | 358930 | 454820 | 806520 | 58921 | 56698 | 370360 | 74405 | 129640 | 0 | 214310 | 41286 | 1631500 | 1877800 | 0 | 125100 |
| RSELEEQQ | DYNC1H1; DYN | Q14204; A0A | DYHC1_HUMA | Unmodified | 0 | 0 | 0 | 0 | 1450200 | 0 | 593410 | 0 | 0 | 1349400 | 0 | 5268500 | 0 | 1101500 | 0 | 0 | 0 | 699760 |
| RSELEEQQ | DYNC1H1; DYN | Q14204; A0A | DYHC1_HUMA | Oxidation (M) | 0 | 0 | 307240 | 0 | 0 | 0 | 149910 | 98731 | 106090 | 179130 | 0 | 1001400 | 179500 | 0 | 0 | 0 | 0 | 414640 |
| RSELVNWY | MCM6 | Q14566 | MCM6_HUMAN | Unmodified | 0 | 0 | 0 | 0 | 0 | 0 | 0 | 0 | 0 | 0 | 0 | 2724200 | 0 | 0 | 0 | 0 | 0 | 0 |
| RSENKNEC | NPTN; NPTN; N | Q9Y639; Q9I | NPTN_HUMAN | Oxidation (M) | 0 | 0 | 0 | 110090 | 875160 | 453970 | 134090 | 148550 | 105470 | 94713 | 163180 | 586640 | 1788000 | 1437100 | 0 | 0 | 672180 | 1630600 |
| RSENKNEC | NPTN; NPTN; N | Q9Y639; Q9I | NPTN_HUMAN | Unmodified | 0 | 0 | 0 | 0 | 10336000 | 0 | 881520 | 775120 | 352370 | 0 | 2013800 | 1381500 | 0 | 11501000 | 0 | 0 | 3284500 | 14168000 |
| RSFITTDVN | LARS1; LARS1; | Q9P2J5; A0A | SYLC_HUMAN | Unmodified | 0 | 0 | 0 | 0 | 0 | 0 | 0 | 24584 | 0 | 230350 | 1593800 | 3068000 | 0 | 0 | 0 | 0 | 0 | 0 |
| RSFLLDLLN | DDX3X; DDX3X | O00571; A0A | DDX3X_HUMAI | Unmodified | 0 | 88073 | 194420 | 0 | 0 | 0 | 0 | 0 | 0 | 0 | 0 | 182250 | 0 | 0 | 239920 | 119360 | 0 | 0 |
| RSFTILLSN | DSC2 | Q02487 | DSC2_HUMAN | Unmodified | 0 | 0 | 0 | 0 | 0 | 0 | 0 | 0 | 1760500 | 752920 | 0 | 0 | 0 | 0 | 0 | 0 | 50343 | 0 |
| RSGNNNG | EFNB2 | P52799 | EFNB2_HUMAN | Unmodified | 0 | 0 | 0 | 0 | 0 | 0 | 0 | 0 | 0 | 0 | 0 | 4896700 | 0 | 0 | 0 | 3099400 | 0 | 0 |
| RSGQVLEV | ATP6V1B2; ATP | P21281; H0\ | VATB2_HUMAN | Unmodified | 182340 | 147720 | 226030 | 536110 | 1036300 | 279680 | 1418500 | 307120 | 1208500 | 509090 | 346480 | 2466000 | 467790 | 999060 | 270810 | 295870 | 0 | 1224600 |
| RSGVAYIA/ | ATIC | P31939 | PUR9_HUMAN | Unmodified | 155800 | 113920 | 505510 | 364150 | 1010100 | 185040 | 0 | 53478 | 1171700 | 554570 | 413040 | 5222000 | 0 | 0 | 339570 | 345480 | 0 | 955780 |
| RSGVAYIA/ | ATIC | P31939 | PUR9_HUMAN | Unmodified | 0 | 0 | 0 | 0 | 0 | 0 | 0 | 0 | 0 | 0 | 0 | 0 | 1872500 | 0 | 0 | 0 | 727010 | 280390 |
| RSHDFSNS | PARP12; PARP1 | Q9H0J9; G5I | PAR12_HUMAN | Unmodified | 195940 | 231450 | 412450 | 288280 | 0 | 0 | 0 | 0 | 0 | 0 | 0 | 0 | 0 | 0 | 315100 | 409210 | 0 | 160090 |
| RSHQLIHA. | ZC3HAV1L | Q96H79 | ZCCHL_HUMAN | Unmodified | 0 | 0 | 0 | 0 | 257280 | 0 | 0 | 0 | 30993 | 41774 | 31498 | 181260 | 55146 | 0 | 121550 | 0 | 74333 | 67073 |
| RSIIKPTVPI | AVIL | O75366 | AVIL_HUMAN A | Unmodified | 0 | 0 | 0 | 0 | 0 | 0 | 0 | 0 | 0 | 0 | 903800 | 6706300 | 0 | 0 | 0 | 0 | 0 | 0 |
| RSIQEELQC | PLEC | Q15149 | PLEC_HUMAN | Unmodified | 0 | 0 | 0 | 0 | 4214800 | 0 | 0 | 0 | 0 | 239560 | 0 | 0 | 704390 | 0 | 0 | 0 | 7322000 | 8985300 |
| RSIQFVDW | TUBA4A; TUBA1 | P68366; P68 | TBA4A_HUMAN | Unmodified | 4637200 | 2844400 | 5435200 | 5912400 | 30553000 | 5009800 | 20834000 | 2488500 | 29981000 | 20828000 | 8894100 | 61699000 | 20182000 | 22708000 | 5567800 | 4871300 | 13836000 | 28710000 |
| RSIQFVDW | TUBA4A; TUBA1 | P68366; P68 | TBA4A_HUMAN | Unmodified | 0 | 0 | 0 | 0 | 0 | 0 | 11698000 | 0 | 0 | 0 | 0 | 0 | 5663000 | 6524100 | 0 | 0 | 8747400 | 0 |
| RSITVTQDE | ACAA1; ACAA1; | P09110; C9J | THIK_HUMAN 3 | Oxidation (M) | 0 | 0 | 0 | 0 | 0 | 0 | 0 | 0 | 2260300 | 477530 | 0 | 0 | 0 | 0 | 0 | 0 | 0 | 0 |
| RSITVTQDE | ACAA1; ACAA1; | P09110; C9J | THIK_HUMAN 3 | Unmodified | 0 | 0 | 0 | 0 | 0 | 0 | 425110 | 0 | 24195000 | 4334500 | 0 | 0 | 148060 | 267030 | 0 | 0 | 0 | 0 |
| RSKAEELGI | ACAA1; ACAA1; | P09110; C9J | THIK_HUMAN 3 | Unmodified | 0 | 0 | 0 | 0 | 0 | 0 | 0 | 0 | 3368000 | 1006500 | 45885 | 0 | 0 | 0 | 0 | 0 | 0 | 0 |
| RSKFTCAC | HABP2 | Q14520 | HABP2_HUMAN | Unmodified | 0 | 0 | 0 | 0 | 0 | 0 | 0 | 0 | 9380900 | 4190300 | 0 | 0 | 0 | 0 | 0 | 0 | 0 | 0 |
| RSLEQARM | CIT; CIT; CIT; CI | O14578; A0A | CTRO_HUMAN | Unmodified | 161320 | 0 | 0 | 80694 | 5277700 | 301240 | 235900 | 24340 | 0 | 106350 | 0 | 0 | 250910 | 128290 | 0 | 0 | 333430 | 339420 |
| RSLFNYHD | SEPTIN11; SEPT | Q9NVA2; D6 | SEP11_HUMAN | Unmodified | 0 | 0 | 0 | 0 | 349670 | 89415 | 0 | 0 | 0 | 190420 | 803030 | 1816600 | 0 | 0 | 154440 | 72869 | 0 | 0 |
| RSLFNYHD | SEPTIN11; SEPT | Q9NVA2; D6 | SEP11_HUMAN | Unmodified | 0 | 0 | 0 | 0 | 2562000 | 0 | 0 | 0 | 0 | 0 | 0 | 0 | 0 | 0 | 0 | 0 | 0 | 0 |
| RSLFSEEA/ | SRA1 | Q9HD15 | SRA1_HUMAN | Unmodified | 60196 | 0 | 0 | 328130 | 522670 | 0 | 0 | 0 | 0 | 0 | 197620 | 655610 | 608920 | 763770 | 0 | 56300 | 309580 | 519150 |
| RSLGYAYVI | PABPC4; PABP( | Q13310; P11 | PABP4_HUMAN | Unmodified | 14850000 | 7050900 | 3477800 | 1677400 | 0 | 0 | 149380 | 63278 | 111480 | 321430 | 0 | 1284100 | 37415 | 0 | 3425900 | 4639900 | 360770 | 0 |
| RSLHDPDC | APMAP; APMAP | Q9HDC9; H( | APMAP_HUMAI | Unmodified | 0 | 0 | 0 | 0 | 0 | 0 | 0 | 0 | 12006000 | 0 | 0 | 0 | 0 | 0 | 0 | 0 | 0 | 0 |
| RSLLSIPNT | PRKRA; PRKRA; | O75569; B4I | PRKRA_HUMAI | Unmodified | 0 | 0 | 0 | 319040 | 0 | 0 | 329440 | 0 | 16533 | 29309 | 0 | 22260 | 369860 | 26193 | 587730 | 0 | 0 | 0 |
| RSLLSMLVI | CPD | O75976 | CBPD_HUMAN | Unmodified | 0 | 0 | 0 | 0 | 0 | 0 | 0 | 0 | 1025700 | 0 | 0 | 0 | 0 | 0 | 0 | 0 | 0 | 0 |
| RSLVESGC | RNF41 | Q9H4P4 | RNF41_HUMAN | Unmodified | 0 | 0 | 0 | 0 | 926930 | 134040 | 0 | 0 | 0 | 0 | 0 | 0 | 0 | 0 | 0 | 0 | 0 | 0 |
| RSLYVRALI | DLG1; DLG1; D | Q12959; A0A | DLG1_HUMAN | Unmodified | 0 | 0 | 0 | 0 | 5097500 | 0 | 180030 | 0 | 0 | 189340 | 0 | 0 | 388280 | 1681900 | 0 | 0 | 3463600 | 1977900 |
| RSNIMWLE | IGSF3 | O75054 | IGSF3_HUMAN | Oxidation (M) | 0 | 0 | 0 | 77033 | 0 | 0 | 0 | 0 | 0 | 0 | 0 | 0 | 0 | 0 | 0 | 126730 | 0 | 0 |
| RSNSQSYI( | PRKD1; PRKD1 | Q15139; F8W | KPCD1_HUMAI | Unmodified | 0 | 0 | 54106 | 0 | 0 | 0 | 0 | 0 | 318250 | 0 | 0 | 0 | 677950 | 484380 | 258510 | 0 | 39890 | 0 |
| RSPFLQVFI | SEC23B; SEC23 | Q15437; A0A | SC23B_HUMAN | Unmodified | 0 | 0 | 0 | 0 | 0 | 0 | 0 | 0 | 0 | 0 | 0 | 1031900 | 0 | 0 | 0 | 0 | 0 | 0 |
| RSPNDFAL | CARS1; CARS1 | P49589; B4E | SYCC_HUMAN | Unmodified | 0 | 0 | 0 | 0 | 0 | 0 | 0 | 9159.3 | 111470 | 75415 | 43347 | 1977400 | 152050 | 114600 | 44376 | 25806 | 69634 | 0 |
| RSPPIPLAK | PDIA4; PDIA4 | P13667; A0A | PDIA4_HUMAN | Unmodified | 0 | 0 | 0 | 0 | 4615700 | 0 | 18260000 | 8423100 | 22661000 | 44674000 | 1845500 | 0 | 4504100 | 7259700 | 0 | 0 | 249590 | 4643100 |
| RSPYNYLQ | P3H1; P3H1 | Q32P28; E2( | P3H1_HUMAN | Unmodified | 0 | 0 | 0 | 0 | 0 | 0 | 0 | 0 | 1474600 | 587610 | 0 | 0 | 0 | 0 | 0 | 0 | 0 | 0 |
| RSQISMEQ | CA14; CA14 | Q9ULX7; U3I | CAH14_HUMAI | Unmodified | 0 | 0 | 0 | 0 | 0 | 0 | 0 | 0 | 0 | 0 | 0 | 793880 | 0 | 0 | 0 | 0 | 0 | 0 |
| RSRLTPVSI | SQSTM1; SQST | Q13501; E7E | SQSTM_HUMAI | Unmodified | 0 | 0 | 0 | 0 | 1094900 | 0 | 416890 | 0 | 0 | 0 | 0 | 0 | 775960 | 459040 | 0 | 0 | 385680 | 953150 |
| RSSGFISEL | CARMIL1; CARI | Q5VZK9; A0A | CARL1_HUMAN | Unmodified | 0 | 0 | 0 | 0 | 2935300 | 292270 | 0 | 0 | 0 | 73781 | 139910 | 1785300 | 1030500 | 0 | 0 | 0 | 332760 | 664190 |
| RSSHSNAY | ZMPSTE24 | O75844 | FACE1_HUMAN | Unmodified | 0 | 0 | 0 | 0 | 0 | 0 | 0 | 0 | 996640 | 80049 | 401300 | 0 | 0 | 0 | 0 | 0 | 0 | 0 |
| RSSHVTIDI | MELTF; MELTF | P08582; C9J | TRFM_HUMAN | Unmodified | 0 | 30816 | 0 | 0 | 738290 | 552710 | 78071 | 0 | 915160 | 488420 | 0 | 164160 | 187460 | 85614 | 0 | 0 | 0 | 49969 |
| RSSITEPEG | TP53BP2 | Q13625 | ASPP2_HUMAN | Unmodified | 0 | 0 | 0 | 0 | 0 | 174390 | 0 | 0 | 0 | 0 | 0 | 0 | 0 | 0 | 0 | 0 | 174240 | 0 |
| RSSNPPAR | EIF4B; EIF4B | P23588; E7E | IF4B_HUMAN E | Unmodified | 0 | 0 | 0 | 91170 | 2236000 | 370250 | 143360 | 0 | 58672 | 412140 | 50960 | 0 | 1934400 | 873230 | 0 | 0 | 988940 | 1723300 |
| RSSSDEQC | IL1RAP | Q9NPH3 | IL1AP_HUMAN | Unmodified | 0 | 0 | 0 | 0 | 828640 | 0 | 0 | 0 | 0 | 0 | 58548 | 0 | 0 | 0 | 0 | 0 | 0 | 566130 |

| | | | | | | | | | | | | | | | | | | | | | | |
|---|---|---|---|---|---|---|---|---|---|---|---|---|---|---|---|---|---|---|---|---|---|---|
| RSTQGVTL | PPP1R12A; PPP | O14974; F8V | MYPT1_HUMAN | Unmodified | 0 | 0 | 0 | 0 | 956800 | 258390 | 0 | 44401 | 0 | 0 | 0 | 0 | 1726700 | 0 | 0 | 0 | 291170 | 621050 |
| RSTTELGEN | UTRN | P46939 | UTRN_HUMAN | Unmodified | 0 | 0 | 653120 | 230980 | 0 | 0 | 0 | 0 | 0 | 0 | 0 | 0 | 0 | 0 | 823030 | 980950 | 0 | 0 |
| RSTVLGLP( | KIAA1522 | Q9P206 | K1522_HUMAN | Unmodified | 0 | 0 | 0 | 0 | 324340 | 0 | 0 | 0 | 0 | 0 | 0 | 0 | 975090 | 0 | 241160 | 36121 | 136520 | 0 |
| RSTYNHLS | RAB14; RAB14 | P61106; X6R | RAB14_HUMAN | Unmodified | 254550 | 436510 | 292540 | 0 | 0 | 0 | 0 | 0 | 497460 | 0 | 0 | 1270900 | 0 | 0 | 0 | 273160 | 0 | 0 |
| RSVGELPS | CKAP4 | Q07065 | CKAP4_HUMAN | Unmodified | 0 | 0 | 0 | 0 | 0 | 0 | 248580 | 198960 | 15249000 | 1185100 | 0 | 0 | 0 | 0 | 0 | 0 | 0 | 1042700 |
| RSVLALTHI | PTGFRN | Q9P2B2 | FPRP_HUMAN | Unmodified | 0 | 2115200 | 17555000 | 4566100 | 0 | 0 | 0 | 0 | 211490 | 0 | 0 | 3574100 | 2158600 | 0 | 0 | 5764000 | 0 | 0 |
| RSVLALTHI | PTGFRN | Q9P2B2 | FPRP_HUMAN | Unmodified | 0 | 0 | 0 | 2950100 | 288010 | 0 | 0 | 0 | 0 | 0 | 0 | 0 | 3958900 | 1418800 | 0 | 0 | 0 | 0 |
| RSVPPATP/ | PPP1R18; PPP1 | Q6NYC8; A0 | PPR18_HUMAN | Unmodified | 0 | 0 | 0 | 0 | 1196700 | 145770 | 0 | 41103 | 0 | 0 | 0 | 0 | 0 | 0 | 0 | 0 | 1062500 | 236410 |
| RSYDVPPP | PGAM1; PGAM4 | P18669; Q8N | PGAM1_HUMA | Unmodified | 0 | 0 | 0 | 0 | 6003600 | 618770 | 0 | 0 | 0 | 0 | 0 | 4053600 | 1387900 | 1560900 | 0 | 316280 | 1741900 | 4957500 |
| RSYDVPPP | PGAM1; PGAM4 | P18669; Q8N | PGAM1_HUMA | Oxidation (M) | 0 | 0 | 0 | 511980 | 4491700 | 1185100 | 338360 | 166680 | 0 | 0 | 1063500 | 7008000 | 3896400 | 726180 | 0 | 411730 | 1080100 | 2585500 |
| RSYSLSELS | SH3BP4 | Q9P0V3 | SH3B4_HUMAN | Unmodified | 0 | 0 | 0 | 0 | 1395600 | 0 | 0 | 0 | 920460 | 617750 | 0 | 357820 | 0 | 393640 | 225670 | 0 | 604380 | 840560 |
| RSYSLSELS | SH3BP4 | Q9P0V3 | SH3B4_HUMAN | Unmodified | 0 | 0 | 2502000 | 0 | 0 | 0 | 0 | 0 | 0 | 532750 | 0 | 0 | 0 | 0 | 108370 | 0 | 2494500 | 0 |
| RSYVTMTA | CHORDC1; CH | Q9UHD1; E9 | CHRD1_HUMA | Unmodified | 0 | 0 | 0 | 0 | 0 | 0 | 0 | 0 | 0 | 0 | 130700 | 1363000 | 0 | 0 | 0 | 0 | 0 | 0 |
| RTAATLATH | LRRC47 | Q8N1G4 | LRC47_HUMAN | Unmodified | 0 | 0 | 0 | 0 | 143970 | 0 | 38526 | 0 | 358560 | 150890 | 91311 | 873530 | 0 | 324160 | 406970 | 355930 | 109560 | 250110 |
| RTAATLATH | LRRC47 | Q8N1G4 | LRC47_HUMAN | Unmodified | 0 | 0 | 0 | 0 | 405420 | 0 | 0 | 76520 | 0 | 155300 | 0 | 0 | 0 | 410570 | 0 | 0 | 0 | 0 |
| RTADSSSS | CBX5; CBX5 | P45973; F8V | CBX5_HUMAN | Unmodified | 0 | 0 | 0 | 0 | 0 | 0 | 0 | 0 | 366830 | 47947 | 0 | 0 | 0 | 0 | 0 | 0 | 0 | 0 |
| RTAEHEAA( | KTN1 | Q86UP2 | KTN1_HUMAN | Unmodified | 0 | 0 | 0 | 0 | 0 | 0 | 0 | 0 | 193730 | 439090 | 43568 | 694170 | 0 | 0 | 0 | 0 | 14799 | 0 |
| RTAEVWMI | GALNT14; GALN | Q96FL9; Q8I | GLT14_HUMAN | Unmodified | 0 | 0 | 0 | 0 | 0 | 0 | 0 | 0 | 0 | 0 | 0 | 0 | 0 | 0 | 0 | 0 | 0 | 0 |
| RTANDAVE | PTGFRN | Q9P2B2 | FPRP_HUMAN | Unmodified | 1116900 | 1338000 | 2938800 | 536730 | 592080 | 65823 | 0 | 0 | 175900 | 14346 | 0 | 1793000 | 5664200 | 3114900 | 3987400 | 3559700 | 0 | 0 |
| RTANDAVE | PTGFRN | Q9P2B2 | FPRP_HUMAN | Unmodified | 0 | 0 | 0 | 0 | 241490 | 0 | 355230 | 56774 | 0 | 0 | 0 | 0 | 3415300 | 1944900 | 0 | 0 | 0 | 0 |
| RTDDIPVW | SKP1; SKP1; SK | P63208; E7E | SKP1_HUMAN | Unmodified | 943460 | 0 | 1400900 | 1088300 | 2198500 | 528720 | 0 | 0 | 2842600 | 285740 | 2118800 | 9927400 | 1837800 | 0 | 1763400 | 1539000 | 209610 | 1224800 |
| RTDKDTEIT | EPCAM; EPCAM | P16422; B5N | EPCAM_HUMA | Unmodified | 1017600 | 3187400 | 3271800 | 0 | 0 | 0 | 0 | 0 | 55185000 | 185860 | 6872900 | 30577000 | 0 | 0 | 1661000 | 25472000 | 0 | 0 |
| RTDLCDHA | CNPY2; CNPY2 | Q9Y2B0; H0 | CNPY2_HUMAI | Unmodified | 0 | 174510 | 0 | 0 | 0 | 0 | 0 | 0 | 93329000 | 28805000 | 1505900 | 304910 | 1083100 | 0 | 0 | 0 | 0 | 0 |
| RTEGYAAF( | RALBP1; RALBP | Q15311; J3Q | RBP1_HUMAN | Unmodified | 0 | 0 | 0 | 0 | 0 | 0 | 0 | 0 | 0 | 0 | 0 | 0 | 0 | 0 | 95263 | 307940 | 0 | 0 |
| RTEITIVKPC | SEPTIN9; SEPTI | Q9UHD8; K7 | SEPT9_HUMAN | Unmodified | 98514 | 124580 | 224810 | 411190 | 640880 | 98131 | 0 | 0 | 433370 | 164320 | 247390 | 1462300 | 349210 | 88131 | 182290 | 203200 | 738330 | 599270 |
| RTELFIAAE( | RPL8; RPL8; RP | P62917; E9P | RL8_HUMAN 6( | Unmodified | 0 | 0 | 0 | 0 | 13451000 | 0 | 3024400 | 0 | 0 | 0 | 0 | 0 | 0 | 2786200 | 0 | 0 | 0 | 3191600 |
| RTELFIAAE( | RPL8; RPL8; RP | P62917; E9P | RL8_HUMAN 6( | Unmodified | 0 | 0 | 0 | 0 | 0 | 0 | 1253100 | 0 | 0 | 0 | 0 | 1759800 | 0 | 0 | 0 | 0 | 0 | 0 |
| RTEPILESPI | TNIK | Q9UKE5 | TNIK_HUMAN T | Unmodified | 0 | 0 | 0 | 0 | 0 | 0 | 32332 | 31080 | 483840 | 0 | 222480 | 744660 | 0 | 0 | 0 | 0 | 0 | 0 |
| RTFLETSTD | EPB41L3; EPB4 | Q9Y2J2; A0A | E41L3_HUMAN | Unmodified | 0 | 0 | 0 | 0 | 0 | 0 | 0 | 0 | 0 | 0 | 0 | 0 | 0 | 259620 | 0 | 0 | 1862900 | 10202000 |
| RTGASFQQ | SCAMP2 | O15127 | SCAM2_HUMA | Unmodified | 0 | 0 | 361670 | 1577800 | 0 | 0 | 0 | 0 | 834020 | 0 | 0 | 0 | 0 | 0 | 0 | 371890 | 0 | 0 |
| RTGETGIEE | USP9X | Q93008 | USP9X_HUMAN | Unmodified | 0 | 0 | 0 | 0 | 0 | 0 | 94140 | 0 | 0 | 0 | 0 | 2557100 | 0 | 0 | 199980 | 0 | 0 | 0 |
| RTGFSTSPI | PRMT1; PRMT1 | Q99873; E9F | ANM1_HUMAN | Unmodified | 0 | 0 | 0 | 0 | 0 | 0 | 0 | 18549 | 0 | 0 | 238330 | 1088400 | 0 | 0 | 0 | 87034 | 52069 | 0 |
| RTGPAATTI | P4HB; P4HB; P4 | P07237; A0A | PDIA1_HUMAN | Unmodified | 0 | 0 | 0 | 497370 | 0 | 0 | 0 | 0 | 0 | 325880 | 0 | 0 | 0 | 0 | 0 | 0 | 0 | 0 |
| RTGPTQQC | MCRIP2; MCRIF | Q9BUT9; I3L | MCRI2_HUMAN | Unmodified | 201460 | 109150 | 22859 | 0 | 0 | 0 | 0 | 0 | 0 | 0 | 0 | 51235 | 0 | 0 | 0 | 0 | 0 | 0 |
| RTHNDIIHN | EZR; EZR | P15311; E7E | EZRI_HUMAN E | Oxidation (M) | 1138200 | 96073 | 336420 | 582580 | 845810 | 446660 | 21046 | 0 | 3425800 | 208230 | 602670 | 7683000 | 493050 | 147570 | 1061600 | 1068400 | 55081 | 208510 |
| RTHNDIIHN | EZR; EZR | P15311; E7E | EZRI_HUMAN E | Unmodified | 3023700 | 0 | 2097000 | 5536300 | 4019300 | 4917100 | 0 | 0 | 26852000 | 0 | 2832900 | 70369000 | 0 | 2156000 | 0 | 4406700 | 140500 | 0 |
| RTHNGESV | PRPS2; PRPS1; | P11908; P60 | PRPS2_HUMAN | Unmodified | 0 | 0 | 0 | 0 | 0 | 0 | 0 | 0 | 0 | 0 | 0 | 1373200 | 0 | 1884400 | 0 | 0 | 0 | 0 |
| RTIAVLLDD | CCDC126; CCI | Q96EE4; C9J | CC126_HUMAI | Unmodified | 0 | 0 | 0 | 0 | 0 | 0 | 0 | 0 | 4211000 | 0 | 210970 | 0 | 0 | 0 | 0 | 0 | 0 | 0 |
| RTIEGLRGF | DCTN1 | E7EX90 | E7EX90_HUMA | Unmodified | 0 | 0 | 0 | 0 | 0 | 0 | 0 | 0 | 0 | 0 | 0 | 0 | 653730 | 479770 | 0 | 0 | 0 | 0 |
| RTIPWLEDI | ACTN4; ACTN4; | O43707; F5C | ACTN4_HUMAI | Unmodified | 0 | 0 | 150470 | 114070 | 928670 | 465600 | 0 | 0 | 540350 | 0 | 343940 | 1194600 | 0 | 0 | 0 | 182890 | 215670 | 0 |
| RTIQFVDW | TUBA3C; TUBA: | P0DPH7; Q7 | TBA3C_HUMAN | Unmodified | 597360 | 343310 | 1180500 | 1794000 | 10244000 | 4084600 | 17088000 | 1166600 | 10725000 | 4908600 | 5811200 | 99571000 | 2366700 | 2366400 | 162310 | 0 | 3030600 | 11656000 |
| RTIQFVDW | TUBA3C; TUBA: | P0DPH7; Q7 | TBA3C_HUMAN | Unmodified | 0 | 0 | 0 | 0 | 0 | 0 | 6040500 | 0 | 0 | 555610 | 0 | 0 | 0 | 0 | 0 | 0 | 0 | 0 |
| RTKAENIPA | SLC30A1 | Q9Y6M5 | ZNT1_HUMAN | Unmodified | 0 | 0 | 0 | 0 | 2219900 | 289220 | 0 | 0 | 0 | 0 | 0 | 0 | 1282500 | 0 | 0 | 0 | 0 | 914610 |
| RTLKPQPQ | C11orf52; C11( | Q96A22; E9F | CK052_HUMAN | Unmodified | 366760 | 442260 | 153490 | 168250 | 0 | 0 | 0 | 0 | 570930 | 0 | 0 | 0 | 1516000 | 689050 | 324230 | 266290 | 23836 | 42989 |
| RTLLFTFNV | CCDC47; CCD( | Q96A33; A0/ | CCD47_HUMA | Unmodified | 19943 | 0 | 173810 | 214570 | 0 | 0 | 745700 | 0 | 424170 | 99659 | 0 | 668490 | 631250 | 1058100 | 124540 | 141880 | 0 | 0 |
| RTLLIDELN | ANLN; ANLN | Q9NQW6; H | ANLN_HUMAN | Unmodified | 0 | 0 | 0 | 0 | 6990500 | 0 | 0 | 0 | 711180 | 0 | 0 | 0 | 0 | 0 | 0 | 0 | 661250 | 385830 |
| RTLQNQLV | MACF1; MACF1 | Q9UPN3; H3 | MACF1_HUMAI | Unmodified | 0 | 0 | 0 | 0 | 464730 | 0 | 104000 | 62802 | 0 | 263410 | 0 | 554320 | 217490 | 253430 | 0 | 88949 | 81324 | 0 |
| RTLTGTAAL | LTA4H; LTA4H | P09960; B4D | LKHA4_HUMAN | Unmodified | 0 | 0 | 0 | 0 | 0 | 0 | 0 | 0 | 233180 | 0 | 109600 | 2140100 | 0 | 0 | 0 | 0 | 0 | 0 |
| RTMMACG( | CCT7 | Q99832 | TCPH_HUMAN 2 | Oxidation (M) | 0 | 0 | 0 | 0 | 0 | 0 | 0 | 0 | 0 | 0 | 0 | 3535100 | 0 | 0 | 0 | 0 | 0 | 0 |
| RTNPPGGK | JPT2; JPT2; JPT2 | Q9H910; H3 | JUPI2_HUMAN | Unmodified | 0 | 0 | 0 | 0 | 1009300 | 0 | 0 | 0 | 0 | 48092 | 0 | 0 | 509740 | 0 | 0 | 0 | 79085 | 0 |
| RTNQRGAE | DDAH1; DDAH1 | O94760; B4D | DDAH1_HUMA | Unmodified | 0 | 0 | 0 | 0 | 3411200 | 193850 | 87358 | 0 | 0 | 533120 | 0 | 0 | 3139800 | 1720700 | 0 | 0 | 1720900 | 2466400 |
| RTNSTFNQ | RPL18; RPL18; I | Q07020; A0/ | RL18_HUMAN ( | Unmodified | 103280 | 0 | 0 | 0 | 134180 | 0 | 0 | 0 | 86534 | 0 | 0 | 2211200 | 0 | 0 | 0 | 0 | 74985 | 0 |
| RTPFGAYG | ACAA2; ACAA2 | P42765; A0A | THIM_HUMAN | Unmodified | 88270 | 0 | 0 | 0 | 0 | 0 | 840570 | 0 | 6737600 | 2872300 | 0 | 0 | 0 | 0 | 0 | 0 | 0 | 0 |
| RTPNGVVL | DYNC1H1; DYN | Q14204; A0/ | DYHC1_HUMA | Unmodified | 0 | 0 | 0 | 0 | 0 | 0 | 0 | 0 | 0 | 0 | 0 | 782990 | 0 | 0 | 0 | 0 | 0 | 0 |
| RTPQPVTIK | CASC3 | O15234 | CASC3_HUMAI | Unmodified | 2165300 | 1599800 | 1122100 | 901440 | 0 | 0 | 0 | 0 | 0 | 0 | 0 | 0 | 0 | 0 | 611140 | 620280 | 0 | 0 |
| RTPSEAER\ | NUMBL; NUMB | Q9Y6R0; A0/ | NUMBL_HUMA | Unmodified | 0 | 0 | 0 | 0 | 199230 | 0 | 0 | 0 | 0 | 0 | 0 | 0 | 0 | 0 | 0 | 0 | 0 | 0 |
| RTPVIDADK | DYNC1H1; DYN | Q14204; A0/ | DYHC1_HUMA | Unmodified | 0 | 0 | 738220 | 271730 | 2331900 | 142490 | 110780 | 18379 | 0 | 1099400 | 715800 | 10085000 | 1279200 | 73029 | 0 | 0 | 329530 | 1070600 |
| RTPVMEGY | LOXL2; LOXL2 | Q9Y4K0; H0 | LOXL2_HUMAN | Unmodified | 0 | 0 | 0 | 0 | 166000 | 0 | 0 | 0 | 675560 | 87765 | 0 | 0 | 0 | 213340 | 0 | 0 | 0 | 0 |
| RTQDPNTII | RASA3 | Q14644 | RASA3_HUMAN | Unmodified | 0 | 0 | 0 | 0 | 0 | 0 | 0 | 0 | 0 | 0 | 0 | 0 | 0 | 0 | 0 | 0 | 171850 | 0 |
| RTQGVSTT | PCYT2; PCYT2; | Q99447; I3L | PCY2_HUMAN | Unmodified | 0 | 0 | 243410 | 0 | 130480 | 0 | 0 | 0 | 129250 | 15360 | 135020 | 2040100 | 0 | 0 | 168770 | 0 | 0 | 0 |
| RTQPCTYC | TRAF4; TRAF4; | Q9BUZ4; K7I | TRAF4_HUMAN | Unmodified | 0 | 0 | 57597 | 0 | 0 | 0 | 0 | 0 | 0 | 0 | 0 | 0 | 0 | 0 | 82334 | 68385 | 0 | 0 |
| RTRDELEVI | RO60 | P10155 | RO60_HUMAN | Unmodified | 13022000 | 14396000 | 5113700 | 4738800 | 0 | 0 | 0 | 35553 | 0 | 0 | 0 | 1365500 | 0 | 0 | 11340000 | 7072200 | 184690 | 0 |
| RTRLGLEDI | STK38 | Q15208 | STK38_HUMAN | Unmodified | 0 | 0 | 0 | 0 | 2991600 | 0 | 0 | 0 | 0 | 0 | 0 | 0 | 0 | 0 | 0 | 0 | 240990 | 719160 |
| RTSMGGT( | CTNNB1; CTNN | P35222; A0A | CTNB1_HUMAI | Unmodified | 65366 | 64863 | 128050 | 0 | 0 | 0 | 0 | 0 | 0 | 0 | 0 | 616530 | 0 | 0 | 372090 | 213080 | 0 | 0 |
| RTSMGGT( | CTNNB1; CTNN | P35222; A0A | CTNB1_HUMAI | Oxidation (M) | 0 | 0 | 0 | 0 | 0 | 0 | 0 | 0 | 0 | 0 | 0 | 219960 | 0 | 0 | 348080 | 259920 | 0 | 0 |
| RTSPARGY | CTNND1; CTNN | O60716; A0/ | CTND1_HUMAI | Unmodified | 0 | 0 | 0 | 0 | 3575800 | 454560 | 0 | 0 | 0 | 0 | 204210 | 0 | 2742000 | 2103900 | 0 | 0 | 783840 | 687550 |
| RTSSAQVE | IMPDH2; IMPDH | P12268; A0A | IMDH2_HUMAN | Unmodified | 76311 | 0 | 512090 | 926210 | 7885100 | 1008100 | 6462900 | 5618200 | 6494400 | 3291600 | 2195200 | 5827200 | 3779200 | 1828600 | 648000 | 1163900 | 10598000 | 7320000 |
| RTSSQMED | N/A | N/A | N/A | Acetyl (Protein N | 0 | 0 | 815240 | 736970 | 0 | 0 | 0 | 0 | 0 | 0 | 0 | 0 | 0 | 182010 | 2090900 | 2224600 | 0 | 0 |
| RTTDVMFG | AHCYL1 | O43865 | SAHH2_HUMAI | Unmodified | 0 | 0 | 0 | 0 | 1129800 | 475210 | 0 | 0 | 0 | 0 | 0 | 181010 | 0 | 0 | 373030 | 215160 | 249340 | 430330 |
| RTTEATLNS | AXL; AXL | P30530; M0I | UFO_HUMAN T | Unmodified | 0 | 0 | 0 | 0 | 1115000 | 0 | 0 | 0 | 0 | 0 | 0 | 0 | 0 | 0 | 0 | 0 | 0 | 0 |
| RTTTGDVQ | IGF2R; IGF2R | P11717; S4F | MPRI_HUMAN ( | Unmodified | 0 | 0 | 761210 | 383150 | 0 | 0 | 0 | 0 | 39847000 | 5781900 | 2169000 | 4146000 | 125410 | 58586 | 0 | 0 | 0 | 0 |
| RTVDLSSH | RPN1; RPN1 | P04843; F8V | RPN1_HUMAN | Unmodified | 0 | 0 | 0 | 28184 | 0 | 0 | 0 | 0 | 59375000 | 5004300 | 1259700 | 3101400 | 0 | 0 | 0 | 0 | 0 | 0 |
| RTVETEAV( | PPP2R5D; PPP2 | Q14738; H0 | 2A5D_HUMAN | Unmodified | 0 | 0 | 0 | 0 | 628560 | 865730 | 0 | 979040 | 272430 | 207590 | 0 | 445310 | 289610 | 171290 | 0 | 0 | 0 | 0 |
| RTVIEQQP\ | GGT1; GGT3P; | P19440; A6N | GGT1_HUMAN | Unmodified | 1158700 | 376910 | 943650 | 964060 | 0 | 0 | 0 | 73128 | 2002800 | 0 | 272030 | 1678100 | 0 | 0 | 2068900 | 1558400 | 25332 | 0 |
| RTVPPAVT( | ALDOA; ALDOA | P04075; J3K | ALDOA_HUMAI | Unmodified | 0 | 0 | 0 | 0 | 5535400 | 769880 | 0 | 0 | 0 | 0 | 914910 | 0 | 0 | 0 | 0 | 0 | 0 | 2181000 |
| RTVQQLRL | GNG12 | Q9UBI6 | GBG12_HUMAI | Unmodified | 0 | 0 | 0 | 1114500 | 3145400 | 0 | 0 | 0 | 532850 | 302200 | 0 | 0 | 1051100 | 653890 | 0 | 0 | 367620 | 897700 |
| RTVQSLEID | KRT18; KRT18 | P05783; F8V | K1C18_HUMAN | Oxidation (M) | 0 | 0 | 0 | 0 | 0 | 0 | 0 | 0 | 0 | 0 | 0 | 1419500 | 0 | 0 | 0 | 21247 | 0 | 0 |
| RTVQSLEID | KRT18; KRT18 | P05783; F8V | K1C18_HUMAN | Unmodified | 0 | 0 | 0 | 0 | 0 | 0 | 0 | 0 | 383530 | 0 | 0 | 668970 | 0 | 0 | 0 | 0 | 0 | 0 |
| RTYEEGLKI | STIP1; STIP1; S | P31948; F5H | STIP1_HUMAN | Unmodified | 0 | 0 | 0 | 0 | 0 | 70494 | 25215 | 0 | 497070 | 291450 | 124400 | 381500 | 2836400 | 294690 | 0 | 22566 | 690570 | 619890 |
| RTYVTPPG | OSGEP; OSGEP | Q9NPF4; G3 | OSGEP_HUMAI | Unmodified | 0 | 38817 | 60609 | 0 | 0 | 0 | 0 | 0 | 0 | 0 | 0 | 2063400 | 0 | 0 | 148860 | 36604 | 0 | 0 |
| RVAAALPG | RAB6A; RAB6A | P20340; F5H | RAB6A_HUMAN | Unmodified | 0 | 220300 | 0 | 865580 | 0 | 0 | 0 | 0 | 1455000 | 109900 | 473260 | 2465400 | 0 | 0 | 0 | 182030 | 0 | 0 |
| RVAAALPG | RAB6A; RAB6A | P20340; F5H | RAB6A_HUMAN | Oxidation (M) | 0 | 0 | 0 | 0 | 0 | 0 | 0 | 0 | 151230 | 0 | 0 | 1151700 | 0 | 0 | 0 | 85449 | 0 | 0 |
| RVAEAHVP | WASHC5; WASI | Q12768; E7E | WASC5_HUMA | Unmodified | 0 | 0 | 0 | 28134 | 120200 | 31510 | 20358 | 0 | 51074 | 0 | 0 | 0 | 41336 | 0 | 0 | 0 | 0 | 0 |
| RVAGGPQI | SELENBP1 | Q13228 | SBP1_HUMAN | Oxidation (M) | 0 | 0 | 0 | 206610 | 0 | 0 | 0 | 0 | 0 | 0 | 0 | 0 | 0 | 0 | 471380 | 350090 | 0 | 0 |
| RVALSPAG | NNT; NNT; NNT | Q13423; A0/ | NNTM_HUMAN | Unmodified | 0 | 0 | 0 | 0 | 202960 | 0 | 770000 | 0 | 8088400 | 1749000 | 0 | 596760 | 0 | 0 | 0 | 81406 | 0 | 0 |
| RVASALPG | RAB6B | Q9NRW1 | RAB6B_HUMAN | Unmodified | 0 | 0 | 0 | 0 | 445150 | 0 | 152610 | 0 | 557830 | 0 | 526370 | 2753100 | 256040 | 0 | 0 | 0 | 0 | 0 |
| RVASALPG | RAB6B | Q9NRW1 | RAB6B_HUMAN | Oxidation (M) | 0 | 0 | 0 | 0 | 0 | 0 | 0 | 0 | 0 | 0 | 171800 | 682870 | 0 | 0 | 0 | 0 | 0 | 0 |
| RVASGPSP | MVP | Q14764 | MVP_HUMAN N | Unmodified | 1598900 | 517360 | 0 | 0 | 2028100 | 0 | 0 | 0 | 392020 | 0 | 0 | 3889700 | 0 | 0 | 0 | 0 | 0 | 0 |
| RVASLLSQI | AVIL; AVIL | O75366; F8V | AVIL_HUMAN A | Unmodified | 0 | 0 | 0 | 0 | 0 | 0 | 0 | 0 | 0 | 0 | 1403800 | 5300500 | 0 | 0 | 0 | 0 | 0 | 0 |
| RVATPVDW | PRDX6 | P30041 | PRDX6_HUMAN | Oxidation (M) | 493340 | 0 | 1568200 | 3515900 | 11907000 | 1822800 | 0 | 378660 | 2875500 | 2297400 | 0 | 9453700 | 22898000 | 18043000 | 3008800 | 3251300 | 27933000 | 44266000 |
| RVATPVDW | PRDX6 | P30041 | PRDX6_HUMAN | Unmodified | 0 | 0 | 0 | 1013100 | 11011000 | 1526600 | 66598 | 0 | 3876100 | 0 | 0 | 1387400 | 15125000 | 13324000 | 0 | 1855300 | 17980000 | 44126000 |
| RVAVILNEF | CBWD2; CBWD | Q8IUF1; Q9E | CBWD2_HUMA | Unmodified | 0 | 0 | 0 | 0 | 0 | 0 | 0 | 0 | 0 | 0 | 0 | 0 | 986560 | 0 | 0 | 0 | 0 | 0 |
| RVAYCFEV | POFUT1 | Q9H488 | OFUT1_HUMAN | Unmodified | 0 | 0 | 0 | 0 | 0 | 0 | 0 | 0 | 1180600 | 0 | 0 | 0 | 0 | 0 | 0 | 0 | 0 | 0 |
| RVCEEIAIP | RPS17; RPS17; | P08708; H0 | RS17_HUMAN | Unmodified | 0 | 0 | 0 | 0 | 0 | 0 | 0 | 1238200 | 0 | 0 | 223290 | 14762000 | 1064100 | 0 | 0 | 0 | 0 | 0 |
| RVCEEIAIP | RPS17; RPS17; | P08708; H0 | RS17_HUMAN | Unmodified | 232240 | 115820 | 300890 | 0 | 0 | 0 | 0 | 0 | 0 | 71778 | 272780 | 2834200 | 0 | 0 | 0 | 158420 | 0 | 0 |
| RVDFHDV( | RAB24; RAB24 | Q969Q5; F8 | RAB24_HUMAN | Unmodified | 399720 | 285350 | 551190 | 352140 | 0 | 0 | 0 | 0 | 0 | 54995 | 0 | 0 | 0 | 0 | 294340 | 0 | 76799 | 0 |
| RVEAALSA( | PLCB2; PLCB2 | Q00722; Q9I | PLCB2_HUMAN | Unmodified | 0 | 0 | 0 | 0 | 0 | 0 | 0 | 0 | 0 | 0 | 0 | 1141900 | 0 | 0 | 0 | 0 | 0 | 0 |
| RVEAAQET | DCTN1; DCTN1 | Q14203; Q6 | DCTN1_HUMAI | Unmodified | 0 | 417720 | 0 | 0 | 0 | 0 | 0 | 0 | 291570 | 145190 | 0 | 1507300 | 1176900 | 508970 | 0 | 0 | 0 | 1025000 |
| RVEAAQET | DCTN1; DCTN1 | Q14203; Q6 | DCTN1_HUMAI | Unmodified | 0 | 0 | 0 | 193540 | 1285000 | 0 | 163910 | 106380 | 197540 | 232160 | 0 | 2636200 | 563330 | 408850 | 460970 | 0 | 0 | 639590 |
| RVEDEVNS | SLC35A4 | L0R6Q1 | S35U4_HUMAN | Unmodified | 0 | 1061300 | 0 | 0 | 0 | 0 | 1094100 | 282200 | 1255500 | 642730 | 0 | 540090 | 0 | 0 | 0 | 0 | 0 | 0 |
| RVEILSEGN | RACGAP1; RAC | Q9H0H5; F8 | RGAP1_HUMAI | Unmodified | 953320 | 0 | 1397700 | 4681300 | 37315000 | 7028000 | 44742 | 0 | 9945100 | 6310500 | 7127000 | 8736100 | 3645600 | 2433700 | 2258600 | 823250 | 2852800 | 2966000 |
| RVEIMEEES | USP14; USP14 | P54578; A6N | UBP14_HUMAN | Unmodified | 0 | 0 | 0 | 874200 | 1462400 | 632790 | 300360 | 165830 | 966530 | 489760 | 1471300 | 7526400 | 0 | 682090 | 477660 | 0 | 748490 | 797360 |
| RVEIMEEES | USP14; USP14 | P54578; A6N | UBP14_HUMAN | Oxidation (M) | 0 | 0 | 0 | 0 | 0 | 0 | 0 | 0 | 0 | 0 | 0 | 0 | 0 | 0 | 0 | 0 | 0 | 0 |
| RVEIQMPKI | SEPTIN9; SEPTI | Q9UHD8; K7 | SEPT9_HUMAN | Unmodified | 0 | 0 | 0 | 0 | 0 | 0 | 0 | 0 | 1484800 | 0 | 0 | 0 | 0 | 0 | 0 | 0 | 1384700 | 1950500 |
| RVEIQMPKI | SEPTIN9; SEPTI | Q9UHD8; K7 | SEPT9_HUMAN | Oxidation (M) | 0 | 0 | 0 | 0 | 0 | 0 | 0 | 0 | 0 | 0 | 0 | 3095000 | 0 | 0 | 0 | 0 | 0 | 1213600 |
| RVEISAPSN | PAK4; PAK4; PA | O96013; M0I | PAK4_HUMAN | Unmodified | 0 | 262650 | 801630 | 0 | 69215 | 0 | 0 | 0 | 0 | 0 | 641780 | 374820 | 0 | 0 | 460240 | 570970 | 1929300 | 0 |
| RVEISAPSN | PAK4; PAK4; PA | O96013; M0I | PAK4_HUMAN | Unmodified | 0 | 0 | 0 | 62781 | 1245700 | 0 | 0 | 0 | 0 | 0 | 0 | 0 | 955110 | 0 | 0 | 0 | 184740 | 692330 |
| RVEPGSPA | SLC9A3R2 | Q15599 | NHRF2_HUMAI | Unmodified | 659690 | 702300 | 312580 | 1245100 | 542750 | 0 | 0 | 0 | 598090 | 0 | 86382 | 2553800 | 32478 | 0 | 1440600 | 1252800 | 89709 | 178580 |
| RVESPVLPI | SMAD1; SMAD5 | Q15797; Q9 | SMAD1_HUMA | Unmodified | 0 | 0 | 0 | 0 | 877390 | 0 | 0 | 117470 | 0 | 0 | 881520 | 2898800 | 1369300 | 0 | 0 | 0 | 457120 | 1121700 |

| | | | | | | | | | | | | | | | | | | | | | | |
|---|---|---|---|---|---|---|---|---|---|---|---|---|---|---|---|---|---|---|---|---|---|---|
| RVEVVEED | ZNF185; ZNF18 | O15231; H7I | ZN185_HUMAN | Unmodified | 0 | 0 | 90109 | 0 | 1689300 | 78196 | 0 | 0 | 414030 | 37012 | 0 | 539520 | 0 | 0 | 0 | 0 | 1569500 | 0 |
| RVEVVEED | ZNF185; ZNF18 | O15231; H7I | ZN185_HUMAN | Unmodified | 0 | 0 | 0 | 0 | 353400 | 0 | 0 | 0 | 0 | 0 | 0 | 0 | 0 | 0 | 0 | 0 | 861660 | 361440 |
| RVFITDDFH | HSP90B1; HSP | P14625; A0A | ENPL_HUMAN | 2 Oxidation (M) | 0 | 0 | 0 | 76944 | 0 | 0 | 1204400 | 81605 | 5506800 | 5673700 | 2871000 | 2533900 | 0 | 0 | 0 | 0 | 0 | 0 |
| RVFITDDFH | HSP90B1; HSP | P14625; A0A | ENPL_HUMAN | Oxidation (M) | 333660 | 165780 | 0 | 503420 | 0 | 0 | 2212800 | 0 | 49775000 | 25190000 | 5968300 | 4285100 | 0 | 146250 | 0 | 30335 | 0 | 0 |
| RVFITDDFH | HSP90B1; HSP | P14625; A0A | ENPL_HUMAN | Unmodified | 0 | 0 | 0 | 0 | 63870 | 0 | 0 | 0 | 46633000 | 8309800 | 227290 | 819520 | 0 | 0 | 0 | 0 | 0 | 0 |
| RVFNDCRD | GNAS; GNAS; G | Q5JWF2; P6 | GNAS1_HUMA | Unmodified | 0 | 0 | 803930 | 1107800 | 2069200 | 0 | 0 | 0 | 0 | 0 | 1006300 | 0 | 3166000 | 1012700 | 0 | 0 | 1196700 | 1797500 |
| RVFNPYTE | EFHD2 | Q96C19 | EFHD2_HUMA | Unmodified | 0 | 0 | 0 | 0 | 2764900 | 0 | 0 | 0 | 474790 | 231890 | 0 | 466130 | 0 | 0 | 0 | 0 | 0 | 261050 |
| RVGFQYEG | G6PD | P11413 | G6PD_HUMAN | Unmodified | 115630 | 0 | 0 | 314350 | 30813000 | 11604000 | 642760 | 591460 | 1882100 | 132470 | 0 | 4855000 | 4552500 | 3517600 | 660680 | 164160 | 2836600 | 4769600 |
| RVGVWNV | PLOD1 | Q02809 | PLOD1_HUMA | Unmodified | 0 | 0 | 35179 | 0 | 0 | 0 | 0 | 0 | 4781200 | 2630000 | 110860 | 0 | 0 | 0 | 0 | 0 | 0 | 0 |
| RVGVWNV | PLOD3; PLOD3 | O60568; H7 | PLOD3_HUMA | Unmodified | 0 | 0 | 0 | 0 | 0 | 0 | 0 | 0 | 3600900 | 279470 | 0 | 0 | 0 | 0 | 0 | 0 | 0 | 0 |
| RVHLAPPL | MGAT1 | P26572 | MGAT1_HUMA | Unmodified | 0 | 0 | 0 | 0 | 0 | 0 | 0 | 174470 | 2787400 | 1376400 | 0 | 0 | 0 | 0 | 0 | 0 | 0 | 69431 |
| RVHLMNPN | YARS1; YARS1 | P54577; A0A | SYYC_HUMAN | Oxidation (M) | 0 | 0 | 0 | 0 | 275180 | 0 | 144210 | 18495 | 235230 | 0 | 0 | 207920 | 552510 | 347450 | 0 | 0 | 240380 | 90748 |
| RVHPVSTM | LDHA | P00338 | LDHA_HUMAN | Unmodified | 0 | 0 | 0 | 0 | 0 | 0 | 0 | 0 | 0 | 0 | 508440 | 9088200 | 150620 | 0 | 0 | 0 | 0 | 751110 |
| RVHVTQED | PSMC5 | P62195 | PRS8_HUMAN | Unmodified | 0 | 0 | 0 | 0 | 878280 | 0 | 237810 | 0 | 1195900 | 532330 | 0 | 1355900 | 1275000 | 322790 | 0 | 0 | 313290 | 729840 |
| RVIAEGDL | BPNT1; BPNT1; | O95861; F8 | BPNT1_HUMA | Unmodified | 0 | 0 | 0 | 0 | 698870 | 0 | 0 | 0 | 329720 | 138420 | 490980 | 1192400 | 522590 | 0 | 0 | 0 | 0 | 406780 |
| RVIESGPD | PDP1 | Q9P0J1 | PDP1_HUMAN | Unmodified | 0 | 10084 | 0 | 0 | 0 | 0 | 0 | 0 | 599150 | 245420 | 0 | 0 | 0 | 0 | 0 | 0 | 0 | 47078 |
| RVIGDNPW | ECPAS; ECPAS; | Q5VYK3; R4 | ECM29_HUMA | Unmodified | 0 | 0 | 0 | 0 | 0 | 0 | 0 | 0 | 0 | 0 | 0 | 0 | 0 | 0 | 0 | 0 | 0 | 0 |
| RVIISAPSA | GAPDH; GAPDH | P04406; E7E | G3P_HUMAN G | 2 Oxidation (M) | 12901000 | 8746100 | 25039000 | 24678000 | 52879000 | 14707000 | 18443000 | 16178000 | 6725300 | 5609700 | 39585000 | 2.3E+08 | 11387000 | 21533000 | 24487000 | 25027000 | 5602900 | 11848000 |
| RVIISAPSA | GAPDH; GAPDH | P04406; E7E | G3P_HUMAN G | Oxidation (M) | 7099200 | 6360300 | 6841600 | 13293000 | 1.41E+08 | 28822000 | 25702000 | 4817100 | 29850000 | 7565800 | 25078000 | 1.96E+08 | 60777000 | 35354000 | 18760000 | 9844800 | 37783000 | 1.04E+08 |
| RVIISAPSA | GAPDH; GAPDH | P04406; E7E | G3P_HUMAN G | Unmodified | 0 | 0 | 0 | 885610 | 83935000 | 1348100 | 1645300 | 0 | 24420000 | 918520 | 1343600 | 8112900 | 8351400 | 7434300 | 0 | 0 | 9793700 | 50509000 |
| RVIISAPSA | GAPDH; GAPDH | P04406; E7E | G3P_HUMAN G | 2 Oxidation (M) | 995790 | 0 | 0 | 2196500 | 84396000 | 10880000 | 29880000 | 7874900 | 282900 | 10012000 | 567340 | 1817400 | 36433000 | 28514000 | 0 | 0 | 64661000 | 47335000 |
| RVIISAPSA | GAPDH; GAPDH | P04406; E7E | G3P_HUMAN G | Oxidation (M) | 0 | 0 | 0 | 943920 | 1.95E+08 | 12169000 | 56205000 | 1485400 | 1100700 | 8924900 | 0 | 1459500 | 81457000 | 39741000 | 0 | 0 | 1.49E+08 | 1.31E+08 |
| RVIISAPSA | GAPDH; GAPDH | P04406; E7E | G3P_HUMAN G | Unmodified | 0 | 0 | 0 | 0 | 1.12E+08 | 3360800 | 20198000 | 131680 | 1580200 | 3340500 | 0 | 0 | 24568000 | 2678500 | 0 | 0 | 48712000 | 86972000 |
| RVIQEGRG | AIP; AIP | O00170; E9F | AIP_HUMAN AH | Unmodified | 0 | 50733 | 1170000 | 0 | 1615100 | 266700 | 0 | 39629 | 299210 | 140590 | 56666 | 1090100 | 2670100 | 167250 | 317230 | 0 | 1146500 | 1492100 |
| RVKDSEDV | KRAS | P01116 | RASK_HUMAN | Unmodified | 0 | 0 | 0 | 0 | 0 | 0 | 0 | 0 | 0 | 0 | 0 | 0 | 0 | 0 | 0 | 0 | 0 | 0 |
| RVLDGLHN | UGDH | O60701 | UGDH_HUMAN | Unmodified | 0 | 0 | 0 | 0 | 2994100 | 1041900 | 0 | 0 | 0 | 0 | 0 | 0 | 0 | 837960 | 0 | 0 | 0 | 0 |
| RVLDGLHN | UGDH | O60701 | UGDH_HUMAN | Unmodified | 0 | 0 | 0 | 0 | 1365800 | 0 | 0 | 0 | 0 | 0 | 0 | 0 | 495130 | 0 | 0 | 0 | 0 | 0 |
| RVLDVEVY | POMGNT1; PON | Q8WZA1; Q6 | PMGT1_HUMA | Unmodified | 0 | 0 | 0 | 0 | 0 | 0 | 0 | 0 | 1436700 | 0 | 0 | 877520 | 653660 | 0 | 0 | 0 | 0 | 0 |
| RVLIAAHG | PGAM1; PGAM2 | P18669; P15 | PGAM1_HUMA | Unmodified | 46393 | 0 | 85284 | 0 | 287140 | 138840 | 0 | 0 | 126200 | 0 | 85430 | 684940 | 242900 | 63953 | 76412 | 69625 | 43763 | 173810 |
| RVLIAAHG | PGAM1; PGAM2 | P18669; P15 | PGAM1_HUMA | Unmodified | 0 | 0 | 0 | 0 | 2029100 | 354130 | 194690 | 0 | 0 | 73117 | 0 | 0 | 508780 | 526650 | 0 | 0 | 373690 | 715910 |
| RVLLIYDGI | YOD1 | Q5VVQ6 | OTU1_HUMAN | Unmodified | 652370 | 1984600 | 73977 | 133810 | 0 | 0 | 0 | 0 | 0 | 0 | 0 | 204110 | 0 | 0 | 420160 | 191060 | 0 | 0 |
| RVLNTEAN | QSOX1 | O00391 | QSOX1_HUMA | Unmodified | 0 | 0 | 0 | 0 | 0 | 0 | 0 | 0 | 1332000 | 0 | 0 | 0 | 0 | 0 | 0 | 0 | 0 | 0 |
| RVLNTEAN | QSOX1 | O00391 | QSOX1_HUMA | Unmodified | 0 | 0 | 0 | 0 | 0 | 0 | 0 | 0 | 496880 | 437710 | 0 | 0 | 0 | 0 | 0 | 0 | 18949 | 0 |
| RVLQALEG | RPS19; RPS19 | P39019; A0A | RS19_HUMAN | Unmodified | 1104500 | 0 | 1266800 | 2503600 | 0 | 0 | 296710 | 0 | 352720 | 361760 | 826690 | 5479700 | 1782600 | 703700 | 1970400 | 841390 | 0 | 0 |
| RVLQESCE | FZD6 | O60353 | FZD6_HUMAN | Unmodified | 0 | 0 | 0 | 0 | 704590 | 0 | 79548 | 0 | 274950 | 274590 | 0 | 0 | 0 | 0 | 0 | 0 | 0 | 127940 |
| RVLQQFAD | HSD17B4; HSD | P51659; E7E | DHB4_HUMAN | Unmodified | 0 | 0 | 0 | 0 | 0 | 0 | 264690 | 0 | 6393600 | 348440 | 167330 | 0 | 0 | 0 | 0 | 0 | 0 | 0 |
| RVLQQFAD | HSD17B4; HSD | P51659; E7E | DHB4_HUMAN | Unmodified | 0 | 0 | 0 | 0 | 145700 | 0 | 1147900 | 0 | 6828700 | 3744300 | 124130 | 0 | 441090 | 104660 | 0 | 0 | 0 | 20277 |
| RVLSGEFQ | RRM1; RRM1 | P23921; E9P | RIR1_HUMAN F | Unmodified | 41203 | 0 | 56626 | 62583 | 0 | 0 | 0 | 0 | 199670 | 0 | 0 | 4405900 | 220400 | 0 | 209820 | 63431 | 0 | 0 |
| RVNAIEEV | GGA1; GGA1 | Q9UJY5; B0 | GGA1_HUMAN | Unmodified | 483740 | 648240 | 0 | 0 | 0 | 0 | 0 | 0 | 0 | 0 | 378300 | 888310 | 392770 | 0 | 171040 | 109960 | 0 | 0 |
| RVNKEESL | KTN1 | Q86UP2 | KTN1_HUMAN | Oxidation (M) | 0 | 0 | 0 | 0 | 0 | 0 | 0 | 0 | 0 | 0 | 0 | 3063800 | 0 | 230370 | 0 | 0 | 0 | 0 |
| RVNLAIWD | RAB21 | Q9UL25 | RAB21_HUMAN | Unmodified | 3761200 | 0 | 0 | 0 | 0 | 0 | 282960 | 0 | 1574200 | 0 | 0 | 5285900 | 0 | 0 | 0 | 3018800 | 0 | 432540 |
| RVNLAIWD | RAB21 | Q9UL25 | RAB21_HUMAN | Unmodified | 0 | 0 | 0 | 534100 | 0 | 0 | 0 | 0 | 277310 | 0 | 0 | 0 | 85820 | 0 | 0 | 0 | 360160 | 0 |
| RVNSASSS | POLDIP3; POLD | Q9BY77; F6 | PDIP3_HUMAN | Unmodified | 0 | 0 | 0 | 0 | 0 | 0 | 0 | 0 | 0 | 0 | 116030 | 959700 | 0 | 0 | 0 | 0 | 0 | 0 |
| RVNSKFDT | TRIM25; TRIM2 | Q14258; A0 | TRI25_HUMAN | Unmodified | 48301 | 0 | 54776 | 291370 | 21389000 | 5690500 | 461070 | 0 | 0 | 282260 | 365190 | 1782800 | 2759900 | 2383100 | 0 | 282810 | 2537400 | 3532800 |
| RVNSLPEV | ITGA6 | P23229 | ITA6_HUMAN I | Unmodified | 0 | 0 | 0 | 0 | 1776300 | 397550 | 0 | 0 | 0 | 0 | 214810 | 1007400 | 0 | 0 | 0 | 0 | 1484300 | 6688000 |
| RVNSLPEV | ITGA6 | P23229 | ITA6_HUMAN I | Unmodified | 0 | 0 | 0 | 0 | 2806500 | 0 | 0 | 0 | 0 | 0 | 0 | 624270 | 3446300 | 0 | 0 | 0 | 4039500 | 15513000 |
| RVPADEAY | FARP2 | O94887 | FARP2_HUMAN | Unmodified | 0 | 0 | 268810 | 252770 | 0 | 0 | 0 | 0 | 0 | 0 | 0 | 0 | 369400 | 841640 | 356740 | 306810 | 0 | 0 |
| RVPVYIILD | FAM83F | Q8NEG4 | FA83F_HUMAN | Unmodified | 0 | 0 | 0 | 0 | 0 | 0 | 0 | 0 | 0 | 0 | 0 | 606290 | 0 | 0 | 908880 | 1832800 | 0 | 0 |
| RVQLAEDL | OPA1; OPA1; O | O60313; E5 | OPA1_HUMAN | Unmodified | 0 | 0 | 0 | 0 | 291350 | 0 | 218580 | 0 | 29578 | 65269 | 0 | 915770 | 0 | 0 | 0 | 0 | 133740 | 0 |
| RVQVYLHE | CUL1; CUL1; C | Q13616; A0 | CUL1_HUMAN | Unmodified | 0 | 0 | 0 | 0 | 160330 | 0 | 86469 | 0 | 0 | 0 | 0 | 304910 | 0 | 0 | 0 | 0 | 0 | 0 |
| RVREAMCF | JUP; JUP; JUP; J | P14923; C9J | PLAK_HUMAN | Oxidation (M) | 0 | 0 | 0 | 0 | 1289000 | 0 | 0 | 0 | 0 | 0 | 0 | 0 | 1008500 | 1025000 | 0 | 0 | 1199100 | 0 |
| RVSEVEAV | CIT; CIT; CIT; C | O14578; A0 | CTRO_HUMAN | Unmodified | 0 | 0 | 0 | 0 | 1490100 | 0 | 0 | 87553 | 0 | 0 | 0 | 0 | 0 | 0 | 0 | 0 | 59550 | 389680 |
| RVSIMSYAL | SNX9; SNX9; SN | Q9Y5X1; A0 | SNX9_HUMAN | Oxidation (M) | 1373100 | 494470 | 677940 | 0 | 0 | 0 | 0 | 0 | 0 | 0 | 0 | 0 | 0 | 0 | 0 | 438860 | 0 | 0 |
| RVSLVGAD | SCRIB; SCRIB; | Q14160; A0 | SCRIB_HUMAN | Unmodified | 106340 | 0 | 224040 | 375870 | 844880 | 0 | 0 | 30969 | 2670600 | 879350 | 189790 | 828690 | 475070 | 0 | 248850 | 158630 | 529170 | 1643500 |
| RVSPVGET | ANTXR2; ANTXR | P58335; A0A | ANTR2_HUMAN | Unmodified | 0 | 0 | 0 | 0 | 492130 | 0 | 0 | 0 | 304610 | 0 | 0 | 0 | 123990 | 0 | 0 | 0 | 226940 | 4425000 |
| RVSVCAET | PRKAR2A | P13861 | KAP2_HUMAN | Unmodified | 0 | 0 | 0 | 0 | 0 | 0 | 0 | 0 | 133950 | 0 | 0 | 0 | 0 | 0 | 470650 | 347610 | 0 | 0 |
| RVSVELTN | JUP | P14923 | PLAK_HUMAN | Unmodified | 0 | 0 | 3153000 | 5039300 | 3611400 | 0 | 0 | 0 | 1878400 | 0 | 2896600 | 8761700 | 1912600 | 0 | 5141200 | 6060900 | 602800 | 928970 |
| RVTAYTVD | AHNAK | Q09666 | AHNK_HUMAN | Unmodified | 0 | 53929 | 0 | 0 | 3193200 | 697300 | 0 | 0 | 0 | 0 | 0 | 0 | 0 | 0 | 0 | 0 | 0 | 2004700 |
| RVTDHLGV | DNAJB12; DNA | Q9NXW2; V | DJB12_HUMAN | Unmodified | 216580 | 416580 | 0 | 0 | 0 | 0 | 0 | 0 | 2472700 | 203010 | 0 | 358770 | 0 | 44896 | 0 | 0 | 0 | 329870 |
| RVTFSLLD | TRAF4; TRAF4 | Q9BUZ4; Q2 | TRAF4_HUMAN | Unmodified | 7540700 | 1838500 | 0 | 0 | 0 | 0 | 0 | 0 | 0 | 0 | 0 | 13689000 | 0 | 0 | 4249000 | 1378700 | 0 | 0 |
| RVTIMPKD | H3-4; H3-3A; H | Q16695; P8 | H31T_HUMAN | Unmodified | 0 | 0 | 0 | 0 | 811640 | 0 | 4649100 | 1684900 | 0 | 0 | 0 | 912670 | 0 | 0 | 0 | 0 | 0 | 2163200 |
| RVTIMPKD | H3-4; H3-3A; H | Q16695; P8 | H31T_HUMAN | Oxidation (M) | 0 | 0 | 0 | 0 | 0 | 0 | 87758 | 2949800 | 0 | 0 | 0 | 0 | 0 | 0 | 0 | 0 | 0 | 0 |
| RVTLELGG | ALDH2; ALDH1 | P05091; P0 | ALDH2_HUMA | Unmodified | 0 | 0 | 0 | 0 | 0 | 0 | 0 | 0 | 0 | 0 | 4673400 | 1.55E+08 | 0 | 0 | 0 | 0 | 0 | 0 |
| RVTNDARE | SNAP25 | P60880 | SNP25_HUMA | Unmodified | 0 | 0 | 0 | 0 | 0 | 0 | 0 | 0 | 2855600 | 0 | 0 | 0 | 0 | 0 | 0 | 0 | 0 | 0 |
| RVTNVGSL | WASL | O00401 | WASL_HUMAN | Unmodified | 1023800 | 3313200 | 153280 | 947210 | 0 | 0 | 153780 | 0 | 370240 | 55977 | 0 | 0 | 324210 | 0 | 3805200 | 317070 | 0 | 0 |
| RVTNVGSL | WASL | O00401 | WASL_HUMAN | Unmodified | 3257100 | 0 | 0 | 5999800 | 0 | 0 | 0 | 0 | 0 | 0 | 0 | 489070 | 0 | 0 | 0 | 4839600 | 0 | 0 |
| RVTSIADRL | PRPS1; PRPS1; | P60891; B1A | PRPS1_HUMAN | Unmodified | 0 | 0 | 138730 | 556740 | 2655500 | 0 | 3382200 | 0 | 634930 | 918230 | 496630 | 1570700 | 6541000 | 2600200 | 0 | 0 | 0 | 2466500 |
| RVTSIADRL | PRPS2 | P11908 | PRPS2_HUMAN | Unmodified | 0 | 0 | 0 | 0 | 0 | 0 | 1563600 | 0 | 0 | 1519200 | 341500 | 0 | 3858000 | 1765100 | 206000 | 0 | 1731400 | 1859400 |
| RVVAEPVE | NDUFS3 | O75489 | NDUS3_HUMA | Unmodified | 206100 | 0 | 0 | 0 | 0 | 0 | 0 | 0 | 3341700 | 180260 | 0 | 509670 | 0 | 0 | 0 | 0 | 0 | 0 |
| RVVALSMS | WDR82 | Q6UXN9 | WDR82_HUMA | Oxidation (M) | 0 | 0 | 0 | 0 | 0 | 0 | 0 | 102960 | 0 | 0 | 0 | 2137000 | 0 | 0 | 0 | 0 | 0 | 0 |
| RVVFMGD | PIGO | Q8TEQ8 | PIGO_HUMAN | Oxidation (M) | 0 | 0 | 0 | 0 | 0 | 0 | 0 | 0 | 870620 | 529420 | 0 | 0 | 0 | 0 | 0 | 0 | 0 | 0 |
| RVVHASGD | EPB41L3; EPB4 | Q9Y2J2; A0A | E41L3_HUMAN | Unmodified | 0 | 0 | 0 | 0 | 0 | 0 | 0 | 0 | 0 | 0 | 0 | 0 | 268930 | 0 | 0 | 0 | 445520 | 2754900 |
| RVVLLAEG | AHCYL1 | O43865 | SAHH2_HUMA | Unmodified | 112120 | 68107 | 832400 | 187520 | 482530 | 89395 | 18607 | 0 | 378060 | 0 | 38302 | 0 | 99615 | 29545 | 633150 | 482270 | 206770 | 0 |
| RVVMRVDF | PGK1 | P00558 | PGK1_HUMAN | Unmodified | 0 | 0 | 0 | 0 | 612320 | 0 | 0 | 0 | 0 | 282790 | 0 | 0 | 448430 | 0 | 0 | 0 | 939290 | 1971300 |
| RVWGSPG | OS9; OS9 | Q13438; B4 | OS9_HUMAN P | Unmodified | 0 | 0 | 0 | 0 | 0 | 0 | 0 | 0 | 1569900 | 154210 | 0 | 0 | 0 | 0 | 0 | 0 | 179990 | 0 |
| RVYATILNA | FASN; FASN | P49327; A0A | FAS_HUMAN F | Unmodified | 640760 | 0 | 1490600 | 1921300 | 1025300 | 0 | 0 | 0 | 1132600 | 0 | 2312700 | 51504000 | 596740 | 0 | 4169400 | 2617900 | 253990 | 0 |
| RVYATILNA | FASN; FASN | P49327; A0A | FAS_HUMAN F | Unmodified | 0 | 0 | 0 | 6200000 | 0 | 0 | 0 | 0 | 991360 | 0 | 0 | 2088400 | 5843500 | 0 | 2022500 | 0 | 0 | 0 |
| RVYKGEFQ | TPST1 | O60507 | TPST1_HUMAN | Unmodified | 0 | 0 | 0 | 0 | 0 | 0 | 0 | 0 | 1006700 | 571760 | 0 | 0 | 0 | 0 | 0 | 0 | 0 | 0 |
| RVYSLFLD | RUVBL2 | Q9Y230 | RUVB2_HUMA | Unmodified | 425970 | 0 | 0 | 0 | 0 | 0 | 31242 | 21826 | 0 | 0 | 0 | 5431200 | 0 | 0 | 171500 | 0 | 0 | 0 |
| RVYVSGLM | APMAP; APMAP | Q9HDC9; H | APMAP_HUMA | Unmodified | 0 | 0 | 0 | 0 | 0 | 0 | 0 | 0 | 1700400 | 823300 | 0 | 0 | 0 | 32707 | 0 | 0 | 0 | 0 |
| RVYVSLPN | SPAST; SPAST; | Q9UBP0; A0 | SPAST_HUMAN | Unmodified | 0 | 0 | 0 | 104680 | 3282500 | 448100 | 0 | 0 | 0 | 588800 | 122450 | 0 | 668790 | 600460 | 0 | 0 | 320020 | 184080 |
| RWDQTAD | SF3B1 | O75533 | SF3B1_HUMAN | Unmodified | 0 | 0 | 0 | 0 | 0 | 0 | 0 | 0 | 0 | 0 | 106110 | 0 | 0 | 0 | 57619 | 0 | 0 | 0 |
| RWEATQQ | KLK14; KLK14 | Q9P0G3; A0 | KLK14_HUMAN | Unmodified | 0 | 0 | 0 | 0 | 0 | 0 | 0 | 0 | 2095600 | 64026 | 0 | 0 | 0 | 0 | 0 | 0 | 0 | 0 |
| RWENMPV | VPS36; VPS36 | Q86VN1; A0 | VPS36_HUMAN | Unmodified | 445960 | 227160 | 178340 | 149620 | 0 | 0 | 0 | 0 | 0 | 0 | 0 | 182440 | 0 | 0 | 710440 | 454040 | 0 | 0 |
| RWENMPV | VPS36; VPS36 | Q86VN1; A0 | VPS36_HUMAN | Oxidation (M) | 517510 | 662180 | 0 | 0 | 0 | 0 | 0 | 0 | 0 | 0 | 0 | 79091 | 0 | 0 | 760570 | 882530 | 0 | 0 |
| RWEVADL | UQCRC2; UQC | P22695; H3 | QCR2_HUMAN | Unmodified | 0 | 0 | 0 | 0 | 0 | 0 | 0 | 0 | 1067300 | 0 | 0 | 0 | 0 | 0 | 0 | 0 | 0 | 0 |
| RWEVADL | UQCRC2; UQC | P22695; H3 | QCR2_HUMAN | Unmodified | 0 | 0 | 0 | 0 | 0 | 0 | 0 | 0 | 1456700 | 286770 | 0 | 0 | 0 | 0 | 0 | 0 | 0 | 0 |
| RWFGFLEA | EPRS1; EPRS1; | P07814; V9 | SYEP_HUMAN | Unmodified | 0 | 0 | 0 | 0 | 559450 | 0 | 257610 | 0 | 0 | 645180 | 32071 | 0 | 0 | 1250800 | 0 | 0 | 1643800 | 767070 |
| RWIGSETD | PPP1R12A; PPP | O14974; F8 | MYPT1_HUMAN | Unmodified | 188990 | 0 | 0 | 0 | 4590400 | 558660 | 0 | 0 | 0 | 223840 | 0 | 0 | 1439100 | 1274200 | 0 | 0 | 795250 | 0 |
| RWIHDVRT | ARRDC1; ARRD | Q8N5I2; Q5 | ARRD1_HUMA | Unmodified | 0 | 0 | 0 | 0 | 2090200 | 0 | 0 | 0 | 0 | 0 | 0 | 113970 | 1167000 | 0 | 0 | 0 | 816530 | 1749400 |
| RWILQPCS | ARL15; ARL15 | Q9NXU5; A0 | ARL15_HUMAN | Oxidation (M) | 0 | 0 | 0 | 0 | 0 | 0 | 0 | 0 | 0 | 0 | 0 | 1813700 | 0 | 0 | 0 | 0 | 0 | 0 |
| RWILQPCS | ARL15; ARL15 | Q9NXU5; A0 | ARL15_HUMAN | Unmodified | 0 | 0 | 0 | 0 | 0 | 0 | 0 | 0 | 0 | 0 | 0 | 169580 | 0 | 0 | 0 | 0 | 0 | 0 |
| RWIYEDVE | CALU | O43852 | CALU_HUMAN | Unmodified | 0 | 0 | 0 | 0 | 0 | 0 | 0 | 0 | 1001000 | 0 | 0 | 0 | 0 | 0 | 0 | 0 | 0 | 0 |
| RWIYEDVE | CALU | O43852 | CALU_HUMAN | Unmodified | 0 | 0 | 364410 | 251260 | 0 | 0 | 415040 | 0 | 3509400 | 1443800 | 631760 | 0 | 245130 | 0 | 0 | 0 | 0 | 398230 |
| RWLHEIN | RAB35; RAB35 | Q15286; F5 | RAB35_HUMAN | Unmodified | 750220 | 455960 | 0 | 0 | 0 | 0 | 0 | 0 | 0 | 0 | 0 | 0 | 0 | 0 | 795640 | 411300 | 0 | 0 |
| RWLLLCNF | PYGB | P11216 | PYGB_HUMAN | Unmodified | 0 | 0 | 0 | 0 | 345250 | 0 | 39416 | 0 | 760660 | 446950 | 120540 | 884870 | 0 | 0 | 84721 | 45246 | 537500 | 3073900 |
| RWLLLCNF | PYGL; PYGL | P06737; E9P | PYGL_HUMAN | Unmodified | 0 | 0 | 0 | 0 | 237910 | 70991 | 0 | 0 | 0 | 0 | 0 | 0 | 0 | 0 | 93074 | 20011 | 89144 | 491560 |
| RWLPAGD | EEF2 | P13639 | EF2_HUMAN E | Oxidation (M) | 154440 | 479360 | 239560 | 612560 | 4240600 | 127290 | 3117500 | 0 | 251220 | 392110 | 165900 | 1161200 | 1836400 | 766050 | 1319700 | 69765 | 993900 | 2581500 |
| RWLPAGD | EEF2 | P13639 | EF2_HUMAN E | Unmodified | 0 | 0 | 0 | 0 | 569930 | 46138 | 203090 | 0 | 70682 | 172800 | 27541 | 107310 | 0 | 125330 | 89680 | 0 | 0 | 0 |
| RWMEACL | IQGAP1; IQGAP | P46940; A0A | IQGA1_HUMAN | Oxidation (M) | 0 | 0 | 0 | 235310 | 11142000 | 94533 | 0 | 0 | 0 | 0 | 0 | 0 | 0 | 0 | 0 | 0 | 1091300 | 0 |
| RWNSLPSE | TOMM34 | Q15785 | TOM34_HUMA | Unmodified | 0 | 0 | 0 | 0 | 233220 | 0 | 48197 | 21089 | 162810 | 0 | 0 | 0 | 364820 | 0 | 0 | 0 | 313610 | 193100 |
| RWQLSVAT | DDX21 | Q9NR30 | DDX21_HUMA | Unmodified | 0 | 0 | 0 | 0 | 0 | 0 | 0 | 0 | 0 | 0 | 0 | 0 | 0 | 0 | 0 | 0 | 0 | 0 |
| RWQTLLSV | GNS; GNS; GN | P15586; H7 | GNS_HUMAN N | Unmodified | 0 | 21148 | 0 | 0 | 1087400 | 92389 | 41892 | 0 | 8915900 | 6023900 | 1291300 | 0 | 2578300 | 2019200 | 0 | 0 | 0 | 874020 |
| RWSSSDW | KYAT3; KYAT3 | Q6YP21; A0 | KAT3_HUMAN I | Unmodified | 0 | 0 | 0 | 0 | 0 | 0 | 0 | 28583 | 200080 | 51765 | 151980 | 87929 | 0 | 0 | 0 | 0 | 0 | 0 |
| RWVDPNS | AKR1C3; AKR1 | P42330; A0A | AK1C3_HUMAN | Unmodified | 106660 | 91314 | 0 | 0 | 3399800 | 293810 | 0 | 0 | 0 | 0 | 0 | 0 | 0 | 0 | 0 | 0 | 358830 | 0 |
| RYADLTED | PGAM1 | P18669 | PGAM1_HUMA | Unmodified | 0 | 0 | 0 | 0 | 0 | 0 | 0 | 0 | 0 | 0 | 0 | 591200 | 0 | 0 | 214160 | 0 | 0 | 0 |
| RYDDPEV | HSPA9; HSPA9; | P38646; H0 | GRP75_HUMA | Unmodified | 0 | 0 | 0 | 0 | 0 | 0 | 0 | 0 | 3403700 | 0 | 0 | 0 | 0 | 0 | 0 | 0 | 0 | 0 |
| RYDDPEV | HSPA9; HSPA9; | P38646; H0 | GRP75_HUMA | Unmodified | 0 | 0 | 0 | 0 | 325190 | 0 | 0 | 29041 | 117080 | 548290 | 9464.5 | 0 | 461740 | 178990 | 0 | 0 | 0 | 250580 |
| RYDEVPSD | MARVELD2; MA | Q8N4S9; A1 | MALD2_HUMA | Unmodified | 474090 | 312610 | 428770 | 177670 | 621690 | 0 | 144230 | 202520 | 0 | 0 | 270980 | 1730400 | 172990 | 0 | 677020 | 532960 | 0 | 857960 |

| | | | | | | | | | | | | | | | | | | | | | | |
|---|---|---|---|---|---|---|---|---|---|---|---|---|---|---|---|---|---|---|---|---|---|---|
| RYDGSQQ | NXF1; NXF1; N | Q9UBU9; E9 | NXF1_HUMAN | Unmodified | 0 | 0 | 0 | 0 | 0 | 0 | 204450 | 0 | 0 | 0 | 0 | 0 | 0 | 0 | 0 | 0 | 0 | 0 |
| RYDLSALVI | IGF2R; IGF2R; I | P11717; A0A | MPRI_HUMAN | Unmodified | 0 | 0 | 0 | 0 | 0 | 823960 | 0 | 0 | 10900000 | 88752 | 0 | 0 | 0 | 0 | 0 | 0 | 0 | 0 |
| RYDLSALVI | IGF2R; IGF2R; I | P11717; A0A | MPRI_HUMAN | Unmodified | 0 | 0 | 0 | 0 | 0 | 0 | 0 | 0 | 0 | 1822700 | 0 | 0 | 399190 | 0 | 0 | 0 | 0 | 0 |
| RYEDQELT | BAZ1B | Q9UIG0 | BAZ1B_HUMAN | Unmodified | 110260 | 0 | 0 | 0 | 119040 | 0 | 0 | 0 | 32550000 | 32900000 | 2.48E+08 | 0 | 0 | 0 | 0 | 0 | 0 | 44302 |
| RYEEIVKEV | EEF1A1P5; EEF | Q5VTE0; P68 | EF1A3_HUMAN | Unmodified | 0 | 0 | 0 | 0 | 2964600 | 0 | 1224700 | 0 | 0 | 0 | 0 | 0 | 2127200 | 2581200 | 0 | 0 | 806950 | 246230 |
| RYEPIQATP | TJP1; TJP1; TJP1 | Q07157; A0A | ZO1_HUMAN T | Unmodified | 0 | 0 | 0 | 0 | 9385700 | 0 | 0 | 0 | 0 | 0 | 0 | 0 | 6661000 | 0 | 0 | 0 | 0 | 0 |
| RYESHPVC | CHCHD3; CHC | Q9NX63; C9 | MIC19_HUMAN | Unmodified | 0 | 0 | 0 | 0 | 0 | 0 | 228240 | 53884 | 254620 | 139880 | 0 | 57565 | 0 | 321480 | 0 | 0 | 0 | 0 |
| RYEVPLETF | GAA; GAA; GAA | P10253; I3L | LYAG_HUMAN | Unmodified | 0 | 0 | 0 | 732860 | 0 | 0 | 0 | 0 | 4763600 | 249340 | 0 | 0 | 0 | 0 | 0 | 0 | 0 | 0 |
| RYEVPVPLI | MGAM2 | Q2M2H8 | MGAL_HUMAN | Unmodified | 0 | 0 | 0 | 0 | 0 | 0 | 0 | 0 | 0 | 0 | 750210 | 6855100 | 0 | 0 | 0 | 0 | 0 | 0 |
| RYFGGTED | FSCN1 | Q16658 | FSCN1_HUMAI | Unmodified | 0 | 0 | 0 | 0 | 0 | 0 | 0 | 0 | 0 | 0 | 0 | 2198700 | 0 | 0 | 0 | 0 | 0 | 0 |
| RYFTWDPS | GANAB; GANAE | Q14697; F5H | GANAB_HUMA | Unmodified | 0 | 0 | 0 | 0 | 0 | 0 | 0 | 0 | 2073500 | 704230 | 0 | 0 | 0 | 0 | 0 | 0 | 0 | 0 |
| RYFYFYNT | PREP; PREP | P48147; A0A | PPCE_HUMAN | Unmodified | 71277 | 0 | 0 | 0 | 0 | 0 | 0 | 0 | 0 | 0 | 689980 | 681690 | 0 | 2990200 | 0 | 0 | 0 | 0 |
| RYGCVFQC | TRAF3; TRAF3 | Q13114; H0Y | TRAF3_HUMAN | Unmodified | 0 | 0 | 0 | 0 | 5309600 | 1315600 | 0 | 0 | 0 | 0 | 0 | 0 | 0 | 43493 | 0 | 0 | 130030 | 0 |
| RYGDEELH | INSR | P06213 | INSR_HUMAN I | Unmodified | 0 | 0 | 0 | 0 | 0 | 0 | 0 | 0 | 372720 | 70689 | 0 | 0 | 0 | 0 | 0 | 0 | 0 | 0 |
| RYGEDSEQ | ATP6AP2; ATP6 | O75787; A0A | RENR_HUMAN | Unmodified | 112030 | 0 | 161600 | 175020 | 0 | 19767 | 0 | 0 | 2724100 | 254010 | 0 | 0 | 0 | 0 | 186310 | 92096 | 0 | 0 |
| RYGPMEEP | GPX4; GPX4; GI | P36969; A0A | GPX4_HUMAN | Unmodified | 0 | 0 | 0 | 0 | 0 | 0 | 0 | 0 | 572510 | 281060 | 0 | 0 | 0 | 0 | 0 | 0 | 303280 | 587960 |
| RYGPMEEP | GPX4; GPX4; GI | P36969; A0A | GPX4_HUMAN | Oxidation (M) | 0 | 525840 | 0 | 0 | 0 | 0 | 632600 | 2889000 | 0 | 988620 | 0 | 0 | 0 | 0 | 0 | 653200 | 0 | 0 |
| RYHPLFGN | ARAP1; ARAP1 | Q96P48; E7E | ARAP1_HUMAN | Unmodified | 0 | 0 | 0 | 0 | 0 | 0 | 0 | 0 | 0 | 0 | 0 | 441580 | 0 | 0 | 0 | 0 | 0 | 75418 |
| RYIFDNVAI | RCN1 | Q15293 | RCN1_HUMAN | Unmodified | 0 | 0 | 0 | 0 | 426180 | 0 | 438830 | 0 | 3586500 | 2597300 | 0 | 0 | 1035000 | 0 | 0 | 0 | 0 | 376630 |
| RYKDLEQQ | CCDC50 | Q8IVM0 | CCD50_HUMA | Unmodified | 0 | 0 | 0 | 0 | 0 | 0 | 0 | 0 | 0 | 0 | 0 | 0 | 1822300 | 2141100 | 0 | 0 | 0 | 0 |
| RYLENGKE | HLA-A; HLA-H; I | P04439; P01 | HLAA_HUMAN | Unmodified | 0 | 0 | 0 | 0 | 0 | 0 | 177900 | 45921 | 3614900 | 361420 | 132720 | 14159000 | 0 | 0 | 0 | 111280 | 0 | 0 |
| RYLKSEPIP | PDIA3 | P30101 | PDIA3_HUMAN | Unmodified | 0 | 0 | 0 | 0 | 714940 | 0 | 1636500 | 1305200 | 442340 | 8741300 | 0 | 0 | 774080 | 276980 | 0 | 0 | 0 | 0 |
| RYLNSGGF | PLOD2; PLOD2; | O00469; C9J | PLOD2_HUMAI | Unmodified | 0 | 0 | 0 | 0 | 0 | 0 | 0 | 0 | 6729800 | 1602400 | 0 | 0 | 0 | 0 | 0 | 0 | 0 | 0 |
| RYNCEYSG | NPC1 | O15118 | NPC1_HUMAN | Unmodified | 184820 | 123830 | 109360 | 0 | 826800 | 0 | 111430 | 24322 | 878170 | 0 | 0 | 147550 | 348250 | 302930 | 180180 | 254050 | 0 | 375230 |
| RYNEDLELI | PSMA2; N/A; PS | P25787; A0A | PSA2_HUMAN | Unmodified | 0 | 317790 | 94126 | 290460 | 6935500 | 0 | 43314 | 0 | 752000 | 3387400 | 617690 | 0 | 3644400 | 859130 | 378200 | 0 | 860490 | 1982700 |
| RYPAYELYL | PGAP1; PGAP1 | Q75T13; B4D | PGAP1_HUMAI | Unmodified | 0 | 0 | 0 | 0 | 0 | 0 | 0 | 0 | 2220300 | 0 | 0 | 0 | 0 | 0 | 0 | 0 | 0 | 0 |
| RYPLSGMG | AGPS; AGPS; A | O00116; A0A | ADAS_HUMAN | Unmodified | 0 | 0 | 0 | 0 | 0 | 0 | 0 | 0 | 6185400 | 495520 | 0 | 0 | 0 | 0 | 0 | 0 | 0 | 0 |
| RYPLSGMG | AGPS; AGPS; A | O00116; A0A | ADAS_HUMAN | Oxidation (M) | 0 | 0 | 0 | 0 | 0 | 0 | 0 | 0 | 2387700 | 959530 | 0 | 0 | 0 | 0 | 0 | 0 | 0 | 0 |
| RYPNVFGI | SQOR; SQOR | Q9Y6N5; H3 | SQOR_HUMAN | Unmodified | 0 | 0 | 0 | 0 | 0 | 0 | 231840 | 646120 | 5214600 | 31743 | 0 | 0 | 102900 | 0 | 0 | 0 | 0 | 0 |
| RYPTPYPDI | ERBIN | Q96RT1 | ERBIN_HUMAN | Unmodified | 0 | 0 | 0 | 0 | 7014200 | 0 | 0 | 145580 | 0 | 0 | 0 | 0 | 162410 | 1266400 | 0 | 0 | 183150 | 1457300 |
| RYPTPYPEI | LRRC7; LRRC7; | Q96NW7; A0 | LRRC7_HUMAI | Unmodified | 0 | 0 | 0 | 0 | 5014800 | 0 | 0 | 0 | 0 | 0 | 0 | 539290 | 492330 | 0 | 0 | 0 | 469370 | 0 |
| RYQEALHL | PSMD11 | O00231 | PSD11_HUMAI | Unmodified | 0 | 0 | 0 | 0 | 0 | 0 | 0 | 0 | 0 | 0 | 152100 | 2376200 | 0 | 0 | 0 | 0 | 0 | 0 |
| RYQEALHL | PSMD11 | O00231 | PSD11_HUMAI | Unmodified | 0 | 0 | 0 | 0 | 615860 | 268450 | 400650 | 0 | 57521 | 305270 | 91146 | 0 | 1336200 | 1006000 | 0 | 0 | 384470 | 545600 |
| RYQQEVDF | KIF5B; KIF5C; K | P33176; O60 | KINH_HUMAN | Unmodified | 0 | 138640 | 0 | 0 | 2909900 | 32127 | 46663 | 128270 | 1161400 | 552350 | 381070 | 4599100 | 0 | 733200 | 433990 | 260970 | 155930 | 1736600 |
| RYRGPGIFI | TMX4 | Q9H1E5 | TMX4_HUMAN | Unmodified | 0 | 0 | 0 | 0 | 0 | 0 | 2085400 | 0 | 3124600 | 105860 | 0 | 0 | 0 | 334640 | 0 | 0 | 560530 | 0 |
| RYSPQFVK | GLT8D1 | Q68CQ7 | GL8D1_HUMAI | Unmodified | 0 | 0 | 0 | 165720 | 0 | 0 | 0 | 0 | 3360200 | 0 | 124960 | 0 | 248010 | 165970 | 0 | 0 | 0 | 20269 |
| RYSTNGKI | SRI; SRI; SRI | P30626; B4D | SORCN_HUMA | Unmodified | 0 | 0 | 0 | 0 | 687880 | 0 | 466080 | 0 | 368440 | 691210 | 0 | 0 | 507300 | 805320 | 0 | 0 | 393770 | 1342000 |
| RYSYLTEP | ANK3; ANK3 | H0Y3A4; H0 | H0Y3A4_HUMA | Oxidation (M) | 0 | 0 | 155030 | 170910 | 0 | 0 | 0 | 0 | 0 | 0 | 0 | 128810 | 0 | 0 | 820410 | 772800 | 0 | 0 |
| RYSYLTEP | ANK3; ANK3 | H0Y3A4; H0 | H0Y3A4_HUMA | Unmodified | 0 | 0 | 0 | 0 | 0 | 0 | 0 | 0 | 0 | 0 | 0 | 0 | 0 | 0 | 0 | 0 | 0 | 0 |
| RYTCGEAP | GSTM3; GSTM3 | P21266; A0A | GSTM3_HUMAI | Unmodified | 0 | 281740 | 0 | 0 | 0 | 0 | 0 | 0 | 0 | 0 | 0 | 0 | 4999500 | 7870900 | 2843600 | 871510 | 1128100 | 1474600 |
| RYTLNVLEI | PLS3; PLS3 | P13797; A0A | PLST_HUMAN I | Unmodified | 0 | 0 | 0 | 0 | 1494300 | 1600500 | 0 | 0 | 277670 | 387540 | 0 | 0 | 0 | 0 | 0 | 0 | 912780 | 0 |
| RYVHTFFG | QSOX2; QSOX2 | Q6ZRP7; H0 | QSOX2_HUMAI | Unmodified | 0 | 0 | 0 | 0 | 0 | 0 | 0 | 0 | 1564900 | 121390 | 0 | 612570 | 0 | 0 | 0 | 389970 | 0 | 0 |
| RYVSSLTEE | LMAN1 | P49257 | LMAN1_HUMAI | Unmodified | 0 | 0 | 0 | 0 | 0 | 0 | 0 | 152070 | 6808600 | 400720 | 0 | 0 | 0 | 0 | 0 | 0 | 29662 | 0 |
| RYYGGAEV | SHMT2; SHMT2 | P34897; G3 | GLYM_HUMAN | Unmodified | 0 | 0 | 0 | 0 | 0 | 0 | 0 | 0 | 4143400 | 0 | 0 | 0 | 0 | 0 | 0 | 0 | 0 | 48574 |
| RYYGGAEV | SHMT2; SHMT2 | P34897; G3 | GLYM_HUMAN | Unmodified | 0 | 916040 | 0 | 0 | 668140 | 0 | 194170 | 0 | 11511000 | 1419700 | 0 | 0 | 1292500 | 449380 | 0 | 0 | 454460 | 149380 |
| RYYVNTTTI | GAS7; GAS7; G | O60861; A0A | GAS7_HUMAN | Unmodified | 0 | 0 | 0 | 0 | 0 | 0 | 0 | 0 | 0 | 0 | 0 | 1667800 | 0 | 0 | 0 | 0 | 0 | 0 |
| SAAAEERS | SEPTIN7; SEPTI | Q16181; A0A | SEPT7_HUMAN | Unmodified | 0 | 0 | 0 | 0 | 563590 | 0 | 58589 | 0 | 0 | 0 | 0 | 0 | 207810 | 164860 | 0 | 0 | 189480 | 624050 |
| SAAAETFV | PSD3 | Q9NYI0 | PSD3_HUMAN | Unmodified | 0 | 0 | 0 | 0 | 1455900 | 0 | 0 | 0 | 0 | 0 | 0 | 0 | 0 | 0 | 0 | 0 | 0 | 0 |
| SAAAIALPP | NBEA; LRBA; LR | Q8NFP9; P5 | NBEA_HUMAN | Unmodified | 0 | 0 | 0 | 0 | 0 | 424910 | 0 | 0 | 0 | 0 | 0 | 2733800 | 0 | 0 | 63699 | 85836 | 490320 | 0 |
| SAAASNLS | PAM16; PAM16; | Q9Y3D7; I3L | TIM16_HUMAN | Unmodified | 0 | 0 | 0 | 0 | 0 | 0 | 0 | 0 | 4050200 | 0 | 0 | 0 | 0 | 0 | 0 | 0 | 0 | 0 |
| SAADAEGY | P4HA2; P4HA2; | O15460; E7E | P4HA2_HUMAI | Unmodified | 0 | 0 | 79212 | 0 | 0 | 0 | 0 | 0 | 7035400 | 3287700 | 0 | 0 | 0 | 0 | 106970 | 0 | 0 | 0 |
| SAADEVDG | IQGAP1; IQGAP | P46940; A0A | IQGA1_HUMAN | Acetyl (Protein N | 0 | 710320 | 0 | 0 | 0 | 0 | 0 | 0 | 1992700 | 0 | 0 | 1573700 | 0 | 0 | 0 | 617430 | 0 | 0 |
| SAADEVDG | IQGAP1; IQGAP | P46940; A0A | IQGA1_HUMAN | Acetyl (Protein N | 0 | 0 | 0 | 3806200 | 0 | 0 | 0 | 111770 | 0 | 0 | 0 | 2571000 | 0 | 0 | 83062 | 3150700 | 0 | 0 |
| SAADIIALAI | CRMP1; CRMP1 | Q14194; E9F | DPYL1_HUMAN | Unmodified | 67429000 | 46136000 | 56670000 | 8346000 | 0 | 0 | 1981600 | 1342200 | 2941800 | 1076400 | 0 | 0 | 248420 | 0 | 96159000 | 67971000 | 0 | 0 |
| SAADIIALAI | CRMP1; CRMP1 | Q14194; E9F | DPYL1_HUMAN | Unmodified | 0 | 0 | 0 | 0 | 0 | 0 | 635350 | 241640 | 0 | 0 | 0 | 0 | 0 | 0 | 0 | 0 | 0 | 0 |
| SAADLISLP | EFCAB14; EFCA | O75071; A0A | EFC14_HUMAN | Unmodified | 0 | 0 | 0 | 0 | 0 | 0 | 45566 | 0 | 12713000 | 8285400 | 275340 | 0 | 0 | 0 | 0 | 0 | 0 | 0 |
| SAADLISQA | DPYSL3 | Q14195 | DPYL3_HUMAN | Unmodified | 0 | 0 | 0 | 0 | 1492400 | 0 | 504110 | 292610 | 0 | 447040 | 29896000 | 71454000 | 0 | 0 | 0 | 0 | 202140 | 0 |
| SAADLISQA | DPYSL3 | Q14195 | DPYL3_HUMAN | Unmodified | 0 | 0 | 0 | 0 | 1013100 | 0 | 1734900 | 505060 | 2339400 | 78443 | 0 | 0 | 79413 | 0 | 0 | 0 | 332800 | 0 |
| SAADSISES | USP5 | P45974 | UBP5_HUMAN | Unmodified | 2913800 | 1429200 | 330790 | 877170 | 0 | 0 | 354850 | 0 | 0 | 0 | 0 | 921250 | 0 | 0 | 1225000 | 1020300 | 0 | 0 |
| SAAEEKAA | SLC19A1 | P41440 | S19A1_HUMAN | Unmodified | 0 | 0 | 0 | 0 | 2864600 | 0 | 59268 | 0 | 93712 | 0 | 0 | 0 | 531820 | 142410 | 0 | 0 | 484850 | 1807500 |
| SAAETVTK | HSPE1; HSPE1- | P61604; S4R | CH10_HUMAN | Oxidation (M) | 0 | 0 | 0 | 0 | 218720 | 82058 | 0 | 0 | 0 | 480720 | 0 | 0 | 0 | 86034 | 0 | 0 | 0 | 0 |
| SAALLLQNI | ANK2; ANK2; AI | Q01484; A0A | ANK2_HUMAN | Unmodified | 0 | 0 | 0 | 0 | 0 | 0 | 0 | 0 | 420190 | 265570 | 0 | 0 | 0 | 0 | 0 | 0 | 0 | 0 |
| SAALPIFSS | RSL1D1; RSL1D | O76021; J3Q | RL1D1_HUMAN | Unmodified | 0 | 0 | 0 | 0 | 0 | 0 | 0 | 0 | 162330 | 0 | 0 | 793980 | 0 | 0 | 0 | 0 | 0 | 0 |
| SAALQPEPI | SORL1 | Q92673 | SORL_HUMAN | Unmodified | 0 | 0 | 0 | 0 | 558640 | 593910 | 0 | 0 | 1296100 | 166320 | 0 | 0 | 0 | 0 | 0 | 34628 | 1570500 | 0 |
| SAALQSTQ | AIDA | Q96BJ3 | AIDA_HUMAN | Unmodified | 2580100 | 2290300 | 6941400 | 6241800 | 1615300 | 1036500 | 899670 | 1181800 | 4135000 | 1096700 | 3316100 | 6018900 | 10772000 | 8215900 | 24210000 | 16000000 | 3131500 | 8607500 |
| SAAMLGNS | SNX2 | O60749 | SNX2_HUMAN | Oxidation (M) | 0 | 0 | 156880 | 161970 | 84862 | 73628 | 0 | 0 | 0 | 32910 | 121900 | 936210 | 320050 | 0 | 10429 | 356150 | 0 | 0 |
| SAAMLGNS | SNX2 | O60749 | SNX2_HUMAN | Unmodified | 25830 | 0 | 0 | 46836 | 0 | 0 | 0 | 0 | 0 | 0 | 0 | 1037600 | 90494 | 0 | 60540 | 0 | 0 | 0 |
| SAAMTLNE | BCLAF1; BCLAF | Q9NYF8; E9I | BCLF1_HUMAN | Unmodified | 0 | 0 | 0 | 186160 | 0 | 52427 | 262760 | 78329 | 0 | 0 | 0 | 0 | 424670 | 181670 | 0 | 0 | 0 | 0 |
| SAAMTLNE | BCLAF1; BCLAF | Q9NYF8; E9I | BCLF1_HUMAN | Oxidation (M) | 0 | 0 | 0 | 0 | 0 | 0 | 1954300 | 154600 | 0 | 0 | 0 | 78235 | 0 | 150760 | 0 | 0 | 58102 | 0 |
| SAAPHPGD | CDH2; CDH2 | P19022; C9J | CADH2_HUMA | Unmodified | 127320 | 0 | 0 | 0 | 0 | 0 | 0 | 0 | 3345200 | 676690 | 4349600 | 7878700 | 0 | 0 | 0 | 0 | 20178000 | 16050000 |
| SAAPSALG | IGSF9 | Q9P2J2 | TUTLA_HUMAN | Unmodified | 0 | 0 | 0 | 0 | 0 | 0 | 0 | 0 | 0 | 0 | 295700 | 534360 | 0 | 0 | 0 | 0 | 0 | 0 |
| SAAPSTLD | SRPRB; SRPRB | Q9Y5M8; H7 | SRPRB_HUMAI | Unmodified | 0 | 68248 | 0 | 0 | 0 | 0 | 0 | 0 | 95750 | 44484 | 0 | 296270 | 562140 | 350320 | 0 | 0 | 204870 | 0 |
| SAAQAAAQ | SERBP1 | Q8NC51 | PAIRB_HUMAN | Unmodified | 383440 | 284750 | 307370 | 227160 | 371730 | 68092 | 0 | 0 | 279310 | 113640 | 347870 | 4281700 | 1638000 | 180430 | 419790 | 297600 | 411210 | 191070 |
| SAARESHP | SMAP; C11orf5 | O00193; E9F | SMAP_HUMAN | Acetyl (Protein N | 0 | 0 | 0 | 0 | 0 | 0 | 0 | 9853.7 | 0 | 53750 | 58237 | 0 | 0 | 0 | 0 | 0 | 0 | 278410 |
| SAARLIQNI | MME; MME; MM | P08473; C9I | NEP_HUMAN N | Oxidation (M) | 0 | 0 | 0 | 0 | 0 | 0 | 0 | 0 | 0 | 0 | 0 | 0 | 1129400 | 0 | 0 | 0 | 0 | 0 |
| SAATLITHP | MTCH2; MTCH2 | Q9Y6C9; E9I | MTCH2_HUMA | Unmodified | 0 | 0 | 0 | 0 | 0 | 0 | 0 | 0 | 4640400 | 1441500 | 0 | 199030 | 0 | 0 | 0 | 0 | 0 | 0 |
| SAATLQQEI | FBLN1 | P23142 | FBLN1_HUMAN | Unmodified | 0 | 0 | 0 | 0 | 0 | 0 | 0 | 0 | 0 | 520100 | 0 | 0 | 0 | 0 | 0 | 0 | 0 | 0 |
| SAATSGAG | HUWE1 | Q7Z6Z7 | HUWE1_HUMA | Unmodified | 0 | 0 | 0 | 0 | 0 | 0 | 0 | 0 | 0 | 0 | 0 | 2913800 | 0 | 0 | 271050 | 0 | 0 | 0 |
| SAATWDEF | SPTBN1; SPTBN | Q01082; A0A | SPTB2_HUMAN | Unmodified | 0 | 0 | 0 | 0 | 0 | 0 | 0 | 0 | 328760 | 0 | 0 | 1583400 | 0 | 0 | 0 | 0 | 0 | 0 |
| SAAYERLEI | LMBRD1; LMBR | Q9NUN5; A0 | LMBD1_HUMAI | Unmodified | 0 | 0 | 0 | 962340 | 0 | 0 | 0 | 0 | 30138000 | 4864600 | 0 | 0 | 0 | 0 | 0 | 0 | 816710 | 0 |
| SACGVCPC | RPL34 | P49207 | RL34_HUMAN | Unmodified | 0 | 0 | 0 | 0 | 0 | 0 | 0 | 0 | 0 | 0 | 0 | 3623200 | 0 | 0 | 36165 | 0 | 0 | 0 |
| SADAAAGA | SLC9A3R1; SLC | O14745; J3Q | NHRF1_HUMAI | Acetyl (Protein N | 0 | 0 | 0 | 821980 | 0 | 0 | 0 | 277190 | 503190 | 242150 | 400480 | 0 | 5725300 | 3641000 | 793290 | 715280 | 583080 | 0 |
| SADAAAGA | SLC9A3R1; SLC | O14745; J3Q | NHRF1_HUMAI | Acetyl (Protein N | 0 | 0 | 0 | 0 | 0 | 0 | 1844400 | 1685900 | 0 | 0 | 18805 | 0 | 4236500 | 13815000 | 0 | 0 | 4291200 | 1625900 |
| SADAAAGA | SLC9A3R1; SLC | O14745; J3Q | NHRF1_HUMAI | Acetyl (Protein N | 0 | 0 | 0 | 0 | 0 | 0 | 0 | 0 | 0 | 0 | 0 | 0 | 0 | 665360 | 0 | 0 | 951690 | 0 |
| SADCYNEI | TMEM87A | Q8NBN3 | TM87A_HUMAN | Unmodified | 0 | 0 | 0 | 0 | 0 | 0 | 0 | 0 | 783040 | 621750 | 0 | 0 | 0 | 0 | 0 | 0 | 0 | 0 |
| SADDFIDK | NCKAP1 | Q9Y2A7 | NCKP1_HUMAI | Unmodified | 0 | 0 | 222820 | 0 | 0 | 0 | 0 | 0 | 128840 | 30108 | 0 | 1875200 | 0 | 0 | 0 | 0 | 0 | 0 |
| SADDSLSG | LLGL1; LLGL1 | Q15334; A0A | L2GL1_HUMAN | Unmodified | 0 | 66462 | 0 | 0 | 777990 | 275320 | 0 | 0 | 525930 | 224970 | 0 | 0 | 591430 | 0 | 414400 | 262230 | 119860 | 0 |
| SADDTYLQ | PLEKHA7; PLEK | Q6IQ23; H0Y | PKHA7_HUMAI | Unmodified | 0 | 0 | 0 | 0 | 0 | 0 | 0 | 0 | 0 | 0 | 0 | 0 | 641960 | 513160 | 138430 | 0 | 0 | 0 |
| SADESGQA | SPTAN1; SPTAN | Q13813; A0A | SPTN1_HUMAN | Unmodified | 0 | 0 | 0 | 0 | 594110 | 348050 | 0 | 0 | 0 | 0 | 0 | 4103200 | 0 | 0 | 0 | 0 | 181130 | 0 |
| SADEVLFT | SNX5; SNX5; SN | Q9Y5X3; Q5 | SNX5_HUMAN | Unmodified | 0 | 0 | 0 | 0 | 0 | 0 | 0 | 0 | 0 | 0 | 0 | 0 | 0 | 0 | 0 | 0 | 0 | 0 |
| SADEVLFT | SNX5; SNX5; SN | Q9Y5X3; Q5 | SNX5_HUMAN | Unmodified | 0 | 0 | 0 | 0 | 0 | 0 | 0 | 0 | 253990 | 0 | 0 | 0 | 1206900 | 1223100 | 0 | 0 | 68609 | 691020 |
| SADGSAPA | SLC7A5 | Q01650 | LAT1_HUMAN I | Unmodified | 7109200 | 8218100 | 6309700 | 2111000 | 1099900 | 685400 | 217700 | 48161 | 5773700 | 1350700 | 8994900 | 1.11E+08 | 0 | 0 | 12559000 | 8913900 | 417050 | 0 |
| SADGVIVS | SNX6; SNX6; SN | Q9UNH7; A0 | SNX6_HUMAN | Unmodified | 0 | 0 | 180810 | 0 | 588100 | 0 | 0 | 234820 | 310670 | 0 | 3189100 | 1125100 | 0 | 0 | 246020 | 0 | 250990 | 0 |
| SADGVIVS | SNX6; SNX6; SN | Q9UNH7; A0 | SNX6_HUMAN | Unmodified | 111660 | 0 | 0 | 0 | 0 | 0 | 0 | 0 | 0 | 0 | 348740 | 1459300 | 0 | 0 | 422620 | 130420 | 0 | 0 |
| SADLEHDQ | CLASP1; CLASF | Q7Z460; F8V | CLAP1_HUMAN | Oxidation (M) | 0 | 0 | 0 | 0 | 0 | 0 | 0 | 0 | 0 | 0 | 36643 | 276400 | 0 | 0 | 0 | 16136 | 0 | 0 |
| SADTLWDI | LDHB; LDHB | P07195; A0A | LDHB_HUMAN | Unmodified | 2199200 | 1237500 | 2150600 | 1655000 | 20656000 | 3192900 | 0 | 0 | 0 | 0 | 18561000 | 1.07E+08 | 10624000 | 4327100 | 2879700 | 1818900 | 5894300 | 7607900 |
| SADTLWDI | LDHB; LDHB | P07195; A0A | LDHB_HUMAN | Unmodified | 383420 | 319350 | 0 | 140260 | 202070 | 0 | 0 | 0 | 0 | 209400 | 0 | 0 | 0 | 0 | 165210 | 0 | 2881400 | 0 |
| SADTLWDI | LDHB; LDHB | P07195; A0A | LDHB_HUMAN | Unmodified | 0 | 0 | 0 | 0 | 2661900 | 363840 | 0 | 0 | 334610 | 287720 | 0 | 0 | 1909200 | 1917700 | 0 | 0 | 11962000 | 1528900 |
| SADTLWGI | LDHA | P00338 | LDHA_HUMAN | Unmodified | 1133600 | 1084600 | 2810600 | 5255400 | 59034000 | 5623700 | 5838200 | 4144100 | 9163600 | 5730200 | 11494000 | 1.11E+08 | 12415000 | 7997500 | 3150900 | 3196900 | 38380000 | 70654000 |
| SADTLWGI | LDHA | P00338 | LDHA_HUMAN | Unmodified | 0 | 0 | 0 | 0 | 460850 | 0 | 475840 | 23684 | 0 | 0 | 0 | 10206 | 329590 | 196850 | 0 | 0 | 12460000 | 1131900 |
| SAEAGIAGI | DHX29; DHX29 | Q7Z478; A0A | DHX29_HUMAI | Unmodified | 0 | 0 | 0 | 0 | 0 | 0 | 0 | 164450 | 104830 | 45749 | 0 | 3157400 | 0 | 0 | 0 | 0 | 84971 | 0 |
| SAEDSFTGI | LLGL2; LLGL2 | Q6P1M3; J3 | L2GL2_HUMAN | Unmodified | 384190 | 389010 | 301970 | 644270 | 500970 | 165460 | 0 | 37389 | 4071900 | 833540 | 595030 | 2390600 | 0 | 0 | 694800 | 783480 | 0 | 0 |
| SAEDYNSS | PAK1; PAK1; PA | Q13153; E9F | PAK1_HUMAN | Unmodified | 0 | 0 | 0 | 0 | 676320 | 0 | 0 | 0 | 136460 | 125910 | 108010 | 191510 | 0 | 0 | 0 | 0 | 0 | 0 |
| SAEEEAADI | PCNP | Q8WW12 | PCNP_HUMAN | Unmodified | 0 | 0 | 0 | 0 | 0 | 0 | 0 | 0 | 274200 | 174280 | 1275800 | 0 | 0 | 0 | 0 | 0 | 0 | 0 |
| SAEEGELAI | TNKS1BP1 | Q9C0C2 | TB182_HUMAN | Unmodified | 0 | 0 | 0 | 0 | 0 | 0 | 0 | 0 | 0 | 0 | 0 | 0 | 340060 | 170240 | 0 | 0 | 0 | 0 |
| SAEETFQLS | STXBP3 | O00186 | STXB3_HUMAN | Unmodified | 0 | 239860 | 0 | 470480 | 0 | 0 | 0 | 0 | 0 | 0 | 0 | 0 | 0 | 0 | 502960 | 6664000 | 0 | 0 |
| SAEFPDFYI | CHGB | P05060 | SCG1_HUMAN | Unmodified | 0 | 0 | 0 | 0 | 0 | 0 | 0 | 0 | 1258100 | 0 | 0 | 0 | 0 | 0 | 0 | 0 | 0 | 0 |
| SAEGPGGL | FAM171A2 | A8MVW0 | F1712_HUMAN | Unmodified | 0 | 0 | 0 | 0 | 0 | 0 | 113680 | 61294 | 445650 | 80854 | 0 | 427720 | 0 | 102810 | 0 | 0 | 89036 | 0 |
| SAEGPGGL | FAM171A2 | A8MVW0 | F1712_HUMAN | Unmodified | 0 | 0 | 0 | 0 | 0 | 0 | 0 | 0 | 0 | 0 | 0 | 0 | 0 | 0 | 0 | 0 | 0 | 1418900 |

| | | | | | | | | | | | | | | | | | | | | | | |
|---|---|---|---|---|---|---|---|---|---|---|---|---|---|---|---|---|---|---|---|---|---|---|
| SAEHYTET/ | SRPK1; SRPK1; | Q96SB4; H3 | SRPK1_HUMAN | Unmodified | 0 | 0 | 0 | 0 | 0 | 0 | 0 | 0 | 0 | 0 | 0 | 6079900 | 0 | 0 | 0 | 0 | 0 | 0 |
| SAEIDSDD | KIF5B | P33176 | KINH_HUMAN | Unmodified | 118800 | 48974 | 0 | 0 | 1753200 | 409540 | 194470 | 184620 | 587840 | 232560 | 449100 | 2127800 | 578570 | 0 | 0 | 156240 | 539030 | 945010 |
| SAEKEISLW | NME2; NME1-N | P22392; Q32 | NDKB_HUMAN | Unmodified | 0 | 0 | 0 | 0 | 10654000 | 838710 | 3252100 | 0 | 151220 | 1399300 | 0 | 0 | 15231000 | 3787000 | 0 | 726410 | 28342000 | 10819000 |
| SAEMEKSK | TRIM16; TRIM16 | O95361; Q3( | TRI16_HUMAN | Unmodified | 1598900 | 0 | 0 | 0 | 0 | 0 | 0 | 0 | 0 | 0 | 0 | 0 | 0 | 0 | 0 | 0 | 0 | 0 |
| SAENGIYS\ | SLC6A8 | P48029 | SC6A8_HUMAN | Unmodified | 0 | 0 | 0 | 209500 | 0 | 0 | 0 | 0 | 0 | 0 | 0 | 1208200 | 309570 | 0 | 0 | 271080 | 0 | 0 |
| SAEQPEQK | AKAP12 | Q02952 | AKA12_HUMAN | Unmodified | 0 | 0 | 0 | 0 | 0 | 25320 | 35798 | 0 | 0 | 0 | 0 | 0 | 113710 | 0 | 0 | 0 | 884920 | 823460 |
| SAESPTSP\ | AKAP12 | Q02952 | AKA12_HUMAN | Unmodified | 0 | 0 | 0 | 0 | 718830 | 86037 | 0 | 0 | 0 | 0 | 0 | 0 | 0 | 508100 | 0 | 0 | 361100 | 853500 |
| SAESPTSP\ | AKAP12 | Q02952 | AKA12_HUMAN | Unmodified | 0 | 0 | 0 | 0 | 1362100 | 0 | 0 | 0 | 0 | 0 | 0 | 0 | 0 | 0 | 0 | 0 | 1340100 | 3114300 |
| SAETNIDQI | DST; DST; DST; | Q03001; A0/ | DYST_HUMAN | Unmodified | 25585 | 206140 | 68154 | 179090 | 1149400 | 0 | 0 | 0 | 0 | 145590 | 0 | 405590 | 108880 | 600110 | 113520 | 59549 | 199930 | 331860 |
| SAEVETSE( | HMOX2; HMOX | P30519; I3L2 | HMOX2_HUMA | Acetyl (Protein N | 0 | 0 | 0 | 0 | 576750 | 0 | 112230 | 0 | 0 | 0 | 0 | 0 | 404980 | 0 | 0 | 0 | 213750 | 222640 |
| SAEVPEAA! | ARPP19; ARPP1 | P56211; H3E | ARP19_HUMAN | Acetyl (Protein N | 0 | 0 | 287630 | 184790 | 0 | 0 | 0 | 0 | 0 | 0 | 255820 | 0 | 371220 | 0 | 0 | 288290 | 0 | 0 |
| SAFEETGK | CNPY3; CNPY3 | Q9BT09; A0/ | CNPY3_HUMAI | Unmodified | 0 | 0 | 0 | 0 | 114110 | 267270 | 124570 | 81883 | 19304000 | 3096100 | 0 | 101150 | 0 | 0 | 0 | 0 | 119730 | 0 |
| SAFFFWGL | FAM234A; FAM2 | Q9H0X4; B4I | F234A_HUMAN | Unmodified | 0 | 798510 | 0 | 0 | 0 | 0 | 0 | 0 | 0 | 0 | 0 | 1017500 | 0 | 0 | 1187900 | 446220 | 0 | 0 |
| SAFLLQNLI | HSPBP1 | Q9NZL4 | HPBP1_HUMAI | Unmodified | 0 | 0 | 0 | 0 | 0 | 0 | 0 | 0 | 0 | 0 | 0 | 1522300 | 0 | 0 | 0 | 0 | 0 | 0 |
| SAFPFSDKI | CLINT1 | Q14677 | EPN4_HUMAN | Unmodified | 0 | 0 | 0 | 0 | 0 | 0 | 0 | 0 | 0 | 0 | 0 | 0 | 377840 | 458790 | 0 | 0 | 0 | 0 |
| SAFPHLQD | XPO1; XPO1; XF | O14980; A0/ | XPO1_HUMAN | Unmodified | 0 | 0 | 0 | 0 | 0 | 0 | 0 | 66090 | 137210 | 160170 | 0 | 3468900 | 0 | 0 | 0 | 0 | 0 | 0 |
| SAFSEEVNI | FLOT2; FLOT2; I | Q14254; J3C | FLOT2_HUMAN | Unmodified | 1139500 | 0 | 0 | 2749500 | 176650 | 3842900 | 4782000 | 1564300 | 9820200 | 5622700 | 2581400 | 8692500 | 4547800 | 8859600 | 4582400 | 3645200 | 0 | 3135900 |
| SAFSEEVNI | FLOT2; FLOT2; I | Q14254; J3C | FLOT2_HUMAN | Unmodified | 0 | 0 | 0 | 0 | 294540 | 0 | 1436100 | 0 | 0 | 824320 | 0 | 0 | 7032100 | 1372100 | 0 | 0 | 0 | 0 |
| SAFSNLFG | TFRC; TFRC | P02786; F8V | TFR1_HUMAN | Unmodified | 7979400 | 9922600 | 16320000 | 29773000 | 5417000 | 1206800 | 315980 | 83380 | 6143900 | 767380 | 539090 | 5310900 | 5230500 | 1066200 | 18570000 | 13620000 | 919520 | 2909100 |
| SAFSNLFG | TFRC; TFRC | P02786; F8V | TFR1_HUMAN | Unmodified | 0 | 0 | 0 | 0 | 20635000 | 2849200 | 168850 | 0 | 152010 | 1291200 | 0 | 0 | 22253000 | 7512700 | 0 | 0 | 3660800 | 8488600 |
| SAGACTAA | LAP3 | P28838 | AMPL_HUMAN | Unmodified | 0 | 0 | 0 | 0 | 0 | 0 | 21099 | 0 | 0 | 0 | 0 | 1976900 | 0 | 0 | 0 | 0 | 0 | 0 |
| SAGAMLKP | ECPAS; ECPAS | Q5VYK3; J3K | ECM29_HUMAI | Unmodified | 0 | 0 | 0 | 0 | 0 | 0 | 89884 | 0 | 0 | 42064 | 0 | 324990 | 0 | 0 | 0 | 0 | 0 | 0 |
| SAGDRWVI | KIF23; KIF23; KI | Q02241; A0/ | KIF23_HUMAN | Oxidation (M) | 0 | 0 | 0 | 0 | 18398000 | 2280000 | 0 | 0 | 1183000 | 816100 | 0 | 0 | 0 | 0 | 0 | 0 | 3975100 | 0 |
| SAGDRWVI | KIF23; KIF23; KI | Q02241; A0/ | KIF23_HUMAN | 2 Oxidation (M) | 0 | 0 | 0 | 1342200 | 6708400 | 1354700 | 0 | 0 | 0 | 665170 | 0 | 0 | 0 | 681780 | 0 | 0 | 1485500 | 0 |
| SAGDRWVI | KIF23; KIF23; KI | Q02241; A0/ | KIF23_HUMAN | Unmodified | 0 | 0 | 0 | 0 | 20234000 | 262840 | 0 | 0 | 771010 | 490630 | 0 | 0 | 0 | 0 | 0 | 0 | 1055300 | 0 |
| SAGDVDTL | AGRN; AGRN; A | O00468; A0/ | AGRIN_HUMAN | Unmodified | 753470 | 0 | 1550200 | 0 | 0 | 0 | 0 | 0 | 0 | 0 | 0 | 0 | 0 | 0 | 751320 | 672190 | 0 | 0 |
| SAGGASVP | MAL2 | Q969L2 | MAL2_HUMAN | Acetyl (Protein N | 2002800 | 3038500 | 0 | 1244700 | 0 | 0 | 0 | 0 | 1109200 | 0 | 0 | 0 | 0 | 0 | 2343000 | 937650 | 0 | 0 |
| SAGGIGVA | RRM1; RRM1; R | P23921; H0Y | RIR1_HUMAN F | Unmodified | 0 | 0 | 0 | 185990 | 0 | 0 | 0 | 0 | 0 | 0 | 0 | 2124600 | 0 | 0 | 137330 | 317220 | 0 | 0 |
| SAGGRPGS | GIPC1; GIPC1; | O14908; K7E | GIPC1_HUMAN | Unmodified | 0 | 0 | 61256 | 0 | 156270 | 0 | 0 | 0 | 167940 | 0 | 85095 | 312630 | 22577 | 0 | 199610 | 0 | 0 | 98370 |
| SAGIKVIMV | ATP1A1; ATP1A | P05023; P50 | AT1A1_HUMAN | Unmodified | 0 | 0 | 0 | 0 | 4706900 | 0 | 458630 | 0 | 0 | 1030900 | 0 | 0 | 5519500 | 2201500 | 0 | 0 | 1778700 | 721990 |
| SAGLAFSL | SERPINH1; SER | P50454; E9P | SERPH_HUMAI | Oxidation (M) | 0 | 0 | 0 | 0 | 0 | 0 | 0 | 0 | 1334900 | 482320 | 0 | 0 | 0 | 0 | 0 | 0 | 0 | 0 |
| SAGLAFSL | SERPINH1; SER | P50454; E9P | SERPH_HUMAI | Unmodified | 0 | 0 | 0 | 0 | 0 | 0 | 0 | 0 | 32383000 | 2252400 | 0 | 0 | 0 | 0 | 0 | 0 | 0 | 0 |
| SAGLFQNP | CCS; CCS; CCS | O14618; J3K | CCS_HUMAN ( | Unmodified | 664380 | 708380 | 0 | 726030 | 1025100 | 0 | 0 | 0 | 373090 | 186640 | 0 | 1632200 | 501280 | 0 | 1532300 | 0 | 443150 | 0 |
| SAGPQSQL | ADD2; ADD2 | P35612; A0A | ADDB_HUMAN | Unmodified | 0 | 0 | 0 | 0 | 2105000 | 620900 | 0 | 305630 | 0 | 104870 | 0 | 0 | 0 | 0 | 0 | 0 | 0 | 0 |
| SAGPSDHI | N4BP1 | O75113 | N4BP1_HUMAI | Unmodified | 0 | 174810 | 0 | 38035 | 0 | 0 | 0 | 0 | 0 | 0 | 0 | 0 | 0 | 0 | 15029 | 11055 | 0 | 0 |
| SAGQGEVL | FLNA; FLNA; FL | P21333; A0A | FLNA_HUMAN | Unmodified | 1827900 | 2398600 | 1489300 | 2551800 | 746460 | 367750 | 388860 | 0 | 641740 | 0 | 0 | 2266200 | 514980 | 0 | 4576100 | 1533000 | 0 | 0 |
| SAGQGEVL | FLNA; FLNA; FL | P21333; A0A | FLNA_HUMAN | Unmodified | 544320 | 0 | 90101 | 1584000 | 2232300 | 547310 | 0 | 0 | 0 | 0 | 940690 | 0 | 0 | 578550 | 0 | 528600 | 1178700 | 0 |
| SAGRPTINI | TATDN3 | Q17R31 | TATD3_HUMAN | Unmodified | 468650 | 0 | 144320 | 0 | 354200 | 0 | 0 | 61485 | 0 | 33535 | 0 | 0 | 133340 | 122230 | 0 | 0 | 189820 | 139850 |
| SAGSALYA | GMPPA; GMPP/ | Q96IJ6; A0A( | GMPPA_HUMA | Unmodified | 0 | 0 | 0 | 459980 | 0 | 108350 | 0 | 0 | 396490 | 0 | 0 | 2107500 | 0 | 0 | 0 | 0 | 0 | 643360 |
| SAGSIPGEI | RELA; RELA; RE | Q04206; A0/ | TF65_HUMAN | Unmodified | 0 | 0 | 25556 | 0 | 0 | 130590 | 1734300 | 0 | 1094900 | 0 | 0 | 253250 | 2845400 | 2658100 | 38374 | 431670 | 124800 | 936030 |
| SAGVLLLV | CEMIP2; CEMIP | Q9UHN6; B4 | CEIP2_HUMAN | Unmodified | 0 | 0 | 0 | 0 | 0 | 0 | 0 | 0 | 0 | 0 | 0 | 0 | 0 | 0 | 0 | 0 | 0 | 0 |
| SAGVQCF( | GART; GART; G/ | P22102; C9J | PUR2_HUMAN | Unmodified | 0 | 0 | 0 | 0 | 0 | 0 | 0 | 286380 | 324040 | 0 | 0 | 2315000 | 864300 | 0 | 0 | 593300 | 0 | 0 |
| SAHAGTYE | SSR4; SSR4 | P51571; A6N | SSRD_HUMAN | Unmodified | 0 | 0 | 0 | 0 | 394360 | 67503 | 65203 | 0 | 1770400 | 2824000 | 261830 | 0 | 0 | 0 | 0 | 0 | 0 | 204150 |
| SAHAGTYE | SSR4; SSR4 | P51571; A6N | SSRD_HUMAN | Unmodified | 388580 | 0 | 0 | 0 | 0 | 0 | 0 | 0 | 9598100 | 13207000 | 650750 | 0 | 0 | 0 | 0 | 0 | 391050 | 0 |
| SAHEEEED | FAM171A1 | Q5VUB5 | F1711_HUMAN | Unmodified | 0 | 0 | 0 | 0 | 0 | 0 | 0 | 0 | 0 | 0 | 0 | 0 | 0 | 0 | 0 | 0 | 0 | 0 |
| SAHLLPEH | ITGA6; ITGA6 | P23229; C9J | ITA6_HUMAN I | Unmodified | 0 | 0 | 0 | 0 | 3096200 | 0 | 620080 | 0 | 0 | 0 | 0 | 0 | 0 | 0 | 972530 | 0 | 2097200 | 6992900 |
| SAHLQWM | RPL28; RPL28; I | P46779; H0Y | RL28_HUMAN ( | Acetyl (Protein N | 0 | 0 | 1623000 | 0 | 0 | 0 | 0 | 0 | 0 | 0 | 0 | 20690000 | 0 | 0 | 0 | 0 | 0 | 0 |
| SAHLQWM | RPL28; RPL28; I | P46779; H0Y | RL28_HUMAN ( | Acetyl (Protein N | 231670 | 0 | 0 | 0 | 0 | 0 | 0 | 0 | 0 | 573300 | 0 | 3565600 | 0 | 0 | 0 | 0 | 0 | 0 |
| SAHLQWM | RPL28; RPL28; I | P46779; H0Y | RL28_HUMAN ( | Acetyl (Protein N | 0 | 0 | 0 | 0 | 0 | 0 | 3069500 | 213030 | 0 | 0 | 0 | 1422000 | 816200 | 628390 | 0 | 0 | 0 | 0 |
| SAHNPDDI | CASC3; CASC3 | O15234; J3K | CASC3_HUMAI | Unmodified | 704100 | 502230 | 0 | 0 | 0 | 0 | 0 | 0 | 0 | 20945 | 0 | 0 | 0 | 0 | 0 | 312000 | 0 | 0 |
| SAIAKQAAI | RDX; RDX; RDX | P35241; A0A | RADI_HUMAN I | Unmodified | 0 | 518530 | 0 | 2533100 | 4039600 | 13363000 | 5821500 | 2906400 | 0 | 108030 | 0 | 0 | 1579300 | 591060 | 82276 | 29272 | 627110 | 9795200 |
| SAICIQSW | MYO1C; MYO1( | O00159; F5H | MYO1C_HUMA | Unmodified | 0 | 731030 | 0 | 1086200 | 0 | 0 | 0 | 0 | 0 | 0 | 0 | 0 | 0 | 0 | 2173900 | 1565700 | 0 | 0 |
| SAICRVTLL | EPB41L1; EPB4 | Q9H4G0; Q4 | E41L1_HUMAN | Unmodified | 0 | 0 | 0 | 0 | 911170 | 0 | 0 | 0 | 0 | 0 | 0 | 0 | 0 | 0 | 0 | 0 | 542550 | 725850 |
| SAIDHAGN | DNAJC13; DNA | O75165; A0/ | DJC13_HUMAN | Unmodified | 0 | 0 | 0 | 0 | 0 | 0 | 0 | 0 | 0 | 0 | 0 | 1678000 | 0 | 0 | 0 | 0 | 0 | 0 |
| SAIGEGMTI | ATP6V1B2 | P21281 | VATB2_HUMAN | Unmodified | 0 | 0 | 0 | 0 | 169320 | 0 | 241750 | 0 | 0 | 0 | 0 | 0 | 0 | 0 | 0 | 0 | 111850 | 133050 |
| SAINEVVTR | RPL31; RPL31; I | P62899; H7( | RL31_HUMAN ( | Unmodified | 8834400 | 9771400 | 605280 | 508720 | 0 | 302390 | 0 | 0 | 0 | 0 | 3704100 | 53619000 | 2030700 | 118230 | 3997000 | 3082900 | 0 | 0 |
| SAINEVVTR | RPL31; RPL31; I | P62899; H7( | RL31_HUMAN ( | Unmodified | 0 | 0 | 0 | 138080 | 4718100 | 851460 | 7034700 | 1437100 | 0 | 737260 | 340670 | 0 | 1714500 | 2165600 | 0 | 0 | 422640 | 2544700 |
| SAIPHPLIM | COPS2; COPS2 | P61201; B4D | CSN2_HUMAN | Oxidation (M) | 0 | 0 | 0 | 0 | 488340 | 0 | 0 | 0 | 0 | 242710 | 0 | 1906000 | 0 | 0 | 0 | 0 | 0 | 0 |
| SAIQNLHSI | EIF1; EIF1 | P41567; K7E | EIF1_HUMAN E | Acetyl (Protein N | 32642 | 136410 | 141190 | 53844 | 0 | 0 | 0 | 0 | 99644 | 0 | 0 | 145190 | 0 | 0 | 798970 | 550500 | 0 | 0 |
| SAITPGGLF | SHMT2; SHMT2 | P34897; H0Y | GLYM_HUMAN | Unmodified | 118560 | 244910 | 61964 | 274580 | 0 | 1755600 | 50634 | 19138 | 30574000 | 882570 | 2458300 | 1199300 | 0 | 0 | 95183 | 66979 | 0 | 0 |
| SAIVHLINY | JUP; JUP; JUP; J | P14923; C9J | PLAK_HUMAN | Unmodified | 1816600 | 2657000 | 20505000 | 33235000 | 33167000 | 7101900 | 441700 | 61507 | 13477000 | 2504300 | 198680 | 0 | 16871000 | 7848900 | 26037000 | 30199000 | 6212500 | 12560000 |
| SAIVHLINY | JUP; JUP; JUP; J | P14923; C9J | PLAK_HUMAN | Unmodified | 0 | 0 | 0 | 974780 | 1.42E+08 | 20926000 | 0 | 0 | 0 | 5912100 | 0 | 0 | 64641000 | 47353000 | 0 | 0 | 24960000 | 38174000 |
| SAKAISEQT | ACLY; ACLY | P53396; K7E | ACLY_HUMAN | Acetyl (Protein N | 0 | 0 | 0 | 0 | 0 | 0 | 368490 | 83837 | 714860 | 0 | 0 | 0 | 0 | 0 | 0 | 0 | 0 | 0 |
| SAKAISEQT | ACLY; ACLY | P53396; K7E | ACLY_HUMAN | Acetyl (Protein N | 0 | 0 | 0 | 0 | 0 | 0 | 1941300 | 0 | 0 | 0 | 0 | 0 | 0 | 0 | 0 | 0 | 0 | 0 |
| SAKMDIDV | AHNAK | Q09666 | AHNK_HUMAN | Oxidation (M) | 0 | 0 | 0 | 0 | 10930000 | 1072300 | 0 | 0 | 0 | 75513 | 0 | 0 | 0 | 0 | 0 | 0 | 1331400 | 1671900 |
| SAKMDIDV | AHNAK | Q09666 | AHNK_HUMAN | Unmodified | 0 | 0 | 0 | 0 | 3313700 | 0 | 0 | 0 | 0 | 0 | 0 | 0 | 0 | 0 | 0 | 0 | 0 | 2512500 |
| SAKPALLKI | ITGB4 | P16144 | ITB4_HUMAN I | Unmodified | 0 | 0 | 0 | 0 | 5085600 | 291340 | 0 | 0 | 0 | 0 | 0 | 0 | 633300 | 292300 | 0 | 0 | 10495000 | 10908000 |
| SAKPTKPA/ | CALD1; CALD1; | Q05682; C9J | CALD1_HUMAN | Unmodified | 0 | 0 | 0 | 0 | 0 | 0 | 0 | 0 | 0 | 0 | 2467600 | 779610 | 0 | 345090 | 299610 | 349290 | 235530 | 291850 |
| SALAAAQT( | GOLIM4 | O00461 | GOLI4_HUMAN | Unmodified | 0 | 0 | 0 | 0 | 214000 | 0 | 0 | 515310 | 8664000 | 2027400 | 0 | 0 | 0 | 0 | 0 | 0 | 0 | 0 |
| SALAFLNR | RACGAP1; RAC | Q9H0H5; F8 | RGAP1_HUMAI | Unmodified | 144170 | 0 | 299090 | 459840 | 3500900 | 473320 | 0 | 0 | 2696600 | 0 | 833310 | 3211500 | 124520 | 0 | 406520 | 0 | 648710 | 0 |
| SALAFLNR( | RACGAP1; RAC | Q9H0H5; F8 | RGAP1_HUMAI | Unmodified | 0 | 0 | 0 | 0 | 0 | 0 | 0 | 0 | 0 | 0 | 0 | 0 | 0 | 0 | 0 | 0 | 0 | 0 |
| SALALAIKV | PSMA4; PSMA4 | P25789; H0Y | PSA4_HUMAN | Unmodified | 0 | 0 | 0 | 0 | 0 | 0 | 0 | 0 | 0 | 0 | 0 | 0 | 0 | 0 | 0 | 0 | 332050 | 0 |
| SALASVIM( | PPP2R1A; PPP2 | P30153; B3K | 2AAA_HUMAN | Oxidation (M) | 0 | 0 | 83822 | 0 | 99927 | 22129 | 81385 | 0 | 27956 | 137300 | 95698 | 604850 | 247610 | 0 | 867690 | 423950 | 168930 | 81132 |
| SALCNADS | PDS5A | Q29RF7 | PDS5A_HUMAI | Unmodified | 0 | 0 | 0 | 0 | 0 | 377370 | 200390 | 0 | 87361 | 0 | 0 | 0 | 0 | 0 | 0 | 0 | 0 | 0 |
| SALDNGP( | ITCH; ITCH; ITC | Q96J02; A0A | ITCH_HUMAN I | Unmodified | 1849100 | 2822900 | 660380 | 552810 | 0 | 0 | 0 | 186490 | 0 | 0 | 0 | 0 | 483130 | 0 | 2800400 | 1988000 | 0 | 0 |
| SALDNGP( | ITCH; ITCH; ITC | Q96J02; A0A | ITCH_HUMAN I | Unmodified | 1081100 | 2160400 | 0 | 1532800 | 0 | 0 | 0 | 0 | 0 | 0 | 0 | 0 | 0 | 0 | 1323100 | 0 | 0 | 0 |
| SALDTAAR | FAM3C; FAM3C | Q92520; C9J | FAM3C_HUMAI | Unmodified | 0 | 0 | 47033 | 42986 | 288800 | 0 | 861420 | 0 | 3372600 | 3415900 | 12759 | 257770 | 338910 | 358290 | 17384 | 13533 | 341660 | 654190 |
| SALEAESES | SYTL4 | Q96C24 | SYTL4_HUMAN | Unmodified | 0 | 0 | 0 | 0 | 2265300 | 0 | 0 | 0 | 0 | 0 | 0 | 0 | 0 | 0 | 0 | 0 | 1174800 | 1162800 |
| SALEMGFN | BIRC2; BIRC2; E | Q13490; E9F | BIRC2_HUMAN | Oxidation (M) | 0 | 0 | 0 | 0 | 1574400 | 0 | 0 | 0 | 0 | 0 | 0 | 0 | 0 | 0 | 0 | 0 | 0 | 0 |
| SALEMGFN | BIRC2; BIRC2; E | Q13490; E9F | BIRC2_HUMAN | Unmodified | 0 | 0 | 0 | 0 | 8390400 | 901310 | 0 | 0 | 0 | 463250 | 0 | 0 | 974490 | 0 | 0 | 0 | 69972 | 0 |
| SALESSPAI | CALCA; CALCA | P01258; P06 | CALC_HUMAN | Unmodified | 0 | 0 | 0 | 0 | 0 | 0 | 0 | 0 | 8995600 | 0 | 0 | 0 | 0 | 0 | 0 | 0 | 0 | 0 |
| SALESSPDI | CALCB; CALCB | P10092; E7E | CALCB_HUMAI | Unmodified | 0 | 0 | 0 | 0 | 0 | 0 | 0 | 0 | 834440 | 93225 | 0 | 0 | 0 | 0 | 0 | 0 | 0 | 0 |
| SALESSPDI | CALCB; CALCB | P10092; E7E | CALCB_HUMAI | Unmodified | 0 | 0 | 0 | 0 | 0 | 0 | 0 | 0 | 1111400 | 331350 | 0 | 0 | 0 | 0 | 0 | 0 | 437030 | 0 |
| SALEWYAH | MBLAC2 | Q68D91 | MBLC2_HUMAI | Acetyl (Protein N | 0 | 0 | 1363600 | 1284000 | 367580 | 0 | 0 | 264220 | 771350 | 186600 | 2674900 | 3888200 | 0 | 719420 | 0 | 1151200 | 899300 | 806480 |
| SALGHALP | RTBDN; RTBDN | Q9BSG5; K7 | RTBDN_HUMAI | Unmodified | 0 | 0 | 0 | 0 | 0 | 0 | 0 | 0 | 2091500 | 0 | 0 | 0 | 0 | 0 | 0 | 0 | 0 | 0 |
| SALIVQGPI | PACC1 | Q9H813 | PACC1_HUMAI | Unmodified | 127260 | 256280 | 317540 | 0 | 0 | 0 | 0 | 0 | 4017900 | 329550 | 358890 | 4605200 | 0 | 0 | 359810 | 220120 | 0 | 0 |
| SALIVQGPI | PACC1 | Q9H813 | PACC1_HUMAI | Unmodified | 0 | 0 | 0 | 0 | 0 | 0 | 0 | 0 | 594140 | 154440 | 0 | 0 | 39100 | 0 | 0 | 0 | 0 | 0 |
| SALLDHIW | PPFIBP2; PPFIB | Q8ND30; E9 | LIPB2_HUMAN | Unmodified | 0 | 0 | 505210 | 374990 | 0 | 0 | 0 | 0 | 0 | 0 | 171010 | 984720 | 0 | 0 | 691790 | 549050 | 0 | 0 |
| SALLPGPLE | INPPL1; INPPL1 | O15357; H0 | SHIP2_HUMAN | Unmodified | 0 | 0 | 0 | 0 | 0 | 0 | 0 | 0 | 0 | 0 | 0 | 692610 | 0 | 0 | 0 | 0 | 0 | 0 |
| SALMEQR | JTB; N/A | O76095; A0/ | JTB_HUMAN Pr | Unmodified | 0 | 0 | 13355 | 0 | 21931 | 0 | 0 | 7252.3 | 5085700 | 349470 | 20937 | 438970 | 0 | 0 | 12119 | 0 | 105800 | 12218 |
| SALPVSPSI | EPB41L5 | Q9HCM4 | E41L5_HUMAN | Unmodified | 6063100 | 0 | 0 | 0 | 0 | 1718600 | 0 | 0 | 20271000 | 6370800 | 0 | 0 | 9237200 | 9738100 | 14409000 | 0 | 1165400 | 6930300 |
| SALQSINE\ | SERPINH1; SER | P50454; E9P | SERPH_HUMAI | Unmodified | 0 | 0 | 0 | 0 | 0 | 0 | 0 | 0 | 21186000 | 3047700 | 0 | 0 | 0 | 0 | 0 | 0 | 0 | 0 |
| SALQSINE\ | SERPINH1; SER | P50454; E9P | SERPH_HUMAI | Unmodified | 0 | 0 | 0 | 0 | 0 | 0 | 0 | 0 | 1719100 | 0 | 0 | 0 | 0 | 0 | 0 | 0 | 0 | 0 |
| SALQVEQK | ASAP2 | O43150 | ASAP2_HUMAN | Unmodified | 0 | 0 | 0 | 0 | 0 | 0 | 0 | 0 | 0 | 0 | 0 | 319610 | 0 | 46691 | 0 | 0 | 0 | 0 |
| SALSGHLE | ANXA2P2; ANX/ | A6NMY6; P0 | AXA2L_HUMAN | Unmodified | 363130 | 1773300 | 32141 | 1419100 | 1.31E+08 | 28119000 | 3388800 | 0 | 11763000 | 43160000 | 55839 | 0 | 976090 | 2016100 | 1396600 | 0 | 35463000 | 19762000 |
| SALSGHLE | ANXA2P2; ANX/ | A6NMY6; P0 | AXA2L_HUMAN | Unmodified | 0 | 0 | 0 | 0 | 2.28E+08 | 23013000 | 9365800 | 0 | 358130 | 49144000 | 0 | 0 | 1370500 | 2440700 | 0 | 0 | 63825000 | 42245000 |
| SALSGHLE | ANXA2P2; ANX/ | A6NMY6; P0 | AXA2L_HUMAN | Oxidation (M) | 0 | 0 | 0 | 0 | 18586000 | 807520 | 2105600 | 0 | 0 | 853020 | 0 | 0 | 0 | 0 | 0 | 0 | 21262000 | 2425700 |
| SALTLQFM | RALA; RALB; RA | P11233; P11 | RALA_HUMAN | Oxidation (M) | 0 | 0 | 0 | 0 | 4588000 | 0 | 0 | 0 | 0 | 4123300 | 260190 | 0 | 9028800 | 5663700 | 0 | 0 | 2668700 | 0 |
| SALTLQFM | RALA; RALB; RA | P11233; P11 | RALA_HUMAN | Unmodified | 0 | 0 | 0 | 387290 | 5782300 | 0 | 2826200 | 0 | 0 | 3373600 | 0 | 0 | 0 | 4303700 | 0 | 0 | 3080600 | 5222300 |
| SALTVQFV( | RAP1B; RAP1A; | P61224; P62 | RAP1B_HUMAN | Unmodified | 409810 | 3126700 | 428250 | 2855200 | 2279900 | 179470 | 944600 | 0 | 0 | 0 | 963400 | 1194600 | 2928900 | 724450 | 4204400 | 605890 | 404760 | 1485000 |
| SALTVQFV( | RAP1B; RAP1A; | P61224; P62 | RAP1B_HUMAN | Unmodified | 908250 | 11673000 | 0 | 19019000 | 70474000 | 9970000 | 92985000 | 0 | 0 | 7462700 | 0 | 0 | 1.08E+08 | 54337000 | 0 | 0 | 27977000 | 45957000 |
| SALTVQFV | RAP2B | P61225 | RAP2B_HUMAN | Unmodified | 0 | 0 | 0 | 0 | 0 | 0 | 0 | 0 | 0 | 0 | 0 | 627690 | 0 | 0 | 1554100 | 0 | 0 | 0 |
| SALTVQFV | RAP2B | P61225 | RAP2B_HUMAN | Unmodified | 0 | 0 | 0 | 859090 | 0 | 0 | 0 | 0 | 0 | 0 | 0 | 0 | 0 | 1504900 | 0 | 0 | 2477000 | 0 |
| SALYSPSDI | QSOX1; QSOX1 | O00391; A8N | QSOX1_HUMAI | Unmodified | 0 | 0 | 296800 | 130540 | 0 | 0 | 0 | 0 | 4915000 | 363090 | 170170 | 0 | 0 | 0 | 251300 | 54822 | 0 | 0 |
| SALYSPSDI | QSOX1; QSOX1 | O00391; A8N | QSOX1_HUMAI | Unmodified | 0 | 0 | 0 | 0 | 0 | 0 | 0 | 0 | 0 | 533910 | 0 | 0 | 0 | 0 | 0 | 0 | 0 | 0 |
| SALYYVDLI | APEH; APEH; AI | P13798; C9J | ACPH_HUMAN | Unmodified | 0 | 0 | 251950 | 0 | 0 | 0 | 0 | 0 | 304730 | 0 | 0 | 2510500 | 0 | 0 | 0 | 267810 | 0 | 0 |
| SAMAHDPI | FAM20B | O75063 | XYLK_HUMAN ( | Oxidation (M) | 0 | 0 | 0 | 0 | 0 | 0 | 0 | 0 | 35421000 | 1381100 | 1112500 | 2612300 | 0 | 0 | 0 | 0 | 0 | 0 |
| SAMAHDPI | FAM20B | O75063 | XYLK_HUMAN ( | Unmodified | 0 | 0 | 0 | 0 | 340330 | 0 | 277980 | 0 | 17958000 | 812200 | 0 | 0 | 0 | 0 | 0 | 0 | 0 | 0 |
| SAMAHDPI | FAM20B | O75063 | XYLK_HUMAN ( | Oxidation (M) | 0 | 0 | 0 | 0 | 257780 | 0 | 3141800 | 0 | 281850 | 13370000 | 0 | 0 | 295610 | 0 | 0 | 0 | 0 | 0 |

| | | | | | | | | | | | | | | | | | | | | | | |
|---|---|---|---|---|---|---|---|---|---|---|---|---|---|---|---|---|---|---|---|---|---|---|
| SAMMQQY | RPS6KA4 | A0A0A6YYC( | A0A0A6YYC0_H | Unmodified | 0 | 0 | 0 | 677840 | 0 | 0 | 0 | 0 | 0 | 0 | 0 | 0 | 1020300 | 0 | 0 | 0 | 0 | 0 |
| SANAEDAQ | S100A14 | Q9HCY8 | S10AE_HUMAN | Unmodified | 0 | 0 | 0 | 0 | 0 | 0 | 0 | 0 | 585440 | 0 | 0 | 0 | 0 | 0 | 0 | 0 | 0 | 0 |
| SANAEDAQ | S100A14 | Q9HCY8 | S10AE_HUMAN | Unmodified | 0 | 0 | 0 | 0 | 0 | 0 | 0 | 0 | 416590 | 1325400 | 0 | 0 | 0 | 0 | 0 | 0 | 0 | 0 |
| SANFLDHL | ITFG1; ITFG1; IT | Q8TB96; F5( | TIP_HUMAN T-( | Unmodified | 160030 | 317390 | 1696500 | 2549200 | 965040 | 399380 | 0 | 0 | 37802000 | 6638100 | 399010 | 1074600 | 790360 | 420230 | 1215200 | 751470 | 0 | 219730 |
| SANPAMQI | PEF1 | Q9UBV8 | PEF1_HUMAN I | Unmodified | 0 | 0 | 0 | 0 | 0 | 0 | 0 | 0 | 4877100 | 0 | 0 | 0 | 0 | 0 | 255660 | 0 | 57853 | 0 |
| SANPQDHI | CHDH | Q8NE62 | CHDH_HUMAN | Unmodified | 0 | 0 | 0 | 0 | 0 | 0 | 0 | 0 | 1058300 | 0 | 0 | 0 | 0 | 0 | 0 | 0 | 0 | 0 |
| SANSELGG | COPS8; COPS8 | Q99627; E9F | CSN8_HUMAN | Unmodified | 0 | 0 | 0 | 0 | 0 | 0 | 0 | 0 | 0 | 0 | 0 | 2910200 | 0 | 0 | 0 | 0 | 0 | 0 |
| SANTQVVR | ARHGAP1; ARH | Q07960; H0 | RHG01_HUMA | Unmodified | 2572400 | 125590 | 31497 | 315140 | 868690 | 267010 | 0 | 28876 | 0 | 243900 | 0 | 0 | 0 | 0 | 0 | 25557 | 135820 | 1920600 |
| SANVNEFP | CSE1L | P55060 | XPO2_HUMAN | Unmodified | 0 | 0 | 0 | 0 | 0 | 0 | 0 | 30152 | 204440 | 0 | 582650 | 13457000 | 0 | 0 | 0 | 0 | 0 | 0 |
| SANYFDSE | NDST1; NDST1 | P52848; E7E | NDST1_HUMA | Unmodified | 0 | 0 | 0 | 0 | 0 | 0 | 0 | 0 | 837240 | 0 | 0 | 0 | 0 | 0 | 0 | 0 | 0 | 0 |
| SAPASPVQ | ADD2 | P35612 | ADDB_HUMAN | Unmodified | 0 | 0 | 0 | 0 | 240150 | 0 | 0 | 0 | 0 | 0 | 0 | 0 | 0 | 0 | 0 | 0 | 0 | 0 |
| SAPDQEVR | ARL3 | P36405 | ARL3_HUMAN | Unmodified | 0 | 69393 | 22231 | 0 | 0 | 291020 | 1898100 | 0 | 0 | 652470 | 0 | 0 | 4267000 | 0 | 105780 | 44030 | 112660 | 240680 |
| SAPDQEVR | ARL3 | P36405 | ARL3_HUMAN | Unmodified | 0 | 0 | 0 | 0 | 2164300 | 271300 | 6061200 | 0 | 0 | 802350 | 0 | 0 | 10161000 | 7648000 | 0 | 0 | 4484800 | 3288100 |
| SAPELKTGI | NCL; NCL; NCL | P19338; C9J | NUCL_HUMAN | Unmodified | 0 | 0 | 0 | 0 | 0 | 0 | 50480 | 30602 | 73045 | 173200 | 0 | 0 | 157040 | 0 | 0 | 0 | 0 | 0 |
| SAPEPKKPI | EIF4B; EIF4B | P23588; E7E | IF4B_HUMAN E | Unmodified | 0 | 0 | 0 | 373130 | 85396 | 0 | 82495 | 9216.8 | 171320 | 0 | 0 | 0 | 291320 | 73492 | 0 | 0 | 23876 | 83226 |
| SAPGEDEE | SORT1; SORT1 | Q99523; A0A | SORT_HUMAN | Unmodified | 0 | 0 | 0 | 0 | 0 | 0 | 0 | 0 | 564090 | 27814 | 0 | 35210 | 0 | 0 | 0 | 0 | 0 | 0 |
| SAPGEDEE | SORT1; SORT1 | Q99523; A0A | SORT_HUMAN | Unmodified | 0 | 0 | 0 | 0 | 73767 | 0 | 239060 | 117710 | 2012200 | 0 | 0 | 0 | 0 | 0 | 0 | 0 | 0 | 0 |
| SAPLDAALI | IFI35 | P80217 | IN35_HUMAN I | Acetyl (Protein N | 0 | 0 | 0 | 0 | 3546000 | 0 | 0 | 0 | 0 | 576210 | 0 | 0 | 605890 | 0 | 0 | 0 | 2488700 | 3004300 |
| SAPLIADQ | NCEH1; NCEH1 | Q6PIU2; A0A | NCEH1_HUMA | Unmodified | 0 | 0 | 9880.5 | 0 | 496960 | 167650 | 23470 | 0 | 4250700 | 760480 | 204440 | 149550 | 0 | 0 | 0 | 0 | 0 | 128840 |
| SAPLPNDS | PIP4K2A | P48426 | PI42A_HUMAN | Unmodified | 233190 | 154070 | 0 | 362750 | 221440 | 141710 | 45000 | 0 | 281750 | 44069 | 421210 | 1428700 | 0 | 0 | 0 | 0 | 0 | 0 |
| SAPSIPKEN | MDH1; MDH1; I | P40925; C9J | MDHC_HUMAN | Unmodified | 0 | 80746 | 0 | 0 | 0 | 0 | 0 | 0 | 0 | 0 | 0 | 0 | 0 | 0 | 0 | 0 | 1141600 | 0 |
| SAPSSFAP | PROSER2; PRO | Q86WR7; D3 | PRSR2_HUMA | Unmodified | 0 | 0 | 0 | 0 | 5509300 | 273390 | 47786 | 0 | 0 | 143530 | 129650 | 0 | 0 | 52275 | 0 | 0 | 291200 | 63522 |
| SAPVSTTET | VPS13A | Q96RL7 | VP13A_HUMAN | Unmodified | 0 | 0 | 0 | 0 | 81775 | 0 | 0 | 0 | 0 | 0 | 0 | 310920 | 0 | 0 | 0 | 0 | 42203 | 0 |
| SAQAQMR | LUC7L2 | Q9Y383 | LC7L2_HUMAN | Acetyl (Protein N | 0 | 0 | 0 | 0 | 0 | 0 | 0 | 0 | 0 | 0 | 4395300 | 0 | 0 | 0 | 0 | 0 | 0 | 0 |
| SAQERSQA | FKBP15 | Q5T1M5 | FKB15_HUMAN | Unmodified | 0 | 0 | 0 | 0 | 1823300 | 0 | 0 | 0 | 70294 | 568690 | 0 | 253720 | 150170 | 0 | 0 | 0 | 0 | 0 |
| SAQGTGFE | ZDHHC5; ZDHH | Q9C0B5; H0 | ZDHC5_HUMA | Unmodified | 0 | 0 | 0 | 706680 | 0 | 0 | 0 | 0 | 0 | 0 | 0 | 0 | 0 | 0 | 625400 | 1251900 | 0 | 0 |
| SAQGTGFE | ZDHHC5; ZDHH | Q9C0B5; H0 | ZDHC5_HUMA | Unmodified | 707610 | 618750 | 754400 | 1453600 | 15876000 | 1398900 | 436890 | 0 | 0 | 2619900 | 0 | 0 | 10079000 | 9889600 | 1378100 | 648610 | 9526300 | 15819000 |
| SAQHYTET | SRPK2; SRPK2; | P78362; C9J | SRPK2_HUMAN | Unmodified | 0 | 385810 | 107060 | 61379 | 0 | 0 | 41627 | 0 | 0 | 0 | 22684 | 418700 | 0 | 0 | 0 | 151640 | 0 | 0 |
| SAQIMESQ | MPP7; MPP7 | Q5T2T1; U5( | MPP7_HUMAN | Unmodified | 0 | 0 | 0 | 0 | 2455100 | 0 | 0 | 0 | 970570 | 198100 | 0 | 151340 | 442210 | 0 | 0 | 0 | 0 | 710730 |
| SAQQESLP | MIA3; MIA3 | Q5JRA6; A0A | TGO1_HUMAN | Unmodified | 0 | 0 | 0 | 0 | 0 | 0 | 0 | 0 | 1396800 | 320690 | 0 | 0 | 0 | 0 | 0 | 0 | 0 | 0 |
| SARDVFTK | VDAC1; VDAC1 | P21796; C9J | VDAC1_HUMA | Unmodified | 0 | 0 | 0 | 0 | 405450 | 0 | 1192900 | 0 | 19572000 | 21352000 | 554400 | 0 | 440150 | 882490 | 0 | 0 | 14610 | 460710 |
| SAREAAASI | SUN2 | Q9UH99 | SUN2_HUMAN | Unmodified | 0 | 0 | 0 | 0 | 0 | 0 | 0 | 0 | 1635000 | 697540 | 0 | 0 | 0 | 0 | 0 | 0 | 0 | 0 |
| SARPLHSL | CC2D1A; CC2D | Q6P1N0; A0 | C2D1A_HUMA | Unmodified | 559750 | 278200 | 0 | 0 | 0 | 0 | 0 | 0 | 699290 | 0 | 0 | 0 | 121770 | 0 | 0 | 497460 | 0 | 0 |
| SARQDLEC | CGN | Q9P2M7 | CING_HUMAN | Unmodified | 0 | 0 | 0 | 0 | 0 | 0 | 0 | 0 | 119710 | 0 | 0 | 355800 | 0 | 0 | 0 | 0 | 0 | 0 |
| SARVINYEE | TPPP3 | Q9BW30 | TPPP3_HUMAN | Unmodified | 0 | 0 | 0 | 0 | 0 | 0 | 994060 | 630060 | 0 | 0 | 0 | 0 | 0 | 0 | 0 | 0 | 0 | 0 |
| SARVINYEE | TPPP3 | Q9BW30 | TPPP3_HUMAN | Unmodified | 0 | 0 | 0 | 0 | 0 | 0 | 3314500 | 1717300 | 0 | 0 | 0 | 0 | 0 | 0 | 0 | 0 | 0 | 0 |
| SARWLSTS | FASN; FASN | P49327; A0A | FAS_HUMAN F | Unmodified | 0 | 0 | 0 | 0 | 600830 | 0 | 0 | 0 | 0 | 0 | 0 | 0 | 632850 | 217020 | 0 | 0 | 323280 | 697370 |
| SASAAQRE | TLN2; TLN2; TLI | Q9Y4G6; H0 | TLN2_HUMAN | Unmodified | 0 | 0 | 0 | 0 | 0 | 0 | 20125 | 183460 | 0 | 30055 | 0 | 0 | 23638 | 18146 | 0 | 0 | 136660 | 302870 |
| SASDNSKA | NHS; NHS; NHS | Q6T4R5; A0A | NHS_HUMAN N | Unmodified | 0 | 0 | 0 | 0 | 811700 | 90899 | 0 | 0 | 78516 | 37945 | 0 | 0 | 0 | 0 | 0 | 0 | 244560 | 652140 |
| SASKQFHN | GLIPR2; GLIPR2 | Q9H4G4; Q5 | GAPR1_HUMA | Unmodified | 0 | 0 | 0 | 0 | 521300 | 95597 | 0 | 0 | 0 | 0 | 0 | 8575.3 | 323930 | 221980 | 0 | 0 | 454420 | 395010 |
| SASLDVEPI | STRN3 | Q13033 | STRN3_HUMAN | Unmodified | 0 | 0 | 0 | 0 | 0 | 0 | 0 | 0 | 0 | 180220 | 0 | 1304200 | 0 | 0 | 0 | 0 | 0 | 0 |
| SASLVTRQI | TAOK3 | Q9H2K8 | TAOK3_HUMAN | Unmodified | 0 | 0 | 0 | 0 | 2895600 | 0 | 1182700 | 0 | 0 | 0 | 0 | 0 | 568240 | 2210500 | 0 | 0 | 1764500 | 793420 |
| SASMDSLT | CRAT; CRAT; CI | P43155; A0A | CACP_HUMAN | Unmodified | 0 | 0 | 0 | 0 | 0 | 123490 | 0 | 0 | 1499100 | 0 | 0 | 302640 | 0 | 0 | 267080 | 0 | 0 | 0 |
| SASPDAAG | ANK3 | Q12955 | ANK3_HUMAN | Unmodified | 0 | 0 | 0 | 0 | 0 | 0 | 0 | 0 | 0 | 0 | 0 | 0 | 709000 | 924020 | 0 | 0 | 0 | 0 |
| SASPDDDL | SMAP; C11orf5 | O00193; E9F | SMAP_HUMAN | Unmodified | 0 | 0 | 0 | 0 | 0 | 0 | 0 | 0 | 368060 | 0 | 371250 | 870880 | 0 | 0 | 0 | 0 | 0 | 0 |
| SASPTIEAQ | IRAG2; IRAG2; I | Q12912; F5H | IRAG2_HUMAN | Unmodified | 0 | 0 | 0 | 0 | 0 | 0 | 0 | 0 | 0 | 0 | 710780 | 2822000 | 0 | 0 | 0 | 0 | 0 | 0 |
| SASPYHGF | DCP1A; DCP1A | Q9NPI6; A0A | DCP1A_HUMA | Unmodified | 103070 | 0 | 0 | 0 | 0 | 0 | 0 | 0 | 0 | 0 | 0 | 604310 | 0 | 0 | 0 | 0 | 0 | 0 |
| SASRPAAEI | FAM171A2 | A8MVW0 | F1712_HUMAN | Unmodified | 0 | 0 | 0 | 0 | 155580 | 2389500 | 390220 | 176970 | 121030 | 0 | 0 | 0 | 180930 | 0 | 26112 | 26582 | 35311 | 1897000 |
| SASRPAAEI | FAM171A2 | A8MVW0 | F1712_HUMAN | Unmodified | 0 | 0 | 0 | 0 | 0 | 0 | 0 | 77042 | 937630 | 899290 | 0 | 0 | 0 | 81807 | 0 | 0 | 4118500 | 7342600 |
| SASSDTSEI | SLC9A3R1; SLC | O14745; J3Q | NHRF1_HUMA | Unmodified | 0 | 0 | 127090 | 0 | 0 | 0 | 0 | 0 | 0 | 0 | 24425 | 0 | 0 | 0 | 432560 | 262380 | 0 | 0 |
| SASSGAEG | DTD1 | Q8TEA8 | DTD1_HUMAN | Unmodified | 0 | 0 | 0 | 0 | 0 | 0 | 0 | 0 | 590310 | 124950 | 394090 | 880800 | 0 | 0 | 0 | 0 | 79187 | 154020 |
| SASSKTEV | LAP3; LAP3 | P28838; H0 | AMPL_HUMAN | Unmodified | 0 | 0 | 35233 | 128150 | 1155000 | 285830 | 1131700 | 717480 | 365130 | 369200 | 185350 | 546410 | 1305400 | 896130 | 70738 | 76078 | 922920 | 781010 |
| SASSLLEQ | YTHDF2; YTHD | Q9Y5A9; S4F | YTHD2_HUMA | Acetyl (Protein N | 0 | 0 | 0 | 0 | 0 | 0 | 58378 | 27512 | 0 | 0 | 0 | 513430 | 370460 | 213240 | 0 | 0 | 0 | 0 |
| SASSLNHT | LPAR1; LPAR1 | Q92633; Q6 | LPAR1_HUMAN | Unmodified | 0 | 0 | 0 | 228830 | 0 | 0 | 0 | 0 | 0 | 0 | 0 | 0 | 0 | 0 | 0 | 0 | 2479000 | 335520 |
| SASSSAAG | VPS37A; VPS37 | Q8NEZ2; E5I | VP37A_HUMAN | Unmodified | 5409000 | 6224000 | 1407900 | 890370 | 1829000 | 0 | 0 | 0 | 0 | 0 | 0 | 0 | 1143400 | 0 | 1195600 | 1374300 | 0 | 0 |
| SASVGPPM | MEGF8 | Q7Z7M0 | MEGF8_HUMA | Unmodified | 0 | 0 | 0 | 0 | 0 | 0 | 0 | 0 | 1328000 | 145270 | 0 | 251770 | 0 | 0 | 0 | 0 | 0 | 0 |
| SASVVSVIS | DAD1; DAD1; D | P61803; F5G | DAD1_HUMAN | Unmodified | 0 | 0 | 0 | 0 | 0 | 0 | 0 | 0 | 1984100 | 0 | 0 | 0 | 0 | 0 | 0 | 0 | 0 | 0 |
| SATAATAPF | VAMP2; N/A; VA | P63027; L7N | VAMP2_HUMA | Acetyl (Protein N | 620930 | 0 | 0 | 1418200 | 0 | 0 | 0 | 0 | 0 | 0 | 445670 | 0 | 0 | 352640 | 701200 | 0 | 0 | 0 |
| SATDVKVT | LAMB1; LAMB1; | P07942; A0A | LAMB1_HUMA | Unmodified | 0 | 0 | 0 | 0 | 472800 | 117130 | 0 | 32740 | 0 | 0 | 0 | 0 | 0 | 0 | 0 | 0 | 105840 | 951430 |
| SATKVTAD | PHGDH; PHGD | O43175; A0A | SERA_HUMAN | Unmodified | 0 | 0 | 0 | 210740 | 427210 | 0 | 346380 | 27818 | 0 | 820060 | 0 | 0 | 5763100 | 1723600 | 0 | 0 | 434170 | 769340 |
| SATLASIDA | KIF5B | P33176 | KINH_HUMAN | Unmodified | 0 | 0 | 0 | 0 | 0 | 648710 | 555710 | 688600 | 1348100 | 909490 | 710550 | 5040200 | 1525700 | 0 | 0 | 0 | 0 | 0 |
| SATLLQLITI | DNM1L | O00429 | DNM1L_HUMA | Unmodified | 0 | 0 | 0 | 0 | 0 | 0 | 130630 | 61244 | 0 | 0 | 173050 | 3647500 | 0 | 0 | 0 | 0 | 0 | 0 |
| SATLQFRP | SDK1; SDK1; SI | Q7Z5N4; F8 | SDK1_HUMAN | Unmodified | 0 | 0 | 0 | 0 | 0 | 0 | 49312 | 17119 | 431950 | 318040 | 0 | 0 | 0 | 0 | 0 | 0 | 0 | 0 |
| SATLSSTES | AKAP12 | Q02952 | AKA12_HUMAN | Unmodified | 0 | 0 | 0 | 0 | 0 | 0 | 0 | 0 | 0 | 0 | 0 | 0 | 0 | 0 | 0 | 0 | 0 | 1025200 |
| SATNLGRQ | RACGAP1 | Q9H0H5 | RGAP1_HUMA | Oxidation (M) | 0 | 0 | 0 | 0 | 1555400 | 0 | 0 | 0 | 0 | 0 | 0 | 0 | 0 | 0 | 0 | 0 | 355720 | 507040 |
| SATNLGRQ | RACGAP1 | Q9H0H5 | RGAP1_HUMA | Unmodified | 0 | 0 | 0 | 0 | 9982700 | 0 | 0 | 0 | 0 | 597340 | 0 | 0 | 170200 | 0 | 0 | 0 | 1398700 | 0 |
| SATNNIAQ | GNG7; GNG7 | O60262; K7E | GBG7_HUMAN | Acetyl (Protein N | 0 | 0 | 0 | 0 | 0 | 0 | 307570 | 171380 | 0 | 0 | 0 | 0 | 0 | 319110 | 0 | 0 | 0 | 0 |
| SATNNIAQ | GNG7; GNG7 | O60262; K7E | GBG7_HUMAN | Acetyl (Protein N | 0 | 0 | 0 | 0 | 763860 | 126900 | 0 | 396470 | 0 | 0 | 0 | 0 | 0 | 977320 | 1543700 | 0 | 581450 | 1935900 |
| SATQFTGR | NAXE; NAXE | Q8NCW5; A( | NNRE_HUMAN | Unmodified | 143120 | 123450 | 0 | 227800 | 0 | 0 | 0 | 0 | 290830 | 0 | 0 | 2413000 | 110210 | 0 | 229720 | 154680 | 0 | 0 |
| SATQFTGR | NAXE; NAXE | Q8NCW5; A( | NNRE_HUMAN | Unmodified | 755180 | 406060 | 0 | 0 | 0 | 0 | 0 | 0 | 0 | 0 | 28809 | 0 | 488660 | 0 | 0 | 0 | 0 | 0 |
| SATQFTGR | NAXE; NAXE | Q8NCW5; A( | NNRE_HUMAN | Unmodified | 0 | 0 | 0 | 0 | 1841100 | 0 | 924640 | 0 | 0 | 1162400 | 0 | 0 | 1561700 | 1380400 | 0 | 0 | 1655700 | 1330000 |
| SATQMFEL | DAAM1 | Q9Y4D1 | DAAM1_HUMA | Unmodified | 0 | 0 | 0 | 85988 | 0 | 0 | 139220 | 0 | 298910 | 0 | 0 | 1679400 | 0 | 0 | 65007 | 55743 | 0 | 0 |
| SATQMFEL | DAAM1 | Q9Y4D1 | DAAM1_HUMA | Unmodified | 0 | 0 | 0 | 0 | 0 | 0 | 0 | 0 | 0 | 0 | 0 | 0 | 0 | 0 | 0 | 0 | 0 | 0 |
| SATSSASRI | FARP1; FARP1 | Q9Y4F1; C9J | FARP1_HUMAN | Unmodified | 0 | 0 | 0 | 0 | 0 | 0 | 0 | 0 | 200110 | 29800 | 70094 | 427760 | 0 | 0 | 0 | 0 | 0 | 0 |
| SATSSSPG | ATP2B1; ATP2B | P20020; E7E | AT2B1_HUMAN | Unmodified | 406540 | 0 | 765040 | 1165500 | 1572900 | 0 | 527840 | 254310 | 0 | 670710 | 0 | 0 | 10246000 | 14338000 | 1876200 | 1146600 | 1947500 | 4526700 |
| SATSVDQR | YTHDF3; YTHD | Q7Z739; S4F | YTHD3_HUMA | Acetyl (Protein N | 552120 | 373940 | 210670 | 401730 | 342100 | 10910 | 0 | 0 | 129300 | 0 | 87221 | 0 | 402030 | 0 | 132590 | 227140 | 0 | 0 |
| SATTTVMN | ITGB2; ITGB2; I | P05107; D3D | ITB2_HUMAN I | Unmodified | 52549 | 0 | 70685 | 50023 | 272220 | 0 | 699190 | 0 | 254000 | 175220 | 25979 | 281560 | 0 | 0 | 143150 | 0 | 103000 | 0 |
| SATVCERG | SDF2; SDF2 | Q99470; K7E | SDF2_HUMAN | Unmodified | 0 | 0 | 20427 | 135020 | 65493 | 323040 | 1024700 | 399930 | 879210 | 6189600 | 177650 | 0 | 0 | 473200 | 0 | 0 | 0 | 476810 |
| SATYFDSE | NDST2; N/A | P52849; S4F | NDST2_HUMA | Unmodified | 0 | 0 | 0 | 0 | 0 | 0 | 0 | 0 | 2448800 | 553050 | 0 | 0 | 0 | 0 | 0 | 0 | 0 | 0 |
| SAVAADEQ | N/A | N/A | N/A | Unmodified | 3370500 | 15008000 | 967440 | 21778000 | 382690 | 39108000 | 30926000 | 13001000 | 0 | 209070 | 8455200 | 15867000 | 287860 | 1.9E+08 | 2098200 | 3460700 | 35881000 | 52475000 |
| SAVADKHE | EIF3H; EIF3H; E | O15372; E5F | EIF3H_HUMAN | Unmodified | 1608500 | 0 | 0 | 1352300 | 2031500 | 0 | 0 | 317400 | 0 | 0 | 0 | 7800800 | 3046200 | 0 | 641480 | 0 | 416600 | 1039000 |
| SAVDLSCS | PKP3; PKP3; PK | Q9Y446; E9F | PKP3_HUMAN | Unmodified | 0 | 0 | 0 | 0 | 536480 | 0 | 0 | 0 | 165520 | 0 | 0 | 504380 | 2428600 | 592800 | 673190 | 622630 | 351490 | 380940 |
| SAVDQAG | RAP1GDS1; RAI | P52306; D6F | GDS1_HUMAN | Unmodified | 0 | 0 | 0 | 0 | 0 | 0 | 0 | 0 | 332610 | 0 | 0 | 2236600 | 0 | 0 | 0 | 0 | 0 | 0 |
| SAVDQAG | RAP1GDS1; RAI | P52306; D6F | GDS1_HUMAN | Unmodified | 0 | 0 | 0 | 0 | 0 | 0 | 393560 | 0 | 0 | 0 | 0 | 0 | 0 | 0 | 0 | 0 | 0 | 0 |
| SAVEAERL | RPN1; RPN1 | P04843; B7Z | RPN1_HUMAN | Unmodified | 0 | 0 | 703040 | 624350 | 1271900 | 262660 | 1075100 | 272480 | 1187700 | 1329500 | 337670 | 1184300 | 1561300 | 1953900 | 433050 | 209760 | 145600 | 643000 |
| SAVEAQNE | LRRFIP1 | Q32MZ4 | LRRF1_HUMAN | Unmodified | 198690 | 0 | 124560 | 0 | 720040 | 29874 | 37235 | 65116 | 404350 | 172110 | 0 | 357530 | 489220 | 0 | 0 | 0 | 173420 | 698650 |
| SAVEDEGL | HSPA1A; HSPA | P0DMV8; P0 | HS71A_HUMA | Unmodified | 15525000 | 15617000 | 3777200 | 9860100 | 63472000 | 8942400 | 16288000 | 6085500 | 173580 | 21840000 | 1364000 | 8351400 | 44170000 | 23016000 | 3694000 | 3693800 | 20024000 | 29931000 |
| SAVEEMEA | DKK3; DKK3; DI | Q9UBP4; F6 | DKK3_HUMAN | Unmodified | 0 | 0 | 0 | 0 | 0 | 0 | 0 | 0 | 0 | 0 | 2452100 | 0 | 0 | 0 | 0 | 0 | 0 | 0 |
| SAVENCQI | PGD | P52209 | 6PGD_HUMAN | Unmodified | 0 | 0 | 0 | 0 | 0 | 0 | 0 | 0 | 195480 | 66138 | 0 | 2601600 | 0 | 0 | 0 | 25273 | 0 | 0 |
| SAVENCQI | PGD | P52209 | 6PGD_HUMAN | Unmodified | 548770 | 343690 | 296410 | 325300 | 497040 | 0 | 121620 | 34903 | 1418800 | 670800 | 0 | 6769600 | 359720 | 196970 | 644650 | 611160 | 145880 | 147310 |
| SAVGFDYQ | CTTN | Q14247 | SRC8_HUMAN | Unmodified | 427000 | 527500 | 0 | 431830 | 1151700 | 328910 | 0 | 0 | 134530 | 0 | 0 | 562500 | 543470 | 241320 | 475340 | 626580 | 169760 | 0 |
| SAVGFEYQ | CTTN | Q14247 | SRC8_HUMAN | Unmodified | 453520 | 0 | 0 | 646180 | 83558 | 0 | 0 | 0 | 71509 | 0 | 0 | 607840 | 194450 | 74583 | 886730 | 500430 | 152740 | 0 |
| SAVGFEYQ | CTTN | Q14247 | SRC8_HUMAN | Unmodified | 962570 | 1127400 | 0 | 1132000 | 0 | 0 | 0 | 0 | 0 | 0 | 0 | 0 | 0 | 0 | 475420 | 0 | 0 | 0 |
| SAVGFEYQ | CTTN | Q14247 | SRC8_HUMAN | Unmodified | 97533 | 516020 | 0 | 83646 | 5939800 | 1493300 | 127840 | 0 | 0 | 0 | 0 | 0 | 872160 | 2259900 | 0 | 0 | 912490 | 2333100 |
| SAVGHEYQ | CTTN | Q14247 | SRC8_HUMAN | Unmodified | 0 | 0 | 25364 | 0 | 752610 | 101130 | 331460 | 46158 | 20657 | 0 | 0 | 0 | 256230 | 198650 | 0 | 0 | 501840 | 330740 |
| SAVLQEAR | COPG1; COPG | Q9Y678; Q9 | COPG1_HUMA | Unmodified | 7392000 | 196890 | 267990 | 3779100 | 628320 | 625010 | 0 | 63616 | 765430 | 316160 | 0 | 2495200 | 528720 | 144010 | 1877700 | 2111000 | 357790 | 698450 |
| SAVLQEAR | COPG1 | Q9Y678 | COPG1_HUMA | Unmodified | 0 | 0 | 0 | 0 | 116690 | 0 | 0 | 0 | 0 | 0 | 0 | 0 | 250180 | 94572 | 0 | 0 | 0 | 159310 |
| SAVLQEAR | COPG1 | Q9Y678 | COPG1_HUMA | Unmodified | 0 | 0 | 0 | 0 | 477700 | 0 | 458470 | 135960 | 0 | 0 | 0 | 0 | 475260 | 468570 | 0 | 0 | 471340 | 932450 |
| SAVLSEPVI | FAHD1 | Q6P587 | FAHD1_HUMA | Oxidation (M) | 0 | 0 | 0 | 0 | 0 | 0 | 0 | 0 | 5369700 | 1180900 | 0 | 0 | 0 | 0 | 0 | 0 | 0 | 0 |
| SAVLSEPVI | FAHD1 | Q6P587 | FAHD1_HUMA | Unmodified | 3209300 | 0 | 0 | 0 | 0 | 0 | 0 | 0 | 2477000 | 0 | 0 | 0 | 0 | 0 | 0 | 0 | 0 | 0 |
| SAVPFNQY | LIMCH1; LIMCH | Q9UPQ0; H( | LIMC1_HUMAN | Unmodified | 0 | 0 | 0 | 0 | 0 | 0 | 275850 | 331920 | 713590 | 0 | 0 | 0 | 0 | 985050 | 0 | 0 | 0 | 0 |
| SAVQKGTS | CAPN5; CAPN5 | O15484; E7E | CAN5_HUMAN | Unmodified | 0 | 0 | 0 | 0 | 327210 | 57136 | 59468 | 37341 | 0 | 100200 | 0 | 0 | 1420200 | 993330 | 0 | 0 | 398420 | 1819300 |
| SAVRTGISH | PFKL | P17858 | PFKAL_HUMAN | Unmodified | 0 | 0 | 0 | 0 | 2338900 | 0 | 470730 | 0 | 0 | 328050 | 0 | 0 | 431760 | 337010 | 0 | 0 | 897370 | 3709600 |
| SAVRVGIAI | PFKP; PFKP; PF | Q01813; A0A | PFKAP_HUMAN | Unmodified | 0 | 0 | 0 | 0 | 321850 | 0 | 0 | 0 | 0 | 0 | 0 | 0 | 0 | 0 | 0 | 0 | 0 | 0 |
| SAVSDKQD | HSPA4L; HSPA | O95757; E9F | HS74L_HUMAN | Unmodified | 0 | 0 | 0 | 0 | 260250 | 35442 | 74626 | 62351 | 0 | 0 | 92545 | 234120 | 936030 | 0 | 0 | 0 | 364990 | 327060 |
| SAVSILER | DNAJC16; DNA | Q9Y2G8; Q5 | DJC16_HUMAN | Unmodified | 0 | 0 | 0 | 0 | 0 | 0 | 0 | 0 | 673180 | 0 | 0 | 0 | 0 | 0 | 0 | 0 | 0 | 0 |
| SAVSPDLH | PKP4; PKP4 | Q99569; A0A | PKP4_HUMAN | Unmodified | 0 | 0 | 623420 | 1044700 | 131630 | 0 | 0 | 0 | 969480 | 0 | 640720 | 1400900 | 98376 | 0 | 717940 | 554160 | 1082800 | 547590 |
| SAVTTVVN | ITGB1; ITGB1; I | P05556; H7( | ITB1_HUMAN I | Unmodified | 5264500 | 3390800 | 9542500 | 5176000 | 31074000 | 9020800 | 1113900 | 285360 | 5798000 | 1109700 | 2616300 | 10509000 | 1605000 | 0 | 8473600 | 8348300 | 4679700 | 36166000 |
| SAVTTVVN | ITGB1; ITGB1; I | P05556; H7( | ITB1_HUMAN I | Unmodified | 2489600 | 2284100 | 128020 | 6457100 | 4.31E+08 | 19425000 | 14359000 | 10670000 | 3230600 | 6839800 | 714110 | 2070800 | 25511000 | 20680000 | 966470 | 1278400 | 1.55E+08 | 5.88E+08 |

| | | | | | | | | | | | | | | | | | | | | | | |
|---|---|---|---|---|---|---|---|---|---|---|---|---|---|---|---|---|---|---|---|---|---|---|
| SAVVEMLIN | ILK; ILK; ILK | Q13418; A0A | ILK_HUMAN Int | Oxidation (M) | 0 | 0 | 0 | 0 | 717820 | 72968 | 299870 | 0 | 0 | 0 | 0 | 0 | 390890 | 225190 | 0 | 0 | 141460 | 159780 |
| SAWLSGYE | P4HA1 | P13674 | P4HA1_HUMAN | Unmodified | 0 | 0 | 0 | 0 | 0 | 0 | 0 | 0 | 9821600 | 1636300 | 1187200 | 205860 | 647570 | 0 | 0 | 0 | 0 | 0 |
| SAWNSDP | USP54; USP54 | Q70EL1; A0A | UBP54_HUMAN | Unmodified | 0 | 0 | 0 | 0 | 0 | 0 | 0 | 0 | 0 | 0 | 0 | 0 | 452770 | 202620 | 0 | 0 | 0 | 0 |
| SAWRVISSI | YWHAQ; YWHA | P27348; E9P | 1433T_HUMAN | Unmodified | 0 | 0 | 0 | 1053200 | 3785100 | 566890 | 649960 | 51358 | 188660 | 528780 | 272830 | 0 | 1987800 | 1300400 | 547980 | 223970 | 1385400 | 2746200 |
| SAWRVISSI | YWHAQ; YWHA | P27348; E9P | 1433T_HUMAN | Unmodified | 0 | 0 | 261590 | 121050 | 19795000 | 7425500 | 678010 | 0 | 389520 | 1713700 | 215190 | 2014000 | 10764000 | 7917200 | 308010 | 260790 | 9536800 | 20159000 |
| SAYALCTFA | ITPA | Q9BY32 | ITPA_HUMAN I | Unmodified | 19605 | 0 | 0 | 0 | 0 | 0 | 0 | 0 | 0 | 0 | 0 | 6051300 | 0 | 0 | 0 | 0 | 0 | 0 |
| SAYALCTFA | ITPA | Q9BY32 | ITPA_HUMAN I | Unmodified | 0 | 0 | 0 | 0 | 0 | 0 | 0 | 0 | 0 | 176600 | 0 | 0 | 0 | 0 | 0 | 0 | 473660 | 1527400 |
| SAYCPYSH | CDA | P32320 | CDD_HUMAN ( | Unmodified | 0 | 0 | 0 | 0 | 0 | 0 | 0 | 43739 | 0 | 0 | 0 | 0 | 0 | 0 | 0 | 0 | 703280 | 0 |
| SAYCPYSH | CDA | P32320 | CDD_HUMAN ( | Unmodified | 0 | 0 | 0 | 0 | 4878700 | 0 | 0 | 0 | 0 | 0 | 0 | 0 | 0 | 0 | 0 | 0 | 1538300 | 0 |
| SAYDDTEN | MIA3; MIA3 | Q5JRA6; A0A | TGO1_HUMAN | Unmodified | 0 | 0 | 0 | 0 | 0 | 0 | 0 | 0 | 725170 | 0 | 0 | 0 | 0 | 0 | 0 | 0 | 0 | 0 |
| SAYDSTME | ACLY | P53396 | ACLY_HUMAN | 2 Oxidation (M) | 0 | 0 | 0 | 0 | 0 | 0 | 0 | 0 | 0 | 0 | 0 | 1917000 | 0 | 0 | 0 | 0 | 0 | 0 |
| SAYDSTME | ACLY | P53396 | ACLY_HUMAN | Oxidation (M) | 0 | 0 | 0 | 0 | 0 | 0 | 0 | 0 | 0 | 0 | 0 | 3151300 | 0 | 0 | 0 | 0 | 0 | 0 |
| SAYEFSETE | LTA4H | P09960 | LKHA4_HUMAN | Unmodified | 0 | 0 | 0 | 0 | 986550 | 0 | 0 | 0 | 492500 | 0 | 411070 | 1802800 | 0 | 0 | 0 | 0 | 0 | 0 |
| SAYLHPQQ | TRIM26 | Q12899 | TRI26_HUMAN | Unmodified | 7824000 | 5208500 | 5763800 | 6672600 | 0 | 0 | 0 | 0 | 0 | 0 | 0 | 0 | 414740 | 375040 | 7773100 | 5940200 | 0 | 0 |
| SAYQEAMD | SFN | P31947 | 1433S_HUMAN | Unmodified | 1619800 | 663080 | 0 | 0 | 0 | 817650 | 0 | 0 | 0 | 1802800 | 0 | 0 | 0 | 0 | 0 | 0 | 736390 | 0 |
| SAYQEAMD | SFN | P31947 | 1433S_HUMAN | Oxidation (M) | 313890 | 0 | 0 | 0 | 221250 | 0 | 0 | 0 | 0 | 0 | 0 | 0 | 0 | 0 | 0 | 0 | 124950 | 0 |
| SAYQTIDSA | AGTRAP | Q6RW13 | ATRAP_HUMAN | Unmodified | 292320 | 0 | 1500300 | 2729200 | 0 | 0 | 0 | 2767400 | 0 | 0 | 0 | 0 | 0 | 0 | 545230 | 524080 | 0 | 0 |
| SAYVSYDV | TMEM106B | Q9NUM4 | T106B_HUMAN | Unmodified | 0 | 0 | 0 | 0 | 128910 | 0 | 0 | 0 | 1250300 | 214800 | 0 | 0 | 0 | 0 | 0 | 0 | 12074 | 260650 |
| SAYVSYDV | TMEM106B | Q9NUM4 | T106B_HUMAN | Unmodified | 0 | 0 | 111950 | 0 | 0 | 0 | 0 | 79209 | 0 | 825560 | 0 | 2046000 | 0 | 201940 | 0 | 0 | 0 | 0 |
| SAYYPEDLI | GPC3; GPC3 | P51654; H0Y | GPC3_HUMAN | Unmodified | 0 | 0 | 0 | 0 | 0 | 0 | 0 | 0 | 8886000 | 2037500 | 0 | 0 | 0 | 0 | 0 | 0 | 0 | 0 |
| SCAEWVSL | ATP5PD | O75947 | ATP5H_HUMAN | Unmodified | 0 | 0 | 0 | 46689 | 0 | 0 | 770800 | 93050 | 5076800 | 1045200 | 0 | 141420 | 0 | 0 | 0 | 0 | 0 | 0 |
| SCAHDWV | NME2P1; NME2 | O60361; P22 | NDK8_HUMAN | Unmodified | 0 | 0 | 0 | 2100300 | 14779000 | 2107100 | 1348800 | 0 | 2210400 | 1186900 | 4820300 | 20045000 | 9176800 | 3805500 | 1687400 | 1066600 | 12219000 | 13869000 |
| SCALDQN | ATRN | O75882 | ATRN_HUMAN | Unmodified | 0 | 0 | 0 | 0 | 0 | 0 | 0 | 0 | 696470 | 0 | 0 | 0 | 0 | 0 | 0 | 0 | 0 | 0 |
| SCASRLAL | GOT1 | P17174 | AATC_HUMAN | Unmodified | 0 | 0 | 0 | 0 | 784700 | 294150 | 44819 | 52139 | 0 | 322730 | 0 | 0 | 55003 | 762990 | 0 | 0 | 203200 | 391860 |
| SCATFSSSI | SEC31A; SEC31 | O94979; D6I | SC31A_HUMAN | Unmodified | 0 | 0 | 403530 | 0 | 0 | 0 | 0 | 85738 | 37711 | 112990 | 39943 | 68570 | 0 | 0 | 0 | 0 | 0 | 192520 |
| SCAVAEYG | HP; HPR; HPR; | P00738; P00 | HPT_HUMAN H | Unmodified | 0 | 0 | 0 | 0 | 211250 | 0 | 794790 | 647690 | 0 | 0 | 0 | 0 | 361840 | 445610 | 0 | 0 | 0 | 405690 |
| SCCDYALH | DPYSL3 | Q14195 | DPYL3_HUMAN | Unmodified | 0 | 0 | 0 | 0 | 0 | 0 | 0 | 0 | 0 | 0 | 0 | 2062000 | 0 | 0 | 0 | 0 | 0 | 0 |
| SCCDYSLH | DPYSL2; DPYSL | Q16555; A0A | DPYL2_HUMAN | Unmodified | 0 | 0 | 0 | 0 | 787460 | 367750 | 0 | 0 | 346590 | 204070 | 0 | 1067500 | 0 | 0 | 0 | 0 | 0 | 0 |
| SCCSCCP | MT1X; MT2A; M | P80297; P02 | MT1X_HUMAN | Unmodified | 0 | 0 | 0 | 0 | 0 | 0 | 0 | 0 | 0 | 0 | 0 | 0 | 0 | 0 | 0 | 0 | 398250 | 0 |
| SCDEDLTS | DLG5; DLG5 | Q8TDM6; A0 | DLG5_HUMAN | Unmodified | 0 | 0 | 0 | 0 | 0 | 0 | 0 | 0 | 0 | 0 | 0 | 0 | 0 | 0 | 0 | 43945 | 801170 | 1107200 |
| SCDFLSSFI | SERPINB6; SER | P35237; A0A | SPB6_HUMAN | Unmodified | 0 | 0 | 0 | 0 | 0 | 0 | 0 | 0 | 0 | 0 | 0 | 0 | 0 | 0 | 0 | 0 | 0 | 0 |
| SCDFLSSFI | SERPINB6; SER | P35237; A0A | SPB6_HUMAN | Unmodified | 0 | 0 | 0 | 0 | 281930 | 0 | 0 | 0 | 0 | 0 | 60556 | 396720 | 636080 | 150230 | 0 | 0 | 870140 | 0 |
| SCDPKGPL | EEF2 | P13639 | EF2_HUMAN El | Oxidation (M) | 0 | 0 | 0 | 0 | 3001800 | 0 | 0 | 229890 | 17825 | 244440 | 0 | 0 | 0 | 0 | 0 | 0 | 709030 | 753090 |
| SCDPKGPL | EEF2 | P13639 | EF2_HUMAN El | Unmodified | 0 | 0 | 0 | 0 | 1089200 | 0 | 353700 | 17745 | 364140 | 168620 | 0 | 154220 | 267550 | 203840 | 0 | 0 | 345270 | 671360 |
| SCEGQNPI | ANLN | Q9NQW6 | ANLN_HUMAN | Unmodified | 0 | 0 | 0 | 347300 | 3435900 | 1195400 | 0 | 101310 | 0 | 0 | 0 | 0 | 159250 | 0 | 0 | 0 | 597590 | 1406200 |
| SCEGQNPI | ANLN | Q9NQW6 | ANLN_HUMAN | Unmodified | 0 | 0 | 0 | 0 | 12967000 | 1584100 | 815290 | 589780 | 0 | 383050 | 0 | 131980 | 275390 | 850970 | 0 | 0 | 14313000 | 4344700 |
| SCENLAPFI | AIMP2; AIMP2 | Q13155; A8I | AIMP2_HUMAN | Unmodified | 0 | 0 | 0 | 0 | 0 | 0 | 0 | 0 | 0 | 0 | 0 | 5215300 | 0 | 0 | 0 | 0 | 0 | 0 |
| SCEVIIGSTI | STXBP3 | O00186 | STXB3_HUMAN | Unmodified | 1042000 | 0 | 0 | 0 | 0 | 0 | 919140 | 783520 | 2456600 | 1656400 | 0 | 0 | 3607100 | 12880000 | 0 | 0 | 4212000 | 4493600 |
| SCEVIIGSTI | STXBP3 | O00186 | STXB3_HUMAN | Unmodified | 0 | 0 | 0 | 0 | 2226400 | 174030 | 0 | 0 | 0 | 1346300 | 0 | 0 | 2344200 | 4507100 | 614830 | 0 | 2841200 | 3642000 |
| SCEVPTRLI | CD55; CD55; C | P08174; H3E | DAF_HUMAN C | Unmodified | 157080 | 181140 | 0 | 0 | 205430 | 0 | 0 | 0 | 0 | 0 | 0 | 726630 | 34400 | 0 | 0 | 80249 | 0 | 0 |
| SCFEYTANI | ITGA6 | P23229 | ITA6_HUMAN I | Unmodified | 0 | 0 | 0 | 789140 | 11876000 | 0 | 0 | 0 | 927050 | 0 | 0 | 0 | 5708200 | 2048300 | 0 | 0 | 12040000 | 40545000 |
| SCFQQNNI | UTRN; UTRN; U | P46939; H0Y | UTRN_HUMAN | Unmodified | 0 | 83114 | 1099400 | 1292400 | 760070 | 225230 | 35392 | 0 | 1068900 | 419580 | 0 | 326290 | 560150 | 203280 | 1108800 | 1579900 | 353640 | 696100 |
| SCGSSSHE | HUWE1; HUWE | Q7Z6Z7; H0Y | HUWE1_HUMA | Unmodified | 0 | 0 | 0 | 0 | 0 | 0 | 0 | 0 | 0 | 53692 | 90025 | 797180 | 0 | 0 | 0 | 0 | 95310 | 179850 |
| SCGSSTPD | EIF2S3; EIF2S3 | P41091; H7E | IF2G_HUMAN E | Unmodified | 63417 | 0 | 70101 | 385370 | 0 | 0 | 0 | 0 | 0 | 0 | 417550 | 2119800 | 0 | 0 | 0 | 0 | 0 | 0 |
| SCHAGFGS | MELTF | P08582 | TRFM_HUMAN | Unmodified | 34878 | 0 | 0 | 0 | 2951200 | 1556400 | 0 | 0 | 3084500 | 644650 | 67616 | 346380 | 0 | 0 | 0 | 0 | 1057900 | 0 |
| SCHAVSQT | PLCB4 | A0A7P0MRI8 | A0A7P0MRI8_H | Unmodified | 0 | 0 | 0 | 0 | 0 | 0 | 0 | 0 | 0 | 0 | 0 | 0 | 0 | 0 | 0 | 178310 | 777540 | 3745400 |
| SCHAVSQT | PLCB4 | A0A7P0MRI8 | A0A7P0MRI8_H | Unmodified | 0 | 0 | 0 | 0 | 0 | 0 | 0 | 0 | 0 | 0 | 0 | 0 | 231970 | 0 | 0 | 0 | 1275300 | 578330 |
| SCHRFEQA | NOMO1; NOMO | Q15155; P69 | NOMO1_HUMA | Unmodified | 0 | 0 | 0 | 0 | 0 | 0 | 0 | 0 | 8876500 | 846430 | 388710 | 0 | 0 | 0 | 0 | 0 | 0 | 0 |
| SCIMNSGA | ADAM9; ADAM9 | Q13443; A0A | ADAM9_HUMA | Unmodified | 92684 | 73039 | 0 | 0 | 0 | 0 | 0 | 0 | 0 | 0 | 0 | 119860 | 0 | 0 | 305590 | 810740 | 0 | 0 |
| SCIYDAKTD | P2RX4; P2RX4; | Q99571; F69 | P2RX4_HUMAN | Unmodified | 0 | 0 | 0 | 0 | 0 | 0 | 0 | 0 | 1036700 | 0 | 0 | 0 | 0 | 0 | 0 | 0 | 0 | 0 |
| SCLIHTSEK | SLC39A6 | Q13433 | S39A6_HUMAN | Unmodified | 0 | 0 | 0 | 0 | 0 | 0 | 0 | 0 | 141700 | 0 | 0 | 756620 | 115830 | 0 | 0 | 58290 | 0 | 0 |
| SCLLHQFIE | RAB4A | P20338 | RAB4A_HUMAN | Unmodified | 0 | 0 | 0 | 905920 | 580780 | 0 | 370090 | 249980 | 0 | 0 | 243100 | 962620 | 1523800 | 472290 | 0 | 708460 | 154940 | 0 |
| SCLLHQFTI | RAB14; RAB14 | P61106; X6R | RAB14_HUMAN | Unmodified | 1123500 | 935890 | 960030 | 1623700 | 7125300 | 0 | 3696900 | 1577500 | 2363800 | 1874500 | 1652000 | 5676400 | 3861400 | 1255600 | 1234100 | 1521300 | 890420 | 0 |
| SCLLHQFTI | RAB14; RAB14 | P61106; X6R | RAB14_HUMAN | Unmodified | 287410 | 210700 | 0 | 736450 | 1396000 | 0 | 454880 | 0 | 0 | 27593 | 0 | 1948100 | 656120 | 274780 | 513440 | 322380 | 0 | 294250 |
| SCLLLQFTD | RAB2A; RAB2B; | P61019; Q8V | RAB2A_HUMAN | Unmodified | 0 | 0 | 0 | 166530 | 0 | 0 | 1389700 | 0 | 1204100 | 1352300 | 306430 | 686000 | 0 | 0 | 0 | 0 | 87873 | 0 |
| SCLLLQFTD | RAB2A; RAB2B; | P61019; Q8V | RAB2A_HUMAN | Unmodified | 131640 | 118230 | 0 | 137750 | 0 | 0 | 111230 | 0 | 854990 | 837960 | 0 | 1041200 | 0 | 0 | 187110 | 317460 | 0 | 0 |
| SCNCLLLK | ENO1; ENO1 | P06733; A0A | ENOA_HUMAN | Unmodified | 0 | 0 | 0 | 0 | 2903300 | 0 | 1746700 | 1154500 | 7001900 | 4006000 | 4098900 | 14090000 | 7718500 | 6111000 | 2145400 | 0 | 5402900 | 0 |
| SCNCLLLK | ENO1; ENO1 | P06733; A0A | ENOA_HUMAN | Unmodified | 0 | 0 | 0 | 0 | 0 | 0 | 0 | 0 | 0 | 0 | 0 | 0 | 0 | 0 | 0 | 0 | 2766500 | 0 |
| SCPDAIKE | CD9; CD9; CD9 | P21926; A6N | CD9_HUMAN ( | Unmodified | 1074800 | 642450 | 0 | 107540 | 0 | 0 | 0 | 0 | 0 | 0 | 0 | 0 | 0 | 0 | 974840 | 653730 | 0 | 0 |
| SCPETLTH | TNS3; TNS3 | Q68CZ2; E9I | TENS3_HUMAN | Oxidation (M) | 0 | 0 | 0 | 0 | 264480 | 0 | 0 | 0 | 0 | 0 | 0 | 0 | 0 | 0 | 0 | 0 | 264530 | 270640 |
| SCPETLTH | TNS3; TNS3 | Q68CZ2; E9I | TENS3_HUMAN | Unmodified | 0 | 0 | 0 | 0 | 301490 | 0 | 0 | 0 | 0 | 0 | 0 | 0 | 0 | 0 | 0 | 0 | 0 | 353280 |
| SCPVVQSS | MARS1 | P56192 | SYMC_HUMAN | Unmodified | 63223 | 0 | 213270 | 0 | 0 | 0 | 0 | 0 | 0 | 0 | 281040 | 1716300 | 0 | 0 | 0 | 0 | 0 | 0 |
| SCPVVQSS | MARS1 | P56192 | SYMC_HUMAN | Unmodified | 56367 | 0 | 22492 | 216450 | 0 | 0 | 142700 | 0 | 0 | 219620 | 256130 | 3003000 | 0 | 0 | 0 | 0 | 0 | 0 |
| SCRTITFEQ | TPPP | O94811 | TPPP_HUMAN | Unmodified | 0 | 0 | 0 | 0 | 0 | 0 | 1311600 | 0 | 0 | 0 | 0 | 0 | 0 | 0 | 0 | 0 | 0 | 0 |
| SCSEVRRL | GPC4 | O75487 | GPC4_HUMAN | Unmodified | 0 | 0 | 0 | 0 | 0 | 0 | 0 | 0 | 0 | 130710 | 54714 | 0 | 577100 | 874900 | 0 | 0 | 1121200 | 1415400 |
| SCSGVEFS | VDAC3; VDAC3 | Q9Y277; E5F | VDAC3_HUMAI | Unmodified | 0 | 0 | 0 | 0 | 0 | 0 | 0 | 0 | 16851000 | 2599200 | 579180 | 122940 | 90710 | 0 | 0 | 0 | 0 | 0 |
| SCSGVEFS | VDAC3; VDAC3 | Q9Y277; E5F | VDAC3_HUMAI | Unmodified | 0 | 0 | 0 | 0 | 0 | 0 | 0 | 0 | 0 | 663700 | 0 | 0 | 0 | 0 | 0 | 0 | 0 | 0 |
| SCSGVEFS | VDAC2; VDAC2 | P45880; Q5J | VDAC2_HUMAI | Unmodified | 0 | 0 | 0 | 0 | 0 | 0 | 0 | 0 | 22913000 | 1802600 | 281050 | 0 | 0 | 0 | 0 | 0 | 104090 | 0 |
| SCSGVEFS | VDAC2; VDAC2 | P45880; Q5J | VDAC2_HUMAI | Unmodified | 0 | 0 | 0 | 0 | 777100 | 0 | 5354600 | 3934400 | 2.84E+08 | 1.04E+08 | 663100 | 0 | 0 | 0 | 0 | 0 | 582960 | 1650500 |
| SCSGVMEF | VDAC3 | E5RFP6 | E5RFP6_HUMA | Unmodified | 0 | 0 | 0 | 0 | 0 | 0 | 0 | 0 | 2707500 | 176700 | 0 | 0 | 0 | 0 | 0 | 0 | 0 | 0 |
| SCSGVMEF | VDAC3 | E5RFP6 | E5RFP6_HUMA | Oxidation (M) | 0 | 0 | 0 | 0 | 0 | 0 | 0 | 0 | 896390 | 0 | 0 | 0 | 0 | 0 | 0 | 0 | 0 | 0 |
| SCSPELQQ | CBR1 | P16152 | CBR1_HUMAN | Unmodified | 72718 | 0 | 0 | 0 | 0 | 0 | 0 | 84737 | 0 | 0 | 0 | 0 | 415720 | 339860 | 235040 | 175620 | 2141200 | 1927900 |
| SCSPLAFS | MVB12A; MVB1 | Q96EY5; E9F | MB12A_HUMAI | Unmodified | 0 | 0 | 0 | 0 | 0 | 0 | 0 | 0 | 61240 | 0 | 0 | 0 | 856100 | 0 | 0 | 0 | 0 | 31730 |
| SCSQEERII | PMM2; PMM2; F | O15305; H3I | PMM2_HUMAN | Unmodified | 0 | 0 | 0 | 0 | 251050 | 0 | 0 | 0 | 439520 | 0 | 0 | 959310 | 0 | 0 | 0 | 0 | 0 | 0 |
| SCSVTDAV | NEDD4L; NEDD | Q96PU5; A0 | NED4L_HUMAN | Unmodified | 1388100 | 356820 | 0 | 328390 | 0 | 0 | 0 | 0 | 0 | 0 | 0 | 0 | 1957900 | 295450 | 1191800 | 843550 | 0 | 0 |
| SCTECVRV | ITGB4 | P16144 | ITB4_HUMAN I | Unmodified | 0 | 0 | 0 | 0 | 238760 | 93559 | 0 | 0 | 0 | 0 | 0 | 0 | 65716 | 0 | 0 | 0 | 662170 | 2026700 |
| SCTEGKGE | GFM1 | Q96RP9 | EFGM_HUMAN | Unmodified | 0 | 0 | 0 | 0 | 0 | 0 | 0 | 0 | 650850 | 0 | 0 | 0 | 0 | 0 | 0 | 0 | 0 | 0 |
| SCTLARDP | PUF60; PUF60; | Q9UHX1; A0 | PUF60_HUMAN | Unmodified | 0 | 0 | 25705 | 84496 | 384370 | 0 | 0 | 0 | 522160 | 616470 | 427460 | 971150 | 148350 | 163070 | 147160 | 22656 | 538330 | 161790 |
| SCTPSPDQ | ZC3HAV1; ZC3I | Q7Z2W4; C9 | ZCCHV_HUMA | Unmodified | 433130 | 0 | 555600 | 868070 | 0 | 0 | 0 | 0 | 0 | 0 | 0 | 0 | 0 | 0 | 653170 | 0 | 0 | 0 |
| SCTPSPDQ | ZC3HAV1; ZC3I | Q7Z2W4; C9 | ZCCHV_HUMA | Unmodified | 0 | 0 | 0 | 215190 | 0 | 0 | 0 | 0 | 0 | 0 | 0 | 0 | 467870 | 111440 | 0 | 0 | 0 | 0 |
| SCTTDVRP | TRIM26; TRIM2 | Q12899; A2A | TRI26_HUMAN | Unmodified | 1237200 | 359540 | 321930 | 1004000 | 0 | 0 | 0 | 0 | 0 | 0 | 0 | 0 | 0 | 0 | 763620 | 484480 | 0 | 0 |
| SCVEEPEPI | HDGF | P51858 | HDGF_HUMAN | Unmodified | 0 | 0 | 0 | 0 | 0 | 0 | 0 | 0 | 113940 | 0 | 0 | 376560 | 0 | 0 | 0 | 0 | 0 | 0 |
| SCVEEPEPI | HDGF | P51858 | HDGF_HUMAN | Unmodified | 0 | 0 | 0 | 0 | 0 | 0 | 0 | 0 | 0 | 0 | 478550 | 748060 | 0 | 0 | 0 | 0 | 0 | 0 |
| SCVEEPEPI | HDGF | P51858 | HDGF_HUMAN | Unmodified | 0 | 0 | 0 | 0 | 0 | 0 | 0 | 0 | 0 | 0 | 0 | 501720 | 0 | 0 | 0 | 0 | 0 | 0 |
| SCVLFNCA | PDCD6IP; PDCI | Q8WUM4; A | PDC6I_HUMAN | Unmodified | 3202600 | 2786000 | 67876 | 1455400 | 0 | 270170 | 0 | 0 | 493520 | 0 | 0 | 0 | 0 | 1787700 | 2607600 | 0 | 429720 | 714990 |
| SCVQCQA | ITGB4 | P16144 | ITB4_HUMAN I | Unmodified | 0 | 0 | 0 | 0 | 0 | 0 | 0 | 0 | 0 | 0 | 0 | 0 | 0 | 0 | 0 | 0 | 187470 | 326490 |
| SCVQCQA | ITGB4 | P16144 | ITB4_HUMAN I | Unmodified | 0 | 500270 | 7975.7 | 34718 | 0 | 0 | 0 | 0 | 0 | 0 | 0 | 0 | 0 | 0 | 0 | 0 | 385580 | 0 |
| SCVSSAW | NOSTRIN | Q8IVI9 | NOSTN_HUMA | Oxidation (M) | 0 | 0 | 397760 | 1433500 | 0 | 0 | 0 | 0 | 0 | 0 | 0 | 0 | 0 | 0 | 0 | 1441200 | 0 | 0 |
| SCYDLSCH | GARS1; GARS1 | P41250; A0A | GARS_HUMAN | Unmodified | 0 | 0 | 0 | 0 | 0 | 0 | 0 | 0 | 222410 | 0 | 0 | 1089300 | 0 | 0 | 0 | 103410 | 0 | 0 |
| SDAAMTVA | KIT | P10721 | KIT_HUMAN Ma | Oxidation (M) | 0 | 0 | 0 | 0 | 0 | 0 | 0 | 0 | 146430 | 0 | 0 | 0 | 0 | 0 | 0 | 0 | 0 | 0 |
| SDAAMTVA | KIT | P10721 | KIT_HUMAN Ma | Unmodified | 0 | 0 | 0 | 0 | 0 | 0 | 0 | 15458 | 709570 | 501140 | 392290 | 3040900 | 0 | 0 | 0 | 0 | 0 | 0 |
| SDAAVDTS | PTMA; PTMA | P06454; B8Z | PTMA_HUMAN | Acetyl (Protein N | 0 | 0 | 0 | 0 | 0 | 0 | 142730 | 100930 | 750000 | 345620 | 95589 | 0 | 0 | 0 | 0 | 0 | 318960 | 95836 |
| SDAAVDTS | PTMA; PTMA | P06454; B8Z | PTMA_HUMAN | Acetyl (Protein N | 0 | 0 | 0 | 0 | 290750 | 654090 | 3056800 | 2939800 | 0 | 0 | 0 | 634340 | 0 | 0 | 0 | 0 | 7446600 | 0 |
| SDAEEDGC | CANX; CANX; C | P27824; A0A | CALX_HUMAN | Unmodified | 0 | 0 | 0 | 0 | 0 | 0 | 0 | 0 | 978500 | 0 | 215520 | 0 | 0 | 0 | 0 | 0 | 0 | 0 |
| SDAGLESD | DDX21 | Q9NR30 | DDX21_HUMAI | Unmodified | 0 | 0 | 0 | 0 | 0 | 0 | 0 | 0 | 0 | 0 | 473930 | 0 | 0 | 0 | 0 | 0 | 0 | 0 |
| SDAIYSAR | LRP1 | Q07954 | LRP1_HUMAN | Unmodified | 0 | 18895 | 91657 | 44276 | 23102 | 0 | 0 | 0 | 850250 | 0 | 122510 | 321520 | 0 | 0 | 154340 | 24687 | 0 | 0 |
| SDALETLGI | HNRNPL; HNRI | P14866; M0 | HNRPL_HUMAI | Oxidation (M) | 0 | 0 | 0 | 0 | 0 | 0 | 0 | 94899 | 921140 | 0 | 1144100 | 7609600 | 0 | 497690 | 0 | 0 | 0 | 0 |
| SDALNSAII | CTNNA1; CTNN | P35221; G3 | CTNA1_HUMAI | Unmodified | 443100 | 0 | 762740 | 2273400 | 1400200 | 970730 | 0 | 0 | 243420 | 75821 | 2805700 | 0 | 0 | 0 | 842110 | 1406400 | 0 | 1446300 |
| SDANRASS | EVL; EVL; EVL; E | Q9UI08; G3 | EVL_HUMAN E | Unmodified | 0 | 0 | 0 | 0 | 0 | 0 | 0 | 1323000 | 0 | 0 | 0 | 0 | 447250 | 0 | 0 | 0 | 0 | 0 |
| SDAPTGDV | GOLPH3 | Q9H4A6 | GOLP3_HUMAI | Unmodified | 0 | 0 | 0 | 0 | 1041400 | 230390 | 0 | 0 | 0 | 0 | 597480 | 1537500 | 804090 | 0 | 0 | 2238600 | 0 | 574050 |
| SDAPTSSSI | SH3BP4 | Q9P0V3 | SH3B4_HUMAN | Unmodified | 0 | 0 | 0 | 317640 | 953370 | 0 | 845850 | 0 | 1002300 | 0 | 103920 | 0 | 0 | 0 | 436490 | 0 | 0 | 293200 |
| SDATADDL | DRG1 | Q9Y295 | DRG1_HUMAN | Unmodified | 48528 | 0 | 74635 | 0 | 0 | 0 | 0 | 0 | 0 | 0 | 0 | 1747600 | 0 | 0 | 134090 | 0 | 0 | 0 |
| SDAVHLFS | ADAM22; ADAM | Q9P0K1; F8 | ADA22_HUMAN | Unmodified | 220090 | 167730 | 0 | 0 | 0 | 0 | 0 | 110490 | 638090 | 0 | 0 | 0 | 0 | 0 | 0 | 0 | 0 | 0 |
| SDAYYCTG | PARP1; PARP1; | P09874; A0A | PARP1_HUMAN | Unmodified | 287860 | 0 | 0 | 0 | 0 | 0 | 0 | 66024 | 488230 | 0 | 374690 | 634080 | 0 | 0 | 0 | 0 | 0 | 0 |
| SDCYTELEI | S100A16 | Q96FQ6 | S10AG_HUMAI | Acetyl (Protein N | 0 | 0 | 356280 | 0 | 3663300 | 892130 | 0 | 0 | 0 | 697310 | 0 | 0 | 235490 | 234610 | 0 | 0 | 7690300 | 6331200 |
| SDDFGQTV | SORL1 | Q92673 | SORL_HUMAN | Unmodified | 0 | 0 | 0 | 0 | 0 | 0 | 0 | 19389 | 1031000 | 0 | 0 | 0 | 0 | 0 | 0 | 35228 | 0 | 0 |
| SDDHPDV | APPL1 | Q9UKG1 | DP13A_HUMAN | Unmodified | 52348 | 225330 | 112590 | 82213 | 0 | 0 | 0 | 0 | 1043100 | 0 | 67616 | 525670 | 0 | 0 | 53681 | 0 | 0 | 0 |
| SDDSNGIE | EMB; EMB | Q6PCB8; D6 | EMB_HUMAN E | Unmodified | 0 | 0 | 0 | 0 | 376380 | 0 | 0 | 0 | 0 | 0 | 0 | 0 | 0 | 0 | 128540 | 77298 | 0 | 0 |
| SDDSVIQLI | CADM1; CADM | Q9BY67; X5I | CADM1_HUMA | Unmodified | 0 | 0 | 45493 | 0 | 55948 | 0 | 0 | 0 | 160030 | 0 | 23578 | 0 | 0 | 0 | 0 | 0 | 0 | 0 |

| | | | | | | | | | | | | | | | | | | | | | | |
|---|---|---|---|---|---|---|---|---|---|---|---|---|---|---|---|---|---|---|---|---|---|---|
| SDEAVKPF | FASN; FASN | P49327; A0A | FAS_HUMAN F | Unmodified | 0 | 0 | 0 | 0 | 0 | 0 | 0 | 0 | 0 | 0 | 0 | 16866000 | 0 | 0 | 1379600 | 485600 | 0 | 0 |
| SDEDYLYR | POR; POR; POR | P16435; H0Y | NCPR_HUMAN | Unmodified | 0 | 0 | 61323 | 0 | 0 | 0 | 0 | 0 | 72199 | 0 | 0 | 539400 | 0 | 0 | 0 | 26056 | 0 | 0 |
| SDEEYRELI | CYFIP2; CYFIP2 | Q96F07; H7C | CYFP2_HUMAN | Unmodified | 0 | 122530 | 0 | 0 | 0 | 0 | 0 | 0 | 0 | 0 | 35017 | 184340 | 0 | 0 | 185930 | 361640 | 0 | 0 |
| SDEFSLAD | MAPK1IP1L | Q8NDC0 | MISSL_HUMAN | Acetyl (Protein N | 902570 | 2723500 | 817640 | 1415000 | 12674000 | 2097100 | 3124300 | 3917300 | 9667100 | 4151600 | 1156700 | 0 | 6243500 | 5667900 | 914420 | 721600 | 2783000 | 17899000 |
| SDEGESMF | CEMIP2; CEMIP | Q9UHN6; F5 | CEIP2_HUMAN | Unmodified | 0 | 0 | 0 | 0 | 265760 | 0 | 0 | 0 | 800570 | 110610 | 247210 | 0 | 0 | 0 | 0 | 0 | 0 | 0 |
| SDEGIYR | NCAM2; NCAM | O15394; H9I | NCAM2_HUMA | Unmodified | 0 | 0 | 1068500 | 0 | 0 | 0 | 0 | 0 | 0 | 0 | 0 | 0 | 0 | 0 | 1068300 | 874940 | 0 | 0 |
| SDEMNVK | MYL6; MYL6; MY | P60660; G8J | MYL6_HUMAN | Unmodified | 0 | 0 | 46864 | 0 | 3022700 | 306270 | 0 | 0 | 52209 | 0 | 0 | 151090 | 305470 | 0 | 0 | 0 | 0 | 284910 |
| SDEMNVK\ | MYL6; MYL6; MY | P60660; G8J | MYL6_HUMAN | 2 Oxidation (M) | 0 | 0 | 0 | 0 | 506980 | 35313 | 1368000 | 0 | 0 | 32350 | 0 | 0 | 908670 | 153640 | 0 | 0 | 7555100 | 0 |
| SDENEDPS | MYOF; MYOF | Q9NZM1; H0 | MYOF_HUMAN | Unmodified | 0 | 0 | 0 | 0 | 1500500 | 0 | 0 | 0 | 0 | 0 | 0 | 0 | 0 | 0 | 0 | 0 | 0 | 129660 |
| SDENIPGG | CPD | O75976 | CBPD_HUMAN | Unmodified | 0 | 0 | 0 | 0 | 0 | 0 | 0 | 0 | 1160800 | 0 | 58073 | 0 | 0 | 0 | 0 | 0 | 0 | 0 |
| SDESIKEED | DNMT1; DNMT | P26358; A0A | DNMT1_HUMA | Unmodified | 0 | 0 | 0 | 0 | 0 | 0 | 0 | 0 | 32094 | 0 | 0 | 75382 | 0 | 0 | 0 | 0 | 0 | 0 |
| SDFDSNPF | SCAMP1 | O15126 | SCAM1_HUMA | Acetyl (Protein N | 2241500 | 3156100 | 91670 | 444740 | 15186000 | 2891600 | 1069100 | 0 | 3391400 | 1263500 | 0 | 0 | 10040000 | 6332200 | 6137100 | 8022300 | 1144800 | 3872000 |
| SDFGKFVL | CALR; CALR; CA | P27797; A0A | CALR_HUMAN | Unmodified | 0 | 0 | 0 | 0 | 165010 | 0 | 838680 | 645520 | 5632200 | 8258000 | 74274 | 0 | 0 | 126160 | 0 | 0 | 0 | 0 |
| SDFGKFVL | CALR; CALR; CA | P27797; A0A | CALR_HUMAN | Unmodified | 0 | 0 | 0 | 0 | 167720 | 0 | 2130000 | 196020 | 2080600 | 18342000 | 0 | 0 | 0 | 320230 | 0 | 0 | 0 | 0 |
| SDFNNYVA | PLPP2; PLPP2; I | O43688; K7E | PLPP2_HUMAN | Unmodified | 0 | 0 | 0 | 0 | 0 | 0 | 0 | 0 | 0 | 0 | 0 | 4638000 | 0 | 0 | 0 | 0 | 0 | 0 |
| SDFQVNLN | CTNND1; CTNN | O60716; H0Y | CTND1_HUMA | Unmodified | 534900 | 587980 | 3695500 | 4300500 | 2469100 | 656730 | 0 | 0 | 0 | 0 | 3776000 | 7528000 | 2032900 | 1247700 | 3755800 | 4998600 | 980950 | 2335600 |
| SDFQVNLN | CTNND1; CTNN | O60716; H0Y | CTND1_HUMA | Unmodified | 0 | 0 | 0 | 0 | 20328000 | 854160 | 0 | 0 | 0 | 0 | 0 | 0 | 9822500 | 8636600 | 0 | 0 | 4627500 | 30448000 |
| SDFSDSIK | MIA3; MIA3 | Q5JRA6; A0A | TGO1_HUMAN | Unmodified | 0 | 0 | 0 | 0 | 0 | 0 | 0 | 55173 | 3477600 | 101160 | 217290 | 94198 | 0 | 0 | 0 | 0 | 0 | 0 |
| SDFYDIVLV | PCYOX1 | Q9UHG3 | PCYOX_HUMAI | Unmodified | 0 | 0 | 48256 | 0 | 0 | 0 | 0 | 18301 | 2048400 | 0 | 0 | 18556 | 0 | 0 | 0 | 0 | 0 | 0 |
| SDGALLLG | WDR77 | Q9BQA1 | MEP50_HUMAN | Unmodified | 0 | 0 | 0 | 0 | 0 | 0 | 0 | 0 | 0 | 0 | 248000 | 5602000 | 0 | 0 | 0 | 0 | 0 | 0 |
| SDGAPASD | BASP1 | P80723 | BASP1_HUMAN | Unmodified | 40044 | 0 | 0 | 0 | 14017000 | 1104500 | 168000 | 743730 | 6490000 | 1672300 | 0 | 0 | 0 | 0 | 0 | 0 | 1549700 | 4179000 |
| SDGGYTYD | RARS1 | P54136 | SYRC_HUMAN | Unmodified | 0 | 0 | 172640 | 186420 | 182880 | 0 | 0 | 0 | 0 | 193820 | 1276800 | 6676800 | 264600 | 0 | 302630 | 29042 | 0 | 0 |
| SDGIYIINLI | RPSA; RPSA; RF | P08865; A0A | RSSA_HUMAN | Unmodified | 0 | 0 | 0 | 0 | 466330 | 249670 | 465520 | 173850 | 214870 | 0 | 1790700 | 6089600 | 281800 | 285980 | 0 | 0 | 0 | 682760 |
| SDGIYIINLI | RPSA; RPSA; RF | P08865; A0A | RSSA_HUMAN | Unmodified | 0 | 0 | 0 | 0 | 0 | 0 | 0 | 0 | 0 | 0 | 0 | 0 | 0 | 0 | 0 | 0 | 0 | 0 |
| SDGSLEDG | SERINC1 | Q9NRX5 | SERC1_HUMAI | Unmodified | 328690 | 1574500 | 1957900 | 834030 | 0 | 0 | 0 | 57718 | 2399000 | 703230 | 101790 | 2729500 | 0 | 0 | 996260 | 1120100 | 0 | 0 |
| SDHGDVSI | STAMBP; STAMI | O95630; C9J | STABP_HUMAN | Acetyl (Protein N | 719840 | 845870 | 0 | 384320 | 1180400 | 652800 | 108580 | 156560 | 1021100 | 552120 | 137800 | 613590 | 845590 | 525560 | 107250 | 219710 | 153740 | 265550 |
| SDHLTVVN | DHX36; DHX36 | Q9H2U1; E7 | DHX36_HUMAN | Unmodified | 0 | 0 | 0 | 0 | 0 | 0 | 0 | 0 | 0 | 0 | 0 | 3604000 | 0 | 0 | 476300 | 249890 | 0 | 0 |
| SDIDEIVLV | HSPA5; HSPA5 | P11021; A0A | BIP_HUMAN Er | Unmodified | 0 | 0 | 0 | 0 | 0 | 0 | 0 | 0 | 24596000 | 764450 | 6824600 | 932890 | 0 | 0 | 328860 | 0 | 0 | 0 |
| SDIDEIVLV | HSPA5; HSPA5 | P11021; A0A | BIP_HUMAN Er | Unmodified | 0 | 0 | 0 | 0 | 1505300 | 0 | 2721800 | 737390 | 57721000 | 26657000 | 12554000 | 620470 | 0 | 674120 | 1901500 | 0 | 0 | 1092800 |
| SDIDEIVLV | HSPA5; HSPA5 | P11021; A0A | BIP_HUMAN Er | Unmodified | 0 | 0 | 0 | 0 | 0 | 0 | 0 | 0 | 0 | 3425200 | 0 | 0 | 0 | 0 | 0 | 0 | 0 | 0 |
| SDIDLLEEH | DAAM1 | Q9Y4D1 | DAAM1_HUMA | Unmodified | 0 | 0 | 266040 | 0 | 0 | 53195 | 0 | 0 | 301520 | 0 | 129380 | 3855700 | 0 | 0 | 618220 | 348330 | 0 | 0 |
| SDIGEVILV | HSPA9; HSPA9; | P38646; H0Y | GRP75_HUMAI | Unmodified | 0 | 0 | 0 | 0 | 0 | 0 | 0 | 0 | 2266000 | 0 | 539730 | 0 | 0 | 0 | 0 | 0 | 399570 | 0 |
| SDIHTEAVC | DIP2B | Q9P265 | DIP2B_HUMAN | Unmodified | 240890 | 173190 | 0 | 0 | 111550 | 0 | 0 | 0 | 841550 | 223150 | 0 | 1768900 | 518920 | 0 | 642700 | 459320 | 420590 | 3035400 |
| SDITDTGLA | FZD6 | O60353 | FZD6_HUMAN | Unmodified | 0 | 0 | 0 | 0 | 3806500 | 0 | 0 | 0 | 2126000 | 0 | 0 | 0 | 0 | 0 | 0 | 0 | 0 | 0 |
| SDIVQLRGI | HDLBP; HDLBP | Q00341; A0A | VIGLN_HUMAN | Unmodified | 0 | 0 | 0 | 0 | 362590 | 0 | 0 | 0 | 0 | 0 | 118710 | 278980 | 837240 | 294090 | 0 | 0 | 0 | 0 |
| SDIYVCMIS | GDI2 | P50395 | GDIB_HUMAN | Unmodified | 0 | 0 | 308960 | 1003500 | 969700 | 0 | 389370 | 0 | 3539300 | 451500 | 1143700 | 401230 | 526010 | 0 | 1824100 | 0 | 0 | 0 |
| SDKLPEEM | MCAM | P43121 | MUC18_HUMA | Unmodified | 0 | 0 | 0 | 0 | 762260 | 0 | 0 | 0 | 2440400 | 0 | 0 | 1748200 | 0 | 0 | 0 | 0 | 0 | 216470 |
| SDKLPEEM | MCAM | P43121 | MUC18_HUMA | Oxidation (M) | 0 | 0 | 0 | 0 | 0 | 0 | 0 | 0 | 524610 | 751680 | 0 | 723140 | 0 | 0 | 0 | 0 | 0 | 0 |
| SDKLPYK | AHCY | P23526 | SAHH_HUMAN | Acetyl (Protein N | 499500 | 402750 | 1323900 | 1477500 | 10916000 | 3527500 | 1762400 | 1110700 | 2033200 | 2174100 | 2737100 | 12962000 | 4391400 | 5059100 | 1551900 | 1307400 | 9279200 | 0 |
| SDKLPYKV | AHCY | P23526 | SAHH_HUMAN | Acetyl (Protein N | 0 | 0 | 0 | 0 | 367510 | 226830 | 285160 | 0 | 0 | 181170 | 0 | 232130 | 397110 | 227660 | 0 | 0 | 1907500 | 719180 |
| SDKPDLSE | TMSB15B; TMSI | P0CG35; P0 | TB15B_HUMAN | Acetyl (Protein N | 126700 | 95222 | 0 | 0 | 0 | 221260 | 0 | 424990 | 4380600 | 1197700 | 110200 | 313100 | 0 | 0 | 0 | 0 | 0 | 285050 |
| SDKPDLSE | TMSB15B; TMSI | P0CG35; P0 | TB15B_HUMAN | Acetyl (Protein N | 220880 | 0 | 0 | 0 | 0 | 0 | 0 | 134690 | 1915500 | 2691300 | 0 | 0 | 1250200 | 0 | 0 | 0 | 0 | 0 |
| SDKPDMAE | TMSB4X | P62328 | TYB4_HUMAN | Acetyl (Protein N | 0 | 0 | 470960 | 0 | 1865100 | 444760 | 1135900 | 64048 | 6971900 | 2946400 | 164640 | 0 | 373330 | 446800 | 191460 | 193240 | 2485400 | 6226200 |
| SDKPDMAE | TMSB4X | P62328 | TYB4_HUMAN | Acetyl (Protein N | 154710 | 0 | 2033200 | 1863500 | 14517000 | 2516800 | 5952900 | 4991700 | 29937000 | 8203000 | 2626800 | 592570 | 1687100 | 4887000 | 687880 | 362070 | 14243000 | 40160000 |
| SDKPDMAE | TMSB4X | P62328 | TYB4_HUMAN | Acetyl (Protein N | 547890 | 477960 | 0 | 1507200 | 20140000 | 5625500 | 22756000 | 29587000 | 12313000 | 30293000 | 1218800 | 677000 | 4647000 | 12181000 | 378170 | 480520 | 46894000 | 67479000 |
| SDKPDMAE | TMSB4X | P62328 | TYB4_HUMAN | Acetyl (Protein N | 56764 | 645320 | 0 | 3505600 | 98224000 | 14666000 | 85029000 | 49404000 | 1.17E+08 | 1.22E+08 | 2044000 | 1209000 | 17156000 | 28062000 | 718960 | 712440 | 2.32E+08 | 3.14E+08 |
| SDKSELKAI | DYNC1I2; DYN | Q13409; E7E | DC1I2_HUMAN | Acetyl (Protein N | 0 | 0 | 0 | 0 | 0 | 68459 | 0 | 0 | 421360 | 0 | 668410 | 279600 | 391190 | 170540 | 0 | 0 | 0 | 0 |
| SDKSPDLA | EIF2A; EIF2A | Q9BY44; F8V | EIF2A_HUMAN | Unmodified | 0 | 0 | 0 | 0 | 0 | 0 | 29845 | 85204 | 0 | 0 | 614980 | 1153000 | 0 | 0 | 0 | 0 | 0 | 0 |
| SDKTEEIAE | DDX21 | Q9NR30 | DDX21_HUMAI | Unmodified | 0 | 0 | 0 | 0 | 0 | 0 | 0 | 0 | 0 | 0 | 560170 | 934800 | 0 | 0 | 0 | 0 | 0 | 0 |
| SDKVPDNK | COPG1; COPG | Q9Y678; H0Y | COPG1_HUMA | Unmodified | 0 | 0 | 0 | 0 | 0 | 0 | 165810 | 0 | 0 | 0 | 0 | 620290 | 0 | 183160 | 0 | 0 | 0 | 0 |
| SDLAVEAG | IGSF8; IGSF8 | Q969P0; C9. | IGSF8_HUMAN | Unmodified | 5618200 | 8635100 | 5709500 | 2131700 | 0 | 0 | 0 | 0 | 0 | 0 | 0 | 1472300 | 0 | 0 | 0 | 0 | 0 | 0 |
| SDLAVEAG | IGSF8; IGSF8 | Q969P0; C9. | IGSF8_HUMAN | Unmodified | 2633000 | 3081400 | 1631200 | 2244400 | 112960 | 0 | 0 | 0 | 0 | 0 | 0 | 0 | 340980 | 0 | 2005200 | 1125600 | 0 | 0 |
| SDLAVPSEI | LGALS3BP | Q08380 | LG3BP_HUMAN | Unmodified | 0 | 0 | 0 | 0 | 0 | 0 | 0 | 0 | 14109000 | 1979300 | 2474700 | 32155000 | 0 | 0 | 0 | 0 | 0 | 616430 |
| SDLCETSAI | CNPY1; CNPY1 | Q3B7I2; A0A | CNPY1_HUMAI | Unmodified | 0 | 0 | 0 | 0 | 0 | 0 | 0 | 0 | 0 | 0 | 1013400 | 351040 | 0 | 0 | 0 | 0 | 0 | 0 |
| SDLEDKRQ | RALA; RALA | P11233; H7C | RALA_HUMAN | Unmodified | 831480 | 972130 | 0 | 1823900 | 293090 | 48647 | 144330 | 133130 | 5519300 | 854740 | 4129900 | 2817100 | 974350 | 269450 | 180390 | 669150 | 295040 | 1101800 |
| SDLLLEGFI | MYH10 | P35580 | MYH10_HUMAI | Unmodified | 0 | 0 | 0 | 61259 | 0 | 0 | 0 | 0 | 0 | 0 | 0 | 1701600 | 0 | 0 | 422140 | 294580 | 0 | 0 |
| SDLLLSGR | PRMT5; PRMT5; | O14744; G3V | ANM5_HUMAN | Unmodified | 0 | 0 | 0 | 0 | 0 | 0 | 0 | 0 | 0 | 0 | 0 | 2570700 | 0 | 0 | 0 | 0 | 0 | 0 |
| SDLLLSGRI | PRMT5; PRMT5; | O14744; G3V | ANM5_HUMAN | Unmodified | 229260 | 214970 | 0 | 576160 | 661870 | 246660 | 255320 | 0 | 1602700 | 289020 | 1203100 | 9281700 | 705680 | 169880 | 852160 | 461420 | 398430 | 926130 |
| SDLLSAIR | WASF2 | Q9Y6W5 | WASF2_HUMAI | Unmodified | 62702 | 0 | 0 | 0 | 0 | 0 | 0 | 0 | 1204600 | 0 | 315560 | 1238000 | 0 | 0 | 0 | 0 | 0 | 0 |
| SDLLSAIRC | WASF2 | Q9Y6W5 | WASF2_HUMAI | Unmodified | 0 | 0 | 23160 | 38816 | 439660 | 141910 | 0 | 0 | 0 | 0 | 0 | 127540 | 0 | 131140 | 0 | 0 | 350060 | 0 |
| SDLPGSDK | SLC9A3R2; SLC | Q15599; H3I | NHRF2_HUMAI | Unmodified | 503840 | 316020 | 0 | 0 | 0 | 0 | 0 | 0 | 465100 | 0 | 0 | 1919600 | 0 | 0 | 645900 | 608960 | 0 | 0 |
| SDLQNKTC | KIF11; KIF11; KI | P52732; A0A | KIF11_HUMAN | Unmodified | 0 | 0 | 0 | 0 | 320100 | 0 | 0 | 0 | 0 | 0 | 0 | 0 | 0 | 0 | 0 | 0 | 0 | 0 |
| SDLQRPNF | MUC13 | Q9H3R2 | MUC13_HUMA | Unmodified | 6187100 | 0 | 0 | 90102 | 0 | 0 | 0 | 0 | 4658700 | 297060 | 0 | 0 | 0 | 0 | 0 | 54933 | 0 | 151780 |
| SDLRLIEVT | CNPY3; CNPY3 | Q9BT09; A0A | CNPY3_HUMAI | Unmodified | 0 | 0 | 0 | 0 | 0 | 0 | 1227500 | 121210 | 0 | 11424000 | 0 | 0 | 0 | 0 | 0 | 0 | 0 | 0 |
| SDLSLER | PTGFRN | Q9P2B2 | FPRP_HUMAN | Unmodified | 8508500 | 3527200 | 20050000 | 4221600 | 0 | 0 | 77171 | 0 | 0 | 0 | 302700 | 15702000 | 0 | 30524 | 24854000 | 35086000 | 1475100 | 0 |
| SDLSLGSR | SNX18 | Q96RF0 | SNX18_HUMAI | Unmodified | 1434900 | 2450200 | 3903000 | 4448500 | 0 | 0 | 0 | 0 | 0 | 0 | 0 | 0 | 425050 | 0 | 6734200 | 4657500 | 0 | 0 |
| SDLVNEEA | CEACAM5; CEA | P06731; A0A | CEAM5_HUMAI | Unmodified | 0 | 0 | 0 | 0 | 0 | 0 | 0 | 73393 | 912640 | 353760 | 0 | 0 | 0 | 0 | 0 | 0 | 0 | 0 |
| SDMDQIVN | NIBAN1 | Q9BZQ8 | NIBA1_HUMAN | Unmodified | 0 | 0 | 0 | 0 | 0 | 0 | 0 | 0 | 1167200 | 247720 | 0 | 0 | 0 | 0 | 0 | 0 | 0 | 0 |
| SDNARQQI | MYH10 | P35580 | MYH10_HUMAI | Unmodified | 0 | 14186 | 0 | 0 | 181790 | 13726 | 0 | 0 | 0 | 0 | 0 | 63509 | 0 | 0 | 0 | 0 | 0 | 0 |
| SDNARQQI | MYH10 | P35580 | MYH10_HUMAI | Unmodified | 0 | 0 | 0 | 0 | 2870700 | 602640 | 476540 | 0 | 0 | 80045 | 0 | 0 | 0 | 0 | 0 | 0 | 0 | 0 |
| SDNGELED | PAK2 | Q13177 | PAK2_HUMAN | Acetyl (Protein N | 0 | 0 | 0 | 29125 | 650070 | 0 | 0 | 0 | 0 | 0 | 2582400 | 2603700 | 398560 | 0 | 162530 | 0 | 183320 | 0 |
| SDNVVTSE | PTGFRN | Q9P2B2 | FPRP_HUMAN | Oxidation (M) | 0 | 0 | 39284 | 106200 | 0 | 0 | 0 | 0 | 0 | 0 | 0 | 0 | 242660 | 0 | 1188200 | 42369 | 0 | 0 |
| SDPDHLAE | MYO6; MYO6; M | Q9UM54; A0 | MYO6_HUMAN | Unmodified | 0 | 0 | 0 | 0 | 0 | 0 | 0 | 0 | 0 | 0 | 0 | 0 | 0 | 0 | 2739200 | 2513300 | 0 | 0 |
| SDPDHLAE | MYO6; MYO6; M | Q9UM54; A0 | MYO6_HUMAN | Unmodified | 51457 | 0 | 261830 | 904810 | 156480 | 452300 | 366830 | 42834 | 2607600 | 860130 | 493340 | 575460 | 0 | 169490 | 774560 | 647600 | 0 | 93149 |
| SDPFLEFFF | CPNE1; CPNE1 | Q99829; Q5. | CPNE1_HUMAI | Unmodified | 486550 | 794560 | 0 | 0 | 0 | 0 | 0 | 0 | 4355500 | 0 | 0 | 3962600 | 0 | 0 | 1549200 | 896420 | 0 | 0 |
| SDPFLEFFF | CPNE1; CPNE1 | Q99829; Q5. | CPNE1_HUMAI | Unmodified | 132280 | 62924 | 296900 | 545120 | 505710 | 303960 | 0 | 0 | 1135300 | 202040 | 430980 | 173900 | 200860 | 133300 | 411830 | 139990 | 76636 | 932200 |
| SDPIQEIR | ERP44 | Q9BS26 | ERP44_HUMAN | Unmodified | 0 | 0 | 0 | 0 | 0 | 0 | 0 | 0 | 19716000 | 184380 | 139560 | 317800 | 0 | 0 | 0 | 0 | 0 | 0 |
| SDPIQEIRD | ERP44 | Q9BS26 | ERP44_HUMAN | Unmodified | 0 | 0 | 0 | 0 | 0 | 0 | 0 | 0 | 1.35E+08 | 6011000 | 780430 | 188700 | 0 | 0 | 0 | 0 | 204060 | 0 |
| SDPIQEIRD | ERP44 | Q9BS26 | ERP44_HUMAN | Unmodified | 229270 | 0 | 180860 | 748060 | 0 | 0 | 10138000 | 0 | 1.37E+08 | 1.3E+08 | 6403000 | 227180 | 2647400 | 0 | 902840 | 142590 | 0 | 0 |
| SDPLCVLFI | CPNE3; CPNE3 | O75131; A0A | CPNE3_HUMAI | Unmodified | 0 | 0 | 0 | 0 | 0 | 0 | 123990 | 0 | 118850 | 88185 | 0 | 0 | 0 | 0 | 0 | 0 | 0 | 0 |
| SDPLCVLFI | CPNE3; CPNE3 | O75131; A0A | CPNE3_HUMAI | Unmodified | 0 | 0 | 198870 | 1284500 | 0 | 0 | 0 | 0 | 13270000 | 557120 | 0 | 160350 | 0 | 950100 | 1981700 | 436570 | 0 | 0 |
| SDPLCVLFI | CPNE3; CPNE3 | O75131; A0A | CPNE3_HUMAI | Unmodified | 0 | 0 | 0 | 0 | 1347000 | 0 | 0 | 0 | 821680 | 2628800 | 0 | 0 | 0 | 0 | 0 | 0 | 1137200 | 3933800 |
| SDPLLIGIP | GNG12 | Q9UBI6 | GBG12_HUMAI | Unmodified | 277400 | 0 | 582890 | 0 | 0 | 76558 | 0 | 0 | 0 | 0 | 0 | 0 | 0 | 0 | 1430600 | 1463700 | 0 | 0 |
| SDPLLIGIP | GNG12 | Q9UBI6 | GBG12_HUMAI | Unmodified | 461850 | 0 | 0 | 336130 | 429980 | 196650 | 96145 | 0 | 730120 | 0 | 0 | 185350 | 0 | 0 | 0 | 630740 | 184890 | 1155100 |
| SDPNQHL1 | AGL | P35573 | GDE_HUMAN ( | Unmodified | 0 | 0 | 0 | 0 | 0 | 0 | 0 | 0 | 0 | 0 | 0 | 971290 | 0 | 0 | 0 | 0 | 0 | 0 |
| SDPNQHL1 | AGL | P35573 | GDE_HUMAN ( | Unmodified | 0 | 0 | 0 | 0 | 0 | 0 | 0 | 0 | 0 | 0 | 789170 | 1079100 | 0 | 0 | 0 | 0 | 0 | 0 |
| SDPNRETD | DDB1; DDB1; D | Q16531; A0A | DDB1_HUMAN | Unmodified | 0 | 0 | 0 | 0 | 0 | 0 | 0 | 0 | 0 | 0 | 0 | 1563700 | 0 | 0 | 0 | 1036200 | 0 | 0 |
| SDPPQEAA | EEF1A2; EEF1A | Q05639; A0A | EF1A2_HUMAN | Unmodified | 17285000 | 5015600 | 2458400 | 14112000 | 2912900 | 0 | 0 | 0 | 1514600 | 0 | 0 | 0 | 0 | 3020700 | 31137000 | 14138000 | 0 | 0 |
| SDPPTLLTF | STK10 | O94804 | STK10_HUMAN | Unmodified | 0 | 0 | 0 | 0 | 1812400 | 23344 | 0 | 0 | 0 | 0 | 0 | 0 | 0 | 0 | 0 | 0 | 1111900 | 1626500 |
| SDPSIVRLL | HPCA; HPCAL1 | P84074; P37 | HPCA_HUMAN | Unmodified | 0 | 0 | 0 | 0 | 2435700 | 177940 | 0 | 0 | 0 | 181580 | 0 | 0 | 0 | 0 | 0 | 0 | 2057700 | 2735300 |
| SDPTSYAG | ATP1B3; ATP1B | P54709; C9J | AT1B3_HUMAN | Unmodified | 0 | 85981 | 943340 | 487060 | 0 | 24475 | 0 | 0 | 1619700 | 39389 | 301190 | 11766000 | 0 | 0 | 3730800 | 2612700 | 0 | 0 |
| SDPTSYAG | ATP1B3; ATP1B | P54709; C9J | AT1B3_HUMAN | Unmodified | 0 | 0 | 0 | 0 | 553780 | 1505600 | 646400 | 179120 | 9564400 | 1666800 | 3032300 | 1.4E+08 | 0 | 1121800 | 1482500 | 0 | 419500 | 1337100 |
| SDPVTLNV | CEACAM6 | P40199 | CEAM6_HUMAN | Unmodified | 0 | 0 | 0 | 0 | 0 | 0 | 0 | 0 | 13104000 | 2171000 | 0 | 0 | 0 | 0 | 0 | 0 | 0 | 652230 |
| SDPVVSYR | EEF2 | P13639 | EF2_HUMAN El | Unmodified | 620280 | 1083500 | 621340 | 982280 | 315370 | 0 | 29270 | 0 | 1703000 | 0 | 2465700 | 36390000 | 0 | 0 | 1158700 | 1217800 | 78006 | 506720 |
| SDPVVSYR | EEF2 | P13639 | EF2_HUMAN El | Unmodified | 907340 | 0 | 0 | 0 | 8133600 | 1409800 | 1382500 | 438290 | 0 | 555110 | 2094700 | 2710100 | 1406700 | 1687500 | 0 | 661510 | 2816600 | 5105900 |
| SDPVVSYR | EEF2 | P13639 | EF2_HUMAN El | Unmodified | 0 | 0 | 0 | 0 | 2846100 | 0 | 1274300 | 423610 | 0 | 0 | 0 | 0 | 294770 | 863940 | 0 | 0 | 3552800 | 0 |
| SDPYLEFH | CPNE3; CPNE3 | O75131; A0A | CPNE3_HUMAI | Unmodified | 128240 | 158920 | 0 | 1254700 | 476490 | 0 | 218480 | 0 | 11994000 | 911370 | 158060 | 210660 | 867720 | 305970 | 0 | 0 | 192430 | 832410 |
| SDPYLEFH | CPNE3; CPNE3 | O75131; A0A | CPNE3_HUMAI | Oxidation (M) | 0 | 0 | 0 | 0 | 1157100 | 0 | 0 | 0 | 0 | 1029800 | 0 | 0 | 624300 | 436460 | 0 | 0 | 0 | 1930300 |
| SDQDYILK | PA2G4; PA2G4; | Q9UQ80; F8 | PA2G4_HUMAN | Unmodified | 0 | 0 | 0 | 0 | 0 | 0 | 129690 | 0 | 704220 | 0 | 1642000 | 12058000 | 0 | 0 | 0 | 0 | 0 | 0 |
| SDQKQEQI | STAT1; STAT1; S | P42224; A0A | STAT1_HUMAN | Unmodified | 183470 | 261510 | 0 | 0 | 4981700 | 2301000 | 43079 | 27194 | 109900 | 143930 | 3821200 | 14741000 | 1848500 | 1276400 | 296480 | 225840 | 2158800 | 2791400 |
| SDQNLQTA | IPO7 | O95373 | IPO7_HUMAN I | Unmodified | 0 | 0 | 0 | 0 | 0 | 0 | 0 | 0 | 0 | 0 | 0 | 697090 | 0 | 0 | 0 | 0 | 0 | 0 |
| SDQQLDC | CAPZB; CAPZB; | P47756; A0A | CAPZB_HUMAI | Acetyl (Protein N | 28192 | 19038 | 0 | 16757 | 264410 | 0 | 0 | 0 | 198280 | 39322 | 19343 | 0 | 75203 | 75558 | 0 | 0 | 0 | 224850 |
| SDQQLDC | CAPZB; CAPZB; | P47756; A0A | CAPZB_HUMAI | Acetyl (Protein N | 0 | 0 | 0 | 0 | 4137400 | 0 | 526290 | 0 | 0 | 0 | 0 | 0 | 366120 | 0 | 0 | 0 | 5240600 | 1316400 |
| SDQSYVISI | ACSL4 | O60488 | ACSL4_HUMAI | Unmodified | 0 | 0 | 0 | 0 | 5124900 | 0 | 0 | 2028700 | 0 | 2986100 | 3181200 | 3987700 | 716690 | 0 | 0 | 0 | 18181000 | 4620700 |
| SDQSYVISI | ACSL4 | O60488 | ACSL4_HUMAI | Unmodified | 0 | 0 | 0 | 0 | 0 | 0 | 0 | 0 | 0 | 0 | 703780 | 10044000 | 0 | 0 | 0 | 0 | 256580 | 0 |
| SDQSYVISI | ACSL4 | O60488 | ACSL4_HUMAI | Unmodified | 0 | 0 | 0 | 0 | 0 | 0 | 0 | 0 | 0 | 0 | 0 | 0 | 0 | 0 | 0 | 0 | 2057500 | 0 |
| SDQYDTAD | MPP7; MPP7; M | Q5T2T1; U5( | MPP7_HUMAN | Unmodified | 0 | 0 | 0 | 0 | 1514300 | 0 | 0 | 0 | 0 | 0 | 0 | 0 | 62009 | 384910 | 0 | 0 | 49490 | 0 |
| SDSDRGGI | ADGRE5 | P48960 | AGRE5_HUMAN | Unmodified | 17265 | 0 | 2586900 | 918830 | 4821600 | 778980 | 0 | 0 | 0 | 0 | 0 | 725680 | 0 | 0 | 0 | 0 | 1024900 | 5546900 |
| SDSEDICLI | RSL1D1; RSL1D | O76021; J3Q | RL1D1_HUMAN | Unmodified | 0 | 0 | 0 | 0 | 0 | 0 | 0 | 0 | 0 | 0 | 0 | 2004200 | 0 | 0 | 0 | 0 | 0 | 0 |

| | | | | | | | | | | | | | | | | | | | | | | |
|---|---|---|---|---|---|---|---|---|---|---|---|---|---|---|---|---|---|---|---|---|---|---|
| SDSFENPV | XRCC6; XRCC6 | P12956; B1A | XRCC6_HUMAI | Unmodified | 755910 | 546510 | 253240 | 0 | 0 | 0 | 0 | 0 | 461750 | 0 | 3182400 | 30753000 | 0 | 0 | 295970 | 316400 | 0 | 0 |
| SDSFYFVDI | UNC119B | A6NIH7 | U119B_HUMAN | Unmodified | 385170 | 309080 | 0 | 0 | 0 | 0 | 0 | 0 | 0 | 0 | 0 | 486020 | 0 | 0 | 0 | 208020 | 0 | 0 |
| SDSGKPYY | PRPF40A; PRPF | O75400; H7( | PR40A_HUMAN | Unmodified | 0 | 0 | 35876 | 0 | 0 | 0 | 0 | 0 | 33610 | 0 | 191210 | 1146200 | 0 | 0 | 0 | 0 | 0 | 0 |
| SDSHGPKE | CKAP4 | Q07065 | CKAP4_HUMAN | Unmodified | 0 | 0 | 0 | 0 | 1551800 | 0 | 184230 | 0 | 6023400 | 1059100 | 254490 | 0 | 754860 | 2008900 | 0 | 28183 | 0 | 334970 |
| SDSHPSDA | ACADVL; ACAD | P49748; K7E | ACADV_HUMAI | Unmodified | 0 | 0 | 44749 | 0 | 0 | 243360 | 0 | 0 | 2828000 | 0 | 63514 | 153030 | 227030 | 0 | 52898 | 71786 | 0 | 0 |
| SDSILLTHD | PAK4 | O96013 | PAK4_HUMAN | Unmodified | 216250 | 158830 | 226590 | 144060 | 0 | 0 | 0 | 0 | 0 | 0 | 0 | 851390 | 0 | 0 | 621070 | 405810 | 0 | 0 |
| SDSSSKKD | PDIA6 | Q15084 | PDIA6_HUMAN | Unmodified | 0 | 949170 | 0 | 0 | 0 | 0 | 0 | 0 | 18502000 | 2095200 | 3295500 | 0 | 0 | 0 | 0 | 0 | 0 | 0 |
| SDSVILNVI | CEACAM5; CEA | P06731; A0A | CEAM5_HUMAI | Unmodified | 0 | 0 | 0 | 0 | 0 | 0 | 0 | 0 | 1113800 | 0 | 0 | 0 | 0 | 0 | 0 | 0 | 0 | 0 |
| SDSVLPASI | MINK1 | Q8N4C8 | MINK1_HUMAN | Unmodified | 0 | 0 | 0 | 178780 | 0 | 0 | 0 | 0 | 288830 | 0 | 0 | 585090 | 0 | 0 | 446530 | 0 | 0 | 0 |
| SDTFINLRE | CAPN2 | P17655 | CAN2_HUMAN | Unmodified | 0 | 0 | 0 | 266120 | 644930 | 0 | 0 | 0 | 359850 | 137930 | 168080 | 0 | 86776 | 0 | 0 | 0 | 270570 | 187110 |
| SDTISSDLT | N/A | N/A | N/A | Unmodified | 0 | 0 | 0 | 0 | 0 | 1211800 | 1545900 | 0 | 0 | 0 | 0 | 0 | 2181300 | 8907300 | 0 | 0 | 0 | 0 |
| SDTITFK | CPT2; CPT2; CF | P23786; A0A | CPT2_HUMAN | Unmodified | 0 | 43913 | 0 | 334650 | 354130 | 0 | 18891 | 28777 | 313620 | 56049 | 0 | 965230 | 0 | 0 | 219330 | 145140 | 0 | 0 |
| SDTQTHTM | PARP12 | Q9H0J9 | PAR12_HUMAN | Unmodified | 0 | 0 | 0 | 302740 | 0 | 0 | 0 | 0 | 0 | 0 | 0 | 0 | 48924 | 0 | 670420 | 561130 | 0 | 0 |
| SDTSESGA | MAGED2; MAGI | Q9UNF1; Q5 | MAGD2_HUMA | Acetyl (Protein N | 440260 | 0 | 0 | 0 | 76204 | 0 | 425300 | 247010 | 342900 | 274510 | 1050100 | 322740 | 0 | 0 | 0 | 0 | 322190 | 401410 |
| SDTSFMFQ | ANXA4; ANXA4; | P09525; B4E | ANXA4_HUMAN | Oxidation (M) | 105370 | 0 | 76417 | 0 | 0 | 0 | 0 | 0 | 881200 | 70459 | 212050 | 2137300 | 0 | 0 | 77710 | 82937 | 0 | 0 |
| SDTSFMFQ | ANXA4; ANXA4; | P09525; B4E | ANXA4_HUMAN | Unmodified | 0 | 0 | 344550 | 328970 | 0 | 426100 | 0 | 0 | 4716000 | 0 | 557120 | 2153400 | 0 | 0 | 281330 | 53124 | 0 | 0 |
| SDTSGDYE | ANXA3; ANXA3 | P12429; D6F | ANXA3_HUMAN | Unmodified | 0 | 213390 | 0 | 0 | 2544100 | 805030 | 0 | 120000 | 584360 | 0 | 0 | 0 | 0 | 0 | 0 | 123680 | 0 | 0 |
| SDTSGHFE | ANXA7 | P20073 | ANXA7_HUMAN | Unmodified | 0 | 0 | 0 | 0 | 0 | 0 | 0 | 0 | 11008000 | 0 | 0 | 0 | 0 | 0 | 233890 | 92378 | 55957 | 0 |
| SDTSGHFQ | ANXA11 | P50995 | ANX11_HUMAN | Unmodified | 0 | 0 | 0 | 0 | 0 | 0 | 0 | 0 | 1097400 | 0 | 0 | 219850 | 0 | 0 | 0 | 0 | 0 | 18889 |
| SDTVPSPR | FN1 | P02751 | FINC_HUMAN I | Unmodified | 0 | 0 | 1264400 | 392390 | 0 | 0 | 0 | 0 | 0 | 0 | 0 | 0 | 0 | 22172 | 0 | 0 | 0 | 0 |
| SDTVPSPRI | FN1 | P02751 | FINC_HUMAN I | Unmodified | 0 | 0 | 375200 | 0 | 0 | 115310 | 0 | 58008 | 0 | 1437700 | 0 | 0 | 86419 | 0 | 0 | 0 | 0 | 0 |
| SDVEAIFSK | HNRNPCL3; HN | B7ZW38; P0I | HNRC3_HUMA | Unmodified | 331770 | 0 | 0 | 0 | 336660 | 0 | 1078400 | 48509 | 0 | 285600 | 361710 | 1969300 | 130890 | 0 | 0 | 0 | 0 | 223070 |
| SDVEAIFSK | HNRNPCL3; HN | B7ZW38; P0I | HNRC3_HUMA | Unmodified | 0 | 0 | 0 | 0 | 0 | 0 | 1317400 | 561220 | 0 | 0 | 0 | 622200 | 0 | 0 | 0 | 0 | 0 | 0 |
| SDVEINYSI | NOMO1; NOMC | Q15155; P69 | NOMO1_HUMA | Unmodified | 0 | 0 | 0 | 0 | 0 | 0 | 0 | 0 | 1857100 | 0 | 0 | 0 | 0 | 0 | 0 | 0 | 0 | 0 |
| SDVHTEAV | DIP2A; DIP2C; I | Q14689; Q9 | DIP2A_HUMAN | Unmodified | 0 | 0 | 0 | 429480 | 0 | 0 | 0 | 0 | 0 | 0 | 0 | 0 | 0 | 0 | 0 | 0 | 0 | 0 |
| SDVKFEDR | TM9SF3; TM9SF | Q9HD45; Q5 | TM9S3_HUMAN | Unmodified | 103100 | 0 | 0 | 0 | 0 | 0 | 0 | 0 | 4093000 | 637460 | 180830 | 579380 | 0 | 0 | 155330 | 0 | 0 | 0 |
| SDVLELTDI | PDIA3; PDIA3; F | P30101; F8V | PDIA3_HUMAN | Unmodified | 0 | 0 | 2367000 | 553430 | 159550 | 0 | 304340 | 397390 | 2.07E+08 | 9257600 | 25925000 | 10895000 | 0 | 0 | 4713900 | 4406600 | 0 | 0 |
| SDVLQPGA | SEC24C; SEC24 | P53992; G5E | SC24C_HUMAI | Unmodified | 0 | 0 | 0 | 0 | 0 | 0 | 0 | 0 | 306680 | 0 | 0 | 633000 | 0 | 0 | 0 | 0 | 0 | 0 |
| SDVLQPGA | SEC24C; SEC24 | P53992; G5E | SC24C_HUMAI | Unmodified | 0 | 0 | 0 | 0 | 0 | 0 | 0 | 0 | 1257000 | 184690 | 0 | 412500 | 0 | 0 | 0 | 0 | 0 | 0 |
| SDVLVEYQ | GNS; GNS; GNS | P15586; H7C | GNS_HUMAN N | Unmodified | 0 | 0 | 0 | 0 | 0 | 0 | 0 | 0 | 612170 | 662930 | 0 | 0 | 0 | 0 | 0 | 0 | 217220 | 0 |
| SDVQIHVP | PRKRA; PRKRA; | O75569; A0A | PRKRA_HUMAN | Unmodified | 0 | 0 | 0 | 0 | 0 | 0 | 0 | 0 | 0 | 0 | 0 | 632150 | 0 | 0 | 0 | 0 | 0 | 0 |
| SDVSGLSD | SYT4; SYT4 | Q9H2B2; M0 | SYT4_HUMAN S | Unmodified | 0 | 0 | 2304800 | 1528200 | 0 | 0 | 0 | 0 | 0 | 0 | 0 | 0 | 0 | 0 | 7515600 | 5587900 | 0 | 0 |
| SDVWSFGI | YES1; YES1 | P07947; J3Q | YES_HUMAN Ty | Unmodified | 0 | 36065 | 50761 | 724900 | 937430 | 276990 | 54204 | 0 | 437740 | 643540 | 51832 | 0 | 796940 | 279630 | 1043500 | 249220 | 1178600 | 781620 |
| SDVYCEVC | PSAP; PSAP; PS | P07602; A0A | SAP_HUMAN P | Unmodified | 0 | 0 | 0 | 0 | 124210 | 34105 | 61679 | 13251 | 6270600 | 8456100 | 538190 | 73453 | 0 | 0 | 0 | 0 | 21892 | 75164 |
| SDYDREAL | RCN2; RCN2 | Q14257; H0 | RCN2_HUMAN | Unmodified | 0 | 0 | 0 | 0 | 0 | 0 | 0 | 0 | 8577600 | 1948200 | 0 | 0 | 0 | 0 | 0 | 0 | 0 | 0 |
| SDYDREAL | RCN2; RCN2 | Q14257; H0 | RCN2_HUMAN | Unmodified | 0 | 0 | 0 | 0 | 0 | 0 | 0 | 0 | 0 | 3070900 | 0 | 0 | 0 | 0 | 0 | 0 | 0 | 0 |
| SEAASVSTI | NCAM1 | A0A0D9SF3( | A0A0D9SF30_H | Oxidation (M) | 0 | 0 | 0 | 0 | 0 | 0 | 0 | 0 | 2305000 | 0 | 0 | 0 | 0 | 0 | 0 | 0 | 0 | 0 |
| SEAASVSTI | NCAM1 | A0A0D9SF3( | A0A0D9SF30_H | Unmodified | 0 | 0 | 0 | 0 | 0 | 0 | 0 | 0 | 3369200 | 408910 | 0 | 0 | 0 | 0 | 0 | 0 | 0 | 0 |
| SEACPCQF | SCRIB; SCRIB; S | Q14160; A0A | SCRIB_HUMAN | Unmodified | 0 | 0 | 0 | 0 | 184560 | 50873 | 0 | 0 | 192150 | 84624 | 0 | 70491 | 0 | 58281 | 0 | 0 | 500340 | 288500 |
| SEAEEAITSI | ELAVL1 | Q15717 | ELAV1_HUMAN | Unmodified | 303150 | 148930 | 313070 | 0 | 0 | 0 | 0 | 0 | 0 | 0 | 0 | 5263300 | 0 | 0 | 229110 | 385390 | 0 | 0 |
| SEAGLAGA | AP3S1 | Q92572 | AP3S1_HUMAN | Unmodified | 54744 | 50023 | 34916 | 0 | 136440 | 0 | 30456 | 19053 | 0 | 27293 | 166510 | 1048500 | 83922 | 0 | 31907 | 0 | 47082 | 47638 |
| SEAGLAGA | AP3S1 | Q92572 | AP3S1_HUMAN | Unmodified | 0 | 0 | 0 | 0 | 579080 | 0 | 0 | 202080 | 0 | 0 | 0 | 0 | 0 | 0 | 0 | 0 | 205280 | 328390 |
| SEAHLTELL | CNPY2; N/A; CI | Q9Y2B0; F8V | CNPY2_HUMAI | Unmodified | 0 | 0 | 0 | 0 | 0 | 0 | 0 | 0 | 60888000 | 388760 | 1214500 | 697780 | 0 | 0 | 63157 | 82809 | 0 | 0 |
| SEAHLTELL | CNPY2; N/A; CI | Q9Y2B0; F8V | CNPY2_HUMAI | Unmodified | 0 | 0 | 0 | 0 | 0 | 0 | 0 | 0 | 24819000 | 0 | 0 | 0 | 0 | 0 | 0 | 0 | 0 | 0 |
| SEAIIEHLC | SFRP1; SFRP1 | Q8N474; Q6 | SFRP1_HUMAN | Unmodified | 0 | 0 | 0 | 0 | 32687 | 0 | 0 | 0 | 529830 | 50011 | 68940 | 0 | 296110 | 0 | 0 | 227450 | 0 | 0 |
| SEAIQTSHF | FOCAD; FOCAD | Q5VW36; S4 | FOCAD_HUMA | Unmodified | 0 | 0 | 0 | 0 | 0 | 38782 | 0 | 0 | 2025000 | 0 | 0 | 759580 | 0 | 0 | 0 | 56555 | 0 | 0 |
| SEAITSVDD | EPS8L2; EPS8L2 | Q9H6S3; E9I | ES8L2_HUMAN | Unmodified | 0 | 0 | 0 | 0 | 220130 | 0 | 0 | 0 | 197780 | 0 | 0 | 68057 | 41204 | 0 | 130120 | 190360 | 28931 | 25158 |
| SEAITSVDD | EPS8L2; EPS8L2 | Q9H6S3; E9I | ES8L2_HUMAN | Unmodified | 0 | 0 | 0 | 0 | 857660 | 138040 | 0 | 0 | 0 | 76610 | 0 | 0 | 344900 | 249780 | 1816100 | 0 | 0 | 97280 |
| SEALAVDG | CHGA; CHGA | P10645; G5E | CMGA_HUMAN | Unmodified | 78493 | 0 | 0 | 208390 | 0 | 0 | 0 | 129110 | 578660 | 1418100 | 0 | 0 | 0 | 0 | 0 | 0 | 0 | 0 |
| SEALAVDG | CHGA; CHGA | P10645; G5E | CMGA_HUMAN | Unmodified | 0 | 0 | 0 | 0 | 0 | 0 | 0 | 0 | 14012000 | 9138000 | 0 | 0 | 0 | 0 | 0 | 0 | 0 | 0 |
| SEALAVDG | CHGA; CHGA | P10645; G5E | CMGA_HUMAN | Unmodified | 0 | 0 | 0 | 0 | 0 | 0 | 0 | 0 | 1533300 | 0 | 0 | 0 | 0 | 0 | 0 | 0 | 0 | 0 |
| SEALGVGD | TES | Q9UGI8 | TES_HUMAN Te | Unmodified | 0 | 0 | 0 | 0 | 1022800 | 0 | 0 | 0 | 0 | 0 | 0 | 212810 | 0 | 0 | 0 | 0 | 221640 | 724100 |
| SEALGVGD | TES | Q9UGI8 | TES_HUMAN Te | Oxidation (M) | 88353 | 0 | 0 | 176600 | 0 | 0 | 0 | 0 | 0 | 0 | 0 | 735750 | 182780 | 0 | 226770 | 237910 | 175690 | 0 |
| SEALGVGE | LMCD1; LMCD1 | Q9NZU5; H7 | LMCD1_HUMA | Unmodified | 0 | 0 | 0 | 0 | 0 | 0 | 0 | 0 | 0 | 0 | 0 | 1248400 | 0 | 0 | 0 | 0 | 0 | 0 |
| SEALGVGE | LMCD1; LMCD1 | Q9NZU5; H7 | LMCD1_HUMA | Unmodified | 0 | 0 | 0 | 0 | 0 | 0 | 0 | 0 | 0 | 0 | 1529800 | 5760600 | 0 | 0 | 0 | 0 | 0 | 0 |
| SEAPNWAT | EIF3E; EIF3E; EI | P60228; B3K | EIF3E_HUMAN | Unmodified | 0 | 0 | 0 | 0 | 0 | 0 | 0 | 0 | 0 | 0 | 0 | 1064000 | 0 | 0 | 313840 | 0 | 0 | 0 |
| SEAPVQNL | USP43; USP43 | Q70EL4; V9C | UBP43_HUMAN | Unmodified | 88461 | 343670 | 68550 | 23881 | 0 | 26245 | 0 | 0 | 200570 | 0 | 0 | 0 | 0 | 0 | 285690 | 390590 | 0 | 0 |
| SEASAFGA | RRAS | P10301 | RRAS_HUMAN | Unmodified | 2705700 | 2678800 | 2242800 | 3269500 | 0 | 732950 | 345550 | 114390 | 0 | 0 | 0 | 1131900 | 202390 | 0 | 7670400 | 3058500 | 490250 | 0 |
| SEASLHPVI | ACTL6A; ACTL6 | O96019; H7( | ACL6A_HUMAN | Oxidation (M) | 0 | 0 | 180560 | 276570 | 445480 | 0 | 0 | 0 | 508140 | 239280 | 678650 | 3312600 | 0 | 0 | 0 | 136100 | 0 | 0 |
| SEASSEDL | TNFAIP2; TNFAI | Q03169; H0 | TNAP2_HUMAN | Acetyl (Protein N | 0 | 0 | 0 | 0 | 1516100 | 1480800 | 397990 | 0 | 0 | 0 | 0 | 431430 | 0 | 0 | 0 | 0 | 486190 | 0 |
| SEASSEDL | TNFAIP2; TNFAI | Q03169; H0 | TNAP2_HUMAN | Acetyl (Protein N | 0 | 0 | 0 | 0 | 1872900 | 0 | 0 | 0 | 0 | 0 | 0 | 0 | 0 | 0 | 0 | 0 | 466430 | 525730 |
| SEAVVEYVI | SND1; SND1 | Q7KZF4; H7( | SND1_HUMAN | Unmodified | 0 | 0 | 617330 | 39571 | 0 | 0 | 0 | 0 | 1802500 | 225760 | 0 | 5994600 | 0 | 0 | 216400 | 151810 | 0 | 0 |
| SECGALEH | CFC1; CFC1B | P0CG37; P0 | CFC1_HUMAN | Unmodified | 1704300 | 782660 | 0 | 0 | 0 | 0 | 0 | 0 | 0 | 0 | 0 | 0 | 0 | 0 | 0 | 0 | 0 | 0 |
| SEDELRPE | REPS1; REPS1; | Q96D71; H0 | REPS1_HUMAN | Unmodified | 0 | 0 | 285580 | 266300 | 409880 | 0 | 63527 | 0 | 0 | 77763 | 0 | 0 | 191030 | 0 | 233440 | 405760 | 135010 | 0 |
| SEDEPTRSI | TNPO2; TNPO1 | O14787; Q9 | TNPO2_HUMAI | Unmodified | 0 | 0 | 30293 | 100020 | 576640 | 88220 | 200450 | 106240 | 458270 | 241350 | 38145 | 0 | 454090 | 185070 | 64273 | 0 | 437270 | 823810 |
| SEDEVGSL | ITGA6; ITGA6 | P23229; H7E | ITA6_HUMAN In | Unmodified | 0 | 0 | 173900 | 0 | 0 | 0 | 0 | 0 | 0 | 0 | 0 | 3796200 | 487530 | 0 | 1439100 | 1864900 | 65524 | 0 |
| SEDFGVNE | ANXA1 | P04083 | ANXA1_HUMAN | Unmodified | 75757 | 0 | 0 | 0 | 0 | 0 | 0 | 0 | 0 | 0 | 28030000 | 26755000 | 0 | 0 | 0 | 0 | 0 | 0 |
| SEDFSLPA | COX4I1; COX4I | P13073; Q86 | COX41_HUMAI | Unmodified | 0 | 0 | 0 | 0 | 0 | 0 | 0 | 0 | 3220100 | 0 | 0 | 0 | 0 | 0 | 0 | 0 | 0 | 0 |
| SEDFSLPA | COX4I1; COX4I | P13073; Q86 | COX41_HUMAI | Unmodified | 0 | 0 | 0 | 0 | 0 | 0 | 288530 | 0 | 1828400 | 3169800 | 0 | 0 | 84007 | 0 | 0 | 0 | 0 | 0 |
| SEDGVEGD | AHNAK | Q09666 | AHNK_HUMAN | Unmodified | 0 | 0 | 0 | 32342 | 7514500 | 376770 | 0 | 0 | 63534 | 0 | 0 | 0 | 0 | 0 | 0 | 0 | 1405100 | 2823400 |
| SEDGVEGD | AHNAK | Q09666 | AHNK_HUMAN | Unmodified | 0 | 0 | 0 | 0 | 5532300 | 1164500 | 0 | 0 | 0 | 0 | 0 | 0 | 0 | 0 | 0 | 0 | 1665500 | 3011200 |
| SEDGVEGD | AHNAK | Q09666 | AHNK_HUMAN | Unmodified | 0 | 0 | 0 | 0 | 3179000 | 1408900 | 0 | 0 | 0 | 0 | 0 | 0 | 0 | 0 | 0 | 0 | 0 | 0 |
| SEDLDNSII | SMC3 | Q9UQE7 | SMC3_HUMAN | Unmodified | 103010 | 85707 | 0 | 0 | 80902 | 0 | 0 | 0 | 0 | 0 | 386250 | 1089600 | 0 | 0 | 65273 | 26357 | 0 | 0 |
| SEDLLDYG | RPN1; RPN1 | P04843; B7Z | RPN1_HUMAN | Unmodified | 0 | 0 | 0 | 0 | 0 | 0 | 0 | 0 | 8871900 | 3682000 | 0 | 1183500 | 0 | 0 | 0 | 0 | 0 | 0 |
| SEDLLDYG | RPN1; RPN1 | P04843; B7Z | RPN1_HUMAN | Unmodified | 3163300 | 7178700 | 320700 | 92106 | 0 | 0 | 0 | 0 | 44763000 | 2507300 | 874860 | 169780 | 0 | 0 | 1031000 | 1833800 | 0 | 0 |
| SEDPDQQ | VPS35 | Q96QK1 | VPS35_HUMAN | Unmodified | 0 | 0 | 0 | 0 | 0 | 0 | 0 | 0 | 378730 | 0 | 0 | 3056800 | 0 | 0 | 135360 | 161500 | 0 | 0 |
| SEDPDQQ | VPS35 | Q96QK1 | VPS35_HUMAN | Unmodified | 0 | 0 | 620060 | 0 | 0 | 0 | 0 | 0 | 1189400 | 0 | 0 | 0 | 0 | 0 | 0 | 797600 | 0 | 0 |
| SEDPPGQE | CLUH; CLUH; C | O75153; I3L | CLU_HUMAN C | Unmodified | 0 | 0 | 0 | 0 | 0 | 0 | 0 | 0 | 0 | 0 | 572150 | 1345300 | 0 | 0 | 0 | 0 | 0 | 0 |
| SEDSIAGS( | AKAP12 | Q02952 | AKA12_HUMAN | Unmodified | 0 | 0 | 0 | 0 | 1351900 | 0 | 0 | 0 | 0 | 0 | 0 | 0 | 0 | 0 | 0 | 0 | 661020 | 2405400 |
| SEDSIAGS( | AKAP12 | Q02952 | AKA12_HUMAN | Unmodified | 0 | 0 | 0 | 0 | 1256700 | 0 | 0 | 0 | 0 | 0 | 0 | 0 | 0 | 0 | 0 | 0 | 0 | 1471600 |
| SEDVPYTA | ATP6AP1; ATP6 | Q15904; A0A | VAS1_HUMAN | Unmodified | 0 | 0 | 0 | 0 | 0 | 0 | 45811 | 0 | 7407800 | 3509900 | 0 | 0 | 0 | 0 | 0 | 0 | 0 | 0 |
| SEDYVDIV( | PLOD1 | Q02809 | PLOD1_HUMAI | Unmodified | 0 | 0 | 0 | 0 | 0 | 0 | 428260 | 0 | 3500100 | 314930 | 147270 | 0 | 0 | 0 | 0 | 0 | 0 | 0 |
| SEDYVDIV( | PLOD1 | Q02809 | PLOD1_HUMAI | Unmodified | 0 | 0 | 0 | 0 | 0 | 0 | 0 | 0 | 2506000 | 0 | 0 | 0 | 0 | 0 | 0 | 0 | 0 | 0 |
| SEDYVELV( | PLOD3; PLOD3; | O60568; H7( | PLOD3_HUMAI | Unmodified | 0 | 0 | 0 | 0 | 0 | 0 | 0 | 0 | 0 | 0 | 0 | 0 | 0 | 0 | 0 | 0 | 0 | 0 |
| SEEACMLR | SPINT2; SPINT2 | O43291; K7E | SPIT2_HUMAN | Oxidation (M) | 72406 | 0 | 0 | 0 | 0 | 0 | 0 | 0 | 2158600 | 500730 | 0 | 242790 | 0 | 0 | 0 | 0 | 0 | 0 |
| SEEACMLR | SPINT2; SPINT2 | O43291; K7E | SPIT2_HUMAN | Unmodified | 17155 | 0 | 0 | 0 | 0 | 0 | 0 | 0 | 16673000 | 1188800 | 0 | 0 | 0 | 0 | 0 | 0 | 0 | 0 |
| SEEAHAED | SERBP1 | Q8NC51 | PAIRB_HUMAN | Unmodified | 167810 | 0 | 146350 | 0 | 0 | 0 | 0 | 0 | 0 | 0 | 785500 | 1229500 | 0 | 0 | 198200 | 157740 | 0 | 0 |
| SEEEFIHIN | KARS1; KARS1; | Q15046; H3I | SYK_HUMAN Ly | Unmodified | 0 | 0 | 0 | 0 | 0 | 0 | 0 | 0 | 0 | 0 | 0 | 0 | 0 | 0 | 0 | 0 | 0 | 0 |
| SEEEIDFLR | EIF3L; EIF3L; EII | Q9Y262; B0( | EIF3L_HUMAN | Unmodified | 0 | 0 | 0 | 0 | 32062 | 1119600 | 0 | 0 | 0 | 0 | 0 | 982620 | 0 | 0 | 0 | 0 | 0 | 0 |
| SEEEQSSS | HNRNPC; HNR | P07910; G3 | HNRPC_HUMA | Unmodified | 0 | 0 | 18656 | 0 | 0 | 0 | 0 | 0 | 21075 | 0 | 48689 | 161440 | 0 | 0 | 0 | 0 | 0 | 0 |
| SEEERVTET | RDX; RDX; RDX; | P35241; A0A | RADI_HUMAN I | Unmodified | 621280 | 303850 | 38290 | 867460 | 118940 | 69940 | 7443.7 | 0 | 1000200 | 0 | 64792 | 9230200 | 81329 | 0 | 2166300 | 1523300 | 46817 | 0 |
| SEEERVTET | RDX; RDX; RDX; | P35241; A0A | RADI_HUMAN I | Unmodified | 207940 | 232600 | 0 | 554150 | 153100 | 129890 | 0 | 0 | 753330 | 67137 | 231710 | 770330 | 127220 | 52301 | 79903 | 0 | 38762 | 52548 |
| SEEETSPLV | SLC1A4 | P43007 | SATT_HUMAN I | Unmodified | 1186700 | 320300 | 2818200 | 0 | 202820 | 0 | 1373000 | 0 | 34925000 | 3330000 | 4053800 | 36371000 | 0 | 0 | 992950 | 237210 | 0 | 0 |
| SEEETSPLV | SLC1A4 | P43007 | SATT_HUMAN I | Unmodified | 0 | 0 | 0 | 0 | 872260 | 0 | 0 | 193370 | 3404800 | 1315000 | 0 | 0 | 1192000 | 0 | 0 | 0 | 0 | 0 |
| SEEFAVCRI | IFNGR1; IFNGR | P15260; A0A | INGR1_HUMAN | Unmodified | 0 | 0 | 0 | 0 | 328900 | 115980 | 0 | 0 | 513580 | 0 | 150720 | 250470 | 0 | 33742 | 0 | 0 | 44306 | 147580 |
| SEEIKDTIL( | SLK | Q9H2G2 | SLK_HUMAN S | Unmodified | 0 | 0 | 59383 | 390420 | 220940 | 0 | 0 | 0 | 3211100 | 231420 | 775480 | 399000 | 1326900 | 610300 | 0 | 175050 | 0 | 232270 |
| SEEMHRLT | BAIAP2; BAIAP2 | Q9UQB8; I3I | BAIP2_HUMAN | Unmodified | 0 | 0 | 0 | 0 | 0 | 138510 | 0 | 0 | 0 | 0 | 48591 | 0 | 0 | 57001 | 403350 | 0 | 0 | 0 |
| SEENEEPM | HSPA4; HSPA4 | P34932; A0A | HSP74_HUMAN | Unmodified | 0 | 0 | 168290 | 0 | 0 | 0 | 0 | 0 | 0 | 0 | 180290 | 1854300 | 0 | 0 | 0 | 0 | 0 | 0 |
| SEEPAGQII | SRRM2 | Q9UQ35 | SRRM2_HUMAI | Unmodified | 0 | 0 | 0 | 0 | 0 | 0 | 0 | 0 | 0 | 0 | 583690 | 2030700 | 261460 | 0 | 0 | 0 | 0 | 0 |
| SEEQLKEE( | DLD; DLD | P09622; E9P | DLDH_HUMAN | Unmodified | 0 | 0 | 0 | 0 | 0 | 226110 | 0 | 0 | 3275800 | 100290 | 364680 | 0 | 65262 | 0 | 0 | 0 | 25648 | 0 |
| SEESVSRLI | CKAP4 | Q07065 | CKAP4_HUMAN | Unmodified | 0 | 0 | 0 | 0 | 0 | 0 | 98938 | 0 | 2306300 | 0 | 379510 | 0 | 0 | 0 | 0 | 0 | 0 | 0 |
| SEETATSDI | SPRED1 | Q7Z699 | SPRE1_HUMAN | Acetyl (Protein N | 0 | 0 | 0 | 0 | 0 | 0 | 0 | 0 | 0 | 0 | 0 | 0 | 0 | 0 | 0 | 0 | 843260 | 118450 |
| SEETLDEGI | FKBP3 | Q00688 | FKBP3_HUMAN | Unmodified | 556860 | 0 | 935550 | 0 | 0 | 0 | 0 | 0 | 351230 | 0 | 0 | 1061900 | 0 | 0 | 1139100 | 434610 | 0 | 0 |
| SEETLDEGI | FKBP3 | Q00688 | FKBP3_HUMAN | Unmodified | 381660 | 208630 | 0 | 128620 | 0 | 0 | 0 | 35419 | 357730 | 0 | 528230 | 378560 | 173300 | 0 | 152490 | 76889 | 0 | 0 |
| SEETNQQE | DNAJC13 | O75165 | DJC13_HUMAN | Unmodified | 0 | 0 | 83402 | 0 | 0 | 0 | 0 | 0 | 0 | 0 | 0 | 2123500 | 0 | 0 | 45469 | 0 | 0 | 0 |
| SEEWADNI | PLIN3; PLIN3 | O60664; K7E | PLIN3_HUMAN | Unmodified | 0 | 1001600 | 362900 | 106790 | 0 | 0 | 0 | 0 | 0 | 0 | 124050 | 0 | 0 | 0 | 397400 | 193490 | 0 | 0 |
| SEFGEVFL | PTK7; PTK7 | Q13308; C9J | PTK7_HUMAN I | Unmodified | 0 | 0 | 952510 | 1240500 | 2039900 | 0 | 355550 | 142690 | 6586800 | 1734000 | 2112400 | 9373700 | 2128300 | 0 | 4586900 | 1678600 | 0 | 0 |
| SEFLLTDYF | NOSTRIN; NOS | Q8IVI9; A0A( | NOSTN_HUMA | Oxidation (M) | 454340 | 463000 | 445060 | 413270 | 0 | 0 | 0 | 0 | 0 | 0 | 0 | 0 | 0 | 0 | 3222400 | 588640 | 0 | 0 |
| SEFPLLR | DOP1B | Q9Y3R5 | DOP2_HUMAN | Unmodified | 0 | 0 | 18290 | 0 | 0 | 0 | 0 | 0 | 749430 | 86500 | 23322 | 552340 | 0 | 18808 | 83969 | 68676 | 0 | 0 |

| | | | | | | | | | | | | | | | | | | | | | | |
|---|---|---|---|---|---|---|---|---|---|---|---|---|---|---|---|---|---|---|---|---|---|---|
| SEFVILRDE | YARS1; YARS1 | P54577; A0A | SYYC_HUMAN | Unmodified | 0 | 0 | 462260 | 501320 | 1097800 | 356850 | 119080 | 95131 | 1857500 | 345070 | 1321500 | 3674000 | 1404800 | 690770 | 820080 | 763190 | 246020 | 90656 |
| SEGAVVNV | IGSF3 | O75054 | IGSF3_HUMAN | Unmodified | 85025 | 74143 | 100760 | 131710 | 0 | 293990 | 0 | 0 | 0 | 260650 | 0 | 316500 | 0 | 0 | 205060 | 131590 | 0 | 0 |
| SEGDSVGE | RER1; RER1; RE | O15258; Q5 | RER1_HUMAN | Acetyl (Protein N | 0 | 0 | 0 | 0 | 0 | 18512 | 0 | 0 | 165460 | 0 | 0 | 0 | 0 | 0 | 0 | 0 | 0 | 0 |
| SEGDSVGE | RER1; RER1; RE | O15258; Q5 | RER1_HUMAN | Acetyl (Protein N | 225620 | 0 | 0 | 0 | 0 | 0 | 0 | 0 | 266010 | 0 | 0 | 425030 | 0 | 0 | 0 | 0 | 0 | 0 |
| SEGEFKQT | F11R; F11R | Q9Y624; A0A | JAM1_HUMAN | Unmodified | 1043600 | 1058000 | 0 | 1393000 | 0 | 0 | 0 | 0 | 1381600 | 1254200 | 517830 | 0 | 3329200 | 1015900 | 0 | 0 | 0 | 0 |
| SEGFDTYR | PCNA | P12004 | PCNA_HUMAN | Unmodified | 0 | 0 | 0 | 0 | 0 | 0 | 0 | 0 | 409070 | 0 | 0 | 4496400 | 0 | 0 | 0 | 0 | 0 | 0 |
| SEGGFIWA | IDH1 | O75874 | IDHC_HUMAN | Unmodified | 0 | 0 | 0 | 0 | 774470 | 0 | 0 | 0 | 982760 | 0 | 0 | 5474000 | 0 | 2517900 | 0 | 0 | 0 | 0 |
| SEGGLGGI | IGF2R; IGF2R | P11717; A0A | MPRI_HUMAN | Oxidation (M) | 0 | 0 | 0 | 0 | 0 | 0 | 0 | 0 | 1885900 | 365160 | 0 | 0 | 0 | 0 | 0 | 0 | 0 | 0 |
| SEGGLGGI | IGF2R; IGF2R | P11717; A0A | MPRI_HUMAN | Oxidation (M) | 0 | 0 | 7394900 | 11367000 | 2225700 | 6624400 | 1084800 | 0 | 23309000 | 12799000 | 26786000 | 6549000 | 1109300 | 1448800 | 3917900 | 0 | 0 | 4223300 |
| SEGGLGGI | IGF2R; IGF2R | P11717; A0A | MPRI_HUMAN | Unmodified | 78738 | 0 | 0 | 0 | 711140 | 0 | 688430 | 0 | 46024000 | 6890600 | 1979200 | 1613200 | 711300 | 650460 | 185970 | 0 | 0 | 0 |
| SEGIASTVT | EPHB3 | P54753 | EPHB3_HUMAN | Unmodified | 256990 | 530970 | 0 | 0 | 0 | 0 | 0 | 0 | 0 | 0 | 0 | 0 | 0 | 0 | 0 | 0 | 0 | 0 |
| SEGLEISK | MGAT4A | Q9UM21 | MGT4A_HUMAN | Unmodified | 0 | 0 | 0 | 0 | 0 | 0 | 0 | 0 | 706430 | 0 | 0 | 0 | 0 | 0 | 0 | 0 | 0 | 0 |
| SEGVVAVLI | FASN; FASN | P49327; A0A | FAS_HUMAN F | Unmodified | 0 | 0 | 661060 | 1808300 | 0 | 0 | 0 | 0 | 466350 | 0 | 3676300 | 34480000 | 0 | 0 | 3621900 | 3134000 | 0 | 0 |
| SEGVVAVLI | FASN; FASN | P49327; A0A | FAS_HUMAN F | Unmodified | 0 | 0 | 0 | 552790 | 152710 | 264050 | 0 | 0 | 0 | 0 | 0 | 0 | 0 | 0 | 1285900 | 1471200 | 0 | 0 |
| SEHISLAQ | ABCA2; ABCA2; | Q9BZC7; A0 | ABCA2_HUMAN | Unmodified | 0 | 0 | 0 | 0 | 0 | 429820 | 292230 | 0 | 162140 | 0 | 0 | 0 | 0 | 0 | 0 | 0 | 0 | 0 |
| SEHPGLSI | HMGB2; HMGB | P26583; D6 | HMGB2_HUMA | Unmodified | 0 | 0 | 0 | 0 | 0 | 0 | 280880 | 0 | 517860 | 430720 | 2213700 | 1416000 | 0 | 0 | 0 | 0 | 0 | 0 |
| SEHSHSTT | ITGB4 | P16144 | ITB4_HUMAN I | Unmodified | 0 | 0 | 0 | 0 | 0 | 0 | 0 | 0 | 0 | 61808 | 0 | 0 | 0 | 0 | 0 | 0 | 569080 | 0 |
| SEIDLLNIR | ANXA6; ANXA6; | P08133; E5R | ANXA6_HUMAN | Unmodified | 0 | 0 | 0 | 0 | 0 | 0 | 0 | 0 | 3684700 | 896120 | 0 | 0 | 0 | 0 | 0 | 0 | 0 | 0 |
| SEIDLLNIR | ANXA6; ANXA6; | P08133; E5R | ANXA6_HUMAN | Unmodified | 3687700 | 4876800 | 13469000 | 8212400 | 0 | 0 | 0 | 0 | 39382000 | 2602700 | 0 | 0 | 0 | 0 | 12834000 | 6953900 | 0 | 0 |
| SEIDLVQIK | ANXA7 | P20073 | ANXA7_HUMAN | Unmodified | 0 | 1993100 | 1284100 | 0 | 0 | 0 | 0 | 534140 | 49532000 | 2711300 | 76044 | 3224600 | 0 | 136580 | 3710000 | 2247300 | 0 | 0 |
| SEIDMNDI | ANXA1 | P04083 | ANXA1_HUMAN | Oxidation (M) | 0 | 317080 | 0 | 0 | 0 | 0 | 0 | 0 | 0 | 0 | 4036100 | 1.06E+08 | 0 | 0 | 0 | 437650 | 0 | 0 |
| SEIDMNDI | ANXA1 | P04083 | ANXA1_HUMAN | Unmodified | 373410 | 437430 | 0 | 415560 | 0 | 0 | 0 | 0 | 0 | 0 | 47476000 | 8.4E+08 | 0 | 0 | 0 | 480810 | 0 | 0 |
| SEIDMNDI | ANXA1 | P04083 | ANXA1_HUMAN | Unmodified | 0 | 0 | 0 | 0 | 0 | 0 | 0 | 0 | 0 | 0 | 0 | 3208000 | 0 | 0 | 0 | 0 | 0 | 0 |
| SEIEYYAML | RPL30; RPL30 | P62888; E5R | RL30_HUMAN | Oxidation (M) | 438650 | 432670 | 640390 | 0 | 0 | 0 | 0 | 0 | 0 | 0 | 0 | 1916000 | 0 | 0 | 595140 | 524770 | 0 | 0 |
| SEIEYYAML | RPL30; RPL30 | P62888; E5R | RL30_HUMAN | Unmodified | 0 | 0 | 0 | 0 | 0 | 0 | 0 | 0 | 0 | 0 | 0 | 764430 | 0 | 0 | 0 | 0 | 0 | 442170 |
| SEIIPMFSN | PPP2R1A; PPP2 | P30153; B3K | 2AAA_HUMAN | Oxidation (M) | 0 | 0 | 200910 | 1438300 | 0 | 0 | 0 | 0 | 0 | 0 | 0 | 9356200 | 0 | 1078400 | 74579 | 49932 | 0 | 0 |
| SEINQQTPI | INPP5D; INPP5I | Q92835; H0 | SHIP1_HUMAN | Unmodified | 0 | 0 | 0 | 0 | 0 | 0 | 0 | 0 | 0 | 0 | 639790 | 1058500 | 0 | 0 | 0 | 0 | 0 | 0 |
| SEIPTKDVP | EPB41; EPB41; | P11171; A0A | EPB41_HUMAN | Unmodified | 0 | 0 | 0 | 0 | 208770 | 0 | 352630 | 1272400 | 0 | 127160 | 0 | 0 | 314960 | 0 | 0 | 0 | 0 | 0 |
| SEIQFLISDI | DSG2 | Q14126 | DSG2_HUMAN | Unmodified | 0 | 0 | 0 | 0 | 142900 | 0 | 0 | 0 | 1773300 | 0 | 0 | 0 | 0 | 0 | 0 | 62785 | 0 | 0 |
| SEISENTDA | LIMA1; LIMA1; L | Q9UHB6; F8 | LIMA1_HUMAN | Unmodified | 876910 | 694270 | 0 | 976150 | 1309500 | 329170 | 0 | 0 | 70888 | 0 | 0 | 381010 | 1110800 | 0 | 1275600 | 970770 | 0 | 0 |
| SEISGDLAF | ANXA6; ANXA6; | P08133; E7E | ANXA6_HUMAN | Unmodified | 593970 | 590000 | 6342700 | 7780100 | 2456300 | 747510 | 768590 | 377650 | 26693000 | 3885300 | 60744 | 707920 | 3161700 | 1717600 | 5916400 | 1754700 | 239040 | 2184300 |
| SEKFAFQAI | HSP90B1; HSP | P14625; A0A | ENPL_HUMAN | Unmodified | 0 | 0 | 0 | 0 | 0 | 0 | 172630 | 0 | 0 | 2037600 | 0 | 0 | 0 | 0 | 0 | 0 | 0 | 0 |
| SEKFAFQAI | HSP90B1; HSP | P14625; A0A | ENPL_HUMAN | Unmodified | 0 | 0 | 0 | 0 | 0 | 0 | 96573 | 0 | 0 | 796950 | 0 | 0 | 0 | 0 | 0 | 0 | 0 | 0 |
| SEKSVEAA | PTMS; PTMS | P20962; F5G | PTMS_HUMAN | Acetyl (Protein N | 0 | 0 | 0 | 0 | 0 | 0 | 262660 | 273320 | 0 | 0 | 0 | 0 | 0 | 0 | 0 | 0 | 464210 | 0 |
| SEKSVEAA | PTMS; PTMS | P20962; F5G | PTMS_HUMAN | Unmodified | 0 | 0 | 0 | 0 | 0 | 0 | 0 | 0 | 1819600 | 0 | 0 | 0 | 0 | 0 | 0 | 0 | 0 | 0 |
| SELDMLDI | ANXA6 | P08133 | ANXA6_HUMAN | Unmodified | 0 | 0 | 0 | 0 | 0 | 0 | 0 | 0 | 4472200 | 905690 | 0 | 0 | 0 | 0 | 0 | 0 | 0 | 0 |
| SELDMLDI | ANXA6 | P08133 | ANXA6_HUMAN | Unmodified | 21538 | 0 | 21315 | 0 | 0 | 0 | 0 | 0 | 7437400 | 480310 | 0 | 0 | 0 | 0 | 0 | 18776 | 0 | 0 |
| SELDMLDI | ANXA6 | P08133 | ANXA6_HUMAN | Oxidation (M) | 255200 | 300310 | 0 | 1004700 | 0 | 0 | 54697 | 0 | 5434900 | 1676700 | 33999 | 0 | 48031 | 0 | 2003900 | 544600 | 0 | 48244 |
| SELDQLRQ | GNB1; GNB1; G | P62873; F6U | GBB1_HUMAN | Acetyl (Protein N | 9031000 | 10794000 | 5200300 | 20244000 | 2.31E+08 | 48972000 | 62518000 | 22931000 | 1.38E+08 | 48881000 | 63055000 | 1.76E+08 | 79732000 | 89029000 | 11961000 | 8115000 | 1.5E+08 | 2.89E+08 |
| SELDQLRQ | GNB1; GNB1; G | P62873; F6U | GBB1_HUMAN | Unmodified | 0 | 0 | 0 | 0 | 0 | 0 | 125340 | 0 | 0 | 193840 | 0 | 237950 | 291520 | 0 | 0 | 0 | 0 | 0 |
| SELDQLRQ | GNB1; GNB1; G | P62873; F6U | GBB1_HUMAN | Acetyl (Protein N | 0 | 0 | 0 | 0 | 0 | 0 | 0 | 0 | 0 | 920010 | 0 | 0 | 373700 | 0 | 0 | 0 | 3783100 | 2121600 |
| SELEAIDVK | DNAH17 | Q9UFH2 | DYH17_HUMAN | Unmodified | 0 | 0 | 0 | 0 | 0 | 0 | 0 | 0 | 0 | 0 | 2314000 | 0 | 0 | 0 | 0 | 0 | 0 | 0 |
| SELEMVSG | TOM1; TOM1 | O60784; B0 | TOM1_HUMAN | Oxidation (M) | 21963 | 0 | 0 | 0 | 0 | 0 | 0 | 0 | 0 | 0 | 0 | 188540 | 0 | 0 | 0 | 0 | 0 | 0 |
| SELEMVSG | TOM1; TOM1 | O60784; B0 | TOM1_HUMAN | Unmodified | 0 | 0 | 0 | 0 | 0 | 63342 | 0 | 0 | 0 | 0 | 0 | 591330 | 0 | 0 | 0 | 0 | 0 | 0 |
| SELEQLR | RUNX1; PCNT; | Q01196; O9 | RUNX1_HUMAN | Acetyl (Protein N | 0 | 0 | 0 | 0 | 0 | 0 | 0 | 223610 | 0 | 0 | 1481900 | 0 | 0 | 121950 | 79739 | 91172 | 0 | 0 |
| SELEQLRQ | GNB2; GNB4; G | P62879; Q9 | GBB2_HUMAN | Acetyl (Protein N | 7578000 | 7206800 | 1941100 | 6717700 | 6122900 | 1925500 | 3253400 | 380890 | 4938000 | 0 | 28656000 | 93745000 | 3856100 | 1951200 | 6352800 | 8821700 | 5991900 | 7407000 |
| SELEQLRQ | GNB4; GNB4; G | Q9HAV0; A0 | GBB4_HUMAN | Acetyl (Protein N | 0 | 0 | 0 | 0 | 3990400 | 0 | 0 | 0 | 0 | 0 | 0 | 0 | 0 | 0 | 0 | 0 | 1838000 | 1665400 |
| SELEQLRQ | GNB2; GNB2; G | P62879; C9J | GBB2_HUMAN | Acetyl (Protein N | 8658700 | 13192000 | 0 | 23417000 | 17640000 | 4019100 | 2404400 | 0 | 0 | 0 | 21721000 | 23966000 | 7333800 | 0 | 17467000 | 5406400 | 8697700 | 14526000 |
| SELFGETAF | AGRN; AGRN; A | O00468; A0 | AGRIN_HUMAN | Unmodified | 0 | 0 | 838230 | 0 | 0 | 0 | 0 | 0 | 182840 | 0 | 446100 | 0 | 0 | 0 | 271630 | 252110 | 0 | 0 |
| SELHIENLN | BSG; BSG; BSG | P35613; A0A | BASI_HUMAN E | Unmodified | 2188600 | 1576400 | 9261100 | 7513700 | 0 | 0 | 0 | 0 | 11843000 | 913470 | 0 | 16876000 | 394320 | 1809300 | 15964000 | 8297600 | 417680 | 583790 |
| SELHIENLN | BSG; BSG; BSG | P35613; A0A | BASI_HUMAN E | Oxidation (M) | 2788700 | 2662300 | 17712000 | 6714800 | 0 | 0 | 69468 | 232180 | 5415200 | 1125100 | 330890 | 47699000 | 445870 | 0 | 27029000 | 28342000 | 641990 | 133940 |
| SELLGCTEI | EDIL3 | O43854 | EDIL3_HUMAN | Unmodified | 116240 | 0 | 0 | 0 | 1319500 | 471100 | 0 | 0 | 1583400 | 0 | 0 | 0 | 0 | 0 | 0 | 0 | 0 | 1676600 |
| SELLSQLQ | MRPS31 | Q92665 | RT31_HUMAN | Unmodified | 0 | 0 | 0 | 0 | 0 | 0 | 1392100 | 49803 | 488210 | 0 | 0 | 0 | 0 | 0 | 0 | 0 | 0 | 0 |
| SELPLDPLF | SLC1A5; SLC1A | Q15758; M0 | AAAT_HUMAN | Unmodified | 5861300 | 4783200 | 9636600 | 21104000 | 90194000 | 20735000 | 6098800 | 1816000 | 1.3E+08 | 26131000 | 81673000 | 2.45E+08 | 49202000 | 23700000 | 30848000 | 34130000 | 19822000 | 11695000 |
| SELPRFQEI | NIBAN2 | Q96TA1 | NIBA2_HUMAN | Unmodified | 649440 | 407520 | 9401300 | 1334400 | 2454000 | 417390 | 0 | 0 | 2553300 | 1035700 | 74690 | 54975 | 489640 | 51025 | 1014800 | 247280 | 33894 | 90967 |
| SELPVVDE | CNNM4 | Q6P4Q7 | CNNM4_HUMA | Unmodified | 0 | 0 | 0 | 0 | 0 | 0 | 0 | 0 | 0 | 0 | 307330 | 1122600 | 0 | 0 | 0 | 0 | 0 | 0 |
| SELRGYPC | QSOX2; QSOX2 | Q6ZRP7; H0 | QSOX2_HUMAN | Unmodified | 0 | 0 | 0 | 0 | 0 | 0 | 0 | 0 | 1710500 | 134470 | 82793 | 0 | 0 | 0 | 0 | 0 | 0 | 0 |
| SELSGNFEI | ANXA13; ANXA | P27216; P50 | ANX13_HUMAN | Unmodified | 1354200 | 753510 | 943560 | 942920 | 6736500 | 1197500 | 286140 | 755620 | 56271000 | 7710200 | 464870 | 0 | 2670600 | 2999200 | 1517500 | 682570 | 899410 | 8440900 |
| SELSSWR | SUN1; SUN1; S | O94901; H0 | SUN1_HUMAN | Unmodified | 0 | 0 | 0 | 0 | 0 | 0 | 0 | 0 | 4779400 | 235830 | 0 | 0 | 0 | 0 | 0 | 0 | 0 | 0 |
| SEMEVQDA | PA2G4 | Q9UQ80 | PA2G4_HUMAN | Unmodified | 0 | 0 | 0 | 0 | 167580 | 0 | 0 | 0 | 235420 | 96422 | 305920 | 4465300 | 0 | 0 | 0 | 0 | 0 | 0 |
| SEMEVQDA | PA2G4 | Q9UQ80 | PA2G4_HUMAN | Oxidation (M) | 179020 | 0 | 371200 | 0 | 0 | 0 | 0 | 0 | 0 | 0 | 0 | 1116700 | 0 | 0 | 0 | 0 | 0 | 0 |
| SEMTPEEL | SNRPD2; SNRP | P62316; K7E | SMD2_HUMAN | Unmodified | 0 | 0 | 53334 | 0 | 0 | 0 | 0 | 0 | 0 | 0 | 384590 | 510460 | 0 | 0 | 0 | 0 | 0 | 0 |
| SEMTPEEL | SNRPD2; SNRP | P62316; K7E | SMD2_HUMAN | Oxidation (M) | 3654000 | 2002200 | 0 | 2249700 | 3852900 | 0 | 0 | 0 | 3736200 | 3020900 | 3599100 | 6717000 | 12295000 | 0 | 0 | 0 | 0 | 0 |
| SENGLEFT | VDAC1; VDAC1 | P21796; C9J | VDAC1_HUMAN | Unmodified | 0 | 0 | 0 | 0 | 0 | 0 | 0 | 0 | 14829000 | 794960 | 0 | 0 | 0 | 0 | 0 | 0 | 0 | 0 |
| SENGLEFT | VDAC1; VDAC1 | P21796; C9J | VDAC1_HUMAN | Unmodified | 0 | 0 | 0 | 0 | 0 | 0 | 0 | 0 | 6830800 | 4666300 | 0 | 0 | 0 | 0 | 0 | 0 | 0 | 0 |
| SENLGQGI | RPA1 | P27694 | RFA1_HUMAN | Unmodified | 0 | 0 | 0 | 0 | 1352100 | 0 | 2146600 | 0 | 0 | 219650 | 0 | 4970800 | 0 | 0 | 0 | 0 | 0 | 0 |
| SENRVVVS | SRSF1; SRSF1; | Q07955; J3K | SRSF1_HUMAN | Unmodified | 774260 | 305180 | 590960 | 443010 | 0 | 0 | 230680 | 0 | 0 | 0 | 0 | 348510 | 98711 | 0 | 453840 | 178810 | 0 | 0 |
| SEPIPESNI | PDIA3 | P30101 | PDIA3_HUMAN | Unmodified | 0 | 0 | 0 | 0 | 4528600 | 0 | 3155700 | 2740700 | 1.66E+08 | 66617000 | 40977000 | 13850000 | 0 | 0 | 0 | 0 | 608180 | 2765100 |
| SEPIPESNI | PDIA3 | P30101 | PDIA3_HUMAN | Unmodified | 0 | 0 | 0 | 0 | 0 | 0 | 0 | 0 | 0 | 2368600 | 0 | 0 | 0 | 0 | 0 | 0 | 0 | 0 |
| SEPPTLAQ | SLK | Q9H2G2 | SLK_HUMAN S | Unmodified | 0 | 0 | 0 | 68898 | 0 | 0 | 0 | 0 | 0 | 0 | 400470 | 0 | 0 | 0 | 57922 | 64105 | 0 | 0 |
| SEPQNLGC | RAB5B | P61020 | RAB5B_HUMAN | Unmodified | 2544200 | 985590 | 642950 | 1744000 | 0 | 0 | 33120 | 19527 | 2741500 | 234800 | 522300 | 1023900 | 185260 | 74703 | 1657900 | 1472300 | 0 | 0 |
| SEPSLEPES | ZDHHC5; ZDHH | Q9C0B5; H0 | ZDHC5_HUMA | Unmodified | 226990 | 340020 | 732880 | 482860 | 767440 | 0 | 0 | 0 | 660040 | 227800 | 473840 | 658000 | 680720 | 842980 | 743960 | 580790 | 1016000 | 937500 |
| SEQAVAQL | KLC2; KLC4; KL | Q9H0B6; Q9 | KLC2_HUMAN | Unmodified | 0 | 0 | 0 | 0 | 0 | 0 | 0 | 0 | 0 | 0 | 0 | 0 | 0 | 0 | 0 | 0 | 0 | 0 |
| SEQEAQAP | TMEM125; TME | Q96AQ2; E5 | TM125_HUMAN | Acetyl (Protein N | 21294 | 236700 | 139190 | 176720 | 0 | 0 | 0 | 0 | 342760 | 0 | 0 | 196270 | 0 | 0 | 79371 | 138410 | 0 | 0 |
| SEQEEYEA | MYO1C; MYO1 | O00159; F5 | MYO1C_HUMA | Unmodified | 0 | 0 | 357730 | 0 | 0 | 0 | 0 | 0 | 0 | 0 | 0 | 0 | 245500 | 0 | 19157000 | 2733100 | 0 | 0 |
| SEQFINLR | CAPN1 | P07384 | CAN1_HUMAN | Unmodified | 0 | 0 | 155560 | 147650 | 0 | 0 | 0 | 0 | 0 | 0 | 145720 | 1002400 | 0 | 0 | 6301700 | 0 | 404880 | 65021 |
| SEQFINLRE | CAPN1 | P07384 | CAN1_HUMAN | Unmodified | 0 | 0 | 0 | 0 | 685240 | 0 | 0 | 0 | 0 | 217490 | 366180 | 0 | 509690 | 0 | 0 | 0 | 254900 | 556740 |
| SEQGTLTS | DLGAP4 | Q9Y2H0 | DLGP4_HUMAN | Unmodified | 0 | 0 | 307400 | 208360 | 346660 | 0 | 0 | 156320 | 280460 | 110500 | 276530 | 251180 | 0 | 0 | 187780 | 159020 | 682530 | 1295700 |
| SEQLGGD | MUC5B | Q9HC84 | MUC5B_HUMA | Unmodified | 0 | 0 | 0 | 752030 | 0 | 0 | 0 | 0 | 0 | 0 | 0 | 0 | 0 | 0 | 0 | 0 | 0 | 550240 |
| SEQLKPLK | EPHA2 | P29317 | EPHA2_HUMAN | Unmodified | 232410 | 99393 | 0 | 0 | 297390 | 0 | 0 | 70184 | 0 | 0 | 0 | 0 | 0 | 0 | 0 | 0 | 2689200 | 4331600 |
| SEQSICQA | ENAH; EVL; EVL | Q8N8S7; Q9 | ENAH_HUMAN | Acetyl (Protein N | 0 | 0 | 0 | 0 | 0 | 0 | 89855 | 98869 | 646050 | 845280 | 0 | 2379900 | 319100 | 235860 | 0 | 329200 | 586170 | 156960 |
| SEQSVAQL | KLC1; KLC1; KL | Q07866; G3 | KLC1_HUMAN | Unmodified | 0 | 0 | 44217 | 0 | 425040 | 135040 | 0 | 0 | 536100 | 0 | 0 | 1066900 | 128980 | 0 | 98634 | 62179 | 173930 | 309230 |
| SEQSVAQL | KLC1; KLC1; KL | Q07866; G3 | KLC1_HUMAN | Unmodified | 0 | 0 | 0 | 216980 | 1546800 | 347940 | 0 | 97500 | 1832700 | 350300 | 438260 | 2310200 | 822060 | 0 | 191800 | 130560 | 0 | 837420 |
| SEQSVAQL | KLC1; KLC1; KL | Q07866; G3 | KLC1_HUMAN | Unmodified | 0 | 0 | 0 | 0 | 1313100 | 0 | 0 | 0 | 0 | 270920 | 0 | 0 | 0 | 0 | 0 | 0 | 57215 | 716160 |
| SERETAVRI | P3H1; P3H1 | Q32P28; H7 | P3H1_HUMAN | Oxidation (M) | 0 | 0 | 0 | 0 | 0 | 0 | 78755 | 139680 | 0 | 0 | 0 | 0 | 1260600 | 1319600 | 0 | 0 | 410320 | 694420 |
| SERPPIFEI | LRP1 | Q07954 | LRP1_HUMAN | Unmodified | 0 | 0 | 0 | 0 | 0 | 0 | 0 | 0 | 1749000 | 0 | 0 | 0 | 0 | 0 | 0 | 0 | 0 | 0 |
| SERPPTILM | CARMIL1; CARI | Q5VZK9; A0 | CARL1_HUMAN | Oxidation (M) | 0 | 0 | 0 | 86140 | 856800 | 0 | 0 | 0 | 0 | 0 | 336400 | 703470 | 213930 | 271760 | 0 | 71857 | 205610 | 505310 |
| SESEVEEA | DBN1; DBN1; D | Q16643; D6 | DREB_HUMAN | Unmodified | 0 | 0 | 0 | 0 | 570230 | 0 | 0 | 0 | 0 | 0 | 1335600 | 2677900 | 0 | 0 | 0 | 0 | 0 | 0 |
| SESGYGFN | SNX27; SNX27 | Q96L92; A0A | SNX27_HUMAN | Unmodified | 0 | 0 | 0 | 0 | 870600 | 238870 | 0 | 0 | 0 | 0 | 0 | 0 | 0 | 0 | 0 | 0 | 0 | 122960 |
| SESLVVCD | GMFB; GMFB; G | P60983; G3 | GMFB_HUMAN | Acetyl (Protein N | 68908 | 39377 | 14115 | 0 | 203000 | 47424 | 0 | 71405 | 86159 | 236340 | 0 | 58140 | 787470 | 552440 | 216290 | 0 | 761990 | 218480 |
| SESLVVCD | GMFB; GMFB; G | P60983; G3 | GMFB_HUMAN | Acetyl (Protein N | 0 | 0 | 0 | 0 | 0 | 254250 | 452490 | 0 | 0 | 325830 | 0 | 0 | 623510 | 557310 | 0 | 0 | 2412500 | 491080 |
| SESPKEPE | HNRNPA1; HNI | P09651; F8V | ROA1_HUMAN | Unmodified | 0 | 0 | 0 | 0 | 0 | 0 | 0 | 11900 | 119560 | 81115 | 503890 | 1282700 | 0 | 0 | 0 | 0 | 0 | 0 |
| SESPKEPE | HNRNPA1; HNI | P09651; F8V | ROA1_HUMAN | Unmodified | 0 | 0 | 0 | 1349900 | 38287 | 0 | 0 | 1227000 | 0 | 1126400 | 6164400 | 0 | 0 | 0 | 0 | 0 | 0 | 0 |
| SESQCRVV | PAICS; PAICS; F | P22234; D6 | PUR6_HUMAN | Unmodified | 0 | 0 | 0 | 0 | 856680 | 0 | 578450 | 0 | 0 | 444240 | 0 | 0 | 1780400 | 1062000 | 0 | 0 | 2033400 | 1620000 |
| SESQMDITI | MME; MME; MM | P08473; Q3 | NEP_HUMAN N | Oxidation (M) | 0 | 0 | 3606700 | 829960 | 0 | 0 | 0 | 0 | 0 | 0 | 0 | 0 | 0 | 0 | 3039300 | 2928700 | 0 | 0 |
| SESQMDITI | MME; MME; MM | P08473; Q3 | NEP_HUMAN N | Unmodified | 0 | 0 | 2846100 | 1774900 | 0 | 0 | 0 | 0 | 0 | 0 | 0 | 0 | 0 | 0 | 3470300 | 2852900 | 0 | 0 |
| SESSPEAYI | PLCG2; PLCG2 | P16885; A0A | PLCG2_HUMAN | Unmodified | 0 | 0 | 0 | 0 | 0 | 413420 | 0 | 20887 | 348310 | 132260 | 1360000 | 13332000 | 184550 | 0 | 0 | 0 | 0 | 155860 |
| SESTQPLLI | IL6ST | P40189 | IL6RB_HUMAN | Unmodified | 0 | 0 | 0 | 0 | 0 | 0 | 0 | 0 | 5466900 | 0 | 0 | 0 | 0 | 0 | 0 | 483200 | 0 | 0 |
| SESVPPVTI | BSG; BSG; BSG | P35613; A0A | BASI_HUMAN E | Unmodified | 0 | 0 | 9063200 | 18871000 | 30729000 | 7579100 | 3010800 | 583660 | 32717000 | 1905600 | 3755500 | 31240000 | 16423000 | 6481200 | 18683000 | 18983000 | 4730800 | 53778000 |
| SESVPPVTI | BSG; BSG; BSG | P35613; A0A | BASI_HUMAN E | Unmodified | 0 | 733130 | 0 | 2187200 | 34306000 | 4576800 | 4970900 | 0 | 518470 | 1747900 | 0 | 0 | 26385000 | 12783000 | 1122500 | 0 | 15141000 | 41221000 |
| SESVVYADI | MPZL1 | O95297 | MPZL1_HUMAN | Unmodified | 1075200 | 605620 | 1585100 | 452650 | 0 | 0 | 0 | 0 | 545250 | 0 | 245410 | 3459100 | 74951 | 0 | 1082000 | 922120 | 0 | 0 |
| SESVVYADI | MPZL1 | O95297 | MPZL1_HUMAN | Unmodified | 0 | 0 | 0 | 0 | 2525400 | 0 | 0 | 0 | 0 | 0 | 0 | 0 | 0 | 0 | 0 | 0 | 0 | 1525600 |
| SETAPAAP | H1-2 | P16403 | H12_HUMAN H | Acetyl (Protein N | 30646000 | 1747700 | 0 | 94959 | 202960 | 0 | 945390 | 2.13E+08 | 848560 | 103930 | 428760 | 2039000 | 0 | 0 | 109140 | 0 | 0 | 0 |
| SETAPAAP | H1-2 | P16403 | H12_HUMAN H | Unmodified | 1151400 | 0 | 0 | 0 | 0 | 0 | 43679 | 644230 | 144490 | 0 | 0 | 0 | 0 | 0 | 0 | 0 | 0 | 94383 |
| SETAPAAP | H1-2 | P16403 | H12_HUMAN H | Acetyl (Protein N | 0 | 0 | 0 | 0 | 0 | 0 | 812990 | 21268000 | 0 | 0 | 0 | 0 | 0 | 0 | 0 | 0 | 0 | 0 |
| SETAPAAP | H1-2 | P16403 | H12_HUMAN H | Acetyl (Protein N | 0 | 0 | 0 | 0 | 1006100 | 236090 | 4929000 | 11496000 | 0 | 0 | 0 | 0 | 0 | 0 | 0 | 0 | 0 | 473070 |
| SETAPAAP | H1-2 | P16403 | H12_HUMAN H | Unmodified | 0 | 0 | 0 | 0 | 0 | 0 | 0 | 810300 | 0 | 0 | 0 | 0 | 0 | 0 | 0 | 0 | 0 | 0 |
| SETAPAAP | H1-4 | P10412 | H14_HUMAN H | Unmodified | 0 | 340930 | 0 | 0 | 0 | 0 | 0 | 1363400 | 0 | 0 | 0 | 0 | 0 | 0 | 0 | 0 | 0 | 0 |
| SETAPAAP | H1-4 | P10412 | H14_HUMAN H | Acetyl (Protein N | 0 | 0 | 0 | 0 | 0 | 0 | 0 | 1.91E+08 | 0 | 0 | 0 | 0 | 0 | 0 | 0 | 0 | 0 | 0 |

| | | | | | | | | | | | | | | | | | | | | | | |
|---|---|---|---|---|---|---|---|---|---|---|---|---|---|---|---|---|---|---|---|---|---|---|
| SETAPAAP | H1-4 | P10412 | H14_HUMAN H | Acetyl (Protein N | 166400 | 1536900 | 0 | 0 | 1587900 | 0 | 4404700 | 5.52E+08 | 443540 | 442130 | 0 | 0 | 422490 | 0 | 0 | 0 | 0 | 1536800 |
| SETAPAAP | H1-4 | P10412 | H14_HUMAN H | Unmodified | 0 | 0 | 0 | 0 | 0 | 0 | 0 | 7516500 | 0 | 0 | 0 | 0 | 0 | 0 | 0 | 0 | 0 | 0 |
| SETAPAAP | H1-4 | P10412 | H14_HUMAN H | Acetyl (Protein N | 67302 | 0 | 0 | 0 | 1020600 | 0 | 5649600 | 3.31E+08 | 0 | 239810 | 0 | 0 | 1509000 | 0 | 0 | 0 | 87210 | 1096400 |
| SETAPAAP | H1-4 | P10412 | H14_HUMAN H | Unmodified | 0 | 0 | 0 | 0 | 0 | 0 | 0 | 2086800 | 0 | 0 | 0 | 0 | 112860 | 20572 | 0 | 0 | 0 | 0 |
| SETAPAETA | H1-5 | P16401 | H15_HUMAN H | Unmodified | 1009800 | 669420 | 0 | 0 | 0 | 0 | 0 | 591710 | 0 | 0 | 271230 | 857750 | 190150 | 0 | 0 | 0 | 0 | 0 |
| SETAPAETA | H1-5 | P16401 | H15_HUMAN H | Acetyl (Protein N | 6423100 | 3350700 | 0 | 0 | 112830 | 0 | 0 | 3979200 | 0 | 0 | 798650 | 0 | 0 | 0 | 0 | 0 | 0 | 0 |
| SETAPAETA | H1-5 | P16401 | H15_HUMAN H | Acetyl (Protein N | 0 | 0 | 0 | 0 | 0 | 0 | 0 | 26845000 | 0 | 0 | 0 | 0 | 0 | 0 | 0 | 0 | 0 | 0 |
| SETAPAETA | H1-5 | P16401 | H15_HUMAN H | Unmodified | 0 | 0 | 0 | 0 | 0 | 0 | 0 | 1084000 | 0 | 0 | 0 | 0 | 0 | 0 | 0 | 0 | 0 | 0 |
| SETAPAETA | H1-5 | P16401 | H15_HUMAN H | Acetyl (Protein N | 0 | 0 | 0 | 0 | 0 | 0 | 164600 | 306650 | 0 | 0 | 0 | 0 | 0 | 0 | 0 | 0 | 0 | 0 |
| SETDLLDIR | ANXA11; ANXA1 | P50995; H0Y | ANX11_HUMAN | Unmodified | 361420 | 0 | 0 | 201890 | 0 | 0 | 0 | 0 | 11185000 | 0 | 784740 | 0 | 0 | 0 | 348060 | 350000 | 0 | 0 |
| SETDLLDIR | ANXA11; ANXA1 | P50995; H0Y | ANX11_HUMAN | Unmodified | 295890 | 332870 | 301420 | 0 | 943160 | 131400 | 0 | 0 | 42734000 | 1591100 | 925310 | 4284400 | 575650 | 294700 | 265010 | 688310 | 357240 | 902170 |
| SETDLLDIR | ANXA11; ANXA1 | P50995; H0Y | ANX11_HUMAN | Unmodified | 310930 | 286770 | 0 | 174840 | 0 | 0 | 0 | 0 | 17631000 | 1106700 | 0 | 1516500 | 0 | 0 | 0 | 198940 | 0 | 0 |
| SETEDSILH | APPL1 | Q9UKG1 | DP13A_HUMAN | Unmodified | 417050 | 769400 | 576570 | 257830 | 0 | 0 | 0 | 0 | 1348600 | 176310 | 39548 | 1465500 | 0 | 0 | 983350 | 600420 | 132080 | 0 |
| SETELKDTY | PSMD2; PSMD2 | Q13200; H7 | PSMD2_HUMA | Unmodified | 86533 | 0 | 0 | 78451 | 34676 | 28728 | 0 | 0 | 101350 | 107100 | 15218 | 1614200 | 112620 | 26803 | 0 | 0 | 102240 | 30663 |
| SETFLLLQP | TMEM132A | Q24JP5 | T132A_HUMAN | Unmodified | 0 | 0 | 0 | 0 | 0 | 0 | 0 | 0 | 334200 | 0 | 0 | 0 | 0 | 0 | 195350 | 51403 | 0 | 0 |
| SETITADVN | UAP1 | Q16222 | UAP1_HUMAN | Unmodified | 0 | 0 | 0 | 74726 | 0 | 79664 | 0 | 0 | 0 | 0 | 0 | 970500 | 0 | 0 | 0 | 0 | 0 | 0 |
| SETSGPQIK | NUDC; NUDC; | Q9Y266; A0A | NUDC_HUMAN | Unmodified | 0 | 0 | 324870 | 494700 | 0 | 0 | 0 | 140520 | 5686200 | 786490 | 119250 | 2707100 | 947130 | 45642 | 0 | 0 | 94097 | 0 |
| SETSGSFEI | ANXA4; ANXA4 | P09525; Q6F | ANXA4_HUMAN | Unmodified | 542880 | 411220 | 316020 | 489150 | 686390 | 174330 | 230880 | 0 | 7478200 | 3048300 | 2532600 | 3995800 | 350590 | 1049200 | 2231700 | 934010 | 67362 | 149990 |
| SETVLTCAT | ITGA3 | P26006 | ITA3_HUMAN I | Unmodified | 0 | 0 | 0 | 0 | 3713200 | 1562200 | 0 | 0 | 484520 | 0 | 0 | 2483500 | 0 | 0 | 0 | 0 | 1695900 | 3356400 |
| SEVAAENH | PDIA5 | Q14554 | PDIA5_HUMAN | Unmodified | 0 | 0 | 0 | 0 | 0 | 0 | 0 | 9985.3 | 4102700 | 655930 | 0 | 0 | 0 | 0 | 0 | 0 | 0 | 0 |
| SEVAAENH | PDIA5 | Q14554 | PDIA5_HUMAN | Unmodified | 0 | 0 | 0 | 0 | 0 | 0 | 64463 | 0 | 0 | 3965400 | 0 | 0 | 0 | 0 | 0 | 0 | 0 | 0 |
| SEVATLTAA | ATP5F1C | P36542 | ATPG_HUMAN | Unmodified | 3265900 | 539370 | 3045600 | 0 | 0 | 0 | 0 | 0 | 8873400 | 1737800 | 158370 | 216240 | 0 | 0 | 0 | 841950 | 0 | 0 |
| SEVDMLK | ANXA2P2; ANX | A6NMY6; P0 | AXA2L_HUMAN | Oxidation (M) | 0 | 191390 | 0 | 0 | 337680 | 76327 | 87338 | 21849 | 920180 | 130200 | 0 | 0 | 37977 | 37587 | 0 | 0 | 31693 | 0 |
| SEVDMLK | ANXA2P2; ANX | A6NMY6; P0 | AXA2L_HUMAN | Unmodified | 425540 | 0 | 0 | 0 | 0 | 0 | 0 | 0 | 15123000 | 1399500 | 0 | 83923 | 0 | 0 | 0 | 467610 | 0 | 0 |
| SEVGLVSEI | PGAP6 | Q9HCN3 | PGAP6_HUMAN | Unmodified | 345690 | 269020 | 741930 | 266430 | 0 | 0 | 0 | 0 | 286050 | 0 | 0 | 1428200 | 0 | 0 | 1080500 | 605790 | 0 | 0 |
| SEVLGTQS | YKT6; YKT6 | O15498; A0A | YKT6_HUMAN | Unmodified | 0 | 0 | 0 | 0 | 0 | 0 | 0 | 0 | 0 | 0 | 0 | 0 | 0 | 0 | 0 | 0 | 0 | 0 |
| SEVLRVTHI | PSMC3; PSMC3 | P17980; E9P | PRS6A_HUMAN | Unmodified | 0 | 0 | 127350 | 0 | 877850 | 0 | 0 | 0 | 0 | 0 | 0 | 0 | 544260 | 0 | 0 | 0 | 0 | 997810 |
| SEVLRVTHI | PSMC3; PSMC3 | P17980; E9P | PRS6A_HUMAN | Oxidation (M) | 0 | 0 | 0 | 82429 | 255870 | 157980 | 0 | 0 | 321440 | 79878 | 95828 | 0 | 248000 | 68403 | 0 | 0 | 80550 | 212880 |
| SEVLRVTHI | PSMC3; PSMC3 | P17980; E9P | PRS6A_HUMAN | Unmodified | 0 | 0 | 0 | 0 | 1103600 | 0 | 13187 | 0 | 0 | 240310 | 0 | 0 | 977410 | 247190 | 0 | 0 | 284840 | 2332300 |
| SEVPLVDV | MYO1B; MYO1B | O43795; H7 | MYO1B_HUMA | Unmodified | 239740 | 559210 | 442160 | 185250 | 6234400 | 1792100 | 376550 | 203840 | 1255900 | 618180 | 4203400 | 37386000 | 228840 | 0 | 1694300 | 1107500 | 263920 | 2022900 |
| SEVQQPVH | LIMA1; LIMA1; L | Q9UHB6; F8 | LIMA1_HUMAN | Unmodified | 1121500 | 890920 | 0 | 995390 | 450110 | 0 | 0 | 0 | 0 | 0 | 0 | 1292100 | 306650 | 0 | 649520 | 889380 | 0 | 0 |
| SEVSSDED | HADHA; HADH | P40939; A0A | ECHA_HUMAN | Unmodified | 0 | 0 | 0 | 267130 | 0 | 121010 | 0 | 0 | 981970 | 0 | 0 | 0 | 0 | 245890 | 0 | 0 | 0 | 0 |
| SEVTRSLLC | AIDA | Q96BJ3 | AIDA_HUMAN | Acetyl (Protein N | 0 | 0 | 0 | 0 | 1609400 | 227040 | 0 | 493130 | 0 | 383180 | 0 | 458950 | 3497600 | 0 | 0 | 0 | 3946200 | 5738500 |
| SEWKFTITF | CAPZA1 | P52907 | CAZA1_HUMAN | Unmodified | 0 | 258020 | 0 | 0 | 5278900 | 0 | 0 | 0 | 0 | 545000 | 0 | 0 | 3883500 | 2039200 | 142340 | 0 | 814270 | 1020500 |
| SEWPREPC | EPS8L2; EPS8L | Q9H6S3; A0A | ES8L2_HUMAN | Unmodified | 0 | 0 | 0 | 0 | 0 | 1660400 | 0 | 0 | 676680 | 0 | 0 | 0 | 0 | 0 | 550430 | 336530 | 0 | 0 |
| SEWQSMT | SUN2 | Q9UH99 | SUN2_HUMAN | Oxidation (M) | 0 | 0 | 0 | 0 | 0 | 0 | 0 | 0 | 5684500 | 0 | 0 | 0 | 0 | 0 | 0 | 0 | 0 | 0 |
| SEWQSMT | SUN2 | Q9UH99 | SUN2_HUMAN | Unmodified | 0 | 0 | 0 | 0 | 0 | 0 | 0 | 0 | 9193400 | 0 | 0 | 0 | 0 | 0 | 0 | 0 | 0 | 0 |
| SEWSDLLS | PFKM; PFKM; PI | P08237; A0A | PFKAM_HUMAN | Unmodified | 0 | 0 | 49097 | 0 | 0 | 211270 | 0 | 0 | 0 | 0 | 0 | 149910 | 0 | 0 | 0 | 0 | 0 | 0 |
| SEYDRGVN | PSMA5 | P28066 | PSA5_HUMAN | Unmodified | 0 | 0 | 0 | 0 | 0 | 0 | 0 | 0 | 205180 | 0 | 73562 | 1221100 | 0 | 0 | 25627 | 0 | 0 | 110200 |
| SFADINLYF | NRAS | P01111 | RASN_HUMAN | Unmodified | 528330 | 454730 | 539170 | 367590 | 0 | 0 | 0 | 361100 | 1784400 | 159820 | 318020 | 5808500 | 0 | 0 | 1004700 | 1434400 | 0 | 0 |
| SFADINLYF | NRAS | P01111 | RASN_HUMAN | Unmodified | 1785600 | 2191000 | 5823200 | 6265700 | 15004000 | 0 | 0 | 1952500 | 9284100 | 4621600 | 1655100 | 4506800 | 17462000 | 12927000 | 5691300 | 4289800 | 5547800 | 6246800 |
| SFADINLYF | NRAS | P01111 | RASN_HUMAN | Unmodified | 0 | 0 | 0 | 1482500 | 0 | 322960 | 722940 | 0 | 0 | 1683100 | 0 | 0 | 2078700 | 1163700 | 0 | 0 | 0 | 0 |
| SFAECAYG | ADAM9; ADAM9 | Q13443; A0A | ADAM9_HUMA | Unmodified | 593990 | 88970 | 0 | 620710 | 1946100 | 0 | 0 | 0 | 0 | 0 | 0 | 0 | 1241400 | 1515800 | 391020 | 306580 | 387800 | 583440 |
| SFAGAVSP | MCM5; MCM5 | P33992; B1A | MCM5_HUMAN | Unmodified | 0 | 0 | 0 | 0 | 0 | 0 | 0 | 0 | 539880 | 0 | 210700 | 1754200 | 0 | 0 | 0 | 0 | 0 | 0 |
| SFAGNLNT | PFKP; PFKP; PF | Q01813; A0A | PFKAP_HUMAN | Unmodified | 0 | 0 | 0 | 0 | 4040700 | 1229300 | 0 | 94453 | 778520 | 394990 | 644890 | 6194700 | 737580 | 0 | 0 | 227200 | 1763900 | 3289700 |
| SFAGNLNT | PFKP; PFKP; PF | Q01813; A0A | PFKAP_HUMAN | Unmodified | 0 | 0 | 0 | 0 | 0 | 0 | 0 | 0 | 0 | 0 | 0 | 1498700 | 0 | 0 | 0 | 0 | 0 | 0 |
| SFAPGSAA | LRATD2 | Q96KN1 | LRAT2_HUMAN | Unmodified | 0 | 0 | 5491400 | 197240 | 735300 | 0 | 195020 | 0 | 2078100 | 454220 | 2405000 | 0 | 2244500 | 0 | 0 | 0 | 3465600 | 0 |
| SFATHTAA | NCOA7 | Q8NI08 | NCOA7_HUMA | Unmodified | 41160 | 0 | 0 | 0 | 263850 | 0 | 0 | 0 | 0 | 47738 | 0 | 0 | 0 | 0 | 0 | 0 | 0 | 0 |
| SFAVGMFK | ACADVL; ACAD | P49748; K7E | ACADV_HUMA | Oxidation (M) | 490260 | 0 | 0 | 0 | 0 | 0 | 0 | 0 | 1522900 | 444980 | 0 | 0 | 0 | 0 | 0 | 0 | 0 | 0 |
| SFCPGGTD | UBE2O; UBE2O | Q9C0C9; K7 | UBE2O_HUMA | Unmodified | 0 | 0 | 0 | 0 | 0 | 0 | 0 | 0 | 0 | 0 | 0 | 0 | 0 | 0 | 0 | 0 | 0 | 0 |
| SFCPGGTD | UBE2O; UBE2O | Q9C0C9; K7 | UBE2O_HUMA | Unmodified | 0 | 0 | 0 | 0 | 0 | 0 | 0 | 0 | 0 | 0 | 0 | 0 | 0 | 0 | 0 | 0 | 924560 | 0 |
| SFCSMAGF | DDAH1; DDAH1 | O94760; B4D | DDAH1_HUMA | Oxidation (M) | 260610 | 0 | 163710 | 0 | 0 | 0 | 0 | 0 | 0 | 566800 | 605470 | 0 | 1338300 | 2484800 | 0 | 0 | 1664200 | 1777200 |
| SFCSMAGF | DDAH1; DDAH1 | O94760; B4D | DDAH1_HUMA | Unmodified | 0 | 0 | 159930 | 0 | 0 | 0 | 0 | 0 | 1234200 | 0 | 0 | 0 | 0 | 0 | 411690 | 108750 | 1285300 | 1105600 |
| SFDAVVFD | RALGPS2 | Q86X27 | RGPS2_HUMA | Unmodified | 0 | 0 | 10597 | 0 | 0 | 0 | 0 | 0 | 0 | 0 | 0 | 154840 | 0 | 0 | 0 | 0 | 0 | 0 |
| SFDDPIVQ | HSPA4L; HSPA4 | O95757; E9F | HS74L_HUMAN | Unmodified | 215710 | 0 | 0 | 0 | 0 | 0 | 0 | 0 | 0 | 0 | 0 | 2812800 | 371220 | 320550 | 0 | 98834 | 0 | 0 |
| SFDDPIVQ | HSPA4L; HSPA4 | O95757; E9F | HS74L_HUMAN | Unmodified | 0 | 0 | 0 | 0 | 0 | 0 | 0 | 0 | 0 | 0 | 0 | 0 | 2702300 | 0 | 0 | 0 | 0 | 0 |
| SFDGFSSA | HEBP2 | Q9Y5Z4 | HEBP2_HUMAN | Unmodified | 52931 | 379130 | 324860 | 254730 | 0 | 0 | 18210 | 0 | 936210 | 793090 | 339940 | 3291200 | 876870 | 368050 | 0 | 625740 | 0 | 156840 |
| SFDLASCD | ST14 | Q9Y5Y6 | ST14_HUMAN | Unmodified | 0 | 430530 | 918080 | 524800 | 0 | 0 | 0 | 0 | 160480 | 0 | 425980 | 0 | 0 | 0 | 0 | 0 | 0 | 0 |
| SFDLLILK | THOC2; THOC2 | Q8NI27; A0A | THOC2_HUMA | Unmodified | 0 | 0 | 59750 | 0 | 0 | 0 | 0 | 16000 | 0 | 0 | 67073 | 813400 | 0 | 0 | 0 | 33149 | 0 | 0 |
| SFDLLVK | CACYBP | Q9HB71 | CYBP_HUMAN | Unmodified | 207580 | 135220 | 341110 | 451940 | 65693 | 0 | 30941 | 110220 | 2260300 | 0 | 4349700 | 22699000 | 1388900 | 82460 | 734760 | 853320 | 104550 | 118620 |
| SFDSNAFT | COMMD3; COM | Q9UBI1; R4 | COMD3_HUMA | Unmodified | 0 | 0 | 0 | 163970 | 37328 | 0 | 0 | 0 | 276020 | 217610 | 45690 | 730630 | 0 | 0 | 0 | 0 | 0 | 1005300 |
| SFDTSLIR | ATP6AP2; ATP6 | O75787; A0A | RENR_HUMAN | Unmodified | 0 | 0 | 293500 | 164890 | 0 | 0 | 0 | 0 | 4713700 | 467340 | 699650 | 0 | 0 | 45925 | 133580 | 109970 | 0 | 0 |
| SFDVNKPG | EIF2S3; EIF2S3 | P41091; Q2 | IF2G_HUMAN | Unmodified | 0 | 0 | 0 | 0 | 0 | 0 | 0 | 0 | 0 | 0 | 0 | 0 | 2747300 | 0 | 0 | 0 | 1387100 | 2410500 |
| SFEALLADL | DCLK1; DCLK1 | O15075; Q5 | DCLK1_HUMAN | Unmodified | 0 | 0 | 0 | 0 | 0 | 0 | 0 | 0 | 0 | 8682.4 | 0 | 262700 | 0 | 0 | 0 | 0 | 0 | 0 |
| SFEAPATIN | STRAP | Q9Y3F4 | STRAP_HUMAN | Unmodified | 848500 | 418420 | 422560 | 431200 | 0 | 360230 | 607190 | 375970 | 530290 | 325620 | 1275300 | 8312400 | 2508700 | 857190 | 470120 | 362410 | 0 | 0 |
| SFEAPIKLV | HM13; HM13; H | Q8TCT9; A0A | HM13_HUMAN | Unmodified | 0 | 0 | 71334 | 0 | 0 | 0 | 0 | 0 | 0 | 977390 | 0 | 0 | 0 | 0 | 0 | 0 | 0 | 0 |
| SFECTCPR | DLL3 | Q9NYJ7 | DLL3_HUMAN | Unmodified | 0 | 0 | 0 | 0 | 0 | 0 | 0 | 0 | 765010 | 27175 | 42288 | 0 | 0 | 0 | 0 | 0 | 0 | 0 |
| SFEDIHHY | KRAS | P01116 | RASK_HUMAN | Unmodified | 905820 | 4152600 | 234420 | 118490 | 0 | 66909 | 36285 | 137550 | 513010 | 0 | 182400 | 882060 | 0 | 0 | 609270 | 1550800 | 15855 | 0 |
| SFEDIHHY | KRAS | P01116 | RASK_HUMAN | Unmodified | 1946100 | 1509600 | 2536300 | 307500 | 7359400 | 1026600 | 587760 | 0 | 6784100 | 1475500 | 1911700 | 6349100 | 7062700 | 5115800 | 1305000 | 2345900 | 903070 | 4348000 |
| SFEDLTDH | ADAM17 | P78536 | ADA17_HUMAN | Unmodified | 0 | 0 | 0 | 459170 | 2012700 | 0 | 0 | 30440 | 1515800 | 356830 | 0 | 475320 | 0 | 0 | 0 | 0 | 1135700 | 2524700 |
| SFEDLTDH | ADAM17 | P78536 | ADA17_HUMAN | Unmodified | 0 | 98892 | 0 | 0 | 7745300 | 1185000 | 0 | 0 | 1073600 | 436630 | 0 | 0 | 0 | 1107900 | 0 | 0 | 5330600 | 12790000 |
| SFEEKVENI | TPD52; N/A; TP | P55327; H0Y | TPD52_HUMAN | Unmodified | 0 | 0 | 0 | 0 | 6856100 | 658330 | 288400 | 721600 | 3849600 | 7472100 | 392580 | 449920 | 0 | 1109300 | 0 | 0 | 2312500 | 1747900 |
| SFEENVND | ZFYVE16 | Q7Z3T8 | ZFY16_HUMAN | Unmodified | 0 | 0 | 0 | 0 | 465760 | 0 | 0 | 0 | 0 | 0 | 0 | 200300 | 0 | 0 | 0 | 0 | 0 | 0 |
| SFEERVETT | TPD52L1; TPD5 | Q16890; E9F | TPD53_HUMAN | Unmodified | 56867 | 72940 | 155110 | 142930 | 519270 | 136280 | 0 | 0 | 0 | 0 | 0 | 0 | 2064100 | 0 | 164810 | 102560 | 0 | 0 |
| SFEEVINLV | ERN1; ERN1 | O75460; A0A | ERN1_HUMAN | Unmodified | 0 | 0 | 0 | 0 | 0 | 0 | 0 | 0 | 3885900 | 0 | 0 | 0 | 0 | 3614700 | 0 | 0 | 0 | 222060 |
| SFEGKTDR | IDE; IDE; IDE; ID | P14735; A0A | IDE_HUMAN In | Unmodified | 0 | 0 | 0 | 0 | 0 | 0 | 0 | 0 | 0 | 0 | 0 | 241720 | 0 | 0 | 0 | 0 | 0 | 0 |
| SFEGMVVA | TPBG | Q13641 | TPBG_HUMAN | Unmodified | 0 | 0 | 0 | 0 | 0 | 0 | 0 | 0 | 91340 | 0 | 0 | 79973 | 0 | 0 | 0 | 0 | 0 | 0 |
| SFEGMVVA | TPBG | Q13641 | TPBG_HUMAN | Oxidation (M) | 0 | 18922 | 0 | 0 | 0 | 0 | 0 | 0 | 35189 | 61917 | 0 | 5345800 | 0 | 0 | 0 | 0 | 0 | 0 |
| SFEGNNN | NRP1; NRP1; N | O14786; E9F | NRP1_HUMAN | Unmodified | 259710 | 138710 | 698680 | 453420 | 1391600 | 450680 | 0 | 0 | 285820 | 0 | 0 | 0 | 233270 | 0 | 491240 | 1525800 | 712690 | 3785600 |
| SFEGNNN | NRP1; NRP1; N | O14786; E9F | NRP1_HUMAN | Unmodified | 0 | 0 | 0 | 60223 | 25881000 | 4007200 | 0 | 0 | 265140 | 158160 | 0 | 0 | 1515300 | 620360 | 0 | 0 | 9673700 | 69058000 |
| SFEGNVFM | TMEM30A; TME | Q9NV96; A0 | CC50A_HUMA | Oxidation (M) | 0 | 0 | 0 | 0 | 0 | 0 | 0 | 0 | 976480 | 876410 | 0 | 0 | 0 | 0 | 0 | 0 | 0 | 0 |
| SFEGNVFM | TMEM30A; TME | Q9NV96; A0 | CC50A_HUMA | Unmodified | 0 | 0 | 0 | 0 | 0 | 0 | 0 | 0 | 3004400 | 386670 | 26113 | 0 | 0 | 0 | 0 | 0 | 0 | 0 |
| SFEGNVFM | TMEM30A; TME | Q9NV96; A0 | CC50A_HUMA | Oxidation (M) | 0 | 0 | 0 | 604610 | 0 | 0 | 0 | 0 | 390500 | 1285400 | 3095200 | 0 | 844400 | 0 | 0 | 0 | 0 | 0 |
| SFEGRELL | CPE | P16870 | CBPE_HUMAN | Unmodified | 0 | 0 | 0 | 0 | 0 | 0 | 0 | 0 | 928670 | 2229400 | 0 | 0 | 0 | 0 | 0 | 0 | 0 | 0 |
| SFENDAQS | INPPL1; INPPL1 | O15357; A0A | SHIP2_HUMAN | Unmodified | 0 | 0 | 0 | 0 | 0 | 0 | 0 | 0 | 178980 | 77107 | 0 | 1488900 | 0 | 0 | 0 | 0 | 0 | 0 |
| SFENIQNW | RAB13; RAB13 | P51153; A0A | RAB13_HUMAN | Unmodified | 0 | 0 | 0 | 0 | 1472800 | 0 | 0 | 0 | 862980 | 0 | 228730 | 0 | 0 | 0 | 0 | 0 | 285260 | 712060 |
| SFENIQNW | RAB13; RAB13 | P51153; A0A | RAB13_HUMAN | Oxidation (M) | 1538900 | 0 | 3904400 | 0 | 551240 | 343230 | 0 | 0 | 0 | 0 | 140910 | 571370 | 0 | 0 | 2965100 | 919830 | 0 | 205660 |
| SFENISK | RAB10 | P61026 | RAB10_HUMAN | Unmodified | 0 | 0 | 0 | 0 | 1535700 | 686080 | 781490 | 417200 | 0 | 437720 | 1319200 | 6187400 | 0 | 0 | 0 | 0 | 0 | 0 |
| SFENKPPA | TJP1; TJP1; TJP1 | Q07157; A0A | ZO1_HUMAN T | Unmodified | 0 | 0 | 571020 | 709970 | 4800300 | 0 | 0 | 189580 | 0 | 311540 | 0 | 0 | 6260400 | 3292100 | 1204400 | 0 | 0 | 0 |
| SFENSLGIN | UGP2; UGP2; U | Q16851; E7E | UGPA_HUMAN | Unmodified | 0 | 0 | 0 | 0 | 2656100 | 0 | 0 | 0 | 2122700 | 1974200 | 0 | 6311900 | 0 | 0 | 0 | 0 | 0 | 0 |
| SFEQISANI | PFKM; PFKM; PI | P08237; A0A | PFKAM_HUMAN | Unmodified | 0 | 0 | 0 | 293140 | 72871 | 0 | 1206500 | 0 | 0 | 0 | 457630 | 2959600 | 0 | 0 | 472840 | 333750 | 566210 | 0 |
| SFESTVGQ | M6PR; M6PR; M | P20645; F5G | MPRD_HUMAN | Unmodified | 0 | 0 | 0 | 0 | 0 | 0 | 0 | 0 | 43015000 | 0 | 0 | 4901600 | 0 | 0 | 0 | 0 | 0 | 0 |
| SFESTVGQ | M6PR; M6PR; M | P20645; F5G | MPRD_HUMAN | Unmodified | 0 | 0 | 0 | 0 | 0 | 0 | 0 | 0 | 4946600 | 1763600 | 3991500 | 0 | 0 | 0 | 0 | 0 | 152680 | 0 |
| SFEVVFND | TXNIP | Q9H3M7 | TXNIP_HUMAN | Unmodified | 0 | 0 | 0 | 0 | 1317200 | 0 | 0 | 152290 | 0 | 0 | 0 | 0 | 298640 | 0 | 0 | 0 | 0 | 0 |
| SFFDNISSE | LSM14B; LSM1 | Q9BX40; Q5 | LS14B_HUMAN | Unmodified | 530420 | 624120 | 508890 | 1253100 | 0 | 0 | 0 | 0 | 114900 | 0 | 0 | 373100 | 2424200 | 904900 | 894290 | 381830 | 223620 | 0 |
| SFFHQHYL | BAG6; BAG6; B | P46379; A0A | BAG6_HUMAN | Unmodified | 0 | 0 | 0 | 0 | 0 | 0 | 0 | 0 | 0 | 0 | 0 | 2921400 | 0 | 0 | 0 | 188110 | 0 | 0 |
| SFGDTKFV | DPP3; DPP3; DI | Q9NY33; E9I | DPP3_HUMAN | Unmodified | 0 | 0 | 0 | 0 | 0 | 98095 | 0 | 223840 | 0 | 441440 | 50012 | 0 | 690690 | 286640 | 0 | 0 | 1009700 | 1679900 |
| SFGMTPVL | NAGLU; NAGLU | P54802; K7E | ANAG_HUMAN | Oxidation (M) | 0 | 0 | 0 | 0 | 0 | 0 | 1464500 | 0 | 1555300 | 31269 | 0 | 0 | 0 | 0 | 0 | 0 | 0 | 0 |
| SFGNPSLD | FAM20B | O75063 | XYLK_HUMAN | Unmodified | 0 | 0 | 0 | 0 | 0 | 0 | 0 | 0 | 10772000 | 0 | 509480 | 0 | 0 | 0 | 0 | 0 | 0 | 0 |
| SFGQTPSQ | LRBA; LRBA; LR | P50851; A0A | LRBA_HUMAN | Unmodified | 0 | 0 | 0 | 0 | 0 | 0 | 0 | 0 | 0 | 0 | 0 | 2962000 | 0 | 0 | 0 | 0 | 0 | 0 |
| SFGSAQEF | COPB2; COPB2 | P35606; H0Y | COPB2_HUMA | Unmodified | 92346 | 86279 | 0 | 0 | 0 | 0 | 0 | 0 | 375660 | 0 | 107940 | 2137400 | 0 | 0 | 389110 | 258490 | 0 | 0 |
| SFGSAQEF | COPB2; COPB2 | P35606; H0Y | COPB2_HUMA | Unmodified | 0 | 0 | 0 | 0 | 2628500 | 0 | 940500 | 0 | 1329400 | 1368600 | 0 | 387570 | 1161300 | 1058000 | 440940 | 249600 | 1089700 | 0 |
| SFGTRPLS | TNKS1BP1 | Q9C0C2 | TB182_HUMAN | Unmodified | 0 | 0 | 0 | 751370 | 0 | 0 | 0 | 0 | 0 | 0 | 0 | 2434100 | 0 | 0 | 2327300 | 1415800 | 0 | 0 |
| SFHFDPLS | ZDHHC5; ZDHH | Q9C0B5; H0 | ZDHC5_HUMA | Unmodified | 0 | 0 | 1918000 | 1512100 | 2277800 | 136660 | 0 | 0 | 962150 | 536880 | 866110 | 2947100 | 1589500 | 891350 | 2207800 | 1630400 | 1527600 | 1469400 |
| SFHFDPLS | ZDHHC5; ZDHH | Q9C0B5; H0 | ZDHC5_HUMA | Unmodified | 0 | 305970 | 0 | 0 | 8854200 | 0 | 159230 | 81772 | 0 | 0 | 0 | 0 | 7959200 | 10657000 | 0 | 0 | 11728000 | 13164000 |
| SFHNELLT | BAIAP2; BAIAP2 | Q9UQB8; I3L | BAIP2_HUMAN | Unmodified | 0 | 295850 | 865620 | 140910 | 0 | 0 | 0 | 0 | 0 | 0 | 0 | 0 | 0 | 0 | 1755000 | 2674100 | 0 | 0 |
| SFHNELLT | BAIAP2; BAIAP2 | Q9UQB8; I3L | BAIP2_HUMAN | Unmodified | 0 | 0 | 471310 | 5313700 | 0 | 0 | 0 | 0 | 0 | 0 | 0 | 0 | 0 | 0 | 0 | 1824200 | 0 | 0 |
| SFHNELLT | BAIAP2; BAIAP2 | Q9UQB8; I3L | BAIP2_HUMAN | Unmodified | 0 | 863330 | 0 | 0 | 2278900 | 0 | 124270 | 0 | 0 | 0 | 0 | 0 | 1590600 | 3704300 | 0 | 0 | 527240 | 882690 |
| SFHSFYQL | MYO1D; MYO1D | O94832; J3Q | MYO1D_HUMA | Unmodified | 3754400 | 1638600 | 49698 | 0 | 0 | 0 | 0 | 148190 | 1546200 | 0 | 0 | 0 | 0 | 0 | 744980 | 0 | 0 | 0 |

| | | | | | | | | | | | | | | | | | | | | | | |
|---|---|---|---|---|---|---|---|---|---|---|---|---|---|---|---|---|---|---|---|---|---|---|
| SFHSFYQLI | MYO1D; MYO1D | O94832; J3Q | MYO1D_HUMA | Oxidation (M) | 13711000 | 10522000 | 1462600 | 3701100 | 0 | 0 | 0 | 0 | 0 | 0 | 0 | 0 | 0 | 0 | 7722900 | 5850200 | 0 | 0 |
| SFHTCPGL | CHST1 | O43916 | CHST1_HUMAI | Unmodified | 0 | 0 | 0 | 0 | 0 | 0 | 0 | 0 | 1513800 | 0 | 0 | 0 | 0 | 0 | 0 | 0 | 0 | 0 |
| SFIDPYQHI | PLA2G4A | P47712 | PA24A_HUMAN | Acetyl (Protein N | 0 | 0 | 0 | 0 | 0 | 0 | 0 | 0 | 0 | 0 | 245530 | 2043300 | 0 | 0 | 0 | 0 | 0 | 0 |
| SFIKDYPVV | ENO1; ENO1 | P06733; A0A | ENOA_HUMAN | Unmodified | 0 | 0 | 0 | 0 | 34597000 | 1244000 | 3137600 | 0 | 0 | 1637100 | 0 | 0 | 14933000 | 9222200 | 0 | 0 | 20107000 | 43577000 |
| SFITTDVNP | LARS1; LARS1; | Q9P2J5; A0A | SYLC_HUMAN | Unmodified | 0 | 0 | 12791 | 0 | 0 | 49336 | 100600 | 0 | 0 | 0 | 329240 | 1896500 | 0 | 0 | 231050 | 0 | 0 | 0 |
| SFITWLR | SEC24B | O95487 | SC24B_HUMAI | Unmodified | 0 | 0 | 0 | 0 | 0 | 0 | 0 | 0 | 0 | 0 | 0 | 1294100 | 0 | 0 | 0 | 0 | 0 | 0 |
| SFKDEWTQ | AASDHPPT; AAS | Q9NRN7; E9 | ADPPT_HUMAI | Oxidation (M) | 42800 | 0 | 0 | 0 | 0 | 0 | 0 | 0 | 159760 | 0 | 0 | 1556200 | 0 | 0 | 121710 | 50710 | 0 | 0 |
| SFKHTHIDI | TM9SF2 | Q99805 | TM9S2_HUMAI | Unmodified | 0 | 0 | 0 | 0 | 2010700 | 0 | 0 | 0 | 813990 | 223070 | 0 | 852570 | 0 | 0 | 0 | 0 | 142180 | 0 |
| SFKLSGFSI | MARCKS | P29966 | MARCS_HUMA | Unmodified | 0 | 0 | 0 | 125390 | 476250 | 0 | 5627800 | 2529500 | 0 | 4326800 | 0 | 445560 | 1104800 | 1290900 | 0 | 0 | 11246000 | 11383000 |
| SFKSDEVT( | MAP4K5 | Q9Y4K4 | M4K5_HUMAN | Unmodified | 0 | 0 | 0 | 67871 | 0 | 0 | 159530 | 63709 | 527670 | 178930 | 158180 | 1105100 | 0 | 117680 | 0 | 0 | 115370 | 0 |
| SFLAFPTLR | AGRN; AGRN; A | O00468; A0A | AGRIN_HUMAN | Unmodified | 591160 | 0 | 1383400 | 0 | 0 | 0 | 0 | 0 | 1362900 | 335410 | 0 | 755060 | 275630 | 0 | 731660 | 2223500 | 0 | 0 |
| SFLDDFESI | WIPF2; WIPF2; | Q8TF74; J3K | WIPF2_HUMAN | Unmodified | 788830 | 796340 | 1429700 | 0 | 0 | 0 | 0 | 0 | 0 | 0 | 0 | 0 | 0 | 0 | 2642900 | 1101800 | 0 | 0 |
| SFLDDFESI | WIPF2; WIPF2 | Q8TF74; A8N | WIPF2_HUMAN | Unmodified | 497840 | 1914900 | 0 | 2121300 | 0 | 0 | 0 | 0 | 0 | 0 | 0 | 0 | 0 | 0 | 0 | 0 | 0 | 0 |
| SFLDDPIW | CYFIP1; CYFIP1 | Q7L576; H0Y | CYFP1_HUMAN | Unmodified | 117610 | 0 | 68943 | 0 | 0 | 0 | 224480 | 0 | 0 | 70914 | 0 | 994480 | 0 | 0 | 0 | 0 | 0 | 0 |
| SFLDSGYR | VCL | P18206 | VINC_HUMAN | Unmodified | 578030 | 833880 | 860810 | 819510 | 1230400 | 1261700 | 187110 | 71704 | 735520 | 323390 | 579460 | 5778800 | 501760 | 0 | 1305200 | 910690 | 1099400 | 190890 |
| SFLDSGYR | VCL | P18206 | VINC_HUMAN | Unmodified | 175410 | 204990 | 190770 | 941690 | 27521000 | 7728500 | 54332000 | 1189500 | 682130 | 12618000 | 1586400 | 745860 | 18050000 | 25742000 | 284820 | 136670 | 2.75E+08 | 21135000 |
| SFLEDIR | PGD; PGD | P52209; K7E | 6PGD_HUMAN | Unmodified | 0 | 0 | 0 | 0 | 0 | 0 | 0 | 0 | 1000000 | 609070 | 89615 | 4976900 | 0 | 0 | 0 | 0 | 0 | 0 |
| SFLEDIRK | PGD; PGD | P52209; K7E | 6PGD_HUMAN | Unmodified | 0 | 0 | 0 | 0 | 0 | 0 | 0 | 83595 | 2948000 | 99768 | 0 | 2782200 | 0 | 0 | 0 | 0 | 0 | 0 |
| SFLEEVLAS | VPS45; VPS45; | Q9NRW7; A0 | VPS45_HUMAN | Unmodified | 2063100 | 1121000 | 478320 | 0 | 0 | 0 | 130030 | 0 | 1689100 | 960270 | 557780 | 4815400 | 0 | 0 | 462360 | 1304300 | 0 | 0 |
| SFLEVLDGI | HOMER2 | Q9NSB8 | HOME2_HUMA | Unmodified | 0 | 0 | 0 | 0 | 0 | 0 | 0 | 0 | 0 | 0 | 0 | 0 | 1942000 | 0 | 0 | 0 | 0 | 0 |
| SFLFEPVVK | PPP4R3A; PPP4 | Q6IN85; G3 | P4R3A_HUMAN | Unmodified | 0 | 0 | 0 | 0 | 0 | 0 | 0 | 0 | 0 | 0 | 0 | 802970 | 0 | 0 | 0 | 0 | 0 | 0 |
| SFLFQLLK | CDK5 | Q00535 | CDK5_HUMAN | Unmodified | 218250 | 0 | 355120 | 148080 | 0 | 0 | 705270 | 0 | 0 | 290570 | 216610 | 590640 | 0 | 0 | 448000 | 416360 | 0 | 0 |
| SFLFSSR | MOB1A | Q9H8S9 | MOB1A_HUMA | Acetyl (Protein N | 174920 | 50174 | 0 | 0 | 2225100 | 580190 | 0 | 0 | 539800 | 85065 | 338190 | 1063100 | 844890 | 878110 | 0 | 0 | 814880 | 1587000 |
| SFLHEQEEI | IQGAP2; IQGAP | Q13576; D6I | IQGA2_HUMAN | Unmodified | 0 | 0 | 1669900 | 868110 | 0 | 0 | 0 | 0 | 0 | 143150 | 0 | 481180 | 232560 | 153280 | 2412700 | 1549300 | 0 | 0 |
| SFLIEGPLIN | PLEKHN1 | Q494U1 | PKHN1_HUMAI | Unmodified | 74129 | 37393 | 42332 | 73021 | 0 | 10614 | 0 | 0 | 0 | 0 | 0 | 0 | 0 | 0 | 185790 | 115340 | 0 | 0 |
| SFLINFIHTI | PLXNB2 | O15031 | PLXB2_HUMAN | Unmodified | 188140 | 16148 | 300050 | 0 | 0 | 0 | 20739 | 0 | 73597 | 136000 | 0 | 687890 | 0 | 0 | 5409000 | 1022100 | 0 | 0 |
| SFLINFIHTI | PLXNB2 | O15031 | PLXB2_HUMAN | Unmodified | 48387 | 0 | 0 | 207770 | 65389 | 36874 | 0 | 0 | 60710 | 86610 | 0 | 75831 | 0 | 0 | 437870 | 61790 | 33370 | 46009 |
| SFLIWVNEI | CKMT1A; CKMT | P12532; F8V | KCRU_HUMAN | Unmodified | 519160 | 288050 | 815380 | 414350 | 0 | 0 | 0 | 0 | 543040 | 0 | 310920 | 7219300 | 175250 | 0 | 464050 | 439210 | 0 | 0 |
| SFLLALPAP | PIK3R2; PIK3R2 | O00459; E9F | P85B_HUMAN | Unmodified | 0 | 0 | 0 | 0 | 0 | 0 | 0 | 0 | 64958 | 0 | 0 | 84299 | 0 | 0 | 82604 | 95490 | 0 | 0 |
| SFLLALPAP | PIK3R2; PIK3R2 | O00459; E9F | P85B_HUMAN | Unmodified | 0 | 0 | 0 | 0 | 108830 | 0 | 0 | 0 | 163300 | 0 | 0 | 338750 | 0 | 0 | 0 | 0 | 0 | 0 |
| SFLLDLLNA | DDX3X; DDX3X; | O00571; A0A | DDX3X_HUMAI | Unmodified | 758220 | 756720 | 1210600 | 357940 | 0 | 0 | 0 | 0 | 303790 | 0 | 193300 | 666840 | 0 | 0 | 2422800 | 1078300 | 108810 | 0 |
| SFLMSLCPI | PLCB2 | Q00722 | PLCB2_HUMAI | Oxidation (M) | 0 | 0 | 0 | 0 | 0 | 0 | 865790 | 0 | 0 | 0 | 0 | 1421600 | 0 | 0 | 0 | 0 | 0 | 0 |
| SFLSQGQV | ATP5PO; ATP5P | P48047; H7C | ATPO_HUMAN | Unmodified | 0 | 0 | 128870 | 0 | 0 | 0 | 0 | 0 | 2866200 | 250420 | 0 | 276500 | 0 | 0 | 0 | 0 | 0 | 0 |
| SFLSQGQV | ATP5PO; ATP5P | P48047; H7C | ATPO_HUMAN | Unmodified | 0 | 0 | 0 | 0 | 0 | 124680 | 0 | 0 | 9004000 | 957590 | 0 | 0 | 0 | 0 | 0 | 0 | 0 | 0 |
| SFMALSQD | CAPRIN1; CAPF | Q14444; G3 | CAPR1_HUMAI | Unmodified | 0 | 0 | 0 | 0 | 0 | 0 | 0 | 0 | 805150 | 0 | 0 | 2156900 | 296000 | 0 | 0 | 0 | 0 | 0 |
| SFNIPLLVL | HDAC3 | O15379 | HDAC3_HUMA | Unmodified | 0 | 0 | 0 | 0 | 0 | 0 | 0 | 0 | 0 | 0 | 0 | 124990 | 0 | 0 | 0 | 0 | 0 | 0 |
| SFNLIIAK | CPT2; CPT2; CF | P23786; A0A | CPT2_HUMAN | Unmodified | 0 | 0 | 0 | 0 | 0 | 0 | 0 | 10982 | 800890 | 477960 | 0 | 46911 | 0 | 0 | 0 | 307900 | 0 | 0 |
| SFNLSALEI | APEH; APEH; AF | P13798; C9J | ACPH_HUMAN | Unmodified | 0 | 0 | 1339800 | 1617700 | 2232800 | 372550 | 1311800 | 278470 | 1888700 | 713270 | 3360800 | 13165000 | 0 | 3114700 | 1825200 | 0 | 3934200 | 4174800 |
| SFNMMSPI | ESPN; ESPN; ES | B1AK53; A0A | ESPN_HUMAN | 2 Oxidation (M) | 0 | 0 | 0 | 0 | 0 | 0 | 0 | 0 | 0 | 0 | 0 | 2047700 | 0 | 0 | 0 | 0 | 0 | 0 |
| SFNRGDVF | VIL1; AVIL | P09327; O75 | VILI_HUMAN Vi | Unmodified | 1026900 | 451240 | 0 | 0 | 0 | 0 | 137340 | 0 | 7898800 | 5922000 | 5781400 | 28062000 | 0 | 0 | 0 | 0 | 416840 | 2345000 |
| SFNWSKLF | DAAM1 | Q9Y4D1 | DAAM1_HUMA | Unmodified | 0 | 0 | 0 | 0 | 4703500 | 0 | 5781800 | 0 | 437620 | 1351400 | 0 | 0 | 1698400 | 1325900 | 0 | 0 | 0 | 0 |
| SFPAAAVLF | VPS37D; VPS37 | Q86XT2; C9J | VP37D_HUMAN | Unmodified | 0 | 722360 | 0 | 0 | 0 | 0 | 0 | 0 | 0 | 0 | 0 | 0 | 0 | 0 | 4710000 | 1907400 | 0 | 0 |
| SFPAAIEHT | UBA6; UBA6 | A0AVT1; H0Y | UBA6_HUMAN | Unmodified | 74594 | 48365 | 75646 | 0 | 80453 | 0 | 0 | 0 | 0 | 243280 | 0 | 4077300 | 0 | 0 | 107150 | 91170 | 0 | 0 |
| SFPCGSEH | PLXNB2; PLXNE | O15031; E2F | PLXB2_HUMAN | Unmodified | 50981 | 0 | 0 | 353640 | 111910 | 64473 | 0 | 0 | 402590 | 144590 | 0 | 228870 | 218310 | 2609600 | 432310 | 472190 | 53053 | 84416 |
| SFPDFPTP( | APRT; APRT; AP | P07741; H3E | APT_HUMAN A | Unmodified | 650920 | 567690 | 679400 | 0 | 2505800 | 0 | 0 | 0 | 517440 | 0 | 64787 | 14030000 | 0 | 0 | 2142400 | 1622500 | 0 | 0 |
| SFPDFPTP( | APRT; APRT; AP | P07741; H3E | APT_HUMAN A | Unmodified | 0 | 29992 | 0 | 349340 | 848830 | 101400 | 0 | 0 | 0 | 164760 | 22366 | 0 | 2067000 | 1052400 | 30748 | 24432 | 513980 | 395600 |
| SFPDKAPV | TJP1; TJP1; TJP1 | Q07157; A0A | ZO1_HUMAN T | Unmodified | 0 | 0 | 0 | 320300 | 2277500 | 558300 | 47148 | 0 | 0 | 53953 | 0 | 0 | 1541900 | 968540 | 0 | 50032 | 551920 | 350650 |
| SFPDQFST | CPD | O75976 | CBPD_HUMAN | Unmodified | 0 | 0 | 328780 | 782830 | 0 | 0 | 0 | 43808 | 70897000 | 6296700 | 536240 | 0 | 0 | 0 | 0 | 0 | 0 | 0 |
| SFPHLSKP | C1orf226; N/A; | A1L170; F8V | CA226_HUMAI | Unmodified | 0 | 0 | 0 | 0 | 1725800 | 5807600 | 0 | 0 | 0 | 0 | 0 | 0 | 2681200 | 2904200 | 0 | 0 | 0 | 0 |
| SFPKDDPS | RPL3; RPL3; RP | P39023; H7C | RL3_HUMAN 6 | Unmodified | 4229200 | 10286000 | 0 | 894320 | 3380900 | 752600 | 5461100 | 440140 | 811630 | 610660 | 2550800 | 17097000 | 1404900 | 938420 | 0 | 0 | 0 | 805790 |
| SFPLHFDEI | ERO1A; ERO1A | Q96HE7; G3 | ERO1A_HUMAI | Unmodified | 0 | 0 | 0 | 0 | 306760 | 0 | 0 | 0 | 8840700 | 1198900 | 0 | 0 | 0 | 0 | 0 | 21940 | 0 | 0 |
| SFPLHFDEI | ERO1A; ERO1A | Q96HE7; G3 | ERO1A_HUMAI | Unmodified | 0 | 0 | 0 | 0 | 0 | 0 | 0 | 0 | 0 | 834390 | 0 | 0 | 0 | 0 | 0 | 0 | 0 | 0 |
| SFPPPGPA | CLNS1A; CLNS | P54105; E9P | ICLN_HUMAN I | Unmodified | 0 | 0 | 0 | 0 | 0 | 0 | 0 | 0 | 0 | 0 | 0 | 2033400 | 0 | 0 | 0 | 0 | 0 | 0 |
| SFPPPGPA | CLNS1A; CLNS | P54105; E9P | ICLN_HUMAN I | Unmodified | 0 | 0 | 0 | 0 | 1365600 | 0 | 0 | 0 | 0 | 0 | 0 | 0 | 0 | 0 | 0 | 0 | 1223700 | 745770 |
| SFPQLRATI | ANXA7 | P20073 | ANXA7_HUMAI | Unmodified | 0 | 0 | 109290 | 0 | 1104100 | 0 | 0 | 0 | 2146800 | 5261300 | 0 | 0 | 886110 | 0 | 0 | 0 | 1115500 | 9066500 |
| SFPQLRATI | ANXA7 | P20073 | ANXA7_HUMAI | Oxidation (M) | 0 | 0 | 0 | 0 | 0 | 93896 | 0 | 0 | 0 | 1727000 | 0 | 0 | 157040 | 80572 | 0 | 0 | 0 | 0 |
| SFPRDELM | CRTAP; CRTAP | O75718; C9J | CRTAP_HUMAI | Unmodified | 0 | 0 | 0 | 0 | 0 | 0 | 0 | 0 | 985900 | 0 | 0 | 0 | 0 | 0 | 0 | 0 | 0 | 0 |
| SFPVNSDV | ARCN1; ARCN1 | P48444; B0Y | COPD_HUMAN | Unmodified | 0 | 0 | 328270 | 0 | 1209800 | 0 | 1140500 | 387170 | 939740 | 716070 | 672390 | 5069900 | 2187900 | 0 | 0 | 324870 | 933020 | 1233400 |
| SFQGDDSI | LIMCH1; LIMCH | Q9UPQ0; HC | LIMC1_HUMAN | Unmodified | 0 | 0 | 0 | 0 | 0 | 0 | 545860 | 994730 | 0 | 260170 | 0 | 572510 | 1400600 | 2019600 | 478750 | 0 | 30276 | 0 |
| SFQNYIRQ | RASA3 | Q14644 | RASA3_HUMAI | Unmodified | 0 | 0 | 0 | 0 | 681450 | 0 | 190960 | 255900 | 0 | 157660 | 0 | 0 | 0 | 0 | 0 | 0 | 3350800 | 3469000 |
| SFQQELDA | TSNAX; TSNAX; | Q99598; Q5 | TSNAX_HUMAI | Unmodified | 0 | 0 | 0 | 0 | 0 | 0 | 0 | 0 | 0 | 0 | 0 | 3061900 | 0 | 0 | 0 | 0 | 0 | 0 |
| SFQRALELI | DNAJC7; DNAJ | Q99615; A0A | DNJC7_HUMAI | Unmodified | 0 | 0 | 0 | 0 | 527190 | 0 | 173100 | 110450 | 1256400 | 0 | 0 | 0 | 716320 | 1598300 | 0 | 0 | 0 | 220290 |
| SFRDFKVQ | ABHD12; ABHD | Q8N2K0; A0A | ABD12_HUMAI | Unmodified | 0 | 0 | 0 | 0 | 0 | 0 | 0 | 0 | 1069700 | 487390 | 0 | 0 | 0 | 0 | 0 | 0 | 0 | 0 |
| SFRILAQIT | PDLIM5; PDLIM | Q96HC4; HC | PDLI5_HUMAN | Unmodified | 0 | 0 | 0 | 43663 | 812910 | 0 | 0 | 0 | 0 | 160420 | 0 | 0 | 0 | 0 | 0 | 0 | 2357100 | 1165300 |
| SFRVTIPTD | DAG1; DAG1; D | Q14118; C9J | DAG1_HUMAN | Unmodified | 0 | 0 | 41193 | 0 | 0 | 125720 | 0 | 0 | 1127100 | 480210 | 131920 | 0 | 121200 | 136460 | 0 | 0 | 0 | 83399 |
| SFRWLHNI | TALDO1 | P37837 | TALDO_HUMAI | Oxidation (M) | 0 | 0 | 0 | 0 | 7889400 | 743200 | 0 | 0 | 0 | 1565800 | 0 | 0 | 3209100 | 1332400 | 0 | 0 | 1649100 | 1711600 |
| SFRWLHNI | TALDO1 | P37837 | TALDO_HUMAI | Unmodified | 0 | 0 | 256910 | 0 | 1766700 | 293070 | 0 | 0 | 0 | 0 | 260500 | 0 | 0 | 0 | 0 | 0 | 494680 | 487400 |
| SFRWYLEN | GALNT16 | Q8N428 | GLT16_HUMAN | Unmodified | 0 | 0 | 0 | 0 | 0 | 0 | 0 | 0 | 1908300 | 1963200 | 0 | 0 | 0 | 0 | 0 | 0 | 0 | 0 |
| SFSCSATLI | ICAM1; ICAM1 | P05362; E7E | ICAM1_HUMAN | Unmodified | 0 | 0 | 382200 | 617990 | 1703700 | 1236700 | 0 | 0 | 0 | 0 | 0 | 0 | 0 | 0 | 773180 | 549250 | 0 | 0 |
| SFSEDAVTI | INF2; INF2; INF | Q27J81; A0A | INF2_HUMAN I | Unmodified | 0 | 0 | 0 | 0 | 0 | 0 | 0 | 0 | 0 | 507330 | 0 | 0 | 0 | 0 | 0 | 0 | 0 | 625270 |
| SFSEFGKLI | HNRNPR; HNRI | O43390; B4I | HNRPR_HUMA | Unmodified | 0 | 0 | 0 | 0 | 1235700 | 229000 | 0 | 0 | 0 | 0 | 0 | 873920 | 149070 | 0 | 0 | 0 | 0 | 2042300 |
| SFSEFGKLI | HNRNPR; HNRI | O43390; B4I | HNRPR_HUMA | Unmodified | 0 | 0 | 0 | 0 | 252750 | 0 | 511440 | 284100 | 0 | 151660 | 576330 | 0 | 368050 | 110340 | 0 | 0 | 0 | 0 |
| SFSELEEK | ULK3 | Q6PHR2 | ULK3_HUMAN | Unmodified | 0 | 0 | 47273 | 0 | 0 | 0 | 0 | 0 | 0 | 0 | 184210 | 308710 | 0 | 0 | 0 | 0 | 0 | 100360 |
| SFSEPFGRI | MLF1; MLF1; MI | P58340; A0A | MLF1_HUMAN | Unmodified | 412680 | 421590 | 348040 | 441990 | 0 | 0 | 31238 | 0 | 0 | 0 | 0 | 0 | 432010 | 0 | 367190 | 258060 | 0 | 1539500 |
| SFSEVELHI | DAG1 | Q14118 | DAG1_HUMAN | Unmodified | 0 | 0 | 171550 | 174240 | 94073 | 106720 | 0 | 0 | 1378000 | 240330 | 132870 | 0 | 0 | 0 | 97832 | 0 | 0 | 0 |
| SFSKESDD | HSP90B1; HSP | P14625; A0A | ENPL_HUMAN | Oxidation (M) | 0 | 0 | 0 | 0 | 0 | 0 | 0 | 0 | 381880 | 3578700 | 0 | 0 | 0 | 0 | 0 | 0 | 0 | 0 |
| SFSKLPIQL | CHRM3 | P20309 | ACM3_HUMAN | Unmodified | 0 | 94091 | 0 | 0 | 0 | 0 | 0 | 0 | 697160 | 65400 | 0 | 0 | 775890 | 0 | 0 | 65912 | 0 | 2364600 |
| SFSKSDLVI | NRDC; NRDC; I | O43847; G3 | NRDC_HUMAN | Unmodified | 0 | 0 | 0 | 0 | 996380 | 0 | 0 | 0 | 0 | 388190 | 0 | 0 | 0 | 0 | 0 | 0 | 664960 | 0 |
| SFSLGDIYF | IGF2R; IGF2R | P11717; S4F | MPRI_HUMAN | Unmodified | 0 | 0 | 1077200 | 0 | 0 | 0 | 729820 | 0 | 57834000 | 15287000 | 5298900 | 7213600 | 955330 | 1140900 | 0 | 717350 | 0 | 0 |
| SFSNLITNL | ABCC4; ABCC4 | O15439; A0A | MRP4_HUMAN | Unmodified | 0 | 0 | 1451100 | 0 | 0 | 0 | 0 | 0 | 0 | 0 | 0 | 0 | 0 | 0 | 1661400 | 0 | 0 | 0 |
| SFSSDRSY | ANK3; ANK3; AI | Q12955; B1A | ANK3_HUMAN | Unmodified | 0 | 0 | 0 | 0 | 0 | 0 | 0 | 0 | 0 | 0 | 0 | 0 | 393220 | 281710 | 0 | 0 | 0 | 0 |
| SFSTALYGI | ACTN4; ACTN1; | O43707; P12 | ACTN4_HUMAI | Unmodified | 0 | 0 | 159750 | 327240 | 8566100 | 1401900 | 957110 | 0 | 1538400 | 1322400 | 600590 | 2335900 | 2658200 | 0 | 0 | 284060 | 4934600 | 1977300 |
| SFSVILEHF | ATP13A3; ATP1 | Q9H7F0; H7 | AT133_HUMAN | Unmodified | 0 | 0 | 0 | 271090 | 541250 | 122860 | 0 | 0 | 1638200 | 305380 | 213040 | 807360 | 153320 | 0 | 0 | 0 | 0 | 122300 |
| SFSWGIDP | SORL1 | Q92673 | SORL_HUMAN | Unmodified | 2718000 | 0 | 0 | 0 | 0 | 0 | 0 | 0 | 1053800 | 94334 | 0 | 0 | 0 | 0 | 0 | 0 | 0 | 0 |
| SFSWGIDP | SORL1 | Q92673 | SORL_HUMAN | Unmodified | 0 | 0 | 0 | 0 | 0 | 0 | 0 | 0 | 2750100 | 0 | 0 | 0 | 0 | 0 | 0 | 0 | 0 | 0 |
| SFTASCPV | FAHD1 | Q6P587 | FAHD1_HUMAI | Unmodified | 0 | 0 | 0 | 0 | 0 | 0 | 0 | 0 | 1896400 | 0 | 0 | 0 | 0 | 0 | 0 | 0 | 103790 | 0 |
| SFTETMSSI | MYG1; MYG1 | Q9HB07; F8 | MYG1_HUMAN | Unmodified | 0 | 0 | 0 | 0 | 0 | 0 | 54917 | 0 | 3695500 | 0 | 438670 | 2072000 | 225930 | 0 | 0 | 0 | 382480 | 145710 |
| SFTIKPVGF | ITGB3; N/A | P05106; H3E | ITB3_HUMAN I | Unmodified | 1296600 | 1636100 | 1609800 | 5666600 | 1639300 | 0 | 3073700 | 1139400 | 0 | 0 | 0 | 0 | 2379200 | 2615000 | 2080500 | 1863000 | 0 | 818360 |
| SFTLDDESI | PROSER2; PRO | Q86WR7; X6 | PRSR2_HUMAN | Unmodified | 0 | 0 | 0 | 0 | 4813300 | 0 | 0 | 0 | 0 | 0 | 0 | 0 | 0 | 0 | 74984 | 0 | 0 | 633150 |
| SFTLDDESI | PROSER2; PRO | Q86WR7; X6 | PRSR2_HUMAN | Unmodified | 0 | 0 | 0 | 0 | 986090 | 0 | 0 | 0 | 0 | 0 | 0 | 0 | 0 | 0 | 0 | 0 | 0 | 0 |
| SFTPDHVV | CAVIN1 | Q6NZI2 | CAVN1_HUMAI | Unmodified | 0 | 0 | 0 | 0 | 10593000 | 0 | 0 | 26879 | 0 | 278580 | 0 | 0 | 0 | 0 | 0 | 0 | 0 | 0 |
| SFTPVFLGS | GFM1; GFM1; G | Q96RP9; C9I | EFGM_HUMAN | Unmodified | 0 | 0 | 0 | 0 | 0 | 0 | 0 | 0 | 1605800 | 0 | 0 | 0 | 0 | 0 | 0 | 0 | 0 | 0 |
| SFTRARDYI | RUVBL2; RUVBI | Q9Y230; M0I | RUVB2_HUMAI | Unmodified | 0 | 0 | 0 | 0 | 0 | 0 | 92927 | 74384 | 0 | 0 | 0 | 0 | 228170 | 175580 | 0 | 0 | 0 | 0 |
| SFTSALTSR | SORCS2; SORC | Q96PQ0; B5 | SORC2_HUMAI | Unmodified | 267210 | 1233300 | 0 | 0 | 0 | 0 | 0 | 0 | 4326500 | 67743 | 0 | 0 | 0 | 0 | 0 | 0 | 49110 | 0 |
| SFTSEDDL | AP3B1 | O00203 | AP3B1_HUMAN | Unmodified | 0 | 0 | 28763 | 0 | 0 | 0 | 0 | 0 | 0 | 0 | 0 | 1644500 | 0 | 0 | 0 | 17050 | 0 | 0 |
| SFTVMELVI | LACTB2 | Q53H82 | LACB2_HUMAN | Unmodified | 0 | 0 | 0 | 0 | 0 | 0 | 0 | 0 | 4327300 | 498300 | 0 | 0 | 0 | 0 | 0 | 0 | 0 | 0 |
| SFVASNDE | PLXNB2; PLXNE | O15031; E2F | PLXB2_HUMAN | Unmodified | 446740 | 301560 | 151640 | 2180600 | 4421800 | 606490 | 0 | 0 | 1217100 | 0 | 0 | 0 | 1421700 | 0 | 1378300 | 870190 | 1520500 | 0 |
| SFVEKVMQ | MYH9 | P35579 | MYH9_HUMAN | Unmodified | 0 | 0 | 0 | 0 | 282010 | 0 | 0 | 0 | 490750 | 220970 | 1296500 | 0 | 0 | 0 | 214760 | 1670500 | 311760 | 0 |
| SFVESVEEI | MICAL3 | Q7RTP6 | MICA3_HUMAN | Unmodified | 0 | 0 | 0 | 0 | 1713000 | 0 | 0 | 0 | 0 | 0 | 0 | 0 | 0 | 0 | 0 | 0 | 0 | 0 |
| SFVETKADS | PLCG2; PLCG2 | P16885; A0A | PLCG2_HUMAI | Unmodified | 0 | 0 | 0 | 0 | 0 | 0 | 0 | 0 | 0 | 0 | 0 | 5784900 | 0 | 0 | 0 | 135010 | 0 | 0 |
| SFVFILEPK | PLCG2; PLCG2 | P16885; A0A | PLCG2_HUMAI | Unmodified | 0 | 0 | 0 | 0 | 0 | 0 | 0 | 0 | 0 | 0 | 7294400 | 30921000 | 0 | 0 | 0 | 0 | 0 | 0 |
| SFVLNLGK | LGALS1 | P09382 | LEG1_HUMAN | Unmodified | 0 | 0 | 0 | 0 | 0 | 0 | 0 | 0 | 415940 | 0 | 0 | 0 | 0 | 0 | 0 | 0 | 30830000 | 0 |
| SFVLNLGKI | LGALS1 | P09382 | LEG1_HUMAN | Unmodified | 0 | 0 | 0 | 0 | 3326300 | 450750 | 538150 | 0 | 640450 | 0 | 0 | 0 | 0 | 0 | 0 | 0 | 9976600 | 4801800 |
| SFVLNLGKI | LGALS1 | P09382 | LEG1_HUMAN | Unmodified | 0 | 0 | 0 | 0 | 1605000 | 0 | 0 | 0 | 0 | 0 | 0 | 0 | 86209 | 0 | 0 | 0 | 22054000 | 0 |
| SFVPNMIG | GSPT1; GSPT2; | P15170; Q8I | ERF3A_HUMAN | Oxidation (M) | 0 | 0 | 0 | 0 | 0 | 0 | 2957000 | 0 | 0 | 422530 | 0 | 0 | 8007200 | 3598300 | 0 | 0 | 4324200 | 4535600 |
| SFVQGLGV | GPC1; GPC1; G | P35052; H7C | GPC1_HUMAN | Unmodified | 0 | 0 | 0 | 2438600 | 1870300 | 0 | 0 | 0 | 505210 | 132000 | 0 | 0 | 581840 | 153080 | 0 | 0 | 0 | 0 |
| SFVVTSVV | ST14 | Q9Y5Y6 | ST14_HUMAN | Unmodified | 1608000 | 1763300 | 1330200 | 2859400 | 0 | 0 | 0 | 0 | 1142200 | 280420 | 0 | 0 | 1902500 | 1060600 | 3030800 | 1412400 | 0 | 0 |
| SFVYGGCL | SPINT1; SPINT1 | O43278; H3I | SPIT1_HUMAN | Unmodified | 840430 | 688530 | 714000 | 552960 | 0 | 0 | 0 | 0 | 0 | 0 | 387040 | 0 | 879050 | 805330 | 959010 | 621040 | 44077 | 0 |
| SFWELIGE | S100A14 | Q9HCY8 | S10AE_HUMAN | Unmodified | 0 | 0 | 0 | 0 | 0 | 0 | 0 | 0 | 2985800 | 602120 | 0 | 0 | 0 | 0 | 0 | 0 | 0 | 0 |

| | | | | | | | | | | | | | | | | | | | | | | |
|---|---|---|---|---|---|---|---|---|---|---|---|---|---|---|---|---|---|---|---|---|---|---|
| SFYGMADK | MGAT2 | Q10469 | MGAT2_HUMAI | Unmodified | 0 | 0 | 0 | 0 | 0 | 0 | 0 | 0 | 16989000 | 993620 | 0 | 0 | 0 | 0 | 0 | 0 | 0 | 0 |
| SFYGMADK | MGAT2 | Q10469 | MGAT2_HUMAI | Oxidation (M) | 0 | 0 | 0 | 28191 | 0 | 0 | 0 | 0 | 3652600 | 524810 | 0 | 714910 | 0 | 0 | 0 | 0 | 56179 | 0 |
| SFYGSTLFL | FASN; FASN | P49327; A0A | FAS_HUMAN F | Unmodified | 0 | 0 | 0 | 0 | 0 | 0 | 0 | 0 | 0 | 0 | 0 | 11248000 | 0 | 0 | 0 | 0 | 0 | 0 |
| SFYPEEVSS | HSPA8; HSPA8; | P11142; E9P | HSP7C_HUMAI | Oxidation (M) | 29987000 | 23716000 | 32530000 | 46402000 | 1.06E+08 | 23333000 | 57333000 | 26012000 | 31035000 | 41463000 | 15768000 | 80502000 | 1.45E+08 | 80281000 | 41058000 | 25604000 | 32213000 | 33844000 |
| SFYPEEVSS | HSPA8; HSPA8; | P11142; E9P | HSP7C_HUMAI | Unmodified | 0 | 8170800 | 2832800 | 10287000 | 1.35E+08 | 14550000 | 40085000 | 5200400 | 1.28E+08 | 38468000 | 5647900 | 29841000 | 96063000 | 72117000 | 10014000 | 3677100 | 25658000 | 79407000 |
| SFYTAYLQF | QSOX1 | O00391 | QSOX1_HUMAI | Unmodified | 0 | 0 | 0 | 219590 | 0 | 0 | 0 | 0 | 5144800 | 0 | 0 | 0 | 0 | 0 | 198180 | 0 | 0 | 204460 |
| SGAEAQTP | TGOLN2; TGOL | O43493; A0/ | TGON2_HUMA | Unmodified | 0 | 0 | 0 | 0 | 0 | 0 | 0 | 0 | 51826000 | 1915900 | 0 | 0 | 0 | 0 | 0 | 0 | 0 | 0 |
| SGAEDQTP | TGOLN2; TGOL | O43493; A0/ | TGON2_HUMA | Unmodified | 0 | 0 | 0 | 0 | 0 | 0 | 0 | 0 | 11231000 | 0 | 124790 | 0 | 0 | 0 | 0 | 0 | 0 | 0 |
| SGAEEQGF | TGOLN2; TGOL | O43493; A0/ | TGON2_HUMA | Unmodified | 0 | 0 | 0 | 20548 | 0 | 30043 | 18626 | 0 | 28375000 | 1389700 | 2118600 | 2393200 | 63917 | 0 | 42953 | 0 | 0 | 0 |
| SGAFGHLF | TUBB3; N/A; TU | Q13509; A0/ | TBB3_HUMAN | Unmodified | 0 | 0 | 0 | 0 | 12111000 | 1116700 | 1740800 | 0 | 10369000 | 2250600 | 740440 | 5753300 | 0 | 0 | 0 | 0 | 751740 | 1572000 |
| SGAGQPLC | PLSCR3; PLSCF | Q9NRY6; I3L | PLS3_HUMAN | Unmodified | 673690 | 0 | 0 | 0 | 0 | 0 | 0 | 0 | 0 | 0 | 0 | 0 | 0 | 0 | 696890 | 0 | 0 | 0 |
| SGAGVISVI | ELAPOR2; ELAP | A8MWY0; H7 | ELAP2_HUMAN | Unmodified | 0 | 399020 | 0 | 0 | 150370 | 143210 | 48494 | 224670 | 4347000 | 285230 | 0 | 1044700 | 0 | 0 | 0 | 0 | 0 | 0 |
| SGAGVPAV | NCSTN; NCSTN | Q92542; H0 | NICA_HUMAN | Unmodified | 0 | 0 | 0 | 0 | 0 | 0 | 0 | 0 | 8420700 | 1201200 | 232450 | 3068500 | 0 | 0 | 0 | 0 | 0 | 0 |
| SGALDVLQ | RPSA; RPSA; RF | P08865; A0A | RSSA_HUMAN | Acetyl (Protein N | 4422200 | 3842700 | 731000 | 3619100 | 7697000 | 650670 | 6036400 | 632620 | 1776500 | 7056000 | 7463200 | 20677000 | 10286000 | 5506900 | 1984300 | 3913800 | 1401600 | 1262100 |
| SGALDVLQ | RPSA; RPSA; RF | P08865; A0A | RSSA_HUMAN | Acetyl (Protein N | 0 | 0 | 0 | 0 | 1393800 | 126100 | 424210 | 0 | 910210 | 196550 | 184110 | 946720 | 811580 | 349590 | 0 | 0 | 0 | 0 |
| SGALLACG | PSMD2 | Q13200 | PSMD2_HUMA | Unmodified | 0 | 0 | 0 | 0 | 0 | 0 | 232190 | 124150 | 0 | 0 | 0 | 827390 | 0 | 0 | 0 | 0 | 0 | 0 |
| SGALQIESS | PTPRF; PTPRF | P10586; A2A | PTPRF_HUMAN | Unmodified | 1349800 | 0 | 0 | 0 | 0 | 0 | 0 | 0 | 1080800 | 0 | 0 | 0 | 0 | 0 | 0 | 0 | 0 | 0 |
| SGAPPPSG | SRPK1; SRPK1 | Q96SB4; H3 | SRPK1_HUMAN | Unmodified | 0 | 0 | 0 | 0 | 0 | 0 | 0 | 0 | 0 | 0 | 0 | 1534100 | 0 | 0 | 0 | 0 | 0 | 0 |
| SGAPVLSLI | PAAF1; PAAF1 | Q9BRP4; F5I | PAAF1_HUMAN | Unmodified | 34345000 | 22144000 | 2089800 | 0 | 0 | 0 | 0 | 0 | 181580 | 0 | 0 | 2064800 | 0 | 0 | 576440 | 15110000 | 0 | 0 |
| SGAQATWI | HPX | P02790 | HEMO_HUMAN | Unmodified | 0 | 0 | 0 | 434160 | 0 | 0 | 0 | 0 | 0 | 0 | 0 | 2997300 | 0 | 0 | 216100 | 0 | 0 | 0 |
| SGASLKLTL | ACLY | P53396 | ACLY_HUMAN | Unmodified | 0 | 0 | 0 | 0 | 2463500 | 0 | 9162500 | 2418900 | 0 | 998500 | 0 | 0 | 881050 | 438770 | 0 | 0 | 1755600 | 213840 |
| SGASVVAIF | HP1BP3; HP1BI | Q5SSJ5; B0C | HP1B3_HUMAN | Unmodified | 1585100 | 468470 | 759110 | 0 | 1961000 | 226600 | 0 | 0 | 1394100 | 260030 | 326830 | 459210 | 990580 | 0 | 847710 | 998300 | 406260 | 474380 |
| SGAVEETFI | HSDL2 | Q6YN16 | HSDL2_HUMAI | Unmodified | 0 | 0 | 0 | 0 | 0 | 0 | 0 | 0 | 2844200 | 220780 | 0 | 0 | 0 | 0 | 0 | 0 | 0 | 0 |
| SGAVEETFI | HSDL2 | Q6YN16 | HSDL2_HUMAI | Unmodified | 0 | 0 | 0 | 0 | 435220 | 1947900 | 1061200 | 1075300 | 0 | 321160 | 0 | 0 | 0 | 0 | 0 | 0 | 241530 | 2888200 |
| SGAVEETFI | HSDL2 | Q6YN16 | HSDL2_HUMAI | Unmodified | 0 | 0 | 0 | 0 | 0 | 0 | 0 | 0 | 618680 | 943610 | 0 | 0 | 0 | 0 | 0 | 0 | 0 | 0 |
| SGAWDESC | COPA; COPA; C | P53621; A0A | COPA_HUMAN | Unmodified | 0 | 0 | 0 | 29155 | 0 | 387730 | 0 | 0 | 472730 | 362580 | 0 | 5722200 | 0 | 0 | 0 | 0 | 0 | 0 |
| SGAWDESC | COPA; COPA; C | P53621; A0A | COPA_HUMAN | Unmodified | 0 | 0 | 0 | 0 | 0 | 0 | 0 | 0 | 0 | 5662800 | 0 | 232230 | 0 | 0 | 0 | 0 | 0 | 0 |
| SGAWGAQ | EHBP1L1 | Q8N3D4 | EH1L1_HUMAN | Oxidation (M) | 0 | 0 | 0 | 0 | 550650 | 0 | 0 | 0 | 0 | 0 | 0 | 53455 | 0 | 0 | 0 | 0 | 0 | 0 |
| SGAWGAQ | EHBP1L1 | Q8N3D4 | EH1L1_HUMAN | Unmodified | 0 | 0 | 0 | 0 | 1049800 | 0 | 0 | 0 | 113750 | 0 | 0 | 0 | 0 | 0 | 0 | 0 | 134570 | 0 |
| SGAYLFLPI | MAN2A1 | Q16706 | MA2A1_HUMAI | Unmodified | 0 | 0 | 0 | 0 | 0 | 0 | 0 | 0 | 1016000 | 676020 | 0 | 668830 | 0 | 0 | 0 | 0 | 0 | 0 |
| SGAYLIPLL | DDX6; DDX6 | P26196; Q8I | DDX6_HUMAN | Unmodified | 0 | 905230 | 387220 | 0 | 0 | 0 | 0 | 0 | 0 | 0 | 264220 | 6853300 | 0 | 0 | 761300 | 713810 | 0 | 0 |
| SGAYLIPLL | DDX6; DDX6 | P26196; Q8I | DDX6_HUMAN | Unmodified | 1728200 | 3058000 | 0 | 2918000 | 3812300 | 704210 | 2515700 | 0 | 1441100 | 3059600 | 1470800 | 7261500 | 6475900 | 2185300 | 2780600 | 2194100 | 1505700 | 683940 |
| SGCIVNNL | FLNB; FLNB | O75369; E7E | FLNB_HUMAN | Unmodified | 0 | 0 | 0 | 0 | 3616800 | 1043600 | 0 | 0 | 213370 | 36954 | 61298 | 1236800 | 0 | 307680 | 0 | 0 | 1286900 | 908500 |
| SGCIVNNL | FLNB; FLNB | O75369; E7E | FLNB_HUMAN | Unmodified | 0 | 0 | 0 | 0 | 8140100 | 0 | 653690 | 164270 | 0 | 0 | 0 | 0 | 0 | 3300700 | 0 | 0 | 10449000 | 1048500 |
| SGCKVDLC | GART | P22102 | PUR2_HUMAN | Unmodified | 0 | 0 | 0 | 0 | 413430 | 0 | 352110 | 0 | 0 | 283280 | 0 | 111050 | 894100 | 388830 | 0 | 0 | 753090 | 630630 |
| SGCTSLTP | BABAM2; BABA | Q9NXR7; F8 | BABA2_HUMAN | Unmodified | 34713 | 253200 | 0 | 80097 | 0 | 35058 | 52145 | 42700 | 0 | 0 | 0 | 771910 | 0 | 39972 | 38037 | 39408 | 0 | 0 |
| SGDAAIVD | EEF1A1P5; EEF | Q5VTE0; P68 | EF1A3_HUMAN | Oxidation (M) | 0 | 0 | 1109500 | 0 | 0 | 0 | 0 | 0 | 0 | 0 | 0 | 0 | 0 | 0 | 1177400 | 0 | 0 | 0 |
| SGDAAIVD | EEF1A1P5; EEF | Q5VTE0; P68 | EF1A3_HUMAN | Unmodified | 0 | 0 | 493340 | 0 | 0 | 0 | 0 | 0 | 0 | 0 | 0 | 0 | 0 | 0 | 351890 | 747720 | 0 | 0 |
| SGDAAIVD | EEF1A1P5; EEF | Q5VTE0; P68 | EF1A3_HUMAN | 2 Oxidation (M) | 1.02E+08 | 81189000 | 21315000 | 39745000 | 30469000 | 11103000 | 10306000 | 1478100 | 3030500 | 9633100 | 12510000 | 21728000 | 54786000 | 30361000 | 58021000 | 54007000 | 26506000 | 47350000 |
| SGDAAIVD | EEF1A1P5; EEF | Q5VTE0; P68 | EF1A3_HUMAN | Oxidation (M) | 16258000 | 29325000 | 1805100 | 23599000 | 25311000 | 0 | 0 | 0 | 13173000 | 2159600 | 958770 | 5083500 | 46739000 | 16482000 | 14165000 | 7856600 | 14208000 | 18880000 |
| SGDAAIVD | EEF1A1P5; EEF | Q5VTE0; P68 | EF1A3_HUMAN | Unmodified | 1153400 | 0 | 0 | 0 | 3311400 | 0 | 0 | 0 | 5990200 | 0 | 0 | 0 | 0 | 0 | 0 | 0 | 994160 | 2718200 |
| SGDAAIVEI | EEF1A2; EEF1A | Q05639; A0/ | EF1A2_HUMAN | 2 Oxidation (M) | 20630000 | 25230000 | 5404300 | 28065000 | 2132100 | 252210 | 4316000 | 0 | 0 | 0 | 0 | 0 | 7845800 | 2821600 | 37328000 | 15274000 | 2426300 | 5506200 |
| SGDAAIVEI | EEF1A2; EEF1A | Q05639; A0/ | EF1A2_HUMAN | Oxidation (M) | 1589300 | 1021200 | 0 | 3516200 | 2588700 | 779890 | 0 | 0 | 0 | 0 | 0 | 0 | 3514300 | 1365700 | 3924400 | 1627500 | 0 | 5960500 |
| SGDFSCG | LDLR; LDLR; LD | P01130; J3K | LDLR_HUMAN | Unmodified | 0 | 0 | 0 | 0 | 0 | 0 | 571330 | 0 | 2764300 | 0 | 0 | 0 | 0 | 0 | 0 | 0 | 0 | 0 |
| SGDGEGTF | CYFIP1 | Q7L576 | CYFP1_HUMAN | Unmodified | 531740 | 170740 | 1291500 | 355350 | 912190 | 142400 | 42085 | 68512 | 410130 | 0 | 249150 | 3752500 | 519050 | 279050 | 1594000 | 688820 | 178120 | 291500 |
| SGDITYGQ | PLCG1 | P19174 | PLCG1_HUMAI | Unmodified | 0 | 0 | 0 | 0 | 0 | 0 | 0 | 0 | 77234 | 0 | 0 | 1445200 | 0 | 0 | 0 | 53432 | 364840 | 0 |
| SGDLGDM | CTNND1; CTNN | O60716; H0 | CTND1_HUMAN | Oxidation (M) | 0 | 0 | 61608 | 0 | 0 | 0 | 0 | 0 | 0 | 0 | 638680 | 192530 | 0 | 0 | 202310 | 1180000 | 0 | 0 |
| SGDLPYDC | LSR; LSR | Q86X29; S4F | LSR_HUMAN Li | Unmodified | 0 | 0 | 375010 | 0 | 136890 | 59669 | 0 | 0 | 12097000 | 1281000 | 1877800 | 319820 | 0 | 0 | 0 | 0 | 0 | 0 |
| SGDLPYDC | LSR; LSR | Q86X29; S4F | LSR_HUMAN Li | Unmodified | 74973 | 71149 | 2730300 | 6361800 | 0 | 0 | 0 | 0 | 24096000 | 289010 | 1110100 | 177230 | 0 | 0 | 145200 | 175680 | 0 | 0 |
| SGDLPYDC | LSR; LSR | Q86X29; S4F | LSR_HUMAN Li | Unmodified | 3324900 | 0 | 4370100 | 6305500 | 1205000 | 1380000 | 0 | 0 | 14369000 | 1292900 | 1699700 | 0 | 4715100 | 1079200 | 4093700 | 3707200 | 163020 | 295550 |
| SGDLYVLA | IARS2 | Q9NSE4 | SYIM_HUMAN | Unmodified | 0 | 0 | 0 | 0 | 0 | 0 | 110080 | 0 | 1752400 | 459830 | 0 | 0 | 0 | 0 | 42063 | 28997 | 0 | 0 |
| SGDNPVYI | ABHD12; ABHD | Q8N2K0; A0/ | ABD12_HUMAN | Unmodified | 0 | 0 | 0 | 0 | 0 | 0 | 0 | 0 | 28419000 | 4022200 | 0 | 0 | 0 | 0 | 0 | 0 | 53520 | 0 |
| SGDNPVYI | ABHD12; ABHD | Q8N2K0; A0/ | ABD12_HUMAN | Unmodified | 0 | 0 | 0 | 0 | 0 | 0 | 0 | 0 | 26426000 | 0 | 0 | 0 | 0 | 0 | 1260700 | 975190 | 0 | 0 |
| SGDPMILS | GLG1; GLG1 | Q92896; H3I | GSLG1_HUMAI | Oxidation (M) | 0 | 0 | 0 | 0 | 0 | 0 | 0 | 99541 | 6925900 | 609150 | 341980 | 250290 | 0 | 0 | 0 | 0 | 0 | 0 |
| SGDPMILS | GLG1; GLG1 | Q92896; H3I | GSLG1_HUMAI | Unmodified | 0 | 0 | 0 | 38061 | 0 | 0 | 0 | 0 | 793930 | 0 | 0 | 0 | 0 | 0 | 0 | 0 | 0 | 13526 |
| SGDPMILS | GLG1; GLG1 | Q92896; H3I | GSLG1_HUMAI | 2 Oxidation (M) | 0 | 0 | 0 | 0 | 0 | 0 | 0 | 0 | 1873700 | 692000 | 0 | 295980 | 0 | 0 | 0 | 0 | 0 | 0 |
| SGDSEVYQ | SSBP1; SSBP1; | Q04837; C9 | SSBP_HUMAN | Unmodified | 118620 | 0 | 0 | 0 | 0 | 0 | 0 | 105670 | 10416000 | 1911800 | 0 | 6420100 | 0 | 0 | 0 | 0 | 0 | 0 |
| SGDTLLLLF | TMEM43; N/A | Q9BTV4; V9 | TMM43_HUMAN | Unmodified | 0 | 0 | 0 | 0 | 0 | 0 | 0 | 0 | 29590000 | 5676900 | 0 | 0 | 0 | 0 | 0 | 0 | 1448900 | 0 |
| SGDTLLLLF | TMEM43; N/A | Q9BTV4; V9 | TMM43_HUMAN | Unmodified | 0 | 0 | 0 | 0 | 0 | 0 | 34001 | 0 | 34795000 | 0 | 1381200 | 0 | 0 | 0 | 0 | 0 | 0 | 0 |
| SGDYWIDF | COL2A1 | P02458 | CO2A1_HUMAI | Oxidation (M) | 0 | 0 | 0 | 0 | 0 | 65115 | 0 | 0 | 0 | 0 | 3967100 | 0 | 0 | 0 | 0 | 0 | 0 | 37977 |
| SGDYWIDF | COL2A1 | P02458 | CO2A1_HUMAI | Unmodified | 0 | 0 | 0 | 0 | 0 | 0 | 0 | 0 | 0 | 0 | 368530 | 0 | 0 | 84820 | 0 | 0 | 0 | 0 |
| SGDYYRFL | DSP | P15924 | DESP_HUMAN | Unmodified | 0 | 0 | 0 | 0 | 0 | 0 | 0 | 0 | 0 | 323140 | 0 | 0 | 30032 | 0 | 0 | 0 | 0 | 0 |
| SGEANVRF | PTGFRN | Q9P2B2 | FPRP_HUMAN | Unmodified | 0 | 0 | 0 | 244340 | 0 | 0 | 178470 | 0 | 0 | 0 | 0 | 0 | 1050300 | 1057200 | 0 | 0 | 0 | 0 |
| SGEATDGA | CHGA | P10645 | CMGA_HUMAN | Oxidation (M) | 0 | 0 | 0 | 0 | 0 | 0 | 0 | 0 | 10471000 | 1431900 | 0 | 0 | 0 | 0 | 0 | 0 | 0 | 0 |
| SGEATDGA | CHGA | P10645 | CMGA_HUMAN | Unmodified | 0 | 0 | 0 | 0 | 0 | 0 | 0 | 0 | 56388000 | 3366300 | 0 | 0 | 0 | 0 | 0 | 0 | 0 | 0 |
| SGEDEQQI | PA2G4; PA2G4 | Q9UQ80; F8 | PA2G4_HUMAI | Acetyl (Protein N | 0 | 0 | 0 | 0 | 3928000 | 3335100 | 0 | 1493100 | 3777800 | 0 | 0 | 13742000 | 4937900 | 5050900 | 0 | 3647200 | 1313600 | 0 |
| SGEDEQQI | PA2G4; PA2G4 | Q9UQ80; F8 | PA2G4_HUMAI | Acetyl (Protein N | 0 | 0 | 0 | 0 | 0 | 0 | 0 | 434680 | 538710 | 635020 | 0 | 0 | 0 | 0 | 118290 | 0 | 0 | 444660 |
| SGEEDFES | PIN1; PIN1; PIN | Q13526; K7E | PIN1_HUMAN I | Unmodified | 6682300 | 4203000 | 3153000 | 2227800 | 0 | 0 | 0 | 136830 | 414140 | 671580 | 572140 | 0 | 1423200 | 697520 | 0 | 2609300 | 547300 | 0 |
| SGEELDGF | CMTM4 | Q8IZR5 | CKLF4_HUMAN | Oxidation (M) | 0 | 926820 | 0 | 0 | 0 | 0 | 0 | 0 | 0 | 0 | 0 | 0 | 0 | 0 | 915720 | 1590500 | 0 | 0 |
| SGEELDGF | CMTM4 | Q8IZR5 | CKLF4_HUMAN | Unmodified | 837060 | 1200000 | 754880 | 833640 | 0 | 0 | 0 | 0 | 0 | 0 | 0 | 0 | 0 | 0 | 1270600 | 0 | 0 | 0 |
| SGEENPAS | PAFAH1B3; PAF | Q15102; M0 | PA1B3_HUMAN | Acetyl (Protein N | 489440 | 261700 | 0 | 264040 | 0 | 0 | 0 | 61230 | 2568200 | 504760 | 2138500 | 17998000 | 0 | 0 | 286790 | 401400 | 0 | 0 |
| SGEENPAS | PAFAH1B3; PAF | Q15102; M0 | PA1B3_HUMAN | Acetyl (Protein N | 0 | 0 | 0 | 0 | 311690 | 0 | 0 | 0 | 1236800 | 0 | 461980 | 0 | 0 | 0 | 0 | 0 | 0 | 0 |
| SGEGSQHI | HIKESHI; HIKES | Q53FT3; E9F | HIKES_HUMAN | Oxidation (M) | 0 | 0 | 0 | 244120 | 0 | 0 | 0 | 0 | 0 | 0 | 33901 | 588220 | 0 | 241810 | 0 | 0 | 0 | 0 |
| SGEILYR | GBE1; GBE1 | Q04446; E9F | GLGB_HUMAN | Unmodified | 99542 | 60640 | 0 | 166160 | 224270 | 33940 | 0 | 0 | 854390 | 378540 | 76353 | 757900 | 90369 | 0 | 0 | 0 | 122870 | 220870 |
| SGEILYRISI | GBE1; GBE1 | Q04446; E9F | GLGB_HUMAN | Unmodified | 0 | 0 | 0 | 0 | 626030 | 0 | 277460 | 28211 | 0 | 671440 | 0 | 0 | 206760 | 222920 | 0 | 0 | 1444600 | 1076600 |
| SGEISLPIKI | MRTFB; MRTFB | Q9ULH7; I3L | MRTFB_HUMAN | Unmodified | 0 | 0 | 0 | 0 | 105490 | 0 | 0 | 0 | 136670 | 124910 | 0 | 0 | 223140 | 238400 | 0 | 0 | 210950 | 135330 |
| SGELAVRAI | ACADVL; ACAD | P49748; G3 | ACADV_HUMAN | Unmodified | 0 | 0 | 0 | 0 | 0 | 0 | 0 | 0 | 0 | 819520 | 0 | 0 | 0 | 0 | 0 | 0 | 0 | 0 |
| SGELEQEE | CHGA; CHGA | P10645; G5E | CMGA_HUMAN | Unmodified | 0 | 0 | 0 | 0 | 0 | 0 | 0 | 0 | 2555000 | 1095200 | 0 | 210890 | 0 | 0 | 0 | 0 | 0 | 0 |
| SGELEQEE | CHGA; CHGA | P10645; G5E | CMGA_HUMAN | Unmodified | 0 | 0 | 0 | 0 | 0 | 0 | 0 | 0 | 2.17E+08 | 41033000 | 0 | 0 | 0 | 0 | 0 | 0 | 0 | 0 |
| SGELEQEE | CHGA; CHGA | P10645; G5E | CMGA_HUMAN | Unmodified | 0 | 0 | 0 | 0 | 0 | 0 | 0 | 0 | 470070 | 8146500 | 0 | 0 | 0 | 0 | 0 | 0 | 0 | 105820 |
| SGELPPNII | SFXN1; SFXN1; | Q9H9B4; D6 | SFXN1_HUMAN | Acetyl (Protein N | 0 | 0 | 0 | 0 | 0 | 0 | 0 | 0 | 0 | 0 | 0 | 0 | 0 | 0 | 0 | 0 | 0 | 0 |
| SGELQGDE | SLCO3A1 | Q9UIG8 | SO3A1_HUMAI | Unmodified | 0 | 0 | 0 | 0 | 0 | 0 | 0 | 0 | 216640 | 0 | 0 | 107830 | 0 | 0 | 0 | 0 | 0 | 0 |
| SGELQGGI | EXOC4 | Q96A65 | EXOC4_HUMAN | Unmodified | 85605 | 146210 | 0 | 0 | 0 | 0 | 0 | 0 | 621400 | 0 | 40336 | 714860 | 0 | 0 | 787190 | 0 | 0 | 0 |
| SGELSNRF | AIF1L; AIF1L | Q9BQI0; Q5J | AIF1L_HUMAN | Acetyl (Protein N | 0 | 0 | 0 | 497660 | 0 | 0 | 0 | 0 | 0 | 387830 | 0 | 0 | 2766300 | 1214800 | 0 | 0 | 0 | 0 |
| SGEPGQTS | MIEN1; MIEN1 | Q9BRT3; J3K | MIEN1_HUMAN | Acetyl (Protein N | 298210 | 216040 | 0 | 0 | 0 | 0 | 0 | 0 | 0 | 180650 | 296870 | 3885900 | 0 | 0 | 0 | 318610 | 0 | 0 |
| SGEPQTFTI | ITGB1; ITGB1; I | P05556; E7E | ITB1_HUMAN I | Unmodified | 4038000 | 3405100 | 7886700 | 8410500 | 1.78E+08 | 33520000 | 3150800 | 240510 | 22312000 | 6848400 | 2212700 | 11324000 | 10086000 | 8633100 | 7781700 | 5792200 | 42938000 | 1.96E+08 |
| SGEPQTFTI | ITGB1; ITGB1; I | P05556; E7E | ITB1_HUMAN I | Unmodified | 0 | 54970 | 0 | 0 | 1460000 | 0 | 395880 | 261520 | 0 | 0 | 0 | 0 | 338020 | 412280 | 0 | 0 | 4212100 | 3231600 |
| SGEQIFTQI | CASKIN2 | Q8WXE0 | CSKI2_HUMAN | Unmodified | 68443 | 52605 | 152550 | 167480 | 0 | 0 | 0 | 0 | 0 | 0 | 254530 | 807640 | 0 | 0 | 221430 | 199760 | 0 | 0 |
| SGERLKEA | KIF20A | O95235 | KI20A_HUMAN | Unmodified | 0 | 0 | 0 | 0 | 619630 | 0 | 0 | 0 | 0 | 0 | 0 | 0 | 539100 | 0 | 0 | 0 | 345230 | 201580 |
| SGERPVTA | CIAO2B; CIAO2 | Q9Y3D0; H3 | CIA2B_HUMAN | Unmodified | 0 | 68734 | 0 | 226570 | 0 | 0 | 0 | 0 | 0 | 0 | 501570 | 1979900 | 0 | 0 | 0 | 182910 | 0 | 0 |
| SGESARSL | GSTO1 | P78417 | GSTO1_HUMAI | Acetyl (Protein N | 0 | 0 | 0 | 0 | 9094500 | 1075200 | 481890 | 1003800 | 0 | 76260 | 0 | 0 | 1837200 | 0 | 0 | 0 | 1859100 | 95366 |
| SGESARSL | GSTO1 | P78417 | GSTO1_HUMAI | Acetyl (Protein N | 0 | 0 | 0 | 0 | 1541800 | 0 | 0 | 217390 | 0 | 0 | 0 | 0 | 434220 | 0 | 0 | 0 | 850380 | 296050 |
| SGESQVLF | HSPA13 | P48723 | HSP13_HUMAN | Unmodified | 0 | 0 | 0 | 0 | 449820 | 0 | 0 | 0 | 12671000 | 552260 | 0 | 466750 | 0 | 0 | 0 | 0 | 0 | 0 |
| SGESVEEV | SIT1; SIT1 | Q9Y3P8; A0/ | SIT1_HUMAN S | Unmodified | 16946 | 0 | 0 | 0 | 0 | 0 | 0 | 0 | 15854000 | 2381000 | 0 | 0 | 0 | 0 | 0 | 0 | 0 | 0 |
| SGESVEEV | SIT1; SIT1 | Q9Y3P8; A0/ | SIT1_HUMAN S | Unmodified | 0 | 0 | 0 | 0 | 0 | 0 | 0 | 0 | 1866800 | 925390 | 0 | 0 | 0 | 0 | 0 | 0 | 0 | 0 |
| SGETEDTFI | ENO1; ENO3; E | P06733; P13 | ENOA_HUMAN | Unmodified | 0 | 0 | 0 | 0 | 346580 | 0 | 0 | 0 | 0 | 69775 | 0 | 0 | 411190 | 0 | 0 | 0 | 0 | 298050 |
| SGETVKHE | CASC3 | O15234 | CASC3_HUMAI | Unmodified | 572990 | 559800 | 0 | 895440 | 0 | 397720 | 0 | 14310 | 0 | 28858 | 0 | 0 | 0 | 0 | 0 | 0 | 0 | 0 |
| SGETWDSH | DPP3; DPP3; DI | Q9NY33; G3 | DPP3_HUMAN | Unmodified | 0 | 0 | 0 | 0 | 0 | 0 | 0 | 0 | 0 | 0 | 0 | 759410 | 0 | 0 | 65712 | 57457 | 0 | 0 |
| SGEVLVNV | EIF3I; EIF3I; EIF | Q13347; A0/ | EIF3I_HUMAN I | Unmodified | 1046100 | 1026800 | 1910200 | 2310800 | 3189700 | 492180 | 59028 | 113410 | 1422100 | 32851 | 6034000 | 9995700 | 4103500 | 5426000 | 1552400 | 1475600 | 2310100 | 2425300 |
| SGEVPPQK | LIN7C; LIN7C | Q9NUP9; G3 | LIN7C_HUMAN | Unmodified | 0 | 0 | 0 | 0 | 1771500 | 1600900 | 2472300 | 1933600 | 145180 | 330870 | 0 | 0 | 881980 | 594710 | 0 | 0 | 0 | 0 |
| SGEYWIDP | COL1A1 | P02452 | CO1A1_HUMAI | Unmodified | 37854 | 0 | 264390 | 261000 | 249440 | 663340 | 109040 | 46048 | 0 | 0 | 0 | 0 | 0 | 465640 | 132480 | 0 | 94050 | 807340 |
| SGFEPASLI | MYH9; MYH9; M | P35579; Q5I | MYH9_HUMAN | Unmodified | 51805 | 0 | 751920 | 40592 | 0 | 0 | 0 | 0 | 0 | 0 | 0 | 967930 | 620540 | 46517 | 1176900 | 443530 | 0 | 0 |
| SGFEPASLI | MYH9; MYH9; M | P35579; Q5I | MYH9_HUMAN | Unmodified | 0 | 0 | 299030 | 0 | 17746000 | 0 | 0 | 0 | 0 | 266510 | 2416900 | 5043100 | 82746 | 0 | 0 | 0 | 4297100 | 6059900 |
| SGFEPPGD | FNBP1L; FNBP1 | Q5T0N5; A0/ | FBP1L_HUMAN | Unmodified | 4000900 | 3784300 | 442950 | 1281300 | 0 | 0 | 0 | 0 | 1329400 | 0 | 0 | 358620 | 747460 | 446080 | 2372900 | 787620 | 0 | 0 |
| SGFGRENC | ALDH9A1 | P49189 | AL9A1_HUMAN | Unmodified | 0 | 0 | 0 | 20524000 | 2415000 | 0 | 1721900 | 0 | 0 | 1286200 | 0 | 0 | 4354700 | 2825100 | 0 | 0 | 3270600 | 2027200 |
| SGFHTVPG | LRP1 | Q07954 | LRP1_HUMAN I | Unmodified | 0 | 0 | 0 | 0 | 0 | 0 | 0 | 0 | 1476400 | 0 | 0 | 0 | 0 | 0 | 0 | 0 | 0 | 0 |
| SGFQINLN | SYPL1; SYPL1 | Q16563; C9J | SYPL1_HUMAN | Acetyl (Protein N | 0 | 0 | 292170 | 0 | 0 | 0 | 0 | 0 | 231890 | 0 | 0 | 0 | 0 | 0 | 366180 | 204720 | 0 | 0 |
| SGFQINLN | SYPL1; SYPL1 | Q16563; C9J | SYPL1_HUMAN | Unmodified | 0 | 0 | 385520 | 107870 | 158210 | 24144 | 11058 | 0 | 184840 | 276430 | 67747 | 17930 | 0 | 0 | 819780 | 344030 | 105300 | 0 |
| SGFQINLN | SYPL1; SYPL1 | Q16563; C9J | SYPL1_HUMAN | Acetyl (Protein N | 0 | 0 | 72978 | 43174 | 50008 | 0 | 0 | 0 | 42503 | 0 | 35982 | 84491 | 0 | 0 | 372320 | 43408 | 0 | 0 |

| | | | | | | | | | | | | | | | | | | | | | | |
|---|---|---|---|---|---|---|---|---|---|---|---|---|---|---|---|---|---|---|---|---|---|---|
| SGFSLDNC | NSFL1C; NSFL1 | Q9UNZ2; F2 | NSF1C_HUMAI | Unmodified | 299180 | 0 | 122120 | 90009 | 0 | 0 | 0 | 0 | 359840 | 0 | 409290 | 1380000 | 0 | 0 | 0 | 390560 | 0 | 0 |
| SGFSTEER/ | PPA1; PPA1 | Q15181; Q5! | IPYR_HUMAN I | Acetyl (Protein N | 0 | 0 | 0 | 0 | 5279500 | 1587000 | 144880 | 135990 | 0 | 0 | 1595900 | 0 | 4859700 | 1764900 | 20836 | 0 | 1993400 | 6722200 |
| SGFSTEER/ | PPA1; PPA1 | Q15181; Q5! | IPYR_HUMAN I | Acetyl (Protein N | 0 | 0 | 0 | 0 | 4879800 | 0 | 0 | 0 | 0 | 0 | 0 | 146010 | 6011900 | 3279500 | 0 | 0 | 5495700 | 0 |
| SGFTPLHIA | ANK3; ANK3; AN | Q12955; A0/ | ANK3_HUMAN | Unmodified | 0 | 0 | 0 | 3284200 | 0 | 0 | 0 | 0 | 0 | 0 | 0 | 0 | 3139400 | 0 | 10585000 | 0 | 0 | 0 |
| SGFWGWF | ANKRD13A; AN | Q8IZ07; H0Y | AN13A_HUMAN | Unmodified | 0 | 0 | 0 | 197700 | 1972600 | 0 | 0 | 0 | 0 | 0 | 0 | 0 | 0 | 79481 | 0 | 0 | 473280 | 1179600 |
| SGGASHSE | PCMT1; PCMT1 | P22061; F8V | PIMT_HUMAN I | Unmodified | 0 | 0 | 0 | 0 | 0 | 0 | 132390 | 0 | 607950 | 0 | 0 | 474980 | 0 | 467560 | 0 | 0 | 0 | 0 |
| SGGASHSE | PCMT1; PCMT1 | P22061; F8V | PIMT_HUMAN I | Unmodified | 1045100 | 0 | 27352 | 0 | 4969000 | 929120 | 2263100 | 1098500 | 6808500 | 5638900 | 2310700 | 11159000 | 0 | 5359000 | 0 | 285720 | 3075600 | 3233800 |
| SGGAVEPL | ZPR1; ZPR1; ZP | O75312; H7( | ZPR1_HUMAN . | Unmodified | 0 | 0 | 0 | 0 | 0 | 0 | 0 | 0 | 423890 | 40543 | 0 | 1772200 | 0 | 0 | 1791900 | 0 | 0 | 0 |
| SGGDLTLH | WDR48 | Q8TAF3 | WDR48_HUMA | Unmodified | 0 | 0 | 0 | 0 | 3271100 | 0 | 730950 | 392570 | 0 | 346430 | 0 | 0 | 649990 | 0 | 0 | 0 | 2345600 | 3184300 |
| SGGDTAPA | FARP2; FARP2 | O94887; H7( | FARP2_HUMAN | Unmodified | 0 | 0 | 0 | 459610 | 646910 | 0 | 0 | 0 | 0 | 0 | 0 | 0 | 0 | 0 | 0 | 437550 | 26897 | 0 |
| SGGEAFVL | SNX18 | Q96RF0 | SNX18_HUMAN | Unmodified | 0 | 0 | 0 | 0 | 0 | 69738 | 0 | 0 | 0 | 0 | 0 | 0 | 3912500 | 0 | 0 | 0 | 0 | 0 |
| SGGEAFVL | SNX18 | Q96RF0 | SNX18_HUMAN | Unmodified | 1126800 | 657520 | 0 | 2157500 | 787150 | 0 | 0 | 0 | 0 | 0 | 0 | 0 | 3464600 | 0 | 920470 | 650990 | 0 | 0 |
| SGGELDIV | PRRC1 | Q96M27 | PRRC1_HUMAI | Unmodified | 0 | 0 | 0 | 0 | 0 | 0 | 0 | 0 | 121880 | 0 | 211800 | 625680 | 598040 | 0 | 0 | 0 | 229170 | 477330 |
| SGGFIKDS | RBM12B; RBM1 | Q8IXT5; B9Z | RB12B_HUMAN | Oxidation (M) | 0 | 0 | 0 | 0 | 2684400 | 9726100 | 6009800 | 420820 | 0 | 0 | 0 | 0 | 3409800 | 3672700 | 0 | 0 | 0 | 0 |
| SGGFIKDS | RBM12B; RBM1 | Q8IXT5; B9Z | RB12B_HUMAN | Unmodified | 0 | 0 | 0 | 0 | 6954900 | 8530000 | 4678300 | 0 | 0 | 0 | 0 | 0 | 0 | 7875700 | 0 | 0 | 375260 | 16021000 |
| SGGGGDIF | SERPINB6; SER | P35237; A0A | SPB6_HUMAN | Unmodified | 664200 | 281820 | 997590 | 1298400 | 652440 | 158600 | 0 | 20939 | 0 | 146130 | 1880700 | 26098000 | 459030 | 356210 | 3185900 | 1765500 | 0 | 1143700 |
| SGGGGLM | VASP; VASP | P50552; K7E | VASP_HUMAN | Oxidation (M) | 134130 | 183620 | 0 | 44727 | 0 | 0 | 0 | 0 | 0 | 0 | 0 | 0 | 250040 | 0 | 0 | 80061 | 0 | 0 |
| SGGGTGEE | PPM1G | O15355 | PPM1G_HUMA | Unmodified | 0 | 0 | 0 | 0 | 0 | 0 | 0 | 0 | 0 | 0 | 455970 | 1008900 | 0 | 0 | 0 | 0 | 0 | 0 |
| SGGGTGEE | PPM1G | O15355 | PPM1G_HUMA | Unmodified | 0 | 0 | 0 | 0 | 329440 | 0 | 0 | 0 | 0 | 0 | 0 | 1405700 | 282090 | 0 | 0 | 0 | 166390 | 0 |
| SGGGVIRG | SRSF1; SRSF1; | Q07955; J3K | SRSF1_HUMAN | Acetyl (Protein N | 657730 | 326960 | 138850 | 314070 | 0 | 0 | 0 | 0 | 0 | 0 | 0 | 233640 | 139120 | 36091 | 255320 | 141420 | 0 | 0 |
| SGGGYIVV | AGL | P35573 | GDE_HUMAN ( | Unmodified | 0 | 0 | 0 | 0 | 0 | 0 | 0 | 0 | 0 | 0 | 0 | 2880600 | 0 | 0 | 0 | 0 | 0 | 0 |
| SGGIETIAN | HSPA4L; HSPA4 | O95757; D6I | HS74L_HUMAN | Unmodified | 0 | 0 | 0 | 0 | 0 | 0 | 0 | 0 | 0 | 0 | 0 | 1418700 | 0 | 0 | 0 | 0 | 0 | 0 |
| SGGIPALVF | JUP; JUP; JUP; J | P14923; C9J | PLAK_HUMAN . | Unmodified | 0 | 0 | 1782500 | 463440 | 0 | 0 | 0 | 0 | 0 | 157230 | 0 | 5722500 | 0 | 0 | 2466400 | 2059100 | 0 | 0 |
| SGGIVISPF | CCDC6 | Q16204 | CCDC6_HUMA | Unmodified | 215040 | 0 | 0 | 497140 | 208000 | 61115 | 0 | 0 | 4705800 | 257060 | 1312000 | 15575000 | 670070 | 0 | 435540 | 316610 | 126250 | 406640 |
| SGGKLGD( | ARHGAP10 | A1A4S6 | RHG10_HUMAI | Unmodified | 0 | 405410 | 0 | 399460 | 500620 | 0 | 893120 | 183870 | 0 | 461760 | 0 | 1171100 | 57849 | 0 | 97861 | 57457 | 577300 | 0 |
| SGGLGGSI | LRPPRC; LRPPF | P42704; B8Z | LPPRC_HUMAN | Unmodified | 0 | 0 | 0 | 0 | 0 | 0 | 0 | 0 | 10576000 | 1271600 | 0 | 648730 | 0 | 161400 | 0 | 0 | 0 | 0 |
| SGGLPLVL! | USP9X | Q93008 | USP9X_HUMAN | Unmodified | 0 | 0 | 0 | 0 | 0 | 0 | 0 | 0 | 0 | 0 | 0 | 67804 | 0 | 0 | 0 | 0 | 0 | 0 |
| SGGLSVEV | GALNT2 | Q10471 | GALT2_HUMAN | Unmodified | 0 | 0 | 0 | 0 | 0 | 0 | 0 | 0 | 2609300 | 712010 | 0 | 0 | 0 | 0 | 0 | 0 | 0 | 0 |
| SGGMERPI | GANAB; GANAE | Q14697; F5H | GANAB_HUMA | Oxidation (M) | 0 | 0 | 0 | 0 | 0 | 0 | 0 | 183930 | 11556000 | 3014200 | 0 | 457360 | 0 | 0 | 0 | 0 | 0 | 0 |
| SGGMERPI | GANAB; GANAE | Q14697; F5H | GANAB_HUMA | Unmodified | 0 | 0 | 0 | 0 | 0 | 0 | 0 | 0 | 39450000 | 5257600 | 0 | 0 | 0 | 0 | 0 | 276210 | 0 | 0 |
| SGGMSNEI | ACLY | P53396 | ACLY_HUMAN | Oxidation (M) | 272520 | 0 | 0 | 0 | 0 | 0 | 0 | 0 | 315870 | 0 | 130020 | 7117100 | 0 | 0 | 296830 | 745500 | 0 | 0 |
| SGGMSNEI | ACLY | P53396 | ACLY_HUMAN | Unmodified | 0 | 0 | 0 | 0 | 0 | 0 | 0 | 0 | 0 | 0 | 0 | 3457100 | 0 | 0 | 0 | 0 | 0 | 0 |
| SGGNLVLV | SORD | Q00796 | DHSO_HUMAN | Oxidation (M) | 0 | 1203200 | 0 | 0 | 0 | 0 | 0 | 0 | 0 | 0 | 0 | 598420 | 0 | 0 | 288520 | 0 | 0 | 0 |
| SGGNSYGS | ILF3; ILF3 | Q12906; K7E | ILF3_HUMAN I | Unmodified | 921950 | 986530 | 0 | 0 | 0 | 0 | 240450 | 0 | 0 | 0 | 0 | 0 | 0 | 0 | 0 | 0 | 0 | 0 |
| SGGQLPSL | FAM171A1 | Q5VUB5 | F1711_HUMAN | Unmodified | 0 | 0 | 114420 | 82405 | 495680 | 0 | 0 | 63365 | 0 | 0 | 240590 | 550450 | 321700 | 83504 | 94496 | 140570 | 0 | 444950 |
| SGGSDRTL | LGALS3BP | Q08380 | LG3BP_HUMAN | Unmodified | 0 | 0 | 84845 | 0 | 2895700 | 0 | 6894200 | 9516600 | 41909000 | 31766000 | 48619000 | 1723500 | 18240000 | 30876000 | 60607 | 69059 | 2440800 | 1.32E+08 |
| SGGSPLSP | SEZ6L2; SEZ6L | Q6UXD5; H3 | SE6L2_HUMAN | Oxidation (M) | 0 | 0 | 0 | 0 | 0 | 0 | 0 | 0 | 929210 | 60676 | 0 | 0 | 0 | 0 | 0 | 0 | 0 | 0 |
| SGGTALHV | PPP1R12A; PPF | O14974; F8V | MYPT1_HUMAN | Unmodified | 0 | 0 | 0 | 0 | 630150 | 543590 | 0 | 0 | 0 | 0 | 26931 | 284860 | 428130 | 238440 | 0 | 0 | 336300 | 210160 |
| SGGTEGLL | SRP68 | Q9UHB9 | SRP68_HUMAN | Unmodified | 0 | 0 | 0 | 0 | 0 | 0 | 0 | 0 | 0 | 0 | 89411 | 641030 | 0 | 0 | 0 | 0 | 0 | 0 |
| SGGTEGLL | SRP68 | Q9UHB9 | SRP68_HUMAN | Unmodified | 0 | 0 | 0 | 0 | 0 | 0 | 0 | 0 | 175670 | 111790 | 394510 | 403080 | 349310 | 92929 | 0 | 0 | 82299 | 23254 |
| SGGVGGSI | RPA1; RPA1 | P27694; I3L | RFA1_HUMAN | Unmodified | 78738 | 0 | 0 | 0 | 459640 | 0 | 40406 | 0 | 83402 | 0 | 0 | 2304200 | 0 | 0 | 0 | 0 | 0 | 0 |
| SGGYGGSI | CIRBP; CIRBP; ( | Q14011; K7E | CIRBP_HUMAN | Unmodified | 2554800 | 2063300 | 1721900 | 2023300 | 0 | 27705 | 40783 | 48738 | 313020 | 43528 | 31961 | 195990 | 600380 | 123890 | 1525500 | 1530400 | 0 | 0 |
| SGHCTACL | PPAT | Q06203 | PUR1_HUMAN | Unmodified | 0 | 0 | 0 | 0 | 0 | 0 | 0 | 0 | 0 | 0 | 421340 | 1157900 | 1101600 | 0 | 0 | 0 | 684510 | 332850 |
| SGHEQVVE | ANK3; ANK3 | Q12955; A0/ | ANK3_HUMAN | Oxidation (M) | 0 | 0 | 727390 | 690810 | 0 | 0 | 0 | 0 | 0 | 0 | 0 | 0 | 159550 | 102670 | 1978100 | 2508000 | 0 | 0 |
| SGHEQVVE | ANK3; ANK3 | Q12955; A0/ | ANK3_HUMAN | Unmodified | 0 | 0 | 0 | 197140 | 0 | 0 | 0 | 0 | 0 | 0 | 0 | 0 | 110210 | 0 | 385660 | 188870 | 0 | 0 |
| SGHEQVVE | ANK3; ANK3 | Q12955; A0/ | ANK3_HUMAN | Oxidation (M) | 0 | 0 | 453340 | 1696000 | 0 | 0 | 0 | 0 | 0 | 0 | 0 | 0 | 3858300 | 5091300 | 2460200 | 873310 | 0 | 0 |
| SGHHPGET | FXYD3; FXYD3 | Q14802; S4F | FXYD3_HUMAN | Unmodified | 5930800 | 2009200 | 9121200 | 4549200 | 0 | 1173800 | 0 | 0 | 1970700 | 1092800 | 0 | 0 | 2630700 | 6089400 | 13816000 | 10217000 | 0 | 0 |
| SGHIQDYQ | EDIL3 | O43854 | EDIL3_HUMAN | Unmodified | 1953400 | 1406700 | 160870 | 405090 | 3.72E+08 | 1.17E+08 | 4642700 | 686130 | 1.16E+08 | 10404000 | 231220 | 212950 | 0 | 0 | 150600 | 101560 | 8685500 | 1.06E+08 |
| SGHIQDYQ | EDIL3 | O43854 | EDIL3_HUMAN | 2 Oxidation (M) | 0 | 0 | 0 | 0 | 1.33E+08 | 11087000 | 0 | 0 | 0 | 3627400 | 0 | 0 | 0 | 0 | 0 | 0 | 7550400 | 36168000 |
| SGHIQDYQ | EDIL3 | O43854 | EDIL3_HUMAN | Oxidation (M) | 0 | 0 | 0 | 0 | 41778000 | 1879300 | 541880 | 0 | 0 | 0 | 0 | 0 | 0 | 0 | 0 | 0 | 1921900 | 23324000 |
| SGHIQDYQ | EDIL3 | O43854 | EDIL3_HUMAN | Unmodified | 0 | 0 | 0 | 0 | 1527000 | 0 | 0 | 0 | 0 | 0 | 0 | 0 | 0 | 0 | 0 | 0 | 0 | 1377500 |
| SGIDFGSLI | ABCC4; ABCC4 | O15439; A0/ | MRP4_HUMAN | Unmodified | 0 | 0 | 3072100 | 1565100 | 0 | 0 | 0 | 0 | 0 | 0 | 0 | 261260 | 0 | 0 | 2994500 | 960690 | 0 | 0 |
| SGILAAESII | ETFDH | Q16134 | ETFD_HUMAN | Unmodified | 0 | 27256 | 159240 | 0 | 36688 | 0 | 61648 | 0 | 510840 | 0 | 0 | 0 | 61247 | 0 | 0 | 0 | 175090 | 0 |
| SGILPILVK | KPNA3 | O00505 | IMA4_HUMAN | Unmodified | 466960 | 74388 | 418110 | 349540 | 1234500 | 457080 | 513490 | 254090 | 624850 | 210890 | 1424300 | 1766900 | 547790 | 307500 | 164440 | 369510 | 481950 | 454710 |
| SGINCPIQI | NPC2; NPC2; N | P61916; J3K | NPC2_HUMAN | Unmodified | 0 | 0 | 445520 | 0 | 1799200 | 1155900 | 1352000 | 843920 | 14580000 | 9952500 | 0 | 222100 | 975040 | 1667000 | 0 | 0 | 93362 | 0 |
| SGIQGKIPA | MYOF; MYOF | Q9NZM1; H( | MYOF_HUMAN | Unmodified | 0 | 0 | 0 | 0 | 4252300 | 1498000 | 0 | 0 | 80545 | 0 | 0 | 0 | 0 | 0 | 0 | 0 | 581470 | 0 |
| SGISTGHW | FUT8; FUT8; FU | Q9BYC5; A0 | FUT8_HUMAN . | Unmodified | 0 | 0 | 44909 | 0 | 0 | 0 | 0 | 0 | 9022200 | 355100 | 52978 | 0 | 0 | 0 | 181500 | 0 | 0 | 0 |
| SGISTGHW | FUT8; FUT8; FU | Q9BYC5; A0 | FUT8_HUMAN . | Unmodified | 0 | 0 | 0 | 0 | 0 | 0 | 14836 | 0 | 1202000 | 0 | 0 | 0 | 0 | 0 | 0 | 0 | 0 | 0 |
| SGISTNHAI | SNX33 | Q8WV41 | SNX33_HUMAN | Unmodified | 2512600 | 2048900 | 10343000 | 16094000 | 0 | 0 | 0 | 0 | 0 | 0 | 0 | 0 | 0 | 0 | 30926000 | 9801600 | 0 | 0 |
| SGKFTQQD | PITRM1; PITRM | Q5JRX3; A0A | PREP_HUMAN | Unmodified | 0 | 0 | 0 | 0 | 276550 | 0 | 38325 | 0 | 560520 | 160540 | 31627 | 0 | 0 | 0 | 0 | 0 | 0 | 142870 |
| SGKPGPFL | PAFAH1B1; PAF | P43034; A0A | LIS1_HUMAN F | Unmodified | 49054 | 35993 | 103370 | 66628 | 0 | 0 | 0 | 0 | 0 | 23368 | 0 | 1214600 | 0 | 0 | 212950 | 181210 | 0 | 0 |
| SGKPGPFL | PAFAH1B1; PAF | P43034; A0A | LIS1_HUMAN F | Unmodified | 0 | 0 | 0 | 0 | 531150 | 378530 | 850760 | 61457 | 0 | 0 | 0 | 659310 | 207070 | 0 | 0 | 0 | 0 | 0 |
| SGKSPILVA | DDX3X; DDX3Y | O00571; O1! | DDX3X_HUMAI | Unmodified | 1102300 | 481450 | 0 | 1354000 | 0 | 0 | 0 | 0 | 0 | 0 | 0 | 0 | 1656500 | 549280 | 325850 | 441340 | 0 | 0 |
| SGKTELAEI | EPRS1 | P07814 | SYEP_HUMAN | Oxidation (M) | 0 | 0 | 0 | 747380 | 0 | 0 | 0 | 444820 | 0 | 482530 | 2741400 | 15591000 | 0 | 0 | 881010 | 971180 | 0 | 0 |
| SGKYDLDF | ENO1; ENO1 | P06733; A0A | ENOA_HUMAN | Unmodified | 0 | 0 | 2594000 | 0 | 0 | 0 | 0 | 0 | 4017200 | 0 | 2556300 | 18974000 | 268200 | 0 | 2271100 | 0 | 163380 | 224870 |
| SGKYDLDF | ENO1; ENO1 | P06733; A0A | ENOA_HUMAN | Unmodified | 994070 | 1105200 | 590030 | 10845000 | 1687600 | 366120 | 0 | 74054 | 37367000 | 1769600 | 22006000 | 1.1E+08 | 877780 | 0 | 7323800 | 11559000 | 355220 | 505090 |
| SGKYVLGY | RPL30; RPL30; I | P62888; A0A | RL30_HUMAN ( | Unmodified | 0 | 0 | 0 | 0 | 444380 | 0 | 2218700 | 1622900 | 0 | 149300 | 0 | 0 | 0 | 2042600 | 0 | 0 | 0 | 0 |
| SGLAYCPN | LPXN; LPXN | O60711; B7Z | LPXN_HUMAN | Unmodified | 0 | 0 | 0 | 0 | 266080 | 0 | 0 | 0 | 0 | 0 | 0 | 0 | 205470 | 129950 | 0 | 0 | 159290 | 0 |
| SGLDHKFG | SCG3 | Q8WXD2 | SCG3_HUMAN | Unmodified | 0 | 0 | 0 | 0 | 0 | 356220 | 0 | 0 | 91984000 | 14078000 | 0 | 0 | 0 | 0 | 0 | 0 | 0 | 0 |
| SGLELEPEI | ARHGEF1; ARH | Q92888; M0 | ARHG1_HUMA | Unmodified | 0 | 0 | 0 | 0 | 0 | 57986 | 0 | 0 | 259680 | 0 | 0 | 98980 | 0 | 0 | 0 | 104240 | 0 | 0 |
| SGLGELILP | FH | P07954 | FUMH_HUMAN | Oxidation (M) | 0 | 0 | 0 | 0 | 0 | 0 | 319260 | 59984 | 6015100 | 928920 | 1284900 | 3481400 | 0 | 0 | 0 | 0 | 89216 | 0 |
| SGLGELILP | FH | P07954 | FUMH_HUMAN | Unmodified | 0 | 0 | 0 | 0 | 0 | 0 | 0 | 0 | 4269600 | 0 | 55612 | 460310 | 57912 | 45727 | 0 | 0 | 0 | 0 |
| SGLGRLFG | CHMP4A; CHM | Q9BY43; E9F | CHM4A_HUMA | Acetyl (Protein N | 0 | 0 | 0 | 0 | 3551300 | 394010 | 0 | 0 | 0 | 1619200 | 0 | 0 | 0 | 1114100 | 0 | 0 | 1192800 | 797870 |
| SGLIFYR | SHMT2; SHMT2 | P34897; H0Y | GLYM_HUMAN | Unmodified | 0 | 0 | 0 | 0 | 0 | 0 | 0 | 0 | 7835300 | 380710 | 0 | 0 | 0 | 0 | 0 | 0 | 0 | 0 |
| SGLKKQEL | XRCC6; XRCC6 | P12956; B1A | XRCC6_HUMAI | Unmodified | 0 | 0 | 0 | 0 | 618240 | 527380 | 1135000 | 872740 | 0 | 415650 | 0 | 0 | 354610 | 180770 | 0 | 0 | 344070 | 216220 |
| SGLLEVLVF | SNTB1; SNTB1 | Q13884; A0/ | SNTB1_HUMAN | Unmodified | 0 | 0 | 0 | 0 | 157710 | 0 | 0 | 0 | 7423300 | 833250 | 0 | 0 | 0 | 0 | 0 | 55931 | 109100 | 218990 |
| SGLLVLTTF | COASY; COASY | Q13057; K7E | COASY_HUMAN | Unmodified | 0 | 0 | 0 | 0 | 0 | 0 | 0 | 0 | 0 | 0 | 0 | 0 | 0 | 0 | 0 | 0 | 0 | 0 |
| SGLNQGAI | HECTD1; HECT | Q9ULT8; H0Y | HECD1_HUMA | Unmodified | 0 | 0 | 0 | 0 | 1762200 | 0 | 0 | 0 | 0 | 0 | 182380 | 0 | 1130500 | 0 | 0 | 0 | 400250 | 0 |
| SGLQVEDL | FPGS; FPGS; FF | Q05932; Q5. | FOLC_HUMAN | Unmodified | 0 | 0 | 132450 | 241610 | 0 | 0 | 0 | 0 | 666660 | 87736 | 0 | 0 | 89509 | 82162 | 303280 | 256410 | 0 | 132390 |
| SGLQVSQL | TUBGCP4; TUB | Q9UGJ1; H3 | GCP4_HUMAN | Unmodified | 0 | 0 | 0 | 0 | 0 | 0 | 0 | 0 | 0 | 0 | 0 | 917280 | 0 | 0 | 0 | 0 | 0 | 0 |
| SGLRAQDF | SLC9A3R2; SLC | Q15599; H3I | NHRF2_HUMAI | Unmodified | 0 | 0 | 0 | 400030 | 0 | 0 | 0 | 0 | 0 | 0 | 0 | 0 | 0 | 0 | 0 | 0 | 0 | 0 |
| SGLRPGTQ | CLIC2; CLIC2 | O15247; A6F | CLIC2_HUMAN | Acetyl (Protein N | 0 | 0 | 0 | 0 | 1167700 | 27047 | 0 | 0 | 73521 | 0 | 0 | 0 | 3314200 | 7235700 | 0 | 0 | 130210 | 146410 |
| SGLRQDPQ | MITD1; MITD1; I | Q8WV92; A0 | MITD1_HUMAN | Unmodified | 89888 | 122680 | 123240 | 65490 | 3427600 | 634430 | 130560 | 50009 | 555420 | 392600 | 433220 | 1364500 | 133140 | 98299 | 59225 | 54582 | 141280 | 0 |
| SGLRQDPQ | MITD1; MITD1; I | Q8WV92; A0 | MITD1_HUMAN | Unmodified | 1260700 | 928990 | 293560 | 1379200 | 1003000 | 1691700 | 0 | 0 | 1599100 | 1441400 | 863720 | 11631000 | 418280 | 66604 | 363480 | 453350 | 305670 | 327240 |
| SGLRVYST! | SH3BGRL3; SH | Q9H299; Q5 | SH3L3_HUMAN | Acetyl (Protein N | 0 | 0 | 0 | 0 | 809690 | 0 | 0 | 86660 | 0 | 172370 | 0 | 0 | 0 | 0 | 0 | 0 | 888820 | 903320 |
| SGLRVYST! | SH3BGRL3; SH | Q9H299; Q5 | SH3L3_HUMAN | Acetyl (Protein N | 243550 | 0 | 230710 | 0 | 41961000 | 8595800 | 29169000 | 16243000 | 415200 | 10716000 | 491460 | 0 | 12343000 | 11862000 | 0 | 0 | 2.08E+08 | 2.2E+08 |
| SGLSSPIYII | UMPS; UMPS; I | P11172; F2Z | UMPS_HUMAN | Unmodified | 0 | 0 | 0 | 0 | 0 | 0 | 968790 | 0 | 195190 | 500930 | 0 | 1169800 | 0 | 0 | 0 | 0 | 569640 | 0 |
| SGLTFLPK | PGAP1; PGAP1 | Q75T13; B4I | PGAP1_HUMAI | Unmodified | 0 | 0 | 0 | 0 | 17947 | 1583500 | 0 | 0 | 2637500 | 0 | 0 | 0 | 0 | 0 | 0 | 0 | 0 | 0 |
| SGLTPLHL/ | ANK3 | Q12955 | ANK3_HUMAN | Unmodified | 0 | 0 | 120480 | 89802 | 0 | 0 | 0 | 0 | 0 | 0 | 0 | 0 | 0 | 0 | 319760 | 211150 | 0 | 0 |
| SGLTPLHL/ | ANK3 | Q12955 | ANK3_HUMAN | Unmodified | 0 | 0 | 0 | 0 | 0 | 0 | 0 | 0 | 436860 | 0 | 0 | 0 | 10584000 | 5018100 | 0 | 0 | 0 | 0 |
| SGLTPLHV/ | ANK3 | Q12955 | ANK3_HUMAN | Unmodified | 0 | 0 | 4855400 | 3385800 | 0 | 0 | 34681 | 208150 | 0 | 0 | 0 | 94806 | 20575000 | 13069000 | 6523800 | 5656000 | 0 | 0 |
| SGMNVARI | PKM; PKM; PKM | P14618; H3E | KPYM_HUMAN | Oxidation (M) | 0 | 0 | 0 | 0 | 1.18E+08 | 4691400 | 15970000 | 4241200 | 0 | 932190 | 0 | 0 | 19535000 | 14257000 | 0 | 0 | 50883000 | 52930000 |
| SGMNVARI | PKM; PKM; PKM | P14618; H3E | KPYM_HUMAN | Unmodified | 0 | 0 | 0 | 0 | 5.23E+08 | 54698000 | 25724000 | 2687000 | 0 | 8081200 | 333460 | 0 | 60413000 | 39242000 | 0 | 0 | 1.46E+08 | 2.41E+08 |
| SGMNVARI | PKM; PKM; PKM | P14618; H3E | KPYM_HUMAN | Unmodified | 0 | 0 | 0 | 0 | 7282900 | 552440 | 1032500 | 0 | 0 | 319900 | 0 | 0 | 947800 | 0 | 0 | 0 | 16393000 | 10240000 |
| SGMNVARI | PKM; PKM; PKM | P14618; H3E | KPYM_HUMAN | Oxidation (M) | 0 | 0 | 0 | 0 | 4703800 | 298310 | 738710 | 0 | 0 | 0 | 0 | 0 | 0 | 0 | 0 | 0 | 0 | 0 |
| SGMRFNEV | PIN4; PIN4; PIN | Q9Y237; H0Y | PIN4_HUMAN I | Oxidation (M) | 0 | 0 | 0 | 0 | 190980 | 0 | 0 | 0 | 0 | 89378 | 0 | 0 | 57100 | 88833 | 0 | 0 | 147880 | 428690 |
| SGMRFNEV | PIN4; PIN4; PIN | Q9Y237; H0Y | PIN4_HUMAN I | Unmodified | 0 | 0 | 0 | 0 | 878010 | 152430 | 71747 | 0 | 0 | 246590 | 0 | 65531 | 0 | 504680 | 0 | 0 | 747160 | 767120 |
| SGMSHEEI | PRPF8 | Q6P2Q9 | PRP8_HUMAN | Oxidation (M) | 0 | 0 | 0 | 0 | 0 | 0 | 0 | 0 | 0 | 0 | 0 | 0 | 0 | 0 | 0 | 0 | 0 | 0 |
| SGMVQTEA | PTPN6 | P29350 | PTN6_HUMAN | Unmodified | 0 | 0 | 0 | 0 | 0 | 0 | 0 | 0 | 0 | 0 | 0 | 486280 | 0 | 0 | 0 | 0 | 0 | 0 |
| SGMVQTEA | PTPN11; PTPN1 | Q06124; A0/ | PTN11_HUMAN | Oxidation (M) | 0 | 0 | 0 | 0 | 0 | 0 | 0 | 0 | 0 | 0 | 0 | 208920 | 0 | 0 | 0 | 0 | 0 | 0 |
| SGMVQTEA | PTPN11; PTPN1 | Q06124; A0/ | PTN11_HUMAN | Unmodified | 0 | 0 | 0 | 0 | 0 | 0 | 0 | 0 | 0 | 0 | 0 | 726210 | 0 | 0 | 0 | 159810 | 68547 | 177580 |
| SGNALFHA | EIF3A | Q14152 | EIF3A_HUMAN | Unmodified | 417930 | 161370 | 103730 | 404410 | 0 | 0 | 0 | 0 | 0 | 0 | 0 | 4853100 | 0 | 0 | 92483 | 0 | 0 | 0 |
| SGNFGGSI | HNRNPA2B1; H | P22626; A0A | ROA2_HUMAN | Unmodified | 0 | 0 | 0 | 0 | 405860 | 0 | 0 | 0 | 0 | 596970 | 0 | 933970 | 1028600 | 0 | 0 | 0 | 0 | 346330 |
| SGNGNKPI | RAB13; RAB13 | P51153; A0A | RAB13_HUMAN | Unmodified | 2559900 | 6828900 | 1101400 | 0 | 0 | 0 | 0 | 0 | 0 | 0 | 0 | 0 | 0 | 16661000 | 861020 | 938290 | 10901 | 707700 |
| SGNIVAGIA | GMPS | P49915 | GUAA_HUMAN | Unmodified | 0 | 0 | 0 | 0 | 1546800 | 196150 | 407090 | 247760 | 0 | 372590 | 850800 | 2446400 | 1576900 | 592880 | 0 | 0 | 0 | 786400 |
| SGNIVAGIA | GMPS | P49915 | GUAA_HUMAN | Unmodified | 129710 | 111650 | 0 | 1133000 | 2079200 | 0 | 478460 | 189040 | 68584 | 247270 | 712530 | 20529000 | 1579800 | 270410 | 2572500 | 2523700 | 700900 | 374050 |
| SGNLTEDD | PLS3; PLS3 | P13797; A0A | PLST_HUMAN | Unmodified | 0 | 0 | 0 | 0 | 0 | 0 | 0 | 0 | 0 | 0 | 0 | 0 | 0 | 0 | 0 | 0 | 327690 | 0 |
| SGNMIQSA | CLASP1; CLASF | Q7Z460; F8V | CLAP1_HUMAN | Unmodified | 0 | 0 | 0 | 0 | 203410 | 0 | 0 | 0 | 0 | 0 | 0 | 647230 | 0 | 0 | 0 | 0 | 0 | 0 |
| SGNWESSI | SNX9; SNX9; SN | Q9Y5X1; A0/ | SNX9_HUMAN | Unmodified | 9699200 | 11437000 | 1774300 | 3333100 | 0 | 0 | 0 | 0 | 515940 | 0 | 0 | 1807200 | 1734000 | 143300 | 2922500 | 4509000 | 0 | 0 |

| | | | | | | | | | | | | | | | | | | | | | | |
|---|---|---|---|---|---|---|---|---|---|---|---|---|---|---|---|---|---|---|---|---|---|---|
| SGNYFASH | MGRN1; MGRN | O60291; B4I | MGRN1_HUMA | Oxidation (M) | 51446 | 20613 | 175670 | 82292 | 46614 | 0 | 0 | 0 | 0 | 0 | 0 | 318700 | 63254 | 0 | 147390 | 220290 | 0 | 0 |
| SGPASTFN | LARS1; LARS1; | Q9P2J5; A0A | SYLC_HUMAN | Unmodified | 0 | 0 | 5525000 | 0 | 0 | 75892 | 0 | 0 | 0 | 0 | 27425 | 1321500 | 0 | 0 | 7607900 | 6306000 | 0 | 0 |
| SGPASTFN | LARS1; LARS1; | Q9P2J5; A0A | SYLC_HUMAN | Unmodified | 0 | 0 | 317820 | 0 | 0 | 0 | 0 | 0 | 571060 | 0 | 0 | 6927200 | 0 | 0 | 0 | 0 | 0 | 0 |
| SGPASTFN | LARS1; LARS1; | Q9P2J5; A0A | SYLC_HUMAN | Unmodified | 0 | 0 | 0 | 0 | 0 | 0 | 0 | 0 | 704510 | 0 | 0 | 8653500 | 0 | 0 | 0 | 0 | 0 | 0 |
| SGPEGDSK | UBAP1; UBAP1 | Q9NZ09; A0/ | UBAP1_HUMAI | Unmodified | 0 | 0 | 0 | 0 | 348400 | 53666 | 0 | 0 | 0 | 6686.6 | 141940 | 784460 | 0 | 47438 | 0 | 0 | 0 | 0 |
| SGPEPLQE | CORO7; CORO | P57737; A0A | CORO7_HUMA | Unmodified | 38411 | 33136 | 39297 | 0 | 0 | 0 | 0 | 0 | 0 | 0 | 659400 | 2610500 | 0 | 0 | 55309 | 47552 | 0 | 0 |
| SGPFAPVLS | UQCRFS1P1; U | P0C7P4; P47 | UCRIL_HUMAN | Unmodified | 0 | 0 | 0 | 0 | 0 | 0 | 0 | 0 | 3933100 | 618900 | 0 | 0 | 267990 | 0 | 0 | 0 | 0 | 0 |
| SGPFGQIFI | TUBB2A; TUBB4 | Q13885; P04 | TBB2A_HUMAN | Unmodified | 22700000 | 21605000 | 11933000 | 35428000 | 45768000 | 10609000 | 70901000 | 0 | 2.28E+08 | 48185000 | 78014000 | 2.45E+08 | 47305000 | 13735000 | 69039000 | 32443000 | 13475000 | 48719000 |
| SGPFGQLF | TUBB6; TUBB6; | Q9BUF5; K7I | TBB6_HUMAN | Unmodified | 0 | 0 | 0 | 0 | 2905400 | 0 | 0 | 0 | 1025200 | 136100 | 0 | 0 | 0 | 0 | 0 | 0 | 0 | 0 |
| SGPGAYES | CACNA2D1 | P54289 | CA2D1_HUMAI | Oxidation (M) | 590510 | 1080300 | 0 | 0 | 0 | 0 | 461300 | 292240 | 848450 | 180190 | 0 | 0 | 0 | 0 | 0 | 0 | 0 | 0 |
| SGPGAYES | CACNA2D1 | P54289 | CA2D1_HUMAI | Unmodified | 214500 | 398850 | 0 | 0 | 0 | 0 | 1397500 | 608560 | 3133900 | 845420 | 0 | 0 | 0 | 0 | 0 | 0 | 0 | 0 |
| SGPHCSES | PRCP; PRCP; PI | P42785; B7Z | PCP_HUMAN L | Unmodified | 0 | 0 | 49833 | 0 | 0 | 0 | 0 | 0 | 524310 | 73116 | 26319 | 115400 | 0 | 0 | 23002 | 217930 | 0 | 0 |
| SGPIFIVVPI | CKAP5 | Q14008 | CKAP5_HUMAI | Unmodified | 0 | 0 | 0 | 0 | 0 | 0 | 0 | 0 | 0 | 0 | 0 | 1079900 | 0 | 0 | 0 | 0 | 0 | 0 |
| SGPKPFSAI | CAP1 | Q01518 | CAP1_HUMAN | Unmodified | 0 | 0 | 0 | 0 | 1455700 | 393340 | 0 | 0 | 797550 | 155060 | 0 | 2596200 | 0 | 0 | 0 | 0 | 440450 | 603590 |
| SGPRLEEFI | N4BP1 | O75113 | N4BP1_HUMAI | Unmodified | 517570 | 636560 | 184290 | 179310 | 72647 | 0 | 0 | 0 | 67226 | 0 | 0 | 0 | 0 | 0 | 146150 | 197290 | 165520 | 0 |
| SGPRPVVLS | GUK1; GUK1; G | Q16774; B1/ | KGUA_HUMAN | Acetyl (Protein N | 0 | 0 | 0 | 41465 | 1469300 | 0 | 870190 | 741840 | 263130 | 219610 | 0 | 0 | 1565500 | 1322300 | 0 | 30842 | 1738400 | 3187800 |
| SGPVGSVV | ERBIN | Q96RT1 | ERBIN_HUMAN | Unmodified | 40792 | 0 | 0 | 45905 | 19462000 | 0 | 527090 | 13603 | 616510 | 179800 | 0 | 1013900 | 0 | 0 | 0 | 99467 | 1350900 | 3257400 |
| SGPVILADE | ARFIP1 | P53367 | ARFP1_HUMAN | Oxidation (M) | 216860 | 0 | 0 | 0 | 0 | 0 | 0 | 0 | 0 | 0 | 0 | 0 | 0 | 0 | 0 | 318510 | 0 | 0 |
| SGQAAASC | CHSY1 | Q86X52 | CHSS1_HUMAI | Unmodified | 0 | 0 | 0 | 0 | 0 | 0 | 0 | 0 | 306400 | 0 | 0 | 0 | 0 | 0 | 0 | 0 | 0 | 0 |
| SGQAEVVR | ANK3 | Q12955 | ANK3_HUMAN | Unmodified | 0 | 0 | 0 | 0 | 0 | 0 | 0 | 0 | 0 | 156240 | 0 | 0 | 1999100 | 467370 | 138830 | 207930 | 0 | 0 |
| SGQCVQAI | GNB5; GNB5 | O14775; H0 | GNB5_HUMAN | Unmodified | 0 | 0 | 0 | 0 | 0 | 0 | 0 | 0 | 519660 | 0 | 0 | 0 | 160390 | 0 | 0 | 0 | 0 | 0 |
| SGQDPYAF | GALNT2 | Q10471 | GALT2_HUMAN | Unmodified | 0 | 0 | 0 | 0 | 0 | 0 | 0 | 0 | 0 | 0 | 0 | 0 | 0 | 0 | 0 | 0 | 0 | 0 |
| SGQFFLESI | DMXL1; DMXL1 | Q9Y485; F5H | DMXL1_HUMAI | Unmodified | 0 | 0 | 0 | 0 | 0 | 0 | 683610 | 0 | 33157 | 0 | 50197 | 207700 | 0 | 0 | 0 | 54173 | 0 | 0 |
| SGQGAFGI | RPL4; RPL4; RP | P36578; H3E | RL4_HUMAN 6 | Oxidation (M) | 1931500 | 430000 | 1774800 | 814110 | 0 | 0 | 0 | 0 | 0 | 0 | 87846 | 2044300 | 0 | 0 | 478900 | 318370 | 0 | 0 |
| SGQGAFGI | RPL4; RPL4; RP | P36578; H3E | RL4_HUMAN 6 | Unmodified | 2792100 | 2236100 | 1086700 | 1939500 | 0 | 0 | 0 | 0 | 355160 | 0 | 0 | 8834100 | 0 | 0 | 0 | 850950 | 0 | 0 |
| SGQIKSEVI | MYO1B; MYO1E | O43795; H7 | MYO1B_HUMA | Unmodified | 0 | 0 | 0 | 0 | 15999000 | 1756600 | 0 | 0 | 468840 | 0 | 645030 | 496610 | 956750 | 555550 | 0 | 0 | 1839400 | 1882000 |
| SGQIKSEVI | MYO1B; MYO1E | O43795; H7 | MYO1B_HUMA | Oxidation (M) | 0 | 0 | 0 | 432530 | 1303900 | 0 | 1243000 | 0 | 0 | 0 | 0 | 0 | 0 | 0 | 0 | 0 | 876330 | 0 |
| SGQIRNLES | PPME1 | Q9Y570 | PPME1_HUMAI | Unmodified | 0 | 0 | 0 | 0 | 712730 | 0 | 289530 | 0 | 0 | 0 | 0 | 0 | 608070 | 510280 | 0 | 0 | 648040 | 485510 |
| SGQLGIQE | SCG2 | P13521 | SCG2_HUMAN | Unmodified | 250540 | 0 | 0 | 0 | 242400 | 0 | 214810 | 534300 | 5142000 | 2779500 | 0 | 282180 | 0 | 0 | 339850 | 0 | 0 | 0 |
| SGQQIVGP | NSFL1C; NSFL1 | Q9UNZ2; F2 | NSF1C_HUMAI | Unmodified | 0 | 0 | 0 | 0 | 341550 | 0 | 0 | 0 | 317820 | 48153 | 504880 | 515450 | 0 | 0 | 0 | 0 | 0 | 0 |
| SGQSLTDR | PICALM; PICALI | Q13492; E9F | PICAL_HUMAN | Acetyl (Protein N | 2133200 | 3189600 | 908180 | 0 | 10708000 | 1736000 | 832550 | 515340 | 2829300 | 3579600 | 871170 | 2614800 | 8487300 | 6716200 | 1108300 | 889520 | 3535900 | 3713900 |
| SGRFFTVKI | RPN1 | P04843 | RPN1_HUMAN | Unmodified | 0 | 0 | 0 | 0 | 251360 | 0 | 0 | 0 | 16187000 | 36462000 | 0 | 0 | 336070 | 180760 | 0 | 0 | 0 | 0 |
| SGRHVVFY | CHID1; CHID1; | Q9BWS9; H | CHID1_HUMAN | Unmodified | 0 | 0 | 0 | 0 | 0 | 0 | 0 | 0 | 0 | 868800 | 0 | 0 | 0 | 0 | 0 | 0 | 0 | 0 |
| SGRLLLAG | GNB1; GNB1 | P62873; B3K | GBB1_HUMAN | Unmodified | 0 | 0 | 0 | 0 | 19777000 | 2337500 | 0 | 0 | 0 | 5699800 | 0 | 0 | 3187700 | 6788200 | 0 | 0 | 25621000 | 18977000 |
| SGRLLLAG | GNB1; GNB1 | P62873; B3K | GBB1_HUMAN | Unmodified | 0 | 0 | 0 | 0 | 0 | 0 | 0 | 0 | 0 | 1961600 | 0 | 0 | 0 | 0 | 0 | 0 | 0 | 0 |
| SGRLLLAG | GNB4; GNB4; G | Q9HAV0; A0 | GBB4_HUMAN | Unmodified | 0 | 0 | 0 | 0 | 733370 | 734320 | 0 | 0 | 0 | 0 | 0 | 0 | 0 | 0 | 0 | 0 | 620890 | 0 |
| SGRLWSK | RPL35A; RPL35 | P18077; F8V | RL35A_HUMAN | Acetyl (Protein N | 0 | 0 | 0 | 189860 | 960200 | 0 | 730950 | 3066200 | 0 | 635080 | 484850 | 0 | 1422700 | 0 | 0 | 0 | 0 | 0 |
| SGRYEAAF | C8orf82; C8orf | Q6P1X6; H0 | CH082_HUMAI | Unmodified | 0 | 0 | 0 | 0 | 0 | 0 | 0 | 0 | 768440 | 0 | 0 | 0 | 0 | 0 | 0 | 0 | 0 | 0 |
| SGSAADTQ | PSMB6; PSMB6 | P28072; A0A | PSB6_HUMAN | Unmodified | 0 | 0 | 0 | 0 | 0 | 0 | 0 | 0 | 0 | 0 | 0 | 0 | 2291300 | 495980 | 0 | 0 | 0 | 0 |
| SGSAHEYS | CPE | P16870 | CBPE_HUMAN | Unmodified | 0 | 0 | 0 | 0 | 0 | 0 | 0 | 0 | 70033000 | 4555700 | 1218100 | 0 | 0 | 0 | 0 | 0 | 0 | 275960 |
| SGSCAHPI | APLP1; APLP1; | P51693; B7Z | APLP1_HUMAN | Unmodified | 0 | 0 | 0 | 0 | 0 | 0 | 0 | 0 | 4856300 | 432210 | 0 | 0 | 0 | 0 | 0 | 0 | 0 | 0 |
| SGSEEVFR | DAAM1 | Q9Y4D1 | DAAM1_HUMA | Unmodified | 0 | 64871 | 0 | 0 | 0 | 0 | 0 | 0 | 249640 | 0 | 263850 | 2138300 | 20902 | 0 | 0 | 52596 | 0 | 0 |
| SGSEEVFR | DAAM1 | Q9Y4D1 | DAAM1_HUMA | Unmodified | 39099 | 34185 | 0 | 194630 | 0 | 0 | 201620 | 0 | 193400 | 100660 | 0 | 0 | 189440 | 0 | 0 | 0 | 0 | 40063 |
| SGSGSRG | ACSS2; ACSS2; | Q9NR19; B4 | ACSA_HUMAN | Unmodified | 3074700 | 209200 | 54367 | 8087400 | 3403800 | 3149000 | 2233600 | 3979400 | 0 | 0 | 0 | 0 | 2454800 | 1108300 | 400680 | 4226300 | 6762900 | 0 |
| SGSGTMNI | CAPZB; CAPZB; | P47756; B1A | CAPZB_HUMAI | Oxidation (M) | 1108800 | 0 | 0 | 937980 | 511630 | 0 | 0 | 0 | 958980 | 196910 | 773310 | 3288400 | 329770 | 0 | 832500 | 678950 | 0 | 0 |
| SGSGTMNI | CAPZB; CAPZB; | P47756; B1A | CAPZB_HUMAI | Unmodified | 2524800 | 0 | 0 | 0 | 0 | 0 | 0 | 0 | 4544300 | 1101000 | 3155600 | 16846000 | 0 | 2308500 | 0 | 0 | 0 | 0 |
| SGSHTVAV | PFKM; PFKM; PI | P08237; A0A | PFKAM_HUMAI | Oxidation (M) | 0 | 0 | 0 | 0 | 0 | 0 | 0 | 0 | 0 | 0 | 0 | 709670 | 0 | 0 | 0 | 0 | 0 | 0 |
| SGSLAVDN | AGL | P35573 | GDE_HUMAN ( | Unmodified | 0 | 0 | 0 | 0 | 0 | 0 | 0 | 0 | 161960 | 88369 | 0 | 732360 | 0 | 0 | 0 | 0 | 0 | 0 |
| SGSLENVP | BLVRA | P53004 | BIEA_HUMAN E | Unmodified | 923320 | 636910 | 646530 | 1083800 | 7010100 | 0 | 1181600 | 1528900 | 539970 | 709750 | 466300 | 1354600 | 6565900 | 5414600 | 1113800 | 1162500 | 2657400 | 4716200 |
| SGSLENVP | BLVRA | P53004 | BIEA_HUMAN E | Unmodified | 0 | 0 | 0 | 0 | 401550 | 0 | 691750 | 413740 | 0 | 99245 | 0 | 0 | 672850 | 768600 | 0 | 0 | 1313000 | 329170 |
| SGSLENVP | BLVRA | P53004 | BIEA_HUMAN E | Unmodified | 0 | 0 | 0 | 0 | 512100 | 0 | 696960 | 50494 | 0 | 0 | 0 | 0 | 578030 | 339430 | 0 | 0 | 3168300 | 0 |
| SGSLRQCF | ADGRG6 | Q86SQ4 | AGRG6_HUMA | Unmodified | 0 | 0 | 0 | 0 | 1936100 | 0 | 0 | 0 | 0 | 0 | 0 | 0 | 794820 | 0 | 0 | 0 | 0 | 33392 |
| SGSMDPS | KHDRBS1 | Q07666 | KHDR1_HUMAI | Unmodified | 145730 | 30606 | 0 | 0 | 240840 | 0 | 0 | 0 | 182670 | 597150 | 646920 | 1570300 | 537740 | 350400 | 20118 | 30372 | 0 | 0 |
| SGSNLPISI | SUCO | Q9UBS9 | SUCO_HUMAN | Unmodified | 0 | 0 | 0 | 0 | 0 | 0 | 0 | 0 | 581050 | 283660 | 0 | 0 | 0 | 0 | 0 | 0 | 0 | 0 |
| SGSPSDNS | AKT1 | P31749 | AKT1_HUMAN I | Unmodified | 0 | 0 | 0 | 0 | 1170400 | 167910 | 0 | 0 | 0 | 108610 | 0 | 1394000 | 221100 | 75599 | 0 | 0 | 147970 | 0 |
| SGSPSDNS | AKT1 | P31749 | AKT1_HUMAN I | Oxidation (M) | 0 | 0 | 0 | 199930 | 0 | 0 | 0 | 0 | 0 | 0 | 271180 | 0 | 0 | 0 | 284370 | 338990 | 0 | 0 |
| SGSSFVHC | RANBP2 | P49792 | RBP2_HUMAN | Unmodified | 0 | 0 | 0 | 0 | 0 | 0 | 0 | 0 | 0 | 0 | 0 | 1131800 | 454680 | 97624 | 0 | 0 | 0 | 101930 |
| SGSSRVLIT | EIF4A1; EIF4A2; | P60842; Q1 | IF4A1_HUMAN | Unmodified | 0 | 0 | 0 | 0 | 244460 | 0 | 0 | 0 | 0 | 0 | 0 | 0 | 126790 | 0 | 0 | 0 | 75655 | 123130 |
| SGSSSVAA | GNG5 | P63218 | GBG5_HUMAN | Acetyl (Protein N | 491820 | 448170 | 0 | 454360 | 5742100 | 0 | 2244500 | 2693700 | 3096200 | 945860 | 1966500 | 8876500 | 3960700 | 0 | 703520 | 782500 | 0 | 8694900 |
| SGSTAVGV | PPM1A; PPM1A | P35813; E9P | PPM1A_HUMAI | Unmodified | 800880 | 0 | 0 | 0 | 0 | 0 | 0 | 0 | 0 | 0 | 0 | 5969900 | 0 | 0 | 758180 | 482870 | 0 | 0 |
| SGSVEEQL | RTN4IP1 | Q8WWV3 | RT4I1_HUMAN | Unmodified | 0 | 35195 | 0 | 0 | 0 | 0 | 25335 | 0 | 661450 | 224990 | 98482 | 0 | 0 | 0 | 30527 | 0 | 0 | 0 |
| SGSYSLPR | SARG | Q9BW04 | SARG_HUMAN | Unmodified | 0 | 0 | 0 | 0 | 0 | 0 | 0 | 0 | 0 | 0 | 0 | 0 | 0 | 542490 | 0 | 0 | 0 | 0 |
| SGTDVDAA | CASP3; CASP3; | P42574; A8N | CASP3_HUMAI | Unmodified | 0 | 0 | 0 | 0 | 0 | 0 | 0 | 0 | 260160 | 0 | 559380 | 2810800 | 0 | 0 | 0 | 0 | 0 | 0 |
| SGTDVDAA | CASP3; CASP3; | P42574; A8N | CASP3_HUMAI | Unmodified | 59018 | 0 | 192830 | 104590 | 0 | 177010 | 0 | 0 | 373930 | 0 | 1018800 | 553680 | 0 | 0 | 60756 | 0 | 293810 | 208300 |
| SGTGIAAM | ATP6V0C | P27449 | VATL_HUMAN | Oxidation (M) | 0 | 28184 | 0 | 0 | 159810 | 0 | 0 | 0 | 2622200 | 843760 | 0 | 560910 | 0 | 0 | 0 | 0 | 0 | 0 |
| SGTGIAAM | ATP6V0C | P27449 | VATL_HUMAN | Unmodified | 0 | 0 | 0 | 0 | 634620 | 0 | 0 | 0 | 1595200 | 164090 | 0 | 0 | 0 | 0 | 0 | 0 | 0 | 0 |
| SGTIFDNFI | CALR; CALR; C/ | P27797; K7E | CALR_HUMAN | Unmodified | 0 | 0 | 0 | 0 | 0 | 0 | 0 | 0 | 0 | 1033200 | 0 | 0 | 0 | 666560 | 0 | 0 | 0 | 890410 |
| SGTNILYW | ELAPOR2; ELAP | A8MWY0; H7 | ELAP2_HUMAN | Unmodified | 0 | 0 | 0 | 0 | 228340 | 0 | 0 | 0 | 1682400 | 1075400 | 0 | 0 | 1059400 | 0 | 0 | 0 | 56504 | 0 |
| SGTNILYW | ELAPOR2; ELAP | A8MWY0; H7 | ELAP2_HUMAN | Oxidation (M) | 0 | 0 | 0 | 0 | 0 | 0 | 294270 | 0 | 0 | 128200 | 0 | 0 | 0 | 0 | 0 | 0 | 0 | 0 |
| SGTQDIQP | ZC3HAV1; ZC3I | Q7Z2W4; C9 | ZCCHV_HUMA | Unmodified | 2029800 | 0 | 0 | 11201000 | 10405000 | 3194000 | 0 | 0 | 1767900 | 0 | 0 | 566560 | 4565000 | 2120800 | 0 | 0 | 0 | 0 |
| SGTQDIQP | ZC3HAV1; ZC3I | Q7Z2W4; C9 | ZCCHV_HUMA | Unmodified | 0 | 0 | 0 | 0 | 16161000 | 2813100 | 0 | 0 | 0 | 0 | 0 | 0 | 2819100 | 4058500 | 0 | 0 | 1547400 | 0 |
| SGTRPGPF | MICALL1; MICA | Q8N3F8; B0 | MILK1_HUMAN | Unmodified | 218010 | 279360 | 130110 | 0 | 1022400 | 338600 | 0 | 19918 | 0 | 87544 | 126490 | 228360 | 855010 | 523070 | 243630 | 0 | 0 | 399880 |
| SGTRVDHC | GPI; GPI; GPI; N | P06744; K7E | G6PI_HUMAN ( | Unmodified | 1666600 | 1078000 | 4087900 | 9143900 | 72164000 | 13625000 | 4645600 | 0 | 34255000 | 7278500 | 9363600 | 1.13E+08 | 21154000 | 11168000 | 4474000 | 5198100 | 14805000 | 25743000 |
| SGTSEFLNI | HSP90B1; HSP | P14625; A0A | ENPL_HUMAN | Unmodified | 720210 | 309440 | 1408800 | 640740 | 108000 | 0 | 5975400 | 3600400 | 3.77E+08 | 93309000 | 50943000 | 15047000 | 1958800 | 1485800 | 1807300 | 1332600 | 171320 | 2656600 |
| SGTVDPQE | SRI; SRI | P30626; C9J | SORCN_HUMA | Unmodified | 788490 | 945560 | 0 | 173320 | 0 | 0 | 136010 | 57274 | 12174000 | 1955400 | 0 | 4014200 | 0 | 0 | 0 | 1070700 | 0 | 0 |
| SGVAYIAAF | ATIC | P31939 | PUR9_HUMAN | Unmodified | 0 | 0 | 0 | 0 | 0 | 0 | 0 | 0 | 0 | 0 | 212510 | 1326700 | 0 | 0 | 0 | 0 | 0 | 0 |
| SGVEAFILG | SNX33 | Q8WV41 | SNX33_HUMAI | 2 Oxidation (M) | 0 | 0 | 0 | 1451900 | 0 | 0 | 0 | 0 | 0 | 0 | 0 | 0 | 0 | 0 | 3248900 | 1133700 | 0 | 0 |
| SGVGLSSY | SARG | Q9BW04 | SARG_HUMAN | Unmodified | 0 | 0 | 536140 | 769150 | 0 | 0 | 0 | 0 | 0 | 0 | 0 | 0 | 1773900 | 2003200 | 469720 | 583000 | 0 | 0 |
| SGVGNIFIK | PABPC5; PABP | Q96DU9; P1 | PABP5_HUMAN | Unmodified | 65626000 | 57380000 | 17297000 | 20438000 | 3269700 | 0 | 3656900 | 1434600 | 236730 | 0 | 919340 | 9958300 | 15537000 | 0 | 8191600 | 19262000 | 2671900 | 0 |
| SGVGNIFIK | PABPC5; PABP | Q96DU9; P1 | PABP5_HUMAN | Unmodified | 0 | 532220 | 0 | 84425 | 231410 | 0 | 678980 | 356950 | 0 | 0 | 0 | 0 | 1263300 | 870070 | 0 | 0 | 301250 | 116750 |
| SGVGNIFIK | PABPC1; PABP | P11940; E5R | PABP1_HUMAN | Unmodified | 0 | 0 | 0 | 0 | 46784 | 0 | 210100 | 13324 | 0 | 0 | 84830 | 0 | 593570 | 434430 | 0 | 0 | 0 | 46061 |
| SGVGNVFI | PABPC4; PABP | Q13310; Q5 | PABP4_HUMAN | Unmodified | 14686000 | 12634000 | 5936000 | 5144100 | 0 | 0 | 0 | 525890 | 0 | 996170 | 0 | 1902300 | 0 | 0 | 3702200 | 2894100 | 185960 | 0 |
| SGVGNVFI | PABPC4; PABP | Q13310; B1/ | PABP4_HUMAN | Unmodified | 50626 | 89653 | 0 | 0 | 0 | 0 | 87967 | 58987 | 0 | 0 | 0 | 0 | 264720 | 0 | 0 | 0 | 54236 | 0 |
| SGVGTALLI | TFRC; TFRC | P02786; G3 | TFR1_HUMAN | Unmodified | 2247600 | 7596900 | 8225400 | 2570200 | 0 | 0 | 0 | 0 | 12430000 | 3007200 | 0 | 6079000 | 0 | 0 | 14579000 | 8720800 | 0 | 0 |
| SGVISDTEL | PDCD6; PDCD6 | O75340; H0 | PDCD6_HUMA | Unmodified | 6629600 | 5904700 | 781050 | 3756900 | 0 | 0 | 0 | 0 | 24278000 | 1659000 | 0 | 666910 | 0 | 0 | 9101300 | 1912200 | 0 | 0 |
| SGVKIHVSI | UBA1; UBA1 | P22314; Q5 | UBA1_HUMAN | Unmodified | 0 | 0 | 0 | 0 | 0 | 0 | 991120 | 0 | 0 | 0 | 0 | 0 | 0 | 0 | 0 | 0 | 1500600 | 0 |
| SGVLDESTI | OXSR1; OXSR1 | O95747; C9 | OXSR1_HUMAI | Unmodified | 0 | 0 | 0 | 0 | 0 | 0 | 0 | 0 | 65663 | 0 | 0 | 1862800 | 0 | 0 | 119510 | 0 | 0 | 0 |
| SGVLDTLTI | MYCBP; N/A; M | Q99417; A0/ | MYCBP_HUMA | Unmodified | 779040 | 538060 | 207810 | 199800 | 3548400 | 1102900 | 1890800 | 1236000 | 0 | 901380 | 355750 | 2175600 | 3040800 | 1327000 | 77842 | 378950 | 1131400 | 1234500 |
| SGVLSVSS | LAMA4; LAMA4; | Q16363; E5F | LAMA4_HUMAI | Unmodified | 0 | 0 | 0 | 0 | 0 | 0 | 0 | 0 | 436800 | 0 | 0 | 0 | 0 | 0 | 0 | 0 | 0 | 0 |
| SGVRAGEA | EHBP1L1; EHBI | Q8N3D4; E9 | EH1L1_HUMAN | Unmodified | 0 | 0 | 0 | 0 | 420390 | 0 | 0 | 0 | 0 | 101030 | 0 | 0 | 0 | 0 | 0 | 0 | 330810 | 0 |
| SGVSLAALI | H1-3; H1-4; H1 | P16402; P10 | H13_HUMAN H | Unmodified | 12125000 | 7267400 | 0 | 125280 | 1106800 | 174880 | 1270800 | 30418000 | 297470 | 267830 | 0 | 254020 | 253140 | 0 | 311150 | 81605 | 51143 | 369730 |
| SGVSLAALI | H1-3; H1-4; H1 | P16402; P10 | H13_HUMAN H | Unmodified | 0 | 3642000 | 0 | 0 | 0 | 0 | 260470 | 10840000 | 0 | 0 | 0 | 0 | 0 | 0 | 0 | 0 | 0 | 0 |
| SGVSLPQR | AHNAK2 | Q8IVF2 | AHNK2_HUMAI | Unmodified | 0 | 0 | 0 | 0 | 4580600 | 1027400 | 0 | 0 | 0 | 0 | 0 | 0 | 0 | 0 | 0 | 0 | 0 | 4016500 |
| SGWADER | SRSF9; SRSF9 | Q13242; S4F | SRSF9_HUMAN | Acetyl (Protein N | 173630 | 666830 | 0 | 0 | 0 | 0 | 0 | 0 | 0 | 0 | 174830 | 0 | 0 | 0 | 75206 | 81203 | 0 | 0 |
| SGWAIDPF | MAN2A1 | Q16706 | MA2A1_HUMAI | Unmodified | 0 | 0 | 0 | 0 | 0 | 0 | 0 | 0 | 620970 | 60604 | 0 | 0 | 0 | 0 | 0 | 0 | 0 | 0 |
| SGWAIDPF | MAN2A1 | Q16706 | MA2A1_HUMAI | Oxidation (M) | 0 | 0 | 0 | 0 | 0 | 1423000 | 122010 | 0 | 1575100 | 515420 | 0 | 2141700 | 0 | 0 | 0 | 204840 | 0 | 0 |
| SGWLLRQS | PLEKHB2; PLEK | Q96CS7; A0/ | PKHB2_HUMAI | Unmodified | 0 | 0 | 0 | 0 | 130960 | 0 | 0 | 0 | 0 | 0 | 0 | 0 | 176170 | 537620 | 0 | 0 | 59879 | 70513 |
| SGYFDERY | CKMT1A; CKMT | P12532; C9J | KCRU_HUMAN | Unmodified | 295760 | 43525 | 909580 | 692210 | 0 | 0 | 0 | 0 | 541830 | 114030 | 765690 | 1196300 | 667930 | 249860 | 195030 | 274020 | 0 | 0 |
| SGYHQSAS | ESD; ESD; ESD; | P10768; X6R | ESTD_HUMAN | Unmodified | 0 | 789170 | 2318200 | 3217400 | 2128800 | 727510 | 0 | 596890 | 6300600 | 651560 | 4955700 | 26794000 | 2541800 | 1430200 | 2440100 | 2885000 | 1480500 | 1792400 |
| SGYLAGDK | MSN | P26038 | MOES_HUMAN | Unmodified | 0 | 0 | 0 | 0 | 0 | 0 | 0 | 32811 | 413260 | 0 | 54537 | 0 | 0 | 1364300 | 0 | 0 | 10707000 | 0 |
| SGYLAGDK | MSN | P26038 | MOES_HUMAN | Unmodified | 0 | 0 | 0 | 0 | 367710 | 159710 | 287610 | 191980 | 0 | 0 | 0 | 0 | 137570 | 308520 | 0 | 0 | 450510 | 293680 |
| SGYLAGDK | MSN | P26038 | MOES_HUMAN | Unmodified | 0 | 0 | 0 | 0 | 3481100 | 0 | 0 | 0 | 0 | 0 | 0 | 0 | 814870 | 754940 | 0 | 0 | 10476000 | 0 |
| SGYLLPDTI | HSP90B1; HSP | P14625; A0A | ENPL_HUMAN | Unmodified | 710680 | 184210 | 2025700 | 781910 | 829850 | 0 | 955590 | 968070 | 2.9E+08 | 51298000 | 38479000 | 25209000 | 301880 | 759870 | 2410700 | 1938300 | 165980 | 1186600 |
| SGYLSSER | EZR; EZR | P15311; E7E | EZRI_HUMAN E | Unmodified | 1857100 | 0 | 927920 | 1513200 | 216040 | 2548900 | 46519 | 0 | 11664000 | 1460400 | 1921600 | 18826000 | 709780 | 447380 | 1817400 | 2672500 | 351550 | 472670 |
| SGYLSSERI | EZR; EZR | P15311; E7E | EZRI_HUMAN E | Unmodified | 0 | 0 | 0 | 0 | 960510 | 742750 | 0 | 0 | 0 | 2418100 | 1940900 | 816050 | 0 | 0 | 0 | 0 | 0 | 0 |
| SGYQALPW | AP1M1; AP1M2 | Q9BXS5; Q9 | AP1M1_HUMAI | Unmodified | 0 | 0 | 1365300 | 0 | 0 | 0 | 0 | 0 | 0 | 0 | 0 | 1803900 | 0 | 0 | 0 | 0 | 0 | 0 |
| SGYRIDFYI | SETSIP; SET; SE | P0DME0; Q0 | SETLP_HUMAN | Unmodified | 0 | 0 | 0 | 0 | 0 | 0 | 0 | 0 | 0 | 808790 | 0 | 0 | 0 | 0 | 0 | 0 | 0 | 0 |
| SGYRIQAS | LSR; LSR | Q86X29; S4F | LSR_HUMAN Li | Unmodified | 0 | 0 | 249590 | 1564700 | 187890 | 109850 | 0 | 0 | 5136700 | 454990 | 489160 | 0 | 0 | 0 | 252960 | 248450 | 39539 | 0 |
| SGYRIQAS | LSR; LSR | Q86X29; S4F | LSR_HUMAN Li | Oxidation (M) | 108410 | 90607 | 161270 | 564720 | 161110 | 0 | 0 | 0 | 1796300 | 251610 | 346320 | 81095 | 215920 | 0 | 142220 | 168910 | 0 | 0 |

| | | | | | | | | | | | | | | | | | | | | | | |
|---|---|---|---|---|---|---|---|---|---|---|---|---|---|---|---|---|---|---|---|---|---|---|
| SGYRIQASG | LSR; LSR | Q86X29; S4F | LSR_HUMAN Li | Unmodified | 0 | 0 | 0 | 0 | 11506000 | 777120 | 0 | 0 | 1871400 | 8050800 | 0 | 0 | 2242200 | 728100 | 0 | 0 | 374120 | 1470700 |
| SGYRIQASG | LSR; LSR | Q86X29; S4F | LSR_HUMAN Li | Oxidation (M) | 0 | 0 | 0 | 476230 | 2993000 | 540850 | 0 | 0 | 536900 | 10408000 | 580880 | 0 | 1888700 | 867780 | 0 | 0 | 256600 | 338250 |
| SGYRIQASG | LSR; LSR | Q86X29; S4F | LSR_HUMAN Li | 2 Oxidation (M) | 0 | 0 | 0 | 119880 | 396400 | 0 | 0 | 0 | 0 | 1622800 | 0 | 0 | 0 | 0 | 0 | 0 | 0 | 0 |
| SHAAEAPG | ENPP1; ENPP1; | P22413; A0A | ENPP1_HUMAN | Unmodified | 0 | 0 | 0 | 0 | 0 | 0 | 0 | 0 | 1417800 | 0 | 0 | 0 | 0 | 0 | 0 | 0 | 825010 | 0 |
| SHAGYQTI | LAMP1 | P11279 | LAMP1_HUMAN | Unmodified | 0 | 237980 | 518950 | 0 | 0 | 0 | 0 | 0 | 1777700 | 175210 | 0 | 757970 | 0 | 0 | 970760 | 759340 | 345770 | 536220 |
| SHAVACVN | TNPO1; TNPO1 | Q92973; A0A | TNPO1_HUMAN | Unmodified | 0 | 0 | 0 | 0 | 0 | 0 | 0 | 0 | 0 | 0 | 0 | 668390 | 0 | 0 | 332620 | 200610 | 0 | 166350 |
| SHAVDKIQ | TSPAN6; TSPAN | O43657; A0A | TSN6_HUMAN | Unmodified | 0 | 0 | 0 | 0 | 0 | 0 | 0 | 0 | 0 | 663350 | 0 | 0 | 0 | 0 | 0 | 0 | 0 | 0 |
| SHDQAVRT | LTA4H | P09960 | LKHA4_HUMAN | Unmodified | 0 | 0 | 0 | 0 | 3028200 | 28738 | 0 | 0 | 0 | 0 | 0 | 0 | 623640 | 403270 | 0 | 0 | 476180 | 1247800 |
| SHDVYWD | PIGS | Q96S52 | PIGS_HUMAN ( | Unmodified | 0 | 0 | 0 | 0 | 0 | 0 | 0 | 0 | 978070 | 0 | 0 | 0 | 0 | 0 | 0 | 0 | 0 | 0 |
| SHEAEVLK | STMN1; STMN1 | P16949; A2A | STMN1_HUMAN | Unmodified | 0 | 99833 | 0 | 0 | 2350500 | 377760 | 829990 | 1159100 | 6983300 | 0 | 1183200 | 5083000 | 1790500 | 369070 | 0 | 0 | 0 | 1500600 |
| SHEETDSTI | SCG3 | Q8WXD2 | SCG3_HUMAN | Unmodified | 142730 | 0 | 0 | 0 | 0 | 0 | 0 | 0 | 3064600 | 0 | 0 | 0 | 0 | 0 | 0 | 0 | 0 | 0 |
| SHEGETAY | SRSF1; SRSF1; | Q07955; J3K | SRSF1_HUMAN | Unmodified | 508020 | 464860 | 137800 | 260300 | 0 | 0 | 0 | 12476 | 0 | 0 | 0 | 301330 | 0 | 0 | 203070 | 333600 | 77679 | 0 |
| SHFEQWG | HNRNPA1L2; H | Q32P51; P09 | RA1L2_HUMAN | Unmodified | 0 | 217900 | 0 | 0 | 0 | 0 | 0 | 0 | 387980 | 0 | 0 | 524950 | 0 | 0 | 0 | 0 | 0 | 0 |
| SHFEQWG | HNRNPA1L2; H | Q32P51; P09 | RA1L2_HUMAN | Oxidation (M) | 86006 | 0 | 0 | 0 | 0 | 0 | 0 | 0 | 810240 | 0 | 200360 | 1949300 | 0 | 0 | 0 | 0 | 0 | 0 |
| SHFEQWG | HNRNPA1L2; H | Q32P51; P09 | RA1L2_HUMAN | Oxidation (M) | 0 | 0 | 0 | 0 | 0 | 0 | 0 | 0 | 0 | 0 | 0 | 524980 | 0 | 0 | 0 | 0 | 0 | 0 |
| SHFGRDLN | ANXA6; ANXA6 | P08133; E5R | ANXA6_HUMAN | Unmodified | 0 | 0 | 0 | 0 | 0 | 0 | 0 | 0 | 731690 | 280040 | 0 | 0 | 146680 | 0 | 0 | 0 | 0 | 0 |
| SHFGRDLN | ANXA6; ANXA6 | P08133; E5R | ANXA6_HUMAN | Oxidation (M) | 0 | 0 | 0 | 430930 | 2534500 | 359560 | 1393100 | 0 | 838820 | 5475800 | 0 | 0 | 1876900 | 1442400 | 0 | 0 | 1321400 | 2944000 |
| SHFGRDLN | ANXA6; ANXA6 | P08133; E5R | ANXA6_HUMAN | Unmodified | 0 | 0 | 0 | 247730 | 7249800 | 865440 | 1773600 | 0 | 4929400 | 9122300 | 0 | 0 | 2579200 | 1216000 | 0 | 0 | 1888800 | 7436100 |
| SHFTPNSR | HGD | Q93099 | HGD_HUMAN I | Unmodified | 0 | 0 | 0 | 0 | 0 | 0 | 0 | 0 | 0 | 0 | 0 | 0 | 2241900 | 1502700 | 0 | 0 | 0 | 0 |
| SHGAGLVF | CMTM7; CMTM | Q96FZ5; F8V | CKLF7_HUMAN | Acetyl (Protein N | 0 | 0 | 135970 | 118820 | 0 | 0 | 0 | 0 | 0 | 0 | 0 | 107320 | 0 | 0 | 243460 | 249180 | 802800 | 978010 |
| SHGLFGLN | ZC3HAV1L | Q96H79 | ZCCHL_HUMAN | Unmodified | 138820 | 165890 | 351990 | 319260 | 254680 | 0 | 0 | 0 | 295960 | 0 | 208300 | 1384200 | 0 | 0 | 505020 | 463650 | 0 | 0 |
| SHGVLGLY | SLC25A1 | P53007 | TXTP_HUMAN T | Unmodified | 0 | 0 | 0 | 0 | 0 | 0 | 0 | 0 | 28701 | 408170 | 0 | 0 | 0 | 0 | 0 | 0 | 0 | 0 |
| SHGVLLEV | MITD1; MITD1; I | Q8WV92; H7 | MITD1_HUMAN | Unmodified | 0 | 0 | 0 | 0 | 0 | 0 | 0 | 0 | 0 | 0 | 0 | 2176200 | 0 | 0 | 0 | 0 | 0 | 0 |
| SHGVQTM | NCAM2; NCAM | O15394; H9I | NCAM2_HUMA | Oxidation (M) | 0 | 0 | 0 | 0 | 0 | 0 | 0 | 0 | 0 | 0 | 0 | 0 | 0 | 0 | 696500 | 302260 | 0 | 0 |
| SHIEIIRGH | GSR | P00390 | GSHR_HUMAN | Unmodified | 0 | 0 | 0 | 0 | 22577000 | 3043000 | 0 | 0 | 0 | 2365200 | 0 | 2970100 | 7391200 | 5044200 | 0 | 0 | 2807500 | 0 |
| SHIGVVPQ | ABCB6 | Q9NP58 | ABCB6_HUMAN | Unmodified | 513810 | 0 | 750960 | 1298400 | 0 | 0 | 0 | 51808 | 0 | 0 | 0 | 2093200 | 0 | 0 | 2538900 | 1278700 | 0 | 0 |
| SHIQQLRV | UGGT1 | Q9NYU2 | UGGG1_HUMA | Oxidation (M) | 0 | 0 | 0 | 0 | 0 | 0 | 0 | 0 | 3272200 | 547700 | 630390 | 0 | 0 | 0 | 0 | 0 | 0 | 69059 |
| SHIQQLRV | UGGT1 | Q9NYU2 | UGGG1_HUMA | Unmodified | 0 | 0 | 0 | 0 | 448630 | 200780 | 0 | 0 | 9282200 | 238950 | 0 | 0 | 0 | 0 | 0 | 0 | 406170 | 0 |
| SHKAESSE | P4HTM; P4HTM | Q9NXG6; V9 | P4HTM_HUMAN | Unmodified | 26086 | 24935 | 34343 | 0 | 0 | 0 | 0 | 0 | 2286700 | 197110 | 0 | 17035 | 22109 | 0 | 0 | 0 | 0 | 0 |
| SHKDEVVK | IQGAP1; IQGAP | P46940; A0A | IQGA1_HUMAN | Unmodified | 0 | 40292 | 0 | 0 | 1274300 | 1006500 | 0 | 0 | 0 | 0 | 0 | 0 | 546660 | 0 | 0 | 0 | 73136 | 0 |
| SHKLESIKE | EEA1 | Q15075 | EEA1_HUMAN I | Unmodified | 0 | 0 | 0 | 0 | 2135500 | 0 | 0 | 0 | 0 | 0 | 0 | 0 | 0 | 0 | 0 | 0 | 0 | 0 |
| SHKSPADI | CAB39; CAB39 | Q9Y376; A0A | CAB39_HUMAN | Unmodified | 0 | 0 | 0 | 0 | 1430000 | 0 | 0 | 0 | 241930 | 136200 | 0 | 0 | 590740 | 515710 | 0 | 0 | 1066800 | 1153400 |
| SHKSPADI | CAB39; CAB39 | Q9Y376; A0A | CAB39_HUMAN | Unmodified | 0 | 0 | 0 | 0 | 80606 | 0 | 0 | 22971 | 0 | 0 | 0 | 0 | 0 | 0 | 0 | 0 | 387130 | 0 |
| SHLAIVQK | CNNM3; CNNM | Q8NE01; Q6 | CNNM3_HUMA | Unmodified | 0 | 0 | 0 | 0 | 0 | 0 | 80693 | 0 | 0 | 158100 | 0 | 1972000 | 0 | 0 | 0 | 0 | 0 | 0 |
| SHLEVPLEI | CCDC124; CCD | Q96CT7; M0 | CC124_HUMAN | Unmodified | 1074800 | 696570 | 0 | 229560 | 0 | 0 | 27475 | 0 | 569370 | 0 | 0 | 0 | 491620 | 0 | 277950 | 431320 | 0 | 0 |
| SHLKITDSA | TMED10; TMED | P49755; G3V | TMEDA_HUMAN | Unmodified | 0 | 0 | 0 | 0 | 0 | 0 | 0 | 0 | 0 | 537390 | 0 | 0 | 0 | 0 | 0 | 0 | 0 | 0 |
| SHLLNCCF | LUC7L2; LUC7L | Q9Y383; A0A | LC7L2_HUMAN | Unmodified | 0 | 62846 | 0 | 0 | 0 | 0 | 0 | 0 | 0 | 0 | 0 | 568050 | 0 | 0 | 0 | 0 | 0 | 0 |
| SHLMSLYS | BAG2 | O95816 | BAG2_HUMAN | Unmodified | 0 | 0 | 0 | 0 | 613150 | 249480 | 338860 | 0 | 346390 | 0 | 0 | 0 | 273240 | 317050 | 0 | 0 | 162190 | 355600 |
| SHLMYAVR | TSC22D1; TSC2 | Q15714; O75 | T22D1_HUMAN | Oxidation (M) | 0 | 0 | 0 | 0 | 1091800 | 0 | 138350 | 0 | 0 | 410140 | 0 | 0 | 1342100 | 0 | 0 | 0 | 0 | 852830 |
| SHLPYDVT | IQGAP3; IQGAP | Q86VI3; F2Z | IQGA3_HUMAN | Unmodified | 0 | 0 | 0 | 0 | 0 | 0 | 0 | 0 | 0 | 266960 | 0 | 601630 | 0 | 180650 | 0 | 0 | 100250 | 0 |
| SHLQYFGR | ITGA2; ITGA2; I | P17301; E7E | ITA2_HUMAN I | Unmodified | 0 | 0 | 1790300 | 265230 | 0 | 0 | 0 | 0 | 0 | 0 | 0 | 0 | 0 | 0 | 572720 | 658580 | 0 | 0 |
| SHMAEESK | TRIP10; TRIP10; | Q15642; W4 | CIP4_HUMAN ( | Unmodified | 622000 | 0 | 0 | 0 | 0 | 0 | 0 | 0 | 334190 | 0 | 0 | 0 | 0 | 0 | 0 | 0 | 0 | 0 |
| SHMMDVQ | PDIA4 | P13667 | PDIA4_HUMAN | Unmodified | 0 | 0 | 132430 | 0 | 0 | 0 | 0 | 0 | 14443000 | 853260 | 1167900 | 1073300 | 0 | 0 | 0 | 0 | 0 | 0 |
| SHMMDVQ | PDIA4 | P13667 | PDIA4_HUMAN | 2 Oxidation (M) | 0 | 0 | 189310 | 0 | 0 | 0 | 0 | 0 | 1627100 | 179240 | 83162 | 0 | 0 | 0 | 186830 | 0 | 0 | 0 |
| SHMMDVQ | PDIA4 | P13667 | PDIA4_HUMAN | Oxidation (M) | 58937 | 0 | 0 | 0 | 0 | 0 | 0 | 0 | 7166300 | 586730 | 887930 | 910800 | 0 | 0 | 12299 | 0 | 0 | 0 |
| SHMMDVQ | PDIA4 | P13667 | PDIA4_HUMAN | 2 Oxidation (M) | 0 | 0 | 0 | 0 | 0 | 0 | 0 | 122270 | 4222300 | 1834900 | 758420 | 309400 | 0 | 0 | 0 | 0 | 0 | 0 |
| SHMMDVQ | PDIA4 | P13667 | PDIA4_HUMAN | Oxidation (M) | 0 | 0 | 0 | 384290 | 1965000 | 7599300 | 121690 | 0 | 17632000 | 5659900 | 182270 | 0 | 0 | 0 | 0 | 0 | 0 | 2967100 |
| SHMMDVQ | PDIA4 | P13667 | PDIA4_HUMAN | Unmodified | 0 | 0 | 0 | 0 | 446530 | 0 | 0 | 385810 | 10285000 | 600410 | 0 | 0 | 0 | 1116200 | 476580 | 0 | 0 | 765690 |
| SHMMDVQ | PDIA4 | A0A499FI48 | A0A499FI48_H | Oxidation (M) | 0 | 0 | 0 | 0 | 0 | 0 | 0 | 0 | 1048500 | 34361 | 101680 | 0 | 0 | 0 | 0 | 0 | 0 | 0 |
| SHMMDVQ | PDIA4 | A0A499FI48 | A0A499FI48_H | Unmodified | 0 | 0 | 0 | 0 | 0 | 0 | 0 | 0 | 1421800 | 73047 | 120250 | 124220 | 0 | 0 | 0 | 0 | 0 | 0 |
| SHPEFSSN | USP13; USP13; | Q92995; A0A | UBP13_HUMAN | Unmodified | 482030 | 0 | 65884 | 622850 | 0 | 0 | 0 | 0 | 0 | 0 | 44072 | 0 | 176520 | 0 | 0 | 0 | 0 | 0 |
| SHPELQTP | TGOLN2; TGOL | O43493; A0A | TGON2_HUMA | Unmodified | 0 | 0 | 0 | 0 | 147600 | 0 | 0 | 0 | 4969600 | 699440 | 0 | 15559 | 0 | 0 | 0 | 0 | 0 | 244760 |
| SHPEPQTP | TGOLN2; TGOL | O43493; A0A | TGON2_HUMA | Unmodified | 0 | 0 | 0 | 142920 | 0 | 0 | 0 | 0 | 1356000 | 0 | 13504 | 43707 | 0 | 0 | 0 | 0 | 0 | 0 |
| SHPEPQTP | TGOLN2; TGOL | O43493; A0A | TGON2_HUMA | Unmodified | 0 | 270530 | 0 | 0 | 127850 | 0 | 0 | 0 | 313160 | 1237700 | 0 | 0 | 16531 | 0 | 0 | 0 | 0 | 30478 |
| SHPEVSIEH | MUC5AC | P98088 | MUC5A_HUMA | Unmodified | 0 | 0 | 0 | 0 | 0 | 0 | 0 | 0 | 0 | 0 | 0 | 0 | 0 | 0 | 0 | 0 | 0 | 2931500 |
| SHPISAKLD | DST; DST; DST; | Q03001; A0A | DYST_HUMAN | Unmodified | 0 | 0 | 0 | 0 | 983240 | 0 | 0 | 0 | 0 | 0 | 0 | 114850 | 0 | 0 | 0 | 0 | 0 | 0 |
| SHPLDLSP | ASAP1; ASAP1; | Q9ULH1; H0 | ASAP1_HUMAN | Unmodified | 0 | 0 | 0 | 479730 | 0 | 0 | 0 | 0 | 0 | 0 | 0 | 0 | 2515800 | 339680 | 0 | 0 | 362320 | 912680 |
| SHPSPQAK | PPID | Q08752 | PPID_HUMAN I | Acetyl (Protein N | 0 | 0 | 0 | 0 | 0 | 0 | 0 | 0 | 1353700 | 0 | 3901900 | 11110000 | 0 | 0 | 0 | 62657 | 0 | 0 |
| SHPSVPYH | ST6GALNAC4; S | Q9H4F1; A6I | SIA7D_HUMAN | Unmodified | 0 | 0 | 0 | 0 | 0 | 0 | 0 | 0 | 1935800 | 0 | 0 | 0 | 0 | 0 | 0 | 0 | 0 | 0 |
| SHQGLDR | FASN; FASN | P49327; A0A | FAS_HUMAN F | Unmodified | 169210 | 496500 | 616780 | 1882800 | 496740 | 146650 | 761200 | 168530 | 172890 | 722590 | 4206800 | 11243000 | 161330 | 68111 | 4405800 | 2820100 | 308800 | 0 |
| SHQTGIQA | TWF1; TWF1 | Q12792; F8V | TWF1_HUMAN | Acetyl (Protein N | 574700 | 724700 | 601540 | 258580 | 744450 | 0 | 0 | 0 | 120060 | 0 | 321890 | 1504500 | 0 | 0 | 989530 | 837230 | 337930 | 0 |
| SHQWFGA | ITGAV | P06756 | ITAV_HUMAN I | Unmodified | 0 | 0 | 0 | 0 | 0 | 0 | 0 | 0 | 0 | 0 | 0 | 0 | 0 | 0 | 0 | 0 | 0 | 0 |
| SHQWFGA | ITGAV | P06756 | ITAV_HUMAN I | Unmodified | 0 | 0 | 0 | 0 | 7933800 | 0 | 0 | 76030 | 0 | 0 | 0 | 0 | 1287700 | 711510 | 0 | 0 | 2165600 | 2701200 |
| SHRDPYAT | TOM1L1; TOM1 | O75674; J3K | TM1L1_HUMAN | Unmodified | 16848000 | 17467000 | 18624000 | 9685800 | 4786100 | 1689300 | 1880200 | 406820 | 3156700 | 2707400 | 0 | 3965700 | 29365000 | 14598000 | 18408000 | 13079000 | 6853000 | 11694000 |
| SHRGMTP | EPB41L3; EPB4 | Q9Y2J2; A0A | E41L3_HUMAN | Unmodified | 0 | 0 | 0 | 0 | 0 | 0 | 133020 | 71969 | 0 | 0 | 0 | 0 | 0 | 0 | 0 | 0 | 362940 | 1239900 |
| SHRGMTP | EPB41L3; EPB4 | Q9Y2J2; A0A | E41L3_HUMAN | Oxidation (M) | 0 | 0 | 0 | 0 | 0 | 0 | 0 | 0 | 0 | 0 | 292170 | 0 | 0 | 31209 | 0 | 0 | 141230 | 919250 |
| SHRTEMD | HNRNPF; HNRI | P52597; A0A | HNRPF_HUMAN | Unmodified | 0 | 0 | 250290 | 0 | 507930 | 106060 | 124980 | 189070 | 686430 | 210530 | 1320900 | 3106300 | 3051500 | 0 | 72215 | 39011 | 420250 | 0 |
| SHSAHFFE | DDT; DDTL; DD | P30046; A6N | DOPD_HUMAN | Unmodified | 407990 | 290440 | 0 | 0 | 0 | 0 | 0 | 0 | 0 | 0 | 0 | 4225600 | 0 | 0 | 4056700 | 1121200 | 18306 | 0 |
| SHSIFLINV | KIF5B | P33176 | KINH_HUMAN | Unmodified | 0 | 0 | 0 | 0 | 0 | 0 | 0 | 0 | 0 | 0 | 0 | 1205000 | 0 | 0 | 0 | 0 | 0 | 0 |
| SHSIFLINV | KIF5B | P33176 | KINH_HUMAN | Unmodified | 85410 | 0 | 0 | 0 | 1443000 | 531280 | 243750 | 0 | 1080100 | 561350 | 1401500 | 2395500 | 1116100 | 1032100 | 0 | 0 | 273450 | 708930 |
| SHSITNME | ERBIN | Q96RT1 | ERBIN_HUMAN | Unmodified | 0 | 0 | 305640 | 77801 | 17542000 | 1536000 | 0 | 98803 | 0 | 0 | 0 | 0 | 1160500 | 1266100 | 542810 | 102190 | 1069300 | 1916700 |
| SHSITNME | ERBIN | Q96RT1 | ERBIN_HUMAN | Oxidation (M) | 0 | 0 | 50555 | 90033 | 4967400 | 791020 | 18590 | 0 | 0 | 0 | 0 | 0 | 502080 | 317550 | 139390 | 0 | 0 | 320730 |
| SHSLAQPP | TJP1; TJP1; TJP1 | Q07157; A0A | ZO1_HUMAN T | Unmodified | 0 | 0 | 0 | 0 | 0 | 0 | 0 | 0 | 0 | 0 | 0 | 0 | 2334700 | 1503700 | 0 | 0 | 0 | 0 |
| SHSLAQPP | TJP1; TJP1; TJP1 | Q07157; A0A | ZO1_HUMAN T | Unmodified | 0 | 0 | 639470 | 1486400 | 3020400 | 0 | 0 | 0 | 0 | 0 | 0 | 222430 | 1336900 | 694140 | 252100 | 318980 | 0 | 0 |
| SHSPSASQ | LMO7; LMO7; LI | Q8WWI1; A0 | LMO7_HUMAN | Unmodified | 0 | 0 | 22488 | 0 | 0 | 139980 | 0 | 0 | 0 | 28437 | 0 | 0 | 0 | 0 | 0 | 0 | 0 | 551260 |
| SHSQEPQD | TRPM5; TRPM5; | Q9NZQ8; A0 | TRPM5_HUMAN | Unmodified | 0 | 0 | 0 | 0 | 0 | 0 | 0 | 0 | 0 | 0 | 628750 | 1335400 | 0 | 0 | 0 | 0 | 0 | 0 |
| SHSSPSLN | PSD3; PSD3; PS | Q9NYI0; E5R | PSD3_HUMAN | Unmodified | 0 | 0 | 0 | 0 | 887920 | 144570 | 55967 | 276520 | 0 | 0 | 0 | 0 | 248310 | 173800 | 0 | 106900 | 2738000 | 0 |
| SHSVFNIK | KIF23; KIF23; KI | Q02241; A0A | KIF23_HUMAN | Unmodified | 0 | 0 | 0 | 0 | 1672300 | 0 | 0 | 0 | 976890 | 0 | 0 | 0 | 0 | 163440 | 211700 | 0 | 239780 | 316320 |
| SHTCFNRL | WWP1; ITCH; W | Q9H0M0; Q9 | WWP1_HUMAN | Unmodified | 1422600 | 1333300 | 484480 | 706870 | 0 | 0 | 0 | 0 | 373140 | 0 | 0 | 0 | 824720 | 137480 | 757810 | 693070 | 0 | 257830 |
| SHTDQLVL | UBQLN1 | Q9UMX0 | UBQL1_HUMAN | Unmodified | 0 | 0 | 0 | 0 | 0 | 0 | 0 | 0 | 0 | 202900 | 0 | 890760 | 0 | 0 | 164220 | 0 | 0 | 0 |
| SHTEEDCT | UQCRH | P07919 | QCR6_HUMAN | Unmodified | 0 | 0 | 0 | 0 | 0 | 0 | 0 | 0 | 49787000 | 877660 | 0 | 219690 | 0 | 0 | 41874 | 0 | 0 | 0 |
| SHTEEDCT | UQCRH | P07919 | QCR6_HUMAN | Unmodified | 0 | 0 | 0 | 0 | 0 | 0 | 0 | 0 | 5747100 | 3528200 | 0 | 0 | 210390 | 0 | 0 | 0 | 0 | 0 |
| SHTFLPK | CNP | P09543 | CN37_HUMAN | Unmodified | 0 | 0 | 0 | 0 | 551900 | 0 | 156510 | 189470 | 663250 | 133130 | 409680 | 1556900 | 0 | 0 | 0 | 23564 | 198360 | 343410 |
| SHTFPELA | FARSB | Q9NSD9 | SYFB_HUMAN | Unmodified | 51838 | 0 | 0 | 0 | 0 | 0 | 64176 | 52321 | 0 | 0 | 188100 | 2403900 | 1002200 | 465320 | 73033 | 0 | 0 | 0 |
| SHTGEEAA | CPNE2 | Q96FN4 | CPNE2_HUMAN | Unmodified | 112610 | 107940 | 0 | 0 | 0 | 543890 | 89945 | 45803 | 5994900 | 166120 | 53911 | 0 | 1419400 | 0 | 190490 | 103230 | 0 | 181570 |
| SHTILLVQP | ERH; ERH | P84090; G3V | ERH_HUMAN E | Acetyl (Protein N | 0 | 0 | 1690700 | 0 | 317460 | 0 | 0 | 0 | 304180 | 0 | 1171800 | 5954600 | 200560 | 0 | 635100 | 732050 | 0 | 0 |
| SHTILLVQP | ERH; ERH | P84090; G3V | ERH_HUMAN E | Acetyl (Protein N | 1443700 | 1778600 | 437020 | 1939100 | 0 | 1413600 | 846870 | 408910 | 3896300 | 2055000 | 8967400 | 10063000 | 4440500 | 1324400 | 0 | 1211300 | 114880 | 0 |
| SHTLEFQSI | LRATD2 | Q96KN1 | LRAT2_HUMAN | Oxidation (M) | 0 | 0 | 0 | 0 | 0 | 0 | 0 | 0 | 2202100 | 914090 | 46181 | 370960 | 0 | 0 | 0 | 0 | 0 | 0 |
| SHTPSPTSI | CAP2; CAP2; CA | P40123; B7Z | CAP2_HUMAN | Unmodified | 0 | 0 | 0 | 0 | 0 | 0 | 0 | 68962 | 0 | 0 | 0 | 0 | 96351 | 44558 | 116360 | 0 | 111710 | 116940 |
| SHTSEGAH | NDRG1; NDRG | Q92597; E7E | NDRG1_HUMA | Unmodified | 354700 | 0 | 948810 | 3075800 | 9669700 | 1975500 | 24983 | 0 | 2072100 | 2395400 | 0 | 934120 | 975050 | 0 | 1021700 | 2281400 | 9154200 | 15951000 |
| SHTSEGTR | NDRG1; NDRG | Q92597; E7E | NDRG1_HUMA | Unmodified | 0 | 0 | 0 | 0 | 5688000 | 487450 | 0 | 0 | 0 | 751130 | 0 | 0 | 83581 | 1114000 | 0 | 0 | 9260600 | 14984000 |
| SHVCWHV | NAA16; NAA15; | Q6N069; Q9 | NAA16_HUMAN | Unmodified | 0 | 0 | 0 | 67878 | 0 | 0 | 0 | 0 | 0 | 0 | 0 | 1770300 | 0 | 0 | 0 | 0 | 0 | 0 |
| SHVEDGDI | RRBP1; RRBP1; | Q9P2E9; A0A | RRBP1_HUMAN | Unmodified | 0 | 0 | 0 | 0 | 0 | 0 | 0 | 0 | 0 | 0 | 0 | 352190 | 779090 | 336540 | 0 | 0 | 0 | 0 |
| SHVEQLVQ | THNSL1 | Q8IYQ7 | THNS1_HUMAN | Unmodified | 1088900 | 1116100 | 0 | 2630300 | 0 | 0 | 0 | 0 | 764560 | 281080 | 0 | 0 | 0 | 0 | 1542000 | 0 | 0 | 0 |
| SHVFLSGM | UBA6 | A0AVT1 | UBA6_HUMAN | Oxidation (M) | 0 | 0 | 0 | 0 | 0 | 0 | 0 | 0 | 0 | 179760 | 0 | 963190 | 0 | 0 | 0 | 0 | 1450000 | 658060 |
| SHYAAEEIS | SPTBN2 | O15020 | SPTN2_HUMAN | Unmodified | 0 | 0 | 147830 | 0 | 0 | 0 | 0 | 0 | 0 | 0 | 0 | 506930 | 158870 | 0 | 186410 | 188560 | 95618 | 0 |
| SHYADVDF | SLC38A2; SLC3 | Q96QD8; F8 | S38A2_HUMAN | Unmodified | 5284100 | 7898000 | 5188700 | 8829000 | 16081000 | 2441600 | 5577500 | 0 | 12008000 | 5262400 | 7445400 | 20390000 | 8539800 | 2676300 | 3422700 | 3890200 | 733180 | 1287000 |
| SHYADVDF | SLC38A2; SLC3 | Q96QD8; F8 | S38A2_HUMAN | Unmodified | 4707300 | 4961900 | 3315700 | 627380 | 4053100 | 750410 | 1481000 | 320190 | 8135000 | 2142300 | 3605300 | 23192000 | 1788200 | 676620 | 2808300 | 1992400 | 1106400 | 4109500 |
| SHYKVGEN | H1-0 | P07305 | H10_HUMAN H | Unmodified | 753370 | 0 | 0 | 0 | 1139200 | 257860 | 0 | 7741100 | 0 | 13517 | 0 | 0 | 0 | 0 | 0 | 0 | 58923 | 1183900 |
| SHYKVGEN | H1-0 | P07305 | H10_HUMAN H | Unmodified | 0 | 0 | 0 | 0 | 250980 | 0 | 0 | 1008200 | 0 | 0 | 0 | 0 | 0 | 0 | 0 | 0 | 284090 | 0 |
| SIAAATSAL | TLN2; TLN1; TLI | Q9Y4G6; Q9 | TLN2_HUMAN | Unmodified | 855360 | 931010 | 1750200 | 4348500 | 9929500 | 2538400 | 10927000 | 4393000 | 1957400 | 1212400 | 566030 | 3503700 | 6620200 | 10554000 | 1861700 | 994970 | 3171200 | 5089200 |
| SIAFHSAVS | STRAP | Q9Y3F4 | STRAP_HUMAN | Unmodified | 0 | 272020 | 22954 | 555660 | 2343800 | 230530 | 803050 | 248430 | 1019600 | 0 | 1307300 | 7229700 | 3766200 | 0 | 312260 | 405760 | 989950 | 926700 |
| SIAFPSIGS | MACROH2A1; N | O75367; B4D | H2AY_HUMAN | Unmodified | 8045100 | 0 | 0 | 0 | 0 | 0 | 0 | 0 | 0 | 0 | 0 | 0 | 0 | 0 | 0 | 0 | 0 | 0 |
| SIAFPSIGS | MACROH2A1; N | O75367; B4D | H2AY_HUMAN | Unmodified | 0 | 0 | 0 | 0 | 0 | 0 | 0 | 477070 | 0 | 0 | 0 | 0 | 0 | 0 | 0 | 0 | 0 | 0 |
| SIAIGSQPV | ERLEC1 | Q96DZ1 | ERLEC_HUMAN | Unmodified | 0 | 0 | 0 | 0 | 0 | 0 | 394740 | 0 | 6997100 | 594380 | 397650 | 0 | 0 | 0 | 0 | 0 | 0 | 0 |
| SIAIGSQPV | ERLEC1 | Q96DZ1 | ERLEC_HUMAN | Unmodified | 0 | 0 | 0 | 0 | 0 | 0 | 0 | 0 | 4780600 | 3607600 | 0 | 0 | 0 | 0 | 0 | 0 | 0 | 0 |
| SIAPSIFGG | MCM5; MCM5 | P33992; B1A | MCM5_HUMAN | Oxidation (M) | 0 | 0 | 0 | 0 | 667150 | 0 | 0 | 0 | 120280 | 0 | 472040 | 1341500 | 259270 | 0 | 0 | 0 | 372870 | 0 |
| SIASEPTEK | ITGA1 | P56199 | ITA1_HUMAN I | Unmodified | 872210 | 643250 | 0 | 0 | 500820 | 148140 | 0 | 17985 | 0 | 0 | 0 | 0 | 0 | 0 | 39447 | 47343 | 1654800 | 4322100 |
| SIATEIILEIK | ECE1; ECE1; EC | P42892; B4D | ECE1_HUMAN | Unmodified | 0 | 0 | 0 | 0 | 723060 | 0 | 0 | 0 | 1894700 | 1656500 | 0 | 0 | 0 | 0 | 0 | 0 | 0 | 0 |

| | | | | | | | | | | | | | | | | | | | | | | |
|---|---|---|---|---|---|---|---|---|---|---|---|---|---|---|---|---|---|---|---|---|---|---|
| SIATLMTDE | SLC4A7; SLC4A | Q9Y6M7; C9 | S4A7_HUMAN | Unmodified | 0 | 0 | 0 | 438760 | 552020 | 156490 | 0 | 0 | 2924200 | 732150 | 578110 | 2168800 | 0 | 0 | 536630 | 278870 | 1014200 | 656750 |
| SIATLMTDE | SLC4A7; SLC4A | Q9Y6M7; C9 | S4A7_HUMAN | Oxidation (M) | 0 | 0 | 0 | 0 | 0 | 0 | 0 | 0 | 0 | 0 | 0 | 1739100 | 0 | 0 | 0 | 0 | 0 | 0 |
| SIAVAEAAC | STK26; STK26; | Q9P289; B4E | STK26_HUMAN | Unmodified | 0 | 0 | 100190 | 581810 | 3571300 | 889210 | 0 | 120760 | 0 | 310380 | 983520 | 3106500 | 5073800 | 4389200 | 95044 | 543240 | 1970800 | 2919200 |
| SIAVAEAAC | STK26; STK26; | Q9P289; B4E | STK26_HUMAN | Unmodified | 0 | 0 | 0 | 0 | 128460 | 0 | 43135 | 44575 | 0 | 42548 | 0 | 0 | 113750 | 0 | 0 | 0 | 829790 | 265320 |
| SICEVLDLE | DEK; DEK; DEK; | P35659; B4D | DEK_HUMAN P | Unmodified | 0 | 0 | 0 | 0 | 0 | 0 | 800650 | 15701 | 0 | 0 | 0 | 0 | 5745600 | 1952000 | 0 | 0 | 0 | 0 |
| SICREMSG | ANXA11; ANXA1 | P50995; H0Y | ANX11_HUMAN | Oxidation (M) | 0 | 0 | 0 | 0 | 2849200 | 0 | 0 | 0 | 1473700 | 1712600 | 0 | 0 | 0 | 859680 | 0 | 0 | 2414500 | 4587800 |
| SICREMSG | ANXA11; ANXA1 | P50995; H0Y | ANX11_HUMAN | Unmodified | 0 | 0 | 0 | 0 | 3101400 | 0 | 0 | 14396 | 4322700 | 0 | 0 | 0 | 1366000 | 1566400 | 0 | 0 | 0 | 10800000 |
| SICTTVLELI | YWHAQ; YWHA | P27348; E9P | 1433T_HUMAN | Unmodified | 218670 | 148650 | 0 | 7778100 | 4328200 | 949450 | 598800 | 0 | 6117700 | 1721700 | 5799000 | 13032000 | 3505000 | 1148400 | 11589000 | 456410 | 1494500 | 3081600 |
| SIDDEITEAI | PRKAA1 | Q13131 | AAPK1_HUMAN | Unmodified | 0 | 274050 | 0 | 0 | 0 | 0 | 0 | 58310 | 1284700 | 123410 | 137160 | 0 | 0 | 0 | 738820 | 0 | 0 | 0 |
| SIDDIGHLI | TTC17 | Q96AE7 | TTC17_HUMAN | Unmodified | 0 | 0 | 0 | 0 | 0 | 0 | 0 | 0 | 0 | 204960 | 0 | 0 | 0 | 0 | 0 | 0 | 0 | 0 |
| SIDDLEEK | TPM1; TPM1 | H7BYY1; Q6 | H7BYY1_HUMA | Unmodified | 17127 | 0 | 0 | 0 | 9688000 | 0 | 0 | 0 | 243270 | 605540 | 228480 | 0 | 2464500 | 1004100 | 0 | 0 | 131670 | 1132300 |
| SIDDTSNFI | STK38 | Q15208 | STK38_HUMAN | Unmodified | 0 | 0 | 0 | 0 | 1638600 | 0 | 0 | 0 | 0 | 0 | 0 | 2618400 | 0 | 0 | 0 | 0 | 0 | 2150600 |
| SIDFPLTK | ZMPSTE24; ZMF | O75844; A0A | FACE1_HUMAN | Unmodified | 0 | 0 | 75305 | 0 | 0 | 0 | 265500 | 0 | 5902200 | 445790 | 0 | 743280 | 34015 | 0 | 52115 | 0 | 0 | 0 |
| SIDLPIQSS | HSPA4L; HSPA4 | O95757; E9F | HS74L_HUMAN | Unmodified | 0 | 0 | 0 | 0 | 0 | 0 | 0 | 0 | 0 | 0 | 0 | 0 | 1041900 | 0 | 0 | 0 | 0 | 0 |
| SIDNGIFVC | SDCBP; SDCBP | O00560; G5I | SDCB1_HUMAI | Unmodified | 0 | 22010000 | 30366 | 31848000 | 8392700 | 1093300 | 274510 | 0 | 8356400 | 14256000 | 137690 | 750650 | 15304000 | 7723000 | 62138000 | 0 | 204520 | 1850500 |
| SIDNKALYD | PABPC4; PABP( | Q13310; P11 | PABP4_HUMAN | Unmodified | 0 | 0 | 0 | 685230 | 0 | 0 | 652600 | 0 | 0 | 0 | 0 | 0 | 2605900 | 0 | 0 | 0 | 466600 | 0 |
| SIDPGLKED | NCAPD2; NCAF | Q15021; F5C | CND1_HUMAN | Unmodified | 0 | 0 | 0 | 0 | 0 | 0 | 0 | 0 | 0 | 0 | 0 | 248260 | 0 | 0 | 0 | 0 | 0 | 0 |
| SIDQDYER | TJP2; TJP2; TJP2 | Q9UDY2; A0 | ZO2_HUMAN T | Unmodified | 0 | 0 | 0 | 0 | 371260 | 0 | 0 | 0 | 2241800 | 257630 | 0 | 65027 | 0 | 0 | 0 | 0 | 0 | 0 |
| SIDRISEVKI | ENTR1; ENTR1 | Q96C92; H7 | ENTR1_HUMAN | Unmodified | 0 | 0 | 217540 | 0 | 614820 | 0 | 0 | 0 | 0 | 0 | 0 | 0 | 0 | 0 | 134160 | 0 | 0 | 0 |
| SIDSEFSNA | DNER | Q8NFT8 | DNER_HUMAN | Unmodified | 0 | 0 | 0 | 0 | 0 | 0 | 0 | 0 | 3229500 | 222390 | 0 | 0 | 0 | 0 | 0 | 0 | 0 | 0 |
| SIDSNPYD | NSF; NSF; NSF | P46459; I3L( | NSF_HUMAN V | Unmodified | 0 | 0 | 688140 | 0 | 0 | 0 | 0 | 0 | 476580 | 0 | 0 | 734420 | 214670 | 212260 | 0 | 1008700 | 86649 | 132980 |
| SIDTGMGL | AARS1; AARS1; | P49588; A0A | SYAC_HUMAN | Unmodified | 0 | 0 | 0 | 295500 | 0 | 0 | 0 | 0 | 0 | 0 | 7936300 | 2876200 | 0 | 0 | 0 | 0 | 0 | 0 |
| SIDTGMGL | AARS1; AARS1; | P49588; A0A | SYAC_HUMAN | Oxidation (M) | 0 | 0 | 0 | 0 | 5364400 | 1174600 | 437800 | 0 | 0 | 2053900 | 0 | 0 | 3099000 | 4184600 | 0 | 0 | 75694 | 967220 |
| SIEALMEMI | DLG3; DLG3 | Q92796; Q5 | DLG3_HUMAN | Unmodified | 0 | 0 | 0 | 0 | 0 | 0 | 0 | 0 | 785460 | 162020 | 194000 | 691430 | 233840 | 0 | 0 | 0 | 185850 | 93877 |
| SIEASADVS | AHNAK2 | Q8IVF2 | AHNK2_HUMAI | Unmodified | 0 | 0 | 0 | 0 | 571500 | 0 | 0 | 0 | 0 | 0 | 148510 | 0 | 0 | 0 | 0 | 0 | 148180 | 289830 |
| SIEAVHEDI | DTYMK; DTYMK | P23919; H7( | KTHY_HUMAN | Unmodified | 166970 | 0 | 0 | 520490 | 1043300 | 0 | 0 | 0 | 635700 | 146620 | 131250 | 0 | 278480 | 0 | 228670 | 90527 | 214780 | 138630 |
| SIEDFAHSS | IDH1 | O75874 | IDHC_HUMAN | Oxidation (M) | 1753200 | 1606100 | 1367200 | 2479700 | 3622900 | 1028200 | 2052600 | 133790 | 2370000 | 2556000 | 1831200 | 9498900 | 14070000 | 7886200 | 3560100 | 4115600 | 1405000 | 651490 |
| SIEDFAHSS | IDH1 | O75874 | IDHC_HUMAN | Unmodified | 0 | 0 | 0 | 32136 | 165330 | 0 | 60540 | 0 | 239700 | 0 | 0 | 0 | 332200 | 190270 | 0 | 0 | 0 | 80410 |
| SIEDLHR | FMNL2 | Q96PY5 | FMNL2_HUMAI | Unmodified | 6078300 | 3355300 | 72999 | 6456300 | 0 | 0 | 136800 | 0 | 0 | 0 | 88533 | 224970 | 108010 | 0 | 399440 | 55072 | 0 | 0 |
| SIEDMTEEA | IDE; IDE; IDE; ID | P14735; A0A | IDE_HUMAN In | Unmodified | 0 | 0 | 0 | 0 | 0 | 0 | 0 | 0 | 0 | 0 | 0 | 779380 | 0 | 0 | 0 | 0 | 0 | 0 |
| SIEEQLGTE | DDX6 | P26196 | DDX6_HUMAN | Unmodified | 3685300 | 3147200 | 4056400 | 3918400 | 0 | 0 | 1533300 | 0 | 2921000 | 1205900 | 3339500 | 15520000 | 2115100 | 770090 | 4269300 | 2745900 | 0 | 851910 |
| SIEEVIK | ALDH1A3; ALDI | P47895; H0Y | AL1A3_HUMAN | Unmodified | 0 | 0 | 366540 | 0 | 1830600 | 525230 | 142580 | 116680 | 570670 | 158810 | 447520 | 0 | 962210 | 343980 | 0 | 0 | 4433400 | 5347800 |
| SIEEVIKRAI | ALDH1A3; ALDI | P47895; H0Y | AL1A3_HUMAN | Unmodified | 0 | 0 | 0 | 0 | 371680 | 0 | 216700 | 0 | 0 | 0 | 0 | 56956 | 0 | 0 | 0 | 0 | 24877000 | 8447800 |
| SIEGRRSEA | SCRIB; SCRIB; | Q14160; A0A | SCRIB_HUMAN | Unmodified | 0 | 0 | 0 | 0 | 0 | 0 | 0 | 128230 | 0 | 79374 | 0 | 0 | 0 | 0 | 0 | 0 | 614390 | 0 |
| SIEIPRPVD | U2AF2; U2AF2 | P26368; K7E | U2AF2_HUMAN | Unmodified | 3796300 | 1730000 | 799410 | 2610200 | 2920300 | 545630 | 2836100 | 2427900 | 1521000 | 1511000 | 1604300 | 11651000 | 7256600 | 2408400 | 829330 | 1129500 | 1730100 | 2570300 |
| SIENAKQD | EXTL3 | O43909 | EXTL3_HUMAN | Unmodified | 0 | 0 | 0 | 0 | 75798 | 0 | 294810 | 0 | 1567800 | 568110 | 94427 | 143060 | 0 | 0 | 0 | 0 | 0 | 91769 |
| SIEQSIEQE | STX1A; STX1A; | Q16623; A8N | STX1A_HUMAN | Unmodified | 272180 | 349010 | 0 | 0 | 0 | 0 | 154740 | 157470 | 0 | 0 | 0 | 0 | 0 | 0 | 0 | 0 | 0 | 0 |
| SIESTLDDL | AGRN; AGRN; A | O00468; A0A | AGRIN_HUMAN | Unmodified | 0 | 0 | 277650 | 284860 | 0 | 0 | 7808 | 0 | 346340 | 0 | 0 | 0 | 0 | 0 | 526370 | 0 | 0 | 200330 |
| SIESTLDDL | AGRN; AGRN; A | O00468; A0A | AGRIN_HUMAN | Unmodified | 545440 | 0 | 1698800 | 537080 | 0 | 581880 | 0 | 0 | 1004700 | 205270 | 0 | 0 | 0 | 0 | 610560 | 169170 | 16390 | 34645 |
| SIEVIENR | NASP; NASP; N | P49321; E9P | NASP_HUMAN | Unmodified | 0 | 0 | 0 | 541100 | 907310 | 411510 | 324770 | 238570 | 2843800 | 1782000 | 1921100 | 5659500 | 1535700 | 241190 | 60198 | 309100 | 546480 | 176290 |
| SIEYSPQLE | HSPG2; HSPG2 | P98160; A0A | PGBM_HUMAN | Unmodified | 0 | 0 | 1111300 | 249710 | 0 | 0 | 0 | 0 | 399290 | 0 | 0 | 0 | 0 | 0 | 0 | 701840 | 0 | 0 |
| SIFGEDALA | COPB1 | P53618 | COPB_HUMAN | Unmodified | 0 | 0 | 0 | 624990 | 0 | 320270 | 0 | 0 | 530740 | 758640 | 1942400 | 6817600 | 3818600 | 482260 | 0 | 0 | 503350 | 1341500 |
| SIFKPFIFVD | SCRN1 | Q12765 | SCRN1_HUMAI | Unmodified | 73387 | 0 | 0 | 0 | 336820 | 103370 | 0 | 0 | 2072800 | 82108 | 0 | 0 | 0 | 0 | 57397 | 0 | 0 | 122200 |
| SIFLFLDR | CUL4B; CUL4A; | Q13620; Q1 | CUL4B_HUMAI | Unmodified | 0 | 0 | 0 | 0 | 0 | 0 | 0 | 0 | 0 | 0 | 0 | 815790 | 0 | 0 | 0 | 0 | 0 | 0 |
| SIFQHIQSA | THRAP3; THRAI | Q9Y2W1; A0 | TR150_HUMAN | Unmodified | 87217 | 0 | 88419 | 59675 | 0 | 0 | 6211200 | 2033700 | 450510 | 227350 | 0 | 4208500 | 2007400 | 0 | 122110 | 93178 | 0 | 0 |
| SIFSPVVKL | MYOF; MYOF | Q9NZM1; H( | MYOF_HUMAN | Oxidation (M) | 0 | 0 | 0 | 0 | 1884100 | 367050 | 0 | 0 | 0 | 0 | 0 | 0 | 0 | 0 | 0 | 0 | 0 | 0 |
| SIFSPVVKL | MYOF; MYOF | Q9NZM1; H( | MYOF_HUMAN | Unmodified | 0 | 0 | 0 | 0 | 2422400 | 60774 | 0 | 0 | 0 | 0 | 0 | 0 | 0 | 0 | 0 | 0 | 30908 | 0 |
| SIFTPTNQII | SBDS; SBDS; SI | Q9Y3A5; A0A | SBDS_HUMAN | Acetyl (Protein N | 749490 | 796110 | 1656500 | 1236200 | 909480 | 0 | 232080 | 0 | 1286900 | 0 | 0 | 2188800 | 909010 | 0 | 1072500 | 0 | 359720 | 713780 |
| SIFVTCGD | ERI3; ERI3; ERI3 | O43414; F6C | ERI3_HUMAN E | Unmodified | 781340 | 0 | 0 | 0 | 0 | 0 | 0 | 0 | 0 | 0 | 0 | 0 | 0 | 0 | 619640 | 453210 | 0 | 0 |
| SIFYEEELSI | CDC42BPA; CD | Q5VT25; A0A | MRCKA_HUMA | Unmodified | 0 | 0 | 0 | 0 | 1329300 | 0 | 0 | 0 | 0 | 0 | 0 | 0 | 1122800 | 0 | 0 | 0 | 1149900 | 6042600 |
| SIFYVPDMI | TRAP1; TRAP1; | Q12931; I3L( | TRAP1_HUMAN | Unmodified | 0 | 0 | 0 | 0 | 220870 | 0 | 460060 | 0 | 796960 | 0 | 0 | 321150 | 678260 | 0 | 0 | 0 | 0 | 0 |
| SIGAAAKS( | HSPA4; HSPA4; | P34932; A0A | HSP74_HUMAI | Unmodified | 0 | 0 | 0 | 0 | 332400 | 0 | 347180 | 40837 | 550460 | 219500 | 0 | 0 | 319620 | 0 | 0 | 0 | 248540 | 0 |
| SIGASVEFI | HSPG2 | P98160 | PGBM_HUMAN | Unmodified | 0 | 0 | 0 | 0 | 0 | 0 | 0 | 0 | 0 | 0 | 0 | 0 | 1400100 | 0 | 0 | 0 | 130270 | 2959600 |
| SIGASVEFI | HSPG2 | P98160 | PGBM_HUMAN | Unmodified | 0 | 0 | 0 | 0 | 0 | 0 | 0 | 0 | 0 | 1449700 | 0 | 0 | 0 | 0 | 0 | 0 | 60596 | 519210 |
| SIGATLTDV | VPS13C; VPS13 | Q709C8; A0 | VP13C_HUMAI | Unmodified | 0 | 0 | 0 | 0 | 0 | 426850 | 0 | 0 | 0 | 1779500 | 0 | 0 | 282820 | 0 | 0 | 0 | 0 | 0 |
| SIGEAIQYL | MAPKAPK2 | P49137 | MAPK2_HUMAI | Unmodified | 123920 | 0 | 0 | 0 | 0 | 0 | 0 | 0 | 364290 | 0 | 0 | 3530000 | 0 | 0 | 0 | 125650 | 0 | 0 |
| SIGEYDVLF | PRKDC | P78527 | PRKDC_HUMAI | Unmodified | 0 | 0 | 0 | 0 | 0 | 0 | 679670 | 0 | 226930 | 0 | 0 | 0 | 542250 | 0 | 0 | 0 | 0 | 924580 |
| SIGNLTQL( | LRSAM1; LRSAI | Q6UWE0; A( | LRSM1_HUMAI | Unmodified | 0 | 0 | 0 | 0 | 753130 | 0 | 0 | 0 | 211920 | 0 | 189420 | 0 | 0 | 0 | 0 | 0 | 0 | 0 |
| SIGSAVDQ | RACGAP1; RAC | Q9H0H5; F8 | RGAP1_HUMAI | Unmodified | 0 | 81947 | 552700 | 2068100 | 18220000 | 2575700 | 503390 | 240770 | 2111300 | 1584700 | 1919000 | 4139800 | 1424200 | 786990 | 580510 | 147350 | 4519500 | 1889000 |
| SIGSIALR | PGAP1; PGAP1 | Q75T13; B4D | PGAP1_HUMAN | Unmodified | 0 | 0 | 0 | 0 | 0 | 0 | 0 | 0 | 5161900 | 586750 | 0 | 0 | 310620 | 0 | 0 | 0 | 0 | 0 |
| SIGVSNFNI | AKR1C2; AKR1( | P52895; B4D | AK1C2_HUMAN | Oxidation (M) | 0 | 0 | 0 | 0 | 582400 | 0 | 0 | 0 | 0 | 0 | 0 | 0 | 0 | 0 | 0 | 0 | 0 | 0 |
| SIGVSNFNI | AKR1C2; AKR1( | P52895; B4D | AK1C2_HUMAN | Unmodified | 0 | 0 | 0 | 0 | 484610 | 0 | 227890 | 0 | 0 | 0 | 0 | 0 | 0 | 0 | 0 | 0 | 271000 | 0 |
| SIHDFCLVS | SPINT2; SPINT2 | O43291; K7E | SPIT2_HUMAN | Unmodified | 397900 | 185330 | 495850 | 602810 | 1641700 | 1018100 | 844520 | 0 | 68134000 | 15918000 | 0 | 66074 | 1062600 | 886320 | 396090 | 249660 | 88126 | 707870 |
| SIHDFCLVS | SPINT2; SPINT2 | O43291; K7E | SPIT2_HUMAN | Unmodified | 0 | 0 | 0 | 0 | 0 | 0 | 0 | 0 | 0 | 1958800 | 0 | 0 | 0 | 0 | 0 | 0 | 0 | 0 |
| SIHIVALGN | RHOBTB3; RHC | O94955; D6I | RHBT3_HUMAI | Acetyl (Protein N | 0 | 0 | 0 | 0 | 951360 | 0 | 0 | 0 | 0 | 186640 | 382830 | 886330 | 0 | 0 | 0 | 0 | 0 | 347520 |
| SIHLFIDSLI | LPL | P06858 | LIPL_HUMAN L | Unmodified | 0 | 20783 | 119750 | 0 | 0 | 0 | 0 | 0 | 1227300 | 382490 | 0 | 0 | 2335500 | 1795000 | 0 | 0 | 0 | 0 |
| SIHSFMTHI | ATP2B4 | P23634 | AT2B4_HUMAN | Oxidation (M) | 0 | 0 | 0 | 0 | 1196800 | 0 | 0 | 0 | 0 | 0 | 0 | 0 | 0 | 0 | 0 | 0 | 0 | 1407600 |
| SIIEKLDEPI | RASA2; RASA2 | Q15283; A0A | RASA2_HUMAN | Unmodified | 0 | 0 | 0 | 0 | 776460 | 108780 | 0 | 0 | 0 | 136300 | 149680 | 0 | 419140 | 0 | 0 | 0 | 0 | 0 |
| SIIGMGTG | NDRG1; NDRG | Q92597; E5F | NDRG1_HUMA | Oxidation (M) | 0 | 0 | 0 | 103660 | 0 | 0 | 0 | 0 | 1512800 | 2105900 | 355130 | 1852300 | 0 | 0 | 326240 | 547450 | 1761200 | 106780 |
| SIIGMGTG | NDRG1; NDRG | Q92597; E5F | NDRG1_HUMA | Unmodified | 0 | 0 | 0 | 0 | 0 | 0 | 0 | 0 | 4099100 | 1592500 | 63276 | 479180 | 0 | 0 | 0 | 0 | 0 | 0 |
| SIIGVIPYFP | PRPSAP2; PRPS | O60256; E7E | KPRB_HUMAN | Unmodified | 0 | 0 | 0 | 0 | 494710 | 0 | 1019300 | 0 | 274750 | 186610 | 267650 | 1203500 | 0 | 0 | 113280 | 161080 | 0 | 0 |
| SIIKPTVPDI | AVIL | O75366 | AVIL_HUMAN A | Unmodified | 0 | 0 | 0 | 0 | 0 | 0 | 0 | 0 | 0 | 0 | 12889000 | 64047000 | 0 | 0 | 0 | 0 | 0 | 0 |
| SIIKPTVPDI | AVIL | O75366 | AVIL_HUMAN A | Unmodified | 0 | 0 | 0 | 0 | 0 | 0 | 405610 | 325940 | 0 | 0 | 0 | 0 | 0 | 0 | 0 | 0 | 0 | 0 |
| SIIQEVAR | TMX3 | Q96JJ7 | TMX3_HUMAN | Unmodified | 0 | 0 | 0 | 0 | 161720 | 0 | 0 | 0 | 6822200 | 501170 | 207160 | 924680 | 310090 | 0 | 777870 | 568220 | 526410 | 620340 |
| SIIQEVARD | TMX3 | Q96JJ7 | TMX3_HUMAN | Unmodified | 0 | 0 | 0 | 0 | 0 | 0 | 0 | 0 | 853400 | 0 | 0 | 0 | 0 | 0 | 0 | 0 | 0 | 0 |
| SIIQQHNLE | PIK3C2A | O00443 | P3C2A_HUMAN | Unmodified | 0 | 57804 | 167690 | 109910 | 0 | 0 | 0 | 0 | 0 | 0 | 0 | 2169600 | 0 | 0 | 222300 | 126000 | 0 | 0 |
| SIIQSAQQD | GCN1 | Q92616 | GCN1_HUMAN | Unmodified | 130040 | 121160 | 149780 | 128120 | 552270 | 79624 | 231660 | 84781 | 185520 | 117210 | 80075 | 640230 | 361600 | 462070 | 149760 | 207060 | 210770 | 365560 |
| SIISMFDRE | PDCD6 | O75340 | PDCD6_HUMA | Oxidation (M) | 4214900 | 4230700 | 4355500 | 3909600 | 537740 | 442220 | 99709 | 30784 | 8540800 | 3700600 | 96972 | 14390000 | 157630 | 89034 | 13022000 | 6629600 | 170310 | 581560 |
| SIISMFDRE | PDCD6 | O75340 | PDCD6_HUMA | Unmodified | 0 | 0 | 0 | 0 | 0 | 928260 | 133930 | 0 | 16262000 | 3032100 | 0 | 1433800 | 582900 | 294680 | 0 | 0 | 384630 | 515270 |
| SIITCRVSLI | EPB41L5 | Q9HCM4 | E41L5_HUMAN | Unmodified | 0 | 0 | 0 | 0 | 251120 | 0 | 0 | 0 | 0 | 437610 | 0 | 0 | 0 | 0 | 0 | 0 | 378050 | 0 |
| SIITYVATYY | SPTBN2; SPTBN | O15020; A4( | SPTN2_HUMAN | Unmodified | 0 | 0 | 46059 | 0 | 0 | 12048 | 71558 | 0 | 0 | 52898 | 0 | 65183 | 0 | 262040 | 281100 | 118270 | 74997 | 0 |
| SIKDTICNQ | CORO1C; COR( | Q9ULV4; H0 | COR1C_HUMA | Unmodified | 304490 | 141870 | 0 | 165270 | 1041000 | 228270 | 0 | 34983 | 805700 | 158320 | 736190 | 4064000 | 154730 | 0 | 53126 | 99273 | 0 | 162760 |
| SIKENASA( | RAB13; RAB13 | P51153; A0A | RAB13_HUMAN | Unmodified | 0 | 0 | 0 | 2586100 | 0 | 0 | 5893.2 | 0 | 46946 | 0 | 0 | 0 | 0 | 0 | 44282 | 132630 | 0 | 0 |
| SILAEYK | DRG1 | Q9Y295 | DRG1_HUMAN | Unmodified | 0 | 0 | 0 | 0 | 104770 | 0 | 0 | 301200 | 223760 | 114660 | 302220 | 2178700 | 0 | 0 | 0 | 0 | 0 | 243470 |
| SILALVDG( | MRPL39; MRPL | Q9NYK5; C9 | RM39_HUMAN | Oxidation (M) | 0 | 0 | 0 | 0 | 1249500 | 171720 | 370210 | 0 | 1504400 | 1274400 | 74909 | 0 | 861780 | 0 | 0 | 0 | 702140 | 305930 |
| SILAPLYQC | FAM20B | O75063 | XYLK_HUMAN ( | Unmodified | 0 | 0 | 0 | 0 | 0 | 0 | 0 | 0 | 3624800 | 130050 | 0 | 0 | 0 | 0 | 0 | 0 | 0 | 0 |
| SILCCLR | PTGES3; PTGES | Q15185; A0A | TEBP_HUMAN | Unmodified | 448090 | 0 | 0 | 0 | 1054000 | 0 | 0 | 0 | 2794200 | 0 | 0 | 2331200 | 1429200 | 0 | 0 | 0 | 429880 | 0 |
| SILDADPEA | MEAK7 | Q6P9B6 | MEAK7_HUMAI | Unmodified | 0 | 0 | 0 | 0 | 0 | 0 | 0 | 0 | 1443500 | 0 | 0 | 0 | 0 | 0 | 0 | 0 | 1873200 | 0 |
| SILDADPEA | MEAK7 | Q6P9B6 | MEAK7_HUMAI | Unmodified | 0 | 0 | 0 | 0 | 21369000 | 1579900 | 0 | 0 | 0 | 1801300 | 0 | 0 | 0 | 0 | 0 | 0 | 14696000 | 19044000 |
| SILDVSMG | TNFAIP2; TNFAI | Q03169; H0Y | TNAP2_HUMAN | Oxidation (M) | 0 | 0 | 0 | 0 | 670510 | 0 | 0 | 0 | 117370 | 0 | 0 | 253890 | 0 | 0 | 0 | 0 | 1018300 | 0 |
| SILEELLLK | BMX | P51813 | BMX_HUMAN ( | Unmodified | 0 | 0 | 0 | 0 | 0 | 0 | 234640 | 0 | 0 | 0 | 2172000 | 9997300 | 0 | 0 | 0 | 0 | 0 | 0 |
| SILEELLLKI | BMX | P51813 | BMX_HUMAN ( | Unmodified | 0 | 0 | 0 | 0 | 0 | 0 | 0 | 0 | 0 | 0 | 1052800 | 1651500 | 0 | 0 | 0 | 15476 | 0 | 0 |
| SILEPDTSN | WWP1 | Q9H0M0 | WWP1_HUMAN | Unmodified | 0 | 346260 | 1046200 | 671100 | 0 | 0 | 0 | 0 | 0 | 0 | 800390 | 0 | 626020 | 366730 | 950020 | 818220 | 0 | 0 |
| SILEPDTSN | WWP1 | Q9H0M0 | WWP1_HUMAN | Unmodified | 0 | 0 | 0 | 0 | 989790 | 290470 | 0 | 0 | 0 | 0 | 0 | 0 | 1409900 | 589960 | 0 | 0 | 584950 | 1175500 |
| SILEYLR | LRRC47 | Q8N1G4 | LRC47_HUMAN | Unmodified | 0 | 0 | 0 | 0 | 116110 | 38315 | 0 | 0 | 393500 | 0 | 200650 | 1519900 | 0 | 0 | 0 | 0 | 0 | 378640 |
| SILFVPTSAI | HSP90B1; HSP9 | P14625; A0A | ENPL_HUMAN | Unmodified | 1409400 | 395020 | 4404000 | 1327300 | 919240 | 46003 | 2499400 | 1354400 | 5.25E+08 | 1.18E+08 | 1.1E+08 | 59588000 | 1007600 | 1068700 | 3091300 | 1956000 | 0 | 1158100 |
| SILFVPTSAI | HSP90B1; HSP9 | P14625; A0A | ENPL_HUMAN | Unmodified | 0 | 0 | 44282 | 109190 | 3796100 | 934380 | 26039000 | 804050 | 8203000 | 3.4E+08 | 5501900 | 0 | 4432900 | 6218300 | 0 | 0 | 1327900 | 3147200 |
| SILFVPTSAI | HSP90B1; HSP9 | P14625; A0A | ENPL_HUMAN | Unmodified | 0 | 0 | 0 | 0 | 0 | 0 | 0 | 0 | 0 | 5127700 | 0 | 0 | 0 | 0 | 0 | 0 | 0 | 0 |
| SILGTLTVE | FLOT2; FLOT2; | Q14254; J3C | FLOT2_HUMAN | Unmodified | 1522000 | 1018100 | 3014100 | 2728800 | 5451000 | 765160 | 373050 | 0 | 21335000 | 6254200 | 2004800 | 1796300 | 672250 | 2188000 | 14882000 | 11968000 | 183760 | 0 |
| SILGVQCE | TENM4 | Q6N022 | TEN4_HUMAN | Unmodified | 0 | 0 | 0 | 0 | 0 | 0 | 0 | 0 | 182700 | 0 | 0 | 0 | 0 | 0 | 0 | 0 | 355640 | 0 |
| SILKIDDVV | CCT4 | P50991 | TCPD_HUMAN | Unmodified | 0 | 0 | 0 | 74067 | 356680 | 0 | 311780 | 290860 | 376470 | 1251000 | 0 | 863110 | 749980 | 334040 | 0 | 0 | 663320 | 133520 |
| SILKPSTPIF | TJP2; TJP2; TJP2 | Q9UDY2; A0 | ZO2_HUMAN T | Unmodified | 0 | 0 | 0 | 0 | 2509900 | 0 | 0 | 0 | 2062800 | 1406700 | 0 | 0 | 0 | 0 | 0 | 0 | 0 | 0 |
| SILMLGAMI | CARD6 | Q9BX69 | CARD6_HUMAI | Oxidation (M) | 0 | 0 | 0 | 0 | 844030 | 0 | 9896100 | 5475700 | 0 | 0 | 0 | 0 | 303960 | 6518700 | 0 | 0 | 0 | 0 |
| SILPTAPR | EIF4B; EIF4B; EI | P23588; F8V | IF4B_HUMAN E | Unmodified | 0 | 0 | 214660 | 0 | 209020 | 0 | 0 | 0 | 689630 | 147580 | 477670 | 3045700 | 101990 | 0 | 0 | 0 | 94162 | 0 |
| SILRCSENI | SLC5A3 | P53794 | SC5A3_HUMAI | Unmodified | 0 | 0 | 0 | 0 | 0 | 0 | 0 | 0 | 0 | 0 | 0 | 0 | 0 | 0 | 0 | 0 | 567590 | 2132100 |
| SILRDALSD | CCT4 | P50991 | TCPD_HUMAN | Unmodified | 0 | 0 | 0 | 0 | 1151200 | 0 | 1243100 | 0 | 73969 | 1301700 | 0 | 0 | 3462900 | 4651800 | 99796 | 0 | 732900 | 4728700 |
| SILTAIPNDI | COG8; COG8; N | Q96MW5; H | COG8_HUMAN | Unmodified | 0 | 21581 | 0 | 0 | 0 | 0 | 303850 | 6963.1 | 0 | 0 | 0 | 793340 | 0 | 0 | 0 | 0 | 0 | 0 |
| SILTQIFGAI | KIDINS220; KID | Q9ULH0; E9 | KDIS_HUMAN | Unmodified | 105060 | 49461 | 152470 | 1051700 | 1112500 | 59232 | 0 | 0 | 897480 | 228680 | 552570 | 3257900 | 530770 | 0 | 277530 | 580820 | 0 | 0 |
| SILYDERSV | PSMA2; N/A; PS | P25787; A0A | PSA2_HUMAN | Unmodified | 0 | 411980 | 0 | 0 | 7047500 | 668090 | 2135900 | 1301300 | 0 | 946490 | 1743400 | 0 | 5480900 | 3162300 | 0 | 0 | 5078200 | 3431200 |

| | | | | | | | | | | | | | | | | | | | | | | |
|---|---|---|---|---|---|---|---|---|---|---|---|---|---|---|---|---|---|---|---|---|---|---|
| SIMDPNQN | RND3; RND3 | P61587; Q5 | RND3_HUMAN | Unmodified | 0 | 0 | 260080 | 0 | 431580 | 84154 | 372580 | 462150 | 0 | 0 | 0 | 111810 | 0 | 0 | 0 | 257210 | 465210 | 1040000 |
| SIMSYNGG | PSMB3; PSMB3 | P49720; A0A | PSB3_HUMAN | Acetyl (Protein N | 0 | 0 | 0 | 0 | 0 | 0 | 0 | 0 | 0 | 0 | 748740 | 0 | 0 | 0 | 0 | 0 | 0 | 0 |
| SIMTDLYYL | FUT8; FUT8; FU | Q9BYC5; A0 | FUT8_HUMAN | Unmodified | 0 | 0 | 0 | 0 | 0 | 0 | 0 | 0 | 5030200 | 0 | 0 | 0 | 0 | 0 | 0 | 0 | 0 | 0 |
| SINAGGHK | VDAC2 | P45880 | VDAC2_HUMAI | Unmodified | 0 | 0 | 0 | 0 | 180010 | 0 | 770420 | 0 | 2368000 | 24575000 | 0 | 972930 | 83052 | 142270 | 27058 | 0 | 234940 | 0 |
| SINGELYAF | SNX27; SNX27 | Q96L92; A0A | SNX27_HUMAN | Unmodified | 781150 | 389360 | 153870 | 201500 | 0 | 0 | 0 | 0 | 851050 | 0 | 126650 | 1941000 | 0 | 0 | 0 | 1120000 | 0 | 0 |
| SINGILFPG | GGH; GGH; GG | Q92820; A0A | GGH_HUMAN ( | Unmodified | 0 | 0 | 0 | 0 | 0 | 0 | 0 | 0 | 64230000 | 8614700 | 519660 | 1076400 | 0 | 0 | 0 | 1385000 | 0 | 0 |
| SINGILFPG | GGH; GGH; GG | Q92820; A0A | GGH_HUMAN ( | Unmodified | 0 | 0 | 0 | 0 | 0 | 0 | 0 | 0 | 5858100 | 1758400 | 240990 | 248150 | 0 | 0 | 0 | 0 | 0 | 0 |
| SINGILFPG | GGH; GGH; GG | Q92820; A0A | GGH_HUMAN ( | Unmodified | 0 | 0 | 0 | 0 | 326770 | 0 | 0 | 0 | 706520 | 8094000 | 0 | 0 | 348030 | 341900 | 0 | 0 | 0 | 0 |
| SINHASLIS | XPO6 | Q96QU8 | XPO6_HUMAN | Unmodified | 8376400 | 4862500 | 51479 | 0 | 0 | 0 | 0 | 0 | 0 | 0 | 0 | 1090700 | 0 | 0 | 0 | 52085 | 0 | 0 |
| SINHQIESF | LIMCH1; LIMCH | Q9UPQ0; HC | LIMC1_HUMAN | Unmodified | 17411 | 0 | 165680 | 299840 | 32465 | 0 | 0 | 0 | 180660 | 0 | 328900 | 0 | 356190 | 197640 | 276950 | 0 | 0 | 0 |
| SINNDTTY( | LACTB2 | Q53H82 | LACB2_HUMAN | Unmodified | 0 | 0 | 0 | 0 | 1382100 | 355000 | 0 | 0 | 3028700 | 791680 | 0 | 0 | 0 | 179100 | 0 | 0 | 960370 | 907640 |
| SINNPDMR | ACLY | P53396 | ACLY_HUMAN | Unmodified | 228500 | 0 | 226940 | 0 | 0 | 0 | 97104 | 0 | 298360 | 104810 | 648880 | 18448000 | 111650 | 0 | 470680 | 279100 | 41881 | 58301 |
| SINNPDMR | ACLY | P53396 | ACLY_HUMAN | Unmodified | 0 | 0 | 0 | 0 | 2642900 | 0 | 4069100 | 679470 | 0 | 567630 | 120060 | 256230 | 874250 | 304870 | 119790 | 0 | 383300 | 0 |
| SINNPDMR | ACLY | P53396 | ACLY_HUMAN | Oxidation (M) | 0 | 0 | 0 | 0 | 0 | 0 | 0 | 120790 | 0 | 0 | 0 | 0 | 0 | 0 | 0 | 0 | 0 | 0 |
| SINPDEAV | HSPA1L; HSPA( | P34931; P17 | HS71L_HUMAN | Unmodified | 0 | 0 | 0 | 0 | 1343900 | 0 | 267550 | 125930 | 0 | 259740 | 298420 | 0 | 888310 | 0 | 0 | 28623 | 1518400 | 764670 |
| SINPDEAV | HSPA1L; HSPA | P34931; P0D | HS71L_HUMAN | Oxidation (M) | 354180 | 1075100 | 214780 | 0 | 0 | 0 | 0 | 0 | 0 | 0 | 0 | 0 | 0 | 16163 | 1392000 | 0 | 0 | 0 |
| SINPDEAV | HSPA8; HSPA8; | P11142; E9P | HSP7C_HUMAI | Unmodified | 3598100 | 3063700 | 8129400 | 3213400 | 2827600 | 847370 | 0 | 0 | 5092600 | 1827300 | 1055000 | 3206100 | 2677500 | 0 | 11196000 | 3084200 | 1139500 | 0 |
| SINPLGGF | RPL3; RPL3 | P39023; G5E | RL3_HUMAN 6 | Oxidation (M) | 11677000 | 27816000 | 1877100 | 5985000 | 6285500 | 2224700 | 31202000 | 111150 | 1907600 | 4230200 | 2009200 | 21008000 | 4604200 | 2893400 | 14123000 | 3960400 | 1632200 | 10494000 |
| SINPLGGF | RPL3; RPL3 | P39023; G5E | RL3_HUMAN 6 | Unmodified | 382790 | 870420 | 0 | 118890 | 1125800 | 0 | 999950 | 0 | 687160 | 393900 | 53867 | 321820 | 0 | 415230 | 0 | 0 | 0 | 927850 |
| SINQPVAF | SLIRP; SLIRP; S | Q9GZT3; H0 | SLIRP_HUMAN | Unmodified | 2498700 | 2059000 | 0 | 3227700 | 0 | 984200 | 225700 | 2372500 | 5151700 | 2649900 | 0 | 0 | 1161200 | 942810 | 0 | 2186000 | 815070 | 512660 |
| SINQQSGA | FUBP3 | Q96I24 | FUBP3_HUMAN | Unmodified | 0 | 0 | 0 | 0 | 99689 | 0 | 0 | 0 | 77946 | 0 | 0 | 288030 | 0 | 0 | 0 | 0 | 0 | 0 |
| SINRSLIVG | TGM2 | P21980 | TGM2_HUMAN | Unmodified | 0 | 0 | 0 | 0 | 1554700 | 324110 | 0 | 0 | 0 | 0 | 0 | 0 | 0 | 0 | 0 | 0 | 0 | 0 |
| SINSILDYIS | COPS2; COPS2 | P61201; B4D | CSN2_HUMAN | Unmodified | 0 | 0 | 0 | 0 | 0 | 0 | 0 | 0 | 0 | 0 | 0 | 2212800 | 0 | 0 | 0 | 1235200 | 0 | 896670 |
| SINTEVVAC | PRDX4; PRDX4; | Q13162; A6N | PRDX4_HUMAN | Unmodified | 0 | 0 | 0 | 0 | 0 | 0 | 0 | 0 | 32389000 | 1449700 | 0 | 0 | 0 | 0 | 0 | 0 | 0 | 0 |
| SINTEVVAC | PRDX4; PRDX4; | Q13162; A6N | PRDX4_HUMAN | Unmodified | 0 | 0 | 0 | 0 | 0 | 0 | 0 | 0 | 24530000 | 0 | 804400 | 576880 | 0 | 0 | 0 | 0 | 0 | 0 |
| SIPAGAEP( | JPT2; JPT2; JPT2 | Q9H910; A6I | JUPI2_HUMAN | Unmodified | 0 | 0 | 0 | 0 | 0 | 4295100 | 0 | 0 | 4533400 | 0 | 0 | 0 | 0 | 0 | 0 | 0 | 0 | 430400 |
| SIPASDLP( | MYO1D; MYO1D | O94832; J3Q | MYO1D_HUMA | Unmodified | 8326000 | 9478300 | 4491700 | 1266000 | 383230 | 0 | 0 | 0 | 1735400 | 433830 | 0 | 1442000 | 0 | 302440 | 6452700 | 3839600 | 619960 | 2863400 |
| SIPASDLP( | MYO1D; MYO1D | O94832; J3Q | MYO1D_HUMA | Unmodified | 0 | 0 | 0 | 0 | 659420 | 0 | 0 | 0 | 0 | 154980 | 0 | 0 | 1033700 | 1015700 | 0 | 0 | 739050 | 3578800 |
| SIPHITSDR | CRMP1 | Q14194 | DPYL1_HUMAN | Unmodified | 0 | 0 | 0 | 0 | 0 | 0 | 2971900 | 1108500 | 0 | 791810 | 0 | 0 | 0 | 0 | 0 | 0 | 0 | 0 |
| SIPICTLK | UBA1 | P22314 | UBA1_HUMAN | Unmodified | 0 | 0 | 569490 | 0 | 1757400 | 262960 | 3490600 | 1847800 | 1786600 | 1663900 | 0 | 5868400 | 2323800 | 0 | 0 | 0 | 6211000 | 3000300 |
| SIPLDEGED | DYNC1H1; DYN | Q14204; A0A | DYHC1_HUMA | Unmodified | 0 | 0 | 0 | 0 | 0 | 0 | 0 | 0 | 0 | 0 | 0 | 2303700 | 0 | 0 | 0 | 0 | 0 | 0 |
| SIPQPTNAL | DAAM1 | Q9Y4D1 | DAAM1_HUMA | Unmodified | 0 | 0 | 0 | 0 | 2202700 | 0 | 356550 | 1002400 | 956620 | 293550 | 0 | 2862900 | 0 | 0 | 0 | 0 | 690740 | 1003300 |
| SIQADGLV | EEF1B2 | P24534 | EF1B_HUMAN | Unmodified | 0 | 0 | 0 | 0 | 2544600 | 0 | 0 | 0 | 2389800 | 325390 | 353920 | 23424000 | 2792200 | 0 | 0 | 0 | 0 | 0 |
| SIQCLTVH | WDR1 | O75083 | WDR1_HUMAN | Unmodified | 0 | 0 | 0 | 0 | 2599900 | 357350 | 174320 | 185050 | 187040 | 177300 | 0 | 1157700 | 1345100 | 0 | 0 | 0 | 1056700 | 2253000 |
| SIQCLTVH | WDR1 | O75083 | WDR1_HUMAN | Unmodified | 0 | 0 | 0 | 0 | 0 | 0 | 44775 | 99282 | 0 | 0 | 0 | 0 | 0 | 0 | 0 | 0 | 250710 | 0 |
| SIQEIQELD | ARHGDIA; ARH | P52565; J3K | GDIR1_HUMAN | Unmodified | 0 | 0 | 0 | 0 | 1959300 | 358910 | 277970 | 417500 | 242230 | 233710 | 19826000 | 1249200 | 0 | 337040 | 0 | 0 | 1172800 | 1097000 |
| SIQEIQELD | ARHGDIA; ARH | P52565; J3K | GDIR1_HUMAN | Unmodified | 4380900 | 4778300 | 6057300 | 14058000 | 83027000 | 13687000 | 13536000 | 9839200 | 13357000 | 6356300 | 26875000 | 1.21E+08 | 58312000 | 28768000 | 17078000 | 18378000 | 51860000 | 1.07E+08 |
| SIQFVDWC | TUBA4A; TUBA1 | P68366; P68 | TBA4A_HUMAN | Unmodified | 153690 | 219470 | 0 | 220300 | 1412300 | 252090 | 3169500 | 954420 | 1818000 | 2378400 | 240400 | 9765200 | 878640 | 805600 | 266930 | 120200 | 617550 | 834740 |
| SIQFVDWC | TUBA4A; TUBA1 | P68366; P68 | TBA4A_HUMAN | Unmodified | 0 | 0 | 0 | 0 | 558580 | 0 | 3050100 | 0 | 0 | 0 | 0 | 0 | 1491000 | 1840100 | 0 | 0 | 848070 | 0 |
| SIQGDGEG | NAPB; NAPB | Q9H115; A0 | SNAB_HUMAN | Unmodified | 144940 | 0 | 0 | 0 | 0 | 0 | 93134 | 64562 | 251380 | 101790 | 0 | 88808 | 52909 | 0 | 0 | 0 | 0 | 0 |
| SIQGSSTS | DIP2B | Q9P265 | DIP2B_HUMAN | Unmodified | 350700 | 0 | 0 | 0 | 0 | 0 | 0 | 0 | 605320 | 89647 | 0 | 2053000 | 0 | 0 | 521300 | 458560 | 424520 | 536040 |
| SIQLDGLV | EEF1D; EEF1D; | P29692; E9P | EF1D_HUMAN | Unmodified | 0 | 0 | 0 | 0 | 1828900 | 318830 | 0 | 0 | 1813000 | 713720 | 2289500 | 15470000 | 0 | 0 | 0 | 0 | 139820 | 0 |
| SIQLDGLV | EEF1D; EEF1D; | P29692; E9P | EF1D_HUMAN | Unmodified | 0 | 0 | 0 | 0 | 0 | 0 | 0 | 0 | 0 | 279210 | 0 | 0 | 0 | 0 | 0 | 0 | 1674400 | 0 |
| SIQMLTLLR | TMED1; TMED1 | Q13445; K7E | TMED1_HUMAI | Unmodified | 0 | 0 | 0 | 0 | 0 | 0 | 0 | 0 | 20432000 | 965700 | 0 | 0 | 0 | 0 | 0 | 0 | 0 | 0 |
| SIQMLTLLR | TMED1; TMED1 | Q13445; K7E | TMED1_HUMAI | Oxidation (M) | 0 | 0 | 0 | 0 | 0 | 0 | 0 | 0 | 5828900 | 939860 | 0 | 0 | 0 | 0 | 0 | 0 | 0 | 0 |
| SIQSLHNQ | TRAF3; TRAF3; | Q13114; A6N | TRAF3_HUMAN | Unmodified | 0 | 0 | 0 | 0 | 12112000 | 0 | 0 | 0 | 0 | 0 | 0 | 0 | 0 | 0 | 0 | 0 | 0 | 0 |
| SIRHVSFQ | SPRED1 | Q7Z699 | SPRE1_HUMAN | Unmodified | 0 | 0 | 0 | 0 | 0 | 0 | 0 | 0 | 0 | 0 | 0 | 0 | 0 | 0 | 0 | 0 | 0 | 1543800 |
| SIRPDNMS | HDAC2; HDAC | Q92769; Q1 | HDAC2_HUMA | Oxidation (M) | 0 | 0 | 0 | 0 | 0 | 0 | 17501 | 0 | 0 | 0 | 0 | 1525900 | 0 | 0 | 0 | 0 | 0 | 0 |
| SIRPDNMS | HDAC2; HDAC | Q92769; Q1 | HDAC2_HUMA | Unmodified | 0 | 0 | 0 | 0 | 0 | 0 | 0 | 21505 | 2489500 | 39401 | 0 | 2765700 | 0 | 0 | 0 | 0 | 245770 | 0 |
| SIRPGLSPY | GANAB | Q14697 | GANAB_HUMA | Unmodified | 0 | 0 | 0 | 0 | 133570 | 0 | 383550 | 0 | 0 | 15435000 | 0 | 0 | 134930 | 0 | 0 | 0 | 0 | 207130 |
| SISEQRPP( | KIAA1522 | Q9P206 | K1522_HUMAN | Unmodified | 0 | 0 | 0 | 54002 | 2188800 | 482420 | 1846300 | 62111000 | 1029300 | 0 | 0 | 900330 | 0 | 869550 | 1007900 | 830260 | 0 | 821570 |
| SISESAFSA | GPC4 | O75487 | GPC4_HUMAN | Unmodified | 2088200 | 2417300 | 0 | 113050 | 294430 | 0 | 0 | 0 | 1821500 | 508840 | 250500 | 0 | 4179100 | 3266800 | 0 | 245590 | 101970 | 250790 |
| SISFHPSGI | CSTF1; CSTF1 | Q05048; A0A | CSTF1_HUMAN | Unmodified | 282510 | 0 | 0 | 0 | 0 | 0 | 0 | 0 | 0 | 0 | 0 | 2829000 | 0 | 0 | 309160 | 352070 | 0 | 0 |
| SISHYHETL | TM9SF3; TM9SF | Q9HD45; Q5 | TM9S3_HUMAN | Unmodified | 0 | 0 | 0 | 0 | 0 | 0 | 0 | 0 | 907120 | 2676100 | 0 | 0 | 0 | 0 | 0 | 0 | 0 | 0 |
| SISIGPISK | LSS; LSS | P48449; A0A | LSS_HUMAN La | Unmodified | 0 | 0 | 0 | 0 | 0 | 0 | 471350 | 0 | 0 | 0 | 0 | 0 | 0 | 0 | 0 | 0 | 0 | 0 |
| SISLYYTGE | NCL; NCL; NCL | P19338; C9J | NUCL_HUMAN | Unmodified | 0 | 0 | 0 | 0 | 1131600 | 0 | 0 | 0 | 2532400 | 861630 | 2286700 | 15466000 | 1954600 | 0 | 0 | 0 | 0 | 0 |
| SISMTTFTQ | AGFG1; AGFG1 | P52594; B8Z | AGFG1_HUMAI | Unmodified | 0 | 0 | 0 | 0 | 4021700 | 272480 | 0 | 0 | 0 | 145760 | 0 | 0 | 348800 | 0 | 0 | 0 | 249740 | 212160 |
| SISMTTFTQ | AGFG1; AGFG1 | P52594; B8Z | AGFG1_HUMAI | Oxidation (M) | 0 | 0 | 0 | 0 | 554030 | 0 | 0 | 0 | 0 | 0 | 0 | 0 | 0 | 0 | 0 | 0 | 179170 | 0 |
| SISNPPGSI | WNK1; WNK1; | Q9H4A3; F5 | WNK1_HUMAN | Unmodified | 0 | 0 | 0 | 0 | 0 | 0 | 0 | 940960 | 0 | 0 | 0 | 0 | 0 | 0 | 0 | 0 | 0 | 0 |
| SISPEWK | MYO1C; MYO1( | O00159; F5H | MYO1C_HUMA | Unmodified | 1855100 | 1700200 | 0 | 3495700 | 1135000 | 901440 | 0 | 54462 | 241620 | 453130 | 0 | 542040 | 894640 | 185300 | 7624200 | 4772600 | 177170 | 302750 |
| SISPEWKQ | MYO1C; MYO1( | O00159; F5H | MYO1C_HUMA | Unmodified | 110040 | 333640 | 0 | 1635300 | 4052000 | 2536000 | 568740 | 186620 | 0 | 136400 | 0 | 0 | 2542700 | 1592100 | 0 | 124900 | 1490000 | 1715600 |
| SISQLESLN | ROCK1; ROCK1 | Q13464; A0A | ROCK1_HUMAI | Unmodified | 0 | 20544 | 0 | 389210 | 36411 | 0 | 0 | 0 | 1087500 | 0 | 0 | 808080 | 0 | 0 | 197580 | 0 | 423570 | 695410 |
| SISQSSTDS | AHCYL1 | O43865 | SAHH2_HUMAI | Unmodified | 0 | 0 | 0 | 0 | 1268900 | 0 | 0 | 0 | 0 | 134180 | 0 | 0 | 786170 | 475690 | 0 | 0 | 333530 | 1697300 |
| SISSSRSEN | PSD2 | Q9BQI7 | PSD2_HUMAN | Unmodified | 0 | 0 | 0 | 0 | 0 | 0 | 0 | 0 | 0 | 0 | 892210 | 0 | 0 | 0 | 0 | 0 | 0 | 0 |
| SISTLMSDK | SLC4A2; SLC4A | P04920; A0A | B3A2_HUMAN | Unmodified | 0 | 0 | 0 | 0 | 0 | 0 | 0 | 0 | 0 | 0 | 0 | 0 | 0 | 0 | 0 | 0 | 0 | 0 |
| SISVIDSPG | EHD4 | Q9H223 | EHD4_HUMAN | Unmodified | 0 | 2386700 | 3333300 | 6388200 | 9573200 | 3038400 | 1041600 | 95517 | 994570 | 592810 | 0 | 4380000 | 5688000 | 5388700 | 7363500 | 6425200 | 5797000 | 10179000 |
| SISVIDSPG | EHD4 | Q9H223 | EHD4_HUMAN | Unmodified | 0 | 0 | 0 | 0 | 0 | 0 | 0 | 0 | 0 | 0 | 0 | 0 | 0 | 0 | 0 | 0 | 627550 | 0 |
| SITGEEMSI | MYOF | Q9NZM1 | MYOF_HUMAN | Unmodified | 130840 | 0 | 0 | 0 | 9909100 | 451510 | 349120 | 0 | 236960 | 0 | 0 | 0 | 0 | 190750 | 27751 | 0 | 0 | 0 |
| SITGEEMSI | MYOF | Q9NZM1 | MYOF_HUMAN | Oxidation (M) | 0 | 0 | 0 | 0 | 3404200 | 478710 | 0 | 0 | 0 | 0 | 0 | 0 | 0 | 0 | 0 | 0 | 0 | 0 |
| SITHDIEEK | SEPTIN9; SEPTI | Q9UHD8; K7 | SEPT9_HUMAN | Unmodified | 36139 | 0 | 0 | 224590 | 3837100 | 900280 | 469010 | 317440 | 921140 | 444140 | 871220 | 3174800 | 932200 | 815720 | 129340 | 149360 | 2596200 | 3445200 |
| SITHDIEEK | SEPTIN9; SEPTI | Q9UHD8; K7 | SEPT9_HUMAN | Unmodified | 0 | 0 | 787270 | 0 | 162090 | 514670 | 0 | 117610 | 4237400 | 362860 | 0 | 0 | 587630 | 0 | 0 | 0 | 996170 | 164640 |
| SITHDIEEK | SEPTIN9; SEPTI | Q9UHD8; K7 | SEPT9_HUMAN | Unmodified | 0 | 0 | 0 | 0 | 536760 | 0 | 332360 | 0 | 0 | 0 | 0 | 0 | 0 | 0 | 166040 | 41210 | 823380 | 282570 |
| SITNTTVCT | SF1; SF1 | Q15637; A0A | SF01_HUMAN | Unmodified | 0 | 0 | 0 | 0 | 0 | 47260 | 36200 | 0 | 165390 | 104960 | 207390 | 382270 | 0 | 0 | 0 | 0 | 0 | 0 |
| SITPSRILQ | AKR1A1 | P14550 | AK1A1_HUMAN | Unmodified | 0 | 0 | 0 | 0 | 2477000 | 349720 | 0 | 0 | 37724 | 1407800 | 0 | 0 | 3654500 | 2663500 | 0 | 0 | 2323200 | 1720700 |
| SITPSRILQ | AKR1A1 | P14550 | AK1A1_HUMAN | Oxidation (M) | 0 | 0 | 0 | 0 | 449660 | 0 | 446610 | 0 | 0 | 238480 | 0 | 0 | 192930 | 309710 | 0 | 0 | 658620 | 0 |
| SITSLSVAD | VPS37A; VPS37 | Q8NEZ2; E5I | VP37A_HUMAN | Unmodified | 1532100 | 1076900 | 822280 | 223060 | 0 | 62773 | 0 | 0 | 0 | 0 | 0 | 0 | 0 | 0 | 1070300 | 520970 | 0 | 0 |
| SITVTQDE( | ACAA1; ACAA1; | P09110; C9J | THIK_HUMAN | Oxidation (M) | 0 | 0 | 0 | 0 | 0 | 0 | 0 | 0 | 1672900 | 1181800 | 0 | 0 | 0 | 0 | 0 | 0 | 0 | 0 |
| SITVTQDE( | ACAA1; ACAA1; | P09110; C9J | THIK_HUMAN | Unmodified | 0 | 0 | 0 | 0 | 0 | 0 | 0 | 0 | 7764700 | 3836700 | 0 | 0 | 131290 | 0 | 0 | 0 | 0 | 0 |
| SIVAFLKDP | PDIA5; PDIA5 | Q14554; C9J | PDIA5_HUMAN | Unmodified | 0 | 0 | 0 | 0 | 0 | 0 | 0 | 0 | 1094200 | 110480 | 0 | 0 | 0 | 0 | 0 | 0 | 0 | 0 |
| SIVAFLKDP | PDIA5; PDIA5 | Q14554; C9J | PDIA5_HUMAN | Unmodified | 0 | 0 | 0 | 0 | 0 | 0 | 0 | 0 | 2351500 | 574130 | 0 | 117860 | 0 | 0 | 0 | 0 | 0 | 0 |
| SIVDLYTGN | ERO1B; ERO1B | Q86YB8; Q5 | ERO1B_HUMAI | Unmodified | 0 | 0 | 0 | 0 | 0 | 0 | 40570 | 174830 | 3383400 | 215930 | 0 | 0 | 0 | 0 | 311070 | 0 | 196470 | 0 |
| SIVDYKPNI | ACTN4; ACTN4 | O43707; F5C | ACTN4_HUMAI | Unmodified | 0 | 0 | 0 | 0 | 0 | 0 | 0 | 0 | 0 | 0 | 0 | 0 | 0 | 0 | 1198500 | 0 | 0 | 0 |
| SIVEEIEDL | RPN2; RPN2; RI | P04844; F2Z | RPN2_HUMAN | Unmodified | 0 | 0 | 0 | 0 | 0 | 0 | 126850 | 0 | 1104200 | 527880 | 14498 | 56441 | 0 | 0 | 8096.6 | 0 | 0 | 0 |
| SIVEGIIEEE | PPME1 | Q9Y570 | PPME1_HUMAI | Unmodified | 0 | 0 | 0 | 0 | 1335900 | 0 | 0 | 0 | 53395 | 910980 | 0 | 1078000 | 2018500 | 887060 | 0 | 0 | 759950 | 1212700 |
| SIVEGIIEEE | PPME1 | Q9Y570 | PPME1_HUMAI | Unmodified | 0 | 0 | 0 | 0 | 0 | 0 | 0 | 0 | 0 | 0 | 1300400 | 0 | 0 | 0 | 0 | 0 | 0 | 0 |
| SIVIRPLEP( | EFTUD2; EFTUD | Q15029; K7E | U5S1_HUMAN | Unmodified | 123400 | 132340 | 91988 | 57032 | 0 | 0 | 0 | 0 | 74207 | 0 | 194160 | 1129300 | 39736 | 0 | 55034 | 0 | 0 | 0 |
| SIVLEPIYW | EFNB2 | P52799 | EFNB2_HUMAN | Unmodified | 0 | 0 | 0 | 0 | 0 | 0 | 0 | 0 | 0 | 0 | 0 | 0 | 0 | 0 | 0 | 472420 | 0 | 0 |
| SIVLLSTRG | CEMIP2; CEMIP | Q9UHN6; B4 | CEIP2_HUMAN | Unmodified | 0 | 0 | 483640 | 444760 | 1572300 | 412070 | 0 | 0 | 1911100 | 704590 | 0 | 153250 | 331600 | 0 | 960360 | 0 | 0 | 0 |
| SIVLQSFYR | PPM1L | Q5SGD2 | PPM1L_HUMAN | Unmodified | 0 | 0 | 0 | 0 | 0 | 0 | 0 | 0 | 24044000 | 2081500 | 0 | 0 | 0 | 0 | 0 | 0 | 0 | 0 |
| SIVLQSFYR | PPM1L | Q5SGD2 | PPM1L_HUMAN | Oxidation (M) | 0 | 0 | 0 | 0 | 0 | 0 | 0 | 0 | 0 | 711140 | 0 | 0 | 0 | 0 | 0 | 0 | 0 | 0 |
| SIVPQGGS | HSPG2 | P98160 | PGBM_HUMAN | Unmodified | 0 | 0 | 6265400 | 2427300 | 245270 | 327580 | 0 | 0 | 5081200 | 0 | 310840 | 0 | 0 | 0 | 2058600 | 1879600 | 0 | 700890 |
| SIVPTMHY( | CPT2; CPT2; CF | P23786; A0A | CPT2_HUMAN | Unmodified | 0 | 0 | 0 | 0 | 0 | 0 | 0 | 0 | 1236100 | 642280 | 0 | 0 | 0 | 0 | 0 | 0 | 0 | 0 |
| SIVTDLVSQ | OPA1; OPA1; OI | O60313; E5K | OPA1_HUMAN | Unmodified | 0 | 0 | 0 | 0 | 0 | 0 | 0 | 0 | 0 | 0 | 0 | 1001200 | 0 | 0 | 0 | 0 | 0 | 0 |
| SIWAFGPD | EFTUD2; EFTUD | Q15029; K7E | U5S1_HUMAN | Unmodified | 147160 | 202920 | 30954 | 118000 | 285960 | 0 | 0 | 0 | 0 | 42939 | 103310 | 355390 | 291210 | 0 | 198100 | 44313 | 0 | 135480 |
| SIWDEFISD | CLDND1; CLDN | Q9NY35; D6 | CLDN1_HUMAI | Unmodified | 2691500 | 1054100 | 0 | 39786 | 207850 | 0 | 0 | 0 | 1718700 | 522850 | 907790 | 1980300 | 0 | 0 | 1143500 | 64724 | 0 | 0 |
| SIYGERFPD | PPIB; PPIB; PPIB | P23284; A0A | PPIB_HUMAN F | Unmodified | 1304400 | 438460 | 4981600 | 2036400 | 871980 | 740760 | 6221600 | 3473300 | 2.77E+08 | 66922000 | 42915000 | 21115000 | 4468000 | 6698900 | 4486900 | 3240800 | 254260 | 1483800 |
| SIYGERFPD | PPIB; PPIB; PPIB | P23284; A0A | PPIB_HUMAN F | Unmodified | 0 | 0 | 0 | 150730 | 206620 | 0 | 1982800 | 0 | 2318500 | 9698500 | 1669100 | 0 | 679170 | 1303000 | 0 | 0 | 127730 | 228970 |
| SIYGSRFPD | PPIF; PPIF; PPIF | P30405; H0 | PPIF_HUMAN F | Unmodified | 0 | 0 | 0 | 0 | 367900 | 0 | 790850 | 111870 | 3648800 | 1780800 | 201090 | 129200 | 353700 | 175050 | 0 | 0 | 674550 | 590320 |
| SIYRYDLAS | SORL1 | Q92673 | SORL_HUMAN | Unmodified | 0 | 0 | 0 | 0 | 0 | 0 | 0 | 0 | 2483300 | 303960 | 0 | 0 | 0 | 0 | 0 | 0 | 0 | 0 |
| SIYTRSVIDI | PAK2 | Q13177 | PAK2_HUMAN | Unmodified | 0 | 0 | 0 | 0 | 1639900 | 0 | 0 | 995320 | 0 | 0 | 0 | 0 | 0 | 1100300 | 0 | 0 | 4477200 | 1354300 |
| SIYTRSVIDI | PAK2 | Q13177 | PAK2_HUMAN | Unmodified | 0 | 0 | 0 | 0 | 321740 | 0 | 0 | 0 | 0 | 0 | 0 | 0 | 0 | 0 | 0 | 0 | 803790 | 0 |
| SIYVGNVD | PABPN1; PABPI | Q86U42; B4I | PABP2_HUMAN | Unmodified | 0 | 0 | 0 | 0 | 0 | 0 | 0 | 0 | 0 | 0 | 585020 | 2875200 | 0 | 0 | 0 | 0 | 0 | 0 |
| SIYVKFPLLI | IARS2 | Q9NSE4 | SYIM_HUMAN I | Unmodified | 0 | 0 | 0 | 0 | 0 | 0 | 0 | 0 | 864910 | 242420 | 0 | 0 | 0 | 0 | 0 | 0 | 0 | 0 |
| SIYYITGESI | HSP90AB1; HSI | P08238; Q5 | HS90B_HUMAN | Unmodified | 4781300 | 2487500 | 0 | 9037800 | 21085000 | 5436100 | 1653600 | 746510 | 17486000 | 4791200 | 15794000 | 1.53E+08 | 10566000 | 6291200 | 10180000 | 9005300 | 5697500 | 6889800 |
| SIYYITGESI | HSP90AB1 | P08238 | HS90B_HUMAN | Unmodified | 453170 | 357320 | 0 | 1814800 | 8925300 | 1585500 | 683470 | 220440 | 2710300 | 3396300 | 825100 | 3588300 | 5379300 | 468940 | 0 | 0 | 3864600 | 4253600 |
| SIYYITGESI | HSP90AB1 | P08238 | HS90B_HUMAN | Unmodified | 0 | 0 | 0 | 319400 | 19655000 | 5038800 | 792130 | 0 | 366330 | 9916600 | 0 | 0 | 21466000 | 11118000 | 0 | 0 | 13331000 | 13976000 |
| SKAEELGLI | ACAA1; ACAA1; | P09110; C9J | THIK_HUMAN | Unmodified | 0 | 0 | 0 | 0 | 0 | 0 | 12195 | 0 | 17161000 | 2164700 | 437970 | 94849 | 0 | 0 | 0 | 0 | 0 | 0 |
| SKAEEWGV | RALB; RALB | P11234; C9J | RALB_HUMAN | Unmodified | 1190000 | 1665200 | 1042700 | 599720 | 3966900 | 1623300 | 0 | 0 | 0 | 0 | 4259200 | 19231000 | 2253700 | 1818200 | 2805000 | 2731800 | 1846800 | 2384700 |
| SKAELMEIS | SSB; SSB; SSB; | P05455; E9P | LA_HUMAN Lu | Unmodified | 0 | 0 | 0 | 0 | 0 | 0 | 108450 | 52167 | 396250 | 0 | 874060 | 3814300 | 149300 | 0 | 251000 | 51320 | 38526 | 0 |

| | | | | | | | | | | | | | | | | | | | | | | |
|---|---|---|---|---|---|---|---|---|---|---|---|---|---|---|---|---|---|---|---|---|---|---|
| SKASGAPP | TPRN; TPRN | Q4KMQ1; H3 | TPRN_HUMAN | Unmodified | 0 | 0 | 0 | 0 | 2920400 | 262800 | 983830 | 69408 | 0 | 422680 | 0 | 0 | 982430 | 448160 | 0 | 0 | 2086500 | 702590 |
| SKATITLYG | CEMIP2; CEMIP | Q9UHN6; F5 | CEIP2_HUMAN | Unmodified | 0 | 0 | 0 | 0 | 851210 | 0 | 0 | 0 | 438780 | 278610 | 0 | 0 | 0 | 0 | 0 | 0 | 0 | 0 |
| SKDDQVTV | TKT; TKT | P29401; A0A | TKT_HUMAN Tr | Unmodified | 0 | 0 | 0 | 0 | 28789000 | 1002900 | 588200 | 0 | 2780100 | 4946600 | 1487700 | 1905600 | 0 | 1442200 | 0 | 0 | 6187800 | 9090600 |
| SKDHFGLE | TLN1 | Q9Y490 | TLN1_HUMAN | Oxidation (M) | 0 | 0 | 334280 | 0 | 946010 | 0 | 614910 | 352320 | 0 | 0 | 0 | 0 | 0 | 0 | 215010 | 229950 | 0 | 0 |
| SKDHFGLE | TLN1 | Q9Y490 | TLN1_HUMAN | Unmodified | 0 | 0 | 0 | 0 | 2495100 | 0 | 0 | 0 | 298740 | 0 | 0 | 0 | 0 | 0 | 0 | 0 | 210860 | 949570 |
| SKDHFGLE | TLN1 | Q9Y490 | TLN1_HUMAN | Unmodified | 0 | 0 | 0 | 0 | 2487800 | 642070 | 896500 | 0 | 0 | 0 | 0 | 0 | 0 | 0 | 0 | 0 | 316050 | 1254400 |
| SKDHFGLE | TLN1 | Q9Y490 | TLN1_HUMAN | Oxidation (M) | 0 | 0 | 0 | 460320 | 0 | 0 | 1091600 | 587140 | 99814 | 0 | 0 | 0 | 493250 | 1782300 | 0 | 0 | 353500 | 0 |
| SKDIVLVAY | AKR1C3; AKR1C | P42330; A0A | AK1C3_HUMAN | Unmodified | 0 | 0 | 0 | 0 | 2191900 | 0 | 0 | 0 | 0 | 0 | 0 | 0 | 0 | 0 | 0 | 0 | 253970 | 347660 |
| SKDKEDQV | NCAM2; NCAM2 | O15394; H9 | NCAM2_HUMA | Unmodified | 0 | 0 | 547980 | 1254000 | 0 | 0 | 0 | 0 | 0 | 0 | 0 | 0 | 0 | 0 | 1421800 | 823820 | 0 | 0 |
| SKDLAEVG | TNKS1BP1 | Q9C0C2 | TB182_HUMAN | Unmodified | 0 | 0 | 0 | 0 | 771650 | 0 | 0 | 0 | 0 | 0 | 0 | 0 | 0 | 5753600 | 0 | 0 | 812140 | 559960 |
| SKDMVCGI | MKRN1 | Q9UHC7 | MKRN1_HUMA | Oxidation (M) | 122100 | 0 | 0 | 0 | 0 | 0 | 0 | 0 | 0 | 0 | 0 | 0 | 0 | 0 | 0 | 0 | 0 | 0 |
| SKDPNSQV | DCTD; DCTD; D | P32321; D6F | DCTD_HUMAN | Unmodified | 0 | 0 | 0 | 0 | 81265 | 0 | 0 | 0 | 0 | 31609 | 212380 | 488760 | 0 | 62637 | 0 | 0 | 137760 | 79210 |
| SKDPSGLL | EXOC4 | Q96A65 | EXOC4_HUMAI | Unmodified | 10594 | 0 | 0 | 0 | 1216300 | 224100 | 1624300 | 18481 | 614580 | 362690 | 0 | 0 | 2450200 | 960640 | 0 | 0 | 764980 | 798110 |
| SKDTCIVIS | MYO1D; MYO1D | O94832; J3Q | MYO1D_HUMA | Unmodified | 351240 | 156740 | 0 | 0 | 0 | 0 | 0 | 0 | 0 | 0 | 0 | 0 | 0 | 0 | 0 | 0 | 0 | 0 |
| SKDTCIVIS | MYO1D; MYO1D | O94832; J3Q | MYO1D_HUMA | Unmodified | 2478100 | 2769300 | 0 | 754090 | 0 | 0 | 200690 | 128220 | 0 | 0 | 0 | 0 | 0 | 256930 | 1291900 | 0 | 0 | 201960 |
| SKDVFSKP | PTGFRN | Q9P2B2 | FPRP_HUMAN | Unmodified | 0 | 21912 | 0 | 2749400 | 1675100 | 93708 | 1019200 | 0 | 0 | 0 | 0 | 0 | 10747000 | 9121800 | 0 | 0 | 0 | 0 |
| SKDVNGPE | CHP1; CHP1; C | Q99653; F5 | CHP1_HUMAN | Unmodified | 0 | 206190 | 0 | 477500 | 654280 | 189800 | 0 | 0 | 177050 | 92446 | 0 | 1213200 | 1776300 | 776270 | 209960 | 351870 | 0 | 267580 |
| SKDVVEPLI | VARS1; VARS1 | P26640; A0A | SYVC_HUMAN | Unmodified | 0 | 0 | 0 | 0 | 0 | 0 | 0 | 0 | 0 | 0 | 0 | 3532800 | 0 | 0 | 0 | 0 | 0 | 0 |
| SKDYEFMV | CKB; CKB; CKB | P12277; H0 | KCRB_HUMAN | Oxidation (M) | 0 | 4452000 | 137070 | 1497400 | 0 | 0 | 3853500 | 0 | 780160 | 0 | 0 | 2278400 | 0 | 0 | 2162700 | 699170 | 228640 | 0 |
| SKEDQGLS | PAIP2B | Q9ULR5 | PAI2B_HUMAN | Unmodified | 18265 | 352120 | 245660 | 0 | 0 | 0 | 0 | 0 | 0 | 0 | 0 | 421060 | 93459 | 0 | 0 | 209660 | 0 | 0 |
| SKEDTIYEA | ITGA1 | P56199 | ITA1_HUMAN I | Unmodified | 1529600 | 1554600 | 0 | 0 | 0 | 0 | 0 | 0 | 0 | 0 | 0 | 0 | 0 | 0 | 0 | 0 | 0 | 0 |
| SKEDTIYEA | ITGA1 | P56199 | ITA1_HUMAN I | Unmodified | 0 | 0 | 0 | 0 | 0 | 0 | 0 | 0 | 0 | 0 | 0 | 0 | 0 | 0 | 0 | 0 | 802390 | 1699700 |
| SKEEACKA | GART; GART; G | P22102; C9 | PUR2_HUMAN | Unmodified | 0 | 0 | 0 | 0 | 119770 | 0 | 347860 | 320790 | 0 | 0 | 0 | 0 | 88159 | 122960 | 0 | 0 | 1849100 | 67664 |
| SKEEAFDAI | TPPP3 | Q9BW30 | TPPP3_HUMAN | Unmodified | 0 | 0 | 0 | 0 | 318310 | 0 | 10939000 | 1340400 | 0 | 0 | 0 | 0 | 0 | 0 | 0 | 0 | 0 | 0 |
| SKEEERLN | MPZL2 | O60487 | MPZL2_HUMA | Unmodified | 310410 | 331500 | 587720 | 1044500 | 42373 | 130830 | 0 | 43643 | 0 | 0 | 75254 | 805640 | 860060 | 276040 | 724430 | 263120 | 0 | 0 |
| SKEEILAINI | CES2; CES2 | O00748; J3Q | EST2_HUMAN | Unmodified | 0 | 0 | 0 | 0 | 0 | 0 | 0 | 0 | 600080 | 0 | 0 | 0 | 0 | 0 | 0 | 0 | 0 | 0 |
| SKEEQELH | GPM6A | P51674 | GPM6A_HUMA | Unmodified | 28510000 | 18028000 | 0 | 9018 | 0 | 25142 | 0 | 38883 | 0 | 0 | 0 | 808260 | 0 | 0 | 0 | 68769 | 14018 | 0 |
| SKEESSGTI | FERMT2; FERM | Q96AC1; A0 | FERM2_HUMAI | Unmodified | 0 | 0 | 0 | 0 | 0 | 191440 | 0 | 0 | 767700 | 0 | 96025 | 0 | 0 | 0 | 50980 | 51983 | 90378 | 426600 |
| SKEESSGTI | FERMT2; FERM | Q96AC1; A0 | FERM2_HUMAI | Unmodified | 0 | 0 | 0 | 0 | 7422300 | 2054500 | 0 | 0 | 0 | 85217 | 0 | 0 | 53058 | 144130 | 0 | 0 | 3288600 | 6478400 |
| SKEESSGTI | FERMT2; FERM | Q96AC1; A0 | FERM2_HUMAI | Oxidation (M) | 0 | 0 | 0 | 0 | 1032300 | 62557 | 0 | 0 | 0 | 0 | 0 | 0 | 0 | 0 | 0 | 0 | 456340 | 971800 |
| SKEEWNEI | PHACTR4 | Q8IZ21 | PHAR4_HUMAI | Unmodified | 0 | 0 | 0 | 0 | 2299900 | 0 | 0 | 0 | 1415600 | 1350100 | 0 | 0 | 2307000 | 1300500 | 0 | 0 | 87915 | 2864100 |
| SKEEYQQT | PRPF4 | O43172 | PRP4_HUMAN | Unmodified | 0 | 0 | 0 | 0 | 332930 | 0 | 0 | 165990 | 0 | 134820 | 0 | 626570 | 156610 | 0 | 0 | 0 | 82830 | 108280 |
| SKEGSIWN | LMAN2; LMAN2 | Q12907; D6I | LMAN2_HUMAI | Unmodified | 0 | 0 | 0 | 0 | 172820 | 0 | 210370 | 0 | 10453000 | 841020 | 129700 | 0 | 0 | 0 | 0 | 0 | 0 | 0 |
| SKEIENGH | CUL3; CUL3 | Q13618; A0 | CUL3_HUMAN | Unmodified | 0 | 0 | 0 | 0 | 1056600 | 0 | 0 | 125980 | 0 | 0 | 0 | 894430 | 363150 | 0 | 0 | 0 | 206650 | 0 |
| SKENPRNF | TAGLN2; TAGLN | P37802; X6 | TAGL2_HUMAN | Unmodified | 115110 | 43188 | 0 | 402380 | 59602000 | 11856000 | 3950400 | 2403700 | 327420 | 3443700 | 382670 | 0 | 9652800 | 6013200 | 0 | 0 | 14970000 | 21119000 |
| SKEPQLIAF | SLC9A1 | P19634 | SL9A1_HUMAN | Unmodified | 341550 | 241330 | 1385400 | 503720 | 0 | 0 | 0 | 0 | 543090 | 0 | 0 | 2540500 | 0 | 0 | 1049300 | 818450 | 0 | 0 |
| SKEQEVNE | RAI14 | Q9P0K7 | RAI14_HUMAN | Unmodified | 0 | 141090 | 110350 | 154130 | 854260 | 402340 | 0 | 59976 | 144180 | 74439 | 125570 | 983860 | 329050 | 296100 | 222660 | 177930 | 378720 | 725650 |
| SKESVPEFI | STMN1; STMN1 | P16949; A2 | STMN1_HUMAI | Unmodified | 0 | 0 | 0 | 0 | 696740 | 175030 | 1724900 | 1677900 | 6741100 | 11659000 | 2768600 | 1844700 | 314080 | 598470 | 0 | 0 | 863840 | 3055600 |
| SKEVGQQL | SRGAP2; SRGA | O75044; P0 | SRGP2_HUMAI | Unmodified | 0 | 0 | 0 | 0 | 3094300 | 0 | 0 | 25961 | 0 | 213010 | 0 | 0 | 665220 | 0 | 0 | 0 | 0 | 1951000 |
| SKFADLSE | VIM; VIM; VIM | P08670; B0 | VIME_HUMAN | Unmodified | 0 | 0 | 0 | 0 | 0 | 0 | 0 | 0 | 0 | 0 | 0 | 0 | 0 | 0 | 0 | 0 | 0 | 472070 |
| SKFALITW | COTL1; COTL1 | Q14019; H3 | COTL1_HUMAN | Unmodified | 0 | 6257600 | 0 | 46496 | 357650 | 60817 | 0 | 0 | 383010 | 362900 | 0 | 54130 | 19706 | 0 | 16162 | 0 | 517830 | 0 |
| SKFDNLYG | AHCY | P23526 | SAHH_HUMAN | Unmodified | 209850 | 0 | 464600 | 0 | 0 | 0 | 0 | 0 | 726740 | 0 | 240970 | 10150000 | 0 | 0 | 652770 | 797040 | 0 | 0 |
| SKFDNLYG | AHCY | P23526 | SAHH_HUMAN | Unmodified | 0 | 0 | 0 | 0 | 3909300 | 0 | 0 | 0 | 0 | 0 | 0 | 0 | 1239500 | 1429400 | 0 | 0 | 2165900 | 4309000 |
| SKFENEEF | CBFB; CBFB; C | Q13951; J3K | PEBB_HUMAN | Unmodified | 0 | 0 | 0 | 0 | 0 | 0 | 0 | 0 | 0 | 0 | 0 | 712760 | 0 | 0 | 0 | 0 | 0 | 0 |
| SKFENEEF | CBFB; CBFB; C | Q13951; J3K | PEBB_HUMAN | Unmodified | 0 | 0 | 112590 | 167900 | 0 | 0 | 0 | 0 | 0 | 0 | 289100 | 1382200 | 301160 | 237510 | 0 | 0 | 0 | 0 |
| SKFFIEEMI | GALE; GALE; G | Q14376; Q5 | GALE_HUMAN | Unmodified | 0 | 0 | 0 | 0 | 165550 | 37898 | 0 | 0 | 0 | 0 | 0 | 0 | 0 | 44494 | 0 | 0 | 203100 | 0 |
| SKFGANAII | ENO1; ENO3; E | P06733; P13 | ENOA_HUMAN | Unmodified | 0 | 0 | 0 | 0 | 5243700 | 680030 | 1001600 | 0 | 828090 | 4054800 | 0 | 548540 | 3955200 | 1567500 | 0 | 0 | 4746800 | 6822500 |
| SKFGANAII | ENO1; ENO1; E | P06733; K7E | ENOA_HUMAN | Unmodified | 0 | 0 | 0 | 0 | 0 | 0 | 0 | 0 | 0 | 236320 | 0 | 0 | 0 | 0 | 0 | 0 | 0 | 0 |
| SKFVLVKF | ERP29; ERP29 | P30040; F8V | ERP29_HUMAN | Unmodified | 0 | 0 | 0 | 0 | 0 | 0 | 0 | 0 | 6823900 | 6451100 | 0 | 0 | 0 | 0 | 0 | 0 | 6197900 | 0 |
| SKFVPYRD | KIF13A; KIF13A | Q9H1H9; A0 | KI13A_HUMAN | Unmodified | 0 | 0 | 0 | 0 | 780680 | 42958 | 54595 | 0 | 0 | 0 | 0 | 0 | 0 | 0 | 0 | 0 | 64322 | 63706 |
| SKGDQPLY | RASA3 | Q14644 | RASA3_HUMAN | Unmodified | 0 | 0 | 0 | 0 | 130610 | 0 | 0 | 0 | 727880 | 84121 | 30720 | 177750 | 0 | 0 | 0 | 0 | 50524 | 252730 |
| SKGFGFVC | PABPC4; PABP | Q13310; P1 | PABP4_HUMAN | Unmodified | 21155000 | 10034000 | 0 | 6525800 | 0 | 0 | 0 | 0 | 0 | 0 | 0 | 0 | 759000 | 251240 | 0 | 991350 | 0 | 0 |
| SKGFGFVS | PABPC1; PABP | P11940; Q9 | PABP1_HUMAN | Unmodified | 7757400 | 5508200 | 0 | 8517900 | 0 | 0 | 555880 | 123210 | 355140 | 714110 | 0 | 0 | 2186300 | 1162900 | 0 | 886540 | 0 | 0 |
| SKGGEGGI | SYTL4 | Q96C24 | SYTL4_HUMAN | Unmodified | 0 | 0 | 0 | 0 | 0 | 0 | 0 | 0 | 0 | 0 | 0 | 0 | 0 | 0 | 0 | 0 | 0 | 0 |
| SKGHYEVT | AHNAK | Q09666 | AHNK_HUMAN | Unmodified | 0 | 0 | 0 | 0 | 2448100 | 217670 | 0 | 0 | 0 | 140190 | 0 | 0 | 0 | 0 | 0 | 0 | 2115800 | 1440400 |
| SKGKDSLS | LYN; LYN | P07948; E5 | LYN_HUMAN T | Unmodified | 0 | 0 | 0 | 46219 | 788020 | 188840 | 149800 | 420390 | 0 | 0 | 0 | 0 | 0 | 0 | 0 | 35182 | 38221 | 129760 |
| SKGPAVGII | HSPA8; HSPA8 | P11142; E9 | HSP7C_HUMA | Acetyl (Protein N | 0 | 431480 | 0 | 0 | 822920 | 0 | 0 | 0 | 799970 | 626330 | 0 | 0 | 3014100 | 0 | 0 | 0 | 0 | 0 |
| SKGPSAAG | METAP2; META | P50579; F8V | MAP2_HUMAN | Unmodified | 132550 | 166040 | 0 | 97472 | 0 | 0 | 0 | 0 | 186360 | 0 | 344560 | 3000100 | 0 | 0 | 51224 | 27985 | 0 | 0 |
| SKGQSSLA | RALB; RALB; RA | P11234; C9 | RALB_HUMAN | Unmodified | 0 | 0 | 0 | 0 | 1126300 | 135750 | 0 | 0 | 0 | 0 | 0 | 0 | 311980 | 0 | 0 | 0 | 1960500 | 869650 |
| SKGSLEITE | ZDHHC5; ZDHH | Q9C0B5; H0 | ZDHC5_HUMA | Unmodified | 0 | 0 | 0 | 833210 | 1403500 | 0 | 0 | 0 | 604240 | 373870 | 1437100 | 0 | 1186100 | 836480 | 0 | 0 | 647920 | 841370 |
| SKGYAFIEF | NCL; NCL; NCL | P19338; C9 | NUCL_HUMAN | Unmodified | 0 | 0 | 0 | 0 | 316810 | 0 | 0 | 0 | 0 | 768000 | 0 | 0 | 2005500 | 0 | 0 | 0 | 391810 | 0 |
| SKHLQEQL | NF2; NF2 | P35240; A0 | MERL_HUMAN | Unmodified | 0 | 0 | 0 | 0 | 827920 | 0 | 0 | 0 | 0 | 272410 | 0 | 0 | 0 | 977800 | 0 | 0 | 1456400 | 2828600 |
| SKIEDYFPE | GNAS; GNAS; G | Q5JWF2; P6 | GNAS1_HUMA | Unmodified | 1822900 | 1636100 | 4592300 | 8875200 | 3118400 | 990090 | 424060 | 24300 | 5931900 | 1390400 | 3427900 | 24902000 | 3962500 | 1552100 | 16576000 | 11584000 | 1705500 | 3675700 |
| SKIEMEETL | TNIP1; TNIP1; T | Q15025; A0 | TNIP1_HUMAN | Unmodified | 0 | 0 | 0 | 0 | 261450 | 0 | 67770 | 0 | 141360 | 211870 | 0 | 354270 | 0 | 70806 | 0 | 0 | 741560 | 478880 |
| SKIETEIK | CACYBP; CACY | Q9HB71; B2 | CYBP_HUMAN | Unmodified | 92218 | 55016 | 0 | 454660 | 2079800 | 0 | 0 | 0 | 1633000 | 705810 | 2119600 | 7818900 | 2715700 | 0 | 0 | 141210 | 0 | 0 |
| SKINVNEIF | RAP1B; RAP1A | P61224; P62 | RAP1B_HUMAN | Unmodified | 28895000 | 35476000 | 586160 | 94752000 | 35935000 | 8511500 | 14474000 | 0 | 12791000 | 7965000 | 11842000 | 13888000 | 67714000 | 36349000 | 16970000 | 10243000 | 5582800 | 20581000 |
| SKINVNEIF | RAP1B; RAP1A | P61224; P62 | RAP1B_HUMAN | Unmodified | 0 | 0 | 0 | 0 | 1904800 | 293610 | 1254400 | 0 | 0 | 0 | 0 | 0 | 3186900 | 3303000 | 0 | 0 | 41403 | 1110600 |
| SKIQKLESD | FGB | P02675 | FIBB_HUMAN F | Oxidation (M) | 0 | 851700 | 0 | 0 | 0 | 0 | 0 | 0 | 0 | 0 | 0 | 0 | 0 | 0 | 0 | 0 | 0 | 0 |
| SKISVNDFI | DLAT; DLAT; DL | P10515; A0 | ODP2_HUMAN | Unmodified | 0 | 0 | 0 | 0 | 0 | 0 | 0 | 0 | 1104500 | 0 | 0 | 0 | 0 | 0 | 0 | 0 | 225020 | 0 |
| SKITVTSEV | RPL22; RPL22; | P35268; K7E | RL22_HUMAN | Unmodified | 398420 | 414570 | 0 | 0 | 2291800 | 0 | 5142300 | 2091300 | 0 | 1076200 | 555030 | 8237000 | 37792 | 0 | 0 | 0 | 0 | 0 |
| SKITVTSEV | RPL22; RPL22; | P35268; K7E | RL22_HUMAN | Unmodified | 0 | 0 | 0 | 0 | 0 | 0 | 0 | 0 | 0 | 0 | 0 | 0 | 0 | 0 | 0 | 0 | 0 | 0 |
| SKKDATLE | PKP1 | Q13835 | PKP1_HUMAN | Unmodified | 0 | 0 | 0 | 0 | 0 | 0 | 0 | 0 | 0 | 0 | 0 | 0 | 687740 | 0 | 0 | 0 | 1148000 | 0 |
| SKKVEEAE | CBX3; CBX3; C | Q13185; S4 | CBX3_HUMAN | Unmodified | 0 | 0 | 0 | 0 | 784220 | 0 | 98717 | 98359 | 134210 | 143180 | 0 | 412170 | 179370 | 0 | 0 | 0 | 204440 | 0 |
| SKLDAEVS | CTNNA1; CTNN | P35221; G3 | CTNA1_HUMAI | Unmodified | 5011300 | 4129200 | 7544600 | 22957000 | 53503000 | 12204000 | 0 | 0 | 2725000 | 1833900 | 7063900 | 13731000 | 64673000 | 41906000 | 25232000 | 18767000 | 0 | 0 |
| SKLESELAN | SMC1A; SMC1A | Q14683; G8 | SMC1A_HUMA | Unmodified | 0 | 0 | 0 | 0 | 377080 | 0 | 0 | 0 | 0 | 0 | 0 | 0 | 212260 | 0 | 0 | 0 | 0 | 0 |
| SKLGVEML | VSIG10; VSIG1 | Q8N0Z9; F5 | VSI10_HUMAN | Unmodified | 0 | 0 | 0 | 0 | 0 | 0 | 199350 | 0 | 881150 | 0 | 0 | 788040 | 0 | 0 | 0 | 0 | 25742 | 0 |
| SKLIRVENF | PTPRJ; PTPRJ; P | Q12913; A0 | PTPRJ_HUMAN | Unmodified | 0 | 0 | 0 | 0 | 627540 | 98182 | 78348 | 0 | 0 | 0 | 0 | 0 | 0 | 0 | 0 | 0 | 120600 | 192180 |
| SKLLSPYSF | DIP2B | Q9P265 | DIP2B_HUMAN | Unmodified | 0 | 0 | 0 | 0 | 0 | 0 | 0 | 0 | 0 | 84569 | 0 | 0 | 225110 | 0 | 0 | 0 | 564020 | 384350 |
| SKLPKPVQ | PARP1 | P09874 | PARP1_HUMAN | Unmodified | 466470 | 325010 | 108810 | 62289 | 403050 | 99253 | 754650 | 1262700 | 1167800 | 1482200 | 569020 | 2635700 | 576600 | 322240 | 138700 | 128210 | 237420 | 155360 |
| SKLPNTVL | EHD1; EHD1 | Q9H4M9; A0 | EHD1_HUMAN | Unmodified | 87754 | 104730 | 0 | 502330 | 8442000 | 2586700 | 580020 | 152010 | 309170 | 322830 | 687230 | 2961200 | 4751700 | 3704700 | 1336000 | 277570 | 7256800 | 13812000 |
| SKLPNTVL | EHD1; EHD1 | Q9H4M9; A0 | EHD1_HUMAN | Unmodified | 0 | 0 | 0 | 0 | 332150 | 0 | 249430 | 0 | 0 | 0 | 0 | 0 | 109930 | 0 | 0 | 0 | 2224300 | 374620 |
| SKLSIIPQE | ABCC5 | O15440 | MRP5_HUMAN | Unmodified | 0 | 0 | 0 | 0 | 0 | 0 | 0 | 0 | 0 | 0 | 0 | 335310 | 0 | 0 | 0 | 0 | 0 | 0 |
| SKLSQNNF | VDAC3 | Q9Y277 | VDAC3_HUMAN | Unmodified | 0 | 0 | 0 | 0 | 0 | 0 | 71462 | 50324 | 3761300 | 1847900 | 23003000 | 0 | 0 | 0 | 0 | 0 | 0 | 0 |
| SKLTFSCL | PTGES3; PTGES | Q15185; A0 | TEBP_HUMAN | Unmodified | 1507200 | 935120 | 764070 | 3939200 | 31281000 | 4123800 | 9904100 | 6739800 | 10781000 | 13214000 | 6120900 | 19392000 | 31060000 | 18372000 | 1910100 | 1853400 | 24938000 | 13169000 |
| SKLVDEEP | ABCF2 | Q9UG63 | ABCF2_HUMAN | Unmodified | 0 | 132040 | 0 | 536310 | 1196800 | 0 | 128300 | 0 | 0 | 93379 | 0 | 627510 | 0 | 381570 | 0 | 162970 | 552890 | 604370 |
| SKMIYASSI | CFL1; DSTN; C | P23528; P60 | COF1_HUMAN | Unmodified | 0 | 0 | 0 | 0 | 32180000 | 0 | 6384700 | 7242900 | 0 | 0 | 0 | 36554 | 7619200 | 7558400 | 0 | 0 | 16650000 | 7818700 |
| SKMIYASSI | CFL1; DSTN; C | P23528; P60 | COF1_HUMAN | Unmodified | 0 | 0 | 0 | 0 | 6997000 | 1246500 | 4858800 | 419100 | 0 | 932990 | 0 | 0 | 1719300 | 4962300 | 0 | 0 | 10174000 | 1790700 |
| SKMIYASSI | CFL1; DSTN; C | P23528; P60 | COF1_HUMAN | Oxidation (M) | 0 | 0 | 0 | 613860 | 1739300 | 0 | 1158400 | 1639700 | 0 | 0 | 0 | 0 | 0 | 561670 | 0 | 0 | 3163000 | 259370 |
| SKNDCDEE | PLA2G12A; PLA | Q9BZM1; A0 | PG12A_HUMAI | Unmodified | 0 | 0 | 0 | 0 | 0 | 0 | 0 | 0 | 0 | 163100 | 0 | 0 | 0 | 0 | 0 | 0 | 0 | 0 |
| SKNNDLQI | LUZP1 | Q86V48 | LUZP1_HUMAN | Unmodified | 0 | 0 | 0 | 0 | 0 | 0 | 0 | 0 | 0 | 24645 | 93065 | 0 | 0 | 0 | 0 | 0 | 663650 | 109380 |
| SKNPDDLC | GGA3; GGA3; G | Q9NZ52; J3 | GGA3_HUMAN | Unmodified | 0 | 0 | 0 | 0 | 376330 | 0 | 507240 | 84239 | 0 | 0 | 0 | 0 | 151070 | 0 | 0 | 0 | 1107700 | 1343800 |
| SKPELPPG | PYM1 | Q9BRP8 | PYM1_HUMAN | Unmodified | 1327900 | 642260 | 0 | 0 | 0 | 0 | 0 | 904780 | 0 | 628240 | 0 | 840020 | 2947400 | 0 | 0 | 558390 | 0 | 0 |
| SKPGAAM | HNRNPL; HNRI | P14866; M0 | HNRPL_HUMAI | Oxidation (M) | 0 | 0 | 0 | 0 | 0 | 0 | 0 | 0 | 54402 | 0 | 0 | 633520 | 0 | 0 | 0 | 0 | 0 | 0 |
| SKPGCSMI | DPY19L1; DPY1 | Q2PZI1; A0A | D19L1_HUMAN | Unmodified | 0 | 0 | 0 | 0 | 0 | 0 | 0 | 0 | 448620 | 0 | 0 | 0 | 0 | 0 | 0 | 0 | 0 | 0 |
| SKPGSTGP | MINK1 | Q8N4C8 | MINK1_HUMAN | Unmodified | 0 | 0 | 0 | 1329900 | 2272900 | 0 | 0 | 0 | 1155400 | 0 | 0 | 0 | 1030700 | 0 | 0 | 0 | 0 | 0 |
| SKPHSEAG | PKM; PKM; PKM | P14618; B4 | KPYM_HUMAN | Acetyl (Protein N | 0 | 0 | 0 | 0 | 0 | 0 | 0 | 0 | 0 | 0 | 0 | 2960700 | 0 | 0 | 0 | 248010 | 0 | 0 |
| SKPIPIMPA | DNM1L | O00429 | DNM1L_HUMA | Unmodified | 0 | 0 | 0 | 0 | 0 | 0 | 0 | 38248 | 0 | 0 | 156510 | 669020 | 335690 | 164510 | 0 | 0 | 0 | 0 |
| SKPLDNAS | STXBP3 | O00186 | STXB3_HUMAN | Unmodified | 0 | 0 | 0 | 0 | 999070 | 0 | 924240 | 1533500 | 841770 | 1092900 | 172810 | 0 | 1259900 | 2593100 | 0 | 0 | 4305000 | 3297200 |
| SKPSEPIQF | DLG1; DLG1; D | Q12959; A0 | DLG1_HUMAN | Unmodified | 0 | 0 | 0 | 0 | 0 | 0 | 0 | 0 | 4572400 | 1205600 | 0 | 1893100 | 0 | 0 | 0 | 0 | 0 | 0 |
| SKPSFHAP | MAP4K4; MAP4 | O95819; C9 | M4K4_HUMAN | Unmodified | 70040 | 0 | 0 | 317860 | 0 | 0 | 0 | 0 | 981330 | 289030 | 0 | 1732900 | 0 | 0 | 0 | 0 | 324750 | 2655400 |
| SKPTVDIV | ASAP1; ASAP1; | Q9ULH1; H0 | ASAP1_HUMAN | Unmodified | 307840 | 274250 | 0 | 99348 | 514980 | 103140 | 0 | 0 | 0 | 294700 | 121210 | 380640 | 0 | 0 | 183650 | 0 | 0 | 568560 |
| SKPVMIVTE | EPHA3; EPHA3 | P29320; C9 | EPHA3_HUMAI | 2 Oxidation (M) | 122970 | 233720 | 0 | 0 | 0 | 0 | 0 | 0 | 0 | 0 | 0 | 0 | 0 | 0 | 0 | 0 | 0 | 0 |
| SKPVMIVTE | EPHA3; EPHA3 | P29320; C9 | EPHA3_HUMAI | Oxidation (M) | 100730 | 525690 | 0 | 0 | 0 | 224330 | 0 | 0 | 70432 | 0 | 0 | 0 | 0 | 0 | 0 | 0 | 978040 | 0 |
| SKPYLTVD | PLCB1; PLCB1; | Q9NQ66; A0 | PLCB1_HUMAN | Unmodified | 0 | 0 | 0 | 0 | 0 | 0 | 0 | 0 | 197450 | 36047 | 240680 | 0 | 0 | 0 | 0 | 0 | 400120 | 50621 |
| SKQEALKN | CAPZB | B1AK88 | B1AK88_HUMA | Unmodified | 3470700 | 3269200 | 0 | 1900900 | 29565000 | 1838700 | 1457200 | 696890 | 3184200 | 3371200 | 1437100 | 14207000 | 15816000 | 7029700 | 8441600 | 1551800 | 5506000 | 12515000 |
| SKQELPPPI | UTRN | P46939 | UTRN_HUMAN | Unmodified | 0 | 0 | 0 | 0 | 0 | 0 | 0 | 0 | 344930 | 0 | 0 | 0 | 0 | 0 | 0 | 405010 | 0 | 1064900 |
| SKQLETKQ | ANLN | Q9NQW6 | ANLN_HUMAN | Unmodified | 0 | 0 | 0 | 0 | 0 | 0 | 0 | 0 | 0 | 0 | 0 | 0 | 0 | 0 | 0 | 0 | 2585800 | 0 |
| SKQPLPSA | AURKA; AURKA | O14965; Q5 | AURKA_HUMAN | Unmodified | 0 | 0 | 0 | 703580 | 0 | 1287500 | 0 | 0 | 1384700 | 597620 | 0 | 0 | 1571000 | 0 | 0 | 0 | 978040 | 0 |
| SKQQPTQF | SEPTIN2; SEPT | Q15019; C9 | SEPT2_HUMAN | Acetyl (Protein N | 0 | 0 | 0 | 0 | 7167500 | 92051 | 5174500 | 1356600 | 0 | 0 | 0 | 0 | 0 | 0 | 0 | 0 | 20408000 | 3506300 |

| | | | | | | | | | | | | | | | | | | | | | | |
|---|---|---|---|---|---|---|---|---|---|---|---|---|---|---|---|---|---|---|---|---|---|---|
| SKQSLIDW | ECPAS; ECPAS; | Q5VYK3; R4( | ECM29_HUMAI | Unmodified | 0 | 0 | 0 | 0 | 671160 | 0 | 217950 | 0 | 0 | 155340 | 0 | 0 | 129470 | 0 | 0 | 0 | 1060400 | 0 |
| SKRALEQQ | MYH9 | P35579 | MYH9_HUMAN | Unmodified | 0 | 0 | 0 | 0 | 510340 | 0 | 0 | 0 | 0 | 0 | 0 | 0 | 0 | 0 | 0 | 0 | 0 | 0 |
| SKRDPSEEI | RPS6KA1; RPS6 | Q15418; Q1! | KS6A1_HUMAN | Unmodified | 0 | 0 | 0 | 0 | 0 | 0 | 0 | 10961 | 518080 | 108900 | 0 | 617520 | 197430 | 69204 | 0 | 0 | 68147 | 183260 |
| SKRPNPCA | PPP1R1B; PPP1 | Q9UD71; J3I | PPR1B_HUMAN | Unmodified | 0 | 0 | 0 | 0 | 0 | 0 | 0 | 44369 | 0 | 0 | 395470 | 1290300 | 0 | 0 | 0 | 0 | 0 | 0 |
| SKSEEAHA | SERBP1 | Q8NC51 | PAIRB_HUMAN | Unmodified | 0 | 423750 | 0 | 370830 | 0 | 0 | 0 | 0 | 0 | 0 | 486750 | 839160 | 0 | 0 | 568620 | 171510 | 0 | 0 |
| SKSESPKEI | HNRNPA1; HNI | P09651; F8V | ROA1_HUMAN | Acetyl (Protein N | 0 | 0 | 0 | 0 | 852050 | 195190 | 234450 | 205520 | 809160 | 552850 | 399080 | 1800800 | 0 | 0 | 0 | 0 | 302700 | 14728 |
| SKSSSTTYF | ADGRG6 | Q86SQ4 | AGRG6_HUMA | Unmodified | 0 | 0 | 0 | 0 | 1247900 | 108450 | 0 | 35174 | 0 | 0 | 0 | 0 | 0 | 0 | 0 | 0 | 0 | 0 |
| SKSVKEDS | IQGAP1; IQGAP | P46940; H0Y | IQGA1_HUMAN | Unmodified | 0 | 0 | 0 | 0 | 1235200 | 368750 | 0 | 0 | 0 | 0 | 0 | 0 | 108640 | 61285 | 0 | 0 | 46789 | 82443 |
| SKTDNSSL | BAZ1B | Q9UIG0 | BAZ1B_HUMAN | Unmodified | 0 | 602540 | 0 | 0 | 0 | 0 | 0 | 594300 | 0 | 0 | 0 | 0 | 0 | 0 | 0 | 0 | 0 | 0 |
| SKTEDHEE | TMEM59; TMEM | Q9BXS4; Q5 | TMM59_HUMAI | Unmodified | 0 | 0 | 0 | 0 | 0 | 0 | 212820 | 23165 | 6679400 | 1171700 | 0 | 0 | 0 | 0 | 0 | 0 | 8432.2 | 206250 |
| SKTEDHEE | TMEM59; TMEM | Q9BXS4; Q5 | TMM59_HUMAI | Unmodified | 0 | 0 | 0 | 0 | 1937000 | 0 | 1358900 | 789710 | 0 | 2339500 | 0 | 0 | 0 | 0 | 0 | 0 | 0 | 493520 |
| SKVAHILNI | HSDL2 | Q6YN16 | HSDL2_HUMAI | Unmodified | 0 | 0 | 0 | 0 | 0 | 0 | 177750 | 0 | 255640 | 5425600 | 0 | 0 | 0 | 0 | 0 | 0 | 0 | 0 |
| SKVDEAVA | PABPC4; PABP( | Q13310; B1/ | PABP4_HUMAN | Unmodified | 5500300 | 3186300 | 1010800 | 169550 | 0 | 0 | 48609 | 36868 | 245770 | 0 | 176710 | 584500 | 0 | 0 | 1575400 | 2259600 | 0 | 0 |
| SKVDEAVA | PABPC4; PABP( | Q13310; B1/ | PABP4_HUMAN | Unmodified | 439320 | 394810 | 0 | 0 | 857890 | 0 | 0 | 0 | 0 | 0 | 0 | 0 | 0 | 0 | 0 | 270620 | 0 | 148470 |
| SKVDEAVA | PABPC1; PABP( | P11940; Q9I | PABP1_HUMAN | Unmodified | 14229000 | 18268000 | 2181200 | 340590 | 0 | 0 | 0 | 0 | 0 | 0 | 0 | 2453700 | 0 | 0 | 2970200 | 5379000 | 123910 | 0 |
| SKVDLDRL | PDGFD | Q9GZP0 | PDGFD_HUMA | Unmodified | 0 | 0 | 0 | 0 | 0 | 498990 | 0 | 0 | 0 | 0 | 0 | 0 | 0 | 0 | 0 | 0 | 0 | 0 |
| SKVDQIQEI | IQGAP1; IQGAP | P46940; A0A | IQGA1_HUMAN | Unmodified | 1420800 | 855780 | 1195800 | 807090 | 7229100 | 2227400 | 726220 | 321890 | 1896600 | 1198200 | 355490 | 9491400 | 613540 | 414540 | 1802500 | 1367000 | 598420 | 0 |
| SKVDQVQI | IQGAP2; IQGAP | Q13576; E7I | IQGA2_HUMAN | Unmodified | 0 | 0 | 1289700 | 646810 | 0 | 0 | 0 | 0 | 0 | 0 | 0 | 0 | 0 | 0 | 1346500 | 1312700 | 0 | 0 |
| SKVEQPQD | IQGAP3; IQGAP | Q86VI3; F2Z | IQGA3_HUMAN | Unmodified | 242580 | 310610 | 0 | 258120 | 276530 | 0 | 0 | 0 | 463500 | 0 | 0 | 486630 | 0 | 0 | 123360 | 100300 | 0 | 0 |
| SKVGGTKP | TPD52; TPD52 | P55327; E5R | TPD52_HUMAN | Unmodified | 0 | 0 | 0 | 0 | 1773700 | 0 | 0 | 0 | 1419500 | 3947800 | 0 | 0 | 2640600 | 645410 | 0 | 0 | 878540 | 0 |
| SKVPAFVRI | PITPNB; PITPNI | P48739; A0A | PIPNB_HUMAN | Oxidation (M) | 0 | 0 | 0 | 0 | 0 | 0 | 251960 | 0 | 0 | 0 | 0 | 0 | 1111400 | 0 | 0 | 0 | 723420 | 0 |
| SKVTVDTG | EIF5B; EIF5B | O60841; A0/ | IF2P_HUMAN E | Unmodified | 265970 | 0 | 0 | 169960 | 56626 | 0 | 0 | 0 | 491650 | 0 | 0 | 616620 | 0 | 0 | 0 | 339840 | 0 | 132180 |
| SKVYEDPAI | EXOC7; EXOC7 | Q9UPT5; B4I | EXOC7_HUMAI | Unmodified | 0 | 0 | 0 | 0 | 5717100 | 0 | 1755600 | 0 | 0 | 411670 | 0 | 0 | 0 | 0 | 0 | 0 | 1674300 | 0 |
| SKWGSKPS | CEP55 | Q53EZ4 | CEP55_HUMAN | Unmodified | 0 | 0 | 0 | 0 | 0 | 0 | 0 | 0 | 550600 | 0 | 0 | 0 | 0 | 25628 | 125630 | 0 | 0 | 0 |
| SKWSFLFS | TBC1D15; TBC1 | Q8TC07; F8\ | TBC15_HUMAN | Unmodified | 0 | 0 | 0 | 0 | 0 | 0 | 551500 | 0 | 17048 | 231340 | 0 | 0 | 170030 | 87280 | 0 | 0 | 0 | 0 |
| SKWTPLET | GALNT6; GALN | Q8NCL4; S4 | GALT6_HUMAN | Unmodified | 0 | 0 | 0 | 0 | 0 | 0 | 0 | 0 | 570960 | 0 | 0 | 0 | 0 | 0 | 0 | 0 | 391550 | 0 |
| SKYDEIFYN | EHD2 | Q9NZN4 | EHD2_HUMAN | Unmodified | 0 | 0 | 0 | 0 | 8719400 | 1911900 | 0 | 36629 | 0 | 0 | 0 | 0 | 0 | 0 | 0 | 0 | 0 | 0 |
| SKYDEIFYN | EHD2 | Q9NZN4 | EHD2_HUMAN | Unmodified | 0 | 0 | 0 | 0 | 21867000 | 4342300 | 0 | 0 | 813710 | 0 | 0 | 0 | 0 | 0 | 0 | 0 | 0 | 0 |
| SLAAAAGK | SST | P61278 | SMS_HUMAN S | Unmodified | 0 | 0 | 0 | 0 | 0 | 0 | 0 | 0 | 0 | 1331800 | 0 | 0 | 0 | 0 | 0 | 0 | 0 | 0 |
| SLAADDEG | ARVCF; ARVCF; | O00192; E9I | ARVC_HUMAN | Unmodified | 0 | 0 | 503740 | 0 | 0 | 0 | 0 | 0 | 158120 | 0 | 265660 | 221610 | 0 | 0 | 1442800 | 2052500 | 0 | 0 |
| SLAAEWGK | DECR1 | Q16698 | DECR_HUMAN | Unmodified | 0 | 0 | 184310 | 359610 | 0 | 0 | 52957 | 0 | 5115300 | 506200 | 0 | 587190 | 0 | 0 | 257400 | 0 | 100180 | 365980 |
| SLAASSSF | MOGS; MOGS; | Q13724; A0/ | MOGS_HUMAN | Unmodified | 0 | 0 | 0 | 0 | 0 | 0 | 0 | 0 | 7417800 | 475860 | 0 | 1021800 | 92884 | 0 | 0 | 0 | 86380 | 0 |
| SLADCNFS | RICTOR | Q6R327 | RICTR_HUMAN | Unmodified | 0 | 0 | 0 | 0 | 1003000 | 0 | 0 | 0 | 0 | 0 | 0 | 0 | 0 | 0 | 0 | 0 | 711480 | 0 |
| SLADEAEVI | GAS7 | O60861 | GAS7_HUMAN | Unmodified | 0 | 0 | 0 | 0 | 0 | 0 | 0 | 0 | 0 | 0 | 2229100 | 270320 | 474040 | 0 | 0 | 0 | 0 | 0 |
| SLADELAL | LDHB; LDHB; LI | P07195; A0A | LDHB_HUMAN | Unmodified | 100520 | 185280 | 119710 | 920900 | 1524600 | 100390 | 0 | 0 | 43893 | 0 | 1801200 | 3684700 | 835490 | 245230 | 1205400 | 198110 | 174020 | 752780 |
| SLADFEK | PSMD11 | O00231 | PSD11_HUMAN | Unmodified | 0 | 0 | 0 | 0 | 0 | 0 | 0 | 0 | 0 | 0 | 343440 | 448170 | 0 | 0 | 0 | 0 | 0 | 0 |
| SLADIAREE | PSMA3; PSMA3 | P25788; G3\ | PSA3_HUMAN | Unmodified | 0 | 0 | 1912000 | 0 | 0 | 0 | 0 | 0 | 2565900 | 167110 | 1055200 | 30431000 | 0 | 0 | 0 | 1971700 | 0 | 0 |
| SLADIEEEY | MVB12A; MVB1 | Q96EY5; E9I | MB12A_HUMAI | Unmodified | 0 | 15872000 | 0 | 9229400 | 4353000 | 712790 | 0 | 17337 | 928850 | 1164300 | 0 | 435570 | 0 | 11306000 | 0 | 4337500 | 1006500 | 5684300 |
| SLADIGKTV | SLC4A4 | Q9Y6R1 | S4A4_HUMAN | Unmodified | 0 | 0 | 0 | 0 | 510920 | 73501 | 0 | 0 | 0 | 0 | 0 | 0 | 243430 | 162470 | 0 | 0 | 151560 | 215030 |
| SLADIKESN | CAMSAP2 | Q08AD1 | CAMP2_HUMA | Unmodified | 0 | 0 | 0 | 0 | 229210 | 71552 | 89520 | 0 | 0 | 119460 | 0 | 0 | 560730 | 208320 | 0 | 0 | 307480 | 260310 |
| SLAEANNL | CCNY | Q8ND76 | CCNY_HUMAN | Unmodified | 96218 | 0 | 0 | 355610 | 0 | 0 | 44345 | 0 | 0 | 0 | 0 | 3784400 | 0 | 0 | 3383000 | 1692800 | 18500 | 0 |
| SLAEDFAE( | HYOU1; HYOU1 | Q9Y4L1; J3Q | HYOU1_HUMA | Unmodified | 0 | 0 | 0 | 0 | 0 | 0 | 237550 | 0 | 11292000 | 964600 | 3247200 | 1595400 | 0 | 0 | 184880 | 55429 | 0 | 0 |
| SLAEDFAE( | HYOU1; HYOU1 | Q9Y4L1; J3Q | HYOU1_HUMA | Unmodified | 0 | 0 | 0 | 0 | 0 | 0 | 0 | 0 | 673040 | 0 | 0 | 0 | 0 | 0 | 0 | 0 | 0 | 0 |
| SLAEDFAE( | HYOU1; HYOU1 | Q9Y4L1; J3Q | HYOU1_HUMA | Unmodified | 0 | 0 | 0 | 872850 | 0 | 0 | 0 | 0 | 23356000 | 8450700 | 712760 | 338500 | 0 | 0 | 0 | 0 | 0 | 0 |
| SLAEGNLL | VPS37B; VPS37 | Q9H9H4; F5 | VP37B_HUMAN | Unmodified | 63624000 | 79001000 | 51754000 | 63891000 | 325850 | 1502700 | 0 | 188060 | 5549100 | 2788900 | 5166500 | 13754000 | 38067000 | 321340 | 1E+08 | 95591000 | 0 | 227230 |
| SLAEGNLL | VPS37B; VPS37 | Q9H9H4; F5 | VP37B_HUMAN | Unmodified | 0 | 0 | 0 | 0 | 0 | 0 | 0 | 0 | 0 | 0 | 0 | 0 | 2521700 | 0 | 0 | 0 | 0 | 0 |
| SLAEGYFD | LANCL1; LANCI | O43813; E9I | LANC1_HUMAI | Unmodified | 849010 | 630730 | 345830 | 0 | 0 | 0 | 0 | 0 | 0 | 0 | 0 | 3821200 | 0 | 0 | 534910 | 446050 | 0 | 0 |
| SLAEHRTPI | STIP1 | P31948 | STIP1_HUMAN | Unmodified | 0 | 0 | 0 | 0 | 633200 | 0 | 0 | 0 | 0 | 485980 | 0 | 377080 | 561000 | 0 | 0 | 0 | 735270 | 314260 |
| SLAEHRTPI | STIP1 | P31948 | STIP1_HUMAN | Unmodified | 160400 | 0 | 95116 | 301200 | 3002900 | 433610 | 471420 | 278030 | 984860 | 1498800 | 924720 | 1429000 | 3397800 | 2067000 | 97118 | 81903 | 3135600 | 1890700 |
| SLAEIEKEYI | MVB12B | Q9H7P6 | MB12B_HUMAI | Unmodified | 269780 | 316560 | 0 | 582760 | 0 | 0 | 0 | 0 | 0 | 219910 | 0 | 0 | 0 | 0 | 0 | 0 | 0 | 0 |
| SLAERNLEI | VPS37C; VPS37 | A5D8V6; F5I | VP37C_HUMAN | Unmodified | 120350 | 0 | 130860 | 517050 | 0 | 0 | 0 | 0 | 228290 | 0 | 0 | 0 | 495970 | 0 | 0 | 51602 | 0 | 0 |
| SLAGPAGA | LFNG | Q8NES3 | LFNG_HUMAN | Unmodified | 0 | 0 | 0 | 0 | 0 | 0 | 0 | 0 | 0 | 0 | 0 | 0 | 0 | 0 | 0 | 0 | 0 | 0 |
| SLAGSSGP | EEF1D; EEF1D; | P29692; E9P | EF1D_HUMAN | Unmodified | 1420500 | 289490 | 188860 | 1603800 | 4176700 | 1479600 | 505630 | 227580 | 3501000 | 919330 | 6578700 | 50638000 | 4140600 | 1340400 | 1551000 | 1419800 | 2103800 | 5155000 |
| SLAINPKDI | TLN2; TLN2; TLI | Q9Y4G6; H0 | TLN2_HUMAN | Unmodified | 0 | 0 | 0 | 0 | 0 | 0 | 94032 | 0 | 0 | 0 | 0 | 431060 | 0 | 0 | 0 | 0 | 412600 | 0 |
| SLALEWAC | PECR | Q9BY49 | PECR_HUMAN | Unmodified | 0 | 0 | 0 | 0 | 0 | 0 | 112610 | 0 | 744800 | 0 | 0 | 0 | 0 | 0 | 0 | 0 | 0 | 0 |
| SLAMEMVL | ECHS1 | P30084 | ECHM_HUMAN | Oxidation (M) | 0 | 0 | 0 | 0 | 0 | 222640 | 3580000 | 0 | 0 | 0 | 0 | 0 | 1707600 | 4037900 | 0 | 467510 | 443610 | 1604100 |
| SLANAESQ | GPN1 | Q9HCN4 | GPN1_HUMAN | Unmodified | 98554 | 0 | 99288 | 124060 | 0 | 0 | 0 | 0 | 0 | 0 | 0 | 0 | 264390 | 0 | 90138 | 51032 | 0 | 0 |
| SLAPQAGT | IGF2R | P11717 | MPRI_HUMAN | Unmodified | 0 | 0 | 0 | 0 | 0 | 0 | 0 | 0 | 11692000 | 801360 | 0 | 776120 | 0 | 0 | 0 | 0 | 0 | 0 |
| SLAPSIHGI | MCM3; MCM3; | P25205; A0A | MCM3_HUMAN | Unmodified | 0 | 0 | 0 | 0 | 0 | 0 | 0 | 126630 | 0 | 0 | 0 | 162110 | 0 | 0 | 0 | 0 | 0 | 0 |
| SLAQAEAEI | PLEC | Q15149 | PLEC_HUMAN | Unmodified | 0 | 0 | 0 | 0 | 0 | 0 | 0 | 0 | 0 | 0 | 0 | 364950 | 0 | 0 | 0 | 0 | 0 | 0 |
| SLASLALR | MBLAC2 | Q68D91 | MBLC2_HUMAI | Unmodified | 71723 | 79456 | 475840 | 240330 | 0 | 0 | 0 | 0 | 0 | 0 | 0 | 622500 | 0 | 0 | 298380 | 445770 | 0 | 0 |
| SLATRTDQI | PSMD10; PSMD | O75832; B1/ | PSD10_HUMAN | Unmodified | 0 | 0 | 0 | 0 | 201510 | 51268 | 12672 | 0 | 835550 | 50356 | 53971 | 104960 | 724020 | 108030 | 0 | 38566 | 455550 | 492600 |
| SLATTRPTV | VPS4A | Q9UN37 | VPS4A_HUMAN | Unmodified | 2988200 | 3237600 | 2343800 | 5992800 | 17112000 | 2895800 | 503240 | 0 | 8902800 | 7420500 | 4986800 | 17199000 | 17038000 | 11307000 | 2420900 | 2524100 | 1082600 | 4042400 |
| SLAVQRPA | PI4KA | P42356 | PI4KA_HUMAN | Unmodified | 105990 | 158210 | 332210 | 0 | 0 | 0 | 28604 | 45763 | 0 | 70570 | 0 | 219420 | 0 | 167870 | 301820 | 50293 | 0 | 0 |
| SLAYLTAAT | COPA; COPA; C | P53621; A0A | COPA_HUMAN | Unmodified | 0 | 0 | 197860 | 0 | 0 | 0 | 0 | 0 | 0 | 0 | 0 | 1350800 | 0 | 0 | 0 | 0 | 0 | 0 |
| SLAYVDNIL | PKN2; PKN2 | Q16513; B1/ | PKN2_HUMAN | Unmodified | 0 | 0 | 0 | 0 | 1584400 | 0 | 0 | 0 | 0 | 0 | 0 | 0 | 0 | 0 | 0 | 0 | 0 | 0 |
| SLCIPFKPL | EPRS1 | P07814 | SYEP_HUMAN | Unmodified | 335370 | 0 | 217100 | 0 | 1441600 | 0 | 1835700 | 272800 | 1606400 | 805190 | 1399900 | 5946900 | 5094600 | 3271200 | 257360 | 214560 | 4020500 | 3379900 |
| SLCPETWP | MYO1C; MYO1( | O00159; F5I | MYO1C_HUMA | Unmodified | 3486100 | 499610 | 1112400 | 7084000 | 1897300 | 865360 | 538250 | 0 | 0 | 0 | 0 | 0 | 0 | 0 | 15626000 | 7088600 | 390520 | 394310 |
| SLDALGWE | PANK4; PANK4; | Q9NVE7; H0 | PANK4_HUMAI | Unmodified | 0 | 0 | 0 | 0 | 0 | 0 | 0 | 0 | 0 | 0 | 0 | 0 | 0 | 0 | 0 | 0 | 867430 | 1148000 |
| SLDAYPILN | ITGA3 | P26006 | ITA3_HUMAN I | Unmodified | 0 | 0 | 0 | 0 | 0 | 0 | 0 | 0 | 0 | 0 | 0 | 0 | 0 | 0 | 0 | 0 | 0 | 3468400 |
| SLDDEVNA | SEPTIN6; SEPTI | Q14141; B1/ | SEPT6_HUMAN | Unmodified | 0 | 0 | 0 | 0 | 2342700 | 0 | 356800 | 0 | 0 | 0 | 0 | 2224700 | 1200900 | 831250 | 0 | 0 | 1935400 | 2416400 |
| SLDDVIKR | ALDH1A1 | P00352 | AL1A1_HUMAN | Unmodified | 0 | 0 | 0 | 0 | 0 | 0 | 0 | 0 | 0 | 314500 | 1987200 | 16573000 | 0 | 0 | 0 | 0 | 0 | 0 |
| SLDDVIKR/ | ALDH1A1 | P00352 | AL1A1_HUMAN | Unmodified | 0 | 0 | 0 | 0 | 0 | 0 | 285450 | 0 | 0 | 546780 | 0 | 0 | 0 | 0 | 0 | 0 | 125550 | 0 |
| SLDEMEK | SFPQ; SFPQ | P23246; H0Y | SFPQ_HUMAN | Unmodified | 0 | 104900 | 0 | 0 | 602590 | 0 | 46052 | 262900 | 1056100 | 325560 | 468430 | 1719000 | 1575700 | 0 | 0 | 0 | 131260 | 0 |
| SLDEMEKQ | SFPQ; SFPQ | P23246; H0Y | SFPQ_HUMAN | Unmodified | 0 | 0 | 0 | 0 | 136960 | 0 | 195440 | 234220 | 0 | 444630 | 0 | 0 | 1132900 | 177580 | 0 | 0 | 138770 | 698810 |
| SLDEQCVV | PLSCR1; PLSCF | O15162; C9J | PLS1_HUMAN I | Unmodified | 952320 | 70992 | 0 | 0 | 0 | 0 | 185590 | 0 | 2093700 | 390930 | 92622 | 603780 | 0 | 0 | 0 | 0 | 0 | 0 |
| SLDETTLRF | TLR9; N/A | Q9NR96; H0 | TLR9_HUMAN T | Unmodified | 0 | 0 | 0 | 0 | 0 | 14636 | 0 | 20129 | 346680 | 377790 | 490010 | 3366900 | 0 | 0 | 0 | 0 | 0 | 0 |
| SLDFYTR | GLO1 | Q04760 | LGUL_HUMAN | Unmodified | 0 | 698460 | 0 | 0 | 1491400 | 0 | 0 | 0 | 2834700 | 1457100 | 0 | 11357000 | 5574400 | 0 | 3561800 | 1674100 | 350060 | 0 |
| SLDFYTRVI | GLO1 | Q04760 | LGUL_HUMAN | Oxidation (M) | 0 | 0 | 0 | 0 | 3300300 | 768300 | 0 | 0 | 0 | 68744 | 0 | 159240 | 10339000 | 5571200 | 0 | 0 | 0 | 0 |
| SLDGAPIG | EPB41L2; EPB4 | O43491; A0/ | E41L2_HUMAN | Oxidation (M) | 0 | 0 | 0 | 0 | 919510 | 0 | 0 | 0 | 0 | 0 | 0 | 0 | 0 | 0 | 0 | 0 | 337930 | 1022700 |
| SLDGASVN | EPB41L3; EPB4 | A0A0A0MRA | A0A0A0MRA8_I | Oxidation (M) | 0 | 75201 | 0 | 0 | 0 | 85603 | 52563 | 0 | 0 | 0 | 0 | 0 | 322090 | 89647 | 0 | 0 | 314280 | 2378000 |
| SLDGASVN | EPB41L3; EPB4 | A0A0A0MRA | A0A0A0MRA8_I | Unmodified | 0 | 0 | 0 | 58480 | 0 | 62996 | 0 | 0 | 227440 | 107320 | 0 | 0 | 328310 | 199730 | 0 | 0 | 1115900 | 5447100 |
| SLDGASVN | EPB41L3; EPB4 | A0A0A0MRA | A0A0A0MRA8_I | Unmodified | 0 | 0 | 0 | 0 | 0 | 0 | 0 | 0 | 0 | 0 | 0 | 0 | 0 | 0 | 0 | 0 | 456480 | 1158500 |
| SLDGIPFTV | LPP; LPP | Q93052; A0/ | LPP_HUMAN Li | Unmodified | 0 | 0 | 0 | 0 | 2241500 | 0 | 0 | 0 | 0 | 0 | 0 | 0 | 0 | 0 | 0 | 0 | 0 | 0 |
| SLDGIPFTV | LPP; LPP | Q93052; A0/ | LPP_HUMAN Li | Unmodified | 0 | 0 | 0 | 0 | 637480 | 0 | 0 | 0 | 0 | 96359 | 0 | 0 | 1469300 | 431440 | 0 | 0 | 173050 | 192590 |
| SLDIQVPNI | QARS1; QARS1 | P47897; H7( | SYQ_HUMAN G | Unmodified | 40467 | 0 | 0 | 0 | 0 | 0 | 81936 | 30660 | 215210 | 0 | 365680 | 2536100 | 0 | 0 | 0 | 0 | 111170 | 0 |
| SLDKDPLLI | ATP2B1 | P20020 | AT2B1_HUMAN | Oxidation (M) | 0 | 0 | 0 | 2172300 | 628500 | 0 | 0 | 0 | 645690 | 582960 | 1288100 | 7909000 | 1446400 | 1513200 | 3381800 | 2920500 | 395260 | 568920 |
| SLDKDPLLI | ATP2B1 | P20020 | AT2B1_HUMAN | Unmodified | 0 | 0 | 0 | 459010 | 877440 | 0 | 0 | 74721 | 1310600 | 0 | 335840 | 2617100 | 2248200 | 794480 | 735050 | 534790 | 0 | 952660 |
| SLDKDPML | ATP2B4 | P23634 | AT2B4_HUMAN | 2 Oxidation (M) | 0 | 0 | 0 | 0 | 0 | 0 | 0 | 0 | 0 | 0 | 571280 | 106520 | 0 | 0 | 0 | 1650500 | 0 | 0 |
| SLDKDPML | ATP2B4 | P23634 | AT2B4_HUMAN | Oxidation (M) | 0 | 0 | 0 | 0 | 0 | 0 | 386870 | 0 | 0 | 0 | 0 | 4470400 | 0 | 0 | 0 | 0 | 0 | 0 |
| SLDLDISK | ACAA2; ACAA2; | P42765; A0A | THIM_HUMAN | Unmodified | 0 | 0 | 0 | 0 | 0 | 0 | 0 | 0 | 8689600 | 1356800 | 0 | 0 | 0 | 0 | 0 | 0 | 0 | 0 |
| SLDLDSIIAI | KRT1 | P04264 | K2C1_HUMAN | Unmodified | 0 | 0 | 0 | 0 | 95106 | 0 | 0 | 0 | 0 | 20423 | 0 | 0 | 86778 | 0 | 0 | 0 | 393080 | 224710 |
| SLDLFNCE | ANP32A; ANP3 | P39687; Q9 | AN32A_HUMAN | Unmodified | 159310 | 205780 | 112430 | 0 | 368600 | 113230 | 183950 | 0 | 420010 | 468910 | 663840 | 888500 | 0 | 0 | 176040 | 126360 | 0 | 143490 |
| SLDLFNCE | ANP32A; ANP3 | P39687; H0Y | AN32A_HUMAN | Unmodified | 0 | 0 | 0 | 0 | 12840000 | 904380 | 0 | 0 | 0 | 2184400 | 2695900 | 2631400 | 11609000 | 2168200 | 0 | 0 | 3086300 | 944340 |
| SLDLFNCE | ANP32B | Q92688 | AN32B_HUMAI | Unmodified | 0 | 0 | 0 | 0 | 1713000 | 0 | 2323900 | 0 | 730780 | 0 | 682720 | 0 | 4535300 | 1545000 | 0 | 0 | 693290 | 0 |
| SLDLIESLLI | CNOT1 | A5YKK6 | CNOT1_HUMA | Unmodified | 0 | 0 | 0 | 0 | 0 | 0 | 0 | 0 | 0 | 0 | 0 | 1295400 | 0 | 0 | 0 | 55078 | 0 | 0 |
| SLDLTGPLI | CELSR3 | Q9NYQ7 | CELR3_HUMAN | Unmodified | 0 | 0 | 0 | 0 | 0 | 0 | 0 | 0 | 565350 | 0 | 0 | 0 | 0 | 0 | 0 | 0 | 0 | 0 |
| SLDLVTMK | SEPTIN8; SEPTI | Q92599; Q1 | SEPT8_HUMAN | Unmodified | 0 | 0 | 0 | 0 | 6603900 | 266120 | 0 | 44638 | 1709700 | 1016400 | 0 | 4324300 | 0 | 0 | 0 | 0 | 3727100 | 1504200 |
| SLDNEVEK | NOSTRIN; NOS | Q8IVI9; E9PI | NOSTN_HUMA | Unmodified | 3304300 | 1455400 | 0 | 5097400 | 0 | 0 | 0 | 0 | 0 | 30457 | 0 | 0 | 0 | 1045800 | 10800000 | 4574900 | 0 | 393840 |
| SLDNIWISI | EEPD1 | Q7L9B9 | EEPD1_HUMAN | Unmodified | 0 | 0 | 165930 | 0 | 406200 | 0 | 0 | 356030 | 0 | 357630 | 72904 | 2753700 | 0 | 929190 | 0 | 0 | 4585100 | 4132900 |
| SLDNNYST | CTNND1; CTNN | O60716; H0Y | CTND1_HUMAI | Unmodified | 0 | 0 | 0 | 742550 | 0 | 0 | 0 | 0 | 0 | 0 | 331770 | 2287600 | 0 | 0 | 1070800 | 1018400 | 88312 | 0 |
| SLDNNYST | CTNND1; CTNN | O60716; H0Y | CTND1_HUMAI | Unmodified | 2419800 | 2407000 | 6809500 | 9737400 | 0 | 0 | 194110 | 0 | 0 | 0 | 0 | 10975000 | 0 | 0 | 11229000 | 9632700 | 0 | 0 |
| SLDPENSE | SHTN1 | A0MZ66 | SHOT1_HUMAI | Unmodified | 0 | 0 | 0 | 0 | 481970 | 0 | 0 | 0 | 0 | 336240 | 0 | 0 | 1071800 | 853260 | 0 | 0 | 130460 | 469540 |
| SLDPFKPFI | LRP1 | Q07954 | LRP1_HUMAN I | Unmodified | 0 | 0 | 98330 | 0 | 0 | 0 | 0 | 0 | 455800 | 224750 | 0 | 29717 | 0 | 0 | 0 | 0 | 0 | 0 |
| SLDPFKPFI | LRP1 | Q07954 | LRP1_HUMAN I | Unmodified | 0 | 0 | 0 | 0 | 0 | 0 | 0 | 0 | 1102200 | 46258 | 0 | 0 | 0 | 0 | 0 | 0 | 0 | 0 |
| SLDPSRPV | GUSB | P08236 | BGLR_HUMAN | Unmodified | 0 | 0 | 0 | 0 | 0 | 0 | 268820 | 0 | 37071000 | 17974000 | 0 | 0 | 91586 | 0 | 0 | 0 | 0 | 0 |
| SLDQAIND | SMC3 | Q9UQE7 | SMC3_HUMAN | Unmodified | 0 | 0 | 0 | 0 | 593100 | 0 | 0 | 0 | 0 | 0 | 0 | 425470 | 0 | 0 | 0 | 0 | 0 | 0 |
| SLDQAISR | CYFIP2; CYFIP2 | Q96F07; H7( | CYFP2_HUMAN | Unmodified | 0 | 0 | 0 | 0 | 0 | 0 | 0 | 0 | 0 | 0 | 0 | 0 | 0 | 0 | 0 | 450960 | 0 | 0 |
| SLDQDPVV | HSPH1; HSPH1 | Q92598; Q5 | HS105_HUMAN | Unmodified | 0 | 4668000 | 0 | 300990 | 0 | 88897 | 177580 | 0 | 1338100 | 0 | 1679000 | 5202500 | 0 | 0 | 907410 | 630330 | 70691 | 469440 |

| | | | | | | | | | | | | | | | | | | | | | | |
|---|---|---|---|---|---|---|---|---|---|---|---|---|---|---|---|---|---|---|---|---|---|---|
| SLDQDPVV | HSPH1; HSPH1 | Q92598; Q5 | HS105_HUMAN | Unmodified | 0 | 0 | 0 | 1367400 | 1565600 | 0 | 1529400 | 713250 | 0 | 0 | 0 | 0 | 2788000 | 0 | 0 | 0 | 2458700 | 4453300 |
| SLDQDPVV | HSPH1; HSPH1 | Q92598; Q5 | HS105_HUMAN | Unmodified | 0 | 0 | 0 | 0 | 396360 | 0 | 616940 | 827030 | 0 | 134930 | 0 | 0 | 257830 | 0 | 0 | 0 | 2505400 | 641880 |
| SLDRNLPS | SRGN | P10124 | SRGN_HUMAN | Oxidation (M) | 0 | 0 | 0 | 0 | 0 | 0 | 0 | 0 | 0 | 0 | 0 | 0 | 0 | 0 | 0 | 0 | 0 | 1642400 |
| SLDSDESE | PDAP1 | Q13442 | HAP28_HUMAN | Unmodified | 0 | 0 | 0 | 0 | 569520 | 32735 | 0 | 0 | 109620 | 123270 | 0 | 0 | 71153 | 0 | 0 | 0 | 69598 | 219650 |
| SLDSYPVTS | MYO6; MYO6; M | A0A1Y0BRN | A0A1Y0BRN3_I | Unmodified | 141110 | 77346 | 589460 | 285120 | 0 | 0 | 0 | 14546 | 269630 | 0 | 479460 | 571960 | 0 | 0 | 577380 | 773630 | 0 | 0 |
| SLDSYPVTS | MYO6; MYO6; M | A0A1Y0BRN | A0A1Y0BRN3_I | Unmodified | 0 | 0 | 0 | 0 | 483050 | 0 | 0 | 0 | 0 | 156480 | 0 | 73548 | 1095800 | 0 | 0 | 0 | 0 | 578620 |
| SLDVNQDS | S100A13 | Q99584 | S10AD_HUMAN | Unmodified | 0 | 0 | 0 | 461470 | 29395000 | 269330 | 0 | 0 | 3695900 | 2264500 | 0 | 1811200 | 0 | 0 | 0 | 0 | 8241700 | 31524000 |
| SLDVNQDS | S100A13 | Q99584 | S10AD_HUMAN | Unmodified | 0 | 0 | 0 | 0 | 3714200 | 569710 | 0 | 0 | 0 | 0 | 0 | 0 | 0 | 0 | 0 | 636460 | 3931900 | 8311800 |
| SLDVNQDS | S100A13 | Q99584 | S10AD_HUMAN | Unmodified | 0 | 0 | 0 | 0 | 1748100 | 0 | 0 | 0 | 0 | 0 | 0 | 0 | 422320 | 0 | 0 | 0 | 4228100 | 6247200 |
| SLDWENPT | AGL | P35573 | GDE_HUMAN ( | Unmodified | 0 | 0 | 0 | 0 | 0 | 0 | 0 | 0 | 0 | 0 | 78276 | 1287600 | 0 | 0 | 0 | 0 | 0 | 0 |
| SLDYEALQ | PCDHB8; PCDH | Q9UN66; Q9 | PCDB8_HUMAN | Unmodified | 80378 | 89546 | 0 | 0 | 0 | 0 | 0 | 0 | 941180 | 33902 | 1461000 | 26929000 | 0 | 0 | 17406 | 0 | 0 | 0 |
| SLEAASEK | TPM4; TPM4; TF | P67936; A0A | TPM4_HUMAN | Unmodified | 0 | 0 | 0 | 0 | 1226000 | 192240 | 21212 | 0 | 0 | 0 | 0 | 0 | 184010 | 0 | 0 | 0 | 37701 | 0 |
| SLEAEILQL | MYH10 | P35580 | MYH10_HUMAN | Unmodified | 0 | 0 | 0 | 525600 | 238940 | 0 | 22035 | 0 | 0 | 0 | 0 | 214200 | 504660 | 0 | 891140 | 30281 | 0 | 63253 |
| SLEAEILQL | MYH10 | P35580 | MYH10_HUMAN | Unmodified | 0 | 0 | 0 | 19240 | 1928900 | 648650 | 112370 | 0 | 0 | 75916 | 0 | 0 | 714370 | 389290 | 0 | 0 | 84608 | 123790 |
| SLEAQADK | TPM2; TPM2; TF | P07951; A0A | TPM2_HUMAN | Unmodified | 0 | 162050 | 393900 | 0 | 201760 | 318320 | 0 | 36923 | 561900 | 0 | 194920 | 633140 | 0 | 0 | 0 | 0 | 113940 | 8953800 |
| SLEAQVAH | NUMA1; NUMA | Q14980; H0 | NUMA1_HUMAN | Unmodified | 0 | 0 | 0 | 0 | 0 | 0 | 0 | 0 | 0 | 0 | 226090 | 841480 | 0 | 0 | 0 | 0 | 0 | 0 |
| SLEDALRQ | IMMT; IMMT; IM | Q16891; B9 | MIC60_HUMAN | Unmodified | 0 | 0 | 0 | 0 | 0 | 0 | 292330 | 0 | 0 | 266150 | 0 | 0 | 0 | 0 | 0 | 0 | 0 | 0 |
| SLEDALSSI | ANXA6; ANXA6; | P08133; E7E | ANXA6_HUMAN | Unmodified | 0 | 0 | 0 | 0 | 3539200 | 619240 | 1647500 | 797780 | 23827000 | 5919600 | 0 | 0 | 0 | 0 | 0 | 0 | 686910 | 2442700 |
| SLEDALSSI | ANXA6; ANXA6; | P08133; E7E | ANXA6_HUMAN | Unmodified | 464470 | 0 | 6316000 | 5017000 | 8303900 | 1219900 | 3684800 | 765620 | 47936000 | 15351000 | 0 | 0 | 6685400 | 6137600 | 4869500 | 2433900 | 2843000 | 8832300 |
| SLEDDIRSI | ANXA4; ANXA4; | P09525; B4E | ANXA4_HUMAN | Unmodified | 856180 | 0 | 92469 | 1340900 | 1341000 | 222050 | 67072 | 12254 | 54656000 | 6397600 | 10394000 | 829180 | 717900 | 225240 | 436580 | 0 | 111780 | 278790 |
| SLEDDIRSI | ANXA4; ANXA4; | P09525; B4E | ANXA4_HUMAN | Oxidation (M) | 349510 | 460530 | 0 | 692290 | 0 | 0 | 0 | 0 | 5191100 | 1629600 | 6595800 | 0 | 0 | 0 | 186910 | 158800 | 0 | 0 |
| SLEDLQDE | STAT1; STAT1; S | P42224; A0A | STAT1_HUMAN | Unmodified | 0 | 0 | 0 | 0 | 1699400 | 0 | 437210 | 51536 | 0 | 0 | 0 | 691820 | 0 | 321760 | 0 | 0 | 0 | 1742800 |
| SLEDQVEM | PRKCSH; PRKC | P14314; K7E | GLU2B_HUMAN | Oxidation (M) | 424000 | 0 | 0 | 493440 | 0 | 0 | 0 | 148780 | 53934000 | 10333000 | 3792900 | 625070 | 0 | 0 | 0 | 116830 | 0 | 0 |
| SLEDQVEM | PRKCSH; PRKC | P14314; K7E | GLU2B_HUMAN | Unmodified | 0 | 0 | 0 | 0 | 0 | 0 | 0 | 0 | 1.19E+08 | 9426300 | 1470100 | 0 | 0 | 0 | 0 | 0 | 0 | 0 |
| SLEDQVEM | PRKCSH; PRKC | P14314; K7E | GLU2B_HUMAN | Oxidation (M) | 0 | 0 | 0 | 0 | 0 | 0 | 0 | 39558 | 0 | 595080 | 0 | 151280 | 0 | 0 | 0 | 0 | 0 | 0 |
| SLEDVTAE | RAI14 | Q9P0K7 | RAI14_HUMAN | Unmodified | 0 | 0 | 0 | 0 | 1386100 | 0 | 0 | 0 | 0 | 0 | 0 | 0 | 0 | 179360 | 0 | 0 | 701690 | 1083800 |
| SLEDVVIDI | MANSC1; MANS | Q9H8J5; F5 | MANS1_HUMAN | Unmodified | 0 | 0 | 0 | 0 | 0 | 0 | 0 | 0 | 0 | 0 | 0 | 0 | 0 | 0 | 0 | 0 | 0 | 0 |
| SLEDVVR | ARHGAP29; EIF | Q52LW3; Q1 | RHG29_HUMAN | Unmodified | 0 | 0 | 0 | 0 | 0 | 0 | 0 | 0 | 0 | 0 | 0 | 1750300 | 0 | 0 | 0 | 0 | 0 | 0 |
| SLEDVVRA | EIF3A | Q14152 | EIF3A_HUMAN | Unmodified | 0 | 0 | 0 | 0 | 1045100 | 199120 | 1748100 | 0 | 0 | 572970 | 0 | 0 | 0 | 628310 | 0 | 0 | 329240 | 250530 |
| SLEEAQEW | ECHDC1 | Q9NTX5 | ECHD1_HUMAN | Unmodified | 0 | 0 | 0 | 0 | 0 | 0 | 0 | 0 | 0 | 135600 | 0 | 1580300 | 0 | 0 | 0 | 0 | 0 | 0 |
| SLEEIYLFSI | RPS2; RPS2; RP | P15880; E9P | RS2_HUMAN 4 | Unmodified | 250950 | 122200 | 196070 | 82672 | 0 | 0 | 28344 | 0 | 123060 | 20761 | 153330 | 1834700 | 0 | 0 | 382390 | 238960 | 0 | 0 |
| SLEELLLDA | SCRIB; LRRC1; | Q14160; Q9 | SCRIB_HUMAN | Unmodified | 255790 | 389250 | 336960 | 1520700 | 0 | 0 | 0 | 0 | 7891400 | 470370 | 1086800 | 3739900 | 0 | 0 | 0 | 548420 | 132410 | 0 |
| SLEELLLDA | SCRIB; SCRIB; S | Q14160; A0 | SCRIB_HUMAN | Unmodified | 0 | 0 | 61505 | 0 | 0 | 0 | 0 | 0 | 812390 | 245380 | 130930 | 0 | 0 | 0 | 35740 | 0 | 0 | 0 |
| SLEELLLDA | SCRIB; SCRIB; S | Q14160; A0 | SCRIB_HUMAN | Unmodified | 0 | 0 | 0 | 528030 | 5426300 | 0 | 0 | 0 | 256820 | 6371900 | 72517 | 0 | 2983400 | 1209200 | 202300 | 0 | 1643100 | 0 |
| SLEELSK | ATP2C1; ATP2C | P98194; B4E | AT2C1_HUMAN | Unmodified | 265340 | 0 | 0 | 0 | 0 | 96204 | 21981 | 9182.4 | 0 | 0 | 0 | 1139900 | 0 | 231780 | 0 | 0 | 0 | 0 |
| SLEFLAHPI | VLDLR; VLDLR; | P98155; A0A | VLDLR_HUMAN | Unmodified | 0 | 0 | 0 | 0 | 0 | 76541 | 0 | 0 | 112500 | 22584 | 0 | 0 | 0 | 0 | 0 | 10686 | 0 | 0 |
| SLEFSYQE | PXDN; PXDN | Q92626; H7 | PXDN_HUMAN | Unmodified | 0 | 0 | 0 | 0 | 0 | 0 | 0 | 0 | 0 | 107410 | 298290 | 351840 | 0 | 0 | 0 | 0 | 0 | 0 |
| SLEGADLP | RFC4; RFC4; RF | P35249; C9 | RFC4_HUMAN | Unmodified | 41121 | 26106 | 0 | 34615 | 1758100 | 204090 | 461970 | 60651 | 1014900 | 279330 | 17076 | 125930 | 244240 | 422070 | 26447 | 23896 | 28487 | 30034 |
| SLEGDLED | LMNB1; LMNB1 | P20700; A0A | LMNB1_HUMAN | Unmodified | 0 | 0 | 0 | 0 | 0 | 0 | 0 | 0 | 0 | 0 | 0 | 501120 | 0 | 0 | 0 | 0 | 0 | 0 |
| SLEGDLED | LMNB1; LMNB1 | P20700; A0A | LMNB1_HUMAN | Unmodified | 0 | 0 | 0 | 0 | 0 | 0 | 0 | 0 | 0 | 0 | 0 | 0 | 894820 | 0 | 0 | 0 | 0 | 0 |
| SLEGGGA | XXYLT1; XXYLT1 | Q8NBI6; H7 | XXLT1_HUMAN | Unmodified | 0 | 0 | 0 | 0 | 0 | 0 | 0 | 0 | 1856500 | 0 | 0 | 0 | 0 | 0 | 0 | 0 | 0 | 0 |
| SLEHSEGP | SLC38A10; SLC | Q9HBR0; H0 | S38AA_HUMAN | Unmodified | 0 | 0 | 0 | 0 | 0 | 0 | 0 | 0 | 9307500 | 210780 | 0 | 0 | 0 | 0 | 0 | 0 | 0 | 0 |
| SLEHSEGP | SLC38A10; SLC | Q9HBR0; H0 | S38AA_HUMAN | Unmodified | 0 | 0 | 0 | 0 | 0 | 0 | 0 | 0 | 382810 | 725020 | 0 | 0 | 0 | 0 | 0 | 0 | 0 | 0 |
| SLEIEEK | PPP2R2C; PPP2 | Q9Y2T4; P6 | 2ABG_HUMAN | Unmodified | 0 | 0 | 0 | 0 | 2409700 | 635260 | 322840 | 148880 | 4575500 | 44812 | 0 | 102710 | 1113200 | 0 | 0 | 0 | 408130 | 730630 |
| SLEIRTLAV | CTNNA1; CTNN | P35221; E5 | CTNA1_HUMAN | Unmodified | 0 | 0 | 0 | 0 | 6063100 | 1455500 | 0 | 0 | 0 | 181700 | 0 | 0 | 5242800 | 6121600 | 0 | 0 | 0 | 0 |
| SLEISKDW | COBLL1; COBL | Q53SF7; A0 | COBL1_HUMAN | Unmodified | 0 | 0 | 0 | 0 | 0 | 0 | 0 | 0 | 0 | 0 | 0 | 0 | 715970 | 616350 | 0 | 0 | 0 | 0 |
| SLEISKDW | COBLL1; COBL | Q53SF7; A0 | COBL1_HUMAN | Oxidation (M) | 0 | 0 | 0 | 0 | 0 | 0 | 0 | 0 | 0 | 0 | 0 | 0 | 596360 | 828710 | 0 | 0 | 0 | 1030000 |
| SLEKSELIQ | EXOC7; EXOC7 | Q9UPT5; B4 | EXOC7_HUMAN | Unmodified | 0 | 0 | 0 | 0 | 1446800 | 0 | 299520 | 42024 | 0 | 0 | 0 | 0 | 0 | 0 | 0 | 0 | 756690 | 0 |
| SLELFSIDP | CELSR3 | Q9NYQ7 | CELR3_HUMAN | Unmodified | 0 | 0 | 0 | 0 | 0 | 0 | 160830 | 0 | 371380 | 0 | 0 | 0 | 0 | 0 | 0 | 0 | 0 | 0 |
| SLELHEEF | ARFIP1 | P53367 | ARFP1_HUMAN | Unmodified | 217670 | 0 | 377970 | 518070 | 417230 | 0 | 444500 | 126650 | 406820 | 0 | 299300 | 0 | 627360 | 979810 | 167590 | 371420 | 127700 | 181150 |
| SLELNPNN | SUGT1 | Q9Y2Z0 | SGT1_HUMAN | Unmodified | 0 | 0 | 0 | 0 | 1783200 | 180770 | 0 | 0 | 306880 | 188970 | 415140 | 1057400 | 415490 | 0 | 0 | 0 | 844750 | 0 |
| SLELYRELD | RNASET2; N/A; | O00584; H0 | RNT2_HUMAN | Unmodified | 24692 | 0 | 57705 | 302240 | 188230 | 126810 | 0 | 0 | 867550 | 3481400 | 1200000 | 0 | 6826000 | 3340000 | 147220 | 32989 | 0 | 205900 |
| SLENAIEW | PPME1 | Q9Y570 | PPME1_HUMAN | Unmodified | 0 | 0 | 0 | 0 | 748440 | 0 | 0 | 116130 | 652560 | 0 | 314980 | 1772200 | 2173000 | 0 | 0 | 0 | 0 | 662560 |
| SLENIFAES | KIF14 | Q15058 | KIF14_HUMAN | Unmodified | 0 | 0 | 0 | 0 | 1833600 | 0 | 0 | 0 | 0 | 0 | 0 | 1139400 | 0 | 0 | 0 | 0 | 0 | 0 |
| SLEPAENVI | SRC | P12931 | SRC_HUMAN F | Unmodified | 0 | 0 | 0 | 0 | 0 | 764970 | 0 | 0 | 1003800 | 894450 | 0 | 0 | 0 | 0 | 0 | 0 | 7152600 | 13285000 |
| SLEPGTPR | GALK1; GALK1; | P51570; K7E | GALK1_HUMAN | Unmodified | 0 | 0 | 18359 | 0 | 20079 | 0 | 0 | 0 | 0 | 0 | 70318 | 502000 | 0 | 0 | 197910 | 146740 | 15016 | 0 |
| SLEPGTPR | GALK1; GALK1; | P51570; K7E | GALK1_HUMAN | Unmodified | 0 | 92158 | 0 | 0 | 0 | 0 | 379900 | 224180 | 342410 | 0 | 0 | 912250 | 0 | 162750 | 0 | 0 | 0 | 128520 |
| SLEPLMEM | DLG2; DLG2; D | Q15700; E9 | DLG2_HUMAN | Oxidation (M) | 0 | 0 | 0 | 0 | 0 | 0 | 0 | 0 | 0 | 0 | 0 | 0 | 438060 | 528410 | 0 | 0 | 0 | 0 |
| SLEPLMEM | DLG2; DLG2; D | Q15700; E9 | DLG2_HUMAN | Unmodified | 0 | 0 | 0 | 0 | 0 | 458500 | 0 | 0 | 0 | 0 | 0 | 0 | 542920 | 386440 | 0 | 0 | 0 | 0 |
| SLEQDALR | SCRIB; SCRIB; S | Q14160; A0 | SCRIB_HUMAN | Unmodified | 394930 | 384690 | 303710 | 353590 | 0 | 0 | 0 | 66650 | 2046400 | 417420 | 288820 | 793170 | 0 | 0 | 0 | 155400 | 691600 | 1286700 |
| SLEQDALR | SCRIB; SCRIB; S | Q14160; A0 | SCRIB_HUMAN | Unmodified | 0 | 0 | 0 | 0 | 0 | 0 | 0 | 0 | 0 | 0 | 0 | 0 | 0 | 0 | 0 | 0 | 376250 | 1926200 |
| SLEQNIQL | MCM7 | P33993 | MCM7_HUMAN | Unmodified | 35447 | 33573 | 0 | 0 | 0 | 0 | 0 | 0 | 657360 | 0 | 120060 | 1507400 | 0 | 0 | 0 | 127100 | 0 | 0 |
| SLEQRIVEL | CIT; CIT; CIT; CI | O14578; A0 | CTRO_HUMAN | Unmodified | 0 | 0 | 0 | 0 | 1381700 | 0 | 0 | 0 | 0 | 0 | 0 | 0 | 142820 | 105120 | 0 | 0 | 411030 | 275490 |
| SLERAEAGI | TUFM | P49411 | EFTU_HUMAN | Unmodified | 0 | 41315 | 0 | 124170 | 1557500 | 0 | 1120600 | 222670 | 16935000 | 4994400 | 1115600 | 211010 | 578760 | 722330 | 0 | 28088 | 653280 | 1480000 |
| SLESINSR | RPL30; RPL30; | P62888; A0 | RL30_HUMAN | Unmodified | 6294400 | 6709100 | 9630200 | 2268700 | 0 | 0 | 0 | 0 | 100420 | 141290 | 1913500 | 29973000 | 676560 | 252110 | 6279600 | 3750500 | 0 | 0 |
| SLESINSRL | RPL30; RPL30; | P62888; A0 | RL30_HUMAN | Oxidation (M) | 0 | 0 | 0 | 218020 | 608050 | 32460 | 1563800 | 561060 | 0 | 360980 | 54225 | 0 | 1173100 | 0 | 0 | 0 | 0 | 154910 |
| SLESINSRL | RPL30; RPL30; | P62888; A0 | RL30_HUMAN | Unmodified | 0 | 0 | 0 | 0 | 0 | 0 | 1734400 | 0 | 0 | 0 | 65453 | 0 | 1446900 | 1058800 | 0 | 0 | 0 | 0 |
| SLESLDTSI | SF3A3 | Q12874 | SF3A3_HUMAN | Unmodified | 2760000 | 0 | 0 | 394470 | 0 | 0 | 0 | 0 | 0 | 0 | 0 | 1254900 | 0 | 0 | 0 | 0 | 0 | 0 |
| SLESLHSF | PLEC; PLEC; PL | Q15149; H0 | PLEC_HUMAN | Unmodified | 0 | 0 | 2566000 | 0 | 251110 | 108900 | 0 | 0 | 253240 | 240290 | 2093700 | 1148700 | 0 | 0 | 0 | 0 | 2339500 | 286450 |
| SLESNPEQ | MOV10; MOV1C | Q9HCE1; Q | MOV10_HUMA | Oxidation (M) | 2359300 | 1570400 | 996200 | 1015300 | 1155700 | 636180 | 0 | 0 | 0 | 138440 | 0 | 0 | 303980 | 0 | 1266900 | 1602800 | 0 | 0 |
| SLESNPEQ | MOV10; MOV1C | Q9HCE1; Q | MOV10_HUMA | Unmodified | 631470 | 2859000 | 0 | 0 | 0 | 0 | 0 | 0 | 0 | 0 | 0 | 0 | 462410 | 0 | 0 | 1898000 | 0 | 0 |
| SLESTTLTE | CSRP2; CSRP2; | Q16527; F8 | CSRP2_HUMAN | Unmodified | 0 | 0 | 0 | 0 | 425330 | 0 | 431430 | 265020 | 0 | 255990 | 0 | 0 | 0 | 0 | 0 | 0 | 1695100 | 287160 |
| SLETCMYD | DYNC1H1; DYN | Q14204; A0 | DYHC1_HUMA | Unmodified | 0 | 0 | 0 | 0 | 0 | 0 | 0 | 0 | 0 | 0 | 0 | 478690 | 0 | 0 | 0 | 0 | 0 | 0 |
| SLETENAG | LMNA; LMNA; L | P02545; A0 | LMNA_HUMAN | Unmodified | 0 | 0 | 0 | 0 | 0 | 0 | 0 | 0 | 415210 | 0 | 0 | 0 | 0 | 0 | 0 | 0 | 0 | 0 |
| SLETENRR | KRT18; KRT18 | P05783; F8 | K1C18_HUMAN | Unmodified | 0 | 0 | 0 | 0 | 0 | 1020800 | 0 | 0 | 347610 | 1509500 | 1005100 | 0 | 2978200 | 119940 | 0 | 0 | 384900 | 0 |
| SLETENSA | LMNB1; LMNB1 | P20700; A0 | LMNB1_HUMAN | Unmodified | 0 | 0 | 0 | 0 | 0 | 0 | 0 | 0 | 284830 | 0 | 884770 | 2689300 | 0 | 0 | 0 | 0 | 0 | 0 |
| SLETSAFVK | SCPEP1; SCPE | Q9HB40; I3 | RISC_HUMAN | Unmodified | 0 | 0 | 0 | 0 | 0 | 0 | 123430 | 215410 | 2950500 | 1557300 | 0 | 165770 | 229370 | 224240 | 54235 | 0 | 95482 | 578720 |
| SLETVYLER | PPP1R7; PPP1R | Q15435; H7 | PP1R7_HUMAN | Unmodified | 0 | 0 | 2828500 | 465170 | 550280 | 0 | 0 | 169550 | 876380 | 0 | 1033500 | 4017400 | 49072 | 0 | 66221 | 93293 | 0 | 0 |
| SLETVYLER | PPP1R7; PPP1R | Q15435; H7 | PP1R7_HUMAN | Unmodified | 0 | 0 | 0 | 0 | 0 | 0 | 0 | 0 | 0 | 1462000 | 0 | 0 | 578840 | 0 | 0 | 0 | 108630 | 0 |
| SLEVSDTRI | IRAK4; IRAK4 | Q9NWZ3; A | IRAK4_HUMAN | Unmodified | 0 | 0 | 218330 | 0 | 334530 | 0 | 669590 | 0 | 247340 | 0 | 0 | 0 | 2818900 | 2170600 | 0 | 0 | 0 | 0 |
| SLEVSER | PRKAR2A; PRK | P13861; H7 | KAP2_HUMAN | Unmodified | 0 | 0 | 0 | 0 | 0 | 364470 | 78069 | 11400 | 0 | 0 | 0 | 0 | 0 | 622180 | 1281000 | 0 | 0 | 0 |
| SLFDYFLTD | TIGAR; TIGAR; | Q9NQ88; A0 | TIGAR_HUMAN | Unmodified | 0 | 0 | 0 | 0 | 0 | 0 | 0 | 0 | 0 | 76692 | 0 | 89501 | 112270 | 0 | 0 | 0 | 0 | 0 |
| SLFFPDEAI | GCLC; GCLC; G | P48506; A0 | GSH1_HUMAN | Unmodified | 0 | 0 | 0 | 0 | 1152400 | 0 | 0 | 0 | 0 | 0 | 0 | 2250600 | 0 | 0 | 0 | 0 | 0 | 0 |
| SLFFPDEAI | GCLC; GCLC; G | P48506; A0 | GSH1_HUMAN | Unmodified | 0 | 0 | 0 | 0 | 4179500 | 1286200 | 0 | 0 | 0 | 183360 | 0 | 555080 | 359250 | 323860 | 0 | 0 | 156730 | 0 |
| SLFFPDEAI | GCLC; GCLC; G | P48506; A0 | GSH1_HUMAN | Unmodified | 0 | 0 | 0 | 0 | 10009000 | 0 | 0 | 0 | 0 | 0 | 0 | 0 | 0 | 1261000 | 0 | 0 | 447710 | 0 |
| SLFFSSKEI | ARHGAP29; AR | Q52LW3; F8 | RHG29_HUMAN | Unmodified | 0 | 0 | 0 | 0 | 473240 | 0 | 0 | 198820 | 0 | 0 | 0 | 0 | 0 | 0 | 0 | 0 | 1513200 | 1928100 |
| SLFGKTEV | EXOC2 | Q96KP1 | EXOC2_HUMAN | Unmodified | 0 | 0 | 0 | 0 | 549310 | 0 | 390640 | 65684 | 265330 | 314630 | 0 | 628700 | 148520 | 0 | 0 | 0 | 184580 | 138060 |
| SLFINHHP | CCNY; CCNY; C | Q8ND76; R4 | CCNY_HUMAN | Unmodified | 0 | 0 | 0 | 0 | 279220 | 81958 | 0 | 0 | 0 | 0 | 0 | 0 | 30416 | 0 | 0 | 0 | 555600 | 1184400 |
| SLFPEGNP | MYO1B; MYO1E | O43795; E9 | MYO1B_HUMA | Unmodified | 399960 | 447020 | 182140 | 223220 | 7194500 | 2788200 | 1414400 | 1183900 | 1695900 | 1110000 | 4689400 | 26527000 | 0 | 99386 | 1019000 | 580390 | 2932000 | 869900 |
| SLFSNVVT | ATIC; ATIC | P31939; H7 | PUR9_HUMAN | Unmodified | 306760 | 573780 | 1618500 | 1362200 | 7826700 | 0 | 0 | 0 | 1645600 | 2252900 | 1681000 | 13981000 | 9617400 | 9299100 | 1579700 | 1154500 | 5113900 | 2417900 |
| SLFSSIGEV | ELAVL1; ELAVL | Q15717; M0 | ELAV1_HUMAN | Unmodified | 162150 | 909970 | 1880700 | 1520700 | 2109800 | 0 | 727250 | 421620 | 274680 | 0 | 458510 | 8133500 | 1099200 | 631690 | 1542700 | 592020 | 1148000 | 3307300 |
| SLGAEPLE | NNT; NNT; NNT | Q13423; E9 | NNTM_HUMAN | Unmodified | 0 | 0 | 0 | 0 | 0 | 0 | 0 | 18670 | 5160900 | 608420 | 0 | 0 | 0 | 0 | 0 | 0 | 0 | 0 |
| SLGAEPLE | NNT; NNT; NNT | Q13423; E9 | NNTM_HUMAN | Unmodified | 0 | 0 | 0 | 0 | 0 | 0 | 1058100 | 379110 | 5444700 | 3332600 | 0 | 0 | 0 | 0 | 0 | 0 | 0 | 0 |
| SLGDDISSI | ANXA3; ANXA3; | P12429; D6 | ANXA3_HUMAN | Unmodified | 19257 | 0 | 0 | 0 | 42155000 | 10973000 | 0 | 0 | 0 | 0 | 0 | 1761000 | 0 | 316950 | 0 | 0 | 9178700 | 0 |
| SLGDGIFW | MBLAC2 | Q68D91 | MBLC2_HUMAN | Unmodified | 0 | 0 | 0 | 0 | 0 | 0 | 0 | 0 | 0 | 0 | 738850 | 1373800 | 0 | 0 | 0 | 0 | 0 | 0 |
| SLGEIPIVE | COPB1 | P53618 | COPB_HUMAN | Unmodified | 0 | 0 | 0 | 0 | 0 | 0 | 0 | 0 | 0 | 0 | 0 | 6148700 | 0 | 0 | 0 | 0 | 0 | 0 |
| SLGELINIC | ABCC5; ABCC5 | O15440; F8 | MRP5_HUMAN | Unmodified | 0 | 0 | 0 | 0 | 0 | 0 | 0 | 0 | 0 | 0 | 0 | 2982400 | 0 | 0 | 0 | 0 | 0 | 0 |
| SLGETQLV | CKAP4 | Q07065 | CKAP4_HUMAN | Unmodified | 0 | 0 | 0 | 898530 | 491950 | 108060 | 465110 | 0 | 458860 | 432750 | 0 | 0 | 78126 | 608920 | 0 | 0 | 0 | 0 |
| SLGGAVGS | ZNRF2 | Q8NHG8 | ZNRF2_HUMAN | Unmodified | 0 | 0 | 0 | 0 | 101290 | 0 | 0 | 0 | 504650 | 0 | 0 | 0 | 0 | 0 | 0 | 0 | 0 | 0 |
| SLGGKVVT | DLG5; DLG5 | Q8TDM6; A0 | DLG5_HUMAN | Unmodified | 0 | 0 | 0 | 0 | 0 | 0 | 0 | 0 | 0 | 0 | 0 | 0 | 0 | 0 | 0 | 0 | 260150 | 0 |
| SLGGLQPE | C13orf46 | A0A1B0GUA | CM046_HUMAN | Unmodified | 1428800 | 1736800 | 3160300 | 4527400 | 0 | 0 | 0 | 0 | 0 | 0 | 55389 | 84023 | 110400 | 0 | 1433200 | 1463400 | 0 | 0 |
| SLGHPEPL | PSEN1; PSEN1; | P49768; G3 | PSN1_HUMAN | Unmodified | 0 | 97164 | 0 | 84152 | 274110 | 0 | 0 | 0 | 284630 | 98095 | 0 | 9816900 | 213750 | 0 | 0 | 0 | 0 | 262360 |
| SLGLNPNH | FURIN; FURIN | P09958; A0 | FURIN_HUMAN | Unmodified | 0 | 0 | 0 | 0 | 0 | 0 | 0 | 0 | 1837300 | 0 | 4337500 | 831690 | 0 | 0 | 0 | 0 | 0 | 0 |
| SLGLSDEEI | LARS1; LARS1; | Q9P2J5; A0 | SYLC_HUMAN | Unmodified | 0 | 0 | 0 | 0 | 0 | 0 | 0 | 0 | 0 | 0 | 336000 | 4393600 | 0 | 0 | 0 | 0 | 0 | 0 |
| SLGNIKFIG | EIF4G1; EIF4G1 | Q04637; C9 | IF4G1_HUMAN | Unmodified | 0 | 0 | 0 | 0 | 259400 | 0 | 356380 | 0 | 0 | 128580 | 0 | 0 | 707220 | 369020 | 0 | 0 | 0 | 558730 |
| SLGPNQEN | COBLL1; COBL | Q53SF7; A0 | COBL1_HUMAN | Oxidation (M) | 1154800 | 2082300 | 0 | 0 | 0 | 0 | 0 | 0 | 0 | 0 | 0 | 0 | 0 | 0 | 0 | 547190 | 0 | 0 |
| SLGPPGPP | MYOF | Q9NZM1 | MYOF_HUMAN | Unmodified | 0 | 0 | 0 | 0 | 4732400 | 0 | 0 | 0 | 1901000 | 0 | 0 | 79792 | 0 | 0 | 0 | 0 | 0 | 0 |
| SLGPPGPP | MYOF | Q9NZM1 | MYOF_HUMAN | Unmodified | 0 | 0 | 0 | 0 | 24010000 | 3649900 | 49810 | 0 | 0 | 360290 | 0 | 0 | 0 | 0 | 0 | 0 | 506180 | 1386600 |
| SLGPPQGE | PRKDC | P78527 | PRKDC_HUMAN | Unmodified | 0 | 0 | 0 | 0 | 0 | 0 | 0 | 0 | 0 | 0 | 0 | 1005300 | 0 | 0 | 0 | 0 | 0 | 0 |

| | | | | | | | | | | | | | | | | | | | | | | |
|---|---|---|---|---|---|---|---|---|---|---|---|---|---|---|---|---|---|---|---|---|---|---|
| SLGPVELLL | PKN1; PKN1; PI | Q16512; K7E | PKN1_HUMAN | Unmodified | 0 | 0 | 0 | 219200 | 224970 | 108380 | 0 | 0 | 1187400 | 298830 | 386240 | 724500 | 0 | 0 | 0 | 0 | 215750 | 155950 |
| SLGRGEFR | MYO1C; MYO1( | O00159; F5H | MYO1C_HUMA | Unmodified | 0 | 0 | 0 | 0 | 5475300 | 0 | 172550 | 0 | 0 | 0 | 0 | 0 | 0 | 39356 | 0 | 0 | 481300 | 0 |
| SLGSASPG | ZDHHC5 | Q9C0B5 | ZDHC5_HUMA | Unmodified | 239190 | 164060 | 667590 | 0 | 0 | 0 | 0 | 0 | 365030 | 58140 | 264300 | 753580 | 0 | 0 | 576550 | 556910 | 145840 | 0 |
| SLGSASPG | ZDHHC5 | Q9C0B5 | ZDHC5_HUMA | Unmodified | 0 | 0 | 999690 | 1313600 | 13846000 | 1417300 | 242480 | 203400 | 1090300 | 1710700 | 623350 | 538880 | 11625000 | 10287000 | 587310 | 561090 | 13430000 | 19241000 |
| SLGSASPG | ZDHHC5 | Q9C0B5 | ZDHC5_HUMA | Unmodified | 0 | 0 | 0 | 619830 | 4540800 | 614580 | 0 | 0 | 132220 | 600860 | 333330 | 0 | 5296400 | 4744500 | 0 | 0 | 5017600 | 6670200 |
| SLGSVQAP | KRT18; KRT18 | P05783; F8V | K1C18_HUMAN | Unmodified | 605750 | 524200 | 0 | 904480 | 1097600 | 0 | 0 | 0 | 877000 | 218580 | 431350 | 1729400 | 996840 | 0 | 0 | 526270 | 0 | 418310 |
| SLGTAGPP | GPRIN1 | Q7Z2K8 | GRIN1_HUMAN | Unmodified | 0 | 0 | 0 | 0 | 0 | 0 | 0 | 0 | 1482600 | 120390 | 781190 | 224200 | 0 | 0 | 0 | 0 | 0 | 0 |
| SLGTAGPP | GPRIN1 | Q7Z2K8 | GRIN1_HUMAN | Unmodified | 0 | 0 | 0 | 0 | 312730 | 0 | 143600 | 0 | 1346500 | 897460 | 0 | 0 | 499520 | 0 | 0 | 0 | 625400 | 1307400 |
| SLGTDLMN | ITGB1; ITGB1; I | P05556; H7( | ITB1_HUMAN I | Oxidation (M) | 332140 | 176490 | 376840 | 0 | 24931000 | 6634100 | 558080 | 89015 | 2852900 | 255930 | 201110 | 511880 | 1654100 | 0 | 322920 | 400600 | 4237900 | 21702000 |
| SLGTDLMN | ITGB1; ITGB1; I | P05556; H7( | ITB1_HUMAN I | Unmodified | 0 | 0 | 0 | 0 | 34322000 | 2983500 | 228900 | 0 | 5568600 | 770670 | 71682 | 1498800 | 4049900 | 910300 | 0 | 0 | 3819800 | 18699000 |
| SLGTDLMN | ITGB1; ITGB1; I | P05556; H7( | ITB1_HUMAN I | 2 Oxidation (M) | 0 | 0 | 0 | 0 | 0 | 0 | 0 | 0 | 0 | 0 | 0 | 125970 | 0 | 0 | 102290 | 123720 | 359960 | 1465200 |
| SLGTDLMN | ITGB1; ITGB1; I | P05556; H7( | ITB1_HUMAN I | Oxidation (M) | 569750 | 379670 | 0 | 1255000 | 13183000 | 723800 | 515560 | 0 | 1462300 | 977320 | 0 | 2404700 | 1100900 | 0 | 519120 | 647850 | 3662700 | 1143400 |
| SLGTDLMN | ITGB1; ITGB1; I | P05556; H7( | ITB1_HUMAN I | Unmodified | 1322400 | 426660 | 0 | 0 | 1.86E+08 | 18544000 | 0 | 0 | 22526000 | 3415900 | 0 | 1888000 | 4477200 | 0 | 1178700 | 655860 | 14851000 | 1.24E+08 |
| SLGTGAPV | CSTF2; CSTF2; | P33240; E7E | CSTF2_HUMAN | Unmodified | 0 | 0 | 0 | 690400 | 0 | 0 | 0 | 208270 | 0 | 0 | 0 | 2755800 | 1553400 | 0 | 0 | 0 | 0 | 0 |
| SLGTIQQC | PRKDC | P78527 | PRKDC_HUMA | Unmodified | 0 | 0 | 0 | 0 | 0 | 201100 | 377300 | 0 | 0 | 0 | 0 | 0 | 0 | 0 | 0 | 0 | 0 | 80499 |
| SLGTRPPH | MYO6; MYO6; M | Q9UM54; A0 | MYO6_HUMAN | Unmodified | 0 | 0 | 3274100 | 672100 | 1868800 | 0 | 300410 | 196980 | 287560 | 651460 | 0 | 1935500 | 2109100 | 2008200 | 6405600 | 4494000 | 369610 | 0 |
| SLGVESDV | STON1; STON1 | Q9Y6Q2; Q5 | STON1_HUMA | Unmodified | 0 | 0 | 0 | 0 | 0 | 0 | 0 | 176290 | 0 | 0 | 269640 | 459460 | 0 | 0 | 0 | 0 | 0 | 0 |
| SLGYAYVN | PABPC4; PABP( | Q13310; P11 | PABP4_HUMAN | Unmodified | 53198000 | 44533000 | 12454000 | 12519000 | 1711700 | 0 | 129080 | 183940 | 1557700 | 68183 | 320580 | 5565700 | 601260 | 153330 | 7372200 | 16499000 | 2994800 | 0 |
| SLGYAYVN | PABPC4; PABP( | Q13310; P11 | PABP4_HUMAN | Oxidation (M) | 4039100 | 11619000 | 480460 | 6556600 | 0 | 0 | 0 | 0 | 0 | 0 | 0 | 0 | 0 | 0 | 2704200 | 0 | 0 | 0 |
| SLGYTGAR | TSTD1; TSTD1 | Q8NFU3; R4 | TSTD1_HUMAN | Unmodified | 607220 | 335280 | 132210 | 440660 | 0 | 0 | 0 | 0 | 2771400 | 442120 | 0 | 368120 | 309920 | 102640 | 537870 | 24368 | 0 | 0 |
| SLHAESPG | FCHSD2; FCHS | O94868; B8Z | FCSD2_HUMA | Unmodified | 0 | 0 | 0 | 0 | 103980 | 0 | 0 | 0 | 0 | 0 | 0 | 0 | 0 | 0 | 0 | 0 | 0 | 416980 |
| SLHDAIMIV | CCT7 | Q99832 | TCPH_HUMAN | Unmodified | 0 | 0 | 0 | 0 | 0 | 0 | 47879 | 0 | 958210 | 0 | 326830 | 4357300 | 0 | 0 | 0 | 0 | 0 | 0 |
| SLHDALCV | CCT2; CCT2; C( | P78371; F5G | TCPB_HUMAN | Unmodified | 0 | 0 | 0 | 0 | 0 | 0 | 0 | 0 | 356060 | 65100 | 323730 | 4670400 | 38582 | 0 | 53599 | 104940 | 0 | 46430 |
| SLHDALCV | TCP1; TCP1; TC | P17987; F5G | TCPA_HUMAN | Unmodified | 0 | 0 | 0 | 0 | 0 | 0 | 0 | 0 | 0 | 0 | 0 | 2755600 | 0 | 0 | 0 | 0 | 0 | 0 |
| SLHDPDGL | APMAP; APMAP | Q9HDC9; H( | APMAP_HUMA | Unmodified | 0 | 0 | 0 | 0 | 0 | 0 | 0 | 0 | 2.03E+08 | 5277500 | 0 | 345490 | 0 | 3445800 | 260810 | 0 | 0 | 0 |
| SLHDPDGL | APMAP; APMAP | Q9HDC9; H( | APMAP_HUMA | Unmodified | 0 | 0 | 0 | 0 | 0 | 0 | 0 | 0 | 7360900 | 8396700 | 0 | 0 | 0 | 0 | 0 | 0 | 0 | 0 |
| SLHINGGG | STK10; STK10 | O94804; H0Y | STK10_HUMAN | Unmodified | 0 | 0 | 0 | 0 | 679400 | 289380 | 0 | 0 | 217890 | 87908 | 0 | 224050 | 0 | 0 | 0 | 20073 | 39003 | 0 |
| SLHNRELS | SCG3 | Q8WXD2 | SCG3_HUMAN | Unmodified | 0 | 0 | 0 | 0 | 0 | 0 | 0 | 0 | 4555200 | 1442400 | 0 | 0 | 0 | 0 | 0 | 0 | 0 | 0 |
| SLHPAQIFI | SEL1L3 | Q68CR1 | SE1L3_HUMAN | Unmodified | 0 | 0 | 0 | 0 | 0 | 0 | 0 | 0 | 456440 | 0 | 0 | 0 | 0 | 0 | 0 | 0 | 0 | 0 |
| SLHPWYGI | MICALL1; MICA | Q8N3F8; B0 | MILK1_HUMAN | Unmodified | 0 | 0 | 0 | 106660 | 0 | 0 | 0 | 0 | 0 | 0 | 0 | 0 | 1269100 | 544040 | 0 | 194370 | 0 | 0 |
| SLHQAIEGI | ANXA6; ANXA6; | P08133; E5R | ANXA6_HUMAN | Unmodified | 1221300 | 912150 | 10675000 | 9591000 | 78779000 | 11924000 | 61319000 | 23239000 | 98711000 | 82534000 | 0 | 0 | 61157000 | 24467000 | 12123000 | 4899200 | 42524000 | 1.13E+08 |
| SLHQFLLEI | ARPC1A; N/A; A | Q92747; F8V | ARC1A_HUMAN | Acetyl (Protein N | 4162400 | 0 | 2489100 | 0 | 297990 | 0 | 0 | 299030 | 0 | 0 | 0 | 8215800 | 145090 | 112640 | 60257 | 1041600 | 0 | 0 |
| SLHQFLLEI | ARPC1A; N/A; A | Q92747; F8V | ARC1A_HUMAN | Acetyl (Protein N | 0 | 0 | 0 | 0 | 0 | 0 | 0 | 0 | 0 | 0 | 0 | 3708800 | 0 | 0 | 0 | 0 | 0 | 0 |
| SLHQQSTC | PROM1 | O43490 | PROM1_HUMA | Unmodified | 0 | 0 | 0 | 0 | 0 | 0 | 0 | 0 | 0 | 0 | 15317000 | 53655000 | 0 | 0 | 0 | 0 | 0 | 0 |
| SLHSIIR | TTLL12 | Q14166 | TTL12_HUMAN | Unmodified | 50590 | 23151000 | 84901 | 144910 | 65161 | 0 | 0 | 0 | 117680 | 48381 | 142180 | 1197500 | 90914 | 31745 | 35789000 | 131110 | 89509 | 0 |
| SLHWSVLL | ENSA | Q5T5H1 | Q5T5H1_HUM | Oxidation (M) | 0 | 379570 | 392130 | 610540 | 1193100 | 3152200 | 2718700 | 1020600 | 0 | 0 | 0 | 0 | 0 | 3107300 | 435810 | 0 | 0 | 5162900 |
| SLIANLAAA | ADK; ADK; ADK; | P55263; A0A | ADK_HUMAN A | Unmodified | 0 | 0 | 0 | 0 | 0 | 1512700 | 0 | 0 | 291190 | 0 | 369000 | 3457600 | 0 | 0 | 0 | 0 | 419160 | 311310 |
| SLICSISNE | PRPF19; PRPF1 | Q9UMS4; F5 | PRP19_HUMAN | Acetyl (Protein N | 0 | 0 | 0 | 0 | 0 | 0 | 0 | 0 | 0 | 0 | 0 | 3913600 | 1392200 | 0 | 0 | 0 | 919540 | 0 |
| SLIDRTIDE | PITRM1; PITRM | Q5JRX3; A0A | PREP_HUMAN | Unmodified | 0 | 0 | 0 | 0 | 0 | 0 | 0 | 0 | 482870 | 0 | 0 | 0 | 0 | 0 | 0 | 0 | 0 | 0 |
| SLIEEGGD | PSMD6; PSMD6 | Q15008; H7( | PSMD6_HUMA | Unmodified | 369330 | 0 | 362610 | 151560 | 0 | 0 | 0 | 0 | 0 | 0 | 0 | 5653000 | 0 | 0 | 585990 | 120380 | 0 | 28265 |
| SLIEVYMRF | NAGA | P17050 | NAGAB_HUMA | Unmodified | 0 | 0 | 0 | 0 | 97059 | 0 | 0 | 0 | 1008600 | 345180 | 342540 | 0 | 212560 | 197060 | 0 | 0 | 0 | 0 |
| SLIFPNFEP | FARP1; FARP1 | Q9Y4F1; C9J | FARP1_HUMAN | Unmodified | 0 | 0 | 0 | 0 | 1249900 | 0 | 0 | 0 | 2054000 | 3500100 | 5165100 | 9194900 | 0 | 0 | 0 | 0 | 2005300 | 966870 |
| SLIGKKGQ | H2AZ2; H2AZ1; | Q71UI9; P0C | H2AV_HUMAN | Unmodified | 0 | 0 | 0 | 0 | 0 | 0 | 1095600 | 9542300 | 0 | 0 | 0 | 0 | 0 | 0 | 0 | 0 | 0 | 31835 |
| SLIGVEYKP | EPRS1; EPRS1 | P07814; V9G | SYEP_HUMAN | Unmodified | 0 | 0 | 1068700 | 493010 | 267540 | 0 | 0 | 0 | 906150 | 0 | 1493900 | 6211600 | 605130 | 519880 | 527590 | 574750 | 58926 | 351300 |
| SLIGVEYKP | EPRS1 | P07814 | SYEP_HUMAN | Unmodified | 0 | 0 | 0 | 0 | 0 | 0 | 0 | 0 | 587230 | 0 | 275320 | 1784800 | 0 | 0 | 0 | 0 | 0 | 450620 |
| SLIGVRNEI | UGP2; UGP2; U | Q16851; E7E | UGPA_HUMAN | Unmodified | 0 | 0 | 0 | 0 | 6071900 | 1067200 | 2196800 | 0 | 1222600 | 4168200 | 334870 | 0 | 3062300 | 0 | 0 | 0 | 2725000 | 3711100 |
| SLIIDEGED | CCS; CCS; CCS | O14618; J3K | CCS_HUMAN ( | Unmodified | 0 | 0 | 0 | 0 | 3132600 | 418870 | 0 | 0 | 1900500 | 535100 | 1263800 | 6223300 | 0 | 0 | 0 | 0 | 751660 | 1553200 |
| SLIINTNPV | IQGAP2; IQGAP | Q13576; E7E | IQGA2_HUMAN | Unmodified | 0 | 0 | 0 | 1381700 | 0 | 0 | 0 | 0 | 0 | 0 | 0 | 0 | 0 | 0 | 1977200 | 1271100 | 0 | 0 |
| SLILTALQR | IPO4; IPO4; IPO | Q8TEX9; H0Y | IPO4_HUMAN I | Unmodified | 0 | 0 | 0 | 0 | 0 | 0 | 0 | 0 | 0 | 0 | 0 | 690840 | 0 | 0 | 0 | 0 | 0 | 0 |
| SLIPSYIR | RAB6A; RAB6B; | P20340; Q9N | RAB6A_HUMAN | Unmodified | 563400 | 588530 | 609770 | 576030 | 0 | 0 | 0 | 0 | 2197800 | 0 | 524980 | 16698000 | 0 | 0 | 846520 | 1250100 | 0 | 0 |
| SLIQFMK | CPD | O75976 | CBPD_HUMAN | Oxidation (M) | 0 | 0 | 0 | 0 | 0 | 0 | 0 | 0 | 3178900 | 1636200 | 0 | 0 | 0 | 0 | 0 | 0 | 0 | 0 |
| SLIQFMK | CPD | O75976 | CBPD_HUMAN | Unmodified | 0 | 0 | 0 | 0 | 1375100 | 218620 | 0 | 0 | 18539000 | 3752500 | 0 | 0 | 0 | 106780 | 0 | 0 | 0 | 0 |
| SLIQRFPVT | YTHDC2; YTHD | Q9H6S0; D6 | YTDC2_HUMAN | Unmodified | 0 | 0 | 0 | 0 | 0 | 0 | 0 | 0 | 0 | 0 | 0 | 0 | 522360 | 0 | 0 | 0 | 0 | 0 |
| SLISWNYLI | MACF1; MACF1 | Q9UPN3; H3 | MACF1_HUMA | Unmodified | 0 | 0 | 0 | 0 | 201910 | 0 | 42728 | 0 | 292420 | 391500 | 0 | 323230 | 0 | 0 | 0 | 0 | 0 | 0 |
| SLKDMEES | PRKCSH; PRKC | P14314; K7E | GLU2B_HUMA | Unmodified | 0 | 0 | 0 | 0 | 183480 | 0 | 0 | 0 | 1206300 | 174420 | 0 | 0 | 0 | 0 | 0 | 0 | 48354 | 0 |
| SLKDQLTD | GOLGB1 | Q14789 | GOGB1_HUMA | Unmodified | 0 | 0 | 0 | 0 | 0 | 0 | 0 | 0 | 0 | 95970 | 0 | 0 | 354440 | 194470 | 0 | 0 | 0 | 0 |
| SLKEEDGE | CHURC1; CHU | Q8WUH1; H | CHUR_HUMAN | Unmodified | 162770 | 0 | 0 | 0 | 714500 | 0 | 0 | 0 | 362860 | 0 | 0 | 1730500 | 0 | 0 | 0 | 0 | 868420 | 902270 |
| SLKEISDGI | EGFR; EGFR; E( | P00533; Q5( | EGFR_HUMAN | Unmodified | 0 | 134820 | 0 | 0 | 6813400 | 0 | 160650 | 0 | 0 | 112600 | 0 | 0 | 0 | 0 | 0 | 0 | 0 | 815130 |
| SLKLQASN | RALY; RALY; RA | Q9UKM9; Q5 | RALY_HUMAN I | Acetyl (Protein N | 0 | 0 | 0 | 0 | 0 | 0 | 232120 | 481070 | 0 | 0 | 0 | 0 | 244880 | 0 | 0 | 0 | 0 | 0 |
| SLKPDSVR | AKR7L; AKR7A2 | Q8NHP1; O4 | ARK74_HUMAN | Unmodified | 0 | 0 | 0 | 0 | 2515200 | 438030 | 5823700 | 1385900 | 0 | 3141500 | 0 | 0 | 3139100 | 4915600 | 0 | 0 | 7545800 | 1974300 |
| SLKPESGG | JAK1; JAK1; JAK | P23458; A0A | JAK1_HUMAN T | Unmodified | 0 | 0 | 0 | 0 | 1835700 | 171430 | 217470 | 0 | 0 | 583740 | 0 | 0 | 0 | 0 | 0 | 0 | 451190 | 825410 |
| SLKPGALSI | STAMBP; STAMI | O95630; C9J | STABP_HUMAN | Unmodified | 3452800 | 127230 | 724060 | 1344100 | 970240 | 304820 | 0 | 0 | 2353200 | 322850 | 1046800 | 1802200 | 1039600 | 350260 | 1017600 | 1268600 | 0 | 245100 |
| SLKPRVVDI | CUL2; CUL2; C | Q13617; Q5 | CUL2_HUMAN | Acetyl (Protein N | 0 | 0 | 0 | 0 | 0 | 0 | 0 | 67398 | 0 | 0 | 0 | 0 | 0 | 0 | 0 | 0 | 881760 | 61737 |
| SLKPSNHA | BMP3 | P12645 | BMP3_HUMAN | Unmodified | 0 | 0 | 0 | 0 | 0 | 0 | 0 | 0 | 297870 | 0 | 0 | 0 | 0 | 0 | 0 | 0 | 0 | 0 |
| SLLALEKEE | DAAM1 | Q9Y4D1 | DAAM1_HUMA | Unmodified | 0 | 0 | 0 | 0 | 0 | 0 | 24959 | 0 | 0 | 62125 | 0 | 905400 | 0 | 0 | 0 | 0 | 0 | 68874 |
| SLLALEKEE | DAAM1 | Q9Y4D1 | DAAM1_HUMA | Unmodified | 0 | 0 | 0 | 673690 | 796290 | 0 | 637170 | 0 | 1296800 | 504840 | 381360 | 0 | 717300 | 752580 | 0 | 631430 | 255590 | 0 |
| SLLDASEE | VCL | P18206 | VINC_HUMAN | Unmodified | 320500 | 752550 | 451370 | 159840 | 2886900 | 1227200 | 627440 | 174560 | 0 | 1391000 | 376040 | 4655900 | 344160 | 0 | 960020 | 693400 | 8252300 | 0 |
| SLLDASEE | VCL | P18206 | VINC_HUMAN | Unmodified | 0 | 0 | 0 | 0 | 526830 | 324530 | 599730 | 0 | 0 | 265970 | 0 | 0 | 0 | 0 | 0 | 0 | 5871600 | 109670 |
| SLLDFLPR | SEC24C; SEC2 | P53992; G5E | SC24C_HUMA | Unmodified | 48486 | 46927 | 20636 | 0 | 0 | 0 | 0 | 0 | 108070 | 74444 | 0 | 736840 | 0 | 0 | 0 | 41185 | 0 | 0 |
| SLLDIISDPI | SCFD1; SCFD1; | Q8WVM8; A( | SCFD1_HUMA | Unmodified | 44845 | 0 | 0 | 25811 | 330210 | 115700 | 0 | 19106 | 188250 | 48568 | 153320 | 663350 | 0 | 0 | 99047 | 0 | 0 | 52400 |
| SLLEAVSFL | PHKG2; PHKG2 | P15735; H3E | PHKG2_HUMA | Unmodified | 0 | 0 | 0 | 0 | 0 | 0 | 0 | 0 | 0 | 0 | 0 | 718420 | 0 | 0 | 0 | 0 | 0 | 0 |
| SLLEPFVYL | NKIRAS2; NKIR | Q9NYR9; K7 | KBRS2_HUMAN | Unmodified | 0 | 0 | 0 | 0 | 1655900 | 218070 | 0 | 0 | 0 | 54134 | 0 | 0 | 820530 | 0 | 0 | 0 | 320330 | 0 |
| SLLEQDLA | CIT; CIT; CIT; CI | O14578; A0A | CTRO_HUMAN | Unmodified | 0 | 0 | 0 | 56770 | 1090100 | 314160 | 0 | 0 | 236970 | 0 | 0 | 0 | 0 | 0 | 0 | 0 | 228130 | 0 |
| SLLEQENH | SLITRK6 | Q9H5Y7 | SLIK6_HUMAN | Oxidation (M) | 1972900 | 1538400 | 0 | 0 | 0 | 0 | 0 | 0 | 1353400 | 1111100 | 0 | 0 | 0 | 0 | 0 | 130330 | 287100 | 338100 |
| SLLEQYHL | GUSB | P08236 | BGLR_HUMAN | Unmodified | 59640 | 0 | 563370 | 424270 | 451570 | 0 | 164720 | 66987 | 58784000 | 26632000 | 1012700 | 540590 | 702260 | 470080 | 1006600 | 221990 | 0 | 0 |
| SLLEQYHL | GUSB | P08236 | BGLR_HUMAN | Unmodified | 0 | 0 | 99224 | 57479 | 0 | 0 | 0 | 0 | 12393000 | 2089700 | 0 | 0 | 0 | 0 | 108180 | 93557 | 0 | 0 |
| SLLFDLLQ | TRPM5; TRPM5; | Q9NZQ8; A0 | TRPM5_HUMA | Unmodified | 17992 | 40184 | 0 | 0 | 0 | 0 | 0 | 109700 | 0 | 0 | 0 | 2884100 | 0 | 0 | 0 | 0 | 0 | 0 |
| SLLGEVEQ | PPL; PPL; PPL | O60437; K7E | PEPL_HUMAN I | Unmodified | 0 | 0 | 0 | 0 | 2046200 | 1370900 | 0 | 0 | 0 | 0 | 0 | 100020 | 12437 | 302090 | 0 | 0 | 31806 | 170380 |
| SLLGKDVLI | PGK1; PGK2 | P00558; P07 | PGK1_HUMAN | Unmodified | 0 | 0 | 0 | 1806100 | 47216000 | 6745700 | 38007000 | 3228500 | 3111600 | 23202000 | 1646200 | 1572500 | 57276000 | 39486000 | 0 | 0 | 1.93E+08 | 2.48E+08 |
| SLLGKDVLI | PGK1 | P00558 | PGK1_HUMAN | Unmodified | 0 | 0 | 0 | 0 | 0 | 0 | 4262500 | 0 | 0 | 0 | 0 | 0 | 1037100 | 2299600 | 0 | 0 | 33692000 | 7397800 |
| SLLHGDFH | SLC3A2; SLC3A | P08195; A0A | 4F2_HUMAN 4I | Unmodified | 9294000 | 11998000 | 608530 | 15446000 | 3039300 | 871560 | 451260 | 0 | 64293000 | 8075900 | 26158000 | 66515000 | 726940 | 1075600 | 30923000 | 424210 | 445310 | 515880 |
| SLLHGDFH | SLC3A2; SLC3A | P08195; A0A | 4F2_HUMAN 4I | Unmodified | 0 | 0 | 0 | 0 | 0 | 1653900 | 0 | 0 | 299530 | 1009800 | 2658000 | 0 | 727320 | 0 | 0 | 0 | 122030 | 0 |
| SLLINAVEA | PYCR1; PYCR2; | P32322; Q96 | P5CR1_HUMAN | Unmodified | 0 | 0 | 0 | 0 | 0 | 0 | 0 | 0 | 5045500 | 297360 | 0 | 0 | 0 | 0 | 0 | 0 | 0 | 0 |
| SLLKPLELE | GXYLT1 | Q4G148 | GXLT1_HUMAN | Unmodified | 0 | 0 | 0 | 0 | 0 | 0 | 0 | 0 | 1469400 | 0 | 0 | 0 | 0 | 0 | 0 | 0 | 0 | 0 |
| SLLLSVNAI | GANAB; GANAE | Q14697; F5H | GANAB_HUMA | Unmodified | 0 | 226550 | 353270 | 375430 | 0 | 0 | 0 | 0 | 2.06E+08 | 6209000 | 4063000 | 2868200 | 257700 | 0 | 0 | 0 | 0 | 0 |
| SLLLTTIPQI | MTAP; MTAP | Q13126; B4I | MTAP_HUMAN | Unmodified | 0 | 0 | 0 | 0 | 768330 | 0 | 0 | 0 | 0 | 288630 | 164920 | 0 | 0 | 0 | 0 | 0 | 1224000 | 1092700 |
| SLLNNSKLI | CCDC88A; CCI | Q3V6T2; A0A | GRDN_HUMAN | Unmodified | 0 | 0 | 0 | 0 | 1250100 | 0 | 0 | 0 | 0 | 0 | 0 | 0 | 0 | 0 | 0 | 0 | 399300 | 679260 |
| SLLNPVLLF | PLCB2; PLCB2 | Q00722; Q9I | PLCB2_HUMAN | Acetyl (Protein N | 0 | 0 | 28428 | 0 | 154550 | 475440 | 402870 | 345180 | 0 | 0 | 1212200 | 3787100 | 75537 | 0 | 0 | 0 | 0 | 0 |
| SLLQALNE | DYNC1H1; DYN | Q14204; A0A | DYHC1_HUMA | Unmodified | 0 | 0 | 0 | 406080 | 248720 | 0 | 0 | 0 | 1313000 | 718460 | 455170 | 11671000 | 0 | 0 | 582300 | 362800 | 266490 | 0 |
| SLLQGTIQI | CEMIP2; CEMIP | Q9UHN6; F5 | CEIP2_HUMAN | Oxidation (M) | 666330 | 304590 | 199760 | 708720 | 116500 | 0 | 0 | 0 | 1271100 | 234860 | 33342 | 401760 | 0 | 0 | 1463800 | 789050 | 0 | 0 |
| SLLQGTIQI | CEMIP2; CEMIP | Q9UHN6; F5 | CEIP2_HUMAN | Unmodified | 0 | 0 | 0 | 0 | 0 | 0 | 0 | 0 | 2644200 | 0 | 0 | 0 | 0 | 0 | 0 | 0 | 0 | 0 |
| SLLRSLLGI | CUL4A; CUL4A | Q13619; A0A | CUL4A_HUMAN | Unmodified | 0 | 0 | 0 | 0 | 0 | 0 | 0 | 0 | 9935400 | 634350 | 227500 | 0 | 0 | 0 | 0 | 0 | 0 | 0 |
| SLLSAEEA | SND1 | Q7KZF4 | SND1_HUMAN | Unmodified | 1005400 | 654340 | 1279000 | 1635400 | 0 | 779080 | 1021200 | 993810 | 703850 | 554240 | 1675700 | 11655000 | 5177300 | 2486000 | 1619900 | 1030700 | 1391300 | 2369200 |
| SLLSAEEA | SND1 | Q7KZF4 | SND1_HUMAN | Unmodified | 0 | 0 | 0 | 0 | 0 | 0 | 202950 | 78204 | 0 | 0 | 0 | 0 | 0 | 0 | 0 | 0 | 614960 | 273400 |
| SLLSGLLK | AKT1; AKT1; AK | P31749; A0A | AKT1_HUMAN I | Unmodified | 0 | 0 | 3065400 | 2700300 | 130560 | 0 | 22108 | 0 | 321980 | 32958 | 4646400 | 8562900 | 2166900 | 0 | 1667200 | 879820 | 177560 | 67463 |
| SLLSHEFQI | CPD | O75976 | CBPD_HUMAN | Unmodified | 116410 | 0 | 0 | 0 | 5688500 | 1316100 | 0 | 0 | 4256600 | 1571700 | 0 | 0 | 0 | 0 | 0 | 131790 | 0 | 0 |
| SLLSILQR | HUWE1; HUWE | Q7Z6Z7; H0Y | HUWE1_HUMA | Unmodified | 0 | 0 | 0 | 0 | 0 | 0 | 0 | 0 | 0 | 0 | 0 | 1924800 | 0 | 0 | 0 | 0 | 0 | 0 |
| SLLSMLVE | CPD | O75976 | CBPD_HUMAN | Oxidation (M) | 0 | 0 | 20072 | 54827 | 0 | 0 | 0 | 0 | 4177300 | 988540 | 0 | 74676 | 0 | 0 | 0 | 17833 | 0 | 0 |
| SLLSMLVE | CPD | O75976 | CBPD_HUMAN | Unmodified | 0 | 0 | 0 | 0 | 0 | 0 | 0 | 0 | 1284900 | 131720 | 0 | 0 | 0 | 0 | 0 | 0 | 0 | 0 |
| SLLSNLDE | PDCD6IP | Q8WUM4 | PDC6I_HUMAN | Unmodified | 7210600 | 0 | 2110300 | 2061900 | 1208100 | 795220 | 0 | 0 | 1647500 | 0 | 0 | 0 | 961770 | 0 | 1642500 | 3760700 | 0 | 1151900 |
| SLLSNLDE | PDCD6IP | Q8WUM4 | PDC6I_HUMAN | Unmodified | 1.05E+08 | 1.16E+08 | 61859000 | 49712000 | 38633000 | 7295000 | 1439300 | 415540 | 60765000 | 12267000 | 7206000 | 2872700 | 36110000 | 14663000 | 73107000 | 69797000 | 2154200 | 12393000 |
| SLLSSASAT | FAM114A1 | Q8IWE2 | NXP20_HUMAN | Unmodified | 0 | 0 | 44036 | 119930 | 0 | 0 | 0 | 0 | 252870 | 116160 | 0 | 0 | 521700 | 1022100 | 0 | 0 | 573620 | 1557200 |
| SLLSSTSTT | PPP1R12A; PPP | O14974; F8V | MYPT1_HUMAN | Unmodified | 0 | 0 | 0 | 145620 | 1058500 | 0 | 0 | 62186 | 0 | 0 | 0 | 81151 | 704020 | 0 | 0 | 95163 | 647850 | 516250 |
| SLLTYGR | UQCRC1 | P31930 | QCR1_HUMAN | Unmodified | 0 | 0 | 0 | 0 | 0 | 0 | 0 | 0 | 12920000 | 322110 | 150550 | 0 | 0 | 0 | 0 | 0 | 0 | 0 |
| SLLVIPNTL | TCP1; TCP1; TC | P17987; F5H | TCPA_HUMAN | Unmodified | 637130 | 562770 | 0 | 1619500 | 1890200 | 378880 | 0 | 0 | 3708600 | 3251600 | 1921300 | 5796700 | 10256000 | 2003600 | 3928500 | 844250 | 1203400 | 1515800 |
| SLLVNPEGI | FASN; FASN | P49327; A0A | FAS_HUMAN Fa | Oxidation (M) | 0 | 0 | 1436500 | 951630 | 0 | 0 | 0 | 0 | 154220 | 0 | 2284100 | 8087400 | 0 | 0 | 1680800 | 1180800 | 0 | 0 |
| SLLVNPEGI | FASN; FASN | P49327; A0A | FAS_HUMAN Fa | Unmodified | 0 | 0 | 0 | 0 | 0 | 0 | 0 | 0 | 127060 | 0 | 0 | 2821000 | 0 | 0 | 151740 | 50710 | 0 | 0 |
| SLLVTELGS | IMPA1; IMPA1; I | P29218; E5R | IMPA1_HUMAN | Unmodified | 238360 | 0 | 0 | 0 | 0 | 0 | 0 | 0 | 2963200 | 1068300 | 706230 | 2390800 | 0 | 0 | 0 | 206220 | 0 | 0 |
| SLLWNVFE | N/A | N/A | N/A | Unmodified | 0 | 0 | 4842300 | 372630 | 0 | 252790 | 0 | 0 | 0 | 0 | 5107300 | 154200 | 0 | 0 | 0 | 0 | 0 | 0 |

| | | | | | | | | | | | | | | | | | | | | | | |
|---|---|---|---|---|---|---|---|---|---|---|---|---|---|---|---|---|---|---|---|---|---|---|
| SLMDETPLI | PPP1R16A | Q96I34 | PP16A_HUMAN | Unmodified | 0 | 680190 | 0 | 0 | 0 | 0 | 0 | 0 | 0 | 0 | 1988200 | 0 | 0 | 0 | 0 | 0 | 1109300 | 0 |
| SLMDHTIPI | SDCBP; SDCBP | O00560; G5I | SDCB1_HUMAI | Unmodified | 8973000 | 14281000 | 6929400 | 16104000 | 2.01E+08 | 31418000 | 18483000 | 4171200 | 2.45E+08 | 1.13E+08 | 5148700 | 24634000 | 90073000 | 62043000 | 21180000 | 8349500 | 5783100 | 32801000 |
| SLMDHTIPI | SDCBP; SDCBP | O00560; G5I | SDCB1_HUMAI | Oxidation (M) | 8958600 | 7544700 | 11298000 | 2880200 | 13018000 | 2687600 | 4193200 | 1161500 | 17199000 | 20894000 | 3486700 | 13247000 | 24078000 | 16939000 | 17019000 | 9197800 | 199540 | 2209700 |
| SLMESGAN | EHBP1L1 | Q8N3D4 | EH1L1_HUMAN | Oxidation (M) | 0 | 0 | 0 | 0 | 887870 | 0 | 0 | 0 | 0 | 33976 | 0 | 0 | 0 | 0 | 0 | 0 | 189460 | 200640 |
| SLMPQIIPG | TMTC3 | Q6ZXV5 | TMTC3_HUMAI | Unmodified | 0 | 0 | 0 | 0 | 0 | 0 | 0 | 0 | 1719400 | 330950 | 0 | 0 | 0 | 0 | 0 | 0 | 0 | 0 |
| SLMPYFLLT | PSMD3 | O43242 | PSMD3_HUMA | Oxidation (M) | 46409 | 0 | 0 | 0 | 0 | 0 | 0 | 0 | 0 | 0 | 74925 | 3248900 | 45603 | 0 | 0 | 266890 | 747330 | 0 |
| SLMPYFLLT | PSMD3 | O43242 | PSMD3_HUMA | Unmodified | 0 | 0 | 0 | 0 | 0 | 0 | 0 | 0 | 32538 | 0 | 0 | 101090 | 0 | 0 | 0 | 0 | 0 | 0 |
| SLMSADNV | DHX15 | O43143 | DHX15_HUMAI | Oxidation (M) | 0 | 0 | 0 | 0 | 0 | 0 | 0 | 0 | 0 | 0 | 0 | 772680 | 0 | 0 | 0 | 0 | 0 | 0 |
| SLMSVSGA | RELL1 | Q8IUW5 | RELL1_HUMAN | Unmodified | 0 | 0 | 0 | 0 | 1418000 | 924440 | 218810 | 0 | 0 | 0 | 0 | 0 | 1825800 | 0 | 0 | 0 | 0 | 0 |
| SLMVDHVT | SRA1 | Q9HD15 | SRA1_HUMAN | Unmodified | 0 | 0 | 0 | 0 | 0 | 0 | 0 | 0 | 0 | 0 | 0 | 0 | 0 | 0 | 0 | 0 | 0 | 2013000 |
| SLNDEKLE | FAM111B | Q6SJ93 | F111B_HUMAN | Unmodified | 632900 | 0 | 0 | 0 | 558890 | 0 | 0 | 0 | 218420 | 0 | 0 | 0 | 0 | 0 | 0 | 0 | 31297 | 0 |
| SLNDLIEEF | PBK; PBK | Q96KB5; E5I | TOPK_HUMAN | Unmodified | 230540 | 114640 | 116120 | 419510 | 879470 | 213970 | 0 | 54514 | 315640 | 289540 | 0 | 0 | 1264500 | 395450 | 589360 | 131830 | 347170 | 438080 |
| SLNECESLI | DLG5; DLG5 | Q8TDM6; A0 | DLG5_HUMAN | Unmodified | 0 | 0 | 0 | 0 | 0 | 0 | 0 | 0 | 0 | 0 | 0 | 0 | 0 | 0 | 0 | 0 | 683650 | 0 |
| SLNGTIENI | UACA; UACA; U | Q9BZF9; H0 | UACA_HUMAN | Unmodified | 0 | 0 | 0 | 0 | 782430 | 0 | 0 | 0 | 67570 | 0 | 0 | 209240 | 685120 | 414650 | 0 | 0 | 0 | 1140700 |
| SLNIKTDPV | IQGAP1; IQGAP | P46940; A0A | IQGA1_HUMAN | Unmodified | 1522300 | 1292300 | 412660 | 1784800 | 38055000 | 7704100 | 2564500 | 1213900 | 6985700 | 3291100 | 1316500 | 9285900 | 5680800 | 985460 | 998390 | 1498900 | 1754400 | 1119400 |
| SLNILTAFQ | ERP29; ERP29 | P30040; F8V | ERP29_HUMAN | Unmodified | 0 | 0 | 619170 | 0 | 835870 | 0 | 3410700 | 1318600 | 3.3E+08 | 1.33E+08 | 7768300 | 5240100 | 0 | 0 | 0 | 589830 | 0 | 0 |
| SLNILTAFQ | ERP29; ERP29 | P30040; F8V | ERP29_HUMAN | Unmodified | 0 | 0 | 0 | 0 | 326400 | 0 | 684900 | 0 | 84769000 | 36089000 | 1944900 | 539270 | 0 | 0 | 0 | 0 | 0 | 0 |
| SLNLSTEFI | SERPINB5; SER | P36952; C9J | SPB5_HUMAN | Unmodified | 0 | 0 | 0 | 0 | 2837200 | 0 | 0 | 0 | 0 | 0 | 0 | 0 | 3561700 | 2955200 | 0 | 0 | 16639000 | 9596700 |
| SLNNFCSC | PRPF4 | O43172 | PRP4_HUMAN | Unmodified | 0 | 0 | 0 | 0 | 0 | 0 | 0 | 0 | 0 | 0 | 0 | 962150 | 0 | 0 | 0 | 0 | 0 | 0 |
| SLNNQIESI | COL2A1 | P02458 | CO2A1_HUMAI | Unmodified | 0 | 0 | 0 | 0 | 0 | 0 | 0 | 0 | 0 | 0 | 1008200 | 0 | 0 | 0 | 0 | 0 | 0 | 0 |
| SLNNQIESI | COL2A1 | P02458 | CO2A1_HUMAI | Unmodified | 0 | 0 | 0 | 0 | 0 | 0 | 0 | 0 | 0 | 0 | 10683000 | 204550 | 0 | 0 | 0 | 0 | 0 | 0 |
| SLNNQIETI | COL1A2; COL1/ | P08123; A0A | CO1A2_HUMAI | Unmodified | 0 | 0 | 312110 | 0 | 0 | 0 | 0 | 0 | 554140 | 242320 | 0 | 0 | 0 | 0 | 0 | 0 | 0 | 0 |
| SLNNQIETI | COL1A2; COL1/ | P08123; A0A | CO1A2_HUMAI | Unmodified | 0 | 0 | 1132500 | 0 | 0 | 0 | 0 | 0 | 0 | 0 | 0 | 0 | 0 | 0 | 0 | 0 | 0 | 0 |
| SLNPATIYP | SLC5A6 | Q9Y289 | SC5A6_HUMAN | Unmodified | 559260 | 0 | 1287100 | 0 | 0 | 0 | 0 | 0 | 823580 | 0 | 2758800 | 14481000 | 0 | 0 | 0 | 1128800 | 0 | 0 |
| SLNPNLLA | EMC1 | Q8N766 | EMC1_HUMAN | Unmodified | 0 | 0 | 0 | 0 | 0 | 0 | 0 | 0 | 4235300 | 440730 | 0 | 290070 | 0 | 0 | 0 | 0 | 0 | 0 |
| SLNQSLAE | ATP1B3; ATP1B | P54709; F8V | AT1B3_HUMAN | Unmodified | 0 | 0 | 4378300 | 664410 | 0 | 0 | 0 | 0 | 0 | 0 | 7558700 | 17707000 | 0 | 0 | 5377800 | 4949600 | 0 | 0 |
| SLNRFCAG | ATP11C | A0A067XG5 | A0A067XG54_I | Unmodified | 0 | 0 | 0 | 0 | 0 | 0 | 0 | 0 | 0 | 0 | 0 | 0 | 138650 | 151630 | 0 | 0 | 0 | 0 |
| SLNRITDIG | MET | P08581 | MET_HUMAN I | Oxidation (M) | 0 | 0 | 0 | 67886 | 896280 | 296170 | 0 | 0 | 0 | 432630 | 0 | 0 | 0 | 0 | 0 | 0 | 1855300 | 1515700 |
| SLNTGIITV | ADAM10; ADAM | O14672; H0 | ADA10_HUMAN | Unmodified | 2896500 | 2968000 | 4586900 | 4974600 | 4649000 | 1521900 | 700530 | 185800 | 3174800 | 0 | 0 | 6698300 | 5301000 | 3129400 | 5515700 | 4249800 | 139330 | 2877700 |
| SLNWEEMI | PSMG2; PSMG2 | Q969U7; K7I | PSMG2_HUMA | Unmodified | 0 | 0 | 0 | 0 | 0 | 0 | 0 | 0 | 0 | 0 | 0 | 898560 | 182130 | 0 | 0 | 0 | 0 | 0 |
| SLNWSSFV | LIMA1; LIMA1; L | Q9UHB6; F8 | LIMA1_HUMAN | Unmodified | 0 | 0 | 0 | 0 | 1489700 | 0 | 0 | 0 | 0 | 0 | 0 | 0 | 0 | 0 | 0 | 0 | 0 | 0 |
| SLNYWCNI | GLOD4; GLOD4 | Q9HC38; F6 | GLOD4_HUMA | Oxidation (M) | 52314 | 0 | 0 | 0 | 299960 | 0 | 0 | 0 | 75259 | 0 | 0 | 256680 | 275760 | 0 | 51215 | 0 | 0 | 0 |
| SLPADILYE | HINT2 | Q9BX68 | HINT2_HUMAN | Unmodified | 0 | 0 | 0 | 0 | 0 | 0 | 0 | 0 | 612120 | 68515 | 0 | 0 | 0 | 0 | 0 | 0 | 0 | 0 |
| SLPADILYE | HINT2 | Q9BX68 | HINT2_HUMAN | Unmodified | 0 | 0 | 0 | 0 | 0 | 0 | 0 | 0 | 0 | 1244400 | 0 | 0 | 0 | 0 | 0 | 0 | 0 | 0 |
| SLPASLSFL | SCRIB; SCRIB; S | Q14160; A0/ | SCRIB_HUMAN | Unmodified | 480970 | 430100 | 14417000 | 2032000 | 0 | 1838000 | 0 | 0 | 4009700 | 409560 | 991900 | 7371700 | 0 | 0 | 5605600 | 1296000 | 99067 | 1011100 |
| SLPDVRLE | GALNT1; GALN | Q10472; Q8I | GALT1_HUMAN | Unmodified | 0 | 0 | 0 | 0 | 0 | 0 | 0 | 0 | 1282100 | 50336 | 231000 | 1040900 | 0 | 0 | 0 | 0 | 0 | 0 |
| SLPEEDVAI | DKC1; DKC1; D | O60832; H7 | DKC1_HUMAN | Unmodified | 1923300 | 0 | 238500 | 0 | 0 | 0 | 0 | 0 | 394450 | 0 | 0 | 925260 | 1871100 | 0 | 0 | 0 | 0 | 315860 |
| SLPERFQAI | ARVCF; ARVCF; | O00192; E9F | ARVC_HUMAN | Unmodified | 0 | 0 | 285480 | 617700 | 0 | 0 | 0 | 94866 | 0 | 0 | 0 | 0 | 0 | 0 | 671330 | 0 | 0 | 68899 |
| SLPERFQAI | ARVCF; ARVCF; | O00192; E9F | ARVC_HUMAN | Unmodified | 0 | 0 | 0 | 0 | 1166600 | 0 | 0 | 0 | 0 | 0 | 0 | 0 | 3530200 | 1676500 | 0 | 0 | 0 | 4982600 |
| SLPEYLYSS | MBLAC2 | Q68D91 | MBLC2_HUMAI | Unmodified | 0 | 0 | 258700 | 0 | 0 | 0 | 0 | 0 | 0 | 0 | 515380 | 1230800 | 123990 | 0 | 0 | 296940 | 0 | 0 |
| SLPEYLYSS | MBLAC2 | Q68D91 | MBLC2_HUMAI | Unmodified | 217690 | 0 | 871180 | 704840 | 0 | 113330 | 198710 | 0 | 0 | 0 | 0 | 2027300 | 0 | 0 | 795760 | 385260 | 0 | 0 |
| SLPFGAQS | DPP7; DPP7; DI | Q9UHL4; R4 | DPP2_HUMAN | Unmodified | 861170 | 0 | 0 | 0 | 0 | 0 | 0 | 0 | 2774000 | 138510 | 152240 | 322260 | 0 | 0 | 1574200 | 58611 | 0 | 0 |
| SLPGAEDY | CRTAP; CRTAP | O75718; C9J | CRTAP_HUMAN | Unmodified | 0 | 0 | 61514 | 0 | 0 | 0 | 0 | 0 | 6378000 | 432410 | 0 | 0 | 0 | 0 | 0 | 0 | 0 | 0 |
| SLPGAEDY | CRTAP; CRTAP | O75718; C9J | CRTAP_HUMAN | Unmodified | 0 | 0 | 0 | 0 | 0 | 0 | 0 | 19592 | 1373000 | 2700800 | 0 | 0 | 0 | 0 | 0 | 0 | 0 | 0 |
| SLPGEQEQ | NOTCH2 | Q04721 | NOTC2_HUMA | Unmodified | 137360 | 0 | 209120 | 235800 | 599710 | 0 | 135790 | 0 | 524190 | 216970 | 0 | 0 | 0 | 0 | 241320 | 276670 | 336540 | 1183400 |
| SLPIYSPEK | MYO1B; MYO1E | O43795; C9I | MYO1B_HUMA | Unmodified | 0 | 0 | 0 | 0 | 448580 | 148200 | 0 | 36358 | 0 | 0 | 3324900 | 8160600 | 0 | 0 | 633890 | 207180 | 0 | 0 |
| SLPIYSPEK | MYO1B; MYO1E | O43795; C9I | MYO1B_HUMA | Unmodified | 483160 | 430950 | 0 | 258550 | 0 | 0 | 0 | 0 | 1483400 | 0 | 6424200 | 16451000 | 0 | 0 | 352760 | 327710 | 0 | 0 |
| SLPLNPKPI | SNRPF; SNRPF; | P62306; A0A | RUXF_HUMAN | Acetyl (Protein N | 15520000 | 15320000 | 0 | 0 | 4583300 | 0 | 2208000 | 409230 | 0 | 0 | 9426000 | 17794000 | 14302000 | 5417000 | 0 | 0 | 0 | 0 |
| SLPLPEKPI | QSOX2; QSOX2 | Q6ZRP7; H0 | QSOX2_HUMAI | Unmodified | 0 | 0 | 0 | 0 | 0 | 0 | 0 | 0 | 6966200 | 1375000 | 139130 | 983750 | 0 | 0 | 0 | 0 | 474330 | 0 |
| SLPPLDSRI | VPS37D; VPS37 | Q86XT2; C9J | VP37D_HUMAN | Unmodified | 505190 | 316580 | 227070 | 0 | 0 | 0 | 0 | 0 | 0 | 0 | 0 | 0 | 0 | 0 | 624930 | 263960 | 0 | 0 |
| SLPQEWLV | UGGT1; UGGT1 | Q9NYU2; H7 | UGGG1_HUMA | Unmodified | 492960 | 0 | 0 | 0 | 0 | 0 | 0 | 0 | 638620 | 441490 | 0 | 0 | 0 | 0 | 0 | 0 | 0 | 0 |
| SLPQSDPV | GLOD4; GLOD4 | Q9HC38; F6 | GLOD4_HUMA | Unmodified | 158300 | 136500 | 0 | 377610 | 665810 | 233320 | 0 | 104790 | 821700 | 803940 | 1109300 | 4188700 | 2679500 | 1346800 | 269160 | 355780 | 212580 | 341690 |
| SLPREESA | PLCB2 | Q00722 | PLCB2_HUMAN | Unmodified | 0 | 212120 | 0 | 0 | 0 | 75244 | 0 | 0 | 0 | 0 | 400960 | 619020 | 0 | 112100 | 0 | 0 | 0 | 155260 |
| SLPSASTSI | CHMP2B; CHM | Q9UQN3; C9 | CHM2B_HUMA | Unmodified | 0 | 0 | 0 | 0 | 0 | 0 | 0 | 0 | 0 | 1082800 | 0 | 0 | 545920 | 0 | 0 | 0 | 0 | 0 |
| SLPSDLAFI | TNKS1BP1 | Q9C0C2 | TB182_HUMAN | Unmodified | 0 | 0 | 0 | 0 | 0 | 0 | 0 | 0 | 0 | 0 | 0 | 715030 | 0 | 0 | 0 | 0 | 0 | 0 |
| SLPSEASE | NRCAM; NRCAI | Q92823; A0/ | NRCAM_HUMA | Unmodified | 0 | 0 | 0 | 0 | 0 | 0 | 0 | 0 | 1243000 | 312680 | 0 | 0 | 0 | 0 | 0 | 0 | 0 | 0 |
| SLPSERILP. | SLC9A1 | P19634 | SL9A1_HUMAN | Unmodified | 0 | 0 | 0 | 0 | 1174700 | 280040 | 0 | 0 | 0 | 284030 | 0 | 0 | 2794400 | 3744600 | 0 | 0 | 117960 | 609540 |
| SLPSGSHQ | MPZL1 | O95297 | MPZL1_HUMAN | Unmodified | 5027200 | 0 | 5494600 | 6073900 | 35817000 | 6112800 | 118240 | 220380 | 5849900 | 1000300 | 0 | 11864000 | 6335400 | 3188700 | 5179900 | 4138700 | 2987600 | 19865000 |
| SLPSLVPLF | TRRAP; TRRAP; | Q9Y4A5; F2 | TRRAP_HUMAN | Unmodified | 0 | 0 | 0 | 0 | 0 | 0 | 0 | 58171 | 454100 | 0 | 0 | 477380 | 0 | 0 | 0 | 0 | 0 | 0 |
| SLPVPQLVI | ACOT7 | O00154 | BACH_HUMAN | Unmodified | 0 | 0 | 0 | 0 | 327080 | 0 | 22430 | 97013 | 1909600 | 694730 | 1104800 | 2909900 | 600970 | 693010 | 0 | 0 | 0 | 561230 |
| SLPYKLISLI | MYOF | Q9NZM1 | MYOF_HUMAN | Unmodified | 0 | 0 | 0 | 0 | 4071600 | 0 | 0 | 0 | 0 | 0 | 0 | 0 | 0 | 0 | 0 | 0 | 65815 | 929380 |
| SLQADTTN | ERC1; ERC1; EF | Q8IUD2; G8 | RB6I2_HUMAN | Unmodified | 0 | 0 | 0 | 0 | 1573600 | 253330 | 0 | 0 | 0 | 0 | 0 | 0 | 0 | 5169800 | 0 | 0 | 0 | 0 |
| SLQALGEV | FARSA; FARSA; | Q9Y285; K7E | SYFA_HUMAN | Unmodified | 49948 | 61614 | 40633 | 141700 | 56971 | 0 | 17356 | 0 | 159530 | 26557 | 59464 | 1575100 | 528140 | 72487 | 146910 | 106840 | 27449 | 42510 |
| SLQALGEV | FARSA; FARSA; | Q9Y285; K7E | SYFA_HUMAN | Unmodified | 0 | 0 | 0 | 0 | 1663300 | 63913 | 1063800 | 148280 | 0 | 2249400 | 598720 | 0 | 3663100 | 2332400 | 0 | 0 | 853220 | 2051400 |
| SLQAWKDI | SULF2 | Q8IWU5 | SULF2_HUMAN | Unmodified | 0 | 0 | 0 | 0 | 0 | 0 | 0 | 0 | 0 | 0 | 0 | 0 | 0 | 0 | 0 | 0 | 0 | 363590 |
| SLQAWKDI | SULF2 | Q8IWU5 | SULF2_HUMAN | Unmodified | 0 | 0 | 0 | 0 | 0 | 0 | 0 | 0 | 0 | 0 | 0 | 0 | 631030 | 0 | 0 | 0 | 0 | 2954600 |
| SLQDASRG | PBXIP1 | Q96AQ6 | PBIP1_HUMAN | Unmodified | 0 | 0 | 0 | 0 | 0 | 0 | 0 | 0 | 1167800 | 134140 | 856730 | 897730 | 0 | 0 | 0 | 0 | 0 | 0 |
| SLQEANAE | DSG2 | Q14126 | DSG2_HUMAN | Unmodified | 0 | 0 | 0 | 0 | 0 | 0 | 0 | 0 | 0 | 0 | 0 | 0 | 0 | 0 | 0 | 0 | 0 | 0 |
| SLQEANAE | DSG2 | Q14126 | DSG2_HUMAN | Unmodified | 0 | 0 | 0 | 0 | 7434200 | 1345900 | 0 | 65072 | 0 | 734910 | 0 | 0 | 4059700 | 2698200 | 0 | 0 | 2024700 | 1330700 |
| SLQEANAE | DSG2 | Q14126 | DSG2_HUMAN | Unmodified | 0 | 0 | 0 | 0 | 25053000 | 2464800 | 0 | 0 | 0 | 0 | 0 | 0 | 5461400 | 5797000 | 0 | 0 | 0 | 4127900 |
| SLQEELSL | GOLGB1; GOLG | Q14789; E7E | GOGB1_HUMA | Unmodified | 0 | 0 | 0 | 0 | 0 | 0 | 0 | 0 | 0 | 0 | 0 | 0 | 430740 | 327320 | 0 | 0 | 0 | 0 |
| SLQELFLAI | PARP1; PARP1 | P09874; A0A | PARP1_HUMAN | Unmodified | 130230 | 125160 | 0 | 0 | 0 | 0 | 0 | 0 | 86209 | 150050 | 107870 | 65831 | 0 | 0 | 0 | 0 | 0 | 0 |
| SLQELFLAI | PARP1; PARP1 | P09874; A0A | PARP1_HUMAN | Unmodified | 0 | 0 | 0 | 0 | 0 | 0 | 511710 | 0 | 0 | 0 | 0 | 0 | 1966000 | 0 | 0 | 0 | 0 | 0 |
| SLQEMKLQ | SERPINF1 | P36955 | PEDF_HUMAN | Oxidation (M) | 0 | 0 | 32497 | 0 | 0 | 0 | 0 | 0 | 0 | 1432600 | 0 | 0 | 0 | 0 | 0 | 0 | 0 | 0 |
| SLQEQADA | TPM1; TPM1; TF | H0YKJ4; H0 | H0YKJ4_HUMA | Unmodified | 0 | 0 | 102530 | 0 | 0 | 0 | 0 | 33019 | 182650 | 0 | 298390 | 1058100 | 0 | 0 | 423450 | 504000 | 0 | 0 |
| SLQESLQK | ROCK1; ROCK1 | Q13464; A0/ | ROCK1_HUMAI | Unmodified | 44451 | 0 | 0 | 0 | 1972400 | 0 | 0 | 39037 | 0 | 420560 | 53199 | 0 | 712830 | 0 | 0 | 0 | 1409300 | 1505000 |
| SLQGDKLE | TGFBI; TGFBI | Q15582; H0 | BGH3_HUMAN | Unmodified | 0 | 0 | 0 | 0 | 1178200 | 365490 | 631020 | 425890 | 0 | 0 | 0 | 0 | 0 | 0 | 0 | 0 | 203270 | 384940 |
| SLQGTSFEI | PTPRK; PTPRK; | Q15262; Q5 | PTPRK_HUMAN | Unmodified | 123250 | 0 | 0 | 0 | 362790 | 0 | 0 | 0 | 607510 | 175050 | 0 | 89264 | 0 | 0 | 0 | 0 | 58769 | 0 |
| SLQHQEQI | PIMREG; PIMRE | Q9BSJ6; I3L | PIMRE_HUMAN | Unmodified | 0 | 74159 | 0 | 0 | 165670 | 0 | 0 | 0 | 143760 | 150650 | 0 | 0 | 0 | 0 | 0 | 0 | 0 | 0 |
| SLQLEAQT | ARIH2 | O95376 | ARI2_HUMAN E | Unmodified | 0 | 0 | 0 | 0 | 0 | 0 | 0 | 1472200 | 0 | 0 | 0 | 821390 | 0 | 0 | 0 | 0 | 0 | 106670 |
| SLQLPASP | PLEKHA6; PLEK | Q9Y2H5; A0 | PKHA6_HUMAI | Unmodified | 153500 | 0 | 133640 | 308680 | 357100 | 0 | 0 | 0 | 579830 | 153360 | 168730 | 544660 | 226410 | 83897 | 362180 | 204930 | 268840 | 0 |
| SLQMYLER | ECI1; ECI1 | P42126; Q9 | ECI1_HUMAN E | Oxidation (M) | 0 | 0 | 0 | 0 | 370650 | 0 | 112530 | 0 | 1371600 | 0 | 0 | 0 | 0 | 0 | 106870 | 88638 | 0 | 88347 |
| SLQMYLER | ECI1; ECI1 | P42126; Q9 | ECI1_HUMAN E | Unmodified | 0 | 0 | 0 | 0 | 123120 | 0 | 0 | 0 | 1327100 | 0 | 0 | 0 | 0 | 0 | 0 | 0 | 0 | 0 |
| SLQPLAPR | MVP | Q14764 | MVP_HUMAN I | Unmodified | 0 | 270680 | 0 | 0 | 1490900 | 413920 | 0 | 10584 | 1504200 | 612250 | 0 | 7327800 | 0 | 0 | 0 | 0 | 0 | 590840 |
| SLQQKLETI | CIT; CIT; CIT; CI | O14578; A0/ | CTRO_HUMAN | Unmodified | 0 | 0 | 0 | 211670 | 5545500 | 764460 | 0 | 0 | 80587 | 0 | 0 | 0 | 205000 | 98452 | 0 | 0 | 0 | 0 |
| SLQQLAEE | SPTAN1; SPTAN | Q13813; A0/ | SPTN1_HUMAN | Unmodified | 0 | 0 | 0 | 0 | 0 | 0 | 0 | 0 | 362940 | 0 | 0 | 2927400 | 0 | 0 | 0 | 0 | 0 | 0 |
| SLQQQQQ | MINK1; MINK1 | Q8N4C8; A0 | MINK1_HUMAN | Unmodified | 0 | 0 | 306210 | 658810 | 968300 | 446470 | 48578 | 64086 | 810160 | 574600 | 185660 | 476490 | 1490800 | 1443900 | 401110 | 380720 | 699200 | 2086000 |
| SLQQRLEQ | ROCK1; ROCK1 | Q13464; A0/ | ROCK1_HUMAI | Unmodified | 0 | 0 | 0 | 0 | 2700700 | 0 | 773820 | 0 | 0 | 676440 | 0 | 0 | 1372200 | 0 | 0 | 0 | 7049300 | 0 |
| SLQSDLEA | UTRN | P46939 | UTRN_HUMAN | Unmodified | 0 | 0 | 687460 | 495470 | 12405000 | 0 | 0 | 0 | 1216600 | 433220 | 0 | 0 | 332520 | 0 | 2289500 | 1285800 | 0 | 0 |
| SLQSELDEI | OCLN; OCLN | Q16625; A0/ | OCLN_HUMAN | Unmodified | 1596300 | 1134300 | 3218400 | 3560600 | 2314100 | 0 | 306910 | 0 | 3180700 | 457430 | 183540 | 641080 | 2942300 | 1944400 | 7229200 | 4783500 | 0 | 0 |
| SLQSELDEI | OCLN; OCLN | Q16625; A0/ | OCLN_HUMAN | Unmodified | 200150 | 965560 | 0 | 4209800 | 2501600 | 0 | 0 | 0 | 0 | 1572100 | 0 | 0 | 5026900 | 2582700 | 0 | 0 | 143670 | 325210 |
| SLQSKDEQ | PTK7; PTK7; PTI | Q13308; H0 | PTK7_HUMAN I | Unmodified | 186680 | 0 | 0 | 0 | 0 | 0 | 0 | 0 | 832580 | 0 | 340230 | 1378100 | 0 | 0 | 211780 | 0 | 0 | 357870 |
| SLQSKDEQ | PTK7; PTK7; PTI | Q13308; H0 | PTK7_HUMAN I | Unmodified | 0 | 0 | 0 | 0 | 452560 | 0 | 0 | 26117 | 2234600 | 158700 | 590790 | 2828600 | 0 | 0 | 0 | 28366 | 0 | 352540 |
| SLQSLLGP | CORO7; CORO | P57737; I3L1 | CORO7_HUMA | Unmodified | 0 | 0 | 0 | 0 | 0 | 0 | 0 | 223520 | 0 | 0 | 0 | 1017800 | 0 | 0 | 0 | 0 | 0 | 0 |
| SLQSVAEEI | RPL15; RPL15; I | P61313; E7E | RL15_HUMAN ( | Unmodified | 6764900 | 5547500 | 3904200 | 4539900 | 204640 | 0 | 0 | 0 | 1783500 | 0 | 2691300 | 78424000 | 967610 | 395240 | 4718700 | 4404700 | 0 | 59339 |
| SLQSVAEEI | RPL15; RPL15; I | P61313; E7E | RL15_HUMAN ( | Unmodified | 0 | 0 | 0 | 0 | 0 | 183990 | 98708 | 0 | 215140 | 160990 | 1193900 | 0 | 0 | 0 | 0 | 0 | 0 | 0 |
| SLQTVSGR | API5; API5; API5 | Q9BZZ5; G3 | API5_HUMAN A | Unmodified | 0 | 0 | 0 | 174390 | 184270 | 47644 | 0 | 0 | 0 | 0 | 0 | 941660 | 775900 | 0 | 0 | 0 | 0 | 0 |
| SLQTYLK | CLASP2; CLASF | O75122; B3I | CLAP2_HUMAN | Unmodified | 108190 | 0 | 153770 | 0 | 1404700 | 1728500 | 335090 | 159660 | 316680 | 195960 | 0 | 3041000 | 0 | 190010 | 0 | 0 | 258600 | 213040 |
| SLQVPSIYY | MGAT4A | Q9UM21 | MGT4A_HUMAI | Unmodified | 0 | 0 | 0 | 0 | 0 | 0 | 0 | 0 | 1977600 | 0 | 0 | 0 | 0 | 0 | 0 | 0 | 0 | 0 |
| SLQWFGAI | ITGA5 | P08648 | ITA5_HUMAN I | Unmodified | 958890 | 163980 | 264390 | 2128800 | 0 | 511050 | 0 | 0 | 0 | 0 | 0 | 503030 | 182760 | 0 | 2210400 | 1216900 | 1429600 | 0 |
| SLRADYNL | VARS1; VARS1 | P26640; A0A | SYVC_HUMAN | Unmodified | 0 | 0 | 0 | 0 | 0 | 0 | 0 | 45482 | 0 | 0 | 0 | 952660 | 0 | 0 | 169180 | 0 | 0 | 0 |
| SLRCADDV | GSKIP; GSKIP; ( | Q9P0R6; G3 | GSKIP_HUMAN | Unmodified | 0 | 0 | 0 | 0 | 0 | 0 | 79568 | 0 | 0 | 0 | 0 | 0 | 134440 | 295850 | 0 | 0 | 677890 | 115850 |
| SLRDLTPEI | DCPS; DCPS | Q96C86; A0 | DCPS_HUMAN | Unmodified | 0 | 0 | 0 | 0 | 0 | 0 | 0 | 0 | 0 | 0 | 45974 | 347220 | 0 | 0 | 0 | 0 | 0 | 0 |
| SLRDQDKD | MYO1B; MYO1E | O43795; C9J | MYO1B_HUMA | Unmodified | 0 | 0 | 0 | 0 | 0 | 0 | 0 | 0 | 0 | 0 | 342100 | 3185900 | 0 | 0 | 0 | 0 | 0 | 0 |
| SLRECELY | PRKAR1A; PRKA | P10644; X6R | KAP0_HUMAN | Unmodified | 0 | 0 | 0 | 0 | 737300 | 0 | 516150 | 182310 | 1631600 | 643480 | 325870 | 714040 | 0 | 0 | 0 | 0 | 1272500 | 1160100 |
| SLRELDSG | KCTD3 | Q9Y597 | KCTD3_HUMAI | Unmodified | 0 | 0 | 0 | 0 | 192230 | 42604 | 0 | 0 | 0 | 0 | 0 | 0 | 48217 | 74022 | 0 | 0 | 139830 | 585820 |
| SLRELDSG | KCTD3 | Q9Y597 | KCTD3_HUMAI | Unmodified | 0 | 0 | 0 | 0 | 0 | 0 | 0 | 0 | 0 | 0 | 0 | 0 | 420530 | 461680 | 0 | 0 | 664260 | 0 |
| SLREPVAQ | TANC1 | Q9C0D5 | TANC1_HUMAI | Unmodified | 0 | 0 | 280860 | 492610 | 676070 | 0 | 0 | 44772 | 0 | 0 | 115030 | 0 | 971790 | 1026000 | 492060 | 378360 | 215250 | 704130 |
| SLRIQEQV | PKP2; PKP2 | Q99959; A0/ | PKP2_HUMAN | Unmodified | 0 | 0 | 0 | 0 | 250970 | 143440 | 0 | 0 | 0 | 40138 | 0 | 0 | 166080 | 249670 | 0 | 0 | 1857000 | 1755200 |

| | | | | | | | | | | | | | | | | | | | | | | |
|---|---|---|---|---|---|---|---|---|---|---|---|---|---|---|---|---|---|---|---|---|---|---|
| SLRIQEQV( | PKP2; PKP2 | Q99959; A0/ | PKP2_HUMAN | Unmodified | 0 | 0 | 0 | 0 | 1011300 | 0 | 0 | 0 | 0 | 0 | 150800 | 0 | 154750 | 275150 | 0 | 0 | 2699900 | 1834600 |
| SLRKQTPSI | LSM6 | P62312 | LSM6_HUMAN | Acetyl (Protein N | 0 | 41039 | 0 | 111170 | 1324800 | 0 | 702040 | 750170 | 0 | 287120 | 0 | 0 | 1452300 | 949190 | 31404 | 0 | 353060 | 702250 |
| SLRLGDRL' | HMCN2 | Q8NDA2 | HMCN2_HUMA | Unmodified | 0 | 0 | 0 | 0 | 0 | 0 | 0 | 93397 | 0 | 0 | 516410 | 1217200 | 0 | 0 | 0 | 0 | 0 | 0 |
| SLRLGPGG | PKP3; PKP3; PK | Q9Y446; E9F | PKP3_HUMAN | Unmodified | 0 | 0 | 0 | 0 | 367710 | 114810 | 1282100 | 0 | 0 | 0 | 427210 | 0 | 2717500 | 552590 | 0 | 0 | 0 | 212810 |
| SLRPEMEN | IGF1R; IGF1R | P08069; C9J | IGF1R_HUMAN | Unmodified | 0 | 0 | 0 | 0 | 419670 | 121020 | 0 | 0 | 0 | 0 | 0 | 0 | 0 | 0 | 0 | 0 | 0 | 40211 |
| SLSAIDRAC | AIFM1; AIFM1; / | O95831; A0/ | AIFM1_HUMAN | Unmodified | 0 | 0 | 38950 | 554190 | 98452 | 0 | 227050 | 119990 | 759110 | 603320 | 224130 | 740880 | 1382500 | 820940 | 0 | 0 | 51883 | 143880 |
| SLSAIYLEK | PLEC; PLEC | Q15149; H0Y | PLEC_HUMAN | Unmodified | 41156 | 0 | 119750 | 38027 | 956720 | 343350 | 0 | 94622 | 216930 | 631680 | 0 | 628080 | 147880 | 0 | 62956 | 82639 | 2453600 | 1985400 |
| SLSALGNV | KIF5B | P33176 | KINH_HUMAN | Unmodified | 0 | 0 | 0 | 0 | 11643000 | 1249700 | 4491800 | 0 | 3019200 | 840570 | 0 | 978290 | 7663500 | 4125500 | 914160 | 0 | 4194600 | 7379200 |
| SLSALGNV | KIF5B | P33176 | KINH_HUMAN | Oxidation (M) | 0 | 0 | 0 | 0 | 427830 | 0 | 552420 | 0 | 0 | 334860 | 0 | 0 | 356910 | 498830 | 0 | 0 | 662560 | 201410 |
| SLSALGNV | KIF5C; KIF5A; K | O60282; Q12 | KIF5C_HUMAN | Unmodified | 0 | 0 | 0 | 0 | 0 | 0 | 157030 | 0 | 578200 | 100290 | 0 | 0 | 492190 | 220120 | 0 | 0 | 0 | 0 |
| SLSDTVEK | DCTN1; DCTN1 | Q14203; A0/ | DCTN1_HUMA | Unmodified | 0 | 0 | 0 | 0 | 153000 | 23533 | 0 | 60318 | 112170 | 25780 | 0 | 1004100 | 714510 | 0 | 0 | 0 | 35470 | 628440 |
| SLSEGHPT. | ACADVL; ACAD | P49748; J3Q | ACADV_HUMAI | Unmodified | 0 | 0 | 0 | 0 | 0 | 0 | 0 | 0 | 4783800 | 1520800 | 0 | 0 | 0 | 0 | 0 | 0 | 0 | 0 |
| SLSELESLK | OSBP | P22059 | OSBP1_HUMAI | Unmodified | 462300 | 0 | 326880 | 1738300 | 2850800 | 1445600 | 95916 | 45755 | 1834200 | 847580 | 946950 | 4183800 | 3199900 | 1250900 | 794620 | 1095100 | 487100 | 1829100 |
| SLSELLGPY | NCKAP1 | Q9Y2A7 | NCKP1_HUMAI | Oxidation (M) | 0 | 0 | 0 | 0 | 0 | 0 | 0 | 0 | 0 | 339570 | 0 | 1505100 | 175790 | 0 | 0 | 349270 | 0 | 0 |
| SLSELVDDI | PSD2; PSD2 | Q9BQI7; A0/ | PSD2_HUMAN | Unmodified | 0 | 0 | 0 | 0 | 0 | 0 | 0 | 0 | 0 | 0 | 3806700 | 0 | 0 | 217550 | 0 | 0 | 0 | 0 |
| SLSELVDDI | PSD2; PSD2 | Q9BQI7; A0/ | PSD2_HUMAN | Unmodified | 0 | 0 | 0 | 0 | 0 | 0 | 0 | 0 | 0 | 0 | 563250 | 378070 | 0 | 0 | 0 | 0 | 0 | 0 |
| SLSETFLPN | STEAP2; STEAP | Q8NFT2; C9. | STEA2_HUMAN | Unmodified | 109170 | 137550 | 6205000 | 5985900 | 106270 | 25929 | 281190 | 0 | 0 | 42886 | 72666 | 413960 | 943210 | 274660 | 5545600 | 8792100 | 0 | 74470 |
| SLSETFLPN | STEAP2; STEAP | Q8NFT2; C9. | STEA2_HUMAN | Unmodified | 0 | 0 | 73350 | 8278600 | 0 | 0 | 0 | 0 | 0 | 168460 | 0 | 0 | 3121800 | 1680600 | 2091000 | 0 | 0 | 0 |
| SLSFPKLDS | UBAP1; UBAP1 | Q9NZ09; A0/ | UBAP1_HUMAI | Unmodified | 1117500 | 1362100 | 37894 | 254960 | 2045700 | 390300 | 153790 | 0 | 0 | 0 | 2329300 | 0 | 704780 | 460570 | 83826 | 146670 | 69104 | 159190 |
| SLSLESTDF | CDC42BPA; CD | Q5VT25; A0/ | MRCKA_HUMA | Unmodified | 0 | 0 | 0 | 0 | 409150 | 0 | 0 | 130420 | 0 | 0 | 0 | 0 | 260040 | 0 | 0 | 0 | 887160 | 0 |
| SLSLGEVLI | TM9SF1; TM9SF | O15321; E9F | TM9S1_HUMAN | Unmodified | 0 | 884820 | 741200 | 539870 | 0 | 0 | 0 | 81278 | 2012400 | 1212100 | 2406100 | 0 | 0 | 0 | 0 | 0 | 0 | 0 |
| SLSLGEVLI | TM9SF1; TM9SF | O15321; E9F | TM9S1_HUMAN | Oxidation (M) | 0 | 0 | 0 | 0 | 0 | 0 | 0 | 0 | 6965500 | 1920800 | 0 | 0 | 0 | 0 | 0 | 0 | 0 | 0 |
| SLSLGEVLI | TM9SF1; TM9SF | O15321; E9F | TM9S1_HUMAN | Unmodified | 0 | 0 | 0 | 0 | 0 | 0 | 682580 | 0 | 2959200 | 0 | 0 | 0 | 343320 | 0 | 0 | 0 | 548980 | 0 |
| SLSLGGHV | SEPTIN8; SEPTI | Q92599; A6N | SEPT8_HUMAN | Unmodified | 0 | 0 | 0 | 0 | 3177200 | 526420 | 914340 | 46418 | 200180 | 0 | 69116 | 398100 | 2346800 | 0 | 0 | 158140 | 828500 | 611500 |
| SLSLPERDI | UTRN | P46939 | UTRN_HUMAN | Unmodified | 0 | 0 | 0 | 1509900 | 0 | 0 | 0 | 0 | 588450 | 200590 | 0 | 0 | 184620 | 0 | 0 | 536060 | 178420 | 0 |
| SLSLSKLEC | TMEM43 | Q9BTV4 | TMM43_HUMAI | Unmodified | 0 | 0 | 0 | 0 | 0 | 0 | 0 | 0 | 29682000 | 1514000 | 0 | 0 | 0 | 0 | 0 | 0 | 153150 | 0 |
| SLSLSKLEC | TMEM43 | Q9BTV4 | TMM43_HUMAI | Unmodified | 0 | 0 | 0 | 0 | 0 | 0 | 0 | 0 | 5729600 | 0 | 0 | 0 | 0 | 0 | 0 | 0 | 0 | 0 |
| SLSLSLADS | PKP3; PKP3; PK | Q9Y446; E9F | PKP3_HUMAN | Unmodified | 0 | 0 | 0 | 0 | 4099300 | 0 | 0 | 0 | 0 | 0 | 0 | 5355600 | 0 | 0 | 0 | 0 | 0 | 0 |
| SLSNKLTLI | PGK1 | P00558 | PGK1_HUMAN | Acetyl (Protein N | 66107 | 0 | 541920 | 2542200 | 9365400 | 1087300 | 2398100 | 1711900 | 6229600 | 2886700 | 7379600 | 35406000 | 6720300 | 0 | 1589500 | 1707900 | 15398000 | 20484000 |
| SLSNKLTLI | PGK1 | P00558 | PGK1_HUMAN | Acetyl (Protein N | 4427900 | 4726500 | 0 | 17189000 | 1.88E+08 | 24907000 | 1.06E+08 | 58968000 | 28023000 | 72116000 | 18482000 | 46267000 | 2.02E+08 | 1.26E+08 | 3414200 | 4855700 | 6.89E+08 | 6.76E+08 |
| SLSNTKPT\ | VPS4B | O75351 | VPS4B_HUMAI | Unmodified | 3825700 | 2713800 | 5850400 | 5384800 | 11454000 | 2981600 | 134350 | 207820 | 16649000 | 13269000 | 4659200 | 3969800 | 7086700 | 4075600 | 3328300 | 2790500 | 1522200 | 3568500 |
| SLSNVIAHE | LTA4H | P09960 | LKHA4_HUMAI | Unmodified | 0 | 0 | 253110 | 0 | 5135400 | 1398600 | 329320 | 0 | 2192400 | 1004400 | 4649700 | 16458000 | 924340 | 452340 | 1242600 | 67912 | 2229300 | 1890300 |
| SLSPIIGK | EPB41L1; EPB4 | Q9H4G0; H7 | E41L1_HUMAN | Unmodified | 0 | 0 | 0 | 0 | 0 | 0 | 0 | 0 | 0 | 0 | 1856900 | 0 | 0 | 0 | 0 | 471090 | 0 | 0 |
| SLSPIIGKD | EPB41L1; EPB4 | Q9H4G0; H7 | E41L1_HUMAN | Unmodified | 0 | 0 | 0 | 0 | 0 | 0 | 1233300 | 0 | 0 | 0 | 0 | 0 | 0 | 4776900 | 0 | 0 | 2302100 | 2354300 |
| SLSPPQSQ | BAIAP2; BAIAP2 | Q9UQB8; I3I | BAIP2_HUMAN | Unmodified | 102560 | 5877600 | 257110 | 734120 | 2192500 | 0 | 104960 | 46435 | 202420 | 81934 | 0 | 0 | 1046900 | 0 | 320930 | 423810 | 460920 | 0 |
| SLSPPQSQ | BAIAP2; BAIAP2 | Q9UQB8; I3I | BAIP2_HUMAN | Unmodified | 0 | 0 | 0 | 0 | 749260 | 0 | 44738 | 431950 | 748230 | 0 | 0 | 0 | 0 | 1319700 | 0 | 0 | 1023600 | 0 |
| SLSQIHEA/ | MYOF; MYOF | Q9NZM1; H( | MYOF_HUMAN | Unmodified | 0 | 0 | 0 | 0 | 6386200 | 2539300 | 0 | 0 | 435060 | 0 | 0 | 0 | 0 | 0 | 0 | 0 | 0 | 0 |
| SLSQQIENI | COL1A1 | P02452 | CO1A1_HUMAI | Unmodified | 0 | 0 | 0 | 699270 | 0 | 0 | 0 | 0 | 0 | 0 | 0 | 157900 | 0 | 0 | 0 | 0 | 0 | 0 |
| SLSQQIENI | COL1A1 | P02452 | CO1A1_HUMAI | Unmodified | 0 | 0 | 825580 | 412620 | 0 | 1128300 | 137420 | 35804 | 0 | 0 | 0 | 0 | 0 | 265150 | 344850 | 0 | 93619 | 0 |
| SLSSPTDN | GIT1; GIT1; GIT | Q9Y2X7; A0/ | GIT1_HUMAN / | Unmodified | 0 | 0 | 0 | 0 | 0 | 0 | 0 | 0 | 0 | 0 | 0 | 201340 | 0 | 0 | 0 | 0 | 0 | 0 |
| SLSSPTVTL | NEDD4L; NEDD | Q96PU5; A0/ | NED4L_HUMAI | Unmodified | 0 | 0 | 0 | 1363000 | 15050 | 0 | 164160 | 0 | 0 | 0 | 0 | 0 | 0 | 0 | 0 | 2162400 | 0 | 257280 |
| SLSSPTVTL | NEDD4L; NEDD | Q96PU5; A0/ | NED4L_HUMAI | Unmodified | 319590 | 449770 | 0 | 598390 | 0 | 0 | 0 | 0 | 0 | 0 | 0 | 0 | 864920 | 0 | 0 | 0 | 0 | 0 |
| SLSSPTVTL | NEDD4L; NEDD | Q96PU5; A0/ | NED4L_HUMAI | Unmodified | 588220 | 953870 | 0 | 1354500 | 0 | 0 | 0 | 0 | 0 | 0 | 0 | 0 | 2250900 | 934380 | 0 | 0 | 571680 | 1352200 |
| SLSSSLDD' | VAPB | O95292 | VAPB_HUMAN | Unmodified | 0 | 0 | 0 | 0 | 634990 | 0 | 0 | 0 | 783560 | 249310 | 154690 | 551080 | 0 | 290220 | 0 | 0 | 0 | 620720 |
| SLSSSIGS | ADGRG6 | Q86SQ4 | AGRG6_HUMA | Unmodified | 0 | 0 | 0 | 3256800 | 16288000 | 2569000 | 290540 | 0 | 0 | 0 | 317330 | 0 | 834190 | 28666 | 2458600 | 4121500 | 0 | 277170 |
| SLSSYNYIF | MYO1D; MYO1D | O94832; J3Q | MYO1D_HUMA | Unmodified | 5479700 | 5188600 | 1636000 | 1176300 | 644970 | 0 | 177030 | 0 | 0 | 734610 | 0 | 740600 | 531580 | 220800 | 2732300 | 2714500 | 1730100 | 3764900 |
| SLSTAPVV( | ARFGEF3 | Q5TH69 | BIG3_HUMAN I | Unmodified | 0 | 0 | 125480 | 0 | 18846 | 0 | 0 | 0 | 183850 | 71014 | 0 | 0 | 0 | 0 | 0 | 0 | 0 | 0 |
| SLSTSGESI | DNAJC5 | Q9H3Z4 | DNJC5_HUMAN | Unmodified | 0 | 0 | 450310 | 1241500 | 0 | 0 | 0 | 0 | 0 | 0 | 0 | 6966600 | 0 | 0 | 0 | 0 | 0 | 0 |
| SLSVQDGL | DST; DST; DST; | Q03001; A0/ | DYST_HUMAN | Oxidation (M) | 0 | 0 | 0 | 0 | 237340 | 0 | 0 | 188710 | 0 | 0 | 0 | 0 | 0 | 2910700 | 0 | 0 | 0 | 1899300 |
| SLSYPFLLP | PLD1 | Q13393 | PLD1_HUMAN | Unmodified | 0 | 0 | 0 | 0 | 0 | 0 | 0 | 0 | 0 | 0 | 0 | 911390 | 0 | 0 | 0 | 0 | 0 | 0 |
| SLTAHSLLF | IQGAP3; IQGAP | Q86VI3; F2Z | IQGA3_HUMAN | Unmodified | 0 | 0 | 0 | 0 | 669890 | 0 | 0 | 0 | 474720 | 0 | 0 | 0 | 0 | 0 | 0 | 0 | 0 | 0 |
| SLTATGGN | FLNC | Q14315 | FLNC_HUMAN | Unmodified | 1614800 | 0 | 0 | 3927400 | 1019600 | 657010 | 242720 | 81646 | 0 | 298840 | 0 | 0 | 0 | 24552 | 0 | 4114200 | 221770 | 345220 |
| SLTEHGVA. | MPDZ; MPDZ; M | O75970; B7Z | MPDZ_HUMAN | Unmodified | 0 | 0 | 0 | 0 | 551450 | 0 | 0 | 13862 | 0 | 0 | 0 | 0 | 0 | 0 | 0 | 0 | 44731 | 0 |
| SLTETELTK | OSMR | Q99650 | OSMR_HUMAN | Unmodified | 0 | 0 | 0 | 0 | 6109200 | 482550 | 0 | 0 | 296820 | 0 | 0 | 0 | 0 | 0 | 0 | 0 | 0 | 0 |
| SLTEYLQN\ | KIF5B | P33176 | KINH_HUMAN | Unmodified | 142830 | 73906 | 204120 | 0 | 2178300 | 329940 | 403280 | 339400 | 989220 | 659680 | 575450 | 2741600 | 702540 | 471250 | 192230 | 150700 | 0 | 663320 |
| SLTEYLQN\ | KIF5B | P33176 | KINH_HUMAN | Unmodified | 0 | 0 | 154090 | 0 | 1225900 | 0 | 460550 | 0 | 653710 | 330030 | 0 | 1540100 | 839590 | 0 | 136460 | 249280 | 0 | 0 |
| SLTGLLSSF | UBR4 | Q5T4S7 | UBR4_HUMAN | Unmodified | 39153 | 0 | 0 | 0 | 0 | 0 | 0 | 0 | 0 | 111560 | 0 | 467520 | 0 | 0 | 48730 | 25326 | 0 | 0 |
| SLTGVASW | PHGDH | A0A286YF78 | A0A286YF78_H | Unmodified | 0 | 0 | 0 | 0 | 0 | 0 | 0 | 0 | 0 | 0 | 0 | 4092800 | 0 | 0 | 0 | 0 | 0 | 54675 |
| SLTGVVNA | PHGDH; PHGD | O43175; A0/ | SERA_HUMAN | Unmodified | 0 | 0 | 0 | 0 | 0 | 0 | 0 | 0 | 269920 | 0 | 0 | 2974100 | 0 | 723810 | 0 | 623080 | 0 | 0 |
| SLTIAEFK | TBCB; TBCB; TE | Q99426; K7E | TBCB_HUMAN | Unmodified | 59250 | 32381 | 102790 | 95619 | 1137100 | 179370 | 668620 | 0 | 1749900 | 1737900 | 580060 | 1775000 | 408010 | 249010 | 246720 | 222460 | 641090 | 159670 |
| SLTLSFVSE | LUZP1; LUZP1 | Q86V48; E5F | LUZP1_HUMAN | Unmodified | 0 | 0 | 0 | 0 | 791020 | 268190 | 0 | 0 | 0 | 0 | 0 | 0 | 0 | 0 | 0 | 0 | 251070 | 193640 |
| SLTNDWEI | HSP90AB1; HSI | P08238; P07 | HS90B_HUMAI | Unmodified | 15091000 | 16616000 | 20141000 | 41309000 | 1.51E+08 | 42693000 | 29396000 | 10680000 | 1.08E+08 | 1.02E+08 | 1.27E+08 | 7.72E+08 | 72651000 | 28800000 | 45497000 | 31644000 | 60999000 | 72625000 |
| SLTNDWEI | HSP90AB1; HSI | P08238; P07 | HS90B_HUMAI | Unmodified | 0 | 0 | 0 | 59022 | 816860 | 0 | 0 | 0 | 0 | 0 | 0 | 0 | 0 | 0 | 0 | 0 | 1191900 | 1071600 |
| SLTPAVPVE | CAST; CAST; C/ | P20810; A0/ | ICAL_HUMAN ( | Unmodified | 50689 | 81784 | 86320 | 97447 | 1357600 | 244450 | 0 | 0 | 304440 | 0 | 310210 | 544620 | 349970 | 194520 | 69123 | 88828 | 426950 | 954510 |
| SLTPDDDV | CPM; CPM; CPI | P14384; F8V | CBPM_HUMAN | Unmodified | 0 | 0 | 0 | 0 | 0 | 0 | 0 | 0 | 3152500 | 786790 | 378380 | 0 | 0 | 0 | 0 | 0 | 0 | 0 |
| SLTPDDDV | CPM; CPM; CPI | P14384; F8V | CBPM_HUMAN | Oxidation (M) | 292530 | 211960 | 97065 | 0 | 5859100 | 6754000 | 0 | 0 | 35967000 | 2571900 | 0 | 0 | 0 | 0 | 0 | 0 | 0 | 0 |
| SLTPDDDV | CPM; CPM; CPI | P14384; F8V | CBPM_HUMAN | Unmodified | 0 | 0 | 0 | 0 | 0 | 0 | 0 | 0 | 10564000 | 0 | 0 | 0 | 0 | 0 | 0 | 0 | 0 | 0 |
| SLTQVRAM | IMPDH2; IMPDI | P12268; A0/ | IMDH2_HUMAI | Oxidation (M) | 0 | 0 | 0 | 0 | 305770 | 0 | 3207400 | 0 | 0 | 176310 | 0 | 0 | 0 | 0 | 99664 | 0 | 1623700 | 0 |
| SLTSWTNI | ZC3HAV1; ZC3I | Q7Z2W4; C9 | ZCCHV_HUMA | Unmodified | 730260 | 0 | 565710 | 1858100 | 6350300 | 1014500 | 0 | 0 | 0 | 435320 | 0 | 0 | 1865200 | 192460 | 477250 | 991830 | 212460 | 522060 |
| SLTSWTNI | ZC3HAV1; ZC3I | Q7Z2W4; C9 | ZCCHV_HUMA | Unmodified | 0 | 0 | 0 | 0 | 2350200 | 32173 | 339580 | 0 | 0 | 78947 | 0 | 0 | 0 | 0 | 0 | 0 | 283730 | 508040 |
| SLTTAFFR | RAB27A; RAB27 | P51159; O0( | RB27A_HUMAN | Unmodified | 183930 | 319690 | 648610 | 0 | 0 | 0 | 0 | 0 | 0 | 0 | 0 | 0 | 0 | 0 | 862540 | 609120 | 0 | 0 |
| SLTTLGLVI | KIF13A; KIF13A | Q9H1H9; A0 | KI13A_HUMAN | Unmodified | 0 | 0 | 0 | 0 | 388470 | 0 | 0 | 0 | 18084 | 0 | 0 | 7380.6 | 22773 | 0 | 38523 | 0 | 71345 | 0 |
| SLTTLGLVI | KIF13A; KIF13A | Q9H1H9; A0 | KI13A_HUMAN | Unmodified | 0 | 0 | 0 | 0 | 4178300 | 0 | 0 | 0 | 0 | 0 | 0 | 0 | 0 | 0 | 0 | 0 | 310310 | 0 |
| SLTTYGLSA | EIF2AK3; EIF2A | Q9NZJ5; A0/ | E2AK3_HUMAN | Unmodified | 0 | 0 | 0 | 0 | 0 | 0 | 0 | 0 | 1093400 | 350890 | 0 | 0 | 0 | 0 | 0 | 0 | 0 | 248710 |
| SLTYLSILR | SNRPA1; SNRP. | P09661; H0\ | RU2A_HUMAN | Unmodified | 0 | 0 | 0 | 0 | 0 | 0 | 0 | 0 | 0 | 0 | 0 | 2253200 | 0 | 0 | 0 | 0 | 0 | 0 |
| SLTYLSILRI | SNRPA1; SNRP. | P09661; H0\ | RU2A_HUMAN | Unmodified | 633060 | 201710 | 71838 | 0 | 0 | 0 | 0 | 36416 | 0 | 94724 | 0 | 3024000 | 0 | 0 | 0 | 0 | 0 | 0 |
| SLVASLAEF | UBA1 | P22314 | UBA1_HUMAN | Unmodified | 0 | 0 | 0 | 520510 | 3059000 | 482430 | 5621800 | 50005 | 2117400 | 2204000 | 1638500 | 3031500 | 4913400 | 248250 | 1003700 | 0 | 4599400 | 3711500 |
| SLVDLTAVI | NDUFS3; NDUF | O75489; E9F | NDUS3_HUMA | Unmodified | 0 | 0 | 0 | 0 | 0 | 0 | 0 | 0 | 1164200 | 0 | 0 | 0 | 0 | 0 | 0 | 0 | 0 | 0 |
| SLVDTVYAI | ARHGEF7; ARH | B1ALK7; A0/ | B1ALK7_HUMA | Unmodified | 0 | 0 | 0 | 0 | 650370 | 123560 | 492630 | 0 | 563670 | 616010 | 0 | 0 | 638560 | 298600 | 0 | 0 | 259450 | 1191600 |
| SLVDYENA | SNX6; SNX6 | Q9UNH7; A( | SNX6_HUMAN | Unmodified | 124890 | 0 | 0 | 0 | 3198500 | 0 | 152710 | 0 | 0 | 76704 | 584320 | 0 | 1448100 | 823420 | 0 | 0 | 176830 | 551060 |
| SLVDYENA | SNX6; SNX6 | Q9UNH7; A( | SNX6_HUMAN | Unmodified | 0 | 0 | 0 | 0 | 0 | 0 | 412450 | 0 | 338210 | 0 | 0 | 0 | 0 | 0 | 0 | 0 | 373650 | 340380 |
| SLVEIADTV | RANBP6; IPO5; | O60518; O0( | RNBP6_HUMAI | Unmodified | 0 | 0 | 686560 | 0 | 240630 | 0 | 0 | 174370 | 0 | 0 | 0 | 0 | 0 | 0 | 853290 | 57834 | 591670 | 221590 |
| SLVEIIEHG | SF3B1 | O75533 | SF3B1_HUMAN | Unmodified | 0 | 0 | 214550 | 0 | 0 | 0 | 0 | 0 | 359610 | 0 | 0 | 16293000 | 0 | 0 | 0 | 0 | 0 | 0 |
| SLVEPKEDI | AKAP12 | Q02952 | AKA12_HUMAN | Unmodified | 0 | 0 | 0 | 0 | 1150200 | 0 | 0 | 0 | 0 | 0 | 0 | 0 | 0 | 0 | 0 | 0 | 812410 | 2176100 |
| SLVESVSSS | PPA2 | Q9H2U2 | IPYR2_HUMAN | Unmodified | 0 | 517410 | 0 | 0 | 0 | 0 | 0 | 0 | 572000 | 0 | 155010 | 512240 | 0 | 0 | 0 | 0 | 0 | 0 |
| SLVESVSSS | PPA2 | Q9H2U2 | IPYR2_HUMAN | Unmodified | 0 | 0 | 0 | 0 | 0 | 0 | 0 | 0 | 2861400 | 296630 | 0 | 645610 | 0 | 0 | 0 | 0 | 0 | 0 |
| SLVGCGHF | CIT; CIT | O14578; H0Y | CTRO_HUMAN | Unmodified | 0 | 0 | 0 | 0 | 0 | 0 | 149380 | 0 | 0 | 0 | 0 | 0 | 0 | 0 | 0 | 0 | 862630 | 0 |
| SLVGTPYW | PAK4; PAK4 | O96013; M0 | PAK4_HUMAN | Oxidation (M) | 0 | 0 | 0 | 0 | 0 | 0 | 0 | 0 | 0 | 0 | 0 | 1363900 | 0 | 0 | 240940 | 221190 | 0 | 0 |
| SLVHAGVP | EIF2B4; EIF2B4 | Q9UI10; A0/ | EI2BD_HUMAN | Unmodified | 0 | 0 | 29521 | 0 | 0 | 0 | 264260 | 0 | 161050 | 39755 | 48340 | 2166500 | 0 | 0 | 0 | 0 | 0 | 0 |
| SLVHDLFQ | SMC4; SMC4 | Q9NTJ3; E9F | SMC4_HUMAN | Unmodified | 0 | 19845 | 0 | 0 | 876170 | 208180 | 943400 | 0 | 308860 | 439000 | 0 | 123880 | 1367300 | 295130 | 0 | 0 | 294090 | 746560 |
| SLVIISTLD( | EIF2AK3; EIF2AI | Q9NZJ5; A0/ | E2AK3_HUMAN | Unmodified | 0 | 0 | 0 | 0 | 0 | 0 | 0 | 0 | 1457100 | 168940 | 0 | 0 | 0 | 0 | 0 | 0 | 0 | 0 |
| SLVKWESE | CRABP2 | P29373 | RABP2_HUMAI | Unmodified | 0 | 0 | 0 | 0 | 1974000 | 601760 | 1211100 | 349560 | 1430900 | 1570800 | 0 | 189220 | 0 | 0 | 0 | 0 | 9841200 | 5317300 |
| SLVKWESE | CRABP2 | P29373 | RABP2_HUMAI | Oxidation (M) | 0 | 0 | 0 | 0 | 0 | 0 | 0 | 0 | 0 | 0 | 0 | 0 | 0 | 88697 | 0 | 0 | 3213900 | 674110 |
| SLVKWESE | CRABP2 | P29373 | RABP2_HUMAI | Unmodified | 0 | 0 | 0 | 0 | 0 | 0 | 213940 | 0 | 0 | 211560 | 0 | 0 | 0 | 0 | 0 | 0 | 9415100 | 46587 |
| SLVLENVTI | CRYBG1; CRYB | Q9Y4K1; A0/ | CRBG1_HUMA | Unmodified | 0 | 0 | 0 | 0 | 1102100 | 412810 | 0 | 0 | 0 | 0 | 510040 | 0 | 32933 | 0 | 0 | 0 | 0 | 486550 |
| SLVLSQSK | SEMA4C | Q9C0C4 | SEM4C_HUMAI | Unmodified | 266730 | 144250 | 0 | 0 | 86439 | 1113700 | 0 | 0 | 1265400 | 0 | 0 | 143090 | 0 | 0 | 0 | 0 | 141790 | 0 |
| SLVQEMVC | SNRNP200; SN | O75643; A0/ | U520_HUMAN | Unmodified | 0 | 0 | 0 | 0 | 0 | 0 | 0 | 0 | 0 | 0 | 0 | 1475200 | 0 | 0 | 0 | 0 | 0 | 0 |
| SLVQEMVC | SNRNP200; SN | O75643; A0/ | U520_HUMAN | Unmodified | 0 | 0 | 0 | 0 | 0 | 0 | 0 | 0 | 0 | 0 | 0 | 2136500 | 0 | 0 | 0 | 0 | 0 | 0 |
| SLVQGELV | ADD1; ADD1; A | P35611; E7E | ADDA_HUMAN | Unmodified | 0 | 0 | 0 | 957310 | 0 | 0 | 224730 | 584030 | 0 | 0 | 0 | 0 | 330330 | 0 | 1402700 | 779400 | 259520 | 0 |
| SLVSDKTSI | LAD1; LAD1 | O00515; E9F | LAD1_HUMAN | Unmodified | 0 | 0 | 0 | 0 | 438290 | 0 | 0 | 0 | 0 | 107800 | 0 | 0 | 1949800 | 631870 | 0 | 0 | 0 | 0 |
| SLVSGLIR | GCN1 | Q92616 | GCN1_HUMAN | Unmodified | 0 | 0 | 82929 | 0 | 0 | 0 | 0 | 0 | 0 | 0 | 0 | 4704600 | 0 | 0 | 0 | 94602 | 0 | 0 |
| SLVSKGTL\ | H1-3; H1-4; H1 | P16402; P10 | H13_HUMAN H | Unmodified | 870040 | 562180 | 0 | 0 | 6735700 | 901130 | 25113000 | 5.94E+08 | 0 | 243810 | 0 | 0 | 1266600 | 422180 | 0 | 0 | 238390 | 2861200 |
| SLVSKGTL\ | H1-3; H1-4; H1 | P16402; P10 | H13_HUMAN H | Unmodified | 0 | 0 | 0 | 0 | 866350 | 0 | 972550 | 1.53E+08 | 0 | 0 | 0 | 0 | 115600 | 0 | 0 | 0 | 0 | 301780 |
| SLVSVTKEC | HSP90AB1; HSI | P08238; Q58 | HS90B_HUMAI | Unmodified | 0 | 0 | 7815300 | 0 | 27282000 | 5852400 | 0 | 1196300 | 5192800 | 4625600 | 3844500 | 5640600 | 17151000 | 11145000 | 2984700 | 0 | 0 | 8999800 |
| SLVSVTKEC | HSP90AB1; HSI | P08238; Q58 | HS90B_HUMAI | Unmodified | 0 | 0 | 0 | 0 | 7102300 | 1665000 | 0 | 0 | 688620 | 1985800 | 0 | 1119300 | 6754500 | 2216200 | 0 | 0 | 896970 | 3923100 |
| SLVTNCKP' | GNPTAB; GNPT | Q3T906; H0\ | GNPTA_HUMAI | Unmodified | 0 | 0 | 0 | 0 | 29325 | 0 | 112050 | 0 | 1359100 | 131890 | 0 | 0 | 0 | 0 | 0 | 0 | 279620 | 68681 |
| SLVTQYLN/ | SLC3A2; SLC3A | P08195; A0/ | 4F2_HUMAN 4I | Unmodified | 0 | 0 | 0 | 0 | 0 | 0 | 0 | 0 | 0 | 0 | 0 | 0 | 0 | 0 | 686890 | 0 | 0 | 0 |
| SLVTVETD/ | NOMO1; NOMC | Q15155; P69 | NOMO1_HUMAI | Unmodified | 0 | 0 | 0 | 0 | 0 | 0 | 0 | 0 | 1712900 | 464850 | 435580 | 0 | 0 | 204800 | 0 | 0 | 0 | 118670 |
| SLVTVLGIP | CADM3; CADM3 | Q8N126; A0/ | CADM3_HUMA | Unmodified | 0 | 0 | 0 | 0 | 0 | 0 | 2741700 | 115900 | 0 | 0 | 0 | 0 | 0 | 0 | 0 | 0 | 0 | 0 |

| | | | | | | | | | | | | | | | | | | | | | | |
|---|---|---|---|---|---|---|---|---|---|---|---|---|---|---|---|---|---|---|---|---|---|---|
| SLVVSWSP | PTPRB; PTPRB; | P23467; F8V | PTPRB_HUMAN | Unmodified | 0 | 349460 | 11915 | 15908 | 0 | 0 | 0 | 0 | 0 | 0 | 0 | 0 | 0 | 0 | 0 | 0 | 294080 | 0 |
| SLVWGPGI | POGLUT3 | Q7Z4H8 | PLGT3_HUMAN | Unmodified | 0 | 0 | 0 | 0 | 0 | 0 | 0 | 0 | 7621600 | 155870 | 0 | 0 | 0 | 0 | 0 | 0 | 0 | 0 |
| SLVYQLNFI | MGAT2 | Q10469 | MGAT2_HUMAI | Unmodified | 0 | 0 | 0 | 0 | 0 | 0 | 0 | 0 | 5850200 | 311300 | 0 | 53030 | 0 | 0 | 187320 | 0 | 0 | 0 |
| SLWNDPGI | GNAQ; GNAQ | P50148; B1A | GNAQ_HUMAN | Unmodified | 138860 | 111780 | 0 | 0 | 0 | 0 | 52268 | 0 | 560930 | 0 | 181740 | 1461000 | 0 | 0 | 0 | 27766 | 0 | 0 |
| SLWSSLCL | ADPGK; ADPGK | Q9BRR6; H3 | ADPGK_HUMAI | Unmodified | 0 | 0 | 0 | 0 | 0 | 0 | 0 | 0 | 0 | 106380 | 0 | 0 | 0 | 0 | 0 | 0 | 0 | 0 |
| SLYDEVAA( | EPRS1; EPRS1 | P07814; V9G | SYEP_HUMAN | Unmodified | 0 | 0 | 0 | 0 | 0 | 0 | 0 | 0 | 47214 | 0 | 254990 | 0 | 0 | 0 | 0 | 0 | 157670 | 41197 |
| SLYDEVAA( | EPRS1; EPRS1 | P07814; V9G | SYEP_HUMAN | Unmodified | 0 | 0 | 1524500 | 0 | 1196900 | 0 | 0 | 182590 | 1746000 | 974920 | 1482100 | 12961000 | 6130500 | 2707100 | 0 | 1315400 | 383250 | 0 |
| SLYESWTK | FOLH1B; FOLH | Q9HBA9; Q0 | FOH1B_HUMAI | Unmodified | 4320100 | 164640 | 19890000 | 27512000 | 0 | 0 | 0 | 0 | 0 | 0 | 0 | 0 | 6659200 | 3538100 | 16134000 | 16611000 | 0 | 0 |
| SLYGKDLIA | ANXA6; ANXA6; | P08133; E5R | ANXA6_HUMAI | Unmodified | 0 | 0 | 0 | 74895 | 341350 | 0 | 176500 | 0 | 528840 | 568600 | 0 | 0 | 146100 | 202380 | 0 | 0 | 0 | 0 |
| SLYHDISGI | ANXA11 | P50995 | ANX11_HUMAI | Unmodified | 96647 | 0 | 50754 | 104150 | 131930 | 40770 | 0 | 0 | 6100200 | 325290 | 230500 | 622190 | 0 | 23068 | 0 | 176070 | 51404 | 328710 |
| SLYHDISGI | ANXA11 | P50995 | ANX11_HUMAI | Unmodified | 2447200 | 812520 | 1713800 | 2130900 | 85381000 | 13416000 | 0 | 0 | 3.04E+08 | 86714000 | 9287300 | 15148000 | 23842000 | 40597000 | 2250100 | 2644100 | 34765000 | 2E+08 |
| SLYHDISGI | ANXA11 | P50995 | ANX11_HUMAI | Unmodified | 0 | 0 | 0 | 0 | 2128000 | 66908 | 0 | 0 | 0 | 804220 | 0 | 0 | 331800 | 215250 | 0 | 0 | 5360000 | 12712000 |
| SLYHSVEW | COLGALT1; CO | Q8NBJ5; M0 | GT251_HUMAI | Unmodified | 0 | 0 | 0 | 288230 | 0 | 0 | 0 | 0 | 29024000 | 3140200 | 528880 | 269170 | 0 | 0 | 0 | 0 | 67545 | 0 |
| SLYHSVEW | COLGALT1; CO | Q8NBJ5; M0 | GT251_HUMAI | Unmodified | 0 | 0 | 0 | 0 | 0 | 0 | 0 | 0 | 10509000 | 4486600 | 178360 | 0 | 0 | 0 | 0 | 0 | 0 | 0 |
| SLYKEELEP | POMGNT1 | Q8WZA1 | PMGT1_HUMAI | Unmodified | 0 | 0 | 0 | 0 | 0 | 0 | 0 | 0 | 2625800 | 118540 | 0 | 425220 | 0 | 0 | 0 | 53140 | 0 | 0 |
| SLYLLINSK | UBE3C | Q15386 | UBE3C_HUMAI | Unmodified | 16511000 | 9335900 | 0 | 1795400 | 0 | 0 | 0 | 0 | 0 | 0 | 0 | 0 | 2302200 | 0 | 14043000 | 1463100 | 0 | 0 |
| SLYNVNGL | ASPH | Q12797 | ASPH_HUMAN | Unmodified | 0 | 0 | 0 | 0 | 0 | 0 | 0 | 0 | 0 | 0 | 0 | 0 | 0 | 0 | 0 | 0 | 0 | 0 |
| SLYPDAED | PSD2 | Q9BQI7 | PSD2_HUMAN | Unmodified | 210030 | 562630 | 0 | 0 | 0 | 0 | 0 | 0 | 0 | 0 | 1080200 | 0 | 0 | 0 | 0 | 42273 | 0 | 0 |
| SLYPLTFIQ | GLB1; GLB1 | P16278; E7E | BGAL_HUMAN | Unmodified | 0 | 0 | 0 | 0 | 0 | 0 | 0 | 0 | 4718400 | 3259500 | 626780 | 0 | 0 | 0 | 0 | 0 | 0 | 472790 |
| SLYPSLEDI | SDCBP2; SDCB | Q9H190; O0 | SDCB2_HUMAI | Acetyl (Protein N | 6947200 | 6262200 | 57851000 | 7055000 | 17491000 | 4454600 | 1220100 | 776840 | 48237000 | 12314000 | 3481300 | 21951000 | 8492800 | 7726800 | 36040000 | 17854000 | 407930 | 2770000 |
| SLYPSLEDI | SDCBP; SDCBP | O00560; G5I | SDCB1_HUMAI | Acetyl (Protein N | 19369000 | 11019000 | 6110600 | 20793000 | 70226000 | 9063100 | 1615700 | 807920 | 60064000 | 19996000 | 3107100 | 12881000 | 28827000 | 10552000 | 16972000 | 20032000 | 1636500 | 8730100 |
| SLYQSAGV | FASN; FASN | P49327; A0A | FAS_HUMAN F | Unmodified | 0 | 0 | 0 | 8010600 | 0 | 0 | 0 | 0 | 0 | 0 | 3524100 | 2.01E+08 | 812300 | 0 | 335370 | 0 | 0 | 0 |
| SLYQSAGV | FASN; FASN | P49327; A0A | FAS_HUMAN F | Unmodified | 0 | 0 | 0 | 1446600 | 0 | 0 | 0 | 0 | 0 | 0 | 2777100 | 0 | 3527600 | 0 | 0 | 0 | 0 | 0 |
| SLYSFIKGD | ANXA4; ANXA4 | P09525; Q6I | ANXA4_HUMAI | Unmodified | 0 | 0 | 0 | 0 | 488040 | 0 | 0 | 147370 | 1832800 | 146340 | 0 | 2452300 | 0 | 0 | 0 | 0 | 0 | 0 |
| SLYSFIKGD | ANXA4; ANXA4 | P09525; Q6I | ANXA4_HUMAI | Unmodified | 0 | 1038200 | 0 | 662890 | 20752000 | 4031700 | 10199000 | 1231000 | 24844000 | 30149000 | 9610400 | 1410100 | 6658700 | 13744000 | 0 | 0 | 2457600 | 5625300 |
| SLYSMIKNI | ANXA6; ANXA6 | P08133; E5R | ANXA6_HUMAI | Unmodified | 0 | 0 | 0 | 72971 | 1154500 | 0 | 262500 | 0 | 276990 | 190870 | 0 | 0 | 98306 | 120010 | 0 | 0 | 185160 | 267990 |
| SLYSMIKNI | ANXA6; ANXA6 | P08133; E5R | ANXA6_HUMAI | Unmodified | 0 | 0 | 0 | 0 | 4244100 | 0 | 10533000 | 0 | 3869000 | 7593500 | 0 | 0 | 1629700 | 4468100 | 0 | 0 | 2805000 | 0 |
| SLYSMIKNI | ANXA6; ANXA6 | P08133; E5R | ANXA6_HUMAI | Oxidation (M) | 0 | 0 | 0 | 0 | 2242700 | 0 | 5541900 | 1596200 | 664770 | 6597900 | 0 | 0 | 2237900 | 2328200 | 0 | 0 | 2632800 | 4825400 |
| SLYTHSYE\ | PECR | Q9BY49 | PECR_HUMAN | Unmodified | 0 | 0 | 0 | 0 | 0 | 0 | 0 | 0 | 1032000 | 0 | 0 | 0 | 0 | 0 | 0 | 0 | 0 | 0 |
| SLYVAEYHS | DDX6 | P26196 | DDX6_HUMAN | Unmodified | 1841200 | 1929400 | 0 | 2930800 | 7186100 | 990150 | 2556100 | 1107000 | 1419500 | 1398700 | 925070 | 10680000 | 8314900 | 3294600 | 1929400 | 2000000 | 2745500 | 6613800 |
| SLYVRALFD | DLG1; DLG1; D | Q12959; A0A | DLG1_HUMAN | Unmodified | 0 | 3581100 | 0 | 0 | 0 | 0 | 0 | 0 | 0 | 0 | 0 | 0 | 0 | 79613 | 0 | 9833.7 | 0 | 0 |
| SLYYYIQQD | ANXA2P2; ANX | A6NMY6; P0 | AXA2L_HUMAN | Unmodified | 539340 | 530970 | 0 | 177590 | 1104600 | 469170 | 0 | 0 | 3086600 | 449220 | 0 | 0 | 0 | 0 | 0 | 352980 | 0 | 714030 |
| SLYYYIQQD | ANXA2P2; ANX | A6NMY6; P0 | AXA2L_HUMAN | Unmodified | 4281100 | 3109900 | 305140 | 3827500 | 3.21E+08 | 97657000 | 15542000 | 3568500 | 67581000 | 34052000 | 691900 | 142450 | 1097900 | 3827900 | 1046100 | 823300 | 71708000 | 60205000 |
| SMAAAAAS | TMEM9; TMEM9 | Q9P0T7; B1A | TMEM9_HUMAI | Unmodified | 0 | 0 | 92362 | 184780 | 0 | 0 | 0 | 0 | 1440500 | 342590 | 0 | 0 | 0 | 0 | 102970 | 79655 | 0 | 0 |
| SMAASGNI | UQCRC2 | P22695 | QCR2_HUMAN | Oxidation (M) | 0 | 0 | 0 | 31495 | 0 | 0 | 327540 | 139860 | 2911900 | 1288000 | 0 | 0 | 0 | 68504 | 0 | 0 | 0 | 0 |
| SMAASGNI | UQCRC2 | P22695 | QCR2_HUMAN | Unmodified | 0 | 0 | 0 | 0 | 0 | 0 | 0 | 0 | 1302900 | 199480 | 0 | 0 | 0 | 0 | 0 | 0 | 0 | 0 |
| SMAEDTIN | GPD2 | P43304 | GPDM_HUMAN | Unmodified | 0 | 0 | 0 | 0 | 0 | 0 | 0 | 0 | 5808900 | 809280 | 0 | 0 | 0 | 0 | 0 | 0 | 0 | 0 |
| SMAEDTIN | GPD2 | P43304 | GPDM_HUMAN | Oxidation (M) | 0 | 0 | 0 | 88984 | 0 | 0 | 0 | 0 | 957620 | 166390 | 0 | 0 | 0 | 0 | 0 | 0 | 0 | 0 |
| SMAGVVKS | CHMP1B | Q7LBR1 | CHM1B_HUMA | Unmodified | 0 | 106440 | 0 | 0 | 2676300 | 0 | 122140 | 44587 | 0 | 130260 | 0 | 0 | 237070 | 262610 | 0 | 0 | 222000 | 370470 |
| SMAGVVKS | CHMP1B | Q7LBR1 | CHM1B_HUMA | Oxidation (M) | 0 | 0 | 0 | 0 | 424660 | 0 | 0 | 0 | 0 | 0 | 0 | 0 | 95598 | 0 | 0 | 0 | 0 | 0 |
| SMALADPE | SNTB1; SNTB1 | Q13884; A0A | SNTB1_HUMAI | Unmodified | 297160 | 66570 | 0 | 0 | 0 | 0 | 0 | 0 | 3137200 | 142230 | 0 | 0 | 0 | 0 | 813240 | 0 | 0 | 0 |
| SMALADPE | SNTB1; SNTB1 | Q13884; A0A | SNTB1_HUMAI | Unmodified | 0 | 0 | 0 | 0 | 0 | 0 | 0 | 0 | 1441600 | 930000 | 0 | 0 | 0 | 0 | 0 | 0 | 0 | 0 |
| SMANPPAA | DYNC1H1; DYN | Q14204; A0A | DYHC1_HUMA | Oxidation (M) | 0 | 0 | 54791 | 70725 | 0 | 0 | 0 | 20434 | 0 | 172720 | 90944 | 756280 | 0 | 0 | 75735 | 76958 | 0 | 0 |
| SMCGYQTI | GNS; GNS; GNS | P15586; H7C | GNS_HUMAN I | Oxidation (M) | 0 | 0 | 0 | 0 | 0 | 0 | 0 | 0 | 754610 | 386970 | 0 | 0 | 201730 | 164880 | 0 | 0 | 0 | 0 |
| SMCGYQTI | GNS; GNS; GNS | P15586; H7C | GNS_HUMAN I | Unmodified | 0 | 0 | 0 | 0 | 0 | 0 | 0 | 0 | 2350700 | 0 | 0 | 0 | 0 | 0 | 0 | 0 | 0 | 0 |
| SMDAETYV | AFDN; AFDN; A | P55196; Q51 | AFAD_HUMAN | Unmodified | 0 | 0 | 0 | 0 | 0 | 0 | 0 | 0 | 0 | 0 | 0 | 319060 | 0 | 0 | 0 | 0 | 0 | 0 |
| SMDATLKTI | CHMP1B | Q7LBR1 | CHM1B_HUMA | Oxidation (M) | 0 | 0 | 0 | 0 | 453340 | 0 | 74365 | 85861 | 0 | 97245 | 0 | 0 | 0 | 222540 | 0 | 0 | 0 | 144210 |
| SMDATLKTI | CHMP1B | Q7LBR1 | CHM1B_HUMA | Unmodified | 0 | 0 | 0 | 0 | 3436300 | 0 | 408410 | 0 | 0 | 410660 | 0 | 0 | 339860 | 664270 | 0 | 0 | 536230 | 0 |
| SMDNMTVI | FDX1 | P10109 | ADX_HUMAN A | Unmodified | 0 | 0 | 0 | 0 | 0 | 0 | 188050 | 0 | 759490 | 0 | 92342 | 0 | 0 | 0 | 0 | 0 | 0 | 0 |
| SMEAEMIQ | MYH9 | P35579 | MYH9_HUMAN | 2 Oxidation (M) | 0 | 0 | 0 | 0 | 0 | 0 | 0 | 0 | 0 | 265540 | 0 | 1064100 | 0 | 0 | 0 | 0 | 0 | 0 |
| SMEAEMIQ | MYH9 | P35579 | MYH9_HUMAN | 2 Oxidation (M) | 0 | 109150 | 0 | 0 | 1550300 | 0 | 0 | 0 | 0 | 430230 | 0 | 262290 | 0 | 0 | 1309200 | 0 | 0 | 0 |
| SMEAEMIQ | MYH9 | P35579 | MYH9_HUMAN | Oxidation (M) | 0 | 0 | 0 | 0 | 2412400 | 550250 | 0 | 0 | 495660 | 353020 | 0 | 0 | 0 | 0 | 0 | 0 | 0 | 0 |
| SMEAEMIQ | MYH9 | P35579 | MYH9_HUMAN | Unmodified | 0 | 0 | 0 | 0 | 2487300 | 0 | 0 | 0 | 0 | 0 | 0 | 0 | 0 | 0 | 0 | 0 | 0 | 0 |
| SMEAEMIQ | MYH9 | P35579 | MYH9_HUMAN | Unmodified | 0 | 0 | 0 | 0 | 903480 | 0 | 0 | 0 | 150390 | 0 | 0 | 0 | 0 | 0 | 0 | 0 | 249430 | 0 |
| SMEAGSST | SLC23A2 | Q9UGH3 | S23A2_HUMAN | Unmodified | 0 | 0 | 20600 | 0 | 262840 | 45810 | 0 | 0 | 0 | 36592 | 276870 | 2125400 | 0 | 0 | 0 | 0 | 0 | 247240 |
| SMEASVDV | AHNAK2 | Q8IVF2 | AHNK2_HUMAI | Oxidation (M) | 0 | 0 | 0 | 0 | 583420 | 0 | 0 | 0 | 0 | 0 | 0 | 0 | 0 | 0 | 0 | 0 | 80720 | 566550 |
| SMEEDPQT | PRKDC | P78527 | PRKDC_HUMAI | Oxidation (M) | 0 | 184080 | 0 | 0 | 0 | 41481 | 512390 | 128800 | 815470 | 494560 | 125040 | 444630 | 0 | 0 | 0 | 0 | 0 | 0 |
| SMEMLVEH | PSMD13; PSMD | Q9UNM6; A0 | PSD13_HUMAI | Oxidation (M) | 0 | 0 | 102710 | 78494 | 2389500 | 0 | 1908900 | 101040 | 679040 | 88305 | 369220 | 1607000 | 1703200 | 862990 | 57320 | 66064 | 24725 | 2414200 |
| SMEMLVEH | PSMD13; PSMD | Q9UNM6; A0 | PSD13_HUMAI | Unmodified | 0 | 0 | 0 | 0 | 1590000 | 0 | 0 | 0 | 0 | 69448 | 34873 | 0 | 651320 | 381780 | 0 | 0 | 181900 | 2827200 |
| SMENIMEM | DLG1; DLG1; D | Q12959; A0A | DLG1_HUMAN | Oxidation (M) | 0 | 0 | 0 | 0 | 847000 | 0 | 0 | 0 | 0 | 0 | 0 | 0 | 0 | 0 | 0 | 0 | 423420 | 351870 |
| SMENIMEM | DLG1; DLG1; D | Q12959; A0A | DLG1_HUMAN | Unmodified | 0 | 0 | 0 | 0 | 4296300 | 183120 | 0 | 0 | 722120 | 455130 | 0 | 0 | 0 | 290090 | 0 | 0 | 671640 | 618740 |
| SMENIMEM | DLG1; DLG1; D | Q12959; A0A | DLG1_HUMAN | Unmodified | 0 | 0 | 40586 | 0 | 617900 | 0 | 0 | 0 | 801640 | 0 | 0 | 0 | 0 | 0 | 0 | 0 | 353000 | 261400 |
| SMEYVQRC | SPRED1 | Q7Z699 | SPRE1_HUMAN | Unmodified | 0 | 0 | 0 | 1433000 | 189790 | 0 | 0 | 0 | 0 | 0 | 0 | 0 | 82614 | 54125 | 0 | 0 | 229600 | 498550 |
| SMGFIGHY | ACLY | P53396 | ACLY_HUMAN | Unmodified | 0 | 0 | 0 | 139000 | 684920 | 0 | 1334600 | 28007 | 0 | 971690 | 272400 | 5132300 | 236240 | 0 | 0 | 0 | 281220 | 202930 |
| SMGFIGHY | ACLY | P53396 | ACLY_HUMAN | Oxidation (M) | 0 | 0 | 0 | 0 | 240710 | 0 | 353830 | 0 | 0 | 144680 | 0 | 2891900 | 163130 | 0 | 186760 | 0 | 0 | 0 |
| SMGFIGHY | ACLY | P53396 | ACLY_HUMAN | Unmodified | 0 | 0 | 0 | 0 | 159180 | 0 | 0 | 0 | 0 | 98813 | 0 | 1109600 | 0 | 0 | 0 | 0 | 0 | 0 |
| SMGLPTSD | NUDC | Q9Y266 | NUDC_HUMAN | Unmodified | 0 | 0 | 0 | 0 | 445020 | 0 | 0 | 0 | 1106300 | 267070 | 470320 | 1588300 | 1404600 | 396440 | 0 | 0 | 453150 | 608830 |
| SMGLPTSD | NUDC | Q9Y266 | NUDC_HUMAN | Oxidation (M) | 0 | 0 | 0 | 0 | 0 | 0 | 0 | 0 | 0 | 0 | 0 | 422700 | 1340300 | 0 | 0 | 0 | 0 | 0 |
| SMGLPTSD | NUDC | Q9Y266 | NUDC_HUMAN | Oxidation (M) | 0 | 0 | 0 | 0 | 0 | 142110 | 260710 | 259580 | 268700 | 0 | 0 | 0 | 0 | 659830 | 0 | 0 | 742860 | 926130 |
| SMGLPTSD | NUDC | Q9Y266 | NUDC_HUMAN | Unmodified | 0 | 0 | 0 | 0 | 1380300 | 226980 | 418900 | 298780 | 0 | 1055900 | 0 | 0 | 3143400 | 1673600 | 0 | 0 | 2535200 | 2113800 |
| SMGSQEDI | COPS3; COPS3 | Q9UNS2; K7 | CSN3_HUMAN | Unmodified | 0 | 0 | 0 | 0 | 331740 | 0 | 0 | 131330 | 80484 | 80735 | 0 | 931820 | 0 | 0 | 0 | 0 | 257970 | 719670 |
| SMGYDDLI | CTNND1; CTNN | O60716; A0A | CTND1_HUMAI | 3 Oxidation (M) | 54879 | 57067 | 138600 | 105310 | 0 | 0 | 0 | 0 | 0 | 0 | 0 | 146850 | 0 | 0 | 478130 | 458280 | 0 | 0 |
| SMHPLLYP | N/A | N/A | N/A | Oxidation (M) | 0 | 0 | 0 | 0 | 0 | 0 | 0 | 0 | 2173800 | 0 | 0 | 0 | 0 | 0 | 0 | 0 | 0 | 0 |
| SMIEISR | CCT3; CCT3; C | P49368; Q5S | TCPG_HUMAN | Unmodified | 0 | 0 | 125780 | 0 | 547750 | 0 | 711880 | 342690 | 0 | 0 | 2022000 | 8673800 | 168870 | 393170 | 0 | 0 | 269350 | 252450 |
| SMIGEPTNI | CDC42SE2 | Q9NRR3 | C42S2_HUMAI | 2 Oxidation (M) | 0 | 0 | 0 | 1986000 | 1651900 | 0 | 0 | 0 | 0 | 0 | 0 | 0 | 0 | 0 | 1484400 | 0 | 0 | 0 |
| SMIGEPTNI | CDC42SE2 | Q9NRR3 | C42S2_HUMAI | 3 Oxidation (M) | 0 | 0 | 0 | 0 | 0 | 0 | 0 | 0 | 0 | 0 | 0 | 0 | 679070 | 0 | 7814800 | 4233600 | 0 | 0 |
| SMIGSTAQ | INPPL1; INPPL1 | O15357; A0A | SHIP2_HUMAN | Oxidation (M) | 0 | 1153800 | 0 | 0 | 0 | 0 | 0 | 0 | 0 | 48203 | 281360 | 1492700 | 0 | 0 | 0 | 0 | 0 | 0 |
| SMKGAGTI | ANXA3; ANXA3; | P12429; D6I | ANXA3_HUMAI | Oxidation (M) | 0 | 0 | 0 | 0 | 272700 | 0 | 0 | 0 | 0 | 0 | 0 | 0 | 0 | 0 | 0 | 0 | 0 | 0 |
| SMLDQLG\ | PRXL2A | Q9BRX8 | PXL2A_HUMAN | Oxidation (M) | 0 | 0 | 0 | 0 | 0 | 0 | 521130 | 0 | 11432000 | 3475500 | 223390 | 0 | 0 | 0 | 0 | 0 | 0 | 0 |
| SMLDQLG\ | PRXL2A | Q9BRX8 | PXL2A_HUMAN | Unmodified | 0 | 0 | 0 | 0 | 0 | 0 | 0 | 0 | 3100700 | 0 | 0 | 0 | 0 | 0 | 0 | 0 | 0 | 0 |
| SMLEVNYP | ACTR2; ACTR2 | P61160; F5H | ARP2_HUMAN | 2 Oxidation (M) | 0 | 75007 | 0 | 0 | 0 | 0 | 0 | 0 | 61173 | 0 | 0 | 2162800 | 0 | 0 | 74070 | 47002 | 0 | 0 |
| SMLEVNYP | ACTR2; ACTR2 | P61160; F5H | ARP2_HUMAN | Oxidation (M) | 48525 | 0 | 0 | 0 | 0 | 0 | 0 | 0 | 0 | 0 | 0 | 444950 | 0 | 0 | 0 | 0 | 0 | 0 |
| SMLQATAE | ATL3; ATL3 | Q6DD88; F5 | ATLA3_HUMAN | Unmodified | 0 | 0 | 0 | 0 | 237530 | 212280 | 0 | 0 | 2471400 | 1623700 | 0 | 0 | 0 | 0 | 99687 | 122300 | 149890 | 685360 |
| SMLRLQK | RPL19; RPL19; | P84098; A0A | RL19_HUMAN | Acetyl (Protein N | 0 | 0 | 0 | 0 | 1164100 | 278760 | 940410 | 526010 | 0 | 256080 | 0 | 0 | 2103700 | 1500600 | 0 | 0 | 363970 | 1150400 |
| SMLRLQK | RPL19; RPL19; | P84098; A0A | RL19_HUMAN | Acetyl (Protein N | 0 | 0 | 146540 | 300770 | 0 | 2064300 | 712980 | 730420 | 0 | 0 | 0 | 0 | 239450 | 0 | 0 | 0 | 0 | 0 |
| SMMEAGES | PLCB4; PLCB4 | Q15147; A0A | PLCB4_HUMAI | 2 Oxidation (M) | 0 | 0 | 0 | 833900 | 0 | 0 | 0 | 0 | 0 | 0 | 0 | 0 | 0 | 0 | 830120 | 775200 | 0 | 0 |
| SMMGGGL | HSD17B4; HSD | P51659; E7E | DHB4_HUMAN | Oxidation (M) | 0 | 0 | 0 | 0 | 0 | 0 | 0 | 0 | 2650400 | 269120 | 0 | 0 | 0 | 0 | 0 | 0 | 0 | 91146 |
| SMMGGGL | HSD17B4; HSD | P51659; E7E | DHB4_HUMAN | 2 Oxidation (M) | 0 | 0 | 60572 | 0 | 0 | 0 | 37581 | 140280 | 0 | 5323800 | 512850 | 47131 | 0 | 0 | 0 | 0 | 178980 | 237250 |
| SMMNLQTI | TOP1; TOP1 | P11387; A0A | TOP1_HUMAN | Unmodified | 0 | 0 | 0 | 0 | 177920 | 0 | 0 | 37672 | 0 | 0 | 0 | 1083000 | 0 | 0 | 0 | 0 | 0 | 0 |
| SMMQDREI | MYH9; MYH9 | P35579; Q5E | MYH9_HUMAN | Unmodified | 0 | 0 | 0 | 310300 | 668430 | 0 | 310610 | 0 | 948230 | 0 | 0 | 2193300 | 268590 | 227390 | 0 | 226500 | 288510 | 476140 |
| SMMQDREI | MYH9; MYH9 | P35579; Q5E | MYH9_HUMAN | 2 Oxidation (M) | 0 | 0 | 0 | 0 | 0 | 0 | 0 | 0 | 0 | 596090 | 0 | 0 | 0 | 0 | 0 | 0 | 0 | 0 |
| SMMQDREI | MYH9; MYH9 | P35579; Q5E | MYH9_HUMAN | Oxidation (M) | 0 | 0 | 0 | 0 | 0 | 0 | 137440 | 0 | 0 | 156240 | 145890 | 1242000 | 0 | 0 | 0 | 0 | 0 | 0 |
| SMNSQYLK | CTSA; CTSA; CT | P10619; X6R | PPGB_HUMAN | Unmodified | 0 | 0 | 0 | 0 | 0 | 0 | 0 | 0 | 612070 | 207040 | 0 | 206310 | 0 | 0 | 0 | 0 | 25986 | 52080 |
| SMNSRVFI | HNRNPCL3; HN | B7ZW38; P0I | HNRC3_HUMA | Oxidation (M) | 0 | 0 | 0 | 0 | 0 | 0 | 621250 | 30495 | 0 | 0 | 0 | 0 | 0 | 0 | 0 | 0 | 0 | 0 |
| SMPIRKDD | RPL26; RPL26L | P61254; Q9U | RL26_HUMAN | Unmodified | 5292200 | 9313800 | 3452200 | 1593100 | 181750 | 0 | 0 | 0 | 495230 | 0 | 2360700 | 26329000 | 1172300 | 0 | 6074600 | 2211500 | 0 | 0 |
| SMPPLDQF | SH3GL1 | Q99961 | SH3G1_HUMAI | Oxidation (M) | 2717600 | 1480200 | 2277800 | 2025500 | 0 | 0 | 0 | 0 | 0 | 0 | 0 | 0 | 649880 | 155800 | 0 | 2711000 | 0 | 0 |
| SMPPLDQF | SH3GL1 | Q99961 | SH3G1_HUMAI | Unmodified | 3654600 | 2339800 | 1458300 | 2114800 | 0 | 0 | 50937 | 44523 | 361900 | 207530 | 134230 | 0 | 3402700 | 1520400 | 2772100 | 1478400 | 0 | 0 |
| SMPTSGAL | PGM1; PGM1 | P36871; A0A | PGM1_HUMAN | Unmodified | 0 | 0 | 0 | 0 | 0 | 0 | 0 | 0 | 0 | 0 | 0 | 0 | 0 | 0 | 0 | 0 | 1119000 | 3663500 |
| SMPWNVD | CDC37; CDC37 | Q16543; K7E | CDC37_HUMA | Unmodified | 0 | 0 | 0 | 0 | 2536700 | 0 | 547390 | 145720 | 0 | 195910 | 0 | 0 | 1220700 | 487420 | 0 | 0 | 1843500 | 2808600 |
| SMPWNVD | CDC37; CDC37 | Q16543; K7E | CDC37_HUMA | Oxidation (M) | 0 | 0 | 0 | 0 | 0 | 0 | 0 | 1572200 | 0 | 0 | 0 | 0 | 98966 | 0 | 0 | 0 | 1440800 | 0 |
| SMPWNVD | CDC37; CDC37 | Q16543; K7E | CDC37_HUMA | Unmodified | 117000 | 539570 | 905190 | 387830 | 0 | 0 | 214860 | 0 | 0 | 0 | 0 | 0 | 0 | 178350 | 0 | 0 | 1346700 | 0 |
| SMQDATQI | CGN | Q9P2M7 | CING_HUMAN | Unmodified | 0 | 0 | 0 | 0 | 218080 | 0 | 0 | 0 | 0 | 0 | 74342 | 402270 | 94616 | 0 | 0 | 0 | 0 | 0 |
| SMQNHAA | SDHA; SDHA | P31040; A0A | SDHA_HUMAN | Unmodified | 0 | 0 | 0 | 0 | 0 | 0 | 0 | 0 | 0 | 0 | 0 | 0 | 0 | 0 | 0 | 0 | 0 | 0 |
| SMQNWH( | CNN2; CNN2; ( | Q99439; H3I | CNN2_HUMAN | Oxidation (M) | 197310 | 244540 | 389060 | 557650 | 6380600 | 403490 | 176820 | 62431 | 526650 | 405980 | 0 | 146100 | 633180 | 473600 | 699840 | 485830 | 630670 | 455340 |
| SMQNWH( | CNN2; CNN2; ( | Q99439; H3I | CNN2_HUMAN | Unmodified | 0 | 0 | 0 | 0 | 1289400 | 81685 | 0 | 0 | 0 | 0 | 0 | 0 | 0 | 0 | 0 | 0 | 0 | 184980 |
| SMQPGTSL | TRMT10C | Q7L0Y3 | TM10C_HUMAI | Unmodified | 0 | 0 | 0 | 0 | 0 | 0 | 0 | 0 | 738460 | 315660 | 0 | 0 | 0 | 0 | 0 | 0 | 0 | 0 |
| SMRGGYIG | UBE2M | P61081 | UBC12_HUMAI | Oxidation (M) | 0 | 0 | 0 | 128770 | 0 | 0 | 100720 | 78042 | 0 | 889200 | 0 | 0 | 1294400 | 0 | 0 | 0 | 645590 | 0 |
| SMSDPDQI | CDC42BPB | Q9Y5S2 | MRCKB_HUMA | Unmodified | 0 | 0 | 0 | 0 | 0 | 0 | 0 | 0 | 166040 | 0 | 0 | 311760 | 0 | 0 | 0 | 0 | 0 | 0 |
| SMSTEGLM | CS; CS; CS | O75390; A0A | CISY_HUMAN ( | Unmodified | 0 | 0 | 0 | 0 | 198760 | 0 | 0 | 0 | 3846200 | 397130 | 0 | 0 | 0 | 0 | 0 | 0 | 0 | 0 |

| | | | | | | | | | | | | | | | | | | | | | | |
|---|---|---|---|---|---|---|---|---|---|---|---|---|---|---|---|---|---|---|---|---|---|---|
| SMSTEGLM | CS; CS; CS | O75390; A0A | CISY_HUMAN ( | Oxidation (M) | 0 | 0 | 0 | 0 | 0 | 1582000 | 82359 | 0 | 2161100 | 175600 | 0 | 0 | 0 | 0 | 0 | 0 | 0 | 0 |
| SMSVDETD | COBLL1; COBL | Q53SF7; A0A | COBL1_HUMA | Unmodified | 285930 | 0 | 0 | 0 | 0 | 0 | 0 | 0 | 0 | 0 | 0 | 0 | 0 | 0 | 0 | 0 | 0 | 0 |
| SMSVYCTP | ATP2A2; ATP2A | P16615; H7C | AT2A2_HUMAN | Unmodified | 0 | 0 | 58861 | 83770 | 52621 | 0 | 271030 | 126160 | 387060 | 0 | 62084 | 259830 | 0 | 0 | 54719 | 27352 | 0 | 0 |
| SMTEAEQC | CKB; CKB | P12277; G3V | KCRB_HUMAN | Oxidation (M) | 0 | 0 | 1600000 | 0 | 0 | 0 | 408900 | 79547 | 0 | 0 | 0 | 1428200 | 0 | 0 | 980690 | 393700 | 0 | 0 |
| SMTEAEQC | CKB; CKB | P12277; G3V | KCRB_HUMAN | 2 Oxidation (M) | 2828500 | 2897100 | 3290400 | 22067000 | 4656100 | 3058900 | 3891000 | 0 | 12375000 | 7420100 | 1034400 | 21078000 | 9845000 | 1519100 | 15183000 | 9667900 | 0 | 0 |
| SMTEAEQC | CKB; CKB | P12277; G3V | KCRB_HUMAN | Oxidation (M) | 219960 | 484240 | 177100 | 2587100 | 0 | 121970 | 983110 | 0 | 22207000 | 0 | 193340 | 2569700 | 0 | 316410 | 3821700 | 525740 | 0 | 0 |
| SMTEAEQC | CKB; CKB | P12277; G3V | KCRB_HUMAN | Unmodified | 0 | 0 | 0 | 0 | 0 | 0 | 0 | 0 | 1952400 | 0 | 0 | 68509 | 92643 | 0 | 0 | 0 | 0 | 0 |
| SMTLEIR | SDHA; SDHA; S | P31040; D6F | SDHA_HUMAN | Unmodified | 0 | 0 | 0 | 0 | 0 | 0 | 0 | 0 | 2248800 | 0 | 0 | 0 | 0 | 0 | 0 | 0 | 0 | 0 |
| SMTPAQAD | EPB41; EPB41; | P11171; A0A | EPB41_HUMAN | Oxidation (M) | 0 | 238210 | 520780 | 854290 | 0 | 0 | 0 | 160100 | 0 | 242990 | 0 | 1877000 | 0 | 400300 | 0 | 897320 | 0 | 0 |
| SMTPAQAD | EPB41; EPB41; | P11171; A0A | EPB41_HUMAN | Unmodified | 0 | 0 | 0 | 666030 | 0 | 0 | 0 | 0 | 833820 | 0 | 0 | 0 | 0 | 0 | 0 | 0 | 0 | 0 |
| SMVAVMDS | CAPNS1; CAPN | P04632; A0A | CPNS1_HUMAI | Unmodified | 924010 | 72184 | 0 | 0 | 0 | 0 | 0 | 0 | 241700 | 0 | 0 | 828620 | 0 | 0 | 0 | 44603 | 0 | 0 |
| SMVAVMDS | CAPNS1; CAPN | P04632; A0A | CPNS1_HUMAI | Oxidation (M) | 517120 | 0 | 49119 | 0 | 0 | 0 | 0 | 0 | 356200 | 0 | 187880 | 702220 | 0 | 0 | 80422 | 106540 | 0 | 0 |
| SMVEDITGI | DSP | P15924 | DESP_HUMAN | Unmodified | 0 | 0 | 0 | 0 | 0 | 0 | 0 | 0 | 689080 | 0 | 0 | 239940 | 0 | 0 | 0 | 0 | 0 | 0 |
| SMVEQLDK | BCCIP | Q9P287 | BCCIP_HUMAN | Oxidation (M) | 19716 | 0 | 0 | 0 | 54386 | 42055 | 32248 | 17249 | 0 | 0 | 0 | 287270 | 54197 | 18248 | 0 | 0 | 0 | 0 |
| SMVSTIMKI | BIRC6; BIRC6 | Q9NR09; A0 | BIRC6_HUMAN | Unmodified | 79106 | 93669 | 101330 | 96341 | 0 | 0 | 171150 | 0 | 0 | 83442 | 205050 | 204100 | 0 | 0 | 278260 | 67381 | 0 | 0 |
| SMYEEEINI | LMNB1; LMNB1 | P20700; A0A | LMNB1_HUMAI | Oxidation (M) | 0 | 0 | 0 | 0 | 0 | 0 | 0 | 0 | 0 | 0 | 0 | 597670 | 0 | 0 | 0 | 0 | 0 | 0 |
| SMYLDIEYA | NCAM2; NCAM | O15394; H9I | NCAM2_HUMA | Oxidation (M) | 393600 | 0 | 0 | 1801100 | 0 | 0 | 0 | 295570 | 0 | 0 | 0 | 0 | 0 | 125560 | 2518100 | 1283300 | 0 | 0 |
| SNAAAYLQ | CTNND1; CTNN | O60716; E9F | CTND1_HUMAI | Unmodified | 518360 | 210730 | 61778 | 0 | 158370 | 473810 | 0 | 0 | 0 | 33156 | 0 | 6145800 | 0 | 0 | 6061700 | 3715500 | 236500 | 166430 |
| SNAAAYLQ | CTNND1; CTNN | O60716; E9F | CTND1_HUMAI | Unmodified | 0 | 0 | 2168300 | 7905000 | 39875000 | 7587900 | 0 | 0 | 0 | 391480 | 4821400 | 7098200 | 31087000 | 30947000 | 5644300 | 3419000 | 26080000 | 48394000 |
| SNAAAYLQ | CTNND1; CTNN | O60716; E9F | CTND1_HUMAI | Unmodified | 0 | 0 | 277880 | 1111300 | 0 | 3251800 | 0 | 0 | 0 | 0 | 0 | 0 | 1768900 | 894310 | 100330 | 0 | 4854200 | 0 |
| SNAEESSL | DCUN1D3 | Q8IWE4 | DCNL3_HUMA | Unmodified | 57748 | 60273 | 196290 | 279750 | 390960 | 0 | 0 | 0 | 329790 | 54768 | 0 | 233260 | 0 | 0 | 385960 | 199580 | 241590 | 493250 |
| SNAGKINY | UGGT1 | Q9NYU2 | UGGG1_HUMA | Unmodified | 0 | 0 | 0 | 0 | 0 | 0 | 0 | 0 | 0 | 602420 | 0 | 0 | 0 | 0 | 0 | 0 | 0 | 0 |
| SNAHYNLC | SPTBN1; SPTBN | Q01082; A0A | SPTB2_HUMAN | Unmodified | 0 | 0 | 0 | 0 | 2016500 | 0 | 0 | 0 | 0 | 453150 | 0 | 5037700 | 0 | 0 | 0 | 0 | 0 | 1258200 |
| SNALDIIFQ | C1S; C1S; C1S; | P09871; A0A | C1S_HUMAN C | Unmodified | 0 | 0 | 0 | 13414 | 34883 | 43969 | 0 | 0 | 0 | 0 | 0 | 0 | 0 | 0 | 0 | 0 | 0 | 112730 |
| SNAQGIDL | CPE; CPE; CPE; | P16870; C9J | CBPE_HUMAN | Unmodified | 0 | 0 | 0 | 0 | 0 | 0 | 0 | 0 | 46403000 | 605660 | 0 | 0 | 425780 | 0 | 0 | 0 | 0 | 0 |
| SNAQGIDL | CPE; CPE; CPE; | P16870; C9J | CBPE_HUMAN | Unmodified | 0 | 0 | 0 | 0 | 0 | 0 | 309070 | 0 | 4967800 | 233260 | 0 | 0 | 0 | 0 | 0 | 0 | 0 | 0 |
| SNAQGIDL | CPE; CPE; CPE; | P16870; C9J | CBPE_HUMAN | Unmodified | 0 | 0 | 0 | 0 | 59620 | 0 | 264430 | 0 | 1.46E+08 | 29961000 | 0 | 0 | 883310 | 0 | 0 | 0 | 0 | 0 |
| SNAQRQDI | ANXA2; ANXA2; | P07355; H0Y | ANXA2_HUMAI | Unmodified | 773430 | 636900 | 71159 | 25593 | 12400000 | 4949600 | 0 | 0 | 40058000 | 4778000 | 871160 | 376450 | 0 | 0 | 62418 | 65374 | 1951500 | 2425700 |
| SNAQRQDI | ANXA2; ANXA2; | P07355; H0Y | ANXA2_HUMAI | Unmodified | 116420 | 82906 | 0 | 0 | 4862100 | 3140300 | 0 | 0 | 6533500 | 2391700 | 0 | 0 | 0 | 0 | 0 | 0 | 972880 | 667840 |
| SNCRYEIE | IGF2R; IGF2R; I | P11717; A0A | MPRI_HUMAN ( | Unmodified | 0 | 0 | 0 | 0 | 0 | 0 | 0 | 170760 | 12066000 | 4692600 | 1005900 | 0 | 0 | 0 | 0 | 0 | 0 | 672700 |
| SNCVLEAF | MYO1D; MYO1D | O94832; J3Q | MYO1D_HUMA | Unmodified | 5530800 | 0 | 1761400 | 1948700 | 941610 | 0 | 0 | 0 | 1212600 | 490480 | 302850 | 976760 | 1063900 | 1100300 | 3020500 | 1807100 | 1207500 | 4590400 |
| SNDEEAFT | CBS; CBSL | P35520; P0D | CBS_HUMAN C | Unmodified | 0 | 0 | 0 | 0 | 163000 | 0 | 0 | 0 | 331380 | 1007500 | 2092400 | 11880000 | 1206200 | 153710 | 0 | 0 | 0 | 0 |
| SNDLFPVH | UBE2O; UBE2O | Q9C0C9; K7 | UBE2O_HUMAI | Unmodified | 0 | 0 | 0 | 0 | 0 | 0 | 0 | 0 | 0 | 0 | 0 | 3324400 | 0 | 0 | 0 | 0 | 0 | 0 |
| SNDRNSMI | KIF23; KIF23; KI | Q02241; A0A | KIF23_HUMAN | Oxidation (M) | 89954 | 0 | 0 | 507430 | 0 | 0 | 2860000 | 0 | 333730 | 0 | 491720 | 0 | 0 | 0 | 0 | 0 | 0 | 0 |
| SNDRNSMI | KIF23; KIF23; KI | Q02241; A0A | KIF23_HUMAN | Unmodified | 0 | 1385100 | 0 | 0 | 180480 | 0 | 0 | 0 | 939530 | 0 | 0 | 0 | 0 | 0 | 0 | 0 | 0 | 0 |
| SNEEGSEE | SFN | P31947 | 1433S_HUMAN | Unmodified | 3148900 | 0 | 0 | 0 | 2645900 | 2356400 | 0 | 0 | 19960000 | 1531300 | 0 | 0 | 78217 | 0 | 0 | 0 | 1143100 | 1273600 |
| SNEHDDC | IGF2R | P11717 | MPRI_HUMAN ( | Unmodified | 0 | 0 | 619730 | 0 | 0 | 0 | 0 | 0 | 55351000 | 3941400 | 3266400 | 5671100 | 0 | 0 | 442370 | 547630 | 0 | 0 |
| SNEHDDC | IGF2R | P11717 | MPRI_HUMAN ( | Unmodified | 0 | 0 | 0 | 0 | 0 | 0 | 0 | 0 | 0 | 3569400 | 0 | 0 | 0 | 0 | 0 | 0 | 0 | 0 |
| SNEILTAIIQ | KPNB1; KPNB1 | Q14974; J3Q | IMB1_HUMAN I | Unmodified | 26797 | 48190 | 0 | 42402 | 14821 | 0 | 0 | 0 | 1516600 | 11208 | 113070 | 1024200 | 0 | 0 | 40618 | 63804 | 0 | 0 |
| SNEILTAIIQ | KPNB1; KPNB1 | Q14974; J3Q | IMB1_HUMAN I | Oxidation (M) | 224230 | 209800 | 160680 | 194400 | 0 | 27414 | 54477 | 0 | 315770 | 76025 | 472520 | 1644300 | 50505 | 0 | 313820 | 386660 | 0 | 78610 |
| SNEILTAIIQ | KPNB1; KPNB1 | Q14974; J3Q | IMB1_HUMAN I | Unmodified | 0 | 0 | 0 | 0 | 5608000 | 1561200 | 881010 | 0 | 1462000 | 1361500 | 1224600 | 0 | 2026000 | 1389000 | 0 | 0 | 1808700 | 1974700 |
| SNELGDVC | RNH1; RNH1; R | P13489; E9P | RINI_HUMAN R | Unmodified | 0 | 0 | 0 | 0 | 0 | 0 | 0 | 0 | 0 | 0 | 300460 | 1905700 | 0 | 0 | 0 | 0 | 0 | 0 |
| SNETNGYL | SLC1A4 | P43007 | SATT_HUMAN I | Unmodified | 0 | 459600 | 0 | 0 | 0 | 0 | 0 | 166860 | 0 | 0 | 0 | 14892000 | 832830 | 0 | 673690 | 299530 | 0 | 0 |
| SNEYQLID | GNAS; GNAS; G | Q5JWF2; P6 | GNAS1_HUMAI | Unmodified | 0 | 92067 | 221040 | 273590 | 0 | 0 | 0 | 0 | 0 | 0 | 0 | 910290 | 0 | 0 | 2529100 | 954160 | 0 | 0 |
| SNFAEALA | P4HB; P4HB; P4 | P07237; A0A | PDIA1_HUMAN | Unmodified | 84069 | 244680 | 0 | 0 | 274750 | 0 | 0 | 0 | 916850 | 807780 | 0 | 0 | 0 | 0 | 0 | 37871 | 0 | 0 |
| SNFDSTLIS | TYRP1 | P17643 | TYRP1_HUMAN | Unmodified | 0 | 0 | 0 | 0 | 0 | 0 | 0 | 0 | 4193400 | 169850 | 102220 | 0 | 0 | 0 | 0 | 0 | 0 | 0 |
| SNFEPFFM | GNS; GNS; GNS | P15586; H7C | GNS_HUMAN N | 2 Oxidation (M) | 0 | 0 | 0 | 0 | 0 | 0 | 0 | 0 | 1106700 | 0 | 0 | 0 | 0 | 0 | 0 | 0 | 0 | 0 |
| SNFEPFFM | GNS; GNS; GNS | P15586; H7C | GNS_HUMAN N | Unmodified | 0 | 0 | 0 | 0 | 0 | 0 | 0 | 0 | 465320 | 0 | 0 | 0 | 0 | 0 | 0 | 0 | 0 | 0 |
| SNFGYNIPI | PSMA3; PSMA3 | P25788; G3V | PSA3_HUMAN | Unmodified | 0 | 0 | 0 | 0 | 0 | 0 | 0 | 54858 | 569880 | 0 | 0 | 13203000 | 0 | 528320 | 0 | 556310 | 0 | 0 |
| SNFKPSLL | AP2M1; AP2M1 | Q96CW1; C9 | AP2M1_HUMAI | Unmodified | 0 | 0 | 0 | 0 | 414760 | 392450 | 95643 | 0 | 0 | 0 | 0 | 2178500 | 0 | 0 | 178430 | 189400 | 250420 | 613340 |
| SNFPARAA | RIDA; RIDA; RID | P52758; H0Y | RIDA_HUMAN 2 | Unmodified | 0 | 0 | 91503 | 0 | 975650 | 184290 | 1082600 | 281440 | 0 | 1576200 | 0 | 0 | 0 | 204140 | 0 | 0 | 1900200 | 749050 |
| SNFSNSAD | CBX5 | P45973 | CBX5_HUMAN | Unmodified | 0 | 0 | 0 | 0 | 0 | 0 | 154180 | 433540 | 328600 | 490650 | 0 | 382790 | 149520 | 0 | 0 | 0 | 0 | 210610 |
| SNFSNSAD | CBX5 | P45973 | CBX5_HUMAN | Unmodified | 0 | 0 | 151460 | 58866 | 0 | 7326.7 | 89920 | 242540 | 99034 | 228890 | 0 | 649850 | 31097 | 0 | 0 | 0 | 63527 | 0 |
| SNFVPTNV | CALCB; CALCB | P10092; E7E | CALCB_HUMAI | Unmodified | 0 | 0 | 0 | 97407 | 541640 | 137860 | 0 | 223070 | 953340 | 1012500 | 0 | 0 | 0 | 0 | 0 | 0 | 0 | 497230 |
| SNGAYNIF | NID1 | P14543 | NID1_HUMAN | Unmodified | 0 | 0 | 0 | 0 | 0 | 250410 | 0 | 76136 | 693170 | 0 | 0 | 39620 | 0 | 0 | 0 | 0 | 0 | 0 |
| SNGDLSPK | MARCKSL1 | P49006 | MRP_HUMAN N | Unmodified | 0 | 366960 | 2864000 | 2246200 | 0 | 0 | 0 | 0 | 0 | 0 | 2951300 | 19214000 | 1477400 | 995650 | 3425500 | 2274200 | 1407000 | 3692600 |
| SNGDLSPK | MARCKSL1 | P49006 | MRP_HUMAN N | Unmodified | 0 | 0 | 0 | 1837100 | 0 | 0 | 0 | 0 | 0 | 0 | 0 | 5509400 | 1227200 | 0 | 1784500 | 0 | 0 | 0 |
| SNGEFWD | CD109 | Q6YHK3 | CD109_HUMAI | Unmodified | 0 | 0 | 0 | 402380 | 295230 | 0 | 0 | 0 | 735380 | 0 | 0 | 0 | 227120 | 228770 | 0 | 97537 | 0 | 0 |
| SNGEFWD | CD109 | Q6YHK3 | CD109_HUMAI | Unmodified | 0 | 0 | 0 | 0 | 0 | 0 | 0 | 0 | 1682600 | 222630 | 0 | 0 | 0 | 43600 | 0 | 0 | 0 | 0 |
| SNGLICGG | ITGB1; ITGB1; I | P05556; H7C | ITB1_HUMAN I | Unmodified | 314170 | 779070 | 368320 | 233970 | 0 | 0 | 0 | 0 | 1093400 | 0 | 72452 | 0 | 176570 | 0 | 377210 | 364030 | 0 | 421870 |
| SNGWILPT | AKR7A2 | O43488 | ARK72_HUMAN | Oxidation (M) | 338410 | 0 | 0 | 0 | 0 | 371450 | 0 | 0 | 187910 | 0 | 0 | 325390 | 0 | 0 | 0 | 0 | 55906 | 0 |
| SNHYDPEE | TJP1; TJP1; TJP1 | Q07157; A0A | ZO1_HUMAN T | Unmodified | 0 | 0 | 57588 | 278610 | 0 | 0 | 0 | 0 | 107540 | 0 | 0 | 0 | 358710 | 0 | 127440 | 110330 | 0 | 0 |
| SNIDALLSF | RALY; RALY; RA | Q9UKM9; Q5 | RALY_HUMAN | Unmodified | 0 | 0 | 0 | 269130 | 0 | 0 | 0 | 0 | 0 | 1090300 | 0 | 0 | 460260 | 0 | 0 | 0 | 0 | 0 |
| SNIHCNTI | HSD17B4; HSD | P51659; A0A | DHB4_HUMAN | Unmodified | 0 | 0 | 0 | 0 | 0 | 0 | 0 | 0 | 1766600 | 82817 | 122370 | 0 | 0 | 0 | 0 | 0 | 0 | 0 |
| SNIHSSVM | ANLN; ANLN; A | Q9NQW6; H | ANLN_HUMAN | Unmodified | 0 | 0 | 0 | 0 | 781850 | 127960 | 0 | 0 | 0 | 0 | 0 | 0 | 0 | 0 | 0 | 0 | 0 | 0 |
| SNILERNK | CAPG; CAPG; C | P40121; E7E | CAPG_HUMAN | Unmodified | 0 | 0 | 0 | 0 | 1857600 | 0 | 0 | 0 | 0 | 0 | 0 | 0 | 0 | 0 | 0 | 0 | 0 | 930010 |
| SNILWDK | OCLN; OCLN; ( | Q16625; A0A | OCLN_HUMAN | Unmodified | 669970 | 1440300 | 0 | 2649800 | 923280 | 0 | 0 | 0 | 2748300 | 0 | 0 | 2505400 | 1020100 | 0 | 6101200 | 3624700 | 81608 | 227670 |
| SNILWDKE | OCLN; OCLN; ( | Q16625; A0A | OCLN_HUMAN | Unmodified | 0 | 0 | 0 | 0 | 2692100 | 0 | 0 | 0 | 489980 | 1688400 | 0 | 0 | 3997400 | 3744000 | 0 | 0 | 0 | 0 |
| SNIMWLDF | IGSF3 | O75054 | IGSF3_HUMAN | Oxidation (M) | 0 | 963280 | 0 | 0 | 0 | 0 | 0 | 0 | 0 | 0 | 0 | 0 | 0 | 0 | 1322700 | 882770 | 0 | 0 |
| SNITIDPDV | FRYL; FRYL; FRY | O94915; F22 | FRYL_HUMAN | Acetyl (Protein N | 0 | 0 | 0 | 0 | 669490 | 83563 | 0 | 125870 | 722650 | 238770 | 134530 | 1018200 | 209990 | 60702 | 0 | 0 | 0 | 140490 |
| SNIVTVGD | UTRN | P46939 | UTRN_HUMAN | Unmodified | 0 | 0 | 751270 | 2045000 | 0 | 0 | 0 | 0 | 0 | 0 | 0 | 0 | 350590 | 0 | 1236700 | 1172600 | 0 | 0 |
| SNIVTVGD | UTRN | P46939 | UTRN_HUMAN | Unmodified | 0 | 0 | 0 | 0 | 1988900 | 0 | 0 | 0 | 0 | 1322800 | 0 | 0 | 1782600 | 0 | 0 | 0 | 1107000 | 1802700 |
| SNIWLAAV | AP2M1; AP2M1 | Q96CW1; C9 | AP2M1_HUMAI | Unmodified | 102510 | 125760 | 95707 | 45243 | 0 | 0 | 0 | 0 | 0 | 0 | 0 | 2025900 | 0 | 0 | 298000 | 313190 | 0 | 0 |
| SNKDGGN | PRKAR2A; PRK | P13861; H7C | KAP2_HUMAN | Unmodified | 0 | 0 | 0 | 135860 | 1134200 | 0 | 34091 | 75731 | 285880 | 197750 | 0 | 107600 | 2907900 | 1625800 | 42908 | 57528 | 171510 | 921890 |
| SNKEGSG | TSNAX; TSNAX- | Q99598; Q9I | TSNAX_HUMAI | Acetyl (Protein N | 0 | 151300 | 344530 | 0 | 839560 | 842030 | 932970 | 4311700 | 0 | 155560 | 11394 | 0 | 579320 | 419150 | 0 | 93347 | 1706800 | 1140400 |
| SNLAQVEH | PLEKHA7; PLEK | Q6IQ23; H0Y | PKHA7_HUMAI | Unmodified | 0 | 0 | 0 | 351500 | 64905 | 0 | 0 | 0 | 0 | 0 | 0 | 163160 | 230430 | 213220 | 247280 | 124740 | 0 | 0 |
| SNLAYDIVC | HSPA4; HSPA4 | P34932; A0A | HSP74_HUMAN | Unmodified | 108630 | 66967 | 59232 | 172080 | 1963800 | 364210 | 617800 | 25158 | 3942000 | 1120600 | 5192500 | 10922000 | 3208200 | 888770 | 261900 | 296440 | 569340 | 584250 |
| SNLEMEIEK | CD2AP | Q9Y5K6 | CD2AP_HUMAI | Unmodified | 5058000 | 4595200 | 3885400 | 5065200 | 0 | 0 | 0 | 0 | 0 | 0 | 0 | 0 | 1317100 | 0 | 7764500 | 4132400 | 0 | 0 |
| SNLEMEIEK | CD2AP | Q9Y5K6 | CD2AP_HUMAI | Oxidation (M) | 4972600 | 2903200 | 3515600 | 3428300 | 0 | 0 | 0 | 0 | 0 | 250150 | 0 | 512430 | 584220 | 133000 | 4901000 | 3632100 | 0 | 0 |
| SNLGSVVL | DHX15 | O43143 | DHX15_HUMAI | Unmodified | 0 | 0 | 0 | 0 | 0 | 0 | 0 | 0 | 74456 | 0 | 0 | 1960700 | 0 | 0 | 0 | 0 | 0 | 0 |
| SNLIQQWL | CD109 | Q6YHK3 | CD109_HUMAI | Unmodified | 497220 | 418330 | 0 | 0 | 140500 | 0 | 0 | 0 | 6002200 | 2568500 | 0 | 0 | 0 | 0 | 0 | 0 | 1210800 | 878740 |
| SNLISGSVI | PCYOX1; PCYO | Q9UHG3; F8 | PCYOX_HUMAI | Unmodified | 0 | 0 | 0 | 0 | 0 | 0 | 0 | 0 | 2819500 | 279370 | 0 | 0 | 36116 | 0 | 0 | 0 | 0 | 0 |
| SNLISGSVI | PCYOX1; PCYO | Q9UHG3; F8 | PCYOX_HUMAI | Oxidation (M) | 0 | 0 | 0 | 0 | 0 | 0 | 0 | 0 | 2728400 | 2296000 | 0 | 0 | 0 | 0 | 0 | 0 | 0 | 0 |
| SNLLSRFTI | RAB11A; RAB11 | P62491; Q15 | RB11A_HUMAN | Unmodified | 0 | 0 | 0 | 180060 | 80545000 | 9137000 | 11374000 | 73379 | 0 | 1377500 | 0 | 11471000 | 7441200 | 0 | 0 | 0 | 3834400 | 8829200 |
| SNLLTQDN | NOMO1; NOMC | Q15155; P69 | NOMO1_HUMA | 2 Oxidation (M) | 0 | 0 | 0 | 0 | 0 | 0 | 0 | 0 | 3115500 | 1691200 | 0 | 0 | 0 | 0 | 0 | 0 | 0 | 0 |
| SNLLWDT( | APOBEC3B; AP( | Q9UH17; Q9 | ABC3B_HUMAI | Unmodified | 0 | 0 | 982160 | 0 | 0 | 0 | 0 | 0 | 0 | 0 | 0 | 0 | 140100 | 0 | 2287000 | 1088100 | 0 | 0 |
| SNLNSLDE | GGCT; N/A; GG | O75223; M0 | GGCT_HUMAN | Unmodified | 492970 | 288140 | 671180 | 634130 | 712380 | 0 | 251270 | 33649 | 1854200 | 1811200 | 889640 | 2988600 | 2874700 | 1595600 | 1201500 | 1137100 | 534550 | 69806 |
| SNLPDNLK | DYNC1H1; DYN | Q14204; A0A | DYHC1_HUMA | Unmodified | 0 | 40668 | 947950 | 53518 | 344510 | 240650 | 1434000 | 623500 | 0 | 129320 | 130160 | 2539100 | 0 | 57314 | 153330 | 139730 | 225710 | 508020 |
| SNLPTDAS | PREX1 | Q8TCU6 | PREX1_HUMAN | Unmodified | 0 | 0 | 0 | 0 | 0 | 0 | 0 | 0 | 0 | 0 | 0 | 548830 | 0 | 0 | 0 | 0 | 0 | 0 |
| SNLSDRYC | VPS37C; VPS37 | A5D8V6; F5H | VP37C_HUMAN | Unmodified | 6737700 | 3432000 | 4347900 | 1725300 | 0 | 0 | 0 | 43451 | 639590 | 860250 | 0 | 0 | 2254100 | 0 | 5717200 | 6328900 | 0 | 0 |
| SNLVDNTN | DDX19B; DDX1 | Q9UMR2; Q9 | DD19B_HUMAN | Unmodified | 860180 | 623470 | 305250 | 447420 | 1028900 | 0 | 0 | 258080 | 375920 | 0 | 806220 | 1724300 | 0 | 0 | 252850 | 419030 | 0 | 787180 |
| SNLVISDPI | BAIAP2; BAIAP2 | Q9UQB8; I3L | BAIP2_HUMAN | Unmodified | 506920 | 532140 | 188090 | 11993000 | 20885000 | 3319700 | 0 | 0 | 6378000 | 1888600 | 0 | 2356500 | 11788000 | 9379900 | 3581100 | 2602500 | 3845800 | 4655400 |
| SNMEKHLF | CHMP1B | Q7LBR1 | CHM1B_HUMA | Acetyl (Protein N | 0 | 0 | 0 | 0 | 1580600 | 0 | 225870 | 54561 | 0 | 0 | 0 | 0 | 0 | 0 | 0 | 0 | 221900 | 0 |
| SNMGHPEI | FASN; FASN | P49327; A0A | FAS_HUMAN F | Unmodified | 0 | 168870 | 2014900 | 1779600 | 1927600 | 495190 | 1697200 | 338930 | 792770 | 699780 | 1764800 | 31009000 | 0 | 1181900 | 1272800 | 1066400 | 1992600 | 520290 |
| SNMGHPEI | FASN; FASN | P49327; A0A | FAS_HUMAN F | Unmodified | 213600 | 0 | 641360 | 860040 | 2025000 | 854720 | 939430 | 83893 | 732790 | 295120 | 989770 | 17609000 | 808290 | 538240 | 618610 | 291060 | 3312600 | 210510 |
| SNMSPDKI | SLC4A7 | H7C3C4 | H7C3C4_HUM | Unmodified | 0 | 0 | 0 | 0 | 1085500 | 0 | 0 | 29879 | 585900 | 122700 | 477070 | 1783800 | 0 | 182850 | 227810 | 360750 | 276380 | 1258600 |
| SNNIINETT | SMC4; SMC4 | Q9NTJ3; E9F | SMC4_HUMAN | Unmodified | 0 | 0 | 0 | 0 | 0 | 0 | 0 | 0 | 0 | 0 | 0 | 464550 | 0 | 0 | 0 | 0 | 0 | 0 |
| SNNSETSS | RGPD3; RGPD8 | A6NKT7; O14 | RGPD3_HUMAI | Unmodified | 0 | 0 | 0 | 0 | 0 | 0 | 0 | 0 | 0 | 0 | 202490 | 1473200 | 364670 | 0 | 0 | 20768 | 0 | 141790 |
| SNNSMAQ | CHMP2A; CHM | O43633; M0 | CHM2A_HUMA | Unmodified | 1741500 | 2277500 | 1860100 | 2506200 | 12337000 | 2236000 | 409180 | 197650 | 12976000 | 3279900 | 2464800 | 13051000 | 4517200 | 2134500 | 1882400 | 1491700 | 557280 | 2679100 |
| SNNSMAQ | CHMP2A; CHM | O43633; M0 | CHM2A_HUMA | Oxidation (M) | 421120 | 128350 | 541210 | 235660 | 820480 | 286500 | 51634 | 9992.5 | 732870 | 608640 | 579540 | 1694300 | 372270 | 318060 | 207610 | 612510 | 52329 | 240720 |
| SNNSMAQ | CHMP2A; CHM | O43633; M0 | CHM2A_HUMA | 2 Oxidation (M) | 0 | 0 | 104680 | 0 | 199610 | 0 | 0 | 0 | 309050 | 72087 | 0 | 415440 | 264580 | 0 | 0 | 0 | 0 | 80427 |
| SNNSMAQ | CHMP2A; CHM | O43633; M0 | CHM2A_HUMA | Unmodified | 0 | 0 | 0 | 0 | 7309000 | 486240 | 998300 | 420430 | 460810 | 1729500 | 88969 | 0 | 3265400 | 6836700 | 0 | 0 | 1346300 | 2668200 |
| SNNSMAQ | CHMP2A; CHM | O43633; M0 | CHM2A_HUMA | Oxidation (M) | 0 | 0 | 0 | 0 | 1041500 | 112020 | 587400 | 449580 | 47904 | 500090 | 0 | 0 | 0 | 986190 | 12540 | 0 | 0 | 0 |
| SNPEDQIL | MLEC; MLEC; M | Q14165; F5C | MLEC_HUMAN | Unmodified | 216990 | 194310 | 0 | 0 | 0 | 0 | 0 | 0 | 11924000 | 1062800 | 0 | 0 | 0 | 0 | 405110 | 0 | 0 | 0 |
| SNPFAHLA | ACO1 | P21399 | ACOC_HUMAN | Acetyl (Protein N | 0 | 0 | 0 | 0 | 1464400 | 0 | 531000 | 0 | 0 | 16330 | 0 | 1163300 | 0 | 0 | 0 | 0 | 428100 | 684340 |
| SNPFAHLA | ACO1 | P21399 | ACOC_HUMAN | Acetyl (Protein N | 0 | 0 | 0 | 0 | 3589300 | 0 | 2751100 | 0 | 0 | 0 | 0 | 0 | 1481200 | 0 | 0 | 0 | 1583300 | 1604700 |
| SNPGDFTL | PTPN11; PTPN1 | Q06124; A0A | PTN11_HUMAN | Unmodified | 206450 | 136660 | 127610 | 0 | 0 | 0 | 0 | 0 | 0 | 0 | 0 | 2175800 | 0 | 0 | 126150 | 220580 | 0 | 0 |
| SNPGHYLC | IDE; IDE; IDE; ID | P14735; A0A | IDE_HUMAN In | Unmodified | 53792 | 30848 | 0 | 458040 | 1332700 | 365060 | 138080 | 0 | 0 | 163250 | 1243700 | 6898600 | 894690 | 0 | 378780 | 0 | 0 | 282120 |
| SNPSAPPP | TMBIM1; TMBIM | Q969X1; C9J | LFG3_HUMAN | Acetyl (Protein N | 0 | 0 | 129990 | 0 | 0 | 0 | 0 | 0 | 0 | 0 | 0 | 0 | 0 | 0 | 0 | 196990 | 0 | 0 |

| | | | | | | | | | | | | | | | | | | | | | | |
|---|---|---|---|---|---|---|---|---|---|---|---|---|---|---|---|---|---|---|---|---|---|---|
| SNPSAVAG | TJP2; TJP2; TJP2 | Q9UDY2; A0 | ZO2_HUMAN T | Unmodified | 0 | 19181 | 0 | 287700 | 4121900 | 1287900 | 0 | 0 | 3639300 | 3193500 | 525640 | 1114900 | 872130 | 375530 | 177790 | 0 | 897360 | 935070 |
| SNPSENEE | DNAJC3 | Q13217 | DNJC3_HUMAI | Unmodified | 0 | 0 | 0 | 0 | 0 | 0 | 234780 | 115300 | 2075900 | 1602600 | 0 | 0 | 0 | 0 | 0 | 0 | 0 | 0 |
| SNPSVFVIF | ZC3HAV1; ZC3I | Q7Z2W4; C9 | ZCCHV_HUMA | Unmodified | 1115400 | 1071300 | 1568700 | 3989100 | 0 | 0 | 0 | 0 | 1542500 | 0 | 0 | 1111900 | 0 | 0 | 1571100 | 2655600 | 0 | 0 |
| SNPTFYLQI | ACAD11; ACAD | Q709F0; F8V | ACD11_HUMAI | Unmodified | 0 | 0 | 0 | 0 | 0 | 0 | 0 | 0 | 1699900 | 378230 | 0 | 319640 | 89354 | 0 | 0 | 0 | 0 | 0 |
| SNQQLENI | IQGAP1; IQGAP | P46940; A0A | IQGA1_HUMAN | Oxidation (M) | 561300 | 388170 | 271280 | 377260 | 964420 | 0 | 0 | 0 | 520620 | 0 | 619190 | 2080500 | 89340 | 27584 | 655150 | 481570 | 0 | 0 |
| SNQQLENI | IQGAP1; IQGAP | P46940; A0A | IQGA1_HUMAN | Unmodified | 113420 | 109460 | 56850 | 335610 | 2176000 | 280240 | 55142 | 0 | 2916400 | 230980 | 85962 | 0 | 189190 | 0 | 148720 | 74555 | 0 | 0 |
| SNQQLVDI | PSME3; PSME3 | P61289; K7E | PSME3_HUMAI | Unmodified | 0 | 0 | 0 | 0 | 642500 | 0 | 386100 | 266320 | 0 | 242940 | 740060 | 4504000 | 703860 | 132370 | 0 | 0 | 527010 | 0 |
| SNQQNLG | RRM1; RRM1 | P23921; E9P | RIR1_HUMAN F | Unmodified | 356220 | 1003900 | 0 | 0 | 450560 | 0 | 0 | 334790 | 735230 | 401440 | 0 | 2960700 | 600460 | 284410 | 1068700 | 538510 | 311740 | 531520 |
| SNQTAYVP | LIMCH1; LIMCH | Q9UPQ0; HC | LIMC1_HUMAN | Unmodified | 0 | 0 | 0 | 0 | 0 | 0 | 194900 | 260200 | 0 | 0 | 0 | 0 | 909830 | 1186200 | 0 | 0 | 0 | 0 |
| SNQTMGW | MAP4K4; MAP4 | O95819; C9J | M4K4_HUMAN | Unmodified | 40740 | 0 | 0 | 0 | 0 | 0 | 0 | 38984 | 408460 | 96553 | 186240 | 825560 | 0 | 0 | 0 | 0 | 0 | 0 |
| SNQTTTDN | CNOT10; CNOT | Q9H9A5; H7 | CNO10_HUMA | Unmodified | 0 | 0 | 0 | 0 | 319140 | 0 | 0 | 0 | 0 | 93736 | 0 | 0 | 286280 | 82461 | 0 | 0 | 0 | 0 |
| SNQVAFQH | EXOC5; EXOC5 | O00471; G3 | EXOC5_HUMAI | Unmodified | 299070 | 0 | 315080 | 778150 | 19004000 | 3935900 | 1917100 | 95274 | 1857000 | 2631500 | 735890 | 3711900 | 3044800 | 2507600 | 907420 | 589250 | 4204000 | 4676400 |
| SNRAASAA | CDV3 | Q9UKY7 | CDV3_HUMAN | Unmodified | 0 | 0 | 0 | 0 | 563300 | 0 | 0 | 0 | 0 | 0 | 0 | 0 | 241260 | 0 | 0 | 0 | 0 | 106700 |
| SNRHVAVT | KIF5B; KIF5A; KI | P33176; Q12 | KINH_HUMAN | Unmodified | 13624 | 7.69E+08 | 0 | 87309 | 424320 | 382090 | 61916 | 0 | 266020 | 141720 | 191580 | 122280 | 1009900 | 596470 | 28733 | 24232 | 65778 | 273120 |
| SNSDSARL | ADGRG1 | Q9Y653 | AGRG1_HUMA | Unmodified | 0 | 0 | 0 | 394850 | 0 | 0 | 0 | 0 | 2211700 | 168070 | 490920 | 222020 | 70486 | 0 | 0 | 887440 | 0 | 0 |
| SNSEALKQ | MINK1; MINK1; | Q8N4C8; A0 | MINK1_HUMAN | Unmodified | 0 | 0 | 0 | 0 | 784100 | 0 | 0 | 0 | 0 | 148140 | 0 | 0 | 0 | 935120 | 0 | 0 | 0 | 761260 |
| SNSFFEGV | STK38 | Q15208 | STK38_HUMAN | Unmodified | 0 | 0 | 0 | 0 | 0 | 0 | 0 | 0 | 0 | 0 | 0 | 1222300 | 0 | 0 | 0 | 190450 | 0 | 0 |
| SNSFSDER | AHNAK | Q09666 | AHNK_HUMAN | Unmodified | 0 | 0 | 0 | 0 | 4119600 | 0 | 0 | 0 | 0 | 0 | 0 | 0 | 0 | 0 | 0 | 0 | 0 | 2725100 |
| SNSFSDER | AHNAK | Q09666 | AHNK_HUMAN | Unmodified | 0 | 0 | 0 | 0 | 4817300 | 0 | 0 | 0 | 0 | 0 | 0 | 0 | 0 | 0 | 0 | 0 | 2412400 | 2033700 |
| SNSLAFIK | DYNC1H1; DYN | Q14204; A0A | DYHC1_HUMA | Unmodified | 0 | 0 | 388620 | 0 | 952650 | 0 | 1006100 | 168160 | 589130 | 0 | 1641800 | 3772300 | 0 | 0 | 0 | 509990 | 511700 | 1387200 |
| SNSNIHPIF | POGLUT2 | Q6UW63 | PLGT2_HUMAN | Unmodified | 0 | 138150 | 0 | 0 | 0 | 0 | 0 | 0 | 284440 | 29999 | 0 | 0 | 0 | 0 | 0 | 0 | 0 | 0 |
| SNSPGTAS | ZC2HC1A; ZC2I | Q96GY0; H0 | ZC21A_HUMAN | Unmodified | 0 | 0 | 0 | 0 | 218090 | 0 | 53561 | 79645 | 280130 | 148060 | 0 | 156630 | 330280 | 173470 | 40615 | 0 | 94249 | 191080 |
| SNSPLPSIC | PCDH1 | Q08174 | PCDH1_HUMA | Unmodified | 0 | 0 | 0 | 226680 | 0 | 0 | 0 | 0 | 309010 | 0 | 3727100 | 2100000 | 236870 | 156870 | 499620 | 268490 | 0 | 0 |
| SNSVSLVPI | RICTOR | Q6R327 | RICTR_HUMAN | Unmodified | 145880 | 0 | 0 | 496310 | 419780 | 0 | 0 | 0 | 408870 | 0 | 0 | 1812100 | 164680 | 0 | 263050 | 168330 | 137250 | 225780 |
| SNTAGSQS | FKBP4 | Q02790 | FKBP4_HUMAN | Unmodified | 0 | 0 | 799310 | 904090 | 0 | 49782 | 0 | 0 | 553030 | 0 | 205860 | 3570900 | 825190 | 130580 | 762030 | 715500 | 103500 | 545360 |
| SNTGGQAI | EEF2 | P13639 | EF2_HUMAN EI | Unmodified | 0 | 0 | 0 | 0 | 0 | 0 | 0 | 0 | 543590 | 0 | 469250 | 1133100 | 0 | 0 | 1261200 | 0 | 0 | 0 |
| SNTGGQAI | EEF2 | P13639 | EF2_HUMAN EI | Unmodified | 2466300 | 0 | 0 | 6038800 | 8255300 | 0 | 0 | 0 | 6154500 | 0 | 12812000 | 13794000 | 3020100 | 0 | 2543300 | 0 | 0 | 0 |
| SNTHEFVN | VPS26A; VPS26 | O75436; S4F | VP26A_HUMAN | Unmodified | 0 | 0 | 214400 | 219210 | 490500 | 293090 | 98775 | 56056 | 1015700 | 239920 | 451370 | 3917900 | 323890 | 260900 | 426000 | 333960 | 223020 | 319230 |
| SNTISKPYI | COBLL1; COBLI | Q53SF7; A0A | COBL1_HUMAI | Unmodified | 3130700 | 2680000 | 0 | 588320 | 187140 | 0 | 0 | 0 | 0 | 0 | 0 | 0 | 2229200 | 1072200 | 278530 | 473670 | 0 | 442620 |
| SNTISKPYI | COBLL1; COBLI | Q53SF7; A0A | COBL1_HUMAI | Unmodified | 0 | 1390600 | 0 | 525140 | 0 | 0 | 0 | 0 | 0 | 0 | 0 | 0 | 1472000 | 0 | 0 | 339500 | 0 | 0 |
| SNTLERSG | SARG | Q9BW04 | SARG_HUMAN | Unmodified | 0 | 0 | 0 | 0 | 0 | 0 | 0 | 0 | 0 | 0 | 0 | 0 | 1939100 | 3135300 | 0 | 0 | 0 | 0 |
| SNTPILVD | GPI; GPI; GPI; G | P06744; K7E | G6PI_HUMAN ( | Unmodified | 80718 | 0 | 0 | 67215 | 812850 | 0 | 28070 | 15224 | 1111600 | 292640 | 0 | 17635000 | 0 | 0 | 67098 | 71284 | 728990 | 0 |
| SNTPILVD | GPI; GPI; GPI; G | P06744; K7E | G6PI_HUMAN ( | Unmodified | 0 | 101540 | 0 | 1315700 | 3903400 | 0 | 0 | 0 | 12347000 | 1577000 | 8013800 | 19584000 | 1335300 | 0 | 1058800 | 563520 | 358710 | 543350 |
| SNTPILVD | GPI; GPI; GPI; G | P06744; K7E | G6PI_HUMAN ( | Oxidation (M) | 479390 | 420160 | 0 | 0 | 43807 | 0 | 0 | 0 | 4772500 | 1952400 | 3428400 | 37244000 | 924140 | 0 | 4049600 | 1455300 | 708570 | 266740 |
| SNTPILVD | GPI; GPI; GPI; G | P06744; K7E | G6PI_HUMAN ( | Unmodified | 0 | 0 | 0 | 0 | 1932600 | 0 | 0 | 0 | 458530 | 350790 | 0 | 0 | 0 | 0 | 0 | 0 | 0 | 520840 |
| SNTPILVD | GPI; GPI; GPI; G | P06744; K7E | G6PI_HUMAN ( | Oxidation (M) | 0 | 0 | 0 | 0 | 0 | 0 | 0 | 0 | 0 | 745260 | 0 | 0 | 0 | 0 | 0 | 0 | 0 | 0 |
| SNTPMGDI | UBR4; UBR4; U | Q5T4S7; A0A | UBR4_HUMAN | Oxidation (M) | 0 | 0 | 0 | 0 | 0 | 0 | 0 | 0 | 0 | 0 | 18333 | 68499 | 0 | 0 | 0 | 0 | 0 | 0 |
| SNTSNEEF | NHS; NHS; NHS | Q6T4R5; A0A | NHS_HUMAN N | Unmodified | 0 | 0 | 0 | 0 | 1044900 | 0 | 0 | 0 | 0 | 0 | 0 | 0 | 0 | 0 | 0 | 0 | 490860 | 0 |
| SNTSPEEL | TLN1 | Q9Y490 | TLN1_HUMAN | Unmodified | 0 | 0 | 1138900 | 319380 | 1390400 | 0 | 0 | 196080 | 1970900 | 0 | 0 | 0 | 0 | 303820 | 2117100 | 2402400 | 256180 | 0 |
| SNVIDSML | SMC4; SMC4; S | Q9NTJ3; C9J | SMC4_HUMAN | Unmodified | 0 | 0 | 0 | 0 | 1094100 | 41445 | 680040 | 0 | 0 | 302360 | 0 | 0 | 180370 | 0 | 103710 | 43682 | 145640 | 0 |
| SNVIVALAF | SORL1 | Q92673 | SORL_HUMAN | Unmodified | 1537300 | 0 | 0 | 0 | 0 | 0 | 0 | 0 | 3694900 | 273150 | 0 | 0 | 210920 | 0 | 0 | 0 | 0 | 0 |
| SNVIVALAF | SORL1 | Q92673 | SORL_HUMAN | Unmodified | 0 | 0 | 0 | 0 | 0 | 0 | 0 | 0 | 0 | 737640 | 0 | 0 | 0 | 0 | 0 | 0 | 0 | 0 |
| SNVQPTAA | AURKB; AURKB | Q96GD4; J3I | AURKB_HUMAI | Unmodified | 64340 | 0 | 389610 | 696530 | 0 | 0 | 0 | 0 | 0 | 0 | 0 | 0 | 0 | 0 | 223390 | 120800 | 0 | 0 |
| SNVSDAVA | TPI1 | P60174 | TPIS_HUMAN T | Unmodified | 6982800 | 4834900 | 9569100 | 9716300 | 25133000 | 2335800 | 595540 | 346850 | 5299000 | 2205700 | 18119000 | 64709000 | 21917000 | 1435800 | 16367000 | 11013000 | 12622000 | 16331000 |
| SNVSDAVA | TPI1 | P60174 | TPIS_HUMAN T | Unmodified | 0 | 0 | 0 | 0 | 12671000 | 1497700 | 1973000 | 0 | 0 | 1582700 | 0 | 0 | 6713400 | 5500600 | 0 | 0 | 20084000 | 9885400 |
| SNYNFEK | RAN; RAN; RAN | P62826; J3K | RAN_HUMAN G | Unmodified | 869370 | 645990 | 1922600 | 0 | 2092300 | 0 | 297510 | 0 | 2237000 | 86555 | 2075800 | 41294000 | 826820 | 136470 | 0 | 628950 | 74689 | 1021000 |
| SNYNFEKP | RAN; RAN; RAN | P62826; J3K | RAN_HUMAN G | Unmodified | 2081900 | 2423800 | 0 | 3217100 | 16459000 | 5096600 | 8795700 | 116240 | 6157800 | 7397800 | 2272100 | 9024800 | 15757000 | 7139700 | 0 | 0 | 6819200 | 16216000 |
| SNYNFEKP | RAN; RAN; RAN | P62826; J3K | RAN_HUMAN G | Unmodified | 0 | 0 | 0 | 0 | 35910000 | 8540100 | 8242800 | 44413 | 2420600 | 5455300 | 2530000 | 2393700 | 18934000 | 15219000 | 0 | 0 | 10075000 | 18561000 |
| SNYNLPMI | RPL4; RPL4 | P36578; H3E | RL4_HUMAN 6 | Unmodified | 272630 | 252040 | 115290 | 0 | 0 | 0 | 0 | 38324 | 0 | 0 | 0 | 1227100 | 0 | 0 | 163670 | 0 | 0 | 0 |
| SPADQDRI | SRP19; SRP19; | P09132; A0A | SRP19_HUMAN | Unmodified | 0 | 0 | 0 | 0 | 0 | 0 | 0 | 0 | 0 | 0 | 0 | 158190 | 0 | 0 | 143580 | 77418 | 0 | 0 |
| SPADQDRI | SRP19; SRP19; | P09132; A0A | SRP19_HUMAN | Unmodified | 159590 | 0 | 0 | 0 | 413770 | 0 | 0 | 0 | 0 | 0 | 0 | 1618000 | 0 | 0 | 0 | 0 | 0 | 287320 |
| SPAEAATAI | NCAM1 | A0A0D9SF3 | A0A0D9SF30_I | Unmodified | 0 | 0 | 0 | 0 | 0 | 0 | 0 | 0 | 1544800 | 267710 | 0 | 0 | 116690 | 0 | 0 | 0 | 0 | 0 |
| SPAECREE | TNKS1BP1; TNI | Q9C0C2; E9 | TB182_HUMAN | Unmodified | 0 | 0 | 0 | 0 | 0 | 0 | 0 | 0 | 23954 | 0 | 0 | 254410 | 0 | 0 | 0 | 0 | 0 | 0 |
| SPAEQKNL | RANBP3; RANB | Q9H6Z4; K7I | RANB3_HUMAI | Unmodified | 487580 | 0 | 21151 | 0 | 955620 | 0 | 0 | 0 | 0 | 0 | 0 | 0 | 0 | 0 | 131540 | 0 | 0 | 210480 |
| SPAGETPFI | POGLUT3 | Q7Z4H8 | PLGT3_HUMAN | Unmodified | 216000 | 0 | 0 | 0 | 75225 | 0 | 0 | 0 | 6226900 | 240420 | 0 | 0 | 0 | 0 | 207860 | 0 | 0 | 0 |
| SPAGLQVL | EEF1B2; EEF1B | P24534; F2Z | EF1B_HUMAN I | Unmodified | 324500 | 401950 | 0 | 3007100 | 14879000 | 5698100 | 1446600 | 548050 | 8616100 | 5925100 | 32486000 | 1.64E+08 | 40912000 | 9657300 | 5369500 | 515640 | 7667800 | 11527000 |
| SPAGSIVHI | NARS1 | O43776 | SYNC_HUMAN | Unmodified | 0 | 582500 | 0 | 0 | 0 | 0 | 0 | 0 | 349500 | 0 | 0 | 1847300 | 0 | 0 | 2110600 | 1940400 | 0 | 0 |
| SPAGSIVHI | NARS1 | O43776 | SYNC_HUMAN | Unmodified | 0 | 0 | 0 | 0 | 0 | 0 | 0 | 0 | 0 | 0 | 0 | 2806200 | 0 | 871190 | 0 | 0 | 0 | 0 |
| SPAGVNLL | PROM1 | O43490 | PROM1_HUMA | Unmodified | 0 | 0 | 0 | 0 | 19792 | 0 | 86672 | 0 | 0 | 0 | 0 | 135080 | 0 | 0 | 0 | 0 | 0 | 0 |
| SPAHLSLIR | SRP72 | O76094 | SRP72_HUMAN | Unmodified | 0 | 0 | 203920 | 0 | 479770 | 70384 | 35663 | 24154 | 0 | 0 | 0 | 444460 | 0 | 0 | 0 | 0 | 14640 | 0 |
| SPAHPTPL | CNNM4 | Q6P4Q7 | CNNM4_HUMA | Unmodified | 0 | 0 | 18555 | 0 | 0 | 0 | 0 | 0 | 0 | 0 | 0 | 588480 | 0 | 0 | 57872 | 0 | 0 | 0 |
| SPAISWVN | UGGT1 | Q9NYU2 | UGGG1_HUMA | Unmodified | 0 | 0 | 0 | 0 | 0 | 0 | 0 | 0 | 1532800 | 25670 | 0 | 0 | 0 | 0 | 0 | 0 | 0 | 0 |
| SPAITATLE | UBA3; UBA3 | Q8TBC4; F8 | UBA3_HUMAN | Unmodified | 0 | 0 | 957990 | 0 | 0 | 0 | 0 | 169680 | 0 | 290910 | 0 | 2690000 | 777010 | 0 | 339890 | 346370 | 183170 | 0 |
| SPALSSWG | POMGNT1 | Q8WZA1 | PMGT1_HUMAI | Unmodified | 0 | 0 | 97281 | 0 | 0 | 0 | 0 | 0 | 8186400 | 309340 | 0 | 0 | 0 | 0 | 0 | 30817 | 0 | 0 |
| SPAPEQFQ | SH3BP4 | Q9P0V3 | SH3B4_HUMAI | Unmodified | 0 | 0 | 0 | 0 | 0 | 0 | 0 | 0 | 406120 | 0 | 0 | 0 | 0 | 0 | 0 | 0 | 0 | 0 |
| SPAPGNVA | SARG | Q9BW04 | SARG_HUMAN | Unmodified | 288660 | 145390 | 361290 | 600630 | 0 | 0 | 49896 | 48641 | 392250 | 12715 | 0 | 320980 | 3386500 | 1990700 | 669430 | 465440 | 26211 | 58552 |
| SPAPKPSD | CALD1; CALD1; | Q05682; E7E | CALD1_HUMAI | Unmodified | 0 | 0 | 961120 | 2145400 | 571400 | 0 | 0 | 28364 | 0 | 0 | 0 | 0 | 390960 | 0 | 3630700 | 1185500 | 0 | 0 |
| SPAPPLLH | CCAR2; CCAR2 | Q8N163; E5I | CCAR2_HUMAI | Unmodified | 130980 | 0 | 0 | 96263 | 0 | 0 | 805310 | 364050 | 0 | 0 | 0 | 375310 | 339660 | 256380 | 0 | 0 | 0 | 245570 |
| SPAPSILK | KIF14 | Q15058 | KIF14_HUMAN | Unmodified | 0 | 0 | 0 | 0 | 0 | 0 | 0 | 0 | 0 | 0 | 0 | 0 | 0 | 17173 | 0 | 0 | 1229900 | 0 |
| SPAQAELS | EPB41L4B | Q9H329 | E41LB_HUMAN | Unmodified | 150970 | 137350 | 2607800 | 195590 | 0 | 0 | 0 | 0 | 839980 | 123360 | 0 | 1070500 | 0 | 0 | 4330500 | 2984300 | 0 | 293490 |
| SPAQILLR | AKR1A1 | P14550 | AK1A1_HUMAN | Unmodified | 121850 | 100590 | 656060 | 392180 | 0 | 0 | 0 | 0 | 1436000 | 130690 | 245520 | 3562900 | 95398 | 0 | 1068600 | 848650 | 0 | 0 |
| SPASDTYIV | NACA; NACA; N | E9PAV3; Q1 | NACAM_HUMA | Unmodified | 597950 | 0 | 211660 | 263780 | 0 | 0 | 0 | 0 | 0 | 0 | 0 | 11324000 | 0 | 0 | 0 | 104250 | 0 | 0 |
| SPAYTLVW | HSPG2 | P98160 | PGBM_HUMAN | Unmodified | 0 | 0 | 2576400 | 0 | 0 | 0 | 0 | 0 | 0 | 0 | 0 | 0 | 0 | 0 | 0 | 677120 | 0 | 0 |
| SPCIVLADA | ALDH1A1 | P00352 | AL1A1_HUMAN | Unmodified | 0 | 0 | 0 | 0 | 0 | 0 | 0 | 0 | 0 | 0 | 0 | 7850800 | 0 | 0 | 0 | 0 | 0 | 0 |
| SPDAAGAT | GGT7 | Q9UJ14 | GGT7_HUMAN | Unmodified | 0 | 0 | 0 | 0 | 0 | 0 | 0 | 0 | 2281600 | 207670 | 0 | 0 | 0 | 0 | 0 | 0 | 0 | 0 |
| SPDDAVFQ | CPD | O75976 | CBPD_HUMAN | Unmodified | 0 | 0 | 0 | 0 | 4602200 | 1638000 | 97432 | 50316 | 51391000 | 11292000 | 295860 | 935840 | 405190 | 0 | 0 | 0 | 0 | 0 |
| SPDDPSRY | ENO1; ENO1 | P06733; A0A | ENOA_HUMAN | Unmodified | 1472200 | 901390 | 4774000 | 7296800 | 48202000 | 11074000 | 1356800 | 307350 | 36973000 | 7924600 | 10959000 | 24390000 | 20449000 | 6728300 | 6885500 | 4218800 | 8799900 | 24302000 |
| SPDDPSRY | ENO1; ENO1 | P06733; A0A | ENOA_HUMAN | Unmodified | 0 | 0 | 0 | 0 | 0 | 0 | 0 | 0 | 0 | 0 | 0 | 0 | 0 | 0 | 0 | 0 | 257430 | 170390 |
| SPDEATAAI | FARP1; FARP1; | Q9Y4F1; C9J | FARP1_HUMAN | Unmodified | 0 | 0 | 0 | 0 | 0 | 0 | 0 | 0 | 91253 | 0 | 157810 | 344560 | 0 | 0 | 0 | 0 | 0 | 0 |
| SPDEAYAIA | SUCLA2; SUCL | Q9P2R7; A0A | SUCB1_HUMAI | Unmodified | 0 | 0 | 0 | 0 | 720950 | 0 | 141590 | 17813 | 0 | 252670 | 0 | 0 | 0 | 0 | 0 | 0 | 0 | 272170 |
| SPDEGFEG | FOLH1B; FOLH | Q9HBA9; Q0 | FOH1B_HUMAI | Unmodified | 424910 | 1022200 | 30945000 | 26252000 | 0 | 0 | 0 | 0 | 0 | 0 | 0 | 0 | 0 | 277430 | 628090 | 30589000 | 0 | 0 |
| SPDELPSA | DIAPH1; DIAPH | O60610; A0A | DIAP1_HUMAN | Unmodified | 0 | 0 | 0 | 0 | 0 | 0 | 0 | 0 | 0 | 0 | 83907 | 128930 | 0 | 0 | 0 | 0 | 312920 | 0 |
| SPDEVLRE | PHLDA2 | Q53GA4 | PHLA2_HUMAN | Unmodified | 789830 | 720840 | 835390 | 350280 | 1531500 | 853870 | 0 | 0 | 0 | 0 | 0 | 0 | 1110900 | 770310 | 1323400 | 838940 | 265750 | 1485200 |
| SPDEVTLTS | PACSIN3; PACS | Q9UKS6; H0 | PACN3_HUMAI | Unmodified | 9186300 | 7271200 | 322560 | 0 | 0 | 0 | 0 | 0 | 0 | 0 | 0 | 40472 | 0 | 0 | 8472200 | 4125200 | 0 | 0 |
| SPDEVTLTS | PACSIN3; PACS | Q9UKS6; A0 | PACN3_HUMAI | Unmodified | 4046900 | 3034100 | 1971600 | 0 | 0 | 0 | 0 | 0 | 0 | 0 | 0 | 0 | 0 | 0 | 0 | 0 | 0 | 0 |
| SPDFTNEN | ATP1A1 | P05023 | AT1A1_HUMAN | Unmodified | 2410300 | 2484300 | 25830000 | 3390400 | 0 | 0 | 0 | 704590 | 16580000 | 4180200 | 12325000 | 1.63E+08 | 230210 | 805510 | 38849000 | 30718000 | 1429800 | 0 |
| SPDGASEY | PTPRJ; PTPRJ; P | Q12913; A0A | PTPRJ_HUMAN | Unmodified | 215440 | 299120 | 0 | 0 | 0 | 245680 | 0 | 0 | 1053300 | 0 | 0 | 4713400 | 0 | 0 | 1574400 | 316230 | 401130 | 0 |
| SPDGYLQI | MGAT4B; MGAT | Q9UQ53; HC | MGT4B_HUMAI | Unmodified | 0 | 0 | 0 | 0 | 0 | 0 | 0 | 0 | 0 | 0 | 0 | 0 | 0 | 0 | 0 | 0 | 0 | 0 |
| SPDMQGSI | LLGL2; LLGL2 | Q6P1M3; J3C | L2GL2_HUMAN | Unmodified | 0 | 0 | 0 | 0 | 0 | 0 | 0 | 0 | 6154700 | 1156900 | 0 | 0 | 0 | 0 | 0 | 0 | 0 | 0 |
| SPDNPMN | CCDC124 | Q96CT7 | CC124_HUMAI | Unmodified | 202450 | 164760 | 275770 | 298740 | 121110 | 0 | 0 | 0 | 0 | 0 | 139980 | 0 | 0 | 0 | 226440 | 205540 | 0 | 0 |
| SPDSDVAA | EIF5B; EIF5B | O60841; A0A | IF2P_HUMAN E | Unmodified | 159350 | 95733 | 134570 | 203190 | 0 | 0 | 0 | 0 | 0 | 0 | 0 | 1028800 | 1137800 | 340550 | 153710 | 126950 | 0 | 0 |
| SPDSDVAA | EIF5B; EIF5B | O60841; A0A | IF2P_HUMAN E | Unmodified | 0 | 0 | 0 | 0 | 348060 | 0 | 0 | 73483 | 0 | 0 | 141140 | 460940 | 1461300 | 552640 | 0 | 0 | 0 | 248800 |
| SPDSTANV | SRGAP2; SRGA | O75044; P0I | SRGP2_HUMAI | Unmodified | 176250 | 0 | 159040 | 1506800 | 1261200 | 222620 | 0 | 0 | 292250 | 94300 | 263570 | 1392200 | 991780 | 263390 | 350210 | 295480 | 114300 | 349510 |
| SPDTILQK | EIF2S2 | P20042 | IF2B_HUMAN E | Unmodified | 256070 | 182400 | 0 | 620180 | 279390 | 0 | 0 | 94643 | 618290 | 55658 | 723820 | 4834200 | 0 | 0 | 0 | 448560 | 0 | 0 |
| SPDTNYLFI | PPP2CB; PPP2C | P62714; P67 | PP2AB_HUMAN | Oxidation (M) | 0 | 0 | 0 | 0 | 0 | 0 | 22379 | 17290 | 0 | 0 | 0 | 241520 | 0 | 0 | 0 | 0 | 0 | 78148 |
| SPDVGLYG | IQGAP1; IQGAP | P46940; A0A | IQGA1_HUMAN | Unmodified | 0 | 0 | 0 | 0 | 0 | 0 | 0 | 0 | 630140 | 0 | 0 | 1076200 | 0 | 0 | 0 | 187210 | 0 | 0 |
| SPEDLER | SIX5; GPI; GPI | Q8N196; P0 | SIX5_HUMAN H | Unmodified | 234900 | 0 | 96108 | 0 | 0 | 0 | 0 | 0 | 2074100 | 0 | 8597.6 | 15469000 | 0 | 0 | 1052200 | 47001 | 0 | 0 |
| SPEDLERLI | GPI; GPI | P06744; A0A | G6PI_HUMAN ( | Unmodified | 0 | 0 | 669990 | 0 | 413240 | 0 | 0 | 33503 | 887530 | 192200 | 578190 | 1436500 | 307630 | 0 | 0 | 0 | 154130 | 0 |
| SPEEGAETI | CBR1 | P16152 | CBR1_HUMAN | Unmodified | 0 | 0 | 2714600 | 0 | 11335000 | 2858700 | 0 | 0 | 7534100 | 0 | 184340 | 2281800 | 2732900 | 220150 | 0 | 1538800 | 1559400 | 8085600 |
| SPEELAALV | RABGGTA; RAB | Q92696; H0 | PGTA_HUMAN | Unmodified | 0 | 0 | 0 | 0 | 0 | 0 | 0 | 0 | 0 | 0 | 84343 | 339180 | 0 | 0 | 0 | 0 | 0 | 0 |
| SPEERLFPE | MICAL3; MICAL | Q7RTP6; J3C | MICA3_HUMAN | Unmodified | 0 | 0 | 0 | 0 | 2619300 | 928730 | 0 | 0 | 0 | 0 | 0 | 0 | 0 | 0 | 0 | 0 | 0 | 0 |
| SPEHSAEP | KIDINS220 | Q9ULH0 | KDIS_HUMAN I | Unmodified | 0 | 0 | 0 | 805620 | 2196600 | 115150 | 0 | 0 | 12565000 | 455210 | 0 | 0 | 810540 | 1315200 | 0 | 0 | 494360 | 433350 |
| SPEKIEEVL | ADD3 | Q9UEY8 | ADDG_HUMAN | Unmodified | 35715 | 0 | 0 | 117490 | 0 | 0 | 0 | 0 | 0 | 0 | 0 | 0 | 0 | 273750 | 406560 | 447710 | 0 | 0 |
| SPELQEEF | ARFIP2; ARFIP2 | P53365; A0A | ARFP2_HUMAN | Unmodified | 0 | 0 | 130530 | 0 | 0 | 0 | 0 | 0 | 0 | 0 | 0 | 1366600 | 0 | 0 | 0 | 0 | 0 | 0 |
| SPELVQAM | ATXN10; ATXN1 | Q9UBB4; A0 | ATX10_HUMAN | Oxidation (M) | 0 | 0 | 0 | 0 | 0 | 0 | 0 | 79098 | 0 | 0 | 0 | 1268400 | 0 | 0 | 0 | 0 | 0 | 0 |
| SPEPEATLT | LIMCH1; LIMCH | Q9UPQ0; HC | LIMC1_HUMAN | Unmodified | 0 | 0 | 118810 | 0 | 0 | 0 | 0 | 0 | 0 | 0 | 294280 | 533570 | 0 | 0 | 0 | 0 | 0 | 0 |
| SPEQQLSA | KPNA1; KPNA1; | P52294; C9J | IMA5_HUMAN I | Unmodified | 0 | 0 | 0 | 134280 | 216390 | 37695 | 0 | 0 | 422790 | 141540 | 205740 | 0 | 559120 | 141340 | 0 | 0 | 57104 | 640130 |
| SPESVALIW | ACSS1; ACSS1 | Q9NUB1; Q5 | ACS2L_HUMAN | Unmodified | 0 | 0 | 0 | 0 | 0 | 0 | 1202100 | 0 | 595930 | 779130 | 0 | 0 | 0 | 0 | 0 | 0 | 0 | 0 |
| SPEVLLGS | CDK1; CDK1; C | P06493; A0A | CDK1_HUMAN | Unmodified | 52132 | 1007300 | 767890 | 0 | 412680 | 0 | 0 | 0 | 534570 | 0 | 0 | 2512300 | 0 | 93891 | 1866500 | 578900 | 0 | 0 |

| | | | | | | | | | | | | | | | | | | | | | | |
|---|---|---|---|---|---|---|---|---|---|---|---|---|---|---|---|---|---|---|---|---|---|---|
| SPFEQHIK | FAM3C | Q92520 | FAM3C_HUMAI | Unmodified | 0 | 0 | 0 | 0 | 0 | 0 | 0 | 27171 | 2545900 | 427420 | 294550 | 734110 | 0 | 0 | 0 | 1003000 | 117390 | 308310 |
| SPFEQHVK | FAM3A; FAM3A; | P98173; D3D | FAM3A_HUMAI | Unmodified | 0 | 0 | 0 | 0 | 0 | 0 | 0 | 0 | 1433100 | 0 | 0 | 169620 | 0 | 0 | 0 | 0 | 0 | 0 |
| SPFEVKVG | FLNA; FLNA; FL | P21333; A0A | FLNA_HUMAN | Unmodified | 0 | 0 | 0 | 463830 | 0 | 0 | 181780 | 160920 | 0 | 0 | 0 | 0 | 555320 | 84458 | 0 | 144400 | 37028 | 0 |
| SPFEVQVG | FLNB; FLNB | O75369; E7E | FLNB_HUMAN | Oxidation (M) | 0 | 0 | 1102500 | 1035200 | 7634900 | 1832200 | 162060 | 71358 | 379040 | 0 | 0 | 3221300 | 1431700 | 630800 | 0 | 0 | 1998100 | 1788900 |
| SPFEVQVG | FLNB; FLNB | O75369; E7E | FLNB_HUMAN | Unmodified | 189240 | 0 | 0 | 0 | 11052000 | 1645800 | 0 | 0 | 208030 | 0 | 0 | 0 | 551470 | 0 | 0 | 0 | 3006700 | 3330100 |
| SPFEVSVDI | FLNB; FLNB | O75369; E7E | FLNB_HUMAN | Unmodified | 0 | 0 | 0 | 0 | 2032000 | 554380 | 85594 | 0 | 226800 | 0 | 0 | 554900 | 0 | 0 | 0 | 0 | 382050 | 331580 |
| SPFEVSVDI | FLNB; FLNB | O75369; E7E | FLNB_HUMAN | Unmodified | 0 | 0 | 0 | 0 | 744030 | 0 | 0 | 0 | 0 | 0 | 0 | 0 | 203970 | 0 | 0 | 0 | 239480 | 0 |
| SPFEVYVDI | FLNA; FLNA; FL | P21333; A0A | FLNA_HUMAN | Unmodified | 1735400 | 2681700 | 2260700 | 4846500 | 4404100 | 1119900 | 3167300 | 1085500 | 524550 | 464940 | 91002 | 2928900 | 2168700 | 1946500 | 4340300 | 3695000 | 1622600 | 651030 |
| SPFEVYVDI | FLNA; FLNA; FL | P21333; A0A | FLNA_HUMAN | Unmodified | 0 | 126680 | 0 | 701770 | 403010 | 185870 | 783970 | 303910 | 0 | 127580 | 0 | 0 | 497530 | 293490 | 0 | 107910 | 356360 | 0 |
| SPFEVYVDI | FLNA; FLNA; FL | P21333; A0A | FLNA_HUMAN | Unmodified | 0 | 0 | 0 | 0 | 0 | 0 | 2510600 | 1171900 | 0 | 0 | 0 | 0 | 611240 | 0 | 0 | 0 | 459590 | 0 |
| SPFIYSPIIA | TRAPPC9; TRAF | Q96Q05; H0 | TPPC9_HUMAN | Unmodified | 0 | 0 | 0 | 0 | 0 | 0 | 0 | 0 | 128560 | 0 | 0 | 382030 | 0 | 0 | 113280 | 263250 | 170470 | 0 |
| SPFLQVFNI | SEC23B; SEC23 | Q15437; A0A | SC23B_HUMAI | Unmodified | 85612 | 110140 | 31471 | 0 | 0 | 0 | 0 | 0 | 155250 | 0 | 16046 | 984000 | 0 | 0 | 63182 | 67383 | 0 | 0 |
| SPFRVNVG | FLNC | Q14315 | FLNC_HUMAN | Unmodified | 0 | 0 | 0 | 0 | 1516000 | 421820 | 0 | 0 | 0 | 0 | 0 | 0 | 0 | 0 | 0 | 0 | 640970 | 0 |
| SPFSVAVSI | FLNA; FLNA; FL | P21333; A0A | FLNA_HUMAN | Unmodified | 30907 | 0 | 291520 | 0 | 8222800 | 1564800 | 0 | 0 | 0 | 1195700 | 1079800 | 4106700 | 173540 | 0 | 1400500 | 476240 | 5956700 | 30225 |
| SPFTVGVA | FLNB; FLNB | O75369; E7E | FLNB_HUMAN | Unmodified | 0 | 0 | 0 | 0 | 9813100 | 2858600 | 0 | 0 | 557890 | 689690 | 1792400 | 6866200 | 0 | 0 | 3000500 | 0 | 3448600 | 0 |
| SPFVVNVAI | FLNC | Q14315 | FLNC_HUMAN | Unmodified | 0 | 0 | 0 | 0 | 106500 | 0 | 0 | 0 | 0 | 0 | 0 | 0 | 0 | 0 | 0 | 0 | 755620 | 0 |
| SPFVVQVG | FLNB; FLNB | O75369; E7E | FLNB_HUMAN | Unmodified | 0 | 0 | 0 | 0 | 981810 | 553480 | 0 | 0 | 0 | 0 | 0 | 0 | 0 | 0 | 0 | 0 | 0 | 133820 |
| SPFVVQVG | FLNB; FLNB | O75369; E7E | FLNB_HUMAN | Unmodified | 0 | 0 | 0 | 0 | 3165100 | 0 | 0 | 0 | 0 | 0 | 0 | 0 | 0 | 1115300 | 0 | 0 | 5070200 | 1557800 |
| SPGADLLQ | TKFC; TKFC; TK | Q3LXA3; I3L | TKFC_HUMAN | Unmodified | 0 | 0 | 674190 | 405990 | 0 | 0 | 0 | 0 | 0 | 0 | 1573900 | 5689900 | 354360 | 112940 | 1997400 | 911000 | 0 | 0 |
| SPGADLSLI | ACOX3 | O15254 | ACOX3_HUMAI | Unmodified | 0 | 0 | 0 | 0 | 0 | 0 | 605660 | 551730 | 420200 | 0 | 35749 | 0 | 0 | 0 | 0 | 0 | 0 | 0 |
| SPGAFDMS | EPN1 | Q9Y6I3 | EPN1_HUMAN | Unmodified | 134550 | 452100 | 0 | 0 | 0 | 0 | 0 | 0 | 0 | 0 | 0 | 0 | 0 | 0 | 0 | 227820 | 0 | 0 |
| SPGALETPS | AKAP2; PALM2A | Q9Y2D5; C9 | AKAP2_HUMAN | Unmodified | 0 | 0 | 0 | 0 | 990810 | 213120 | 0 | 0 | 341610 | 0 | 0 | 0 | 0 | 0 | 0 | 0 | 103900 | 314160 |
| SPGALETPS | AKAP2; PALM2A | Q9Y2D5; C9 | AKAP2_HUMAN | Unmodified | 0 | 0 | 0 | 0 | 3603500 | 0 | 146040 | 111260 | 0 | 0 | 0 | 0 | 0 | 0 | 0 | 0 | 367410 | 336430 |
| SPGAPGPL | ZYX; ZYX; ZYX | Q15942; H0 | ZYX_HUMAN Zy | Unmodified | 74225 | 0 | 239740 | 0 | 371090 | 0 | 0 | 0 | 0 | 0 | 0 | 0 | 66335 | 0 | 247620 | 351990 | 174360 | 708450 |
| SPGASVSS | KIAA1522 | Q9P206 | K1522_HUMAN | Unmodified | 0 | 0 | 184960 | 0 | 0 | 0 | 0 | 0 | 0 | 0 | 0 | 655060 | 0 | 0 | 384300 | 519810 | 0 | 0 |
| SPGLTPEPI | SELPLG; SELPL | Q14242; A0A | SELPL_HUMAN | Unmodified | 37334 | 693410 | 0 | 0 | 0 | 0 | 0 | 0 | 0 | 0 | 506340 | 1885000 | 471880 | 0 | 0 | 35032 | 0 | 0 |
| SPGLVLCH | TYSND1 | Q2T9J0 | TYSD1_HUMAN | Unmodified | 0 | 65074 | 0 | 42069 | 0 | 16277 | 0 | 0 | 3283800 | 116480 | 0 | 0 | 0 | 0 | 0 | 0 | 0 | 0 |
| SPGSQGVI | NDUFA2 | O43678 | NDUA2_HUMA | Unmodified | 121410 | 0 | 0 | 0 | 546250 | 0 | 0 | 0 | 1845400 | 314530 | 0 | 0 | 0 | 0 | 0 | 0 | 0 | 0 |
| SPGSVVFR | ATP6AP2; ATP6 | O75787; A0A | RENR_HUMAN | Unmodified | 0 | 0 | 0 | 0 | 0 | 4041700 | 1907000 | 513460 | 3262100 | 203420 | 267970 | 0 | 0 | 385450 | 0 | 0 | 0 | 0 |
| SPGTKDVV | SLC12A2; SLC1 | P55011; G3 | S12A2_HUMAN | Unmodified | 0 | 0 | 0 | 254120 | 0 | 0 | 0 | 0 | 0 | 0 | 0 | 0 | 0 | 0 | 0 | 0 | 1035500 | 700300 |
| SPHFQVVN | RABGAP1 | Q9Y3P9 | RBGP1_HUMAI | Unmodified | 0 | 0 | 71476 | 0 | 971720 | 197790 | 336480 | 454890 | 0 | 426480 | 154490 | 631770 | 480440 | 120500 | 0 | 0 | 0 | 0 |
| SPHPAIQPI | PABPC4; PABP | Q13310; B1 | PABP4_HUMAN | Oxidation (M) | 4393400 | 4143200 | 0 | 0 | 0 | 0 | 0 | 0 | 0 | 0 | 0 | 0 | 0 | 0 | 3985800 | 0 | 0 | 0 |
| SPHPAIQPI | PABPC4; PABP | Q13310; B1 | PABP4_HUMAN | Unmodified | 1273800 | 1113200 | 0 | 0 | 0 | 0 | 0 | 0 | 0 | 0 | 0 | 0 | 0 | 0 | 1098000 | 454250 | 0 | 0 |
| SPHQGFMI | NUDT19 | A8MXV4 | NUD19_HUMA | Unmodified | 0 | 0 | 0 | 0 | 0 | 0 | 0 | 0 | 1200800 | 0 | 0 | 0 | 0 | 0 | 0 | 0 | 0 | 0 |
| SPHSDVGS | ROR2; ROR2 | Q01974; B1 | ROR2_HUMAN | Unmodified | 0 | 0 | 0 | 0 | 0 | 0 | 0 | 0 | 1753500 | 175510 | 0 | 0 | 0 | 0 | 0 | 0 | 0 | 0 |
| SPIIFIGTGE | SRP54; SRP54; | P61011; G3 | SRP54_HUMAN | Unmodified | 0 | 0 | 0 | 134290 | 0 | 0 | 0 | 0 | 0 | 0 | 186150 | 603450 | 196040 | 0 | 32168 | 0 | 0 | 0 |
| SPIIFIGTGE | SRP54; SRP54; | P61011; G3 | SRP54_HUMAN | Unmodified | 35165 | 0 | 0 | 0 | 2250100 | 0 | 1063900 | 0 | 0 | 374070 | 0 | 0 | 2299600 | 1313000 | 0 | 0 | 0 | 2254200 |
| SPILLGSLA | HIBADH | P31937 | 3HIDH_HUMAN | Unmodified | 40003 | 0 | 0 | 0 | 481290 | 117520 | 60213 | 0 | 5909200 | 1255400 | 276560 | 87482 | 221150 | 94363 | 113360 | 0 | 21451 | 270940 |
| SPILVATAV | DDX3X; DDX3Y | O00571; O1 | DDX3X_HUMAI | Unmodified | 945860 | 849220 | 113100 | 684710 | 0 | 0 | 0 | 0 | 212770 | 0 | 0 | 884150 | 0 | 0 | 1233000 | 1009000 | 0 | 0 |
| SPISFLSK | MICAL3; MICAL | Q7RTP6; A0 | MICA3_HUMAN | Unmodified | 0 | 0 | 0 | 0 | 363250 | 0 | 0 | 41480 | 0 | 0 | 0 | 0 | 0 | 257460 | 0 | 0 | 297000 | 103150 |
| SPITGNDLS | GARS1; GARS1 | P41250; A0A | GARS_HUMAN | Oxidation (M) | 0 | 0 | 361030 | 491670 | 974370 | 408620 | 1264200 | 207200 | 1623800 | 2682700 | 1462900 | 3750900 | 3816700 | 1935400 | 501180 | 507110 | 889730 | 664810 |
| SPITGNDLS | GARS1; GARS1 | P41250; A0A | GARS_HUMAN | Unmodified | 0 | 0 | 129210 | 0 | 105490 | 0 | 0 | 0 | 338320 | 143990 | 0 | 61429 | 132480 | 0 | 0 | 0 | 127330 | 102020 |
| SPKDNVDI | LMAN2 | Q12907 | LMAN2_HUMAI | Unmodified | 0 | 0 | 0 | 0 | 218750 | 107380 | 0 | 0 | 6507900 | 1408900 | 0 | 0 | 0 | 0 | 0 | 0 | 0 | 407420 |
| SPKGEFKD | CLINT1 | Q14677 | EPN4_HUMAN | Unmodified | 0 | 0 | 98668 | 0 | 479410 | 0 | 36305 | 0 | 0 | 0 | 45090 | 319530 | 0 | 0 | 0 | 0 | 23915 | 231750 |
| SPKPLLTGI | MOGS; MOGS; | Q13724; A0 | MOGS_HUMAN | Oxidation (M) | 0 | 0 | 0 | 0 | 0 | 0 | 0 | 0 | 3199800 | 0 | 0 | 0 | 0 | 0 | 0 | 0 | 0 | 0 |
| SPKPLLTGI | MOGS; MOGS; | Q13724; A0 | MOGS_HUMAN | Unmodified | 30642 | 0 | 0 | 2015600 | 0 | 0 | 0 | 0 | 1725300 | 0 | 0 | 0 | 0 | 0 | 0 | 0 | 0 | 0 |
| SPKPVKPVI | PCDH1 | Q08174 | PCDH1_HUMA | Unmodified | 285840 | 364690 | 499770 | 857050 | 0 | 0 | 0 | 0 | 1924900 | 222280 | 8782600 | 19569000 | 1679500 | 203400 | 551140 | 1737300 | 79625 | 0 |
| SPLAQMEE | SEPTIN7; SEPTI | Q16181; E7E | SEPT7_HUMAN | Unmodified | 0 | 0 | 0 | 0 | 0 | 0 | 0 | 0 | 745370 | 167440 | 659780 | 6384100 | 0 | 0 | 0 | 409910 | 0 | 0 |
| SPLAQMEE | SEPTIN7; SEPTI | Q16181; E7E | SEPT7_HUMAN | Unmodified | 0 | 0 | 0 | 0 | 31372000 | 5635500 | 1765100 | 746050 | 0 | 52429 | 77869 | 6176900 | 5939000 | 5007500 | 0 | 0 | 6224100 | 8100400 |
| SPLAQMEE | SEPTIN7; SEPTI | Q16181; E7E | SEPT7_HUMAN | Oxidation (M) | 0 | 0 | 0 | 0 | 2764300 | 0 | 324740 | 0 | 0 | 0 | 0 | 0 | 0 | 0 | 0 | 0 | 569360 | 642420 |
| SPLFNQILT | PATJ; PATJ; PAT | Q8NI35; A0A | INADL_HUMAN | Unmodified | 0 | 0 | 0 | 769860 | 3750500 | 508250 | 0 | 0 | 646370 | 535640 | 68954 | 0 | 2090200 | 1745400 | 242400 | 195760 | 1064600 | 1434500 |
| SPLGEVAIF | CPNE3 | O75131 | CPNE3_HUMAI | Unmodified | 561610 | 0 | 2704400 | 298560 | 0 | 0 | 81675 | 67100 | 6008200 | 458990 | 69411 | 1819900 | 0 | 0 | 2414400 | 2384200 | 0 | 0 |
| SPLGEVAIF | CPNE3 | O75131 | CPNE3_HUMAI | Unmodified | 0 | 0 | 0 | 57170 | 0 | 0 | 0 | 0 | 249980 | 0 | 0 | 0 | 0 | 0 | 430240 | 0 | 0 | 15901 |
| SPLGEVAIF | CPNE3 | O75131 | CPNE3_HUMAI | Unmodified | 0 | 0 | 0 | 661990 | 0 | 0 | 795470 | 0 | 3495400 | 2105900 | 0 | 0 | 0 | 0 | 0 | 0 | 0 | 0 |
| SPLIIFSDCI | ALDH9A1 | P49189 | AL9A1_HUMAN | Oxidation (M) | 98144 | 0 | 0 | 60846 | 0 | 0 | 0 | 0 | 0 | 0 | 130160 | 1005000 | 165360 | 0 | 174010 | 173830 | 0 | 0 |
| SPLLYVTRI | NDST1; NDST1 | P52848; E7E | NDST1_HUMAI | Unmodified | 0 | 0 | 0 | 0 | 0 | 0 | 0 | 0 | 1825000 | 198290 | 0 | 553720 | 0 | 0 | 0 | 0 | 0 | 0 |
| SPLPLGFSI | MVB12A; MVB1 | Q96EY5; E9F | MB12A_HUMAI | Oxidation (M) | 0 | 0 | 0 | 0 | 0 | 0 | 0 | 0 | 0 | 0 | 0 | 0 | 494440 | 529730 | 0 | 0 | 0 | 0 |
| SPLPWQHI | HSPG2 | P98160 | PGBM_HUMAN | Unmodified | 0 | 0 | 358310 | 740580 | 0 | 0 | 523060 | 0 | 0 | 0 | 0 | 0 | 0 | 0 | 0 | 0 | 0 | 258880 |
| SPLSSPQQ | CNOT11; CNOT | Q9UKZ1; H7 | CNO11_HUMA | Unmodified | 0 | 0 | 0 | 0 | 2224700 | 0 | 0 | 15686 | 354520 | 0 | 0 | 354470 | 0 | 0 | 0 | 0 | 44695 | 0 |
| SPLTILQDI | CDCA3; CDCA3 | Q99618; J3K | CDCA3_HUMA | Unmodified | 276830 | 342860 | 477110 | 1196300 | 304480 | 114110 | 0 | 44529 | 651030 | 0 | 65652 | 261960 | 1034000 | 155020 | 786970 | 385560 | 281390 | 173010 |
| SPLTPLPR | ATP6V0A4; ATP | Q9HBG4; A0 | VPP4_HUMAN | Unmodified | 0 | 169120 | 0 | 0 | 0 | 0 | 0 | 0 | 0 | 0 | 0 | 1887500 | 0 | 0 | 0 | 0 | 0 | 0 |
| SPLTQEQLI | STUB1; STUB1 | Q9UNE7; H3 | CHIP_HUMAN | Oxidation (M) | 396680 | 0 | 0 | 0 | 0 | 0 | 0 | 0 | 0 | 198810 | 0 | 1406000 | 0 | 0 | 0 | 0 | 0 | 0 |
| SPLVAAMQ | IPO7; IPO7 | O95373; E9F | IPO7_HUMAN I | Unmodified | 211270 | 0 | 62218 | 0 | 0 | 0 | 0 | 0 | 752430 | 96393 | 0 | 4473800 | 0 | 0 | 403060 | 140420 | 0 | 0 |
| SPLVAAMQ | IPO7; IPO7 | O95373; E9F | IPO7_HUMAN I | Oxidation (M) | 0 | 0 | 0 | 0 | 0 | 0 | 0 | 0 | 0 | 0 | 0 | 1062800 | 0 | 0 | 0 | 0 | 0 | 0 |
| SPLVFVLCI | C2orf72 | A6NCS6 | CB072_HUMAN | Unmodified | 254940 | 0 | 0 | 0 | 0 | 0 | 0 | 0 | 0 | 0 | 0 | 0 | 0 | 0 | 728710 | 393350 | 0 | 0 |
| SPLVMDVL | DYNC1H1; DYN | Q14204; A0 | DYHC1_HUMA | Oxidation (M) | 0 | 0 | 0 | 0 | 0 | 0 | 0 | 0 | 0 | 0 | 0 | 1025500 | 0 | 0 | 0 | 0 | 0 | 0 |
| SPLVMDVL | DYNC1H1; DYN | Q14204; A0 | DYHC1_HUMA | Unmodified | 0 | 0 | 0 | 0 | 0 | 0 | 0 | 0 | 20127 | 0 | 10046 | 568880 | 0 | 0 | 20003 | 0 | 0 | 0 |
| SPMEFVEYI | PLCB3 | Q01970 | PLCB3_HUMAN | Unmodified | 0 | 0 | 0 | 0 | 1545900 | 435070 | 210650 | 129410 | 0 | 0 | 0 | 0 | 130230 | 0 | 139250 | 0 | 397030 | 0 |
| SPNGTIRNI | IDH1; IDH1 | O75874; C9J | IDHC_HUMAN | Unmodified | 0 | 0 | 0 | 0 | 0 | 0 | 0 | 0 | 0 | 0 | 0 | 0 | 0 | 2471600 | 0 | 0 | 100100 | 0 |
| SPNHGTVE | PKP4; PKP4 | Q99569; A0 | PKP4_HUMAN | Unmodified | 0 | 0 | 191770 | 160470 | 0 | 0 | 0 | 0 | 0 | 0 | 0 | 1018000 | 0 | 0 | 108270 | 150580 | 0 | 0 |
| SPNKHNRI | EEF2; EEF2 | P13639; M0 | EF2_HUMAN El | Unmodified | 0 | 33179 | 38275 | 0 | 1587000 | 0 | 11870 | 0 | 0 | 237700 | 0 | 0 | 0 | 605410 | 13911 | 0 | 3555300 | 660790 |
| SPPEAGTQ | GPR107; GPR1 | Q5VW38; G | GP107_HUMAN | Unmodified | 49062 | 0 | 80527 | 0 | 1472600 | 0 | 122330 | 292170 | 17656000 | 4123700 | 550110 | 513870 | 319920 | 263570 | 79134 | 99926 | 0 | 0 |
| SPPFFENL | ACO1 | P21399 | ACOC_HUMAN | Unmodified | 0 | 0 | 0 | 35412 | 526820 | 0 | 0 | 0 | 0 | 309560 | 231860 | 598670 | 77863 | 112320 | 0 | 28113 | 0 | 0 |
| SPPGPHGI | NOTCH3 | Q9UM47 | NOTC3_HUMA | Unmodified | 99086 | 65950 | 0 | 0 | 0 | 0 | 0 | 0 | 0 | 0 | 0 | 0 | 0 | 0 | 0 | 0 | 0 | 0 |
| SPPGQVTE | AKR1B1; AKR1E | P15121; E9P | ALDR_HUMAN | Unmodified | 0 | 0 | 0 | 0 | 3387500 | 0 | 0 | 0 | 136530 | 0 | 3790300 | 22169000 | 0 | 0 | 0 | 0 | 35458 | 0 |
| SPPILPPHI | PRKAB2 | O43741 | AAKB2_HUMAN | Unmodified | 114590 | 0 | 129040 | 214040 | 262850 | 62862 | 10501 | 0 | 746890 | 27008 | 408430 | 275950 | 172580 | 0 | 528900 | 261200 | 58292 | 306830 |
| SPPIPLAK | PDIA4; PDIA4 | P13667; A0A | PDIA4_HUMAN | Unmodified | 0 | 0 | 0 | 0 | 0 | 0 | 0 | 0 | 1640600 | 0 | 555620 | 0 | 0 | 0 | 0 | 0 | 0 | 0 |
| SPPIPLAKV | PDIA4; PDIA4 | P13667; A0A | PDIA4_HUMAN | Unmodified | 0 | 0 | 0 | 0 | 0 | 0 | 126180 | 133050 | 4067700 | 1374400 | 107050 | 0 | 0 | 30155 | 0 | 0 | 0 | 0 |
| SPPLLESPI | SNX17 | Q15036 | SNX17_HUMAN | Unmodified | 0 | 62937 | 0 | 143860 | 3043100 | 0 | 0 | 0 | 320620 | 0 | 0 | 0 | 334490 | 0 | 0 | 0 | 202580 | 1195100 |
| SPPLNFLS | XPO4 | Q9C0E2 | XPO4_HUMAN | Unmodified | 0 | 0 | 0 | 0 | 0 | 0 | 0 | 0 | 0 | 0 | 0 | 868450 | 0 | 0 | 0 | 0 | 0 | 0 |
| SPPNVQFT | PLXNB2 | O15031 | PLXB2_HUMAN | Unmodified | 0 | 0 | 376790 | 64984 | 0 | 0 | 0 | 0 | 0 | 0 | 0 | 0 | 0 | 0 | 932250 | 543870 | 0 | 0 |
| SPPREGSQ | GPS1; GPS1; G | Q13098; A8 | CSN1_HUMAN | Unmodified | 0 | 100870 | 0 | 0 | 0 | 0 | 0 | 0 | 0 | 0 | 0 | 257260 | 0 | 113150 | 0 | 0 | 0 | 0 |
| SPPTPTGL | KIAA2013 | Q8IYS2 | K2013_HUMAN | Unmodified | 0 | 0 | 0 | 0 | 0 | 0 | 0 | 0 | 15152000 | 1501300 | 190500 | 1263500 | 0 | 0 | 0 | 0 | 0 | 0 |
| SPPTPTGL | KIAA2013 | Q8IYS2 | K2013_HUMAN | Unmodified | 0 | 0 | 29526 | 0 | 0 | 0 | 0 | 0 | 4130800 | 179950 | 72739 | 339170 | 0 | 0 | 79813 | 44315 | 0 | 0 |
| SPPTVLVIC | NAXE; NAXE; N | Q8NCW5; A | NNRE_HUMAN | Unmodified | 0 | 0 | 0 | 0 | 0 | 0 | 0 | 0 | 992790 | 0 | 0 | 1155400 | 0 | 0 | 0 | 0 | 0 | 0 |
| SPPVMVAG | FLOT1; FLOT1; | O75955; A2 | FLOT1_HUMAN | Unmodified | 0 | 0 | 848940 | 416830 | 200760 | 48816 | 0 | 0 | 3962300 | 0 | 258490 | 3476300 | 273310 | 30420 | 2273700 | 932030 | 55158 | 61384 |
| SPPYIFSPI | CYP51A1; N/A; | Q16850; C9 | CP51A_HUMAN | Unmodified | 0 | 0 | 0 | 0 | 0 | 0 | 0 | 0 | 0 | 0 | 0 | 114320 | 0 | 0 | 0 | 0 | 0 | 0 |
| SPPYQLDS | BCAM; BCAM; E | P50895; A0A | BCAM_HUMAN | Unmodified | 456630 | 452500 | 0 | 0 | 0 | 0 | 0 | 0 | 122800 | 96690 | 196550 | 0 | 0 | 0 | 0 | 0 | 0 | 0 |
| SPPYTAFL | EIF4B; EIF4B; E | P23588; F8V | IF4B_HUMAN E | Unmodified | 289780 | 0 | 0 | 0 | 129230 | 0 | 0 | 0 | 397160 | 239630 | 0 | 661600 | 0 | 0 | 588920 | 332620 | 0 | 0 |
| SPQDEGLI | MICAL3 | Q7RTP6 | MICA3_HUMAN | Unmodified | 0 | 0 | 0 | 0 | 2982300 | 362070 | 0 | 0 | 0 | 87413 | 0 | 0 | 0 | 0 | 0 | 0 | 0 | 0 |
| SPQEEALQ | ARHGEF2; ARH | Q92974; V9 | ARHG2_HUMA | Unmodified | 0 | 0 | 0 | 0 | 0 | 0 | 0 | 0 | 295860 | 195730 | 354180 | 1309600 | 0 | 0 | 0 | 58849 | 241050 | 0 |
| SPQIFDDEI | MYO1D; MYO1 | O94832; J3Q | MYO1D_HUMA | Unmodified | 6435100 | 2767600 | 1022600 | 1376700 | 0 | 0 | 0 | 0 | 990920 | 0 | 0 | 478160 | 0 | 0 | 1906600 | 738270 | 0 | 0 |
| SPQISMSD | AHNAK | Q09666 | AHNK_HUMAN | Unmodified | 29480 | 19612 | 0 | 43058 | 930230 | 94303 | 0 | 0 | 0 | 75790 | 0 | 0 | 0 | 0 | 617360 | 27853 | 90237 | 481030 |
| SPQISMSD | AHNAK | Q09666 | AHNK_HUMAN | Unmodified | 0 | 0 | 0 | 0 | 1203300 | 252520 | 0 | 0 | 0 | 0 | 0 | 65347 | 0 | 0 | 0 | 0 | 613570 | 854460 |
| SPQISMSD | AHNAK | Q09666 | AHNK_HUMAN | Oxidation (M) | 0 | 0 | 0 | 0 | 1067600 | 0 | 0 | 0 | 0 | 141450 | 0 | 0 | 0 | 0 | 0 | 0 | 0 | 0 |
| SPQMVSAI | CTNNB1; CTNN | P35222; A0A | CTNB1_HUMAI | Unmodified | 810830 | 0 | 2164900 | 2700400 | 0 | 0 | 0 | 0 | 2659100 | 0 | 0 | 6235100 | 0 | 0 | 3158200 | 2445200 | 0 | 0 |
| SPQMVSAI | CTNNB1; CTNN | P35222; A0A | CTNB1_HUMAI | Oxidation (M) | 0 | 0 | 262230 | 266260 | 0 | 0 | 0 | 0 | 0 | 0 | 0 | 1607400 | 0 | 0 | 452780 | 540630 | 0 | 0 |
| SPQPPVEEI | HNRNPU; HNR | Q00839; A0 | HNRPU_HUMA | Unmodified | 0 | 0 | 0 | 0 | 0 | 0 | 1011500 | 0 | 0 | 0 | 0 | 372980 | 0 | 0 | 0 | 0 | 0 | 0 |
| SPQPSSPA | TRPM4 | Q8TD43 | TRPM4_HUMAI | Unmodified | 637040 | 1007300 | 585770 | 306640 | 0 | 0 | 0 | 0 | 0 | 0 | 0 | 0 | 0 | 0 | 721910 | 705750 | 0 | 0 |
| SPRDYSNF | PRKCD | Q05655 | KPCD_HUMAN | Unmodified | 0 | 0 | 0 | 370140 | 654290 | 0 | 0 | 0 | 932230 | 216640 | 271070 | 3065700 | 382270 | 0 | 567910 | 0 | 820190 | 2157000 |
| SPSAQELK | STUB1; STUB1 | Q9UNE7; H3 | CHIP_HUMAN | Unmodified | 0 | 0 | 315090 | 0 | 0 | 0 | 0 | 0 | 0 | 0 | 0 | 0 | 0 | 0 | 0 | 0 | 0 | 0 |
| SPSDSSTA | DBN1; DBN1 | Q16643; D6 | DREB_HUMAN | Unmodified | 0 | 0 | 0 | 0 | 0 | 0 | 0 | 0 | 0 | 0 | 911570 | 1988400 | 0 | 0 | 0 | 0 | 0 | 0 |
| SPSDTEGL | MPZL1 | O95297 | MPZL1_HUMAN | Unmodified | 588330 | 448880 | 2743100 | 0 | 1580300 | 476000 | 352240 | 265440 | 1125200 | 1119100 | 1713700 | 5586800 | 262490 | 380050 | 1797700 | 1412800 | 1496100 | 792390 |
| SPSELFAQI | THRAP3; THRAI | Q9Y2W1; A0 | TR150_HUMAN | Unmodified | 0 | 0 | 154960 | 0 | 0 | 0 | 300730 | 0 | 0 | 0 | 0 | 1029900 | 0 | 0 | 0 | 0 | 0 | 0 |
| SPSESFIGF | PRKD1; PRKD1 | Q15139; F8 | KPCD1_HUMAI | Unmodified | 0 | 0 | 0 | 0 | 0 | 0 | 0 | 0 | 456530 | 0 | 0 | 0 | 400090 | 0 | 0 | 0 | 0 | 0 |
| SPSGGAAC | KCTD12 | Q96CX2 | KCD12_HUMAI | Unmodified | 0 | 0 | 0 | 0 | 0 | 0 | 0 | 0 | 1507500 | 189820 | 0 | 1195400 | 0 | 0 | 0 | 0 | 0 | 0 |
| SPSKEPHF | SLC38A3; SLC3 | Q99624; A0 | S38A3_HUMAN | Unmodified | 0 | 0 | 0 | 0 | 0 | 0 | 0 | 0 | 611380 | 139300 | 0 | 0 | 0 | 0 | 0 | 0 | 0 | 0 |
| SPSKPLPE | SSB; SSB; SSB; | P05455; E9P | LA_HUMAN Lu | Unmodified | 775020 | 0 | 0 | 0 | 0 | 0 | 0 | 0 | 141900 | 0 | 1379500 | 4612400 | 0 | 0 | 0 | 86285 | 0 | 0 |
| SPSLGSDL | SNTB2 | Q13425 | SNTB2_HUMAN | Unmodified | 159120 | 149160 | 1272800 | 0 | 0 | 0 | 0 | 0 | 592400 | 0 | 0 | 800640 | 0 | 0 | 2171400 | 1503300 | 0 | 0 |

| | | | | | | | | | | | | | | | | | | | | | | |
|---|---|---|---|---|---|---|---|---|---|---|---|---|---|---|---|---|---|---|---|---|---|---|
| SPSPEFSGI | FOLH1B; FOLH | Q9HBA9; Q0 | FOH1B_HUMAI | Unmodified | 0 | 0 | 1817700 | 5297000 | 0 | 0 | 0 | 0 | 0 | 0 | 0 | 0 | 0 | 0 | 1834200 | 0 | 0 | 0 |
| SPSPEFSGI | FOLH1B; FOLH | Q9HBA9; Q0 | FOH1B_HUMAI | Oxidation (M) | 74475 | 150500 | 1291200 | 790130 | 0 | 0 | 0 | 0 | 0 | 0 | 0 | 0 | 0 | 0 | 339970 | 1623900 | 0 | 0 |
| SPSQDNTD | RASAL2; RASAL | Q9UJF2; H0Y | NGAP_HUMAN | Unmodified | 0 | 0 | 0 | 0 | 1469700 | 154010 | 41559 | 61797 | 0 | 72112 | 0 | 0 | 0 | 0 | 0 | 0 | 294630 | 100670 |
| SPSSALSS\ | FGD6; FGD6 | Q6ZV73; H0Y | FGD6_HUMAN | Unmodified | 0 | 0 | 0 | 0 | 1025300 | 264300 | 0 | 0 | 0 | 0 | 0 | 0 | 0 | 0 | 0 | 0 | 0 | 0 |
| SPSSLLIAN | FAM91A1 | Q658Y4 | F91A1_HUMAN | Unmodified | 0 | 0 | 0 | 0 | 0 | 0 | 0 | 0 | 0 | 0 | 0 | 655150 | 0 | 0 | 0 | 0 | 0 | 0 |
| SPSTDIFDN | GULP1; GULP1; | Q9UBP9; H0 | GULP1_HUMAI | 2 Oxidation (M) | 0 | 363090 | 138700 | 1108300 | 0 | 0 | 0 | 0 | 0 | 0 | 0 | 0 | 0 | 0 | 0 | 0 | 0 | 0 |
| SPSWTAPP | ZFAND2B; ZFAN | Q8WV99; A0 | ZFN2B_HUMAN | Unmodified | 2616300 | 4075000 | 233760 | 898220 | 0 | 0 | 0 | 0 | 0 | 0 | 0 | 0 | 0 | 0 | 1592200 | 1576500 | 0 | 0 |
| SPSYIVCN1 | PLXNA4 | Q9HCM2 | PLXA4_HUMAN | Unmodified | 0 | 2215500 | 0 | 0 | 0 | 0 | 0 | 0 | 0 | 0 | 0 | 0 | 0 | 0 | 1926400 | 2570500 | 0 | 0 |
| SPTAALNES | CEP55 | Q53EZ4 | CEP55_HUMAN | Unmodified | 342470 | 0 | 0 | 0 | 0 | 1060000 | 0 | 0 | 0 | 740780 | 0 | 0 | 0 | 454190 | 0 | 0 | 0 | 0 |
| SPTDWALF | DBNL; DBNL; D | Q9UJU6; F8\ | DBNL_HUMAN | Unmodified | 884040 | 932610 | 347460 | 548040 | 462870 | 487130 | 0 | 0 | 0 | 0 | 768350 | 3460500 | 0 | 0 | 1510700 | 567630 | 0 | 2598200 |
| SPTGEWLP | SFTPB | P07988 | PSPB_HUMAN | Unmodified | 0 | 0 | 0 | 0 | 0 | 0 | 62477 | 0 | 961060 | 90318 | 0 | 0 | 122040 | 81725 | 0 | 0 | 0 | 0 |
| SPTGSWQI | PTGFRN | Q9P2B2 | FPRP_HUMAN | Unmodified | 4306700 | 3982400 | 14463000 | 11314000 | 0 | 0 | 229460 | 22915 | 627740 | 0 | 1150500 | 20413000 | 4727300 | 1571300 | 15261000 | 16071000 | 0 | 0 |
| SPTITGFFA | NOXO1 | Q8NFA2 | NOXO1_HUMA | Unmodified | 0 | 0 | 0 | 0 | 0 | 0 | 0 | 0 | 0 | 0 | 356810 | 0 | 0 | 459400 | 0 | 0 | 0 | 0 |
| SPTKAPHL( | EPB41L2; EPB4 | O43491; A0/ | E41L2_HUMAN | Unmodified | 0 | 0 | 0 | 0 | 352570 | 0 | 0 | 0 | 0 | 0 | 0 | 0 | 0 | 0 | 0 | 0 | 166310 | 232910 |
| SPTKAPHL( | EPB41L2; EPB4 | O43491; H0Y | E41L2_HUMAN | Unmodified | 0 | 0 | 0 | 0 | 1423600 | 234140 | 0 | 0 | 0 | 0 | 0 | 66484 | 0 | 0 | 0 | 0 | 440230 | 1220900 |
| SPTLGIAR | CDCA3; CDCA3 | Q99618; F8V | CDCA3_HUMA | Unmodified | 2091600 | 1608100 | 1574600 | 102020 | 0 | 0 | 0 | 240970 | 81972 | 28180 | 91040 | 343130 | 379010 | 63205 | 116000 | 1343400 | 60883 | 74348 |
| SPTNTISVY | SEZ6L; SEZ6L; S | Q9BYH1; B0 | SE6L1_HUMAN | Unmodified | 0 | 0 | 0 | 0 | 0 | 0 | 0 | 0 | 1583800 | 0 | 0 | 0 | 0 | 0 | 0 | 0 | 0 | 0 |
| SPVAKDVD | VCP; VCP; VCP; | P55072; C9J | TERA_HUMAN | Unmodified | 0 | 0 | 0 | 0 | 12050000 | 1628900 | 1443300 | 982960 | 0 | 5098200 | 0 | 0 | 13487000 | 9053900 | 0 | 0 | 2094300 | 1664900 |
| SPVAVQSTI | ECH1; ECH1 | Q13011; M0I | ECH1_HUMAN | Unmodified | 0 | 0 | 0 | 0 | 0 | 0 | 0 | 0 | 1683100 | 0 | 0 | 0 | 0 | 0 | 0 | 0 | 0 | 0 |
| SPVDIVTG( | LRBA; LRBA; LR | P50851; A0A | LRBA_HUMAN | Unmodified | 0 | 39786 | 0 | 0 | 0 | 0 | 0 | 0 | 0 | 0 | 227080 | 2556500 | 0 | 0 | 32827 | 55825 | 0 | 0 |
| SPVDVLQIF | PARP4 | Q9UKK3 | PARP4_HUMAN | Unmodified | 29714000 | 34534000 | 30150 | 31172000 | 0 | 0 | 0 | 196190 | 292560 | 0 | 0 | 3285600 | 0 | 0 | 22616000 | 18029000 | 0 | 0 |
| SPVEFVEYN | PLCB1; PLCB1; | Q9NQ66; A0 | PLCB1_HUMAN | Unmodified | 0 | 0 | 0 | 0 | 0 | 0 | 0 | 279800 | 1093500 | 301500 | 0 | 0 | 0 | 0 | 0 | 0 | 0 | 0 |
| SPVESTTEF | SEC24C; SEC24 | P53992; G5E | SC24C_HUMAI | Unmodified | 0 | 0 | 0 | 0 | 0 | 0 | 0 | 0 | 1280700 | 565390 | 171730 | 1227800 | 0 | 0 | 0 | 0 | 0 | 332730 |
| SPVFIAQV( | GRK5 | P34947 | GRK5_HUMAN | Unmodified | 0 | 0 | 225740 | 0 | 315900 | 0 | 168690 | 0 | 0 | 185420 | 6777400 | 6354600 | 0 | 0 | 0 | 36882 | 663850 | 518800 |
| SPVFLYEDI | COL5A1 | P20908 | CO5A1_HUMAI | Unmodified | 0 | 0 | 651800 | 649810 | 0 | 0 | 0 | 0 | 0 | 0 | 0 | 408520 | 0 | 0 | 0 | 0 | 0 | 0 |
| SPVIEFWEI | SLC6A8; SLC6A | P48029; H7( | SC6A8_HUMAN | Unmodified | 0 | 0 | 710410 | 0 | 0 | 0 | 0 | 0 | 0 | 0 | 0 | 2632600 | 0 | 0 | 0 | 713920 | 0 | 0 |
| SPVISIDPP! | HSPG2 | P98160 | PGBM_HUMAN | Unmodified | 0 | 0 | 2803900 | 1708100 | 0 | 0 | 0 | 0 | 25441000 | 41950 | 642750 | 0 | 0 | 930550 | 0 | 1032500 | 0 | 2330500 |
| SPVPSPGS | MICALL1 | B0QY86 | B0QY86_HUM/ | Unmodified | 0 | 0 | 0 | 225330 | 5726700 | 7350600 | 1576900 | 43337 | 0 | 0 | 0 | 0 | 777430 | 5366000 | 0 | 0 | 944780 | 11665000 |
| SPVQENSS | CLDND1; CLDN | Q9NY35; D6 | CLDN1_HUMAI | Unmodified | 187730 | 155090 | 266070 | 0 | 0 | 0 | 0 | 63154 | 0 | 146460 | 0 | 1128000 | 0 | 0 | 538190 | 470510 | 300070 | 0 |
| SPVYLTVLK | NPEPPS; NPEPI | P55786; A0A | PSA_HUMAN P | Unmodified | 0 | 0 | 999050 | 689670 | 473950 | 330560 | 0 | 0 | 1640100 | 277810 | 1343300 | 10931000 | 0 | 0 | 1160200 | 1275800 | 43892 | 0 |
| SPVYSHFN | ABCC1; ABCC1 | P33527; I3L4 | MRP1_HUMAN | Unmodified | 0 | 0 | 801440 | 167850 | 0 | 0 | 0 | 0 | 0 | 0 | 987340 | 8401300 | 0 | 0 | 1188500 | 995620 | 0 | 0 |
| SPWSNKYI | CAPZB; CAPZB; | P47756; A0A | CAPZB_HUMAN | Oxidation (M) | 470900 | 227470 | 182210 | 215650 | 0 | 0 | 0 | 0 | 0 | 0 | 162320 | 2889500 | 0 | 0 | 415700 | 677710 | 0 | 0 |
| SPWSNKYI | CAPZB; CAPZB; | P47756; A0A | CAPZB_HUMAN | Unmodified | 131090 | 209420 | 0 | 0 | 0 | 0 | 0 | 0 | 0 | 0 | 0 | 3210400 | 0 | 0 | 0 | 95339 | 0 | 0 |
| SPYLYPLY( | GDI1; GDI2; GD | P31150; P50 | GDIA_HUMAN | Unmodified | 2001500 | 1384300 | 918530 | 4845800 | 7392800 | 0 | 0 | 0 | 21125000 | 10033000 | 5964000 | 13406000 | 6108000 | 3314700 | 7770900 | 2248200 | 0 | 4437200 |
| SPYNYLQV | P3H1; P3H1 | Q32P28; E2( | P3H1_HUMAN | Unmodified | 0 | 0 | 0 | 0 | 0 | 2366000 | 0 | 0 | 1602600 | 0 | 0 | 0 | 0 | 0 | 0 | 0 | 0 | 1461600 |
| SPYQEFTDI | RPS2; RPS2; RP | P15880; E9P | RS2_HUMAN 4 | Unmodified | 7643500 | 6814400 | 4850400 | 4875300 | 4421900 | 980350 | 5190700 | 2375100 | 4892200 | 1506900 | 7906700 | 50980000 | 5608300 | 2955800 | 4492500 | 3719900 | 1439400 | 3350800 |
| SPYTVTVG( | FLNA; FLNA; FL | P21333; F8V | FLNA_HUMAN | Unmodified | 0 | 197940 | 0 | 0 | 0 | 0 | 0 | 0 | 0 | 373700 | 0 | 434900 | 0 | 0 | 0 | 84979 | 0 | 0 |
| SQAEFEK | DBI | P07108 | ACBP_HUMAN | Acetyl (Protein N | 867230 | 281950 | 973660 | 479160 | 504100 | 39502 | 37225 | 49292 | 337510 | 216580 | 498050 | 1189600 | 192090 | 144650 | 1288700 | 767480 | 310520 | 602160 |
| SQAEFEKA | DBI | P07108 | ACBP_HUMAN | Acetyl (Protein N | 0 | 0 | 0 | 236750 | 1301100 | 0 | 1141600 | 0 | 706060 | 5619900 | 0 | 952840 | 1242300 | 0 | 0 | 0 | 142150 | 1009400 |
| SQAKPGTP | EPHB4; EPHB4 | P54760; Q9( | EPHB4_HUMAN | Unmodified | 359700 | 0 | 0 | 0 | 811860 | 155150 | 0 | 0 | 0 | 0 | 0 | 0 | 238840 | 105670 | 0 | 142070 | 340720 | 1072200 |
| SQALNAFG | DNAJC3 | Q13217 | DNJC3_HUMAN | Unmodified | 0 | 0 | 0 | 0 | 0 | 43131 | 0 | 0 | 3832300 | 894500 | 86899 | 0 | 0 | 0 | 0 | 0 | 0 | 0 |
| SQAPGQP( | DAZAP1; DAZAF | Q96EP5; K7E | DAZP1_HUMAN | Unmodified | 0 | 0 | 0 | 0 | 0 | 0 | 0 | 0 | 412290 | 0 | 0 | 751730 | 164180 | 0 | 0 | 0 | 0 | 0 |
| SQASGPEP | SIT1; SIT1 | Q9Y3P8; A0/ | SIT1_HUMAN S | Unmodified | 0 | 0 | 0 | 0 | 0 | 0 | 0 | 0 | 1810700 | 1450100 | 0 | 0 | 0 | 0 | 0 | 0 | 0 | 0 |
| SQAVKDRA | ARPC5; ARPC5 | O15511; B1/ | ARPC5_HUMAN | Unmodified | 0 | 0 | 0 | 0 | 551680 | 0 | 827030 | 771080 | 0 | 65572 | 0 | 0 | 532460 | 642490 | 0 | 0 | 893220 | 217350 |
| SQCLQVPE | PDCD2L; N/A; N | Q9BRP1; U3 | PDD2L_HUMAN | Unmodified | 90521 | 0 | 0 | 0 | 0 | 0 | 0 | 0 | 75484 | 0 | 103380 | 620200 | 0 | 0 | 84352 | 63827 | 0 | 0 |
| SQCLSTLVI | STK25 | O00506 | STK25_HUMAN | Unmodified | 124320 | 0 | 131430 | 467860 | 6227500 | 770240 | 265260 | 13171 | 203910 | 178070 | 456200 | 688320 | 4024000 | 1889300 | 535130 | 609300 | 1266400 | 5070800 |
| SQDADSP( | AFDN; AFDN; A | P55196; H0Y | AFAD_HUMAN | Unmodified | 0 | 0 | 0 | 0 | 1058400 | 0 | 0 | 93496 | 97471 | 0 | 221560 | 1206400 | 1453300 | 1084700 | 0 | 0 | 0 | 418230 |
| SQDAEVGI | CCT7; CCT7 | Q99832; F8V | TCPH_HUMAN | Unmodified | 96047 | 127380 | 68465 | 296750 | 2220000 | 229130 | 1063900 | 88348 | 350350 | 620480 | 541390 | 1376200 | 5638600 | 3500300 | 535950 | 142280 | 1208900 | 2134600 |
| SQDAQLVL | SLC12A7; SLC1 | Q9Y666; A0/ | S12A7_HUMAN | Oxidation (M) | 189320 | 0 | 203470 | 219960 | 0 | 0 | 0 | 587530 | 0 | 0 | 0 | 1121500 | 593510 | 527550 | 563970 | 393620 | 0 | 0 |
| SQDAQLVL | SLC12A7; SLC1 | Q9Y666; A0/ | S12A7_HUMAN | Unmodified | 0 | 0 | 0 | 0 | 0 | 0 | 602540 | 148990 | 840600 | 125890 | 0 | 586640 | 1290500 | 0 | 250940 | 205330 | 0 | 0 |
| SQDDRDKI | ATP1B3; ATP1B | P54709; C9J | AT1B3_HUMAN | Unmodified | 0 | 18956 | 0 | 348500 | 5075100 | 246420 | 0 | 0 | 131620 | 669510 | 136830 | 223890 | 3226400 | 2797800 | 0 | 0 | 255980 | 0 |
| SQDHFHVI | TIA1; TIA1; TIA1 | P31483; E5R | TIA1_HUMAN N | Unmodified | 0 | 0 | 0 | 0 | 146750 | 0 | 0 | 0 | 221880 | 0 | 0 | 830710 | 0 | 0 | 0 | 0 | 0 | 0 |
| SQDIPFVS1 | COBLL1; COBLI | Q53SF7; A0/ | COBL1_HUMAI | Unmodified | 658280 | 543370 | 0 | 406710 | 0 | 0 | 0 | 0 | 0 | 0 | 0 | 0 | 0 | 0 | 439140 | 419020 | 0 | 0 |
| SQDIYLR | RPL18; RPL18; I | Q07020; F8V | RL18_HUMAN ( | Unmodified | 0 | 1964900 | 597380 | 0 | 59334 | 0 | 0 | 0 | 0 | 469130 | 66966 | 5313000 | 0 | 0 | 1982800 | 0 | 0 | 80287 |
| SQDLHCG/ | CNPY2; N/A; CI | Q9Y2B0; F8V | CNPY2_HUMAI | Unmodified | 0 | 0 | 25727 | 0 | 0 | 0 | 0 | 0 | 2777300 | 93852 | 0 | 79455 | 0 | 0 | 133970 | 10752 | 0 | 0 |
| SQDPPALA | ZNF185; ZNF18 | O15231; H7I | ZN185_HUMAN | Unmodified | 0 | 0 | 0 | 0 | 160550 | 0 | 0 | 177520 | 0 | 0 | 29691 | 891230 | 0 | 0 | 0 | 68518 | 357200 | 0 |
| SQDPPALA | ZNF185; ZNF18 | O15231; H7I | ZN185_HUMAN | Unmodified | 0 | 0 | 0 | 0 | 0 | 0 | 0 | 0 | 0 | 0 | 0 | 0 | 0 | 0 | 0 | 0 | 929670 | 0 |
| SQDPRNPI | TKT; TKT; TKT | P29401; F8V | TKT_HUMAN Tr | Unmodified | 0 | 0 | 0 | 0 | 3211700 | 511980 | 208020 | 55563 | 206740 | 685130 | 203970 | 0 | 937400 | 184060 | 0 | 0 | 796400 | 788700 |
| SQDSGIAEI | PHYHIPL | Q96FC7 | PHIPL_HUMAN | Unmodified | 0 | 0 | 0 | 0 | 0 | 0 | 289990 | 179430 | 586800 | 190240 | 1046200 | 0 | 0 | 0 | 0 | 0 | 0 | 182940 |
| SQDTEVDN | SLC3A2; SLC3A | P08195; A0A | 4F2_HUMAN 4I | Acetyl (Protein N | 106540 | 147600 | 207350 | 0 | 599110 | 0 | 0 | 0 | 863870 | 0 | 586790 | 262520 | 0 | 0 | 220100 | 116700 | 0 | 138590 |
| SQDTEVDN | SLC3A2; SLC3A | P08195; A0A | 4F2_HUMAN 4I | Acetyl (Protein N | 0 | 0 | 661900 | 0 | 2843400 | 0 | 0 | 547890 | 0 | 0 | 0 | 0 | 853570 | 1060700 | 912450 | 822940 | 2629500 | 0 |
| SQDTEVDN | SLC3A2; SLC3A | P08195; A0A | 4F2_HUMAN 4I | Acetyl (Protein N | 0 | 0 | 0 | 0 | 3868200 | 0 | 0 | 225690 | 1074500 | 0 | 0 | 469080 | 0 | 0 | 1528200 | 0 | 2040800 | 0 |
| SQDTEVDN | SLC3A2; SLC3A | P08195; A0A | 4F2_HUMAN 4I | Acetyl (Protein N | 0 | 0 | 0 | 0 | 6950200 | 0 | 0 | 0 | 0 | 0 | 0 | 0 | 0 | 0 | 991870 | 480310 | 1087300 | 0 |
| SQDTEVDN | SLC3A2; SLC3A | P08195; A0A | 4F2_HUMAN 4I | Acetyl (Protein N | 0 | 0 | 0 | 0 | 13452000 | 0 | 0 | 0 | 0 | 0 | 0 | 0 | 0 | 0 | 0 | 0 | 0 | 2873300 |
| SQDTEVDN | SLC3A2; SLC3A | P08195; A0A | 4F2_HUMAN 4I | Acetyl (Protein N | 0 | 0 | 0 | 0 | 3557300 | 0 | 0 | 0 | 744220 | 0 | 0 | 0 | 0 | 0 | 0 | 0 | 0 | 0 |
| SQEAGGR\ | CD151; CD151; | P48509; E9P | CD151_HUMAI | Unmodified | 1681100 | 284130 | 266820 | 0 | 519780 | 297350 | 0 | 0 | 287590 | 0 | 0 | 0 | 0 | 0 | 324430 | 190060 | 777550 | 762910 |
| SQEATEAAI | SEPTIN9; SEPTI | Q9UHD8; K7 | SEPT9_HUMAN | Unmodified | 0 | 0 | 0 | 232010 | 2366600 | 0 | 0 | 0 | 473510 | 0 | 0 | 0 | 229640 | 0 | 0 | 97479 | 1898300 | 2465600 |
| SQEEPKDT | BCLAF1; BCLAF | Q9NYF8; E9I | BCLF1_HUMAN | Unmodified | 0 | 0 | 534380 | 427670 | 0 | 986790 | 0 | 0 | 0 | 0 | 3611800 | 2458700 | 242740 | 0 | 273710 | 0 | 0 | 0 |
| SQEESEEG | CHGB; CHGB | P05060; A0A | SCG1_HUMAN | Unmodified | 0 | 0 | 0 | 0 | 0 | 0 | 0 | 0 | 660160 | 0 | 0 | 0 | 0 | 0 | 0 | 0 | 200740 | 0 |
| SQEESEEG | CHGB; CHGB | P05060; A0A | SCG1_HUMAN | Unmodified | 0 | 0 | 0 | 0 | 0 | 0 | 0 | 0 | 433830 | 73939 | 0 | 0 | 0 | 0 | 0 | 0 | 0 | 214250 |
| SQEESEEG | CHGB; CHGB | P05060; A0A | SCG1_HUMAN | Unmodified | 0 | 0 | 0 | 0 | 0 | 0 | 0 | 0 | 657270 | 0 | 0 | 0 | 0 | 0 | 0 | 0 | 0 | 0 |
| SQEGDSIS' | CPD | O75976 | CBPD_HUMAN | Unmodified | 0 | 0 | 0 | 0 | 0 | 0 | 0 | 0 | 8087400 | 811530 | 0 | 174430 | 0 | 0 | 0 | 0 | 0 | 0 |
| SQEGENEE | INPP5D; INPP5I | Q92835; H0Y | SHIP1_HUMAN | Unmodified | 0 | 0 | 0 | 0 | 0 | 0 | 0 | 0 | 0 | 0 | 300880 | 943780 | 0 | 0 | 0 | 0 | 0 | 0 |
| SQEGESVT | SEL1L; SEL1L | Q9UBV2; G3 | SE1L1_HUMAN | Unmodified | 0 | 0 | 136370 | 0 | 0 | 0 | 0 | 0 | 4883500 | 0 | 0 | 0 | 0 | 0 | 0 | 0 | 0 | 0 |
| SQEGESVT | SEL1L; SEL1L | Q9UBV2; G3 | SE1L1_HUMAN | Unmodified | 0 | 0 | 0 | 0 | 0 | 0 | 0 | 0 | 4412400 | 0 | 0 | 0 | 0 | 0 | 0 | 0 | 0 | 0 |
| SQEGRPVQ | COPS6; COPS6 | Q7L5N1; E7I | CSN6_HUMAN | Unmodified | 0 | 0 | 150660 | 0 | 0 | 0 | 0 | 0 | 2424100 | 410960 | 143770 | 2237000 | 0 | 0 | 426340 | 306040 | 0 | 0 |
| SQEGRPVQ | COPS6; COPS6 | Q7L5N1; E7I | CSN6_HUMAN | Unmodified | 0 | 126770 | 0 | 155120 | 0 | 0 | 0 | 39212 | 0 | 0 | 669770 | 2443400 | 224790 | 0 | 174930 | 0 | 225810 | 0 |
| SQEGSQF\ | ETF1; ETF1 | P62495; B7Z | ERF1_HUMAN I | Unmodified | 58639 | 58620 | 694430 | 388110 | 662730 | 249530 | 23780 | 0 | 956880 | 71435 | 2292300 | 2407200 | 0 | 0 | 922040 | 809860 | 165520 | 691830 |
| SQEHFSHN | PPP3CB; PPP3( | P16298; Q5F | PP2BB_HUMAN | Unmodified | 2294800 | 1739600 | 24984 | 0 | 799600 | 0 | 184540 | 54841 | 1600300 | 379360 | 58229 | 4922800 | 0 | 168010 | 0 | 3409200 | 247020 | 99601 |
| SQEHFTAL( | INPP5A; INPP5/ | Q14642; H0Y | I5P1_HUMAN I | Unmodified | 0 | 0 | 0 | 2181700 | 1060900 | 0 | 0 | 0 | 1251000 | 0 | 0 | 0 | 374820 | 0 | 0 | 4241300 | 283470 | 0 |
| SQEKADLY | BPNT2 | Q9NX62 | IMPA3_HUMAN | Unmodified | 0 | 0 | 0 | 0 | 0 | 0 | 0 | 0 | 402040 | 2021500 | 0 | 0 | 0 | 0 | 0 | 0 | 320460 | 0 |
| SQEKPREIN | DARS1; DARS1 | P14868; H7( | SYDC_HUMAN | Unmodified | 82534 | 42704 | 0 | 0 | 1663500 | 0 | 299300 | 0 | 321790 | 0 | 0 | 738830 | 1326100 | 0 | 37887 | 51044 | 454300 | 1409400 |
| SQELEVKN | DYNC1H1; DYN | Q14204; A0/ | DYHC1_HUMA | Unmodified | 0 | 0 | 0 | 0 | 301170 | 0 | 0 | 39064 | 509010 | 34038 | 0 | 0 | 0 | 0 | 0 | 0 | 198000 | 155930 |
| SQELSGPA | ANKRD13A | Q8IZ07 | AN13A_HUMAN | Unmodified | 917010 | 573300 | 0 | 1577700 | 0 | 0 | 0 | 0 | 0 | 0 | 0 | 0 | 0 | 0 | 693040 | 0 | 0 | 0 |
| SQEMVHLV | CD44; CD44; C | P16070; H0Y | CD44_HUMAN | Unmodified | 0 | 0 | 0 | 0 | 3201000 | 0 | 0 | 0 | 0 | 0 | 0 | 0 | 0 | 0 | 0 | 0 | 0 | 2436100 |
| SQEMVHLV | CD44; CD44; C | P16070; H0Y | CD44_HUMAN | Oxidation (M) | 134070 | 129430 | 0 | 0 | 0 | 241660 | 0 | 0 | 0 | 0 | 0 | 0 | 0 | 0 | 0 | 0 | 293280 | 565150 |
| SQEMVHLV | CD44; CD44; C | P16070; H0Y | CD44_HUMAN | Unmodified | 76948 | 0 | 0 | 0 | 3543900 | 0 | 0 | 0 | 0 | 0 | 0 | 0 | 0 | 0 | 0 | 0 | 405680 | 2601000 |
| SQEMVHLV | CD44; CD44; C | P16070; H0Y | CD44_HUMAN | Oxidation (M) | 0 | 0 | 0 | 0 | 9542300 | 0 | 0 | 0 | 0 | 0 | 0 | 0 | 0 | 0 | 0 | 0 | 0 | 7693400 |
| SQEMVHLV | CD44; CD44; C | P16070; H0Y | CD44_HUMAN | 2 Oxidation (M) | 0 | 0 | 0 | 0 | 3148600 | 0 | 0 | 0 | 0 | 0 | 0 | 0 | 0 | 0 | 0 | 0 | 0 | 579400 |
| SQEMVHLV | CD44; CD44; C | P16070; H0Y | CD44_HUMAN | Unmodified | 0 | 0 | 0 | 0 | 4489400 | 0 | 0 | 0 | 0 | 0 | 0 | 0 | 0 | 0 | 0 | 0 | 0 | 6815600 |
| SQEPIPDD( | MTDH; MTDH; N | Q86UE4; H0 | LYRIC_HUMAN | Unmodified | 3204200 | 1184400 | 3626100 | 380290 | 0 | 0 | 380960 | 0 | 124020 | 0 | 53153 | 341310 | 0 | 0 | 5199100 | 4351900 | 0 | 552180 |
| SQEPIPDD( | MTDH; MTDH; N | Q86UE4; H0 | LYRIC_HUMAN | Unmodified | 0 | 0 | 0 | 0 | 0 | 0 | 0 | 0 | 0 | 0 | 0 | 0 | 0 | 0 | 0 | 0 | 0 | 0 |
| SQEQLAAE | EZR; EZR | P15311; E7E | EZRI_HUMAN E | Unmodified | 3526800 | 1227400 | 548480 | 4580000 | 45197000 | 13782000 | 186280 | 462220 | 39955000 | 9483500 | 5562700 | 40130000 | 16417000 | 9834900 | 6208500 | 5071100 | 2215500 | 7947900 |
| SQERPTFYI | MAPK14; MAPK | Q16539; B5I | MK14_HUMAN | Acetyl (Protein N | 0 | 1120700 | 269500 | 450150 | 2121800 | 87033 | 0 | 0 | 778710 | 59794 | 0 | 1073500 | 1154500 | 1099800 | 407100 | 373790 | 1528200 | 1525600 |
| SQERVEGK | ASS1 | P00966 | ASSY_HUMAN . | Unmodified | 0 | 0 | 0 | 0 | 77681 | 0 | 6525800 | 2930800 | 56540 | 634330 | 0 | 0 | 671030 | 721900 | 0 | 0 | 68392 | 225170 |
| SQESGGTE | GPRIN3 | Q6ZVF9 | GRIN3_HUMAN | Unmodified | 0 | 0 | 0 | 0 | 0 | 0 | 235060 | 0 | 0 | 0 | 0 | 0 | 0 | 0 | 0 | 0 | 0 | 0 |
| SQETECTYI | MDH2; MDH2 | P40926; G3X | MDHM_HUMAN | Unmodified | 46143 | 0 | 118760 | 176670 | 1472700 | 174670 | 3340800 | 223170 | 15682000 | 5286000 | 185040 | 397070 | 2055200 | 1158000 | 0 | 0 | 366350 | 488260 |
| SQETECTYI | MDH2; MDH2 | P40926; G3X | MDHM_HUMAN | Unmodified | 0 | 0 | 0 | 0 | 0 | 136920 | 2438400 | 323470 | 10842000 | 3401400 | 0 | 74671 | 1172500 | 1384800 | 0 | 0 | 125570 | 1440900 |
| SQETECTYI | MDH2; MDH2 | P40926; G3X | MDHM_HUMAN | Unmodified | 0 | 0 | 0 | 0 | 0 | 0 | 2199000 | 296380 | 0 | 0 | 170940 | 0 | 0 | 274280 | 0 | 0 | 0 | 0 |
| SQETGDLD | FBLN1; FBLN1; | P23142; B1A | FBLN1_HUMAN | Unmodified | 0 | 0 | 0 | 0 | 0 | 0 | 0 | 0 | 0 | 832140 | 782620 | 0 | 69335 | 0 | 0 | 0 | 0 | 0 |
| SQETGDLD | FBLN1; FBLN1; | P23142; B1A | FBLN1_HUMAN | Unmodified | 0 | 0 | 0 | 0 | 0 | 0 | 0 | 0 | 3576100 | 862900 | 0 | 0 | 0 | 0 | 0 | 0 | 0 | 0 |
| SQEVEDKT | KIF5B | P33176 | KINH_HUMAN | Unmodified | 0 | 0 | 0 | 0 | 3759800 | 0 | 925740 | 343020 | 0 | 1438700 | 0 | 0 | 1660700 | 0 | 0 | 0 | 800700 | 1378300 |
| SQFLINGKI | GUSB | P08236 | BGLR_HUMAN | Unmodified | 0 | 0 | 110440 | 0 | 0 | 0 | 0 | 0 | 1113600 | 342850 | 0 | 164220 | 0 | 0 | 208860 | 0 | 0 | 0 |
| SQFLINGKI | GUSB | P08236 | BGLR_HUMAN | Unmodified | 0 | 0 | 0 | 0 | 0 | 0 | 926760 | 0 | 8177800 | 0 | 0 | 0 | 0 | 0 | 1281700 | 0 | 0 | 0 |
| SQFSDKPV | CHID1; CHID1; | Q9BWS9; E9 | CHID1_HUMAN | Unmodified | 0 | 22867 | 137650 | 326920 | 0 | 0 | 0 | 0 | 28452000 | 1116000 | 826040 | 9782700 | 0 | 0 | 317410 | 69723 | 0 | 0 |
| SQFSPAVF | BTN2A2; BTN2/ | Q8WVV5; Q9 | BT2A2_HUMAN | Unmodified | 0 | 0 | 0 | 0 | 848080 | 0 | 0 | 0 | 1255600 | 68359 | 0 | 2312000 | 2880100 | 0 | 0 | 458560 | 0 | 0 |
| SQFTITPGS | ACO2; ACO2; A | Q99798; A2/ | ACON_HUMAN | Unmodified | 0 | 0 | 0 | 0 | 0 | 0 | 289780 | 0 | 6369500 | 0 | 225410 | 0 | 0 | 96435 | 0 | 0 | 0 | 0 |
| SQFVVTSN | ST14 | Q9Y5Y6 | ST14_HUMAN S | Unmodified | 1627100 | 1209500 | 5493100 | 3520600 | 0 | 0 | 46544 | 143740 | 1845300 | 0 | 150730 | 297900 | 0 | 0 | 4541300 | 4252500 | 156840 | 0 |
| SQGAALDK | MDH1; MDH1; I | P40925; C9J | MDHC_HUMAN | Unmodified | 0 | 0 | 0 | 0 | 71966 | 0 | 0 | 2272500 | 0 | 103560 | 0 | 627480 | 0 | 56596 | 0 | 118280 | 223480 | 228000 |

| | | | | | | | | | | | | | | | | | | | | | | |
|---|---|---|---|---|---|---|---|---|---|---|---|---|---|---|---|---|---|---|---|---|---|---|
| SQGAALDK | MDH1; MDH1; I | P40925; C9J | MDHC_HUMAN | Unmodified | 0 | 0 | 0 | 0 | 0 | 0 | 947340 | 0 | 67964 | 0 | 0 | 0 | 0 | 482350 | 205470 | 0 | 1795600 | 0 |
| SQGCSEQ\ | PRKDC | P78527 | PRKDC_HUMAI | Unmodified | 0 | 0 | 0 | 0 | 0 | 0 | 0 | 0 | 857510 | 0 | 0 | 734550 | 0 | 0 | 0 | 0 | 0 | 0 |
| SQGDASK\ | FLNA; FLNA; FL | P21333; A0A | FLNA_HUMAN | Unmodified | 228080 | 486360 | 0 | 1311600 | 2679300 | 1605400 | 3021400 | 1127400 | 0 | 348610 | 0 | 0 | 1289600 | 1260400 | 0 | 0 | 1074500 | 0 |
| SQGDSNP/ | PAFAH1B2 | P68402 | PA1B2_HUMAN | Acetyl (Protein N | 0 | 0 | 0 | 0 | 0 | 0 | 0 | 0 | 0 | 0 | 435170 | 416960 | 0 | 0 | 0 | 0 | 0 | 0 |
| SQGDSNP/ | PAFAH1B2 | P68402 | PA1B2_HUMAN | Acetyl (Protein N | 278360 | 255660 | 662660 | 0 | 0 | 0 | 0 | 0 | 0 | 0 | 0 | 14298000 | 0 | 0 | 0 | 1303600 | 0 | 0 |
| SQGDSNP/ | PAFAH1B2 | P68402 | PA1B2_HUMAN | Acetyl (Protein N | 0 | 0 | 0 | 286250 | 0 | 116350 | 0 | 0 | 1295400 | 0 | 705680 | 10338000 | 0 | 0 | 527660 | 0 | 0 | 0 |
| SQGGEPTY | PFN2; PFN2; PF | P35080; G5E | PROF2_HUMAN | Unmodified | 4233300 | 3676100 | 1152800 | 957040 | 10069000 | 2212100 | 0 | 272890 | 4446800 | 4139300 | 2905100 | 29140000 | 3238800 | 2126000 | 2110300 | 2589300 | 3928600 | 4663100 |
| SQGGEPTY | PFN2; PFN2; PF | P35080; G5E | PROF2_HUMAN | Unmodified | 0 | 0 | 0 | 0 | 28149000 | 4727800 | 1471200 | 953430 | 0 | 4289700 | 418710 | 163550 | 12105000 | 8551600 | 0 | 0 | 22569000 | 17035000 |
| SQGKPIELT | PPP2R5E | Q16537 | 2A5E_HUMAN ! | Unmodified | 235920 | 0 | 1453700 | 825670 | 551360 | 0 | 0 | 0 | 2120600 | 180580 | 0 | 11600000 | 0 | 0 | 1511900 | 0 | 367770 | 559220 |
| SQGLDSRL | CKAP4 | Q07065 | CKAP4_HUMAN | Unmodified | 0 | 0 | 0 | 0 | 0 | 0 | 876430 | 808500 | 370480 | 450060 | 0 | 0 | 0 | 0 | 0 | 0 | 0 | 0 |
| SQGMASLN | CTNNA1; CTNN | P35221; E5R | CTNA1_HUMAI | Oxidation (M) | 1005100 | 1562100 | 3968000 | 7290900 | 26671000 | 4591100 | 70800 | 253710 | 0 | 351340 | 2736500 | 8869700 | 19782000 | 18517000 | 7204000 | 8291600 | 0 | 298630 |
| SQGMASLN | CTNNA1; CTNN | P35221; E5R | CTNA1_HUMAI | Unmodified | 40066 | 0 | 0 | 857570 | 12586000 | 1059600 | 331410 | 0 | 126990 | 0 | 33506 | 733290 | 6060800 | 6796300 | 694870 | 71340 | 0 | 0 |
| SQGPALTA | TNIK | Q9UKE5 | TNIK_HUMAN T | Unmodified | 0 | 0 | 0 | 0 | 739750 | 157820 | 453000 | 206560 | 491410 | 786950 | 0 | 1535300 | 239130 | 697560 | 0 | 0 | 298320 | 1615000 |
| SQGPLEVA | MACROH2A2; N | Q9P0M6; A0 | H2AW_HUMAN | Unmodified | 196130 | 0 | 0 | 0 | 550310 | 41891 | 0 | 180220 | 0 | 0 | 257550 | 1167500 | 0 | 0 | 0 | 0 | 0 | 0 |
| SQGPSLDL | FMNL1; FMNL1 | O95466; A0/ | FMNL1_HUMAI | Unmodified | 0 | 0 | 0 | 0 | 5740900 | 1740500 | 0 | 0 | 0 | 0 | 0 | 0 | 0 | 0 | 0 | 0 | 320920 | 1158800 |
| SQGQDVTS | UCHL3; UCHL3 | P15374; Q5T | UCHL3_HUMAI | Unmodified | 0 | 0 | 0 | 0 | 0 | 0 | 0 | 0 | 0 | 0 | 0 | 999840 | 745900 | 528650 | 132590 | 0 | 0 | 0 |
| SQGQLNEI | ITCH; ITCH; ITC | Q96J02; A0A | ITCH_HUMAN I | Oxidation (M) | 954600 | 1114600 | 493570 | 0 | 0 | 0 | 0 | 0 | 0 | 0 | 0 | 364160 | 0 | 0 | 776740 | 730490 | 0 | 0 |
| SQGQLNEI | ITCH; ITCH; ITC | Q96J02; A0A | ITCH_HUMAN I | Unmodified | 439790 | 438180 | 78237 | 65237 | 0 | 0 | 95946 | 0 | 554040 | 312250 | 0 | 0 | 91786 | 0 | 287960 | 192600 | 0 | 0 |
| SQGSRNAI | BAIAP2L1 | Q9UHR4 | BI2L1_HUMAN | Unmodified | 2612200 | 0 | 1737400 | 0 | 1124900 | 186390 | 0 | 0 | 0 | 324440 | 0 | 0 | 49271 | 500770 | 0 | 0 | 641360 | 59431 |
| SQGYDWS | DPP9 | Q86TI2 | DPP9_HUMAN | Unmodified | 0 | 0 | 0 | 0 | 0 | 0 | 0 | 0 | 0 | 0 | 0 | 328910 | 0 | 0 | 0 | 0 | 0 | 0 |
| SQHLGEEL | ABCC4; ABCC4 | O15439; A0/ | MRP4_HUMAN | Unmodified | 26119 | 0 | 5031600 | 706010 | 0 | 0 | 39333 | 0 | 0 | 0 | 0 | 794720 | 0 | 0 | 3098400 | 3076500 | 0 | 0 |
| SQIDVALS( | UPF1 | Q92900 | RENT1_HUMAN | Unmodified | 0 | 1500600 | 0 | 222430 | 0 | 0 | 0 | 0 | 0 | 0 | 0 | 0 | 0 | 0 | 0 | 0 | 0 | 0 |
| SQIFSTASD | HSPA5; HSPA5; | P11021; A0A | BIP_HUMAN Er | Unmodified | 323590 | 210290 | 1121300 | 504680 | 2505300 | 2010500 | 19776000 | 96916 | 79189000 | 1.14E+08 | 4981400 | 6124900 | 806840 | 1573800 | 2039700 | 563880 | 0 | 0 |
| SQIFSTASD | HSPA5; HSPA5 | P11021; A0A | BIP_HUMAN Er | Unmodified | 0 | 0 | 0 | 0 | 0 | 0 | 12178000 | 0 | 1588300 | 7332900 | 0 | 0 | 0 | 0 | 0 | 0 | 389540 | 0 |
| SQIHDIVLV | HSPA8; HSPA8; | P11142; E9P | HSP7C_HUMAI | Unmodified | 69013000 | 75412000 | 77028000 | 76644000 | 16130000 | 3299900 | 1900500 | 593510 | 1.28E+08 | 8822200 | 34573000 | 2.34E+08 | 19890000 | 6302600 | 80637000 | 64596000 | 1042900 | 8453300 |
| SQIHDIVLV | HSPA8; HSPA8; | P11142; E9P | HSP7C_HUMAI | Unmodified | 8715400 | 5422600 | 9301600 | 40631000 | 5.69E+08 | 61271000 | 1.6E+08 | 19031000 | 36429000 | 1.42E+08 | 22456000 | 2715300 | 6.26E+08 | 2.35E+08 | 3657900 | 2791200 | 45354000 | 1.6E+08 |
| SQIHDIVLV | HSPA8; HSPA8; | P11142; E9P | HSP7C_HUMAI | Unmodified | 0 | 0 | 0 | 0 | 56115000 | 3200700 | 60097000 | 4932100 | 0 | 18126000 | 0 | 0 | 61203000 | 60835000 | 0 | 0 | 22965000 | 21992000 |
| SQIKNEINII | EMC1; EMC1 | Q8N766; A0 | EMC1_HUMAN | Unmodified | 0 | 0 | 0 | 0 | 0 | 31191 | 46110 | 0 | 6157700 | 1104100 | 82547 | 0 | 0 | 0 | 0 | 0 | 0 | 0 |
| SQISFDNL1 | FAM171A1 | Q5VUB5 | F1711_HUMAN | Unmodified | 0 | 0 | 0 | 0 | 0 | 0 | 0 | 0 | 0 | 0 | 0 | 239370 | 0 | 0 | 0 | 0 | 0 | 0 |
| SQISSVNSI | MPRIP; MPRIP; | Q6WCQ1; H | MPRIP_HUMAN | Unmodified | 0 | 0 | 0 | 0 | 0 | 0 | 0 | 0 | 303800 | 0 | 0 | 1553900 | 0 | 0 | 0 | 0 | 0 | 0 |
| SQISTESFL | OPLAH | O14841 | OPLA_HUMAN | Unmodified | 0 | 0 | 0 | 0 | 0 | 0 | 0 | 0 | 0 | 0 | 0 | 952730 | 0 | 0 | 0 | 0 | 0 | 0 |
| SQIVTNVR | HSPA4L; HSPA4 | O95757; E9F | HS74L_HUMAN | Unmodified | 0 | 0 | 0 | 62306 | 158110 | 0 | 59518 | 0 | 315740 | 20494 | 158160 | 2379900 | 1305300 | 127450 | 50031 | 42523 | 37830 | 0 |
| SQIVTNVRI | HSPA4L; HSPA4 | O95757; E9F | HS74L_HUMAN | Unmodified | 0 | 0 | 0 | 0 | 0 | 0 | 0 | 0 | 0 | 0 | 0 | 0 | 641040 | 0 | 0 | 0 | 0 | 0 |
| SQKPEAGG | TBC1D4 | O60343 | TBCD4_HUMAI | Unmodified | 0 | 0 | 0 | 0 | 167940 | 0 | 0 | 0 | 0 | 0 | 0 | 76932 | 0 | 0 | 0 | 0 | 0 | 0 |
| SQKQEEEN | ENSA; ENSA; EI | O43768; A0/ | ENSA_HUMAN | Acetyl (Protein N | 181580 | 102020 | 97169 | 129930 | 332700 | 0 | 0 | 20076 | 464990 | 0 | 396630 | 1314600 | 305510 | 93560 | 215660 | 219860 | 0 | 304680 |
| SQKQEEEN | ENSA; ENSA; EI | O43768; A0/ | ENSA_HUMAN | Acetyl (Protein N | 485650 | 0 | 0 | 0 | 4993500 | 1279000 | 0 | 321890 | 0 | 679170 | 885940 | 0 | 5027500 | 0 | 0 | 270460 | 1699300 | 5222300 |
| SQLDIIIHSI | XRCC5 | P13010 | XRCC5_HUMAI | Unmodified | 0 | 0 | 0 | 0 | 218550 | 215550 | 1998300 | 402540 | 336680 | 565900 | 997400 | 7364500 | 0 | 0 | 0 | 0 | 0 | 270660 |
| SQLDIIIHSI | XRCC5 | P13010 | XRCC5_HUMAI | Unmodified | 0 | 0 | 0 | 0 | 187660 | 0 | 594850 | 0 | 0 | 199490 | 0 | 3508600 | 0 | 22099 | 0 | 0 | 0 | 129970 |
| SQLEGLEE! | PLXNB2 | O15031 | PLXB2_HUMAN | Unmodified | 0 | 0 | 1070600 | 0 | 0 | 0 | 0 | 0 | 0 | 0 | 0 | 0 | 0 | 0 | 0 | 0 | 0 | 0 |
| SQLEGLEE! | PLXNB2 | O15031 | PLXB2_HUMAN | Unmodified | 0 | 0 | 2379000 | 1962500 | 0 | 0 | 0 | 0 | 0 | 0 | 0 | 10090000 | 0 | 0 | 7489100 | 6128700 | 0 | 0 |
| SQLETSLK | AKR7L; AKR7A2 | Q8NHP1; O4 | ARK74_HUMAN | Unmodified | 0 | 0 | 0 | 0 | 79188 | 48877 | 192240 | 0 | 2705700 | 107930 | 127100 | 1961500 | 0 | 0 | 0 | 0 | 0 | 0 |
| SQLIMQAE | FLOT1; FLOT1; I | O75955; A0/ | FLOT1_HUMAN | Unmodified | 0 | 0 | 178710 | 297550 | 8753500 | 750600 | 0 | 13874 | 16304000 | 1733700 | 477400 | 1199300 | 1942900 | 0 | 339200 | 55949 | 296700 | 552440 |
| SQLIMQAE | FLOT1; FLOT1; I | O75955; A0/ | FLOT1_HUMAN | Oxidation (M) | 0 | 0 | 2166500 | 2667200 | 6911100 | 2218100 | 592010 | 678640 | 4994900 | 4258000 | 2395100 | 7633200 | 3281200 | 0 | 4410800 | 2436200 | 149940 | 550670 |
| SQLLGSAH | SPTAN1; SPTAN | Q13813; A0/ | SPTN1_HUMAN | Unmodified | 0 | 0 | 428930 | 98282 | 0 | 0 | 0 | 0 | 474110 | 28612 | 325780 | 3956500 | 0 | 0 | 458920 | 332680 | 0 | 0 |
| SQLLILDR | STXBP1; STXBP | P61764; A0A | STXB1_HUMAN | Unmodified | 439110 | 112450 | 0 | 0 | 0 | 0 | 0 | 0 | 0 | 0 | 0 | 5293800 | 0 | 0 | 0 | 89221 | 0 | 0 |
| SQLLQYVYI | MCM4; MCM4; | P33991; A0A | MCM4_HUMAN | Unmodified | 0 | 0 | 0 | 0 | 2255800 | 0 | 966950 | 0 | 27205 | 0 | 0 | 0 | 0 | 0 | 0 | 0 | 112960 | 0 |
| SQLLSYIDF | MCM7 | P33993 | MCM7_HUMAN | Unmodified | 139440 | 64384 | 248420 | 130290 | 662310 | 109770 | 1474300 | 0 | 646060 | 351360 | 459940 | 941630 | 205450 | 0 | 75504 | 65988 | 126550 | 153530 |
| SQLNSQSV | USO1 | O60763 | USO1_HUMAN | Unmodified | 0 | 0 | 0 | 0 | 273200 | 0 | 97713 | 0 | 320150 | 116950 | 166870 | 1772200 | 661920 | 237020 | 0 | 0 | 207100 | 690940 |
| SQLNSQSV | USO1 | O60763 | USO1_HUMAN | Unmodified | 0 | 0 | 0 | 0 | 308980 | 81768 | 216970 | 0 | 0 | 194880 | 0 | 0 | 1031100 | 474240 | 0 | 0 | 251390 | 769880 |
| SQLQGAM( | ITCH; ITCH; ITC | Q96J02; A0A | ITCH_HUMAN I | Unmodified | 2911300 | 4186600 | 1674700 | 1812500 | 216380 | 0 | 0 | 0 | 778910 | 67262 | 91769 | 208640 | 142750 | 0 | 3690200 | 3515300 | 0 | 0 |
| SQLQGAM( | ITCH; ITCH; ITC | Q96J02; A0A | ITCH_HUMAN I | Oxidation (M) | 1665500 | 345730 | 880450 | 621680 | 0 | 0 | 0 | 0 | 0 | 175020 | 0 | 56981 | 0 | 0 | 1171800 | 596780 | 0 | 86943 |
| SQLQITVIS | ITCH; ITCH; ITC | Q96J02; A0A | ITCH_HUMAN I | Unmodified | 7859300 | 7879100 | 7070000 | 5339700 | 8925900 | 0 | 0 | 252460 | 2153000 | 285900 | 0 | 0 | 728120 | 0 | 7152100 | 4614500 | 2747300 | 9179600 |
| SQLVYQSR | LGALS3BP | Q08380 | LG3BP_HUMAN | Unmodified | 0 | 0 | 0 | 0 | 37152 | 0 | 0 | 0 | 844050 | 1151000 | 1065800 | 311290 | 2114200 | 1013700 | 0 | 0 | 0 | 49920 |
| SQMAAVEF | ATP2A3 | Q93084 | AT2A3_HUMAN | Oxidation (M) | 0 | 0 | 0 | 0 | 0 | 0 | 0 | 0 | 0 | 0 | 82725 | 704670 | 0 | 0 | 0 | 0 | 0 | 0 |
| SQMAAVEF | ATP2A3 | Q93084 | AT2A3_HUMAN | Unmodified | 0 | 0 | 0 | 0 | 0 | 0 | 0 | 0 | 0 | 0 | 0 | 1487900 | 0 | 0 | 0 | 0 | 0 | 0 |
| SQMEEVQI | EPS8; EPS8; EP | Q12929; A0/ | EPS8_HUMAN | Unmodified | 0 | 0 | 0 | 0 | 1265000 | 291810 | 0 | 0 | 0 | 0 | 0 | 0 | 145800 | 0 | 0 | 0 | 0 | 0 |
| SQMLLSR | NIPSNAP1; NIP | Q9BPW8; C9 | NIPS1_HUMAN | Unmodified | 0 | 0 | 0 | 0 | 0 | 0 | 0 | 0 | 584380 | 120320 | 0 | 0 | 0 | 0 | 0 | 0 | 0 | 0 |
| SQMSGVE/ | FKBP15 | Q5T1M5 | FKB15_HUMAN | Unmodified | 0 | 0 | 0 | 0 | 540590 | 0 | 0 | 0 | 0 | 0 | 0 | 205790 | 0 | 0 | 0 | 0 | 0 | 0 |
| SQMSPQGI | ITGB4 | P16144 | ITB4_HUMAN I | Unmodified | 0 | 0 | 61787 | 0 | 864580 | 0 | 0 | 0 | 0 | 0 | 0 | 0 | 0 | 0 | 319850 | 395000 | 2341700 | 1764800 |
| SQNEQEAF | HOMER1; HOM | Q86YM7; Q8 | HOME1_HUMA | Unmodified | 0 | 0 | 0 | 0 | 0 | 0 | 0 | 0 | 0 | 0 | 0 | 0 | 2118800 | 0 | 0 | 0 | 0 | 0 |
| SQNIVTDS | SPTBN1; SPTBN | Q01082; A0/ | SPTB2_HUMAN | Unmodified | 0 | 765850 | 24066 | 0 | 0 | 0 | 0 | 0 | 0 | 0 | 0 | 1962900 | 0 | 0 | 0 | 0 | 0 | 0 |
| SQNKEDYA | GSTO1; GSTO1 | P78417; Q5T | GSTO1_HUMAI | Unmodified | 0 | 0 | 0 | 0 | 0 | 0 | 82512 | 0 | 0 | 299690 | 70608 | 1783100 | 0 | 0 | 0 | 0 | 0 | 0 |
| SQNRFYTD | MAN2A1 | Q16706 | MA2A1_HUMAI | Unmodified | 0 | 0 | 0 | 0 | 349280 | 0 | 0 | 0 | 2433800 | 1430500 | 803530 | 0 | 0 | 0 | 0 | 0 | 347680 | 0 |
| SQNVMAA/ | TCP1; TCP1; TC | P17987; F5G | TCPA_HUMAN | Oxidation (M) | 0 | 0 | 0 | 0 | 0 | 0 | 0 | 0 | 0 | 0 | 0 | 3222100 | 0 | 0 | 0 | 0 | 0 | 0 |
| SQNVMAA/ | TCP1; TCP1; TC | P17987; F5G | TCPA_HUMAN | Unmodified | 31933 | 31897 | 0 | 0 | 170900 | 0 | 0 | 0 | 596740 | 212460 | 2384800 | 1388400 | 0 | 0 | 76262 | 51302 | 0 | 0 |
| SQPAILLLT | TOR1AIP1; TOR | Q5JTV8; A0A | TOIP1_HUMAN | Unmodified | 0 | 0 | 0 | 0 | 0 | 0 | 0 | 0 | 16883000 | 0 | 653230 | 0 | 0 | 0 | 0 | 0 | 0 | 0 |
| SQPAILLLT | TOR1AIP1; TOR | Q5JTV8; A0A | TOIP1_HUMAN | Unmodified | 0 | 0 | 0 | 0 | 0 | 0 | 0 | 0 | 365990 | 0 | 0 | 0 | 0 | 0 | 0 | 0 | 0 | 0 |
| SQPAPAEFI | MPST; MPST | P25325; B1A | THTM_HUMAN | Unmodified | 4264100 | 1913500 | 0 | 0 | 200870 | 0 | 0 | 139160 | 0 | 716040 | 0 | 0 | 1210100 | 560680 | 0 | 0 | 369790 | 385590 |
| SQPELIEAH | TBCD; TBCD; TI | Q9BTW9; A0 | TBCD_HUMAN | Unmodified | 0 | 0 | 0 | 0 | 64853 | 0 | 0 | 26569 | 0 | 0 | 0 | 569890 | 146180 | 0 | 0 | 0 | 24727 | 0 |
| SQPEPLVVI | SELENBP1 | Q13228 | SBP1_HUMAN | Unmodified | 0 | 0 | 0 | 1553200 | 0 | 0 | 0 | 0 | 272260 | 126210 | 0 | 0 | 2386600 | 0 | 0 | 3487800 | 0 | 2998600 |
| SQPERDW | DDX17; DDX17; | Q92841; A0/ | DDX17_HUMAI | Unmodified | 0 | 0 | 906980 | 0 | 0 | 0 | 0 | 0 | 536180 | 0 | 626560 | 7811600 | 0 | 0 | 0 | 0 | 0 | 0 |
| SQPLTSPE | UTRN | P46939 | UTRN_HUMAN | Unmodified | 0 | 0 | 473200 | 601020 | 0 | 344970 | 0 | 0 | 897890 | 565840 | 0 | 453000 | 0 | 0 | 1568100 | 0 | 0 | 202240 |
| SQPNLDTC | CST4; CST1; CS | P01036; P01 | CYTS_HUMAN | Unmodified | 0 | 0 | 0 | 0 | 0 | 162860 | 0 | 0 | 1664200 | 0 | 0 | 0 | 0 | 0 | 0 | 0 | 390420 | 0 |
| SQPNLDTC | CST4; CST1; CS | P01036; P01 | CYTS_HUMAN | Unmodified | 0 | 0 | 0 | 0 | 0 | 0 | 0 | 0 | 942330 | 0 | 0 | 0 | 0 | 0 | 0 | 0 | 0 | 0 |
| SQPSETERI | ITGA3 | P26006 | ITA3_HUMAN I | Unmodified | 581450 | 680300 | 0 | 301370 | 2.23E+08 | 31739000 | 0 | 244720 | 852080 | 549060 | 318320 | 4325900 | 872350 | 707030 | 565340 | 489150 | 30898000 | 1.38E+08 |
| SQPVSQPL | EPS8L2; EPS8L | Q9H6S3; A0 | ES8L2_HUMAN | Unmodified | 0 | 0 | 0 | 0 | 2499900 | 1289500 | 0 | 0 | 560120 | 0 | 436350 | 275550 | 0 | 0 | 818630 | 0 | 0 | 0 |
| SQPWDLTF | EPN3 | Q9H201 | EPN3_HUMAN | Oxidation (M) | 283080 | 485300 | 113530 | 0 | 0 | 0 | 0 | 0 | 0 | 0 | 0 | 0 | 0 | 0 | 0 | 182000 | 0 | 0 |
| SQQCNGK | ST14 | Q9Y5Y6 | ST14_HUMAN ! | Unmodified | 691810 | 0 | 346280 | 403500 | 0 | 0 | 0 | 0 | 180760 | 14056 | 14305 | 0 | 599620 | 189720 | 380070 | 760640 | 0 | 0 |
| SQQERDW | DDX5; DDX5; D | P17844; A0A | DDX5_HUMAN | Unmodified | 0 | 0 | 0 | 0 | 284910 | 0 | 181080 | 50412 | 0 | 375500 | 1446900 | 2304300 | 0 | 0 | 0 | 0 | 0 | 0 |
| SQQHHGL | RTBDN; RTBDN | Q9BSG5; K7 | RTBDN_HUMAI | Unmodified | 0 | 0 | 0 | 0 | 0 | 0 | 0 | 0 | 977630 | 666590 | 42851 | 520240 | 0 | 0 | 0 | 0 | 0 | 0 |
| SQQIQQM/ | CIT; CIT; CIT; CI | A0A7I2V4H7 | A0A7I2V4H7_H | Unmodified | 0 | 0 | 61128 | 134930 | 1713400 | 243420 | 433480 | 0 | 0 | 0 | 33204 | 78550 | 0 | 0 | 0 | 0 | 103890 | 109740 |
| SQQPFVVT | BMP8B; BMP8A | P34820; Q7Z | BMP8B_HUMAI | Unmodified | 0 | 0 | 0 | 0 | 0 | 0 | 0 | 0 | 1360500 | 0 | 0 | 0 | 0 | 0 | 0 | 0 | 0 | 0 |
| SQQSEVTR | SH3BGRL3; SH | Q9H299; Q5 | SH3L3_HUMAN | Unmodified | 108800 | 40572 | 0 | 156920 | 28254000 | 4865800 | 9046800 | 4646100 | 270740 | 3840700 | 455380 | 0 | 6987200 | 6680600 | 149300 | 37374 | 48682000 | 1.29E+08 |
| SQQWLLR | GALNT1; GALN | Q10472; Q8I | GALT1_HUMAN | Unmodified | 691290 | 513900 | 1293500 | 2101300 | 0 | 0 | 0 | 0 | 1802200 | 0 | 516840 | 1626100 | 0 | 0 | 1436200 | 1585100 | 0 | 0 |
| SQRDPLAA | LTV1 | Q96GA3 | LTV1_HUMAN I | Unmodified | 0 | 0 | 0 | 0 | 0 | 0 | 0 | 0 | 0 | 0 | 0 | 0 | 0 | 0 | 0 | 0 | 0 | 0 |
| SQRLAASA | KIF20A | O95235 | KI20A_HUMAN | Unmodified | 0 | 0 | 0 | 0 | 0 | 0 | 0 | 0 | 0 | 0 | 0 | 0 | 0 | 0 | 0 | 0 | 191670 | 333340 |
| SQRYAYVV | MUC5B | Q9HC84 | MUC5B_HUMA | Unmodified | 0 | 0 | 0 | 0 | 0 | 0 | 0 | 0 | 0 | 0 | 0 | 0 | 0 | 2218900 | 0 | 0 | 39153 | 811360 |
| SQRYESLK | RPL29; RPL29 | P47914; A0A | RL29_HUMAN ( | Unmodified | 0 | 0 | 0 | 0 | 0 | 0 | 0 | 0 | 0 | 116070 | 0 | 38312 | 816400 | 274710 | 0 | 0 | 0 | 187630 |
| SQSAAVTP | ADRM1; ADRM1 | Q16186; A0/ | ADRM1_HUMA | Unmodified | 89787 | 0 | 0 | 0 | 0 | 0 | 0 | 0 | 634160 | 0 | 0 | 479850 | 0 | 0 | 103730 | 92591 | 0 | 95382 |
| SQSEQLLR | BMX | P51813 | BMX_HUMAN ( | Unmodified | 0 | 0 | 0 | 0 | 0 | 0 | 30978 | 0 | 0 | 0 | 217200 | 0 | 17691 | 0 | 0 | 0 | 0 | 0 |
| SQSESSDE | C2CD5 | Q86YS7 | C2CD5_HUMA | Unmodified | 0 | 0 | 0 | 0 | 276970 | 0 | 0 | 0 | 1404100 | 101270 | 103500 | 457550 | 0 | 0 | 0 | 0 | 0 | 0 |
| SQSFAGFS | RIPOR2; RIPOR | Q9Y4F9; B7Z | RIPR2_HUMAN | Unmodified | 0 | 358610 | 0 | 0 | 0 | 0 | 0 | 0 | 0 | 183340 | 0 | 842970 | 0 | 0 | 0 | 0 | 0 | 0 |
| SQSFAGFS | RIPOR2; RIPOR | Q9Y4F9; B7Z | RIPR2_HUMAN | Unmodified | 0 | 0 | 0 | 0 | 0 | 0 | 0 | 181390 | 527860 | 155610 | 0 | 1165300 | 1247700 | 488700 | 0 | 15994 | 0 | 0 |
| SQSLQDQI | MINK1 | Q8N4C8 | MINK1_HUMAN | Unmodified | 0 | 0 | 0 | 0 | 1866700 | 0 | 0 | 0 | 0 | 1445800 | 0 | 0 | 0 | 0 | 0 | 0 | 0 | 2855600 |
| SQSLSLESI | GPR39 | O43194 | GPR39_HUMAI | Unmodified | 0 | 0 | 0 | 0 | 726460 | 0 | 0 | 0 | 0 | 0 | 0 | 0 | 0 | 0 | 0 | 0 | 517680 | 1401400 |
| SQSPADQI | B4GALNT1; B4( | Q00973; H0\ | B4GN1_HUMAI | Unmodified | 0 | 0 | 0 | 0 | 0 | 0 | 0 | 0 | 15970000 | 570450 | 0 | 0 | 0 | 0 | 297600 | 0 | 0 | 0 |
| SQSPTCQI | TSPAN31; TSPA | Q12999; F8\ | TSN31_HUMAN | Unmodified | 0 | 0 | 0 | 0 | 0 | 0 | 0 | 0 | 1384500 | 314360 | 0 | 0 | 0 | 0 | 0 | 0 | 0 | 0 |
| SQSRPNDI | PRKAA1 | Q13131 | AAPK1_HUMAN | Unmodified | 0 | 0 | 0 | 168790 | 478420 | 0 | 55740 | 0 | 885260 | 0 | 0 | 272540 | 0 | 114680 | 145390 | 128430 | 0 | 607050 |
| SQSSHSYD | CTNND1; CTNN | O60716; H0\ | CTND1_HUMAI | Unmodified | 27812 | 0 | 230210 | 245260 | 0 | 0 | 0 | 36288 | 0 | 0 | 673610 | 1491800 | 60997 | 0 | 588750 | 721730 | 58041 | 0 |
| SQSSHSYD | CTNND1; CTNN | O60716; H0\ | CTND1_HUMAI | Unmodified | 0 | 3302000 | 7080800 | 10714000 | 10103000 | 4620500 | 0 | 0 | 0 | 99673 | 5182700 | 16896000 | 8417400 | 3759700 | 22038000 | 16916000 | 7166200 | 19340000 |
| SQSVRPGA | HSPG2 | P98160 | PGBM_HUMAN | Unmodified | 0 | 0 | 4276900 | 2039000 | 0 | 364020 | 0 | 0 | 1750200 | 204830 | 408470 | 165860 | 0 | 353270 | 0 | 2076500 | 0 | 461490 |
| SQTEAVTFI | POLDIP2; POLD | Q9Y2S7; B4I | PDIP2_HUMAN | Unmodified | 102900 | 161600 | 0 | 0 | 0 | 0 | 0 | 61129 | 1016800 | 0 | 0 | 0 | 0 | 0 | 0 | 0 | 0 | 0 |
| SQTGVGEL | EXOC4 | Q96A65 | EXOC4_HUMAI | Unmodified | 0 | 0 | 0 | 0 | 0 | 0 | 0 | 0 | 221820 | 0 | 0 | 858850 | 0 | 0 | 361870 | 172120 | 0 | 0 |
| SQTIYEIIDN | PSAT1 | Q9Y617 | SERC_HUMAN | Unmodified | 0 | 0 | 0 | 0 | 0 | 0 | 0 | 0 | 710070 | 0 | 703210 | 1531900 | 0 | 0 | 0 | 0 | 0 | 0 |
| SQTIYEIIDN | PSAT1 | Q9Y617 | SERC_HUMAN | Unmodified | 0 | 0 | 0 | 776700 | 3262600 | 0 | 276970 | 0 | 0 | 239860 | 0 | 0 | 3793200 | 4442700 | 0 | 0 | 677970 | 534260 |
| SQTPSERP | POMK; POMK; F | Q9H5K3; A0 | SG196_HUMAN | Unmodified | 0 | 0 | 67776 | 0 | 0 | 0 | 236880 | 150260 | 14262000 | 2875500 | 530420 | 318480 | 0 | 177000 | 0 | 0 | 0 | 0 |
| SQTSQNSH | IGSF3 | O75054 | IGSF3_HUMAN | Unmodified | 2082300 | 2123800 | 3350100 | 545040 | 0 | 0 | 0 | 0 | 343280 | 0 | 0 | 4640100 | 113560 | 0 | 5263200 | 3014500 | 0 | 0 |
| SQTSQNSH | IGSF3 | O75054 | IGSF3_HUMAN | Unmodified | 0 | 0 | 0 | 883520 | 1006700 | 0 | 708430 | 0 | 0 | 0 | 0 | 0 | 16328000 | 0 | 0 | 0 | 0 | 0 |
| SQTTLQEQ | ARHGEF39 | Q8N4T4 | ARG39_HUMAI | Unmodified | 305340 | 446610 | 380950 | 531690 | 0 | 0 | 58095 | 93472 | 377660 | 0 | 0 | 0 | 0 | 25583 | 558970 | 0 | 77926 | 66254 |

| | | | | | | | | | | | | | | | | | | | | | | |
|---|---|---|---|---|---|---|---|---|---|---|---|---|---|---|---|---|---|---|---|---|---|---|
| SQTYESDG | ADAM9; ADAM9 | Q13443; A0A | ADAM9_HUMA | Unmodified | 1225200 | 1314300 | 0 | 4004000 | 1262500 | 0 | 0 | 0 | 0 | 50767 | 0 | 0 | 0 | 0 | 0 | 32468 | 0 | 0 |
| SQVDHVIW | AHCYL1; AHCY | O43865; Q9( | SAHH2_HUMAI | Unmodified | 210610 | 0 | 0 | 0 | 1241600 | 0 | 0 | 0 | 257800 | 2238200 | 0 | 2207100 | 0 | 0 | 3192000 | 1318400 | 120700 | 110250 |
| SQVDHVIW | AHCYL1; AHCY | O43865; Q9( | SAHH2_HUMAI | Unmodified | 0 | 0 | 374320 | 0 | 0 | 0 | 0 | 0 | 0 | 0 | 0 | 0 | 0 | 0 | 809730 | 342130 | 0 | 0 |
| SQVEPADY | PITPNB; PITPNE | P48739; A0A | PIPNB_HUMAN | Unmodified | 1229400 | 611800 | 373660 | 774070 | 0 | 0 | 0 | 0 | 1290600 | 0 | 995310 | 3609200 | 0 | 0 | 2135200 | 1235100 | 0 | 141610 |
| SQVFSTAAI | HSPA9; HSPA9; | P38646; A0A | GRP75_HUMAN | Unmodified | 0 | 26037 | 0 | 162090 | 732950 | 0 | 662120 | 151800 | 12407000 | 1642200 | 613180 | 0 | 300770 | 0 | 0 | 0 | 358710 | 354410 |
| SQVFSTAAI | HSPA9; HSPA9; | P38646; A0A | GRP75_HUMAN | Unmodified | 0 | 0 | 0 | 0 | 0 | 0 | 0 | 0 | 530310 | 980860 | 0 | 0 | 0 | 0 | 0 | 0 | 0 | 0 |
| SQVMTHLH | APLP2 | Q06481 | APLP2_HUMAN | Unmodified | 0 | 0 | 0 | 47164 | 0 | 0 | 0 | 0 | 765220 | 60767 | 0 | 0 | 0 | 0 | 0 | 0 | 0 | 0 |
| SQVMTHLF | APP; APP; APP; | P05067; H7( | A4_HUMAN Am | Unmodified | 964350 | 127260 | 0 | 0 | 0 | 0 | 0 | 0 | 798480 | 13302 | 0 | 0 | 0 | 102670 | 0 | 0 | 0 | 0 |
| SQVSYRTQ | ITGB4 | P16144 | ITB4_HUMAN In | Unmodified | 0 | 0 | 0 | 0 | 5281500 | 0 | 0 | 0 | 0 | 0 | 0 | 0 | 0 | 0 | 0 | 0 | 10033000 | 20681000 |
| SQVVAGTN | CSTB; CSTB | P04080; A0A | CYTB_HUMAN | Unmodified | 0 | 300950 | 0 | 484500 | 0 | 0 | 0 | 0 | 0 | 0 | 0 | 0 | 0 | 0 | 2198200 | 1440200 | 0 | 0 |
| SQVVFSAEI | PLOD1; PLOD1; | Q02809; Q5. | PLOD1_HUMAI | Unmodified | 0 | 0 | 0 | 0 | 2481000 | 0 | 0 | 0 | 7468100 | 317270 | 166730 | 0 | 0 | 0 | 0 | 0 | 0 | 0 |
| SQVVFSAEI | PLOD1; PLOD1; | Q02809; Q5. | PLOD1_HUMAI | Unmodified | 0 | 0 | 0 | 0 | 0 | 0 | 0 | 0 | 1013700 | 391610 | 0 | 0 | 0 | 0 | 0 | 0 | 0 | 0 |
| SQWSPALT | UBE2D3; UBE2I | P61077; P62 | UB2D3_HUMAI | Unmodified | 3997400 | 4903100 | 0 | 1169200 | 0 | 0 | 0 | 197060 | 1051400 | 0 | 105910 | 10883000 | 0 | 0 | 8034800 | 6070100 | 0 | 0 |
| SQYDYILP( | NOMO1; NOMC | Q15155; P69 | NOMO1_HUMA | Unmodified | 0 | 0 | 0 | 0 | 0 | 297600 | 0 | 0 | 2186000 | 0 | 269420 | 210740 | 0 | 0 | 0 | 0 | 0 | 0 |
| SQYHDLQ/ | SH2D4A; SH2D | Q9H788; H0 | SH24A_HUMAN | Unmodified | 115430 | 61561 | 0 | 202910 | 1640900 | 254690 | 0 | 0 | 69195 | 0 | 0 | 0 | 1117300 | 617340 | 270110 | 246230 | 333920 | 596320 |
| SQYNFIAD\ | HTRA2; HTRA2 | O43464; A0/ | HTRA2_HUMAN | Unmodified | 0 | 0 | 0 | 0 | 0 | 0 | 0 | 0 | 818360 | 0 | 0 | 698770 | 0 | 0 | 0 | 0 | 0 | 0 |
| SRAAPSPQ | CHMP4C | Q96CF2 | CHM4C_HUMA | Unmodified | 0 | 0 | 0 | 305430 | 1898100 | 0 | 0 | 0 | 2260500 | 1296200 | 0 | 0 | 1177200 | 424390 | 0 | 0 | 0 | 0 |
| SRAASSQR | CHMP4C | Q96CF2 | CHM4C_HUMA | Unmodified | 0 | 0 | 0 | 0 | 0 | 0 | 0 | 0 | 0 | 230850 | 0 | 0 | 0 | 0 | 0 | 0 | 0 | 0 |
| SRAEQMAS | TJP2; TJP2; TJP2 | Q9UDY2; A0. | ZO2_HUMAN T | Unmodified | 0 | 0 | 0 | 0 | 0 | 0 | 0 | 24996 | 2144700 | 0 | 0 | 838650 | 0 | 0 | 0 | 0 | 0 | 0 |
| SRAHLVAVI | ANXA11; ANXA1 | P50995; H0Y | ANX11_HUMAN | Unmodified | 0 | 0 | 0 | 1486500 | 2696700 | 0 | 122830 | 0 | 22911000 | 5099500 | 1285800 | 1738800 | 1842400 | 1378300 | 287500 | 0 | 615880 | 3676100 |
| SRALEHFPI | GPD2; GPD2 | P43304; F5G | GPDM_HUMAN | Unmodified | 40637 | 0 | 0 | 110150 | 0 | 0 | 169720 | 0 | 758660 | 266320 | 0 | 0 | 70044 | 0 | 0 | 0 | 0 | 0 |
| SRANEEYG | TJP2; TJP2; TJP2 | Q9UDY2; A0. | ZO2_HUMAN T | Unmodified | 0 | 0 | 0 | 0 | 438810 | 386540 | 0 | 0 | 1440300 | 155160 | 0 | 2524900 | 0 | 0 | 0 | 0 | 0 | 0 |
| SRANEEYG | TJP2; TJP2; TJP2 | Q9UDY2; A0. | ZO2_HUMAN T | Unmodified | 0 | 0 | 0 | 0 | 5195200 | 0 | 0 | 0 | 0 | 3150400 | 0 | 0 | 0 | 0 | 0 | 0 | 962080 | 0 |
| SRAPMITPE | PLXNA2 | O75051 | PLXA2_HUMAN | Unmodified | 0 | 0 | 0 | 0 | 0 | 0 | 0 | 0 | 1701100 | 27967 | 0 | 0 | 0 | 0 | 0 | 0 | 0 | 0 |
| SRATLYALS | APOB; APOB | P04114; A8N | APOB_HUMAN | Unmodified | 0 | 0 | 0 | 0 | 0 | 2243400 | 0 | 0 | 0 | 0 | 0 | 0 | 0 | 0 | 0 | 0 | 0 | 0 |
| SRDALTQIN | ATP2B4 | P23634 | AT2B4_HUMAN | Unmodified | 343950 | 0 | 911070 | 1320400 | 6668500 | 1146600 | 1207500 | 167840 | 2080900 | 1220900 | 5290300 | 29921000 | 588540 | 0 | 1586600 | 1677800 | 2891700 | 3135100 |
| SRDALTQIN | ATP2B4 | P23634 | AT2B4_HUMAN | Unmodified | 0 | 0 | 0 | 0 | 4546300 | 0 | 1773400 | 0 | 0 | 0 | 0 | 327730 | 0 | 0 | 0 | 0 | 579250 | 3643700 |
| SRDDLYDC | LSR; LSR | Q86X29; S4F | LSR_HUMAN Li | Unmodified | 5860500 | 4617400 | 11220000 | 8034800 | 2685200 | 1718000 | 0 | 0 | 56533000 | 10815000 | 4212400 | 11188000 | 5571000 | 2679800 | 7001300 | 12388000 | 264150 | 566100 |
| SRDDSQLN | TMEM30A; TME | Q9NV96; E5I | CC50A_HUMAI | Unmodified | 208960 | 258000 | 2156000 | 2464200 | 4922500 | 0 | 867430 | 0 | 15286000 | 1914200 | 6616000 | 6515800 | 1500700 | 481390 | 1844700 | 1038000 | 56218 | 0 |
| SRDGLLTVI | PLEKHB1; PLEK | Q9UF11; F5H | PKHB1_HUMAI | Unmodified | 0 | 0 | 413230 | 0 | 0 | 0 | 0 | 0 | 160720 | 102460 | 0 | 939940 | 0 | 0 | 390670 | 323420 | 0 | 0 |
| SRDLLVQQ | HNRNPUL2; HN | Q1KMD3; H3 | HNRL2_HUMAI | Unmodified | 111210 | 0 | 0 | 0 | 0 | 74054 | 984120 | 438820 | 519340 | 271870 | 222970 | 0 | 1000800 | 392570 | 0 | 0 | 136680 | 678440 |
| SRDPHYDE | LSR; LSR | Q86X29; S4F | LSR_HUMAN Li | Unmodified | 1271700 | 821730 | 2052100 | 2209200 | 0 | 0 | 0 | 0 | 8063800 | 764640 | 817540 | 2744900 | 0 | 0 | 3171000 | 3099300 | 0 | 0 |
| SREAIDSPV | UPF1 | Q92900 | RENT1_HUMAN | Unmodified | 7831000 | 12489000 | 5099000 | 4893500 | 566340 | 0 | 0 | 0 | 0 | 399820 | 0 | 97888 | 0 | 235500 | 1081700 | 4122600 | 0 | 0 |
| SREDLSAQ | TJP1; TJP1; TJP1 | Q07157; A0/ | ZO1_HUMAN T | Unmodified | 0 | 0 | 706190 | 0 | 0 | 0 | 0 | 0 | 0 | 0 | 0 | 836330 | 0 | 86397 | 701070 | 504930 | 0 | 0 |
| SREDLSAQ | TJP1; TJP1; TJP1 | Q07157; A0/ | ZO1_HUMAN T | Unmodified | 0 | 0 | 0 | 1687100 | 2658800 | 250270 | 0 | 0 | 0 | 0 | 0 | 241470 | 2638100 | 225910 | 137830 | 890790 | 0 | 0 |
| SREDLTAV\ | TJP2; TJP2; TJP2 | Q9UDY2; A0. | ZO2_HUMAN T | Unmodified | 0 | 0 | 0 | 0 | 0 | 0 | 0 | 0 | 0 | 1976300 | 0 | 0 | 0 | 0 | 0 | 0 | 0 | 0 |
| SREGSRTD | HSP90B1; HSP9 | P14625; A0A | ENPL_HUMAN | Unmodified | 0 | 0 | 0 | 0 | 0 | 0 | 0 | 0 | 275340 | 37497 | 0 | 280350 | 0 | 0 | 22711 | 0 | 0 | 0 |
| SREIFLSQP | PPP1CA; PPP1( | P62136; P62 | PP1A_HUMAN | Unmodified | 1013900 | 2112700 | 770410 | 1527500 | 9479900 | 3372700 | 3710800 | 0 | 1514200 | 5552600 | 1263100 | 2868600 | 3334300 | 12679000 | 4930100 | 1095600 | 8362000 | 6365100 |
| SREMQDVI | NDRG1; NDRG | Q92597; A0/ | NDRG1_HUMA | Acetyl (Protein N | 0 | 0 | 930610 | 391740 | 0 | 0 | 0 | 0 | 1698100 | 0 | 0 | 356810 | 0 | 0 | 429030 | 467680 | 881230 | 5137000 |
| SREMQDVI | NDRG1; NDRG | Q92597; A0/ | NDRG1_HUMA | Acetyl (Protein N | 0 | 0 | 0 | 0 | 2287100 | 0 | 0 | 0 | 4038900 | 922230 | 0 | 0 | 0 | 0 | 0 | 0 | 1778600 | 1660600 |
| SREPVTLG/ | AGRN; AGRN; A | O00468; A0/ | AGRIN_HUMAN | Unmodified | 1030500 | 0 | 1216500 | 651680 | 0 | 0 | 0 | 0 | 902750 | 0 | 0 | 0 | 58562 | 0 | 631600 | 706760 | 0 | 0 |
| SREVLELG( | CNPY4; CNPY4 | Q8N129; A0/ | CNPY4_HUMAN | Unmodified | 0 | 67500 | 0 | 0 | 0 | 0 | 0 | 0 | 9598700 | 1982800 | 0 | 0 | 0 | 0 | 0 | 0 | 0 | 0 |
| SREWDME/ | CKAP4 | Q07065 | CKAP4_HUMAI | Unmodified | 0 | 0 | 0 | 0 | 0 | 0 | 0 | 0 | 1482500 | 0 | 0 | 0 | 0 | 0 | 0 | 0 | 0 | 0 |
| SREYQLND | GNAI1; GNAI1 | P63096; A0A | GNAI1_HUMAN | Unmodified | 0 | 0 | 0 | 0 | 0 | 0 | 0 | 0 | 292380 | 0 | 340750 | 8309700 | 5646400 | 1540300 | 0 | 0 | 378600 | 0 |
| SREYQLND | GNAI2 | P04899 | GNAI2_HUMAN | Unmodified | 2340700 | 0 | 897220 | 2448500 | 0 | 0 | 0 | 0 | 5555700 | 0 | 4147600 | 38040000 | 0 | 0 | 2868000 | 1538400 | 571530 | 752080 |
| SREYQLND | GNAO1; GNAO | P09471; H3E | GNAO_HUMAN | Unmodified | 57146 | 0 | 121470 | 0 | 2218600 | 880060 | 892290 | 406950 | 1472900 | 370380 | 106600 | 0 | 0 | 0 | 22828 | 20778 | 121730 | 614090 |
| SREYQLND | GNAI3 | P08754 | GNAI3_HUMAN | Unmodified | 570230 | 321110 | 623060 | 0 | 0 | 85940 | 0 | 0 | 703000 | 374430 | 331950 | 6467900 | 0 | 0 | 1051600 | 1342900 | 0 | 0 |
| SRFVQDLS | UGP2; UGP2 | Q16851; C9J | UGPA_HUMAN | Acetyl (Protein N | 92271 | 0 | 0 | 426670 | 5065500 | 1408800 | 0 | 123520 | 0 | 868400 | 187490 | 1772700 | 790940 | 222250 | 0 | 247340 | 1592900 | 1360500 |
| SRGDNVY\ | GSDMD; GSDM | P57764; G3\ | GSDMD_HUMA | Unmodified | 0 | 0 | 0 | 0 | 655760 | 193680 | 501700 | 88511 | 1446700 | 456390 | 152870 | 335040 | 0 | 1150100 | 0 | 0 | 1238100 | 2412600 |
| SRGEGPEA | C1orf198 | Q9H425 | CA198_HUMAN | Unmodified | 313710 | 0 | 691010 | 492450 | 561830 | 380250 | 1747000 | 179240 | 1110000 | 285790 | 769190 | 2610600 | 1274000 | 378590 | 396180 | 349850 | 528800 | 1097500 |
| SRGFAFVTI | RBMXL3; RBMX | Q8N7X1; O7 | RMXL3_HUMAN | Unmodified | 0 | 0 | 0 | 0 | 0 | 0 | 606070 | 77497 | 1024300 | 963890 | 2595100 | 3039300 | 9790400 | 0 | 0 | 1224300 | 0 | 0 |
| SRGFAFVYI | TRA2B; TRA2B | P62995; H7E | TRA2B_HUMAN | Unmodified | 0 | 0 | 0 | 0 | 0 | 0 | 285310 | 0 | 0 | 0 | 0 | 872810 | 0 | 0 | 0 | 0 | 0 | 0 |
| SRGFGFITF | RBM3 | P98179 | RBM3_HUMAN | Oxidation (M) | 7018600 | 3882700 | 2638000 | 3750700 | 0 | 0 | 0 | 0 | 0 | 0 | 200590 | 0 | 2346600 | 727410 | 2175000 | 3374300 | 0 | 0 |
| SRGFGFVL | HNRNPD; HNR | Q14103; H0' | HNRPD_HUMA | Unmodified | 0 | 0 | 0 | 0 | 372510 | 0 | 263760 | 0 | 89445 | 491970 | 81911 | 252480 | 318460 | 107310 | 0 | 0 | 0 | 128750 |
| SRGFGFVT | CIRBP; CIRBP; ( | Q14011; K7E | CIRBP_HUMAN | Unmodified | 1100400 | 990290 | 1074800 | 0 | 0 | 0 | 0 | 0 | 0 | 0 | 0 | 8129300 | 0 | 0 | 2738100 | 1601000 | 0 | 0 |
| SRGGLTFIC | MAN1A2; MAN1 | O60476; H0' | MA1A2_HUMAI | Unmodified | 0 | 0 | 0 | 0 | 0 | 0 | 0 | 0 | 2009200 | 572460 | 0 | 73685 | 0 | 0 | 0 | 0 | 0 | 0 |
| SRGLFSAN | ZRANB2 | O95218 | ZRAB2_HUMAN | Unmodified | 0 | 0 | 0 | 0 | 150930 | 0 | 0 | 57101 | 0 | 27073 | 0 | 0 | 112130 | 0 | 0 | 0 | 155470 | 0 |
| SRGLPCTE | LAD1; LAD1 | O00515; E9F | LAD1_HUMAN | Unmodified | 48696 | 0 | 124810 | 131510 | 0 | 0 | 0 | 0 | 0 | 0 | 161690 | 1528900 | 3884600 | 2101000 | 96680 | 130570 | 0 | 0 |
| SRGPATVE | GIPC1; GIPC1 | O14908; K7E | GIPC1_HUMAN | Unmodified | 982680 | 0 | 2848600 | 2694200 | 11637000 | 3875700 | 0 | 373420 | 2535200 | 1210600 | 0 | 4957600 | 5231600 | 3130500 | 2597400 | 2304700 | 179900 | 20592000 |
| SRGPEEVN | BAIAP2L1 | Q9UHR4 | BI2L1_HUMAN | Acetyl (Protein N | 269120 | 0 | 2137900 | 4103100 | 6005300 | 2105600 | 0 | 0 | 3071800 | 341680 | 0 | 768310 | 6344500 | 2290100 | 1699600 | 0 | 252570 | 736920 |
| SRGSLGGL | FARP1 | A0A1B0GV6 | A0A1B0GV68_I | Unmodified | 0 | 0 | 0 | 0 | 0 | 0 | 0 | 0 | 0 | 358970 | 0 | 0 | 0 | 0 | 0 | 0 | 537500 | 1002600 |
| SRGSPSSY | EPN3; EPN3 | Q9H201; I6L | EPN3_HUMAN | Unmodified | 299280 | 66272 | 0 | 0 | 0 | 0 | 702100 | 258940 | 0 | 0 | 0 | 16910 | 88666 | 0 | 184720 | 125220 | 0 | 0 |
| SRGVGELA | TRIM26; TRIM2( | Q12899; A2/ | TRI26_HUMAN | Unmodified | 0 | 0 | 0 | 0 | 32282 | 0 | 47983 | 0 | 0 | 0 | 0 | 0 | 317970 | 387130 | 22049 | 0 | 25722 | 0 |
| SRGVGGA\ | PKP3; PKP3; PK | Q9Y446; E9F | PKP3_HUMAN | Unmodified | 0 | 0 | 827430 | 1018000 | 0 | 0 | 0 | 0 | 748680 | 0 | 0 | 1239900 | 192040 | 0 | 1324500 | 1064500 | 0 | 0 |
| SRHPSSSS | KIAA1522 | Q9P206 | K1522_HUMAN | Unmodified | 0 | 0 | 0 | 0 | 0 | 0 | 0 | 0 | 0 | 0 | 0 | 0 | 0 | 0 | 430880 | 311990 | 0 | 0 |
| SRIFQNLN | PLOD3; PLOD3 | O60568; H7( | PLOD3_HUMAI | Unmodified | 0 | 0 | 0 | 0 | 0 | 0 | 24173 | 0 | 8075400 | 2903500 | 0 | 0 | 0 | 0 | 0 | 0 | 0 | 0 |
| SRKESYSIY | H2BC19P; H2B | Q6DRA6; P0 | H2B2D_HUMAI | Unmodified | 0 | 0 | 0 | 0 | 0 | 0 | 8321800 | 24351000 | 0 | 0 | 0 | 1948800 | 499780 | 0 | 0 | 0 | 0 | 0 |
| SRKESYSV\ | H2BC4; H2BC1 | P62807; O6( | H2B1C_HUMAI | Unmodified | 5254300 | 4286600 | 0 | 0 | 7367000 | 804760 | 1168200 | 71312000 | 50599 | 314880 | 0 | 4784200 | 973640 | 886180 | 0 | 0 | 52782 | 0 |
| SRKGNIEL/ | LIMCH1; LIMCH | Q9UPQ0; HC | LIMC1_HUMAN | Unmodified | 0 | 0 | 0 | 0 | 0 | 0 | 0 | 0 | 0 | 489360 | 0 | 0 | 761690 | 0 | 0 | 0 | 0 | 0 |
| SRKLAAVT\ | SEPTIN7; SEPTI | Q16181; E7E | SEPT7_HUMAN | Unmodified | 0 | 716980 | 0 | 73317 | 246570 | 22314 | 0 | 0 | 0 | 0 | 0 | 0 | 48948 | 0 | 0 | 0 | 242360 | 183850 |
| SRLEVTDD | PPM1A | P35813 | PPM1A_HUMAI | Unmodified | 368600 | 435620 | 0 | 0 | 918620 | 0 | 0 | 0 | 181970 | 39321 | 293590 | 850330 | 207520 | 0 | 820530 | 665590 | 27367 | 115180 |
| SRLLDSVE( | TWF2; N/A; TWI | Q6IBS0; H7C | TWF2_HUMAN | Unmodified | 0 | 0 | 0 | 0 | 863100 | 465740 | 0 | 0 | 0 | 775620 | 1359500 | 1806000 | 0 | 0 | 0 | 0 | 1818300 | 4602800 |
| SRLNQSQF | PRPF8; PRPF8 | Q6P2Q9; I3L | PRP8_HUMAN | Unmodified | 191750 | 236580 | 119510 | 0 | 0 | 0 | 0 | 0 | 0 | 0 | 0 | 2815200 | 0 | 0 | 0 | 63256 | 0 | 0 |
| SRLQGGVL | ACP3 | P15309 | PPAP_HUMAN | Unmodified | 0 | 0 | 0 | 0 | 0 | 0 | 0 | 0 | 2331400 | 1142600 | 0 | 0 | 0 | 0 | 0 | 0 | 0 | 0 |
| SRLSAIEIDI | CLDN12 | P56749 | CLD12_HUMAN | Unmodified | 1031700 | 1341700 | 2052700 | 3956700 | 3113300 | 1190000 | 245690 | 41031 | 2327400 | 2841000 | 0 | 1026400 | 5095600 | 3421800 | 6264300 | 5128500 | 858800 | 732500 |
| SRLTIYNAN | NCAM2; NCAM | O15394; H9I | NCAM2_HUMA | Unmodified | 0 | 47710 | 0 | 839420 | 213270 | 0 | 292420 | 0 | 0 | 0 | 0 | 434350 | 0 | 0 | 564550 | 390940 | 647210 | 337350 |
| SRLTPVSPE | SQSTM1; SQST | Q13501; E7E | SQSTM_HUMAN | Unmodified | 0 | 0 | 0 | 0 | 1229400 | 111230 | 311220 | 58227 | 255630 | 364860 | 0 | 52604 | 703310 | 402160 | 0 | 0 | 461180 | 1169700 |
| SRLVNTSEI | VTI1B | Q9UEU0 | VTI1B_HUMAN | Unmodified | 0 | 0 | 0 | 0 | 878570 | 0 | 0 | 21761 | 0 | 0 | 0 | 95322 | 0 | 0 | 0 | 66877 | 0 | 921520 |
| SRMGPSG( | LASP1; LASP1 | Q14847; C9J | LASP1_HUMAN | Unmodified | 1551900 | 1107700 | 0 | 535380 | 432270 | 0 | 0 | 0 | 0 | 0 | 0 | 201590 | 0 | 0 | 971340 | 702310 | 0 | 0 |
| SRNITYLPA | PHB; PHB | P35232; C9J | PHB_HUMAN F | Unmodified | 0 | 0 | 0 | 0 | 0 | 0 | 0 | 110120 | 2602900 | 2286400 | 0 | 0 | 593310 | 0 | 0 | 0 | 0 | 0 |
| SRNTDEM\ | HNRNPK; HNRI | P61978; Q5I | HNRPK_HUMA | Oxidation (M) | 0 | 0 | 0 | 0 | 0 | 0 | 0 | 0 | 0 | 0 | 0 | 998780 | 0 | 0 | 0 | 0 | 0 | 0 |
| SRNTDEM\ | HNRNPK; HNRI | P61978; Q5I | HNRPK_HUMA | Unmodified | 0 | 0 | 0 | 0 | 97289 | 0 | 111210 | 0 | 0 | 0 | 0 | 0 | 48073 | 0 | 0 | 0 | 0 | 32625 |
| SRPELLEY/ | TRIM25 | Q14258 | TRI25_HUMAN | Unmodified | 0 | 0 | 0 | 0 | 3125900 | 0 | 0 | 0 | 0 | 0 | 0 | 670580 | 114130 | 0 | 0 | 0 | 0 | 0 |
| SRPFTVAAS | LIMA1; LIMA1; L | Q9UHB6; F8 | LIMA1_HUMAN | Unmodified | 526580 | 0 | 231850 | 0 | 0 | 0 | 0 | 0 | 0 | 0 | 0 | 325890 | 0 | 0 | 407170 | 446870 | 0 | 0 |
| SRPPMPVV | NEO1; NEO1 | Q92859; Q5I | NEO1_HUMAN | Oxidation (M) | 393610 | 231470 | 513710 | 0 | 0 | 0 | 0 | 29679 | 0 | 0 | 0 | 0 | 0 | 0 | 956250 | 417190 | 0 | 0 |
| SRPPMPVV | NEO1; NEO1 | Q92859; Q5I | NEO1_HUMAN | Unmodified | 0 | 0 | 117280 | 212730 | 0 | 0 | 0 | 0 | 412170 | 0 | 0 | 0 | 0 | 0 | 667110 | 0 | 0 | 0 |
| SRPVSMDA | STK10 | O94804 | STK10_HUMAN | Unmodified | 0 | 0 | 0 | 0 | 934250 | 87188 | 0 | 0 | 0 | 0 | 0 | 0 | 0 | 78860 | 0 | 0 | 1226500 | 1482800 |
| SRQELEQH | DYNC1H1; DYN | Q14204; A0/ | DYHC1_HUMAN | Unmodified | 0 | 0 | 289170 | 0 | 5075700 | 0 | 2509100 | 0 | 2754700 | 3234800 | 1177500 | 3006700 | 0 | 1581100 | 0 | 0 | 1272200 | 2015500 |
| SRQELIALY | CASC3; CASC3 | O15234; J3K | CASC3_HUMAI | Unmodified | 1147200 | 1700800 | 856250 | 0 | 0 | 0 | 0 | 0 | 0 | 0 | 0 | 0 | 0 | 163230 | 0 | 1426900 | 0 | 0 |
| SRQGSLAN | SPN | P16150 | LEUK_HUMAN | Unmodified | 0 | 0 | 0 | 341280 | 527800 | 0 | 1876900 | 0 | 0 | 0 | 0 | 3411400 | 1018700 | 918240 | 0 | 0 | 28292 | 237790 |
| SRQGSLAN | SPN | P16150 | LEUK_HUMAN | Oxidation (M) | 256210 | 352550 | 0 | 368100 | 978630 | 0 | 1801100 | 83553 | 442070 | 0 | 0 | 1109900 | 1223900 | 705760 | 287910 | 333090 | 117060 | 216560 |
| SRQLLLTPI | MYO1C; MYO1( | O00159; F5H | MYO1C_HUMA | Unmodified | 187510 | 0 | 133450 | 415990 | 24192000 | 5426800 | 78223 | 0 | 1055200 | 0 | 191040 | 0 | 11598000 | 10526000 | 12277000 | 6089900 | 4531700 | 10910000 |
| SRQLLLTPI | MYO1C; MYO1( | O00159; F5H | MYO1C_HUMA | Unmodified | 0 | 0 | 0 | 963020 | 27696000 | 2644600 | 1634400 | 0 | 0 | 183550 | 0 | 0 | 6487400 | 4542800 | 0 | 0 | 9008500 | 5790100 |
| SRQPPLVT( | EXOC2; EXOC2 | Q96KP1; Q2I | EXOC2_HUMAI | Unmodified | 74610 | 72505 | 226950 | 374320 | 2183100 | 418710 | 239210 | 91089 | 1168500 | 559770 | 231970 | 1257200 | 762960 | 463110 | 194920 | 175480 | 409510 | 579820 |
| SRQPTKTV/ | N/A | N/A | N/A | Unmodified | 0 | 0 | 0 | 0 | 5136000 | 0 | 71875 | 1056400 | 329260 | 0 | 604270 | 0 | 0 | 1101000 | 0 | 0 | 3690300 | 2542400 |
| SRRNLDFQ | RTCB | Q9Y3I0 | RTCB_HUMAN | Unmodified | 0 | 0 | 0 | 0 | 0 | 16731 | 0 | 0 | 0 | 0 | 0 | 0 | 0 | 0 | 0 | 0 | 5537500 | 0 |
| SRRVDFETI | MYL6B; MYL6B | P14649; F8V | MYL6B_HUMAI | Oxidation (M) | 0 | 0 | 0 | 0 | 660680 | 0 | 15339 | 0 | 0 | 0 | 0 | 0 | 0 | 0 | 0 | 0 | 511750 | 89794 |
| SRRVDFETI | MYL6B; MYL6B | P14649; F8V | MYL6B_HUMAI | Unmodified | 0 | 0 | 0 | 0 | 426800 | 19997 | 0 | 0 | 40381 | 19133 | 0 | 0 | 0 | 0 | 0 | 0 | 62889 | 0 |
| SRSCGEVR | GPC1 | P35052 | GPC1_HUMAN | Unmodified | 0 | 0 | 0 | 0 | 6669500 | 2080500 | 1894000 | 865900 | 0 | 248290 | 0 | 0 | 1661000 | 2859000 | 0 | 0 | 318500 | 0 |
| SRSDIDVN. | CLASP2; CLASF | O75122; B3I | CLAP2_HUMAN | Unmodified | 132500 | 113160 | 0 | 67451 | 200230 | 0 | 0 | 0 | 96334 | 68219 | 24254 | 0 | 0 | 0 | 0 | 49759 | 95166 | 191110 |
| SRSDIDVN. | CLASP1; CLASF | Q7Z460; F8V | CLAP1_HUMAN | Unmodified | 0 | 0 | 0 | 0 | 175850 | 0 | 0 | 27546 | 0 | 48824 | 65319 | 126040 | 142640 | 80353 | 0 | 0 | 148710 | 156640 |
| SRSFTLDDI | PROSER2; PRO | Q86WR7; X6 | PRSR2_HUMAN | Unmodified | 0 | 0 | 0 | 0 | 3970000 | 0 | 0 | 0 | 280660 | 0 | 0 | 0 | 109110 | 0 | 0 | 0 | 233240 | 0 |
| SRSHTSEG | NDRG1; NDRG | Q92597; E7E | NDRG1_HUMA | Unmodified | 134660 | 118330 | 346390 | 473750 | 10332000 | 2559900 | 0 | 0 | 953540 | 1391500 | 0 | 169810 | 1260400 | 0 | 240080 | 1153900 | 10141000 | 17902000 |
| SRSLSQIHI | MYOF; MYOF | Q9NZM1; HC | MYOF_HUMAN | Unmodified | 0 | 0 | 0 | 0 | 5846200 | 0 | 0 | 0 | 0 | 0 | 19162 | 0 | 0 | 0 | 0 | 0 | 324110 | 0 |
| SRSLVDYEI | SNX6; SNX6 | Q9UNH7; AC | SNX6_HUMAN | Unmodified | 0 | 0 | 0 | 0 | 2764000 | 375450 | 0 | 0 | 0 | 0 | 32664 | 146900 | 0 | 0 | 0 | 0 | 0 | 80698 |
| SRSNVQPT | AURKB; AURKB | Q96GD4; J3I | AURKB_HUMAI | Acetyl (Protein N | 0 | 0 | 0 | 0 | 844530 | 0 | 0 | 0 | 0 | 0 | 0 | 0 | 0 | 82708 | 0 | 0 | 250250 | 224380 |
| SRSPGSPV | ADD1; ADD1; A | P35611; E7E | ADDA_HUMAN | Unmodified | 0 | 0 | 262690 | 309870 | 12937000 | 1300400 | 110700 | 68089 | 435560 | 67832 | 0 | 0 | 254730 | 219120 | 405540 | 333950 | 1079600 | 7327100 |
| SRSQGCSE | PRKDC | P78527 | PRKDC_HUMAN | Unmodified | 0 | 0 | 0 | 0 | 82304 | 0 | 293520 | 182110 | 0 | 0 | 0 | 824520 | 0 | 0 | 0 | 0 | 0 | 0 |
| SRSTTELDI | LMO7; LMO7; LI | Q8WWI1; A0 | LMO7_HUMAN | Unmodified | 0 | 0 | 0 | 0 | 693010 | 0 | 0 | 0 | 0 | 0 | 0 | 0 | 0 | 0 | 0 | 0 | 0 | 359000 |

| | | | | | | | | | | | | | | | | | | | | | | |
|---|---|---|---|---|---|---|---|---|---|---|---|---|---|---|---|---|---|---|---|---|---|---|
| SRSVIEQG( | PDCD6IP | Q8WUM4 | PDC6I_HUMAN | Unmodified | 1.02E+08 | 91469000 | 19289000 | 31820000 | 1.38E+08 | 19822000 | 10291000 | 1162300 | 77005000 | 53702000 | 9797200 | 12824000 | 1.32E+08 | 55956000 | 40005000 | 35192000 | 14109000 | 47805000 |
| SRSVIEQG( | PDCD6IP | Q8WUM4 | PDC6I_HUMAN | Unmodified | 0 | 0 | 0 | 0 | 1089100 | 0 | 253610 | 0 | 0 | 352330 | 0 | 0 | 2478000 | 1141900 | 0 | 0 | 0 | 603160 |
| SRSVLQPS | NCKAP1 | Q9Y2A7 | NCKP1_HUMAI | Acetyl (Protein N | 0 | 0 | 0 | 0 | 2857000 | 380250 | 685920 | 456770 | 0 | 356990 | 0 | 0 | 1526800 | 1438900 | 0 | 0 | 1519300 | 2008500 |
| SRSVLQPS | NCKAP1 | Q9Y2A7 | NCKP1_HUMAI | Acetyl (Protein N | 0 | 0 | 0 | 0 | 0 | 0 | 249420 | 404890 | 0 | 0 | 0 | 0 | 492170 | 0 | 0 | 0 | 1299500 | 0 |
| SRSYNDEL | RTCB | Q9Y3I0 | RTCB_HUMAN | Acetyl (Protein N | 0 | 0 | 0 | 0 | 850140 | 0 | 1543600 | 446780 | 539860 | 0 | 97223 | 0 | 1649100 | 878350 | 0 | 0 | 773320 | 696430 |
| SRSYNDEL | RTCB | Q9Y3I0 | RTCB_HUMAN | Acetyl (Protein N | 0 | 0 | 0 | 0 | 0 | 0 | 0 | 350300 | 0 | 0 | 0 | 0 | 0 | 0 | 0 | 0 | 758380 | 0 |
| SRTASGSS | NDRG1; NDRG | Q92597; E7E | NDRG1_HUMA | Unmodified | 122460 | 57566 | 198550 | 307670 | 4524800 | 1209700 | 0 | 0 | 2623200 | 1013000 | 0 | 0 | 271760 | 197870 | 216290 | 417320 | 5704700 | 4487300 |
| SRTHSTSS | NDRG3; NDRG | Q9UGV2; F8 | NDRG3_HUMA | Unmodified | 0 | 0 | 0 | 0 | 746780 | 228080 | 0 | 0 | 0 | 0 | 807820 | 5098800 | 788440 | 299120 | 458990 | 286220 | 262890 | 0 |
| SRTLTAVHI | RPS7; RPS7; RP | P62081; A0A | RS7_HUMAN 4 | Unmodified | 0 | 0 | 220040 | 2146400 | 5489100 | 467460 | 2764700 | 0 | 266910 | 820850 | 365460 | 1031400 | 7289500 | 1022000 | 3004300 | 389260 | 229300 | 2525200 |
| SRTLTAVHI | RPS7; RPS7; RP | P62081; A0A | RS7_HUMAN 4 | Unmodified | 0 | 0 | 0 | 0 | 1569200 | 0 | 0 | 0 | 0 | 0 | 0 | 580650 | 3825500 | 425750 | 0 | 770020 | 0 | 738570 |
| SRTSTARS | CXADR | P78310 | CXAR_HUMAN | Unmodified | 0 | 0 | 0 | 0 | 4637700 | 0 | 608520 | 0 | 0 | 0 | 0 | 0 | 0 | 0 | 0 | 0 | 115570 | 0 |
| SRTSVQTEI | CTNNA1; CTNN | P35221; P26 | CTNA1_HUMAI | Unmodified | 390870 | 385450 | 673570 | 1985800 | 3416200 | 1238000 | 0 | 0 | 716000 | 0 | 669510 | 2419200 | 4756600 | 1524800 | 1499600 | 1488000 | 0 | 0 |
| SRVAPAEP( | TKFC; TKFC; TK | Q3LXA3; I3L | TKFC_HUMAN | Unmodified | 0 | 0 | 0 | 0 | 342980 | 0 | 84907 | 27364 | 0 | 75673 | 0 | 0 | 340320 | 468230 | 25591 | 57720 | 113060 | 183980 |
| SRVITVDGI | NDUFA10; NDU | O95299; Q8I | NDUAA_HUMA | Unmodified | 0 | 0 | 0 | 0 | 0 | 0 | 95511 | 0 | 551270 | 88071 | 0 | 0 | 0 | 0 | 0 | 0 | 0 | 179540 |
| SRVIVQQP( | MYO1D; MYO1E | O94832; J3Q | MYO1D_HUMA | Unmodified | 9004200 | 7821600 | 226250 | 684170 | 360970 | 0 | 0 | 0 | 143690 | 0 | 0 | 227420 | 0 | 68768 | 6796000 | 840540 | 433750 | 86822 |
| SRVTQSNF | VDAC1; VDAC1 | P21796; C9J | VDAC1_HUMAI | Unmodified | 0 | 0 | 0 | 0 | 0 | 0 | 0 | 0 | 3091600 | 214450 | 0 | 0 | 0 | 0 | 0 | 0 | 0 | 0 |
| SRVTQSNF | VDAC1 | P21796 | VDAC1_HUMAI | Unmodified | 0 | 0 | 0 | 0 | 1668400 | 0 | 11444000 | 0 | 55882000 | 42523000 | 1845500 | 0 | 3479900 | 4919000 | 0 | 0 | 829080 | 1573000 |
| SRVVAGEV | IGSF8; IGSF8 | Q969P0; C9J | IGSF8_HUMAN | Unmodified | 27828000 | 34962000 | 17174000 | 14765000 | 789620 | 0 | 0 | 0 | 1945100 | 281890 | 0 | 0 | 2135100 | 881980 | 15632000 | 15468000 | 88489 | 430430 |
| SRVVAGEV | IGSF8; IGSF8 | Q969P0; C9J | IGSF8_HUMAN | Unmodified | 4259700 | 2071900 | 1200500 | 10300000 | 24777000 | 3859700 | 5501900 | 734670 | 0 | 1963600 | 221030 | 0 | 67908000 | 24106000 | 608970 | 632820 | 8067700 | 22857000 |
| SRWDETPA | SF3B1 | O75533 | SF3B1_HUMAN | Oxidation (M) | 281470 | 411520 | 0 | 0 | 0 | 0 | 0 | 0 | 0 | 0 | 0 | 173640 | 0 | 0 | 201240 | 130790 | 0 | 0 |
| SRWETGEV | LSP1; LSP1; LSI | P33241; E9P | LSP1_HUMAN I | Unmodified | 0 | 0 | 0 | 0 | 114970 | 0 | 386650 | 142860 | 314420 | 149230 | 0 | 0 | 1031300 | 0 | 0 | 0 | 0 | 0 |
| SRYDSIDS | GCLC; GCLC; G | P48506; A0A | GSH1_HUMAN | Unmodified | 0 | 0 | 0 | 112830 | 4476700 | 585140 | 44983 | 36750 | 0 | 177750 | 675050 | 998870 | 372290 | 279790 | 112490 | 0 | 115150 | 199760 |
| SRYDVFCF | HAPLN1 | P10915 | HPLN1_HUMAI | Unmodified | 0 | 0 | 281440 | 0 | 0 | 0 | 0 | 0 | 2273600 | 0 | 0 | 0 | 0 | 0 | 0 | 0 | 0 | 0 |
| SRYEQVDL | DDX50 | Q9BQ39 | DDX50_HUMAI | Unmodified | 0 | 0 | 0 | 0 | 0 | 0 | 325650 | 553950 | 0 | 112290 | 46176 | 119930 | 0 | 0 | 0 | 0 | 0 | 0 |
| SRYETYIHL | PSD2 | Q9BQI7 | PSD2_HUMAN | Oxidation (M) | 0 | 0 | 0 | 0 | 0 | 0 | 0 | 0 | 0 | 0 | 532330 | 0 | 0 | 0 | 0 | 0 | 0 | 0 |
| SSAAAAAA | SPTLC2 | O15270 | SPTC2_HUMAN | Unmodified | 0 | 0 | 0 | 0 | 67008 | 0 | 0 | 26314 | 0 | 0 | 0 | 793610 | 0 | 0 | 0 | 0 | 0 | 0 |
| SSACDAGI | ANKRD13A | Q8IZ07 | AN13A_HUMAI | Acetyl (Protein N | 1826400 | 964640 | 0 | 0 | 0 | 0 | 0 | 65259 | 0 | 0 | 174070 | 289740 | 0 | 0 | 0 | 1334500 | 0 | 0 |
| SSACFTLQI | RANGAP1; RAN | P46060; H0Y | RAGP1_HUMAI | Unmodified | 0 | 0 | 0 | 0 | 0 | 0 | 0 | 0 | 0 | 223090 | 0 | 0 | 696680 | 0 | 0 | 0 | 608600 | 0 |
| SSAEAQTPI | TGOLN2; TGOL | O43493; A0A | TGON2_HUMA | Unmodified | 0 | 0 | 2489800 | 0 | 0 | 0 | 0 | 0 | 17256000 | 1051000 | 0 | 0 | 0 | 0 | 215080 | 0 | 0 | 0 |
| SSAETNELI | N4BP1 | O75113 | N4BP1_HUMAI | Unmodified | 850500 | 795950 | 92142 | 309180 | 25865000 | 3790700 | 0 | 2136600 | 1238500 | 1715000 | 0 | 2182600 | 6382300 | 3877000 | 1008800 | 369220 | 4169700 | 10629000 |
| SSAEVIAQA | DPYSL2; DPYSL | Q16555; A0A | DPYL2_HUMAN | Unmodified | 0 | 0 | 0 | 0 | 9989000 | 2000500 | 19710 | 0 | 41898 | 1519200 | 366890 | 4724000 | 0 | 0 | 0 | 0 | 678680 | 1315700 |
| SSAEVIAQA | DPYSL2; DPYSL | Q16555; A0A | DPYL2_HUMAN | Unmodified | 0 | 0 | 0 | 0 | 3604300 | 0 | 0 | 31589 | 0 | 462460 | 0 | 0 | 0 | 0 | 0 | 0 | 297570 | 0 |
| SSAFGTLN | ANO6; ANO6; A | Q4KMQ2; A0 | ANO6_HUMAN | Unmodified | 0 | 0 | 0 | 0 | 0 | 417870 | 0 | 0 | 0 | 0 | 465160 | 1922900 | 0 | 22205 | 0 | 960650 | 0 | 0 |
| SSAGGGGI | ANKRD26; ANK | Q9UPS8; A0A | ANR26_HUMAI | Unmodified | 0 | 366560 | 0 | 0 | 0 | 0 | 0 | 0 | 0 | 0 | 176040 | 0 | 0 | 0 | 0 | 1162000 | 0 | 0 |
| SSAIPDFRI | AGFG1; AGFG1 | P52594; B8Z | AGFG1_HUMAI | Unmodified | 0 | 0 | 0 | 0 | 0 | 213240 | 0 | 0 | 253520 | 0 | 0 | 584090 | 0 | 0 | 0 | 0 | 54891 | 0 |
| SSAIVEIFSI | LIMA1; LIMA1; L | Q9UHB6; F8 | LIMA1_HUMAN | Unmodified | 545920 | 644590 | 689810 | 571340 | 1594900 | 0 | 0 | 0 | 313890 | 0 | 0 | 931380 | 2408200 | 1193500 | 643170 | 774460 | 0 | 0 |
| SSALVIQSY | MYO1B; MYO1E | O43795; E9F | MYO1B_HUMA | Unmodified | 0 | 0 | 0 | 0 | 367960 | 0 | 0 | 0 | 332890 | 0 | 941910 | 7959000 | 0 | 0 | 0 | 0 | 0 | 0 |
| SSALVIQSY | MYO1B; MYO1E | O43795; E9F | MYO1B_HUMA | Unmodified | 0 | 0 | 0 | 0 | 1783900 | 694380 | 528520 | 0 | 0 | 229160 | 242920 | 80541 | 0 | 0 | 0 | 0 | 0 | 525040 |
| SSANVEEA | RAB8B; RAB8B | Q92930; H0Y | RAB8B_HUMAI | Unmodified | 0 | 0 | 1637600 | 666660 | 149220 | 237510 | 0 | 0 | 1144200 | 67630 | 151150 | 4314000 | 468950 | 0 | 1186000 | 2351700 | 0 | 0 |
| SSANVEEA | RAB8B; RAB8B | Q92930; H0Y | RAB8B_HUMAI | Unmodified | 0 | 0 | 0 | 0 | 557250 | 31760 | 0 | 0 | 0 | 0 | 0 | 0 | 0 | 0 | 0 | 0 | 0 | 82843 |
| SSANVEEA | RAB8B; RAB8B | Q92930; H0Y | RAB8B_HUMAI | Oxidation (M) | 0 | 0 | 0 | 0 | 1477800 | 0 | 0 | 0 | 0 | 0 | 0 | 0 | 1329000 | 0 | 0 | 0 | 364110 | 918710 |
| SSAPPITPE | CHGB; CHGB | P05060; A0A | SCG1_HUMAN | Unmodified | 0 | 0 | 0 | 0 | 0 | 0 | 0 | 0 | 1107200 | 150690 | 0 | 0 | 133430 | 0 | 0 | 0 | 0 | 0 |
| SSAPPITPE | CHGB; CHGB | P05060; A0A | SCG1_HUMAN | Unmodified | 0 | 0 | 0 | 0 | 848240 | 0 | 19676 | 113970 | 0 | 1025900 | 0 | 0 | 0 | 0 | 0 | 0 | 266160 | 206580 |
| SSAPSRVPI | PARP12; PARP1 | Q9H0J9; H7( | PAR12_HUMAN | Unmodified | 0 | 546590 | 693290 | 933970 | 0 | 0 | 0 | 0 | 0 | 0 | 0 | 0 | 659060 | 0 | 556000 | 589190 | 0 | 0 |
| SSAPTTPPS | PPP2R5E | Q16537 | 2A5E_HUMAN | Acetyl (Protein N | 0 | 0 | 0 | 0 | 0 | 0 | 1313500 | 1596200 | 0 | 0 | 0 | 108150 | 0 | 0 | 0 | 0 | 3698900 | 0 |
| SSAQDPQA | VARS1; VARS1; | P26640; A2A | SYVC_HUMAN | Unmodified | 0 | 0 | 0 | 0 | 0 | 0 | 0 | 0 | 0 | 0 | 205010 | 11942000 | 0 | 0 | 343570 | 195220 | 345290 | 0 |
| SSAQGKPL | RANGAP1 | P46060 | RAGP1_HUMAI | Unmodified | 0 | 0 | 0 | 0 | 0 | 23653 | 55558 | 0 | 0 | 0 | 30678 | 419760 | 244650 | 99852 | 0 | 0 | 156840 | 99381 |
| SSASFNVIE | ITGAV; ITGAV | P06756; H7E | ITAV_HUMAN I | Unmodified | 3590200 | 1647900 | 2781600 | 1753300 | 3318800 | 0 | 0 | 0 | 5037400 | 0 | 0 | 0 | 1994700 | 0 | 6985600 | 4710200 | 0 | 731680 |
| SSASFSTTA | STXBP1; STXBP | P61764; A0A | STXB1_HUMAN | Unmodified | 560900 | 531190 | 364610 | 512860 | 338350 | 0 | 374250 | 0 | 1209600 | 646050 | 0 | 1648400 | 530990 | 321690 | 370910 | 338410 | 0 | 515970 |
| SSASSGPQ | ITGA5 | P08648 | ITA5_HUMAN I | Unmodified | 0 | 0 | 0 | 499600 | 2264400 | 436760 | 0 | 0 | 0 | 0 | 0 | 0 | 0 | 0 | 1030600 | 1185700 | 146660 | 262360 |
| SSATGSVF | L3HYPDH; L3H | Q96EM0; E9I | T3HPD_HUMAI | Unmodified | 0 | 0 | 0 | 0 | 383210 | 1573000 | 0 | 0 | 114510 | 0 | 0 | 0 | 0 | 274410 | 0 | 0 | 522510 | 258900 |
| SSAVDPEP | HP1BP3; HP1BI | Q5SSJ5; B0C | HP1B3_HUMAI | Unmodified | 17688000 | 7904000 | 0 | 0 | 243480 | 0 | 1085800 | 101590 | 179510 | 50822 | 255320 | 1157100 | 0 | 68008 | 0 | 0 | 0 | 1622800 |
| SSCSLIAFN | CORO7; CORO | P57737; I3L1 | CORO7_HUMA | Unmodified | 0 | 0 | 0 | 0 | 0 | 0 | 0 | 0 | 0 | 0 | 1121200 | 0 | 0 | 863920 | 0 | 0 | 0 | 0 |
| SSCTLFQD | MINPP1 | Q9UNW1 | MINP1_HUMAN | Unmodified | 25017 | 0 | 0 | 0 | 26278 | 73966 | 0 | 0 | 3621900 | 1113400 | 908260 | 190980 | 41983 | 59101 | 0 | 38433 | 71815 | 0 |
| SSCTLPEG | TM9SF4; TM9SF | Q92544; A0A | TM9S4_HUMAN | Unmodified | 0 | 0 | 0 | 0 | 0 | 0 | 0 | 0 | 1331000 | 137230 | 0 | 0 | 0 | 0 | 0 | 0 | 0 | 0 |
| SSDANLYR | CAMP; CAMP | P49913; J3KI | CAMP_HUMAN | Unmodified | 0 | 66972 | 83021 | 0 | 0 | 0 | 0 | 0 | 5546800 | 88364 | 0 | 0 | 0 | 0 | 0 | 0 | 0 | 0 |
| SSDCASGL | DKK1 | O94907 | DKK1_HUMAN | Unmodified | 0 | 0 | 0 | 0 | 0 | 0 | 0 | 0 | 283510 | 0 | 0 | 0 | 0 | 0 | 0 | 0 | 84761 | 146250 |
| SSDDQKIIS | KIAA0319L; KIA | Q8IZA0; C9J | K319L_HUMAN | Unmodified | 0 | 0 | 0 | 0 | 0 | 0 | 0 | 0 | 635230 | 1030700 | 0 | 0 | 0 | 0 | 0 | 0 | 0 | 0 |
| SSDDRNPI | GPRIN1 | Q7Z2K8 | GRIN1_HUMAN | Unmodified | 0 | 0 | 0 | 76355 | 430540 | 0 | 292210 | 138760 | 2148100 | 404360 | 0 | 0 | 236440 | 0 | 0 | 0 | 870190 | 1998300 |
| SSDDRNPI | GPRIN1 | Q7Z2K8 | GRIN1_HUMAN | Oxidation (M) | 0 | 0 | 0 | 0 | 0 | 0 | 157070 | 214080 | 218290 | 195880 | 0 | 0 | 230700 | 41397 | 0 | 0 | 434580 | 0 |
| SSDEAVILC | PSMD13; PSMD | Q9UNM6; A0 | PSD13_HUMAN | Unmodified | 0 | 0 | 47245 | 0 | 0 | 0 | 265030 | 287170 | 0 | 0 | 0 | 1370900 | 387940 | 295470 | 0 | 0 | 0 | 412690 |
| SSDFGTSY | SORCS2; SORC | Q96PQ0; B5 | SORC2_HUMAI | Unmodified | 0 | 312920 | 0 | 0 | 229170 | 0 | 0 | 0 | 1606900 | 361340 | 0 | 180450 | 0 | 0 | 0 | 0 | 127960 | 959950 |
| SSDGRPDS | AFDN; AFDN; A | P55196; Q5T | AFAD_HUMAN | Unmodified | 0 | 0 | 0 | 0 | 0 | 0 | 0 | 0 | 0 | 0 | 1101300 | 4080900 | 0 | 12939 | 0 | 0 | 0 | 0 |
| SSDGSLSH | PPFIA1; PPFIA1 | Q13136; E9F | LIPA1_HUMAN | Unmodified | 0 | 0 | 0 | 115510 | 248070 | 0 | 0 | 0 | 317630 | 124640 | 125860 | 76683 | 183570 | 69597 | 0 | 0 | 21227 | 80490 |
| SSDILSVLK | IQGAP2; IQGAP | Q13576; D6I | IQGA2_HUMAN | Unmodified | 0 | 0 | 2538600 | 1492900 | 0 | 0 | 0 | 0 | 0 | 0 | 0 | 0 | 2432400 | 1914600 | 908000 | 1269200 | 0 | 0 |
| SSDLINKMI | KIAA1549 | Q9HCM3 | K1549_HUMAN | Unmodified | 0 | 0 | 0 | 0 | 0 | 0 | 0 | 0 | 494410 | 0 | 0 | 0 | 0 | 0 | 0 | 0 | 0 | 0 |
| SSDNVSVT | LRP11; LRP11 | Q86VZ4; Q5 | LRP11_HUMAN | Unmodified | 0 | 0 | 0 | 0 | 0 | 0 | 0 | 0 | 1317000 | 0 | 0 | 0 | 0 | 0 | 0 | 0 | 0 | 0 |
| SSDSYPLAI | ARHGAP29 | Q52LW3 | RHG29_HUMAI | Unmodified | 0 | 0 | 0 | 0 | 482170 | 0 | 0 | 0 | 0 | 0 | 0 | 0 | 0 | 0 | 0 | 0 | 517420 | 900120 |
| SSDVYVSYI | SORL1 | Q92673 | SORL_HUMAN | Unmodified | 0 | 0 | 0 | 0 | 0 | 0 | 0 | 0 | 2266500 | 395290 | 0 | 0 | 0 | 0 | 0 | 0 | 0 | 0 |
| SSEEIESAF | SPTAN1; SPTAN | Q13813; A0A | SPTN1_HUMAN | Unmodified | 17396 | 0 | 82620 | 0 | 0 | 0 | 0 | 0 | 0 | 0 | 173500 | 5427000 | 0 | 0 | 87613 | 0 | 0 | 0 |
| SSEELLSAL | MYO6; MYO6; N | Q9UM54; A0 | MYO6_HUMAN | Unmodified | 0 | 0 | 3168000 | 2508100 | 0 | 0 | 0 | 0 | 1317700 | 0 | 0 | 1602600 | 1124800 | 0 | 3332000 | 4659500 | 0 | 0 |
| SSEELLSAL | MYO6; MYO6; N | Q9UM54; A0 | MYO6_HUMAN | Unmodified | 0 | 0 | 965460 | 914110 | 0 | 0 | 0 | 0 | 0 | 0 | 0 | 0 | 0 | 0 | 383250 | 1748000 | 0 | 0 |
| SSEHENAY | PDZK1IP1 | Q13113 | PDZ1I_HUMAN | Unmodified | 0 | 0 | 126610 | 212780 | 0 | 0 | 0 | 0 | 0 | 0 | 47969 | 0 | 0 | 0 | 159020 | 165760 | 0 | 0 |
| SSEHINEGI | BSG; BSG; BSG | P35613; A0A | BASI_HUMAN E | Oxidation (M) | 510630 | 387880 | 3533200 | 3596500 | 5528100 | 663370 | 528760 | 473150 | 1000600 | 302650 | 424890 | 6817200 | 5819800 | 3732200 | 4042200 | 5650600 | 4257100 | 12059000 |
| SSEHINEGI | BSG; BSG; BSG | P35613; A0A | BASI_HUMAN E | Unmodified | 483820 | 304010 | 1940800 | 5535400 | 19751000 | 4179100 | 2787700 | 626550 | 5565800 | 527640 | 781380 | 6804100 | 10434000 | 7016100 | 2815200 | 2767700 | 8967700 | 23932000 |
| SSEHINEGI | BSG; BSG; BSG | P35613; A0A | BASI_HUMAN E | Oxidation (M) | 0 | 0 | 0 | 0 | 5606000 | 831780 | 0 | 0 | 0 | 0 | 0 | 0 | 4490800 | 3943600 | 0 | 0 | 4136700 | 8558200 |
| SSEHINEGI | BSG; BSG; BSG | P35613; A0A | BASI_HUMAN E | Oxidation (M) | 0 | 0 | 0 | 0 | 8223900 | 0 | 0 | 0 | 0 | 0 | 0 | 0 | 6985800 | 9605900 | 0 | 0 | 4626900 | 13589000 |
| SSEIEQAV( | ARPC5L | Q9BPX5 | ARP5L_HUMAN | Oxidation (M) | 0 | 0 | 71241 | 0 | 3805000 | 690230 | 42310 | 0 | 2104400 | 0 | 0 | 2235200 | 4783100 | 1941400 | 0 | 514570 | 227060 | 2088800 |
| SSEKNEDF | AKAP12 | Q02952 | AKA12_HUMAN | Unmodified | 0 | 0 | 0 | 0 | 2837800 | 0 | 0 | 0 | 0 | 0 | 0 | 0 | 0 | 0 | 0 | 0 | 1958600 | 3620400 |
| SSELEGDTI | MARK2; MARK2 | Q7KZI7; E7E | MARK2_HUMAI | Unmodified | 0 | 0 | 0 | 0 | 0 | 0 | 0 | 0 | 0 | 0 | 567840 | 0 | 0 | 0 | 854710 | 1332800 | 0 | 0 |
| SSELEPECI | MEGF9 | Q9H1U4 | MEGF9_HUMAI | Unmodified | 68253 | 0 | 0 | 0 | 80923 | 0 | 0 | 0 | 592880 | 68478 | 57808 | 120860 | 0 | 0 | 0 | 0 | 0 | 0 |
| SSELNTIVC | MGAT4A | Q9UM21 | MGT4A_HUMAI | Unmodified | 0 | 0 | 0 | 0 | 0 | 0 | 0 | 0 | 826540 | 0 | 0 | 0 | 0 | 0 | 0 | 0 | 0 | 0 |
| SSELNTIVC | MGAT4A | Q9UM21 | MGT4A_HUMAI | Unmodified | 0 | 0 | 0 | 0 | 0 | 0 | 100570 | 0 | 1907200 | 0 | 0 | 0 | 0 | 0 | 0 | 0 | 0 | 0 |
| SSELQAIKT | RALY; RALY; RA | Q9UKM9; Q5 | RALY_HUMAN I | Unmodified | 0 | 0 | 0 | 0 | 0 | 0 | 542410 | 102140 | 0 | 1406000 | 0 | 0 | 707180 | 680080 | 0 | 0 | 0 | 498220 |
| SSELRPGEI | HTRA1; HTRA1; | Q92743; A0A | HTRA1_HUMAN | Unmodified | 0 | 0 | 0 | 0 | 0 | 0 | 0 | 0 | 0 | 731310 | 0 | 0 | 0 | 0 | 0 | 0 | 997880 | 9135400 |
| SSEMNVLII | CAP1 | Q01518 | CAP1_HUMAN | Oxidation (M) | 1042000 | 0 | 257710 | 4914600 | 81938000 | 9670800 | 41850000 | 4389400 | 10113000 | 5119800 | 1973200 | 5913100 | 30456000 | 26351000 | 5059200 | 1205800 | 22178000 | 30029000 |
| SSEMNVLII | CAP1 | Q01518 | CAP1_HUMAN | Unmodified | 0 | 0 | 0 | 0 | 6785800 | 780630 | 2801400 | 48527 | 4359300 | 579500 | 78493 | 0 | 1649100 | 1873100 | 0 | 0 | 2136300 | 4987900 |
| SSENFDELI | CRABP1 | P29762 | RABP1_HUMAN | Unmodified | 385050 | 910410 | 0 | 0 | 0 | 353300 | 0 | 0 | 1345000 | 377670 | 20335 | 0 | 0 | 0 | 0 | 0 | 145930 | 0 |
| SSENPNEV | USP5 | P45974 | UBP5_HUMAN | Unmodified | 1513600 | 1282100 | 735260 | 864860 | 142400 | 0 | 0 | 32244 | 609190 | 0 | 593780 | 2868200 | 0 | 0 | 811080 | 837190 | 0 | 0 |
| SSENPNEV | USP5 | P45974 | UBP5_HUMAN | Unmodified | 0 | 0 | 0 | 289040 | 1487300 | 0 | 0 | 0 | 0 | 1172000 | 533560 | 0 | 4928600 | 2338000 | 0 | 0 | 971130 | 1311700 |
| SSEPPPPP( | CARHSP1; CAR | Q9Y2V2; H3I | CHSP1_HUMAI | Acetyl (Protein N | 0 | 0 | 0 | 0 | 0 | 0 | 0 | 311520 | 2609400 | 681890 | 0 | 1002900 | 0 | 0 | 0 | 0 | 0 | 0 |
| SSEPPPPP( | CARHSP1; CAR | Q9Y2V2; H3I | CHSP1_HUMAI | Acetyl (Protein N | 0 | 0 | 0 | 0 | 0 | 0 | 0 | 819660 | 0 | 1337700 | 0 | 0 | 0 | 0 | 0 | 0 | 586290 | 0 |
| SSEPPPPYI | CDIP1; CDIP1; ( | Q9H305; K7I | CDIP1_HUMAN | Acetyl (Protein N | 0 | 462010 | 0 | 62174 | 0 | 0 | 4027100 | 750800 | 2367800 | 0 | 0 | 0 | 0 | 0 | 45748 | 52647 | 0 | 0 |
| SSEPTEDVI | TGOLN2; TGOL | O43493; A0A | TGON2_HUMA | Unmodified | 0 | 0 | 0 | 0 | 0 | 0 | 0 | 0 | 1090800 | 265660 | 168150 | 84836 | 0 | 0 | 0 | 0 | 0 | 0 |
| SSEPTEDVI | TGOLN2; TGOL | O43493; A0A | TGON2_HUMA | Unmodified | 0 | 0 | 0 | 0 | 170780 | 0 | 0 | 0 | 18212000 | 731740 | 1108900 | 1447300 | 0 | 0 | 0 | 0 | 0 | 0 |
| SSEPVQHE | TJP2; TJP2; TJP2 | Q9UDY2; A0 | ZO2_HUMAN T | Unmodified | 0 | 0 | 0 | 0 | 2765200 | 6496900 | 0 | 0 | 7933300 | 2686700 | 0 | 0 | 396190 | 0 | 0 | 0 | 347500 | 0 |
| SSEQGLDE | SCEL | O95171 | SCEL_HUMAN | Unmodified | 0 | 0 | 0 | 0 | 1164600 | 255500 | 0 | 0 | 0 | 0 | 0 | 0 | 0 | 0 | 0 | 0 | 0 | 0 |
| SSEQILATL | PRPF19 | Q9UMS4 | PRP19_HUMAN | Unmodified | 0 | 0 | 78737 | 0 | 1503100 | 0 | 1572300 | 860640 | 1256900 | 1676600 | 1229100 | 7331800 | 1343100 | 738940 | 235770 | 152000 | 2089000 | 1721600 |
| SSERSLPD( | DLGAP4; DLGA | Q9Y2H0; A0A | DLGP4_HUMAI | Unmodified | 0 | 0 | 0 | 0 | 0 | 0 | 0 | 0 | 0 | 0 | 0 | 0 | 0 | 0 | 0 | 0 | 851840 | 0 |
| SSERSSLF( | KIDINS220 | Q9ULH0 | KDIS_HUMAN I | Unmodified | 0 | 0 | 0 | 0 | 3805400 | 363880 | 354130 | 126680 | 0 | 0 | 0 | 337910 | 720210 | 648290 | 0 | 80511 | 114360 | 369760 |
| SSESPSSSI | DOP1B | Q9Y3R5 | DOP2_HUMAN | Unmodified | 0 | 0 | 17060 | 0 | 0 | 16474 | 0 | 0 | 100510 | 0 | 18152 | 202950 | 0 | 0 | 0 | 30935 | 10048 | 80236 |
| SSETMQLK | SLC7A6; SLC7A | Q92536; H3I | YLAT2_HUMAN | Unmodified | 0 | 0 | 0 | 0 | 0 | 0 | 0 | 0 | 0 | 0 | 0 | 2208300 | 0 | 0 | 0 | 0 | 0 | 0 |
| SSETQQPL | COBL; COBL; C | O75128; C9J | COBL_HUMAN | Unmodified | 1462200 | 3095400 | 0 | 0 | 0 | 0 | 0 | 0 | 0 | 0 | 0 | 451620 | 0 | 0 | 0 | 0 | 0 | 0 |
| SSEVDVSD | SIGIRR; SIGIRR | Q6IA17; H0Y | SIGIR_HUMAN | Unmodified | 0 | 0 | 846360 | 704030 | 0 | 151620 | 0 | 0 | 0 | 0 | 0 | 0 | 0 | 0 | 1270400 | 681290 | 0 | 0 |
| SSEVFTTFK | TXLNA | P40222 | TXLNA_HUMAN | Unmodified | 0 | 0 | 625860 | 0 | 489580 | 198760 | 0 | 0 | 0 | 0 | 0 | 623120 | 569570 | 0 | 0 | 0 | 0 | 1141200 |
| SSEYERIEII | BRCC3 | A0A087WZR | A0A087WZR3_I | Unmodified | 0 | 0 | 0 | 88235 | 0 | 0 | 0 | 0 | 118000 | 0 | 0 | 384060 | 304520 | 0 | 0 | 0 | 0 | 294500 |
| SSEYHLEDI | GNAZ | P19086 | GNAZ_HUMAN | Unmodified | 236610 | 254490 | 0 | 239460 | 0 | 0 | 732700 | 0 | 780680 | 0 | 0 | 0 | 0 | 0 | 0 | 246820 | 0 | 0 |
| SSFADISNI | TOMM34 | Q15785 | TOM34_HUMAI | Unmodified | 77805 | 307890 | 40912 | 457940 | 1094300 | 258400 | 141850 | 16936 | 99381 | 105280 | 153450 | 831100 | 1230000 | 535380 | 1060200 | 402420 | 506800 | 1432400 |
| SSFADISNI | TOMM34 | Q15785 | TOM34_HUMAI | Unmodified | 0 | 0 | 0 | 0 | 174880 | 0 | 0 | 0 | 3359600 | 187110 | 0 | 0 | 194550 | 102560 | 0 | 0 | 643070 | 488360 |

| | | | | | | | | | | | | | | | | | | | | | | |
|---|---|---|---|---|---|---|---|---|---|---|---|---|---|---|---|---|---|---|---|---|---|---|
| SSFDEMLP | OGDH; OGDH; | Q02218; E9F | ODO1_HUMAN | Oxidation (M) | 0 | 0 | 0 | 0 | 0 | 0 | 0 | 0 | 1801000 | 88049 | 444990 | 0 | 0 | 0 | 0 | 0 | 0 | 0 |
| SSFGGVQN | IGSF3 | O75054 | IGSF3_HUMAN | Oxidation (M) | 203780 | 0 | 0 | 0 | 0 | 0 | 0 | 0 | 787040 | 0 | 0 | 0 | 0 | 0 | 2677300 | 0 | 0 | 0 |
| SSFHSDPQ | SARG | Q9BW04 | SARG_HUMAN | Unmodified | 0 | 0 | 0 | 0 | 0 | 0 | 0 | 0 | 0 | 0 | 0 | 0 | 2864200 | 3967100 | 0 | 0 | 0 | 0 |
| SSFLIMWR | VPS25; VPS25 | Q9BRG1; K7 | VPS25_HUMAN | Oxidation (M) | 2158900 | 0 | 0 | 227950 | 0 | 0 | 0 | 0 | 0 | 0 | 0 | 732840 | 0 | 0 | 5915800 | 4443100 | 0 | 0 |
| SSFLQVFN | SEC23A; SEC23 | Q15436; F5H | SC23A_HUMAN | Unmodified | 44379 | 0 | 0 | 0 | 0 | 0 | 0 | 0 | 0 | 0 | 0 | 605510 | 0 | 0 | 0 | 0 | 0 | 15640 |
| SSFLVDCS | FLNB; FLNB; FL | O75369; A0A | FLNB_HUMAN | Unmodified | 1335500 | 331060 | 0 | 0 | 6606800 | 1157000 | 0 | 0 | 149270 | 0 | 0 | 796090 | 0 | 608880 | 0 | 268360 | 5308600 | 2027400 |
| SSFQEIPKL | EFCAB14; EFCA | O75071; A0A | EFC14_HUMAN | Unmodified | 0 | 0 | 0 | 0 | 881960 | 0 | 569090 | 66216 | 1024000 | 7764700 | 0 | 0 | 223850 | 204530 | 0 | 0 | 216720 | 636780 |
| SSFREEDN | AP1G1; AP1G1; | O43747; H3I | AP1G1_HUMAN | Unmodified | 0 | 0 | 0 | 1028800 | 0 | 0 | 0 | 0 | 282320 | 0 | 234410 | 2216400 | 0 | 0 | 454270 | 688620 | 0 | 0 |
| SSFSGISR | FAM131B | Q86XD5 | F131B_HUMAN | Unmodified | 0 | 0 | 0 | 0 | 0 | 0 | 0 | 0 | 0 | 0 | 0 | 0 | 0 | 0 | 0 | 0 | 405750 | 0 |
| SSFTVDCS | FLNA; FLNA; FL | P21333; A0A | FLNA_HUMAN | Unmodified | 0 | 0 | 0 | 0 | 11412000 | 1097600 | 16318000 | 15255000 | 840910 | 850800 | 0 | 1723700 | 5812400 | 10041000 | 1576400 | 62554 | 10434000 | 7674200 |
| SSFTVDCS | FLNA; FLNA; FL | P21333; A0A | FLNA_HUMAN | Oxidation (M) | 0 | 0 | 0 | 0 | 0 | 0 | 1161000 | 0 | 0 | 0 | 0 | 0 | 0 | 0 | 0 | 0 | 1017500 | 0 |
| SSFTVDCS | FLNA; FLNA; FL | P21333; A0A | FLNA_HUMAN | Unmodified | 0 | 0 | 0 | 0 | 0 | 0 | 891540 | 0 | 0 | 0 | 0 | 0 | 0 | 0 | 0 | 0 | 2327200 | 0 |
| SSFTVQDLI | IL6ST; IL6ST; IL6 | P40189; A0A | IL6RB_HUMAN | Unmodified | 0 | 0 | 191000 | 424160 | 0 | 0 | 0 | 0 | 7942900 | 424470 | 0 | 0 | 0 | 0 | 228280 | 126920 | 0 | 32717 |
| SSFVAPLEK | SERPINF1 | P36955 | PEDF_HUMAN | Unmodified | 0 | 0 | 0 | 0 | 0 | 0 | 0 | 0 | 2454600 | 1093000 | 13610000 | 0 | 0 | 0 | 0 | 0 | 0 | 0 |
| SSFYVNGL | PFN1; PFN1; PF | P07737; I3L3 | PROF1_HUMAN | Unmodified | 0 | 0 | 289360 | 2324300 | 3080100 | 0 | 1319000 | 636690 | 4842900 | 849180 | 2692800 | 35172000 | 0 | 367200 | 275320 | 114900 | 1422000 | 0 |
| SSGAGSTS | RAB11FIP5; RAI | Q9BXF6; A0A | RFIP5_HUMAN | Unmodified | 0 | 0 | 0 | 0 | 4191500 | 0 | 0 | 111500 | 0 | 0 | 0 | 0 | 3168700 | 0 | 0 | 0 | 643040 | 0 |
| SSGASASA | GNG10 | P50151 | GBG10_HUMA | Acetyl (Protein N | 0 | 0 | 0 | 0 | 4792300 | 1036600 | 627750 | 0 | 0 | 566140 | 0 | 0 | 197520 | 679790 | 0 | 0 | 1910600 | 2875900 |
| SSGASASA | GNG10 | P50151 | GBG10_HUMA | Acetyl (Protein N | 0 | 0 | 0 | 0 | 1268600 | 0 | 0 | 0 | 0 | 562910 | 0 | 0 | 0 | 0 | 0 | 0 | 0 | 934370 |
| SSGATPVS | SRCIN1; SRCIN | Q9C0H9; A0 | SRCN1_HUMA | Unmodified | 0 | 0 | 0 | 0 | 0 | 0 | 0 | 0 | 0 | 0 | 123330 | 829620 | 0 | 0 | 0 | 0 | 0 | 0 |
| SSGCDVNI | AHNAK | Q09666 | AHNK_HUMAN | Unmodified | 0 | 0 | 0 | 0 | 4019900 | 0 | 0 | 0 | 0 | 0 | 0 | 0 | 0 | 1659200 | 0 | 0 | 1256500 | 2309200 |
| SSGCDVNI | AHNAK | Q09666 | AHNK_HUMAN | Unmodified | 0 | 0 | 0 | 0 | 5694800 | 1669700 | 0 | 0 | 0 | 0 | 0 | 0 | 0 | 0 | 0 | 0 | 2990200 | 4471500 |
| SSGEIVYC | RPL18A; RPL18 | Q02543; M0 | RL18A_HUMAN | Unmodified | 3272300 | 1429300 | 1114300 | 918680 | 458600 | 0 | 1494800 | 1299800 | 305150 | 223810 | 690320 | 24197000 | 43091 | 0 | 1369700 | 1013100 | 0 | 249510 |
| SSGFLNLIK | CARMIL1; CARN | Q5VZK9; A0A | CARL1_HUMAN | Unmodified | 0 | 0 | 0 | 0 | 2286700 | 208540 | 0 | 44249 | 0 | 0 | 0 | 1672000 | 574610 | 435010 | 0 | 0 | 949280 | 1035200 |
| SSGGFVW | IDH2; IDH2 | P48735; H0Y | IDHP_HUMAN | Unmodified | 0 | 0 | 0 | 0 | 0 | 0 | 0 | 200270 | 5701200 | 3220100 | 162350 | 0 | 0 | 477400 | 0 | 0 | 0 | 0 |
| SSGGGYG | DHX36 | Q9H2U1 | DHX36_HUMA | Unmodified | 0 | 48953 | 111110 | 0 | 0 | 0 | 0 | 0 | 0 | 0 | 13038 | 205860 | 0 | 0 | 133860 | 149460 | 0 | 0 |
| SSGGKGG | HLA-C; HLA-C; | Q5QT33; Q5 | Q5QT33_HUM | Unmodified | 0 | 0 | 0 | 0 | 1542700 | 0 | 0 | 0 | 0 | 0 | 0 | 0 | 0 | 0 | 0 | 0 | 523690 | 1994000 |
| SSGGKGG | HLA-B; HLA-B; | P01889; D5H | HLAB_HUMAN | Unmodified | 0 | 0 | 0 | 0 | 1526400 | 0 | 0 | 0 | 0 | 0 | 0 | 0 | 0 | 0 | 0 | 0 | 0 | 1419200 |
| SSGGKGG | HLA-A; HLA-B; | A0A1W2PRT | A0A1W2PRT9_ | Unmodified | 0 | 0 | 0 | 0 | 11049000 | 1363500 | 0 | 0 | 247440 | 0 | 0 | 0 | 0 | 0 | 0 | 0 | 683070 | 6519500 |
| SSGGPGA | PLXNB2; PLXNE | O15031; E2F | PLXB2_HUMAN | Unmodified | 0 | 0 | 0 | 123090 | 460410 | 129530 | 34760 | 16473 | 0 | 68148 | 0 | 44456 | 190250 | 0 | 245490 | 338280 | 264140 | 277070 |
| SSGGREDL | JPT1; JPT1; JPT1 | Q9UK76; J3K | JUPI1_HUMAN | Unmodified | 0 | 0 | 1632900 | 3204100 | 4328300 | 2216100 | 0 | 45044 | 683370 | 440040 | 388260 | 2081000 | 5673700 | 3055200 | 1889100 | 2209100 | 1595800 | 2220300 |
| SSGGSEHS | LARP4; LARP4; | Q71RC2; X6I | LARP4_HUMAN | Unmodified | 190000 | 148460 | 119920 | 197850 | 0 | 0 | 0 | 0 | 0 | 0 | 0 | 0 | 0 | 0 | 72370 | 76775 | 0 | 0 |
| SSGGSYRD | CIRBP | Q14011 | CIRBP_HUMAN | Unmodified | 2662200 | 2452200 | 1736100 | 1441900 | 0 | 0 | 535810 | 58466 | 138220 | 25368 | 0 | 0 | 0 | 0 | 1188800 | 2155000 | 0 | 86702 |
| SSGHSSSE | SRRM2 | Q9UQ35 | SRRM2_HUMA | Unmodified | 0 | 0 | 0 | 0 | 0 | 0 | 0 | 0 | 188380 | 36081 | 384300 | 0 | 0 | 0 | 111600 | 0 | 0 | 0 |
| SSGIHVAL | CBR1; CBR1; C | P16152; E9P | CBR1_HUMAN | Acetyl (Protein N | 113740 | 0 | 351710 | 1157400 | 7675100 | 1684500 | 418150 | 323250 | 1898800 | 1050700 | 473760 | 4137200 | 929710 | 984930 | 1103200 | 618740 | 4475500 | 3550000 |
| SSGLNLCT | NADK2; NADK2 | Q4G0N4; H0 | NAKD2_HUMA | Unmodified | 0 | 0 | 0 | 0 | 0 | 0 | 0 | 0 | 410610 | 0 | 0 | 0 | 0 | 0 | 0 | 0 | 0 | 0 |
| SSGLPNIP | TFRC; TFRC | P02786; G3 | TFR1_HUMAN | Unmodified | 7645000 | 4845700 | 15058000 | 11932000 | 1236700 | 918210 | 123380 | 0 | 1.07E+08 | 6632300 | 5419100 | 15149000 | 1520000 | 253420 | 7137400 | 6820600 | 466490 | 988460 |
| SSGLPNIP | TFRC; TFRC | P02786; G3 | TFR1_HUMAN | Unmodified | 0 | 0 | 0 | 0 | 2698100 | 854940 | 310360 | 173170 | 0 | 1531100 | 0 | 0 | 934870 | 958700 | 0 | 0 | 218810 | 2302200 |
| SSGLTAVW | COPA; COPA; C | P53621; A0A | COPA_HUMAN | Unmodified | 0 | 0 | 189420 | 0 | 0 | 0 | 0 | 199490 | 0 | 0 | 0 | 3000700 | 0 | 0 | 0 | 202350 | 0 | 0 |
| SSGPPPPS | CAPRIN1; CAPF | Q14444; E9F | CAPR1_HUMA | Unmodified | 0 | 0 | 0 | 0 | 1388600 | 0 | 0 | 0 | 0 | 0 | 1569500 | 4467200 | 1168700 | 0 | 0 | 0 | 0 | 0 |
| SSGPPSYF | EMC7; EMC7; E | Q9NPA0; H0 | EMC7_HUMAN | Unmodified | 0 | 0 | 0 | 0 | 426980 | 0 | 0 | 0 | 771170 | 0 | 0 | 0 | 0 | 0 | 0 | 0 | 0 | 0 |
| SSGPYGG | HNRNPA1L2; H | Q32P51; P09 | RA1L2_HUMAN | Unmodified | 429580 | 268660 | 0 | 0 | 865640 | 329350 | 0 | 0 | 868540 | 156160 | 0 | 3143500 | 435610 | 0 | 0 | 0 | 0 | 382200 |
| SSGPYGG | HNRNPA1; HNF | P09651; A0A | ROA1_HUMAN | Unmodified | 0 | 0 | 0 | 0 | 1411500 | 290330 | 0 | 260280 | 0 | 630670 | 0 | 0 | 2031400 | 363770 | 0 | 30252 | 143870 | 630520 |
| SSGQSNNI | ITGB1BP1; ITGE | O14713; C9J | ITBP1_HUMAN | Unmodified | 0 | 0 | 0 | 0 | 0 | 0 | 0 | 128630 | 0 | 134300 | 0 | 0 | 395900 | 324870 | 0 | 0 | 358750 | 927790 |
| SSGRSDDN | CMPK1; CMPK1 | P30085; A0A | KCY_HUMAN U | Unmodified | 0 | 0 | 77284 | 0 | 0 | 0 | 460210 | 430060 | 574860 | 455840 | 92133 | 78585 | 3679400 | 0 | 0 | 0 | 0 | 0 |
| SSGRVYYF | PIN1; PIN1 | Q13526; K7E | PIN1_HUMAN I | Unmodified | 0 | 0 | 0 | 1812000 | 0 | 0 | 0 | 0 | 0 | 0 | 0 | 0 | 461510 | 0 | 0 | 0 | 0 | 0 |
| SSGSEGSS | TNKS1BP1 | Q9C0C2 | TB182_HUMAN | Unmodified | 0 | 0 | 0 | 92380 | 693830 | 0 | 0 | 10345 | 0 | 0 | 0 | 986300 | 1792200 | 1709600 | 0 | 440880 | 206630 | 246840 |
| SSGSNQPF | ZDHHC20; ZDH | Q5W0Z9; B4 | ZDH20_HUMA | Unmodified | 317180 | 393090 | 650170 | 826250 | 2221000 | 0 | 0 | 0 | 1136600 | 112490 | 301860 | 1228400 | 182690 | 0 | 847450 | 810980 | 0 | 0 |
| SSGSPYGG | HNRNPA3; HNF | P51991; A0A | ROA3_HUMAN | Unmodified | 0 | 0 | 0 | 0 | 723450 | 0 | 320810 | 699250 | 0 | 662250 | 0 | 0 | 1967100 | 445270 | 0 | 0 | 169520 | 976000 |
| SSGTSSGV | CSNK1G2; CSN | P78368; K7E | KC1G2_HUMA | Oxidation (M) | 19542 | 91265 | 304090 | 262930 | 0 | 0 | 0 | 0 | 0 | 0 | 0 | 112520 | 0 | 0 | 185600 | 239510 | 0 | 0 |
| SSGTVYPK | CD2AP | Q9Y5K6 | CD2AP_HUMA | Unmodified | 0 | 3339000 | 3746700 | 180760 | 0 | 0 | 46604 | 37455 | 701620 | 52933 | 291240 | 658480 | 1098200 | 0 | 5880200 | 2931800 | 0 | 0 |
| SSGVPVDG | NTPCR; NTPCR | Q9BSD7; Q5 | NTPCR_HUMA | Unmodified | 0 | 0 | 0 | 0 | 0 | 0 | 0 | 0 | 0 | 0 | 0 | 744690 | 0 | 0 | 381350 | 354720 | 0 | 408540 |
| SSGVPVDG | NTPCR; NTPCR | Q9BSD7; Q5 | NTPCR_HUMA | Unmodified | 312890 | 138500 | 380240 | 273320 | 0 | 0 | 0 | 0 | 0 | 0 | 0 | 420020 | 1424200 | 652310 | 292510 | 146320 | 0 | 0 |
| SSHETLNI | EPB41L2; EPB4 | O43491; E9F | E41L2_HUMAN | Unmodified | 128950 | 56777 | 0 | 0 | 11713000 | 0 | 152660 | 204550 | 0 | 0 | 0 | 0 | 319050 | 444580 | 0 | 0 | 2190900 | 2326300 |
| SSHNLAVH | ZFP91; ZFP91-C | Q96JP5; H3E | ZFP91_HUMAN | Oxidation (M) | 0 | 0 | 0 | 0 | 0 | 0 | 249400 | 0 | 0 | 0 | 0 | 0 | 0 | 0 | 0 | 2.23E+08 | 0 | 0 |
| SSHSNAYF | ZMPSTE24 | O75844 | FACE1_HUMAN | Unmodified | 0 | 0 | 0 | 0 | 0 | 0 | 0 | 0 | 1440900 | 155390 | 269700 | 0 | 0 | 0 | 0 | 0 | 0 | 0 |
| SSHVTIDTL | MELTF; MELTF | P08582; C9J | TRFM_HUMAN | Unmodified | 0 | 0 | 0 | 0 | 1016600 | 0 | 0 | 0 | 952680 | 484500 | 0 | 383870 | 128050 | 0 | 0 | 0 | 428570 | 877840 |
| SSHVTIDTL | MELTF; MELTF | P08582; C9J | TRFM_HUMAN | Unmodified | 0 | 81722 | 0 | 0 | 1968500 | 81910 | 0 | 0 | 0 | 536500 | 0 | 0 | 470060 | 0 | 0 | 0 | 566920 | 51654 |
| SSHYDELL | SND1; SND1 | Q7KZF4; H7 | SND1_HUMAN | Unmodified | 671030 | 618220 | 1859200 | 1563500 | 0 | 23299 | 0 | 0 | 0 | 0 | 649590 | 8390300 | 50427 | 63595 | 1863700 | 1183000 | 0 | 0 |
| SSIAGLLLK | PLEC; PLEC | Q15149; H0 | PLEC_HUMAN | Unmodified | 0 | 0 | 0 | 0 | 2314800 | 0 | 0 | 0 | 0 | 0 | 0 | 0 | 0 | 0 | 0 | 0 | 3686600 | 2967400 |
| SSIAGLLLK | PLEC; PLEC | Q15149; H0 | PLEC_HUMAN | Unmodified | 0 | 0 | 0 | 0 | 0 | 0 | 0 | 0 | 0 | 0 | 0 | 0 | 0 | 0 | 0 | 0 | 1503200 | 0 |
| SSIDSEPAL | NOMO1; NOMC | Q15155; P69 | NOMO1_HUMA | Unmodified | 0 | 0 | 0 | 0 | 0 | 0 | 0 | 0 | 6464800 | 938260 | 1340200 | 284320 | 0 | 0 | 0 | 0 | 0 | 0 |
| SSIGTGYDI | PSMA3; PSMA3 | P25788; G3 | PSA3_HUMAN | Acetyl (Protein N | 22647 | 48234 | 0 | 0 | 20633 | 0 | 0 | 0 | 0 | 421260 | 0 | 1882800 | 0 | 0 | 0 | 0 | 0 | 0 |
| SSIHNFMTI | ATP2B1; ATP2B | P20020; E7E | AT2B1_HUMAN | Oxidation (M) | 0 | 0 | 0 | 0 | 0 | 0 | 0 | 0 | 0 | 0 | 0 | 0 | 0 | 0 | 479610 | 327180 | 0 | 0 |
| SSIHNFMTI | ATP2B1; ATP2B | P20020; E7E | AT2B1_HUMAN | Oxidation (M) | 0 | 0 | 0 | 6058800 | 0 | 0 | 0 | 0 | 0 | 0 | 0 | 0 | 2695300 | 0 | 0 | 0 | 0 | 0 |
| SSIIVAADE | RCC2 | Q9P258 | RCC2_HUMAN | Unmodified | 0 | 0 | 0 | 0 | 2432600 | 0 | 1651600 | 0 | 0 | 2308200 | 1306800 | 2555000 | 0 | 0 | 0 | 0 | 0 | 0 |
| SSILLDVKP | EEF1B2 | P24534 | EF1B_HUMAN I | Oxidation (M) | 266710 | 0 | 1199200 | 174490 | 0 | 0 | 0 | 0 | 0 | 0 | 3421400 | 10730000 | 307890 | 0 | 1014100 | 543350 | 0 | 0 |
| SSILLDVKP | EEF1B2 | P24534 | EF1B_HUMAN I | Unmodified | 0 | 0 | 0 | 0 | 317380 | 0 | 0 | 0 | 0 | 0 | 0 | 641570 | 0 | 0 | 0 | 0 | 0 | 0 |
| SSILPFTPP | SMAD2; SMAD2 | Q15796; K7E | SMAD2_HUMA | Acetyl (Protein N | 0 | 0 | 0 | 0 | 0 | 0 | 0 | 0 | 65094 | 92297 | 61612 | 121330 | 0 | 278510 | 0 | 0 | 0 | 0 |
| SSILQAESK | APPL1 | Q9UKG1 | DP13A_HUMAN | Unmodified | 160940 | 85210 | 0 | 0 | 3080000 | 720430 | 448800 | 0 | 0 | 1087000 | 0 | 187870 | 622650 | 1162300 | 0 | 21416 | 0 | 0 |
| SSIMDVR | NCLN; NCLN; N | Q969V3; K7E | NCLN_HUMAN | Unmodified | 0 | 0 | 0 | 0 | 0 | 0 | 0 | 0 | 5053300 | 218410 | 192890 | 0 | 0 | 0 | 0 | 0 | 0 | 0 |
| SSINDAAEI | MYO1D; MYO1D | O94832; J3Q | MYO1D_HUMA | Unmodified | 3075800 | 3573200 | 1157700 | 0 | 0 | 0 | 0 | 0 | 0 | 47004 | 0 | 696900 | 0 | 0 | 3342700 | 2221400 | 0 | 0 |
| SSIVCRFVC | RAB31; RAB31 | Q13636; J3Q | RAB31_HUMAN | Oxidation (M) | 0 | 0 | 0 | 0 | 1614400 | 0 | 0 | 0 | 0 | 0 | 0 | 0 | 0 | 0 | 0 | 0 | 0 | 0 |
| SSIWQFFSI | SPAG9 | O60271 | JIP4_HUMAN C | Unmodified | 0 | 0 | 0 | 0 | 0 | 0 | 0 | 0 | 0 | 0 | 0 | 217780 | 0 | 0 | 0 | 0 | 0 | 0 |
| SSKASLGSI | AHNAK | Q09666 | AHNK_HUMAN | Unmodified | 0 | 0 | 0 | 0 | 0 | 0 | 0 | 0 | 0 | 0 | 0 | 0 | 0 | 0 | 0 | 0 | 1188800 | 0 |
| SSKGGPGS | ANXA1; ANXA1; | P04083; Q5T | ANXA1_HUMAN | Unmodified | 0 | 0 | 0 | 0 | 12436000 | 962810 | 919140 | 0 | 165440 | 0 | 0 | 1593100 | 0 | 0 | 0 | 0 | 7194900 | 1593800 |
| SSKTASTNI | GNG12 | Q9UBI6 | GBG12_HUMA | Acetyl (Protein N | 0 | 0 | 0 | 0 | 616980 | 0 | 0 | 72100 | 0 | 0 | 0 | 0 | 206230 | 304470 | 0 | 0 | 1402200 | 410120 |
| SSLAASSSI | MKRN1; MKRN1 | Q9UHC7; C9 | MKRN1_HUMA | Oxidation (M) | 1246100 | 3813600 | 0 | 0 | 0 | 0 | 0 | 0 | 0 | 0 | 0 | 0 | 0 | 0 | 2518700 | 0 | 0 | 0 |
| SSLDGPIVL | CYFIP2; CYFIP2 | Q96F07; H7 | CYFP2_HUMAN | Unmodified | 0 | 0 | 0 | 0 | 0 | 0 | 0 | 0 | 0 | 0 | 0 | 1077700 | 0 | 0 | 0 | 545020 | 0 | 0 |
| SSLEGPTIL | CYFIP1; CYFIP1 | Q7L576; A0A | CYFP1_HUMAN | Unmodified | 342860 | 0 | 0 | 0 | 0 | 0 | 0 | 0 | 0 | 0 | 0 | 2678200 | 0 | 0 | 1033400 | 604000 | 92369 | 0 |
| SSLFELAK | SLC38A1; SLC3 | Q9H2H9; F8 | S38A1_HUMAN | Unmodified | 293240 | 277820 | 0 | 0 | 0 | 0 | 0 | 0 | 0 | 0 | 0 | 4958500 | 0 | 0 | 525030 | 419800 | 0 | 0 |
| SSLGPVGL | TCP1; TCP1; TC | P17987; F5H | TCPA_HUMAN | Unmodified | 299540 | 856440 | 0 | 772620 | 3264300 | 799480 | 2481800 | 1304300 | 5289300 | 3816900 | 2759700 | 15879000 | 7579600 | 8746400 | 1521600 | 610820 | 4155900 | 4007800 |
| SSLGQSAS | SF3B2; SF3B2; | Q13435; E9F | SF3B2_HUMAN | Unmodified | 2116700 | 0 | 0 | 0 | 0 | 0 | 0 | 0 | 0 | 0 | 200690 | 332170 | 0 | 0 | 0 | 0 | 0 | 0 |
| SSLGSLQTI | ZC3HAV1; ZC3H | Q7Z2W4; C9 | ZCCHV_HUMA | Unmodified | 0 | 0 | 0 | 385390 | 3647800 | 617380 | 0 | 0 | 0 | 0 | 0 | 0 | 1386200 | 246180 | 0 | 0 | 322520 | 0 |
| SSLGSLQTI | ZC3HAV1; ZC3H | Q7Z2W4; C9 | ZCCHV_HUMA | Unmodified | 729390 | 837210 | 2361000 | 2252700 | 8645100 | 2519300 | 139060 | 41887 | 898290 | 0 | 0 | 0 | 2365300 | 1168500 | 1367900 | 1758300 | 172190 | 981840 |
| SSLHYKPTI | ADGRG1; ADGF | Q9Y653; H3I | AGRG1_HUMA | Unmodified | 349410 | 0 | 187000 | 130750 | 0 | 0 | 0 | 0 | 1697700 | 0 | 284140 | 75675 | 0 | 0 | 0 | 123470 | 66324 | 0 |
| SSLISALFR | ABCC4 | O15439 | MRP4_HUMAN | Unmodified | 0 | 0 | 5174200 | 4499000 | 257940 | 0 | 0 | 0 | 0 | 0 | 0 | 978880 | 0 | 0 | 3740700 | 4377900 | 0 | 0 |
| SSLISALFRI | ABCC4 | O15439 | MRP4_HUMAN | Unmodified | 0 | 0 | 0 | 123600 | 868740 | 0 | 774360 | 0 | 0 | 361990 | 0 | 0 | 0 | 0 | 0 | 0 | 173690 | 0 |
| SSLLSAVLG | ABCC4; ABCC4 | O15439; A0A | MRP4_HUMAN | Unmodified | 0 | 0 | 1124200 | 1652900 | 0 | 0 | 643740 | 0 | 0 | 0 | 0 | 0 | 0 | 0 | 3294500 | 917760 | 0 | 0 |
| SSLLTELSN | UXT | Q9UBK9 | UXT_HUMAN P | Unmodified | 0 | 0 | 0 | 0 | 0 | 0 | 0 | 0 | 0 | 0 | 0 | 806400 | 0 | 0 | 0 | 0 | 0 | 0 |
| SSLMDTAD | DIP2B | Q9P265 | DIP2B_HUMAN | Unmodified | 0 | 0 | 161770 | 262640 | 0 | 0 | 0 | 0 | 886770 | 217850 | 729450 | 0 | 0 | 0 | 211490 | 192210 | 0 | 0 |
| SSLMDTAD | DIP2B | Q9P265 | DIP2B_HUMAN | Oxidation (M) | 0 | 0 | 0 | 0 | 0 | 0 | 0 | 0 | 266540 | 0 | 215410 | 917880 | 0 | 0 | 0 | 0 | 0 | 0 |
| SSLMNRYV | RAB9A | P51151 | RAB9A_HUMAN | Oxidation (M) | 0 | 0 | 0 | 0 | 0 | 0 | 171260 | 0 | 0 | 157890 | 0 | 0 | 1210100 | 351660 | 0 | 0 | 0 | 0 |
| SSLNDPLC | PROM1 | O43490 | PROM1_HUMA | Unmodified | 0 | 0 | 0 | 0 | 695450 | 0 | 0 | 0 | 0 | 0 | 1190700 | 10995000 | 0 | 0 | 0 | 0 | 625660 | 649750 |
| SSLNPILFR | DDOST; DDOST | P39656; U3K | OST48_HUMAN | Unmodified | 49160 | 0 | 0 | 0 | 0 | 0 | 0 | 0 | 3.31E+08 | 22185000 | 6779100 | 3363600 | 219070 | 994140 | 0 | 0 | 0 | 0 |
| SSLPAVSD | WASF2 | Q9Y6W5 | WASF2_HUMA | Unmodified | 0 | 0 | 0 | 0 | 1701600 | 0 | 0 | 0 | 547480 | 273310 | 0 | 0 | 0 | 0 | 0 | 0 | 509000 | 1311900 |
| SSLPNGEG | PPM1H | Q9ULR3 | PPM1H_HUMA | Unmodified | 0 | 0 | 0 | 0 | 0 | 0 | 0 | 0 | 0 | 0 | 0 | 0 | 0 | 0 | 0 | 0 | 0 | 0 |
| SSLQPITTL | PTK7; PTK7 | Q13308; C9J | PTK7_HUMAN I | Unmodified | 0 | 0 | 0 | 0 | 769270 | 0 | 0 | 0 | 5524700 | 700970 | 1890700 | 8678400 | 921520 | 228810 | 6047100 | 3147800 | 0 | 0 |
| SSLQPITTL | PTK7; PTK7 | Q13308; C9J | PTK7_HUMAN I | Unmodified | 0 | 0 | 0 | 4022800 | 18536000 | 2874700 | 2587000 | 99823 | 9612700 | 5629600 | 3122700 | 2454000 | 12526000 | 4788600 | 0 | 0 | 6308400 | 19021000 |
| SSLQSQCL | DYNC1H1; DYN | Q14204; A0A | DYHC1_HUMA | Unmodified | 3052900 | 2466600 | 1801200 | 1331100 | 0 | 1555500 | 387780 | 230590 | 62561 | 0 | 265090 | 597120 | 0 | 0 | 0 | 2016600 | 277840 | 208060 |
| SSLSHAMV | SAMM50 | Q9Y512 | SAM50_HUMA | Oxidation (M) | 0 | 0 | 0 | 0 | 0 | 165720 | 0 | 47814 | 969840 | 134820 | 90964 | 0 | 0 | 249910 | 0 | 0 | 0 | 0 |
| SSLSHAMV | SAMM50 | Q9Y512 | SAM50_HUMA | Unmodified | 0 | 0 | 0 | 0 | 0 | 0 | 0 | 0 | 8903100 | 563170 | 84894 | 0 | 0 | 0 | 0 | 0 | 0 | 0 |
| SSLSHAMV | SAMM50 | Q9Y512 | SAM50_HUMA | Unmodified | 0 | 0 | 0 | 0 | 3241600 | 0 | 0 | 0 | 0 | 1144300 | 0 | 0 | 0 | 0 | 0 | 0 | 0 | 991470 |
| SSLSNEVN | UACA; UACA; U | Q9BZF9; H0 | UACA_HUMAN | Unmodified | 0 | 0 | 0 | 0 | 489040 | 44455 | 0 | 25548 | 0 | 0 | 23833 | 932760 | 400730 | 152020 | 0 | 36641 | 506720 | 265620 |
| SSLSSAQA | SPTAN1; SPTAN | Q13813; A0A | SPTN1_HUMAN | Unmodified | 0 | 0 | 0 | 0 | 0 | 0 | 0 | 0 | 1016200 | 0 | 96427 | 3079000 | 0 | 0 | 486110 | 218760 | 0 | 0 |
| SSLTLGLFF | ABCC1; ABCC1 | P33527; I3L4 | MRP1_HUMAN | Unmodified | 0 | 0 | 266940 | 0 | 0 | 0 | 0 | 0 | 0 | 0 | 0 | 3867400 | 0 | 0 | 521120 | 707760 | 0 | 0 |
| SSLVLSML | RELCH; RELCH | Q9P260; A0A | RELCH_HUMA | Unmodified | 0 | 0 | 0 | 1343900 | 0 | 0 | 0 | 0 | 0 | 0 | 0 | 0 | 2565600 | 3936000 | 0 | 0 | 0 | 0 |
| SSLVSALLG | ABCC3 | O15438 | MRP3_HUMAN | Unmodified | 0 | 0 | 0 | 0 | 250090 | 0 | 0 | 0 | 0 | 0 | 0 | 0 | 0 | 0 | 0 | 0 | 0 | 0 |
| SSLYEGLEI | ATP2B2; ATP2B | Q01814; P20 | AT2B2_HUMAN | Unmodified | 2538400 | 1463600 | 4118300 | 3709300 | 567290 | 0 | 0 | 0 | 1360400 | 1154300 | 398820 | 3183300 | 2778900 | 1067500 | 548340 | 8986800 | 298830 | 1384600 |
| SSMAPSQF | N/A; AAK1; AAK | Q6ZSR9; A0A | YJ005_HUMAN | Oxidation (M) | 0 | 0 | 0 | 0 | 0 | 0 | 0 | 0 | 0 | 0 | 0 | 0 | 1188500 | 2418500 | 0 | 0 | 0 | 0 |

| | | | | | | | | | | | | | | | | | | | | | | |
|---|---|---|---|---|---|---|---|---|---|---|---|---|---|---|---|---|---|---|---|---|---|---|
| SSMAVHSL | CD109 | Q6YHK3 | CD109_HUMAI | Unmodified | 0 | 0 | 0 | 0 | 0 | 0 | 0 | 0 | 841800 | 235370 | 0 | 0 | 0 | 0 | 0 | 0 | 0 | 0 |
| SSMAVHSL | CD109 | Q6YHK3 | CD109_HUMAI | Unmodified | 0 | 0 | 0 | 0 | 0 | 0 | 0 | 13290 | 0 | 977010 | 0 | 0 | 0 | 0 | 0 | 0 | 0 | 0 |
| SSMFRAEE | ARHGAP29; ARI | Q52LW3; F8' | RHG29_HUMAI | Unmodified | 0 | 0 | 0 | 0 | 0 | 167660 | 0 | 0 | 0 | 0 | 0 | 0 | 0 | 0 | 0 | 0 | 2315900 | 2710200 |
| SSMFRAEE | ARHGAP29; ARI | Q52LW3; F8' | RHG29_HUMAI | Oxidation (M) | 0 | 0 | 0 | 0 | 0 | 0 | 0 | 0 | 0 | 0 | 0 | 0 | 0 | 0 | 0 | 0 | 234360 | 394410 |
| SSMNPEYD | RAB1A; RAB1A | P62820; E7E | RAB1A_HUMAI | Acetyl (Protein N | 0 | 0 | 0 | 0 | 0 | 0 | 0 | 439280 | 0 | 269950 | 1634800 | 3678500 | 1021800 | 0 | 0 | 0 | 0 | 0 |
| SSMNVDE/ | RAB13; RAB13 | P51153; A0A | RAB13_HUMAI | Oxidation (M) | 5313500 | 4731700 | 5422900 | 3981700 | 3665100 | 0 | 0 | 0 | 3636100 | 501180 | 813000 | 1359900 | 998120 | 0 | 3762100 | 2869100 | 250380 | 885050 |
| SSMNVDE/ | RAB13; RAB13 | P51153; A0A | RAB13_HUMAI | Unmodified | 1426900 | 2117700 | 87222 | 1384100 | 2067900 | 34143 | 0 | 0 | 5710200 | 436920 | 0 | 610200 | 369040 | 140440 | 1417600 | 574100 | 150190 | 2044600 |
| SSMNVDE/ | RAB13; RAB13 | P51153; A0A | RAB13_HUMAI | Oxidation (M) | 498930 | 0 | 0 | 2659500 | 78228000 | 6412600 | 3600600 | 0 | 423510 | 7730100 | 307450 | 0 | 29055000 | 14881000 | 0 | 0 | 10385000 | 16472000 |
| SSMNVDE/ | RAB13; RAB13 | P51153; A0A | RAB13_HUMAI | Unmodified | 0 | 0 | 0 | 108540 | 8060600 | 207250 | 0 | 10005 | 0 | 1370800 | 0 | 0 | 2694700 | 2571400 | 0 | 0 | 838620 | 1680500 |
| SSMNVDE/ | RAB13; RAB13 | P51153; A0A | RAB13_HUMAI | Oxidation (M) | 0 | 0 | 0 | 0 | 480660 | 58397 | 0 | 0 | 0 | 209120 | 0 | 0 | 0 | 0 | 0 | 0 | 907250 | 353660 |
| SSMSVTSLI | SFXN1 | Q9H9B4 | SFXN1_HUMAI | Unmodified | 0 | 0 | 0 | 0 | 0 | 0 | 0 | 0 | 639710 | 0 | 0 | 429270 | 0 | 0 | 0 | 0 | 0 | 0 |
| SSMTFLTR | RTCB | Q9Y3I0 | RTCB_HUMAN | Unmodified | 0 | 0 | 0 | 0 | 0 | 0 | 0 | 0 | 0 | 0 | 0 | 2761200 | 0 | 0 | 0 | 0 | 0 | 0 |
| SSMTFLTR | RTCB | Q9Y3I0 | RTCB_HUMAN | Oxidation (M) | 71793 | 0 | 108970 | 0 | 0 | 0 | 0 | 0 | 0 | 0 | 176630 | 2613400 | 0 | 0 | 225410 | 156670 | 0 | 0 |
| SSNEATNII | FOLH1; FOLH1 | Q04609; E9F | FOLH1_HUMAI | Unmodified | 0 | 0 | 0 | 0 | 26072 | 0 | 0 | 0 | 0 | 0 | 0 | 0 | 0 | 0 | 0 | 1114800 | 0 | 0 |
| SSNECFK | CNBP; CNBP | P62633; D6F | CNBP_HUMAN | Acetyl (Protein N | 239880 | 137990 | 98812 | 0 | 571660 | 0 | 51946 | 53669 | 0 | 54493 | 57573 | 951100 | 277280 | 0 | 211450 | 102650 | 355620 | 699030 |
| SSNFGYITS | PAN3; PAN3 | Q58A45; A0/ | PAN3_HUMAN | Unmodified | 330450 | 234710 | 0 | 0 | 0 | 0 | 0 | 0 | 0 | 0 | 134290 | 0 | 0 | 0 | 0 | 0 | 0 | 0 |
| SSNFLYLQ' | HTATIP2 | Q9BUP3 | HTAI2_HUMAN | Unmodified | 0 | 0 | 0 | 0 | 0 | 0 | 0 | 0 | 1500200 | 0 | 0 | 0 | 0 | 0 | 0 | 0 | 0 | 0 |
| SSNIQVGD | ATP9A | O75110 | ATP9A_HUMAI | Unmodified | 0 | 0 | 0 | 0 | 0 | 0 | 0 | 0 | 2803000 | 0 | 0 | 0 | 0 | 0 | 0 | 0 | 0 | 0 |
| SSNIRPSPI | SPRED1 | Q7Z699 | SPRE1_HUMAI | Unmodified | 220130 | 0 | 0 | 0 | 0 | 0 | 0 | 0 | 0 | 0 | 0 | 0 | 0 | 0 | 0 | 78101 | 397750 | 0 |
| SSNLRGEC | NUP210 | Q8TEM1 | PO210_HUMAI | Unmodified | 0 | 0 | 0 | 0 | 0 | 0 | 0 | 45797 | 8803200 | 1186100 | 0 | 0 | 0 | 0 | 0 | 0 | 0 | 0 |
| SSNTYTLTI | ALCAM; ALCAM | Q13740; F5C | CD166_HUMAI | Unmodified | 0 | 1322600 | 0 | 0 | 0 | 0 | 0 | 0 | 426680 | 0 | 0 | 0 | 0 | 0 | 0 | 0 | 0 | 0 |
| SSNTYTLTI | ALCAM; ALCAM | Q13740; F5C | CD166_HUMAI | Unmodified | 4395900 | 2974100 | 618330 | 0 | 0 | 0 | 0 | 0 | 2985600 | 133920 | 0 | 542160 | 0 | 0 | 0 | 1442500 | 0 | 0 |
| SSNVVIKGI | PBK; PBK | Q96KB5; E5F | TOPK_HUMAN | Unmodified | 355670 | 344130 | 0 | 0 | 1550200 | 0 | 844230 | 424700 | 791680 | 0 | 0 | 0 | 1567500 | 0 | 0 | 0 | 678690 | 0 |
| SSPAAKPG | MAPRE2; MAPR | Q15555; K7E | MARE2_HUMAI | Unmodified | 0 | 0 | 0 | 0 | 222000 | 120340 | 0 | 0 | 210100 | 447020 | 50524 | 105220 | 103260 | 105120 | 0 | 0 | 48459 | 85710 |
| SSPCIHYFT | SCRN1 | Q12765 | SCRN1_HUMAI | Unmodified | 0 | 87156 | 0 | 0 | 261200 | 0 | 0 | 0 | 2162500 | 196010 | 0 | 0 | 0 | 0 | 0 | 0 | 300740 | 136190 |
| SSPCIHYFT | SCRN1 | Q12765 | SCRN1_HUMAI | Unmodified | 0 | 0 | 0 | 0 | 0 | 94990 | 0 | 0 | 0 | 107680 | 0 | 0 | 0 | 0 | 0 | 0 | 242390 | 276400 |
| SSPDLTGV' | TOM1L2 | Q6ZVM7 | TM1L2_HUMAI | Unmodified | 873420 | 1423300 | 387940 | 0 | 1082900 | 0 | 161460 | 19959 | 0 | 0 | 0 | 444810 | 0 | 250470 | 3567600 | 2276700 | 1413000 | 0 |
| SSPEIQEPII | RELCH; RELCH | Q9P260; A0/ | RELCH_HUMAI | Unmodified | 19581 | 133740 | 41171 | 59941 | 579470 | 42084 | 294020 | 65305 | 51369 | 71384 | 45119 | 183430 | 870420 | 0 | 36705 | 37525 | 408810 | 602140 |
| SSPEPVALT | COPG1 | Q9Y678 | COPG1_HUMA | Unmodified | 0 | 0 | 0 | 0 | 0 | 33817 | 0 | 0 | 707240 | 0 | 0 | 5650000 | 0 | 0 | 0 | 0 | 0 | 0 |
| SSPEQSYQ | GPRC5C; GPRC | Q9NQ84; A8 | GPC5C_HUMA | Oxidation (M) | 0 | 0 | 1060400 | 1387200 | 0 | 596350 | 0 | 27634 | 1414900 | 239760 | 0 | 93461 | 0 | 0 | 1962700 | 1594700 | 0 | 114860 |
| SSPEQSYQ | GPRC5C; GPRC | Q9NQ84; A8 | GPC5C_HUMA | Unmodified | 187160 | 0 | 546050 | 1158500 | 0 | 0 | 0 | 0 | 3737500 | 297340 | 0 | 0 | 0 | 0 | 1861100 | 1251200 | 219210 | 1073200 |
| SSPEQSYQ | GPRC5C; GPRC | Q9NQ84; A8 | GPC5C_HUMA | Unmodified | 0 | 0 | 0 | 1133600 | 1078500 | 0 | 0 | 0 | 4515900 | 0 | 0 | 0 | 1185500 | 0 | 300700 | 0 | 0 | 987800 |
| SSPEQSYQ | GPRC5C; GPRC | Q9NQ84; A8 | GPC5C_HUMA | Oxidation (M) | 0 | 0 | 1029000 | 5233300 | 0 | 0 | 0 | 0 | 2752900 | 2828700 | 0 | 0 | 5849000 | 0 | 3067200 | 1259900 | 0 | 0 |
| SSPGLEEQ | IGSF3 | O75054 | IGSF3_HUMAN | Unmodified | 684300 | 711620 | 0 | 290620 | 0 | 0 | 0 | 0 | 0 | 0 | 0 | 1182800 | 0 | 0 | 1669900 | 1364800 | 0 | 0 |
| SSPGLSDTI | MINDY3 | Q9H8M7 | MINY3_HUMAI | Unmodified | 1655900 | 881460 | 0 | 429960 | 0 | 0 | 0 | 0 | 0 | 0 | 0 | 443210 | 0 | 0 | 411370 | 388230 | 0 | 0 |
| SSPGQTPE | TTLL12 | Q14166 | TTL12_HUMAN | Unmodified | 0 | 0 | 0 | 0 | 0 | 0 | 0 | 0 | 0 | 0 | 0 | 3676400 | 0 | 0 | 0 | 0 | 0 | 0 |
| SSPHSTYYI | KIDINS220 | Q9ULH0 | KDIS_HUMAN I | Unmodified | 0 | 0 | 0 | 0 | 1255600 | 0 | 0 | 0 | 0 | 0 | 0 | 0 | 453800 | 0 | 0 | 0 | 0 | 0 |
| SSPLDNKLI | SCG3 | Q8WXD2 | SCG3_HUMAN | Unmodified | 6988000 | 9439700 | 0 | 4774700 | 0 | 0 | 393890 | 1377500 | 63267000 | 42051000 | 0 | 0 | 0 | 0 | 4890100 | 0 | 86387 | 0 |
| SSPLDNKLI | SCG3 | Q8WXD2 | SCG3_HUMAN | Unmodified | 0 | 0 | 0 | 0 | 0 | 0 | 0 | 0 | 0 | 997030 | 0 | 0 | 0 | 0 | 0 | 0 | 0 | 0 |
| SSPMVQVI | LARP4; LARP4; | Q71RC2; A0. | LARP4_HUMAI | Unmodified | 110750 | 162440 | 0 | 0 | 0 | 0 | 0 | 0 | 0 | 0 | 0 | 0 | 0 | 0 | 41885 | 50256 | 0 | 0 |
| SSPNVAN( | AFDN; AFDN; A | P55196; Q5T | AFAD_HUMAN | Unmodified | 0 | 0 | 0 | 0 | 121630 | 0 | 0 | 0 | 0 | 0 | 224600 | 490480 | 215850 | 147760 | 0 | 32713 | 0 | 0 |
| SSPPAPPLI | SH3BP1; SH3BI | Q9Y3L3; A0/ | 3BP1_HUMAN | Unmodified | 355180 | 0 | 0 | 0 | 0 | 0 | 0 | 0 | 0 | 0 | 0 | 0 | 0 | 0 | 0 | 0 | 0 | 0 |
| SSPPEGKLI | DAP; DAP | P51397; B4D | DAP1_HUMAN | Acetyl (Protein N | 230580 | 0 | 0 | 0 | 2288300 | 1047000 | 597260 | 0 | 133070 | 105750 | 0 | 652690 | 2952500 | 2878500 | 654080 | 0 | 4097600 | 6876400 |
| SSPPEGKLI | DAP; DAP | P51397; B4D | DAP1_HUMAN | Acetyl (Protein N | 0 | 0 | 0 | 0 | 658270 | 0 | 685090 | 0 | 0 | 0 | 0 | 0 | 0 | 1116100 | 0 | 0 | 4910700 | 904670 |
| SSPQAPED | ACOT8; ACOT8; | O14734; E9F | ACOT8_HUMAI | Acetyl (Protein N | 0 | 0 | 0 | 0 | 0 | 0 | 0 | 0 | 6329600 | 394290 | 0 | 0 | 0 | 0 | 0 | 0 | 0 | 0 |
| SSPQFGVT | SLC25A13 | Q9UJS0 | CMC2_HUMAN | Unmodified | 0 | 0 | 0 | 0 | 0 | 0 | 0 | 0 | 135850 | 42209 | 0 | 0 | 0 | 0 | 0 | 0 | 0 | 0 |
| SSPQGSLIS | N/A | N/A | N/A | Unmodified | 0 | 0 | 0 | 0 | 57447 | 0 | 0 | 19462 | 4.55E+08 | 14970000 | 0 | 6444300 | 0 | 0 | 0 | 0 | 0 | 0 |
| SSPQIRNPI | CASK; CASK; C/ | O14936; A0/ | CSKP_HUMAN | Unmodified | 503670 | 0 | 2170400 | 3726000 | 4813700 | 903400 | 175070 | 149440 | 0 | 2816100 | 881630 | 0 | 2410300 | 1652500 | 2959600 | 2473500 | 750510 | 2275300 |
| SSPQIRNPI | CASK; CASK; C/ | O14936; A0/ | CSKP_HUMAN | Unmodified | 0 | 0 | 0 | 0 | 10731000 | 3696400 | 1970300 | 932850 | 335330 | 5212400 | 0 | 4632200 | 12091000 | 10170000 | 0 | 0 | 4172100 | 8359500 |
| SSPSFSSLI | ALCAM; ALCAM | Q13740; F5C | CD166_HUMAI | Unmodified | 2519500 | 1621100 | 100290 | 1385300 | 8177800 | 238490 | 0 | 0 | 3043500 | 1340600 | 505750 | 0 | 0 | 0 | 954520 | 0 | 3789200 | 9678000 |
| SSPSGPSN | BRI3BP; BRI3BI | Q8WY22; A0 | BRI3B_HUMAN | Unmodified | 0 | 0 | 0 | 0 | 0 | 0 | 0 | 0 | 2129000 | 181360 | 74643 | 112650 | 0 | 0 | 0 | 0 | 0 | 0 |
| SSPSVKPA' | CIAPIN1; CIAPII | Q6FI81; H3B | CPIN1_HUMAI | Unmodified | 0 | 0 | 0 | 0 | 975230 | 0 | 0 | 59487 | 0 | 0 | 563330 | 1030400 | 549080 | 514350 | 0 | 0 | 450790 | 855530 |
| SSPSVLIHS | RAI14 | Q9P0K7 | RAI14_HUMAN | Unmodified | 0 | 0 | 0 | 120680 | 507580 | 99139 | 29878 | 91985 | 0 | 0 | 0 | 1504100 | 207590 | 228090 | 0 | 112540 | 265790 | 384900 |
| SSPTDKHT | ATP2B2; ATP2B | Q01814; Q1( | AT2B2_HUMAI | Unmodified | 0 | 61357 | 0 | 0 | 34225000 | 0 | 0 | 0 | 0 | 0 | 0 | 1191900 | 0 | 28021000 | 0 | 0 | 3143900 | 0 |
| SSPTDKHT | ATP2B1; ATP2B | P20020; E7E | AT2B1_HUMAI | Unmodified | 0 | 0 | 0 | 0 | 183360 | 0 | 0 | 0 | 0 | 0 | 0 | 0 | 478270 | 262100 | 0 | 0 | 512580 | 0 |
| SSPVAHSIL | TTC17 | Q96AE7 | TTC17_HUMAI | Unmodified | 0 | 0 | 0 | 46203 | 0 | 0 | 0 | 61914 | 1774700 | 53766 | 0 | 0 | 0 | 0 | 0 | 0 | 0 | 0 |
| SSPVDLVT/ | IMPA1; IMPA1; I | P29218; E5R | IMPA1_HUMAI | Unmodified | 0 | 0 | 0 | 0 | 1461700 | 0 | 1059200 | 1169500 | 1572000 | 1683200 | 170080 | 0 | 3933900 | 1374000 | 0 | 0 | 1884200 | 2352000 |
| SSPVEFECI | CPNE3; CPNE3 | O75131; H0' | CPNE3_HUMAI | Unmodified | 1044900 | 0 | 1744600 | 2038800 | 368430 | 0 | 480150 | 417450 | 19407000 | 1595500 | 460460 | 1476500 | 734670 | 208100 | 1583500 | 1899700 | 219360 | 717240 |
| SSPVEFECI | CPNE3; CPNE3 | O75131; H0' | CPNE3_HUMAI | Unmodified | 0 | 0 | 0 | 0 | 0 | 0 | 0 | 0 | 0 | 0 | 0 | 0 | 0 | 0 | 0 | 0 | 0 | 0 |
| SSPVVIDAS | FN1; FN1; FN1 | P02751; H0' | FINC_HUMAN I | Unmodified | 717060 | 3925800 | 3022600 | 1432000 | 0 | 69575 | 0 | 0 | 382660 | 593010 | 246810 | 2348500 | 0 | 0 | 896710 | 3966100 | 0 | 0 |
| SSPVVNDG | GRSF1; GRSF1; | Q12849; H0' | GRSF1_HUMAI | Unmodified | 170590 | 0 | 8106300 | 422910 | 0 | 0 | 0 | 0 | 1086800 | 0 | 0 | 0 | 172850 | 0 | 1030200 | 595070 | 0 | 0 |
| SSQAQAQE | SLC29A2; SLC2 | Q14542; G5I | S29A2_HUMAI | Unmodified | 300250 | 415020 | 1078300 | 63363 | 0 | 0 | 0 | 0 | 181090 | 0 | 121830 | 2968600 | 0 | 0 | 735100 | 445180 | 0 | 12060 |
| SSQAQAQE | SLC29A2 | Q14542 | S29A2_HUMAI | Unmodified | 0 | 0 | 0 | 0 | 0 | 0 | 0 | 0 | 0 | 0 | 0 | 1342200 | 0 | 0 | 0 | 478310 | 0 | 0 |
| SSQEEVRS | HNRNPF | P52597 | HNRPF_HUMAI | Unmodified | 0 | 0 | 195460 | 0 | 1291700 | 301970 | 1699100 | 1109200 | 958040 | 1375900 | 532500 | 294860 | 3967800 | 1113000 | 0 | 0 | 552980 | 79202 |
| SSQEVRDE | CCER2 | I3L3R5 | CCER2_HUMAI | Unmodified | 0 | 0 | 0 | 0 | 0 | 0 | 0 | 67152 | 0 | 0 | 0 | 0 | 0 | 0 | 0 | 0 | 0 | 0 |
| SSQGGSLF | CHGB; CHGB | P05060; A0A | SCG1_HUMAN | Unmodified | 0 | 0 | 836110 | 0 | 0 | 0 | 0 | 0 | 2462400 | 0 | 0 | 0 | 0 | 0 | 0 | 0 | 0 | 0 |
| SSQIGAVV! | PNPO; PNPO; P | Q9NVS9; J3( | PNPO_HUMAN | Unmodified | 259580 | 0 | 1771700 | 2381500 | 4041900 | 0 | 0 | 45400 | 1514300 | 326130 | 1477400 | 6818200 | 2721600 | 1593200 | 3020100 | 899560 | 912640 | 6749900 |
| SSQLGDTT | FGD6; FGD6 | Q6ZV73; F8\ | FGD6_HUMAN | Unmodified | 0 | 0 | 0 | 0 | 799580 | 136320 | 0 | 0 | 0 | 24714 | 0 | 0 | 0 | 0 | 0 | 0 | 0 | 156670 |
| SSQLNNP( | USP24 | Q9UPU5 | UBP24_HUMAI | Unmodified | 0 | 0 | 0 | 0 | 0 | 0 | 0 | 0 | 378110 | 0 | 0 | 874370 | 0 | 0 | 0 | 0 | 0 | 0 |
| SSQPWQA | KLK14; KLK14 | Q9P0G3; A0. | KLK14_HUMAI | Unmodified | 0 | 0 | 0 | 0 | 0 | 0 | 0 | 0 | 5656300 | 1291600 | 0 | 0 | 0 | 0 | 0 | 0 | 0 | 0 |
| SSQTSGTN | USO1 | O60763 | USO1_HUMAN | Unmodified | 0 | 0 | 0 | 0 | 0 | 0 | 0 | 0 | 226300 | 0 | 209080 | 1310000 | 0 | 0 | 0 | 0 | 0 | 0 |
| SSQTSGTN | USO1 | O60763 | USO1_HUMAN | Unmodified | 0 | 0 | 0 | 0 | 0 | 0 | 0 | 0 | 0 | 1376500 | 0 | 0 | 2743800 | 0 | 0 | 0 | 1949800 | 0 |
| SSQTVAVH | NECTIN3; NECT | Q9NQS3; H7 | NECT3_HUMAI | Unmodified | 386430 | 293530 | 418510 | 370940 | 0 | 0 | 0 | 0 | 1422300 | 0 | 0 | 0 | 0 | 0 | 297240 | 270540 | 0 | 0 |
| SSQVSSYP | ILDR1 | Q86SU0 | ILDR1_HUMAN | Unmodified | 95411 | 144740 | 133830 | 0 | 0 | 0 | 0 | 0 | 416120 | 131620 | 0 | 575670 | 0 | 0 | 362350 | 0 | 0 | 0 |
| SSQVSSYP | ILDR1 | Q86SU0 | ILDR1_HUMAN | Oxidation (M) | 138850 | 0 | 0 | 0 | 0 | 0 | 0 | 0 | 0 | 0 | 0 | 0 | 0 | 0 | 0 | 0 | 0 | 0 |
| SSREFGSV | EEF1AKNMT | Q8N6R0 | EFNMT_HUMAI | Unmodified | 0 | 0 | 0 | 0 | 290700 | 0 | 307970 | 0 | 0 | 137660 | 0 | 0 | 216220 | 642990 | 0 | 0 | 0 | 0 |
| SSRGFGFT | MAGI1; MAGI1; | Q96QZ7; H7 | MAGI1_HUMAI | Unmodified | 0 | 0 | 0 | 0 | 0 | 0 | 34530 | 0 | 0 | 905360 | 0 | 0 | 310350 | 354050 | 0 | 0 | 204970 | 129530 |
| SSSADFGT | AGFG1; AGFG1 | P52594; B8Z | AGFG1_HUMAI | Unmodified | 549440 | 0 | 0 | 0 | 5664700 | 567970 | 482900 | 199990 | 542960 | 309070 | 0 | 0 | 1004800 | 850350 | 0 | 0 | 868380 | 2013300 |
| SSSAFEAAI | WWP1 | Q9H0M0 | WWP1_HUMAI | Unmodified | 0 | 260950 | 505620 | 643540 | 0 | 0 | 0 | 0 | 145100 | 46529 | 0 | 0 | 240120 | 75988 | 1039400 | 598120 | 0 | 0 |
| SSSAGGQ( | LSR; LSR | Q86X29; S4F | LSR_HUMAN Li | Unmodified | 0 | 0 | 4126900 | 2067500 | 0 | 0 | 0 | 0 | 12750000 | 7051200 | 1200900 | 7085700 | 0 | 0 | 8730400 | 6055000 | 29662 | 1721500 |
| SSSAGGQ( | LSR; LSR | Q86X29; S4F | LSR_HUMAN Li | Unmodified | 722830 | 720650 | 1669200 | 6940400 | 0 | 0 | 0 | 0 | 17720000 | 3154800 | 782510 | 2584100 | 0 | 0 | 1789300 | 3570500 | 0 | 0 |
| SSSDPQA( | HSPA14; HSPA: | Q0VDF9; H7 | HSP7E_HUMAI | Unmodified | 0 | 0 | 0 | 0 | 0 | 0 | 118390 | 105400 | 0 | 0 | 0 | 0 | 213060 | 103060 | 0 | 0 | 0 | 0 |
| SSSGAKDK | CNP; CNP; CNI | P09543; K7E | CN37_HUMAN | Acetyl (Protein N | 0 | 0 | 0 | 0 | 3330000 | 0 | 802830 | 0 | 0 | 825820 | 0 | 0 | 1839900 | 4781500 | 0 | 0 | 4809800 | 3138600 |
| SSSGFRVD | GLCE | O94923 | GLCE_HUMAN | Unmodified | 0 | 0 | 807650 | 0 | 0 | 0 | 0 | 0 | 417870 | 56799 | 0 | 398730 | 0 | 0 | 0 | 0 | 0 | 0 |
| SSSGILSM\ | NRP1; NRP1; N | O14786; Q5. | NRP1_HUMAN | Unmodified | 0 | 0 | 0 | 35222 | 244170 | 0 | 0 | 0 | 73250 | 0 | 0 | 0 | 0 | 0 | 0 | 0 | 0 | 347970 |
| SSSGLLEW | HNRNPL; HNRI | P14866; M0( | HNRPL_HUMAI | Unmodified | 120680 | 0 | 178410 | 0 | 0 | 0 | 0 | 0 | 445940 | 0 | 613790 | 4342600 | 0 | 0 | 0 | 0 | 0 | 189250 |
| SSSGLTYIA | MAN1A1 | P33908 | MA1A1_HUMAI | Unmodified | 0 | 0 | 0 | 0 | 0 | 0 | 0 | 0 | 0 | 0 | 405300 | 1048600 | 0 | 0 | 0 | 0 | 194070 | 0 |
| SSSGLTYIA | MAN1A1 | P33908 | MA1A1_HUMAI | Unmodified | 0 | 0 | 0 | 0 | 0 | 0 | 0 | 0 | 0 | 884510 | 262580 | 390980 | 0 | 0 | 0 | 0 | 0 | 0 |
| SSSGTPDLI | ALDH1A1; ALDI | P00352; Q5S | AL1A1_HUMAI | Acetyl (Protein N | 0 | 0 | 0 | 0 | 0 | 0 | 557590 | 0 | 3743100 | 2000700 | 36738000 | 5.22E+08 | 0 | 0 | 0 | 0 | 0 | 0 |
| SSSHSRAG | FLNA; FLNA; FL | P21333; A0A | FLNA_HUMAN | Acetyl (Protein N | 0 | 0 | 0 | 0 | 429760 | 0 | 0 | 0 | 0 | 0 | 0 | 0 | 0 | 204670 | 0 | 0 | 187350 | 466570 |
| SSSHSRAG | FLNA; FLNA; FL | P21333; A0A | FLNA_HUMAN | Acetyl (Protein N | 0 | 0 | 0 | 0 | 3728800 | 0 | 0 | 0 | 0 | 0 | 0 | 0 | 1604000 | 1041800 | 0 | 0 | 760540 | 833540 |
| SSSISEEKG | ACIN1; ACIN1; . | Q9UKV3; E7I | ACINU_HUMAI | Unmodified | 0 | 0 | 0 | 0 | 0 | 0 | 0 | 0 | 26881 | 0 | 53611 | 142480 | 0 | 0 | 0 | 0 | 0 | 0 |
| SSSLGSFYI | HTT; HTT; HTT; | P42858; A0A | HD_HUMAN Hu | Unmodified | 0 | 0 | 0 | 0 | 0 | 0 | 0 | 0 | 0 | 0 | 0 | 2419800 | 0 | 0 | 0 | 0 | 0 | 0 |
| SSSNLALAI | CPD; CPD | O75976; J3C | CBPD_HUMAN | Unmodified | 3280000 | 3373300 | 2089400 | 4962800 | 16394000 | 6276100 | 0 | 1074600 | 9020100 | 1890400 | 13083000 | 0 | 0 | 1367900 | 3025200 | 655790 | 0 | 0 |
| SSSNLALAI | CPD; CPD | O75976; J3C | CBPD_HUMAN | Unmodified | 0 | 0 | 0 | 0 | 0 | 0 | 0 | 0 | 1343000 | 3752800 | 0 | 0 | 0 | 0 | 0 | 0 | 0 | 0 |
| SSSPAPAD | G3BP1; G3BP1; | Q13283; A0/ | G3BP1_HUMAI | Unmodified | 70442 | 66444 | 60315 | 87125 | 180860 | 106710 | 0 | 0 | 0 | 348630 | 277530 | 4336500 | 758210 | 0 | 0 | 0 | 0 | 119760 |
| SSSPAPEFI | FARP1; FARP1; | Q9Y4F1; C9. | FARP1_HUMAI | Unmodified | 115560 | 0 | 0 | 0 | 0 | 0 | 0 | 0 | 1531200 | 0 | 0 | 1382600 | 0 | 0 | 0 | 197910 | 0 | 0 |
| SSSPGAGC | IQSEC2; IQSEC | Q5JU85; A0/ | IQEC2_HUMAI | Unmodified | 206100 | 105930 | 0 | 108280 | 0 | 0 | 0 | 0 | 0 | 0 | 0 | 0 | 0 | 0 | 64070 | 0 | 0 | 0 |
| SSSPVNVK | HNRNPU; HNR | Q00839; A0/ | HNRPU_HUMA | Acetyl (Protein N | 0 | 0 | 0 | 0 | 0 | 1244100 | 0 | 246230 | 0 | 150080 | 1280300 | 7053500 | 0 | 362560 | 0 | 0 | 0 | 122400 |
| SSSQIPAQI | PRRC2C; PRRC | Q9Y520; A0/ | PRC2C_HUMAI | Unmodified | 0 | 0 | 0 | 0 | 0 | 0 | 0 | 0 | 0 | 0 | 243830 | 1152200 | 0 | 0 | 0 | 0 | 0 | 0 |
| SSSQPSSC | SQSTM1; SQSTI | Q13501; E7E | SQSTM_HUMAI | Unmodified | 0 | 0 | 0 | 0 | 7265100 | 0 | 0 | 0 | 0 | 0 | 0 | 0 | 4284600 | 2534200 | 0 | 0 | 2634500 | 5184100 |
| SSSRKYTEI | DCTPP1; DCTPI | Q9H773; H3 | DCTP1_HUMAI | Unmodified | 0 | 0 | 0 | 0 | 0 | 0 | 0 | 0 | 0 | 0 | 0 | 0 | 2197800 | 0 | 0 | 0 | 1353000 | 773040 |
| SSSSEDGS | EMC1 | Q8N766 | EMC1_HUMAN | Unmodified | 0 | 0 | 0 | 0 | 0 | 0 | 0 | 23879 | 7185400 | 1175100 | 126910 | 0 | 0 | 0 | 0 | 0 | 0 | 0 |
| SSSSEDSS | NOLC1; NOLC1 | Q14978; A0/ | NOLC1_HUMA | Unmodified | 0 | 0 | 0 | 0 | 0 | 0 | 0 | 0 | 0 | 0 | 0 | 247540 | 142510 | 0 | 0 | 0 | 0 | 0 |
| SSSSFHPAI | IL4R | P24394 | IL4RA_HUMAN | Unmodified | 0 | 0 | 0 | 0 | 591600 | 82785 | 0 | 0 | 0 | 0 | 0 | 0 | 0 | 0 | 0 | 0 | 0 | 66218 |
| SSSSFTPFI | MAP4K4; MAP4 | O95819; C9. | M4K4_HUMAN | Unmodified | 0 | 0 | 0 | 439460 | 465340 | 0 | 0 | 23078 | 2188200 | 450740 | 1429900 | 3009500 | 382480 | 0 | 1067500 | 0 | 523050 | 1692100 |
| SSSSGHYV | USP14; USP14 | P54578; A6N | UBP14_HUMAI | Unmodified | 0 | 0 | 489710 | 0 | 533010 | 0 | 0 | 0 | 0 | 0 | 0 | 1677300 | 835860 | 0 | 0 | 0 | 0 | 0 |
| SSSSNFLN | MUC13 | Q9H3R2 | MUC13_HUMA | Unmodified | 324050 | 221410 | 0 | 0 | 0 | 0 | 0 | 0 | 519150 | 0 | 0 | 0 | 0 | 0 | 0 | 0 | 0 | 0 |
| SSSSRGSS | FLNC | Q14315 | FLNC_HUMAN | Unmodified | 0 | 0 | 0 | 0 | 3825300 | 355460 | 40896 | 0 | 0 | 0 | 0 | 0 | 0 | 32697 | 0 | 0 | 1549000 | 679420 |

| | | | | | | | | | | | | | | | | | | | | | | |
|---|---|---|---|---|---|---|---|---|---|---|---|---|---|---|---|---|---|---|---|---|---|---|
| SSSSSGVP | EXOC7; EXOC7 | Q9UPT5; B4I | EXOC7_HUMAI | Unmodified | 0 | 0 | 104580 | 280930 | 2557800 | 440130 | 0 | 95850 | 451430 | 364760 | 0 | 838870 | 915570 | 509030 | 106180 | 171770 | 414630 | 1004800 |
| SSSSSLLR | TMEM237; TMEI | Q96Q45; A0/ | TM237_HUMAN | Unmodified | 35663 | 322090 | 497520 | 62564 | 0 | 280320 | 0 | 0 | 0 | 0 | 0 | 0 | 104100 | 0 | 0 | 323310 | 145620 | 171820 |
| SSSSVTTSE | VASP; VASP | P50552; K7E | VASP_HUMAN | Unmodified | 0 | 0 | 0 | 303070 | 0 | 0 | 0 | 0 | 435580 | 0 | 0 | 2474900 | 0 | 0 | 0 | 0 | 0 | 0 |
| SSSSYHTW | GEMIN5 | Q8TEQ6 | GEMI5_HUMAN | Unmodified | 0 | 0 | 0 | 0 | 0 | 0 | 0 | 0 | 0 | 0 | 0 | 1410900 | 0 | 0 | 0 | 0 | 0 | 0 |
| SSSTEPPPI | CPSF7; CPSF7; | Q8N684; C9 | CPSF7_HUMAN | Unmodified | 494450 | 344920 | 0 | 0 | 0 | 0 | 0 | 0 | 8284300 | 1206700 | 370580 | 0 | 0 | 0 | 0 | 0 | 0 | 0 |
| SSSTEPSLL | CNKSR3; CNKS | Q6P9H4; A0 | CNKR3_HUMAI | Unmodified | 0 | 0 | 10965 | 0 | 0 | 0 | 0 | 0 | 306190 | 148220 | 169790 | 95996 | 0 | 0 | 0 | 0 | 0 | 0 |
| SSSTEPSLL | CNKSR3; CNKS | Q6P9H4; A0 | CNKR3_HUMAI | Unmodified | 0 | 0 | 0 | 0 | 986850 | 0 | 0 | 0 | 0 | 1031700 | 0 | 0 | 517700 | 403880 | 0 | 0 | 258460 | 614300 |
| SSSTGNLLI | BAIAP2; BAIAP2 | Q9UQB8; I3I | BAIP2_HUMAN | Unmodified | 0 | 0 | 1023600 | 989150 | 2837400 | 0 | 0 | 173540 | 1792200 | 577820 | 754570 | 0 | 2782200 | 0 | 0 | 3953700 | 0 | 2067500 |
| SSSTGNLLI | BAIAP2; BAIAP2 | Q9UQB8; I3I | BAIP2_HUMAN | Unmodified | 0 | 0 | 0 | 0 | 7311900 | 0 | 0 | 0 | 0 | 0 | 0 | 0 | 2044600 | 3082900 | 0 | 0 | 0 | 2962600 |
| SSSVPHPFI | ARHGEF16 | Q5VV41 | ARHGG_HUMA | Unmodified | 0 | 0 | 0 | 0 | 0 | 0 | 0 | 0 | 1190500 | 266500 | 0 | 0 | 0 | 0 | 0 | 630630 | 0 | 0 |
| SSSYFKDSI | SNX9; SNX9; SI | Q9Y5X1; A0/ | SNX9_HUMAN | Unmodified | 0 | 0 | 0 | 0 | 0 | 72840 | 0 | 0 | 0 | 0 | 0 | 0 | 1843800 | 401190 | 0 | 0 | 0 | 0 |
| SSTATTSCI | ELAPOR2; ELAP | A8MWY0; H7 | ELAP2_HUMAN | Unmodified | 0 | 0 | 0 | 0 | 0 | 0 | 165160 | 114280 | 0 | 0 | 0 | 0 | 0 | 29458 | 0 | 0 | 0 | 0 |
| SSTDALIEN | HERC1; HERC1 | Q15751; H0 | HERC1_HUMAI | Unmodified | 1708400 | 0 | 0 | 0 | 0 | 64978 | 0 | 0 | 35086 | 38476 | 0 | 28772 | 55010 | 0 | 0 | 0 | 0 | 0 |
| SSTDILRCI | AHNAK2 | Q8IVF2 | AHNK2_HUMAI | Unmodified | 0 | 0 | 0 | 0 | 0 | 0 | 0 | 0 | 0 | 0 | 0 | 0 | 0 | 134700 | 0 | 0 | 172320 | 355730 |
| SSTIQDQQ | WASF1; WASF1 | Q92558; Q5 | WASF1_HUMAI | Unmodified | 0 | 0 | 0 | 0 | 0 | 0 | 0 | 0 | 661500 | 160160 | 609520 | 4283200 | 0 | 0 | 0 | 0 | 260670 | 0 |
| SSTKPAAG | ZYX; ZYX; ZYX; Z | Q15942; H0 | ZYX_HUMAN Zy | Unmodified | 87933 | 0 | 0 | 0 | 0 | 0 | 0 | 0 | 0 | 0 | 176440 | 0 | 0 | 0 | 289910 | 0 | 876440 | 1458100 |
| SSTLEPEDI | EVA1B | Q9NVM1 | EVA1B_HUMAN | Unmodified | 37418 | 0 | 98235 | 0 | 0 | 0 | 0 | 0 | 0 | 0 | 0 | 0 | 0 | 0 | 638010 | 408350 | 0 | 0 |
| SSTPEEVKI | CFL1; CFL1; CF | P23528; G3 | COF1_HUMAN | Unmodified | 0 | 0 | 0 | 0 | 628000 | 0 | 0 | 151250 | 0 | 0 | 0 | 108600 | 0 | 70774 | 0 | 0 | 215950 | 212340 |
| SSTQFNKG | CNN2; CNN2; ( | Q99439; H3I | CNN2_HUMAN | Acetyl (Protein N | 0 | 0 | 0 | 0 | 7100600 | 837120 | 450650 | 262750 | 0 | 0 | 0 | 0 | 899710 | 666960 | 0 | 0 | 1220300 | 916300 |
| SSTRSQNP | CUL1; CUL1; CI | Q13616; A0/ | CUL1_HUMAN | Acetyl (Protein N | 0 | 0 | 0 | 0 | 642950 | 132550 | 108530 | 93459 | 0 | 91479 | 0 | 0 | 0 | 0 | 0 | 0 | 0 | 0 |
| SSTSFANIC | OSTM1 | Q86WC4 | OSTM1_HUMAI | Unmodified | 0 | 0 | 0 | 0 | 483330 | 0 | 188440 | 114710 | 171400 | 0 | 0 | 0 | 289890 | 315980 | 0 | 0 | 0 | 0 |
| SSTSLTSEE | SYT4 | Q9H2B2 | SYT4_HUMAN S | Unmodified | 0 | 0 | 0 | 4240300 | 0 | 0 | 0 | 0 | 0 | 0 | 0 | 21559 | 0 | 0 | 0 | 0 | 0 | 205270 |
| SSTSNAND | IQGAP2; IQGAP | Q13576; D6I | IQGA2_HUMAN | Unmodified | 0 | 0 | 0 | 394200 | 0 | 0 | 0 | 0 | 1792800 | 873010 | 0 | 0 | 1395700 | 1386600 | 0 | 0 | 0 | 0 |
| SSTSPNLQ | VPS4B; VPS4B; | O75351; K7E | VPS4B_HUMAN | Acetyl (Protein N | 0 | 602530 | 0 | 270990 | 0 | 191540 | 0 | 0 | 3362700 | 170680 | 0 | 1675300 | 0 | 0 | 583710 | 501010 | 298810 | 194920 |
| SSTTSMTS | PSMD2; PSMD2 | Q13200; H7 | PSMD2_HUMA | Unmodified | 0 | 0 | 0 | 350030 | 346730 | 256040 | 176720 | 0 | 0 | 285310 | 346610 | 3650800 | 571560 | 343660 | 0 | 126010 | 496940 | 561480 |
| SSTTVSSFA | CUL4B; CUL4B; | Q13620; A0/ | CUL4B_HUMAI | Unmodified | 0 | 0 | 0 | 86970 | 1255100 | 0 | 0 | 0 | 76463 | 139110 | 57876 | 559900 | 0 | 0 | 0 | 0 | 488610 | 749760 |
| SSTVAPAQI | KIF23; KIF23; KI | Q02241; A0/ | KIF23_HUMAN | Unmodified | 0 | 0 | 592640 | 1332500 | 2034000 | 130170 | 0 | 0 | 3001800 | 438670 | 218450 | 1704100 | 0 | 0 | 741910 | 362450 | 676250 | 0 |
| SSTVQDQC | WASF3; WASF3 | Q9UPY6; A0 | WASF3_HUMAI | Unmodified | 50525 | 0 | 135950 | 183580 | 0 | 0 | 0 | 0 | 0 | 0 | 0 | 268540 | 0 | 242960 | 156990 | 183760 | 0 | 127480 |
| SSTYEVRLT | ISG15; ISG15; I | P05161; A0A | ISG15_HUMAN | Unmodified | 157270 | 0 | 368380 | 0 | 185360 | 0 | 0 | 0 | 0 | 150400 | 0 | 0 | 0 | 155460 | 54521 | 0 | 433740 | 814260 |
| SSVAVLTQI | HDLBP; HDLBP | Q00341; A0/ | VIGLN_HUMAN | Acetyl (Protein N | 0 | 0 | 0 | 0 | 1367800 | 0 | 125240 | 175640 | 0 | 0 | 0 | 0 | 1252400 | 3118700 | 0 | 0 | 780150 | 1172100 |
| SSVESLQN | TRAF3; TRAF3; | Q13114; A6I | TRAF3_HUMAN | Unmodified | 0 | 0 | 0 | 0 | 22366000 | 4893800 | 0 | 0 | 0 | 0 | 0 | 0 | 0 | 0 | 0 | 0 | 1672100 | 318250 |
| SSVGAPVE | UPF1 | Q92900 | RENT1_HUMAN | Unmodified | 926500 | 3085700 | 277700 | 966380 | 0 | 0 | 0 | 0 | 0 | 0 | 0 | 0 | 0 | 0 | 1014000 | 413320 | 0 | 0 |
| SSVLENFV | DNM3; DNM2; I | Q9UQ16; P5 | DYN3_HUMAN | Unmodified | 0 | 492250 | 0 | 0 | 0 | 0 | 0 | 0 | 0 | 0 | 0 | 0 | 0 | 0 | 0 | 0 | 0 | 0 |
| SSVLENFV | DNM3; DNM2; I | Q9UQ16; P5 | DYN3_HUMAN | Unmodified | 6516700 | 7497900 | 2367900 | 5293800 | 6719500 | 1222900 | 1741400 | 27657 | 14005000 | 7203600 | 949310 | 5909400 | 5089000 | 2222100 | 5235800 | 3045400 | 1863100 | 2064000 |
| SSVLESLV | DNM1L; DNM1I | O00429; F8 | DNM1L_HUMA | Unmodified | 0 | 0 | 0 | 0 | 0 | 0 | 0 | 0 | 2044800 | 0 | 267410 | 3253300 | 0 | 0 | 0 | 0 | 0 | 0 |
| SSVLESLV | DNM1L; DNM1I | O00429; F8 | DNM1L_HUMA | Unmodified | 0 | 0 | 0 | 0 | 0 | 0 | 0 | 0 | 0 | 0 | 0 | 1221000 | 1417200 | 0 | 0 | 0 | 0 | 0 |
| SSVLIAQQI | RANGAP1 | P46060 | RAGP1_HUMAI | Unmodified | 0 | 0 | 0 | 0 | 0 | 0 | 155670 | 0 | 0 | 0 | 0 | 817600 | 0 | 0 | 0 | 0 | 0 | 0 |
| SSVLQLR | NPTX1 | Q15818 | NPTX1_HUMAN | Unmodified | 0 | 0 | 0 | 0 | 0 | 0 | 0 | 0 | 66715000 | 0 | 0 | 166210 | 0 | 0 | 0 | 0 | 0 | 0 |
| SSVLQLRE | NPTX1 | Q15818 | NPTX1_HUMAN | Unmodified | 0 | 0 | 0 | 0 | 0 | 0 | 0 | 0 | 2179800 | 3448100 | 0 | 0 | 0 | 0 | 0 | 0 | 0 | 0 |
| SSVMDGAI | WDR20; WDR2 | Q8TBZ3; A0/ | WDR20_HUMA | Unmodified | 5306300 | 4910900 | 0 | 0 | 1310800 | 0 | 0 | 959790 | 0 | 247580 | 0 | 0 | 0 | 0 | 0 | 0 | 491290 | 1618300 |
| SSVNCPFS | DHX9 | Q08211 | DHX9_HUMAN | Unmodified | 0 | 0 | 0 | 0 | 0 | 0 | 0 | 0 | 0 | 0 | 0 | 1040800 | 0 | 0 | 0 | 0 | 0 | 0 |
| SSVNCPFS | DHX9 | Q08211 | DHX9_HUMAN | Oxidation (M) | 0 | 0 | 0 | 0 | 0 | 0 | 0 | 0 | 0 | 0 | 0 | 1074200 | 0 | 0 | 0 | 0 | 0 | 0 |
| SSVNELTQ | EEA1 | Q15075 | EEA1_HUMAN | Unmodified | 0 | 0 | 0 | 0 | 759270 | 0 | 96868 | 64930 | 89234 | 82935 | 0 | 160010 | 0 | 0 | 0 | 0 | 0 | 0 |
| SSVQQQPI | WASL | O00401 | WASL_HUMAN | Acetyl (Protein N | 485950 | 0 | 395300 | 614200 | 168570 | 0 | 0 | 158640 | 0 | 0 | 85964 | 382580 | 206500 | 0 | 982620 | 349750 | 0 | 94037 |
| SSVQQQPI | WASL | O00401 | WASL_HUMAN | Acetyl (Protein N | 0 | 0 | 0 | 0 | 2497100 | 104160 | 0 | 0 | 0 | 202380 | 0 | 0 | 2459800 | 1820000 | 0 | 0 | 791490 | 510550 |
| SSVVGGFV | MYO6; MYO6; M | Q9UM54; A0 | MYO6_HUMAN | Unmodified | 779920 | 0 | 2602000 | 5913800 | 2124400 | 478980 | 1276800 | 51347 | 3791500 | 2596700 | 988640 | 6722400 | 3461600 | 4169200 | 7290600 | 5815000 | 934820 | 350110 |
| SSVVLDDG | LNPEP | Q9UIQ6 | LCAP_HUMAN | Unmodified | 0 | 0 | 0 | 0 | 0 | 0 | 0 | 22371 | 0 | 591290 | 0 | 226640 | 247920 | 0 | 0 | 0 | 81057 | 42953 |
| SSVVPCEL | SLC30A1 | Q9Y6M5 | ZNT1_HUMAN | Unmodified | 0 | 0 | 0 | 0 | 2266400 | 381060 | 0 | 0 | 0 | 0 | 0 | 0 | 1833600 | 2080600 | 0 | 0 | 1343600 | 2926100 |
| SSWLEEDI | P4HA2 | O15460 | P4HA2_HUMAI | Unmodified | 0 | 0 | 0 | 0 | 0 | 0 | 0 | 0 | 0 | 2202200 | 0 | 0 | 0 | 0 | 0 | 0 | 0 | 0 |
| SSWRDCSI | MINDY3 | Q9H8M7 | MINY3_HUMAN | Unmodified | 958650 | 554480 | 190530 | 418420 | 0 | 0 | 145900 | 0 | 0 | 0 | 0 | 0 | 0 | 0 | 368420 | 432080 | 0 | 0 |
| SSWRVISSI | YWHAB; YWHA | P31946; Q0 | 1433B_HUMAN | Unmodified | 0 | 0 | 113730 | 243770 | 17727000 | 3033200 | 2759500 | 791440 | 0 | 3036600 | 921240 | 0 | 8377900 | 12061000 | 76359 | 81023 | 9644900 | 10973000 |
| SSWRVISSI | YWHAG | P61981 | 1433G_HUMAN | Unmodified | 0 | 0 | 0 | 0 | 1996600 | 0 | 2818700 | 0 | 0 | 1072400 | 312110 | 0 | 2153000 | 5678500 | 0 | 0 | 7238700 | 3062000 |
| SSWRVVSS | YWHAZ; YWHA | P63104; E5R | 1433Z_HUMAN | Unmodified | 0 | 0 | 0 | 317340 | 3015600 | 700230 | 879030 | 1052000 | 320020 | 1196600 | 234070 | 0 | 1194700 | 1210500 | 51108 | 0 | 1142600 | 4965300 |
| SSWRVVSS | YWHAZ; YWHA | P63104; E5R | 1433Z_HUMAN | Unmodified | 0 | 0 | 0 | 0 | 43294000 | 12808000 | 14250000 | 1416900 | 0 | 20446000 | 1478300 | 0 | 29793000 | 32803000 | 0 | 0 | 56493000 | 66939000 |
| SSWVFIAAI | FAM3B; FAM3B | P58499; A8 | FAM3B_HUMAI | Unmodified | 0 | 0 | 0 | 0 | 0 | 0 | 0 | 0 | 2696400 | 120080 | 37570 | 896270 | 0 | 0 | 0 | 0 | 0 | 240030 |
| SSYASQHS | PKP4; PKP4 | Q99569; A0/ | PKP4_HUMAN | Unmodified | 81276000 | 49615000 | 7.38E+09 | 23374000 | 0 | 0 | 35695000 | 12476000 | 322580 | 24222 | 0 | 504450 | 1.09E+10 | 25236000 | 6.91E+09 | 7.16E+09 | 190420 | 290280 |
| SSYDVFQL | FSCN1 | Q16658 | FSCN1_HUMAI | Unmodified | 0 | 0 | 0 | 0 | 2772600 | 17181000 | 3674700 | 821820 | 0 | 746780 | 3556900 | 0 | 0 | 6454800 | 0 | 0 | 1077000 | 6799400 |
| SSYNLVPR | GNPAT; GNPAT | O15228; Q5 | GNPAT_HUMAI | Unmodified | 1907000 | 2112500 | 0 | 1869100 | 11066000 | 54874 | 1539900 | 0 | 4436700 | 1551800 | 342090 | 1452600 | 32636 | 6332700 | 0 | 0 | 16006000 | 35739000 |
| SSYPIWEDI | MTSS2; MTSS2 | Q765P7; A0/ | MTSS2_HUMAN | Unmodified | 0 | 0 | 0 | 0 | 0 | 0 | 0 | 0 | 0 | 0 | 0 | 1183500 | 0 | 0 | 0 | 0 | 389600 | 0 |
| SSYYMIGE | NNMT | P40261 | NNMT_HUMAN | Oxidation (M) | 0 | 0 | 0 | 0 | 536550 | 0 | 0 | 0 | 0 | 0 | 0 | 0 | 0 | 0 | 0 | 0 | 0 | 0 |
| SSYYVVSG | MAP1B | P46821 | MAP1B_HUMAI | Unmodified | 0 | 0 | 0 | 0 | 0 | 0 | 1364500 | 0 | 0 | 810330 | 266820 | 0 | 3935200 | 0 | 0 | 0 | 0 | 975070 |
| STAALEED | ARHGEF6; ARH | Q15052; Q1 | ARHG6_HUMA | Unmodified | 0 | 0 | 0 | 0 | 0 | 0 | 0 | 219600 | 0 | 314200 | 0 | 0 | 0 | 0 | 0 | 0 | 247300 | 349450 |
| STAAPLTM | POFUT1 | Q9H488 | OFUT1_HUMAN | Oxidation (M) | 0 | 0 | 0 | 0 | 0 | 0 | 0 | 0 | 494790 | 0 | 0 | 0 | 0 | 0 | 0 | 0 | 0 | 0 |
| STAGALDK | LRRC8C | Q8TDW0 | LRC8C_HUMAI | Unmodified | 0 | 0 | 0 | 0 | 594170 | 95095 | 14810 | 0 | 373740 | 0 | 22175 | 57277 | 0 | 0 | 0 | 0 | 0 | 151120 |
| STAGDTHL | HSPA8; HSPA2; | P11142; P54 | HSP7C_HUMAI | Unmodified | 2505600 | 3113000 | 2924300 | 4156000 | 847960 | 255280 | 77612 | 40540 | 6781900 | 390400 | 2174500 | 14835000 | 1116700 | 387880 | 2658200 | 3261300 | 0 | 264360 |
| STAGDTHL | HSPA8; HSPA8; | P11142; E9P | HSP7C_HUMAI | Oxidation (M) | 1108900 | 0 | 1002600 | 14080000 | 1.05E+08 | 20380000 | 20192000 | 0 | 7248500 | 48447000 | 11722000 | 0 | 1.44E+08 | 63905000 | 1122300 | 524120 | 11145000 | 40287000 |
| STAGDTHL | HSPA8; HSPA8; | P11142; E9P | HSP7C_HUMAI | Unmodified | 0 | 0 | 230160 | 10029000 | 1.68E+08 | 18843000 | 25255000 | 0 | 46556000 | 68911000 | 3816700 | 0 | 1.33E+08 | 73799000 | 0 | 0 | 11859000 | 64466000 |
| STAGDTHL | HSPA2 | P54652 | HSP72_HUMAN | Unmodified | 0 | 0 | 0 | 0 | 3915600 | 1098000 | 0 | 0 | 0 | 72688 | 0 | 0 | 0 | 0 | 0 | 0 | 0 | 0 |
| STAGNFLV | SRPK1; SRPK1 | Q96SB4; H3 | SRPK1_HUMAN | Unmodified | 0 | 0 | 0 | 1075900 | 481890 | 0 | 0 | 0 | 370810 | 519250 | 402210 | 4062300 | 0 | 493730 | 0 | 0 | 0 | 0 |
| STAGNFLV | SRPK1; SRPK1 | Q96SB4; H3 | SRPK1_HUMAN | Unmodified | 0 | 0 | 0 | 0 | 0 | 0 | 0 | 0 | 0 | 0 | 0 | 0 | 0 | 0 | 0 | 0 | 240480 | 0 |
| STAIFFYCD | IGF2R; IGF2R | P11717; A0A | MPRI_HUMAN | Unmodified | 0 | 0 | 0 | 0 | 0 | 0 | 0 | 0 | 10855000 | 1778100 | 295670 | 2375100 | 0 | 0 | 0 | 0 | 0 | 0 |
| STAIFFYCD | IGF2R; IGF2R | P11717; A0A | MPRI_HUMAN | Unmodified | 0 | 0 | 0 | 734940 | 964140 | 0 | 0 | 0 | 29985000 | 10170000 | 3818700 | 0 | 988940 | 0 | 0 | 0 | 0 | 0 |
| STAISCIGA | THSD4 | Q6ZMP0 | THSD4_HUMAI | Unmodified | 0 | 0 | 0 | 153240 | 799270 | 131160 | 0 | 0 | 0 | 157210 | 133060 | 0 | 0 | 0 | 0 | 0 | 0 | 0 |
| STAISLFYEI | EEF2 | P13639 | EF2_HUMAN El | Unmodified | 0 | 0 | 0 | 0 | 453640 | 0 | 0 | 0 | 0 | 0 | 0 | 0 | 0 | 0 | 0 | 0 | 257220 | 0 |
| STAISLFYEI | EEF2 | P13639 | EF2_HUMAN El | Unmodified | 0 | 0 | 0 | 0 | 226720 | 0 | 0 | 0 | 0 | 0 | 0 | 0 | 82675 | 0 | 59745 | 0 | 383920 | 0 |
| STAIVHIQLI | SLC30A4 | O14863 | ZNT4_HUMAN | Unmodified | 0 | 0 | 380350 | 0 | 0 | 0 | 0 | 0 | 0 | 0 | 0 | 0 | 0 | 0 | 585080 | 0 | 0 | 0 |
| STANKYQV | HDGF | P51858 | HDGF_HUMAN | Unmodified | 0 | 0 | 0 | 540520 | 2521600 | 0 | 0 | 0 | 0 | 959750 | 0 | 0 | 1240500 | 0 | 0 | 0 | 0 | 313050 |
| STCIYGGAI | DDX5; DDX17; I | P17844; Q9 | DDX5_HUMAN | Unmodified | 0 | 137780 | 0 | 0 | 0 | 0 | 0 | 365040 | 0 | 0 | 0 | 3854600 | 0 | 0 | 139920 | 0 | 0 | 0 |
| STDADLAM | BIN3; BIN3; BIN | Q9NQY0; E5 | BIN3_HUMAN | Unmodified | 0 | 0 | 16229 | 0 | 758490 | 0 | 0 | 0 | 44366 | 0 | 0 | 0 | 145220 | 0 | 0 | 0 | 250280 | 0 |
| STDALRKIC | ATP2B1; ATP2B | P20020; F8 | AT2B1_HUMAN | Unmodified | 0 | 0 | 0 | 0 | 369960 | 0 | 0 | 0 | 0 | 0 | 0 | 0 | 2548200 | 2134200 | 0 | 0 | 468060 | 0 |
| STDASLDS | FAM171A2 | A8MVW0 | F1712_HUMAN | Unmodified | 0 | 0 | 0 | 0 | 112160 | 0 | 0 | 0 | 276690 | 0 | 0 | 0 | 0 | 0 | 0 | 0 | 0 | 416840 |
| STDRKQTLI | FMNL1; FMNL1 | O95466; A0/ | FMNL1_HUMAI | Unmodified | 0 | 0 | 0 | 0 | 1100800 | 330810 | 182860 | 0 | 0 | 0 | 0 | 0 | 0 | 0 | 0 | 0 | 0 | 0 |
| STDSPVEC | F3 | P13726 | TF_HUMAN Tis | Unmodified | 105290 | 44792 | 0 | 0 | 0 | 0 | 0 | 0 | 0 | 0 | 0 | 0 | 0 | 0 | 0 | 0 | 0 | 0 |
| STDVPNTG | PKP4; PKP4; PK | Q99569; H0 | PKP4_HUMAN | Unmodified | 0 | 0 | 187810 | 768400 | 23517 | 133020 | 32064 | 0 | 0 | 66569 | 0 | 271700 | 0 | 0 | 246100 | 216550 | 0 | 0 |
| STEATLSFG | CNTN1 | Q12860 | CNTN1_HUMA | Unmodified | 1208700 | 1612800 | 0 | 0 | 0 | 0 | 0 | 0 | 3727900 | 0 | 0 | 0 | 0 | 0 | 0 | 0 | 0 | 0 |
| STEDLSPQI | ERBIN | Q96RT1 | ERBIN_HUMAN | Unmodified | 0 | 0 | 0 | 0 | 7061100 | 905410 | 0 | 0 | 0 | 0 | 0 | 0 | 783620 | 0 | 0 | 0 | 526140 | 1365100 |
| STEEAFEVI | KIF23; KIF23; KI | Q02241; A0/ | KIF23_HUMAN | Unmodified | 0 | 0 | 0 | 1830600 | 0 | 0 | 0 | 0 | 107290 | 0 | 283180 | 1165300 | 0 | 0 | 0 | 0 | 0 | 0 |
| STEEAFEVI | KIF23; KIF23; KI | Q02241; A0/ | KIF23_HUMAN | Unmodified | 0 | 0 | 0 | 733960 | 11004000 | 3026100 | 652420 | 0 | 1214400 | 1496500 | 0 | 0 | 434130 | 0 | 0 | 79922 | 1977400 | 0 |
| STEEAFEVI | KIF23; KIF23; KI | Q02241; A0/ | KIF23_HUMAN | Unmodified | 0 | 0 | 56944 | 452400 | 9028500 | 1174700 | 513500 | 101600 | 509840 | 1274800 | 672250 | 0 | 206550 | 0 | 0 | 0 | 1227100 | 214840 |
| STEGGQSL | SCEL | O95171 | SCEL_HUMAN | Unmodified | 0 | 0 | 0 | 0 | 1562800 | 0 | 0 | 0 | 0 | 0 | 0 | 0 | 0 | 0 | 0 | 0 | 0 | 0 |
| STEHNMGI | MGAT2 | Q10469 | MGAT2_HUMAI | Oxidation (M) | 0 | 0 | 0 | 0 | 0 | 0 | 0 | 0 | 454150 | 0 | 0 | 0 | 0 | 0 | 0 | 0 | 0 | 0 |
| STEHNMGI | MGAT2 | Q10469 | MGAT2_HUMAI | Unmodified | 0 | 0 | 0 | 0 | 609370 | 0 | 0 | 0 | 10291000 | 0 | 151700 | 0 | 0 | 0 | 0 | 0 | 0 | 0 |
| STEHNMGI | MGAT2 | Q10469 | MGAT2_HUMAI | Unmodified | 0 | 0 | 0 | 0 | 0 | 0 | 0 | 0 | 0 | 526990 | 0 | 0 | 0 | 0 | 0 | 0 | 0 | 0 |
| STEKLEKEI | NOSTRIN; NOS | Q8IVI9; A0A | NOSTN_HUMA | Unmodified | 4479100 | 4155100 | 0 | 10505000 | 0 | 0 | 0 | 0 | 0 | 117890 | 0 | 0 | 8678500 | 4356300 | 3028400 | 2659900 | 0 | 0 |
| STELLIR | H3-2; H3-4; H3 | Q5TEC6; Q1 | H3PS2_HUMAN | Unmodified | 5.06E+08 | 2.81E+08 | 5421900 | 2774900 | 41731000 | 16513000 | 41903000 | 1.63E+08 | 42477000 | 35933000 | 39510000 | 3.07E+09 | 8457000 | 5966800 | 2049300 | 3307500 | 845740 | 8406800 |
| STELLIRK | H3-2; H3-4; H3 | Q5TEC6; Q1 | H3PS2_HUMAN | Unmodified | 1278100 | 1041500 | 0 | 0 | 251400 | 0 | 307700 | 11926 | 1375500 | 115330 | 0 | 13135000 | 0 | 0 | 0 | 0 | 0 | 0 |
| STELLIRKLI | H3-2; H3-4; H3 | Q5TEC6; Q1 | H3PS2_HUMAN | Unmodified | 0 | 0 | 0 | 0 | 5921100 | 0 | 5342900 | 3521600 | 0 | 2367400 | 0 | 1473700 | 0 | 1360800 | 0 | 0 | 0 | 1225600 |
| STELPGKNI | PTPN12 | Q05209 | PTN12_HUMAN | Unmodified | 0 | 0 | 0 | 68913 | 222340 | 0 | 0 | 0 | 0 | 0 | 0 | 0 | 0 | 0 | 0 | 0 | 0 | 833670 |
| STEPELIQV | SLC1A5; SLC1A | Q15758; M0 | AAAT_HUMAN | Unmodified | 4104300 | 2914500 | 6399300 | 7957300 | 969250 | 2472900 | 413750 | 306310 | 29174000 | 4817000 | 47127000 | 2.22E+08 | 557870 | 558630 | 8940200 | 10854000 | 1452000 | 0 |
| STEPELIQV | SLC1A5; SLC1A | Q15758; M0 | AAAT_HUMAN | Unmodified | 0 | 0 | 0 | 2650100 | 20819000 | 4623200 | 0 | 0 | 1243900 | 3567600 | 591540 | 0 | 14012000 | 3466400 | 0 | 0 | 7506900 | 4161000 |
| STESLQAN | RPL13; RPL13 | P26373; J3Q | RL13_HUMAN | Unmodified | 1380600 | 5195300 | 0 | 0 | 1492100 | 0 | 192980 | 372810 | 1384500 | 248850 | 2992100 | 39531000 | 0 | 0 | 0 | 0 | 0 | 434810 |
| STESLQAN | RPL13; RPL13 | P26373; J3Q | RL13_HUMAN | Unmodified | 0 | 0 | 0 | 0 | 1460700 | 0 | 4847100 | 4023000 | 0 | 405600 | 0 | 0 | 75572 | 0 | 0 | 0 | 316930 | 1842400 |
| STEVLPEK | UTRN | P46939 | UTRN_HUMAN | Unmodified | 0 | 0 | 377110 | 0 | 0 | 0 | 0 | 0 | 0 | 0 | 0 | 0 | 0 | 0 | 0 | 0 | 0 | 0 |
| STFDAHIYE | PRKCD; PRKCD | Q05655; C9 | KPCD_HUMAN | Unmodified | 0 | 0 | 0 | 0 | 0 | 0 | 0 | 0 | 1081600 | 0 | 560500 | 3877000 | 0 | 0 | 247750 | 345820 | 0 | 0 |
| STFSTLQEI | HCK; HCK; HCI | P08631; J3K | HCK_HUMAN T | Unmodified | 0 | 0 | 0 | 0 | 0 | 0 | 0 | 0 | 0 | 0 | 2211200 | 3886200 | 0 | 0 | 0 | 0 | 0 | 0 |
| STFSTLQEI | HCK; HCK; HCI | P08631; J3K | HCK_HUMAN T | Unmodified | 0 | 94163 | 0 | 0 | 0 | 0 | 0 | 0 | 0 | 0 | 4363500 | 3470900 | 0 | 164170 | 0 | 0 | 0 | 0 |
| STFVLDEFI | EEF1G | P26641 | EF1G_HUMAN | Unmodified | 0 | 0 | 0 | 0 | 2061700 | 465230 | 0 | 96616 | 1508400 | 1001700 | 3068900 | 23697000 | 780660 | 839590 | 1076600 | 1055000 | 1061500 | 1894600 |
| STFVLDEFI | EEF1G | P26641 | EF1G_HUMAN | Unmodified | 583000 | 0 | 494290 | 698280 | 981470 | 0 | 0 | 0 | 2193200 | 373700 | 0 | 31146000 | 0 | 0 | 477820 | 618040 | 360040 | 256700 |
| STGASFIR | MYO6; MYO6; M | Q9UM54; A0 | MYO6_HUMAN | Unmodified | 0 | 105530 | 123280 | 0 | 120040 | 0 | 0 | 0 | 286270 | 0 | 0 | 1872700 | 0 | 0 | 804200 | 851740 | 0 | 0 |

| | | | | | | | | | | | | | | | | | | | | | | |
|---|---|---|---|---|---|---|---|---|---|---|---|---|---|---|---|---|---|---|---|---|---|---|
| STGCDFAV | OXCT1; OXCT1 | P55809; E9P | SCOT1_HUMAI | Unmodified | 0 | 0 | 0 | 0 | 0 | 0 | 243490 | 85272 | 2213100 | 401910 | 0 | 1518200 | 0 | 231650 | 557850 | 568490 | 0 | 0 |
| STGDIAGT\ | TJP2; TJP2; TJP2 | Q9UDY2; A0 | ZO2_HUMAN T | Unmodified | 93804 | 0 | 362840 | 1250000 | 1842000 | 0 | 102110 | 25026 | 6856800 | 1650000 | 1042600 | 1234800 | 58093 | 0 | 264030 | 376720 | 0 | 0 |
| STGDSQNI | GAS7; GAS7; G | O60861; A0/ | GAS7_HUMAN | Unmodified | 0 | 0 | 0 | 0 | 0 | 0 | 0 | 0 | 0 | 0 | 0 | 186390 | 0 | 0 | 0 | 0 | 0 | 0 |
| STGDVPHT | USP8 | P40818 | UBP8_HUMAN | Unmodified | 0 | 0 | 0 | 0 | 2773000 | 0 | 0 | 0 | 250400 | 93197 | 0 | 156050 | 0 | 0 | 0 | 0 | 104230 | 273020 |
| STGEAFVQ | HNRNPH3 | P31942 | HNRH3_HUMA | Unmodified | 642230 | 637260 | 0 | 397930 | 0 | 0 | 0 | 0 | 0 | 0 | 0 | 3085100 | 0 | 0 | 418630 | 180780 | 0 | 0 |
| STGEAFVQ | HNRNPH1; HNI | P31943; P55 | HNRH1_HUMA | Unmodified | 0 | 0 | 0 | 0 | 503660 | 215640 | 1995300 | 90945 | 1425100 | 891940 | 1766500 | 9954000 | 0 | 0 | 0 | 0 | 281800 | 750340 |
| STGEGVIR | CLSTN1; CLSTN | O94985; A0/ | CSTN1_HUMAI | Unmodified | 0 | 0 | 1307000 | 0 | 34318 | 317300 | 29388 | 0 | 4578300 | 631470 | 3236200 | 475870 | 0 | 0 | 0 | 0 | 96455 | 75832 |
| STGEPLGH | LPP; LPP; LPP; l | Q93052; C9J | LPP_HUMAN Li | Unmodified | 0 | 0 | 0 | 0 | 0 | 0 | 170260 | 0 | 0 | 123430 | 0 | 0 | 0 | 0 | 0 | 206410 | 0 | 0 |
| STGEPLGH | LPP; LPP; LPP; l | Q93052; C9J | LPP_HUMAN Li | Unmodified | 0 | 0 | 0 | 0 | 2482600 | 0 | 0 | 0 | 0 | 0 | 0 | 0 | 0 | 0 | 0 | 0 | 0 | 0 |
| STGFETLVV | ECI2; ECI2; ECI | O75521; F8V | ECI2_HUMAN E | Unmodified | 0 | 0 | 0 | 0 | 0 | 0 | 0 | 29447 | 9262700 | 1234600 | 268990 | 0 | 0 | 0 | 0 | 0 | 0 | 0 |
| STGFSNPA | CCDC28A; CCI | Q8IWP9; B4I | CC28A_HUMAI | Unmodified | 718140 | 767090 | 0 | 0 | 213390 | 39456 | 424050 | 48364 | 0 | 0 | 0 | 311300 | 27513 | 93164 | 0 | 0 | 120450 | 235900 |
| STGGAPTFI | PFN1; PFN1 | P07737; K7E | PROF1_HUMAI | Unmodified | 33703000 | 27492000 | 33070000 | 34525000 | 89640000 | 15778000 | 8773400 | 7168200 | 79075000 | 16246000 | 45300000 | 1.38E+08 | 71227000 | 17373000 | 42269000 | 36668000 | 16785000 | 22239000 |
| STGGAPTFI | PFN1; PFN1 | P07737; K7E | PROF1_HUMAI | Unmodified | 32906000 | 28184000 | 3360200 | 48403000 | 6.92E+08 | 1.15E+08 | 1.54E+08 | 96224000 | 63272000 | 1.39E+08 | 44087000 | 72666000 | 3.63E+08 | 1.36E+08 | 15016000 | 12730000 | 2.75E+08 | 2.75E+08 |
| STGGAPTFI | PFN1; PFN1 | P07737; K7E | PROF1_HUMAI | Oxidation (M) | 0 | 0 | 0 | 80046 | 16931000 | 468480 | 11664000 | 344060 | 0 | 2053100 | 0 | 0 | 2972000 | 5023100 | 0 | 0 | 28347000 | 1981800 |
| STGGAPTFI | PFN1; PFN1 | P07737; K7E | PROF1_HUMAI | Unmodified | 0 | 0 | 0 | 0 | 0 | 0 | 12866000 | 0 | 0 | 3018800 | 0 | 0 | 2759600 | 0 | 0 | 0 | 17771000 | 4184300 |
| STGGDFGI | RAB6A; RAB6A; | P20340; F5G | RAB6A_HUMAI | Acetyl (Protein N | 97888 | 0 | 0 | 160570 | 0 | 0 | 0 | 0 | 0 | 0 | 0 | 858740 | 0 | 0 | 0 | 0 | 0 | 0 |
| STGKENKIT | HSPA8; HSPA2; | P11142; P54 | HSP7C_HUMAI | Unmodified | 0 | 0 | 0 | 0 | 0 | 0 | 0 | 0 | 0 | 2041900 | 0 | 0 | 0 | 0 | 0 | 0 | 1275700 | 0 |
| STGLILDSG | ACTL6A | O96019 | ACL6A_HUMAI | Unmodified | 1438300 | 0 | 0 | 0 | 0 | 0 | 0 | 0 | 8728200 | 1287900 | 4158200 | 13197000 | 0 | 0 | 0 | 1217400 | 0 | 0 |
| STGLNAVP | ITGAV | P06756 | ITAV_HUMAN II | Unmodified | 2846800 | 2414400 | 3482700 | 4444600 | 4893200 | 2714700 | 0 | 19750 | 12412000 | 2540400 | 1706100 | 14029000 | 2084700 | 496000 | 5572900 | 5992200 | 2144000 | 986080 |
| STGLNAVP | ITGAV | P06756 | ITAV_HUMAN II | 2 Oxidation (M) | 0 | 0 | 0 | 0 | 329750 | 0 | 0 | 0 | 0 | 0 | 0 | 0 | 0 | 0 | 0 | 0 | 233210 | 1082800 |
| STGNFLTLT | LARS1; LARS1; | Q9P2J5; A0A | SYLC_HUMAN | Unmodified | 0 | 0 | 0 | 0 | 0 | 0 | 0 | 30315 | 801360 | 0 | 1004300 | 19195000 | 70327 | 0 | 0 | 0 | 0 | 0 |
| STGPGASL | CLDN3 | O15551 | CLD3_HUMAN | Unmodified | 0 | 0 | 303530 | 0 | 138440 | 0 | 0 | 0 | 698260 | 662790 | 0 | 0 | 0 | 0 | 0 | 542310 | 0 | 0 |
| STGPGASL | CLDN3 | O15551 | CLD3_HUMAN | Unmodified | 8238200 | 6659800 | 26872000 | 14625000 | 4087400 | 1292900 | 0 | 0 | 29280000 | 2775400 | 856600 | 2895100 | 2488600 | 0 | 39690000 | 23922000 | 576630 | 1515400 |
| STGPGASL | CLDN3 | O15551 | CLD3_HUMAN | Unmodified | 3028300 | 2693400 | 0 | 11279000 | 22793000 | 5994400 | 0 | 0 | 8484100 | 19162000 | 0 | 0 | 11636000 | 5984500 | 915990 | 1430500 | 13548000 | 34806000 |
| STGSIVGQ | ALDH4A1 | P30038 | AL4A1_HUMAI | Unmodified | 0 | 0 | 0 | 0 | 0 | 396310 | 297910 | 240700 | 898520 | 0 | 0 | 0 | 0 | 246080 | 0 | 0 | 0 | 322000 |
| STGSPTQE | ZNF185 | H0Y704 | H0Y704_HUMA | Unmodified | 0 | 0 | 0 | 0 | 0 | 0 | 0 | 0 | 1248200 | 295800 | 0 | 0 | 0 | 0 | 0 | 0 | 502830 | 0 |
| STGTFVVS | ALDH9A1 | P49189 | AL9A1_HUMAI | Acetyl (Protein N | 0 | 0 | 0 | 759050 | 8260900 | 577880 | 1297100 | 1563500 | 0 | 2332200 | 455930 | 0 | 14064000 | 10167000 | 0 | 0 | 8002100 | 6565100 |
| STGTRAPPI | KIAA1522 | Q9P206 | K1522_HUMAI | 2 Oxidation (M) | 0 | 0 | 0 | 2686800 | 0 | 0 | 0 | 0 | 0 | 0 | 0 | 0 | 3302400 | 0 | 0 | 582170 | 0 | 0 |
| STHGLAILG | PITRM1; PITRM | Q5JRX3; A0A | PREP_HUMAN | Unmodified | 0 | 0 | 0 | 0 | 0 | 0 | 0 | 0 | 1085100 | 372360 | 0 | 0 | 50523 | 0 | 0 | 0 | 0 | 0 |
| STHGPGM | PACSIN3; PACS | Q9UKS6; H0 | PACN3_HUMAI | Oxidation (M) | 2589600 | 2950200 | 0 | 0 | 0 | 0 | 0 | 0 | 0 | 0 | 0 | 0 | 0 | 0 | 0 | 0 | 0 | 0 |
| STHGPGM | PACSIN3; PACS | Q9UKS6; H0 | PACN3_HUMAI | 2 Oxidation (M) | 6806200 | 14425000 | 358600 | 8031800 | 0 | 0 | 0 | 0 | 0 | 0 | 0 | 0 | 1566900 | 0 | 8973900 | 922280 | 0 | 0 |
| STHNEDQ | PPM1H | Q9ULR3 | PPM1H_HUMA | Unmodified | 0 | 0 | 0 | 0 | 0 | 0 | 0 | 36076 | 151870 | 150500 | 146040 | 1458100 | 113630 | 0 | 0 | 0 | 0 | 0 |
| STHQAAIV | CALD1; CALD1; | Q05682; C9J | CALD1_HUMAI | Unmodified | 0 | 0 | 0 | 0 | 768560 | 0 | 1027800 | 0 | 0 | 610280 | 0 | 0 | 0 | 928410 | 0 | 0 | 9302700 | 256860 |
| STHSELLEI | ABCD3; ABCD3 | P28288; A0A | ABCD3_HUMAI | Unmodified | 0 | 0 | 0 | 0 | 0 | 0 | 0 | 0 | 1803100 | 0 | 0 | 0 | 0 | 0 | 0 | 0 | 0 | 0 |
| STHTDGG | PAPLN; PAPLN; | O95428; H0 | PPN_HUMAN P | Unmodified | 0 | 0 | 0 | 0 | 0 | 0 | 0 | 0 | 195210 | 0 | 0 | 0 | 0 | 0 | 0 | 0 | 0 | 0 |
| STHTLDLSI | LGALS3BP; LG/ | Q08380; K7E | LG3BP_HUMAI | Unmodified | 0 | 0 | 0 | 0 | 0 | 0 | 0 | 0 | 1669800 | 0 | 0 | 2722700 | 0 | 0 | 0 | 0 | 0 | 0 |
| STIANLMGI | DST; DST; DST; | Q03001; A0/ | DYST_HUMAN | Unmodified | 0 | 0 | 0 | 0 | 898630 | 0 | 0 | 9593 | 0 | 0 | 0 | 0 | 0 | 0 | 0 | 0 | 0 | 0 |
| STIGGQIM | GSPT1; GSPT1; | P15170; H3E | ERF3A_HUMAI | Oxidation (M) | 0 | 0 | 0 | 0 | 138100 | 0 | 0 | 0 | 0 | 0 | 0 | 0 | 172860 | 78163 | 0 | 0 | 57622 | 0 |
| STIGVEFAT | RAB11A; RAB11 | P62491; Q15 | RB11A_HUMAI | Unmodified | 6860300 | 8842400 | 9535500 | 21705000 | 3449900 | 2883100 | 0 | 1037500 | 6280600 | 101800 | 3458300 | 193370 | 10741000 | 713040 | 11313000 | 14405000 | 747910 | 535510 |
| STIGVEFAT | RAB11A; RAB11 | P62491; Q15 | RB11A_HUMAI | Unmodified | 0 | 0 | 0 | 0 | 10542000 | 1778000 | 4283400 | 1517700 | 178220 | 927680 | 0 | 0 | 3059200 | 4214600 | 0 | 0 | 2183400 | 3038700 |
| STIGVEFAT | RAB11A; RAB11 | P62491; Q15 | RB11A_HUMAI | Unmodified | 0 | 0 | 0 | 74613 | 11794000 | 0 | 5236100 | 357650 | 0 | 376710 | 0 | 0 | 2826900 | 1011700 | 0 | 0 | 925370 | 489800 |
| STIIGESISR | RARS1 | P54136 | SYRC_HUMAN | Unmodified | 0 | 0 | 0 | 0 | 0 | 0 | 198500 | 0 | 0 | 0 | 0 | 10819000 | 0 | 0 | 0 | 0 | 0 | 0 |
| STIKPSQM | PAN2 | Q504Q3 | PAN2_HUMAN | Unmodified | 388840 | 487300 | 0 | 0 | 0 | 0 | 0 | 0 | 0 | 0 | 0 | 0 | 226580 | 0 | 161260 | 138610 | 0 | 0 |
| STIQRQSS | ERBIN | Q96RT1 | ERBIN_HUMAI | Unmodified | 0 | 0 | 0 | 0 | 1475800 | 414910 | 0 | 0 | 0 | 0 | 0 | 0 | 240160 | 0 | 0 | 0 | 147030 | 0 |
| STISTELAL | NUBP2; NUBP2 | Q9Y5Y2; H3I | NUBP2_HUMAI | Unmodified | 0 | 0 | 0 | 0 | 704650 | 0 | 199650 | 0 | 0 | 123070 | 0 | 0 | 0 | 415840 | 0 | 0 | 0 | 558170 |
| STITEIKEC | GLG1; GLG1; G | Q92896; H3I | GSLG1_HUMAI | Unmodified | 0 | 80852 | 0 | 263660 | 1456000 | 873780 | 0 | 164060 | 8060400 | 5025100 | 1404700 | 655510 | 307200 | 419640 | 0 | 0 | 49040 | 0 |
| STITSREIQ | H2BC4; H2BC1 | P62807; P06 | H2B1C_HUMAI | Unmodified | 3459600 | 642900 | 0 | 0 | 1776800 | 0 | 1136700 | 14396000 | 0 | 860360 | 276470 | 17059000 | 0 | 0 | 0 | 0 | 0 | 5725700 |
| STITSREIQ | H2BC4; H2BC1 | P62807; P06 | H2B1C_HUMAI | Unmodified | 0 | 1214100 | 0 | 0 | 42612000 | 16846000 | 25318000 | 0 | 0 | 0 | 0 | 5344000 | 0 | 0 | 0 | 0 | 0 | 14592000 |
| STITSREVQ | H2BU1 | Q8N257 | H2B3B_HUMAI | Unmodified | 28723 | 38076 | 0 | 0 | 0 | 0 | 0 | 556260 | 0 | 0 | 0 | 480610 | 0 | 0 | 0 | 0 | 0 | 0 |
| STIYLVR | ITFG1; ITFG1 | Q8TB96; F5C | TIP_HUMAN T- | Unmodified | 32793 | 37085 | 0 | 21516 | 686740 | 0 | 0 | 0 | 7938200 | 3275500 | 0 | 325970 | 0 | 0 | 407370 | 195120 | 0 | 0 |
| STIYLVRSG | ITFG1; ITFG1 | Q8TB96; F5C | TIP_HUMAN T- | Unmodified | 0 | 0 | 0 | 0 | 223220 | 0 | 0 | 0 | 0 | 0 | 0 | 0 | 0 | 0 | 0 | 0 | 0 | 0 |
| STKVPIYLK | CDC42EP2 | O14613 | BORG1_HUMA | Acetyl (Protein N | 0 | 0 | 0 | 0 | 269860 | 0 | 0 | 67496 | 0 | 0 | 0 | 0 | 0 | 0 | 0 | 0 | 1927900 | 1518400 |
| STLADRLLE | GUF1 | Q8N442 | GUF1_HUMAN | Unmodified | 0 | 0 | 0 | 15118 | 95049 | 0 | 32279 | 0 | 384740 | 0 | 0 | 0 | 0 | 0 | 0 | 0 | 51859 | 34836 |
| STLAEIEDV | MYOF; MYOF | Q9NZM1; H0 | MYOF_HUMAN | Unmodified | 0 | 0 | 0 | 0 | 108420 | 40974 | 0 | 0 | 0 | 0 | 0 | 0 | 0 | 0 | 0 | 0 | 0 | 0 |
| STLDIVLAN | RRM1 | P23921 | RIR1_HUMAN F | Unmodified | 231490 | 0 | 0 | 0 | 531650 | 0 | 308480 | 279570 | 118180 | 298070 | 1406600 | 4761400 | 0 | 0 | 505530 | 792390 | 271190 | 0 |
| STLEPVEK | HSPA6; HSPA1/ | P17066; P0E | HSP76_HUMAI | Unmodified | 8621200 | 6969000 | 6896400 | 8754200 | 3586900 | 164800 | 404720 | 192920 | 17788000 | 2915500 | 2503600 | 14351000 | 2753400 | 1234300 | 5014400 | 4234400 | 616090 | 2162700 |
| STLEQKPSI | CD2AP | Q9Y5K6 | CD2AP_HUMAI | Unmodified | 0 | 323620 | 0 | 0 | 0 | 0 | 0 | 263380 | 0 | 0 | 0 | 0 | 0 | 0 | 1445400 | 1704800 | 0 | 0 |
| STLGDLDT | PDE4DIP; PDE4 | Q5VU43; E9I | MYOME_HUMA | Unmodified | 0 | 0 | 0 | 0 | 0 | 0 | 0 | 0 | 3336200 | 357700 | 0 | 0 | 289750 | 0 | 0 | 0 | 0 | 0 |
| STLIDTLFN | SEPTIN10; SEP | Q9P0V9; B5I | SEP10_HUMAI | Unmodified | 0 | 0 | 0 | 0 | 3568900 | 0 | 0 | 0 | 0 | 0 | 0 | 0 | 0 | 0 | 0 | 0 | 2372600 | 0 |
| STLINSLFL | SEPTIN2; SEPT | Q15019; C9I | SEPT2_HUMAI | Unmodified | 0 | 7328.7 | 0 | 275250 | 0 | 0 | 0 | 0 | 9933.6 | 73068 | 0 | 462590 | 0 | 0 | 0 | 0 | 0 | 0 |
| STLINSLFL | SEPTIN2; SEPT | Q15019; C9I | SEPT2_HUMAI | Unmodified | 146640 | 197700 | 150890 | 2061400 | 18464000 | 3367800 | 1057200 | 0 | 433100 | 1190200 | 155180 | 934310 | 1446500 | 1573100 | 2305600 | 295720 | 3630600 | 11406000 |
| STLINSLFL | SEPTIN2; SEPT | Q15019; C9I | SEPT2_HUMAI | Unmodified | 0 | 0 | 0 | 705980 | 2662500 | 1191700 | 0 | 0 | 0 | 1249200 | 0 | 0 | 848760 | 0 | 0 | 0 | 2043300 | 2340100 |
| STLINSLFL | SEPTIN7; SEPT | Q16181; A0/ | SEPT7_HUMAI | Unmodified | 1009000 | 922540 | 628850 | 3012200 | 36270000 | 6378700 | 2939500 | 0 | 8578900 | 4309200 | 2559100 | 8592300 | 10618000 | 5136300 | 3410300 | 695990 | 8224700 | 7000300 |
| STLINSLFL | SEPTIN7; SEPT | Q16181; A0/ | SEPT7_HUMAI | Unmodified | 0 | 0 | 0 | 0 | 18498000 | 7935200 | 3461500 | 0 | 0 | 979170 | 0 | 0 | 0 | 0 | 0 | 0 | 5603200 | 3682400 |
| STLINSLFL | SEPTIN7; SEPT | Q16181; A0/ | SEPT7_HUMAI | Unmodified | 0 | 0 | 0 | 0 | 17714000 | 1119500 | 0 | 0 | 0 | 0 | 0 | 0 | 0 | 998730 | 0 | 0 | 8011400 | 3565100 |
| STLINTLFK | SEPTIN9; SEPT | Q9UHD8; K7 | SEPT9_HUMAI | Unmodified | 0 | 0 | 522880 | 1178100 | 18531000 | 4798800 | 3414700 | 1552900 | 1659900 | 2111600 | 1326600 | 11435000 | 4241100 | 4308000 | 0 | 815480 | 18009000 | 18858000 |
| STLKEDVGI | DYNC1H1; DYN | Q14204; A0/ | DYHC1_HUMA | Unmodified | 147320 | 86637 | 99953 | 325310 | 0 | 0 | 0 | 0 | 0 | 0 | 711720 | 5525600 | 0 | 0 | 289800 | 0 | 0 | 97931 |
| STLLFGQR | KIF5B | P33176 | KINH_HUMAN | Unmodified | 0 | 0 | 230590 | 275680 | 1088800 | 0 | 107090 | 0 | 943400 | 359480 | 441440 | 3068000 | 542420 | 0 | 0 | 223830 | 0 | 0 |
| STLLLLLTG | ABCF1; ABCF1; | Q8NE71; H0 | ABCF1_HUMAI | Unmodified | 0 | 0 | 137180 | 0 | 0 | 0 | 0 | 0 | 292520 | 0 | 408590 | 3803000 | 0 | 0 | 478900 | 713850 | 0 | 0 |
| STLMDTLFI | SEPTIN11; SEP | Q9NVA2; D6 | SEP11_HUMAI | Oxidation (M) | 0 | 0 | 0 | 481410 | 0 | 0 | 0 | 0 | 0 | 0 | 0 | 1611400 | 0 | 0 | 0 | 0 | 0 | 0 |
| STLMDTLFI | SEPTIN11; SEP | Q9NVA2; D6 | SEP11_HUMAI | Unmodified | 616480 | 1550300 | 0 | 0 | 0 | 0 | 0 | 0 | 0 | 0 | 0 | 2626400 | 0 | 0 | 98753 | 0 | 0 | 0 |
| STLMDTLFI | SEPTIN11; SEP | Q9NVA2; D6 | SEP11_HUMAI | Oxidation (M) | 0 | 0 | 0 | 179660 | 7314100 | 2279900 | 1336100 | 33484 | 0 | 537580 | 39474 | 0 | 2110600 | 0 | 0 | 0 | 6431000 | 0 |
| STLMDTLFI | SEPTIN11; SEP | Q9NVA2; D6 | SEP11_HUMAI | Unmodified | 0 | 0 | 0 | 0 | 24836000 | 2022400 | 493120 | 0 | 0 | 1395200 | 0 | 0 | 2699600 | 2067100 | 0 | 0 | 7149200 | 2827600 |
| STLNEIYFG | CAPZB; CAPZB; | P47756; B1A | CAPZB_HUMAI | Unmodified | 0 | 8836600 | 0 | 1237100 | 1421100 | 137300 | 0 | 196320 | 2303200 | 383040 | 0 | 11594000 | 1328300 | 0 | 0 | 2650800 | 532070 | 892800 |
| STLNPQWI | PRKCA; PRKCA | P17252; J3K | KPCA_HUMAN | Unmodified | 0 | 0 | 0 | 45018 | 1546400 | 41577 | 1298100 | 417000 | 0 | 34801 | 722900 | 258720 | 0 | 143580 | 0 | 0 | 551090 | 214400 |
| STLPDADR | ACTN4; ACTN4; | O43707; F5C | ACTN4_HUMAI | Unmodified | 0 | 0 | 0 | 0 | 0 | 0 | 0 | 0 | 0 | 51275 | 0 | 2765000 | 0 | 0 | 281350 | 0 | 0 | 0 |
| STLPDADR | ACTN4; ACTN4; | O43707; F5C | ACTN4_HUMAI | Unmodified | 132390 | 0 | 0 | 257290 | 4390400 | 2285600 | 137620 | 0 | 1815400 | 842780 | 0 | 2078900 | 254820 | 0 | 296010 | 226280 | 919370 | 0 |
| STLQEVVG | IGSF8; IGSF8 | Q969P0; C9J | IGSF8_HUMAN | Unmodified | 10533000 | 9707600 | 5968100 | 3540400 | 0 | 0 | 0 | 0 | 777240 | 0 | 0 | 1960300 | 373510 | 0 | 7882300 | 9513600 | 0 | 0 |
| STLQEVVG | IGSF8; IGSF8 | Q969P0; C9J | IGSF8_HUMAN | Unmodified | 0 | 1420300 | 0 | 639760 | 0 | 0 | 0 | 0 | 0 | 0 | 0 | 0 | 518420 | 0 | 920900 | 0 | 0 | 0 |
| STLQTMESI | CKAP4 | Q07065 | CKAP4_HUMAI | Unmodified | 0 | 0 | 0 | 48417 | 0 | 0 | 0 | 0 | 884280 | 265560 | 91903 | 386050 | 0 | 0 | 61475 | 0 | 0 | 0 |
| STLSQTVPS | ANLN | Q9NQW6 | ANLN_HUMAN | Unmodified | 155650 | 192940 | 149390 | 776570 | 1672100 | 0 | 0 | 0 | 404220 | 0 | 51680 | 295110 | 0 | 0 | 178080 | 182970 | 781860 | 845790 |
| STLSQTVPS | ANLN | Q9NQW6 | ANLN_HUMAN | Unmodified | 0 | 0 | 0 | 0 | 6544600 | 1030600 | 571160 | 452810 | 0 | 0 | 0 | 0 | 885470 | 805260 | 0 | 0 | 7311100 | 3060700 |
| STLTDSLV | EEF2 | P13639 | EF2_HUMAN El | Unmodified | 1986000 | 0 | 1173200 | 2885500 | 14505000 | 3428300 | 3295500 | 2881500 | 2516000 | 2408300 | 7059100 | 34842000 | 7271200 | 4230000 | 2776300 | 0 | 5296300 | 8442800 |
| STLVHSLFI | SEPTIN5; SEPT | Q99719; F8V | SEPT5_HUMAI | Unmodified | 42469 | 143160 | 301670 | 943440 | 2476600 | 1226400 | 2481300 | 0 | 391670 | 374050 | 0 | 303600 | 4178900 | 3016700 | 739470 | 372960 | 1984200 | 1765800 |
| STLVNTLFK | SEPTIN3; SEPT | Q9UH03; B1 | SEPT3_HUMAI | Unmodified | 0 | 0 | 0 | 0 | 0 | 0 | 1476100 | 1154900 | 0 | 0 | 12215000 | 0 | 0 | 0 | 0 | 0 | 0 | 0 |
| STLYVSPHI | VARS1; VARS1; | P26640; A2A | SYVC_HUMAN | Acetyl (Protein N | 0 | 0 | 0 | 0 | 0 | 0 | 0 | 0 | 0 | 0 | 0 | 3947600 | 0 | 0 | 0 | 0 | 0 | 0 |
| STMDNPTT | DYNLRB1; DYN | Q9NP97; B1 | DLRB1_HUMAI | Oxidation (M) | 267700 | 0 | 0 | 0 | 1920000 | 0 | 616060 | 0 | 1413300 | 850030 | 0 | 0 | 2201200 | 1065800 | 0 | 0 | 806420 | 3866900 |
| STMDNPTT | DYNLRB1; DYN | Q9NP97; B1 | DLRB1_HUMAI | 2 Oxidation (M) | 0 | 0 | 0 | 0 | 0 | 0 | 0 | 40358 | 182380 | 0 | 0 | 1439100 | 0 | 0 | 0 | 0 | 428180 | 1377900 |
| STMEGGTL | PLCG2; PLCG2 | P16885; A0A | PLCG2_HUMAI | Oxidation (M) | 0 | 0 | 0 | 0 | 0 | 0 | 0 | 0 | 0 | 0 | 0 | 1517700 | 0 | 0 | 0 | 0 | 0 | 0 |
| STMEGGTL | PLCG2; PLCG2 | P16885; A0A | PLCG2_HUMAI | Unmodified | 0 | 0 | 0 | 0 | 0 | 0 | 0 | 0 | 0 | 0 | 0 | 6036200 | 0 | 0 | 0 | 0 | 0 | 0 |
| STMGSSFS | KDF1 | Q8NAX2 | KDF1_HUMAN | Unmodified | 0 | 0 | 0 | 178190 | 0 | 0 | 0 | 0 | 770650 | 270290 | 0 | 155120 | 666530 | 700310 | 0 | 0 | 0 | 0 |
| STMGSSFS | KDF1 | Q8NAX2 | KDF1_HUMAN | Oxidation (M) | 0 | 0 | 0 | 0 | 0 | 0 | 0 | 0 | 320500 | 188640 | 143110 | 0 | 625030 | 602210 | 0 | 0 | 0 | 0 |
| STMMTQE | MYO6; MYO6; N | Q9UM54; A0 | MYO6_HUMAN | Unmodified | 61550 | 0 | 304570 | 504930 | 0 | 0 | 0 | 0 | 501840 | 0 | 0 | 427100 | 156550 | 0 | 512090 | 523910 | 0 | 0 |
| STMMTQE | MYO6; MYO6; N | Q9UM54; A0 | MYO6_HUMAN | Oxidation (M) | 0 | 0 | 1893500 | 228330 | 0 | 0 | 0 | 0 | 0 | 0 | 0 | 103960 | 0 | 68301 | 370050 | 495710 | 0 | 0 |
| STMMTQE | MYO6; MYO6; N | Q9UM54; A0 | MYO6_HUMAN | Oxidation (M) | 0 | 0 | 92957 | 187320 | 0 | 0 | 0 | 0 | 0 | 0 | 0 | 0 | 170240 | 186970 | 0 | 0 | 0 | 0 |
| STNEAMEV | HSPA4 | P34932 | HSP74_HUMAI | Unmodified | 0 | 0 | 0 | 0 | 531990 | 0 | 0 | 0 | 0 | 0 | 0 | 1334300 | 0 | 0 | 0 | 0 | 221500 | 354500 |
| STNENANT | KPNA2; KPNA2; | P52292; J3Q | IMA1_HUMAN I | Acetyl (Protein N | 0 | 0 | 0 | 0 | 591340 | 0 | 235640 | 91098 | 0 | 0 | 0 | 0 | 0 | 241910 | 0 | 0 | 895440 | 326730 |
| STNGDTFL | HSPA9; HSPA9; | P38646; H0 | GRP75_HUMAI | Unmodified | 0 | 0 | 0 | 0 | 1047200 | 111560 | 80682 | 54406 | 26854000 | 1376600 | 1262500 | 394310 | 0 | 76469 | 0 | 0 | 71397 | 97508 |
| STNGDTFL | HSPA9; HSPA9; | P38646; H0 | GRP75_HUMAI | Unmodified | 0 | 0 | 0 | 0 | 0 | 357090 | 0 | 0 | 354410 | 0 | 0 | 0 | 954980 | 0 | 0 | 0 | 0 | 45538 |
| STNINFYEI | ITGA2 | P17301 | ITA2_HUMAN II | Unmodified | 53734 | 305500 | 0 | 212400 | 0 | 67243 | 0 | 0 | 0 | 0 | 0 | 0 | 0 | 0 | 3280200 | 37560 | 0 | 0 |
| STNLHDYG | APP; APP; APP; | P05067; H7C | A4_HUMAN Am | Oxidation (M) | 0 | 0 | 153980 | 161820 | 457890 | 819900 | 94850 | 43539 | 3149700 | 1716400 | 716920 | 67276 | 0 | 0 | 0 | 29870 | 170250 | 862930 |
| STNLHDYG | APP; APP; APP; | P05067; H7C | A4_HUMAN Am | Unmodified | 210870 | 114040 | 0 | 0 | 410360 | 162760 | 0 | 0 | 8768500 | 972120 | 54012 | 0 | 46191 | 0 | 0 | 0 | 0 | 402630 |
| STNPGISIG | HMGB3; HMGB | O15347; E7E | HMGB3_HUMA | Unmodified | 0 | 0 | 0 | 0 | 604910 | 0 | 0 | 0 | 2547600 | 2427600 | 1277300 | 1755500 | 0 | 0 | 139760 | 62972 | 0 | 0 |
| STNPGISIG | HMGB3; HMGB | O15347; E7E | HMGB3_HUMA | Unmodified | 0 | 0 | 0 | 0 | 285320 | 0 | 0 | 0 | 1478400 | 322590 | 0 | 834220 | 0 | 0 | 0 | 0 | 107630 | 136330 |
| STNPSVATI | GRPR | P30550 | GRPR_HUMAN | Unmodified | 0 | 0 | 0 | 0 | 0 | 0 | 0 | 0 | 0 | 227510 | 0 | 0 | 1274800 | 0 | 0 | 0 | 2566200 | 5557800 |
| STPAITLESI | CYB5R3; CYB5I | P00387; B1A | NB5R3_HUMAI | Unmodified | 101410 | 0 | 0 | 0 | 0 | 0 | 0 | 0 | 0 | 0 | 0 | 1236900 | 0 | 0 | 0 | 0 | 0 | 0 |
| STPAVIQGI | GPR108; GPR1( | Q9NPR9; MC | GP108_HUMAI | Unmodified | 0 | 0 | 0 | 0 | 258480 | 0 | 0 | 0 | 448890 | 0 | 106550 | 0 | 0 | 0 | 0 | 0 | 0 | 176690 |
| STPEYFAEF | ANXA6 | P08133 | ANXA6_HUMAI | Unmodified | 184760 | 473270 | 2421300 | 1302500 | 0 | 0 | 139470 | 0 | 7328300 | 1202400 | 0 | 119380 | 0 | 0 | 3112400 | 275490 | 39247 | 0 |
| STPEYFAEF | ANXA6 | P08133 | ANXA6_HUMAI | Unmodified | 0 | 0 | 461410 | 890210 | 1814500 | 1199000 | 1973000 | 133410 | 2698900 | 13786000 | 0 | 0 | 3114800 | 1978400 | 425730 | 0 | 1628700 | 1996900 |

| | | | | | | | | | | | | | | | | | | | | | | |
|---|---|---|---|---|---|---|---|---|---|---|---|---|---|---|---|---|---|---|---|---|---|---|
| STPGVANN | SYTL2 | A0A0U1RR0 | A0A0U1RR07_H | Unmodified | 204880 | 1553600 | 982880 | 57687000 | 0 | 0 | 0 | 0 | 0 | 0 | 0 | 0 | 0 | 0 | 274500 | 0 | 0 | 40879 |
| STPHRTPII | ANLN; ANLN | Q9NQW6; H | ANLN_HUMAN | Unmodified | 0 | 0 | 0 | 0 | 7314200 | 808410 | 0 | 0 | 148970 | 0 | 0 | 0 | 265210 | 0 | 0 | 0 | 0 | 1258000 |
| STPLSQPS | ENAH; ENAH | Q8N8S7; A0 | ENAH_HUMAN | Unmodified | 0 | 0 | 0 | 0 | 0 | 0 | 0 | 0 | 0 | 0 | 464250 | 3276500 | 0 | 0 | 0 | 160520 | 0 | 0 |
| STPSHGSV | EPS15L1; EPS1 | Q9UBC2; M | EP15R_HUMAN | Unmodified | 263230 | 367470 | 0 | 0 | 0 | 0 | 0 | 0 | 0 | 0 | 0 | 867250 | 487240 | 0 | 0 | 209510 | 0 | 0 |
| STPTQGQT | SDK2; SDK2 | Q58EX2; H7 | SDK2_HUMAN | Unmodified | 33876 | 0 | 0 | 0 | 0 | 0 | 0 | 0 | 671300 | 0 | 0 | 0 | 0 | 0 | 0 | 0 | 0 | 0 |
| STPVIVSAT | AKAP12 | Q02952 | AKA12_HUMAN | Unmodified | 0 | 0 | 0 | 0 | 2104200 | 175410 | 0 | 0 | 0 | 0 | 0 | 0 | 0 | 0 | 0 | 0 | 2428800 | 4142000 |
| STPVIVSAT | AKAP12 | Q02952 | AKA12_HUMAN | Unmodified | 0 | 0 | 0 | 0 | 1864700 | 261610 | 0 | 0 | 0 | 0 | 0 | 0 | 0 | 0 | 0 | 0 | 1460600 | 2719500 |
| STPVNVPIS | PDCD6IP | Q8WUM4 | PDC6I_HUMAN | Unmodified | 84252000 | 78614000 | 0 | 30066000 | 16447000 | 4609300 | 936040 | 720720 | 30170000 | 4771000 | 5748900 | 17164000 | 15362000 | 6971300 | 46391000 | 38826000 | 1298800 | 6704800 |
| STPVNVPIS | PDCD6IP | Q8WUM4 | PDC6I_HUMAN | Unmodified | 12768000 | 12470000 | 108450 | 21096000 | 2.09E+08 | 22983000 | 40704000 | 3838000 | 6875800 | 1.25E+08 | 2503500 | 283850 | 3.09E+08 | 2.13E+08 | 0 | 1471000 | 49006000 | 1.01E+08 |
| STPVPEVV | OCLN; OCLN | Q16625; A0 | OCLN_HUMAN | Unmodified | 0 | 0 | 3684700 | 6873600 | 15843000 | 4133400 | 17289 | 0 | 19772000 | 8366700 | 0 | 1573900 | 17826000 | 16530000 | 18945000 | 6315700 | 0 | 1388600 |
| STPVPEVV | OCLN; OCLN | Q16625; A0 | OCLN_HUMAN | Unmodified | 0 | 0 | 0 | 0 | 20414000 | 2599000 | 0 | 0 | 0 | 2454800 | 0 | 0 | 14689000 | 24075000 | 0 | 152230 | 0 | 6643800 |
| STPVTVVLF | EFTUD2 | Q15029 | U5S1_HUMAN | Unmodified | 0 | 0 | 0 | 0 | 485350 | 0 | 0 | 0 | 0 | 0 | 0 | 2172900 | 515520 | 0 | 0 | 0 | 0 | 381300 |
| STQAATQV | PMPCB; PMPCE | O75439; G3 | MPPB_HUMAN | Unmodified | 0 | 0 | 82464 | 0 | 0 | 0 | 0 | 0 | 1273000 | 0 | 0 | 0 | 0 | 0 | 0 | 142860 | 0 | 0 |
| STQAGDDL | GAS7 | O60861 | GAS7_HUMAN | Oxidation (M) | 0 | 0 | 0 | 0 | 0 | 0 | 0 | 0 | 0 | 0 | 0 | 804980 | 0 | 0 | 0 | 0 | 0 | 0 |
| STQAGDDL | GAS7 | O60861 | GAS7_HUMAN | Unmodified | 0 | 46191 | 0 | 0 | 0 | 0 | 0 | 0 | 0 | 0 | 0 | 1832300 | 0 | 0 | 0 | 0 | 0 | 0 |
| STQAPLIIRI | NAMPT; NAMPT | P43490; A0A | NAMPT_HUMA | Unmodified | 705280 | 0 | 2283500 | 3881500 | 42400000 | 6127500 | 956950 | 273800 | 3364600 | 3026000 | 6797400 | 28157000 | 14949000 | 2809300 | 2794700 | 3791600 | 2795400 | 3252800 |
| STQDEINQ | EPS15L1; EPS1 | Q9UBC2; M | EP15R_HUMAN | Unmodified | 0 | 0 | 0 | 0 | 0 | 0 | 0 | 0 | 121230 | 0 | 0 | 11538 | 2375900 | 1121300 | 10360 | 0 | 738000 | 0 |
| STQDLSEG | AKAP5 | P24588 | AKAP5_HUMAN | Unmodified | 0 | 0 | 0 | 0 | 0 | 0 | 0 | 32572 | 0 | 0 | 0 | 0 | 0 | 0 | 165200 | 244050 | 0 | 0 |
| STQLPGGE | PAAT | Q9H8K7 | PAAT_HUMAN | Unmodified | 1684700 | 187800 | 0 | 42811 | 0 | 0 | 83558 | 0 | 0 | 0 | 0 | 184900 | 0 | 0 | 0 | 37684 | 0 | 0 |
| STQYQELL | MACF1; MACF1 | Q9UPN3; H | MACF1_HUMA | Unmodified | 0 | 0 | 0 | 0 | 0 | 0 | 0 | 0 | 94452 | 204000 | 0 | 0 | 30491 | 39650 | 0 | 19699 | 67780 | 0 |
| STRESFNPI | SEPHS1; SEPHS | P49903; Q5 | SPS1_HUMAN | Acetyl (Protein N | 0 | 0 | 0 | 0 | 0 | 0 | 0 | 0 | 0 | 0 | 0 | 0 | 592700 | 0 | 0 | 0 | 0 | 0 |
| STSAGSTP | DTNB; DTNB; D | O60941; E7 | DTNB_HUMAN | Unmodified | 0 | 0 | 300860 | 365780 | 0 | 0 | 0 | 0 | 0 | 0 | 0 | 0 | 401650 | 0 | 0 | 431900 | 0 | 0 |
| STSAPQMS | UBAP2L; UBAP | Q14157; F8 | UBP2L_HUMAN | Unmodified | 0 | 0 | 0 | 0 | 350880 | 0 | 0 | 0 | 154950 | 59006 | 0 | 1255900 | 0 | 139390 | 0 | 0 | 242660 | 0 |
| STSCLELTK | CNNM4 | Q6P4Q7 | CNNM4_HUMA | Unmodified | 0 | 0 | 0 | 0 | 0 | 0 | 0 | 0 | 367340 | 150810 | 742250 | 969610 | 0 | 288740 | 0 | 0 | 0 | 0 |
| STSEGLLGI | UTRN | P46939 | UTRN_HUMAN | Unmodified | 0 | 0 | 24610 | 192200 | 492430 | 0 | 0 | 0 | 0 | 336960 | 0 | 0 | 498110 | 232890 | 298190 | 66407 | 439780 | 257540 |
| STSEPAVPI | ASPH; ASPH; A | Q12797; A0 | ASPH_HUMAN | Unmodified | 0 | 0 | 0 | 0 | 0 | 0 | 0 | 0 | 4184900 | 735460 | 0 | 0 | 0 | 0 | 0 | 0 | 0 | 0 |
| STSESFIQH | BCLAF1; BCLAF | Q9NYF8; E9 | BCLF1_HUMAN | Unmodified | 0 | 48513 | 0 | 114300 | 347760 | 85696 | 387560 | 0 | 604320 | 279030 | 380380 | 1326000 | 933170 | 338500 | 0 | 0 | 126950 | 333220 |
| STSFQTQL | BTBD17 | A6NE02 | BTBDH_HUMA | Unmodified | 0 | 0 | 0 | 0 | 0 | 0 | 0 | 0 | 2667700 | 0 | 0 | 0 | 0 | 0 | 0 | 0 | 0 | 0 |
| STSGPGMF | PACSIN1; PACS | Q9BY11; F6 | PACN1_HUMA | 2 Oxidation (M) | 21337000 | 11110000 | 2052300 | 1990900 | 0 | 0 | 0 | 0 | 0 | 0 | 0 | 0 | 1514500 | 0 | 5449000 | 759450 | 0 | 0 |
| STSGPRPG | TRAFD1 | O14545 | TRAD1_HUMAN | Unmodified | 484670 | 153960 | 24019 | 0 | 0 | 0 | 0 | 0 | 0 | 0 | 0 | 0 | 209050 | 96686 | 32318 | 25342 | 0 | 0 |
| STSIQSFK | PLS3; PLS3 | P13797; A0A | PLST_HUMAN | Unmodified | 203350 | 189230 | 0 | 0 | 52499 | 505450 | 0 | 39828 | 0 | 0 | 0 | 0 | 291130 | 210790 | 60774 | 97642 | 848200 | 460560 |
| STSKEPVDI | HSPB11; HSPB | Q9Y547; X6 | IFT25_HUMAN | Unmodified | 311900 | 0 | 350150 | 403810 | 732930 | 283920 | 559200 | 0 | 601900 | 646680 | 287530 | 1082300 | 787810 | 446940 | 720840 | 609220 | 1186600 | 576550 |
| STSLDVEPI | STRN | O43815 | STRN_HUMAN | Unmodified | 0 | 0 | 0 | 0 | 0 | 0 | 0 | 0 | 0 | 0 | 0 | 659200 | 0 | 0 | 0 | 0 | 0 | 0 |
| STSQSIVSL | USP43; USP43; | Q70EL4; V9 | UBP43_HUMAN | Unmodified | 0 | 0 | 0 | 645670 | 0 | 769190 | 62495 | 44944 | 0 | 214590 | 0 | 120360 | 1374100 | 0 | 0 | 0 | 0 | 0 |
| STSRIFLLTI | MYO1B; MYO1E | O43795; H7 | MYO1B_HUMA | Unmodified | 0 | 0 | 0 | 0 | 47394000 | 2382800 | 5055900 | 364360 | 0 | 1356700 | 0 | 0 | 666890 | 953310 | 0 | 0 | 10181000 | 6751800 |
| STSRQFVD | RACGAP1; RAC | Q9H0H5; F8 | RGAP1_HUMA | Unmodified | 0 | 0 | 0 | 0 | 2325300 | 304170 | 182890 | 73546 | 0 | 439290 | 0 | 0 | 0 | 190220 | 0 | 0 | 803880 | 272400 |
| STSRQFVD | RACGAP1; RAC | Q9H0H5; F8 | RGAP1_HUMA | Unmodified | 0 | 0 | 0 | 0 | 5789700 | 608570 | 0 | 0 | 0 | 88706 | 0 | 0 | 1506200 | 0 | 0 | 0 | 3452200 | 887380 |
| STSRVTAFL | NLN; NLN; NLN | Q9BYT8; E9 | NEUL_HUMAN | Unmodified | 0 | 0 | 0 | 0 | 1310200 | 244050 | 242250 | 0 | 162390 | 579660 | 55578 | 0 | 374750 | 233980 | 0 | 0 | 0 | 0 |
| STSSFPCP | LGALS3BP | Q08380 | LG3BP_HUMAN | Unmodified | 0 | 0 | 196180 | 0 | 304090 | 345160 | 200600 | 148180 | 33829000 | 3255800 | 16021000 | 10194000 | 6907300 | 3572800 | 102780 | 141310 | 204980 | 18747000 |
| STSSGHRE | ZC3HAV1; ZC3 | Q7Z2W4; C | ZCCHV_HUMA | Unmodified | 54903 | 0 | 0 | 0 | 354050 | 0 | 0 | 0 | 0 | 0 | 0 | 0 | 0 | 0 | 0 | 0 | 0 | 0 |
| STSTPNVH | RAF1; RAF1 | P04049; H7 | RAF1_HUMAN | Unmodified | 0 | 0 | 0 | 0 | 181210 | 0 | 0 | 0 | 251460 | 130260 | 0 | 0 | 295170 | 0 | 0 | 0 | 0 | 160000 |
| STSTQGLEI | DST; DST; DST; | Q03001; A0 | DYST_HUMAN | Unmodified | 24817 | 0 | 0 | 0 | 0 | 0 | 0 | 0 | 501750 | 0 | 0 | 606780 | 0 | 0 | 0 | 86022 | 0 | 0 |
| STSTQGLEI | DST; DST; DST; | Q03001; A0 | DYST_HUMAN | Unmodified | 0 | 0 | 0 | 0 | 1170000 | 254260 | 0 | 0 | 0 | 434370 | 0 | 0 | 848330 | 441400 | 75495 | 0 | 711890 | 774820 |
| STTDNDVR | RAI14 | Q9P0K7 | RAI14_HUMAN | Unmodified | 0 | 0 | 0 | 1209300 | 994130 | 0 | 0 | 0 | 0 | 0 | 0 | 0 | 0 | 0 | 0 | 0 | 1200500 | 1281600 |
| STTELPLTV | CLPTM1L; CLPT | Q96KA5; G5 | CLP1L_HUMAN | Unmodified | 0 | 0 | 0 | 0 | 0 | 0 | 0 | 0 | 1644500 | 180610 | 0 | 433710 | 0 | 0 | 0 | 0 | 0 | 0 |
| STTGPFDR | TMOD3 | Q9NYL9 | TMOD3_HUMA | Unmodified | 0 | 0 | 0 | 0 | 11037000 | 2851800 | 209760 | 0 | 1762600 | 668240 | 1288800 | 2460400 | 3868800 | 1594500 | 0 | 0 | 1215000 | 1861300 |
| STTLFSR | ACLY | P53396 | ACLY_HUMAN | Unmodified | 0 | 0 | 0 | 0 | 491410 | 134870 | 405860 | 0 | 1296100 | 84369 | 320360 | 7959200 | 0 | 0 | 327040 | 0 | 0 | 0 |
| STTLFSRHT | ACLY | P53396 | ACLY_HUMAN | Unmodified | 0 | 0 | 0 | 0 | 401160 | 0 | 1624800 | 504810 | 0 | 224050 | 0 | 0 | 0 | 0 | 0 | 0 | 209960 | 0 |
| STTLRVDQ | ATP2A1; ATP2A | O14983; P1 | AT2A1_HUMAN | Unmodified | 0 | 0 | 0 | 0 | 904530 | 0 | 1277500 | 0 | 0 | 0 | 0 | 0 | 0 | 0 | 0 | 0 | 0 | 0 |
| STTPPPAEF | G3BP2 | A0A7I2V647 | A0A7I2V647_H | Unmodified | 0 | 0 | 0 | 0 | 0 | 0 | 0 | 0 | 0 | 0 | 0 | 1735100 | 0 | 0 | 0 | 0 | 0 | 0 |
| STTQASSH | SARG | Q9BW04 | SARG_HUMAN | Unmodified | 0 | 0 | 121810 | 0 | 0 | 0 | 0 | 0 | 0 | 0 | 0 | 0 | 0 | 0 | 88489 | 162450 | 0 | 0 |
| STTQASSH | SARG | Q9BW04 | SARG_HUMAN | Unmodified | 30126 | 0 | 62757 | 163060 | 0 | 0 | 0 | 0 | 0 | 0 | 0 | 0 | 1433900 | 1049700 | 0 | 0 | 0 | 0 |
| STTQASSH | SARG | Q9BW04 | SARG_HUMAN | Unmodified | 0 | 0 | 0 | 0 | 0 | 0 | 0 | 0 | 0 | 0 | 0 | 0 | 7348100 | 3670000 | 0 | 0 | 0 | 0 |
| STTQVADS | EXOC4 | Q96A65 | EXOC4_HUMA | Unmodified | 0 | 0 | 0 | 0 | 0 | 0 | 0 | 0 | 0 | 0 | 0 | 387890 | 0 | 0 | 0 | 0 | 96403 | 0 |
| STTSDIPH | FAM98B; FAM9 | Q52LJ0; H0 | FA98B_HUMAN | Unmodified | 0 | 0 | 0 | 0 | 549470 | 174170 | 125620 | 0 | 0 | 0 | 226200 | 565280 | 519540 | 260880 | 0 | 0 | 132700 | 279390 |
| STTSITDAK | ANLN | Q9NQW6 | ANLN_HUMAN | Unmodified | 0 | 0 | 0 | 0 | 0 | 178870 | 0 | 0 | 0 | 0 | 0 | 0 | 0 | 0 | 0 | 0 | 796750 | 0 |
| STTSITDAK | ANLN | Q9NQW6 | ANLN_HUMAN | Unmodified | 0 | 0 | 0 | 0 | 385640 | 0 | 286580 | 186580 | 0 | 0 | 0 | 0 | 0 | 0 | 0 | 0 | 5833800 | 183650 |
| STTSVSEEI | PAWR | Q96IZ0 | PAWR_HUMAN | Unmodified | 0 | 277560 | 0 | 147590 | 0 | 0 | 0 | 0 | 0 | 0 | 58813 | 316220 | 0 | 0 | 0 | 151500 | 0 | 0 |
| STTTGHLIY | EEF1A2; EEF1A | Q05639; Q5 | EF1A2_HUMAN | Unmodified | 1.55E+08 | 1.19E+08 | 69594000 | 71446000 | 1.06E+08 | 6540900 | 20846000 | 9775600 | 22607000 | 11002000 | 11606000 | 83313000 | 77326000 | 33987000 | 1.29E+08 | 94322000 | 25616000 | 57925000 |
| STVAQLVK | ATP5F1A; ATP5 | P25705; K7E | ATPA_HUMAN | Unmodified | 0 | 0 | 0 | 0 | 0 | 0 | 387230 | 23536 | 7946800 | 1792900 | 0 | 499210 | 0 | 23757 | 0 | 0 | 0 | 0 |
| STVAQLVK | ATP5F1A; ATP5 | P25705; K7E | ATPA_HUMAN | Unmodified | 0 | 0 | 0 | 0 | 73542 | 0 | 524710 | 64600 | 9564500 | 2592600 | 125170 | 95129 | 0 | 177270 | 0 | 0 | 0 | 0 |
| STVDAPTA | CCT7 | Q99832 | TCPH_HUMAN | Unmodified | 0 | 0 | 54899 | 33692 | 0 | 49419 | 56977 | 0 | 410790 | 30023 | 27610 | 695330 | 80523 | 0 | 28858 | 49521 | 0 | 0 |
| STVDAVQG | RNF31; RNF31; | Q96EP0; H0 | RNF31_HUMAN | Unmodified | 0 | 730010 | 0 | 0 | 0 | 0 | 0 | 0 | 0 | 0 | 0 | 0 | 0 | 0 | 0 | 0 | 0 | 0 |
| STVEGIQAS | VCL | P18206 | VINC_HUMAN | Unmodified | 0 | 0 | 0 | 0 | 3031500 | 0 | 1943900 | 514510 | 394810 | 1783700 | 198480 | 2488700 | 0 | 0 | 0 | 307550 | 9336700 | 0 |
| STVEGIQAS | VCL | P18206 | VINC_HUMAN | Unmodified | 0 | 0 | 0 | 0 | 0 | 0 | 570390 | 0 | 0 | 0 | 0 | 0 | 0 | 0 | 0 | 0 | 2319200 | 0 |
| STVFGTALI | LSS; LSS | P48449; C9 | LSS_HUMAN La | Unmodified | 52193 | 0 | 0 | 0 | 0 | 0 | 0 | 0 | 45203 | 24368 | 0 | 546240 | 0 | 0 | 0 | 0 | 0 | 0 |
| STVFTIFGS | SORL1 | Q92673 | SORL_HUMAN | Unmodified | 0 | 0 | 0 | 0 | 0 | 0 | 0 | 0 | 3100400 | 425540 | 0 | 0 | 0 | 0 | 0 | 0 | 0 | 0 |
| STVGLQCL | SH3BP4 | Q9P0V3 | SH3B4_HUMA | Unmodified | 0 | 0 | 0 | 0 | 292370 | 181460 | 213700 | 24940 | 0 | 396760 | 0 | 359340 | 0 | 0 | 143780 | 0 | 0 | 0 |
| STVGVDFK | RAB12; RAB12 | Q6IQ22; A0A | RAB12_HUMAN | Unmodified | 0 | 0 | 125710 | 655130 | 0 | 0 | 0 | 0 | 0 | 0 | 0 | 0 | 495770 | 0 | 0 | 0 | 0 | 0 |
| STVHEILCK | ANXA2P2; ANX | A6NMY6; P0 | AXA2L_HUMAN | Acetyl (Protein N | 2157000 | 884610 | 0 | 1147000 | 8.81E+08 | 99173000 | 6839700 | 5504000 | 13619000 | 10601000 | 494100 | 0 | 2309500 | 4772100 | 338700 | 665230 | 3.59E+08 | 3.85E+08 |
| STVHEILCK | ANXA2P2; ANX | A6NMY6; P0 | AXA2L_HUMAN | Acetyl (Protein N | 0 | 0 | 0 | 0 | 1.53E+08 | 8921800 | 14422000 | 174320 | 0 | 4806100 | 0 | 0 | 470130 | 1312000 | 0 | 0 | 8.45E+08 | 77112000 |
| STVISGPG | HSH2D | Q96JZ2 | HSH2D_HUMA | Unmodified | 0 | 0 | 0 | 0 | 88137 | 0 | 0 | 0 | 143510 | 0 | 33032 | 0 | 0 | 0 | 0 | 278660 | 0 | 51627 |
| STVLGLPQ | KIAA1522 | Q9P206 | K1522_HUMAN | Unmodified | 0 | 0 | 1048300 | 1781600 | 1880300 | 0 | 0 | 0 | 736420 | 0 | 0 | 932280 | 4548900 | 2590700 | 2351500 | 1579600 | 747800 | 1852500 |
| STVLSLDW | ARPC1B; ARPC | O15143; Q9 | ARC1B_HUMA | Unmodified | 0 | 0 | 0 | 0 | 3003000 | 0 | 0 | 0 | 0 | 0 | 0 | 787610 | 0 | 0 | 0 | 0 | 0 | 0 |
| STVPTDFS | CD47; CD47 | Q08722; A0 | CD47_HUMAN | Unmodified | 2202800 | 1374300 | 5180500 | 2532100 | 1464500 | 110220 | 0 | 3154800 | 45989000 | 6256700 | 18307000 | 1.36E+08 | 1176200 | 0 | 5042000 | 5195400 | 0 | 1910700 |
| STVPTDFS | CD47; CD47 | Q08722; A0 | CD47_HUMAN | Unmodified | 2377100 | 6519600 | 43403 | 11837000 | 9547000 | 1218600 | 764590 | 40942 | 70905000 | 12779000 | 38477000 | 44907000 | 11657000 | 3976700 | 2746600 | 3893800 | 1169500 | 10699000 |
| STVPTDFS | CD47; CD47 | Q08722; A0 | CD47_HUMAN | Unmodified | 239900 | 0 | 0 | 1302500 | 8206800 | 220220 | 0 | 0 | 4342100 | 3209600 | 1700300 | 241720 | 3501600 | 1710400 | 0 | 0 | 347290 | 2680100 |
| STVPVRAA | UGDH; UGDH; | O60701; E7 | UGDH_HUMAN | Unmodified | 0 | 0 | 0 | 40948 | 15824000 | 64458 | 327600 | 128440 | 0 | 769440 | 0 | 0 | 0 | 1714300 | 0 | 0 | 897120 | 1487100 |
| STVRDIDP | DYNLRB2; DYN | Q8TF09; Q9 | DLRB2_HUMAN | Unmodified | 0 | 0 | 0 | 563030 | 448290 | 0 | 181770 | 0 | 1997000 | 466120 | 730690 | 1798300 | 603660 | 194750 | 0 | 0 | 1084000 | 867880 |
| STVRDIDP | DYNLRB2; DYN | Q8TF09; Q9 | DLRB2_HUMAN | Unmodified | 0 | 0 | 0 | 0 | 194190 | 85184 | 71700 | 0 | 0 | 120940 | 0 | 0 | 440060 | 136790 | 0 | 0 | 439250 | 188760 |
| STVSETFM | SRGAP2; SRGA | O75044; P0 | SRGP2_HUMA | Unmodified | 100560 | 0 | 0 | 0 | 2659100 | 0 | 0 | 0 | 646570 | 291060 | 0 | 1111300 | 0 | 0 | 326940 | 100010 | 682720 | 921530 |
| STVSETFM | SRGAP2; SRGA | O75044; P0 | SRGP2_HUMA | Oxidation (M) | 0 | 0 | 0 | 0 | 0 | 10451 | 0 | 0 | 0 | 0 | 0 | 209070 | 1103500 | 0 | 0 | 0 | 443470 | 0 |
| STVSETYLS | SRGAP1; SRGA | Q7Z6B7; G5 | SRGP1_HUMA | Unmodified | 0 | 0 | 0 | 0 | 886200 | 0 | 215920 | 0 | 0 | 0 | 0 | 0 | 0 | 0 | 0 | 0 | 289350 | 0 |
| STVVPVPY | GTF2I; GTF2I | P78347; A0A | GTF2I_HUMAN | Unmodified | 0 | 0 | 84454 | 0 | 276150 | 0 | 314630 | 203710 | 0 | 0 | 173140 | 2994100 | 0 | 0 | 61328 | 0 | 479680 | 0 |
| STVYIIENA | ATP11C; ATP11 | Q8NB49; A0 | AT11C_HUMAN | Unmodified | 0 | 0 | 376690 | 0 | 0 | 0 | 0 | 56067 | 501720 | 68061 | 266140 | 3716600 | 0 | 0 | 557880 | 537550 | 0 | 0 |
| STYDGLHV | CNKSR2; CNK3 | Q8WXI2; G9 | CNKR2_HUMA | Unmodified | 0 | 0 | 0 | 0 | 0 | 0 | 0 | 0 | 1634600 | 0 | 0 | 1165900 | 0 | 0 | 0 | 0 | 0 | 0 |
| STYDGLHV | CNK3/IPCEF1; | G9CGD6; Q | CNIPF_HUMAN | Unmodified | 0 | 0 | 0 | 0 | 0 | 0 | 0 | 0 | 1886200 | 1039500 | 1529300 | 442560 | 0 | 0 | 0 | 0 | 0 | 0 |
| STYEQVDLI | DDX21 | Q9NR30 | DDX21_HUMA | Unmodified | 0 | 0 | 0 | 0 | 0 | 0 | 756320 | 781930 | 0 | 0 | 0 | 0 | 0 | 0 | 0 | 0 | 0 | 0 |
| STYNHLSS | RAB14; RAB14 | P61106; X6 | RAB14_HUMAN | Unmodified | 785620 | 511630 | 959740 | 1125400 | 143990 | 0 | 0 | 0 | 1046700 | 190330 | 0 | 7052100 | 260650 | 0 | 806690 | 514690 | 0 | 0 |
| STYPRPHE | CTSZ; CTSZ; CT | Q9UBR2; A0 | CATZ_HUMAN | Unmodified | 0 | 0 | 401760 | 690770 | 434850 | 0 | 0 | 0 | 2342100 | 955850 | 0 | 0 | 1378500 | 0 | 1145200 | 1165600 | 0 | 0 |
| STYTSYPK | HGS; HGS; HG | O14964; A0 | HGS_HUMAN | Unmodified | 436130 | 0 | 129730 | 0 | 0 | 0 | 284120 | 80439 | 495050 | 247930 | 0 | 158010 | 0 | 0 | 124180 | 0 | 482580 | 1387400 |
| SVAAIHPSL | PRPSAP2; PRPS | O60256; E7 | KPRB_HUMAN | Oxidation (M) | 0 | 0 | 0 | 0 | 1022200 | 387830 | 0 | 318210 | 1430700 | 711680 | 1636600 | 0 | 1151800 | 0 | 0 | 268120 | 528540 | 518970 |
| SVACDVGY | ITGA2; ITGA2 | P17301; E7 | ITA2_HUMAN | Unmodified | 210830 | 115470 | 267060 | 0 | 230030 | 101830 | 0 | 0 | 0 | 189750 | 217710 | 2212100 | 0 | 0 | 446760 | 278400 | 0 | 0 |
| SVACDVGY | ITGA2; ITGA2 | P17301; E7 | ITA2_HUMAN | Unmodified | 481450 | 0 | 276270 | 501600 | 0 | 0 | 0 | 0 | 0 | 0 | 0 | 0 | 0 | 0 | 1338500 | 1216200 | 0 | 0 |
| SVAGGEIR | DCTPP1 | Q9H773 | DCTP1_HUMA | Acetyl (Protein N | 0 | 0 | 0 | 0 | 0 | 0 | 0 | 0 | 73721 | 30421 | 0 | 2267200 | 0 | 0 | 0 | 0 | 0 | 127620 |
| SVAGIGQT | SCARB1; SCAR | Q8WTV0; F5 | SCRB1_HUMA | Unmodified | 499570 | 230500 | 328220 | 0 | 433780 | 118670 | 0 | 0 | 144030 | 100020 | 0 | 1665300 | 0 | 0 | 628300 | 543420 | 42948 | 0 |
| SVALCVAA | SLC44A1 | Q8WWI5 | CTL1_HUMAN | Unmodified | 0 | 0 | 0 | 0 | 0 | 0 | 0 | 0 | 1350300 | 0 | 0 | 1772900 | 0 | 0 | 0 | 0 | 0 | 374460 |
| SVALCVAA | SLC44A1 | Q8WWI5 | CTL1_HUMAN | Unmodified | 1489700 | 9955.5 | 0 | 1065700 | 7717900 | 0 | 1602100 | 0 | 940480 | 4056500 | 0 | 0 | 19393000 | 6363800 | 0 | 0 | 1348400 | 2861800 |
| SVAQQASL | RAP1GDS1 | P52306 | GDS1_HUMAN | Unmodified | 0 | 0 | 0 | 0 | 0 | 0 | 0 | 0 | 0 | 0 | 0 | 2098600 | 0 | 0 | 0 | 0 | 0 | 0 |
| SVAQQASL | RAP1GDS1 | P52306 | GDS1_HUMAN | Unmodified | 0 | 0 | 0 | 0 | 106980 | 0 | 950050 | 0 | 0 | 0 | 0 | 0 | 0 | 0 | 0 | 0 | 0 | 0 |
| SVASGGW | DCTN4; DCTN4 | Q9UJW0; E5 | DCTN4_HUMA | Unmodified | 55194 | 61970 | 0 | 151520 | 0 | 0 | 0 | 0 | 0 | 0 | 0 | 1962600 | 0 | 0 | 259150 | 128430 | 0 | 0 |
| SVASIGNTI | EFCAB14; EFCA | O75071; A0 | EFC14_HUMAN | Unmodified | 49204 | 0 | 0 | 0 | 0 | 0 | 0 | 0 | 20865000 | 8214200 | 0 | 0 | 0 | 0 | 0 | 0 | 481060 | 0 |
| SVASSLRD | SCNN1A; SCNN | P37088; C5 | SCNNA_HUMA | Unmodified | 0 | 0 | 0 | 0 | 168340 | 0 | 0 | 0 | 1040500 | 67124 | 0 | 0 | 0 | 0 | 0 | 0 | 0 | 0 |
| SVASSQPA | TJP1; TJP1; TJP1 | Q07157; A0 | ZO1_HUMAN T | Unmodified | 0 | 0 | 0 | 0 | 1848400 | 564090 | 0 | 0 | 0 | 101280 | 0 | 0 | 0 | 0 | 0 | 0 | 203310 | 0 |
| SVATGPMT | GPRIN1 | Q7Z2K8 | GRIN1_HUMAN | Unmodified | 0 | 0 | 0 | 0 | 0 | 0 | 0 | 50864 | 1001000 | 0 | 105260 | 0 | 777970 | 0 | 0 | 0 | 0 | 0 |
| SVDALDDL | LSR; LSR | Q86X29; S4 | LSR_HUMAN Li | Unmodified | 1375900 | 0 | 2498900 | 1739200 | 299200 | 0 | 0 | 189620 | 9435100 | 1854800 | 1502900 | 2720100 | 323040 | 273960 | 3218600 | 3679900 | 80270 | 0 |
| SVDALDDL | LSR; LSR | Q86X29; S4 | LSR_HUMAN Li | Unmodified | 0 | 0 | 0 | 486130 | 3049800 | 614380 | 0 | 0 | 369830 | 0 | 0 | 0 | 1247800 | 554580 | 0 | 0 | 0 | 0 |
| SVDCFLHD | STXBP3 | O00186 | STXB3_HUMAN | Unmodified | 0 | 0 | 0 | 0 | 523680 | 0 | 84622 | 15047 | 0 | 0 | 0 | 0 | 509290 | 525660 | 0 | 0 | 282980 | 1026800 |
| SVDCFLHD | STXBP3 | O00186 | STXB3_HUMAN | Unmodified | 0 | 0 | 0 | 0 | 4037200 | 557670 | 608540 | 0 | 279230 | 0 | 0 | 152670 | 4333600 | 6336700 | 0 | 0 | 8595400 | 7322100 |

| | | | | | | | | | | | | | | | | | | | | | | |
|---|---|---|---|---|---|---|---|---|---|---|---|---|---|---|---|---|---|---|---|---|---|---|
| SVDCFLHD | STXBP3 | O00186 | STXB3_HUMAN | Unmodified | 0 | 0 | 0 | 0 | 365800 | 0 | 466540 | 37698 | 0 | 0 | 0 | 0 | 0 | 2003900 | 0 | 0 | 4161500 | 1931900 |
| SVDDTSQA | COLEC12 | Q5KU26 | COL12_HUMAN | Unmodified | 0 | 0 | 0 | 670240 | 84088 | 0 | 72495 | 0 | 0 | 48210 | 0 | 0 | 0 | 0 | 0 | 224840 | 66663 | 362580 |
| SVDETLRLV | PRDX4; PRDX4 | Q13162; H7 | PRDX4_HUMAN | Unmodified | 0 | 0 | 0 | 0 | 0 | 0 | 0 | 0 | 25969000 | 20931000 | 13061000 | 0 | 342900 | 0 | 0 | 0 | 0 | 0 |
| SVDEVFDE | CMPK1; CMPK1 | P30085; A0A | KCY_HUMAN U | Unmodified | 0 | 0 | 0 | 0 | 55440 | 0 | 0 | 0 | 0 | 16176 | 0 | 0 | 128660 | 79393 | 0 | 0 | 0 | 15762 |
| SVDFDSLTY | CD2AP | Q9Y5K6 | CD2AP_HUMAN | Unmodified | 2197000 | 3251400 | 11669000 | 9330100 | 0 | 290010 | 0 | 26623 | 1662300 | 0 | 754680 | 15440000 | 2589300 | 0 | 16013000 | 10971000 | 100390 | 0 |
| SVDGPIRLF | GSPT1; GSPT1 | P15170; H3 | ERF3A_HUMAN | Unmodified | 0 | 0 | 119050 | 0 | 0 | 0 | 0 | 0 | 0 | 0 | 258390 | 1506800 | 0 | 0 | 270740 | 0 | 0 | 0 |
| SVDIWLR | PVR; PVR; PVR | P15151; A0A | PVR_HUMAN P | Unmodified | 0 | 0 | 0 | 0 | 0 | 0 | 0 | 0 | 0 | 0 | 3601400 | 11099000 | 0 | 0 | 0 | 0 | 0 | 0 |
| SVDLAEYAI | PPFIBP2; PPFIB | Q8ND30; Q8 | LIPB2_HUMAN | Unmodified | 0 | 0 | 0 | 686020 | 1402200 | 0 | 946710 | 0 | 0 | 0 | 0 | 1278500 | 0 | 0 | 0 | 0 | 2004300 | 878390 |
| SVDLQTR | GOLM1; GOLM1 | Q8NBJ4; F8 | GOLM1_HUMA | Unmodified | 1448200 | 1557300 | 0 | 0 | 0 | 0 | 0 | 0 | 5095500 | 372700 | 17400 | 644280 | 77432 | 0 | 21663 | 0 | 0 | 0 |
| SVDNRPQA | ARFGEF1; ARFG | Q9Y6D6; E5 | BIG1_HUMAN I | Unmodified | 0 | 0 | 0 | 0 | 0 | 0 | 0 | 0 | 755810 | 0 | 0 | 0 | 0 | 0 | 0 | 0 | 0 | 0 |
| SVDPATFH | RTN4R; RTN4R; | Q9BZR6; H7 | RTN4R_HUMAN | Unmodified | 368700 | 0 | 0 | 305600 | 0 | 0 | 0 | 0 | 732860 | 0 | 0 | 0 | 0 | 0 | 164120 | 96692 | 0 | 0 |
| SVDPDSPA | SLC9A3R1 | O14745 | NHRF1_HUMAN | Unmodified | 0 | 0 | 761260 | 772530 | 0 | 0 | 0 | 0 | 0 | 69661 | 529380 | 8114600 | 837610 | 271140 | 1879800 | 2091200 | 89312 | 0 |
| SVDPDSPA | SLC9A3R1 | O14745 | NHRF1_HUMAN | Unmodified | 0 | 0 | 757630 | 827480 | 0 | 0 | 0 | 0 | 741410 | 0 | 231280 | 3705100 | 739990 | 0 | 1680400 | 1631300 | 0 | 0 |
| SVDPDSPA | SLC9A3R1 | O14745 | NHRF1_HUMAN | Unmodified | 0 | 0 | 0 | 705690 | 3608500 | 0 | 0 | 0 | 0 | 0 | 0 | 0 | 10961000 | 6410400 | 389080 | 95445 | 1079100 | 0 |
| SVDPGSPA | SLC9A3R2; SLC | Q15599; H3 | NHRF2_HUMAN | Unmodified | 269000 | 264140 | 690550 | 1241300 | 2299100 | 648800 | 0 | 0 | 845910 | 307340 | 0 | 1947300 | 3167300 | 1723100 | 1086800 | 836430 | 1249500 | 2410900 |
| SVDRTSLH | ASAP2 | O43150 | ASAP2_HUMAN | Unmodified | 79122 | 0 | 0 | 0 | 839770 | 438430 | 0 | 0 | 0 | 187200 | 129620 | 0 | 469190 | 0 | 0 | 57227 | 471550 | 468420 |
| SVDSSGET | PIP5K1A; PIP5K | Q99755; A2 | PI51A_HUMAN | Unmodified | 0 | 0 | 0 | 0 | 561730 | 0 | 0 | 42957 | 0 | 20791 | 16056 | 541630 | 0 | 0 | 81151 | 0 | 0 | 1086000 |
| SVDSTATAI | ERBIN | Q96RT1 | ERBIN_HUMAN | Unmodified | 0 | 0 | 240870 | 957490 | 21332000 | 3284800 | 0 | 87301 | 1154600 | 243380 | 386660 | 733800 | 2636900 | 2911600 | 410770 | 265380 | 1565100 | 4387600 |
| SVDSTATAI | ERBIN | Q96RT1 | ERBIN_HUMAN | Oxidation (M) | 0 | 0 | 0 | 0 | 4811800 | 371470 | 0 | 0 | 0 | 0 | 0 | 0 | 525080 | 0 | 0 | 25880 | 0 | 1418500 |
| SVEAAAELS | PTMS; PTMS | P20962; F5G | PTMS_HUMAN | Unmodified | 1270000 | 216810 | 0 | 484830 | 6673500 | 1258400 | 907680 | 2896100 | 1903600 | 7805800 | 7347200 | 28465000 | 18332000 | 7221200 | 32913 | 504950 | 2516000 | 1258000 |
| SVEAAAELS | PTMS; PTMS | P20962; F5G | PTMS_HUMAN | Unmodified | 0 | 80661 | 0 | 0 | 0 | 0 | 950980 | 797110 | 0 | 0 | 1170400 | 960430 | 1301000 | 1947500 | 0 | 0 | 0 | 0 |
| SVEDLLDRI | SASH1; SASH1 | O94885; A0 | SASH1_HUMAN | Unmodified | 0 | 0 | 0 | 0 | 376300 | 0 | 0 | 0 | 236810 | 207390 | 0 | 0 | 210510 | 0 | 0 | 0 | 244820 | 232950 |
| SVEDRFDQ | DSP | P15924 | DESP_HUMAN | Unmodified | 0 | 0 | 0 | 0 | 446780 | 0 | 0 | 72264 | 348780 | 1306000 | 0 | 0 | 1089100 | 1635100 | 0 | 0 | 2184500 | 1223600 |
| SVEEISTLV | MRPL43; MRPL4 | Q8N983; H0 | RM43_HUMAN | Unmodified | 0 | 0 | 0 | 0 | 0 | 0 | 0 | 0 | 793020 | 173550 | 0 | 0 | 0 | 0 | 0 | 0 | 0 | 0 |
| SVEETLR | PRDX3 | P30048 | PRDX3_HUMAN | Unmodified | 0 | 0 | 0 | 0 | 0 | 0 | 0 | 0 | 14392000 | 415550 | 0 | 143070 | 0 | 0 | 0 | 0 | 122410 | 0 |
| SVEEVASEI | DSP | P15924 | DESP_HUMAN | Unmodified | 0 | 0 | 0 | 0 | 124780 | 0 | 0 | 0 | 1762300 | 253950 | 0 | 215330 | 0 | 0 | 71587 | 0 | 0 | 0 |
| SVEGLQEG | CLUH; CLUH; C | O75153; I3L | CLU_HUMAN C | Unmodified | 0 | 0 | 0 | 0 | 0 | 0 | 0 | 0 | 130250 | 0 | 248240 | 3485900 | 0 | 0 | 0 | 0 | 0 | 0 |
| SVEGRPLW | CPD | O75976 | CBPD_HUMAN | Unmodified | 0 | 0 | 0 | 360980 | 0 | 0 | 0 | 0 | 67670000 | 4905700 | 493030 | 354590 | 0 | 0 | 0 | 0 | 0 | 0 |
| SVELEDVK | AP1M1; AP1M2 | Q9BXS5; Q9 | AP1M1_HUMAN | Unmodified | 101340 | 0 | 404070 | 701020 | 71573 | 0 | 0 | 0 | 247720 | 0 | 166450 | 1126400 | 0 | 25505 | 113950 | 583860 | 0 | 79217 |
| SVELEEALF | H1-10 | Q92522 | H1X_HUMAN H | Acetyl (Protein N | 0 | 0 | 0 | 0 | 0 | 0 | 0 | 1283900 | 0 | 0 | 0 | 0 | 0 | 0 | 0 | 0 | 0 | 0 |
| SVELSWQE | EPHA7; EPHA7; | Q15375; A0 | EPHA7_HUMAN | Unmodified | 98262 | 0 | 0 | 90411 | 0 | 0 | 0 | 0 | 1766400 | 0 | 0 | 0 | 0 | 0 | 0 | 0 | 0 | 0 |
| SVEMHHEA | EEF1A2; EEF1A | Q05639; P6 | EF1A2_HUMAN | Oxidation (M) | 16885000 | 17414000 | 999750 | 2553000 | 8847200 | 1069000 | 1231000 | 637860 | 3897000 | 2150900 | 0 | 477950 | 4763400 | 1611800 | 1149200 | 1153700 | 4855200 | 6922100 |
| SVEMHHEA | EEF1A2; EEF1A | Q05639; P6 | EF1A2_HUMAN | Unmodified | 3078800 | 2852600 | 0 | 0 | 10547000 | 1856900 | 440520 | 99883 | 5186100 | 561130 | 0 | 0 | 1642000 | 577130 | 0 | 68555 | 3033300 | 7459000 |
| SVENLATAT | PIGS | Q96S52 | PIGS_HUMAN ( | Unmodified | 0 | 0 | 0 | 0 | 0 | 0 | 0 | 0 | 103880 | 177410 | 0 | 0 | 0 | 0 | 0 | 0 | 0 | 0 |
| SVEQQVIG | BAG5 | Q9UL15 | BAG5_HUMAN | Unmodified | 0 | 245950 | 151670 | 226570 | 430700 | 111010 | 234200 | 87713 | 4465700 | 2183100 | 102370 | 0 | 1212700 | 0 | 196340 | 0 | 0 | 0 |
| SVEQQVIG | BAG5 | Q9UL15 | BAG5_HUMAN | Unmodified | 443160 | 1037200 | 0 | 0 | 2287900 | 277050 | 0 | 136360 | 0 | 3671000 | 0 | 0 | 2144400 | 1850200 | 251630 | 321500 | 524610 | 0 |
| SVESRELYV | CPD | O75976 | CBPD_HUMAN | Unmodified | 0 | 0 | 0 | 0 | 1963200 | 0 | 0 | 0 | 0 | 3397200 | 0 | 0 | 0 | 0 | 0 | 0 | 0 | 0 |
| SVESRELYV | CPD | O75976 | CBPD_HUMAN | Oxidation (M) | 0 | 0 | 229670 | 642870 | 1035300 | 0 | 0 | 0 | 306650 | 5289300 | 0 | 0 | 220100 | 0 | 567800 | 0 | 0 | 0 |
| SVESVPKLV | ADH5 | P11766 | ADHX_HUMAN | Unmodified | 0 | 0 | 0 | 0 | 113820 | 0 | 210400 | 0 | 473430 | 0 | 0 | 0 | 0 | 0 | 0 | 0 | 439220 | 232510 |
| SVETDSSTV | CYFIP2; CYFIP2 | Q96F07; H7 | CYFP2_HUMAN | Unmodified | 0 | 0 | 605960 | 1878100 | 0 | 0 | 144680 | 75043 | 1725100 | 464500 | 757210 | 0 | 1160700 | 546540 | 1618400 | 1081800 | 0 | 99163 |
| SVETGHLD | TNIK; MINK1; M | Q9UKE5; Q8 | TNIK_HUMAN T | Unmodified | 0 | 0 | 0 | 0 | 0 | 0 | 117830 | 0 | 0 | 0 | 0 | 599590 | 137710 | 0 | 0 | 0 | 0 | 0 |
| SVETLKEMI | PKM; PKM; PKM | P14618; H3 | KPYM_HUMAN | Unmodified | 0 | 0 | 0 | 0 | 1257200 | 0 | 0 | 102820 | 0 | 0 | 0 | 0 | 0 | 0 | 0 | 0 | 1632400 | 1312900 |
| SVEVHGAV | LAMA5 | O15230 | LAMA5_HUMAN | Unmodified | 68212 | 0 | 0 | 0 | 0 | 79077 | 0 | 0 | 0 | 0 | 0 | 121850 | 0 | 0 | 0 | 0 | 90276 | 622580 |
| SVEVQGET | USO1 | O60763 | USO1_HUMAN | Unmodified | 0 | 151160 | 0 | 0 | 0 | 0 | 0 | 82037 | 1094900 | 0 | 567930 | 4537900 | 689350 | 0 | 0 | 0 | 202220 | 640000 |
| SVFALTNGI | RPL6 | Q02878 | RL6_HUMAN 6 | Unmodified | 1691600 | 693120 | 411930 | 542150 | 95094 | 0 | 33728 | 21694 | 104980 | 45468 | 385010 | 13696000 | 320530 | 104140 | 465680 | 455850 | 0 | 141870 |
| SVFALTNGI | RPL6 | Q02878 | RL6_HUMAN 6 | Unmodified | 0 | 0 | 0 | 0 | 805930 | 0 | 332210 | 0 | 0 | 285680 | 19985 | 0 | 4264400 | 0 | 0 | 0 | 260960 | 0 |
| SVFGKLFG. | CHMP4B | Q9H444 | CHM4B_HUMA | Acetyl (Protein N | 0 | 0 | 0 | 0 | 0 | 0 | 644340 | 323030 | 0 | 340400 | 0 | 578540 | 0 | 0 | 0 | 0 | 1384800 | 542360 |
| SVFIIFFQGI | ATP6V0A1; ATP | Q93050; B7 | VPP1_HUMAN | Unmodified | 0 | 0 | 0 | 0 | 0 | 0 | 43244 | 0 | 123880 | 536200 | 0 | 16644 | 0 | 0 | 36237 | 0 | 0 | 0 |
| SVFILGASG | HTATIP2; HTATI | Q9BUP3; E9 | HTAI2_HUMAN | Unmodified | 0 | 0 | 0 | 0 | 0 | 0 | 0 | 0 | 4759300 | 0 | 0 | 572270 | 0 | 0 | 0 | 0 | 0 | 0 |
| SVFILGASG | HTATIP2; HTATI | Q9BUP3; E9 | HTAI2_HUMAN | Unmodified | 0 | 0 | 0 | 0 | 6201300 | 1745500 | 2666300 | 310840 | 3426000 | 6321900 | 0 | 0 | 4449900 | 1724700 | 0 | 0 | 3109200 | 3330800 |
| SVFLRTDQI | CAPN5; CAPN5 | O15484; E7 | CAN5_HUMAN | Unmodified | 0 | 0 | 227410 | 456970 | 295340 | 0 | 0 | 0 | 0 | 150040 | 0 | 466740 | 748210 | 480100 | 428960 | 461530 | 159580 | 0 |
| SVFPEQAN | PSMD3 | O43242 | PSMD3_HUMA | Unmodified | 0 | 0 | 322480 | 0 | 0 | 0 | 0 | 32952 | 0 | 0 | 0 | 5001200 | 0 | 0 | 274720 | 145760 | 0 | 0 |
| SVFVGNIPY | CSTF2T; CSTF2 | Q9H0L4; P3 | CSTFT_HUMAN | Unmodified | 0 | 0 | 0 | 0 | 0 | 0 | 0 | 0 | 0 | 0 | 375550 | 1218600 | 0 | 0 | 0 | 0 | 2208200 | 0 |
| SVFVGNLP | RBM34; RBM34 | P42696; A2A | RBM34_HUMAN | Unmodified | 0 | 0 | 0 | 0 | 0 | 0 | 765350 | 98407 | 0 | 0 | 0 | 310070 | 0 | 0 | 0 | 0 | 0 | 0 |
| SVGAPGGA | KIAA1522 | Q9P206 | K1522_HUMAN | Unmodified | 0 | 0 | 0 | 174340 | 0 | 0 | 0 | 0 | 0 | 0 | 0 | 0 | 402330 | 327710 | 436930 | 420820 | 245120 | 0 |
| SVGDGETV | YBX1; YBX2; YB | P67809; Q9 | YBOX1_HUMAN | Unmodified | 3083700 | 1741000 | 1298600 | 3085300 | 0 | 0 | 0 | 373470 | 0 | 0 | 2906900 | 11475000 | 0 | 0 | 981160 | 1463000 | 0 | 0 |
| SVGDGETV | YBX1; YBX1 | P67809; H0 | YBOX1_HUMAN | Unmodified | 0 | 0 | 0 | 958440 | 0 | 0 | 0 | 0 | 0 | 0 | 0 | 0 | 0 | 0 | 0 | 0 | 0 | 0 |
| SVGDLAPP | CTSF; CTSF; CT | Q9UBX1; A0 | CATF_HUMAN | Unmodified | 0 | 0 | 0 | 0 | 0 | 0 | 0 | 0 | 1732400 | 0 | 0 | 0 | 0 | 0 | 0 | 0 | 0 | 0 |
| SVGFIGAG | PYCR2; PYCR2; | Q96C36; A0 | P5CR2_HUMAN | Acetyl (Protein N | 0 | 0 | 0 | 0 | 0 | 0 | 0 | 0 | 0 | 110740 | 0 | 0 | 0 | 0 | 0 | 0 | 0 | 0 |
| SVGGACVL | RPL27A; RPL27 | P46776; E9P | RL27A_HUMAN | Unmodified | 438630 | 0 | 291470 | 0 | 0 | 0 | 849480 | 503730 | 0 | 0 | 921970 | 4956900 | 589190 | 0 | 0 | 156350 | 0 | 0 |
| SVGGIRASI | PSAT1 | Q9Y617 | SERC_HUMAN | Unmodified | 0 | 0 | 0 | 0 | 1614400 | 130620 | 0 | 0 | 0 | 167010 | 64812 | 0 | 1290100 | 792210 | 0 | 0 | 0 | 617750 |
| SVGLANQE | ARG2 | P78540 | ARGI2_HUMAN | Unmodified | 0 | 0 | 0 | 0 | 0 | 0 | 0 | 0 | 874490 | 0 | 0 | 0 | 0 | 0 | 0 | 0 | 0 | 0 |
| SVGLEVYT | GPAA1; GPAA1; | O43292; E9 | GPAA1_HUMAN | Unmodified | 0 | 0 | 0 | 0 | 0 | 422590 | 0 | 0 | 9562700 | 2951700 | 113350 | 0 | 0 | 0 | 0 | 0 | 0 | 0 |
| SVGMIAGG | CYB5R3 | P00387 | NB5R3_HUMAN | 2 Oxidation (M) | 219010 | 85970 | 298970 | 445650 | 1004000 | 326890 | 871580 | 393890 | 188640 | 582870 | 510580 | 822050 | 377010 | 54318 | 178760 | 445900 | 319750 | 455800 |
| SVGNTIDPY | MAP1B; MAP1B | P46821; D6 | MAP1B_HUMAN | Unmodified | 0 | 0 | 0 | 0 | 0 | 0 | 521720 | 406720 | 960150 | 745320 | 186400 | 3274400 | 0 | 0 | 0 | 0 | 0 | 0 |
| SVGPLPQA | MLPH; MLPH; M | Q9BV36; H7 | MELPH_HUMAN | Unmodified | 0 | 0 | 0 | 0 | 0 | 0 | 0 | 0 | 0 | 0 | 0 | 0 | 542110 | 0 | 0 | 129030 | 267400 | 614720 |
| SVGYKPDF | HPRT1 | P00492 | HPRT_HUMAN | Unmodified | 430260 | 0 | 0 | 0 | 464330 | 0 | 0 | 71932 | 2457000 | 0 | 1759800 | 6354700 | 0 | 0 | 0 | 0 | 366030 | 588930 |
| SVGYPHPD | NPTN; NPTN; N | Q9Y639; Q9 | NPTN_HUMAN | Unmodified | 0 | 0 | 0 | 0 | 2572300 | 0 | 0 | 0 | 4582500 | 884690 | 8906300 | 16344000 | 0 | 0 | 0 | 0 | 0 | 1972600 |
| SVGYPHPD | NPTN; NPTN; N | Q9Y639; Q9 | NPTN_HUMAN | Unmodified | 0 | 0 | 367060 | 0 | 0 | 0 | 0 | 0 | 0 | 0 | 0 | 1650100 | 588570 | 355010 | 0 | 546930 | 0 | 0 |
| SVGYPHPD | NPTN; NPTN; N | Q9Y639; Q9 | NPTN_HUMAN | Unmodified | 0 | 0 | 0 | 0 | 9705400 | 521610 | 1163100 | 0 | 0 | 936160 | 875320 | 272960 | 4251500 | 4421000 | 0 | 0 | 3370600 | 10192000 |
| SVHAWEIS | TNPO3; TNPO3 | Q9Y5L0; C9 | TNPO3_HUMAN | Unmodified | 448560 | 59074 | 215190 | 0 | 0 | 0 | 0 | 0 | 1290800 | 236110 | 0 | 1587300 | 15606 | 0 | 0 | 0 | 0 | 0 |
| SVHEGALK | EPB41L2; EPB4 | O43491; E9 | E41L2_HUMAN | Unmodified | 0 | 0 | 0 | 0 | 814070 | 0 | 0 | 0 | 0 | 0 | 0 | 0 | 0 | 0 | 0 | 0 | 155150 | 336210 |
| SVHFPGQA | GFPT1; GFPT1; | Q06210; A0 | GFPT1_HUMAN | Unmodified | 0 | 0 | 0 | 0 | 739000 | 139330 | 0 | 32942 | 1015400 | 0 | 311560 | 1594200 | 2653700 | 923270 | 0 | 0 | 0 | 0 |
| SVHKVEPIT | PSMA2; N/A; PS | P25787; A0A | PSA2_HUMAN | Unmodified | 0 | 0 | 0 | 0 | 0 | 0 | 0 | 0 | 0 | 0 | 0 | 1476000 | 0 | 0 | 0 | 0 | 0 | 0 |
| SVHSLISDF | STXBP1; STXBP | P61764; A0A | STXB1_HUMAN | Unmodified | 1636700 | 1936200 | 0 | 475060 | 7452800 | 605650 | 18085000 | 10063000 | 7007000 | 2966200 | 831160 | 12990000 | 3173800 | 3966900 | 155300 | 1136000 | 1592000 | 5996800 |
| SVIDPVPAF | PAK2 | Q13177 | PAK2_HUMAN | Unmodified | 0 | 0 | 0 | 0 | 0 | 0 | 0 | 0 | 0 | 0 | 0 | 0 | 251540 | 363850 | 0 | 0 | 2721900 | 0 |
| SVIDYQTHI | TMED7; TMED7 | Q9Y3B3; A0 | TMED7_HUMAN | Unmodified | 179890 | 0 | 0 | 670130 | 0 | 0 | 0 | 0 | 23885000 | 1219200 | 1520200 | 1716500 | 133430 | 0 | 840620 | 810740 | 0 | 0 |
| SVIEQGGIC | PDCD6IP | Q8WUM4 | PDC6I_HUMAN | Unmodified | 62861000 | 65678000 | 23616000 | 30605000 | 5721900 | 1044700 | 565330 | 60778 | 22125000 | 2794500 | 4538200 | 18276000 | 4293700 | 2724500 | 58556000 | 29990000 | 1138400 | 4520100 |
| SVILDCTPT | BCAM; BCAM | P50895; A0A | BCAM_HUMAN | Oxidation (M) | 1564500 | 843460 | 0 | 0 | 0 | 0 | 0 | 0 | 61243 | 0 | 0 | 0 | 0 | 0 | 0 | 0 | 0 | 0 |
| SVILEAFSS | CAND1; CAND1 | Q86VP6; A0 | CAND1_HUMA | Unmodified | 0 | 0 | 404790 | 410180 | 0 | 0 | 1739100 | 0 | 899390 | 456630 | 1585200 | 7141900 | 0 | 0 | 626380 | 610330 | 0 | 0 |
| SVILISTVR | MOV10; MOV10 | Q9HCE1; Q5 | MOV10_HUMA | Unmodified | 4452400 | 5635300 | 1475600 | 1176200 | 0 | 0 | 0 | 499340 | 984770 | 0 | 0 | 0 | 0 | 0 | 3457700 | 4574500 | 0 | 0 |
| SVINLLR | RPEL1; RPE; RP | Q2QD12; Q9 | RPEL1_HUMAN | Unmodified | 77275 | 0 | 0 | 136650 | 407830 | 0 | 0 | 0 | 656390 | 0 | 753200 | 2178500 | 0 | 0 | 0 | 117550 | 0 | 104690 |
| SVISGGLDA | FAM114A2; FAM | Q9NRY5; E5 | F1142_HUMAN | Unmodified | 0 | 0 | 109110 | 0 | 0 | 0 | 457380 | 0 | 0 | 0 | 0 | 1429100 | 0 | 0 | 146290 | 145320 | 0 | 0 |
| SVITFDKGC | SORL1 | Q92673 | SORL_HUMAN | Unmodified | 0 | 0 | 0 | 0 | 0 | 0 | 0 | 0 | 384880 | 284340 | 0 | 0 | 0 | 0 | 0 | 0 | 0 | 0 |
| SVKEDSNL | IQGAP1; IQGAP | P46940; H0 | IQGA1_HUMAN | Unmodified | 1217600 | 1057000 | 463910 | 899260 | 13060000 | 3157900 | 496880 | 131440 | 2199000 | 545020 | 702670 | 4017400 | 1856400 | 471790 | 1253000 | 991480 | 305010 | 710300 |
| SVKEDSNL | IQGAP1; IQGAP | P46940; H0 | IQGA1_HUMAN | Unmodified | 597010 | 481110 | 0 | 904560 | 24348000 | 5276200 | 0 | 231880 | 3088700 | 1675000 | 136310 | 1566700 | 3284500 | 1027900 | 335940 | 433460 | 465620 | 949630 |
| SVKIEEREG | HEBP1; HEBP1 | Q9NRV9; A0 | HEBP1_HUMAN | Unmodified | 0 | 0 | 0 | 0 | 2991100 | 0 | 0 | 0 | 0 | 0 | 0 | 0 | 1261900 | 933630 | 0 | 0 | 0 | 1755600 |
| SVKIEEREG | HEBP1; HEBP1 | Q9NRV9; A0 | HEBP1_HUMAN | Oxidation (M) | 0 | 0 | 0 | 0 | 0 | 0 | 0 | 0 | 0 | 0 | 0 | 0 | 2210800 | 0 | 0 | 0 | 0 | 0 |
| SVKVIVVGN | MDH1; MDH1; I | P40925; C9 | MDHC_HUMAN | Unmodified | 0 | 0 | 0 | 0 | 231420 | 0 | 1827000 | 828680 | 0 | 0 | 0 | 0 | 145210 | 335610 | 0 | 0 | 3620300 | 319060 |
| SVLAETEGI | KTN1; KTN1; KT | Q86UP2; G3 | KTN1_HUMAN | Unmodified | 0 | 0 | 0 | 0 | 0 | 0 | 0 | 0 | 120990 | 0 | 0 | 1612600 | 462280 | 0 | 0 | 0 | 0 | 0 |
| SVLALTHEC | PTGFRN | Q9P2B2 | FPRP_HUMAN | Unmodified | 4188000 | 4203100 | 26198000 | 8813000 | 311880 | 0 | 0 | 0 | 0 | 0 | 0 | 7134100 | 0 | 0 | 7938300 | 11019000 | 0 | 0 |
| SVLALTHEC | PTGFRN | Q9P2B2 | FPRP_HUMAN | Unmodified | 1538300 | 933120 | 2812100 | 16254000 | 1503000 | 1192700 | 161280 | 0 | 0 | 0 | 0 | 0 | 11779000 | 2459400 | 1708100 | 804350 | 0 | 326740 |
| SVLCSTPTI | PBK; PBK | Q96KB5; E5 | TOPK_HUMAN | Oxidation (M) | 0 | 510700 | 163340 | 417400 | 654150 | 0 | 0 | 341630 | 0 | 369970 | 0 | 0 | 2335200 | 1029900 | 253770 | 194450 | 1156700 | 0 |
| SVLDAAQIV | HSPH1 | Q92598 | HS105_HUMAN | Unmodified | 0 | 0 | 0 | 164670 | 0 | 0 | 124470 | 0 | 0 | 0 | 236940 | 2036800 | 0 | 0 | 369410 | 271040 | 0 | 0 |
| SVLDKLSAI | TMEM33; TMEM | P57088; H0 | TMM33_HUMAN | Unmodified | 0 | 0 | 0 | 0 | 0 | 0 | 0 | 0 | 0 | 336780 | 0 | 0 | 0 | 0 | 0 | 0 | 35027 | 0 |
| SVLEAQR | TOLLIP; TOLLIP; | Q9H0E2; F2 | TOLIP_HUMAN | Unmodified | 4095500 | 4029100 | 0 | 0 | 0 | 0 | 40193 | 0 | 0 | 107010 | 485650 | 99271 | 0 | 0 | 5029700 | 0 | 0 | 0 |
| SVLELIPELI | LRPPRC; LRPPR | P42704; H7 | LPPRC_HUMAN | Unmodified | 0 | 0 | 0 | 0 | 0 | 0 | 0 | 0 | 651730 | 0 | 229210 | 0 | 0 | 0 | 0 | 0 | 0 | 0 |
| SVLEQTK | HSPA4; HSPA4; | P34932; A0A | HSP74_HUMAN | Unmodified | 0 | 45738 | 193420 | 194830 | 172970 | 28314 | 29335 | 0 | 615160 | 51054 | 155890 | 5719600 | 414530 | 0 | 303070 | 154540 | 79133 | 114940 |
| SVLGEADQ | AARS1; AARS1; | P49588; A0A | SYAC_HUMAN | Unmodified | 756230 | 0 | 0 | 0 | 0 | 0 | 0 | 0 | 1167100 | 189570 | 274930 | 5589500 | 558400 | 0 | 1216800 | 359850 | 0 | 0 |
| SVLHEVME | MCM4; MCM4; | P33991; A0A | MCM4_HUMAN | Unmodified | 0 | 0 | 0 | 238470 | 273890 | 0 | 242910 | 35548 | 408700 | 0 | 0 | 1157000 | 103230 | 0 | 0 | 281230 | 52981 | 121550 |
| SVLHEVME | MCM4; MCM4; | P33991; A0A | MCM4_HUMAN | Oxidation (M) | 0 | 0 | 0 | 0 | 0 | 0 | 0 | 0 | 0 | 0 | 0 | 0 | 0 | 0 | 0 | 0 | 0 | 0 |
| SVLIGEFLE | RNMT; RNMT | O43148; K7 | MCES_HUMAN | Unmodified | 0 | 146840 | 0 | 0 | 0 | 0 | 0 | 149130 | 0 | 0 | 349370 | 1472900 | 0 | 0 | 0 | 0 | 35120 | 0 |
| SVLISLKQA | CLTA; CLTA | P09496; C9 | CLCA_HUMAN | Unmodified | 0 | 0 | 0 | 0 | 0 | 0 | 737110 | 335310 | 0 | 0 | 0 | 0 | 0 | 0 | 0 | 0 | 806050 | 0 |
| SVLLLAYKP | DDB1; DDB1; D | Q16531; A0 | DDB1_HUMAN | Oxidation (M) | 0 | 0 | 0 | 0 | 0 | 0 | 0 | 0 | 0 | 0 | 0 | 1337200 | 0 | 0 | 0 | 82079 | 0 | 0 |
| SVLQPSQC | NCKAP1 | Q9Y2A7 | NCKP1_HUMAN | Unmodified | 0 | 0 | 0 | 2009400 | 3486400 | 555250 | 0 | 211230 | 2280900 | 1120600 | 407590 | 3603800 | 3371400 | 1285700 | 1905700 | 1931800 | 617980 | 1790400 |
| SVLQSPLFS | MPDZ; MPDZ | O75970; F5 | MPDZ_HUMAN | Unmodified | 0 | 0 | 0 | 0 | 366350 | 0 | 0 | 0 | 0 | 0 | 0 | 0 | 0 | 0 | 0 | 0 | 0 | 0 |
| SVLRPAVL | RANBP3; RANB | Q9H6Z4; B7 | RANB3_HUMAN | Unmodified | 0 | 0 | 0 | 0 | 0 | 0 | 0 | 0 | 0 | 0 | 0 | 0 | 0 | 0 | 0 | 0 | 0 | 0 |
| SVLTGGLD. | FAM114A1 | Q8IWE2 | NXP20_HUMAN | Unmodified | 0 | 0 | 0 | 0 | 0 | 0 | 254080 | 0 | 0 | 0 | 0 | 0 | 549280 | 977020 | 0 | 0 | 0 | 1935000 |
| SVLVVTYSE | PAN3; PAN3 | Q58A45; A0 | PAN3_HUMAN | Unmodified | 945340 | 0 | 0 | 0 | 0 | 0 | 0 | 0 | 0 | 0 | 0 | 0 | 0 | 0 | 0 | 0 | 0 | 0 |

| | | | | | | | | | | | | | | | | | | | | | | |
|---|---|---|---|---|---|---|---|---|---|---|---|---|---|---|---|---|---|---|---|---|---|---|
| SVLVVTYSD | PAN3; PAN3 | Q58A45; A0A | PAN3_HUMAN | Unmodified | 630440 | 340040 | 89861 | 152040 | 0 | 0 | 0 | 0 | 0 | 0 | 0 | 0 | 0 | 0 | 0 | 81059 | 0 | 0 |
| SVLWVSAD | NUMBL; NUMB | Q9Y6R0; A0A | NUMBL_HUMA | Unmodified | 0 | 0 | 0 | 0 | 1077500 | 0 | 0 | 0 | 0 | 151960 | 0 | 0 | 0 | 0 | 0 | 0 | 0 | 0 |
| SVLWWLPV | SUMF2; SUMF2 | Q8NBJ7; C9J | SUMF2_HUMA | Unmodified | 0 | 0 | 0 | 0 | 393200 | 92426 | 1491900 | 0 | 60562000 | 32245000 | 922650 | 420010 | 337740 | 513860 | 0 | 0 | 0 | 0 |
| SVMAAAQV | HSPA4L; HSPA4 | O95757; E9F | HS74L_HUMAN | Oxidation (M) | 0 | 0 | 0 | 859940 | 0 | 0 | 0 | 0 | 0 | 0 | 0 | 0 | 0 | 0 | 0 | 0 | 0 | 0 |
| SVMDATQL | HSPA4; HSPA4; | P34932; A0A | HSP74_HUMAN | Oxidation (M) | 0 | 0 | 0 | 0 | 0 | 0 | 0 | 0 | 0 | 0 | 611000 | 1833000 | 0 | 0 | 0 | 0 | 0 | 0 |
| SVMLDRAE | POFUT2; POFU | Q9Y2G5; H7 | OFUT2_HUMAN | Unmodified | 0 | 0 | 0 | 0 | 0 | 0 | 0 | 0 | 2755300 | 0 | 342370 | 0 | 0 | 0 | 0 | 0 | 0 | 0 |
| SVMSLAEA | CALB2; CALB2 | P22676; H3E | CALB2_HUMAN | Oxidation (M) | 44947 | 0 | 68153 | 0 | 0 | 0 | 322980 | 117060 | 0 | 0 | 73231 | 0 | 0 | 0 | 128690 | 81576 | 0 | 0 |
| SVMSLAEA | CALB2; CALB2 | P22676; H3E | CALB2_HUMAN | Unmodified | 0 | 0 | 0 | 0 | 0 | 0 | 0 | 542550 | 0 | 0 | 0 | 0 | 0 | 0 | 0 | 0 | 0 | 0 |
| SVMSLAEA | CALB2; CALB2 | P22676; H3E | CALB2_HUMAN | Unmodified | 0 | 0 | 0 | 0 | 0 | 0 | 710910 | 111860 | 0 | 0 | 0 | 666830 | 0 | 0 | 0 | 0 | 0 | 0 |
| SVMTEEYK | FUBP1; FUBP1; | Q96AE4; C9J | FUBP1_HUMAN | 2 Oxidation (M) | 0 | 0 | 0 | 0 | 0 | 0 | 0 | 0 | 0 | 0 | 0 | 262650 | 0 | 0 | 0 | 0 | 0 | 0 |
| SVNDLRQE | GALNT7; GALN | Q86SF2; E9F | GALT7_HUMAN | Unmodified | 0 | 0 | 32404 | 0 | 0 | 0 | 0 | 0 | 443430 | 0 | 57377 | 322750 | 0 | 0 | 106270 | 61844 | 0 | 0 |
| SVNDQPS | SF3B1 | O75533 | SF3B1_HUMAN | Unmodified | 0 | 1040400 | 442500 | 0 | 0 | 2147300 | 0 | 0 | 3172900 | 0 | 1287600 | 26604000 | 672000 | 0 | 0 | 0 | 0 | 0 |
| SVNELIYKR | RPL7; RPL7 | P18124; A8M | RL7_HUMAN 6 | Unmodified | 648490 | 1103300 | 1022300 | 0 | 189820 | 0 | 0 | 0 | 0 | 0 | 0 | 5524700 | 65499 | 0 | 730400 | 0 | 97179 | 124800 |
| SVNESLNN | CLTC; CLTC; CL | Q00610; K7E | CLH1_HUMAN | Unmodified | 14159000 | 12367000 | 1229300 | 0 | 0 | 53121 | 0 | 0 | 1846600 | 753650 | 0 | 24819000 | 0 | 0 | 6828200 | 5422800 | 0 | 0 |
| SVNFDMTS | PDCD6IP | Q8WUM4 | PDC6I_HUMAN | Oxidation (M) | 6219600 | 6474700 | 0 | 2785100 | 5500100 | 2444300 | 0 | 249070 | 1782600 | 1454200 | 0 | 0 | 5302500 | 0 | 4583400 | 3291600 | 1239900 | 202200 |
| SVNFDMTS | PDCD6IP | Q8WUM4 | PDC6I_HUMAN | Unmodified | 31782000 | 40905000 | 12901000 | 18330000 | 45302000 | 11011000 | 0 | 0 | 14703000 | 13098000 | 2487800 | 8921700 | 40107000 | 25038000 | 20819000 | 16365000 | 4737800 | 19004000 |
| SVNGLAFY | COPB2; COPB2 | P35606; H0Y | COPB2_HUMA | Unmodified | 50087 | 26409 | 0 | 53953 | 0 | 0 | 150640 | 0 | 116550 | 44516 | 28866 | 449800 | 0 | 0 | 0 | 34985 | 0 | 0 |
| SVNVEQTD | DHX36; DHX36 | Q9H2U1; A0 | DHX36_HUMA | Unmodified | 0 | 0 | 0 | 0 | 0 | 0 | 0 | 0 | 0 | 0 | 0 | 1873800 | 0 | 0 | 726330 | 543150 | 0 | 0 |
| SVNVKPLV | SORD | Q00796 | DHSO_HUMAN | Unmodified | 0 | 0 | 0 | 0 | 0 | 0 | 0 | 0 | 0 | 0 | 0 | 1345200 | 2951800 | 1117500 | 0 | 0 | 0 | 0 |
| SVNVNRSV | EIF5; EIF5; EIF5 | P55010; H0Y | IF5_HUMAN Eu | Acetyl (Protein N | 0 | 0 | 0 | 0 | 1666800 | 0 | 0 | 0 | 0 | 131920 | 0 | 282740 | 0 | 0 | 0 | 0 | 414360 | 907170 |
| SVNVNRSV | EIF5; EIF5; EIF5 | P55010; H0Y | IF5_HUMAN Eu | Acetyl (Protein N | 0 | 71989 | 0 | 0 | 2050500 | 302350 | 292760 | 358040 | 0 | 0 | 0 | 0 | 0 | 0 | 0 | 0 | 1579400 | 1943000 |
| SVPAFIDISI | EIF3M; EIF3M; E | Q7L2H7; E9I | EIF3M_HUMAN | Acetyl (Protein N | 222410 | 178780 | 0 | 273150 | 0 | 0 | 0 | 0 | 127600 | 0 | 263410 | 0 | 0 | 0 | 0 | 121800 | 0 | 0 |
| SVPASDCV | HEPACAM2 | A8MVW5 | HECA2_HUMA | Unmodified | 88084 | 0 | 0 | 0 | 0 | 0 | 0 | 0 | 0 | 0 | 0 | 8488500 | 0 | 0 | 42808 | 1265800 | 0 | 0 |
| SVPDEIILLF | LRRC40 | Q9H9A6 | LRC40_HUMAN | Unmodified | 0 | 0 | 0 | 0 | 0 | 0 | 0 | 0 | 0 | 0 | 0 | 0 | 0 | 0 | 0 | 0 | 0 | 0 |
| SVPEAPVG | IGSF8; IGSF8 | Q969P0; C9J | IGSF8_HUMAN | Unmodified | 16909000 | 9799200 | 12678000 | 16231000 | 442360 | 831270 | 231920 | 27952 | 1598000 | 269800 | 398190 | 2926100 | 26631000 | 844990 | 10907000 | 10659000 | 884210 | 3060400 |
| SVPEDDGH | AKAP12 | Q02952 | AKA12_HUMAN | Unmodified | 0 | 0 | 0 | 0 | 4538100 | 491290 | 0 | 0 | 0 | 0 | 0 | 539460 | 307570 | 0 | 0 | 0 | 2660700 | 5922600 |
| SVPEVDPA | TOM1L2 | Q6ZVM7 | TM1L2_HUMAN | Oxidation (M) | 1570600 | 0 | 1154400 | 1157100 | 0 | 0 | 0 | 0 | 2332400 | 0 | 0 | 0 | 393380 | 0 | 1188500 | 1709300 | 0 | 0 |
| SVPEVDPA | TOM1L2 | Q6ZVM7 | TM1L2_HUMAN | Unmodified | 777350 | 934310 | 902960 | 1325600 | 0 | 0 | 0 | 0 | 0 | 0 | 0 | 0 | 439110 | 0 | 1524700 | 1009000 | 0 | 0 |
| SVPHLQKV | ANXA2P2; ANX | A6NMY6; P0 | AXA2L_HUMAN | Unmodified | 0 | 0 | 0 | 0 | 0 | 0 | 0 | 0 | 0 | 0 | 0 | 0 | 0 | 0 | 0 | 14996 | 587160 | 0 |
| SVPHYAAE | NUDCD1; NUD | Q96RS6; A0 | NUDC1_HUMA | Oxidation (M) | 0 | 0 | 0 | 0 | 632870 | 0 | 0 | 87986 | 288000 | 289510 | 0 | 0 | 0 | 0 | 0 | 0 | 1093300 | 341760 |
| SVPLAATSN | OCIAD1; OCIAD | Q9NX40; D6 | OCAD1_HUMA | Unmodified | 0 | 0 | 0 | 0 | 0 | 0 | 0 | 0 | 802380 | 0 | 0 | 0 | 0 | 0 | 0 | 0 | 0 | 0 |
| SVPLAATSN | OCIAD1; OCIAD | Q9NX40; D6 | OCAD1_HUMA | Oxidation (M) | 0 | 0 | 0 | 0 | 0 | 0 | 0 | 0 | 936440 | 321990 | 0 | 0 | 0 | 0 | 0 | 0 | 0 | 0 |
| SVPNQPST | ECPAS; ECPAS; | Q5VYK3; J3K | ECM29_HUMA | Unmodified | 40324 | 0 | 0 | 0 | 0 | 0 | 479590 | 275790 | 381930 | 103710 | 375850 | 2290600 | 56904 | 0 | 164990 | 70160 | 0 | 0 |
| SVPRGEAA | PCSK1N | Q9UHG2 | PCS1N_HUMA | Unmodified | 0 | 0 | 0 | 0 | 0 | 0 | 162220 | 79942 | 13008000 | 8719000 | 0 | 0 | 0 | 0 | 0 | 0 | 0 | 0 |
| SVPTPLVM | PROSER2 | Q86WR7 | PRSR2_HUMAN | Unmodified | 0 | 0 | 0 | 0 | 2378700 | 439390 | 0 | 0 | 0 | 0 | 0 | 164510 | 0 | 0 | 0 | 53188 | 158890 | 0 |
| SVPTSTVFY | TKT; TKT | P29401; A0A | TKT_HUMAN Tr | Unmodified | 0 | 0 | 0 | 0 | 8520300 | 3391400 | 0 | 45851 | 4324600 | 2095300 | 5308700 | 18637000 | 1163900 | 0 | 0 | 0 | 1353500 | 2746600 |
| SVPVCESQ | CHAMP1 | Q96JM3 | CHAP1_HUMA | Unmodified | 0 | 0 | 0 | 0 | 0 | 0 | 0 | 0 | 0 | 0 | 0 | 0 | 0 | 0 | 0 | 0 | 1048500 | 0 |
| SVPVTPARI | CDCA3; CDCA3 | Q99618; F8W | CDCA3_HUMA | Unmodified | 0 | 90297 | 150710 | 0 | 1434200 | 354800 | 404120 | 0 | 0 | 0 | 29653 | 0 | 519930 | 0 | 110890 | 57191 | 325040 | 446450 |
| SVQALIKDF | STXBP2; STXBP | Q15833; M0 | STXB2_HUMAN | Unmodified | 0 | 0 | 0 | 0 | 555980 | 0 | 0 | 0 | 0 | 0 | 0 | 0 | 0 | 665550 | 0 | 0 | 1638500 | 355990 |
| SVQANLDC | INF2; INF2; INF | Q27J81; A0A | INF2_HUMAN I | Unmodified | 0 | 0 | 0 | 0 | 0 | 0 | 53300 | 0 | 432810 | 0 | 0 | 377030 | 0 | 0 | 1237500 | 0 | 0 | 0 |
| SVQELRRE | NOMO1; NOMO | Q15155; P69 | NOMO1_HUMA | Unmodified | 0 | 0 | 0 | 0 | 221170 | 0 | 0 | 0 | 545850 | 551730 | 277530 | 0 | 0 | 0 | 0 | 0 | 0 | 0 |
| SVQGSDQ | SEMA3C | Q99985 | SEM3C_HUMA | Unmodified | 124670 | 0 | 714190 | 567370 | 0 | 76573 | 0 | 0 | 0 | 0 | 0 | 0 | 0 | 0 | 733110 | 476940 | 0 | 0 |
| SVQLSWTP | NRCAM; NRCAM | Q92823; A0A | NRCAM_HUMA | Unmodified | 0 | 0 | 0 | 0 | 0 | 0 | 0 | 0 | 1016700 | 0 | 0 | 0 | 0 | 0 | 0 | 0 | 0 | 0 |
| SVQLYGPT | CPNE8; CPNE8 | Q86YQ8; E7I | CPNE8_HUMA | Unmodified | 0 | 0 | 0 | 0 | 2887500 | 0 | 0 | 0 | 0 | 0 | 0 | 0 | 0 | 0 | 0 | 0 | 0 | 0 |
| SVQNDSQ | DSP | P15924 | DESP_HUMAN | Unmodified | 0 | 0 | 0 | 0 | 0 | 0 | 0 | 0 | 1111500 | 782610 | 0 | 55597 | 0 | 0 | 0 | 0 | 0 | 0 |
| SVQPTSEEI | SEPTIN9; SEPTI | Q9UHD8; K7 | SEPT9_HUMAN | Unmodified | 0 | 0 | 0 | 0 | 6669300 | 735720 | 430190 | 537630 | 239030 | 212060 | 195280 | 945400 | 0 | 575120 | 83882 | 0 | 2590800 | 293450 |
| SVQPTSEEI | SEPTIN9; SEPTI | Q9UHD8; K7 | SEPT9_HUMAN | Unmodified | 0 | 0 | 0 | 0 | 0 | 0 | 732200 | 0 | 50321 | 0 | 0 | 0 | 0 | 0 | 0 | 0 | 3518200 | 508600 |
| SVQSHKDL | AP3D1; AP3D1; | O14617; A0A | AP3D1_HUMAN | Unmodified | 0 | 0 | 0 | 0 | 0 | 0 | 0 | 0 | 0 | 0 | 4392900 | 0 | 0 | 0 | 0 | 0 | 0 | 0 |
| SVQSQGFK | ARL3 | P36405 | ARL3_HUMAN | Unmodified | 179200 | 0 | 160720 | 466560 | 0 | 91110 | 0 | 0 | 390280 | 67475 | 212370 | 2037100 | 2153400 | 1443400 | 623980 | 66272 | 0 | 1223400 |
| SVQSQGFK | ARL3 | P36405 | ARL3_HUMAN | Unmodified | 0 | 0 | 0 | 0 | 1283300 | 132780 | 3936700 | 417200 | 0 | 739220 | 0 | 0 | 5042300 | 3466200 | 0 | 0 | 396380 | 2128300 |
| SVQTFADK | CAPZB | B1AK88 | B1AK88_HUMA | Unmodified | 3703100 | 2997800 | 2203200 | 2117400 | 16561000 | 1206900 | 280900 | 154610 | 6143500 | 1352000 | 1309000 | 27728000 | 7442200 | 3475700 | 1886400 | 0 | 657310 | 5822900 |
| SVQTTLQT | ERLIN2; ERLIN1 | O94905; O75 | ERLN2_HUMAN | Unmodified | 0 | 0 | 0 | 0 | 0 | 0 | 0 | 0 | 1813800 | 147950 | 0 | 0 | 0 | 0 | 0 | 0 | 0 | 0 |
| SVQVSTLK | ARPC1A; ARPC | Q92747; E9F | ARC1A_HUMAN | Unmodified | 332330 | 367660 | 1046600 | 312400 | 124200 | 55953 | 83076 | 34215 | 492110 | 0 | 376220 | 6044900 | 183080 | 0 | 672470 | 820250 | 28381 | 71268 |
| SVQVVSAA | AHCYL2 | Q96HN2 | SAHH3_HUMA | Acetyl (Protein N | 0 | 90827 | 0 | 0 | 5937400 | 409310 | 0 | 32101 | 0 | 0 | 0 | 0 | 0 | 0 | 0 | 0 | 635040 | 931300 |
| SVQYDDVF | ALCAM; ALCAM | Q13740; F5G | CD166_HUMA | Unmodified | 5363700 | 2467200 | 2731600 | 2046400 | 890940 | 0 | 163000 | 0 | 3369100 | 317980 | 0 | 1954000 | 217160 | 0 | 2343700 | 3256700 | 240600 | 1342200 |
| SVRFDSWH | VWDE; VWDE; V | Q8N2E2; E5I | VWDE_HUMAN | Unmodified | 0 | 0 | 0 | 0 | 6865900 | 0 | 0 | 0 | 0 | 0 | 0 | 0 | 2201200 | 0 | 0 | 0 | 0 | 0 |
| SVRPGVAI | HGD; HGD | Q93099; H7C | HGD_HUMAN H | Unmodified | 0 | 0 | 0 | 312470 | 0 | 0 | 189770 | 0 | 0 | 0 | 0 | 0 | 369150 | 1202000 | 0 | 0 | 0 | 361100 |
| SVSAFAPIC | ESD; ESD; ESD; | P10768; X6R | ESTD_HUMAN | Unmodified | 0 | 0 | 0 | 0 | 650490 | 382570 | 0 | 122620 | 0 | 0 | 0 | 797140 | 646550 | 529690 | 0 | 0 | 872710 | 554170 |
| SVSAFAPIC | ESD; ESD; ESD; | P10768; X6R | ESTD_HUMAN | Unmodified | 0 | 0 | 0 | 0 | 6174100 | 0 | 1495500 | 338730 | 1023500 | 662600 | 1087900 | 3201200 | 1947900 | 1576400 | 0 | 0 | 6893700 | 3393900 |
| SVSARSAA | SEPTIN7; SEPTI | Q16181; A0A | SEPT7_HUMAN | Acetyl (Protein N | 0 | 0 | 0 | 0 | 605630 | 0 | 263820 | 108130 | 0 | 0 | 0 | 0 | 0 | 292930 | 0 | 0 | 824200 | 313120 |
| SVSDNDIR | VCP; VCP; VCP; | P55072; C9J | TERA_HUMAN | Oxidation (M) | 0 | 0 | 0 | 324440 | 8520000 | 3025500 | 768520 | 25114 | 0 | 4615200 | 487710 | 0 | 17398000 | 4871200 | 0 | 0 | 2515300 | 0 |
| SVSDNDIR | VCP; VCP; VCP; | P55072; C9J | TERA_HUMAN | Unmodified | 0 | 0 | 0 | 0 | 4440900 | 568380 | 0 | 0 | 685010 | 1502300 | 0 | 0 | 3784800 | 2049800 | 0 | 0 | 806450 | 1662200 |
| SVSDQFYR | EIF5; EIF5; EIF5 | P55010; H0Y | IF5_HUMAN Eu | Unmodified | 0 | 0 | 0 | 0 | 825750 | 9800300 | 0 | 451570 | 0 | 111860 | 84351 | 471720 | 629510 | 0 | 264690 | 0 | 688640 | 1107900 |
| SVSDYDGK | P4HB; P4HB; P4 | P07237; A0A | PDIA1_HUMAN | Unmodified | 0 | 0 | 0 | 63077 | 2395300 | 541540 | 5798700 | 3190900 | 11484000 | 75593000 | 344930 | 0 | 1888600 | 4070200 | 0 | 0 | 697870 | 1655400 |
| SVSEDVHV | ACOT8; ACOT8; | O14734; E9F | ACOT8_HUMA | Unmodified | 0 | 0 | 0 | 0 | 0 | 0 | 0 | 0 | 10143000 | 1095500 | 0 | 0 | 0 | 0 | 116050 | 0 | 0 | 0 |
| SVSHQTVQ | TACC2; TACC2; | O95359; Q4V | TACC2_HUMA | Unmodified | 0 | 0 | 0 | 0 | 615140 | 104230 | 0 | 0 | 0 | 151120 | 0 | 0 | 297950 | 184540 | 46141 | 0 | 0 | 0 |
| SVSIPAPAY | AGO3; AGO2 | Q9H9G7; Q9 | AGO3_HUMAN | Unmodified | 0 | 0 | 0 | 543640 | 0 | 0 | 0 | 0 | 176090 | 285560 | 0 | 1252700 | 24452 | 0 | 52598 | 0 | 0 | 0 |
| SVSLTGAPI | KHSRP; KHSRP | Q92945; M0 | FUBP2_HUMAN | Unmodified | 0 | 0 | 0 | 0 | 0 | 0 | 0 | 0 | 1856400 | 126110 | 1034800 | 3160700 | 0 | 0 | 0 | 0 | 0 | 0 |
| SVSSPSSF | PTPRM; PTPRM; | P28827; E7E | PTPRM_HUMA | Unmodified | 0 | 0 | 0 | 0 | 1304200 | 0 | 0 | 0 | 0 | 0 | 0 | 0 | 169400 | 0 | 0 | 0 | 141220 | 804740 |
| SVSTPSEA | DYNC1I2; DYN | Q13409; E7E | DC1I2_HUMAN | Unmodified | 0 | 0 | 0 | 0 | 310290 | 35113 | 56062 | 0 | 630500 | 165380 | 154820 | 444740 | 0 | 180600 | 0 | 232670 | 290940 | 261780 |
| SVSTPSEA | DYNC1I2; DYN | E7EUM4; E7I | E7EUM4_HUM | Unmodified | 0 | 0 | 0 | 0 | 2846600 | 0 | 159310 | 54030 | 0 | 687030 | 0 | 0 | 1832800 | 1880600 | 0 | 0 | 4400300 | 6014000 |
| SVSTSTTFV | DNAJB2; DNAJE | P25686; C9J | DNJB2_HUMAN | Unmodified | 1591900 | 1564300 | 930810 | 917900 | 0 | 0 | 0 | 0 | 195860 | 0 | 0 | 351540 | 218600 | 0 | 993830 | 974490 | 0 | 0 |
| SVSTTNIAG | ARHGEF2; ARH | Q92974; V9C | ARHG2_HUMA | Unmodified | 41052 | 0 | 40270 | 71300 | 358420 | 0 | 97040 | 13195 | 527390 | 222210 | 464740 | 1671200 | 220340 | 112980 | 41944 | 40720 | 87412 | 0 |
| SVSTTNIAG | ARHGEF2; ARH | Q92974; V9C | ARHG2_HUMA | Unmodified | 0 | 0 | 0 | 0 | 0 | 0 | 0 | 0 | 947710 | 0 | 631570 | 2185200 | 0 | 0 | 0 | 0 | 0 | 0 |
| SVSYPSLLS | MAN2A1 | Q16706 | MA2A1_HUMA | 2 Oxidation (M) | 0 | 0 | 0 | 0 | 0 | 0 | 0 | 0 | 1896900 | 368120 | 1917300 | 1565200 | 0 | 0 | 0 | 0 | 0 | 0 |
| SVSYQLSPI | PPP1R16A | Q96I34 | PP16A_HUMAN | Unmodified | 0 | 0 | 0 | 0 | 0 | 0 | 0 | 0 | 0 | 0 | 0 | 0 | 0 | 1031000 | 0 | 0 | 1058900 | 0 |
| SVTCTYSP | TP53; TP53; TP5 | P04637; E7E | P53_HUMAN C | Unmodified | 0 | 0 | 0 | 0 | 0 | 0 | 388740 | 294800 | 0 | 0 | 0 | 0 | 452730 | 0 | 0 | 0 | 0 | 0 |
| SVTDSIRDE | DNM1L; DNM1L | O00429; F8V | DNM1L_HUMA | Unmodified | 153950 | 0 | 225640 | 0 | 0 | 0 | 1799700 | 163330 | 0 | 0 | 0 | 6439200 | 879080 | 681990 | 201390 | 127460 | 0 | 0 |
| SVTDYAQQ | MYO6; MYO6 | A0A0A0MRM | A0A0A0MRM8_ | Unmodified | 0 | 400100 | 1339100 | 1649300 | 0 | 0 | 0 | 0 | 0 | 0 | 0 | 0 | 726330 | 402220 | 1677000 | 1168400 | 0 | 0 |
| SVTDYAQQ | MYO6; MYO6 | A0A0A0MRM | A0A0A0MRM8_ | Unmodified | 0 | 0 | 0 | 627030 | 0 | 0 | 0 | 0 | 0 | 0 | 0 | 0 | 2102100 | 1490600 | 0 | 0 | 0 | 0 |
| SVTEQGAE | YWHAZ; YWHA | P63104; E5R | 1433Z_HUMAN | Unmodified | 1290700 | 1543800 | 1682500 | 4439900 | 1614500 | 0 | 188650 | 172300 | 15592000 | 2812700 | 14419000 | 38594000 | 1018600 | 253650 | 5294100 | 6259100 | 890540 | 2274200 |
| SVTEQGAE | YWHAZ; YWHA | P63104; E5R | 1433Z_HUMAN | Unmodified | 11473000 | 14087000 | 15006000 | 34861000 | 3.26E+08 | 71894000 | 99134000 | 6948300 | 1.3E+08 | 1.38E+08 | 48049000 | 73728000 | 1.04E+08 | 1.32E+08 | 18674000 | 22672000 | 3.42E+08 | 4.38E+08 |
| SVTEQGAE | YWHAZ; YWHA | P63104; E5R | 1433Z_HUMAN | Unmodified | 0 | 0 | 0 | 0 | 966050 | 1053100 | 15827000 | 0 | 0 | 5345100 | 0 | 0 | 13010000 | 17731000 | 0 | 0 | 91504000 | 39347000 |
| SVTGTDVD | TPPP3 | Q9BW30 | TPPP3_HUMAN | Unmodified | 0 | 0 | 0 | 0 | 0 | 0 | 4334800 | 2844600 | 0 | 0 | 0 | 0 | 0 | 0 | 0 | 0 | 0 | 0 |
| SVTGTDVD | TPPP3 | Q9BW30 | TPPP3_HUMAN | Unmodified | 0 | 0 | 0 | 0 | 0 | 0 | 0 | 154990 | 0 | 0 | 0 | 0 | 0 | 0 | 0 | 0 | 0 | 0 |
| SVTHANAL | SLC4A2 | P04920 | B3A2_HUMAN | Unmodified | 182670 | 0 | 678160 | 197760 | 0 | 0 | 0 | 33043 | 0 | 0 | 0 | 0 | 0 | 0 | 1406300 | 750200 | 0 | 0 |
| SVTHANAL | SLC4A2 | P04920 | B3A2_HUMAN | Oxidation (M) | 0 | 0 | 234190 | 80917 | 132950 | 0 | 0 | 9620.6 | 198510 | 0 | 0 | 0 | 0 | 0 | 331580 | 309120 | 0 | 0 |
| SVTHFDSL | ACSL4; ACSL4; | O60488; D6I | ACSL4_HUMAN | Unmodified | 0 | 0 | 0 | 0 | 6058700 | 0 | 0 | 0 | 0 | 0 | 2892900 | 4956600 | 0 | 0 | 0 | 0 | 1119700 | 2405700 |
| SVTIQPPTG | VANGL1 | Q8TAA9 | VANG1_HUMA | Unmodified | 0 | 0 | 0 | 0 | 0 | 0 | 0 | 0 | 0 | 0 | 0 | 0 | 0 | 0 | 0 | 0 | 0 | 0 |
| SVTLGYLFS | ADAR; ADAR; AI | P55265; A0A | DSRAD_HUMA | Unmodified | 0 | 0 | 0 | 0 | 0 | 0 | 0 | 0 | 0 | 25045 | 0 | 3974600 | 0 | 0 | 0 | 0 | 0 | 0 |
| SVTLHTDV | PPIL3; PPIL3; PI | Q9H2H8; F8 | PPIL3_HUMAN | Acetyl (Protein N | 0 | 595610 | 0 | 0 | 0 | 0 | 0 | 0 | 0 | 204050 | 0 | 0 | 0 | 0 | 0 | 0 | 0 | 0 |
| SVTLLIKGP | CCT6A | P40227 | TCPZ_HUMAN | Unmodified | 0 | 0 | 0 | 0 | 2612200 | 68822 | 563680 | 0 | 0 | 1164800 | 0 | 0 | 4920100 | 4793500 | 0 | 0 | 1182100 | 2122800 |
| SVTLQFRP | SDK2; SDK2 | Q58EX2; H7C | SDK2_HUMAN | Unmodified | 0 | 0 | 0 | 0 | 0 | 0 | 0 | 0 | 0 | 299860 | 0 | 0 | 0 | 0 | 0 | 0 | 0 | 0 |
| SVTNEDVT | PCCB; PCCB; P | P05166; C9J | PCCB_HUMAN | Unmodified | 671010 | 2232200 | 0 | 0 | 2537300 | 0 | 0 | 408040 | 1243000 | 300780 | 0 | 0 | 0 | 0 | 1865300 | 50265 | 2440200 | 0 |
| SVTPERIAE | AKR1B1 | P15121 | ALDR_HUMAN | Unmodified | 0 | 0 | 0 | 0 | 14879000 | 2310800 | 0 | 0 | 0 | 0 | 0 | 0 | 0 | 0 | 0 | 0 | 0 | 0 |
| SVTQADAG | KIRREL1; KIRRE | Q96J84; Q5V | KIRR1_HUMAN | Unmodified | 0 | 0 | 0 | 0 | 891940 | 0 | 0 | 0 | 0 | 0 | 0 | 0 | 0 | 0 | 0 | 0 | 362660 | 0 |
| SVTQADAG | KIRREL1; KIRRE | Q96J84; Q5V | KIRR1_HUMAN | Unmodified | 0 | 0 | 0 | 0 | 1551000 | 0 | 0 | 0 | 0 | 0 | 0 | 0 | 0 | 0 | 0 | 0 | 690610 | 605260 |
| SVTREDTG | F11R; F11R | Q9Y624; A0A | JAM1_HUMAN | Unmodified | 2895700 | 1748300 | 1108300 | 2294800 | 8426800 | 1119300 | 0 | 0 | 4728700 | 657270 | 277880 | 1926200 | 7088700 | 6501000 | 1977700 | 1206700 | 1920200 | 2148400 |
| SVTREDTG | F11R; F11R | Q9Y624; A0A | JAM1_HUMAN | Oxidation (M) | 1282700 | 886590 | 831250 | 1269300 | 2395000 | 884230 | 0 | 0 | 264110 | 488710 | 0 | 1285200 | 3458500 | 3086900 | 1373600 | 1786900 | 0 | 1104200 |
| SVTREDTG | F11R; F11R | Q9Y624; A0A | JAM1_HUMAN | Oxidation (M) | 0 | 0 | 0 | 0 | 427440 | 0 | 0 | 0 | 0 | 0 | 0 | 0 | 436350 | 424420 | 0 | 0 | 274830 | 318510 |
| SVTREDTG | F11R; F11R | Q9Y624; A0A | JAM1_HUMAN | Unmodified | 0 | 0 | 0 | 0 | 2292500 | 0 | 0 | 0 | 0 | 0 | 0 | 0 | 2510700 | 2387900 | 0 | 0 | 1422400 | 1156500 |
| SVVALHNL | PSMD7 | P51665 | PSMD7_HUMA | Unmodified | 33970 | 36726 | 198940 | 97359 | 430770 | 0 | 363050 | 0 | 0 | 0 | 472450 | 5777000 | 1250900 | 538180 | 118260 | 88152 | 0 | 443040 |
| SVVAPPGA | GSPT1; GSPT1; | P15170; H3E | ERF3A_HUMAN | Unmodified | 0 | 0 | 0 | 0 | 114910 | 0 | 50177 | 38309 | 0 | 0 | 76081 | 408040 | 0 | 121270 | 0 | 0 | 57387 | 0 |
| SVVAPPGA | GSPT1; GSPT1; | P15170; H3E | ERF3A_HUMAN | Unmodified | 0 | 0 | 0 | 0 | 867770 | 0 | 3442500 | 245290 | 0 | 182580 | 0 | 0 | 5848700 | 0 | 0 | 0 | 3830100 | 1534200 |
| SVVFTSAQ | CA14 | Q9ULX7 | CAH14_HUMA | Unmodified | 0 | 0 | 0 | 0 | 560450 | 0 | 0 | 0 | 0 | 0 | 311550 | 5698900 | 0 | 0 | 0 | 0 | 0 | 0 |
| SVVGDALE | MUC5B | Q9HC84 | MUC5B_HUMA | Unmodified | 0 | 0 | 0 | 0 | 0 | 0 | 0 | 0 | 0 | 0 | 0 | 0 | 0 | 0 | 0 | 0 | 0 | 2224800 |
| SVVLMSHL | PGK1 | P00558 | PGK1_HUMAN | Unmodified | 0 | 0 | 80578 | 0 | 2071600 | 358530 | 0 | 0 | 5729300 | 198420 | 0 | 3903100 | 289880 | 0 | 378450 | 0 | 1175500 | 6456200 |
| SVVLMSHL | PGK1 | P00558 | PGK1_HUMAN | Oxidation (M) | 74916 | 29230 | 345500 | 131380 | 399010 | 0 | 0 | 0 | 1213000 | 74116 | 802190 | 10118000 | 118290 | 0 | 533210 | 417090 | 539990 | 2845400 |
| SVVLMSHL | PGK1 | P00558 | PGK1_HUMAN | 2 Oxidation (M) | 0 | 0 | 2265700 | 0 | 435180 | 0 | 0 | 37946 | 0 | 0 | 1204400 | 17788000 | 0 | 0 | 1094300 | 384780 | 0 | 2007700 |

| | | | | | | | | | | | | | | | | | | | | | | |
|---|---|---|---|---|---|---|---|---|---|---|---|---|---|---|---|---|---|---|---|---|---|---|
| SVVLMSHL | PGK1 | P00558 | PGK1_HUMAN | 2 Oxidation (M) | 0 | 0 | 0 | 2791600 | 45632000 | 5449400 | 17864000 | 331820 | 1159500 | 14317000 | 5733800 | 16071000 | 13091000 | 33822000 | 0 | 0 | 1.67E+08 | 1.48E+08 |
| SVVLMSHL | PGK1 | P00558 | PGK1_HUMAN | Oxidation (M) | 0 | 0 | 0 | 1461700 | 30182000 | 3154800 | 9821600 | 0 | 5149700 | 7634000 | 1227700 | 1230400 | 25798000 | 7229500 | 0 | 0 | 79388000 | 1.07E+08 |
| SVVLMSHL | PGK1 | P00558 | PGK1_HUMAN | Unmodified | 0 | 0 | 0 | 0 | 5037700 | 0 | 0 | 0 | 2194600 | 1771300 | 0 | 0 | 0 | 0 | 0 | 0 | 10492000 | 19067000 |
| SVVNMEWI | EPRS1; EPRS1 | P07814; V9G | SYEP_HUMAN | Oxidation (M) | 247510 | 335810 | 14856000 | 2415100 | 0 | 2250300 | 0 | 0 | 457080 | 117380 | 8797900 | 1672400 | 0 | 0 | 2632000 | 586290 | 0 | 602270 |
| SVVQQQQ | RNF41; RNF41; | Q9H4P4; F8 | RNF41_HUMAN | Unmodified | 0 | 0 | 0 | 0 | 1560300 | 248510 | 44201 | 0 | 0 | 0 | 0 | 0 | 125050 | 85469 | 0 | 0 | 22068 | 0 |
| SVVSWHYL | DST; DST; DST; | Q03001; A0 | DYST_HUMAN | Unmodified | 0 | 0 | 0 | 0 | 1851800 | 0 | 0 | 0 | 0 | 0 | 0 | 0 | 0 | 0 | 0 | 0 | 236300 | 0 |
| SVVTPARV | LYPLA2; LYPLA | O95372; Q5 | LYPA2_HUMAN | Unmodified | 0 | 0 | 0 | 0 | 464820 | 0 | 0 | 0 | 0 | 0 | 0 | 0 | 93834 | 0 | 0 | 0 | 673140 | 561230 |
| SVVTSIFGV | IARS1; IARS1 | P41252; A0A | SYIC_HUMAN I | Unmodified | 0 | 0 | 760250 | 0 | 2204400 | 460300 | 1689100 | 357030 | 1275500 | 1893100 | 1503300 | 18091000 | 952900 | 0 | 0 | 0 | 677690 | 476910 |
| SVVYQETN | PRPSAP1; PRPS | Q14558; C9J | KPRA_HUMAN | Unmodified | 14348 | 0 | 0 | 0 | 0 | 0 | 0 | 0 | 0 | 0 | 0 | 864280 | 0 | 0 | 0 | 0 | 0 | 0 |
| SVWEYVDF | SBF1; SBF1 | O95248; H0 | MTMR5_HUMA | Unmodified | 0 | 0 | 0 | 0 | 282350 | 0 | 0 | 0 | 0 | 0 | 0 | 935610 | 0 | 0 | 0 | 0 | 0 | 0 |
| SVWGIQVT | NDST2; N/A | P52849; S4R | NDST2_HUMAI | Unmodified | 0 | 0 | 0 | 0 | 0 | 0 | 0 | 0 | 2985000 | 0 | 0 | 0 | 0 | 0 | 0 | 0 | 0 | 0 |
| SVYDGPEQ | RASA3 | Q14644 | RASA3_HUMAN | Unmodified | 0 | 0 | 0 | 0 | 0 | 0 | 0 | 0 | 0 | 0 | 0 | 0 | 0 | 0 | 0 | 0 | 0 | 1943900 |
| SVYQLLSEI | THOC1; THOC1 | Q96FV9; R4 | THOC1_HUMA | Unmodified | 0 | 0 | 0 | 0 | 0 | 0 | 0 | 0 | 0 | 0 | 0 | 0 | 293690 | 124900 | 0 | 0 | 0 | 0 |
| SVYSWDIV | EIF3D; EIF3D | O15371; B0 | EIF3D_HUMAN | Unmodified | 0 | 262090 | 0 | 478300 | 0 | 0 | 0 | 0 | 0 | 0 | 0 | 2472200 | 0 | 0 | 264070 | 288230 | 0 | 0 |
| SVYSWDIV | EIF3D; EIF3D | O15371; B0 | EIF3D_HUMAN | Unmodified | 0 | 0 | 0 | 0 | 1937100 | 0 | 0 | 0 | 332590 | 0 | 0 | 0 | 1156200 | 0 | 0 | 0 | 0 | 207340 |
| SWANHEN | UEVLD | Q8IX04 | UEVLD_HUMAI | Unmodified | 0 | 0 | 0 | 0 | 344540 | 106060 | 0 | 0 | 492420 | 0 | 0 | 0 | 0 | 33693 | 0 | 0 | 0 | 36622 |
| SWANKME | P4HA2; P4HA2; | O15460; E7E | P4HA2_HUMAI | Unmodified | 0 | 0 | 0 | 0 | 0 | 0 | 0 | 0 | 1613000 | 1228100 | 0 | 0 | 0 | 0 | 0 | 0 | 0 | 0 |
| SWCPDCV | TXNDC17; TXNI | Q9BRA2; I3L | TXD17_HUMAN | Unmodified | 0 | 0 | 0 | 0 | 5177700 | 1001900 | 253080 | 118190 | 0 | 1574600 | 0 | 0 | 5816200 | 4937500 | 0 | 0 | 4149800 | 4862700 |
| SWDIFFR | OGDH; OGDH; | Q02218; E9F | ODO1_HUMAN | Unmodified | 0 | 0 | 0 | 0 | 0 | 0 | 0 | 0 | 667110 | 83125 | 0 | 0 | 0 | 0 | 0 | 0 | 0 | 0 |
| SWDWRNV | CTSZ; CTSZ; CT | Q9UBR2; A0 | CATZ_HUMAN | Unmodified | 0 | 0 | 0 | 0 | 0 | 0 | 0 | 0 | 0 | 418010 | 0 | 0 | 0 | 0 | 0 | 0 | 0 | 0 |
| SWEEQMIE | TFAM | Q00059 | TFAM_HUMAN | Unmodified | 384570 | 179570 | 1348500 | 1438200 | 302940 | 0 | 65505 | 191380 | 4074100 | 150510 | 693710 | 594280 | 0 | 0 | 1019400 | 778320 | 549970 | 341950 |
| SWEQKLEE | CDC37; CDC37 | Q16543; K7E | CDC37_HUMA | Unmodified | 82418 | 0 | 0 | 247710 | 0 | 0 | 0 | 0 | 515680 | 35934 | 410450 | 1268800 | 26514 | 0 | 144760 | 0 | 0 | 0 |
| SWETKPGL | GGT7 | Q9UJ14 | GGT7_HUMAN | Oxidation (M) | 0 | 0 | 0 | 0 | 0 | 0 | 0 | 0 | 2246300 | 385820 | 0 | 0 | 0 | 0 | 0 | 0 | 0 | 0 |
| SWETKPGL | GGT7 | Q9UJ14 | GGT7_HUMAN | Unmodified | 0 | 0 | 0 | 0 | 0 | 0 | 0 | 0 | 7643500 | 626800 | 0 | 0 | 0 | 0 | 0 | 0 | 0 | 0 |
| SWGQQAQ | CARMIL1 | Q5VZK9 | CARL1_HUMAI | Unmodified | 0 | 0 | 1358900 | 0 | 240840 | 0 | 0 | 0 | 0 | 0 | 0 | 0 | 0 | 0 | 184190 | 0 | 0 | 0 |
| SWHDMQT | TPD52L1; TPD5 | Q16890; E9F | TPD53_HUMAI | Unmodified | 0 | 0 | 0 | 0 | 0 | 0 | 0 | 0 | 0 | 0 | 0 | 0 | 260950 | 0 | 0 | 0 | 0 | 0 |
| SWHDVQV | TPD52L2; TPD5 | O43399; A0 | TPD54_HUMAI | Unmodified | 63734 | 0 | 0 | 0 | 1967800 | 0 | 0 | 0 | 0 | 259960 | 0 | 307740 | 1031900 | 0 | 118050 | 537060 | 28935 | 723570 |
| SWHPEEFN | PDLIM5; PDLIM | Q96HC4; H0 | PDLI5_HUMAN | Unmodified | 0 | 0 | 0 | 0 | 0 | 0 | 125690 | 0 | 0 | 0 | 0 | 622580 | 0 | 88340 | 0 | 0 | 389120 | 253960 |
| SWHPVCQ | TMPRSS2; TMPI | O15393; C9J | TMPS2_HUMAI | Unmodified | 39720 | 0 | 1383200 | 2242400 | 0 | 0 | 0 | 0 | 0 | 0 | 0 | 0 | 224450 | 0 | 2129900 | 1879900 | 0 | 0 |
| SWIEEQAM | LAP3; LAP3 | P28838; H0 | AMPL_HUMAN | Unmodified | 0 | 0 | 0 | 0 | 1086500 | 109480 | 197480 | 0 | 0 | 0 | 46739 | 0 | 403310 | 144880 | 0 | 0 | 0 | 0 |
| SWIEETMR | KLK14; KLK14 | Q9P0G3; A0 | KLK14_HUMAN | Unmodified | 0 | 0 | 0 | 0 | 0 | 152330 | 0 | 0 | 1303400 | 0 | 0 | 0 | 0 | 0 | 0 | 0 | 0 | 0 |
| SWITKQDE | ZNF185; ZNF18 | O15231; H7I | ZN185_HUMAI | Unmodified | 0 | 0 | 0 | 0 | 779620 | 0 | 0 | 0 | 0 | 0 | 183370 | 0 | 0 | 0 | 0 | 0 | 1213300 | 0 |
| SWKEDSEV | SH2D4A | Q9H788 | SH24A_HUMAI | Unmodified | 0 | 0 | 0 | 0 | 2003900 | 0 | 0 | 0 | 0 | 0 | 0 | 0 | 1149700 | 0 | 159860 | 0 | 610390 | 0 |
| SWLIIQR | AOC1; AOC1; A | P19801; C9J | AOC1_HUMAN | Unmodified | 0 | 0 | 0 | 0 | 0 | 0 | 0 | 0 | 1405400 | 0 | 0 | 0 | 0 | 0 | 0 | 0 | 0 | 0 |
| SWLKPRLG | ACLY; ACLY | P53396; K7E | ACLY_HUMAN | Unmodified | 0 | 0 | 0 | 0 | 0 | 0 | 635880 | 0 | 0 | 0 | 0 | 0 | 352300 | 0 | 0 | 0 | 0 | 0 |
| SWLPQNC | PAICS; PAICS; | P22234; D6F | PUR6_HUMAN | Oxidation (M) | 0 | 0 | 0 | 0 | 0 | 0 | 0 | 0 | 0 | 0 | 0 | 912510 | 0 | 0 | 0 | 0 | 27267 | 0 |
| SWNCAPG | EXTL3; EXTL3 | O43909; E7E | EXTL3_HUMAN | Unmodified | 0 | 2285500 | 0 | 0 | 0 | 0 | 125570 | 0 | 1598400 | 0 | 0 | 0 | 230940 | 0 | 0 | 0 | 0 | 81947 |
| SWNETLTS | ATP5PD; ATP5P | O75947; F5H | ATP5H_HUMAI | Unmodified | 0 | 0 | 0 | 0 | 0 | 0 | 0 | 0 | 7592500 | 0 | 0 | 310170 | 0 | 0 | 0 | 55491 | 0 | 0 |
| SWNWLNG | GAS6 | Q14393 | GAS6_HUMAN | Unmodified | 0 | 0 | 0 | 0 | 0 | 0 | 0 | 0 | 3229800 | 163690 | 0 | 0 | 0 | 0 | 0 | 0 | 0 | 0 |
| SWQDELA | MTDH | Q86UE4 | LYRIC_HUMAN | Unmodified | 0 | 0 | 0 | 0 | 0 | 0 | 0 | 0 | 12304000 | 1470200 | 1233100 | 414590 | 0 | 0 | 0 | 0 | 0 | 0 |
| SWRESCD | CAPZA1 | P52907 | CAZA1_HUMAI | Unmodified | 595840 | 396410 | 0 | 0 | 0 | 0 | 0 | 0 | 0 | 137450 | 0 | 2425000 | 0 | 0 | 781560 | 0 | 0 | 0 |
| SWRESCD | CAPZA1 | P52907 | CAZA1_HUMAI | Unmodified | 0 | 0 | 0 | 0 | 2577400 | 359920 | 1212600 | 135670 | 0 | 1359700 | 0 | 0 | 5193100 | 3976700 | 0 | 0 | 4079800 | 3100700 |
| SWTAADM | HLA-A; HLA-H; | P04439; P01 | HLAA_HUMAN | Unmodified | 0 | 0 | 0 | 2096100 | 357660 | 0 | 0 | 0 | 14013000 | 284180 | 0 | 2974700 | 0 | 0 | 0 | 0 | 358950 | 1655900 |
| SWTAADM | HLA-A; HLA-H; | P04439; P01 | HLAA_HUMAN | Oxidation (M) | 0 | 0 | 0 | 0 | 0 | 0 | 788770 | 0 | 366410 | 0 | 0 | 1820900 | 0 | 0 | 0 | 0 | 0 | 0 |
| SWTAADM | HLA-A; HLA-A; | Q53Z42; A0 | Q53Z42_HUMA | Oxidation (M) | 0 | 0 | 0 | 0 | 0 | 0 | 0 | 0 | 2040000 | 0 | 1791300 | 3617100 | 0 | 0 | 0 | 0 | 0 | 0 |
| SWTAADM | HLA-A; HLA-A; | Q53Z42; A0 | Q53Z42_HUMA | Unmodified | 0 | 0 | 0 | 0 | 0 | 0 | 0 | 0 | 10400000 | 831810 | 395960 | 24720000 | 0 | 0 | 0 | 0 | 0 | 0 |
| SWTAADTA | HLA-B; HLA-C; | P01889; P10 | HLAB_HUMAN | Unmodified | 193460 | 1157600 | 0 | 1025700 | 147870 | 0 | 0 | 0 | 35754000 | 2632500 | 722450 | 13045000 | 0 | 0 | 0 | 0 | 0 | 0 |
| SWTAADTA | HLA-B; HLA-C; | P01889; P10 | HLAB_HUMAN | Unmodified | 0 | 0 | 0 | 0 | 237710 | 28768 | 0 | 0 | 2095700 | 39590 | 0 | 1363800 | 0 | 0 | 0 | 0 | 0 | 0 |
| SWTPFPTQ | CHMP2A | M0R1T5 | M0R1T5_HUM | Unmodified | 0 | 0 | 0 | 0 | 0 | 0 | 0 | 0 | 68864 | 40977 | 0 | 0 | 261940 | 0 | 0 | 0 | 0 | 0 |
| SWVDKMQ | CACNA2D1; CA | P54289; E7E | CA2D1_HUMAI | Unmodified | 0 | 0 | 0 | 0 | 0 | 0 | 20439000 | 0 | 0 | 2129100 | 0 | 0 | 0 | 0 | 0 | 0 | 0 | 0 |
| SWVDKMQ | CACNA2D1; CA | P54289; E7E | CA2D1_HUMAI | Oxidation (M) | 0 | 0 | 0 | 0 | 0 | 0 | 2330200 | 515070 | 0 | 525240 | 0 | 0 | 0 | 0 | 0 | 0 | 0 | 0 |
| SWVIDQK | OLA1; OLA1; OL | Q9NTK5; J3K | OLA1_HUMAN | Unmodified | 0 | 0 | 0 | 0 | 536010 | 0 | 0 | 49729 | 254930 | 0 | 576630 | 8550000 | 0 | 0 | 0 | 0 | 335030 | 0 |
| SWVIDQKK | OLA1; OLA1; OL | Q9NTK5; J3K | OLA1_HUMAN | Unmodified | 48082 | 0 | 0 | 269720 | 0 | 0 | 0 | 0 | 251580 | 54620 | 191050 | 879960 | 100860 | 31807 | 136750 | 89005 | 0 | 0 |
| SWVNQME | IQGAP1; IQGAP | P46940; A0A | IQGA1_HUMAI | Oxidation (M) | 1285200 | 1547700 | 0 | 648640 | 5503600 | 0 | 0 | 0 | 0 | 525310 | 0 | 0 | 665760 | 0 | 976060 | 1546600 | 723360 | 0 |
| SWVNQME | IQGAP1; IQGAP | P46940; A0A | IQGA1_HUMAI | Unmodified | 0 | 0 | 0 | 0 | 12771000 | 3842900 | 1664200 | 0 | 3360200 | 526810 | 0 | 1741300 | 0 | 0 | 0 | 0 | 0 | 0 |
| SYAEELAK | HSD17B12; HS | Q53GQ0; A0 | DHB12_HUMAI | Unmodified | 0 | 0 | 0 | 0 | 1205500 | 620490 | 0 | 0 | 2107500 | 0 | 93380 | 859220 | 0 | 0 | 0 | 0 | 1192500 | 1412100 |
| SYAVVGVP | ACAA1; ACAA1; | P09110; C9J | THIK_HUMAN | Oxidation (M) | 0 | 0 | 0 | 0 | 0 | 0 | 0 | 0 | 3764200 | 1114200 | 0 | 0 | 0 | 0 | 0 | 0 | 0 | 0 |
| SYAVVGVP | ACAA1; ACAA1; | P09110; C9J | THIK_HUMAN | Unmodified | 0 | 0 | 0 | 0 | 0 | 0 | 0 | 0 | 187750 | 0 | 0 | 0 | 0 | 0 | 0 | 0 | 0 | 0 |
| SYCAEIAH | RPL32; RPL32; | P62910; D3Y | RL32_HUMAN | Unmodified | 2622000 | 1548000 | 1150500 | 662600 | 1805900 | 206700 | 2965600 | 2311900 | 769290 | 81171 | 515080 | 14346000 | 2593800 | 278190 | 1099500 | 846820 | 522880 | 1729600 |
| SYCAEIAH | RPL32; RPL32; | P62910; D3Y | RL32_HUMAN | Unmodified | 0 | 0 | 0 | 0 | 0 | 0 | 345790 | 117150 | 0 | 0 | 0 | 0 | 163610 | 64587 | 0 | 0 | 0 | 0 |
| SYCNDQS | HPRT1 | P00492 | HPRT_HUMAN | Unmodified | 0 | 0 | 0 | 0 | 257620 | 0 | 161840 | 108700 | 372950 | 105550 | 136410 | 974660 | 0 | 0 | 0 | 0 | 217470 | 316300 |
| SYCNDQS | HPRT1 | P00492 | HPRT_HUMAN | Unmodified | 0 | 0 | 0 | 0 | 0 | 0 | 0 | 188420 | 0 | 0 | 0 | 0 | 0 | 0 | 0 | 0 | 539670 | 0 |
| SYCYVSKE | IDI1 | Q13907 | IDI1_HUMAN Is | Unmodified | 33024 | 0 | 0 | 0 | 0 | 0 | 47908 | 0 | 0 | 0 | 0 | 480150 | 0 | 0 | 0 | 0 | 0 | 0 |
| SYDHSRH | NBR1 | Q14596 | NBR1_HUMAN | Unmodified | 0 | 0 | 0 | 62428000 | 0 | 0 | 483220 | 0 | 0 | 0 | 0 | 0 | 0 | 0 | 0 | 0 | 0 | 0 |
| SYDLTPVD | SCFD1; SCFD1 | Q8WVM8; A | SCFD1_HUMAI | Unmodified | 0 | 0 | 0 | 0 | 0 | 0 | 0 | 0 | 251470 | 86300 | 0 | 782420 | 0 | 0 | 0 | 0 | 0 | 0 |
| SYDNACQI | TMEFF2 | Q9UIK5 | TEFF2_HUMAN | Unmodified | 755410 | 0 | 0 | 0 | 0 | 0 | 0 | 0 | 0 | 0 | 0 | 0 | 393400 | 346680 | 0 | 173650 | 0 | 0 |
| SYDRAITVF | PSMA7 | O14818 | PSA7_HUMAN | Acetyl (Protein N | 0 | 0 | 0 | 0 | 4504100 | 0 | 0 | 0 | 0 | 1077300 | 0 | 0 | 9174400 | 2655900 | 0 | 0 | 3546600 | 4666700 |
| SYDRAITVF | PSMA7 | O14818 | PSA7_HUMAN | Acetyl (Protein N | 0 | 0 | 0 | 0 | 18867000 | 3392100 | 0 | 0 | 0 | 9830000 | 491920 | 0 | 20984000 | 20511000 | 0 | 0 | 11987000 | 12902000 |
| SYDTDQGF | FN3KRP; FN3KF | Q9HA64; I3L | KT3K_HUMAN | Unmodified | 109540 | 48607 | 52618 | 225920 | 357270 | 0 | 52893 | 46718 | 0 | 285340 | 0 | 0 | 0 | 230530 | 0 | 0 | 559550 | 1208800 |
| SYDVPPPP | PGAM1; PGAM | P18669; Q8 | PGAM1_HUMA | Oxidation (M) | 1590100 | 975930 | 3837000 | 6423600 | 42383000 | 10941000 | 1609500 | 2384200 | 3099700 | 3438700 | 14643000 | 70945000 | 16631000 | 9180400 | 5576400 | 4609000 | 10602000 | 21321000 |
| SYDVPPPP | PGAM1; PGAM | P18669; Q8 | PGAM1_HUMA | Unmodified | 353970 | 251650 | 597190 | 1399400 | 35847000 | 6257000 | 951840 | 568350 | 4866300 | 779230 | 3203600 | 12186000 | 14896000 | 3119800 | 1341100 | 860500 | 7891200 | 32254000 |
| SYDYDLIIG | TXNRD1; TXNRI | Q16881; A0 | TRXR1_HUMAN | Unmodified | 0 | 0 | 0 | 0 | 17081000 | 4475400 | 0 | 181990 | 2661400 | 3877600 | 4020800 | 3082600 | 2817100 | 1762400 | 0 | 0 | 2498400 | 9613.7 |
| SYDYDLIIG | TXNRD1; TXNRI | Q16881; A0 | TRXR1_HUMAN | Unmodified | 0 | 0 | 0 | 0 | 0 | 0 | 0 | 0 | 0 | 0 | 0 | 0 | 0 | 0 | 0 | 0 | 1833400 | 631870 |
| SYDYQSVC | CACNA2D1 | P54289 | CA2D1_HUMAI | Unmodified | 365950 | 0 | 0 | 0 | 0 | 0 | 0 | 58611 | 374870 | 0 | 0 | 0 | 0 | 0 | 0 | 0 | 0 | 0 |
| SYEAQDPE | C12orf57; C12 | Q99622; F5 | C10_HUMAN P | Unmodified | 398350 | 598760 | 0 | 856290 | 2397700 | 562860 | 782590 | 735610 | 869930 | 426360 | 0 | 1352500 | 2176000 | 1625500 | 0 | 288630 | 1124900 | 1369900 |
| SYEAQDPE | C12orf57; C12 | Q99622; F5 | C10_HUMAN P | Unmodified | 0 | 153280 | 0 | 0 | 0 | 0 | 974620 | 952770 | 0 | 0 | 0 | 0 | 0 | 0 | 0 | 58895 | 1514100 | 837350 |
| SYEAYVLNI | ATP1B1; ATP1B | P05026; V9 | AT1B1_HUMAN | Unmodified | 2772300 | 0 | 11115000 | 11973000 | 651460 | 380830 | 0 | 0 | 33408000 | 4022800 | 6326600 | 43468000 | 605730 | 0 | 17080000 | 17645000 | 0 | 0 |
| SYEAYVLNI | ATP1B1; ATP1B | P05026; V9 | AT1B1_HUMAN | Unmodified | 0 | 0 | 17434 | 0 | 1197100 | 103410 | 729370 | 0 | 0 | 1137600 | 0 | 0 | 1305200 | 1350400 | 0 | 0 | 293130 | 265490 |
| SYEDMIGE | CTNND1; CTNN | O60716; A0 | CTND1_HUMAI | Oxidation (M) | 0 | 0 | 0 | 3083300 | 0 | 0 | 0 | 0 | 0 | 0 | 0 | 0 | 0 | 0 | 0 | 0 | 0 | 0 |
| SYEDMIGE | CTNND1; CTNN | O60716; A0 | CTND1_HUMAI | Oxidation (M) | 0 | 0 | 0 | 0 | 0 | 0 | 0 | 0 | 0 | 0 | 0 | 0 | 0 | 9019500 | 0 | 0 | 1729200 | 3861700 |
| SYEECIIESI | IGF2R | P11717 | MPRI_HUMAN | Unmodified | 0 | 0 | 0 | 0 | 0 | 238140 | 0 | 0 | 11870000 | 1988600 | 818530 | 1904200 | 0 | 0 | 0 | 0 | 0 | 0 |
| SYEILLHEV | KRIT1; KRIT1; K | O00522; C9J | KRIT1_HUMAN | Unmodified | 0 | 0 | 0 | 0 | 0 | 0 | 244080 | 141170 | 0 | 143240 | 0 | 0 | 172790 | 503430 | 0 | 0 | 0 | 469690 |
| SYELPDGQ | POTEE; POTEF; | Q6S8J3; A5A | POTEE_HUMAN | Unmodified | 1.18E+08 | 91276000 | 1.16E+08 | 2.25E+08 | 3.31E+08 | 1.25E+08 | 18183000 | 18003000 | 2.86E+08 | 41825000 | 1.62E+08 | 1.01E+09 | 1.68E+08 | 57050000 | 2.15E+08 | 1.91E+08 | 39969000 | 99455000 |
| SYELPDGQ | POTEE; POTEF; | Q6S8J3; A5A | POTEE_HUMAN | Unmodified | 7211900 | 2391100 | 4805000 | 34884000 | 2.58E+08 | 1.28E+08 | 15710000 | 2384600 | 31560000 | 20024000 | 31157000 | 2053100 | 1.33E+08 | 45419000 | 6869600 | 3140700 | 44106000 | 70281000 |
| SYELQESN | SEPTIN11; SEP | Q9NVA2; D6 | SEP11_HUMAN | Unmodified | 100660 | 87880 | 0 | 248460 | 0 | 0 | 0 | 0 | 0 | 41592 | 1073000 | 2795400 | 0 | 0 | 0 | 0 | 0 | 0 |
| SYEPLEDP | TALDO1; TALDC | P37837; F2Z | TALDO_HUMAN | Unmodified | 0 | 0 | 510040 | 686910 | 6804600 | 1032200 | 0 | 0 | 0 | 0 | 5421400 | 15653000 | 3127300 | 0 | 0 | 0 | 1094800 | 1557600 |
| SYEQLREK | WWP2 | O00308 | WWP2_HUMAN | Unmodified | 330830 | 0 | 404170 | 0 | 0 | 0 | 0 | 0 | 251890 | 0 | 0 | 0 | 0 | 0 | 630110 | 716570 | 0 | 0 |
| SYESLFIR | CRTAP; CRTAP | O75718; C9J | CRTAP_HUMAN | Unmodified | 0 | 0 | 27091 | 21406 | 0 | 0 | 0 | 0 | 5363600 | 571750 | 45546 | 0 | 0 | 0 | 20557 | 0 | 94112 | 0 |
| SYESMCEY | SMOC1 | Q9H4F8 | SMOC1_HUMA | Unmodified | 0 | 0 | 0 | 0 | 0 | 0 | 0 | 0 | 451690 | 0 | 0 | 0 | 0 | 0 | 0 | 0 | 0 | 0 |
| SYEVTVENI | PIGK; PIGK | Q92643; A6 | GPI8_HUMAN | Unmodified | 0 | 0 | 0 | 0 | 0 | 0 | 0 | 0 | 671720 | 0 | 0 | 0 | 0 | 0 | 0 | 0 | 0 | 0 |
| SYEYKSNP | TJP2; TJP2; TJP2 | Q9UDY2; A0 | ZO2_HUMAN T | Unmodified | 0 | 0 | 0 | 0 | 90729 | 0 | 0 | 0 | 0 | 0 | 0 | 0 | 0 | 0 | 0 | 0 | 162490 | 0 |
| SYGATATSI | SLC35B2 | Q8TB61 | S35B2_HUMAN | Unmodified | 0 | 0 | 0 | 0 | 0 | 0 | 0 | 0 | 1608100 | 303490 | 23654 | 1737800 | 99920 | 0 | 0 | 0 | 0 | 0 |
| SYGEEDFE | SLC4A2; SLC4A | P04920; C9J | B3A2_HUMAN | Unmodified | 80480 | 54318 | 226640 | 76418 | 0 | 0 | 0 | 0 | 0 | 0 | 0 | 471670 | 0 | 0 | 472110 | 461820 | 0 | 0 |
| SYGFNAAR | MSH6; MSH6; | P52701; A0A | MSH6_HUMAN | Unmodified | 0 | 0 | 0 | 0 | 0 | 0 | 4616500 | 106070 | 0 | 76055 | 0 | 0 | 0 | 0 | 0 | 0 | 0 | 0 |
| SYGIPFIETS | NRAS; KRAS | P01111; P01 | RASN_HUMAN | Unmodified | 0 | 9088100 | 0 | 7624300 | 11485000 | 8460100 | 0 | 5743900 | 10708000 | 4322400 | 19072000 | 18596000 | 16973000 | 14595000 | 7738300 | 7758700 | 2128800 | 5166700 |
| SYGPAPGA | AIMP2 | Q13155 | AIMP2_HUMAN | Unmodified | 0 | 0 | 0 | 0 | 0 | 0 | 0 | 0 | 0 | 0 | 0 | 5590100 | 0 | 0 | 0 | 0 | 0 | 0 |
| SYGPAPGA | AIMP2 | Q13155 | AIMP2_HUMAN | Unmodified | 0 | 0 | 0 | 0 | 1999300 | 0 | 0 | 0 | 0 | 1511600 | 0 | 0 | 4982000 | 1277000 | 0 | 0 | 0 | 1351500 |
| SYGPLDMY | STX12; STX12 | Q86Y82; B1 | STX12_HUMAN | Acetyl (Protein N | 0 | 0 | 0 | 786650 | 0 | 0 | 0 | 0 | 1142600 | 145940 | 168340 | 0 | 249710 | 0 | 0 | 0 | 0 | 0 |
| SYGPLDMY | STX12; STX12 | Q86Y82; B1 | STX12_HUMAN | Acetyl (Protein N | 0 | 0 | 0 | 0 | 0 | 0 | 0 | 0 | 847670 | 0 | 0 | 524560 | 0 | 0 | 117790 | 0 | 0 | 0 |
| SYGQDHLI | SLC5A6 | Q9Y289 | SC5A6_HUMAI | Unmodified | 0 | 0 | 670300 | 785300 | 0 | 0 | 0 | 0 | 0 | 0 | 1301900 | 2200100 | 0 | 132050 | 477280 | 357210 | 0 | 0 |
| SYGTCVNT | ADGRE2; ADGR | Q9UHX3; B4 | AGRE2_HUMAI | Unmodified | 0 | 0 | 0 | 0 | 0 | 0 | 426510 | 0 | 0 | 0 | 1705700 | 0 | 0 | 0 | 0 | 0 | 0 | 0 |
| SYGVENR | GPRC5A; GPRC | Q8NFJ5; A0 | RAI3_HUMAN F | Unmodified | 0 | 0 | 0 | 148560 | 39942000 | 16127000 | 0 | 0 | 0 | 0 | 0 | 0 | 0 | 0 | 0 | 0 | 0 | 0 |
| SYHEEFNP | AK2; AK2; AK2; | P54819; A0A | KAD2_HUMAN | Oxidation (M) | 0 | 0 | 0 | 0 | 0 | 0 | 0 | 0 | 671740 | 0 | 639770 | 2691900 | 0 | 0 | 0 | 0 | 0 | 0 |
| SYHEEFNP | AK2; AK2; AK2; | P54819; A0A | KAD2_HUMAN | Unmodified | 0 | 0 | 0 | 0 | 0 | 0 | 0 | 0 | 4458600 | 0 | 0 | 757700 | 0 | 0 | 0 | 0 | 0 | 0 |
| SYHLLVR | PTGFRN | Q9P2B2 | FPRP_HUMAN | Unmodified | 1773700 | 1527300 | 9247300 | 2584100 | 0 | 0 | 0 | 0 | 0 | 0 | 0 | 5604100 | 0 | 0 | 5038700 | 4921800 | 0 | 0 |
| SYHLLVRD | PTGFRN | Q9P2B2 | FPRP_HUMAN | Unmodified | 84501 | 32277 | 534930 | 2328400 | 1318800 | 857890 | 197920 | 0 | 0 | 0 | 0 | 0 | 4451600 | 1414200 | 88156 | 55866 | 31894 | 74087 |
| SYIATQGCI | PTPN11 | Q06124 | PTN11_HUMAI | Unmodified | 0 | 0 | 0 | 0 | 395460 | 0 | 0 | 0 | 0 | 205330 | 131430 | 1434700 | 0 | 0 | 0 | 0 | 175060 | 1784100 |
| SYIEYQLTP | SNX9; SNX9; SI | Q9Y5X1; A0 | SNX9_HUMAN | Unmodified | 8587300 | 7678700 | 929800 | 771120 | 416770 | 55290 | 0 | 0 | 175970 | 120020 | 402990 | 2524500 | 836170 | 322590 | 1606900 | 2768600 | 0 | 91106 |
| SYIEYQLTP | SNX9; SNX9; SI | Q9Y5X1; A0 | SNX9_HUMAN | Unmodified | 0 | 0 | 0 | 391030 | 495630 | 0 | 44521 | 0 | 0 | 0 | 0 | 0 | 1709600 | 1271400 | 0 | 0 | 0 | 0 |

| | | | | | | | | | | | | | | | | | | | | | | |
|---|---|---|---|---|---|---|---|---|---|---|---|---|---|---|---|---|---|---|---|---|---|---|
| SYIGSNHS | CXADR | P78310 | CXAR_HUMAN | Unmodified | 1047000 | 1157400 | 0 | 0 | 24494000 | 2558300 | 320880 | 0 | 3024000 | 764620 | 323770 | 946860 | 924430 | 0 | 781730 | 0 | 682710 | 880190 |
| SYIGSNHS | CXADR | P78310 | CXAR_HUMAN | 2 Oxidation (M) | 298560 | 149220 | 0 | 0 | 1994700 | 568400 | 110960 | 0 | 1210300 | 0 | 0 | 0 | 0 | 0 | 239400 | 176870 | 231800 | 0 |
| SYIGSNHS | CXADR | P78310 | CXAR_HUMAN | Oxidation (M) | 470770 | 318320 | 0 | 98292 | 8409700 | 2234900 | 75608 | 0 | 848400 | 224600 | 0 | 0 | 132220 | 0 | 106910 | 320030 | 237990 | 65538 |
| SYIKSNLEL | C3orf20 | Q8ND61 | CC020_HUMAN | Acetyl (Protein N | 0 | 0 | 0 | 0 | 0 | 9883000 | 0 | 0 | 0 | 0 | 0 | 0 | 0 | 0 | 0 | 0 | 0 | 0 |
| SYIKSNLEL | C3orf20 | Q8ND61 | CC020_HUMAN | Acetyl (Protein N | 827990 | 1482600 | 0 | 912690 | 0 | 0 | 330930 | 0 | 0 | 0 | 324020 | 0 | 0 | 2091600 | 842350 | 115270 | 0 | 0 |
| SYIPGQPVT | TAX1BP3; TAX1 | O14907; A0A | TX1B3_HUMAN | Acetyl (Protein N | 0 | 0 | 166030 | 0 | 0 | 0 | 0 | 0 | 0 | 0 | 223190 | 890370 | 0 | 0 | 0 | 0 | 0 | 0 |
| SYIPGQPVT | TAX1BP3; TAX1 | O14907; A0A | TX1B3_HUMAN | Acetyl (Protein N | 0 | 962560 | 1156600 | 3914400 | 26584000 | 11032000 | 1267500 | 0 | 0 | 525500 | 2879000 | 2657800 | 6467600 | 9807400 | 4561800 | 2440300 | 9782500 | 8892800 |
| SYIQSLPQT | MAPK13 | O15264 | MK13_HUMAN | Unmodified | 0 | 0 | 43005 | 0 | 0 | 0 | 0 | 0 | 2587500 | 662270 | 195220 | 1949500 | 0 | 0 | 0 | 0 | 0 | 0 |
| SYISFDILR | CARS1; CARS1 | P49589; B4D | SYCC_HUMAN | Unmodified | 0 | 0 | 0 | 0 | 0 | 0 | 0 | 0 | 0 | 0 | 0 | 630520 | 0 | 0 | 772210 | 0 | 434450 | 0 |
| SYISFDILRI | CARS1; CARS1 | P49589; B4D | SYCC_HUMAN | Unmodified | 0 | 0 | 0 | 0 | 0 | 0 | 0 | 0 | 0 | 0 | 0 | 778570 | 0 | 0 | 68490 | 35587 | 0 | 0 |
| SYISYKLTP | SNX33; SNX33 | Q8WV41; H3 | SNX33_HUMAN | Unmodified | 0 | 0 | 0 | 1847100 | 0 | 0 | 0 | 0 | 0 | 0 | 0 | 0 | 0 | 0 | 0 | 0 | 0 | 0 |
| SYISYKLVP | SNX18 | Q96RF0 | SNX18_HUMAN | Unmodified | 0 | 0 | 0 | 0 | 0 | 0 | 0 | 0 | 0 | 0 | 0 | 0 | 1152700 | 256220 | 0 | 0 | 0 | 0 |
| SYISYKLVP | SNX18 | Q96RF0 | SNX18_HUMAN | Unmodified | 0 | 0 | 0 | 0 | 0 | 0 | 0 | 0 | 0 | 0 | 0 | 0 | 240080 | 0 | 0 | 0 | 0 | 0 |
| SYIYSGSHI | WDR1 | O75083 | WDR1_HUMAN | Unmodified | 1594000 | 1412700 | 0 | 0 | 17203000 | 3691500 | 0 | 116820 | 2059400 | 0 | 0 | 11088000 | 3046800 | 3814200 | 2119400 | 1702400 | 1351600 | 2575900 |
| SYIYSGSHI | WDR1 | O75083 | WDR1_HUMAN | Unmodified | 0 | 0 | 0 | 0 | 19602000 | 2711300 | 569920 | 0 | 0 | 1496300 | 0 | 0 | 10731000 | 4998800 | 0 | 0 | 4419600 | 4066200 |
| SYKGEGRQ | ZC3HAV1; ZC3I | Q7Z2W4; C9 | ZCCHV_HUMA | Unmodified | 0 | 0 | 0 | 129490 | 5146900 | 1216100 | 212970 | 83061 | 0 | 210570 | 0 | 0 | 1732600 | 1838200 | 0 | 0 | 388530 | 689730 |
| SYKIAYSNE | EDIL3 | O43854 | EDIL3_HUMAN | Unmodified | 0 | 0 | 0 | 0 | 1430200 | 76659 | 98839 | 74883 | 0 | 0 | 0 | 0 | 0 | 0 | 0 | 0 | 65802 | 448750 |
| SYKIAYSNE | EDIL3 | O43854 | EDIL3_HUMAN | Unmodified | 0 | 0 | 0 | 0 | 5759300 | 0 | 676370 | 73419 | 0 | 0 | 0 | 0 | 0 | 0 | 0 | 0 | 481230 | 3226200 |
| SYKIAYSNE | EDIL3 | O43854 | EDIL3_HUMAN | Oxidation (M) | 0 | 0 | 0 | 0 | 2372700 | 0 | 883990 | 174780 | 79189 | 0 | 0 | 0 | 64166 | 0 | 0 | 0 | 336280 | 1354400 |
| SYKPVFWS | IARS2 | Q9NSE4 | SYIM_HUMAN | Unmodified | 0 | 0 | 0 | 0 | 0 | 0 | 0 | 11575 | 1541800 | 0 | 0 | 0 | 0 | 0 | 0 | 0 | 0 | 0 |
| SYLENPAFI | DBT; DBT; DBT | P11182; A0A | ODB2_HUMAN | Oxidation (M) | 0 | 0 | 0 | 0 | 0 | 0 | 64155 | 0 | 471860 | 216080 | 0 | 0 | 37322 | 38431 | 20808 | 0 | 0 | 0 |
| SYLLSMGIA | VPS36; VPS36 | Q86VN1; A0A | VPS36_HUMAN | Oxidation (M) | 2237900 | 1665200 | 880810 | 514880 | 0 | 0 | 0 | 0 | 0 | 0 | 0 | 187000 | 0 | 0 | 1967600 | 1441700 | 0 | 0 |
| SYLPNAHL | TAX1BP1; TAX1 | Q86VP1; B8A | TAXB1_HUMAN | Unmodified | 381990 | 0 | 0 | 0 | 0 | 0 | 701420 | 0 | 0 | 248470 | 0 | 0 | 0 | 0 | 0 | 0 | 385630 | 0 |
| SYLPQTVR | IL6ST | P40189 | IL6RB_HUMAN | Oxidation (M) | 0 | 0 | 0 | 0 | 926340 | 89848 | 141040 | 0 | 0 | 0 | 0 | 689570 | 0 | 0 | 0 | 0 | 0 | 188410 |
| SYLPQTVR | IL6ST | P40189 | IL6RB_HUMAN | Unmodified | 140610 | 0 | 0 | 0 | 1426700 | 0 | 297810 | 0 | 0 | 71503 | 0 | 0 | 93413 | 0 | 0 | 104470 | 122450 | 435020 |
| SYLPVQFQ | CEMIP2; CEMIP | Q9UHN6; B4 | CEIP2_HUMAN | Unmodified | 522100 | 215730 | 0 | 454930 | 0 | 0 | 0 | 0 | 2115100 | 192110 | 0 | 487910 | 0 | 0 | 731480 | 630920 | 0 | 0 |
| SYLTEQVN | PCK2; PCK2 | Q16822; H0 | PCKGM_HUMA | Unmodified | 37881 | 0 | 0 | 531230 | 436850 | 0 | 0 | 103050 | 2323600 | 0 | 0 | 0 | 180890 | 0 | 133810 | 89603 | 0 | 0 |
| SYLTPVRD | PPP1R12A; PPP | O14974; F8 | MYPT1_HUMAN | Unmodified | 0 | 0 | 0 | 0 | 734270 | 0 | 28530 | 28827 | 0 | 0 | 0 | 0 | 395470 | 341180 | 0 | 90681 | 216140 | 329600 |
| SYLVQYMQ | UPF1 | Q92900 | RENT1_HUMAN | Oxidation (M) | 548340 | 484840 | 210700 | 0 | 0 | 0 | 774060 | 0 | 152360 | 0 | 0 | 0 | 0 | 0 | 186120 | 187570 | 0 | 0 |
| SYLYFTQFK | MYDGF; MYDGI | Q969H8; M0 | MYDGF_HUMA | Unmodified | 0 | 0 | 0 | 0 | 712800 | 0 | 0 | 284910 | 71150000 | 13848000 | 5563000 | 1521300 | 700810 | 642310 | 0 | 0 | 284570 | 1146600 |
| SYLYFTQFK | MYDGF; MYDGI | Q969H8; M0 | MYDGF_HUMA | Unmodified | 0 | 0 | 0 | 0 | 0 | 0 | 0 | 0 | 7790600 | 15721000 | 0 | 692000 | 0 | 0 | 0 | 0 | 3015500 | 0 |
| SYNDELQF | RTCB | Q9Y3I0 | RTCB_HUMAN | Unmodified | 0 | 0 | 1268200 | 0 | 0 | 0 | 0 | 159410 | 0 | 0 | 244920 | 7358800 | 0 | 0 | 0 | 0 | 0 | 0 |
| SYNEQRIR | AKR1C2; AKR1 | P52895; Q0 | AK1C2_HUMAN | Oxidation (M) | 0 | 0 | 0 | 0 | 3263900 | 0 | 0 | 0 | 0 | 0 | 0 | 0 | 0 | 0 | 0 | 0 | 0 | 0 |
| SYNEQRIR | AKR1C2; AKR1 | P52895; Q0 | AK1C2_HUMAN | Unmodified | 0 | 0 | 0 | 0 | 635680 | 0 | 0 | 0 | 0 | 0 | 0 | 0 | 0 | 0 | 0 | 0 | 0 | 0 |
| SYNRFQIA | PSMB4 | P28070 | PSB4_HUMAN | Unmodified | 0 | 0 | 0 | 0 | 2229900 | 683260 | 0 | 0 | 4503300 | 2554500 | 1998500 | 13288000 | 1676100 | 191910 | 0 | 0 | 678500 | 937240 |
| SYPDGWN | FOLH1 | Q04609 | FOLH1_HUMAN | Unmodified | 6952600 | 5537100 | 27499000 | 35047000 | 0 | 0 | 0 | 0 | 107470 | 0 | 0 | 0 | 1622600 | 703570 | 22309000 | 15488000 | 0 | 0 |
| SYPGLLIVP | SBF1 | O95248 | MTMR5_HUMA | Unmodified | 0 | 0 | 0 | 0 | 0 | 0 | 0 | 0 | 0 | 0 | 0 | 1376600 | 0 | 0 | 0 | 0 | 0 | 0 |
| SYPNEEKD | ADAM17 | P78536 | ADA17_HUMAN | Unmodified | 0 | 0 | 0 | 0 | 705180 | 84076 | 62674 | 0 | 0 | 117310 | 0 | 0 | 160920 | 283140 | 0 | 0 | 736750 | 1925600 |
| SYPSSTNV | AVIL | O75366 | AVIL_HUMAN A | Unmodified | 0 | 0 | 0 | 0 | 0 | 0 | 76693 | 42898 | 0 | 77223 | 849290 | 7295000 | 0 | 0 | 0 | 0 | 174730 | 0 |
| SYPSSTNV | AVIL | O75366 | AVIL_HUMAN A | Oxidation (M) | 0 | 0 | 0 | 0 | 0 | 54233 | 0 | 0 | 0 | 0 | 0 | 8385400 | 0 | 0 | 0 | 0 | 0 | 0 |
| SYQDAVLE | FKBP7 | Q9Y680 | FKBP7_HUMAN | Unmodified | 0 | 0 | 0 | 0 | 0 | 0 | 0 | 0 | 0 | 0 | 519970 | 0 | 0 | 0 | 0 | 0 | 0 | 0 |
| SYQDAVLE | FKBP7 | Q9Y680 | FKBP7_HUMAN | Unmodified | 0 | 0 | 0 | 0 | 0 | 0 | 0 | 0 | 2526300 | 1419300 | 0 | 0 | 0 | 0 | 0 | 0 | 0 | 0 |
| SYQDFFEA | ATF6; ATF6; ATF | P18850; A0A | ATF6A_HUMAN | Unmodified | 0 | 0 | 668320 | 13418 | 0 | 0 | 0 | 0 | 1741700 | 27992 | 0 | 0 | 0 | 0 | 0 | 0 | 0 | 0 |
| SYQDPSNA | NSFL1C; NSFL1 | Q9UNZ2; F2 | NSF1C_HUMAN | Unmodified | 309450 | 190140 | 0 | 0 | 0 | 0 | 0 | 0 | 0 | 0 | 0 | 797040 | 0 | 0 | 119410 | 193420 | 0 | 0 |
| SYQFGIIGN | TNKS1BP1 | Q9C0C2 | TB182_HUMAN | Unmodified | 2094300 | 273940 | 0 | 401460 | 0 | 0 | 0 | 0 | 85700 | 0 | 89573 | 2453200 | 149100 | 0 | 0 | 0 | 0 | 0 |
| SYQLRPGT | NIPSNAP2; NIP | O75323; C9 | NIPS2_HUMAN | Oxidation (M) | 0 | 0 | 0 | 0 | 0 | 0 | 0 | 0 | 939600 | 0 | 0 | 0 | 0 | 0 | 0 | 0 | 0 | 0 |
| SYQLRPGT | NIPSNAP2; NIP | O75323; C9 | NIPS2_HUMAN | Unmodified | 0 | 0 | 0 | 0 | 0 | 671260 | 0 | 0 | 2101600 | 0 | 0 | 0 | 0 | 0 | 0 | 0 | 0 | 0 |
| SYQTNTAD | B4GALNT1 | Q00973 | B4GN1_HUMAN | Unmodified | 0 | 0 | 0 | 0 | 0 | 0 | 0 | 0 | 779340 | 0 | 0 | 0 | 0 | 0 | 0 | 0 | 0 | 0 |
| SYQVICRD | PLAT; PLAT; PL | P00750; E5R | TPA_HUMAN Ti | Unmodified | 0 | 0 | 0 | 0 | 0 | 0 | 0 | 0 | 0 | 0 | 0 | 0 | 0 | 0 | 0 | 0 | 0 | 620680 |
| SYQVPMLA | SERPINE2; SER | P07093; C9J | GDN_HUMAN ( | Unmodified | 43914 | 0 | 18078 | 0 | 0 | 0 | 0 | 0 | 0 | 17650 | 0 | 0 | 0 | 0 | 0 | 61904 | 0 | 0 |
| SYRDDIDL | TSPAN14; TSPA | Q8NG11; A6 | TSN14_HUMAN | Unmodified | 11391000 | 7022100 | 6501300 | 11802000 | 402170 | 319520 | 0 | 0 | 4063800 | 2639700 | 1330900 | 7504400 | 526290 | 163780 | 30284000 | 10471000 | 0 | 0 |
| SYRGDREA | PNPO; PNPO; P | Q9NVS9; J3 | PNPO_HUMAN | Unmodified | 527030 | 400910 | 861840 | 1587500 | 7120900 | 2188500 | 1106800 | 0 | 761800 | 0 | 432390 | 3219600 | 10258000 | 5200600 | 2083500 | 1446600 | 3762000 | 11478000 |
| SYRSMTPA | EPB41; EPB41; | P11171; A0A | EPB41_HUMAN | Oxidation (M) | 0 | 0 | 0 | 0 | 1660900 | 0 | 4637800 | 1290300 | 0 | 369640 | 0 | 0 | 1050500 | 1996700 | 0 | 0 | 612950 | 1451600 |
| SYRSMTPA | EPB41; EPB41; | P11171; A0A | EPB41_HUMAN | Unmodified | 0 | 0 | 0 | 0 | 0 | 0 | 1879000 | 121610 | 0 | 0 | 0 | 0 | 0 | 0 | 0 | 0 | 0 | 0 |
| SYRYWDAE | GOT1 | P17174 | AATC_HUMAN | Unmodified | 0 | 0 | 0 | 0 | 155900 | 0 | 0 | 0 | 0 | 155360 | 291610 | 0 | 0 | 68894 | 0 | 0 | 0 | 0 |
| SYSEDDIHI | YTHDF1; YTHDI | Q9BYJ9; Q7Z | YTHD1_HUMAN | Unmodified | 491270 | 0 | 0 | 0 | 0 | 0 | 0 | 0 | 73157 | 0 | 0 | 489260 | 0 | 0 | 166420 | 183620 | 0 | 0 |
| SYSFDEIRK | NAMPT; NAMPT | P43490; A0A | NAMPT_HUMAN | Unmodified | 0 | 0 | 0 | 0 | 22289000 | 552530 | 180440 | 44017 | 0 | 84690 | 0 | 556670 | 694920 | 408880 | 0 | 0 | 345080 | 445230 |
| SYSKPPQC | SFRP5 | Q5T4F7 | SFRP5_HUMAN | Unmodified | 5864700 | 1865900 | 0 | 0 | 0 | 0 | 0 | 0 | 1084400 | 0 | 0 | 0 | 0 | 0 | 0 | 0 | 0 | 0 |
| SYSLQVPSI | CDCP1 | Q9H5V8 | CDCP1_HUMA | Unmodified | 156220 | 0 | 461520 | 2930100 | 47377000 | 2126300 | 0 | 0 | 418350 | 2450900 | 664180 | 789010 | 10833000 | 5269400 | 6207300 | 652500 | 60665000 | 31301000 |
| SYSLQVPSI | CDCP1 | Q9H5V8 | CDCP1_HUMA | Unmodified | 0 | 0 | 0 | 0 | 130880 | 0 | 0 | 0 | 0 | 0 | 0 | 0 | 97981 | 48559 | 0 | 0 | 505040 | 354830 |
| SYSMIVNN | CACYBP | Q9HB71 | CYBP_HUMAN | Oxidation (M) | 108220 | 199910 | 95701 | 0 | 874010 | 0 | 1876100 | 162670 | 1338000 | 2106600 | 3399800 | 23712000 | 6253400 | 4023300 | 247550 | 195800 | 0 | 0 |
| SYSMIVNN | CACYBP | Q9HB71 | CYBP_HUMAN | Oxidation (M) | 0 | 0 | 188450 | 329250 | 2550100 | 235600 | 993950 | 94836 | 728760 | 1667400 | 893790 | 7504000 | 2867500 | 2390300 | 0 | 0 | 625220 | 642480 |
| SYSPPVPC | SORL1 | Q92673 | SORL_HUMAN | Unmodified | 0 | 0 | 0 | 45396 | 243990 | 0 | 78449 | 0 | 1888400 | 269230 | 111560 | 115410 | 83945 | 0 | 19507 | 0 | 177100 | 0 |
| SYSPSDLVI | ELP3; ELP3 | Q9H9T3; B4I | ELP3_HUMAN | Unmodified | 0 | 0 | 847290 | 0 | 0 | 0 | 0 | 0 | 396460 | 0 | 0 | 894770 | 0 | 0 | 0 | 0 | 0 | 0 |
| SYSPYDML | ANXA2P2; ANX | A6NMY6; P0 | AXA2L_HUMAN | Oxidation (M) | 0 | 0 | 0 | 0 | 13762000 | 6897200 | 497650 | 232280 | 113170 | 2004900 | 0 | 0 | 0 | 0 | 0 | 0 | 4835900 | 2532600 |
| SYSPYDML | ANXA2P2; ANX | A6NMY6; P0 | AXA2L_HUMAN | Unmodified | 0 | 0 | 0 | 0 | 67969000 | 21647000 | 1791500 | 94102 | 4566100 | 3756900 | 0 | 0 | 0 | 0 | 0 | 0 | 17877000 | 12500000 |
| SYSQSILLD | CD109 | Q6YHK3 | CD109_HUMAN | Unmodified | 0 | 0 | 0 | 0 | 0 | 0 | 0 | 0 | 11085000 | 17670000 | 0 | 0 | 0 | 0 | 0 | 0 | 16338000 | 0 |
| SYSRPSPL | USP6NL; USP6I | Q92738; X6F | US6NL_HUMAN | Unmodified | 0 | 0 | 0 | 0 | 0 | 0 | 0 | 0 | 0 | 0 | 0 | 0 | 0 | 0 | 280800 | 0 | 0 | 0 |
| SYSRQSSS | C2CD5; C2CD5 | Q86YS7; H0 | C2CD5_HUMA | Unmodified | 0 | 0 | 0 | 0 | 202180 | 0 | 88521 | 63982 | 0 | 57134 | 0 | 0 | 117540 | 100390 | 0 | 0 | 185650 | 178730 |
| SYSRRTTE | AXL; AXL | P30530; M0 | UFO_HUMAN T | Unmodified | 0 | 0 | 0 | 0 | 2854900 | 0 | 431560 | 0 | 0 | 0 | 0 | 0 | 589360 | 245070 | 0 | 0 | 0 | 54558 |
| SYSSGGED | EIF3I; EIF3I; EIF | Q13347; A0A | EIF3I_HUMAN | Unmodified | 51288 | 51639 | 280210 | 296890 | 79155 | 0 | 0 | 0 | 172400 | 0 | 274930 | 3883000 | 0 | 0 | 179470 | 289350 | 0 | 0 |
| SYSTTYEE | SLC1A5; SLC1A | Q15758; H0 | AAAT_HUMAN | Unmodified | 0 | 0 | 0 | 0 | 0 | 0 | 0 | 0 | 59250 | 0 | 49943 | 9112500 | 0 | 0 | 0 | 0 | 0 | 0 |
| SYSVGASG | CPSF7; CPSF7 | Q8N684; F5I | CPSF7_HUMAN | Unmodified | 261670 | 247370 | 74874 | 147600 | 0 | 0 | 0 | 0 | 0 | 0 | 0 | 188690 | 224950 | 0 | 61149 | 57053 | 0 | 0 |
| SYSVGASG | CPSF7; CPSF7 | Q8N684; F5I | CPSF7_HUMAN | Unmodified | 755660 | 600520 | 0 | 0 | 143810 | 0 | 36411 | 17284 | 0 | 25287 | 0 | 0 | 0 | 25070 | 0 | 0 | 0 | 0 |
| SYTAADAT | UBA6; UBA6 | A0AVT1; H0 | UBA6_HUMAN | Unmodified | 0 | 0 | 0 | 0 | 762570 | 0 | 0 | 289940 | 347190 | 224900 | 0 | 4049400 | 670480 | 257860 | 0 | 0 | 3025000 | 5065500 |
| SYTITGLQP | FN1; FN1 | P02751; H0 | FINC_HUMAN | Unmodified | 564930 | 1957100 | 8417200 | 1455700 | 0 | 924490 | 0 | 0 | 0 | 737220 | 0 | 1455000 | 161910 | 0 | 1044500 | 4014400 | 191400 | 0 |
| SYTLVVAK | EPB41L2; EPB4 | O43491; E9F | E41L2_HUMAN | Unmodified | 0 | 0 | 0 | 150850 | 7660100 | 1182900 | 338130 | 851580 | 0 | 0 | 0 | 0 | 659020 | 461300 | 147810 | 101000 | 7040200 | 2763900 |
| SYTVGVMN | SERPINH1; SER | P50454; E9P | SERPH_HUMAN | Oxidation (M) | 0 | 0 | 0 | 0 | 0 | 0 | 0 | 0 | 766950 | 354110 | 0 | 0 | 0 | 0 | 0 | 0 | 0 | 0 |
| SYTVGVMN | SERPINH1; SER | P50454; E9P | SERPH_HUMAN | Unmodified | 0 | 0 | 0 | 0 | 0 | 0 | 0 | 0 | 3119400 | 0 | 0 | 0 | 0 | 0 | 0 | 0 | 0 | 0 |
| SYVDLLKLE | NAA15; NAA15 | Q9BXJ9; A0A | NAA15_HUMAN | Unmodified | 0 | 0 | 0 | 0 | 0 | 0 | 0 | 0 | 0 | 22231 | 32674 | 971950 | 0 | 0 | 0 | 0 | 64330 | 0 |
| SYVDTGGV | FMNL1 | O95466 | FMNL1_HUMAN | Unmodified | 0 | 99116 | 0 | 0 | 1496400 | 700110 | 724360 | 0 | 0 | 0 | 24573 | 0 | 0 | 0 | 0 | 0 | 0 | 0 |
| SYVHSALK | USP8; USP8; U | P40818; H0 | UBP8_HUMAN | Unmodified | 0 | 0 | 0 | 0 | 623490 | 257640 | 0 | 0 | 0 | 0 | 0 | 0 | 0 | 0 | 0 | 0 | 0 | 136700 |
| SYVISSFTE | PLCB2 | Q00722 | PLCB2_HUMAN | Unmodified | 0 | 0 | 0 | 0 | 0 | 0 | 0 | 0 | 0 | 0 | 0 | 3650100 | 330570 | 0 | 0 | 0 | 0 | 2507600 |
| SYVITGSW | RABGAP1 | Q9Y3P9 | RBGP1_HUMAN | Unmodified | 0 | 0 | 0 | 0 | 1084000 | 0 | 0 | 0 | 139850 | 0 | 0 | 890310 | 0 | 0 | 0 | 0 | 0 | 0 |
| SYVSEKDV | LRPPRC | P42704 | LPPRC_HUMAN | Unmodified | 0 | 0 | 0 | 0 | 0 | 0 | 142270 | 0 | 79688 | 201020 | 0 | 0 | 0 | 0 | 0 | 0 | 0 | 0 |
| SYYRRTHS | ARHGAP5; ARH | Q13017; H0 | RHG05_HUMAN | Unmodified | 0 | 0 | 0 | 0 | 296140 | 0 | 0 | 0 | 0 | 0 | 0 | 0 | 0 | 0 | 0 | 61249 | 0 | 0 |
| SYYSVQTA | PCYOX1L; PCY | Q8NBM8; E7 | PCYXL_HUMAN | Unmodified | 0 | 0 | 0 | 0 | 0 | 0 | 0 | 0 | 5093700 | 544280 | 0 | 0 | 0 | 0 | 0 | 0 | 0 | 0 |
| SYYYAISDF | LAMC1 | P11047 | LAMC1_HUMAN | Unmodified | 0 | 0 | 0 | 683830 | 0 | 0 | 0 | 0 | 0 | 0 | 0 | 0 | 0 | 324730 | 0 | 203440 | 0 | 913610 |
| TAAAAAEH | PSMD3 | O43242 | PSMD3_HUMA | Unmodified | 155790 | 150810 | 1041600 | 484260 | 0 | 0 | 283220 | 101900 | 2189200 | 0 | 594680 | 10605000 | 2570200 | 289700 | 591860 | 1013400 | 0 | 698200 |
| TAAANAAA | SCAMP3 | O14828 | SCAM3_HUMA | Unmodified | 0 | 458650 | 795550 | 217190 | 1147400 | 0 | 0 | 0 | 608710 | 0 | 0 | 914390 | 1590200 | 0 | 0 | 343290 | 439100 | 0 |
| TAAARLPPS | MVB12A; MVB1 | Q96EY5; M0 | MB12A_HUMAN | Unmodified | 7140900 | 2850700 | 5191400 | 6554200 | 2568900 | 6251300 | 560250 | 411780 | 588120 | 707090 | 605570 | 808140 | 9250700 | 5622700 | 5140300 | 4039700 | 865570 | 2930100 |
| TAAATATVL | PHLDA3 | Q9Y5J5 | PHLA3_HUMAN | Acetyl (Protein N | 95013 | 0 | 0 | 0 | 0 | 0 | 0 | 113960 | 0 | 0 | 0 | 0 | 700670 | 392210 | 152710 | 177380 | 197150 | 0 |
| TAAAVAAQ | SQOR; SQOR | Q9Y6N5; H3 | SQOR_HUMAN | Unmodified | 0 | 0 | 0 | 0 | 0 | 0 | 0 | 0 | 3428900 | 0 | 0 | 0 | 68166 | 0 | 0 | 0 | 0 | 0 |
| TAAAVAAQ | SQOR; SQOR | Q9Y6N5; H3 | SQOR_HUMAN | Unmodified | 0 | 0 | 0 | 0 | 0 | 0 | 0 | 0 | 318530 | 677120 | 0 | 0 | 0 | 0 | 0 | 0 | 0 | 0 |
| TAAEDAIR | RBM4; RBM4B; | Q9BWF3; Q | RBM4_HUMAN | Unmodified | 207650 | 211750 | 1431300 | 0 | 0 | 20880 | 49018 | 0 | 57102 | 0 | 1448300 | 573860 | 126370 | 6521100 | 209160 | 0 | 33183 | 0 |
| TAAELLQS | SEPTIN6; SEPT | Q14141; B1 | SEPT6_HUMAN | Unmodified | 0 | 0 | 0 | 0 | 1561000 | 0 | 0 | 0 | 0 | 0 | 0 | 2202300 | 0 | 0 | 0 | 0 | 2180600 | 3081300 |
| TAAELLQS | SEPTIN6; SEPT | Q14141; B1 | SEPT6_HUMAN | Unmodified | 0 | 0 | 0 | 0 | 569720 | 0 | 0 | 0 | 0 | 0 | 0 | 1191800 | 321500 | 0 | 59823 | 0 | 0 | 0 |
| TAAELLQS | SEPTIN6; SEPT | Q14141; B1 | SEPT6_HUMAN | Unmodified | 0 | 0 | 0 | 0 | 2765200 | 0 | 290830 | 291090 | 0 | 0 | 0 | 0 | 966020 | 0 | 0 | 0 | 2054500 | 3047800 |
| TAAENAEK | LRRFIP2 | Q9Y608 | LRRF2_HUMAN | Unmodified | 0 | 0 | 0 | 0 | 2816100 | 0 | 87090 | 167100 | 265450 | 289470 | 0 | 0 | 818030 | 634890 | 0 | 0 | 868590 | 1520400 |
| TAAESFKGI | P4HB; P4HB; P4 | P07237; A0A | PDIA1_HUMAN | Unmodified | 15029 | 0 | 0 | 0 | 833070 | 347280 | 0 | 0 | 115730 | 379510 | 0 | 0 | 0 | 206790 | 0 | 0 | 333470 | 455910 |
| TAAFLLPIL | DDX3X; DDX3X | O00571; A0A | DDX3X_HUMAN | Oxidation (M) | 0 | 0 | 0 | 0 | 123460 | 0 | 41986 | 0 | 0 | 24178 | 0 | 0 | 80820 | 178770 | 0 | 0 | 61779 | 0 |
| TAAFVLAMI | DDX19A; DDX2 | Q9NUU7; Q5 | DD19A_HUMAN | Unmodified | 0 | 0 | 0 | 0 | 0 | 0 | 0 | 0 | 593960 | 0 | 221160 | 167280 | 0 | 0 | 0 | 0 | 0 | 0 |
| TAAKGEAA | MARCKS | P29966 | MARCS_HUMA | Unmodified | 6443300 | 5055700 | 1954800 | 14933000 | 3592400 | 0 | 9310500 | 12488000 | 1.38E+08 | 42871000 | 22596000 | 62218000 | 31680000 | 12190000 | 7611100 | 7575100 | 68300000 | 75795000 |
| TAALPTFR | TMEM30A; TME | Q9NV96; Q3 | CC50A_HUMAN | Unmodified | 751010 | 806320 | 231890 | 0 | 790790 | 304940 | 0 | 0 | 0 | 0 | 0 | 5325300 | 0 | 88242 | 1224700 | 20278 | 580030 | 0 |
| TAANKSPC | TANC1 | Q9C0D5 | TANC1_HUMAN | Unmodified | 0 | 0 | 0 | 0 | 0 | 0 | 0 | 0 | 0 | 0 | 0 | 0 | 193510 | 0 | 0 | 0 | 46182 | 0 |
| TAAQVLIR | AKR1B10 | O60218 | AK1BA_HUMAN | Unmodified | 0 | 0 | 0 | 0 | 454530 | 0 | 0 | 0 | 0 | 0 | 0 | 0 | 0 | 0 | 0 | 0 | 0 | 0 |
| TAASGIPYI | SPTBN1; SPTBN | Q01082; A0A | SPTB2_HUMAN | Unmodified | 415520 | 0 | 0 | 0 | 6028300 | 1685900 | 0 | 497010 | 4665400 | 0 | 2230600 | 7594100 | 0 | 0 | 0 | 0 | 2907500 | 1980000 |
| TAASGVEAI | PDLIM1 | O00151 | PDLI1_HUMAN | Unmodified | 0 | 0 | 0 | 0 | 5484400 | 1569400 | 0 | 0 | 0 | 0 | 0 | 0 | 0 | 377390 | 0 | 0 | 3921300 | 3017600 |
| TAAVANSM | UGGT1 | Q9NYU2 | UGGG1_HUMA | Unmodified | 0 | 0 | 51240 | 0 | 0 | 0 | 0 | 0 | 10725000 | 1861200 | 1897700 | 0 | 0 | 0 | 0 | 0 | 62072 | 0 |

| | | | | | | | | | | | | | | | | | | | | | | |
|---|---|---|---|---|---|---|---|---|---|---|---|---|---|---|---|---|---|---|---|---|---|---|
| TAAVANSM | UGGT1 | Q9NYU2 | UGGG1_HUMA | Oxidation (M) | 0 | 0 | 0 | 0 | 0 | 0 | 0 | 0 | 848800 | 753440 | 0 | 0 | 0 | 0 | 0 | 0 | 0 | 0 |
| TAAVANSM | UGGT1 | Q9NYU2 | UGGG1_HUMA | Unmodified | 0 | 0 | 0 | 0 | 0 | 0 | 0 | 22490 | 4259100 | 1643500 | 276660 | 0 | 40119 | 0 | 0 | 0 | 0 | 0 |
| TAAVANSM | UGGT1 | Q9NYU2 | UGGG1_HUMA | Oxidation (M) | 0 | 0 | 0 | 139000 | 0 | 0 | 0 | 0 | 455670 | 218980 | 64929 | 0 | 0 | 0 | 0 | 0 | 0 | 0 |
| TAAYGHFG | MAT2A | P31153 | METK2_HUMAN | Unmodified | 0 | 0 | 0 | 0 | 974540 | 0 | 0 | 0 | 0 | 0 | 0 | 692140 | 0 | 0 | 0 | 236490 | 449820 | 1319400 |
| TACGAPSG | ITGA6 | P23229 | ITA6_HUMAN I | Unmodified | 0 | 0 | 0 | 531070 | 9104400 | 1133200 | 0 | 0 | 0 | 0 | 0 | 2411400 | 5498900 | 2918300 | 883240 | 155930 | 29746000 | 44402000 |
| TADDPSLS | DDOST; DDOST | P39656; U3K | OST48_HUMAN | Unmodified | 0 | 0 | 846920 | 0 | 0 | 0 | 0 | 0 | 75339000 | 7528400 | 0 | 2171500 | 0 | 0 | 0 | 438250 | 0 | 0 |
| TADEIPMEF | FUT8; FUT8 | Q9BYC5; G3 | FUT8_HUMAN | Oxidation (M) | 0 | 0 | 0 | 0 | 0 | 0 | 0 | 0 | 4399600 | 869730 | 0 | 0 | 0 | 0 | 0 | 0 | 0 | 0 |
| TADEIPMEF | FUT8; FUT8 | Q9BYC5; G3 | FUT8_HUMAN | Unmodified | 0 | 0 | 0 | 0 | 0 | 0 | 0 | 0 | 16289000 | 0 | 0 | 0 | 0 | 0 | 0 | 0 | 0 | 0 |
| TADGIVSHI | PDIA3 | P30101 | PDIA3_HUMAN | Unmodified | 0 | 0 | 0 | 0 | 0 | 0 | 0 | 0 | 3849500 | 390510 | 799310 | 601650 | 0 | 0 | 0 | 0 | 0 | 0 |
| TADGLFHT | PELI2 | Q9HAT8 | PELI2_HUMAN | Unmodified | 0 | 0 | 0 | 55165 | 0 | 0 | 22167 | 0 | 1218000 | 662070 | 0 | 575640 | 0 | 31366 | 0 | 24938 | 0 | 0 |
| TADKIEGFL | PLCG2; PLCG2 | P16885; A0A | PLCG2_HUMA | Oxidation (M) | 0 | 0 | 0 | 0 | 0 | 0 | 0 | 0 | 0 | 0 | 2446300 | 5771900 | 0 | 0 | 0 | 0 | 0 | 0 |
| TADKIEGFL | PLCG2; PLCG2 | P16885; A0A | PLCG2_HUMA | Unmodified | 0 | 0 | 0 | 0 | 0 | 0 | 0 | 0 | 0 | 89962 | 667560 | 1985600 | 0 | 0 | 0 | 104450 | 0 | 0 |
| TADRQYME | CDC37; CDC37 | Q16543; K7E | CDC37_HUMA | Oxidation (M) | 0 | 0 | 0 | 0 | 120730 | 0 | 0 | 0 | 0 | 0 | 249140 | 1701000 | 170480 | 0 | 0 | 0 | 101950 | 174280 |
| TADSVFCP | EFNB2 | P52799 | EFNB2_HUMAN | Unmodified | 108660 | 0 | 0 | 0 | 0 | 0 | 0 | 0 | 236120 | 55856 | 0 | 1483500 | 0 | 0 | 0 | 1560900 | 0 | 0 |
| TAEAGGVT | PARP1; PARP1; | P09874; A0A | PARP1_HUMAN | Unmodified | 0 | 110480 | 0 | 0 | 902480 | 0 | 898510 | 3109500 | 0 | 0 | 0 | 0 | 0 | 0 | 0 | 0 | 0 | 564060 |
| TAEAQLAYE | FLOT2; FLOT2; | Q14254; J3Q | FLOT2_HUMAN | Unmodified | 134560 | 194440 | 1661700 | 664810 | 270000 | 0 | 452950 | 168480 | 4871800 | 1732800 | 979250 | 11546000 | 96428 | 0 | 2142100 | 1550400 | 0 | 0 |
| TAEAQLAYE | FLOT2; FLOT2; | Q14254; J3Q | FLOT2_HUMAN | Unmodified | 775390 | 0 | 6582700 | 4806000 | 35201000 | 3246800 | 7110400 | 1875000 | 14455000 | 9298800 | 3230700 | 7849700 | 14941000 | 10822000 | 5364200 | 3225100 | 1368900 | 2937200 |
| TAEDDETP | PIK3C2A | O00443 | P3C2A_HUMAN | Unmodified | 0 | 0 | 0 | 0 | 0 | 0 | 0 | 0 | 0 | 0 | 533070 | 726890 | 0 | 0 | 0 | 0 | 0 | 0 |
| TAEDYQALI | H6PD | O95479 | G6PE_HUMAN | Unmodified | 0 | 0 | 0 | 0 | 0 | 0 | 0 | 45569 | 2664900 | 866640 | 0 | 0 | 0 | 0 | 0 | 0 | 0 | 0 |
| TAEDYSVD | ACSL4; ACSL4 | O60488; H0Y | ACSL4_HUMAN | Unmodified | 0 | 54151000 | 0 | 0 | 0 | 0 | 0 | 0 | 140310 | 0 | 371930 | 5775800 | 0 | 0 | 37801 | 0 | 0 | 0 |
| TAEEFLQQI | SCIN | Q9Y6U3 | ADSV_HUMAN | Unmodified | 0 | 0 | 0 | 52205 | 0 | 0 | 0 | 0 | 1034600 | 646120 | 0 | 0 | 0 | 0 | 0 | 0 | 0 | 0 |
| TAEEFLQQI | SCIN | Q9Y6U3 | ADSV_HUMAN | Oxidation (M) | 0 | 0 | 0 | 0 | 0 | 0 | 0 | 0 | 133240 | 174870 | 0 | 0 | 0 | 0 | 83667 | 0 | 0 | 0 |
| TAEELMNF | CCT8; CCT8 | P50990; H7C | TCPQ_HUMAN | Unmodified | 0 | 0 | 0 | 0 | 0 | 0 | 0 | 0 | 546400 | 0 | 554950 | 2855400 | 0 | 0 | 0 | 0 | 0 | 0 |
| TAEELMNF | CCT8; CCT8 | P50990; H7C | TCPQ_HUMAN | Oxidation (M) | 0 | 0 | 0 | 0 | 1483900 | 0 | 500250 | 21352 | 0 | 1192200 | 1155100 | 2560000 | 2482000 | 1665100 | 0 | 0 | 1517200 | 231360 |
| TAEELMNF | CCT8; CCT8 | P50990; H7C | TCPQ_HUMAN | Unmodified | 0 | 0 | 0 | 0 | 2246700 | 0 | 0 | 0 | 1158900 | 0 | 0 | 0 | 21782000 | 0 | 0 | 0 | 0 | 1703200 |
| TAEELMNF | CCT8; CCT8 | P50990; H7C | TCPQ_HUMAN | 2 Oxidation (M) | 0 | 0 | 0 | 0 | 0 | 0 | 0 | 407070 | 0 | 0 | 349370 | 0 | 1243100 | 0 | 0 | 0 | 0 | 0 |
| TAEEMKATI | FKBP4; FKBP4 | Q02790; F5H | FKBP4_HUMAN | 2 Oxidation (M) | 0 | 0 | 0 | 0 | 0 | 0 | 0 | 350870 | 0 | 0 | 0 | 0 | 2114800 | 925180 | 0 | 0 | 0 | 0 |
| TAEEMKATI | FKBP4; FKBP4 | Q02790; F5H | FKBP4_HUMAN | Oxidation (M) | 0 | 0 | 0 | 0 | 0 | 0 | 0 | 0 | 0 | 0 | 0 | 0 | 2073200 | 1191000 | 0 | 0 | 0 | 0 |
| TAEEVAALII | PRPF8 | Q6P2Q9 | PRP8_HUMAN | Unmodified | 278020 | 93348 | 153530 | 73531 | 139730 | 0 | 0 | 240780 | 111460 | 57700 | 634320 | 2376300 | 0 | 0 | 124190 | 31500 | 0 | 0 |
| TAEFAPFV | CASK; CASK; C | O14936; A0A | CSKP_HUMAN | Unmodified | 0 | 0 | 0 | 107360 | 0 | 0 | 0 | 0 | 0 | 0 | 0 | 0 | 0 | 0 | 145410 | 0 | 0 | 0 |
| TAEFAPFV | CASK; CASK; C | O14936; A0A | CSKP_HUMAN | Unmodified | 0 | 0 | 0 | 124840 | 281720 | 82744 | 0 | 0 | 151680 | 513060 | 0 | 0 | 163310 | 121300 | 327950 | 0 | 0 | 0 |
| TAEFQVAR | FARSB | Q9NSD9 | SYFB_HUMAN | Unmodified | 0 | 0 | 0 | 0 | 0 | 0 | 0 | 0 | 0 | 0 | 0 | 3766400 | 0 | 349820 | 576930 | 0 | 0 | 0 |
| TAEFQVAR | FARSB | Q9NSD9 | SYFB_HUMAN | Unmodified | 0 | 0 | 0 | 0 | 898360 | 0 | 2499000 | 0 | 0 | 194220 | 0 | 0 | 1172200 | 1293100 | 0 | 0 | 509780 | 861520 |
| TAEHFQEA | SDF4; SDF4; SD | Q9BRK5; H0 | CAB45_HUMA | Unmodified | 186280 | 0 | 862400 | 659860 | 674560 | 179310 | 265590 | 434970 | 58669000 | 9039800 | 3632300 | 1726300 | 2302500 | 1892000 | 499590 | 453670 | 80536 | 1387000 |
| TAEHFQEA | SDF4; SDF4; SD | Q9BRK5; H0 | CAB45_HUMA | Oxidation (M) | 0 | 0 | 0 | 0 | 0 | 0 | 107850 | 115510 | 7679400 | 2834100 | 515510 | 373560 | 803690 | 317000 | 0 | 258320 | 27834 | 534250 |
| TAEHFQEA | SDF4; SDF4; SD | Q9BRK5; H0 | CAB45_HUMA | Unmodified | 0 | 0 | 0 | 0 | 0 | 0 | 0 | 0 | 0 | 1303000 | 0 | 0 | 0 | 501110 | 0 | 0 | 0 | 0 |
| TAEIDSVK | KIF14 | Q15058 | KIF14_HUMAN | Unmodified | 0 | 0 | 0 | 0 | 1693600 | 0 | 0 | 0 | 0 | 0 | 0 | 0 | 0 | 0 | 0 | 0 | 0 | 0 |
| TAEIEPKAD | BCAM; BCAM | P50895; A0A | BCAM_HUMAN | Unmodified | 1478200 | 514130 | 0 | 0 | 711800 | 0 | 0 | 0 | 580050 | 0 | 0 | 0 | 0 | 0 | 0 | 0 | 0 | 355290 |
| TAELLQVA | P4HA2 | O15460 | P4HA2_HUMA | Unmodified | 0 | 0 | 0 | 0 | 0 | 0 | 0 | 0 | 1222500 | 0 | 0 | 0 | 0 | 518190 | 0 | 0 | 0 | 0 |
| TAELLQVA | P4HA2 | O15460 | P4HA2_HUMA | Unmodified | 0 | 0 | 0 | 0 | 0 | 0 | 0 | 0 | 3366400 | 1860900 | 0 | 0 | 0 | 0 | 0 | 0 | 0 | 0 |
| TAELNVQV | MIA3; MIA3 | Q5JRA6; A0A | TGO1_HUMAN | Unmodified | 0 | 0 | 0 | 0 | 0 | 0 | 0 | 0 | 14908000 | 891710 | 2715600 | 0 | 0 | 0 | 145540 | 125790 | 0 | 0 |
| TAELSPFIVI | MPP1 | Q00013 | EM55_HUMAN | Unmodified | 0 | 0 | 0 | 0 | 412460 | 0 | 0 | 0 | 183580 | 0 | 0 | 0 | 0 | 0 | 0 | 0 | 0 | 0 |
| TAENATSG | PA2G4 | Q9UQ80 | PA2G4_HUMA | Unmodified | 1713400 | 1438700 | 861900 | 823320 | 1731200 | 0 | 169060 | 450790 | 0 | 128210 | 1355900 | 5182500 | 2168600 | 240910 | 840950 | 660460 | 379170 | 425650 |
| TAENFRAL | PPID; PPIF; RA | Q08752; P30 | PPID_HUMAN | Unmodified | 0 | 0 | 0 | 0 | 1019500 | 153120 | 338780 | 101740 | 0 | 477500 | 0 | 0 | 334620 | 258550 | 0 | 0 | 652860 | 0 |
| TAENFRQF | PPIH; PPIH; PP | O43447; A0A | PPIH_HUMAN | Unmodified | 0 | 0 | 0 | 0 | 627870 | 0 | 381400 | 0 | 0 | 0 | 0 | 0 | 5368900 | 1002600 | 0 | 0 | 430510 | 558290 |
| TAERSYRE | EXOC7; EXOC7 | Q9UPT5; B4I | EXOC7_HUMA | Unmodified | 0 | 0 | 0 | 0 | 1653300 | 441400 | 111970 | 0 | 0 | 397640 | 0 | 0 | 676390 | 395790 | 0 | 0 | 430400 | 690040 |
| TAESQTPTF | ADAR; ADAR; A | P55265; A0A | DSRAD_HUMA | Unmodified | 0 | 0 | 0 | 0 | 0 | 0 | 0 | 0 | 87638 | 0 | 0 | 1092400 | 384590 | 0 | 0 | 0 | 0 | 0 |
| TAEVLANK | DYNC1H1; DYN | Q14204; A0A | DYHC1_HUMA | Unmodified | 0 | 0 | 380840 | 498390 | 34018 | 438430 | 232500 | 31445 | 357010 | 142770 | 95608 | 2129200 | 688650 | 0 | 1159700 | 815730 | 0 | 0 |
| TAEVWMD | GALNT14; GAL | Q96FL9; Q8 | GLT14_HUMA | Unmodified | 25962 | 0 | 0 | 0 | 0 | 0 | 0 | 0 | 2490200 | 148810 | 0 | 0 | 0 | 0 | 0 | 0 | 0 | 0 |
| TAFDDAIAE | YWHAG | P61981 | 1433G_HUMA | Unmodified | 126240 | 875140 | 1073400 | 1588300 | 1336900 | 200470 | 286840 | 34712 | 856760 | 386780 | 878500 | 2868600 | 774820 | 1093400 | 7086700 | 859070 | 625220 | 626170 |
| TAFDEAIAE | YWHAQ | P27348 | 1433T_HUMAN | Unmodified | 321540 | 344140 | 1300400 | 2202900 | 1111100 | 0 | 217470 | 0 | 1469800 | 343540 | 1486400 | 4532000 | 0 | 0 | 4458400 | 765480 | 484490 | 390690 |
| TAFDEAIAE | YWHAB | P31946 | 1433B_HUMA | Unmodified | 0 | 497290 | 558570 | 749940 | 956010 | 0 | 0 | 0 | 1817900 | 346430 | 481230 | 1515900 | 0 | 0 | 5283000 | 0 | 567360 | 830160 |
| TAFDEAIAE | YWHAZ; YWHA | P63104; B7Z | 1433Z_HUMAN | Unmodified | 0 | 853440 | 959870 | 584470 | 722950 | 0 | 0 | 0 | 1652400 | 1405900 | 280430 | 903070 | 513690 | 0 | 2787100 | 400620 | 659120 | 854200 |
| TAFHQEQG | TPMT | P51580 | TPMT_HUMAN | Unmodified | 0 | 0 | 0 | 0 | 0 | 0 | 33213 | 0 | 0 | 0 | 0 | 0 | 135870 | 0 | 0 | 0 | 123080 | 182620 |
| TAFLASLVS | TEP1; TEP1; TE | Q99973; G3 | TEP1_HUMAN | Unmodified | 0 | 0 | 0 | 0 | 0 | 0 | 0 | 0 | 0 | 0 | 0 | 134160 | 23108 | 0 | 0 | 0 | 0 | 0 |
| TAFPSLPD | NUDT19 | A8MXV4 | NUD19_HUMA | Unmodified | 0 | 0 | 117020 | 0 | 0 | 0 | 0 | 0 | 15416000 | 0 | 557780 | 0 | 0 | 0 | 0 | 0 | 0 | 2626500 |
| TAFTDLPE | SNTB1; SNTB1 | Q13884; A0A | SNTB1_HUMA | Unmodified | 130290 | 53236 | 0 | 0 | 170340 | 66444 | 0 | 0 | 10453000 | 1563400 | 81139 | 664520 | 0 | 0 | 0 | 0 | 73944 | 0 |
| TAFYLAEFF | GGH | Q92820 | GGH_HUMAN | Unmodified | 0 | 0 | 0 | 0 | 0 | 11326 | 0 | 0 | 1778600 | 1671700 | 35860 | 0 | 79188 | 0 | 0 | 0 | 0 | 0 |
| TAFYLAEFF | GGH | Q92820 | GGH_HUMAN | Unmodified | 240920 | 0 | 183680 | 466680 | 1696300 | 886700 | 0 | 0 | 1.19E+08 | 68655000 | 6725500 | 1814200 | 2516400 | 1590800 | 503080 | 153180 | 0 | 511290 |
| TAGDHPGF | GFPT1; GFPT1; | Q06210; A0A | GFPT1_HUMA | Oxidation (M) | 0 | 0 | 0 | 0 | 136850 | 0 | 0 | 0 | 0 | 106150 | 0 | 0 | 652060 | 451910 | 0 | 0 | 0 | 0 |
| TAGGWAA | GGT1; GGTLC3 | P19440; B5N | GGT1_HUMAN | Unmodified | 0 | 2405700 | 0 | 2146400 | 0 | 0 | 0 | 0 | 1437500 | 203110 | 0 | 672210 | 0 | 0 | 2140500 | 0 | 0 | 0 |
| TAGGWAA | GGT1; GGTLC3 | P19440; B5N | GGT1_HUMAN | Unmodified | 125750 | 179690 | 528230 | 808250 | 0 | 0 | 0 | 0 | 474270 | 0 | 0 | 450110 | 0 | 63315 | 1425700 | 441780 | 204820 | 59903 |
| TAGNSEFL | EIF4G2; EIF4G2 | P78344; H0Y | IF4G2_HUMAN | Unmodified | 0 | 0 | 0 | 521590 | 0 | 0 | 0 | 0 | 0 | 0 | 0 | 0 | 0 | 0 | 0 | 0 | 0 | 0 |
| TAGNSEFL | EIF4G2; EIF4G2 | P78344; H0Y | IF4G2_HUMAN | Unmodified | 0 | 51079 | 0 | 1981100 | 738840 | 141900 | 0 | 0 | 87839 | 0 | 63724 | 0 | 948900 | 499870 | 0 | 0 | 385120 | 885620 |
| TAGPIASAC | MAP4; MAP4 | P27816; E7E | MAP4_HUMAN | Unmodified | 0 | 0 | 0 | 0 | 504210 | 0 | 0 | 292930 | 0 | 0 | 0 | 0 | 447100 | 0 | 0 | 0 | 886910 | 746260 |
| TAGTICLET | GSPT1; GSPT2; | P15170; Q8I | ERF3A_HUMA | Unmodified | 0 | 0 | 0 | 0 | 0 | 0 | 0 | 42248 | 0 | 0 | 0 | 3432000 | 0 | 0 | 76146 | 0 | 0 | 0 |
| TAGVKVET | ETFB | P38117 | ETFB_HUMAN | Unmodified | 89325 | 81155 | 0 | 0 | 0 | 0 | 0 | 0 | 3105900 | 150600 | 0 | 0 | 0 | 0 | 85436 | 88871 | 0 | 0 |
| TAGYPNVN | SPTBN1; SPTBN | Q01082; A0A | SPTB2_HUMA | Unmodified | 0 | 0 | 0 | 0 | 0 | 0 | 0 | 0 | 0 | 0 | 0 | 507790 | 0 | 0 | 0 | 0 | 0 | 0 |
| TAHDEYW | OGFOD3; OGF | Q6PK18; J3Q | OGFD3_HUMA | Unmodified | 0 | 17261 | 0 | 0 | 0 | 55939 | 0 | 0 | 1538200 | 0 | 0 | 0 | 0 | 0 | 0 | 0 | 0 | 38965 |
| TAHIDVHF | ITGA6 | P23229 | ITA6_HUMAN I | Unmodified | 0 | 0 | 249150 | 0 | 658510 | 359540 | 0 | 0 | 0 | 0 | 0 | 877290 | 510690 | 178240 | 384330 | 0 | 241860 | 2168500 |
| TAHIDVHF | ITGA6 | P23229 | ITA6_HUMAN I | Unmodified | 0 | 0 | 0 | 352170 | 367090 | 0 | 0 | 0 | 0 | 0 | 0 | 0 | 0 | 0 | 0 | 0 | 1978200 | 3326200 |
| TAHLDEEV | MTHFD1; MTHF | P11586; V9G | C1TC_HUMAN | Oxidation (M) | 463810 | 245280 | 593580 | 0 | 0 | 0 | 3110200 | 0 | 2390300 | 0 | 0 | 11555000 | 0 | 0 | 778770 | 507380 | 1326300 | 0 |
| TAIEEVQAE | P3H1 | Q32P28 | P3H1_HUMAN | Unmodified | 0 | 0 | 0 | 0 | 0 | 0 | 0 | 0 | 1259300 | 126160 | 71189 | 0 | 0 | 13475 | 0 | 0 | 0 | 0 |
| TAIEEVQAE | P3H1 | Q32P28 | P3H1_HUMAN | Unmodified | 0 | 0 | 0 | 0 | 0 | 0 | 0 | 0 | 286820 | 0 | 0 | 0 | 0 | 0 | 0 | 0 | 0 | 0 |
| TAIGKAGY | ENO1; ENO1 | P06733; A0A | ENOA_HUMAN | Unmodified | 0 | 0 | 0 | 0 | 2384700 | 0 | 0 | 28535 | 0 | 505840 | 0 | 0 | 1656800 | 1002500 | 0 | 0 | 3549200 | 2148900 |
| TAIHTAAM | HSDL2 | Q6YN16 | HSDL2_HUMA | Oxidation (M) | 0 | 0 | 0 | 0 | 0 | 0 | 0 | 0 | 3036000 | 254000 | 0 | 0 | 0 | 0 | 0 | 0 | 0 | 0 |
| TAIHTAAM | HSDL2 | Q6YN16 | HSDL2_HUMA | Unmodified | 0 | 0 | 0 | 0 | 0 | 0 | 0 | 0 | 9496900 | 338700 | 0 | 0 | 2466500 | 0 | 189210 | 76247 | 0 | 0 |
| TAIIANVHP | KIF15; KIF15 | Q9NS87; D6 | KIF15_HUMAN | Unmodified | 0 | 0 | 0 | 0 | 73144 | 0 | 0 | 0 | 299600 | 97078 | 146540 | 183700 | 60219 | 14590 | 15754 | 0 | 33190 | 58129 |
| TAIPFTQEP | CYFIP2; CYFIP2 | Q96F07; H7C | CYFP2_HUMA | Unmodified | 0 | 0 | 0 | 0 | 0 | 0 | 0 | 0 | 0 | 0 | 0 | 2689500 | 0 | 0 | 1780200 | 0 | 0 | 0 |
| TAIQAAGY | ENO3; ENO3 | P13929; E5R | ENOB_HUMAN | Unmodified | 0 | 0 | 0 | 0 | 0 | 0 | 0 | 0 | 880360 | 224650 | 228870 | 927590 | 0 | 0 | 0 | 0 | 0 | 0 |
| TAIQTTEIK | UTRN | P46939 | UTRN_HUMAN | Unmodified | 147420 | 0 | 887770 | 702370 | 2414700 | 0 | 0 | 0 | 1160700 | 1122700 | 684700 | 103220 | 385290 | 0 | 1318900 | 1164000 | 45971 | 0 |
| TAIQTTEIKE | UTRN | P46939 | UTRN_HUMAN | Unmodified | 0 | 0 | 0 | 0 | 684180 | 0 | 0 | 55412 | 0 | 0 | 0 | 0 | 0 | 0 | 0 | 0 | 1062600 | 647650 |
| TAISTNIK | OPLAH | O14841 | OPLA_HUMAN | Unmodified | 0 | 0 | 0 | 0 | 182880 | 0 | 0 | 0 | 303830 | 17824 | 0 | 853460 | 0 | 0 | 0 | 0 | 0 | 25306 |
| TAITVEHLA | DDX21 | Q9NR30 | DDX21_HUMA | Unmodified | 0 | 0 | 0 | 0 | 0 | 0 | 412130 | 517670 | 0 | 0 | 0 | 380060 | 0 | 0 | 0 | 0 | 0 | 0 |
| TAKEDFSE | TXNDC16 | Q9P2K2 | TXD16_HUMAN | Unmodified | 0 | 0 | 0 | 0 | 0 | 0 | 0 | 0 | 1388700 | 610300 | 0 | 0 | 0 | 0 | 0 | 0 | 0 | 0 |
| TALAAKIAE | NSF; NSF | P46459; I3L | NSF_HUMAN V | Unmodified | 0 | 0 | 0 | 0 | 0 | 0 | 1889000 | 0 | 0 | 0 | 0 | 0 | 1057800 | 582930 | 0 | 0 | 1727400 | 887220 |
| TALALAIAQ | RUVBL1; RUVB | Q9Y265; E7E | RUVB1_HUMA | Unmodified | 105730 | 0 | 189740 | 134160 | 0 | 0 | 0 | 0 | 283710 | 0 | 401210 | 5181200 | 0 | 0 | 184320 | 119620 | 0 | 0 |
| TALALEVGI | CRKL | P46109 | CRKL_HUMAN | Unmodified | 0 | 0 | 0 | 0 | 1462500 | 0 | 0 | 0 | 0 | 0 | 0 | 0 | 0 | 0 | 0 | 0 | 0 | 0 |
| TALALEVGI | CRKL | P46109 | CRKL_HUMAN | Unmodified | 0 | 0 | 0 | 0 | 710250 | 0 | 0 | 0 | 0 | 138230 | 0 | 0 | 0 | 0 | 0 | 0 | 0 | 0 |
| TALAMGAD | ETFB | P38117 | ETFB_HUMAN | Unmodified | 0 | 0 | 0 | 0 | 0 | 0 | 0 | 0 | 1537500 | 0 | 0 | 0 | 0 | 0 | 0 | 0 | 0 | 0 |
| TALASGGV | B4GAT1 | O43505 | B4GA1_HUMA | Unmodified | 0 | 0 | 0 | 0 | 0 | 0 | 0 | 142900 | 1567700 | 0 | 0 | 699910 | 0 | 0 | 99443 | 53548 | 0 | 0 |
| TALEEEIK | IQGAP2; IQGAP | Q13576; E9F | IQGA2_HUMAN | Unmodified | 0 | 380750 | 1491800 | 461130 | 1162400 | 0 | 124280 | 73758 | 0 | 445940 | 385540 | 1663800 | 0 | 0 | 1078900 | 1015300 | 140510 | 0 |
| TALHHAAF | ANKRD28; ANK | O15084; B4D | ANR28_HUMA | Unmodified | 0 | 0 | 0 | 0 | 0 | 0 | 0 | 0 | 234880 | 69560 | 0 | 1436600 | 0 | 0 | 0 | 0 | 0 | 0 |
| TALIHDGL | RPS12 | P25398 | RS12_HUMAN | Unmodified | 0 | 0 | 0 | 0 | 5130500 | 553700 | 131730 | 41866 | 0 | 748980 | 0 | 0 | 3892800 | 0 | 0 | 0 | 2684900 | 3417800 |
| TALLDISCV | ACSL4 | O60488 | ACSL4_HUMAN | Unmodified | 0 | 0 | 0 | 0 | 0 | 513180 | 190420 | 166360 | 0 | 0 | 0 | 1960600 | 130410 | 0 | 0 | 0 | 1393100 | 244530 |
| TALLEANST | PLEKHB1; PLEK | F5H3M1; F5I | F5H3M1_HUM | Unmodified | 0 | 0 | 0 | 0 | 0 | 0 | 0 | 0 | 0 | 0 | 0 | 0 | 0 | 0 | 0 | 687190 | 0 | 0 |
| TALLGLTK | DHRS4 | Q9BTZ2 | DHRS4_HUMA | Unmodified | 0 | 0 | 0 | 0 | 0 | 0 | 1660900 | 634900 | 12932000 | 3838500 | 0 | 0 | 0 | 0 | 0 | 0 | 44778 | 0 |
| TALLHVFAK | RND3; RND3 | P61587; Q53 | RND3_HUMAN | Unmodified | 0 | 0 | 0 | 0 | 377380 | 0 | 0 | 31984 | 0 | 0 | 0 | 0 | 0 | 0 | 0 | 0 | 172900 | 577190 |
| TALLSLFGII | SUN2; SUN2 | Q9UH99; B0 | SUN2_HUMAN | Unmodified | 0 | 0 | 0 | 0 | 0 | 0 | 0 | 0 | 9868800 | 971260 | 0 | 0 | 0 | 0 | 0 | 0 | 0 | 0 |
| TALLSVGP | IGSF3 | O75054 | IGSF3_HUMAN | Unmodified | 4549800 | 3338900 | 521930 | 1344800 | 0 | 0 | 0 | 0 | 591420 | 106310 | 35568 | 2186400 | 0 | 0 | 7803800 | 2847400 | 0 | 0 |
| TALNHCNL | OPA1; OPA1; O | O60313; E5K | OPA1_HUMAN | Unmodified | 0 | 0 | 0 | 0 | 0 | 0 | 0 | 0 | 46422 | 0 | 49716 | 738140 | 0 | 0 | 0 | 0 | 0 | 0 |
| TALPAQSA | SPTBN1; SPTBN | Q01082; A0A | SPTB2_HUMA | Unmodified | 0 | 0 | 0 | 0 | 0 | 0 | 0 | 0 | 0 | 0 | 637450 | 3224300 | 0 | 0 | 0 | 0 | 0 | 0 |
| TALPFNRFL | LMO7; LMO7; L | Q8WWI1; E9 | LMO7_HUMAN | Unmodified | 0 | 0 | 0 | 0 | 1083000 | 97636 | 55808 | 42528 | 0 | 0 | 0 | 0 | 0 | 0 | 0 | 0 | 1010500 | 947570 |
| TALPTSGS | EPN1 | Q9Y6I3 | EPN1_HUMAN | Unmodified | 1541300 | 2280900 | 331710 | 0 | 0 | 0 | 23556 | 18147 | 172390 | 0 | 600860 | 0 | 0 | 0 | 0 | 1156300 | 0 | 0 |
| TALQEEIK | IQGAP3; IQGAP | Q86VI3; P46 | IQGA3_HUMAN | Unmodified | 268110 | 666530 | 540890 | 607050 | 2164200 | 822110 | 0 | 18513 | 0 | 146870 | 0 | 5653900 | 0 | 0 | 839630 | 853970 | 0 | 0 |
| TALQPLKH | CD276; CD276 | Q5ZPR3; A0A | CD276_HUMA | Unmodified | 0 | 0 | 0 | 0 | 6711500 | 910460 | 0 | 0 | 0 | 804250 | 0 | 0 | 6185200 | 8127700 | 0 | 0 | 1584800 | 2398600 |
| TALRHPPF | TNS3; TNS3 | Q68CZ2; E9I | TENS3_HUMA | Unmodified | 0 | 0 | 0 | 0 | 2933300 | 0 | 0 | 0 | 0 | 252490 | 0 | 0 | 0 | 0 | 0 | 0 | 999790 | 3273700 |
| TALVANTS | ATP6V1A | P38606 | VATA_HUMAN | Unmodified | 0 | 0 | 0 | 0 | 0 | 0 | 0 | 0 | 0 | 0 | 0 | 1202300 | 0 | 0 | 0 | 0 | 0 | 0 |
| TALVERDD | GPD2; GPD2 | P43304; F5G | GPDM_HUMAN | Unmodified | 0 | 0 | 0 | 0 | 0 | 0 | 0 | 20630 | 0 | 2187200 | 0 | 0 | 0 | 0 | 0 | 0 | 0 | 0 |

| | | | | | | | | | | | | | | | | | | | | | | |
|---|---|---|---|---|---|---|---|---|---|---|---|---|---|---|---|---|---|---|---|---|---|---|
| TALYWAVE | KIDINS220; KID | Q9ULH0; E9 | KDIS_HUMAN I | Unmodified | 0 | 0 | 0 | 0 | 2765400 | 0 | 0 | 0 | 0 | 0 | 0 | 977860 | 0 | 0 | 0 | 0 | 0 | 0 |
| TAMIATISP/ | KIF14 | Q15058 | KIF14_HUMAN | Oxidation (M) | 0 | 0 | 0 | 632940 | 577050 | 153850 | 0 | 21000 | 0 | 0 | 134850 | 138620 | 0 | 0 | 326620 | 0 | 0 | 0 |
| TAMLLALQI | ABHD11 | Q8NFV4 | ABHDB_HUMA | Oxidation (M) | 0 | 0 | 0 | 0 | 0 | 0 | 0 | 0 | 1156300 | 153340 | 0 | 0 | 0 | 0 | 0 | 0 | 0 | 0 |
| TAMLLALQI | ABHD11 | Q8NFV4 | ABHDB_HUMA | Unmodified | 0 | 0 | 0 | 0 | 0 | 0 | 0 | 23112 | 2117000 | 0 | 0 | 0 | 0 | 0 | 0 | 0 | 0 | 0 |
| TAMNVNDI | RAB5B | P61020 | RAB5B_HUMAN | Oxidation (M) | 6442600 | 5218200 | 943060 | 3409700 | 6346800 | 1511600 | 1367100 | 64802 | 778340 | 1460700 | 1140400 | 7553100 | 1087400 | 1383500 | 4708000 | 3957400 | 0 | 0 |
| TAMNVNDI | RAB5B | P61020 | RAB5B_HUMAN | Unmodified | 523490 | 310700 | 18789 | 125650 | 1631500 | 98752 | 59457 | 0 | 4849200 | 1413400 | 25515 | 1073400 | 633230 | 152300 | 102600 | 33995 | 339540 | 358830 |
| TAMNVNEII | RAB5C | P51148 | RAB5C_HUMAI | 2 Oxidation (M) | 5121000 | 6546700 | 1702000 | 13893000 | 3612600 | 296620 | 1356500 | 528680 | 1015400 | 1237500 | 4237600 | 12159000 | 1764600 | 1212200 | 8331500 | 6307500 | 0 | 0 |
| TAMNVNEII | RAB5C | P51148 | RAB5C_HUMAI | Oxidation (M) | 310300 | 459360 | 172330 | 661970 | 2405600 | 244160 | 426370 | 24625 | 5988300 | 2224700 | 332610 | 1440400 | 2578600 | 1632200 | 122420 | 297160 | 0 | 0 |
| TAMNVNEII | RAB5C | P51148 | RAB5C_HUMAI | Unmodified | 0 | 0 | 0 | 0 | 0 | 0 | 0 | 0 | 1396300 | 0 | 0 | 0 | 0 | 0 | 0 | 0 | 0 | 0 |
| TAMSTPHV | MSN | P26038 | MOES_HUMAN | 2 Oxidation (M) | 0 | 0 | 0 | 0 | 25152000 | 8116700 | 0 | 0 | 0 | 0 | 0 | 0 | 5487400 | 5805700 | 0 | 575080 | 9983000 | 11683000 |
| TAMSTPHV | MSN | P26038 | MOES_HUMAN | Oxidation (M) | 0 | 0 | 0 | 0 | 10056000 | 19993000 | 0 | 0 | 0 | 0 | 0 | 0 | 11912000 | 6539100 | 0 | 0 | 31406000 | 37012000 |
| TAMSTPHV | MSN | P26038 | MOES_HUMAN | Unmodified | 0 | 0 | 0 | 0 | 55607000 | 9967700 | 0 | 0 | 0 | 0 | 0 | 0 | 3633700 | 3772900 | 0 | 0 | 18530000 | 27837000 |
| TANDAVELI | PTGFRN | Q9P2B2 | FPRP_HUMAN | Unmodified | 5420400 | 4380700 | 15030000 | 19648000 | 6106400 | 2000000 | 4317100 | 474520 | 170170 | 64580 | 0 | 7928100 | 0 | 29090000 | 19851000 | 14990000 | 0 | 1878500 |
| TANDAVELI | PTGFRN | Q9P2B2 | FPRP_HUMAN | Unmodified | 0 | 0 | 0 | 0 | 0 | 0 | 0 | 0 | 0 | 0 | 0 | 0 | 2751400 | 2707300 | 0 | 0 | 0 | 0 |
| TANDMIHA | MSN | P26038 | MOES_HUMAN | Unmodified | 175920 | 0 | 253000 | 1314100 | 14215000 | 4747100 | 0 | 87867 | 0 | 0 | 0 | 0 | 2059500 | 1241900 | 432940 | 0 | 3510400 | 6593300 |
| TANDMIHA | MSN | P26038 | MOES_HUMAN | Oxidation (M) | 0 | 0 | 332070 | 798790 | 8377700 | 1566800 | 69441 | 59840 | 0 | 0 | 0 | 0 | 860220 | 657130 | 813810 | 459270 | 2662600 | 3790700 |
| TANDMIHA | MSN | P26038 | MOES_HUMAN | Unmodified | 0 | 0 | 0 | 0 | 42999000 | 0 | 825850 | 0 | 0 | 0 | 0 | 0 | 0 | 2524200 | 0 | 0 | 1197200 | 3615600 |
| TANEGGSL | SLC38A2 | Q96QD8 | S38A2_HUMAN | Unmodified | 394580 | 0 | 0 | 0 | 0 | 0 | 0 | 0 | 0 | 0 | 0 | 0 | 0 | 0 | 0 | 0 | 0 | 0 |
| TANKDHLV | FKBP3 | Q00688 | FKBP3_HUMAN | Unmodified | 0 | 0 | 0 | 0 | 4082100 | 0 | 0 | 0 | 0 | 2183200 | 0 | 578280 | 4125700 | 0 | 0 | 0 | 0 | 0 |
| TANLVISNV | NOSTRIN; NOS | Q8IVI9; A0A( | NOSTN_HUMA | Unmodified | 3789000 | 14063000 | 3944300 | 14979000 | 0 | 0 | 0 | 0 | 0 | 0 | 0 | 0 | 3484900 | 2865300 | 36725000 | 18044000 | 0 | 0 |
| TANQASDT | ADIRF | Q15847 | ADIRF_HUMAN | Unmodified | 3647500 | 1753700 | 0 | 0 | 0 | 0 | 0 | 353860 | 771360 | 432870 | 0 | 0 | 0 | 0 | 0 | 0 | 3294700 | 15668000 |
| TANQASDT | ADIRF | Q15847 | ADIRF_HUMAN | Unmodified | 0 | 0 | 0 | 0 | 0 | 0 | 428520 | 1461300 | 0 | 0 | 0 | 0 | 0 | 0 | 0 | 0 | 560010 | 2985700 |
| TANQASDT | ADIRF | Q15847 | ADIRF_HUMAN | Unmodified | 0 | 0 | 0 | 0 | 0 | 0 | 0 | 0 | 0 | 0 | 295800 | 0 | 0 | 0 | 0 | 0 | 1642700 | 0 |
| TAPAAPAEI | PHLDA2 | Q53GA4 | PHLA2_HUMAN | Unmodified | 0 | 1961900 | 587810 | 407410 | 0 | 275780 | 78584 | 0 | 0 | 0 | 0 | 0 | 0 | 0 | 939400 | 773930 | 0 | 0 |
| TAPAAPAEI | PHLDA2 | Q53GA4 | PHLA2_HUMAN | Unmodified | 0 | 0 | 0 | 2025400 | 1032000 | 0 | 0 | 0 | 0 | 0 | 0 | 753130 | 1035500 | 0 | 0 | 0 | 0 | 0 |
| TAPGALTAI | TBC1D10B | Q4KMP7 | TB10B_HUMAN | Unmodified | 0 | 0 | 0 | 0 | 0 | 0 | 0 | 0 | 1175200 | 0 | 195270 | 1473400 | 0 | 0 | 390120 | 861500 | 0 | 0 |
| TAPGPATT( | SYNGR3 | O43761 | SNG3_HUMAN | Unmodified | 0 | 0 | 0 | 0 | 0 | 0 | 129270 | 0 | 6872800 | 0 | 0 | 421470 | 0 | 0 | 0 | 0 | 0 | 0 |
| TAPLIATR | SNTA1 | Q13424 | SNTA1_HUMAN | Unmodified | 0 | 0 | 462640 | 244160 | 677900 | 266420 | 0 | 144390 | 234580 | 0 | 71124 | 2175500 | 0 | 0 | 195050 | 231870 | 263950 | 112850 |
| TAPSCTEAI | MKRN1; MKRN1 | Q9UHC7; C9 | MKRN1_HUMA | Unmodified | 1068000 | 153490 | 1308400 | 516230 | 261070 | 0 | 0 | 0 | 0 | 126420 | 0 | 0 | 0 | 0 | 252770 | 158040 | 114730 | 0 |
| TAPSCTEAI | MKRN1; MKRN1 | Q9UHC7; C9 | MKRN1_HUMA | Unmodified | 6806100 | 4308100 | 306010 | 2482000 | 0 | 0 | 0 | 0 | 0 | 0 | 0 | 0 | 0 | 0 | 2282900 | 2136500 | 0 | 0 |
| TAPSCTEAI | MKRN1; MKRN1 | Q9UHC7; C9 | MKRN1_HUMA | Unmodified | 2609800 | 1888600 | 0 | 0 | 0 | 0 | 0 | 0 | 0 | 0 | 0 | 0 | 0 | 0 | 0 | 0 | 0 | 0 |
| TAPYVVTG! | PAFAH1B1; PAF | P43034; I3L3 | LIS1_HUMAN P | Unmodified | 89073 | 0 | 296960 | 1222400 | 5310600 | 1066400 | 2007900 | 1254500 | 0 | 857110 | 1926800 | 7409200 | 2980900 | 3775900 | 1641700 | 969980 | 760550 | 1858500 |
| TAQAAAALI | GALK1; GALK1 | P51570; K7E | GALK1_HUMAN | Unmodified | 0 | 0 | 0 | 0 | 666450 | 160250 | 0 | 385120 | 179850 | 0 | 0 | 809970 | 0 | 0 | 309070 | 258130 | 981370 | 295110 |
| TAQEAFES\ | FMNL1; FMNL1 | O95466; K7E | FMNL1_HUMAI | Unmodified | 0 | 0 | 0 | 131560 | 1986300 | 5102000 | 0 | 0 | 0 | 0 | 0 | 0 | 0 | 0 | 0 | 0 | 0 | 265560 |
| TAQEHDKR | VTA1; VTA1 | Q9NP79; A0, | VTA1_HUMAN \ | Unmodified | 1495900 | 871160 | 470770 | 3036100 | 0 | 48776 | 0 | 0 | 5257800 | 843340 | 996980 | 7337800 | 893950 | 457400 | 1189100 | 977130 | 0 | 0 |
| TAQELSFR | ARHGAP4; ARH | P98171; E9P | RHG04_HUMAI | Unmodified | 0 | 0 | 0 | 0 | 0 | 0 | 0 | 0 | 0 | 0 | 0 | 282800 | 0 | 0 | 0 | 0 | 0 | 0 |
| TAQGSLSLI | EIF3H; EIF3H; E | O15372; E5F | EIF3H_HUMAN | Unmodified | 0 | 0 | 0 | 0 | 30720 | 1034200 | 28223 | 0 | 1074000 | 499490 | 487880 | 10722000 | 0 | 0 | 0 | 0 | 26051 | 0 |
| TAQNHPMI | LSM4; LSM4; LS | Q9Y4Z0; M0( | LSM4_HUMAN | Unmodified | 145240 | 124260 | 0 | 0 | 0 | 0 | 0 | 0 | 0 | 0 | 199530 | 1309900 | 853630 | 520440 | 0 | 0 | 0 | 0 |
| TAQNLSIFL | DIAPH1; DIAPH | O60610; A0/ | DIAP1_HUMAN | Unmodified | 0 | 0 | 32871 | 504210 | 31475 | 0 | 0 | 14280 | 1613700 | 200560 | 359530 | 3358200 | 0 | 0 | 444950 | 352930 | 0 | 0 |
| TAQRADLQ | EEA1 | Q15075 | EEA1_HUMAN I | Unmodified | 0 | 0 | 0 | 0 | 1081500 | 0 | 0 | 0 | 0 | 0 | 0 | 0 | 471250 | 0 | 0 | 0 | 0 | 0 |
| TARATGASI | DSG2 | Q14126 | DSG2_HUMAN | Oxidation (M) | 0 | 0 | 0 | 0 | 2619100 | 0 | 0 | 0 | 0 | 0 | 428150 | 0 | 0 | 23484 | 0 | 0 | 0 | 0 |
| TARELTPQ\ | VCL | P18206 | VINC_HUMAN ' | Unmodified | 0 | 107460 | 0 | 0 | 1084900 | 0 | 0 | 74932 | 0 | 364290 | 0 | 0 | 84103 | 0 | 0 | 0 | 2147800 | 0 |
| TARSIASQV | ATP7A | Q04656 | ATP7A_HUMAN | Unmodified | 0 | 0 | 0 | 0 | 0 | 0 | 265400 | 107790 | 0 | 0 | 0 | 0 | 100980 | 257270 | 0 | 0 | 0 | 51535 |
| TASDAGFP | SEZ6L2; SEZ6L2 | Q6UXD5; A0 | SE6L2_HUMAN | Unmodified | 0 | 0 | 0 | 0 | 0 | 0 | 100200 | 0 | 9992700 | 474940 | 0 | 0 | 0 | 0 | 555240 | 0 | 0 | 0 |
| TASDCGAV | PSAP; PSAP; PS | P07602; Q5E | SAP_HUMAN P | Unmodified | 0 | 0 | 0 | 0 | 0 | 0 | 1704600 | 0 | 0 | 395020 | 0 | 0 | 0 | 0 | 0 | 0 | 52279 | 0 |
| TASDMVST | PSMD12 | O00232 | PSD12_HUMAN | Unmodified | 36779 | 0 | 163180 | 143320 | 0 | 0 | 0 | 0 | 0 | 0 | 0 | 3119600 | 295430 | 85355 | 214220 | 197760 | 0 | 0 |
| TASDMVST | PSMD12 | O00232 | PSD12_HUMAN | Oxidation (M) | 0 | 0 | 0 | 0 | 0 | 3409300 | 604630 | 344950 | 0 | 0 | 0 | 0 | 0 | 68765 | 0 | 0 | 599250 | 0 |
| TASDVVQP | LSM14B; LSM14 | Q9BX40; Q5 | LS14B_HUMAN | Unmodified | 0 | 0 | 1643200 | 0 | 0 | 0 | 0 | 0 | 0 | 0 | 0 | 0 | 484100 | 0 | 0 | 320890 | 0 | 0 |
| TASFSESRA | ACLY | P53396 | ACLY_HUMAN | Unmodified | 225890 | 128160 | 519120 | 668810 | 444200 | 160180 | 233590 | 0 | 1291500 | 114000 | 284060 | 5669500 | 0 | 0 | 557550 | 796090 | 108710 | 262180 |
| TASFSESRA | ACLY | P53396 | ACLY_HUMAN | Unmodified | 576360 | 180940 | 62235 | 0 | 623320 | 310850 | 379180 | 213810 | 1234300 | 317470 | 1190700 | 8148600 | 1215000 | 171790 | 763300 | 769850 | 0 | 384550 |
| TASGSSVT! | NDRG1; NDRG1 | Q92597; E7E | NDRG1_HUMA | Unmodified | 673810 | 132790 | 469790 | 49895 | 0 | 0 | 0 | 0 | 483780 | 386600 | 0 | 225940 | 144240 | 41632 | 510250 | 1081900 | 1278400 | 154490 |
| TASGSSVT! | NDRG1; NDRG1 | Q92597; E7E | NDRG1_HUMA | Unmodified | 0 | 0 | 0 | 0 | 2472700 | 0 | 0 | 0 | 0 | 0 | 0 | 0 | 0 | 0 | 0 | 0 | 0 | 0 |
| TASGVNQL | CACNA2D1; CA | P54289; E7E | CA2D1_HUMAI | Unmodified | 3330200 | 1477100 | 0 | 0 | 0 | 0 | 6458900 | 4806000 | 7684000 | 2075800 | 0 | 465480 | 0 | 0 | 0 | 0 | 0 | 0 |
| TASHGDLII | SEZ6L2; SEZ6L2 | Q6UXD5; H3 | SE6L2_HUMAN | Unmodified | 2014600 | 2239100 | 1416400 | 755340 | 0 | 0 | 0 | 10212 | 2471000 | 0 | 13356 | 801260 | 0 | 0 | 1803800 | 2047200 | 0 | 0 |
| TASIGTLGR | NEO1; NEO1 | Q92859; Q5 | NEO1_HUMAN | Unmodified | 0 | 0 | 0 | 571090 | 2483300 | 34325 | 285920 | 110690 | 133880 | 70330 | 0 | 0 | 539580 | 332610 | 41131 | 820910 | 0 | 0 |
| TASISIADPI | BTN3A2; BTN3A | P78410; O0( | BT3A2_HUMAN | Unmodified | 0 | 0 | 0 | 0 | 0 | 0 | 0 | 0 | 0 | 0 | 0 | 1024200 | 0 | 0 | 0 | 0 | 0 | 0 |
| TASISSSPS | LARP1 | Q6PKG0 | LARP1_HUMAN | Unmodified | 0 | 0 | 0 | 0 | 0 | 0 | 0 | 0 | 0 | 0 | 0 | 0 | 0 | 0 | 0 | 0 | 0 | 0 |
| TASLEELRE | PLCB2 | Q00722 | PLCB2_HUMAN | Unmodified | 0 | 14338 | 0 | 0 | 0 | 0 | 0 | 0 | 0 | 0 | 0 | 1199000 | 0 | 0 | 0 | 0 | 0 | 0 |
| TASLFEQK | MYOF | Q9NZM1 | MYOF_HUMAN | Unmodified | 0 | 0 | 0 | 0 | 25576000 | 3091600 | 98916 | 30271 | 2582400 | 555450 | 3741700 | 4238700 | 0 | 0 | 0 | 0 | 401930 | 3265400 |
| TASLTSAAS | NDRG2; NDRG2 | Q9UN36; G3 | NDRG2_HUMA | Unmodified | 0 | 0 | 0 | 0 | 435500 | 0 | 143760 | 0 | 0 | 0 | 0 | 0 | 954130 | 330210 | 196910 | 83849 | 0 | 0 |
| TASNIGGFI | RAB3GAP2 | Q9H2M9 | RBGPR_HUMAI | Unmodified | 0 | 0 | 0 | 0 | 483820 | 0 | 0 | 182420 | 0 | 0 | 2512800 | 2024700 | 176070 | 0 | 0 | 0 | 0 | 0 |
| TASNVEEAI | RAB2A; RAB2A | P61019; H7( | RAB2A_HUMAN | Unmodified | 757930 | 1959700 | 3259500 | 4085600 | 3305600 | 289670 | 2359900 | 1286100 | 39816000 | 9847600 | 2758200 | 8863000 | 2907300 | 0 | 1134500 | 2079400 | 0 | 338400 |
| TASNVEEAI | RAB2A; RAB2A | P61019; H7( | RAB2A_HUMAN | Unmodified | 178670 | 0 | 0 | 0 | 0 | 0 | 3057400 | 0 | 3728900 | 12798000 | 50008 | 1238400 | 210120 | 8960000 | 0 | 0 | 0 | 0 |
| TASRPDDII | CARMIL1; CARN | Q5VZK9; A0/ | CARL1_HUMAN | Unmodified | 0 | 0 | 124590 | 0 | 1837600 | 0 | 0 | 0 | 0 | 0 | 232240 | 0 | 1241700 | 223660 | 0 | 71317 | 0 | 0 |
| TASSPDSLI | PLXNA1 | Q9UIW2 | PLXA1_HUMAN | Unmodified | 0 | 0 | 0 | 0 | 386500 | 0 | 0 | 0 | 0 | 0 | 0 | 0 | 575290 | 349990 | 0 | 0 | 461140 | 854590 |
| TASSVIELT( | IGF2R; IGF2R | P11717; S4R | MPRI_HUMAN ( | Unmodified | 197190 | 0 | 268330 | 232480 | 0 | 0 | 0 | 0 | 19340000 | 2457500 | 1484800 | 2698300 | 0 | 0 | 0 | 364470 | 52942 | 0 |
| TASTNNIA( | GNG12 | Q9UBI6 | GBG12_HUMAI | Unmodified | 1154900 | 81026 | 1635900 | 1238200 | 17574000 | 4071800 | 560840 | 504470 | 2230000 | 0 | 0 | 0 | 7188700 | 6138600 | 2906800 | 2727000 | 7214300 | 19401000 |
| TASTNNIA( | GNG12 | Q9UBI6 | GBG12_HUMAI | Unmodified | 0 | 36645 | 0 | 0 | 0 | 0 | 0 | 0 | 0 | 0 | 0 | 0 | 0 | 5964200 | 0 | 0 | 3328700 | 22972 |
| TASTPTPP( | EIF4G1; EIF4G1 | Q04637; C9J | IF4G1_HUMAN | Unmodified | 0 | 0 | 0 | 0 | 0 | 0 | 0 | 0 | 0 | 0 | 0 | 1807800 | 0 | 0 | 0 | 0 | 0 | 0 |
| TASTSFTNI | HS2ST1; HS2ST | Q7LGA3; K7I | HS2ST_HUMAN | Unmodified | 0 | 0 | 0 | 0 | 63112 | 0 | 228770 | 156410 | 9482200 | 2991600 | 151430 | 406100 | 112950 | 0 | 0 | 0 | 0 | 0 |
| TASVFLLEK | THOC3; THOC3 | Q96J01; D6F | THOC3_HUMA | Unmodified | 274430 | 170150 | 0 | 101490 | 913080 | 0 | 438120 | 818950 | 769610 | 648790 | 621530 | 2829900 | 5617800 | 1654500 | 0 | 0 | 1900000 | 3356700 |
| TASVSINQI | EPHA2 | P29317 | EPHA2_HUMAN | Unmodified | 0 | 57103 | 89509 | 0 | 0 | 66754 | 0 | 0 | 0 | 0 | 0 | 452830 | 0 | 0 | 105600 | 0 | 0 | 0 |
| TASWLAAE | DDX19B; DDX1 | Q9UMR2; Q | DD19B_HUMAI | Unmodified | 0 | 0 | 0 | 0 | 0 | 0 | 0 | 0 | 0 | 0 | 0 | 0 | 0 | 0 | 0 | 0 | 0 | 0 |
| TASYTDVW | GLT8D1 | Q68CQ7 | GL8D1_HUMAN | Unmodified | 0 | 0 | 0 | 0 | 0 | 0 | 0 | 0 | 19493000 | 2095300 | 0 | 0 | 0 | 0 | 0 | 0 | 0 | 0 |
| TATDEAYKI | SPTAN1; SPTAN | Q13813; A0/ | SPTN1_HUMAN | Unmodified | 0 | 0 | 0 | 0 | 216290 | 78357 | 0 | 0 | 0 | 81675 | 80193 | 2137100 | 0 | 0 | 139450 | 70576 | 148120 | 0 |
| TATESFASI | PKM; PKM; PKM | P14618; H3E | KPYM_HUMAN | Unmodified | 3975900 | 3637500 | 14443000 | 4968800 | 11441000 | 4934700 | 2601500 | 1247300 | 15352000 | 3475300 | 29059000 | 2.04E+08 | 4031600 | 608180 | 19027000 | 12188000 | 5252300 | 7690400 |
| TATESFASI | PKM; PKM; PKM | P14618; H3E | KPYM_HUMAN | Unmodified | 16061000 | 18347000 | 6322200 | 69244000 | 1.36E+08 | 29241000 | 3575300 | 55370 | 1.24E+08 | 12393000 | 1.51E+08 | 7.07E+08 | 41786000 | 3691200 | 77437000 | 37179000 | 29621000 | 66372000 |
| TATGFGFAI | PIK3R2 | O00459 | P85B_HUMAN | Unmodified | 0 | 94708 | 10220 | 0 | 243800 | 0 | 0 | 0 | 1065400 | 65140 | 0 | 216520 | 129990 | 0 | 0 | 0 | 0 | 0 |
| TATGYGFAI | PIK3R1 | P27986 | P85A_HUMAN | Unmodified | 0 | 0 | 63723 | 0 | 0 | 0 | 168650 | 0 | 140170 | 0 | 73042 | 131690 | 2133300 | 461840 | 0 | 113200 | 0 | 0 |
| TATITEPRG | EFTUD2 | Q15029 | U5S1_HUMAN | Unmodified | 0 | 0 | 0 | 0 | 1781200 | 0 | 696790 | 212740 | 0 | 0 | 0 | 0 | 1745300 | 745700 | 0 | 0 | 1070000 | 0 |
| TATLRHDAI | PSMD3 | O43242 | PSMD3_HUMA | Unmodified | 0 | 0 | 0 | 0 | 0 | 0 | 0 | 0 | 0 | 0 | 0 | 3670200 | 0 | 0 | 0 | 0 | 0 | 0 |
| TATLRPYLS | ARPC4; ARPC4 | P59998; A0A | ARPC4_HUMAI | Acetyl (Protein N | 1169300 | 2566800 | 0 | 4204300 | 5838100 | 1487200 | 529600 | 179100 | 1362100 | 950080 | 1935400 | 10171000 | 4909000 | 0 | 2352900 | 1666700 | 719030 | 3960500 |
| TATMLCAA( | PTPRF; PTPRF | P10586; A2A | PTPRF_HUMAN | Unmodified | 0 | 0 | 0 | 0 | 0 | 0 | 0 | 0 | 143180 | 0 | 0 | 0 | 0 | 0 | 0 | 0 | 0 | 0 |
| TATNIIKKS! | ADGRG6 | Q86SQ4 | AGRG6_HUMA | Unmodified | 0 | 0 | 0 | 0 | 479550 | 59489 | 0 | 0 | 0 | 0 | 0 | 0 | 0 | 0 | 0 | 0 | 0 | 0 |
| TATPEIVDN | FLNB; FLNB | O75369; E7E | FLNB_HUMAN | Unmodified | 190790 | 154240 | 431960 | 318240 | 458730 | 0 | 0 | 0 | 0 | 0 | 0 | 2628900 | 0 | 0 | 1143900 | 691770 | 209720 | 0 |
| TATPQQAQ | TPI1; TPI1 | P60174; U3K | TPIS_HUMAN T | Unmodified | 0 | 0 | 0 | 0 | 24234000 | 0 | 13913000 | 0 | 0 | 9841700 | 4675800 | 18734000 | 0 | 69287000 | 9990100 | 3823900 | 1.04E+08 | 21101000 |
| TATPQQAQ | TPI1; TPI1 | P60174; U3K | TPIS_HUMAN T | Unmodified | 200770 | 0 | 0 | 110680 | 422790 | 0 | 235830 | 31561 | 0 | 2438000 | 0 | 0 | 1677600 | 869140 | 0 | 0 | 1432900 | 243120 |
| TATPQQAQ | TPI1; TPI1 | P60174; U3K | TPIS_HUMAN T | Unmodified | 0 | 0 | 0 | 0 | 1344300 | 0 | 3833600 | 316130 | 0 | 0 | 0 | 0 | 659980 | 0 | 0 | 0 | 17016000 | 0 |
| TATQLAVNI | CCT7; CCT7 | Q99832; F8V | TCPH_HUMAN | Unmodified | 776500 | 1108400 | 390120 | 321080 | 737080 | 152120 | 220380 | 30590 | 563030 | 316390 | 1159200 | 10584000 | 1671700 | 559690 | 561400 | 537970 | 289160 | 1071800 |
| TATSLAEGI | TMEM43 | Q9BTV4 | TMM43_HUMAI | Unmodified | 0 | 0 | 0 | 0 | 0 | 0 | 0 | 0 | 10323000 | 1853700 | 0 | 409910 | 0 | 0 | 0 | 0 | 667810 | 0 |
| TATVCGREI | CDC42EP2 | O14613 | BORG1_HUMA | Unmodified | 0 | 0 | 0 | 0 | 0 | 0 | 0 | 0 | 0 | 0 | 0 | 0 | 0 | 0 | 0 | 0 | 1016600 | 902370 |
| TATVYPEP( | ME1 | P48163 | MAOX_HUMAN | Unmodified | 0 | 0 | 0 | 224990 | 1909000 | 851770 | 143600 | 98207 | 132200 | 118800 | 560400 | 3134600 | 1235300 | 558970 | 0 | 0 | 1329800 | 1784500 |
| TATYNGPP, | MARK3; MARK3 | P27448; A0A | MARK3_HUMAI | Unmodified | 0 | 0 | 0 | 0 | 654450 | 0 | 0 | 0 | 427380 | 0 | 0 | 388940 | 502300 | 0 | 287790 | 511300 | 0 | 0 |
| TAVAHRPG | MYDGF | Q969H8 | MYDGF_HUMA | Unmodified | 0 | 0 | 0 | 0 | 0 | 0 | 0 | 0 | 7588300 | 318870 | 0 | 0 | 0 | 0 | 12849 | 0 | 0 | 0 |
| TAVAHRPG | MYDGF | Q969H8 | MYDGF_HUMA | Unmodified | 28890 | 0 | 0 | 128770 | 1554200 | 145040 | 1012700 | 685770 | 41638000 | 38822000 | 1478600 | 415130 | 443800 | 1304400 | 0 | 103900 | 654980 | 1463900 |
| TAVAPIER | SLC25A4; SLC2 | P12235; P12 | ADT1_HUMAN | Unmodified | 538960 | 380850 | 201680 | 194350 | 307170 | 60674 | 143140 | 119060 | 46106000 | 8169500 | 2293400 | 1166500 | 0 | 78736 | 552900 | 227980 | 13772 | 25023 |
| TAVCDIPPI | TUBB2A; TUBB | Q13885; A6N | TBB2A_HUMAN | Unmodified | 2477600 | 3197300 | 0 | 0 | 2451000 | 0 | 946860 | 0 | 0 | 0 | 0 | 12314000 | 0 | 0 | 61055 | 4261400 | 1894100 | 0 |
| TAVCDIPPI | TUBB2A; TUBB | Q13885; A6N | TBB2A_HUMAN | Unmodified | 0 | 116800 | 0 | 0 | 3485600 | 74315 | 9058100 | 5885000 | 1022300 | 2006600 | 2133800 | 12378000 | 4744100 | 5138400 | 1087300 | 515390 | 1528300 | 2980000 |
| TAVDSLVA\ | CKAP4 | Q07065 | CKAP4_HUMAN | Unmodified | 0 | 0 | 0 | 0 | 0 | 0 | 0 | 0 | 4391100 | 229450 | 0 | 0 | 0 | 0 | 0 | 0 | 0 | 0 |
| TAVEHATDI | LRBA; LRBA; LR | A0A494C1L5 | A0A494C1L5_H | Unmodified | 0 | 0 | 0 | 0 | 0 | 0 | 57942 | 0 | 0 | 0 | 0 | 647330 | 461380 | 0 | 0 | 56159 | 0 | 0 |
| TAVETAVLL | CCT3; CCT3 | P49368; B4D | TCPG_HUMAN | Unmodified | 0 | 0 | 571290 | 0 | 0 | 129180 | 15920 | 18446 | 2455600 | 775630 | 2567400 | 34994000 | 0 | 14648 | 869170 | 1049300 | 0 | 0 |
| TAVETAVLL | CCT3; CCT3 | P49368; B4D | TCPG_HUMAN | Unmodified | 47146 | 19818 | 0 | 271210 | 8208100 | 436600 | 1887300 | 0 | 254320 | 2676900 | 147070 | 92388 | 19485000 | 8772100 | 0 | 0 | 4237000 | 6347900 |
| TAVETAVLL | CCT3; CCT3 | P49368; B4D | TCPG_HUMAN | Unmodified | 0 | 0 | 0 | 0 | 428920 | 85266 | 0 | 0 | 179510 | 0 | 340650 | 0 | 611110 | 446180 | 0 | 0 | 260640 | 403220 |
| TAVFGFETS | MVP | Q14764 | MVP_HUMAN I | Unmodified | 668470 | 410370 | 0 | 0 | 0 | 0 | 0 | 0 | 0 | 216740 | 0 | 17802000 | 0 | 0 | 0 | 0 | 0 | 0 |
| TAVFGFETS | MVP | Q14764 | MVP_HUMAN I | Oxidation (M) | 907930 | 499730 | 0 | 0 | 0 | 0 | 120320 | 0 | 0 | 0 | 0 | 554810 | 0 | 0 | 321700 | 0 | 0 | 0 |
| TAVHAGNII | CTNNA1; CTNN | P35221; E5R | CTNA1_HUMAI | Acetyl (Protein N | 775950 | 587300 | 6751900 | 2056400 | 4501100 | 459650 | 0 | 0 | 986260 | 0 | 1525700 | 9654900 | 729840 | 0 | 6340600 | 7548200 | 0 | 0 |
| TAVHAGNII | CTNNA1; CTNN | P35221; E5R | CTNA1_HUMAI | Acetyl (Protein N | 2670600 | 3368100 | 4210700 | 17893000 | 1.85E+08 | 29809000 | 0 | 164790 | 2401000 | 2776300 | 6250100 | 13158000 | 67552000 | 41645000 | 16891000 | 14927000 | 0 | 0 |
| TAVHAGVIS | CRISPLD2; CRI | Q9H0B8; H3 | CRLD2_HUMAI | Oxidation (M) | 0 | 0 | 0 | 0 | 0 | 0 | 0 | 87132 | 1007700 | 221790 | 0 | 0 | 0 | 0 | 0 | 0 | 0 | 0 |
| TAVHAGVIS | CRISPLD2; CRI | Q9H0B8; H3 | CRLD2_HUMAI | Unmodified | 0 | 0 | 0 | 0 | 0 | 0 | 0 | 37570 | 1602000 | 310850 | 0 | 0 | 0 | 0 | 0 | 0 | 0 | 0 |

| | | | | | | | | | | | | | | | | | | | | |
|---|---|---|---|---|---|---|---|---|---|---|---|---|---|---|---|---|---|---|---|---|
| TAVIDHHN | DDOST; DDOST | P39656; U3K | OST48_HUMAN | Unmodified | 0 | 0 | 0 | 0 | 0 | 0 | 0 | 0 | 4.24E+08 | 51770000 | 27424000 | 1551700 | 0 | 0 | 0 | 0 | 0 | 0 |
| TAVIDHHN | DDOST; DDOST | P39656; U3K | OST48_HUMAN | Unmodified | 0 | 0 | 0 | 0 | 0 | 0 | 257000 | 0 | 5110500 | 23701000 | 0 | 0 | 0 | 0 | 0 | 0 | 0 | 0 |
| TAVLTAFAN | ACTL6A; ACTL6 | O96019; O9 | ACL6A_HUMAN | Unmodified | 182030 | 103700 | 0 | 0 | 123810 | 0 | 0 | 0 | 337830 | 172200 | 476450 | 1635800 | 0 | 0 | 0 | 69893 | 0 | 0 |
| TAVSSELR | UGGT1 | Q9NYU2 | UGGG1_HUMA | Unmodified | 0 | 108260 | 0 | 0 | 0 | 0 | 0 | 0 | 489460 | 0 | 0 | 0 | 0 | 0 | 27617 | 0 | 0 | 0 |
| TAVSSELRT | UGGT1 | Q9NYU2 | UGGG1_HUMA | Unmodified | 0 | 0 | 0 | 65058 | 609840 | 0 | 0 | 328790 | 48847000 | 19747000 | 3638500 | 1221900 | 390330 | 398050 | 232560 | 190570 | 0 | 0 |
| TAVVVGTIT | RPL18; RPL18; | Q07020; F8V | RL18_HUMAN | Unmodified | 1662000 | 1451600 | 554490 | 705410 | 0 | 0 | 0 | 0 | 413550 | 0 | 3616200 | 27052000 | 508410 | 0 | 645080 | 634150 | 0 | 340860 |
| TAVVVGTIT | RPL18; RPL18; | Q07020; F8V | RL18_HUMAN | Unmodified | 10067000 | 9172400 | 6694400 | 3270900 | 3218500 | 639650 | 9966500 | 4400200 | 1689000 | 709730 | 4893800 | 69954000 | 2411600 | 0 | 7361700 | 4634500 | 431990 | 2275400 |
| TAVVVGTIT | RPL18; RPL18; | Q07020; F8V | RL18_HUMAN | Unmodified | 0 | 0 | 0 | 0 | 671630 | 0 | 5328900 | 673700 | 0 | 0 | 0 | 0 | 1926200 | 1015300 | 0 | 0 | 493850 | 983860 |
| TAVYKVPPF | CAVIN1 | Q6NZI2 | CAVN1_HUMA | Unmodified | 0 | 0 | 0 | 0 | 1107600 | 0 | 0 | 0 | 0 | 0 | 0 | 0 | 0 | 0 | 0 | 0 | 0 | 0 |
| TAVYKVPPF | CAVIN1 | Q6NZI2 | CAVN1_HUMA | Unmodified | 0 | 0 | 0 | 0 | 1290900 | 0 | 0 | 0 | 0 | 0 | 0 | 0 | 0 | 0 | 0 | 0 | 0 | 0 |
| TAWFNGVE | TUBA1C; TUBA | F8VRK1; F5H | F8VRK1_HUMA | Acetyl (Protein N | 0 | 0 | 0 | 0 | 0 | 38039 | 0 | 0 | 159890 | 905190 | 38880 | 0 | 0 | 0 | 0 | 0 | 88535 | 0 |
| TAWGQQP | PSMD13; PSMD | Q9UNM6; A0 | PSD13_HUMAN | Unmodified | 0 | 0 | 0 | 0 | 0 | 0 | 695470 | 199030 | 0 | 230330 | 0 | 3330300 | 0 | 0 | 0 | 295410 | 0 | 0 |
| TAWLDGKI | PPIB; PPIB; PPI | P23284; A0A | PPIB_HUMAN | Unmodified | 0 | 0 | 0 | 0 | 537700 | 217450 | 593380 | 262270 | 0 | 2517900 | 0 | 0 | 0 | 177610 | 0 | 0 | 437700 | 0 |
| TAWRLDCA | CNP; CNP; CNI | P09543; K7E | CN37_HUMAN | Unmodified | 1328400 | 550390 | 2173400 | 3742400 | 17738000 | 3570500 | 5296600 | 2641500 | 3928000 | 3725700 | 13356000 | 23590000 | 15785000 | 11688000 | 3124200 | 3167000 | 8823800 | 16833000 |
| TAWRLDCA | CNP; CNP; CNI | P09543; K7E | CN37_HUMAN | Unmodified | 0 | 0 | 0 | 0 | 1007600 | 0 | 517650 | 566380 | 0 | 71295 | 0 | 0 | 628760 | 1007800 | 0 | 0 | 428340 | 1228000 |
| TAYFSLDTF | CDH1; CDH1; ( | P12830; H3E | CADH1_HUMA | Unmodified | 0 | 520370 | 1229200 | 0 | 0 | 0 | 0 | 0 | 1304100 | 0 | 299540 | 0 | 0 | 0 | 1795300 | 724630 | 0 | 0 |
| TAYGPNGN | CCT8; CCT8 | P50990; H7C | TCPQ_HUMAN | Unmodified | 0 | 0 | 0 | 0 | 0 | 0 | 0 | 0 | 491330 | 216350 | 475450 | 2338300 | 3296100 | 559020 | 0 | 0 | 153850 | 0 |
| TAYLFSR | XPOT | O43592 | XPOT_HUMAN | Unmodified | 245810 | 259460 | 0 | 0 | 0 | 0 | 0 | 0 | 0 | 0 | 0 | 1891300 | 0 | 0 | 0 | 51399 | 0 | 0 |
| TCATDLQTI | BPNT1; BPNT1; | O95861; F8V | BPNT1_HUMAN | Unmodified | 78096 | 0 | 0 | 0 | 0 | 23294 | 0 | 168250 | 224690 | 21412 | 0 | 542900 | 0 | 0 | 0 | 54942 | 0 | 80376 |
| TCDGVQCA | RAB18; RAB18; | Q9NP72; H0 | RAB18_HUMAN | Unmodified | 0 | 0 | 75750 | 0 | 296140 | 0 | 445740 | 46706 | 582700 | 465030 | 366700 | 1968900 | 340270 | 0 | 0 | 228930 | 0 | 0 |
| TCDISFSDF | UBE2M | P61081 | UBC12_HUMA | Unmodified | 0 | 0 | 0 | 338470 | 0 | 0 | 0 | 51305 | 1208900 | 0 | 0 | 484300 | 567660 | 0 | 373460 | 156820 | 505490 | 542770 |
| TCDLVGEK | M6PR; M6PR; M | P20645; F5G | MPRD_HUMAN | Unmodified | 0 | 184090 | 0 | 407060 | 0 | 0 | 2467100 | 1045600 | 39140000 | 9326500 | 2400500 | 6738200 | 8120200 | 2133700 | 207630 | 167720 | 158690 | 176530 |
| TCDLVGEK | M6PR; M6PR; M | P20645; F5G | MPRD_HUMAN | Unmodified | 0 | 0 | 0 | 0 | 0 | 0 | 0 | 501540 | 161380 | 0 | 0 | 0 | 0 | 0 | 0 | 0 | 0 | 0 |
| TCEECNFK | ITGB4 | P16144 | ITB4_HUMAN I | Unmodified | 0 | 0 | 0 | 0 | 0 | 0 | 0 | 0 | 0 | 0 | 0 | 0 | 117360 | 0 | 0 | 0 | 864730 | 2625500 |
| TCEECNFK | ITGB4 | P16144 | ITB4_HUMAN I | Unmodified | 0 | 0 | 0 | 0 | 316640 | 304540 | 0 | 0 | 0 | 0 | 0 | 0 | 0 | 25122 | 0 | 0 | 2849200 | 1088600 |
| TCEESSFCI | GANAB | Q14697 | GANAB_HUMA | Unmodified | 0 | 0 | 118820 | 112090 | 274690 | 105550 | 392630 | 393220 | 30432000 | 15021000 | 798650 | 776610 | 146730 | 626770 | 0 | 0 | 158100 | 333630 |
| TCEESSFCI | GANAB | Q14697 | GANAB_HUMA | Unmodified | 0 | 0 | 0 | 0 | 0 | 0 | 0 | 0 | 8945900 | 2174200 | 0 | 0 | 0 | 0 | 0 | 0 | 0 | 0 |
| TCESDTLE | UGGT1 | Q9NYU2 | UGGG1_HUMA | Unmodified | 0 | 0 | 0 | 0 | 0 | 0 | 0 | 0 | 5140400 | 0 | 0 | 0 | 0 | 0 | 0 | 0 | 0 | 0 |
| TCESLGAG | HSPG2 | P98160 | PGBM_HUMAN | Unmodified | 0 | 0 | 0 | 0 | 0 | 0 | 0 | 0 | 538330 | 0 | 167140 | 266200 | 0 | 0 | 0 | 0 | 0 | 0 |
| TCEVAAWC | P2RX4; P2RX4 | Q99571; F6S | P2RX4_HUMAN | Unmodified | 0 | 0 | 0 | 0 | 0 | 0 | 0 | 0 | 3737300 | 0 | 0 | 552910 | 0 | 0 | 0 | 0 | 0 | 0 |
| TCFSMVPA | ISOC2; ISOC2; | Q96AB3; K7I | ISOC2_HUMAN | Unmodified | 0 | 0 | 0 | 0 | 0 | 0 | 0 | 0 | 1089700 | 214560 | 0 | 0 | 0 | 0 | 0 | 0 | 0 | 0 |
| TCFSPNRV | EIF3F; EIF3F | O00303; H0Y | EIF3F_HUMAN | Unmodified | 0 | 0 | 0 | 0 | 0 | 0 | 0 | 0 | 0 | 0 | 296560 | 4290700 | 0 | 0 | 362890 | 0 | 299940 | 0 |
| TCHESYDR | MAN1A2; MAN1 | O60476; H0Y | MA1A2_HUMA | Unmodified | 282860 | 150770 | 0 | 0 | 0 | 68024 | 150090 | 0 | 2856700 | 0 | 23634 | 0 | 0 | 551360 | 0 | 0 | 0 | 0 |
| TCILDVNP | PALS2; PALS2 | Q9NZW5; B8 | PALS2_HUMAN | Unmodified | 0 | 0 | 0 | 0 | 0 | 0 | 0 | 0 | 827410 | 0 | 955360 | 2008600 | 0 | 0 | 0 | 0 | 0 | 0 |
| TCLDNLAS | IQGAP1; IQGAP | P46940; H0Y | IQGA1_HUMAN | Unmodified | 885270 | 0 | 751030 | 873630 | 16922000 | 2742000 | 0 | 65304 | 2440800 | 1095000 | 392600 | 3723500 | 3137800 | 2277800 | 715990 | 567380 | 0 | 1940900 |
| TCLDNLAS | IQGAP1; IQGAP | P46940; H0Y | IQGA1_HUMAN | Unmodified | 0 | 0 | 437460 | 0 | 825130 | 225550 | 247780 | 96392 | 0 | 0 | 0 | 0 | 799630 | 438420 | 0 | 0 | 0 | 167250 |
| TCLLISYTTI | CDC42; CDC42 | P60953; Q5J | CDC42_HUMA | Unmodified | 268360 | 351860 | 0 | 0 | 0 | 0 | 0 | 0 | 0 | 0 | 0 | 0 | 0 | 0 | 775130 | 447470 | 0 | 0 |
| TCLLIVFSK | RHOA; RHOC; I | P61586; P08 | RHOA_HUMAN | Unmodified | 3392800 | 3333800 | 0 | 10264000 | 6794000 | 998330 | 0 | 0 | 10197000 | 0 | 8640900 | 11811000 | 6292900 | 2652000 | 10969000 | 4984700 | 2489700 | 5325600 |
| TCLLNETGI | GLUL; GLUL | P15104; A0A | GLNA_HUMAN | Unmodified | 0 | 0 | 0 | 0 | 0 | 0 | 0 | 196380 | 0 | 0 | 0 | 615630 | 0 | 0 | 0 | 0 | 0 | 0 |
| TCMDVLRE | KIF23; KIF23; KI | Q02241; A0A | KIF23_HUMAN | Oxidation (M) | 0 | 0 | 0 | 112370 | 4629000 | 1278800 | 26749 | 0 | 675800 | 428030 | 47993 | 604260 | 86848 | 0 | 0 | 0 | 517900 | 0 |
| TCMDVLRE | KIF23; KIF23; KI | Q02241; A0A | KIF23_HUMAN | Unmodified | 0 | 0 | 0 | 0 | 5278700 | 0 | 0 | 0 | 1708000 | 525490 | 434960 | 424440 | 209910 | 393240 | 85930 | 0 | 683920 | 263550 |
| TCMDVLRE | KIF23; KIF23; KI | Q02241; A0A | KIF23_HUMAN | 2 Oxidation (M) | 0 | 0 | 170930 | 0 | 653510 | 256740 | 0 | 0 | 0 | 76981 | 74968 | 0 | 0 | 0 | 0 | 0 | 0 | 0 |
| TCNTMNQI | COPS6; COPS6 | Q7L5N1; E7I | CSN6_HUMAN | Unmodified | 0 | 0 | 0 | 0 | 0 | 0 | 0 | 0 | 0 | 0 | 159380 | 783510 | 0 | 0 | 0 | 0 | 125760 | 0 |
| TCPLSWVC | DIP2B | Q9P265 | DIP2B_HUMAN | Unmodified | 217320 | 0 | 0 | 0 | 0 | 0 | 0 | 0 | 0 | 0 | 0 | 1312100 | 659430 | 0 | 0 | 223000 | 0 | 0 |
| TCPVIISSTA | THNSL1 | Q8IYQ7 | THNS1_HUMA | Unmodified | 0 | 0 | 0 | 0 | 0 | 0 | 0 | 0 | 604510 | 148890 | 0 | 294320 | 0 | 0 | 0 | 0 | 266730 | 0 |
| TCPYVSFR | PTGS1; PTGS1 | P23219; A0A | PGH1_HUMAN | Unmodified | 0 | 0 | 0 | 0 | 0 | 0 | 0 | 416490 | 0 | 0 | 11033000 | 2802300 | 0 | 0 | 0 | 0 | 0 | 0 |
| TCQEVPSV | PAQR7 | Q86WK9 | PAQR7_HUMA | Unmodified | 0 | 0 | 0 | 0 | 0 | 283870 | 0 | 0 | 278060 | 0 | 0 | 0 | 0 | 0 | 0 | 0 | 0 | 0 |
| TCQQTWEI | ATP6V1C1 | P21283 | VATC1_HUMAN | Unmodified | 0 | 0 | 0 | 0 | 0 | 0 | 0 | 0 | 0 | 0 | 0 | 257370 | 0 | 0 | 0 | 0 | 634610 | 0 |
| TCRNPSGI | MUC5B | Q9HC84 | MUC5B_HUMA | Unmodified | 0 | 0 | 0 | 0 | 0 | 0 | 0 | 0 | 0 | 0 | 0 | 0 | 0 | 0 | 0 | 0 | 226740 | 3705200 |
| TCRPSTQS | MGAT2 | Q10469 | MGAT2_HUMA | Unmodified | 0 | 0 | 0 | 0 | 0 | 0 | 218440 | 0 | 11079000 | 2449200 | 0 | 0 | 0 | 0 | 0 | 0 | 0 | 0 |
| TCRVECSD | MASP1; MASP1 | P48740; F8V | MASP1_HUMA | Unmodified | 0 | 0 | 0 | 0 | 0 | 0 | 0 | 0 | 0 | 0 | 0 | 0 | 0 | 0 | 419710 | 381120 | 0 | 0 |
| TCSPASLS | TRIM25; TRIM25 | Q14258; A0A | TRI25_HUMAN | Unmodified | 0 | 0 | 0 | 0 | 3887400 | 0 | 0 | 0 | 0 | 0 | 0 | 0 | 0 | 0 | 0 | 0 | 0 | 1309200 |
| TCSVSSSG | ELFN2 | Q5R3F8 | PPR29_HUMAN | Unmodified | 0 | 0 | 0 | 0 | 0 | 0 | 0 | 0 | 0 | 0 | 0 | 0 | 0 | 0 | 0 | 0 | 0 | 0 |
| TCTDIKPEV | DHX15 | O43143 | DHX15_HUMA | Unmodified | 53892 | 0 | 0 | 0 | 0 | 0 | 0 | 0 | 0 | 0 | 0 | 3843500 | 0 | 0 | 0 | 0 | 0 | 0 |
| TCTFILR | CCT7 | Q99832 | TCPH_HUMAN | Unmodified | 155910 | 0 | 0 | 154810 | 0 | 0 | 0 | 0 | 0 | 0 | 542400 | 10064000 | 0 | 0 | 0 | 390920 | 0 | 0 |
| TCTLLEKFP | ASPH | Q12797 | ASPH_HUMAN | Unmodified | 0 | 0 | 0 | 0 | 0 | 0 | 0 | 0 | 1395600 | 299460 | 0 | 0 | 78114 | 0 | 0 | 0 | 0 | 0 |
| TCTTLSGLA | DIPK1B | Q5VUD6 | DIK1B_HUMAN | Unmodified | 0 | 0 | 0 | 0 | 0 | 0 | 0 | 0 | 2629600 | 122680 | 0 | 2576500 | 0 | 0 | 0 | 0 | 0 | 0 |
| TCTTVAFTC | RPL7A; RPL7A | P62424; Q5T | RL7A_HUMAN | Unmodified | 1151100 | 736240 | 773900 | 521890 | 0 | 0 | 0 | 188750 | 0 | 235890 | 308770 | 18333000 | 0 | 219940 | 941510 | 610080 | 0 | 0 |
| TCTTVAFTC | RPL7A; RPL7A | P62424; Q5T | RL7A_HUMAN | Unmodified | 2882200 | 2197600 | 0 | 0 | 1477000 | 0 | 4190900 | 2456800 | 0 | 0 | 0 | 2441200 | 805320 | 1067200 | 386760 | 771060 | 0 | 0 |
| TCVVVAEEI | DIP2B; DIP2C; I | Q9P265; Q9 | DIP2B_HUMAN | Unmodified | 0 | 0 | 125930 | 0 | 0 | 0 | 661520 | 0 | 294880 | 472430 | 169540 | 1705800 | 1151700 | 462760 | 564750 | 289630 | 26575 | 0 |
| TCWDADPI | KIT | P10721 | KIT_HUMAN M | Unmodified | 0 | 0 | 0 | 0 | 0 | 0 | 0 | 0 | 0 | 329720 | 0 | 1764900 | 0 | 0 | 0 | 0 | 0 | 0 |
| TCYEGNGI | PLAU; PLAU; PL | P00749; E7E | UROK_HUMAN | Unmodified | 0 | 0 | 0 | 0 | 565620 | 465310 | 0 | 0 | 0 | 0 | 0 | 0 | 0 | 0 | 0 | 0 | 391700 | 821540 |
| TDAAVEMK | DBN1; DBN1 | Q16643; D6I | DREB_HUMAN | Unmodified | 0 | 0 | 1166300 | 692720 | 1610700 | 483130 | 54658 | 61857 | 5155400 | 0 | 19357 | 2571000 | 1330800 | 728050 | 671360 | 121850 | 946050 | 1262500 |
| TDAAVEMK | DBN1; DBN1 | Q16643; D6I | DREB_HUMAN | Oxidation (M) | 0 | 0 | 23462 | 0 | 170900 | 0 | 0 | 0 | 42308 | 0 | 0 | 177580 | 0 | 0 | 0 | 0 | 52959 | 0 |
| TDAAVSFAI | SLC25A5 | P05141 | ADT2_HUMAN | Acetyl (Protein N | 0 | 0 | 0 | 141280 | 607640 | 0 | 6557100 | 2008200 | 0 | 6551300 | 935320 | 0 | 0 | 0 | 0 | 24190 | 199250 | 255490 |
| TDAETLPAL | PAPSS1 | O43252 | PAPS1_HUMAN | Unmodified | 0 | 0 | 0 | 0 | 2994200 | 1885600 | 161680 | 51692 | 0 | 0 | 587860 | 1428800 | 125340 | 0 | 0 | 0 | 0 | 0 |
| TDAGFTLR | VCL | P18206 | VINC_HUMAN | Unmodified | 0 | 0 | 0 | 0 | 0 | 0 | 0 | 0 | 459300 | 0 | 0 | 0 | 0 | 0 | 0 | 0 | 219240 | 0 |
| TDAPQPDV | CANX; CANX; C | P27824; A0A | CALX_HUMAN | Unmodified | 151560 | 0 | 0 | 0 | 0 | 0 | 0 | 0 | 1864700 | 0 | 173250 | 250150 | 0 | 0 | 0 | 0 | 0 | 47583 |
| TDAVEALTA | HNRNPLL; HNI | Q8WVV9; C9 | HNRLL_HUMA | Unmodified | 0 | 254810 | 0 | 0 | 0 | 0 | 0 | 0 | 0 | 0 | 0 | 531040 | 0 | 0 | 0 | 0 | 0 | 56168 |
| TDAVNEALI | UTRN | P46939 | UTRN_HUMAN | Unmodified | 0 | 0 | 338230 | 0 | 0 | 0 | 0 | 0 | 273510 | 0 | 0 | 0 | 0 | 0 | 1627700 | 678210 | 0 | 0 |
| TDDCHPW | GOT1 | P17174 | AATC_HUMAN | Unmodified | 0 | 0 | 0 | 0 | 340250 | 0 | 134880 | 0 | 0 | 0 | 262150 | 5021400 | 0 | 0 | 0 | 0 | 0 | 0 |
| TDDEVVQR | HSP90B1; HSP | P14625; A0A | ENPL_HUMAN | Unmodified | 0 | 0 | 0 | 0 | 0 | 0 | 0 | 0 | 1024800 | 132760 | 223040 | 262620 | 97979 | 79187 | 0 | 0 | 0 | 0 |
| TDDEVVQR | HSP90B1; HSP | P14625; A0A | ENPL_HUMAN | Unmodified | 0 | 0 | 0 | 0 | 0 | 0 | 0 | 0 | 33686000 | 653830 | 1355900 | 3914200 | 0 | 0 | 0 | 0 | 0 | 0 |
| TDDEVVQR | HSP90B1; HSP | P14625; A0A | ENPL_HUMAN | Unmodified | 0 | 0 | 0 | 0 | 548960 | 0 | 0 | 0 | 16433000 | 1421800 | 1564600 | 0 | 0 | 0 | 0 | 0 | 0 | 0 |
| TDDIPVWD | SKP1; SKP1; SK | P63208; E7E | SKP1_HUMAN | Unmodified | 0 | 0 | 0 | 0 | 0 | 0 | 13024 | 0 | 0 | 0 | 0 | 468160 | 0 | 9639.7 | 0 | 0 | 0 | 0 |
| TDDYGRDI | SPTAN1; SPTAN | Q13813; A0A | SPTN1_HUMAN | Unmodified | 23528 | 0 | 70470 | 0 | 0 | 0 | 0 | 0 | 0 | 0 | 0 | 936340 | 0 | 0 | 189200 | 46188 | 0 | 0 |
| TDDYLDQP | GDI2; GDI2; GD | P50395; Q5S | GDIB_HUMAN | Unmodified | 0 | 224020 | 0 | 249720 | 0 | 0 | 72918 | 0 | 651780 | 157100 | 159110 | 5210900 | 0 | 0 | 975220 | 849650 | 0 | 0 |
| TDDYLDQP | GDI2; GDI2; GD | P50395; Q5S | GDIB_HUMAN | Unmodified | 0 | 0 | 0 | 0 | 0 | 0 | 140500 | 67362 | 950700 | 0 | 0 | 1637700 | 0 | 0 | 0 | 394860 | 0 | 0 |
| TDEEGDFR | JAM3 | Q9BX67 | JAM3_HUMAN | Unmodified | 0 | 0 | 0 | 0 | 0 | 0 | 0 | 0 | 277190 | 0 | 0 | 1238800 | 0 | 0 | 47010 | 23959 | 0 | 0 |
| TDEFCQAK | LEMD2 | Q8NC56 | LEMD2_HUMA | Unmodified | 0 | 0 | 0 | 0 | 0 | 0 | 0 | 0 | 0 | 0 | 0 | 0 | 0 | 0 | 0 | 0 | 0 | 0 |
| TDEFPRHG | ACAT2 | Q9BWD1 | THIC_HUMAN | Unmodified | 0 | 0 | 0 | 108560 | 408180 | 0 | 0 | 0 | 849750 | 214550 | 692520 | 0 | 186730 | 0 | 0 | 0 | 77946 | 116530 |
| TDEFQLHTI | VDAC1 | P21796 | VDAC1_HUMA | Unmodified | 0 | 0 | 0 | 0 | 0 | 0 | 0 | 0 | 29938000 | 5443500 | 479340 | 0 | 0 | 0 | 0 | 0 | 0 | 0 |
| TDEGIAYR | PRDX2 | P32119 | PRDX2_HUMAN | Unmodified | 1464700 | 1281300 | 4362300 | 2095700 | 0 | 0 | 0 | 0 | 0 | 0 | 6210300 | 80458000 | 0 | 0 | 2176200 | 3577400 | 0 | 0 |
| TDEGIAYR | PRDX2 | P32119 | PRDX2_HUMAN | Unmodified | 0 | 0 | 0 | 0 | 0 | 54126 | 0 | 0 | 0 | 0 | 536200 | 0 | 595690 | 0 | 0 | 0 | 171030 | 234120 |
| TDEGIAYR | PRDX2 | P32119 | PRDX2_HUMAN | Unmodified | 0 | 0 | 0 | 0 | 150160 | 0 | 72598 | 0 | 0 | 586510 | 0 | 0 | 1328200 | 100340 | 0 | 0 | 272950 | 0 |
| TDEIGEICV | DIP2B | Q9P265 | DIP2B_HUMAN | Unmodified | 0 | 0 | 0 | 0 | 0 | 0 | 0 | 0 | 0 | 0 | 0 | 1931700 | 0 | 143390 | 0 | 0 | 0 | 0 |
| TDELIQK | ABCC4 | O15439 | MRP4_HUMAN | Unmodified | 0 | 278640 | 2145000 | 256360 | 0 | 0 | 73716 | 0 | 480850 | 0 | 0 | 284520 | 0 | 0 | 2556900 | 1498800 | 0 | 0 |
| TDELPQLV | FN1; FN1 | P02751; H0Y | FINC_HUMAN | Unmodified | 0 | 1261500 | 1182100 | 424330 | 0 | 881490 | 0 | 0 | 0 | 0 | 0 | 0 | 0 | 0 | 0 | 0 | 0 | 0 |
| TDEQALLS | LAMTOR1; LAM | Q6IAA8; H0Y | LTOR1_HUMAN | Unmodified | 0 | 0 | 834700 | 758690 | 0 | 0 | 0 | 0 | 3630200 | 0 | 1564800 | 3829000 | 0 | 0 | 0 | 631970 | 0 | 0 |
| TDFFIGGEE | NUDC; NUDC | Q9Y266; A0A | NUDC_HUMAN | Oxidation (M) | 0 | 0 | 0 | 0 | 0 | 0 | 0 | 0 | 0 | 0 | 0 | 1405500 | 0 | 0 | 0 | 0 | 0 | 0 |
| TDFIKIEEFL | CLIC2 | O15247 | CLIC2_HUMAN | Unmodified | 0 | 0 | 0 | 0 | 0 | 0 | 0 | 0 | 0 | 0 | 0 | 0 | 210930 | 587030 | 0 | 0 | 0 | 0 |
| TDGAAPNV | CNTN1 | Q12860 | CNTN1_HUMA | Unmodified | 157080 | 166930 | 0 | 0 | 0 | 0 | 0 | 0 | 0 | 0 | 0 | 0 | 0 | 0 | 0 | 0 | 0 | 0 |
| TDGALLVN | SERPINH1; SER | P50454; E9P | SERPH_HUMA | Oxidation (M) | 0 | 0 | 0 | 0 | 0 | 0 | 0 | 0 | 812890 | 0 | 0 | 0 | 0 | 0 | 0 | 0 | 0 | 0 |
| TDGALLVN | SERPINH1; SER | P50454; E9P | SERPH_HUMA | Unmodified | 0 | 0 | 0 | 0 | 0 | 0 | 0 | 0 | 1446600 | 0 | 0 | 0 | 0 | 0 | 0 | 0 | 0 | 0 |
| TDGDYDYL | RAB27B; RAB27 | O00194; K7E | RB27B_HUMAN | Acetyl (Protein N | 0 | 0 | 0 | 455440 | 732460 | 0 | 887090 | 1125300 | 0 | 0 | 0 | 0 | 0 | 1732200 | 255830 | 157320 | 0 | 555850 |
| TDGDYDYL | RAB27B; RAB27 | O00194; K7E | RB27B_HUMAN | Unmodified | 0 | 0 | 0 | 0 | 0 | 0 | 0 | 302570 | 0 | 0 | 0 | 0 | 0 | 0 | 0 | 0 | 0 | 0 |
| TDGEPGPC | DCTPP1 | Q9H773 | DCTP1_HUMA | Unmodified | 0 | 0 | 0 | 0 | 0 | 0 | 0 | 0 | 0 | 0 | 0 | 2982000 | 0 | 0 | 0 | 0 | 0 | 0 |
| TDGGDALS | C1QL1 | O75973 | C1QRF_HUMA | Unmodified | 0 | 0 | 0 | 0 | 0 | 0 | 0 | 0 | 10834000 | 417730 | 0 | 0 | 0 | 0 | 0 | 0 | 0 | 0 |
| TDGKGDF | AGRN; AGRN; A | O00468; A0A | AGRIN_HUMAN | Unmodified | 0 | 0 | 1590800 | 78357 | 69218 | 410360 | 200890 | 202180 | 0 | 0 | 0 | 0 | 0 | 0 | 981300 | 133380 | 0 | 187470 |
| TDGKVFQF | RPL24; RPL24; | P83731; C9J | RL24_HUMAN | Unmodified | 0 | 0 | 0 | 0 | 0 | 23424 | 306370 | 0 | 816000 | 0 | 0 | 0 | 563580 | 198270 | 0 | 0 | 24495 | 40195 |
| TDIARTEYL | DYNC1H1; DYN | Q14204; A0A | DYHC1_HUMA | Unmodified | 0 | 0 | 0 | 0 | 0 | 0 | 0 | 0 | 791660 | 114470 | 205530 | 3604300 | 0 | 0 | 293470 | 196160 | 0 | 0 |
| TDIEDLFK | PLCB4; PLCB4 | Q15147; A0A | PLCB4_HUMAN | Unmodified | 0 | 0 | 492020 | 0 | 0 | 0 | 0 | 0 | 0 | 0 | 0 | 0 | 0 | 0 | 488810 | 603490 | 0 | 0 |
| TDIGDVQIL | PIPOX | Q9P0Z9 | SOX_HUMAN P | Unmodified | 0 | 0 | 0 | 0 | 0 | 0 | 0 | 0 | 4001200 | 0 | 83221 | 0 | 0 | 0 | 0 | 0 | 0 | 0 |
| TDIQIALPS | DUT; DUT; DUT | P33316; H0Y | DUT_HUMAN D | Unmodified | 0 | 0 | 0 | 0 | 0 | 0 | 0 | 0 | 0 | 0 | 0 | 2285100 | 0 | 0 | 0 | 0 | 0 | 0 |
| TDITYPAGF | RPS4X | P62701 | RS4X_HUMAN | Oxidation (M) | 506050 | 425880 | 106710 | 212850 | 0 | 0 | 0 | 0 | 0 | 0 | 252720 | 3526400 | 54827 | 0 | 537330 | 429560 | 0 | 0 |
| TDKDTEITC | EPCAM; EPCAM | P16422; B5N | EPCAM_HUMA | Unmodified | 4735600 | 0 | 0 | 0 | 0 | 0 | 0 | 0 | 1390100 | 1114600 | 964670 | 7754700 | 0 | 0 | 7590600 | 5440300 | 0 | 0 |
| TDKDTEITC | EPCAM; EPCAM | P16422; B5N | EPCAM_HUMA | Unmodified | 2587300 | 0 | 881880 | 1105500 | 0 | 0 | 0 | 0 | 1268500 | 0 | 521030 | 0 | 441580 | 199770 | 3810000 | 1735700 | 0 | 0 |
| TDKQPLQK | LARS1; LARS1; | Q9P2J5; A0A | SYLC_HUMAN | Unmodified | 0 | 0 | 0 | 0 | 0 | 0 | 303740 | 202830 | 0 | 0 | 0 | 1136500 | 0 | 0 | 0 | 0 | 0 | 0 |
| TDKTLVLLN | PFN1; PFN1 | P07737; K7E | PROF1_HUMAN | Unmodified | 0 | 0 | 0 | 0 | 1383300 | 0 | 530280 | 202450 | 0 | 0 | 0 | 0 | 279530 | 472810 | 0 | 0 | 1923800 | 0 |

| | | | | | | | | | | | | | | | | | | | | | | |
|---|---|---|---|---|---|---|---|---|---|---|---|---|---|---|---|---|---|---|---|---|---|---|
| TDKTLVLLM | PFN1; PFN1 | P07737; K7E | PROF1_HUMAN | Unmodified | 0 | 0 | 0 | 0 | 395210 | 0 | 0 | 0 | 0 | 0 | 0 | 0 | 1265900 | 0 | 0 | 0 | 636920 | 0 |
| TDLDGLVQ | PROM1 | O43490 | PROM1_HUMA | Unmodified | 0 | 0 | 0 | 0 | 0 | 0 | 0 | 0 | 0 | 0 | 0 | 5562500 | 0 | 0 | 0 | 0 | 0 | 0 |
| TDLEKDIISI | ANXA2P2; ANX/ | A6NMY6; P0 | AXA2L_HUMAN | Unmodified | 38869 | 0 | 0 | 0 | 1861300 | 207530 | 182320 | 30176 | 822550 | 304990 | 0 | 446680 | 0 | 0 | 0 | 0 | 128580 | 666750 |
| TDLEKDIISI | ANXA2P2; ANX/ | A6NMY6; P0 | AXA2L_HUMAN | Unmodified | 261000 | 0 | 35850 | 58127 | 80058000 | 12751000 | 1167200 | 267010 | 3464900 | 4570300 | 0 | 291940 | 0 | 0 | 58962 | 0 | 16042000 | 13830000 |
| TDLGDSPL | SORL1; SORL1; | Q92673; E9F | SORL_HUMAN | Unmodified | 0 | 0 | 0 | 0 | 0 | 0 | 0 | 0 | 1914100 | 147790 | 0 | 0 | 139160 | 0 | 0 | 0 | 0 | 0 |
| TDLGHILDI | NISCH; NISCH | Q9Y2I1; C9J | NISCH_HUMAN | Unmodified | 0 | 0 | 0 | 0 | 0 | 10630 | 0 | 0 | 0 | 0 | 0 | 606860 | 0 | 0 | 0 | 0 | 0 | 0 |
| TDLIELLAR | FRYL; FRYL; FR | O94915; F22 | FRYL_HUMAN I | Unmodified | 0 | 46781 | 0 | 0 | 0 | 0 | 0 | 0 | 0 | 0 | 0 | 309910 | 0 | 0 | 0 | 0 | 0 | 0 |
| TDLKGDDL | TOM1L2; TOM1 | Q6ZVM7; F5I | TM1L2_HUMAN | Unmodified | 3514200 | 3144500 | 1050100 | 1196800 | 0 | 0 | 0 | 0 | 0 | 0 | 0 | 0 | 77109 | 81879 | 3391500 | 3582600 | 0 | 0 |
| TDLLLEPYN | MYH9 | P35579 | MYH9_HUMAN | Unmodified | 0 | 0 | 1239600 | 385610 | 0 | 0 | 0 | 0 | 0 | 0 | 369920 | 8374500 | 0 | 0 | 1650100 | 945150 | 0 | 0 |
| TDLNPDNL | CKB; CKB; CKB | P12277; G3\ | KCRB_HUMAN | Unmodified | 663050 | 1065100 | 5472100 | 2751800 | 1348900 | 382820 | 4216100 | 2133200 | 2833800 | 680160 | 1420400 | 26643000 | 1710300 | 921170 | 12686000 | 8492400 | 0 | 0 |
| TDLNPDNL | CKB; CKB; CKB | P12277; G3\ | KCRB_HUMAN | Unmodified | 0 | 0 | 0 | 0 | 0 | 0 | 3088000 | 1184100 | 0 | 0 | 0 | 0 | 0 | 0 | 0 | 0 | 0 | 0 |
| TDLNRTFPI | GRTP1; GRTP1 | Q5TC63; A0/ | GRTP1_HUMAN | Unmodified | 0 | 0 | 156160 | 246350 | 62073 | 0 | 105250 | 0 | 95435 | 0 | 0 | 0 | 0 | 0 | 375600 | 182830 | 82902 | 0 |
| TDLSNVQE | POGLUT1 | Q8NBL1 | PGLT1_HUMAN | Unmodified | 0 | 0 | 0 | 0 | 0 | 0 | 0 | 0 | 841290 | 360690 | 70578 | 44485 | 0 | 0 | 0 | 0 | 0 | 0 |
| TDLTGAFRI | DDC; DDC; DD( | P20711; A0A | DDC_HUMAN / | Unmodified | 201680 | 256510 | 0 | 0 | 0 | 113350 | 0 | 0 | 2136800 | 356570 | 30239 | 1107800 | 0 | 0 | 0 | 277870 | 0 | 0 |
| TDMDQIITS | NIBAN2 | Q96TA1 | NIBA2_HUMAN | Unmodified | 0 | 0 | 0 | 0 | 0 | 0 | 139450 | 0 | 0 | 0 | 0 | 1132100 | 294350 | 0 | 0 | 0 | 0 | 0 |
| TDNAGDQI | HNRNPL | P14866 | HNRPL_HUMAI | Unmodified | 629900 | 386220 | 0 | 0 | 0 | 0 | 0 | 0 | 873420 | 0 | 759660 | 13626000 | 0 | 0 | 0 | 0 | 0 | 0 |
| TDPHLCDF | FTL | P02792 | FRIL_HUMAN F | Unmodified | 0 | 0 | 0 | 0 | 0 | 0 | 0 | 0 | 12988000 | 334640 | 292900 | 0 | 0 | 1069600 | 0 | 456950 | 0 | 0 |
| TDPPKTHM | HLA-A; HLA-A; I | P04439; A0A | HLAA_HUMAN | Unmodified | 0 | 0 | 0 | 0 | 0 | 0 | 0 | 0 | 1288100 | 384890 | 0 | 0 | 0 | 0 | 0 | 0 | 0 | 770840 |
| TDPRPDAA | TPP2; TPP2; TPI | P29144; A0A | TPP2_HUMAN | Unmodified | 0 | 0 | 0 | 0 | 0 | 0 | 0 | 181230 | 1016300 | 53340 | 436400 | 7355300 | 0 | 0 | 85758 | 170480 | 275050 | 708550 |
| TDPSILGGI | ATP5PO | P48047 | ATPO_HUMAN | Oxidation (M) | 0 | 0 | 0 | 938460 | 0 | 0 | 0 | 0 | 1322400 | 109480 | 0 | 0 | 0 | 0 | 222290 | 0 | 0 | 0 |
| TDPSILGGI | ATP5PO | P48047 | ATPO_HUMAN | Unmodified | 0 | 0 | 75849 | 0 | 1802400 | 614670 | 0 | 52316 | 6984800 | 205780 | 7321000 | 623970 | 1017400 | 1127900 | 0 | 206950 | 1140600 | 317840 |
| TDPTTLTDE | MTHFD1; MTHF | P11586; V9G | C1TC_HUMAN | Unmodified | 0 | 0 | 0 | 0 | 0 | 0 | 0 | 0 | 0 | 0 | 320310 | 4130800 | 0 | 0 | 0 | 239020 | 0 | 0 |
| TDPTTLTDE | MTHFD1; MTHF | P11586; V9G | C1TC_HUMAN | Unmodified | 0 | 0 | 0 | 0 | 0 | 0 | 0 | 0 | 3349300 | 0 | 0 | 6273600 | 0 | 0 | 0 | 0 | 0 | 0 |
| TDQEVLGE | GRTP1 | Q5TC63 | GRTP1_HUMAN | Unmodified | 0 | 0 | 0 | 516240 | 0 | 0 | 0 | 0 | 136800 | 0 | 0 | 0 | 0 | 189720 | 1105400 | 948290 | 0 | 0 |
| TDQVIQSLI | UBE2L5; UBE2L | A0A1B0GUS | UB2L5_HUMAN | Unmodified | 0 | 0 | 41692 | 0 | 0 | 0 | 0 | 0 | 0 | 0 | 303170 | 699360 | 0 | 0 | 2062900 | 192670 | 0 | 0 |
| TDQVIQSLI | UBE2L3 | P68036 | UB2L3_HUMAN | Unmodified | 0 | 315440 | 0 | 306000 | 291110 | 0 | 0 | 0 | 0 | 0 | 0 | 0 | 0 | 0 | 3832200 | 0 | 0 | 0 |
| TDRYTIHSC | SF3B5 | Q9BWJ5 | SF3B5_HUMAN | Acetyl (Protein N | 0 | 0 | 0 | 0 | 818500 | 109690 | 0 | 153940 | 0 | 189560 | 191750 | 1382500 | 2301800 | 741300 | 0 | 0 | 264720 | 321350 |
| TDSAATRIP | ZPR1; ZPR1; ZP | O75312; H7( | ZPR1_HUMAN | Unmodified | 0 | 0 | 0 | 0 | 0 | 0 | 0 | 0 | 165270 | 0 | 435940 | 1295100 | 0 | 0 | 0 | 54607 | 0 | 0 |
| TDSDIIAK | SNRPA; SNRPA | P09012; M0I | SNRPA_HUMAI | Unmodified | 0 | 0 | 43164 | 0 | 0 | 0 | 0 | 0 | 0 | 0 | 0 | 1629900 | 0 | 0 | 0 | 0 | 0 | 0 |
| TDSDIISK | SNRPB2 | P08579 | RU2B_HUMAN | Unmodified | 355790 | 219940 | 0 | 0 | 0 | 0 | 0 | 0 | 0 | 0 | 0 | 532390 | 0 | 23939 | 93714 | 0 | 0 | 0 |
| TDTCMSSN | ITGB3; N/A | P05106; H3E | ITB3_HUMAN I | Unmodified | 0 | 0 | 0 | 0 | 0 | 0 | 0 | 0 | 0 | 0 | 0 | 0 | 0 | 0 | 0 | 0 | 0 | 0 |
| TDTESELDI | MTHFD1; MTHF | P11586; V9G | C1TC_HUMAN | Unmodified | 0 | 0 | 0 | 0 | 0 | 0 | 0 | 0 | 0 | 408380 | 1133400 | 1736900 | 0 | 0 | 0 | 141490 | 0 | 0 |
| TDTGEPMG | HSP90AA1 | P07900 | HS90A_HUMAI | Oxidation (M) | 0 | 0 | 281150 | 0 | 0 | 0 | 0 | 0 | 289900 | 0 | 192790 | 4876600 | 0 | 0 | 178640 | 293670 | 0 | 0 |
| TDTGEPMG | HSP90AA1 | P07900 | HS90A_HUMAI | Unmodified | 979670 | 1639300 | 2777700 | 767650 | 181590 | 0 | 41074 | 0 | 9556700 | 55296 | 397560 | 47237000 | 0 | 0 | 2112800 | 2289400 | 19472 | 156380 |
| TDTGEPMG | HSP90AA1 | P07900 | HS90A_HUMAI | Unmodified | 33037 | 0 | 0 | 0 | 170700 | 118780 | 0 | 0 | 604970 | 155650 | 624300 | 851180 | 0 | 0 | 0 | 62126 | 80416 | 0 |
| TDTGVSLQ | GDF15 | Q99988 | GDF15_HUMAI | Unmodified | 56814 | 0 | 862530 | 436700 | 0 | 0 | 0 | 0 | 6092400 | 0 | 0 | 0 | 234050 | 0 | 2381700 | 345360 | 0 | 0 |
| TDTLEDLFF | XRCC5 | P13010 | XRCC5_HUMAI | Unmodified | 0 | 0 | 492070 | 0 | 0 | 0 | 0 | 0 | 0 | 0 | 1965200 | 0 | 0 | 0 | 0 | 0 | 0 | 0 |
| TDTLEDLFF | XRCC5 | P13010 | XRCC5_HUMAI | Unmodified | 481810 | 0 | 0 | 0 | 0 | 0 | 0 | 0 | 0 | 197820 | 1030200 | 8720200 | 0 | 0 | 0 | 0 | 0 | 0 |
| TDTSDKKL( | YWHAQ; YWHA | P27348; E9P | 1433T_HUMAN | Unmodified | 0 | 0 | 0 | 83380 | 638300 | 154390 | 116900 | 81949 | 0 | 179500 | 0 | 0 | 456710 | 298980 | 0 | 0 | 507670 | 473910 |
| TDTVLILCR | CACYBP | Q9HB71 | CYBP_HUMAN | Unmodified | 0 | 0 | 0 | 0 | 0 | 0 | 31752 | 0 | 374860 | 0 | 0 | 3498300 | 0 | 0 | 0 | 0 | 0 | 0 |
| TDTVLILCR | CACYBP | Q9HB71 | CYBP_HUMAN | Unmodified | 0 | 0 | 0 | 0 | 0 | 0 | 0 | 0 | 0 | 0 | 0 | 5106700 | 0 | 0 | 0 | 38727 | 0 | 0 |
| TDVFQSLR | CYFIP2; CYFIP2 | Q96F07; H7( | CYFP2_HUMAN | Unmodified | 0 | 0 | 0 | 0 | 881030 | 0 | 0 | 0 | 697800 | 0 | 0 | 838990 | 0 | 0 | 0 | 0 | 0 | 0 |
| TDVHSPGN | MME; MME; MM | P08473; A0A | NEP_HUMAN N | Unmodified | 0 | 298230 | 9301000 | 5689800 | 0 | 0 | 0 | 0 | 1239400 | 91683 | 0 | 172260 | 0 | 0 | 7806900 | 70751 | 0 | 0 |
| TDVHSPGN | MME; MME; MM | P08473; A0A | NEP_HUMAN N | Unmodified | 0 | 0 | 822920 | 864080 | 0 | 0 | 0 | 0 | 0 | 1301800 | 0 | 0 | 11101000 | 4670300 | 0 | 0 | 0 | 62279 |
| TDVPLSSAI | POMGNT1 | Q8WZA1 | PMGT1_HUMAI | Unmodified | 0 | 0 | 0 | 0 | 0 | 0 | 0 | 40795 | 2259700 | 0 | 0 | 0 | 0 | 0 | 0 | 0 | 0 | 0 |
| TDWLDGK | PPIF; PPIE; PPIE | P30405; Q9U | PPIF_HUMAN F | Unmodified | 93599 | 0 | 0 | 0 | 210670 | 0 | 91401 | 0 | 6230500 | 783650 | 0 | 218430 | 0 | 0 | 804600 | 0 | 0 | 182860 |
| TDYMVGSY | ARHGDIA; ARH( | P52565; J3K | GDIR1_HUMAN | Oxidation (M) | 0 | 0 | 283640 | 0 | 0 | 0 | 0 | 0 | 0 | 0 | 0 | 531310 | 0 | 0 | 260560 | 204160 | 0 | 0 |
| TDYMVGSY | ARHGDIA; ARH( | P52565; J3K | GDIR1_HUMAN | Unmodified | 0 | 0 | 0 | 0 | 0 | 0 | 0 | 0 | 0 | 0 | 978590 | 1094800 | 154840 | 0 | 141680 | 0 | 0 | 0 |
| TDYMVGSY | ARHGDIA; ARH( | P52565; J3K | GDIR1_HUMAN | Oxidation (M) | 0 | 0 | 550620 | 0 | 0 | 0 | 0 | 0 | 0 | 211810 | 0 | 0 | 0 | 0 | 0 | 0 | 0 | 0 |
| TDYNASVS | HNRNPK; HNRI | P61978; Q5T | HNRPK_HUMA | Unmodified | 427010 | 363040 | 0 | 101190 | 0 | 0 | 0 | 0 | 244030 | 35298 | 897900 | 10675000 | 0 | 0 | 118320 | 238030 | 0 | 0 |
| TEAAETTPN | SLC35B2 | Q8TB61 | S35B2_HUMAN | Unmodified | 0 | 0 | 0 | 0 | 0 | 0 | 0 | 0 | 1487100 | 0 | 0 | 0 | 0 | 0 | 0 | 0 | 0 | 0 |
| TEAEIAHIAI | FLOT1; FLOT1; I | O75955; A0/ | FLOT1_HUMAN | Unmodified | 0 | 0 | 1515100 | 0 | 0 | 0 | 254720 | 0 | 4663500 | 1117700 | 1195800 | 5697700 | 0 | 0 | 3335200 | 1427600 | 0 | 0 |
| TEAPAAPA/ | BASP1 | P80723 | BASP1_HUMAN | Unmodified | 0 | 0 | 0 | 0 | 866350 | 0 | 48503 | 0 | 2056800 | 170950 | 0 | 0 | 0 | 0 | 0 | 0 | 0 | 0 |
| TEATQGLVI | AGRN; AGRN; A | O00468; A0/ | AGRIN_HUMAN | Unmodified | 0 | 0 | 529360 | 0 | 0 | 0 | 0 | 0 | 0 | 0 | 0 | 0 | 0 | 0 | 335290 | 470090 | 0 | 0 |
| TEAYLEAIRI | SCG3 | Q8WXD2 | SCG3_HUMAN | Unmodified | 0 | 0 | 0 | 0 | 0 | 0 | 0 | 0 | 465580 | 0 | 0 | 0 | 0 | 0 | 0 | 0 | 0 | 0 |
| TECALLGLI | ATP2B1; ATP2B | P20020; E7E | AT2B1_HUMAN | Unmodified | 0 | 0 | 0 | 0 | 0 | 0 | 0 | 0 | 0 | 0 | 0 | 93700 | 0 | 0 | 1046600 | 0 | 0 | 26447 |
| TECYGYAL( | PPP5C; PPP5C | P53041; H0Y | PPP5_HUMAN | Unmodified | 0 | 0 | 0 | 0 | 0 | 0 | 0 | 0 | 505130 | 0 | 0 | 1608100 | 0 | 0 | 0 | 0 | 0 | 0 |
| TEDERVTNI | NCAM2; NCAM | O15394; H9I | NCAM2_HUMA | Unmodified | 0 | 0 | 4354900 | 0 | 0 | 0 | 0 | 0 | 0 | 0 | 0 | 0 | 0 | 0 | 5347700 | 1290900 | 0 | 0 |
| TEDFFKLEF | FOLH1; NAALAI | Q04609; Q9 | FOLH1_HUMAI | Unmodified | 1099300 | 0 | 1617400 | 5361500 | 0 | 0 | 0 | 0 | 1255900 | 0 | 0 | 0 | 0 | 0 | 2761400 | 3238300 | 0 | 0 |
| TEDHEEAG | TMEM59; TMEM | Q9BXS4; Q5 | TMM59_HUMAI | Unmodified | 193290 | 71227 | 0 | 0 | 0 | 0 | 206620 | 0 | 265950 | 73724 | 0 | 0 | 0 | 0 | 213610 | 235600 | 0 | 0 |
| TEDIVAVQI | MME; MME; MM | P08473; A0A | NEP_HUMAN N | Unmodified | 143220 | 242570 | 10158000 | 5360300 | 0 | 103300 | 643650 | 119590 | 2729400 | 842840 | 0 | 358280 | 978250 | 0 | 17108000 | 13823000 | 151150 | 44780 |
| TEDLEATSE | VAMP8; VAMP8 | Q9BV40; C9 | VAMP8_HUMAI | Unmodified | 138120 | 96846 | 605310 | 239970 | 0 | 0 | 40126 | 0 | 448890 | 0 | 495180 | 1757000 | 0 | 0 | 312520 | 292550 | 0 | 0 |
| TEDPDLPAI | PRPF8; PRPF8 | Q6P2Q9; I3L | PRP8_HUMAN | Unmodified | 208330 | 364680 | 0 | 0 | 0 | 461570 | 0 | 0 | 0 | 0 | 0 | 369330 | 0 | 0 | 0 | 109480 | 0 | 0 |
| TEDVMFISI | ERBIN | Q96RT1 | ERBIN_HUMAN | Oxidation (M) | 0 | 0 | 0 | 701230 | 2454300 | 322090 | 0 | 0 | 0 | 0 | 0 | 1208400 | 0 | 0 | 271760 | 0 | 0 | 0 |
| TEEALQLYI | SRP72; SRP72 | O76094; D6I | SRP72_HUMAN | Unmodified | 185730 | 0 | 0 | 0 | 0 | 0 | 0 | 0 | 0 | 0 | 56493 | 5615400 | 0 | 0 | 0 | 0 | 0 | 0 |
| TEEARPSP/ | TBC1D10B | Q4KMP7 | TB10B_HUMAN | Unmodified | 83662 | 0 | 318190 | 0 | 0 | 0 | 0 | 0 | 0 | 0 | 0 | 719940 | 0 | 0 | 237090 | 237160 | 0 | 0 |
| TEEERTPNI | NCAM1; NCAM | P13591; A0A | NCAM1_HUMA | Unmodified | 0 | 0 | 0 | 0 | 0 | 0 | 0 | 0 | 226770 | 0 | 0 | 1329200 | 0 | 0 | 0 | 0 | 0 | 0 |
| TEEIHENVI | LRPAP1; LRPAP | P30533; A0A | AMRP_HUMAN | Unmodified | 24343 | 0 | 0 | 0 | 0 | 0 | 0 | 0 | 0 | 0 | 469550 | 0 | 0 | 0 | 0 | 0 | 0 | 0 |
| TEEIYGFLG | CHST1 | O43916 | CHST1_HUMAI | Unmodified | 0 | 0 | 0 | 0 | 0 | 0 | 0 | 0 | 2899500 | 0 | 613740 | 0 | 0 | 0 | 0 | 0 | 0 | 0 |
| TEEQMEEY | BTN2A2; BTN2/ | Q8WVV5; Q7 | BT2A2_HUMAN | Unmodified | 0 | 0 | 0 | 0 | 0 | 0 | 0 | 0 | 351550 | 0 | 0 | 883000 | 0 | 0 | 0 | 0 | 0 | 0 |
| TEETHPDD | SPRED2; SPREI | Q7Z698; C9J | SPRE2_HUMAN | Acetyl (Protein N | 0 | 0 | 0 | 0 | 943670 | 0 | 0 | 0 | 0 | 0 | 0 | 0 | 0 | 1001200 | 0 | 0 | 3860900 | 3191700 |
| TEEVSNLK | LARP1; LARP1; | Q6PKG0; H0 | LARP1_HUMAN | Unmodified | 0 | 0 | 847170 | 0 | 0 | 0 | 0 | 0 | 0 | 0 | 0 | 138010 | 124600 | 0 | 137670 | 102430 | 0 | 0 |
| TEFGMAIGI | SORT1; SORT1; | Q99523; A0/ | SORT_HUMAN | Unmodified | 0 | 183600 | 0 | 0 | 0 | 1544500 | 0 | 0 | 2512400 | 322370 | 0 | 879820 | 312500 | 0 | 0 | 0 | 0 | 0 |
| TEFGMAIGI | SORT1; SORT1; | Q99523; A0/ | SORT_HUMAN | Oxidation (M) | 0 | 0 | 0 | 0 | 0 | 0 | 0 | 0 | 989120 | 0 | 0 | 0 | 0 | 0 | 0 | 0 | 0 | 0 |
| TEFKVDSDI | BSG; BSG; BSG | P35613; A0A | BASI_HUMAN E | Oxidation (M) | 0 | 0 | 0 | 916410 | 0 | 4354900 | 0 | 0 | 0 | 0 | 0 | 0 | 2936300 | 0 | 0 | 0 | 1888600 | 7023800 |
| TEFLSFMN | S100A11 | P31949 | S10AB_HUMAN | Unmodified | 46653 | 104740 | 18294 | 184220 | 3167900 | 455350 | 0 | 0 | 1075800 | 521500 | 0 | 0 | 774150 | 421370 | 278480 | 0 | 208210 | 1368200 |
| TEFLSFMN | S100A11 | P31949 | S10AB_HUMAN | Oxidation (M) | 163460 | 119030 | 50001 | 402270 | 3890100 | 532000 | 48834 | 24804 | 899220 | 278660 | 43355 | 38186 | 603140 | 432470 | 429120 | 115360 | 290060 | 1132800 |
| TEFLSLLAK | MYO1E | Q12965 | MYO1E_HUMAI | Unmodified | 0 | 0 | 0 | 0 | 0 | 0 | 0 | 0 | 0 | 0 | 0 | 489990 | 0 | 0 | 0 | 0 | 0 | 0 |
| TEFTQGMIL | AHSA1; AHSA1; | O95433; H0 | AHSA1_HUMAI | 2 Oxidation (M) | 0 | 0 | 0 | 0 | 0 | 0 | 0 | 0 | 0 | 0 | 0 | 0 | 0 | 0 | 0 | 0 | 0 | 0 |
| TEGAAEAK/ | BASP1 | P80723 | BASP1_HUMAN | Unmodified | 0 | 0 | 0 | 0 | 5321300 | 1249200 | 0 | 161960 | 5746200 | 851600 | 0 | 0 | 0 | 0 | 0 | 0 | 424760 | 1991200 |
| TEGDEEAEI | XRCC6 | P12956 | XRCC6_HUMAI | Unmodified | 0 | 0 | 0 | 0 | 0 | 0 | 0 | 0 | 0 | 0 | 0 | 4014200 | 0 | 0 | 0 | 0 | 0 | 0 |
| TEGLSVLS( | EIF2S1; EIF2S1; | P05198; H0 | IF2A_HUMAN E | Unmodified | 16492 | 12291 | 0 | 0 | 12923 | 0 | 0 | 0 | 55569 | 0 | 0 | 118220 | 0 | 0 | 19268 | 0 | 0 | 0 |
| TEGQELITV | IFNGR1; IFNGR | P15260; A0A | INGR1_HUMAN | Unmodified | 0 | 0 | 0 | 0 | 0 | 0 | 0 | 0 | 555330 | 0 | 0 | 0 | 0 | 0 | 0 | 0 | 0 | 0 |
| TEHAEFLH( | DNM2 | P50570 | DYN2_HUMAN | Unmodified | 0 | 180810 | 63140 | 0 | 0 | 0 | 0 | 0 | 0 | 0 | 0 | 0 | 49328 | 25353 | 75034 | 0 | 0 | 0 |
| TEHLGLNW | PSME4; PSME4 | Q14997; F8V | PSME4_HUMAI | Unmodified | 0 | 0 | 0 | 0 | 0 | 0 | 0 | 0 | 0 | 0 | 0 | 636730 | 0 | 0 | 0 | 0 | 0 | 0 |
| TEHQVPSS | COBLL1; COBL | Q53SF7; A0/ | COBL1_HUMAI | Unmodified | 783970 | 1012900 | 412270 | 0 | 0 | 0 | 0 | 0 | 0 | 0 | 0 | 0 | 478330 | 0 | 442420 | 0 | 0 | 0 |
| TEHQVPSS | COBLL1; COBL | Q53SF7; A0/ | COBL1_HUMAI | Oxidation (M) | 738650 | 1791500 | 0 | 338110 | 0 | 302360 | 0 | 0 | 0 | 0 | 0 | 0 | 0 | 0 | 406610 | 480520 | 0 | 0 |
| TEIDEVVLV | HSPA13 | P48723 | HSP13_HUMAN | Unmodified | 0 | 0 | 0 | 0 | 0 | 0 | 0 | 0 | 3429000 | 0 | 0 | 0 | 0 | 0 | 0 | 0 | 0 | 0 |
| TEIDKPSQI | FN1; FN1 | P02751; H0 | FINC_HUMAN I | Oxidation (M) | 0 | 0 | 0 | 0 | 0 | 0 | 0 | 0 | 0 | 0 | 0 | 0 | 0 | 0 | 0 | 1049800 | 0 | 0 |
| TEIIILATR | RPS3; RPS3; RP | P23396; H0 | RS3_HUMAN 4 | Unmodified | 0 | 0 | 0 | 0 | 0 | 0 | 0 | 0 | 0 | 305500 | 763300 | 1050600 | 0 | 0 | 0 | 704400 | 404780 | 0 |
| TEIIPDRDG | PCYT2; PCYT2; | Q99447; I3L | PCY2_HUMAN | Unmodified | 0 | 0 | 0 | 0 | 0 | 0 | 0 | 0 | 0 | 0 | 0 | 1526400 | 0 | 0 | 0 | 0 | 0 | 0 |
| TEILPPFFK | SF3B1 | O75533 | SF3B1_HUMAN | Unmodified | 828450 | 325030 | 628920 | 0 | 0 | 0 | 0 | 0 | 128070 | 0 | 0 | 11265000 | 0 | 0 | 0 | 0 | 0 | 0 |
| TEISDKITSI | CKAP5 | Q14008 | CKAP5_HUMAN | Unmodified | 157570 | 0 | 0 | 209080 | 47456 | 51364 | 0 | 0 | 486900 | 61966 | 188430 | 829000 | 270820 | 215200 | 254270 | 299350 | 39973 | 219710 |
| TEISFRPNI | ATP1B1; ATP1B | P05026; V9G | AT1B1_HUMAN | Unmodified | 214630 | 0 | 3545500 | 858890 | 212600 | 492320 | 0 | 0 | 0 | 286460 | 962590 | 7051600 | 468650 | 167090 | 2699800 | 3162900 | 29196 | 0 |
| TEISVESK | TWF2; TWF2; T\ | Q6IBS0; D6F | TWF2_HUMAN | Unmodified | 53992 | 44749 | 72032 | 68811 | 1140000 | 0 | 0 | 50734 | 0 | 73212 | 0 | 0 | 60969 | 63419 | 21708 | 74480 | 612180 | 1359800 |
| TEKQLEAID | ACTN4; ACTN4 | O43707; F5C | ACTN4_HUMAI | Unmodified | 0 | 0 | 0 | 0 | 1005600 | 511090 | 0 | 0 | 0 | 312110 | 0 | 0 | 0 | 0 | 0 | 0 | 843560 | 0 |
| TEKYDPMP | PLCG2; PLCG2 | P16885; A0A | PLCG2_HUMAI | Unmodified | 0 | 0 | 0 | 0 | 0 | 0 | 0 | 0 | 0 | 0 | 613510 | 2518800 | 0 | 0 | 0 | 0 | 0 | 0 |
| TELAEPIAIF | EPRS1 | P07814 | SYEP_HUMAN | Oxidation (M) | 0 | 0 | 375800 | 0 | 0 | 0 | 0 | 111050 | 0 | 0 | 1267300 | 3994100 | 0 | 0 | 346340 | 388870 | 0 | 0 |
| TELDAHLEI | SH3GLB1; SH3( | Q9Y371; A0/ | SHLB1_HUMAN | Unmodified | 0 | 0 | 0 | 88978 | 0 | 0 | 365360 | 0 | 534900 | 0 | 111610 | 0 | 368380 | 0 | 266950 | 123670 | 0 | 0 |
| TELEDTLDS | MYH9 | P35579 | MYH9_HUMAN | Unmodified | 0 | 0 | 451560 | 0 | 0 | 0 | 0 | 0 | 213510 | 0 | 0 | 0 | 0 | 0 | 336090 | 160320 | 0 | 0 |
| TELISVSEVI | STAT1; STAT1; S | P42224; A0A | STAT1_HUMAN | Unmodified | 0 | 0 | 0 | 0 | 0 | 0 | 0 | 0 | 0 | 0 | 0 | 0 | 0 | 0 | 0 | 114050 | 0 | 0 |
| TELLMLQDI | N/A | N/A | N/A | Oxidation (M) | 0 | 0 | 0 | 0 | 16585000 | 418070 | 11156000 | 911600 | 0 | 42825000 | 0 | 0 | 3211600 | 0 | 0 | 0 | 2607800 | 10958000 |
| TELPQFVSY | TTLL12 | Q14166 | TTL12_HUMAN | Unmodified | 0 | 0 | 0 | 0 | 0 | 0 | 0 | 0 | 0 | 0 | 101200 | 5123200 | 0 | 0 | 0 | 0 | 0 | 0 |
| TELQGLIG( | BAG5 | Q9UL15 | BAG5_HUMAN | Unmodified | 118900 | 157830 | 271410 | 322790 | 121560 | 33149 | 0 | 0 | 1293200 | 247700 | 74941 | 229450 | 0 | 0 | 1013500 | 344750 | 0 | 56652 |
| TELSLLLQN | RNF31; RNF31; | Q96EP0; H0 | RNF31_HUMAN | Unmodified | 1281600 | 0 | 0 | 0 | 0 | 0 | 0 | 0 | 0 | 0 | 0 | 0 | 0 | 0 | 0 | 0 | 0 | 0 |
| TELTPEER | UTRN; UTRN | P46939; Q5T | UTRN_HUMAN | Unmodified | 0 | 0 | 713500 | 680500 | 92143 | 0 | 0 | 76729 | 821520 | 212270 | 547650 | 0 | 0 | 0 | 663740 | 926330 | 0 | 0 |
| TEMEDLMS | MYH9 | P35579 | MYH9_HUMAN | Unmodified | 60611 | 0 | 125450 | 143260 | 585590 | 0 | 134450 | 0 | 866660 | 0 | 296450 | 2643700 | 0 | 0 | 229620 | 218270 | 0 | 120360 |
| TEMEDLMS | MYH9 | P35579 | MYH9_HUMAN | 2 Oxidation (M) | 0 | 0 | 0 | 0 | 0 | 0 | 0 | 0 | 0 | 87364 | 0 | 474030 | 0 | 0 | 48407 | 65726 | 0 | 0 |
| TEMEDLMS | MYH9 | P35579 | MYH9_HUMAN | Oxidation (M) | 0 | 30277 | 42723 | 39937 | 0 | 0 | 0 | 0 | 0 | 0 | 445180 | 425530 | 0 | 0 | 123910 | 69006 | 0 | 0 |

| | | | | | | | | | | | | | | | | | | | | | | |
|---|---|---|---|---|---|---|---|---|---|---|---|---|---|---|---|---|---|---|---|---|---|---|
| TEMKQERE | ITGAV | P06756 | ITAV_HUMAN I | Unmodified | 382470 | 0 | 0 | 408150 | 426310 | 0 | 0 | 0 | 1780300 | 0 | 0 | 1117600 | 399280 | 0 | 0 | 0 | 159160 | 155610 |
| TEMQDNTY | DHX15 | O43143 | DHX15_HUMA | Oxidation (M) | 0 | 0 | 0 | 0 | 0 | 0 | 0 | 0 | 0 | 0 | 0 | 724770 | 0 | 0 | 0 | 0 | 0 | 0 |
| TEMVTISDA | EPB41L2; EPB4 | O43491; Q6 | E41L2_HUMAN | Unmodified | 0 | 0 | 0 | 0 | 391720 | 0 | 0 | 0 | 1095500 | 241550 | 0 | 0 | 0 | 0 | 0 | 0 | 0 | 0 |
| TEMVTISDA | EPB41L2; EPB4 | O43491; Q6 | E41L2_HUMAN | Oxidation (M) | 0 | 0 | 0 | 239110 | 268000 | 256760 | 0 | 43217 | 0 | 0 | 0 | 0 | 0 | 0 | 341260 | 231800 | 0 | 291280 |
| TEMVTISDA | EPB41L2; EPB4 | O43491; Q6 | E41L2_HUMAN | Unmodified | 0 | 0 | 0 | 0 | 4833000 | 415070 | 0 | 0 | 0 | 0 | 0 | 0 | 0 | 0 | 354050 | 133800 | 0 | 0 |
| TENLLGSYI | PSME1 | Q06323 | PSME1_HUMA | Unmodified | 0 | 0 | 0 | 0 | 866100 | 0 | 0 | 0 | 1495100 | 470690 | 1937000 | 4000700 | 74502 | 0 | 0 | 0 | 0 | 460690 |
| TENNDHIN | SUMO2 | P61956 | SUMO2_HUMA | Unmodified | 59068 | 0 | 239460 | 0 | 0 | 0 | 0 | 0 | 0 | 0 | 366860 | 943120 | 0 | 0 | 251650 | 169300 | 0 | 0 |
| TENPLILIDI | LONP1; LONP1 | P36776; K7E | LONM_HUMAN | Unmodified | 0 | 17193 | 0 | 34277 | 0 | 0 | 0 | 0 | 2049700 | 0 | 52375 | 78961 | 119870 | 23774 | 25801 | 0 | 0 | 46482 |
| TENYLGGE | AMPD3; AMPD3 | Q01432; E9F | AMPD3_HUMA | Unmodified | 0 | 0 | 0 | 0 | 0 | 0 | 0 | 0 | 0 | 0 | 0 | 2318500 | 0 | 0 | 0 | 0 | 0 | 0 |
| TEPFTQGK | AKAP12 | Q02952 | AKA12_HUMAN | Unmodified | 0 | 0 | 5502000 | 0 | 123960 | 0 | 0 | 0 | 0 | 0 | 0 | 142240 | 0 | 0 | 0 | 0 | 496510 | 143340 |
| TEPGFHW | DNASE1L1 | P49184 | DNSL1_HUMA | Unmodified | 1463400 | 1538500 | 1175500 | 767960 | 0 | 0 | 0 | 0 | 866540 | 0 | 103730 | 1463000 | 0 | 213480 | 1378600 | 992170 | 0 | 0 |
| TEPSIAEYT | DNAJC3; DNAJ | Q13217; X6F | DNJC3_HUMA | Unmodified | 0 | 0 | 0 | 0 | 0 | 106490 | 0 | 66215 | 88556 | 555300 | 0 | 0 | 0 | 0 | 35421 | 0 | 0 | 0 |
| TEPTAQQN | EIF3CL; EIF3C | B5ME19; Q9 | EIFCL_HUMAN | Unmodified | 0 | 0 | 0 | 0 | 0 | 0 | 0 | 0 | 0 | 0 | 0 | 5539300 | 0 | 0 | 0 | 0 | 0 | 0 |
| TEQAISFAK | SLC25A6 | P12236 | ADT3_HUMAN | Acetyl (Protein N | 0 | 362740 | 377740 | 0 | 1853200 | 179030 | 1326700 | 422970 | 15163000 | 3137300 | 1155700 | 2208800 | 0 | 0 | 0 | 249150 | 0 | 0 |
| TEQDLALG | STXBP3 | O00186 | STXB3_HUMA | Unmodified | 269240 | 0 | 883820 | 0 | 0 | 0 | 0 | 0 | 340900 | 0 | 0 | 2963600 | 211590 | 112980 | 926480 | 711010 | 0 | 238660 |
| TEQGGAHI | XRCC5 | P13010 | XRCC5_HUMA | Unmodified | 0 | 0 | 0 | 0 | 0 | 0 | 0 | 0 | 529890 | 0 | 0 | 1675800 | 0 | 0 | 0 | 762290 | 0 | 0 |
| TEQMTLR | RACK1; RACK1; | P63244; D6F | RACK1_HUMA | Acetyl (Protein N | 0 | 0 | 0 | 0 | 0 | 0 | 0 | 0 | 0 | 0 | 0 | 539430 | 0 | 54752 | 0 | 0 | 0 | 0 |
| TEQMTLR | RACK1; RACK1; | P63244; D6F | RACK1_HUMA | Acetyl (Protein N | 499980 | 14111 | 825590 | 267440 | 0 | 0 | 0 | 0 | 353720 | 0 | 0 | 5031700 | 367230 | 0 | 241460 | 515050 | 0 | 0 |
| TEQMTLRG | RACK1; RACK1; | P63244; D6F | RACK1_HUMA | Acetyl (Protein N | 0 | 1075200 | 21428000 | 7034500 | 3811600 | 717180 | 3249000 | 1710200 | 0 | 1045600 | 31005000 | 0 | 4127700 | 9445900 | 4112400 | 0 | 1574400 | 15890000 |
| TEQMTLRG | RACK1; RACK1; | P63244; D6F | RACK1_HUMA | Acetyl (Protein N | 0 | 0 | 0 | 0 | 969040 | 0 | 1256400 | 886640 | 113910 | 307780 | 976040 | 0 | 2273700 | 0 | 0 | 0 | 558920 | 1275600 |
| TEREQFVEI | CALU; CALU | O43852; H0 | CALU_HUMAN | Unmodified | 0 | 0 | 0 | 0 | 250980 | 0 | 0 | 0 | 6560900 | 389480 | 0 | 1111100 | 0 | 0 | 279990 | 130650 | 0 | 178270 |
| TERFDCHH | FHL2; FHL2; FH | Q14192; F8V | FHL2_HUMAN | Acetyl (Protein N | 0 | 0 | 0 | 0 | 238430 | 0 | 0 | 0 | 0 | 0 | 0 | 0 | 0 | 0 | 0 | 0 | 594800 | 248020 |
| TERFDCHH | FHL2; FHL2; FH | Q14192; F8V | FHL2_HUMAN | Acetyl (Protein N | 0 | 0 | 0 | 0 | 0 | 0 | 0 | 0 | 0 | 0 | 178000 | 0 | 0 | 0 | 0 | 0 | 510700 | 614180 |
| TERSDLEY | CIT; CIT; CIT; CI | O14578; A0 | CTRO_HUMAN | Unmodified | 0 | 34142 | 0 | 0 | 7577600 | 0 | 644250 | 0 | 718550 | 0 | 0 | 730500 | 350200 | 0 | 0 | 0 | 0 | 0 |
| TESAVSQM | IARS1; IARS1; I | P41252; A0A | SYIC_HUMAN I | Unmodified | 0 | 0 | 0 | 0 | 0 | 0 | 0 | 0 | 140760 | 0 | 0 | 1470400 | 0 | 0 | 0 | 0 | 0 | 0 |
| TESFDVVTI | CLDND1; CLDN | Q9NY35; D6 | CLDN1_HUMA | Unmodified | 412320 | 0 | 639380 | 0 | 2217500 | 0 | 243050 | 55422 | 3838600 | 1349000 | 2449400 | 4521900 | 189820 | 260950 | 679390 | 440770 | 0 | 0 |
| TESGEETDI | TGOLN2; TGOL | O43493; A0 | TGON2_HUMA | Unmodified | 0 | 0 | 0 | 0 | 0 | 0 | 0 | 0 | 880710 | 112650 | 128540 | 0 | 435660 | 0 | 0 | 0 | 0 | 0 |
| TESNQEVA | EIF4E; EIF4E; EI | P06730; D6F | IF4E_HUMAN E | Unmodified | 0 | 0 | 0 | 0 | 0 | 0 | 0 | 0 | 431820 | 0 | 387660 | 5710500 | 0 | 36204 | 0 | 0 | 48771 | 43777 |
| TESPRTSG | MAP2K7 | O14733 | MP2K7_HUMA | Unmodified | 0 | 0 | 0 | 0 | 300200 | 0 | 0 | 0 | 0 | 202750 | 0 | 0 | 206640 | 592700 | 0 | 0 | 286080 | 0 |
| TESTLNALL | NPTX2 | P47972 | NPTX2_HUMA | Unmodified | 0 | 0 | 0 | 0 | 0 | 0 | 0 | 0 | 4981400 | 0 | 0 | 0 | 0 | 0 | 0 | 0 | 0 | 0 |
| TESTPITAVI | COPG1 | Q9Y678 | COPG1_HUMA | Unmodified | 0 | 0 | 0 | 0 | 0 | 0 | 0 | 0 | 0 | 0 | 215700 | 637480 | 0 | 0 | 0 | 0 | 0 | 0 |
| TETAAHSLI | CEP55; CEP55 | Q53EZ4; H0 | CEP55_HUMA | Unmodified | 0 | 0 | 0 | 0 | 900840 | 0 | 0 | 0 | 0 | 0 | 0 | 0 | 0 | 0 | 0 | 0 | 0 | 0 |
| TETDSVGTI | RACGAP1; RAC | Q9H0H5; H0 | RGAP1_HUMA | Unmodified | 0 | 0 | 235400 | 305700 | 0 | 0 | 0 | 0 | 357610 | 0 | 258730 | 992640 | 0 | 0 | 113510 | 0 | 0 | 0 |
| TETGQELE | GLG1; GLG1 | Q92896; H3 | GSLG1_HUMA | Unmodified | 0 | 0 | 0 | 0 | 0 | 0 | 0 | 0 | 2757300 | 0 | 0 | 0 | 0 | 0 | 0 | 0 | 0 | 0 |
| TETQEKNP | TMSB4X | P62328 | TYB4_HUMAN | Unmodified | 0 | 0 | 0 | 0 | 2122800 | 539070 | 332130 | 500120 | 0 | 4634800 | 0 | 0 | 112580 | 0 | 0 | 0 | 2644200 | 8340000 |
| TETSASPH | HCK; HCK; HCK | P08631; J3K | HCK_HUMAN T | Unmodified | 0 | 0 | 0 | 0 | 0 | 0 | 0 | 0 | 0 | 0 | 0 | 5069700 | 0 | 0 | 0 | 0 | 0 | 0 |
| TETVEEPMI | HSP90B1; HSP | P14625; A0A | ENPL_HUMAN | Oxidation (M) | 0 | 0 | 0 | 0 | 0 | 0 | 0 | 26266 | 3827200 | 1095600 | 653950 | 0 | 0 | 0 | 0 | 0 | 0 | 0 |
| TETVEEPMI | HSP90B1; HSP | P14625; A0A | ENPL_HUMAN | Unmodified | 0 | 0 | 0 | 0 | 0 | 0 | 95774 | 112020 | 11076000 | 2679900 | 1962000 | 239510 | 0 | 0 | 0 | 0 | 0 | 0 |
| TETVEEPMI | HSP90B1; HSP | P14625; A0A | ENPL_HUMAN | Oxidation (M) | 0 | 0 | 0 | 0 | 0 | 0 | 0 | 0 | 1079400 | 248360 | 419820 | 0 | 0 | 0 | 0 | 0 | 0 | 0 |
| TETVEEPMI | HSP90B1; HSP | P14625; A0A | ENPL_HUMAN | Unmodified | 0 | 0 | 174050 | 0 | 0 | 0 | 0 | 0 | 6355400 | 1008000 | 2060500 | 707400 | 0 | 0 | 368060 | 116460 | 0 | 0 |
| TETVEEPMI | HSP90B1; HSP | P14625; A0A | ENPL_HUMAN | Oxidation (M) | 0 | 0 | 601620 | 0 | 0 | 0 | 0 | 0 | 43475000 | 793400 | 12431000 | 14734000 | 0 | 0 | 1862500 | 421830 | 0 | 0 |
| TETVEEPMI | HSP90B1; HSP | P14625; A0A | ENPL_HUMAN | Unmodified | 0 | 0 | 433650 | 0 | 0 | 0 | 0 | 0 | 1.56E+08 | 1439900 | 18420000 | 20188000 | 0 | 0 | 0 | 173410 | 0 | 0 |
| TEVEQQRD | NIBAN1 | Q9BZQ8 | NIBA1_HUMA | Unmodified | 435710 | 89357 | 585960 | 446210 | 351690 | 0 | 0 | 0 | 9252500 | 848970 | 0 | 0 | 0 | 0 | 215700 | 239140 | 0 | 0 |
| TEVLSEDLL | ARHGAP17; AR | Q68EM7; C9 | RHG17_HUMA | Unmodified | 0 | 12001 | 0 | 0 | 0 | 0 | 0 | 0 | 104240 | 0 | 76197 | 304990 | 0 | 0 | 0 | 0 | 0 | 0 |
| TEVNGFDP | SMC2 | O95347 | SMC2_HUMAN | Unmodified | 0 | 0 | 0 | 0 | 0 | 0 | 0 | 0 | 0 | 0 | 0 | 140960 | 0 | 0 | 0 | 0 | 0 | 0 |
| TEVNSGFF | CCT6A; CCT6B | P40227; Q9 | TCPZ_HUMAN | Unmodified | 369050 | 0 | 1226700 | 1039600 | 4366000 | 682640 | 1132800 | 748770 | 888450 | 552130 | 3490900 | 24885000 | 8149300 | 2912300 | 769680 | 1084100 | 1061200 | 3041600 |
| TEVNYTQL | GPX4; GPX4; GI | P36969; A0A | GPX4_HUMAN | Unmodified | 3035900 | 3584300 | 1250900 | 1213300 | 103860 | 419460 | 0 | 0 | 1478300 | 26572 | 189790 | 1770500 | 0 | 0 | 4554400 | 0 | 0 | 0 |
| TEVVDSDG | ACAD9; ACAD9 | Q9H845; A0 | ACAD9_HUMA | Unmodified | 0 | 0 | 0 | 0 | 123720 | 0 | 0 | 22639 | 2245700 | 374580 | 0 | 77862 | 0 | 0 | 0 | 0 | 0 | 0 |
| TEWETAAP | RPS5; RPS5; RP | P46782; M0 | RS5_HUMAN 4 | Acetyl (Protein N | 1466300 | 1685100 | 553760 | 740410 | 1233400 | 0 | 748330 | 519720 | 683550 | 326440 | 2376000 | 10651000 | 209900 | 730940 | 615000 | 0 | 0 | 414750 |
| TEYLSNADI | DYNC1H1; DYN | Q14204; A0 | DYHC1_HUMA | Unmodified | 0 | 0 | 0 | 0 | 0 | 0 | 0 | 0 | 0 | 0 | 0 | 1621500 | 0 | 0 | 0 | 0 | 0 | 0 |
| TEYLSNADI | DYNC1H1; DYN | Q14204; A0 | DYHC1_HUMA | Unmodified | 0 | 0 | 81101 | 138870 | 0 | 1607900 | 0 | 0 | 378070 | 66784 | 160250 | 1445700 | 0 | 0 | 0 | 0 | 0 | 0 |
| TFAADHNY | ERBIN; ERBIN | Q96RT1; B4 | ERBIN_HUMA | Unmodified | 0 | 0 | 582330 | 1876400 | 10312000 | 1687000 | 0 | 554130 | 3399600 | 192980 | 957770 | 4538100 | 4533600 | 1412200 | 1177200 | 864890 | 908160 | 591330 |
| TFAILER | B3GLCT | Q6Y288 | B3GLT_HUMA | Unmodified | 0 | 0 | 0 | 0 | 0 | 89093 | 35127 | 34402 | 1497200 | 79263 | 0 | 0 | 0 | 0 | 0 | 0 | 0 | 0 |
| TFAPEEISA | HSPA5; HSPA5 | P11021; A0A | BIP_HUMAN Er | Unmodified | 0 | 0 | 0 | 0 | 5308000 | 110420 | 11840000 | 2247500 | 3.3E+08 | 54484000 | 4890900 | 848360 | 538100 | 2978700 | 1393800 | 0 | 979920 | 5524800 |
| TFAPEEISA | HSPA5; HSPA5 | P11021; A0A | BIP_HUMAN Er | Oxidation (M) | 0 | 0 | 5720100 | 0 | 4525000 | 215420 | 42051000 | 26260000 | 1.04E+08 | 1.77E+08 | 28988000 | 7378100 | 4825600 | 7733300 | 3478400 | 0 | 0 | 5862700 |
| TFAPSFQQ | CDCP1 | Q9H5V8 | CDCP1_HUMA | Unmodified | 0 | 0 | 0 | 923890 | 0 | 0 | 0 | 0 | 0 | 0 | 0 | 11597000 | 0 | 0 | 1452000 | 2065700 | 971580 | 0 |
| TFAPSFQQ | CDCP1 | Q9H5V8 | CDCP1_HUMA | Unmodified | 0 | 0 | 0 | 0 | 0 | 0 | 0 | 0 | 0 | 0 | 0 | 0 | 0 | 0 | 0 | 0 | 1363600 | 0 |
| TFAQGLAV | GPC4 | O75487 | GPC4_HUMAN | Unmodified | 950750 | 0 | 0 | 0 | 0 | 0 | 0 | 0 | 630660 | 466420 | 0 | 0 | 0 | 0 | 0 | 0 | 0 | 0 |
| TFAVDETS | UPF1 | Q92900 | RENT1_HUMA | Unmodified | 4885700 | 5005000 | 2830600 | 4397100 | 1742300 | 0 | 2947500 | 905580 | 0 | 1044200 | 796020 | 4064000 | 5184000 | 0 | 3664900 | 3358100 | 984040 | 880760 |
| TFAVSSKVI | CHMP2B; CHM | Q9UQN3; A0 | CHM2B_HUMA | Unmodified | 0 | 0 | 0 | 0 | 600840 | 0 | 94907 | 25585 | 0 | 247120 | 0 | 0 | 264770 | 290200 | 0 | 0 | 161910 | 194350 |
| TFCGTPDY | PRKCB; PRKCA | P05771; P17 | KPCB_HUMAN | Unmodified | 0 | 0 | 0 | 0 | 0 | 0 | 3019200 | 0 | 0 | 0 | 0 | 1426900 | 0 | 1338500 | 889220 | 0 | 0 | 0 |
| TFCGTPEYI | AKT3; AKT2; AK | Q9Y243; P31 | AKT3_HUMAN | Unmodified | 0 | 0 | 235770 | 0 | 0 | 0 | 0 | 0 | 0 | 0 | 0 | 2099400 | 0 | 0 | 1789900 | 0 | 0 | 0 |
| TFCHWEHI | NRP1; NRP1; N | O14786; E9F | NRP1_HUMAN | Unmodified | 0 | 0 | 0 | 0 | 1310600 | 151270 | 0 | 0 | 0 | 0 | 0 | 0 | 0 | 71470 | 0 | 484690 | 292160 | 3042100 |
| TFCQLILDF | EEF2 | P13639 | EF2_HUMAN E | Unmodified | 593250 | 96587 | 566480 | 814030 | 4288200 | 910860 | 1866100 | 28644 | 1897200 | 1648700 | 2744200 | 11040000 | 2030000 | 229210 | 1690800 | 1021300 | 1179800 | 3071500 |
| TFDAQIVIIE | GSPT1 | P15170 | ERF3A_HUMA | Unmodified | 0 | 0 | 1009500 | 0 | 0 | 0 | 0 | 0 | 844640 | 0 | 910680 | 6083300 | 0 | 0 | 651250 | 621900 | 0 | 0 |
| TFDEFQELI | PIK3C2A | O00443 | P3C2A_HUMA | Unmodified | 0 | 15050 | 0 | 0 | 0 | 0 | 0 | 0 | 0 | 0 | 0 | 2629500 | 0 | 0 | 0 | 0 | 0 | 0 |
| TFDEIASGF | SLC2A1 | P11166 | GTR1_HUMAN | Unmodified | 5302500 | 6552100 | 14417000 | 10337000 | 259550 | 0 | 857440 | 1232000 | 1367800 | 1709900 | 1043500 | 36783000 | 32654 | 0 | 13747000 | 11483000 | 3290600 | 0 |
| TFDEIASGF | SLC2A1 | P11166 | GTR1_HUMAN | Unmodified | 0 | 0 | 0 | 6253100 | 15920000 | 8890800 | 0 | 0 | 4477800 | 0 | 0 | 0 | 0 | 0 | 0 | 0 | 6073300 | 0 |
| TFDQLTPDI | RCN1 | Q15293 | RCN1_HUMAN | Unmodified | 0 | 0 | 0 | 0 | 288400 | 0 | 120930 | 0 | 18014000 | 2441100 | 2508100 | 1138600 | 349600 | 0 | 0 | 0 | 0 | 0 |
| TFDQLTPDI | RCN1 | Q15293 | RCN1_HUMAN | Unmodified | 0 | 0 | 0 | 0 | 74666 | 0 | 130460 | 21023 | 0 | 583160 | 0 | 0 | 46449 | 168990 | 0 | 0 | 0 | 245120 |
| TFDQLTPEI | CALU | O43852 | CALU_HUMAN | Unmodified | 0 | 0 | 0 | 0 | 744250 | 0 | 0 | 0 | 15621000 | 1986700 | 2767100 | 1602400 | 0 | 0 | 0 | 0 | 0 | 980520 |
| TFDQLTPEI | CALU | O43852 | CALU_HUMAN | Unmodified | 0 | 0 | 0 | 0 | 383620 | 0 | 1121300 | 630580 | 0 | 2119300 | 0 | 521350 | 331430 | 460680 | 0 | 0 | 62449 | 488380 |
| TFDSIVMDI | PDIA4; PDIA4 | P13667; A0A | PDIA4_HUMAN | Oxidation (M) | 0 | 0 | 0 | 0 | 0 | 0 | 0 | 0 | 895280 | 0 | 0 | 0 | 0 | 0 | 0 | 0 | 0 | 0 |
| TFDSIVMDI | PDIA4; PDIA4 | P13667; A0A | PDIA4_HUMAN | Unmodified | 0 | 0 | 0 | 0 | 0 | 0 | 0 | 0 | 1857700 | 154280 | 187120 | 0 | 0 | 0 | 0 | 0 | 0 | 0 |
| TFDSIVMDI | PDIA4; PDIA4 | P13667; A0A | PDIA4_HUMAN | Unmodified | 0 | 0 | 236690 | 164290 | 0 | 0 | 358800 | 110790 | 14640000 | 2202800 | 2503300 | 920120 | 198240 | 208580 | 397670 | 0 | 0 | 151490 |
| TFDSIVMDI | PDIA4; PDIA4 | P13667; A0A | PDIA4_HUMAN | Oxidation (M) | 0 | 0 | 69832 | 30356 | 81358 | 0 | 54165 | 140320 | 4737100 | 1211400 | 1381600 | 418330 | 90913 | 48115 | 228700 | 148190 | 0 | 114760 |
| TFDVQIVIIE | GSPT2 | Q8IYD1 | ERF3B_HUMA | Unmodified | 3824000 | 8302900 | 495960 | 5041600 | 0 | 0 | 2414300 | 0 | 0 | 0 | 0 | 0 | 13279000 | 4627000 | 2074100 | 1336100 | 0 | 0 |
| TFDWAEFR | MPRIP; MPRIP; | Q6WCQ1; J3 | MPRIP_HUMA | Unmodified | 0 | 0 | 0 | 133390 | 97699 | 0 | 482380 | 52228 | 0 | 0 | 0 | 762690 | 205110 | 0 | 919220 | 71505 | 98746 | 137950 |
| TFDYKIPYT | SSRP1; SSRP1 | Q08945; E9F | SSRP1_HUMA | Unmodified | 0 | 0 | 0 | 0 | 0 | 0 | 0 | 0 | 0 | 0 | 0 | 3594500 | 0 | 0 | 0 | 0 | 0 | 0 |
| TFEDISEK | GSS; GSS; GSS | P48637; A0A | GSHB_HUMAN | Unmodified | 0 | 0 | 143470 | 346760 | 0 | 0 | 0 | 0 | 800190 | 255570 | 562880 | 2295500 | 255350 | 0 | 0 | 421200 | 0 | 0 |
| TFEEFQELI | PIK3C2B; PIK3C | O00750; F5C | P3C2B_HUMA | Unmodified | 307670 | 0 | 0 | 263030 | 0 | 0 | 0 | 0 | 0 | 0 | 0 | 0 | 0 | 0 | 558980 | 840520 | 0 | 0 |
| TFEGVDPQ | GDI1 | P31150 | GDIA_HUMAN | Oxidation (M) | 0 | 0 | 0 | 0 | 0 | 0 | 0 | 0 | 0 | 0 | 0 | 899740 | 0 | 0 | 0 | 0 | 0 | 0 |
| TFEHVTSEI | PSMD7; PSMD7 | P51665; H3E | PSMD7_HUMA | Unmodified | 155570 | 3008600 | 0 | 1360400 | 577680 | 0 | 0 | 0 | 720500 | 407280 | 1648100 | 4366900 | 0 | 0 | 645400 | 0 | 0 | 0 |
| TFEINPR | HSP90B2P; HSI | Q58FF3; P14 | ENPLL_HUMA | Unmodified | 292140 | 1006700 | 1570100 | 628530 | 546360 | 412700 | 1405300 | 1025300 | 2.38E+08 | 27864000 | 37086000 | 25689000 | 1526300 | 2021300 | 0 | 833430 | 0 | 947770 |
| TFEINPRHI | HSP90B2P; HSI | Q58FF3; P14 | ENPLL_HUMA | Unmodified | 0 | 0 | 0 | 0 | 0 | 0 | 298730 | 0 | 890330 | 8807000 | 0 | 0 | 0 | 0 | 0 | 0 | 0 | 0 |
| TFEINPRHI | HSP90B2P; HSI | Q58FF3; P14 | ENPLL_HUMA | Unmodified | 0 | 0 | 0 | 0 | 0 | 0 | 1079300 | 0 | 0 | 7768000 | 0 | 0 | 298850 | 0 | 0 | 0 | 0 | 0 |
| TFEISASDK | SWAP70; SWAF | Q9UH65; E7 | SWP70_HUMA | Unmodified | 0 | 421810 | 0 | 0 | 928020 | 0 | 64888 | 0 | 104830 | 0 | 338300 | 0 | 0 | 0 | 0 | 0 | 0 | 211250 |
| TFEMSDFIV | FLNB; FLNB | O75369; E7E | FLNB_HUMAN | Oxidation (M) | 0 | 0 | 0 | 0 | 0 | 0 | 0 | 0 | 0 | 0 | 0 | 1295000 | 0 | 0 | 0 | 0 | 0 | 0 |
| TFEMSHQS | TBC1D8; TBC1 | O95759; J3K | TBCD8_HUMA | Unmodified | 0 | 0 | 0 | 0 | 50330 | 0 | 0 | 0 | 0 | 0 | 0 | 0 | 0 | 0 | 0 | 0 | 41291 | 140760 |
| TFESLVDFS | HADH; HADH; | Q16836; A0 | HCDH_HUMAN | Unmodified | 0 | 0 | 0 | 0 | 0 | 0 | 3109500 | 1068800 | 8468100 | 4241600 | 0 | 1187800 | 738030 | 0 | 0 | 0 | 0 | 0 |
| TFEVCDLP | COPB2; COPB2 | P35606; D6F | COPB2_HUMA | Unmodified | 0 | 0 | 0 | 0 | 0 | 0 | 0 | 0 | 0 | 0 | 0 | 0 | 0 | 0 | 0 | 0 | 0 | 0 |
| TFEVCDLP | COPB2; COPB2 | P35606; D6F | COPB2_HUMA | Unmodified | 0 | 0 | 105960 | 0 | 0 | 0 | 175370 | 0 | 0 | 442360 | 0 | 0 | 295030 | 0 | 0 | 0 | 210380 | 197650 |
| TFFKLHDVI | NUCB2; NUCB2 | P80303; Q2L | NUCB2_HUMA | Unmodified | 0 | 0 | 0 | 0 | 0 | 0 | 0 | 0 | 0 | 1243900 | 0 | 0 | 372540 | 1301400 | 0 | 0 | 0 | 98699 |
| TFFLEALR | HDHD2; HDHD | Q9H0R4; K7 | HDHD2_HUMA | Unmodified | 0 | 0 | 0 | 0 | 0 | 0 | 0 | 0 | 174100 | 0 | 0 | 1804400 | 0 | 0 | 0 | 44077 | 0 | 0 |
| TFFPEEISSI | HSPA2 | P54652 | HSP72_HUMA | Oxidation (M) | 183750 | 649910 | 0 | 0 | 951780 | 230650 | 0 | 0 | 0 | 0 | 0 | 470110 | 0 | 1246600 | 0 | 0 | 0 | 0 |
| TFFPEEISSI | HSPA2 | P54652 | HSP72_HUMA | Unmodified | 0 | 0 | 0 | 0 | 356200 | 0 | 0 | 0 | 0 | 0 | 0 | 0 | 0 | 0 | 0 | 0 | 64257 | 0 |
| TFFPEIFQTI | SHCBP1 | Q8NEM2 | SHCBP_HUMA | Unmodified | 0 | 0 | 0 | 0 | 115760 | 0 | 0 | 0 | 0 | 0 | 0 | 0 | 0 | 0 | 0 | 0 | 0 | 0 |
| TFFSFPAV | GARS1; GARS1 | P41250; A0A | GARS_HUMAN | Unmodified | 410820 | 6138.4 | 371540 | 531580 | 578870 | 136200 | 683160 | 0 | 7911100 | 731110 | 1400900 | 20549000 | 481220 | 0 | 1522700 | 1809800 | 216030 | 0 |
| TFGAPRPY | COBLL1; COBL | Q53SF7; A0 | COBL1_HUMA | Unmodified | 1181500 | 889310 | 0 | 0 | 0 | 0 | 353120 | 114080 | 106540 | 0 | 0 | 0 | 1868400 | 921700 | 0 | 0 | 276050 | 0 |
| TFGETHPF | LTA4H | P09960 | LKHA4_HUMA | Unmodified | 0 | 0 | 0 | 0 | 0 | 0 | 0 | 0 | 0 | 0 | 0 | 1281700 | 0 | 0 | 0 | 0 | 0 | 0 |
| TFGHNTMI | SLC12A2; SLC1 | P55011; G3 | S12A2_HUMA | Unmodified | 0 | 62659 | 0 | 847090 | 0 | 0 | 38332 | 0 | 0 | 140860 | 0 | 0 | 0 | 0 | 0 | 0 | 150130 | 1997000 |
| TFGHNTMI | SLC12A2; SLC1 | P55011; G3 | S12A2_HUMA | Oxidation (M) | 0 | 0 | 0 | 0 | 0 | 0 | 0 | 0 | 0 | 0 | 0 | 0 | 0 | 0 | 623080 | 460980 | 0 | 0 |
| TFGLTLGH | EHHADH | Q08426 | ECHP_HUMAN | Unmodified | 41082 | 0 | 0 | 0 | 0 | 0 | 0 | 0 | 1566100 | 0 | 0 | 0 | 0 | 0 | 0 | 0 | 0 | 0 |
| TFGTAIVINI | CUL4A; CUL4A | Q13619; A0 | CUL4A_HUMA | Unmodified | 0 | 0 | 0 | 0 | 0 | 0 | 0 | 0 | 0 | 0 | 0 | 1958400 | 0 | 0 | 0 | 0 | 0 | 0 |
| TFHDLEGN | ERMP1; ERMP1 | Q7Z2K6; E7E | ERMP1_HUMA | Unmodified | 33448 | 0 | 0 | 0 | 16376 | 0 | 185110 | 0 | 3379700 | 219700 | 131090 | 0 | 0 | 30793 | 0 | 0 | 0 | 0 |
| TFHETLDC | CD81; CD81; C | P60033; A6 | CD81_HUMAN | Unmodified | 1.79E+08 | 4015200 | 37735000 | 1.68E+08 | 42417000 | 10285000 | 3032400 | 0 | 23130000 | 8903400 | 9904800 | 1.05E+08 | 2.52E+08 | 1.54E+08 | 35372000 | 1.2E+08 | 5740100 | 42771000 |
| TFHFNTVEI | SERPINB1 | P30740 | ILEU_HUMAN L | Unmodified | 440350 | 0 | 0 | 0 | 0 | 0 | 0 | 0 | 539270 | 95648 | 0 | 0 | 0 | 0 | 0 | 0 | 0 | 0 |
| TFHFNTVEI | SERPINB1 | P30740 | ILEU_HUMAN L | Unmodified | 0 | 0 | 0 | 0 | 0 | 0 | 0 | 0 | 0 | 1741300 | 0 | 0 | 0 | 0 | 0 | 0 | 0 | 0 |
| TFHIFYYLL | MYH9 | P35579 | MYH9_HUMAN | Unmodified | 0 | 0 | 414400 | 1052500 | 0 | 0 | 0 | 0 | 445430 | 0 | 0 | 6380400 | 0 | 0 | 4836100 | 1111800 | 0 | 0 |

| | | | | | | | | | | | | | | | | | | | | | | |
|---|---|---|---|---|---|---|---|---|---|---|---|---|---|---|---|---|---|---|---|---|---|---|
| TFHRAASS | SCAMP2 | O15127 | SCAM2_HUMA | Unmodified | 0 | 0 | 0 | 0 | 1153000 | 365290 | 214230 | 0 | 0 | 0 | 0 | 0 | 167900 | 28007 | 0 | 0 | 0 | 347300 |
| TFHTIGFCF | ZFP36L1; ZFP36 | Q07352; P47 | TISB_HUMAN n | Unmodified | 1843100 | 299080 | 0 | 0 | 0 | 0 | 0 | 0 | 0 | 0 | 238230 | 0 | 0 | 0 | 0 | 0 | 0 | 0 |
| TFHTSSAAI | RABEPK; RABEF | Q7Z6M1; Q5 | RABEK_HUMAN | Unmodified | 0 | 0 | 0 | 0 | 0 | 0 | 0 | 0 | 0 | 0 | 0 | 649100 | 0 | 0 | 0 | 0 | 0 | 0 |
| TFHVETPEE | AKT1 | P31749 | AKT1_HUMAN I | Unmodified | 0 | 0 | 0 | 0 | 0 | 0 | 0 | 0 | 0 | 8147.1 | 109000 | 291240 | 0 | 0 | 0 | 0 | 0 | 0 |
| TFHVETPEE | AKT1 | P31749 | AKT1_HUMAN I | Unmodified | 269060 | 0 | 467000 | 2710500 | 0 | 0 | 0 | 0 | 282880 | 0 | 0 | 0 | 0 | 0 | 7849500 | 3895500 | 0 | 0 |
| TFIAIKPDG | NME1; NME2; N | P15531; P22 | NDKA_HUMAN | Unmodified | 2517400 | 2058800 | 10606000 | 20229000 | 10334000 | 3158300 | 310270 | 161470 | 13965000 | 1986600 | 77141000 | 3.13E+08 | 12113000 | 5761100 | 9882100 | 7828200 | 3688800 | 7694300 |
| TFIAIKPDG | NME1; NME2; N | P15531; P22 | NDKA_HUMAN | Unmodified | 0 | 0 | 0 | 205850 | 97849000 | 8552500 | 3883100 | 70551 | 0 | 6518300 | 3610000 | 249840 | 94754000 | 16935000 | 0 | 0 | 4023000 | 61938000 |
| TFIEINQIFT | SLC2A5 | P22732 | GTR5_HUMAN | Unmodified | 0 | 0 | 0 | 0 | 0 | 0 | 0 | 0 | 0 | 0 | 0 | 43464 | 0 | 0 | 0 | 2379800 | 0 | 0 |
| TFIFSDLDY | WARS1; WARS1 | P23381; G3V | SYWC_HUMAN | 2 Oxidation (M) | 0 | 0 | 0 | 48253 | 153840 | 42991 | 67128 | 0 | 0 | 409360 | 0 | 87562 | 59094 | 65150 | 179520 | 59877 | 151390 | 110540 |
| TFIFSEKPVI | SORL1 | Q92673 | SORL_HUMAN | Unmodified | 0 | 0 | 0 | 0 | 17651 | 0 | 0 | 0 | 5499500 | 1219700 | 295960 | 0 | 0 | 0 | 0 | 0 | 189110 | 0 |
| TFIGPGGN | GARS1; GARS1 | P41250; A0A | GARS_HUMAN | Oxidation (M) | 0 | 0 | 0 | 0 | 11160000 | 1938300 | 9247100 | 0 | 4662000 | 7283500 | 11847000 | 1882800 | 3813700 | 8229800 | 3081700 | 1945300 | 7845300 | 4111200 |
| TFIGPGGN | GARS1; GARS1 | P41250; A0A | GARS_HUMAN | Unmodified | 0 | 0 | 0 | 0 | 1118200 | 0 | 0 | 0 | 2162700 | 365970 | 0 | 0 | 309990 | 0 | 0 | 0 | 526100 | 0 |
| TFIIDYYFEV | CPNE3; CPNE3 | O75131; A0A | CPNE3_HUMA | Unmodified | 0 | 0 | 0 | 0 | 165780 | 0 | 192880 | 0 | 0 | 397570 | 0 | 0 | 164690 | 347440 | 69280 | 0 | 186780 | 982550 |
| TFITQQGIK | AP2M1; AP2M1 | Q96CW1; C9 | AP2M1_HUMA | Unmodified | 0 | 0 | 1198800 | 1650700 | 2814600 | 492870 | 719210 | 203310 | 2724300 | 788940 | 671740 | 5109100 | 720520 | 401880 | 977670 | 1364300 | 1255400 | 1794100 |
| TFITQQGIK | AP2M1 | A0A087WY7 | A0A087WY71_ | Unmodified | 0 | 0 | 0 | 0 | 0 | 0 | 0 | 0 | 0 | 355390 | 0 | 0 | 0 | 0 | 0 | 0 | 0 | 0 |
| TFIWDVK | CDCP1 | Q9H5V8 | CDCP1_HUMA | Unmodified | 139710 | 336710 | 3249100 | 486420 | 0 | 0 | 0 | 0 | 119070 | 0 | 345320 | 16385000 | 33684 | 0 | 3916100 | 4044000 | 368600 | 97637 |
| TFKVDFLTG | PCDHB8 | Q9UN66 | PCDB8_HUMA | Unmodified | 201070 | 144820 | 0 | 0 | 0 | 0 | 0 | 0 | 548730 | 57457 | 0 | 0 | 0 | 0 | 0 | 0 | 0 | 0 |
| TFLALINQV | MYO6; MYO6; N | Q9UM54; A0 | MYO6_HUMAN | Unmodified | 0 | 0 | 0 | 61708 | 0 | 0 | 0 | 0 | 0 | 12110 | 0 | 34951 | 0 | 0 | 162870 | 0 | 0 | 0 |
| TFLALINQV | MYO6; MYO6; N | Q9UM54; A0 | MYO6_HUMAN | Unmodified | 0 | 0 | 1989300 | 955650 | 44452 | 0 | 0 | 0 | 847330 | 19235 | 393140 | 1158900 | 27045 | 0 | 5151700 | 4846900 | 0 | 0 |
| TFLASVQW | MAN2B1; N/A | O00754; M0 | MA2B1_HUMA | Unmodified | 0 | 0 | 0 | 0 | 0 | 0 | 0 | 0 | 813630 | 0 | 103170 | 0 | 0 | 0 | 0 | 0 | 0 | 0 |
| TFLHAVDV | BTN2A1 | Q7KYR7 | BT2A1_HUMAN | Unmodified | 64495 | 0 | 0 | 0 | 198300 | 0 | 0 | 0 | 598940 | 0 | 882500 | 4159900 | 0 | 0 | 333110 | 138800 | 0 | 202520 |
| TFLLTTSSK | TAX1BP1; TAX1 | Q86VP1; B8 | TAXB1_HUMAN | Unmodified | 0 | 0 | 0 | 0 | 3492900 | 1989300 | 1078600 | 0 | 2129200 | 1246300 | 368020 | 662330 | 0 | 0 | 0 | 0 | 1252000 | 3160000 |
| TFLLTYVLQ | XPO4 | Q9C0E2 | XPO4_HUMAN | Unmodified | 0 | 0 | 0 | 0 | 0 | 0 | 853210 | 0 | 0 | 155070 | 0 | 172730 | 0 | 579450 | 0 | 0 | 530710 | 0 |
| TFLQREHQ | ITGA5 | P08648 | ITA5_HUMAN I | Unmodified | 0 | 0 | 0 | 0 | 4054300 | 686450 | 265030 | 114250 | 0 | 0 | 0 | 0 | 0 | 0 | 0 | 0 | 1758500 | 924480 |
| TFLQTVSSE | MRPL53 | Q96EL3 | RM53_HUMAN | Unmodified | 0 | 0 | 0 | 0 | 0 | 0 | 0 | 0 | 772490 | 0 | 0 | 0 | 0 | 0 | 0 | 0 | 0 | 0 |
| TFLRPSPED | TJP2; TJP2; TJP2 | Q9UDY2; A0 | ZO2_HUMAN T | Unmodified | 0 | 0 | 0 | 952940 | 6768800 | 1513500 | 0 | 0 | 6344200 | 6080600 | 797050 | 2564600 | 1795400 | 1186700 | 0 | 0 | 898370 | 161480 |
| TFLTVYWTI | ICAM1 | P05362 | ICAM1_HUMAN | Unmodified | 0 | 0 | 0 | 108920 | 15623000 | 689610 | 0 | 0 | 0 | 0 | 0 | 0 | 0 | 0 | 0 | 0 | 608280 | 6416000 |
| TFLVGER | EEF1G | P26641 | EF1G_HUMAN | Unmodified | 3586000 | 1073900 | 1842700 | 1287200 | 0 | 0 | 0 | 0 | 0 | 0 | 0 | 15910000 | 0 | 0 | 188310 | 203040 | 0 | 0 |
| TFLVWVNE | CKB; CKB | P12277; H0Y | KCRB_HUMAN | Unmodified | 1944800 | 1351000 | 1762600 | 4333700 | 184760 | 0 | 521200 | 0 | 6213400 | 391500 | 2297800 | 36009000 | 1136500 | 452930 | 8614500 | 8492900 | 0 | 0 |
| TFLVWVNE | CKB; CKB | P12277; H0Y | KCRB_HUMAN | Oxidation (M) | 0 | 0 | 160580 | 2503800 | 13401000 | 1525500 | 31941000 | 0 | 771970 | 6534400 | 1183100 | 360540 | 14295000 | 7908000 | 924720 | 273630 | 173510 | 0 |
| TFLVWVNE | CKB; CKB | P12277; H0Y | KCRB_HUMAN | Unmodified | 0 | 0 | 0 | 0 | 2266400 | 0 | 3735400 | 0 | 550010 | 0 | 0 | 0 | 1420300 | 737300 | 0 | 0 | 0 | 0 |
| TFNDYFR | MAN2A1 | Q16706 | MA2A1_HUMA | Unmodified | 0 | 338100 | 0 | 343480 | 0 | 0 | 0 | 0 | 0 | 0 | 0 | 1130000 | 341700 | 0 | 682080 | 274280 | 0 | 0 |
| TFNIKNDF | SKP1; SKP1 | P63208; E5R | SKP1_HUMAN | Unmodified | 1060400 | 0 | 0 | 0 | 4297300 | 0 | 239070 | 0 | 235190 | 0 | 0 | 13340000 | 1461800 | 0 | 3286300 | 1593800 | 0 | 0 |
| TFNIRHPEE | FERMT2; FERM | Q96AC1; G3 | FERM2_HUMA | Unmodified | 0 | 0 | 0 | 0 | 14706000 | 4090100 | 0 | 0 | 0 | 0 | 0 | 0 | 0 | 0 | 0 | 0 | 2100700 | 10464000 |
| TFNLPLLMI | HDAC2 | Q92769 | HDAC2_HUMA | Oxidation (M) | 60425 | 0 | 0 | 0 | 61816 | 15731 | 0 | 0 | 185660 | 116430 | 235290 | 1452100 | 54808 | 0 | 0 | 0 | 0 | 0 |
| TFNLRTELF | TTLL12 | Q14166 | TTL12_HUMAN | Unmodified | 0 | 0 | 465090 | 0 | 0 | 348240 | 79054 | 0 | 0 | 0 | 459410 | 53455 | 0 | 57271 | 0 | 0 | 0 | 245980 |
| TFNPGAGL | SNRPA1; SNRP | P09661; H0Y | RU2A_HUMAN | Unmodified | 900740 | 618460 | 436310 | 701230 | 1502000 | 0 | 0 | 0 | 0 | 105010 | 1270100 | 1444400 | 58631 | 0 | 314610 | 271500 | 0 | 0 |
| TFNQVEIKF | RPS15; RPS15; | P62841; K7E | RS15_HUMAN | Oxidation (M) | 0 | 0 | 0 | 5654700 | 0 | 268000 | 0 | 0 | 731030 | 0 | 0 | 14795000 | 3291500 | 0 | 0 | 0 | 221380 | 2251300 |
| TFNTSTGG | XRCC6; XRCC6 | P12956; B1A | XRCC6_HUMA | Unmodified | 79602 | 0 | 0 | 0 | 0 | 0 | 44742 | 106010 | 0 | 0 | 168220 | 1055800 | 373760 | 0 | 0 | 0 | 0 | 0 |
| TFNTSTGG | XRCC6; XRCC6 | P12956; B1A | XRCC6_HUMA | Unmodified | 0 | 0 | 0 | 0 | 0 | 0 | 0 | 166970 | 0 | 0 | 896550 | 1821900 | 629870 | 0 | 0 | 0 | 0 | 0 |
| TFPACDGS | CISD2; CISD2 | Q8N5K1; D6 | CISD2_HUMAN | Unmodified | 0 | 0 | 0 | 0 | 0 | 0 | 0 | 135060 | 2070300 | 0 | 0 | 176300 | 0 | 0 | 0 | 0 | 0 | 0 |
| TFPALSTR | NRP1; NRP1; N | O14786; E9F | NRP1_HUMAN | Unmodified | 267420 | 209290 | 564970 | 402550 | 981350 | 0 | 0 | 0 | 358480 | 0 | 0 | 0 | 580620 | 0 | 228260 | 1087000 | 236840 | 2014600 |
| TFPGFFSPI | FGA | P02671 | FIBA_HUMAN F | Oxidation (M) | 0 | 1175400 | 0 | 0 | 0 | 0 | 0 | 0 | 0 | 0 | 0 | 0 | 0 | 0 | 0 | 0 | 0 | 0 |
| TFPIRGFQI | CEMIP2 | Q9UHN6 | CEIP2_HUMAN | Unmodified | 0 | 0 | 0 | 0 | 0 | 0 | 0 | 0 | 1595100 | 0 | 0 | 0 | 0 | 0 | 0 | 0 | 0 | 0 |
| TFPLDVGSI | PREP; PREP | P48147; A0A | PPCE_HUMAN | Unmodified | 0 | 606660 | 0 | 0 | 0 | 0 | 0 | 0 | 595480 | 368900 | 757840 | 1664600 | 0 | 0 | 337000 | 0 | 0 | 0 |
| TFPNHYSIV | ENPP4 | Q9Y6X5 | ENPP4_HUMA | Unmodified | 0 | 0 | 0 | 0 | 0 | 0 | 0 | 0 | 2451800 | 0 | 0 | 0 | 0 | 928380 | 0 | 0 | 0 | 0 |
| TFPNHYSIV | ENPP4 | Q9Y6X5 | ENPP4_HUMA | Oxidation (M) | 0 | 0 | 0 | 0 | 0 | 0 | 0 | 0 | 0 | 3811500 | 0 | 0 | 0 | 0 | 0 | 0 | 0 | 0 |
| TFPNHYSIV | ENPP4 | Q9Y6X5 | ENPP4_HUMA | Oxidation (M) | 0 | 0 | 0 | 0 | 0 | 0 | 0 | 0 | 1235000 | 2997900 | 0 | 496430 | 0 | 0 | 0 | 0 | 0 | 0 |
| TFPNHYSIV | ENPP4 | Q9Y6X5 | ENPP4_HUMA | Unmodified | 0 | 0 | 0 | 0 | 0 | 0 | 0 | 0 | 8678600 | 934600 | 286140 | 0 | 0 | 0 | 0 | 0 | 0 | 0 |
| TFPNHYSIV | ENPP1; ENPP1 | P22413; E9P | ENPP1_HUMA | Unmodified | 606730 | 529740 | 417980 | 0 | 0 | 0 | 66756 | 52753 | 1009900 | 0 | 0 | 2176600 | 1910500 | 2066100 | 0 | 3450200 | 0 | 4859900 |
| TFPPDLNFI | CLUH; CLUH; C | O75153; I3L | CLU_HUMAN C | Unmodified | 0 | 0 | 0 | 0 | 0 | 0 | 0 | 0 | 0 | 0 | 50719 | 298340 | 0 | 0 | 0 | 0 | 0 | 0 |
| TFPVLLR | PRKDC | P78527 | PRKDC_HUMA | Unmodified | 0 | 67260 | 162510 | 0 | 0 | 0 | 0 | 0 | 0 | 0 | 0 | 653140 | 0 | 0 | 98430 | 60913 | 0 | 0 |
| TFQEDSSP | PTPN14 | Q15678 | PTN14_HUMAN | Unmodified | 0 | 0 | 0 | 0 | 2325000 | 359720 | 0 | 0 | 0 | 0 | 0 | 0 | 0 | 0 | 0 | 0 | 986700 | 1201800 |
| TFQFQPSK | SCARB1; SCAR | Q8WTV0; A0 | SCRB1_HUMA | Unmodified | 3194200 | 2162400 | 2434600 | 1430800 | 265120 | 0 | 0 | 0 | 897990 | 0 | 0 | 14916000 | 0 | 0 | 3161300 | 2847900 | 117060 | 0 |
| TFQGHTNE | TBL1Y; TBL1XR | Q9BQ87; Q9 | TBL1Y_HUMAN | Unmodified | 0 | 0 | 0 | 0 | 0 | 0 | 0 | 0 | 0 | 0 | 0 | 1418400 | 0 | 0 | 0 | 0 | 0 | 0 |
| TFQNTVELI | ERBIN; ERBIN | Q96RT1; B4 | ERBIN_HUMAN | Unmodified | 0 | 0 | 0 | 0 | 1573500 | 0 | 17812 | 0 | 0 | 83797 | 0 | 0 | 0 | 0 | 0 | 0 | 0 | 0 |
| TFQVVPEIE | PHACTR4 | Q8IZ21 | PHAR4_HUMA | Unmodified | 0 | 0 | 0 | 0 | 3301300 | 0 | 0 | 0 | 815580 | 239140 | 0 | 0 | 0 | 0 | 0 | 0 | 0 | 3985400 |
| TFREEQCE | ADAMTS1; ADA | Q9UHI8; E5F | ATS1_HUMAN | Unmodified | 0 | 0 | 1396000 | 0 | 0 | 0 | 0 | 0 | 0 | 156850 | 0 | 0 | 0 | 0 | 0 | 0 | 0 | 2108100 |
| TFRTERPVI | EIF3I; EIF3I; EIF | Q13347; A0 | EIF3I_HUMAN I | Oxidation (M) | 1347100 | 655390 | 1141700 | 0 | 846990 | 0 | 0 | 0 | 0 | 0 | 652430 | 6187200 | 0 | 360690 | 482130 | 1494900 | 405740 | 0 |
| TFRTERPVI | EIF3I; EIF3I; EIF | Q13347; A0 | EIF3I_HUMAN I | Unmodified | 375860 | 1565200 | 416730 | 651570 | 1968500 | 0 | 0 | 0 | 1148500 | 0 | 831920 | 2723000 | 5003300 | 1096000 | 0 | 250170 | 832630 | 706410 |
| TFSATVRAS | CARHSP1; CAR | Q9Y2V2; H3 | CHSP1_HUMA | Unmodified | 0 | 0 | 0 | 0 | 0 | 0 | 678960 | 747710 | 0 | 0 | 0 | 0 | 32298 | 0 | 0 | 0 | 0 | 1124600 |
| TFSATVRAS | CARHSP1; CAR | Q9Y2V2; H3 | CHSP1_HUMA | Unmodified | 0 | 0 | 0 | 0 | 0 | 0 | 1069100 | 0 | 0 | 0 | 0 | 0 | 0 | 0 | 0 | 0 | 894180 | 303060 |
| TFSAYQR | DAAM1 | Q9Y4D1 | DAAM1_HUMA | Unmodified | 0 | 0 | 0 | 0 | 0 | 0 | 0 | 0 | 0 | 0 | 0 | 0 | 0 | 0 | 0 | 0 | 0 | 0 |
| TFSEILNRV | DYNC1H1; DYN | Q14204; A0 | DYHC1_HUMA | Unmodified | 0 | 0 | 0 | 0 | 0 | 0 | 192830 | 0 | 555480 | 376190 | 0 | 2328000 | 0 | 0 | 0 | 0 | 0 | 0 |
| TFSESSVW | ABCC4; ABCC4 | O15439; A0 | MRP4_HUMAN | Unmodified | 75592 | 0 | 2401000 | 904390 | 0 | 0 | 47246 | 0 | 0 | 0 | 0 | 0 | 0 | 0 | 2270500 | 2853600 | 0 | 0 |
| TFSFAIPLIE | DDX21 | Q9NR30 | DDX21_HUMA | Unmodified | 0 | 0 | 0 | 0 | 0 | 0 | 444390 | 38526 | 0 | 102780 | 358080 | 1231900 | 332380 | 0 | 0 | 0 | 0 | 0 |
| TFSGQTHG | CMBL | Q96DG6 | CMBL_HUMAN | Unmodified | 162590 | 113840 | 750720 | 591320 | 388340 | 0 | 62468 | 0 | 0 | 0 | 57329 | 0 | 0 | 193360 | 690210 | 460030 | 280790 | 490650 |
| TFSHELSDI | PDIA3 | P30101 | PDIA3_HUMAN | Unmodified | 728380 | 1317500 | 994920 | 0 | 2751800 | 841660 | 1117200 | 0 | 2.63E+08 | 57197000 | 34981000 | 4996400 | 5096700 | 868230 | 5633700 | 1150100 | 4058200 | 2633300 |
| TFSHELSDI | PDIA3 | P30101 | PDIA3_HUMAN | Unmodified | 0 | 0 | 0 | 0 | 861530 | 0 | 34652 | 0 | 0 | 6410300 | 0 | 0 | 614250 | 306670 | 0 | 0 | 0 | 0 |
| TFSHELSDI | PDIA3 | P30101 | PDIA3_HUMAN | Unmodified | 0 | 0 | 0 | 0 | 7514300 | 0 | 14616000 | 0 | 0 | 2.13E+08 | 3763700 | 0 | 10796000 | 7534800 | 0 | 0 | 0 | 10929000 |
| TFSLTEVRG | KIF4A | O95239 | KIF4A_HUMAN | Unmodified | 0 | 0 | 0 | 0 | 932550 | 229180 | 234760 | 0 | 0 | 0 | 0 | 0 | 1284200 | 600450 | 0 | 0 | 587990 | 1060600 |
| TFSSIPVSR | DYNC1H1; DYN | Q14204; A0 | DYHC1_HUMA | Unmodified | 0 | 0 | 146040 | 0 | 452770 | 0 | 0 | 26038 | 1088200 | 1046200 | 663630 | 3128500 | 0 | 0 | 0 | 0 | 0 | 0 |
| TFSVSLQK | EXOC6; EXOC6 | Q8TAG9; E7I | EXOC6_HUMA | Unmodified | 0 | 0 | 0 | 0 | 1045000 | 0 | 63809 | 52581 | 213430 | 158870 | 0 | 784790 | 211950 | 0 | 0 | 0 | 0 | 0 |
| TFSVWYVP | FLNA; FLNA; FL | P21333; A0A | FLNA_HUMAN | Unmodified | 1394000 | 4177200 | 943780 | 1789400 | 0 | 0 | 0 | 33565 | 0 | 0 | 0 | 191420 | 0 | 353820 | 2640800 | 1998500 | 0 | 0 |
| TFSYAGFEI | CCT7 | Q99832 | TCPH_HUMAN | Unmodified | 0 | 0 | 0 | 0 | 0 | 0 | 0 | 98155 | 1118400 | 400400 | 0 | 2264500 | 692300 | 0 | 0 | 0 | 296500 | 0 |
| TFSYAGFEI | CCT7 | Q99832 | TCPH_HUMAN | Oxidation (M) | 189960 | 0 | 0 | 0 | 624180 | 0 | 556440 | 428980 | 642120 | 668000 | 1191300 | 4916000 | 1100800 | 789370 | 231850 | 229190 | 698090 | 539660 |
| TFTAWCNS | ACTN2; ACTN4 | P35609; O4 | ACTN2_HUMA | Unmodified | 0 | 0 | 166640 | 371180 | 1375200 | 369120 | 0 | 0 | 701000 | 147830 | 215420 | 1798400 | 610930 | 267830 | 293150 | 369840 | 1167000 | 594530 |
| TFTAWCNS | ACTN2; ACTN4 | P35609; O4 | ACTN2_HUMA | Unmodified | 0 | 0 | 0 | 0 | 14571000 | 209290 | 231310 | 191400 | 1373000 | 949450 | 773470 | 9461600 | 3872800 | 6142600 | 280730 | 0 | 6607600 | 4600000 |
| TFTDCFNC | PPP1CA; PPP1C | P62136; P62 | PP1A_HUMAN | Unmodified | 297160 | 15455 | 0 | 261050 | 699560 | 411050 | 803610 | 0 | 673040 | 0 | 106330 | 990220 | 0 | 401340 | 0 | 0 | 1201000 | 276280 |
| TFTEMDSH | ARCN1; ARCN1 | P48444; Q6 | COPD_HUMAN | Unmodified | 0 | 0 | 0 | 0 | 231920 | 0 | 0 | 0 | 442780 | 70824 | 596900 | 1428400 | 0 | 42663 | 0 | 0 | 0 | 0 |
| TFTINPER | FRYL; FRYL; FR | O94915; F2 | FRYL_HUMAN I | Unmodified | 0 | 0 | 0 | 0 | 616020 | 0 | 0 | 0 | 4215200 | 1027200 | 2618000 | 707190 | 0 | 0 | 0 | 0 | 474980 | 0 |
| TFTPCERLE | MYO10; MYO10 | Q9HD67; A0 | MYO10_HUMA | Unmodified | 0 | 0 | 0 | 0 | 189270 | 0 | 80103 | 32814 | 0 | 0 | 0 | 0 | 0 | 0 | 0 | 0 | 213900 | 0 |
| TFTTQETIT | GPI; GPI; GPI; G | P06744; K7E | G6PI_HUMAN | Unmodified | 0 | 0 | 0 | 0 | 16009000 | 2488000 | 4925300 | 0 | 0 | 0 | 0 | 0 | 0 | 0 | 0 | 0 | 0 | 93661 |
| TFTWLVGK | MYO1C; MYO1C | O00159; F5 | MYO1C_HUMA | Unmodified | 2494800 | 3241100 | 2988900 | 9104400 | 3053300 | 2517800 | 119330 | 167740 | 585070 | 0 | 0 | 0 | 0 | 0 | 16559000 | 8690000 | 1459300 | 324200 |
| TFVEESIYN | ALDH1B1 | P30837 | AL1B1_HUMAN | Unmodified | 0 | 0 | 0 | 0 | 0 | 0 | 0 | 0 | 350810 | 0 | 0 | 0 | 0 | 0 | 0 | 0 | 0 | 0 |
| TFVEYLNA | AGRN | A0A494C0G | A0A494C0G5_ | Unmodified | 0 | 0 | 978830 | 712920 | 0 | 0 | 0 | 0 | 0 | 0 | 0 | 0 | 0 | 0 | 0 | 593830 | 0 | 0 |
| TFVGAGLR | AGRN; AGRN; A | O00468; A0 | AGRIN_HUMA | Unmodified | 1057100 | 0 | 2108900 | 1164200 | 0 | 0 | 0 | 0 | 0 | 0 | 0 | 0 | 0 | 0 | 944140 | 730840 | 0 | 0 |
| TFVNITPAE | PFN1; PFN1 | P07737; I3L | PROF1_HUMA | Unmodified | 34975000 | 98349000 | 30442000 | 59592000 | 3.28E+08 | 50476000 | 76194000 | 17079000 | 1.56E+08 | 1.36E+08 | 1.03E+08 | 1.41E+08 | 2.43E+08 | 1.47E+08 | 71597000 | 85110000 | 1.46E+08 | 96544000 |
| TFVNITPAE | PFN1; PFN1 | P07737; I3L | PROF1_HUMA | Unmodified | 12600000 | 10577000 | 0 | 9296700 | 5830400 | 1401100 | 2015000 | 264800 | 10563000 | 2848200 | 2601800 | 18921000 | 2745400 | 1060800 | 5007200 | 8352800 | 5360100 | 1376400 |
| TFVNITPAE | PFN1; PFN1 | P07737; I3L | PROF1_HUMA | Unmodified | 0 | 0 | 0 | 0 | 0 | 0 | 1697300 | 0 | 0 | 0 | 0 | 0 | 1564600 | 1424100 | 0 | 0 | 6108900 | 746610 |
| TFVPQFQE | BABAM2 | Q9NXR7 | BABA2_HUMA | Unmodified | 0 | 0 | 0 | 0 | 250330 | 0 | 0 | 27599 | 0 | 68219 | 0 | 623200 | 128420 | 0 | 0 | 0 | 0 | 418440 |
| TFVQGLTV | GPC6 | Q9Y625 | GPC6_HUMAN | Unmodified | 1827400 | 1543800 | 0 | 0 | 0 | 387810 | 129860 | 0 | 0 | 0 | 0 | 0 | 0 | 0 | 0 | 0 | 0 | 0 |
| TFVSGACD | GNB2; GNB2; G | P62879; E7E | GBB2_HUMAN | Unmodified | 5960700 | 2294400 | 4635500 | 4036300 | 974210 | 1182700 | 937650 | 393280 | 5043200 | 360840 | 12162000 | 87178000 | 382460 | 518780 | 7991400 | 5179500 | 833040 | 804980 |
| TFVSGACD | GNB4; GNB4; G | Q9HAV0; A0 | GBB4_HUMAN | Unmodified | 0 | 0 | 0 | 948270 | 530800 | 0 | 0 | 0 | 503050 | 57617 | 536260 | 2049400 | 0 | 0 | 0 | 410400 | 374020 | 404210 |
| TFVTEFK | HEG1 | Q9ULI3 | HEG1_HUMAN | Unmodified | 453020 | 446850 | 572720 | 949790 | 2151200 | 0 | 0 | 0 | 0 | 0 | 0 | 0 | 1100100 | 1425400 | 338550 | 586400 | 0 | 990290 |
| TFYSCTTEC | FN1 | P02751 | FINC_HUMAN I | Unmodified | 0 | 246600 | 1707100 | 487690 | 0 | 0 | 0 | 0 | 531020 | 198130 | 139930 | 0 | 0 | 0 | 250120 | 766080 | 0 | 0 |
| TFYSCTTEC | FN1 | P02751 | FINC_HUMAN I | Unmodified | 0 | 0 | 2664100 | 0 | 0 | 1832000 | 0 | 0 | 0 | 535540 | 0 | 0 | 0 | 0 | 0 | 0 | 0 | 1755300 |
| TGAAPIIDV | RPL27A; RPL27 | P46776; E9P | RL27A_HUMAN | Unmodified | 8157900 | 11325000 | 6054700 | 5295200 | 370100 | 3677200 | 132940 | 173960 | 1848600 | 867000 | 5059600 | 69960000 | 705880 | 217850 | 5367400 | 4895800 | 0 | 363780 |
| TGAAPIIDV | RPL27A; RPL27 | P46776; E9P | RL27A_HUMAN | Unmodified | 0 | 0 | 0 | 0 | 2261700 | 404860 | 2553800 | 1599400 | 412570 | 725660 | 0 | 0 | 1207900 | 0 | 0 | 0 | 430230 | 1405200 |
| TGAEGAVL | KIF5B; KIF5C; K | P33176; O6 | KINH_HUMAN | Unmodified | 0 | 0 | 0 | 431720 | 3647000 | 501450 | 309920 | 219380 | 1458100 | 977440 | 1781400 | 4579900 | 1597800 | 694360 | 579380 | 323080 | 409510 | 1082900 |
| TGAEGAVL | KIF5B; KIF5C; K | P33176; O6 | KINH_HUMAN | Unmodified | 0 | 0 | 0 | 0 | 875830 | 1339700 | 526690 | 396970 | 0 | 256990 | 0 | 0 | 265850 | 287770 | 0 | 0 | 613660 | 622640 |
| TGAEIEPSA | GPRASP2 | Q96D09 | GASP2_HUMA | Unmodified | 444240 | 459340 | 1002000 | 299040 | 0 | 0 | 0 | 0 | 0 | 0 | 0 | 0 | 580780 | 177560 | 566090 | 472280 | 0 | 0 |
| TGAELVTC | SDF2L1 | Q9HCN8 | SDF2L_HUMA | Unmodified | 0 | 0 | 489870 | 0 | 0 | 0 | 0 | 54232 | 4403900 | 651910 | 804960 | 510290 | 0 | 0 | 0 | 0 | 0 | 0 |
| TGAFEPAE | NLE1; NLE1; NL | Q9NVX2; A0 | NLE1_HUMAN | Unmodified | 0 | 0 | 0 | 0 | 0 | 0 | 0 | 0 | 0 | 0 | 0 | 1074100 | 0 | 0 | 237170 | 0 | 0 | 0 |
| TGAFSIPVI | DDX1; DDX1; D | Q92499; A0 | DDX1_HUMAN | Unmodified | 0 | 0 | 0 | 0 | 1196600 | 0 | 0 | 0 | 0 | 0 | 0 | 0 | 2109000 | 1139500 | 0 | 0 | 347180 | 926740 |
| TGAIIPELEF | NT5DC2; NT5D | Q9H857; H7 | NT5D2_HUMA | Unmodified | 0 | 0 | 0 | 0 | 0 | 0 | 0 | 0 | 199220 | 0 | 0 | 0 | 0 | 0 | 0 | 0 | 0 | 0 |
| TGAIVDVP | ATP5F1A; ATP5 | P25705; K7E | ATPA_HUMAN | Unmodified | 0 | 0 | 0 | 0 | 0 | 0 | 0 | 0 | 2483400 | 0 | 2324900 | 0 | 0 | 0 | 0 | 0 | 0 | 0 |
| TGALLLQG | BAX | Q07812 | BAX_HUMAN A | Unmodified | 0 | 0 | 283210 | 0 | 0 | 0 | 0 | 0 | 0 | 0 | 0 | 694010 | 0 | 0 | 0 | 0 | 0 | 0 |
| TGALLLQG | BAX | Q07812 | BAX_HUMAN A | Unmodified | 0 | 0 | 0 | 0 | 0 | 0 | 170800 | 10210 | 0 | 235840 | 0 | 0 | 0 | 187710 | 0 | 0 | 205840 | 256070 |

| | | | | | | | | | | | | | | | | | | | | | | |
|---|---|---|---|---|---|---|---|---|---|---|---|---|---|---|---|---|---|---|---|---|---|---|
| TGALSTGAI | LTBP3; LTBP3 | Q9NS15; E9I | LTBP3_HUMAN | Unmodified | 0 | 0 | 0 | 0 | 0 | 0 | 190260 | 0 | 205120 | 47801 | 0 | 0 | 0 | 0 | 0 | 0 | 0 | 0 |
| TGALVLSR | SPN; SPN | P16150; C9J | LEUK_HUMAN | Unmodified | 0 | 0 | 0 | 0 | 0 | 0 | 0 | 21355 | 0 | 0 | 238280 | 16354000 | 0 | 0 | 15528 | 0 | 0 | 0 |
| TGAPCRSE | ENO1; ENO3; E | P06733; P13 | ENOA_HUMAN | Unmodified | 0 | 0 | 0 | 0 | 182980 | 14324000 | 18264 | 5361200 | 0 | 22743000 | 40019 | 46812 | 37056000 | 37563000 | 0 | 0 | 66387000 | 87208000 |
| TGASFQQA | SCAMP2 | O15127 | SCAM2_HUMA | Unmodified | 2023700 | 0 | 5552400 | 4439100 | 550420 | 0 | 0 | 0 | 1494100 | 0 | 0 | 1982800 | 0 | 0 | 0 | 3473900 | 0 | 0 |
| TGAYQYTIF | NUP210 | Q8TEM1 | PO210_HUMAN | Unmodified | 0 | 0 | 0 | 0 | 972570 | 0 | 0 | 0 | 17253000 | 1396100 | 397650 | 403220 | 0 | 0 | 0 | 0 | 0 | 0 |
| TGCELGDC | PAGE1 | O75459 | PAGE1_HUMAN | Unmodified | 0 | 0 | 162340 | 910810 | 0 | 0 | 0 | 0 | 0 | 0 | 0 | 0 | 273920 | 65750 | 227790 | 89283 | 0 | 0 |
| TGCETVDA | SUN1; SUN1; S | O94901; H0Y | SUN1_HUMAN | Unmodified | 41805 | 0 | 0 | 0 | 1208900 | 0 | 0 | 0 | 2981200 | 0 | 0 | 0 | 0 | 0 | 0 | 0 | 0 | 0 |
| TGCNVLLIC | CCT4 | P50991 | TCPD_HUMAN | Unmodified | 0 | 0 | 0 | 0 | 0 | 0 | 566710 | 262150 | 0 | 422310 | 0 | 7341100 | 633040 | 720960 | 0 | 0 | 0 | 204400 |
| TGCTFPEKI | ADK; ADK | P55263; A0A | ADK_HUMAN A | Unmodified | 80112 | 0 | 0 | 0 | 829370 | 0 | 0 | 127930 | 606780 | 0 | 0 | 4748700 | 0 | 0 | 509080 | 367570 | 776820 | 2481800 |
| TGDARIGF\ | DRG1 | Q9Y295 | DRG1_HUMAN | Unmodified | 0 | 0 | 0 | 0 | 0 | 277500 | 222520 | 229730 | 0 | 189370 | 0 | 0 | 0 | 241880 | 0 | 0 | 176300 | 542140 |
| TGDFQLHT | VDAC2; VDAC2 | P45880; A0A | VDAC2_HUMAI | Unmodified | 0 | 0 | 0 | 0 | 597110 | 0 | 1749600 | 1490100 | 4.1E+08 | 24399000 | 0 | 648130 | 0 | 0 | 0 | 0 | 0 | 0 |
| TGDIFTTET | PCDH1; PCDH1 | Q08174; D6I | PCDH1_HUMA | Unmodified | 392020 | 261510 | 344090 | 217080 | 0 | 0 | 0 | 0 | 572500 | 0 | 0 | 1435400 | 0 | 91486 | 486900 | 305790 | 0 | 0 |
| TGDLLEVQ | OPLAH | O14841 | OPLA_HUMAN | Unmodified | 0 | 0 | 0 | 0 | 0 | 0 | 0 | 0 | 0 | 0 | 57945 | 426040 | 0 | 0 | 31110 | 52391 | 0 | 0 |
| TGDPNMGI | CEL | P19835 | CEL_HUMAN B | Oxidation (M) | 0 | 0 | 0 | 0 | 0 | 0 | 0 | 0 | 12357000 | 3952100 | 13610000 | 0 | 0 | 0 | 0 | 0 | 0 | 0 |
| TGDPNMGI | CEL | P19835 | CEL_HUMAN B | Unmodified | 0 | 0 | 0 | 0 | 0 | 0 | 0 | 0 | 7508600 | 317130 | 996120 | 0 | 0 | 0 | 0 | 0 | 0 | 0 |
| TGDPNMGI | CEL | P19835 | CEL_HUMAN B | Oxidation (M) | 0 | 0 | 0 | 0 | 0 | 0 | 0 | 0 | 5603700 | 2052500 | 8622800 | 0 | 0 | 0 | 0 | 0 | 0 | 0 |
| TGDPNMGI | CEL | P19835 | CEL_HUMAN B | Unmodified | 0 | 0 | 0 | 0 | 0 | 0 | 0 | 0 | 8203200 | 671640 | 1037200 | 0 | 0 | 0 | 0 | 0 | 0 | 0 |
| TGDQEERC | MLF2 | Q15773 | MLF2_HUMAN | Unmodified | 11056000 | 21450000 | 3100500 | 5042000 | 0 | 0 | 0 | 0 | 0 | 0 | 0 | 0 | 0 | 0 | 5035100 | 2248700 | 0 | 0 |
| TGDTGMLF | LASP1 | Q14847 | LASP1_HUMAN | Unmodified | 0 | 0 | 0 | 0 | 0 | 0 | 0 | 0 | 806200 | 0 | 0 | 0 | 0 | 0 | 0 | 0 | 0 | 0 |
| TGDTGMLF | LASP1 | Q14847 | LASP1_HUMAN | Oxidation (M) | 2223900 | 2920400 | 1161400 | 0 | 0 | 0 | 0 | 0 | 0 | 0 | 0 | 0 | 0 | 0 | 0 | 0 | 0 | 0 |
| TGDVEDST | FAF1; FAF1 | Q9UNN5; B1 | FAF1_HUMAN I | Unmodified | 791460 | 0 | 319600 | 243920 | 0 | 0 | 111940 | 0 | 535520 | 0 | 0 | 1818400 | 0 | 0 | 0 | 0 | 143380 | 0 |
| TGDVVETFI | TRIO; TRIO; TRI | O75962; E7E | TRIO_HUMAN T | Unmodified | 0 | 944340 | 75836 | 110600 | 0 | 0 | 0 | 0 | 0 | 0 | 0 | 255540 | 0 | 199040 | 0 | 77694 | 0 | 0 |
| TGEAIVDA | PDIA6 | Q15084 | PDIA6_HUMAN | Unmodified | 186850 | 257130 | 116740 | 132390 | 0 | 0 | 0 | 28664 | 1.14E+08 | 1412100 | 5163400 | 2715000 | 55540 | 0 | 726670 | 296850 | 60966 | 0 |
| TGEAQIHAI | POFUT1 | Q9H488 | OFUT1_HUMAN | Unmodified | 0 | 0 | 0 | 0 | 0 | 0 | 0 | 0 | 19832000 | 6973200 | 1550200 | 249810 | 144250 | 0 | 0 | 0 | 0 | 0 |
| TGEEIFGTI | PRMT1; PRMT1 | Q99873; E9F | ANM1_HUMAN | Oxidation (M) | 172740 | 0 | 0 | 92406 | 0 | 0 | 0 | 56968 | 0 | 390480 | 541420 | 5418600 | 0 | 0 | 132120 | 167230 | 146560 | 0 |
| TGEEINFGI | MRE11; MRE11 | P49959; F8V | MRE11_HUMAI | Unmodified | 0 | 297890 | 0 | 0 | 289820 | 0 | 17024 | 0 | 0 | 0 | 102550 | 215000 | 0 | 0 | 0 | 210160 | 112280 | 95932 |
| TGEEVGFV | FLNB; FLNB | O75369; E7E | FLNB_HUMAN | Unmodified | 0 | 0 | 0 | 0 | 4691000 | 2855300 | 0 | 0 | 912090 | 0 | 0 | 3321300 | 601780 | 601380 | 0 | 0 | 1996200 | 1211700 |
| TGEEVGFV | FLNB; FLNB | O75369; E7E | FLNB_HUMAN | Unmodified | 0 | 0 | 0 | 0 | 0 | 0 | 0 | 0 | 0 | 0 | 0 | 205420 | 0 | 0 | 0 | 0 | 768950 | 0 |
| TGEGFIDFI | PTPRF; PTPRF; I | P10586; H0Y | PTPRF_HUMAN | Unmodified | 4117200 | 2462200 | 1978900 | 2132000 | 5256000 | 2620000 | 0 | 26842 | 4663900 | 1307400 | 1588400 | 2341000 | 1444400 | 927900 | 6803500 | 4095700 | 0 | 0 |
| TGEGFLCV | NRAS | P01111 | RASN_HUMAN | Unmodified | 0 | 0 | 0 | 466680 | 706010 | 0 | 0 | 57846 | 1236900 | 613380 | 505420 | 0 | 0 | 677330 | 0 | 636620 | 221760 | 0 |
| TGEILWR | EMC1 | Q8N766 | EMC1_HUMAN | Unmodified | 697580 | 2191600 | 0 | 0 | 0 | 183020 | 0 | 0 | 8992600 | 723360 | 509110 | 412170 | 0 | 0 | 0 | 0 | 0 | 0 |
| TGEMSGPV | PIN1; PIN1 | Q13526; K7E | PIN1_HUMAN I | Oxidation (M) | 1593200 | 4241600 | 355960 | 505120 | 0 | 0 | 0 | 0 | 469130 | 0 | 422690 | 0 | 0 | 0 | 914890 | 1776000 | 0 | 0 |
| TGEMSGPV | PIN1; PIN1 | Q13526; K7E | PIN1_HUMAN I | Oxidation (M) | 3423000 | 8758200 | 929220 | 2266400 | 397330 | 0 | 0 | 0 | 1253300 | 166040 | 270940 | 277610 | 629910 | 0 | 2886400 | 0 | 201900 | 250030 |
| TGEMSGPV | PIN1; PIN1 | Q13526; K7E | PIN1_HUMAN I | Unmodified | 62924 | 429390 | 0 | 0 | 0 | 0 | 0 | 0 | 0 | 0 | 0 | 0 | 14912000 | 4550400 | 0 | 0 | 0 | 90249 |
| TGENFRLIY | RPS4X | P62701 | RS4X_HUMAN | Unmodified | 0 | 0 | 188600 | 974130 | 1675300 | 604690 | 5388600 | 4421800 | 600830 | 1570400 | 1801600 | 0 | 996370 | 904660 | 0 | 0 | 768120 | 1811100 |
| TGENVEDA | RAB14; RAB14 | P61106; X6R | RAB14_HUMAN | Unmodified | 0 | 0 | 300660 | 3171800 | 8448200 | 1137600 | 866630 | 639970 | 5950100 | 1961900 | 2692100 | 14203000 | 2477400 | 980210 | 2691000 | 3267600 | 0 | 3226300 |
| TGENVEDA | RAB14; RAB14 | P61106; X6R | RAB14_HUMAN | Unmodified | 5147000 | 0 | 6718800 | 8775100 | 14119000 | 0 | 5107100 | 1883100 | 11746000 | 11225000 | 10010000 | 39967000 | 6354500 | 0 | 5000600 | 3826000 | 675090 | 3841100 |
| TGEPHCPC | CPD | O75976 | CBPD_HUMAN | Unmodified | 0 | 0 | 48937 | 0 | 0 | 0 | 0 | 0 | 1202500 | 287390 | 0 | 0 | 0 | 0 | 0 | 21541 | 0 | 0 |
| TGETSRALS | SUB1; SUB1; SU | P53999; D6F | TCP4_HUMAN | Unmodified | 0 | 44895 | 0 | 0 | 434250 | 85183 | 70076 | 0 | 0 | 38204 | 0 | 0 | 251320 | 0 | 0 | 0 | 61606 | 458340 |
| TGEVLDTK | FAM3C; FAM3C | Q92520; C9J | FAM3C_HUMAI | Unmodified | 0 | 0 | 0 | 0 | 51954 | 0 | 0 | 0 | 33694000 | 1507000 | 661450 | 0 | 0 | 0 | 0 | 0 | 0 | 20587 |
| TGEYPVPLI | ACADM; ACADM | P11310; Q5T | ACADM_HUMA | Unmodified | 0 | 0 | 0 | 0 | 0 | 0 | 0 | 0 | 1935600 | 0 | 0 | 0 | 0 | 0 | 0 | 0 | 0 | 0 |
| TGFFEFQA | LARS1; LARS1; | Q9P2J5; A0A | SYLC_HUMAN | Unmodified | 0 | 0 | 0 | 0 | 0 | 0 | 0 | 0 | 0 | 0 | 0 | 8708400 | 0 | 0 | 0 | 0 | 0 | 0 |
| TGFINLDKI | DKC1; DKC1; D | O60832; H7C | DKC1_HUMAN | Unmodified | 358090 | 0 | 0 | 0 | 0 | 0 | 0 | 0 | 0 | 0 | 0 | 4119700 | 0 | 0 | 0 | 0 | 0 | 0 |
| TGFQAVTG | EIF2S2 | P20042 | IF2B_HUMAN E | Unmodified | 116600 | 0 | 0 | 0 | 3682300 | 258830 | 922840 | 0 | 1118700 | 406850 | 646860 | 4046900 | 2353600 | 2102000 | 0 | 1565900 | 2657700 | 11247000 |
| TGFSFPTQI | SORBS3; SORB | O60504; H0Y | VINEX_HUMAN | Unmodified | 0 | 918740 | 0 | 0 | 0 | 0 | 0 | 0 | 0 | 0 | 0 | 0 | 0 | 0 | 0 | 0 | 0 | 0 |
| TGFSTSPES | PRMT1; PRMT1 | Q99873; E9F | ANM1_HUMAN | Unmodified | 0 | 0 | 515200 | 0 | 50247 | 106730 | 138420 | 0 | 0 | 0 | 780270 | 17692000 | 0 | 0 | 269860 | 0 | 31488 | 155730 |
| TGFVPSNY | NCK1; NCK1; N | P16333; C9J | NCK1_HUMAN | Unmodified | 326520 | 422050 | 1175000 | 271420 | 427100 | 0 | 0 | 0 | 328050 | 0 | 35713 | 0 | 211450 | 270950 | 588150 | 458160 | 0 | 0 |
| TGGADQSI | SRP19; SRP19 | P09132; A0A | SRP19_HUMAN | Unmodified | 0 | 0 | 0 | 0 | 0 | 0 | 0 | 0 | 0 | 0 | 0 | 0 | 0 | 0 | 0 | 0 | 0 | 0 |
| TGGADQSI | SRP19; SRP19 | P09132; A0A | SRP19_HUMAN | Unmodified | 101250 | 222450 | 0 | 0 | 1169700 | 0 | 0 | 0 | 131530 | 54409 | 0 | 1663900 | 260590 | 0 | 1587400 | 745980 | 52818 | 0 |
| TGGAYGED | FASN; FASN | P49327; A0A | FAS_HUMAN F | Unmodified | 0 | 0 | 190780 | 0 | 0 | 0 | 0 | 0 | 0 | 0 | 0 | 31464000 | 601250 | 0 | 0 | 0 | 278790 | 0 |
| TGGLYSCD | ITGA6 | P23229 | ITA6_HUMAN I | Unmodified | 0 | 0 | 844520 | 0 | 181200 | 0 | 0 | 0 | 0 | 0 | 0 | 5483800 | 0 | 0 | 1715700 | 1530100 | 0 | 352270 |
| TGGMMYF | DIP2B | Q9P265 | DIP2B_HUMAN | 2 Oxidation (M) | 0 | 0 | 0 | 0 | 0 | 0 | 0 | 0 | 0 | 0 | 0 | 751270 | 0 | 0 | 0 | 0 | 0 | 0 |
| TGGTQTDL | TCEA1; TCEA2; | P23193; Q15 | TCEA1_HUMAN | Unmodified | 0 | 0 | 0 | 0 | 423700 | 0 | 95821 | 123220 | 438940 | 443450 | 0 | 974590 | 0 | 0 | 0 | 0 | 71200 | 0 |
| TGGVEAVK | WASL | O00401 | WASL_HUMAN | Unmodified | 119630 | 149390 | 49114 | 0 | 49843 | 0 | 0 | 0 | 71977 | 0 | 0 | 249580 | 0 | 0 | 275270 | 224640 | 44402 | 0 |
| TGHIAAGT | AGA | P20933 | ASPG_HUMAN | Unmodified | 0 | 0 | 0 | 0 | 0 | 0 | 25060 | 14616 | 582350 | 235160 | 0 | 51578 | 0 | 0 | 0 | 0 | 0 | 0 |
| TGHIAAGT | AGA | P20933 | ASPG_HUMAN | Unmodified | 0 | 0 | 0 | 0 | 0 | 0 | 110320 | 45723 | 0 | 99845 | 0 | 0 | 0 | 0 | 0 | 0 | 0 | 0 |
| TGHSLLHT | SDHA; SDHA; N | P31040; D6F | SDHA_HUMAN | Unmodified | 0 | 0 | 0 | 0 | 0 | 0 | 0 | 0 | 865130 | 0 | 0 | 0 | 0 | 0 | 0 | 0 | 0 | 0 |
| TGIANTLGI | INPP5D; INPPL1 | Q92835; O15 | SHIP1_HUMAN | Unmodified | 0 | 0 | 0 | 0 | 878180 | 0 | 38073 | 166700 | 329240 | 0 | 840480 | 1090800 | 0 | 0 | 0 | 0 | 0 | 0 |
| TGIEQGSD | HADHA; HADHA | P40939; A0A | ECHA_HUMAN | Unmodified | 0 | 0 | 0 | 0 | 0 | 0 | 0 | 0 | 806270 | 0 | 0 | 0 | 0 | 0 | 0 | 0 | 0 | 0 |
| TGIQVLLK | CLUH; CLUH; C | O75153; I3L | CLU_HUMAN C | Unmodified | 0 | 0 | 0 | 0 | 0 | 0 | 0 | 0 | 0 | 0 | 0 | 2078900 | 0 | 0 | 0 | 0 | 0 | 0 |
| TGISDVFAI | NCL; NCL; NCL | P19338; C9J | NUCL_HUMAN | Unmodified | 0 | 0 | 0 | 256500 | 455970 | 0 | 0 | 0 | 1294800 | 925280 | 1709400 | 3977500 | 867150 | 0 | 12440 | 0 | 0 | 0 |
| TGISHGHT | PFKL | P17858 | PFKAL_HUMAN | Unmodified | 0 | 0 | 0 | 0 | 573240 | 0 | 69272 | 117220 | 2044400 | 203300 | 506040 | 3430500 | 893990 | 239030 | 0 | 0 | 227110 | 767670 |
| TGIVSGFGI | F10 | P00742 | FA10_HUMAN | Unmodified | 410200 | 2252400 | 2821900 | 6543500 | 988070 | 501520 | 1184600 | 1215100 | 192820 | 0 | 0 | 0 | 2026900 | 876200 | 2442500 | 1335100 | 0 | 564010 |
| TGIVSGFGI | F10 | P00742 | FA10_HUMAN | Unmodified | 0 | 0 | 0 | 46662 | 6473400 | 176330 | 18215000 | 10991000 | 74160 | 12075 | 0 | 0 | 12818000 | 11460000 | 0 | 64764 | 938700 | 3643400 |
| TGIYEEK | CANX; CANX; C | P27824; A0A | CALX_HUMAN | Unmodified | 0 | 0 | 0 | 0 | 3155100 | 365380 | 0 | 1393500 | 3.76E+08 | 66226000 | 6479000 | 0 | 0 | 1067700 | 0 | 0 | 0 | 0 |
| TGKTVSYL | ATOX1; ATOX1; | O00244; E5F | ATOX1_HUMAN | Unmodified | 0 | 0 | 0 | 0 | 248370 | 1604500 | 0 | 0 | 0 | 0 | 0 | 235200 | 614420 | 618140 | 0 | 0 | 814470 | 368460 |
| TGKVDNIQ | SSRP1 | Q08945 | SSRP1_HUMAN | Unmodified | 22206000 | 0 | 0 | 285350 | 0 | 0 | 1820200 | 493420 | 0 | 0 | 0 | 897850 | 104350 | 0 | 0 | 0 | 0 | 0 |
| TGLATKDG | TJP2; TJP2; TJP2 | Q9UDY2; A0 | ZO2_HUMAN T | Unmodified | 0 | 0 | 0 | 63678 | 0 | 0 | 0 | 0 | 630940 | 252900 | 0 | 72714 | 0 | 0 | 0 | 0 | 0 | 0 |
| TGLDSPTG | FN1; FN1 | P02751; H0Y | FINC_HUMAN I | Unmodified | 0 | 2018000 | 0 | 842770 | 0 | 657700 | 818880 | 0 | 0 | 1195300 | 0 | 0 | 0 | 0 | 393040 | 0 | 320940 | 0 |
| TGLEDPER | MYH10; MYH10 | P35580; E7E | MYH10_HUMAI | Unmodified | 0 | 0 | 0 | 0 | 19290000 | 0 | 790710 | 0 | 0 | 0 | 0 | 0 | 1023400 | 0 | 0 | 0 | 0 | 0 |
| TGLFQDVG | ADGRG6 | Q86SQ4 | AGRG6_HUMA | Unmodified | 19876 | 0 | 1765200 | 3127500 | 10872000 | 3813900 | 0 | 0 | 0 | 238860 | 345640 | 0 | 1125700 | 592370 | 3654500 | 2110500 | 129190 | 2064200 |
| TGLFTPDL | DNM2 | P50570 | DYN2_HUMAN | Unmodified | 0 | 0 | 0 | 0 | 0 | 0 | 0 | 0 | 659460 | 31261 | 459590 | 1489400 | 0 | 0 | 0 | 0 | 0 | 0 |
| TGLIDYNQI | SHMT2; SHMT2 | P34897; G3V | GLYM_HUMAN | Unmodified | 0 | 0 | 0 | 0 | 0 | 130400 | 0 | 24084 | 37346000 | 6573300 | 0 | 1310300 | 0 | 0 | 0 | 0 | 0 | 0 |
| TGLIIGKGC | FUBP1; FUBP1 | Q96AE4; E9F | FUBP1_HUMAN | Unmodified | 0 | 0 | 0 | 0 | 0 | 0 | 0 | 0 | 0 | 212100 | 0 | 1058800 | 0 | 0 | 0 | 0 | 111510 | 0 |
| TGLIKGSG | PKM; PKM; PKM | P14618; H3E | KPYM_HUMAN | Unmodified | 0 | 0 | 0 | 0 | 1988800 | 138500 | 178110 | 74740 | 0 | 124410 | 0 | 0 | 161590 | 122700 | 0 | 0 | 783870 | 1015900 |
| TGLIKGSG | PKM; PKM; PKM | P14618; H3E | KPYM_HUMAN | Unmodified | 0 | 0 | 0 | 179570 | 17325000 | 3097700 | 4694500 | 2199900 | 196880 | 2552800 | 0 | 0 | 5616900 | 2081600 | 0 | 0 | 12194000 | 12971000 |
| TGLIPSNY\ | OSTF1 | Q92882 | OSTF1_HUMAN | Unmodified | 0 | 0 | 0 | 0 | 0 | 0 | 0 | 0 | 2328800 | 0 | 0 | 5928600 | 0 | 0 | 2400100 | 2271400 | 0 | 0 |
| TGLIPSNY\ | OSTF1 | Q92882 | OSTF1_HUMAN | Unmodified | 0 | 0 | 0 | 0 | 0 | 0 | 0 | 0 | 0 | 0 | 0 | 848700 | 0 | 0 | 549870 | 175250 | 0 | 0 |
| TGLKAHEP | FLNA; FLNA; FL | P21333; A0A | FLNA_HUMAN | Unmodified | 818890 | 0 | 0 | 0 | 0 | 0 | 0 | 0 | 0 | 0 | 0 | 0 | 0 | 0 | 0 | 994020 | 1436800 | 0 |
| TGLLELR | SNTA1 | Q13424 | SNTA1_HUMAN | Unmodified | 0 | 0 | 164330 | 132820 | 0 | 0 | 0 | 0 | 1013800 | 362300 | 397060 | 3231800 | 0 | 0 | 200370 | 330540 | 0 | 72857 |
| TGLPSTLLA | EPS15; EPS15 | P42566; B1A | EPS15_HUMAN | Unmodified | 535810 | 338030 | 0 | 198340 | 914900 | 144760 | 158950 | 0 | 211760 | 191250 | 0 | 23436 | 2070100 | 0 | 306360 | 32901 | 346450 | 835660 |
| TGLRSPDIF | TMEM219; TMEI | Q86XT9; I3L | TM219_HUMAN | Unmodified | 0 | 0 | 0 | 0 | 0 | 0 | 0 | 0 | 179140 | 35027 | 18616 | 0 | 0 | 0 | 0 | 0 | 0 | 0 |
| TGLSQLHN | EXOC3; EXOC3 | O60645; D6I | EXOC3_HUMAI | Unmodified | 0 | 0 | 217940 | 198700 | 0 | 0 | 0 | 0 | 670780 | 0 | 289490 | 2328000 | 0 | 0 | 165640 | 191710 | 0 | 0 |
| TGLTSGNV | PAAT | Q9H8K7 | PAAT_HUMAN | Unmodified | 1179600 | 554600 | 47283 | 0 | 0 | 0 | 0 | 0 | 0 | 0 | 0 | 0 | 0 | 0 | 0 | 101930 | 0 | 0 |
| TGLVEFAR | ATIC; ATIC | P31939; F8W | PUR9_HUMAN | Unmodified | 0 | 0 | 0 | 0 | 595790 | 0 | 0 | 0 | 0 | 0 | 0 | 5629200 | 1204200 | 0 | 772190 | 0 | 0 | 0 |
| TGLYNYYD | SERPINH1; SER | P50454; E9P | SERPH_HUMAI | Unmodified | 0 | 0 | 0 | 0 | 0 | 0 | 0 | 0 | 2173800 | 245800 | 0 | 0 | 0 | 0 | 0 | 0 | 0 | 0 |
| TGLYNYYD | SERPINH1; SER | P50454; E9P | SERPH_HUMAI | Unmodified | 0 | 0 | 0 | 0 | 0 | 0 | 0 | 0 | 5901400 | 1670600 | 0 | 0 | 0 | 0 | 0 | 156030 | 0 | 0 |
| TGLYNYYD | SERPINH1; SER | P50454; E9P | SERPH_HUMAI | Unmodified | 0 | 0 | 0 | 0 | 0 | 0 | 0 | 0 | 2254100 | 1001200 | 0 | 0 | 0 | 0 | 0 | 0 | 0 | 0 |
| TGLYPSYK | FUT8; FUT8 | Q9BYC5; G3 | FUT8_HUMAN | Unmodified | 0 | 0 | 0 | 0 | 0 | 0 | 0 | 0 | 5640900 | 0 | 0 | 0 | 0 | 0 | 0 | 0 | 0 | 0 |
| TGMGSGS | C2CD5; C2CD5 | Q86YS7; H0Y | C2CD5_HUMA | Unmodified | 0 | 0 | 0 | 0 | 576630 | 0 | 146790 | 0 | 0 | 106560 | 0 | 0 | 0 | 148130 | 0 | 0 | 343690 | 260380 |
| TGMIESNR | DLG3; DLG3 | Q92796; Q5J | DLG3_HUMAN | Unmodified | 0 | 0 | 386780 | 394380 | 0 | 0 | 0 | 0 | 683590 | 0 | 139700 | 798160 | 277700 | 50809 | 355720 | 115790 | 0 | 220100 |
| TGMSYHEL | NPRL2 | Q8WTW4 | NPRL2_HUMAN | Oxidation (M) | 0 | 0 | 0 | 0 | 0 | 0 | 0 | 0 | 0 | 0 | 0 | 1885400 | 0 | 0 | 6722100 | 0 | 0 | 0 |
| TGMYYLR | RRM1; RRM1 | P23921; E9P | RIR1_HUMAN I | Unmodified | 0 | 0 | 0 | 0 | 0 | 0 | 0 | 0 | 0 | 0 | 0 | 1423600 | 168260 | 0 | 0 | 0 | 0 | 0 |
| TGNALPMF | MANEA | Q5SRI9 | MANEA_HUMA | Unmodified | 0 | 151520 | 0 | 1456200 | 0 | 630950 | 106610 | 46974 | 0 | 0 | 0 | 0 | 0 | 0 | 0 | 2746000 | 0 | 296950 |
| TGNALPMF | MANEA | Q5SRI9 | MANEA_HUMA | Oxidation (M) | 0 | 0 | 0 | 0 | 0 | 0 | 0 | 0 | 3157000 | 0 | 0 | 402700 | 0 | 0 | 0 | 0 | 0 | 0 |
| TGNALPMF | MANEA | Q5SRI9 | MANEA_HUMA | Unmodified | 0 | 0 | 0 | 0 | 0 | 0 | 0 | 0 | 4334800 | 0 | 0 | 0 | 0 | 0 | 0 | 0 | 0 | 0 |
| TGNIPALVF | CSE1L | P55060 | XPO2_HUMAN | Unmodified | 26845 | 0 | 0 | 0 | 0 | 0 | 464430 | 340050 | 0 | 34008 | 15007 | 6758200 | 0 | 0 | 104470 | 0 | 0 | 0 |
| TGNLGNV\ | RPL22L1; RPL2 | Q6P5R6; H0 | RL22L_HUMAN | Unmodified | 5441500 | 6083500 | 6802100 | 2721800 | 0 | 0 | 0 | 0 | 0 | 32090 | 352170 | 5177300 | 0 | 89804 | 7829000 | 5874200 | 0 | 0 |
| TGNLGNV\ | RPL22L1; RPL2 | Q6P5R6; H0 | RL22L_HUMAN | Unmodified | 0 | 0 | 0 | 727010 | 0 | 0 | 382580 | 170760 | 0 | 0 | 0 | 0 | 417810 | 214160 | 0 | 0 | 0 | 171410 |
| TGNQHFG\ | NPLOC4 | Q8TAT6 | NPL4_HUMAN | Unmodified | 0 | 0 | 239620 | 201530 | 480910 | 148550 | 0 | 28820 | 1110400 | 161060 | 463500 | 1818700 | 359000 | 229850 | 277040 | 0 | 175610 | 153880 |
| TGNQHFG\ | NPLOC4 | Q8TAT6 | NPL4_HUMAN | Unmodified | 0 | 0 | 0 | 0 | 640010 | 0 | 0 | 162220 | 0 | 107280 | 0 | 0 | 305350 | 180570 | 0 | 0 | 296400 | 0 |
| TGPFAEHS | PTPA; PTPA; N/ | Q15257; A6F | PTPA_HUMAN | Unmodified | 123530 | 325810 | 257120 | 251380 | 263810 | 864190 | 203100 | 0 | 0 | 267100 | 359750 | 2391400 | 225380 | 164610 | 1898000 | 328430 | 1375600 | 0 |
| TGPFVVHW | DMXL1; DMXL1 | Q9Y485; F5H | DMXL1_HUMAI | Unmodified | 0 | 0 | 0 | 0 | 0 | 0 | 0 | 0 | 0 | 0 | 37634 | 312740 | 0 | 0 | 0 | 0 | 0 | 0 |
| TGPIQDHT | NRP1; NRP1; N | O14786; E9F | NRP1_HUMAN | Unmodified | 336800 | 0 | 1819500 | 1508200 | 11545000 | 3527600 | 0 | 0 | 0 | 0 | 0 | 0 | 1425200 | 533770 | 841830 | 3482700 | 2961000 | 25079000 |
| TGPIQDHT | NRP1; NRP1; N | O14786; E9F | NRP1_HUMAN | Unmodified | 0 | 0 | 0 | 0 | 0 | 0 | 0 | 0 | 0 | 115490 | 0 | 0 | 0 | 0 | 0 | 0 | 0 | 2273400 |
| TGPLLIELR | CAPN7 | Q9Y6W3 | CAN7_HUMAN | Unmodified | 0 | 0 | 0 | 0 | 0 | 0 | 0 | 108400 | 1522800 | 140310 | 0 | 7996500 | 0 | 0 | 0 | 0 | 0 | 0 |
| TGPLLIELR | CAPN7 | Q9Y6W3 | CAN7_HUMAN | Unmodified | 0 | 122430 | 0 | 286540 | 387850 | 0 | 0 | 9208 | 2047800 | 706490 | 505110 | 1048900 | 0 | 0 | 0 | 0 | 0 | 0 |
| TGPLPPGP | PTGR1; PTGR1 | Q14914; F2Z | PTGR1_HUMAN | Unmodified | 119590 | 0 | 60025 | 0 | 0 | 0 | 0 | 0 | 0 | 0 | 0 | 0 | 0 | 0 | 189270 | 169200 | 0 | 0 |
| TGPVTLHT | RTCA | O00442 | RTCA_HUMAN | Unmodified | 90244 | 161220 | 215740 | 286120 | 1757600 | 508290 | 762260 | 0 | 732630 | 265040 | 220700 | 0 | 6852800 | 3255500 | 461270 | 757400 | 551890 | 1995800 |

| | | | | | | | | | | | | | | | | | | | | | | |
|---|---|---|---|---|---|---|---|---|---|---|---|---|---|---|---|---|---|---|---|---|---|---|
| TGQAAELG | PSMD11; PSMD | O00231; J3Q | PSD11_HUMAN | Unmodified | 0 | 0 | 0 | 0 | 0 | 213130 | 139220 | 171360 | 0 | 97730 | 579000 | 4827900 | 0 | 201220 | 0 | 0 | 0 | 0 |
| TGQATVAS | PGK1 | P00558 | PGK1_HUMAN | Oxidation (M) | 0 | 0 | 0 | 215100 | 1310500 | 0 | 973620 | 619680 | 2851300 | 804430 | 1043000 | 3666800 | 5153200 | 2594500 | 1212700 | 0 | 4661200 | 6967400 |
| TGQATVAS | PGK1 | P00558 | PGK1_HUMAN | Unmodified | 0 | 0 | 0 | 0 | 1302200 | 0 | 0 | 0 | 4018200 | 1403400 | 0 | 1184200 | 0 | 0 | 0 | 0 | 1830100 | 5651000 |
| TGQATVAS | PGK1 | P00558 | PGK1_HUMAN | Oxidation (M) | 0 | 0 | 0 | 824710 | 1475900 | 786410 | 0 | 470900 | 1292300 | 1153700 | 1945600 | 5742300 | 2124300 | 0 | 972390 | 954770 | 5286500 | 5511400 |
| TGQATVAS | PGK1 | P00558 | PGK1_HUMAN | Unmodified | 0 | 0 | 0 | 0 | 1462500 | 0 | 0 | 0 | 0 | 0 | 0 | 1090400 | 2574000 | 1884200 | 0 | 0 | 2696800 | 5767300 |
| TGQATVAS | PGK1 | P00558 | PGK1_HUMAN | Oxidation (M) | 0 | 0 | 0 | 0 | 0 | 0 | 0 | 0 | 0 | 457690 | 0 | 0 | 0 | 0 | 0 | 0 | 5032500 | 0 |
| TGQATVAS | PGK1 | P00558 | PGK1_HUMAN | Unmodified | 0 | 0 | 0 | 0 | 921130 | 0 | 0 | 0 | 0 | 0 | 0 | 0 | 0 | 0 | 0 | 0 | 1714600 | 1604700 |
| TGQECRDT | PLEKHB2; PLEK | Q96CS7; A0 | PKHB2_HUMAN | Unmodified | 312220 | 0 | 236280 | 333390 | 0 | 0 | 0 | 0 | 224250 | 0 | 18255 | 75056 | 189700 | 159430 | 233310 | 177910 | 0 | 0 |
| TGQEIPVN | CNDP2; CNDP2 | Q96KP4; J3Q | CNDP2_HUMA | Unmodified | 336990 | 0 | 1034300 | 2270900 | 175120 | 0 | 0 | 0 | 3709600 | 165750 | 4325900 | 28660000 | 1974700 | 0 | 4899400 | 4018900 | 0 | 0 |
| TGQEVVFV. | RPN2 | P04844 | RPN2_HUMAN | Unmodified | 0 | 739340 | 0 | 0 | 153070 | 0 | 199300 | 0 | 29385000 | 976350 | 0 | 1009500 | 0 | 0 | 0 | 0 | 0 | 0 |
| TGQEVVFV. | RPN2 | P04844 | RPN2_HUMAN | Unmodified | 0 | 0 | 0 | 0 | 0 | 0 | 0 | 0 | 0 | 1884900 | 0 | 0 | 0 | 0 | 0 | 0 | 136760 | 0 |
| TGQEVVFV. | RPN2 | P04844 | RPN2_HUMAN | Unmodified | 0 | 0 | 0 | 0 | 0 | 0 | 0 | 0 | 2586400 | 0 | 0 | 0 | 0 | 0 | 0 | 0 | 0 | 0 |
| TGQEYKPG | EPRS1; EPRS1 | P07814; V9G | SYEP_HUMAN | Unmodified | 0 | 0 | 650360 | 0 | 0 | 0 | 0 | 0 | 650260 | 264980 | 1576600 | 9851700 | 955900 | 0 | 430220 | 0 | 0 | 0 |
| TGQLAAIK | TNIK; MINK1; M | Q9UKE5; Q8 | TNIK_HUMAN T | Unmodified | 27887 | 0 | 736390 | 932580 | 3090400 | 41411 | 446330 | 35383 | 2493100 | 1207600 | 1078900 | 4360200 | 19045 | 1474500 | 386680 | 130330 | 13036 | 3284000 |
| TGQLDELE | SMAD3 | P84022 | SMAD3_HUMA | Unmodified | 0 | 0 | 0 | 0 | 0 | 0 | 0 | 0 | 0 | 0 | 0 | 0 | 0 | 0 | 0 | 0 | 0 | 0 |
| TGQPDLLG | EVA1B | Q9NVM1 | EVA1B_HUMAN | Unmodified | 976280 | 1390400 | 3113500 | 3842200 | 0 | 0 | 0 | 0 | 0 | 0 | 0 | 88627 | 193410 | 0 | 5700400 | 2833800 | 1169700 | 808000 |
| TGQPMIHY | EWSR1; EWSR1 | Q01844; H7I | EWS_HUMAN F | Oxidation (M) | 0 | 0 | 0 | 0 | 1736200 | 250890 | 0 | 0 | 0 | 0 | 0 | 1438800 | 0 | 0 | 0 | 0 | 0 | 651620 |
| TGQPMIHY | EWSR1; EWSR1 | Q01844; H7I | EWS_HUMAN F | Unmodified | 66307 | 54615 | 0 | 124950 | 0 | 0 | 0 | 0 | 113570 | 0 | 110320 | 1019400 | 0 | 0 | 0 | 116970 | 0 | 0 |
| TGQPMINL | FUS; FUS | P35637; H3E | FUS_HUMAN R | Unmodified | 1242700 | 0 | 103470 | 0 | 2450300 | 0 | 1296200 | 44861 | 1384500 | 955430 | 870350 | 1408500 | 0 | 0 | 0 | 0 | 2033900 | 0 |
| TGQQLTSD | ABI3BP; ABI3BF | Q7Z7G0; D3 | TARSH_HUMAN | Unmodified | 0 | 0 | 0 | 0 | 0 | 0 | 0 | 0 | 1515600 | 0 | 0 | 0 | 0 | 0 | 0 | 0 | 0 | 0 |
| TGQSLATLI | WDR33 | Q9C0J8 | WDR33_HUMA | Unmodified | 0 | 0 | 38809 | 0 | 0 | 0 | 359660 | 0 | 42642 | 0 | 12386 | 723160 | 250150 | 220390 | 0 | 64483 | 92214 | 737950 |
| TGQYSGIYI | SLC25A24 | Q6NUK1 | SCMC1_HUMA | Unmodified | 0 | 0 | 0 | 0 | 0 | 0 | 0 | 0 | 595080 | 154000 | 0 | 0 | 0 | 0 | 0 | 0 | 0 | 0 |
| TGQYSGIYI | SLC25A24 | Q6NUK1 | SCMC1_HUMA | Unmodified | 0 | 0 | 0 | 0 | 0 | 0 | 0 | 0 | 1423200 | 609450 | 0 | 0 | 0 | 0 | 0 | 0 | 0 | 0 |
| TGRDASVH | TNFRSF10B | O14763 | TR10B_HUMAN | Unmodified | 70369 | 14435000 | 283590 | 2218700 | 1149000 | 178350 | 0 | 0 | 87535 | 0 | 1003900 | 506690 | 144160 | 44339 | 1126300 | 311220 | 80179 | 74282 |
| TGRDASVH | TNFRSF10B | O14763 | TR10B_HUMAN | Unmodified | 0 | 0 | 0 | 311330 | 6250500 | 821940 | 0 | 0 | 0 | 51374 | 96424 | 0 | 3074100 | 754700 | 0 | 0 | 937040 | 1488100 |
| TGRETAALI | IL17C | Q9P0M4 | IL17C_HUMAN | Unmodified | 0 | 0 | 0 | 50983000 | 0 | 0 | 77144 | 0 | 648390 | 0 | 0 | 0 | 0 | 0 | 28098000 | 48751000 | 0 | 0 |
| TGRPAGKG | CPNE3; CPNE3 | O75131; H0 | CPNE3_HUMAI | Unmodified | 0 | 0 | 0 | 0 | 155120 | 39693 | 159520 | 88806 | 271150 | 333650 | 0 | 0 | 228180 | 0 | 0 | 0 | 124590 | 796970 |
| TGRVMLGE | NME2P1; NME1 | O60361; P15 | NDK8_HUMAN | Unmodified | 38137 | 0 | 0 | 336620 | 985180 | 165060 | 39304 | 0 | 442170 | 65913 | 1867800 | 338930 | 1719900 | 286170 | 0 | 0 | 226190 | 1681700 |
| TGRVMLGE | NME2P1; NME1 | O60361; P15 | NDK8_HUMAN | Oxidation (M) | 0 | 0 | 0 | 214190 | 0 | 193740 | 0 | 0 | 166120 | 0 | 3255800 | 230780 | 1331600 | 336880 | 0 | 0 | 0 | 0 |
| TGRVYNVT | RPL21; RPL21; I | P46778; G3\ | RL21_HUMAN ( | Unmodified | 63268 | 0 | 0 | 216960 | 9540200 | 729540 | 13132000 | 734100 | 0 | 1419200 | 1494300 | 0 | 6845500 | 3186000 | 0 | 19875 | 328500 | 8061600 |
| TGRVYNVT | RPL21; RPL21 | P46778; M0I | RL21_HUMAN ( | Unmodified | 0 | 0 | 0 | 0 | 150530 | 0 | 8510600 | 0 | 0 | 0 | 0 | 0 | 1257000 | 0 | 0 | 0 | 0 | 199760 |
| TGRWEPDI | APLP1; APLP1; / | P51693; B7Z | APLP1_HUMAN | Unmodified | 0 | 0 | 0 | 0 | 151010 | 9207 | 0 | 0 | 3735400 | 333480 | 16890 | 0 | 0 | 0 | 0 | 0 | 0 | 0 |
| TGRWLAVD | OAT | P04181 | OAT_HUMAN C | Unmodified | 0 | 0 | 0 | 0 | 0 | 0 | 0 | 0 | 0 | 749260 | 0 | 0 | 0 | 0 | 0 | 0 | 0 | 0 |
| TGSDADLV | OAS2; OAS2; O | P29728; F8V | OAS2_HUMAN | Unmodified | 0 | 0 | 0 | 0 | 0 | 0 | 0 | 0 | 0 | 0 | 0 | 0 | 152110 | 177620 | 0 | 0 | 0 | 3148200 |
| TGSESSIDF | COPG1 | Q9Y678 | COPG1_HUMA | Unmodified | 0 | 0 | 0 | 0 | 0 | 0 | 0 | 0 | 0 | 0 | 0 | 718050 | 0 | 0 | 0 | 0 | 0 | 0 |
| TGSNEFKLI | BUB3; BUB3; BI | O43684; J3Q | BUB3_HUMAN | Unmodified | 0 | 0 | 0 | 0 | 0 | 625930 | 0 | 0 | 0 | 0 | 0 | 0 | 693910 | 0 | 0 | 0 | 811900 | 0 |
| TGSPGSEV | CCDC88A; CCI | Q3V6T2; A0A | GRDN_HUMAN | Unmodified | 0 | 0 | 0 | 0 | 29658 | 0 | 0 | 0 | 0 | 107180 | 0 | 375490 | 0 | 0 | 0 | 0 | 0 | 16674 |
| TGSQGQC | RPS28 | P62857 | RS28_HUMAN | Unmodified | 2290100 | 2193800 | 1010300 | 2288700 | 927370 | 0 | 0 | 0 | 2367700 | 88973 | 536060 | 7114200 | 545110 | 0 | 1449100 | 1116600 | 216500 | 280620 |
| TGSQGQC | RPS28 | P62857 | RS28_HUMAN | Oxidation (M) | 2264300 | 507120 | 0 | 1658000 | 0 | 0 | 0 | 0 | 0 | 0 | 270290 | 2300200 | 469840 | 0 | 0 | 0 | 0 | 0 |
| TGSSFSVR | ACOT8; ACOT8 | O14734; E9F | ACOT8_HUMAI | Unmodified | 0 | 0 | 0 | 132260 | 123980 | 1092600 | 82128 | 0 | 5727700 | 498650 | 0 | 0 | 0 | 0 | 0 | 23771 | 35460 | 1012800 |
| TGSVEAQT. | SYT13; SYT13 | Q7L8C5; H0 | SYT13_HUMAN | Unmodified | 0 | 0 | 0 | 0 | 0 | 0 | 0 | 0 | 533520 | 0 | 0 | 0 | 0 | 0 | 0 | 0 | 0 | 0 |
| TGSVVLTAS | ARRDC1 | Q8N5I2 | ARRD1_HUMAI | Unmodified | 2432500 | 1710400 | 1532200 | 2137300 | 0 | 0 | 0 | 0 | 731840 | 0 | 0 | 1197400 | 0 | 0 | 3752800 | 3924600 | 0 | 0 |
| TGSYGALA | PPP1R12A; PPP | O14974; F8V | MYPT1_HUMAI | Unmodified | 0 | 0 | 0 | 0 | 996730 | 0 | 0 | 0 | 0 | 0 | 0 | 0 | 0 | 989530 | 0 | 0 | 447500 | 0 |
| TGTAEMSS | ATP5F1A; ATP5I | P25705; K7E | ATPA_HUMAN | Oxidation (M) | 0 | 0 | 0 | 0 | 0 | 0 | 0 | 0 | 1700300 | 0 | 104730 | 0 | 0 | 0 | 0 | 0 | 0 | 0 |
| TGTAEMSS | ATP5F1A; ATP5I | P25705; K7E | ATPA_HUMAN | Unmodified | 0 | 184730 | 0 | 0 | 0 | 0 | 0 | 0 | 17105000 | 240460 | 536310 | 0 | 435890 | 0 | 0 | 0 | 0 | 0 |
| TGTAYTFFT | DDX5; DDX5; D | P17844; A0A | DDX5_HUMAN | Unmodified | 0 | 0 | 0 | 0 | 0 | 0 | 0 | 610740 | 1153800 | 1062400 | 0 | 3235400 | 571860 | 0 | 0 | 0 | 0 | 0 |
| TGTDKTLVI | COTL1; COTL1 | Q14019; H3I | COTL1_HUMAI | Unmodified | 0 | 0 | 0 | 0 | 0 | 0 | 418150 | 0 | 0 | 0 | 0 | 0 | 0 | 0 | 0 | 0 | 1436100 | 0 |
| TGTFIALDR | PTPRO; PTPRB; | Q16827; P23 | PTPRO_HUMAI | Unmodified | 124430 | 0 | 0 | 0 | 0 | 0 | 0 | 0 | 365380 | 0 | 0 | 4301400 | 0 | 64707 | 0 | 146760 | 0 | 0 |
| TGTITHGTF | ATP7A | Q04656 | ATP7A_HUMAN | Unmodified | 0 | 0 | 850610 | 973280 | 0 | 16547 | 282610 | 750150 | 220940 | 47425 | 169150 | 767810 | 1186300 | 894010 | 1066800 | 1092200 | 0 | 0 |
| TGTITTFEH. | EEF2 | P13639 | EF2_HUMAN El | Unmodified | 1848800 | 3196600 | 1988200 | 5612000 | 8044400 | 2350400 | 597580 | 0 | 8480200 | 2101400 | 13105000 | 62940000 | 2191700 | 774320 | 2971800 | 2720500 | 998560 | 0 |
| TGTITTFEH. | EEF2 | P13639 | EF2_HUMAN El | Oxidation (M) | 690270 | 387940 | 224510 | 1576700 | 2475200 | 95195 | 184460 | 0 | 1436400 | 704490 | 2632500 | 26650000 | 783380 | 1320600 | 1867400 | 515230 | 210890 | 727750 |
| TGTITTFEH. | EEF2 | P13639 | EF2_HUMAN El | Unmodified | 0 | 0 | 0 | 33997 | 8136200 | 841390 | 3366100 | 201290 | 49771 | 1027100 | 139340 | 152270 | 1429700 | 949410 | 0 | 0 | 3464800 | 1788600 |
| TGTITTFEH. | EEF2 | P13639 | EF2_HUMAN El | 2 Oxidation (M) | 0 | 0 | 0 | 303930 | 695710 | 0 | 0 | 332670 | 0 | 0 | 74230 | 0 | 250020 | 0 | 0 | 0 | 913640 | 0 |
| TGTITTFEH. | EEF2 | P13639 | EF2_HUMAN El | Oxidation (M) | 0 | 0 | 0 | 0 | 2929100 | 614130 | 1433500 | 551570 | 0 | 312080 | 0 | 0 | 636580 | 570200 | 0 | 0 | 1007400 | 0 |
| TGTIVVEGI | IARS1; IARS1; I | P41252; A0A | SYIC_HUMAN I | Unmodified | 175370 | 0 | 550210 | 289160 | 0 | 0 | 0 | 0 | 514670 | 112350 | 2063600 | 13021000 | 0 | 0 | 524830 | 0 | 0 | 0 |
| TGTLQPWI | RACGAP1; RAC | Q9H0H5; H0 | RGAP1_HUMAI | Unmodified | 0 | 0 | 320330 | 0 | 6146300 | 1532500 | 0 | 0 | 1008600 | 286410 | 0 | 1566800 | 218590 | 0 | 370520 | 135570 | 422130 | 0 |
| TGTLQPWI | RACGAP1; RAC | Q9H0H5; H0 | RGAP1_HUMAI | Unmodified | 0 | 0 | 0 | 0 | 1761900 | 0 | 0 | 161360 | 0 | 0 | 0 | 0 | 0 | 0 | 0 | 0 | 10791000 | 6740900 |
| TGTLTMNR | ATP2B4; ATP2B | P23634; P20 | AT2B4_HUMAN | Unmodified | 440970 | 351950 | 1765700 | 2044600 | 271080 | 0 | 0 | 0 | 765840 | 274540 | 1140100 | 11230000 | 223900 | 0 | 4082000 | 2439800 | 0 | 0 |
| TGTLTQNE | ATP9A; ATP9B; / | O75110; O4 | ATP9A_HUMAN | Unmodified | 0 | 0 | 0 | 0 | 0 | 0 | 0 | 0 | 326540 | 0 | 0 | 0 | 95452 | 0 | 77044 | 0 | 0 | 0 |
| TGTLTQNII | ATP8B1; ATP8B | O43520; A0A | AT8B1_HUMAN | Unmodified | 112060 | 0 | 0 | 0 | 0 | 0 | 0 | 0 | 191690 | 0 | 0 | 472200 | 0 | 0 | 92268 | 122130 | 0 | 0 |
| TGTLTTNQI | ATP2A2; ATP2A | P16615; Q93 | AT2A2_HUMAN | Unmodified | 0 | 0 | 0 | 0 | 0 | 0 | 0 | 0 | 249410 | 0 | 276800 | 654640 | 116110 | 0 | 0 | 0 | 0 | 0 |
| TGTQYLLR | SERPINB6; SER | P35237; P50 | SPB6_HUMAN | Unmodified | 0 | 745170 | 130490 | 827050 | 199740 | 0 | 0 | 0 | 0 | 0 | 0 | 0 | 0 | 0 | 1102700 | 888400 | 153510 | 208410 |
| TGTTVPESI | MTOR; MTOR; N | P42345; B1A | MTOR_HUMAN | Unmodified | 0 | 0 | 0 | 184300 | 547910 | 0 | 86754 | 0 | 0 | 0 | 0 | 0 | 202800 | 110650 | 151500 | 109050 | 86327 | 230880 |
| TGTVSLEVI | FASN; FASN | P49327; A0A | FAS_HUMAN F | Unmodified | 426140 | 494400 | 2660100 | 2271200 | 0 | 399730 | 0 | 0 | 892700 | 243670 | 5933100 | 50463000 | 0 | 224570 | 3806100 | 1848700 | 127870 | 0 |
| TGVAPDVF. | HMGCS1 | Q01581 | HMCS1_HUMA | Unmodified | 0 | 0 | 0 | 0 | 1390300 | 0 | 0 | 0 | 0 | 0 | 354730 | 3640200 | 206980 | 0 | 0 | 0 | 0 | 0 |
| TGVAPDVF. | HMGCS1 | Q01581 | HMCS1_HUMA | Oxidation (M) | 0 | 0 | 0 | 0 | 0 | 266110 | 0 | 0 | 0 | 350950 | 1038000 | 4557800 | 205670 | 0 | 0 | 0 | 0 | 0 |
| TGVAVNKP | FLNA; FLNA; FL | P21333; A0A | FLNA_HUMAN | Unmodified | 207850 | 600500 | 800720 | 985880 | 72700000 | 11993000 | 790570 | 1014400 | 5852300 | 3384800 | 1117700 | 11850000 | 14771000 | 11151000 | 1015700 | 1136200 | 28665000 | 25460000 |
| TGVAVNKP | FLNA; FLNA; FL | P21333; A0A | FLNA_HUMAN | Unmodified | 0 | 0 | 0 | 0 | 614920 | 110520 | 0 | 28111 | 0 | 0 | 0 | 0 | 221790 | 302970 | 0 | 0 | 1517200 | 267480 |
| TGVELGKP | FLNA; FLNA; FL | P21333; A0A | FLNA_HUMAN | Unmodified | 0 | 0 | 0 | 0 | 1198800 | 0 | 0 | 0 | 336270 | 178540 | 0 | 1934700 | 104240 | 0 | 223280 | 169580 | 786790 | 0 |
| TGVFEEK | CLGN | O14967 | CLGN_HUMAN | Unmodified | 0 | 0 | 0 | 0 | 0 | 0 | 0 | 0 | 12471000 | 3084100 | 1867100 | 576350 | 0 | 166820 | 0 | 0 | 0 | 0 |
| TGVHHYSG | RPL30; RPL30; I | P62888; A0A | RL30_HUMAN ( | Unmodified | 7665700 | 3436500 | 3739700 | 2860300 | 2067000 | 68490 | 583440 | 1079200 | 109820 | 117770 | 1083900 | 17842000 | 2034300 | 428500 | 5600700 | 3361200 | 52918 | 188860 |
| TGVLNENT | EHBP1 | Q8NDI1 | EHBP1_HUMAI | Unmodified | 0 | 199890 | 0 | 0 | 722050 | 194830 | 90385 | 0 | 699950 | 203670 | 164530 | 0 | 0 | 0 | 0 | 0 | 168610 | 300210 |
| TGVLTVAS | P4HA2 | O15460 | P4HA2_HUMAI | Unmodified | 0 | 0 | 0 | 0 | 0 | 228950 | 0 | 0 | 4761500 | 0 | 0 | 0 | 0 | 0 | 0 | 0 | 0 | 216690 |
| TGVQFTVK | STAT1; STAT1; S | P42224; A0A | STAT1_HUMAN | Unmodified | 0 | 199060 | 249130 | 0 | 612190 | 0 | 166130 | 182460 | 0 | 0 | 0 | 2677900 | 282770 | 0 | 0 | 66302 | 0 | 675850 |
| TGVQTFR | COPA; COPA; C | P53621; A0A | COPA_HUMAN | Unmodified | 94208 | 101750 | 0 | 0 | 0 | 0 | 0 | 57170 | 5325200 | 120400 | 0 | 254220 | 0 | 0 | 0 | 18239 | 0 | 0 |
| TGVVPQLV | KPNA2; KPNA2; | P52292; A0A | IMA1_HUMAN I | Unmodified | 0 | 320830 | 581560 | 750660 | 2366000 | 647310 | 594200 | 415880 | 821310 | 688890 | 403250 | 2668800 | 1470900 | 676700 | 656530 | 484590 | 1694000 | 1636600 |
| TGVVTDGV | ETFB; ETFB | P38117; M0 | ETFB_HUMAN I | Unmodified | 3650800 | 13579000 | 59512 | 498390 | 0 | 0 | 86167 | 8121.9 | 1199900 | 34046 | 0 | 694050 | 197690 | 76678 | 2979500 | 3087900 | 0 | 0 |
| TGVVYQIE | CANT1; CANT1 | Q8WVQ1; K7 | CANT1_HUMAI | Unmodified | 0 | 92205 | 0 | 0 | 0 | 407470 | 0 | 0 | 4845500 | 426140 | 0 | 0 | 0 | 0 | 0 | 0 | 0 | 0 |
| TGYAFVDC | IGF2BP3; IGF2E | O00425; F8V | IF2B3_HUMAN | Unmodified | 103280 | 0 | 0 | 0 | 0 | 0 | 0 | 14659 | 0 | 0 | 0 | 155580 | 0 | 0 | 0 | 0 | 0 | 0 |
| TGYKREHII | VDAC1; VDAC1 | P21796; C9J | VDAC1_HUMAI | Unmodified | 266430 | 0 | 0 | 0 | 0 | 886520 | 0 | 0 | 1664700 | 575140 | 0 | 0 | 0 | 0 | 0 | 383860 | 0 | 466920 |
| TGYKREHII | VDAC1; VDAC1 | P21796; C9J | VDAC1_HUMAI | Oxidation (M) | 0 | 0 | 0 | 0 | 0 | 0 | 0 | 0 | 0 | 989540 | 0 | 370470 | 0 | 0 | 0 | 0 | 0 | 0 |
| TGYLGFVR | DIP2B | Q9P265 | DIP2B_HUMAN | Unmodified | 230810 | 249260 | 0 | 0 | 0 | 0 | 0 | 34255 | 0 | 0 | 0 | 3715200 | 0 | 0 | 759770 | 0 | 0 | 0 |
| TGYRVFSAI | PRPSAP1 | Q14558 | KPRA_HUMAN | Unmodified | 0 | 0 | 0 | 0 | 0 | 0 | 817330 | 0 | 0 | 302390 | 0 | 983650 | 734030 | 465530 | 0 | 0 | 296270 | 353810 |
| TGYSFVNC | TSFM; TSFM; TS | P43897; F8V | EFTS_HUMAN I | Unmodified | 0 | 0 | 0 | 0 | 82910 | 0 | 0 | 0 | 582000 | 104250 | 0 | 0 | 0 | 0 | 0 | 0 | 0 | 0 |
| TGYTLDVTI | HNRNPR; SYNC | O43390; O6 | HNRPR_HUMA | Unmodified | 0 | 0 | 0 | 0 | 0 | 0 | 0 | 0 | 0 | 0 | 0 | 6498800 | 0 | 505820 | 0 | 0 | 0 | 0 |
| TGYTLDVTI | HNRNPR; SYNC | O43390; O6 | HNRPR_HUMA | Unmodified | 79411 | 362500 | 219390 | 0 | 0 | 0 | 180710 | 92247 | 952080 | 681050 | 1285300 | 9390100 | 0 | 0 | 274070 | 284820 | 84089 | 0 |
| TGYVLQPE | PLCG2; PLCG2 | P16885; A0A | PLCG2_HUMAI | Unmodified | 0 | 0 | 0 | 0 | 416620 | 0 | 801730 | 56402 | 0 | 0 | 584640 | 6786200 | 0 | 0 | 0 | 0 | 154560 | 0 |
| TGYVLQPE | PLCG2; PLCG2 | P16885; A0A | PLCG2_HUMAI | Oxidation (M) | 0 | 0 | 0 | 0 | 0 | 0 | 0 | 0 | 0 | 0 | 0 | 4595900 | 0 | 0 | 0 | 0 | 0 | 0 |
| TGYYFDGIS | FBLN1; FBLN1 | P23142; B1A | FBLN1_HUMAN | Unmodified | 0 | 0 | 207870 | 0 | 0 | 0 | 0 | 0 | 14522000 | 6063700 | 0 | 0 | 0 | 0 | 0 | 0 | 0 | 0 |
| THAIEFVNY | PLCB4; PLCB4 | Q15147; A0A | PLCB4_HUMAN | Unmodified | 0 | 0 | 0 | 0 | 241540 | 0 | 0 | 67595 | 0 | 0 | 0 | 0 | 583780 | 0 | 343850 | 608570 | 2468800 | 0 |
| THAIEFVNY | PLCB4; PLCB4 | Q15147; A0A | PLCB4_HUMAN | Unmodified | 0 | 0 | 0 | 408120 | 0 | 0 | 90640 | 0 | 116000 | 18684 | 119810 | 0 | 0 | 923860 | 291750 | 0 | 0 | 0 |
| THAVLVALI | PSMA1; N/A; PS | P25786; B4D | PSA1_HUMAN | Unmodified | 0 | 0 | 0 | 0 | 491780 | 125710 | 100890 | 0 | 553470 | 433480 | 0 | 2225100 | 334070 | 0 | 0 | 0 | 217550 | 257260 |
| THDNDLMI | KLK14; KLK14 | Q9P0G3; A0 | KLK14_HUMAN | Unmodified | 0 | 0 | 0 | 0 | 0 | 0 | 142310 | 0 | 7634300 | 102730 | 0 | 0 | 114350 | 523130 | 0 | 0 | 461500 | 845200 |
| THDNDLMI | KLK14; KLK14 | Q9P0G3; A0 | KLK14_HUMAN | Oxidation (M) | 0 | 0 | 0 | 0 | 0 | 0 | 508280 | 266030 | 8142000 | 187590 | 0 | 349990 | 343120 | 0 | 0 | 0 | 0 | 53034 |
| THEAQIQEI | MYH9 | P35579 | MYH9_HUMAN | Unmodified | 0 | 0 | 418880 | 620310 | 878310 | 216140 | 0 | 0 | 959050 | 17417 | 636590 | 1540600 | 0 | 130500 | 821960 | 439470 | 74396 | 120370 |
| THEAQIQEI | MYH9 | P35579 | MYH9_HUMAN | Oxidation (M) | 39174 | 0 | 1112800 | 534090 | 214860 | 0 | 0 | 0 | 0 | 0 | 253930 | 607420 | 0 | 0 | 369790 | 490990 | 0 | 0 |
| THEAQIQEI | MYH9 | P35579 | MYH9_HUMAN | Unmodified | 0 | 0 | 0 | 0 | 0 | 90945 | 0 | 0 | 0 | 0 | 0 | 0 | 0 | 3864400 | 0 | 0 | 2548300 | 0 |
| THEAQIQEI | MYH9 | P35579 | MYH9_HUMAN | Oxidation (M) | 0 | 0 | 0 | 0 | 0 | 526030 | 31977 | 0 | 0 | 0 | 0 | 0 | 0 | 0 | 0 | 0 | 0 | 0 |
| THEDIEAQI | SF3B1; SF3B1; | O75533; B4I | SF3B1_HUMAN | Unmodified | 175260 | 0 | 85844 | 0 | 0 | 0 | 0 | 0 | 0 | 0 | 0 | 0 | 452630 | 175110 | 0 | 0 | 0 | 0 |
| THEEVKMT | UACA; UACA; U | Q9BZF9; H0 | UACA_HUMAN | Unmodified | 0 | 0 | 0 | 0 | 209410 | 0 | 0 | 0 | 0 | 0 | 0 | 0 | 123130 | 101180 | 0 | 0 | 492400 | 351640 |
| THFFLNAG | SAMM50 | Q9Y512 | SAM50_HUMAI | Unmodified | 0 | 0 | 0 | 0 | 0 | 0 | 0 | 0 | 1670300 | 426920 | 0 | 0 | 0 | 0 | 0 | 0 | 0 | 0 |
| THFLMPFP | PITRM1; PITRM | Q5JRX3; A0A | PREP_HUMAN | Unmodified | 0 | 0 | 0 | 0 | 0 | 0 | 0 | 0 | 221270 | 0 | 0 | 186350 | 0 | 0 | 0 | 36802 | 0 | 0 |
| THFNKGPS | CNN3 | Q15417 | CNN3_HUMAN | Acetyl (Protein N | 0 | 46130 | 0 | 114760 | 14958000 | 2090900 | 0 | 0 | 0 | 2532000 | 105480 | 326090 | 0 | 0 | 99078 | 37723 | 4631300 | 3518200 |
| THFNKGPS | CNN3 | Q15417 | CNN3_HUMAN | Acetyl (Protein N | 0 | 0 | 0 | 0 | 199210 | 0 | 0 | 0 | 0 | 76547 | 0 | 0 | 0 | 0 | 0 | 0 | 673810 | 0 |
| THFPQFSY | AKT2 | P31751 | AKT2_HUMAN I | Unmodified | 0 | 0 | 0 | 443500 | 0 | 0 | 0 | 0 | 0 | 0 | 0 | 0 | 435590 | 0 | 403790 | 367830 | 0 | 0 |
| THFRAVDP | SDF4; SDF4; SD | Q9BRK5; H0 | CAB45_HUMAI | Unmodified | 0 | 0 | 0 | 0 | 527040 | 165050 | 0 | 0 | 354420 | 3392300 | 0 | 0 | 936500 | 1082500 | 0 | 0 | 0 | 523680 |
| THFRAVDP | SDF4; SDF4; SD | Q9BRK5; H0 | CAB45_HUMAI | Unmodified | 0 | 0 | 0 | 0 | 0 | 0 | 0 | 0 | 0 | 1541300 | 0 | 0 | 594870 | 1159000 | 0 | 0 | 0 | 232690 |
| THHANHAI | HYAL2 | Q12891 | HYAL2_HUMAN | Unmodified | 781090 | 597000 | 804490 | 0 | 0 | 0 | 0 | 0 | 0 | 0 | 0 | 2113000 | 0 | 0 | 1652500 | 777820 | 0 | 0 |
| THHEMKEY | BCLAF1; BCLAF | Q9NYF8; E9I | BCLF1_HUMAN | Unmodified | 0 | 5851000 | 0 | 0 | 0 | 0 | 0 | 0 | 0 | 0 | 0 | 0 | 3514600 | 0 | 2608700 | 889080 | 0 | 0 |

| | | | | | | | | | | | | | | | | | | | | | | |
|---|---|---|---|---|---|---|---|---|---|---|---|---|---|---|---|---|---|---|---|---|---|---|
| THHNDTEL | GMPS | P49915 | GUAA_HUMAN | Unmodified | 50733 | 18490 | 0 | 0 | 584700 | 225170 | 0 | 76096 | 478580 | 365300 | 92290 | 811760 | 656410 | 0 | 0 | 0 | 136650 | 430680 |
| THIALDGR | ITGB3; N/A; ITG | P05106; H3E | ITB3_HUMAN I | Unmodified | 800030 | 2035600 | 0 | 3542800 | 0 | 0 | 0 | 0 | 0 | 0 | 0 | 0 | 0 | 163640 | 1796800 | 1831000 | 28790 | 0 |
| THIDTVINA | AP2A1 | O95782 | AP2A1_HUMAN | Unmodified | 0 | 0 | 188170 | 0 | 0 | 0 | 0 | 0 | 0 | 0 | 143920 | 1280100 | 0 | 0 | 156150 | 0 | 0 | 0 |
| THIDVIHYR | MCM4; MCM4; | P33991; A0A | MCM4_HUMAN | Unmodified | 0 | 0 | 73527 | 99642 | 0 | 0 | 0 | 38744 | 0 | 0 | 0 | 466510 | 0 | 0 | 0 | 0 | 0 | 0 |
| THIETVINA | AP2A2; AP2A2; | O94973; A0A | AP2A2_HUMAN | Unmodified | 0 | 0 | 0 | 0 | 0 | 0 | 0 | 0 | 0 | 0 | 0 | 1015100 | 0 | 0 | 48690 | 97357 | 0 | 0 |
| THIETVINA | AP2A2; AP2A2; | O94973; A0A | AP2A2_HUMAN | Unmodified | 0 | 0 | 0 | 0 | 0 | 0 | 0 | 0 | 295140 | 0 | 0 | 3502000 | 0 | 0 | 0 | 0 | 0 | 0 |
| THIETVINA | AP2A2; AP2A2; | O94973; A0A | AP2A2_HUMAN | Unmodified | 0 | 0 | 0 | 238030 | 0 | 0 | 0 | 0 | 1262100 | 0 | 0 | 572800 | 0 | 0 | 0 | 141480 | 0 | 0 |
| THILLFLPK | P4HB; P4HB; P4 | P07237; A0A | PDIA1_HUMAN | Unmodified | 0 | 0 | 0 | 0 | 0 | 0 | 0 | 0 | 5564100 | 4433900 | 0 | 0 | 0 | 0 | 0 | 87074 | 44101 | 0 |
| THINIVVIGI | EEF1A2; EEF1A | Q05639; Q5 | EF1A2_HUMAN | Unmodified | 78222000 | 71205000 | 58244000 | 56014000 | 1.85E+08 | 16610000 | 52240000 | 2369500 | 33969000 | 23529000 | 29624000 | 1.07E+08 | 93690000 | 46718000 | 66105000 | 82392000 | 67243000 | 1.05E+08 |
| THINIVVIGI | EEF1A2; EEF1A | Q05639; Q5 | EF1A2_HUMAN | Unmodified | 1622800 | 2193800 | 0 | 1403100 | 60901000 | 994260 | 87127000 | 264910 | 0 | 9486000 | 0 | 0 | 37040000 | 39028000 | 0 | 138430 | 74820000 | 40449000 |
| THIQDNHD | FLNA; FLNA | P21333; Q60 | FLNA_HUMAN | Unmodified | 121680 | 249050 | 0 | 0 | 0 | 0 | 0 | 0 | 0 | 0 | 0 | 239470 | 0 | 0 | 373080 | 267630 | 0 | 0 |
| THIQDNHD | FLNA; FLNA | P21333; Q60 | FLNA_HUMAN | Unmodified | 0 | 0 | 0 | 0 | 1262400 | 0 | 952050 | 0 | 0 | 0 | 0 | 0 | 560740 | 1080100 | 0 | 0 | 631760 | 0 |
| THKPDPGT | TJP2; TJP2; TJP2 | Q9UDY2; A0 | ZO2_HUMAN T | Unmodified | 0 | 0 | 0 | 0 | 0 | 134430 | 0 | 0 | 5668900 | 412260 | 871800 | 0 | 0 | 0 | 0 | 0 | 0 | 0 |
| THLENNRF | PSMD2 | Q13200 | PSMD2_HUMA | Unmodified | 198970 | 0 | 0 | 444680 | 171290 | 0 | 0 | 0 | 468310 | 0 | 195240 | 1586400 | 269680 | 0 | 248300 | 237920 | 0 | 0 |
| THLGEALAI | VGF; VGF | O15240; Q0 | VGF_HUMAN N | Unmodified | 631140 | 1380300 | 0 | 0 | 438230 | 0 | 1809100 | 1989900 | 24145000 | 10490000 | 0 | 0 | 225190 | 0 | 0 | 0 | 166060 | 0 |
| THLGLATN | ZDHHC5; ZDHH | Q9C0B5; H0 | ZDHC5_HUMA | Unmodified | 0 | 0 | 0 | 1754200 | 0 | 0 | 40840 | 0 | 0 | 0 | 0 | 0 | 1334200 | 0 | 3017400 | 1721300 | 0 | 1872900 |
| THLMSESE | CKS1B; CKS1B; | P61024; Q9E | CKS1_HUMAN | Unmodified | 227280 | 236800 | 0 | 105020 | 0 | 0 | 0 | 0 | 547980 | 0 | 0 | 110650 | 0 | 0 | 0 | 0 | 0 | 0 |
| THLPGFVE | PRDX5 | P30044 | PRDX5_HUMAN | Unmodified | 1617900 | 108960 | 1659200 | 2061300 | 22923000 | 3218200 | 5535200 | 1733100 | 32087000 | 16980000 | 3972700 | 20496000 | 24824000 | 15728000 | 3522000 | 3855600 | 18938000 | 20629000 |
| THLPGFVE | PRDX5 | P30044 | PRDX5_HUMAN | Unmodified | 0 | 0 | 0 | 0 | 0 | 0 | 1516800 | 238910 | 0 | 286450 | 0 | 0 | 847230 | 867080 | 0 | 0 | 4608200 | 0 |
| THLPTFSQ | ENPP1 | P22413 | ENPP1_HUMAN | Unmodified | 7869200 | 262480 | 487160 | 0 | 0 | 0 | 0 | 0 | 543570 | 398800 | 0 | 0 | 254530 | 244060 | 6284900 | 1226000 | 2490000 | 3385100 |
| THLSEVQA | LNPEP | Q9UIQ6 | LCAP_HUMAN | Unmodified | 125230 | 0 | 251230 | 196720 | 0 | 0 | 0 | 0 | 0 | 315150 | 106200 | 634860 | 0 | 0 | 1357800 | 648190 | 0 | 0 |
| THLSLSHN | MTHFD1; MTHF | P11586; V9G | C1TC_HUMAN | Unmodified | 0 | 4322200 | 871810 | 1030000 | 5413400 | 977350 | 5366400 | 2610700 | 2872200 | 1692800 | 1111700 | 16956000 | 3163800 | 1420800 | 93089 | 932890 | 4019100 | 4960400 |
| THLSLSHN | MTHFD1; MTHF | P11586; V9G | C1TC_HUMAN | Unmodified | 0 | 0 | 0 | 0 | 916500 | 0 | 670170 | 0 | 0 | 0 | 0 | 0 | 0 | 0 | 0 | 0 | 1748800 | 1344800 |
| THMADENI | FNBP1L; FNBP1 | Q5T0N5; A0A | FBP1L_HUMAN | Oxidation (M) | 1105800 | 912600 | 0 | 0 | 0 | 0 | 0 | 0 | 0 | 0 | 0 | 0 | 0 | 0 | 240150 | 0 | 0 | 0 |
| THMADENI | FNBP1L; FNBP1 | Q5T0N5; A0A | FBP1L_HUMAN | Unmodified | 0 | 575680 | 0 | 0 | 0 | 0 | 0 | 0 | 0 | 0 | 0 | 0 | 0 | 86372 | 0 | 0 | 0 | 0 |
| THMQDLKI | SEPTIN7; SEPTI | Q16181; E7E | SEPT7_HUMAN | Unmodified | 0 | 0 | 0 | 0 | 1400600 | 307610 | 0 | 0 | 0 | 229770 | 0 | 0 | 0 | 0 | 0 | 0 | 0 | 638180 |
| THMQNIKD | SEPTIN9 | Q9UHD8 | SEPT9_HUMAN | Unmodified | 0 | 0 | 0 | 0 | 256470 | 0 | 0 | 0 | 0 | 0 | 0 | 0 | 0 | 0 | 0 | 0 | 369930 | 162390 |
| THMTHHA | HLA-A; HLA-A; H | Q9GJ45; Q53 | Q9GJ45_HUMA | Oxidation (M) | 233550 | 0 | 98119 | 70308 | 2078600 | 856160 | 0 | 0 | 1980400 | 425010 | 3227100 | 13919000 | 0 | 0 | 238510 | 219200 | 0 | 0 |
| THMTHHA | HLA-A; HLA-A; H | Q9GJ45; Q53 | Q9GJ45_HUMA | Unmodified | 0 | 0 | 0 | 0 | 1873400 | 6531700 | 0 | 0 | 53856000 | 186280 | 7140800 | 1.65E+08 | 0 | 0 | 0 | 0 | 0 | 0 |
| THMTHHPI | HLA-A; HLA-H; | P04439; P01 | HLAA_HUMAN | Unmodified | 0 | 0 | 0 | 0 | 0 | 0 | 0 | 0 | 40636000 | 790870 | 754240 | 25144000 | 0 | 0 | 0 | 0 | 0 | 10088000 |
| THMTHHPI | HLA-A; HLA-H; | P04439; P01 | HLAA_HUMAN | Oxidation (M) | 0 | 0 | 0 | 0 | 0 | 0 | 0 | 0 | 0 | 0 | 2163900 | 0 | 0 | 0 | 0 | 0 | 0 | 0 |
| THMVVANT | EPRS1 | P07814 | SYEP_HUMAN | Unmodified | 0 | 0 | 0 | 0 | 1433500 | 0 | 0 | 0 | 1259700 | 456880 | 0 | 20141000 | 871280 | 0 | 0 | 174370 | 0 | 0 |
| THMVVANT | EPRS1 | P07814 | SYEP_HUMAN | Oxidation (M) | 0 | 0 | 71350 | 66429 | 75957 | 33090 | 44045 | 44930 | 0 | 73024 | 286600 | 5325700 | 1306600 | 102300 | 60836 | 0 | 126340 | 77522 |
| THNDIIHNI | EZR; EZR | P15311; E7E | EZRI_HUMAN E | Unmodified | 106840 | 130520 | 12159 | 266850 | 376090 | 70136 | 27427 | 0 | 1666400 | 82918 | 1504400 | 759360 | 0 | 0 | 0 | 27648 | 0 | 0 |
| THNGESVS | PRPS2; PRPS1; | P11908; P60 | PRPS2_HUMAN | Unmodified | 0 | 0 | 0 | 0 | 0 | 0 | 273960 | 0 | 2448100 | 659520 | 777150 | 5829700 | 0 | 0 | 0 | 1040700 | 0 | 0 |
| THNSSLEY | DPYSL2; DPYSL | Q16555; A0A | DPYL2_HUMAN | Unmodified | 0 | 0 | 0 | 0 | 644990 | 247910 | 0 | 0 | 1473100 | 0 | 0 | 1679300 | 0 | 0 | 0 | 0 | 0 | 75227 |
| THNSSLEY | DPYSL2; DPYSL | Q16555; A0A | DPYL2_HUMAN | Oxidation (M) | 0 | 161150 | 0 | 0 | 0 | 0 | 0 | 0 | 378590 | 0 | 0 | 3551400 | 0 | 0 | 0 | 0 | 0 | 0 |
| THNSSLEY | DPYSL2; DPYSL | Q16555; A0A | DPYL2_HUMAN | Unmodified | 0 | 0 | 0 | 0 | 7762500 | 1056600 | 0 | 0 | 0 | 0 | 0 | 0 | 0 | 0 | 0 | 0 | 346860 | 0 |
| THPEVCGR | TRAFD1; TRAFD | O14545; F8V | TRAD1_HUMAN | Unmodified | 886110 | 409080 | 107220 | 0 | 0 | 12334 | 0 | 0 | 0 | 0 | 8132 | 0 | 349640 | 218410 | 41816 | 0 | 59512 | 166840 |
| THPHFVIPY | APP; APP; APP; | P05067; H7C | A4_HUMAN Am | Unmodified | 0 | 0 | 0 | 0 | 79495 | 466760 | 0 | 0 | 3488600 | 725690 | 0 | 0 | 0 | 0 | 0 | 0 | 0 | 382990 |
| THPQTALLI | MATN2 | O00339 | MATN2_HUMAI | Unmodified | 0 | 0 | 0 | 0 | 0 | 0 | 0 | 0 | 484260 | 0 | 0 | 0 | 0 | 0 | 0 | 0 | 0 | 0 |
| THPSGRDF | VIL1; VIL1 | P09327; B4D | VILI_HUMAN Vi | Unmodified | 172610 | 107890 | 0 | 242700 | 0 | 0 | 0 | 61106 | 2616500 | 862230 | 4452400 | 13869000 | 0 | 0 | 0 | 0 | 0 | 0 |
| THPSIFGIF | PPM1L | Q5SGD2 | PPM1L_HUMAI | Unmodified | 0 | 0 | 0 | 0 | 0 | 0 | 0 | 0 | 2286800 | 0 | 0 | 0 | 0 | 0 | 0 | 0 | 0 | 0 |
| THPSVVPG | NSF; NSF; NSF | P46459; I3L0 | NSF_HUMAN V | Unmodified | 0 | 300120 | 412330 | 985320 | 1189400 | 312210 | 827660 | 356860 | 810190 | 314090 | 392360 | 578580 | 6497600 | 3233900 | 0 | 1044900 | 2337600 | 2380400 |
| THPVEIFYT | DHX15 | O43143 | DHX15_HUMAI | Unmodified | 0 | 0 | 0 | 0 | 0 | 0 | 0 | 0 | 0 | 0 | 0 | 3866800 | 0 | 0 | 0 | 0 | 0 | 0 |
| THQHIPYRI | CSNK1D; CSNK | P48730; P49 | KC1D_HUMAN | Unmodified | 215880 | 156380 | 165200 | 433020 | 0 | 0 | 0 | 0 | 0 | 0 | 0 | 231380 | 0 | 0 | 240070 | 214870 | 0 | 0 |
| THRGLSPA | EPB41L2; EPB4 | O43491; A0A | E41L2_HUMAN | Unmodified | 0 | 0 | 0 | 127880 | 26963000 | 4574300 | 985680 | 2235000 | 0 | 0 | 0 | 0 | 1470700 | 1750700 | 0 | 0 | 5943400 | 10270000 |
| THRGLSPA | EPB41L2; EPB4 | O43491; A0A | E41L2_HUMAN | Unmodified | 0 | 0 | 0 | 0 | 600930 | 0 | 46249 | 0 | 0 | 0 | 0 | 0 | 0 | 0 | 0 | 0 | 435670 | 0 |
| THSAVTLGI | RASAL1 | O95294 | RASL1_HUMAN | Unmodified | 0 | 0 | 0 | 0 | 0 | 0 | 0 | 0 | 3474300 | 0 | 0 | 0 | 0 | 0 | 0 | 0 | 0 | 0 |
| THSDQFLV | ATP5F1C | P36542 | ATPG_HUMAN | Unmodified | 0 | 0 | 0 | 0 | 0 | 0 | 0 | 0 | 8975600 | 841950 | 0 | 0 | 0 | 0 | 0 | 0 | 0 | 0 |
| THSDSKPY | GANAB; GANAE | Q14697; F5H | GANAB_HUMA | 2 Oxidation (M) | 0 | 0 | 0 | 0 | 0 | 0 | 0 | 0 | 15665000 | 1023500 | 0 | 406270 | 0 | 0 | 0 | 0 | 0 | 0 |
| THSDSKPY | GANAB; GANAE | Q14697; F5H | GANAB_HUMA | Oxidation (M) | 0 | 0 | 0 | 0 | 0 | 0 | 0 | 0 | 86662000 | 0 | 0 | 0 | 0 | 0 | 0 | 0 | 0 | 0 |
| THSDSKPY | GANAB; GANAE | Q14697; F5H | GANAB_HUMA | Unmodified | 0 | 0 | 0 | 0 | 0 | 0 | 0 | 0 | 26904000 | 0 | 0 | 0 | 0 | 0 | 0 | 0 | 0 | 0 |
| THSEASGM | HPN; HPN | P05981; M0 | HEPS_HUMAN | Oxidation (M) | 0 | 0 | 210070 | 0 | 0 | 143840 | 0 | 0 | 0 | 0 | 0 | 0 | 0 | 0 | 467810 | 268580 | 0 | 0 |
| THSPDAPD | CHD6 | Q8TD26 | CHD6_HUMAN | Oxidation (M) | 0 | 0 | 0 | 0 | 0 | 3266700 | 368970 | 0 | 0 | 0 | 0 | 0 | 0 | 0 | 0 | 0 | 0 | 0 |
| THSPSSKD | PTPRF; PTPRF; | P10586; H0Y | PTPRF_HUMAN | Unmodified | 284350 | 0 | 189820 | 376140 | 2027000 | 752590 | 0 | 0 | 1086200 | 0 | 217000 | 174280 | 0 | 467560 | 687540 | 854590 | 0 | 0 |
| THSQLLIID | STXBP3 | O00186 | STXB3_HUMAN | Unmodified | 0 | 0 | 0 | 0 | 0 | 0 | 0 | 0 | 0 | 0 | 0 | 1721800 | 0 | 0 | 253300 | 312380 | 16796 | 0 |
| THSSPALS | PSD2 | Q9BQI7 | PSD2_HUMAN | Unmodified | 0 | 0 | 0 | 0 | 0 | 0 | 0 | 0 | 0 | 0 | 220650 | 0 | 232240 | 0 | 0 | 0 | 0 | 0 |
| THSTATMP | UBAP1; UBAP1 | Q9NZ09; A0A | UBAP1_HUMAI | Oxidation (M) | 0 | 0 | 0 | 0 | 8719700 | 3025400 | 0 | 0 | 0 | 0 | 0 | 0 | 0 | 8148500 | 0 | 0 | 0 | 0 |
| THSTSSSL | NDRG3; NDRG3 | Q9UGV2; F8 | NDRG3_HUMA | Unmodified | 388520 | 69868 | 129980 | 183330 | 529020 | 138430 | 0 | 0 | 243250 | 0 | 1416400 | 8028900 | 525440 | 159230 | 322380 | 349680 | 57706 | 140930 |
| THSVNGITI | CSDE1; CSDE1 | O75534; E9F | CSDE1_HUMAI | Unmodified | 0 | 183460 | 0 | 303260 | 0 | 0 | 0 | 0 | 0 | 0 | 0 | 2299400 | 1066100 | 652200 | 191160 | 0 | 0 | 0 |
| THTIQGTIK | KIF20A; KIF20A | O95235; D6I | KI20A_HUMAN | Unmodified | 0 | 0 | 0 | 0 | 770350 | 0 | 0 | 0 | 0 | 0 | 0 | 0 | 0 | 0 | 0 | 0 | 212550 | 0 |
| THTMTGSP | KIF23; KIF23; KI | Q02241; A0A | KIF23_HUMAN | Unmodified | 0 | 0 | 0 | 1018200 | 11134000 | 1930400 | 0 | 0 | 3135400 | 0 | 967250 | 2808700 | 257680 | 0 | 1121800 | 0 | 753590 | 0 |
| THTMTGSP | KIF23; KIF23; KI | Q02241; A0A | KIF23_HUMAN | Oxidation (M) | 0 | 0 | 979540 | 960830 | 0 | 1295200 | 0 | 0 | 1414600 | 631920 | 674610 | 2290600 | 0 | 0 | 0 | 0 | 0 | 0 |
| THTMTGSP | KIF23; KIF23; KI | Q02241; A0A | KIF23_HUMAN | 2 Oxidation (M) | 0 | 0 | 0 | 0 | 2497500 | 0 | 0 | 0 | 0 | 552050 | 0 | 0 | 0 | 0 | 0 | 0 | 0 | 0 |
| THTQDAVP | FH | P07954 | FUMH_HUMAN | Unmodified | 0 | 0 | 0 | 0 | 0 | 602340 | 0 | 0 | 1112300 | 3253700 | 590580 | 11877000 | 950150 | 0 | 0 | 616550 | 0 | 0 |
| THTSQAIL | CFAP36; CFAP3 | Q96G28; B5I | CFA36_HUMAN | Unmodified | 300050 | 0 | 0 | 0 | 147290 | 0 | 219380 | 0 | 0 | 0 | 0 | 0 | 126180 | 71550 | 0 | 0 | 148490 | 112610 |
| THTVGSVA | VIL1; VIL1 | P09327; B4D | VILI_HUMAN Vi | Unmodified | 0 | 0 | 0 | 0 | 0 | 0 | 0 | 0 | 252570 | 127680 | 80407 | 431960 | 0 | 0 | 0 | 0 | 0 | 0 |
| THVADFAP | EPRS1 | P07814 | SYEP_HUMAN | Unmodified | 220100 | 191530 | 538030 | 523980 | 0 | 0 | 0 | 0 | 0 | 0 | 805930 | 6301700 | 477580 | 0 | 1289100 | 520360 | 0 | 0 |
| THVTHHPI | HLA-B; HLA-B; | D5H3J5; A0A | D5H3J5_HUMA | Unmodified | 0 | 0 | 0 | 0 | 0 | 0 | 22760 | 0 | 5474100 | 158550 | 0 | 0 | 0 | 0 | 0 | 0 | 0 | 0 |
| THVTHHPL | HLA-C; HLA-C; | P10321; A0A | HLAC_HUMAN | Unmodified | 0 | 24231 | 3671400 | 0 | 2280000 | 1069200 | 108940 | 0 | 44953000 | 1118200 | 26702000 | 1.92E+08 | 0 | 0 | 0 | 91763 | 0 | 57869 |
| THVTHHPV | HLA-C; HLA-C; | Q5QT33; A0A | Q5QT33_HUMA | Unmodified | 0 | 0 | 0 | 0 | 0 | 0 | 0 | 0 | 11130000 | 0 | 7986900 | 4104900 | 0 | 0 | 0 | 0 | 0 | 186420 |
| THWFHRNI | BROX; BROX; B | Q5VW32; F5 | BROX_HUMAN | Acetyl (Protein N | 1765200 | 2545900 | 1488800 | 0 | 4122800 | 922120 | 589630 | 0 | 4980800 | 2358000 | 608680 | 2887400 | 0 | 0 | 1877200 | 1884900 | 571930 | 625450 |
| THYILIQDG | CEMIP2; CEMIP | Q9UHN6; F5 | CEIP2_HUMAN | Unmodified | 963010 | 0 | 1031400 | 823980 | 0 | 0 | 0 | 0 | 10671000 | 1054000 | 0 | 0 | 0 | 0 | 1522000 | 1463000 | 0 | 0 |
| THYIVGYNI | RPN1; RPN1 | P04843; B7Z | RPN1_HUMAN | Unmodified | 0 | 0 | 0 | 0 | 0 | 0 | 0 | 0 | 3681200 | 2475600 | 0 | 0 | 0 | 0 | 0 | 0 | 0 | 0 |
| THYPAQQG | BAG3; BAG3 | O95817; C9J | BAG3_HUMAN | Unmodified | 0 | 0 | 0 | 164490 | 1844300 | 228570 | 0 | 0 | 245510 | 53564 | 0 | 0 | 1322200 | 1430900 | 0 | 0 | 1091800 | 2021600 |
| THYSNIEAI | CAPNS1; CAPN | P04632; A0A | CPNS1_HUMAI | Unmodified | 444710 | 361680 | 0 | 0 | 0 | 0 | 0 | 0 | 1339600 | 0 | 0 | 3781000 | 0 | 0 | 939320 | 698150 | 0 | 0 |
| TIAALLPCLI | ECPAS; ECPAS | Q5VYK3; J3K | ECM29_HUMAI | Unmodified | 0 | 0 | 0 | 0 | 0 | 0 | 0 | 0 | 0 | 0 | 73384 | 541330 | 0 | 0 | 0 | 0 | 0 | 0 |
| TIADHCPD | CTNNA1; CTNN | P35221; G3X | CTNA1_HUMAI | Unmodified | 897190 | 0 | 0 | 1834200 | 119530 | 1063700 | 0 | 0 | 0 | 0 | 0 | 0 | 1200900 | 4975500 | 0 | 0 | 0 | 0 |
| TIADHCPD | CTNNA1; CTNN | P35221; G3X | CTNA1_HUMAI | Unmodified | 0 | 0 | 0 | 0 | 0 | 0 | 0 | 0 | 0 | 0 | 0 | 0 | 0 | 5355700 | 0 | 0 | 0 | 0 |
| TIADIIRTCL | CCT3; CCT3; C | P49368; Q5S | TCPG_HUMAN | Unmodified | 0 | 0 | 53851 | 0 | 5568700 | 1252500 | 8020400 | 577200 | 0 | 4181300 | 55103 | 0 | 11080000 | 8952400 | 0 | 0 | 5633700 | 5833700 |
| TIAECLADE | RPS5; RPS5; RP | P46782; M0F | RS5_HUMAN 4 | Unmodified | 1149900 | 1028500 | 308720 | 1108000 | 863270 | 258950 | 1373500 | 75450 | 1010400 | 884820 | 2290800 | 7536300 | 1126500 | 517610 | 1088900 | 647030 | 340290 | 692470 |
| TIAEIFGNP | PSMD5 | Q16401 | PSMD5_HUMA | Unmodified | 0 | 0 | 77896 | 94421 | 610260 | 0 | 0 | 14135 | 0 | 167360 | 505400 | 1764100 | 0 | 0 | 0 | 0 | 0 | 0 |
| TIAEIFGNP | PSMD5 | Q16401 | PSMD5_HUMA | Unmodified | 13367 | 0 | 0 | 0 | 0 | 408040 | 475360 | 0 | 0 | 499030 | 0 | 626620 | 0 | 0 | 0 | 0 | 278560 | 0 |
| TIAEVEGAG | ARRDC1; ARRD | Q8N5I2; Q5T | ARRD1_HUMAI | Unmodified | 7561300 | 7919300 | 7179300 | 9189500 | 2967200 | 511130 | 258110 | 293290 | 6917600 | 2006200 | 2269500 | 6418700 | 2950000 | 1674900 | 14080000 | 11896000 | 1397300 | 1080800 |
| TIAIIAEGIPE | ACLY | P53396 | ACLY_HUMAN . | Unmodified | 0 | 0 | 0 | 241420 | 261930 | 145380 | 469850 | 0 | 797500 | 329790 | 787480 | 4527500 | 144480 | 120800 | 0 | 250890 | 0 | 0 |
| TIAIIAEGIPE | ACLY | P53396 | ACLY_HUMAN . | Unmodified | 0 | 0 | 0 | 585060 | 2913600 | 778140 | 1741900 | 0 | 1750900 | 988740 | 1826700 | 18178000 | 0 | 0 | 861050 | 665210 | 347100 | 0 |
| TIAMDGTE | ATP5F1B; ATP5I | P06576; H0Y | ATPB_HUMAN . | Oxidation (M) | 0 | 0 | 0 | 0 | 0 | 0 | 0 | 0 | 5467800 | 0 | 133380 | 0 | 0 | 0 | 0 | 0 | 0 | 0 |
| TIAMDGTE | ATP5F1B; ATP5I | P06576; H0Y | ATPB_HUMAN . | Unmodified | 99608 | 118290 | 0 | 0 | 0 | 0 | 0 | 0 | 18893000 | 856730 | 0 | 0 | 0 | 1157900 | 310390 | 0 | 0 | 0 |
| TIAMDGTE | ATP5F1B; ATP5I | P06576; H0Y | ATPB_HUMAN . | Oxidation (M) | 0 | 0 | 0 | 0 | 0 | 650800 | 0 | 0 | 2672200 | 2709800 | 0 | 1175700 | 0 | 0 | 0 | 0 | 0 | 0 |
| TIAMDGTE | ATP5F1B; ATP5I | P06576; H0Y | ATPB_HUMAN . | Unmodified | 0 | 0 | 0 | 0 | 0 | 0 | 413980 | 243210 | 9581200 | 3718100 | 0 | 0 | 0 | 0 | 0 | 0 | 0 | 0 |
| TIAPALVSK | ENO1; ENO1; E | P06733; K7E | ENOA_HUMAN | Unmodified | 351090 | 0 | 159900 | 18769000 | 5.7E+08 | 77194000 | 24386 | 35121000 | 1.12E+08 | 1.4E+08 | 74805000 | 2.24E+08 | 0 | 0 | 212330 | 18756000 | 3.35E+08 | 382740 |
| TIAPALVSK | ENO1; ENO1; E | P06733; A0A | ENOA_HUMAN | Unmodified | 0 | 0 | 0 | 0 | 0 | 0 | 116910 | 113070 | 0 | 0 | 0 | 0 | 0 | 0 | 0 | 0 | 1939800 | 115060 |
| TIAQAVYG | MTHFD1L; MTH | Q6UB35; A0 | C1TM_HUMAN | Unmodified | 171530 | 0 | 0 | 0 | 338370 | 0 | 0 | 121180 | 657290 | 0 | 325760 | 805590 | 0 | 0 | 0 | 0 | 16779 | 0 |
| TIAQGNLSI | UQCRC2; UQC | P22695; H3E | QCR2_HUMAN | Unmodified | 0 | 0 | 0 | 0 | 785870 | 0 | 1198600 | 467280 | 38591000 | 17235000 | 1313600 | 375220 | 0 | 0 | 0 | 60264 | 20369 | 455370 |
| TIAQGNLSI | UQCRC2 | P22695 | QCR2_HUMAN | Unmodified | 0 | 0 | 0 | 0 | 0 | 0 | 378920 | 0 | 0 | 403390 | 0 | 0 | 0 | 0 | 0 | 0 | 0 | 0 |
| TIAQVLVHL | NAPG | Q99747 | SNAG_HUMAN | Unmodified | 211200 | 285740 | 923000 | 1139000 | 5976500 | 447530 | 197680 | 0 | 740710 | 681990 | 1027500 | 1969000 | 1506800 | 697640 | 668350 | 972000 | 1677900 | 3759000 |
| TIATALEYV | BLOC1S1; BLO | P78537; G8J | BL1S1_HUMAN | Unmodified | 0 | 0 | 0 | 0 | 0 | 0 | 0 | 0 | 0 | 0 | 0 | 1061300 | 0 | 0 | 0 | 0 | 0 | 0 |
| TIAVDFASE | HSD17B12; HS | Q53GQ0; E9 | DHB12_HUMAI | Unmodified | 0 | 0 | 0 | 0 | 0 | 0 | 0 | 0 | 1991200 | 0 | 0 | 1057100 | 0 | 0 | 0 | 0 | 0 | 0 |
| TIAVLLDDI | CCDC126; CCD | Q96EE4; C9J | CC126_HUMAI | Unmodified | 0 | 0 | 0 | 0 | 0 | 0 | 0 | 0 | 597060 | 246860 | 8613.4 | 0 | 0 | 0 | 0 | 0 | 0 | 0 |
| TICAILENY | SARS1; SARS1 | P49591; Q5T | SYSC_HUMAN | Unmodified | 0 | 0 | 0 | 0 | 833190 | 0 | 141340 | 74933 | 1193600 | 290060 | 168610 | 2184800 | 0 | 0 | 0 | 330190 | 58141 | 292710 |
| TICSHVQN | RPL9; RPL9; RP | P32969; A0A | RL9_HUMAN 6 | Unmodified | 3001400 | 1999500 | 574260 | 541450 | 223880 | 0 | 88597 | 797000 | 0 | 0 | 967960 | 27969000 | 0 | 0 | 1813200 | 1148300 | 65735 | 250660 |
| TICSHVQN | RPL9; RPL9; RP | P32969; A0A | RL9_HUMAN 6 | Oxidation (M) | 847980 | 0 | 0 | 0 | 44644 | 20440 | 693570 | 0 | 0 | 130470 | 275130 | 4529100 | 0 | 17640 | 0 | 328120 | 176090 | 22213 |
| TIDDLEDK | TPM3; TPM3 | J3KN67; A0A | J3KN67_HUMA | Unmodified | 440040 | 317890 | 1328200 | 1008500 | 22857000 | 1790900 | 485350 | 256040 | 2673700 | 380440 | 3351600 | 8934600 | 5276900 | 2364100 | 4483600 | 1475700 | 1535200 | 1135800 |
| TIDDLEEK | TPM4; TPM4; TF | P67936; A0A | TPM4_HUMAN | Unmodified | 0 | 0 | 0 | 0 | 16657000 | 4567000 | 262310 | 0 | 39870 | 0 | 517090 | 176290 | 5137200 | 0 | 0 | 0 | 809620 | 0 |
| TIDDLEEKL | TPM4; TPM4; TF | P67936; A0A | TPM4_HUMAN | Unmodified | 0 | 0 | 0 | 0 | 888790 | 0 | 0 | 0 | 0 | 0 | 0 | 0 | 153860 | 0 | 0 | 0 | 0 | 0 |
| TIDDRIVHE | MIX23; MIX23; | Q4VC31; C9 | MIX23_HUMAN | Unmodified | 0 | 0 | 0 | 0 | 0 | 0 | 0 | 0 | 0 | 0 | 1436300 | 876970 | 0 | 0 | 0 | 0 | 0 | 0 |
| TIDELDAEV | N/A | N/A | N/A | Oxidation (M) | 0 | 273010 | 0 | 0 | 0 | 0 | 0 | 0 | 0 | 0 | 0 | 0 | 0 | 0 | 0 | 0 | 0 | 0 |
| TIDEMEREA | ABCA12 | Q86UK0 | ABCAC_HUMAI | Unmodified | 0 | 0 | 0 | 0 | 0 | 0 | 0 | 0 | 233590 | 151960 | 0 | 0 | 0 | 0 | 0 | 0 | 0 | 0 |

| | | | | | | | | | | | | | | | | | | | | | | |
|---|---|---|---|---|---|---|---|---|---|---|---|---|---|---|---|---|---|---|---|---|---|---|
| TIDGILLLIE | CUL4B; CUL4A; | Q13620; Q1 | CUL4B_HUMAI | Unmodified | 0 | 0 | 0 | 0 | 0 | 0 | 0 | 0 | 0 | 0 | 0 | 123720 | 0 | 0 | 0 | 0 | 0 | 0 |
| TIDGILLLIE | CUL4B; CUL4A; | Q13620; Q1 | CUL4B_HUMAI | Unmodified | 0 | 0 | 19272 | 46371 | 0 | 0 | 0 | 0 | 0 | 0 | 0 | 1369200 | 0 | 0 | 0 | 0 | 0 | 0 |
| TIDGQQTII/ | CAPZA1 | P52907 | CAZA1_HUMAN | Unmodified | 0 | 0 | 0 | 0 | 2725100 | 0 | 1289300 | 252830 | 0 | 420810 | 0 | 731250 | 3807600 | 3581200 | 0 | 0 | 835260 | 1521300 |
| TIDLCNNPI | UGGT1; UGGT1 | Q9NYU2; H7 | UGGG1_HUMA | Oxidation (M) | 0 | 0 | 0 | 0 | 0 | 0 | 0 | 0 | 655920 | 0 | 0 | 180980 | 0 | 0 | 0 | 0 | 0 | 0 |
| TIDLCNNPI | UGGT1; UGGT1 | Q9NYU2; H7 | UGGG1_HUMA | Unmodified | 0 | 0 | 0 | 0 | 0 | 160210 | 0 | 0 | 3410000 | 0 | 169080 | 0 | 0 | 0 | 0 | 0 | 0 | 0 |
| TIDLCNNPI | UGGT1; UGGT1 | Q9NYU2; H7 | UGGG1_HUMA | Unmodified | 0 | 0 | 0 | 0 | 0 | 0 | 0 | 0 | 1121800 | 0 | 0 | 0 | 0 | 0 | 0 | 0 | 0 | 0 |
| TIDLGAAAH | CLINT1 | Q14677 | EPN4_HUMAN | Unmodified | 253160 | 211040 | 572310 | 621090 | 0 | 0 | 0 | 77687 | 1272000 | 0 | 1692600 | 6275800 | 568560 | 333460 | 334200 | 537260 | 0 | 0 |
| TIDLSNNKI | LRRC57; N/A; L | Q8N9N7; A0 | LRC57_HUMAN | Unmodified | 0 | 1457600 | 407180 | 1730900 | 1302500 | 0 | 0 | 0 | 0 | 0 | 1425400 | 3980300 | 0 | 0 | 1349100 | 1432900 | 0 | 0 |
| TIDLSNNKI | LRRC57; N/A; L | Q8N9N7; A0 | LRC57_HUMAN | Unmodified | 0 | 0 | 0 | 347230 | 5477200 | 0 | 0 | 0 | 0 | 1130200 | 0 | 0 | 1608800 | 1466000 | 0 | 0 | 2232700 | 3665500 |
| TIDNADSS1 | NUP43 | Q8NFH3 | NUP43_HUMAI | Unmodified | 0 | 94513 | 38134 | 0 | 0 | 0 | 0 | 0 | 551030 | 0 | 0 | 1224400 | 0 | 0 | 0 | 32236 | 0 | 0 |
| TIDPELLGK | GNS; GNS; GN | P15586; H7( | GNS_HUMAN N | Unmodified | 243850 | 130320 | 363030 | 804330 | 2200400 | 408170 | 463900 | 253980 | 15816000 | 7283800 | 1485400 | 1950500 | 3266000 | 1108200 | 384330 | 248190 | 0 | 1203200 |
| TIDSFTQN( | KIDINS220; KID | Q9ULH0; E9 | KDIS_HUMAN I | Unmodified | 0 | 0 | 157670 | 0 | 0 | 0 | 0 | 0 | 41038 | 0 | 0 | 486870 | 0 | 0 | 0 | 0 | 0 | 0 |
| TIDWVAFAI | ATP5PD; ATP5P | O75947; F5I | ATP5H_HUMAN | Unmodified | 0 | 0 | 0 | 0 | 0 | 0 | 1292600 | 0 | 3998000 | 2215900 | 65348 | 0 | 571240 | 0 | 0 | 0 | 225420 | 0 |
| TIEAEAAHG | IDH2; IDH2 | P48735; H0Y | IDHP_HUMAN | Unmodified | 0 | 0 | 0 | 0 | 0 | 0 | 0 | 0 | 20430000 | 493530 | 816220 | 426120 | 0 | 0 | 0 | 0 | 0 | 0 |
| TIEDAIAVLS | CCDC124 | Q96CT7 | CC124_HUMAI | Unmodified | 0 | 0 | 0 | 0 | 0 | 0 | 0 | 0 | 557330 | 0 | 420450 | 0 | 0 | 0 | 0 | 0 | 0 | 0 |
| TIEDEDLKF | PPM1H | Q9ULR3 | PPM1H_HUMA | Unmodified | 0 | 0 | 0 | 0 | 0 | 0 | 0 | 0 | 655030 | 331000 | 709890 | 5405800 | 0 | 0 | 0 | 0 | 292060 | 810170 |
| TIEDEDLKF | PPM1H | Q9ULR3 | PPM1H_HUMA | Unmodified | 0 | 0 | 0 | 0 | 0 | 0 | 0 | 0 | 0 | 0 | 412070 | 1468500 | 0 | 0 | 0 | 0 | 0 | 0 |
| TIEDTLMTV | ARFIP1 | P53367 | ARFP1_HUMAN | Unmodified | 0 | 0 | 0 | 0 | 0 | 0 | 845350 | 11368 | 110660 | 0 | 0 | 940520 | 382660 | 0 | 0 | 0 | 0 | 0 |
| TIEEQLDEE | AKAP2; PALM2/ | Q9Y2D5; C9 | AKAP2_HUMAN | Unmodified | 0 | 0 | 0 | 0 | 736890 | 0 | 0 | 0 | 0 | 0 | 0 | 0 | 0 | 0 | 0 | 0 | 0 | 487540 |
| TIEEQLDEE | AKAP2; PALM2/ | Q9Y2D5; C9 | AKAP2_HUMAN | Unmodified | 0 | 0 | 0 | 0 | 1733900 | 377370 | 0 | 0 | 0 | 0 | 130780 | 0 | 0 | 0 | 0 | 0 | 191280 | 146770 |
| TIEESEETLI | SMC2 | O95347 | SMC2_HUMAN | Unmodified | 16529 | 0 | 0 | 30740 | 368810 | 0 | 122550 | 71481 | 700340 | 118550 | 68324 | 988460 | 0 | 42859 | 0 | 30310 | 0 | 0 |
| TIEIKSITHD | SEPTIN9; SEPTI | Q9UHD8; K7 | SEPT9_HUMAN | Unmodified | 0 | 0 | 0 | 0 | 642920 | 0 | 726050 | 149170 | 0 | 0 | 0 | 0 | 0 | 0 | 0 | 0 | 2156300 | 712090 |
| TIEKFEK | EEF1A2; EEF1A | Q05639; Q5 | EF1A2_HUMAN | Unmodified | 23246000 | 30536000 | 0 | 21378000 | 19396000 | 5633300 | 0 | 114440 | 3658900 | 2725000 | 1219900 | 8387300 | 26190000 | 8406100 | 2783200 | 5258900 | 6782100 | 13978000 |
| TIEKFEKEA | EEF1A2; EEF1A | Q05639; Q5 | EF1A2_HUMAN | Oxidation (M) | 0 | 0 | 0 | 153290 | 2158100 | 194090 | 0 | 328670 | 0 | 311270 | 0 | 23544 | 1309300 | 1314700 | 0 | 34342 | 1532100 | 1323000 |
| TIEKFEKEA | EEF1A2; EEF1A | Q05639; Q5 | EF1A2_HUMAN | Unmodified | 0 | 102570 | 0 | 606190 | 8792100 | 1555100 | 873200 | 366730 | 0 | 1365000 | 162100 | 0 | 8352300 | 5018500 | 0 | 0 | 5600300 | 4404600 |
| TIESANQQ | SRP72; SRP72; | O76094; D6I | SRP72_HUMAN | Unmodified | 0 | 0 | 0 | 0 | 321610 | 0 | 335560 | 173290 | 0 | 85414 | 0 | 0 | 0 | 525890 | 0 | 0 | 185550 | 243090 |
| TIESILEPVA | VCL; VCL | P18206; A0A | VINC_HUMAN | Oxidation (M) | 0 | 260310 | 0 | 192550 | 1224100 | 114430 | 0 | 0 | 0 | 123050 | 0 | 0 | 872230 | 692970 | 0 | 0 | 1900100 | 0 |
| TIESKAQEC | ADAM17 | P78536 | ADA17_HUMAN | Unmodified | 0 | 0 | 0 | 0 | 0 | 0 | 0 | 0 | 0 | 0 | 0 | 0 | 0 | 41240 | 0 | 0 | 0 | 587510 |
| TIEYLQPNF | SH3GL2; SH3G | Q99962; Q9 | SH3G2_HUMAI | Unmodified | 40666000 | 37072000 | 28242000 | 36817000 | 2295400 | 395900 | 566950 | 151670 | 5509100 | 3185300 | 358660 | 1015700 | 22422000 | 4175600 | 34591000 | 35013000 | 242180 | 782810 |
| TIEYLQPNF | SH3GL2; SH3G | Q99962; Q9 | SH3G2_HUMAI | Unmodified | 0 | 0 | 0 | 450350 | 3793200 | 927180 | 2218800 | 2419900 | 0 | 1725200 | 0 | 410420 | 19350000 | 11802000 | 387780 | 0 | 803000 | 2083100 |
| TIFAYFTGS | TXNDC17; TXN | Q9BRA2; I3L | TXD17_HUMAN | Unmodified | 2749300 | 1708300 | 3855800 | 6538200 | 25017000 | 5973700 | 2112400 | 712860 | 12047000 | 9289400 | 6391000 | 8361900 | 23577000 | 12544000 | 6365400 | 4488000 | 10524000 | 14256000 |
| TIFAYFTGS | TXNDC17; TXN | Q9BRA2; I3L | TXD17_HUMAN | Unmodified | 0 | 0 | 0 | 0 | 896020 | 0 | 670260 | 257300 | 375670 | 431370 | 0 | 163810 | 992340 | 260330 | 0 | 102290 | 4323600 | 501180 |
| TIFGDMLLT | PLCB2 | Q00722 | PLCB2_HUMAN | Oxidation (M) | 0 | 0 | 0 | 0 | 0 | 0 | 0 | 0 | 0 | 0 | 838090 | 3100100 | 0 | 0 | 0 | 10380 | 0 | 0 |
| TIFINQPQL | ATP8A1; ATP8A | Q9Y2Q0; Q4 | AT8A1_HUMAN | Unmodified | 0 | 0 | 0 | 0 | 0 | 0 | 0 | 0 | 871610 | 0 | 0 | 0 | 0 | 0 | 0 | 0 | 0 | 0 |
| TIFQGIAAK | PGD; PGD; PGD | P52209; K7E | 6PGD_HUMAN | Unmodified | 866520 | 0 | 1593100 | 2984500 | 42073000 | 6350100 | 3990200 | 2546200 | 10056000 | 6608600 | 8669200 | 35319000 | 12293000 | 4240200 | 1422900 | 1600100 | 6358000 | 7629700 |
| TIFSALEND | ACOX3 | O15254 | ACOX3_HUMAI | Unmodified | 0 | 0 | 0 | 0 | 0 | 0 | 0 | 0 | 905670 | 79251 | 0 | 0 | 0 | 0 | 0 | 0 | 0 | 0 |
| TIFVLTK | OPA1; OPA1; OI | O60313; E5I | OPA1_HUMAN | Unmodified | 0 | 0 | 0 | 0 | 330560 | 55275 | 31849 | 28843 | 403210 | 0 | 300550 | 2178700 | 94348 | 42989 | 0 | 0 | 77160 | 156970 |
| TIGAQVYE( | B4GALT5 | O43286 | B4GT5_HUMAN | Unmodified | 0 | 0 | 0 | 0 | 0 | 0 | 0 | 0 | 1658700 | 0 | 268200 | 0 | 0 | 0 | 0 | 0 | 0 | 0 |
| TIGDMFQN | FKBP10; FKBP1 | Q96AY3; H0 | FKB10_HUMAN | Unmodified | 0 | 0 | 0 | 0 | 0 | 82914 | 0 | 127360 | 0 | 298360 | 744570 | 0 | 188380 | 0 | 0 | 0 | 0 | 568720 |
| TIGDMFQN | FKBP10; FKBP1 | Q96AY3; H0 | FKB10_HUMAN | Unmodified | 0 | 0 | 0 | 0 | 0 | 0 | 0 | 0 | 0 | 0 | 0 | 657020 | 0 | 0 | 0 | 37922 | 0 | 119350 |
| TIGGGDDS | TUBA3C; TUBA | P0DPH7; Q7 | TBA3C_HUMAN | Unmodified | 6918700 | 7457600 | 14577000 | 0 | 18817000 | 11571000 | 6374900 | 2736700 | 90297000 | 9533800 | 22404000 | 1.97E+08 | 9596300 | 8347300 | 10994000 | 10064000 | 1571200 | 13654000 |
| TIGGGDDS | TUBA3C; TUBA | P0DPH7; Q7 | TBA3C_HUMAN | Unmodified | 1914600 | 0 | 0 | 6013900 | 31252000 | 9450100 | 19024000 | 1664800 | 20507000 | 21232000 | 5487000 | 4193700 | 33625000 | 5957000 | 0 | 0 | 8541100 | 25335000 |
| TIGGGDDS | TUBA1A; TUBA1 | Q71U36; Q9 | TBA1A_HUMAN | Unmodified | 0 | 0 | 0 | 0 | 0 | 0 | 2286900 | 0 | 0 | 1019400 | 0 | 0 | 1802500 | 0 | 0 | 0 | 1462600 | 2634400 |
| TIGGGDDS | TUBA4A; TUBA4 | P68366; C9J | TBA4A_HUMAN | Unmodified | 1019500 | 2343900 | 1989400 | 8952000 | 682250 | 486200 | 1271300 | 711250 | 0 | 1775900 | 247230 | 5046900 | 1666800 | 966640 | 6901600 | 2035600 | 294430 | 2385900 |
| TIGGGDDS | TUBA4A; TUBA4 | P68366; C9J | TBA4A_HUMAN | Unmodified | 0 | 282440 | 0 | 0 | 0 | 91929 | 1888100 | 123840 | 0 | 2443500 | 60097 | 0 | 2961700 | 1792400 | 0 | 0 | 0 | 0 |
| TIGISVDPR | RPL13; RPL13 | P26373; J3Q | RL13_HUMAN ( | Unmodified | 6186900 | 754580 | 8964000 | 3674400 | 2939600 | 0 | 1651500 | 881200 | 3626700 | 1030400 | 9774700 | 35053000 | 5021400 | 3898900 | 4821200 | 3647600 | 307470 | 3037500 |
| TIGLREVW | EZR; EZR | P15311; E7E | EZRI_HUMAN E | Unmodified | 0 | 250940 | 0 | 0 | 5602100 | 773850 | 0 | 0 | 317520 | 3405400 | 0 | 0 | 2105700 | 1406100 | 0 | 0 | 517300 | 648940 |
| TIGLREVW | EZR; EZR | P15311; E7E | EZRI_HUMAN E | Unmodified | 0 | 0 | 0 | 0 | 1073000 | 0 | 298130 | 0 | 0 | 476180 | 0 | 0 | 728450 | 1026000 | 0 | 0 | 286880 | 0 |
| TIGTGLVTN | TUFM | P49411 | EFTU_HUMAN | Unmodified | 0 | 169390 | 0 | 0 | 0 | 0 | 0 | 0 | 15003000 | 0 | 0 | 83156 | 0 | 0 | 0 | 0 | 0 | 0 |
| TIGVDFLER | RAB23 | Q9ULC3 | RAB23_HUMAN | Unmodified | 0 | 475820 | 922840 | 0 | 0 | 0 | 0 | 67171 | 1727000 | 700770 | 0 | 1637100 | 0 | 0 | 1073800 | 2285500 | 0 | 0 |
| TIGVMTMVI | EPRS1 | P07814 | SYEP_HUMAN | 3 Oxidation (M) | 0 | 0 | 0 | 0 | 0 | 0 | 0 | 0 | 0 | 0 | 0 | 2320300 | 0 | 0 | 0 | 0 | 0 | 0 |
| TIHLLTSLDI | MITD1; MITD1 | Q8WV92; H7 | MITD1_HUMAN | Unmodified | 2352300 | 2513400 | 589250 | 0 | 4760000 | 1192100 | 146570 | 0 | 11529000 | 1747300 | 4788600 | 15973000 | 0 | 0 | 0 | 0 | 0 | 0 |
| TIHLSSIRPI | SND1 | Q7KZF4 | SND1_HUMAN | Unmodified | 506780 | 410250 | 1398900 | 1769500 | 1197200 | 282070 | 467490 | 78140 | 520000 | 225750 | 1874500 | 14330000 | 2092000 | 1266700 | 1266600 | 1201800 | 0 | 645520 |
| TIHSEQAVF | ALCAM; ALCAM | Q13740; F5C | CD166_HUMAI | Unmodified | 0 | 1381600 | 0 | 0 | 0 | 0 | 0 | 0 | 0 | 0 | 0 | 0 | 0 | 0 | 1015600 | 0 | 0 | 0 |
| TIICHVEGF | ALCAM; ALCAM | Q13740; F5C | CD166_HUMAI | Unmodified | 286110 | 0 | 1100200 | 0 | 0 | 0 | 0 | 0 | 0 | 0 | 0 | 0 | 0 | 192130 | 543050 | 433580 | 767300 | 0 |
| TIIFVETK | DDX17; DDX17 | Q92841; A0/ | DDX17_HUMAI | Unmodified | 94195 | 302030 | 120990 | 0 | 1033500 | 102850 | 1267400 | 728500 | 0 | 372390 | 1457800 | 9810400 | 0 | 0 | 0 | 0 | 0 | 209680 |
| TIIHHETIFI | SRGAP1; SRGA | Q7Z6B7; G5I | SRGP1_HUMAI | Unmodified | 0 | 0 | 0 | 0 | 0 | 0 | 0 | 0 | 0 | 0 | 0 | 0 | 0 | 0 | 0 | 0 | 0 | 0 |
| TIINADKDG | PPP3R1; PPP3R | P63098; F6U | CANB1_HUMAI | Unmodified | 0 | 0 | 0 | 0 | 0 | 0 | 0 | 0 | 0 | 0 | 0 | 277340 | 0 | 0 | 0 | 0 | 0 | 0 |
| TIINADKDG | PPP3R1; PPP3R | P63098; F6U | CANB1_HUMAI | Unmodified | 0 | 0 | 0 | 1007400 | 0 | 0 | 0 | 0 | 0 | 861440 | 0 | 0 | 1648100 | 1271300 | 0 | 0 | 0 | 0 |
| TIIPLATAPN | LIN54 | Q6MZP7 | LIN54_HUMAN | Unmodified | 0 | 0 | 366070 | 0 | 0 | 0 | 259190 | 224090 | 0 | 0 | 0 | 0 | 162770 | 46001 | 0 | 0 | 0 | 0 |
| TIIPLISQCT | MDH2; MDH2 | P40926; G3 | MDHM_HUMAI | Unmodified | 0 | 0 | 0 | 276190 | 0 | 0 | 0 | 0 | 16976000 | 781810 | 1280200 | 0 | 0 | 0 | 0 | 0 | 0 | 0 |
| TIIPLISQCT | MDH2; MDH2 | P40926; G3 | MDHM_HUMAI | Unmodified | 0 | 0 | 0 | 0 | 235250 | 0 | 0 | 0 | 14589000 | 2471700 | 176010 | 0 | 476190 | 0 | 0 | 0 | 0 | 76265 |
| TIIPLISQCT | MDH2; MDH2 | P40926; G3 | MDHM_HUMAI | Unmodified | 0 | 0 | 0 | 0 | 0 | 0 | 0 | 0 | 0 | 931510 | 0 | 0 | 0 | 0 | 0 | 0 | 0 | 0 |
| TIIQKEPIDL | L1CAM | P32004 | L1CAM_HUMAI | Unmodified | 0 | 0 | 0 | 0 | 340020 | 0 | 0 | 0 | 1264000 | 107470 | 0 | 0 | 0 | 0 | 0 | 0 | 0 | 0 |
| TIIQNPTDQ | PA2G4; PA2G4 | Q9UQ80; F8 | PA2G4_HUMAN | Unmodified | 950240 | 607570 | 535550 | 528510 | 550180 | 0 | 118440 | 36318 | 607940 | 230660 | 2555200 | 9409800 | 576800 | 0 | 622120 | 605820 | 0 | 0 |
| TIIQNPTDQ | PA2G4; PA2G4 | Q9UQ80; F8 | PA2G4_HUMAN | Unmodified | 2099800 | 1437700 | 689580 | 1712700 | 0 | 0 | 233590 | 265260 | 1805000 | 382680 | 2119800 | 10377000 | 1822000 | 899970 | 1097900 | 0 | 0 | 801050 |
| TIIQSPYEK | SNTB1 | Q13884 | SNTB1_HUMAN | Unmodified | 0 | 0 | 0 | 0 | 1264500 | 0 | 0 | 52271 | 4111300 | 3084200 | 0 | 0 | 0 | 0 | 0 | 0 | 1113700 | 1015500 |
| TIISLDTSQI | AGPS; AGPS; A( | O00116; A0/ | ADAS_HUMAN | Oxidation (M) | 0 | 0 | 0 | 325220 | 1066100 | 0 | 464830 | 0 | 3718700 | 0 | 0 | 0 | 0 | 0 | 163850 | 59751 | 0 | 0 |
| TIISLDTSQI | AGPS; AGPS; A( | O00116; A0/ | ADAS_HUMAN | Unmodified | 0 | 0 | 0 | 0 | 0 | 0 | 0 | 0 | 8946700 | 0 | 0 | 0 | 0 | 0 | 0 | 0 | 0 | 0 |
| TIISYIDEQF | SEPTIN2; SEPTI | Q15019; C9I | SEPT2_HUMAN | Unmodified | 83669 | 0 | 126730 | 182890 | 111540 | 0 | 0 | 0 | 1369200 | 113740 | 553360 | 3796000 | 0 | 0 | 0 | 167230 | 0 | 0 |
| TIITYPWFQ | GNAQ; GNA11; | P50148; P29 | GNAQ_HUMAN | Unmodified | 508240 | 1237400 | 0 | 0 | 733580 | 0 | 573570 | 0 | 343870 | 418570 | 105120 | 1429100 | 0 | 0 | 27015 | 15343 | 0 | 193010 |
| TIIVTELSPV | CEACAM1 | P13688 | CEAM1_HUMAI | Unmodified | 0 | 0 | 0 | 0 | 0 | 0 | 0 | 0 | 0 | 0 | 429440 | 670250 | 0 | 0 | 0 | 0 | 0 | 0 |
| TIIVTELSPV | CEACAM1 | P13688 | CEAM1_HUMAI | Unmodified | 0 | 0 | 0 | 0 | 580960 | 0 | 0 | 0 | 3908200 | 1951300 | 9541100 | 19024000 | 0 | 0 | 0 | 0 | 1194000 | 2084500 |
| TIKAELDAA | ASPH; ASPH; A! | Q12797; A0/ | ASPH_HUMAN | Unmodified | 0 | 0 | 0 | 0 | 0 | 0 | 0 | 0 | 1436900 | 0 | 0 | 783730 | 0 | 0 | 0 | 0 | 181680 | 0 |
| TIKDVFHNI | SLC30A1 | Q9Y6M5 | ZNT1_HUMAN | Unmodified | 0 | 0 | 0 | 0 | 1082000 | 0 | 665140 | 0 | 0 | 0 | 0 | 0 | 2492100 | 0 | 0 | 0 | 2126800 | 0 |
| TIKLWNTL( | RACK1; RACK1; | P63244; D6I | RACK1_HUMAI | Unmodified | 0 | 0 | 0 | 0 | 0 | 0 | 145390 | 12015 | 0 | 0 | 0 | 0 | 0 | 0 | 0 | 0 | 35247 | 0 |
| TIKPDVRSY | FN1; FN1 | P02751; H0Y | FINC_HUMAN I | Unmodified | 0 | 0 | 624410 | 56768 | 134250 | 247360 | 0 | 45718 | 0 | 583130 | 0 | 0 | 186270 | 0 | 0 | 0 | 280010 | 193370 |
| TIKVECYDY | CPNE3; CPNE3 | O75131; H0 | CPNE3_HUMAI | Oxidation (M) | 0 | 0 | 0 | 2814300 | 0 | 0 | 2535900 | 346200 | 3848600 | 3130400 | 0 | 0 | 2437100 | 7175400 | 0 | 0 | 1247400 | 2389100 |
| TIKVECYDY | CPNE3; CPNE3 | O75131; H0 | CPNE3_HUMAI | Unmodified | 0 | 0 | 0 | 0 | 0 | 0 | 3198000 | 0 | 15006000 | 4650900 | 0 | 0 | 3444300 | 3001200 | 0 | 0 | 3542100 | 10202000 |
| TILEDEK | LDLR; LDLR; LD | P01130; J3K | LDLR_HUMAN | Unmodified | 0 | 0 | 0 | 0 | 0 | 0 | 0 | 0 | 0 | 717370 | 0 | 0 | 0 | 0 | 0 | 0 | 0 | 0 |
| TILERNNFT | HINT3 | Q9NQE9 | HINT3_HUMAN | Unmodified | 0 | 0 | 0 | 332710 | 449850 | 273180 | 0 | 16217 | 0 | 516920 | 0 | 0 | 799900 | 1751500 | 0 | 0 | 0 | 0 |
| TILEWEPKF | F3 | P13726 | TF_HUMAN Tis | Unmodified | 5642500 | 5088900 | 0 | 0 | 37847000 | 5123400 | 0 | 0 | 0 | 0 | 0 | 0 | 0 | 0 | 0 | 0 | 0 | 0 |
| TILGPTENII | PMPCB; PMPCE | O75439; G3 | MPPB_HUMAN | Unmodified | 0 | 0 | 400660 | 424660 | 0 | 0 | 172400 | 98153 | 1374600 | 0 | 2387600 | 0 | 0 | 0 | 637810 | 744550 | 0 | 0 |
| TILLNTK | IQGAP1; IQGAP | P46940; H0Y | IQGA1_HUMAN | Unmodified | 1223200 | 0 | 0 | 0 | 7261000 | 0 | 190410 | 19083 | 2145900 | 466690 | 42781 | 0 | 1301600 | 0 | 1238300 | 948540 | 350320 | 571080 |
| TILLSTTDP/ | AFDN; AFDN; A | P55196; Q5T | AFAD_HUMAN | Unmodified | 701040 | 1402600 | 0 | 472290 | 3564700 | 614530 | 1125700 | 0 | 675750 | 458520 | 1222800 | 0 | 0 | 3275800 | 0 | 227840 | 739200 | 362910 |
| TILLSTTDP/ | AFDN; AFDN; A | P55196; Q5T | AFAD_HUMAN | Unmodified | 0 | 0 | 0 | 0 | 745260 | 0 | 0 | 0 | 0 | 0 | 0 | 0 | 992650 | 889480 | 0 | 0 | 0 | 0 |
| TILMMGR | EEF2 | P13639 | EF2_HUMAN EI | Unmodified | 0 | 162960 | 56673 | 0 | 817030 | 0 | 34432 | 0 | 1021600 | 92430 | 1178100 | 11313000 | 0 | 37850 | 0 | 71391 | 206070 | 145420 |
| TILMMGR | EEF2 | P13639 | EF2_HUMAN EI | Oxidation (M) | 30014 | 0 | 0 | 0 | 229190 | 71793 | 0 | 0 | 0 | 101120 | 0 | 7669400 | 0 | 0 | 0 | 0 | 77003 | 114000 |
| TILNLEAKE | IRAG2; IRAG2; I | Q12912; F5I | IRAG2_HUMAN | Unmodified | 0 | 0 | 0 | 0 | 0 | 0 | 0 | 0 | 0 | 0 | 0 | 1415600 | 0 | 0 | 0 | 0 | 0 | 0 |
| TILPLINFK | PLCG2; PLCG2 | P16885; A0A | PLCG2_HUMAI | Unmodified | 0 | 0 | 0 | 0 | 0 | 0 | 0 | 38796 | 648770 | 142500 | 5350900 | 69125000 | 0 | 0 | 0 | 0 | 0 | 0 |
| TILQCALNF | ANXA7 | P20073 | ANXA7_HUMAI | Unmodified | 1963800 | 1235200 | 1334200 | 1675900 | 0 | 0 | 0 | 0 | 77698000 | 4615100 | 399290 | 6388900 | 454360 | 0 | 4377200 | 2958800 | 858380 | 547390 |
| TILSDDWK | CNTN1 | Q12860 | CNTN1_HUMA | Unmodified | 0 | 0 | 0 | 0 | 0 | 0 | 0 | 41753 | 431410 | 0 | 0 | 0 | 0 | 0 | 0 | 0 | 0 | 0 |
| TILSLMTR | ACLY | P53396 | ACLY_HUMAN | Unmodified | 0 | 0 | 77618 | 0 | 0 | 0 | 157260 | 34579 | 657960 | 98524 | 0 | 9721400 | 0 | 0 | 0 | 0 | 0 | 0 |
| TILSNQTVD | RPL9; RPL9 | P32969; D6I | RL9_HUMAN 6 | Unmodified | 1983000 | 7878800 | 0 | 3453400 | 2639700 | 705510 | 2018200 | 857920 | 837270 | 1198400 | 0 | 5831300 | 5995900 | 949800 | 0 | 1201400 | 461240 | 2099800 |
| TILTLTGVS1 | PGP; PGP | A6NDG6; H3 | PGP_HUMAN G | Unmodified | 0 | 0 | 0 | 0 | 0 | 0 | 0 | 0 | 0 | 0 | 1310200 | 1786100 | 254310 | 0 | 0 | 0 | 0 | 0 |
| TILTTSHAFI | DOCK7; DOCK7 | Q96N67; A0/ | DOCK7_HUMA | Unmodified | 0 | 0 | 0 | 0 | 2281100 | 0 | 354080 | 0 | 0 | 0 | 258040 | 2009400 | 573600 | 0 | 0 | 0 | 1209400 | 1881800 |
| TILTTSHLFI | DOCK10; DOCK | Q96BY6; H0 | DOC10_HUMA | Unmodified | 0 | 0 | 0 | 0 | 1084100 | 168000 | 12364 | 0 | 0 | 0 | 0 | 0 | 0 | 0 | 0 | 0 | 208730 | 135340 |
| TIMEQFNP! | BAIAP2; BAIAP2 | Q9UQB8; I3I | BAIP2_HUMAN | Unmodified | 0 | 0 | 0 | 0 | 833130 | 0 | 0 | 0 | 0 | 0 | 0 | 0 | 0 | 0 | 0 | 0 | 0 | 0 |
| TIMHLMINI | DNM2; DNM1; I | P50570; Q0 | DYN2_HUMAN | Unmodified | 94586 | 0 | 0 | 0 | 753710 | 169910 | 321840 | 0 | 331180 | 0 | 0 | 272510 | 441310 | 158860 | 59809 | 0 | 302690 | 387140 |
| TIMVDDSK | TLN1 | Q9Y490 | TLN1_HUMAN | Unmodified | 0 | 0 | 0 | 0 | 2612100 | 529720 | 62606 | 67117 | 524670 | 198250 | 235840 | 917950 | 88520 | 183450 | 0 | 0 | 262850 | 1281000 |
| TINEVENQI | ACTN4; ACTN1 | O43707; P12 | ACTN4_HUMAI | Unmodified | 848770 | 1323600 | 1400700 | 412430 | 2346200 | 344460 | 0 | 0 | 1179000 | 1242900 | 1962500 | 34447000 | 0 | 31383 | 3142200 | 2994600 | 593680 | 0 |
| TINEVENQI | ACTN4; ACTN1 | O43707; P12 | ACTN4_HUMAI | Unmodified | 0 | 0 | 0 | 0 | 0 | 0 | 0 | 0 | 0 | 0 | 0 | 0 | 0 | 0 | 0 | 0 | 0 | 0 |
| TINLYPLTN | NUDT21 | O43809 | CPSF5_HUMAN | Unmodified | 1278600 | 2395900 | 2179600 | 1339300 | 0 | 0 | 0 | 0 | 168420 | 19888 | 158140 | 1016200 | 49127 | 0 | 793980 | 908410 | 0 | 0 |
| TINLYPLTN | NUDT21 | O43809 | CPSF5_HUMAN | Unmodified | 13460000 | 11496000 | 377530 | 8921000 | 1527300 | 506000 | 0 | 0 | 2689500 | 1280700 | 1431800 | 2957500 | 4597400 | 826480 | 1510400 | 3391800 | 0 | 0 |
| TINMLVR | LSS; LSS | P48449; A0A | LSS_HUMAN La | Unmodified | 0 | 0 | 0 | 0 | 0 | 0 | 0 | 10156 | 0 | 48208 | 0 | 680640 | 0 | 0 | 0 | 0 | 0 | 0 |
| TINQIRQD/ | EIF4G2; EIF4G2 | P78344; H0Y | IF4G2_HUMAN | Unmodified | 207430 | 0 | 0 | 0 | 1497800 | 0 | 0 | 0 | 75663 | 36875 | 0 | 0 | 1376700 | 1227600 | 0 | 206550 | 0 | 0 |
| TINWTLLEK | PRKCD | Q05655 | KPCD_HUMAN | Unmodified | 0 | 0 | 77985 | 0 | 660600 | 0 | 0 | 0 | 587560 | 0 | 0 | 920270 | 0 | 0 | 0 | 0 | 1171200 | 1292800 |
| TIPDSDISA: | DDR1; DDR1; D | Q08345; A0/ | DDR1_HUMAN | Unmodified | 198360 | 185710 | 0 | 0 | 0 | 648580 | 0 | 82923 | 503010 | 82660 | 0 | 0 | 0 | 0 | 0 | 0 | 0 | 257080 |
| TIPEASALG | PAN2 | Q504Q3 | PAN2_HUMAN | Unmodified | 431720 | 551000 | 0 | 0 | 0 | 0 | 258210 | 0 | 0 | 0 | 0 | 79491 | 0 | 0 | 0 | 0 | 0 | 0 |

| | | | | | | | | | | | | | | | | | | | | | | |
|---|---|---|---|---|---|---|---|---|---|---|---|---|---|---|---|---|---|---|---|---|---|---|
| TIPGKQQLI | CCT7 | Q99832 | TCPH_HUMAN | Unmodified | 0 | 0 | 0 | 149330 | 1101500 | 273150 | 887690 | 541570 | 293160 | 1806000 | 0 | 266560 | 2428900 | 1491800 | 0 | 0 | 1558000 | 1327000 |
| TIPIDGNFF | ALDH1A1; ALDH | P00352; Q5S | AL1A1_HUMAN | Unmodified | 0 | 0 | 0 | 0 | 0 | 0 | 0 | 0 | 392540 | 132530 | 1047500 | 5.75E+08 | 0 | 0 | 0 | 85313 | 0 | 0 |
| TIPLYNLTG | MYO1D; MYO1D | O94832; J3Q | MYO1D_HUMA | Unmodified | 388920 | 619910 | 1070400 | 111270 | 0 | 0 | 0 | 0 | 132810 | 0 | 107670 | 314410 | 0 | 0 | 1346700 | 457910 | 69725 | 0 |
| TIPLYNLTG | MYO1D; MYO1D | O94832; J3Q | MYO1D_HUMA | Unmodified | 0 | 0 | 0 | 0 | 0 | 0 | 0 | 0 | 2382600 | 1377700 | 809750 | 0 | 0 | 0 | 0 | 0 | 0 | 0 |
| TIPSWATLS | HP1BP3; HP1BI | Q5SSJ5; B0C | HP1B3_HUMAN | Unmodified | 0 | 0 | 0 | 0 | 0 | 0 | 0 | 370980 | 442750 | 0 | 0 | 0 | 0 | 0 | 0 | 0 | 0 | 468260 |
| TIPWLEDR | ACTN4; ACTN4 | O43707; F5C | ACTN4_HUMAN | Unmodified | 1542400 | 0 | 19889 | 1349100 | 4923400 | 3266200 | 0 | 0 | 4629600 | 1799600 | 4615900 | 20052000 | 1181200 | 1086500 | 1909400 | 1697000 | 1465700 | 193340 |
| TIPWLENR | ACTN1; ACTN1 | P12814; A0A | ACTN1_HUMAN | Unmodified | 0 | 508420 | 0 | 0 | 0 | 0 | 0 | 0 | 0 | 0 | 0 | 317650 | 0 | 0 | 0 | 0 | 0 | 0 |
| TIQEMQQK | ACTN4; ACTN4 | O43707; F5C | ACTN4_HUMAN | Unmodified | 0 | 69174 | 519380 | 0 | 298920 | 185910 | 263890 | 0 | 695920 | 49994 | 51899 | 5797900 | 0 | 0 | 807770 | 536690 | 0 | 98144 |
| TIQEMQQK | ACTN4; ACTN4 | O43707; F5C | ACTN4_HUMAN | Unmodified | 133830 | 150450 | 0 | 345770 | 0 | 0 | 0 | 0 | 1449900 | 0 | 0 | 0 | 0 | 0 | 208800 | 206920 | 0 | 0 |
| TIQEMQQK | ACTN4; ACTN4 | O43707; F5C | ACTN4_HUMAN | Oxidation (M) | 0 | 0 | 0 | 126960 | 585440 | 0 | 0 | 0 | 85825 | 0 | 0 | 998700 | 0 | 0 | 92159 | 94825 | 0 | 0 |
| TIQEMQQK | ACTN4; ACTN4 | O43707; F5C | ACTN4_HUMAN | Unmodified | 157370 | 0 | 0 | 399480 | 6099900 | 7432900 | 0 | 93950 | 1844300 | 642220 | 0 | 0 | 0 | 0 | 0 | 0 | 575030 | 0 |
| TIQFVDWC | TUBA3C; TUBA | P0DPH7; Q7 | TBA3C_HUMAN | Unmodified | 104100 | 0 | 65275 | 225830 | 3116500 | 435890 | 5608500 | 1503900 | 4554700 | 4542400 | 2276700 | 21739000 | 1865100 | 1529700 | 0 | 48245 | 485090 | 3629800 |
| TIQFVDWC | TUBA3C; TUBA | P0DPH7; Q7 | TBA3C_HUMAN | Unmodified | 0 | 0 | 0 | 0 | 0 | 0 | 0 | 2005300 | 0 | 0 | 260550 | 0 | 0 | 0 | 0 | 0 | 0 | 0 |
| TIQGDEEDI | NAPA; NAPA; N | P54920; M0F | SNAA_HUMAN | Unmodified | 2424500 | 0 | 0 | 0 | 2065500 | 404580 | 2350800 | 1619700 | 3000800 | 1163200 | 0 | 2247000 | 0 | 0 | 0 | 0 | 1600700 | 2140700 |
| TIQGELGW | EPHA3; EPHA3 | P29320; C9J | EPHA3_HUMAN | Unmodified | 0 | 2620500 | 0 | 0 | 0 | 415060 | 1201400 | 0 | 0 | 0 | 0 | 0 | 0 | 852610 | 0 | 0 | 282140 | 0 |
| TIQIPEHLM | NCOA4; NCOA4 | Q13772; B4D | NCOA4_HUMA | Oxidation (M) | 0 | 0 | 0 | 0 | 0 | 2410700 | 0 | 410020 | 0 | 0 | 0 | 0 | 0 | 301780 | 0 | 0 | 866520 | 795850 |
| TIQMGSFR | CNPY2; N/A; CI | Q9Y2B0; F8V | CNPY2_HUMAI | Unmodified | 0 | 0 | 0 | 53807 | 0 | 0 | 0 | 0 | 5048800 | 1303400 | 712370 | 563790 | 59931 | 0 | 82196 | 77699 | 0 | 0 |
| TIQMGSFR | CNPY2; N/A; CI | Q9Y2B0; F8V | CNPY2_HUMAI | Oxidation (M) | 0 | 67025 | 343670 | 198520 | 0 | 0 | 0 | 0 | 590320 | 0 | 1101200 | 0 | 0 | 0 | 0 | 85911 | 0 | 0 |
| TIQMGSFRI | CNPY2; N/A; CI | Q9Y2B0; F8V | CNPY2_HUMAI | Unmodified | 0 | 0 | 0 | 0 | 0 | 0 | 0 | 0 | 3203200 | 1464500 | 0 | 0 | 0 | 0 | 0 | 0 | 0 | 0 |
| TIQMGSFRI | CNPY2; N/A; CI | Q9Y2B0; F8V | CNPY2_HUMAI | Oxidation (M) | 0 | 0 | 0 | 0 | 0 | 0 | 0 | 0 | 758000 | 1529300 | 0 | 0 | 0 | 0 | 0 | 0 | 0 | 0 |
| TIQVDNTD | LAP3 | P28838 | AMPL_HUMAN | Unmodified | 0 | 0 | 411680 | 0 | 146530 | 0 | 55949 | 38033 | 1345400 | 88783 | 422360 | 3846600 | 0 | 98941 | 192350 | 0 | 0 | 46617 |
| TIQVDNTD | LAP3 | P28838 | AMPL_HUMAN | Unmodified | 0 | 0 | 0 | 0 | 2651200 | 147440 | 1515600 | 0 | 0 | 429330 | 0 | 0 | 2366400 | 2147900 | 0 | 0 | 599370 | 1788100 |
| TIQVSDFVF | FKBP9 | O95302 | FKBP9_HUMAN | Unmodified | 0 | 0 | 0 | 0 | 246510 | 0 | 0 | 0 | 31165000 | 164600 | 3512000 | 0 | 0 | 0 | 0 | 0 | 0 | 163690 |
| TIRDIIALNF | LONP1; LONP1 | P36776; K7E | LONM_HUMAN | Unmodified | 2034100 | 2209000 | 0 | 0 | 0 | 0 | 0 | 0 | 5513100 | 343910 | 0 | 0 | 0 | 0 | 0 | 317460 | 0 | 0 |
| TIRDNFGG | CRELD1; CRELI | Q96HD1; A0 | CREL1_HUMAN | Unmodified | 0 | 0 | 0 | 0 | 0 | 0 | 0 | 0 | 13757000 | 2863900 | 0 | 126380 | 110960 | 0 | 0 | 0 | 0 | 0 |
| TIRPMDME | PDHB; PDHB | P11177; C9J | ODPB_HUMAN | Unmodified | 0 | 0 | 16265 | 0 | 0 | 386500 | 0 | 0 | 905060 | 116010 | 338350 | 0 | 0 | 0 | 0 | 0 | 0 | 0 |
| TIRQFQFTD | PTPRF; PTPRF; I | P10586; H0Y | PTPRF_HUMAN | Unmodified | 0 | 0 | 0 | 0 | 4001000 | 0 | 0 | 217000 | 1166600 | 934460 | 0 | 579650 | 1706500 | 0 | 0 | 0 | 0 | 541560 |
| TIRSTLNPQ | PRKCA; PRKCA | P17252; J3KI | KPCA_HUMAN | Unmodified | 0 | 0 | 0 | 0 | 0 | 0 | 6210300 | 637460 | 0 | 0 | 0 | 0 | 0 | 0 | 0 | 0 | 0 | 0 |
| TIRYPDPLII | RPS4Y2; RPS4X | Q8TD47; P62 | RS4Y2_HUMAN | Unmodified | 639550 | 252620 | 297620 | 423010 | 216500 | 0 | 230830 | 87342 | 4013400 | 7338900 | 434190 | 6774100 | 0 | 0 | 230840 | 284020 | 0 | 185410 |
| TIRYPDPVII | RPS4Y1; RPS4Y | P22090; C9J | RS4Y1_HUMAN | Unmodified | 0 | 0 | 0 | 0 | 0 | 0 | 0 | 0 | 0 | 0 | 806140 | 1895700 | 0 | 0 | 0 | 0 | 61786 | 0 |
| TISDGTISA | FNBP1L; FNBP1 | Q5T0N5; A0/ | FBP1L_HUMAN | Unmodified | 0 | 1268500 | 0 | 0 | 0 | 0 | 0 | 0 | 248410 | 49573 | 0 | 81211 | 0 | 0 | 402580 | 0 | 0 | 0 |
| TISDGTISA | FNBP1L; FNBP1 | Q5T0N5; A0/ | FBP1L_HUMAN | Unmodified | 234170 | 175890 | 0 | 0 | 0 | 0 | 0 | 0 | 0 | 0 | 0 | 0 | 0 | 0 | 0 | 0 | 0 | 0 |
| TISDGTISA | FNBP1L; FNBP1 | Q5T0N5; A0/ | FBP1L_HUMAN | Unmodified | 762290 | 0 | 0 | 505670 | 0 | 0 | 0 | 0 | 136530 | 161260 | 0 | 0 | 458220 | 0 | 0 | 0 | 0 | 209010 |
| TISETIER | TARS1 | P26639 | SYTC_HUMAN | Unmodified | 0 | 0 | 0 | 0 | 186810 | 0 | 189810 | 87709 | 1825800 | 0 | 1641900 | 13992000 | 27801 | 0 | 0 | 601090 | 455470 | 0 |
| TISETIERLC | TARS1 | P26639 | SYTC_HUMAN | Unmodified | 0 | 0 | 636560 | 2200400 | 3244400 | 1874200 | 0 | 0 | 1239500 | 1017600 | 2327500 | 2945900 | 3259700 | 1670000 | 1025600 | 670340 | 584440 | 1659000 |
| TISFHYLSLI | FBLN7; FBLN7; | Q53RD9; H7 | FBLN7_HUMAN | Unmodified | 0 | 0 | 143720 | 0 | 0 | 0 | 0 | 0 | 879460 | 708480 | 0 | 0 | 0 | 242340 | 0 | 0 | 0 | 0 |
| TISHVIIGLK | RUVBL1; RUVBI | Q9Y265; E7E | RUVB1_HUMAI | Unmodified | 0 | 0 | 0 | 0 | 0 | 0 | 0 | 0 | 0 | 0 | 0 | 0 | 0 | 640160 | 0 | 0 | 0 | 0 |
| TISLVIRGT | PLEC; PLEC | Q15149; H0Y | PLEC_HUMAN | Unmodified | 0 | 0 | 0 | 0 | 4975400 | 0 | 0 | 0 | 0 | 447430 | 0 | 0 | 1149200 | 0 | 0 | 0 | 11820000 | 6532200 |
| TISNPEVVN | AFDN; AFDN; A | P55196; Q5T | AFAD_HUMAN | Unmodified | 0 | 0 | 0 | 0 | 0 | 0 | 66818 | 0 | 256360 | 102460 | 489450 | 1141400 | 1753400 | 676970 | 0 | 0 | 0 | 0 |
| TISPMVMD | VCL | P18206 | VINC_HUMAN | Oxidation (M) | 0 | 52107 | 0 | 0 | 265680 | 0 | 0 | 0 | 0 | 609870 | 0 | 265890 | 0 | 380100 | 63000 | 0 | 5913500 | 938450 |
| TISPMVMD | VCL | P18206 | VINC_HUMAN | Unmodified | 0 | 0 | 0 | 0 | 0 | 0 | 413940 | 0 | 732310 | 0 | 0 | 111780 | 0 | 0 | 0 | 70715 | 9662400 | 1625800 |
| TISPMVMD | VCL | P18206 | VINC_HUMAN | 2 Oxidation (M) | 0 | 0 | 0 | 0 | 0 | 0 | 0 | 0 | 1440100 | 0 | 0 | 0 | 0 | 0 | 0 | 0 | 744840 | 0 |
| TISPMVMD | VCL | P18206 | VINC_HUMAN | Oxidation (M) | 0 | 0 | 0 | 0 | 0 | 0 | 0 | 0 | 0 | 0 | 0 | 0 | 0 | 0 | 0 | 0 | 2661200 | 0 |
| TISPMVMD | VCL | P18206 | VINC_HUMAN | Unmodified | 63170 | 0 | 0 | 0 | 0 | 0 | 0 | 0 | 0 | 0 | 0 | 0 | 0 | 0 | 0 | 0 | 1303800 | 0 |
| TISQDEILEI | EHHADH | Q08426 | ECHP_HUMAN | Unmodified | 0 | 0 | 0 | 0 | 0 | 0 | 162070 | 0 | 2494600 | 140780 | 0 | 0 | 0 | 0 | 0 | 0 | 0 | 0 |
| TISQTSAPV | ESYT1 | Q9BSJ8 | ESYT1_HUMAN | Unmodified | 258990 | 279750 | 63726 | 0 | 0 | 0 | 26278 | 0 | 702680 | 189530 | 113960 | 413660 | 0 | 0 | 115190 | 33423 | 0 | 0 |
| TISRIGSTT | PSD3; PSD3; PS | Q9NYI0; E5R | PSD3_HUMAN | Unmodified | 0 | 0 | 0 | 0 | 3013400 | 434780 | 0 | 0 | 0 | 0 | 0 | 0 | 343790 | 192050 | 0 | 0 | 0 | 1479500 |
| TISSDAVLQ | LLGL2; LLGL2 | Q6P1M3; J3C | L2GL2_HUMAN | Unmodified | 752510 | 610970 | 736070 | 116760 | 0 | 0 | 0 | 0 | 781840 | 57878 | 134170 | 982340 | 0 | 0 | 1228400 | 294770 | 0 | 0 |
| TISSDAVLQ | LLGL2; LLGL2 | Q6P1M3; J3C | L2GL2_HUMAN | Unmodified | 1454000 | 1118000 | 742820 | 1949600 | 0 | 188700 | 0 | 0 | 8098900 | 234340 | 818170 | 7283200 | 0 | 0 | 2737100 | 2463800 | 547360 | 0 |
| TISSSLAVV | PLS3; PLS3 | P13797; A0A | PLST_HUMAN I | Unmodified | 0 | 0 | 0 | 0 | 0 | 0 | 0 | 0 | 0 | 0 | 0 | 0 | 0 | 0 | 0 | 0 | 1658400 | 0 |
| TISVATLDG | VLDLR; VLDLR; | P98155; A0A | VLDLR_HUMAN | Unmodified | 0 | 0 | 0 | 0 | 766510 | 0 | 0 | 0 | 2841100 | 633130 | 0 | 0 | 0 | 0 | 109100 | 0 | 0 | 0 |
| TISVATVDG | LRP8; LRP8; LRI | Q14114; A0/ | LRP8_HUMAN I | Unmodified | 0 | 0 | 0 | 0 | 0 | 0 | 0 | 0 | 1590200 | 12841 | 0 | 186260 | 0 | 0 | 0 | 0 | 0 | 0 |
| TISVILFLNK | GNAS; GNAS; G | Q5JWF2; P63 | GNAS1_HUMAI | Unmodified | 0 | 0 | 0 | 0 | 0 | 0 | 0 | 0 | 0 | 0 | 0 | 0 | 139460 | 0 | 553780 | 34098 | 87706 | 0 |
| TITELGEK | DLST | P36957 | ODO2_HUMAN | Unmodified | 8036300 | 10003000 | 247760 | 82902 | 0 | 0 | 537020 | 195100 | 8754100 | 259050 | 63372 | 431120 | 4989200 | 2241000 | 844530 | 303020 | 0 | 0 |
| TITGFQTHT | PSMD2 | Q13200 | PSMD2_HUMA | Unmodified | 0 | 0 | 0 | 0 | 0 | 0 | 191990 | 0 | 0 | 0 | 226010 | 3208400 | 0 | 0 | 0 | 0 | 421840 | 0 |
| TITKQVPNE | NAP1L4; NAP1L | Q99733; H0Y | NP1L4_HUMAN | Unmodified | 0 | 0 | 0 | 0 | 232830 | 959540 | 259390 | 0 | 0 | 0 | 0 | 0 | 480250 | 429320 | 0 | 0 | 604670 | 0 |
| TITLQPGSP | ADAM10 | O14672 | ADA10_HUMAN | Unmodified | 0 | 171630 | 484860 | 546800 | 0 | 0 | 0 | 0 | 0 | 0 | 0 | 0 | 74667 | 0 | 0 | 802220 | 0 | 0 |
| TITLQPGSP | ADAM10 | O14672 | ADA10_HUMAN | Oxidation (M) | 1689100 | 353630 | 0 | 4897500 | 543650 | 0 | 0 | 0 | 0 | 0 | 0 | 0 | 6658800 | 0 | 2888200 | 727560 | 482220 | 0 |
| TITMHKDST | SDCBP; SDCBP | O00560; G5I | SDCB1_HUMAI | Unmodified | 0 | 0 | 0 | 0 | 2594900 | 0 | 944950 | 0 | 2482800 | 6066200 | 0 | 0 | 2417200 | 2155800 | 0 | 0 | 164780 | 721950 |
| TITMHKDST | SDCBP; SDCBP | O00560; G5I | SDCB1_HUMAI | Oxidation (M) | 74845 | 0 | 0 | 166810 | 112810 | 0 | 0 | 0 | 0 | 2534600 | 0 | 0 | 738610 | 108130 | 91602 | 0 | 0 | 0 |
| TITMLDEQ | SNAP23; SNAP2 | O00161; H3I | SNP23_HUMAN | Oxidation (M) | 0 | 0 | 0 | 0 | 563970 | 516680 | 0 | 0 | 0 | 0 | 238900 | 1869100 | 414800 | 259240 | 0 | 0 | 361250 | 1238000 |
| TITMLDEQ | SNAP23; SNAP2 | O00161; H3I | SNP23_HUMAN | Unmodified | 0 | 435080 | 570110 | 516070 | 2900900 | 438150 | 0 | 0 | 457300 | 96264 | 1133100 | 9215300 | 2097400 | 0 | 1049400 | 868470 | 1027700 | 2665000 |
| TITMLDEQ | SNAP23; SNAP2 | O00161; H3I | SNP23_HUMAN | Oxidation (M) | 0 | 0 | 0 | 0 | 2466600 | 0 | 0 | 0 | 0 | 0 | 0 | 0 | 0 | 0 | 0 | 0 | 0 | 0 |
| TITMLDEQ | SNAP23; SNAP2 | O00161; H3I | SNP23_HUMAN | Unmodified | 0 | 0 | 0 | 0 | 4359400 | 0 | 0 | 168250 | 0 | 0 | 0 | 0 | 205930 | 857300 | 0 | 0 | 526270 | 0 |
| TITPGMVC | KLK14; KLK14 | Q9P0G3; A0 | KLK14_HUMAN | Unmodified | 0 | 0 | 0 | 0 | 0 | 140260 | 397070 | 0 | 1161600 | 0 | 0 | 0 | 0 | 71798 | 0 | 0 | 0 | 0 |
| TITSSYYR | RAB1C; RAB1A; | Q92928; P62 | RAB1C_HUMAI | Unmodified | 6655300 | 6373700 | 4344800 | 7580000 | 0 | 0 | 0 | 0 | 8494500 | 445250 | 5449400 | 18490000 | 815670 | 0 | 10723000 | 10440000 | 0 | 0 |
| TITSSYYRG | RAB1A; RAB1A | P62820; E7E | RAB1A_HUMAI | Unmodified | 0 | 0 | 0 | 0 | 0 | 1599100 | 2499000 | 224880 | 0 | 1058300 | 0 | 0 | 0 | 0 | 0 | 0 | 831070 | 907870 |
| TITSSYYRG | RAB1B | Q9H0U4 | RAB1B_HUMAN | Unmodified | 0 | 0 | 0 | 1639400 | 1360300 | 0 | 2011100 | 0 | 0 | 1262400 | 0 | 0 | 0 | 0 | 0 | 0 | 530740 | 1263400 |
| TITSTYYR | RAB35 | Q15286 | RAB35_HUMAN | Unmodified | 7597400 | 9280400 | 3428600 | 1044400 | 0 | 0 | 0 | 0 | 845250 | 0 | 1216400 | 3472500 | 0 | 0 | 10380000 | 7910900 | 0 | 0 |
| TITTAYYR | RAB3D; RAB8B; | O95716; Q9 | RAB3D_HUMAI | Unmodified | 11879000 | 10705000 | 0 | 7006800 | 1372300 | 0 | 0 | 59229 | 2102000 | 364210 | 1019700 | 7017400 | 0 | 0 | 13558000 | 10109000 | 0 | 0 |
| TITVALADG | AARS1; AARS1; | P49588; A0A | SYAC_HUMAN | Unmodified | 324130 | 61555 | 0 | 0 | 0 | 0 | 0 | 0 | 0 | 51435 | 958310 | 14720000 | 0 | 0 | 427590 | 642580 | 0 | 0 |
| TITVTPNR | ITGA6 | P23229 | ITA6_HUMAN I | Unmodified | 0 | 0 | 2315200 | 212750 | 445620 | 70625 | 0 | 379860 | 279950 | 11091 | 236000 | 1249500 | 0 | 350900 | 2357300 | 70021 | 183330 | 678860 |
| TITVTPNRI | ITGA6 | P23229 | ITA6_HUMAN I | Unmodified | 305260 | 232800 | 2830000 | 6674300 | 26951000 | 4170100 | 0 | 57676 | 711880 | 481550 | 2068800 | 12767000 | 13716000 | 5406100 | 5177900 | 5780300 | 26113000 | 75433000 |
| TITVTPNRI | ITGA6 | P23229 | ITA6_HUMAN I | Unmodified | 0 | 0 | 0 | 0 | 12434000 | 1530400 | 0 | 0 | 0 | 0 | 0 | 0 | 4280400 | 4849800 | 0 | 0 | 25807000 | 52129000 |
| TITYQAVPS | EIF2A; EIF2A; EI | Q9BY44; F8V | EIF2A_HUMAN | Unmodified | 0 | 0 | 0 | 0 | 0 | 0 | 0 | 0 | 0 | 0 | 0 | 3109900 | 0 | 0 | 0 | 283210 | 0 | 0 |
| TIVFVETK | DDX5; DDX5; D | P17844; A0A | DDX5_HUMAN | Unmodified | 176790 | 0 | 0 | 0 | 1167800 | 0 | 916350 | 1465700 | 854320 | 654430 | 677660 | 8087000 | 0 | 0 | 0 | 0 | 226020 | 756430 |
| TIVHESERL | COPS4; COPS4 | Q9BT78; D6I | CSN4_HUMAN | Unmodified | 0 | 0 | 0 | 0 | 1370700 | 0 | 0 | 0 | 0 | 0 | 0 | 0 | 0 | 675330 | 0 | 0 | 574330 | 871490 |
| TIVHYSPNL | ITGA1 | P56199 | ITA1_HUMAN I | Unmodified | 0 | 0 | 0 | 0 | 531450 | 0 | 0 | 0 | 0 | 353770 | 339250 | 0 | 0 | 0 | 0 | 0 | 2641000 | 11923000 |
| TIVIRPPFLF | ZDHHC5; ZDHH | Q9C0B5; H0 | ZDHC5_HUMA | Unmodified | 323890 | 275380 | 3633900 | 1366800 | 14050000 | 2435300 | 539490 | 0 | 6503500 | 5152500 | 5183000 | 8918800 | 15512000 | 5983800 | 7207500 | 6816300 | 6940800 | 9665900 |
| TIVITSHPG | DNAJA1 | P31689 | DNJA1_HUMAN | Unmodified | 7879100 | 9211300 | 6819900 | 10972000 | 13596000 | 2903400 | 291490 | 203810 | 48516000 | 8918300 | 2894500 | 17342000 | 13905000 | 4343500 | 11286000 | 11036000 | 591810 | 2287000 |
| TIVITSHPG | DNAJA1 | P31689 | DNJA1_HUMAN | Unmodified | 0 | 0 | 0 | 0 | 401430 | 0 | 0 | 0 | 0 | 300930 | 0 | 0 | 793130 | 812350 | 0 | 0 | 0 | 0 |
| TIVKETYGA | EXOC7; EXOC7 | Q9UPT5; B4I | EXOC7_HUMAI | Unmodified | 0 | 0 | 0 | 0 | 0 | 0 | 137250 | 87644 | 0 | 0 | 0 | 0 | 0 | 0 | 0 | 0 | 0 | 0 |
| TIVMGASFI | TALDO1; TALDC | P37837; F2Z | TALDO_HUMAI | Unmodified | 0 | 0 | 0 | 551070 | 800490 | 126020 | 0 | 0 | 0 | 0 | 3006100 | 4799600 | 812660 | 79751 | 0 | 0 | 108740 | 94010 |
| TIVMGASFI | TALDO1; TALDC | P37837; F2Z | TALDO_HUMAI | Oxidation (M) | 0 | 0 | 106290 | 88367 | 177550 | 68177 | 0 | 106650 | 452820 | 97369 | 2616200 | 2413600 | 0 | 0 | 144320 | 98901 | 0 | 0 |
| TIVMGASFI | TALDO1; TALDC | P37837; F2Z | TALDO_HUMAI | Unmodified | 0 | 0 | 0 | 0 | 7081900 | 0 | 1034000 | 285960 | 0 | 1294500 | 531780 | 0 | 3367700 | 1397500 | 0 | 0 | 2387000 | 1477200 |
| TIVMGASFI | TALDO1; TALDC | P37837; F2Z | TALDO_HUMAI | Oxidation (M) | 0 | 0 | 0 | 0 | 2661300 | 374320 | 385650 | 0 | 0 | 841750 | 401680 | 0 | 1525700 | 886200 | 0 | 0 | 0 | 562370 |
| TIVQSGGIK | RAP1GDS1 | P52306 | GDS1_HUMAN | Unmodified | 0 | 0 | 66553 | 0 | 0 | 0 | 0 | 0 | 0 | 0 | 0 | 4641300 | 199730 | 126070 | 110130 | 105030 | 0 | 0 |
| TIVRVSPHA | COBLL1; COBLI | Q53SF7; A0/ | COBL1_HUMAI | Unmodified | 0 | 0 | 0 | 0 | 0 | 0 | 0 | 0 | 0 | 0 | 0 | 0 | 0 | 0 | 0 | 0 | 0 | 0 |
| TIVTNEYMK | PITPNB; PITPNE | P48739; B3K | PIPNB_HUMAN | Oxidation (M) | 300430 | 218910 | 225590 | 318720 | 0 | 0 | 397770 | 0 | 0 | 318610 | 1117100 | 2039200 | 486050 | 0 | 845530 | 363240 | 0 | 0 |
| TIVVAASTR | FARP2; FARP2 | O94887; H7C | FARP2_HUMAN | Unmodified | 0 | 0 | 0 | 0 | 0 | 0 | 0 | 0 | 0 | 31430 | 0 | 0 | 0 | 0 | 333900 | 233310 | 85159 | 0 |
| TIVWEGKN | GMDS | O60547 | GMDS_HUMAN | Unmodified | 0 | 85187 | 0 | 0 | 0 | 0 | 80390 | 0 | 351840 | 0 | 0 | 1556200 | 0 | 0 | 0 | 182200 | 0 | 214240 |
| TIVYLDGSS | NT5E; NT5E; NT | P21589; H0Y | 5NTD_HUMAN | Unmodified | 0 | 0 | 0 | 0 | 1732000 | 2138900 | 0 | 0 | 0 | 0 | 0 | 0 | 0 | 0 | 0 | 0 | 488200 | 1062400 |
| TIYAGNALC | ETFA; ETFA; ETF | P13804; H0Y | ETFA_HUMAN I | Unmodified | 0 | 0 | 0 | 0 | 0 | 0 | 0 | 0 | 1240500 | 239120 | 0 | 0 | 0 | 0 | 0 | 0 | 0 | 0 |
| TIYKLEEQLI | ROCK1; ROCK1 | Q13464; A0/ | ROCK1_HUMAI | Unmodified | 0 | 0 | 0 | 0 | 3033200 | 0 | 0 | 0 | 0 | 0 | 0 | 0 | 0 | 0 | 0 | 0 | 0 | 757300 |
| TIYVRDPTS | LYN; LYN | P07948; E5R | LYN_HUMAN T | Unmodified | 544500 | 449110 | 645700 | 653400 | 0 | 420590 | 0 | 0 | 1821300 | 0 | 672700 | 0 | 908860 | 271780 | 835490 | 982910 | 85929 | 0 |
| TIYWTDAAS | VLDLR; VLDLR; | P98155; A0A | VLDLR_HUMAN | Unmodified | 0 | 0 | 0 | 0 | 849180 | 0 | 340950 | 0 | 1966100 | 555990 | 0 | 0 | 0 | 0 | 0 | 0 | 0 | 0 |
| TKAEPAVP | MYL6B; MYL6B | P14649; F8V | MYL6B_HUMAI | Unmodified | 0 | 0 | 0 | 0 | 1122300 | 0 | 527770 | 0 | 0 | 0 | 0 | 0 | 372680 | 0 | 0 | 0 | 1644100 | 487270 |
| TKAEQFYC | OGDH; OGDH; | Q02218; E9F | ODO1_HUMAN | Unmodified | 0 | 0 | 0 | 0 | 0 | 0 | 0 | 0 | 470880 | 70964 | 0 | 0 | 0 | 0 | 0 | 0 | 0 | 0 |
| TKAEQFYC | OGDH; OGDH; | Q02218; E9F | ODO1_HUMAN | Unmodified | 215230 | 0 | 0 | 0 | 0 | 0 | 0 | 0 | 905300 | 535340 | 0 | 0 | 0 | 0 | 0 | 0 | 0 | 0 |
| TKAQYEQT | PACSIN3; PACS | Q9UKS6; E9I | PACN3_HUMAI | Unmodified | 1215100 | 1127300 | 0 | 920130 | 0 | 0 | 0 | 0 | 0 | 139140 | 0 | 0 | 216320 | 54432 | 45710 | 116800 | 0 | 0 |
| TKAQYEQT | PACSIN3; PACS | Q9UKS6; E9I | PACN3_HUMAI | Unmodified | 1023700 | 1419200 | 0 | 2593100 | 0 | 0 | 0 | 0 | 0 | 235150 | 0 | 0 | 6205600 | 1816500 | 0 | 0 | 422820 | 348350 |
| TKDDIIEFAI | TMX3 | Q96JJ7 | TMX3_HUMAN | Unmodified | 0 | 0 | 69440 | 0 | 356840 | 0 | 0 | 0 | 4817800 | 530790 | 212700 | 557630 | 0 | 0 | 0 | 0 | 67333 | 0 |
| TKDDIIICEI | SAMM50 | Q9Y512 | SAM50_HUMAI | Unmodified | 0 | 0 | 0 | 0 | 0 | 0 | 0 | 0 | 3594100 | 76350 | 0 | 0 | 0 | 0 | 0 | 0 | 0 | 0 |
| TKDDLEQL | STX3; STX3; ST | Q13277; A0/ | STX3_HUMAN | Unmodified | 413780 | 228380 | 1658700 | 247610 | 0 | 0 | 0 | 0 | 0 | 0 | 268370 | 1438800 | 0 | 0 | 1413400 | 1036800 | 70350 | 0 |
| TKDDSEPS | IRAG2; IRAG2; I | Q12912; F5H | IRAG2_HUMAN | Unmodified | 0 | 0 | 0 | 0 | 0 | 0 | 0 | 0 | 0 | 0 | 434160 | 156930 | 0 | 0 | 0 | 0 | 0 | 0 |
| TKDEPQRE | CD9; CD9; CD9 | P21926; A0A | CD9_HUMAN C | Unmodified | 1.6E+08 | 1.87E+08 | 3.6E+08 | 2.51E+08 | 93468000 | 14965000 | 2354200 | 1370700 | 70387000 | 24892000 | 6705700 | 97701000 | 1.5E+08 | 44736000 | 5.37E+08 | 4.21E+08 | 10936000 | 26272000 |
| TKDEYLINS | IARS2 | Q9NSE4 | SYIM_HUMAN I | Unmodified | 0 | 0 | 0 | 0 | 0 | 0 | 0 | 0 | 3059600 | 228910 | 0 | 0 | 0 | 0 | 0 | 0 | 0 | 0 |
| TKDGQILP | RCC2 | Q9P258 | RCC2_HUMAN | Unmodified | 0 | 0 | 0 | 0 | 0 | 0 | 0 | 0 | 0 | 0 | 0 | 1254500 | 0 | 0 | 0 | 0 | 0 | 0 |
| TKDGVREV | RHOA; RHOA | P61586; A0A | RHOA_HUMAN | Unmodified | 0 | 0 | 0 | 558460 | 12010000 | 4143400 | 1690500 | 0 | 1284100 | 14221000 | 421630 | 0 | 7257700 | 4346800 | 0 | 0 | 3023700 | 14587000 |

| | | | | | | | | | | | | | | | | | | | | | | |
|---|---|---|---|---|---|---|---|---|---|---|---|---|---|---|---|---|---|---|---|---|---|---|
| TKDGVREV | RHOA; RHOA | P61586; A0A | RHOA_HUMAN | Oxidation (M) | 0 | 0 | 0 | 0 | 3106700 | 761320 | 687460 | 80301 | 102920 | 3720500 | 146780 | 0 | 3011300 | 750710 | 0 | 0 | 316960 | 2444000 |
| TKDGVVEII | HSPB1; HSPB1 | P04792; A0A | HSPB1_HUMAI | Unmodified | 116170 | 0 | 781740 | 733490 | 0 | 0 | 0 | 0 | 1039800 | 0 | 0 | 0 | 0 | 0 | 676600 | 0 | 0 | 0 |
| TKDIEDVFY | SRSF1; SRSF1; | Q07955; J3K | SRSF1_HUMAN | Unmodified | 1135700 | 1683700 | 0 | 0 | 0 | 0 | 0 | 31684 | 0 | 0 | 0 | 812130 | 0 | 0 | 0 | 0 | 80936 | 0 |
| TKDIVNGLI | CAPZB | B1AK88 | B1AK88_HUMA | Unmodified | 0 | 0 | 0 | 0 | 11103000 | 1024700 | 2039300 | 611430 | 0 | 1019400 | 0 | 0 | 4639500 | 3451300 | 0 | 0 | 5041900 | 4469300 |
| TKDLLVDQ | NUMBL; NUMB | Q9Y6R0; A0A | NUMBL_HUMA | Unmodified | 0 | 0 | 0 | 0 | 5055200 | 143990 | 63658 | 0 | 94692 | 309710 | 0 | 0 | 764460 | 880990 | 90317 | 0 | 0 | 1619700 |
| TKDNLENP | PLCG2; PLCG2 | P16885; A0A | PLCG2_HUMAI | Unmodified | 0 | 0 | 0 | 0 | 38956 | 0 | 0 | 0 | 0 | 11777 | 1370000 | 3444700 | 0 | 0 | 0 | 0 | 23259 | 0 |
| TKDNLENP | PLCG2; PLCG2 | P16885; A0A | PLCG2_HUMAI | Unmodified | 0 | 0 | 0 | 0 | 0 | 0 | 0 | 0 | 0 | 0 | 768570 | 689460 | 192570 | 0 | 0 | 0 | 0 | 0 |
| TKDSGLPS | DLG1; DLG1; D | Q12959; A0A | DLG1_HUMAN | Unmodified | 0 | 0 | 2912200 | 0 | 1674700 | 419210 | 0 | 81966 | 512030 | 375310 | 0 | 0 | 0 | 0 | 23025 | 0 | 231600 | 884500 |
| TKDSGSISL | SLK | Q9H2G2 | SLK_HUMAN S | Unmodified | 0 | 0 | 0 | 0 | 0 | 0 | 0 | 141870 | 0 | 0 | 0 | 0 | 353200 | 0 | 0 | 0 | 0 | 0 |
| TKDVWEIPI | FYN | P06241 | FYN_HUMAN T | Unmodified | 0 | 0 | 0 | 0 | 0 | 0 | 1344500 | 0 | 2510300 | 2180000 | 119700 | 0 | 0 | 0 | 0 | 0 | 352810 | 425000 |
| TKEAGMQF | HYOU1; HYOU | Q9Y4L1; E9F | HYOU1_HUMA | Unmodified | 0 | 0 | 0 | 0 | 69243 | 0 | 0 | 0 | 2415000 | 2961100 | 0 | 0 | 0 | 0 | 0 | 0 | 0 | 0 |
| TKEAQPGQ | LTBP1; LTBP1; | Q14766; E7E | LTBP1_HUMAN | Unmodified | 0 | 0 | 0 | 0 | 0 | 0 | 0 | 0 | 0 | 0 | 0 | 0 | 0 | 0 | 0 | 0 | 697620 | 1083000 |
| TKEEALELII | PIN1; PIN1; PIN | Q13526; K7E | PIN1_HUMAN I | Unmodified | 1451200 | 1198600 | 488000 | 309440 | 0 | 0 | 0 | 0 | 0 | 0 | 0 | 0 | 0 | 0 | 499540 | 513540 | 0 | 0 |
| TKEEMIVAT | ITGA2; ITGA2; I | P17301; E9P | ITA2_HUMAN I | Oxidation (M) | 0 | 0 | 0 | 512850 | 3401300 | 0 | 0 | 0 | 0 | 0 | 0 | 0 | 0 | 0 | 1070300 | 317660 | 3055100 | 4083700 |
| TKEEMIVAT | ITGA2; ITGA2; I | P17301; E9P | ITA2_HUMAN I | Unmodified | 0 | 0 | 0 | 0 | 6347000 | 3011600 | 0 | 0 | 0 | 0 | 0 | 0 | 0 | 0 | 513650 | 0 | 6474400 | 0 |
| TKEGVREVI | RHOC; RHOC | P08134; Q5J | RHOC_HUMAN | Unmodified | 0 | 0 | 0 | 0 | 2917800 | 0 | 0 | 130690 | 0 | 0 | 0 | 0 | 2366900 | 0 | 0 | 0 | 1111300 | 14744000 |
| TKEGVREVI | RHOC; RHOC | P08134; Q5J | RHOC_HUMAN | Oxidation (M) | 0 | 0 | 0 | 0 | 477820 | 114540 | 0 | 0 | 0 | 0 | 0 | 0 | 896160 | 573270 | 0 | 0 | 357460 | 925190 |
| TKEGVVHG | SNCA; SNCA; S | P37840; D6F | SYUA_HUMAN | Unmodified | 0 | 0 | 0 | 0 | 0 | 0 | 221870 | 167060 | 0 | 0 | 0 | 0 | 0 | 0 | 0 | 0 | 21357 | 0 |
| TKENRQSII | NSF; NSF | P46459; I3L( | NSF_HUMAN V | Unmodified | 0 | 0 | 0 | 0 | 1586000 | 833530 | 560420 | 265310 | 75684 | 0 | 0 | 0 | 1760800 | 2365900 | 0 | 0 | 407700 | 293290 |
| TKEQILEEF | SYNCRIP; SYNC | O60506; B7Z | HNRPQ_HUMA | Unmodified | 0 | 0 | 0 | 23556 | 0 | 0 | 0 | 134820 | 381900 | 173650 | 992380 | 4520300 | 929220 | 0 | 894650 | 0 | 0 | 0 |
| TKEQLGYIV | IDE; IDE; IDE; ID | P14735; A0A | IDE_HUMAN In | Unmodified | 0 | 0 | 0 | 0 | 0 | 0 | 0 | 0 | 0 | 0 | 0 | 1127300 | 0 | 0 | 0 | 0 | 0 | 0 |
| TKEQVTNV | SNCA; SNCA; S | P37840; A0A | SYUA_HUMAN | Unmodified | 0 | 0 | 0 | 0 | 0 | 0 | 460040 | 290140 | 0 | 0 | 0 | 0 | 0 | 0 | 0 | 0 | 0 | 0 |
| TKEVIQEWI | TLN1 | Q9Y490 | TLN1_HUMAN | Unmodified | 0 | 0 | 0 | 875470 | 3030700 | 0 | 1807400 | 765610 | 0 | 274600 | 0 | 0 | 0 | 1402600 | 0 | 0 | 756280 | 0 |
| TKEVIQEWI | TLN1 | Q9Y490 | TLN1_HUMAN | Unmodified | 445210 | 380520 | 0 | 2037900 | 0 | 0 | 0 | 34048 | 300530 | 0 | 0 | 558380 | 0 | 0 | 221620 | 431010 | 0 | 0 |
| TKEVYELLD | PAICS; PAICS | P22234; E9P | PUR6_HUMAN | Unmodified | 0 | 111980 | 0 | 0 | 1046600 | 0 | 1268000 | 1010500 | 1223800 | 1364100 | 0 | 1487600 | 4049800 | 8545900 | 87114 | 293270 | 1840600 | 3748800 |
| TKFENLCK | HSP90AA5P; H | Q58FG0; P0 | HS905_HUMAI | Unmodified | 0 | 0 | 0 | 0 | 1743200 | 695050 | 386020 | 279130 | 0 | 40725 | 177190 | 1339800 | 0 | 356890 | 0 | 0 | 935310 | 2366500 |
| TKFGSTAD | MIA3 | Q5JRA6 | TGO1_HUMAN | Unmodified | 0 | 0 | 0 | 0 | 0 | 0 | 0 | 0 | 1249400 | 76155 | 0 | 0 | 0 | 0 | 48650 | 0 | 0 | 0 |
| TKFGYHIIM | PIN4 | Q9Y237 | PIN4_HUMAN I | Unmodified | 0 | 0 | 0 | 171340 | 0 | 0 | 0 | 0 | 0 | 0 | 0 | 0 | 576750 | 0 | 0 | 0 | 459570 | 0 |
| TKFPAEQY | TSN; TSN; TSN | Q15631; H7( | TSN_HUMAN T | Unmodified | 466270 | 41515 | 0 | 0 | 0 | 0 | 0 | 0 | 0 | 0 | 0 | 4190600 | 0 | 0 | 188100 | 0 | 0 | 0 |
| TKGDPLGT | PTPRJ; PTPRJ; P | Q12913; A0A | PTPRJ_HUMAN | Unmodified | 182900 | 147380 | 0 | 0 | 0 | 0 | 0 | 0 | 1699400 | 0 | 1804900 | 0 | 0 | 0 | 0 | 361730 | 0 | 47014 |
| TKGFHTTID | CNN3; CNN3 | Q15417; E9F | CNN3_HUMAN | Unmodified | 0 | 0 | 0 | 0 | 354340 | 0 | 0 | 0 | 0 | 190660 | 0 | 0 | 0 | 0 | 0 | 0 | 183780 | 0 |
| TKGPDAAS | PSMC1 | P62191 | PRS4_HUMAN | Unmodified | 175710 | 0 | 0 | 219440 | 0 | 0 | 0 | 0 | 0 | 0 | 352000 | 2022000 | 993020 | 0 | 0 | 331270 | 0 | 0 |
| TKGVDEVT | ANXA2; ANXA2; | P07355; H0Y | ANXA2_HUMAI | Unmodified | 863550 | 0 | 0 | 1109900 | 16754000 | 6603700 | 0 | 0 | 43389000 | 5431300 | 491720 | 0 | 0 | 0 | 0 | 0 | 3846400 | 5124400 |
| TKHDVKFP | UBFD1; UBFD1 | O14562; H3I | UBFD1_HUMAI | Unmodified | 34218 | 0 | 0 | 0 | 183100 | 34482 | 250990 | 48220 | 0 | 352760 | 0 | 970030 | 3541900 | 646980 | 0 | 0 | 1451700 | 2836600 |
| TKHKLDVT | CCT5; CCT5; C | P48643; B7Z | TCPE_HUMAN | Unmodified | 0 | 0 | 0 | 0 | 418640 | 0 | 189290 | 114000 | 0 | 189820 | 0 | 0 | 621990 | 504590 | 0 | 0 | 588680 | 391460 |
| TKIEGLDIH | EPHX1 | P07099 | HYEP_HUMAN | Unmodified | 0 | 0 | 725680 | 201420 | 0 | 0 | 0 | 0 | 0 | 0 | 0 | 0 | 0 | 0 | 858110 | 247140 | 0 | 0 |
| TKIIFVVGG | AK1; AK1 | P00568; Q5T | KAD1_HUMAN | Unmodified | 0 | 0 | 0 | 0 | 0 | 0 | 1697100 | 346990 | 0 | 843070 | 0 | 0 | 999830 | 1016900 | 0 | 0 | 1361700 | 459880 |
| TKIISNRGE | EGFR; EGFR; E | P00533; Q5( | EGFR_HUMAN | Unmodified | 0 | 0 | 0 | 0 | 5047600 | 621760 | 0 | 0 | 0 | 0 | 0 | 0 | 587820 | 414130 | 0 | 0 | 628300 | 843200 |
| TKKAEEMA | RBCK1; RBCK1 | Q9BYM8; G3 | HOIL1_HUMAN | Unmodified | 0 | 0 | 0 | 0 | 595020 | 0 | 0 | 18204 | 0 | 0 | 0 | 0 | 0 | 0 | 0 | 0 | 0 | 0 |
| TKLEQLHI | SCYL2; SCYL2; | Q6P3W7; A0 | SCYL2_HUMAN | Oxidation (M) | 0 | 0 | 0 | 0 | 171270 | 0 | 0 | 0 | 0 | 0 | 0 | 0 | 771100 | 461940 | 0 | 0 | 274420 | 326200 |
| TKLGYSPLI | ANK1; ANK1 | P16157; H0Y | ANK1_HUMAN | Unmodified | 0 | 0 | 0 | 0 | 0 | 0 | 1523500 | 0 | 0 | 0 | 0 | 0 | 0 | 0 | 0 | 0 | 0 | 0 |
| TKMTDEEIN | PAK2 | Q13177 | PAK2_HUMAN | Unmodified | 0 | 0 | 0 | 0 | 614520 | 73279 | 60896 | 30227 | 0 | 218910 | 109600 | 151970 | 573850 | 364030 | 0 | 0 | 306760 | 311140 |
| TKNDGFSS | GPR107; GPR1 | Q5VW38; U3 | GP107_HUMAN | Unmodified | 0 | 0 | 0 | 0 | 0 | 0 | 0 | 0 | 945320 | 0 | 0 | 0 | 0 | 0 | 0 | 0 | 0 | 0 |
| TKNEACSS | NAALADL2 | Q58DX5 | NADL2_HUMAI | Unmodified | 0 | 0 | 66505 | 140380 | 0 | 0 | 0 | 0 | 0 | 0 | 0 | 0 | 0 | 327200 | 258380 | 128300 | 0 | 0 |
| TKPADMVII | PLPBP; PLPBP | O94903; E5F | PLPHP_HUMAI | Oxidation (M) | 0 | 0 | 0 | 0 | 2688700 | 0 | 493460 | 0 | 0 | 232860 | 0 | 0 | 196950 | 0 | 0 | 0 | 1201600 | 0 |
| TKPADMVII | PLPBP; PLPBP | O94903; E5F | PLPHP_HUMAI | Unmodified | 0 | 0 | 0 | 0 | 1084800 | 0 | 0 | 0 | 0 | 0 | 0 | 0 | 0 | 0 | 0 | 0 | 0 | 0 |
| TKPIVKPQT | STAU1 | O95793 | STAU1_HUMAI | Unmodified | 416600 | 0 | 109820 | 0 | 0 | 0 | 0 | 0 | 172330 | 41115 | 94871 | 466050 | 0 | 0 | 0 | 0 | 0 | 0 |
| TKPIWTRNI | HSP90AA1 | P07900 | HS90A_HUMAI | Unmodified | 222410 | 0 | 0 | 10143000 | 79542000 | 20456000 | 0 | 0 | 75628000 | 58759000 | 52963000 | 1.91E+08 | 20541000 | 0 | 8123500 | 9417700 | 26260000 | 48188000 |
| TKPIWTRNI | HSP90AB1 | P08238 | HS90B_HUMAI | Unmodified | 0 | 0 | 0 | 45883000 | 1.2E+08 | 18898000 | 6283200 | 189220 | 13062000 | 5013200 | 47968000 | 1.94E+08 | 89803000 | 30745000 | 15705000 | 0 | 37371000 | 55677000 |
| TKPPLAPG | NIBAN1 | Q9BZQ8 | NIBA1_HUMAN | Unmodified | 0 | 0 | 240300 | 2121000 | 1320000 | 2568700 | 0 | 0 | 6191700 | 3142100 | 0 | 0 | 0 | 0 | 960100 | 796260 | 0 | 2904900 |
| TKPPLAPG | NIBAN1 | Q9BZQ8 | NIBA1_HUMAN | Unmodified | 0 | 0 | 0 | 0 | 2070100 | 859440 | 0 | 0 | 1731100 | 0 | 0 | 0 | 0 | 0 | 0 | 0 | 0 | 0 |
| TKPSDEEM | DBI; DBI; DBI | P07108; B8Z | ACBP_HUMAN | Unmodified | 0 | 599790 | 2596600 | 143540 | 0 | 0 | 0 | 0 | 2434100 | 871060 | 511290 | 2321100 | 246690 | 0 | 9185800 | 453720 | 0 | 0 |
| TKPSDEEM | DBI; DBI; DBI | P07108; B8Z | ACBP_HUMAN | Oxidation (M) | 0 | 4075100 | 6698300 | 1227100 | 0 | 0 | 0 | 0 | 0 | 501540 | 1308100 | 4109800 | 0 | 0 | 6052400 | 8141200 | 90162 | 0 |
| TKPSPSQFI | PNPT1; PNPT1 | Q8TCS8; H7 | PNPT1_HUMAI | Unmodified | 0 | 0 | 0 | 0 | 0 | 0 | 0 | 0 | 406440 | 0 | 0 | 0 | 0 | 0 | 0 | 0 | 0 | 0 |
| TKPVTGNLI | DYNC1H1; DYN | Q14204; A0A | DYHC1_HUMA | Unmodified | 0 | 0 | 993820 | 891480 | 3770500 | 355820 | 1002500 | 0 | 2710500 | 1150400 | 1320000 | 7098800 | 2164800 | 1159700 | 958740 | 648710 | 301360 | 2640700 |
| TKPYIQVDI | HSPA5; HSPA5; | P11021; A0A | BIP_HUMAN Er | Unmodified | 0 | 0 | 32385 | 0 | 0 | 0 | 161400 | 162790 | 1196300 | 427790 | 102440 | 343150 | 0 | 22972 | 38817 | 0 | 0 | 0 |
| TKQDEVNA | SPTAN1; SPTAN | Q13813; A0A | SPTN1_HUMAI | Unmodified | 0 | 0 | 0 | 0 | 0 | 102930 | 0 | 0 | 81905 | 0 | 30829 | 1371500 | 0 | 0 | 0 | 0 | 0 | 0 |
| TKQDEVNA | SPTAN1; SPTAN | Q13813; A0A | SPTN1_HUMAI | Unmodified | 0 | 0 | 0 | 0 | 0 | 0 | 0 | 0 | 0 | 0 | 0 | 0 | 0 | 0 | 0 | 0 | 387370 | 0 |
| TKQEQQAS | CASK; CASK; C | O14936; A0A | CSKP_HUMAN | Unmodified | 203950 | 0 | 0 | 492840 | 3539100 | 575680 | 361720 | 370460 | 1215500 | 984990 | 0 | 217220 | 1935300 | 2158600 | 424430 | 357760 | 1084400 | 3839700 |
| TKQEQQAS | CASK; CASK; C | O14936; A0A | CSKP_HUMAN | Unmodified | 0 | 0 | 0 | 0 | 1704400 | 0 | 213590 | 144770 | 430830 | 535340 | 0 | 0 | 442100 | 592330 | 0 | 138650 | 321490 | 127610 |
| TKQQIEEQI | FLOT1; FLOT1; | O75955; A0A | FLOT1_HUMAN | Unmodified | 0 | 0 | 0 | 277180 | 1559000 | 455200 | 490300 | 151470 | 874870 | 1586300 | 278390 | 0 | 1035600 | 476140 | 0 | 0 | 0 | 0 |
| TKQVIQGLI | KLC1; KLC1; KL | Q07866; G3 | KLC1_HUMAN | Unmodified | 0 | 0 | 0 | 0 | 955630 | 0 | 1040300 | 0 | 0 | 402290 | 0 | 0 | 406030 | 328100 | 0 | 0 | 563590 | 648180 |
| TKRSIQFVD | TUBA4A; TUBA1 | P68366; P68 | TBA4A_HUMAN | Unmodified | 0 | 96339 | 0 | 0 | 177790 | 0 | 5748600 | 322100 | 0 | 270320 | 0 | 0 | 0 | 135260 | 0 | 0 | 1479500 | 110340 |
| TKRTIQFVD | TUBA3C; TUBA | P0DPH7; Q7 | TBA3C_HUMAN | Unmodified | 0 | 0 | 0 | 0 | 0 | 0 | 1455500 | 0 | 0 | 0 | 0 | 0 | 51209 | 0 | 0 | 0 | 621490 | 685610 |
| TKSCSGVE | VDAC2; VDAC2 | P45880; Q5J | VDAC2_HUMAI | Unmodified | 0 | 0 | 0 | 0 | 0 | 0 | 0 | 0 | 0 | 212030 | 0 | 0 | 0 | 0 | 0 | 0 | 0 | 0 |
| TKSCSGVE | VDAC2; VDAC2 | P45880; Q5J | VDAC2_HUMAI | Unmodified | 0 | 0 | 0 | 0 | 0 | 0 | 278650 | 0 | 603100 | 3065900 | 0 | 0 | 0 | 0 | 0 | 0 | 0 | 0 |
| TKSENGLEI | VDAC1; VDAC1 | P21796; C9J | VDAC1_HUMAI | Unmodified | 0 | 0 | 386460 | 319730 | 211430 | 0 | 229670 | 116990 | 1.23E+08 | 9867600 | 2725000 | 1162200 | 0 | 672090 | 0 | 808100 | 0 | 182240 |
| TKSENGLEI | VDAC1; VDAC1 | P21796; C9J | VDAC1_HUMAI | Unmodified | 0 | 0 | 0 | 0 | 1046900 | 0 | 2004200 | 514390 | 47968000 | 46556000 | 208160 | 0 | 689670 | 2057600 | 0 | 0 | 0 | 909460 |
| TKSPFEQH | FAM3C | Q92520 | FAM3C_HUMAI | Unmodified | 0 | 0 | 0 | 0 | 399000 | 0 | 0 | 0 | 13155000 | 3736900 | 0 | 967870 | 0 | 0 | 0 | 0 | 61077 | 0 |
| TKSQGGEF | PFN2; PFN2; PF | P35080; G5E | PROF2_HUMAI | Unmodified | 192380 | 205530 | 97250 | 390240 | 0 | 793290 | 62722 | 14318 | 526940 | 148090 | 0 | 0 | 876340 | 563840 | 113300 | 0 | 478050 | 1313100 |
| TKSQGPSL | FMNL1; FMNL1 | O95466; A0A | FMNL1_HUMAI | Unmodified | 151130 | 0 | 0 | 0 | 566460 | 39539 | 0 | 0 | 0 | 0 | 0 | 0 | 0 | 0 | 0 | 0 | 0 | 157880 |
| TKSTGGAP | PFN1; PFN1 | P07737; K7E | PROF1_HUMAI | Unmodified | 0 | 0 | 0 | 0 | 1498100 | 0 | 453760 | 520200 | 0 | 298050 | 0 | 0 | 420030 | 0 | 0 | 0 | 1480600 | 94261 |
| TKTENPLILI | LONP1; LONP1 | P36776; K7E | LONM_HUMAN | Unmodified | 0 | 0 | 0 | 0 | 0 | 0 | 0 | 0 | 1515800 | 0 | 0 | 0 | 0 | 0 | 0 | 0 | 0 | 0 |
| TKTENPLILI | LONP1; LONP1 | P36776; K7E | LONM_HUMAN | Unmodified | 0 | 0 | 0 | 0 | 0 | 0 | 0 | 0 | 2080600 | 0 | 0 | 0 | 0 | 0 | 0 | 0 | 0 | 0 |
| TKTETITGF | FN1; FN1 | P02751; H0Y | FINC_HUMAN I | Unmodified | 0 | 0 | 19914000 | 7916600 | 0 | 0 | 0 | 0 | 0 | 451150 | 1247400 | 1561700 | 0 | 0 | 0 | 15511000 | 0 | 0 |
| TKTPGPGA | RPS14; RPS14; | P62263; E5R | RS14_HUMAN | Unmodified | 324560 | 511850 | 0 | 244830 | 135890 | 0 | 0 | 0 | 172420 | 95733 | 46837 | 821710 | 215490 | 62785 | 0 | 0 | 56664 | 40125 |
| TKTVPGYLI | MVB12A; MVB1 | Q96EY5; E9F | MB12A_HUMAI | Unmodified | 0 | 0 | 0 | 0 | 0 | 0 | 0 | 0 | 0 | 0 | 0 | 0 | 0 | 381980 | 0 | 0 | 0 | 284250 |
| TKTYSEDNI | SCG3; SCG3 | Q8WXD2; H( | SCG3_HUMAN | Unmodified | 0 | 0 | 0 | 0 | 0 | 0 | 0 | 0 | 34139 | 823280 | 0 | 0 | 0 | 0 | 0 | 0 | 0 | 0 |
| TKVEAFQT | PSME2; PSME2 | Q9UL46; A0A | PSME2_HUMAI | Unmodified | 0 | 0 | 0 | 0 | 0 | 0 | 195780 | 0 | 0 | 142580 | 0 | 1775800 | 0 | 0 | 0 | 0 | 0 | 0 |
| TKVEATKNI | EPRS1; EPRS1 | P07814; V9C | SYEP_HUMAN | Unmodified | 54750 | 0 | 0 | 315690 | 1522800 | 98772 | 0 | 298890 | 375510 | 482220 | 106630 | 1847900 | 5291900 | 1784200 | 0 | 0 | 663820 | 1774700 |
| TKVNKDNN | ABR; ABR; ABR; | Q12979; A0A | ABR_HUMAN A | Unmodified | 0 | 0 | 0 | 0 | 981880 | 0 | 0 | 0 | 0 | 0 | 0 | 0 | 0 | 0 | 0 | 0 | 0 | 627990 |
| TKVQNIHP | NUCB2; NUCB2 | P80303; E9P | NUCB2_HUMA | Unmodified | 0 | 0 | 0 | 0 | 0 | 0 | 389930 | 0 | 11344000 | 5502600 | 0 | 0 | 0 | 0 | 0 | 0 | 0 | 0 |
| TKVVVTMEI | OXCT1; OXCT1 | P55809; E9P | SCOT1_HUMAI | Unmodified | 0 | 0 | 0 | 0 | 0 | 0 | 436730 | 0 | 231670 | 407470 | 0 | 118230 | 0 | 0 | 0 | 0 | 0 | 75233 |
| TKYFFFDD | SPCS3 | P61009 | SPCS3_HUMAI | Unmodified | 0 | 0 | 0 | 0 | 0 | 0 | 0 | 0 | 2587400 | 952650 | 0 | 0 | 0 | 0 | 0 | 0 | 37378 | 0 |
| TKYPNYEFI | FUT8; FUT8 | Q9BYC5; G3 | FUT8_HUMAN | Unmodified | 0 | 0 | 0 | 0 | 0 | 0 | 0 | 0 | 1516400 | 0 | 427080 | 0 | 0 | 0 | 0 | 0 | 0 | 0 |
| TKYYLQDG | IGF2R | P11717 | MPRI_HUMAN | Unmodified | 273530 | 724920 | 0 | 0 | 0 | 0 | 0 | 0 | 0 | 2319200 | 0 | 0 | 913840 | 0 | 0 | 0 | 0 | 0 |
| TLAALVDH | TUBGCP3; TUB | Q96CW5; A( | GCP3_HUMAN | Unmodified | 0 | 0 | 0 | 0 | 88068 | 0 | 86162 | 0 | 0 | 0 | 0 | 518280 | 265900 | 21540 | 22504 | 18528 | 0 | 0 |
| TLAEGFPGI | PROM2 | Q8N271 | PROM2_HUMA | Unmodified | 0 | 0 | 0 | 0 | 393660 | 0 | 0 | 0 | 0 | 0 | 196490 | 2561400 | 0 | 0 | 0 | 1562600 | 0 | 0 |
| TLAEHQQL | UCHL5; UCHL5 | Q9Y5K5; H0 | UCHL5_HUMAI | Unmodified | 0 | 0 | 306320 | 220690 | 1065000 | 0 | 526170 | 519520 | 1360400 | 1330700 | 629210 | 2863800 | 1558700 | 773970 | 0 | 0 | 524850 | 464180 |
| TLAEIAKAEI | PSPC1; PSPC1 | Q8WXF1; X6 | PSPC1_HUMAI | Unmodified | 0 | 0 | 0 | 0 | 592430 | 0 | 391400 | 0 | 0 | 33523 | 427780 | 2046000 | 0 | 0 | 0 | 0 | 0 | 0 |
| TLAEIAKVEI | NONO; NONO; | Q15233; C9I | NONO_HUMAN | Unmodified | 0 | 0 | 0 | 0 | 0 | 0 | 726290 | 0 | 0 | 963460 | 681670 | 1535500 | 0 | 0 | 0 | 0 | 0 | 0 |
| TLAEINANF | DYNC1I2 | Q13409 | DC1I2_HUMAN | Unmodified | 0 | 0 | 0 | 0 | 0 | 0 | 0 | 0 | 96050 | 38557 | 738570 | 532680 | 0 | 0 | 0 | 0 | 0 | 0 |
| TLAEINANF | DYNC1I2 | Q13409 | DC1I2_HUMAN | Unmodified | 192030 | 218940 | 0 | 829310 | 1385000 | 0 | 0 | 0 | 2025300 | 1276800 | 1326100 | 7218800 | 1725700 | 516470 | 1067300 | 337170 | 0 | 1461100 |
| TLAESALQI | TLN1 | Q9Y490 | TLN1_HUMAN | Unmodified | 1144400 | 1363700 | 938400 | 3130100 | 22156000 | 3580000 | 30206000 | 937860 | 1513100 | 1394900 | 545820 | 2911300 | 14846000 | 14723000 | 2391100 | 1416000 | 8534100 | 10289000 |
| TLAESALQI | TLN1 | Q9Y490 | TLN1_HUMAN | Unmodified | 0 | 0 | 0 | 0 | 0 | 546290 | 2598800 | 0 | 0 | 0 | 131560 | 0 | 0 | 0 | 0 | 0 | 2670700 | 0 |
| TLAESALQ | TLN2; TLN2; TLI | Q9Y4G6; H0 | TLN2_HUMAN | Unmodified | 0 | 0 | 0 | 0 | 0 | 0 | 0 | 0 | 0 | 0 | 0 | 0 | 0 | 0 | 0 | 0 | 233470 | 87907 |
| TLAEVCLG | DSG2 | Q14126 | DSG2_HUMAN | Unmodified | 0 | 0 | 1210500 | 2083400 | 5487300 | 1263900 | 0 | 0 | 1663900 | 120010 | 1650600 | 3063500 | 2526800 | 1429900 | 1789200 | 1447700 | 446850 | 879380 |
| TLAEVCLG | DSG2 | Q14126 | DSG2_HUMAN | Unmodified | 0 | 0 | 0 | 0 | 3912600 | 0 | 0 | 0 | 344760 | 0 | 0 | 0 | 0 | 0 | 0 | 0 | 642130 | 960950 |
| TLAEVCLG | DSG2 | Q14126 | DSG2_HUMAN | Unmodified | 0 | 0 | 0 | 0 | 8815200 | 918810 | 0 | 0 | 104150 | 108710 | 0 | 0 | 4162100 | 503580 | 0 | 0 | 2589100 | 1156300 |
| TLAFTSVDL | ACOT13 | Q9NPJ3 | ACO13_HUMAI | Unmodified | 0 | 0 | 0 | 0 | 915120 | 113000 | 0 | 0 | 3308600 | 563470 | 0 | 0 | 0 | 0 | 0 | 0 | 571020 | 0 |
| TLAFTSVDL | ACOT13 | Q9NPJ3 | ACO13_HUMAI | Unmodified | 0 | 0 | 0 | 0 | 1032800 | 0 | 284410 | 95679 | 151710 | 651860 | 0 | 0 | 237340 | 260710 | 0 | 0 | 221180 | 204090 |
| TLAGDVHI | VPS29; VPS29 | Q9UBQ0; F8 | VPS29_HUMAN | Unmodified | 1060600 | 973260 | 1393700 | 1508400 | 7990200 | 1993000 | 1039600 | 0 | 5425700 | 2002000 | 2441800 | 9468300 | 1502400 | 4405600 | 2722800 | 3012100 | 2392500 | 2115900 |
| TLAGDVHI | VPS29; VPS29 | Q9UBQ0; F8 | VPS29_HUMAN | Unmodified | 0 | 0 | 0 | 0 | 1098900 | 0 | 661870 | 0 | 0 | 0 | 0 | 0 | 0 | 0 | 0 | 0 | 669240 | 0 |
| TLAGLPLLV | EIF5B; EIF5B | O60841; A0A | IF2P_HUMAN E | Unmodified | 0 | 892200 | 0 | 522170 | 0 | 0 | 0 | 0 | 0 | 422830 | 0 | 25108 | 3420800 | 1105400 | 576620 | 0 | 324950 | 639320 |
| TLAHFRPIE | CHP1; CHP1; C | Q99653; F5( | CHP1_HUMAN | Unmodified | 1711200 | 837110 | 1379200 | 2136700 | 1509700 | 0 | 0 | 0 | 0 | 257140 | 572470 | 3361200 | 1711200 | 0 | 1941000 | 1935800 | 367390 | 281330 |
| TLAIELAPR | DHRS4L1; DHR | P0CG22; Q6 | DR4L1_HUMAN | Unmodified | 0 | 0 | 0 | 0 | 0 | 0 | 634980 | 220420 | 12674000 | 4721000 | 0 | 0 | 0 | 0 | 0 | 0 | 0 | 0 |
| TLAMDTILA | GNPDA1; GNPI | P46926; D6F | GNPI1_HUMAN | Unmodified | 0 | 0 | 0 | 0 | 0 | 0 | 0 | 0 | 0 | 0 | 0 | 845020 | 0 | 0 | 0 | 0 | 0 | 0 |
| TLAMLPGM | POMGNT2 | Q8NAT1 | PMGT2_HUMAI | Oxidation (M) | 0 | 0 | 0 | 0 | 0 | 0 | 0 | 374550 | 0 | 0 | 0 | 0 | 0 | 0 | 0 | 0 | 0 | 0 |
| TLAPTWEEI | TXNDC5 | Q8NBS9 | TXND5_HUMAI | Unmodified | 0 | 477910 | 0 | 0 | 0 | 0 | 0 | 38778 | 4221000 | 1159700 | 353040 | 0 | 0 | 0 | 0 | 0 | 707790 | 0 |

| | | | | | | | | | | | | | | | | | | | | | | |
|---|---|---|---|---|---|---|---|---|---|---|---|---|---|---|---|---|---|---|---|---|---|---|
| TLAPTWEEI | TXNDC5 | Q8NBS9 | TXND5_HUMAI | Unmodified | 0 | 0 | 0 | 0 | 2755100 | 0 | 1338200 | 1086400 | 65418000 | 25818000 | 3571600 | 831380 | 829510 | 1497200 | 0 | 480590 | 483730 | 1632800 |
| TLAPTWEEI | TXNDC5 | Q8NBS9 | TXND5_HUMAI | Unmodified | 0 | 0 | 0 | 0 | 655180 | 0 | 1336100 | 52522 | 1277000 | 5931700 | 0 | 0 | 0 | 0 | 0 | 0 | 197700 | 1157000 |
| TLAQLNPES | GPI; GPI; GPI; G | P06744; A0A | G6PI_HUMAN | Unmodified | 2857900 | 1631000 | 2871300 | 7115900 | 19547000 | 6556300 | 5383900 | 657230 | 25225000 | 15683000 | 14731000 | 76197000 | 11053000 | 6983800 | 7561500 | 8305800 | 6264600 | 5101100 |
| TLASAGGP | ADD1; ADD1; A | P35611; E7E | ADDA_HUMAN | Unmodified | 0 | 0 | 98864 | 872620 | 495210 | 0 | 0 | 0 | 0 | 0 | 0 | 0 | 349420 | 526750 | 620890 | 617680 | 0 | 0 |
| TLATDILMC | DHX15 | O43143 | DHX15_HUMAI | Oxidation (M) | 0 | 0 | 0 | 0 | 0 | 0 | 0 | 0 | 183410 | 0 | 442200 | 4904000 | 0 | 0 | 0 | 0 | 0 | 0 |
| TLATIKEQN | PSMD12 | O00232 | PSD12_HUMAI | Unmodified | 0 | 0 | 0 | 0 | 408700 | 80734 | 199650 | 82226 | 0 | 42891 | 0 | 0 | 300170 | 90901 | 0 | 0 | 239930 | 291740 |
| TLATLSENN | MRPL46 | Q9H2W6 | RM46_HUMAN | Unmodified | 0 | 0 | 0 | 0 | 0 | 0 | 0 | 0 | 481860 | 0 | 0 | 0 | 0 | 0 | 0 | 0 | 0 | 0 |
| TLATQLNQ | DST; DST; DST; | Q03001; A0A | DYST_HUMAN | Unmodified | 0 | 64083 | 0 | 0 | 1084400 | 685970 | 104530 | 53585 | 0 | 0 | 0 | 225440 | 0 | 0 | 0 | 0 | 73636 | 142470 |
| TLATWATK | KPNB1; KPNB1 | Q14974; J3K | IMB1_HUMAN | Unmodified | 588850 | 479890 | 962460 | 1175200 | 7372600 | 256820 | 2721400 | 1356300 | 5395000 | 1919200 | 411550 | 638970 | 3374900 | 2733200 | 1067900 | 956780 | 6823800 | 7067900 |
| TLAVERLLE | CTNNA1; CTNN | P35221; E5R | CTNA1_HUMAI | Unmodified | 0 | 0 | 0 | 0 | 0 | 0 | 0 | 0 | 0 | 0 | 0 | 0 | 185320 | 50305 | 0 | 0 | 0 | 0 |
| TLAVERLLE | CTNNA1; CTNN | P35221; E5R | CTNA1_HUMAI | Unmodified | 0 | 0 | 0 | 162750 | 12489000 | 125920 | 0 | 0 | 0 | 192390 | 0 | 0 | 9519900 | 994020 | 0 | 0 | 0 | 0 |
| TLAVLLDNI | MGAT5 | Q09328 | MGT5A_HUMAI | Unmodified | 0 | 0 | 0 | 0 | 0 | 0 | 0 | 0 | 2258000 | 389810 | 149420 | 203280 | 0 | 0 | 0 | 6960.1 | 0 | 12005 |
| TLAVPPWIE | POFUT1 | Q9H488 | OFUT1_HUMAI | Unmodified | 0 | 0 | 0 | 0 | 0 | 0 | 318270 | 0 | 16430000 | 636840 | 1097100 | 0 | 0 | 0 | 0 | 0 | 0 | 0 |
| TLAYLLPAI | DDX17; DDX17 | Q92841; A0A | DDX17_HUMAI | Unmodified | 135890 | 82213 | 0 | 0 | 0 | 0 | 0 | 0 | 212880 | 0 | 241900 | 6366600 | 0 | 0 | 262350 | 0 | 0 | 0 |
| TLCTLGPNI | NBEAL2; NBEAI | Q6ZNJ1; H0 | NBEL2_HUMAI | Unmodified | 247110 | 229360 | 122850 | 542470 | 0 | 0 | 0 | 0 | 0 | 0 | 0 | 0 | 527260 | 1143300 | 176420 | 268190 | 0 | 0 |
| TLDAQDRS | TNKS1BP1 | Q9C0C2 | TB182_HUMAI | Unmodified | 0 | 0 | 0 | 0 | 0 | 0 | 0 | 0 | 0 | 0 | 0 | 0 | 0 | 2328700 | 0 | 0 | 0 | 0 |
| TLDDIVGR | DST; DST; DST; | Q03001; A0A | DYST_HUMAN | Unmodified | 108090 | 101320 | 146260 | 255460 | 488920 | 0 | 176060 | 0 | 1006500 | 155880 | 335250 | 1568300 | 0 | 410890 | 272290 | 293740 | 125170 | 307510 |
| TLDDIVGR | DST; DST; DST; | Q03001; A0A | DYST_HUMAN | Unmodified | 0 | 0 | 0 | 0 | 1033500 | 238920 | 0 | 0 | 0 | 0 | 0 | 0 | 0 | 0 | 0 | 0 | 0 | 0 |
| TLDDVPGP | SDK2; SDK2 | Q58EX2; H7 | SDK2_HUMAN | Oxidation (M) | 0 | 442710 | 0 | 0 | 0 | 0 | 68651 | 0 | 610180 | 89300 | 0 | 0 | 0 | 1099300 | 0 | 0 | 39520 | 0 |
| TLDEFTIIQ | C1R; C1R; C1R | P00736; B4E | C1R_HUMAN C | Unmodified | 0 | 0 | 0 | 64003 | 1496400 | 181270 | 43441 | 0 | 1265200 | 48316 | 0 | 0 | 131160 | 0 | 0 | 0 | 0 | 228010 |
| TLDEILENQ | UTRN | P46939 | UTRN_HUMAN | Unmodified | 0 | 0 | 0 | 594280 | 0 | 569990 | 0 | 0 | 0 | 403850 | 0 | 0 | 258610 | 992070 | 947150 | 674530 | 446290 | 473210 |
| TLDELGIHL | EFTUD2 | Q15029 | U5S1_HUMAN | Unmodified | 0 | 0 | 0 | 0 | 0 | 0 | 0 | 0 | 0 | 0 | 0 | 655330 | 0 | 0 | 0 | 0 | 0 | 0 |
| TLDGELDEI | LAMB1; LAMB1 | P07942; G3 | LAMB1_HUMAI | Unmodified | 0 | 0 | 0 | 0 | 0 | 0 | 0 | 0 | 0 | 0 | 0 | 0 | 0 | 0 | 0 | 0 | 0 | 409120 |
| TLDGGLNV | UGP2; UGP2; U | Q16851; E7E | UGPA_HUMAN | Unmodified | 0 | 0 | 0 | 0 | 945070 | 0 | 0 | 0 | 0 | 182500 | 0 | 0 | 697570 | 337590 | 0 | 0 | 401040 | 0 |
| TLDGIFSEQ | RCC2 | Q9P258 | RCC2_HUMAN | Oxidation (M) | 0 | 0 | 0 | 0 | 0 | 0 | 0 | 0 | 698700 | 0 | 1265400 | 2466300 | 928560 | 0 | 0 | 0 | 0 | 0 |
| TLDGPESN | RFTN1; RFTN1 | Q14699; G3 | RFTN1_HUMAI | Unmodified | 0 | 0 | 0 | 0 | 0 | 0 | 0 | 0 | 0 | 0 | 0 | 1427300 | 0 | 0 | 0 | 0 | 0 | 0 |
| TLDGPESN | RFTN1; RFTN1 | Q14699; G3 | RFTN1_HUMAI | Oxidation (M) | 0 | 0 | 0 | 0 | 7804900 | 2486200 | 0 | 0 | 0 | 0 | 0 | 511880 | 0 | 0 | 0 | 0 | 0 | 0 |
| TLDGPESN | RFTN1; RFTN1 | Q14699; G3 | RFTN1_HUMAI | Unmodified | 0 | 0 | 0 | 0 | 4070700 | 0 | 0 | 0 | 0 | 0 | 0 | 0 | 0 | 0 | 0 | 0 | 0 | 0 |
| TLDINVK | DHRS4L2; DHR | Q6PKH6; Q9 | DR4L2_HUMAI | Unmodified | 520850 | 711020 | 926670 | 2466500 | 582160 | 0 | 654960 | 145150 | 13726000 | 109830 | 0 | 0 | 0 | 425640 | 724880 | 1049600 | 0 | 0 |
| TLDINVKAF | DHRS4L2; DHR | Q6PKH6; Q9 | DR4L2_HUMAI | Unmodified | 0 | 0 | 0 | 0 | 0 | 0 | 0 | 0 | 773750 | 759650 | 0 | 0 | 0 | 0 | 0 | 0 | 22912 | 0 |
| TLDLLTMK | SEPTIN10; SEPT | Q9P0V9; B5 | SEP10_HUMAI | Unmodified | 0 | 0 | 0 | 0 | 3427400 | 264210 | 146890 | 0 | 0 | 0 | 0 | 0 | 82394 | 0 | 0 | 0 | 1531600 | 882820 |
| TLDLLTMK | SEPTIN10; SEPT | Q9P0V9; B5 | SEP10_HUMAI | Oxidation (M) | 0 | 0 | 0 | 0 | 751360 | 257540 | 0 | 0 | 0 | 0 | 0 | 0 | 0 | 0 | 0 | 0 | 484560 | 0 |
| TLDLPIYVTI | COPA; COPA; C | P53621; A0A | COPA_HUMAN | Unmodified | 0 | 0 | 372680 | 475740 | 0 | 0 | 0 | 0 | 1265800 | 0 | 1368500 | 23506000 | 0 | 0 | 502180 | 899640 | 0 | 0 |
| TLDNGGFY | HCK; HCK; HCK | P08631; J3K | HCK_HUMAN T | Unmodified | 0 | 0 | 0 | 0 | 0 | 0 | 0 | 0 | 0 | 0 | 0 | 0 | 0 | 0 | 0 | 0 | 0 | 0 |
| TLDNLGYR | ADD3 | Q9UEY8 | ADDG_HUMAN | Unmodified | 0 | 0 | 417760 | 271120 | 0 | 0 | 0 | 1063400 | 0 | 0 | 505420 | 172870 | 110800 | 0 | 495090 | 477390 | 0 | 0 |
| TLDPDPAIF | CSE1L | P55060 | XPO2_HUMAN | Unmodified | 0 | 0 | 0 | 0 | 399840 | 0 | 208970 | 61055 | 325400 | 215130 | 206380 | 1147900 | 34552 | 0 | 0 | 99907 | 315040 | 300470 |
| TLDQAIMK | RAD50; N/A; RA | Q92878; A0A | RAD50_HUMAI | Unmodified | 0 | 0 | 0 | 0 | 135670 | 0 | 312190 | 31678 | 0 | 189820 | 0 | 1191800 | 271570 | 0 | 0 | 0 | 0 | 0 |
| TLDRSGDL | CTNND1; CTNN | O60716; H0 | CTND1_HUMAI | Unmodified | 0 | 0 | 984860 | 1131000 | 1196100 | 0 | 0 | 16740 | 0 | 0 | 0 | 2070900 | 0 | 0 | 2092300 | 960200 | 361060 | 729660 |
| TLDRSGDL | CTNND1; CTNN | O60716; H0 | CTND1_HUMAI | Oxidation (M) | 880750 | 111770 | 1334200 | 1500600 | 0 | 0 | 0 | 0 | 0 | 0 | 0 | 3655800 | 0 | 0 | 3744500 | 2682500 | 0 | 0 |
| TLDRSGDL | CTNND1; CTNN | A0A669KB62 | A0A669KB62_H | Oxidation (M) | 0 | 0 | 0 | 0 | 0 | 0 | 3161400 | 0 | 0 | 0 | 0 | 0 | 0 | 6081200 | 0 | 0 | 0 | 0 |
| TLDSWRDE | RAB7A; RAB7A | P51149; A0A | RAB7A_HUMAI | Unmodified | 0 | 0 | 0 | 0 | 0 | 0 | 83988 | 55975 | 0 | 1380700 | 2935100 | 12495000 | 0 | 173800 | 80678 | 51984 | 0 | 0 |
| TLDSWRDE | RAB7A; RAB7A | P51149; A0A | RAB7A_HUMAI | Unmodified | 0 | 0 | 0 | 0 | 0 | 0 | 0 | 0 | 0 | 0 | 0 | 0 | 0 | 0 | 512600 | 0 | 0 | 0 |
| TLDTGETPS | CNOT11 | Q9UKZ1 | CNO11_HUMA | Unmodified | 0 | 0 | 0 | 0 | 0 | 0 | 0 | 0 | 0 | 22948 | 56186 | 551410 | 0 | 0 | 0 | 0 | 142090 | 480150 |
| TLDVFNIIL | VPS13C | Q709C8 | VP13C_HUMAI | Unmodified | 27433 | 19320 | 32997 | 21701 | 0 | 0 | 0 | 0 | 155910 | 56501 | 106020 | 512750 | 0 | 0 | 62109 | 34305 | 0 | 0 |
| TLEAEFNSF | SHTN1 | A0MZ66 | SHOT1_HUMAI | Unmodified | 0 | 0 | 0 | 0 | 0 | 0 | 0 | 0 | 314440 | 168320 | 0 | 691450 | 86826 | 0 | 144110 | 0 | 0 | 0 |
| TLEDGKKE | TXNDC12 | O95881 | TXD12_HUMAI | Oxidation (M) | 0 | 0 | 0 | 0 | 0 | 0 | 0 | 0 | 0 | 828040 | 0 | 0 | 0 | 0 | 0 | 0 | 0 | 0 |
| TLEDGKKE | TXNDC12 | O95881 | TXD12_HUMAI | Unmodified | 0 | 0 | 0 | 0 | 0 | 0 | 0 | 0 | 564220 | 811720 | 0 | 0 | 0 | 0 | 0 | 0 | 150930 | 0 |
| TLEDPDLN | CAND1; CAND1 | Q86VP6; A0 | CAND1_HUMA | Unmodified | 0 | 0 | 0 | 0 | 0 | 0 | 0 | 0 | 0 | 0 | 0 | 0 | 0 | 0 | 0 | 0 | 0 | 0 |
| TLEDPDLN | CAND1; CAND1 | Q86VP6; A0 | CAND1_HUMA | Unmodified | 183850 | 0 | 0 | 0 | 0 | 0 | 0 | 0 | 2288200 | 693080 | 813800 | 6608400 | 0 | 0 | 0 | 0 | 0 | 0 |
| TLEDRFFEH | ATP6V0D1; ATP | P61421; R4C | VA0D1_HUMAI | Unmodified | 0 | 0 | 0 | 343230 | 0 | 0 | 210970 | 0 | 1835900 | 565070 | 644210 | 1678400 | 373120 | 277820 | 463470 | 107640 | 0 | 44461 |
| TLEEAIR | ANXA11 | P50995 | ANX11_HUMAI | Unmodified | 0 | 1278700 | 544820 | 1025700 | 406320 | 0 | 0 | 0 | 34880000 | 1414000 | 2443800 | 4975600 | 0 | 0 | 76625 | 1305300 | 111440 | 587700 |
| TLEEAIRSD | ANXA11 | P50995 | ANX11_HUMAI | Unmodified | 1416700 | 0 | 0 | 2745900 | 10280000 | 5680000 | 1001600 | 0 | 1.1E+08 | 16457000 | 8715900 | 4100100 | 10524000 | 6792800 | 627460 | 720760 | 2710300 | 10329000 |
| TLEEAVGN | COPG1; COPG | Q9Y678; H0 | COPG1_HUMAI | Unmodified | 0 | 0 | 0 | 0 | 0 | 0 | 0 | 0 | 0 | 355490 | 361850 | 5983400 | 600260 | 0 | 0 | 0 | 409900 | 0 |
| TLEEDEEEI | RANBP1; RANB | P43487; C9J | RANG_HUMAN | Unmodified | 0 | 0 | 0 | 0 | 921420 | 0 | 0 | 0 | 0 | 0 | 0 | 0 | 0 | 0 | 0 | 0 | 619290 | 239940 |
| TLEEEAKTH | MYH9 | P35579 | MYH9_HUMAN | Unmodified | 0 | 0 | 0 | 0 | 1326800 | 122060 | 0 | 0 | 0 | 50775 | 0 | 0 | 518780 | 117100 | 0 | 0 | 368550 | 231690 |
| TLEEEAKTH | MYH9 | P35579 | MYH9_HUMAN | Unmodified | 0 | 0 | 0 | 0 | 27219000 | 3800700 | 1702800 | 291510 | 0 | 718640 | 0 | 0 | 4430200 | 3646600 | 0 | 0 | 12072000 | 9141100 |
| TLEEEAKTH | MYH9 | P35579 | MYH9_HUMAN | Oxidation (M) | 0 | 0 | 0 | 0 | 2686400 | 610560 | 0 | 91534 | 0 | 197970 | 0 | 0 | 0 | 0 | 0 | 0 | 944690 | 808070 |
| TLEEEFARK | SLK | Q9H2G2 | SLK_HUMAN S | Unmodified | 0 | 0 | 0 | 0 | 1777000 | 0 | 0 | 0 | 0 | 1011500 | 0 | 0 | 5487300 | 5002900 | 0 | 0 | 863070 | 0 |
| TLEEGHDF | NUBP2; NUBP2 | Q9Y5Y2; H3 | NUBP2_HUMAI | Unmodified | 1103600 | 584980 | 0 | 453970 | 0 | 0 | 1288800 | 0 | 742180 | 0 | 0 | 0 | 0 | 0 | 1000100 | 0 | 0 | 0 |
| TLEELHLTC | LRRC8A | Q8IWT6 | LRC8A_HUMAI | Unmodified | 209300 | 59255 | 0 | 0 | 0 | 0 | 0 | 0 | 0 | 0 | 0 | 965410 | 0 | 0 | 0 | 0 | 0 | 0 |
| TLEELYLDA | ERBIN; LRRC7; | Q96RT1; Q9 | ERBIN_HUMAI | Unmodified | 0 | 0 | 128030 | 536290 | 27948000 | 3580900 | 0 | 65039 | 425720 | 938460 | 0 | 42078 | 5122500 | 3527600 | 616320 | 215150 | 2003900 | 4317300 |
| TLEEMELRI | ERC2; ERC1; EF | O15083; Q8 | ERC2_HUMAN | Unmodified | 0 | 0 | 0 | 0 | 644290 | 0 | 0 | 0 | 0 | 0 | 0 | 0 | 0 | 0 | 0 | 0 | 0 | 0 |
| TLEFAGQD | ACTL8 | Q9H568 | ACTL8_HUMAI | Unmodified | 0 | 0 | 0 | 0 | 0 | 0 | 0 | 0 | 0 | 0 | 1034200 | 2082200 | 0 | 0 | 0 | 0 | 0 | 0 |
| TLEFFWR | MAN2A1 | Q16706 | MA2A1_HUMAI | Unmodified | 0 | 0 | 0 | 0 | 0 | 0 | 0 | 0 | 0 | 0 | 0 | 779300 | 0 | 0 | 0 | 0 | 0 | 0 |
| TLEFKPYVII | MPP7; MPP7 | Q5T2T1; U5 | MPP7_HUMAN | Unmodified | 0 | 0 | 307230 | 0 | 848200 | 0 | 0 | 0 | 2162700 | 0 | 214020 | 845750 | 0 | 0 | 0 | 404030 | 0 | 0 |
| TLEGDGDC | AHNAK2 | Q8IVF2 | AHNK2_HUMA | Unmodified | 0 | 0 | 0 | 0 | 4706000 | 0 | 0 | 0 | 0 | 0 | 0 | 0 | 0 | 0 | 0 | 0 | 1227100 | 7073100 |
| TLEGFRLLV | SBF1; SBF1 | O95248; H0 | MTMR5_HUMA | Unmodified | 0 | 0 | 0 | 0 | 0 | 0 | 172270 | 124420 | 0 | 88847 | 0 | 0 | 252250 | 276060 | 0 | 0 | 1388800 | 0 |
| TLEGPENC | LRBA; LRBA; LR | P50851; A0A | LRBA_HUMAN | Unmodified | 0 | 0 | 0 | 0 | 0 | 0 | 0 | 0 | 169630 | 0 | 91081 | 1329500 | 0 | 14872 | 0 | 0 | 0 | 0 |
| TLEGPSFLS | NAALADL2 | Q58DX5 | NADL2_HUMAI | Unmodified | 0 | 0 | 1280100 | 2332000 | 0 | 0 | 0 | 0 | 0 | 0 | 0 | 1012400 | 1294700 | 0 | 3920700 | 0 | 159910 | 0 |
| TLEGPSFLS | NAALADL2 | Q58DX5 | NADL2_HUMAI | Unmodified | 0 | 173310 | 0 | 0 | 847990 | 0 | 0 | 124700 | 0 | 253700 | 0 | 0 | 1142800 | 0 | 0 | 0 | 0 | 0 |
| TLEGQAQE | AHNAK2 | Q8IVF2 | AHNK2_HUMA | Unmodified | 0 | 0 | 0 | 0 | 4206700 | 1331900 | 159160 | 0 | 0 | 0 | 0 | 0 | 0 | 0 | 0 | 0 | 1151200 | 9463300 |
| TLEHLQLSI | HK2; HK2 | P52789; E9P | HXK2_HUMAN | Unmodified | 23434 | 46920 | 0 | 0 | 85551 | 0 | 0 | 62999 | 0 | 0 | 0 | 2486000 | 166060 | 0 | 0 | 22214 | 201190 | 2580500 |
| TLEKEPRTF | PRXL2A | Q9BRX8 | PXL2A_HUMAI | Unmodified | 0 | 0 | 0 | 0 | 0 | 0 | 203840 | 888120 | 0 | 608010 | 0 | 0 | 0 | 0 | 0 | 0 | 338340 | 0 |
| TLEKYEREA | GSPT1; GSPT2; | P15170; Q8 | ERF3A_HUMAI | Unmodified | 0 | 0 | 0 | 0 | 0 | 0 | 0 | 0 | 0 | 0 | 0 | 0 | 0 | 0 | 0 | 0 | 0 | 0 |
| TLELGADFS | PROM2 | Q8N271 | PROM2_HUMA | Unmodified | 1259200 | 0 | 601000 | 5331700 | 0 | 0 | 0 | 0 | 0 | 0 | 0 | 0 | 0 | 0 | 2427000 | 1641200 | 0 | 0 |
| TLELQASS | FGD4; FGD4; F | Q96M96; B7 | FGD4_HUMAN | Unmodified | 0 | 0 | 0 | 0 | 0 | 0 | 0 | 0 | 0 | 0 | 0 | 287030 | 0 | 0 | 0 | 0 | 0 | 0 |
| TLELQGLIN | DSP | P15924 | DESP_HUMAN | Unmodified | 32253 | 0 | 0 | 24589 | 0 | 0 | 0 | 0 | 294870 | 0 | 7930.2 | 125420 | 0 | 0 | 0 | 28535 | 0 | 0 |
| TLELRVAYL | BCAM; BCAM | P50895; A0A | BCAM_HUMAN | Unmodified | 0 | 0 | 0 | 0 | 457560 | 39453 | 0 | 0 | 0 | 90467 | 0 | 0 | 0 | 0 | 0 | 0 | 41854 | 219630 |
| TLEMLSLD | LRSAM1; LRSA | Q6UWE0; A | LRSM1_HUMAI | Oxidation (M) | 0 | 0 | 0 | 0 | 384120 | 0 | 179620 | 0 | 0 | 0 | 0 | 0 | 520800 | 241770 | 0 | 0 | 0 | 0 |
| TLEPVKPPT | ABI1; ABI1; ABI | Q8IZP0; A0A | ABI1_HUMAN | Oxidation (M) | 594400 | 429390 | 318660 | 550940 | 539820 | 174940 | 0 | 82352 | 405790 | 149480 | 426100 | 2001600 | 410490 | 0 | 567200 | 508930 | 204580 | 167270 |
| TLEPVRPP | ABI2; ABI2; ABI | Q9NYB9; H7 | ABI2_HUMAN | Oxidation (M) | 710610 | 393140 | 0 | 212570 | 334850 | 0 | 0 | 0 | 352260 | 25077 | 1008500 | 3684000 | 0 | 0 | 252680 | 255610 | 158350 | 251180 |
| TLEQHDNI | USO1 | O60763 | USO1_HUMAN | Unmodified | 0 | 0 | 0 | 0 | 0 | 0 | 0 | 70850 | 0 | 0 | 167640 | 1837200 | 807460 | 295950 | 0 | 0 | 0 | 324120 |
| TLEQQENF | ANK3; ANK3; AN | Q12955; A0A | ANK3_HUMAN | Unmodified | 0 | 0 | 1634400 | 1503100 | 0 | 0 | 0 | 0 | 0 | 0 | 0 | 675210 | 0 | 0 | 2844700 | 2851700 | 0 | 0 |
| TLEQTGHV | IQGAP2; IQGAP | Q13576; E7E | IQGA2_HUMAI | Unmodified | 0 | 0 | 0 | 0 | 0 | 0 | 0 | 0 | 0 | 0 | 0 | 0 | 0 | 0 | 0 | 0 | 0 | 0 |
| TLESQAVEI | NLE1; NLE1; NL | Q9NVX2; A0 | NLE1_HUMAN | Unmodified | 0 | 0 | 0 | 0 | 0 | 0 | 0 | 0 | 0 | 68694 | 81281 | 577770 | 0 | 0 | 0 | 0 | 0 | 0 |
| TLESSIQGL | ARPC1A | Q92747 | ARC1A_HUMAI | Unmodified | 1003700 | 1069500 | 1434000 | 1349100 | 0 | 0 | 0 | 0 | 1431600 | 0 | 560430 | 7813200 | 589470 | 0 | 991300 | 1685400 | 0 | 0 |
| TLETLIR | MCM3; MCM3; | P25205; Q7 | MCM3_HUMAN | Unmodified | 0 | 17504 | 143000 | 75139 | 430560 | 0 | 63644 | 51325 | 398140 | 175120 | 240530 | 1338800 | 298750 | 30608 | 38369 | 0 | 28488 | 155910 |
| TLETLLLPT | IQGAP2; IQGAP | Q13576; D6 | IQGA2_HUMAI | Unmodified | 0 | 0 | 945080 | 1281500 | 1436800 | 0 | 0 | 0 | 0 | 0 | 0 | 0 | 0 | 0 | 0 | 1124800 | 0 | 0 |
| TLETLPEK | HMBS; HMBS; | P08397; A0A | HEM3_HUMAN | Unmodified | 0 | 724720 | 392060 | 0 | 0 | 0 | 428220 | 0 | 145440 | 153750 | 556160 | 986010 | 854950 | 460120 | 88301 | 137960 | 0 | 0 |
| TLETQSALC | SEC24A | O95486 | SC24A_HUMAI | Unmodified | 0 | 0 | 0 | 0 | 0 | 0 | 0 | 0 | 0 | 0 | 0 | 1036300 | 0 | 0 | 0 | 0 | 0 | 0 |
| TLEVEIEPG | DNAJB11; N/A | Q9UBS4; H7 | DJB11_HUMAI | Unmodified | 0 | 0 | 0 | 0 | 0 | 0 | 0 | 0 | 1365100 | 0 | 742880 | 359940 | 0 | 0 | 0 | 171500 | 0 | 0 |
| TLEVEIEPG | DNAJB11; N/A | Q9UBS4; H7 | DJB11_HUMAI | Oxidation (M) | 0 | 0 | 0 | 0 | 0 | 0 | 0 | 0 | 5936600 | 0 | 0 | 0 | 0 | 0 | 0 | 0 | 0 | 0 |
| TLEVEIEPG | DNAJB11; N/A | Q9UBS4; H7 | DJB11_HUMAI | Unmodified | 0 | 0 | 0 | 0 | 0 | 0 | 0 | 0 | 2108700 | 0 | 0 | 0 | 0 | 0 | 0 | 0 | 0 | 0 |
| TLEVVSPSC | MAP1B | P46821 | MAP1B_HUMAI | Unmodified | 0 | 0 | 0 | 0 | 0 | 0 | 0 | 139810 | 306040 | 0 | 0 | 1090000 | 0 | 0 | 0 | 0 | 0 | 0 |
| TLEYKIEILE | KIF23; KIF23; KI | Q02241; A0A | KIF23_HUMAI | Unmodified | 0 | 0 | 0 | 663370 | 26176000 | 4077800 | 1496700 | 91175 | 1926600 | 3628300 | 755090 | 470230 | 1040900 | 1673200 | 0 | 0 | 6464400 | 1544200 |
| TLFALGTPN | B3GNT5 | Q9BYG0 | B3GN5_HUMAI | Unmodified | 0 | 0 | 0 | 0 | 0 | 0 | 0 | 0 | 1048200 | 0 | 491240 | 0 | 0 | 0 | 0 | 0 | 0 | 0 |
| TLFALGTPN | B3GNT5 | Q9BYG0 | B3GN5_HUMAI | Unmodified | 0 | 0 | 0 | 0 | 0 | 0 | 0 | 0 | 1152000 | 0 | 438720 | 0 | 0 | 0 | 0 | 0 | 0 | 0 |
| TLFANTGG | PTBP2; PTBP2; | Q9UKA9; A0 | PTBP2_HUMAI | Unmodified | 0 | 0 | 0 | 0 | 0 | 0 | 0 | 0 | 0 | 0 | 0 | 857230 | 0 | 0 | 0 | 0 | 0 | 0 |
| TLFATEDAL | MYO1C; MYO1C | O00159; F5H | MYO1C_HUMA | Unmodified | 0 | 0 | 0 | 0 | 0 | 0 | 0 | 0 | 0 | 0 | 0 | 0 | 0 | 0 | 23969000 | 0 | 0 | 0 |
| TLFATEDAL | MYO1C; MYO1C | O00159; F5H | MYO1C_HUMA | Unmodified | 0 | 0 | 433000 | 0 | 0 | 0 | 0 | 0 | 0 | 0 | 0 | 0 | 0 | 0 | 273580 | 248720 | 0 | 0 |
| TLFAYEGTN | CD2AP | Q9Y5K6 | CD2AP_HUMAI | Unmodified | 1066800 | 0 | 4809900 | 0 | 0 | 23463 | 0 | 0 | 0 | 0 | 0 | 0 | 0 | 0 | 4870500 | 2037300 | 0 | 0 |
| TLFAYEGTN | CD2AP | Q9Y5K6 | CD2AP_HUMAI | Unmodified | 0 | 14223000 | 0 | 3209500 | 0 | 375890 | 0 | 0 | 0 | 0 | 0 | 1354100 | 0 | 0 | 7819400 | 0 | 0 | 0 |
| TLFDEGWI | COL21A1; COL | Q96P44; F5 | COLA1_HUMAN | Unmodified | 0 | 0 | 0 | 0 | 0 | 0 | 0 | 0 | 6078700 | 1334700 | 0 | 0 | 0 | 0 | 0 | 0 | 23225 | 0 |
| TLFEDAGYI | GNPTG | Q9UJJ9 | GNPTG_HUMA | Unmodified | 0 | 0 | 0 | 0 | 0 | 0 | 0 | 193560 | 5665400 | 0 | 0 | 0 | 0 | 0 | 0 | 0 | 0 | 207740 |
| TLFEDSFQ | ITCH; ITCH; ITC | Q96J02; A0A | ITCH_HUMAN | Oxidation (M) | 675030 | 571480 | 36956 | 136100 | 0 | 0 | 0 | 0 | 0 | 0 | 0 | 21020 | 0 | 0 | 438570 | 40653 | 0 | 0 |
| TLFEDSFQ | ITCH; ITCH; ITC | Q96J02; A0A | ITCH_HUMAN | Oxidation (M) | 72352 | 236170 | 0 | 0 | 0 | 0 | 0 | 0 | 0 | 0 | 0 | 0 | 0 | 0 | 153630 | 0 | 0 | 0 |
| TLFGLHLS | ATIC; ATIC | P31939; H7 | PUR9_HUMAN | Unmodified | 0 | 0 | 0 | 0 | 0 | 0 | 0 | 0 | 590060 | 0 | 0 | 1276000 | 0 | 0 | 0 | 0 | 0 | 0 |
| TLFIFPGLL | GPR108; GPR1 | Q9NPR9; M | GP108_HUMAI | Unmodified | 0 | 0 | 0 | 0 | 0 | 0 | 0 | 0 | 38587000 | 3315900 | 847590 | 0 | 0 | 0 | 0 | 0 | 0 | 0 |
| TLFILFR | ZCCHC3 | Q9NUD5 | ZCHC3_HUMA | Unmodified | 0 | 0 | 0 | 0 | 0 | 0 | 0 | 0 | 0 | 0 | 0 | 109240 | 0 | 0 | 0 | 0 | 0 | 0 |

| | | | | | | | | | | | | | | | | | | | | | | |
|---|---|---|---|---|---|---|---|---|---|---|---|---|---|---|---|---|---|---|---|---|---|---|
| TLFILRGLP( | CNP; CNP; CNI | P09543; K7E | CN37_HUMAN | Unmodified | 0 | 0 | 0 | 0 | 4535100 | 1451000 | 1749100 | 267850 | 0 | 707250 | 95515 | 0 | 2681000 | 4503200 | 0 | 0 | 2321600 | 4639000 |
| TLFKLEDLR | MYO1B; MYO1E | O43795; E9F | MYO1B_HUMA | Unmodified | 0 | 247480 | 0 | 0 | 0 | 474230 | 182660 | 20400 | 0 | 473470 | 773490 | 11662000 | 0 | 0 | 0 | 95362 | 0 | 0 |
| TLFPLIEAK | XRCC5 | P13010 | XRCC5_HUMAI | Unmodified | 1188900 | 443540 | 735550 | 451130 | 5366000 | 769890 | 2874300 | 3928900 | 0 | 1380400 | 5471600 | 35407000 | 1985100 | 993150 | 401570 | 266820 | 793150 | 2535700 |
| TLFPVVLEÇ | LAMA4; LAMA4; | Q16363; A0/ | LAMA4_HUMAI | Unmodified | 0 | 0 | 0 | 0 | 0 | 0 | 0 | 0 | 539560 | 0 | 0 | 0 | 0 | 0 | 0 | 0 | 0 | 0 |
| TLFQIQAEF | UAP1 | Q16222 | UAP1_HUMAN | Unmodified | 0 | 0 | 0 | 0 | 6608600 | 0 | 362740 | 0 | 0 | 199180 | 0 | 0 | 2293700 | 2159100 | 0 | 0 | 646240 | 1236700 |
| TLFQIQAEF | UAP1 | Q16222 | UAP1_HUMAN | Unmodified | 0 | 0 | 0 | 0 | 3085700 | 0 | 0 | 0 | 0 | 0 | 0 | 0 | 691810 | 1159600 | 0 | 0 | 0 | 442190 |
| TLFQNDFW | TMTC3; TMTC3; | Q6ZXV5; F8V | TMTC3_HUMAI | Oxidation (M) | 0 | 0 | 0 | 0 | 35390 | 0 | 0 | 0 | 892030 | 248510 | 0 | 0 | 0 | 0 | 0 | 0 | 0 | 0 |
| TLFQNDFW | TMTC3; TMTC3; | Q6ZXV5; F8V | TMTC3_HUMAI | Unmodified | 19777 | 0 | 0 | 0 | 0 | 0 | 0 | 0 | 1127600 | 0 | 0 | 0 | 0 | 0 | 0 | 0 | 0 | 0 |
| TLFQPQTG. | SEC24C; SEC2 | P53992; G5E | SC24C_HUMAI | Unmodified | 0 | 0 | 0 | 0 | 669820 | 0 | 304760 | 182280 | 637150 | 408870 | 0 | 2689500 | 427540 | 0 | 0 | 0 | 2917300 | 5396600 |
| TLFSNIVLS | ACTR1A; ACTR1 | P61163; R4C | ACTZ_HUMAN . | Unmodified | 0 | 0 | 0 | 0 | 0 | 0 | 0 | 0 | 0 | 88536 | 0 | 631480 | 56187 | 0 | 192080 | 18929 | 0 | 0 |
| TLFTDLFEK | CSNK1G1; CSN | Q9HCP0; U3 | KC1G1_HUMAI | Unmodified | 0 | 669100 | 0 | 0 | 0 | 0 | 0 | 0 | 0 | 0 | 357110 | 874510 | 0 | 0 | 88877 | 509110 | 0 | 0 |
| TLFTDLFEK | CSNK1G1; CSN | Q9HCP0; U3 | KC1G1_HUMAI | Unmodified | 0 | 0 | 0 | 0 | 0 | 0 | 0 | 0 | 0 | 0 | 0 | 778470 | 0 | 0 | 234720 | 385430 | 0 | 0 |
| TLGAGAFG | KIT; CSF1R; CSI | P10721; P07 | KIT_HUMAN M | Unmodified | 0 | 0 | 1452900 | 1415400 | 0 | 0 | 72969 | 0 | 3293500 | 390290 | 807440 | 2940200 | 498450 | 0 | 0 | 1914600 | 31429 | 0 |
| TLGDFAAE | PARP1; PARP1; | P09874; A0A | PARP1_HUMAN | Unmodified | 1282600 | 664320 | 0 | 0 | 1596500 | 569400 | 0 | 3810300 | 0 | 0 | 1352100 | 2643000 | 1829400 | 0 | 0 | 0 | 946340 | 540370 |
| TLGECGFT | ELOB; ELOB; EL | Q15370; B82 | ELOB_HUMAN | Unmodified | 0 | 0 | 0 | 0 | 0 | 0 | 0 | 0 | 1407300 | 0 | 208470 | 8156500 | 0 | 0 | 0 | 0 | 0 | 0 |
| TLGEGEFG. | AXL; AXL | P30530; M0F | UFO_HUMAN T | Oxidation (M) | 0 | 0 | 0 | 0 | 1072500 | 0 | 0 | 0 | 0 | 0 | 0 | 0 | 0 | 0 | 0 | 0 | 0 | 0 |
| TLGGKQPV | MACF1; MACF1 | Q9UPN3; H3 | MACF1_HUMAI | Unmodified | 0 | 0 | 0 | 0 | 157620 | 0 | 0 | 0 | 446130 | 107340 | 145160 | 849130 | 0 | 0 | 0 | 0 | 89628 | 0 |
| TLGGKQPV | MACF1; MACF1 | Q9UPN3; H3 | MACF1_HUMAI | Unmodified | 38963 | 0 | 209940 | 248150 | 0 | 0 | 0 | 945510 | 0 | 0 | 0 | 0 | 1120500 | 3594200 | 0 | 0 | 0 | 828530 |
| TLGGLEME | HYOU1; HYOU | Q9Y4L1; E9F | HYOU1_HUMA | Oxidation (M) | 0 | 0 | 276680 | 0 | 0 | 0 | 0 | 0 | 5193400 | 0 | 849360 | 858950 | 633460 | 248480 | 316680 | 179700 | 0 | 0 |
| TLGGLEME | HYOU1; HYOU | Q9Y4L1; E9F | HYOU1_HUMA | Unmodified | 0 | 0 | 111620 | 0 | 0 | 0 | 0 | 0 | 16126000 | 0 | 756610 | 0 | 145290 | 0 | 0 | 0 | 66970 | 0 |
| TLGGYLVEI | UNC5B | Q8IZJ1 | UNC5B_HUMA | Oxidation (M) | 0 | 0 | 0 | 0 | 0 | 491760 | 0 | 0 | 545240 | 0 | 0 | 2235500 | 0 | 0 | 0 | 0 | 0 | 0 |
| TLGHNLLV | CUL4A; CUL4A | Q13619; A0/ | CUL4A_HUMAI | Unmodified | 0 | 0 | 0 | 0 | 0 | 0 | 0 | 138060 | 0 | 68006 | 0 | 1435600 | 0 | 0 | 0 | 0 | 0 | 0 |
| TLGILGLGF | PHGDH; PHGD | O43175; A0/ | SERA_HUMAN | Unmodified | 0 | 0 | 0 | 10468000 | 3272200 | 1318400 | 2920800 | 1116800 | 4754400 | 3221800 | 3183900 | 16250000 | 38487000 | 11503000 | 0 | 0 | 2293900 | 3585000 |
| TLGKDHPA | KLC1; KLC2; KL | Q07866; Q9I | KLC1_HUMAN | Unmodified | 0 | 0 | 0 | 0 | 7346400 | 0 | 2446500 | 18624 | 225720 | 1712800 | 0 | 0 | 4323900 | 2835800 | 0 | 0 | 8272200 | 4481400 |
| TLGLGEEW | PLOD3; PLOD3 | O60568; C9J | PLOD3_HUMAI | Unmodified | 0 | 0 | 0 | 0 | 0 | 0 | 0 | 0 | 1160600 | 0 | 0 | 0 | 0 | 0 | 0 | 0 | 0 | 0 |
| TLGLGEEW | PLOD3; PLOD3 | O60568; C9J | PLOD3_HUMAI | Unmodified | 0 | 0 | 65761 | 0 | 0 | 0 | 0 | 0 | 33319000 | 2241500 | 495990 | 321700 | 0 | 0 | 0 | 0 | 0 | 0 |
| TLGMVYIYI | ITGAV | P06756 | ITAV_HUMAN I | Oxidation (M) | 224010 | 0 | 0 | 0 | 0 | 0 | 0 | 0 | 0 | 0 | 0 | 1090200 | 0 | 0 | 0 | 277890 | 0 | 0 |
| TLGMVYIYI | ITGAV | P06756 | ITAV_HUMAN I | Unmodified | 0 | 0 | 0 | 0 | 0 | 0 | 0 | 0 | 0 | 0 | 0 | 1459800 | 0 | 0 | 0 | 0 | 0 | 0 |
| TLGSAFLNI | ITGA7; ITGA7 | Q13683; J3K | ITA7_HUMAN I | Unmodified | 0 | 0 | 0 | 0 | 0 | 1656900 | 0 | 0 | 608590 | 156500 | 71152 | 0 | 0 | 0 | 0 | 0 | 0 | 0 |
| TLGSFEFE( | MTOR | P42345 | MTOR_HUMAN | Unmodified | 0 | 0 | 25568 | 0 | 14165 | 819480 | 0 | 0 | 403640 | 74050 | 0 | 846920 | 97604 | 1163900 | 0 | 0 | 627200 | 0 |
| TLGTGSFG | PRKACA; PRKA( | P17612; P22 | KAPCA_HUMAI | Unmodified | 0 | 0 | 0 | 0 | 865040 | 199200 | 540130 | 31650 | 4307100 | 1442600 | 0 | 0 | 475430 | 0 | 0 | 0 | 267510 | 1899600 |
| TLGTGSFG | PRKACA; PRKA( | P17612; P22 | KAPCA_HUMAI | Oxidation (M) | 0 | 0 | 0 | 0 | 1300600 | 0 | 2130900 | 155660 | 191340 | 774560 | 0 | 0 | 1954800 | 244880 | 0 | 0 | 394130 | 794720 |
| TLGTGSFG | PRKACA; PRKA( | P17612; P22 | KAPCA_HUMAI | Unmodified | 0 | 0 | 0 | 0 | 2076300 | 0 | 1093800 | 0 | 0 | 0 | 0 | 0 | 660130 | 1104300 | 0 | 0 | 0 | 0 |
| TLGTMIAGI | ANXA7 | P20073 | ANXA7_HUMAI | Oxidation (M) | 0 | 0 | 0 | 0 | 0 | 0 | 0 | 0 | 5063500 | 142830 | 0 | 151620 | 0 | 0 | 79422 | 44949 | 0 | 0 |
| TLGTMIAGI | ANXA7 | P20073 | ANXA7_HUMAI | Unmodified | 0 | 0 | 0 | 0 | 0 | 0 | 0 | 0 | 10281000 | 647880 | 0 | 684730 | 0 | 0 | 0 | 0 | 0 | 0 |
| TLGTMIAGI | ANXA7 | P20073 | ANXA7_HUMAI | Oxidation (M) | 1192700 | 1119100 | 307590 | 1720400 | 0 | 0 | 0 | 0 | 25412000 | 4068800 | 315060 | 951510 | 0 | 564100 | 2508700 | 754060 | 0 | 1443500 |
| TLGTMIAGI | ANXA7 | P20073 | ANXA7_HUMAI | Unmodified | 0 | 0 | 0 | 0 | 1244500 | 0 | 0 | 0 | 51691000 | 4225000 | 0 | 0 | 456740 | 0 | 664450 | 0 | 0 | 0 |
| TLGTMIAGI | ANXA7 | P20073 | ANXA7_HUMAI | Oxidation (M) | 0 | 0 | 0 | 36356 | 383370 | 0 | 262450 | 0 | 0 | 2568400 | 0 | 0 | 419830 | 246780 | 214090 | 0 | 2219500 | 1974800 |
| TLGTPEFVT | IGF2R; IGF2R; I | P11717; A0A | MPRI_HUMAN | Unmodified | 0 | 0 | 0 | 0 | 0 | 0 | 0 | 0 | 23734000 | 7343000 | 0 | 774800 | 0 | 0 | 0 | 0 | 0 | 0 |
| TLGTPEFVT | IGF2R; IGF2R; I | P11717; A0A | MPRI_HUMAN | Unmodified | 0 | 0 | 0 | 0 | 0 | 0 | 0 | 0 | 0 | 5801100 | 0 | 0 | 620720 | 0 | 0 | 0 | 0 | 0 |
| TLGVDLVAI | CAD; CAD | P27708; F8V | PYR1_HUMAN | Unmodified | 0 | 429960 | 689650 | 6500400 | 0 | 0 | 0 | 0 | 0 | 0 | 12901000 | 1214200 | 1353300 | 7362000 | 0 | 0 | 0 | 0 |
| TLGVERFEE | SLC4A2; SLC4A | P04920; C9J | B3A2_HUMAN . | Unmodified | 294600 | 250940 | 397320 | 241520 | 0 | 0 | 45350 | 0 | 0 | 0 | 27942 | 293590 | 0 | 0 | 2348100 | 1103800 | 0 | 0 |
| TLGVHPDD | PLXNA4 | Q9HCM2 | PLXA4_HUMAN | Unmodified | 0 | 0 | 0 | 0 | 0 | 0 | 262840 | 0 | 0 | 0 | 0 | 0 | 0 | 0 | 0 | 0 | 0 | 132760 |
| TLHADPGD | GMIP | Q9P107 | GMIP_HUMAN | Unmodified | 0 | 0 | 0 | 0 | 0 | 0 | 0 | 0 | 0 | 239560 | 0 | 434220 | 0 | 0 | 0 | 0 | 0 | 0 |
| TLHCTWDF | OSMR | Q99650 | OSMR_HUMAN | Unmodified | 0 | 0 | 0 | 0 | 1057600 | 255840 | 0 | 0 | 1157700 | 0 | 0 | 0 | 0 | 0 | 0 | 0 | 0 | 0 |
| TLHEWLQ( | CAD; CAD | P27708; F8V | PYR1_HUMAN | Unmodified | 71710 | 0 | 0 | 0 | 0 | 35384 | 0 | 0 | 188120 | 0 | 0 | 661530 | 0 | 0 | 98769 | 0 | 0 | 0 |
| TLHLQTGN | GCLM | P48507 | GSH0_HUMAN | Unmodified | 176860 | 152630 | 172320 | 189320 | 4787000 | 737150 | 94254 | 0 | 1291300 | 726560 | 158320 | 1209200 | 3231200 | 1151400 | 422850 | 345180 | 1909700 | 1133100 |
| TLHPAPNA | PLCD3 | Q8N3E9 | PLCD3_HUMAI | Unmodified | 0 | 0 | 0 | 0 | 0 | 0 | 0 | 0 | 0 | 0 | 0 | 0 | 0 | 484810 | 0 | 0 | 950760 | 1281600 |
| TLHPAPNA | PLCD3 | Q8N3E9 | PLCD3_HUMAI | Unmodified | 0 | 0 | 156680 | 264220 | 822110 | 0 | 0 | 0 | 0 | 0 | 0 | 0 | 0 | 143460 | 0 | 0 | 1338900 | 2865700 |
| TLHPAVHA | ATIC; ATIC | P31939; C9J | PUR9_HUMAN | Unmodified | 0 | 0 | 167970 | 0 | 3239100 | 0 | 0 | 0 | 194200 | 216760 | 90452 | 1991800 | 701620 | 441300 | 249750 | 0 | 475820 | 622590 |
| TLHPDLGTI | LDHA | P00338 | LDHA_HUMAN | Unmodified | 0 | 0 | 711340 | 0 | 0 | 0 | 0 | 0 | 0 | 0 | 1726700 | 8791300 | 0 | 0 | 701340 | 366160 | 0 | 0 |
| TLHPDLGTI | LDHA | P00338 | LDHA_HUMAN | Unmodified | 0 | 72859 | 285460 | 0 | 461270 | 121990 | 0 | 0 | 325460 | 417010 | 1429500 | 11707000 | 89058 | 36776 | 373930 | 558570 | 135150 | 494850 |
| TLHPDLGTI | LDHA | P00338 | LDHA_HUMAN | Unmodified | 2014100 | 2253100 | 0 | 5604900 | 19445000 | 3087700 | 464670 | 264450 | 5501600 | 2892200 | 20551000 | 1.42E+08 | 6056900 | 2040800 | 4203300 | 3624900 | 10468000 | 29604000 |
| TLHSTFQPI | MYOF | Q9NZM1 | MYOF_HUMAN | Unmodified | 105500 | 0 | 0 | 0 | 1252400 | 0 | 0 | 0 | 0 | 0 | 0 | 412630 | 0 | 0 | 0 | 54372 | 0 | 307130 |
| TLHSTFQPI | MYOF | Q9NZM1 | MYOF_HUMAN | Oxidation (M) | 526220 | 0 | 0 | 0 | 2466100 | 0 | 0 | 0 | 0 | 0 | 0 | 1035400 | 0 | 0 | 0 | 0 | 0 | 352440 |
| TLHTTASN | DYNC1H1; DYN | Q14204; A0/ | DYHC1_HUMA | Unmodified | 0 | 0 | 40948 | 0 | 0 | 0 | 0 | 0 | 0 | 0 | 72958 | 390680 | 0 | 0 | 129860 | 73679 | 0 | 0 |
| TLHTTASN | DYNC1H1; DYN | Q14204; A0/ | DYHC1_HUMA | Unmodified | 0 | 0 | 0 | 0 | 0 | 0 | 0 | 0 | 0 | 0 | 0 | 277020 | 0 | 0 | 0 | 0 | 0 | 0 |
| TLHWDTDI | DYNC1I2; DYN( | Q13409; E7E | DC1I2_HUMAN | Unmodified | 90941 | 0 | 70077 | 120030 | 0 | 0 | 235840 | 0 | 252130 | 421000 | 0 | 0 | 722300 | 411970 | 0 | 95127 | 0 | 0 |
| TLHYECIVL | NUDT5; NUDT5 | Q9UKK9; H0 | NUDT5_HUMAI | Unmodified | 94578 | 54543 | 0 | 261990 | 2329500 | 843240 | 723480 | 185480 | 2259500 | 1695200 | 2757300 | 4526700 | 1610800 | 0 | 301960 | 190970 | 986290 | 657740 |
| TLIDFVK | SCG3 | Q8WXD2 | SCG3_HUMAN | Unmodified | 0 | 0 | 0 | 0 | 0 | 0 | 0 | 0 | 46847000 | 22188000 | 0 | 0 | 0 | 0 | 0 | 0 | 0 | 0 |
| TLIEAGLPQ | HNRNPR; HNRI | O43390; A0/ | HNRPR_HUMA | Unmodified | 0 | 55071 | 0 | 0 | 0 | 0 | 0 | 0 | 0 | 0 | 0 | 6779700 | 0 | 0 | 0 | 0 | 0 | 0 |
| TLIEDEIATI | STK3 | Q13188 | STK3_HUMAN | Unmodified | 0 | 0 | 168920 | 122030 | 0 | 0 | 0 | 0 | 432250 | 0 | 0 | 828190 | 0 | 0 | 259220 | 0 | 0 | 0 |
| TLIEFLLR | LAP3 | P28838 | AMPL_HUMAN | Unmodified | 0 | 0 | 0 | 0 | 152530 | 32674 | 0 | 0 | 1491800 | 133230 | 360360 | 1113800 | 0 | 56899 | 0 | 349180 | 0 | 85656 |
| TLIEPFTLLA | NKIRAS1; NKIR/ | Q9NYS0; G5 | KBRS1_HUMAN | Unmodified | 0 | 0 | 0 | 0 | 345640 | 69012 | 0 | 0 | 58879 | 33339 | 31790 | 75106 | 246070 | 271930 | 0 | 0 | 0 | 284390 |
| TLIGHVPD( | PCK2; PCK2; P( | Q16822; H0 | PCKGM_HUMA | Unmodified | 0 | 0 | 0 | 0 | 0 | 0 | 0 | 0 | 182050 | 2166100 | 0 | 0 | 0 | 0 | 0 | 0 | 1104000 | 0 |
| TLIGHVPD( | PCK2; PCK2; P( | Q16822; H0 | PCKGM_HUMA | Unmodified | 0 | 0 | 0 | 0 | 0 | 0 | 0 | 0 | 3214800 | 1953900 | 0 | 0 | 0 | 0 | 0 | 0 | 0 | 0 |
| TLILNCVNF | PLXNA2 | O75051 | PLXA2_HUMAN | Unmodified | 129740 | 48579 | 35791 | 123480 | 0 | 0 | 0 | 0 | 1935000 | 999750 | 0 | 0 | 363620 | 323080 | 189320 | 44680 | 265980 | 581350 |
| TLINAEDPF | IQGAP1; IQGAP | P46940; A0A | IQGA1_HUMAN | Oxidation (M) | 768440 | 404870 | 533650 | 545950 | 8432400 | 0 | 343620 | 298240 | 1289900 | 0 | 0 | 2572000 | 846230 | 0 | 663850 | 1043500 | 447310 | 3281200 |
| TLINAEDPF | IQGAP1; IQGAP | P46940; A0A | IQGA1_HUMAN | Unmodified | 0 | 0 | 0 | 0 | 9812800 | 0 | 554260 | 0 | 1136500 | 419820 | 0 | 554620 | 0 | 0 | 0 | 0 | 0 | 0 |
| TLINELVK | DYNC1H1; DYN | Q14204; A0/ | DYHC1_HUMA | Unmodified | 0 | 1140500 | 0 | 0 | 0 | 0 | 0 | 0 | 0 | 0 | 0 | 2867100 | 0 | 264380 | 0 | 162500 | 0 | 0 |
| TLIQNCGA | CCT3; CCT3 | P49368; B4D | TCPG_HUMAN | Unmodified | 0 | 0 | 0 | 0 | 0 | 0 | 112720 | 103740 | 1306500 | 473240 | 2512200 | 27848000 | 4435700 | 286820 | 0 | 0 | 400330 | 691150 |
| TLISASSDT | WDR48; WDR4 | Q8TAF3; C9J | WDR48_HUMA | Unmodified | 0 | 0 | 0 | 0 | 929920 | 145820 | 32486 | 0 | 494720 | 536960 | 0 | 1863800 | 0 | 137360 | 0 | 239350 | 568350 | 458860 |
| TLISSTDFL | LRP8; LRP8; LRI | Q14114; A0/ | LRP8_HUMAN | Unmodified | 0 | 0 | 0 | 0 | 318520 | 760440 | 39097 | 0 | 1600000 | 1116200 | 352160 | 0 | 0 | 0 | 0 | 0 | 1304200 | 2592500 |
| TLIVTHSN( | AQR | O60306 | AQR_HUMAN F | Unmodified | 0 | 227810 | 0 | 0 | 0 | 0 | 0 | 0 | 82467 | 60717 | 0 | 283290 | 0 | 0 | 0 | 0 | 42548 | 0 |
| TLKAPDGLI | MANEAL | Q5VSG8 | MANEL_HUMAI | Unmodified | 0 | 0 | 0 | 0 | 0 | 0 | 0 | 0 | 82130 | 37095 | 0 | 0 | 0 | 0 | 0 | 0 | 0 | 0 |
| TLKDVLND | UTRN | P46939 | UTRN_HUMAN | Unmodified | 0 | 0 | 0 | 102700 | 445050 | 18329 | 0 | 0 | 0 | 286720 | 0 | 0 | 323060 | 208650 | 0 | 231670 | 535620 | 293500 |
| TLKIPAMTI/ | HSPD1; HSPD1 | P10809; A0A | CH60_HUMAN | Oxidation (M) | 0 | 0 | 0 | 0 | 108900 | 0 | 276280 | 0 | 145370 | 689730 | 0 | 0 | 1021600 | 511550 | 0 | 0 | 70721 | 0 |
| TLKPGTMM | YES1; YES1 | P07947; J3Q | YES_HUMAN T | 2 Oxidation (M) | 0 | 0 | 0 | 0 | 2960500 | 0 | 0 | 0 | 0 | 340420 | 0 | 0 | 481230 | 0 | 0 | 0 | 2227300 | 2137200 |
| TLKPGTMS | SRC | P12931 | SRC_HUMAN P | 2 Oxidation (M) | 0 | 0 | 0 | 0 | 0 | 0 | 0 | 0 | 0 | 0 | 0 | 0 | 0 | 0 | 0 | 0 | 756950 | 0 |
| TLKPGTMS | SRC | P12931 | SRC_HUMAN P | Oxidation (M) | 0 | 0 | 0 | 0 | 329430 | 0 | 0 | 0 | 0 | 0 | 0 | 0 | 288520 | 0 | 0 | 0 | 1434800 | 121000 |
| TLKPGTMS | FYN | P06241 | FYN_HUMAN T | Unmodified | 0 | 0 | 0 | 0 | 0 | 0 | 0 | 0 | 985720 | 0 | 0 | 0 | 0 | 0 | 0 | 0 | 0 | 0 |
| TLKPGTMS | LYN | P07948 | LYN_HUMAN T | 2 Oxidation (M) | 0 | 774840 | 0 | 0 | 3386500 | 0 | 0 | 0 | 0 | 1559200 | 0 | 303670 | 0 | 532780 | 2835400 | 0 | 0 | 0 |
| TLKPQPQQ | C11orf52; C11( | Q96A22; E9F | CK052_HUMAN | Unmodified | 5171400 | 5414000 | 2336200 | 2371700 | 0 | 0 | 0 | 0 | 5856700 | 1080200 | 0 | 856840 | 4983700 | 1235700 | 4067100 | 3280400 | 152580 | 2084100 |
| TLLADQGE | MTA2 | O94776 | MTA2_HUMAN | Unmodified | 0 | 0 | 0 | 0 | 0 | 0 | 0 | 0 | 7067300 | 0 | 462940 | 1358000 | 0 | 0 | 0 | 0 | 0 | 0 |
| TLLAQANSI | LAMB1; LAMB1; | P07942; G3 | LAMB1_HUMAN | Unmodified | 0 | 0 | 0 | 0 | 0 | 233220 | 0 | 129550 | 0 | 0 | 0 | 167670 | 0 | 0 | 139360 | 0 | 201310 | 2578000 |
| TLLAQANSI | LAMB1; LAMB1; | P07942; G3 | LAMB1_HUMAN | Unmodified | 0 | 0 | 0 | 0 | 0 | 0 | 0 | 0 | 0 | 0 | 0 | 0 | 0 | 0 | 0 | 0 | 251400 | 1889900 |
| TLLARACA/ | PSMC3; PSMC3 | P17980; E9P | PRS6A_HUMAN | Unmodified | 0 | 0 | 0 | 0 | 1007900 | 0 | 2166400 | 850980 | 0 | 130190 | 0 | 0 | 0 | 531900 | 0 | 0 | 3767700 | 0 |
| TLLARAVAS | PSMC6; PSMC6 | P62333; A0A | PRS10_HUMAN | Unmodified | 0 | 0 | 0 | 0 | 0 | 0 | 3386400 | 0 | 0 | 0 | 0 | 0 | 0 | 0 | 0 | 0 | 3447700 | 297290 |
| TLLDRVKLE | PYCR2; PYCR2; | Q96C36; A0/ | P5CR2_HUMAN | Unmodified | 0 | 0 | 0 | 0 | 0 | 0 | 1267400 | 95670 | 2445400 | 1020800 | 0 | 0 | 0 | 0 | 0 | 0 | 0 | 0 |
| TLLEALEQF | CC2D1A; CC2D | Q6P1N0; K7I | C2D1A_HUMAI | Unmodified | 500970 | 852900 | 496020 | 0 | 0 | 0 | 0 | 0 | 0 | 0 | 0 | 911050 | 0 | 0 | 714360 | 578750 | 0 | 0 |
| TLLEGSGLI | FASN; FASN | P49327; A0A | FAS_HUMAN F | Unmodified | 0 | 0 | 0 | 192830 | 0 | 0 | 0 | 0 | 0 | 0 | 0 | 0 | 88184 | 0 | 397440 | 0 | 0 | 0 |
| TLLEGSGLI | FASN; FASN | P49327; A0A | FAS_HUMAN F | Unmodified | 0 | 0 | 0 | 0 | 58711 | 0 | 0 | 0 | 0 | 0 | 0 | 0 | 172810 | 0 | 0 | 0 | 0 | 61466 |
| TLLEILDGK | HOMER1; HOM | Q86YM7; Q8 | HOME1_HUMA | Unmodified | 48990 | 0 | 62103 | 454600 | 113810 | 0 | 0 | 0 | 69052 | 0 | 115180 | 1712900 | 269860 | 141760 | 58965 | 98758 | 0 | 0 |
| TLLEILDGK | HOMER1; HOM | Q86YM7; Q8 | HOME1_HUMA | Unmodified | 0 | 0 | 0 | 0 | 0 | 0 | 0 | 0 | 0 | 0 | 0 | 0 | 864950 | 1101000 | 0 | 0 | 0 | 0 |
| TLLENTAITI | TNPO1; TNPO1 | Q92973; A0/ | TNPO1_HUMAI | Unmodified | 195560 | 0 | 390080 | 231070 | 0 | 0 | 0 | 0 | 0 | 0 | 0 | 1827100 | 0 | 0 | 0 | 267440 | 0 | 0 |
| TLLETLQK | PCYOX1; PCYO | Q9UHG3; F8 | PCYOX_HUMAI | Unmodified | 0 | 0 | 0 | 0 | 940780 | 21427000 | 3419400 | 231630 | 1654200 | 2543400 | 228840 | 66549 | 162760 | 0 | 0 | 0 | 390320 | 60338 |
| TLLGTKLEA | TRAF2; TRAF2; | Q12933; B1/ | TRAF2_HUMAN | Unmodified | 0 | 0 | 0 | 0 | 5108900 | 1407100 | 0 | 45385 | 0 | 0 | 0 | 0 | 0 | 143510 | 0 | 0 | 0 | 0 |
| TLLHLAVSL | ANKRD13A | Q8IZ07 | AN13A_HUMAN | Unmodified | 140550 | 206870 | 27983 | 118610 | 0 | 0 | 188620 | 0 | 0 | 0 | 0 | 0 | 0 | 0 | 212240 | 63241 | 0 | 0 |
| TLLIDELNK | ANLN; ANLN | Q9NQW6; H | ANLN_HUMAN | Unmodified | 0 | 0 | 0 | 284050 | 8822900 | 2119600 | 240450 | 0 | 196230 | 0 | 107650 | 0 | 381510 | 288210 | 0 | 0 | 1031800 | 813190 |
| TLLIKTVETF | VIM | P08670 | VIME_HUMAN | Unmodified | 0 | 0 | 0 | 0 | 0 | 0 | 0 | 0 | 0 | 0 | 0 | 0 | 385270 | 0 | 0 | 0 | 733770 | 0 |
| TLLLDGNPI | LRRC40 | Q9H9A6 | LRC40_HUMAN | Unmodified | 0 | 0 | 0 | 0 | 0 | 0 | 0 | 0 | 1078000 | 269170 | 0 | 2712400 | 0 | 0 | 0 | 0 | 0 | 0 |
| TLLLEPEGF | ANO9 | A1A5B4 | ANO9_HUMAN | Unmodified | 0 | 0 | 0 | 0 | 0 | 0 | 0 | 0 | 0 | 0 | 0 | 1922900 | 0 | 0 | 0 | 0 | 0 | 0 |
| TLLLNIFQD | ERO1B; ERO1B | Q86YB8; Q5 | ERO1B_HUMAI | Unmodified | 0 | 0 | 0 | 0 | 0 | 0 | 0 | 0 | 4804200 | 1488300 | 0 | 0 | 0 | 0 | 0 | 0 | 0 | 0 |
| TLLNETPE( | PROM1 | O43490 | PROM1_HUMA | Unmodified | 0 | 0 | 0 | 0 | 0 | 0 | 0 | 0 | 0 | 0 | 5708000 | 18034000 | 0 | 0 | 0 | 0 | 0 | 0 |
| TLLPEDSQI | MACF1; MACF1 | Q9UPN3; H3 | MACF1_HUMAI | Unmodified | 0 | 0 | 0 | 0 | 0 | 0 | 0 | 0 | 380450 | 0 | 0 | 575420 | 0 | 0 | 0 | 0 | 0 | 0 |
| TLLQQNQ( | GOLGA3; GOL( | Q08378; A0/ | GOGA3_HUMA | Unmodified | 0 | 0 | 0 | 0 | 0 | 0 | 0 | 0 | 0 | 0 | 0 | 215730 | 0 | 573430 | 0 | 0 | 0 | 197900 |
| TLLSDPTYF | STIP1; STIP1 | P31948; F5H | STIP1_HUMAN | Unmodified | 0 | 0 | 287960 | 0 | 0 | 0 | 101760 | 251310 | 425640 | 0 | 0 | 6344700 | 0 | 0 | 746790 | 319380 | 0 | 1875700 |
| TLLSDPTYF | STIP1; STIP1 | P31948; F5H | STIP1_HUMAN | Unmodified | 0 | 732690 | 0 | 667580 | 0 | 0 | 0 | 0 | 0 | 0 | 2940900 | 12681000 | 548490 | 0 | 1990400 | 866700 | 0 | 0 |
| TLLSESSS( | USP24 | Q9UPU5 | UBP24_HUMAI | Unmodified | 0 | 589240 | 0 | 0 | 0 | 0 | 0 | 0 | 0 | 0 | 0 | 339040 | 0 | 0 | 0 | 0 | 0 | 0 |

| | | | | | | | | | | | | | | | | | | | | | | |
|---|---|---|---|---|---|---|---|---|---|---|---|---|---|---|---|---|---|---|---|---|---|---|
| TLLSNLEEA | CLU; CLU; CLU | P10909; H0Y | CLUS_HUMAN | Unmodified | 0 | 0 | 0 | 0 | 330020 | 0 | 409020 | 706970 | 3465700 | 1430500 | 0 | 531620 | 611060 | 476030 | 0 | 242230 | 796550 | 0 |
| TLLSNLEEA | CLU; CLU; CLU | P10909; H0Y | CLUS_HUMAN | Unmodified | 0 | 0 | 0 | 0 | 2303200 | 2172000 | 139330 | 21803 | 3171200 | 1397900 | 0 | 0 | 0 | 0 | 0 | 0 | 24381 | 6059800 |
| TLLSTDHAI | DOCK6; DOCK6 | Q96HP0; K7I | DOCK6_HUMA | Unmodified | 0 | 0 | 0 | 0 | 235710 | 27123 | 0 | 0 | 0 | 137710 | 0 | 81404 | 0 | 0 | 0 | 68892 | 314450 | 152950 |
| TLLVNNNR | SNRPA1; SNRP | P09661; H0Y | RU2A_HUMAN | Unmodified | 175490 | 136200 | 260940 | 0 | 0 | 0 | 0 | 0 | 0 | 0 | 0 | 2341400 | 0 | 100350 | 0 | 141210 | 0 | 80510 |
| TLLWTELFF | NDUFS8; NDUF | O00217; F8W | NDUS8_HUMA | Unmodified | 0 | 0 | 0 | 0 | 0 | 0 | 0 | 0 | 1209200 | 251390 | 0 | 0 | 0 | 0 | 0 | 0 | 0 | 0 |
| TLMNLGGL | TAGLN2; TAGLN | P37802; X6R | TAGL2_HUMAN | Unmodified | 0 | 0 | 0 | 0 | 0 | 0 | 0 | 0 | 1098100 | 0 | 364220 | 2242400 | 0 | 0 | 0 | 0 | 0 | 0 |
| TLMNLGGL | TAGLN2; TAGLN | P37802; X6R | TAGL2_HUMAN | Oxidation (M) | 0 | 0 | 0 | 0 | 0 | 0 | 0 | 0 | 0 | 0 | 0 | 2710400 | 0 | 0 | 0 | 0 | 0 | 0 |
| TLMNLGGL | TAGLN2; TAGLN | P37802; X6R | TAGL2_HUMAN | Unmodified | 0 | 0 | 0 | 0 | 2375600 | 316740 | 0 | 0 | 395250 | 0 | 0 | 0 | 122770 | 87303 | 0 | 0 | 643250 | 579090 |
| TLMNLGGL | TAGLN2; TAGLN | P37802; X6R | TAGL2_HUMAN | Oxidation (M) | 474470 | 513820 | 0 | 363850 | 19910000 | 3192400 | 1962500 | 0 | 2105600 | 2400900 | 0 | 0 | 0 | 1653600 | 1153300 | 0 | 5365000 | 8452000 |
| TLMNLGGL | TAGLN2; TAGLN | P37802; X6R | TAGL2_HUMAN | Oxidation (M) | 0 | 0 | 0 | 11778000 | 51824000 | 25871000 | 8112200 | 0 | 4099200 | 8842900 | 2975200 | 4506100 | 16096000 | 8658900 | 4585000 | 0 | 23754000 | 18232000 |
| TLMNLGGL | TAGLN2; TAGLN | P37802; X6R | TAGL2_HUMAN | Unmodified | 0 | 0 | 0 | 497500 | 24772000 | 0 | 0 | 0 | 5270000 | 166310 | 448680 | 0 | 0 | 1620700 | 484670 | 0 | 3143900 | 9733900 |
| TLNAHEHF | PAFAH1B1; PAF | P43034; I3L | LIS1_HUMAN P | Unmodified | 0 | 0 | 0 | 0 | 2069200 | 0 | 1146700 | 0 | 0 | 492160 | 1617400 | 3101100 | 1254400 | 1277500 | 0 | 123410 | 1097700 | 284620 |
| TLNDELEIIE | HSPD1; HSPD1 | P10809; A0A | CH60_HUMAN | Oxidation (M) | 0 | 0 | 0 | 0 | 0 | 0 | 0 | 81329 | 3522800 | 308860 | 773410 | 0 | 190570 | 272710 | 122220 | 152770 | 0 | 0 |
| TLNDELEIIE | HSPD1; HSPD1 | P10809; A0A | CH60_HUMAN | Unmodified | 0 | 0 | 0 | 0 | 1010200 | 0 | 490720 | 0 | 10256000 | 113050 | 0 | 0 | 0 | 0 | 0 | 0 | 749950 | 0 |
| TLNDELEIIE | HSPD1; HSPD1 | P10809; A0A | CH60_HUMAN | Oxidation (M) | 0 | 0 | 0 | 0 | 0 | 0 | 0 | 0 | 1753900 | 0 | 0 | 0 | 0 | 0 | 0 | 0 | 0 | 182340 |
| TLNDELEIIE | HSPD1; HSPD1 | P10809; A0A | CH60_HUMAN | Unmodified | 0 | 0 | 0 | 0 | 0 | 0 | 0 | 0 | 4450100 | 0 | 0 | 0 | 0 | 0 | 0 | 0 | 0 | 0 |
| TLNDELEIIE | HSPD1; HSPD1 | P10809; A0A | CH60_HUMAN | Oxidation (M) | 0 | 0 | 0 | 0 | 0 | 0 | 490540 | 0 | 0 | 1816400 | 0 | 0 | 1033000 | 1589600 | 0 | 0 | 0 | 0 |
| TLNDELEIIE | HSPD1; HSPD1 | P10809; A0A | CH60_HUMAN | Unmodified | 0 | 0 | 0 | 0 | 0 | 0 | 0 | 0 | 0 | 606970 | 0 | 0 | 614460 | 784980 | 0 | 0 | 0 | 0 |
| TLNEWSSC | GCLM | P48507 | GSH0_HUMAN | Unmodified | 0 | 0 | 0 | 0 | 3553200 | 0 | 0 | 0 | 0 | 0 | 0 | 0 | 1467400 | 0 | 0 | 0 | 0 | 0 |
| TLNHLISGF | DICER1; DICER | Q9UPY3; H0 | DICER_HUMAN | Unmodified | 0 | 0 | 0 | 0 | 0 | 0 | 12014 | 0 | 0 | 0 | 0 | 344010 | 0 | 0 | 0 | 0 | 0 | 0 |
| TLNLAGNL | NISCH; NISCH | Q9Y2I1; C9J | NISCH_HUMAN | Unmodified | 30123 | 0 | 0 | 0 | 1081400 | 328040 | 343920 | 0 | 721560 | 876020 | 154950 | 869160 | 316170 | 131870 | 0 | 0 | 266900 | 179860 |
| TLNMDMFT | EDIL3 | O43854 | EDIL3_HUMAN | 2 Oxidation (M) | 0 | 0 | 0 | 0 | 0 | 0 | 0 | 0 | 592930 | 0 | 0 | 0 | 0 | 0 | 0 | 0 | 0 | 0 |
| TLNMDMFT | EDIL3 | O43854 | EDIL3_HUMAN | Unmodified | 0 | 0 | 0 | 0 | 457820 | 0 | 0 | 0 | 1704300 | 0 | 0 | 0 | 0 | 0 | 0 | 0 | 0 | 0 |
| TLNMDMFT | EDIL3 | O43854 | EDIL3_HUMAN | Oxidation (M) | 109280 | 0 | 0 | 0 | 51973000 | 25750000 | 0 | 0 | 15918000 | 1535600 | 0 | 0 | 0 | 0 | 0 | 0 | 1970100 | 18313000 |
| TLNMDMFT | EDIL3 | O43854 | EDIL3_HUMAN | Unmodified | 0 | 0 | 0 | 0 | 31880000 | 5696800 | 0 | 0 | 0 | 0 | 0 | 0 | 0 | 0 | 0 | 0 | 659500 | 9303200 |
| TLNMDMFT | EDIL3 | O43854 | EDIL3_HUMAN | 2 Oxidation (M) | 0 | 0 | 0 | 0 | 10690000 | 3856500 | 0 | 82631 | 1422600 | 163220 | 0 | 289320 | 0 | 0 | 0 | 30190 | 215270 | 2401000 |
| TLNMTTSPI | GMPS | P49915 | GUAA_HUMAN | Unmodified | 0 | 0 | 62696 | 160000 | 1832500 | 0 | 273200 | 247000 | 327960 | 336900 | 393280 | 1084900 | 1056300 | 100370 | 0 | 0 | 0 | 2140000 |
| TLNNLYAE | GOLGA7 | Q7Z5G4 | GOGA7_HUMA | Unmodified | 0 | 0 | 0 | 0 | 0 | 0 | 457320 | 0 | 152550 | 977980 | 199710 | 4403700 | 0 | 700180 | 1046900 | 0 | 0 | 0 |
| TLNPAFNE | SYT2 | Q8N9I0 | SYT2_HUMAN | Unmodified | 0 | 0 | 0 | 0 | 0 | 0 | 9995700 | 22872 | 0 | 0 | 0 | 0 | 0 | 0 | 0 | 0 | 0 | 0 |
| TLNPKAEV | CCT6A | P40227 | TCPZ_HUMAN | Unmodified | 0 | 0 | 0 | 0 | 0 | 0 | 110910 | 0 | 0 | 485470 | 0 | 0 | 1037500 | 294240 | 0 | 0 | 389160 | 141150 |
| TLNPKWNE | NEDD4L; NEDD | Q96PU5; A0 | NED4L_HUMAN | Unmodified | 0 | 0 | 0 | 0 | 1924100 | 0 | 99899 | 0 | 677470 | 4974800 | 0 | 0 | 1706200 | 1022700 | 0 | 0 | 0 | 1160600 |
| TLNPVFNE | SYT1; SYT1 | P21579; J3K | SYT1_HUMAN | Unmodified | 0 | 0 | 0 | 0 | 2824400 | 0 | 610400 | 0 | 3579100 | 4572300 | 0 | 0 | 0 | 0 | 0 | 0 | 176700 | 359270 |
| TLNQLGTP | STX7 | O15400 | STX7_HUMAN | Unmodified | 110000 | 0 | 401320 | 198710 | 0 | 0 | 0 | 147790 | 0 | 0 | 145340 | 890780 | 0 | 86827 | 343320 | 309210 | 0 | 0 |
| TLNQLGTP | STX7 | O15400 | STX7_HUMAN | Unmodified | 0 | 0 | 871840 | 2674000 | 744840 | 0 | 205360 | 79855 | 3159200 | 307690 | 693690 | 0 | 433040 | 255930 | 0 | 0 | 0 | 0 |
| TLNQQLTN | DNM2; DNM2 | P50570; K7E | DYN2_HUMAN | Unmodified | 2464700 | 3518100 | 1676700 | 1227900 | 0 | 0 | 0 | 0 | 1120400 | 0 | 287890 | 1910300 | 0 | 0 | 1740400 | 1585300 | 0 | 0 |
| TLNQQLTN | DNM2; DNM2 | P50570; K7E | DYN2_HUMAN | Unmodified | 415830 | 466890 | 0 | 865940 | 1318200 | 0 | 174700 | 0 | 888160 | 838400 | 0 | 0 | 3107100 | 1457400 | 0 | 0 | 744300 | 941670 |
| TLNVLFDSI | KIF20B; KIF20B | Q96Q89; A0 | KI20B_HUMAN | Unmodified | 24756 | 26849 | 0 | 0 | 1831000 | 375400 | 117220 | 0 | 47541 | 89321 | 32142 | 62319 | 103600 | 47692 | 0 | 0 | 767350 | 298050 |
| TLNVNKIIA | NT5E; NT5E | P21589; Q9 | 5NTD_HUMAN | Unmodified | 0 | 0 | 0 | 0 | 1115500 | 0 | 0 | 0 | 0 | 0 | 0 | 0 | 0 | 0 | 0 | 0 | 486380 | 0 |
| TLNVNKIIA | NT5E; NT5E | P21589; Q9 | 5NTD_HUMAN | Unmodified | 0 | 0 | 0 | 0 | 7156700 | 0 | 0 | 0 | 0 | 0 | 0 | 0 | 1036700 | 0 | 0 | 0 | 6651400 | 2424900 |
| TLNVNKIIA | NT5E; NT5E | P21589; Q9 | 5NTD_HUMAN | Oxidation (M) | 0 | 0 | 0 | 0 | 1692200 | 223810 | 0 | 0 | 0 | 0 | 0 | 228920 | 0 | 0 | 0 | 0 | 4141800 | 1978800 |
| TLPAAPAST | PHACTR4 | Q8IZ21 | PHAR4_HUMAN | Unmodified | 0 | 0 | 0 | 0 | 0 | 0 | 0 | 0 | 1234500 | 0 | 0 | 0 | 0 | 0 | 0 | 0 | 0 | 0 |
| TLPDEVLTK | API5; API5; API5 | Q9BZZ5; G3V | API5_HUMAN | Unmodified | 0 | 0 | 0 | 0 | 0 | 0 | 0 | 0 | 0 | 0 | 1043200 | 3140100 | 0 | 0 | 0 | 0 | 0 | 0 |
| TLPDEVLTK | API5; API5; API5 | Q9BZZ5; G3V | API5_HUMAN | Unmodified | 0 | 0 | 0 | 0 | 0 | 0 | 0 | 0 | 0 | 382190 | 0 | 289350 | 0 | 0 | 0 | 0 | 0 | 0 |
| TLPETLDPA | NDUFB4; NDUF | O95168; F22 | NDUB4_HUMA | Unmodified | 0 | 14851 | 0 | 0 | 594230 | 0 | 354370 | 217030 | 2053200 | 0 | 0 | 515280 | 0 | 332020 | 0 | 0 | 0 | 33777 |
| TLPETSLPN | FAM126A; FAM | Q9BYI3; A0A | HYCCI_HUMAN | Unmodified | 0 | 0 | 158370 | 0 | 1673900 | 0 | 154760 | 226870 | 137410 | 0 | 0 | 0 | 337210 | 289550 | 246760 | 0 | 319710 | 1968100 |
| TLPGSDWT | MARS1 | P56192 | SYMC_HUMAN | Unmodified | 0 | 0 | 0 | 0 | 0 | 0 | 0 | 0 | 0 | 0 | 795960 | 2400000 | 134820 | 0 | 231060 | 0 | 0 | 0 |
| TLPGWNTD | ADSS2 | P30520 | PURA2_HUMAN | Unmodified | 0 | 0 | 0 | 0 | 1229300 | 0 | 0 | 0 | 117160 | 331920 | 2749800 | 11206000 | 394190 | 261400 | 63318 | 144640 | 31711 | 0 |
| TLPGWNTD | ADSS2 | P30520 | PURA2_HUMAN | Unmodified | 0 | 0 | 0 | 0 | 2533600 | 266030 | 1716100 | 343540 | 0 | 249630 | 0 | 0 | 1010200 | 0 | 0 | 0 | 2658600 | 1868100 |
| TLPLTGSTF | ADAR; ADAR; AI | P55265; A0A | DSRAD_HUMAN | Oxidation (M) | 0 | 0 | 0 | 0 | 0 | 0 | 0 | 0 | 0 | 0 | 0 | 1567500 | 0 | 0 | 0 | 0 | 0 | 0 |
| TLPNIQEV | MTMR10; MTMR | Q9NXD2; H3 | MTMRA_HUMA | Unmodified | 81575 | 0 | 38023 | 56732 | 337290 | 71193 | 0 | 27594 | 0 | 0 | 32348 | 233190 | 63029 | 98414 | 0 | 45071 | 0 | 296260 |
| TLPQAEALI | AK3 | Q9UIJ7 | KAD3_HUMAN | Unmodified | 0 | 0 | 0 | 0 | 0 | 0 | 0 | 0 | 2597000 | 0 | 0 | 0 | 0 | 0 | 0 | 0 | 0 | 0 |
| TLPQQESLI | CARMIL1; CARM | Q5VZK9; A0 | CARL1_HUMAN | Unmodified | 0 | 0 | 0 | 0 | 1942100 | 0 | 0 | 0 | 0 | 0 | 0 | 1674600 | 0 | 0 | 181420 | 0 | 607570 | 1222300 |
| TLPQQESLI | CARMIL1; CARM | Q5VZK9; A0 | CARL1_HUMAN | Unmodified | 0 | 0 | 0 | 0 | 3753400 | 0 | 0 | 0 | 0 | 0 | 542750 | 0 | 723280 | 645150 | 0 | 0 | 0 | 0 |
| TLPSGLDD | MOGS; MOGS; | Q13724; A0 | MOGS_HUMAN | Unmodified | 0 | 0 | 0 | 0 | 0 | 0 | 0 | 193030 | 8996500 | 0 | 0 | 0 | 0 | 0 | 0 | 0 | 0 | 0 |
| TLQAGLSS | ELP1; ELP1; ELF | O95163; A0 | ELP1_HUMAN | Unmodified | 0 | 0 | 0 | 0 | 204120 | 0 | 0 | 0 | 90917 | 251860 | 0 | 413350 | 0 | 0 | 0 | 0 | 226120 | 207280 |
| TLQALEFH | LGALS3BP | Q08380 | LG3BP_HUMAN | Unmodified | 0 | 0 | 478960 | 350780 | 608770 | 717480 | 1518200 | 0 | 1.28E+08 | 26694000 | 1.18E+08 | 17335000 | 10112000 | 7044600 | 893310 | 535330 | 539170 | 19294000 |
| TLQALEFH | LGALS3BP | Q08380 | LG3BP_HUMAN | Unmodified | 0 | 0 | 0 | 0 | 372310 | 375470 | 4135700 | 0 | 0 | 3415800 | 1919100 | 0 | 7203300 | 7706400 | 0 | 0 | 981630 | 14135000 |
| TLQALQIPA | IQGAP1; IQGAP | P46940; A0A | IQGA1_HUMAN | Unmodified | 0 | 0 | 287790 | 0 | 0 | 0 | 0 | 0 | 0 | 0 | 0 | 1595500 | 0 | 0 | 0 | 0 | 0 | 0 |
| TLQALQIPA | IQGAP1; IQGAP | P46940; A0A | IQGA1_HUMAN | Unmodified | 0 | 127450 | 0 | 51395 | 1792000 | 0 | 0 | 0 | 0 | 69377 | 0 | 0 | 86885 | 0 | 88854 | 0 | 221650 | 0 |
| TLQALQIPA | IQGAP1; IQGAP | P46940; A0A | IQGA1_HUMAN | Unmodified | 0 | 0 | 0 | 0 | 2945800 | 1055900 | 0 | 0 | 0 | 885830 | 0 | 0 | 218920 | 0 | 0 | 0 | 0 | 0 |
| TLQELRAHI | DCN; DCN; DC | P07585; F8V | PGS2_HUMAN | Unmodified | 0 | 0 | 0 | 0 | 0 | 0 | 484440 | 0 | 0 | 0 | 0 | 0 | 0 | 0 | 0 | 0 | 0 | 0 |
| TLQEVTQLS | HADHA; HADHA | P40939; A0A | ECHA_HUMAN | Unmodified | 0 | 0 | 0 | 0 | 0 | 0 | 0 | 0 | 8347400 | 388820 | 0 | 255230 | 0 | 0 | 0 | 0 | 0 | 0 |
| TLQEVTQLS | HADHA; HADHA | P40939; A0A | ECHA_HUMAN | Unmodified | 0 | 0 | 0 | 0 | 0 | 0 | 312950 | 0 | 111060 | 469040 | 0 | 0 | 0 | 0 | 0 | 0 | 0 | 0 |
| TLQEVTQLS | HADHA; HADHA | P40939; A0A | ECHA_HUMAN | Unmodified | 0 | 0 | 0 | 0 | 279140 | 0 | 1532100 | 0 | 0 | 2167200 | 0 | 0 | 277230 | 0 | 0 | 0 | 137870 | 0 |
| TLQIFNIEM | CLTCL1; CLTC; | P53675; Q0 | CLH2_HUMAN | Oxidation (M) | 10428000 | 8339100 | 1874200 | 4419800 | 8588100 | 1067200 | 4658100 | 1585900 | 574110 | 838750 | 0 | 7503400 | 8160000 | 2508300 | 5476600 | 7572500 | 1924200 | 2727600 |
| TLQIFNIEM | CLTCL1; CLTC; | P53675; Q0 | CLH2_HUMAN | Unmodified | 0 | 0 | 0 | 0 | 3447600 | 838790 | 2483800 | 0 | 1273000 | 539290 | 106270 | 1965800 | 2518900 | 0 | 0 | 0 | 1555700 | 0 |
| TLQIFNIEM | CLTCL1; CLTC; | P53675; Q0 | CLH2_HUMAN | Oxidation (M) | 0 | 0 | 0 | 0 | 0 | 0 | 576410 | 262780 | 0 | 0 | 0 | 0 | 128860 | 0 | 0 | 0 | 325190 | 0 |
| TLQISYDAL | ENTR1; ENTR1 | Q96C92; H7 | ENTR1_HUMAN | Unmodified | 0 | 378440 | 0 | 371680 | 360960 | 0 | 0 | 117850 | 0 | 0 | 0 | 0 | 286710 | 110180 | 221560 | 160150 | 0 | 0 |
| TLQKGEYF | PRKG2 | Q13237 | KGP2_HUMAN | Unmodified | 0 | 0 | 0 | 0 | 0 | 0 | 1423700 | 988120 | 0 | 0 | 0 | 0 | 0 | 0 | 0 | 0 | 0 | 0 |
| TLQKLDEY | CLIC4 | Q9Y696 | CLIC4_HUMAN | Oxidation (M) | 0 | 0 | 0 | 0 | 3680500 | 0 | 0 | 0 | 0 | 0 | 0 | 0 | 928800 | 0 | 721490 | 0 | 1379600 | 2286000 |
| TLQLDNNF | PRPF19 | Q9UMS4 | PRP19_HUMAN | Unmodified | 138120 | 0 | 0 | 70331 | 539610 | 1264000 | 0 | 489730 | 137430 | 745970 | 0 | 4482900 | 480950 | 0 | 205590 | 0 | 225950 | 487810 |
| TLQLEYR | CPNE1; CPNE1 | Q99829; Q5 | CPNE1_HUMAN | Unmodified | 0 | 0 | 108480 | 547960 | 0 | 2552700 | 88476 | 46613 | 608450 | 448490 | 4347000 | 702150 | 0 | 818630 | 123920 | 0 | 205680 | 963370 |
| TLQLEYRF | CPNE1; CPNE1 | Q99829; Q5 | CPNE1_HUMAN | Unmodified | 143700 | 90724 | 262640 | 310340 | 3239300 | 816280 | 546500 | 0 | 2605100 | 1632600 | 1032000 | 2063600 | 853790 | 1246200 | 323380 | 201990 | 4650800 | 4836100 |
| TLQNREHE | STAT1; STAT1; S | P42224; A0A | STAT1_HUMAN | Unmodified | 0 | 0 | 0 | 0 | 228540 | 0 | 0 | 0 | 0 | 0 | 1107000 | 118340 | 0 | 0 | 0 | 0 | 61994 | 446910 |
| TLQPGDGE | MEGF8 | Q7Z7M0 | MEGF8_HUMA | Unmodified | 0 | 0 | 0 | 0 | 339340 | 0 | 0 | 0 | 517560 | 90697 | 0 | 0 | 0 | 0 | 0 | 0 | 163050 | 132920 |
| TLQQERED | CDC42BPA; CD | Q5VT25; A0 | MRCKA_HUMAN | Unmodified | 0 | 0 | 336370 | 426490 | 1073000 | 401710 | 0 | 93165 | 1455300 | 406190 | 504260 | 1104900 | 1468200 | 2484900 | 547510 | 781470 | 1202100 | 6335500 |
| TLQQLRQD | CGN | Q9P2M7 | CING_HUMAN | Unmodified | 0 | 0 | 0 | 0 | 387620 | 81351 | 0 | 0 | 0 | 0 | 0 | 596130 | 0 | 104880 | 1407800 | 0 | 48618 | 0 |
| TLQQNAES | BAG2 | O95816 | BAG2_HUMAN | Unmodified | 0 | 0 | 265540 | 0 | 4359300 | 1138000 | 370650 | 281000 | 0 | 1859200 | 0 | 170270 | 1618300 | 924880 | 208460 | 136450 | 2904700 | 1373200 |
| TLQQQTFK | RAD23B; RAD2 | P54727; P54 | RD23B_HUMAN | Unmodified | 3687000 | 3789800 | 1012400 | 692290 | 406100 | 456360 | 748060 | 179360 | 213960 | 1122800 | 248380 | 1324300 | 715340 | 265690 | 1048700 | 1758500 | 972260 | 259400 |
| TLQQQTFK | RAD23B; RAD2 | P54727; Q5 | RD23B_HUMAN | Unmodified | 9101100 | 8242200 | 1371100 | 1669100 | 2950500 | 0 | 689920 | 836530 | 2073200 | 2486800 | 1300800 | 0 | 3222400 | 0 | 4545600 | 4675800 | 1987700 | 109560 |
| TLQRAPTIS | OXSR1; OXSR1 | O95747; C9 | OXSR1_HUMAN | Unmodified | 0 | 0 | 0 | 0 | 626900 | 0 | 0 | 34332 | 393510 | 101710 | 0 | 229680 | 723880 | 319520 | 0 | 101830 | 0 | 592510 |
| TLQRDCLV | PIP5K1A; PIPSL | Q99755; A2 | PI51A_HUMAN | Unmodified | 0 | 0 | 0 | 0 | 1086100 | 0 | 369330 | 128330 | 299250 | 436770 | 0 | 0 | 484290 | 0 | 0 | 0 | 1289100 | 1168900 |
| TLQRLSSGI | PKP3; PKP3 | Q9Y446; E9 | PKP3_HUMAN | Unmodified | 0 | 0 | 0 | 0 | 916960 | 0 | 0 | 0 | 0 | 354920 | 0 | 0 | 1846500 | 1063700 | 0 | 0 | 0 | 0 |
| TLQRNYGR | GOLM1 | Q8NBJ4 | GOLM1_HUMA | Unmodified | 0 | 0 | 0 | 0 | 0 | 0 | 0 | 0 | 0 | 1562800 | 0 | 0 | 0 | 0 | 0 | 0 | 0 | 0 |
| TLQTMQNF | CHMP2B; CHM | Q9UQN3; C9 | CHM2B_HUMA | Oxidation (M) | 1265900 | 0 | 0 | 0 | 2582500 | 0 | 156250 | 88071 | 570470 | 646510 | 208920 | 476170 | 1741900 | 947860 | 0 | 69703 | 405030 | 2289400 |
| TLQTMQNF | CHMP2B; CHM | Q9UQN3; C9 | CHM2B_HUMA | Unmodified | 2600400 | 0 | 0 | 1706400 | 7275700 | 1280600 | 683170 | 293190 | 3060000 | 2399800 | 786370 | 2544000 | 7880200 | 4145600 | 788360 | 0 | 899080 | 4569800 |
| TLQVQAAQ | BLOC1S1; BLO | P78537; G8J | BL1S1_HUMAN | Unmodified | 0 | 0 | 0 | 0 | 824630 | 0 | 0 | 117300 | 161610 | 0 | 0 | 1801000 | 515850 | 345590 | 0 | 0 | 0 | 223390 |
| TLQVSPLDI | PTPN6 | P29350 | PTN6_HUMAN | Unmodified | 1726300 | 0 | 0 | 0 | 737910 | 0 | 0 | 0 | 524150 | 0 | 0 | 1338600 | 178580 | 0 | 77396 | 0 | 0 | 0 |
| TLQYLSQG | MRPS25; MRPS | P82663; E7E | RT25_HUMAN | Unmodified | 0 | 0 | 0 | 0 | 0 | 0 | 0 | 0 | 337080 | 0 | 0 | 0 | 0 | 0 | 0 | 0 | 0 | 0 |
| TLREQDDE | PKP3 | Q9Y446 | PKP3_HUMAN | Unmodified | 0 | 0 | 0 | 0 | 882240 | 82016 | 20115 | 0 | 0 | 0 | 0 | 6929800 | 358520 | 42494 | 2251700 | 1914700 | 425280 | 210280 |
| TLREQGVE | TLN1 | Q9Y490 | TLN1_HUMAN | Unmodified | 1349000 | 1461100 | 2297600 | 1484100 | 0 | 0 | 0 | 156370 | 1320200 | 0 | 1064100 | 1533200 | 0 | 0 | 2981000 | 2130400 | 0 | 295940 |
| TLRGSFSS | GPR180 | Q86V85 | GP180_HUMAN | Unmodified | 0 | 0 | 0 | 0 | 0 | 0 | 0 | 0 | 536820 | 0 | 0 | 0 | 0 | 0 | 0 | 0 | 0 | 0 |
| TLRHPNILA | SCYL1; SCYL1; | Q96KG9; E9 | SCYL1_HUMAN | Unmodified | 0 | 0 | 71041 | 0 | 362700 | 47533 | 130520 | 0 | 0 | 0 | 0 | 337900 | 0 | 0 | 207790 | 0 | 0 | 432350 |
| TLRLAGTQI | UBA1; UBA1 | P22314; Q5J | UBA1_HUMAN | Unmodified | 0 | 0 | 0 | 0 | 0 | 0 | 0 | 0 | 0 | 4202100 | 200130 | 0 | 986890 | 0 | 0 | 0 | 643120 | 622480 |
| TLRQKVEIL | MIA3 | Q5JRA6 | TGO1_HUMAN | Unmodified | 0 | 0 | 0 | 0 | 0 | 0 | 0 | 0 | 0 | 0 | 0 | 0 | 341560 | 370810 | 0 | 0 | 0 | 86923 |
| TLRQVEVIN | RANGAP1; RAN | P46060; H0Y | RAGP1_HUMAN | Unmodified | 0 | 0 | 0 | 0 | 532120 | 0 | 0 | 0 | 0 | 164260 | 0 | 0 | 1258800 | 409010 | 0 | 0 | 364800 | 0 |
| TLRQVEVIN | RANGAP1; RAN | P46060; H0Y | RAGP1_HUMAN | Unmodified | 0 | 0 | 0 | 0 | 0 | 0 | 659130 | 0 | 0 | 223690 | 0 | 0 | 0 | 866030 | 0 | 0 | 1517200 | 1365800 |
| TLSALLPA | IQGAP3; IQGAP | Q86VI3; F2Z | IQGA3_HUMAN | Unmodified | 360330 | 604680 | 0 | 399390 | 5026400 | 904920 | 217930 | 0 | 1339400 | 1116700 | 163800 | 370740 | 239610 | 0 | 165070 | 0 | 716710 | 330600 |
| TLSALLPA | IQGAP3; IQGAP | Q86VI3; F2Z | IQGA3_HUMAN | Unmodified | 0 | 0 | 0 | 0 | 967770 | 0 | 0 | 0 | 0 | 0 | 0 | 0 | 0 | 0 | 0 | 0 | 0 | 0 |
| TLSALRSPL | IQGAP1; IQGAP | P46940; A0A | IQGA1_HUMAN | Unmodified | 0 | 0 | 0 | 0 | 11376000 | 3903700 | 0 | 0 | 1336700 | 3200000 | 0 | 0 | 0 | 0 | 0 | 0 | 2301000 | 0 |
| TLSCPTKEF | POGLUT3 | Q7Z4H8 | PLGT3_HUMAN | Unmodified | 0 | 0 | 0 | 0 | 0 | 0 | 0 | 0 | 888870 | 511470 | 0 | 0 | 0 | 0 | 0 | 0 | 0 | 125870 |
| TLSDATTKF | CYRIB; CYRIB; | Q9NUQ9; A0 | CYRIB_HUMAN | Unmodified | 0 | 0 | 0 | 0 | 7713600 | 802940 | 515720 | 323910 | 0 | 1143900 | 28113 | 0 | 1005100 | 1150800 | 0 | 0 | 2665400 | 4575600 |
| TLSDLKLD | AKR1B1; AKR1B | P15121; E9P | ALDR_HUMAN | Unmodified | 0 | 0 | 0 | 0 | 8977900 | 1349100 | 0 | 0 | 0 | 158350 | 252090 | 1618400 | 0 | 0 | 0 | 0 | 1184500 | 0 |
| TLSEVDYAI | INPPL1; INPPL1 | O15357; H0Y | SHIP2_HUMAN | Unmodified | 0 | 0 | 0 | 0 | 0 | 0 | 0 | 0 | 0 | 0 | 0 | 991330 | 0 | 0 | 0 | 0 | 0 | 0 |
| TLSEVKLEV | UTRN | P46939 | UTRN_HUMAN | Unmodified | 0 | 0 | 0 | 326440 | 588990 | 0 | 0 | 0 | 129830 | 275080 | 0 | 0 | 434030 | 189270 | 305750 | 0 | 906200 | 508430 |
| TLSFGSDLI | EIF3A | Q14152 | EIF3A_HUMAN | Unmodified | 0 | 0 | 0 | 0 | 0 | 0 | 0 | 0 | 0 | 0 | 0 | 2123800 | 0 | 0 | 0 | 0 | 0 | 0 |
| TLSGMESY | CCT4 | P50991 | TCPD_HUMAN | Unmodified | 155450 | 0 | 0 | 105610 | 0 | 0 | 0 | 0 | 0 | 0 | 0 | 2218400 | 0 | 0 | 0 | 0 | 0 | 0 |
| TLSGMESY | CCT4 | P50991 | TCPD_HUMAN | Oxidation (M) | 0 | 0 | 0 | 0 | 0 | 0 | 0 | 0 | 130310 | 0 | 169130 | 1316300 | 0 | 0 | 200720 | 0 | 0 | 0 |
| TLSHNLLV | CUL4B; CUL4B | Q13620; A0 | CUL4B_HUMAN | Unmodified | 0 | 0 | 0 | 0 | 0 | 0 | 0 | 0 | 0 | 0 | 0 | 1251000 | 0 | 0 | 0 | 0 | 0 | 0 |

| | | | | | | | | | | | | | | | | | | | | | | |
|---|---|---|---|---|---|---|---|---|---|---|---|---|---|---|---|---|---|---|---|---|---|---|
| TLSHPQQN | TLN1 | Q9Y490 | TLN1_HUMAN | Unmodified | 0 | 683540 | 809270 | 1920100 | 15190000 | 3085200 | 14403000 | 5083700 | 1332100 | 685080 | 0 | 3940700 | 7562400 | 8693300 | 1103300 | 1291600 | 3439800 | 6187100 |
| TLSHPQQN | TLN1 | Q9Y490 | TLN1_HUMAN | Oxidation (M) | 0 | 1014200 | 0 | 0 | 0 | 0 | 1818400 | 0 | 0 | 0 | 0 | 0 | 60056 | 0 | 0 | 0 | 1132200 | 0 |
| TLSINEEEP | PPFIBP2; PPFIB | Q8ND30; HC | LIPB2_HUMAN | Unmodified | 0 | 0 | 0 | 368220 | 0 | 0 | 0 | 0 | 0 | 0 | 0 | 0 | 0 | 0 | 377350 | 295420 | 0 | 0 |
| TLSKYESSE | CRELD2; CRELI | Q6UXH1; A6 | CREL2_HUMAN | Unmodified | 0 | 0 | 0 | 0 | 0 | 0 | 0 | 18633 | 2811300 | 64117 | 0 | 0 | 52002 | 0 | 0 | 0 | 0 | 0 |
| TLSLDEVYL | XPNPEP1; XPNI | Q9NQW7; Q | XPP1_HUMAN | Unmodified | 0 | 0 | 0 | 0 | 0 | 0 | 0 | 0 | 0 | 0 | 0 | 1140200 | 0 | 0 | 0 | 0 | 0 | 0 |
| TLSLEAPAR | ABCC5 | O15440 | MRP5_HUMAN | Unmodified | 0 | 0 | 0 | 0 | 0 | 0 | 0 | 0 | 0 | 0 | 221200 | 4360200 | 0 | 0 | 0 | 0 | 0 | 0 |
| TLSLSGNQ | LRRC57; N/A; L | Q8N9N7; A0 | LRC57_HUMAN | Unmodified | 0 | 0 | 0 | 3097900 | 1339700 | 0 | 0 | 0 | 0 | 0 | 0 | 1199000 | 49769 | 0 | 85205 | 0 | 0 | 0 |
| TLSNAEDYI | OSTF1 | Q92882 | OSTF1_HUMAN | Unmodified | 0 | 0 | 0 | 0 | 0 | 0 | 0 | 0 | 0 | 0 | 0 | 1026500 | 0 | 0 | 0 | 0 | 0 | 0 |
| TLSNATMH | CYRIA | Q9H0Q0 | CYRIA_HUMAN | Oxidation (M) | 0 | 0 | 0 | 0 | 0 | 0 | 0 | 0 | 338420 | 0 | 105780 | 471630 | 0 | 0 | 0 | 0 | 0 | 0 |
| TLSNATMH | CYRIA | Q9H0Q0 | CYRIA_HUMAN | Unmodified | 0 | 0 | 0 | 0 | 0 | 0 | 0 | 0 | 1533500 | 0 | 205510 | 4123400 | 426220 | 296930 | 0 | 0 | 0 | 0 |
| TLSPTGNIS | SUN1; SUN1; S | O94901; H0 | SUN1_HUMAN | Unmodified | 1866500 | 1881200 | 980310 | 0 | 0 | 0 | 267470 | 107360 | 3697000 | 143790 | 0 | 0 | 0 | 0 | 1171900 | 719770 | 0 | 0 |
| TLSPTPASA | INCENP | Q9NQS7 | INCE_HUMAN | Unmodified | 0 | 0 | 0 | 0 | 650680 | 0 | 0 | 0 | 0 | 0 | 0 | 0 | 709960 | 400910 | 0 | 0 | 162140 | 0 |
| TLSQWIAEI | MPI; MPI; MPI; I | P34949; H3E | MPI_HUMAN M | Unmodified | 0 | 0 | 245660 | 225250 | 1683900 | 260950 | 338610 | 0 | 281840 | 413110 | 473950 | 1086500 | 1118100 | 984790 | 315420 | 382690 | 490670 | 604690 |
| TLSSKVEDI | OSBP | P22059 | OSBP1_HUMAI | Unmodified | 300180 | 0 | 132950 | 398090 | 456010 | 0 | 0 | 0 | 940260 | 132450 | 252390 | 2131500 | 1360500 | 273020 | 474110 | 359410 | 102580 | 549140 |
| TLSSLYSEN | OSTM1; OSTM1 | Q86WC4; AC | OSTM1_HUMAI | Unmodified | 0 | 0 | 0 | 0 | 0 | 0 | 0 | 299310 | 5208300 | 0 | 230350 | 0 | 0 | 0 | 0 | 0 | 0 | 0 |
| TLSSPSNRI | ATXN2L | Q8WWM7 | ATX2L_HUMAN | Unmodified | 686580 | 408140 | 0 | 276090 | 0 | 0 | 0 | 0 | 0 | 319000 | 0 | 0 | 381400 | 0 | 0 | 160830 | 0 | 0 |
| TLSTDAAP( | CLMP | Q9H6B4 | CLMP_HUMAN | Unmodified | 0 | 0 | 0 | 0 | 0 | 0 | 0 | 0 | 0 | 0 | 0 | 0 | 0 | 0 | 0 | 0 | 1265200 | 6046200 |
| TLSTDAAP( | CLMP | Q9H6B4 | CLMP_HUMAN | Unmodified | 0 | 0 | 0 | 0 | 0 | 0 | 0 | 0 | 0 | 0 | 0 | 0 | 0 | 0 | 0 | 0 | 0 | 1579400 |
| TLSTSDDVI | EXOC4 | Q96A65 | EXOC4_HUMAI | Unmodified | 124680 | 87738 | 302010 | 302390 | 1025900 | 347200 | 0 | 35645 | 0 | 358540 | 385030 | 2203700 | 285050 | 109070 | 325200 | 399750 | 66193 | 274860 |
| TLSVQEAA/ | LEMD3 | Q9Y2U8 | MAN1_HUMAN | Unmodified | 0 | 0 | 0 | 0 | 0 | 0 | 0 | 0 | 8504100 | 0 | 0 | 121130 | 0 | 0 | 0 | 0 | 0 | 0 |
| TLSVSPSVE | MTR; MTR; MTR | Q99707; A0/ | METH_HUMAN | Unmodified | 0 | 0 | 0 | 0 | 0 | 0 | 0 | 0 | 697630 | 0 | 0 | 0 | 0 | 0 | 0 | 0 | 0 | 0 |
| TLSYLLPAI\ | DDX5; DDX5; D | P17844; A0A | DDX5_HUMAN | Unmodified | 139900 | 0 | 0 | 0 | 0 | 0 | 0 | 0 | 0 | 0 | 0 | 3989300 | 0 | 0 | 0 | 0 | 0 | 0 |
| TLTALPGLF | AP4E1; AP4E1; | Q9UPM8; HC | AP4E1_HUMAN | Unmodified | 31197 | 19332 | 0 | 53174 | 146120 | 0 | 72002 | 23088 | 52494 | 0 | 27163 | 112040 | 31435 | 41163 | 0 | 0 | 30610 | 113860 |
| TLTAQSHM | DST; DST; DST; | Q03001; A0/ | DYST_HUMAN | Unmodified | 0 | 0 | 1079200 | 1517100 | 316740 | 122970 | 74560 | 9568.4 | 0 | 0 | 0 | 0 | 0 | 0 | 973600 | 1048500 | 0 | 0 |
| TLTAVHDAI | RPS7; RPS7; RP | P62081; A0A | RS7_HUMAN 4 | Unmodified | 76639 | 2337600 | 0 | 1127700 | 1777300 | 0 | 1396100 | 0 | 61949 | 851330 | 898500 | 898590 | 3843600 | 0 | 3804500 | 707090 | 0 | 1580400 |
| TLTAVHDAI | RPS7; RPS7; RP | P62081; A0A | RS7_HUMAN 4 | Unmodified | 2224400 | 3017800 | 168810 | 2976800 | 1450300 | 298910 | 1044300 | 0 | 1302000 | 497460 | 1976200 | 3056300 | 8583100 | 1477700 | 1126900 | 413700 | 267920 | 1981700 |
| TLTDELAAL | HSD17B11; HS | Q8NBQ5; D6 | DHB11_HUMAI | Unmodified | 0 | 0 | 0 | 0 | 146410 | 0 | 0 | 45844 | 1073100 | 346650 | 0 | 0 | 0 | 0 | 0 | 0 | 0 | 0 |
| TLTDGVLDI | PPFIA1; PPFIA1 | Q13136; E9F | LIPA1_HUMAN | Unmodified | 0 | 0 | 275240 | 199590 | 1599500 | 2725300 | 976590 | 329610 | 324480 | 608170 | 761000 | 0 | 268550 | 170730 | 423700 | 313720 | 97845 | 2806900 |
| TLTEGDSP( | CYB561; CYB56 | P49447; J3K | CY561_HUMAN | Unmodified | 0 | 0 | 505250 | 1710500 | 0 | 0 | 0 | 0 | 173800 | 31460 | 0 | 118700 | 0 | 0 | 0 | 409490 | 0 | 0 |
| TLTELNK | SNAP23; SNAP2 | O00161; H3I | SNP23_HUMAN | Unmodified | 0 | 0 | 0 | 87031 | 12192000 | 3214000 | 0 | 0 | 278860 | 50189 | 0 | 233560 | 0 | 50749 | 48634 | 25415 | 3768700 | 7660900 |
| TLTEPCPLA | PIGT; PIGT; PIG | Q969N2; A0 | PIGT_HUMAN ( | Unmodified | 0 | 0 | 0 | 0 | 0 | 0 | 0 | 0 | 11793000 | 1094800 | 0 | 522000 | 0 | 0 | 0 | 0 | 0 | 0 |
| TLTEPSVDF | RICTOR | Q6R327 | RICTR_HUMAN | Unmodified | 0 | 0 | 0 | 281840 | 0 | 0 | 0 | 0 | 0 | 0 | 0 | 1373400 | 0 | 1081900 | 0 | 0 | 0 | 0 |
| TLTETDLYC | PLXNB2; PLXNE | O15031; H0 | PLXB2_HUMAN | Unmodified | 0 | 0 | 0 | 0 | 0 | 0 | 0 | 256430 | 0 | 0 | 0 | 0 | 1809100 | 1258600 | 727350 | 0 | 2205000 | 2382200 |
| TLTFALR | DCPS; DCPS | Q96C86; A0 | DCPS_HUMAN | Unmodified | 0 | 0 | 0 | 0 | 0 | 0 | 0 | 0 | 0 | 0 | 0 | 1482000 | 0 | 0 | 0 | 0 | 0 | 0 |
| TLTGLINVG | DNAJC10; DNA | Q8IXB1; A0A | DJC10_HUMAN | Unmodified | 0 | 0 | 0 | 0 | 0 | 0 | 0 | 0 | 7454200 | 671170 | 0 | 0 | 0 | 0 | 0 | 0 | 0 | 0 |
| TLTGTAALT | LTA4H; LTA4H | P09960; B4D | LKHA4_HUMAN | Unmodified | 177860 | 0 | 0 | 0 | 484800 | 0 | 0 | 0 | 1492000 | 62206 | 2230700 | 11157000 | 0 | 0 | 0 | 122030 | 0 | 0 |
| TLTGVLPEF | UBLCP1 | Q8WVY7 | UBCP1_HUMAI | Unmodified | 0 | 0 | 0 | 0 | 0 | 0 | 0 | 0 | 0 | 439050 | 0 | 447180 | 669540 | 0 | 0 | 0 | 0 | 0 |
| TLTIDCVQF | CPT2; CPT2; CF | P23786; A0A | CPT2_HUMAN | Unmodified | 0 | 0 | 0 | 0 | 0 | 0 | 296400 | 0 | 1252800 | 0 | 0 | 0 | 0 | 0 | 0 | 0 | 0 | 0 |
| TLTKIDNSL | ACVR1 | Q04771 | ACVR1_HUMAI | Unmodified | 0 | 0 | 0 | 134950 | 1552700 | 234040 | 87555 | 39040 | 0 | 0 | 0 | 0 | 532960 | 142750 | 0 | 0 | 466130 | 945360 |
| TLTLLSVTR | CEACAM5; CEA | P06731; P31 | CEAM5_HUMAI | Unmodified | 0 | 0 | 0 | 0 | 0 | 0 | 0 | 28121 | 23532000 | 531170 | 0 | 0 | 0 | 0 | 0 | 0 | 0 | 0 |
| TLTLLSVTR | CEACAM5 | P06731 | CEAM5_HUMAI | Unmodified | 0 | 0 | 0 | 0 | 103170 | 0 | 0 | 0 | 10919000 | 0 | 91283 | 572850 | 0 | 0 | 0 | 0 | 0 | 0 |
| TLTLNCVNI | PLXNA1 | Q9UIW2 | PLXA1_HUMAN | Unmodified | 53277 | 0 | 63299 | 0 | 954070 | 0 | 0 | 30407 | 0 | 97573 | 0 | 0 | 659390 | 532420 | 103390 | 187340 | 445570 | 1409500 |
| TLTLPSLPLI | CD109 | Q6YHK3 | CD109_HUMAI | Unmodified | 157990 | 602040 | 0 | 0 | 0 | 0 | 0 | 0 | 3990500 | 460900 | 0 | 0 | 0 | 0 | 0 | 0 | 179890 | 590330 |
| TLTLVDTGI | HSP90AB1; HSI | P08238; Q5 | HS90B_HUMAI | Unmodified | 265360 | 0 | 687390 | 746300 | 1370100 | 0 | 0 | 0 | 22642000 | 918140 | 4269700 | 23742000 | 1010900 | 0 | 2090300 | 383600 | 0 | 0 |
| TLTLVDTGI | HSP90AB1; HSI | P08238; Q5 | HS90B_HUMAI | Unmodified | 163620 | 0 | 0 | 309580 | 606830 | 157610 | 40610 | 0 | 580260 | 843730 | 0 | 176760 | 1385300 | 330840 | 2306500 | 0 | 535320 | 386470 |
| TLTPDQWA | BOLA2; BOLA2E | Q9H3K6; A0 | BOLA2_HUMAN | Unmodified | 0 | 0 | 0 | 0 | 0 | 0 | 0 | 245390 | 0 | 0 | 0 | 0 | 584740 | 509380 | 0 | 0 | 0 | 0 |
| TLTPGGHA | CRIP2 | P52943 | CRIP2_HUMAN | Unmodified | 0 | 0 | 0 | 0 | 0 | 0 | 5511600 | 0 | 1773600 | 3398100 | 0 | 0 | 0 | 353710 | 0 | 0 | 19201000 | 13080000 |
| TLTPGGHA | CRIP2; CRIP2; ( | P52943; H0 | CRIP2_HUMAN | Unmodified | 0 | 0 | 0 | 0 | 688370 | 0 | 14314000 | 351150 | 4206200 | 5220200 | 0 | 0 | 0 | 0 | 0 | 0 | 18014000 | 16432000 |
| TLTPISAAY/ | ATIC | P31939 | PUR9_HUMAN | Unmodified | 200900 | 0 | 380140 | 185870 | 7254600 | 1168700 | 410470 | 144090 | 1836800 | 208720 | 1127500 | 14248000 | 434380 | 672190 | 0 | 102760 | 4855500 | 4331800 |
| TLTPSYYR | RAB18; RAB18; | Q9NP72; H0 | RAB18_HUMAN | Unmodified | 238000 | 513030 | 0 | 517840 | 0 | 1132100 | 0 | 0 | 808300 | 386930 | 823250 | 7145100 | 0 | 0 | 0 | 399600 | 0 | 461380 |
| TLTPTPDGI | ARHGEF1; ARH | Q92888; M0 | ARHG1_HUMA | Unmodified | 0 | 0 | 42364 | 0 | 635270 | 145180 | 0 | 110690 | 101430 | 0 | 28004 | 672530 | 0 | 0 | 70739 | 57682 | 3254200 | 140880 |
| TLTSGGHA | CRIP1; N/A; N/ | P50238; K4D | CRIP1_HUMAN | Unmodified | 0 | 0 | 0 | 0 | 11903000 | 3569800 | 41921000 | 2308900 | 1049600 | 5631500 | 1166600 | 0 | 0 | 450290 | 0 | 0 | 1906700 | 4308100 |
| TLTSGGHA | CRIP1; N/A; N/ | P50238; K4D | CRIP1_HUMAN | Oxidation (M) | 0 | 0 | 0 | 0 | 2619500 | 951560 | 7505800 | 1437200 | 214280 | 1943200 | 0 | 0 | 0 | 0 | 0 | 0 | 199420 | 547440 |
| TLTSSQAC\ | FURIN; FURIN | P09958; A0A | FURIN_HUMAN | Unmodified | 0 | 0 | 0 | 0 | 0 | 0 | 0 | 117570 | 0 | 0 | 744240 | 0 | 0 | 0 | 0 | 0 | 0 | 0 |
| TLTTLREDL | CCDC88C | Q9P219 | DAPLE_HUMAN | Unmodified | 0 | 0 | 0 | 119300 | 0 | 0 | 0 | 0 | 0 | 0 | 0 | 0 | 419830 | 295240 | 37320 | 52538 | 45713 | 79887 |
| TLTTSGTLS | ITGB4 | P16144 | ITB4_HUMAN I | Oxidation (M) | 0 | 0 | 0 | 0 | 1535900 | 131880 | 0 | 0 | 0 | 0 | 0 | 0 | 185050 | 0 | 0 | 0 | 8281400 | 7877900 |
| TLTTSGTLS | ITGB4 | P16144 | ITB4_HUMAN I | Unmodified | 0 | 0 | 0 | 0 | 1768200 | 0 | 0 | 0 | 0 | 61172 | 0 | 0 | 0 | 0 | 419730 | 0 | 3411600 | 6443600 |
| TLTTVQGIA | EIF1B; EIF1; EIF | O60739; P41 | EIF1B_HUMAN | Unmodified | 1830500 | 1474700 | 1298900 | 2215000 | 1236000 | 311120 | 287710 | 318030 | 604050 | 294610 | 457950 | 1076400 | 4197800 | 3227900 | 2765700 | 2171600 | 0 | 1023800 |
| TLTTVQGIA | EIF1B; EIF1; EIF | O60739; P41 | EIF1B_HUMAN | Unmodified | 737520 | 764310 | 1168200 | 1441200 | 3287500 | 0 | 834380 | 573340 | 539910 | 0 | 0 | 1148600 | 3503100 | 2471700 | 1429800 | 1236400 | 1389300 | 4757100 |
| TLTTVQGIA | EIF1B; EIF1; EIF | O60739; P41 | EIF1B_HUMAN | Unmodified | 0 | 0 | 0 | 0 | 1618000 | 243430 | 1175300 | 155340 | 0 | 128120 | 0 | 0 | 2697300 | 3681900 | 0 | 0 | 2074400 | 1300200 |
| TLVAHYPP\ | MSH6; MSH6; I | P52701; A0A | MSH6_HUMAN | Unmodified | 0 | 0 | 0 | 0 | 16090 | 0 | 0 | 214480 | 64871 | 194610 | 0 | 641530 | 0 | 0 | 0 | 0 | 0 | 0 |
| TLVEDLFAD | SNRNP200; SN | O75643; A0/ | U520_HUMAN | Unmodified | 0 | 0 | 0 | 0 | 0 | 0 | 0 | 0 | 0 | 0 | 0 | 5434900 | 0 | 0 | 0 | 0 | 0 | 0 |
| TLVFEQR | TMED5; TMED5 | Q9Y3A6; B1/ | TMED5_HUMAI | Unmodified | 0 | 0 | 0 | 0 | 0 | 0 | 0 | 0 | 53496000 | 0 | 0 | 0 | 0 | 0 | 0 | 0 | 0 | 0 |
| TLVGSENP | IQGAP2; IQGAP | Q13576; D6I | IQGA2_HUMAN | Unmodified | 0 | 0 | 1798400 | 1639500 | 311000 | 0 | 0 | 0 | 0 | 0 | 0 | 0 | 613790 | 400540 | 1725000 | 1207200 | 0 | 0 |
| TLVKEVVQI | COTL1; COTL1 | Q14019; H3I | COTL1_HUMAN | Unmodified | 0 | 0 | 0 | 0 | 853950 | 0 | 819390 | 0 | 0 | 0 | 0 | 0 | 0 | 189020 | 0 | 0 | 5555100 | 569750 |
| TLVLLDNLI | DDOST; DDOST | P39656; U3K | OST48_HUMAN | Unmodified | 0 | 0 | 0 | 0 | 0 | 0 | 0 | 0 | 34264000 | 4176600 | 0 | 3134700 | 0 | 0 | 0 | 0 | 0 | 0 |
| TLVLLDNLI | DDOST; DDOST | P39656; U3K | OST48_HUMAN | Unmodified | 0 | 0 | 0 | 0 | 0 | 0 | 45491 | 0 | 3867900 | 7367900 | 92746 | 0 | 0 | 0 | 0 | 0 | 0 | 0 |
| TLVLLDNLI | DDOST; DDOST | P39656; U3K | OST48_HUMAN | Unmodified | 0 | 0 | 42149 | 0 | 0 | 0 | 0 | 0 | 0 | 3835300 | 0 | 0 | 0 | 0 | 0 | 0 | 0 | 0 |
| TLVLLMGKI | PFN1; PFN1 | P07737; K7E | PROF1_HUMAN | Unmodified | 0 | 0 | 0 | 0 | 14724000 | 1304100 | 7018600 | 0 | 0 | 599280 | 0 | 0 | 6245400 | 824060 | 0 | 0 | 11402000 | 4259800 |
| TLVLLMGKI | PFN1; PFN1 | P07737; K7E | PROF1_HUMAN | Oxidation (M) | 0 | 0 | 0 | 0 | 6681400 | 0 | 3981500 | 832820 | 0 | 2096000 | 0 | 0 | 4184400 | 1678200 | 0 | 0 | 6381300 | 1026900 |
| TLVLLMGKI | PFN1; PFN1 | P07737; K7E | PROF1_HUMAN | Oxidation (M) | 0 | 0 | 0 | 0 | 8014700 | 1227200 | 5445900 | 623830 | 0 | 3102000 | 0 | 0 | 4209300 | 3642700 | 0 | 0 | 11460000 | 5075100 |
| TLVLLMGKI | PFN1; PFN1 | P07737; K7E | PROF1_HUMAN | Unmodified | 0 | 0 | 0 | 0 | 43658000 | 1613300 | 16538000 | 716020 | 237980 | 6110900 | 0 | 0 | 10563000 | 6626600 | 0 | 0 | 35074000 | 10810000 |
| TLVLSCVSI | PLXNA4 | Q9HCM2 | PLXA4_HUMAN | Unmodified | 0 | 0 | 0 | 0 | 0 | 0 | 132490 | 0 | 0 | 283890 | 0 | 451030 | 0 | 0 | 0 | 0 | 0 | 0 |
| TLVLSNLSY | NCL; NCL; NCL | P19338; C9J | NUCL_HUMAN | Unmodified | 0 | 0 | 0 | 0 | 0 | 0 | 0 | 0 | 0 | 632290 | 0 | 0 | 5871900 | 2080700 | 0 | 0 | 506780 | 0 |
| TLVNPANV | LYPLA1; LYPLA | O75608; B4D | LYPA1_HUMAN | Unmodified | 0 | 0 | 0 | 0 | 3758100 | 1311400 | 701370 | 728610 | 3437500 | 3210800 | 2924100 | 17602000 | 2364900 | 2668400 | 0 | 0 | 0 | 0 |
| TLVPHYCEI | SCYL1; SCYL1; | Q96KG9; E9I | SCYL1_HUMAN | Unmodified | 52324 | 0 | 0 | 884030 | 513110 | 0 | 244310 | 100720 | 46243 | 122270 | 180840 | 847630 | 350960 | 182790 | 0 | 0 | 567540 | 0 |
| TLVQEDSAI | SOGA1; SOGA1 | O94964; X6F | SOGA1_HUMAI | Unmodified | 0 | 0 | 0 | 0 | 129110 | 0 | 0 | 0 | 0 | 0 | 0 | 398900 | 1206100 | 1704400 | 0 | 98116 | 233060 | 277430 |
| TLVQQLYTI | MCU; MCU | Q8NE86; S4I | MCU_HUMAN ( | Unmodified | 0 | 0 | 0 | 0 | 0 | 0 | 0 | 0 | 1520300 | 0 | 0 | 0 | 0 | 0 | 0 | 0 | 0 | 0 |
| TLVRPSEH/ | TACSTD2 | P09758 | TACD2_HUMAN | Unmodified | 0 | 0 | 0 | 0 | 4809900 | 1315000 | 0 | 0 | 0 | 0 | 0 | 0 | 0 | 0 | 0 | 0 | 0 | 0 |
| TLVSQILDR | RIPOR2; RIPOR | Q9Y4F9; F5C | RIPR2_HUMAN | Unmodified | 0 | 0 | 0 | 0 | 0 | 0 | 0 | 0 | 0 | 0 | 0 | 814050 | 0 | 0 | 0 | 0 | 0 | 0 |
| TLVSVTKEC | HSP90AA1 | P07900 | HS90A_HUMAN | Unmodified | 1110700 | 0 | 0 | 3400900 | 11903000 | 3347400 | 2942800 | 1263100 | 10712000 | 7057400 | 7231500 | 22451000 | 4351900 | 3099200 | 0 | 1187800 | 4157800 | 5212500 |
| TLVTFSDEF | INSR; INSR | P06213; M0F | INSR_HUMAN I | Unmodified | 153800 | 0 | 373580 | 194850 | 0 | 0 | 0 | 0 | 797490 | 0 | 0 | 0 | 0 | 0 | 244130 | 272490 | 0 | 0 |
| TLVTLGKEI | PDK3 | Q15120 | PDK3_HUMAN | Unmodified | 0 | 0 | 0 | 0 | 0 | 0 | 0 | 0 | 837040 | 0 | 0 | 0 | 0 | 0 | 0 | 0 | 0 | 0 |
| TLVTLREDL | CCDC88A; CCI | Q3V6T2; A0/ | GRDN_HUMAN | Unmodified | 0 | 0 | 0 | 0 | 742550 | 0 | 211600 | 95400 | 370170 | 116650 | 0 | 256350 | 0 | 117210 | 0 | 0 | 375170 | 377420 |
| TLVTQNSG | JUP | P14923 | PLAK_HUMAN | Unmodified | 0 | 0 | 2296200 | 4463900 | 972810 | 332780 | 0 | 0 | 4549200 | 1093200 | 1481800 | 4474900 | 1013900 | 0 | 33556000 | 5758900 | 247080 | 406860 |
| TLVTQNSG | JUP | P14923 | PLAK_HUMAN | Unmodified | 0 | 0 | 0 | 6350000 | 8826500 | 0 | 0 | 0 | 0 | 375440 | 0 | 0 | 5109700 | 4616100 | 2029800 | 0 | 1508700 | 0 |
| TLVWSEK | LTA4H | P09960 | LKHA4_HUMAN | Unmodified | 13982 | 0 | 24704 | 24304 | 444570 | 0 | 147340 | 172090 | 434920 | 0 | 1161200 | 6568000 | 118300 | 0 | 64877 | 157430 | 98647 | 0 |
| TLVWSEKE | LTA4H | P09960 | LKHA4_HUMAN | Unmodified | 0 | 0 | 0 | 0 | 2938300 | 0 | 0 | 46419 | 0 | 0 | 0 | 0 | 0 | 0 | 0 | 0 | 546060 | 0 |
| TLWEDPGI | GNA11 | P29992 | GNA11_HUMAI | Unmodified | 137260 | 78100 | 0 | 0 | 1109900 | 0 | 0 | 61963 | 0 | 110900 | 72592 | 812560 | 0 | 0 | 0 | 0 | 0 | 848230 |
| TLWIWECG | RNH1; RNH1 | P13489; H0 | RINI_HUMAN R | Unmodified | 48933 | 0 | 0 | 46953 | 145360 | 0 | 45388 | 0 | 69426 | 47375 | 68282 | 1055700 | 73969 | 57286 | 0 | 33822 | 1285800 | 304670 |
| TLWNGQK | CAP1 | Q01518 | CAP1_HUMAN | Unmodified | 261260 | 0 | 0 | 606380 | 11137000 | 16305000 | 2063000 | 5025200 | 1460600 | 2206000 | 954890 | 4423200 | 197980 | 11217000 | 0 | 373200 | 5412900 | 21568000 |
| TLWNGQKI | CAP1 | Q01518 | CAP1_HUMAN | Unmodified | 1793400 | 0 | 0 | 705480 | 944190 | 0 | 2098100 | 0 | 0 | 14566 | 0 | 1202900 | 0 | 72566 | 0 | 0 | 0 | 0 |
| TLYDESCSI | PSMG2; PSMG2 | Q969U7; K7I | PSMG2_HUMA | Unmodified | 0 | 0 | 0 | 0 | 0 | 0 | 0 | 171720 | 0 | 0 | 0 | 456260 | 0 | 0 | 0 | 0 | 0 | 0 |
| TLYFADTYL | LLGL2; LLGL2 | Q6P1M3; J3C | L2GL2_HUMAN | Unmodified | 0 | 0 | 0 | 0 | 675040 | 0 | 0 | 204300 | 4994500 | 1797500 | 923840 | 2952700 | 0 | 0 | 0 | 0 | 590790 | 633630 |
| TLYLLAPSF | CIT; CIT; CIT; CI | O14578; A0/ | CTRO_HUMAN | Unmodified | 0 | 0 | 0 | 0 | 0 | 0 | 0 | 0 | 0 | 0 | 0 | 999750 | 0 | 0 | 0 | 0 | 0 | 0 |
| TLYNNQPII | BLMH; BLMH; B | Q13867; K7E | BLMH_HUMAN | Unmodified | 0 | 0 | 0 | 34368 | 0 | 0 | 587010 | 599210 | 630510 | 637680 | 0 | 2192300 | 0 | 564960 | 0 | 98305 | 0 | 0 |
| TLYNNQPII | BLMH; BLMH; B | Q13867; K7E | BLMH_HUMAN | Unmodified | 0 | 0 | 0 | 0 | 669660 | 0 | 1837400 | 1290200 | 0 | 3007100 | 0 | 0 | 1568300 | 1461800 | 0 | 0 | 1163800 | 883600 |
| TLYVEEVVF | GARS1; GARS1 | P41250; A0A | GARS_HUMAN | Unmodified | 97760 | 155680 | 112150 | 863100 | 1451500 | 611030 | 1489700 | 0 | 2403500 | 4134100 | 1909600 | 3842400 | 5207700 | 1597500 | 2621900 | 233590 | 909010 | 1283900 |
| TMAAEVLSI | NCLN | Q969V3 | NCLN_HUMAN | Unmodified | 0 | 0 | 0 | 0 | 0 | 0 | 0 | 0 | 1937000 | 2770900 | 0 | 0 | 1124800 | 0 | 0 | 0 | 0 | 0 |
| TMAAGDPA | DOP1B | Q9Y3R5 | DOP2_HUMAN | Unmodified | 0 | 0 | 0 | 0 | 0 | 0 | 0 | 0 | 1214800 | 178110 | 0 | 0 | 0 | 0 | 0 | 186660 | 0 | 0 |
| TMADGNEI | YWHAH; YWHA | Q04917; A2I | 1433F_HUMAN | Unmodified | 0 | 0 | 0 | 0 | 0 | 0 | 707890 | 0 | 0 | 0 | 0 | 0 | 0 | 0 | 0 | 0 | 3352000 | 470240 |
| TMADLHTA | TAX1BP1; TAX1 | Q86VP1; B8 | TAXB1_HUMAN | Unmodified | 0 | 0 | 0 | 0 | 0 | 0 | 0 | 0 | 252610 | 38639 | 0 | 0 | 0 | 0 | 0 | 0 | 0 | 0 |
| TMADSSYC | FERMT1; FERM | Q9BQL6; G3 | FERM1_HUMAI | Oxidation (M) | 0 | 0 | 0 | 0 | 83534 | 0 | 11439 | 0 | 0 | 0 | 0 | 0 | 76172 | 29945 | 0 | 0 | 0 | 0 |
| TMAFHGLL | ZYG11B; ZYG11 | Q9C0D3; A8 | ZY11B_HUMAN | Oxidation (M) | 0 | 0 | 0 | 66528 | 0 | 0 | 0 | 0 | 0 | 0 | 0 | 228470 | 0 | 0 | 0 | 28614 | 0 | 0 |
| TMASQVSII | MRPL45; MRPL | Q9BRJ2; A0A | RM45_HUMAN | Unmodified | 0 | 0 | 0 | 0 | 820480 | 999750 | 0 | 0 | 329820 | 0 | 0 | 0 | 0 | 0 | 0 | 0 | 0 | 0 |
| TMDASER | ALDH1A1; ALDI | P00352; Q5 | AL1A1_HUMAN | Unmodified | 0 | 0 | 0 | 0 | 0 | 0 | 0 | 0 | 0 | 0 | 0 | 706880 | 0 | 0 | 0 | 0 | 0 | 0 |
| TMDKSELV | YWHAB; YWHA | P31946; Q4 | 1433B_HUMAN | Acetyl (Protein N | 1278800 | 709970 | 111710 | 423770 | 4802400 | 1404400 | 1445300 | 1221800 | 200780 | 493100 | 557990 | 0 | 3595200 | 1580500 | 125970 | 172360 | 7019700 | 5594700 |
| TMDKSELV | YWHAB; YWHA | P31946; Q4 | 1433B_HUMAN | Acetyl (Protein N | 154380 | 210920 | 0 | 774090 | 15246000 | 2173900 | 3466700 | 2739700 | 1298600 | 1768700 | 1150100 | 2245300 | 4931200 | 4560200 | 0 | 0 | 20040000 | 16244000 |
| TMDKSELV | YWHAB; YWHA | P31946; Q4 | 1433B_HUMAN | Acetyl (Protein N | 0 | 0 | 0 | 0 | 0 | 0 | 290330 | 313910 | 121330 | 135330 | 227320 | 1340500 | 200150 | 136240 | 0 | 70199 | 1055500 | 1005900 |

| | | | | | | | | | | | | | | | | | | | | | | |
|---|---|---|---|---|---|---|---|---|---|---|---|---|---|---|---|---|---|---|---|---|---|---|
| TMDKSELV | YWHAB; YWHA | P31946; Q4\ | 1433B_HUMAN | Acetyl (Protein N | 0 | 0 | 0 | 0 | 944030 | 0 | 974240 | 966920 | 0 | 0 | 0 | 0 | 340040 | 306610 | 0 | 0 | 3336400 | 0 |
| TMDLPFLE/ | PLCG1 | P19174 | PLCG1_HUMAI | Oxidation (M) | 0 | 0 | 0 | 0 | 0 | 0 | 0 | 0 | 0 | 0 | 0 | 1363200 | 0 | 0 | 0 | 0 | 0 | 0 |
| TMDYETSE | DAAM1 | Q9Y4D1 | DAAM1_HUMA | Unmodified | 0 | 0 | 0 | 0 | 0 | 0 | 0 | 0 | 0 | 0 | 0 | 630250 | 0 | 0 | 90947 | 0 | 0 | 0 |
| TMELAAQS | CHMP4A; CHM | Q9BY43; E9I | CHM4A_HUMA | Unmodified | 0 | 910810 | 0 | 104140 | 0 | 0 | 0 | 0 | 955600 | 0 | 66541 | 427660 | 0 | 0 | 0 | 0 | 0 | 0 |
| TMELLQSD | EFCAB14; EFCA | O75071; A0/ | EFC14_HUMAN | Oxidation (M) | 0 | 0 | 0 | 0 | 0 | 0 | 0 | 23251 | 2381700 | 611260 | 0 | 0 | 0 | 0 | 0 | 0 | 0 | 0 |
| TMELLQSD | EFCAB14; EFCA | O75071; A0/ | EFC14_HUMAN | Unmodified | 0 | 0 | 0 | 0 | 0 | 0 | 139680 | 0 | 6093500 | 458230 | 0 | 0 | 0 | 0 | 0 | 0 | 0 | 0 |
| TMGCLATT | ABAT; ABAT; AB | P80404; H3E | GABT_HUMAN | Unmodified | 28177 | 0 | 0 | 0 | 0 | 0 | 441670 | 0 | 861960 | 207350 | 0 | 0 | 0 | 0 | 0 | 0 | 0 | 0 |
| TMGFCYQI | IPO7 | O95373 | IPO7_HUMAN I | Oxidation (M) | 0 | 0 | 0 | 0 | 0 | 0 | 0 | 0 | 0 | 0 | 0 | 1067400 | 0 | 0 | 0 | 0 | 0 | 0 |
| TMGYSVTA | PMM2; PMM2; F | O15305; H3I | PMM2_HUMAN | Unmodified | 0 | 0 | 0 | 0 | 0 | 0 | 0 | 0 | 539150 | 0 | 419030 | 1120700 | 0 | 0 | 0 | 0 | 0 | 0 |
| TMIVHDDV | MAP4K4; MAP4 | O95819; C9J | M4K4_HUMAN | Unmodified | 0 | 0 | 0 | 0 | 179910 | 0 | 0 | 0 | 613220 | 103400 | 186500 | 436880 | 143240 | 0 | 0 | 35906 | 119150 | 1264400 |
| TMIVHDDV | MAP4K4 | E7EN19 | E7EN19_HUMA | Oxidation (M) | 0 | 0 | 0 | 0 | 0 | 0 | 0 | 0 | 0 | 0 | 0 | 0 | 0 | 0 | 0 | 0 | 0 | 914580 |
| TMKEKEELI | TAX1BP1; TAX1I | Q86VP1; B8Z | TAXB1_HUMAN | Unmodified | 0 | 0 | 0 | 0 | 0 | 0 | 617960 | 0 | 0 | 292140 | 0 | 0 | 0 | 0 | 0 | 0 | 362650 | 566560 |
| TMLDFPQH | NLN | Q9BYT8 | NEUL_HUMAN | Unmodified | 0 | 0 | 0 | 0 | 0 | 0 | 0 | 0 | 682160 | 355750 | 138330 | 845440 | 0 | 0 | 0 | 0 | 0 | 0 |
| TMLEDLGM | SKP1; N/A; SKP | P63208; E5R | SKP1_HUMAN | 2 Oxidation (M) | 0 | 0 | 0 | 0 | 1889400 | 0 | 0 | 0 | 0 | 0 | 122280 | 0 | 2228000 | 0 | 0 | 0 | 0 | 0 |
| TMLELLNQ | PSMC3; PSMC3 | P17980; E9P | PRS6A_HUMAN | Oxidation (M) | 0 | 0 | 0 | 0 | 0 | 0 | 0 | 0 | 566160 | 0 | 0 | 1094800 | 0 | 0 | 0 | 0 | 353790 | 32348 |
| TMLELLNQ | PSMC3; PSMC3 | P17980; E9P | PRS6A_HUMAN | Unmodified | 0 | 0 | 17599 | 99329 | 0 | 0 | 0 | 0 | 1121200 | 0 | 0 | 0 | 0 | 0 | 0 | 0 | 0 | 0 |
| TMLERLASI | GANAB; GANAE | Q14697; F5H | GANAB_HUMA | Unmodified | 0 | 0 | 0 | 0 | 0 | 0 | 0 | 0 | 0 | 6881600 | 0 | 0 | 0 | 0 | 0 | 0 | 0 | 0 |
| TMLESAGG | TLN1 | Q9Y490 | TLN1_HUMAN | Unmodified | 0 | 0 | 0 | 804890 | 8543500 | 1713800 | 2733900 | 627350 | 751720 | 325110 | 0 | 1115600 | 2857200 | 3440800 | 152970 | 66010 | 1402600 | 5434400 |
| TMLESAGG | TLN1 | Q9Y490 | TLN1_HUMAN | Oxidation (M) | 0 | 0 | 0 | 0 | 2170100 | 227010 | 1290500 | 1441600 | 0 | 33355 | 0 | 548980 | 1666100 | 1831300 | 0 | 222050 | 911040 | 2281900 |
| TMLESLIAD | CYFIP1; CYFIP2 | Q7L576; Q9 | CYFP1_HUMAN | Oxidation (M) | 145190 | 220000 | 367050 | 1099400 | 189490 | 0 | 0 | 0 | 322300 | 97651 | 657690 | 4614700 | 0 | 0 | 1109000 | 979690 | 475040 | 0 |
| TMLYTPQEI | RUVBL1; RUVBI | Q9Y265; E7E | RUVB1_HUMAI | 2 Oxidation (M) | 0 | 0 | 0 | 0 | 0 | 0 | 0 | 0 | 0 | 0 | 0 | 868160 | 0 | 0 | 0 | 0 | 0 | 0 |
| TMLYTPQEI | RUVBL1; RUVBI | Q9Y265; E7E | RUVB1_HUMAI | Unmodified | 0 | 0 | 0 | 0 | 0 | 0 | 0 | 0 | 0 | 0 | 0 | 1428200 | 0 | 0 | 0 | 0 | 0 | 0 |
| TMLYTPQEI | RUVBL1; RUVBI | Q9Y265; E7E | RUVB1_HUMAI | Oxidation (M) | 0 | 0 | 0 | 0 | 0 | 0 | 0 | 0 | 0 | 0 | 0 | 2556600 | 0 | 0 | 0 | 0 | 0 | 0 |
| TMNISPEQI | SEC23IP | Q9Y6Y8 | S23IP_HUMAN | Unmodified | 0 | 0 | 0 | 0 | 0 | 0 | 0 | 0 | 0 | 0 | 0 | 647210 | 0 | 0 | 0 | 0 | 0 | 0 |
| TMNRLTQL | ERBIN; ERBIN | Q96RT1; B4I | ERBIN_HUMAN | Oxidation (M) | 0 | 0 | 0 | 0 | 3162400 | 0 | 0 | 0 | 0 | 0 | 0 | 0 | 2421500 | 0 | 0 | 0 | 0 | 0 |
| TMQALEK | QSER1; VPS28; | Q2KHR3; Q9 | QSER1_HUMAI | Unmodified | 21349000 | 28159000 | 21402000 | 22313000 | 10984000 | 2947800 | 3449800 | 2432700 | 0 | 3127400 | 0 | 0 | 36720000 | 5483700 | 43887000 | 24773000 | 2297900 | 8814800 |
| TMQALEK | QSER1; VPS28; | Q2KHR3; Q9 | QSER1_HUMAI | Oxidation (M) | 10842000 | 45480 | 72471 | 196840 | 1398900 | 254750 | 0 | 0 | 59952 | 0 | 0 | 0 | 657940 | 45009 | 0 | 0 | 10545 | 0 |
| TMQGSEVV | PDCD6IP | Q8WUM4 | PDC6I_HUMAN | Oxidation (M) | 15983000 | 12040000 | 11516000 | 12055000 | 16811000 | 4637700 | 1510900 | 920690 | 5684300 | 6583300 | 262770 | 5739900 | 18582000 | 12929000 | 12746000 | 16237000 | 3428700 | 12308000 |
| TMQGSEVV | PDCD6IP | Q8WUM4 | PDC6I_HUMAN | Unmodified | 4115600 | 4157800 | 2987600 | 5711600 | 25113000 | 0 | 0 | 300520 | 13743000 | 8768600 | 1162400 | 4723800 | 15901000 | 8970100 | 5067700 | 2989200 | 3856900 | 13129000 |
| TMQNTND | CTNNB1; CTNN | P35222; A0A | CTNB1_HUMAI | Unmodified | 0 | 0 | 2584400 | 3689300 | 290800 | 295030 | 0 | 0 | 750200 | 48135 | 516650 | 4297400 | 0 | 0 | 5799600 | 5110900 | 371710 | 406720 |
| TMQNTND | CTNNB1; CTNN | P35222; A0A | CTNB1_HUMAI | Oxidation (M) | 0 | 0 | 0 | 0 | 0 | 0 | 0 | 0 | 0 | 0 | 0 | 0 | 0 | 0 | 0 | 0 | 0 | 0 |
| TMQNTSDI | JUP; JUP; JUP; J | P14923; C9J | PLAK_HUMAN | Oxidation (M) | 0 | 0 | 1980800 | 2967100 | 0 | 0 | 0 | 0 | 330720 | 0 | 188190 | 1486300 | 0 | 0 | 2975900 | 3279400 | 0 | 0 |
| TMQNTSDI | JUP; JUP; JUP; J | P14923; C9J | PLAK_HUMAN | Unmodified | 0 | 0 | 130830 | 3560500 | 240010 | 0 | 0 | 0 | 880510 | 66158 | 367130 | 2923000 | 0 | 0 | 4905500 | 3237100 | 0 | 0 |
| TMQNVKHI | TFRC; TFRC | P02786; G3\ | TFR1_HUMAN | Unmodified | 0 | 0 | 0 | 0 | 0 | 0 | 0 | 0 | 3014600 | 0 | 0 | 0 | 0 | 0 | 0 | 0 | 0 | 0 |
| TMQTLLSL | HPRT1 | P00492 | HPRT_HUMAN | Unmodified | 0 | 0 | 0 | 0 | 0 | 0 | 0 | 0 | 1226400 | 219620 | 0 | 642170 | 0 | 0 | 0 | 0 | 0 | 0 |
| TMQTLLSL | HPRT1 | P00492 | HPRT_HUMAN | Oxidation (M) | 98484 | 95716 | 0 | 0 | 1015200 | 0 | 0 | 0 | 309910 | 652560 | 974140 | 4165500 | 0 | 0 | 0 | 51435 | 392840 | 0 |
| TMRSNLEM | CD2AP | Q9Y5K6 | CD2AP_HUMAI | Oxidation (M) | 0 | 0 | 0 | 0 | 436380 | 0 | 0 | 51021 | 0 | 0 | 0 | 0 | 673500 | 510700 | 0 | 0 | 38720 | 0 |
| TMRSNLEM | CD2AP | Q9Y5K6 | CD2AP_HUMAI | Unmodified | 0 | 0 | 0 | 0 | 0 | 0 | 0 | 0 | 0 | 0 | 0 | 0 | 3935900 | 1523200 | 0 | 0 | 0 | 0 |
| TMTIHNGM | FGB; FGB | P02675; D6F | FIBB_HUMAN F | 2 Oxidation (M) | 0 | 3237200 | 0 | 0 | 0 | 0 | 0 | 0 | 0 | 0 | 0 | 0 | 0 | 0 | 0 | 0 | 0 | 0 |
| TMTLFSALF | DYNC1H1; DYN | Q14204; A0/ | DYHC1_HUMA | Unmodified | 0 | 0 | 0 | 0 | 0 | 0 | 0 | 0 | 91894 | 0 | 0 | 654240 | 0 | 0 | 0 | 0 | 0 | 0 |
| TMTNALR | PSMD3 | O43242 | PSMD3_HUMA | Unmodified | 0 | 0 | 36505 | 0 | 0 | 0 | 0 | 0 | 254400 | 13648 | 29675 | 677990 | 0 | 0 | 46080 | 0 | 183390 | 10189 |
| TMVDELFAI | RAP2A | P10114 | RAP2A_HUMAN | Oxidation (M) | 0 | 0 | 0 | 0 | 122220 | 0 | 0 | 0 | 164550 | 0 | 56862 | 896600 | 0 | 0 | 0 | 0 | 0 | 337850 |
| TMWSPPGI | CDON; CDON | Q4KMG0; E9 | CDON_HUMAN | Oxidation (M) | 0 | 0 | 0 | 0 | 1088400 | 0 | 0 | 0 | 114500 | 0 | 0 | 0 | 0 | 0 | 0 | 0 | 530360 | 416920 |
| TMYLEDR | RAB6A; RAB6B; | P20340; Q9I | RAB6A_HUMAN | Oxidation (M) | 0 | 207310 | 0 | 0 | 0 | 0 | 0 | 0 | 0 | 0 | 0 | 2516200 | 76389 | 0 | 427760 | 323600 | 0 | 0 |
| TMYVLPFSI | PCK2; PCK2; P( | Q16822; H0' | PCKGM_HUMA | Oxidation (M) | 0 | 0 | 0 | 0 | 0 | 12670 | 315570 | 0 | 2591700 | 152200 | 64650 | 0 | 0 | 0 | 0 | 0 | 0 | 0 |
| TMYVLPFSI | PCK2; PCK2; P( | Q16822; H0' | PCKGM_HUMA | Unmodified | 0 | 0 | 0 | 0 | 187900 | 0 | 0 | 0 | 143870 | 17737 | 0 | 0 | 207540 | 58581 | 20641 | 0 | 178790 | 251340 |
| TMYVLPFSI | PCK2; PCK2; P( | Q16822; H0' | PCKGM_HUMA | 2 Oxidation (M) | 0 | 0 | 0 | 0 | 0 | 0 | 0 | 0 | 5218600 | 1234800 | 0 | 0 | 40379 | 20228 | 0 | 0 | 0 | 0 |
| TNAAAMNI | LARP4; LARP4; | Q71RC2; X6I | LARP4_HUMAN | Oxidation (M) | 83846 | 215720 | 0 | 74146 | 0 | 0 | 0 | 0 | 246070 | 0 | 0 | 0 | 269130 | 70546 | 141710 | 39626 | 0 | 0 |
| TNADTDGI | DLD; DLD | P09622; E9P | DLDH_HUMAN | Unmodified | 0 | 0 | 44278 | 0 | 164610 | 0 | 643550 | 188100 | 5482700 | 1108700 | 89918 | 0 | 310620 | 221050 | 0 | 0 | 0 | 256580 |
| TNADTDGI | DLD; DLD | P09622; E9P | DLDH_HUMAN | Oxidation (M) | 0 | 0 | 0 | 0 | 0 | 0 | 0 | 0 | 202410 | 14415 | 0 | 0 | 0 | 0 | 0 | 0 | 0 | 0 |
| TNADVMTA | LYN | P07948 | LYN_HUMAN T | Oxidation (M) | 0 | 0 | 0 | 0 | 0 | 0 | 0 | 0 | 0 | 0 | 0 | 0 | 0 | 0 | 121730 | 206210 | 0 | 0 |
| TNAEIRAIN | ANXA6; ANXA6; | P08133; E7E | ANXA6_HUMAI | Unmodified | 0 | 0 | 100220 | 0 | 827780 | 0 | 0 | 0 | 0 | 491480 | 0 | 0 | 234030 | 0 | 0 | 0 | 0 | 254120 |
| TNAITLGSA | RAB25 | P57735 | RAB25_HUMAN | Unmodified | 0 | 0 | 0 | 0 | 0 | 0 | 0 | 0 | 0 | 0 | 0 | 0 | 0 | 0 | 0 | 0 | 0 | 0 |
| TNAMGLIR | KIF14 | Q15058 | KIF14_HUMAN | Oxidation (M) | 0 | 0 | 0 | 0 | 388630 | 32106 | 60005 | 0 | 0 | 0 | 0 | 0 | 81801 | 34282 | 0 | 0 | 283740 | 0 |
| TNAMGLIR | KIF14 | Q15058 | KIF14_HUMAN | Oxidation (M) | 0 | 0 | 0 | 0 | 282030 | 27738 | 70546 | 0 | 0 | 0 | 0 | 0 | 35839 | 49542 | 0 | 0 | 145560 | 30876 |
| TNATSPGV | SRGAP2; SRGA | O75044; B7Z | SRGP2_HUMAI | Unmodified | 0 | 681900 | 0 | 0 | 274460 | 0 | 224300 | 21321 | 0 | 0 | 0 | 213540 | 0 | 0 | 37509 | 42959 | 0 | 125070 |
| TNAVSEIK | DBNL; DBNL; D | Q9UJU6; B4I | DBNL_HUMAN | Unmodified | 0 | 0 | 444600 | 27215 | 1384100 | 677080 | 1410400 | 206120 | 0 | 0 | 151290 | 53163 | 7220600 | 1684500 | 553330 | 612000 | 270470 | 3305800 |
| TNDDHVS( | IQSEC2; IQSEC | Q5JU85; A0/ | IQEC2_HUMAN | Unmodified | 0 | 98385 | 0 | 0 | 0 | 0 | 0 | 0 | 0 | 0 | 0 | 0 | 0 | 0 | 40981 | 0 | 0 | 0 |
| TNEAQAIET | UBE2NL; UBE2I | Q5JXB2; P61 | UE2NL_HUMAI | Unmodified | 2282900 | 3914700 | 1782200 | 628200 | 0 | 0 | 0 | 0 | 2515400 | 148470 | 2095300 | 16444000 | 0 | 0 | 14092000 | 2938200 | 0 | 0 |
| TNEAQAIET | UBE2NL; UBE2I | Q5JXB2; P61 | UE2NL_HUMAI | Unmodified | 0 | 0 | 0 | 0 | 0 | 0 | 0 | 0 | 0 | 838510 | 0 | 443410 | 0 | 0 | 0 | 0 | 0 | 0 |
| TNEGVLCV | DSC2; DSC2 | Q02487; A0/ | DSC2_HUMAN | Unmodified | 0 | 0 | 575370 | 0 | 1595400 | 0 | 0 | 0 | 2892500 | 795700 | 956120 | 3132400 | 0 | 0 | 0 | 0 | 328090 | 346410 |
| TNEKVELQI | VIM; VIM; VIM | P08670; B0Y | VIME_HUMAN \ | Unmodified | 0 | 0 | 0 | 0 | 3236100 | 1547900 | 0 | 0 | 0 | 0 | 0 | 0 | 245880 | 0 | 0 | 0 | 0 | 1998800 |
| TNEPVWEE | ESYT2; ESYT2; E | A0FGR8; A0/ | ESYT2_HUMAN | Unmodified | 0 | 0 | 0 | 0 | 2566000 | 0 | 0 | 0 | 0 | 0 | 126190 | 0 | 0 | 0 | 0 | 0 | 0 | 0 |
| TNEQMHQ | ANXA6; ANXA6; | P08133; E5R | ANXA6_HUMAI | Oxidation (M) | 65671 | 0 | 161230 | 0 | 0 | 0 | 0 | 0 | 5070000 | 0 | 0 | 0 | 0 | 0 | 0 | 0 | 0 | 0 |
| TNEQMHQ | ANXA6; ANXA6; | P08133; E5R | ANXA6_HUMAI | Unmodified | 338050 | 0 | 2577700 | 1545400 | 0 | 101950 | 195190 | 297850 | 41305000 | 0 | 0 | 0 | 0 | 0 | 2917500 | 1637200 | 0 | 0 |
| TNEQMHQ | ANXA6; ANXA6; | P08133; E5R | ANXA6_HUMAI | Oxidation (M) | 0 | 0 | 0 | 1094000 | 0 | 0 | 0 | 0 | 3408300 | 0 | 0 | 0 | 0 | 0 | 0 | 0 | 0 | 0 |
| TNEQMHQ | ANXA6; ANXA6; | P08133; E5R | ANXA6_HUMAI | Unmodified | 0 | 0 | 0 | 0 | 0 | 0 | 0 | 0 | 10688000 | 5752800 | 0 | 0 | 0 | 0 | 0 | 0 | 0 | 0 |
| TNFFGTR | CBR1; CBR1; C | P16152; E9P | CBR1_HUMAN | Unmodified | 163650 | 202680 | 518610 | 0 | 356820 | 0 | 0 | 9093.9 | 372090 | 52112 | 0 | 3852000 | 0 | 36420 | 892400 | 450800 | 90997 | 93844 |
| TNFFGTRD | CBR1 | P16152 | CBR1_HUMAN | Unmodified | 0 | 0 | 0 | 0 | 9396000 | 4172500 | 1171100 | 0 | 0 | 1760200 | 0 | 0 | 3142500 | 2965300 | 0 | 0 | 6847100 | 7628600 |
| TNFFIQLVR | MRPL23; MRPL: | Q16540; A8I | RM23_HUMAN | Unmodified | 0 | 0 | 0 | 0 | 0 | 0 | 0 | 0 | 950170 | 0 | 0 | 0 | 0 | 0 | 0 | 0 | 0 | 0 |
| TNFFVNGL | TRIM56; TRIM5( | Q9BRZ2; C9. | TRI56_HUMAN | Unmodified | 0 | 0 | 0 | 0 | 194190 | 28932 | 0 | 0 | 0 | 0 | 0 | 0 | 0 | 0 | 0 | 0 | 0 | 0 |
| TNGDNYLY | IGF2R; IGF2R | P11717; S4F | MPRI_HUMAN ( | Unmodified | 0 | 0 | 0 | 0 | 0 | 0 | 0 | 0 | 5027100 | 0 | 0 | 0 | 0 | 0 | 0 | 0 | 0 | 0 |
| TNGDQAS( | INSR | P06213 | INSR_HUMAN I | Unmodified | 0 | 0 | 0 | 0 | 0 | 0 | 0 | 0 | 490120 | 97276 | 0 | 0 | 0 | 0 | 0 | 0 | 0 | 0 |
| TNGGFVEL | GALNT7; GALN | Q86SF2; H0' | GALT7_HUMAN | Unmodified | 0 | 0 | 0 | 0 | 0 | 0 | 0 | 0 | 1469600 | 146150 | 0 | 582010 | 0 | 0 | 0 | 0 | 0 | 0 |
| TNGKEPELI | RPL21 | P46778 | RL21_HUMAN ( | Oxidation (M) | 2688400 | 2712600 | 680290 | 0 | 0 | 0 | 0 | 0 | 0 | 0 | 65837 | 2539000 | 0 | 0 | 0 | 824680 | 0 | 0 |
| TNHIGHTG | RACK1; RACK1; | P63244; D6F | RACK1_HUMAI | Unmodified | 2241100 | 440530 | 348940 | 202040 | 0 | 0 | 0 | 0 | 283840 | 0 | 520400 | 23450000 | 0 | 0 | 0 | 244750 | 0 | 0 |
| TNHIGHTG | RACK1; RACK1; | P63244; D6F | RACK1_HUMAI | Oxidation (M) | 935740 | 0 | 0 | 0 | 0 | 0 | 366390 | 0 | 0 | 0 | 0 | 3012900 | 0 | 0 | 0 | 0 | 0 | 0 |
| TNHSNIMD | COPS2; COPS2 | P61201; H0Y | CSN2_HUMAN | Oxidation (M) | 0 | 0 | 0 | 0 | 0 | 0 | 0 | 0 | 0 | 0 | 0 | 1347900 | 0 | 0 | 0 | 0 | 0 | 0 |
| TNIAIFCETI | ACSL3; ACSL3; | O95573; A0/ | ACSL3_HUMAN | Unmodified | 0 | 0 | 0 | 0 | 0 | 0 | 0 | 0 | 0 | 0 | 0 | 2099400 | 0 | 0 | 0 | 0 | 0 | 0 |
| TNIFQIQR | HUWE1; HUWE | Q7Z6Z7; H0' | HUWE1_HUMA | Unmodified | 750410 | 0 | 214680 | 0 | 0 | 0 | 0 | 0 | 142480 | 0 | 106550 | 1473100 | 0 | 0 | 561750 | 522760 | 0 | 0 |
| TNIIFVAIHF | STEAP2; STEAP | Q8NFT2; C9. | STEA2_HUMAN | Unmodified | 311670 | 135350 | 2505200 | 654520 | 443180 | 75404 | 0 | 0 | 0 | 92285 | 0 | 0 | 0 | 664890 | 7257900 | 1093400 | 222250 | 515750 |
| TNIIPVIEDA | FBL; FBL; FBL; F | P22087; M0F | FBRL_HUMAN | Unmodified | 0 | 0 | 0 | 0 | 0 | 0 | 521360 | 441310 | 0 | 0 | 0 | 0 | 0 | 111180 | 0 | 0 | 0 | 124270 |
| TNIQQAVA | SUMF1; SUMF1 | Q8NBK3; F5 | SUMF1_HUMAI | Unmodified | 0 | 0 | 0 | 0 | 111520 | 0 | 44266 | 0 | 2232600 | 602000 | 0 | 0 | 0 | 0 | 0 | 0 | 0 | 0 |
| TNIQVTVT | CD109 | Q6YHK3 | CD109_HUMAI | Unmodified | 179470 | 94665 | 0 | 0 | 0 | 0 | 0 | 0 | 1254200 | 123320 | 0 | 0 | 0 | 0 | 0 | 0 | 50955 | 0 |
| TNISTASAT | PTPN12; PTPN1 | Q05209; H3I | PTN12_HUMAN | Unmodified | 0 | 0 | 0 | 0 | 367770 | 0 | 0 | 0 | 0 | 0 | 0 | 0 | 0 | 0 | 0 | 0 | 0 | 429490 |
| TNIVTASVD | ACOT7; ACOT7 | O00154; K7E | BACH_HUMAN | Unmodified | 0 | 0 | 0 | 0 | 3400600 | 795150 | 3135600 | 1164600 | 5719400 | 4818000 | 1506800 | 18682000 | 4016000 | 3601500 | 0 | 0 | 5852500 | 7264500 |
| TNIVTASVD | ACOT7; ACOT7 | O00154; K7E | BACH_HUMAN | Unmodified | 0 | 0 | 0 | 0 | 0 | 0 | 158350 | 0 | 0 | 0 | 0 | 0 | 0 | 0 | 0 | 0 | 540050 | 0 |
| TNKEVPGL | C1orf21; C1orf: | Q9H246; A0/ | CA021_HUMAN | Unmodified | 0 | 0 | 0 | 0 | 0 | 0 | 0 | 0 | 0 | 0 | 0 | 0 | 870570 | 403500 | 0 | 0 | 65427 | 0 |
| TNLATENQ | CCPG1 | Q9ULG6 | CCPG1_HUMA | Unmodified | 0 | 0 | 0 | 0 | 435680 | 0 | 0 | 0 | 764610 | 0 | 0 | 267320 | 0 | 0 | 0 | 0 | 0 | 129070 |
| TNLDESDV | RTCB | Q9Y3I0 | RTCB_HUMAN | Unmodified | 0 | 308970 | 719600 | 403990 | 0 | 0 | 0 | 0 | 145100 | 0 | 1282500 | 6339600 | 0 | 0 | 460120 | 530010 | 0 | 339600 |
| TNLEEFASI | SNF8; SNF8; SI | Q96H20; D6 | SNF8_HUMAN | Unmodified | 4692800 | 5162500 | 4307800 | 4877600 | 1742700 | 356190 | 588460 | 0 | 821700 | 254530 | 407190 | 1816300 | 5728200 | 3225500 | 7754000 | 5667700 | 160070 | 1231000 |
| TNLIVNYLP | ELAVL1; ELAVL | Q15717; M0 | ELAV1_HUMAN | Unmodified | 0 | 0 | 0 | 0 | 0 | 0 | 0 | 0 | 110730 | 0 | 43477 | 1110300 | 0 | 0 | 0 | 0 | 0 | 0 |
| TNLIVNYLP | ELAVL1; ELAVL | Q15717; M0 | ELAV1_HUMAN | Oxidation (M) | 187200 | 0 | 92090 | 79275 | 0 | 0 | 0 | 0 | 0 | 0 | 200260 | 4799800 | 0 | 0 | 163970 | 242510 | 0 | 0 |
| TNLKAEAE | CHDH | Q8NE62 | CHDH_HUMAN | Unmodified | 373540 | 0 | 0 | 0 | 0 | 0 | 0 | 0 | 978110 | 246000 | 0 | 0 | 0 | 0 | 0 | 0 | 0 | 0 |
| TNLKPGSP | SYT4 | Q9H2B2 | SYT4_HUMAN | Unmodified | 615630 | 419190 | 10943000 | 11669000 | 0 | 0 | 1101500 | 131500 | 155820 | 118530 | 0 | 0 | 557730 | 0 | 28901000 | 18560000 | 0 | 0 |
| TNLLQVCE | VCL | P18206 | VINC_HUMAN | Unmodified | 646690 | 2251300 | 327170 | 3069500 | 5506000 | 944670 | 3948300 | 340660 | 1802700 | 2168700 | 843420 | 8408800 | 4418700 | 6660300 | 2263000 | 1831500 | 16313000 | 6429600 |
| TNLPIFK | SNX12; SNX3; S | Q9UMY4; O6 | SNX12_HUMAN | Unmodified | 0 | 0 | 0 | 0 | 8740400 | 0 | 5208100 | 65890 | 2595000 | 1314700 | 1635500 | 2566900 | 3180600 | 2938700 | 0 | 0 | 2224500 | 2670600 |
| TNLSAVVA | SRP72; SRP72; | O76094; D6I | SRP72_HUMAN | Unmodified | 0 | 0 | 0 | 0 | 0 | 0 | 0 | 228430 | 0 | 0 | 0 | 2081900 | 0 | 0 | 0 | 0 | 38595 | 395140 |
| TNLSVHED | KIF5B | P33176 | KINH_HUMAN | Unmodified | 0 | 0 | 0 | 0 | 0 | 0 | 176210 | 109530 | 0 | 99931 | 0 | 0 | 111650 | 514360 | 0 | 0 | 283250 | 0 |
| TNLSYYEYI | BMX | P51813 | BMX_HUMAN ( | Unmodified | 0 | 0 | 0 | 0 | 0 | 0 | 0 | 0 | 0 | 0 | 0 | 9861400 | 0 | 0 | 0 | 0 | 0 | 0 |
| TNLYIRGL | RBMS2; RBMS2 | Q15434; F8V | RBMS2_HUMAI | Unmodified | 0 | 0 | 0 | 0 | 2194900 | 0 | 0 | 0 | 0 | 0 | 0 | 0 | 0 | 0 | 0 | 0 | 0 | 0 |
| TNNFIIAVK | GGT7 | Q9UJ14 | GGT7_HUMAN | Unmodified | 349470 | 631630 | 555150 | 61230 | 233330 | 0 | 0 | 0 | 3788200 | 312950 | 0 | 133130 | 0 | 0 | 254010 | 112250 | 0 | 0 |
| TNNFIIAVK | GGT7 | Q9UJ14 | GGT7_HUMAN | Unmodified | 0 | 0 | 0 | 0 | 0 | 0 | 0 | 0 | 624780 | 1500800 | 0 | 0 | 0 | 0 | 0 | 0 | 0 | 0 |
| TNNFNVET | GYS2; GYS1 | P54840; P13 | GYS2_HUMAN | Unmodified | 0 | 0 | 0 | 0 | 0 | 0 | 0 | 0 | 0 | 0 | 143650 | 5816800 | 0 | 0 | 0 | 0 | 0 | 0 |
| TNNLETGP | VIL1; VIL1; VIL1 | P09327; H7( | VILI_HUMAN Vi | Unmodified | 0 | 0 | 0 | 0 | 0 | 0 | 0 | 0 | 945140 | 100480 | 133710 | 5918200 | 0 | 0 | 0 | 0 | 23343 | 0 |
| TNNPLPLE( | XPO7; XPO7; XF | Q9UIA9; H0' | XPO7_HUMAN | Unmodified | 0 | 0 | 0 | 0 | 0 | 0 | 0 | 0 | 0 | 0 | 94848 | 1216100 | 0 | 0 | 0 | 0 | 0 | 0 |
| TNPAASME | NDST2; N/A; N/ | P52849; S4F | NDST2_HUMAI | Unmodified | 0 | 0 | 0 | 0 | 0 | 0 | 0 | 0 | 991830 | 0 | 0 | 0 | 0 | 0 | 0 | 0 | 0 | 0 |
| TNPEKTED/ | CARMIL1; CARI | Q5VZK9; A0/ | CARL1_HUMAN | Unmodified | 0 | 0 | 0 | 121880 | 0 | 0 | 2021500 | 13884 | 231350 | 0 | 272240 | 1623200 | 0 | 0 | 0 | 0 | 0 | 0 |
| TNPEVHNY | EPB41L4B | Q9H329 | E41LB_HUMAN | Unmodified | 3243800 | 3310400 | 147740 | 3500700 | 214830 | 0 | 0 | 0 | 1556600 | 239500 | 167400 | 239490 | 0 | 0 | 8403800 | 6363600 | 94926 | 426290 |

| | | | | | | | | | | | | | | | | | | | | | | |
|---|---|---|---|---|---|---|---|---|---|---|---|---|---|---|---|---|---|---|---|---|---|---|
| TNPGAFRE | FRYL; FRYL; FRY | O94915; F2Z | FRYL_HUMAN I | Unmodified | 57310 | 0 | 0 | 86876 | 0 | 0 | 0 | 0 | 331820 | 0 | 0 | 491330 | 0 | 0 | 0 | 0 | 0 | 0 |
| TNPGYLLSI | PIK3C2A; PIK3C | O00443; A0A | P3C2A_HUMAN | Unmodified | 0 | 0 | 0 | 0 | 0 | 0 | 0 | 0 | 348140 | 0 | 0 | 1172200 | 0 | 0 | 0 | 0 | 0 | 0 |
| TNPPLIQEK | EIF4G2; EIF4G2 | P78344; H0Y | IF4G2_HUMAN | Unmodified | 111590 | 0 | 0 | 277860 | 847250 | 0 | 0 | 0 | 0 | 0 | 264800 | 547670 | 688710 | 0 | 0 | 88331 | 0 | 0 |
| TNPSDRVL | ATP2B4 | P23634 | AT2B4_HUMAN | Acetyl (Protein N | 0 | 0 | 110020 | 0 | 0 | 0 | 33888 | 0 | 0 | 43199 | 0 | 3957000 | 0 | 0 | 0 | 0 | 79740 | 112160 |
| TNPSDRVL | ATP2B4 | P23634 | AT2B4_HUMAN | Acetyl (Protein N | 0 | 0 | 0 | 0 | 0 | 0 | 0 | 0 | 0 | 161420 | 0 | 4497700 | 0 | 0 | 0 | 280120 | 0 | 0 |
| TNPVSSGP | GPRIN1 | Q7Z2K8 | GRIN1_HUMAN | Unmodified | 0 | 0 | 0 | 0 | 0 | 0 | 0 | 0 | 1475700 | 92784 | 0 | 0 | 0 | 0 | 0 | 0 | 0 | 122770 |
| TNQCLQG( | USP54; USP54; | Q70EL1; A0A | UBP54_HUMAN | Unmodified | 0 | 0 | 0 | 0 | 0 | 0 | 0 | 0 | 275660 | 229640 | 0 | 0 | 0 | 906950 | 0 | 0 | 0 | 0 |
| TNQDAMET | PARD3; PARD3; | Q8TEW0; Q5 | PARD3_HUMAN | Unmodified | 0 | 0 | 0 | 0 | 0 | 0 | 0 | 0 | 470400 | 0 | 0 | 0 | 0 | 0 | 0 | 0 | 617850 | 508540 |
| TNQELQEII | ANXA2P2; ANX | A6NMY6; P0 | AXA2L_HUMAN | Unmodified | 645740 | 471190 | 317720 | 280510 | 2015800 | 56927 | 359990 | 142120 | 19980000 | 1117300 | 257870 | 1624700 | 0 | 0 | 228390 | 25260 | 277670 | 768700 |
| TNQELQEII | ANXA2P2; ANX | A6NMY6; P0 | AXA2L_HUMAN | Unmodified | 1953700 | 0 | 0 | 2062800 | 50009000 | 29957000 | 434250 | 194010 | 82532000 | 8319600 | 0 | 485800 | 0 | 0 | 693560 | 450330 | 7264600 | 11929000 |
| TNQELQEII | ANXA2P2; ANX | A6NMY6; P0 | AXA2L_HUMAN | Unmodified | 0 | 0 | 0 | 0 | 4336700 | 0 | 0 | 0 | 0 | 996150 | 0 | 0 | 106630 | 0 | 0 | 0 | 1030500 | 1890200 |
| TNQFSVTR | ERGIC3; ERGIC | Q9Y282; H0Y | ERGI3_HUMAN | Unmodified | 0 | 0 | 0 | 0 | 3274100 | 0 | 315220 | 129050 | 3201200 | 105570 | 1298700 | 2121200 | 3668300 | 1604400 | 0 | 0 | 512160 | 3723700 |
| TNQIGTVNI | IST1; IST1; IST1 | P53990; H3E | IST1_HUMAN IS | Unmodified | 1410500 | 1356400 | 923700 | 855070 | 177740 | 0 | 0 | 0 | 4762700 | 131790 | 850260 | 17081000 | 0 | 0 | 1235400 | 645110 | 0 | 202630 |
| TNQIGTVNI | IST1; IST1; IST1 | P53990; H3E | IST1_HUMAN IS | Unmodified | 455250 | 224480 | 1800000 | 1790300 | 7073400 | 1027100 | 47198 | 18972 | 15336000 | 1436100 | 5799700 | 5735200 | 4406200 | 1251700 | 237060 | 174850 | 0 | 1123300 |
| TNQIGTVNI | IST1; IST1; IST1 | P53990; H3E | IST1_HUMAN IS | Oxidation (M) | 262470 | 122190 | 0 | 416600 | 1564500 | 540330 | 0 | 0 | 944110 | 545930 | 826670 | 1505800 | 0 | 0 | 0 | 0 | 132250 | 0 |
| TNRLAELEE | CLTC; CLTC; CL | Q00610; J3K | CLH1_HUMAN | Unmodified | 2575100 | 1161300 | 1411300 | 2896500 | 0 | 0 | 0 | 0 | 630690 | 35212 | 0 | 2026900 | 0 | 0 | 1041000 | 877940 | 131910 | 0 |
| TNRLAELEE | CLTC; CLTC; CL | Q00610; J3K | CLH1_HUMAN | Unmodified | 972020 | 0 | 0 | 5275800 | 4768200 | 759110 | 2976700 | 500370 | 1025800 | 0 | 8123600 | 0 | 18572000 | 11471000 | 0 | 0 | 6834600 | 12946000 |
| TNRNDNSS | MYO1D; MYO1E | O94832; J3Q | MYO1D_HUMA | Unmodified | 0 | 145860 | 0 | 0 | 72908 | 0 | 0 | 0 | 0 | 0 | 0 | 0 | 0 | 0 | 387060 | 203220 | 0 | 2189700 |
| TNSAVRTC | EIF3I; EIF3I; EIF | Q13347; A0A | EIF3I_HUMAN I | Oxidation (M) | 0 | 0 | 0 | 0 | 0 | 0 | 0 | 0 | 0 | 0 | 0 | 0 | 1718800 | 0 | 0 | 0 | 2607200 | 0 |
| TNSTFNQV | RPL18; RPL18; I | Q07020; A0A | RL18_HUMAN ( | Unmodified | 1649800 | 1689500 | 1041400 | 252620 | 0 | 0 | 0 | 196400 | 0 | 0 | 0 | 10898000 | 0 | 0 | 1529100 | 744780 | 0 | 0 |
| TNSTFNQV | RPL18; RPL18; I | Q07020; A0A | RL18_HUMAN ( | Unmodified | 2555600 | 4340100 | 1045800 | 0 | 0 | 0 | 0 | 0 | 0 | 0 | 0 | 16108000 | 0 | 0 | 4378200 | 2304800 | 0 | 0 |
| TNTEEKNTI | TMSB15B; TMSI | P0CG35; P0 | TB15B_HUMAN | Unmodified | 0 | 0 | 0 | 0 | 0 | 0 | 264690 | 330200 | 0 | 418720 | 0 | 0 | 0 | 110280 | 0 | 0 | 0 | 0 |
| TNTIGTLNI | UXS1; UXS1; U) | Q8NBZ7; C9 | UXS1_HUMAN | Unmodified | 0 | 0 | 0 | 0 | 0 | 0 | 0 | 0 | 3959200 | 563680 | 0 | 0 | 0 | 0 | 0 | 0 | 0 | 750450 |
| TNTLAVTG( | AAMP; AAMP; A | Q13685; C9J | AAMP_HUMAN | Unmodified | 0 | 0 | 0 | 0 | 604770 | 0 | 0 | 0 | 0 | 0 | 128600 | 672100 | 0 | 0 | 0 | 0 | 0 | 0 |
| TNTLRLSQ1 | CEP55 | Q53EZ4 | CEP55_HUMAN | Unmodified | 0 | 0 | 0 | 0 | 2367800 | 0 | 0 | 0 | 0 | 0 | 0 | 0 | 0 | 0 | 2547800 | 0 | 0 | 0 |
| TNTPADVFI | RO60; RO60 | P10155; G5E | RO60_HUMAN | Unmodified | 301080 | 8844500 | 53545 | 616450 | 0 | 0 | 0 | 0 | 0 | 0 | 0 | 0 | 0 | 0 | 2552100 | 52189 | 0 | 0 |
| TNTPADVFI | RO60; RO60 | P10155; G5E | RO60_HUMAN | Unmodified | 0 | 0 | 0 | 0 | 0 | 0 | 695710 | 0 | 0 | 0 | 0 | 0 | 3362800 | 2614500 | 0 | 0 | 0 | 0 |
| TNTQTYREI | HLA-B; HLA-B; I | D5H3J5; A0A | D5H3J5_HUMA | Unmodified | 36724 | 0 | 0 | 0 | 0 | 0 | 136580 | 85112 | 3186300 | 474320 | 229180 | 21274 | 0 | 42972 | 0 | 0 | 0 | 3178600 |
| TNTVPFDL\ | MUL1 | Q969V5 | MUL1_HUMAN | Unmodified | 193000 | 0 | 0 | 0 | 0 | 0 | 0 | 0 | 3931700 | 216440 | 68511 | 0 | 0 | 0 | 0 | 136890 | 0 | 0 |
| TNVLGHLQ | PFKL | P17858 | PFKAL_HUMAN | Unmodified | 0 | 0 | 746690 | 644820 | 0 | 0 | 182680 | 89710 | 532120 | 0 | 280490 | 3710800 | 0 | 0 | 1435800 | 1207000 | 0 | 457500 |
| TNVNGGAI | ACAA2; ACAA2; | P42765; A0A | THIM_HUMAN | Unmodified | 352330 | 0 | 0 | 0 | 0 | 0 | 0 | 0 | 8528700 | 912650 | 0 | 0 | 0 | 0 | 0 | 0 | 0 | 0 |
| TNVPSWCI | INPPL1; INPPL1 | O15357; A0A | SHIP2_HUMAN | Unmodified | 0 | 0 | 0 | 0 | 0 | 0 | 0 | 0 | 0 | 0 | 121410 | 0 | 0 | 0 | 0 | 0 | 0 | 0 |
| TNVYISSSA | SORL1 | Q92673 | SORL_HUMAN | Unmodified | 0 | 0 | 0 | 0 | 0 | 0 | 57573 | 0 | 631550 | 133000 | 0 | 342210 | 98725 | 0 | 0 | 33670 | 0 | 0 |
| TNYNDRYD | RPL7A; RPL7A | P62424; Q5T | RL7A_HUMAN ( | Unmodified | 1877200 | 170440 | 912060 | 404470 | 64637 | 0 | 0 | 0 | 0 | 31843 | 576940 | 30056000 | 0 | 0 | 1099200 | 843390 | 0 | 14726 |
| TPAAETLSC | NPTX1 | Q15818 | NPTX1_HUMAN | Unmodified | 29579 | 0 | 0 | 0 | 0 | 0 | 0 | 0 | 4.41E+08 | 19970000 | 0 | 0 | 0 | 0 | 796560 | 0 | 0 | 0 |
| TPADCPVIA | ERP44 | Q9BS26 | ERP44_HUMAN | Unmodified | 0 | 0 | 0 | 0 | 0 | 0 | 0 | 0 | 58567000 | 2818200 | 1527500 | 2303300 | 0 | 37650 | 0 | 0 | 0 | 0 |
| TPADCPVIA | ERP44 | Q9BS26 | ERP44_HUMAN | Oxidation (M) | 0 | 0 | 0 | 0 | 0 | 0 | 0 | 0 | 0 | 6305700 | 0 | 0 | 0 | 130980 | 0 | 0 | 0 | 0 |
| TPAFAESVI | DLST; DLST; DL | P36957; Q8( | ODO2_HUMAN | Unmodified | 0 | 0 | 0 | 0 | 0 | 0 | 1034900 | 299480 | 13667000 | 2983700 | 0 | 0 | 0 | 0 | 885700 | 0 | 54631 | 272480 |
| TPAGNFVTI | NAMPT; NAMPT | P43490; A0A | NAMPT_HUMA | Unmodified | 0 | 86732 | 0 | 0 | 0 | 0 | 0 | 0 | 0 | 0 | 0 | 0 | 293750 | 0 | 670960 | 0 | 0 | 0 |
| TPAILEASA | CTNND1; CTNN | O60716; A0A | CTND1_HUMA | Unmodified | 1515400 | 924750 | 3476500 | 4899300 | 0 | 0 | 0 | 0 | 0 | 0 | 1368700 | 8351000 | 407230 | 442610 | 11926000 | 6119300 | 0 | 0 |
| TPAILEASA | CTNND1; CTNN | O60716; A0A | CTND1_HUMA | Unmodified | 0 | 0 | 0 | 452360 | 2916200 | 341890 | 0 | 0 | 0 | 0 | 563070 | 0 | 5038700 | 3948800 | 0 | 0 | 791630 | 3330600 |
| TPALIVYGD | ABHD14B; ABH | Q96IU4; F8W | ABHEB_HUMA | Oxidation (M) | 0 | 0 | 0 | 0 | 4179900 | 0 | 0 | 0 | 0 | 430680 | 1397800 | 1727100 | 0 | 0 | 0 | 0 | 815600 | 0 |
| TPALIVYGD | ABHD14B; ABH | Q96IU4; F8W | ABHEB_HUMA | Unmodified | 0 | 0 | 0 | 0 | 2567400 | 0 | 0 | 0 | 595100 | 169810 | 0 | 105500 | 0 | 0 | 0 | 0 | 0 | 0 |
| TPALVFEH\ | CSNK2A3; CSN | Q8NEV1; P6 | CSK23_HUMAN | Unmodified | 0 | 0 | 0 | 0 | 0 | 0 | 0 | 0 | 708460 | 0 | 0 | 9158800 | 0 | 0 | 0 | 0 | 0 | 0 |
| TPALVNAA\ | ATP5MG; N/A; / | O75964; A0A | ATP5L_HUMAN | Unmodified | 0 | 0 | 0 | 0 | 0 | 0 | 0 | 0 | 849820 | 152480 | 0 | 0 | 0 | 0 | 0 | 0 | 0 | 0 |
| TPALVNAA\ | ATP5MG; N/A; / | O75964; A0A | ATP5L_HUMAN | Unmodified | 0 | 0 | 0 | 0 | 0 | 0 | 0 | 0 | 1457100 | 0 | 0 | 0 | 0 | 0 | 0 | 0 | 0 | 0 |
| TPAQFDAD | ANXA1; ANXA1 | P04083; Q5T | ANXA1_HUMAN | Unmodified | 455240 | 0 | 0 | 0 | 754950 | 0 | 0 | 0 | 469680 | 0 | 95160000 | 2.63E+09 | 0 | 0 | 0 | 750940 | 0 | 0 |
| TPAQFDAD | ANXA1; ANXA1 | P04083; Q5T | ANXA1_HUMAN | Oxidation (M) | 40568 | 81611 | 0 | 0 | 8406200 | 2880900 | 350040 | 11797 | 117540 | 265820 | 70092000 | 1.81E+08 | 320030 | 335400 | 13983 | 254400 | 968740 | 1706900 |
| TPAQFDAD | ANXA1; ANXA1 | P04083; Q5T | ANXA1_HUMAN | Unmodified | 76482 | 0 | 0 | 0 | 42735000 | 10009000 | 1386100 | 0 | 48165 | 1255400 | 1.82E+08 | 5.89E+08 | 0 | 0 | 0 | 33568 | 2310300 | 3520100 |
| TPAQYDAS | ANXA2P2; ANX | A6NMY6; P0 | AXA2L_HUMAN | Unmodified | 2168400 | 1638500 | 0 | 1519800 | 99922000 | 33609000 | 2223600 | 1068700 | 51581000 | 11913000 | 607940 | 0 | 483450 | 827030 | 0 | 0 | 24286000 | 14556000 |
| TPAQYDAS | ANXA2P2; ANX | A6NMY6; P0 | AXA2L_HUMAN | Oxidation (M) | 0 | 0 | 0 | 0 | 3714400 | 798820 | 211400 | 0 | 0 | 407850 | 0 | 0 | 0 | 59954 | 0 | 0 | 3749300 | 955070 |
| TPAQYDAS | ANXA2P2; ANX | A6NMY6; P0 | AXA2L_HUMAN | Unmodified | 89784 | 0 | 0 | 0 | 19057000 | 3744600 | 1394900 | 776570 | 0 | 3040700 | 0 | 0 | 0 | 227080 | 0 | 0 | 18188000 | 5216800 |
| TPAVSISCL | SLC30A1 | Q9Y6M5 | ZNT1_HUMAN | Unmodified | 0 | 0 | 0 | 0 | 12578000 | 1292900 | 0 | 0 | 2242600 | 1660100 | 1615800 | 3947100 | 803840 | 400520 | 0 | 0 | 2077500 | 2600000 |
| TPCEEILVK | FLNA; FLNA; FL | P21333; A0A | FLNA_HUMAN | Unmodified | 1166600 | 1173000 | 812320 | 2382000 | 0 | 0 | 0 | 0 | 0 | 191950 | 288730 | 0 | 0 | 0 | 1825100 | 1521200 | 0 | 0 |
| TPCNAGTF | BUB3; BUB3; BI | O43684; J3Q | BUB3_HUMAN | Unmodified | 0 | 0 | 0 | 0 | 0 | 0 | 0 | 0 | 0 | 0 | 0 | 3879100 | 0 | 0 | 0 | 0 | 0 | 0 |
| TPCQQELD | IGFBP2; IGFBP2 | P18065; C9J | IBP2_HUMAN I | Unmodified | 74228 | 0 | 701930 | 134520 | 0 | 0 | 0 | 0 | 56668000 | 1745300 | 0 | 0 | 0 | 0 | 175260 | 73706 | 0 | 0 |
| TPCSSLLPI | SPATS2L; SPAT | Q9NUQ6; B8 | SPS2L_HUMAN | Unmodified | 77065 | 0 | 0 | 0 | 0 | 0 | 0 | 0 | 0 | 0 | 0 | 1316800 | 0 | 0 | 0 | 0 | 0 | 0 |
| TPDGNLDC | SLC7A1 | P30825 | SL7A1_HUMAN | Unmodified | 0 | 0 | 1314000 | 1963000 | 681180 | 0 | 0 | 0 | 495320 | 104670 | 260580 | 1863500 | 202620 | 0 | 1107900 | 982180 | 593800 | 123190 |
| TPDGQGLS | ECPAS; ECPAS | Q5VYK3; J3K | ECM29_HUMA | Unmodified | 0 | 0 | 0 | 0 | 0 | 0 | 0 | 0 | 60428 | 0 | 0 | 385620 | 0 | 0 | 0 | 0 | 0 | 69307 |
| TPDQAILW | QSOX2; QSOX2 | Q6ZRP7; H0 | QSOX2_HUMA | Unmodified | 0 | 0 | 48314 | 0 | 0 | 0 | 0 | 0 | 6728500 | 1281400 | 0 | 507290 | 0 | 0 | 0 | 37934 | 0 | 0 |
| TPDQAILW | QSOX2; QSOX2 | Q6ZRP7; H0 | QSOX2_HUMA | Unmodified | 0 | 0 | 0 | 0 | 0 | 0 | 0 | 0 | 6815400 | 0 | 0 | 0 | 0 | 0 | 0 | 0 | 0 | 0 |
| TPDTPNND | PKN2; PKN2 | Q16513; B1A | PKN2_HUMAN | Unmodified | 19544000 | 18019000 | 0 | 7100100 | 0 | 0 | 1432600 | 0 | 19324000 | 3297400 | 8435200 | 9910700 | 5583500 | 2081300 | 0 | 488580 | 0 | 0 |
| TPEELDDSI | CTNNA1; CTNN | P35221; G3 | CTNA1_HUMA | Unmodified | 0 | 0 | 0 | 196890 | 0 | 0 | 0 | 0 | 295420 | 0 | 747200 | 3435500 | 0 | 0 | 1775300 | 1858800 | 0 | 482480 |
| TPEELFHPI | SLC2A1 | P11166 | GTR1_HUMAN | Unmodified | 43544 | 45592 | 871110 | 0 | 315740 | 0 | 105750 | 65629 | 715010 | 56669 | 827450 | 2832700 | 0 | 0 | 0 | 221000 | 0 | 0 |
| TPEENEPT( | GNPTG | Q9UJJ9 | GNPTG_HUMA | Unmodified | 0 | 0 | 0 | 0 | 0 | 0 | 0 | 0 | 2420500 | 0 | 0 | 0 | 0 | 0 | 0 | 0 | 0 | 0 |
| TPEEYPESA | DDAH1; DDAH1 | O94760; B4D | DDAH1_HUMA | Unmodified | 141090 | 111980 | 0 | 206610 | 0 | 0 | 0 | 0 | 520260 | 0 | 404580 | 784450 | 0 | 0 | 566510 | 372820 | 0 | 0 |
| TPEFEEFN( | ANO6; ANO6; A | Q4KMQ2; A0 | ANO6_HUMAN | Unmodified | 138900 | 0 | 652460 | 0 | 0 | 0 | 0 | 0 | 814080 | 0 | 221870 | 4822000 | 0 | 0 | 1606300 | 655510 | 0 | 0 |
| TPEIVAPQS | SGPL1 | O95470 | SGPL1_HUMAN | Unmodified | 0 | 0 | 0 | 0 | 0 | 0 | 0 | 0 | 1451500 | 0 | 0 | 522950 | 0 | 0 | 0 | 0 | 0 | 0 |
| TPELAEEFK | RGPD3; RGPD2 | A6NKT7; P0I | RGPD3_HUMA | Unmodified | 150700 | 384310 | 0 | 201140 | 0 | 0 | 0 | 0 | 0 | 0 | 0 | 799070 | 0 | 0 | 0 | 112430 | 0 | 0 |
| TPELNLDQ | CANX; CANX; C | P27824; A0A | CALX_HUMAN | Unmodified | 1429100 | 0 | 2373500 | 940600 | 976900 | 0 | 1153100 | 0 | 8.67E+08 | 12847000 | 27724000 | 24038000 | 0 | 0 | 2202500 | 1581700 | 0 | 0 |
| TPELNLDQ | CANX; CANX; C | P27824; A0A | CALX_HUMAN | Oxidation (M) | 419570 | 234910 | 0 | 1020400 | 1747600 | 441120 | 1062600 | 71255 | 6.55E+08 | 69858000 | 35369000 | 3756400 | 397090 | 301790 | 0 | 261500 | 0 | 398530 |
| TPELNLDQ | CANX; CANX; C | P27824; A0A | CALX_HUMAN | Unmodified | 112140 | 0 | 0 | 125120 | 938510 | 0 | 482000 | 0 | 7.21E+08 | 24772000 | 7271600 | 414090 | 99198 | 0 | 0 | 0 | 0 | 326570 |
| TPELNLDQ | CANX; CANX; C | P27824; A0A | CALX_HUMAN | Oxidation (M) | 0 | 0 | 0 | 0 | 0 | 0 | 1166600 | 0 | 27446000 | 13042000 | 563540 | 0 | 0 | 469960 | 0 | 0 | 0 | 1356100 |
| TPELNLDQ | CANX; CANX; C | P27824; A0A | CALX_HUMAN | Unmodified | 0 | 0 | 0 | 0 | 0 | 0 | 0 | 0 | 47388000 | 4213700 | 0 | 0 | 0 | 0 | 0 | 0 | 0 | 0 |
| TPELTWER | AHCYL1; AHCY | O43865; Q9( | SAHH2_HUMA | Unmodified | 770010 | 692830 | 711030 | 0 | 0 | 0 | 0 | 0 | 262690 | 0 | 0 | 2382300 | 0 | 0 | 2859300 | 1872800 | 0 | 0 |
| TPESFLGPI | EPN1 | Q9Y6I3 | EPN1_HUMAN | Unmodified | 1745900 | 1202000 | 555910 | 397440 | 0 | 0 | 0 | 0 | 373350 | 0 | 0 | 0 | 0 | 0 | 0 | 946250 | 0 | 962930 |
| TPESFLGPS | EPN3 | Q9H201 | EPN3_HUMAN | Unmodified | 429720 | 371840 | 91311 | 227500 | 0 | 0 | 0 | 0 | 0 | 0 | 0 | 0 | 289870 | 0 | 761940 | 310960 | 0 | 0 |
| TPESSHEG | ECE1; ECE1; EC | P42892; B4D | ECE1_HUMAN | Unmodified | 0 | 0 | 0 | 0 | 0 | 0 | 0 | 0 | 14189000 | 213440 | 0 | 0 | 0 | 0 | 0 | 0 | 0 | 0 |
| TPETAEFLG | SH3BP1 | Q9Y3L3 | 3BP1_HUMAN | Unmodified | 185480 | 386600 | 0 | 0 | 0 | 0 | 0 | 0 | 0 | 168730 | 0 | 0 | 0 | 0 | 0 | 0 | 0 | 0 |
| TPFFIFNLA | TMEM260 | Q9NX78 | TM260_HUMAN | Unmodified | 0 | 0 | 0 | 0 | 0 | 31058 | 0 | 0 | 283150 | 10242 | 0 | 0 | 0 | 0 | 0 | 0 | 0 | 0 |
| TPFFIFNLA | TMEM260 | Q9NX78 | TM260_HUMAN | Oxidation (M) | 0 | 412100 | 0 | 0 | 0 | 0 | 0 | 0 | 0 | 385170 | 0 | 0 | 0 | 0 | 0 | 0 | 0 | 0 |
| TPFLLSGTS | HADHB; HADHI | P55084; C9J | ECHB_HUMAN | Unmodified | 0 | 0 | 0 | 0 | 0 | 0 | 0 | 0 | 2659600 | 0 | 0 | 0 | 0 | 0 | 0 | 0 | 0 | 0 |
| TPFLLVGT( | CDC42; CDC42 | P60953; Q5J | CDC42_HUMA | Unmodified | 0 | 10220000 | 0 | 0 | 17667000 | 0 | 514720 | 449030 | 572120 | 4044200 | 296530 | 559590 | 13115000 | 0 | 18253 | 0 | 2607600 | 27532000 |
| TPFLLVGT( | CDC42; CDC42 | P60953; Q5J | CDC42_HUMA | Unmodified | 0 | 0 | 0 | 0 | 859580 | 0 | 35748 | 0 | 0 | 0 | 0 | 0 | 0 | 1297000 | 0 | 0 | 2147800 | 0 |
| TPFVTHPG' | FN1; FN1; FN1 | P02751; H0Y | FINC_HUMAN I | Oxidation (M) | 0 | 0 | 5061700 | 0 | 0 | 0 | 0 | 0 | 0 | 0 | 0 | 0 | 0 | 0 | 0 | 0 | 0 | 0 |
| TPGGGSR( | EXOC5; EXOC5 | O00471; G3 | EXOC5_HUMA | Unmodified | 0 | 0 | 0 | 0 | 245700 | 0 | 0 | 0 | 0 | 136710 | 0 | 0 | 0 | 21772 | 0 | 0 | 30952 | 69940 |
| TPGNRIVYI | RPL34 | P49207 | RL34_HUMAN ( | Unmodified | 0 | 0 | 0 | 0 | 0 | 0 | 0 | 179130 | 0 | 0 | 0 | 0 | 2195300 | 0 | 0 | 0 | 0 | 0 |
| TPGPGAQS | RPS14; RPS14; | P62263; E5R | RS14_HUMAN | Unmodified | 6081900 | 7382400 | 5676600 | 4552700 | 4640300 | 486070 | 472100 | 207410 | 2004200 | 884980 | 3337400 | 29868000 | 5877200 | 1069100 | 3631900 | 4207100 | 155620 | 1053900 |
| TPGTGSLA | PDXP | Q96GD0 | PLPP_HUMAN ( | Unmodified | 0 | 0 | 0 | 0 | 0 | 0 | 0 | 0 | 0 | 0 | 0 | 821170 | 0 | 0 | 0 | 0 | 0 | 0 |
| TPGTWNHI | GOT1 | P17174 | AATC_HUMAN | Oxidation (M) | 0 | 0 | 0 | 0 | 0 | 0 | 0 | 0 | 0 | 149560 | 976940 | 1940400 | 0 | 0 | 0 | 0 | 0 | 0 |
| TPGVFDPG | GNS; GNS; GNS | P15586; H7( | GNS_HUMAN I | Unmodified | 0 | 0 | 0 | 0 | 0 | 0 | 0 | 0 | 2064400 | 0 | 0 | 446880 | 0 | 0 | 0 | 0 | 0 | 0 |
| TPGVFDPG | GNS; GNS; GNS | P15586; H7( | GNS_HUMAN I | Unmodified | 0 | 0 | 0 | 0 | 0 | 0 | 138010 | 83258 | 3883600 | 303930 | 0 | 0 | 0 | 172130 | 0 | 0 | 0 | 0 |
| TPGYVVTPI | QARS1; QARS1 | P47897; A0A | SYQ_HUMAN G | Oxidation (M) | 0 | 0 | 0 | 0 | 0 | 0 | 0 | 0 | 0 | 0 | 610200 | 5222600 | 0 | 0 | 0 | 0 | 0 | 0 |
| TPIAAGHPS | IDH3A; IDH3A; | P50213; H0Y | IDH3A_HUMAN | Unmodified | 0 | 0 | 0 | 0 | 0 | 0 | 0 | 0 | 3313700 | 354820 | 0 | 0 | 0 | 0 | 0 | 0 | 0 | 221850 |
| TPIESLFIEA | MYO9B; MYO9E | Q13459; M0I | MYO9B_HUMA | Unmodified | 0 | 0 | 0 | 0 | 463860 | 0 | 0 | 45913 | 0 | 0 | 0 | 0 | 0 | 0 | 0 | 0 | 0 | 0 |
| TPIGKPATA | GNPAT | O15228 | GNPAT_HUMA | Unmodified | 0 | 0 | 0 | 0 | 0 | 0 | 0 | 0 | 1538000 | 0 | 0 | 0 | 0 | 0 | 0 | 0 | 0 | 0 |
| TPIGSFLGS | ACAT1; ACAT1; | P24752; E9P | THIL_HUMAN A | Unmodified | 0 | 0 | 0 | 0 | 0 | 0 | 0 | 0 | 1492600 | 50427 | 0 | 20502 | 0 | 0 | 0 | 0 | 0 | 0 |
| TPIITPNTK | ANLN; ANLN | Q9NQW6; H | ANLN_HUMAN | Unmodified | 0 | 0 | 0 | 93147 | 773160 | 90026 | 22993 | 0 | 917130 | 0 | 0 | 0 | 0 | 0 | 0 | 0 | 144840 | 89750 |
| TPIQVESSP | CDCA3; CDCA3 | Q99618; F8V | CDCA3_HUMA | Unmodified | 1007200 | 0 | 1183000 | 2009500 | 1776700 | 0 | 0 | 0 | 1494200 | 0 | 820920 | 2026000 | 2944900 | 0 | 1113400 | 0 | 173330 | 543830 |
| TPISPLKTG | ANLN | Q9NQW6 | ANLN_HUMAN | Unmodified | 0 | 0 | 0 | 0 | 0 | 0 | 0 | 0 | 0 | 0 | 0 | 0 | 0 | 0 | 0 | 0 | 634470 | 0 |
| TPKGPSSVI | NPM1; NPM1; N | P06748; A0A | NPM_HUMAN I | Unmodified | 0 | 0 | 0 | 0 | 2623200 | 0 | 0 | 180200 | 227010 | 988600 | 0 | 151720 | 4825200 | 0 | 0 | 0 | 470710 | 131370 |
| TPKGPSSVI | NPM1; NPM1; N | P06748; A0A | NPM_HUMAN I | Unmodified | 0 | 0 | 0 | 0 | 468370 | 0 | 0 | 0 | 0 | 244910 | 0 | 0 | 851710 | 365470 | 0 | 0 | 0 | 0 |
| TPKTPKGPS | NPM1; NPM1; N | P06748; A0A | NPM_HUMAN I | Unmodified | 0 | 0 | 0 | 0 | 1160300 | 169040 | 138490 | 0 | 0 | 716480 | 0 | 0 | 1846300 | 631150 | 0 | 0 | 501140 | 445810 |
| TPLAVELEV | ACOT2; ACOT1 | P49753; Q8( | ACOT2_HUMA | Unmodified | 0 | 0 | 0 | 0 | 0 | 0 | 0 | 0 | 361700 | 0 | 0 | 0 | 0 | 0 | 0 | 0 | 204290 | 0 |
| TPLDQHTL | SEL1L3 | Q68CR1 | SE1L3_HUMAN | Unmodified | 0 | 0 | 0 | 391340 | 0 | 0 | 0 | 183780 | 1448900 | 230500 | 0 | 0 | 0 | 0 | 0 | 0 | 957800 | 0 |
| TPLGGIHIS | DIP2B | Q9P265 | DIP2B_HUMAN | Unmodified | 346820 | 0 | 645580 | 0 | 0 | 0 | 74279 | 40095 | 734030 | 572900 | 739440 | 5874200 | 86660 | 0 | 1533000 | 1082700 | 0 | 0 |
| TPLGGIHLS | DIP2C; DIP2C; I | Q9Y2E4; A0A | DIP2C_HUMAN | Unmodified | 0 | 87311 | 593440 | 0 | 0 | 435170 | 0 | 0 | 1511600 | 192270 | 0 | 0 | 0 | 0 | 525110 | 299610 | 0 | 0 |
| TPLGSISAE | TANC1 | Q9C0D5 | TANC1_HUMA | Unmodified | 0 | 0 | 515620 | 617850 | 0 | 0 | 0 | 0 | 0 | 132550 | 0 | 0 | 147310 | 0 | 406480 | 296080 | 0 | 0 |
| TPLHEIALS | DHX9 | Q08211 | DHX9_HUMAN | Unmodified | 194900 | 0 | 0 | 0 | 0 | 0 | 0 | 0 | 203020 | 0 | 0 | 7173300 | 0 | 0 | 444100 | 0 | 0 | 0 |

| | | | | | | | | | | | | | | | | | | | | | | |
|---|---|---|---|---|---|---|---|---|---|---|---|---|---|---|---|---|---|---|---|---|---|---|
| TPLHMTALI | ANKRD28; ANK | O15084; B4[ | ANR28_HUMAN | Unmodified | 0 | 0 | 0 | 0 | 0 | 0 | 0 | 41038 | 0 | 46235 | 0 | 164650 | 0 | 105150 | 0 | 0 | 0 | 0 |
| TPLIADVYY | NAAA; NAAA; N. | Q02083; B4[ | NAAA_HUMAN | Unmodified | 0 | 0 | 38366 | 58224 | 0 | 0 | 0 | 0 | 891320 | 104260 | 41027 | 39620 | 0 | 0 | 59175 | 34919 | 0 | 0 |
| TPLLDDPVI | DHX37; DHX37 | Q8IY37; F5H | DHX37_HUMAI | Unmodified | 38826 | 0 | 17032 | 0 | 1440400 | 0 | 154030 | 0 | 47601 | 0 | 16394 | 185230 | 0 | 0 | 34281 | 20985 | 0 | 811590 |
| TPLLDEEEE | ATP2B4 | P23634 | AT2B4_HUMAN | Unmodified | 0 | 217870 | 0 | 0 | 814650 | 0 | 928060 | 1083300 | 234760 | 303260 | 195840 | 0 | 0 | 0 | 0 | 0 | 0 | 756610 |
| TPLLLMLG( | APEH; APEH | P13798; C9J | ACPH_HUMAN | Oxidation (M) | 40026 | 0 | 90627 | 85542 | 0 | 0 | 0 | 0 | 0 | 0 | 63250 | 1034400 | 0 | 0 | 87274 | 152570 | 0 | 0 |
| TPLPPELAD | ZNF706; ZNF70 | Q9Y5V0; H0' | ZN706_HUMAN | Unmodified | 0 | 0 | 0 | 0 | 0 | 0 | 0 | 0 | 0 | 0 | 0 | 0 | 0 | 0 | 0 | 0 | 0 | 1265300 |
| TPLPTLNEF | MARK2; MARK2 | Q7KZI7; E7E | MARK2_HUMAI | Unmodified | 0 | 0 | 658770 | 0 | 0 | 0 | 0 | 0 | 0 | 591760 | 1040700 | 5744800 | 122910 | 0 | 412680 | 561370 | 0 | 251560 |
| TPLQGILQL | HIP1R | O75146 | HIP1R_HUMAN | Unmodified | 0 | 173680 | 213000 | 350350 | 0 | 0 | 0 | 0 | 269320 | 0 | 500170 | 1934800 | 0 | 0 | 490130 | 229300 | 0 | 0 |
| TPLQVAKG | PSMD10; PSMD | O75832; B1/ | PSD10_HUMAN | Unmodified | 0 | 0 | 0 | 0 | 0 | 0 | 124360 | 27662 | 0 | 0 | 0 | 0 | 0 | 121400 | 0 | 0 | 1070500 | 0 |
| TPLSAAAHT | DYNC1I2 | Q13409 | DC1I2_HUMAN | Unmodified | 0 | 0 | 0 | 0 | 0 | 0 | 0 | 40976 | 0 | 0 | 0 | 1978600 | 0 | 0 | 0 | 0 | 0 | 0 |
| TPLSEAEFE | CPSF6; CPSF6 | Q16630; F8V | CPSF6_HUMAN | Oxidation (M) | 561860 | 612190 | 0 | 0 | 0 | 0 | 0 | 0 | 0 | 0 | 0 | 0 | 0 | 0 | 0 | 547540 | 0 | 0 |
| TPLSQSMS | RAB11FIP1; RAI | Q6WKZ4; E7 | RFIP1_HUMAN | Unmodified | 0 | 0 | 0 | 0 | 3685500 | 452320 | 0 | 0 | 0 | 0 | 0 | 412590 | 238410 | 450000 | 0 | 0 | 0 | 465700 |
| TPLTHALP( | GPC1; GPC1; G | P35052; H7( | GPC1_HUMAN | Unmodified | 739480 | 1016900 | 970980 | 667700 | 0 | 0 | 55053 | 0 | 487090 | 0 | 0 | 0 | 0 | 0 | 651440 | 778050 | 0 | 0 |
| TPLYDLR | MEMO1 | Q9Y316 | MEMO1_HUMA | Unmodified | 0 | 34238 | 130510 | 36245 | 676070 | 139700 | 0 | 23794 | 0 | 0 | 0 | 1057600 | 0 | 0 | 0 | 42386 | 61557 | 0 |
| TPMGIVLD/ | ARL6IP5 | O75915 | PRAF3_HUMAN | Unmodified | 0 | 63240 | 27127 | 0 | 100200 | 58292 | 0 | 0 | 0 | 150980 | 165190 | 0 | 0 | 0 | 0 | 25993 | 291230 | 76664 |
| TPMIAGGLI | GALNT2 | Q10471 | GALT2_HUMAN | 2 Oxidation (M) | 0 | 0 | 0 | 0 | 0 | 0 | 0 | 0 | 848220 | 0 | 0 | 649540 | 0 | 0 | 0 | 0 | 0 | 0 |
| TPNFPSSVI | MYOF; MYOF | Q9NZM1; H0 | MYOF_HUMAN | Oxidation (M) | 0 | 0 | 0 | 0 | 3029000 | 749030 | 0 | 0 | 0 | 0 | 0 | 0 | 0 | 0 | 0 | 0 | 0 | 0 |
| TPNFPSSVI | MYOF; MYOF | Q9NZM1; H0 | MYOF_HUMAN | Unmodified | 0 | 0 | 0 | 0 | 2975000 | 169680 | 75732 | 0 | 0 | 0 | 0 | 0 | 0 | 0 | 61887 | 0 | 0 | 0 |
| TPNGPNID | HSPH1; HSPH1 | Q92598; Q5' | HS105_HUMAN | Unmodified | 0 | 0 | 49305 | 0 | 233640 | 39939 | 0 | 0 | 0 | 0 | 79197 | 3320000 | 0 | 0 | 64980 | 56458 | 0 | 0 |
| TPNGVVLA | DYNC1H1; DYN | Q14204; A0/ | DYHC1_HUMA | Unmodified | 0 | 0 | 0 | 0 | 0 | 0 | 0 | 0 | 0 | 0 | 0 | 3543600 | 0 | 0 | 0 | 0 | 0 | 0 |
| TPNKIDLYD | CACNA2D1; CA | P54289; E7E | CA2D1_HUMAI | Unmodified | 462520 | 249950 | 0 | 0 | 361100 | 110100 | 0 | 0 | 332080 | 0 | 0 | 0 | 0 | 0 | 0 | 0 | 404300 | 0 |
| TPNNPSCN | CACNA2D1; CA | P54289; H0' | CA2D1_HUMAI | Unmodified | 585840 | 0 | 0 | 0 | 0 | 0 | 0 | 0 | 411490 | 0 | 0 | 0 | 0 | 0 | 0 | 0 | 0 | 0 |
| TPNPTASEI | PAN3; PAN3 | Q58A45; A0/ | PAN3_HUMAN | Unmodified | 727580 | 631800 | 0 | 266910 | 0 | 0 | 0 | 0 | 0 | 0 | 0 | 0 | 0 | 0 | 0 | 238260 | 0 | 0 |
| TPNQDTEF | PNMA2 | Q9UL42 | PNMA2_HUMA | Unmodified | 1960300 | 2533700 | 0 | 0 | 0 | 0 | 0 | 0 | 0 | 0 | 0 | 0 | 0 | 0 | 0 | 0 | 0 | 0 |
| TPNQDTEF | PNMA2 | Q9UL42 | PNMA2_HUMA | Unmodified | 1811800 | 3739800 | 0 | 0 | 633960 | 0 | 0 | 0 | 1423300 | 0 | 0 | 0 | 0 | 0 | 0 | 0 | 0 | 364200 |
| TPPDKKLEI | MYOF | Q9NZM1 | MYOF_HUMAN | Unmodified | 0 | 0 | 0 | 0 | 1766100 | 118440 | 81651 | 0 | 0 | 0 | 0 | 0 | 0 | 0 | 0 | 0 | 177930 | 696520 |
| TPPGLVGV. | TBC1D2 | Q9BYX2 | TBD2A_HUMAN | Unmodified | 0 | 0 | 0 | 0 | 476760 | 0 | 19287 | 0 | 0 | 0 | 0 | 0 | 0 | 39426 | 0 | 0 | 0 | 0 |
| TPPLLENSI | SLC30A7; SLC3 | Q8NEW0; H( | ZNT7_HUMAN | Unmodified | 33887 | 0 | 72963 | 48296 | 0 | 0 | 119900 | 38298 | 1578700 | 299890 | 0 | 128420 | 46974 | 0 | 82352 | 37479 | 0 | 0 |
| TPPPELLQT | HTT; HTT; HTT; | P42858; A0/ | HD_HUMAN Hu | Unmodified | 0 | 0 | 0 | 0 | 0 | 0 | 0 | 0 | 186760 | 0 | 0 | 711340 | 0 | 0 | 0 | 0 | 0 | 0 |
| TPPPPAMIF | LSR; LSR | Q86X29; S4F | LSR_HUMAN Li | 2 Oxidation (M) | 343030 | 0 | 582110 | 458420 | 0 | 0 | 0 | 0 | 0 | 1048200 | 0 | 5338900 | 0 | 0 | 932860 | 899220 | 0 | 0 |
| TPPPPAMIF | LSR; LSR | Q86X29; S4F | LSR_HUMAN Li | 2 Oxidation (M) | 0 | 0 | 636760 | 0 | 0 | 0 | 0 | 0 | 1802900 | 0 | 0 | 0 | 0 | 0 | 1852400 | 1777300 | 0 | 0 |
| TPPPRPPLF | ARHGAP8; PRR | P85298; H0' | RHG08_HUMAI | Unmodified | 0 | 0 | 160550 | 0 | 0 | 0 | 0 | 0 | 0 | 0 | 133680 | 781550 | 0 | 0 | 219740 | 0 | 0 | 0 |
| TPPRDLPTI | EIF4EBP1 | Q13541 | 4EBP1_HUMAN | Oxidation (M) | 0 | 0 | 0 | 422520 | 0 | 0 | 0 | 0 | 0 | 0 | 424360 | 4574700 | 0 | 0 | 382950 | 460480 | 0 | 0 |
| TPPTAVTEI | TUBGCP4; TUB | Q9UGJ1; H3 | GCP4_HUMAN | Unmodified | 0 | 0 | 0 | 0 | 0 | 0 | 0 | 0 | 0 | 0 | 0 | 1894100 | 0 | 0 | 0 | 0 | 0 | 0 |
| TPPYADPSI | RELA; RELA; RE | Q04206; A0/ | TF65_HUMAN 1 | Unmodified | 0 | 0 | 0 | 0 | 0 | 0 | 0 | 0 | 0 | 0 | 0 | 697910 | 0 | 0 | 0 | 0 | 0 | 0 |
| TPQGPPEI | CDV3; CDV3; C | Q9UKY7; D6 | CDV3_HUMAN | Unmodified | 653020 | 0 | 0 | 0 | 0 | 0 | 0 | 0 | 0 | 0 | 0 | 0 | 576400 | 0 | 0 | 369320 | 0 | 0 |
| TPQHVKITI | EGFR; EGFR; E( | P00533; Q5( | EGFR_HUMAN | Unmodified | 0 | 0 | 0 | 0 | 2969800 | 459500 | 0 | 28030 | 0 | 0 | 0 | 0 | 479050 | 291900 | 0 | 0 | 0 | 0 |
| TPQPIGEV | CLGN; CLGN | O14967; D6I | CLGN_HUMAN | Unmodified | 0 | 0 | 0 | 0 | 0 | 0 | 0 | 0 | 3017300 | 0 | 52888 | 0 | 0 | 0 | 0 | 0 | 0 | 0 |
| TPSALAILEI | ALDOC; ALDOC | P09972; J3Q | ALDOC_HUMA | Unmodified | 382930 | 83038 | 253900 | 227370 | 0 | 0 | 0 | 0 | 0 | 0 | 341660 | 14056000 | 52034 | 0 | 777620 | 539090 | 0 | 0 |
| TPSAYELPY | ST3GAL4; ST3G | Q11206; E9F | SIA4C_HUMAN | Unmodified | 795150 | 0 | 0 | 0 | 0 | 0 | 0 | 0 | 920400 | 0 | 0 | 0 | 0 | 0 | 0 | 0 | 0 | 0 |
| TPSEGGKL | HSD17B4; HSD | P51659; E7E | DHB4_HUMAN | Unmodified | 0 | 0 | 0 | 0 | 0 | 0 | 0 | 0 | 959750 | 0 | 0 | 0 | 0 | 0 | 0 | 0 | 0 | 0 |
| TPSFGISAP | AHNAK | Q09666 | AHNK_HUMAN | Unmodified | 46654 | 0 | 0 | 69249 | 1743500 | 0 | 0 | 0 | 0 | 0 | 0 | 0 | 0 | 109350 | 0 | 26380 | 0 | 1563500 |
| TPSFGISAP | AHNAK | Q09666 | AHNK_HUMAN | Unmodified | 0 | 0 | 0 | 0 | 3159700 | 0 | 0 | 0 | 0 | 0 | 0 | 0 | 0 | 0 | 0 | 0 | 2133200 | 2991400 |
| TPSGLYLGT | HSPG2; HSPG2 | P98160; H0' | PGBM_HUMAN | Unmodified | 0 | 0 | 2149200 | 379730 | 0 | 0 | 0 | 0 | 502650 | 0 | 220490 | 0 | 0 | 0 | 0 | 629550 | 0 | 0 |
| TPSGNLVN | ABCC1; ABCC1 | P33527; I3L4 | MRP1_HUMAN | Unmodified | 0 | 0 | 636110 | 102210 | 0 | 0 | 49759 | 0 | 47478 | 0 | 268360 | 3642100 | 0 | 0 | 782410 | 840200 | 0 | 0 |
| TPSKIPYSD | MAN1B1; MAN1 | Q9UKM7; H( | MA1B1_HUMAI | Unmodified | 0 | 0 | 0 | 0 | 0 | 0 | 0 | 0 | 9073800 | 1223600 | 0 | 574160 | 0 | 0 | 0 | 0 | 0 | 0 |
| TPSLDAEG | AGRN; AGRN; A | O00468; A0/ | AGRIN_HUMAN | Unmodified | 0 | 0 | 314780 | 0 | 0 | 0 | 0 | 0 | 0 | 50596 | 0 | 0 | 0 | 0 | 0 | 0 | 0 | 0 |
| TPSNELYKI | PDCD6IP | Q8WUM4 | PDC6I_HUMAN | Unmodified | 5190300 | 5218000 | 1160700 | 0 | 141050 | 0 | 77264 | 85477 | 1170700 | 142160 | 0 | 1634800 | 105950 | 0 | 3156200 | 3880200 | 0 | 0 |
| TPSNELYKI | PDCD6IP | Q8WUM4 | PDC6I_HUMAN | Unmodified | 34940000 | 30846000 | 9373400 | 18918000 | 1608400 | 769230 | 0 | 0 | 17980000 | 1274800 | 2808200 | 8097900 | 3169700 | 1104500 | 12526000 | 15559000 | 110270 | 968330 |
| TPSPKEEDI | NUCKS1 | Q9H1E3 | NUCKS_HUMA | Unmodified | 0 | 0 | 0 | 0 | 0 | 0 | 192190 | 19634 | 473760 | 0 | 0 | 1949400 | 779070 | 183010 | 0 | 0 | 0 | 0 |
| TPSSDGTV | IGSF21 | Q96ID5 | IGS21_HUMAN | Unmodified | 0 | 0 | 124610 | 0 | 0 | 0 | 0 | 0 | 957380 | 0 | 22378 | 691380 | 0 | 0 | 0 | 0 | 0 | 0 |
| TPSSDVLVI | RBBP7; RBBP4; | Q16576; Q0( | RBBP7_HUMAN | Unmodified | 0 | 0 | 0 | 0 | 0 | 0 | 0 | 0 | 2397900 | 0 | 0 | 9899800 | 0 | 0 | 0 | 334680 | 0 | 0 |
| TPSSSSLS( | RACGAP1 | Q9H0H5 | RGAP1_HUMAI | Unmodified | 0 | 0 | 0 | 1878900 | 4227000 | 0 | 0 | 0 | 0 | 0 | 0 | 0 | 0 | 0 | 401920 | 0 | 594080 | 0 |
| TPSSSSLS( | RACGAP1 | Q9H0H5 | RGAP1_HUMAI | Unmodified | 2104600 | 831140 | 0 | 0 | 1113800 | 116270 | 18814 | 0 | 0 | 0 | 0 | 0 | 0 | 0 | 0 | 0 | 65821 | 47782 |
| TPSSTGPSI | PTPRJ; PTPRJ; P | Q12913; A0/ | PTPRJ_HUMAN | Unmodified | 0 | 0 | 0 | 496020 | 624420 | 0 | 0 | 0 | 2221900 | 0 | 1300700 | 3690100 | 0 | 0 | 0 | 0 | 0 | 0 |
| TPTAVVAP\ | CLIP1; CLIP1; C | P30622; F5F | CLIP1_HUMAN | Unmodified | 0 | 0 | 0 | 0 | 439880 | 87422 | 43786 | 0 | 0 | 0 | 228590 | 381120 | 856980 | 303510 | 0 | 0 | 134590 | 178710 |
| TPTEALASF | PTGES2; PTGES | Q9H7Z7; A0/ | PGES2_HUMAN | Unmodified | 0 | 0 | 0 | 0 | 0 | 610410 | 0 | 0 | 0 | 0 | 0 | 1521200 | 0 | 0 | 444660 | 0 | 0 | 0 |
| TPTFAGGLI | GALNT3; GALN' | Q14435; E7E | GALT3_HUMAN | Unmodified | 0 | 0 | 0 | 0 | 0 | 0 | 0 | 0 | 1483000 | 436980 | 654450 | 0 | 0 | 0 | 0 | 0 | 0 | 0 |
| TPTIIALVVL | ENTPD3; ENTPI | O75355; C9J | ENTP3_HUMAN | Unmodified | 0 | 0 | 99355 | 0 | 0 | 0 | 0 | 0 | 0 | 0 | 0 | 0 | 0 | 0 | 0 | 0 | 330090 | 0 |
| TPTLILYGEI | ABHD14A; ABH | Q9BUJ0; K7I | ABHEA_HUMAN | Unmodified | 0 | 0 | 0 | 0 | 0 | 0 | 0 | 0 | 260430 | 42132 | 46892 | 0 | 0 | 0 | 0 | 0 | 135370 | 0 |
| TPTLYLDFK | PIR | O00625 | PIR_HUMAN Pi | Unmodified | 75165 | 0 | 97846 | 0 | 841190 | 0 | 538800 | 0 | 0 | 369930 | 588630 | 5550700 | 0 | 0 | 170560 | 149000 | 0 | 0 |
| TPTPEPAEV | NRBP1; NRBP1 | Q9UHY1; F8' | NRBP_HUMAN | Unmodified | 0 | 0 | 0 | 29235 | 173160 | 0 | 0 | 0 | 0 | 0 | 0 | 300930 | 0 | 0 | 40724 | 0 | 0 | 259410 |
| TPTPSTPG( | CHD4; CHD4; ( | Q14839; A0/ | CHD4_HUMAN | Unmodified | 0 | 0 | 0 | 0 | 0 | 0 | 0 | 0 | 0 | 0 | 604010 | 0 | 0 | 0 | 0 | 0 | 0 | 0 |
| TPTQLEGA | MTA2 | O94776 | MTA2_HUMAN | Unmodified | 0 | 0 | 0 | 0 | 0 | 0 | 0 | 0 | 0 | 0 | 196240 | 0 | 0 | 0 | 0 | 0 | 0 | 0 |
| TPTQTNGS | GTF2I; GTF2I | P78347; A0/ | GTF2I_HUMAN | Unmodified | 0 | 0 | 0 | 0 | 0 | 17794 | 0 | 0 | 0 | 27262 | 0 | 319980 | 0 | 0 | 0 | 0 | 0 | 112880 |
| TPTSVNQTI | KIAA1109; KIAA | Q2LD37; A0/ | K1109_HUMAN | Unmodified | 0 | 0 | 0 | 0 | 0 | 0 | 0 | 0 | 0 | 0 | 0 | 1210800 | 0 | 0 | 0 | 0 | 0 | 0 |
| TPTVEYICT( | CAB39; CAB39 | Q9Y376; A0/ | CAB39_HUMAN | Oxidation (M) | 0 | 24108 | 0 | 17790 | 0 | 0 | 0 | 0 | 0 | 0 | 0 | 0 | 0 | 0 | 88988 | 0 | 0 | 0 |
| TPVEEVPA/ | SF3B3 | Q15393 | SF3B3_HUMAN | Unmodified | 578120 | 0 | 0 | 0 | 0 | 0 | 0 | 0 | 0 | 0 | 0 | 3959100 | 0 | 0 | 0 | 0 | 0 | 0 |
| TPVEPEVAI | RPS20; RPS20; | P60866; A0A | RS20_HUMAN | Unmodified | 1581100 | 1299400 | 2423100 | 486060 | 188130 | 420180 | 0 | 0 | 1120200 | 103440 | 5031400 | 30775000 | 0 | 0 | 2841900 | 1732200 | 22352 | 0 |
| TPVEVPVG( | PLEC; PLEC | Q15149; H0' | PLEC_HUMAN | Unmodified | 215830 | 0 | 0 | 0 | 1014400 | 0 | 0 | 0 | 63695 | 44240 | 0 | 0 | 0 | 0 | 39832 | 30366 | 2724500 | 1202600 |
| TPVFVSPTF | WASF1 | Q92558 | WASF1_HUMAI | Unmodified | 0 | 0 | 0 | 0 | 0 | 0 | 0 | 0 | 299200 | 0 | 281750 | 1342900 | 0 | 0 | 0 | 0 | 0 | 0 |
| TPVIDADKF | DYNC1H1; DYN | Q14204; A0/ | DYHC1_HUMA | Unmodified | 49139 | 0 | 0 | 220560 | 47922 | 0 | 0 | 417690 | 298240 | 475040 | 1178000 | 967800 | 92268 | 0 | 92680 | 94140 | 255380 | 91471 |
| TPVIVTLK | HYOU1; HYOU: | Q9Y4L1; J3Q | HYOU1_HUMA | Unmodified | 0 | 0 | 1013600 | 419050 | 0 | 0 | 0 | 94169 | 38784000 | 6652000 | 0 | 0 | 0 | 1094700 | 0 | 485970 | 0 | 0 |
| TPVKEPNSI | RAB10 | P61026 | RAB10_HUMAN | Unmodified | 1161500 | 1664400 | 1964400 | 387420 | 450370 | 0 | 0 | 0 | 0 | 0 | 0 | 5416000 | 0 | 0 | 3467500 | 3389800 | 0 | 0 |
| TPVLFDIYE | ANXA11 | P50995 | ANX11_HUMAN | Unmodified | 2006700 | 1188700 | 707850 | 1028900 | 15996000 | 4358800 | 0 | 0 | 2.19E+08 | 30034000 | 17102000 | 18290000 | 4101300 | 10152000 | 1620700 | 1639000 | 4819200 | 24666000 |
| TPVLFDIYE | ANXA11 | P50995 | ANX11_HUMAN | Unmodified | 0 | 72355 | 0 | 148520 | 16649000 | 1695300 | 0 | 0 | 20242000 | 22787000 | 497980 | 136660 | 5482400 | 6821600 | 0 | 0 | 3673500 | 22282000 |
| TPVMEGYV | LOXL2; LOXL2 | Q9Y4K0; H0' | LOXL2_HUMAN | Unmodified | 0 | 0 | 0 | 0 | 0 | 0 | 0 | 0 | 714240 | 187670 | 0 | 0 | 0 | 0 | 0 | 0 | 0 | 0 |
| TPVPSDIDI | MTHFD1; MTHF | P11586; V9C | C1TC_HUMAN | Unmodified | 388930 | 257440 | 1369500 | 1157800 | 3687300 | 0 | 1073500 | 0 | 4890400 | 2246600 | 4346000 | 14222000 | 1756900 | 2650500 | 1236200 | 1428200 | 1606600 | 4029300 |
| TPVQYLLPI | CTDSP1; CTDSI | Q9GZU7; H7 | CTDS1_HUMAI | Unmodified | 0 | 0 | 5457200 | 0 | 0 | 0 | 0 | 0 | 0 | 0 | 0 | 1398800 | 9751300 | 0 | 0 | 0 | 0 | 523550 |
| TPVSEDML | ATP6V1B2; ATP | P21281; P15 | VATB2_HUMAN | Unmodified | 66896 | 0 | 0 | 0 | 0 | 0 | 0 | 0 | 2041700 | 93951 | 914810 | 2861500 | 0 | 0 | 0 | 0 | 0 | 0 |
| TPVSITEHP | DYNC1H1; DYN | Q14204; A0/ | DYHC1_HUMA | Unmodified | 0 | 0 | 47979 | 0 | 0 | 0 | 0 | 0 | 0 | 0 | 0 | 608420 | 0 | 0 | 34430 | 49623 | 0 | 0 |
| TPVSSQAP | INPP5D; INPP5 | Q92835; H0' | SHIP1_HUMAN | Unmodified | 0 | 0 | 0 | 0 | 0 | 0 | 0 | 0 | 0 | 0 | 295910 | 772200 | 0 | 0 | 0 | 0 | 0 | 0 |
| TPVTDPAT( | ACADVL | P49748 | ACADV_HUMAI | Unmodified | 307860 | 179190 | 0 | 0 | 0 | 0 | 0 | 0 | 1127100 | 246690 | 151630 | 150830 | 0 | 0 | 0 | 96919 | 0 | 0 |
| TPVTDPAT( | ACADVL | P49748 | ACADV_HUMAI | Unmodified | 0 | 0 | 0 | 0 | 0 | 0 | 0 | 0 | 0 | 998700 | 0 | 0 | 56571 | 55221 | 21890 | 31157 | 50411 | 0 |
| TPWRGILLI | VPS4B; VPS4A; | O75351; Q9I | VPS4B_HUMAN | Unmodified | 0 | 0 | 0 | 0 | 225590 | 48290 | 0 | 0 | 0 | 183970 | 0 | 0 | 259160 | 151420 | 0 | 0 | 0 | 0 |
| TPYLGDVA' | DPYSL5 | Q9BPU6 | DPYL5_HUMAN | Unmodified | 0 | 0 | 0 | 0 | 0 | 0 | 0 | 0 | 0 | 0 | 401570 | 2110700 | 0 | 0 | 0 | 0 | 0 | 0 |
| TPYSDGVL' | SGCE; SGCE; S | O43556; A0/ | SGCE_HUMAN | Unmodified | 1218800 | 1795100 | 0 | 0 | 0 | 0 | 0 | 0 | 535600 | 0 | 0 | 0 | 0 | 0 | 0 | 0 | 0 | 0 |
| TPYTIMFGF | CANX; CANX; C | P27824; A0A | CALX_HUMAN | Oxidation (M) | 0 | 832880 | 0 | 149140 | 1519800 | 0 | 596130 | 0 | 88791000 | 1928400 | 2031300 | 4052700 | 0 | 0 | 488270 | 503810 | 0 | 0 |
| TPYTIMFGF | CANX; CANX; C | P27824; A0A | CALX_HUMAN | Unmodified | 0 | 0 | 0 | 0 | 0 | 47191 | 0 | 0 | 1.32E+08 | 0 | 1385100 | 0 | 0 | 0 | 0 | 0 | 11074 | 0 |
| TPYTIMFGF | CANX; CANX; C | P27824; A0A | CALX_HUMAN | Oxidation (M) | 0 | 0 | 0 | 0 | 0 | 0 | 0 | 15013 | 11918000 | 1198100 | 718720 | 0 | 0 | 0 | 0 | 0 | 0 | 0 |
| TPYTIMFGF | CANX; CANX; C | P27824; A0A | CALX_HUMAN | Unmodified | 0 | 0 | 0 | 0 | 0 | 0 | 0 | 0 | 27770000 | 803070 | 228400 | 0 | 0 | 0 | 0 | 0 | 0 | 0 |
| TQADLDSL' | DCXR; DCXR; D | Q7Z4W1; J3I | DCXR_HUMAN | Unmodified | 108180 | 115970 | 0 | 483420 | 0 | 0 | 0 | 0 | 1377700 | 0 | 0 | 1867300 | 0 | 0 | 986140 | 588850 | 0 | 0 |
| TQAEHYLW | AVIL | O75366 | AVIL_HUMAN A | Unmodified | 0 | 0 | 0 | 0 | 0 | 0 | 0 | 0 | 0 | 0 | 1854200 | 6652500 | 0 | 0 | 0 | 0 | 0 | 0 |
| TQAFEADN | ROCK1; ROCK1 | Q13464; A0/ | ROCK1_HUMAI | Unmodified | 0 | 0 | 0 | 280410 | 2828000 | 657500 | 190400 | 101400 | 507150 | 586700 | 95501 | 843510 | 1143000 | 416180 | 0 | 0 | 1609400 | 2386600 |
| TQAFEADN | ROCK1; ROCK1 | Q13464; A0/ | ROCK1_HUMAI | Unmodified | 38450 | 765410 | 1062000 | 111150 | 0 | 0 | 218630 | 216770 | 0 | 0 | 0 | 0 | 0 | 0 | 578090 | 86395 | 1081900 | 281270 |
| TQAIVCQQ | ABCE1; ABCE1; | P61221; D6F | ABCE1_HUMAN | Unmodified | 417870 | 0 | 0 | 450010 | 2040500 | 0 | 0 | 0 | 0 | 0 | 1655600 | 7139400 | 1635700 | 0 | 0 | 506720 | 1099000 | 0 |
| TQAPPSVIL | DENND6A; DEN | Q8IWF6; H7( | DEN6A_HUMAN | Unmodified | 0 | 0 | 0 | 0 | 0 | 0 | 0 | 0 | 0 | 0 | 215080 | 753770 | 0 | 0 | 0 | 0 | 0 | 0 |
| TQAPPTPD | SLC12A7; SLC1 | Q9Y666; A0/ | S12A7_HUMAN | Unmodified | 0 | 0 | 0 | 0 | 0 | 0 | 0 | 163940 | 0 | 149550 | 0 | 0 | 1035500 | 285050 | 0 | 0 | 0 | 509320 |
| TQAVFNHT | CSTF1 | Q05048 | CSTF1_HUMAN | Unmodified | 0 | 0 | 0 | 0 | 0 | 0 | 0 | 0 | 0 | 0 | 0 | 1812900 | 3555400 | 0 | 0 | 0 | 0 | 0 |
| TQAVLDQS | RMC1; RMC1; F | Q96DM3; K7 | RMC1_HUMAN | Unmodified | 0 | 256270 | 0 | 0 | 523270 | 0 | 0 | 0 | 0 | 0 | 0 | 276720 | 0 | 0 | 0 | 0 | 0 | 0 |
| TQAYQDQI | PGM1; PGM1 | P36871; A0A | PGM1_HUMAN | Unmodified | 0 | 0 | 0 | 245870 | 0 | 152940 | 691800 | 330410 | 357640 | 139300 | 0 | 0 | 0 | 994330 | 0 | 0 | 974550 | 5258800 |
| TQDASGPE | AFDN; AFDN; A | P55196; Q5T | AFAD_HUMAN | Unmodified | 0 | 0 | 0 | 0 | 0 | 0 | 0 | 0 | 0 | 183020 | 507930 | 1626600 | 0 | 0 | 21352 | 0 | 0 | 0 |
| TQDDIETVL | KANK2; KANK2 | Q63ZY3; K7E | KANK2_HUMAN | Unmodified | 0 | 0 | 0 | 0 | 644480 | 68677 | 0 | 0 | 0 | 0 | 0 | 0 | 0 | 0 | 0 | 0 | 198620 | 0 |
| TQDGTAQ( | ITGB4 | P16144 | ITB4_HUMAN I | Unmodified | 0 | 0 | 0 | 0 | 0 | 0 | 0 | 0 | 0 | 0 | 0 | 0 | 0 | 0 | 0 | 0 | 1111800 | 2122700 |
| TQDGVALE | ATP2B4 | P23634 | AT2B4_HUMAN | Unmodified | 0 | 0 | 1058400 | 0 | 4115900 | 0 | 1172500 | 127860 | 0 | 0 | 4071800 | 13371000 | 0 | 0 | 763000 | 1888200 | 0 | 0 |
| TQDNSCSF | ST14 | Q9Y5Y6 | ST14_HUMAN | Unmodified | 3200800 | 1832600 | 4183500 | 6462000 | 0 | 0 | 0 | 0 | 2909500 | 0 | 0 | 609100 | 586610 | 0 | 7744500 | 5322300 | 0 | 0 |
| TQDPAKAP | ASS1 | P00966 | ASSY_HUMAN | Unmodified | 0 | 0 | 0 | 0 | 176560 | 0 | 0 | 302150 | 415300 | 238800 | 0 | 0 | 576470 | 0 | 0 | 0 | 0 | 0 |
| TQDPAKAP | ASS1 | P00966 | ASSY_HUMAN | Unmodified | 0 | 0 | 0 | 0 | 787820 | 0 | 2646300 | 1342400 | 1017900 | 1085100 | 0 | 0 | 2423900 | 1249100 | 0 | 0 | 0 | 0 |
| TQDQISNII | LASP1; LASP1 | Q14847; C9J | LASP1_HUMAN | Unmodified | 3023200 | 2478700 | 540170 | 749840 | 1569800 | 0 | 0 | 0 | 0 | 369510 | 455110 | 529670 | 652770 | 0 | 0 | 1507500 | 0 | 726690 |

| | | | | | | | | | | | | | | | | | | | | | | |
|---|---|---|---|---|---|---|---|---|---|---|---|---|---|---|---|---|---|---|---|---|---|---|
| TQDQISNIK | LASP1; LASP1 | Q14847; C9J | LASP1_HUMAN | Unmodified | 0 | 0 | 0 | 144100 | 6447400 | 370570 | 1722200 | 1590600 | 86604 | 0 | 0 | 0 | 3119900 | 989630 | 0 | 0 | 2226400 | 3962500 |
| TQEEGEER | DIABLO; DIABL | Q9NR28; A0 | DBLOH_HUMA | Unmodified | 0 | 0 | 0 | 0 | 0 | 0 | 0 | 0 | 3907000 | 878310 | 266480 | 1722500 | 0 | 0 | 0 | 0 | 0 | 0 |
| TQEEIVAK | PDIA4; PDIA4 | P13667; A0A | PDIA4_HUMAN | Unmodified | 0 | 0 | 0 | 0 | 0 | 0 | 0 | 0 | 1365300 | 0 | 0 | 0 | 0 | 0 | 0 | 0 | 0 | 0 |
| TQEHFTHN | PPP3CA; PPP3C | Q08209; E7E | PP2BA_HUMAN | Unmodified | 0 | 0 | 546520 | 0 | 864790 | 56070 | 79307 | 0 | 0 | 119370 | 42844 | 5484700 | 0 | 0 | 0 | 904360 | 663360 | 1628200 |
| TQEKEEMN | DAAM1 | Q9Y4D1 | DAAM1_HUMA | Unmodified | 118230 | 53624 | 0 | 185970 | 373850 | 68251 | 238180 | 0 | 394990 | 124870 | 0 | 453920 | 266120 | 216340 | 22934 | 0 | 155210 | 213050 |
| TQEKPPKEI | CHMP3 | Q9Y3E7 | CHMP3_HUMA | Unmodified | 469620 | 112580 | 0 | 492580 | 2568200 | 264420 | 469260 | 346660 | 1320900 | 1880500 | 0 | 0 | 1833700 | 2163900 | 0 | 0 | 0 | 751350 |
| TQEQLALEI | MSN | P26038 | MOES_HUMAN | Unmodified | 0 | 0 | 0 | 0 | 7359500 | 297730 | 0 | 0 | 0 | 0 | 0 | 247770 | 386690 | 0 | 0 | 0 | 1587900 | 2781400 |
| TQESAGLA | HYAL2 | Q12891 | HYAL2_HUMAN | Unmodified | 0 | 651730 | 61840 | 248640 | 0 | 0 | 0 | 0 | 140350 | 0 | 0 | 0 | 0 | 0 | 877250 | 52923 | 0 | 0 |
| TQETLSQA | TPD52L2; TPD5 | O43399; A0A | TPD54_HUMAN | Unmodified | 0 | 0 | 0 | 0 | 1121900 | 0 | 0 | 228630 | 0 | 232370 | 0 | 0 | 0 | 0 | 0 | 0 | 303880 | 908490 |
| TQETPSHLI | TC2N | Q8N9U0 | TAC2N_HUMAN | Unmodified | 64982 | 73587 | 521530 | 1161500 | 0 | 0 | 0 | 0 | 0 | 0 | 0 | 0 | 0 | 0 | 585220 | 566090 | 0 | 0 |
| TQEVHHPI | ADGRG6 | Q86SQ4 | AGRG6_HUMA | Unmodified | 81652 | 0 | 541030 | 723320 | 1601400 | 174490 | 0 | 0 | 0 | 0 | 0 | 257490 | 53220 | 0 | 879790 | 795480 | 0 | 0 |
| TQEWDNSI | TENM4 | Q6N022 | TEN4_HUMAN | Unmodified | 0 | 0 | 0 | 48357 | 0 | 0 | 0 | 0 | 0 | 0 | 24000 | 32287 | 0 | 0 | 0 | 0 | 143910 | 474160 |
| TQFEDCTV | ABCC1; ABCC1 | P33527; I3L | MRP1_HUMAN | Unmodified | 0 | 0 | 0 | 0 | 0 | 0 | 0 | 0 | 0 | 0 | 0 | 1910400 | 0 | 0 | 0 | 328250 | 0 | 0 |
| TQFNSLQC | SRC | P12931 | SRC_HUMAN P | Unmodified | 354410 | 0 | 0 | 0 | 0 | 164020 | 0 | 0 | 968650 | 240160 | 145400 | 363010 | 0 | 0 | 0 | 29361 | 290710 | 290810 |
| TQFTTREGI | WDR20; WDR2 | Q8TBZ3; H0 | WDR20_HUMA | Unmodified | 0 | 0 | 0 | 0 | 2510900 | 424740 | 220390 | 156330 | 488330 | 182830 | 0 | 108040 | 1284800 | 672830 | 0 | 0 | 952490 | 1689200 |
| TQGFGGS | SPINT1; SPINT1 | O43278; H3I | SPIT1_HUMAN | Unmodified | 0 | 0 | 0 | 0 | 0 | 0 | 0 | 0 | 1444200 | 61400 | 476830 | 0 | 0 | 0 | 1731000 | 181770 | 0 | 0 |
| TQGFLALFS | RUVBL2; RUVBI | Q9Y230; M0I | RUVB2_HUMAN | Unmodified | 177620 | 0 | 770920 | 299150 | 0 | 0 | 0 | 0 | 338350 | 10174 | 734630 | 9735200 | 36299 | 0 | 146270 | 453730 | 11129 | 0 |
| TQGFLALFS | RUVBL2; RUVBI | Q9Y230; M0I | RUVB2_HUMAN | Unmodified | 0 | 0 | 0 | 0 | 0 | 1831700 | 0 | 0 | 463630 | 0 | 257380 | 5775300 | 0 | 0 | 0 | 0 | 547380 | 0 |
| TQGLVPEH | MYOF | Q9NZM1 | MYOF_HUMAN | Unmodified | 0 | 0 | 0 | 0 | 4226800 | 0 | 0 | 0 | 0 | 0 | 0 | 0 | 0 | 0 | 0 | 0 | 0 | 0 |
| TQGNVFAT | EIF3D; EIF3D; E | O15371; B0 | EIF3D_HUMAN | Oxidation (M) | 71086 | 176110 | 32457 | 600160 | 284020 | 21989 | 199160 | 0 | 16078 | 245830 | 111480 | 315610 | 0 | 0 | 840270 | 58025 | 139350 | 162160 |
| TQGSSTSW | ARHGAP12; AR | Q8IWW6; H | RHG12_HUMA | Unmodified | 0 | 0 | 0 | 0 | 2702000 | 0 | 183990 | 51158 | 1786900 | 0 | 177830 | 0 | 212310 | 182650 | 0 | 0 | 0 | 828740 |
| TQGTKIASD | RPS3A; RPS3A; | P61247; D6F | RS3A_HUMAN | Unmodified | 0 | 0 | 0 | 0 | 0 | 0 | 170010 | 155300 | 0 | 0 | 0 | 0 | 0 | 79758 | 0 | 0 | 57581 | 59089 |
| TQHEILRQ | LMAN1 | P49257 | LMAN1_HUMA | Unmodified | 0 | 0 | 0 | 79279 | 988450 | 64173 | 326400 | 86518 | 6126200 | 7519500 | 1346300 | 0 | 0 | 707640 | 0 | 0 | 53035 | 2176400 |
| TQHGVLSC | STX4 | Q12846 | STX4_HUMAN | Unmodified | 392560 | 384040 | 672220 | 768580 | 3317400 | 0 | 0 | 27747 | 1785500 | 1257700 | 779770 | 3109200 | 1340300 | 803220 | 717440 | 666790 | 1508400 | 2552300 |
| TQHLSNDI | MAPK1 | P28482 | MK01_HUMAN | Unmodified | 86727 | 0 | 0 | 80803 | 0 | 0 | 0 | 0 | 26038 | 0 | 0 | 0 | 0 | 0 | 252660 | 80146 | 0 | 0 |
| TQHNGMN | HUWE1 | Q7Z6Z7 | HUWE1_HUMA | Unmodified | 142420 | 0 | 0 | 0 | 0 | 0 | 0 | 0 | 0 | 10408 | 0 | 354250 | 0 | 0 | 0 | 0 | 0 | 0 |
| TQHWPIRL | MYORG | Q6NSJ0 | MYORG_HUMA | Unmodified | 48766 | 0 | 0 | 0 | 0 | 0 | 0 | 0 | 302120 | 0 | 0 | 0 | 0 | 0 | 0 | 0 | 0 | 0 |
| TQIALSPN | ARPC1A; ARPC | Q92747; E9F | ARC1A_HUMAN | Unmodified | 0 | 205630 | 159320 | 274190 | 0 | 0 | 0 | 0 | 0 | 0 | 0 | 1034700 | 0 | 0 | 238870 | 136690 | 0 | 0 |
| TQIALSPN | ARPC1A; ARPC | Q92747; E9F | ARC1A_HUMAN | Unmodified | 590040 | 0 | 661720 | 285180 | 0 | 0 | 0 | 0 | 202780 | 0 | 0 | 5101100 | 0 | 0 | 467410 | 949270 | 0 | 0 |
| TQIDHYVG | RTN3 | O95197 | RTN3_HUMAN | Unmodified | 0 | 0 | 352920 | 113890 | 0 | 0 | 0 | 0 | 0 | 75227 | 0 | 2616600 | 0 | 0 | 0 | 198120 | 0 | 0 |
| TQIDHYVG | RTN3 | O95197 | RTN3_HUMAN | Unmodified | 0 | 0 | 421330 | 0 | 326300 | 0 | 686690 | 0 | 546180 | 775460 | 225300 | 699110 | 0 | 55829 | 0 | 305300 | 49861 | 736840 |
| TQILAASYE | SPTBN1; SPTBN | Q01082; A0 | SPTB2_HUMAN | Unmodified | 232460 | 0 | 0 | 0 | 0 | 0 | 0 | 0 | 251750 | 0 | 0 | 6239600 | 0 | 0 | 0 | 0 | 0 | 0 |
| TQIPLQARE | BIRC2; BIRC2; | Q13490; E9F | BIRC2_HUMAN | Unmodified | 0 | 0 | 0 | 0 | 954930 | 521670 | 0 | 0 | 0 | 57983 | 0 | 0 | 0 | 0 | 0 | 0 | 0 | 0 |
| TQIPTQR | PSMA1; PSMA1 | P25786; F5G | PSA1_HUMAN | Unmodified | 0 | 0 | 0 | 124780 | 799910 | 0 | 0 | 0 | 1395900 | 0 | 0 | 5974900 | 131660 | 0 | 122160 | 106040 | 112230 | 1035400 |
| TQIQSQESI | EPS15L1; EPS1 | Q9UBC2; M | EP15R_HUMAN | Unmodified | 454340 | 435580 | 223520 | 165160 | 383160 | 0 | 79023 | 0 | 273840 | 78899 | 627310 | 1160100 | 0 | 0 | 162440 | 266870 | 201080 | 0 |
| TQIQSQESI | EPS15L1; EPS1 | Q9UBC2; M | EP15R_HUMAN | Unmodified | 0 | 159350 | 0 | 0 | 0 | 0 | 0 | 0 | 0 | 257880 | 0 | 0 | 1126700 | 471170 | 0 | 0 | 0 | 981830 |
| TQLAVCQC | FKBP4; FKBP4 | Q02790; H0 | FKBP4_HUMAN | Unmodified | 0 | 0 | 0 | 1382200 | 2061700 | 0 | 964450 | 488130 | 873340 | 425590 | 1759400 | 16669000 | 11361000 | 6696700 | 974450 | 1245500 | 1216700 | 2431700 |
| TQLEELEDI | MYH9; MYH10; | P35579; P35 | MYH9_HUMAN | Unmodified | 0 | 0 | 910310 | 1859400 | 3354100 | 1749200 | 0 | 0 | 3128200 | 889840 | 1733200 | 5968300 | 0 | 0 | 7150000 | 2662600 | 101990 | 0 |
| TQLEINGR | MBNL1; MBNL3 | Q9NR56; Q9 | MBNL1_HUMA | Unmodified | 1661400 | 0 | 503270 | 355340 | 686690 | 0 | 0 | 0 | 365870 | 201630 | 0 | 0 | 0 | 0 | 0 | 0 | 0 | 0 |
| TQLEINGRI | MBNL1; MBNL3 | Q9NR56; Q9 | MBNL1_HUMA | Unmodified | 288920 | 145760 | 0 | 622230 | 0 | 234210 | 224070 | 176140 | 0 | 0 | 0 | 0 | 4204800 | 1445000 | 0 | 76277 | 243870 | 635420 |
| TQLEVRPV | ROBO1; ROBO1 | Q9Y6N7; A0 | ROBO1_HUMA | Unmodified | 0 | 122280 | 0 | 0 | 0 | 0 | 0 | 0 | 2127800 | 276380 | 387990 | 960870 | 0 | 0 | 0 | 0 | 208790 | 966940 |
| TQLGRCFC | LTBP1; LTBP1; | Q14766; E7E | LTBP1_HUMAN | Unmodified | 0 | 0 | 0 | 0 | 0 | 0 | 0 | 0 | 66273 | 0 | 0 | 0 | 0 | 0 | 0 | 0 | 924400 | 1112300 |
| TQLLQDVQ | KTN1; KTN1; KT | Q86UP2; H0 | KTN1_HUMAN | Unmodified | 0 | 0 | 0 | 0 | 0 | 0 | 0 | 0 | 96473 | 0 | 0 | 1181400 | 450880 | 0 | 0 | 0 | 0 | 0 |
| TQLNLLLDI | MYO6; MYO6; M | Q9UM54; A0 | MYO6_HUMAN | Unmodified | 0 | 0 | 1396600 | 0 | 0 | 0 | 0 | 0 | 0 | 0 | 0 | 0 | 0 | 0 | 568130 | 0 | 0 | 0 |
| TQLNPTSL | MARK3; MARK1 | P27448; Q9F | MARK3_HUMA | Unmodified | 2162300 | 4589900 | 103890 | 113430 | 0 | 0 | 134490 | 145740 | 0 | 0 | 124160 | 463980 | 905860 | 0 | 0 | 302000 | 0 | 0 |
| TQLNSSSL | MARK2; MARK2 | Q7KZI7; E7E | MARK2_HUMA | Unmodified | 0 | 0 | 369320 | 0 | 247210 | 0 | 0 | 0 | 163800 | 106200 | 0 | 556510 | 0 | 0 | 1207000 | 1522200 | 0 | 0 |
| TQLPYEYYS | TM9SF4; TM9SF | Q92544; F22 | TM9S4_HUMAN | Unmodified | 0 | 0 | 0 | 0 | 0 | 0 | 0 | 35667 | 49533000 | 1580400 | 0 | 1174800 | 0 | 0 | 0 | 0 | 0 | 0 |
| TQLPYEYYS | TM9SF4; TM9SF | Q92544; F22 | TM9S4_HUMAN | Unmodified | 0 | 0 | 0 | 0 | 0 | 0 | 0 | 0 | 2527500 | 357700 | 0 | 0 | 0 | 0 | 0 | 0 | 0 | 0 |
| TQMELEED | ITM2C; ITM2C; | Q9NQX7; E7 | ITM2C_HUMAN | Oxidation (M) | 0 | 0 | 0 | 0 | 0 | 0 | 0 | 0 | 474420 | 48569 | 0 | 0 | 0 | 25841 | 0 | 0 | 0 | 0 |
| TQMELEED | ITM2C; ITM2C; | Q9NQX7; E7 | ITM2C_HUMAN | Unmodified | 0 | 0 | 0 | 0 | 0 | 0 | 0 | 0 | 2340700 | 0 | 0 | 0 | 0 | 0 | 0 | 0 | 0 | 0 |
| TQMLDQEE | KIF5B | P33176 | KINH_HUMAN | Oxidation (M) | 0 | 0 | 0 | 0 | 0 | 0 | 25417 | 0 | 133540 | 0 | 0 | 890410 | 0 | 0 | 0 | 0 | 0 | 114660 |
| TQMLDQEE | KIF5B | P33176 | KINH_HUMAN | Unmodified | 0 | 0 | 0 | 0 | 362490 | 0 | 0 | 0 | 0 | 150270 | 0 | 1322200 | 34277 | 82724 | 0 | 0 | 44352 | 0 |
| TQNDVDIA | EPCAM; EPCAM | P16422; B5N | EPCAM_HUMA | Unmodified | 24417000 | 24226000 | 16288000 | 22230000 | 0 | 1845700 | 0 | 24100 | 1.5E+08 | 23741000 | 12725000 | 19036000 | 15690000 | 6240200 | 49336000 | 28409000 | 0 | 1858900 |
| TQNDVDIA | EPCAM; EPCAM | P16422; B5N | EPCAM_HUMA | Unmodified | 0 | 81360 | 0 | 0 | 0 | 0 | 0 | 0 | 0 | 1415200 | 0 | 0 | 0 | 0 | 0 | 0 | 0 | 0 |
| TQNDVDIA | EPCAM; EPCAM | P16422; B5N | EPCAM_HUMA | Unmodified | 0 | 0 | 0 | 0 | 993100 | 0 | 0 | 0 | 0 | 8045500 | 0 | 0 | 2312300 | 1311400 | 0 | 63859 | 632510 | 0 |
| TQNDVLHA | RDX; RDX; RDX | P35241; A0A | RADI_HUMAN | Unmodified | 779620 | 641500 | 1636400 | 0 | 12000000 | 3125200 | 409740 | 401530 | 3108900 | 930360 | 1309200 | 5042500 | 9286700 | 3454600 | 1739600 | 0 | 1648600 | 2722100 |
| TQNFLIQKI | BCL10; BCL10 | O95999; A0 | BCL10_HUMAN | Unmodified | 0 | 0 | 0 | 0 | 263240 | 0 | 167800 | 0 | 0 | 146870 | 0 | 0 | 297180 | 326420 | 0 | 0 | 1348800 | 364560 |
| TQNGEIVQ | DIP2B; DIP2B | Q9P265; H0 | DIP2B_HUMAN | Unmodified | 21194000 | 3290000 | 0 | 2455300 | 2559800 | 0 | 661280 | 609520 | 669620 | 0 | 456820 | 3046500 | 1631000 | 0 | 2343100 | 2318400 | 13090000 | 31981000 |
| TQNQPRFS | MIA3; MIA3 | Q5JRA6; A0A | TGO1_HUMAN | Unmodified | 0 | 0 | 0 | 0 | 0 | 0 | 0 | 0 | 0 | 1221800 | 0 | 0 | 0 | 0 | 0 | 0 | 0 | 0 |
| TQNSWTQI | B4GALT3 | O60512 | B4GT3_HUMAN | Unmodified | 0 | 0 | 0 | 0 | 0 | 0 | 0 | 0 | 3009400 | 0 | 0 | 0 | 0 | 0 | 0 | 0 | 0 | 0 |
| TQNVLGEK | RPS3; RPS3; RP | P23396; H0 | RS3_HUMAN 4 | Unmodified | 2386900 | 2203600 | 1568800 | 0 | 0 | 0 | 0 | 0 | 0 | 0 | 0 | 24277000 | 612890 | 0 | 1272900 | 1547000 | 0 | 0 |
| TQPESSSM | MGAT5 | Q09328 | MGT5A_HUMA | Unmodified | 0 | 0 | 213780 | 0 | 1879300 | 0 | 0 | 324380 | 481010 | 0 | 0 | 0 | 351660 | 0 | 196640 | 103480 | 0 | 1155900 |
| TQPGEGLE | ERICH5 | Q6P6B1 | ERIC5_HUMAN | Unmodified | 0 | 0 | 0 | 0 | 0 | 0 | 0 | 0 | 0 | 0 | 0 | 0 | 0 | 0 | 0 | 0 | 618620 | 2467200 |
| TQPILDILLK | CAB39; CAB39 | Q9Y376; A0 | CAB39_HUMAN | Unmodified | 366710 | 347890 | 0 | 0 | 392330 | 296740 | 138460 | 0 | 453980 | 0 | 905650 | 3993300 | 0 | 0 | 1736600 | 1276500 | 303090 | 80145 |
| TQPNLDNC | CST3 | P01034 | CYTC_HUMAN | Unmodified | 20063 | 0 | 803700 | 67921 | 0 | 399740 | 0 | 0 | 21625000 | 8132600 | 0 | 0 | 0 | 701230 | 510070 | 68724 | 41452 | 831840 |
| TQPPEDISC | SEC31A; SEC31 | O94979; D6I | SC31A_HUMAN | Unmodified | 0 | 0 | 0 | 0 | 0 | 0 | 0 | 0 | 139600 | 0 | 0 | 685770 | 46588 | 0 | 0 | 0 | 0 | 47290 |
| TQPVQGEF | NCAM1; NCAM | P13591; A0A | NCAM1_HUMA | Unmodified | 0 | 0 | 0 | 0 | 0 | 0 | 0 | 0 | 2189000 | 0 | 0 | 637130 | 0 | 0 | 0 | 0 | 0 | 0 |
| TQPVREPS | NCAM1; NCAM | H7BYX6; A0 | H7BYX6_HUMA | Unmodified | 0 | 0 | 0 | 0 | 0 | 0 | 0 | 0 | 1386400 | 0 | 0 | 0 | 0 | 0 | 0 | 0 | 0 | 0 |
| TQQAAQAN | SF3A1; SF3A1 | Q15459; H7 | SF3A1_HUMAN | Unmodified | 0 | 3173300 | 266120 | 414610 | 0 | 0 | 0 | 0 | 0 | 223210 | 759490 | 0 | 1716800 | 0 | 0 | 170160 | 0 | 0 |
| TQQDRPEI | ATP2C1; ATP2C | P98194; B4E | AT2C1_HUMAN | Unmodified | 0 | 0 | 0 | 0 | 0 | 126290 | 0 | 183100 | 1874300 | 146920 | 0 | 0 | 0 | 0 | 0 | 0 | 0 | 0 |
| TQQQVEAE | COPS7B; COPS | Q9H9Q2; J3I | CSN7B_HUMA | Unmodified | 0 | 0 | 0 | 0 | 0 | 0 | 0 | 0 | 0 | 0 | 329000 | 1484700 | 0 | 0 | 0 | 0 | 0 | 0 |
| TQQVIHIID | CSNK1G3 | Q9Y6M4 | KC1G3_HUMA | Unmodified | 0 | 6482000 | 1296400 | 0 | 541750 | 0 | 0 | 0 | 0 | 0 | 0 | 3197000 | 0 | 0 | 1956600 | 1883200 | 0 | 0 |
| TQRPADVIF | MCM7 | P33993 | MCM7_HUMAN | Unmodified | 0 | 0 | 0 | 0 | 0 | 0 | 0 | 0 | 495300 | 0 | 0 | 1379900 | 0 | 0 | 0 | 0 | 0 | 0 |
| TQRPAMVC | PTPRF; PTPRF; | P10586; H0 | PTPRF_HUMAN | Unmodified | 412440 | 405350 | 0 | 346940 | 0 | 0 | 0 | 0 | 1258200 | 0 | 0 | 0 | 0 | 0 | 907230 | 393380 | 0 | 0 |
| TQRVDDPC | SUN1; SUN1; S | O94901; H0 | SUN1_HUMAN | Unmodified | 113980 | 0 | 0 | 0 | 177910 | 0 | 0 | 0 | 4260100 | 546840 | 0 | 0 | 39753 | 45987 | 0 | 0 | 0 | 0 |
| TQSCCYLW | VIL1; VIL1; VIL1 | P09327; H7 | VILI_HUMAN Vi | Unmodified | 0 | 0 | 0 | 0 | 0 | 0 | 0 | 0 | 0 | 0 | 0 | 810910 | 0 | 0 | 0 | 0 | 0 | 0 |
| TQSISYLAY | PLCH1; PLCH1 | Q4KWH8; A | PLCH1_HUMA | Unmodified | 0 | 0 | 0 | 296720 | 199990 | 52296 | 0 | 0 | 549330 | 116800 | 183840 | 0 | 1052800 | 0 | 0 | 92983 | 269610 | 216560 |
| TQSLPVTEK | ANLN | Q9NQW6 | ANLN_HUMAN | Unmodified | 98792 | 183010 | 0 | 56484 | 7250500 | 1041200 | 37158 | 120770 | 0 | 77634 | 0 | 232850 | 555050 | 361610 | 151990 | 67052 | 3627400 | 1919200 |
| TQSLPVTEK | ANLN | Q9NQW6 | ANLN_HUMAN | Unmodified | 0 | 0 | 0 | 0 | 934690 | 117890 | 87502 | 98195 | 0 | 0 | 0 | 0 | 0 | 170370 | 0 | 0 | 614090 | 187540 |
| TQSPCFGD | SCRN1 | Q12765 | SCRN1_HUMA | Unmodified | 0 | 0 | 0 | 0 | 557470 | 144170 | 94975 | 25811 | 316250 | 77060 | 0 | 0 | 0 | 0 | 28655 | 0 | 238850 | 334880 |
| TQSSLVPAI | CUTA; CUTA | O60888; C9I | CUTA_HUMAN | Unmodified | 0 | 0 | 132920 | 9122500 | 9302100 | 0 | 0 | 0 | 15422000 | 5498400 | 3687400 | 17647000 | 5154500 | 0 | 2481800 | 106430 | 2503900 | 6776400 |
| TQSVEITKT | CD2AP | Q9Y5K6 | CD2AP_HUMA | Unmodified | 23059000 | 17128000 | 11370000 | 20700000 | 3655800 | 829670 | 114740 | 38208 | 1554300 | 740150 | 802380 | 2341300 | 10564000 | 2821900 | 31225000 | 19618000 | 0 | 290940 |
| TQSVEITKT | CD2AP | Q9Y5K6 | CD2AP_HUMA | Unmodified | 0 | 0 | 0 | 0 | 1753900 | 0 | 0 | 0 | 0 | 0 | 0 | 0 | 3061000 | 0 | 0 | 0 | 0 | 0 |
| TQSYPTDW | PACSIN2; PACS | Q9UNF0; A0 | PACN2_HUMA | Unmodified | 4922500 | 904500 | 109510 | 1019100 | 0 | 0 | 0 | 0 | 0 | 0 | 0 | 0 | 434750 | 0 | 6879900 | 1939000 | 0 | 0 |
| TQSYPTDW | PACSIN2; PACS | Q9UNF0; A0 | PACN2_HUMA | Unmodified | 0 | 0 | 0 | 1162300 | 0 | 0 | 0 | 0 | 0 | 0 | 0 | 0 | 3262600 | 0 | 0 | 0 | 0 | 0 |
| TQTAIASED | SPTBN1; SPTBN | Q01082; A0 | SPTB2_HUMAN | Unmodified | 0 | 0 | 0 | 0 | 0 | 0 | 0 | 0 | 0 | 0 | 0 | 0 | 0 | 0 | 0 | 0 | 0 | 0 |
| TQTAVASE | SPTBN2 | O15020 | SPTN2_HUMAN | Unmodified | 0 | 0 | 0 | 0 | 0 | 0 | 0 | 0 | 0 | 0 | 0 | 1289400 | 0 | 0 | 1917800 | 0 | 0 | 0 |
| TQTESSTPI | PDPK2P; PDPK | Q6A1A2; O1 | PDPK2_HUMAN | Unmodified | 0 | 0 | 0 | 0 | 0 | 0 | 0 | 0 | 0 | 0 | 0 | 0 | 0 | 0 | 233900 | 509710 | 61390 | 0 |
| TQTFSGDF | TMEM87A | Q8NBN3 | TM87A_HUMAN | Unmodified | 0 | 0 | 0 | 0 | 0 | 0 | 0 | 0 | 642150 | 27606 | 119940 | 295870 | 0 | 0 | 0 | 17401 | 0 | 0 |
| TQTFSGDF | TMEM87A | Q8NBN3 | TM87A_HUMAN | Oxidation (M) | 0 | 0 | 106580 | 77155 | 0 | 0 | 0 | 0 | 803260 | 171150 | 393050 | 3548900 | 0 | 0 | 196020 | 0 | 0 | 0 |
| TQTIFLKYP | MYOF; MYOF | Q9NZM1; H | MYOF_HUMAN | Unmodified | 0 | 49337 | 0 | 0 | 4351800 | 945040 | 0 | 0 | 0 | 0 | 0 | 0 | 0 | 0 | 74355 | 79794 | 0 | 581180 |
| TQTIHSTYS | LTBP1; LTBP1; | Q14766; E7E | LTBP1_HUMAN | Unmodified | 0 | 0 | 0 | 0 | 0 | 0 | 671980 | 0 | 0 | 0 | 0 | 0 | 0 | 0 | 0 | 0 | 1402100 | 5251500 |
| TQTPPVSP | CTTN; CTTN; C | Q14247; H0 | SRC8_HUMAN | Unmodified | 1217400 | 997850 | 2446700 | 786820 | 1741800 | 0 | 0 | 0 | 0 | 0 | 0 | 6273000 | 2112100 | 514570 | 3812900 | 2275200 | 0 | 0 |
| TQTVCNFT | FABP5P3; FABP | A8MUU1; Q | FB5L3_HUMAN | Unmodified | 1961700 | 763250 | 204730 | 0 | 4837900 | 1232700 | 0 | 0 | 1665600 | 0 | 8514400 | 28882000 | 216900 | 0 | 0 | 0 | 4430700 | 12792000 |
| TQTVCNFT | FABP5P3; FABP | A8MUU1; Q | FB5L3_HUMAN | Unmodified | 0 | 0 | 0 | 0 | 1174600 | 0 | 590840 | 0 | 0 | 0 | 0 | 0 | 0 | 0 | 0 | 0 | 4590800 | 6840900 |
| TQTVCNFT | FABP5P3; FABP | A8MUU1; Q | FB5L3_HUMAN | Unmodified | 0 | 0 | 0 | 0 | 18304000 | 2331600 | 362650 | 0 | 0 | 0 | 2844300 | 0 | 0 | 0 | 0 | 0 | 28164000 | 65357000 |
| TQTVTISDN | EPB41; EPB41; | P11171; A0A | EPB41_HUMAN | Unmodified | 0 | 0 | 772290 | 810530 | 0 | 0 | 0 | 172860 | 935610 | 151160 | 1622600 | 1028400 | 548470 | 563230 | 764460 | 605830 | 305680 | 473050 |
| TQTVTISDN | EPB41; EPB41; | P11171; A0A | EPB41_HUMAN | Unmodified | 0 | 0 | 0 | 1820800 | 1990600 | 295230 | 231130 | 291380 | 3085300 | 1646800 | 571440 | 0 | 3129600 | 3064000 | 0 | 0 | 1102800 | 4646300 |
| TQTVTISDN | EPB41; EPB41; | P11171; A0A | EPB41_HUMAN | Unmodified | 0 | 0 | 0 | 0 | 1280200 | 0 | 0 | 3096800 | 0 | 2550200 | 0 | 0 | 3869600 | 7131700 | 0 | 0 | 5718800 | 11220000 |
| TQVAEYK | GOLIM4; GOLIM | O00461; F8 | GOLI4_HUMAN | Unmodified | 82175 | 0 | 0 | 0 | 755310 | 6332 | 85096 | 24248 | 1453900 | 0 | 0 | 1836600 | 0 | 0 | 0 | 0 | 394060 | 330920 |
| TQVAGGQI | RANGAP1; RAN | P46060; B0 | RAGP1_HUMA | Unmodified | 0 | 0 | 0 | 0 | 0 | 0 | 0 | 261440 | 0 | 0 | 0 | 0 | 0 | 381760 | 0 | 0 | 0 | 0 |
| TQVELLEAF | GYS1 | P13807 | GYS1_HUMAN | Unmodified | 0 | 0 | 0 | 0 | 0 | 0 | 0 | 0 | 0 | 0 | 0 | 5571400 | 0 | 0 | 0 | 0 | 0 | 0 |
| TQVELLEAF | GYS1 | P13807 | GYS1_HUMAN | Unmodified | 0 | 0 | 0 | 716670 | 0 | 1870600 | 0 | 0 | 0 | 0 | 3118500 | 5138100 | 0 | 0 | 0 | 0 | 0 | 174240 |
| TQVGLIQY | ITGA2; ITGA2; I | P17301; E9P | ITA2_HUMAN I | Unmodified | 1877600 | 1103000 | 2448400 | 2744900 | 2511100 | 1684100 | 0 | 0 | 0 | 0 | 0 | 0 | 498820 | 52553 | 2816500 | 3238100 | 1177600 | 1093300 |
| TQVGLIQY | ITGA2; ITGA2; I | P17301; E9P | ITA2_HUMAN I | Unmodified | 0 | 0 | 0 | 0 | 27493000 | 3893700 | 0 | 0 | 0 | 0 | 0 | 0 | 1800000 | 869980 | 0 | 304320 | 19923000 | 9932000 |
| TQVKELKEI | ROCK1; ROCK1 | Q13464; A0 | ROCK1_HUMA | Unmodified | 0 | 0 | 0 | 0 | 138080 | 0 | 27500 | 22992 | 0 | 0 | 0 | 0 | 0 | 18897 | 0 | 0 | 482230 | 96759 |
| TQVLSPDS | AFDN; AFDN; A | P55196; H0 | AFAD_HUMAN | Unmodified | 0 | 0 | 0 | 0 | 3518800 | 0 | 953470 | 826400 | 0 | 172230 | 1247900 | 1111100 | 3677600 | 3838800 | 0 | 0 | 1129200 | 0 |
| TQVTVQYM | MAT2A | P31153 | METK2_HUMAN | Oxidation (M) | 0 | 0 | 0 | 0 | 0 | 0 | 0 | 0 | 0 | 0 | 0 | 569230 | 0 | 0 | 0 | 0 | 0 | 0 |
| TQVTVQYM | MAT2A | P31153 | METK2_HUMAN | Unmodified | 0 | 577230 | 322800 | 0 | 0 | 0 | 0 | 0 | 0 | 0 | 0 | 2214200 | 0 | 0 | 99439 | 0 | 0 | 0 |

| | | | | | | | | | | | | | | | | | | | | | | |
|---|---|---|---|---|---|---|---|---|---|---|---|---|---|---|---|---|---|---|---|---|---|---|
| TQYACPPF | IGF2R; IGF2R | P11717; A0A | MPRI_HUMAN | Unmodified | 53110 | 0 | 0 | 0 | 0 | 0 | 0 | 0 | 3085000 | 212740 | 0 | 633540 | 0 | 0 | 0 | 52932 | 73138 | 0 |
| TQYIFNNM | MAN2A1 | Q16706 | MA2A1_HUMAI | Unmodified | 0 | 0 | 0 | 0 | 0 | 0 | 541000 | 8822.2 | 931110 | 0 | 0 | 157330 | 328920 | 0 | 0 | 0 | 0 | 0 |
| TQYLQSYG | VPS33B; VPS33 | Q9H267; F5I | VP33B_HUMAN | Unmodified | 51049 | 0 | 33137 | 43770 | 707840 | 190470 | 0 | 0 | 191770 | 0 | 79525 | 495540 | 0 | 0 | 56050 | 56633 | 0 | 0 |
| TQYNQVPS | CXADR | P78310 | CXAR_HUMAN | Unmodified | 1121900 | 1066900 | 271090 | 512860 | 42555000 | 2480200 | 1708900 | 1188600 | 6295800 | 592460 | 286520 | 7133100 | 0 | 0 | 367590 | 273510 | 3931100 | 524760 |
| TQYSSAML | NPEPPS; NPEPI | P55786; A0A | PSA_HUMAN P | Oxidation (M) | 0 | 0 | 0 | 0 | 0 | 0 | 0 | 0 | 0 | 0 | 0 | 178280 | 0 | 0 | 0 | 0 | 0 | 0 |
| TRADVKGG | UGP2; UGP2; U | Q16851; E7E | UGPA_HUMAN | Unmodified | 0 | 0 | 0 | 0 | 1782600 | 443760 | 1167700 | 280760 | 0 | 1323000 | 0 | 0 | 1034100 | 536810 | 0 | 0 | 1592500 | 1035700 |
| TRAELDTV | HLA-DQB1; HL/ | P01920; A0A | DQB1_HUMAN | Unmodified | 0 | 0 | 0 | 0 | 0 | 0 | 0 | 0 | 1011300 | 0 | 0 | 0 | 0 | 0 | 0 | 0 | 53150 | 0 |
| TRANEQLTI | KIF5C; KIF5C; K | O60282; A0/ | KIF5C_HUMAN | Unmodified | 0 | 0 | 0 | 0 | 0 | 0 | 0 | 0 | 0 | 0 | 0 | 0 | 110300 | 0 | 0 | 0 | 0 | 0 |
| TRAQAVHP | PCCA; PCCA; P | P05165; A0A | PCCA_HUMAN | Unmodified | 268110 | 0 | 0 | 0 | 0 | 0 | 0 | 0 | 693020 | 176640 | 0 | 0 | 0 | 0 | 0 | 0 | 0 | 0 |
| TRDDEPVC | APMAP; APMAP | Q9HDC9; H( | APMAP_HUMAI | Unmodified | 0 | 0 | 181780 | 0 | 128570 | 0 | 0 | 21235 | 1.59E+08 | 6019700 | 1045200 | 403950 | 0 | 0 | 0 | 115470 | 0 | 0 |
| TRDELEVIH | RO60 | P10155 | RO60_HUMAN | Unmodified | 42852000 | 43854000 | 7811500 | 30342000 | 0 | 0 | 0 | 0 | 1889900 | 0 | 1098200 | 4117100 | 1056500 | 0 | 18464000 | 15286000 | 0 | 0 |
| TRDENIKV | CD109 | Q6YHK3 | CD109_HUMAI | Unmodified | 0 | 0 | 0 | 0 | 1665200 | 0 | 0 | 0 | 0 | 973500 | 888540 | 0 | 0 | 0 | 0 | 0 | 4281000 | 2827800 |
| TRDFKPGD | PSIP1 | O75475 | PSIP1_HUMAN | Acetyl (Protein N | 0 | 0 | 0 | 0 | 0 | 0 | 120580 | 649330 | 0 | 157060 | 0 | 0 | 0 | 0 | 0 | 0 | 0 | 0 |
| TRDGGSVV | SPAG9; SPAG9 | O60271; A0/ | JIP4_HUMAN C | Unmodified | 0 | 0 | 0 | 0 | 0 | 0 | 0 | 394100 | 913930 | 0 | 0 | 539150 | 108940 | 0 | 0 | 0 | 272490 | 0 |
| TRDGLTPLI | ANK3; ANK3 | Q12955; A0/ | ANK3_HUMAN | Unmodified | 0 | 0 | 374670 | 928520 | 0 | 0 | 0 | 0 | 0 | 50993 | 0 | 33474 | 1649700 | 1321400 | 1159800 | 1163600 | 0 | 0 |
| TRDGSDYE | GANAB; GANAE | Q14697; F5H | GANAB_HUMA | Oxidation (M) | 0 | 0 | 0 | 0 | 0 | 0 | 0 | 0 | 27296000 | 4511200 | 0 | 0 | 0 | 0 | 0 | 0 | 0 | 0 |
| TRDGSDYE | GANAB; GANAE | Q14697; F5H | GANAB_HUMA | Unmodified | 0 | 0 | 0 | 0 | 0 | 0 | 0 | 0 | 10918000 | 0 | 0 | 0 | 0 | 0 | 0 | 0 | 0 | 0 |
| TRDKGEIPI | DLG1; DLG1; D | Q12959; A0/ | DLG1_HUMAN | Unmodified | 0 | 240660 | 0 | 0 | 2001400 | 0 | 162490 | 107590 | 172470 | 1279900 | 31547 | 0 | 1531000 | 1663100 | 0 | 0 | 1030400 | 1265200 |
| TREAQQAL | LAMC1 | P11047 | LAMC1_HUMAI | Unmodified | 829370 | 355050 | 279510 | 0 | 1508200 | 1009600 | 0 | 0 | 1664400 | 952910 | 269820 | 288690 | 1540400 | 1822700 | 350450 | 555160 | 238970 | 7940100 |
| TREEECHF | PRDX4; PRDX4 | Q13162; A6I | PRDX4_HUMAI | Unmodified | 0 | 0 | 0 | 0 | 0 | 0 | 0 | 0 | 45780000 | 787260 | 2761900 | 4635700 | 0 | 0 | 0 | 0 | 0 | 0 |
| TREEECHF | PRDX4; PRDX4 | Q13162; A6I | PRDX4_HUMAI | Unmodified | 0 | 0 | 0 | 0 | 0 | 0 | 0 | 0 | 0 | 2660000 | 0 | 0 | 0 | 0 | 0 | 0 | 0 | 0 |
| TREGAEDK | BPNT2; BPNT2 | Q9NX62; H0 | IMPA3_HUMAN | Unmodified | 0 | 0 | 0 | 0 | 0 | 0 | 0 | 0 | 0 | 165820 | 0 | 0 | 0 | 0 | 0 | 0 | 0 | 0 |
| TREGNDLY | ATP5F1B; ATP5I | P06576; H0\ | ATPB_HUMAN | Unmodified | 0 | 0 | 0 | 0 | 66533 | 1718700 | 918600 | 0 | 34584000 | 1866600 | 727910 | 0 | 0 | 0 | 0 | 0 | 0 | 1466200 |
| TREGVVQG | SNCB | Q16143 | SYUB_HUMAN | Unmodified | 0 | 0 | 0 | 0 | 0 | 0 | 2448200 | 1465200 | 0 | 0 | 0 | 0 | 0 | 0 | 0 | 0 | 0 | 0 |
| TREIFTNLA | DENND10; DEN | Q8TCE6; Q6 | DEN10_HUMAI | Unmodified | 0 | 15448 | 0 | 0 | 1951000 | 565890 | 0 | 0 | 0 | 166170 | 0 | 1473000 | 1086900 | 649880 | 0 | 0 | 0 | 888190 |
| TREKLMNV | FRYL; FRYL; FR | O94915; H0 | FRYL_HUMAN | Unmodified | 0 | 0 | 0 | 0 | 944540 | 484770 | 566100 | 121440 | 0 | 162620 | 0 | 0 | 2112500 | 0 | 0 | 0 | 2330600 | 1674200 |
| TREVQERLI | CENPE; CENPE | Q02224; A0/ | CENPE_HUMAI | Unmodified | 198880 | 0 | 0 | 0 | 0 | 1240900 | 1171600 | 0 | 566690 | 0 | 185070 | 115930 | 0 | 282190 | 0 | 0 | 0 | 3713100 |
| TREYFYYVI | C8orf82; C8orf | Q6P1X6; J3K | CH082_HUMAI | Unmodified | 0 | 0 | 0 | 0 | 0 | 0 | 134190 | 0 | 2522700 | 1005300 | 0 | 0 | 0 | 1587200 | 0 | 0 | 0 | 1012400 |
| TRGGGQS\ | KIF23; KIF23; KI | Q02241; A0/ | KIF23_HUMAN | Unmodified | 79155 | 0 | 1308600 | 0 | 2952300 | 621280 | 1139200 | 55212 | 550670 | 229390 | 268870 | 961070 | 362830 | 240760 | 384050 | 152960 | 600900 | 381660 |
| TRGGGQS\ | KIF23; KIF23; KI | Q02241; A0/ | KIF23_HUMAN | Unmodified | 0 | 0 | 0 | 0 | 1420100 | 0 | 0 | 0 | 342120 | 0 | 0 | 0 | 684890 | 0 | 0 | 0 | 634940 | 0 |
| TRHLALNL | SMC1A; SMC1A | Q14683; G8. | SMC1A_HUMA | Unmodified | 24648 | 37637 | 0 | 0 | 194740 | 0 | 194880 | 70800 | 121780 | 253580 | 0 | 0 | 194450 | 0 | 0 | 0 | 61965 | 28409 |
| TRIAEVGG\ | C11orf54; C11c | Q9H0W9; AC | CK054_HUMAI | Unmodified | 63288 | 0 | 36533 | 97398 | 246460 | 52283 | 128630 | 0 | 136970 | 186910 | 79938 | 115170 | 0 | 306290 | 69206 | 0 | 1508700 | 1254200 |
| TRIAEVGG\ | C11orf54; C11c | Q9H0W9; AC | CK054_HUMAI | Unmodified | 0 | 0 | 0 | 0 | 0 | 0 | 0 | 0 | 0 | 2800400 | 0 | 0 | 1420500 | 0 | 0 | 0 | 1578400 | 5031500 |
| TRIDEANQI | SNAP25 | P60880 | SNP25_HUMAI | Unmodified | 0 | 0 | 0 | 0 | 154850 | 0 | 1365400 | 1160100 | 0 | 720240 | 0 | 0 | 0 | 114280 | 0 | 0 | 0 | 0 |
| TRIEGLLAA | ARCN1; ARCN1 | P48444; E9P | COPD_HUMAN | Unmodified | 0 | 0 | 139350 | 0 | 0 | 0 | 0 | 0 | 293120 | 249820 | 277780 | 4746000 | 0 | 0 | 182590 | 183290 | 0 | 0 |
| TRIIDVVYN | RPS8; RPS8 | P62241; Q5J | RS8_HUMAN 4 | Unmodified | 3072100 | 3500800 | 1753500 | 1515200 | 0 | 0 | 2008200 | 0 | 2393000 | 872390 | 3943600 | 13627000 | 0 | 0 | 3130800 | 1752800 | 0 | 0 |
| TRIIDVVYN | RPS8; RPS8 | P62241; Q5J | RS8_HUMAN 4 | Unmodified | 0 | 0 | 0 | 0 | 6471200 | 2453300 | 22283000 | 2575300 | 0 | 1999500 | 0 | 0 | 2487900 | 3528800 | 0 | 0 | 1960300 | 5982000 |
| TRIVIVPSL | CPD | O75976 | CBPD_HUMAN | Unmodified | 0 | 0 | 0 | 0 | 0 | 0 | 0 | 0 | 1158700 | 0 | 0 | 0 | 0 | 0 | 0 | 0 | 0 | 0 |
| TRKPLEALY | BPHL | Q86WA6 | BPHL_HUMAN | Unmodified | 0 | 0 | 0 | 0 | 0 | 0 | 0 | 0 | 975510 | 0 | 0 | 0 | 0 | 0 | 0 | 0 | 0 | 0 |
| TRLAFLNV | GPD2 | P43304 | GPDM_HUMAN | Unmodified | 0 | 0 | 0 | 0 | 0 | 0 | 0 | 0 | 4054400 | 838870 | 0 | 0 | 0 | 0 | 0 | 0 | 0 | 0 |
| TRLENLEQ | NPTX1 | Q15818 | NPTX1_HUMAI | Unmodified | 0 | 0 | 0 | 0 | 0 | 79281 | 0 | 0 | 13505000 | 553170 | 0 | 0 | 0 | 0 | 0 | 0 | 0 | 0 |
| TRLIGPNCI | SUCLG1; SUCL | P53597; A0A | SUCA_HUMAN | Unmodified | 0 | 0 | 0 | 0 | 113110 | 0 | 422680 | 195330 | 2514500 | 1359200 | 0 | 0 | 401470 | 346860 | 0 | 0 | 153940 | 708530 |
| TRLPLELET | TMEM237; TMEI | Q96Q45; A0 | TM237_HUMAI | Unmodified | 0 | 0 | 339980 | 0 | 7887400 | 299830 | 0 | 0 | 2927400 | 0 | 0 | 0 | 1734000 | 0 | 0 | 216670 | 428160 | 1363200 |
| TRLPLELET | TMEM237; TMEI | Q96Q45; A0 | TM237_HUMAI | Unmodified | 0 | 0 | 144570 | 374980 | 6723800 | 504510 | 0 | 352170 | 200850 | 104740 | 200060 | 1031700 | 1304400 | 2091800 | 135280 | 0 | 1118500 | 2265000 |
| TRLQQELD | MYH9 | P35579 | MYH9_HUMAN | Unmodified | 141030 | 1569000 | 289830 | 3307500 | 972330 | 244110 | 0 | 0 | 0 | 1383200 | 117530 | 921100 | 1382100 | 0 | 6283600 | 813160 | 90260 | 0 |
| TRLQQELD | MYH9 | P35579 | MYH9_HUMAN | Unmodified | 0 | 0 | 0 | 0 | 36337000 | 0 | 2503900 | 0 | 0 | 964660 | 0 | 0 | 14712000 | 14031000 | 0 | 0 | 26279000 | 19302000 |
| TRPAANPIC | RRM1; RRM1 | P23921; E9P | RIR1_HUMAN F | Unmodified | 326260 | 237700 | 1237500 | 1718500 | 1704200 | 222350 | 188650 | 218510 | 865770 | 652600 | 488460 | 7097000 | 1163700 | 629460 | 1324700 | 823700 | 392700 | 279890 |
| TRPDGNCI | OTUB1; OTUB1 | Q96FW1; J3I | OTUB1_HUMAI | Unmodified | 0 | 0 | 0 | 0 | 267100 | 286560 | 0 | 0 | 143400 | 0 | 0 | 2394500 | 0 | 0 | 0 | 0 | 323610 | 593490 |
| TRPDGNCI | OTUB1; OTUB1 | Q96FW1; J3I | OTUB1_HUMAI | Unmodified | 0 | 0 | 0 | 0 | 1564100 | 760170 | 0 | 0 | 0 | 351600 | 0 | 0 | 789270 | 985010 | 0 | 0 | 1938400 | 1650100 |
| TRPEGEPS | GINS4; GINS4 | Q9BRT9; E5I | SLD5_HUMAN | Unmodified | 0 | 0 | 0 | 0 | 0 | 0 | 0 | 0 | 0 | 0 | 0 | 523020 | 0 | 0 | 0 | 0 | 0 | 0 |
| TRQVCPLD | RBX1 | P62877 | RBX1_HUMAN | Unmodified | 0 | 0 | 0 | 557670 | 2232100 | 0 | 1325900 | 673690 | 0 | 0 | 0 | 686740 | 1953900 | 3712700 | 0 | 0 | 5080300 | 1122200 |
| TRSIGSAVI | RACGAP1; RAC | Q9H0H5; F8 | RGAP1_HUMAI | Unmodified | 0 | 0 | 0 | 106150 | 10246000 | 1750600 | 414650 | 52235 | 242960 | 732410 | 119310 | 0 | 266140 | 442680 | 0 | 0 | 2162800 | 977970 |
| TRSLISMDE | TSNAX; TSNAX; | Q99598; Q5\ | TSNAX_HUMAI | Unmodified | 0 | 0 | 0 | 68605 | 525940 | 21363 | 0 | 155100 | 1256500 | 276860 | 0 | 387750 | 196230 | 0 | 0 | 0 | 190240 | 0 |
| TRSNPENN | PSMD4; PSMD4 | P55036; A6P | PSMD4_HUMA | Unmodified | 0 | 0 | 0 | 0 | 3097900 | 0 | 1524800 | 0 | 0 | 455110 | 0 | 0 | 3952100 | 2155600 | 0 | 0 | 1105600 | 1682700 |
| TRSPVTFLS | CNOT1 | A5YKK6 | CNOT1_HUMA | Unmodified | 0 | 0 | 0 | 0 | 0 | 0 | 0 | 0 | 0 | 0 | 0 | 402000 | 0 | 0 | 0 | 0 | 0 | 0 |
| TRSTAEGE | PPL; PPL | O60437; K7E | PEPL_HUMAN | Unmodified | 0 | 0 | 0 | 0 | 3815400 | 592660 | 0 | 0 | 0 | 0 | 0 | 0 | 476440 | 281970 | 0 | 0 | 0 | 0 |
| TRTEELIVQ | NLN | Q9BYT8 | NEUL_HUMAN | Unmodified | 0 | 0 | 0 | 0 | 1375000 | 0 | 0 | 0 | 1233000 | 329510 | 0 | 2256900 | 0 | 0 | 0 | 0 | 0 | 0 |
| TRTFAVSSK | CHMP2B; CHM | Q9UQN3; AC | CHM2B_HUMA | Unmodified | 319830 | 238450 | 181140 | 189680 | 0 | 0 | 0 | 0 | 991870 | 620060 | 284830 | 710480 | 1976900 | 990760 | 187360 | 184200 | 0 | 0 |
| TRTFAVSSK | CHMP2B; CHM | Q9UQN3; AC | CHM2B_HUMA | Unmodified | 42352 | 0 | 0 | 0 | 1423100 | 0 | 80529 | 150510 | 0 | 558130 | 0 | 0 | 0 | 1153000 | 0 | 0 | 0 | 297060 |
| TRTFAVSSK | CHMP2B; CHM | Q9UQN3; AC | CHM2B_HUMA | Oxidation (M) | 0 | 151460 | 0 | 314540 | 2123700 | 428270 | 0 | 0 | 0 | 0 | 0 | 0 | 568510 | 95640 | 0 | 0 | 147350 | 882660 |
| TRTFNTSTC | XRCC6; XRCC6 | P12956; B1A | XRCC6_HUMAI | Unmodified | 805020 | 0 | 329790 | 92395 | 1028700 | 0 | 731990 | 857660 | 358800 | 248280 | 1125000 | 8756600 | 770160 | 0 | 0 | 270000 | 0 | 635630 |
| TRTFNTSTC | XRCC6; XRCC6 | P12956; B1A | XRCC6_HUMAI | Unmodified | 0 | 0 | 0 | 0 | 0 | 0 | 0 | 0 | 0 | 80018 | 0 | 1492200 | 258600 | 0 | 0 | 0 | 0 | 1326700 |
| TRTFYGLH | LAP3 | P28838 | AMPL_HUMAN | Unmodified | 0 | 0 | 0 | 0 | 5508000 | 157160 | 1743400 | 0 | 1403400 | 1304600 | 794430 | 2070400 | 14277000 | 5211400 | 0 | 0 | 2441500 | 3065000 |
| TRTFYGLH | LAP3 | P28838 | AMPL_HUMAN | Unmodified | 0 | 0 | 0 | 0 | 4477200 | 247310 | 1120900 | 0 | 660510 | 862890 | 381340 | 497400 | 6666500 | 2628200 | 0 | 0 | 1903800 | 2260100 |
| TRTYHSVG | IGF2R; IGF2R; I | P11717; A0A | MPRI_HUMAN | Unmodified | 0 | 0 | 0 | 0 | 0 | 0 | 0 | 0 | 1996300 | 287940 | 188100 | 244020 | 0 | 0 | 0 | 0 | 0 | 0 |
| TRVEGGQL | SHC1; SHC1; S | P29353; Q5T | SHC1_HUMAN | Unmodified | 798260 | 0 | 0 | 91212 | 291990 | 0 | 0 | 0 | 1016000 | 0 | 201550 | 1067900 | 46152 | 154990 | 118200 | 0 | 0 | 66690 |
| TRVVSYRVI | MVP | Q14764 | MVP_HUMAN N | Unmodified | 0 | 625320 | 0 | 249650 | 0 | 0 | 0 | 0 | 0 | 0 | 0 | 3824000 | 0 | 0 | 0 | 0 | 0 | 0 |
| TRWDSELC | PIGT; PIGT; PIG | Q969N2; A0 | PIGT_HUMAN | Unmodified | 0 | 0 | 0 | 0 | 0 | 0 | 0 | 0 | 2979100 | 532120 | 0 | 0 | 116310 | 0 | 0 | 0 | 0 | 0 |
| TRWLNSPN | ADD3 | Q9UEY8 | ADDG_HUMAN | Unmodified | 0 | 0 | 0 | 0 | 271570 | 0 | 0 | 0 | 384320 | 0 | 0 | 0 | 620490 | 551840 | 93679 | 0 | 504320 | 449140 |
| TRWLNSPN | ADD3 | Q9UEY8 | ADDG_HUMAN | Oxidation (M) | 0 | 0 | 0 | 82674 | 206960 | 0 | 0 | 38701 | 0 | 0 | 0 | 0 | 214390 | 219810 | 0 | 0 | 17811 | 124830 |
| TRYIFDLFY | BUD31 | P41223 | BUD31_HUMAI | Unmodified | 0 | 0 | 0 | 0 | 0 | 0 | 895820 | 29296 | 0 | 0 | 0 | 94203 | 289190 | 229910 | 0 | 0 | 0 | 0 |
| TSAAAQLD | LPXN; LPXN; LP | O60711; E9I | LPXN_HUMAN | Oxidation (M) | 0 | 0 | 0 | 0 | 0 | 193350 | 0 | 0 | 0 | 0 | 0 | 0 | 0 | 3386000 | 0 | 0 | 508130 | 619160 |
| TSAALSTVC | TPD52L2; TPD5 | O43399; A0/ | TPD54_HUMAI | Unmodified | 0 | 0 | 0 | 0 | 913550 | 0 | 0 | 219290 | 0 | 395920 | 83638 | 0 | 1339600 | 545980 | 0 | 0 | 0 | 682710 |
| TSAALSTVC | TPD52L2; TPD5 | O43399; A0/ | TPD54_HUMAI | Unmodified | 0 | 0 | 0 | 0 | 887910 | 395990 | 183860 | 0 | 430860 | 0 | 0 | 0 | 746090 | 0 | 0 | 0 | 866250 | 397710 |
| TSACGLFS | PPP2R1A; PPP2 | P30153; P30 | 2AAA_HUMAN | Unmodified | 0 | 0 | 0 | 0 | 0 | 0 | 0 | 0 | 181220 | 0 | 293770 | 1312900 | 0 | 0 | 0 | 0 | 0 | 0 |
| TSAEFLMK | SHCBP1 | Q8NEM2 | SHCBP_HUMA | Unmodified | 0 | 0 | 0 | 0 | 911100 | 297940 | 0 | 35917 | 389440 | 0 | 0 | 0 | 0 | 0 | 62444 | 0 | 0 | 0 |
| TSAKEEDAI | UCHL5; UCHL5 | Q9Y5K5; Q5I | UCHL5_HUMAI | Unmodified | 0 | 0 | 0 | 0 | 0 | 230080 | 0 | 0 | 513550 | 0 | 277470 | 5384900 | 0 | 0 | 0 | 0 | 0 | 0 |
| TSALENYIN | LSM8; LSM8; LS | O95777; C9. | LSM8_HUMAN | Acetyl (Protein N | 0 | 0 | 0 | 0 | 0 | 0 | 0 | 0 | 0 | 0 | 0 | 966370 | 0 | 0 | 0 | 0 | 0 | 0 |
| TSAPITCEL | DYNC1H1; DYN | Q14204; A0/ | DYHC1_HUMA | Unmodified | 230060 | 0 | 379020 | 0 | 668350 | 0 | 662690 | 203300 | 762520 | 477320 | 713450 | 3590800 | 0 | 0 | 234690 | 0 | 0 | 736130 |
| TSAQGILQ | MFSD6; MFSD6 | Q6ZSS7; C9I | MFSD6_HUMAI | Unmodified | 72880 | 46247 | 57004 | 37014 | 0 | 0 | 0 | 0 | 1831700 | 0 | 0 | 699370 | 0 | 0 | 245960 | 113940 | 0 | 0 |
| TSAQHALT | CXCR4 | P61073 | CXCR4_HUMAI | Unmodified | 568120 | 1071500 | 45576 | 0 | 39786 | 0 | 0 | 0 | 1843000 | 683200 | 0 | 0 | 51178 | 0 | 38452 | 0 | 28499 | 0 |
| TSAQVVLD | CKAP5 | Q14008 | CKAP5_HUMAI | Unmodified | 0 | 0 | 0 | 0 | 0 | 0 | 0 | 0 | 0 | 0 | 0 | 1127000 | 0 | 0 | 0 | 0 | 0 | 0 |
| TSAQVVLD | CKAP5 | Q14008 | CKAP5_HUMAI | Unmodified | 0 | 0 | 0 | 0 | 800460 | 0 | 456560 | 0 | 0 | 720500 | 248650 | 0 | 2251100 | 713220 | 0 | 0 | 482080 | 517790 |
| TSASIILR | TCP1; TCP1; TC | P17987; F5G | TCPA_HUMAN | Unmodified | 158420 | 98812 | 327170 | 229490 | 0 | 0 | 0 | 0 | 1123600 | 76184 | 1742200 | 18147000 | 0 | 0 | 363920 | 402530 | 0 | 0 |
| TSATVGPK | H2AX | P16104 | H2AX_HUMAN | Unmodified | 0 | 214680 | 0 | 0 | 0 | 81776 | 0 | 28481 | 0 | 0 | 0 | 0 | 136430 | 40003 | 0 | 0 | 20057 | 1878000 |
| TSATWLAL | ATP1A1 | P05023 | AT1A1_HUMAI | Unmodified | 3325700 | 1236000 | 34223000 | 25937000 | 16220000 | 2384400 | 1056700 | 0 | 22358000 | 5339500 | 27175000 | 1.55E+08 | 15657000 | 4337300 | 51213000 | 42087000 | 1270900 | 0 |
| TSAVPSPC | ACADVL | P49748 | ACADV_HUMAI | Unmodified | 0 | 0 | 0 | 0 | 0 | 0 | 0 | 0 | 2564100 | 174480 | 0 | 0 | 79256 | 62156 | 0 | 0 | 0 | 0 |
| TSCALTIHA | ECPAS; ECPAS | Q5VYK3; J3K | ECM29_HUMAI | Unmodified | 0 | 0 | 0 | 57659 | 0 | 0 | 0 | 0 | 258850 | 70881 | 84081 | 1145200 | 249360 | 259980 | 0 | 61087 | 146400 | 0 |
| TSCEFTGD | ATP6V1B2; ATP | P21281; H0 | VATB2_HUMAI | Unmodified | 0 | 0 | 0 | 0 | 0 | 414640 | 0 | 0 | 0 | 0 | 506810 | 836790 | 0 | 0 | 0 | 0 | 0 | 0 |
| TSCFNVGN | ELAPOR2; ELAP | A8MWY0; H7 | ELAP2_HUMAI | Unmodified | 0 | 0 | 0 | 0 | 0 | 0 | 0 | 0 | 1195800 | 112820 | 134440 | 0 | 0 | 0 | 0 | 0 | 0 | 0 |
| TSDANETEI | NCBP1; NCBP1 | Q09161; F22 | NCBP1_HUMAI | Unmodified | 0 | 0 | 0 | 0 | 370360 | 0 | 0 | 0 | 0 | 0 | 333180 | 3122800 | 0 | 0 | 0 | 59434 | 0 | 46086 |
| TSDDEVFK | CPD | O75976 | CBPD_HUMAN | Unmodified | 0 | 0 | 0 | 0 | 0 | 0 | 0 | 0 | 21130000 | 3848400 | 0 | 198770 | 0 | 0 | 0 | 0 | 0 | 0 |
| TSDDEVFK | CPD | O75976 | CBPD_HUMAN | Unmodified | 0 | 0 | 0 | 0 | 1988200 | 166860 | 143170 | 85428 | 0 | 7700900 | 0 | 0 | 173280 | 102050 | 0 | 0 | 0 | 57950 |
| TSDFNTFL | CHORDC1; CH | Q9UHD1; E9 | CHRD1_HUMA | Unmodified | 0 | 0 | 0 | 0 | 734510 | 0 | 402430 | 289450 | 282130 | 663510 | 393380 | 1670700 | 2219800 | 741530 | 0 | 0 | 859040 | 1581900 |
| TSDHKPVS | OCRL; OCRL; O | Q01968; A0/ | OCRL_HUMAN | Unmodified | 154420 | 148450 | 108600 | 147380 | 299520 | 0 | 102200 | 0 | 168740 | 132380 | 79781 | 1069500 | 203980 | 167830 | 182300 | 242230 | 198100 | 253360 |
| TSDIEANQI | UMAD1 | C9J7I0 | UMAD1_HUMA | Unmodified | 6452500 | 4469900 | 6861700 | 4867300 | 0 | 0 | 163470 | 106040 | 977700 | 58512 | 0 | 0 | 671120 | 0 | 8508200 | 6721800 | 19278 | 0 |
| TSDIEANQI | UMAD1 | C9J7I0 | UMAD1_HUMA | Unmodified | 1084400 | 3504600 | 0 | 1943900 | 0 | 0 | 0 | 0 | 0 | 0 | 714030 | 0 | 1312400 | 0 | 0 | 0 | 0 | 0 |
| TSDIFGSP\ | JPT2; JPT2; JPT2 | Q9H910; H3 | JUPI2_HUMAN | Unmodified | 449030 | 680220 | 548180 | 847720 | 474360 | 0 | 0 | 0 | 574140 | 0 | 750310 | 1227800 | 1011500 | 0 | 827030 | 736840 | 0 | 402000 |
| TSDLIVLGL | TARDBP; TARDI | Q13148; A0/ | TADBP_HUMAI | Unmodified | 501400 | 425700 | 199620 | 191010 | 0 | 0 | 0 | 0 | 303420 | 0 | 171630 | 1692600 | 339050 | 0 | 0 | 270320 | 0 | 312450 |
| TSDLIVLGL | TARDBP; TARDI | Q13148; A0/ | TADBP_HUMAI | Unmodified | 0 | 0 | 0 | 0 | 1573700 | 0 | 2076400 | 0 | 0 | 0 | 0 | 0 | 0 | 0 | 0 | 0 | 172650 | 0 |
| TSDQTWV | DYNC1H1; DYN | Q14204; A0/ | DYHC1_HUMA | Unmodified | 0 | 0 | 0 | 0 | 0 | 0 | 0 | 0 | 119550 | 50798 | 85118 | 959680 | 0 | 0 | 0 | 0 | 0 | 0 |
| TSDSYGFK | PTPRK; PTPRK; | Q15262; H0 | PTPRK_HUMAI | Unmodified | 0 | 1127700 | 0 | 0 | 630910 | 0 | 0 | 0 | 1139300 | 0 | 0 | 0 | 0 | 0 | 0 | 0 | 1756100 | 0 |
| TSDTIAPWI | SH2D4A | Q9H788 | SH24A_HUMAI | Unmodified | 87594 | 18784 | 43224 | 31822 | 541270 | 0 | 0 | 0 | 305190 | 222750 | 109110 | 1091100 | 261240 | 410250 | 487970 | 111930 | 2336900 | 0 |
| TSDVTSTK | DDX6; DDX6 | P26196; Q8I | DDX6_HUMAN | Unmodified | 264230 | 0 | 0 | 0 | 0 | 0 | 0 | 0 | 0 | 48387 | 3019600 | 0 | 0 | 0 | 57395 | 0 | 50543 | 0 |
| TSDVYWVL | CASD1 | Q96PB1 | CASD1_HUMAI | Unmodified | 0 | 0 | 0 | 0 | 0 | 0 | 0 | 0 | 1473800 | 733480 | 0 | 0 | 0 | 0 | 0 | 0 | 0 | 0 |
| TSEDASEYI | PARP4 | Q9UKK3 | PARP4_HUMAI | Unmodified | 0 | 0 | 0 | 0 | 0 | 0 | 0 | 0 | 0 | 0 | 0 | 1205900 | 0 | 0 | 0 | 0 | 0 | 0 |
| TSEFTAAFV | IPO9 | Q96P70 | IPO9_HUMAN I | Unmodified | 0 | 0 | 0 | 71566 | 0 | 0 | 0 | 0 | 0 | 0 | 0 | 1045800 | 0 | 0 | 0 | 0 | 0 | 492170 |
| TSEGVTNEI | FKBP5 | Q13451 | FKBP5_HUMAI | Unmodified | 0 | 0 | 0 | 0 | 471200 | 266640 | 108700 | 0 | 0 | 45666 | 0 | 0 | 875300 | 689850 | 0 | 0 | 0 | 0 |
| TSEISIPADI | UTRN | P46939 | UTRN_HUMAN | Unmodified | 0 | 0 | 1227800 | 1318800 | 0 | 0 | 0 | 0 | 1625700 | 0 | 0 | 448080 | 315380 | 44249 | 1905500 | 1187300 | 0 | 0 |

| | | | | | | | | | | | | | | | | | | | | | | |
|---|---|---|---|---|---|---|---|---|---|---|---|---|---|---|---|---|---|---|---|---|---|---|
| TSEKLGEW | TPD52L2; TPD5 | O43399; A0A | TPD54_HUMAN | Unmodified | 0 | 0 | 0 | 0 | 650410 | 0 | 0 | 0 | 0 | 0 | 0 | 0 | 0 | 398600 | 0 | 0 | 467810 | 355240 |
| TSESLCQN | XPO1; XPO1; XP | O14980; A0A | XPO1_HUMAN | Unmodified | 0 | 0 | 0 | 0 | 0 | 0 | 0 | 0 | 0 | 0 | 0 | 960990 | 0 | 0 | 0 | 0 | 0 | 0 |
| TSETLSQA | TPD52; N/A | P55327; H0Y | TPD52_HUMAN | Unmodified | 0 | 0 | 0 | 0 | 1459200 | 206100 | 604080 | 0 | 1265600 | 0 | 0 | 1477200 | 1041800 | 1088100 | 0 | 0 | 1051500 | 26792 |
| TSEVDLAK | PDCD6IP; PDCI | Q8WUM4; F8 | PDC6I_HUMAN | Unmodified | 13824000 | 13695000 | 11919000 | 6093600 | 4240700 | 168460 | 339160 | 0 | 4318000 | 0 | 247560 | 4774100 | 5174100 | 4836800 | 10892000 | 10690000 | 500280 | 2926100 |
| TSEVDLAKI | PDCD6IP; PDCI | Q8WUM4; F8 | PDC6I_HUMAN | Unmodified | 14965000 | 15147000 | 0 | 5125400 | 17202000 | 2549600 | 2496100 | 1709400 | 4848300 | 9386000 | 640470 | 995190 | 23456000 | 11870000 | 1659700 | 1839100 | 382760 | 7967200 |
| TSEVLNAAI | TMPRSS2; TMPI | O15393; C9J | TMPS2_HUMAN | Unmodified | 101140 | 0 | 2399200 | 2233800 | 131120 | 0 | 20344 | 95923 | 0 | 37465 | 0 | 0 | 0 | 118650 | 2059900 | 1840500 | 183410 | 92225 |
| TSEVNCYR | CNBP | P62633 | CNBP_HUMAN | Unmodified | 792370 | 0 | 417380 | 0 | 0 | 0 | 0 | 0 | 304860 | 0 | 0 | 4441200 | 0 | 0 | 501400 | 586680 | 0 | 0 |
| TSEVNCYR | CNBP | P62633 | CNBP_HUMAN | Unmodified | 0 | 0 | 0 | 0 | 175840 | 0 | 0 | 0 | 0 | 0 | 73313 | 97168 | 873050 | 0 | 0 | 0 | 59627 | 0 |
| TSEVNCYR | CNBP | P62633 | CNBP_HUMAN | Unmodified | 0 | 0 | 0 | 0 | 521270 | 141060 | 211340 | 151410 | 0 | 241540 | 0 | 0 | 788500 | 1201600 | 0 | 0 | 1819100 | 867210 |
| TSEVPYAGI | EIF5B; EIF5B | O60841; A0A | IF2P_HUMAN E | Unmodified | 51320 | 0 | 0 | 0 | 0 | 0 | 0 | 0 | 0 | 0 | 0 | 1125800 | 0 | 0 | 0 | 0 | 0 | 0 |
| TSEVPYAGI | EIF5B; EIF5B | O60841; A0A | IF2P_HUMAN E | Unmodified | 0 | 0 | 446920 | 332930 | 535550 | 0 | 30962 | 216440 | 222480 | 104500 | 363580 | 7365800 | 700650 | 244750 | 0 | 0 | 0 | 0 |
| TSEVVRLPY | EMC7; EMC7; E | Q9NPA0; H0 | EMC7_HUMAN | Unmodified | 0 | 0 | 0 | 0 | 0 | 0 | 0 | 0 | 2761200 | 233110 | 0 | 0 | 0 | 0 | 0 | 0 | 0 | 0 |
| TSEYFSLSH | SEMA3C | Q99985 | SEM3C_HUMA | Unmodified | 355230 | 0 | 1457700 | 1706500 | 0 | 0 | 0 | 0 | 0 | 0 | 0 | 0 | 0 | 0 | 1589900 | 672350 | 0 | 0 |
| TSFFQALGI | RPLP0; RPLP0P | P05388; Q8N | RLA0_HUMAN | Unmodified | 2507300 | 22383000 | 1389100 | 1032700 | 3367200 | 973710 | 3064800 | 756960 | 233250 | 1942400 | 4346700 | 65521000 | 0 | 2940000 | 24791000 | 23609000 | 1171800 | 1961800 |
| TSFHALTSI | SLC25A11; SLC | Q02978; I3L | M2OM_HUMAN | Unmodified | 0 | 0 | 0 | 0 | 0 | 0 | 0 | 0 | 1354600 | 217480 | 0 | 0 | 0 | 0 | 0 | 0 | 0 | 0 |
| TSFIQYLLE | EHD2 | Q9NZN4 | EHD2_HUMAN | Unmodified | 0 | 0 | 0 | 0 | 750850 | 284490 | 109480 | 0 | 0 | 0 | 0 | 0 | 0 | 0 | 0 | 0 | 0 | 0 |
| TSFLLNLR | MYO1C; MYO1C | O00159; F5H | MYO1C_HUMA | Unmodified | 2251000 | 2483200 | 2615800 | 2724000 | 0 | 0 | 0 | 0 | 0 | 0 | 0 | 813020 | 0 | 0 | 9092400 | 4238300 | 0 | 0 |
| TSFLTELIDI | STK25; STK25 | O00506; H7C | STK25_HUMAN | Unmodified | 0 | 0 | 0 | 0 | 1970700 | 264880 | 0 | 0 | 0 | 0 | 357010 | 0 | 909240 | 840250 | 0 | 0 | 731960 | 903320 |
| TSFPEDTVI | CD55; CD55; C | P08174; B1A | DAF_HUMAN C | Unmodified | 0 | 0 | 0 | 0 | 0 | 0 | 0 | 0 | 1545000 | 0 | 0 | 14726000 | 0 | 0 | 0 | 0 | 0 | 0 |
| TSFPEDTVI | CD55; CD55; C | P08174; B1A | DAF_HUMAN C | Unmodified | 4666500 | 0 | 0 | 0 | 0 | 0 | 0 | 0 | 866470 | 0 | 1116300 | 2411600 | 0 | 0 | 0 | 0 | 0 | 1262500 |
| TSFPEDTVI | CD55; CD55; C | P08174; B1A | DAF_HUMAN C | Unmodified | 388590 | 599900 | 0 | 317480 | 0 | 553220 | 0 | 0 | 0 | 0 | 722410 | 311240 | 0 | 0 | 0 | 0 | 535640 | 5154300 |
| TSFPMTVK | TMEM181 | Q9P2C4 | TM181_HUMAN | Unmodified | 67768 | 0 | 0 | 38685 | 0 | 0 | 0 | 0 | 5791500 | 655880 | 173850 | 792750 | 0 | 0 | 0 | 0 | 0 | 0 |
| TSFPMTVK | TMEM181 | Q9P2C4 | TM181_HUMAN | Oxidation (M) | 0 | 0 | 0 | 0 | 0 | 0 | 0 | 0 | 2810300 | 0 | 0 | 0 | 0 | 0 | 0 | 0 | 0 | 0 |
| TSFPMTVK | TMEM181 | Q9P2C4 | TM181_HUMAN | Unmodified | 0 | 0 | 0 | 0 | 1090400 | 0 | 0 | 0 | 4143600 | 51281 | 0 | 0 | 0 | 0 | 0 | 0 | 871020 | 841650 |
| TSFQEVPL | TAX1BP1; TAX1 | Q86VP1; B8 | TAXB1_HUMAN | Acetyl (Protein N | 0 | 0 | 0 | 0 | 574080 | 0 | 4273200 | 0 | 472510 | 1288500 | 0 | 0 | 0 | 0 | 0 | 0 | 2452500 | 0 |
| TSFTPVGD | RPN2 | P04844 | RPN2_HUMAN | Oxidation (M) | 0 | 0 | 0 | 0 | 178530 | 0 | 0 | 0 | 12227000 | 10740000 | 0 | 0 | 0 | 0 | 34039 | 0 | 0 | 0 |
| TSFTPVGD | RPN2 | P04844 | RPN2_HUMAN | Unmodified | 0 | 0 | 0 | 0 | 0 | 0 | 0 | 0 | 3694700 | 1301000 | 0 | 0 | 0 | 0 | 0 | 0 | 0 | 0 |
| TSFVNFTDI | EIF2S2 | P20042 | IF2B_HUMAN E | Unmodified | 0 | 0 | 0 | 0 | 0 | 0 | 841590 | 266250 | 256150 | 0 | 0 | 1887800 | 0 | 0 | 0 | 0 | 0 | 0 |
| TSGEGGCI | IGF2R; IGF2R; I | P11717; A0A | MPRI_HUMAN | Unmodified | 0 | 0 | 0 | 0 | 0 | 0 | 0 | 0 | 7665600 | 969400 | 0 | 441540 | 0 | 0 | 0 | 0 | 0 | 0 |
| TSGFLCND | ITGB4 | P16144 | ITB4_HUMAN I | Unmodified | 0 | 0 | 0 | 0 | 1255700 | 0 | 0 | 0 | 0 | 0 | 0 | 0 | 0 | 0 | 0 | 0 | 4403300 | 4885000 |
| TSGHDHPI | KLC1; KLC2; KL | Q07866; Q9I | KLC1_HUMAN | Unmodified | 0 | 0 | 0 | 0 | 1870500 | 429880 | 0 | 0 | 1600800 | 912950 | 331900 | 829170 | 959960 | 1706900 | 188400 | 0 | 1124700 | 1638500 |
| TSGHVDKF | GARS1; GARS1 | P41250; F8W | GARS_HUMAN | Unmodified | 0 | 0 | 0 | 0 | 1237400 | 191120 | 497420 | 121820 | 0 | 274360 | 0 | 0 | 1247700 | 1065100 | 0 | 0 | 1087400 | 721210 |
| TSGIFVKPV | EXTL2 | Q9UBQ6 | EXTL2_HUMAN | Oxidation (M) | 0 | 0 | 0 | 0 | 0 | 0 | 0 | 0 | 1099300 | 0 | 666850 | 112660 | 0 | 276570 | 0 | 0 | 0 | 0 |
| TSGIFVKPV | EXTL2 | Q9UBQ6 | EXTL2_HUMAN | Unmodified | 22745 | 0 | 0 | 0 | 0 | 0 | 0 | 0 | 9532900 | 695720 | 500130 | 350370 | 0 | 0 | 0 | 0 | 0 | 0 |
| TSGKFSGE | LRPAP1; LRPAP | P30533; A0A | AMRP_HUMAN | Unmodified | 0 | 0 | 0 | 0 | 0 | 0 | 0 | 0 | 817630 | 229480 | 0 | 0 | 0 | 0 | 0 | 0 | 0 | 0 |
| TSGLAGEP | MIA3; MIA3 | Q5JRA6; A0A | TGO1_HUMAN | Unmodified | 0 | 0 | 0 | 0 | 0 | 0 | 273520 | 78042 | 6968000 | 885660 | 0 | 213660 | 0 | 0 | 0 | 0 | 188960 | 0 |
| TSGLCVPL | CD320; CD320 | Q9NPF0; M0 | CD320_HUMA | Unmodified | 0 | 0 | 0 | 0 | 0 | 0 | 0 | 0 | 4116600 | 0 | 0 | 0 | 0 | 0 | 0 | 0 | 0 | 0 |
| TSGSVYITL | SRP14; SRP14 | P37108; H0Y | SRP14_HUMAN | Unmodified | 179020 | 139320 | 0 | 118370 | 0 | 0 | 0 | 0 | 0 | 0 | 0 | 2927000 | 0 | 0 | 407980 | 9074000 | 0 | 0 |
| TSGTLISFIY | NNT; NNT; NNT | Q13423; E9P | NNTM_HUMAN | Unmodified | 0 | 0 | 0 | 0 | 0 | 126540 | 243430 | 0 | 13520000 | 0 | 228470 | 0 | 0 | 59433 | 0 | 0 | 0 | 0 |
| TSGVLSQP | MAP2K7 | O14733 | MP2K7_HUMA | Unmodified | 224390 | 255300 | 0 | 0 | 50584 | 0 | 0 | 0 | 365100 | 369280 | 0 | 356250 | 86677 | 69701 | 0 | 204340 | 773750 | 119160 |
| TSHVENDY | SPRY4; SPRY4 | Q9C004; A0A | SPY4_HUMAN | Unmodified | 0 | 0 | 0 | 0 | 0 | 0 | 0 | 0 | 0 | 0 | 0 | 0 | 0 | 0 | 0 | 0 | 0 | 3944500 |
| TSHWEEAI | NANP | Q8TBE9 | NANP_HUMAN | Unmodified | 0 | 0 | 0 | 26527 | 0 | 0 | 0 | 0 | 0 | 0 | 0 | 1239400 | 0 | 0 | 0 | 0 | 34975 | 0 |
| TSHYLTPQ | RICTOR | Q6R327 | RICTR_HUMAN | Unmodified | 0 | 0 | 0 | 0 | 314560 | 0 | 0 | 0 | 0 | 0 | 0 | 0 | 0 | 0 | 0 | 0 | 614490 | 0 |
| TSIAIDTIIN | ATP5F1A; ATP5I | P25705; K7E | ATPA_HUMAN | Unmodified | 0 | 0 | 0 | 0 | 0 | 0 | 838830 | 194870 | 11112000 | 2473500 | 429770 | 0 | 0 | 0 | 0 | 0 | 0 | 0 |
| TSIAIDTIIN | ATP5F1A; ATP5I | P25705; K7E | ATPA_HUMAN | Unmodified | 0 | 0 | 0 | 0 | 0 | 0 | 0 | 0 | 3334100 | 244150 | 0 | 0 | 0 | 0 | 0 | 0 | 0 | 0 |
| TSIDAYDNI | CLTC; CLTC; CI | Q00610; K7E | CLH1_HUMAN | Unmodified | 8208500 | 5764200 | 1433500 | 444190 | 0 | 0 | 0 | 0 | 0 | 0 | 0 | 29214000 | 0 | 0 | 3365600 | 4985600 | 0 | 0 |
| TSIDAYDNI | CLTC; CLTC; CI | Q00610; K7E | CLH1_HUMAN | Unmodified | 15918000 | 17087000 | 3072000 | 4951600 | 104040 | 72283 | 0 | 0 | 3164500 | 0 | 711540 | 33844000 | 0 | 0 | 8806600 | 10325000 | 0 | 0 |
| TSIEDQDEI | EIF6; N/A | P56537; A0A | IF6_HUMAN Eu | Unmodified | 0 | 2438700 | 69568 | 549120 | 0 | 0 | 0 | 0 | 0 | 0 | 0 | 0 | 0 | 103110 | 2264300 | 0 | 0 | 0 |
| TSIFWNDV | EIF3B; EIF3B; EI | P55884; C9J | EIF3B_HUMAN | Unmodified | 0 | 242640 | 276560 | 118410 | 0 | 0 | 0 | 0 | 101780 | 0 | 0 | 4695200 | 0 | 0 | 379360 | 533880 | 0 | 0 |
| TSILAAANP | MCM6 | Q14566 | MCM6_HUMAN | Unmodified | 0 | 0 | 0 | 0 | 0 | 0 | 0 | 0 | 862060 | 300200 | 333810 | 1675500 | 0 | 0 | 0 | 0 | 0 | 0 |
| TSILAAANP | MCM6 | Q14566 | MCM6_HUMAN | Unmodified | 0 | 0 | 0 | 0 | 2113400 | 0 | 308820 | 207540 | 0 | 463690 | 0 | 0 | 964910 | 0 | 0 | 0 | 284700 | 530200 |
| TSILQTLCE | TNIP1; TNIP1; T | Q15025; A0A | TNIP1_HUMAN | Unmodified | 0 | 0 | 0 | 0 | 237970 | 96942 | 0 | 0 | 0 | 156030 | 0 | 0 | 221850 | 64988 | 0 | 0 | 0 | 774410 |
| TSILRDGIT | BTN3A2; BTN3A | P78410; O00 | BT3A2_HUMAN | Unmodified | 0 | 0 | 0 | 47862 | 697830 | 233410 | 0 | 0 | 671440 | 188150 | 266350 | 364940 | 410530 | 166740 | 0 | 54699 | 279480 | 1419200 |
| TSIPRAPSS | GPSM1; GPSM1 | Q86YR5; A0A | GPSM1_HUMA | Unmodified | 0 | 0 | 0 | 0 | 1091900 | 26055 | 0 | 0 | 564950 | 87394 | 0 | 0 | 89390 | 50182 | 0 | 0 | 50193 | 90886 |
| TSIRDPCA | CACNA2D1 | P54289 | CA2D1_HUMA | Unmodified | 318390 | 0 | 0 | 0 | 0 | 0 | 182090 | 85435 | 297060 | 129650 | 0 | 0 | 0 | 0 | 0 | 0 | 0 | 0 |
| TSISRYDSS | PLXNA2 | O75051 | PLXA2_HUMAN | Unmodified | 0 | 0 | 0 | 0 | 0 | 0 | 58678 | 0 | 1937800 | 188760 | 0 | 0 | 0 | 0 | 2391700 | 0 | 0 | 0 |
| TSIYPFLDFI | BTD; BTD; BTD; | P43251; A0A | BTD_HUMAN B | Unmodified | 0 | 0 | 0 | 240700 | 0 | 0 | 0 | 0 | 232140 | 0 | 0 | 0 | 0 | 0 | 0 | 0 | 14343 | 19674 |
| TSIYPFLDFI | BTD; BTD; BTD; | P43251; A0A | BTD_HUMAN B | Oxidation (M) | 0 | 0 | 0 | 0 | 101380 | 72239 | 0 | 0 | 188000 | 40911 | 0 | 0 | 0 | 0 | 0 | 0 | 0 | 0 |
| TSKAEELLA | DNM1L | O00429 | DNM1L_HUMA | Unmodified | 0 | 136230 | 0 | 232790 | 257400 | 79373 | 1402600 | 432670 | 31349 | 194260 | 373780 | 1593600 | 950600 | 701760 | 0 | 0 | 141710 | 389840 |
| TSKETDVVI | CD2AP | Q9Y5K6 | CD2AP_HUMA | Unmodified | 0 | 4725800 | 0 | 4831900 | 0 | 0 | 0 | 0 | 0 | 0 | 0 | 0 | 0 | 0 | 2500800 | 2108600 | 0 | 0 |
| TSKGPEEEI | ACY1; ACY1; AE | Q03154; F8V | ACY1_HUMAN | Acetyl (Protein N | 0 | 0 | 0 | 918310 | 0 | 0 | 0 | 46257 | 246920 | 0 | 516260 | 7556900 | 272710 | 0 | 0 | 0 | 195760 | 0 |
| TSKPFNPLI | OSBP; OSBP | P22059; H0Y | OSBP1_HUMA | Unmodified | 201250 | 0 | 0 | 0 | 0 | 0 | 115360 | 0 | 973280 | 0 | 0 | 2410700 | 0 | 0 | 0 | 0 | 0 | 0 |
| TSLADQEE | ATP8A1; ATP8A | Q9Y2Q0; Q4 | AT8A1_HUMAN | Unmodified | 0 | 0 | 0 | 0 | 0 | 0 | 0 | 0 | 393360 | 0 | 0 | 0 | 0 | 0 | 0 | 0 | 0 | 0 |
| TSLALDESL | PABPN1; PABPI | Q86U42; B4I | PABP2_HUMAN | Unmodified | 0 | 0 | 0 | 567820 | 0 | 0 | 0 | 0 | 0 | 178300 | 0 | 2537100 | 0 | 1145800 | 0 | 0 | 0 | 0 |
| TSLDPSLEI | CCDC134; CCI | Q9H6E4; B0 | CC134_HUMA | Unmodified | 0 | 0 | 0 | 0 | 0 | 0 | 2731100 | 0 | 3178800 | 3996300 | 0 | 0 | 0 | 0 | 0 | 0 | 0 | 0 |
| TSLDWWIT | SCARB2; SCARI | Q14108; A0A | SCRB2_HUMA | Unmodified | 416830 | 0 | 0 | 0 | 149810 | 13638 | 378840 | 28215 | 10219000 | 1797700 | 0 | 1747500 | 0 | 0 | 1742900 | 840850 | 135140 | 0 |
| TSLEDFYLD | SERPINF1; SER | P36955; I3L1 | PEDF_HUMAN | 2 Oxidation (M) | 0 | 0 | 0 | 0 | 0 | 0 | 0 | 0 | 0 | 0 | 2977000 | 276390 | 0 | 0 | 0 | 0 | 0 | 0 |
| TSLETQKLI | PPFIBP2; PPFIB | Q8ND30; E9 | LIPB2_HUMAN | Oxidation (M) | 0 | 0 | 74732 | 379880 | 0 | 0 | 0 | 0 | 0 | 0 | 144940 | 166250 | 0 | 0 | 287650 | 350400 | 0 | 0 |
| TSLETRFM1 | ANXA6; ANXA6; | P08133; E7E | ANXA6_HUMA | Unmodified | 0 | 316310 | 151340 | 0 | 680850 | 0 | 503440 | 0 | 511420 | 1121900 | 68040 | 0 | 323640 | 1296400 | 724110 | 0 | 164930 | 1492600 |
| TSLEVPEIN | THOC6 | Q86W42 | THOC6_HUMA | Unmodified | 0 | 0 | 0 | 0 | 0 | 0 | 0 | 0 | 0 | 0 | 0 | 100540 | 0 | 0 | 0 | 0 | 0 | 0 |
| TSLFEEDKE | WASHC2C; WA | Q9Y4E1; A0A | WAC2C_HUMA | Unmodified | 0 | 0 | 0 | 0 | 733470 | 108760 | 0 | 0 | 0 | 0 | 0 | 0 | 0 | 0 | 0 | 0 | 349950 | 0 |
| TSLFENDKI | LUZP1 | Q86V48 | LUZP1_HUMAN | Unmodified | 0 | 0 | 0 | 0 | 643520 | 0 | 0 | 0 | 0 | 0 | 0 | 0 | 0 | 0 | 0 | 0 | 92786 | 0 |
| TSLGPNGL | CCT5; CCT5; C | P48643; B7Z | TCPE_HUMAN | Unmodified | 0 | 0 | 0 | 343960 | 1044000 | 0 | 0 | 0 | 0 | 243080 | 671090 | 4340700 | 4670800 | 1625500 | 335390 | 322770 | 547470 | 1427900 |
| TSLLMVYS | RHOF; RHOF | Q9HBH0; F5 | RHOF_HUMAN | Oxidation (M) | 1814800 | 0 | 0 | 0 | 3990600 | 539450 | 0 | 0 | 0 | 0 | 0 | 0 | 0 | 0 | 0 | 0 | 966300 | 1836900 |
| TSLLTLEDV | DHX29; DHX29 | Q7Z478; A0A | DHX29_HUMA | Unmodified | 0 | 0 | 0 | 0 | 0 | 0 | 0 | 46314 | 0 | 0 | 459550 | 2018900 | 0 | 0 | 0 | 0 | 0 | 0 |
| TSLMERFTI | RAB12; RAB12 | Q6IQ22; A0A | RAB12_HUMAN | Unmodified | 0 | 0 | 0 | 0 | 352040 | 0 | 38738 | 0 | 0 | 0 | 0 | 0 | 0 | 0 | 0 | 0 | 0 | 191250 |
| TSLMNQYV | RAB7A; RAB7A; | P51149; C9J | RAB7A_HUMAN | Oxidation (M) | 0 | 0 | 0 | 0 | 730490 | 240990 | 351960 | 99175 | 327920 | 262240 | 32905 | 1824800 | 2075300 | 486130 | 0 | 0 | 164570 | 585070 |
| TSLMNQYV | RAB7A; RAB7A; | P51149; C9J | RAB7A_HUMAN | Unmodified | 1255800 | 1265300 | 926580 | 1839300 | 6658700 | 1370800 | 1747100 | 585260 | 5568700 | 3267300 | 3673200 | 12595000 | 3380500 | 2734000 | 1213700 | 951840 | 826850 | 3464700 |
| TSLMNQYV | RAB7A; RAB7A; | P51149; C9J | RAB7A_HUMAN | Unmodified | 618580 | 706850 | 596320 | 809430 | 2378200 | 0 | 629390 | 121730 | 1566000 | 304050 | 584260 | 2997700 | 0 | 441440 | 911170 | 643520 | 293040 | 972490 |
| TSLMNQYV | RAB7A; RAB7A; | P51149; C9J | RAB7A_HUMAN | Oxidation (M) | 767090 | 410060 | 787530 | 960720 | 83288 | 351510 | 268310 | 118150 | 398560 | 0 | 792670 | 4005400 | 1320900 | 399770 | 1013300 | 1279200 | 144240 | 318330 |
| TSLNILIKDI | CD109 | Q6YHK3 | CD109_HUMA | Unmodified | 96183 | 0 | 0 | 0 | 0 | 0 | 0 | 0 | 2538800 | 859040 | 0 | 0 | 0 | 0 | 0 | 0 | 252600 | 0 |
| TSLNPNDL | NEDD4L; NEDD | Q96PU5; A0A | NED4L_HUMA | Unmodified | 5177400 | 6346600 | 4011400 | 1018000 | 0 | 0 | 0 | 0 | 0 | 0 | 0 | 0 | 0 | 0 | 7923100 | 4597600 | 0 | 0 |
| TSLPWQGL | CSNK1A1; CSN | P48729; E7E | KC1A_HUMAN | Unmodified | 343170 | 0 | 0 | 0 | 321260 | 0 | 0 | 0 | 2227000 | 269070 | 0 | 1487500 | 0 | 0 | 752680 | 0 | 0 | 0 |
| TSLSALTQL | STXBP3 | O00186 | STXB3_HUMAN | Unmodified | 0 | 0 | 0 | 0 | 6986100 | 862050 | 607950 | 145740 | 0 | 0 | 564500 | 1348700 | 0 | 2232000 | 540580 | 0 | 894340 | 7093700 |
| TSLSALTQL | STXBP3 | O00186 | STXB3_HUMAN | Oxidation (M) | 0 | 0 | 0 | 0 | 1086500 | 0 | 0 | 0 | 0 | 0 | 0 | 1429800 | 929710 | 0 | 0 | 0 | 618800 | 1090000 |
| TSLSMFSK | SNX2; SNX2 | O60749; D6I | SNX2_HUMAN | Unmodified | 0 | 34431 | 0 | 0 | 1104000 | 146630 | 740160 | 90720 | 0 | 0 | 0 | 0 | 394640 | 405200 | 0 | 0 | 522520 | 425560 |
| TSLSMFSK | SNX2; SNX2 | O60749; D6I | SNX2_HUMAN | Oxidation (M) | 0 | 0 | 0 | 0 | 364610 | 0 | 673270 | 555520 | 666290 | 0 | 0 | 0 | 303820 | 173740 | 0 | 0 | 503760 | 203210 |
| TSMESLIHI | NDUFS2; NDUF | O75306; A0A | NDUS2_HUMA | Unmodified | 0 | 0 | 0 | 0 | 0 | 0 | 0 | 0 | 1478800 | 640280 | 0 | 0 | 0 | 0 | 0 | 0 | 0 | 0 |
| TSMGGTQ | CTNNB1; CTNN | P35222; A0A | CTNB1_HUMA | Oxidation (M) | 379900 | 129740 | 151090 | 202800 | 0 | 0 | 0 | 0 | 0 | 0 | 0 | 607910 | 0 | 0 | 811210 | 912900 | 0 | 0 |
| TSMGGTQ | CTNNB1; CTNN | P35222; A0A | CTNB1_HUMA | Unmodified | 63059 | 79230 | 0 | 183890 | 347830 | 0 | 0 | 0 | 0 | 0 | 0 | 2216500 | 165760 | 0 | 891460 | 962450 | 0 | 0 |
| TSMNVNEI | RAB5A | P20339 | RAB5A_HUMAN | Oxidation (M) | 0 | 0 | 0 | 0 | 0 | 0 | 0 | 0 | 1882600 | 1522600 | 141160 | 441590 | 0 | 0 | 0 | 0 | 0 | 0 |
| TSMNVNEI | RAB5A | P20339 | RAB5A_HUMAN | 2 Oxidation (M) | 522550 | 562180 | 318100 | 738510 | 0 | 23281 | 78363 | 43607 | 400940 | 405990 | 62689 | 1338500 | 124370 | 0 | 105880 | 380380 | 0 | 107030 |
| TSMTSQKS | TACC1; TACC1; | O75410; E7E | TACC1_HUMA | 2 Oxidation (M) | 0 | 0 | 0 | 0 | 0 | 1496000 | 0 | 0 | 0 | 0 | 0 | 0 | 0 | 0 | 0 | 0 | 128630 | 1564000 |
| TSNHAIVLA | ACOX1 | Q15067 | ACOX1_HUMA | Unmodified | 0 | 0 | 0 | 0 | 0 | 0 | 66108 | 73023 | 8661800 | 2700100 | 1829400 | 289210 | 0 | 0 | 0 | 0 | 44248 | 0 |
| TSNIRANFE | CTTN; CTTN | Q14247; H0Y | SRC8_HUMAN | Unmodified | 0 | 0 | 0 | 0 | 14267000 | 0 | 0 | 244040 | 0 | 927710 | 230700 | 6954500 | 4847700 | 5098200 | 0 | 0 | 1915100 | 5811500 |
| TSNLLEIR | IGSF3 | O75054 | IGSF3_HUMAN | Unmodified | 1461100 | 2061900 | 0 | 0 | 0 | 0 | 0 | 0 | 593570 | 0 | 0 | 7175000 | 0 | 0 | 2954300 | 1585500 | 0 | 0 |
| TSNLLEIRV | IGSF3 | O75054 | IGSF3_HUMAN | Unmodified | 0 | 0 | 0 | 0 | 567920 | 117310 | 48121 | 0 | 0 | 0 | 0 | 0 | 496380 | 94226 | 0 | 0 | 0 | 0 |
| TSNLLLSH | CDK11A; CDK1 | Q9UQ88; P2 | CD11A_HUMA | Unmodified | 0 | 0 | 126760 | 0 | 85444 | 0 | 211670 | 114020 | 0 | 61301 | 0 | 548520 | 0 | 0 | 0 | 0 | 0 | 0 |
| TSNLNEEL | ATP8A2; ATP8A | Q9NTI2; Q9Y | AT8A2_HUMAN | Unmodified | 0 | 0 | 0 | 0 | 713780 | 0 | 0 | 0 | 589040 | 753060 | 0 | 0 | 0 | 0 | 0 | 0 | 0 | 0 |
| TSPADGKP | ANK3; ANK3; AI | Q12955; H0Y | ANK3_HUMAN | Unmodified | 0 | 0 | 0 | 0 | 0 | 0 | 0 | 0 | 0 | 0 | 0 | 0 | 741770 | 710510 | 0 | 0 | 0 | 0 |
| TSPAPGGP | TNK2; TNK2; TN | Q07912; A0A | ACK1_HUMAN | Unmodified | 0 | 0 | 0 | 0 | 0 | 0 | 0 | 0 | 0 | 0 | 0 | 0 | 603700 | 0 | 0 | 0 | 0 | 0 |
| TSPENVQD | RANBP2 | P49792 | RBP2_HUMAN | Unmodified | 0 | 0 | 0 | 0 | 0 | 0 | 0 | 0 | 0 | 0 | 0 | 991000 | 225890 | 0 | 0 | 0 | 0 | 0 |
| TSPERDPLI | POGLUT1 | Q8NBL1 | PGLT1_HUMAN | Unmodified | 0 | 0 | 0 | 0 | 0 | 0 | 0 | 0 | 3156400 | 294240 | 208620 | 201660 | 0 | 0 | 0 | 0 | 0 | 0 |
| TSPFPILLSI | PLCB1; PLCB1; | Q9NQ66; A0 | PLCB1_HUMAN | Unmodified | 0 | 0 | 0 | 0 | 0 | 0 | 0 | 0 | 1004800 | 0 | 0 | 0 | 0 | 0 | 0 | 0 | 0 | 0 |
| TSPKPAVVI | MARS1; MARS1 | P56192; H0Y | SYMC_HUMAN | Unmodified | 0 | 0 | 0 | 0 | 0 | 0 | 0 | 0 | 1381800 | 0 | 0 | 1810500 | 0 | 0 | 0 | 0 | 0 | 0 |
| TSPNEGLS | ATP2B1 | P20020 | AT2B1_HUMAN | Unmodified | 0 | 0 | 348940 | 0 | 0 | 0 | 0 | 0 | 260330 | 197860 | 861700 | 1146000 | 0 | 0 | 392740 | 0 | 0 | 0 |
| TSPNEGLS | ATP2B1 | P20020 | AT2B1_HUMAN | Unmodified | 1872700 | 2067100 | 6388900 | 4912000 | 0 | 0 | 125740 | 0 | 4288400 | 0 | 4218500 | 42824000 | 4816500 | 5396900 | 16848000 | 10765000 | 874440 | 0 |
| TSPNEGLS | ATP2B1 | P20020 | AT2B1_HUMAN | Unmodified | 0 | 0 | 0 | 260780 | 5924800 | 704250 | 2004500 | 235010 | 0 | 759810 | 670570 | 0 | 15969000 | 10039000 | 0 | 0 | 1734800 | 2963200 |
| TSPPGPAP | NUDT19 | A8MXV4 | NUD19_HUMA | Unmodified | 0 | 0 | 0 | 0 | 0 | 0 | 81080 | 0 | 2641300 | 0 | 0 | 0 | 0 | 89213 | 0 | 0 | 0 | 0 |

| | | | | | | | | | | | | | | | | | | | | | | |
|---|---|---|---|---|---|---|---|---|---|---|---|---|---|---|---|---|---|---|---|---|---|---|
| TSPPGPAP | NUDT19 | A8MXV4 | NUD19_HUMA | Unmodified | 0 | 0 | 0 | 0 | 0 | 0 | 0 | 0 | 3733900 | 0 | 0 | 0 | 0 | 0 | 0 | 0 | 0 | 0 |
| TSPQEGER | PRKDC | P78527 | PRKDC_HUMA | Unmodified | 60997 | 0 | 0 | 0 | 0 | 0 | 499940 | 965440 | 0 | 0 | 0 | 644020 | 0 | 0 | 0 | 0 | 0 | 0 |
| TSPQELSEI | NEDD4L; NEDD | Q96PU5; A0A | NED4L_HUMA | Unmodified | 1279900 | 1560000 | 1238500 | 2503800 | 0 | 0 | 0 | 0 | 1317000 | 0 | 0 | 1395700 | 0 | 0 | 1769400 | 1582800 | 0 | 0 |
| TSPQELSEI | NEDD4L; NEDD | Q96PU5; A0A | NED4L_HUMA | Unmodified | 702500 | 848070 | 0 | 0 | 0 | 0 | 0 | 0 | 0 | 0 | 0 | 608880 | 0 | 0 | 1640900 | 1224800 | 0 | 0 |
| TSPQQKLI | ATP1A1; ATP1A | P05023; P50 | AT1A1_HUMAN | Unmodified | 0 | 0 | 0 | 0 | 311670 | 0 | 0 | 0 | 0 | 0 | 0 | 0 | 825590 | 1101000 | 0 | 0 | 961830 | 0 |
| TSPSGKPA | APEX1; APEX1; | P27695; A0A | APEX1_HUMAN | Unmodified | 0 | 161140 | 509960 | 0 | 226320 | 208030 | 533440 | 304170 | 0 | 0 | 0 | 3592600 | 0 | 0 | 193320 | 328940 | 0 | 234770 |
| TSPSPERSI | TRIM14 | Q14142 | TRI14_HUMAN | Unmodified | 0 | 0 | 0 | 738650 | 494100 | 0 | 517300 | 459340 | 0 | 0 | 0 | 562870 | 0 | 257320 | 0 | 0 | 321130 | 478360 |
| TSPSSPAPI | SLC4A2; SLC4A | P04920; A0A | B3A2_HUMAN | Unmodified | 926950 | 905250 | 2069200 | 1576200 | 0 | 0 | 43787 | 0 | 0 | 0 | 519040 | 2064600 | 383440 | 110940 | 3808200 | 2785000 | 0 | 29297 |
| TSPVADAA | SQOR | Q9Y6N5 | SQOR_HUMAN | Unmodified | 0 | 0 | 0 | 0 | 0 | 134690 | 0 | 0 | 12308000 | 0 | 0 | 0 | 0 | 0 | 0 | 0 | 0 | 0 |
| TSPVAIFNG | DTL; DTL | Q9NZJ0; F5G | DTL_HUMAN D | Unmodified | 0 | 0 | 0 | 0 | 0 | 53066 | 101900 | 81753 | 0 | 0 | 0 | 207860 | 0 | 0 | 0 | 16357 | 0 | 0 |
| TSPVEGLS | ATP2B4 | P23634 | AT2B4_HUMAN | Unmodified | 0 | 0 | 2184700 | 0 | 6866500 | 1487300 | 3415700 | 1197300 | 725990 | 1434400 | 6503600 | 38226000 | 82138 | 0 | 613390 | 2899500 | 818460 | 2267100 |
| TSPVVAPTS | SRGAP2; SRGA | O75044; B7Z | SRGP2_HUMA | Unmodified | 0 | 0 | 0 | 0 | 992840 | 0 | 0 | 0 | 2126400 | 203190 | 297430 | 1495600 | 950300 | 0 | 0 | 0 | 0 | 0 |
| TSPYDHML | MPST; MPST | P25325; B1A | THTM_HUMAN | Unmodified | 0 | 0 | 0 | 0 | 0 | 0 | 0 | 0 | 0 | 0 | 0 | 853690 | 0 | 0 | 0 | 0 | 0 | 0 |
| TSPYDSGP | FAM20B | O75063 | XYLK_HUMAN | Unmodified | 0 | 3075700 | 8928800 | 6609000 | 0 | 28441000 | 0 | 0 | 7120900 | 488390 | 3720000 | 0 | 0 | 0 | 3564100 | 321900 | 0 | 16422000 |
| TSPYPIILSF | PLCB2 | Q00722 | PLCB2_HUMAN | Unmodified | 616950 | 0 | 0 | 0 | 0 | 0 | 0 | 0 | 0 | 0 | 0 | 7672500 | 0 | 0 | 0 | 0 | 0 | 0 |
| TSPYPVILSI | PLCB3 | Q01970 | PLCB3_HUMAN | Unmodified | 12371 | 0 | 0 | 0 | 1043500 | 0 | 0 | 0 | 742720 | 0 | 0 | 1438000 | 0 | 0 | 0 | 0 | 0 | 0 |
| TSQGNSFN | PLCB3 | Q01970 | PLCB3_HUMAN | Unmodified | 130450 | 129050 | 0 | 0 | 276280 | 126290 | 0 | 0 | 359800 | 0 | 240900 | 563040 | 0 | 0 | 0 | 0 | 160850 | 0 |
| TSQIFLAK | MACF1; MACF1 | Q9UPN3; H3 | MACF1_HUMA | Unmodified | 16424 | 0 | 29994 | 57977 | 153800 | 0 | 212170 | 11439 | 275940 | 201970 | 0 | 292370 | 0 | 194150 | 63219 | 0 | 78676 | 571510 |
| TSQLLETLN | DCTN1; DCTN1 | Q14203; A0A | DCTN1_HUMA | Unmodified | 0 | 0 | 0 | 0 | 0 | 0 | 0 | 0 | 0 | 0 | 0 | 3662100 | 0 | 0 | 0 | 0 | 0 | 0 |
| TSQLLETLN | DCTN1; DCTN1 | Q14203; A0A | DCTN1_HUMA | Unmodified | 0 | 0 | 0 | 0 | 1589300 | 0 | 206910 | 0 | 2908200 | 120830 | 0 | 0 | 979100 | 669590 | 0 | 0 | 1647500 | 0 |
| TSQTVATFL | THOP1; THOP1 | P52888; K7E | THOP1_HUMA | Unmodified | 125570 | 0 | 116720 | 228030 | 736870 | 222990 | 0 | 0 | 0 | 503290 | 716050 | 2932000 | 180740 | 0 | 267040 | 229620 | 362710 | 51878 |
| TSQTWHG | GGCT; GGCT; G | O75223; H7I | GGCT_HUMAN | Unmodified | 0 | 0 | 0 | 0 | 0 | 0 | 0 | 0 | 0 | 482880 | 0 | 0 | 1849400 | 439080 | 0 | 0 | 0 | 0 |
| TSRFASFPE | USP13; USP13; | Q92995; A0A | UBP13_HUMA | Unmodified | 0 | 0 | 0 | 0 | 846130 | 0 | 169610 | 0 | 0 | 0 | 187690 | 568020 | 594810 | 0 | 0 | 0 | 1546000 | 0 |
| TSRPENAII | TKT; TKT | P29401; A0A | TKT_HUMAN Tr | Unmodified | 1019100 | 600940 | 674130 | 1026300 | 27603000 | 5274900 | 1232100 | 636710 | 15755000 | 5655700 | 7785600 | 43602000 | 5022900 | 1720500 | 605060 | 906050 | 4564300 | 10118000 |
| TSSAETPTI | IMMT; IMMT; IM | Q16891; B9A | MIC60_HUMAN | Unmodified | 0 | 0 | 0 | 0 | 0 | 0 | 0 | 0 | 1432900 | 0 | 0 | 0 | 0 | 0 | 0 | 0 | 0 | 0 |
| TSSAEVTIQ | ADGRE5 | P48960 | AGRE5_HUMA | Unmodified | 0 | 0 | 0 | 0 | 6023000 | 612310 | 0 | 118680 | 0 | 0 | 0 | 0 | 0 | 0 | 0 | 0 | 869490 | 6438700 |
| TSSFAEPG | GSK3A; GSK3A; | P49840; A0A | GSK3A_HUMA | Unmodified | 0 | 0 | 0 | 0 | 861710 | 71734 | 211770 | 107320 | 0 | 0 | 221720 | 605040 | 844160 | 0 | 0 | 0 | 0 | 114580 |
| TSSFLDQR | ECE1; ECE1; EC | P42892; B4D | ECE1_HUMAN | Unmodified | 285380 | 1075400 | 98623 | 0 | 459840 | 61401 | 0 | 455330 | 7105600 | 0 | 250260 | 1376600 | 290490 | 0 | 68351 | 133310 | 65883 | 139460 |
| TSSFLDQRI | ECE1; ECE1; EC | P42892; B4D | ECE1_HUMAN | Unmodified | 50396 | 0 | 61931 | 274600 | 1517400 | 0 | 0 | 0 | 5198200 | 1569600 | 0 | 0 | 0 | 0 | 0 | 63484 | 0 | 0 |
| TSSGDPPS | CDCA3; CDCA3 | Q99618; F8V | CDCA3_HUMA | Unmodified | 0 | 0 | 0 | 0 | 4213100 | 518470 | 462800 | 160380 | 353750 | 329880 | 0 | 176980 | 2118700 | 1107500 | 0 | 0 | 1022100 | 1420500 |
| TSSGKKEV | EPHA2 | P29317 | EPHA2_HUMA | Unmodified | 0 | 0 | 0 | 0 | 12331000 | 2261500 | 0 | 15549 | 0 | 0 | 0 | 0 | 525240 | 311120 | 0 | 0 | 30609000 | 35041000 |
| TSSGLGGS | PAWR; PAWR; F | Q96IZ0; F8W | PAWR_HUMAN | Unmodified | 218920 | 89538 | 0 | 0 | 0 | 0 | 0 | 0 | 0 | 0 | 51315 | 154020 | 0 | 0 | 0 | 861710 | 0 | 0 |
| TSSGTSIAG | SMARCAL1; SM | Q9NZC9; C9 | SMAL1_HUMA | Unmodified | 0 | 0 | 0 | 0 | 525160 | 0 | 0 | 0 | 0 | 0 | 0 | 0 | 0 | 0 | 0 | 0 | 681180 | 0 |
| TSSIIFKCDI | IGF2R | P11717 | MPRI_HUMAN | Unmodified | 353360 | 0 | 0 | 676170 | 772610 | 171170 | 0 | 0 | 28605000 | 615390 | 898270 | 5593700 | 0 | 0 | 0 | 587950 | 0 | 0 |
| TSSIIIALTD | ANTXR2; ANTXF | P58335; A0A | ANTR2_HUMA | Unmodified | 0 | 0 | 0 | 0 | 597930 | 0 | 0 | 0 | 41810 | 0 | 0 | 0 | 0 | 0 | 0 | 0 | 0 | 2591100 |
| TSSPIPAQH | RNF31; RNF31 | Q96EP0; H0 | RNF31_HUMA | Unmodified | 1892200 | 1892900 | 0 | 0 | 0 | 0 | 0 | 0 | 0 | 0 | 0 | 0 | 0 | 0 | 0 | 0 | 0 | 0 |
| TSSQYVAS | COBL; COBL; C | O75128; J3K | COBL_HUMAN | Unmodified | 350710 | 647280 | 0 | 0 | 0 | 0 | 0 | 0 | 0 | 0 | 0 | 0 | 277560 | 0 | 0 | 65633 | 0 | 0 |
| TSSTVNTK | SLC4A7; SLC4A | Q9Y6M7; C9 | S4A7_HUMAN | Unmodified | 0 | 0 | 451660 | 367800 | 0 | 97254 | 0 | 0 | 381710 | 59707 | 456920 | 2121400 | 104170 | 85791 | 453420 | 618740 | 166930 | 670950 |
| TSSTVNTK | SLC4A7; SLC4A | Q9Y6M7; C9 | S4A7_HUMAN | Unmodified | 0 | 0 | 0 | 614740 | 771530 | 0 | 0 | 0 | 0 | 293720 | 201430 | 0 | 258760 | 45039 | 0 | 0 | 477260 | 1250700 |
| TSSVPEYV | PKP2 | Q99959 | PKP2_HUMAN | Unmodified | 0 | 0 | 0 | 0 | 0 | 0 | 0 | 0 | 0 | 0 | 803450 | 1454000 | 0 | 0 | 0 | 0 | 1905200 | 2179300 |
| TSSVPVVSI | GPSM2; GPSM2 | P81274; A0A | GPSM2_HUMA | Unmodified | 0 | 0 | 0 | 0 | 0 | 0 | 0 | 0 | 853090 | 390590 | 0 | 0 | 858840 | 0 | 57665 | 0 | 0 | 0 |
| TSSVSNPQ | PXN; PXN; PXN; | P49023; F5H | PAXI_HUMAN F | Unmodified | 0 | 0 | 0 | 0 | 1316900 | 0 | 0 | 0 | 0 | 0 | 0 | 0 | 354600 | 438860 | 0 | 0 | 5487200 | 4803100 |
| TSSVSSISQ | PI4KA | P42356 | PI4KA_HUMAN | Unmodified | 0 | 0 | 461790 | 453640 | 0 | 0 | 0 | 0 | 0 | 0 | 0 | 0 | 0 | 0 | 400910 | 239370 | 0 | 0 |
| TSSVVTLEV | AFDN; AFDN; A | P55196; Q5T | AFAD_HUMAN | Unmodified | 0 | 0 | 0 | 220570 | 346290 | 0 | 0 | 0 | 0 | 362780 | 586520 | 1692300 | 0 | 0 | 0 | 231410 | 0 | 0 |
| TSTADYAM | FGG; FGG; FGG | P02679; C9J | FIBG_HUMAN | Oxidation (M) | 0 | 1879500 | 0 | 0 | 0 | 0 | 0 | 0 | 0 | 0 | 0 | 0 | 0 | 0 | 0 | 0 | 0 | 0 |
| TSTDRHSL | FRMD6 | Q96NE9 | FRMD6_HUMA | Unmodified | 0 | 0 | 0 | 0 | 1409700 | 261630 | 0 | 0 | 86098 | 0 | 0 | 0 | 0 | 0 | 0 | 0 | 893240 | 1681700 |
| TSTILAAAR | RFC4; RFC4; RF | P35249; C9J | RFC4_HUMAN | Unmodified | 263870 | 153640 | 44198 | 279390 | 136920 | 39605 | 0 | 0 | 439030 | 62584 | 0 | 212450 | 84420 | 0 | 0 | 136710 | 0 | 0 |
| TSTILAAARI | RFC4; RFC4; RF | P35249; C9J | RFC4_HUMAN | Unmodified | 0 | 0 | 0 | 0 | 70832 | 59226 | 76836 | 0 | 0 | 264310 | 0 | 0 | 287400 | 112090 | 0 | 0 | 67962 | 0 |
| TSTILGDITS | FERMT2; FERM | Q96AC1; A0A | FERM2_HUMA | Unmodified | 0 | 87600 | 0 | 13569 | 1230600 | 0 | 41129 | 0 | 27457 | 113920 | 0 | 0 | 33976 | 0 | 82249 | 0 | 303600 | 382330 |
| TSTPEDFIR | TLN1 | Q9Y490 | TLN1_HUMAN | Unmodified | 640100 | 966340 | 2059800 | 2983000 | 4451200 | 182380 | 1413700 | 523620 | 1640300 | 655570 | 458390 | 3167800 | 1831700 | 1870000 | 1244300 | 1081000 | 344880 | 2349600 |
| TSTPEDFIR | TLN1 | Q9Y490 | TLN1_HUMAN | Unmodified | 0 | 0 | 0 | 0 | 7043800 | 481380 | 2951700 | 724460 | 147280 | 326730 | 0 | 514900 | 1216600 | 0 | 0 | 0 | 1767900 | 2299400 |
| TSTPEDFIR | TLN1 | Q9Y490 | TLN1_HUMAN | Oxidation (M) | 0 | 0 | 0 | 0 | 1356300 | 400540 | 1263600 | 369290 | 0 | 0 | 0 | 0 | 592030 | 1001700 | 0 | 0 | 667390 | 732300 |
| TSTPLAPLP | MICAL3 | Q7RTP6 | MICA3_HUMA | Unmodified | 0 | 0 | 0 | 0 | 1983300 | 662600 | 0 | 0 | 0 | 0 | 191350 | 0 | 184490 | 0 | 0 | 0 | 0 | 0 |
| TSTPLDREI | FAT1; FAT1 | Q14517; A0A | FAT1_HUMAN | Unmodified | 139600 | 0 | 0 | 0 | 0 | 0 | 0 | 0 | 0 | 0 | 77529 | 0 | 0 | 0 | 0 | 595410 | 0 | 552310 |
| TSTQKHTH | IGSF8; IGSF8 | Q969P0; C9J | IGSF8_HUMAN | Unmodified | 0 | 0 | 0 | 0 | 0 | 0 | 0 | 0 | 0 | 0 | 229940 | 0 | 5158800 | 2439200 | 0 | 0 | 555380 | 0 |
| TSTRQTYFL | PSMC3; PSMC3 | P17980; E9P | PRS6A_HUMA | Unmodified | 0 | 0 | 0 | 0 | 5556900 | 0 | 2148000 | 0 | 0 | 2008300 | 0 | 0 | 0 | 0 | 0 | 0 | 3639700 | 2487100 |
| TSTSLGKG | SARG | Q9BW04 | SARG_HUMAN | Unmodified | 0 | 0 | 0 | 0 | 457940 | 123640 | 0 | 0 | 0 | 0 | 0 | 0 | 3870400 | 3563900 | 0 | 0 | 268720 | 55117 |
| TSTSQAVF | DARS1; DARS1; | P14868; C9J | SYDC_HUMAN | Unmodified | 0 | 0 | 0 | 0 | 0 | 19710 | 0 | 0 | 691000 | 0 | 344620 | 3965800 | 0 | 0 | 264590 | 307510 | 0 | 0 |
| TSTTGVATT | LMO7; LMO7; L | Q8WWI1; A0 | LMO7_HUMAN | Unmodified | 0 | 0 | 0 | 0 | 456890 | 111060 | 51452 | 26638 | 119150 | 0 | 0 | 0 | 0 | 0 | 0 | 0 | 237150 | 446620 |
| TSTTSSMV | PPP1R2; PPP1R | P41236; E7E | IPP2_HUMAN F | Unmodified | 0 | 0 | 0 | 0 | 584930 | 0 | 0 | 0 | 0 | 191400 | 0 | 0 | 367760 | 178090 | 74189 | 0 | 265640 | 606360 |
| TSTTVRTTE | CRABP1; CRAB | P29762; P29 | RABP1_HUMA | Unmodified | 0 | 0 | 0 | 0 | 322750 | 131070 | 0 | 49871 | 0 | 361680 | 0 | 0 | 0 | 0 | 0 | 0 | 799670 | 1393100 |
| TSVDEITSG | CEP55 | Q53EZ4 | CEP55_HUMA | Unmodified | 1599100 | 1425200 | 2938000 | 4292900 | 18569000 | 2846100 | 249710 | 130260 | 10533000 | 5145300 | 954520 | 2534600 | 2349400 | 677500 | 2770800 | 1479500 | 1140900 | 1314700 |
| TSVDEITSG | CEP55 | Q53EZ4 | CEP55_HUMA | Unmodified | 0 | 136940 | 106950 | 0 | 804410 | 229440 | 0 | 0 | 0 | 399720 | 0 | 0 | 0 | 0 | 0 | 68974 | 0 | 0 |
| TSVDEITSG | CEP55 | Q53EZ4 | CEP55_HUMA | Unmodified | 154430 | 0 | 0 | 0 | 7556400 | 1771300 | 76784 | 303160 | 345200 | 1729300 | 0 | 0 | 633880 | 142620 | 0 | 0 | 283240 | 976220 |
| TSVETALR | CAPZA2; CAPZA | P47755; A0A | CAZA2_HUMA | Unmodified | 1613400 | 957860 | 474010 | 398350 | 121740 | 43900 | 269240 | 0 | 930350 | 102460 | 530820 | 9612800 | 68953 | 0 | 774750 | 670880 | 0 | 0 |
| TSVFLVR | GEMIN5 | Q8TEQ6 | GEMI5_HUMA | Unmodified | 0 | 67457 | 0 | 85006 | 0 | 0 | 0 | 0 | 162910 | 0 | 215140 | 3663400 | 470660 | 488480 | 0 | 0 | 0 | 0 |
| TSVIQGIHT | LYRM7 | Q5U5X0 | LYRM7_HUMA | Unmodified | 0 | 0 | 0 | 0 | 0 | 0 | 0 | 18805 | 2592900 | 404910 | 0 | 0 | 130360 | 0 | 0 | 0 | 214600 | 26177 |
| TSVKEDLN | RAB23 | Q9ULC3 | RAB23_HUMA | Unmodified | 0 | 837270 | 0 | 2346100 | 2286200 | 730250 | 58458 | 0 | 3491300 | 1322800 | 677330 | 0 | 1865400 | 403400 | 1326000 | 2089400 | 0 | 1548000 |
| TSVLEMIAC | OPA1; OPA1; OI | O60313; E5K | OPA1_HUMAN | Oxidation (M) | 0 | 0 | 0 | 0 | 0 | 0 | 0 | 0 | 0 | 0 | 0 | 1525000 | 0 | 0 | 0 | 0 | 29273 | 355070 |
| TSVLYQYTI | RAB27A; RAB27 | P51159; H3E | RB27A_HUMA | Unmodified | 26478 | 287140 | 0 | 737340 | 106260 | 0 | 288300 | 104050 | 0 | 0 | 0 | 0 | 0 | 0 | 0 | 101850 | 0 | 0 |
| TSVPVDSFI | ACAD10; ACAD | Q6JQN1; F8V | ACD10_HUMA | Unmodified | 0 | 0 | 0 | 0 | 0 | 0 | 0 | 0 | 105820 | 0 | 0 | 0 | 0 | 0 | 0 | 0 | 0 | 0 |
| TSVQFQNF | AP2A1 | O95782 | AP2A1_HUMA | Unmodified | 0 | 0 | 0 | 0 | 1577100 | 0 | 1139000 | 68361 | 0 | 0 | 0 | 2177600 | 0 | 0 | 0 | 0 | 3658900 | 2569100 |
| TSVQPTFT | UBR4 | Q5T4S7 | UBR4_HUMAN | Unmodified | 0 | 0 | 0 | 0 | 0 | 0 | 0 | 0 | 159080 | 0 | 0 | 652720 | 0 | 0 | 0 | 0 | 0 | 0 |
| TSVQTEDD | CTNNA1; CTNN | P35221; P26 | CTNA1_HUMA | Unmodified | 1585300 | 2151300 | 8539900 | 11161000 | 6735900 | 3577300 | 0 | 0 | 1232800 | 797610 | 6280600 | 20527000 | 14517000 | 4905400 | 14701000 | 11530000 | 0 | 0 |
| TSVSLAVSI | AP2A2; AP2A2; | O94973; A0A | AP2A2_HUMA | Unmodified | 60461 | 73444 | 263670 | 0 | 0 | 0 | 0 | 0 | 0 | 0 | 125030 | 2963700 | 0 | 0 | 381170 | 368180 | 0 | 0 |
| TSVVVGTW | ERLEC1 | Q96DZ1 | ERLEC_HUMA | Unmodified | 0 | 0 | 0 | 0 | 0 | 0 | 0 | 0 | 1609500 | 411680 | 0 | 0 | 0 | 0 | 0 | 0 | 0 | 0 |
| TSWAGAP | NES | P48681 | NEST_HUMAN | Unmodified | 0 | 0 | 0 | 0 | 0 | 92833 | 0 | 0 | 145850 | 0 | 0 | 0 | 0 | 0 | 0 | 0 | 974010 | 1971800 |
| TSWINLK | UTRN | P46939 | UTRN_HUMAN | Unmodified | 0 | 0 | 0 | 0 | 1396900 | 2433500 | 0 | 25899 | 1025300 | 1612000 | 180910 | 0 | 768750 | 0 | 1162000 | 729780 | 1170800 | 808230 |
| TSWSTVTR | GLCE; GLCE | O94923; H0 | GLCE_HUMAN | Unmodified | 0 | 286830 | 0 | 0 | 604480 | 169890 | 0 | 56586 | 2870700 | 134920 | 0 | 1302200 | 0 | 0 | 0 | 0 | 56471 | 0 |
| TSWVQPLL | GGCX | P38435 | VKGC_HUMAN | Oxidation (M) | 0 | 0 | 0 | 0 | 0 | 0 | 0 | 0 | 91551 | 42880 | 0 | 0 | 0 | 0 | 0 | 0 | 0 | 0 |
| TSYAQHQ | RPS3A; RPS3A; | P61247; D6F | RS3A_HUMAN | Unmodified | 353840 | 955990 | 1311600 | 836920 | 1636900 | 261000 | 77858 | 534660 | 534200 | 392940 | 136340 | 11759000 | 728730 | 555580 | 1185700 | 929580 | 91135 | 606260 |
| TSYETADK | CAPN7 | Q9Y6W3 | CAN7_HUMAN | Unmodified | 0 | 0 | 0 | 0 | 788660 | 0 | 201640 | 80947 | 0 | 366410 | 0 | 0 | 0 | 249940 | 0 | 0 | 119450 | 251580 |
| TSYIIMFGP | CLGN | O14967 | CLGN_HUMAN | Oxidation (M) | 0 | 0 | 0 | 0 | 0 | 0 | 0 | 0 | 5017200 | 866450 | 1216100 | 0 | 0 | 384360 | 0 | 0 | 0 | 0 |
| TSYIIMFGP | CLGN | O14967 | CLGN_HUMAN | Unmodified | 0 | 0 | 0 | 0 | 76313 | 0 | 0 | 0 | 4968700 | 0 | 0 | 0 | 0 | 0 | 194450 | 0 | 177180 | 0 |
| TSYIIMFGP | CLGN | O14967 | CLGN_HUMAN | Oxidation (M) | 0 | 0 | 0 | 0 | 0 | 0 | 0 | 0 | 560880 | 381190 | 188620 | 0 | 0 | 0 | 0 | 0 | 0 | 0 |
| TSYLTELIDI | STK26; STK26; | Q9P289; B4E | STK26_HUMA | Unmodified | 221270 | 0 | 201240 | 388340 | 5105400 | 1711300 | 202700 | 0 | 697910 | 500850 | 1065500 | 1302500 | 5605500 | 2982200 | 589280 | 269580 | 1041500 | 2340400 |
| TSYLTELIDI | STK24; STK24; | Q9Y6E0; H0 | STK24_HUMA | Unmodified | 17468 | 0 | 0 | 0 | 4640600 | 0 | 484370 | 0 | 705320 | 0 | 0 | 1313100 | 0 | 1290600 | 775810 | 215210 | 797640 | 1379700 |
| TTAFSVWT | ELAPOR1; ELAP | Q6UXG2; E9 | ELAP1_HUMA | Unmodified | 804640 | 213220 | 113820 | 0 | 0 | 0 | 0 | 0 | 1458900 | 118090 | 0 | 0 | 0 | 0 | 0 | 0 | 0 | 0 |
| TTAHFPHP | EMC1 | Q8N766 | EMC1_HUMAN | Unmodified | 0 | 0 | 0 | 0 | 0 | 0 | 0 | 0 | 3454500 | 386720 | 0 | 0 | 0 | 0 | 0 | 88790 | 0 | 0 |
| TTAHYDPA | ACOX3; ACOX3 | O15254; D6I | ACOX3_HUMA | Unmodified | 0 | 0 | 0 | 0 | 0 | 0 | 0 | 0 | 1281600 | 110180 | 0 | 0 | 0 | 0 | 0 | 0 | 0 | 0 |
| TTANAIYCF | PSMD11; PSMD | O00231; J3Q | PSD11_HUMA | Unmodified | 0 | 0 | 0 | 0 | 0 | 0 | 0 | 0 | 0 | 0 | 0 | 520940 | 0 | 0 | 0 | 0 | 0 | 0 |
| TTANRNHT | KIAA1217; KIAA | Q5T5P2; Q5 | SKT_HUMAN S | Unmodified | 0 | 0 | 0 | 0 | 0 | 0 | 38213 | 0 | 0 | 0 | 0 | 0 | 0 | 308440 | 0 | 0 | 1175900 | 328250 |
| TTAQVLIR | AKR1B15; AKR1 | C9JRZ8; P15 | AK1BF_HUMA | Unmodified | 142910 | 0 | 0 | 297270 | 84762 | 477060 | 102410 | 128800 | 0 | 0 | 277210 | 7544400 | 145210 | 140900 | 0 | 0 | 0 | 132910 |
| TTARDQDL | EPRS1 | P07814 | SYEP_HUMAN | Unmodified | 0 | 0 | 0 | 354570 | 1891100 | 335990 | 1054000 | 484740 | 730510 | 639950 | 686940 | 4318400 | 3653700 | 1677800 | 0 | 0 | 1092000 | 1539000 |
| TTARDQDL | EPRS1 | P07814 | SYEP_HUMAN | Oxidation (M) | 0 | 0 | 0 | 0 | 331120 | 0 | 179680 | 117640 | 140050 | 236420 | 183300 | 848990 | 1336200 | 400700 | 0 | 0 | 236760 | 392030 |
| TTASDGEQ | MPPE1; MPPE1 | Q53F39; K7E | MPPE1_HUMA | Unmodified | 0 | 0 | 0 | 0 | 0 | 0 | 0 | 0 | 834400 | 0 | 0 | 0 | 118790 | 0 | 0 | 0 | 0 | 0 |
| TTASTSQV | FTH1; FTH1; FT | P02794; E9P | FRIH_HUMAN | Acetyl (Protein N | 0 | 639410 | 376890 | 0 | 0 | 0 | 0 | 56095 | 21166000 | 0 | 75100 | 0 | 0 | 0 | 449900 | 123140 | 0 | 0 |
| TTASTSQV | FTH1; FTH1; FT | P02794; E9P | FRIH_HUMAN | Acetyl (Protein N | 0 | 0 | 0 | 0 | 784300 | 0 | 1115500 | 0 | 17226000 | 27779000 | 302230 | 0 | 570590 | 315850 | 0 | 0 | 1472000 | 1042400 |
| TTAVEIDYD | FYB1 | O15117 | FYB1_HUMAN | Unmodified | 752370 | 833590 | 348180 | 529320 | 0 | 0 | 0 | 0 | 0 | 0 | 0 | 721730 | 0 | 0 | 343900 | 493400 | 0 | 32902 |
| TTCSCSLA | GET3 | O43681 | GET3_HUMAN | Unmodified | 0 | 0 | 0 | 0 | 0 | 0 | 0 | 0 | 0 | 0 | 0 | 0 | 475340 | 0 | 0 | 0 | 755940 | 1008100 |
| TTCSSGSA | CMTM7; CMTM | Q96FZ5; F8V | CKLF7_HUMA | Unmodified | 0 | 0 | 0 | 0 | 0 | 0 | 0 | 0 | 910050 | 329850 | 0 | 0 | 0 | 0 | 0 | 128620 | 0 | 0 |
| TTDDIVKVE | RABL6; RABL6; | Q3YEC7; F2Z | RABL6_HUMA | Unmodified | 0 | 0 | 0 | 0 | 0 | 0 | 1050200 | 114330 | 0 | 0 | 148130 | 0 | 0 | 0 | 0 | 0 | 0 | 0 |
| TTDEGAKN | FKBP5 | Q13451 | FKBP5_HUMA | Unmodified | 0 | 0 | 0 | 0 | 1061300 | 0 | 0 | 243560 | 0 | 0 | 0 | 0 | 1875200 | 0 | 0 | 0 | 0 | 0 |
| TTDFSDFLS | CHORDC1; CH | Q9UHD1; E9 | CHRD1_HUMA | Unmodified | 0 | 0 | 0 | 21428 | 0 | 0 | 0 | 0 | 329630 | 59519 | 112060 | 252720 | 105240 | 240820 | 0 | 0 | 230310 | 0 |
| TTDGVYEG | ACLY | P53396 | ACLY_HUMAN | Unmodified | 0 | 0 | 0 | 172920 | 3718100 | 0 | 9287500 | 0 | 0 | 422120 | 0 | 0 | 583150 | 316850 | 0 | 0 | 0 | 0 |
| TTDGYLLR | RPS3A; RPS3A; | P61247; D6F | RS3A_HUMAN | Unmodified | 6570100 | 3053800 | 2986300 | 1442500 | 562410 | 0 | 57352 | 158460 | 922480 | 433620 | 5163000 | 50830000 | 309460 | 245690 | 3895900 | 4954900 | 0 | 0 |
| TTDGYLLRI | RPS3A; RPS3A; | P61247; D6F | RS3A_HUMAN | Unmodified | 0 | 0 | 0 | 69815 | 1322900 | 116380 | 4369400 | 0 | 0 | 624750 | 43867 | 0 | 453270 | 981940 | 0 | 0 | 0 | 1149400 |
| TTDGYLLRI | RPS3A; RPS3A; | P61247; D6F | RS3A_HUMAN | Unmodified | 0 | 0 | 115810 | 0 | 202080 | 31804 | 704240 | 0 | 0 | 122930 | 20868 | 33954 | 127260 | 0 | 0 | 0 | 46790 | 0 |
| TTDIVIR | UBE2O; UBE2O | Q9C0C9; K7 | UBE2O_HUMA | Unmodified | 0 | 0 | 46560 | 91510 | 0 | 0 | 0 | 0 | 0 | 0 | 0 | 1670600 | 58794 | 0 | 63892 | 87389 | 0 | 0 |

| | | | | | | | | | | | | | | | | | | | | | | |
|---|---|---|---|---|---|---|---|---|---|---|---|---|---|---|---|---|---|---|---|---|---|---|
| TTDLLTDW | DYNC1H1; DYN | Q14204; A0A | DYHC1_HUMA | Unmodified | 0 | 0 | 0 | 0 | 0 | 0 | 0 | 0 | 745970 | 0 | 201830 | 4911400 | 0 | 0 | 0 | 0 | 0 | 0 |
| TTDLSIQPA | AHNAK2 | Q8IVF2 | AHNK2_HUMA | Unmodified | 0 | 0 | 0 | 0 | 601680 | 0 | 0 | 0 | 0 | 0 | 0 | 0 | 0 | 0 | 0 | 0 | 0 | 706470 |
| TTDLSIQPA | AHNAK2 | Q8IVF2 | AHNK2_HUMA | Unmodified | 0 | 0 | 0 | 0 | 0 | 0 | 0 | 0 | 0 | 0 | 0 | 0 | 0 | 0 | 0 | 0 | 0 | 7862000 |
| TTDLSIQPH | AHNAK2 | Q8IVF2 | AHNK2_HUMA | Unmodified | 0 | 0 | 0 | 0 | 1513800 | 225450 | 0 | 0 | 0 | 0 | 0 | 0 | 0 | 0 | 0 | 0 | 0 | 886110 |
| TTDLSIQPL | AHNAK2 | Q8IVF2 | AHNK2_HUMA | Unmodified | 0 | 0 | 0 | 0 | 951910 | 0 | 0 | 0 | 0 | 0 | 0 | 0 | 0 | 0 | 0 | 0 | 0 | 1376600 |
| TTDMAPSK | MAP4; MAP4; M | P27816; E7E | MAP4_HUMAN | Unmodified | 0 | 0 | 244810 | 21259 | 535530 | 0 | 246910 | 111260 | 0 | 151590 | 0 | 298770 | 426880 | 315410 | 0 | 127290 | 0 | 458140 |
| TTDTASVQI | CORO1C; COR | Q9ULV4; H0 | COR1C_HUMA | Unmodified | 0 | 0 | 0 | 260130 | 23024000 | 5107500 | 1213200 | 1030100 | 2316700 | 1565400 | 2277600 | 0 | 1872700 | 1072400 | 0 | 262260 | 2493900 | 2535600 |
| TTDTASVQI | CORO1C; COR | Q9ULV4; H0 | COR1C_HUMA | Unmodified | 0 | 0 | 0 | 0 | 0 | 0 | 0 | 0 | 0 | 0 | 0 | 0 | 0 | 0 | 0 | 0 | 1962300 | 1079300 |
| TTDVEGRLS | USO1 | O60763 | USO1_HUMAN | Unmodified | 0 | 0 | 0 | 0 | 1576100 | 0 | 0 | 0 | 0 | 0 | 0 | 0 | 2106700 | 918240 | 0 | 0 | 335850 | 1831800 |
| TTDVMFGC | AHCYL1 | O43865 | SAHH2_HUMA | Unmodified | 0 | 0 | 206360 | 203210 | 497930 | 166040 | 65015 | 41436 | 172510 | 69319 | 0 | 547000 | 156950 | 93910 | 327820 | 173170 | 227570 | 214540 |
| TTEADDPFI | SMOC1 | Q9H4F8 | SMOC1_HUMA | Unmodified | 0 | 0 | 0 | 0 | 0 | 0 | 0 | 0 | 833020 | 0 | 0 | 0 | 0 | 0 | 0 | 0 | 0 | 0 |
| TTEADDPFI | SMOC1 | Q9H4F8 | SMOC1_HUMA | Unmodified | 0 | 0 | 0 | 0 | 0 | 0 | 0 | 0 | 1528400 | 1419400 | 0 | 0 | 0 | 0 | 0 | 0 | 0 | 0 |
| TTEDEVHIC | LDLR; LDLR | P01130; J3K | LDLR_HUMAN | Unmodified | 37926 | 0 | 0 | 0 | 0 | 0 | 0 | 0 | 1555400 | 161720 | 0 | 700530 | 0 | 0 | 0 | 0 | 0 | 0 |
| TTEDLSFK | YES1; YES1 | P07947; J3Q | YES_HUMAN Ty | Unmodified | 912250 | 571140 | 1081900 | 346940 | 0 | 0 | 0 | 0 | 0 | 0 | 0 | 1816700 | 0 | 0 | 2417000 | 2430700 | 31665 | 43656 |
| TTEEDRGN | PTGFRN | Q9P2B2 | FPRP_HUMAN | Unmodified | 8590100 | 5082200 | 10394000 | 14502000 | 813760 | 1105000 | 156480 | 37265 | 0 | 1148100 | 0 | 18246000 | 4819700 | 1297200 | 12921000 | 16547000 | 0 | 0 |
| TTEENPIFV | TBK1; TBK1; TB | Q9UHD2; AC | TBK1_HUMAN S | Unmodified | 0 | 0 | 0 | 0 | 521520 | 0 | 0 | 0 | 766110 | 76985 | 0 | 0 | 0 | 0 | 0 | 0 | 0 | 0 |
| TTEGFGPEI | ITGB4; ITGB4 | P16144; J3K | ITB4_HUMAN In | Unmodified | 0 | 0 | 0 | 0 | 9190100 | 2792600 | 0 | 0 | 0 | 0 | 0 | 0 | 574820 | 0 | 0 | 0 | 23788000 | 35324000 |
| TTEINFK | CRABP1; CRAB | P29762; P29 | RABP1_HUMAN | Unmodified | 2368000 | 0 | 2871800 | 2032600 | 176450 | 0 | 0 | 58643 | 2581100 | 1015000 | 2263900 | 0 | 0 | 1854100 | 2001300 | 2773200 | 0 | 0 |
| TTEINFKVG | CRABP2 | P29373 | RABP2_HUMAN | Unmodified | 0 | 0 | 0 | 0 | 0 | 0 | 0 | 0 | 0 | 0 | 0 | 0 | 0 | 0 | 0 | 0 | 4014600 | 0 |
| TTEINFKVG | CRABP1; CRAB | P29762; B5N | RABP1_HUMAN | Unmodified | 0 | 0 | 0 | 0 | 0 | 0 | 0 | 0 | 817370 | 0 | 0 | 0 | 0 | 0 | 0 | 0 | 0 | 1299200 |
| TTEMETIYD | RUVBL2; RUVBI | Q9Y230; M0 | RUVB2_HUMAN | Unmodified | 0 | 0 | 0 | 0 | 2162400 | 116800 | 211750 | 154720 | 135850 | 0 | 102610 | 2042200 | 693240 | 741520 | 0 | 0 | 248820 | 237960 |
| TTEMETIYD | RUVBL2; RUVBI | Q9Y230; M0 | RUVB2_HUMAN | Oxidation (M) | 0 | 0 | 0 | 0 | 436170 | 0 | 0 | 73655 | 0 | 0 | 0 | 3314800 | 392170 | 0 | 0 | 0 | 0 | 0 |
| TTENIDVSN | TMEM237; TME | Q96Q45; A0A | TM237_HUMAN | Oxidation (M) | 0 | 0 | 69546 | 0 | 0 | 0 | 0 | 0 | 0 | 0 | 0 | 886160 | 0 | 0 | 199270 | 960730 | 0 | 0 |
| TTENNYCP | EFNB1 | P98172 | EFNB1_HUMAN | Unmodified | 133580 | 0 | 79811 | 66098 | 0 | 0 | 0 | 0 | 0 | 0 | 44478 | 434170 | 0 | 0 | 141470 | 176890 | 93127 | 112340 |
| TTEVGSVSE | EPB41L2; EPB4 | O43491; E9F | E41L2_HUMAN | Acetyl (Protein N | 0 | 0 | 0 | 0 | 21453000 | 0 | 1731100 | 3167200 | 0 | 0 | 0 | 0 | 3172400 | 3349800 | 0 | 0 | 11778000 | 13536000 |
| TTEVGSVSE | EPB41L2; EPB4 | O43491; E9F | E41L2_HUMAN | Unmodified | 0 | 0 | 0 | 0 | 0 | 0 | 0 | 0 | 0 | 0 | 0 | 0 | 0 | 0 | 0 | 0 | 312680 | 730180 |
| TTEYLQPNI | SH3GL3 | Q99963 | SH3G3_HUMA | Unmodified | 0 | 0 | 0 | 0 | 0 | 0 | 0 | 0 | 0 | 0 | 0 | 0 | 1558500 | 1408500 | 0 | 0 | 182500 | 40403 |
| TTFDKLETE | MPP7 | Q5T2T1 | MPP7_HUMAN | Unmodified | 0 | 0 | 228180 | 832830 | 12337000 | 508600 | 0 | 0 | 2168500 | 2651400 | 0 | 0 | 3307500 | 1155300 | 649580 | 346600 | 364420 | 1657500 |
| TTFDVYTES | CEL | P19835 | CEL_HUMAN B | Unmodified | 0 | 0 | 0 | 0 | 0 | 0 | 0 | 0 | 11224000 | 2107500 | 14301000 | 1247000 | 0 | 0 | 0 | 0 | 0 | 0 |
| TTFDVYTES | CEL | P19835 | CEL_HUMAN B | Unmodified | 0 | 0 | 0 | 0 | 0 | 0 | 0 | 0 | 2863500 | 167310 | 0 | 0 | 0 | 0 | 0 | 0 | 0 | 0 |
| TTFDVYTES | CEL | P19835 | CEL_HUMAN B | Unmodified | 0 | 0 | 0 | 0 | 0 | 0 | 0 | 0 | 353820 | 2294800 | 0 | 0 | 405860 | 390330 | 0 | 0 | 0 | 0 |
| TTFEDLIQR | SEC31A; SEC31 | O94979; H0Y | SC31A_HUMAN | Unmodified | 0 | 0 | 0 | 0 | 0 | 0 | 458580 | 0 | 0 | 259160 | 0 | 0 | 1689300 | 1979200 | 0 | 0 | 412390 | 1095800 |
| TTFFHHEYI | COPA; COPA; C | P53621; A0A | COPA_HUMAN | Unmodified | 0 | 0 | 0 | 0 | 0 | 0 | 0 | 0 | 0 | 0 | 0 | 2763800 | 0 | 0 | 0 | 0 | 0 | 0 |
| TTFHEASRI | ELAPOR1; ELAP | Q6UXG2; E9 | ELAP1_HUMAN | Unmodified | 0 | 0 | 0 | 0 | 0 | 0 | 212160 | 0 | 0 | 468370 | 0 | 0 | 0 | 0 | 0 | 0 | 0 | 0 |
| TTFIRHLIEC | EHD1; EHD1; E | Q9H4M9; C9 | EHD1_HUMAN | Oxidation (M) | 0 | 0 | 0 | 0 | 1592500 | 596780 | 0 | 0 | 0 | 170490 | 98029 | 229950 | 1350200 | 412190 | 0 | 0 | 1253300 | 1396800 |
| TTFIRHLIEC | EHD1; EHD1; E | Q9H4M9; C9 | EHD1_HUMAN | Unmodified | 0 | 0 | 0 | 0 | 1282700 | 0 | 0 | 0 | 0 | 0 | 0 | 0 | 0 | 0 | 0 | 0 | 460740 | 2556000 |
| TTFIRYLLEC | EHD4; EHD3 | Q9H223; Q9 | EHD4_HUMAN | Oxidation (M) | 0 | 0 | 605060 | 1260700 | 2277800 | 375950 | 1175200 | 0 | 0 | 269290 | 0 | 0 | 1710300 | 9203200 | 510170 | 0 | 62748 | 3057500 |
| TTFIRYLLEC | EHD4; EHD3 | Q9H223; Q9 | EHD4_HUMAN | Unmodified | 32397 | 1387800 | 0 | 0 | 194370 | 0 | 0 | 0 | 0 | 99860 | 0 | 0 | 0 | 83816 | 142730 | 13736 | 0 | 0 |
| TTFLYLVNF | CEMIP2; CEMIP | Q9UHN6; H3 | CEIP2_HUMAN | Unmodified | 0 | 614790 | 1841100 | 0 | 0 | 615350 | 0 | 0 | 2071700 | 0 | 144560 | 422170 | 0 | 0 | 1249400 | 1351600 | 0 | 0 |
| TTFLYLVNF | CEMIP2; CEMIP | Q9UHN6; H3 | CEIP2_HUMAN | Unmodified | 0 | 0 | 0 | 0 | 0 | 0 | 0 | 0 | 1777600 | 404680 | 821110 | 1035700 | 0 | 0 | 0 | 0 | 0 | 0 |
| TTFLYRYTD | RAB27B; RAB27 | O00194; K7E | RB27B_HUMAN | Unmodified | 0 | 0 | 0 | 0 | 2688400 | 0 | 947970 | 222350 | 0 | 0 | 0 | 0 | 444140 | 1136000 | 0 | 0 | 0 | 0 |
| TTFQLQVR | ITGB5; ITGB5; I | P18084; V9C | ITB5_HUMAN In | Unmodified | 0 | 0 | 0 | 0 | 0 | 0 | 0 | 0 | 0 | 0 | 294960 | 4638300 | 0 | 0 | 0 | 0 | 0 | 0 |
| TTFTVAQNI | SULT1A3; SULT | P0DMM9; P5 | ST1A3_HUMAN | Unmodified | 418650 | 465630 | 0 | 724420 | 1699100 | 0 | 0 | 0 | 0 | 0 | 0 | 0 | 497600 | 0 | 429910 | 241320 | 0 | 0 |
| TTFYTALGR | XPO7; XPO7 | Q9UIA9; E7E | XPO7_HUMAN | Unmodified | 51316 | 0 | 0 | 0 | 0 | 0 | 0 | 0 | 0 | 0 | 0 | 1654800 | 0 | 0 | 36638 | 26395 | 0 | 0 |
| TTGARVELS | CNP | P09543 | CN37_HUMAN | Unmodified | 0 | 0 | 0 | 446550 | 0 | 0 | 0 | 0 | 0 | 0 | 9111900 | 4524100 | 0 | 0 | 2327400 | 0 | 0 | 0 |
| TTGARVELS | CNP | P09543 | CN37_HUMAN | Unmodified | 0 | 0 | 0 | 4790400 | 3645500 | 821320 | 0 | 0 | 1351400 | 1555800 | 6714200 | 768110 | 6857200 | 0 | 0 | 0 | 0 | 3692400 |
| TTGASFEK | SCAMP1; SCAM | O15126; A0A | SCAM1_HUMA | Unmodified | 0 | 0 | 773440 | 0 | 107550 | 13976 | 0 | 0 | 0 | 0 | 0 | 2386200 | 0 | 0 | 422850 | 375200 | 0 | 0 |
| TTGDWLNC | EPHA3 | P29320 | EPHA3_HUMAN | Unmodified | 832990 | 0 | 0 | 0 | 0 | 0 | 313600 | 0 | 0 | 0 | 0 | 0 | 18164 | 0 | 0 | 119270 | 0 | 0 |
| TTGEENGV | DNM1L; DNM1L | O00429; F8V | DNM1L_HUMA | Unmodified | 0 | 0 | 0 | 0 | 0 | 0 | 0 | 0 | 0 | 0 | 0 | 677220 | 0 | 0 | 0 | 0 | 0 | 0 |
| TTGEVVSGY | CKAP5; CKAP5 | Q14008; E9F | CKAP5_HUMAN | Unmodified | 61070 | 0 | 0 | 148500 | 0 | 0 | 83807 | 0 | 509350 | 172650 | 299940 | 2673200 | 1029500 | 236780 | 0 | 240900 | 0 | 249980 |
| TTGFGMIYI | RPS24; RPS24; | P62847; A0A | RS24_HUMAN | Oxidation (M) | 1339500 | 1137300 | 667720 | 819670 | 534470 | 60161 | 722120 | 168140 | 0 | 92600 | 788130 | 5683100 | 511090 | 155860 | 1073200 | 1149000 | 67617 | 106270 |
| TTGFGMIYI | RPS24; RPS24; | P62847; A0A | RS24_HUMAN | Oxidation (M) | 0 | 0 | 0 | 0 | 0 | 0 | 0 | 0 | 0 | 0 | 0 | 2002300 | 0 | 0 | 0 | 0 | 0 | 0 |
| TTGFGMIYI | RPS24; RPS24; | P62847; A0A | RS24_HUMAN | Unmodified | 40137 | 56189 | 0 | 118240 | 0 | 0 | 56212 | 0 | 376940 | 0 | 53840 | 310480 | 0 | 0 | 115000 | 73239 | 0 | 0 |
| TTGIIETQFS | GNAT1; GNAT3 | P11488; A8M | GNAT1_HUMA | Unmodified | 0 | 0 | 0 | 163700 | 0 | 306190 | 0 | 0 | 0 | 89442 | 0 | 767890 | 2169200 | 0 | 0 | 0 | 0 | 0 |
| TTGIVETHF | GNAI3; GNAI2; | P08754; P04 | GNAI3_HUMAN | Unmodified | 8526100 | 11345000 | 4736000 | 9813600 | 11030000 | 2545000 | 7351200 | 2231200 | 24155000 | 10278000 | 22567000 | 1.52E+08 | 9460500 | 7069900 | 10173000 | 8926800 | 7291300 | 10057000 |
| TTGIVETHF | GNAI2; GNAI1; | P04899; P63 | GNAI2_HUMAN | Unmodified | 0 | 0 | 0 | 0 | 1850300 | 0 | 2944800 | 94606 | 0 | 1063300 | 0 | 0 | 928520 | 1913200 | 0 | 0 | 6019800 | 3091900 |
| TTGIVETHF | GNAI3 | P08754 | GNAI3_HUMAN | Unmodified | 0 | 0 | 0 | 0 | 1081100 | 0 | 695050 | 73850 | 226890 | 955270 | 0 | 0 | 2111000 | 0 | 0 | 0 | 1940400 | 0 |
| TTGIVMDS | POTEJ; POTEE; | P0CG39; Q6 | POTEJ_HUMAN | Oxidation (M) | 0 | 0 | 0 | 0 | 0 | 0 | 0 | 0 | 0 | 0 | 0 | 1446400 | 0 | 0 | 11671 | 18625 | 0 | 0 |
| TTGIVMDS | ACTG1; ACTB; | P63261; P60 | ACTG_HUMAN | Oxidation (M) | 74135000 | 1.12E+08 | 35407000 | 35950000 | 0 | 0 | 0 | 0 | 24242000 | 3407700 | 12794000 | 4.43E+08 | 0 | 0 | 2.78E+08 | 2.17E+08 | 0 | 0 |
| TTGIVMDS | ACTG1; ACTB; | P63261; P60 | ACTG_HUMAN | Unmodified | 5774500 | 7110100 | 1892300 | 2931200 | 0 | 0 | 0 | 0 | 23410000 | 598330 | 0 | 25769000 | 0 | 0 | 13767000 | 8239400 | 0 | 0 |
| TTGIVMDS | ACTG1; ACTB; | P63261; P60 | ACTG_HUMAN | Oxidation (M) | 0 | 0 | 0 | 0 | 0 | 0 | 0 | 0 | 0 | 0 | 0 | 0 | 0 | 0 | 18974000 | 0 | 0 | 0 |
| TTGKPIEAS | ANXA6 | P08133 | ANXA6_HUMAN | Unmodified | 0 | 0 | 0 | 0 | 0 | 0 | 69511 | 22329 | 936270 | 179830 | 0 | 0 | 0 | 0 | 254740 | 0 | 0 | 0 |
| TTGKPIEAS | ANXA6 | P08133 | ANXA6_HUMAN | Unmodified | 0 | 0 | 8416700 | 1177300 | 7034100 | 927690 | 8979100 | 423170 | 23679000 | 22844000 | 0 | 0 | 2253500 | 9858000 | 1744200 | 832260 | 1940500 | 4641300 |
| TTGPPSGQ | CYB5D2 | Q8WUJ1 | NEUFC_HUMA | Unmodified | 0 | 0 | 0 | 0 | 0 | 0 | 0 | 0 | 749510 | 29680 | 33542 | 0 | 0 | 0 | 0 | 0 | 0 | 0 |
| TTGPSSEG | FAM234B | A2RU67 | F234B_HUMAN | Unmodified | 188940 | 0 | 0 | 0 | 0 | 0 | 0 | 0 | 7757600 | 1310200 | 510230 | 1005200 | 0 | 0 | 0 | 149140 | 0 | 0 |
| TTGQVVAM | CDK1; CDK1; C | P06493; A0A | CDK1_HUMAN | Unmodified | 0 | 0 | 0 | 2478300 | 0 | 784340 | 0 | 462670 | 0 | 0 | 0 | 34517000 | 2005400 | 984120 | 1828900 | 0 | 0 | 1240400 |
| TTGSPENEI | ERMP1; ERMP1 | Q7Z2K6; E7E | ERMP1_HUMA | Unmodified | 0 | 183140 | 0 | 0 | 105900 | 0 | 66715 | 0 | 6888500 | 0 | 0 | 0 | 0 | 0 | 0 | 23237 | 0 | 0 |
| TTGTPPDS | DCTN2; DCTN2 | Q13561; A8K | DCTN2_HUMA | Unmodified | 266020 | 0 | 0 | 0 | 1587500 | 0 | 0 | 0 | 0 | 0 | 0 | 4379900 | 1256300 | 0 | 0 | 491640 | 0 | 0 |
| TTGTPPDS | DCTN2; DCTN2 | Q13561; A8K | DCTN2_HUMA | Unmodified | 0 | 0 | 0 | 0 | 0 | 0 | 902310 | 0 | 0 | 0 | 0 | 0 | 2181400 | 856040 | 0 | 0 | 0 | 0 |
| TTGVVLDS | ACTR1A; ACTR1 | P61163; P42 | ACTZ_HUMAN | 2 Oxidation (M) | 0 | 0 | 0 | 0 | 0 | 0 | 0 | 0 | 0 | 0 | 0 | 8839800 | 0 | 0 | 0 | 0 | 0 | 0 |
| TTHFVEGG | RPL7; RPL7 | P18124; A8M | RL7_HUMAN 60 | Unmodified | 14412000 | 15440000 | 8032500 | 6576600 | 280180 | 161420 | 147550 | 188730 | 2052300 | 267680 | 8099300 | 85898000 | 636890 | 132240 | 9313300 | 6991500 | 348310 | 0 |
| TTHFVEGG | RPL7; RPL7 | P18124; A8M | RL7_HUMAN 60 | Unmodified | 174580 | 58826 | 117750 | 498980 | 3409500 | 373750 | 353220 | 182920 | 0 | 365310 | 226180 | 0 | 3563000 | 619480 | 85856 | 0 | 569670 | 704390 |
| TTHHTTERI | SLITRK6 | Q9H5Y7 | SLIK6_HUMAN | Unmodified | 1286700 | 0 | 0 | 0 | 0 | 0 | 0 | 0 | 1432800 | 0 | 0 | 0 | 0 | 0 | 0 | 0 | 0 | 0 |
| TTHLFVVTT | VPS11; VPS11; | Q9H270; B7. | VPS11_HUMAN | Unmodified | 0 | 31656 | 0 | 39348 | 0 | 0 | 0 | 0 | 0 | 0 | 0 | 370430 | 0 | 0 | 74398 | 56537 | 0 | 0 |
| TTHLHNDL | PLOD1 | Q02809 | PLOD1_HUMA | Unmodified | 0 | 756870 | 0 | 0 | 0 | 0 | 0 | 0 | 1713500 | 0 | 0 | 0 | 0 | 0 | 0 | 0 | 0 | 0 |
| TTHLHNDL | PLOD1 | Q02809 | PLOD1_HUMA | Unmodified | 0 | 0 | 0 | 0 | 0 | 113200 | 0 | 0 | 3423400 | 290610 | 0 | 0 | 0 | 0 | 0 | 0 | 0 | 0 |
| TTHLIAKEE | PSMD12 | O00232 | PSD12_HUMAN | Unmodified | 0 | 0 | 0 | 0 | 1443900 | 66418 | 926580 | 0 | 0 | 0 | 0 | 4185100 | 0 | 179310 | 0 | 0 | 186140 | 742590 |
| TTHNSAFE | IGSF3 | O75054 | IGSF3_HUMAN | Unmodified | 0 | 1731000 | 0 | 0 | 0 | 0 | 0 | 0 | 0 | 0 | 0 | 1559400 | 0 | 0 | 0 | 0 | 0 | 0 |
| TTHNSAFE | IGSF3 | O75054 | IGSF3_HUMAN | Unmodified | 1416800 | 1442500 | 507030 | 956790 | 149130 | 0 | 0 | 0 | 0 | 0 | 0 | 1680000 | 300880 | 0 | 2364100 | 0 | 0 | 0 |
| TTHVVNLG | NUDCD1 | Q96RS6 | NUDC1_HUMA | Unmodified | 0 | 0 | 0 | 0 | 0 | 0 | 0 | 0 | 0 | 0 | 0 | 0 | 0 | 0 | 0 | 0 | 0 | 0 |
| TTHYTPLAC | DKC1; DKC1; D | O60832; H7 | DKC1_HUMAN | Unmodified | 0 | 0 | 0 | 0 | 0 | 0 | 0 | 0 | 0 | 0 | 0 | 519010 | 0 | 0 | 0 | 0 | 0 | 0 |
| TTIEAIHGLI | COPS5 | Q92905 | CSN5_HUMAN | Unmodified | 0 | 0 | 106930 | 226460 | 706480 | 92040 | 94041 | 0 | 944100 | 259120 | 820590 | 2111800 | 1080600 | 273600 | 236990 | 339080 | 0 | 1066900 |
| TTIEAIHGLI | COPS5 | Q92905 | CSN5_HUMAN | Unmodified | 0 | 0 | 0 | 0 | 0 | 0 | 639170 | 0 | 0 | 0 | 0 | 0 | 0 | 0 | 0 | 0 | 538780 | 3072900 |
| TTIENIQLPI | MCM4; MCM4; | P33991; A0A | MCM4_HUMAN | Unmodified | 64039 | 0 | 0 | 0 | 0 | 0 | 0 | 34400 | 0 | 0 | 0 | 931890 | 0 | 0 | 78575 | 0 | 0 | 0 |
| TTIFSPEGR | PSMA4; PSMA4 | P25789; H0Y | PSA4_HUMAN | Unmodified | 0 | 0 | 796210 | 2213800 | 1430800 | 222400 | 46173 | 37129 | 5296800 | 1226200 | 4073400 | 12510000 | 779150 | 0 | 733690 | 623090 | 420860 | 0 |
| TTIFTDAK | ELOB; ELOB; EL | Q15370; A0A | ELOB_HUMAN | Unmodified | 219060 | 334230 | 679320 | 470810 | 5589600 | 1097800 | 0 | 393680 | 1780600 | 567380 | 1632600 | 7264200 | 4063200 | 1401200 | 557110 | 970600 | 1084000 | 2562400 |
| TTIFTDAKE | ELOB; ELOB; EL | Q15370; A0A | ELOB_HUMAN | Unmodified | 0 | 0 | 0 | 0 | 6618500 | 1964600 | 94587 | 1112600 | 0 | 1414100 | 0 | 0 | 7506800 | 6638900 | 0 | 0 | 5613400 | 4325700 |
| TTIFTDAKE | ELOB; ELOB; EL | Q15370; A0A | ELOB_HUMAN | Unmodified | 0 | 0 | 0 | 322940 | 883980 | 360120 | 410340 | 0 | 0 | 0 | 0 | 0 | 381320 | 423050 | 0 | 0 | 0 | 688950 |
| TTIIWDAHT | TBL1Y; TBL1XR | Q9BQ87; Q9 | TBL1Y_HUMAN | Unmodified | 0 | 0 | 0 | 0 | 0 | 0 | 0 | 368000 | 214030 | 137750 | 400160 | 1014600 | 0 | 0 | 0 | 0 | 0 | 0 |
| TTINNEYNY | IGF1R; IGF1R; I | P08069; H0Y | IGF1R_HUMAN | Unmodified | 0 | 0 | 71038 | 0 | 1582700 | 0 | 0 | 0 | 0 | 0 | 0 | 0 | 0 | 0 | 0 | 0 | 0 | 0 |
| TTISVAHLL | FAM98A; FAM98 | Q8NCA5; E9 | FA98A_HUMAN | Unmodified | 0 | 0 | 0 | 0 | 0 | 0 | 0 | 0 | 0 | 0 | 0 | 1401500 | 0 | 0 | 0 | 0 | 0 | 0 |
| TTISVAHLL | FAM98A; FAM98 | Q8NCA5; E9 | FA98A_HUMAN | Unmodified | 0 | 0 | 0 | 0 | 3569000 | 295330 | 2632600 | 138220 | 0 | 0 | 464550 | 622070 | 5501200 | 1024900 | 0 | 0 | 251500 | 2293800 |
| TTITMAHLL | FAM98B | Q52LJ0 | FA98B_HUMAN | Unmodified | 0 | 0 | 0 | 0 | 1099000 | 0 | 490440 | 0 | 0 | 229430 | 0 | 0 | 1188800 | 628780 | 0 | 0 | 391560 | 988430 |
| TTITTTTSS | MYADM; MYADI | Q96S97; A0A | MYADM_HUMA | Oxidation (M) | 7037600 | 9760600 | 10713000 | 8958100 | 0 | 0 | 0 | 322350 | 794410 | 340350 | 0 | 0 | 470620 | 0 | 9407000 | 13639000 | 0 | 0 |
| TTKADIQMI | ALCAM; ALCAM | Q13740; F5C | CD166_HUMAN | Unmodified | 0 | 0 | 0 | 0 | 3992600 | 0 | 581800 | 270200 | 0 | 737790 | 0 | 0 | 0 | 605240 | 0 | 0 | 884700 | 6374600 |
| TTKADIQMI | ALCAM; ALCAM | Q13740; F5C | CD166_HUMAN | Oxidation (M) | 182040 | 381480 | 0 | 481080 | 833220 | 0 | 0 | 0 | 0 | 399830 | 0 | 0 | 0 | 0 | 0 | 0 | 536840 | 1421000 |
| TTKPYPADI | CNTN1 | Q12860 | CNTN1_HUMA | Unmodified | 426300 | 461250 | 0 | 0 | 0 | 0 | 0 | 0 | 822470 | 0 | 0 | 0 | 0 | 0 | 0 | 0 | 0 | 0 |
| TTKQEPMLI | RO60; RO60 | P10155; H0Y | RO60_HUMAN | Oxidation (M) | 2645800 | 0 | 0 | 0 | 0 | 0 | 0 | 0 | 0 | 0 | 0 | 0 | 0 | 0 | 0 | 0 | 0 | 0 |
| TTLKEDTVT | UEVLD | Q8IX04 | UEVLD_HUMAN | Unmodified | 38681 | 0 | 0 | 239000 | 4666600 | 286610 | 0 | 0 | 1286100 | 686930 | 200680 | 986650 | 2457500 | 456990 | 23532 | 31415 | 39998 | 0 |
| TTLLHMLKI | SAR1B; SAR1A; | Q9Y6B6; Q9 | SAR1B_HUMAN | Unmodified | 0 | 0 | 0 | 0 | 0 | 0 | 0 | 0 | 0 | 0 | 0 | 817410 | 0 | 0 | 0 | 0 | 0 | 0 |
| TTLLKMAD | NDRG1; NDRG | Q92597; E7E | NDRG1_HUMA | Unmodified | 6918900 | 8193700 | 12194000 | 12260000 | 0 | 0 | 0 | 0 | 0 | 0 | 0 | 0 | 1151500 | 249460 | 7284900 | 3486800 | 0 | 0 |
| TTLLNYILTE | CBWD2; CBWD | Q8IUF1; Q4V | CBWD2_HUMA | Unmodified | 0 | 0 | 0 | 0 | 0 | 0 | 0 | 0 | 167970 | 275640 | 0 | 1068500 | 1530100 | 1074500 | 0 | 0 | 0 | 0 |
| TTLLPGLLK | FARSB | Q9NSD9 | SYFB_HUMAN | Unmodified | 76735 | 89764 | 229300 | 131630 | 0 | 0 | 220780 | 273810 | 0 | 584680 | 1129600 | 11127000 | 1667900 | 318190 | 150370 | 599530 | 0 | 0 |
| TTLPGVVN | SMPDL3B; SMP | Q92485; F8V | ASM3B_HUMA | Unmodified | 3020800 | 1726300 | 9612600 | 8061400 | 0 | 0 | 0 | 0 | 3809700 | 0 | 738800 | 6580600 | 276200 | 0 | 11219000 | 10480000 | 0 | 0 |
| TTLPTFQSF | SNX5; SNX5 | Q9Y5X3; Q5 | SNX5_HUMAN | Unmodified | 87195 | 0 | 68748 | 384870 | 0 | 0 | 0 | 0 | 139260 | 0 | 0 | 2398600 | 0 | 48909 | 188020 | 129280 | 0 | 0 |
| TTLQQEKPI | TFPI | P10646 | TFPI1_HUMAN | Unmodified | 0 | 261000 | 966560 | 2518400 | 0 | 0 | 0 | 0 | 325810 | 0 | 0 | 0 | 0 | 0 | 0 | 576040 | 0 | 0 |
| TTLQVDTLN | MYH10 | P35580 | MYH10_HUMA | Unmodified | 0 | 0 | 193630 | 410780 | 0 | 0 | 198400 | 0 | 0 | 0 | 153940 | 4671700 | 0 | 0 | 3229500 | 746820 | 0 | 0 |
| TTLQVDTLN | MYH10 | P35580 | MYH10_HUMA | Unmodified | 0 | 0 | 0 | 433700 | 11678000 | 2043300 | 1184500 | 426250 | 0 | 0 | 0 | 0 | 3928900 | 2794900 | 0 | 0 | 0 | 0 |
| TTLRYSVQL | RUVBL1; RUVBI | Q9Y265; H7 | RUVB1_HUMAN | Unmodified | 0 | 0 | 0 | 0 | 1012500 | 0 | 535240 | 0 | 0 | 196840 | 0 | 0 | 1824000 | 0 | 0 | 0 | 1353000 | 1114100 |

| | | | | | | | | | | | | | | | | | | | | | | |
|---|---|---|---|---|---|---|---|---|---|---|---|---|---|---|---|---|---|---|---|---|---|---|
| TTLSLPAPA | PLPP3; N/A; N/ | O14495; A0A | PLPP3_HUMAN | Unmodified | 0 | 0 | 0 | 117100 | 546240 | 0 | 0 | 0 | 0 | 0 | 0 | 331440 | 0 | 0 | 0 | 0 | 0 | 0 |
| TTLTAAITK | TUFM; TUFM | P49411; H3E | EFTU_HUMAN | Unmodified | 160230 | 0 | 0 | 0 | 0 | 0 | 673720 | 215960 | 6775500 | 910200 | 167300 | 0 | 0 | 250530 | 0 | 0 | 90289 | 412630 |
| TTLTSDESV | SEL1L; SEL1L | Q9UBV2; G3 | SE1L1_HUMAN | Unmodified | 0 | 0 | 0 | 0 | 0 | 0 | 89720 | 18131 | 12300000 | 176570 | 447200 | 307390 | 0 | 0 | 0 | 0 | 0 | 0 |
| TTLVYGIFT | SEMA3C | Q99985 | SEM3C_HUMAI | Unmodified | 0 | 0 | 217900 | 226020 | 0 | 0 | 0 | 0 | 0 | 0 | 0 | 0 | 0 | 0 | 0 | 125600 | 0 | 0 |
| TTMRDLSQ | STXBP1; STXBP | P61764; A0A | STXB1_HUMAN | Oxidation (M) | 0 | 0 | 0 | 0 | 670550 | 0 | 482870 | 758590 | 0 | 119280 | 0 | 0 | 410410 | 237840 | 0 | 0 | 102600 | 203400 |
| TTMRDLSQ | STXBP1; STXBP | P61764; A0A | STXB1_HUMAN | Unmodified | 0 | 0 | 0 | 0 | 1887900 | 141620 | 5252100 | 8764.3 | 713220 | 953800 | 0 | 0 | 690870 | 575770 | 0 | 0 | 579140 | 1100500 |
| TTNAGPLH | EBP; EBP; N/A; | Q15125; C9J | EBP_HUMAN 3 | Unmodified | 0 | 0 | 0 | 0 | 0 | 0 | 0 | 0 | 3174800 | 591250 | 0 | 0 | 0 | 0 | 0 | 0 | 0 | 0 |
| TTNAGPLH | EBP; EBP; N/A; | Q15125; C9J | EBP_HUMAN 3 | Acetyl (Protein N | 310230 | 0 | 644390 | 157100 | 5796700 | 642060 | 0 | 0 | 72469000 | 11411000 | 18926000 | 70547000 | 0 | 0 | 392920 | 225070 | 0 | 815240 |
| TTNFAGILS | PARP1; PARP1 | P09874; A0A | PARP1_HUMAN | Unmodified | 2299800 | 1225200 | 0 | 169150 | 0 | 0 | 0 | 0 | 1878400 | 981460 | 1741900 | 7291200 | 0 | 0 | 0 | 0 | 0 | 0 |
| TTNKFEGLS | ASAP1; ASAP1; | Q9ULH1; H0 | ASAP1_HUMAN | Unmodified | 0 | 0 | 0 | 52549 | 161910 | 0 | 0 | 0 | 50268 | 48257 | 0 | 0 | 0 | 0 | 0 | 0 | 297650 | 982890 |
| TTNLSVEEC | GALNT3; GALN | Q14435; C9J | GALT3_HUMAN | Unmodified | 195250 | 63194 | 0 | 0 | 0 | 0 | 0 | 0 | 1019800 | 67566 | 179430 | 463090 | 261260 | 0 | 0 | 0 | 0 | 0 |
| TTNQSTEFI | SLITRK6 | Q9H5Y7 | SLIK6_HUMAN | Unmodified | 4814200 | 0 | 0 | 0 | 0 | 0 | 0 | 0 | 4541100 | 1439000 | 0 | 0 | 0 | 0 | 0 | 0 | 0 | 0 |
| TTNQSTEFI | SLITRK6 | Q9H5Y7 | SLIK6_HUMAN | Unmodified | 0 | 0 | 0 | 0 | 0 | 0 | 0 | 0 | 0 | 1934400 | 0 | 0 | 0 | 0 | 0 | 0 | 0 | 752700 |
| TTNSSLEEI | VPS13C | Q709C8 | VP13C_HUMAN | Unmodified | 0 | 0 | 0 | 0 | 0 | 936190 | 0 | 48929 | 0 | 0 | 0 | 1269500 | 0 | 0 | 0 | 0 | 0 | 0 |
| TTNTVEPLF | LY6D | Q14210 | LY6D_HUMAN | Unmodified | 0 | 0 | 0 | 0 | 0 | 308750 | 1095400 | 431660 | 0 | 0 | 0 | 0 | 0 | 0 | 0 | 0 | 48920 | 0 |
| TTNTVEPLF | LY6D | Q14210 | LY6D_HUMAN | Unmodified | 0 | 0 | 0 | 0 | 0 | 0 | 6725200 | 3891100 | 0 | 0 | 0 | 0 | 0 | 0 | 0 | 0 | 0 | 0 |
| TTNVLGAVI | PDS5B; PDS5B | Q9NTI5; A9I | PDS5B_HUMAI | Unmodified | 0 | 0 | 0 | 0 | 0 | 0 | 0 | 633210 | 0 | 0 | 0 | 0 | 0 | 0 | 0 | 0 | 0 | 0 |
| TTPDVIFVF | RPS24; RPS24; | P62847; A0A | RS24_HUMAN | Unmodified | 198630 | 144350 | 0 | 171570 | 0 | 0 | 0 | 0 | 0 | 0 | 0 | 1991500 | 178910 | 0 | 0 | 107750 | 0 | 0 |
| TTPDVIFVF | RPS24; RPS24; | P62847; A0A | RS24_HUMAN | Unmodified | 0 | 0 | 0 | 0 | 0 | 0 | 809970 | 0 | 0 | 0 | 0 | 0 | 0 | 0 | 0 | 0 | 0 | 225880 |
| TTPECGPT | JTB | O76095 | JTB_HUMAN Pr | Unmodified | 63631 | 0 | 0 | 0 | 1246400 | 119690 | 141530 | 164390 | 7228800 | 1035400 | 278430 | 590900 | 226140 | 0 | 0 | 0 | 0 | 390740 |
| TTPEYVFH | DYNC1I2 | Q13409 | DC1I2_HUMAN | Oxidation (M) | 0 | 0 | 0 | 0 | 0 | 108540 | 308110 | 77542 | 0 | 388280 | 133680 | 2252600 | 0 | 0 | 0 | 0 | 146260 | 76662 |
| TTPGKENFI | KPNA1; KPNA1; | P52294; C9J | IMA5_HUMAN I | Acetyl (Protein N | 0 | 116840 | 0 | 45614 | 952740 | 663760 | 459430 | 465740 | 0 | 158910 | 0 | 0 | 1114000 | 1457000 | 0 | 0 | 2814800 | 1919100 |
| TTPLKPSPL | HUWE1 | Q7Z6Z7 | HUWE1_HUMA | Unmodified | 0 | 0 | 0 | 0 | 0 | 0 | 0 | 0 | 346390 | 0 | 0 | 1601900 | 0 | 0 | 0 | 0 | 0 | 0 |
| TTPPPGRPI | LRATD2 | Q96KN1 | LRAT2_HUMAN | Unmodified | 0 | 0 | 137150 | 305240 | 308920 | 0 | 0 | 0 | 812990 | 363210 | 0 | 0 | 389650 | 181690 | 198380 | 0 | 791810 | 1013500 |
| TTPPSVFFP | FMNL2; FMNL3 | Q96PY5; Q8I | FMNL2_HUMAI | Unmodified | 0 | 0 | 115130 | 0 | 2623000 | 0 | 0 | 0 | 1410200 | 745970 | 1739100 | 4760800 | 0 | 0 | 0 | 0 | 0 | 1371700 |
| TTPPSVFFP | FMNL2; FMNL2 | Q96PY5; C9I | FMNL2_HUMAI | Unmodified | 0 | 0 | 0 | 0 | 896370 | 335290 | 0 | 0 | 0 | 67421 | 0 | 0 | 0 | 0 | 0 | 0 | 613090 | 473480 |
| TTPPVLDSI | IGHG1; IGHG1; | P01857; A0A | IGHG1_HUMAI | Unmodified | 2061900 | 4979600 | 4884100 | 2909500 | 0 | 0 | 0 | 0 | 0 | 0 | 0 | 0 | 749410 | 0 | 3340900 | 3147700 | 0 | 0 |
| TTPSVVAFT | HSPA9; HSPA9; | P38646; H0Y | GRP75_HUMAI | Unmodified | 60546 | 23431 | 0 | 0 | 0 | 0 | 0 | 90959 | 4992100 | 0 | 0 | 195740 | 0 | 0 | 0 | 0 | 0 | 0 |
| TTPSVVAFT | HSPA9; HSPA9; | P38646; H0Y | GRP75_HUMAI | Unmodified | 0 | 0 | 0 | 0 | 107370 | 0 | 0 | 0 | 15138000 | 677190 | 0 | 770070 | 0 | 0 | 0 | 0 | 0 | 0 |
| TTPSVVAFT | HSPA9; HSPA9; | P38646; H0Y | GRP75_HUMAI | Oxidation (M) | 0 | 0 | 0 | 0 | 96544 | 0 | 0 | 0 | 6954300 | 1757900 | 225430 | 0 | 376720 | 0 | 55214 | 0 | 0 | 0 |
| TTPSYVAFT | HSPA1L; HSPA( | P34931; P17 | HS71L_HUMAN | Unmodified | 4217200 | 21457000 | 13341000 | 8793800 | 0 | 0 | 0 | 0 | 20841000 | 324300 | 0 | 1.02E+08 | 193140 | 68439 | 21994000 | 20360000 | 0 | 0 |
| TTPSYVAFT | HSPA1L; HSPA1 | P34931; P0D | HS71L_HUMAN | Unmodified | 78545000 | 86104000 | 65945000 | 77356000 | 23572000 | 3841200 | 2048300 | 0 | 1.43E+08 | 11957000 | 20938000 | 1.09E+08 | 30642000 | 9567900 | 53666000 | 49080000 | 2490400 | 11105000 |
| TTPTPSTNS | MARK2; MARK2 | Q7KZI7; E7E | MARK2_HUMAI | Unmodified | 3175900 | 0 | 203930 | 0 | 151100 | 0 | 0 | 0 | 327470 | 0 | 739970 | 4333800 | 413320 | 0 | 301410 | 208420 | 0 | 152410 |
| TTPYQIACC | TARS1; TARS1 | P26639; D6F | SYTC_HUMAN | Unmodified | 0 | 0 | 0 | 32633 | 2963600 | 163790 | 164750 | 55894 | 1531200 | 0 | 1826500 | 4911800 | 905060 | 0 | 0 | 1012000 | 0 | 1002700 |
| TTQAAAIIDI | CDC42BPA; CD | Q5VT25; A0A | MRCKA_HUMA | Unmodified | 0 | 0 | 0 | 0 | 93777 | 0 | 0 | 0 | 0 | 0 | 0 | 1672500 | 0 | 0 | 440600 | 419690 | 14507 | 0 |
| TTQDQLEK | PKIB; PKIB | Q9C010; Q5 | IPKB_HUMAN ( | Unmodified | 0 | 0 | 0 | 0 | 0 | 0 | 0 | 100410 | 0 | 0 | 0 | 0 | 1181800 | 1388100 | 0 | 0 | 429760 | 3256200 |
| TTQETIDKT | ADIRF | Q15847 | ADIRF_HUMAN | Unmodified | 0 | 0 | 0 | 0 | 0 | 0 | 1638000 | 87226 | 0 | 550610 | 0 | 0 | 0 | 0 | 0 | 0 | 8077000 | 11225000 |
| TTQETIDKT | ADIRF | Q15847 | ADIRF_HUMAN | Unmodified | 0 | 0 | 0 | 0 | 0 | 0 | 0 | 0 | 0 | 0 | 0 | 0 | 0 | 0 | 0 | 0 | 1417400 | 1454100 |
| TTQFSCTL( | FABP5; FABP5 | Q01469; I6L | FABP5_HUMAN | Unmodified | 0 | 0 | 0 | 0 | 332550 | 0 | 0 | 0 | 0 | 0 | 890920 | 4792600 | 0 | 0 | 0 | 0 | 0 | 587980 |
| TTQFSCTL( | FABP5; FABP5 | Q01469; I6L | FABP5_HUMAN | Unmodified | 0 | 0 | 0 | 0 | 0 | 0 | 57075 | 53147 | 0 | 0 | 4977700 | 1699700 | 0 | 0 | 0 | 0 | 1205400 | 733130 |
| TTQFSCTL( | FABP5; FABP5 | Q01469; I6L | FABP5_HUMAN | Unmodified | 2733700 | 0 | 0 | 0 | 23909000 | 7854900 | 813360 | 672360 | 1123600 | 672980 | 18531000 | 43314000 | 0 | 0 | 0 | 0 | 44447000 | 94137000 |
| TTQIGCLLF | AP2A2; AP2A2 | O94973; A0A | AP2A2_HUMAN | Unmodified | 0 | 0 | 0 | 0 | 0 | 0 | 0 | 0 | 398070 | 0 | 0 | 1734700 | 0 | 0 | 141410 | 127590 | 0 | 0 |
| TTQKWEMI | GALNT7; GALN | Q86SF2; H0Y | GALT7_HUMAN | Unmodified | 0 | 0 | 0 | 140580 | 0 | 155370 | 119140 | 82892 | 0 | 1288300 | 332360 | 518630 | 256200 | 134910 | 43077 | 0 | 0 | 262900 |
| TTQLGPGR | SKOR1; DNAJB1 | P84550; Q9U | SKOR1_HUMAI | Unmodified | 0 | 0 | 0 | 0 | 0 | 0 | 0 | 0 | 8790000 | 51431 | 93508 | 1697500 | 0 | 0 | 0 | 162410 | 0 | 0 |
| TTQLGPGR | DNAJB11 | Q9UBS4 | DJB11_HUMAN | Oxidation (M) | 0 | 0 | 0 | 0 | 0 | 0 | 0 | 0 | 1398600 | 491200 | 0 | 0 | 0 | 0 | 0 | 0 | 0 | 0 |
| TTQLGPGR | DNAJB11 | Q9UBS4 | DJB11_HUMAN | Unmodified | 0 | 0 | 0 | 0 | 0 | 0 | 0 | 0 | 15834000 | 815530 | 0 | 0 | 0 | 0 | 0 | 0 | 0 | 0 |
| TTQLLFLK | UFL1 | O94874 | UFL1_HUMAN | Unmodified | 0 | 0 | 0 | 0 | 313190 | 0 | 0 | 95447 | 0 | 121760 | 0 | 377560 | 0 | 0 | 0 | 0 | 0 | 0 |
| TTQNRYSF | PKP1 | Q13835 | PKP1_HUMAN | Unmodified | 0 | 0 | 0 | 0 | 0 | 0 | 0 | 0 | 0 | 0 | 0 | 0 | 0 | 0 | 0 | 0 | 1581700 | 1650300 |
| TTQQIDLQ( | PDLIM1 | O00151 | PDLI1_HUMAN | Acetyl (Protein N | 34199 | 0 | 39092 | 137400 | 398370 | 128490 | 0 | 0 | 699280 | 144320 | 68440 | 0 | 67771 | 59778 | 0 | 80046 | 29208 | 494120 |
| TTQQIDLQ( | PDLIM1 | O00151 | PDLI1_HUMAN | Acetyl (Protein N | 0 | 0 | 0 | 0 | 485840 | 0 | 0 | 0 | 0 | 102310 | 0 | 0 | 0 | 0 | 0 | 0 | 667840 | 631880 |
| TTQQSPED | CAMP; CAMP | P49913; J3K | CAMP_HUMAN | Unmodified | 0 | 0 | 0 | 0 | 0 | 0 | 0 | 0 | 4941800 | 287900 | 0 | 0 | 0 | 0 | 0 | 0 | 0 | 0 |
| TTQVPQFIL | DHX9 | Q08211 | DHX9_HUMAN | Unmodified | 346890 | 162410 | 72539 | 67709 | 687320 | 0 | 647640 | 77436 | 416180 | 383660 | 384590 | 2397700 | 0 | 0 | 186370 | 73926 | 0 | 610050 |
| TTQVTQFIL | DHX36; DHX36 | Q9H2U1; E7 | DHX36_HUMAI | Unmodified | 1693400 | 1358100 | 0 | 253630 | 0 | 0 | 0 | 0 | 0 | 0 | 0 | 1053100 | 0 | 0 | 952590 | 0 | 0 | 0 |
| TTQWHRPS | NEDD4L; NEDD | Q96PU5; A0 | NED4L_HUMAI | Unmodified | 1190300 | 2721000 | 583550 | 933390 | 0 | 0 | 0 | 0 | 271950 | 0 | 0 | 719160 | 0 | 0 | 1808300 | 845150 | 0 | 0 |
| TTQWHRPS | NEDD4L; NEDD | Q96PU5; A0 | NED4L_HUMAI | Oxidation (M) | 1599200 | 0 | 1592000 | 1034300 | 0 | 0 | 0 | 0 | 0 | 0 | 0 | 643730 | 0 | 0 | 2540800 | 0 | 0 | 0 |
| TTRFASFPD | USP5 | P45974 | UBP5_HUMAN | Unmodified | 0 | 0 | 0 | 0 | 2626500 | 487210 | 1836900 | 0 | 0 | 457870 | 0 | 0 | 3662700 | 2766400 | 0 | 0 | 4031000 | 1864700 |
| TTRFLAEEG | STT3A; STT3A; S | P46977; E9P | STT3A_HUMAN | Unmodified | 0 | 0 | 0 | 0 | 0 | 0 | 0 | 0 | 1374800 | 0 | 0 | 0 | 0 | 0 | 0 | 0 | 0 | 0 |
| TTRTPDTST | MAP1B | P46821 | MAP1B_HUMAI | Unmodified | 0 | 0 | 0 | 0 | 135470 | 0 | 287110 | 185560 | 0 | 112820 | 0 | 0 | 270950 | 202420 | 0 | 0 | 133540 | 196760 |
| TTRTPEEG( | MAP1B | P46821 | MAP1B_HUMAI | Unmodified | 0 | 0 | 0 | 0 | 94062 | 0 | 221820 | 101550 | 0 | 0 | 0 | 0 | 193750 | 221930 | 0 | 0 | 58454 | 133090 |
| TTRTSDVG | MAP1B | P46821 | MAP1B_HUMAI | Unmodified | 58332 | 0 | 0 | 0 | 106880 | 0 | 313620 | 165200 | 40386 | 72106 | 0 | 0 | 193270 | 144430 | 0 | 0 | 130230 | 159640 |
| TTRYEFEQ( | NADK2 | Q4G0N4 | NAKD2_HUMAI | Unmodified | 0 | 0 | 0 | 0 | 0 | 0 | 0 | 0 | 901520 | 176560 | 0 | 0 | 0 | 0 | 0 | 0 | 0 | 0 |
| TTSAEKNT( | ADAM10 | O14672 | ADA10_HUMAN | Unmodified | 0 | 0 | 0 | 0 | 1238900 | 0 | 1439400 | 0 | 0 | 0 | 0 | 0 | 950310 | 275320 | 0 | 0 | 1058500 | 496460 |
| TTSASSHL | GLUL; GLUL | P15104; A0A | GLNA_HUMAN | Acetyl (Protein N | 0 | 0 | 0 | 0 | 100780 | 0 | 0 | 154520 | 0 | 58945 | 0 | 0 | 0 | 18481 | 0 | 0 | 282400 | 127720 |
| TTSFAESC | GSK3B; GSK3B; | P49841; A0A | GSK3B_HUMAI | Unmodified | 0 | 0 | 0 | 0 | 0 | 0 | 62637 | 0 | 0 | 0 | 0 | 0 | 0 | 0 | 0 | 0 | 0 | 435640 |
| TTSGALFPS | ADRM1; ADRM1 | Q16186; A0A | ADRM1_HUMA | Acetyl (Protein N | 0 | 0 | 0 | 0 | 2250200 | 293600 | 0 | 904980 | 0 | 50880 | 0 | 0 | 4644800 | 50573 | 0 | 0 | 1292900 | 1647200 |
| TTSHMNGI | PALS1; PALS1 | Q8N3R9; G3 | PALS1_HUMAN | Acetyl (Protein N | 0 | 0 | 0 | 0 | 393960 | 92526 | 0 | 0 | 0 | 0 | 0 | 0 | 0 | 0 | 0 | 0 | 138140 | 290530 |
| TTSHTELVF | SEZ6L; SEZ6L; S | Q9BYH1; B0 | SE6L1_HUMAN | Unmodified | 0 | 0 | 0 | 0 | 0 | 0 | 0 | 0 | 1062400 | 0 | 0 | 0 | 0 | 0 | 0 | 0 | 0 | 0 |
| TTSNTLHYI | HUWE1 | Q7Z6Z7 | HUWE1_HUMA | Unmodified | 0 | 0 | 0 | 0 | 0 | 0 | 0 | 0 | 0 | 0 | 0 | 4795500 | 0 | 0 | 0 | 0 | 0 | 0 |
| TTSPGLFFS | FMNL1; FMNL1 | O95466; K7E | FMNL1_HUMAI | Unmodified | 0 | 0 | 0 | 0 | 580700 | 84729 | 0 | 0 | 0 | 0 | 0 | 0 | 0 | 0 | 0 | 0 | 0 | 38017 |
| TTSSISPHF | PHLDB2; PHLD | Q86SQ0; A0 | PHLB2_HUMAN | Unmodified | 0 | 0 | 0 | 0 | 570980 | 0 | 0 | 31998 | 0 | 0 | 0 | 0 | 0 | 76679 | 0 | 0 | 131810 | 284760 |
| TTSSTDGPI | CNOT2; CNOT2 | Q9NZN8; H0 | CNOT2_HUMA | Unmodified | 0 | 0 | 0 | 67882 | 647090 | 0 | 0 | 0 | 0 | 68537 | 0 | 1066200 | 147630 | 63870 | 0 | 0 | 44376 | 181130 |
| TTSTSLHV | PTPRB; PTPRB; | P23467; F8V | PTPRB_HUMAN | Unmodified | 508030 | 0 | 0 | 0 | 0 | 0 | 0 | 0 | 0 | 0 | 0 | 0 | 0 | 0 | 0 | 0 | 0 | 0 |
| TTTESEVMI | MTHFD1; MTHF | P11586; V9C | C1TC_HUMAN | Unmodified | 101200 | 0 | 294980 | 586050 | 2455900 | 0 | 0 | 75888 | 3865000 | 532650 | 647600 | 6338900 | 477760 | 108500 | 620370 | 400870 | 0 | 558410 |
| TTTESEVMI | MTHFD1; MTHF | P11586; V9C | C1TC_HUMAN | Oxidation (M) | 16646 | 0 | 0 | 0 | 58768 | 0 | 16875 | 0 | 69041 | 0 | 83921 | 328850 | 105810 | 0 | 0 | 40838 | 20562 | 56774 |
| TTTFKDPRF | WWP2 | O00308 | WWP2_HUMAN | Unmodified | 1372100 | 1467100 | 667400 | 481370 | 0 | 0 | 0 | 0 | 101780 | 0 | 0 | 167440 | 0 | 0 | 1384400 | 1390300 | 0 | 0 |
| TTTGDVQV | IGF2R; IGF2R | P11717; S4F | MPRI_HUMAN | Unmodified | 130740 | 0 | 558210 | 251000 | 0 | 0 | 0 | 0 | 16638000 | 2013400 | 1712300 | 2419700 | 200850 | 0 | 343940 | 278430 | 0 | 0 |
| TTTGDVQV | IGF2R; IGF2R | P11717; S4F | MPRI_HUMAN | Unmodified | 0 | 0 | 0 | 0 | 0 | 0 | 0 | 0 | 278710 | 396500 | 0 | 0 | 0 | 0 | 0 | 0 | 0 | 0 |
| TTTGNKVF | RPL5; RPL5; RP | P46777; Q5T | RL5_HUMAN 60 | Unmodified | 0 | 0 | 0 | 0 | 251620 | 0 | 347380 | 184330 | 0 | 215850 | 0 | 0 | 2026900 | 0 | 0 | 0 | 503700 | 713020 |
| TTTGSYIAN | PSMB6; PSMB6 | P28072; I3L3 | PSB6_HUMAN | Unmodified | 84849 | 80578 | 385820 | 420060 | 0 | 73858 | 207980 | 216180 | 924390 | 294520 | 361350 | 4743300 | 0 | 0 | 295070 | 457130 | 0 | 0 |
| TTTGSYIAN | PSMB6; PSMB6 | P28072; I3L3 | PSB6_HUMAN | Unmodified | 0 | 0 | 718880 | 0 | 0 | 0 | 156780 | 52146 | 1088300 | 0 | 634560 | 2983100 | 148840 | 0 | 0 | 593150 | 0 | 0 |
| TTTGSYIAN | PSMB6; PSMB6 | P28072; I3L3 | PSB6_HUMAN | Unmodified | 0 | 0 | 0 | 0 | 0 | 0 | 0 | 0 | 1778600 | 92671 | 0 | 4953000 | 0 | 0 | 0 | 0 | 0 | 0 |
| TTTHVPPEL | ZMPSTE24; ZMF | O75844; A0A | FACE1_HUMAN | Unmodified | 0 | 0 | 0 | 0 | 0 | 0 | 0 | 0 | 9494000 | 0 | 0 | 241380 | 0 | 0 | 0 | 0 | 0 | 0 |
| TTTHVPPEL | ZMPSTE24; ZMF | O75844; A0A | FACE1_HUMAN | Oxidation (M) | 0 | 0 | 0 | 0 | 0 | 0 | 0 | 0 | 697120 | 0 | 0 | 274990 | 0 | 0 | 0 | 24082 | 0 | 0 |
| TTTIAVEVK | HDLBP; HDLBP | Q00341; A0A | VIGLN_HUMAN | Unmodified | 113010 | 99663 | 0 | 143220 | 465870 | 0 | 121860 | 50771 | 65955 | 162010 | 97103 | 195380 | 666710 | 218150 | 0 | 0 | 101930 | 140300 |
| TTTIYEEDK | KIF23; KIF23; KI | Q02241; A0A | KIF23_HUMAN | Unmodified | 0 | 0 | 367230 | 0 | 1563500 | 0 | 0 | 0 | 591770 | 0 | 0 | 522200 | 0 | 0 | 461950 | 155460 | 108130 | 0 |
| TTTIYEEDKI | KIF23; KIF23; KI | Q02241; A0A | KIF23_HUMAN | Unmodified | 0 | 0 | 0 | 0 | 105570 | 541220 | 0 | 0 | 2583800 | 0 | 661850 | 1498000 | 0 | 0 | 0 | 0 | 302240 | 0 |
| TTTIYEEDKI | KIF23; KIF23; KI | Q02241; A0A | KIF23_HUMAN | Unmodified | 0 | 0 | 0 | 241670 | 66780000 | 17330000 | 2424400 | 732350 | 2318500 | 14254000 | 0 | 0 | 3283700 | 2620400 | 0 | 0 | 10385000 | 4433100 |
| TTTLNEQL( | ATP8B1; ATP8B | O43520; A0A | AT8B1_HUMAN | Unmodified | 0 | 0 | 0 | 0 | 0 | 0 | 0 | 0 | 0 | 0 | 0 | 4483300 | 0 | 0 | 0 | 0 | 0 | 0 |
| TTTLSGTAP | MAP4; MAP4; M | P27816; H7C | MAP4_HUMAN | Unmodified | 0 | 0 | 0 | 0 | 546610 | 0 | 103330 | 239050 | 0 | 0 | 0 | 428360 | 1318500 | 372000 | 0 | 0 | 482890 | 955390 |
| TTTLTTTQR | KIF5C; KIF5C; K | O60282; A0A | KIF5C_HUMAN | Unmodified | 0 | 0 | 0 | 0 | 0 | 0 | 161440 | 0 | 104450 | 0 | 0 | 0 | 868780 | 423570 | 0 | 0 | 0 | 0 |
| TTTTAVISK | MVD; MVD; MVI | P53602; H3E | MVD1_HUMAN | Unmodified | 0 | 0 | 0 | 0 | 0 | 0 | 0 | 0 | 335490 | 0 | 0 | 0 | 0 | 40940 | 0 | 0 | 246300 | 44641 |
| TTTTTFKGV | JPT1; JPT1; JPT1 | Q9UK76; J3K | JUPI1_HUMAN | Acetyl (Protein N | 0 | 0 | 0 | 0 | 3694300 | 372540 | 371710 | 790770 | 0 | 223010 | 0 | 0 | 7191500 | 5155100 | 0 | 0 | 4288400 | 4102100 |
| TTTVVNVE( | SLC1A4 | P43007 | SATT_HUMAN I | Unmodified | 943560 | 581430 | 414450 | 163080 | 0 | 0 | 0 | 0 | 7610000 | 4859900 | 0 | 13828000 | 0 | 0 | 1187900 | 0 | 505960 | 0 |
| TTTWEDPR | NEDD4; NEDD4 | P46934; Q96 | NEDD4_HUMA | Unmodified | 1797200 | 1663300 | 23792 | 1042000 | 301540 | 554370 | 182370 | 96253 | 2679200 | 187510 | 728520 | 1261400 | 317670 | 251770 | 3150600 | 1681400 | 162860 | 264540 |
| TTTWERPLI | WWP2 | O00308 | WWP2_HUMAN | Unmodified | 0 | 958580 | 0 | 0 | 0 | 0 | 0 | 0 | 0 | 0 | 0 | 0 | 0 | 0 | 1407400 | 742840 | 0 | 0 |
| TTTWERPQ | WWP1 | Q9H0M0 | WWP1_HUMAN | Unmodified | 369720 | 0 | 376570 | 746540 | 0 | 0 | 0 | 0 | 0 | 0 | 0 | 0 | 0 | 0 | 612750 | 565890 | 0 | 0 |
| TTTWQRPT | WWP2 | O00308 | WWP2_HUMAN | Unmodified | 0 | 0 | 163040 | 2570400 | 0 | 0 | 0 | 0 | 0 | 0 | 0 | 0 | 0 | 0 | 2764000 | 2026400 | 0 | 0 |
| TTTWQRPT | ITCH; ITCH; ITC | Q96J02; A0A | ITCH_HUMAN I | Unmodified | 9510200 | 8844500 | 3364700 | 1571000 | 0 | 0 | 0 | 0 | 1002000 | 0 | 0 | 991710 | 0 | 0 | 5495300 | 5536300 | 0 | 0 |
| TTTWQRPT | ITCH; ITCH; ITC | Q96J02; A0A | ITCH_HUMAN I | Unmodified | 0 | 0 | 0 | 943120 | 0 | 0 | 0 | 0 | 0 | 0 | 0 | 0 | 0 | 0 | 0 | 0 | 0 | 0 |
| TTTWQRPT | WWP1 | Q9H0M0 | WWP1_HUMAN | Unmodified | 195160 | 0 | 0 | 384540 | 0 | 136720 | 0 | 384590 | 0 | 0 | 0 | 0 | 0 | 4105900 | 930310 | 219540 | 0 | 380890 |
| TTTYTQGVI | JUP; JUP; JUP; J | P14923; C9J | PLAK_HUMAN | Unmodified | 0 | 0 | 0 | 0 | 1013900 | 0 | 0 | 0 | 0 | 0 | 0 | 327600 | 514510 | 0 | 0 | 0 | 0 | 0 |
| TTTYTQGVI | JUP; JUP; JUP; J | P14923; C9J | PLAK_HUMAN | Unmodified | 0 | 0 | 0 | 0 | 2693000 | 200790 | 0 | 0 | 0 | 0 | 0 | 0 | 783260 | 952750 | 0 | 0 | 907800 | 0 |
| TTVDAMKL | CHMP5 | Q9NZZ3 | CHMP5_HUMA | Unmodified | 0 | 0 | 0 | 0 | 2852700 | 0 | 187590 | 130200 | 0 | 638430 | 0 | 0 | 605490 | 803960 | 0 | 0 | 261660 | 406100 |
| TTVLAMDQ | NNT; NNT; NNT | Q13423; E9F | NNTM_HUMAN | Oxidation (M) | 0 | 0 | 0 | 0 | 0 | 0 | 0 | 0 | 479510 | 114820 | 81389 | 0 | 0 | 0 | 0 | 0 | 0 | 0 |
| TTVLAMDQ | NNT; NNT; NNT | Q13423; E9F | NNTM_HUMAN | Unmodified | 0 | 0 | 0 | 0 | 0 | 0 | 0 | 0 | 2035600 | 0 | 0 | 0 | 0 | 0 | 0 | 0 | 0 | 0 |
| TTVLLADIN | RIDA; RIDA; RID | P52758; H0Y | RIDA_HUMAN | Unmodified | 0 | 0 | 0 | 0 | 1369200 | 0 | 190690 | 0 | 817090 | 1078900 | 0 | 0 | 219440 | 197580 | 0 | 0 | 153550 | 299800 |
| TTVPEVEPF | PRPH; PRPH | P41219; H7C | PERI_HUMAN F | Unmodified | 0 | 0 | 0 | 0 | 47781 | 0 | 139610 | 403280 | 0 | 0 | 84763 | 0 | 0 | 0 | 0 | 0 | 0 | 0 |
| TTVPRPAPY | RAB3GAP1 | Q15042 | RB3GP_HUMAI | Unmodified | 0 | 0 | 0 | 0 | 279160 | 41174 | 0 | 0 | 0 | 0 | 159650 | 921150 | 0 | 0 | 0 | 0 | 123920 | 345680 |
| TTVTVPND( | RACGAP1; RAC | Q9H0H5; H0 | RGAP1_HUMAI | Unmodified | 0 | 0 | 0 | 1870300 | 41086000 | 3948600 | 0 | 0 | 3909500 | 3926600 | 546200 | 533860 | 3791200 | 1864500 | 0 | 0 | 4005900 | 3429000 |

| | | | | | | | | | | | | | | | | | | | | | | |
|---|---|---|---|---|---|---|---|---|---|---|---|---|---|---|---|---|---|---|---|---|---|---|
| TTVTVPND( | RACGAP1; RAC | Q9H0H5; H( | RGAP1_HUMAI | Unmodified | 0 | 0 | 0 | 0 | 29664000 | 2875400 | 0 | 0 | 0 | 2663000 | 0 | 0 | 0 | 0 | 0 | 0 | 5239300 | 3942900 |
| TTVVNDNG | PLCG2; PLCG2 | P16885; A0A | PLCG2_HUMAI | Unmodified | 44150 | 0 | 0 | 0 | 471560 | 439720 | 0 | 0 | 0 | 0 | 3546300 | 25071000 | 0 | 0 | 0 | 0 | 267760 | 819580 |
| TTVVYPATE | DCPS; DCPS | Q96C86; A0/ | DCPS_HUMAN | Unmodified | 0 | 0 | 0 | 0 | 167680 | 0 | 11010 | 0 | 1073300 | 246390 | 908910 | 2405000 | 665120 | 0 | 0 | 0 | 354650 | 249450 |
| TTVYVVED( | CYSTM1 | Q9H1C7 | CYTM1_HUMAI | Unmodified | 652990 | 378610 | 776000 | 0 | 0 | 0 | 0 | 0 | 322850 | 44658 | 136490 | 4137300 | 0 | 0 | 1079400 | 1297900 | 0 | 0 |
| TTVYVVED( | CYSTM1 | Q9H1C7 | CYTM1_HUMAI | Unmodified | 1260500 | 0 | 1629800 | 0 | 362650 | 0 | 0 | 0 | 0 | 0 | 0 | 2184700 | 0 | 0 | 1079700 | 0 | 0 | 0 |
| TTWDRPEP | ITCH; ITCH; ITC | Q96J02; A0A | ITCH_HUMAN I | Unmodified | 848920 | 45753 | 0 | 0 | 0 | 0 | 0 | 0 | 1096200 | 375110 | 0 | 1118300 | 0 | 0 | 0 | 0 | 0 | 0 |
| TTWGDGG | SNAP23; SNAP2 | O00161; H3I | SNP23_HUMAI | Unmodified | 634010 | 247270 | 682590 | 692610 | 586010 | 0 | 0 | 0 | 192000 | 0 | 650470 | 2920800 | 430120 | 172070 | 1697700 | 1290000 | 0 | 359720 |
| TTWGDGG | SNAP23; SNAP2 | O00161; H3I | SNP23_HUMAI | Unmodified | 0 | 0 | 0 | 0 | 0 | 1013600 | 0 | 0 | 0 | 0 | 0 | 0 | 1450900 | 0 | 0 | 0 | 1445500 | 2389000 |
| TTWMKADI | ADD2 | P35612 | ADDB_HUMAN | Unmodified | 0 | 0 | 0 | 0 | 280480 | 0 | 0 | 0 | 0 | 0 | 0 | 0 | 0 | 0 | 0 | 0 | 0 | 0 |
| TTYMDPTG | NUDT5; NUDT5 | Q9UKK9; A6I | NUDT5_HUMAI | Unmodified | 61969 | 75273 | 202350 | 0 | 4600100 | 582950 | 1041600 | 132510 | 1248800 | 851150 | 630500 | 5001900 | 2766600 | 1875900 | 0 | 372230 | 1197400 | 909780 |
| TTYPIDETM | GFUS; GFUS; G | Q13630; E9F | FCL_HUMAN G | Unmodified | 0 | 0 | 0 | 78237 | 0 | 0 | 0 | 0 | 398780 | 0 | 0 | 363310 | 157620 | 0 | 192130 | 0 | 344910 | 1122800 |
| TVAAESHEI | MYO9B; MYO9E | Q13459; M0I | MYO9B_HUMA | Unmodified | 0 | 269060 | 0 | 0 | 247280 | 0 | 0 | 35382 | 0 | 19553 | 0 | 0 | 0 | 0 | 0 | 0 | 52442 | 60306 |
| TVAATNMN | KIF1B; KIF1C; K | O60333; O4: | KIF1B_HUMAN | Unmodified | 0 | 0 | 0 | 0 | 0 | 0 | 0 | 0 | 92777 | 0 | 0 | 460470 | 0 | 0 | 0 | 0 | 0 | 0 |
| TVAAVMAC | EMC1 | Q8N766 | EMC1_HUMAN | Unmodified | 0 | 0 | 0 | 0 | 0 | 0 | 0 | 0 | 969390 | 0 | 0 | 0 | 0 | 0 | 0 | 0 | 0 | 0 |
| TVAAVMAC | EMC1 | Q8N766 | EMC1_HUMAN | Unmodified | 0 | 0 | 0 | 0 | 23820 | 0 | 103210 | 87117 | 5154600 | 1149800 | 656820 | 337640 | 36944 | 54425 | 0 | 0 | 0 | 70628 |
| TVAEGHGI | EML2; EML2; EI | O95834; A0/ | EMAL2_HUMAI | Unmodified | 59734 | 83014 | 22773 | 0 | 0 | 0 | 0 | 0 | 145320 | 0 | 86643 | 1272100 | 0 | 0 | 49416 | 75035 | 0 | 0 |
| TVAELEAEK | NNT; NNT; NNT | Q13423; E9F | NNTM_HUMAN | Unmodified | 868600 | 3964500 | 0 | 0 | 0 | 0 | 0 | 0 | 4860900 | 0 | 38422 | 81165 | 0 | 0 | 0 | 0 | 0 | 0 |
| TVAGGAW1 | RPL37A; RPL37 | P61513; C9J | RL37A_HUMAN | Unmodified | 13074000 | 11187000 | 3650500 | 3632700 | 2294300 | 0 | 4544000 | 4077100 | 2203900 | 882100 | 4598700 | 42833000 | 1063800 | 1090200 | 6305100 | 4861500 | 664440 | 3463200 |
| TVAGIIVEPI | ABAT; ABAT; AB | P80404; H3E | GABT_HUMAN | Unmodified | 0 | 0 | 0 | 0 | 0 | 0 | 0 | 0 | 9573500 | 1202600 | 0 | 0 | 0 | 0 | 0 | 0 | 0 | 0 |
| TVAGQDAV | BLVRB; BLVRB; | P30043; M0( | BLVRB_HUMAI | Unmodified | 188030 | 116240 | 0 | 72921 | 0 | 0 | 0 | 0 | 0 | 0 | 0 | 0 | 0 | 0 | 1072300 | 46526 | 0 | 0 |
| TVAGQDAV | BLVRB; BLVRB; | P30043; M0( | BLVRB_HUMAI | Oxidation (M) | 720840 | 836610 | 547400 | 1408100 | 0 | 0 | 0 | 0 | 0 | 0 | 0 | 0 | 1411200 | 0 | 3050800 | 1271700 | 0 | 0 |
| TVAGYGQY | EPHB3 | P54753 | EPHB3_HUMAI | Unmodified | 929120 | 1320800 | 513830 | 0 | 0 | 0 | 0 | 0 | 119810 | 0 | 0 | 646620 | 0 | 0 | 264870 | 139840 | 26412 | 0 |
| TVAIHSDVI | PCCA; PCCA; P | P05165; A0A | PCCA_HUMAN | Unmodified | 0 | 0 | 0 | 0 | 0 | 0 | 0 | 0 | 2631500 | 414280 | 0 | 0 | 0 | 0 | 0 | 0 | 0 | 0 |
| TVALLTEK | AFG3L2 | Q9Y4W6 | AFG32_HUMAI | Unmodified | 0 | 0 | 0 | 0 | 0 | 0 | 131030 | 62572 | 2060600 | 382720 | 0 | 0 | 0 | 0 | 0 | 0 | 0 | 0 |
| TVALWDLR | RBBP7; RBBP4; | Q16576; Q0! | RBBP7_HUMAI | Unmodified | 0 | 0 | 36735 | 58750 | 615310 | 263620 | 120560 | 0 | 2111000 | 728000 | 1156400 | 6928400 | 2511700 | 306420 | 0 | 31692 | 352860 | 0 |
| TVALWDLR | RBBP7; RBBP4; | Q16576; Q0! | RBBP7_HUMAI | Unmodified | 0 | 0 | 0 | 0 | 2342100 | 0 | 1546500 | 0 | 0 | 2091500 | 0 | 1533400 | 2117300 | 1614500 | 0 | 0 | 2106800 | 928920 |
| TVAMHEVF | SNX6; SNX6; SI | Q9UNH7; A( | SNX6_HUMAN | Oxidation (M) | 0 | 173900 | 0 | 0 | 459470 | 0 | 0 | 20910 | 414290 | 60759 | 0 | 297880 | 25249 | 0 | 0 | 0 | 177530 | 30124 |
| TVAPMPPA | GART; GART; G/ | P22102; C9J | PUR2_HUMAN | Unmodified | 86268 | 0 | 0 | 0 | 1055400 | 0 | 0 | 0 | 0 | 0 | 0 | 0 | 628820 | 0 | 0 | 232690 | 250750 | 625760 |
| TVASPGVT\ | ATIC; ATIC | P31939; C9J | PUR9_HUMAN | Unmodified | 0 | 0 | 0 | 0 | 212650 | 0 | 0 | 0 | 0 | 100990 | 0 | 0 | 341410 | 239480 | 83840 | 0 | 367570 | 134780 |
| TVASPGVT\ | ATIC; ATIC | P31939; C9J | PUR9_HUMAN | Unmodified | 0 | 0 | 0 | 0 | 532740 | 0 | 147090 | 0 | 0 | 0 | 0 | 0 | 353410 | 0 | 0 | 0 | 1884800 | 264850 |
| TVASRYDE\ | PDCD6IP | Q8WUM4 | PDC6I_HUMAN | Unmodified | 1.19E+08 | 1.39E+08 | 55159000 | 82673000 | 68365000 | 15078000 | 4326300 | 2844200 | 54681000 | 41888000 | 10926000 | 20484000 | 91478000 | 37698000 | 53733000 | 65516000 | 5815300 | 29310000 |
| TVASRYDE\ | PDCD6IP | Q8WUM4 | PDC6I_HUMAN | Unmodified | 0 | 1822100 | 0 | 260780 | 6206300 | 280690 | 826640 | 0 | 0 | 4419900 | 0 | 0 | 12403000 | 7837800 | 0 | 0 | 289820 | 2797200 |
| TVASVLLR | FRYL; FRYL; FR\ | O94915; H0\ | FRYL_HUMAN I | Unmodified | 0 | 0 | 0 | 0 | 0 | 0 | 0 | 0 | 0 | 0 | 0 | 1281200 | 0 | 0 | 0 | 0 | 0 | 0 |
| TVATGRDIII | MYO1D; MYO1I | O94832; J3C | MYO1D_HUMA | Unmodified | 652690 | 907170 | 555830 | 0 | 0 | 0 | 0 | 0 | 133320 | 0 | 0 | 0 | 0 | 0 | 953520 | 268110 | 0 | 0 |
| TVATPLNQ\ | EIF4H; EIF4H; E | Q15056; A0/ | IF4H_HUMAN E | Unmodified | 4814600 | 3675000 | 3964800 | 11482000 | 0 | 0 | 0 | 0 | 840060 | 0 | 472730 | 1135200 | 2490500 | 667900 | 15109000 | 7922700 | 0 | 0 |
| TVAVITSDG | LSM8; LSM8; LS | O95777; C9J | LSM8_HUMAN | Unmodified | 75147 | 51016 | 0 | 23941 | 0 | 0 | 0 | 0 | 0 | 0 | 0 | 1015500 | 70345 | 0 | 35773 | 21471 | 0 | 0 |
| TVAVTIIPSI | ATP7A | Q04656 | ATP7A_HUMAN | Unmodified | 0 | 0 | 0 | 0 | 0 | 0 | 0 | 0 | 0 | 0 | 35504 | 0 | 0 | 0 | 130530 | 182230 | 0 | 0 |
| TVAVWDSE | SNRNP40 | Q96DI7 | SNR40_HUMAI | Unmodified | 0 | 0 | 0 | 0 | 0 | 0 | 0 | 0 | 0 | 0 | 0 | 713350 | 0 | 0 | 0 | 0 | 0 | 0 |
| TVAVWDSE | SNRNP40 | Q96DI7 | SNR40_HUMAI | Unmodified | 0 | 0 | 0 | 0 | 0 | 0 | 0 | 0 | 0 | 0 | 0 | 2176300 | 368600 | 0 | 0 | 0 | 0 | 0 |
| TVAYHEAG | AFG3L2 | Q9Y4W6 | AFG32_HUMAI | Unmodified | 0 | 0 | 0 | 0 | 0 | 0 | 0 | 0 | 1366400 | 570210 | 0 | 0 | 0 | 0 | 0 | 0 | 327350 | 0 |
| TVAYTEQK | PDIA3 | P30101 | PDIA3_HUMAN | Unmodified | 2091800 | 574630 | 0 | 0 | 22691 | 0 | 314070 | 476550 | 30931000 | 3842700 | 0 | 1361300 | 0 | 1314600 | 1570800 | 1710600 | 0 | 0 |
| TVAYTEQKI | PDIA3 | P30101 | PDIA3_HUMAN | Unmodified | 0 | 0 | 0 | 0 | 0 | 0 | 626370 | 748670 | 0 | 2923700 | 0 | 0 | 0 | 0 | 0 | 0 | 0 | 0 |
| TVCAHEELI | MFAP2 | P55001 | MFAP2_HUMAI | Unmodified | 0 | 0 | 45532 | 0 | 0 | 0 | 0 | 0 | 2104800 | 0 | 0 | 0 | 0 | 0 | 0 | 58802 | 0 | 0 |
| TVDDVIKE( | CHMP2B; CHM | Q9UQN3; C! | CHM2B_HUMA | Unmodified | 989770 | 1239300 | 0 | 273840 | 99944 | 29201 | 0 | 0 | 2297300 | 347770 | 0 | 740760 | 0 | 56395 | 0 | 332440 | 27801 | 99211 |
| TVDDVIKE( | CHMP2B; CHM | Q9UQN3; C! | CHM2B_HUMA | Unmodified | 726100 | 1547300 | 0 | 751640 | 791630 | 0 | 0 | 0 | 2071000 | 380680 | 0 | 125840 | 481240 | 0 | 98284 | 148580 | 510050 | 628780 |
| TVDFTQDS | STRAP | Q9Y3F4 | STRAP_HUMAN | Unmodified | 491490 | 0 | 0 | 0 | 3029500 | 983120 | 1665800 | 1874600 | 901630 | 1204000 | 0 | 9937200 | 9632100 | 4129800 | 0 | 0 | 2127300 | 3073800 |
| TVDGPSGK | GAPDH; GAPDI | P04406; E7E | G3P_HUMAN G | Unmodified | 0 | 0 | 0 | 0 | 551700 | 0 | 560310 | 251890 | 0 | 172390 | 0 | 0 | 0 | 296680 | 0 | 0 | 1963000 | 346270 |
| TVDIEEADF | TMEM168; TMEI | Q9H0V1; H7 | TM168_HUMAN | Unmodified | 0 | 0 | 0 | 151630 | 0 | 0 | 0 | 32413 | 38532 | 48585 | 0 | 0 | 121140 | 0 | 0 | 85663 | 0 | 0 |
| TVDIYGHV | PTPRF; PTPRF; I | P10586; H0\ | PTPRF_HUMAN | Oxidation (M) | 0 | 0 | 0 | 349320 | 0 | 2355800 | 0 | 24776 | 164910 | 0 | 0 | 0 | 0 | 0 | 638190 | 929470 | 0 | 0 |
| TVDLKPDW | STIP1; STIP1; S1 | P31948; F5H | STIP1_HUMAN | Unmodified | 0 | 346480 | 7290700 | 2337400 | 1091900 | 278390 | 414350 | 0 | 904200 | 545410 | 788840 | 2880800 | 1590700 | 981280 | 0 | 0 | 3109600 | 782500 |
| TVDLQDAE | MCM3; MCM3; | P25205; Q7Z | MCM3_HUMAN | Unmodified | 0 | 0 | 0 | 1046600 | 1833500 | 218030 | 1815900 | 0 | 0 | 736060 | 0 | 0 | 332620 | 0 | 0 | 0 | 0 | 0 |
| TVDLSSHL/ | RPN1; RPN1 | P04843; F8V | RPN1_HUMAN | Unmodified | 0 | 0 | 0 | 0 | 0 | 0 | 0 | 0 | 2548700 | 4651100 | 0 | 0 | 0 | 0 | 0 | 0 | 0 | 0 |
| TVDNFVAL/ | PPIB; PPIB; PPII | P23284; A0A | PPIB_HUMAN F | Unmodified | 0 | 0 | 0 | 0 | 1122100 | 0 | 13118000 | 0 | 1.22E+08 | 95974000 | 34627000 | 16826000 | 0 | 15345000 | 0 | 941890 | 0 | 0 |
| TVDNFVAL/ | PPIB; PPIB; PPII | P23284; A0A | PPIB_HUMAN F | Unmodified | 0 | 0 | 0 | 0 | 1033700 | 483450 | 1122000 | 415970 | 405450 | 2334600 | 0 | 0 | 734850 | 0 | 0 | 24164 | 904240 | 1073200 |
| TVDNFVAL/ | PPIB; PPIB; PPII | P23284; A0A | PPIB_HUMAN F | Unmodified | 0 | 0 | 0 | 0 | 0 | 0 | 0 | 193840 | 0 | 1216800 | 0 | 0 | 0 | 0 | 0 | 0 | 0 | 0 |
| TVDTKQAQ | KRAS | P01116 | RASK_HUMAN | Unmodified | 0 | 0 | 0 | 0 | 0 | 110370 | 0 | 54237 | 27376 | 243900 | 0 | 0 | 571660 | 163880 | 0 | 0 | 0 | 0 |
| TVDWALAE | OGDHL; OGDH | Q9ULD0; Q0 | OGDHL_HUMA | Oxidation (M) | 0 | 416720 | 790050 | 0 | 177060 | 0 | 0 | 0 | 386990 | 0 | 371490 | 5771000 | 406250 | 0 | 925170 | 0 | 95852 | 197910 |
| TVEACPVVI | IGF2R; IGF2R | P11717; A0A | MPRI_HUMAN ( | Unmodified | 0 | 0 | 0 | 0 | 0 | 0 | 0 | 0 | 2059100 | 1427200 | 0 | 0 | 0 | 0 | 0 | 0 | 0 | 0 |
| TVEACPVVI | IGF2R; IGF2R | P11717; A0A | MPRI_HUMAN ( | Unmodified | 0 | 0 | 0 | 0 | 0 | 0 | 0 | 0 | 6782300 | 624170 | 312930 | 1309600 | 0 | 0 | 186600 | 124860 | 0 | 0 |
| TVEAEAAH( | IDH1 | O75874 | IDHC_HUMAN | Unmodified | 573380 | 412070 | 436810 | 0 | 0 | 0 | 0 | 0 | 1392700 | 0 | 1475800 | 4775100 | 845420 | 523630 | 1415200 | 886710 | 0 | 0 |
| TVEAEAAH( | IDH1 | O75874 | IDHC_HUMAN | Unmodified | 0 | 0 | 0 | 0 | 5707300 | 1106400 | 0 | 17368 | 198010 | 901600 | 0 | 0 | 18628000 | 8018400 | 0 | 0 | 899750 | 46858 |
| TVEDKYKC | TRAF3; TRAF3; 1 | Q13114; A6I | TRAF3_HUMAN | Unmodified | 0 | 0 | 0 | 0 | 837180 | 44533 | 0 | 0 | 0 | 0 | 0 | 0 | 0 | 0 | 0 | 0 | 0 | 0 |
| TVEDVGSP | ITGA3; ITGA3 | P26006; H0\ | ITA3_HUMAN I | Unmodified | 705460 | 123010 | 614840 | 23944 | 1812800 | 2663900 | 101220 | 126250 | 128340 | 88734 | 474240 | 4559000 | 155550 | 0 | 544670 | 265680 | 475920 | 3309600 |
| TVEEAADL! | ASRGL1 | Q7L266 | ASGL1_HUMAI | Oxidation (M) | 0 | 0 | 0 | 0 | 0 | 0 | 102260 | 0 | 0 | 577910 | 597630 | 0 | 523240 | 269690 | 344600 | 0 | 0 | 0 |
| TVEEVTVEF | TJP1; TJP1; TJP1 | Q07157; A0/ | ZO1_HUMAN T | Unmodified | 0 | 0 | 0 | 0 | 340920 | 0 | 106620 | 77160 | 2099500 | 0 | 97462 | 57051 | 580780 | 344070 | 43048 | 46947 | 124000 | 0 |
| TVEEVTVEF | TJP1; TJP1; TJP1 | Q07157; A0/ | ZO1_HUMAN T | Unmodified | 0 | 116540 | 721040 | 0 | 3761300 | 1076100 | 106760 | 65908 | 417730 | 248470 | 0 | 948100 | 3259400 | 1947500 | 796430 | 872200 | 367040 | 0 |
| TVEGAGSI/ | SNCA; SNCA; S | P37840; A0A | SYUA_HUMAN | Unmodified | 0 | 0 | 0 | 0 | 2200900 | 0 | 199040 | 162880 | 0 | 0 | 297960 | 1687300 | 0 | 0 | 0 | 0 | 0 | 1765800 |
| TVEGVLIVH | NUDT21; NUDT | O43809; H3I | CPSF5_HUMAI | Unmodified | 23699 | 0 | 0 | 0 | 499830 | 0 | 174240 | 0 | 0 | 139760 | 0 | 2118600 | 291760 | 0 | 0 | 0 | 0 | 0 |
| TVEHGFPH | LLGL2; LLGL2; I | Q6P1M3; J3( | L2GL2_HUMAN | Unmodified | 447120 | 788840 | 0 | 0 | 928060 | 0 | 0 | 0 | 4382900 | 2117400 | 0 | 2607200 | 0 | 0 | 0 | 0 | 520820 | 0 |
| TVEHGFPN | LLGL1; LLGL1 | Q15334; A0/ | L2GL1_HUMAN | Unmodified | 494100 | 556340 | 596860 | 1500200 | 0 | 0 | 0 | 0 | 2383700 | 240950 | 377230 | 3004600 | 164990 | 68400 | 1974900 | 1270900 | 0 | 176140 |
| TVEINGEK | RAB35; RAB30; | Q15286; Q1! | RAB35_HUMAN | Unmodified | 893540 | 757810 | 488020 | 452770 | 5207000 | 0 | 183390 | 60041 | 578080 | 16669 | 889230 | 371280 | 1888700 | 1098000 | 938390 | 500770 | 1354900 | 3446800 |
| TVEIPDPVE | TGM2 | P21980 | TGM2_HUMAN | Unmodified | 0 | 0 | 0 | 0 | 9505800 | 1681700 | 0 | 0 | 342080 | 198220 | 0 | 0 | 752480 | 0 | 48442 | 0 | 0 | 601330 |
| TVEIPDPVE | TGM2 | P21980 | TGM2_HUMAN | Unmodified | 0 | 0 | 0 | 0 | 2179400 | 0 | 50480 | 0 | 0 | 0 | 0 | 0 | 0 | 0 | 0 | 0 | 0 | 0 |
| TVEIVHIDI/ | PITPNB; PITPNI | P48739; A0A | PIPNB_HUMAN | Unmodified | 0 | 0 | 0 | 0 | 2033400 | 0 | 0 | 0 | 0 | 0 | 0 | 0 | 1970200 | 0 | 0 | 0 | 0 | 1240000 |
| TVELDVESI | N/A | N/A | N/A | Unmodified | 0 | 0 | 0 | 0 | 765610 | 55626 | 209230 | 65600 | 0 | 679360 | 885670 | 0 | 0 | 0 | 0 | 0 | 0 | 726370 |
| TVELVIVAD | ADAM15 | Q13444 | ADA15_HUMAN | Unmodified | 4396800 | 445110 | 0 | 0 | 272830 | 106290 | 0 | 0 | 0 | 0 | 0 | 0 | 0 | 0 | 530880 | 3705000 | 0 | 0 |
| TVENFVAL/ | PPIC | P45877 | PPIC_HUMAN I | Unmodified | 0 | 0 | 188980 | 0 | 0 | 0 | 1335700 | 793320 | 1967800 | 1370700 | 0 | 0 | 0 | 414260 | 628430 | 0 | 0 | 0 |
| TVENIKDPI | DYNC1H1; DYN | Q14204; A0/ | DYHC1_HUMA | Unmodified | 0 | 46777 | 1257900 | 484240 | 223150 | 2237300 | 297270 | 0 | 115770 | 0 | 1189900 | 3614000 | 0 | 510940 | 0 | 0 | 0 | 325230 |
| TVENVTVF( | VAT1; VAT1; VA1 | Q99536; K7E | VAT1_HUMAN ! | Unmodified | 0 | 0 | 0 | 349270 | 2285400 | 2508000 | 0 | 0 | 1005900 | 0 | 0 | 0 | 441670 | 0 | 0 | 0 | 0 | 2177800 |
| TVERNVED | DNAJB12; DNA. | Q9NXW2; V! | DJB12_HUMAN | Unmodified | 0 | 0 | 0 | 0 | 0 | 0 | 251630 | 0 | 3281400 | 0 | 0 | 0 | 0 | 0 | 0 | 0 | 0 | 0 |
| TVESEAASY | PSME3; PSME3; | P61289; A0A | PSME3_HUMAI | Unmodified | 0 | 0 | 0 | 0 | 0 | 0 | 0 | 0 | 0 | 0 | 748730 | 3182300 | 0 | 0 | 0 | 0 | 0 | 0 |
| TVESITDIR | LAMP1 | P11279 | LAMP1_HUMAI | Unmodified | 339910 | 298310 | 749080 | 866280 | 0 | 0 | 0 | 0 | 55725000 | 2868100 | 869060 | 4634200 | 0 | 0 | 427040 | 345430 | 21815 | 0 |
| TVESITDIR/ | LAMP1 | P11279 | LAMP1_HUMAI | Unmodified | 1866000 | 1285200 | 3101900 | 4029500 | 18648000 | 2548700 | 4782300 | 2441200 | 3.21E+08 | 68554000 | 17005000 | 5202100 | 7868000 | 6838000 | 2399900 | 2024400 | 642770 | 7221600 |
| TVESITDIR/ | LAMP1 | P11279 | LAMP1_HUMAI | Unmodified | 2604800 | 1088800 | 6113300 | 6115400 | 25432000 | 4657400 | 8239600 | 2211900 | 3.83E+08 | 83285000 | 32064000 | 3710900 | 12266000 | 14807000 | 2869400 | 1567600 | 398310 | 14835000 |
| TVESLEETL | PTGR1; PTGR1 | Q14914; F2Z | PTGR1_HUMAN | Unmodified | 197320 | 0 | 0 | 43600 | 4674200 | 1216500 | 0 | 0 | 0 | 0 | 0 | 0 | 0 | 111610 | 0 | 0 | 282470 | 0 |
| TVESLEETL | PTGR1; PTGR1 | Q14914; F2Z | PTGR1_HUMAN | Unmodified | 0 | 0 | 134110 | 0 | 3234800 | 921530 | 150820 | 0 | 0 | 569170 | 0 | 0 | 0 | 0 | 341840 | 0 | 0 | 0 |
| TVETLLLR | ACAD9; ACAD9 | Q9H845; H0 | ACAD9_HUMAI | Unmodified | 0 | 0 | 0 | 0 | 0 | 0 | 0 | 0 | 1356100 | 0 | 12120 | 0 | 0 | 0 | 0 | 0 | 0 | 0 |
| TVETRDGQ | VIM | P08670 | VIME_HUMAN \ | Unmodified | 0 | 0 | 0 | 0 | 5810100 | 1037200 | 2237300 | 2167500 | 370500 | 0 | 0 | 0 | 0 | 0 | 0 | 0 | 11809000 | 21561000 |
| TVETVPQKI | N/A | N/A | N/A | Unmodified | 0 | 0 | 0 | 0 | 1174200 | 90571 | 134730 | 0 | 0 | 639990 | 0 | 0 | 341360 | 520020 | 0 | 0 | 134390 | 491010 |
| TVEVGRAYI | GSPT1; GSPT1; | P15170; H3E | ERF3A_HUMAN | Unmodified | 0 | 0 | 0 | 148640 | 3893700 | 0 | 1507800 | 749770 | 0 | 0 | 504110 | 0 | 4108900 | 2347100 | 0 | 0 | 1263300 | 7508500 |
| TVEVGRAYI | GSPT1; GSPT1; | P15170; H3E | ERF3A_HUMAN | Unmodified | 0 | 0 | 0 | 0 | 0 | 0 | 0 | 0 | 0 | 0 | 0 | 0 | 2404100 | 1493700 | 0 | 0 | 0 | 0 |
| TVEVLEPEV | CYFIP1; CYFIP2 | Q7L576; Q9( | CYFP1_HUMAN | Unmodified | 0 | 0 | 0 | 0 | 949410 | 0 | 402540 | 477010 | 1459300 | 639110 | 1252700 | 4386800 | 0 | 0 | 0 | 0 | 461410 | 0 |
| TVEVPFKGI | GMPR2; GMPR2 | Q9P2T1; H0\ | GMPR2_HUMA | Unmodified | 0 | 0 | 0 | 0 | 0 | 0 | 0 | 0 | 339410 | 0 | 315860 | 3042100 | 0 | 0 | 0 | 0 | 0 | 0 |
| TVEVPIHQL | TMPPE | Q6ZT21 | TMPPE_HUMAI | Oxidation (M) | 0 | 0 | 0 | 0 | 605980 | 0 | 0 | 0 | 242880 | 150890 | 48949 | 0 | 0 | 0 | 0 | 0 | 0 | 0 |
| TVEVPIHQL | TMPPE | Q6ZT21 | TMPPE_HUMAI | Unmodified | 0 | 0 | 0 | 0 | 0 | 0 | 0 | 0 | 2110500 | 416900 | 59743 | 0 | 0 | 0 | 0 | 0 | 0 | 0 |
| TVEYAPCR | ITGAV | P06756 | ITAV_HUMAN I | Unmodified | 0 | 0 | 527160 | 0 | 0 | 0 | 0 | 0 | 0 | 63479 | 56134 | 3364700 | 0 | 50062 | 722590 | 653700 | 44153 | 35097 |
| TVEYHRLD( | KCT2 | Q8NC54 | KCT2_HUMAN | Unmodified | 0 | 0 | 0 | 0 | 968350 | 0 | 0 | 0 | 654080 | 0 | 0 | 0 | 0 | 0 | 0 | 0 | 0 | 0 |
| TVEYHRLD( | KCT2 | Q8NC54 | KCT2_HUMAN | Oxidation (M) | 0 | 0 | 0 | 0 | 0 | 0 | 847350 | 0 | 0 | 0 | 0 | 0 | 0 | 0 | 0 | 0 | 0 | 235900 |
| TVEYLHSQ | RPS6KA1; RPS6 | Q15418; Q5! | KS6A1_HUMAN | Unmodified | 0 | 0 | 0 | 0 | 0 | 0 | 0 | 0 | 206720 | 157770 | 0 | 665690 | 0 | 0 | 0 | 0 | 0 | 0 |
| TVFAEHISD | RPL3; RPL3; RP | P39023; B5I | RL3_HUMAN 6 | Unmodified | 6357100 | 3009300 | 2515700 | 1492800 | 843620 | 344290 | 2837800 | 1377600 | 448380 | 336430 | 2853800 | 59419000 | 484450 | 1057500 | 2601600 | 2459500 | 188370 | 1245800 |
| TVFAEHISD | RPL3; RPL3; RP | P39023; B5I | RL3_HUMAN 6 | Unmodified | 0 | 0 | 0 | 0 | 0 | 0 | 139850 | 82053 | 0 | 0 | 0 | 2368300 | 0 | 0 | 0 | 0 | 0 | 0 |
| TVFDEAIIAI | RHOQ | P17081 | RHOQ_HUMAN | Unmodified | 0 | 0 | 23192 | 0 | 118620 | 0 | 78280 | 0 | 62375 | 59795 | 32300 | 312740 | 115420 | 149230 | 0 | 38122 | 74517 | 79255 |
| TVFDEAIR | RAC2; RAC3; R/ | P15153; P60 | RAC2_HUMAN | Unmodified | 48767000 | 58313000 | 40178000 | 40861000 | 13028000 | 4731600 | 875920 | 406640 | 9519000 | 1390600 | 13357000 | 55973000 | 13830000 | 8733100 | 49110000 | 51966000 | 3887800 | 17540000 |
| TVFDEAIRA | RAC3; RAC1 | P60763; P63 | RAC3_HUMAN | Unmodified | 0 | 0 | 0 | 0 | 4872500 | 644010 | 1136900 | 35615 | 226810 | 527370 | 84571 | 0 | 3562900 | 1969000 | 0 | 0 | 3223100 | 5242400 |
| TVFDEAIRA | RAC3; RAC1 | P60763; P63 | RAC3_HUMAN | Unmodified | 0 | 0 | 0 | 0 | 2642200 | 0 | 0 | 0 | 0 | 336670 | 0 | 0 | 2818400 | 2500100 | 0 | 0 | 1613300 | 1967500 |
| TVFEALQAI | AP2A2; AP2A2; | O94973; A0/ | AP2A2_HUMAN | Unmodified | 0 | 251270 | 395860 | 0 | 180490 | 0 | 0 | 0 | 0 | 0 | 0 | 3645000 | 181990 | 0 | 739890 | 521980 | 176970 | 0 |
| TVFEALQAI | AP2A1 | O95782 | AP2A1_HUMAN | Oxidation (M) | 198640 | 0 | 179290 | 186740 | 0 | 0 | 0 | 31544 | 0 | 0 | 0 | 1741100 | 0 | 0 | 0 | 450530 | 0 | 0 |
| TVFEALQAI | AP2A1 | O95782 | AP2A1_HUMAN | Unmodified | 0 | 0 | 0 | 0 | 254200 | 0 | 0 | 0 | 963870 | 0 | 0 | 718040 | 263460 | 848890 | 0 | 0 | 482620 | 435480 |
| TVFFESLER | UBA6 | A0AVT1 | UBA6_HUMAN | Unmodified | 0 | 0 | 0 | 0 | 0 | 0 | 0 | 0 | 0 | 0 | 0 | 4643500 | 0 | 0 | 0 | 0 | 0 | 0 |

| | | | | | | | | | | | | | | | | | | | | | | |
|---|---|---|---|---|---|---|---|---|---|---|---|---|---|---|---|---|---|---|---|---|---|---|
| TVFFESLER | UBA6 | A0AVT1 | UBA6_HUMAN | Unmodified | 0 | 0 | 0 | 0 | 990620 | 0 | 419710 | 40565 | 0 | 271990 | 0 | 0 | 502570 | 792090 | 0 | 0 | 1797800 | 4622200 |
| TVFFWAPIN | MPC2; MPC2 | O95563; Q5I | MPC2_HUMAN | Unmodified | 0 | 0 | 0 | 0 | 0 | 143490 | 0 | 0 | 399120 | 0 | 0 | 0 | 0 | 0 | 0 | 0 | 0 | 0 |
| TVFGVEPDI | CNDP2 | Q96KP4 | CNDP2_HUMA | Unmodified | 382210 | 285360 | 979430 | 958610 | 653410 | 419650 | 0 | 0 | 1846800 | 104370 | 2436900 | 21418000 | 636710 | 416650 | 2053700 | 1994600 | 414490 | 0 |
| TVFGVEPDI | CNDP2 | Q96KP4 | CNDP2_HUMA | Unmodified | 0 | 0 | 0 | 746170 | 1925200 | 1256900 | 0 | 0 | 0 | 666140 | 948250 | 0 | 29905000 | 11922000 | 723500 | 85069 | 0 | 1956100 |
| TVFLSGLPF | MRPS30 | Q9NP92 | RT30_HUMAN | Unmodified | 0 | 0 | 0 | 0 | 0 | 0 | 0 | 0 | 1097200 | 285130 | 0 | 0 | 262610 | 0 | 0 | 0 | 0 | 0 |
| TVFPGAVP\ | EPS15L1; EPS1 | Q9UBC2; M( | EP15R_HUMAN | Unmodified | 0 | 0 | 0 | 0 | 0 | 0 | 0 | 0 | 0 | 0 | 541770 | 2226500 | 0 | 0 | 0 | 0 | 0 | 0 |
| TVFPLADVS | CEMIP2; CEMIP | Q9UHN6; B4 | CEIP2_HUMAN | Unmodified | 756000 | 775510 | 338460 | 0 | 16023000 | 1715600 | 0 | 0 | 4467200 | 6889200 | 0 | 0 | 1604200 | 1963900 | 4678400 | 796590 | 0 | 320010 |
| TVFRPPLDI | NOL6; NOL6 | Q9H6R4; A0. | NOL6_HUMAN | Unmodified | 36406 | 0 | 0 | 0 | 0 | 0 | 0 | 0 | 0 | 0 | 0 | 482100 | 0 | 0 | 0 | 0 | 0 | 0 |
| TVFSPTLPA | ZC3HAV1; ZC3H | Q7Z2W4; C9 | ZCCHV_HUMA | Unmodified | 1555700 | 1286400 | 2625400 | 3262000 | 19262000 | 4908900 | 22180 | 18072 | 786190 | 514410 | 0 | 0 | 5228300 | 3207600 | 1601800 | 1784300 | 0 | 3175500 |
| TVFSPTLPA | ZC3HAV1; ZC3H | Q7Z2W4; C9 | ZCCHV_HUMA | Unmodified | 0 | 0 | 0 | 0 | 2124600 | 0 | 0 | 0 | 0 | 0 | 0 | 0 | 0 | 0 | 0 | 0 | 0 | 0 |
| TVFSPTLPA | ZC3HAV1; ZC3H | Q7Z2W4; C9 | ZCCHV_HUMA | Unmodified | 0 | 0 | 0 | 0 | 5033700 | 0 | 0 | 0 | 0 | 0 | 0 | 0 | 0 | 830920 | 0 | 0 | 0 | 890370 |
| TVFTKLDYY | TOR1A | O14656 | TOR1A_HUMAN | Unmodified | 0 | 0 | 0 | 0 | 0 | 0 | 522910 | 63469 | 7499600 | 2303300 | 0 | 0 | 0 | 0 | 0 | 0 | 0 | 0 |
| TVFVGNHP | ATP11C; ATP11 | Q8NB49; A0. | AT11C_HUMAN | Unmodified | 0 | 0 | 1346200 | 0 | 0 | 0 | 0 | 0 | 0 | 0 | 0 | 2970300 | 0 | 0 | 1661700 | 750640 | 0 | 0 |
| TVGAALDIL | GLB1; GLB1 | P16278; E7E | BGAL_HUMAN | Unmodified | 0 | 0 | 0 | 0 | 0 | 0 | 0 | 80916 | 1312600 | 868220 | 47787 | 121560 | 62481 | 54144 | 0 | 0 | 0 | 93267 |
| TVGALQVL( | PRKDC | P78527 | PRKDC_HUMA | Unmodified | 184390 | 188550 | 343540 | 0 | 0 | 0 | 243690 | 126100 | 158360 | 90607 | 0 | 4610400 | 0 | 0 | 171830 | 0 | 0 | 0 |
| TVGATALPF | CCT8; CCT8 | P50990; H7( | TCPQ_HUMAN | Unmodified | 589400 | 0 | 1054900 | 1166000 | 0 | 0 | 0 | 0 | 3773100 | 2221500 | 4900000 | 14664000 | 7594700 | 2058900 | 0 | 1009300 | 0 | 0 |
| TVGATVEFT | UNC119B | A6NIH7 | U119B_HUMAN | Unmodified | 402100 | 566710 | 0 | 81967 | 697220 | 0 | 0 | 0 | 0 | 363890 | 0 | 483300 | 1477000 | 0 | 0 | 0 | 708640 | 1006800 |
| TVGEIMWG | SCARB1; SCARI | Q8WTV0; A0 | SCRB1_HUMAI | Oxidation (M) | 104840 | 212360 | 837220 | 36898 | 0 | 0 | 0 | 0 | 0 | 0 | 55541 | 5244000 | 0 | 0 | 1693500 | 272670 | 0 | 0 |
| TVGIDDLT( | AK3 | Q9UIJ7 | KAD3_HUMAN | Unmodified | 0 | 0 | 0 | 0 | 0 | 0 | 0 | 0 | 11508000 | 2782900 | 0 | 0 | 0 | 0 | 0 | 0 | 0 | 0 |
| TVGLFLQG | GPD2 | P43304 | GPDM_HUMAN | Unmodified | 0 | 0 | 0 | 0 | 0 | 0 | 0 | 0 | 7791200 | 1097600 | 0 | 0 | 0 | 0 | 0 | 0 | 0 | 0 |
| TVGLFLQG | GPD2 | P43304 | GPDM_HUMAN | Unmodified | 0 | 0 | 0 | 0 | 0 | 0 | 0 | 0 | 0 | 348510 | 0 | 0 | 0 | 0 | 0 | 0 | 0 | 0 |
| TVGLGDAIS | ADPGK | Q9BRR6 | ADPGK_HUMAI | Unmodified | 0 | 0 | 0 | 0 | 0 | 0 | 0 | 0 | 7888500 | 630580 | 63432 | 0 | 0 | 0 | 0 | 0 | 0 | 0 |
| TVGLREVW | RDX; RDX; RDX; | P35241; E9P | RADI_HUMAN I | Unmodified | 0 | 0 | 0 | 112090 | 458160 | 0 | 0 | 0 | 0 | 416470 | 0 | 0 | 212380 | 305300 | 71621 | 0 | 232700 | 0 |
| TVGQLYKE( | MYH9 | P35579 | MYH9_HUMAN | Unmodified | 0 | 83124 | 0 | 0 | 1843300 | 238560 | 1047500 | 375950 | 0 | 1099500 | 0 | 0 | 640190 | 0 | 0 | 0 | 481740 | 464970 |
| TVGQYEYYI | EFNB2 | P52799 | EFNB2_HUMAN | Unmodified | 202880 | 0 | 0 | 0 | 0 | 0 | 0 | 0 | 0 | 0 | 0 | 0 | 0 | 0 | 0 | 1776900 | 0 | 0 |
| TVGTPIASV | PSMD1; PSMD1 | Q99460; A0/ | PSMD1_HUMA | Unmodified | 0 | 0 | 956160 | 0 | 0 | 0 | 0 | 0 | 0 | 0 | 0 | 3353100 | 0 | 0 | 0 | 0 | 0 | 0 |
| TVGVEPAAI | RPL28; RPL28; I | P46779; H0Y | RL28_HUMAN ( | Unmodified | 2979500 | 2370100 | 2784300 | 1132900 | 2816900 | 639050 | 0 | 709290 | 160280 | 53808 | 431680 | 23635000 | 0 | 0 | 2595300 | 2525300 | 288640 | 484870 |
| TVGVEPAAI | RPL28; RPL28; I | P46779; H0Y | RL28_HUMAN ( | Unmodified | 3180600 | 2478200 | 0 | 1447500 | 3287500 | 0 | 2500500 | 1965200 | 363230 | 363760 | 706030 | 0 | 6006500 | 1113300 | 0 | 446470 | 250540 | 1972300 |
| TVGVEPAAI | RPL28; RPL28; I | P46779; H0Y | RL28_HUMAN ( | Unmodified | 220510 | 296920 | 0 | 220780 | 79346 | 0 | 0 | 114880 | 0 | 59489 | 0 | 0 | 348440 | 172770 | 0 | 0 | 0 | 0 |
| TVGVNNPV | LONP2; LONP2 | Q86WA8; H3 | LONP2_HUMAI | Unmodified | 0 | 0 | 0 | 0 | 0 | 0 | 0 | 0 | 5190600 | 1981100 | 444090 | 0 | 0 | 0 | 0 | 0 | 0 | 0 |
| TVGWNVP\ | MELTF; MELTF | P08582; C9J | TRFM_HUMAN | Unmodified | 0 | 0 | 0 | 0 | 0 | 0 | 0 | 0 | 8995800 | 0 | 0 | 2618600 | 0 | 0 | 0 | 0 | 0 | 0 |
| TVHAGEVG | ADA; ADA | P00813; F5G | ADA_HUMAN A | Unmodified | 0 | 0 | 0 | 0 | 0 | 0 | 0 | 0 | 0 | 0 | 750900 | 2012900 | 0 | 0 | 0 | 124480 | 0 | 0 |
| TVHLYDNW | ELOVL7; ELOVL | A1L3X0; D6F | ELOV7_HUMAN | Unmodified | 0 | 0 | 0 | 0 | 0 | 0 | 0 | 0 | 2754600 | 531890 | 0 | 0 | 0 | 0 | 0 | 0 | 0 | 0 |
| TVHSLACLI | CAD; CAD; CAD | P27708; F8V | PYR1_HUMAN | Unmodified | 0 | 259470 | 0 | 0 | 438640 | 0 | 0 | 0 | 81883 | 24002 | 0 | 1046300 | 0 | 0 | 124740 | 0 | 0 | 0 |
| TVHTEWTQ | APEH; APEH; AI | P13798; C9J | ACPH_HUMAN | Unmodified | 0 | 0 | 0 | 0 | 0 | 0 | 0 | 0 | 0 | 0 | 66754 | 1263900 | 0 | 0 | 0 | 0 | 0 | 139620 |
| TVHYLPILFI | CLPTM1L; CLPT | Q96KA5; G5I | CLP1L_HUMAN | Unmodified | 0 | 0 | 0 | 0 | 0 | 0 | 0 | 0 | 6494200 | 961900 | 147510 | 0 | 0 | 0 | 0 | 0 | 0 | 0 |
| TVHYLPILFI | CLPTM1L; CLPT | Q96KA5; G5I | CLP1L_HUMAN | Unmodified | 0 | 0 | 0 | 0 | 0 | 0 | 0 | 0 | 13424000 | 1451700 | 320040 | 0 | 3664400 | 0 | 12592000 | 0 | 0 | 0 |
| TVIDSQTHY | TMED3 | Q9Y3Q3 | TMED3_HUMAI | Unmodified | 0 | 0 | 0 | 0 | 186990 | 0 | 11653 | 23379 | 4523000 | 384300 | 99563 | 1270800 | 0 | 0 | 0 | 0 | 0 | 0 |
| TVIDYNGEI | P4HB; P4HB; P4 | P07237; A0A | PDIA1_HUMAN | Unmodified | 0 | 0 | 0 | 0 | 0 | 175810 | 15296 | 0 | 6182700 | 66299 | 0 | 1055400 | 0 | 0 | 0 | 0 | 0 | 0 |
| TVIDYNGEI | P4HB; P4HB; P4 | P07237; A0A | PDIA1_HUMAN | Unmodified | 0 | 0 | 0 | 0 | 0 | 0 | 0 | 0 | 3072200 | 0 | 0 | 0 | 62106 | 0 | 0 | 0 | 0 | 0 |
| TVIDYNGEI | P4HB; P4HB; P4 | P07237; A0A | PDIA1_HUMAN | Unmodified | 0 | 0 | 132200 | 0 | 0 | 0 | 95292 | 0 | 8165000 | 743040 | 0 | 0 | 0 | 0 | 159160 | 128960 | 0 | 0 |
| TVIEPMACI | ATP2B4 | P23634 | AT2B4_HUMAN | Unmodified | 0 | 0 | 0 | 0 | 0 | 0 | 0 | 0 | 0 | 0 | 658000 | 3316700 | 0 | 0 | 0 | 0 | 399050 | 275740 |
| TVIEPMACI | ATP2B4 | P23634 | AT2B4_HUMAN | Oxidation (M) | 0 | 0 | 0 | 445770 | 0 | 0 | 0 | 0 | 0 | 0 | 589080 | 2918600 | 0 | 0 | 0 | 0 | 0 | 0 |
| TVIEPMASE | ATP2B1; ATP2B | P20020; E7E | AT2B1_HUMAN | Unmodified | 294940 | 0 | 807980 | 1244100 | 1096200 | 0 | 0 | 0 | 4504500 | 670840 | 1546200 | 38726000 | 5095100 | 1708100 | 2541400 | 826790 | 0 | 770940 |
| TVIEPMASE | ATP2B1; ATP2B | P20020; E7E | AT2B1_HUMAN | Oxidation (M) | 1025200 | 515710 | 3136700 | 2138700 | 0 | 0 | 0 | 0 | 0 | 0 | 0 | 6635000 | 1881800 | 972820 | 5057300 | 3928500 | 0 | 0 |
| TVIEQQPVL | GGT1; GGT3P; ( | P19440; A6N | GGT1_HUMAN | Unmodified | 2729800 | 0 | 0 | 0 | 0 | 0 | 157310 | 0 | 1531100 | 0 | 268910 | 235660 | 0 | 0 | 532940 | 0 | 0 | 0 |
| TVIESSPES | GEMIN5 | Q8TEQ6 | GEMI5_HUMAN | Unmodified | 0 | 0 | 0 | 0 | 0 | 0 | 0 | 37948 | 0 | 0 | 388950 | 2825800 | 0 | 0 | 0 | 0 | 0 | 0 |
| TVIFVATGE | PGLS; PGLS | O95336; M0I | 6PGL_HUMAN | Unmodified | 579030 | 375060 | 0 | 173820 | 2001400 | 412820 | 934050 | 242650 | 2635600 | 3050300 | 697080 | 3204300 | 626480 | 419240 | 167650 | 404850 | 1168900 | 2239200 |
| TVIGSVLLR | AP3B1 | O00203 | AP3B1_HUMAN | Unmodified | 0 | 0 | 0 | 0 | 0 | 0 | 0 | 0 | 0 | 0 | 314540 | 1943600 | 0 | 0 | 0 | 0 | 0 | 0 |
| TVIIDHGSC | ACTL8 | Q9H568 | ACTL8_HUMAN | Unmodified | 0 | 0 | 0 | 0 | 0 | 0 | 0 | 0 | 616410 | 0 | 1720800 | 5508100 | 0 | 0 | 0 | 111860 | 0 | 0 |
| TVIIEQSWG | HSPD1; HSPD1 | P10809; A0A | CH60_HUMAN | Unmodified | 149330 | 0 | 385310 | 0 | 0 | 0 | 275210 | 134310 | 9650000 | 1823100 | 1116800 | 520190 | 0 | 0 | 184630 | 215360 | 0 | 0 |
| TVIIEQSWG | HSPD1; HSPD1 | P10809; A0A | CH60_HUMAN | Unmodified | 0 | 0 | 0 | 0 | 0 | 0 | 317040 | 0 | 0 | 628130 | 136260 | 0 | 0 | 0 | 0 | 0 | 0 | 0 |
| TVIIEQSWG | HSPD1; HSPD1 | P10809; A0A | CH60_HUMAN | Unmodified | 0 | 0 | 0 | 0 | 0 | 0 | 1331700 | 175890 | 0 | 1664300 | 0 | 0 | 0 | 167300 | 0 | 0 | 241270 | 0 |
| TVIKEGEEQ | POLR2B; POLR2 | P30876; C9J | RPB2_HUMAN | Unmodified | 0 | 0 | 0 | 74443 | 361730 | 0 | 527540 | 0 | 0 | 356350 | 0 | 349990 | 110360 | 0 | 0 | 0 | 218980 | 404650 |
| TVIKKDEQE | PSMC4 | P43686 | PRS6B_HUMAN | Unmodified | 0 | 0 | 0 | 0 | 1825200 | 0 | 138240 | 889560 | 895190 | 326610 | 527540 | 1212600 | 8989700 | 2108000 | 0 | 0 | 4565200 | 1582300 |
| TVIKPESCV | RACGAP1; RAC | Q9H0H5; F8 | RGAP1_HUMAI | Unmodified | 0 | 0 | 0 | 0 | 6831900 | 1312700 | 476000 | 155230 | 641160 | 904530 | 246720 | 650440 | 631620 | 524080 | 0 | 0 | 3247800 | 1304400 |
| TVIKPESCV | RACGAP1; RAC | Q9H0H5; F8 | RGAP1_HUMAI | Unmodified | 0 | 0 | 0 | 0 | 541530 | 127390 | 0 | 0 | 0 | 83598 | 0 | 0 | 0 | 0 | 0 | 0 | 686780 | 0 |
| TVIRLPSGS | AHNAK | Q09666 | AHNK_HUMAN | Unmodified | 282410 | 210160 | 562980 | 618230 | 8252100 | 1663500 | 0 | 0 | 254810 | 0 | 0 | 0 | 0 | 0 | 405050 | 318160 | 1653800 | 3073600 |
| TVIRLPSGS | AHNAK | Q09666 | AHNK_HUMAN | Unmodified | 0 | 0 | 0 | 0 | 5902100 | 0 | 0 | 0 | 0 | 0 | 0 | 0 | 0 | 0 | 0 | 0 | 959850 | 2445100 |
| TVISQSLSK | ATP6V1A | P38606 | VATA_HUMAN ' | Unmodified | 890470 | 1087700 | 1370900 | 1123700 | 4275200 | 845520 | 4815200 | 1637900 | 3422000 | 1732100 | 1900700 | 10109000 | 6150300 | 5759000 | 2098600 | 1691100 | 1769900 | 2524900 |
| TVITEEFKVI | FUBP3 | Q96I24 | FUBP3_HUMAN | Unmodified | 669600 | 462120 | 0 | 33963 | 0 | 380890 | 0 | 75237 | 369580 | 0 | 357540 | 1872700 | 0 | 0 | 0 | 122410 | 0 | 0 |
| TVIVHGFTL | IARS2 | Q9NSE4 | SYIM_HUMAN I | Unmodified | 0 | 0 | 0 | 0 | 0 | 0 | 0 | 0 | 1164700 | 0 | 0 | 0 | 0 | 0 | 0 | 0 | 0 | 0 |
| TVIVHGFTL | IARS2 | Q9NSE4 | SYIM_HUMAN I | Unmodified | 0 | 0 | 0 | 0 | 0 | 0 | 0 | 0 | 2197900 | 596740 | 0 | 0 | 0 | 0 | 0 | 0 | 0 | 216360 |
| TVIVNMVD' | EIF5; EIF5; EIF5 | P55010; H0Y | IF5_HUMAN Eu | Oxidation (M) | 0 | 0 | 0 | 0 | 437900 | 0 | 79942 | 0 | 0 | 0 | 107880 | 336030 | 0 | 0 | 0 | 0 | 435220 | 0 |
| TVIVNMVD' | EIF5; EIF5; EIF5 | P55010; H0Y | IF5_HUMAN Eu | Unmodified | 0 | 0 | 0 | 0 | 4032500 | 0 | 310670 | 0 | 0 | 0 | 0 | 3938800 | 1064900 | 0 | 0 | 0 | 2239300 | 2432400 |
| TVIVSKDEC | SORL1; SORL1; | Q92673; E9F | SORL_HUMAN | Unmodified | 0 | 0 | 0 | 0 | 0 | 0 | 0 | 0 | 2030000 | 0 | 196430 | 0 | 0 | 0 | 0 | 0 | 0 | 0 |
| TVIVTPSQR | PCK2; PCK2 | Q16822; H0' | PCKGM_HUMA | Unmodified | 14545000 | 48609000 | 0 | 692440 | 0 | 0 | 90998 | 68133 | 4155500 | 0 | 0 | 261750 | 413810 | 0 | 884860 | 7035900 | 138510 | 0 |
| TVKEEAEKF | PRKCSH; PRKC | P14314; K7E | GLU2B_HUMAI | Unmodified | 0 | 0 | 0 | 0 | 0 | 0 | 0 | 21125 | 13995000 | 401080 | 0 | 511200 | 0 | 0 | 0 | 0 | 0 | 0 |
| TVKGGISET | EPB41L2; EPB4 | O43491; P11 | E41L2_HUMAN | Unmodified | 0 | 0 | 0 | 0 | 0 | 0 | 837230 | 1271300 | 0 | 234730 | 0 | 0 | 0 | 1196400 | 0 | 0 | 4554300 | 2719300 |
| TVKGPDGL | MTPN; MTPN | P58546; C9J | MTPN_HUMAN | Unmodified | 0 | 937050 | 0 | 3116700 | 9012000 | 3893700 | 5933500 | 5336000 | 1933900 | 2859300 | 7349300 | 16050000 | 12208000 | 12911000 | 2143000 | 1880300 | 7824800 | 9765700 |
| TVKGPDGL | MTPN; MTPN | P58546; C9J | MTPN_HUMAN | Unmodified | 0 | 0 | 0 | 0 | 1835200 | 0 | 0 | 0 | 0 | 0 | 0 | 0 | 0 | 0 | 0 | 0 | 4035500 | 2280900 |
| TVKHELLPS | PON2; PON2; P | Q15165; G3 | PON2_HUMAN | Unmodified | 0 | 0 | 0 | 0 | 0 | 0 | 0 | 0 | 6761200 | 0 | 0 | 0 | 0 | 0 | 0 | 0 | 0 | 0 |
| TVKHGAGA | PIP4K2C | Q8TBX8 | PI42C_HUMAN | Unmodified | 0 | 0 | 0 | 0 | 164330 | 0 | 0 | 172730 | 2595700 | 630250 | 452940 | 71130 | 0 | 508050 | 0 | 0 | 972330 | 250220 |
| TVKHGAGA | PIP4K2B; PIP4K | P78356; P48 | PI42B_HUMAN | Unmodified | 0 | 0 | 0 | 0 | 0 | 0 | 1702500 | 8338100 | 0 | 0 | 0 | 0 | 311830 | 959870 | 0 | 0 | 1581800 | 0 |
| TVKNDNSS | MYH14; MYH9; | Q7Z406; P35 | MYH14_HUMAI | Unmodified | 0 | 0 | 0 | 0 | 1751200 | 0 | 1082100 | 326110 | 0 | 446950 | 0 | 0 | 397090 | 497980 | 0 | 0 | 759120 | 0 |
| TVLDKAVQ. | PDCD6IP | Q8WUM4 | PDC6I_HUMAN | Unmodified | 164840 | 254440 | 0 | 0 | 668540 | 0 | 0 | 0 | 0 | 158190 | 0 | 0 | 887620 | 0 | 0 | 0 | 208100 | 739920 |
| TVLDPVTGI | SF3B3 | Q15393 | SF3B3_HUMAN | Unmodified | 0 | 0 | 0 | 0 | 0 | 0 | 0 | 0 | 0 | 0 | 496330 | 6670200 | 0 | 0 | 0 | 0 | 0 | 0 |
| TVLDQQQT | LRPPRC | P42704 | LPPRC_HUMAN | Unmodified | 439520 | 0 | 0 | 0 | 12047 | 0 | 0 | 0 | 3067100 | 233460 | 0 | 0 | 320560 | 0 | 0 | 0 | 0 | 0 |
| TVLDQQQT | LRPPRC | P42704 | LPPRC_HUMAN | Unmodified | 0 | 0 | 0 | 0 | 0 | 0 | 133280 | 0 | 2236600 | 1929800 | 0 | 0 | 0 | 0 | 0 | 0 | 0 | 205790 |
| TVLELMNPI | IQGAP1; IQGAP | P46940; A0A | IQGA1_HUMAN | Oxidation (M) | 0 | 214880 | 0 | 0 | 3645400 | 650500 | 0 | 0 | 0 | 473440 | 0 | 0 | 0 | 0 | 0 | 0 | 0 | 0 |
| TVLELQYVL | CAPRIN1; CAPF | Q14444; G3\ | CAPR1_HUMAI | Unmodified | 0 | 0 | 0 | 0 | 0 | 0 | 0 | 0 | 0 | 0 | 0 | 1527000 | 0 | 0 | 0 | 0 | 0 | 0 |
| TVLELQYVL | CAPRIN1; CAPF | Q14444; G3\ | CAPR1_HUMAI | Unmodified | 0 | 0 | 0 | 0 | 2670400 | 0 | 0 | 0 | 0 | 0 | 0 | 115010 | 0 | 0 | 0 | 0 | 0 | 0 |
| TVLEQVLPI | DNAJB1; DNAJE | P25685; A0A | DNJB1_HUMAN | Unmodified | 789720 | 585000 | 226660 | 405680 | 2823600 | 781990 | 362700 | 164320 | 2326500 | 1299100 | 385420 | 859710 | 2429500 | 540220 | 391260 | 308080 | 1441600 | 2144300 |
| TVLGQQIL( | PKP2 | Q99959 | PKP2_HUMAN | Unmodified | 0 | 0 | 0 | 0 | 0 | 0 | 0 | 0 | 143450 | 0 | 646040 | 665820 | 0 | 0 | 0 | 0 | 505940 | 0 |
| TVLGTPEVL | HADHA; HADH/ | P40939; H0Y | ECHA_HUMAN | Unmodified | 0 | 0 | 0 | 0 | 0 | 0 | 0 | 0 | 1910400 | 543280 | 412400 | 0 | 0 | 0 | 0 | 0 | 0 | 0 |
| TVLGTPEVL | HADHA; HADH/ | P40939; H0Y | ECHA_HUMAN | Unmodified | 0 | 0 | 0 | 0 | 0 | 0 | 721610 | 0 | 3174100 | 652150 | 0 | 0 | 0 | 0 | 0 | 0 | 0 | 0 |
| TVLIMELINI | ATP5F1B; ATP5I | P06576; H0Y | ATPB_HUMAN . | Oxidation (M) | 113960 | 35116 | 23492 | 110110 | 109150 | 36596 | 1075200 | 72338 | 2989600 | 3539700 | 284390 | 174420 | 110510 | 327620 | 76930 | 23497 | 44752 | 118720 |
| TVLIMELINI | ATP5F1B; ATP5I | P06576; H0Y | ATPB_HUMAN . | Unmodified | 0 | 0 | 0 | 0 | 79167 | 0 | 318150 | 0 | 7314800 | 1254400 | 41703 | 17746 | 50203 | 88704 | 0 | 0 | 0 | 0 |
| TVLMLADQ | CSNK1A1L; CSI | Q8N752; P4 | KC1AL_HUMAN | Oxidation (M) | 323900 | 73126 | 60042 | 279220 | 0 | 0 | 289180 | 0 | 0 | 0 | 0 | 136280 | 0 | 0 | 1192700 | 274090 | 0 | 0 |
| TVLNSEVLE | SHTN1 | A0MZ66 | SHOT1_HUMAI | Unmodified | 0 | 0 | 275880 | 473210 | 2502900 | 0 | 0 | 683370 | 0 | 1212300 | 314210 | 0 | 0 | 4503700 | 486130 | 304210 | 1294200 | 348580 |
| TVLPFSQEF | CYFIP1 | Q7L576 | CYFP1_HUMAN | Unmodified | 1014400 | 0 | 696030 | 1359800 | 0 | 0 | 0 | 0 | 888060 | 294670 | 737620 | 7676200 | 0 | 0 | 1734500 | 2076700 | 0 | 0 |
| TVLPFSQEF | CYFIP1 | Q7L576 | CYFP1_HUMAN | Unmodified | 0 | 0 | 258530 | 599820 | 0 | 0 | 0 | 0 | 0 | 0 | 0 | 747040 | 0 | 0 | 398150 | 350290 | 0 | 0 |
| TVLQQVIED | RARS2; RARS2 | Q5T160; H0Y | SYRM_HUMAN | Unmodified | 0 | 0 | 0 | 0 | 4078200 | 1046200 | 0 | 0 | 1242800 | 0 | 0 | 0 | 0 | 0 | 0 | 0 | 1713600 | 3013200 |
| TVLQRPLSI | UPF1 | Q92900 | RENT1_HUMAN | Unmodified | 4091100 | 4821800 | 4963800 | 3893700 | 1605700 | 183980 | 2838600 | 645320 | 1657800 | 995220 | 595560 | 2555500 | 3388400 | 3098500 | 2196800 | 2190500 | 932600 | 1440400 |
| TVLRAGDR | CTNNB1; CTNN | P35222; A0A | CTNB1_HUMAI | Unmodified | 0 | 0 | 0 | 1249100 | 266210 | 215750 | 0 | 0 | 0 | 0 | 0 | 0 | 0 | 338510 | 65509 | 0 | 1111800 | 299350 |
| TVLSANADI | HYOU1; HYOU1 | Q9Y4L1; E9F | HYOU1_HUMA | Unmodified | 0 | 0 | 0 | 0 | 0 | 0 | 0 | 0 | 2526500 | 590300 | 0 | 0 | 0 | 0 | 0 | 0 | 0 | 0 |
| TVLSANADI | HYOU1; HYOU1 | Q9Y4L1; E9F | HYOU1_HUMA | Oxidation (M) | 0 | 0 | 0 | 0 | 0 | 0 | 0 | 0 | 818820 | 418050 | 0 | 0 | 0 | 0 | 0 | 0 | 0 | 0 |
| TVLSANADI | HYOU1; HYOU1 | Q9Y4L1; E9F | HYOU1_HUMA | 2 Oxidation (M) | 0 | 0 | 0 | 253660 | 0 | 0 | 1208500 | 0 | 1330300 | 1404500 | 396220 | 0 | 0 | 44150 | 0 | 0 | 0 | 391590 |
| TVLSANADI | HYOU1; HYOU1 | Q9Y4L1; E9F | HYOU1_HUMA | Oxidation (M) | 0 | 0 | 0 | 0 | 0 | 0 | 0 | 0 | 0 | 2406400 | 0 | 0 | 0 | 0 | 0 | 0 | 0 | 0 |
| TVLTAADVL | CCDC134 | Q9H6E4 | CC134_HUMAI | Unmodified | 0 | 0 | 0 | 0 | 0 | 0 | 0 | 0 | 4393900 | 1134700 | 638550 | 0 | 0 | 0 | 90189 | 0 | 0 | 0 |
| TVLVSSGLF | LRP1 | Q07954 | LRP1_HUMAN I | Unmodified | 0 | 0 | 599820 | 0 | 0 | 0 | 67831 | 19886 | 4707000 | 200540 | 0 | 0 | 0 | 0 | 0 | 110290 | 0 | 0 |
| TVMLGTAA' | RAB25 | P57735 | RAB25_HUMAN | Oxidation (M) | 0 | 0 | 0 | 494960 | 0 | 0 | 0 | 0 | 0 | 0 | 0 | 397850 | 0 | 0 | 262170 | 407750 | 0 | 0 |
| TVMLIPGDI | BCCIP | Q9P287 | BCCIP_HUMAN | 2 Oxidation (M) | 1281100 | 0 | 146450 | 381620 | 0 | 0 | 0 | 19315 | 0 | 0 | 0 | 1424600 | 266560 | 85971 | 453290 | 368000 | 0 | 0 |
| TVMNGALV | SLC6A6; SLC6A | P31641; A0A | SC6A6_HUMAN | 2 Oxidation (M) | 0 | 0 | 0 | 0 | 0 | 0 | 0 | 0 | 0 | 0 | 0 | 949290 | 0 | 0 | 0 | 0 | 0 | 0 |
| TVMPYISTT | ITGB1; ITGB1; I | P05556; H7( | ITB1_HUMAN Ii | Oxidation (M) | 944030 | 503900 | 1039200 | 948730 | 45322000 | 13126000 | 3022900 | 1904400 | 2508100 | 957600 | 770360 | 1555300 | 3300700 | 204430 | 1176700 | 68739 | 17786000 | 50740000 |
| TVMPYISTT | ITGB1; ITGB1; I | P05556; H7( | ITB1_HUMAN Ii | Unmodified | 2682700 | 0 | 2638100 | 4289800 | 3.72E+08 | 59672000 | 17462000 | 7915200 | 22294000 | 6102900 | 2313300 | 8929300 | 18620000 | 14399000 | 4074300 | 2187100 | 1.26E+08 | 10744000 |
| TVMVDDSK | TLN2 | Q9Y4G6 | TLN2_HUMAN | Unmodified | 149300 | 502060 | 533030 | 27132 | 1350500 | 162220 | 706500 | 1301200 | 0 | 0 | 23363 | 0 | 1227800 | 1353700 | 245060 | 671230 | 0 | 1282400 |
| TVMVDDSK | TLN2 | Q9Y4G6 | TLN2_HUMAN | Oxidation (M) | 0 | 0 | 52582 | 0 | 0 | 0 | 0 | 421070 | 0 | 0 | 0 | 0 | 13158 | 0 | 0 | 0 | 0 | 0 |
| TVNDDQSF | PIK3C2B; PIK3( | O00750; F5( | P3C2B_HUMAI | Unmodified | 897030 | 740980 | 0 | 634100 | 0 | 0 | 0 | 0 | 0 | 0 | 0 | 817530 | 0 | 0 | 2702700 | 1849700 | 0 | 0 |

| | | | | | | | | | | | | | | | | | | | | | | |
|---|---|---|---|---|---|---|---|---|---|---|---|---|---|---|---|---|---|---|---|---|---|---|
| TVNDIVSAL | BIRC2; BIRC2; E | Q13490; E9F | BIRC2_HUMAN | Unmodified | 0 | 0 | 0 | 0 | 0 | 1367100 | 0 | 0 | 472180 | 0 | 0 | 0 | 0 | 0 | 0 | 0 | 0 | 0 |
| TVNDLSSQ | UTRN; UTRN | P46939; Q5T | UTRN_HUMAN | Unmodified | 0 | 0 | 1674600 | 1606000 | 1346800 | 64625 | 0 | 0 | 2718100 | 1355500 | 0 | 531440 | 0 | 0 | 6139500 | 4612200 | 146510 | 0 |
| TVNELQNL | IGF2BP3 | O00425 | IF2B3_HUMAN | Unmodified | 583410 | 580350 | 0 | 165240 | 0 | 0 | 0 | 0 | 0 | 0 | 0 | 878560 | 0 | 0 | 0 | 0 | 0 | 0 |
| TVNELQNL | IGF2BP3 | O00425 | IF2B3_HUMAN | Unmodified | 1279400 | 0 | 0 | 0 | 0 | 0 | 0 | 0 | 0 | 0 | 0 | 0 | 1686800 | 625220 | 0 | 0 | 0 | 0 |
| TVNESASLI | INSR | P06213 | INSR_HUMAN I | Unmodified | 0 | 0 | 0 | 0 | 0 | 0 | 0 | 0 | 764740 | 0 | 0 | 0 | 0 | 0 | 0 | 189100 | 0 | 0 |
| TVNLNLWD | RHOG | P84095 | RHOG_HUMAN | Unmodified | 6352100 | 394060 | 2200000 | 8697400 | 1037900 | 0 | 0 | 0 | 882270 | 0 | 4915300 | 13681000 | 0 | 0 | 5097000 | 4285500 | 54377 | 942570 |
| TVNLSVTPS | PHACTR4 | Q8IZ21 | PHAR4_HUMAI | Unmodified | 0 | 0 | 0 | 0 | 1694600 | 260140 | 0 | 138920 | 1071100 | 770330 | 0 | 0 | 1160000 | 980300 | 0 | 0 | 903630 | 2173900 |
| TVNTFSQS | BRI3BP; BRI3BF | Q8WY22; A0 | BRI3B_HUMAN | Unmodified | 0 | 0 | 0 | 0 | 0 | 0 | 1327700 | 0 | 5540500 | 1255800 | 0 | 0 | 0 | 0 | 0 | 0 | 0 | 0 |
| TVNTMALL | DOP1B | Q9Y3R5 | DOP2_HUMAN | Unmodified | 0 | 0 | 0 | 0 | 0 | 0 | 0 | 0 | 327650 | 0 | 0 | 0 | 0 | 0 | 0 | 0 | 0 | 0 |
| TVNVPFHL | RAB3GAP2 | Q9H2M9 | RBGPR_HUMAI | Unmodified | 25086 | 18661 | 127930 | 0 | 0 | 25782 | 50651 | 52560 | 250830 | 58928 | 254140 | 1377100 | 0 | 0 | 118000 | 104320 | 0 | 34079 |
| TVNVVQFE | GARS1; GARS1 | P41250; A0A | GARS_HUMAN | Unmodified | 0 | 232560 | 329950 | 1551300 | 2703300 | 1054700 | 839480 | 815100 | 1176400 | 1500000 | 3262300 | 15811000 | 4391000 | 1806100 | 1411000 | 1263800 | 591950 | 2323400 |
| TVNVVQFE | GARS1; GARS1 | P41250; A0A | GARS_HUMAN | Unmodified | 0 | 0 | 0 | 0 | 1475200 | 131770 | 2093600 | 1301600 | 0 | 683050 | 0 | 0 | 2873100 | 2196800 | 0 | 0 | 2132600 | 807990 |
| TVPEELVKF | PRPF19; PRPF1 | Q9UMS4; F5 | PRP19_HUMAN | Unmodified | 0 | 0 | 1501700 | 0 | 0 | 0 | 1458100 | 369190 | 1555900 | 798930 | 2478900 | 9403000 | 0 | 0 | 0 | 0 | 1029400 | 1599300 |
| TVPFCSTF | TKT; TKT | P29401; A0A | TKT_HUMAN Tr | Unmodified | 144590 | 0 | 0 | 150530 | 1928700 | 180840 | 0 | 0 | 1907600 | 1518500 | 256010 | 4041800 | 833110 | 0 | 68887 | 199230 | 1134800 | 975770 |
| TVPGYLRIG | MVB12A; MVB1 | Q96EY5; E9F | MB12A_HUMAI | Oxidation (M) | 17627 | 0 | 0 | 0 | 892520 | 0 | 0 | 0 | 0 | 0 | 0 | 0 | 3985500 | 2753000 | 65845 | 0 | 0 | 622420 |
| TVPKTVDN | PPIB; PPIB; PPIE | P23284; A0A | PPIB_HUMAN F | Unmodified | 0 | 0 | 0 | 0 | 0 | 0 | 594800 | 375910 | 319400 | 2314900 | 0 | 0 | 110950 | 0 | 0 | 0 | 96643 | 0 |
| TVPLAGHV | SEPTIN6; SEPTI | Q14141; B1A | SEPT6_HUMAN | Unmodified | 0 | 0 | 0 | 229010 | 329640 | 0 | 0 | 24760 | 0 | 0 | 0 | 5956200 | 698620 | 1031800 | 382520 | 140780 | 0 | 0 |
| TVPLLSKVD | RACGAP1; RAC | Q9H0H5; H0 | RGAP1_HUMAI | Unmodified | 0 | 0 | 0 | 0 | 2241300 | 436100 | 350730 | 0 | 0 | 0 | 0 | 446430 | 1000400 | 0 | 0 | 0 | 2067300 | 0 |
| TVPLNGVIF | GNG2 | G3V415 | G3V415_HUMA | Acetyl (Protein N | 0 | 0 | 0 | 0 | 5748700 | 4681300 | 0 | 0 | 0 | 0 | 0 | 0 | 528000 | 612710 | 0 | 0 | 0 | 1101500 |
| TVPNDATQ | NCAM1; NCAM | P13591; A0A | NCAM1_HUMA | Unmodified | 0 | 0 | 0 | 0 | 0 | 0 | 2369500 | 4981400 | 0 | 16002000 | 0 | 0 | 68117 | 0 | 0 | 0 | 0 | 0 |
| TVPPAVTGI | ALDOA; ALDOA | P04075; J3KI | ALDOA_HUMAI | Unmodified | 0 | 0 | 0 | 0 | 0 | 531140 | 0 | 0 | 0 | 1124700 | 0 | 218140 | 0 | 774980 | 320350 | 0 | 2069500 | 0 |
| TVPVEAVTS | CTTN | Q14247 | SRC8_HUMAN | Unmodified | 2581900 | 2047600 | 1942000 | 3047200 | 18296000 | 1832500 | 308360 | 448620 | 683430 | 524570 | 476140 | 0 | 5081300 | 4606200 | 2259900 | 2531800 | 5383000 | 7320100 |
| TVPVEAVTS | CTTN | Q14247 | SRC8_HUMAN | Unmodified | 0 | 0 | 0 | 0 | 161340 | 0 | 33954 | 12899 | 0 | 0 | 0 | 0 | 0 | 173120 | 0 | 0 | 573930 | 0 |
| TVPVEAVTS | CTTN | Q14247 | SRC8_HUMAN | Unmodified | 0 | 0 | 0 | 0 | 898160 | 58022 | 467270 | 71912 | 0 | 0 | 0 | 0 | 0 | 945360 | 0 | 0 | 2462600 | 318470 |
| TVPVPSGK | GPRIN1 | Q7Z2K8 | GRIN1_HUMAN | Unmodified | 0 | 0 | 0 | 0 | 0 | 0 | 0 | 115870 | 2416800 | 211980 | 0 | 0 | 0 | 147920 | 0 | 0 | 0 | 0 |
| TVPVPSGK | GPRIN1 | Q7Z2K8 | GRIN1_HUMAN | Unmodified | 0 | 0 | 0 | 0 | 0 | 0 | 0 | 0 | 2045700 | 0 | 0 | 0 | 583240 | 0 | 0 | 0 | 1030200 | 6363400 |
| TVPWEQPA | TTYH3 | Q9C0H2 | TTYH3_HUMAN | Unmodified | 1113800 | 1256500 | 1564900 | 1567800 | 0 | 0 | 0 | 160980 | 6964200 | 3899200 | 423720 | 8716100 | 0 | 0 | 2810300 | 1769700 | 0 | 0 |
| TVQAVLTVF | SERPINF1; SER | P36955; I3L4 | PEDF_HUMAN | Unmodified | 0 | 0 | 0 | 0 | 278210 | 0 | 0 | 0 | 1561700 | 534950 | 16241000 | 895560 | 0 | 0 | 0 | 0 | 0 | 0 |
| TVQCKVTLI | EPB41L2; EPB4 | O43491; A0A | E41L2_HUMAN | Unmodified | 0 | 0 | 0 | 0 | 1029200 | 0 | 0 | 0 | 0 | 0 | 0 | 0 | 0 | 0 | 0 | 0 | 1426700 | 694770 |
| TVQCKVTLI | EPB41L2; EPB4 | O43491; A0A | E41L2_HUMAN | Unmodified | 0 | 0 | 0 | 0 | 0 | 0 | 0 | 0 | 0 | 0 | 0 | 0 | 0 | 0 | 0 | 0 | 624420 | 0 |
| TVQGPPTS | OAT | P04181 | OAT_HUMAN C | Unmodified | 0 | 0 | 0 | 0 | 0 | 0 | 0 | 36313 | 4529600 | 480670 | 0 | 0 | 0 | 0 | 0 | 0 | 0 | 0 |
| TVQIEASTV | SEPTIN2; SEPTI | Q15019; C9I | SEPT2_HUMAN | Unmodified | 185090 | 474190 | 162020 | 837230 | 0 | 0 | 0 | 0 | 633790 | 0 | 1579000 | 7245500 | 0 | 0 | 568880 | 476030 | 0 | 0 |
| TVQIEASTV | SEPTIN2; SEPTI | Q15019; C9I | SEPT2_HUMAN | Unmodified | 0 | 0 | 0 | 0 | 2299200 | 0 | 393680 | 413080 | 0 | 595310 | 0 | 0 | 0 | 0 | 0 | 0 | 1242900 | 1379500 |
| TVQLTSSEL | LRPPRC | P42704 | LPPRC_HUMAN | Unmodified | 36670 | 61390 | 54039 | 31013 | 0 | 0 | 0 | 0 | 3174500 | 187510 | 236490 | 0 | 0 | 0 | 3641300 | 80623 | 0 | 0 |
| TVQLTSSEL | LRPPRC | P42704 | LPPRC_HUMAN | Unmodified | 0 | 0 | 0 | 0 | 620100 | 0 | 1455900 | 0 | 2937100 | 7294200 | 0 | 0 | 0 | 0 | 0 | 0 | 122150 | 0 |
| TVQLVWK | IL6ST; IL6ST | P40189; A0A | IL6RB_HUMAN | Unmodified | 0 | 0 | 0 | 0 | 0 | 0 | 0 | 0 | 636910 | 0 | 0 | 0 | 0 | 0 | 0 | 0 | 0 | 0 |
| TVQPKPDY | ACY1; ACY1; AE | Q03154; F8V | ACY1_HUMAN | Unmodified | 0 | 0 | 0 | 0 | 0 | 0 | 0 | 107620 | 0 | 0 | 1049800 | 4646500 | 0 | 0 | 0 | 0 | 0 | 0 |
| TVQPVAMG | CTNND1; CTNN | O60716; E9F | CTND1_HUMAI | Oxidation (M) | 0 | 5010300 | 0 | 11460000 | 0 | 1661900 | 0 | 0 | 0 | 0 | 2878000 | 5548700 | 0 | 0 | 18467000 | 10037000 | 1618200 | 3516700 |
| TVQPVAMG | CTNND1; CTNN | O60716; A0A | CTND1_HUMAI | Oxidation (M) | 0 | 0 | 0 | 0 | 23327000 | 0 | 399610 | 0 | 0 | 0 | 0 | 0 | 0 | 4076400 | 0 | 0 | 23555000 | 45669000 |
| TVQQHAGI | HAGH; HAGH; I | Q16775; H3I | GLO2_HUMAN | Unmodified | 0 | 0 | 0 | 0 | 396120 | 0 | 0 | 0 | 0 | 0 | 0 | 0 | 262240 | 0 | 0 | 0 | 292810 | 425670 |
| TVQQHAGI | HAGH; HAGH; I | Q16775; H3I | GLO2_HUMAN | Oxidation (M) | 71398 | 43571 | 96584 | 93774 | 0 | 21002 | 0 | 0 | 0 | 56246 | 66013 | 0 | 248840 | 0 | 0 | 100520 | 133610 | 161430 |
| TVQQLRLE | GNG12 | Q9UBI6 | GBG12_HUMAI | Unmodified | 241600 | 344940 | 0 | 1466900 | 3958100 | 0 | 0 | 0 | 2282200 | 1998400 | 0 | 0 | 3110800 | 2104700 | 833750 | 454590 | 1194900 | 5882600 |
| TVQQLRLE | GNG12 | Q9UBI6 | GBG12_HUMAI | Unmodified | 0 | 0 | 0 | 0 | 41511000 | 0 | 550340 | 0 | 0 | 3121900 | 0 | 0 | 35465000 | 18812000 | 0 | 0 | 6669300 | 56006000 |
| TVQRTQDN | ST14 | Q9Y5Y6 | ST14_HUMAN S | Unmodified | 0 | 0 | 0 | 765970 | 0 | 0 | 0 | 0 | 0 | 0 | 0 | 0 | 1665500 | 0 | 0 | 0 | 0 | 0 |
| TVQRVEDG | PACSIN3; PACS | Q9UKS6; E9I | PACN3_HUMAI | Unmodified | 3331400 | 1063200 | 298570 | 0 | 0 | 0 | 0 | 0 | 0 | 0 | 0 | 0 | 2133900 | 645400 | 0 | 571750 | 0 | 0 |
| TVQTAAAN | SCAMP1; SCAM | O15126; A0A | SCAM1_HUMA | Unmodified | 3836700 | 1728200 | 8735200 | 9768400 | 2268300 | 0 | 1168300 | 154550 | 5018900 | 853050 | 0 | 4570600 | 3297900 | 0 | 2863800 | 6326200 | 0 | 0 |
| TVQTAAAN | SCAMP1; SCAM | O15126; A0A | SCAM1_HUMA | Unmodified | 0 | 0 | 0 | 583300 | 12588000 | 403960 | 0 | 863200 | 0 | 621380 | 0 | 0 | 12701000 | 3997000 | 0 | 0 | 277820 | 1946100 |
| TVQTDHNA | AP1G1 | O43747 | AP1G1_HUMAN | Unmodified | 36759 | 0 | 294420 | 108270 | 0 | 14359 | 13488 | 0 | 195550 | 20849 | 10686 | 876390 | 0 | 0 | 178660 | 144780 | 0 | 0 |
| TVQVEQSK | SEPTIN7; SEPTI | Q16181; A0A | SEPT7_HUMAN | Unmodified | 0 | 0 | 0 | 0 | 508900 | 0 | 0 | 0 | 0 | 0 | 0 | 0 | 0 | 0 | 0 | 0 | 769120 | 506040 |
| TVQVIMETF | PLCG2; PLCG2 | P16885; A0A | PLCG2_HUMAI | Oxidation (M) | 0 | 0 | 0 | 0 | 0 | 0 | 0 | 0 | 0 | 0 | 1405800 | 1549500 | 0 | 0 | 0 | 0 | 0 | 0 |
| TVQVIMETF | PLCG2; PLCG2 | P16885; A0A | PLCG2_HUMAI | Unmodified | 0 | 0 | 0 | 0 | 0 | 0 | 0 | 0 | 0 | 0 | 2219200 | 12301000 | 0 | 0 | 0 | 0 | 0 | 0 |
| TVQVVPHIT | CTPS1; CTPS1; | P17812; A0A | PYRG1_HUMAI | Oxidation (M) | 209450 | 0 | 0 | 0 | 0 | 0 | 0 | 0 | 1158700 | 396540 | 0 | 1883500 | 0 | 0 | 0 | 0 | 546720 | 0 |
| TVQYQNEL | RAB22A | Q9UL26 | RB22A_HUMAN | Unmodified | 1855900 | 1731500 | 1325200 | 2723200 | 127560 | 143770 | 418670 | 247260 | 5669500 | 1371700 | 498210 | 3207900 | 1512100 | 815950 | 2371600 | 2327800 | 1237700 | 1013700 |
| TVRDSDSG | MLF2 | Q15773 | MLF2_HUMAN | Unmodified | 0 | 926080 | 0 | 0 | 556390 | 0 | 0 | 0 | 0 | 0 | 0 | 0 | 0 | 0 | 0 | 0 | 0 | 0 |
| TVRDSDSG | MLF2 | Q15773 | MLF2_HUMAN | Unmodified | 2203100 | 2435900 | 0 | 1383400 | 0 | 0 | 109070 | 0 | 0 | 0 | 0 | 0 | 0 | 0 | 503780 | 274190 | 0 | 0 |
| TVRGSLER | CAVIN1 | Q6NZI2 | CAVN1_HUMAI | Unmodified | 0 | 0 | 0 | 0 | 24254000 | 2311600 | 0 | 81736 | 0 | 0 | 0 | 0 | 0 | 0 | 0 | 0 | 27779 | 0 |
| TVRLWDHA | STRAP | Q9Y3F4 | STRAP_HUMAN | Unmodified | 0 | 0 | 0 | 0 | 686580 | 73536 | 416170 | 0 | 4760200 | 72143 | 0 | 0 | 1409400 | 694130 | 0 | 0 | 677150 | 335000 |
| TVRNDNSS | MYH3; MYH4; M | P11055; Q9Y | MYH3_HUMAN | Unmodified | 0 | 0 | 28782 | 0 | 6096500 | 1785000 | 390700 | 0 | 197050 | 322270 | 0 | 0 | 182540 | 205510 | 0 | 0 | 2215700 | 683730 |
| TVRQAEML | AK2; AK2; AK2; | P54819; F8V | KAD2_HUMAN | Unmodified | 0 | 0 | 0 | 0 | 259830 | 0 | 51370 | 0 | 0 | 377110 | 0 | 126000 | 0 | 0 | 0 | 0 | 0 | 27874 |
| TVRVVATK | LSR; LSR; LSR; I | Q86X29; M0 | LSR_HUMAN Li | Unmodified | 0 | 0 | 0 | 0 | 19291000 | 0 | 1005000 | 0 | 0 | 1202400 | 0 | 0 | 135630 | 0 | 0 | 0 | 3136300 | 0 |
| TVSDLIDQ | EIF3L; EIF3L; EI | Q9Y262; B0 | EIF3L_HUMAN | Unmodified | 0 | 0 | 578310 | 534020 | 0 | 0 | 0 | 0 | 0 | 0 | 0 | 7490100 | 0 | 0 | 489570 | 617830 | 0 | 0 |
| TVSEWLES | EPHA2 | P29317 | EPHA2_HUMAN | Unmodified | 0 | 0 | 0 | 0 | 0 | 1687600 | 0 | 0 | 0 | 0 | 576070 | 3596500 | 0 | 0 | 0 | 0 | 0 | 0 |
| TVSFASSQ | ENTPD4 | Q9Y227 | ENTP4_HUMAN | Unmodified | 0 | 0 | 0 | 0 | 0 | 815940 | 91288 | 0 | 775170 | 0 | 0 | 0 | 0 | 0 | 0 | 0 | 0 | 0 |
| TVSGVNGF | ATP6V1B2; ATP | P21281; E5R | VATB2_HUMAN | Unmodified | 208370 | 0 | 471290 | 0 | 0 | 0 | 0 | 0 | 626900 | 0 | 0 | 2915300 | 0 | 0 | 395110 | 438130 | 0 | 0 |
| TVSGVNGF | ATP6V1B2; ATP | P21281; H0Y | VATB2_HUMAN | Unmodified | 0 | 0 | 0 | 0 | 1591500 | 0 | 0 | 0 | 457800 | 881650 | 0 | 0 | 0 | 0 | 0 | 0 | 0 | 944300 |
| TVSHEAEVI | GOLGB1; GOLG | Q14789; E7E | GOGB1_HUMA | Unmodified | 0 | 0 | 69797 | 0 | 0 | 0 | 0 | 0 | 29932 | 0 | 0 | 361060 | 143090 | 0 | 0 | 0 | 0 | 25002 |
| TVSHLYQE | TANC1 | Q9C0D5 | TANC1_HUMAI | Unmodified | 0 | 0 | 0 | 317170 | 243930 | 132100 | 0 | 0 | 0 | 64395 | 0 | 42823 | 430280 | 369020 | 298870 | 261040 | 29427 | 349710 |
| TVSKVDDF | PGD; PGD; PGD | P52209; K7E | 6PGD_HUMAN | Unmodified | 825030 | 1173200 | 780600 | 2372600 | 10132000 | 2446700 | 487910 | 280950 | 9187500 | 3412000 | 8775800 | 23010000 | 4399600 | 2324000 | 1624000 | 1291900 | 644060 | 2292000 |
| TVSLGAGA | NPM1; NPM1; N | P06748; A0A | NPM_HUMAN N | Oxidation (M) | 1381400 | 2970800 | 1541100 | 1269200 | 2221100 | 287620 | 124910 | 0 | 11217000 | 3170700 | 30288000 | 13989000 | 3084700 | 6975300 | 96786 | 0 | 1865900 | 14145000 |
| TVSLGAGA | NPM1; NPM1; N | P06748; A0A | NPM_HUMAN N | Unmodified | 0 | 0 | 0 | 0 | 676410 | 0 | 0 | 0 | 7790700 | 323950 | 1803900 | 0 | 999410 | 0 | 0 | 0 | 920280 | 0 |
| TVSLGAGA | NPM1; NPM1; N | P06748; A0A | NPM_HUMAN N | Oxidation (M) | 0 | 0 | 0 | 0 | 874960 | 0 | 426850 | 0 | 0 | 785250 | 0 | 0 | 3651700 | 995260 | 0 | 0 | 0 | 0 |
| TVSLLDENI | PRKDC | P78527 | PRKDC_HUMAI | Unmodified | 0 | 0 | 221720 | 0 | 0 | 0 | 3791200 | 2671300 | 514370 | 227870 | 0 | 2783000 | 96557 | 0 | 0 | 0 | 0 | 334840 |
| TVSPALISR | CAND1; CAND1 | Q86VP6; A0A | CAND1_HUMA | Unmodified | 269200 | 254570 | 664640 | 727270 | 510730 | 0 | 106420 | 0 | 3600100 | 1484100 | 2136000 | 9598300 | 1144600 | 1230100 | 985290 | 748480 | 752710 | 599450 |
| TVSPALISR | CAND1; CAND1 | Q86VP6; A0A | CAND1_HUMA | Unmodified | 0 | 0 | 0 | 209020 | 1952500 | 290760 | 476790 | 182470 | 0 | 477970 | 0 | 0 | 1275600 | 1155100 | 0 | 0 | 745950 | 1080600 |
| TVSPDRLEI | KPNB1; KPNB1 | Q14974; J3Q | IMB1_HUMAN I | Unmodified | 2473900 | 2311800 | 155100 | 5152300 | 20024000 | 3517500 | 7457800 | 2431600 | 465600 | 6307800 | 7251100 | 25085000 | 15497000 | 464220 | 4310300 | 4407300 | 12541000 | 11112000 |
| TVSPDRLEI | KPNB1; KPNB1 | Q14974; J3Q | IMB1_HUMAN I | Unmodified | 0 | 0 | 0 | 0 | 0 | 0 | 214140 | 0 | 0 | 258530 | 0 | 0 | 228700 | 298030 | 0 | 0 | 446450 | 357070 |
| TVSQIISLQ | RSL1D1; RSL1D | O76021; J3Q | RL1D1_HUMAN | Unmodified | 459740 | 232140 | 0 | 0 | 0 | 0 | 730400 | 567490 | 0 | 0 | 0 | 6640400 | 0 | 0 | 0 | 0 | 0 | 0 |
| TVSSEDVAI | PIP4K2B | P78356 | PI42B_HUMAN | Unmodified | 0 | 0 | 0 | 0 | 0 | 0 | 0 | 52450 | 535160 | 40883 | 0 | 603810 | 0 | 0 | 192620 | 118430 | 74931 | 0 |
| TVSSGELAT | SEMA4C; SEMA | Q9C0C4; C9 | SEM4C_HUMAI | Unmodified | 0 | 0 | 0 | 0 | 0 | 0 | 0 | 0 | 713610 | 0 | 0 | 0 | 0 | 0 | 0 | 0 | 0 | 0 |
| TVSSGELAT | SEMA4C; SEMA | Q9C0C4; C9 | SEM4C_HUMAI | Unmodified | 0 | 0 | 0 | 0 | 0 | 0 | 0 | 0 | 1730600 | 286690 | 0 | 0 | 0 | 0 | 0 | 0 | 0 | 0 |
| TVSSHEVFI | SNX5; SNX5; SN | Q9Y5X3; Q5 | SNX5_HUMAN | Unmodified | 311830 | 0 | 133080 | 134990 | 0 | 32883 | 50130 | 0 | 349990 | 202760 | 299290 | 1672800 | 241770 | 114980 | 379070 | 326780 | 74319 | 0 |
| TVSSPIPYT | CCDC6 | Q16204 | CCDC6_HUMA | Unmodified | 0 | 0 | 0 | 0 | 0 | 0 | 0 | 0 | 0 | 0 | 0 | 2475400 | 4068800 | 0 | 0 | 0 | 0 | 2225100 |
| TVSSTKLVS | IGF2R; IGF2R | P11717; A0A | MPRI_HUMAN ( | Unmodified | 0 | 0 | 0 | 0 | 0 | 0 | 0 | 0 | 0 | 222580 | 0 | 0 | 386620 | 512030 | 0 | 0 | 118510 | 76055 |
| TVSTLHHVI | PDXK; PDXK | O00764; F2Z | PDXK_HUMAN | Unmodified | 302510 | 0 | 0 | 321550 | 0 | 0 | 0 | 0 | 290620 | 194230 | 0 | 712830 | 0 | 727290 | 0 | 1229200 | 0 | 0 |
| TVSTLHHVI | PDXK; PDXK | O00764; F2Z | PDXK_HUMAN | Unmodified | 0 | 0 | 0 | 0 | 0 | 0 | 0 | 0 | 0 | 0 | 0 | 0 | 0 | 0 | 0 | 0 | 874370 | 1052500 |
| TVSYNGILC | TGM2 | P21980 | TGM2_HUMAN | Unmodified | 0 | 0 | 0 | 0 | 2528000 | 647370 | 0 | 0 | 0 | 0 | 0 | 0 | 0 | 0 | 0 | 0 | 0 | 0 |
| TVSYTQFLL | CABLES1 | Q8TDN4 | CABL1_HUMAN | Unmodified | 0 | 0 | 29458 | 0 | 0 | 0 | 0 | 0 | 228730 | 0 | 82969 | 177230 | 0 | 0 | 17620 | 109550 | 0 | 0 |
| TVTAEVQP | P3H1 | Q32P28 | P3H1_HUMAN | Unmodified | 0 | 0 | 0 | 0 | 0 | 0 | 0 | 0 | 1649100 | 408900 | 0 | 0 | 0 | 0 | 0 | 0 | 0 | 0 |
| TVTAEVQP | P3H1 | Q32P28 | P3H1_HUMAN | Unmodified | 0 | 0 | 0 | 0 | 0 | 0 | 0 | 0 | 0 | 647930 | 0 | 0 | 0 | 0 | 0 | 0 | 0 | 0 |
| TVTAMDVV | H4C1 | P62805 | H4_HUMAN Hi | Oxidation (M) | 97677000 | 58028000 | 3764800 | 1862300 | 1834500 | 3298900 | 17821000 | 55324000 | 974900 | 1480200 | 1705000 | 7.05E+08 | 395740 | 406500 | 370050 | 2516200 | 0 | 2208100 |
| TVTAMDVV | H4C1 | P62805 | H4_HUMAN Hi | Unmodified | 32178000 | 848620 | 0 | 0 | 5482100 | 1801700 | 16388000 | 26160000 | 3752700 | 2487700 | 0 | 4E+08 | 0 | 1115300 | 0 | 0 | 0 | 2650300 |
| TVTAMDVV | H4C1 | P62805 | H4_HUMAN Hi | Oxidation (M) | 2.78E+08 | 2.47E+08 | 2557700 | 1045600 | 569560 | 144280 | 1414500 | 2431900 | 1513000 | 341560 | 216480 | 2.06E+09 | 481300 | 87585 | 700270 | 2388700 | 101160 | 210040 |
| TVTAMDVV | H4C1 | P62805 | H4_HUMAN Hi | Unmodified | 1.76E+08 | 83554000 | 545860 | 169850 | 432610 | 0 | 1057000 | 500220 | 2401200 | 189870 | 0 | 1.03E+09 | 292500 | 0 | 184350 | 189050 | 0 | 0 |
| TVTAMDVV | H4C1 | P62805 | H4_HUMAN Hi | Unmodified | 0 | 0 | 0 | 0 | 0 | 0 | 2569300 | 509240 | 0 | 0 | 0 | 0 | 0 | 0 | 0 | 0 | 0 | 0 |
| TVTAMDVV | H4C1 | P62805 | H4_HUMAN Hi | Oxidation (M) | 55403 | 29544 | 0 | 0 | 35633 | 0 | 348560 | 619090 | 0 | 0 | 0 | 429310 | 69835 | 0 | 0 | 0 | 0 | 49338 |
| TVTATFGYF | SYPL1; SYPL1; S | Q16563; A0A | SYPL1_HUMAN | Unmodified | 0 | 0 | 0 | 0 | 0 | 0 | 108970 | 0 | 4692400 | 1174800 | 187020 | 19856000 | 0 | 0 | 18479000 | 9556300 | 0 | 0 |
| TVTEYKIDE | EIF3G; EIF3G; E | O75821; K7E | EIF3G_HUMAN | Unmodified | 280130 | 186940 | 0 | 451180 | 870500 | 431630 | 0 | 0 | 377120 | 145020 | 0 | 2516100 | 532370 | 0 | 0 | 284500 | 228090 | 1006100 |
| TVTEYKIDE | EIF3G; EIF3G; E | O75821; K7E | EIF3G_HUMAN | Unmodified | 197000 | 137730 | 127470 | 411790 | 0 | 0 | 0 | 55502 | 0 | 0 | 0 | 1843900 | 0 | 0 | 167380 | 168210 | 0 | 715430 |
| TVTGEFIDK | IGSF3 | O75054 | IGSF3_HUMAN | Unmodified | 1240500 | 2226400 | 1053800 | 1496000 | 4947500 | 1498700 | 626920 | 251500 | 1222700 | 179640 | 0 | 4053700 | 2790700 | 958400 | 1840200 | 1365500 | 0 | 0 |
| TVTISDHG | LAMP2 | P13473 | LAMP2_HUMAN | Unmodified | 0 | 0 | 2387100 | 1347100 | 0 | 0 | 0 | 0 | 0 | 0 | 0 | 516730 | 0 | 0 | 2946100 | 0 | 0 | 0 |
| TVTISQVSD | SH3KBP1; SH3I | Q96B97; Q5 | SH3K1_HUMAN | Unmodified | 264490 | 0 | 401580 | 539670 | 2499000 | 320930 | 0 | 0 | 115920 | 0 | 0 | 0 | 1305800 | 0 | 334380 | 391110 | 88803 | 0 |
| TVTLGGDV | SMC2 | O95347 | SMC2_HUMAN | Unmodified | 0 | 0 | 0 | 0 | 369160 | 0 | 0 | 42853 | 670080 | 0 | 370230 | 1567900 | 0 | 0 | 0 | 0 | 0 | 0 |
| TVTLVEAIK | MOV10; MOV10 | Q9HCE1; Q5 | MOV10_HUMA | Unmodified | 7148100 | 2217300 | 1969800 | 466090 | 0 | 0 | 0 | 0 | 0 | 0 | 0 | 0 | 0 | 0 | 550650 | 597580 | 0 | 0 |
| TVTNAVVT | HSPA8; HSPA8; | P11142; E9P | HSP7C_HUMAI | Unmodified | 53836000 | 61860000 | 87825000 | 1.03E+08 | 50807000 | 11541000 | 5991200 | 2086600 | 1.19E+08 | 24804000 | 49659000 | 2.28E+08 | 58833000 | 31971000 | 75385000 | 46576000 | 6763100 | 25118000 |
| TVTNAVVT | HSPA8; HSPA8; | P11142; E9P | HSP7C_HUMAI | Unmodified | 1919700 | 0 | 0 | 25765000 | 6.85E+08 | 80078000 | 2.75E+08 | 98260000 | 18199000 | 1.94E+08 | 10342000 | 0 | 5.79E+08 | 3.72E+08 | 0 | 0 | 1.2E+08 | 2.3E+08 |
| TVTNAVVT | HSPA8; HSPA8; | P11142; E9P | HSP7C_HUMAI | Unmodified | 0 | 0 | 0 | 0 | 18342000 | 1073300 | 21406000 | 0 | 0 | 8004500 | 0 | 0 | 9601700 | 25690000 | 1216000 | 0 | 19561000 | 8669200 |
| TVTPDASV | TBC1D24; TBC1 | Q9ULP9; A0A | TBC24_HUMAN | Unmodified | 0 | 0 | 0 | 0 | 0 | 0 | 0 | 0 | 386160 | 0 | 0 | 2879600 | 0 | 0 | 473350 | 0 | 253430 | 0 |
| TVTPVSPLG | RAET1L; ULBP2 | Q5VY80; Q9I | ULBP6_HUMAN | Unmodified | 80037 | 122000 | 649970 | 494750 | 0 | 0 | 0 | 0 | 1752400 | 238580 | 0 | 0 | 0 | 0 | 968310 | 554610 | 0 | 0 |
| TVTSATIVYI | UPF1 | Q92900 | RENT1_HUMAN | Unmodified | 4572100 | 3906600 | 4851900 | 6473300 | 742710 | 253600 | 532170 | 170470 | 1440200 | 388350 | 0 | 2871700 | 2065600 | 820140 | 3380700 | 2447500 | 288640 | 663240 |

| | | | | | | | | | | | | | | | | | | | | | | |
|---|---|---|---|---|---|---|---|---|---|---|---|---|---|---|---|---|---|---|---|---|---|---|
| TVTTRTVQF | CTNND1; CTNN | O60716; E9F | CTND1_HUMAI | Oxidation (M) | 0 | 0 | 0 | 0 | 0 | 0 | 0 | 0 | 0 | 236820 | 0 | 0 | 0 | 0 | 0 | 0 | 1107000 | 1759400 |
| TVTTRTVQF | CTNND1; CTNN | O60716; A0A | CTND1_HUMAI | Oxidation (M) | 0 | 0 | 0 | 0 | 0 | 0 | 0 | 0 | 0 | 0 | 0 | 0 | 0 | 0 | 0 | 0 | 2937700 | 0 |
| TVTYEDPQ | TOM1L2; TOM1 | Q6ZVM7; F5I | TM1L2_HUMAN | Unmodified | 7643600 | 126840 | 2802500 | 5132000 | 720720 | 51503 | 0 | 0 | 1260300 | 309250 | 0 | 0 | 7868600 | 0 | 8562500 | 5527100 | 0 | 2690800 |
| TVVALCGQ | CD151; CD151 | P48509; E9P | CD151_HUMAI | Unmodified | 40333000 | 19903000 | 6695900 | 9428600 | 0 | 0 | 198470 | 220320 | 4661800 | 0 | 0 | 17025000 | 0 | 0 | 26579000 | 20447000 | 264170 | 462530 |
| TVVALCGQ | CD151; CD151 | P48509; E9P | CD151_HUMAI | Unmodified | 2432000 | 1679000 | 779480 | 247930 | 1608900 | 0 | 0 | 0 | 249440 | 0 | 0 | 0 | 0 | 0 | 634570 | 1271100 | 467890 | 2998400 |
| TVVAPMLD | COLGALT1; CO | Q8NBJ5; M0 | GT251_HUMAN | Unmodified | 0 | 0 | 0 | 0 | 0 | 0 | 0 | 0 | 10096000 | 237990 | 0 | 0 | 0 | 0 | 0 | 0 | 0 | 0 |
| TVVFAGTR | PLXNA1 | Q9UIW2 | PLXA1_HUMAN | Unmodified | 0 | 0 | 0 | 158480 | 277830 | 0 | 0 | 0 | 328470 | 14740 | 0 | 72783 | 112780 | 0 | 0 | 175630 | 104920 | 89267 |
| TVVNKDVF | RPL27; RPL27 | P61353; K7E | RL27_HUMAN ( | Unmodified | 423200 | 1457600 | 0 | 780300 | 1745500 | 139710 | 210350 | 439110 | 0 | 121500 | 0 | 0 | 1048700 | 1157000 | 0 | 0 | 0 | 850910 |
| TVVPFSVP( | ABCA3; ABCA3 | Q99758; H0Y | ABCA3_HUMAI | Unmodified | 0 | 0 | 0 | 0 | 0 | 0 | 0 | 0 | 2634000 | 0 | 0 | 0 | 0 | 0 | 0 | 0 | 0 | 0 |
| TVVSGLVN | AIMP1; AIMP1 | Q12904; A0A | AIMP1_HUMAN | Oxidation (M) | 0 | 0 | 0 | 357780 | 0 | 0 | 0 | 0 | 0 | 0 | 1177200 | 3815100 | 2615500 | 0 | 0 | 472290 | 0 | 0 |
| TVVSGLVN | AIMP1; AIMP1 | Q12904; A0A | AIMP1_HUMAN | Unmodified | 0 | 0 | 0 | 0 | 854320 | 0 | 0 | 0 | 1033600 | 185360 | 707960 | 1748400 | 1112800 | 0 | 0 | 0 | 0 | 0 |
| TVVSGLVQ | YARS1; YARS1 | P54577; A0A | SYYC_HUMAN | Unmodified | 0 | 0 | 399000 | 242470 | 0 | 0 | 0 | 113730 | 0 | 0 | 528300 | 5074200 | 0 | 0 | 0 | 560190 | 0 | 0 |
| TVVSGLVQ | YARS1; YARS1 | P54577; A0A | SYYC_HUMAN | Unmodified | 0 | 0 | 0 | 0 | 0 | 0 | 0 | 0 | 0 | 0 | 0 | 3328700 | 0 | 0 | 0 | 380860 | 0 | 0 |
| TVVSGSCA | RCC2 | Q9P258 | RCC2_HUMAN | Unmodified | 10995000 | 0 | 0 | 0 | 698810 | 0 | 265510 | 0 | 0 | 0 | 0 | 5712800 | 0 | 0 | 0 | 0 | 0 | 422830 |
| TVVSHSLTI | TMPO | P42166 | LAP2A_HUMAN | Unmodified | 260440 | 0 | 509090 | 82544 | 159290 | 0 | 754670 | 123210 | 269770 | 305770 | 0 | 442580 | 276290 | 0 | 0 | 0 | 576640 | 0 |
| TVVTEHALI | GEMIN5 | Q8TEQ6 | GEMI5_HUMAN | Unmodified | 0 | 0 | 0 | 0 | 0 | 0 | 0 | 0 | 0 | 196040 | 649870 | 6290000 | 0 | 0 | 0 | 0 | 0 | 0 |
| TVVTGEQIV | GALNT3; GALN | Q14435; E7E | GALT3_HUMAN | Unmodified | 0 | 0 | 0 | 0 | 0 | 0 | 0 | 0 | 1881300 | 0 | 0 | 493000 | 0 | 0 | 0 | 0 | 0 | 0 |
| TVVTGEQIV | GALNT3; GALN | Q14435; E7E | GALT3_HUMAN | Unmodified | 0 | 0 | 0 | 0 | 0 | 0 | 0 | 0 | 131470 | 454970 | 0 | 0 | 237640 | 463310 | 0 | 0 | 0 | 0 |
| TVVTGIEMF | TUFM | P49411 | EFTU_HUMAN | Oxidation (M) | 0 | 0 | 525990 | 0 | 2626100 | 541510 | 2167900 | 954820 | 1321100 | 361510 | 0 | 158300 | 728820 | 2470800 | 291040 | 0 | 194360 | 0 |
| TVVTGIEMF | TUFM | P49411 | EFTU_HUMAN | Unmodified | 90388 | 0 | 199560 | 0 | 284430 | 0 | 0 | 0 | 10994000 | 794030 | 0 | 0 | 0 | 0 | 0 | 0 | 0 | 217880 |
| TVWDWELI | HSP90B1; HSP9 | P14625; A0A | ENPL_HUMAN | Unmodified | 0 | 239720 | 53695 | 267800 | 0 | 61674 | 0 | 0 | 556060 | 0 | 205880 | 0 | 0 | 158320 | 0 | 75190 | 0 | 0 |
| TVWDWELI | HSP90B1; HSP9 | P14625; A0A | ENPL_HUMAN | Unmodified | 0 | 0 | 0 | 0 | 0 | 0 | 0 | 0 | 17441000 | 1893800 | 245680 | 0 | 0 | 0 | 0 | 0 | 0 | 0 |
| TVWDWELI | HSP90B1; HSP9 | P14625; A0A | ENPL_HUMAN | Oxidation (M) | 256130 | 0 | 0 | 0 | 0 | 0 | 2553200 | 0 | 11301000 | 3136000 | 1498700 | 1633300 | 0 | 0 | 496950 | 0 | 0 | 0 |
| TVWEISQK | RRM1; RRM1 | P23921; E9P | RIR1_HUMAN F | Unmodified | 0 | 0 | 0 | 331440 | 31356000 | 1676800 | 490420 | 409720 | 0 | 0 | 645810 | 4310100 | 4380400 | 1224300 | 3936100 | 0 | 20399000 | 29357000 |
| TVWSSGDI | SPINT2; SPINT2 | O43291; K7E | SPIT2_HUMAN | Unmodified | 0 | 0 | 363640 | 49405 | 0 | 0 | 0 | 0 | 4357700 | 131650 | 0 | 0 | 0 | 0 | 60997 | 65729 | 0 | 0 |
| TVWSSGDI | SPINT2; SPINT2 | O43291; K7E | SPIT2_HUMAN | Unmodified | 1837900 | 1095800 | 1414100 | 3438400 | 6032100 | 2057300 | 0 | 0 | 1.65E+08 | 18838000 | 937190 | 0 | 1962200 | 306990 | 774840 | 1380700 | 0 | 1320700 |
| TVYFAEEV( | CSRP1; CSRP1 | P21291; E9P | CSRP1_HUMAI | Unmodified | 4030900 | 1584500 | 1176100 | 1596200 | 16839000 | 4760400 | 2961600 | 1989800 | 14010000 | 6552600 | 1021400 | 3169700 | 15881000 | 8620300 | 2588500 | 1941600 | 8520900 | 4298600 |
| TVYFAEK | CRIP2; CRIP2 | P52943; H0Y | CRIP2_HUMAN | Unmodified | 456390 | 369100 | 0 | 102620 | 1503500 | 0 | 0 | 4693900 | 5559900 | 3421500 | 0 | 0 | 0 | 0 | 0 | 0 | 5878900 | 7508000 |
| TVYGGGCS | CCT2; CCT2 | P78371; F5G | TCPB_HUMAN | Oxidation (M) | 0 | 0 | 0 | 0 | 0 | 0 | 0 | 0 | 0 | 0 | 0 | 1999400 | 0 | 0 | 0 | 0 | 0 | 0 |
| TVYGGGCS | CCT2; CCT2 | P78371; F5G | TCPB_HUMAN | Oxidation (M) | 0 | 0 | 0 | 437980 | 0 | 0 | 202050 | 0 | 1238600 | 290930 | 0 | 10472000 | 2105600 | 498310 | 338140 | 219660 | 0 | 0 |
| TVYGGGCS | CCT2; CCT2 | P78371; F5G | TCPB_HUMAN | Unmodified | 0 | 0 | 0 | 0 | 0 | 0 | 0 | 125220 | 4639800 | 0 | 0 | 785950 | 0 | 0 | 0 | 0 | 0 | 0 |
| TVYGGGCS | CCT2; CCT2 | P78371; F5G | TCPB_HUMAN | 2 Oxidation (M) | 75996 | 0 | 178110 | 124940 | 0 | 0 | 0 | 0 | 0 | 0 | 443830 | 2604500 | 576440 | 76624 | 222280 | 100620 | 0 | 0 |
| TVYHAEEV( | CSRP2; CSRP2; | Q16527; F8V | CSRP2_HUMAI | Unmodified | 541070 | 185280 | 0 | 0 | 178400 | 0 | 0 | 0 | 0 | 0 | 535630 | 3398700 | 0 | 0 | 0 | 495460 | 24160 | 0 |
| TVYSHLFDI | MYO6; MYO6; M | Q9UM54; A0 | MYO6_HUMAN | Unmodified | 0 | 0 | 1502700 | 0 | 0 | 797740 | 0 | 0 | 0 | 0 | 0 | 2090900 | 0 | 0 | 1436000 | 1512400 | 0 | 0 |
| TVYSILR | EIF2S1; EIF2S1; | P05198; H0Y | IF2A_HUMAN E | Unmodified | 2572800 | 0 | 827180 | 923830 | 0 | 0 | 0 | 61103 | 0 | 22938 | 994320 | 9180800 | 4253400 | 3016400 | 557090 | 3238400 | 0 | 0 |
| TVYVSVLPT | IARS1 | P41252 | SYIC_HUMAN I | Unmodified | 0 | 0 | 113240 | 0 | 1015200 | 0 | 796310 | 0 | 0 | 765540 | 813890 | 2530100 | 860560 | 479140 | 0 | 0 | 1135800 | 0 |
| TWDHFWL | LTA4H | P09960 | LKHA4_HUMAN | Unmodified | 37711 | 449850 | 0 | 112450 | 0 | 0 | 0 | 0 | 0 | 241580 | 299690 | 1705300 | 0 | 0 | 530880 | 314840 | 0 | 0 |
| TWEQQQE | DBNL; DBNL; D | Q9UJU6; B4I | DBNL_HUMAN | Unmodified | 0 | 0 | 0 | 0 | 0 | 0 | 0 | 0 | 666830 | 0 | 242500 | 1859300 | 0 | 0 | 676900 | 0 | 29361 | 0 |
| TWFSIPEK | DHFR2; DHFR; | Q86XF0; P0( | DYR2_HUMAN | Unmodified | 0 | 0 | 0 | 0 | 0 | 0 | 0 | 6762.3 | 0 | 0 | 0 | 0 | 1200200 | 0 | 0 | 0 | 0 | 0 |
| TWGADVIN | MTAP; MTAP; M | Q13126; J3K | MTAP_HUMAN | Unmodified | 0 | 0 | 0 | 0 | 0 | 0 | 0 | 0 | 0 | 0 | 0 | 0 | 0 | 0 | 0 | 0 | 0 | 0 |
| TWGLHLFS | MFGE8 | Q08431 | MFGM_HUMAN | Unmodified | 3045400 | 3064300 | 4714500 | 14344000 | 7261200 | 2550000 | 0 | 0 | 6247200 | 0 | 0 | 3083700 | 805730 | 0 | 15934000 | 11171000 | 0 | 1360900 |
| TWGLHLFS | MFGE8 | Q08431 | MFGM_HUMAN | Unmodified | 0 | 0 | 136170 | 20756000 | 3.99E+08 | 65738000 | 12299000 | 61824 | 8984400 | 9383200 | 3490700 | 2009700 | 30385000 | 10897000 | 73825 | 0 | 32977000 | 76430000 |
| TWKEVCFA | CLTC; CLTC; CI | Q00610; J3K | CLH1_HUMAN | Unmodified | 113710 | 103890 | 0 | 0 | 0 | 0 | 0 | 0 | 33796 | 0 | 0 | 0 | 0 | 0 | 0 | 0 | 0 | 0 |
| TWKFVEGL | MDH1; MDH1; I | P40925; B8Z | MDHC_HUMAN | Unmodified | 0 | 0 | 0 | 0 | 0 | 0 | 858440 | 0 | 0 | 414640 | 0 | 0 | 534190 | 134470 | 32818 | 0 | 188610 | 279510 |
| TWKFVEGL | MDH1; MDH1; I | P40925; B8Z | MDHC_HUMAN | Unmodified | 0 | 0 | 0 | 0 | 17125000 | 0 | 7962800 | 0 | 363070 | 5939400 | 0 | 0 | 6091400 | 9779700 | 0 | 0 | 9454500 | 13087000 |
| TWKGSDH | NANS; NANS; N | Q9NR45; Q5 | SIAS_HUMAN S | Unmodified | 0 | 0 | 0 | 0 | 1182700 | 0 | 118250 | 0 | 0 | 368910 | 0 | 0 | 3939900 | 1676100 | 0 | 0 | 244060 | 0 |
| TWKGTAFG | ADH5 | P11766 | ADHX_HUMAN | Unmodified | 0 | 0 | 0 | 160320 | 587370 | 215790 | 0 | 0 | 0 | 125620 | 0 | 715610 | 345150 | 0 | 0 | 0 | 0 | 830500 |
| TWKPTLVIL | ARPC1B; ARPC | O15143; C9J | ARC1B_HUMAI | Unmodified | 0 | 0 | 0 | 0 | 0 | 0 | 0 | 0 | 450200 | 0 | 0 | 10521000 | 0 | 0 | 0 | 0 | 0 | 0 |
| TWMHTPE | GULP1; GULP1 | Q9UBP9; B8 | GULP1_HUMAI | Unmodified | 0 | 0 | 0 | 0 | 283630 | 0 | 0 | 0 | 0 | 0 | 0 | 0 | 468060 | 612060 | 114810 | 0 | 0 | 313350 |
| TWMVGTKI | EHD2 | Q9NZN4 | EHD2_HUMAN | Oxidation (M) | 22068 | 0 | 0 | 86371 | 1543500 | 673290 | 46060 | 0 | 0 | 0 | 0 | 0 | 39943 | 0 | 0 | 0 | 0 | 0 |
| TWMVGTKI | EHD2 | Q9NZN4 | EHD2_HUMAN | Unmodified | 0 | 0 | 0 | 0 | 1927600 | 250850 | 0 | 0 | 0 | 0 | 0 | 0 | 0 | 0 | 0 | 0 | 0 | 0 |
| TWNAVLLR | GALE; GALE; G | Q14376; Q5 | GALE_HUMAN | Unmodified | 83273 | 75732 | 224550 | 122360 | 450810 | 98487 | 0 | 0 | 123070 | 40172 | 79859 | 327020 | 97175 | 0 | 343490 | 524570 | 531900 | 0 |
| TWNDPSV( | HSPA5; HSPA5; | P11021; A0A | BIP_HUMAN Er | Unmodified | 0 | 0 | 720680 | 0 | 0 | 0 | 415260 | 246700 | 56784000 | 7346900 | 7408400 | 2276600 | 336520 | 418720 | 1245200 | 334630 | 0 | 403140 |
| TWNDPSV( | HSPA5; HSPA5; | P11021; A0A | BIP_HUMAN Er | Unmodified | 0 | 0 | 0 | 0 | 0 | 0 | 0 | 0 | 407280 | 8024800 | 0 | 0 | 0 | 0 | 0 | 0 | 0 | 0 |
| TWNDPSV( | HSPA5; HSPA5; | P11021; A0A | BIP_HUMAN Er | Unmodified | 0 | 0 | 0 | 0 | 0 | 0 | 3507500 | 0 | 0 | 6290700 | 0 | 0 | 0 | 0 | 0 | 0 | 0 | 0 |
| TWNLGLSN | IGF2R; IGF2R | P11717; A0A | MPRI_HUMAN ( | Unmodified | 234320 | 1430700 | 700650 | 1599700 | 364850 | 27192 | 0 | 41087 | 35419000 | 9626800 | 3775100 | 6641300 | 998610 | 1808300 | 1225700 | 1209300 | 0 | 0 |
| TWPSSHN | CREG1; CREG1 | O75629; A0A | CREG1_HUMAI | Unmodified | 0 | 0 | 0 | 0 | 0 | 0 | 0 | 0 | 4164700 | 2216600 | 1428200 | 14008 | 0 | 0 | 303640 | 0 | 0 | 0 |
| TWPTIIDTD | DIP2B | Q9P265 | DIP2B_HUMAN | Unmodified | 0 | 448760 | 642840 | 821290 | 0 | 0 | 0 | 0 | 1244400 | 319200 | 1104600 | 3280600 | 679080 | 110350 | 1740900 | 1112700 | 234900 | 0 |
| TWQDSDT | TOR1AIP2 | Q8NFQ8 | TOIP2_HUMAN | Unmodified | 0 | 0 | 0 | 0 | 0 | 0 | 0 | 0 | 3707700 | 130730 | 173640 | 0 | 102720 | 0 | 0 | 0 | 0 | 0 |
| TWSQGFP( | TOM1L1; TOM1 | O75674; J3K | TM1L1_HUMAN | Unmodified | 2189600 | 2503900 | 5335300 | 1316400 | 0 | 0 | 0 | 0 | 167700 | 0 | 0 | 0 | 0 | 0 | 5771000 | 3844700 | 0 | 0 |
| TWSQGFP( | TOM1L1; TOM1 | O75674; J3K | TM1L1_HUMAN | Unmodified | 0 | 0 | 0 | 0 | 0 | 0 | 0 | 0 | 0 | 0 | 0 | 0 | 1048700 | 1075000 | 0 | 0 | 0 | 0 |
| TWSQGFP( | TOM1L1; TOM1 | O75674; J3K | TM1L1_HUMAN | Unmodified | 0 | 0 | 0 | 0 | 0 | 0 | 0 | 0 | 0 | 489060 | 0 | 0 | 2675000 | 0 | 0 | 0 | 18492 | 0 |
| TWTDFEHN | ZC3HAV1; ZC3I | Q7Z2W4; C9 | ZCCHV_HUMA | Unmodified | 0 | 0 | 478020 | 0 | 4838000 | 1199200 | 419220 | 0 | 0 | 206970 | 0 | 0 | 0 | 0 | 0 | 0 | 423540 | 0 |
| TWTDFEHN | ZC3HAV1; ZC3I | Q7Z2W4; C9 | ZCCHV_HUMA | Oxidation (M) | 485850 | 0 | 970830 | 0 | 250800 | 669510 | 0 | 0 | 131120 | 155200 | 0 | 0 | 0 | 0 | 512820 | 433560 | 0 | 0 |
| TWTVVDAK | XRCC5 | P13010 | XRCC5_HUMAI | Unmodified | 212300 | 0 | 0 | 0 | 330210 | 0 | 1035200 | 0 | 491450 | 251480 | 517760 | 7882000 | 0 | 0 | 125320 | 0 | 0 | 547680 |
| TWVQTRLG | UTRN | P46939 | UTRN_HUMAN | Unmodified | 0 | 0 | 0 | 0 | 3131200 | 0 | 0 | 0 | 0 | 0 | 0 | 0 | 934340 | 1624300 | 0 | 0 | 4112900 | 2664900 |
| TYAEVDRQ | FNBP1; FNBP1; | Q96RU3; H0 | FNBP1_HUMAN | Unmodified | 962160 | 980900 | 94211 | 154900 | 6425800 | 1111100 | 731300 | 518330 | 61071 | 0 | 0 | 0 | 1408700 | 697440 | 0 | 122560 | 1216400 | 2592900 |
| TYAICGAIR | RPS21; RPS21; | P63220; Q9I | RS21_HUMAN | Unmodified | 1417300 | 925430 | 796080 | 939570 | 180340 | 733300 | 998130 | 1027100 | 420440 | 510850 | 590610 | 6114500 | 2126200 | 485730 | 657010 | 563080 | 412260 | 949700 |
| TYAPVAFR | PIP5K1A; PIP5K | Q99755; O6I | PI51A_HUMAN | Unmodified | 268900 | 0 | 0 | 52780 | 0 | 0 | 38981 | 0 | 313790 | 0 | 143920 | 3126400 | 0 | 0 | 551750 | 431480 | 96394 | 24675 |
| TYARRPTPI | HMGCS1 | Q01581 | HMCS1_HUMA | Unmodified | 0 | 0 | 0 | 0 | 2943900 | 0 | 0 | 0 | 539540 | 0 | 0 | 0 | 1392700 | 0 | 0 | 0 | 0 | 0 |
| TYASTEQE( | ABCC1; ABCC1 | P33527; A0A | MRP1_HUMAN | Unmodified | 0 | 0 | 0 | 0 | 0 | 0 | 0 | 0 | 0 | 0 | 0 | 4906300 | 0 | 0 | 0 | 789880 | 0 | 0 |
| TYATDYK | SLC1A4 | P43007 | SATT_HUMAN I | Unmodified | 196720 | 189980 | 215560 | 0 | 0 | 0 | 0 | 0 | 2172300 | 390460 | 0 | 5452100 | 0 | 213780 | 308510 | 0 | 0 | 0 |
| TYAYLFSHI | CEL | P19835 | CEL_HUMAN B | Unmodified | 0 | 0 | 0 | 0 | 0 | 0 | 0 | 25397 | 15365000 | 0 | 9989700 | 2360800 | 0 | 0 | 0 | 0 | 101500 | 0 |
| TYAYLFSHI | CEL | P19835 | CEL_HUMAN B | Unmodified | 0 | 0 | 0 | 0 | 0 | 0 | 0 | 0 | 3858600 | 1006300 | 1784300 | 0 | 0 | 0 | 0 | 0 | 0 | 0 |
| TYAYLFSHI | CEL | P19835 | CEL_HUMAN B | Oxidation (M) | 135130 | 0 | 0 | 0 | 2683300 | 0 | 240230 | 223450 | 1405000 | 1709900 | 3129500 | 0 | 0 | 223380 | 0 | 0 | 0 | 0 |
| TYDATTHFI | GDI2 | P50395 | GDIB_HUMAN | Unmodified | 1722200 | 803230 | 3098400 | 4581300 | 1865900 | 1062500 | 242590 | 217130 | 19458000 | 3010700 | 9980200 | 29850000 | 1677600 | 954540 | 4552500 | 5227000 | 590790 | 1353700 |
| TYDATTHFI | GDI2 | P50395 | GDIB_HUMAN | Unmodified | 0 | 0 | 0 | 0 | 2602100 | 1196100 | 0 | 0 | 395980 | 4960100 | 0 | 0 | 1112200 | 2589000 | 0 | 30434 | 3996700 | 0 |
| TYDLNKPPI | PRKG2 | Q13237 | KGP2_HUMAN | Unmodified | 0 | 0 | 0 | 0 | 0 | 0 | 0 | 345510 | 0 | 0 | 0 | 165390 | 0 | 0 | 0 | 0 | 0 | 0 |
| TYDRDNSG | PDCD6; PDCD6 | O75340; A0A | PDCD6_HUMA | Oxidation (M) | 2080600 | 2088100 | 1074700 | 850980 | 134170 | 90241 | 0 | 0 | 4204500 | 429140 | 0 | 1339000 | 56092 | 0 | 2942600 | 2478600 | 0 | 0 |
| TYDRDNSG | PDCD6; PDCD6 | O75340; A0A | PDCD6_HUMA | Unmodified | 3573500 | 0 | 0 | 0 | 448040 | 0 | 0 | 0 | 7898300 | 753240 | 0 | 2632300 | 0 | 0 | 4334300 | 2653600 | 0 | 0 |
| TYDTYVGIC | FKBP9 | O95302 | FKBP9_HUMAN | Oxidation (M) | 0 | 0 | 0 | 0 | 0 | 0 | 0 | 0 | 281540 | 434310 | 0 | 0 | 0 | 0 | 0 | 0 | 0 | 0 |
| TYEAIGEMI | SNX33 | Q8WV41 | SNX33_HUMAN | Oxidation (M) | 51484 | 0 | 2273800 | 31527 | 0 | 0 | 0 | 0 | 0 | 0 | 0 | 0 | 0 | 0 | 2646100 | 1581400 | 0 | 0 |
| TYEDIKENL | MPST; MPST | P25325; B1A | THTM_HUMAN | Unmodified | 0 | 0 | 0 | 0 | 0 | 0 | 0 | 0 | 272880 | 215610 | 319540 | 0 | 313750 | 0 | 0 | 0 | 0 | 665650 |
| TYEEGLK | STIP1; STIP1; S | P31948; F5H | STIP1_HUMAN | Unmodified | 0 | 0 | 0 | 0 | 0 | 0 | 0 | 0 | 212240 | 0 | 0 | 3032800 | 0 | 0 | 0 | 0 | 0 | 147850 |
| TYEEGLKHI | STIP1; STIP1; S | P31948; F5H | STIP1_HUMAN | Unmodified | 0 | 0 | 0 | 157910 | 135010 | 0 | 0 | 0 | 823070 | 267490 | 601010 | 2664400 | 0 | 0 | 0 | 0 | 201770 | 96292 |
| TYEEIASLV | SNX9; SNX9; SN | Q9Y5X1; A0A | SNX9_HUMAN | Unmodified | 10618000 | 8355800 | 1464300 | 1722600 | 0 | 0 | 0 | 0 | 1034200 | 0 | 797260 | 5109900 | 826110 | 0 | 2389800 | 2905100 | 0 | 0 |
| TYEGMMH | LYPLA1; LYPLA | O75608; B4I | LYPA1_HUMAN | Unmodified | 0 | 0 | 0 | 0 | 372870 | 0 | 0 | 0 | 793890 | 157850 | 204910 | 0 | 0 | 0 | 0 | 0 | 0 | 193860 |
| TYEGMMH | LYPLA1; LYPLA | O75608; B4I | LYPA1_HUMAN | Oxidation (M) | 0 | 0 | 0 | 0 | 129080 | 0 | 0 | 0 | 158090 | 52613 | 119370 | 1052000 | 0 | 68327 | 0 | 0 | 58503 | 0 |
| TYEQLKEEL | AVIL | O75366 | AVIL_HUMAN A | Oxidation (M) | 0 | 0 | 0 | 0 | 0 | 0 | 0 | 0 | 0 | 0 | 783720 | 4225600 | 0 | 0 | 0 | 0 | 0 | 0 |
| TYEQLKEEL | AVIL | O75366 | AVIL_HUMAN A | Unmodified | 0 | 0 | 0 | 0 | 0 | 0 | 0 | 0 | 0 | 0 | 0 | 2070200 | 100830 | 517280 | 0 | 0 | 0 | 0 |
| TYEQVLENI | TST | Q16762 | THTR_HUMAN | Unmodified | 0 | 0 | 0 | 0 | 0 | 349610 | 0 | 0 | 3734500 | 547170 | 0 | 37763 | 0 | 0 | 0 | 0 | 0 | 0 |
| TYEVPNTPI | KCNJ2 | P63252 | KCNJ2_HUMAN | Unmodified | 0 | 0 | 0 | 0 | 0 | 0 | 439950 | 99398 | 0 | 0 | 0 | 0 | 0 | 0 | 0 | 0 | 0 | 0 |
| TYFSHLDLS | HBQ1 | P09105 | HBAT_HUMAN | Unmodified | 159030 | 288580 | 0 | 151590 | 0 | 0 | 0 | 0 | 0 | 0 | 0 | 0 | 0 | 0 | 122260 | 96244 | 0 | 0 |
| TYGADLAS | SERPINB1 | P30740 | ILEU_HUMAN L | Unmodified | 663700 | 0 | 163220 | 0 | 0 | 714020 | 1764300 | 1165100 | 412980 | 0 | 0 | 0 | 0 | 0 | 0 | 0 | 0 | 0 |
| TYGADLAS | SERPINB1 | P30740 | ILEU_HUMAN L | Unmodified | 0 | 0 | 0 | 620920 | 1157100 | 401940 | 1105200 | 290190 | 3783900 | 3619600 | 3823300 | 0 | 400750 | 1818400 | 0 | 0 | 578590 | 6925500 |
| TYGFHFTTI | DYNC1LI2; DYN | O43237; B4E | DC1L2_HUMAN | Unmodified | 221290 | 3250400 | 370930 | 765200 | 2580000 | 542290 | 1566300 | 72979 | 945710 | 1119500 | 2622400 | 6546900 | 2253100 | 1832200 | 324380 | 323020 | 1497200 | 1530700 |
| TYGGCEGI | ELOC; ELOC; EI | Q15369; R4( | ELOC_HUMAN | Oxidation (M) | 0 | 0 | 0 | 0 | 1235400 | 217720 | 148950 | 132890 | 84659 | 153910 | 263610 | 1142100 | 359730 | 48307 | 0 | 0 | 560380 | 694330 |
| TYGGCEGI | ELOC; ELOC; EI | Q15369; R4( | ELOC_HUMAN | Unmodified | 0 | 0 | 0 | 0 | 1777500 | 196320 | 39470 | 0 | 482920 | 283880 | 0 | 1303000 | 1201100 | 889510 | 0 | 0 | 747890 | 1139300 |
| TYGGYQGI | HAPLN1; HAPLI | P10915; D6F | HPLN1_HUMAI | Unmodified | 0 | 0 | 0 | 0 | 991480 | 80422 | 9436.3 | 0 | 2922100 | 402080 | 55375 | 0 | 22914 | 0 | 15763 | 30625 | 5981.9 | 0 |
| TYGGYQGI | HAPLN1; HAPLI | P10915; D6F | HPLN1_HUMAI | Unmodified | 0 | 0 | 0 | 0 | 0 | 0 | 0 | 0 | 79654 | 3643200 | 0 | 0 | 0 | 0 | 0 | 0 | 0 | 0 |
| TYGHLYMC | GPC4 | O75487 | GPC4_HUMAN | Oxidation (M) | 354700 | 389640 | 270540 | 0 | 0 | 0 | 0 | 0 | 0 | 0 | 0 | 0 | 0 | 0 | 0 | 191440 | 0 | 0 |
| TYGLMDSH | HSPG2 | P98160 | PGBM_HUMAN | Oxidation (M) | 0 | 0 | 246760 | 0 | 0 | 0 | 0 | 0 | 0 | 0 | 0 | 0 | 0 | 0 | 0 | 0 | 0 | 0 |
| TYGLVPAA( | MELTF | P08582 | TRFM_HUMAN | Unmodified | 0 | 0 | 0 | 0 | 18602000 | 7650800 | 0 | 0 | 14601000 | 3756700 | 1654000 | 1712200 | 0 | 0 | 0 | 108890 | 2362100 | 0 |
| TYGQNGH | SLC39A8 | Q9C0K1 | S39A8_HUMAN | Unmodified | 0 | 0 | 110020 | 196860 | 0 | 0 | 0 | 0 | 0 | 0 | 0 | 0 | 0 | 0 | 0 | 95013 | 0 | 0 |
| TYGQSTYSI | PSMC2; PSMC2 | P35998; C9J | PRS7_HUMAN | Unmodified | 0 | 54003 | 0 | 249700 | 2007200 | 0 | 134860 | 0 | 0 | 0 | 0 | 689300 | 2081500 | 612590 | 0 | 172360 | 840090 | 596210 |
| TYGQSTYSI | PSMC2; PSMC2 | P35998; C9J | PRS7_HUMAN | Unmodified | 0 | 0 | 0 | 0 | 1019100 | 418680 | 677170 | 293610 | 0 | 257970 | 0 | 0 | 910130 | 862020 | 0 | 0 | 776010 | 1364300 |
| TYGQSTYSI | PSMC2; PSMC2 | P35998; C9J | PRS7_HUMAN | Unmodified | 0 | 0 | 0 | 0 | 107440 | 0 | 0 | 0 | 0 | 0 | 0 | 0 | 442710 | 0 | 0 | 0 | 2887400 | 585990 |
| TYGSEFVYS | DNAJC13 | O75165 | DJC13_HUMAN | Unmodified | 0 | 0 | 0 | 0 | 0 | 0 | 0 | 0 | 0 | 0 | 0 | 1883500 | 0 | 16890 | 0 | 0 | 0 | 0 |
| TYGVSFFLV | TLN2; TLN1 | Q9Y4G6; Q9 | TLN2_HUMAN | Unmodified | 959720 | 4347800 | 1699600 | 4669000 | 4070800 | 0 | 6800800 | 0 | 1468200 | 0 | 533390 | 5301600 | 4322400 | 2114700 | 2810500 | 3101100 | 563840 | 1091000 |

| | | | | | | | | | | | | | | | | | | | | | | |
|---|---|---|---|---|---|---|---|---|---|---|---|---|---|---|---|---|---|---|---|---|---|---|
| TYHALSNLI | PSMD11; PSMD | O00231; J3Q | PSD11_HUMAN | Unmodified | 0 | 0 | 0 | 0 | 0 | 0 | 0 | 0 | 0 | 0 | 0 | 1128400 | 0 | 0 | 0 | 0 | 0 | 0 |
| TYHSVGDS | IGF2R; IGF2R; I | P11717; A0A | MPRI_HUMAN | Unmodified | 0 | 0 | 0 | 0 | 0 | 11004 | 0 | 0 | 18694000 | 1173100 | 2568400 | 1907300 | 0 | 54808 | 0 | 86400 | 0 | 0 |
| TYHSVGDS | IGF2R; IGF2R; I | P11717; A0A | MPRI_HUMAN | Unmodified | 0 | 0 | 0 | 0 | 0 | 0 | 0 | 0 | 0 | 729870 | 0 | 0 | 0 | 0 | 0 | 0 | 0 | 0 |
| TYHVGEQV | FN1; FN1 | P02751; H0Y | FINC_HUMAN | Unmodified | 1065900 | 7958200 | 45795000 | 23104000 | 3932800 | 16455000 | 1905900 | 672520 | 2620200 | 2199700 | 1130500 | 1421300 | 20784000 | 24906000 | 3222300 | 12772000 | 2513800 | 22677000 |
| TYIASFQNC | CRISPLD1; CRI | Q9H336; B7 | CRLD1_HUMAN | Unmodified | 0 | 0 | 0 | 0 | 0 | 0 | 0 | 0 | 826930 | 0 | 0 | 0 | 0 | 0 | 0 | 0 | 0 | 0 |
| TYIDPETYE | EPHA7; EPHA7; | Q15375; A0A | EPHA7_HUMAN | Unmodified | 0 | 0 | 0 | 0 | 0 | 0 | 0 | 218340 | 2902300 | 915760 | 1154800 | 0 | 0 | 0 | 0 | 0 | 393720 | 659770 |
| TYINPFVSF | SEC24A | O95486 | SC24A_HUMAN | Unmodified | 0 | 0 | 0 | 0 | 0 | 61102 | 0 | 0 | 0 | 0 | 0 | 381490 | 0 | 0 | 0 | 0 | 0 | 0 |
| TYINPFVSF | SEC24A | O95486 | SC24A_HUMAN | Unmodified | 0 | 0 | 0 | 0 | 0 | 0 | 0 | 0 | 0 | 0 | 0 | 557610 | 0 | 0 | 0 | 0 | 0 | 0 |
| TYIPPKGET | HMGB1P1; HM | B2RPK0; P09 | HGB1A_HUMA | Unmodified | 0 | 0 | 0 | 0 | 0 | 0 | 0 | 0 | 0 | 0 | 0 | 1263200 | 0 | 0 | 0 | 0 | 0 | 0 |
| TYIRQDNEI | MVP; MVP; MVP | Q14764; H3I | MVP_HUMAN M | Unmodified | 0 | 0 | 0 | 0 | 1107900 | 860830 | 0 | 0 | 675800 | 574820 | 0 | 0 | 0 | 0 | 0 | 0 | 0 | 0 |
| TYITDPVSA | LPP; LPP; LPP; L | Q93052; C9J | LPP_HUMAN Li | Unmodified | 640300 | 0 | 0 | 0 | 2159500 | 0 | 198770 | 0 | 0 | 0 | 0 | 0 | 2733900 | 2422800 | 0 | 0 | 699120 | 929630 |
| TYKNPSLR | CAP2; CAP2; C | P40123; B7Z | CAP2_HUMAN | Unmodified | 0 | 0 | 0 | 0 | 1141100 | 0 | 221870 | 326750 | 0 | 0 | 0 | 43644 | 0 | 0 | 0 | 0 | 0 | 707340 |
| TYLARLPW | PBXIP1 | Q96AQ6 | PBIP1_HUMAN | Unmodified | 2877900 | 0 | 0 | 0 | 0 | 0 | 0 | 0 | 2900700 | 1505000 | 1205500 | 32744 | 0 | 0 | 0 | 0 | 0 | 0 |
| TYLGNALV | FN1 | P02751 | FINC_HUMAN | Unmodified | 0 | 0 | 0 | 1837700 | 0 | 0 | 0 | 0 | 1196200 | 317350 | 440580 | 1344000 | 0 | 0 | 0 | 0 | 0 | 31416 |
| TYLLDFRSI | PRKAA1 | Q13131 | AAPK1_HUMAN | Unmodified | 0 | 0 | 0 | 51447 | 0 | 0 | 0 | 0 | 141860 | 561110 | 52012 | 0 | 0 | 0 | 0 | 0 | 0 | 0 |
| TYLLPDRSS | SYTL3 | Q4VX76 | SYTL3_HUMAN | Unmodified | 0 | 86592 | 98235 | 375250 | 363430 | 60600 | 828340 | 716080 | 119630 | 95938 | 303600 | 0 | 374200 | 661660 | 149680 | 231070 | 0 | 194490 |
| TYLPAVDEI | APOC2; APOC4 | P02655; K7E | APOC2_HUMAI | Unmodified | 0 | 0 | 0 | 0 | 0 | 0 | 0 | 0 | 0 | 0 | 0 | 0 | 0 | 0 | 0 | 0 | 822460 | 0 |
| TYLPSQVSI | COPB2; COPB2 | P35606; H0Y | COPB2_HUMAI | Unmodified | 0 | 0 | 0 | 0 | 0 | 0 | 0 | 0 | 0 | 0 | 0 | 3849600 | 0 | 0 | 0 | 0 | 0 | 0 |
| TYLVSGQP | PTGES2; PTGES | Q9H7Z7; A0 | PGES2_HUMAN | Oxidation (M) | 0 | 0 | 0 | 0 | 0 | 0 | 0 | 0 | 1497600 | 167650 | 88104 | 231570 | 174060 | 0 | 0 | 0 | 0 | 0 |
| TYNASITLQ | TOP1; TOP1 | P11387; A0A | TOP1_HUMAN | Unmodified | 733420 | 0 | 0 | 0 | 0 | 0 | 0 | 0 | 0 | 0 | 100380 | 3677800 | 0 | 0 | 0 | 0 | 0 | 0 |
| TYNDALFR | CLDND1; CLDN | Q9NY35; D6 | CLDN1_HUMAI | Unmodified | 3653000 | 2023200 | 0 | 0 | 0 | 0 | 0 | 0 | 964120 | 0 | 0 | 15556000 | 0 | 0 | 4218100 | 2117800 | 0 | 0 |
| TYNFLPEFL | SERPINB1 | P30740 | ILEU_HUMAN L | Unmodified | 190430 | 142250 | 0 | 0 | 338650 | 135520 | 107640 | 0 | 1660200 | 1718200 | 0 | 110050 | 0 | 0 | 0 | 0 | 291660 | 466630 |
| TYNNLDVS | PRKAG1; PRKA | P54619; H0Y | AAKG1_HUMAN | Unmodified | 202710 | 0 | 0 | 0 | 388690 | 0 | 198480 | 165890 | 620920 | 0 | 0 | 438240 | 0 | 912900 | 143650 | 3199300 | 260460 | 1192900 |
| TYNTDVPL | UGP2; UGP2; U | Q16851; E7E | UGPA_HUMAN | Unmodified | 0 | 0 | 0 | 0 | 0 | 0 | 0 | 0 | 2715100 | 2393700 | 0 | 1146700 | 0 | 0 | 0 | 0 | 0 | 0 |
| TYNTNAQV | ALPL | P05186 | PPBT_HUMAN | Unmodified | 617040 | 0 | 0 | 0 | 0 | 0 | 0 | 336580 | 0 | 0 | 0 | 0 | 0 | 0 | 0 | 0 | 0 | 0 |
| TYPGVMHS | LYPLA2; LYPLA | O95372; Q5 | LYPA2_HUMAN | Oxidation (M) | 0 | 0 | 0 | 58906 | 0 | 0 | 277580 | 57635 | 0 | 53488 | 0 | 820010 | 0 | 0 | 61416 | 0 | 248120 | 404940 |
| TYPGVMHS | LYPLA2; LYPLA | O95372; Q5 | LYPA2_HUMAN | Unmodified | 32914 | 0 | 0 | 0 | 314880 | 152650 | 0 | 0 | 87489 | 0 | 0 | 335090 | 0 | 0 | 0 | 0 | 391930 | 621510 |
| TYPGVMHS | LYPLA2; LYPLA | O95372; Q5 | LYPA2_HUMAN | 2 Oxidation (M) | 0 | 0 | 0 | 0 | 112820 | 0 | 335630 | 45466 | 88724 | 0 | 11768 | 461290 | 0 | 0 | 0 | 0 | 130750 | 262520 |
| TYPGVMHS | LYPLA2; LYPLA | O95372; Q5 | LYPA2_HUMAN | Oxidation (M) | 0 | 0 | 0 | 0 | 0 | 0 | 0 | 0 | 0 | 0 | 0 | 0 | 0 | 0 | 0 | 0 | 1613300 | 0 |
| TYPKDPYQ | ACP2; ACP2; A | P11117; B7Z | PPAL_HUMAN | Unmodified | 0 | 0 | 0 | 0 | 881530 | 0 | 0 | 0 | 5765400 | 2621000 | 458830 | 0 | 0 | 0 | 0 | 0 | 0 | 245130 |
| TYQASSAAI | FAM168A | Q92567 | F168A_HUMAN | Unmodified | 60174 | 149780 | 141540 | 0 | 381120 | 0 | 0 | 0 | 774950 | 142490 | 0 | 0 | 0 | 0 | 217780 | 239510 | 0 | 700130 |
| TYQAYHME | ARHGAP4; ARH | P98171; C9J | RHG04_HUMAI | Unmodified | 11455000 | 11618000 | 2334900 | 226020 | 0 | 0 | 92525 | 36340 | 0 | 0 | 0 | 280920 | 0 | 0 | 4886500 | 3963800 | 0 | 0 |
| TYQGLESLI | CHORDC1 | Q9UHD1 | CHRD1_HUMA | Oxidation (M) | 0 | 0 | 0 | 0 | 1132100 | 0 | 796430 | 0 | 684760 | 963930 | 1983700 | 4163000 | 0 | 2332700 | 0 | 0 | 1898600 | 0 |
| TYQGSYGF | TP53; TP53; TP | P04637; E7E | P53_HUMAN C | Unmodified | 0 | 0 | 0 | 0 | 0 | 0 | 1731300 | 77265 | 0 | 0 | 0 | 0 | 0 | 0 | 0 | 0 | 43798 | 0 |
| TYQLVYFLI | RNPEP; RNPEP | Q9H4A4; A6 | AMPB_HUMAN | Unmodified | 0 | 0 | 0 | 0 | 1253400 | 0 | 0 | 0 | 868790 | 627140 | 0 | 9397100 | 0 | 0 | 0 | 0 | 0 | 0 |
| TYQSITER | EXOC4 | Q96A65 | EXOC4_HUMAI | Unmodified | 0 | 0 | 0 | 0 | 0 | 0 | 0 | 0 | 0 | 0 | 0 | 536990 | 0 | 0 | 0 | 0 | 0 | 0 |
| TYQVCNVN | EPHA3; EPHA3 | P29320; C9J | EPHA3_HUMAN | Unmodified | 0 | 605190 | 806550 | 92773 | 0 | 0 | 0 | 0 | 0 | 0 | 0 | 0 | 0 | 258290 | 0 | 22588 | 0 | 0 |
| TYQVCNVN | EPHA3; EPHA3 | P29320; C9J | EPHA3_HUMAN | Oxidation (M) | 773710 | 635040 | 0 | 0 | 0 | 0 | 0 | 0 | 0 | 0 | 0 | 0 | 0 | 0 | 0 | 0 | 0 | 0 |
| TYRPLNDV | PPIL3; PPIL3; P | Q9H2H8; B8 | PPIL3_HUMAN | Unmodified | 0 | 0 | 0 | 0 | 4971000 | 0 | 580740 | 0 | 0 | 467960 | 0 | 0 | 809340 | 1215100 | 0 | 0 | 392600 | 0 |
| TYSKPIFGS | PTPRB; PTPRB; | P23467; F8V | PTPRB_HUMAN | Unmodified | 289570 | 0 | 0 | 0 | 0 | 0 | 205390 | 99549 | 0 | 1518100 | 0 | 0 | 0 | 0 | 0 | 0 | 0 | 0 |
| TYSLGSALF | VIM; VIM; VIM; | P08670; B0Y | VIME_HUMAN | Unmodified | 24191 | 0 | 0 | 0 | 335690 | 0 | 0 | 0 | 62896 | 0 | 0 | 0 | 49016 | 0 | 0 | 0 | 252020 | 288220 |
| TYSSTSSFS | NOSTRIN; NOS | Q8IVI9; A0A | NOSTN_HUMA | Unmodified | 2704500 | 2072800 | 4534800 | 5996400 | 0 | 0 | 0 | 0 | 0 | 0 | 0 | 0 | 2667700 | 2091500 | 8676600 | 4452400 | 0 | 0 |
| TYSTSFTPN | GSR | P00390 | GSHR_HUMAN | Oxidation (M) | 363250 | 0 | 333810 | 509970 | 141820 | 0 | 542450 | 0 | 663560 | 896990 | 1150000 | 1435300 | 2878900 | 1752500 | 187910 | 653230 | 1171400 | 0 |
| TYSTSFTPN | GSR | P00390 | GSHR_HUMAN | Unmodified | 274690 | 146160 | 278380 | 0 | 22479000 | 3313700 | 791990 | 509910 | 3299100 | 1178500 | 1117700 | 5922800 | 5984000 | 4225500 | 584520 | 269200 | 3151800 | 2517300 |
| TYSVEYLPK | FLNB; FLNB | O75369; E7E | FLNB_HUMAN | Unmodified | 0 | 0 | 788290 | 572560 | 4538800 | 1273100 | 0 | 78886 | 429890 | 242210 | 688960 | 6798100 | 644170 | 0 | 626390 | 577470 | 1920400 | 629460 |
| TYSVEYLPK | FLNB; FLNB | O75369; E7E | FLNB_HUMAN | Unmodified | 0 | 0 | 0 | 0 | 397010 | 44389 | 20574 | 12386 | 0 | 0 | 0 | 0 | 0 | 30109 | 0 | 0 | 187130 | 112690 |
| TYSVGVCTI | IGF2R | P11717 | MPRI_HUMAN | Unmodified | 0 | 0 | 0 | 0 | 0 | 732760 | 0 | 0 | 9030800 | 828710 | 663560 | 1773100 | 0 | 0 | 0 | 0 | 0 | 0 |
| TYSVSFDSI | FOLH1 | Q04609 | FOLH1_HUMAI | Unmodified | 8818900 | 11151000 | 11589000 | 17536000 | 0 | 0 | 0 | 0 | 0 | 0 | 0 | 0 | 27454 | 493040 | 44995000 | 20987000 | 0 | 0 |
| TYSWDNA | RAB3D | O95716 | RAB3D_HUMAI | Unmodified | 0 | 0 | 0 | 2636700 | 0 | 0 | 0 | 0 | 0 | 0 | 0 | 0 | 0 | 0 | 0 | 0 | 0 | 0 |
| TYSWDNA | RAB3A; RAB3A | P20336; S4F | RAB3A_HUMAN | Unmodified | 0 | 0 | 2532200 | 874650 | 33659 | 0 | 197020 | 32335 | 0 | 0 | 0 | 0 | 0 | 201210 | 1266400 | 571490 | 631990 | 17706 |
| TYSYLNK | NPC2; NPC2; N | P61916; J3K | NPC2_HUMAN | Unmodified | 0 | 0 | 177810 | 70561 | 54084 | 43358 | 0 | 0 | 16200000 | 518900 | 106750 | 0 | 158020 | 105660 | 236210 | 83817 | 0 | 0 |
| TYSYLNKLF | NPC2; NPC2; N | P61916; J3K | NPC2_HUMAN | Unmodified | 0 | 0 | 0 | 0 | 0 | 0 | 0 | 0 | 2591600 | 421550 | 0 | 0 | 0 | 0 | 0 | 0 | 0 | 0 |
| TYSYLNKLF | NPC2; NPC2; N | P61916; J3K | NPC2_HUMAN | Unmodified | 0 | 0 | 0 | 0 | 0 | 1620800 | 0 | 0 | 1145900 | 3129900 | 0 | 0 | 130790 | 0 | 0 | 0 | 0 | 0 |
| TYSYLTPDL | RPS2; RPS2; RP | P15880; E9P | RS2_HUMAN 4 | Unmodified | 4342600 | 3248700 | 3504500 | 2641300 | 0 | 0 | 1292200 | 407760 | 826310 | 1144500 | 3186500 | 55563000 | 718810 | 131660 | 4041100 | 2545900 | 0 | 736420 |
| TYSYLTPDL | RPS2; RPS2; RP | P15880; E9P | RS2_HUMAN 4 | Unmodified | 0 | 0 | 0 | 0 | 0 | 2073500 | 6516800 | 42328 | 1429000 | 2971700 | 1989500 | 2102800 | 6454500 | 6814500 | 0 | 0 | 1624200 | 10887000 |
| TYSYLTPDL | RPS2; RPS2; RP | P15880; E9P | RS2_HUMAN 4 | Unmodified | 0 | 0 | 0 | 0 | 0 | 0 | 5993600 | 0 | 0 | 0 | 0 | 0 | 2025900 | 0 | 0 | 0 | 0 | 0 |
| TYTAYVDLE | YARS1; YARS1 | P54577; A0A | SYYC_HUMAN | Unmodified | 0 | 133760 | 0 | 885810 | 3005300 | 0 | 0 | 403680 | 0 | 0 | 613920 | 2179300 | 1687500 | 5609400 | 650380 | 586930 | 486320 | 1244300 |
| TYTAYVDLE | YARS1; YARS1 | P54577; A0A | SYYC_HUMAN | Unmodified | 0 | 0 | 0 | 0 | 3800500 | 0 | 1766600 | 0 | 0 | 981000 | 0 | 0 | 3991900 | 0 | 0 | 0 | 1318200 | 1541100 |
| TYTDAMQT | TSPAN7; TSPAN | P41732; B4D | TSN7_HUMAN | Unmodified | 0 | 0 | 0 | 0 | 0 | 0 | 0 | 0 | 4715800 | 667740 | 0 | 0 | 0 | 0 | 192720 | 0 | 0 | 0 |
| TYTDAMQT | TSPAN7; TSPAN | P41732; B4D | TSN7_HUMAN | Oxidation (M) | 0 | 0 | 0 | 0 | 0 | 0 | 0 | 0 | 338940 | 136200 | 0 | 0 | 0 | 0 | 0 | 0 | 0 | 0 |
| TYTDAMQT | TSPAN7; TSPAN | P41732; B4D | TSN7_HUMAN | Unmodified | 0 | 0 | 0 | 0 | 0 | 0 | 0 | 0 | 1665300 | 115820 | 0 | 0 | 0 | 0 | 0 | 0 | 0 | 0 |
| TYTDELTPI | NDUFS3 | O75489 | NDUS3_HUMA | Unmodified | 0 | 0 | 0 | 0 | 0 | 0 | 0 | 0 | 799590 | 204020 | 0 | 158710 | 0 | 0 | 0 | 0 | 0 | 367450 |
| TYTDRVFFL | PLXNB2 | O15031 | PLXB2_HUMAN | Unmodified | 240800 | 121430 | 1805500 | 325380 | 265290 | 0 | 166180 | 0 | 951640 | 91819 | 363540 | 2189900 | 0 | 108030 | 2054000 | 1199600 | 0 | 0 |
| TYTEEELNA | MICAL3; MICAL | Q7RTP6; J3C | MICA3_HUMAN | Unmodified | 0 | 0 | 0 | 0 | 2814200 | 0 | 0 | 0 | 0 | 0 | 103630 | 0 | 0 | 0 | 0 | 0 | 0 | 0 |
| TYTLTDYLK | DPP4; DPP4; DI | P27487; A0A | DPP4_HUMAN | Unmodified | 383110 | 244490 | 8919400 | 3969900 | 0 | 0 | 0 | 0 | 0 | 0 | 0 | 0 | 0 | 0 | 8423700 | 4597600 | 0 | 0 |
| TYVDPHTYI | EPHA2 | P29317 | EPHA2_HUMAN | Unmodified | 1499300 | 1320900 | 1125600 | 1358600 | 90944000 | 12920000 | 183120 | 0 | 419370 | 158750 | 4041000 | 13598000 | 3654100 | 3308400 | 1476300 | 993360 | 57210000 | 1.84E+08 |
| TYVDPHTYI | EPHA2 | P29317 | EPHA2_HUMAN | Unmodified | 0 | 0 | 0 | 1792000 | 1208300 | 0 | 1146500 | 0 | 1403100 | 0 | 10658000 | 0 | 717460 | 925840 | 0 | 0 | 8060100 | 5438500 |
| TYVDPHTYI | EPHA2 | P29317 | EPHA2_HUMAN | Unmodified | 0 | 0 | 0 | 0 | 1302400 | 0 | 0 | 0 | 0 | 0 | 0 | 0 | 0 | 0 | 0 | 0 | 45859000 | 21152000 |
| TYVDPHTYI | EPHA3; EPHA3 | P29320; C9J | EPHA3_HUMAN | Unmodified | 3036300 | 2369700 | 0 | 658630 | 0 | 0 | 0 | 0 | 0 | 0 | 0 | 0 | 0 | 0 | 0 | 0 | 0 | 893810 |
| TYVGPMTE | PLOD3 | O60568 | PLOD3_HUMAI | Oxidation (M) | 0 | 0 | 0 | 0 | 0 | 0 | 0 | 0 | 3887700 | 1670000 | 399540 | 0 | 0 | 0 | 0 | 0 | 0 | 0 |
| TYVGPMTE | PLOD3 | O60568 | PLOD3_HUMAI | Unmodified | 151280 | 945140 | 0 | 0 | 0 | 0 | 0 | 0 | 9653100 | 1387900 | 0 | 85334 | 0 | 0 | 37175 | 0 | 1074800 | 260210 |
| TYVGVVDG | VIL1 | P09327 | VILI_HUMAN Vi | Unmodified | 501580 | 171640 | 443760 | 492000 | 0 | 0 | 173700 | 346930 | 2533800 | 352530 | 1950700 | 14918000 | 0 | 0 | 315990 | 301880 | 0 | 0 |
| TYVPDGFG | PLCB4; PLCB4 | Q15147; A0 | PLCB4_HUMAN | Unmodified | 0 | 0 | 0 | 0 | 0 | 0 | 0 | 0 | 0 | 0 | 0 | 0 | 67256 | 0 | 0 | 0 | 433800 | 74524 |
| TYVRDKPH | TUFM; TUFM | P49411; H3E | EFTU_HUMAN | Unmodified | 0 | 0 | 0 | 0 | 393530 | 161770 | 756470 | 0 | 2064300 | 1301800 | 0 | 0 | 0 | 360260 | 0 | 0 | 49507 | 244060 |
| TYVSPTHV | MAPK13 | O15264 | MK13_HUMAN | Unmodified | 0 | 0 | 0 | 0 | 0 | 0 | 0 | 0 | 1155000 | 676630 | 0 | 956120 | 0 | 0 | 0 | 0 | 0 | 0 |
| TYWEDNK | CPE | P16870 | CBPE_HUMAN | Unmodified | 0 | 0 | 0 | 0 | 0 | 0 | 0 | 0 | 32557000 | 2343300 | 0 | 0 | 0 | 0 | 0 | 0 | 0 | 0 |
| TYWEDNK | CPE | P16870 | CBPE_HUMAN | Unmodified | 0 | 0 | 0 | 0 | 0 | 0 | 0 | 0 | 5840700 | 680110 | 0 | 0 | 0 | 0 | 0 | 0 | 0 | 0 |
| TYWEDNK | CPE | P16870 | CBPE_HUMAN | Unmodified | 0 | 0 | 0 | 0 | 0 | 0 | 0 | 0 | 1032300 | 31799000 | 0 | 0 | 0 | 0 | 0 | 0 | 148810 | 0 |
| TYYLMDPS | PDPK1; PDPK1 | O15530; E9F | PDPK1_HUMAN | Oxidation (M) | 0 | 0 | 0 | 81947 | 0 | 0 | 0 | 0 | 0 | 0 | 0 | 31468 | 0 | 0 | 311240 | 222840 | 0 | 0 |
| TYYVFGRH | KIF14 | Q15058 | KIF14_HUMAN | Unmodified | 83109 | 0 | 0 | 75775 | 915700 | 0 | 71280 | 0 | 103930 | 0 | 0 | 93320 | 57948 | 0 | 46586 | 29304 | 102130 | 66853 |
| TYYYNTETK | PRPF40A; PRPF | O75400; H0Y | PR40A_HUMAN | Unmodified | 0 | 0 | 0 | 0 | 0 | 0 | 0 | 0 | 0 | 24726 | 0 | 480720 | 0 | 0 | 0 | 0 | 0 | 0 |
| VAAALPGM | RAB6A; RAB6A | P20340; F5H | RAB6A_HUMAN | Unmodified | 0 | 0 | 346180 | 308390 | 0 | 0 | 0 | 0 | 1305000 | 187820 | 0 | 2611700 | 0 | 0 | 0 | 0 | 0 | 0 |
| VAAALPGM | RAB6A; RAB6A | P20340; F5H | RAB6A_HUMAN | Oxidation (M) | 0 | 0 | 159740 | 208060 | 0 | 0 | 0 | 0 | 124710 | 0 | 0 | 611650 | 0 | 0 | 0 | 0 | 0 | 0 |
| VAAALTAMI | TYMP; TYMP; TY | P19971; C9J | TYPH_HUMAN | Oxidation (M) | 0 | 0 | 0 | 0 | 710270 | 0 | 0 | 0 | 0 | 0 | 0 | 0 | 0 | 0 | 0 | 0 | 0 | 0 |
| VAAAVDLIII | FASN; FASN | P49327; A0A | FAS_HUMAN F | Unmodified | 144060 | 65785 | 106220 | 112400 | 0 | 0 | 0 | 18549 | 0 | 0 | 403420 | 4760900 | 0 | 0 | 532780 | 600150 | 0 | 0 |
| VAAERAES | DHX36; DHX36 | Q9H2U1; E7 | DHX36_HUMAI | Unmodified | 0 | 0 | 82157 | 0 | 0 | 0 | 0 | 0 | 0 | 0 | 31719 | 322750 | 0 | 0 | 116880 | 172540 | 0 | 0 |
| VAAETQSP | NUP50 | Q9UKX7 | NUP50_HUMAI | Unmodified | 0 | 0 | 0 | 0 | 0 | 0 | 0 | 0 | 0 | 0 | 0 | 517310 | 0 | 834870 | 0 | 0 | 0 | 0 |
| VAAGAQIH | EIF2B5; EIF2B5 | Q13144; A0 | EI2BE_HUMAN | Unmodified | 0 | 0 | 0 | 0 | 0 | 0 | 0 | 0 | 0 | 0 | 0 | 1062400 | 0 | 0 | 46780 | 0 | 0 | 30484 |
| VAAGLGLS | ZC3H10 | Q96K80 | ZC3HA_HUMAI | Unmodified | 4153000 | 1242200 | 713570 | 4654700 | 0 | 0 | 0 | 0 | 0 | 0 | 0 | 0 | 0 | 0 | 454450 | 3461200 | 0 | 0 |
| VAAGLPPIL | SCLY; SCLY; SC | Q96I15; H7C | SCLY_HUMAN | Unmodified | 0 | 0 | 0 | 0 | 0 | 0 | 0 | 0 | 0 | 0 | 0 | 0 | 0 | 0 | 0 | 0 | 0 | 0 |
| VAAGTLDA | NCAPH; NCAPH | Q15003; E9F | CND2_HUMAN | Unmodified | 0 | 0 | 0 | 0 | 524130 | 0 | 887140 | 520460 | 141130 | 110460 | 0 | 569820 | 96399 | 0 | 359330 | 40696 | 243460 | 781710 |
| VAAGVLCE | JUP; CTNNB1; | P14923; P35 | PLAK_HUMAN | Unmodified | 2605200 | 266010 | 4030300 | 1772200 | 0 | 0 | 0 | 0 | 0 | 0 | 1111600 | 8204800 | 423360 | 0 | 7227100 | 8585700 | 0 | 0 |
| VAAHEAEE | VPS37A; VPS37 | Q8NEZ2; H0 | VP37A_HUMAN | Unmodified | 3823800 | 0 | 0 | 0 | 0 | 0 | 0 | 0 | 3896100 | 194420 | 225620 | 647830 | 0 | 0 | 1106700 | 1260900 | 0 | 0 |
| VAAIEALND | ST13P4; ST13; | Q8IZP2; P50 | ST134_HUMAN | Unmodified | 1564300 | 1018600 | 2364100 | 2757300 | 1000600 | 0 | 0 | 11632 | 6838600 | 1662000 | 6388500 | 19499000 | 9606200 | 969100 | 3445000 | 3041700 | 452850 | 568010 |
| VAAISESPV | PTGFR | P43088 | PF2R_HUMAN | Unmodified | 0 | 0 | 547880 | 622800 | 0 | 0 | 114130 | 45786 | 0 | 0 | 0 | 0 | 0 | 0 | 273580 | 0 | 0 | 0 |
| VAALDCME | QSOX2 | Q6ZRP7 | QSOX2_HUMAI | Oxidation (M) | 0 | 0 | 0 | 0 | 0 | 0 | 0 | 0 | 1234700 | 0 | 0 | 0 | 0 | 0 | 0 | 0 | 0 | 0 |
| VAALDCME | QSOX2 | Q6ZRP7 | QSOX2_HUMAI | Unmodified | 0 | 0 | 0 | 0 | 0 | 0 | 0 | 0 | 5770700 | 203640 | 0 | 366750 | 0 | 0 | 0 | 0 | 0 | 198970 |
| VAALEASG | MMS19; MMS1 | Q96T76; H0Y | MMS19_HUMA | Unmodified | 0 | 0 | 0 | 0 | 0 | 0 | 0 | 0 | 0 | 0 | 0 | 649840 | 0 | 0 | 0 | 0 | 0 | 0 |
| VAALGSDL | BTN3A3; BTN3 | O00478; C9J | BT3A3_HUMAN | Unmodified | 0 | 0 | 0 | 0 | 377950 | 0 | 160600 | 0 | 0 | 0 | 0 | 690970 | 0 | 0 | 0 | 0 | 167330 | 227480 |
| VAALGSDL | BTN3A1 | O00481 | BT3A1_HUMAN | Unmodified | 0 | 0 | 0 | 0 | 217980 | 159810 | 0 | 0 | 1277200 | 57677 | 135400 | 1520600 | 0 | 0 | 120330 | 0 | 0 | 215970 |
| VAALIDRLI | CEMIP2 | Q9UHN6 | CEIP2_HUMAN | Unmodified | 368960 | 293790 | 831730 | 2033000 | 6112800 | 1594200 | 0 | 0 | 10559000 | 1828300 | 271400 | 0 | 926860 | 237790 | 569640 | 466270 | 63206 | 62641 |
| VAALLERT | PTPN23 | Q9H3S7 | PTN23_HUMAN | Unmodified | 111900 | 0 | 0 | 114550 | 0 | 605680 | 0 | 0 | 0 | 0 | 0 | 0 | 0 | 0 | 0 | 0 | 58108 | 0 |
| VAALQNLV | KPNB1; KPNB1 | Q14974; J3K | IMB1_HUMAN | Unmodified | 313470 | 282670 | 257060 | 366500 | 471850 | 243980 | 609880 | 345140 | 765520 | 1360300 | 1709700 | 8742500 | 0 | 0 | 476150 | 1127200 | 701230 | 0 |
| VAALQTVG | KIAA2013 | Q8IYS2 | K2013_HUMAN | Unmodified | 0 | 0 | 0 | 0 | 0 | 0 | 0 | 0 | 2202800 | 0 | 0 | 0 | 0 | 0 | 0 | 0 | 0 | 0 |
| VAANAVDP | TRIP10; TRIP10 | Q15642; W4 | CIP4_HUMAN | Unmodified | 0 | 190600 | 0 | 0 | 12845000 | 1004300 | 0 | 0 | 0 | 673550 | 0 | 0 | 576610 | 0 | 0 | 0 | 1390600 | 3541500 |
| VAAPDVVV | DYNC1H1; DYN | Q14204; A0 | DYHC1_HUMA | Unmodified | 0 | 0 | 0 | 0 | 0 | 0 | 0 | 0 | 0 | 0 | 0 | 981750 | 0 | 0 | 0 | 0 | 0 | 0 |
| VAAPNLSR | CXADR | P78310 | CXAR_HUMAN | Unmodified | 0 | 0 | 0 | 0 | 40173000 | 0 | 3067200 | 51315 | 2657800 | 1686700 | 1845500 | 5109400 | 93019 | 1405400 | 1191800 | 806030 | 1398500 | 1210200 |
| VAASIGNA | PDLIM1 | O00151 | PDLI1_HUMAN | Oxidation (M) | 0 | 0 | 0 | 105880 | 1152700 | 0 | 487090 | 291410 | 0 | 187510 | 0 | 0 | 511700 | 979540 | 0 | 0 | 1968600 | 1794800 |
| VAASIGNA | PDLIM1 | O00151 | PDLI1_HUMAN | Unmodified | 0 | 0 | 0 | 0 | 354700 | 0 | 0 | 105150 | 0 | 53182 | 73845 | 129700 | 0 | 0 | 0 | 0 | 503890 | 551410 |

| | | | | | | | | | | | | | | | | | | | | | | |
|---|---|---|---|---|---|---|---|---|---|---|---|---|---|---|---|---|---|---|---|---|---|---|
| VAASNIVQI | PSMB2 | P49721 | PSB2_HUMAN | Unmodified | 0 | 0 | 0 | 0 | 0 | 0 | 0 | 73040 | 0 | 0 | 0 | 652930 | 0 | 0 | 0 | 0 | 0 | 0 |
| VAASNIVQI | PSMB2 | P49721 | PSB2_HUMAN | Unmodified | 116480 | 219150 | 115540 | 80220 | 0 | 0 | 51359 | 16698 | 1019500 | 45173 | 145320 | 6965700 | 35772 | 0 | 112310 | 134210 | 0 | 0 |
| VAASPEDIK | MRPS27; MRPS | Q92552; D6I | RT27_HUMAN : | Unmodified | 0 | 0 | 0 | 0 | 194930 | 0 | 107750 | 10281 | 1925300 | 278130 | 339210 | 139220 | 0 | 0 | 0 | 0 | 67294 | 0 |
| VAASPEDIK | MRPS27; MRPS | Q92552; D6I | RT27_HUMAN : | Unmodified | 0 | 0 | 0 | 0 | 0 | 0 | 64332 | 0 | 0 | 168940 | 0 | 0 | 0 | 0 | 0 | 0 | 0 | 0 |
| VAASVLNP' | ATP5PO; ATP5P | P48047; A0A | ATPO_HUMAN | Unmodified | 0 | 0 | 0 | 0 | 0 | 0 | 481990 | 119890 | 4606100 | 1289000 | 0 | 0 | 0 | 0 | 0 | 0 | 0 | 0 |
| VAASVLNP' | ATP5PO; ATP5P | P48047; A0A | ATPO_HUMAN | Unmodified | 0 | 0 | 0 | 0 | 0 | 0 | 0 | 0 | 2118200 | 0 | 0 | 174650 | 0 | 0 | 0 | 0 | 0 | 0 |
| VAATRQEIF | COPG1 | Q9Y678 | COPG1_HUMA | Unmodified | 0 | 0 | 0 | 0 | 1871700 | 0 | 758140 | 0 | 0 | 1229000 | 0 | 0 | 4255900 | 423370 | 0 | 0 | 1647100 | 6271000 |
| VAAVEMLK | MYO1D; MYO1I | O94832; J3Q | MYO1D_HUMA | Unmodified | 7280700 | 7685800 | 2449700 | 2167000 | 2071100 | 294490 | 0 | 0 | 1432400 | 593790 | 524570 | 0 | 4867900 | 3715600 | 2951700 | 2435100 | 1805200 | 14805000 |
| VAAVEMLK | MYO1D; MYO1I | O94832; J3Q | MYO1D_HUMA | Oxidation (M) | 0 | 0 | 1165600 | 0 | 0 | 0 | 0 | 0 | 0 | 0 | 0 | 0 | 0 | 0 | 0 | 380050 | 0 | 1547500 |
| VAAVQPSD | PLEKHA6; PLEK | Q9Y2H5; A0 | PKHA6_HUMAI | Unmodified | 0 | 0 | 0 | 0 | 0 | 0 | 0 | 0 | 0 | 0 | 0 | 613830 | 0 | 0 | 143370 | 78546 | 0 | 0 |
| VAAVSTIVN | PNPLA8; PNPL/ | Q9NP80; A0 | PLPL8_HUMAN | Unmodified | 73215 | 376500 | 0 | 0 | 408350 | 83577 | 0 | 0 | 3572900 | 197700 | 0 | 0 | 0 | 0 | 0 | 40629 | 332960 | 0 |
| VAAVSTIVN | PNPLA8; PNPL/ | Q9NP80; A0 | PLPL8_HUMAN | Unmodified | 0 | 0 | 0 | 0 | 0 | 0 | 0 | 0 | 0 | 933510 | 0 | 0 | 0 | 0 | 0 | 0 | 0 | 0 |
| VAAYAAQLI | NAPB; NAPB | Q9H115; A0 | SNAB_HUMAN | Unmodified | 0 | 0 | 0 | 0 | 0 | 0 | 472670 | 279790 | 0 | 354430 | 0 | 0 | 0 | 0 | 0 | 0 | 0 | 0 |
| VACAEEWC | TIPRL | O75663 | TIPRL_HUMAN | Unmodified | 0 | 0 | 0 | 390090 | 777740 | 184930 | 108000 | 69186 | 429940 | 144800 | 241410 | 2399200 | 1039800 | 0 | 0 | 0 | 986820 | 0 |
| VACGDSVL | SLC19A1; SLC1 | P41440; H3I | S19A1_HUMAN | Unmodified | 465950 | 0 | 737300 | 693120 | 0 | 0 | 0 | 50721 | 654140 | 0 | 0 | 0 | 0 | 0 | 1711700 | 282710 | 266930 | 142060 |
| VACIGAWH | RPL3L; RPL3; RI | Q92901; P39 | RL3L_HUMAN ( | Unmodified | 2675900 | 35120 | 1083200 | 0 | 0 | 0 | 65290 | 108820 | 466950 | 145820 | 1381800 | 19867000 | 292030 | 132840 | 0 | 982480 | 0 | 0 |
| VACITEQVL | RPN1; RPN1 | P04843; B7Z | RPN1_HUMAN | Unmodified | 0 | 0 | 0 | 0 | 0 | 0 | 0 | 0 | 0 | 0 | 12200 | 988950 | 0 | 0 | 0 | 0 | 0 | 0 |
| VADAMKQI | VAT1; VAT1 | Q99536; K7E | VAT1_HUMAN : | Unmodified | 0 | 0 | 0 | 0 | 1391900 | 205360 | 66295 | 11121 | 0 | 0 | 0 | 34369 | 0 | 0 | 0 | 0 | 594720 | 303420 |
| VADEGSFT | CD276; CD276 | Q5ZPR3; H0 | CD276_HUMAI | Unmodified | 88565 | 0 | 871370 | 128010 | 0 | 540150 | 199870 | 0 | 31848 | 865950 | 0 | 1029200 | 0 | 0 | 1549200 | 0 | 0 | 0 |
| VADFGLAR | SRMS; MST1R; | Q9H3Y6; Q0 | SRMS_HUMAN | Unmodified | 836180 | 586750 | 415410 | 1010400 | 0 | 0 | 0 | 0 | 3902200 | 1079300 | 0 | 2411300 | 0 | 0 | 895830 | 672150 | 1594300 | 1179400 |
| VADFGLAR | MET | P08581 | MET_HUMAN H | Unmodified | 14051 | 0 | 0 | 0 | 1722500 | 454250 | 0 | 0 | 799200 | 102180 | 90115 | 0 | 0 | 0 | 0 | 0 | 813090 | 3238500 |
| VADIGLAA\ | AHCY | P23526 | SAHH_HUMAN | Unmodified | 0 | 0 | 0 | 0 | 0 | 0 | 0 | 0 | 865480 | 315130 | 1241200 | 10399000 | 0 | 0 | 0 | 0 | 0 | 0 |
| VADIGLAA\ | AHCY | P23526 | SAHH_HUMAN | Unmodified | 552630 | 598210 | 1159500 | 1345100 | 16739000 | 2078700 | 1162200 | 760630 | 2514800 | 2900600 | 3302700 | 19291000 | 5320700 | 2169500 | 1744600 | 1473500 | 8604900 | 9160600 |
| VADISGDT | SND1; SND1 | Q7KZF4; H7 | SND1_HUMAN | Unmodified | 447870 | 0 | 517880 | 0 | 0 | 17266 | 225330 | 0 | 377370 | 537820 | 239150 | 3336000 | 0 | 0 | 922100 | 490190 | 49885 | 0 |
| VADKIQLIN | PGK1 | P00558 | PGK1_HUMAN | Oxidation (M) | 0 | 0 | 0 | 105840 | 1444700 | 2882500 | 859690 | 280530 | 0 | 2664300 | 122720 | 0 | 2122000 | 4985900 | 0 | 0 | 3427400 | 6303600 |
| VADKIQLIN | PGK1 | P00558 | PGK1_HUMAN | Unmodified | 89621 | 0 | 0 | 485090 | 9915200 | 1152300 | 1965600 | 304440 | 4059300 | 4783700 | 6566600 | 1852500 | 7071100 | 4529800 | 42163 | 0 | 20737000 | 28255000 |
| VADLLQHII | PTPRK; PTPRK; | Q15262; H0 | PTPRK_HUMAN | Oxidation (M) | 130560 | 0 | 0 | 172070 | 0 | 0 | 0 | 0 | 1176200 | 1763500 | 3869800 | 0 | 0 | 0 | 193130 | 0 | 773550 | 234520 |
| VADLLQHII | PTPRK; PTPRK; | Q15262; H0 | PTPRK_HUMAN | Unmodified | 0 | 0 | 0 | 0 | 0 | 0 | 0 | 0 | 1412500 | 0 | 0 | 0 | 0 | 0 | 0 | 0 | 565860 | 0 |
| VADMALHY | CCT8; CCT8 | P50990; H7 | TCPQ_HUMAN | Unmodified | 0 | 0 | 791440 | 0 | 6222600 | 2596000 | 1230900 | 457390 | 3003000 | 1604700 | 3070300 | 18599000 | 6673600 | 1506900 | 616610 | 825830 | 1473700 | 5510700 |
| VADMALHY | CCT8; CCT8 | P50990; H7 | TCPQ_HUMAN | Oxidation (M) | 89639 | 0 | 209160 | 0 | 1374500 | 194250 | 233570 | 44151 | 446870 | 270720 | 716690 | 362820 | 2203500 | 616170 | 0 | 0 | 280240 | 1150200 |
| VADPAYLP | GNAQ; GNAQ | P50148; B1A | GNAQ_HUMAN | Unmodified | 0 | 0 | 0 | 0 | 0 | 0 | 0 | 0 | 0 | 0 | 0 | 90458000 | 0 | 0 | 0 | 0 | 0 | 0 |
| VADPDHDI | MAPK1 | P28482 | MK01_HUMAN | Unmodified | 965280 | 382740 | 960590 | 1063500 | 0 | 0 | 0 | 0 | 2637200 | 0 | 539850 | 3465500 | 76590 | 0 | 1400600 | 1246200 | 0 | 51028 |
| VADPRSPS | CDCA3; CDCA3 | Q99618; F8 | CDCA3_HUMA | Unmodified | 0 | 0 | 0 | 0 | 12872000 | 1609300 | 0 | 0 | 0 | 1496400 | 0 | 0 | 6336800 | 3957800 | 0 | 0 | 811450 | 3374700 |
| VADWTGAI | EIF3D; EIF3D; E | O15371; B0 | EIF3D_HUMAN | Unmodified | 0 | 257340 | 515040 | 476610 | 1215900 | 214200 | 211660 | 206860 | 0 | 200400 | 175480 | 4042900 | 1271900 | 544240 | 350570 | 441490 | 244270 | 356650 |
| VADWTGAI | EIF3D; EIF3D; E | O15371; B0 | EIF3D_HUMAN | Unmodified | 718470 | 102590 | 0 | 0 | 0 | 0 | 0 | 0 | 0 | 0 | 0 | 850800 | 209590 | 0 | 0 | 135720 | 0 | 0 |
| VADYCENN | ABI1; ABI1; ABI | Q8IZP0; A0A | ABI1_HUMAN A | Unmodified | 0 | 0 | 0 | 205930 | 669560 | 0 | 0 | 0 | 525270 | 230040 | 0 | 427640 | 1119700 | 418000 | 325320 | 0 | 0 | 0 |
| VADYCENN | ABI1; ABI1; ABI | Q8IZP0; A0A | ABI1_HUMAN A | Unmodified | 0 | 0 | 0 | 0 | 0 | 0 | 79801 | 56812 | 0 | 261600 | 0 | 0 | 1112100 | 632630 | 0 | 0 | 452620 | 0 |
| VADYCENN | ABI2; ABI2; ABI2 | Q9NYB9; E7 | ABI2_HUMAN A | Unmodified | 0 | 0 | 0 | 0 | 0 | 0 | 0 | 0 | 0 | 0 | 0 | 1141600 | 0 | 0 | 0 | 0 | 0 | 0 |
| VAEAHENII | CRYZ | Q08257 | QOR_HUMAN ( | Unmodified | 0 | 0 | 114080 | 0 | 3105200 | 0 | 298190 | 0 | 2132500 | 773870 | 136370 | 1439500 | 863230 | 1109600 | 50899 | 93333 | 0 | 1046700 |
| VAEAVATFL | PPME1 | Q9Y570 | PPME1_HUMAI | Unmodified | 0 | 540190 | 260950 | 0 | 0 | 0 | 0 | 0 | 6135500 | 122400 | 544900 | 2425600 | 0 | 0 | 0 | 0 | 0 | 0 |
| VAEDEAEA | SLC3A2; SLC3A | P08195; A0A | 4F2_HUMAN 4I | Unmodified | 665680 | 2881000 | 5278100 | 2982200 | 4481300 | 1149200 | 531720 | 124990 | 5001300 | 1110000 | 6714100 | 18092000 | 260100 | 266080 | 5239200 | 777640 | 269240 | 1702500 |
| VAEDEAEA | SLC3A2; SLC3A | P08195; A0A | 4F2_HUMAN 4I | Unmodified | 0 | 0 | 0 | 2138600 | 27508000 | 0 | 2153000 | 0 | 1347600 | 2222400 | 1587800 | 0 | 2745200 | 0 | 0 | 0 | 2082600 | 5704200 |
| VAEEHAPS | PSMC1 | P62191 | PRS4_HUMAN | Unmodified | 445910 | 0 | 1952100 | 2004700 | 2592200 | 979570 | 590990 | 0 | 3714800 | 0 | 0 | 2313900 | 1312000 | 0 | 0 | 503680 | 0 | 0 |
| VAEEMTFFI | TOR1A | O14656 | TOR1A_HUMAI | Oxidation (M) | 0 | 0 | 0 | 0 | 0 | 0 | 90419 | 0 | 1464700 | 0 | 0 | 0 | 0 | 0 | 0 | 0 | 0 | 0 |
| VAEEMTFFI | TOR1A | O14656 | TOR1A_HUMAI | Unmodified | 0 | 0 | 0 | 0 | 0 | 0 | 0 | 0 | 4979100 | 0 | 0 | 0 | 199360 | 0 | 0 | 0 | 0 | 0 |
| VAEEMTFFI | TOR1A | O14656 | TOR1A_HUMAI | Oxidation (M) | 0 | 0 | 0 | 0 | 0 | 0 | 0 | 0 | 1068800 | 0 | 0 | 0 | 0 | 0 | 0 | 0 | 0 | 0 |
| VAEEMTFFI | TOR1A | O14656 | TOR1A_HUMAI | Unmodified | 118080 | 0 | 374580 | 102460 | 0 | 0 | 0 | 0 | 4933100 | 250180 | 0 | 106220 | 0 | 0 | 0 | 0 | 0 | 0 |
| VAEEMTFFI | TOR1B | O14657 | TOR1B_HUMAI | Oxidation (M) | 0 | 0 | 0 | 0 | 0 | 0 | 0 | 0 | 688700 | 0 | 0 | 0 | 0 | 0 | 0 | 0 | 0 | 0 |
| VAEEMTFFI | TOR1B | O14657 | TOR1B_HUMAI | Unmodified | 0 | 0 | 0 | 0 | 0 | 0 | 0 | 0 | 5677600 | 0 | 0 | 0 | 0 | 0 | 0 | 0 | 0 | 0 |
| VAEFHTELE | PSMD8; PSMD8 | P48556; K7E | PSMD8_HUMA | Unmodified | 0 | 0 | 0 | 0 | 0 | 0 | 0 | 0 | 0 | 0 | 0 | 1680500 | 0 | 0 | 0 | 0 | 0 | 0 |
| VAEGQTLD | HSPG2 | P98160 | PGBM_HUMAN | Unmodified | 114400 | 0 | 3872200 | 1070200 | 0 | 598540 | 0 | 0 | 737710 | 310890 | 510220 | 883170 | 356170 | 360730 | 294760 | 719200 | 0 | 704930 |
| VAEIEHAEK | CCT2; CCT2; C( | P78371; F5G | TCPB_HUMAN | Unmodified | 0 | 0 | 0 | 0 | 0 | 0 | 0 | 0 | 131140 | 0 | 0 | 1805200 | 117960 | 46709 | 0 | 0 | 0 | 13112 |
| VAEIEHAEK | CCT2; CCT2; C( | P78371; F5G | TCPB_HUMAN | Unmodified | 0 | 0 | 0 | 0 | 0 | 0 | 0 | 281480 | 28614 | 1211500 | 0 | 0 | 917990 | 0 | 0 | 0 | 0 | 0 |
| VAEIMQND | PPM1L | Q5SGD2 | PPM1L_HUMAI | Oxidation (M) | 0 | 431920 | 0 | 0 | 5658400 | 0 | 0 | 0 | 3170200 | 0 | 0 | 0 | 0 | 0 | 0 | 0 | 545440 | 0 |
| VAEITELILK | EXOSC9; EXOS( | Q06265; D6I | EXOS9_HUMAI | Unmodified | 0 | 0 | 0 | 0 | 0 | 0 | 0 | 0 | 0 | 109000 | 267380 | 1257800 | 0 | 0 | 0 | 0 | 0 | 0 |
| VAEKASDK | CCER2; CCER2 | I3L3R5; I3L3 | CCER2_HUMAI | Unmodified | 0 | 0 | 0 | 0 | 0 | 0 | 656070 | 2250000 | 0 | 0 | 0 | 0 | 0 | 0 | 0 | 0 | 0 | 0 |
| VAEKEELE( | SH2D4A; SH2D | Q9H788; H0 | SH24A_HUMAI | Unmodified | 0 | 211070 | 0 | 276900 | 2947400 | 0 | 0 | 0 | 0 | 0 | 0 | 0 | 1286200 | 1340400 | 313970 | 245150 | 250060 | 1155200 |
| VAEKLDEIY | SYNCRIP; SYN( | O60506; A0/ | HNRPQ_HUMA | Unmodified | 0 | 0 | 0 | 0 | 0 | 0 | 0 | 0 | 0 | 0 | 1187100 | 10213000 | 0 | 0 | 0 | 0 | 0 | 0 |
| VAEKLDEIY | SYNCRIP; SYN( | O60506; A0/ | HNRPQ_HUMA | Unmodified | 0 | 0 | 0 | 47892 | 326440 | 0 | 0 | 0 | 0 | 261940 | 420640 | 316160 | 598960 | 264500 | 0 | 0 | 173810 | 120500 |
| VAELEDEK | NPTX2 | P47972 | NPTX2_HUMAI | Unmodified | 0 | 0 | 0 | 0 | 0 | 0 | 0 | 0 | 5070200 | 558800 | 0 | 0 | 0 | 0 | 0 | 0 | 0 | 0 |
| VAELEHGS | NPTXR; NPTXR | O95502; A0/ | NPTXR_HUMAI | Unmodified | 0 | 0 | 0 | 0 | 0 | 0 | 0 | 0 | 10487000 | 1416900 | 54686 | 0 | 0 | 0 | 0 | 0 | 0 | 0 |
| VAELENSEI | CD58; CD58; C | P19256; H0 | LFA3_HUMAN I | Unmodified | 0 | 0 | 626650 | 0 | 559620 | 0 | 0 | 113570 | 0 | 12097 | 0 | 0 | 2373300 | 94574 | 0 | 0 | 0 | 5723700 |
| VAELFEK | SNX5; SNX5; SI | Q9Y5X3; Q5 | SNX5_HUMAN | Unmodified | 72742 | 32295 | 140270 | 109480 | 0 | 0 | 0 | 0 | 0 | 0 | 210060 | 2544600 | 511690 | 234250 | 122790 | 158220 | 0 | 0 |
| VAELHKTH | EPB41L2; EPB4 | O43491; A0/ | E41L2_HUMAN | Unmodified | 0 | 0 | 0 | 0 | 383700 | 0 | 0 | 0 | 0 | 0 | 0 | 0 | 0 | 85687 | 0 | 0 | 461620 | 0 |
| VAELILHG( | FAM114A1 | Q8IWE2 | NXP20_HUMAI | Unmodified | 54615 | 0 | 0 | 0 | 738400 | 191230 | 473960 | 273200 | 284700 | 309060 | 132840 | 0 | 1423900 | 1108400 | 0 | 213730 | 964560 | 885380 |
| VAELLLAK | C9orf64 | Q5T6V5 | QSPP_HUMAN | Unmodified | 158770 | 228880 | 0 | 150850 | 1075600 | 0 | 412070 | 30869 | 447540 | 221600 | 543420 | 1170600 | 0 | 0 | 289600 | 73562 | 0 | 0 |
| VAELNDVT | RNF141; RNF14 | Q8WVD5; E | RN141_HUMAI | Unmodified | 0 | 538730 | 0 | 0 | 0 | 0 | 0 | 0 | 0 | 0 | 0 | 0 | 0 | 0 | 0 | 0 | 0 | 0 |
| VAELSSDD | GLG1; GLG1 | Q92896; H3I | GSLG1_HUMAI | Unmodified | 0 | 0 | 0 | 0 | 0 | 0 | 0 | 36610 | 17753000 | 529320 | 1039100 | 6696800 | 0 | 0 | 0 | 0 | 0 | 0 |
| VAENVADK | GOLM2; GOLM2 | Q6P4E1; A0 | GOLM2_HUMA | Unmodified | 0 | 0 | 0 | 0 | 0 | 0 | 0 | 0 | 4581400 | 220700 | 0 | 0 | 0 | 138200 | 0 | 0 | 0 | 0 |
| VAEQDRAY | HLA-B; HLA-B; I | D5H3J5; A0/ | D5H3J5_HUMA | Unmodified | 0 | 0 | 0 | 0 | 0 | 0 | 0 | 0 | 1616300 | 0 | 0 | 0 | 0 | 0 | 0 | 0 | 0 | 0 |
| VAEQDRAY | HLA-B; HLA-B; I | D5H3J5; A0/ | D5H3J5_HUMA | Unmodified | 0 | 0 | 0 | 0 | 0 | 0 | 0 | 0 | 10239000 | 807760 | 0 | 0 | 0 | 0 | 0 | 0 | 105180 | 0 |
| VAEQDRAY | HLA-B; HLA-B; I | A0A140T951 | A0A140T951_H | Unmodified | 0 | 0 | 0 | 0 | 0 | 0 | 0 | 0 | 0 | 0 | 0 | 0 | 0 | 0 | 0 | 0 | 0 | 0 |
| VAEQDRAY | HLA-B; HLA-B; I | A0A140T951 | A0A140T951_H | Unmodified | 0 | 0 | 0 | 64219 | 0 | 0 | 0 | 0 | 0 | 0 | 0 | 6319700 | 0 | 0 | 133510 | 0 | 0 | 0 |
| VAEQTPLS/ | ALDH1B1 | P30837 | AL1B1_HUMAN | Unmodified | 0 | 0 | 0 | 0 | 133280 | 0 | 151140 | 0 | 757720 | 118680 | 182540 | 0 | 0 | 82386 | 0 | 0 | 0 | 0 |
| VAEQVLNA | SCPEP1 | Q9HB40 | RISC_HUMAN I | Unmodified | 0 | 1824000 | 0 | 533300 | 0 | 0 | 0 | 0 | 1791100 | 480420 | 0 | 521450 | 0 | 0 | 314330 | 0 | 0 | 0 |
| VAETANEEI | ABCE1; ABCE1 | P61221; D6F | ABCE1_HUMAI | Unmodified | 40720 | 0 | 107120 | 0 | 203660 | 0 | 0 | 0 | 66032 | 0 | 84193 | 2588800 | 54251 | 0 | 196360 | 115020 | 92625 | 0 |
| VAETAVQLI | ETF1; ETF1; ETF | P62495; B7Z | ERF1_HUMAN I | Unmodified | 78662 | 0 | 138150 | 0 | 0 | 0 | 55098 | 57154 | 0 | 35493 | 104870 | 849070 | 121950 | 24809 | 42077 | 84459 | 0 | 0 |
| VAEVEGEQ | LETM1 | O95202 | LETM1_HUMAI | Unmodified | 0 | 0 | 0 | 0 | 0 | 0 | 40633 | 0 | 1023600 | 68589 | 111450 | 124300 | 0 | 0 | 0 | 0 | 192770 | 0 |
| VAEVLEGK | AARSD1; PTGES | Q9BTE6; B3I | AASD1_HUMAI | Unmodified | 0 | 0 | 0 | 0 | 0 | 0 | 0 | 176410 | 319840 | 198340 | 449120 | 1059700 | 0 | 35991 | 0 | 0 | 0 | 0 |
| VAFERGEEI | DHX9 | Q08211 | DHX9_HUMAN | Unmodified | 168610 | 0 | 0 | 0 | 92513 | 33844 | 0 | 30332 | 119190 | 762330 | 392100 | 7853400 | 0 | 0 | 0 | 103260 | 14573 | 228750 |
| VAFITGGG! | DECR2; DECR2 | Q9NUI1; Q4 | DECR2_HUMAI | Unmodified | 0 | 0 | 0 | 0 | 0 | 0 | 0 | 0 | 3022600 | 666280 | 0 | 0 | 0 | 0 | 0 | 0 | 0 | 0 |
| VAFITGGGT | DECR1; DECR1 | Q16698; E5F | DECR_HUMAN | Unmodified | 0 | 0 | 0 | 0 | 0 | 0 | 0 | 415970 | 8194900 | 2576300 | 0 | 0 | 0 | 0 | 0 | 0 | 0 | 3858400 |
| VAFPLEVV( | IL1RAP; IL1RAP | Q9NPH3; C9 | IL1AP_HUMAN | Unmodified | 0 | 125700 | 1262500 | 1312200 | 2158400 | 871920 | 305320 | 133420 | 504940 | 162320 | 0 | 1094600 | 702960 | 618070 | 1138200 | 682050 | 0 | 709670 |
| VAFRDCEG | FSCN1 | Q16658 | FSCN1_HUMAI | Unmodified | 0 | 0 | 0 | 0 | 14325000 | 2507600 | 1444800 | 1234100 | 0 | 2385900 | 812080 | 0 | 128760 | 206610 | 0 | 0 | 12070000 | 8333300 |
| VAFTGSTE\ | ALDH1A1 | P00352 | AL1A1_HUMAN | Unmodified | 47759 | 0 | 210390 | 0 | 0 | 0 | 347620 | 0 | 1979200 | 132680 | 13334000 | 7.85E+08 | 0 | 0 | 0 | 286380 | 358950 | 160290 |
| VAFTGSTE\ | ALDH1A1 | P00352 | AL1A1_HUMAN | Unmodified | 0 | 0 | 0 | 0 | 0 | 0 | 0 | 0 | 0 | 564060 | 1795100 | 3009400 | 0 | 0 | 0 | 0 | 0 | 0 |
| VAFTLLEK | YKT6; YKT6; YK | O15498; A0/ | YKT6_HUMAN : | Unmodified | 2585300 | 2743200 | 1139300 | 429610 | 65941 | 0 | 0 | 0 | 1579100 | 284410 | 272280 | 2066000 | 0 | 0 | 1428300 | 2141700 | 0 | 0 |
| VAFVNWIN | PLS1; PLS1 | Q14651; C9J | PLSI_HUMAN F | Unmodified | 0 | 0 | 0 | 0 | 122820 | 748850 | 64364 | 104680 | 1644900 | 453310 | 1336100 | 4838300 | 2772000 | 0 | 0 | 0 | 275670 | 275620 |
| VAGALAEA( | TKFC; TKFC | Q3LXA3; H0 | TKFC_HUMAN | Unmodified | 0 | 0 | 0 | 0 | 0 | 0 | 0 | 25020 | 0 | 0 | 18874 | 3889000 | 1062900 | 515570 | 0 | 0 | 0 | 0 |
| VAGAQIQG | NNT; NNT; NNT | Q13423; A0/ | NNTM_HUMAN | Unmodified | 0 | 298630 | 0 | 0 | 0 | 0 | 0 | 0 | 3066100 | 0 | 0 | 0 | 0 | 0 | 0 | 0 | 105250 | 0 |
| VAGASRIIG | ADH5; ADH5 | P11766; H0 | ADHX_HUMAN | Unmodified | 0 | 0 | 0 | 0 | 803750 | 0 | 0 | 237090 | 0 | 110160 | 0 | 0 | 170820 | 429590 | 0 | 0 | 849390 | 410950 |
| VAGDCLDE | LRRC59 | Q96AG4 | LRC59_HUMAN | Unmodified | 0 | 0 | 0 | 0 | 473720 | 46613 | 0 | 0 | 0 | 0 | 0 | 96327 | 0 | 222380 | 22633 | 91580 | 64257 | 0 |
| VAGDIESLI | CACNA2D2; CA | Q9NY47; C9 | CA2D2_HUMAI | Unmodified | 0 | 0 | 0 | 0 | 0 | 0 | 0 | 0 | 2694400 | 472570 | 0 | 0 | 0 | 0 | 0 | 0 | 0 | 0 |
| VAGDIESLI | CACNA2D2; CA | Q9NY47; C9 | CA2D2_HUMAI | Unmodified | 49459 | 54116 | 0 | 0 | 0 | 0 | 430450 | 0 | 8649700 | 665600 | 719160 | 1858800 | 0 | 0 | 0 | 0 | 0 | 0 |
| VAGDIESLI | CACNA2D2; CA | Q9NY47; C9 | CA2D2_HUMAI | Unmodified | 0 | 0 | 0 | 0 | 1663800 | 0 | 5068400 | 101160 | 2356400 | 8788500 | 242670 | 0 | 1232600 | 612650 | 0 | 0 | 0 | 920680 |
| VAGFTGAS | ITCH | Q96J02 | ITCH_HUMAN I | Unmodified | 320960 | 187610 | 0 | 0 | 0 | 0 | 0 | 0 | 0 | 0 | 0 | 0 | 0 | 0 | 46163 | 0 | 27465 | 75629 |
| VAGGAAPS | NOLC1; NOLC1 | Q14978; A0/ | NOLC1_HUMA | Unmodified | 0 | 0 | 0 | 0 | 138300 | 0 | 0 | 0 | 0 | 23981 | 81058 | 250530 | 237380 | 73038 | 0 | 12124 | 99144 | 135350 |
| VAGHPLAQ | SNX3 | O60493 | SNX3_HUMAN | Oxidation (M) | 0 | 0 | 0 | 0 | 1369400 | 0 | 2323700 | 0 | 0 | 0 | 0 | 0 | 1080700 | 1496900 | 0 | 0 | 330580 | 0 |
| VAGHPLAQ | SNX3 | O60493 | SNX3_HUMAN | Unmodified | 0 | 0 | 0 | 0 | 1051800 | 0 | 475310 | 0 | 0 | 111440 | 0 | 816530 | 468450 | 228430 | 0 | 0 | 0 | 249070 |
| VAGHPNIV | DECR1; DECR1 | Q16698; E5F | DECR_HUMAN | Unmodified | 0 | 0 | 0 | 0 | 0 | 0 | 0 | 32261 | 22161000 | 1807500 | 0 | 0 | 0 | 0 | 0 | 0 | 0 | 0 |
| VAGHPNIV | DECR1; DECR1 | Q16698; E5F | DECR_HUMAN | Unmodified | 0 | 0 | 0 | 0 | 0 | 0 | 602570 | 0 | 1682000 | 2476000 | 0 | 0 | 0 | 0 | 0 | 0 | 836240 | 0 |
| VAGHSLGY | ELAVL1; ELAVL | Q15717; M0 | ELAV1_HUMAN | Unmodified | 282280 | 293780 | 1144600 | 830940 | 0 | 0 | 303350 | 38069 | 1195400 | 308990 | 964760 | 13656000 | 491080 | 223690 | 480270 | 832860 | 0 | 235290 |
| VAGIESHSI | NUMA1; NUMA | Q14980; A0/ | NUMA1_HUMA | Unmodified | 14380 | 0 | 0 | 0 | 0 | 0 | 0 | 0 | 0 | 0 | 0 | 266180 | 0 | 0 | 0 | 0 | 0 | 0 |
| VAGKEIYSI | INPP5A; INPP5/ | Q14642; Q5 | I5P1_HUMAN I | Unmodified | 0 | 0 | 0 | 0 | 488230 | 0 | 151450 | 0 | 0 | 401500 | 0 | 0 | 273350 | 255260 | 0 | 0 | 147890 | 1190100 |
| VAGLLTQK | IGF2R; IGF2R | P11717; A0A | MPRI_HUMAN ( | Unmodified | 715600 | 547460 | 1022300 | 730850 | 469970 | 0 | 838620 | 1159600 | 41303000 | 7951500 | 3618300 | 7965700 | 616590 | 794150 | 686890 | 1307000 | 116290 | 204120 |
| VAGLLTQKI | IGF2R; IGF2R | P11717; A0A | MPRI_HUMAN ( | Unmodified | 0 | 0 | 0 | 106070 | 259200 | 0 | 0 | 20957 | 4922800 | 5808900 | 353800 | 0 | 577690 | 448890 | 0 | 0 | 0 | 127140 |
| VAGMDVEI | YWHAE; YWHAI | P62258; I3L | 1433E_HUMAN | Oxidation (M) | 137460 | 0 | 0 | 0 | 363510 | 0 | 217710 | 0 | 0 | 0 | 0 | 4923200 | 244340 | 263780 | 0 | 0 | 0 | 0 |
| VAGMLQRF | EXOC3; EXOC3 | O60645; D6I | EXOC3_HUMAI | Unmodified | 0 | 0 | 0 | 0 | 0 | 0 | 0 | 0 | 1250600 | 919580 | 0 | 782190 | 0 | 0 | 0 | 0 | 0 | 0 |
| VAGMLQRF | EXOC3; EXOC3 | O60645; D6I | EXOC3_HUMAI | Oxidation (M) | 0 | 0 | 0 | 316060 | 0 | 0 | 0 | 0 | 364060 | 0 | 381350 | 763580 | 0 | 0 | 0 | 0 | 0 | 0 |

| | | | | | | | | | | | | | | | | | | | | | | |
|---|---|---|---|---|---|---|---|---|---|---|---|---|---|---|---|---|---|---|---|---|---|---|
| VAGNFHFA | ERGIC3; ERGIC | Q9Y282; A0A | ERGI3_HUMAN | Unmodified | 0 | 0 | 0 | 0 | 0 | 0 | 0 | 0 | 405040 | 180750 | 80060 | 0 | 0 | 0 | 0 | 0 | 0 | 0 |
| VAGNFHIT | ERGIC2; ERGIC | Q96RQ1; A0. | ERGI2_HUMAN | Unmodified | 0 | 0 | 0 | 0 | 0 | 0 | 0 | 0 | 745110 | 0 | 0 | 247570 | 0 | 0 | 0 | 0 | 0 | 0 |
| VAGQDGS | SUMO3; SUMO | P55854; P61 | SUMO3_HUMA | Unmodified | 2090200 | 2159300 | 2395200 | 1861600 | 11585000 | 2803900 | 3353800 | 896320 | 16195000 | 22764000 | 0 | 18846000 | 2929200 | 2155200 | 3576300 | 2000200 | 9814900 | 8162400 |
| VAGQDGS | SUMO3; SUMO | P55854; P61 | SUMO3_HUMA | Unmodified | 0 | 0 | 0 | 0 | 1889900 | 0 | 562920 | 538060 | 0 | 535400 | 0 | 0 | 371840 | 0 | 0 | 0 | 5625000 | 2012800 |
| VAGSPGW | SLC3A2; SLC3A | P08195; A0A | 4F2_HUMAN 4I | Unmodified | 2162000 | 2409200 | 1184600 | 432090 | 628900 | 473300 | 0 | 0 | 1034300 | 516830 | 261380 | 15341000 | 0 | 0 | 2950500 | 1881000 | 0 | 0 |
| VAGTGEGC | DBNL; DBNL; D | Q9UJU6; B4I | DBNL_HUMAN | Oxidation (M) | 0 | 121400 | 0 | 0 | 0 | 0 | 0 | 0 | 2849400 | 226330 | 0 | 2060000 | 0 | 0 | 1463400 | 0 | 0 | 0 |
| VAGTQACA | ADPGK | Q9BRR6 | ADPGK_HUMA | Unmodified | 0 | 0 | 0 | 0 | 0 | 0 | 0 | 0 | 0 | 0 | 0 | 0 | 0 | 0 | 0 | 0 | 0 | 0 |
| VAGVVAPT | RECQL | P46063 | RECQ1_HUMA | Unmodified | 0 | 0 | 76590 | 0 | 0 | 0 | 1633500 | 584160 | 0 | 270370 | 0 | 0 | 0 | 0 | 0 | 0 | 0 | 0 |
| VAGYAALL | NAPA; NAPA; N. | P54920; M0 | SNAA_HUMAN | Unmodified | 294840 | 0 | 0 | 0 | 972820 | 0 | 3670100 | 2427700 | 1153400 | 1000200 | 0 | 1537900 | 4858600 | 2165800 | 2079000 | 1472700 | 2067700 | 4005900 |
| VAHALAEG | TPI1; TPI1; TPI1 | P60174; U3 | TPIS_HUMAN T | Unmodified | 35296 | 0 | 0 | 149990 | 0 | 0 | 0 | 0 | 1248500 | 442980 | 4133800 | 5741700 | 4447200 | 0 | 0 | 32675 | 0 | 0 |
| VAHALAEG | TPI1; TPI1; TPI1 | P60174; U3 | TPIS_HUMAN T | Unmodified | 9920300 | 0 | 148320 | 7066300 | 1.43E+08 | 16903000 | 4502100 | 0 | 5553500 | 13203000 | 4777600 | 7143200 | 1.53E+08 | 50874000 | 19119000 | 673150 | 62516000 | 1.03E+08 |
| VAHILNISP | HSDL2 | Q6YN16 | HSDL2_HUMA | Unmodified | 0 | 922410 | 0 | 0 | 0 | 0 | 1821300 | 0 | 34175000 | 21773000 | 0 | 0 | 512920 | 488710 | 0 | 0 | 110640 | 0 |
| VAHPIRPK | PRPF19; PRPF1 | Q9UMS4; F5 | PRP19_HUMA | Unmodified | 0 | 0 | 0 | 0 | 806450 | 0 | 0 | 0 | 0 | 801660 | 0 | 1808800 | 1244300 | 341150 | 0 | 0 | 320300 | 0 |
| VAHQLQAL | CHGA; CHGA | P10645; G5 | CMGA_HUMAN | Unmodified | 0 | 0 | 0 | 0 | 0 | 0 | 0 | 0 | 4990100 | 1113700 | 0 | 0 | 0 | 0 | 0 | 0 | 0 | 0 |
| VAHSFNCT | PA2G4; PA2G4; | Q9UQ80; F8 | PA2G4_HUMA | Unmodified | 0 | 0 | 0 | 0 | 2638500 | 0 | 3903400 | 0 | 0 | 770700 | 0 | 9437900 | 0 | 952770 | 1276200 | 0 | 1180200 | 1048900 |
| VAHSFNCT | PA2G4; PA2G4; | Q9UQ80; F8 | PA2G4_HUMA | Oxidation (M) | 643500 | 0 | 0 | 297930 | 745070 | 0 | 291800 | 0 | 324800 | 419870 | 0 | 7192400 | 735350 | 1001100 | 374840 | 217800 | 237700 | 65936 |
| VAHTSLDA | HSH2D; HSH2D | Q96JZ2; K7E | HSH2D_HUMA | Unmodified | 939170 | 766840 | 2970900 | 3146500 | 0 | 0 | 0 | 0 | 0 | 0 | 0 | 0 | 0 | 0 | 4274900 | 3498100 | 0 | 0 |
| VAHVEHEE | GPC3; GPC3 | P51654; H0 | GPC3_HUMAN | Unmodified | 0 | 0 | 0 | 4920200 | 0 | 1302000 | 0 | 0 | 9605400 | 3830600 | 1155700 | 257550 | 210030 | 208270 | 0 | 0 | 293740 | 51186 |
| VAHVEVAP | XPR1 | Q9UBH6 | XPR1_HUMAN | Unmodified | 0 | 0 | 0 | 0 | 0 | 0 | 0 | 0 | 587590 | 283160 | 0 | 2467200 | 0 | 0 | 0 | 99876 | 0 | 0 |
| VAIAALEVL | OAT | P04181 | OAT_HUMAN C | Unmodified | 0 | 0 | 0 | 0 | 757360 | 0 | 0 | 0 | 3898400 | 284210 | 0 | 0 | 0 | 0 | 0 | 0 | 0 | 0 |
| VAIDVGYR | AKR1B1; AKR1B | P15121; E9P | ALDR_HUMAN | Unmodified | 157440 | 116380 | 0 | 490530 | 5488900 | 383700 | 0 | 0 | 59152 | 4133200 | 8811000 | 51524000 | 289640 | 21368 | 25250 | 266440 | 222910 | 0 |
| VAIDVGYR | AKR1B1; AKR1B | P15121; E9P | ALDR_HUMAN | Unmodified | 0 | 0 | 0 | 0 | 1982800 | 0 | 0 | 0 | 0 | 0 | 0 | 0 | 0 | 0 | 0 | 0 | 0 | 0 |
| VAIEHLDK | CCT5; CCT5; C | P48643; B7Z | TCPE_HUMAN | Unmodified | 0 | 0 | 0 | 0 | 0 | 0 | 0 | 0 | 697470 | 68686 | 0 | 3180100 | 0 | 0 | 0 | 0 | 0 | 45549 |
| VAIEPGAP | SORD; SORD | Q00796; A0A | DHSO_HUMAN | Unmodified | 0 | 87287 | 643780 | 830440 | 1635100 | 0 | 0 | 188060 | 1256500 | 949500 | 857510 | 6763800 | 2633600 | 0 | 1109300 | 1252600 | 0 | 0 |
| VAIEPGAP | SORD; SORD | Q00796; A0A | DHSO_HUMAN | Unmodified | 173140 | 0 | 3206900 | 1741400 | 0 | 0 | 0 | 0 | 612960 | 76804 | 0 | 6789900 | 77980 | 0 | 2456000 | 2346000 | 0 | 0 |
| VAIHNADL | DLG5 | Q8TDM6 | DLG5_HUMAN | Unmodified | 0 | 0 | 0 | 0 | 3726700 | 427530 | 0 | 0 | 0 | 265480 | 0 | 0 | 0 | 0 | 0 | 0 | 5400800 | 5752500 |
| VAILIPFR | B4GALT5 | O43286 | B4GT5_HUMA | Unmodified | 0 | 0 | 791530 | 0 | 0 | 0 | 0 | 0 | 7476900 | 737270 | 736290 | 5906300 | 0 | 0 | 0 | 0 | 0 | 0 |
| VAIPNSAV | ELAPOR2; ELAP | A8MWY0; C | ELAP2_HUMA | Unmodified | 0 | 193420 | 0 | 0 | 0 | 0 | 0 | 0 | 7374000 | 2766400 | 0 | 0 | 0 | 0 | 0 | 0 | 0 | 0 |
| VAIVKPGVF | HYOU1; HYOU1 | Q9Y4L1; J3Q | HYOU1_HUMA | Oxidation (M) | 342110 | 0 | 2527900 | 1574800 | 806730 | 0 | 1423400 | 821350 | 48336000 | 52054000 | 13981000 | 5334300 | 3390300 | 4536300 | 2739800 | 1390800 | 144130 | 1472100 |
| VAIVKPGVF | HYOU1; HYOU1 | Q9Y4L1; J3Q | HYOU1_HUMA | Unmodified | 0 | 0 | 0 | 0 | 316400 | 0 | 388780 | 0 | 21804000 | 6297800 | 853550 | 0 | 285560 | 497480 | 0 | 0 | 0 | 0 |
| VAIVKPGVF | HYOU1; HYOU1 | Q9Y4L1; J3Q | HYOU1_HUMA | Unmodified | 0 | 0 | 0 | 0 | 0 | 0 | 0 | 0 | 0 | 612100 | 0 | 0 | 0 | 0 | 0 | 0 | 0 | 0 |
| VAIVKPGVF | HYOU1; HYOU1 | Q9Y4L1; J3Q | HYOU1_HUMA | Oxidation (M) | 0 | 0 | 0 | 0 | 0 | 0 | 0 | 0 | 0 | 676370 | 0 | 0 | 0 | 0 | 0 | 0 | 0 | 0 |
| VAKAEAAG | TNFRSF10B | O14763 | TR10B_HUMA | Unmodified | 0 | 0 | 0 | 0 | 1449400 | 0 | 0 | 0 | 0 | 0 | 0 | 0 | 0 | 0 | 0 | 0 | 0 | 0 |
| VAKEIARAV | CUTA; CUTA; C | O60888; H0 | CUTA_HUMAN | Unmodified | 0 | 0 | 0 | 0 | 0 | 0 | 395640 | 177930 | 0 | 0 | 0 | 0 | 0 | 0 | 0 | 0 | 958540 | 0 |
| VAKLPFVIN | NPTX1 | Q15818 | NPTX1_HUMA | Unmodified | 0 | 0 | 0 | 0 | 0 | 0 | 0 | 0 | 18168000 | 2488000 | 0 | 0 | 0 | 0 | 0 | 0 | 0 | 0 |
| VAKLPFVIN | NPTX1 | Q15818 | NPTX1_HUMA | Unmodified | 0 | 0 | 0 | 0 | 0 | 0 | 0 | 0 | 1937600 | 0 | 0 | 0 | 0 | 104020 | 0 | 0 | 0 | 0 |
| VAKPVVEM | IDH2; IDH2 | P48735; H0 | IDHP_HUMAN | Unmodified | 5728300 | 2782400 | 7427300 | 918240 | 0 | 0 | 0 | 0 | 18778000 | 388730 | 186580 | 0 | 3341800 | 2171700 | 1696700 | 1338400 | 18358 | 0 |
| VAKPVVEM | IDH2; IDH2 | P48735; H0 | IDHP_HUMAN | Oxidation (M) | 445580 | 0 | 0 | 0 | 0 | 0 | 35924 | 0 | 7248900 | 91869 | 118010 | 142140 | 508530 | 0 | 0 | 0 | 0 | 0 |
| VAKPVVEM | IDH2; IDH2 | P48735; H0 | IDHP_HUMAN | 2 Oxidation (M) | 0 | 0 | 0 | 0 | 0 | 0 | 0 | 0 | 1779100 | 57923 | 75827 | 103230 | 0 | 0 | 22093 | 0 | 0 | 0 |
| VALAEACR | TPP2; TPP2; TP | P29144; A0A | TPP2_HUMAN | Unmodified | 0 | 0 | 0 | 0 | 0 | 0 | 0 | 0 | 2798900 | 0 | 0 | 3439600 | 0 | 0 | 0 | 0 | 0 | 0 |
| VALCTQVA | PTK7; PTK7; PT | Q13308; H0 | PTK7_HUMAN | Unmodified | 0 | 0 | 0 | 0 | 0 | 0 | 0 | 0 | 1367700 | 265250 | 0 | 621770 | 0 | 0 | 0 | 0 | 0 | 0 |
| VALCTQVA | PTK7; PTK7; PT | Q13308; H0 | PTK7_HUMAN | Unmodified | 0 | 0 | 0 | 0 | 5004400 | 0 | 3339000 | 0 | 790730 | 1642400 | 841300 | 0 | 4983400 | 2966500 | 0 | 0 | 2811100 | 4890400 |
| VALEGLRP | ILK; ILK | Q13418; A0A | ILK_HUMAN Int | Unmodified | 0 | 0 | 197250 | 0 | 1637900 | 0 | 2012600 | 564310 | 0 | 0 | 0 | 0 | 0 | 857230 | 0 | 0 | 0 | 2027000 |
| VALETAEK | ABR; ABR; ABR; | Q12979; A0A | ABR_HUMAN A | Unmodified | 0 | 0 | 0 | 0 | 114020 | 0 | 0 | 0 | 0 | 0 | 0 | 0 | 0 | 0 | 0 | 0 | 325960 | 150850 |
| VALIGFPSV | DRG2; DRG2; D | P55039; J3Q | DRG2_HUMAN | Unmodified | 0 | 0 | 0 | 0 | 0 | 0 | 0 | 0 | 331220 | 0 | 0 | 1548100 | 383820 | 0 | 0 | 358440 | 0 | 0 |
| VALILQNV | GBE1; GBE1 | Q04446; E9F | GLGB_HUMAN | Unmodified | 0 | 0 | 79133 | 0 | 0 | 0 | 0 | 0 | 1433600 | 112800 | 109970 | 1279700 | 0 | 0 | 108770 | 115610 | 0 | 0 |
| VALLEQQM | CEP55 | Q53EZ4 | CEP55_HUMA | Oxidation (M) | 0 | 0 | 202200 | 0 | 4327900 | 0 | 0 | 0 | 1126500 | 269540 | 0 | 833650 | 0 | 0 | 0 | 0 | 0 | 0 |
| VALLEQQM | CEP55 | Q53EZ4 | CEP55_HUMA | Unmodified | 0 | 0 | 0 | 0 | 1863400 | 0 | 0 | 0 | 2163900 | 0 | 0 | 0 | 0 | 0 | 0 | 0 | 159770 | 66894 |
| VALLEQQM | CEP55 | Q53EZ4 | CEP55_HUMA | Unmodified | 0 | 0 | 0 | 0 | 3103300 | 0 | 0 | 0 | 0 | 0 | 0 | 0 | 0 | 0 | 0 | 0 | 0 | 0 |
| VALLLLDQ | ANK3 | Q12955 | ANK3_HUMAN | Unmodified | 0 | 0 | 0 | 2474000 | 0 | 0 | 0 | 0 | 0 | 0 | 0 | 0 | 14418000 | 13640000 | 1360300 | 10120000 | 0 | 0 |
| VALLSGGG | TKFC; TKFC; TK | Q3LXA3; H0 | TKFC_HUMAN | Unmodified | 0 | 0 | 0 | 0 | 0 | 0 | 0 | 0 | 0 | 0 | 0 | 1573800 | 0 | 0 | 0 | 222890 | 0 | 0 |
| VALNTLAR | QPRT | Q15274 | NADC_HUMAN | Unmodified | 0 | 0 | 0 | 0 | 0 | 0 | 0 | 0 | 0 | 0 | 0 | 9296700 | 0 | 0 | 0 | 0 | 0 | 0 |
| VALNVSCA | CPNE3; CPNE3 | O75131; A0A | CPNE3_HUMA | Unmodified | 1342500 | 0 | 408920 | 5525300 | 6750500 | 707780 | 3591500 | 2121600 | 58754000 | 16752000 | 1025500 | 2956000 | 5699700 | 6495500 | 549140 | 4489200 | 4718400 | 11869000 |
| VALNVSCA | CPNE3; CPNE3 | O75131; A0A | CPNE3_HUMA | Unmodified | 0 | 0 | 0 | 0 | 889060 | 290190 | 2702400 | 783890 | 309600 | 3708400 | 253890 | 2729000 | 1849400 | 982910 | 0 | 0 | 2213800 | 1927400 |
| VALQAEKL | PCBD1 | P61457 | PHS_HUMAN P | Unmodified | 318600 | 529120 | 0 | 1275600 | 5353800 | 797580 | 391100 | 0 | 10714000 | 14683000 | 2406100 | 6476600 | 15510000 | 6729700 | 400890 | 540130 | 5137400 | 8346000 |
| VALQQSFS | DIP2B | Q9P265 | DIP2B_HUMA | Unmodified | 272520 | 178890 | 300590 | 306930 | 1151400 | 0 | 302050 | 140850 | 1731700 | 651550 | 243570 | 8165200 | 0 | 44000 | 1014000 | 846180 | 0 | 1016200 |
| VALRGEDV | COPZ1; COPZ1 | P61923; F8V | COPZ1_HUMA | Unmodified | 0 | 0 | 0 | 0 | 0 | 0 | 0 | 0 | 0 | 0 | 0 | 0 | 0 | 0 | 0 | 0 | 0 | 0 |
| VALSEPQP | SRRT; SRRT; SR | Q9BXP5; H7 | SRRT_HUMAN | Unmodified | 0 | 0 | 0 | 0 | 501840 | 126350 | 118890 | 106700 | 222820 | 193100 | 371710 | 3728500 | 245120 | 0 | 0 | 0 | 64616 | 109520 |
| VALVGLEA | ADAM15 | Q13444 | ADA15_HUMA | Unmodified | 1750700 | 3261200 | 318440 | 0 | 0 | 0 | 0 | 0 | 0 | 0 | 0 | 3593900 | 0 | 0 | 1400600 | 922260 | 0 | 0 |
| VALVTASTD | DHRS4L2; DHR | Q6PKH6; Q9 | DR4L2_HUMA | Unmodified | 0 | 0 | 0 | 0 | 0 | 0 | 0 | 0 | 2695600 | 1836800 | 0 | 0 | 0 | 0 | 0 | 0 | 0 | 0 |
| VALVTASTD | DHRS4L2; DHR | Q6PKH6; Q9 | DR4L2_HUMA | Unmodified | 0 | 0 | 0 | 0 | 0 | 0 | 0 | 0 | 4557100 | 1327200 | 0 | 0 | 0 | 0 | 0 | 0 | 0 | 0 |
| VALVTFNS | CAND1; CAND1 | Q86VP6; A0A | CAND1_HUMA | Unmodified | 0 | 0 | 0 | 0 | 0 | 0 | 534210 | 0 | 0 | 0 | 0 | 1162500 | 0 | 0 | 0 | 0 | 0 | 0 |
| VALVTGAA | HPGD; HPGD; H | P15428; E9P | PGDH_HUMAN | Unmodified | 0 | 0 | 0 | 0 | 0 | 0 | 0 | 0 | 0 | 0 | 0 | 0 | 348580 | 0 | 0 | 0 | 0 | 1372600 |
| VALVTGAA | HPGD; HPGD; H | P15428; E9P | PGDH_HUMAN | Unmodified | 0 | 0 | 0 | 0 | 0 | 0 | 0 | 0 | 0 | 0 | 0 | 0 | 0 | 0 | 0 | 0 | 0 | 1463000 |
| VALVYGQM | ATP5F1B; ATP5 | P06576; H0 | ATPB_HUMAN | Oxidation (M) | 0 | 0 | 0 | 0 | 0 | 0 | 0 | 0 | 5241100 | 1387100 | 0 | 0 | 0 | 232640 | 0 | 0 | 0 | 0 |
| VALVYGQM | ATP5F1B; ATP5 | P06576; H0 | ATPB_HUMAN | Unmodified | 0 | 0 | 0 | 0 | 0 | 0 | 0 | 0 | 11340000 | 890090 | 0 | 0 | 0 | 0 | 0 | 0 | 0 | 0 |
| VALVYGQM | ATP5F1B; ATP5 | P06576; H0 | ATPB_HUMAN | Oxidation (M) | 0 | 0 | 0 | 0 | 0 | 0 | 200130 | 0 | 0 | 882800 | 0 | 1150100 | 0 | 0 | 0 | 0 | 0 | 0 |
| VALVYGQM | ATP5F1B; ATP5 | P06576; H0 | ATPB_HUMAN | Unmodified | 77824 | 0 | 61355 | 246330 | 469210 | 0 | 79853 | 0 | 0 | 1597800 | 168330 | 589840 | 0 | 0 | 110720 | 0 | 0 | 0 |
| VAMANIQP | VCL | P18206 | VINC_HUMAN | 2 Oxidation (M) | 0 | 0 | 0 | 0 | 3090200 | 0 | 0 | 0 | 0 | 605430 | 429060 | 0 | 0 | 0 | 0 | 0 | 4086100 | 209930 |
| VAMANIQP | VCL | P18206 | VINC_HUMAN | Oxidation (M) | 0 | 0 | 0 | 0 | 1256600 | 246210 | 0 | 86243 | 861300 | 0 | 0 | 926410 | 0 | 0 | 0 | 0 | 1938500 | 375820 |
| VAMFLTDS | KARS1 | Q15046 | SYK_HUMAN L | Oxidation (M) | 0 | 0 | 0 | 0 | 275450 | 0 | 0 | 58283 | 0 | 0 | 0 | 1041200 | 0 | 0 | 0 | 0 | 0 | 0 |
| VAMFLTDS | KARS1 | Q15046 | SYK_HUMAN L | Unmodified | 0 | 0 | 0 | 0 | 282660 | 525810 | 0 | 117390 | 565740 | 150210 | 0 | 1695900 | 415380 | 225100 | 0 | 0 | 516940 | 576670 |
| VAMQDATA | ACO2; ACO2; A | Q99798; A2A | ACON_HUMAN | 2 Oxidation (M) | 0 | 0 | 0 | 0 | 0 | 0 | 0 | 0 | 625070 | 0 | 0 | 0 | 0 | 0 | 0 | 0 | 0 | 0 |
| VAMQDATA | ACO2; ACO2; A | Q99798; A2A | ACON_HUMAN | Oxidation (M) | 0 | 0 | 0 | 0 | 0 | 0 | 0 | 0 | 499840 | 0 | 0 | 0 | 0 | 0 | 0 | 0 | 0 | 0 |
| VAMQDATA | ACO2; ACO2; A | Q99798; A2A | ACON_HUMAN | Unmodified | 0 | 0 | 0 | 0 | 0 | 0 | 0 | 0 | 102690 | 0 | 0 | 0 | 0 | 0 | 0 | 0 | 0 | 0 |
| VAMSHFEP | ACO2; ACO2; A | Q99798; A2A | ACON_HUMAN | Unmodified | 0 | 0 | 0 | 0 | 0 | 0 | 472020 | 0 | 5997200 | 691240 | 0 | 0 | 0 | 0 | 0 | 0 | 0 | 0 |
| VAMSHFEP | ACO2; ACO2; A | Q99798; A2A | ACON_HUMAN | Oxidation (M) | 0 | 0 | 0 | 0 | 1254100 | 0 | 2256600 | 0 | 7428400 | 2475500 | 0 | 0 | 477550 | 0 | 0 | 0 | 0 | 0 |
| VANDNAPE | LAMTOR3 | Q9UHA4 | LTOR3_HUMA | Unmodified | 0 | 0 | 0 | 3289100 | 4259100 | 930550 | 1107100 | 0 | 6416000 | 2339100 | 3775300 | 7321600 | 5358700 | 2832800 | 0 | 0 | 0 | 1767600 |
| VANDNAPE | LAMTOR3 | Q9UHA4 | LTOR3_HUMA | Unmodified | 0 | 0 | 0 | 0 | 6049900 | 0 | 290640 | 0 | 0 | 1973400 | 0 | 0 | 5899600 | 4913000 | 0 | 0 | 0 | 3931700 |
| VANEAEFIL | LRBA; LRBA; LR | P50851; A0A | LRBA_HUMAN | Unmodified | 0 | 0 | 0 | 0 | 0 | 0 | 0 | 0 | 963440 | 127260 | 0 | 3094300 | 0 | 0 | 0 | 0 | 0 | 0 |
| VANIIAEVA | RAP1GDS1; RA | P52306; D6 | GDS1_HUMAN | Unmodified | 0 | 0 | 0 | 0 | 0 | 0 | 0 | 0 | 0 | 386290 | 0 | 1236500 | 0 | 0 | 0 | 0 | 0 | 0 |
| VANIIAEVA | RAP1GDS1 | P52306 | GDS1_HUMAN | Oxidation (M) | 0 | 0 | 0 | 0 | 0 | 0 | 0 | 0 | 0 | 0 | 0 | 1075100 | 0 | 0 | 0 | 0 | 0 | 0 |
| VANILHDD | ERP44 | Q9BS26 | ERP44_HUMA | Unmodified | 0 | 0 | 0 | 0 | 0 | 0 | 0 | 0 | 78393000 | 943970 | 713060 | 1181300 | 0 | 0 | 0 | 0 | 0 | 0 |
| VANLCGIN | PRKCD; PRKCQ | Q05655; Q0 | KPCD_HUMAN | Unmodified | 99601 | 0 | 0 | 0 | 483340 | 173930 | 649320 | 137120 | 677070 | 278740 | 0 | 1353900 | 615000 | 0 | 138260 | 125050 | 19352 | 1588900 |
| VANVELYY | CLTC; CLTC; C | Q00610; J3K | CLH1_HUMAN | Unmodified | 4581300 | 3333300 | 1266000 | 271580 | 0 | 182920 | 734500 | 147090 | 197830 | 139270 | 0 | 7454000 | 0 | 0 | 3023100 | 2853100 | 62709 | 0 |
| VANVIVDH | MIF4GD; MIF4G | A9UHW6; J3 | MI4GD_HUMA | Unmodified | 0 | 0 | 0 | 0 | 0 | 0 | 0 | 137120 | 0 | 0 | 0 | 763360 | 2830200 | 0 | 0 | 0 | 0 | 0 |
| VANVSLLA | RPS23 | P62266 | RS23_HUMAN | Unmodified | 2487600 | 2508100 | 2940800 | 2062900 | 7873000 | 870050 | 12868000 | 8695400 | 2434300 | 2520800 | 5653100 | 28657000 | 9171500 | 4391400 | 2759300 | 1608400 | 0 | 7219300 |
| VAPAEPQE | TKFC; TKFC; TK | Q3LXA3; I3L | TKFC_HUMAN | Unmodified | 190810 | 0 | 0 | 0 | 0 | 0 | 0 | 0 | 268400 | 0 | 147270 | 1302400 | 458540 | 110000 | 328460 | 430830 | 104900 | 0 |
| VAPAPAAV | DLAT; DLAT; DL | P10515; A0A | ODP2_HUMAN | Oxidation (M) | 0 | 0 | 0 | 217780 | 0 | 0 | 0 | 0 | 3986400 | 469750 | 214440 | 0 | 0 | 0 | 0 | 0 | 0 | 0 |
| VAPAPAAV | DLAT; DLAT; DL | P10515; A0A | ODP2_HUMAN | Unmodified | 0 | 0 | 0 | 0 | 0 | 0 | 0 | 0 | 861820 | 0 | 0 | 0 | 0 | 0 | 0 | 0 | 0 | 0 |
| VAPAPAVV | RPL7A; RPL7A | P62424; Q5 | RL7A_HUMAN | Unmodified | 2544300 | 2633200 | 1178800 | 1320900 | 0 | 0 | 0 | 0 | 0 | 172690 | 1477800 | 4648600 | 2495100 | 0 | 1691400 | 1357800 | 163810 | 295660 |
| VAPDEHPI | ACTBL2 | Q562R1 | ACTBL_HUMA | Unmodified | 5.42E+08 | 3.1E+08 | 7.11E+08 | 7.94E+08 | 7.22E+08 | 2.19E+08 | 2068700 | 2231600 | 0 | 1895100 | 22289000 | 1.53E+08 | 18565000 | 4055100 | 1.08E+09 | 33673000 | 3886300 | 2.02E+08 |
| VAPEEHPT | ACTC1; ACTA1; | P68032; P68 | ACTC_HUMAN | Unmodified | 380260 | 369190 | 975990 | 473900 | 0 | 0 | 212540 | 173230 | 0 | 0 | 0 | 353630 | 544790 | 0 | 441260 | 334580 | 0 | 0 |
| VAPEEHPV | ACTG1; ACTB; | P63261; P60 | ACTG_HUMAN | Unmodified | 5.66E+08 | 3.22E+08 | 7.42E+08 | 8.4E+08 | 7.46E+08 | 2.36E+08 | 69085000 | 66207000 | 1.04E+09 | 55571000 | 6.12E+08 | 3.23E+09 | 4.7E+08 | 1.22E+08 | 1.17E+09 | 9.06E+08 | 1.21E+08 | 2.06E+08 |
| VAPEEHPV | ACTG1; ACTB; | P63261; P60 | ACTG_HUMAN | Unmodified | 0 | 531490 | 0 | 0 | 2901600 | 0 | 8305700 | 4403600 | 0 | 0 | 0 | 0 | 1059400 | 1322200 | 441640 | 123650 | 13973000 | 2498400 |
| VAPEMFFS | ACTL8 | Q9H568 | ACTL8_HUMA | Oxidation (M) | 0 | 0 | 0 | 0 | 2475200 | 0 | 0 | 0 | 0 | 0 | 2763100 | 2202700 | 0 | 188710 | 0 | 0 | 0 | 2036900 |
| VAPGYYTL | ITGB4 | P16144 | ITB4_HUMAN | Unmodified | 0 | 0 | 0 | 0 | 0 | 0 | 0 | 0 | 0 | 0 | 0 | 192650 | 0 | 0 | 0 | 0 | 0 | 942840 |
| VAPHALSE | IPO11; IPO11; I | Q9UI26; D6 | IPO11_HUMAN | Unmodified | 0 | 62167 | 0 | 0 | 0 | 0 | 0 | 0 | 91091 | 294160 | 80380 | 1691100 | 0 | 0 | 0 | 0 | 0 | 0 |
| VAPPGLTQ | ATP1B1; ATP1B | P05026; V9C | AT1B1_HUMA | Unmodified | 3650000 | 2111000 | 33353000 | 12358000 | 0 | 0 | 0 | 48612 | 28420000 | 5011200 | 4206800 | 1.18E+08 | 0 | 0 | 31045000 | 33373000 | 0 | 0 |
| VAPPGLTQ | ATP1B1; ATP1B | P05026; V9C | AT1B1_HUMA | Unmodified | 0 | 1340100 | 0 | 30647000 | 2082800 | 0 | 0 | 0 | 50389000 | 793140 | 6474900 | 33726000 | 1144200 | 0 | 2847100 | 2623400 | 0 | 0 |
| VAPSAVLG | GMPPA; GMPPA | Q96IJ6; A0A | GMPPA_HUMA | Unmodified | 0 | 0 | 0 | 0 | 432440 | 208470 | 31243 | 121690 | 1294500 | 1324600 | 412880 | 2363700 | 571390 | 345180 | 0 | 341900 | 563840 | 1108500 |
| VAPTVETS | SUOX | P51687 | SUOX_HUMAN | Unmodified | 108470 | 0 | 0 | 0 | 0 | 0 | 0 | 0 | 0 | 0 | 0 | 3237600 | 0 | 0 | 0 | 136900 | 0 | 0 |
| VAQGVSGA | IMPDH2; IMPDH | P12268; A0A | IMDH2_HUMA | Unmodified | 553840 | 274720 | 854400 | 1608600 | 1843000 | 311230 | 1134800 | 518660 | 319130 | 0 | 825790 | 2291500 | 594190 | 210580 | 754220 | 772840 | 635780 | 2143900 |
| VAQGVSGA | IMPDH2; IMPDH | P12268; A0A | IMDH2_HUMA | Unmodified | 0 | 0 | 0 | 0 | 376490 | 0 | 436510 | 994030 | 0 | 509300 | 0 | 0 | 2244500 | 0 | 0 | 0 | 1085500 | 550310 |
| VAQLCDFN | CLTB; CLTB | P09497; D6F | CLCB_HUMAN | Unmodified | 4366100 | 3962700 | 2994600 | 3965400 | 3225100 | 177670 | 2952100 | 1954700 | 402810 | 403490 | 0 | 3023900 | 10658000 | 5862700 | 2952300 | 3189000 | 2689600 | 2470000 |
| VAQLEAQC | FGG; FGG; FGG | P02679; C9J | FIBG_HUMAN | Unmodified | 0 | 828810 | 0 | 0 | 0 | 0 | 0 | 0 | 0 | 0 | 0 | 0 | 0 | 0 | 0 | 0 | 0 | 0 |
| VAQLEQVY | SNRPD3 | P62318 | SMD3_HUMAN | Unmodified | 1260100 | 1372100 | 849310 | 462690 | 0 | 0 | 0 | 0 | 1929300 | 0 | 1357300 | 12555000 | 125930 | 0 | 836460 | 830460 | 125590 | 0 |
| VAQLEQVY | SNRPD3 | P62318 | SMD3_HUMAN | Unmodified | 972740 | 590270 | 569290 | 1597800 | 1594500 | 245590 | 0 | 14149000 | 973200 | 869160 | 4351300 | 2346700 | 3328500 | 612560 | 321500 | 348280 | 0 | 0 |
| VAQLPLSL | NPTXR; NPTXR | O95502; A0A | NPTXR_HUMA | Unmodified | 0 | 0 | 0 | 0 | 0 | 0 | 0 | 0 | 18622000 | 2649100 | 1378800 | 811670 | 0 | 0 | 0 | 0 | 0 | 0 |

| | | | | | | | | | | | | | | | | | | | | | | |
|---|---|---|---|---|---|---|---|---|---|---|---|---|---|---|---|---|---|---|---|---|---|---|
| VAQPTITDN | FLNA; FLNA; FL | P21333; H0Y | FLNA_HUMAN | Unmodified | 62989 | 0 | 0 | 0 | 285670 | 207660 | 0 | 0 | 0 | 163220 | 0 | 1832600 | 0 | 0 | 429460 | 330740 | 273670 | 0 |
| VAQSPHTK | RACGAP1; RAC | Q9H0H5; H0 | RGAP1_HUMAN | Unmodified | 0 | 0 | 0 | 0 | 2481800 | 1210200 | 25063 | 0 | 1363700 | 1991500 | 0 | 0 | 0 | 0 | 0 | 0 | 1632600 | 0 |
| VAQTEFDR | SH3GLB2; SH3 | Q9NR46; B7 | SHLB2_HUMAN | Unmodified | 0 | 0 | 0 | 0 | 393510 | 35744 | 380920 | 219730 | 428030 | 155910 | 0 | 449990 | 0 | 0 | 0 | 281410 | 173980 | 163540 |
| VAQVAEITY | ERLIN2; ERLIN2 | O94905; E5F | ERLN2_HUMAN | Unmodified | 0 | 0 | 0 | 0 | 808670 | 48710 | 1001600 | 560900 | 36338000 | 12008000 | 1282500 | 403120 | 1014000 | 1197600 | 0 | 0 | 82587 | 377940 |
| VAQWMIQ( | ADK; ADK; ADK; | P55263; A0A | ADK_HUMAN A | Oxidation (M) | 0 | 0 | 0 | 0 | 3638000 | 0 | 1339400 | 0 | 85569 | 0 | 0 | 0 | 0 | 0 | 0 | 346580 | 1477000 | 0 |
| VAQWMIQ( | ADK; ADK; ADK; | P55263; A0A | ADK_HUMAN A | Unmodified | 73744 | 0 | 148280 | 520580 | 5207000 | 1516900 | 3600700 | 1144400 | 2539500 | 870480 | 1890200 | 4401200 | 2355400 | 1069900 | 0 | 112390 | 5926700 | 14484000 |
| VARAAAEV | TFRC; TFRC | P02786; G3\ | TFR1_HUMAN | Unmodified | 0 | 0 | 0 | 0 | 331510 | 394640 | 297950 | 18564 | 0 | 578810 | 0 | 0 | 83772 | 57111 | 0 | 0 | 130630 | 170760 |
| VARLCDFN | CLTA; CLTA | P09496; C9J | CLCA_HUMAN | Unmodified | 0 | 0 | 0 | 0 | 822530 | 0 | 0 | 20459 | 0 | 0 | 0 | 0 | 865910 | 858000 | 0 | 0 | 370520 | 843790 |
| VASAGSAIS | MIB1 | Q86YT6 | MIB1_HUMAN I | Unmodified | 0 | 0 | 42785 | 0 | 1511600 | 0 | 592450 | 0 | 0 | 0 | 0 | 0 | 0 | 0 | 0 | 0 | 356820 | 0 |
| VASALPGM | RAB6B | Q9NRW1 | RAB6B_HUMAN | Unmodified | 0 | 0 | 0 | 428660 | 0 | 0 | 147190 | 0 | 427110 | 0 | 0 | 1969400 | 0 | 0 | 0 | 0 | 0 | 0 |
| VASEAPLEI | NEK9; NEK9; NI | Q8TD19; A0 | NEK9_HUMAN | Unmodified | 0 | 0 | 0 | 0 | 0 | 0 | 0 | 0 | 0 | 0 | 0 | 763280 | 0 | 0 | 0 | 0 | 0 | 0 |
| VASGPSPG | MVP | Q14764 | MVP_HUMAN N | Unmodified | 8736300 | 1162400 | 0 | 0 | 4868000 | 0 | 0 | 0 | 2506400 | 925680 | 0 | 13123000 | 0 | 0 | 0 | 0 | 0 | 2944300 |
| VASIFDFK | ENTPD3 | O75355 | ENTP3_HUMAN | Unmodified | 0 | 0 | 0 | 0 | 0 | 0 | 0 | 7857.9 | 1794400 | 1027100 | 0 | 0 | 0 | 0 | 0 | 0 | 0 | 133890 |
| VASISFDAS | TMEM131 | Q92545 | TM131_HUMAN | Unmodified | 0 | 0 | 78403 | 0 | 0 | 0 | 0 | 47116 | 2327200 | 647170 | 0 | 0 | 0 | 0 | 0 | 0 | 0 | 0 |
| VASLEESE( | CKAP4 | Q07065 | CKAP4_HUMAN | Unmodified | 0 | 0 | 298630 | 155590 | 817470 | 62424 | 927940 | 773870 | 8517700 | 2875500 | 508700 | 157660 | 894220 | 1798000 | 117760 | 94671 | 179030 | 1277400 |
| VASLLSQD | AVIL; AVIL | O75366; F8V | AVIL_HUMAN A | Unmodified | 43924 | 0 | 42601 | 28259 | 0 | 0 | 0 | 0 | 0 | 0 | 703080 | 1438400 | 0 | 0 | 112110 | 62289 | 0 | 0 |
| VASSDLVN | FLOT1; FLOT1; I | O75955; A0 | FLOT1_HUMAN | Unmodified | 0 | 0 | 0 | 48721 | 336100 | 0 | 0 | 0 | 293040 | 221840 | 0 | 12994 | 0 | 0 | 120350 | 0 | 63958 | 0 |
| VASSDLVN | FLOT1; FLOT1; I | O75955; A0 | FLOT1_HUMAN | Oxidation (M) | 0 | 0 | 0 | 0 | 2151100 | 222710 | 571250 | 0 | 0 | 1004900 | 158760 | 0 | 0 | 0 | 1089100 | 35992 | 0 | 0 |
| VASVEMVK | ASNS; ASNS; AS | P08243; F8V | ASNS_HUMAN | Unmodified | 0 | 0 | 212740 | 453520 | 1090200 | 0 | 165050 | 159510 | 1667500 | 310260 | 1483500 | 14943000 | 0 | 662440 | 343520 | 431740 | 0 | 0 |
| VASVEMVK | ASNS; ASNS; AS | P08243; F8V | ASNS_HUMAN | Oxidation (M) | 0 | 0 | 0 | 0 | 0 | 8656.2 | 42081 | 38012 | 0 | 0 | 179650 | 1663200 | 301990 | 66961 | 0 | 0 | 55082 | 0 |
| VATAQDDI | CCT6A | P40227 | TCPZ_HUMAN | Unmodified | 0 | 0 | 0 | 460780 | 4590500 | 0 | 2157100 | 0 | 449070 | 1654700 | 172820 | 635530 | 11498000 | 4509300 | 729940 | 53955 | 4825500 | 6003600 |
| VATGQELS | MICU2 | Q8IYU8 | MICU2_HUMAN | Unmodified | 0 | 0 | 0 | 54197 | 0 | 0 | 13207 | 0 | 205800 | 0 | 0 | 88351 | 0 | 0 | 0 | 0 | 0 | 0 |
| VATPLPDPI | DSG2 | Q14126 | DSG2_HUMAN | Unmodified | 0 | 0 | 553720 | 1035800 | 4005500 | 0 | 0 | 0 | 2852600 | 0 | 1840600 | 1264500 | 1178600 | 775400 | 0 | 441550 | 149330 | 1356700 |
| VATPPPSVI | TNS3 | Q68CZ2 | TENS3_HUMAN | Unmodified | 0 | 0 | 0 | 0 | 1011900 | 0 | 0 | 0 | 0 | 0 | 0 | 0 | 0 | 0 | 0 | 0 | 692670 | 0 |
| VATPPPSVI | TNS3 | Q68CZ2 | TENS3_HUMAN | Unmodified | 0 | 0 | 0 | 0 | 572310 | 0 | 0 | 0 | 0 | 0 | 0 | 0 | 0 | 0 | 0 | 0 | 405430 | 364070 |
| VATQAVED | NADK2; NADK2 | Q4G0N4; A0 | NAKD2_HUMAN | Unmodified | 0 | 0 | 0 | 0 | 0 | 0 | 0 | 0 | 475000 | 0 | 0 | 0 | 80073 | 0 | 0 | 0 | 0 | 80563 |
| VATQEGKE | GGCT; N/A; GG | O75223; M0 | GGCT_HUMAN | Unmodified | 0 | 0 | 0 | 0 | 286450 | 29647 | 0 | 0 | 426550 | 1837100 | 19876 | 0 | 2252900 | 1109900 | 0 | 0 | 0 | 0 |
| VATRGLCA | RPS3; RPS3; RP | P23396; H0Y | RS3_HUMAN 4 | Unmodified | 0 | 0 | 0 | 0 | 0 | 0 | 103900 | 0 | 0 | 0 | 0 | 0 | 0 | 0 | 0 | 0 | 0 | 0 |
| VATRGLCA | RPS3; RPS3; RP | P23396; H0Y | RS3_HUMAN 4 | Unmodified | 0 | 0 | 0 | 0 | 732060 | 0 | 2690600 | 265950 | 0 | 446430 | 0 | 0 | 0 | 656590 | 0 | 0 | 0 | 0 |
| VATWFNQI | RPL13; RPL13; I | P26373; H3E | RL13_HUMAN ( | Unmodified | 2136500 | 1654200 | 1095000 | 896440 | 0 | 0 | 0 | 104340 | 0 | 0 | 0 | 0 | 824860 | 0 | 934030 | 812750 | 0 | 0 |
| VATWFNQI | RPL13; RPL13; I | P26373; H3E | RL13_HUMAN ( | Unmodified | 870330 | 784980 | 567850 | 384700 | 758700 | 0 | 103190 | 143580 | 0 | 35241 | 308360 | 3054600 | 668840 | 263060 | 632010 | 504450 | 0 | 0 |
| VAVAAASKI | CRABP1; CRAB | P29762; B5N | RABP1_HUMAN | Unmodified | 0 | 0 | 0 | 0 | 0 | 0 | 0 | 0 | 3633500 | 2953800 | 0 | 0 | 0 | 0 | 0 | 0 | 6591700 | 13217000 |
| VAVAIPNRI | HSD17B4; HSD | P51659; E7E | DHB4_HUMAN | Unmodified | 0 | 0 | 0 | 0 | 0 | 0 | 0 | 0 | 1.83E+08 | 2913300 | 3494300 | 1426900 | 0 | 0 | 0 | 0 | 0 | 0 |
| VAVCDIPPI | TUBB6; TUBB3; | Q9BUF5; Q1 | TBB6_HUMAN | Unmodified | 0 | 0 | 0 | 0 | 662260 | 223010 | 561570 | 343930 | 477840 | 233900 | 0 | 55480 | 0 | 48141 | 0 | 0 | 161830 | 625930 |
| VAVEAKNP | EEF2 | P13639 | EF2_HUMAN El | Unmodified | 1431800 | 0 | 0 | 3071500 | 12976000 | 2812900 | 2249300 | 1002500 | 4905300 | 2158000 | 2463200 | 4577200 | 5089300 | 1355300 | 0 | 613460 | 1539800 | 3091600 |
| VAVEWDK | CANT1; CANT1 | Q8WVQ1; K7 | CANT1_HUMAN | Unmodified | 0 | 0 | 4209800 | 0 | 0 | 0 | 0 | 55909 | 10431000 | 0 | 0 | 213240 | 0 | 0 | 0 | 0 | 0 | 0 |
| VAVEWDKI | CANT1; CANT1 | Q8WVQ1; K7 | CANT1_HUMAN | Unmodified | 0 | 0 | 0 | 0 | 0 | 0 | 0 | 0 | 8741300 | 1248700 | 0 | 0 | 0 | 0 | 0 | 0 | 0 | 0 |
| VAVEYLDP | BUB3; BUB3 | O43684; J3C | BUB3_HUMAN | Unmodified | 0 | 0 | 0 | 0 | 0 | 0 | 0 | 0 | 0 | 0 | 0 | 3239900 | 0 | 0 | 0 | 0 | 0 | 0 |
| VAVEYLDP | BUB3; BUB3 | O43684; J3C | BUB3_HUMAN | Unmodified | 85272 | 133030 | 0 | 0 | 0 | 0 | 0 | 0 | 0 | 0 | 1015500 | 4410800 | 123200 | 0 | 176880 | 167340 | 0 | 0 |
| VAVFDTGL | MBTPS1 | Q14703 | MBTP1_HUMAN | Unmodified | 0 | 0 | 0 | 0 | 0 | 0 | 0 | 0 | 1698900 | 0 | 254730 | 0 | 0 | 0 | 110040 | 85089 | 0 | 253820 |
| VAVFFGGL | DDX39B; DDX3 | Q13838; F6T | DX39B_HUMAN | Unmodified | 0 | 0 | 0 | 0 | 0 | 0 | 0 | 0 | 410960 | 0 | 0 | 3726500 | 0 | 0 | 0 | 0 | 0 | 0 |
| VAVFFGGL | DDX39B; DDX3 | Q13838; F6T | DX39B_HUMAN | Unmodified | 0 | 0 | 0 | 0 | 0 | 0 | 0 | 0 | 0 | 0 | 0 | 1212800 | 0 | 0 | 0 | 0 | 0 | 0 |
| VAVFFGGL | DDX39B; DDX3 | Q13838; F6T | DX39B_HUMAN | Unmodified | 0 | 0 | 0 | 0 | 0 | 0 | 541330 | 0 | 0 | 517220 | 387390 | 7408400 | 0 | 0 | 0 | 53100 | 0 | 0 |
| VAVFTENA | MRPL1; MRPL1 | Q9BYD6; H0 | RM01_HUMAN | Unmodified | 0 | 0 | 0 | 0 | 0 | 0 | 0 | 0 | 4043600 | 0 | 167470 | 564450 | 0 | 0 | 0 | 0 | 0 | 0 |
| VAVGNQP | MX1; MX1; MX1 | P20591; A0A | MX1_HUMAN I | Unmodified | 0 | 0 | 0 | 0 | 0 | 0 | 0 | 0 | 0 | 0 | 0 | 0 | 0 | 0 | 0 | 0 | 0 | 909220 |
| VAVGQEQ | FLNC | Q14315 | FLNC_HUMAN | Unmodified | 0 | 0 | 0 | 0 | 552360 | 0 | 0 | 0 | 0 | 0 | 0 | 0 | 0 | 0 | 0 | 0 | 178840 | 0 |
| VAVGQEQ | FLNC | Q14315 | FLNC_HUMAN | Unmodified | 0 | 0 | 0 | 0 | 0 | 0 | 0 | 0 | 0 | 0 | 0 | 0 | 0 | 0 | 0 | 0 | 649560 | 0 |
| VAVHHAEA | MBLAC2 | Q68D91 | MBLC2_HUMAN | Unmodified | 202890 | 0 | 456190 | 0 | 0 | 0 | 0 | 0 | 0 | 0 | 84018 | 3846700 | 0 | 0 | 1094000 | 1486100 | 43131 | 0 |
| VAVLETQL | KANK2; KANK2 | Q63ZY3; K7E | KANK2_HUMAN | Unmodified | 0 | 0 | 0 | 0 | 507580 | 0 | 0 | 0 | 0 | 0 | 0 | 0 | 0 | 0 | 0 | 0 | 360280 | 233330 |
| VAVLGASG | MDH2 | P40926 | MDHM_HUMAN | Unmodified | 367790 | 207570 | 287550 | 347220 | 433180 | 371880 | 3032400 | 0 | 61880000 | 22328000 | 881530 | 1371000 | 3959500 | 2126300 | 443870 | 580070 | 0 | 441940 |
| VAVLLLAG( | UAP1 | Q16222 | UAP1_HUMAN | Unmodified | 0 | 0 | 0 | 0 | 3388100 | 0 | 0 | 0 | 0 | 218310 | 0 | 0 | 2751800 | 0 | 0 | 0 | 640690 | 900740 |
| VAVLSQNR | ACAT2 | Q9BWD1 | THIC_HUMAN / | Unmodified | 0 | 0 | 0 | 0 | 94849 | 0 | 0 | 0 | 283930 | 221720 | 0 | 2030600 | 0 | 0 | 0 | 0 | 67982 | 0 |
| VAVLSQNR | ACAT2 | Q9BWD1 | THIC_HUMAN / | Unmodified | 412510 | 266740 | 208100 | 466910 | 5041500 | 1193600 | 299800 | 162870 | 3891800 | 1386700 | 956470 | 10900000 | 2447700 | 0 | 320320 | 347910 | 624090 | 1074000 |
| VAVLVKPEI | INPPL1; INPPL1 | O15357; A0 | SHIP2_HUMAN | Unmodified | 0 | 0 | 0 | 0 | 684660 | 0 | 0 | 16612 | 0 | 18924 | 0 | 954520 | 0 | 0 | 0 | 0 | 0 | 0 |
| VAVMEEGN | PLCB2 | Q00722 | PLCB2_HUMAN | Unmodified | 0 | 64382 | 0 | 0 | 0 | 0 | 0 | 95290 | 0 | 142260 | 0 | 3189100 | 0 | 0 | 0 | 0 | 286640 | 251600 |
| VAVNDAHL | LGALS3 | P17931 | LEG3_HUMAN | Unmodified | 0 | 0 | 0 | 0 | 0 | 0 | 0 | 51631 | 0 | 0 | 0 | 9679600 | 0 | 0 | 1846400 | 0 | 0 | 0 |
| VAVNDAHL | LGALS3 | P17931 | LEG3_HUMAN | Unmodified | 0 | 0 | 0 | 0 | 1288700 | 472660 | 0 | 0 | 0 | 0 | 0 | 219720 | 0 | 0 | 0 | 0 | 1011000 | 523540 |
| VAVNNFGV | MYO10; MYO10 | Q9HD67; D6 | MYO10_HUMAN | Unmodified | 0 | 0 | 0 | 0 | 404520 | 0 | 78070 | 0 | 104860 | 126270 | 0 | 0 | 0 | 0 | 388780 | 0 | 214270 | 103360 |
| VAVPFYIPA | SCAMP1 | A0A087WZA | A0A087WZA6_I | Unmodified | 0 | 9047900 | 0 | 14706000 | 34770000 | 0 | 33765000 | 0 | 759390 | 4673200 | 4726900 | 0 | 11682000 | 91999000 | 0 | 0 | 0 | 706550 |
| VAVPLLSS( | CORO7; CORO | P57737; A0A | CORO7_HUMA | Unmodified | 0 | 0 | 0 | 0 | 0 | 0 | 0 | 0 | 0 | 0 | 0 | 901940 | 0 | 0 | 0 | 31742 | 0 | 0 |
| VAVPSTIHC | ACO2; ACO2; A | Q99798; A2A | ACON_HUMAN | Unmodified | 0 | 0 | 0 | 0 | 0 | 0 | 325180 | 95328 | 2593200 | 444040 | 0 | 0 | 75892 | 0 | 0 | 0 | 0 | 0 |
| VAVQAVQA | GUK1; GUK1; G | Q16774; B1A | KGUA_HUMAN | Unmodified | 0 | 0 | 0 | 0 | 0 | 0 | 0 | 0 | 422700 | 0 | 0 | 0 | 0 | 0 | 0 | 0 | 0 | 0 |
| VAVQGDV\ | EPRS1; EPRS1 | P07814; V9G | SYEP_HUMAN | Unmodified | 167400 | 93912 | 404540 | 0 | 0 | 0 | 0 | 0 | 788100 | 191970 | 307520 | 9312700 | 0 | 0 | 343280 | 290310 | 26742 | 0 |
| VAVQGDV\ | EPRS1; EPRS1 | P07814; V9G | SYEP_HUMAN | Unmodified | 0 | 0 | 0 | 0 | 0 | 0 | 0 | 0 | 0 | 0 | 1480100 | 991050 | 0 | 0 | 0 | 0 | 0 | 0 |
| VAVQRGVC | TNS3; TNS3 | Q68CZ2; E9I | TENS3_HUMAN | Unmodified | 0 | 0 | 0 | 0 | 0 | 0 | 0 | 0 | 0 | 0 | 0 | 0 | 0 | 0 | 0 | 0 | 474370 | 2253600 |
| VAVTEGCQ | FLNB; FLNB | O75369; E7E | FLNB_HUMAN | Unmodified | 0 | 0 | 0 | 0 | 0 | 0 | 0 | 24663 | 158550 | 0 | 0 | 737040 | 0 | 0 | 0 | 0 | 0 | 0 |
| VAVTEGCQ | FLNB; FLNB | O75369; E7E | FLNB_HUMAN | Unmodified | 138760 | 59914 | 757020 | 858770 | 15363000 | 4538700 | 327090 | 226340 | 354270 | 663930 | 0 | 3134300 | 2334100 | 1857500 | 383300 | 348380 | 6405000 | 3565700 |
| VAVVAGYG | AHCY | P23526 | SAHH_HUMAN | Unmodified | 1149100 | 847430 | 3483200 | 2651400 | 6085500 | 486730 | 803060 | 323390 | 853970 | 0 | 11215000 | 45406000 | 1338300 | 1516600 | 5604100 | 4092800 | 1208000 | 5155700 |
| VAVVAGYG | AHCY | P23526 | SAHH_HUMAN | Unmodified | 0 | 0 | 0 | 0 | 5908700 | 1280000 | 0 | 0 | 442610 | 4084000 | 0 | 0 | 4571600 | 2234000 | 0 | 0 | 5029900 | 7057900 |
| VAVVDYVEI | TXNRD2; TXNRI | Q9NNW7; A( | TRXR2_HUMAN | Unmodified | 0 | 0 | 0 | 0 | 0 | 0 | 0 | 0 | 0 | 0 | 0 | 0 | 0 | 0 | 0 | 0 | 0 | 0 |
| VAVVGNVD | GTPBP1; GTPBI | O00178; F5I | GTPB1_HUMAN | Unmodified | 529510 | 373510 | 0 | 497170 | 1372400 | 0 | 0 | 0 | 1396400 | 0 | 161350 | 2460800 | 3352800 | 730620 | 1408900 | 881480 | 573650 | 664040 |
| VAVVNQIAI | SPTBN1; SPTBN | Q01082; A0A | SPTB2_HUMAN | Unmodified | 0 | 0 | 0 | 0 | 358440 | 101440 | 0 | 0 | 274160 | 119830 | 186370 | 6998000 | 0 | 0 | 0 | 0 | 0 | 0 |
| VAVVQYSG | COL6A1; COL6 | P12109; A0A | CO6A1_HUMAN | Unmodified | 328600 | 0 | 4027000 | 0 | 0 | 171370 | 54113 | 0 | 3357400 | 153040 | 0 | 1903000 | 0 | 0 | 1004900 | 498770 | 0 | 0 |
| VAVYVPGS | EIF2A; EIF2A; EI | Q9BY44; C9I | EIF2A_HUMAN | Unmodified | 177840 | 220000 | 189080 | 0 | 0 | 0 | 0 | 143370 | 0 | 0 | 516590 | 3707700 | 0 | 0 | 0 | 222410 | 0 | 0 |
| VAWVSHDS | ARPC1B; ARPC | O15143; C9J | ARC1B_HUMAN | Unmodified | 0 | 0 | 0 | 0 | 0 | 0 | 0 | 0 | 0 | 0 | 0 | 941740 | 92209 | 0 | 0 | 0 | 0 | 0 |
| VAWVSHDS | ARPC1B; ARPC | O15143; C9J | ARC1B_HUMAN | Unmodified | 1136900 | 479050 | 0 | 845830 | 453430 | 0 | 263520 | 103300 | 0 | 46518 | 0 | 8789400 | 0 | 109290 | 0 | 772090 | 2047200 | 0 |
| VAYCFEVA | POFUT1 | Q9H488 | OFUT1_HUMAN | Unmodified | 0 | 0 | 0 | 0 | 0 | 0 | 0 | 0 | 874160 | 0 | 0 | 0 | 0 | 0 | 0 | 0 | 0 | 0 |
| VAYEELLDI | SEZ6L2 | A0A087WYL | A0A087WYL5_I | Unmodified | 0 | 0 | 0 | 0 | 0 | 0 | 0 | 51682 | 1628500 | 0 | 0 | 0 | 0 | 0 | 0 | 0 | 0 | 0 |
| VAYFQSALI | PTPN23 | Q9H3S7 | PTN23_HUMAN | Unmodified | 862940 | 850540 | 0 | 0 | 0 | 0 | 0 | 0 | 1297300 | 0 | 0 | 408730 | 0 | 0 | 0 | 146940 | 0 | 0 |
| VAYFQSALI | PTPN23 | Q9H3S7 | PTN23_HUMAN | Unmodified | 149690 | 0 | 0 | 0 | 0 | 110710 | 0 | 0 | 618110 | 194360 | 0 | 0 | 0 | 0 | 0 | 0 | 0 | 141460 |
| VAYIPDEM | IGF2BP3 | O00425 | IF2B3_HUMAN | Oxidation (M) | 0 | 331580 | 0 | 345820 | 0 | 0 | 0 | 0 | 0 | 0 | 0 | 2517700 | 0 | 0 | 0 | 0 | 0 | 0 |
| VAYLDPLEI | BCAM; BCAM | P50895; A0A | BCAM_HUMAN | Unmodified | 0 | 7539000 | 0 | 0 | 1136000 | 0 | 0 | 0 | 4362600 | 0 | 343540 | 0 | 0 | 0 | 0 | 0 | 0 | 662860 |
| VAYSLNGH | MFGE8; MFGE8 | Q08431; H0Y | MFGM_HUMAN | Unmodified | 0 | 0 | 2430100 | 3172100 | 13559000 | 3916800 | 0 | 0 | 16398000 | 6043300 | 861930 | 3857500 | 885630 | 659350 | 4935400 | 0 | 979450 | 1300500 |
| VAYSLNGH | MFGE8; MFGE8 | Q08431; H0Y | MFGM_HUMAN | Unmodified | 5726800 | 1562200 | 13949000 | 31573000 | 1.13E+08 | 40448000 | 4221800 | 0 | 4415900 | 1096500 | 0 | 14927000 | 12679000 | 9058500 | 34454000 | 16776000 | 10707000 | 13661000 |
| VAYSLNGH | MFGE8; MFGE8 | Q08431; H0Y | MFGM_HUMAN | Unmodified | 0 | 0 | 0 | 0 | 5959800 | 317550 | 0 | 0 | 0 | 0 | 0 | 0 | 0 | 553670 | 0 | 0 | 0 | 0 |
| VAYSNDSA | MFGE8 | Q08431 | MFGM_HUMAN | Unmodified | 207870 | 389330 | 2432600 | 3706000 | 243650 | 0 | 0 | 0 | 356970 | 0 | 0 | 966740 | 0 | 0 | 5093100 | 2270700 | 0 | 69436 |
| VAYSNDSA | MFGE8 | Q08431 | MFGM_HUMAN | Unmodified | 0 | 0 | 0 | 287410 | 20574000 | 4613300 | 0 | 0 | 0 | 316910 | 0 | 0 | 2691700 | 0 | 0 | 0 | 917830 | 745010 |
| VAYVSFGPI | RPL14; RPL14 | P50914; E7E | RL14_HUMAN ( | Unmodified | 9237100 | 606840 | 6454900 | 2972800 | 301810 | 0 | 395780 | 154830 | 3184000 | 223310 | 4601300 | 91829000 | 202060 | 0 | 5886300 | 5649300 | 105440 | 319110 |
| VCAHITNIP | RBM12 | Q9NTZ6 | RBM12_HUMAN | Unmodified | 0 | 0 | 0 | 0 | 0 | 0 | 130200 | 67110 | 0 | 0 | 347600 | 1243600 | 0 | 0 | 0 | 71850 | 40232 | 0 |
| VCALLSCTS | AKR1B1 | P15121 | ALDR_HUMAN | Unmodified | 0 | 0 | 0 | 0 | 225480 | 0 | 0 | 0 | 0 | 0 | 1026100 | 1736000 | 0 | 0 | 0 | 0 | 0 | 0 |
| VCALLSCTS | AKR1B1 | P15121 | ALDR_HUMAN | Unmodified | 0 | 0 | 0 | 0 | 2004700 | 652530 | 0 | 0 | 0 | 0 | 0 | 643460 | 0 | 0 | 0 | 0 | 0 | 0 |
| VCALYIMA( | MLEC; MLEC | Q14165; F5I | MLEC_HUMAN | Unmodified | 0 | 0 | 28987 | 0 | 0 | 0 | 23468 | 84098 | 1165800 | 0 | 0 | 0 | 0 | 0 | 0 | 38868 | 0 | 24292 |
| VCANGEVT | N/A | N/A | N/A | Unmodified | 0 | 0 | 0 | 0 | 496600 | 166890 | 0 | 25076 | 0 | 0 | 490950 | 3579900 | 239430 | 0 | 87679 | 0 | 295520 | 789310 |
| VCARGEIIG | PPA1; PPA1 | Q15181; Q5 | IPYR_HUMAN I | Unmodified | 0 | 0 | 56035 | 517610 | 3611100 | 0 | 635270 | 0 | 0 | 0 | 1942600 | 760810 | 8030100 | 3569600 | 0 | 55752 | 5212100 | 6139800 |
| VCASVTFK | CSE1L | P55060 | XPO2_HUMAN | Unmodified | 55026 | 30673 | 45257 | 83295 | 0 | 0 | 427950 | 0 | 0 | 224410 | 311910 | 2120500 | 0 | 0 | 119310 | 98121 | 0 | 0 |
| VCDGQEH | GAS6 | Q14393 | GAS6_HUMAN | Unmodified | 0 | 0 | 0 | 0 | 0 | 0 | 449220 | 441300 | 1690500 | 0 | 0 | 0 | 0 | 0 | 0 | 0 | 0 | 0 |
| VCEAGGLF | EPHX2; EPHX2; | P34913; E5R | HYES_HUMAN | Unmodified | 87931 | 0 | 0 | 72528 | 0 | 0 | 0 | 45431 | 463700 | 0 | 0 | 0 | 743720 | 241070 | 0 | 0 | 494760 | 120170 |
| VCEATYDT | PTP4A1; PTP4A | Q93096; A0 | TP4A1_HUMAN | Unmodified | 0 | 0 | 0 | 0 | 0 | 0 | 0 | 0 | 235930 | 0 | 0 | 608190 | 0 | 0 | 0 | 0 | 0 | 0 |
| VCEDLDTS | VDAC2; VDAC2 | P45880; A0A | VDAC2_HUMAN | Unmodified | 0 | 0 | 0 | 0 | 0 | 0 | 0 | 0 | 2468000 | 0 | 0 | 0 | 0 | 0 | 0 | 0 | 0 | 0 |
| VCEEIAIIPS | RPS17; RPS17; | P08708; H0Y | RS17_HUMAN | Unmodified | 950040 | 0 | 0 | 0 | 0 | 0 | 0 | 0 | 0 | 0 | 878020 | 2270100 | 0 | 0 | 0 | 545530 | 0 | 0 |
| VCEEIAIIPS | RPS17; RPS17; | P08708; H0Y | RS17_HUMAN | Unmodified | 916980 | 0 | 0 | 503010 | 0 | 0 | 0 | 0 | 0 | 0 | 0 | 0 | 0 | 0 | 242610 | 438720 | 0 | 0 |
| VCEHLNLL | EPB41L2; EPB4 | O43491; Q9 | E41L2_HUMAN | Unmodified | 0 | 0 | 0 | 0 | 822640 | 501130 | 69671 | 51686 | 0 | 0 | 0 | 0 | 0 | 0 | 210790 | 0 | 497940 | 0 |
| VCENIPIVL | RAN; RAN; RAN | P62826; J3K | RAN_HUMAN C | Unmodified | 0 | 0 | 0 | 418710 | 0 | 0 | 54074 | 0 | 114150 | 0 | 0 | 11321000 | 0 | 0 | 0 | 0 | 0 | 0 |
| VCENIPIVL | RAN; RAN; RAN | P62826; J3K | RAN_HUMAN C | Unmodified | 2609400 | 666010 | 0 | 0 | 0 | 0 | 0 | 45449 | 3262000 | 0 | 3604200 | 49908000 | 0 | 0 | 0 | 1533900 | 0 | 951630 |
| VCENIPIVL | RAN; RAN; RAN | P62826; J3K | RAN_HUMAN C | Unmodified | 0 | 0 | 0 | 0 | 0 | 0 | 0 | 0 | 0 | 0 | 395150 | 5949700 | 195370 | 0 | 0 | 0 | 0 | 0 |
| VCEVTYDK | PTP4A3; PTP4A | O75365; E5F | TP4A3_HUMAN | Unmodified | 0 | 0 | 0 | 0 | 102060 | 0 | 0 | 0 | 155180 | 0 | 0 | 55905 | 0 | 20804 | 0 | 0 | 0 | 0 |
| VCFAQHTP | CLCN3; CLCN3 | P51790; A0A | CLCN3_HUMAN | Unmodified | 0 | 0 | 948710 | 0 | 318470 | 0 | 183970 | 0 | 773490 | 0 | 0 | 1627400 | 0 | 0 | 0 | 682070 | 0 | 0 |
| VCGDSDK( | CCT6A | P40227 | TCPZ_HUMAN | Unmodified | 0 | 0 | 0 | 91165 | 1228600 | 0 | 726810 | 460270 | 0 | 726350 | 0 | 0 | 3265900 | 2314200 | 112520 | 101210 | 0 | 1680200 |
| VCGKLSSD | IGF2R; IGF2R | P11717; A0A | MPRI_HUMAN | Unmodified | 0 | 0 | 0 | 0 | 0 | 0 | 0 | 0 | 0 | 164780 | 31779 | 0 | 0 | 0 | 0 | 0 | 0 | 0 |

| | | | | | | | | | | | | | | | | | | | | | | |
|---|---|---|---|---|---|---|---|---|---|---|---|---|---|---|---|---|---|---|---|---|---|---|
| VCGSDGV1 | AGRN; AGRN; A | O00468; A0A | AGRIN_HUMAN | Unmodified | 0 | 0 | 0 | 0 | 0 | 0 | 0 | 0 | 0 | 0 | 0 | 0 | 1169400 | 591610 | 0 | 0 | 0 | 424290 |
| VCGYDTPF | BCKDHB | P21953 | ODBB_HUMAN | Unmodified | 0 | 0 | 0 | 0 | 0 | 0 | 0 | 0 | 736720 | 0 | 0 | 0 | 0 | 0 | 0 | 0 | 0 | 0 |
| VCHAHPTL | DLD; DLD | P09622; E9P | DLDH_HUMAN | Unmodified | 0 | 0 | 0 | 0 | 0 | 0 | 0 | 0 | 3417800 | 656560 | 0 | 0 | 0 | 0 | 0 | 0 | 0 | 0 |
| VCHLGDQI | EXOC5; EXOC5 | O00471; A0A | EXOC5_HUMAI | Unmodified | 0 | 0 | 0 | 292010 | 1020600 | 59442 | 0 | 0 | 2445400 | 0 | 0 | 0 | 64093 | 0 | 0 | 180420 | 0 | 0 |
| VCIEHHTFF | EPB41L1; EPB4 | Q9H4G0; A0 | E41L1_HUMAN | Unmodified | 0 | 0 | 0 | 0 | 0 | 0 | 0 | 0 | 0 | 0 | 0 | 1230300 | 0 | 0 | 418660 | 307120 | 0 | 0 |
| VCIESEHSI | ATOX1; ATOX1; | O00244; E5F | ATOX1_HUMAN | Oxidation (M) | 415470 | 0 | 106160 | 0 | 388160 | 0 | 0 | 0 | 0 | 0 | 232420 | 0 | 2037900 | 423510 | 444260 | 585820 | 176050 | 0 |
| VCLESAVEI | BRCC3; BRCC3 | P46736; A0A | BRCC3_HUMAI | Unmodified | 0 | 0 | 0 | 0 | 0 | 0 | 0 | 0 | 0 | 0 | 0 | 778600 | 0 | 0 | 0 | 0 | 0 | 0 |
| VCLKPGLS | DLL3; DLL3 | Q9NYJ7; M0 | DLL3_HUMAN | Unmodified | 0 | 0 | 0 | 0 | 0 | 0 | 0 | 0 | 4308400 | 0 | 0 | 0 | 0 | 0 | 0 | 0 | 0 | 0 |
| VCLLDVQP | MPP7; MPP7 | Q5T2T1; U5C | MPP7_HUMAN | Unmodified | 0 | 0 | 0 | 0 | 1552100 | 733570 | 333300 | 157410 | 0 | 0 | 0 | 0 | 0 | 0 | 57228 | 0 | 133280 | 97820 |
| VCLLGCGI | ADH5; ADH5 | P11766; H0Y | ADHX_HUMAN | Unmodified | 0 | 0 | 0 | 0 | 0 | 0 | 111780 | 0 | 0 | 0 | 0 | 3240300 | 0 | 0 | 0 | 0 | 0 | 0 |
| VCLLGDTG | RAB22A; RAB31 | Q9UL26; Q1 | RB22A_HUMAN | Unmodified | 1204300 | 1182400 | 927750 | 3065400 | 9259500 | 1170100 | 12879000 | 429200 | 2716100 | 845640 | 0 | 1244000 | 7052100 | 9752300 | 515040 | 2812000 | 1074600 | 3337500 |
| VCLQPIR | PTPRD; PTPRF; | P23468; P10 | PTPRD_HUMAN | Unmodified | 679000 | 612820 | 429400 | 373190 | 0 | 0 | 0 | 0 | 469010 | 107760 | 0 | 714250 | 0 | 0 | 775020 | 1128700 | 0 | 0 |
| VCMDFNIIF | IARS1; IARS1; I | P41252; A0A | SYIC_HUMAN I | Unmodified | 0 | 0 | 0 | 0 | 0 | 0 | 0 | 0 | 0 | 0 | 0 | 1242200 | 0 | 0 | 0 | 0 | 0 | 0 |
| VCMEETLW | ITGA6 | P23229 | ITA6_HUMAN I | Oxidation (M) | 0 | 0 | 0 | 169440 | 2634600 | 0 | 0 | 0 | 0 | 0 | 0 | 678320 | 1242100 | 0 | 0 | 0 | 2839900 | 9708800 |
| VCMEETLW | ITGA6 | P23229 | ITA6_HUMAN I | Unmodified | 0 | 0 | 0 | 0 | 1313500 | 0 | 0 | 0 | 0 | 0 | 0 | 0 | 676010 | 0 | 0 | 0 | 0 | 5245100 |
| VCMYFTYK | ELOC; ELOC | Q15369; E5F | ELOC_HUMAN | Unmodified | 36719 | 0 | 0 | 138930 | 1733100 | 269530 | 811510 | 225790 | 283500 | 316460 | 0 | 762760 | 1135200 | 966260 | 167580 | 72595 | 2101300 | 1749300 |
| VCNFLASQ | HSD17B10 | Q99714 | HCD2_HUMAN | Unmodified | 0 | 0 | 0 | 0 | 0 | 0 | 0 | 27289 | 4401800 | 387680 | 0 | 0 | 0 | 0 | 0 | 0 | 0 | 0 |
| VCNPIITK | HSPA8; HSPA8; | P11142; E9P | HSP7C_HUMAI | Unmodified | 31469000 | 16949000 | 22392000 | 30066000 | 99521000 | 28654000 | 68892000 | 22369000 | 44426000 | 62070000 | 20556000 | 73316000 | 1.69E+08 | 1.16E+08 | 21396000 | 12706000 | 68656000 | 67769000 |
| VCNYGLTF | VDAC3; VDAC3 | Q9Y277; E5F | VDAC3_HUMAI | Unmodified | 0 | 0 | 0 | 0 | 0 | 0 | 0 | 0 | 12658000 | 1311500 | 0 | 0 | 0 | 0 | 0 | 0 | 0 | 0 |
| VCQGEREI | HSPA9; HSPA9; | P38646; H0Y | GRP75_HUMAN | Unmodified | 0 | 0 | 0 | 0 | 34761 | 0 | 26302 | 0 | 422360 | 130990 | 0 | 0 | 0 | 0 | 0 | 0 | 0 | 18966 |
| VCQGIGM | PPIL1 | Q9Y3C6 | PPIL1_HUMAN | Unmodified | 438750 | 372830 | 233800 | 208280 | 0 | 0 | 0 | 0 | 0 | 0 | 0 | 922690 | 0 | 0 | 396330 | 247140 | 0 | 0 |
| VCQGIGM | PPIL1 | Q9Y3C6 | PPIL1_HUMAN | Oxidation (M) | 228290 | 0 | 0 | 0 | 0 | 0 | 0 | 0 | 0 | 0 | 0 | 0 | 0 | 0 | 225030 | 204450 | 0 | 0 |
| VCQGTSN | EGFR; EGFR; E | P00533; Q5 | EGFR_HUMAN | Unmodified | 85883 | 274600 | 0 | 0 | 701810 | 0 | 0 | 0 | 0 | 0 | 0 | 0 | 0 | 0 | 549690 | 245280 | 0 | 0 |
| VCQSLINVI | PIP4P1; PIP4P1 | Q86T03; G3 | PP4P1_HUMAN | Unmodified | 107450 | 0 | 0 | 0 | 632510 | 0 | 0 | 0 | 1475000 | 0 | 0 | 0 | 0 | 0 | 0 | 0 | 0 | 296520 |
| VCREFTERI | UCHL1; UCHL1 | P09936; D6F | UCHL1_HUMAI | Unmodified | 1148600 | 13285 | 0 | 0 | 9347100 | 2683900 | 470680 | 293190 | 10164000 | 2379500 | 8713800 | 8182800 | 26084 | 171140 | 0 | 0 | 0 | 0 |
| VCRGNPAI | PLPP2; PLPP2; I | O43688; K7E | PLPP2_HUMAN | Unmodified | 0 | 0 | 0 | 0 | 0 | 0 | 0 | 6396.1 | 11577000 | 627570 | 867130 | 2279200 | 0 | 0 | 0 | 0 | 0 | 0 |
| VCSEVLQM | DNAJC3; DNAJ | Q13217; X6F | DNJC3_HUMAI | Oxidation (M) | 0 | 34077 | 0 | 0 | 0 | 0 | 0 | 0 | 626280 | 0 | 0 | 0 | 0 | 0 | 0 | 0 | 114480 | 0 |
| VCSTLLGLI | TKFC; TKFC; TK | Q3LXA3; I3L | TKFC_HUMAN | Unmodified | 323920 | 0 | 0 | 0 | 0 | 811860 | 0 | 0 | 0 | 0 | 0 | 1026300 | 0 | 0 | 310790 | 257540 | 96524 | 133310 |
| VCTLAIIDP | RPL30; RPL30; I | P62888; A0A | RL30_HUMAN | Unmodified | 4606800 | 4053100 | 3109300 | 5338300 | 0 | 0 | 0 | 21443 | 300060 | 0 | 1301800 | 21931000 | 60420 | 0 | 5516900 | 14691000 | 30362 | 253090 |
| VCTNIIEK | MYOF | Q9NZM1 | MYOF_HUMAN | Unmodified | 0 | 0 | 0 | 0 | 14938000 | 2957400 | 0 | 0 | 0 | 0 | 157530 | 0 | 0 | 0 | 0 | 0 | 0 | 0 |
| VCVEHHTF | EPB41; EPB41L | P11171; Q9 | EPB41_HUMAN | Unmodified | 260040 | 336890 | 207570 | 0 | 0 | 0 | 0 | 0 | 0 | 0 | 0 | 795360 | 0 | 0 | 451740 | 474810 | 0 | 0 |
| VCVEHHTF | EPB41L2; EPB4 | O43491; A0A | E41L2_HUMAN | Unmodified | 0 | 0 | 196680 | 107740 | 0 | 0 | 0 | 0 | 0 | 0 | 0 | 296980 | 0 | 0 | 0 | 0 | 0 | 0 |
| VCVETVES | IDH2 | P48735 | IDHP_HUMAN | Oxidation (M) | 0 | 0 | 0 | 0 | 0 | 0 | 210690 | 0 | 1799800 | 486420 | 0 | 0 | 0 | 0 | 0 | 0 | 0 | 0 |
| VCVETVES | IDH2 | P48735 | IDHP_HUMAN | Unmodified | 0 | 0 | 0 | 0 | 0 | 0 | 395000 | 0 | 6867200 | 1725300 | 0 | 0 | 0 | 0 | 0 | 0 | 0 | 0 |
| VCVGSDVT | MYORG | Q6NSJ0 | MYORG_HUMA | Unmodified | 79042 | 0 | 0 | 0 | 0 | 0 | 0 | 0 | 442130 | 0 | 0 | 304710 | 0 | 0 | 0 | 0 | 0 | 0 |
| VCVKFIQGI | MYO1B; MYO1E | O43795; E9F | MYO1B_HUMA | Unmodified | 0 | 0 | 0 | 0 | 0 | 0 | 228090 | 290760 | 0 | 0 | 0 | 0 | 0 | 0 | 0 | 0 | 415380 | 0 |
| VCVSLLGT | UBE2O; UBE2O | Q9C0C9; K7 | UBE2O_HUMAI | Unmodified | 0 | 0 | 0 | 0 | 0 | 0 | 0 | 0 | 0 | 0 | 0 | 733400 | 0 | 0 | 0 | 0 | 0 | 0 |
| VDALLSAQ | UGGT1 | Q9NYU2 | UGGG1_HUMA | Unmodified | 0 | 0 | 0 | 0 | 115620 | 0 | 0 | 44085 | 7122000 | 253840 | 1133600 | 154600 | 0 | 0 | 0 | 0 | 0 | 0 |
| VDAQEAKT | AKAP12 | Q02952 | AKA12_HUMAN | Unmodified | 34588 | 0 | 0 | 0 | 287080 | 104590 | 18284 | 0 | 0 | 0 | 0 | 199570 | 0 | 0 | 0 | 0 | 178790 | 549040 |
| VDAQEENF | SULF2 | Q8IWU5 | SULF2_HUMAN | Unmodified | 0 | 0 | 0 | 0 | 761930 | 609900 | 0 | 250340 | 0 | 0 | 179540 | 194610 | 866830 | 0 | 0 | 0 | 0 | 0 |
| VDAQFGGI | YARS1; YARS1; | P54577; A0A | SYYC_HUMAN | Unmodified | 183680 | 0 | 402450 | 341400 | 23949 | 0 | 0 | 119730 | 836650 | 0 | 260950 | 1083300 | 526910 | 631130 | 1300300 | 532530 | 0 | 0 |
| VDASGPDV | GPC2 | Q8N158 | GPC2_HUMAN | Unmodified | 0 | 0 | 11804 | 0 | 0 | 0 | 0 | 0 | 162230 | 19791 | 0 | 0 | 0 | 0 | 0 | 0 | 0 | 0 |
| VDATADYIC | ACLY | P53396 | ACLY_HUMAN | Unmodified | 0 | 0 | 151600 | 113580 | 0 | 71626 | 167750 | 0 | 186000 | 0 | 0 | 3996900 | 0 | 0 | 2105600 | 193590 | 82053 | 0 |
| VDATAETDI | PDIA4; PDIA4 | P13667; A0A | PDIA4_HUMAN | Unmodified | 544580 | 1066700 | 673910 | 0 | 0 | 202840 | 3111300 | 0 | 47794000 | 7804400 | 6011700 | 3499000 | 0 | 0 | 2160600 | 1531500 | 0 | 0 |
| VDATAETDI | PDIA4; PDIA4 | P13667; A0A | PDIA4_HUMAN | Unmodified | 0 | 0 | 0 | 0 | 0 | 0 | 0 | 0 | 441530 | 52221 | 0 | 0 | 0 | 0 | 0 | 0 | 0 | 0 |
| VDATEESDI | P4HB; P4HB; P | P07237; A0A | PDIA1_HUMAN | Unmodified | 0 | 0 | 0 | 0 | 0 | 0 | 0 | 0 | 69926000 | 1297900 | 4195000 | 3607100 | 0 | 0 | 1424100 | 907320 | 0 | 0 |
| VDATEESDI | P4HB; P4HB; P | P07237; A0A | PDIA1_HUMAN | Unmodified | 0 | 0 | 0 | 0 | 386300 | 0 | 0 | 736580 | 15240000 | 12782000 | 2304700 | 0 | 923460 | 1122500 | 0 | 0 | 0 | 0 |
| VDAVAARV | CHMP1B | Q7LBR1 | CHM1B_HUMA | Unmodified | 0 | 0 | 0 | 0 | 1460400 | 283620 | 104120 | 115200 | 0 | 505380 | 0 | 0 | 246550 | 0 | 0 | 0 | 0 | 216680 |
| VDAVAARV | CHMP1B | Q7LBR1 | CHM1B_HUMA | Oxidation (M) | 0 | 0 | 0 | 0 | 1042700 | 0 | 151650 | 0 | 0 | 453420 | 0 | 0 | 629090 | 0 | 0 | 0 | 26065 | 331490 |
| VDAVAARV | CHMP1B | Q7LBR1 | CHM1B_HUMA | Oxidation (M) | 0 | 0 | 0 | 0 | 989640 | 0 | 0 | 0 | 0 | 677280 | 0 | 1959000 | 474840 | 556130 | 0 | 0 | 0 | 254880 |
| VDAVAARV | CHMP1B | Q7LBR1 | CHM1B_HUMA | Unmodified | 0 | 0 | 0 | 0 | 5381900 | 0 | 0 | 0 | 0 | 2297500 | 0 | 0 | 1702100 | 0 | 0 | 0 | 239690 | 1163600 |
| VDCDQHSI | ERP44 | Q9BS26 | ERP44_HUMAN | Unmodified | 0 | 0 | 52833 | 0 | 0 | 0 | 0 | 0 | 32492000 | 0 | 0 | 11565 | 0 | 0 | 0 | 0 | 0 | 0 |
| VDCDQHSI | ERP44 | Q9BS26 | ERP44_HUMAN | Unmodified | 0 | 0 | 0 | 0 | 0 | 0 | 0 | 0 | 3188900 | 455050 | 0 | 0 | 0 | 0 | 0 | 0 | 0 | 0 |
| VDCQAYAC | DNAJC10; DNA | Q8IXB1; A0A | DJC10_HUMAN | Unmodified | 0 | 0 | 0 | 0 | 0 | 0 | 0 | 0 | 180100 | 0 | 0 | 0 | 0 | 0 | 0 | 0 | 0 | 0 |
| VDCTAHSD | TXNDC5 | Q8NBS9 | TXND5_HUMAI | Unmodified | 0 | 0 | 0 | 0 | 0 | 0 | 0 | 0 | 960800 | 0 | 0 | 0 | 0 | 0 | 0 | 0 | 0 | 0 |
| VDCTAHSD | TXNDC5 | Q8NBS9 | TXND5_HUMAI | Unmodified | 0 | 0 | 0 | 0 | 0 | 0 | 0 | 84161 | 4046500 | 4719000 | 295390 | 0 | 0 | 0 | 0 | 0 | 0 | 83844 |
| VDCTANTN | PDIA3; PDIA3 | P30101; H7E | PDIA3_HUMAN | Unmodified | 0 | 0 | 0 | 0 | 0 | 0 | 0 | 0 | 71314 | 0 | 0 | 321780 | 0 | 0 | 0 | 0 | 0 | 0 |
| VDCTANTN | PDIA3; PDIA3 | P30101; H7E | PDIA3_HUMAN | Unmodified | 0 | 0 | 0 | 0 | 0 | 0 | 0 | 475200 | 0 | 11199000 | 0 | 0 | 443650 | 0 | 0 | 0 | 0 | 0 |
| VDCTQHYE | TXNDC5 | Q8NBS9 | TXND5_HUMAI | Unmodified | 0 | 0 | 0 | 0 | 0 | 0 | 0 | 0 | 1082800 | 59573 | 199750 | 0 | 0 | 0 | 0 | 0 | 0 | 0 |
| VDDEVVAR | RASAL1 | O95294 | RASL1_HUMAN | Unmodified | 0 | 0 | 0 | 0 | 0 | 0 | 0 | 0 | 0 | 0 | 0 | 0 | 0 | 0 | 0 | 0 | 0 | 0 |
| VDDFLANE | PGD; PGD; PGD | P52209; K7E | 6PGD_HUMAN | Unmodified | 0 | 34198 | 0 | 0 | 693070 | 0 | 0 | 0 | 1621800 | 0 | 1977100 | 8835200 | 294630 | 0 | 903640 | 525390 | 0 | 0 |
| VDDLLIIEI | DYNC1H1; DYN | Q14204; A0A | DYHC1_HUMA | Unmodified | 0 | 0 | 0 | 0 | 0 | 0 | 0 | 0 | 0 | 0 | 0 | 643840 | 0 | 0 | 0 | 0 | 0 | 0 |
| VDDSSEDK | AARS1; AARS1; | P49588; A0A | SYAC_HUMAN | Unmodified | 0 | 104800 | 626410 | 240460 | 887390 | 134110 | 210760 | 116040 | 1531200 | 256340 | 527990 | 5828400 | 843580 | 460400 | 565150 | 769040 | 0 | 394770 |
| VDDVYSVL | CTDSP1; CTDSI | Q9GZU7; H7 | CTDS1_HUMAN | Unmodified | 115350 | 781850 | 473510 | 0 | 744320 | 0 | 0 | 0 | 0 | 0 | 0 | 320790 | 0 | 0 | 1550700 | 1216100 | 107420 | 0 |
| VDEAVAVL | PABPC4; PABP | Q13310; B1 | PABP4_HUMAN | Unmodified | 445860 | 0 | 712260 | 72631 | 0 | 0 | 0 | 0 | 0 | 42800 | 0 | 0 | 0 | 0 | 0 | 0 | 0 | 0 |
| VDEAVAVL | PABPC4; PABP | Q13310; B1 | PABP4_HUMAN | Unmodified | 633390 | 862560 | 1029500 | 0 | 0 | 0 | 125630 | 0 | 0 | 0 | 0 | 0 | 0 | 0 | 140880 | 269950 | 68970 | 0 |
| VDEAVAVL | PABPC1; PABP | P11940; Q9 | PABP1_HUMAN | Unmodified | 5048900 | 6718100 | 2519700 | 315240 | 0 | 0 | 112810 | 0 | 0 | 0 | 0 | 0 | 0 | 0 | 1279600 | 1698900 | 0 | 0 |
| VDEDEEAL | CHMP4A; CHM | Q9BY43; H0 | CHM4A_HUMA | Unmodified | 416680 | 381590 | 0 | 0 | 342320 | 0 | 0 | 0 | 520510 | 0 | 0 | 0 | 0 | 0 | 0 | 0 | 0 | 0 |
| VDEFVTHN | ADH5 | P11766 | ADHX_HUMAN | Unmodified | 169600 | 0 | 224880 | 144430 | 762950 | 157570 | 61314 | 45687 | 813660 | 0 | 2781100 | 14977000 | 167480 | 105730 | 0 | 334290 | 0 | 409230 |
| VDEIVVLSG | FLOT2; FLOT2; | Q14254; J3Q | FLOT2_HUMAN | Unmodified | 771370 | 614580 | 1836600 | 1870100 | 0 | 468600 | 0 | 0 | 0 | 0 | 1195800 | 2412200 | 120060 | 0 | 0 | 0 | 0 | 0 |
| VDFADIGW | BMP3 | P12645 | BMP3_HUMAN | Unmodified | 113230 | 0 | 0 | 0 | 2223300 | 1196800 | 2295200 | 2969700 | 3918100 | 1062300 | 666880 | 8100200 | 1309900 | 0 | 641710 | 162430 | 353540 | 1438800 |
| VDFNVPMK | PGK1; PGK2 | P00558; P07 | PGK1_HUMAN | Unmodified | 0 | 303300 | 699990 | 1257600 | 3971800 | 628350 | 228270 | 0 | 10232000 | 647650 | 4706400 | 31334000 | 1285700 | 448760 | 1785800 | 876550 | 1814600 | 3966100 |
| VDFNVPMK | PGK1; PGK2 | P00558; P07 | PGK1_HUMAN | Oxidation (M) | 0 | 271390 | 1493900 | 950520 | 0 | 1743400 | 0 | 97634 | 1627200 | 0 | 287330 | 10810000 | 144030 | 0 | 0 | 322460 | 796930 | 0 |
| VDFNVPMK | PGK1 | P00558 | PGK1_HUMAN | Unmodified | 0 | 0 | 0 | 211200 | 0 | 0 | 0 | 91861 | 0 | 0 | 0 | 0 | 0 | 0 | 0 | 0 | 314880 | 0 |
| VDFNVPMK | PGK1 | P00558 | PGK1_HUMAN | Unmodified | 0 | 0 | 0 | 0 | 0 | 0 | 0 | 0 | 0 | 0 | 0 | 0 | 0 | 0 | 0 | 0 | 1263300 | 0 |
| VDFPQDQI | MDH2; MDH2 | P40926; G3 | MDHM_HUMAN | Unmodified | 0 | 0 | 0 | 0 | 0 | 0 | 0 | 0 | 14839000 | 284890 | 0 | 1123600 | 0 | 0 | 0 | 0 | 0 | 0 |
| VDFPQDQI | MDH2; MDH2 | P40926; G3 | MDHM_HUMAN | Unmodified | 0 | 0 | 0 | 0 | 0 | 0 | 1598700 | 0 | 2759400 | 3329200 | 0 | 0 | 0 | 0 | 0 | 0 | 0 | 0 |
| VDGEVLRT | ERGIC3; ERGIC | Q9Y282; A0A | ERGI3_HUMAN | Unmodified | 0 | 0 | 0 | 0 | 0 | 0 | 0 | 0 | 813910 | 178500 | 0 | 0 | 0 | 0 | 0 | 0 | 0 | 0 |
| VDGVAQD | TMEM181 | Q9P2C4 | TM181_HUMAN | Unmodified | 0 | 0 | 0 | 0 | 0 | 0 | 0 | 0 | 5872700 | 1015200 | 0 | 1085700 | 0 | 0 | 0 | 0 | 0 | 0 |
| VDHQTGPI | GPI; GPI; GPI; N | P06744; K7E | G6PI_HUMAN | Unmodified | 0 | 0 | 0 | 0 | 0 | 0 | 0 | 0 | 4163800 | 0 | 2536000 | 36211000 | 0 | 0 | 0 | 0 | 0 | 0 |
| VDHVQMIV | DDX6 | P26196 | DDX6_HUMAN | Unmodified | 85014 | 162560 | 150660 | 0 | 0 | 0 | 0 | 0 | 62876 | 0 | 0 | 1232300 | 0 | 0 | 223980 | 94900 | 0 | 0 |
| VDIDAPDVI | AHNAK | Q09666 | AHNK_HUMAN | Unmodified | 0 | 191620 | 332230 | 0 | 2885200 | 301840 | 0 | 0 | 0 | 0 | 0 | 0 | 0 | 0 | 311530 | 0 | 399720 | 867310 |
| VDIKAPDVE | AHNAK | Q09666 | AHNK_HUMAN | Unmodified | 0 | 0 | 0 | 0 | 364590 | 129260 | 0 | 0 | 0 | 0 | 0 | 0 | 0 | 0 | 0 | 272610 | 0 | 0 |
| VDIKAPDVE | AHNAK | Q09666 | AHNK_HUMAN | Unmodified | 0 | 0 | 0 | 0 | 1480000 | 0 | 0 | 0 | 0 | 0 | 0 | 0 | 0 | 0 | 0 | 0 | 0 | 0 |
| VDILDPALL | PSMC3; PSMC3 | P17980; E9P | PRS6A_HUMAN | Unmodified | 128360 | 0 | 0 | 0 | 0 | 0 | 0 | 0 | 0 | 91794 | 162790 | 575500 | 0 | 0 | 0 | 0 | 0 | 0 |
| VDISAPDVI | AHNAK | Q09666 | AHNK_HUMAN | Unmodified | 0 | 0 | 0 | 0 | 1367300 | 0 | 0 | 0 | 0 | 0 | 0 | 0 | 0 | 0 | 0 | 0 | 58272 | 448510 |
| VDIVQAAW | GGCX | P38435 | VKGC_HUMAN | Unmodified | 0 | 0 | 116720 | 0 | 0 | 0 | 0 | 0 | 786190 | 37045 | 0 | 0 | 0 | 0 | 0 | 0 | 0 | 0 |
| VDKDEWIS | MME | P08473 | NEP_HUMAN N | Unmodified | 3000900 | 0 | 1504700 | 3887500 | 0 | 0 | 0 | 0 | 417600 | 0 | 0 | 0 | 591760 | 0 | 9415600 | 4182700 | 0 | 0 |
| VDKDRSG | PDCD6; PDCD6 | O75340; H0 | PDCD6_HUMA | Unmodified | 0 | 0 | 0 | 0 | 0 | 0 | 0 | 0 | 5348000 | 0 | 0 | 0 | 0 | 0 | 0 | 0 | 0 | 0 |
| VDKLDASE | CLTC; CLTC; CI | Q00610; K7E | CLH1_HUMAN | Unmodified | 149710 | 250590 | 4586900 | 0 | 0 | 0 | 0 | 0 | 379270 | 180820 | 0 | 40715000 | 0 | 0 | 2083600 | 0 | 0 | 0 |
| VDKLDASE | CLTC; CLTC; CI | Q00610; K7E | CLH1_HUMAN | Unmodified | 292890 | 172390 | 793510 | 0 | 0 | 0 | 88717 | 147140 | 0 | 0 | 0 | 1717700 | 0 | 0 | 484480 | 423680 | 0 | 0 |
| VDLEGERE | RAP2B | P61225 | RAP2B_HUMAN | Unmodified | 21338 | 13637 | 1733600 | 169530 | 102700 | 0 | 0 | 0 | 0 | 0 | 0 | 324410 | 0 | 0 | 461210 | 186120 | 0 | 57531 |
| VDLENRCC | LMNB1; LMNB1 | P20700; A0A | LMNB1_HUMAI | Unmodified | 0 | 0 | 0 | 0 | 0 | 0 | 0 | 0 | 739850 | 0 | 0 | 0 | 564170 | 0 | 0 | 0 | 0 | 0 |
| VDLEPERE | RAP2C; RAP2C | Q9Y3L5; A0A | RAP2C_HUMAI | Unmodified | 12968 | 35246 | 0 | 0 | 0 | 0 | 0 | 0 | 0 | 0 | 0 | 301900 | 0 | 0 | 0 | 123500 | 0 | 866130 |
| VDLFYLHAI | AKR7L; AKR7L | Q8NHP1; A0 | ARK74_HUMAN | Unmodified | 70514 | 0 | 0 | 355290 | 0 | 0 | 0 | 0 | 0 | 0 | 0 | 0 | 0 | 0 | 224030 | 207700 | 0 | 0 |
| VDLGGFAG | GART | P22102 | PUR2_HUMAN | Unmodified | 0 | 0 | 0 | 133210 | 0 | 0 | 0 | 0 | 393700 | 160540 | 232740 | 1016700 | 0 | 0 | 280720 | 145070 | 0 | 0 |
| VDLKGPQII | AHNAK2 | Q8IVF2 | AHNK2_HUMAI | Unmodified | 0 | 0 | 0 | 0 | 2495100 | 316150 | 0 | 0 | 0 | 0 | 0 | 0 | 0 | 0 | 0 | 0 | 0 | 2355000 |
| VDLPATSV | GALNT2 | Q10471 | GALT2_HUMAN | Unmodified | 0 | 0 | 0 | 0 | 0 | 0 | 0 | 0 | 2281100 | 0 | 0 | 0 | 0 | 0 | 0 | 0 | 0 | 0 |
| VDLPLAVLS | DCTPP1; DCTPI | Q9H773; H3 | DCTP1_HUMAN | Unmodified | 0 | 0 | 0 | 0 | 0 | 0 | 0 | 0 | 0 | 0 | 1483200 | 1824500 | 0 | 0 | 0 | 50816 | 0 | 0 |
| VDLSTSTD | MACROD1; MA | Q9BQ69; A0 | MACD1_HUMA | Unmodified | 0 | 56676000 | 780160 | 551420 | 5212900 | 1486800 | 4039800 | 0 | 3236300 | 574230 | 1010700 | 0 | 502620 | 359100 | 0 | 0 | 0 | 1942800 |
| VDLVDENF | UBE2K; UBE2K | P61086; D6F | UBE2K_HUMAN | Unmodified | 0 | 0 | 0 | 0 | 0 | 0 | 0 | 0 | 0 | 0 | 0 | 411960 | 0 | 0 | 0 | 0 | 0 | 0 |
| VDMNFMR | SLC4A7; SLC4A | Q9Y6M7; C9 | S4A7_HUMAN | Unmodified | 0 | 0 | 0 | 0 | 83235 | 0 | 0 | 0 | 0 | 0 | 0 | 633720 | 0 | 0 | 0 | 0 | 46588 | 0 |
| VDNAYWL | EIF3B | P55884 | EIF3B_HUMAN | Unmodified | 0 | 0 | 0 | 0 | 0 | 0 | 0 | 0 | 0 | 0 | 0 | 595550 | 0 | 0 | 0 | 10317 | 0 | 0 |
| VDNDENEI | NPM1; NPM1; N | P06748; A0A | NPM_HUMAN I | Unmodified | 0 | 0 | 1492300 | 0 | 0 | 0 | 0 | 0 | 1769800 | 0 | 2845000 | 7243000 | 0 | 0 | 766540 | 0 | 0 | 0 |
| VDNEELLPI | ADAM17; ADAM | P78536; A6H | ADA17_HUMAN | Unmodified | 0 | 0 | 470960 | 212330 | 0 | 0 | 0 | 0 | 464210 | 0 | 421900 | 161220 | 0 | 435700 | 0 | 0 | 0 | 0 |
| VDNSSLTG | ATP1A1; ATP1A | P05023; P50 | AT1A1_HUMAN | Unmodified | 1504000 | 1609600 | 25313000 | 5880400 | 133530 | 107720 | 231840 | 73324 | 6071800 | 1470300 | 6526900 | 76080000 | 225270 | 0 | 19449000 | 20578000 | 0 | 28023 |
| VDNSSLTG | ATP1A1 | P05023 | AT1A1_HUMAN | Unmodified | 217660 | 0 | 9712000 | 8230900 | 0 | 0 | 0 | 0 | 6844000 | 0 | 8177300 | 4467500 | 1152600 | 0 | 8496900 | 3496500 | 0 | 0 |
| VDNVEVLD | PIK3C2A; PIK3C | O00443; A0A | P3C2A_HUMAN | Unmodified | 0 | 0 | 0 | 0 | 0 | 0 | 0 | 0 | 700700 | 257810 | 0 | 1074100 | 0 | 0 | 0 | 0 | 0 | 0 |
| VDPALFPP | MOGS; MOGS; | Q13724; A0A | MOGS_HUMAN | Unmodified | 0 | 0 | 0 | 0 | 0 | 0 | 0 | 0 | 1281300 | 60617 | 119500 | 0 | 0 | 0 | 0 | 10441 | 0 | 0 |

| | | | | | | | | | | | | | | | | | | | | | | |
|---|---|---|---|---|---|---|---|---|---|---|---|---|---|---|---|---|---|---|---|---|---|---|
| VDPVSLGK | GPRIN1 | Q7Z2K8 | GRIN1_HUMAN | Unmodified | 771000 | 624340 | 0 | 0 | 0 | 0 | 0 | 0 | 682050 | 29100 | 0 | 0 | 0 | 292780 | 1042700 | 0 | 0 | 0 |
| VDQAIQAQ | SDCBP2 | Q9H190 | SDCB2_HUMAI | Unmodified | 277580 | 0 | 0 | 0 | 0 | 0 | 0 | 0 | 249790 | 0 | 77796 | 0 | 0 | 0 | 258700 | 244910 | 0 | 0 |
| VDQALHTC | THOP1; THOP1 | P52888; K7E | THOP1_HUMAI | Unmodified | 0 | 0 | 0 | 0 | 0 | 0 | 0 | 0 | 564170 | 1644100 | 1345600 | 5433700 | 0 | 0 | 207880 | 269250 | 0 | 0 |
| VDQGLFVC | SDCBP2 | Q9H190 | SDCB2_HUMAI | Unmodified | 0 | 68433 | 0 | 0 | 0 | 0 | 0 | 0 | 0 | 0 | 0 | 0 | 74008 | 0 | 116500 | 0 | 0 | 0 |
| VDQIQEIVT | IQGAP1; IQGAP | P46940; A0A | IQGA1_HUMAN | Unmodified | 0 | 0 | 0 | 0 | 0 | 0 | 0 | 0 | 252760 | 0 | 0 | 1498500 | 0 | 0 | 0 | 0 | 0 | 0 |
| VDQLNLER | INPPL1; INPPL1 | O15357; A0A | SHIP2_HUMAN | Unmodified | 0 | 0 | 0 | 0 | 0 | 0 | 0 | 0 | 0 | 0 | 644310 | 1031800 | 0 | 0 | 0 | 0 | 0 | 0 |
| VDQLNLER | INPPL1; INPPL1 | O15357; A0A | SHIP2_HUMAN | Unmodified | 0 | 0 | 0 | 0 | 162260 | 148620 | 0 | 0 | 0 | 0 | 24453 | 1706200 | 0 | 0 | 0 | 0 | 89935 | 1106200 |
| VDQMEGG | SUN1; SUN1; S | O94901; H0Y | SUN1_HUMAN | Unmodified | 0 | 0 | 0 | 0 | 0 | 0 | 0 | 0 | 4141300 | 0 | 0 | 0 | 0 | 0 | 0 | 0 | 242650 | 0 |
| VDQSLHTN | NLN; NLN; NLN | Q9BYT8; H0Y | NEUL_HUMAN | Unmodified | 0 | 0 | 0 | 0 | 0 | 0 | 0 | 0 | 1501900 | 0 | 672400 | 4032400 | 0 | 0 | 0 | 0 | 0 | 0 |
| VDQVIILYS | CXADR | P78310 | CXAR_HUMAN | Unmodified | 460170 | 0 | 650780 | 0 | 0 | 0 | 0 | 0 | 521040 | 0 | 0 | 11654000 | 0 | 0 | 1618500 | 611190 | 0 | 0 |
| VDREGSPV | STT3A | P46977 | STT3A_HUMAN | Unmodified | 0 | 0 | 0 | 0 | 0 | 0 | 0 | 0 | 749910 | 0 | 0 | 0 | 0 | 0 | 0 | 0 | 0 | 0 |
| VDREQLVQ | YWHAG | P61981 | 1433G_HUMAN | Acetyl (Protein N | 3330900 | 0 | 0 | 8549400 | 52351000 | 16505000 | 23084000 | 21004000 | 8671700 | 4891400 | 13885000 | 64166000 | 28044000 | 62492000 | 17437000 | 0 | 1.06E+08 | 54987000 |
| VDREQLVQ | YWHAG | P61981 | 1433G_HUMAN | Acetyl (Protein N | 0 | 0 | 0 | 0 | 2817200 | 0 | 2604000 | 0 | 0 | 292660 | 0 | 0 | 0 | 3427200 | 0 | 0 | 11548000 | 3041600 |
| VDRGSEET | USP54; USP54; | Q70EL1; A0A | UBP54_HUMAN | Unmodified | 0 | 28950 | 223780 | 180160 | 0 | 0 | 0 | 0 | 0 | 0 | 0 | 0 | 649890 | 435330 | 107380 | 0 | 0 | 0 |
| VDRTLVEDI | SNRNP200; SN | O75643; A0A | U520_HUMAN | Unmodified | 0 | 0 | 0 | 0 | 0 | 69117 | 0 | 12997 | 0 | 0 | 0 | 766080 | 0 | 0 | 0 | 0 | 0 | 0 |
| VDSDKEDE | RAP1GDS1; RAI | P52306; H0Y | GDS1_HUMAN | Unmodified | 0 | 0 | 0 | 0 | 0 | 0 | 0 | 0 | 0 | 0 | 0 | 1220400 | 0 | 0 | 0 | 75166 | 40999 | 0 |
| VDSGYRVH | GANAB; GANAE | Q14697; F5H | GANAB_HUMA | Unmodified | 0 | 0 | 0 | 0 | 1598600 | 210850 | 0 | 0 | 51316000 | 35712000 | 279020 | 0 | 2063400 | 1018600 | 0 | 0 | 214640 | 0 |
| VDSLETEEV | HCK; HCK; HCK | P08631; J3KI | HCK_HUMAN T | Unmodified | 0 | 0 | 0 | 0 | 0 | 0 | 0 | 0 | 0 | 0 | 39353 | 2633300 | 0 | 0 | 0 | 0 | 0 | 48951 |
| VDSLIPWIK | MACF1; MACF1 | Q9UPN3; H3 | MACF1_HUMAI | Unmodified | 0 | 0 | 0 | 0 | 0 | 0 | 0 | 0 | 0 | 0 | 0 | 433500 | 0 | 0 | 0 | 0 | 0 | 0 |
| VDSLLENLI | HNRNPCL3; HN | B7ZW38; P0I | HNRC3_HUMA | Unmodified | 0 | 0 | 0 | 0 | 0 | 0 | 0 | 0 | 0 | 0 | 1058300 | 0 | 0 | 0 | 0 | 0 | 0 | 0 |
| VDSPTVTTI | KLC1; KLC1; KL | Q07866; G3Y | KLC1_HUMAN | Unmodified | 0 | 0 | 0 | 0 | 0 | 0 | 0 | 0 | 292640 | 61687 | 149270 | 1787700 | 0 | 0 | 0 | 0 | 0 | 0 |
| VDSSNYDF | PLCG2; PLCG2 | P16885; A0A | PLCG2_HUMAI | Unmodified | 0 | 0 | 0 | 0 | 0 | 0 | 0 | 0 | 0 | 0 | 0 | 2410100 | 0 | 0 | 0 | 0 | 0 | 0 |
| VDTDDRTE | SLC39A6; SLC3 | Q13433; K7E | S39A6_HUMAN | Unmodified | 0 | 88709 | 642390 | 0 | 72766 | 0 | 0 | 0 | 0 | 0 | 376030 | 1365900 | 0 | 0 | 723420 | 785550 | 18799 | 0 |
| VDTIAPDES | EPHB3 | P54753 | EPHB3_HUMAI | Unmodified | 0 | 0 | 0 | 0 | 0 | 0 | 0 | 0 | 0 | 0 | 0 | 0 | 0 | 0 | 0 | 0 | 0 | 0 |
| VDTQSLDL | PROM2 | Q8N271 | PROM2_HUMA | Unmodified | 0 | 0 | 0 | 0 | 0 | 0 | 0 | 0 | 0 | 0 | 33508 | 484840 | 0 | 0 | 0 | 0 | 0 | 0 |
| VDTQSLDL | PROM2 | Q8N271 | PROM2_HUMA | Unmodified | 2166300 | 1137500 | 3387000 | 786990 | 0 | 0 | 0 | 0 | 0 | 0 | 0 | 270610 | 0 | 0 | 2641100 | 2217700 | 0 | 0 |
| VDTVAAEH | EPHB4; EPHB4 | P54760; Q96 | EPHB4_HUMAI | Unmodified | 0 | 342850 | 267360 | 0 | 0 | 0 | 0 | 0 | 0 | 0 | 0 | 0 | 0 | 0 | 397770 | 11907 | 0 | 0 |
| VDVAVNCA | HSD17B10; HS | Q99714; Q5I | HCD2_HUMAN | Unmodified | 0 | 0 | 0 | 0 | 0 | 0 | 0 | 0 | 614250 | 131710 | 0 | 0 | 0 | 0 | 0 | 0 | 0 | 0 |
| VDVGKDQI | FLNA; FLNA; FL | P21333; A0A | FLNA_HUMAN | Unmodified | 0 | 801260 | 2104900 | 0 | 2610500 | 0 | 579570 | 593330 | 0 | 0 | 0 | 568140 | 0 | 0 | 696630 | 0 | 251020 | 0 |
| VDVIPVNLF | FN1 | P02751 | FINC_HUMAN I | Unmodified | 0 | 0 | 1019600 | 187200 | 0 | 0 | 140460 | 0 | 0 | 0 | 0 | 776340 | 0 | 0 | 242240 | 804300 | 0 | 0 |
| VDVIPVNLF | FN1 | P02751 | FINC_HUMAN I | Unmodified | 0 | 0 | 3454100 | 636890 | 0 | 0 | 0 | 0 | 0 | 0 | 0 | 0 | 0 | 0 | 0 | 1212600 | 0 | 0 |
| VDVIQEPGI | TMX4 | Q9H1E5 | TMX4_HUMAN | Unmodified | 1237200 | 517650 | 0 | 1223000 | 0 | 0 | 0 | 0 | 531570 | 0 | 0 | 0 | 0 | 191440 | 827300 | 751200 | 0 | 457520 |
| VDVLSWVH | GGT7 | Q9UJ14 | GGT7_HUMAN | Unmodified | 0 | 0 | 0 | 0 | 0 | 0 | 0 | 0 | 2862500 | 194420 | 0 | 0 | 0 | 0 | 0 | 0 | 0 | 0 |
| VDVLSWVH | GGT7 | Q9UJ14 | GGT7_HUMAN | Unmodified | 0 | 0 | 0 | 0 | 0 | 0 | 0 | 0 | 1719700 | 0 | 0 | 0 | 51307 | 0 | 0 | 0 | 0 | 0 |
| VDVNTDRK | SDF4; SDF4; SD | Q9BRK5; H0 | CAB45_HUMAI | Unmodified | 0 | 0 | 0 | 0 | 0 | 0 | 74380 | 0 | 0 | 565190 | 0 | 1965500 | 0 | 0 | 0 | 0 | 0 | 0 |
| VDVSAPDV | AHNAK | Q09666 | AHNK_HUMAN | Unmodified | 0 | 0 | 0 | 0 | 0 | 0 | 0 | 0 | 0 | 0 | 0 | 0 | 0 | 0 | 0 | 0 | 154870 | 508580 |
| VDVSAPDV | AHNAK | Q09666 | AHNK_HUMAN | Oxidation (M) | 0 | 0 | 0 | 0 | 490830 | 0 | 0 | 0 | 0 | 0 | 0 | 0 | 0 | 0 | 65373 | 43753 | 0 | 0 |
| VDVTEQPG | TMX1; TMX1 | Q9H3N1; G3 | TMX1_HUMAN | Unmodified | 0 | 0 | 0 | 0 | 0 | 0 | 0 | 0 | 22504000 | 5094000 | 2744000 | 1358800 | 0 | 0 | 0 | 0 | 0 | 0 |
| VDWLTEK | EIF4A3; EIF4A1; | P38919; P60 | IF4A3_HUMAN | Unmodified | 0 | 0 | 0 | 0 | 0 | 0 | 0 | 49691 | 0 | 0 | 0 | 1980000 | 0 | 0 | 1536300 | 1544700 | 0 | 0 |
| VDYIAPWV | TTYH2 | Q9BSA4 | TTYH2_HUMAN | Unmodified | 0 | 0 | 26541 | 0 | 0 | 0 | 0 | 0 | 49093 | 89855 | 0 | 0 | 0 | 0 | 209520 | 0 | 0 | 0 |
| VDYIVEYDY | CD2AP | Q9Y5K6 | CD2AP_HUMAI | Acetyl (Protein N | 0 | 375460 | 0 | 0 | 0 | 0 | 0 | 0 | 0 | 0 | 0 | 0 | 0 | 0 | 379680 | 0 | 0 | 0 |
| VDYKADEV | MYH9; MYH10 | P35579; P35 | MYH9_HUMAN | Oxidation (M) | 0 | 0 | 1050600 | 0 | 0 | 0 | 79766 | 0 | 0 | 0 | 270050 | 5552300 | 0 | 0 | 1070000 | 807210 | 0 | 0 |
| VDYKADEV | MYH9; MYH10 | P35579; P35 | MYH9_HUMAN | Unmodified | 0 | 81399 | 0 | 0 | 502940 | 0 | 0 | 0 | 386930 | 112460 | 0 | 2263100 | 0 | 0 | 378010 | 196910 | 70589 | 0 |
| VDYLSSLGI | FLOT2; FLOT2; I | Q14254; J3Q | FLOT2_HUMAN | Unmodified | 0 | 226530 | 2388800 | 0 | 0 | 0 | 0 | 0 | 1988900 | 0 | 0 | 4307200 | 0 | 0 | 2631800 | 820010 | 20189 | 0 |
| VDYLVTEEE | SYNJ2BP-COX1 | A0A087WTA | A0A087WTA2_I | Unmodified | 0 | 0 | 0 | 0 | 0 | 0 | 0 | 0 | 0 | 0 | 0 | 0 | 0 | 0 | 0 | 0 | 976550 | 0 |
| VDYYEVLG | DNAJB6; DNAJE | O75190; A0A | DNJB6_HUMAN | Unmodified | 2963400 | 6374000 | 3044700 | 0 | 293690 | 0 | 0 | 261540 | 0 | 0 | 0 | 569530 | 0 | 0 | 4420900 | 2372200 | 314530 | 189750 |
| VEADRPGK | RBMX; RBMXL1 | P38159; Q96 | RBMX_HUMAN | Acetyl (Protein N | 0 | 0 | 0 | 0 | 1141500 | 61499 | 1111500 | 1332400 | 0 | 2131700 | 0 | 0 | 1903200 | 875760 | 0 | 0 | 109460 | 104230 |
| VEAKPEVQ | G3BP2; G3BP2; | Q9UN86; A0 | G3BP2_HUMAI | Unmodified | 0 | 0 | 0 | 0 | 0 | 0 | 0 | 0 | 0 | 0 | 53493 | 521560 | 0 | 0 | 0 | 0 | 0 | 0 |
| VEAMLNDF | APLP2; APP; AP | Q06481; P05 | APLP2_HUMAN | Unmodified | 0 | 0 | 271410 | 0 | 0 | 351360 | 0 | 0 | 2755300 | 0 | 127070 | 237840 | 0 | 0 | 61462 | 0 | 26884 | 170870 |
| VEAPQALR | ADK; ADK; ADK | P55263; A0A | ADK_HUMAN A | Unmodified | 142910 | 419170 | 221780 | 443950 | 0 | 0 | 0 | 0 | 473300 | 237610 | 0 | 119600 | 0 | 0 | 1075900 | 827700 | 0 | 0 |
| VEAQLQEL | MYH9 | P35579 | MYH9_HUMAN | Unmodified | 642270 | 0 | 0 | 0 | 1419100 | 727480 | 0 | 0 | 0 | 0 | 0 | 0 | 0 | 0 | 0 | 0 | 0 | 0 |
| VEAQVYILS | TUFM | P49411 | EFTU_HUMAN | Unmodified | 0 | 0 | 0 | 0 | 0 | 0 | 0 | 0 | 318850 | 0 | 0 | 0 | 0 | 0 | 0 | 0 | 0 | 0 |
| VEAYQWSV | PLAA; PLAA; PL | Q9Y263; H0Y | PLAP_HUMAN I | Unmodified | 0 | 0 | 0 | 0 | 0 | 0 | 0 | 0 | 0 | 0 | 0 | 2149200 | 0 | 0 | 0 | 439590 | 0 | 0 |
| VECSDNLF | MASP1; MASP1 | P48740; F8W | MASP1_HUMAI | Unmodified | 597030 | 615230 | 3153100 | 0 | 0 | 0 | 0 | 0 | 452960 | 0 | 0 | 1918100 | 0 | 0 | 1231500 | 384800 | 0 | 0 |
| VECYDYDN | CPNE3; CPNE3 | O75131; H0Y | CPNE3_HUMAI | Unmodified | 0 | 0 | 0 | 0 | 0 | 0 | 0 | 0 | 3718700 | 0 | 0 | 0 | 0 | 0 | 0 | 0 | 0 | 0 |
| VEDEVNSG | SLC35A4 | L0R6Q1 | S35U4_HUMAN | Unmodified | 6397600 | 0 | 0 | 0 | 0 | 0 | 0 | 0 | 2682100 | 1448800 | 0 | 0 | 0 | 0 | 0 | 0 | 0 | 0 |
| VEDGHRLC | PACSIN3; PACS | Q9UKS6; E9I | PACN3_HUMAI | Unmodified | 13248000 | 943960 | 0 | 657750 | 0 | 0 | 0 | 0 | 0 | 0 | 0 | 0 | 55987 | 0 | 3514900 | 2573800 | 0 | 0 |
| VEDLGKVD | WDR48 | Q8TAF3 | WDR48_HUMA | Unmodified | 0 | 172000 | 1290100 | 40641 | 0 | 106440 | 0 | 30100 | 84815 | 0 | 0 | 886120 | 0 | 0 | 272500 | 439780 | 0 | 0 |
| VEDMAELT | MYH9; MYH10; | P35579; P35 | MYH9_HUMAN | Oxidation (M) | 0 | 0 | 0 | 0 | 0 | 0 | 0 | 298910 | 0 | 674290 | 0 | 1862100 | 845820 | 0 | 0 | 597670 | 0 | 0 |
| VEDVSAVEI | HSPH1; HSPH1 | Q92598; A0A | HS105_HUMAN | Unmodified | 0 | 0 | 1643300 | 366140 | 0 | 262220 | 0 | 0 | 0 | 0 | 0 | 3111300 | 0 | 0 | 783810 | 429500 | 0 | 0 |
| VEDYDAAD | BCAM; BCAM | P50895; A0A | BCAM_HUMAN | Unmodified | 0 | 0 | 0 | 0 | 0 | 0 | 0 | 0 | 0 | 0 | 0 | 0 | 0 | 0 | 0 | 0 | 0 | 0 |
| VEEDDYPS | MIA3; MIA3 | Q5JRA6; A0A | TGO1_HUMAN | Unmodified | 0 | 0 | 0 | 0 | 0 | 0 | 0 | 0 | 0 | 0 | 675240 | 0 | 0 | 0 | 0 | 0 | 0 | 0 |
| VEEEIQTLS | TPD52; TPD52; | P55327; H0Y | TPD52_HUMAN | Unmodified | 0 | 0 | 261580 | 0 | 0 | 0 | 0 | 0 | 1693400 | 384510 | 570270 | 1821400 | 54467 | 0 | 0 | 0 | 0 | 0 |
| VEEISPNIR | SRP68 | Q9UHB9 | SRP68_HUMAN | Unmodified | 444750 | 0 | 362270 | 131400 | 0 | 157280 | 0 | 0 | 216310 | 0 | 270800 | 932940 | 0 | 0 | 0 | 29381 | 0 | 166410 |
| VEELTFHLL | PNPLA8; PNPL/ | Q9NP80; A0 | PLPL8_HUMAN | Unmodified | 79992 | 0 | 0 | 0 | 260640 | 0 | 0 | 0 | 154530 | 43952 | 0 | 1639200 | 0 | 0 | 0 | 0 | 0 | 0 |
| VEEQEPELT | C1QBP; C1QBP | Q07021; I3L | C1QBP_HUMAI | Unmodified | 0 | 0 | 0 | 0 | 0 | 0 | 610820 | 0 | 1273300 | 0 | 713030 | 0 | 0 | 0 | 0 | 0 | 0 | 0 |
| VEEQEPELT | C1QBP; C1QBP | Q07021; I3L | C1QBP_HUMAI | Unmodified | 0 | 0 | 0 | 0 | 0 | 0 | 0 | 0 | 2673100 | 0 | 0 | 0 | 0 | 0 | 0 | 573080 | 0 | 0 |
| VEETVKHD | TXNRD1; TXNRI | Q16881; A0A | TRXR1_HUMAN | Unmodified | 0 | 0 | 0 | 0 | 0 | 0 | 0 | 0 | 65433 | 0 | 0 | 1030900 | 0 | 0 | 0 | 0 | 0 | 0 |
| VEEVELPVE | PRMT1; PRMT8; | Q99873; Q9I | ANM1_HUMAN | Unmodified | 0 | 0 | 0 | 0 | 0 | 0 | 0 | 0 | 0 | 109160 | 131940 | 1214800 | 0 | 0 | 0 | 0 | 0 | 0 |
| VEEVGPYT | SCARB2; SCARI | Q14108; A0A | SCRB2_HUMAI | Unmodified | 0 | 0 | 0 | 0 | 0 | 0 | 0 | 0 | 415640 | 738860 | 0 | 0 | 0 | 0 | 90795 | 0 | 0 | 0 |
| VEFATLQEA | NUMA1; NUMA | Q14980; A0A | NUMA1_HUMA | Unmodified | 0 | 62835 | 0 | 54267 | 246620 | 0 | 0 | 0 | 118840 | 0 | 97175 | 0 | 314380 | 0 | 0 | 0 | 185430 | 0 |
| VEFLRPSFT | NECTIN1 | Q15223 | NECT1_HUMAN | Unmodified | 1012100 | 1217400 | 461700 | 271190 | 0 | 0 | 0 | 0 | 701640 | 0 | 0 | 759350 | 0 | 0 | 369360 | 207650 | 0 | 0 |
| VEGCFNQL | TSPAN1 | O60635 | TSN1_HUMAN | Unmodified | 64992 | 0 | 379090 | 0 | 0 | 0 | 0 | 0 | 388000 | 0 | 0 | 0 | 0 | 0 | 755180 | 0 | 0 | 0 |
| VEGDLKGP | AHNAK | Q09666 | AHNK_HUMAN | Unmodified | 0 | 0 | 0 | 0 | 778310 | 0 | 0 | 89783 | 0 | 0 | 0 | 0 | 0 | 0 | 0 | 0 | 2101400 | 0 |
| VEGELEEM | MYH10 | P35580 | MYH10_HUMAI | Unmodified | 0 | 0 | 317180 | 147300 | 0 | 0 | 0 | 0 | 0 | 0 | 0 | 1292200 | 0 | 0 | 363170 | 94545 | 0 | 0 |
| VEGFPTIYF | PDIA4; PDIA4 | P13667; A0A | PDIA4_HUMAN | Unmodified | 0 | 0 | 69393 | 0 | 0 | 0 | 0 | 0 | 9799200 | 0 | 1812800 | 1162300 | 0 | 0 | 0 | 0 | 0 | 142930 |
| VEGFPTIYF | PDIA4; PDIA4 | P13667; A0A | PDIA4_HUMAN | Unmodified | 0 | 0 | 0 | 0 | 0 | 0 | 0 | 0 | 1266100 | 0 | 0 | 0 | 0 | 0 | 121310 | 0 | 0 | 0 |
| VEGGCITKI | CD151; CD151 | P48509; K4D | CD151_HUMAI | Unmodified | 1689500 | 1785100 | 0 | 0 | 0 | 0 | 0 | 0 | 0 | 0 | 0 | 0 | 0 | 174760 | 0 | 0 | 0 | 0 |
| VEGGTPLF | DLST; DLST | P36957; Q86 | ODO2_HUMAN | Unmodified | 0 | 0 | 0 | 0 | 0 | 0 | 0 | 62352 | 1013500 | 0 | 0 | 0 | 0 | 0 | 72719 | 93806 | 0 | 0 |
| VEGGVKG( | AHNAK | Q09666 | AHNK_HUMAN | Unmodified | 0 | 0 | 0 | 0 | 1798000 | 0 | 0 | 0 | 0 | 0 | 0 | 0 | 0 | 0 | 0 | 0 | 0 | 1536000 |
| VEGTEPTTA | NCL; NCL; NCL | P19338; C9J | NUCL_HUMAN | Unmodified | 127270 | 86125 | 75285 | 0 | 42134 | 72501 | 43747 | 0 | 995440 | 349380 | 2507300 | 2045900 | 120700 | 0 | 185210 | 22969 | 0 | 0 |
| VEGTPSDP | GMPPA; GMPP/ | Q96IJ6; A0A( | GMPPA_HUMA | Unmodified | 0 | 0 | 0 | 0 | 0 | 0 | 0 | 0 | 0 | 0 | 115350 | 507980 | 0 | 0 | 0 | 0 | 0 | 0 |
| VEIAGPGFI | RARS1 | P54136 | SYRC_HUMAN | Unmodified | 0 | 0 | 0 | 0 | 0 | 0 | 0 | 0 | 0 | 0 | 0 | 804770 | 0 | 0 | 0 | 0 | 0 | 0 |
| VEIAGPGFI | RARS1 | P54136 | SYRC_HUMAN | Unmodified | 0 | 0 | 0 | 0 | 1371400 | 0 | 0 | 0 | 1045600 | 255120 | 637190 | 7794200 | 1325600 | 0 | 0 | 0 | 0 | 463240 |
| VEIATLTR | PSMA4; PSMA4 | P25789; H0Y | PSA4_HUMAN | Unmodified | 1511400 | 1165000 | 826840 | 0 | 506260 | 0 | 504220 | 778980 | 0 | 201320 | 390360 | 11379000 | 256900 | 0 | 579840 | 0 | 0 | 131750 |
| VEIATLTREI | PSMA4; PSMA4 | P25789; H0Y | PSA4_HUMAN | Unmodified | 0 | 0 | 0 | 0 | 0 | 0 | 0 | 0 | 141900 | 0 | 110160 | 369690 | 0 | 0 | 0 | 0 | 0 | 0 |
| VEIIANDQC | HSPA5; HSPA5; | P11021; A0A | BIP_HUMAN Er | Unmodified | 0 | 0 | 0 | 0 | 0 | 0 | 0 | 0 | 14727000 | 1074500 | 2072900 | 0 | 0 | 0 | 0 | 0 | 0 | 0 |
| VEIIANDQC | HSPA1L; HSPA | P34931; P0D | HS71L_HUMAN | Unmodified | 1183600 | 0 | 0 | 6214400 | 9742500 | 728410 | 1362000 | 0 | 10881000 | 2618500 | 1933400 | 726460 | 9471500 | 0 | 0 | 0 | 0 | 3571600 |
| VEILANDQ( | HSPA6; HSPA7 | P17066; P48 | HSP76_HUMAN | Unmodified | 2590600 | 3208300 | 12395000 | 1165000 | 119660 | 0 | 0 | 67605 | 10495000 | 707160 | 5758200 | 21012000 | 182960 | 0 | 5345800 | 3694800 | 49735 | 0 |
| VEILANDQ( | HSPA6; HSPA7 | P17066; P48 | HSP76_HUMAN | Unmodified | 0 | 1760800 | 5828000 | 1682900 | 0 | 0 | 0 | 0 | 9081600 | 0 | 955950 | 2739700 | 0 | 0 | 3229500 | 1591400 | 0 | 0 |
| VEITTETIK | PIGK; PIGK; PIG | Q92643; B1A | GPI8_HUMAN ( | Unmodified | 0 | 83546 | 0 | 0 | 861570 | 0 | 0 | 0 | 702460 | 0 | 0 | 0 | 165620 | 0 | 0 | 0 | 670330 | 0 |
| VEKEEAGG | SAE1; SAE1; SA | Q9UBE0; B3 | SAE1_HUMAN | Unmodified | 0 | 33732 | 0 | 0 | 0 | 0 | 0 | 0 | 205980 | 79495 | 374550 | 798000 | 0 | 0 | 0 | 20981 | 0 | 0 |
| VEKIPGGIII | CCT3; CCT3; C( | P49368; E9P | TCPG_HUMAN | Unmodified | 51610 | 0 | 58197 | 96161 | 701640 | 0 | 0 | 0 | 476110 | 0 | 255510 | 552790 | 1366800 | 0 | 61809 | 0 | 643660 | 0 |
| VELDNMPL | NONO; NONO; | Q15233; C9I | NONO_HUMAN | Unmodified | 0 | 0 | 0 | 0 | 0 | 0 | 236140 | 173180 | 0 | 0 | 0 | 373070 | 0 | 0 | 0 | 0 | 0 | 0 |
| VELFFKDD/ | PAICS; PAICS; F | P22234; D6F | PUR6_HUMAN | Unmodified | 0 | 0 | 0 | 0 | 0 | 0 | 0 | 0 | 0 | 0 | 1049300 | 2353300 | 0 | 0 | 0 | 35783000 | 0 | 0 |
| VELKGTVRI | ANXA6; ANXA6 | P08133; E5R | ANXA6_HUMAN | Unmodified | 0 | 0 | 0 | 0 | 0 | 0 | 263490 | 398820 | 0 | 2159800 | 0 | 0 | 0 | 0 | 0 | 0 | 88662 | 289140 |
| VELPGTAVI | SLC12A2; SLC1 | P55011; G3X | S12A2_HUMAN | Unmodified | 121440 | 0 | 164240 | 104790 | 0 | 0 | 0 | 0 | 220080 | 0 | 316200 | 503470 | 0 | 0 | 308160 | 364290 | 0 | 0 |
| VELQELND | VIM; VIM; VIM | P08670; B0Y | VIME_HUMAN | Unmodified | 0 | 0 | 0 | 392670 | 14348000 | 4710700 | 227250 | 0 | 0 | 0 | 0 | 0 | 363930 | 275760 | 0 | 0 | 2416600 | 7261500 |
| VELSEQQL | CNP | P09543 | CN37_HUMAN | Unmodified | 242480 | 215400 | 2700300 | 311790 | 0 | 0 | 0 | 0 | 0 | 0 | 3268600 | 28502000 | 0 | 0 | 6290800 | 2287700 | 0 | 0 |
| VELSEQQL | CNP | P09543 | CN37_HUMAN | Unmodified | 1479100 | 0 | 0 | 2054600 | 0 | 0 | 0 | 0 | 5475300 | 0 | 4470000 | 10624000 | 0 | 0 | 2369200 | 5065300 | 0 | 0 |
| VELSQLR | RPL35; UQCRF | P42766; P0C | RL35_HUMAN ( | Unmodified | 0 | 0 | 1416100 | 0 | 0 | 0 | 0 | 0 | 0 | 0 | 2008200 | 27479000 | 0 | 0 | 0 | 686590 | 0 | 0 |
| VELSVGESI | NCAM2; NCAM | O15394; H9I | NCAM2_HUMA | Unmodified | 0 | 0 | 2986200 | 1159400 | 0 | 0 | 0 | 0 | 0 | 0 | 0 | 548180 | 0 | 0 | 4224800 | 2723500 | 0 | 0 |
| VELSVSGQ | CPNE2; CPNE2 | Q96FN4; H3 | CPNE2_HUMAI | Unmodified | 0 | 0 | 178280 | 646030 | 2133200 | 817440 | 322000 | 0 | 4530300 | 337810 | 0 | 0 | 285470 | 0 | 0 | 0 | 0 | 371770 |
| VELVPPTPA | ATP5MG; ATP5N | O75964; Q7 | ATP5L_HUMAN | Unmodified | 0 | 0 | 0 | 0 | 0 | 0 | 0 | 0 | 0 | 864310 | 0 | 0 | 0 | 0 | 0 | 0 | 0 | 0 |
| VELVTGEEI | RANBP2 | P49792 | RBP2_HUMAN | Unmodified | 0 | 0 | 0 | 0 | 0 | 0 | 0 | 0 | 0 | 0 | 0 | 522480 | 0 | 0 | 0 | 0 | 0 | 0 |
| VENCPDEL | LYN | P07948 | LYN_HUMAN T | Unmodified | 0 | 0 | 0 | 0 | 0 | 0 | 0 | 0 | 1144000 | 0 | 0 | 0 | 0 | 0 | 0 | 0 | 0 | 0 |
| VENFEAAVI | SGTA; SGTA; SC | O43765; K7E | SGTA_HUMAN | Unmodified | 0 | 0 | 309780 | 0 | 238810 | 2192500 | 197400 | 0 | 444930 | 0 | 3001900 | 2074400 | 0 | 0 | 0 | 0 | 57354 | 0 |
| VENFEAYFI | PTPRJ; PTPRJ; P | Q12913; A0A | PTPRJ_HUMAN | Unmodified | 0 | 0 | 0 | 0 | 0 | 0 | 0 | 0 | 0 | 0 | 0 | 751170 | 0 | 0 | 0 | 0 | 0 | 0 |
| VENFIQK | PITPNB; PITPNE | P48739; A0A | PIPNB_HUMAN | Unmodified | 87331 | 0 | 0 | 0 | 0 | 0 | 0 | 0 | 244830 | 43283 | 0 | 834660 | 0 | 0 | 382040 | 9665 | 0 | 0 |

| | | | | | | | | | | | | | | | | | | | | | | |
|---|---|---|---|---|---|---|---|---|---|---|---|---|---|---|---|---|---|---|---|---|---|---|
| VENFSVAG | STON1; STON1 | Q9Y6Q2; Q5 | STON1_HUMAI | Unmodified | 510730 | 344950 | 0 | 0 | 0 | 0 | 19271 | 0 | 708550 | 0 | 0 | 380760 | 431760 | 0 | 1270900 | 2116500 | 0 | 0 |
| VENGEGTII | ADAR; ADAR; AI | P55265; A0A | DSRAD_HUMAI | Unmodified | 0 | 463280 | 1128500 | 0 | 0 | 372670 | 0 | 0 | 73999 | 0 | 0 | 682410 | 0 | 0 | 824610 | 0 | 0 | 2599700 |
| VENMVDQI | IARS1; IARS1; I/ | P41252; A0A | SYIC_HUMAN I | Unmodified | 0 | 0 | 0 | 0 | 0 | 0 | 0 | 0 | 0 | 0 | 0 | 1788700 | 0 | 0 | 0 | 0 | 0 | 0 |
| VENMVDQI | IARS1; IARS1; I/ | P41252; A0A | SYIC_HUMAN I | Oxidation (M) | 0 | 0 | 0 | 0 | 0 | 0 | 0 | 0 | 0 | 0 | 0 | 1887900 | 0 | 0 | 0 | 0 | 0 | 0 |
| VENQENVS | CAP1 | Q01518 | CAP1_HUMAN | Unmodified | 224820 | 0 | 0 | 0 | 0 | 0 | 0 | 238480 | 0 | 364900 | 0 | 6289900 | 0 | 0 | 294540 | 206370 | 0 | 0 |
| VENQRKEII | ATP11C; ATP11 | Q8NB49; A0 | AT11C_HUMAN | Unmodified | 0 | 0 | 0 | 182540 | 336550 | 0 | 0 | 0 | 0 | 0 | 83094 | 0 | 25231 | 0 | 0 | 0 | 764090 | 0 |
| VENQYQLL | IKBIP | Q70UQ0 | IKIP_HUMAN In | Unmodified | 0 | 0 | 0 | 0 | 0 | 0 | 0 | 0 | 504920 | 0 | 0 | 0 | 0 | 0 | 0 | 0 | 0 | 0 |
| VENTRWD\ | CACYBP | Q9HB71 | CYBP_HUMAN | Unmodified | 0 | 0 | 0 | 0 | 657920 | 0 | 244760 | 0 | 1436400 | 0 | 538040 | 3875500 | 613190 | 348290 | 0 | 101950 | 98047 | 139990 |
| VEPAVDTSI | FLNB; FLNB | O75369; E7E | FLNB_HUMAN | Unmodified | 6519700 | 0 | 295550 | 157110 | 1561300 | 526170 | 0 | 21279 | 0 | 0 | 28413 | 257460 | 282180 | 0 | 157130 | 55452 | 354390 | 225180 |
| VEPFLPGH | ASNS; ASNS; AS | P08243; F8V | ASNS_HUMAN | Unmodified | 0 | 0 | 0 | 0 | 0 | 0 | 0 | 0 | 0 | 0 | 1638200 | 12624000 | 0 | 0 | 3564900 | 0 | 0 | 0 |
| VEPGLGAD | FLNA; FLNA; FL | P21333; A0A | FLNA_HUMAN | Unmodified | 0 | 0 | 1838200 | 1518000 | 199300 | 0 | 0 | 0 | 534850 | 0 | 777210 | 2351000 | 139970 | 0 | 2549700 | 2369700 | 0 | 0 |
| VEPLDFGG | HEXB; HEXB; HI | P07686; H0 | HEXB_HUMAN | Unmodified | 0 | 0 | 389920 | 0 | 0 | 0 | 0 | 0 | 569940 | 0 | 0 | 0 | 0 | 0 | 176600 | 0 | 0 | 0 |
| VEQATKPSI | RBMX; RBMX; R | P38159; H3E | RBMX_HUMAN | Unmodified | 0 | 0 | 0 | 0 | 0 | 0 | 0 | 0 | 434070 | 26753 | 1076700 | 3878400 | 0 | 0 | 0 | 0 | 21318 | 0 |
| VEQATQAIF | APLP1; APLP1; / | P51693; B7Z | APLP1_HUMAN | Unmodified | 0 | 0 | 0 | 0 | 0 | 0 | 0 | 0 | 2187100 | 0 | 0 | 0 | 0 | 0 | 0 | 0 | 0 | 0 |
| VEQDPLEL | IGSF3 | O75054 | IGSF3_HUMAN | Unmodified | 0 | 0 | 0 | 0 | 0 | 74376 | 0 | 0 | 0 | 0 | 0 | 0 | 0 | 0 | 199460 | 0 | 0 | 0 |
| VEQEKMLL | CCDC88A; CCI | Q3V6T2; A0/ | GRDN_HUMAN | Unmodified | 0 | 0 | 0 | 0 | 0 | 0 | 0 | 0 | 15594000 | 3454200 | 13684000 | 0 | 0 | 0 | 550460 | 366180 | 0 | 0 |
| VEQELSGE | N/A | N/A | N/A | Unmodified | 0 | 0 | 0 | 0 | 360710 | 0 | 0 | 0 | 0 | 0 | 1432800 | 0 | 0 | 0 | 0 | 0 | 0 | 0 |
| VEQHVVD( | RTCB | Q9Y3I0 | RTCB_HUMAN | Unmodified | 0 | 0 | 0 | 0 | 0 | 0 | 0 | 0 | 103180 | 0 | 0 | 284440 | 0 | 0 | 0 | 37473 | 0 | 0 |
| VEQIEAGTF | TXNRD1; TXNRI | Q16881; A0/ | TRXR1_HUMAN | Unmodified | 89189 | 93028 | 0 | 0 | 413620 | 545340 | 10246 | 0 | 0 | 0 | 145850 | 1349200 | 0 | 0 | 0 | 117270 | 0 | 0 |
| VEQLGAEG | LUC7L2; LUC7L | Q9Y383; A0/ | LC7L2_HUMAN | Unmodified | 0 | 0 | 0 | 0 | 41583 | 0 | 0 | 0 | 354490 | 29102 | 1322600 | 1830300 | 0 | 0 | 100410 | 0 | 0 | 0 |
| VEQLMEWS | TUSC3; TUSC3; | Q13454; D6I | TUSC3_HUMAI | Unmodified | 130780 | 121760 | 83887 | 0 | 144730 | 0 | 0 | 0 | 1142800 | 0 | 0 | 0 | 0 | 0 | 0 | 140710 | 0 | 58730 |
| VEQVKFDA | VIL1; VIL1 | P09327; B4E | VILI_HUMAN Vi | Unmodified | 0 | 0 | 0 | 0 | 0 | 0 | 0 | 0 | 1703000 | 274610 | 868260 | 2505800 | 0 | 0 | 0 | 0 | 0 | 0 |
| VEQVLSLEI | VAPB; VAPB | O95292; E5F | VAPB_HUMAN | Unmodified | 0 | 0 | 0 | 0 | 0 | 0 | 0 | 0 | 543970 | 301880 | 0 | 2127200 | 0 | 0 | 0 | 0 | 0 | 0 |
| VERADGYE | RPS3A; RPS3A; | P61247; D6F | RS3A_HUMAN | Unmodified | 846830 | 0 | 1882700 | 410840 | 0 | 0 | 14152 | 0 | 0 | 0 | 0 | 935050 | 0 | 0 | 1811000 | 0 | 0 | 0 |
| VESDLKGP | AHNAK | Q09666 | AHNK_HUMAN | Unmodified | 0 | 0 | 0 | 0 | 1552000 | 0 | 0 | 0 | 0 | 200080 | 0 | 0 | 0 | 0 | 0 | 0 | 141020 | 709830 |
| VESELRSIC | YWHAQ; YWHA | P27348; E9P | 1433T_HUMAN | Unmodified | 0 | 0 | 0 | 0 | 2372500 | 102960 | 0 | 0 | 0 | 341280 | 0 | 0 | 1667400 | 968620 | 0 | 0 | 1188700 | 1212100 |
| VESLEQEA/ | APP; APP; APP; | P05067; H7C | A4_HUMAN Am | Unmodified | 0 | 0 | 0 | 0 | 0 | 0 | 0 | 0 | 1856200 | 0 | 591270 | 182170 | 0 | 0 | 61957 | 0 | 0 | 0 |
| VESLEQEA/ | APP; APP; APP; | P05067; H7C | A4_HUMAN Am | Unmodified | 0 | 0 | 0 | 0 | 0 | 0 | 0 | 0 | 6607800 | 0 | 0 | 0 | 0 | 0 | 0 | 0 | 0 | 0 |
| VESRDKLPI | ITGB1; ITGB1; I | P05556; H7C | ITB1_HUMAN In | Unmodified | 0 | 0 | 0 | 143160 | 6712300 | 1857100 | 0 | 0 | 0 | 0 | 487940 | 0 | 0 | 0 | 0 | 0 | 1114800 | 7371700 |
| VESTDVSD | PSMB8; PSMB8 | P28062; Q5J | PSB8_HUMAN | Unmodified | 0 | 0 | 0 | 0 | 0 | 0 | 0 | 0 | 0 | 0 | 811000 | 4571700 | 0 | 0 | 0 | 0 | 0 | 0 |
| VESTDVSD | PSMB8; PSMB8 | P28062; Q5J | PSB8_HUMAN | Unmodified | 0 | 0 | 0 | 0 | 0 | 0 | 0 | 0 | 1140500 | 0 | 0 | 2808500 | 0 | 0 | 0 | 0 | 0 | 362440 |
| VESVFETLV | HYOU1; HYOU1 | Q9Y4L1; E9F | HYOU1_HUMA | Unmodified | 0 | 0 | 0 | 0 | 0 | 0 | 0 | 0 | 0 | 261240 | 143270 | 0 | 0 | 0 | 0 | 0 | 252580 | 0 |
| VETEDQVL/ | MASP1; MASP1 | P48740; F8V | MASP1_HUMAI | Unmodified | 85300 | 0 | 1594700 | 0 | 0 | 0 | 0 | 0 | 0 | 0 | 0 | 0 | 0 | 0 | 1415200 | 904340 | 0 | 0 |
| VETEQAEIC | CENPF | P49454 | CENPF_HUMAI | Unmodified | 0 | 0 | 0 | 0 | 5214000 | 14490000 | 6501500 | 2028100 | 1280100 | 118140 | 13614000 | 0 | 4003100 | 10130000 | 0 | 0 | 39184 | 23927000 |
| VETFLGWE | PLSCR3; PLSCF | Q9NRY6; I3L | PLS3_HUMAN | Unmodified | 1531900 | 1224800 | 0 | 81152 | 0 | 0 | 0 | 0 | 0 | 0 | 0 | 302450 | 0 | 53638 | 641250 | 523950 | 0 | 0 |
| VETFSGVYI | RPS7; RPS7 | P62081; A0A | RS7_HUMAN 4 | Unmodified | 0 | 0 | 0 | 0 | 0 | 0 | 0 | 0 | 0 | 0 | 0 | 3336100 | 0 | 0 | 0 | 0 | 0 | 0 |
| VETGILRPG | EEF1A2; EEF1A | Q05639; A0/ | EF1A2_HUMAN | Oxidation (M) | 25376000 | 72770000 | 718950 | 43982000 | 0 | 0 | 0 | 0 | 744150 | 323940 | 0 | 0 | 0 | 0 | 2.23E+08 | 43717000 | 244560 | 333890 |
| VETGILRPG | EEF1A2; EEF1A | Q05639; A0/ | EF1A2_HUMAN | Unmodified | 519300 | 1191600 | 161820 | 1667400 | 0 | 0 | 0 | 0 | 424220 | 0 | 0 | 0 | 0 | 0 | 4463900 | 441090 | 0 | 0 |
| VETGVLKP( | EEF1A1P5; EEF | Q5VTE0; P6 | EF1A3_HUMAN | Oxidation (M) | 1.6E+08 | 1.84E+08 | 61514000 | 65389000 | 7024500 | 2291300 | 4406700 | 196960 | 15338000 | 3977500 | 49719000 | 1.25E+08 | 5057200 | 3244300 | 1.79E+08 | 1.54E+08 | 2487500 | 8695400 |
| VETGVLKP( | EEF1A1P5; EEF | Q5VTE0; P6 | EF1A3_HUMAN | Unmodified | 1178800 | 2201500 | 76336 | 404960 | 0 | 0 | 0 | 0 | 6167200 | 0 | 1170200 | 1901600 | 0 | 0 | 1475500 | 440180 | 0 | 0 |
| VETPLEEAI | NAA15; NAA15 | Q9BXJ9; A0/ | NAA15_HUMAI | Unmodified | 0 | 83692 | 0 | 0 | 0 | 0 | 0 | 0 | 0 | 0 | 459700 | 939780 | 0 | 0 | 0 | 0 | 591610 | 0 |
| VETPVLPP\ | SMAD9; SMAD3 | O15198; P8 | SMAD9_HUMA | Unmodified | 0 | 0 | 0 | 0 | 0 | 0 | 372720 | 0 | 284750 | 0 | 313660 | 534970 | 0 | 0 | 0 | 262620 | 0 | 0 |
| VETSDEEIH | EPHX1; EPHX1 | P07099; B1A | HYEP_HUMAN | Unmodified | 0 | 0 | 183530 | 0 | 0 | 0 | 0 | 0 | 832180 | 252060 | 0 | 0 | 0 | 0 | 0 | 34791 | 0 | 0 |
| VEWKFDQ( | F11R | Q9Y624 | JAM1_HUMAN | Unmodified | 677360 | 815680 | 205530 | 313130 | 0 | 0 | 0 | 0 | 507320 | 0 | 130020 | 1319000 | 0 | 0 | 703720 | 573440 | 0 | 0 |
| VEYHAGGS | ADK; ADK; ADK; | P55263; A0/ | ADK_HUMAN A | Unmodified | 0 | 0 | 268400 | 107190 | 399680 | 0 | 645060 | 207790 | 205070 | 41221 | 179730 | 5310400 | 288460 | 0 | 0 | 244620 | 139290 | 470630 |
| VEYHFLSP\ | TFRC; TFRC | P02786; G3\ | TFR1_HUMAN | Unmodified | 4983400 | 5343600 | 5767600 | 5868500 | 687830 | 1680300 | 1201100 | 148190 | 30085000 | 4750500 | 1083600 | 0 | 417720 | 0 | 9462800 | 0 | 0 | 0 |
| VEYSEEELK | XRCC6; XRCC6 | P12956; B1/ | XRCC6_HUMAI | Unmodified | 451500 | 156820 | 255980 | 0 | 0 | 0 | 0 | 0 | 0 | 0 | 482530 | 2435900 | 0 | 0 | 100790 | 0 | 0 | 0 |
| VEYTLGEE | EIF4G1; EIF4G1 | Q04637; E7E | IF4G1_HUMAN | Unmodified | 0 | 0 | 0 | 0 | 0 | 0 | 0 | 0 | 0 | 0 | 0 | 0 | 0 | 0 | 0 | 0 | 0 | 0 |
| VEYYIPGST | CDCP1 | Q9H5V8 | CDCP1_HUMA | Unmodified | 0 | 0 | 479390 | 0 | 0 | 0 | 44684 | 0 | 0 | 0 | 0 | 1459200 | 0 | 0 | 0 | 408650 | 0 | 0 |
| VFADYEAYI | PYGB; PYGB | P11216; H0 | PYGB_HUMAN | Oxidation (M) | 0 | 0 | 0 | 0 | 0 | 0 | 0 | 0 | 0 | 0 | 0 | 0 | 0 | 0 | 0 | 0 | 0 | 1419800 |
| VFADYEAYI | PYGB; PYGB | P11216; H0 | PYGB_HUMAN | Unmodified | 0 | 0 | 0 | 0 | 0 | 0 | 0 | 0 | 0 | 0 | 0 | 0 | 0 | 0 | 0 | 0 | 475900 | 2398400 |
| VFADYEAY\ | PYGL; PYGL | P06737; E9P | PYGL_HUMAN | Unmodified | 0 | 0 | 0 | 184740 | 3202100 | 0 | 0 | 0 | 0 | 0 | 0 | 0 | 0 | 0 | 0 | 104720 | 541540 | 2551900 |
| VFAFDAVSI | IRF2BPL | Q9H1B7 | I2BPL_HUMAN | Oxidation (M) | 0 | 0 | 0 | 0 | 0 | 0 | 625600 | 23812 | 0 | 93261 | 0 | 481430 | 128900 | 0 | 0 | 0 | 0 | 0 |
| VFASELNA( | LARS1; LARS1; | Q9P2J5; A0A | SYLC_HUMAN | Unmodified | 0 | 0 | 0 | 0 | 0 | 0 | 3459800 | 1705300 | 660220 | 0 | 949180 | 2608500 | 1274100 | 0 | 0 | 0 | 172660 | 0 |
| VFASELNA( | LARS1; LARS1; | Q9P2J5; A0A | SYLC_HUMAN | Unmodified | 0 | 0 | 0 | 0 | 0 | 0 | 0 | 0 | 0 | 0 | 0 | 1695200 | 0 | 0 | 0 | 0 | 0 | 0 |
| VFASLPQVI | WDR1; WDR1 | O75083; D6I | WDR1_HUMAN | Unmodified | 2031300 | 1638700 | 1938100 | 1877700 | 3071700 | 1036300 | 0 | 0 | 4045900 | 0 | 1530700 | 10958000 | 1914000 | 768220 | 4784100 | 2017500 | 0 | 0 |
| VFASLPQVI | WDR1; WDR1 | O75083; D6I | WDR1_HUMAN | Unmodified | 0 | 364810 | 0 | 0 | 70539000 | 6656900 | 8160700 | 8475900 | 0 | 4529900 | 491970 | 0 | 33432000 | 14609000 | 0 | 0 | 15922000 | 11895000 |
| VFDADVVP | RET; RET | P07949; Q9E | RET_HUMAN P | Unmodified | 0 | 0 | 0 | 0 | 0 | 0 | 0 | 0 | 1293800 | 0 | 0 | 0 | 0 | 0 | 0 | 0 | 0 | 0 |
| VFDADVVP | RET; RET | P07949; Q9E | RET_HUMAN P | Unmodified | 0 | 0 | 0 | 0 | 356650 | 0 | 0 | 0 | 3484800 | 0 | 0 | 0 | 0 | 0 | 0 | 0 | 0 | 0 |
| VFDAIMNFI | EEF2 | P13639 | EF2_HUMAN El | Oxidation (M) | 0 | 0 | 648970 | 0 | 0 | 0 | 145890 | 0 | 0 | 0 | 1005500 | 2947400 | 0 | 0 | 0 | 0 | 0 | 341240 |
| VFDAIMNFI | EEF2 | P13639 | EF2_HUMAN El | Unmodified | 0 | 0 | 0 | 0 | 0 | 0 | 0 | 0 | 83914 | 0 | 0 | 1259400 | 0 | 0 | 0 | 0 | 0 | 0 |
| VFDAIMNFI | EEF2 | P13639 | EF2_HUMAN El | Unmodified | 0 | 0 | 0 | 0 | 0 | 0 | 0 | 0 | 0 | 0 | 0 | 2484800 | 0 | 0 | 0 | 0 | 0 | 0 |
| VFDAIMNFI | EEF2 | P13639 | EF2_HUMAN El | Unmodified | 186510 | 320000 | 0 | 364200 | 2442000 | 1190300 | 850250 | 72740 | 1420200 | 303590 | 3157100 | 8854800 | 1386600 | 1030800 | 303550 | 263000 | 792950 | 1075300 |
| VFDAIMNFI | EEF2 | P13639 | EF2_HUMAN El | Oxidation (M) | 0 | 0 | 0 | 0 | 450700 | 0 | 70906 | 0 | 0 | 0 | 1406700 | 7469200 | 533810 | 334100 | 291730 | 647500 | 0 | 0 |
| VFDDESDE | HTATSF1 | O43719 | HTSF1_HUMAN | Unmodified | 0 | 0 | 0 | 0 | 0 | 0 | 0 | 0 | 159620 | 0 | 388990 | 1319500 | 0 | 0 | 0 | 0 | 0 | 0 |
| VFDFLVDSI | MYO1E; MYO1E | Q12965; H0 | MYO1E_HUMAI | Unmodified | 0 | 0 | 0 | 0 | 0 | 0 | 0 | 0 | 0 | 0 | 0 | 0 | 0 | 0 | 0 | 0 | 0 | 0 |
| VFDFTFSPE | AKR1A1 | P14550 | AK1A1_HUMAN | Oxidation (M) | 0 | 0 | 0 | 0 | 881910 | 0 | 0 | 0 | 825440 | 1011100 | 69240 | 2115500 | 1562100 | 0 | 973750 | 1069700 | 309540 | 0 |
| VFDFTFSPE | AKR1A1 | P14550 | AK1A1_HUMAN | Oxidation (M) | 0 | 0 | 0 | 0 | 0 | 0 | 0 | 0 | 4436600 | 0 | 0 | 0 | 0 | 0 | 0 | 0 | 663270 | 0 |
| VFDFTFSPE | AKR1A1 | P14550 | AK1A1_HUMAN | Unmodified | 0 | 0 | 0 | 0 | 0 | 0 | 0 | 0 | 0 | 438940 | 0 | 0 | 0 | 647500 | 0 | 0 | 0 | 0 |
| VFDGGVVD | PDE2A; PDE2A; | O00408; H0 | PDE2A_HUMAI | Unmodified | 0 | 0 | 0 | 0 | 0 | 0 | 0 | 0 | 0 | 0 | 0 | 2882200 | 0 | 0 | 0 | 0 | 0 | 0 |
| VFDGIPPP\ | RPL13AP3; RPL | Q6NVV1; P4 | R13P3_HUMAN | Unmodified | 1811100 | 1510100 | 988840 | 888490 | 996140 | 0 | 413280 | 291160 | 1287400 | 298890 | 1163000 | 11743000 | 544130 | 418220 | 0 | 1174700 | 116320 | 450420 |
| VFDGIPPP\ | RPL13AP3; RPL | Q6NVV1; P4 | R13P3_HUMAN | Unmodified | 884680 | 674580 | 0 | 0 | 0 | 0 | 0 | 166940 | 0 | 0 | 1561500 | 4168400 | 0 | 0 | 0 | 0 | 0 | 0 |
| VFDGSITK | ACE; ACE; ACE; | P12821; J3K | ACE_HUMAN A | Unmodified | 710730 | 489070 | 0 | 0 | 0 | 0 | 0 | 0 | 366410 | 0 | 156120 | 0 | 0 | 0 | 0 | 0 | 97565 | 238660 |
| VFDKDGN( | CALML3 | P27482 | CALL3_HUMAN | Unmodified | 0 | 0 | 0 | 0 | 0 | 0 | 0 | 0 | 2472800 | 0 | 811990 | 0 | 0 | 0 | 0 | 0 | 0 | 0 |
| VFDKDGN( | CALM2; CALM3 | P0DP24; P0I | CALM2_HUMAI | Unmodified | 948840 | 2813700 | 0 | 3296300 | 2183600 | 3392800 | 1189400 | 418430 | 2743000 | 493410 | 235040 | 0 | 1368400 | 424230 | 67569 | 152200 | 779510 | 1820000 |
| VFDMLNR | EIF4A1; EIF4A2; | P60842; Q14 | IF4A1_HUMAN | Unmodified | 222990 | 0 | 0 | 0 | 0 | 0 | 0 | 0 | 0 | 0 | 0 | 2751500 | 0 | 0 | 418940 | 162490 | 54641 | 0 |
| VFDPQND | ARPC3 | O15145 | ARPC3_HUMAI | Unmodified | 3926800 | 2717000 | 2645500 | 4142800 | 6379300 | 1795300 | 667500 | 476900 | 2375400 | 733670 | 1811200 | 17897000 | 4796700 | 0 | 5103900 | 3555100 | 879950 | 2773200 |
| VFDPQND | ARPC3 | O15145 | ARPC3_HUMAI | Unmodified | 0 | 0 | 0 | 0 | 4644600 | 809290 | 2710300 | 0 | 0 | 828510 | 0 | 0 | 9076000 | 5000700 | 0 | 0 | 4406000 | 3048100 |
| VFDPVPVG | DHX9 | Q08211 | DHX9_HUMAN | Unmodified | 273950 | 235370 | 220030 | 0 | 594520 | 133620 | 1307500 | 775240 | 451200 | 296240 | 1298100 | 5969900 | 183330 | 192910 | 0 | 156340 | 0 | 321990 |
| VFDQQSSF | INPPL1; INPPL1 | O15357; A0/ | SHIP2_HUMAN | Unmodified | 0 | 0 | 0 | 0 | 0 | 0 | 0 | 0 | 0 | 0 | 281710 | 939010 | 0 | 0 | 0 | 0 | 0 | 0 |
| VFEEDALS\ | DYNC1H1; DYN | Q14204; A0/ | DYHC1_HUMA | Unmodified | 0 | 0 | 0 | 0 | 0 | 0 | 0 | 0 | 0 | 0 | 0 | 776160 | 0 | 0 | 0 | 0 | 0 | 0 |
| VFEEDALS\ | DYNC1H1; DYN | Q14204; A0/ | DYHC1_HUMA | Unmodified | 0 | 0 | 48161 | 0 | 0 | 0 | 0 | 0 | 0 | 0 | 0 | 899120 | 0 | 79204 | 0 | 0 | 0 | 0 |
| VFEFFGDH | NCR3LG1 | Q68D85 | NR3L1_HUMAN | Unmodified | 0 | 0 | 0 | 0 | 0 | 0 | 0 | 0 | 1677700 | 0 | 0 | 0 | 0 | 0 | 0 | 0 | 0 | 0 |
| VFEFGGPE | CRYZ; CRYZ; CF | Q08257; A6I | QOR_HUMAN ( | Unmodified | 0 | 0 | 0 | 0 | 0 | 0 | 0 | 190430 | 1924000 | 437250 | 0 | 1542700 | 1006000 | 0 | 0 | 0 | 0 | 0 |
| VFEGPPKP | NCEH1; NCEH1 | Q6PIU2; H7C | NCEH1_HUMA | Unmodified | 0 | 0 | 0 | 0 | 0 | 0 | 0 | 0 | 1119100 | 403640 | 0 | 0 | 0 | 0 | 0 | 0 | 0 | 0 |
| VFEGQTNII | SH3D19; SH3D | Q5HYK7; A0 | SH319_HUMAN | Unmodified | 0 | 0 | 0 | 0 | 0 | 0 | 0 | 0 | 0 | 0 | 0 | 0 | 559220 | 0 | 0 | 0 | 658690 | 419310 |
| VFEHDSVE | ESD; ESD | P10768; U3K | ESTD_HUMAN | Unmodified | 399350 | 0 | 0 | 912700 | 0 | 0 | 2111100 | 1535900 | 914700 | 1057900 | 875160 | 5938900 | 2256700 | 1739700 | 970190 | 886130 | 5553700 | 2338900 |
| VFEIRNTEC | GMFB; GMFB | P60983; G3\ | GMFB_HUMAN | Unmodified | 1198400 | 0 | 0 | 0 | 8714200 | 2022300 | 3125300 | 0 | 0 | 3751200 | 911690 | 118290 | 7950300 | 10315000 | 0 | 0 | 6608100 | 7193000 |
| VFEMSQD | MCM6 | Q14566 | MCM6_HUMAN | Unmodified | 0 | 0 | 0 | 0 | 443520 | 0 | 336290 | 0 | 230730 | 185090 | 0 | 623060 | 364860 | 0 | 0 | 0 | 151560 | 0 |
| VFEMVEAL( | LLGL2; LLGL2 | Q6P1M3; J3( | L2GL2_HUMAN | Oxidation (M) | 249370 | 339840 | 0 | 424190 | 0 | 0 | 0 | 0 | 902660 | 0 | 0 | 0 | 0 | 0 | 524990 | 476660 | 0 | 0 |
| VFEVMLATI | PCMT1; PCMT1 | P22061; F8V | PIMT_HUMAN I | Unmodified | 90450 | 0 | 0 | 0 | 1664300 | 203880 | 492310 | 0 | 4527400 | 2307000 | 549070 | 2455500 | 2199400 | 3096900 | 0 | 0 | 839540 | 1067700 |
| VFEVMLATI | PCMT1; PCMT1 | P22061; F8V | PIMT_HUMAN I | Oxidation (M) | 424530 | 266520 | 338980 | 401070 | 0 | 325210 | 0 | 0 | 1515300 | 2158400 | 852420 | 2305100 | 0 | 0 | 306580 | 291090 | 0 | 0 |
| VFEVNASN | LARS1; LARS1; | Q9P2J5; B4E | SYLC_HUMAN | Unmodified | 0 | 0 | 155470 | 0 | 0 | 0 | 0 | 0 | 0 | 0 | 776390 | 9789800 | 0 | 0 | 144820 | 127330 | 0 | 0 |
| VFEVSLADI | RPS3A; RPS3A; | P61247; D6F | RS3A_HUMAN | Unmodified | 532450 | 424860 | 0 | 326520 | 0 | 0 | 64184 | 0 | 22997 | 0 | 210010 | 139050 | 54564 | 62788 | 84759 | 51312 | 0 | 0 |
| VFEVSLADI | RPS3A; RPS3A; | P61247; D6F | RS3A_HUMAN | Unmodified | 0 | 5119700 | 1144800 | 2800100 | 0 | 0 | 1720500 | 0 | 0 | 939620 | 3326500 | 24492000 | 0 | 0 | 3378300 | 1615900 | 0 | 0 |
| VFEYDRATI | SMPDL3B; SMP | Q92485; F8\ | ASM3B_HUMAI | Unmodified | 4183300 | 3118800 | 10775000 | 14350000 | 0 | 174830 | 0 | 0 | 11787000 | 502070 | 586560 | 10111000 | 595430 | 208730 | 19704000 | 14424000 | 0 | 0 |
| VFFASVR | TNIK; MINK1; M | Q9UKE5; Q8 | TNIK_HUMAN T | Unmodified | 1883200 | 0 | 282490 | 1086100 | 479540 | 0 | 119830 | 95368 | 1554800 | 1534000 | 0 | 2210200 | 374620 | 0 | 1567100 | 1104200 | 231420 | 464170 |
| VFFDLMR | RALA; RALB | P11233; P11 | RALA_HUMAN | Unmodified | 959720 | 0 | 0 | 2166900 | 3508700 | 0 | 0 | 0 | 13631000 | 501070 | 2050200 | 8153300 | 0 | 0 | 0 | 977830 | 0 | 1053800 |
| VFFDLMR | RALA; RALB | P11233; P11 | RALA_HUMAN | Oxidation (M) | 2876500 | 3333700 | 5397000 | 6285400 | 1161000 | 0 | 0 | 0 | 5770000 | 972210 | 7130000 | 22952000 | 473620 | 913190 | 8678200 | 5830500 | 126750 | 753350 |
| VFFDVDIG( | PPID | Q08752 | PPID_HUMAN I | Unmodified | 0 | 0 | 0 | 186180 | 0 | 0 | 0 | 0 | 0 | 0 | 822320 | 1045600 | 0 | 0 | 0 | 86530 | 0 | 0 |
| VFFRAGVL/ | MYH9 | P35579 | MYH9_HUMAN | Unmodified | 0 | 0 | 0 | 0 | 913470 | 0 | 1341200 | 0 | 0 | 906000 | 0 | 0 | 126630 | 265950 | 0 | 352500 | 0 | 0 |
| VFFTDYGQ | LRP1; LRP1 | Q07954; Q6I | LRP1_HUMAN | Unmodified | 0 | 0 | 0 | 0 | 0 | 0 | 0 | 0 | 2718600 | 0 | 214640 | 0 | 0 | 0 | 0 | 0 | 0 | 0 |
| VFFVYVGG | TMX3; TMX3 | Q96JJ7; B4D | TMX3_HUMAN | Unmodified | 0 | 0 | 0 | 0 | 0 | 0 | 0 | 0 | 2710100 | 0 | 0 | 85600 | 0 | 0 | 0 | 0 | 0 | 0 |
| VFGALKGA | RPL5; RPL5; RP | P46777; Q5 | RL5_HUMAN 6 | Unmodified | 0 | 0 | 0 | 0 | 0 | 0 | 0 | 0 | 0 | 0 | 0 | 0 | 869230 | 0 | 0 | 0 | 0 | 0 |
| VFGAPEVLI | NUP210 | Q8TEM1 | PO210_HUMAI | Unmodified | 0 | 0 | 0 | 0 | 0 | 0 | 0 | 0 | 31081000 | 2347800 | 588760 | 84334 | 0 | 0 | 0 | 0 | 0 | 0 |
| VFGAPNVV | NPLOC4; NPLO | Q8TAT6; A0/ | NPL4_HUMAN | Unmodified | 227430 | 284360 | 83877 | 245210 | 503910 | 110740 | 0 | 0 | 0 | 142390 | 0 | 0 | 425860 | 204470 | 347550 | 59761 | 0 | 396640 |
| VFGAPNVV | NPLOC4; NPLO | Q8TAT6; A0/ | NPL4_HUMAN | Unmodified | 15249000 | 11038000 | 20994000 | 46072000 | 10589000 | 2915300 | 0 | 0 | 1088500 | 1759700 | 0 | 12218000 | 41480000 | 2193400 | 41610000 | 60383000 | 495460 | 1028900 |
| VFGERFGG | HADHA; HADH/ | P40939; A0/ | ECHA_HUMAN | Unmodified | 0 | 0 | 0 | 0 | 0 | 0 | 399750 | 0 | 1661200 | 754810 | 0 | 0 | 0 | 201250 | 0 | 0 | 0 | 1452100 |
| VFGFSLITN | PNP | P00491 | PNPH_HUMAN | Unmodified | 0 | 0 | 0 | 852140 | 0 | 522170 | 0 | 0 | 3195600 | 512560 | 4559900 | 27680000 | 0 | 0 | 0 | 315980 | 0 | 0 |

| | | | | | | | | | | | | | | | | | | | | | | |
|---|---|---|---|---|---|---|---|---|---|---|---|---|---|---|---|---|---|---|---|---|---|---|
| VFGFVARK | TNS3 | Q68CZ2 | TENS3_HUMAN | Unmodified | 0 | 3295400 | 0 | 0 | 1471400 | 0 | 0 | 0 | 0 | 0 | 0 | 0 | 0 | 0 | 0 | 0 | 6366400 | 0 |
| VFGLQGVS | CRMP1; CRMP1 | Q14194; E9F | DPYL1_HUMAN | Unmodified | 0 | 0 | 0 | 0 | 0 | 0 | 844700 | 0 | 0 | 0 | 0 | 0 | 0 | 0 | 0 | 0 | 0 | 0 |
| VFGLQGVS | CRMP1; CRMP1 | Q14194; E9F | DPYL1_HUMAN | Oxidation (M) | 0 | 0 | 0 | 0 | 0 | 0 | 0 | 939650 | 0 | 362960 | 0 | 0 | 0 | 0 | 0 | 0 | 0 | 0 |
| VFGNEIKLE | NCL; NCL; NCL | P19338; C9J | NUCL_HUMAN | Unmodified | 0 | 0 | 0 | 0 | 0 | 0 | 0 | 0 | 0 | 0 | 3185600 | 7669600 | 719940 | 95297 | 0 | 0 | 0 | 0 |
| VFGPGIEGI | FLNB; FLNB | O75369; E7E | FLNB_HUMAN | Unmodified | 47044 | 0 | 1606200 | 221820 | 0 | 912990 | 0 | 0 | 364330 | 57757 | 275260 | 3404500 | 423040 | 0 | 702370 | 638760 | 213090 | 897120 |
| VFGPGIEGI | FLNB; FLNB | O75369; E7E | FLNB_HUMAN | Unmodified | 0 | 0 | 31984 | 0 | 0 | 773880 | 0 | 0 | 0 | 0 | 52059 | 1499900 | 0 | 0 | 0 | 0 | 496370 | 0 |
| VFGPGVER | FLNB; FLNB | O75369; E7E | FLNB_HUMAN | Unmodified | 0 | 0 | 0 | 0 | 0 | 0 | 0 | 0 | 0 | 0 | 0 | 4746900 | 0 | 0 | 0 | 0 | 0 | 0 |
| VFGPGVER | FLNB; FLNB | O75369; E7E | FLNB_HUMAN | Unmodified | 0 | 38675 | 82822 | 0 | 8922400 | 2408000 | 391600 | 45517 | 295330 | 344520 | 98201 | 0 | 1165400 | 571120 | 0 | 0 | 3080200 | 1665700 |
| VFGPTIVAH | RACGAP1; RAC | Q9H0H5; HC | RGAP1_HUMAI | Oxidation (M) | 0 | 0 | 514440 | 0 | 2176300 | 1561900 | 0 | 0 | 2578800 | 505520 | 2193300 | 2024100 | 0 | 0 | 1593200 | 0 | 0 | 0 |
| VFGPTIVAH | RACGAP1; RAC | Q9H0H5; HC | RGAP1_HUMAI | Unmodified | 245710 | 0 | 0 | 0 | 1687800 | 584780 | 0 | 0 | 8938000 | 570990 | 432220 | 0 | 0 | 0 | 0 | 0 | 0 | 0 |
| VFGPTIVAH | RACGAP1; RAC | Q9H0H5; HC | RGAP1_HUMAI | Unmodified | 0 | 0 | 0 | 0 | 4287400 | 811700 | 0 | 0 | 2718900 | 0 | 0 | 0 | 0 | 0 | 0 | 0 | 2473500 | 35909 |
| VFGWVHR | NARS1; NARS1; | O43776; K7E | SYNC_HUMAN | Unmodified | 0 | 0 | 0 | 0 | 0 | 0 | 0 | 0 | 0 | 0 | 0 | 659860 | 0 | 0 | 0 | 0 | 0 | 0 |
| VFHLLGVD | PNP; PNP; PNP | P00491; G3\ | PNPH_HUMAN | Unmodified | 0 | 0 | 0 | 0 | 0 | 0 | 0 | 0 | 0 | 0 | 0 | 2260600 | 0 | 0 | 0 | 100100 | 0 | 0 |
| VFHLPTTTF | OGDH; OGDH; | Q02218; E9F | ODO1_HUMAN | Unmodified | 0 | 0 | 0 | 0 | 0 | 0 | 0 | 0 | 6658900 | 109720 | 0 | 0 | 0 | 0 | 0 | 0 | 0 | 0 |
| VFIDKQTNI | CELF1; CELF2; | Q92879; O9! | CELF1_HUMAN | Unmodified | 0 | 0 | 0 | 0 | 0 | 0 | 944560 | 249710 | 0 | 0 | 0 | 0 | 0 | 0 | 0 | 0 | 0 | 0 |
| VFIGKDCIG | GLRX | P35754 | GLRX1_HUMAI | Unmodified | 0 | 0 | 0 | 0 | 0 | 4144100 | 0 | 0 | 0 | 0 | 0 | 0 | 0 | 0 | 0 | 0 | 1875200 | 0 |
| VFIGNLNT/ | RALY; RALY; RA | Q9UKM9; Q5 | RALY_HUMAN | Unmodified | 0 | 0 | 118960 | 208700 | 0 | 0 | 649560 | 546990 | 509620 | 398380 | 0 | 1691300 | 320820 | 264080 | 0 | 0 | 0 | 537980 |
| VFIGNLNTI | HNRNPCL3; HN | B7ZW38; P0I | HNRC3_HUMA | Unmodified | 1678400 | 1126200 | 1431100 | 946880 | 3993100 | 2171800 | 7485500 | 4085100 | 6202400 | 4032800 | 7227500 | 24492000 | 4235500 | 1408300 | 127560 | 90447 | 76520 | 3133900 |
| VFIGNLNTI | HNRNPCL3; HN | B7ZW38; P0I | HNRC3_HUMA | Unmodified | 0 | 0 | 130970 | 0 | 1547500 | 323580 | 1856000 | 708810 | 1346500 | 1518800 | 2040600 | 9014900 | 1272100 | 0 | 0 | 0 | 0 | 1182700 |
| VFIGNLNTI | HNRNPCL3; HN | B7ZW38; P0I | HNRC3_HUMA | Unmodified | 0 | 0 | 0 | 0 | 0 | 0 | 8258600 | 0 | 0 | 0 | 0 | 0 | 934560 | 0 | 0 | 0 | 0 | 0 |
| VFIPHGLIM | HPRT1 | P00492 | HPRT_HUMAN | Oxidation (M) | 22564 | 35924 | 0 | 0 | 0 | 0 | 0 | 0 | 0 | 0 | 0 | 481930 | 0 | 0 | 0 | 0 | 0 | 0 |
| VFIQRDYS | GOLGA7 | Q7Z5G4 | GOGA7_HUMA | Unmodified | 109070 | 185660 | 0 | 0 | 1010800 | 0 | 38184 | 49992 | 618780 | 163080 | 27836 | 0 | 0 | 0 | 0 | 0 | 772900 | 0 |
| VFITDDFHI | HSP90B1; HSP! | P14625; A0A | ENPL_HUMAN | Unmodified | 0 | 0 | 0 | 0 | 0 | 0 | 170180 | 0 | 39706000 | 3558100 | 559990 | 340560 | 0 | 0 | 0 | 0 | 0 | 0 |
| VFITDDFHI | HSP90B1; HSP! | P14625; A0A | ENPL_HUMAN | 2 Oxidation (M) | 0 | 0 | 0 | 0 | 0 | 0 | 431120 | 0 | 7299100 | 4712400 | 6754400 | 4862700 | 0 | 0 | 0 | 0 | 0 | 0 |
| VFITDDFHI | HSP90B1; HSP! | P14625; A0A | ENPL_HUMAN | Oxidation (M) | 0 | 0 | 0 | 0 | 0 | 0 | 376680 | 41788 | 43333000 | 16913000 | 11587000 | 3600200 | 0 | 0 | 0 | 0 | 0 | 0 |
| VFIYHGSAI | ITGA6 | P23229 | ITA6_HUMAN I | Unmodified | 0 | 0 | 0 | 0 | 0 | 0 | 0 | 0 | 0 | 0 | 0 | 329860 | 0 | 0 | 137330 | 126360 | 240450 | 0 |
| VFIYHGSAI | ITGA6 | P23229 | ITA6_HUMAN I | Unmodified | 493080 | 0 | 3527900 | 5581800 | 39342000 | 9105900 | 0 | 0 | 0 | 494990 | 1081800 | 18316000 | 22994000 | 14447000 | 6882600 | 7796300 | 52425000 | 2.64E+08 |
| VFIYHGSSI | ITGA7; ITGA7 | Q13683; J3K | ITA7_HUMAN I | Unmodified | 0 | 0 | 0 | 0 | 0 | 0 | 1427300 | 0 | 2982000 | 2020700 | 0 | 0 | 0 | 0 | 0 | 0 | 0 | 0 |
| VFKDFSELE | PYGL; PYGL | P06737; E9P | PYGL_HUMAN | Unmodified | 0 | 0 | 0 | 0 | 1825000 | 610080 | 0 | 0 | 0 | 0 | 0 | 0 | 0 | 0 | 0 | 0 | 655160 | 3658800 |
| VFKEAVQG | CACNA2D2; CA | Q9NY47; C9 | CA2D2_HUMAI | Unmodified | 0 | 0 | 0 | 0 | 0 | 0 | 0 | 81103 | 0 | 123760 | 0 | 0 | 0 | 0 | 0 | 0 | 0 | 0 |
| VFKQPSKPI | ALCAM; ALCAM | Q13740; F5C | CD166_HUMAI | Unmodified | 2390800 | 2060500 | 0 | 858920 | 5603800 | 946270 | 1405200 | 605780 | 3789700 | 742470 | 0 | 0 | 827600 | 1738400 | 504850 | 347610 | 3155300 | 5925200 |
| VFLADQPIE | UTRN; UTRN | P46939; Q5T | UTRN_HUMAN | Unmodified | 0 | 178770 | 985670 | 1716000 | 112930 | 0 | 0 | 0 | 2721200 | 429110 | 0 | 0 | 0 | 0 | 2513700 | 2731200 | 238030 | 0 |
| VFLATNSD | NT5C2; NT5C2; | P49902; A0A | 5NTC_HUMAN | Unmodified | 0 | 0 | 0 | 291850 | 366960 | 0 | 54046 | 0 | 115660 | 256190 | 0 | 1095100 | 0 | 0 | 132560 | 148120 | 377130 | 425430 |
| VFLDPNILS | PREP; PREP | P48147; A0A | PPCE_HUMAN | Unmodified | 0 | 0 | 0 | 0 | 0 | 0 | 0 | 0 | 213810 | 0 | 0 | 735630 | 0 | 0 | 0 | 0 | 0 | 0 |
| VFLEFCK | MOV10; MOV10 | Q9HCE1; Q5 | MOV10_HUMA | Unmodified | 496500 | 396370 | 307020 | 182400 | 0 | 0 | 0 | 0 | 0 | 0 | 0 | 0 | 0 | 0 | 417800 | 937070 | 0 | 0 |
| VFLENVIR | H4C1 | P62805 | H4_HUMAN Hi | Unmodified | 5.28E+08 | 3.34E+08 | 1662800 | 631140 | 1494000 | 1646400 | 1545100 | 3240700 | 12240000 | 4115400 | 11436000 | 3.88E+09 | 374450 | 1007900 | 0 | 0 | 1096800 | 1470400 |
| VFLENVIRD | H4C1 | P62805 | H4_HUMAN Hi | Unmodified | 1.31E+09 | 1.33E+09 | 4895500 | 3141800 | 94340000 | 8294400 | 3.69E+08 | 643460 | 26186000 | 61720000 | 12065000 | 2.83E+09 | 9191200 | 12543000 | 0 | 2514400 | 1488700 | 28873000 |
| VFLENVIRD | H4C1 | P62805 | H4_HUMAN Hi | Unmodified | 1.21E+08 | 1.73E+08 | 0 | 0 | 0 | 0 | 1081800 | 0 | 4824100 | 0 | 766000 | 5.12E+08 | 0 | 0 | 0 | 148510 | 0 | 0 |
| VFLGNVETI | ELP1; ELP1; ELF | O95163; A0A | ELP1_HUMAN | Unmodified | 0 | 0 | 0 | 0 | 0 | 0 | 0 | 0 | 0 | 0 | 0 | 2889000 | 0 | 0 | 0 | 0 | 0 | 0 |
| VFLISASSK | RANBP3; RANB | Q9H6Z4; B7 | RANB3_HUMAI | Unmodified | 0 | 0 | 0 | 0 | 114230 | 0 | 2085000 | 0 | 185610 | 237910 | 136140 | 3856200 | 70792 | 0 | 0 | 0 | 0 | 943740 |
| VFLLGEEV/ | PDHB; PDHB; P | P11177; F8V | ODPB_HUMAN | Unmodified | 0 | 0 | 0 | 0 | 0 | 0 | 0 | 0 | 3109000 | 969740 | 0 | 0 | 0 | 0 | 0 | 0 | 0 | 0 |
| VFLLGQTPI | B3GNT2 | Q9NY97 | B3GN2_HUMAI | Oxidation (M) | 0 | 0 | 0 | 0 | 0 | 0 | 0 | 0 | 1283000 | 0 | 506090 | 350360 | 0 | 0 | 0 | 0 | 0 | 0 |
| VFLLGQTPI | B3GNT2 | Q9NY97 | B3GN2_HUMAI | Unmodified | 0 | 0 | 0 | 0 | 50654 | 0 | 1483100 | 0 | 3341900 | 475780 | 0 | 78657 | 0 | 0 | 0 | 0 | 0 | 1202000 |
| VFLLPDK | SYT2; SYT1; SY | Q8N9I0; P21 | SYT2_HUMAN | Unmodified | 332010 | 0 | 169840 | 204850 | 0 | 0 | 0 | 0 | 2521900 | 665780 | 0 | 0 | 0 | 0 | 216490 | 340790 | 0 | 0 |
| VFLNNLCP | PLCB1; PLCB1; | Q9NQ66; A0 | PLCB1_HUMAN | Unmodified | 1312900 | 310050 | 0 | 0 | 0 | 0 | 0 | 0 | 311010 | 0 | 0 | 0 | 0 | 0 | 143930 | 77495 | 0 | 0 |
| VFLPKEELY | MYOF; MYOF | Q9NZM1; HC | MYOF_HUMAN | Oxidation (M) | 0 | 0 | 0 | 0 | 58371000 | 7478700 | 0 | 0 | 0 | 0 | 0 | 0 | 0 | 0 | 0 | 0 | 267850 | 4282100 |
| VFLPKEELY | MYOF; MYOF | Q9NZM1; HC | MYOF_HUMAN | Unmodified | 0 | 0 | 0 | 0 | 24696000 | 0 | 0 | 0 | 0 | 0 | 0 | 0 | 0 | 0 | 0 | 0 | 357360 | 2019900 |
| VFLQNLLS | FAH; FAH | P16930; H0Y | FAAA_HUMAN | Unmodified | 0 | 0 | 25250 | 0 | 0 | 0 | 28950 | 0 | 0 | 0 | 0 | 2138100 | 0 | 0 | 0 | 0 | 0 | 0 |
| VFLQYLPAI | GNA13 | Q14344 | GNA13_HUMAI | Unmodified | 165260 | 0 | 0 | 104270 | 0 | 0 | 0 | 0 | 5952600 | 954240 | 3674500 | 15977000 | 0 | 0 | 0 | 0 | 0 | 0 |
| VFLSDEND | DCHS1 | Q96JQ0 | PCD16_HUMAI | Unmodified | 0 | 0 | 0 | 0 | 0 | 0 | 0 | 0 | 1333300 | 0 | 0 | 0 | 0 | 0 | 0 | 0 | 0 | 0 |
| VFMQEFK | ARPC2 | O15144 | ARPC2_HUMAI | Unmodified | 764730 | 403880 | 363580 | 440600 | 0 | 368170 | 110330 | 40045 | 1396900 | 184920 | 490990 | 12738000 | 290630 | 141950 | 0 | 492710 | 0 | 0 |
| VFMQEFK | ARPC2 | O15144 | ARPC2_HUMAI | Oxidation (M) | 357130 | 407150 | 808360 | 417890 | 0 | 191500 | 0 | 21451 | 252390 | 0 | 96225 | 2559200 | 0 | 0 | 1372300 | 710070 | 0 | 0 |
| VFNAGDDI | PPP1R3D | O95685 | PPR3D_HUMAI | Unmodified | 0 | 0 | 0 | 0 | 0 | 0 | 0 | 0 | 463900 | 0 | 0 | 391830 | 0 | 0 | 0 | 0 | 78871 | 0 |
| VFNDCRDI | GNAS; GNAS; G | Q5JWF2; P6 | GNAS1_HUMAI | Unmodified | 199150 | 105690 | 696210 | 1048900 | 1326000 | 1015600 | 89271 | 0 | 444260 | 133820 | 824400 | 2136700 | 1153600 | 523820 | 874290 | 1050300 | 770670 | 1762300 |
| VFNDYTAVI | ECE1; ECE1; EC | P42892; B4D | ECE1_HUMAN | Oxidation (M) | 0 | 0 | 0 | 0 | 0 | 0 | 0 | 0 | 5045500 | 0 | 0 | 0 | 0 | 20114 | 0 | 0 | 0 | 0 |
| VFNDYTAVI | ECE1; ECE1; EC | P42892; B4D | ECE1_HUMAN | Unmodified | 0 | 0 | 0 | 0 | 0 | 0 | 0 | 0 | 1978900 | 0 | 0 | 0 | 0 | 0 | 23254 | 0 | 0 | 0 |
| VFNETPINF | COPG1 | Q9Y678 | COPG1_HUMA | Unmodified | 0 | 0 | 0 | 0 | 0 | 0 | 0 | 0 | 0 | 0 | 298560 | 2456100 | 0 | 0 | 0 | 0 | 0 | 0 |
| VFNFSTSH | PSME4 | Q14997 | PSME4_HUMAI | Unmodified | 0 | 0 | 0 | 0 | 0 | 0 | 0 | 0 | 0 | 0 | 0 | 599480 | 0 | 0 | 0 | 0 | 0 | 0 |
| VFNGYMR | ST14 | Q9Y5Y6 | ST14_HUMAN | Unmodified | 354040 | 215330 | 1026000 | 527770 | 0 | 0 | 0 | 0 | 259880 | 0 | 0 | 0 | 126060 | 0 | 2624600 | 632620 | 0 | 0 |
| VFNLYPR | DPYSL2; DPYSL | Q16555; A0/ | DPYL2_HUMAN | Unmodified | 130500 | 0 | 0 | 0 | 0 | 0 | 0 | 178210 | 418930 | 0 | 0 | 4357500 | 0 | 0 | 131080 | 84255 | 0 | 0 |
| VFNNISFLF | TBK1; TBK1; TBI | Q9UHD2; AC | TBK1_HUMAN | Oxidation (M) | 0 | 0 | 0 | 0 | 0 | 0 | 0 | 0 | 215020 | 0 | 0 | 0 | 0 | 0 | 0 | 126230 | 0 | 0 |
| VFNPYTEFI | EFHD2 | Q96C19 | EFHD2_HUMAI | Unmodified | 0 | 0 | 0 | 0 | 5013200 | 2306900 | 0 | 0 | 1686200 | 0 | 197030 | 0 | 497020 | 0 | 0 | 0 | 0 | 0 |
| VFNTGGAP | CAD; CAD | P27708; F8V | PYR1_HUMAN | Unmodified | 0 | 0 | 0 | 0 | 0 | 66015 | 0 | 0 | 0 | 73620 | 0 | 750810 | 0 | 0 | 50070 | 0 | 291090 | 186800 |
| VFNTTPDD | RTCB | Q9Y3I0 | RTCB_HUMAN | Unmodified | 0 | 0 | 1311100 | 0 | 772770 | 0 | 0 | 0 | 0 | 164630 | 2876700 | 8636300 | 1987900 | 0 | 0 | 0 | 0 | 1170800 |
| VFNTTPDD | RTCB | Q9Y3I0 | RTCB_HUMAN | Unmodified | 0 | 82818 | 0 | 0 | 957330 | 48211 | 804490 | 0 | 0 | 149460 | 0 | 0 | 1385000 | 1186400 | 75564 | 0 | 501280 | 669960 |
| VFNTTPDD | RTCB | Q9Y3I0 | RTCB_HUMAN | Unmodified | 62190 | 162930 | 0 | 842400 | 1045500 | 133930 | 3207900 | 0 | 0 | 354950 | 0 | 0 | 1318100 | 1098700 | 0 | 0 | 1381900 | 1393000 |
| VFNVFCLY | HNRNPL; HNRI | P14866; M0 | HNRPL_HUMAI | Unmodified | 0 | 0 | 0 | 0 | 0 | 0 | 0 | 0 | 0 | 0 | 77749 | 1211700 | 0 | 0 | 0 | 0 | 0 | 0 |
| VFNYNTLEI | COPB2; COPB2 | P35606; D6F | COPB2_HUMAI | Unmodified | 0 | 0 | 0 | 0 | 0 | 0 | 0 | 0 | 0 | 0 | 0 | 0 | 0 | 0 | 0 | 0 | 0 | 0 |
| VFPLSCAV | MPI; MPI; MPI; I | P34949; H3E | MPI_HUMAN M | Unmodified | 0 | 0 | 0 | 120610 | 1258700 | 0 | 0 | 0 | 104150 | 101090 | 266020 | 948760 | 578120 | 184240 | 266230 | 219180 | 133590 | 228870 |
| VFPPFHLVI | PTPRB; PTPRB; | P23467; F8V | PTPRB_HUMAN | Unmodified | 2386600 | 2620100 | 0 | 0 | 36176 | 0 | 0 | 0 | 0 | 0 | 0 | 0 | 0 | 0 | 0 | 0 | 0 | 34140 |
| VFQASPED | MRPL30; MRPL | Q8TCC3; B8 | RM30_HUMAN | Unmodified | 0 | 0 | 0 | 0 | 0 | 0 | 0 | 0 | 558690 | 67767 | 0 | 0 | 0 | 0 | 0 | 0 | 0 | 0 |
| VFQDKFDV | AVIL | O75366 | AVIL_HUMAN A | Unmodified | 0 | 0 | 0 | 0 | 0 | 0 | 0 | 0 | 0 | 0 | 6750400 | 77545000 | 0 | 0 | 0 | 0 | 0 | 0 |
| VFQEALR | UNC45A; UNC4 | Q9H3U1; A0 | UN45A_HUMAI | Unmodified | 0 | 0 | 0 | 0 | 0 | 0 | 0 | 0 | 0 | 0 | 0 | 0 | 0 | 0 | 0 | 0 | 0 | 0 |
| VFQEALRN | UNC45A; UNC4 | Q9H3U1; A0 | UN45A_HUMAI | Unmodified | 513240 | 0 | 0 | 0 | 5952200 | 402930 | 1056500 | 949080 | 0 | 281060 | 0 | 853270 | 2335200 | 1928600 | 0 | 0 | 1902500 | 2963500 |
| VFQERINCI | UBXN6; UBXN6 | Q9BZV1; K7E | UBXN6_HUMAI | Unmodified | 0 | 0 | 0 | 0 | 0 | 0 | 0 | 0 | 0 | 0 | 0 | 0 | 0 | 2188500 | 0 | 0 | 0 | 0 |
| VFQFLNAK | RPL24; RPL24; I | P83731; C9J | RL24_HUMAN | Unmodified | 6890400 | 7057800 | 4510300 | 3110200 | 2550600 | 449880 | 2525300 | 2081900 | 2590300 | 1393600 | 3769200 | 63914000 | 2304000 | 1361100 | 5603000 | 4085700 | 0 | 1623000 |
| VFQFLNAK | RPL24; RPL24; I | P83731; C9J | RL24_HUMAN | Unmodified | 0 | 0 | 0 | 0 | 405710 | 100360 | 1725900 | 232320 | 0 | 0 | 0 | 0 | 1267300 | 620690 | 0 | 0 | 154660 | 419490 |
| VFQGCGPI | GPC4 | O75487 | GPC4_HUMAN | Unmodified | 755960 | 626320 | 249750 | 266370 | 0 | 0 | 0 | 0 | 0 | 259180 | 0 | 0 | 849140 | 342640 | 0 | 138570 | 0 | 0 |
| VFQGNFDI | EDIL3 | O43854 | EDIL3_HUMAN | Unmodified | 140050 | 125010 | 0 | 0 | 1836600 | 847710 | 19676 | 17025 | 1326000 | 76732 | 0 | 0 | 0 | 0 | 0 | 0 | 58180 | 936500 |
| VFQGNFDI | EDIL3 | O43854 | EDIL3_HUMAN | Unmodified | 839970 | 862240 | 0 | 0 | 1.91E+08 | 71240000 | 4869400 | 1636800 | 29562000 | 11486000 | 0 | 102560 | 0 | 0 | 0 | 0 | 5664700 | 90992000 |
| VFQGNFDI | EDIL3 | O43854 | EDIL3_HUMAN | Unmodified | 0 | 0 | 0 | 0 | 45010000 | 7757400 | 2536900 | 0 | 0 | 1706300 | 0 | 0 | 0 | 0 | 0 | 0 | 7486600 | 25524000 |
| VFQGTDEC | RAD50; N/A; RA | Q92878; A0A | RAD50_HUMAN | Unmodified | 0 | 0 | 0 | 0 | 0 | 0 | 0 | 0 | 0 | 0 | 0 | 1742200 | 0 | 0 | 0 | 0 | 0 | 0 |
| VFQSLPHE | CSTB; CSTB | P04080; A0A | CYTB_HUMAN | Unmodified | 0 | 0 | 0 | 0 | 0 | 0 | 0 | 0 | 1564000 | 0 | 0 | 6002400 | 0 | 0 | 6841900 | 3780600 | 802120 | 0 |
| VFQSSTSQ | KIF5B | P33176 | KINH_HUMAN | Unmodified | 0 | 0 | 0 | 0 | 0 | 0 | 0 | 0 | 0 | 0 | 0 | 1356300 | 0 | 0 | 0 | 0 | 0 | 0 |
| VFQSSTSQ | KIF5B | P33176 | KINH_HUMAN | Unmodified | 0 | 0 | 0 | 0 | 0 | 0 | 0 | 0 | 0 | 0 | 0 | 498440 | 0 | 0 | 0 | 0 | 0 | 0 |
| VFQVEYAM | PSMA3; PSMA3 | P25788; G3\ | PSA3_HUMAN | Unmodified | 153060 | 307240 | 0 | 0 | 0 | 0 | 907270 | 419470 | 1512700 | 92729 | 0 | 5835100 | 0 | 0 | 0 | 481330 | 0 | 0 |
| VFQVEYAM | PSMA3; PSMA3 | P25788; G3\ | PSA3_HUMAN | Oxidation (M) | 0 | 0 | 0 | 0 | 0 | 0 | 0 | 0 | 284930 | 275130 | 389160 | 2319500 | 0 | 0 | 0 | 0 | 0 | 0 |
| VFRASESQ | MIA3; MIA3 | Q5JRA6; A0A | TGO1_HUMAN | Unmodified | 317060 | 0 | 49831 | 0 | 69490 | 0 | 323730 | 49764 | 1952600 | 3068100 | 526510 | 694360 | 0 | 150840 | 0 | 0 | 0 | 201790 |
| VFSANSTA | PRPSAP1 | Q14558 | KPRA_HUMAN | Unmodified | 0 | 0 | 315200 | 0 | 341490 | 41021 | 13597 | 59207 | 15224000 | 743460 | 0 | 0 | 0 | 107350 | 0 | 0 | 485970 | 435360 |
| VFSATLGL | PARP1; PARP1 | P09874; A0A | PARP1_HUMAN | Unmodified | 318970 | 0 | 0 | 0 | 0 | 0 | 397100 | 0 | 0 | 628820 | 0 | 0 | 0 | 0 | 0 | 0 | 0 | 0 |
| VFSDEVQC | EIF3L; EIF3L; EI | Q9Y262; B0 | EIF3L_HUMAN | Unmodified | 445610 | 374510 | 1097800 | 1146900 | 0 | 0 | 0 | 0 | 0 | 0 | 0 | 16240000 | 381850 | 0 | 597380 | 716730 | 0 | 0 |
| VFSGLVST | EEF2 | P13639 | EF2_HUMAN E | Unmodified | 4655400 | 5947100 | 8222900 | 5856200 | 5157500 | 2027100 | 1402100 | 712620 | 3361200 | 0 | 17930000 | 98509000 | 98182 | 0 | 6707500 | 7282600 | 1556800 | 1335300 |
| VFSIGPVFR | DARS1 | P14868 | SYDC_HUMAN | Unmodified | 0 | 0 | 0 | 0 | 0 | 0 | 0 | 0 | 0 | 0 | 0 | 9324800 | 0 | 0 | 0 | 0 | 0 | 0 |
| VFSIGPVFR | DARS1 | P14868 | SYDC_HUMAN | Unmodified | 0 | 57751 | 473250 | 181220 | 0 | 0 | 0 | 0 | 446520 | 92377 | 280210 | 3203300 | 83867 | 0 | 171000 | 126650 | 0 | 0 |
| VFSLDVPD | ELFN2 | Q5R3F8 | PPR29_HUMAN | Unmodified | 0 | 0 | 0 | 0 | 0 | 0 | 0 | 0 | 0 | 0 | 0 | 0 | 0 | 0 | 0 | 0 | 0 | 0 |
| VFSLDVPD | ELFN2 | Q5R3F8 | PPR29_HUMAN | Unmodified | 0 | 206630 | 0 | 382280 | 3499700 | 0 | 0 | 0 | 0 | 257850 | 0 | 0 | 0 | 0 | 174530 | 108700 | 766770 | 0 |
| VFSLIQQA | PARD3; PARD3; | Q8TEW0; Q5 | PARD3_HUMAI | Unmodified | 0 | 0 | 0 | 0 | 211170 | 0 | 0 | 0 | 0 | 44016 | 0 | 0 | 0 | 0 | 0 | 0 | 140760 | 0 |
| VFSLSSEFK | SNRNP200; SN | O75643; A0/ | U520_HUMAN | Unmodified | 322800 | 0 | 0 | 0 | 0 | 0 | 0 | 0 | 0 | 0 | 0 | 4229700 | 0 | 0 | 0 | 396140 | 88473 | 0 |
| VFSSAEAL | UBE3A; UBE3A; | Q05086; A0/ | UBE3A_HUMAN | Unmodified | 314880 | 346480 | 0 | 177370 | 646030 | 222470 | 0 | 0 | 961690 | 305230 | 393010 | 1788000 | 916010 | 0 | 178550 | 382510 | 4709500 | 331160 |
| VFSVAITPD | COL6A1; COL6 | P12109; A0A | CO6A1_HUMAI | Unmodified | 0 | 835980 | 1541000 | 1110000 | 742060 | 1166000 | 980420 | 360340 | 7469700 | 2655900 | 85237 | 874960 | 1276900 | 1334500 | 902430 | 709420 | 305430 | 6201000 |
| VFSVLREE | PPP3CA; PPP3C | Q08209; P1 | PP2BA_HUMAN | Unmodified | 0 | 0 | 0 | 501540 | 437670 | 0 | 153610 | 0 | 0 | 136880 | 276780 | 1186100 | 2655600 | 1334400 | 411270 | 277570 | 486720 | 217390 |
| VFSWGFG | RCC2 | Q9P258 | RCC2_HUMAN | Unmodified | 0 | 0 | 0 | 0 | 0 | 0 | 0 | 0 | 0 | 0 | 0 | 4631400 | 0 | 0 | 0 | 0 | 0 | 0 |
| VFSWLQQI | GART | P22102 | PUR2_HUMAN | Oxidation (M) | 0 | 0 | 0 | 0 | 0 | 0 | 0 | 0 | 0 | 0 | 20206 | 3439300 | 0 | 0 | 336610 | 0 | 0 | 0 |
| VFTAGSEEI | PTK7; PTK7; PTI | Q13308; Q8 | PTK7_HUMAN | Unmodified | 0 | 91223 | 157440 | 120030 | 0 | 0 | 56340 | 0 | 2123100 | 0 | 0 | 0 | 0 | 0 | 192720 | 301640 | 0 | 0 |
| VFTAIANQF | PSMD5 | Q16401 | PSMD5_HUMA | Unmodified | 0 | 0 | 0 | 0 | 159780 | 0 | 0 | 0 | 0 | 0 | 0 | 672940 | 0 | 0 | 0 | 0 | 75703 | 51109 |
| VFTASCDK | RAE1; RAE1; RA | P78406; E9P | RAE1L_HUMAN | Unmodified | 148930 | 0 | 0 | 0 | 326040 | 60905 | 104300 | 103290 | 0 | 33316 | 0 | 946370 | 82391 | 97197 | 0 | 0 | 296450 | 261080 |
| VFTDVPSN | CAPN5; CAPN5 | O15484; E7E | CAN5_HUMAN | Unmodified | 300460 | 0 | 370990 | 457350 | 0 | 117450 | 77253 | 68144 | 0 | 0 | 0 | 0 | 0 | 284660 | 317270 | 486660 | 0 | 0 |
| VFTESEDR | PLAA; PLAA | Q9Y263; E5F | PLAP_HUMAN | Unmodified | 0 | 0 | 0 | 578240 | 0 | 0 | 0 | 0 | 1149500 | 33251 | 179780 | 4032000 | 0 | 0 | 366870 | 389280 | 0 | 0 |
| VFTFSVGQ | CACNA2D2; CA | Q9NY47; C9 | CA2D2_HUMAI | Unmodified | 0 | 0 | 0 | 0 | 0 | 0 | 0 | 0 | 2194200 | 0 | 0 | 0 | 0 | 0 | 0 | 0 | 0 | 0 |

| | | | | | | | | | | | | | | | | | | | | | | |
|---|---|---|---|---|---|---|---|---|---|---|---|---|---|---|---|---|---|---|---|---|---|---|
| VFTTGFSR | CORO1C; COR( | Q9ULV4; Q9 | COR1C_HUMA | Unmodified | 0 | 2715200 | 0 | 1457300 | 3258700 | 0 | 0 | 31677 | 430490 | 622120 | 1220300 | 7898600 | 2898300 | 3222800 | 0 | 83571 | 1240000 | 277560 |
| VFTTQELV( | AHSA1; AHSA1; | O95433; H0Y | AHSA1_HUMAI | Unmodified | 135160 | 77344 | 55136 | 0 | 0 | 0 | 0 | 0 | 0 | 0 | 351490 | 3305300 | 0 | 0 | 498880 | 384470 | 0 | 0 |
| VFTTQELV( | AHSA1; AHSA1; | O95433; H0Y | AHSA1_HUMAI | Unmodified | 0 | 0 | 0 | 0 | 0 | 0 | 0 | 0 | 2274400 | 83854 | 1698400 | 12269000 | 0 | 0 | 0 | 4803200 | 0 | 0 |
| VFTTVGSAI | FASN; FASN | P49327; A0A | FAS_HUMAN Fa | Unmodified | 0 | 0 | 519510 | 0 | 2794900 | 0 | 0 | 0 | 0 | 196300 | 553800 | 14560000 | 639250 | 0 | 1247600 | 1800200 | 0 | 0 |
| VFVANPNK | CAB39; CAB39 | Q9Y376; A0A | CAB39_HUMAI | Unmodified | 0 | 0 | 4944300 | 3687000 | 0 | 0 | 60232 | 0 | 2548500 | 74626 | 4622800 | 35915000 | 0 | 0 | 6326300 | 7338100 | 0 | 0 |
| VFVANPNK | CAB39; CAB39 | Q9Y376; A0A | CAB39_HUMAI | Unmodified | 0 | 68926 | 0 | 0 | 647490 | 208570 | 493810 | 18973 | 104690 | 399050 | 79909 | 202730 | 161000 | 156070 | 0 | 0 | 1175800 | 98074 |
| VFVEEQVY | ALDH1A3; ALDI | P47895; H0Y | AL1A3_HUMAN | Unmodified | 0 | 0 | 0 | 0 | 0 | 0 | 0 | 0 | 0 | 0 | 0 | 0 | 0 | 0 | 0 | 0 | 3748700 | 3690400 |
| VFVGGLSP | HNRNPDL; HNI | O14979; A0A | HNRDL_HUMA | Unmodified | 0 | 0 | 0 | 0 | 656930 | 125690 | 0 | 294150 | 2106900 | 1064000 | 4268400 | 14930000 | 0 | 0 | 0 | 0 | 58697 | 339480 |
| VFVGYNST | SYT1; SYT1 | P21579; J3K | SYT1_HUMAN S | Unmodified | 0 | 0 | 0 | 0 | 0 | 0 | 0 | 0 | 3663100 | 772930 | 0 | 0 | 0 | 0 | 0 | 0 | 0 | 0 |
| VFVHYTGW | FKBP4; FKBP4 | Q02790; F5H | FKBP4_HUMAN | Unmodified | 0 | 0 | 0 | 0 | 0 | 0 | 0 | 0 | 0 | 0 | 0 | 1807600 | 0 | 0 | 0 | 0 | 440700 | 0 |
| VFVLPCIQ( | FLOT1; FLOT1; I | O75955; A2A | FLOT1_HUMAN | Unmodified | 199000 | 0 | 0 | 783550 | 238900 | 328600 | 223020 | 0 | 854330 | 199170 | 0 | 934530 | 343830 | 0 | 1424000 | 1105000 | 0 | 0 |
| VFVSPLAK | DLAT; DLAT; DL | P10515; A0A | ODP2_HUMAN | Unmodified | 0 | 234340 | 0 | 0 | 0 | 0 | 49323 | 254910 | 1678100 | 84471 | 0 | 17932000 | 0 | 0 | 0 | 0 | 0 | 0 |
| VFVVGESY | ITPK1; ITPK1; IT | Q13572; G3V | ITPK1_HUMAN | Unmodified | 0 | 0 | 0 | 0 | 2070200 | 249550 | 253570 | 60826 | 0 | 154280 | 231200 | 1245400 | 1335600 | 860640 | 74697 | 67052 | 1014500 | 0 |
| VFVVGVGN | SCP2; SCP2 | P22307; E9P | SCP2_HUMAN | Unmodified | 0 | 0 | 0 | 0 | 178480 | 0 | 0 | 0 | 3021700 | 437900 | 0 | 0 | 0 | 0 | 0 | 0 | 361460 | 61467 |
| VFWDPSVI | KDM1A; KDM1A | O60341; R4( | KDM1A_HUMA | Unmodified | 0 | 0 | 0 | 141530 | 0 | 0 | 0 | 0 | 0 | 0 | 0 | 2493300 | 0 | 0 | 0 | 77039 | 0 | 0 |
| VFWGSQSI | RASAL1 | O95294 | RASL1_HUMAN | Unmodified | 0 | 0 | 0 | 0 | 0 | 0 | 0 | 0 | 3854400 | 711770 | 0 | 107750 | 0 | 0 | 0 | 0 | 0 | 0 |
| VFWREMVE | ECH1; ECH1; N | Q13011; M0I | ECH1_HUMAN | Unmodified | 0 | 0 | 0 | 0 | 0 | 0 | 518790 | 0 | 0 | 1082200 | 0 | 0 | 0 | 204720 | 0 | 0 | 0 | 0 |
| VFYELQHS | USP7; USP7; US | Q93009; A0A | UBP7_HUMAN | Unmodified | 41367 | 0 | 88418 | 70194 | 120080 | 0 | 26423 | 0 | 0 | 94878 | 215250 | 1375600 | 191860 | 64313 | 0 | 0 | 0 | 79389 |
| VFYPVVFSL | CSGALNACT2 | Q8N6G5 | CGAT2_HUMAI | Unmodified | 0 | 0 | 0 | 0 | 0 | 0 | 0 | 0 | 713010 | 0 | 0 | 0 | 0 | 0 | 0 | 0 | 0 | 0 |
| VGAAEEEL( | RRBP1; RRBP1; | Q9P2E9; A0A | RRBP1_HUMAI | Unmodified | 0 | 0 | 0 | 0 | 0 | 0 | 0 | 0 | 0 | 0 | 0 | 128070 | 2390100 | 0 | 0 | 0 | 0 | 0 |
| VGAAEIISR | SF3B1 | O75533 | SF3B1_HUMAI | Unmodified | 410980 | 352370 | 435840 | 254490 | 341270 | 0 | 0 | 0 | 194820 | 0 | 567320 | 13601000 | 243390 | 0 | 728040 | 314020 | 0 | 0 |
| VGADNHVI | AGL | P35573 | GDE_HUMAN ( | Unmodified | 0 | 0 | 22487 | 0 | 0 | 0 | 90107 | 0 | 219530 | 0 | 73902 | 1433400 | 0 | 0 | 0 | 0 | 0 | 0 |
| VGAEDAD( | AARS1; AARS1; | P49588; A0A | SYAC_HUMAN | Unmodified | 0 | 0 | 0 | 188830 | 0 | 0 | 0 | 0 | 890040 | 0 | 438760 | 2383300 | 0 | 0 | 198140 | 219790 | 0 | 0 |
| VGAEDAD( | AARS1; AARS1; | P49588; A0A | SYAC_HUMAN | Oxidation (M) | 0 | 0 | 0 | 0 | 0 | 0 | 0 | 0 | 148150 | 0 | 0 | 347920 | 0 | 0 | 98619 | 0 | 0 | 0 |
| VGAEDAD( | AARS1; AARS1; | P49588; A0A | SYAC_HUMAN | Unmodified | 0 | 0 | 0 | 0 | 0 | 0 | 0 | 0 | 2468200 | 171240 | 517820 | 0 | 995220 | 0 | 0 | 0 | 0 | 0 |
| VGAENVAI | SQOR; SQOR; S | Q9Y6N5; H3 | SQOR_HUMAN | Unmodified | 0 | 0 | 0 | 0 | 0 | 0 | 89184 | 0 | 2456600 | 0 | 0 | 0 | 0 | 0 | 0 | 0 | 0 | 0 |
| VGAENVAI | SQOR; SQOR; S | Q9Y6N5; H3 | SQOR_HUMAN | Unmodified | 0 | 0 | 0 | 0 | 0 | 0 | 0 | 0 | 1273400 | 1467600 | 0 | 0 | 0 | 0 | 0 | 0 | 0 | 0 |
| VGAFQFSS | VWA2 | Q5GFL6 | VWA2_HUMAN | Unmodified | 0 | 0 | 0 | 0 | 0 | 0 | 0 | 0 | 769190 | 0 | 0 | 0 | 0 | 0 | 0 | 0 | 0 | 0 |
| VGAIPANAI | TLN1 | Q9Y490 | TLN1_HUMAN | Unmodified | 0 | 0 | 0 | 0 | 0 | 2227700 | 0 | 0 | 0 | 1184800 | 860300 | 3998500 | 0 | 0 | 0 | 0 | 50475 | 0 |
| VGAIQRWL | ERP29; ERP29 | P30040; F8V | ERP29_HUMAN | Unmodified | 0 | 0 | 0 | 0 | 0 | 0 | 0 | 0 | 104430 | 588970 | 0 | 0 | 0 | 0 | 0 | 0 | 0 | 0 |
| VGALRIIDS | SUSD2 | Q9UGT4 | SUSD2_HUMAI | Unmodified | 0 | 0 | 0 | 0 | 736890 | 141660 | 0 | 0 | 0 | 393250 | 0 | 0 | 0 | 0 | 0 | 0 | 0 | 0 |
| VGALRIIDS | SUSD2 | Q9UGT4 | SUSD2_HUMAI | Unmodified | 0 | 0 | 0 | 0 | 2024300 | 0 | 77701 | 0 | 411100 | 665730 | 0 | 0 | 0 | 0 | 0 | 0 | 0 | 0 |
| VGASASDG | TBL1XR1; TBL1) | Q9BZK7; O6( | TBL1R_HUMAN | Unmodified | 0 | 0 | 0 | 0 | 0 | 0 | 0 | 0 | 933380 | 0 | 444280 | 0 | 0 | 0 | 0 | 69744 | 0 | 0 |
| VGASEAALI | EPS15L1; EPS1 | Q9UBC2; K7 | EP15R_HUMAN | Unmodified | 0 | 0 | 0 | 0 | 0 | 0 | 0 | 0 | 0 | 0 | 0 | 0 | 0 | 0 | 0 | 0 | 0 | 0 |
| VGASEAALI | EPS15L1; EPS1 | Q9UBC2; K7 | EP15R_HUMAN | Unmodified | 1437700 | 317400 | 80413 | 0 | 87844 | 0 | 0 | 0 | 0 | 169140 | 0 | 1725000 | 0 | 0 | 118880 | 285460 | 0 | 0 |
| VGAVDADK | PDIA6 | Q15084 | PDIA6_HUMAN | Unmodified | 0 | 0 | 0 | 0 | 0 | 323830 | 1347000 | 899230 | 50771000 | 76209000 | 5155300 | 1070500 | 0 | 2605200 | 0 | 0 | 567070 | 2154200 |
| VGCDRLLS | DPP7; DPP7 | Q9UHL4; R4 | DPP2_HUMAN | Unmodified | 0 | 0 | 0 | 0 | 0 | 1152300 | 84052 | 0 | 1370400 | 0 | 0 | 0 | 0 | 0 | 0 | 0 | 178230 | 0 |
| VGCFDPYS | LLGL1; LLGL1 | Q15334; A0A | L2GL1_HUMAN | Unmodified | 0 | 0 | 0 | 0 | 792930 | 286940 | 0 | 0 | 0 | 0 | 0 | 0 | 0 | 0 | 0 | 0 | 787150 | 331810 |
| VGCPVGD/ | CEL | P19835 | CEL_HUMAN B | Unmodified | 0 | 0 | 0 | 0 | 0 | 0 | 0 | 0 | 5937600 | 1780600 | 5195000 | 1193100 | 0 | 0 | 0 | 0 | 0 | 0 |
| VGCPVGD/ | CEL | P19835 | CEL_HUMAN B | Unmodified | 0 | 0 | 93611 | 0 | 0 | 0 | 0 | 232720 | 0 | 617550 | 0 | 0 | 0 | 0 | 0 | 0 | 0 | 0 |
| VGDAIPAVI | PRDX5 | P30044 | PRDX5_HUMAI | Unmodified | 0 | 0 | 0 | 1869500 | 1943200 | 0 | 138000 | 0 | 13247000 | 1187400 | 0 | 8643500 | 1152500 | 0 | 4725400 | 2330700 | 72896 | 0 |
| VGDAIPAVI | PRDX5 | P30044 | PRDX5_HUMAI | Unmodified | 0 | 0 | 0 | 1419100 | 11834000 | 1957000 | 0 | 0 | 12397000 | 9557200 | 840120 | 0 | 16321000 | 8912400 | 463100 | 0 | 8107700 | 11559000 |
| VGDDPAVV | TLN1 | Q9Y490 | TLN1_HUMAN | Unmodified | 959950 | 0 | 1786500 | 953810 | 1702700 | 0 | 0 | 217790 | 0 | 0 | 0 | 3274600 | 0 | 0 | 1342700 | 1753100 | 0 | 0 |
| VGDDVEFE | CSDE1; CSDE1 | O75534; E9F | CSDE1_HUMAI | Unmodified | 157000 | 134240 | 0 | 0 | 0 | 0 | 0 | 0 | 0 | 0 | 0 | 1862400 | 0 | 0 | 424570 | 300700 | 0 | 0 |
| VGDFGDAI | LARP1; LARP1; | Q6PKG0; H0 | LARP1_HUMAN | Unmodified | 416710 | 685730 | 740500 | 0 | 0 | 0 | 0 | 0 | 0 | 0 | 0 | 0 | 300730 | 0 | 611710 | 349940 | 0 | 0 |
| VGDHQFLL | KIAA2013 | Q8IYS2 | K2013_HUMAN | Unmodified | 0 | 0 | 0 | 0 | 0 | 0 | 0 | 0 | 22804000 | 1976800 | 0 | 1998000 | 0 | 0 | 0 | 0 | 0 | 0 |
| VGDHQFLL | KIAA2013 | Q8IYS2 | K2013_HUMAN | Unmodified | 0 | 0 | 0 | 0 | 0 | 0 | 0 | 0 | 2875200 | 4130800 | 0 | 0 | 0 | 0 | 0 | 0 | 0 | 0 |
| VGDHQFLL | KIAA2013 | Q8IYS2 | K2013_HUMAN | Unmodified | 0 | 0 | 0 | 0 | 0 | 0 | 0 | 0 | 0 | 1689600 | 0 | 0 | 0 | 0 | 0 | 0 | 0 | 0 |
| VGDIIQIISK | CASK; CASK; C/ | O14936; A0A | CSKP_HUMAN | Unmodified | 0 | 0 | 3191100 | 0 | 0 | 0 | 0 | 0 | 1329900 | 0 | 178920 | 1049100 | 0 | 557250 | 0 | 0 | 0 | 0 |
| VGDIIQIISK | CASK; CASK; C/ | O14936; A0A | CSKP_HUMAN | Unmodified | 282570 | 205540 | 858120 | 3421900 | 1476700 | 0 | 0 | 0 | 6729700 | 893930 | 0 | 1089800 | 978620 | 0 | 3505700 | 1755400 | 659260 | 0 |
| VGDIIQIISK | CASK; CASK; C/ | O14936; A0A | CSKP_HUMAN | Unmodified | 44976 | 0 | 0 | 2097700 | 10041000 | 0 | 0 | 0 | 990730 | 1893000 | 0 | 0 | 4270500 | 3656100 | 0 | 63694 | 275830 | 6031500 |
| VGDKVLLP | HSPE1; HSPE1; | P61604; B8Z | CH10_HUMAN | Unmodified | 0 | 0 | 0 | 0 | 0 | 0 | 385040 | 0 | 0 | 990660 | 0 | 0 | 0 | 229190 | 0 | 0 | 0 | 0 |
| VGDPQELN | FASN; FASN | P49327; A0A | FAS_HUMAN Fa | Unmodified | 514330 | 207550 | 0 | 1471100 | 1093700 | 1885200 | 0 | 0 | 1031700 | 685150 | 5148400 | 64409000 | 704370 | 0 | 3117700 | 3280800 | 749980 | 349220 |
| VGDPQELN | FASN; FASN | P49327; A0A | FAS_HUMAN Fa | Unmodified | 0 | 0 | 0 | 0 | 0 | 0 | 163200 | 0 | 1038600 | 3970900 | 0 | 0 | 0 | 101530 | 0 | 0 | 0 | 0 |
| VGDQILLAI | MRPL14 | Q6P1L8 | RM14_HUMAN | Unmodified | 0 | 0 | 0 | 0 | 0 | 100050 | 0 | 368770 | 970920 | 446020 | 747830 | 4112100 | 0 | 0 | 236820 | 0 | 376350 | 0 |
| VGDVVEFR | CYB5R1; CYB5I | Q9UHQ9; H7 | NB5R1_HUMAI | Unmodified | 0 | 0 | 0 | 38362 | 0 | 0 | 0 | 0 | 225790 | 109360 | 0 | 0 | 0 | 0 | 0 | 0 | 0 | 0 |
| VGDYGSLS | CTSZ; CTSZ; CT | Q9UBR2; A0. | CATZ_HUMAN | Unmodified | 0 | 0 | 0 | 0 | 295660 | 66646 | 82111 | 0 | 1077400 | 0 | 0 | 0 | 0 | 0 | 0 | 0 | 311620 | 44961 |
| VGDYGSLS | CTSZ; CTSZ; CT | Q9UBR2; A0. | CATZ_HUMAN | Unmodified | 18175 | 0 | 537320 | 438070 | 0 | 615330 | 22280 | 0 | 2651300 | 986130 | 18754 | 0 | 362210 | 191710 | 194390 | 0 | 0 | 0 |
| VGEAFHAS | SMAD1; SMAD5 | Q15797; Q9 | SMAD1_HUMA | Unmodified | 0 | 0 | 269310 | 437060 | 0 | 0 | 0 | 0 | 797080 | 0 | 299710 | 1227800 | 0 | 0 | 819390 | 940990 | 0 | 0 |
| VGEAQETTI | CCNDBP1; CCI | O95273; H3I | CCDB1_HUMA | Unmodified | 434180 | 103080 | 106600 | 0 | 0 | 0 | 0 | 0 | 0 | 0 | 0 | 0 | 0 | 0 | 207300 | 187350 | 55779 | 0 |
| VGEAQETTI | CCNDBP1; CCI | O95273; H3I | CCDB1_HUMA | Oxidation (M) | 934930 | 1227900 | 485450 | 327250 | 0 | 0 | 0 | 0 | 0 | 0 | 0 | 0 | 74265 | 0 | 0 | 1577100 | 0 | 235260 |
| VGEATETAI | ATP2A2; ATP2A | P16615; Q93 | AT2A2_HUMAN | Unmodified | 0 | 0 | 0 | 0 | 0 | 0 | 0 | 1185200 | 901690 | 0 | 0 | 294620 | 0 | 0 | 0 | 45115 | 0 | 76321 |
| VGEEDDGI | L1CAM | P32004 | L1CAM_HUMAI | Unmodified | 0 | 0 | 0 | 0 | 0 | 0 | 0 | 0 | 668670 | 0 | 0 | 0 | 0 | 0 | 0 | 0 | 0 | 0 |
| VGEEDDGI | L1CAM | P32004 | L1CAM_HUMAI | Unmodified | 0 | 0 | 0 | 0 | 0 | 0 | 0 | 0 | 423630 | 128240 | 0 | 0 | 0 | 0 | 0 | 0 | 0 | 0 |
| VGEEDEEA | VGF | O15240 | VGF_HUMAN N | Unmodified | 0 | 0 | 0 | 0 | 0 | 0 | 0 | 0 | 5164000 | 1620200 | 1655700 | 0 | 0 | 0 | 0 | 0 | 0 | 0 |
| VGEEFEEQ | CRABP2 | P29373 | RABP2_HUMAN | Unmodified | 0 | 0 | 0 | 0 | 0 | 0 | 0 | 0 | 775140 | 176850 | 0 | 255820 | 0 | 0 | 0 | 0 | 0 | 551670 |
| VGEGFEEE | CRABP1; CRAB | P29762; B5N | RABP1_HUMAN | Unmodified | 0 | 255060 | 0 | 0 | 0 | 0 | 0 | 0 | 1873100 | 0 | 0 | 0 | 0 | 0 | 0 | 148500 | 0 | 864680 |
| VGEGPGV( | RNPEP; RNPEP; | Q9H4A4; A6I | AMPB_HUMAN | Unmodified | 126940 | 0 | 0 | 0 | 0 | 0 | 0 | 0 | 500230 | 33707 | 207540 | 3519500 | 0 | 0 | 28280 | 0 | 0 | 301090 |
| VGEIVVFRI | SEC11A; SEC11 | P67812; H0Y | SC11A_HUMAN | Unmodified | 0 | 0 | 0 | 0 | 0 | 0 | 0 | 0 | 362050 | 50380 | 0 | 195010 | 0 | 0 | 0 | 0 | 0 | 0 |
| VGEKPVEV | IGSF3 | O75054 | IGSF3_HUMAN | Unmodified | 777160 | 611680 | 0 | 337270 | 0 | 0 | 0 | 0 | 0 | 0 | 0 | 1516900 | 0 | 0 | 1067800 | 1314800 | 56486 | 0 |
| VGELKDDD | MAP2K1 | Q02750 | MP2K1_HUMAI | Unmodified | 476180 | 0 | 607630 | 439100 | 493410 | 169070 | 13044 | 12293 | 439370 | 72746 | 497660 | 1537400 | 576920 | 0 | 757730 | 122440 | 188380 | 734420 |
| VGELKDDD | MAP2K2; N/A | P36507; A0A | MP2K2_HUMAI | Unmodified | 0 | 0 | 0 | 0 | 0 | 0 | 0 | 0 | 0 | 0 | 201720 | 40118 | 0 | 0 | 0 | 0 | 0 | 0 |
| VGELKDDD | MAP2K2 | P36507 | MP2K2_HUMAI | Unmodified | 0 | 0 | 133730 | 603100 | 322680 | 88367 | 684130 | 0 | 0 | 0 | 152800 | 478830 | 0 | 0 | 347170 | 195630 | 0 | 367110 |
| VGELTNLT( | LRRC8A | Q8IWT6 | LRC8A_HUMAN | Unmodified | 53478 | 0 | 0 | 0 | 0 | 0 | 0 | 0 | 0 | 41655 | 0 | 378040 | 0 | 0 | 0 | 23987 | 0 | 0 |
| VGENADS( | H1-0 | P07305 | H10_HUMAN H | Unmodified | 1981900 | 1624200 | 0 | 0 | 0 | 0 | 0 | 947890 | 0 | 0 | 0 | 0 | 0 | 0 | 0 | 0 | 0 | 0 |
| VGENADS( | H1-0 | P07305 | H10_HUMAN H | Unmodified | 0 | 0 | 0 | 0 | 0 | 0 | 0 | 641670 | 0 | 0 | 0 | 0 | 0 | 0 | 0 | 0 | 0 | 104250 |
| VGEPVALSI | SF3B2; SF3B2; | Q13435; E9F | SF3B2_HUMAN | Unmodified | 0 | 0 | 0 | 503460 | 3220900 | 0 | 3405300 | 641210 | 0 | 0 | 0 | 3100300 | 0 | 1202700 | 322590 | 0 | 367010 | 2483300 |
| VGEQAQV\ | CLTC; CLTC; CI | Q00610; J3K | CLH1_HUMAN | Oxidation (M) | 1318300 | 878400 | 472410 | 0 | 0 | 0 | 0 | 0 | 0 | 0 | 0 | 1764200 | 0 | 0 | 746740 | 0 | 0 | 0 |
| VGEQAQV\ | CLTC; CLTC; CI | Q00610; J3K | CLH1_HUMAN | 2 Oxidation (M) | 18219000 | 17195000 | 7665100 | 17341000 | 1547800 | 0 | 595290 | 0 | 0 | 3368100 | 0 | 6895600 | 22252000 | 646210 | 13440000 | 23168000 | 0 | 0 |
| VGEQAQV\ | CLTC; CLTC; CI | Q00610; J3K | CLH1_HUMAN | Oxidation (M) | 0 | 0 | 506760 | 5414100 | 0 | 0 | 0 | 0 | 1733100 | 0 | 0 | 0 | 8888100 | 0 | 0 | 0 | 0 | 0 |
| VGERQPLL | EIF3CL; EIF3C | B5ME19; Q9 | EIFCL_HUMAN | Oxidation (M) | 0 | 0 | 0 | 0 | 0 | 0 | 0 | 0 | 0 | 0 | 0 | 1152900 | 0 | 0 | 0 | 0 | 0 | 0 |
| VGERQPLL | EIF3CL; EIF3C | B5ME19; Q9 | EIFCL_HUMAN | Unmodified | 0 | 0 | 0 | 0 | 0 | 0 | 304880 | 0 | 419610 | 0 | 0 | 2953400 | 727740 | 0 | 303010 | 211890 | 0 | 0 |
| VGETETRLI | DCAF7; DCAF7 | P61962; A0A | DCAF7_HUMAI | Unmodified | 0 | 0 | 0 | 0 | 0 | 0 | 0 | 0 | 0 | 0 | 0 | 1391900 | 0 | 0 | 158240 | 0 | 0 | 0 |
| VGETFHAS | SMAD2; SMAD2 | Q15796; B72 | SMAD2_HUMA | Unmodified | 0 | 270620 | 0 | 284460 | 0 | 0 | 0 | 0 | 0 | 0 | 0 | 1873100 | 0 | 0 | 0 | 0 | 0 | 0 |
| VGETIIDLEI | MYOF; MYOF | Q9NZM1; H( | MYOF_HUMAN | Unmodified | 92794 | 106530 | 0 | 26711 | 12686000 | 2271600 | 0 | 0 | 422620 | 127800 | 40254 | 0 | 0 | 0 | 0 | 0 | 182760 | 593430 |
| VGEVCHIT( | FKBP4; FKBP4 | Q02790; F5H | FKBP4_HUMAN | Unmodified | 158250 | 0 | 287410 | 0 | 0 | 0 | 0 | 0 | 0 | 26629 | 369300 | 6102200 | 164850 | 0 | 597920 | 329170 | 0 | 21442 |
| VGEVEQEA | PPFIA2; PPFIA2 | O75334; G3 | LIPA2_HUMAN | Unmodified | 0 | 0 | 0 | 0 | 0 | 0 | 0 | 0 | 0 | 0 | 0 | 0 | 0 | 0 | 586510 | 0 | 0 | 0 |
| VGEVIVTK | HSPD1; HSPD1 | P10809; A0A | CH60_HUMAN | Unmodified | 74925 | 0 | 0 | 51732 | 1960700 | 579720 | 0 | 0 | 4413100 | 289310 | 1643300 | 439680 | 0 | 756050 | 69195 | 81897 | 0 | 0 |
| VGEVIVTKD | HSPD1; HSPD1 | P10809; A0A | CH60_HUMAN | Oxidation (M) | 164790 | 129800 | 357550 | 280790 | 1013800 | 79585 | 1692700 | 618260 | 33286000 | 5200200 | 1918000 | 682320 | 4078300 | 2380800 | 249560 | 450130 | 283130 | 224440 |
| VGEVIVTKD | HSPD1; HSPD1 | P10809; A0A | CH60_HUMAN | Unmodified | 0 | 0 | 0 | 0 | 0 | 0 | 709350 | 0 | 62038000 | 3572300 | 302250 | 0 | 2490200 | 1651900 | 63125 | 13339 | 0 | 0 |
| VGEYATYE/ | MMP14; MMP14 | P50281; A0A | MMP14_HUMA | Unmodified | 0 | 0 | 0 | 0 | 1748900 | 921440 | 0 | 0 | 0 | 0 | 0 | 0 | 0 | 0 | 0 | 0 | 978570 | 593580 |
| VGFAGFPS | AHNAK2 | Q8IVF2 | AHNK2_HUMAI | Unmodified | 0 | 0 | 0 | 0 | 0 | 0 | 0 | 0 | 0 | 0 | 0 | 0 | 0 | 0 | 0 | 0 | 0 | 1221900 |
| VGFLKPVG | FYB1; FYB1 | O15117; D6I | FYB1_HUMAN | Unmodified | 0 | 0 | 0 | 0 | 0 | 0 | 0 | 0 | 0 | 0 | 0 | 1091500 | 0 | 0 | 0 | 0 | 0 | 0 |
| VGFLPSAG | APEH; APEH | P13798; C9J | ACPH_HUMAN | Unmodified | 147120 | 0 | 434730 | 263990 | 0 | 34360 | 61844 | 0 | 241230 | 60204 | 556680 | 12911000 | 0 | 67608 | 688630 | 908830 | 100970 | 120600 |
| VGGAEGTK | SH3GL2; SH3G | Q99962; Q9 | SH3G2_HUMAI | Unmodified | 14555000 | 15454000 | 27300000 | 9692900 | 1349200 | 401490 | 266280 | 102170 | 4313600 | 832380 | 232250 | 896960 | 5765300 | 1418900 | 26537000 | 25605000 | 52528 | 270790 |
| VGGAEGTK | SH3GL1 | Q99961 | SH3G1_HUMAI | Unmodified | 32980000 | 36878000 | 4310900 | 19426000 | 8418100 | 1003200 | 2639000 | 891170 | 2465500 | 3365200 | 0 | 357810 | 66412000 | 19482000 | 22223000 | 25037000 | 1204100 | 2754100 |
| VGGAEGTK | SH3GL1 | Q99961 | SH3G1_HUMAI | Oxidation (M) | 10071000 | 12865000 | 2298000 | 7915400 | 2129700 | 206680 | 214310 | 222910 | 0 | 319560 | 0 | 75566 | 18553000 | 3964200 | 7456900 | 7962800 | 135600 | 0 |
| VGGAEGTK | SH3GL2 | Q99962 | SH3G2_HUMAI | Unmodified | 1070500 | 339440 | 0 | 0 | 0 | 0 | 0 | 0 | 2000200 | 645080 | 0 | 0 | 0 | 0 | 0 | 0 | 0 | 0 |
| VGGAGNKI | BPNT1; BPNT1 | O95861; A6N | BPNT1_HUMAN | Unmodified | 0 | 0 | 0 | 0 | 95020 | 18001 | 0 | 0 | 0 | 118430 | 0 | 0 | 250440 | 0 | 0 | 0 | 112220 | 129240 |
| VGGASSLE | IRF3; IRF3 | Q14653; M0 | IRF3_HUMAN I | Unmodified | 300810 | 565660 | 0 | 532570 | 292330 | 271300 | 0 | 0 | 547680 | 396130 | 286580 | 1582600 | 0 | 0 | 0 | 0 | 380500 | 418690 |
| VGGEVKLS | PIK3C2B; PIK3C | O00750; F5C | P3C2B_HUMAN | Unmodified | 0 | 0 | 0 | 0 | 112200 | 0 | 122240 | 0 | 0 | 0 | 0 | 0 | 0 | 0 | 0 | 0 | 346210 | 0 |
| VGGFPNFL | NUDT16L1; NU | Q9BRJ7; K7E | TIRR_HUMAN T | Unmodified | 82249 | 119430 | 62358 | 72998 | 0 | 0 | 0 | 0 | 0 | 0 | 0 | 0 | 0 | 0 | 187220 | 163860 | 0 | 0 |
| VGGGWM/ | MACF1; DST; D | Q9UPN3; Q( | MACF1_HUMAI | Unmodified | 0 | 0 | 0 | 0 | 0 | 0 | 0 | 0 | 0 | 0 | 353070 | 0 | 0 | 0 | 0 | 0 | 0 | 0 |
| VGGIYTVL( | GYS1 | P13807 | GYS1_HUMAN | Unmodified | 0 | 0 | 0 | 0 | 0 | 0 | 0 | 0 | 0 | 0 | 0 | 3553900 | 0 | 0 | 212430 | 0 | 0 | 0 |
| VGGLEDQL | CRLF1; CRLF1 | O75462; M0 | CRLF1_HUMAN | Unmodified | 0 | 0 | 0 | 0 | 0 | 0 | 0 | 0 | 4635800 | 0 | 0 | 0 | 0 | 0 | 0 | 0 | 0 | 0 |
| VGGNIEVL( | CHD4; CHD5; ( | Q14839; Q8 | CHD4_HUMAN | Unmodified | 0 | 0 | 0 | 0 | 0 | 0 | 0 | 0 | 0 | 0 | 146180 | 1486400 | 0 | 0 | 0 | 46029 | 0 | 0 |
| VGGPPPEG | XXYLT1; XXYLT1 | Q8NBI6; A0A | XXLT1_HUMAN | 2 Oxidation (M) | 0 | 0 | 0 | 0 | 0 | 0 | 0 | 0 | 1114000 | 236790 | 0 | 0 | 0 | 0 | 0 | 0 | 0 | 0 |
| VGGPPPEG | XXYLT1; XXYLT1 | Q8NBI6; A0A | XXLT1_HUMAN | Oxidation (M) | 0 | 0 | 0 | 0 | 0 | 0 | 0 | 0 | 444570 | 0 | 0 | 0 | 0 | 0 | 0 | 0 | 0 | 0 |
| VGGRHGTI | RIPOR2; RIPOR | Q9Y4F9; F5C | RIPR2_HUMAN | Unmodified | 0 | 0 | 0 | 0 | 0 | 0 | 0 | 0 | 0 | 0 | 0 | 0 | 1068100 | 811400 | 0 | 0 | 0 | 0 |

| | | | | | | | | | | | | | | | | | | | | | | |
|---|---|---|---|---|---|---|---|---|---|---|---|---|---|---|---|---|---|---|---|---|---|---|
| VGGRLEDT | CCT5; CCT5; C | P48643; B7Z | TCPE_HUMAN | Unmodified | 0 | 248250 | 21780 | 0 | 1559800 | 0 | 0 | 0 | 62833 | 0 | 0 | 4399400 | 1178500 | 0 | 66810 | 43239 | 103190 | 0 |
| VGGRYLKG | FSCN1 | Q16658 | FSCN1_HUMAI | Unmodified | 0 | 0 | 0 | 0 | 3206300 | 483340 | 255100 | 144640 | 0 | 252110 | 25485 | 0 | 0 | 0 | 0 | 0 | 1556700 | 1597400 |
| VGGSGVN | AHNAK | Q09666 | AHNK_HUMAN | Unmodified | 78688 | 0 | 0 | 0 | 7569500 | 1021600 | 0 | 0 | 0 | 0 | 0 | 0 | 0 | 191670 | 0 | 0 | 3156300 | 4031100 |
| VGGSSVDL | CTNND1; CTNN | O60716; A0A | CTND1_HUMAI | Unmodified | 138690 | 150890 | 352330 | 404850 | 65492 | 0 | 0 | 0 | 0 | 0 | 129340 | 605390 | 0 | 0 | 684050 | 612520 | 42561 | 82645 |
| VGGSSVDL | CTNND1; CTNN | O60716; A0A | CTND1_HUMAI | Unmodified | 527970 | 0 | 912370 | 2302300 | 0 | 0 | 0 | 0 | 0 | 0 | 795030 | 313470 | 829010 | 0 | 2185900 | 1216500 | 0 | 0 |
| VGGTKPAG | TPD52; TPD52 | P55327; E5R | TPD52_HUMAN | Unmodified | 313000 | 0 | 0 | 0 | 723660 | 0 | 0 | 0 | 6025400 | 1914300 | 1965900 | 7568600 | 0 | 0 | 705520 | 457250 | 0 | 0 |
| VGGTKPAG | TPD52; TPD52 | P55327; E5R | TPD52_HUMAN | Unmodified | 0 | 0 | 0 | 0 | 0 | 0 | 0 | 0 | 0 | 1314900 | 0 | 0 | 0 | 0 | 0 | 0 | 0 | 0 |
| VGGTSDVE | HSPD1; HSPD1 | P10809; A0A | CH60_HUMAN | Unmodified | 8357.7 | 0 | 0 | 0 | 530290 | 0 | 317140 | 110510 | 7166800 | 935540 | 2275000 | 341710 | 0 | 814360 | 0 | 0 | 114880 | 1528300 |
| VGGTSDVE | HSPD1; HSPD1 | P10809; A0A | CH60_HUMAN | Unmodified | 23148 | 634940 | 1422000 | 0 | 1309400 | 202810 | 1836300 | 770290 | 33707000 | 3309900 | 0 | 0 | 5133500 | 3482400 | 159700 | 167670 | 328230 | 2423000 |
| VGGVQSLG | GOT1 | P17174 | AATC_HUMAN | Unmodified | 0 | 0 | 0 | 0 | 0 | 0 | 0 | 0 | 0 | 0 | 1286900 | 3721000 | 0 | 0 | 0 | 90223 | 0 | 0 |
| VGGYILGEI | AP2A2; AP2A1; | O94973; O9 | AP2A2_HUMAN | Unmodified | 133770 | 332790 | 87451 | 0 | 0 | 0 | 0 | 0 | 0 | 0 | 186220 | 880300 | 0 | 0 | 919630 | 205870 | 27173 | 0 |
| VGHIYRLE | GALNT7; GALN | Q86SF2; H0 | GALT7_HUMAN | Unmodified | 0 | 0 | 0 | 0 | 0 | 0 | 0 | 0 | 2973200 | 0 | 0 | 0 | 0 | 0 | 0 | 0 | 0 | 0 |
| VGHSELVG | ATP6V1A; ATP6 | P38606; C9J | VATA_HUMAN | Unmodified | 0 | 0 | 0 | 0 | 0 | 0 | 0 | 0 | 0 | 50090 | 490470 | 2293500 | 0 | 0 | 0 | 0 | 0 | 0 |
| VGHSIRHP | PARP1; PARP1; | P09874; A0A | PARP1_HUMAN | Unmodified | 577260 | 477760 | 0 | 0 | 757800 | 0 | 0 | 0 | 0 | 0 | 0 | 0 | 0 | 0 | 0 | 0 | 0 | 0 |
| VGIDTPDID | AHNAK | Q09666 | AHNK_HUMAN | Unmodified | 314640 | 283450 | 249090 | 289870 | 8597600 | 1894000 | 0 | 0 | 96630 | 87882 | 0 | 0 | 310500 | 319910 | 388330 | 316700 | 1467000 | 3857800 |
| VGIDTPDID | AHNAK | Q09666 | AHNK_HUMAN | Unmodified | 0 | 0 | 0 | 0 | 4029100 | 689450 | 0 | 0 | 0 | 0 | 0 | 0 | 0 | 0 | 0 | 0 | 633030 | 1819600 |
| VGIDTPDID | AHNAK | Q09666 | AHNK_HUMAN | Unmodified | 0 | 0 | 0 | 129280 | 1259900 | 0 | 0 | 0 | 0 | 0 | 0 | 0 | 0 | 0 | 0 | 0 | 1016800 | 0 |
| VGIFNCHG | GALNT13 | Q8IUC8 | GLT13_HUMAN | Unmodified | 0 | 0 | 0 | 0 | 0 | 0 | 0 | 0 | 402040 | 0 | 0 | 0 | 0 | 0 | 1795200 | 0 | 0 | 0 |
| VGIFNCHG | GALNT1 | Q10472 | GALT1_HUMAN | Oxidation (M) | 0 | 0 | 0 | 0 | 0 | 0 | 0 | 0 | 5146700 | 615560 | 686000 | 3061400 | 0 | 0 | 0 | 0 | 0 | 0 |
| VGIFNCHG | GALNT1 | Q10472 | GALT1_HUMAN | Unmodified | 0 | 0 | 0 | 671880 | 429240 | 0 | 0 | 0 | 5044600 | 0 | 669720 | 0 | 0 | 0 | 0 | 0 | 0 | 0 |
| VGIFTGAR | PFKM; PFKM; PI | P08237; F8V | PFKAM_HUMAI | Unmodified | 0 | 377250 | 0 | 0 | 0 | 273240 | 831220 | 250670 | 232140 | 102150 | 147890 | 1343700 | 1147500 | 2477700 | 119930 | 83161 | 149530 | 0 |
| VGIGAFPTE | ADSS2 | P30520 | PURA2_HUMAI | Unmodified | 0 | 0 | 196040 | 1539100 | 0 | 0 | 75068 | 0 | 1925200 | 13434 | 2661900 | 12200000 | 0 | 0 | 0 | 0 | 0 | 0 |
| VGIIAWHP | CORO1B; COR | Q9BR76; F5I | COR1B_HUMA | Unmodified | 368030 | 309860 | 266080 | 288260 | 0 | 0 | 123240 | 51340 | 0 | 0 | 0 | 1869800 | 0 | 0 | 568520 | 519500 | 0 | 0 |
| VGIIPPKGC | PSMC6; PSMC6 | P62333; A0A | PRS10_HUMAN | Unmodified | 0 | 0 | 0 | 0 | 0 | 0 | 0 | 0 | 0 | 0 | 0 | 0 | 0 | 0 | 0 | 0 | 308120 | 0 |
| VGILGSGD | STEAP3; STEAP | Q658P3; B8 | STEA3_HUMAN | Unmodified | 0 | 6597500 | 1693700 | 1317100 | 0 | 0 | 0 | 367000 | 1948800 | 270320 | 2113300 | 2410800 | 73437 | 0 | 2388400 | 1603500 | 0 | 0 |
| VGINYQPP | TUBA3C; TUBA | P0DPH7; Q7 | TBA3C_HUMAN | Unmodified | 17686000 | 10105000 | 20105000 | 21127000 | 86859000 | 32818000 | 93734000 | 1.44E+08 | 1.31E+08 | 57279000 | 37071000 | 3.55E+08 | 59274000 | 45042000 | 19229000 | 16963000 | 26447000 | 78132000 |
| VGINYQPP | TUBA3C; TUBA | P0DPH7; Q7 | TBA3C_HUMAN | Unmodified | 0 | 0 | 0 | 0 | 1719900 | 0 | 3253300 | 4778400 | 0 | 986330 | 0 | 646380 | 1307900 | 1037800 | 0 | 0 | 1967000 | 1760700 |
| VGIPVTDEI | SFXN1 | Q9H9B4 | SFXN1_HUMAN | Unmodified | 0 | 0 | 0 | 0 | 0 | 0 | 169420 | 0 | 11389000 | 1674300 | 0 | 313400 | 0 | 0 | 73295 | 30021 | 0 | 0 |
| VGIRETPEI | FARSB | Q9NSD9 | SYFB_HUMAN | Unmodified | 0 | 0 | 0 | 0 | 829640 | 0 | 457090 | 206770 | 0 | 179980 | 210820 | 0 | 1276000 | 464630 | 0 | 0 | 245500 | 605800 |
| VGIVGNLS | AGAP3; AGAP3; | Q96P47; E7E | AGAP3_HUMAI | Unmodified | 156160 | 0 | 0 | 209460 | 0 | 0 | 0 | 0 | 271990 | 0 | 241130 | 1767500 | 0 | 0 | 0 | 0 | 0 | 0 |
| VGIYSPHTA | NIF3L1; NIF3L1 | Q9GZT8; Q6 | NIF3L_HUMAN | Unmodified | 0 | 0 | 0 | 0 | 0 | 2985100 | 0 | 0 | 4514500 | 0 | 892740 | 4736700 | 0 | 0 | 0 | 0 | 0 | 0 |
| VGKDELFA | FSCN1 | Q16658 | FSCN1_HUMAI | Unmodified | 0 | 0 | 0 | 0 | 0 | 0 | 0 | 0 | 2367100 | 632710 | 4402700 | 3055000 | 0 | 0 | 0 | 0 | 312960 | 684850 |
| VGKDELFA | FSCN1 | Q16658 | FSCN1_HUMAI | Unmodified | 0 | 0 | 0 | 0 | 6178200 | 2060200 | 237950 | 0 | 1078800 | 538370 | 5762600 | 0 | 0 | 0 | 0 | 0 | 2011000 | 7245000 |
| VGKETVQT | VCL | P18206 | VINC_HUMAN | Unmodified | 848830 | 980830 | 211970 | 2443900 | 1588100 | 458500 | 145860 | 0 | 2401400 | 793250 | 871800 | 4429100 | 2072300 | 314640 | 909940 | 709380 | 2374200 | 1968000 |
| VGKGEPG | GCN1 | Q92616 | GCN1_HUMAN | Unmodified | 0 | 197230 | 20847 | 0 | 0 | 0 | 283800 | 0 | 0 | 55349 | 0 | 136040 | 0 | 0 | 128210 | 18621 | 0 | 0 |
| VGKMDATS | TMX3; TMX3; TI | Q96JJ7; B4D | TMX3_HUMAN | Oxidation (M) | 0 | 0 | 0 | 0 | 0 | 0 | 0 | 0 | 3529600 | 158690 | 334900 | 190450 | 0 | 0 | 0 | 0 | 0 | 94965 |
| VGKMDATS | TMX3; TMX3; TI | Q96JJ7; B4D | TMX3_HUMAN | Unmodified | 176330 | 297540 | 0 | 0 | 0 | 0 | 0 | 0 | 2200500 | 0 | 0 | 0 | 0 | 0 | 0 | 0 | 0 | 0 |
| VGKMDATS | TMX3; TMX3; TI | Q96JJ7; B4D | TMX3_HUMAN | Oxidation (M) | 0 | 0 | 0 | 0 | 0 | 0 | 0 | 0 | 2327900 | 2381100 | 578770 | 841280 | 4185700 | 3368200 | 0 | 222100 | 2619400 | 0 |
| VGKMDATS | TMX3; TMX3; TI | Q96JJ7; B4D | TMX3_HUMAN | Unmodified | 0 | 0 | 0 | 0 | 0 | 0 | 0 | 0 | 2407800 | 461170 | 0 | 688950 | 1413400 | 632980 | 0 | 0 | 278910 | 532700 |
| VGLDATNC | BABAM2; BABA | Q9NXR7; F8 | BABA2_HUMAN | Unmodified | 0 | 0 | 0 | 0 | 0 | 0 | 0 | 283260 | 0 | 0 | 66440 | 660040 | 0 | 0 | 0 | 0 | 0 | 0 |
| VGLGLGYL | FLII | Q13045 | FLII_HUMAN Pr | Unmodified | 0 | 0 | 36736 | 94959 | 1083800 | 108670 | 295570 | 0 | 512610 | 301730 | 27069 | 397440 | 877830 | 171810 | 359220 | 215540 | 158150 | 767400 |
| VGLIGSCTI | ACO2; ACO2; A | Q99798; A2A | ACON_HUMAN | Unmodified | 0 | 0 | 0 | 0 | 0 | 0 | 0 | 0 | 972970 | 0 | 0 | 0 | 0 | 0 | 0 | 0 | 0 | 0 |
| VGLLQREE | SUN2 | Q9UH99 | SUN2_HUMAN | Oxidation (M) | 0 | 0 | 0 | 0 | 0 | 0 | 0 | 0 | 1826300 | 0 | 0 | 0 | 0 | 0 | 0 | 0 | 0 | 0 |
| VGLLQREE | SUN2 | Q9UH99 | SUN2_HUMAN | Unmodified | 0 | 0 | 0 | 0 | 0 | 0 | 0 | 636160 | 7156800 | 0 | 0 | 0 | 0 | 731250 | 0 | 0 | 0 | 0 |
| VGLLQREE | SUN2 | Q9UH99 | SUN2_HUMAN | Unmodified | 0 | 0 | 0 | 0 | 0 | 0 | 0 | 0 | 40067000 | 195570 | 0 | 0 | 0 | 0 | 0 | 0 | 0 | 0 |
| VGLLQREE | SUN2 | Q9UH99 | SUN2_HUMAN | Oxidation (M) | 0 | 0 | 0 | 0 | 0 | 0 | 0 | 0 | 4838100 | 0 | 0 | 0 | 0 | 0 | 0 | 0 | 0 | 0 |
| VGLLQYGS | MATN2 | O00339 | MATN2_HUMAI | Unmodified | 0 | 0 | 0 | 0 | 0 | 0 | 0 | 0 | 771060 | 0 | 0 | 0 | 0 | 0 | 0 | 0 | 0 | 0 |
| VGLLQYGS | MATN2 | O00339 | MATN2_HUMAI | Unmodified | 0 | 0 | 0 | 0 | 0 | 0 | 0 | 0 | 1389800 | 0 | 0 | 0 | 0 | 0 | 0 | 0 | 0 | 0 |
| VGLPIGFSL | UBAP1 | Q9NZ09 | UBAP1_HUMAN | Unmodified | 3468200 | 3545500 | 203830 | 126390 | 289830 | 68312 | 0 | 0 | 211600 | 17035 | 0 | 0 | 137960 | 0 | 1096800 | 328160 | 0 | 0 |
| VGLPTQQG | VPS37B | Q9H9H4 | VP37B_HUMAN | Unmodified | 59766000 | 21597000 | 0 | 0 | 0 | 0 | 0 | 0 | 0 | 0 | 0 | 898800 | 0 | 0 | 0 | 16031000 | 299900 | 0 |
| VGLQAQPV | ANK3; ANK3; AI | Q12955; A0A | ANK3_HUMAN | Unmodified | 0 | 0 | 755690 | 1181300 | 0 | 0 | 0 | 0 | 0 | 0 | 0 | 570670 | 928040 | 1021000 | 0 | 2235700 | 0 | 0 |
| VGLQAQPV | ANK3; ANK3; AI | Q12955; A0A | ANK3_HUMAN | Unmodified | 85676 | 0 | 1603300 | 3029000 | 0 | 0 | 0 | 0 | 0 | 0 | 501990 | 1686800 | 3721500 | 7635800 | 4182300 | 3841600 | 0 | 0 |
| VGLQAQPV | ANK3; ANK3; AI | Q12955; A0A | ANK3_HUMAN | Unmodified | 0 | 0 | 0 | 0 | 1055400 | 275060 | 76646 | 0 | 98559 | 0 | 0 | 0 | 240360 | 247910 | 0 | 0 | 290640 | 776510 |
| VGLQVVAV | HSPD1; HSPD1 | P10809; A0A | CH60_HUMAN | Unmodified | 368650 | 181020 | 434070 | 407560 | 1174100 | 133810 | 1879500 | 358170 | 47920000 | 5934500 | 920370 | 884340 | 2307200 | 2075800 | 409590 | 338020 | 317380 | 781320 |
| VGLQVVAV | HSPD1; HSPD1 | P10809; A0A | CH60_HUMAN | Unmodified | 0 | 0 | 0 | 0 | 58911 | 0 | 189160 | 265960 | 702380 | 5644400 | 0 | 0 | 0 | 297760 | 0 | 0 | 48331 | 0 |
| VGLQVVAV | HSPD1; HSPD1 | P10809; A0A | CH60_HUMAN | Unmodified | 0 | 0 | 0 | 0 | 6046100 | 469230 | 12151000 | 4154600 | 5176700 | 28573000 | 0 | 0 | 9938200 | 14429000 | 0 | 0 | 3126100 | 5495700 |
| VGLRLLEVI | SCRIB; SCRIB; | Q14160; A0A | SCRIB_HUMAN | Unmodified | 0 | 0 | 0 | 0 | 0 | 0 | 0 | 0 | 0 | 1484800 | 0 | 0 | 0 | 0 | 0 | 0 | 0 | 0 |
| VGLSPDSP | CASKIN2 | Q8WXE0 | CSKI2_HUMAN | Unmodified | 0 | 0 | 0 | 0 | 0 | 0 | 0 | 0 | 0 | 0 | 0 | 0 | 0 | 0 | 0 | 0 | 0 | 0 |
| VGLTNYAA | RPL5; RPL5; RP | P46777; Q5T | RL5_HUMAN 6 | Unmodified | 4395100 | 0 | 0 | 837050 | 0 | 0 | 2479000 | 0 | 0 | 0 | 889360 | 14415000 | 0 | 0 | 0 | 5615600 | 0 | 0 |
| VGLTNYAA | RPL5; RPL5; RP | P46777; Q5T | RL5_HUMAN 6 | Unmodified | 0 | 0 | 0 | 0 | 0 | 1736900 | 16913 | 0 | 0 | 0 | 422330 | 4860400 | 0 | 0 | 0 | 0 | 0 | 738680 |
| VGLVHGLP | UGCG | Q16739 | CEGT_HUMAN | Unmodified | 0 | 0 | 0 | 0 | 0 | 0 | 0 | 0 | 0 | 0 | 0 | 693180 | 0 | 0 | 0 | 0 | 0 | 0 |
| VGLYDTYSI | GEMIN5 | Q8TEQ6 | GEMI5_HUMAN | Unmodified | 0 | 0 | 0 | 0 | 0 | 0 | 0 | 0 | 0 | 0 | 0 | 1635100 | 0 | 0 | 0 | 0 | 0 | 0 |
| VGMDSTTV | SNX27; SNX27; | Q96L92; A0A | SNX27_HUMAN | Oxidation (M) | 0 | 154300 | 0 | 0 | 250620 | 92649 | 0 | 0 | 0 | 0 | 0 | 0 | 0 | 0 | 84870 | 19535 | 126290 | 0 |
| VGMGQKD | ACTG1; ACTC1; | P63261; P68 | ACTG_HUMAN | Oxidation (M) | 0 | 0 | 0 | 0 | 884110 | 0 | 0 | 0 | 0 | 29197 | 0 | 0 | 0 | 0 | 0 | 0 | 430390 | 0 |
| VGMGQKD | ACTG1; ACTC1; | P63261; P68 | ACTG_HUMAN | Unmodified | 0 | 0 | 0 | 0 | 2391400 | 385960 | 374720 | 128310 | 49729 | 255630 | 0 | 0 | 745020 | 344810 | 0 | 0 | 1645200 | 477590 |
| VGMVETNS | PPIL1 | Q9Y3C6 | PPIL1_HUMAN | Unmodified | 1869500 | 2252000 | 1294200 | 1204700 | 128160 | 31310 | 0 | 0 | 811000 | 0 | 531260 | 7111800 | 167020 | 0 | 3199900 | 1965200 | 20337 | 67624 |
| VGMVETNS | PPIL1 | Q9Y3C6 | PPIL1_HUMAN | Oxidation (M) | 1634400 | 1072900 | 912340 | 572110 | 0 | 0 | 0 | 0 | 0 | 0 | 132310 | 2085500 | 0 | 0 | 1834100 | 1656200 | 155790 | 0 |
| VGNELCEV | EXTL3 | O43909 | EXTL3_HUMAN | Unmodified | 0 | 0 | 0 | 0 | 0 | 0 | 0 | 0 | 336070 | 0 | 0 | 0 | 0 | 0 | 0 | 0 | 0 | 0 |
| VGNGHSN | MAN2A1 | Q16706 | MA2A1_HUMAI | Unmodified | 0 | 0 | 0 | 0 | 0 | 0 | 0 | 0 | 1361400 | 0 | 658480 | 0 | 0 | 0 | 0 | 0 | 120650 | 0 |
| VGNIEIKDL | ACTR2 | P61160 | ARP2_HUMAN | Oxidation (M) | 385980 | 0 | 0 | 305120 | 577970 | 0 | 110020 | 0 | 0 | 0 | 116810 | 771590 | 225750 | 98260 | 0 | 0 | 0 | 0 |
| VGNLGLAT | DDX3X; DDX3X; | O00571; A0A | DDX3X_HUMAI | Unmodified | 901670 | 1169700 | 3148000 | 824640 | 0 | 20326 | 731350 | 0 | 903620 | 575490 | 504480 | 4918900 | 90628 | 0 | 1597000 | 954420 | 0 | 0 |
| VGNLGLAT | DDX3X; DDX3X; | O00571; A0A | DDX3X_HUMAI | Unmodified | 0 | 0 | 514040 | 1497200 | 1059400 | 473510 | 0 | 0 | 0 | 783150 | 374700 | 0 | 757230 | 0 | 0 | 0 | 692370 | 0 |
| VGNLTVVG | GOT1 | P17174 | AATC_HUMAN | Unmodified | 0 | 0 | 0 | 0 | 0 | 0 | 0 | 0 | 0 | 0 | 327170 | 1848700 | 0 | 0 | 0 | 51926 | 0 | 0 |
| VGNNFHN | ANAPC10; ANA | Q9UM13; D6 | APC10_HUMAN | Unmodified | 154830 | 0 | 0 | 0 | 0 | 0 | 0 | 0 | 0 | 0 | 0 | 849410 | 0 | 0 | 0 | 0 | 0 | 0 |
| VGNNLQRI | NDC80 | O14777 | NDC80_HUMA | Oxidation (M) | 0 | 0 | 0 | 0 | 530020 | 0 | 164580 | 0 | 0 | 0 | 0 | 0 | 100390 | 195830 | 0 | 0 | 162220 | 148270 |
| VGNPWDP | ALDH7A1; ALD | P49419; A0A | AL7A1_HUMAN | Unmodified | 0 | 0 | 0 | 0 | 0 | 0 | 0 | 0 | 645710 | 0 | 1110100 | 6110400 | 0 | 1946000 | 0 | 0 | 0 | 0 |
| VGNVEYFG | ATP1B1 | P05026 | AT1B1_HUMAN | Unmodified | 0 | 564290 | 3710500 | 6134700 | 7427900 | 373650 | 1857700 | 0 | 15863000 | 1719200 | 1796800 | 9389500 | 2526300 | 3625200 | 21112000 | 961990 | 782340 | 513870 |
| VGNYETDT | EHBP1 | Q8NDI1 | EHBP1_HUMAI | Unmodified | 0 | 0 | 0 | 77512 | 420760 | 0 | 0 | 0 | 366990 | 60246 | 0 | 0 | 56423 | 0 | 0 | 0 | 21755 | 0 |
| VGPGAGES | GEMIN5 | Q8TEQ6 | GEMI5_HUMAN | Unmodified | 0 | 0 | 0 | 0 | 0 | 0 | 0 | 0 | 0 | 0 | 0 | 1581100 | 0 | 0 | 0 | 0 | 0 | 0 |
| VGPGGSS | TRIM41 | Q8WV44 | TRI41_HUMAN | Unmodified | 348910 | 665470 | 174260 | 96425 | 0 | 0 | 0 | 0 | 0 | 0 | 0 | 0 | 0 | 0 | 262480 | 232510 | 0 | 0 |
| VGPLGFLS | CEL | P19835 | CEL_HUMAN B | Unmodified | 0 | 0 | 0 | 0 | 0 | 0 | 0 | 0 | 718510 | 0 | 281190 | 669060 | 0 | 88564 | 0 | 0 | 0 | 117430 |
| VGPSGAPG | COL2A1 | P02458 | CO2A1_HUMAI | Unmodified | 0 | 0 | 0 | 0 | 0 | 0 | 0 | 0 | 0 | 0 | 799060 | 0 | 0 | 0 | 0 | 0 | 0 | 0 |
| VGPYHNP | TM9SF1; TM9SF | O15321; E9F | TM9S1_HUMAN | Unmodified | 0 | 0 | 0 | 0 | 2066200 | 0 | 0 | 0 | 3051900 | 701730 | 0 | 0 | 0 | 0 | 0 | 0 | 0 | 0 |
| VGQALELP | NUP210 | Q8TEM1 | PO210_HUMAN | Unmodified | 0 | 0 | 0 | 0 | 0 | 0 | 0 | 0 | 51373000 | 2869500 | 454000 | 390320 | 0 | 0 | 0 | 0 | 0 | 0 |
| VGQAMAS | STT3A | P46977 | STT3A_HUMAN | Oxidation (M) | 0 | 0 | 0 | 0 | 0 | 0 | 0 | 0 | 207750 | 54260 | 98988 | 59737 | 0 | 0 | 0 | 0 | 0 | 48741 |
| VGQAMAS | STT3A | P46977 | STT3A_HUMAN | Unmodified | 0 | 0 | 0 | 0 | 0 | 0 | 0 | 0 | 30917000 | 960650 | 1673000 | 728640 | 0 | 0 | 26793 | 0 | 0 | 0 |
| VGQAVALR | CTBP1; CTBP1; | Q13363; H0 | CTBP1_HUMAN | Unmodified | 86098000 | 52255000 | 42190000 | 31504000 | 0 | 9900.6 | 0 | 0 | 1128900 | 477940 | 42014 | 2153700 | 6379500 | 915930 | 39971000 | 45446000 | 0 | 0 |
| VGQAVDV | PSMD2 | Q13200 | PSMD2_HUMA | Unmodified | 1100200 | 305680 | 1103600 | 1132500 | 1673200 | 0 | 130590 | 132230 | 4773300 | 0 | 3133300 | 0 | 1802900 | 634460 | 866480 | 794200 | 1212600 | 1538900 |
| VGQDPVLF | SSBP1; SSBP1; | Q04837; C9 | SSBP_HUMAN | Unmodified | 63032 | 49567 | 70143 | 0 | 56577 | 0 | 0 | 0 | 13322000 | 0 | 491540 | 7560500 | 0 | 0 | 0 | 14642 | 144500 | 137130 |
| VGQDPVLF | SSBP1; SSBP1; | Q04837; C9 | SSBP_HUMAN | Unmodified | 0 | 0 | 0 | 0 | 26906 | 0 | 0 | 0 | 284020 | 104690 | 57191 | 0 | 0 | 0 | 0 | 0 | 0 | 30013 |
| VGQDPVLF | SSBP1; SSBP1; | Q04837; C9 | SSBP_HUMAN | Oxidation (M) | 0 | 0 | 0 | 0 | 0 | 0 | 0 | 0 | 0 | 2865000 | 0 | 0 | 0 | 0 | 0 | 0 | 0 | 0 |
| VGQEIEVR | EIF2S3 | P41091 | IF2G_HUMAN | Unmodified | 230350 | 134220 | 1102900 | 1003300 | 2499100 | 0 | 0 | 204600 | 1831000 | 458320 | 3827800 | 9442700 | 1758400 | 0 | 436320 | 162100 | 20136 | 0 |
| VGQEIEVR | EIF2S3 | P41091 | IF2G_HUMAN | Unmodified | 0 | 0 | 0 | 31448 | 989690 | 0 | 420710 | 40274 | 0 | 639490 | 0 | 0 | 2108900 | 0 | 0 | 0 | 543220 | 1077000 |
| VGQISFDL | AFG3L2 | Q9Y4W6 | AFG32_HUMAN | Unmodified | 0 | 0 | 0 | 0 | 1212800 | 320860 | 661330 | 414510 | 2349300 | 0 | 0 | 0 | 513140 | 0 | 0 | 0 | 407420 | 378600 |
| VGQLSQLT | LRRC8D | Q7L1W4 | LRC8D_HUMAI | Unmodified | 0 | 0 | 0 | 51194 | 59472 | 14482 | 31202 | 0 | 221870 | 45917 | 93570 | 706100 | 59736 | 73063 | 0 | 75110 | 0 | 0 |
| VGQYVSR | ULK3 | Q6PHR2 | ULK3_HUMAN | Unmodified | 0 | 34435 | 0 | 0 | 0 | 0 | 0 | 0 | 0 | 122640 | 0 | 971030 | 0 | 0 | 112480 | 32670 | 0 | 0 |
| VGQYVSRA | ULK3 | Q6PHR2 | ULK3_HUMAN | Unmodified | 0 | 0 | 0 | 322090 | 0 | 0 | 0 | 0 | 0 | 243420 | 222930 | 612680 | 0 | 45736 | 0 | 0 | 0 | 0 |
| VGRPSNIG | PUF60; PUF60; | Q9UHX1; E9 | PUF60_HUMAN | Unmodified | 0 | 0 | 0 | 0 | 0 | 0 | 0 | 0 | 3927100 | 1118900 | 0 | 7165900 | 0 | 0 | 0 | 0 | 26394 | 0 |
| VGRSFCSC | MEAK7; MEAK7 | Q6P9B6; H3 | MEAK7_HUMAI | Unmodified | 0 | 0 | 0 | 0 | 558220 | 0 | 0 | 0 | 0 | 0 | 0 | 0 | 0 | 0 | 0 | 0 | 516540 | 398010 |
| VGRVNETT | PARP4 | Q9UKK3 | PARP4_HUMAN | Unmodified | 0 | 27555 | 0 | 0 | 0 | 0 | 56429 | 0 | 0 | 1682900 | 0 | 1134400 | 0 | 0 | 0 | 0 | 0 | 0 |
| VGSAADIPI | FLNA; FLNA; FL | P21333; H0 | FLNA_HUMAN | Unmodified | 0 | 108100 | 0 | 0 | 5590100 | 0 | 3612700 | 0 | 0 | 47910 | 0 | 0 | 9938200 | 9504100 | 0 | 0 | 3362200 | 1357600 |
| VGSAADIPI | FLNA; FLNA; FL | P21333; H0 | FLNA_HUMAN | Unmodified | 0 | 0 | 0 | 2375000 | 1426400 | 0 | 0 | 0 | 0 | 0 | 0 | 0 | 0 | 0 | 2178500 | 0 | 1387600 | 0 |
| VGSCSDV | B4GALNT1 | Q00973 | B4GN1_HUMAI | Unmodified | 0 | 0 | 0 | 0 | 0 | 0 | 0 | 0 | 1635300 | 62476 | 0 | 0 | 0 | 0 | 0 | 0 | 0 | 0 |
| VGSDQCLL | CXADR | P78310 | CXAR_HUMAN | Unmodified | 0 | 0 | 0 | 0 | 0 | 0 | 0 | 0 | 0 | 0 | 0 | 5965100 | 0 | 0 | 0 | 0 | 0 | 0 |
| VGSEHSLL | ERBIN; ERBIN | Q96RT1; H0 | ERBIN_HUMAN | Unmodified | 0 | 0 | 308270 | 364360 | 6439500 | 1662300 | 64881 | 61145 | 345400 | 0 | 514500 | 678160 | 1134400 | 593130 | 241760 | 189620 | 370890 | 980660 |
| VGSFDPYS | LLGL2; LLGL2 | Q6P1M3; J3 | L2GL2_HUMAN | Unmodified | 472260 | 887890 | 589770 | 1545400 | 2250400 | 698810 | 0 | 0 | 2192900 | 2022700 | 0 | 0 | 0 | 0 | 1949900 | 855540 | 1687500 | 853660 |
| VGSGDTNI | NANS | Q9NR45 | SIAS_HUMAN | Unmodified | 0 | 257430 | 568220 | 491040 | 3749700 | 0 | 860810 | 444780 | 0 | 0 | 1128300 | 0 | 2948400 | 1756200 | 1001500 | 248110 | 0 | 0 |
| VGSHDLTL | EEPD1 | Q7L9B9 | EEPD1_HUMAN | Unmodified | 0 | 0 | 0 | 0 | 0 | 0 | 0 | 0 | 0 | 0 | 0 | 0 | 0 | 0 | 103540 | 0 | 559600 | 2507100 |
| VGSHEQPS | MTSS2; MTSS2 | Q765P7; A0A | MTSS2_HUMAN | Unmodified | 0 | 0 | 0 | 0 | 415810 | 0 | 0 | 0 | 90166 | 0 | 0 | 409680 | 0 | 0 | 0 | 0 | 0 | 0 |

| | | | | | | | | | | | | | | | | | | | | | | |
|---|---|---|---|---|---|---|---|---|---|---|---|---|---|---|---|---|---|---|---|---|---|---|
| VGSHITGG | ATP6V1A; ATP6' | P38606; C9J | VATA_HUMAN ' | Unmodified | 0 | 0 | 0 | 0 | 0 | 0 | 0 | 0 | 0 | 0 | 696350 | 2817800 | 187280 | 0 | 0 | 0 | 0 | 0 |
| VGSLDNAH | MAP2; MAP2 | P11137; A0A | MTAP2_HUMAN | Unmodified | 0 | 0 | 0 | 0 | 0 | 0 | 0 | 0 | 0 | 0 | 0 | 0 | 187920 | 0 | 0 | 0 | 0 | 0 |
| VGSLDNVG | MAP4; MAP4; M | P27816; H7C | MAP4_HUMAN | Unmodified | 174940 | 221990 | 0 | 0 | 0 | 0 | 0 | 0 | 513380 | 0 | 0 | 4932600 | 81273 | 0 | 0 | 0 | 0 | 0 |
| VGSLPLEK | GPRIN1 | Q7Z2K8 | GRIN1_HUMAN | Unmodified | 768720 | 124090 | 1094200 | 0 | 163540 | 0 | 129060 | 246990 | 2761300 | 385350 | 1205500 | 1063400 | 206300 | 228220 | 127230 | 335100 | 428850 | 998240 |
| VGSPFELV | CD109 | Q6YHK3 | CD109_HUMAI | Unmodified | 0 | 0 | 0 | 174830 | 0 | 0 | 0 | 196680 | 11519000 | 3245200 | 0 | 0 | 0 | 0 | 0 | 1572600 | 1284900 | 1278400 |
| VGSPFELV | CD109 | Q6YHK3 | CD109_HUMAI | Unmodified | 0 | 0 | 0 | 0 | 0 | 0 | 0 | 29749 | 4093000 | 222910 | 0 | 0 | 0 | 0 | 0 | 0 | 0 | 0 |
| VGSTKVPM | ATP2A2; ATP2A | P16615; H7C | AT2A2_HUMAN | Unmodified | 0 | 0 | 0 | 374320 | 0 | 0 | 662020 | 345050 | 417420 | 0 | 0 | 921100 | 188860 | 0 | 0 | 87873 | 0 | 0 |
| VGSTSENIT | DDX3X; DDX3Y | O00571; O15 | DDX3X_HUMAN | Unmodified | 1238700 | 1525400 | 1459300 | 802980 | 0 | 109560 | 0 | 0 | 475620 | 0 | 71383 | 2322400 | 0 | 0 | 2083800 | 1430000 | 0 | 0 |
| VGSVEEFQ | MOV10; MOV10 | Q9HCE1; Q5 | MOV10_HUMA | Unmodified | 3591600 | 4810200 | 1125300 | 1653000 | 0 | 0 | 0 | 0 | 0 | 0 | 0 | 1129400 | 0 | 0 | 2626600 | 3291400 | 0 | 0 |
| VGSVLQEG | SDHA; SDHA | P31040; A0A | SDHA_HUMAN | Unmodified | 0 | 0 | 0 | 0 | 0 | 0 | 0 | 0 | 2250000 | 225750 | 0 | 0 | 103690 | 0 | 0 | 0 | 0 | 0 |
| VGSVLQEG | SDHA; SDHA | P31040; A0A | SDHA_HUMAN | Unmodified | 0 | 0 | 0 | 0 | 0 | 0 | 0 | 0 | 0 | 166760 | 0 | 0 | 0 | 0 | 0 | 0 | 0 | 0 |
| VGSVVSVG | SLC20A1 | Q8WUM9 | S20A1_HUMAN | Unmodified | 0 | 0 | 0 | 0 | 0 | 0 | 0 | 0 | 0 | 0 | 0 | 758410 | 0 | 0 | 0 | 0 | 0 | 0 |
| VGTAQLAL' | EIF2B4; EIF2B4 | Q9UI10; A0A | EI2BD_HUMAN | Unmodified | 0 | 0 | 0 | 0 | 0 | 0 | 0 | 0 | 0 | 0 | 0 | 2294100 | 0 | 0 | 174390 | 0 | 0 | 0 |
| VGTDGVIT' | CDH1; CDH1; C | P12830; H3E | CADH1_HUMA | Unmodified | 0 | 420560 | 1059300 | 0 | 0 | 0 | 0 | 0 | 814040 | 156000 | 0 | 0 | 0 | 0 | 902120 | 592210 | 0 | 0 |
| VGTGEPCC | PGD; PGD; PGD | P52209; K7E | 6PGD_HUMAN | Unmodified | 259670 | 0 | 0 | 605940 | 4995100 | 1649100 | 1002400 | 973970 | 1595400 | 1725900 | 942620 | 5744100 | 3434600 | 2733800 | 485510 | 438800 | 1476800 | 2134700 |
| VGTPHFMA | CASK; CASK; C | O14936; A0A | CSKP_HUMAN | Oxidation (M) | 0 | 0 | 298160 | 119790 | 0 | 0 | 0 | 0 | 0 | 0 | 0 | 149200 | 0 | 0 | 321260 | 642890 | 0 | 0 |
| VGTPHFMA | CASK; CASK; C | O14936; A0A | CSKP_HUMAN | Unmodified | 0 | 0 | 0 | 0 | 0 | 0 | 0 | 0 | 0 | 0 | 0 | 0 | 0 | 0 | 0 | 0 | 0 | 0 |
| VGTSFSIPV | EMG1; EMG1; E | Q92979; V9C | NEP1_HUMAN | Unmodified | 0 | 0 | 0 | 0 | 0 | 0 | 0 | 0 | 0 | 0 | 0 | 1215000 | 0 | 0 | 146450 | 0 | 0 | 0 |
| VGTVGYMA | GRK5 | P34947 | GRK5_HUMAN | Oxidation (M) | 0 | 0 | 0 | 0 | 0 | 0 | 0 | 0 | 0 | 0 | 0 | 576050 | 0 | 0 | 0 | 0 | 0 | 0 |
| VGTVIGSNI | PARP1; PARP1 | P09874; A0A | PARP1_HUMAN | Unmodified | 0 | 0 | 92967 | 121030 | 0 | 2193900 | 47419 | 976360 | 325160 | 2505300 | 0 | 472130 | 1864400 | 2397700 | 0 | 126790 | 26044 | 60652 |
| VGVASSEE | SNAP29 | O95721 | SNP29_HUMAN | Unmodified | 0 | 46256 | 0 | 0 | 455710 | 0 | 0 | 0 | 1177000 | 193150 | 184760 | 466200 | 0 | 0 | 0 | 193060 | 0 | 564390 |
| VGVASSEE | SNAP29 | O95721 | SNP29_HUMAN | Unmodified | 0 | 0 | 0 | 27080 | 24585 | 756640 | 20777 | 0 | 0 | 284440 | 0 | 0 | 162170 | 103940 | 0 | 0 | 62560 | 606810 |
| VGVAYKLD | ADSS2 | P30520 | PURA2_HUMAI | Unmodified | 0 | 0 | 0 | 0 | 0 | 0 | 0 | 0 | 0 | 0 | 0 | 1830800 | 0 | 0 | 0 | 0 | 0 | 0 |
| VGVDPLIIP | XPNPEP1; XPNI | Q9NQW7; Q | XPP1_HUMAN | Unmodified | 0 | 0 | 0 | 0 | 0 | 0 | 0 | 0 | 296580 | 0 | 426830 | 2575800 | 0 | 0 | 905180 | 367390 | 0 | 0 |
| VGVDYEGC | PLOD1 | Q02809 | PLOD1_HUMAI | Unmodified | 1926900 | 0 | 0 | 0 | 0 | 0 | 0 | 0 | 803050 | 0 | 0 | 0 | 0 | 0 | 1367200 | 0 | 0 | 0 |
| VGVEVPDV | AHNAK | Q09666 | AHNK_HUMAN | Unmodified | 121860 | 0 | 0 | 0 | 6263900 | 1329400 | 0 | 0 | 0 | 0 | 0 | 0 | 0 | 0 | 0 | 0 | 1293700 | 1836100 |
| VGVEVPDV | AHNAK | Q09666 | AHNK_HUMAN | Unmodified | 0 | 0 | 0 | 0 | 2519300 | 0 | 0 | 167270 | 0 | 71681 | 0 | 0 | 0 | 0 | 0 | 0 | 0 | 1631800 |
| VGVEVPDV | AHNAK | Q09666 | AHNK_HUMAN | Unmodified | 0 | 0 | 0 | 0 | 627940 | 0 | 0 | 0 | 0 | 0 | 0 | 0 | 0 | 0 | 0 | 0 | 807710 | 847980 |
| VGVHIADV | DIS3L2 | Q8IYB7 | DI3L2_HUMAN | Unmodified | 240300 | 328970 | 0 | 0 | 0 | 150080 | 0 | 0 | 140140 | 86954 | 273240 | 2898000 | 0 | 0 | 0 | 0 | 190570 | 80748 |
| VGVIGSICI | MEGF9 | Q9H1U4 | MEGF9_HUMA | Unmodified | 0 | 0 | 0 | 0 | 0 | 0 | 401470 | 0 | 1717600 | 0 | 0 | 369650 | 0 | 0 | 0 | 0 | 31266 | 0 |
| VGVIGSICI | MEGF9 | Q9H1U4 | MEGF9_HUMAI | Unmodified | 0 | 0 | 0 | 0 | 1152300 | 260120 | 232320 | 40289 | 2400800 | 1416400 | 277840 | 0 | 0 | 0 | 0 | 0 | 0 | 0 |
| VGVKPVGS | TXNL1; TXNL1; | O43396; K7E | TXNL1_HUMAN | Unmodified | 2940700 | 553330 | 2839100 | 4351100 | 2182900 | 322710 | 264940 | 254460 | 4593900 | 935160 | 2814700 | 10567000 | 3515200 | 1197700 | 4453000 | 3744900 | 2252600 | 1203400 |
| VGVKPVGS | TXNL1; TXNL1 | O43396; K7E | TXNL1_HUMAN | Unmodified | 0 | 0 | 0 | 0 | 12949000 | 0 | 0 | 184930 | 0 | 5863100 | 0 | 0 | 11030000 | 453360 | 0 | 0 | 18970000 | 582450 |
| VGVLAASM | LIMA1; LIMA1; L | Q9UHB6; F8 | LIMA1_HUMAN | Unmodified | 0 | 0 | 0 | 0 | 1733700 | 197750 | 0 | 0 | 0 | 0 | 0 | 547160 | 0 | 717970 | 0 | 105140 | 188350 | 0 |
| VGVNGFGI | GAPDH | P04406 | G3P_HUMAN ( | Unmodified | 11423000 | 7610300 | 11649000 | 13304000 | 68058000 | 4621800 | 8656300 | 7900200 | 26767000 | 8900500 | 26899000 | 3.51E+08 | 25384000 | 20283000 | 20321000 | 20373000 | 17995000 | 36434000 |
| VGVSEAGL | LRSAM1; LRSAI | Q6UWE0; A( | LRSM1_HUMA | Unmodified | 0 | 79883 | 249530 | 101870 | 0 | 0 | 0 | 0 | 0 | 0 | 67357 | 0 | 0 | 0 | 171300 | 231370 | 0 | 0 |
| VGVSQQPE | PLXNB2; PLXNE | O15031; A6( | PLXB2_HUMAN | Unmodified | 0 | 0 | 1877900 | 1726100 | 0 | 0 | 173650 | 344640 | 3603800 | 751500 | 1290800 | 8999600 | 257770 | 206220 | 4358500 | 3810200 | 368320 | 194080 |
| VGVTVVGP | EPHA3 | P29320 | EPHA3_HUMAI | Unmodified | 575550 | 688350 | 0 | 0 | 704870 | 61419 | 150950 | 69912 | 0 | 141460 | 188440 | 0 | 0 | 0 | 0 | 0 | 280560 | 227470 |
| VGVWNVP' | PLOD1 | Q02809 | PLOD1_HUMAI | Unmodified | 0 | 0 | 0 | 0 | 0 | 0 | 0 | 0 | 675930 | 289190 | 0 | 0 | 0 | 0 | 22199 | 0 | 0 | 0 |
| VGVWNVP' | PLOD3; PLOD3 | O60568; H7( | PLOD3_HUMAI | Unmodified | 0 | 0 | 0 | 0 | 74130 | 0 | 0 | 0 | 3289600 | 93493 | 0 | 0 | 0 | 0 | 0 | 130980 | 31850 | 0 |
| VGVWNVP' | PLOD3; PLOD3 | O60568; H7( | PLOD3_HUMAI | Unmodified | 0 | 0 | 0 | 0 | 0 | 0 | 0 | 0 | 2562400 | 0 | 0 | 0 | 0 | 0 | 0 | 0 | 0 | 0 |
| VGVWNVP' | PLOD2; PLOD2 | O00469; E7E | PLOD2_HUMAI | Unmodified | 0 | 0 | 0 | 0 | 0 | 18286 | 0 | 0 | 478560 | 0 | 0 | 0 | 14013 | 0 | 13775 | 0 | 0 | 0 |
| VGVWNVP' | PLOD2; PLOD2 | O00469; E7E | PLOD2_HUMAI | Oxidation (M) | 0 | 0 | 0 | 0 | 0 | 0 | 6440.5 | 0 | 670780 | 240360 | 0 | 0 | 0 | 0 | 0 | 0 | 0 | 20938 |
| VGWEQLLT | ACTN4; ACTN1 | O43707; P12 | ACTN4_HUMAI | Unmodified | 929970 | 1304000 | 283540 | 0 | 223700 | 261550 | 0 | 0 | 1534900 | 168410 | 0 | 23111000 | 0 | 0 | 2009500 | 2378200 | 0 | 0 |
| VGWIFTDL' | NPLOC4 | Q8TAT6 | NPL4_HUMAN | Unmodified | 582300 | 680230 | 0 | 560010 | 0 | 140590 | 0 | 0 | 2341000 | 3041300 | 830870 | 5618600 | 817640 | 452180 | 714420 | 468220 | 156440 | 404230 |
| VGYPFLR | ITGA1 | P56199 | ITA1_HUMAN I | Unmodified | 1903900 | 1420700 | 0 | 30173 | 130490 | 0 | 0 | 0 | 251190 | 0 | 0 | 301470 | 0 | 0 | 17006 | 0 | 598270 | 188030 |
| VGYTPDWI | CLTCL1; CLTC; | P53675; Q0( | CLH2_HUMAN | Unmodified | 8304400 | 4669700 | 437710 | 964080 | 0 | 21493 | 0 | 0 | 0 | 267620 | 0 | 3283200 | 0 | 0 | 5109300 | 2426200 | 0 | 0 |
| VGYYVNNE | ASF1A | Q9Y294 | ASF1A_HUMAN | Unmodified | 0 | 0 | 0 | 0 | 0 | 0 | 0 | 0 | 0 | 0 | 0 | 1831100 | 0 | 0 | 0 | 0 | 0 | 0 |
| VHACGVNI | CRYZ; CRYZ; CF | Q08257; A6I | QOR_HUMAN ( | Unmodified | 0 | 0 | 0 | 0 | 0 | 0 | 0 | 446930 | 859580 | 0 | 0 | 0 | 116010 | 0 | 0 | 0 | 0 | 0 |
| VHAENGEL | DPYSL5 | Q9BPU6 | DPYL5_HUMAN | Unmodified | 22076 | 0 | 0 | 0 | 0 | 0 | 0 | 134720 | 0 | 0 | 748710 | 3518100 | 0 | 0 | 24933 | 19061 | 104160 | 0 |
| VHAIPLR | GNB2; GNB1; G | P62879; P62 | GBB2_HUMAN | Unmodified | 1874800 | 0 | 983130 | 695150 | 316740 | 149280 | 164350 | 0 | 3706000 | 717080 | 2712500 | 43486000 | 109610 | 0 | 2513600 | 2189900 | 183170 | 184250 |
| VHAIPLRSS | GNB1; GNB1; G | P62873; F6U | GBB1_HUMAN | Oxidation (M) | 0 | 0 | 0 | 0 | 0 | 0 | 0 | 0 | 0 | 602400 | 0 | 0 | 0 | 11023000 | 0 | 0 | 3638500 | 0 |
| VHAIQCDV | DECR1; DECR1 | Q16698; E5F | DECR_HUMAN | Unmodified | 0 | 0 | 0 | 0 | 0 | 0 | 0 | 0 | 2570900 | 259120 | 0 | 0 | 0 | 0 | 0 | 0 | 0 | 0 |
| VHAPGLNL | AHNAK | Q09666 | AHNK_HUMAN | Unmodified | 0 | 0 | 0 | 0 | 2754700 | 270040 | 0 | 0 | 0 | 0 | 0 | 0 | 312110 | 0 | 0 | 0 | 465350 | 2459100 |
| VHAYIISSLI | EHD1; EHD3; E | Q9H4M9; Q9 | EHD1_HUMAN | Unmodified | 586940 | 714190 | 1788500 | 1131500 | 10285000 | 1494700 | 2454100 | 962350 | 362400 | 433830 | 674720 | 10586000 | 2532600 | 3118500 | 1110400 | 1007400 | 4562600 | 3813400 |
| VHAYIISYLI | EHD4; EHD2 | Q9H223; Q9 | EHD4_HUMAN | Unmodified | 305480 | 364370 | 616010 | 175940 | 688900 | 191690 | 0 | 0 | 0 | 128340 | 0 | 1292900 | 189500 | 280880 | 1119400 | 851440 | 0 | 0 |
| VHCCLYFIF | SEPTIN9; SEPTI | Q9UHD8; K7 | SEPT9_HUMAN | Oxidation (M) | 0 | 0 | 107050 | 135740 | 0 | 0 | 0 | 0 | 0 | 0 | 0 | 817800 | 160010 | 0 | 142620 | 0 | 0 | 0 |
| VHCDVHF( | MUC5B | Q9HC84 | MUC5B_HUMA | Unmodified | 0 | 0 | 0 | 0 | 0 | 0 | 0 | 0 | 0 | 0 | 0 | 0 | 0 | 0 | 0 | 0 | 0 | 2581200 |
| VHDSNIYIK | PPM1H | Q9ULR3 | PPM1H_HUMA | Unmodified | 0 | 0 | 0 | 34504 | 0 | 0 | 0 | 0 | 0 | 0 | 0 | 1579800 | 227070 | 0 | 0 | 0 | 0 | 0 |
| VHEELRNL | GANAB; GANAE | Q14697; F5I | GANAB_HUMAI | Unmodified | 0 | 0 | 0 | 0 | 0 | 0 | 0 | 0 | 7130100 | 0 | 0 | 0 | 0 | 0 | 0 | 0 | 0 | 0 |
| VHELNEEI( | LUC7L2; LUC7L | Q9Y383; Q9I | LC7L2_HUMAN | Unmodified | 0 | 0 | 0 | 0 | 1054800 | 0 | 0 | 57044 | 105600 | 420890 | 455800 | 410980 | 0 | 0 | 0 | 0 | 2164100 | 0 |
| VHELQGN/ | ELP1; ELP1; ELF | O95163; A0/ | ELP1_HUMAN I | Unmodified | 0 | 70466 | 0 | 0 | 439170 | 122520 | 176390 | 85254 | 255850 | 0 | 1015900 | 3454100 | 83318 | 0 | 0 | 217500 | 0 | 181890 |
| VHEYNVLLI | LRPAP1; LRPAP | P30533; A0A | AMRP_HUMAN | Unmodified | 0 | 0 | 0 | 0 | 0 | 0 | 0 | 0 | 3289700 | 44122 | 141260 | 74713 | 0 | 0 | 0 | 0 | 0 | 0 |
| VHFEESSKI | XRCC6; XRCC6 | P12956; B1A | XRCC6_HUMAI | Unmodified | 0 | 0 | 0 | 0 | 0 | 0 | 0 | 0 | 0 | 0 | 315410 | 1252200 | 0 | 0 | 0 | 0 | 0 | 0 |
| VHFEESSKI | XRCC6; XRCC6 | P12956; B1A | XRCC6_HUMAI | Unmodified | 85035 | 0 | 0 | 0 | 0 | 0 | 0 | 0 | 0 | 0 | 0 | 1451500 | 0 | 0 | 97384 | 0 | 0 | 0 |
| VHFFTQDH | PPM1A; PPM1A | P35813; E9P | PPM1A_HUMAI | Unmodified | 2322500 | 700550 | 1470600 | 1590600 | 1328000 | 617610 | 790660 | 257290 | 963420 | 414350 | 441080 | 3069500 | 1468900 | 1094900 | 2849100 | 2381500 | 1859400 | 3231900 |
| VHFTAERS | SND1 | Q7KZF4 | SND1_HUMAN | Unmodified | 0 | 0 | 0 | 0 | 0 | 289510 | 421180 | 216350 | 0 | 0 | 0 | 0 | 0 | 933090 | 0 | 0 | 826340 | 0 |
| VHGDPAVV | FUT8; FUT8; FU | Q9BYC5; A0, | FUT8_HUMAN , | Unmodified | 0 | 0 | 303150 | 303970 | 0 | 0 | 0 | 0 | 5671200 | 478330 | 0 | 0 | 0 | 0 | 1558600 | 882940 | 0 | 0 |
| VHGERTISE | TARS1 | P26639 | SYTC_HUMAN | Unmodified | 0 | 0 | 0 | 0 | 8178800 | 1725200 | 630790 | 0 | 0 | 596280 | 0 | 0 | 2848200 | 1891100 | 0 | 0 | 0 | 2067200 |
| VHGLRNV( | ATP5F1A; ATP5I | P25705; K7E | ATPA_HUMAN , | Unmodified | 0 | 0 | 0 | 0 | 0 | 0 | 0 | 0 | 0 | 711680 | 0 | 0 | 0 | 0 | 0 | 0 | 0 | 0 |
| VHGPGIQS | FLNA; FLNA | P21333; Q6( | FLNA_HUMAN | Unmodified | 0 | 2367200 | 1543400 | 0 | 5517500 | 213090 | 1336200 | 1395900 | 2880700 | 102820 | 0 | 3976700 | 1720100 | 654690 | 3242600 | 3759600 | 0 | 794010 |
| VHHFSPVL | TOR2A; TOR2A | Q8N2E6; Q5 | TOR2X_HUMAN | Unmodified | 0 | 0 | 0 | 0 | 0 | 0 | 0 | 0 | 843620 | 0 | 0 | 0 | 0 | 0 | 0 | 0 | 0 | 0 |
| VHIGQVIM: | RPL10; RPL10L; | P27635; Q9( | RL10_HUMAN ( | Oxidation (M) | 2442800 | 1774300 | 1445700 | 710120 | 0 | 0 | 0 | 0 | 0 | 0 | 901010 | 20560000 | 0 | 0 | 1009200 | 1415400 | 0 | 0 |
| VHIGQVIM: | RPL10; RPL10L; | P27635; Q9( | RL10_HUMAN ( | Unmodified | 0 | 0 | 0 | 0 | 0 | 0 | 0 | 0 | 0 | 0 | 0 | 4823400 | 0 | 0 | 0 | 0 | 0 | 0 |
| VHIHHPPE. | LTBP2; LTBP2; | Q14767; G3' | LTBP2_HUMAN | Unmodified | 0 | 0 | 0 | 0 | 0 | 0 | 0 | 0 | 335340 | 0 | 0 | 0 | 0 | 0 | 0 | 0 | 0 | 0 |
| VHKVEDLL | TTC13; TTC13 | Q8NBP0; H0 | TTC13_HUMAN | Unmodified | 0 | 0 | 0 | 0 | 0 | 0 | 0 | 0 | 1016500 | 90745 | 0 | 0 | 0 | 0 | 0 | 0 | 0 | 0 |
| VHLDIQVG | TMED9 | Q9BVK6 | TMED9_HUMAI | Unmodified | 0 | 0 | 0 | 0 | 0 | 0 | 0 | 0 | 2444700 | 238360 | 0 | 0 | 237810 | 0 | 0 | 0 | 0 | 0 |
| VHLDIQVG | TMED9 | Q9BVK6 | TMED9_HUMAI | Unmodified | 0 | 0 | 0 | 0 | 0 | 0 | 0 | 0 | 1737500 | 0 | 0 | 0 | 0 | 0 | 0 | 0 | 0 | 0 |
| VHLDIQVG | TMED4 | Q7Z7H5 | TMED4_HUMAI | Unmodified | 0 | 0 | 0 | 0 | 0 | 0 | 0 | 0 | 9821600 | 775730 | 0 | 0 | 0 | 0 | 0 | 0 | 0 | 0 |
| VHLDKAQ( | RPS7; RPS7 | P62081; A0A | RS7_HUMAN 4 | Unmodified | 0 | 0 | 0 | 0 | 0 | 0 | 444320 | 0 | 0 | 0 | 0 | 0 | 1238900 | 0 | 0 | 0 | 0 | 426630 |
| VHLNTFGF | GPR107; GPR1( | Q5VW38; U3 | GP107_HUMAI | Unmodified | 45104 | 0 | 0 | 0 | 703330 | 0 | 0 | 0 | 13863000 | 731510 | 199060 | 494210 | 340850 | 0 | 0 | 0 | 0 | 0 |
| VHLQLWDT | RAB27B; RAB27 | O00194; K7E | RB27B_HUMAN | Unmodified | 223300 | 320460 | 517240 | 199590 | 0 | 0 | 0 | 0 | 0 | 695430 | 469990 | 329260 | 0 | 0 | 651770 | 673270 | 0 | 0 |
| VHLRPLKLI | AFG3L2 | Q9Y4W6 | AFG32_HUMAN | Unmodified | 0 | 0 | 0 | 0 | 0 | 0 | 228710 | 0 | 2040800 | 0 | 0 | 0 | 0 | 0 | 0 | 0 | 0 | 0 |
| VHLSDSER | SDC2; SDC2; S | P34741; E9P | SDC2_HUMAN | Unmodified | 188720 | 0 | 0 | 0 | 0 | 0 | 130270 | 0 | 0 | 0 | 0 | 0 | 605870 | 0 | 0 | 0 | 0 | 0 |
| VHLVGIDIF | EIF5A; EIF5AL1; | P63241; Q6I | IF5A1_HUMAN | Unmodified | 24591000 | 13220000 | 5558800 | 9722500 | 82143000 | 15485000 | 14978000 | 753570 | 21033000 | 11235000 | 19948000 | 58039000 | 53548000 | 27967000 | 24414000 | 20413000 | 18492000 | 32299000 |
| VHLVGIDIF | EIF5A; EIF5AL1; | P63241; Q6I | IF5A1_HUMAN | Oxidation (M) | 0 | 0 | 0 | 0 | 16047000 | 0 | 1723600 | 0 | 0 | 1764400 | 0 | 0 | 19083000 | 0 | 0 | 0 | 7065700 | 5158800 |
| VHMPMDC | PLEKHB2; PLEK | Q96CS7; A0, | PKHB2_HUMAI | Unmodified | 236890 | 189790 | 84593 | 233760 | 0 | 0 | 0 | 0 | 1619500 | 0 | 0 | 0 | 0 | 0 | 250640 | 80605 | 0 | 0 |
| VHNDAQSI | CALU | O43852 | CALU_HUMAN | Unmodified | 0 | 0 | 5790900 | 109390 | 0 | 0 | 54708 | 0 | 26645000 | 1492800 | 14614000 | 1148700 | 89109 | 0 | 0 | 98049 | 0 | 0 |
| VHPEGKFV | PSMC5; PSMC5 | P62195; J3Q | PRS8_HUMAN | Unmodified | 0 | 0 | 0 | 0 | 1003000 | 0 | 409570 | 0 | 0 | 0 | 0 | 0 | 1236000 | 657940 | 0 | 0 | 590780 | 2574800 |
| VHPEIINEN | TXNDC12 | O95881 | TXD12_HUMAN | Unmodified | 0 | 0 | 239360 | 0 | 190710 | 0 | 0 | 68431 | 32159000 | 1446300 | 2419700 | 849550 | 160710 | 0 | 109880 | 274450 | 0 | 49109 |
| VHPVSTMII | LDHA | P00338 | LDHA_HUMAN | Unmodified | 20735 | 0 | 0 | 0 | 0 | 0 | 0 | 0 | 0 | 0 | 891700 | 5159700 | 0 | 0 | 88008 | 56789 | 0 | 0 |
| VHQLYETI( | DCTN2; DCTN2 | Q13561; A8I | DCTN2_HUMAI | Unmodified | 0 | 0 | 0 | 0 | 0 | 0 | 0 | 0 | 285630 | 0 | 76119 | 3525900 | 0 | 0 | 0 | 0 | 0 | 0 |
| VHQLYETI( | DCTN2; DCTN2 | Q13561; A8I | DCTN2_HUMAI | Unmodified | 0 | 0 | 0 | 0 | 2404500 | 0 | 3527800 | 0 | 0 | 657800 | 0 | 0 | 0 | 0 | 0 | 467650 | 0 | 9531500 |
| VHQVLYR | TST; TST | Q16762; B1/ | THTR_HUMAN | Acetyl (Protein N | 12936000 | 10937000 | 2836500 | 1097700 | 0 | 0 | 0 | 0 | 0 | 0 | 0 | 0 | 0 | 0 | 0 | 5947600 | 0 | 0 |
| VHQVLYR | TST; TST | Q16762; B1/ | THTR_HUMAN | Unmodified | 834110 | 1393900 | 0 | 63430 | 0 | 0 | 0 | 0 | 0 | 0 | 0 | 0 | 0 | 0 | 0 | 595950 | 0 | 0 |
| VHSAVITVF | HSPA2 | P54652 | HSP72_HUMAI | Unmodified | 1963800 | 1258500 | 0 | 0 | 1423300 | 0 | 0 | 0 | 0 | 0 | 74707 | 3002500 | 0 | 0 | 0 | 0 | 199360 | 0 |
| VHSAVITVF | HSPA2 | P54652 | HSP72_HUMAI | Unmodified | 0 | 0 | 0 | 0 | 10216000 | 0 | 0 | 0 | 0 | 0 | 0 | 0 | 0 | 0 | 0 | 0 | 1343000 | 3144100 |
| VHSFPTLK | P4HB; P4HB; P4 | P07237; A0A | PDIA1_HUMAN | Unmodified | 72785 | 0 | 870330 | 0 | 1448900 | 0 | 2237700 | 2203800 | 39233000 | 12402000 | 3049900 | 3393100 | 1688700 | 1153900 | 415480 | 370610 | 128550 | 564480 |
| VHSPSGAL | FLNA; FLNA; FL | P21333; A0A | FLNA_HUMAN | Unmodified | 456620 | 831930 | 488410 | 657390 | 0 | 0 | 0 | 0 | 0 | 0 | 0 | 0 | 240910 | 0 | 1302800 | 947760 | 0 | 0 |
| VHSPSGAV | FLNB; FLNB | O75369; E7E | FLNB_HUMAN | Unmodified | 0 | 0 | 661830 | 0 | 0 | 0 | 0 | 0 | 0 | 0 | 0 | 1283900 | 0 | 0 | 564770 | 0 | 28319 | 0 |
| VHSPSGAV | FLNB; FLNB | O75369; E7E | FLNB_HUMAN | Unmodified | 0 | 0 | 0 | 486800 | 1805800 | 0 | 0 | 0 | 0 | 0 | 208940 | 1250500 | 151340 | 0 | 0 | 174760 | 526630 | 214850 |
| VHSRGGLI | CAPN5; CAPN5 | O15484; E7E | CAN5_HUMAN | Unmodified | 0 | 0 | 0 | 0 | 813980 | 226470 | 433630 | 0 | 0 | 185200 | 0 | 0 | 0 | 1645600 | 0 | 0 | 405820 | 272010 |
| VHSVSALA | UEVLD | Q8IX04 | UEVLD_HUMAN | Unmodified | 0 | 0 | 0 | 0 | 1366900 | 0 | 0 | 0 | 381650 | 157640 | 0 | 289290 | 175110 | 141930 | 0 | 0 | 0 | 0 |
| VHTFVVDC | HSD17B11; HS | Q8NBQ5; DE | DHB11_HUMAI | Unmodified | 0 | 0 | 0 | 0 | 238830 | 113510 | 0 | 0 | 638040 | 0 | 0 | 0 | 0 | 0 | 0 | 0 | 0 | 0 |
| VHVGDEDI | CSTB; CSTB | P04080; A0A | CYTB_HUMAN | Unmodified | 282830 | 233390 | 0 | 110020 | 0 | 0 | 0 | 0 | 0 | 87166 | 0 | 1725800 | 0 | 0 | 1015800 | 592210 | 0 | 0 |
| VHVGDEDI | CSTB; CSTB | P04080; A0A | CYTB_HUMAN | Unmodified | 0 | 0 | 0 | 0 | 3079900 | 0 | 968620 | 0 | 0 | 2607500 | 0 | 0 | 0 | 439640 | 1543100 | 440060 | 4518300 | 0 |
| VHVIFNYK | CALR; CALR; C/ | P27797; A0A | CALR_HUMAN | Unmodified | 190240 | 0 | 250780 | 133300 | 771880 | 122300 | 4039500 | 2025800 | 1E+08 | 57402000 | 5719300 | 1318900 | 167220 | 604740 | 380880 | 367350 | 476680 | 889340 |
| VHVIFNYK( | CALR; CALR; C/ | P27797; A0A | CALR_HUMAN | Unmodified | 0 | 0 | 0 | 0 | 0 | 0 | 2821500 | 1282000 | 349110 | 3468400 | 0 | 0 | 0 | 0 | 0 | 0 | 0 | 0 |
| VHVLWWN | NIPSNAP3B; NI | Q9BS92; Q9 | NPS3B_HUMAI | Unmodified | 0 | 0 | 0 | 0 | 0 | 0 | 0 | 0 | 4513000 | 185890 | 0 | 0 | 0 | 0 | 0 | 0 | 0 | 0 |

| | | | | | | | | | | | | | | | | | | | | | | |
|---|---|---|---|---|---|---|---|---|---|---|---|---|---|---|---|---|---|---|---|---|---|---|
| VHYENNSF | RPN1; RPN1 | P04843; B7Z | RPN1_HUMAN | Oxidation (M) | 0 | 0 | 0 | 0 | 0 | 0 | 0 | 0 | 26316000 | 2264000 | 2239300 | 1316300 | 0 | 0 | 0 | 0 | 0 | 0 |
| VHYENNSF | RPN1; RPN1 | P04843; B7Z | RPN1_HUMAN | Unmodified | 0 | 0 | 0 | 67233 | 0 | 0 | 0 | 0 | 1.35E+08 | 3052700 | 1439600 | 410710 | 0 | 0 | 0 | 0 | 321510 | 0 |
| VIAAEGEMI | STOM | P27105 | STOM_HUMAN | Unmodified | 1006200 | 1078200 | 955500 | 2711500 | 14780000 | 2429600 | 0 | 709870 | 0 | 0 | 0 | 0 | 556590 | 0 | 1203200 | 1004000 | 0 | 650030 |
| VIAAEGEMI | STOM | P27105 | STOM_HUMAN | Oxidation (M) | 349530 | 471450 | 378030 | 917290 | 3246300 | 888250 | 0 | 0 | 0 | 104160 | 102800 | 0 | 394000 | 0 | 434620 | 303510 | 0 | 0 |
| VIAAEGEMI | STOM | P27105 | STOM_HUMAN | Unmodified | 0 | 0 | 0 | 0 | 72874000 | 4916800 | 9065700 | 2768500 | 0 | 0 | 0 | 0 | 3330700 | 4461200 | 0 | 0 | 0 | 0 |
| VIAAEGEMI | STOM | P27105 | STOM_HUMAN | Oxidation (M) | 0 | 0 | 0 | 0 | 12190000 | 1271300 | 2486100 | 1296300 | 0 | 43537 | 0 | 0 | 0 | 0 | 0 | 0 | 97882 | 335070 |
| VIAALLQTM | IPO5; IPO5 | O00410; H0Y | IPO5_HUMAN I | Unmodified | 0 | 0 | 0 | 0 | 322740 | 0 | 123080 | 0 | 0 | 182750 | 0 | 4457400 | 0 | 0 | 0 | 0 | 0 | 0 |
| VIAALLQTM | IPO5; IPO5 | O00410; H0Y | IPO5_HUMAN I | Oxidation (M) | 30065 | 0 | 0 | 86434 | 0 | 0 | 0 | 0 | 0 | 0 | 0 | 2497000 | 0 | 0 | 0 | 56757 | 0 | 0 |
| VIAAMTVGI | AP2B1; AP2B1; | P63010; K7E | AP2B1_HUMAN | Unmodified | 92572 | 138310 | 172040 | 201240 | 1910100 | 371500 | 227840 | 65564 | 819710 | 452540 | 298560 | 3580400 | 986570 | 772570 | 169350 | 193170 | 616770 | 1369500 |
| VIAATNRVD | PSMC3; PSMC3 | P17980; E9P | PRS6A_HUMAN | Unmodified | 621620 | 511580 | 854370 | 1959800 | 6264400 | 812730 | 1532900 | 268450 | 6408500 | 3391300 | 1828500 | 10368000 | 7036300 | 2244900 | 1413500 | 1461900 | 5490100 | 7785900 |
| VIACFDSLK | MBNL1; MBNL2 | Q9NR56; Q5 | MBNL1_HUMA | Unmodified | 1993800 | 1467000 | 702400 | 1064800 | 0 | 0 | 219160 | 0 | 0 | 0 | 0 | 373980 | 0 | 0 | 411060 | 803400 | 0 | 989450 |
| VIADNVKD' | TPI1; TPI1; TPI1 | P60174; U3K | TPIS_HUMAN T | Unmodified | 0 | 54775 | 0 | 136730 | 18342000 | 2054400 | 3362600 | 2633300 | 0 | 3178100 | 0 | 0 | 14068000 | 10961000 | 0 | 0 | 31769000 | 14891000 |
| VIADNVKD' | TPI1; TPI1; TPI1 | P60174; U3K | TPIS_HUMAN T | Unmodified | 0 | 0 | 0 | 0 | 0 | 0 | 0 | 0 | 0 | 0 | 0 | 0 | 0 | 0 | 0 | 0 | 2433600 | 923850 |
| VIAEAYATK | TTC7B; TTC7B; | Q86TV6; G3' | TTC7B_HUMAN | Unmodified | 0 | 0 | 537760 | 188070 | 876780 | 111350 | 141350 | 55909 | 408480 | 61071 | 0 | 288100 | 240000 | 149710 | 325230 | 238430 | 227320 | 599260 |
| VIAGQFFGI | SMPDL3B; SMP | Q92485; F8\ | ASM3B_HUMA | Unmodified | 3626700 | 3487900 | 8058800 | 6419400 | 0 | 0 | 0 | 0 | 3532000 | 0 | 301280 | 8597800 | 0 | 0 | 13377000 | 9811600 | 0 | 0 |
| VIAINVDDF | PPA1 | Q15181 | IPYR_HUMAN I | Unmodified | 3269500 | 0 | 0 | 0 | 9547100 | 0 | 0 | 230300 | 1831400 | 301810 | 6033700 | 24695000 | 0 | 1765500 | 4205300 | 3005700 | 0 | 3614200 |
| VIAINVDDF | PPA1 | Q15181 | IPYR_HUMAN I | Unmodified | 0 | 0 | 0 | 0 | 0 | 0 | 0 | 0 | 0 | 0 | 1405400 | 3973500 | 0 | 0 | 0 | 0 | 0 | 0 |
| VIALINDQR | APLP1; APLP1; | P51693; B7Z | APLP1_HUMAN | Unmodified | 154400 | 1034400 | 162360 | 0 | 281660 | 720840 | 0 | 171920 | 8434200 | 1650200 | 953000 | 0 | 0 | 557840 | 200710 | 185980 | 252330 | 0 |
| VIAPMLSR | PLOD3; PLOD3 | O60568; H7C | PLOD3_HUMAI | Unmodified | 0 | 0 | 0 | 0 | 0 | 0 | 0 | 0 | 11569000 | 1237900 | 0 | 0 | 0 | 0 | 0 | 0 | 0 | 0 |
| VIASAQSGI | EPHB3 | P54753 | EPHB3_HUMAI | Unmodified | 0 | 0 | 0 | 0 | 1283100 | 645950 | 0 | 0 | 0 | 831610 | 0 | 1092500 | 0 | 51701 | 0 | 131730 | 0 | 0 |
| VIASAQSGI | EPHB3 | P54753 | EPHB3_HUMAI | Oxidation (M) | 17230 | 0 | 0 | 0 | 0 | 0 | 65050 | 0 | 0 | 155420 | 0 | 2832200 | 0 | 0 | 0 | 0 | 0 | 0 |
| VIASELGSN | LARS1; LARS1; | Q9P2J5; A0A | SYLC_HUMAN | Oxidation (M) | 0 | 0 | 0 | 0 | 0 | 0 | 0 | 0 | 0 | 0 | 0 | 1258100 | 0 | 0 | 0 | 0 | 0 | 0 |
| VIASELGSN | LARS1; LARS1; | Q9P2J5; A0A | SYLC_HUMAN | Oxidation (M) | 0 | 0 | 0 | 0 | 0 | 0 | 0 | 0 | 0 | 0 | 0 | 1073900 | 0 | 0 | 0 | 0 | 0 | 0 |
| VIASELGSN | LARS1; LARS1; | Q9P2J5; A0A | SYLC_HUMAN | Unmodified | 0 | 0 | 0 | 0 | 0 | 0 | 225370 | 0 | 0 | 0 | 451190 | 2905800 | 0 | 0 | 0 | 0 | 0 | 0 |
| VIASMTVGI | AP1B1; AP1B1 | Q10567; C9J | AP1B1_HUMAN | Unmodified | 0 | 0 | 138850 | 0 | 310570 | 157260 | 0 | 82681 | 304980 | 149940 | 254770 | 1945600 | 428190 | 249900 | 79941 | 123430 | 24666 | 0 |
| VIATDINES | BDH2; BDH2; B | Q9BUT1; D6 | BDH2_HUMAN | Unmodified | 0 | 0 | 0 | 0 | 0 | 0 | 0 | 0 | 0 | 0 | 0 | 0 | 0 | 0 | 0 | 0 | 0 | 0 |
| VIATFTCSG | ALDH9A1 | P49189 | AL9A1_HUMAN | Unmodified | 0 | 0 | 339820 | 498010 | 228060 | 0 | 0 | 0 | 917940 | 243990 | 240970 | 1287700 | 740390 | 400100 | 558630 | 392990 | 157420 | 180900 |
| VIDDTNITR | KRT18; KRT18 | P05783; F8V | K1C18_HUMAN | Unmodified | 0 | 0 | 354640 | 211060 | 0 | 0 | 0 | 0 | 5233100 | 656360 | 7957300 | 29977000 | 0 | 0 | 0 | 468330 | 0 | 0 |
| VIDFDENT/ | RCN2; RCN2 | Q14257; H0Y | RCN2_HUMAN | Unmodified | 0 | 0 | 0 | 0 | 0 | 0 | 0 | 0 | 0 | 0 | 1601200 | 0 | 0 | 0 | 0 | 0 | 0 | 0 |
| VIDFDENT/ | RCN2; RCN2 | Q14257; H0Y | RCN2_HUMAN | Unmodified | 0 | 0 | 0 | 0 | 0 | 0 | 0 | 0 | 4632800 | 385040 | 1943700 | 0 | 0 | 0 | 0 | 0 | 0 | 0 |
| VIDGLETLD | PPIL3; PPIL3; PI | Q9H2H8; B8 | PPIL3_HUMAN | Unmodified | 988750 | 1639400 | 242020 | 990520 | 7421400 | 980730 | 57457 | 547720 | 1588000 | 1512000 | 864380 | 1261200 | 3509300 | 3143000 | 1084300 | 811560 | 1341800 | 2501800 |
| VIDGTSGA' | BIRC6; BIRC6 | Q9NR09; A0 | BIRC6_HUMAN | Unmodified | 0 | 0 | 0 | 0 | 478990 | 0 | 0 | 0 | 0 | 0 | 0 | 1063200 | 0 | 0 | 0 | 0 | 170450 | 0 |
| VIDIFDTDG | PPP3R1; PPP3R | P63098; F6U | CANB1_HUMA | Unmodified | 126360 | 0 | 92127 | 146340 | 0 | 0 | 0 | 0 | 0 | 0 | 271420 | 1060400 | 0 | 0 | 403770 | 152310 | 0 | 0 |
| VIDPATATS | CCT4 | P50991 | TCPD_HUMAN | Unmodified | 566230 | 0 | 1974200 | 1274400 | 788740 | 0 | 696970 | 0 | 5795200 | 0 | 7769500 | 30574000 | 9117700 | 0 | 1066700 | 1509100 | 0 | 416880 |
| VIDPATATS | CCT4 | P50991 | TCPD_HUMAN | Unmodified | 0 | 0 | 0 | 558650 | 13327000 | 1698600 | 3216200 | 1992800 | 1994600 | 5483100 | 1867200 | 0 | 18620000 | 9377900 | 0 | 0 | 4425900 | 8453800 |
| VIDPATATS | CCT4 | P50991 | TCPD_HUMAN | Unmodified | 0 | 0 | 0 | 0 | 4720100 | 0 | 6581300 | 1703900 | 0 | 2713900 | 0 | 0 | 5179100 | 5945900 | 0 | 0 | 4476900 | 3401100 |
| VIDPPDTNI | NELL1; NELL1; | Q92832; J3K | NELL1_HUMAN | Unmodified | 0 | 0 | 0 | 0 | 0 | 0 | 0 | 0 | 811910 | 0 | 0 | 0 | 0 | 0 | 0 | 0 | 0 | 0 |
| VIDPVAPR | EPRS1; EPRS1 | P07814; V9C | SYEP_HUMAN | Unmodified | 0 | 0 | 690920 | 388270 | 0 | 0 | 0 | 0 | 0 | 0 | 1056000 | 5012000 | 842250 | 413100 | 347720 | 263360 | 0 | 0 |
| VIDPVAPRY | EPRS1; EPRS1 | P07814; V9C | SYEP_HUMAN | Unmodified | 0 | 0 | 0 | 0 | 507880 | 393120 | 459200 | 144910 | 0 | 0 | 0 | 0 | 772180 | 1170400 | 0 | 0 | 411760 | 345160 |
| VIDQQNGL | RPA1 | P27694 | RFA1_HUMAN | Unmodified | 0 | 252660 | 0 | 1176400 | 0 | 0 | 0 | 0 | 0 | 0 | 0 | 1167800 | 0 | 0 | 0 | 0 | 0 | 0 |
| VIEDDPEA\ | EPHA7; EPHA7; | Q15375; A0/ | EPHA7_HUMAN | Unmodified | 206480 | 219970 | 0 | 0 | 0 | 0 | 0 | 0 | 950810 | 0 | 0 | 687290 | 0 | 0 | 0 | 208210 | 0 | 0 |
| VIEDNEYTA | HCK; LYN; HCK | P08631; P07 | HCK_HUMAN T | Unmodified | 264460 | 506990 | 247710 | 611360 | 233350 | 0 | 0 | 0 | 735580 | 127670 | 269940 | 1924900 | 267060 | 53468 | 470470 | 635900 | 141980 | 0 |
| VIEDNEYTA | HCK; LYN; HCK | P08631; P07 | HCK_HUMAN T | Unmodified | 0 | 271880 | 0 | 0 | 126150 | 0 | 0 | 0 | 534400 | 0 | 0 | 0 | 0 | 0 | 0 | 393490 | 27038 | 0 |
| VIEDRDHYI | MYOF; MYOF | Q9NZM1; HC | MYOF_HUMAN | Unmodified | 0 | 0 | 0 | 0 | 5680800 | 1134000 | 0 | 0 | 0 | 0 | 0 | 0 | 0 | 0 | 0 | 0 | 350100 | 0 |
| VIEDRDSQI | NUP93 | Q8N1F7 | NUP93_HUMAI | Unmodified | 0 | 96143 | 0 | 0 | 0 | 0 | 44794 | 0 | 0 | 22642 | 211140 | 1461800 | 0 | 0 | 537230 | 0 | 0 | 0 |
| VIEEQLEPA | LRPPRC | P42704 | LPPRC_HUMAN | Unmodified | 0 | 0 | 0 | 0 | 0 | 0 | 0 | 0 | 755140 | 257310 | 0 | 0 | 0 | 0 | 0 | 0 | 0 | 0 |
| VIEGDVVS/ | LIMS2; LIMS1; L | Q7Z4I7; P48 | LIMS2_HUMAN | Unmodified | 403860 | 852320 | 0 | 0 | 0 | 1078700 | 0 | 56042 | 244250 | 389380 | 121750 | 1001500 | 0 | 0 | 0 | 0 | 0 | 0 |
| VIEGRQLS( | MYOF | Q9NZM1 | MYOF_HUMAN | Unmodified | 0 | 0 | 0 | 0 | 3145400 | 476880 | 0 | 0 | 0 | 0 | 0 | 0 | 0 | 0 | 0 | 0 | 0 | 0 |
| VIEHIMEDL | S100A9 | P06702 | S10A9_HUMAN | Oxidation (M) | 0 | 0 | 0 | 2610900 | 187290 | 0 | 0 | 0 | 0 | 0 | 0 | 0 | 0 | 0 | 0 | 0 | 0 | 0 |
| VIEITIYEDR | GOLGA7 | Q7Z5G4 | GOGA7_HUMA | Unmodified | 0 | 0 | 0 | 0 | 828360 | 0 | 45194 | 0 | 437190 | 0 | 171350 | 0 | 0 | 0 | 0 | 0 | 162550 | 237980 |
| VIEITIYEDR | GOLGA7 | Q7Z5G4 | GOGA7_HUMA | Oxidation (M) | 243450 | 236620 | 817920 | 525940 | 0 | 0 | 0 | 0 | 0 | 0 | 0 | 0 | 376210 | 0 | 1945200 | 1060200 | 135030 | 215970 |
| VIELENWTI | ECHDC1; ECHI | Q9NTX5; J3K | ECHD1_HUMA | Unmodified | 0 | 0 | 0 | 0 | 0 | 0 | 0 | 0 | 0 | 0 | 1838500 | 2606200 | 0 | 0 | 0 | 0 | 0 | 0 |
| VIELLPEK | RAB3GAP2 | Q9H2M9 | RBGPR_HUMAI | Unmodified | 0 | 0 | 0 | 0 | 411110 | 0 | 102880 | 60546 | 0 | 0 | 130330 | 0 | 109620 | 0 | 0 | 0 | 73752 | 206750 |
| VIELQEIISK | PPFIA1; PPFIA1 | Q13136; E9F | LIPA1_HUMAN | Unmodified | 0 | 0 | 0 | 0 | 1880800 | 0 | 0 | 105820 | 1073900 | 297710 | 0 | 0 | 666350 | 0 | 0 | 0 | 526740 | 0 |
| VIENADGSI | MYH11 | P35749 | MYH11_HUMAI | Unmodified | 0 | 0 | 0 | 0 | 0 | 2894600 | 0 | 0 | 0 | 0 | 0 | 0 | 0 | 0 | 0 | 0 | 0 | 0 |
| VIENRALKD | TPM3; TPM3; TF | P06753; J3K | TPM3_HUMAN | Unmodified | 0 | 0 | 0 | 0 | 0 | 0 | 0 | 0 | 0 | 0 | 0 | 0 | 0 | 0 | 0 | 0 | 0 | 0 |
| VIEPFSR | PSMD11 | O00231 | PSD11_HUMAN | Unmodified | 0 | 0 | 0 | 0 | 0 | 0 | 0 | 0 | 1241200 | 49287 | 97100 | 9999200 | 0 | 0 | 0 | 0 | 0 | 0 |
| VIEPFSRVQ | PSMD11 | O00231 | PSD11_HUMAN | Unmodified | 0 | 0 | 0 | 1162400 | 0 | 0 | 0 | 0 | 0 | 275380 | 777600 | 0 | 607870 | 0 | 0 | 0 | 24036 | 0 |
| VIEPGCVR | DNAJA2 | O60884 | DNJA2_HUMAN | Unmodified | 3652100 | 1855100 | 1468100 | 1687800 | 1253000 | 603330 | 0 | 0 | 2091800 | 0 | 311960 | 3488800 | 1207500 | 800950 | 0 | 1248800 | 0 | 0 |
| VIEPQYFGL | LRPPRC | P42704 | LPPRC_HUMAN | Unmodified | 0 | 0 | 0 | 0 | 0 | 0 | 0 | 0 | 93970 | 33825 | 0 | 0 | 0 | 0 | 0 | 0 | 0 | 0 |
| VIEPQYFGL | LRPPRC | P42704 | LPPRC_HUMAN | Unmodified | 284390 | 0 | 0 | 0 | 252230 | 0 | 0 | 0 | 1581800 | 552290 | 0 | 0 | 0 | 0 | 0 | 0 | 80733 | 352330 |
| VIEQLLVGL | TBCD; TBCD | Q9BTW9; J3I | TBCD_HUMAN | Unmodified | 0 | 0 | 0 | 0 | 0 | 0 | 0 | 0 | 0 | 0 | 0 | 2487700 | 0 | 0 | 0 | 0 | 0 | 0 |
| VIEQMGK | DDX19B; DDX1 | Q9UMR2; Q5 | DD19B_HUMAI | Unmodified | 150530 | 105660 | 0 | 0 | 287150 | 59248 | 37415 | 11961 | 619800 | 44329 | 0 | 1552600 | 0 | 0 | 0 | 87399 | 0 | 0 |
| VIEQRPDPI | NPRL2 | Q8WTW4 | NPRL2_HUMAI | Unmodified | 164110 | 91942 | 0 | 0 | 0 | 0 | 0 | 0 | 0 | 0 | 0 | 337470 | 0 | 0 | 0 | 0 | 372630 | 0 |
| VIESTQDLC | SPTBN1; SPTBN | Q01082; A0/ | SPTB2_HUMAN | Oxidation (M) | 0 | 0 | 0 | 0 | 1087000 | 0 | 0 | 0 | 0 | 0 | 0 | 1505200 | 0 | 0 | 0 | 0 | 0 | 0 |
| VIEVSHWG | RPN1; RPN1 | P04843; B7Z | RPN1_HUMAN | Unmodified | 0 | 0 | 0 | 0 | 0 | 0 | 0 | 0 | 6052900 | 1854400 | 0 | 0 | 0 | 0 | 0 | 0 | 0 | 0 |
| VIEVSHWG | RPN1; RPN1 | P04843; B7Z | RPN1_HUMAN | Unmodified | 0 | 0 | 0 | 0 | 0 | 0 | 0 | 0 | 2923200 | 0 | 0 | 0 | 1082700 | 0 | 0 | 0 | 119870 | 0 |
| VIEYTDLEK | ABCC4 | O15439 | MRP4_HUMAN | Unmodified | 0 | 236310 | 2854100 | 645220 | 0 | 0 | 0 | 0 | 0 | 0 | 0 | 283520 | 0 | 0 | 3963000 | 3289200 | 0 | 0 |
| VIFFLPWQI | GPD2; GPD2 | P43304; F5G | GPDM_HUMAN | Unmodified | 0 | 0 | 701260 | 0 | 0 | 963760 | 0 | 3017900 | 6571000 | 0 | 888120 | 0 | 0 | 0 | 0 | 0 | 0 | 0 |
| VIFHPEFLS | GYS1 | P13807 | GYS1_HUMAN | Unmodified | 0 | 0 | 0 | 0 | 0 | 0 | 0 | 0 | 0 | 0 | 0 | 832190 | 0 | 0 | 0 | 0 | 0 | 0 |
| VIFKTPNQI | PNMA2 | Q9UL42 | PNMA2_HUMA | Unmodified | 7666900 | 8529700 | 0 | 0 | 0 | 0 | 0 | 0 | 3170700 | 0 | 0 | 0 | 0 | 0 | 0 | 0 | 0 | 0 |
| VIFKTPNQI | PNMA2 | Q9UL42 | PNMA2_HUMA | Unmodified | 494420 | 1441800 | 0 | 0 | 291200 | 0 | 0 | 0 | 0 | 0 | 0 | 0 | 0 | 0 | 0 | 0 | 0 | 0 |
| VIFLENYR | PYGB; PYGM; P | P11216; P11 | PYGB_HUMAN | Unmodified | 0 | 0 | 201300 | 643220 | 159220 | 0 | 0 | 0 | 356360 | 333000 | 0 | 3624500 | 0 | 0 | 471890 | 738400 | 202560 | 868260 |
| VIFLENYRV | PYGB; PYGM; P | P11216; P11 | PYGB_HUMAN | Unmodified | 0 | 0 | 0 | 0 | 2226400 | 0 | 1197400 | 35618 | 0 | 960080 | 0 | 0 | 0 | 0 | 0 | 0 | 3370700 | 8176600 |
| VIFLQGGG | PSAT1 | Q9Y617 | SERC_HUMAN | Unmodified | 0 | 0 | 0 | 0 | 332950 | 0 | 0 | 0 | 0 | 0 | 0 | 0 | 361390 | 0 | 0 | 0 | 0 | 0 |
| VIFPAAEDK | HDLBP; HDLBP | Q00341; A0/ | VIGLN_HUMAN | Unmodified | 0 | 96649 | 242540 | 0 | 0 | 0 | 0 | 0 | 0 | 0 | 549980 | 388820 | 0 | 0 | 331600 | 159590 | 0 | 0 |
| VIFPAAEDK | HDLBP; HDLBP | Q00341; A0/ | VIGLN_HUMAN | Unmodified | 848860 | 1050700 | 282580 | 2524200 | 0 | 0 | 0 | 0 | 525280 | 0 | 0 | 3176700 | 0 | 0 | 0 | 778700 | 0 | 0 |
| VIFPAPTPK | NNT; NNT; NNT | Q13423; E9F | NNTM_HUMAN | Unmodified | 0 | 0 | 0 | 0 | 0 | 0 | 0 | 0 | 3345400 | 481560 | 0 | 0 | 0 | 0 | 0 | 0 | 0 | 0 |
| VIFSGSLDF | DDOST; DDOST | P39656; A0A | OST48_HUMAN | Unmodified | 0 | 0 | 0 | 0 | 0 | 0 | 0 | 0 | 31914 | 140140 | 0 | 0 | 0 | 17700 | 0 | 0 | 0 | 0 |
| VIGAGEFGI | EPHB1; EPHA2; | P54762; P29 | EPHB1_HUMAI | Unmodified | 13105 | 701210 | 396980 | 0 | 17042000 | 3425800 | 0 | 0 | 0 | 0 | 2353000 | 4163600 | 954490 | 310530 | 527880 | 373140 | 13864000 | 54382000 |
| VIGAGEFGI | EPHA2 | P29317 | EPHA2_HUMAN | Oxidation (M) | 0 | 0 | 0 | 0 | 8445300 | 1902900 | 48957 | 0 | 1380900 | 0 | 0 | 0 | 91912 | 46062 | 0 | 0 | 23992000 | 22149000 |
| VIGAGEFGI | EPHA2 | P29317 | EPHA2_HUMAN | Unmodified | 0 | 0 | 0 | 0 | 13894000 | 1532600 | 0 | 0 | 0 | 0 | 0 | 0 | 878180 | 1859600 | 0 | 0 | 47940000 | 61044000 |
| VIGFKPEEI | MYO1D; MYO1E | O94832; J3Q | MYO1D_HUMA | Unmodified | 4835400 | 6525200 | 3824800 | 794520 | 0 | 241410 | 0 | 0 | 1157700 | 665530 | 484050 | 2967000 | 0 | 0 | 8680300 | 4790900 | 365270 | 636800 |
| VIGGDDLS | HPRT1 | P00492 | HPRT_HUMAN | Unmodified | 452540 | 500790 | 0 | 0 | 1979600 | 482140 | 502410 | 259070 | 4403600 | 1351100 | 3563200 | 10667000 | 377210 | 0 | 0 | 0 | 390300 | 516770 |
| VIGHYAGE | FADS2 | O95864 | FADS2_HUMAN | Unmodified | 0 | 0 | 318860 | 167570 | 0 | 0 | 0 | 0 | 717070 | 0 | 338480 | 999670 | 0 | 0 | 421470 | 0 | 0 | 0 |
| VIGIDHIK | PCMT1; PCMT1 | P22061; F6S | PIMT_HUMAN I | Unmodified | 0 | 0 | 0 | 0 | 2123100 | 0 | 0 | 0 | 2561800 | 0 | 0 | 0 | 0 | 0 | 0 | 0 | 0 | 0 |
| VIGKGSEK: | SARS1; SARS1 | P49591; Q5T | SYSC_HUMAN | Unmodified | 39009 | 0 | 0 | 0 | 627150 | 0 | 417450 | 182630 | 19275 | 190860 | 0 | 0 | 3494000 | 420000 | 0 | 0 | 517830 | 178180 |
| VIGLSSDL( | EIF3F; EIF3F | O00303; H0Y | EIF3F_HUMAN | Unmodified | 0 | 0 | 0 | 0 | 627250 | 0 | 170550 | 0 | 0 | 316550 | 1694700 | 15047000 | 0 | 0 | 0 | 0 | 0 | 0 |
| VIGNGSFG | GSK3B; GSK3B | P49841; A0A | GSK3B_HUMAI | Unmodified | 0 | 0 | 0 | 0 | 0 | 0 | 0 | 63643 | 222910 | 0 | 0 | 638300 | 133830 | 0 | 0 | 0 | 0 | 0 |
| VIGNQSLVI | ECH1 | Q13011 | ECH1_HUMAN | Unmodified | 0 | 0 | 0 | 0 | 0 | 0 | 185670 | 28855 | 8293500 | 1427800 | 0 | 0 | 0 | 0 | 0 | 0 | 0 | 0 |
| VIGNQSLVI | ECH1 | Q13011 | ECH1_HUMAN | Unmodified | 0 | 0 | 204250 | 0 | 0 | 0 | 6000800 | 526340 | 34511000 | 19995000 | 940840 | 0 | 1824800 | 1465600 | 558740 | 359260 | 0 | 0 |
| VIGRGAFG | CDC42BPA; CD | Q5VT25; Q9' | MRCKA_HUMA | Unmodified | 0 | 0 | 0 | 0 | 533210 | 262600 | 157260 | 0 | 0 | 344360 | 0 | 0 | 0 | 0 | 0 | 0 | 481640 | 738680 |
| VIGSELVQI | PSMC2; PSMC2 | P35998; C9J | PRS7_HUMAN | Unmodified | 0 | 264350 | 0 | 0 | 0 | 0 | 0 | 0 | 390840 | 0 | 360910 | 8916400 | 0 | 0 | 733730 | 0 | 31898 | 0 |
| VIGSGCNL | LDHAL6A; LDH | Q6ZMR3; P0 | LDH6A_HUMAI | Unmodified | 2492100 | 1174400 | 56455 | 2582200 | 322490 | 0 | 0 | 0 | 1657400 | 319230 | 14776000 | 2.07E+08 | 0 | 0 | 2610900 | 3035000 | 416630 | 503470 |
| VIGSLNSN | ECE1; ECE1; EC | P42892; B4E | ECE1_HUMAN | Unmodified | 0 | 0 | 620970 | 917310 | 4210500 | 1602600 | 96349 | 0 | 22477000 | 1522600 | 51934 | 454660 | 104210 | 0 | 464990 | 0 | 583280 | 813840 |
| VIGSNFAP | TUBGCP3; TUB | Q96CW5; A0 | GCP3_HUMAN | Unmodified | 241590 | 0 | 365160 | 0 | 0 | 0 | 0 | 16749 | 369970 | 0 | 539390 | 2635500 | 215300 | 190790 | 286720 | 243170 | 0 | 0 |
| VIGTLLGTV | EIF3F; EIF3F; EI | O00303; A0/ | EIF3F_HUMAN | Unmodified | 786680 | 706730 | 921460 | 1044000 | 0 | 0 | 0 | 0 | 653940 | 0 | 1723900 | 18169000 | 0 | 0 | 820390 | 975370 | 0 | 0 |
| VIGVKSEGI | GDI2; GDI2 | P50395; Q5S | GDIB_HUMAN | Unmodified | 146520 | 587920 | 0 | 1912800 | 3741500 | 131150 | 811000 | 146290 | 3952000 | 1362000 | 1690000 | 1168500 | 1992800 | 511500 | 0 | 0 | 526390 | 1213100 |
| VIGVKSEGI | GDI2; GDI2 | P50395; Q5S | GDIB_HUMAN | Unmodified | 0 | 0 | 0 | 0 | 2106300 | 0 | 687930 | 488550 | 0 | 0 | 61883 | 0 | 2007000 | 1901200 | 0 | 0 | 1234700 | 2210000 |
| VIHALSENE | DNAJC13 | O75165 | DJC13_HUMAN | Unmodified | 108830 | 0 | 207760 | 483240 | 0 | 0 | 0 | 0 | 0 | 454770 | 0 | 1320900 | 0 | 1784300 | 64857 | 332520 | 0 | 0 |
| VIHCTVPP( | TOLLIP; TOLLIP | Q9H0E2; E9I | TOLIP_HUMAN | Unmodified | 918660 | 1496100 | 110750 | 458830 | 0 | 0 | 0 | 0 | 0 | 0 | 0 | 0 | 0 | 0 | 2090800 | 262130 | 0 | 0 |
| VIHDNFGI\ | GAPDH; GAPDH | P04406; E7E | G3P_HUMAN C | Oxidation (M) | 0 | 19233000 | 0 | 28321000 | 3.65E+08 | 27802000 | 1.32E+08 | 0 | 12117000 | 49232000 | 42079000 | 59130000 | 2.1E+08 | 1.09E+08 | 62919000 | 8323200 | 4.09E+08 | 3.31E+08 |
| VIHDNFGI\ | GAPDH; GAPDH | P04406; E7E | G3P_HUMAN C | Unmodified | 719560 | 3309600 | 491590 | 4495800 | 1.61E+08 | 238100 | 41191000 | 0 | 7480100 | 23676000 | 2230300 | 4345800 | 64592000 | 30709000 | 7816900 | 120640 | 77962000 | 2.29E+08 |
| VIHDNFGI\ | GAPDH; GAPDH | P04406; E7E | G3P_HUMAN C | Oxidation (M) | 0 | 0 | 0 | 0 | 12066000 | 0 | 29364000 | 0 | 0 | 1398300 | 0 | 0 | 4372100 | 5080600 | 0 | 0 | 69368000 | 19937000 |
| VIHDNFGI\ | GAPDH; GAPDH | P04406; E7E | G3P_HUMAN C | Unmodified | 0 | 0 | 0 | 0 | 2289300 | 283160 | 7369900 | 0 | 0 | 231320 | 0 | 0 | 935430 | 812590 | 0 | 0 | 18932000 | 3462300 |
| VIHGVINSF | CUL2; CUL2; C | Q13617; Q5' | CUL2_HUMAN | Unmodified | 106450 | 0 | 0 | 73483 | 0 | 0 | 138390 | 0 | 0 | 0 | 174120 | 984500 | 0 | 0 | 181590 | 89925 | 0 | 368760 |
| VIHRDIKG( | TNIK; MINK1; M | Q9UKE5; Q8 | TNIK_HUMAN T | Unmodified | 0 | 0 | 0 | 0 | 1852700 | 0 | 0 | 0 | 0 | 0 | 0 | 0 | 0 | 0 | 0 | 0 | 126440 | 0 |
| VIHSILNSP | TUBG1; TUBG2 | P23258; Q9I | TBG1_HUMAN | Unmodified | 235640 | 198370 | 467640 | 570920 | 595460 | 681950 | 172250 | 85805 | 531720 | 103920 | 361370 | 3912700 | 550880 | 217870 | 420040 | 371460 | 193200 | 123130 |
| VIHVVTSEN | CTNNA1; CTNN | P35221; G3V | CTNA1_HUMAI | Unmodified | 1659900 | 1518800 | 4604800 | 7846800 | 42902000 | 9932300 | 214480 | 0 | 4571200 | 1808600 | 4002200 | 18979000 | 25454000 | 28125000 | 9299200 | 6627300 | 0 | 0 |

| | | | | | | | | | | | | | | | | | | | | | | |
|---|---|---|---|---|---|---|---|---|---|---|---|---|---|---|---|---|---|---|---|---|---|---|
| VIHVVTSEM | CTNNA1; CTNN | P35221; G3X | CTNA1_HUMAI | Oxidation (M) | 2116900 | 1246500 | 5067400 | 0 | 13605000 | 12799000 | 0 | 0 | 884520 | 545350 | 2661200 | 16790000 | 9497700 | 0 | 8544400 | 9381500 | 0 | 0 |
| VIILGDSGV | RAB7A; RAB7A; | P51149; C9J | RAB7A_HUMAN | Unmodified | 11587000 | 15521000 | 12654000 | 7922600 | 39320000 | 8844000 | 9740200 | 6049700 | 33043000 | 19115000 | 44921000 | 1.09E+08 | 23858000 | 5748000 | 20555000 | 15000000 | 4205000 | 16685000 |
| VIILGDSGV | RAB7A; RAB7A; | P51149; C9J | RAB7A_HUMAN | Unmodified | 0 | 0 | 0 | 0 | 23131000 | 0 | 6482800 | 598390 | 0 | 2996600 | 0 | 0 | 4168400 | 4427300 | 0 | 0 | 3723300 | 4905900 |
| VIILGDSGV | RAB7A; RAB7A; | P51149; C9J | RAB7A_HUMAN | Oxidation (M) | 0 | 0 | 0 | 0 | 1580700 | 0 | 2041200 | 1218600 | 0 | 441420 | 0 | 0 | 2477300 | 228950 | 0 | 0 | 864340 | 554450 |
| VIILGDSGV | RAB7A; RAB7A; | P51149; C9J | RAB7A_HUMAN | Oxidation (M) | 0 | 0 | 0 | 0 | 1799000 | 419670 | 2113900 | 758500 | 0 | 705960 | 0 | 0 | 1214600 | 1303100 | 0 | 0 | 483070 | 713690 |
| VIILGDSGV | RAB7A; RAB7A; | P51149; C9J | RAB7A_HUMAN | Unmodified | 0 | 0 | 0 | 0 | 2608500 | 0 | 1385200 | 174600 | 0 | 605040 | 0 | 0 | 0 | 679980 | 0 | 0 | 696630 | 1131600 |
| VIILSLDRSI | EIF4G2; EIF4G2 | P78344; H0Y | IF4G2_HUMAN | Unmodified | 138520 | 0 | 636640 | 0 | 0 | 0 | 0 | 0 | 0 | 0 | 0 | 0 | 0 | 0 | 355970 | 154990 | 0 | 0 |
| VIILTAAAQ( | BDH2; BDH2; B | Q9BUT1; D6 | BDH2_HUMAN | Unmodified | 0 | 0 | 0 | 0 | 322090 | 0 | 0 | 0 | 0 | 63209 | 0 | 0 | 102270 | 0 | 0 | 0 | 0 | 0 |
| VIIQWNGP | AVIL | O75366 | AVIL_HUMAN A | Unmodified | 0 | 0 | 0 | 0 | 0 | 0 | 0 | 0 | 0 | 339960 | 5594900 | 31016000 | 0 | 0 | 0 | 0 | 551240 | 53138 |
| VIIQWNGP | AVIL | O75366 | AVIL_HUMAN A | Unmodified | 0 | 0 | 0 | 0 | 0 | 0 | 0 | 0 | 0 | 256000 | 1174400 | 14444000 | 0 | 440660 | 0 | 0 | 0 | 0 |
| VIISAPSAD, | GAPDH; GAPDH | P04406; E7E | G3P_HUMAN G | 2 Oxidation (M) | 4250900 | 2590200 | 5621500 | 6930400 | 2430600 | 1249300 | 0 | 2680700 | 1407700 | 1549300 | 39016000 | 90851000 | 1602900 | 1077400 | 6797700 | 8919800 | 2194700 | 1598600 |
| VIISAPSAD, | GAPDH; GAPDH | P04406; E7E | G3P_HUMAN G | Oxidation (M) | 221670 | 0 | 271640 | 511040 | 1162200 | 0 | 2284000 | 205740 | 3210200 | 484020 | 14916000 | 18598000 | 0 | 0 | 559770 | 286960 | 1147000 | 1048600 |
| VIISAPSAD, | GAPDH; GAPDH | P04406; E7E | G3P_HUMAN G | Unmodified | 0 | 0 | 0 | 0 | 0 | 0 | 0 | 0 | 1601400 | 0 | 819580 | 0 | 0 | 0 | 0 | 0 | 0 | 0 |
| VIISAPSAD, | GAPDH; GAPDH | P04406; E7E | G3P_HUMAN G | 2 Oxidation (M) | 1452400 | 521330 | 0 | 8675700 | 2603400 | 0 | 1716400 | 808950 | 0 | 2184800 | 1077900 | 9362600 | 765900 | 538830 | 0 | 413980 | 7995300 | 2162000 |
| VIISAPSAD, | GAPDH; GAPDH | P04406; E7E | G3P_HUMAN G | Oxidation (M) | 0 | 0 | 0 | 1735300 | 6098100 | 0 | 4174400 | 135850 | 2670800 | 891080 | 0 | 1617200 | 0 | 1596600 | 0 | 0 | 5349500 | 5495000 |
| VIISAPSAD, | GAPDH; GAPDH | P04406; E7E | G3P_HUMAN G | Unmodified | 0 | 0 | 0 | 0 | 0 | 0 | 0 | 0 | 0 | 0 | 0 | 0 | 0 | 0 | 0 | 0 | 0 | 1105800 |
| VIITNFPAAI | QARS1; QARS1; | P47897; H7( | SYQ_HUMAN G | Unmodified | 0 | 0 | 0 | 0 | 0 | 0 | 0 | 0 | 211880 | 0 | 307170 | 3442200 | 0 | 0 | 0 | 0 | 0 | 0 |
| VIIVDAENV | DIPK2A; DIPK2A | Q8NDZ4; E7 | DIK2A_HUMAN | Unmodified | 0 | 0 | 0 | 0 | 0 | 0 | 0 | 0 | 12867000 | 671890 | 0 | 0 | 0 | 0 | 0 | 0 | 0 | 0 |
| VIIVDAENV | DIPK2A; DIPK2A | Q8NDZ4; E7 | DIK2A_HUMAN | Unmodified | 0 | 0 | 0 | 0 | 0 | 0 | 0 | 0 | 23029000 | 1996000 | 0 | 0 | 0 | 0 | 0 | 0 | 0 | 0 |
| VIIVVKDGP | HADHA; HADHA | P40939; A0A | ECHA_HUMAN | Unmodified | 0 | 0 | 0 | 0 | 126340 | 0 | 280860 | 0 | 225730 | 1289900 | 0 | 0 | 72806 | 0 | 0 | 0 | 0 | 219600 |
| VIIWNTK | MYOF | Q9NZM1 | MYOF_HUMAN | Unmodified | 70302 | 0 | 25269 | 0 | 5809400 | 947440 | 0 | 0 | 410030 | 121460 | 0 | 81632 | 0 | 0 | 31760 | 29215 | 0 | 0 |
| VIIWNTKD\ | MYOF | Q9NZM1 | MYOF_HUMAN | Unmodified | 0 | 0 | 0 | 0 | 15799000 | 1351500 | 0 | 0 | 0 | 295430 | 0 | 0 | 0 | 0 | 0 | 0 | 143270 | 492870 |
| VIIWREEN( | SEC13; SEC13 | P55735; A8M | SEC13_HUMAN | Unmodified | 0 | 0 | 0 | 0 | 1270000 | 467620 | 205230 | 0 | 584400 | 244850 | 0 | 0 | 1365600 | 673530 | 0 | 0 | 0 | 591380 |
| VIIYQEK | PUF60; PUF60; | Q9UHX1; A0 | PUF60_HUMAN | Unmodified | 0 | 0 | 0 | 0 | 0 | 0 | 0 | 0 | 362270 | 0 | 131980 | 1691900 | 0 | 0 | 0 | 0 | 0 | 0 |
| VIKDFMIQ( | PPIB; PPIB; PPIE | P23284; A0A | PPIB_HUMAN F | Oxidation (M) | 98072 | 0 | 0 | 263220 | 0 | 0 | 1580900 | 143240 | 5400700 | 3765400 | 830760 | 0 | 295090 | 361950 | 0 | 0 | 0 | 0 |
| VIKDFMIQ( | PPIB; PPIB; PPIE | P23284; A0A | PPIB_HUMAN F | Unmodified | 0 | 0 | 0 | 207200 | 36889 | 0 | 346260 | 0 | 24048000 | 5549100 | 752160 | 0 | 0 | 289960 | 0 | 0 | 0 | 0 |
| VIKTDELPA | CLUH; CLUH | I3L2B0; A0A | I3L2B0_HUMAI | Unmodified | 0 | 0 | 0 | 43581 | 94140 | 0 | 26886 | 47063 | 0 | 0 | 0 | 551260 | 410250 | 198050 | 0 | 0 | 0 | 62942 |
| VILDLTPNY | SLC3A2; SLC3A | P08195; A0A | 4F2_HUMAN 4I | Unmodified | 6826000 | 5392000 | 3558100 | 0 | 0 | 0 | 0 | 0 | 3811600 | 1411800 | 2165700 | 80966000 | 0 | 0 | 8027100 | 5492100 | 0 | 0 |
| VILDLTPNY | SLC3A2; SLC3A | P08195; A0A | 4F2_HUMAN 4I | Unmodified | 27612000 | 21142000 | 8184400 | 30956000 | 23027000 | 9596100 | 711650 | 0 | 73753000 | 9005700 | 56908000 | 68799000 | 5883000 | 0 | 36239000 | 10754000 | 0 | 2962900 |
| VILDLTPNY | SLC3A2; SLC3A | P08195; A0A | 4F2_HUMAN 4I | Unmodified | 0 | 0 | 0 | 0 | 5636200 | 1328400 | 0 | 0 | 0 | 0 | 0 | 0 | 0 | 0 | 0 | 0 | 0 | 0 |
| VILDYNTAH | TRIM25 | Q14258 | TRI25_HUMAN | Unmodified | 633930 | 389380 | 0 | 893100 | 2339500 | 947090 | 461510 | 62884 | 0 | 102070 | 0 | 1720900 | 1970500 | 1657800 | 664620 | 246420 | 0 | 341840 |
| VILGSEAAC | DPP3; DPP3; DI | Q9NY33; E9I | DPP3_HUMAN | Unmodified | 269310 | 154300 | 0 | 0 | 0 | 0 | 0 | 0 | 799230 | 0 | 32529 | 12639000 | 0 | 0 | 1299100 | 601280 | 0 | 0 |
| VILHLKEDC | HSP90AB4P; HS | Q58FF6; P08 | H90B4_HUMAI | Unmodified | 12763000 | 10154000 | 5389600 | 19452000 | 24405000 | 13263000 | 1967400 | 359590 | 87109000 | 21114000 | 80323000 | 3.57E+08 | 28033000 | 5597000 | 13924000 | 18099000 | 10122000 | 18585000 |
| VILHLKEDC | HSP90AB1; HSI | P08238; Q58 | HS90B_HUMAI | Unmodified | 1009100 | 503320 | 0 | 2542900 | 7003900 | 1752200 | 779040 | 178770 | 17523000 | 8596800 | 9874600 | 51598000 | 4443200 | 1518400 | 1969400 | 1880000 | 2745900 | 2452500 |
| VILIECSELF | IL1RAPL1; IL1R | Q9NZN1; A0 | IRPL1_HUMAN | Oxidation (M) | 0 | 0 | 0 | 0 | 8463100 | 1593700 | 298380 | 0 | 0 | 0 | 0 | 1545400 | 0 | 1450900 | 0 | 0 | 0 | 3230300 |
| VILLAVTGE | FGD4; FGD4; F( | Q96M96; B7 | FGD4_HUMAN | Unmodified | 0 | 0 | 0 | 0 | 0 | 0 | 0 | 0 | 0 | 0 | 0 | 1320400 | 342290 | 0 | 0 | 0 | 0 | 0 |
| VILLDGSEY | EPB41L3; EPB4 | Q9Y2J2; A0A | E41L3_HUMAN | Unmodified | 0 | 137970 | 0 | 0 | 0 | 62200 | 0 | 0 | 0 | 67024 | 0 | 0 | 0 | 0 | 0 | 0 | 462630 | 2918300 |
| VILLGDGG' | RAB9B; RAB9A; | Q9NP90; P5 | RAB9B_HUMAN | Unmodified | 329950 | 419640 | 601440 | 577270 | 1160900 | 0 | 2353600 | 1574600 | 2808300 | 1651000 | 461000 | 1099000 | 2590400 | 728690 | 1246900 | 824270 | 785290 | 1790000 |
| VILPDLAVL | PLCB4; PLCB4 | Q15147; A0A | PLCB4_HUMAN | Unmodified | 232090 | 196640 | 930180 | 1020400 | 109020 | 43553 | 0 | 0 | 253660 | 257420 | 0 | 0 | 282700 | 180920 | 1552600 | 2162200 | 1086400 | 244120 |
| VILPQTSDA | ADGRG6; ADGF | Q86SQ4; F5I | AGRG6_HUMA | Unmodified | 0 | 0 | 415050 | 683610 | 4535100 | 566210 | 168510 | 282790 | 0 | 0 | 605690 | 0 | 0 | 424770 | 629320 | 627630 | 0 | 0 |
| VILPRGGS\ | ICAM1; ICAM1 | P05362; K7E | ICAM1_HUMAN | Unmodified | 0 | 0 | 0 | 0 | 19467000 | 1745800 | 0 | 0 | 0 | 0 | 0 | 0 | 21637 | 0 | 0 | 0 | 9281500 | 5497300 |
| VILPTPEEQ | NOXO1 | Q8NFA2 | NOXO1_HUMA | Unmodified | 0 | 0 | 0 | 0 | 0 | 0 | 0 | 0 | 0 | 0 | 0 | 1221100 | 0 | 0 | 0 | 0 | 0 | 0 |
| VILQDFTG\ | ACO1 | P21399 | ACOC_HUMAN | Oxidation (M) | 0 | 0 | 0 | 0 | 0 | 0 | 0 | 0 | 0 | 0 | 0 | 808960 | 0 | 0 | 0 | 0 | 0 | 0 |
| VILQPDVHI | SUN2; SUN2 | Q9UH99; B0 | SUN2_HUMAN | Unmodified | 0 | 0 | 0 | 0 | 0 | 723080 | 1014900 | 0 | 37857000 | 1966800 | 0 | 0 | 2102700 | 2888600 | 0 | 0 | 0 | 0 |
| VILTSIDGVI | PTPRF; PTPRF; I | P10586; H0Y | PTPRF_HUMAN | Unmodified | 3929000 | 4327500 | 0 | 3881500 | 1487800 | 0 | 0 | 0 | 8957800 | 0 | 1062500 | 0 | 182510 | 0 | 10292000 | 4084400 | 0 | 486900 |
| VILVQEAGI | ECT2; ECT2; EC | Q9H8V3; C9 | ECT2_HUMAN | Unmodified | 0 | 0 | 0 | 0 | 725100 | 0 | 0 | 0 | 1223500 | 366520 | 0 | 0 | 0 | 0 | 0 | 0 | 0 | 0 |
| VIMDYESLE | PNP | P00491 | PNPH_HUMAN | Unmodified | 0 | 0 | 0 | 0 | 2654200 | 420680 | 0 | 0 | 742140 | 489500 | 0 | 0 | 0 | 0 | 0 | 0 | 534140 | 0 |
| VIMDYESLE | PNP | P00491 | PNPH_HUMAN | Oxidation (M) | 0 | 0 | 0 | 0 | 0 | 0 | 0 | 0 | 0 | 0 | 0 | 4154500 | 0 | 0 | 0 | 0 | 0 | 0 |
| VIMELFDNI | MYOF; MYOF | Q9NZM1; H( | MYOF_HUMAN | Unmodified | 0 | 0 | 0 | 0 | 10087000 | 1931500 | 0 | 0 | 0 | 0 | 0 | 0 | 0 | 0 | 0 | 0 | 748380 | 0 |
| VIMELFDNI | MYOF; MYOF | Q9NZM1; H( | MYOF_HUMAN | Oxidation (M) | 0 | 0 | 0 | 0 | 8515100 | 2095800 | 0 | 0 | 0 | 0 | 0 | 37507 | 0 | 0 | 0 | 0 | 0 | 628750 |
| VIMELFDNI | MYOF; MYOF | Q9NZM1; H( | MYOF_HUMAN | Oxidation (M) | 0 | 0 | 0 | 0 | 16629000 | 1074100 | 0 | 0 | 0 | 0 | 0 | 0 | 0 | 0 | 0 | 0 | 256200 | 2414900 |
| VIMERDTYI | DKC1; DKC1 | O60832; H7( | DKC1_HUMAN | Unmodified | 0 | 0 | 0 | 0 | 0 | 0 | 0 | 0 | 0 | 0 | 0 | 274290 | 0 | 0 | 0 | 0 | 0 | 0 |
| VIMIVTDGF | MATN2 | O00339 | MATN2_HUMAI | Unmodified | 0 | 0 | 0 | 0 | 0 | 0 | 1416900 | 428700 | 2635100 | 0 | 0 | 0 | 260080 | 0 | 0 | 0 | 0 | 0 |
| VIMVGSGG | RALA; RALB; RA | P11233; P11 | RALA_HUMAN | Oxidation (M) | 3955100 | 2782100 | 3999400 | 3827600 | 8603900 | 2029400 | 3962300 | 2439900 | 3692200 | 3051300 | 6029400 | 18286000 | 7963800 | 8139900 | 5842400 | 6149800 | 4220800 | 6058200 |
| VIMVGSGG | RALA; RALB; RA | P11233; P11 | RALA_HUMAN | Unmodified | 3403700 | 5263600 | 3981900 | 6293500 | 23162000 | 5232500 | 8094800 | 8682400 | 26100000 | 11514000 | 12233000 | 49383000 | 16522000 | 22294000 | 6255700 | 4374000 | 13136000 | 21175000 |
| VIMVTGDH | ATP1A1; ATP1A | P05023; P50 | AT1A1_HUMAN | Unmodified | 935980 | 1328600 | 12892000 | 13226000 | 42680000 | 3028000 | 5831800 | 662610 | 15113000 | 7738400 | 14073000 | 97284000 | 28421000 | 19100000 | 13524000 | 8739200 | 4268500 | 4609500 |
| VIMVTGDH | ATP1A1; ATP1A | P05023; P50 | AT1A1_HUMAN | Oxidation (M) | 1198500 | 647300 | 18278000 | 13439000 | 26022000 | 2964300 | 7181200 | 2521900 | 5694900 | 4740500 | 13553000 | 80940000 | 23683000 | 15845000 | 22568000 | 22363000 | 7515400 | 3412100 |
| VIMVTGDH | ATP1A1; ATP1A | P05023; P50 | AT1A1_HUMAN | Unmodified | 0 | 0 | 0 | 0 | 1282400 | 0 | 837370 | 0 | 0 | 82673 | 0 | 0 | 1141500 | 1212900 | 0 | 0 | 1070200 | 0 |
| VINAAHSF\ | GMPS | P49915 | GUAA_HUMAN | Unmodified | 0 | 0 | 0 | 0 | 0 | 0 | 98856 | 0 | 0 | 0 | 932550 | 2060600 | 589790 | 0 | 0 | 0 | 0 | 0 |
| VINAAHSF\ | GMPS | P49915 | GUAA_HUMAN | Unmodified | 0 | 0 | 0 | 0 | 2068100 | 279590 | 0 | 0 | 0 | 232570 | 1533000 | 2318200 | 1440100 | 0 | 0 | 0 | 577060 | 585150 |
| VINAENAAI | SIPA1L1; SIPA1 | O43166; Q9I | SI1L1_HUMAN | Unmodified | 0 | 0 | 0 | 0 | 0 | 0 | 0 | 0 | 0 | 0 | 0 | 0 | 0 | 0 | 0 | 0 | 0 | 0 |
| VINAINVNI | MGAT3 | Q09327 | MGAT3_HUMAI | Unmodified | 0 | 0 | 0 | 0 | 0 | 0 | 0 | 0 | 560230 | 0 | 0 | 133130 | 95845 | 0 | 0 | 0 | 0 | 0 |
| VINDKHDD | PSMD1; PSMD1 | Q99460; A0A | PSMD1_HUMA | Unmodified | 0 | 0 | 0 | 0 | 315610 | 1181600 | 1923000 | 685650 | 0 | 0 | 0 | 4068300 | 0 | 938850 | 0 | 0 | 210300 | 0 |
| VINEEYK | RBBP7; RBBP4; | Q16576; Q0! | RBBP7_HUMAN | Unmodified | 0 | 0 | 0 | 0 | 0 | 0 | 0 | 0 | 420380 | 0 | 0 | 655750 | 0 | 20974 | 0 | 0 | 0 | 0 |
| VINEPTAAA | HSPA9; HSPA9; | P38646; H0Y | GRP75_HUMAN | Unmodified | 0 | 0 | 0 | 0 | 111670 | 0 | 0 | 0 | 5607200 | 99117 | 0 | 505710 | 0 | 0 | 0 | 0 | 130930 | 0 |
| VINEPTAAA | HSPA9; HSPA9; | P38646; H0Y | GRP75_HUMAN | Unmodified | 0 | 0 | 0 | 338610 | 788440 | 0 | 0 | 0 | 29718000 | 2030100 | 1327900 | 0 | 0 | 0 | 0 | 0 | 0 | 0 |
| VINEPTAAA | HSPA13 | P48723 | HSP13_HUMAN | Unmodified | 0 | 0 | 0 | 0 | 0 | 0 | 0 | 0 | 5046900 | 0 | 193960 | 0 | 0 | 0 | 188690 | 0 | 0 | 0 |
| VINLNDNTI | RANGAP1; RAN | P46060; H0Y | RAGP1_HUMAI | Unmodified | 147210 | 0 | 0 | 0 | 286380 | 0 | 0 | 0 | 654310 | 481300 | 0 | 0 | 2900100 | 1048100 | 0 | 0 | 0 | 688860 |
| VINNPSYR | UGT8 | Q16880 | CGT_HUMAN 2 | Unmodified | 2586300 | 1825500 | 0 | 0 | 548370 | 0 | 0 | 0 | 937290 | 51146 | 299260 | 0 | 363570 | 0 | 184840 | 295940 | 0 | 0 |
| VINSVDIK | WDR1; WDR1 | O75083; D6I | WDR1_HUMAN | Unmodified | 3991300 | 2953100 | 3351500 | 4048400 | 62311 | 5475700 | 1669900 | 2189300 | 5959700 | 2124700 | 2875500 | 19601000 | 14500000 | 6851500 | 5310300 | 4294500 | 2780200 | 6181500 |
| VINTPEVLR | EHD4 | Q9H223 | EHD4_HUMAN | Unmodified | 593320 | 0 | 982490 | 320470 | 0 | 174360 | 0 | 0 | 0 | 0 | 0 | 4426100 | 0 | 0 | 7295900 | 6288500 | 0 | 0 |
| VINYEEFKK | TPPP3 | Q9BW30 | TPPP3_HUMAN | Unmodified | 0 | 0 | 0 | 0 | 0 | 0 | 1668100 | 0 | 174600 | 0 | 0 | 0 | 0 | 0 | 0 | 0 | 0 | 0 |
| VIPDDLAQ( | DSC2; DSC2 | Q02487; A0A | DSC2_HUMAN | Unmodified | 0 | 0 | 0 | 0 | 864210 | 0 | 0 | 0 | 3671200 | 0 | 0 | 0 | 1641700 | 0 | 0 | 0 | 731310 | 3342100 |
| VIPEDGPA/ | OGDH; OGDH; | Q02218; E9F | ODO1_HUMAN | Unmodified | 0 | 0 | 0 | 0 | 0 | 0 | 0 | 24897 | 1824100 | 224590 | 0 | 0 | 0 | 0 | 0 | 0 | 0 | 0 |
| VIPEDGPA/ | OGDH; OGDH; | Q02218; E9F | ODO1_HUMAN | Unmodified | 0 | 0 | 0 | 0 | 0 | 0 | 0 | 0 | 834100 | 0 | 0 | 0 | 0 | 0 | 0 | 0 | 0 | 0 |
| VIPELNGK | GAPDH; GAPDH | P04406; E7E | G3P_HUMAN G | Unmodified | 16638000 | 9723400 | 22030000 | 13995000 | 54381000 | 10835000 | 2650900 | 2431300 | 20505000 | 3223800 | 73658000 | 2.8E+08 | 26147000 | 7309500 | 22576000 | 18992000 | 8630900 | 31345000 |
| VIPELNGKI | GAPDH; GAPDH | P04406; E7E | G3P_HUMAN G | Unmodified | 0 | 0 | 0 | 0 | 338680 | 31576 | 0 | 0 | 211850 | 52371 | 0 | 0 | 79287 | 0 | 0 | 0 | 215420 | 164040 |
| VIPHFTATA | PDIA5; PDIA5 | Q14554; H7( | PDIA5_HUMAN | Unmodified | 0 | 0 | 0 | 0 | 0 | 0 | 0 | 0 | 8680500 | 2413000 | 0 | 0 | 0 | 0 | 0 | 0 | 0 | 0 |
| VIPIQCNIRI | PECR; PECR | Q9BY49; B4I | PECR_HUMAN | Unmodified | 0 | 0 | 0 | 0 | 0 | 0 | 0 | 0 | 3138600 | 557770 | 0 | 0 | 0 | 0 | 0 | 0 | 0 | 0 |
| VIPLFTPQC | ADH1B; ADH1C | P00325; P00 | ADH1B_HUMAI | Unmodified | 0 | 0 | 0 | 0 | 0 | 0 | 0 | 0 | 0 | 0 | 1060700 | 858140 | 0 | 0 | 0 | 0 | 0 | 0 |
| VIPPAGQTI | HYOU1; HYOU1 | Q9Y4L1; A0A | HYOU1_HUMA | Unmodified | 0 | 34885 | 0 | 0 | 248230 | 0 | 0 | 0 | 2216600 | 582410 | 0 | 0 | 0 | 0 | 0 | 0 | 0 | 0 |
| VIPSQEHLI | SLC9A3R1 | O14745 | NHRF1_HUMAI | Unmodified | 0 | 0 | 0 | 0 | 0 | 0 | 0 | 0 | 0 | 0 | 0 | 1476100 | 3502300 | 2295700 | 0 | 0 | 0 | 0 |
| VIPSQESV( | OVCA2 | Q8WZ82 | OVCA2_HUMAI | Unmodified | 0 | 0 | 0 | 0 | 0 | 0 | 0 | 0 | 0 | 0 | 0 | 1880500 | 0 | 0 | 0 | 0 | 0 | 0 |
| VIPTLALLK | TXNDC9 | O14530 | TXND9_HUMAI | Unmodified | 49564 | 0 | 0 | 60695 | 751270 | 102270 | 122620 | 134720 | 206900 | 134750 | 461780 | 1695300 | 0 | 0 | 162370 | 108720 | 718340 | 521940 |
| VIQAGQKII | STS; STS; STS; S | P08842; A0A | STS_HUMAN S | Oxidation (M) | 0 | 0 | 0 | 0 | 0 | 0 | 0 | 0 | 969000 | 360300 | 0 | 0 | 0 | 0 | 0 | 0 | 0 | 0 |
| VIQAGQKII | STS; STS; STS; S | P08842; A0A | STS_HUMAN S | Unmodified | 0 | 0 | 0 | 0 | 0 | 0 | 0 | 0 | 2183800 | 161270 | 0 | 0 | 0 | 0 | 0 | 0 | 0 | 0 |
| VIQALAMK( | LRPPRC | P42704 | LPPRC_HUMAN | Unmodified | 0 | 0 | 0 | 0 | 0 | 0 | 0 | 0 | 2847900 | 0 | 0 | 0 | 0 | 0 | 0 | 0 | 0 | 0 |
| VIQAQTAFS | SDCBP; SDCBP | O00560; G5I | SDCB1_HUMAI | Unmodified | 1.25E+08 | 2.02E+08 | 22087000 | 2.18E+08 | 19061000 | 7143200 | 710720 | 0 | 6.53E+08 | 32505000 | 31084000 | 90305000 | 30190000 | 3602300 | 1.48E+08 | 1.32E+08 | 0 | 2515900 |
| VIQATGGG | PANK4 | Q9NVE7 | PANK4_HUMAN | Unmodified | 0 | 0 | 59364 | 59861 | 3196300 | 112460 | 204940 | 0 | 404650 | 249420 | 0 | 0 | 225680 | 0 | 115500 | 66570 | 907370 | 1777500 |
| VIQCFAET( | CLTC; CLTC | Q00610; A0A | CLH1_HUMAN | Unmodified | 9605400 | 5610000 | 1927200 | 3482900 | 949530 | 1090800 | 779950 | 748830 | 0 | 509620 | 227260 | 16861000 | 561460 | 378000 | 4811300 | 6775900 | 674480 | 736930 |
| VIQDCEDE | ITGA1 | P56199 | ITA1_HUMAN I | Unmodified | 0 | 0 | 0 | 0 | 0 | 0 | 0 | 0 | 0 | 0 | 0 | 0 | 0 | 0 | 0 | 0 | 0 | 0 |
| VIQDVLEDI | IQGAP3; IQGAP | Q86VI3; F2Z | IQGA3_HUMAN | Unmodified | 0 | 0 | 366660 | 0 | 230270 | 0 | 0 | 166360 | 380540 | 0 | 172690 | 746830 | 43414 | 0 | 0 | 0 | 0 | 107790 |
| VIQEIVDK | FXR2; FXR1; FXI | P51116; P51 | FXR2_HUMAN I | Unmodified | 358580 | 326940 | 271830 | 331760 | 0 | 0 | 72618 | 0 | 385800 | 0 | 315510 | 2024500 | 0 | 497760 | 839430 | 1254800 | 0 | 0 |
| VIQEIWPVL | TNPO3; TNPO3 | Q9Y5L0; C9J | TNPO3_HUMAI | Unmodified | 0 | 0 | 0 | 341120 | 0 | 0 | 0 | 11839 | 0 | 67102 | 77663 | 173410 | 75346 | 34613 | 0 | 0 | 0 | 74783 |
| VIQENEHAI | SMC4; SMC4 | Q9NTJ3; E9F | SMC4_HUMAN | Unmodified | 0 | 0 | 0 | 0 | 871240 | 226300 | 598340 | 383360 | 0 | 351090 | 0 | 0 | 610280 | 787470 | 0 | 223580 | 0 | 416650 |
| VIQGCGNF | GPC1; GPC1; G | P35052; H7( | GPC1_HUMAN | Unmodified | 0 | 0 | 0 | 0 | 725240 | 221130 | 155650 | 173740 | 0 | 39002 | 0 | 0 | 0 | 304410 | 0 | 0 | 0 | 397270 |
| VIQGDGVD | NAMPT; NAMPT | P43490; A0A | NAMPT_HUMAI | Oxidation (M) | 0 | 0 | 0 | 700570 | 1230300 | 1132100 | 370600 | 0 | 0 | 0 | 330690 | 1056700 | 0 | 0 | 0 | 0 | 0 | 0 |
| VIQGDGVD | NAMPT; NAMPT | P43490; A0A | NAMPT_HUMAI | Unmodified | 0 | 0 | 0 | 0 | 354480 | 0 | 0 | 0 | 35918 | 22970 | 31696 | 0 | 204820 | 0 | 0 | 0 | 0 | 0 |
| VIQHNALEI | OPA1; OPA1; OI | O60313; E5K | OPA1_HUMAN | Unmodified | 0 | 0 | 0 | 0 | 0 | 0 | 0 | 0 | 36022 | 0 | 0 | 1753400 | 143520 | 25711 | 0 | 0 | 13043 | 0 |
| VIQIATSSS' | DHX30; DHX30 | Q7L2E3; H7I | DHX30_HUMAI | Unmodified | 0 | 0 | 0 | 0 | 0 | 0 | 154130 | 85028 | 746050 | 230640 | 0 | 634170 | 0 | 0 | 0 | 0 | 0 | 0 |
| VIQIVINRDI | AP2A2; AP2A2; | O94973; A0A | AP2A2_HUMAN | Unmodified | 0 | 113630 | 0 | 0 | 0 | 0 | 0 | 0 | 0 | 0 | 0 | 712680 | 0 | 0 | 150710 | 0 | 0 | 0 |
| VIQIVLMR | PRKCD; PRKCD | Q05655; C9J | KPCD_HUMAN | Oxidation (M) | 0 | 0 | 0 | 0 | 0 | 0 | 0 | 0 | 0 | 0 | 325300 | 1020600 | 0 | 0 | 0 | 0 | 0 | 0 |
| VIQPHGGG | DSG2 | Q14126 | DSG2_HUMAN | Oxidation (M) | 0 | 0 | 3707900 | 8620700 | 691840 | 0 | 0 | 0 | 4351800 | 0 | 3476000 | 9122400 | 0 | 0 | 4930300 | 10786000 | 0 | 0 |
| VIQPHGGG | DSG2 | Q14126 | DSG2_HUMAN | Unmodified | 0 | 0 | 0 | 0 | 929080 | 0 | 0 | 0 | 4814700 | 0 | 400810 | 1696200 | 0 | 0 | 3420000 | 1134800 | 0 | 0 |
| VIQQSLEQI | ROCK2; ROCK2 | O75116; Q1 | ROCK2_HUMAI | Unmodified | 0 | 0 | 0 | 0 | 1357600 | 0 | 166680 | 121030 | 913230 | 222230 | 356320 | 1629600 | 0 | 0 | 0 | 0 | 0 | 1762700 |
| VIQQSLEQI | ROCK2; ROCK2 | O75116; Q1 | ROCK2_HUMAI | Unmodified | 0 | 0 | 0 | 0 | 1858800 | 0 | 0 | 724990 | 0 | 537760 | 0 | 0 | 1320300 | 3507100 | 0 | 0 | 2907600 | 5151500 |
| VIQSVANY/ | GAK | O14976 | GAK_HUMAN C | Unmodified | 0 | 0 | 0 | 0 | 0 | 0 | 0 | 0 | 0 | 0 | 0 | 859410 | 0 | 0 | 0 | 0 | 0 | 0 |
| VIQVDEMI\ | CDCA8 | Q53HL2 | BOREA_HUMAI | Unmodified | 0 | 0 | 0 | 0 | 2562100 | 0 | 0 | 0 | 0 | 0 | 0 | 0 | 1893800 | 0 | 0 | 0 | 0 | 0 |
| VIQYLAHV/ | MYH10 | P35580 | MYH10_HUMAI | Unmodified | 0 | 0 | 0 | 0 | 1257900 | 0 | 1071200 | 207860 | 0 | 0 | 0 | 646440 | 504980 | 909270 | 217560 | 90645 | 0 | 277110 |

| | | | | | | | | | | | | | | | | | | | | | | |
|---|---|---|---|---|---|---|---|---|---|---|---|---|---|---|---|---|---|---|---|---|---|---|
| VIQYLAHVA | MYH14; MYH14 | Q7Z406; M0 | MYH14_HUMA | Unmodified | 0 | 0 | 60236 | 0 | 438860 | 41626 | 0 | 0 | 193250 | 109760 | 0 | 0 | 196350 | 181200 | 0 | 49859 | 0 | 0 |
| VIQYLAYVA | MYH9; MYH9 | P35579; Q5 | MYH9_HUMAN | Unmodified | 0 | 0 | 0 | 14828000 | 15811000 | 2951200 | 1465300 | 412630 | 1865800 | 1398200 | 1908300 | 10884000 | 3865100 | 1851900 | 0 | 0 | 9865500 | 2873700 |
| VIREEEEEE | PNMA2 | Q9UL42 | PNMA2_HUMA | Unmodified | 12369000 | 11166000 | 0 | 0 | 0 | 0 | 0 | 0 | 0 | 0 | 0 | 0 | 0 | 0 | 0 | 0 | 0 | 0 |
| VIREEEEEE | PNMA2 | Q9UL42 | PNMA2_HUMA | Unmodified | 14418000 | 9414100 | 0 | 0 | 0 | 0 | 0 | 0 | 0 | 0 | 0 | 0 | 0 | 0 | 0 | 0 | 0 | 0 |
| VIRGWDEA | FKBP3 | Q00688 | FKBP3_HUMAN | Unmodified | 0 | 0 | 0 | 0 | 3266100 | 313040 | 324360 | 0 | 0 | 938800 | 0 | 0 | 1519500 | 517160 | 0 | 0 | 483890 | 2846700 |
| VIRPLDQPS | CAND1 | Q86VP6 | CAND1_HUMA | Unmodified | 0 | 268980 | 0 | 216290 | 0 | 0 | 1264900 | 506570 | 4195200 | 1532700 | 0 | 19985000 | 0 | 0 | 0 | 3155600 | 1295900 | 0 |
| VIRSPADLII | OS9; OS9 | Q13438; B4E | OS9_HUMAN P | Unmodified | 0 | 0 | 0 | 0 | 0 | 0 | 0 | 0 | 0 | 439380 | 0 | 0 | 0 | 0 | 0 | 0 | 0 | 0 |
| VIRVYIASS | SH3BGRL | O75368 | SH3L1_HUMAN | Unmodified | 0 | 0 | 0 | 0 | 986420 | 0 | 107520 | 89171 | 0 | 944100 | 0 | 0 | 0 | 0 | 0 | 0 | 368030 | 1204700 |
| VIRVYIASS | SH3BGRL | O75368 | SH3L1_HUMAN | Unmodified | 0 | 0 | 0 | 0 | 425680 | 0 | 0 | 0 | 0 | 179440 | 0 | 0 | 0 | 0 | 0 | 0 | 0 | 338500 |
| VISALSEQA | KIF14 | Q15058 | KIF14_HUMAN | Unmodified | 0 | 0 | 352160 | 0 | 1739900 | 0 | 0 | 0 | 0 | 0 | 0 | 499850 | 0 | 10736 | 411230 | 540300 | 414170 | 0 |
| VISDGEQG | SLC38A10 | Q9HBR0 | S38AA_HUMAN | Unmodified | 0 | 0 | 0 | 0 | 0 | 0 | 0 | 0 | 798070 | 0 | 0 | 0 | 0 | 0 | 0 | 0 | 0 | 0 |
| VISDGEQG | SLC38A10 | Q9HBR0 | S38AA_HUMAN | Unmodified | 0 | 0 | 0 | 0 | 0 | 0 | 0 | 0 | 653750 | 0 | 0 | 0 | 0 | 0 | 0 | 0 | 0 | 0 |
| VISEFVIPS\ | MKRN2; MKRN2 | Q9H000; C9 | MKRN2_HUMA | Unmodified | 120010 | 0 | 0 | 247720 | 0 | 0 | 0 | 0 | 0 | 0 | 0 | 0 | 508230 | 0 | 0 | 0 | 0 | 0 |
| VISELNGK | RPLP2; RPLP2 | P05387; H0 | RLA2_HUMAN ( | Unmodified | 51406000 | 36852000 | 16946000 | 17327000 | 2316300 | 477390 | 436100 | 440020 | 5780200 | 794790 | 5157400 | 78579000 | 9974200 | 2281400 | 41015000 | 27432000 | 746680 | 3794400 |
| VISELNGKI | RPLP2; RPLP2 | P05387; H0 | RLA2_HUMAN ( | Unmodified | 0 | 0 | 0 | 0 | 0 | 0 | 972150 | 0 | 0 | 1986400 | 0 | 0 | 17262000 | 0 | 0 | 0 | 1173800 | 2779400 |
| VISESPPD( | GCN1 | Q92616 | GCN1_HUMAN | Unmodified | 0 | 0 | 0 | 0 | 0 | 0 | 0 | 0 | 0 | 0 | 0 | 1034500 | 0 | 0 | 0 | 0 | 0 | 0 |
| VISFEEQVA | COPS4; COPS4 | Q9BT78; D6 | CSN4_HUMAN | Unmodified | 436960 | 0 | 0 | 0 | 0 | 0 | 0 | 0 | 0 | 0 | 0 | 794230 | 0 | 0 | 0 | 0 | 0 | 0 |
| VISGQFLSI | PLCB3 | Q01970 | PLCB3_HUMAN | Unmodified | 0 | 0 | 0 | 0 | 0 | 0 | 0 | 0 | 157750 | 0 | 0 | 672540 | 0 | 0 | 0 | 0 | 48825 | 0 |
| VISGVLQL( | MYH9 | P35579 | MYH9_HUMAN | Unmodified | 0 | 0 | 42236 | 25262 | 0 | 0 | 0 | 1462100 | 0 | 0 | 202600 | 1313100 | 0 | 0 | 399310 | 60407 | 0 | 0 |
| VISGVLQL( | MYH9 | P35579 | MYH9_HUMAN | Unmodified | 0 | 0 | 0 | 0 | 0 | 0 | 0 | 0 | 0 | 0 | 0 | 19847000 | 0 | 0 | 0 | 4070400 | 0 | 0 |
| VISHFLPNI | DCAF11; DCAF | Q8TEB1; H0 | DCA11_HUMAI | Unmodified | 0 | 0 | 0 | 193990 | 1858000 | 0 | 0 | 0 | 0 | 0 | 0 | 1916800 | 0 | 0 | 0 | 0 | 347500 | 383530 |
| VISHPNYD: | TMPRSS2; TMPI | O15393; C9 | TMPS2_HUMAN | Unmodified | 0 | 0 | 393640 | 612650 | 0 | 0 | 0 | 0 | 0 | 0 | 0 | 0 | 0 | 0 | 521880 | 594870 | 0 | 0 |
| VISHPTLPI: | STRN | O43815 | STRN_HUMAN | Unmodified | 0 | 0 | 0 | 78159 | 0 | 736900 | 0 | 171270 | 373940 | 566010 | 452090 | 3119800 | 0 | 0 | 111140 | 103940 | 477760 | 686130 |
| VISLEDFME | POFUT1 | Q9H488 | OFUT1_HUMAI | Oxidation (M) | 0 | 0 | 0 | 0 | 0 | 0 | 0 | 0 | 2282500 | 607470 | 0 | 0 | 0 | 0 | 0 | 0 | 0 | 0 |
| VISLEDFME | POFUT1 | Q9H488 | OFUT1_HUMAI | Unmodified | 0 | 0 | 0 | 0 | 0 | 0 | 0 | 0 | 7037400 | 406890 | 0 | 0 | 0 | 0 | 0 | 0 | 0 | 0 |
| VISLPELK | SMOC1 | Q9H4F8 | SMOC1_HUMA | Unmodified | 0 | 0 | 0 | 0 | 0 | 0 | 0 | 17408 | 6227000 | 1737300 | 0 | 0 | 0 | 0 | 0 | 0 | 0 | 0 |
| VISNEFNSI | UBE3A; UBE3A; | Q05086; A0 | UBE3A_HUMAN | Unmodified | 203150 | 376990 | 107710 | 131940 | 1206600 | 712620 | 300880 | 0 | 0 | 447440 | 141450 | 839170 | 0 | 0 | 177520 | 0 | 0 | 155060 |
| VISQRYPDI | SELENOT; SELE | P62341; A0 | SELT_HUMAN ' | Unmodified | 0 | 0 | 0 | 0 | 0 | 0 | 0 | 0 | 2808900 | 0 | 0 | 0 | 92781 | 0 | 0 | 0 | 0 | 0 |
| VISSGSVAS | HSPG2; HSPG2 | P98160; A0 | PGBM_HUMAN | Unmodified | 115900 | 0 | 2652200 | 2782200 | 0 | 110370 | 187120 | 595910 | 181110 | 0 | 0 | 359820 | 0 | 0 | 478210 | 1595800 | 0 | 0 |
| VISSIEQKTI | YWHAQ; YWHA | P27348; E9 | 1433T_HUMAN | Unmodified | 1311900 | 1330000 | 571230 | 6561300 | 16855000 | 5520700 | 1137900 | 953270 | 10222000 | 2689600 | 9822300 | 24119000 | 14782000 | 5504000 | 3446200 | 3221700 | 5446600 | 12049000 |
| VISSIEQKTI | YWHAQ; YWHA | P27348; E9 | 1433T_HUMAN | Unmodified | 0 | 0 | 0 | 0 | 2746000 | 556430 | 0 | 0 | 1123400 | 385730 | 1053700 | 2351500 | 1280000 | 497470 | 0 | 135610 | 1080700 | 1659700 |
| VISSIEQKTI | YWHAB; YWHA | P31946; Q4 | 1433B_HUMAN | Unmodified | 0 | 0 | 0 | 0 | 252380 | 0 | 0 | 0 | 0 | 35667 | 0 | 0 | 31441 | 0 | 269130 | 454460 | 60144 | 370970 |
| VISSIEQKTI | YWHAB; YWHA | P31946; A2 | 1433B_HUMAN | Unmodified | 649470 | 1575400 | 0 | 8786000 | 52564000 | 27503000 | 4099700 | 7729700 | 9730600 | 10420000 | 6920500 | 5632900 | 88164000 | 37381000 | 1935600 | 1966900 | 39040000 | 1.32E+08 |
| VISSIEQKTI | YWHAH; YWHA | Q04917; A2 | 1433F_HUMAN | Unmodified | 0 | 0 | 0 | 0 | 602370 | 0 | 0 | 154410 | 0 | 0 | 0 | 0 | 0 | 270710 | 0 | 0 | 899460 | 0 |
| VISSIEQKTI | YWHAH; YWHA | Q04917; A2 | 1433F_HUMAN | Oxidation (M) | 0 | 0 | 0 | 0 | 225200 | 59870 | 110100 | 115920 | 0 | 63375 | 0 | 0 | 1133000 | 187020 | 0 | 0 | 316380 | 148510 |
| VISSIEQKT: | YWHAG | P61981 | 1433G_HUMAN | Unmodified | 0 | 0 | 0 | 1458700 | 7787600 | 1695700 | 2004400 | 1367400 | 399530 | 2214100 | 413890 | 547480 | 12357000 | 13139000 | 0 | 0 | 12532000 | 5875800 |
| VISTMSVGI | GRHPR; GRHPF | Q9UBQ7; U | GRHPR_HUMA | Oxidation (M) | 0 | 0 | 492620 | 872230 | 0 | 0 | 0 | 0 | 653290 | 0 | 423630 | 0 | 0 | 0 | 1152300 | 559940 | 0 | 0 |
| VISYEEGK | RHEB; RHEB | Q15382; C9 | RHEB_HUMAN | Unmodified | 1651500 | 1725500 | 1129400 | 1470700 | 102750 | 253560 | 0 | 45575 | 1672500 | 132800 | 379760 | 1604400 | 68597 | 0 | 1162100 | 1252100 | 26761 | 557190 |
| VITDENWR | TMX1; TMX1 | Q9H3N1; G3 | TMX1_HUMAN | Unmodified | 28463 | 0 | 2981700 | 4152100 | 0 | 309230 | 247390 | 109750 | 26621000 | 1272300 | 438630 | 0 | 13302000 | 1631200 | 0 | 2616500 | 0 | 0 |
| VITEEMNGI | EPB41L2; EPB4 | O43491; E9 | E41L2_HUMAN | Unmodified | 262510 | 187360 | 354870 | 159020 | 142010 | 0 | 0 | 0 | 263600 | 0 | 124620 | 721120 | 0 | 0 | 917660 | 1288700 | 57954 | 266570 |
| VITEPIPVSI | PTPRJ; PTPRJ; P | Q12913; A0 | PTPRJ_HUMAN | Unmodified | 0 | 0 | 0 | 531680 | 1711100 | 0 | 0 | 0 | 5877200 | 809720 | 8998500 | 12285000 | 0 | 0 | 0 | 0 | 0 | 0 |
| VITEPIPVSI | PTPRJ; PTPRJ; P | Q12913; A0 | PTPRJ_HUMAN | Unmodified | 0 | 0 | 0 | 0 | 6344200 | 842660 | 0 | 0 | 3495900 | 364810 | 0 | 0 | 1428300 | 0 | 0 | 0 | 0 | 1610000 |
| VITESTVHL | TRIM16; TRIM1 | O95361; Q3 | TRI16_HUMAN | Unmodified | 0 | 0 | 0 | 0 | 1592800 | 175090 | 165320 | 0 | 305150 | 0 | 0 | 0 | 557970 | 200240 | 0 | 0 | 0 | 0 |
| VITEYLNAC | SND1 | Q7KZF4 | SND1_HUMAN | Unmodified | 581830 | 879500 | 776370 | 917780 | 2551000 | 0 | 2052200 | 1087400 | 0 | 175340 | 476510 | 3210400 | 2143200 | 3766300 | 872730 | 1322600 | 1889100 | 1894800 |
| VITFIDLAGI | GTPBP1 | O00178 | GTPB1_HUMAN | Unmodified | 0 | 0 | 0 | 605110 | 0 | 0 | 0 | 123540 | 0 | 0 | 181330 | 6274200 | 0 | 0 | 0 | 0 | 0 | 0 |
| VITIMQNPF | RPS18 | P62269 | RS18_HUMAN | Oxidation (M) | 327810 | 516650 | 366810 | 0 | 0 | 55868 | 0 | 0 | 0 | 0 | 159570 | 5561800 | 0 | 0 | 414380 | 442660 | 0 | 0 |
| VITIMQNPF | RPS18 | P62269 | RS18_HUMAN | Unmodified | 0 | 0 | 0 | 0 | 0 | 0 | 0 | 0 | 0 | 0 | 0 | 31652000 | 634300 | 0 | 0 | 518470 | 0 | 159920 |
| VITKDDTH\ | PLEK2; PLEK2 | Q9NYT0; H0 | PLEK2_HUMAN | Unmodified | 67057 | 0 | 0 | 0 | 0 | 0 | 0 | 0 | 2610700 | 92675 | 0 | 0 | 0 | 0 | 283990 | 0 | 0 | 54044 |
| VITKDDTH\ | PLEK2; PLEK2 | Q9NYT0; H0 | PLEK2_HUMAN | Unmodified | 0 | 0 | 0 | 63654 | 0 | 0 | 0 | 0 | 313700 | 26663 | 0 | 0 | 0 | 0 | 0 | 0 | 0 | 0 |
| VITLEPDRP | TTC13 | Q8NBP0 | TTC13_HUMAN | Unmodified | 0 | 0 | 0 | 0 | 234700 | 70930 | 0 | 0 | 999360 | 60430 | 88299 | 355540 | 0 | 0 | 0 | 0 | 0 | 0 |
| VITMFVQR | XRCC5; XRCC5 | P13010; C9 | XRCC5_HUMAI | Unmodified | 0 | 0 | 0 | 0 | 0 | 0 | 0 | 0 | 0 | 0 | 75027 | 3105400 | 0 | 0 | 0 | 0 | 0 | 0 |
| VITNQYNN | PDLIM1 | O00151 | PDLI1_HUMAN | Unmodified | 0 | 0 | 0 | 0 | 2352900 | 0 | 0 | 0 | 0 | 0 | 0 | 0 | 0 | 0 | 0 | 0 | 0 | 11066000 |
| VIVAVGDR | NUP210 | Q8TEM1 | PO210_HUMAI | Unmodified | 462130 | 96454 | 0 | 1086500 | 275200 | 175360 | 81693 | 0 | 46215000 | 2686300 | 2492100 | 463160 | 0 | 0 | 0 | 0 | 140790 | 0 |
| VIVAVGDR: | NUP210 | Q8TEM1 | PO210_HUMAI | Unmodified | 1851000 | 1177200 | 823220 | 2556100 | 0 | 0 | 211310 | 0 | 3517600 | 0 | 73596 | 0 | 1344900 | 0 | 1572300 | 958980 | 0 | 21222 |
| VIVAVGDR: | NUP210 | Q8TEM1 | PO210_HUMAI | Unmodified | 0 | 0 | 0 | 339520 | 0 | 0 | 0 | 0 | 8751300 | 0 | 0 | 0 | 0 | 0 | 0 | 0 | 0 | 0 |
| VIVCNTK | LGALS3; LGALS | P17931; G3 | LEG3_HUMAN | Unmodified | 0 | 38267 | 0 | 0 | 0 | 0 | 0 | 0 | 0 | 0 | 0 | 1126000 | 0 | 0 | 131140 | 0 | 0 | 0 |
| VIVDFSSPN | RARS1 | P54136 | SYRC_HUMAN | Unmodified | 0 | 0 | 326320 | 0 | 0 | 0 | 269730 | 154240 | 545500 | 359620 | 1135200 | 6825800 | 1856200 | 0 | 0 | 0 | 0 | 0 |
| VIVDKYRD( | CNP; CNP; CNI | P09543; K7 | CN37_HUMAN | Unmodified | 0 | 0 | 0 | 119150 | 8335200 | 184240 | 2897500 | 2189400 | 0 | 2236400 | 38519 | 0 | 6324500 | 5205600 | 0 | 0 | 4394300 | 8657600 |
| VIVESASNI | MYOF | Q9NZM1 | MYOF_HUMAN | Unmodified | 0 | 0 | 0 | 0 | 8131700 | 1205900 | 0 | 0 | 303370 | 749840 | 404350 | 0 | 0 | 0 | 0 | 0 | 305950 | 1470000 |
| VIVFDGVPS | ABCE1 | P61221 | ABCE1_HUMAN | Unmodified | 115400 | 95416 | 0 | 0 | 0 | 0 | 0 | 0 | 0 | 0 | 551670 | 5307900 | 0 | 0 | 287240 | 164540 | 0 | 0 |
| VIVGGSSE\ | SMC1A; SMC1A | Q14683; G8 | SMC1A_HUMA | Unmodified | 0 | 0 | 0 | 0 | 2417400 | 964570 | 0 | 19035 | 0 | 0 | 0 | 745920 | 0 | 76533 | 0 | 19426 | 44679 | 2473700 |
| VIVLDKGEI | ABCC1; ABCC1 | P33527; I3L | MRP1_HUMAN | Unmodified | 0 | 0 | 0 | 0 | 5794700 | 0 | 0 | 0 | 0 | 364460 | 2654900 | 8461700 | 541330 | 0 | 1459500 | 1137200 | 1708800 | 3763600 |
| VIVLSGLCS | ITFG1; ITFG1; IT | Q8TB96; F5C | TIP_HUMAN T-( | Unmodified | 0 | 0 | 0 | 0 | 0 | 0 | 0 | 0 | 573990 | 301910 | 0 | 0 | 0 | 0 | 0 | 0 | 0 | 0 |
| VIVLSGLCS | ITFG1; ITFG1; IT | Q8TB96; F5C | TIP_HUMAN T-( | Unmodified | 18956 | 0 | 0 | 0 | 0 | 0 | 0 | 0 | 860300 | 0 | 0 | 0 | 0 | 0 | 0 | 0 | 0 | 0 |
| VIVLSSSHS | PSMG2; PSMG2 | Q969U7; K7 | PSMG2_HUMA | Unmodified | 0 | 0 | 142370 | 0 | 0 | 0 | 284710 | 126210 | 0 | 0 | 143800 | 2955000 | 0 | 0 | 0 | 0 | 0 | 0 |
| VIVMPTTK | IARS2 | Q9NSE4 | SYIM_HUMAN I | Unmodified | 0 | 0 | 0 | 0 | 0 | 0 | 67800 | 0 | 1886300 | 0 | 0 | 0 | 0 | 0 | 0 | 0 | 0 | 0 |
| VIVPNMEFI | CSE1L | P55060 | XPO2_HUMAN | Unmodified | 0 | 0 | 0 | 0 | 0 | 0 | 0 | 0 | 981990 | 0 | 417400 | 3433300 | 0 | 0 | 0 | 0 | 0 | 0 |
| VIVQQPGE | MYO1D; MYO1I | O94832; J3Q | MYO1D_HUMA | Unmodified | 7375100 | 4346300 | 1992800 | 2279800 | 0 | 0 | 0 | 0 | 1036300 | 173910 | 300150 | 1208100 | 0 | 0 | 0 | 2855300 | 0 | 0 |
| VIVVGDLS\ | RAB34; RAB34; | Q9BZG1; C9 | RAB34_HUMAN | Unmodified | 0 | 0 | 0 | 0 | 2101000 | 705610 | 0 | 0 | 686220 | 819690 | 0 | 0 | 0 | 0 | 0 | 0 | 0 | 0 |
| VIVVGNPAI | MDH1; MDH1; I | P40925; C9 | MDHC_HUMAN | Unmodified | 724300 | 236910 | 386180 | 992880 | 20141000 | 1902200 | 3529100 | 2022300 | 4014400 | 2244400 | 3291900 | 24532000 | 6451100 | 4807100 | 975300 | 898970 | 11678000 | 10888000 |
| VIVVWNNI( | EXTL2; EXTL2; E | Q9UBQ6; C9 | EXTL2_HUMAN | Unmodified | 0 | 0 | 0 | 0 | 0 | 0 | 0 | 0 | 2262000 | 262700 | 481760 | 0 | 0 | 0 | 0 | 0 | 0 | 0 |
| VIWTAGDV | RABGGTA; RAB | Q92696; H0 | PGTA_HUMAN | Unmodified | 0 | 0 | 0 | 0 | 0 | 0 | 0 | 0 | 0 | 0 | 0 | 1867000 | 0 | 0 | 0 | 0 | 0 | 0 |
| VIYAMAED( | SLC7A1; SLC7A | P30825; P52 | SL7A1_HUMAN | Unmodified | 0 | 0 | 0 | 0 | 0 | 0 | 0 | 0 | 0 | 0 | 0 | 948550 | 0 | 0 | 0 | 0 | 0 | 0 |
| VIYAMAED( | SLC7A1; SLC7A | P30825; P52 | SL7A1_HUMAN | Oxidation (M) | 0 | 0 | 148180 | 0 | 0 | 0 | 0 | 0 | 0 | 0 | 0 | 1400500 | 515850 | 0 | 292200 | 353740 | 0 | 0 |
| VIYDFIEK | WASL | O00401 | WASL_HUMAN | Unmodified | 1524800 | 1156700 | 1469800 | 1136500 | 0 | 0 | 0 | 0 | 0 | 0 | 0 | 0 | 600760 | 555650 | 2254300 | 1412500 | 0 | 0 |
| VIYGPAAR | ENPP1 | P22413 | ENPP1_HUMAN | Unmodified | 176640 | 172440 | 421230 | 21881 | 0 | 34910 | 0 | 0 | 0 | 0 | 0 | 1421000 | 540520 | 292850 | 459600 | 444630 | 420170 | 972910 |
| VIYPAVEGF | NT5E | P21589 | 5NTD_HUMAN | Unmodified | 0 | 0 | 0 | 0 | 826810 | 0 | 0 | 0 | 0 | 0 | 0 | 0 | 0 | 0 | 0 | 0 | 105330 | 243650 |
| VIYPGLPSH | CTH | P32929 | CGL_HUMAN ( | Unmodified | 0 | 0 | 0 | 0 | 0 | 0 | 0 | 99532 | 0 | 238610 | 0 | 3076200 | 2574900 | 1349200 | 0 | 0 | 0 | 0 |
| VIYPTDSR( | SLC44A2; SLC4 | Q8IWA5; K7 | CTL2_HUMAN ( | Unmodified | 233010 | 0 | 72370 | 290560 | 0 | 0 | 0 | 0 | 595300 | 0 | 0 | 301920 | 0 | 0 | 251010 | 151520 | 0 | 0 |
| VIYSQPSAF | F11R; F11R | Q9Y624; A0 | JAM1_HUMAN . | Unmodified | 7043000 | 4114500 | 2604100 | 2630300 | 459710 | 0 | 0 | 0 | 4332600 | 552630 | 952760 | 4559600 | 1120100 | 125890 | 2213600 | 4567600 | 242210 | 160320 |
| VIYSQPSAF | F11R; F11R | Q9Y624; A0 | JAM1_HUMAN . | Unmodified | 780000 | 0 | 571750 | 717260 | 3631800 | 739120 | 0 | 0 | 706790 | 344580 | 454370 | 0 | 2245700 | 795810 | 324970 | 0 | 237350 | 1570400 |
| VIYSQPSAF | F11R; F11R | Q9Y624; A0 | JAM1_HUMAN . | Unmodified | 0 | 0 | 194750 | 0 | 10022000 | 0 | 679870 | 0 | 0 | 1574400 | 0 | 0 | 4998000 | 1566200 | 0 | 0 | 609770 | 4879800 |
| VIYTLSALS\ | VAV1; VAV1; VA | P15498; Q9 | VAV_HUMAN P | Unmodified | 0 | 0 | 0 | 0 | 0 | 0 | 0 | 0 | 0 | 351340 | 0 | 338640 | 0 | 0 | 0 | 58354 | 0 | 0 |
| VIYVLPMLT | LARS1; LARS1; | Q9P2J5; A0 | SYLC_HUMAN | Oxidation (M) | 0 | 0 | 0 | 0 | 0 | 0 | 0 | 0 | 214610 | 0 | 417140 | 2748200 | 0 | 0 | 0 | 0 | 0 | 0 |
| VIYVLPMLT | LARS1; LARS1; | Q9P2J5; A0 | SYLC_HUMAN | Oxidation (M) | 0 | 0 | 0 | 0 | 0 | 0 | 0 | 0 | 0 | 0 | 0 | 2523500 | 0 | 0 | 0 | 0 | 0 | 0 |
| VKADQDSE | TLN1 | Q9Y490 | TLN1_HUMAN ' | Unmodified | 51327 | 0 | 99254 | 252650 | 0 | 0 | 470510 | 233500 | 0 | 0 | 0 | 0 | 0 | 0 | 0 | 0 | 0 | 0 |
| VKADRDES | RPS14; RPS14; | P62263; E5 | RS14_HUMAN . | Unmodified | 1865200 | 2847800 | 1043300 | 2674900 | 2082900 | 0 | 1227200 | 1203600 | 2865200 | 0 | 3056800 | 20829000 | 3869800 | 2202300 | 2072200 | 1074900 | 0 | 834930 |
| VKADRDES | RPS14; RPS14; | P62263; E5 | RS14_HUMAN . | Oxidation (M) | 1586600 | 932700 | 412050 | 994280 | 0 | 0 | 0 | 0 | 0 | 0 | 1813400 | 7783100 | 0 | 0 | 113270 | 0 | 0 | 0 |
| VKAEEDKIF | RALB; RALB; RA | P11234; C9 | RALB_HUMAN | Unmodified | 322890 | 291980 | 505920 | 710700 | 0 | 0 | 0 | 0 | 0 | 0 | 2722200 | 9756600 | 0 | 0 | 1026700 | 898690 | 0 | 0 |
| VKAEGILD\ | PTPRA | P18433 | PTPRA_HUMAN | Unmodified | 224690 | 0 | 0 | 471610 | 0 | 0 | 0 | 0 | 0 | 154500 | 806960 | 2239200 | 253130 | 0 | 0 | 749020 | 0 | 0 |
| VKAEPAKIE | PGK1 | P00558 | PGK1_HUMAN | Unmodified | 875140 | 523260 | 280280 | 1984900 | 1175400 | 0 | 91707 | 0 | 6134100 | 864680 | 3086100 | 35014000 | 3133900 | 0 | 1515000 | 1175000 | 124090 | 219290 |
| VKAEVQNL | CTNNA1 | P35221 | CTNA1_HUMAI | Oxidation (M) | 0 | 0 | 0 | 0 | 1209000 | 0 | 0 | 0 | 0 | 0 | 0 | 0 | 0 | 1981500 | 1372700 | 0 | 0 | 0 |
| VKAEVQNL | CTNNA1 | P35221 | CTNA1_HUMAI | Unmodified | 0 | 0 | 0 | 0 | 743230 | 0 | 0 | 0 | 0 | 0 | 0 | 0 | 0 | 0 | 129520 | 0 | 0 | 0 |
| VKAFGPGL | FLNA; FLNA | P21333; Q6 | FLNA_HUMAN | Unmodified | 0 | 0 | 0 | 0 | 0 | 0 | 1577800 | 1163100 | 0 | 0 | 0 | 0 | 840620 | 0 | 0 | 0 | 1209000 | 0 |
| VKAHGPGL | FLNB; FLNB | O75369; E7 | FLNB_HUMAN | Unmodified | 0 | 0 | 0 | 0 | 891950 | 0 | 0 | 0 | 0 | 0 | 0 | 0 | 0 | 0 | 0 | 0 | 827230 | 0 |
| VKASGPGL | FLNA; FLNA | P21333; Q6 | FLNA_HUMAN | Unmodified | 0 | 0 | 0 | 0 | 1752800 | 0 | 2569200 | 43838 | 0 | 0 | 0 | 1329000 | 0 | 2339200 | 0 | 0 | 1613100 | 307090 |
| VKCSGPGL | FLNA; FLNA | P21333; Q6 | FLNA_HUMAN | Oxidation (M) | 0 | 0 | 0 | 0 | 0 | 0 | 966950 | 539560 | 0 | 0 | 0 | 0 | 0 | 0 | 0 | 0 | 0 | 0 |
| VKDATQIT( | L1CAM | P32004 | L1CAM_HUMAN | Unmodified | 0 | 0 | 0 | 0 | 0 | 0 | 234570 | 112150 | 0 | 0 | 0 | 0 | 0 | 0 | 0 | 0 | 204380 | 0 |
| VKDDIESLI | DYNC1H1; DYI | Q14204; A0 | DYHC1_HUMA | Unmodified | 0 | 0 | 325080 | 169930 | 0 | 0 | 0 | 0 | 0 | 0 | 536260 | 3421000 | 0 | 0 | 0 | 162060 | 0 | 0 |
| VKDGPGGI | SCP2; SCP2 | P22307; H0 | SCP2_HUMAN | Unmodified | 0 | 0 | 0 | 0 | 0 | 0 | 0 | 0 | 498200 | 1559700 | 0 | 0 | 0 | 0 | 0 | 0 | 0 | 0 |
| VKDLLDIAI | DPY19L4 | Q7Z388 | D19L4_HUMAN | Unmodified | 0 | 0 | 0 | 0 | 0 | 0 | 0 | 0 | 468980 | 0 | 0 | 0 | 0 | 0 | 0 | 0 | 0 | 0 |
| VKDSEDVP | KRAS | P01116 | RASK_HUMAN | Unmodified | 0 | 0 | 0 | 0 | 154690 | 65064 | 116870 | 30391 | 474240 | 92435 | 357880 | 0 | 0 | 0 | 0 | 134220 | 0 | 66400 |
| VKDSEDVP | KRAS | P01116 | RASK_HUMAN | Oxidation (M) | 0 | 0 | 10715 | 0 | 0 | 0 | 0 | 0 | 0 | 0 | 381950 | 244260 | 0 | 0 | 0 | 51344 | 0 | 0 |
| VKDSMRVL | STXBP3 | O00186 | STXB3_HUMAN | Unmodified | 0 | 0 | 0 | 0 | 556140 | 45865 | 0 | 60120 | 0 | 0 | 0 | 0 | 171610 | 191670 | 0 | 0 | 148180 | 499230 |
| VKDTDDVP | RAP1B; N/A; RA | P61224; A6 | RAP1B_HUMAI | Unmodified | 841180 | 873030 | 1132600 | 2005800 | 0 | 0 | 0 | 0 | 917100 | 0 | 890140 | 5376500 | 0 | 0 | 1450900 | 670390 | 0 | 0 |
| VKDTDDVP | RAP1B; N/A; RA | P61224; A6 | RAP1B_HUMAI | Oxidation (M) | 1177400 | 1588500 | 5451400 | 1221700 | 0 | 0 | 0 | 0 | 0 | 0 | 61182 | 903790 | 0 | 0 | 2323000 | 3391400 | 0 | 0 |
| VKDTDDVP | RAP1B; N/A; RA | P61224; A6 | RAP1B_HUMAI | Oxidation (M) | 1086600 | 0 | 157620 | 272710 | 0 | 0 | 0 | 0 | 0 | 0 | 0 | 732210 | 0 | 0 | 0 | 319330 | 0 | 16788000 |
| VKDTEDVP | RAP1A; RAP1A | P62834; A0 | RAP1A_HUMAI | Unmodified | 317180 | 300130 | 1032200 | 1392500 | 0 | 0 | 0 | 0 | 779140 | 0 | 718170 | 2728600 | 367790 | 0 | 786690 | 239180 | 0 | 0 |
| VKDTEDVP | RAP1A; RAP1A | P62834; A0 | RAP1A_HUMAI | Oxidation (M) | 165870 | 327500 | 3195800 | 423840 | 0 | 0 | 0 | 0 | 0 | 0 | 411560 | 2013000 | 0 | 0 | 1306400 | 1659100 | 0 | 0 |

| | | | | | | | | | | | | | | | | | | | | | | |
|---|---|---|---|---|---|---|---|---|---|---|---|---|---|---|---|---|---|---|---|---|---|---|
| VKDTFGN | MPI; MPI; MPI; N | P34949; H3E | MPI_HUMAN M | Unmodified | 0 | 0 | 0 | 0 | 1519900 | 0 | 0 | 0 | 0 | 141120 | 35079 | 847400 | 1727100 | 832260 | 0 | 0 | 764260 | 0 |
| VKEDENVP | RALA; RALA | P11233; H7C | RALA_HUMAN | Unmodified | 4511100 | 4492000 | 13244000 | 4671700 | 0 | 0 | 0 | 0 | 14179000 | 539980 | 7909100 | 22813000 | 0 | 575320 | 12702000 | 10225000 | 87390 | 351490 |
| VKEDENVP | RALA; RALA | P11233; H7C | RALA_HUMAN | Unmodified | 12435000 | 1162900 | 0 | 0 | 0 | 0 | 0 | 0 | 31018000 | 1020500 | 16108000 | 49764000 | 0 | 0 | 12044000 | 12356000 | 0 | 0 |
| VKEDPDQC | GLT8D1; GLT8D | Q68CQ7; C9 | GL8D1_HUMAN | Unmodified | 131370 | 0 | 0 | 0 | 206780 | 0 | 0 | 0 | 32936000 | 2481000 | 0 | 0 | 0 | 0 | 0 | 0 | 1114100 | 254490 |
| VKEDPDQC | GLT8D1; GLT8D | Q68CQ7; C9 | GL8D1_HUMAN | Oxidation (M) | 0 | 0 | 0 | 0 | 291110 | 0 | 0 | 0 | 4086800 | 230230 | 0 | 0 | 0 | 0 | 0 | 0 | 4295700 | 0 |
| VKEEESLEN | RAB3GAP1 | Q15042 | RB3GP_HUMAN | Unmodified | 0 | 0 | 0 | 0 | 0 | 0 | 0 | 95341 | 0 | 0 | 0 | 1226100 | 0 | 0 | 0 | 0 | 0 | 0 |
| VKEEIIEAFV | VASP | P50552 | VASP_HUMAN | Unmodified | 0 | 0 | 67105 | 0 | 4378500 | 0 | 0 | 0 | 131380 | 514600 | 1750400 | 796630 | 1526200 | 3079400 | 0 | 0 | 1480200 | 0 |
| VKEEIIEAFV | VASP | P50552 | VASP_HUMAN | Unmodified | 190470 | 0 | 217020 | 0 | 3851900 | 0 | 1134300 | 0 | 703860 | 745250 | 682300 | 2922400 | 809690 | 3674000 | 0 | 0 | 605510 | 1115500 |
| VKEFLQEK | AGFG1; AGFG2 | P52594; O9 | AGFG1_HUMAN | Unmodified | 0 | 0 | 0 | 0 | 488980 | 0 | 0 | 0 | 0 | 0 | 0 | 0 | 0 | 0 | 0 | 0 | 142880 | 0 |
| VKEQKDYW | CALCOCO2; CA | Q13137; D6I | CACO2_HUMA | Unmodified | 0 | 0 | 0 | 0 | 651440 | 0 | 3853900 | 0 | 0 | 147670 | 0 | 0 | 45875 | 0 | 0 | 0 | 1285900 | 2388800 |
| VKEQYQDV | PKP2 | Q99959 | PKP2_HUMAN | Oxidation (M) | 0 | 0 | 0 | 0 | 0 | 0 | 0 | 0 | 0 | 0 | 0 | 179030 | 0 | 0 | 0 | 0 | 74513 | 170710 |
| VKESDKLV | SNX9; SNX9; SN | Q9Y5X1; A0A | SNX9_HUMAN | Unmodified | 2328000 | 5054800 | 0 | 2063200 | 0 | 0 | 0 | 0 | 0 | 0 | 0 | 0 | 0 | 0 | 0 | 0 | 0 | 0 |
| VKETADFK | FLNA; FLNA; FL | P21333; F8W | FLNA_HUMAN | Unmodified | 0 | 0 | 0 | 377550 | 0 | 0 | 287310 | 0 | 0 | 0 | 0 | 0 | 0 | 0 | 0 | 0 | 454620 | 0 |
| VKFDDGAV | PPP1R12A; PPP | O14974; F8W | MYPT1_HUMAN | Unmodified | 27531 | 1492200 | 0 | 363230 | 2190500 | 258860 | 0 | 0 | 297020 | 0 | 0 | 0 | 1146300 | 798490 | 0 | 95082 | 1024500 | 1075600 |
| VKFGESIED | PIGX; PIGX; PIG | Q8TBF5; C9J | PIGX_HUMAN | Unmodified | 0 | 0 | 0 | 0 | 0 | 0 | 0 | 0 | 5244700 | 856220 | 0 | 0 | 30076 | 0 | 0 | 0 | 0 | 0 |
| VKFIHDQTS | SLC25A1; SLC2 | P53007; B4D | TXTP_HUMAN | Unmodified | 0 | 0 | 0 | 0 | 0 | 0 | 0 | 0 | 0 | 116000 | 0 | 0 | 0 | 0 | 0 | 0 | 0 | 0 |
| VKFQAKLEI | MAPRE3; MAPR | Q9UPY8; Q1 | MARE3_HUMAN | Unmodified | 0 | 0 | 0 | 0 | 0 | 0 | 1427400 | 0 | 0 | 483010 | 0 | 0 | 0 | 789510 | 0 | 0 | 0 | 582120 |
| VKFQAKLEI | MAPRE1 | Q15691 | MARE1_HUMAN | Unmodified | 0 | 0 | 0 | 0 | 1654400 | 0 | 1484600 | 0 | 0 | 1211000 | 0 | 0 | 1307100 | 0 | 0 | 0 | 1522200 | 2007800 |
| VKFSGFEA | PTPRJ; PTPRJ; P | Q12913; A0A | PTPRJ_HUMAN | Unmodified | 0 | 0 | 94389 | 0 | 649610 | 0 | 58691 | 944280 | 0 | 365800 | 0 | 0 | 0 | 208240 | 0 | 44528 | 390500 | 183100 |
| VKFTAEELS | IQGAP3; IQGAP | Q86VI3; F2Z | IQGA3_HUMAN | Unmodified | 59190 | 106170 | 39259 | 0 | 469720 | 185230 | 0 | 0 | 0 | 0 | 44548 | 166910 | 0 | 0 | 58157 | 0 | 113610 | 0 |
| VKGDVDVS | AHNAK | Q09666 | AHNK_HUMAN | Unmodified | 0 | 0 | 0 | 0 | 1564000 | 0 | 0 | 0 | 0 | 0 | 0 | 0 | 0 | 0 | 0 | 0 | 1198700 | 0 |
| VKGEYDMT | AHNAK | Q09666 | AHNK_HUMAN | Unmodified | 0 | 0 | 0 | 0 | 3823300 | 0 | 0 | 0 | 0 | 0 | 0 | 0 | 0 | 0 | 0 | 0 | 1336200 | 2187800 |
| VKGEYDMT | AHNAK | Q09666 | AHNK_HUMAN | Oxidation (M) | 0 | 0 | 0 | 0 | 1027500 | 0 | 0 | 0 | 0 | 0 | 0 | 0 | 0 | 0 | 0 | 0 | 440820 | 260140 |
| VKGEYDVT | AHNAK | Q09666 | AHNK_HUMAN | Unmodified | 0 | 0 | 0 | 0 | 9103500 | 1081800 | 0 | 0 | 0 | 0 | 0 | 0 | 0 | 0 | 0 | 0 | 2685200 | 3724800 |
| VKGFGGAN | GBA; GBA; GBA | P04062; A0A | GLCM_HUMAN | Oxidation (M) | 0 | 0 | 0 | 0 | 0 | 0 | 0 | 0 | 0 | 1113400 | 0 | 0 | 0 | 0 | 0 | 0 | 0 | 0 |
| VKGGASEL | LLGL2; LLGL2; I | Q6P1M3; J3C | L2GL2_HUMAN | Unmodified | 0 | 0 | 0 | 0 | 154380 | 0 | 0 | 0 | 0 | 0 | 0 | 421540 | 0 | 0 | 0 | 382610 | 301850 | 63843 |
| VKGGGHV | RPS16; RPS16; | P62249; M0F | RS16_HUMAN | Unmodified | 1174600 | 1118200 | 0 | 363580 | 0 | 0 | 0 | 0 | 0 | 0 | 0 | 7210200 | 0 | 0 | 216450 | 401620 | 0 | 0 |
| VKGGGHV | RPS16; RPS16; | P62249; M0F | RS16_HUMAN | Unmodified | 0 | 0 | 0 | 65795 | 0 | 0 | 448620 | 44052 | 322200 | 54876 | 0 | 0 | 0 | 0 | 0 | 0 | 38106 | 0 |
| VKGGVDVT | AHNAK | Q09666 | AHNK_HUMAN | Unmodified | 0 | 0 | 0 | 0 | 4233600 | 0 | 0 | 0 | 0 | 0 | 0 | 0 | 0 | 0 | 0 | 0 | 1595700 | 2610600 |
| VKGTNEDN | EDIL3 | O43854 | EDIL3_HUMAN | Unmodified | 209530 | 0 | 0 | 0 | 3704100 | 2182800 | 0 | 0 | 20321000 | 1243800 | 0 | 0 | 112570 | 0 | 0 | 0 | 90296 | 1616700 |
| VKGTNEDN | EDIL3 | O43854 | EDIL3_HUMAN | Oxidation (M) | 0 | 0 | 0 | 0 | 548160 | 672760 | 0 | 0 | 3663000 | 0 | 0 | 21707 | 0 | 102290 | 0 | 0 | 451050 | 218260 |
| VKGTNEDN | EDIL3 | O43854 | EDIL3_HUMAN | Oxidation (M) | 0 | 0 | 0 | 0 | 2E+08 | 66739000 | 14475000 | 5519900 | 12574000 | 10420000 | 0 | 0 | 0 | 0 | 0 | 0 | 11699000 | 70909000 |
| VKGTNEDN | EDIL3 | O43854 | EDIL3_HUMAN | Unmodified | 0 | 0 | 0 | 0 | 1.09E+08 | 21052000 | 4827500 | 488650 | 12012000 | 2541700 | 0 | 0 | 0 | 0 | 0 | 0 | 4866500 | 39006000 |
| VKIEPGVDI | RNPEP; RNPEP | Q9H4A4; A6I | AMPB_HUMAN | Unmodified | 0 | 0 | 0 | 0 | 0 | 0 | 526700 | 112070 | 981500 | 160290 | 3460500 | 5077800 | 0 | 0 | 0 | 0 | 0 | 0 |
| VKIETALTSI | NPTX1 | Q15818 | NPTX1_HUMAN | Unmodified | 0 | 0 | 0 | 0 | 0 | 0 | 0 | 0 | 14593000 | 454360 | 0 | 0 | 0 | 0 | 0 | 0 | 62917 | 0 |
| VKIPEGTILT | NANS | Q9NR45 | SIAS_HUMAN S | 2 Oxidation (M) | 0 | 0 | 0 | 234020 | 692500 | 0 | 0 | 0 | 75380 | 126370 | 256520 | 910100 | 761830 | 0 | 117030 | 360220 | 197240 | 297730 |
| VKIPEGTILT | NANS | Q9NR45 | SIAS_HUMAN S | Oxidation (M) | 0 | 1107600 | 233720 | 176270 | 1916900 | 0 | 304510 | 0 | 0 | 234690 | 334320 | 551420 | 1234300 | 1364100 | 148280 | 65568 | 317710 | 634300 |
| VKIPEPHTC | AHNAK2 | Q8IVF2 | AHNK2_HUMAN | Unmodified | 0 | 0 | 0 | 0 | 153740 | 0 | 0 | 0 | 0 | 0 | 0 | 0 | 0 | 0 | 0 | 0 | 0 | 109030 |
| VKIQDAHR | WDR1 | O75083 | WDR1_HUMAN | Unmodified | 0 | 0 | 0 | 0 | 1115200 | 0 | 0 | 0 | 0 | 153170 | 0 | 0 | 480300 | 0 | 0 | 0 | 0 | 0 |
| VKISTEVGI | RPN2 | P04844 | RPN2_HUMAN | Unmodified | 0 | 0 | 0 | 0 | 0 | 0 | 0 | 0 | 2344300 | 1686500 | 0 | 0 | 0 | 0 | 0 | 0 | 0 | 0 |
| VKKYEQGF | IMPDH2; IMPDH | P12268; A0A | IMDH2_HUMAN | Unmodified | 0 | 0 | 0 | 0 | 1304200 | 0 | 877990 | 0 | 0 | 308210 | 0 | 0 | 324710 | 0 | 0 | 0 | 2086800 | 5816100 |
| VKLAADED | NPM1; NPM1; N | P06748; A0A | NPM_HUMAN N | Unmodified | 0 | 0 | 0 | 0 | 317310 | 0 | 0 | 286420 | 0 | 0 | 0 | 741140 | 1248500 | 0 | 0 | 0 | 0 | 352570 |
| VKLAAVDA | PDIA6 | Q15084 | PDIA6_HUMAN | Unmodified | 0 | 0 | 0 | 0 | 0 | 0 | 33124 | 0 | 1401100 | 2024900 | 0 | 0 | 0 | 0 | 0 | 0 | 0 | 0 |
| VKLAAVDA | PDIA6 | Q15084 | PDIA6_HUMAN | Unmodified | 0 | 0 | 0 | 0 | 193810 | 0 | 0 | 0 | 0 | 1020300 | 0 | 0 | 0 | 0 | 0 | 0 | 0 | 0 |
| VKLADIQIE | PFDN1; PFDN1 | O60925; E5F | PFD1_HUMAN | Unmodified | 0 | 0 | 0 | 0 | 1401300 | 0 | 0 | 48465 | 0 | 0 | 0 | 0 | 1291300 | 403540 | 0 | 0 | 4356600 | 1970700 |
| VKLDSPAG | PYCR1; PYCR1 | P32322; E2C | P5CR1_HUMAN | Unmodified | 0 | 0 | 0 | 0 | 0 | 0 | 159510 | 0 | 1791700 | 278640 | 0 | 0 | 0 | 0 | 0 | 0 | 0 | 0 |
| VKLEAEIAT | KRT18; KRT18 | P05783; F8V | K1C18_HUMAN | Unmodified | 0 | 0 | 0 | 0 | 224960 | 0 | 0 | 0 | 0 | 0 | 1071800 | 10689000 | 0 | 0 | 0 | 0 | 0 | 0 |
| VKLEAEIAT | KRT18; KRT18 | P05783; F8V | K1C18_HUMAN | Unmodified | 0 | 0 | 0 | 0 | 0 | 0 | 0 | 0 | 0 | 0 | 0 | 10814000 | 0 | 0 | 0 | 917590 | 0 | 0 |
| VKLEEQLKE | GRPEL1 | Q9HAV7 | GRPE1_HUMAN | Unmodified | 0 | 0 | 0 | 0 | 541700 | 0 | 72882 | 0 | 0 | 492100 | 0 | 0 | 277880 | 290830 | 0 | 0 | 55180 | 0 |
| VKLEGHELI | EHD1; EHD1 | Q9H4M9; A0 | EHD1_HUMAN | Unmodified | 903170 | 1228200 | 699730 | 1589400 | 21732000 | 7061300 | 1699800 | 464990 | 0 | 2207000 | 3942900 | 7443500 | 7767000 | 6686400 | 1923600 | 1482700 | 15503000 | 23698000 |
| VKLEGHELI | EHD3 | Q9NZN3 | EHD3_HUMAN | Unmodified | 0 | 0 | 360730 | 484570 | 0 | 0 | 0 | 0 | 0 | 0 | 0 | 0 | 1286800 | 1030400 | 0 | 578760 | 0 | 0 |
| VKLGNNFA | NSG1 | P42857 | NSG1_HUMAN | Unmodified | 0 | 0 | 0 | 0 | 0 | 0 | 631800 | 0 | 0 | 0 | 0 | 0 | 0 | 0 | 0 | 0 | 0 | 0 |
| VKLIVLVPP | F11R; F11R | Q9Y624; A0A | JAM1_HUMAN | Unmodified | 0 | 0 | 0 | 0 | 0 | 0 | 0 | 0 | 0 | 0 | 0 | 0 | 2019400 | 0 | 0 | 0 | 0 | 0 |
| VKLLLQVQI | SLC25A4; SLC2 | P12235; P12 | ADT1_HUMAN | Unmodified | 0 | 0 | 0 | 0 | 0 | 0 | 25326 | 0 | 0 | 779260 | 0 | 0 | 0 | 0 | 0 | 0 | 0 | 0 |
| VKLPIIGVVI | NUBP1; NUBP1 | P53384; I3L3 | NUBP1_HUMAN | Unmodified | 0 | 0 | 0 | 0 | 359860 | 0 | 72223 | 0 | 52427 | 37047 | 0 | 10735 | 193100 | 515430 | 18873 | 0 | 37636 | 0 |
| VKLPIIGVVI | NUBP1; NUBP1 | P53384; I3L3 | NUBP1_HUMAN | Oxidation (M) | 0 | 0 | 0 | 0 | 0 | 26386 | 121120 | 0 | 0 | 0 | 0 | 1028500 | 203210 | 265060 | 0 | 0 | 85314 | 2595100 |
| VKLQEMEG | MYH9 | P35579 | MYH9_HUMAN | Oxidation (M) | 0 | 54122 | 0 | 173700 | 3667800 | 440730 | 345870 | 159450 | 0 | 923200 | 0 | 649880 | 1069900 | 1142300 | 0 | 90331 | 1285700 | 754520 |
| VKLQEMEG | MYH9 | P35579 | MYH9_HUMAN | Unmodified | 0 | 365730 | 0 | 1113900 | 34404000 | 3973500 | 7536000 | 1486500 | 1226600 | 2265800 | 563970 | 4031600 | 10075000 | 11584000 | 682780 | 525640 | 10687000 | 10411000 |
| VKLQEMEG | MYH9 | P35579 | MYH9_HUMAN | Unmodified | 0 | 0 | 0 | 0 | 174410 | 0 | 487150 | 148610 | 0 | 0 | 0 | 0 | 188820 | 130330 | 0 | 0 | 2853800 | 160560 |
| VKLQIWDT | RAB35; RAB4B; | Q15286; P61 | RAB35_HUMAN | Unmodified | 3532900 | 3539400 | 39894 | 2014100 | 168070 | 48674 | 0 | 0 | 0 | 145980 | 191660 | 1042000 | 70055 | 294430 | 1117300 | 1842900 | 0 | 0 |
| VKLQSLCR | TTC7B; TTC7B; | Q86TV6; Q6I | TTC7B_HUMAN | Unmodified | 2696300 | 1115500 | 137060 | 525300 | 0 | 0 | 0 | 0 | 0 | 0 | 0 | 0 | 0 | 0 | 572060 | 933270 | 0 | 0 |
| VKLTAELIE | SNRPA1; SNRP | P09661; H0Y | RU2A_HUMAN | Unmodified | 728250 | 0 | 0 | 0 | 0 | 0 | 0 | 0 | 56855 | 0 | 0 | 393330 | 0 | 0 | 0 | 23904 | 0 | 0 |
| VKMDCQET | FLNB; FLNB | O75369; E7E | FLNB_HUMAN | Unmodified | 0 | 0 | 0 | 0 | 477500 | 182570 | 0 | 0 | 154120 | 0 | 0 | 248080 | 48365 | 42515 | 58944 | 0 | 356090 | 93995 |
| VKMEGTIS | CIT; CIT; CIT; CI | O14578; A0A | CTRO_HUMAN | Unmodified | 0 | 0 | 0 | 0 | 1105000 | 160550 | 220500 | 0 | 0 | 0 | 0 | 160860 | 80454 | 0 | 0 | 0 | 425880 | 139310 |
| VKMPDVDI | AHNAK | Q09666 | AHNK_HUMAN | Unmodified | 0 | 0 | 0 | 0 | 1195000 | 0 | 0 | 0 | 0 | 0 | 0 | 0 | 0 | 0 | 0 | 0 | 0 | 1131000 |
| VKMPDVDI | AHNAK | Q09666 | AHNK_HUMAN | Unmodified | 0 | 0 | 0 | 0 | 1571400 | 0 | 0 | 0 | 0 | 0 | 0 | 0 | 0 | 0 | 0 | 0 | 724110 | 0 |
| VKMPEQEY | ANTXR1 | Q9H6X2 | ANTR1_HUMAN | Unmodified | 0 | 0 | 56904 | 0 | 6833000 | 510960 | 0 | 0 | 0 | 0 | 0 | 0 | 0 | 0 | 0 | 0 | 607990 | 437660 |
| VKNENVAF | PRKG2; PRKG2 | Q13237; B7Z | KGP2_HUMAN | Unmodified | 0 | 0 | 0 | 0 | 0 | 0 | 513910 | 379620 | 0 | 0 | 0 | 0 | 0 | 0 | 0 | 0 | 0 | 0 |
| VKPAPDETS | ILF2; ILF2; ILF2; | Q12905; B4D | ILF2_HUMAN In | Unmodified | 869680 | 0 | 0 | 245600 | 0 | 0 | 0 | 0 | 0 | 0 | 0 | 1231000 | 326230 | 0 | 0 | 0 | 0 | 0 |
| VKPAPDETS | ILF2; ILF2; ILF2; | Q12905; B4D | ILF2_HUMAN In | Unmodified | 796790 | 0 | 0 | 1586100 | 0 | 0 | 0 | 0 | 165760 | 0 | 780920 | 1446500 | 0 | 0 | 0 | 0 | 0 | 0 |
| VKPFMTGA | N/A; TPT1; TPT1 | Q56UQ5; P1 | TPT1L_HUMAN | Unmodified | 2634400 | 2899100 | 3705300 | 3748600 | 1238500 | 722480 | 241560 | 81562 | 2672900 | 412110 | 3644700 | 19153000 | 2300500 | 738070 | 6604400 | 5197400 | 337620 | 1050400 |
| VKPFMTGA | N/A; TPT1; TPT1 | Q56UQ5; P1 | TPT1L_HUMAN | Oxidation (M) | 427970 | 1303800 | 3917700 | 2512200 | 0 | 0 | 0 | 12877 | 647430 | 253540 | 1103900 | 4452500 | 0 | 0 | 3403700 | 3441700 | 132660 | 248220 |
| VKPFVVRD | HYOU1; HYOU1 | Q9Y4L1; E9F | HYOU1_HUMA | Unmodified | 0 | 0 | 0 | 0 | 0 | 0 | 0 | 0 | 3254400 | 1505400 | 305070 | 0 | 0 | 0 | 0 | 0 | 0 | 0 |
| VKPLQDQN | SF3A3 | Q12874 | SF3A3_HUMAN | Unmodified | 150610 | 116550 | 0 | 0 | 0 | 0 | 0 | 0 | 0 | 0 | 0 | 551010 | 0 | 0 | 148870 | 120990 | 0 | 0 |
| VKPPPSPT | AKAP2 | Q9Y2D5 | AKAP2_HUMAN | Unmodified | 0 | 0 | 0 | 0 | 1913900 | 0 | 0 | 0 | 0 | 0 | 0 | 0 | 0 | 0 | 0 | 0 | 951350 | 2522700 |
| VKPYLPQIC | SF3B1 | O75533 | SF3B1_HUMAN | Unmodified | 234140 | 89879 | 0 | 0 | 150860 | 0 | 0 | 0 | 0 | 0 | 64574 | 477190 | 0 | 0 | 0 | 0 | 0 | 122850 |
| VKQELLEE | VASP | P50552 | VASP_HUMAN | Unmodified | 0 | 0 | 0 | 0 | 372440 | 0 | 0 | 0 | 0 | 0 | 868760 | 2908300 | 0 | 0 | 0 | 0 | 71098 | 169130 |
| VKQENLNL | RPIA | P49247 | RPIA_HUMAN F | Unmodified | 0 | 0 | 0 | 0 | 0 | 0 | 404030 | 44376 | 250470 | 121190 | 728010 | 1604000 | 0 | 126740 | 0 | 0 | 0 | 0 |
| VKQSLAQS | CIT; CIT; CIT; CI | O14578; A0A | CTRO_HUMAN | Unmodified | 0 | 0 | 0 | 0 | 1591100 | 0 | 0 | 0 | 0 | 0 | 257250 | 0 | 0 | 0 | 0 | 0 | 664620 | 0 |
| VKQVERED | FGFRL1; FGFRL | Q8N441; D6 | FGRL1_HUMAN | Unmodified | 0 | 0 | 0 | 0 | 80361 | 0 | 0 | 0 | 0 | 0 | 0 | 0 | 273020 | 275760 | 0 | 0 | 0 | 0 |
| VKQVLLAQ | FLOT2; FLOT2; | Q14254; J3Q | FLOT2_HUMAN | Unmodified | 0 | 0 | 0 | 0 | 5955300 | 880770 | 2020700 | 1022000 | 0 | 1664600 | 0 | 0 | 1826500 | 2226600 | 0 | 0 | 456330 | 611740 |
| VKRENFCN | CTPS1; CTPS1; | P17812; B4E | PYRG1_HUMAN | Unmodified | 0 | 0 | 0 | 0 | 0 | 0 | 574140 | 0 | 0 | 197800 | 0 | 0 | 171160 | 0 | 0 | 0 | 565780 | 1444000 |
| VKSPRDYS | PRKCD | Q05655 | KPCD_HUMAN | Unmodified | 0 | 748120 | 0 | 0 | 0 | 0 | 0 | 0 | 0 | 421140 | 0 | 0 | 265170 | 861640 | 0 | 0 | 3034600 | 3861900 |
| VKSSDEAV | PSMD13; PSMD | Q9UNM6; A0 | PSD13_HUMAN | Unmodified | 0 | 0 | 0 | 0 | 0 | 0 | 730370 | 295310 | 0 | 0 | 0 | 0 | 1853900 | 662890 | 0 | 0 | 308830 | 0 |
| VKTDTVLIL | CACYBP | Q9HB71 | CYBP_HUMAN | Unmodified | 0 | 0 | 0 | 0 | 0 | 0 | 0 | 0 | 0 | 0 | 0 | 1382100 | 0 | 0 | 0 | 0 | 0 | 0 |
| VKTPEMIIQ | AHNAK | Q09666 | AHNK_HUMAN | Unmodified | 0 | 0 | 0 | 0 | 3013100 | 0 | 0 | 0 | 0 | 0 | 0 | 0 | 114300 | 140490 | 0 | 0 | 1781000 | 1485200 |
| VKTPEMIIQ | AHNAK | Q09666 | AHNK_HUMAN | Oxidation (M) | 0 | 0 | 0 | 0 | 991680 | 0 | 0 | 0 | 0 | 0 | 0 | 0 | 0 | 0 | 0 | 0 | 559880 | 0 |
| VKTPSFGIS | AHNAK | Q09666 | AHNK_HUMAN | Unmodified | 0 | 0 | 0 | 0 | 13366000 | 417090 | 0 | 0 | 0 | 0 | 0 | 0 | 119380 | 119780 | 0 | 0 | 5617200 | 9159300 |
| VKTPSFGIS | AHNAK | Q09666 | AHNK_HUMAN | Unmodified | 0 | 0 | 0 | 0 | 3553200 | 0 | 0 | 0 | 0 | 0 | 0 | 0 | 0 | 0 | 0 | 0 | 3232400 | 0 |
| VKTTGIVET | GNAI3; GNAI2; | P08754; P04 | GNAI3_HUMAN | Unmodified | 0 | 0 | 0 | 724590 | 612480 | 341710 | 635590 | 0 | 933100 | 0 | 0 | 0 | 775670 | 0 | 0 | 0 | 792650 | 928090 |
| VKVEHVVS | UGGT1 | Q9NYU2 | UGGG1_HUMA | Unmodified | 0 | 0 | 0 | 0 | 0 | 0 | 0 | 0 | 8398300 | 2814600 | 1192400 | 0 | 0 | 0 | 0 | 0 | 0 | 0 |
| VKVETYND | RPA1; RPA1 | P27694; I3L4 | RFA1_HUMAN | Unmodified | 72688 | 47713 | 0 | 0 | 0 | 0 | 0 | 0 | 1393200 | 531270 | 0 | 1969000 | 0 | 0 | 0 | 16425 | 0 | 0 |
| VKVFGPGV | FLNB; FLNB | O75369; E7E | FLNB_HUMAN | Unmodified | 0 | 0 | 0 | 0 | 430740 | 167520 | 73195 | 55111 | 0 | 0 | 0 | 0 | 0 | 170540 | 0 | 0 | 657900 | 0 |
| VKVGKEDS | SNX1; SNX1 | Q13596; H0Y | SNX1_HUMAN | Unmodified | 0 | 0 | 0 | 0 | 0 | 0 | 0 | 0 | 96705 | 353990 | 0 | 0 | 325090 | 0 | 0 | 0 | 0 | 0 |
| VKVGKEDS | SNX2; SNX2 | O60749; D6I | SNX2_HUMAN | Unmodified | 0 | 0 | 0 | 0 | 0 | 0 | 263820 | 116660 | 0 | 0 | 0 | 0 | 1916900 | 676680 | 0 | 0 | 222160 | 0 |
| VKVTFNSAI | ITM2B; ITM2B; I | Q9Y287; A0A | ITM2B_HUMAN | Unmodified | 0 | 0 | 0 | 536400 | 14391000 | 1719500 | 8023900 | 991630 | 5165600 | 4945000 | 0 | 0 | 2576600 | 1529600 | 0 | 0 | 206060 | 3174100 |
| VKVTQSTE | PRKG2 | Q13237 | KGP2_HUMAN | Unmodified | 0 | 0 | 0 | 0 | 0 | 0 | 2016500 | 1568400 | 0 | 229280 | 0 | 0 | 0 | 0 | 0 | 0 | 0 | 0 |
| VKVYGPGV | FLNA; FLNA; FL | P21333; A0A | FLNA_HUMAN | Unmodified | 0 | 0 | 0 | 0 | 74239 | 0 | 254460 | 171640 | 0 | 0 | 0 | 0 | 41989 | 0 | 0 | 0 | 151970 | 0 |
| VKWGDIEF | ACLY | P53396 | ACLY_HUMAN | Unmodified | 0 | 0 | 0 | 0 | 0 | 0 | 17984 | 0 | 456510 | 0 | 268470 | 2231400 | 0 | 0 | 0 | 197220 | 0 | 0 |
| VKWGDIEF | ACLY | P53396 | ACLY_HUMAN | Unmodified | 0 | 0 | 0 | 0 | 564630 | 0 | 517330 | 0 | 0 | 0 | 0 | 0 | 0 | 0 | 0 | 0 | 0 | 0 |
| VKWGETMI | TC2N; TC2N | Q8N9U0; HC | TAC2N_HUMAN | Oxidation (M) | 0 | 0 | 0 | 151200 | 0 | 0 | 0 | 0 | 0 | 0 | 0 | 0 | 0 | 0 | 314820 | 262440 | 0 | 0 |
| VKYEETVFY | NAMPT; NAMPT | P43490; A0A | NAMPT_HUMAN | Unmodified | 0 | 37394 | 0 | 374100 | 18429000 | 1315100 | 1566500 | 0 | 0 | 208840 | 0 | 86046 | 1957400 | 991180 | 390030 | 0 | 647210 | 3085200 |
| VKYETELAN | KRT18; KRT18 | P05783; F8V | K1C18_HUMAN | Unmodified | 0 | 0 | 0 | 142970 | 0 | 0 | 0 | 0 | 3309600 | 0 | 2437700 | 24195000 | 0 | 0 | 0 | 349180 | 0 | 0 |
| VKYETELAN | KRT18; KRT18 | P05783; F8V | K1C18_HUMAN | Oxidation (M) | 0 | 0 | 0 | 0 | 0 | 0 | 0 | 16497 | 1442700 | 140140 | 781320 | 4075100 | 0 | 0 | 0 | 0 | 0 | 0 |
| VKYEVTED | APPL1 | Q9UKG1 | DP13A_HUMAN | Unmodified | 390370 | 364750 | 0 | 0 | 0 | 0 | 0 | 0 | 852210 | 0 | 0 | 825660 | 0 | 0 | 0 | 496570 | 0 | 0 |
| VKYEVTVSN | ITGA7; ITGA7 | Q13683; J3K | ITA7_HUMAN In | Unmodified | 0 | 0 | 0 | 0 | 0 | 0 | 660920 | 0 | 2002700 | 252610 | 0 | 0 | 0 | 0 | 0 | 0 | 0 | 0 |
| VLAALQER | NAP1L4; NAP1L | Q99733; C9J | NP1L4_HUMAN | Unmodified | 187370 | 0 | 0 | 0 | 0 | 0 | 0 | 0 | 920150 | 39344 | 0 | 7377200 | 0 | 0 | 92152 | 69744 | 121780 | 0 |
| VLADSLHV | PTGFRN | Q9P2B2 | FPRP_HUMAN | Unmodified | 21288000 | 9585300 | 72144000 | 48670000 | 431090 | 0 | 0 | 0 | 282220 | 0 | 0 | 51292000 | 7856600 | 1148300 | 1.01E+08 | 69784000 | 0 | 295770 |
| VLAEGQRIL | NELL1; NELL1; | Q92832; J3K | NELL1_HUMAN | Unmodified | 0 | 0 | 0 | 0 | 0 | 0 | 0 | 0 | 0 | 345910 | 0 | 0 | 0 | 0 | 0 | 0 | 0 | 0 |

| | | | | | | | | | | | | | | | | | | | | | | |
|---|---|---|---|---|---|---|---|---|---|---|---|---|---|---|---|---|---|---|---|---|---|---|
| VLAELPQCI | RAB34; RAB34; | Q9BZG1; C9 | RAB34_HUMAN | Unmodified | 0 | 0 | 0 | 0 | 0 | 0 | 0 | 0 | 858970 | 0 | 0 | 0 | 0 | 0 | 0 | 0 | 0 | 0 |
| VLAELYVSD | NARS1; NARS1; | O43776; K7E | SYNC_HUMAN | Unmodified | 667250 | 452990 | 525480 | 1098000 | 3261200 | 0 | 489220 | 285630 | 3294400 | 1034000 | 3586500 | 16871000 | 3042200 | 2444600 | 1646100 | 1737000 | 764480 | 2028000 |
| VLAESLGPI | TPRN | Q4KMQ1 | TPRN_HUMAN | Oxidation (M) | 0 | 0 | 0 | 117990 | 0 | 0 | 0 | 0 | 0 | 0 | 0 | 0 | 0 | 0 | 229760 | 0 | 0 | 0 |
| VLAFLSSVA | GCN1 | Q92616 | GCN1_HUMAN | Unmodified | 0 | 0 | 0 | 0 | 0 | 0 | 0 | 0 | 0 | 0 | 0 | 576090 | 0 | 0 | 0 | 0 | 0 | 0 |
| VLAFTAVGI | PTPRF; PTPRF; | P10586; H0Y | PTPRF_HUMAN | Unmodified | 1359700 | 1171100 | 189170 | 0 | 0 | 0 | 0 | 292360 | 2779700 | 558870 | 0 | 0 | 0 | 0 | 0 | 803460 | 0 | 0 |
| VLAGDKNF | ACTN4; ACTN4; | O43707; F5C | ACTN4_HUMAI | Unmodified | 0 | 0 | 0 | 0 | 3645900 | 0 | 0 | 131530 | 0 | 611740 | 0 | 0 | 1082800 | 939090 | 0 | 0 | 1161500 | 0 |
| VLAGETLS\ | ILF3; ILF3; ILF3 | Q12906; K7E | ILF3_HUMAN Ir | Unmodified | 759170 | 535480 | 0 | 255560 | 0 | 0 | 0 | 0 | 0 | 0 | 452490 | 1308900 | 0 | 0 | 188580 | 278450 | 0 | 0 |
| VLAGETLS\ | ILF3; ILF3; ILF3 | Q12906; K7E | ILF3_HUMAN Ir | Unmodified | 11755000 | 6171500 | 5348600 | 5649500 | 1349800 | 0 | 1759800 | 1044400 | 0 | 1889700 | 0 | 9166200 | 8103600 | 1533300 | 3220800 | 3450600 | 786100 | 0 |
| VLAHIRDDI | ADAM17; ADAM | P78536; A0A | ADA17_HUMAN | Unmodified | 0 | 0 | 0 | 0 | 0 | 225330 | 0 | 0 | 1048600 | 0 | 0 | 0 | 0 | 0 | 310960 | 0 | 0 | 0 |
| VLAIMREH( | ZC3HAV1; ZC3I | Q7Z2W4; C9 | ZCCHV_HUMA | Oxidation (M) | 627920 | 0 | 236420 | 0 | 8747900 | 2467100 | 0 | 0 | 0 | 241670 | 0 | 0 | 3064200 | 0 | 0 | 0 | 990870 | 1198800 |
| VLAIMREH( | ZC3HAV1; ZC3I | Q7Z2W4; C9 | ZCCHV_HUMA | Unmodified | 0 | 0 | 0 | 0 | 7719400 | 0 | 0 | 0 | 0 | 0 | 0 | 0 | 0 | 1544000 | 0 | 0 | 468820 | 1366100 |
| VLAITSSSIF | ST3GAL4; ST3G | Q11206; E9F | SIA4C_HUMAN | Unmodified | 0 | 0 | 0 | 0 | 0 | 0 | 0 | 0 | 912550 | 0 | 183820 | 675120 | 0 | 0 | 0 | 0 | 0 | 0 |
| VLAKPQNT. | PVR; PVR; PVR | P15151; A0A | PVR_HUMAN P | Unmodified | 5892600 | 4226400 | 2569800 | 1471800 | 305910 | 523420 | 0 | 189540 | 599720 | 85477 | 3840900 | 36913000 | 0 | 0 | 4873500 | 5073000 | 0 | 79465 |
| VLALIQAW/ | TOM1L2 | Q6ZVM7 | TM1L2_HUMAN | Unmodified | 17466 | 151320 | 0 | 0 | 0 | 0 | 0 | 0 | 0 | 0 | 0 | 0 | 0 | 0 | 64082 | 0 | 0 | 0 |
| VLALLDVPI | BPNT2 | Q9NX62 | IMPA3_HUMAN | Unmodified | 0 | 0 | 506000 | 0 | 137050 | 0 | 0 | 0 | 74941000 | 5134600 | 2284500 | 4065400 | 0 | 0 | 0 | 185570 | 0 | 0 |
| VLALLDVPI | BPNT2 | Q9NX62 | IMPA3_HUMAN | Unmodified | 0 | 0 | 0 | 0 | 0 | 0 | 397640 | 402660 | 40992000 | 16645000 | 942410 | 1575500 | 0 | 477100 | 0 | 0 | 0 | 0 |
| VLALLDVPI | BPNT2 | Q9NX62 | IMPA3_HUMAN | Unmodified | 0 | 0 | 0 | 0 | 0 | 0 | 0 | 0 | 0 | 3514300 | 0 | 0 | 0 | 0 | 0 | 0 | 51098 | 0 |
| VLALPEPSF | PLEC; PLEC | Q15149; H0Y | PLEC_HUMAN | Unmodified | 0 | 0 | 0 | 0 | 2935100 | 270470 | 0 | 0 | 0 | 395980 | 0 | 0 | 0 | 0 | 0 | 0 | 6458600 | 4421900 |
| VLALSVETL | RPLP0; RPLP0; | P05388; F8V | RLA0_HUMAN ( | Unmodified | 0 | 537530 | 0 | 155730 | 0 | 0 | 0 | 0 | 0 | 0 | 0 | 0 | 0 | 0 | 521620 | 0 | 0 | 0 |
| VLAMSGDF | PPP2R1A; PPP2 | P30153; B3K | 2AAA_HUMAN | Oxidation (M) | 691610 | 166900 | 635180 | 580490 | 0 | 53347 | 0 | 0 | 0 | 0 | 0 | 3153600 | 457110 | 0 | 856160 | 1176400 | 148970 | 390770 |
| VLAMSGDF | PPP2R1A; PPP2 | P30153; B3K | 2AAA_HUMAN | Unmodified | 202020 | 160540 | 471170 | 521260 | 952880 | 150360 | 182300 | 57435 | 838540 | 291400 | 0 | 7537100 | 647660 | 342090 | 669900 | 523180 | 384110 | 825110 |
| VLANNSLS | SRP72; SRP72; | O76094; D6I | SRP72_HUMAN | Unmodified | 150150 | 0 | 0 | 184720 | 1142800 | 0 | 1418200 | 930390 | 195140 | 338300 | 0 | 3740200 | 1125400 | 1071500 | 104090 | 128940 | 0 | 1195200 |
| VLANPGNS | KPNB1; KPNB1 | Q14974; J3Q | IMB1_HUMAN I | Unmodified | 828550 | 863110 | 3245200 | 1923500 | 3208500 | 0 | 124890 | 24320 | 4865200 | 1451100 | 4155700 | 17097000 | 2427700 | 331360 | 1590000 | 1553700 | 1143500 | 888000 |
| VLANPGNS | KPNB1; KPNB1 | Q14974; J3Q | IMB1_HUMAN I | Unmodified | 0 | 0 | 0 | 0 | 1867700 | 0 | 416810 | 60666 | 0 | 595180 | 0 | 0 | 338580 | 862510 | 0 | 0 | 978770 | 0 |
| VLANQYK | ADSS2 | P30520 | PURA2_HUMAI | Unmodified | 11117 | 109340 | 82670 | 83162 | 269440 | 0 | 493350 | 152500 | 0 | 50884 | 53059 | 9204300 | 552270 | 67644 | 99268 | 0 | 507920 | 303470 |
| VLAPASTLC | DSG2 | Q14126 | DSG2_HUMAN | Oxidation (M) | 73807 | 74737 | 3792200 | 3429500 | 1200800 | 0 | 3904000 | 0 | 1613500 | 738900 | 1579000 | 5371900 | 0 | 0 | 2915800 | 3929900 | 569650 | 0 |
| VLAPASTLC | DSG2 | Q14126 | DSG2_HUMAN | Unmodified | 0 | 0 | 0 | 0 | 2811700 | 350010 | 0 | 0 | 3090500 | 0 | 0 | 0 | 0 | 0 | 0 | 0 | 0 | 0 |
| VLAPELIPSI | KNTC1; KNTC1 | P50748; E7E | KNTC1_HUMAI | Unmodified | 0 | 0 | 0 | 0 | 0 | 0 | 0 | 0 | 0 | 0 | 0 | 930480 | 0 | 0 | 0 | 0 | 0 | 0 |
| VLAPGASH | RAB11FIP5; RAI | Q9BXF6; A0/ | RFIP5_HUMAN | Unmodified | 0 | 0 | 154460 | 394710 | 1209800 | 2029100 | 0 | 0 | 0 | 0 | 0 | 0 | 2133700 | 0 | 0 | 0 | 0 | 2131800 |
| VLAPILPDN | INCENP | Q9NQS7 | INCE_HUMAN I | Unmodified | 0 | 0 | 157260 | 358530 | 0 | 0 | 0 | 68343 | 187720 | 87482 | 0 | 182820 | 434890 | 416520 | 167950 | 60251 | 0 | 0 |
| VLAPILPDN | INCENP | Q9NQS7 | INCE_HUMAN I | Unmodified | 0 | 0 | 483450 | 1861000 | 0 | 0 | 0 | 0 | 0 | 0 | 0 | 0 | 0 | 0 | 0 | 0 | 0 | 0 |
| VLAPILPDN | INCENP | Q9NQS7 | INCE_HUMAN I | Unmodified | 0 | 0 | 0 | 0 | 2473200 | 0 | 0 | 0 | 0 | 0 | 0 | 0 | 2086700 | 0 | 0 | 0 | 0 | 0 |
| VLAQCNID | KIDINS220; KID | Q9ULH0; E9 | KDIS_HUMAN I | Unmodified | 0 | 107150 | 0 | 0 | 2084600 | 0 | 0 | 0 | 0 | 0 | 0 | 0 | 1068700 | 0 | 0 | 0 | 0 | 0 |
| VLAQDVYA | GPHN; GPHN | Q9NQX3; F5 | GEPH_HUMAN | Unmodified | 0 | 0 | 0 | 0 | 0 | 0 | 0 | 0 | 337370 | 0 | 0 | 1911100 | 0 | 0 | 0 | 0 | 0 | 0 |
| VLAQLLR | PCSK1N; CD93 | Q9UHG2; Q9 | PCS1N_HUMAI | Unmodified | 0 | 78872 | 344010 | 0 | 0 | 0 | 0 | 0 | 5344900 | 331340 | 0 | 0 | 0 | 0 | 0 | 774680 | 0 | 0 |
| VLAQNSGF | CCT6A | P40227 | TCPZ_HUMAN | Unmodified | 0 | 0 | 908250 | 652020 | 5453000 | 242050 | 2210500 | 1619600 | 4424500 | 1685000 | 4161200 | 22951000 | 11281000 | 5167100 | 700630 | 1158600 | 2759300 | 5695300 |
| VLARLDRD | PTGFRN | Q9P2B2 | FPRP_HUMAN | Unmodified | 2393300 | 3379700 | 9835300 | 49984000 | 0 | 4542200 | 0 | 0 | 0 | 0 | 0 | 0 | 51814000 | 14084000 | 14781000 | 5969700 | 323880 | 198700 |
| VLATAFDTT | HSPA4; HSPA4; | P34932; A0A | HSP74_HUMAI | Unmodified | 0 | 0 | 299030 | 1740400 | 250280 | 0 | 0 | 61426 | 0 | 174300 | 1869900 | 27936000 | 256200 | 0 | 1002900 | 839030 | 0 | 0 |
| VLATTFDPY | HSPA4L; HSPA4 | O95757; E9F | HS74L_HUMAI | Unmodified | 1137000 | 0 | 0 | 0 | 0 | 0 | 0 | 0 | 511370 | 0 | 0 | 4356700 | 0 | 0 | 0 | 248540 | 0 | 0 |
| VLATVTKPV | RPL6; RPL6; RP | Q02878; F8V | RL6_HUMAN 6( | Unmodified | 588390 | 362910 | 376950 | 169360 | 0 | 0 | 0 | 0 | 0 | 0 | 495780 | 5071600 | 140530 | 0 | 261480 | 298220 | 0 | 0 |
| VLATVTKPV | RPL6; RPL6; RP | Q02878; F8V | RL6_HUMAN 6( | Unmodified | 0 | 0 | 0 | 0 | 0 | 0 | 14665 | 257560 | 0 | 0 | 0 | 0 | 927120 | 0 | 0 | 0 | 0 | 0 |
| VLAVNQEN | ACTN4; ACTN4; | O43707; F5C | ACTN4_HUMAI | Oxidation (M) | 252990 | 0 | 1389400 | 785370 | 6051200 | 1870400 | 0 | 192260 | 1345800 | 938410 | 1032400 | 15175000 | 651890 | 1455300 | 1160600 | 1480600 | 1493900 | 0 |
| VLAVNQEN | ACTN4; ACTN4; | O43707; F5C | ACTN4_HUMAI | Unmodified | 697530 | 439420 | 746180 | 1018900 | 14800000 | 7374800 | 0 | 0 | 6942800 | 2034500 | 1751800 | 17055000 | 2932300 | 0 | 1691500 | 1059600 | 3224700 | 0 |
| VLAVNQEN | ACTN1; ACTN1; | P12814; G3\ | ACTN1_HUMAI | Oxidation (M) | 161890 | 0 | 0 | 0 | 1168300 | 2306300 | 319550 | 331860 | 1297200 | 0 | 0 | 4927400 | 0 | 0 | 834020 | 586880 | 926810 | 0 |
| VLAVNQEN | ACTN1; ACTN1; | P12814; G3\ | ACTN1_HUMAI | Unmodified | 0 | 0 | 0 | 0 | 2520800 | 1556000 | 0 | 0 | 0 | 0 | 0 | 2926600 | 0 | 0 | 294080 | 0 | 1005400 | 214690 |
| VLAVNSIGI | PTPRF; PTPRF; | P10586; H0Y | PTPRF_HUMAN | Unmodified | 149890 | 115030 | 0 | 64697 | 0 | 0 | 368970 | 0 | 0 | 0 | 0 | 0 | 275400 | 267900 | 0 | 0 | 932330 | 646840 |
| VLAVNSIGI | PTPRF; PTPRF; | P10586; H0Y | PTPRF_HUMAN | Unmodified | 855030 | 1495300 | 0 | 0 | 0 | 0 | 0 | 0 | 770630 | 0 | 0 | 0 | 106790 | 0 | 0 | 0 | 0 | 0 |
| VLAVTAIR | GART | P22102 | PUR2_HUMAN | Unmodified | 0 | 0 | 0 | 0 | 0 | 0 | 0 | 0 | 0 | 0 | 0 | 3018900 | 0 | 0 | 0 | 151970 | 0 | 0 |
| VLAVTAIRE | GART | P22102 | PUR2_HUMAN | Unmodified | 0 | 0 | 450180 | 1434200 | 1312800 | 367720 | 469830 | 0 | 2957500 | 730280 | 3387500 | 9786700 | 3004000 | 461670 | 1523600 | 754510 | 0 | 971430 |
| VLCALTEAN | PTPN23 | Q9H3S7 | PTN23_HUMAN | Unmodified | 116870 | 0 | 0 | 0 | 385620 | 0 | 0 | 44363 | 117360 | 0 | 0 | 0 | 0 | 0 | 0 | 0 | 0 | 0 |
| VLCELADL( | TCP1; TCP1; TC | P17987; F5H | TCPA_HUMAN | Unmodified | 0 | 0 | 0 | 0 | 0 | 0 | 0 | 0 | 0 | 0 | 0 | 8685000 | 0 | 0 | 0 | 0 | 0 | 0 |
| VLCGGDIY | ALDH9A1 | P49189 | AL9A1_HUMAN | Unmodified | 0 | 0 | 0 | 0 | 2161000 | 0 | 174600 | 192530 | 0 | 0 | 0 | 0 | 2052400 | 1395900 | 0 | 0 | 1396000 | 1124800 |
| VLCLAGFR | OVCA2 | Q8WZ82 | OVCA2_HUMAI | Unmodified | 47581 | 53951 | 0 | 0 | 0 | 0 | 0 | 0 | 0 | 0 | 0 | 381630 | 0 | 0 | 53472 | 81704 | 0 | 0 |
| VLCPSNSS | AURKA | O14965 | AURKA_HUMAI | Unmodified | 0 | 253380 | 0 | 476770 | 142220 | 46395 | 0 | 0 | 127130 | 0 | 0 | 0 | 214260 | 104450 | 241350 | 224130 | 57630 | 0 |
| VLCQFCD( | MID1; MID1; MI | O15344; A0/ | TRI18_HUMAN | Unmodified | 0 | 0 | 157270 | 0 | 0 | 0 | 0 | 0 | 0 | 0 | 0 | 0 | 0 | 0 | 0 | 0 | 1544800 | 489530 |
| VLCSSPAPI | ZDHHC5; ZDHH | Q9C0B5; H0 | ZDHC5_HUMA | Unmodified | 447810 | 50979 | 258670 | 277910 | 403800 | 0 | 57331 | 0 | 668120 | 109000 | 226700 | 10644 | 732790 | 89771 | 281450 | 0 | 344490 | 160320 |
| VLCTPPPK | CD46 | P15529 | MCP_HUMAN I | Unmodified | 5593100 | 2074900 | 1984500 | 3800400 | 1452900 | 0 | 152060 | 0 | 1623600 | 640450 | 319260 | 2244900 | 906820 | 846340 | 1662100 | 1603700 | 298160 | 721750 |
| VLDAIIQEK | SMC4; SMC4 | Q9NTJ3; E9F | SMC4_HUMAN | Unmodified | 693790 | 50681 | 0 | 0 | 551810 | 0 | 975170 | 742180 | 0 | 0 | 0 | 344380 | 0 | 500340 | 0 | 992450 | 229890 | 0 |
| VLDANSC( | MARK2; MARK2 | Q7KZI7; E7E | MARK2_HUMAI | Unmodified | 0 | 335870 | 389350 | 0 | 0 | 197510 | 223370 | 137280 | 333110 | 386480 | 487510 | 761530 | 0 | 0 | 700810 | 423540 | 550110 | 294230 |
| VLDASWYS | TST; TST | Q16762; B1/ | THTR_HUMAN | Unmodified | 0 | 0 | 0 | 0 | 0 | 0 | 0 | 0 | 1218000 | 0 | 0 | 0 | 0 | 0 | 0 | 0 | 0 | 0 |
| VLDDGELL' | NSF; NSF | P46459; I3L( | NSF_HUMAN V | Unmodified | 0 | 0 | 0 | 0 | 0 | 0 | 0 | 56973 | 347920 | 0 | 0 | 819480 | 253190 | 350930 | 0 | 0 | 0 | 0 |
| VLDEFSK | YKT6; YKT6; YK | O15498; A0/ | YKT6_HUMAN ! | Unmodified | 1524700 | 1689400 | 520770 | 213070 | 222870 | 0 | 0 | 0 | 1538400 | 1014400 | 0 | 4671800 | 340490 | 384040 | 2529900 | 1277900 | 619590 | 0 |
| VLDFEHFLI | MYL6; MYL6; M' | P60660; G3\ | MYL6_HUMAN | Oxidation (M) | 5705000 | 6342300 | 2641900 | 11994000 | 1.18E+08 | 70131000 | 45154000 | 373080 | 18224000 | 32860000 | 9805200 | 27154000 | 1.16E+08 | 72811000 | 12049000 | 5265900 | 1.2E+08 | 80906000 |
| VLDFEHFLI | MYL6; MYL6; M' | P60660; G3\ | MYL6_HUMAN | Unmodified | 192750 | 309160 | 79975 | 551320 | 69444000 | 7713400 | 1875900 | 0 | 12846000 | 3314300 | 519850 | 540040 | 13854000 | 4962400 | 1203700 | 756480 | 11876000 | 12128000 |
| VLDFHNLP | COL5A1 | P20908 | CO5A1_HUMAI | Unmodified | 17242 | 94050 | 568000 | 397330 | 712550 | 1417900 | 573690 | 281310 | 0 | 142990 | 0 | 78121 | 0 | 1039600 | 207070 | 34876 | 244760 | 1278400 |
| VLDGGKS( | ITGB4 | P16144 | ITB4_HUMAN I | Unmodified | 0 | 0 | 0 | 0 | 1874700 | 0 | 258790 | 0 | 0 | 0 | 0 | 0 | 0 | 0 | 0 | 0 | 758280 | 170350 |
| VLDGLHNE | UGDH | O60701 | UGDH_HUMAN | Unmodified | 0 | 0 | 0 | 0 | 3408500 | 246590 | 0 | 0 | 0 | 0 | 0 | 0 | 1766400 | 121240 | 0 | 0 | 0 | 0 |
| VLDGLHNE | UGDH | O60701 | UGDH_HUMAN | Unmodified | 0 | 0 | 0 | 0 | 3102300 | 0 | 0 | 0 | 0 | 50795 | 0 | 0 | 0 | 28253 | 0 | 0 | 69188 | 0 |
| VLDGPEKV | NUP210 | Q8TEM1 | PO210_HUMAI | Unmodified | 0 | 0 | 0 | 0 | 0 | 0 | 0 | 0 | 39700000 | 1419300 | 711100 | 0 | 0 | 0 | 0 | 0 | 65631 | 0 |
| VLDHDVSK | NF2; NF2 | P35240; A0A | MERL_HUMAN | Unmodified | 0 | 264530 | 0 | 167100 | 735310 | 260050 | 0 | 123050 | 0 | 0 | 0 | 0 | 260590 | 0 | 0 | 451430 | 0 | 983500 |
| VLDIIATINK | HEXB; HEXB; HI | P07686; H0Y | HEXB_HUMAN | Unmodified | 551600 | 2682600 | 330220 | 504280 | 0 | 0 | 0 | 0 | 2575000 | 852100 | 479570 | 119170 | 1057600 | 1998800 | 663190 | 0 | 0 | 0 |
| VLDIIATINK | HEXB; HEXB; HI | P07686; H0Y | HEXB_HUMAN | Unmodified | 0 | 0 | 0 | 0 | 0 | 0 | 0 | 0 | 0 | 0 | 0 | 0 | 0 | 155930 | 0 | 0 | 0 | 0 |
| VLDIIYQPQ | NLE1; NLE1; NL | Q9NVX2; A0, | NLE1_HUMAN | Unmodified | 0 | 0 | 22202 | 0 | 0 | 0 | 59467 | 0 | 0 | 0 | 0 | 373910 | 0 | 0 | 0 | 0 | 0 | 0 |
| VLDKYEDV | PPL; PPL | O60437; K7E | PEPL_HUMAN I | Unmodified | 0 | 0 | 0 | 0 | 1419100 | 2125600 | 0 | 0 | 0 | 0 | 0 | 0 | 0 | 0 | 0 | 0 | 0 | 0 |
| VLDLDLFR\ | SARS1; SARS1 | P49591; Q5I | SYSC_HUMAN | Unmodified | 484900 | 24778 | 986060 | 748860 | 2328000 | 117720 | 0 | 41070 | 1363500 | 355660 | 234930 | 4020200 | 1296500 | 825240 | 1062500 | 665770 | 0 | 0 |
| VLDLELK | ANXA1 | P04083 | ANXA1_HUMAI | Unmodified | 0 | 0 | 764950 | 575130 | 1766900 | 0 | 490260 | 183690 | 0 | 0 | 49292000 | 1.92E+08 | 0 | 0 | 1037900 | 235140 | 258060 | 0 |
| VLDLELKGI | ANXA1 | P04083 | ANXA1_HUMAI | Unmodified | 467290 | 890770 | 235180 | 1226600 | 26908000 | 2212500 | 6764300 | 1127600 | 0 | 546710 | 54566000 | 2.57E+08 | 2847400 | 5365500 | 418960 | 557650 | 1748600 | 3264500 |
| VLDLELKGI | ANXA1 | P04083 | ANXA1_HUMAI | Unmodified | 0 | 0 | 0 | 0 | 752350 | 0 | 0 | 0 | 0 | 0 | 0 | 105300 | 0 | 0 | 0 | 0 | 353360 | 138540 |
| VLDLIVNGI | ATP2B4 | P23634 | AT2B4_HUMAN | Unmodified | 0 | 0 | 78037 | 332010 | 1119000 | 0 | 0 | 0 | 0 | 74566 | 398750 | 1866100 | 0 | 0 | 484280 | 295300 | 0 | 0 |
| VLDLQQIK | PSMD13; PSMD | Q9UNM6; J3 | PSD13_HUMAN | Unmodified | 163890 | 86944 | 1131300 | 810550 | 1244900 | 561020 | 647640 | 445370 | 664110 | 229060 | 522740 | 9149500 | 2672000 | 1645800 | 472660 | 425060 | 587510 | 2398800 |
| VLDMENSF | MACF1; MACF1 | Q9UPN3; H3 | MACF1_HUMAI | Oxidation (M) | 0 | 0 | 0 | 0 | 0 | 0 | 809380 | 282990 | 0 | 0 | 0 | 0 | 301170 | 461270 | 0 | 0 | 0 | 0 |
| VLDNQIKK | CIT; CIT; CIT; CI | O14578; A0/ | CTRO_HUMAN | Unmodified | 0 | 0 | 0 | 0 | 1788500 | 0 | 0 | 0 | 0 | 0 | 0 | 0 | 0 | 0 | 0 | 0 | 0 | 0 |
| VLDNYLTSI | CLIC1 | O00299 | CLIC1_HUMAN | Unmodified | 467190 | 0 | 0 | 734920 | 2028400 | 0 | 0 | 0 | 2485100 | 1179000 | 0 | 1445500 | 0 | 0 | 863330 | 0 | 1849600 | 0 |
| VLDNYLTSI | CLIC1 | O00299 | CLIC1_HUMAN | Unmodified | 6638700 | 6980100 | 0 | 6937900 | 1.6E+08 | 30399000 | 11178000 | 6444000 | 36763000 | 10514000 | 1074800 | 28365000 | 77454000 | 76676000 | 7345300 | 11246000 | 34968000 | 58585000 |
| VLDNYPMF | MMP15 | P51511 | MMP15_HUMA | Oxidation (M) | 423000 | 955920 | 709930 | 0 | 0 | 0 | 0 | 0 | 80661 | 0 | 0 | 0 | 0 | 0 | 1796300 | 1621100 | 0 | 0 |
| VLDPFTIK | TKT; TKT | P29401; A0A | TKT_HUMAN Tr | Unmodified | 0 | 0 | 176920 | 0 | 322530 | 0 | 336590 | 139420 | 0 | 0 | 576420 | 2547700 | 0 | 0 | 0 | 81306 | 0 | 0 |
| VLDPFTIKP | TKT; TKT | P29401; A0A | TKT_HUMAN Tr | Unmodified | 0 | 0 | 0 | 0 | 4643100 | 968380 | 118130 | 0 | 2370100 | 2071600 | 5775600 | 5032000 | 2576000 | 230630 | 0 | 0 | 1033300 | 1136600 |
| VLDPFTIKP | TKT; TKT | P29401; A0A | TKT_HUMAN Tr | Unmodified | 0 | 0 | 0 | 390620 | 7089700 | 1531700 | 0 | 0 | 5365800 | 1541800 | 4189300 | 16722000 | 753330 | 0 | 153480 | 141340 | 444560 | 913360 |
| VLDQDAVS | PSD3; PSD3 | Q9NYI0; E5F | PSD3_HUMAN | Unmodified | 0 | 0 | 0 | 0 | 1099100 | 0 | 662990 | 193700 | 0 | 0 | 0 | 0 | 0 | 0 | 0 | 0 | 337080 | 0 |
| VLDQFQVN | SORCS2; SORC | Q96PQ0; B5 | SORC2_HUMAI | Unmodified | 0 | 0 | 0 | 0 | 0 | 0 | 0 | 0 | 2285800 | 0 | 0 | 0 | 0 | 109650 | 0 | 0 | 491810 | 0 |
| VLDQFQVN | SORCS2; SORC | Q96PQ0; B5 | SORC2_HUMAI | Oxidation (M) | 0 | 0 | 0 | 0 | 0 | 0 | 0 | 118000 | 3720300 | 277330 | 0 | 0 | 153600 | 0 | 0 | 0 | 0 | 0 |
| VLDQYERE | ATG7; ATG7; AT | O95352; H7I | ATG7_HUMAN | Unmodified | 0 | 0 | 24300 | 89602 | 570260 | 70993 | 0 | 39341 | 0 | 187840 | 130500 | 607380 | 489420 | 238190 | 94899 | 59408 | 913650 | 1516900 |
| VLDSDGSD | BAIAP2; BAIAP2 | Q9UQB8; I3I | BAIP2_HUMAN | Unmodified | 158390 | 0 | 653840 | 461060 | 851390 | 0 | 0 | 0 | 1532100 | 99215 | 0 | 2913400 | 0 | 57336 | 3905300 | 450160 | 82229 | 0 |
| VLDSDGSD | BAIAP2; BAIAP2 | Q9UQB8; I3I | BAIP2_HUMAN | Oxidation (M) | 0 | 0 | 0 | 0 | 0 | 0 | 0 | 0 | 0 | 0 | 0 | 0 | 0 | 0 | 0 | 376810 | 0 | 0 |
| VLDSEMVA | SEMA3C | Q99985 | SEM3C_HUMAI | Unmodified | 0 | 0 | 236140 | 0 | 1358900 | 0 | 0 | 115960 | 0 | 0 | 0 | 0 | 0 | 0 | 0 | 0 | 0 | 389830 |
| VLDSFGTEI | MGAT5 | Q09328 | MGT5A_HUMAI | Unmodified | 0 | 0 | 105470 | 0 | 0 | 0 | 0 | 0 | 4095200 | 715840 | 270200 | 1949200 | 0 | 0 | 0 | 0 | 0 | 0 |
| VLDSGAPIk | ATP5F1B; ATP5I | P06576; H0Y | ATPB_HUMAN . | Unmodified | 0 | 0 | 0 | 0 | 0 | 0 | 64655 | 0 | 8915600 | 620280 | 0 | 423170 | 0 | 0 | 0 | 0 | 0 | 0 |
| VLDSGAPIk | ATP5F1B; ATP5I | P06576; H0Y | ATPB_HUMAN . | Unmodified | 0 | 0 | 0 | 0 | 432920 | 63258 | 572310 | 192410 | 43325000 | 7578000 | 1324800 | 795110 | 957770 | 561840 | 0 | 0 | 95447 | 345890 |
| VLDTDTITQ | PLXNB3; PLXNE | Q9ULL4; H7( | PLXB3_HUMAN | Unmodified | 59804 | 0 | 236300 | 0 | 0 | 0 | 0 | 0 | 0 | 0 | 0 | 0 | 0 | 0 | 500250 | 339220 | 0 | 0 |
| VLDTLRDAI | POMK; POMK; F | Q9H5K3; A0, | SG196_HUMAN | Unmodified | 0 | 0 | 0 | 0 | 0 | 0 | 0 | 0 | 2666900 | 0 | 0 | 0 | 0 | 0 | 0 | 0 | 0 | 0 |
| VLDTNDRF | MTHFD1L; MTH | Q6UB35; P1 | C1TM_HUMAN | Unmodified | 0 | 0 | 0 | 0 | 0 | 0 | 0 | 0 | 0 | 0 | 0 | 618250 | 0 | 0 | 0 | 0 | 0 | 0 |
| VLDTPYLLN | PROM1 | O43490 | PROM1_HUMA | Unmodified | 0 | 0 | 0 | 0 | 0 | 0 | 0 | 0 | 0 | 0 | 15714 | 212770 | 14900 | 0 | 0 | 0 | 0 | 0 |
| VLDVEPDN | SPAG1 | Q07617 | SPAG1_HUMAI | Unmodified | 0 | 0 | 0 | 0 | 390980 | 0 | 0 | 221360 | 0 | 245160 | 0 | 0 | 518200 | 287620 | 0 | 0 | 363320 | 170160 |
| VLEAAVAAI | EIF2B1; EIF2B1 | Q14232; F5H | EI2BA_HUMAN | Unmodified | 0 | 0 | 0 | 0 | 0 | 0 | 0 | 0 | 0 | 0 | 106110 | 472810 | 80878 | 0 | 0 | 0 | 0 | 0 |
| VLEACSIAC | SHMT1 | P34896 | GLYC_HUMAN | Unmodified | 0 | 0 | 0 | 0 | 0 | 0 | 0 | 0 | 0 | 0 | 0 | 1638200 | 0 | 0 | 0 | 0 | 0 | 0 |
| VLEAEVLAL | NF2; NF2 | P35240; A0A | MERL_HUMAN | Unmodified | 0 | 0 | 0 | 685110 | 3108600 | 807800 | 285740 | 800530 | 0 | 191560 | 0 | 0 | 1298800 | 1261300 | 0 | 0 | 1243200 | 6915800 |
| VLEALELYN | STAM2 | O75886 | STAM2_HUMAI | Unmodified | 803380 | 0 | 0 | 0 | 2717400 | 383210 | 651740 | 314220 | 1314600 | 838900 | 169820 | 0 | 0 | 848960 | 1051100 | 443650 | 622950 | 2632000 |
| VLEALELYN | STAM2 | O75886 | STAM2_HUMAI | Unmodified | 197010 | 0 | 0 | 0 | 502930 | 0 | 1030500 | 0 | 0 | 156310 | 0 | 0 | 518490 | 373070 | 0 | 0 | 570050 | 1155500 |
| VLEALLPLK | FASN; FASN | P49327; A0A | FAS_HUMAN F | Unmodified | 193490 | 104490 | 635400 | 356200 | 346210 | 0 | 480590 | 267170 | 87715 | 197400 | 1061400 | 57981000 | 0 | 0 | 2066000 | 2135500 | 111740 | 0 |

| | | | | | | | | | | | | | | | | | | | | | | |
|---|---|---|---|---|---|---|---|---|---|---|---|---|---|---|---|---|---|---|---|---|---|---|
| VLEALQVLF | WDR26; WDR2( | Q9H7D7; C9 | WDR26_HUMA | Unmodified | 0 | 0 | 0 | 0 | 0 | 0 | 0 | 0 | 0 | 0 | 0 | 1126000 | 0 | 0 | 0 | 0 | 0 | 0 |
| VLEANDGS | ATXN3; ATXN3; | P54252; A0A | ATX3_HUMAN / | Oxidation (M) | 0 | 716900 | 0 | 41087 | 0 | 0 | 82632 | 43874 | 31785 | 0 | 0 | 0 | 0 | 0 | 0 | 0 | 0 | 0 |
| VLEDDPEA | EPHA4; EPHA5; | P54764; P54 | EPHA4_HUMAI | Unmodified | 1520700 | 1966000 | 204580 | 212190 | 0 | 0 | 114660 | 0 | 0 | 0 | 0 | 0 | 0 | 0 | 0 | 226630 | 0 | 0 |
| VLEDDPEA | EPHA2 | P29317 | EPHA2_HUMAI | Unmodified | 0 | 0 | 0 | 0 | 0 | 0 | 0 | 0 | 0 | 0 | 0 | 610920 | 0 | 0 | 0 | 0 | 0 | 0 |
| VLEDDPEA | EPHA2 | P29317 | EPHA2_HUMAI | Unmodified | 0 | 0 | 0 | 0 | 0 | 0 | 0 | 0 | 0 | 0 | 0 | 3360700 | 0 | 0 | 0 | 0 | 0 | 0 |
| VLEDNSALI | DPP4; DPP4; DI | P27487; A0A | DPP4_HUMAN | Unmodified | 996490 | 503100 | 14869000 | 17356000 | 0 | 253230 | 0 | 52504 | 581220 | 0 | 338840 | 220110 | 0 | 0 | 11540000 | 7978800 | 330980 | 816990 |
| VLEDSDLK | HSPA5; HSPA5 | P11021; A0A | BIP_HUMAN Er | Unmodified | 260310 | 78750 | 4936400 | 562500 | 2154200 | 624490 | 4851300 | 0 | 2.69E+08 | 34154000 | 23714000 | 6861900 | 543190 | 0 | 4616300 | 466560 | 292410 | 1973800 |
| VLEDSDLKI | HSPA5; HSPA5 | P11021; A0A | BIP_HUMAN Er | Unmodified | 0 | 0 | 0 | 0 | 0 | 0 | 0 | 0 | 0 | 775510 | 0 | 198440 | 0 | 0 | 0 | 0 | 0 | 0 |
| VLEDSDLKI | HSPA5; HSPA5 | P11021; A0A | BIP_HUMAN Er | Unmodified | 0 | 82614 | 0 | 422280 | 0 | 0 | 1720700 | 0 | 54023000 | 18632000 | 884150 | 0 | 493580 | 0 | 0 | 0 | 0 | 0 |
| VLEEANQA | DDX17; DDX17; | Q92841; A0/ | DDX17_HUMAI | Unmodified | 0 | 0 | 0 | 0 | 1141600 | 198700 | 454490 | 462200 | 540790 | 556310 | 1204100 | 5067700 | 1363600 | 366290 | 0 | 76427 | 130770 | 233600 |
| VLEEGGFFI | PDLIM7 | Q9NR12 | PDLI7_HUMAN | Unmodified | 151220 | 228790 | 0 | 0 | 2316600 | 887380 | 0 | 0 | 0 | 163890 | 0 | 0 | 461020 | 0 | 0 | 332580 | 543520 | 0 |
| VLEEHGEW | LYN; LYN | P07948; E5R | LYN_HUMAN T | Unmodified | 648380 | 579670 | 592430 | 0 | 1395200 | 389130 | 0 | 0 | 1564900 | 529190 | 0 | 0 | 0 | 0 | 770590 | 1216200 | 0 | 0 |
| VLEELEGVI | PI4KA | P42356 | PI4KA_HUMAN | Unmodified | 0 | 0 | 442810 | 512740 | 0 | 0 | 0 | 0 | 0 | 0 | 0 | 806360 | 0 | 0 | 509080 | 761130 | 0 | 0 |
| VLEEMEAR | LIN7A; LIN7B | O14910; Q9I | LIN7A_HUMAN | Unmodified | 0 | 0 | 0 | 0 | 0 | 0 | 0 | 0 | 769580 | 0 | 0 | 0 | 0 | 0 | 0 | 369890 | 0 | 0 |
| VLEEMEARI | LIN7A; LIN7B | O14910; Q9I | LIN7A_HUMAN | Unmodified | 0 | 0 | 485750 | 296630 | 0 | 0 | 570010 | 392690 | 2041300 | 2039400 | 0 | 0 | 0 | 0 | 380810 | 0 | 0 | 0 |
| VLEEMESRI | LIN7C; LIN7C | Q9NUP9; G3 | LIN7C_HUMAN | Unmodified | 200020 | 410640 | 1486600 | 326360 | 2845200 | 1126300 | 328270 | 0 | 1565000 | 2348600 | 0 | 1782500 | 996280 | 1024000 | 1434900 | 899200 | 7613500 | 883100 |
| VLEEMESRI | LIN7C; LIN7C | Q9NUP9; G3 | LIN7C_HUMAN | Oxidation (M) | 0 | 0 | 183100 | 38086 | 225360 | 116210 | 26547 | 0 | 0 | 327150 | 24805 | 450550 | 253930 | 106130 | 304060 | 0 | 423300 | 0 |
| VLEEPKDFS | OSMR | Q99650 | OSMR_HUMAN | Unmodified | 0 | 0 | 0 | 0 | 1827300 | 372000 | 0 | 0 | 0 | 0 | 0 | 0 | 0 | 0 | 0 | 0 | 0 | 0 |
| VLEEQINEC | SNX4 | O95219 | SNX4_HUMAN | Unmodified | 0 | 0 | 52213 | 0 | 1249400 | 0 | 452040 | 75325 | 844020 | 0 | 0 | 0 | 188110 | 142060 | 0 | 51023 | 0 | 359440 |
| VLEEQLVK | FUT8; FUT8 | Q9BYC5; A0/ | FUT8_HUMAN / | Unmodified | 59742 | 87385 | 0 | 0 | 0 | 0 | 30693 | 17106 | 4148700 | 1337400 | 0 | 0 | 0 | 0 | 0 | 0 | 0 | 0 |
| VLEEYSVP( | WDR6; WDR6; \ | Q9NNW5; Q | WDR6_HUMAN | Unmodified | 150500 | 109360 | 41626 | 0 | 0 | 0 | 0 | 0 | 0 | 0 | 0 | 304890 | 0 | 0 | 29090 | 46313 | 0 | 0 |
| VLEFNCR | GART; GART | P22102; F8V | PUR2_HUMAN | Unmodified | 62844 | 0 | 0 | 75405 | 0 | 0 | 0 | 0 | 0 | 0 | 0 | 2703300 | 0 | 0 | 195350 | 0 | 0 | 0 |
| VLEFNCRF( | GART | P22102 | PUR2_HUMAN | Unmodified | 470140 | 0 | 0 | 1272700 | 975250 | 0 | 0 | 0 | 0 | 93549 | 0 | 0 | 1771500 | 0 | 0 | 0 | 817320 | 1341300 |
| VLEGHTDF | NUP37; NUP37 | Q8NFH4; F8 | NUP37_HUMAI | Unmodified | 57809 | 60561 | 0 | 0 | 0 | 0 | 0 | 0 | 0 | 0 | 27788 | 282100 | 0 | 0 | 36278 | 22385 | 0 | 0 |
| VLEGMEVV | PPIB; PPIB; PPII | P23284; A0A | PPIB_HUMAN F | Unmodified | 0 | 0 | 0 | 0 | 0 | 0 | 0 | 0 | 1.2E+08 | 8461200 | 10377000 | 15798000 | 0 | 0 | 0 | 0 | 0 | 0 |
| VLEGMEVV | PPIB; PPIB; PPII | P23284; A0A | PPIB_HUMAN F | Oxidation (M) | 0 | 63065 | 0 | 0 | 155570 | 39209 | 0 | 46576 | 36442000 | 2784000 | 3334200 | 2712400 | 0 | 0 | 1203000 | 1231600 | 0 | 58164 |
| VLEGMEVV | PPIB; PPIB; PPII | P23284; A0A | PPIB_HUMAN F | Unmodified | 0 | 0 | 0 | 0 | 0 | 0 | 0 | 0 | 2327300 | 0 | 269310 | 0 | 0 | 0 | 0 | 0 | 0 | 0 |
| VLEGMEVV | PPIB; PPIB; PPII | P23284; A0A | PPIB_HUMAN F | Unmodified | 0 | 0 | 0 | 0 | 0 | 0 | 276250 | 44778 | 3574400 | 5031100 | 1251800 | 0 | 0 | 0 | 0 | 0 | 0 | 0 |
| VLEGMVEE | DSG2 | Q14126 | DSG2_HUMAN | Unmodified | 0 | 0 | 0 | 0 | 0 | 0 | 0 | 0 | 0 | 0 | 0 | 0 | 0 | 0 | 0 | 0 | 0 | 0 |
| VLEGMVEE | DSG2 | Q14126 | DSG2_HUMAN | Unmodified | 0 | 0 | 0 | 0 | 3147100 | 0 | 0 | 0 | 618100 | 0 | 0 | 0 | 0 | 0 | 0 | 0 | 0 | 0 |
| VLEKVDVE( | CHP1; CHP1 | Q99653; H0Y | CHP1_HUMAN | Unmodified | 0 | 26903 | 0 | 0 | 0 | 0 | 0 | 0 | 0 | 0 | 0 | 0 | 826920 | 874520 | 0 | 0 | 0 | 0 |
| VLELDNVK | PPP2R1A; PPP2 | P30153; B3K | 2AAA_HUMAN | Unmodified | 0 | 153220 | 995560 | 1228500 | 880010 | 107960 | 0 | 141950 | 1310900 | 283220 | 1798000 | 10314000 | 859430 | 387520 | 1706400 | 1807200 | 264180 | 688140 |
| VLELDPAL/ | AIP | O00170 | AIP_HUMAN AH | Unmodified | 0 | 0 | 0 | 0 | 0 | 0 | 0 | 0 | 1511200 | 268130 | 862170 | 2503200 | 0 | 0 | 0 | 1760600 | 0 | 0 |
| VLELEGEK( | COPS2; COPS2 | P61201; H0Y | CSN2_HUMAN | Unmodified | 47934 | 0 | 0 | 0 | 227890 | 0 | 545680 | 0 | 0 | 162000 | 0 | 4130600 | 170300 | 209260 | 0 | 81191 | 891000 | 121800 |
| VLELEPNNI | RPAP3 | Q9H6T3 | RPAP3_HUMAI | Unmodified | 0 | 0 | 0 | 0 | 0 | 0 | 0 | 0 | 0 | 0 | 0 | 12359000 | 0 | 0 | 0 | 0 | 300750 | 0 |
| VLELQTELK | BICD2 | Q8TD16 | BICD2_HUMAI | Unmodified | 0 | 0 | 0 | 0 | 1003200 | 0 | 0 | 140780 | 0 | 0 | 0 | 103570 | 386950 | 0 | 0 | 0 | 328770 | 248630 |
| VLELVLR | CLUH; PCDHA( | O75153; Q9 | CLU_HUMAN C | Unmodified | 0 | 0 | 8683.3 | 0 | 383850 | 0 | 9074.2 | 0 | 0 | 0 | 50853 | 2285300 | 0 | 0 | 26724 | 27291 | 74533 | 0 |
| VLELVSITAI | SHMT2; SHMT2 | P34897; H0Y | GLYM_HUMAN | Unmodified | 0 | 0 | 0 | 0 | 0 | 0 | 0 | 0 | 36133000 | 0 | 949360 | 975770 | 0 | 207050 | 0 | 0 | 0 | 0 |
| VLELYVSDI | NARS1 | K7ENF0 | K7ENF0_HUMA | Unmodified | 0 | 0 | 342610 | 141740 | 172120 | 178660 | 0 | 34746 | 962970 | 393120 | 774810 | 3160700 | 644310 | 291000 | 276600 | 0 | 143510 | 153840 |
| VLENAEGA | HSPA9; HSPA9; | P38646; H0Y | GRP75_HUMAI | Unmodified | 0 | 0 | 130240 | 298010 | 227300 | 0 | 159520 | 0 | 24322000 | 2566300 | 728410 | 0 | 239090 | 187600 | 149830 | 22939 | 0 | 205120 |
| VLENAEGA | HSPA9; HSPA9; | P38646; H0Y | GRP75_HUMAI | Oxidation (M) | 0 | 0 | 0 | 0 | 2546100 | 0 | 0 | 127430 | 0 | 2481900 | 0 | 0 | 0 | 0 | 0 | 0 | 0 | 0 |
| VLENAEGA | HSPA9; HSPA9; | P38646; H0Y | GRP75_HUMAI | Unmodified | 0 | 0 | 0 | 0 | 1317000 | 0 | 2780600 | 0 | 0 | 834660 | 0 | 0 | 0 | 0 | 0 | 0 | 0 | 0 |
| VLENIELNK | EIF3L; EIF3L; EII | Q9Y262; B0( | EIF3L_HUMAN | Unmodified | 24048 | 1088100 | 445930 | 328150 | 0 | 0 | 0 | 0 | 0 | 0 | 0 | 1779200 | 0 | 0 | 571220 | 0 | 0 | 0 |
| VLENIELNK | EIF3L; EIF3L; EII | Q9Y262; B0( | EIF3L_HUMAN | Unmodified | 0 | 0 | 0 | 0 | 0 | 0 | 0 | 0 | 0 | 0 | 0 | 1161100 | 0 | 0 | 0 | 0 | 0 | 0 |
| VLEPMPST/ | CNTN1 | Q12860 | CNTN1_HUMA | Oxidation (M) | 1059100 | 718230 | 0 | 0 | 0 | 0 | 0 | 104020 | 0 | 0 | 0 | 0 | 0 | 0 | 0 | 0 | 0 | 0 |
| VLEPSGWF | SHCBP1 | Q8NEM2 | SHCBP_HUMA | Unmodified | 0 | 0 | 0 | 0 | 327410 | 0 | 0 | 0 | 0 | 0 | 0 | 0 | 0 | 0 | 0 | 0 | 139020 | 0 |
| VLEPYPSK | LARS1; LARS1; | Q9P2J5; A0A | SYLC_HUMAN | Unmodified | 0 | 0 | 180940 | 0 | 0 | 0 | 0 | 0 | 0 | 0 | 0 | 0 | 0 | 0 | 0 | 407160 | 0 | 0 |
| VLEPYPSKI | LARS1; LARS1; | Q9P2J5; A0A | SYLC_HUMAN | Unmodified | 0 | 0 | 0 | 0 | 0 | 0 | 888580 | 860090 | 0 | 95445 | 0 | 0 | 72010 | 91137 | 0 | 0 | 0 | 0 |
| VLEQLTGQ | RPL11; RPL11 | P62913; Q5\ | RL11_HUMAN ( | Unmodified | 6887500 | 5800300 | 2282500 | 3316400 | 6612300 | 1104400 | 11360000 | 8367300 | 862170 | 1284700 | 6625300 | 41195000 | 4006900 | 3381100 | 5150000 | 3646200 | 948950 | 4023100 |
| VLERLHNE | UBE2Z | Q9H832 | UBE2Z_HUMAI | Oxidation (M) | 0 | 0 | 0 | 0 | 0 | 0 | 0 | 0 | 0 | 0 | 0 | 0 | 0 | 0 | 0 | 0 | 881080 | 0 |
| VLERLHNE | UBE2Z | Q9H832 | UBE2Z_HUMAI | Unmodified | 0 | 0 | 0 | 0 | 0 | 0 | 155840 | 0 | 0 | 0 | 0 | 0 | 0 | 0 | 0 | 0 | 1567000 | 355680 |
| VLESPENK | EHBP1L1 | Q8N3D4 | EH1L1_HUMAI | Unmodified | 0 | 0 | 0 | 0 | 1374200 | 231850 | 0 | 0 | 17387 | 84911 | 0 | 0 | 0 | 0 | 0 | 0 | 269960 | 342850 |
| VLESVVADI | VPS13C | Q709C8 | VP13C_HUMAI | Unmodified | 0 | 0 | 0 | 0 | 0 | 0 | 0 | 0 | 0 | 0 | 0 | 319750 | 0 | 0 | 0 | 0 | 0 | 0 |
| VLETAEDIQ | SPTAN1; SPTAN | Q13813; A0/ | SPTN1_HUMAI | Unmodified | 0 | 0 | 0 | 0 | 239100 | 0 | 0 | 0 | 0 | 0 | 735230 | 0 | 0 | 0 | 0 | 0 | 0 | 0 |
| VLETEEVYS | GRK5 | P34947 | GRK5_HUMAN | Unmodified | 0 | 0 | 0 | 0 | 0 | 0 | 0 | 0 | 0 | 0 | 972370 | 2998900 | 0 | 0 | 0 | 0 | 0 | 0 |
| VLETFLAK | TRIM25; TRIM25 | Q14258; A0/ | TRI25_HUMAN | Unmodified | 241550 | 0 | 281190 | 210730 | 4164100 | 7040900 | 136970 | 0 | 0 | 127430 | 348030 | 6629300 | 230930 | 418970 | 225340 | 266090 | 659170 | 711480 |
| VLETLQEK | GCLC; GCLC; C | P48506; A0A | GSH1_HUMAN | Unmodified | 550920 | 498930 | 468200 | 492370 | 3271800 | 637440 | 0 | 41985 | 186820 | 0 | 552100 | 3739300 | 0 | 0 | 117650 | 143200 | 0 | 147170 |
| VLETLQEK( | GCLC; GCLC; C | P48506; A0A | GSH1_HUMAN | Unmodified | 0 | 0 | 0 | 0 | 560920 | 0 | 39349 | 0 | 0 | 0 | 135270 | 645430 | 0 | 0 | 0 | 0 | 19547 | 0 |
| VLETPVIDN | TMEM94 | Q12767 | TMM94_HUMAI | Unmodified | 0 | 0 | 0 | 0 | 1295700 | 0 | 361440 | 0 | 0 | 150560 | 0 | 1328600 | 0 | 562710 | 117870 | 137160 | 1301500 | 830400 |
| VLETTVEIFI | VPS35 | Q96QK1 | VPS35_HUMAI | Unmodified | 0 | 139670 | 0 | 0 | 0 | 0 | 0 | 0 | 0 | 0 | 0 | 2621800 | 0 | 0 | 0 | 100160 | 0 | 0 |
| VLEVAKDFI | PDIA4; PDIA4 | P13667; A0A | PDIA4_HUMAN | Unmodified | 0 | 0 | 0 | 0 | 0 | 0 | 345400 | 0 | 0 | 2281900 | 0 | 172780 | 0 | 0 | 0 | 0 | 0 | 0 |
| VLEVASGS( | METTL26; METT | Q96S19; H3I | MTL26_HUMAI | Unmodified | 0 | 0 | 1801500 | 1101700 | 0 | 0 | 0 | 0 | 198690 | 0 | 0 | 1599000 | 0 | 0 | 2314600 | 1394000 | 0 | 0 |
| VLEVNPQN | FKBP5 | Q13451 | FKBP5_HUMAI | Unmodified | 218890 | 0 | 139370 | 96691 | 1251700 | 243170 | 108720 | 268280 | 121670 | 86006 | 331680 | 1744600 | 3093800 | 1302500 | 0 | 0 | 198300 | 197930 |
| VLEYLNQEI | SCG2 | P13521 | SCG2_HUMAN | Unmodified | 0 | 0 | 0 | 0 | 0 | 50025 | 119080 | 338290 | 3681600 | 2316100 | 0 | 0 | 0 | 0 | 0 | 0 | 837780 | 0 |
| VLEYLNQEI | SCG2 | P13521 | SCG2_HUMAN | Unmodified | 0 | 0 | 0 | 0 | 0 | 0 | 214570 | 522400 | 119230 | 1263700 | 0 | 0 | 0 | 0 | 0 | 0 | 48660 | 0 |
| VLFAPMR | MVP; MVP; MVP | Q14764; H3I | MVP_HUMAN N | Unmodified | 0 | 0 | 0 | 0 | 0 | 0 | 0 | 0 | 318440 | 0 | 0 | 6796100 | 0 | 0 | 0 | 0 | 0 | 0 |
| VLFASQEIP | FLNB; FLNB | O75369; E7E | FLNB_HUMAN | Unmodified | 0 | 0 | 1357500 | 1624800 | 556890 | 0 | 0 | 0 | 1247500 | 0 | 695170 | 11217000 | 0 | 0 | 1319800 | 1202700 | 339100 | 0 |
| VLFASQEIP | FLNB; FLNB | O75369; E7E | FLNB_HUMAN | Unmodified | 0 | 0 | 0 | 0 | 6311300 | 0 | 0 | 0 | 188690 | 0 | 0 | 1224300 | 0 | 0 | 0 | 0 | 1265600 | 0 |
| VLFASQEIP | FLNB; FLNB | O75369; E7E | FLNB_HUMAN | Unmodified | 0 | 0 | 0 | 0 | 7866700 | 2264500 | 0 | 0 | 0 | 476680 | 0 | 0 | 0 | 0 | 0 | 0 | 6780800 | 856960 |
| VLFDKDVN | STAT1; STAT1; S | P42224; A0A | STAT1_HUMAN | Unmodified | 0 | 0 | 0 | 0 | 312030 | 0 | 0 | 59144 | 0 | 0 | 0 | 0 | 106190 | 0 | 0 | 0 | 0 | 492300 |
| VLFEDPLFF | CAPN5; CAPN5 | O15484; E7E | CAN5_HUMAN | Unmodified | 0 | 0 | 0 | 0 | 2312000 | 592380 | 0 | 0 | 1339300 | 578010 | 0 | 272370 | 4294300 | 741170 | 682710 | 331630 | 1129900 | 2733500 |
| VLFEDPLFF | CAPN5; CAPN5 | O15484; E7E | CAN5_HUMAN | Unmodified | 0 | 0 | 0 | 0 | 2068300 | 446930 | 0 | 0 | 0 | 712780 | 0 | 0 | 0 | 6304200 | 0 | 0 | 1099900 | 2399900 |
| VLFEGTR | CHDH | Q8NE62 | CHDH_HUMAI | Unmodified | 46823 | 0 | 0 | 125590 | 0 | 0 | 0 | 0 | 3584100 | 701030 | 0 | 9079000 | 0 | 0 | 23496 | 0 | 0 | 0 |
| VLFELTGR | AP1M2; AP1M2; | Q9Y6Q5; K7I | AP1M2_HUMAI | Unmodified | 0 | 73147 | 160640 | 0 | 0 | 0 | 0 | 0 | 0 | 0 | 0 | 1618100 | 0 | 0 | 121390 | 109540 | 0 | 0 |
| VLFEYIPQN | CD2AP | Q9Y5K6 | CD2AP_HUMAI | Unmodified | 5803300 | 3650800 | 2675400 | 2230200 | 0 | 0 | 724350 | 0 | 0 | 0 | 464110 | 0 | 1812400 | 649980 | 8456800 | 4053500 | 0 | 0 |
| VLFICTANV | LONP1; LONP1 | P36776; K7E | LONM_HUMAN | Unmodified | 0 | 0 | 0 | 0 | 0 | 0 | 0 | 0 | 2481500 | 0 | 0 | 67358 | 0 | 0 | 0 | 0 | 0 | 1725800 |
| VLFMSSEEI | C1GALT1; C1G/ | Q9NS00; C9 | C1GLT_HUMAI | Oxidation (M) | 0 | 0 | 0 | 0 | 11885000 | 3864500 | 0 | 0 | 279990 | 59432 | 0 | 0 | 0 | 0 | 0 | 0 | 1814800 | 2470400 |
| VLFMSSEEI | C1GALT1; C1G/ | Q9NS00; C9 | C1GLT_HUMAI | Unmodified | 0 | 0 | 0 | 0 | 0 | 0 | 568120 | 454530 | 3139700 | 0 | 123090 | 0 | 0 | 546230 | 212170 | 205680 | 0 | 0 |
| VLFMSSEEI | C1GALT1 | Q9NS00 | C1GLT_HUMAI | Unmodified | 0 | 0 | 0 | 0 | 0 | 0 | 0 | 0 | 3170500 | 498900 | 0 | 0 | 0 | 0 | 0 | 0 | 0 | 0 |
| VLFMSSEEI | C1GALT1 | Q9NS00 | C1GLT_HUMAI | Oxidation (M) | 0 | 0 | 0 | 0 | 0 | 0 | 0 | 24347 | 6059000 | 618620 | 37002 | 0 | 0 | 0 | 0 | 0 | 0 | 0 |
| VLFPGIEDI | PLXNA1 | Q9UIW2 | PLXA1_HUMAI | Unmodified | 0 | 0 | 0 | 0 | 0 | 0 | 0 | 0 | 0 | 0 | 0 | 2166500 | 0 | 0 | 584270 | 504460 | 0 | 0 |
| VLFPGIEDI | PLXNA2; PLXNA | O75051; Q9I | PLXA2_HUMAI | Unmodified | 253710 | 138290 | 199530 | 0 | 0 | 0 | 0 | 0 | 2558300 | 441560 | 0 | 3181200 | 0 | 0 | 231300 | 143810 | 0 | 0 |
| VLFPGNST | MAP1B; MAP1B | P46821; D6F | MAP1B_HUMAI | Unmodified | 0 | 0 | 0 | 0 | 0 | 0 | 335520 | 113340 | 918090 | 0 | 370560 | 1810500 | 0 | 0 | 0 | 0 | 0 | 274990 |
| VLFSGSLR | CTHRC1; ABCC | Q96CG8; P3 | CTHR1_HUMAI | Unmodified | 28604 | 0 | 0 | 50219 | 43810 | 0 | 0 | 6058.2 | 5483000 | 881420 | 0 | 54702 | 0 | 0 | 20269 | 0 | 0 | 0 |
| VLFSNIEDII | PREX1 | Q8TCU6 | PREX1_HUMAI | Unmodified | 0 | 0 | 0 | 0 | 0 | 0 | 0 | 0 | 0 | 0 | 0 | 717410 | 0 | 0 | 0 | 0 | 0 | 0 |
| VLFSSNGG | PTBP1; PTBP1; | P26599; A0A | PTBP1_HUMAI | Unmodified | 0 | 217460 | 91012 | 173360 | 297560 | 0 | 0 | 99185 | 1626500 | 645010 | 1791500 | 12404000 | 0 | 0 | 75784 | 262360 | 0 | 0 |
| VLFVRNLAI | SYNCRIP; SYN( | O60506; B7Z | HNRPQ_HUMA | Unmodified | 0 | 0 | 0 | 0 | 59860 | 0 | 2190500 | 0 | 0 | 1332700 | 0 | 0 | 3036700 | 1204300 | 0 | 0 | 533870 | 0 |
| VLFVRNLAT | HNRNPR; HNRI | O43390; B4I | HNRPR_HUMA | Unmodified | 0 | 0 | 0 | 0 | 0 | 0 | 3031100 | 0 | 0 | 801810 | 0 | 261500 | 2229400 | 1347600 | 23941 | 0 | 124030 | 931670 |
| VLFYEEELV | CDC42BPB | Q9Y5S2 | MRCKB_HUMA | Unmodified | 0 | 0 | 0 | 0 | 1367300 | 0 | 0 | 0 | 0 | 335410 | 0 | 0 | 0 | 689960 | 0 | 0 | 0 | 1680200 |
| VLGAPVPAI | AGRN; AGRN; A | O00468; A0/ | AGRIN_HUMAI | Unmodified | 1677100 | 701320 | 3102000 | 1071400 | 0 | 0 | 0 | 0 | 1964100 | 0 | 747240 | 337760 | 751860 | 404530 | 1629300 | 1414300 | 0 | 0 |
| VLGAPVPAI | AGRN; AGRN; A | O00468; A0/ | AGRIN_HUMAI | Unmodified | 0 | 0 | 0 | 0 | 0 | 0 | 0 | 0 | 0 | 177910 | 0 | 0 | 76018 | 44133 | 0 | 0 | 0 | 0 |
| VLGATLLPI | NPEPPS; NPEPI | P55786; A0A | PSA_HUMAN P | Unmodified | 0 | 0 | 739160 | 0 | 0 | 0 | 0 | 0 | 3140000 | 643070 | 1772300 | 15629000 | 0 | 0 | 1446000 | 1722700 | 0 | 0 |
| VLGCNPGI | LACTB2 | Q53H82 | LACB2_HUMAI | Oxidation (M) | 0 | 0 | 873240 | 0 | 0 | 1980900 | 282930 | 128110 | 1323400 | 0 | 0 | 0 | 0 | 0 | 0 | 0 | 0 | 657860 |
| VLGCNPGI | LACTB2 | Q53H82 | LACB2_HUMAI | Unmodified | 0 | 0 | 0 | 0 | 0 | 0 | 0 | 0 | 1589800 | 0 | 0 | 166140 | 0 | 0 | 0 | 0 | 0 | 0 |
| VLGDTLLTI | PLCG1 | P19174 | PLCG1_HUMAI | Unmodified | 0 | 0 | 0 | 0 | 0 | 0 | 0 | 0 | 0 | 0 | 0 | 1403800 | 0 | 0 | 0 | 0 | 3898500 | 2217300 |
| VLGEAMTG | TLN1 | Q9Y490 | TLN1_HUMAN | Unmodified | 0 | 211420 | 379830 | 659170 | 8936300 | 1437500 | 2583000 | 679410 | 357700 | 0 | 0 | 2854400 | 2161100 | 670690 | 441700 | 0 | 1793100 | 3417600 |
| VLGEAMTG | TLN1 | Q9Y490 | TLN1_HUMAN | Oxidation (M) | 76995 | 0 | 0 | 217410 | 737960 | 0 | 393390 | 246200 | 0 | 0 | 62755 | 329960 | 354010 | 393990 | 205400 | 118100 | 0 | 460610 |
| VLGEEVSFI | PDE2A; PDE2A; | O00408; H0Y | PDE2A_HUMAI | Unmodified | 0 | 0 | 0 | 0 | 0 | 0 | 0 | 0 | 0 | 0 | 603650 | 2244600 | 0 | 0 | 800040 | 0 | 0 | 0 |
| VLGEKPELI | SCIN; SCIN; SC | Q9Y6U3; E5I | ADSV_HUMAN | Unmodified | 0 | 0 | 0 | 146970 | 0 | 0 | 0 | 0 | 4996100 | 1102000 | 0 | 0 | 0 | 0 | 0 | 0 | 0 | 0 |
| VLGESPVPI | AGRN | A0A494C0G | A0A494C0G5_I | Unmodified | 607200 | 0 | 6539400 | 387670 | 0 | 0 | 0 | 0 | 511820 | 0 | 0 | 0 | 0 | 0 | 886410 | 783990 | 0 | 0 |
| VLGESPVPI | AGRN | A0A494C0G | A0A494C0G5_I | Unmodified | 2659500 | 0 | 0 | 960880 | 0 | 0 | 0 | 0 | 1814700 | 0 | 0 | 0 | 2301500 | 0 | 0 | 0 | 0 | 0 |
| VLGETANIL | AKAP12 | Q02952 | AKA12_HUMAI | Unmodified | 0 | 0 | 0 | 0 | 0 | 0 | 0 | 0 | 0 | 0 | 0 | 0 | 0 | 0 | 0 | 0 | 0 | 0 |
| VLGFCHYY | ATP1A3; N/A; A | P13637; A0A | AT1A3_HUMAI | Unmodified | 0 | 0 | 188910 | 0 | 0 | 93694 | 256890 | 0 | 1085700 | 186970 | 0 | 69064 | 0 | 0 | 0 | 0 | 0 | 0 |
| VLGILAMID | PPA1; PPA1 | Q15181; Q5 | IPYR_HUMAN I | Oxidation (M) | 0 | 46023 | 14797 | 216530 | 566330 | 0 | 0 | 0 | 15807 | 0 | 85966 | 305770 | 554410 | 77701 | 532330 | 0 | 150710 | 307480 |
| VLGIVVQN | ADGRG1; ADGI | Q9Y653; A0/ | AGRG1_HUMA | Unmodified | 41552 | 0 | 272730 | 417870 | 0 | 0 | 0 | 0 | 2030900 | 1047400 | 242070 | 0 | 296540 | 235540 | 247800 | 253170 | 487080 | 1241400 |
| VLGKFHPD | KLC2; KLC2 | Q9H0B6; A8 | KLC2_HUMAN | Unmodified | 0 | 0 | 0 | 0 | 194220 | 0 | 278480 | 137870 | 0 | 80413 | 0 | 0 | 0 | 0 | 0 | 0 | 246230 | 212860 |
| VLGLLGALI | MTOR | P42345 | MTOR_HUMAN | Unmodified | 0 | 0 | 0 | 0 | 0 | 0 | 0 | 0 | 0 | 0 | 0 | 534100 | 0 | 0 | 0 | 0 | 0 | 0 |
| VLGLVLLR | SNRPB; SNRPN | P14678; P63 | RSMB_HUMAN | Unmodified | 0 | 0 | 0 | 0 | 0 | 0 | 0 | 0 | 0 | 0 | 262380 | 444570 | 0 | 0 | 0 | 0 | 0 | 0 |
| VLGMDPLP | ILF3 | Q12906 | ILF3_HUMAN Ir | Oxidation (M) | 0 | 0 | 0 | 0 | 0 | 0 | 0 | 0 | 0 | 45120 | 0 | 1976300 | 0 | 0 | 0 | 0 | 0 | 0 |

| | | | | | | | | | | | | | | | | | | | | | | |
|---|---|---|---|---|---|---|---|---|---|---|---|---|---|---|---|---|---|---|---|---|---|---|
| VLGMDPLP | ILF3 | Q12906 | ILF3_HUMAN Ir | Unmodified | 933280 | 743700 | 0 | 569340 | 0 | 0 | 0 | 0 | 399210 | 0 | 497270 | 2330800 | 364310 | 0 | 0 | 500620 | 0 | 0 |
| VLGMTLIQI | GLO1 | Q04760 | LGUL_HUMAN | Unmodified | 612100 | 0 | 0 | 0 | 1611500 | 71431 | 0 | 55678 | 4386400 | 358720 | 2320400 | 6946500 | 3124200 | 0 | 2257000 | 0 | 0 | 528910 |
| VLGMTLIQI | GLO1 | Q04760 | LGUL_HUMAN | Oxidation (M) | 0 | 0 | 0 | 0 | 0 | 0 | 0 | 0 | 508410 | 0 | 966490 | 3581900 | 1089100 | 0 | 1306300 | 0 | 0 | 0 |
| VLGMTLIQI | GLO1 | Q04760 | LGUL_HUMAN | 2 Oxidation (M) | 0 | 0 | 0 | 0 | 0 | 0 | 0 | 0 | 0 | 0 | 0 | 0 | 368340 | 132140 | 0 | 0 | 0 | 0 |
| VLGNLQLN | EXOC7; EXOC7 | Q9UPT5; B4I | EXOC7_HUMAI | Unmodified | 0 | 0 | 0 | 0 | 2467400 | 381920 | 0 | 0 | 0 | 974950 | 0 | 0 | 0 | 0 | 0 | 0 | 0 | 0 |
| VLGNPKSD | MYL6; MYL6; MY | P60660; G8J | MYL6_HUMAN | Unmodified | 0 | 93823 | 0 | 134360 | 1.16E+08 | 14498000 | 10797000 | 6805700 | 331400 | 6701400 | 0 | 0 | 20342000 | 17752000 | 0 | 0 | 41575000 | 21386000 |
| VLGNPKSD | MYL6; MYL6; MY | P60660; G8J | MYL6_HUMAN | Oxidation (M) | 0 | 0 | 0 | 0 | 20752000 | 3148600 | 2530000 | 1435100 | 0 | 190390 | 72230 | 0 | 6604800 | 2754400 | 0 | 237480 | 5122200 | 6426800 |
| VLGNPKSD | MYL6; MYL6; MY | P60660; G8J | MYL6_HUMAN | 2 Oxidation (M) | 0 | 0 | 0 | 0 | 17914000 | 367090 | 5388600 | 0 | 0 | 0 | 0 | 0 | 0 | 5098000 | 0 | 0 | 9143600 | 0 |
| VLGNPKSD | MYL6; MYL6; MY | P60660; G8J | MYL6_HUMAN | Oxidation (M) | 0 | 0 | 0 | 0 | 2209300 | 46375 | 392440 | 0 | 0 | 89645 | 0 | 0 | 221700 | 0 | 0 | 0 | 9804400 | 0 |
| VLGNPKSD | MYL6; MYL6; MY | P60660; G8J | MYL6_HUMAN | Unmodified | 0 | 0 | 0 | 0 | 95626 | 0 | 0 | 0 | 0 | 0 | 0 | 0 | 0 | 0 | 0 | 0 | 125460 | 0 |
| VLGPCSDII | PRKAR1A | P10644 | KAP0_HUMAN | Unmodified | 0 | 0 | 0 | 175120 | 649810 | 321240 | 166720 | 0 | 932110 | 0 | 0 | 1205300 | 0 | 0 | 0 | 0 | 567790 | 1139200 |
| VLGPCSDII | PRKAR1A | P10644 | KAP0_HUMAN | Unmodified | 0 | 0 | 0 | 0 | 0 | 80942 | 231660 | 19849 | 771660 | 102600 | 134760 | 168110 | 0 | 0 | 0 | 0 | 522030 | 379830 |
| VLGQGEEV | PLOD2; PLOD2; | O00469; C9J | PLOD2_HUMAI | Unmodified | 0 | 0 | 0 | 0 | 1865100 | 1276100 | 0 | 0 | 24246000 | 12448000 | 249810 | 553120 | 0 | 0 | 0 | 0 | 0 | 708580 |
| VLGQGSFG | RPS6KA6; RPS6 | Q9UK32; Q1 | KS6A6_HUMAN | Unmodified | 0 | 108110 | 0 | 0 | 0 | 0 | 0 | 97729 | 0 | 489980 | 136590 | 1238800 | 899040 | 0 | 0 | 0 | 0 | 0 |
| VLGQLTET( | GNPNAT1; GNF | Q96EK6; G3Y | GNA1_HUMAN | Oxidation (M) | 216400 | 0 | 521460 | 535500 | 1184600 | 0 | 0 | 0 | 0 | 212490 | 0 | 889110 | 873130 | 843880 | 573830 | 0 | 1389500 | 0 |
| VLGQLTET( | GNPNAT1; GNF | Q96EK6; G3Y | GNA1_HUMAN | Unmodified | 0 | 0 | 0 | 0 | 0 | 0 | 0 | 0 | 0 | 0 | 0 | 0 | 0 | 0 | 0 | 0 | 0 | 0 |
| VLGQYSGF | SEZ6; SEZ6; SEZ | Q53EL9; C9J | SEZ6_HUMAN | Unmodified | 0 | 0 | 0 | 0 | 0 | 0 | 0 | 0 | 2842500 | 0 | 0 | 0 | 0 | 0 | 0 | 0 | 0 | 0 |
| VLGRSPLTI | CDCA3; CDCA3 | Q99618; J3K | CDCA3_HUMA | Unmodified | 0 | 0 | 0 | 0 | 3251100 | 422660 | 58689 | 23236 | 0 | 899480 | 0 | 0 | 3176400 | 749880 | 0 | 0 | 393520 | 1044900 |
| VLGRTGSQ | RPS28 | P62857 | RS28_HUMAN | Unmodified | 0 | 0 | 0 | 0 | 0 | 0 | 0 | 0 | 0 | 476350 | 0 | 0 | 0 | 0 | 0 | 0 | 731520 | 0 |
| VLGSEPAP | PLEKHO2 | Q8TD55 | PKHO2_HUMAI | Unmodified | 0 | 0 | 0 | 0 | 0 | 43079 | 285790 | 0 | 0 | 0 | 126700 | 0 | 0 | 227430 | 0 | 0 | 0 | 722890 |
| VLGSGAFG | ERBB2; ERBB4; | P04626; Q15 | ERBB2_HUMAN | Unmodified | 374600 | 538790 | 2120800 | 4342900 | 31695000 | 5631600 | 0 | 0 | 961050 | 0 | 0 | 0 | 2953000 | 2352100 | 4800800 | 3038300 | 344960 | 4982200 |
| VLGSGAFG | EGFR; EGFR; EC | P00533; Q5C | EGFR_HUMAN | Unmodified | 0 | 0 | 0 | 0 | 2874000 | 0 | 0 | 0 | 0 | 0 | 0 | 0 | 497040 | 363810 | 0 | 0 | 0 | 853740 |
| VLGSQEAL | MAOA | P21397 | AOFA_HUMAN | Unmodified | 0 | 0 | 1887300 | 0 | 0 | 0 | 0 | 0 | 173000 | 74571 | 0 | 0 | 0 | 797750 | 3082800 | 671460 | 0 | 0 |
| VLGTAFDPI | HSPH1; HSPH1 | Q92598; A0A | HS105_HUMAN | Unmodified | 0 | 0 | 1350400 | 2967800 | 2674900 | 0 | 4426000 | 1389100 | 2335300 | 1459000 | 0 | 10410000 | 2378500 | 2117800 | 0 | 1519400 | 1809500 | 2452300 |
| VLGTAFDPI | HSPH1; HSPH1 | Q92598; A0A | HS105_HUMAN | Unmodified | 0 | 0 | 0 | 0 | 0 | 0 | 1864200 | 230140 | 0 | 0 | 0 | 0 | 578420 | 0 | 0 | 0 | 0 | 439790 |
| VLGTEDLYI | CSNK2A1; CSN | P68400; A0A | CSK21_HUMAN | Unmodified | 40178 | 0 | 277340 | 0 | 0 | 0 | 0 | 0 | 0 | 287620 | 0 | 2126600 | 0 | 640860 | 0 | 0 | 0 | 0 |
| VLGTEELY( | CSNK2A2; CSN | P19784; H3E | CSK22_HUMAN | Unmodified | 0 | 0 | 0 | 0 | 0 | 0 | 0 | 0 | 0 | 0 | 0 | 1174200 | 0 | 0 | 0 | 0 | 311110 | 0 |
| VLGTGAYG | RPS6KA5; RPS6 | O75582; O75 | KS6A5_HUMAN | Unmodified | 0 | 0 | 0 | 0 | 0 | 86084 | 849050 | 89869 | 62378 | 64178 | 0 | 0 | 0 | 0 | 0 | 0 | 9016.1 | 0 |
| VLGTSPEAI | CAD; CAD | P27708; F8V | PYR1_HUMAN | Unmodified | 129110 | 0 | 191980 | 306340 | 0 | 0 | 0 | 0 | 0 | 0 | 0 | 2073000 | 0 | 0 | 264720 | 193340 | 0 | 0 |
| VLGVENLR | CHDH | Q8NE62 | CHDH_HUMAN | Unmodified | 0 | 0 | 0 | 0 | 0 | 0 | 0 | 0 | 886870 | 0 | 0 | 0 | 0 | 0 | 0 | 0 | 0 | 0 |
| VLGVPIIVQ | RBM39; RBM39 | Q14498; G3X | RBM39_HUMAI | Unmodified | 0 | 0 | 0 | 0 | 0 | 0 | 0 | 0 | 377300 | 0 | 344810 | 1297800 | 0 | 0 | 0 | 0 | 23324 | 0 |
| VLHDEILNI | SLC35A3; SLC3 | Q9Y2D2; A0A | S35A3_HUMAN | Unmodified | 0 | 0 | 0 | 0 | 0 | 0 | 0 | 0 | 859060 | 0 | 0 | 0 | 0 | 0 | 0 | 0 | 0 | 0 |
| VLHEAEGH | SNRPD3; SNRP | P62318; H3E | SMD3_HUMAN | Unmodified | 722080 | 0 | 0 | 0 | 198940 | 0 | 33835 | 63475 | 283740 | 0 | 0 | 2855300 | 0 | 0 | 0 | 0 | 62499 | 0 |
| VLHEAEGH | SNRPD3; SNRP | P62318; H3E | SMD3_HUMAN | Unmodified | 689800 | 0 | 198280 | 1277900 | 2737000 | 0 | 1240100 | 560440 | 2149700 | 1836300 | 2499400 | 2606500 | 12759000 | 3046900 | 0 | 199120 | 1692800 | 451040 |
| VLHGEQYL | HSD17B4; HSD | P51659; E7E | DHB4_HUMAN | Unmodified | 0 | 0 | 0 | 0 | 0 | 0 | 0 | 0 | 590990 | 0 | 0 | 0 | 0 | 0 | 0 | 0 | 0 | 0 |
| VLHGEQYL | HSD17B4; HSD | P51659; E7E | DHB4_HUMAN | Unmodified | 200810 | 0 | 0 | 0 | 992550 | 0 | 1410200 | 0 | 32648000 | 9064100 | 2454400 | 753750 | 0 | 0 | 0 | 0 | 265520 | 0 |
| VLHGEQYL | HSD17B4; HSD | P51659; E7E | DHB4_HUMAN | Unmodified | 0 | 0 | 0 | 0 | 565320 | 0 | 1764200 | 0 | 0 | 3278600 | 0 | 0 | 0 | 0 | 0 | 0 | 0 | 0 |
| VLHMDRNI | GDI2; GDI2; GD | P50395; V9G | GDIB_HUMAN | Oxidation (M) | 0 | 0 | 0 | 0 | 3420700 | 0 | 1060500 | 0 | 2099000 | 0 | 1233400 | 3252800 | 0 | 0 | 0 | 0 | 1018400 | 1310900 |
| VLHMDRNI | GDI2; GDI2; GD | P50395; V9G | GDIB_HUMAN | Unmodified | 0 | 0 | 0 | 0 | 24796000 | 0 | 1790600 | 0 | 6703200 | 359980 | 0 | 2494500 | 4087500 | 2307400 | 0 | 0 | 4742600 | 9426700 |
| VLHMDRNI | GDI2; GDI2; GD | P50395; V9G | GDIB_HUMAN | Oxidation (M) | 0 | 388000 | 0 | 1268700 | 892760 | 0 | 105440 | 0 | 1030900 | 0 | 0 | 5815200 | 0 | 0 | 0 | 656520 | 1069500 | 0 |
| VLHMDRNI | GDI1; GDI1 | P31150; G5E | GDIA_HUMAN | Oxidation (M) | 0 | 0 | 0 | 0 | 1289300 | 0 | 1354200 | 0 | 2807500 | 3270300 | 0 | 4036700 | 3754400 | 2288200 | 0 | 0 | 2801600 | 2542200 |
| VLHMDRNI | GDI1; GDI1 | P31150; G5E | GDIA_HUMAN | Unmodified | 0 | 0 | 0 | 0 | 0 | 0 | 0 | 0 | 3725900 | 1158800 | 0 | 327180 | 819550 | 935930 | 0 | 0 | 746270 | 784690 |
| VLHNQLVL | ARFIP1 | P53367 | ARFP1_HUMAN | Unmodified | 0 | 0 | 0 | 3139500 | 0 | 0 | 747600 | 0 | 0 | 431970 | 0 | 0 | 1921000 | 374950 | 0 | 0 | 0 | 0 |
| VLHPLEGA | ALCAM; ALCAM | Q13740; F5C | CD166_HUMAI | Unmodified | 16864000 | 13118000 | 3670000 | 7200400 | 14642000 | 1083200 | 3567100 | 0 | 5924200 | 4183300 | 0 | 1104900 | 4488200 | 2457100 | 4676500 | 3045200 | 4442200 | 12551000 |
| VLHTNVVFI | NOTCH1; NOTC | P46531; A0A | NOTC1_HUMA | Unmodified | 0 | 0 | 0 | 0 | 0 | 0 | 0 | 0 | 892920 | 0 | 703020 | 2914100 | 0 | 0 | 0 | 0 | 0 | 0 |
| VLHTYHDS | SEPTIN6; SEPTI | Q14141; B1A | SEPT6_HUMAN | Unmodified | 0 | 0 | 0 | 0 | 0 | 0 | 0 | 0 | 0 | 0 | 0 | 112500 | 0 | 0 | 0 | 0 | 0 | 0 |
| VLHVSENP | LRRC47 | Q8N1G4 | LRC47_HUMAN | Unmodified | 204050 | 0 | 408420 | 478000 | 0 | 0 | 0 | 0 | 1523900 | 523860 | 793510 | 4375200 | 486620 | 0 | 266330 | 328670 | 0 | 0 |
| VLHVSENP | LRRC47 | Q8N1G4 | LRC47_HUMAN | Unmodified | 0 | 0 | 0 | 0 | 0 | 0 | 0 | 0 | 0 | 0 | 0 | 0 | 1185200 | 946690 | 0 | 0 | 0 | 0 |
| VLHYGDLD | ELMO2; ELMO2 | Q96JJ3; H0Y | ELMO2_HUMAI | Unmodified | 0 | 0 | 0 | 0 | 3991500 | 342650 | 0 | 0 | 0 | 0 | 0 | 0 | 726100 | 620480 | 0 | 0 | 1468400 | 1478000 |
| VLHYLAIQK | RAB3GAP1 | Q15042 | RB3GP_HUMAI | Unmodified | 0 | 0 | 0 | 0 | 936720 | 101020 | 0 | 0 | 437390 | 159790 | 218240 | 1529700 | 351630 | 244410 | 0 | 0 | 89151 | 0 |
| VLIAAHGN | PGAM1; PGAM2 | P18669; P15 | PGAM1_HUMA | Unmodified | 0 | 0 | 0 | 0 | 334290 | 181790 | 0 | 0 | 0 | 0 | 0 | 776740 | 238900 | 0 | 123030 | 0 | 305590 | 218580 |
| VLIAAHGN | PGAM1; PGAM2 | P18669; P15 | PGAM1_HUMA | Unmodified | 0 | 0 | 0 | 0 | 4881800 | 0 | 298020 | 0 | 0 | 0 | 0 | 0 | 1398100 | 1506300 | 0 | 0 | 2086500 | 3695000 |
| VLIALLAR | COLGALT1 | Q8NBJ5 | GT251_HUMAN | Unmodified | 0 | 0 | 0 | 0 | 0 | 0 | 0 | 0 | 27079000 | 6942700 | 479990 | 0 | 0 | 0 | 0 | 0 | 0 | 0 |
| VLIANNGIA | ACACB; ACACA | O00763; Q13 | ACACB_HUMAI | Unmodified | 0 | 0 | 0 | 0 | 162920 | 0 | 125030 | 82108 | 0 | 0 | 0 | 1117700 | 0 | 0 | 131280 | 61307 | 0 | 0 |
| VLIDPVSV( | IARS1; IARS1; I/ | P41252; A0A | SYIC_HUMAN I | Unmodified | 0 | 0 | 0 | 0 | 0 | 0 | 0 | 0 | 0 | 0 | 0 | 11316000 | 0 | 0 | 0 | 0 | 0 | 0 |
| VLIEDTDDE | XPR1 | Q9UBH6 | XPR1_HUMAN | Unmodified | 760740 | 0 | 0 | 285260 | 3345300 | 0 | 274450 | 137350 | 0 | 0 | 222100 | 331740 | 725330 | 521150 | 0 | 0 | 0 | 0 |
| VLIEGSINS | ARPC4; ARPC4- | P59998; A0A | ARPC4_HUMAI | Unmodified | 2022400 | 2433800 | 899340 | 2211400 | 397270 | 0 | 0 | 0 | 758660 | 166550 | 1038100 | 7324600 | 51299000 | 0 | 2206900 | 1738400 | 138050 | 153540 |
| VLIEILCTR | ANXA7 | P20073 | ANXA7_HUMAN | Unmodified | 0 | 0 | 1040600 | 4156100 | 0 | 0 | 0 | 0 | 1.6E+08 | 2317300 | 3238300 | 860520 | 0 | 0 | 4693600 | 1583200 | 0 | 0 |
| VLIEILCTRT | ANXA7 | P20073 | ANXA7_HUMAN | Unmodified | 0 | 0 | 0 | 678530 | 0 | 0 | 0 | 0 | 4797100 | 1999000 | 524580 | 0 | 0 | 0 | 0 | 0 | 0 | 0 |
| VLIEILCTRT | ANXA7 | P20073 | ANXA7_HUMAN | Unmodified | 0 | 0 | 0 | 0 | 238800 | 164740 | 0 | 0 | 1304300 | 3051000 | 83444 | 0 | 0 | 0 | 0 | 0 | 337060 | 446180 |
| VLIELDK | RHOU | Q7L0Q8 | RHOU_HUMAN | Unmodified | 707420 | 387940 | 293860 | 460410 | 0 | 0 | 265240 | 77421 | 232650 | 96127 | 593830 | 0 | 755460 | 784290 | 275120 | 190150 | 0 | 0 |
| VLIFDDSFE | ASPH | Q12797 | ASPH_HUMAN | Unmodified | 0 | 0 | 0 | 0 | 83535 | 0 | 0 | 0 | 594530 | 0 | 0 | 0 | 0 | 0 | 0 | 0 | 0 | 0 |
| VLIIGGGD( | SRM; SRM | P19623; K7E | SPEE_HUMAN | Unmodified | 828170 | 542200 | 1275400 | 999300 | 0 | 0 | 0 | 0 | 5751500 | 279200 | 806670 | 8015200 | 0 | 0 | 619530 | 1207100 | 0 | 0 |
| VLIIGGGD( | SRM; SRM | P19623; K7E | SPEE_HUMAN | Unmodified | 0 | 0 | 0 | 0 | 0 | 0 | 0 | 0 | 1179300 | 196110 | 178830 | 0 | 995600 | 0 | 0 | 0 | 0 | 0 |
| VLIIGSGGR | GART; GART; G/ | P22102; C9J | PUR2_HUMAN | Unmodified | 0 | 213160 | 203690 | 757370 | 7705700 | 1375200 | 872570 | 102050 | 620880 | 968180 | 616430 | 5453700 | 4136500 | 3451200 | 706970 | 274030 | 1310800 | 3626800 |
| VLILFCNLV | USP43; USP43 | Q70EL4; V9C | UBP43_HUMAN | Unmodified | 0 | 0 | 0 | 113140 | 0 | 0 | 0 | 0 | 0 | 0 | 0 | 0 | 0 | 0 | 170300 | 146120 | 0 | 0 |
| VLIMDHPSI | STXBP2; STXBP | Q15833; M0I | STXB2_HUMAN | Unmodified | 0 | 0 | 0 | 0 | 0 | 0 | 0 | 0 | 0 | 0 | 0 | 0 | 0 | 0 | 0 | 0 | 648560 | 491850 |
| VLINLFVGT | MME; MME; MM | P08473; A0A | NEP_HUMAN N | Unmodified | 0 | 0 | 935960 | 0 | 0 | 0 | 0 | 0 | 0 | 0 | 0 | 0 | 0 | 0 | 6477900 | 1938500 | 0 | 0 |
| VLINLFVGT | MME; MME; MM | P08473; A0A | NEP_HUMAN N | Unmodified | 0 | 0 | 0 | 10916000 | 0 | 0 | 0 | 0 | 1145000 | 0 | 0 | 0 | 7882500 | 1306700 | 0 | 0 | 0 | 0 |
| VLIQRGYD | TOP2A | P11388 | TOP2A_HUMAN | Unmodified | 0 | 0 | 85939 | 0 | 2014800 | 0 | 2146400 | 533930 | 195230 | 0 | 0 | 369020 | 0 | 0 | 0 | 0 | 0 | 737450 |
| VLISDSLDF | PHGDH; PHGD | O43175; A0A | SERA_HUMAN | Unmodified | 327950 | 0 | 0 | 3461900 | 2399000 | 0 | 6986300 | 2676300 | 5373600 | 2803000 | 0 | 7156800 | 23819000 | 35240000 | 1574700 | 1848300 | 5115600 | 4911700 |
| VLISDSLDF | PHGDH; PHGD | O43175; A0A | SERA_HUMAN | Unmodified | 0 | 0 | 0 | 0 | 0 | 0 | 480330 | 0 | 0 | 51128 | 0 | 0 | 0 | 1330000 | 0 | 0 | 361990 | 0 |
| VLISFKAND | ARPC5; ARPC5 | O15511; B1A | ARPC5_HUMAN | Unmodified | 0 | 0 | 0 | 0 | 2351000 | 0 | 2529700 | 1993400 | 0 | 618130 | 0 | 0 | 8713000 | 4501000 | 0 | 0 | 2474000 | 1505300 |
| VLISSLQDC | CPSF6; CPSF6 | Q16630; F8W | CPSF6_HUMAN | Unmodified | 2875100 | 2317700 | 0 | 1284100 | 0 | 0 | 0 | 0 | 0 | 0 | 0 | 0 | 0 | 1479100 | 417730 | 1012300 | 0 | 0 |
| VLISTDVW | EIF4A3; EIF4A3 | P38919; I3L3 | IF4A3_HUMAN | Unmodified | 1024700 | 1325500 | 647410 | 0 | 0 | 0 | 0 | 0 | 0 | 0 | 297080 | 1906000 | 0 | 0 | 0 | 288100 | 0 | 0 |
| VLITTDLLAI | EIF4A1; EIF4A2; | P60842; Q14 | IF4A1_HUMAN | Unmodified | 6957500 | 4927600 | 2590900 | 5657700 | 3093600 | 751410 | 451030 | 349850 | 6793100 | 2135200 | 6140000 | 29632000 | 4509800 | 809050 | 12972000 | 10149000 | 0 | 1535900 |
| VLITTLITAG | MOV10; MOV10 | Q9HCE1; Q5 | MOV10_HUMA | Unmodified | 0 | 3861100 | 0 | 0 | 0 | 0 | 0 | 0 | 0 | 0 | 0 | 516860 | 0 | 0 | 0 | 0 | 0 | 0 |
| VLIVSEDPE | AIFM1; AIFM1; A | O95831; A0A | AIFM1_HUMAN | Unmodified | 0 | 0 | 0 | 0 | 0 | 0 | 0 | 0 | 938670 | 0 | 933440 | 0 | 0 | 0 | 0 | 0 | 0 | 0 |
| VLIVSEDPE | AIFM1; AIFM1; A | O95831; A0A | AIFM1_HUMAN | Oxidation (M) | 0 | 0 | 0 | 0 | 0 | 0 | 0 | 0 | 0 | 0 | 0 | 1614800 | 1134200 | 0 | 0 | 0 | 0 | 0 |
| VLIWNLDV | CORO2B | Q9UQ03 | COR2B_HUMA | Unmodified | 0 | 0 | 0 | 0 | 0 | 0 | 0 | 0 | 0 | 0 | 382740 | 3117000 | 0 | 0 | 0 | 0 | 0 | 0 |
| VLIYDRFG( | SCFD1; SCFD1; | Q8WVM8; A( | SCFD1_HUMAI | Unmodified | 0 | 0 | 0 | 0 | 0 | 0 | 571660 | 0 | 0 | 156840 | 0 | 43160 | 0 | 0 | 19143 | 0 | 0 | 0 |
| VLKDEFLG( | CAPN5; CAPN5 | O15484; E7E | CAN5_HUMAN | Unmodified | 0 | 0 | 0 | 380780 | 0 | 0 | 0 | 0 | 0 | 0 | 0 | 782080 | 460590 | 0 | 0 | 438560 | 0 | 0 |
| VLKDQPPN | CYRIB | Q9NUQ9 | CYRIB_HUMAN | Unmodified | 0 | 307370 | 45325 | 702390 | 3285100 | 565010 | 0 | 0 | 846590 | 1537900 | 0 | 448510 | 566830 | 389260 | 0 | 23410 | 2242700 | 2528000 |
| VLKDQPPN | CYRIB | Q9NUQ9 | CYRIB_HUMAN | Unmodified | 0 | 0 | 0 | 0 | 5197400 | 964980 | 0 | 0 | 0 | 0 | 0 | 0 | 0 | 0 | 0 | 0 | 7044000 | 11813000 |
| VLKEEGNE | TOMM34 | Q15785 | TOM34_HUMAI | Unmodified | 0 | 0 | 0 | 0 | 0 | 0 | 17912 | 13995 | 0 | 0 | 207220 | 0 | 0 | 0 | 0 | 0 | 245890 | 0 |
| VLKGEIPW | PDPK1; PDPK1 | O15530; E9F | PDPK1_HUMAN | Unmodified | 0 | 0 | 0 | 1339400 | 322570 | 0 | 0 | 0 | 0 | 0 | 0 | 0 | 1648800 | 2621300 | 495760 | 514130 | 0 | 0 |
| VLKIELETY | CCDC6 | Q16204 | CCDC6_HUMA | Unmodified | 0 | 0 | 0 | 0 | 1055800 | 0 | 0 | 0 | 449340 | 405620 | 0 | 512470 | 743240 | 275340 | 0 | 0 | 1052200 | 787830 |
| VLKISEEDE | PEA15; PEA15 | Q15121; B1A | PEA15_HUMAN | Unmodified | 0 | 0 | 0 | 79049 | 181730 | 0 | 0 | 0 | 160370 | 312600 | 0 | 0 | 707620 | 292680 | 0 | 0 | 223490 | 0 |
| VLKPDTTY( | SORL1; SORL1; | Q92673; E9F | SORL_HUMAN | Unmodified | 0 | 0 | 0 | 0 | 0 | 0 | 24527 | 0 | 1481700 | 260080 | 0 | 0 | 0 | 0 | 0 | 0 | 0 | 0 |
| VLKQELGG | SNTB1; SNTB1 | Q13884; A0A | SNTB1_HUMAN | Unmodified | 0 | 0 | 0 | 0 | 441960 | 0 | 0 | 0 | 327660 | 1937100 | 0 | 0 | 0 | 0 | 0 | 0 | 424830 | 657030 |
| VLKQVHPD | H2BC4; H2BC1 | P62807; P06 | H2B1C_HUMAI | Unmodified | 692730 | 422990 | 0 | 0 | 2554200 | 370850 | 11126000 | 1.11E+08 | 0 | 71520 | 0 | 3841900 | 442280 | 220210 | 0 | 0 | 0 | 701860 |
| VLLAEYKN | PKN2 | Q16513 | PKN2_HUMAN | Unmodified | 0 | 0 | 0 | 0 | 738440 | 0 | 112830 | 0 | 0 | 0 | 0 | 681420 | 118950 | 0 | 0 | 0 | 220880 | 0 |
| VLLATLSIPI | EIF3A | Q14152 | EIF3A_HUMAN | Unmodified | 0 | 0 | 73451 | 0 | 0 | 0 | 0 | 0 | 0 | 0 | 140450 | 1742100 | 0 | 0 | 0 | 0 | 0 | 0 |
| VLLATLSIPI | EIF3A | Q14152 | EIF3A_HUMAN | Unmodified | 2741500 | 2358500 | 856350 | 2549200 | 0 | 0 | 0 | 0 | 1117500 | 0 | 1051700 | 28178000 | 0 | 0 | 2389700 | 2649700 | 0 | 0 |
| VLLDAGFTI | CACNA2D1; CA | P54289; H0Y | CA2D1_HUMAI | Unmodified | 0 | 0 | 0 | 0 | 0 | 0 | 0 | 0 | 420210 | 0 | 0 | 0 | 0 | 0 | 0 | 0 | 0 | 0 |
| VLLDNVEN | USP7; USP7; US | Q93009; A0A | UBP7_HUMAN | Unmodified | 71674 | 0 | 0 | 0 | 82311 | 103230 | 0 | 0 | 0 | 0 | 2274700 | 0 | 0 | 0 | 0 | 0 | 0 | 18631 |
| VLLDQLR | APMAP; APMAP | Q9HDC9; H( | APMAP_HUMAI | Unmodified | 125700 | 391390 | 683450 | 182170 | 0 | 0 | 0 | 0 | 62568000 | 3893800 | 0 | 667210 | 0 | 0 | 1227000 | 220590 | 0 | 0 |
| VLLEAAVR | IREB2; IREB2 | P48200; H0Y | IREB2_HUMAN | Unmodified | 0 | 0 | 0 | 20348 | 0 | 0 | 0 | 0 | 0 | 0 | 0 | 1135400 | 0 | 0 | 0 | 0 | 0 | 0 |
| VLLEAGEGI | DPP3; DPP3; DI | Q9NY33; G3 | DPP3_HUMAN | Unmodified | 0 | 0 | 0 | 0 | 0 | 0 | 0 | 0 | 0 | 0 | 0 | 2205900 | 0 | 0 | 0 | 375580 | 0 | 0 |
| VLLESEQFI | SRP14; SRP14 | P37108; H0Y | SRP14_HUMAN | Unmodified | 20245000 | 26675000 | 6094900 | 33072000 | 3568000 | 1003000 | 2104500 | 80362 | 2060700 | 1180300 | 5367800 | 12429000 | 3461700 | 737820 | 76686000 | 46346000 | 207220 | 2214400 |
| VLLESEQFI | SRP14; SRP14 | P37108; H0Y | SRP14_HUMAN | Unmodified | 0 | 0 | 0 | 80871 | 325790 | 0 | 0 | 0 | 0 | 0 | 0 | 0 | 498340 | 330990 | 0 | 0 | 194320 | 152820 |
| VLLESIGSH | PTPRJ; PTPRJ; P | Q12913; A0A | PTPRJ_HUMAN | Unmodified | 777540 | 1193300 | 0 | 418310 | 0 | 0 | 0 | 0 | 5934300 | 123200 | 4771000 | 24676000 | 0 | 0 | 0 | 909130 | 0 | 0 |
| VLLFNATGI | FPGS | Q05932 | FOLC_HUMAN | Unmodified | 50158 | 0 | 0 | 0 | 47367 | 0 | 0 | 0 | 753260 | 58897 | 44159 | 0 | 0 | 0 | 0 | 0 | 0 | 68360 |
| VLLGAHALI | EIF2B4; EIF2B4; | Q9UI10; A0A | EI2BD_HUMAN | Oxidation (M) | 0 | 0 | 0 | 0 | 0 | 0 | 0 | 0 | 0 | 0 | 0 | 0 | 0 | 0 | 0 | 0 | 0 | 0 |
| VLLGETGK | RPS11; RPS11; | P62280; M0I | RS11_HUMAN | Unmodified | 881280 | 664860 | 1018000 | 400090 | 0 | 0 | 86000 | 97455 | 825720 | 0 | 2257500 | 11248000 | 437850 | 24467 | 841010 | 10968 | 76788 | 0 |
| VLLGETGKI | RPS11; RPS11; | P62280; M0I | RS11_HUMAN | Unmodified | 0 | 0 | 0 | 0 | 0 | 0 | 0 | 30414 | 0 | 0 | 0 | 0 | 546310 | 0 | 0 | 0 | 0 | 0 |
| VLLGQDEP | PSMG3 | Q9BT73 | PSMG3_HUMA | Unmodified | 153410 | 0 | 0 | 0 | 2217500 | 930870 | 826490 | 118520 | 0 | 545400 | 714410 | 4531200 | 2263800 | 1734400 | 389090 | 0 | 1289700 | 1198900 |
| VLLGSVSG | SLC25A10; SLC | Q9UBX3; F6I | DIC_HUMAN M | Unmodified | 0 | 0 | 0 | 0 | 0 | 0 | 0 | 0 | 2257800 | 697690 | 0 | 0 | 0 | 0 | 0 | 0 | 0 | 0 |
| VLLIETQR | TMPRSS2; TMPI | O15393; C9J | TMPS2_HUMAN | Unmodified | 28542000 | 59446000 | 2103700 | 2189900 | 0 | 0 | 92332 | 0 | 0 | 1335400 | 0 | 0 | 0 | 0 | 1737600 | 1761400 | 0 | 0 |

| | | | | | | | | | | | | | | | | | | | | | | |
|---|---|---|---|---|---|---|---|---|---|---|---|---|---|---|---|---|---|---|---|---|---|---|
| VLLISDPTTI | IDE; IDE; IDE; ID | P14735; A0A | IDE_HUMAN In | Unmodified | 0 | 0 | 0 | 218920 | 2667300 | 0 | 0 | 0 | 459290 | 0 | 1937200 | 6932300 | 704660 | 176180 | 373390 | 357070 | 382920 | 0 |
| VLLITPGFL( | TIRAP | P58753 | TIRAP_HUMAN | Unmodified | 180900 | 0 | 0 | 0 | 0 | 0 | 7414.2 | 0 | 0 | 0 | 0 | 0 | 0 | 0 | 0 | 0 | 0 | 0 |
| VLLIYDGIH | YOD1 | Q5VVQ6 | OTU1_HUMAN | Unmodified | 1746800 | 1574600 | 0 | 527650 | 0 | 0 | 0 | 0 | 0 | 0 | 0 | 332890 | 0 | 0 | 535500 | 1145700 | 0 | 0 |
| VLLLSSPGL | CCAR2; CCAR2 | Q8N163; E5I | CCAR2_HUMAI | Unmodified | 73774 | 0 | 0 | 0 | 0 | 165370 | 179260 | 0 | 191030 | 25173 | 282530 | 454860 | 0 | 0 | 0 | 0 | 0 | 0 |
| VLLNAEDK | VPS33B; VPS33 | Q9H267; F5I | VP33B_HUMAN | Unmodified | 0 | 0 | 0 | 0 | 534350 | 0 | 0 | 0 | 0 | 0 | 0 | 0 | 0 | 0 | 0 | 0 | 0 | 0 |
| VLLNIGQQ | SLC7A1; SLC7A | P30825; Q5J | SL7A1_HUMAN | Unmodified | 0 | 0 | 464020 | 840810 | 0 | 0 | 0 | 0 | 2424700 | 63621 | 0 | 645720 | 173680 | 0 | 1355600 | 568810 | 0 | 0 |
| VLLNIGQQ | SLC7A1; SLC7A | P30825; Q5J | SL7A1_HUMAN | Oxidation (M) | 0 | 0 | 1112300 | 4261200 | 0 | 0 | 0 | 219470 | 508080 | 0 | 167310 | 3146000 | 177950 | 73328 | 4104200 | 2272600 | 0 | 0 |
| VLLPEYGG | HSPE1; HSPE1; | P61604; B8Z | CH10_HUMAN | Unmodified | 0 | 0 | 1556800 | 0 | 0 | 0 | 3285000 | 0 | 76967000 | 22190000 | 8473200 | 3381400 | 10574000 | 8804300 | 0 | 0 | 575530 | 2800900 |
| VLLPFTR | NUP210 | Q8TEM1 | PO210_HUMAI | Unmodified | 0 | 0 | 0 | 0 | 0 | 0 | 0 | 0 | 66564000 | 8207100 | 1790900 | 852440 | 0 | 0 | 84260 | 62003 | 0 | 0 |
| VLLPTQKID | LTV1 | Q96GA3 | LTV1_HUMAN I | Unmodified | 61826 | 144360 | 0 | 143520 | 0 | 0 | 0 | 0 | 0 | 0 | 0 | 0 | 0 | 0 | 0 | 43803 | 0 | 0 |
| VLLPVLLNK | STXBP3 | O00186 | STXB3_HUMAN | Unmodified | 0 | 0 | 0 | 0 | 10249000 | 2585500 | 403590 | 551300 | 7680700 | 2293900 | 4662100 | 28744000 | 14666000 | 5385500 | 0 | 0 | 3022200 | 9623300 |
| VLLQGKGD | MYH9 | P35579 | MYH9_HUMAN | Unmodified | 0 | 0 | 0 | 1998800 | 24075000 | 0 | 0 | 1468800 | 0 | 802060 | 0 | 4599300 | 0 | 32130 | 0 | 0 | 2934500 | 5444400 |
| VLLQHVHE | ARIH1; ARIH1 | Q9Y4X5; A0A | ARI1_HUMAN E | Unmodified | 0 | 0 | 0 | 0 | 185830 | 0 | 0 | 0 | 0 | 0 | 12337 | 1183500 | 0 | 0 | 0 | 0 | 0 | 58688 |
| VLLQSKDQ | PAICS; PAICS | P22234; E9P | PUR6_HUMAN | Unmodified | 0 | 0 | 0 | 78331 | 166560 | 0 | 159690 | 0 | 0 | 298360 | 0 | 0 | 1286800 | 370180 | 0 | 0 | 554170 | 368370 |
| VLLQSKDQ | PAICS; PAICS | P22234; E9P | PUR6_HUMAN | Unmodified | 0 | 0 | 0 | 0 | 2475400 | 349070 | 1147600 | 1004700 | 0 | 899120 | 0 | 0 | 11223000 | 5032300 | 0 | 0 | 3213500 | 4268000 |
| VLLRPGDF | RAB11FIP1; RAI | Q6WKZ4; E7 | RFIP1_HUMAN | Unmodified | 0 | 0 | 0 | 0 | 2201300 | 0 | 0 | 0 | 0 | 0 | 0 | 0 | 0 | 0 | 0 | 0 | 322200 | 0 |
| VLLSALER | MTHFD1; MTHF | P11586; V9C | C1TC_HUMAN | Unmodified | 0 | 98700 | 903010 | 1196300 | 662180 | 172390 | 0 | 327310 | 5805100 | 0 | 962500 | 37615000 | 671500 | 339690 | 1399700 | 604670 | 474000 | 431180 |
| VLLSDSNLI | CIAPIN1; CIAPI | Q6FI81; H3B | CPIN1_HUMAI | Unmodified | 132840 | 0 | 0 | 248630 | 0 | 0 | 0 | 509300 | 0 | 0 | 577810 | 3121300 | 0 | 0 | 456490 | 201010 | 0 | 0 |
| VLLSEFRPS | PKN1; PKN1 | Q16512; K7E | PKN1_HUMAN | Unmodified | 205670 | 218780 | 507120 | 1166600 | 3409900 | 949050 | 1213300 | 0 | 2701600 | 1500400 | 370140 | 1048000 | 1672700 | 1247200 | 991520 | 1026100 | 2272500 | 2149200 |
| VLLSLEHGI | FASN; FASN | P49327; A0A | FAS_HUMAN F | Unmodified | 0 | 0 | 0 | 340300 | 0 | 0 | 0 | 0 | 0 | 0 | 0 | 432230 | 0 | 0 | 0 | 0 | 0 | 0 |
| VLLTEENK | CDC42BPA; CD | Q5VT25; A0A | MRCKA_HUMA | Unmodified | 0 | 37794 | 180150 | 254090 | 819850 | 38461 | 142400 | 113320 | 3033300 | 456000 | 295490 | 2325800 | 917590 | 975530 | 191670 | 209950 | 371810 | 2435700 |
| VLLTELQPD | EPHA1 | P21709 | EPHA1_HUMAI | Unmodified | 458350 | 349780 | 476890 | 0 | 474810 | 0 | 2488900 | 1383400 | 0 | 0 | 0 | 0 | 0 | 603720 | 2137400 | 556780 | 0 | 0 |
| VLLTMIAR | SEC22B | O75396 | SC22B_HUMAI | Unmodified | 0 | 0 | 0 | 0 | 0 | 0 | 0 | 0 | 0 | 0 | 0 | 0 | 873840 | 346320 | 0 | 0 | 0 | 0 |
| VLLTRPGE( | PTPRK; PTPRK; | Q15262; Q5 | PTPRK_HUMAI | Unmodified | 850020 | 441140 | 202920 | 156170 | 0 | 0 | 0 | 0 | 4477500 | 132780 | 374350 | 273660 | 0 | 0 | 98848 | 87312 | 0 | 0 |
| VLLTTQGVI | DYNC1H1; DYN | Q14204; A0A | DYHC1_HUMA | Unmodified | 0 | 0 | 0 | 0 | 0 | 0 | 0 | 0 | 1523800 | 0 | 0 | 688580 | 0 | 0 | 0 | 0 | 0 | 0 |
| VLLVGADN | MRPL21; MRPL | Q7Z2W9; H3 | RM21_HUMAN | Unmodified | 0 | 0 | 0 | 0 | 0 | 0 | 0 | 0 | 1461200 | 0 | 0 | 0 | 0 | 0 | 0 | 0 | 0 | 0 |
| VLLVLCGG | ANXA4; ANXA4 | P09525; Q6I | ANXA4_HUMAI | Unmodified | 0 | 0 | 0 | 0 | 542630 | 143120 | 0 | 240140 | 2390900 | 1169800 | 2201200 | 0 | 0 | 0 | 204500 | 0 | 0 | 485660 |
| VLLVLELQC | GANAB | Q14697 | GANAB_HUMA | Unmodified | 0 | 0 | 0 | 0 | 0 | 0 | 0 | 0 | 3072800 | 18707000 | 249500 | 0 | 0 | 0 | 0 | 0 | 0 | 0 |
| VLLVPGPEI | VAT1; VAT1 | Q99536; K7E | VAT1_HUMAN | Unmodified | 0 | 66991 | 264980 | 0 | 17361000 | 4794000 | 0 | 0 | 6756100 | 1690000 | 2152900 | 1125000 | 2038400 | 1703900 | 90230 | 0 | 1380800 | 5869900 |
| VLLVPGPEI | VAT1; VAT1 | Q99536; K7E | VAT1_HUMAN | Unmodified | 0 | 0 | 0 | 0 | 75428000 | 9276300 | 3537600 | 4183400 | 1059600 | 4066800 | 0 | 204810 | 0 | 6324100 | 0 | 0 | 16445000 | 15514000 |
| VLMEFVEN | SLC44A1 | Q8WWI5 | CTL1_HUMAN | Unmodified | 740130 | 764980 | 575440 | 662280 | 1309400 | 0 | 0 | 0 | 0 | 1195900 | 244110 | 3421600 | 852250 | 0 | 1032100 | 807380 | 0 | 0 |
| VLMEFVEN | SLC44A1 | Q8WWI5 | CTL1_HUMAN | Oxidation (M) | 262870 | 519950 | 155120 | 0 | 0 | 0 | 0 | 0 | 0 | 166200 | 0 | 112990 | 0 | 0 | 647500 | 313010 | 0 | 0 |
| VLMEKEFP | S100A10 | P60903 | S10AA_HUMAI | Oxidation (M) | 0 | 0 | 0 | 0 | 1178200 | 0 | 0 | 0 | 0 | 0 | 0 | 0 | 258870 | 0 | 0 | 0 | 0 | 0 |
| VLMLTLQN | OXSR1; OXSR1 | O95747; C9J | OXSR1_HUMAI | Oxidation (M) | 0 | 0 | 0 | 0 | 618690 | 0 | 0 | 156810 | 578630 | 0 | 235950 | 1523900 | 710930 | 701780 | 0 | 0 | 0 | 0 |
| VLMYGQPY | BSCL2; BSCL2; | Q96G97; E9I | BSCL2_HUMAI | Unmodified | 0 | 0 | 0 | 0 | 835470 | 0 | 0 | 0 | 1189700 | 0 | 0 | 0 | 0 | 0 | 0 | 0 | 119790 | 0 |
| VLNCDTIS( | PLXNB2 | O15031 | PLXB2_HUMAN | Unmodified | 410000 | 0 | 3201000 | 1221300 | 2951200 | 0 | 764490 | 0 | 1180300 | 1062100 | 2539700 | 2942700 | 1743900 | 1301000 | 2084600 | 1239900 | 1240000 | 3175500 |
| VLNCDTIS( | PLXNB2 | O15031 | PLXB2_HUMAN | Unmodified | 0 | 0 | 0 | 0 | 823500 | 0 | 336800 | 100780 | 0 | 98865 | 0 | 0 | 446370 | 0 | 0 | 0 | 578150 | 663320 |
| VLNCDTIT( | PLXNA2 | O75051 | PLXA2_HUMAN | Unmodified | 0 | 0 | 0 | 0 | 0 | 0 | 0 | 0 | 1175300 | 400050 | 0 | 598070 | 0 | 0 | 0 | 0 | 123050 | 0 |
| VLNDYLEE | IQGAP3; IQGAP | Q86VI3; F2Z | IQGA3_HUMAN | Unmodified | 561440 | 502460 | 0 | 541390 | 0 | 364320 | 0 | 0 | 1407200 | 360180 | 192130 | 1081500 | 0 | 0 | 0 | 156960 | 0 | 0 |
| VLNEECDQ | GRB2; GRB2; G | P62993; J3Q | GRB2_HUMAN | Unmodified | 1124300 | 798130 | 1612300 | 0 | 0 | 0 | 0 | 0 | 256430 | 0 | 0 | 917590 | 0 | 0 | 1850200 | 1004600 | 0 | 0 |
| VLNEECDQ | GRB2; GRB2; G | P62993; J3Q | GRB2_HUMAN | Unmodified | 861410 | 466140 | 0 | 0 | 0 | 0 | 0 | 0 | 0 | 0 | 0 | 80866 | 0 | 0 | 293410 | 0 | 0 | 0 |
| VLNEECDQ | GRB2; GRB2; G | P62993; J3Q | GRB2_HUMAN | Unmodified | 1602000 | 1089700 | 0 | 0 | 843510 | 0 | 614960 | 0 | 613970 | 0 | 0 | 589240 | 1404000 | 0 | 0 | 0 | 2611000 | 0 |
| VLNELTQLI | RAI14 | Q9P0K7 | RAI14_HUMAN | Unmodified | 0 | 0 | 0 | 0 | 27025000 | 0 | 0 | 0 | 0 | 0 | 0 | 0 | 0 | 0 | 0 | 0 | 2773800 | 0 |
| VLNELYSVI | SRGAP2; SRGA | O75044; P0I | SRGP2_HUMAI | Unmodified | 0 | 0 | 0 | 0 | 3616800 | 0 | 0 | 0 | 0 | 0 | 0 | 435930 | 0 | 0 | 0 | 0 | 0 | 422980 |
| VLNGPEGD | NOMO1; NOMC | Q15155; P6 | NOMO1_HUMA | Unmodified | 0 | 0 | 0 | 0 | 0 | 0 | 0 | 0 | 5870100 | 153550 | 131530 | 0 | 0 | 0 | 0 | 0 | 0 | 0 |
| VLNLIQSW | TOM1; TOM1; T | O60784; V9C | TOM1_HUMAN | Unmodified | 1862300 | 2021600 | 115360 | 1099500 | 0 | 0 | 0 | 0 | 553810 | 202880 | 0 | 209750 | 311170 | 275360 | 2444600 | 951570 | 0 | 0 |
| VLNLSDNR | LRRC7; LRRC7; | Q96NW7; A0 | LRRC7_HUMAI | Unmodified | 0 | 0 | 708060 | 1314900 | 0 | 0 | 0 | 0 | 0 | 0 | 0 | 6477500 | 0 | 0 | 682920 | 0 | 0 | 0 |
| VLNLSDNR | LRRC7; LRRC7; | Q96NW7; A0 | LRRC7_HUMAI | Unmodified | 0 | 0 | 0 | 0 | 23664000 | 1354800 | 141790 | 129140 | 0 | 0 | 0 | 0 | 0 | 0 | 0 | 0 | 490160 | 494670 |
| VLNNMEIG | PGK1 | P00558 | PGK1_HUMAN | Oxidation (M) | 0 | 0 | 2814100 | 3576300 | 7090700 | 0 | 2018300 | 2065900 | 4568000 | 4982200 | 6330900 | 24870000 | 7012200 | 4619100 | 4108800 | 3765100 | 13516000 | 12668000 |
| VLNNMEIG | PGK1 | P00558 | PGK1_HUMAN | Unmodified | 0 | 0 | 0 | 377600 | 3587700 | 2512100 | 0 | 244500 | 4930500 | 2134800 | 970470 | 3436800 | 0 | 2028600 | 503210 | 610370 | 6960500 | 10502000 |
| VLNNMEIG | PGK1 | P00558 | PGK1_HUMAN | Oxidation (M) | 0 | 0 | 0 | 0 | 0 | 0 | 0 | 0 | 0 | 0 | 0 | 0 | 0 | 0 | 0 | 0 | 861680 | 0 |
| VLNNMEIG | PGK1 | P00558 | PGK1_HUMAN | 2 Oxidation (M) | 0 | 0 | 0 | 0 | 184180 | 0 | 0 | 0 | 0 | 0 | 35064 | 0 | 0 | 0 | 0 | 0 | 8594000 | 0 |
| VLNQELR | APOD; APOD; A | P05090; C9J | APOD_HUMAN | Unmodified | 0 | 0 | 0 | 0 | 0 | 0 | 0 | 0 | 1.05E+08 | 1605400 | 0 | 0 | 0 | 0 | 0 | 0 | 0 | 0 |
| VLNQQLTN | DNM1; DNM1; I | Q05193; A0A | DYN1_HUMAN | Unmodified | 0 | 0 | 0 | 0 | 93960 | 0 | 0 | 0 | 914540 | 360770 | 0 | 0 | 0 | 0 | 0 | 0 | 0 | 0 |
| VLNSYWV( | RPL15; RPL15; I | P61313; E7E | RL15_HUMAN | Unmodified | 0 | 0 | 0 | 0 | 0 | 0 | 0 | 107410 | 1692900 | 0 | 0 | 45267000 | 0 | 0 | 0 | 0 | 0 | 0 |
| VLNTEANV | QSOX1 | O00391 | QSOX1_HUMAI | Unmodified | 0 | 0 | 0 | 0 | 102790 | 0 | 0 | 0 | 2875500 | 226830 | 157540 | 0 | 176960 | 0 | 0 | 0 | 0 | 0 |
| VLNTEANV | QSOX1 | O00391 | QSOX1_HUMAI | Unmodified | 0 | 0 | 0 | 0 | 0 | 0 | 0 | 238920 | 1200100 | 44859 | 0 | 0 | 0 | 0 | 0 | 0 | 0 | 0 |
| VLNTNIDGI | RPS18; RPS18; | P62269; A0A | RS18_HUMAN | Unmodified | 3977400 | 3055000 | 1297700 | 99002 | 0 | 0 | 433260 | 445350 | 81331 | 436890 | 1816300 | 66444000 | 419610 | 0 | 2637100 | 2365200 | 222510 | 166990 |
| VLNYVDWI | C1R; C1R; C1R; | P00736; B4I | C1R_HUMAN C | Unmodified | 0 | 0 | 0 | 0 | 0 | 0 | 0 | 0 | 812820 | 194790 | 0 | 0 | 0 | 0 | 0 | 0 | 0 | 292290 |
| VLPANSMA | ATP2B4 | P23634 | AT2B4_HUMAN | Unmodified | 0 | 0 | 0 | 0 | 0 | 0 | 0 | 0 | 69580 | 67525 | 1124500 | 2864200 | 0 | 0 | 0 | 0 | 0 | 0 |
| VLPDVLQV | IGSF8; IGSF8 | Q969P0; C9J | IGSF8_HUMAN | Unmodified | 24393000 | 18667000 | 0 | 10935000 | 1189700 | 452320 | 0 | 0 | 1660400 | 201230 | 1082800 | 592280 | 4076800 | 0 | 12285000 | 16696000 | 0 | 578620 |
| VLPDVLQV | IGSF8; IGSF8 | Q969P0; C9J | IGSF8_HUMAN | Unmodified | 0 | 0 | 0 | 0 | 723750 | 0 | 0 | 0 | 0 | 0 | 0 | 0 | 0 | 0 | 0 | 0 | 178560 | 1025200 |
| VLPEFDTP( | HEXB; HEXB | P07686; Q5U | HEXB_HUMAN | Unmodified | 875980 | 650620 | 1832200 | 3450900 | 322720 | 259050 | 0 | 37619 | 10496000 | 1316600 | 1410200 | 888800 | 3838100 | 1313600 | 2879900 | 0 | 0 | 98505 |
| VLPEGGAQ | AGRN; AGRN; A | O00468; A0A | AGRIN_HUMAI | Unmodified | 187020 | 61073 | 0 | 242910 | 0 | 0 | 0 | 0 | 0 | 0 | 0 | 0 | 0 | 0 | 0 | 93237 | 0 | 0 |
| VLPFFERPI | ERO1A | Q96HE7 | ERO1A_HUMAI | Unmodified | 30034 | 0 | 88890 | 0 | 0 | 0 | 0 | 0 | 4009400 | 1498800 | 1836300 | 580480 | 0 | 0 | 246280 | 209920 | 0 | 0 |
| VLPFFERPI | ERO1A | Q96HE7 | ERO1A_HUMAI | Unmodified | 0 | 0 | 244590 | 0 | 0 | 0 | 0 | 0 | 3218000 | 5118800 | 1893900 | 0 | 0 | 0 | 0 | 0 | 0 | 512870 |
| VLPGDYEIL | NOMO1; NOMC | Q15155; P6 | NOMO1_HUMA | Unmodified | 0 | 0 | 0 | 0 | 0 | 0 | 0 | 0 | 32582000 | 10072000 | 4110900 | 0 | 0 | 0 | 0 | 0 | 0 | 0 |
| VLPGHFNT | MBLAC2 | Q68D91 | MBLC2_HUMAI | Unmodified | 284570 | 292290 | 1509600 | 529710 | 0 | 0 | 0 | 0 | 0 | 0 | 0 | 4388200 | 0 | 0 | 843480 | 957610 | 0 | 0 |
| VLPGMHHI | CELF1; CELF1 | Q92879; G5I | CELF1_HUMAN | Unmodified | 615670 | 523040 | 314050 | 641580 | 0 | 0 | 333750 | 0 | 0 | 0 | 0 | 698890 | 516940 | 0 | 527980 | 500360 | 0 | 0 |
| VLPGMHHI | CELF1; CELF1 | Q92879; G5I | CELF1_HUMAN | Oxidation (M) | 1338500 | 2544300 | 56206 | 126910 | 0 | 0 | 0 | 0 | 0 | 0 | 0 | 0 | 371060 | 0 | 0 | 106200 | 0 | 0 |
| VLPGVDAL | PGK1 | P00558 | PGK1_HUMAN | Unmodified | 705950 | 2432400 | 5886100 | 12435000 | 92826000 | 13326000 | 22413000 | 13439000 | 47919000 | 50532000 | 30111000 | 1.2E+08 | 62299000 | 47046000 | 12881000 | 9628000 | 2.69E+08 | 3.3E+08 |
| VLPIEQSLS | PROM1 | O43490 | PROM1_HUMA | Unmodified | 0 | 0 | 68584 | 0 | 129930 | 0 | 60087 | 71592 | 0 | 0 | 297140 | 5181200 | 247540 | 188450 | 0 | 0 | 65109 | 293540 |
| VLPIMFPAL | PPP2R5D; PPP2 | Q14738; H0 | 2A5D_HUMAN | Oxidation (M) | 0 | 0 | 173480 | 0 | 0 | 0 | 0 | 0 | 0 | 0 | 0 | 8133100 | 0 | 0 | 504070 | 320040 | 0 | 0 |
| VLPLEALVT | GCN1 | Q92616 | GCN1_HUMAN | Unmodified | 152910 | 118660 | 0 | 141990 | 0 | 119470 | 346450 | 0 | 492040 | 0 | 93154 | 1101200 | 265380 | 0 | 328820 | 0 | 0 | 277350 |
| VLPLEEAYF | EHD4 | Q9H223 | EHD4_HUMAN | Unmodified | 0 | 1164200 | 0 | 0 | 0 | 0 | 0 | 0 | 0 | 0 | 0 | 0 | 0 | 0 | 0 | 0 | 0 | 0 |
| VLPLSVTHF | DHRS11; DHRS | Q6UWP2; A0 | DHR11_HUMAI | Unmodified | 0 | 0 | 0 | 0 | 0 | 0 | 0 | 0 | 0 | 0 | 0 | 1148600 | 0 | 0 | 0 | 0 | 0 | 0 |
| VLPNGEVS | C1orf226 | A1L170 | CA226_HUMAN | Unmodified | 0 | 0 | 0 | 435460 | 1461500 | 299640 | 1134300 | 0 | 0 | 0 | 0 | 0 | 2260100 | 2422900 | 0 | 0 | 0 | 1055800 |
| VLPPVPTE( | AGPAT1 | Q99943 | PLCA_HUMAN | Unmodified | 0 | 0 | 0 | 0 | 0 | 0 | 0 | 0 | 1050800 | 35571 | 0 | 0 | 0 | 0 | 0 | 0 | 0 | 0 |
| VLPPVPTE( | AGPAT1 | Q99943 | PLCA_HUMAN | Unmodified | 0 | 0 | 0 | 0 | 0 | 0 | 0 | 0 | 2427700 | 0 | 0 | 0 | 1835000 | 0 | 0 | 0 | 0 | 0 |
| VLPSDLDLI | THEM6 | Q8WUY1 | THEM6_HUMAI | Unmodified | 0 | 0 | 0 | 0 | 1309200 | 0 | 859650 | 0 | 5268500 | 195000 | 0 | 0 | 0 | 0 | 0 | 0 | 1021100 | 569550 |
| VLPSDLDLI | THEM6 | Q8WUY1 | THEM6_HUMAI | Oxidation (M) | 0 | 0 | 0 | 0 | 0 | 0 | 0 | 0 | 4255900 | 759180 | 77069 | 1394200 | 0 | 0 | 0 | 0 | 0 | 0 |
| VLPSITTEIL | PHB; PHB; PHB | P35232; C9J | PHB_HUMAN P | Unmodified | 0 | 0 | 0 | 0 | 0 | 0 | 0 | 0 | 1464000 | 0 | 0 | 0 | 0 | 0 | 0 | 0 | 0 | 0 |
| VLPSIVNEV | PHB2; PHB2; PI | Q99623; F5C | PHB2_HUMAN | Unmodified | 0 | 0 | 0 | 0 | 0 | 0 | 0 | 0 | 1155400 | 0 | 0 | 557790 | 0 | 0 | 0 | 0 | 0 | 0 |
| VLPTQPNP | EMC1 | Q8N766 | EMC1_HUMAN | Unmodified | 0 | 0 | 0 | 0 | 0 | 0 | 0 | 0 | 7710600 | 539740 | 559620 | 0 | 0 | 0 | 0 | 0 | 0 | 0 |
| VLPVGDEV | NT5E; NT5E | P21589; Q9 | 5NTD_HUMAN | Unmodified | 424340 | 0 | 0 | 0 | 75353000 | 22221000 | 0 | 0 | 0 | 0 | 126260 | 0 | 0 | 0 | 0 | 0 | 7432100 | 33136000 |
| VLPWLSPE | STRADA; STRAD | Q7RTN6; A0 | STRAA_HUMAN | Unmodified | 0 | 0 | 24687 | 0 | 0 | 0 | 0 | 0 | 280540 | 0 | 73979 | 0 | 0 | 0 | 123110 | 50957 | 0 | 0 |
| VLPYHPAQ | CHPF; CHPF | Q8IZ52; F8W | CHSS2_HUMAI | Unmodified | 0 | 0 | 0 | 0 | 0 | 0 | 0 | 0 | 524530 | 0 | 254940 | 0 | 0 | 0 | 0 | 0 | 0 | 0 |
| VLQAGSSR | ACE; ACE | P12821; A0A | ACE_HUMAN A | Unmodified | 2899700 | 1249200 | 0 | 0 | 0 | 0 | 0 | 0 | 4139900 | 640030 | 0 | 321360 | 0 | 0 | 0 | 0 | 0 | 1899200 |
| VLQALEGLI | RPS19; RPS19 | P39019; A0A | RS19_HUMAN | Unmodified | 2532000 | 1986600 | 1052500 | 951760 | 421340 | 0 | 265330 | 160600 | 725560 | 134300 | 2486700 | 12303000 | 896200 | 712690 | 1272800 | 1073000 | 0 | 20217 |
| VLQALGSE | MYO1C; MYO1( | O00159; F5I | MYO1C_HUMA | Unmodified | 9593900 | 0 | 10337000 | 9628600 | 6961300 | 0 | 0 | 0 | 1305300 | 0 | 0 | 3807600 | 0 | 0 | 19158000 | 9039600 | 1781600 | 0 |
| VLQALGSE | MYO1C; MYO1( | O00159; F5I | MYO1C_HUMA | Unmodified | 2642900 | 4020900 | 0 | 6012300 | 1333900 | 0 | 0 | 0 | 756070 | 0 | 0 | 497810 | 1309600 | 0 | 5567200 | 3247800 | 0 | 0 |
| VLQALGSE | MYO1C; MYO1( | O00159; F5I | MYO1C_HUMA | Unmodified | 814410 | 925190 | 0 | 10525000 | 33490000 | 13424000 | 0 | 0 | 0 | 552810 | 0 | 0 | 8023600 | 7050800 | 0 | 747760 | 5298500 | 9305500 |
| VLQAQGSS | TOMM34 | Q15785 | TOM34_HUMAI | Unmodified | 0 | 110080 | 262150 | 410370 | 0 | 0 | 0 | 0 | 0 | 0 | 0 | 2328800 | 0 | 0 | 425930 | 342940 | 0 | 0 |
| VLQATVVA | HSPE1; HSPE1- | P61604; S4F | CH10_HUMAN | Unmodified | 0 | 0 | 426910 | 153420 | 693650 | 0 | 2287600 | 591750 | 13065000 | 3892600 | 541800 | 364310 | 2458500 | 1727700 | 166490 | 0 | 251520 | 0 |
| VLQATVVA | HSPE1; HSPE1- | P61604; S4F | CH10_HUMAN | Unmodified | 0 | 0 | 0 | 0 | 0 | 0 | 253990 | 0 | 0 | 303540 | 0 | 0 | 0 | 0 | 0 | 0 | 0 | 0 |
| VLQATVVA | HSPE1; HSPE1 | P61604; B8Z | CH10_HUMAN | Unmodified | 0 | 0 | 0 | 0 | 0 | 0 | 1689100 | 597540 | 0 | 1854200 | 0 | 0 | 0 | 854030 | 0 | 0 | 457020 | 0 |
| VLQCHKPV | NUDT4B; NUDT | A0A024RBG | NUD4B_HUMA | Unmodified | 0 | 0 | 0 | 0 | 125370 | 0 | 63788 | 27482 | 0 | 0 | 0 | 0 | 0 | 477950 | 0 | 22882 | 160200 | 0 |
| VLQDMGLI | ILF3 | Q12906 | ILF3_HUMAN I | Oxidation (M) | 763110 | 455170 | 272900 | 260460 | 0 | 0 | 0 | 0 | 0 | 0 | 148890 | 258710 | 0 | 0 | 0 | 273660 | 0 | 0 |
| VLQDMGLI | ILF3 | Q12906 | ILF3_HUMAN I | Unmodified | 582680 | 787160 | 148440 | 0 | 0 | 0 | 0 | 0 | 0 | 0 | 208030 | 384920 | 0 | 0 | 0 | 207230 | 0 | 0 |
| VLQDQLK | GOLM1 | Q8NBJ4 | GOLM1_HUMA | Unmodified | 0 | 0 | 0 | 0 | 95274 | 0 | 83969 | 33643 | 41750000 | 2867400 | 287820 | 399400 | 163140 | 124550 | 261430 | 77517 | 0 | 37449 |
| VLQDTLGR | AVIL | O75366 | AVIL_HUMAN A | Unmodified | 0 | 0 | 0 | 0 | 0 | 0 | 0 | 0 | 0 | 0 | 1051600 | 4268600 | 0 | 0 | 0 | 0 | 0 | 0 |
| VLQDTLGR | AVIL | O75366 | AVIL_HUMAN A | Unmodified | 0 | 0 | 0 | 0 | 0 | 0 | 0 | 0 | 0 | 0 | 0 | 5670500 | 0 | 0 | 0 | 0 | 0 | 0 |
| VLQELDMS | FMNL1; FMNL1 | O95466; A0A | FMNL1_HUMAI | Unmodified | 0 | 0 | 0 | 0 | 2514300 | 47274 | 20383 | 0 | 0 | 0 | 0 | 0 | 0 | 0 | 0 | 0 | 0 | 0 |
| VLQGDLVM | FASN; FASN | P49327; A0A | FAS_HUMAN F | Unmodified | 0 | 0 | 0 | 0 | 0 | 0 | 0 | 0 | 0 | 0 | 0 | 5882100 | 0 | 0 | 0 | 0 | 0 | 0 |
| VLQGDLVM | FASN; FASN | P49327; A0A | FAS_HUMAN F | Oxidation (M) | 0 | 0 | 0 | 0 | 0 | 0 | 0 | 0 | 0 | 0 | 0 | 15184000 | 0 | 0 | 897800 | 565360 | 0 | 0 |
| VLQHILNP | TRRAP; TRRAP; | Q9Y4A5; F2Z | TRRAP_HUMAN | Unmodified | 139040 | 199790 | 0 | 0 | 0 | 0 | 0 | 0 | 0 | 0 | 0 | 499500 | 0 | 0 | 52021 | 22015 | 0 | 0 |
| VLQHYQES | MTCH2 | Q9Y6C9 | MTCH2_HUMA | Unmodified | 0 | 0 | 0 | 0 | 0 | 0 | 208140 | 0 | 1654800 | 538730 | 461130 | 0 | 0 | 78346 | 0 | 0 | 0 | 0 |
| VLQIIPYEFI | PTPRA | P18433 | PTPRA_HUMAN | Unmodified | 0 | 0 | 0 | 0 | 0 | 0 | 0 | 0 | 1007200 | 0 | 471170 | 4968700 | 0 | 0 | 0 | 0 | 0 | 0 |
| VLQIVTNRI | AP2A1 | O95782 | AP2A1_HUMAN | Unmodified | 148640 | 212620 | 383110 | 0 | 0 | 0 | 0 | 0 | 295430 | 0 | 0 | 2958400 | 0 | 0 | 529860 | 370610 | 0 | 0 |

| | | | | | | | | | | | | | | | | | | | | | | |
|---|---|---|---|---|---|---|---|---|---|---|---|---|---|---|---|---|---|---|---|---|---|---|
| VLQLINDN | HYOU1; HYOU1 | Q9Y4L1; J3Q | HYOU1_HUMA | Unmodified | 0 | 0 | 0 | 0 | 0 | 0 | 0 | 0 | 2008100 | 0 | 1048900 | 0 | 0 | 0 | 0 | 0 | 0 | 0 |
| VLQLINDN | HYOU1; HYOU1 | Q9Y4L1; E9F | HYOU1_HUMA | Unmodified | 0 | 0 | 593680 | 0 | 0 | 0 | 0 | 0 | 29428000 | 17698000 | 2620500 | 747600 | 0 | 769270 | 0 | 0 | 0 | 0 |
| VLQLINDN | HYOU1; HYOU1 | Q9Y4L1; E9F | HYOU1_HUMA | Unmodified | 0 | 0 | 0 | 0 | 0 | 0 | 882230 | 0 | 0 | 18982000 | 0 | 0 | 705050 | 1060000 | 0 | 0 | 0 | 303980 |
| VLQLSDTAI | CORO7; CORO | P57737; I3L1 | CORO7_HUMA | Unmodified | 693580 | 0 | 0 | 0 | 35051 | 0 | 0 | 113200 | 0 | 286420 | 0 | 1746100 | 57846 | 234660 | 0 | 0 | 0 | 0 |
| VLQLSDTAI | CORO7; CORO | P57737; I3L1 | CORO7_HUMA | Unmodified | 0 | 0 | 0 | 0 | 0 | 0 | 0 | 0 | 511290 | 0 | 301420 | 2950500 | 0 | 0 | 0 | 0 | 0 | 0 |
| VLQLTSWD | VCL | P18206 | VINC_HUMAN | Unmodified | 95890 | 190250 | 0 | 0 | 0 | 0 | 0 | 0 | 0 | 0 | 0 | 2849800 | 0 | 0 | 1165500 | 237430 | 63517 | 0 |
| VLQLTSWD | VCL | P18206 | VINC_HUMAN | Oxidation (M) | 437570 | 180440 | 0 | 0 | 0 | 0 | 0 | 0 | 0 | 0 | 0 | 1985800 | 0 | 0 | 452280 | 387620 | 0 | 0 |
| VLQLYPNN | FKBP4; FKBP4 | Q02790; H0 | FKBP4_HUMAN | Unmodified | 592900 | 436100 | 2353200 | 2397800 | 3698800 | 925450 | 3515500 | 2558100 | 2170200 | 1152000 | 8164700 | 30908000 | 20730000 | 13284000 | 2220500 | 1972700 | 5497500 | 3415600 |
| VLQNIIK | MON2; MON2 | Q7Z3U7; A0 | MON2_HUMAN | Unmodified | 0 | 0 | 0 | 0 | 0 | 0 | 156030 | 85009 | 577040 | 0 | 250040 | 995210 | 0 | 0 | 0 | 0 | 0 | 0 |
| VLQQFADN | HSD17B4; HSD | P51659; E7E | DHB4_HUMAN | Unmodified | 0 | 0 | 254550 | 3149400 | 282690 | 2071400 | 0 | 0 | 2679300 | 102340 | 897720 | 47780 | 0 | 0 | 844950 | 0 | 0 | 0 |
| VLQQFADN | HSD17B4; HSD | P51659; E7E | DHB4_HUMAN | Unmodified | 541940 | 0 | 0 | 0 | 762500 | 0 | 253250 | 0 | 7412100 | 1811700 | 452680 | 294330 | 0 | 0 | 0 | 0 | 0 | 0 |
| VLQQKLEA | USP14; USP14 | P54578; A6N | UBP14_HUMAN | Unmodified | 0 | 0 | 0 | 0 | 0 | 0 | 0 | 0 | 0 | 0 | 618930 | 1510900 | 0 | 0 | 0 | 169210 | 0 | 0 |
| VLQQKLEA | USP14; USP14 | P54578; A6N | UBP14_HUMAN | Unmodified | 529030 | 414480 | 0 | 1511200 | 1332700 | 0 | 0 | 0 | 2351200 | 749060 | 2990600 | 16386000 | 1809000 | 778470 | 618670 | 559010 | 499690 | 1241400 |
| VLQQLLVL | CDS1 | Q92903 | CDS1_HUMAN | Unmodified | 0 | 0 | 0 | 0 | 0 | 0 | 0 | 0 | 3650800 | 1019400 | 4650200 | 470140 | 0 | 0 | 0 | 0 | 0 | 0 |
| VLQQLRSR | GSDMD; GSDM | P57764; G3 | GSDMD_HUMA | Unmodified | 0 | 0 | 0 | 0 | 0 | 0 | 0 | 0 | 0 | 251650 | 0 | 0 | 0 | 0 | 0 | 0 | 1033000 | 0 |
| VLQQQGS | C11orf52 | Q96A22 | CK052_HUMAN | Unmodified | 226220 | 0 | 61607 | 0 | 0 | 0 | 0 | 0 | 269730 | 0 | 0 | 94026 | 0 | 0 | 357890 | 207720 | 0 | 0 |
| VLQQVKAE | MYH10 | P35580 | MYH10_HUMA | Unmodified | 0 | 0 | 0 | 0 | 0 | 35710 | 297570 | 73229 | 0 | 0 | 0 | 0 | 0 | 757990 | 0 | 0 | 0 | 0 |
| VLQRNCA | MYH9; MYH10 | P35579; P35 | MYH9_HUMAN | Unmodified | 0 | 0 | 0 | 0 | 2935900 | 406120 | 2902900 | 1017900 | 0 | 939400 | 0 | 0 | 0 | 0 | 0 | 0 | 990100 | 852720 |
| VLQSALAA | ILF2; ILF2 | Q12905; B4I | ILF2_HUMAN In | Unmodified | 5104900 | 2829600 | 1192300 | 354060 | 0 | 0 | 0 | 0 | 191080 | 70146 | 0 | 20991000 | 211130 | 0 | 896020 | 1385900 | 0 | 0 |
| VLQSEFCN | LIN7C; LIN7C | Q9NUP9; G3 | LIN7C_HUMAN | Unmodified | 0 | 0 | 0 | 1425200 | 0 | 0 | 0 | 0 | 3888900 | 0 | 0 | 2015600 | 0 | 0 | 2466600 | 3170100 | 0 | 0 |
| VLQSEFCT | LIN7A; LIN7A; L | O14910; H0 | LIN7A_HUMAN | Unmodified | 0 | 0 | 0 | 0 | 1151600 | 0 | 0 | 60830 | 612630 | 0 | 0 | 0 | 0 | 0 | 524590 | 500420 | 0 | 0 |
| VLQTAGMF | FARP2; FARP2; | O94887; C9J | FARP2_HUMAN | Unmodified | 0 | 107840 | 35426 | 505510 | 0 | 0 | 0 | 0 | 420350 | 24901 | 31907 | 0 | 0 | 0 | 594420 | 159970 | 0 | 0 |
| VLQTEQAV | AAMDC; AAMD | Q9H7C9; E9 | AAMDC_HUMA | Unmodified | 70778 | 0 | 0 | 303840 | 223730 | 303900 | 68454 | 18683 | 0 | 0 | 0 | 0 | 298460 | 90135 | 344840 | 289340 | 0 | 1014300 |
| VLQTLGVLI | SIL1; SIL1 | Q9H173; D6 | SIL1_HUMAN N | Unmodified | 0 | 0 | 0 | 0 | 0 | 0 | 0 | 0 | 366890 | 0 | 0 | 0 | 0 | 0 | 0 | 0 | 0 | 0 |
| VLQTSLPEF | ADAM15 | Q13444 | ADA15_HUMAN | Unmodified | 0 | 0 | 0 | 0 | 0 | 0 | 0 | 0 | 1007800 | 0 | 0 | 0 | 0 | 0 | 0 | 0 | 0 | 0 |
| VLQYLQQE | PDE2A; PDE2A; | O00408; H0 | PDE2A_HUMAN | Unmodified | 0 | 0 | 0 | 0 | 0 | 0 | 0 | 0 | 0 | 0 | 0 | 4697800 | 0 | 0 | 0 | 0 | 0 | 0 |
| VLRARDDL | ATP6V1E1; ATP | P36543; C9J | VATE1_HUMAN | Unmodified | 0 | 0 | 0 | 0 | 3189600 | 230390 | 12386000 | 23813 | 0 | 699950 | 0 | 0 | 3553400 | 4792500 | 0 | 0 | 695810 | 1408400 |
| VLRDNIQG | H4C1 | P62805 | H4_HUMAN Hi | Unmodified | 0 | 143570 | 25202 | 0 | 444840 | 78567 | 214090 | 133530 | 0 | 86071 | 0 | 9190900 | 0 | 0 | 0 | 0 | 166500 | 0 |
| VLRDNIQG | H4C1 | P62805 | H4_HUMAN Hi | Unmodified | 23202000 | 15874000 | 135890 | 0 | 1194100 | 3381400 | 5100900 | 14266000 | 1576000 | 1431500 | 3220100 | 2.22E+08 | 234720 | 0 | 0 | 64280 | 82880 | 1193100 |
| VLRGASWI | SUMF2; SUMF2 | Q8NBJ7; J3Q | SUMF2_HUMA | Unmodified | 0 | 0 | 0 | 0 | 0 | 0 | 0 | 0 | 9818200 | 1769900 | 0 | 402580 | 0 | 0 | 0 | 143090 | 0 | 0 |
| VLRGASWI | SUMF2; SUMF2 | Q8NBJ7; J3Q | SUMF2_HUMA | Unmodified | 0 | 0 | 0 | 0 | 0 | 0 | 0 | 0 | 0 | 1575100 | 0 | 0 | 0 | 0 | 0 | 0 | 0 | 0 |
| VLRGVEVT | GATD3B; GATD | A0A0B4J2D5 | GAL3B_HUMAN | Unmodified | 266880 | 0 | 0 | 0 | 0 | 0 | 2180100 | 0 | 3515100 | 2393500 | 0 | 0 | 2320800 | 383270 | 0 | 0 | 0 | 662090 |
| VLRHEEFE | GLOD4; GLOD4 | I3L1F4; F6TL | I3L1F4_HUMAN | Unmodified | 171270 | 0 | 94516 | 471930 | 1803200 | 256050 | 328920 | 262010 | 353280 | 479070 | 663280 | 2073700 | 3131400 | 1793000 | 242220 | 177220 | 0 | 744670 |
| VLRNESPP | CASP8AP2; CA | Q9UKL3; A0 | C8AP2_HUMAN | Unmodified | 0 | 303770 | 0 | 0 | 0 | 0 | 0 | 0 | 0 | 0 | 0 | 0 | 874310 | 0 | 0 | 0 | 0 | 0 |
| VLRNVSGQ | LRRC20; LRRC2 | Q8TCA0; X6 | LRC20_HUMAN | Unmodified | 0 | 0 | 0 | 0 | 8012000 | 0 | 1101600 | 125820 | 0 | 681860 | 0 | 0 | 668170 | 4992700 | 0 | 0 | 5729800 | 0 |
| VLRPPGGG | JPT1; JPT1 | Q9UK76; J3K | JUPI1_HUMAN | Unmodified | 0 | 0 | 0 | 7827900 | 13582000 | 0 | 498750 | 337410 | 829250 | 513290 | 814900 | 3035700 | 9153500 | 7082300 | 18507000 | 7933000 | 4800300 | 8583700 |
| VLRPQVTA | DYNC1H1; DYN | Q14204; A0 | DYHC1_HUMA | Oxidation (M) | 0 | 0 | 0 | 0 | 0 | 0 | 0 | 0 | 276010 | 0 | 615940 | 4283600 | 126270 | 0 | 451650 | 0 | 0 | 0 |
| VLRPQVTA | DYNC1H1; DYN | Q14204; A0 | DYHC1_HUMA | Unmodified | 0 | 0 | 0 | 0 | 0 | 0 | 0 | 0 | 0 | 0 | 0 | 2118600 | 0 | 0 | 0 | 0 | 0 | 0 |
| VLRSFDSM | GSR | P00390 | GSHR_HUMAN | Oxidation (M) | 0 | 0 | 0 | 0 | 1566000 | 0 | 0 | 0 | 0 | 0 | 0 | 0 | 1474500 | 0 | 0 | 0 | 0 | 0 |
| VLRTPGNN | EIF1AD; EIF1AD | Q8N9N8; E9 | EIF1A_HUMAN | Unmodified | 0 | 0 | 0 | 0 | 0 | 0 | 0 | 0 | 0 | 0 | 90863 | 274410 | 0 | 0 | 0 | 0 | 0 | 0 |
| VLRVLDQF | SORCS2; SORC | Q96PQ0; B5 | SORC2_HUMA | Oxidation (M) | 0 | 0 | 0 | 0 | 0 | 0 | 0 | 0 | 107630 | 466800 | 0 | 0 | 0 | 0 | 0 | 0 | 0 | 0 |
| VLRYEGFF | SLC25A12; SLC | O75746; Q9 | CMC1_HUMAN | Unmodified | 0 | 0 | 0 | 0 | 0 | 0 | 36968 | 0 | 2570100 | 683880 | 0 | 0 | 0 | 0 | 0 | 0 | 0 | 0 |
| VLRYETFISI | UXT; UXT | Q9UBK9; S4 | UXT_HUMAN P | Unmodified | 0 | 0 | 0 | 0 | 0 | 0 | 0 | 0 | 0 | 0 | 0 | 0 | 0 | 0 | 0 | 0 | 0 | 0 |
| VLSAPPHFI | EEF1G | P26641 | EF1G_HUMAN | Unmodified | 0 | 28848 | 77132 | 66605 | 0 | 0 | 0 | 0 | 342550 | 207990 | 641210 | 16576000 | 0 | 0 | 1269800 | 165430 | 0 | 0 |
| VLSAPPHFI | EEF1G | P26641 | EF1G_HUMAN | Unmodified | 0 | 0 | 0 | 558380 | 725280 | 0 | 0 | 0 | 1410400 | 0 | 5298000 | 6089500 | 373020 | 148160 | 198750 | 0 | 0 | 0 |
| VLSAVEDRI | TRIM47 | Q96LD4 | TRI47_HUMAN | Unmodified | 0 | 0 | 0 | 0 | 1440900 | 440290 | 0 | 0 | 0 | 105570 | 0 | 0 | 0 | 0 | 0 | 0 | 0 | 0 |
| VLSAVGSA | DLGAP4; DLGA | Q9Y2H0; F8 | DLGP4_HUMA | Unmodified | 0 | 0 | 0 | 0 | 89994 | 0 | 34537 | 0 | 0 | 0 | 0 | 223620 | 0 | 69122 | 0 | 0 | 227360 | 285770 |
| VLSEAAISA | ACSL3; ACSL3; | O95573; A0 | ACSL3_HUMAN | Unmodified | 0 | 0 | 0 | 0 | 0 | 0 | 0 | 173020 | 0 | 0 | 0 | 1146200 | 124340 | 0 | 0 | 0 | 0 | 0 |
| VLSEAAISA | ACSL3; ACSL3; | O95573; A0 | ACSL3_HUMAN | Unmodified | 0 | 0 | 0 | 186950 | 1381000 | 0 | 1703800 | 28978 | 0 | 417940 | 308910 | 455470 | 3924100 | 1102100 | 0 | 0 | 230450 | 1574700 |
| VLSGDLGQ | PCK2; PCK2 | Q16822; H0 | PCKGM_HUMA | Unmodified | 0 | 0 | 0 | 438120 | 0 | 0 | 0 | 0 | 21234000 | 1440000 | 263830 | 0 | 0 | 0 | 0 | 0 | 0 | 0 |
| VLSGEFQI | RRM1; RRM1 | P23921; E9P | RIR1_HUMAN F | Unmodified | 0 | 0 | 0 | 0 | 0 | 0 | 0 | 41373 | 340560 | 215070 | 722790 | 6090400 | 0 | 0 | 493810 | 652320 | 0 | 0 |
| VLSGLEILS | INPPL1; INPPL1 | O15357; A0 | SHIP2_HUMAN | Unmodified | 0 | 0 | 0 | 0 | 0 | 0 | 0 | 0 | 0 | 0 | 1840000 | 3915200 | 0 | 0 | 0 | 0 | 0 | 0 |
| VLSGQDTE | SFT2D2; SFT2D | O95562; Q5 | SFT2B_HUMAN | Unmodified | 506100 | 255860 | 1236800 | 467460 | 0 | 2024400 | 1165000 | 682160 | 384660 | 99622 | 0 | 2732800 | 0 | 0 | 540820 | 329060 | 766410 | 7108300 |
| VLSGTIHA | EFTUD2 | Q15029 | U5S1_HUMAN | Unmodified | 0 | 0 | 0 | 0 | 0 | 0 | 0 | 18926 | 0 | 0 | 170800 | 536640 | 0 | 0 | 552880 | 0 | 270070 | 0 |
| VLSHQDD | CUL5 | Q93034 | CUL5_HUMAN | Unmodified | 0 | 0 | 0 | 0 | 0 | 0 | 0 | 0 | 0 | 0 | 0 | 1101500 | 0 | 0 | 0 | 0 | 0 | 0 |
| VLSIGDGIA | ATP5F1A; ATP5 | P25705; K7E | ATPA_HUMAN | Unmodified | 0 | 0 | 0 | 0 | 2204000 | 691830 | 0 | 0 | 9621900 | 1125900 | 0 | 915740 | 0 | 0 | 0 | 0 | 118090 | 127300 |
| VLSIGLPVA | MON2; MON2 | Q7Z3U7; A0 | MON2_HUMAN | Unmodified | 0 | 0 | 0 | 0 | 0 | 0 | 0 | 0 | 95981 | 0 | 0 | 451730 | 0 | 0 | 0 | 0 | 0 | 0 |
| VLSKGQPL | NOMO1; NOMO | Q15155; P6 | NOMO1_HUMA | Unmodified | 0 | 0 | 0 | 0 | 0 | 0 | 0 | 0 | 0 | 816950 | 0 | 0 | 0 | 0 | 0 | 0 | 0 | 0 |
| VLSLLALVK | PSME2; PSME2 | Q9UL46; A0 | PSME2_HUMA | Unmodified | 144660 | 137250 | 0 | 47372 | 708990 | 204490 | 3272800 | 0 | 210620 | 771410 | 141120 | 720800 | 1893800 | 0 | 158820 | 116990 | 1384100 | 0 |
| VLSLLALVK | PSME2; PSME2 | Q9UL46; A0 | PSME2_HUMA | Unmodified | 0 | 0 | 0 | 0 | 0 | 0 | 332280 | 0 | 0 | 37391 | 0 | 0 | 63031 | 73485 | 0 | 0 | 303860 | 0 |
| VLSLNFSE | G3BP2; G3BP2; | Q9UN86; D6 | G3BP2_HUMA | Unmodified | 0 | 0 | 0 | 0 | 25498 | 70501 | 450280 | 0 | 0 | 296520 | 388300 | 3721800 | 3287000 | 1264400 | 0 | 0 | 414460 | 0 |
| VLSMAPGL | PAPSS2 | O95340 | PAPS2_HUMAN | Oxidation (M) | 0 | 0 | 0 | 0 | 0 | 0 | 0 | 0 | 0 | 0 | 0 | 0 | 0 | 0 | 0 | 0 | 2425700 | 3823900 |
| VLSPSGLLI | SLITRK6 | Q9H5Y7 | SLIK6_HUMAN | Unmodified | 4508600 | 2354800 | 0 | 0 | 66508 | 0 | 0 | 0 | 2999200 | 0 | 519050 | 0 | 0 | 0 | 0 | 0 | 0 | 0 |
| VLSQEEPD | SLC16A10; SLC | Q8TF71; A0 | MOT10_HUMA | Unmodified | 0 | 0 | 646260 | 483530 | 0 | 0 | 0 | 0 | 0 | 0 | 0 | 642830 | 0 | 0 | 720340 | 727800 | 0 | 0 |
| VLSQLTSD | ITGB4; ITGB4 | P16144; J3Q | ITB4_HUMAN I | Unmodified | 0 | 0 | 0 | 0 | 652040 | 592940 | 0 | 0 | 0 | 327160 | 0 | 0 | 0 | 0 | 0 | 0 | 3647300 | 3162900 |
| VLSQLTSD | ITGB4; ITGB4 | P16144; J3Q | ITB4_HUMAN I | Unmodified | 0 | 0 | 0 | 33691 | 4130900 | 0 | 0 | 0 | 0 | 113670 | 0 | 0 | 0 | 0 | 0 | 0 | 6657700 | 8670000 |
| VLSQQAAS | ADSL; ADSL; AD | P30566; A0A | PUR8_HUMAN | Unmodified | 121040 | 0 | 0 | 237250 | 457030 | 0 | 400500 | 168930 | 0 | 175740 | 259730 | 2899100 | 2309200 | 1182000 | 432760 | 485400 | 190180 | 338770 |
| VLSRLGVA | UCHL1; UCHL1 | P09936; D6F | UCHL1_HUMA | Unmodified | 0 | 0 | 0 | 0 | 727770 | 0 | 0 | 0 | 0 | 88203 | 0 | 0 | 0 | 0 | 0 | 0 | 0 | 0 |
| VLSRPNAQ | PHB2; PHB2; PI | Q99623; F5C | PHB2_HUMAN | Oxidation (M) | 65564 | 138430 | 114690 | 251610 | 0 | 0 | 0 | 21992 | 724730 | 102940 | 0 | 369370 | 0 | 0 | 107250 | 34982 | 0 | 0 |
| VLSSIEQK | SFN | P31947 | 1433S_HUMAN | Unmodified | 7619300 | 2.78E+08 | 18821000 | 24232000 | 44632000 | 12877000 | 2233200 | 1292400 | 34154000 | 7991100 | 20103000 | 2.08E+08 | 17547000 | 9258200 | 36038000 | 28248000 | 11150000 | 25391000 |
| VLSSIEQKS | SFN | P31947 | 1433S_HUMAN | Unmodified | 0 | 0 | 0 | 0 | 655690 | 322620 | 0 | 0 | 0 | 72753 | 0 | 0 | 0 | 0 | 0 | 0 | 164830 | 292850 |
| VLSSIEQKS | SFN | P31947 | 1433S_HUMAN | Unmodified | 0 | 0 | 0 | 0 | 2848400 | 0 | 0 | 0 | 461630 | 508820 | 0 | 0 | 0 | 0 | 0 | 0 | 685720 | 809260 |
| VLSSLGYH | ABHD12; ABHD | Q8N2K0; A0 | ABD12_HUMAN | Unmodified | 0 | 0 | 0 | 0 | 0 | 0 | 0 | 0 | 5208200 | 432630 | 0 | 412380 | 0 | 0 | 0 | 26923 | 0 | 0 |
| VLSSLGYH | ABHD12; ABHD | Q8N2K0; A0 | ABD12_HUMAN | Unmodified | 0 | 0 | 0 | 0 | 0 | 0 | 0 | 0 | 1328500 | 0 | 0 | 0 | 0 | 0 | 0 | 0 | 0 | 0 |
| VLSSSEISV | CNTN1 | Q12860 | CNTN1_HUMA | Unmodified | 0 | 1774400 | 0 | 0 | 0 | 0 | 167340 | 0 | 1485200 | 492130 | 54920 | 0 | 0 | 0 | 0 | 0 | 0 | 0 |
| VLSTVHTH | AIMP2; AIMP2 | Q13155; A8 | AIMP2_HUMAN | Unmodified | 0 | 0 | 0 | 0 | 0 | 0 | 0 | 0 | 0 | 0 | 0 | 13258000 | 0 | 0 | 0 | 0 | 0 | 0 |
| VLSVCPSN | JUP | P14923 | PLAK_HUMAN | Oxidation (M) | 228300 | 0 | 669270 | 1064700 | 0 | 0 | 0 | 0 | 0 | 295850 | 491660 | 1677300 | 968700 | 1321500 | 2675300 | 2638200 | 804610 | 458500 |
| VLSVCSSN | CTNNB1; CTNN | P35222; A0A | CTNB1_HUMA | Oxidation (M) | 760400 | 0 | 380320 | 614340 | 0 | 0 | 0 | 0 | 0 | 0 | 0 | 0 | 0 | 0 | 5223600 | 4813300 | 0 | 0 |
| VLSVCSSN | CTNNB1; CTNN | P35222; A0A | CTNB1_HUMA | Unmodified | 0 | 0 | 0 | 0 | 0 | 0 | 0 | 0 | 0 | 0 | 0 | 0 | 0 | 0 | 1539400 | 722750 | 0 | 0 |
| VLSVDESII | ANO6; ANO6; A | Q4KMQ2; A0 | ANO6_HUMAN | Unmodified | 341050 | 339000 | 171640 | 856400 | 847250 | 118420 | 1320500 | 0 | 1342800 | 837990 | 946270 | 2001200 | 532190 | 350800 | 859960 | 347540 | 0 | 0 |
| VLSVETPLS | MPI; MPI; MPI; | P34949; H3 | MPI_HUMAN M | Unmodified | 194840 | 183160 | 407660 | 632500 | 2715700 | 541050 | 149130 | 97927 | 0 | 1058200 | 616040 | 3081000 | 1962800 | 960550 | 671650 | 585390 | 651850 | 1062200 |
| VLSVHTDP | IQGAP3; IQGAP | Q86VI3; F2Z | IQGA3_HUMAN | Unmodified | 0 | 0 | 119400 | 237980 | 447450 | 88446 | 46942 | 0 | 418160 | 0 | 66342 | 367820 | 0 | 0 | 71035 | 0 | 0 | 0 |
| VLSVPESTF | UFM1; UFM1 | P61960; H0 | UFM1_HUMAN | Unmodified | 2384900 | 1792100 | 10537000 | 3912700 | 1864700 | 0 | 781960 | 437190 | 1777900 | 1602200 | 3939800 | 8385500 | 1314800 | 926090 | 15618000 | 9616200 | 2447100 | 2336300 |
| VLTAEELN | RARS1 | P54136 | SYRC_HUMAN | Unmodified | 0 | 0 | 0 | 0 | 0 | 0 | 0 | 0 | 0 | 0 | 0 | 12457000 | 530000 | 0 | 0 | 0 | 0 | 0 |
| VLTEDEMG | FDPS; FDPS; FD | P14324; A0A | FPPS_HUMAN | Oxidation (M) | 224800 | 168500 | 0 | 0 | 0 | 0 | 0 | 0 | 925680 | 72150 | 787890 | 6004700 | 0 | 0 | 196400 | 222060 | 0 | 0 |
| VLTEDEMG | FDPS; FDPS; FD | P14324; A0A | FPPS_HUMAN | Unmodified | 232830 | 0 | 0 | 0 | 0 | 0 | 0 | 0 | 3225900 | 304690 | 1044700 | 10774000 | 0 | 0 | 0 | 0 | 0 | 0 |
| VLTEFTR | IPO9 | Q96P70 | IPO9_HUMAN I | Unmodified | 0 | 0 | 0 | 0 | 0 | 0 | 0 | 0 | 105850 | 26330 | 0 | 4857700 | 0 | 0 | 56409 | 193150 | 84553 | 80460 |
| VLTFLDSH | GALNT2 | Q10471 | GALT2_HUMAN | Unmodified | 0 | 0 | 0 | 0 | 0 | 0 | 0 | 0 | 13286000 | 564240 | 0 | 146960 | 0 | 0 | 0 | 0 | 0 | 85911 |
| VLTFLRRG | HSPH1; HSPH1 | Q92598; A0 | HS105_HUMAN | Unmodified | 0 | 0 | 0 | 0 | 0 | 0 | 2307700 | 0 | 0 | 0 | 0 | 0 | 2844500 | 1012000 | 0 | 0 | 0 | 1119500 |
| VLTGVAGE | IPO7 | O95373 | IPO7_HUMAN I | Unmodified | 0 | 0 | 0 | 0 | 0 | 0 | 0 | 0 | 0 | 0 | 0 | 1785400 | 0 | 0 | 0 | 0 | 0 | 0 |
| VLTGYLNY | ECE1; ECE1; EC | P42892; B4 | ECE1_HUMAN | Unmodified | 0 | 0 | 0 | 0 | 226300 | 0 | 0 | 0 | 169660 | 186210 | 0 | 0 | 0 | 0 | 0 | 0 | 0 | 20763 |
| VLTIDPTEF | COPA; COPA; C | P53621; A0A | COPA_HUMAN | Unmodified | 0 | 0 | 0 | 0 | 0 | 0 | 380320 | 0 | 695850 | 334360 | 0 | 7103300 | 0 | 0 | 0 | 400360 | 0 | 0 |
| VLTILINPA | NDST1 | P52848 | NDST1_HUMA | Unmodified | 0 | 0 | 0 | 0 | 0 | 0 | 0 | 0 | 1647200 | 0 | 0 | 381460 | 0 | 0 | 0 | 0 | 0 | 0 |
| VLTLQNNQ | LGR4 | Q9BXB1 | LGR4_HUMAN | Unmodified | 0 | 0 | 0 | 0 | 3131500 | 0 | 0 | 0 | 700330 | 116790 | 0 | 0 | 0 | 0 | 0 | 0 | 0 | 0 |
| VLTLSDDLE | EIF5 | P55010 | IF5_HUMAN Eu | Unmodified | 0 | 0 | 188600 | 0 | 0 | 606860 | 0 | 0 | 0 | 0 | 729880 | 2269600 | 0 | 0 | 0 | 148660 | 0 | 0 |
| VLTLSDDLE | EIF5 | P55010 | IF5_HUMAN Eu | Unmodified | 70655 | 0 | 0 | 1122200 | 0 | 0 | 0 | 44168 | 1404100 | 0 | 0 | 6734900 | 0 | 0 | 0 | 0 | 0 | 0 |
| VLTMSGW | CLUH; CLUH; C | O75153; I3L | CLU_HUMAN C | Unmodified | 0 | 0 | 0 | 0 | 0 | 0 | 0 | 0 | 3058400 | 505450 | 0 | 376320 | 0 | 0 | 0 | 0 | 0 | 0 |
| VLTNFKSS | ARPC5L | Q9BPX5 | ARP5L_HUMAN | Oxidation (M) | 0 | 0 | 0 | 0 | 0 | 0 | 1340800 | 0 | 0 | 0 | 0 | 0 | 0 | 0 | 0 | 0 | 857850 | 0 |
| VLTPEEQL | EIF3J | O75822 | EIF3J_HUMAN | Unmodified | 0 | 0 | 0 | 0 | 0 | 0 | 69228 | 0 | 0 | 0 | 1665100 | 4052100 | 0 | 0 | 0 | 0 | 0 | 0 |
| VLTPEEQL | EIF3J | O75822 | EIF3J_HUMAN | Unmodified | 0 | 0 | 0 | 0 | 0 | 0 | 0 | 0 | 0 | 0 | 0 | 0 | 748700 | 0 | 0 | 0 | 0 | 0 |
| VLTPELYAE | CKB; CKB; CKB | P12277; G3 | KCRB_HUMAN | Unmodified | 0 | 699960 | 11042000 | 36052000 | 7498400 | 1817800 | 4811600 | 2574900 | 20806000 | 2088200 | 1983900 | 53913000 | 8811300 | 2420000 | 30170000 | 19199000 | 0 | 0 |
| VLTPELYAE | CKB; CKB; CKB | P12277; G3 | KCRB_HUMAN | Unmodified | 247300 | 0 | 0 | 8881800 | 1.53E+08 | 11503000 | 2.16E+08 | 74735000 | 2231200 | 13819000 | 0 | 0 | 1.04E+08 | 69317000 | 1630700 | 508570 | 803940 | 648260 |
| VLTPTQVK | PEBP1 | P30086 | PEBP1_HUMAN | Unmodified | 11786000 | 8425700 | 7530500 | 11085000 | 18609000 | 5282800 | 4371800 | 2216600 | 14235000 | 12910000 | 10782000 | 27571000 | 44562000 | 30557000 | 11291000 | 9534400 | 26404000 | 26660000 |
| VLTQASNP | TMPRSS2; TMP | O15393; C9J | TMPS2_HUMAN | Unmodified | 0 | 0 | 2231400 | 1981300 | 0 | 0 | 0 | 0 | 0 | 0 | 0 | 0 | 0 | 0 | 2598900 | 1148600 | 0 | 0 |
| VLTSDIEK | ME1 | P48163 | MAOX_HUMAN | Unmodified | 0 | 0 | 138050 | 96626 | 0 | 0 | 0 | 0 | 0 | 0 | 127280 | 3397600 | 45112 | 0 | 184780 | 177200 | 0 | 0 |
| VLTSEDEY | NIBAN1 | Q9BZQ8 | NIBA1_HUMAN | Unmodified | 0 | 0 | 0 | 0 | 0 | 0 | 0 | 0 | 1696700 | 456550 | 0 | 0 | 0 | 0 | 0 | 59443 | 0 | 0 |
| VLTSEDEY | NIBAN1 | Q9BZQ8 | NIBA1_HUMAN | Unmodified | 0 | 0 | 0 | 0 | 11626000 | 5538700 | 3558100 | 110400 | 40917000 | 12448000 | 0 | 0 | 1599600 | 914580 | 0 | 0 | 957420 | 2574700 |
| VLTSEDEY | NIBAN1 | Q9BZQ8 | NIBA1_HUMAN | Unmodified | 0 | 0 | 0 | 0 | 0 | 3302600 | 3798500 | 0 | 0 | 10569000 | 0 | 0 | 0 | 0 | 0 | 0 | 0 | 0 |
| VLTSGIFET | GNAS; GNAS; G | Q5JWF2; P6 | GNAS1_HUMA | Unmodified | 1044500 | 825430 | 3149800 | 2424200 | 1072500 | 520200 | 0 | 80170 | 1197500 | 138670 | 2494800 | 19263000 | 592090 | 0 | 7023000 | 6482200 | 556250 | 0 |

| | | | | | | | | | | | | | | | | | | | | | |
|---|---|---|---|---|---|---|---|---|---|---|---|---|---|---|---|---|---|---|---|---|---|
| VLTSGIFETI | GNAS; GNAS; G | Q5JWF2; P63 | GNAS1_HUMAI | Unmodified | 163550 | 0 | 0 | 492450 | 462960 | 166080 | 0 | 0 | 0 | 0 | 212090 | 380950 | 374700 | 221410 | 0 | 500570 | 594760 | 389270 |
| VLTTGYWP | CUL3; CUL3 | Q13618; A0A | CUL3_HUMAN | Unmodified | 0 | 0 | 0 | 0 | 0 | 0 | 0 | 0 | 0 | 0 | 0 | 1421300 | 0 | 0 | 0 | 0 | 0 | 0 |
| VLTVINQTC | RPL35; RPL35 | P42766; F2Z | RL35_HUMAN ( | Unmodified | 2247700 | 3520800 | 4236400 | 1660800 | 3150500 | 0 | 0 | 0 | 1695900 | 0 | 4855400 | 79425000 | 0 | 1271000 | 2718100 | 2237900 | 0 | 1430700 |
| VLTVINQTC | RPL35; RPL35 | P42766; F2Z | RL35_HUMAN ( | Unmodified | 109370 | 151930 | 0 | 0 | 0 | 0 | 156350 | 137810 | 0 | 0 | 0 | 0 | 741530 | 0 | 0 | 0 | 112000 | 0 |
| VLTVINQTC | RPL35; RPL35 | P42766; F2Z | RL35_HUMAN ( | Unmodified | 0 | 0 | 0 | 0 | 583300 | 0 | 368810 | 380940 | 0 | 0 | 0 | 0 | 1553800 | 0 | 0 | 0 | 0 | 0 |
| VLTVPCQK | SYK | P43405 | KSYK_HUMAN | Unmodified | 0 | 0 | 0 | 0 | 0 | 0 | 0 | 0 | 0 | 0 | 0 | 0 | 0 | 0 | 0 | 0 | 0 | 0 |
| VLTWDVGK | AP3M1 | Q9Y2T2 | AP3M1_HUMAI | Unmodified | 120120 | 295180 | 0 | 61619 | 1001100 | 0 | 764880 | 784990 | 36750000 | 17168000 | 150900 | 2319500 | 90091 | 0 | 131390 | 0 | 0 | 382230 |
| VLTWSGQE | UEVLD | Q8IX04 | UEVLD_HUMAI | Unmodified | 228170 | 0 | 0 | 0 | 764510 | 0 | 0 | 91739 | 1956700 | 358050 | 710300 | 0 | 0 | 0 | 0 | 783670 | 0 | 0 |
| VLVALASEE | ESYT2; ESYT2; E | A0FGR8; A0A | ESYT2_HUMAN | Unmodified | 0 | 0 | 0 | 0 | 1284100 | 0 | 0 | 217620 | 196520 | 137260 | 86544 | 0 | 0 | 255110 | 0 | 0 | 0 | 0 |
| VLVALYEEF | MYCBP; N/A | Q99417; A0A | MYCBP_HUMA | Unmodified | 744240 | 975940 | 0 | 4025700 | 15740000 | 728150 | 6868000 | 203900 | 7377200 | 2499300 | 1405300 | 3179500 | 11882000 | 4435300 | 554610 | 309020 | 4316500 | 7341500 |
| VLVAQHDV | ETFA; ETFA; ETF | P13804; H0Y | ETFA_HUMAN E | Unmodified | 0 | 0 | 0 | 0 | 0 | 0 | 4006000 | 0 | 0 | 1408400 | 0 | 0 | 0 | 0 | 0 | 0 | 0 | 0 |
| VLVATCTSA | NECTIN1 | Q15223 | NECT1_HUMAI | Unmodified | 1459600 | 527630 | 0 | 170590 | 0 | 0 | 0 | 0 | 0 | 0 | 0 | 416870 | 0 | 0 | 168010 | 168510 | 0 | 0 |
| VLVCAHR | ITGA3 | P26006 | ITA3_HUMAN I | Unmodified | 179210 | 0 | 509370 | 0 | 932930 | 728640 | 0 | 0 | 171820 | 0 | 0 | 1091100 | 0 | 0 | 64320 | 0 | 0 | 0 |
| VLVCAHRY | ITGA3 | P26006 | ITA3_HUMAN I | Unmodified | 0 | 0 | 0 | 0 | 3246600 | 1265700 | 0 | 0 | 0 | 0 | 1904900 | 0 | 0 | 0 | 0 | 0 | 1966800 | 2342400 |
| VLVCAHRY | ITGA3 | P26006 | ITA3_HUMAN I | Unmodified | 0 | 0 | 0 | 0 | 20619000 | 14450000 | 0 | 0 | 0 | 707050 | 0 | 0 | 0 | 0 | 0 | 0 | 13329000 | 19037000 |
| VLVCGAGF | SORD; SORD | Q00796; A0A | DHSO_HUMAN | Oxidation (M) | 457030 | 617440 | 0 | 196280 | 0 | 0 | 0 | 0 | 0 | 0 | 0 | 0 | 454880 | 0 | 354110 | 328330 | 0 | 0 |
| VLVDGEEH | EMC7; EMC7 | Q9NPA0; H0 | EMC7_HUMAN | Unmodified | 0 | 0 | 222640 | 0 | 0 | 0 | 0 | 0 | 6869700 | 506540 | 1725500 | 475150 | 0 | 0 | 0 | 0 | 0 | 0 |
| VLVDMSR | CCT2; CCT2; C( | P78371; F5G | TCPB_HUMAN | Unmodified | 0 | 0 | 0 | 0 | 0 | 0 | 0 | 0 | 269150 | 0 | 299380 | 10201000 | 0 | 0 | 0 | 0 | 0 | 0 |
| VLVDQTTG | ELAVL1 | Q15717 | ELAV1_HUMAN | Unmodified | 0 | 0 | 558370 | 0 | 322570 | 0 | 0 | 0 | 578840 | 156790 | 809000 | 10352000 | 133550 | 0 | 674480 | 1001800 | 332550 | 0 |
| VLVDSSFG | CPSF1 | Q10570 | CPSF1_HUMAN | Unmodified | 2357200 | 0 | 0 | 1561200 | 0 | 0 | 28831 | 26950 | 0 | 0 | 0 | 435750 | 0 | 0 | 3983400 | 4010400 | 0 | 0 |
| VLVEDTK | TLN1 | Q9Y490 | TLN1_HUMAN | Unmodified | 603290 | 399380 | 264370 | 961180 | 2432400 | 1248800 | 6279000 | 2126100 | 741640 | 888750 | 220150 | 1408500 | 557580 | 3608100 | 998230 | 934920 | 1488200 | 861490 |
| VLVEDTKVI | TLN1 | Q9Y490 | TLN1_HUMAN | Unmodified | 0 | 0 | 0 | 0 | 0 | 0 | 3442700 | 1412900 | 0 | 0 | 0 | 0 | 383890 | 966680 | 0 | 0 | 1305300 | 0 |
| VLVEFSHK | COMMD10; CO | Q9Y6G5; D6 | COMDA_HUMA | Unmodified | 0 | 0 | 0 | 0 | 1035400 | 0 | 0 | 0 | 0 | 0 | 0 | 66151 | 0 | 0 | 0 | 0 | 0 | 0 |
| VLVEGPLN | UPF1 | Q92900 | RENT1_HUMAN | Unmodified | 1539500 | 1617900 | 1298900 | 864140 | 0 | 0 | 0 | 0 | 152030 | 0 | 0 | 2695500 | 0 | 0 | 879340 | 1064000 | 0 | 0 |
| VLVEGPLN | UPF1 | Q92900 | RENT1_HUMAN | Oxidation (M) | 816400 | 1370600 | 0 | 0 | 0 | 0 | 184770 | 0 | 346720 | 89231 | 0 | 0 | 148900 | 58566 | 0 | 0 | 0 | 114910 |
| VLVEPDAG | ECI1 | P42126 | ECI1_HUMAN E | Oxidation (M) | 0 | 0 | 0 | 0 | 0 | 0 | 0 | 0 | 1181500 | 0 | 0 | 0 | 0 | 0 | 0 | 0 | 0 | 0 |
| VLVEPDAG | ECI1 | P42126 | ECI1_HUMAN E | Unmodified | 0 | 0 | 0 | 0 | 0 | 0 | 0 | 0 | 6377700 | 0 | 0 | 0 | 0 | 0 | 0 | 0 | 0 | 0 |
| VLVEQQQD | DDI2; DDI2; DD | Q5TDH0; H0 | DDI2_HUMAN I | Unmodified | 2543900 | 164170 | 1335000 | 3223700 | 0 | 0 | 0 | 0 | 0 | 0 | 0 | 468570 | 1900800 | 239660 | 2086000 | 192100 | 0 | 0 |
| VLVEQTK | SLITRK6 | Q9H5Y7 | SLIK6_HUMAN | Unmodified | 0 | 0 | 0 | 0 | 218900 | 0 | 0 | 16467 | 0 | 183030 | 0 | 0 | 0 | 0 | 0 | 34998 | 0 | 0 |
| VLVEQTKN | SLITRK6 | Q9H5Y7 | SLIK6_HUMAN | Unmodified | 1382400 | 1428800 | 0 | 0 | 0 | 0 | 0 | 0 | 9885500 | 8274800 | 0 | 0 | 557860 | 318770 | 0 | 0 | 887780 | 1225500 |
| VLVERSAAI | HSPE1; HSPE1- | P61604; S4R | CH10_HUMAN | Unmodified | 0 | 0 | 0 | 0 | 0 | 0 | 33070 | 55090 | 466920 | 930810 | 89884 | 0 | 200640 | 89738 | 0 | 0 | 0 | 0 |
| VLVFVMGK | PFN2; PFN2; PF | P35080; C9J | PROF2_HUMAN | Unmodified | 939020 | 670090 | 165500 | 94113 | 1148900 | 45155 | 0 | 0 | 3013500 | 566210 | 0 | 3800000 | 177120 | 100410 | 467050 | 47824 | 899080 | 1275100 |
| VLVFVMGK | PFN2; PFN2; PF | P35080; C9J | PROF2_HUMAN | Oxidation (M) | 2150600 | 1372200 | 33144000 | 0 | 0 | 0 | 0 | 47061 | 663940 | 536800 | 614620 | 4413800 | 0 | 0 | 0 | 0 | 103080 | 0 |
| VLVFVMGK | PFN2; PFN2; PF | P35080; C9J | PROF2_HUMAN | Unmodified | 0 | 0 | 0 | 0 | 0 | 0 | 673760 | 0 | 0 | 490270 | 0 | 0 | 0 | 0 | 0 | 0 | 2526400 | 476420 |
| VLVFVMGK | PFN2; PFN2; PF | P35080; C9J | PROF2_HUMAN | Unmodified | 0 | 0 | 0 | 0 | 486160 | 0 | 0 | 0 | 0 | 504860 | 0 | 0 | 413730 | 0 | 0 | 0 | 1450000 | 363510 |
| VLVGKNFE | P4HB; P4HB; P4 | P07237; A0A | PDIA1_HUMAN | Unmodified | 0 | 0 | 0 | 0 | 0 | 0 | 2409400 | 1017000 | 0 | 5824300 | 0 | 0 | 284290 | 615230 | 0 | 0 | 370740 | 0 |
| VLVGKNFE | P4HB; P4HB; P4 | P07237; A0A | PDIA1_HUMAN | Unmodified | 0 | 0 | 0 | 0 | 0 | 0 | 1235100 | 224650 | 0 | 1188800 | 0 | 0 | 0 | 239040 | 0 | 0 | 0 | 0 |
| VLVHPPQD | CLPP; CLPP | Q16740; M0 | CLPP_HUMAN | Unmodified | 0 | 0 | 0 | 0 | 0 | 0 | 0 | 212140 | 5970600 | 1148900 | 0 | 0 | 0 | 0 | 0 | 0 | 0 | 0 |
| VLVIGAGG | UBA3; UBA3 | Q8TBC4; F8 | UBA3_HUMAN | Unmodified | 0 | 0 | 0 | 0 | 0 | 0 | 0 | 0 | 0 | 0 | 0 | 1393200 | 0 | 0 | 0 | 0 | 0 | 0 |
| VLVLAPTRE | DDX50 | Q9BQ39 | DDX50_HUMAI | Unmodified | 0 | 0 | 0 | 0 | 0 | 0 | 948650 | 1374600 | 0 | 0 | 0 | 0 | 0 | 0 | 0 | 0 | 0 | 0 |
| VLVLSQDY | SPTBN1; SPTBN | Q01082; A0A | SPTB2_HUMAN | Unmodified | 0 | 0 | 0 | 0 | 0 | 0 | 0 | 0 | 0 | 0 | 0 | 2475300 | 0 | 0 | 0 | 0 | 0 | 0 |
| VLVMANDF | PPP2R1B | P30154 | 2AAB_HUMAN | Unmodified | 0 | 0 | 0 | 40169 | 0 | 0 | 0 | 0 | 0 | 0 | 60152 | 531920 | 243240 | 0 | 72684 | 0 | 0 | 2181200 |
| VLVPATDR | EIF3E; EIF3E; EI | P60228; B3K | EIF3E_HUMAN | Unmodified | 88444 | 0 | 0 | 0 | 107250 | 0 | 252460 | 0 | 1154800 | 0 | 142850 | 3360000 | 0 | 0 | 128980 | 156660 | 0 | 0 |
| VLVPATDRI | EIF3E; EIF3E; EI | P60228; B3K | EIF3E_HUMAN | Unmodified | 539440 | 0 | 275480 | 0 | 0 | 0 | 0 | 0 | 0 | 0 | 392470 | 5921000 | 0 | 0 | 0 | 408540 | 0 | 0 |
| VLVPGIPGI | WDR43 | Q15061 | WDR43_HUMA | Unmodified | 0 | 0 | 0 | 0 | 326120 | 0 | 0 | 55285 | 0 | 0 | 0 | 0 | 623190 | 0 | 0 | 0 | 81310 | 0 |
| VLVPQIPR | MGAT2 | Q10469 | MGAT2_HUMAI | Unmodified | 0 | 0 | 0 | 0 | 0 | 0 | 0 | 0 | 6676700 | 1025100 | 0 | 352060 | 0 | 0 | 0 | 0 | 0 | 462020 |
| VLVQNAAG | TLN1 | Q9Y490 | TLN1_HUMAN | Unmodified | 85582 | 77689 | 0 | 88766 | 4378900 | 1823300 | 5263900 | 1878800 | 793470 | 822130 | 352210 | 4570900 | 187350 | 2401500 | 90637 | 158800 | 2415300 | 1009900 |
| VLVTGATGI | MAT2B; MAT2B | Q9NZL9; E5I | MAT2B_HUMAI | Unmodified | 0 | 0 | 0 | 0 | 755280 | 0 | 0 | 0 | 606240 | 170850 | 451610 | 3790500 | 947120 | 0 | 0 | 0 | 0 | 388710 |
| VLVTHETGI | PLXNA4 | Q9HCM2 | PLXA4_HUMAN | Unmodified | 0 | 0 | 0 | 0 | 0 | 0 | 965670 | 410620 | 67427 | 22277 | 17821 | 0 | 39065 | 1678100 | 0 | 0 | 0 | 725830 |
| VLVTTNVC | DDX19B; DDX1 | Q9UMR2; Q9 | DD19B_HUMAI | Unmodified | 0 | 0 | 0 | 0 | 1045100 | 0 | 144650 | 0 | 329630 | 0 | 0 | 2028500 | 1989000 | 0 | 0 | 0 | 358480 | 0 |
| VLVTVFR | ELP1; ELP1; ELF | O95163; A0A | ELP1_HUMAN I | Unmodified | 0 | 0 | 0 | 0 | 0 | 0 | 0 | 0 | 0 | 0 | 236850 | 639450 | 0 | 0 | 0 | 0 | 0 | 0 |
| VLVVDQLS | STXBP1; STXBP | P61764; A0A | STXB1_HUMAN | Unmodified | 0 | 0 | 0 | 0 | 871700 | 0 | 538340 | 139500 | 2309600 | 621520 | 0 | 0 | 0 | 0 | 0 | 0 | 0 | 789630 |
| VLVVDQLS | STXBP1; STXBP | P61764; A0A | STXB1_HUMAN | Oxidation (M) | 388750 | 490250 | 208370 | 275130 | 1078800 | 0 | 1139500 | 1485500 | 699030 | 0 | 0 | 2425800 | 0 | 0 | 266080 | 267380 | 206100 | 618390 |
| VLVVGAGG | UBA2; UBA2; UI | Q9UBT2; K7I | SAE2_HUMAN | Unmodified | 0 | 0 | 0 | 0 | 0 | 0 | 0 | 0 | 493420 | 0 | 575010 | 1499600 | 0 | 0 | 0 | 0 | 0 | 0 |
| VLVVHDGF | PFKM; PFKM; PI | P08237; A0A | PFKAM_HUMAI | Unmodified | 301410 | 0 | 472420 | 676460 | 0 | 0 | 0 | 0 | 0 | 0 | 0 | 7816600 | 183370 | 0 | 1058900 | 850290 | 0 | 0 |
| VLVVVTDG | COL12A1; COL | Q99715; D6I | COCA1_HUMA | Unmodified | 0 | 0 | 0 | 0 | 102900 | 216860 | 0 | 0 | 289150 | 260030 | 50693 | 0 | 0 | 0 | 0 | 0 | 0 | 0 |
| VLVWDLR | BUB3; BUB3 | O43684; J3Q | BUB3_HUMAN | Unmodified | 0 | 0 | 0 | 0 | 0 | 0 | 0 | 0 | 0 | 0 | 0 | 5118900 | 0 | 0 | 0 | 0 | 0 | 0 |
| VLVWVTDG | VWA1; VWA1 | Q6PCB0; J3C | VWA1_HUMAN | Oxidation (M) | 0 | 0 | 1437000 | 684040 | 0 | 0 | 0 | 0 | 0 | 0 | 0 | 0 | 0 | 0 | 292540 | 447540 | 0 | 0 |
| VLVYNNTS | POLR1C; POLR | O15160; A0A | RPAC1_HUMAI | Unmodified | 278290 | 201930 | 234700 | 345860 | 0 | 0 | 0 | 0 | 432030 | 0 | 0 | 2121200 | 0 | 0 | 363220 | 493660 | 0 | 0 |
| VLYAVFSR | PLXNB2; PLXNE | O15031; E2F | PLXB2_HUMAN | Unmodified | 958540 | 1442800 | 1378500 | 179510 | 0 | 0 | 0 | 0 | 1954300 | 0 | 0 | 0 | 0 | 0 | 371640 | 2976500 | 0 | 0 |
| VLYEIPDTY | PRR15L | Q9BU68 | PR15L_HUMAN | Unmodified | 1168300 | 0 | 0 | 0 | 0 | 0 | 0 | 0 | 0 | 693580 | 0 | 0 | 10178000 | 7670600 | 2391100 | 2573300 | 0 | 0 |
| VLYHFLSS( | LLGL2; LLGL2 | Q6P1M3; J3C | L2GL2_HUMAN | Unmodified | 0 | 0 | 0 | 0 | 0 | 0 | 0 | 0 | 0 | 0 | 0 | 0 | 0 | 0 | 673290 | 85293 | 243140 | 0 |
| VLYLLSQL( | KIF4A | O95239 | KIF4A_HUMAN | Oxidation (M) | 0 | 0 | 0 | 0 | 2259000 | 0 | 504160 | 90395 | 0 | 142030 | 0 | 0 | 1189200 | 1267800 | 0 | 260060 | 1342600 | 1818900 |
| VLYLNNNL | SLITRK6 | Q9H5Y7 | SLIK6_HUMAN | Unmodified | 543340 | 2659800 | 0 | 0 | 0 | 0 | 0 | 0 | 145600 | 586120 | 0 | 0 | 0 | 0 | 0 | 0 | 0 | 0 |
| VLYNDLGA | UTRN | P46939 | UTRN_HUMAN | Unmodified | 0 | 0 | 148520 | 0 | 760460 | 832750 | 507520 | 235030 | 0 | 167980 | 0 | 95531 | 957380 | 469320 | 0 | 120730 | 0 | 1032300 |
| VLYNDLGA | UTRN | P46939 | UTRN_HUMAN | Unmodified | 0 | 0 | 1371000 | 2418600 | 0 | 0 | 0 | 0 | 1572000 | 0 | 0 | 0 | 0 | 0 | 2992100 | 2646700 | 523330 | 0 |
| VLYNDLGA | UTRN | P46939 | UTRN_HUMAN | Unmodified | 0 | 0 | 0 | 0 | 2590600 | 0 | 0 | 0 | 0 | 0 | 0 | 0 | 1727700 | 973100 | 0 | 0 | 928400 | 2650500 |
| VLYNLFTK | PARVB; PARVA; | Q9HBI1; Q9I | PARVB_HUMAI | Unmodified | 313580 | 745030 | 1030500 | 1452900 | 0 | 0 | 0 | 0 | 0 | 0 | 0 | 774980 | 0 | 0 | 0 | 0 | 0 | 0 |
| VLYPNDNF | PYGB; PYGM; P | P11216; P11 | PYGB_HUMAN | Unmodified | 0 | 0 | 1041000 | 0 | 0 | 0 | 0 | 0 | 0 | 0 | 332570 | 5576000 | 0 | 0 | 800400 | 764230 | 0 | 0 |
| VLYQVEGF | MYO1B; MYO1E | O43795; E9F | MYO1B_HUMA | Unmodified | 0 | 0 | 0 | 0 | 4546000 | 0 | 542050 | 0 | 1316400 | 398960 | 3943500 | 15169000 | 0 | 0 | 0 | 0 | 0 | 379040 |
| VLYQVEGF | MYO1B; MYO1E | O43795; E9F | MYO1B_HUMA | Unmodified | 114430 | 0 | 0 | 39881 | 260220 | 71686 | 851750 | 0 | 0 | 197020 | 0 | 814980 | 0 | 0 | 0 | 0 | 361640 | 0 |
| VLYVQDSLI | PREP; PREP | P48147; A0A | PPCE_HUMAN | Unmodified | 0 | 0 | 119430 | 0 | 0 | 0 | 0 | 89355 | 0 | 0 | 417520 | 2430400 | 0 | 0 | 0 | 0 | 0 | 0 |
| VLYYMEK | LSR; LSR | Q86X29; S4F | LSR_HUMAN Li | Oxidation (M) | 518070 | 572010 | 208880 | 155460 | 0 | 0 | 0 | 0 | 5130000 | 522690 | 0 | 0 | 0 | 0 | 5216700 | 228710 | 0 | 0 |
| VMAAENIPI | STAT1; STAT1; S | P42224; A0A | STAT1_HUMAN | Unmodified | 0 | 0 | 0 | 0 | 1857300 | 0 | 0 | 0 | 0 | 0 | 0 | 0 | 0 | 0 | 0 | 0 | 0 | 2355800 |
| VMAAENIPI | STAT1; STAT1; S | P42224; A0A | STAT1_HUMAN | Oxidation (M) | 0 | 0 | 0 | 0 | 1015500 | 0 | 198430 | 163900 | 0 | 0 | 0 | 960380 | 0 | 440120 | 0 | 60712 | 664650 | 1635100 |
| VMCLFEEQ | N/A | N/A | N/A | Oxidation (M) | 0 | 0 | 0 | 0 | 0 | 0 | 0 | 0 | 0 | 4531800 | 0 | 8797500 | 0 | 0 | 0 | 0 | 0 | 0 |
| VMDDLDR | AARS1; AARS1; | P49588; A0A | SYAC_HUMAN | Oxidation (M) | 128370 | 0 | 0 | 0 | 604340 | 0 | 88032 | 65182 | 0 | 124150 | 0 | 0 | 0 | 735830 | 0 | 0 | 127640 | 140070 |
| VMDDLDR | AARS1; AARS1; | P49588; A0A | SYAC_HUMAN | Unmodified | 0 | 0 | 0 | 0 | 1025200 | 0 | 0 | 0 | 0 | 606250 | 0 | 0 | 767340 | 0 | 0 | 0 | 228450 | 0 |
| VMDLETK | NHLRC2 | Q8NBF2 | NHLC2_HUMA | Unmodified | 0 | 0 | 0 | 0 | 832980 | 63909 | 0 | 0 | 482400 | 1377200 | 244360 | 677540 | 369150 | 254790 | 0 | 0 | 79204 | 310810 |
| VMDRPGN | ALDH7A1; ALDI | P49419; A0A | AL7A1_HUMAN | Oxidation (M) | 0 | 0 | 0 | 0 | 4001000 | 0 | 0 | 0 | 550860 | 381800 | 359100 | 1705800 | 0 | 0 | 0 | 0 | 6884000 | 30430000 |
| VMEALSLY | STAM | Q92783 | STAM1_HUMAI | Unmodified | 0 | 0 | 0 | 0 | 3799400 | 0 | 132920 | 0 | 2767000 | 1063800 | 0 | 0 | 0 | 0 | 0 | 0 | 550450 | 1236900 |
| VMEALSLY | STAM | Q92783 | STAM1_HUMAI | Oxidation (M) | 780510 | 688740 | 277740 | 340750 | 2400800 | 1303600 | 1146900 | 584580 | 914980 | 1193000 | 0 | 0 | 901330 | 717000 | 284880 | 161300 | 353200 | 760200 |
| VMEEEGLK | STK38 | Q15208 | STK38_HUMAN | Unmodified | 0 | 0 | 0 | 85885 | 1059600 | 172670 | 99311 | 65926 | 392190 | 180090 | 164610 | 834050 | 194300 | 107640 | 0 | 0 | 197240 | 791460 |
| VMEGTVAA | CSK | P41240 | CSK_HUMAN T | Oxidation (M) | 0 | 0 | 0 | 0 | 0 | 0 | 0 | 0 | 0 | 0 | 0 | 902960 | 0 | 0 | 0 | 0 | 0 | 0 |
| VMEGTVAA | CSK | P41240 | CSK_HUMAN T | Unmodified | 0 | 0 | 0 | 0 | 0 | 0 | 0 | 18134 | 2366600 | 0 | 83429 | 1569300 | 0 | 0 | 0 | 0 | 0 | 0 |
| VMEHFIK | HSPA5; HSPA5; | P11021; A0A | BIP_HUMAN Er | Oxidation (M) | 0 | 0 | 0 | 0 | 0 | 0 | 0 | 0 | 1851500 | 67146 | 83033 | 112540 | 0 | 0 | 441350 | 0 | 0 | 0 |
| VMEHFIK | HSPA5; HSPA5; | P11021; A0A | BIP_HUMAN Er | Unmodified | 0 | 0 | 0 | 0 | 0 | 0 | 0 | 0 | 12184000 | 0 | 610500 | 1296300 | 0 | 0 | 386900 | 85692 | 0 | 0 |
| VMELLLK | ANK3; ANK1; AI | Q12955; P16 | ANK3_HUMAN | Oxidation (M) | 0 | 0 | 169870 | 148560 | 0 | 0 | 0 | 0 | 0 | 0 | 0 | 0 | 518260 | 386850 | 359370 | 367770 | 0 | 0 |
| VMELLLK | ANK3; ANK1; AI | Q12955; P16 | ANK3_HUMAN | Unmodified | 0 | 0 | 0 | 0 | 0 | 0 | 0 | 0 | 0 | 0 | 0 | 0 | 0 | 1293200 | 539070 | 0 | 0 | 0 |
| VMETQAQV | VAMP7 | P51809 | VAMP7_HUMAI | Unmodified | 0 | 0 | 118390 | 0 | 0 | 0 | 484400 | 31738 | 1404000 | 337190 | 0 | 181400 | 0 | 0 | 138080 | 110590 | 0 | 271510 |
| VMETQAQV | VAMP7 | P51809 | VAMP7_HUMAI | Oxidation (M) | 0 | 0 | 0 | 149560 | 396360 | 0 | 78824 | 0 | 618740 | 78807 | 252280 | 189410 | 224970 | 0 | 242510 | 114790 | 0 | 0 |
| VMEYINR | CLTC; CLTC | Q00610; A0A | CLH1_HUMAN | Unmodified | 2157000 | 949040 | 430340 | 0 | 0 | 1285100 | 0 | 0 | 581970 | 129690 | 3830400 | 13775000 | 0 | 0 | 1300400 | 553350 | 0 | 548410 |
| VMEYINR | CLTC; CLTC | Q00610; A0A | CLH1_HUMAN | Oxidation (M) | 444880 | 0 | 46619 | 0 | 0 | 0 | 0 | 0 | 0 | 0 | 0 | 797590 | 0 | 0 | 314640 | 190460 | 0 | 0 |
| VMFSSFGC | CELF1; CELF1 | Q92879; G5I | CELF1_HUMAN | Oxidation (M) | 494440 | 424110 | 0 | 0 | 0 | 0 | 0 | 0 | 214130 | 0 | 0 | 181400 | 0 | 0 | 106090 | 105990 | 0 | 0 |
| VMGLVPAK | FASN; FASN | P49327; A0A | FAS_HUMAN F | Unmodified | 0 | 32688 | 294550 | 581760 | 0 | 0 | 243140 | 0 | 396790 | 73870 | 0 | 17715000 | 135980 | 64904 | 325290 | 155220 | 0 | 0 |
| VMIPQDEY | SF1; SF1 | Q15637; A0A | SF01_HUMAN | Oxidation (M) | 0 | 0 | 0 | 287950 | 730840 | 0 | 0 | 0 | 0 | 225420 | 0 | 0 | 1577800 | 0 | 0 | 0 | 0 | 0 |
| VMITHCNL | GALNT7; GALN | Q86SF2; H0Y | GALT7_HUMAN | Unmodified | 0 | 0 | 0 | 0 | 0 | 0 | 305750 | 619890 | 2574000 | 460610 | 0 | 383320 | 0 | 0 | 0 | 0 | 0 | 0 |
| VMITHCNL | GALNT7; GALN | Q86SF2; H0Y | GALT7_HUMAN | Oxidation (M) | 0 | 0 | 0 | 0 | 0 | 0 | 0 | 0 | 0 | 581750 | 0 | 0 | 0 | 0 | 0 | 0 | 21111 | 0 |
| VMIWNLDT | CORO2A | Q92828 | COR2A_HUMAI | Oxidation (M) | 0 | 0 | 0 | 0 | 0 | 0 | 0 | 0 | 0 | 0 | 0 | 723940 | 0 | 0 | 0 | 0 | 0 | 0 |
| VMIYQDEV | CAVIN1 | Q6NZI2 | CAVN1_HUMAI | Unmodified | 0 | 0 | 0 | 0 | 24766000 | 1989000 | 0 | 69373 | 0 | 0 | 0 | 0 | 0 | 0 | 0 | 0 | 0 | 0 |
| VMIYQDEV | CAVIN1 | Q6NZI2 | CAVN1_HUMAI | Oxidation (M) | 0 | 0 | 0 | 0 | 12179000 | 1790600 | 0 | 256160 | 0 | 0 | 0 | 0 | 0 | 0 | 0 | 0 | 0 | 0 |
| VMLETPEYI | CYRIA; CYRIB; ( | Q9H0Q0; Q9 | CYRIA_HUMAN | Oxidation (M) | 0 | 0 | 0 | 0 | 0 | 0 | 0 | 0 | 280370 | 0 | 0 | 2571000 | 0 | 0 | 0 | 0 | 0 | 0 |
| VMLGETNP | NME2P1; NME1 | O60361; P15 | NDK8_HUMAN | Oxidation (M) | 2651600 | 1357000 | 4199600 | 3807700 | 205480 | 118420 | 84303 | 0 | 3571900 | 160970 | 6111700 | 1.91E+08 | 425510 | 111070 | 8125600 | 7589200 | 194180 | 662840 |
| VMLGETNP | NME2P1; NME1 | O60361; P15 | NDK8_HUMAN | Unmodified | 1957500 | 1175900 | 1870000 | 2421200 | 0 | 0 | 0 | 19749 | 8361700 | 0 | 7608000 | 2.3E+08 | 0 | 0 | 4592700 | 3402400 | 0 | 0 |
| VMLGETNP | NME2P1; NME1 | O60361; P15 | NDK8_HUMAN | Oxidation (M) | 262420 | 0 | 0 | 863220 | 1866700 | 0 | 0 | 0 | 987480 | 0 | 0 | 9093900 | 0 | 0 | 327180 | 0 | 2510000 | 1143500 |
| VMLNDPLP | TTC13 | Q8NBP0 | TTC13_HUMAN | Unmodified | 0 | 0 | 0 | 0 | 0 | 0 | 0 | 0 | 751370 | 162610 | 0 | 0 | 0 | 0 | 0 | 0 | 0 | 0 |
| VMLPGQC( | ERI3; ERI3; ERI3 | O43414; F6C | ERI3_HUMAN E | Oxidation (M) | 2004600 | 686130 | 79147 | 369800 | 57848 | 0 | 22501 | 0 | 0 | 310740 | 0 | 103680 | 412400 | 0 | 821020 | 779050 | 0 | 0 |
| VMLPPGAC | AFDN; AFDN; A | P55196; Q5T | AFAD_HUMAN | Unmodified | 0 | 0 | 0 | 0 | 598300 | 0 | 0 | 22466 | 0 | 116480 | 450520 | 1157900 | 0 | 908220 | 0 | 0 | 0 | 132170 |
| VMLSERKG | PRKCB; PRKCB | P05771; H3E | KPCB_HUMAN | Unmodified | 0 | 0 | 0 | 0 | 477410 | 56933 | 1353700 | 279820 | 0 | 0 | 0 | 0 | 545920 | 706510 | 0 | 0 | 79045 | 307870 |

| | | | | | | | | | | | | | | | | | | | | | | |
|---|---|---|---|---|---|---|---|---|---|---|---|---|---|---|---|---|---|---|---|---|---|---|
| VMLSERKG | PRKCB; PRKCB | P05771; H3E | KPCB_HUMAN | Oxidation (M) | 0 | 0 | 0 | 0 | 0 | 0 | 1800900 | 1617900 | 0 | 0 | 0 | 0 | 0 | 0 | 0 | 0 | 0 | 0 |
| VMLTTAGG | MYO6; MYO6; M | Q9UM54; A0 | MYO6_HUMAN | Unmodified | 0 | 124260 | 1277500 | 0 | 0 | 0 | 0 | 0 | 616120 | 0 | 0 | 2545600 | 0 | 0 | 2356000 | 913840 | 0 | 0 |
| VMLVNSMI | VCL | P18206 | VINC_HUMAN | Unmodified | 0 | 0 | 0 | 76455 | 288880 | 0 | 0 | 0 | 533670 | 0 | 0 | 1239900 | 0 | 0 | 0 | 0 | 367440 | 0 |
| VMLVNSMI | VCL | P18206 | VINC_HUMAN | Oxidation (M) | 0 | 107210 | 0 | 0 | 0 | 0 | 0 | 0 | 0 | 0 | 0 | 1293600 | 0 | 0 | 397760 | 149680 | 0 | 0 |
| VMNLISK | ST13; ST13P5 | P50502; Q8N | F10A1_HUMAN | Oxidation (M) | 0 | 0 | 59465 | 0 | 0 | 0 | 134040 | 0 | 400360 | 377910 | 110690 | 1728200 | 1956400 | 1028000 | 0 | 0 | 170800 | 0 |
| VMPAPPPK | EIF4B; EIF4B | P23588; E7E | IF4B_HUMAN E | Oxidation (M) | 0 | 0 | 0 | 0 | 749810 | 0 | 0 | 0 | 0 | 0 | 0 | 0 | 0 | 0 | 0 | 0 | 474660 | 647030 |
| VMPAPPPK | EIF4B; EIF4B | P23588; E7E | IF4B_HUMAN E | Unmodified | 0 | 0 | 0 | 0 | 861810 | 199430 | 0 | 0 | 0 | 169360 | 0 | 0 | 585300 | 330610 | 0 | 0 | 1539100 | 581440 |
| VMPFSTAC | IARS1; IARS1; IA | P41252; A0A | SYIC_HUMAN I | Oxidation (M) | 0 | 0 | 0 | 0 | 0 | 0 | 0 | 0 | 0 | 0 | 0 | 3149800 | 0 | 0 | 0 | 0 | 0 | 0 |
| VMPQNHSI | PAAT | Q9H8K7 | PAAT_HUMAN A | Oxidation (M) | 521770 | 0 | 0 | 0 | 0 | 0 | 0 | 0 | 0 | 0 | 0 | 0 | 0 | 0 | 0 | 81630 | 0 | 0 |
| VMPQNHSI | PAAT | Q9H8K7 | PAAT_HUMAN A | Unmodified | 512790 | 0 | 0 | 0 | 0 | 0 | 0 | 0 | 0 | 0 | 0 | 0 | 0 | 0 | 0 | 0 | 0 | 0 |
| VMQEEIFGI | ALDH3A2; ALDH | P51648; A0A | AL3A2_HUMAN | Oxidation (M) | 805730 | 0 | 111450 | 0 | 0 | 0 | 0 | 0 | 17455000 | 3390300 | 207570 | 368030 | 0 | 0 | 0 | 0 | 0 | 0 |
| VMQEEIFGI | ALDH3A2; ALDH | P51648; A0A | AL3A2_HUMAN | Unmodified | 0 | 0 | 0 | 0 | 0 | 0 | 0 | 0 | 2906100 | 0 | 0 | 0 | 0 | 0 | 0 | 0 | 0 | 0 |
| VMQEQGTI | MYH9 | P35579 | MYH9_HUMAN | Unmodified | 0 | 0 | 0 | 0 | 990820 | 0 | 1575900 | 0 | 0 | 0 | 0 | 0 | 0 | 554610 | 0 | 0 | 2042600 | 520100 |
| VMQPQILE | TTLL12; TTLL12 | Q14166; V9C | TTL12_HUMAN | Oxidation (M) | 0 | 0 | 0 | 0 | 0 | 0 | 0 | 0 | 279510 | 0 | 0 | 1734700 | 0 | 0 | 0 | 0 | 0 | 0 |
| VMQQQQC | SF3A1 | Q15459 | SF3A1_HUMAN | Oxidation (M) | 322850 | 304970 | 0 | 325330 | 0 | 0 | 0 | 0 | 0 | 0 | 0 | 110890 | 136450 | 0 | 0 | 115490 | 0 | 0 |
| VMQQQQC | SF3A1 | Q15459 | SF3A1_HUMAN | Unmodified | 519970 | 386810 | 228580 | 530360 | 0 | 0 | 0 | 0 | 0 | 0 | 0 | 292350 | 0 | 0 | 99403 | 114680 | 0 | 0 |
| VMQVLNAI | SND1 | Q7KZF4 | SND1_HUMAN | Oxidation (M) | 0 | 0 | 0 | 1341500 | 0 | 0 | 0 | 244970 | 311930 | 0 | 957640 | 6465900 | 0 | 447290 | 632320 | 0 | 144370 | 378520 |
| VMQVLNAI | SND1 | Q7KZF4 | SND1_HUMAN | Unmodified | 0 | 0 | 0 | 0 | 376500 | 0 | 0 | 0 | 0 | 0 | 0 | 714860 | 0 | 0 | 0 | 0 | 0 | 0 |
| VMRVEYHF | TFRC; TFRC | P02786; G3V | TFR1_HUMAN | Oxidation (M) | 769600 | 530000 | 3943300 | 4211100 | 0 | 0 | 0 | 0 | 8223500 | 470350 | 248380 | 745940 | 232240 | 0 | 2167000 | 1329000 | 64364 | 0 |
| VMRVEYHF | TFRC; TFRC | P02786; G3V | TFR1_HUMAN | Unmodified | 89518 | 116740 | 129370 | 191150 | 78890 | 0 | 0 | 0 | 6701000 | 0 | 0 | 157060 | 0 | 0 | 93730 | 0 | 0 | 0 |
| VMSAILMEI | TOM1L1; TOM1 | O75674; J3K | TM1L1_HUMAN | 2 Oxidation (M) | 4627200 | 5747900 | 2722000 | 5145800 | 0 | 0 | 0 | 0 | 0 | 0 | 0 | 0 | 0 | 0 | 8694100 | 6479000 | 0 | 0 |
| VMSEFNNI | CD63; CD63; C | P08962; F8V | CD63_HUMAN | Unmodified | 2658800 | 1012000 | 1532100 | 458730 | 0 | 0 | 0 | 0 | 4894400 | 1107900 | 596700 | 1157700 | 0 | 0 | 2750600 | 1590600 | 0 | 0 |
| VMSEFNNI | CD63; CD63; C | P08962; F8V | CD63_HUMAN | Oxidation (M) | 4342400 | 4662200 | 4429000 | 263440 | 0 | 0 | 0 | 0 | 1693900 | 118670 | 75097 | 1105900 | 0 | 0 | 6878700 | 3413600 | 0 | 0 |
| VMSEFNNI | CD63; CD63; C | P08962; F8V | CD63_HUMAN | Unmodified | 0 | 0 | 0 | 0 | 0 | 0 | 327870 | 0 | 2542900 | 0 | 0 | 0 | 0 | 0 | 0 | 0 | 0 | 0 |
| VMSEFNNI | CD63; CD63; C | P08962; F8V | CD63_HUMAN | Oxidation (M) | 0 | 0 | 0 | 0 | 0 | 0 | 271960 | 0 | 1086200 | 127760 | 0 | 0 | 0 | 0 | 0 | 0 | 0 | 0 |
| VMSQEIQE | DYNC1H1; DYN | Q14204; A0A | DYHC1_HUMA | Unmodified | 0 | 0 | 0 | 0 | 372890 | 0 | 85604 | 0 | 1332000 | 233730 | 733720 | 8886400 | 313680 | 0 | 0 | 0 | 0 | 481920 |
| VMSQEIQE | DYNC1H1; DYN | Q14204; A0A | DYHC1_HUMA | Oxidation (M) | 0 | 0 | 0 | 0 | 229070 | 0 | 0 | 20408 | 187850 | 0 | 274150 | 1236600 | 0 | 0 | 0 | 0 | 0 | 0 |
| VMSQEIQE | DYNC1H1; DYN | Q14204; A0A | DYHC1_HUMA | Unmodified | 0 | 0 | 0 | 0 | 1201400 | 0 | 279010 | 38819 | 555530 | 630690 | 0 | 0 | 656980 | 1118400 | 0 | 0 | 217610 | 1446200 |
| VMSQNFTI | G3BP1; G3BP1; | Q13283; A0A | G3BP1_HUMAN | Unmodified | 42009 | 0 | 0 | 0 | 676320 | 0 | 162000 | 88142 | 522650 | 345570 | 749610 | 7150700 | 735380 | 444820 | 0 | 0 | 477280 | 604050 |
| VMSTSSLEI | CASP8AP2; CAS | Q9UKL3; A0A | C8AP2_HUMAN | Unmodified | 0 | 0 | 0 | 0 | 0 | 0 | 0 | 0 | 0 | 0 | 0 | 0 | 0 | 0 | 0 | 0 | 0 | 0 |
| VMTIAPGLI | HSD17B10 | Q99714 | HCD2_HUMAN | Unmodified | 0 | 0 | 0 | 0 | 42924 | 0 | 47834 | 0 | 2643400 | 153800 | 0 | 0 | 0 | 0 | 0 | 0 | 31942 | 0 |
| VMTIPYQPI | PCBP1 | Q15365 | PCBP1_HUMAN | 2 Oxidation (M) | 8801300 | 3827000 | 3830400 | 634140 | 0 | 0 | 0 | 0 | 0 | 0 | 0 | 7289900 | 0 | 0 | 0 | 1481800 | 0 | 0 |
| VMTLITYNK | SORCS2; SORC | Q96PQ0; B5 | SORC2_HUMAN | Unmodified | 0 | 0 | 0 | 0 | 0 | 0 | 0 | 0 | 1999900 | 0 | 0 | 0 | 0 | 0 | 0 | 0 | 0 | 0 |
| VMTVNYNT | MRPS25; MRPS | P82663; E7E | RT25_HUMAN | Unmodified | 0 | 0 | 0 | 0 | 0 | 0 | 0 | 0 | 843300 | 76924 | 0 | 0 | 0 | 0 | 0 | 162390 | 0 | 0 |
| VMTVSFHK | HDAC3; HDAC3 | O15379; Q9 | HDAC3_HUMA | Unmodified | 0 | 131740 | 0 | 0 | 0 | 0 | 0 | 0 | 0 | 0 | 0 | 610740 | 0 | 0 | 0 | 0 | 0 | 0 |
| VMVEGVGA | MIB1 | Q86YT6 | MIB1_HUMAN I | Unmodified | 18323 | 332820 | 0 | 0 | 213560 | 1284900 | 0 | 0 | 3013500 | 0 | 0 | 334340 | 0 | 0 | 0 | 0 | 0 | 0 |
| VMVIVTDGI | ITGA1 | P56199 | ITA1_HUMAN I | Unmodified | 324800 | 296210 | 0 | 0 | 186340 | 0 | 0 | 0 | 125040 | 0 | 0 | 0 | 0 | 0 | 0 | 0 | 0 | 781070 |
| VMVIVTDGI | ITGA1 | P56199 | ITA1_HUMAN I | Unmodified | 0 | 0 | 0 | 0 | 642380 | 0 | 0 | 0 | 0 | 147730 | 0 | 0 | 0 | 0 | 0 | 0 | 3392100 | 7534500 |
| VMVIVTDGI | ITGA1 | P56199 | ITA1_HUMAN I | Oxidation (M) | 0 | 0 | 0 | 0 | 0 | 0 | 18175 | 17530 | 0 | 0 | 0 | 0 | 0 | 0 | 0 | 0 | 922310 | 1040600 |
| VMVIVTDGI | ITGA1 | P56199 | ITA1_HUMAN I | Unmodified | 0 | 0 | 0 | 0 | 0 | 0 | 0 | 0 | 294790 | 412160 | 0 | 0 | 0 | 0 | 0 | 0 | 2347800 | 4150100 |
| VMVIVTDGI | ITGA1 | P56199 | ITA1_HUMAN I | Oxidation (M) | 0 | 0 | 0 | 0 | 0 | 0 | 0 | 0 | 0 | 0 | 0 | 0 | 0 | 0 | 0 | 0 | 0 | 545170 |
| VMVLDFVTI | TXNRD1; TXNRI | Q16881; A0A | TRXR1_HUMAN | Oxidation (M) | 0 | 0 | 94846 | 191670 | 1554600 | 19940 | 0 | 0 | 810650 | 274720 | 1482500 | 7188800 | 0 | 136430 | 200800 | 236350 | 0 | 0 |
| VMVQPINLI | SNRPE | P62304 | RUXE_HUMAN | Oxidation (M) | 6420500 | 5813200 | 1523100 | 790460 | 0 | 0 | 0 | 0 | 955620 | 0 | 499280 | 17260000 | 718000 | 0 | 2748800 | 1506000 | 0 | 0 |
| VMVQPINLI | SNRPE | P62304 | RUXE_HUMAN | Unmodified | 0 | 0 | 0 | 0 | 0 | 0 | 0 | 0 | 168620 | 0 | 0 | 88426 | 0 | 0 | 0 | 0 | 0 | 0 |
| VMVSRAEII | ANXA4; ANXA4 | P09525; Q6F | ANXA4_HUMAN | Oxidation (M) | 60747 | 0 | 0 | 0 | 0 | 14540 | 0 | 42975 | 5081800 | 0 | 1342600 | 0 | 0 | 0 | 38682 | 0 | 0 | 0 |
| VMVSRAEII | ANXA4; ANXA4 | P09525; Q6F | ANXA4_HUMAN | Unmodified | 0 | 0 | 0 | 670610 | 130880 | 6061700 | 0 | 218270 | 2415200 | 366840 | 323000 | 0 | 0 | 0 | 0 | 0 | 273740 | 0 |
| VMVSRAEII | ANXA4; ANXA4 | P09525; Q6F | ANXA4_HUMAN | 2 Oxidation (M) | 290590 | 0 | 190190 | 447000 | 0 | 0 | 0 | 0 | 1636400 | 0 | 3413400 | 1269300 | 0 | 0 | 320120 | 167680 | 0 | 0 |
| VMVSRAEII | ANXA4; ANXA4 | P09525; Q6F | ANXA4_HUMAN | Oxidation (M) | 0 | 0 | 0 | 0 | 248040 | 0 | 0 | 0 | 208130 | 484290 | 144850 | 0 | 0 | 0 | 0 | 0 | 0 | 0 |
| VMVTNVTS | TLN2; TLN1; TLI | Q9Y4G6; Q9 | TLN2_HUMAN | Unmodified | 0 | 0 | 0 | 0 | 2129400 | 142180 | 2240800 | 0 | 221550 | 0 | 0 | 0 | 0 | 0 | 0 | 0 | 1449300 | 308850 |
| VMVTNVTS | TLN2; TLN1; TLI | Q9Y4G6; Q9 | TLN2_HUMAN | Oxidation (M) | 589840 | 480920 | 1193000 | 857650 | 3319200 | 1499700 | 4216000 | 3019500 | 345580 | 881130 | 301040 | 3372800 | 1244900 | 3092400 | 1248500 | 1124100 | 2748100 | 628030 |
| VMVTNVTS | TLN1 | Q9Y490 | TLN1_HUMAN | Oxidation (M) | 0 | 0 | 0 | 0 | 0 | 0 | 257530 | 0 | 0 | 0 | 0 | 0 | 27928 | 401280 | 0 | 0 | 0 | 0 |
| VMVVVTDG | ITGA2; ITGA2; I | P17301; E9P | ITA2_HUMAN I | 2 Oxidation (M) | 698020 | 681230 | 0 | 0 | 1281100 | 784960 | 0 | 0 | 0 | 0 | 393700 | 757840 | 0 | 0 | 2918600 | 766060 | 1729400 | 1258600 |
| VMVVVTDG | ITGA2; ITGA2; I | P17301; E9P | ITA2_HUMAN I | Oxidation (M) | 720810 | 860080 | 735480 | 2123500 | 6395200 | 2921000 | 0 | 0 | 0 | 59979 | 0 | 204430 | 1070200 | 281020 | 2754400 | 1421800 | 7016200 | 5037300 |
| VMVVVTDG | ITGA2; ITGA2; I | P17301; E9P | ITA2_HUMAN I | Unmodified | 187040 | 150760 | 0 | 528220 | 10747000 | 2840800 | 0 | 0 | 0 | 0 | 0 | 0 | 341370 | 0 | 460750 | 220260 | 5600200 | 0 |
| VMYGFCSV | DBN1; DBN1; D | Q16643; D6I | DREB_HUMAN | Oxidation (M) | 0 | 0 | 0 | 223180 | 1150100 | 0 | 141170 | 380880 | 0 | 0 | 0 | 2830300 | 613640 | 517190 | 0 | 208330 | 2325700 | 1092700 |
| VMYGFCSV | DBN1; DBN1; D | Q16643; D6I | DREB_HUMAN | Oxidation (M) | 82500 | 0 | 0 | 0 | 0 | 0 | 0 | 0 | 0 | 0 | 0 | 0 | 0 | 0 | 0 | 0 | 2007900 | 0 |
| VMYGFCSV | DBN1; DBN1; D | Q16643; D6I | DREB_HUMAN | Unmodified | 0 | 0 | 0 | 0 | 1370300 | 1125200 | 0 | 0 | 0 | 0 | 0 | 0 | 107500 | 0 | 0 | 0 | 2694800 | 136050 |
| VNADIEKVI | NSDHL; NSDHL | Q15738; C9J | NSDHL_HUMA | Unmodified | 0 | 0 | 0 | 0 | 0 | 0 | 181540 | 122870 | 0 | 0 | 0 | 0 | 0 | 0 | 0 | 0 | 0 | 0 |
| VNAEGSVD | AK1; AK1 | P00568; Q5T | KAD1_HUMAN | Unmodified | 663200 | 854180 | 0 | 1395200 | 9443400 | 1587000 | 2272100 | 46824 | 2686300 | 4849600 | 939360 | 922720 | 0 | 4728100 | 1857100 | 556210 | 3799800 | 4682600 |
| VNAIEHVIII | ATP6V1D; ATP6 | Q9Y5K8; G3 | VATD_HUMAN | Unmodified | 0 | 0 | 0 | 240690 | 0 | 0 | 0 | 7549.2 | 379140 | 28935 | 305690 | 1361200 | 35735 | 0 | 0 | 0 | 0 | 0 |
| VNAMTFTFI | NOMO1; NOMO | Q15155; P69 | NOMO1_HUMA | Oxidation (M) | 0 | 0 | 0 | 0 | 0 | 0 | 0 | 0 | 0 | 941980 | 696430 | 0 | 0 | 0 | 0 | 0 | 0 | 0 |
| VNAMTFTFI | NOMO1; NOMO | Q15155; P69 | NOMO1_HUMA | Unmodified | 0 | 0 | 0 | 0 | 0 | 0 | 0 | 0 | 1898900 | 203340 | 0 | 98613 | 0 | 0 | 0 | 0 | 0 | 0 |
| VNAQAAAY | COPS5; COPS5 | Q92905; E5F | CSN5_HUMAN | Unmodified | 0 | 0 | 0 | 0 | 0 | 0 | 0 | 0 | 0 | 0 | 0 | 1450400 | 0 | 0 | 0 | 0 | 0 | 0 |
| VNASASSLI | FSCN1 | Q16658 | FSCN1_HUMAI | Unmodified | 0 | 0 | 0 | 69422 | 12229000 | 3003500 | 56828 | 1214600 | 1687000 | 416030 | 332060 | 6044600 | 192400 | 134520 | 46878 | 0 | 5350800 | 6906600 |
| VNAVFEWI | HSD17B4; HSD | P51659; E7E | DHB4_HUMAN | Unmodified | 0 | 0 | 0 | 0 | 0 | 0 | 0 | 0 | 5407100 | 0 | 0 | 0 | 0 | 0 | 0 | 0 | 0 | 0 |
| VNAWTSGI | EDIL3 | O43854 | EDIL3_HUMAN | Unmodified | 2047300 | 280420 | 0 | 75054 | 1.63E+08 | 38478000 | 1014500 | 0 | 17085000 | 6682800 | 0 | 0 | 0 | 0 | 0 | 12223 | 3009200 | 76490000 |
| VNAWTSGI | EDIL3 | O43854 | EDIL3_HUMAN | Unmodified | 0 | 0 | 0 | 0 | 32603000 | 1503300 | 395060 | 0 | 0 | 0 | 0 | 0 | 0 | 0 | 0 | 0 | 623870 | 13360000 |
| VNCLAPGL | DHRS4L1; DHR | P0CG22; Q9 | DR4L1_HUMAN | Unmodified | 0 | 0 | 0 | 0 | 0 | 0 | 0 | 0 | 7829900 | 439520 | 0 | 0 | 0 | 0 | 0 | 0 | 0 | 0 |
| VNDASVNT | TAX1BP1; TAX1I | Q86VP1; B8 | TAXB1_HUMAN | Unmodified | 0 | 0 | 0 | 0 | 0 | 0 | 1873200 | 202590 | 0 | 0 | 0 | 0 | 0 | 0 | 0 | 0 | 1302700 | 2861500 |
| VNDFLAEIF | CKAP5; CKAP5 | Q14008; H0 | CKAP5_HUMAN | Unmodified | 0 | 316140 | 40556 | 42210 | 88140 | 0 | 0 | 0 | 0 | 0 | 136870 | 560830 | 144110 | 0 | 89258 | 66833 | 63831 | 0 |
| VNDGVCDI | PRKCSH; PRKC | P14314; K7E | GLU2B_HUMAI | Unmodified | 0 | 0 | 0 | 0 | 0 | 0 | 0 | 0 | 1333200 | 0 | 0 | 0 | 0 | 0 | 0 | 0 | 0 | 0 |
| VNDKNPH | HGS; HGS; HGS | O14964; A0A | HGS_HUMAN H | Unmodified | 574730 | 462290 | 0 | 0 | 0 | 73902 | 227180 | 0 | 461740 | 499830 | 0 | 0 | 0 | 0 | 0 | 0 | 0 | 84150 |
| VNDTIQIDI | RPS4X | P62701 | RS4X_HUMAN | Unmodified | 0 | 0 | 1166500 | 360550 | 0 | 0 | 189970 | 118970 | 0 | 168200 | 597630 | 3146700 | 0 | 0 | 0 | 0 | 0 | 0 |
| VNDTVQID | RPS4Y2; RPS4Y | Q8TD47; P2 | RS4Y2_HUMAN | Unmodified | 128880 | 171950 | 0 | 0 | 0 | 0 | 0 | 0 | 0 | 0 | 387240 | 0 | 0 | 0 | 0 | 0 | 0 | 0 |
| VNEAAPEK | UBR4 | Q5T4S7 | UBR4_HUMAN | Unmodified | 0 | 0 | 0 | 0 | 0 | 0 | 0 | 0 | 0 | 0 | 0 | 1062200 | 0 | 0 | 0 | 0 | 0 | 0 |
| VNEAREELI | ARF3; ARF1; N/ | P61204; P84 | ARF3_HUMAN | Unmodified | 6777500 | 5916300 | 7225100 | 3072100 | 0 | 0 | 0 | 25019 | 4161300 | 750070 | 2124300 | 58015000 | 0 | 0 | 10891000 | 8055800 | 435450 | 47828 |
| VNEASGDO | DCPS | Q96C86 | DCPS_HUMAN | Unmodified | 0 | 0 | 0 | 0 | 0 | 0 | 0 | 0 | 0 | 0 | 845710 | 1663800 | 0 | 0 | 0 | 0 | 0 | 0 |
| VNEESLDR | ABRAXAS2 | Q15018 | ABRX2_HUMAN | Unmodified | 0 | 121040 | 0 | 0 | 89077 | 0 | 0 | 0 | 0 | 0 | 49045 | 429930 | 0 | 0 | 0 | 30249 | 22639 | 0 |
| VNEIVETNI | PSMD6 | Q15008 | PSMD6_HUMA | Unmodified | 66073 | 0 | 275610 | 164860 | 271280 | 0 | 0 | 0 | 341700 | 44657 | 300410 | 5126500 | 200980 | 0 | 231000 | 198270 | 150840 | 0 |
| VNELKLDD | SRP54; SRP54; | P61011; G3 | SRP54_HUMAN | Unmodified | 96732 | 0 | 154460 | 0 | 0 | 0 | 0 | 0 | 0 | 0 | 492770 | 3037900 | 0 | 0 | 536110 | 467470 | 0 | 0 |
| VNESSHYD | TPP2; TPP2; TPI | P29144; A0A | TPP2_HUMAN | Unmodified | 0 | 0 | 0 | 0 | 0 | 0 | 0 | 0 | 672000 | 0 | 0 | 8255800 | 0 | 0 | 0 | 0 | 0 | 0 |
| VNESSHYD | TPP2; TPP2; TPI | P29144; A0A | TPP2_HUMAN | Unmodified | 0 | 0 | 0 | 0 | 0 | 10944000 | 2109000 | 0 | 0 | 1015700 | 0 | 2023100 | 0 | 0 | 0 | 0 | 0 | 0 |
| VNESSLNW | CNN3; CNN3 | Q15417; E9F | CNN3_HUMAN | Unmodified | 71976 | 143870 | 0 | 0 | 1155900 | 301800 | 0 | 0 | 318700 | 341780 | 394230 | 661680 | 0 | 16665 | 0 | 0 | 248400 | 377840 |
| VNETTEFLS | PARP4 | Q9UKK3 | PARP4_HUMAN | Unmodified | 395980 | 479980 | 0 | 0 | 0 | 0 | 0 | 0 | 0 | 0 | 0 | 1416200 | 0 | 0 | 221390 | 156920 | 0 | 0 |
| VNEYLPGQ | FARP2 | O94887 | FARP2_HUMAN | Unmodified | 177190 | 0 | 0 | 0 | 700590 | 0 | 0 | 0 | 1331500 | 198160 | 0 | 0 | 607470 | 640830 | 0 | 1145400 | 0 | 0 |
| VNFAMNV | PKM; PKM; PKM | P14618; B4D | KPYM_HUMAN | Unmodified | 444940 | 0 | 842220 | 0 | 582450 | 0 | 0 | 0 | 1341300 | 0 | 0 | 67382000 | 0 | 0 | 2226300 | 1610300 | 65440 | 0 |
| VNFAMNV | PKM; PKM; PKM | P14618; B4D | KPYM_HUMAN | Oxidation (M) | 0 | 252670 | 353530 | 0 | 0 | 151410 | 0 | 0 | 352490 | 0 | 299050 | 15905000 | 0 | 0 | 1611700 | 578520 | 75597 | 0 |
| VNFEDCTG | CDH1; CDH1; C | P12830; H3E | CADH1_HUMA | Unmodified | 0 | 0 | 0 | 0 | 0 | 0 | 0 | 0 | 0 | 0 | 0 | 0 | 0 | 0 | 0 | 0 | 0 | 0 |
| VNFEDDSR | SEC31A; SEC31 | O94979; H7I | SC31A_HUMAN | Unmodified | 0 | 0 | 0 | 0 | 0 | 0 | 0 | 0 | 0 | 0 | 0 | 598200 | 0 | 0 | 0 | 0 | 0 | 0 |
| VNFGTDFL | TBCE; TBCE; TB | Q15813; A0A | TBCE_HUMAN | Unmodified | 0 | 0 | 0 | 0 | 0 | 0 | 0 | 0 | 0 | 0 | 478980 | 3413200 | 420540 | 0 | 0 | 0 | 0 | 0 |
| VNFHFILFI | UCHL1; UCHL1 | P09936; D6F | UCHL1_HUMAI | Oxidation (M) | 0 | 0 | 0 | 0 | 2626000 | 0 | 0 | 0 | 0 | 0 | 0 | 0 | 0 | 0 | 0 | 0 | 0 | 0 |
| VNFHMFD | GNAS; GNAS; G | Q5JWF2; P63 | GNAS1_HUMAI | Oxidation (M) | 217960 | 0 | 369980 | 0 | 0 | 0 | 0 | 0 | 0 | 0 | 279290 | 2083100 | 0 | 0 | 165070 | 980260 | 0 | 0 |
| VNFLPEIITI | DYNC1H1; DYN | Q14204; A0A | DYHC1_HUMA | Unmodified | 0 | 0 | 833810 | 200220 | 0 | 0 | 420200 | 0 | 933910 | 669850 | 1633000 | 18510000 | 0 | 0 | 2735500 | 716290 | 0 | 0 |
| VNFLVNAA | CBR4; CBR4 | Q8N4T8; H0 | CBR4_HUMAN | Unmodified | 791670 | 0 | 0 | 0 | 0 | 0 | 0 | 0 | 1067900 | 0 | 0 | 0 | 0 | 0 | 0 | 596550 | 0 | 0 |
| VNFPENGF | DNAJA1 | P31689 | DNJA1_HUMAN | Unmodified | 0 | 0 | 0 | 0 | 0 | 0 | 0 | 0 | 6037800 | 228950 | 256990 | 5123800 | 1613700 | 0 | 10402000 | 1736100 | 0 | 0 |
| VNFPENGF | DNAJA1 | P31689 | DNJA1_HUMAN | Unmodified | 2585200 | 4109900 | 0 | 7330900 | 23967000 | 3682300 | 2434500 | 61347 | 17778000 | 19448000 | 1080300 | 2383000 | 16412000 | 19087000 | 1279500 | 1881200 | 766820 | 2445900 |
| VNFSDRVF | FERMT2; FERM | Q96AC1; G3 | FERM2_HUMAN | Unmodified | 0 | 0 | 0 | 0 | 1064600 | 123940 | 0 | 0 | 0 | 0 | 0 | 0 | 0 | 0 | 0 | 0 | 0 | 305720 |
| VNFTFNR | MOV10; MOV10 | Q9HCE1; Q5 | MOV10_HUMA | Unmodified | 0 | 0 | 0 | 0 | 0 | 0 | 0 | 0 | 0 | 0 | 0 | 0 | 0 | 0 | 446320 | 488880 | 0 | 0 |
| VNFTVDQII | EEF2 | P13639 | EF2_HUMAN EI | Unmodified | 3800500 | 4325100 | 5588100 | 7766600 | 23585000 | 4369300 | 2121000 | 1675300 | 7321900 | 2715000 | 10880000 | 1.09E+08 | 8432500 | 3220600 | 8480500 | 5813900 | 4386500 | 9866800 |
| VNFTVDQII | EEF2 | P13639 | EF2_HUMAN EI | Oxidation (M) | 0 | 0 | 0 | 0 | 18189000 | 2949600 | 5928400 | 2496700 | 0 | 1382300 | 0 | 0 | 4218300 | 3870500 | 0 | 0 | 10102000 | 4613500 |
| VNFTVDQII | EEF2 | P13639 | EF2_HUMAN EI | Unmodified | 0 | 0 | 0 | 31936 | 1.04E+08 | 11183000 | 24552000 | 1215500 | 0 | 6514100 | 92862 | 0 | 18368000 | 14574000 | 0 | 0 | 38365000 | 33285000 |
| VNFTVDQII | EEF2 | P13639 | EF2_HUMAN EI | Unmodified | 0 | 0 | 0 | 0 | 20915000 | 2142200 | 4082600 | 184850 | 0 | 953270 | 33790 | 0 | 3283500 | 4180000 | 0 | 0 | 10461000 | 8526800 |
| VNGDASPA | MARCKS | P29966 | MARCS_HUMA | Unmodified | 0 | 0 | 0 | 0 | 1977800 | 0 | 0 | 0 | 0 | 0 | 2361100 | 3906700 | 0 | 0 | 862970 | 401940 | 571110 | 353630 |
| VNGDASPA | MARCKS | P29966 | MARCS_HUMA | Unmodified | 0 | 0 | 0 | 0 | 0 | 0 | 0 | 0 | 0 | 0 | 6491400 | 24939000 | 0 | 0 | 1652500 | 2135800 | 0 | 2141800 |
| VNGLEEGV | ST14 | Q9Y5Y6 | ST14_HUMAN S | Unmodified | 1644500 | 952990 | 7544700 | 0 | 9368.2 | 0 | 0 | 0 | 414580 | 0 | 1355200 | 2756200 | 0 | 0 | 6249100 | 3071500 | 56326 | 0 |
| VNGRPLEN | RPS16; RPS16; | P62249; M0F | RS16_HUMAN | Unmodified | 1028800 | 1043800 | 688930 | 787000 | 474380 | 187230 | 320210 | 137700 | 655300 | 0 | 1122100 | 18313000 | 471130 | 488400 | 1340900 | 807170 | 80372 | 303820 |
| VNGRPLEN | RPS16; RPS16; | P62249; M0F | RS16_HUMAN | Oxidation (M) | 1203700 | 317310 | 0 | 0 | 0 | 0 | 0 | 0 | 0 | 0 | 0 | 6341300 | 211600 | 0 | 0 | 0 | 0 | 0 |
| VNGSLAVS | PPM1A; PPM1B | P35813; O75 | PPM1A_HUMAI | Unmodified | 0 | 0 | 628110 | 0 | 0 | 0 | 0 | 37879 | 141160 | 0 | 54294 | 840670 | 0 | 0 | 4203900 | 970000 | 0 | 0 |
| VNGVDDA | MYO1B; MYO1E | O43795; E9F | MYO1B_HUMA | Unmodified | 0 | 0 | 179640 | 0 | 0 | 0 | 0 | 209040 | 396200 | 0 | 1064900 | 12872000 | 0 | 0 | 273510 | 330050 | 0 | 0 |

| | | | | | | | | | | | | | | | | | | | | | |
|---|---|---|---|---|---|---|---|---|---|---|---|---|---|---|---|---|---|---|---|---|---|---|
| VNGVDDA | MYO1B; MYO1E | O43795; E9F | MYO1B_HUMA | Unmodified | 0 | 0 | 0 | 27241 | 2287000 | 1610500 | 0 | 48817 | 43055 | 138900 | 306560 | 0 | 87720 | 35465 | 0 | 0 | 507860 | 0 |
| VNGWATLF | ITGA3 | P26006 | ITA3_HUMAN I | Unmodified | 0 | 0 | 479510 | 0 | 0 | 0 | 0 | 0 | 190360 | 0 | 12681 | 2258000 | 0 | 0 | 217710 | 133150 | 0 | 0 |
| VNHEPEPA | BIN1 | O00499 | BIN1_HUMAN I | Unmodified | 0 | 0 | 443030 | 472180 | 0 | 0 | 0 | 0 | 89759 | 0 | 0 | 0 | 0 | 0 | 913720 | 1002000 | 0 | 0 |
| VNHPGSKI | NUCB2; NUCB2 | P80303; Q2L | NUCB2_HUMA | Unmodified | 89991 | 223440 | 0 | 0 | 0 | 0 | 0 | 139850 | 27358000 | 484160 | 473860 | 72302 | 0 | 0 | 20687 | 26306 | 0 | 122550 |
| VNHPGSKI | NUCB2; NUCB2 | P80303; Q2L | NUCB2_HUMA | Unmodified | 0 | 0 | 0 | 0 | 0 | 268900 | 377730 | 0 | 0 | 6048900 | 0 | 0 | 1041200 | 0 | 0 | 0 | 0 | 0 |
| VNIAFNYDI | DDX39B; DDX3 | Q13838; Q5 | DX39B_HUMAN | Oxidation (M) | 210530 | 0 | 0 | 0 | 0 | 0 | 0 | 61477 | 0 | 0 | 0 | 5784200 | 0 | 0 | 354120 | 1109000 | 0 | 0 |
| VNIEGGAIA | ACAT2 | Q9BWD1 | THIC_HUMAN | Unmodified | 0 | 0 | 0 | 0 | 231730 | 0 | 0 | 429650 | 0 | 740250 | 620170 | 10654000 | 282210 | 0 | 0 | 0 | 0 | 0 |
| VNIHGIFSV | HSPA4L; HSPA4 | O95757; E9F | HS74L_HUMAN | Unmodified | 0 | 0 | 0 | 0 | 0 | 0 | 198160 | 0 | 359370 | 33589 | 0 | 478070 | 109300 | 41178 | 27338 | 21843 | 0 | 0 |
| VNIIGEVVD | ADD3 | Q9UEY8 | ADDG_HUMAN | Unmodified | 0 | 0 | 663150 | 654820 | 0 | 0 | 0 | 0 | 0 | 683740 | 0 | 0 | 295620 | 212780 | 1305600 | 1049500 | 0 | 0 |
| VNIIPIIAKAI | SEPTIN11; SEPT | Q9NVA2; D6 | SEP11_HUMAN | Unmodified | 0 | 0 | 0 | 0 | 2156300 | 328400 | 0 | 0 | 0 | 470720 | 0 | 0 | 1581900 | 0 | 0 | 0 | 1084800 | 857810 |
| VNIIPLIAK | SEPTIN14; SEPT | Q6ZU15; Q1 | SEP14_HUMAN | Unmodified | 0 | 0 | 803880 | 731030 | 4514800 | 0 | 0 | 207860 | 0 | 0 | 2149600 | 6884700 | 0 | 0 | 0 | 800490 | 0 | 0 |
| VNIIPLIAKA | SEPTIN7; SEPTI | Q16181; E7E | SEPT7_HUMAN | Unmodified | 18293 | 20856 | 0 | 0 | 2767100 | 0 | 0 | 0 | 50400 | 34418 | 0 | 0 | 0 | 26633 | 0 | 20000 | 264340 | 0 |
| VNIIPLIAKA | SEPTIN7; SEPTI | Q16181; E7E | SEPT7_HUMAN | Unmodified | 0 | 0 | 0 | 0 | 21728000 | 6612000 | 0 | 0 | 0 | 1215200 | 0 | 0 | 6102900 | 8469500 | 0 | 0 | 12681000 | 5031000 |
| VNIIPVIAKA | SEPTIN10; SEPT | Q9P0V9; B5I | SEP10_HUMAN | Unmodified | 0 | 0 | 0 | 0 | 1480700 | 281620 | 0 | 0 | 0 | 0 | 0 | 0 | 394170 | 0 | 0 | 0 | 575070 | 466830 |
| VNIIQFGTG | PARP4 | Q9UKK3 | PARP4_HUMAN | Unmodified | 0 | 0 | 0 | 0 | 0 | 0 | 0 | 0 | 0 | 0 | 0 | 2679400 | 0 | 0 | 0 | 0 | 0 | 0 |
| VNIISVTNK | EIF2A; EIF2A; EI | Q9BY44; H7 | EIF2A_HUMAN | Unmodified | 0 | 54477 | 473120 | 71933 | 794840 | 131630 | 473650 | 1416600 | 0 | 191870 | 711440 | 7086300 | 849500 | 507920 | 0 | 368130 | 733360 | 869180 |
| VNIKPQVDI | AHCY | P23526 | SAHH_HUMAN | Unmodified | 157550 | 120210 | 488160 | 490270 | 1216800 | 0 | 37918 | 18588 | 1576700 | 436560 | 0 | 0 | 0 | 154620 | 0 | 0 | 278450 | 484510 |
| VNILFDFVK | EIF5 | P55010 | IF5_HUMAN Eu | Unmodified | 148300 | 188630 | 166050 | 332550 | 0 | 0 | 0 | 0 | 0 | 0 | 318510 | 3528400 | 138800 | 0 | 420350 | 279820 | 0 | 0 |
| VNILFDFVK | EIF5 | P55010 | IF5_HUMAN Eu | Unmodified | 0 | 0 | 0 | 0 | 0 | 0 | 0 | 0 | 264330 | 0 | 159870 | 2206700 | 0 | 0 | 0 | 0 | 0 | 0 |
| VNINGGAV | ACAT1; ACAT1; | P24752; A0A | THIL_HUMAN A | Unmodified | 0 | 0 | 0 | 0 | 0 | 0 | 0 | 0 | 1288100 | 0 | 0 | 0 | 0 | 0 | 0 | 0 | 0 | 0 |
| VNINTATPA | EEPD1 | Q7L9B9 | EEPD1_HUMAN | Oxidation (M) | 0 | 0 | 82266 | 0 | 0 | 0 | 0 | 0 | 0 | 0 | 0 | 157320 | 0 | 0 | 165510 | 144490 | 0 | 0 |
| VNIVPVIAK | SEPTIN2; SEPTI | Q15019; H7 | SEPT2_HUMAN | Unmodified | 0 | 0 | 0 | 0 | 5935000 | 1546600 | 252930 | 83992 | 0 | 110470 | 0 | 0 | 562870 | 1287200 | 0 | 0 | 521370 | 463620 |
| VNKSPASN | N/A | N/A | N/A | Acetyl (Protein N | 0 | 0 | 0 | 0 | 997090 | 0 | 0 | 612390 | 0 | 0 | 5761600 | 35754000 | 0 | 0 | 0 | 0 | 0 | 0 |
| VNLAELFK | PRDX5 | P30044 | PRDX5_HUMAN | Unmodified | 794160 | 0 | 3567700 | 2904500 | 3605200 | 827040 | 720740 | 233690 | 22262000 | 3126700 | 4415200 | 20856000 | 5426100 | 2165400 | 6705400 | 3885700 | 1785400 | 4532100 |
| VNLAHSEI | TMEM59; TMEM | Q9BXS4; Q5 | TMM59_HUMA | Unmodified | 0 | 307380 | 0 | 0 | 7966600 | 663430 | 11122000 | 1584300 | 9349200 | 3219300 | 0 | 0 | 0 | 868970 | 359830 | 202940 | 0 | 1198300 |
| VNLAIWDT | RAB21 | Q9UL25 | RAB21_HUMAN | Unmodified | 1369300 | 1320800 | 0 | 0 | 0 | 0 | 0 | 176560 | 0 | 0 | 0 | 0 | 0 | 0 | 830090 | 0 | 0 | 0 |
| VNLDALTQ | ABHD11 | Q8NFV4 | ABHDB_HUMA | Unmodified | 0 | 0 | 0 | 1003400 | 0 | 0 | 0 | 0 | 2380600 | 0 | 0 | 0 | 0 | 0 | 0 | 0 | 0 | 0 |
| VNLEEQTP | BMX | P51813 | BMX_HUMAN C | Unmodified | 0 | 115700 | 0 | 0 | 0 | 0 | 0 | 0 | 0 | 0 | 0 | 8193300 | 0 | 0 | 0 | 314510 | 0 | 0 |
| VNLEEQTP | BMX | P51813 | BMX_HUMAN C | Unmodified | 0 | 0 | 0 | 0 | 0 | 0 | 0 | 0 | 0 | 0 | 3210500 | 0 | 0 | 0 | 0 | 0 | 0 | 0 |
| VNLEESSG | SCFD1; SCFD1; | Q8WVM8; A0 | SCFD1_HUMA | Unmodified | 0 | 0 | 0 | 0 | 0 | 0 | 0 | 0 | 0 | 0 | 310340 | 1294000 | 0 | 0 | 0 | 0 | 0 | 0 |
| VNLESPGP | VGF; VGF | O15240; Q0 | VGF_HUMAN N | Unmodified | 1124500 | 1156200 | 0 | 0 | 444450 | 0 | 0 | 24849 | 4304900 | 119800 | 0 | 0 | 0 | 0 | 0 | 0 | 0 | 0 |
| VNLFILGK | ARHGAP5; ARH | Q13017; A0A | RHG05_HUMA | Unmodified | 0 | 0 | 0 | 0 | 0 | 0 | 0 | 0 | 0 | 0 | 0 | 0 | 0 | 0 | 0 | 70599 | 0 | 0 |
| VNLFPGLSI | TNKS1BP1 | Q9C0C2 | TB182_HUMAN | Unmodified | 0 | 0 | 0 | 0 | 1265700 | 0 | 0 | 23622 | 0 | 0 | 222570 | 870690 | 6252400 | 4044700 | 0 | 562060 | 1418100 | 1153400 |
| VNLFTDFDI | EIF3B; EIF3B | P55884; C9J | EIF3B_HUMAN | Unmodified | 0 | 576370 | 792860 | 0 | 239760 | 0 | 0 | 79369 | 0 | 0 | 933410 | 13828000 | 0 | 418250 | 850410 | 1147700 | 274190 | 0 |
| VNLGVGAY | GOT1 | P17174 | AATC_HUMAN | Unmodified | 157890 | 172400 | 285500 | 271950 | 0 | 0 | 0 | 0 | 561420 | 0 | 549640 | 8292800 | 39206 | 0 | 1782700 | 468550 | 0 | 0 |
| VNLGVGAY | GOT1 | P17174 | AATC_HUMAN | Unmodified | 0 | 0 | 127820 | 566410 | 3917900 | 446430 | 0 | 0 | 474820 | 657090 | 1594500 | 2031800 | 4932500 | 2279100 | 0 | 0 | 0 | 1202500 |
| VNLIESDVV | EFCAB14; EFCA | O75071; A0A | EFC14_HUMAN | Unmodified | 0 | 0 | 0 | 0 | 0 | 0 | 0 | 0 | 2151000 | 0 | 0 | 0 | 0 | 0 | 0 | 0 | 0 | 0 |
| VNLIESDVV | EFCAB14; EFCA | O75071; A0A | EFC14_HUMAN | Oxidation (M) | 0 | 0 | 0 | 0 | 0 | 0 | 0 | 0 | 1242400 | 72830 | 0 | 0 | 0 | 0 | 0 | 0 | 0 | 0 |
| VNLIKQDD | NCAM1; NCAM | P13591; A0A | NCAM1_HUMA | Unmodified | 0 | 177720 | 0 | 0 | 0 | 0 | 0 | 0 | 1068700 | 30738 | 0 | 1112200 | 0 | 0 | 0 | 0 | 68659 | 0 |
| VNLLEQEL | LZTS1 | Q9Y250 | LZTS1_HUMAN | Unmodified | 0 | 0 | 0 | 0 | 0 | 0 | 0 | 0 | 0 | 0 | 0 | 256780 | 0 | 0 | 0 | 0 | 0 | 0 |
| VNLLSFTG | ALDH7A1; ALDH | P49419; A0A | AL7A1_HUMAN | Unmodified | 0 | 0 | 0 | 0 | 0 | 0 | 0 | 0 | 0 | 0 | 0 | 2533700 | 0 | 0 | 0 | 0 | 0 | 0 |
| VNLLSVLE | PRXL2A | Q9BRX8 | PXL2A_HUMAN | Unmodified | 0 | 0 | 0 | 0 | 0 | 0 | 534610 | 0 | 57531000 | 2270200 | 1147500 | 1880500 | 0 | 0 | 0 | 0 | 0 | 0 |
| VNLLYSR | ECH1; ECH1 | Q13011; M0 | ECH1_HUMAN | Unmodified | 0 | 0 | 0 | 34343 | 0 | 0 | 0 | 0 | 12807000 | 191260 | 0 | 161460 | 0 | 0 | 0 | 112710 | 0 | 0 |
| VNLNIFLLG | PROM1 | O43490 | PROM1_HUMA | Unmodified | 0 | 0 | 0 | 0 | 0 | 0 | 0 | 0 | 0 | 0 | 1010200 | 2954300 | 0 | 0 | 0 | 0 | 0 | 0 |
| VNLQNNP | BCAP31; BCAP3 | P51572; C9J | BAP31_HUMAN | Oxidation (M) | 0 | 0 | 0 | 0 | 0 | 0 | 0 | 0 | 1099700 | 300560 | 0 | 0 | 0 | 0 | 0 | 0 | 0 | 0 |
| VNLQNNP | BCAP31; BCAP3 | P51572; C9J | BAP31_HUMAN | Unmodified | 0 | 0 | 0 | 62524 | 0 | 0 | 0 | 0 | 7202000 | 600580 | 0 | 785700 | 0 | 0 | 0 | 0 | 0 | 92440 |
| VNLSAAQT | PDCD10; PDCD | Q9BUL8; H7 | PDC10_HUMA | Unmodified | 0 | 0 | 0 | 687600 | 199570 | 0 | 0 | 0 | 898110 | 541160 | 252070 | 4146300 | 0 | 0 | 0 | 1104700 | 302860 | 0 |
| VNLTLGSIV | GANAB; GANAB | E9PKU7; E9F | E9PKU7_HUMA | Unmodified | 0 | 0 | 0 | 0 | 0 | 0 | 0 | 0 | 4653400 | 0 | 285260 | 0 | 0 | 0 | 0 | 0 | 0 | 0 |
| VNMEDLR | SEPTIN8; SEPTI | Q92599; Q1 | SEPT8_HUMAN | Unmodified | 0 | 153990 | 0 | 0 | 0 | 0 | 0 | 0 | 1832200 | 0 | 110920 | 7989500 | 0 | 0 | 95048 | 163930 | 0 | 0 |
| VNMPAKPA | CKAP5 | Q14008 | CKAP5_HUMA | Unmodified | 0 | 0 | 0 | 0 | 0 | 0 | 0 | 0 | 0 | 28221 | 0 | 985360 | 0 | 0 | 0 | 0 | 41028 | 0 |
| VNMPPAVD | MRPS26 | Q9BYN8 | RT26_HUMAN | Oxidation (M) | 0 | 0 | 0 | 0 | 0 | 0 | 0 | 0 | 232230 | 0 | 0 | 0 | 0 | 0 | 0 | 0 | 0 | 0 |
| VNMTELDK | DAAM1 | Q9Y4D1 | DAAM1_HUMA | Unmodified | 298130 | 0 | 0 | 130980 | 593880 | 55014 | 0 | 0 | 1187300 | 0 | 0 | 4227900 | 311070 | 0 | 0 | 52033 | 84190 | 79697 |
| VNMTELDK | DAAM1 | Q9Y4D1 | DAAM1_HUMA | Unmodified | 0 | 0 | 0 | 0 | 0 | 113390 | 745050 | 0 | 0 | 424550 | 0 | 0 | 633770 | 441320 | 0 | 0 | 100330 | 334140 |
| VNMTELDK | DAAM1 | Q9Y4D1 | DAAM1_HUMA | Oxidation (M) | 0 | 0 | 0 | 0 | 338160 | 651730 | 375500 | 0 | 0 | 137850 | 0 | 0 | 293580 | 315160 | 0 | 0 | 62865 | 244130 |
| VNNASLIGI | VDAC3 | Q9Y277 | VDAC3_HUMA | Unmodified | 0 | 0 | 0 | 0 | 0 | 0 | 0 | 0 | 0 | 0 | 673050 | 0 | 0 | 0 | 0 | 0 | 0 | 0 |
| VNNASLIGI | VDAC3 | Q9Y277 | VDAC3_HUMA | Unmodified | 0 | 0 | 0 | 0 | 0 | 0 | 0 | 0 | 0 | 1020600 | 921330 | 0 | 0 | 0 | 0 | 0 | 0 | 0 |
| VNNFGSGI | ACSL4; ACSL4 | O60488; D6 | ACSL4_HUMAN | Unmodified | 0 | 0 | 0 | 0 | 0 | 0 | 0 | 0 | 0 | 0 | 875110 | 5247300 | 0 | 0 | 0 | 0 | 0 | 0 |
| VNNPDNH | MAP4K5 | Q9Y4K4 | M4K5_HUMAN | Unmodified | 0 | 0 | 0 | 0 | 0 | 0 | 0 | 0 | 0 | 0 | 342920 | 1210700 | 0 | 0 | 0 | 0 | 0 | 0 |
| VNNRFELIN | TCF25; TCF25; | Q9BQ70; H3 | TCF25_HUMAN | Unmodified | 296240 | 2240100 | 80197 | 182200 | 0 | 0 | 0 | 0 | 0 | 0 | 0 | 0 | 0 | 0 | 993990 | 333080 | 0 | 0 |
| VNNSGISL | DST; DST; DST; | Q03001; A0A | DYST_HUMAN | Unmodified | 0 | 0 | 0 | 0 | 1121800 | 0 | 0 | 0 | 0 | 0 | 0 | 60508 | 0 | 0 | 0 | 0 | 0 | 0 |
| VNNSSLIGI | VDAC1 | P21796 | VDAC1_HUMA | Unmodified | 0 | 0 | 0 | 0 | 0 | 0 | 1808200 | 1319400 | 5.2E+08 | 1.4E+08 | 11685000 | 4505100 | 3614800 | 6730800 | 1338000 | 0 | 0 | 2083600 |
| VNNSSLIG | VDAC2; VDAC2 | P45880; A0A | VDAC2_HUMA | Unmodified | 507610 | 409770 | 962670 | 1665000 | 3151200 | 8404400 | 3848800 | 1589200 | 2.13E+09 | 2.65E+08 | 9924900 | 4898000 | 1671100 | 2362300 | 791540 | 538750 | 502320 | 5313600 |
| VNNSSLIG | VDAC2; VDAC2 | P45880; A0A | VDAC2_HUMA | Unmodified | 0 | 0 | 0 | 0 | 0 | 0 | 0 | 0 | 0 | 8479800 | 0 | 0 | 0 | 0 | 0 | 0 | 0 | 0 |
| VNNSTGTS | BAIAP2L1 | Q9UHR4 | BI2L1_HUMAN | Unmodified | 0 | 0 | 0 | 216980 | 86132 | 0 | 0 | 0 | 852580 | 0 | 45007 | 512260 | 0 | 0 | 176450 | 0 | 0 | 0 |
| VNNSTMLG | PSMB4 | P28070 | PSB4_HUMAN | Oxidation (M) | 0 | 0 | 0 | 0 | 0 | 0 | 0 | 0 | 2309600 | 0 | 0 | 4277600 | 0 | 0 | 0 | 0 | 0 | 0 |
| VNPALAELI | RNH1; RNH1; R | P13489; E9P | RINI_HUMAN R | Unmodified | 218430 | 245930 | 712260 | 365620 | 0 | 0 | 0 | 0 | 1131700 | 0 | 681230 | 8140700 | 0 | 0 | 544730 | 498810 | 0 | 0 |
| VNPAVIRNI | SCEL | O95171 | SCEL_HUMAN | Unmodified | 0 | 0 | 0 | 0 | 3468300 | 0 | 0 | 0 | 0 | 0 | 0 | 0 | 0 | 0 | 0 | 0 | 0 | 0 |
| VNPDTGYII | SHMT1 | P34896 | GLYC_HUMAN | Unmodified | 0 | 0 | 0 | 0 | 0 | 0 | 0 | 0 | 745470 | 0 | 624350 | 4090400 | 282040 | 0 | 0 | 0 | 0 | 0 |
| VNPESSGK | GPRIN1 | Q7Z2K8 | GRIN1_HUMAN | Unmodified | 0 | 0 | 0 | 0 | 263210 | 0 | 0 | 0 | 0 | 483990 | 0 | 0 | 0 | 0 | 0 | 0 | 302140 | 918380 |
| VNPFRPGD | ZYX; ZYX; ZYX | Q15942; H0 | ZYX_HUMAN Zy | Unmodified | 0 | 0 | 0 | 0 | 0 | 0 | 0 | 0 | 0 | 0 | 0 | 0 | 0 | 0 | 0 | 0 | 0 | 810340 |
| VNPFRPGD | ZYX; ZYX; ZYX | Q15942; H0 | ZYX_HUMAN Zy | Unmodified | 0 | 0 | 0 | 0 | 3372600 | 0 | 0 | 0 | 0 | 0 | 0 | 0 | 684070 | 0 | 0 | 0 | 0 | 7363300 |
| VNPIQGLA | COX4I1 | P13073 | COX41_HUMA | Unmodified | 0 | 0 | 0 | 0 | 0 | 0 | 0 | 0 | 14850000 | 1726200 | 323560 | 397000 | 0 | 0 | 0 | 0 | 0 | 0 |
| VNPLGGAV | ACAA1; ACAA1; | P09110; C9J | THIK_HUMAN | Unmodified | 0 | 0 | 0 | 135330 | 0 | 0 | 0 | 0 | 13644000 | 3491700 | 0 | 0 | 0 | 0 | 0 | 0 | 0 | 0 |
| VNPNTALTI | EXOC6; EXOC6 | Q8TAG9; E7 | EXOC6_HUMA | Unmodified | 0 | 0 | 0 | 0 | 0 | 0 | 0 | 0 | 0 | 0 | 0 | 700000 | 0 | 0 | 179560 | 0 | 0 | 0 |
| VNPQGPGI | GPC1; GPC1; G | P35052; H7 | GPC1_HUMAN | Unmodified | 1112700 | 0 | 821530 | 1083600 | 4491000 | 1858700 | 59717 | 1491300 | 3069100 | 0 | 11326000 | 900060 | 0 | 0 | 528540 | 637370 | 0 | 0 |
| VNPREIAEN | RARS1 | P54136 | SYRC_HUMAN | Unmodified | 0 | 0 | 0 | 0 | 1750900 | 0 | 0 | 0 | 0 | 358430 | 0 | 2872900 | 2354500 | 1065000 | 0 | 0 | 188730 | 1030200 |
| VNPSRLPV | CLTC; CLTC | Q00610; A0A | CLH1_HUMAN | Unmodified | 1617100 | 1748300 | 167860 | 1302400 | 591210 | 581190 | 0 | 0 | 0 | 0 | 0 | 1057600 | 1406400 | 0 | 2578100 | 743670 | 0 | 0 |
| VNPVTALTL | EXOC6B; EXOC | Q9Y2D4; A0A | EXC6B_HUMAN | Unmodified | 0 | 0 | 33305 | 0 | 0 | 0 | 0 | 0 | 0 | 0 | 0 | 311580 | 0 | 0 | 0 | 46198 | 0 | 0 |
| VNQAIWLL | RPS5; RPS5; RP | P46782; M0 | RS5_HUMAN 4 | Unmodified | 339030 | 0 | 97538 | 136190 | 0 | 0 | 0 | 0 | 0 | 0 | 437950 | 2670100 | 89027 | 0 | 173780 | 214070 | 0 | 0 |
| VNQFRQLY | PSMD1; PSMD1 | Q99460; A0A | PSMD1_HUMA | Unmodified | 0 | 0 | 0 | 0 | 195280 | 48573 | 114900 | 80186 | 0 | 0 | 0 | 0 | 241490 | 202960 | 0 | 0 | 241280 | 235960 |
| VNQIGSVTI | ENO2; ENO2 | P09104; F5H | ENOG_HUMAN | Unmodified | 167620 | 0 | 0 | 0 | 2256700 | 275170 | 3193500 | 2037800 | 2820800 | 2010600 | 0 | 528670 | 305550 | 315420 | 71164 | 0 | 6430300 | 6813400 |
| VNQIGSVTI | ENO1; ENO1 | P06733; A0A | ENOA_HUMAN | Unmodified | 7841500 | 2096300 | 8344800 | 13432000 | 2.47E+08 | 32346000 | 25153000 | 16631000 | 69116000 | 76119000 | 25907000 | 1.75E+08 | 1.49E+08 | 91295000 | 9659200 | 10012000 | 2.05E+08 | 1.98E+08 |
| VNQIGSVTI | ENO1; ENO1 | P06733; A0A | ENOA_HUMAN | Oxidation (M) | 0 | 0 | 0 | 0 | 0 | 0 | 0 | 0 | 0 | 0 | 0 | 0 | 0 | 0 | 0 | 0 | 1987000 | 0 |
| VNQLQRQI | ROCK2; ROCK2 | O75116; E9F | ROCK2_HUMA | Unmodified | 0 | 0 | 0 | 0 | 2066200 | 0 | 0 | 0 | 0 | 0 | 0 | 0 | 1099800 | 428050 | 0 | 0 | 1666200 | 1933900 |
| VNQPASFA | FLNB; FLNB | O75369; E7E | FLNB_HUMAN | Unmodified | 145610 | 0 | 0 | 374470 | 577770 | 0 | 0 | 0 | 196920 | 0 | 0 | 5476100 | 112770 | 0 | 754300 | 724070 | 0 | 0 |
| VNQPASFA | FLNB; FLNB | O75369; E7E | FLNB_HUMAN | Unmodified | 0 | 0 | 0 | 0 | 7008400 | 1174400 | 0 | 0 | 0 | 0 | 0 | 0 | 0 | 624590 | 0 | 0 | 4328200 | 0 |
| VNQPASFA | FLNA; FLNA; FL | P21333; A0A | FLNA_HUMAN | Unmodified | 709320 | 1909800 | 1175900 | 622210 | 2608600 | 1265100 | 1776500 | 991720 | 0 | 275020 | 0 | 0 | 609610 | 920000 | 1976300 | 2168200 | 2253700 | 143900 |
| VNQSALEA | APPL1 | Q9UKG1 | DP13A_HUMAN | Unmodified | 147720 | 256580 | 236510 | 0 | 0 | 0 | 0 | 0 | 0 | 0 | 0 | 0 | 0 | 0 | 279820 | 100680 | 0 | 0 |
| VNQWTTN | DYNLT1 | P63172 | DYLT1_HUMAN | Unmodified | 0 | 0 | 0 | 0 | 370650 | 70502 | 480560 | 0 | 217080 | 605750 | 214800 | 1377400 | 0 | 372920 | 0 | 0 | 0 | 0 |
| VNRLDMY | AP3M2; AP3M1; | P53677; Q9Y | AP3M2_HUMA | Unmodified | 0 | 0 | 0 | 0 | 1102300 | 147660 | 284630 | 113430 | 264260 | 197570 | 246000 | 952970 | 204340 | 222760 | 0 | 0 | 119300 | 391890 |
| VNRLDMY | AP3M2; AP3M1; | P53677; Q9Y | AP3M2_HUMA | Oxidation (M) | 0 | 0 | 21549 | 0 | 297530 | 48735 | 164690 | 37874 | 36660 | 242190 | 0 | 195670 | 80366 | 61427 | 16189 | 0 | 17965 | 266830 |
| VNRLLDSLI | TFG; TFG; TFG; | Q92734; A0A | TFG_HUMAN P | Unmodified | 0 | 0 | 0 | 0 | 23187000 | 0 | 975980 | 401870 | 6543100 | 3476100 | 582120 | 0 | 5252300 | 3545700 | 0 | 0 | 2682500 | 15909000 |
| VNSININQ | MYO1C; MYO1C | O00159; F5H | MYO1C_HUMA | Unmodified | 712490 | 619240 | 723490 | 646270 | 0 | 0 | 0 | 0 | 0 | 0 | 0 | 425620 | 26600 | 0 | 2172300 | 2252600 | 132450 | 0 |
| VNSKFDTIY | TRIM25; TRIM25 | Q14258; A0A | TRI25_HUMAN | Unmodified | 0 | 0 | 0 | 0 | 1590800 | 0 | 0 | 0 | 0 | 0 | 0 | 0 | 0 | 0 | 0 | 0 | 0 | 0 |
| VNSLAPGP | DECR2; DECR2 | Q9NUI1; Q4 | DECR2_HUMA | Unmodified | 0 | 53259 | 0 | 0 | 0 | 0 | 209420 | 0 | 3555000 | 83617 | 0 | 0 | 0 | 0 | 0 | 52681 | 0 | 0 |
| VNSLGETA | SPTAN1; SPTAN | Q13813; A0A | SPTN1_HUMAN | Unmodified | 340560 | 0 | 131510 | 0 | 223690 | 0 | 0 | 0 | 259070 | 216680 | 0 | 1584400 | 385560 | 0 | 130110 | 105570 | 0 | 0 |
| VNSLPEVLI | ITGA6 | P23229 | ITA6_HUMAN I | Unmodified | 0 | 0 | 127710 | 0 | 2432600 | 0 | 10930 | 0 | 0 | 144200 | 887520 | 3682500 | 5080000 | 1499900 | 0 | 0 | 2141400 | 9732900 |
| VNSLPEVLI | ITGA6 | P23229 | ITA6_HUMAN I | Unmodified | 0 | 0 | 0 | 0 | 1965300 | 0 | 0 | 0 | 0 | 0 | 0 | 0 | 2599600 | 0 | 0 | 0 | 9074400 | 29260000 |
| VNSVYVLVI | FAR1; FAR1 | Q8WVX9; E9 | FACR1_HUMAN | Unmodified | 0 | 0 | 0 | 0 | 0 | 0 | 0 | 0 | 0 | 0 | 0 | 0 | 0 | 0 | 0 | 0 | 0 | 0 |
| VNTFSALLL | GET3 | O43681 | GET3_HUMAN | Unmodified | 0 | 0 | 0 | 0 | 0 | 0 | 0 | 0 | 0 | 63189 | 0 | 2553000 | 0 | 0 | 0 | 0 | 0 | 2220200 |
| VNTLEEGK | NPTX1 | Q15818 | NPTX1_HUMAN | Unmodified | 0 | 0 | 142030 | 0 | 0 | 0 | 0 | 0 | 5776900 | 0 | 0 | 0 | 0 | 0 | 0 | 42800 | 0 | 0 |
| VNTLIRPD | RPL23A; RPL23 | P62750; K7E | RL23A_HUMAN | Unmodified | 0 | 0 | 0 | 0 | 0 | 0 | 133610 | 0 | 0 | 0 | 516860 | 3000000 | 390530 | 0 | 0 | 0 | 0 | 0 |
| VNTLIRPD | RPL23A; RPL23 | P62750; K7E | RL23A_HUMAN | Unmodified | 0 | 0 | 0 | 0 | 0 | 0 | 0 | 0 | 0 | 0 | 1072300 | 0 | 0 | 0 | 0 | 0 | 287710 | 0 |
| VNTPTTTVY | TARS1 | P26639 | SYTC_HUMAN | Unmodified | 0 | 0 | 1306500 | 0 | 3356400 | 1988300 | 130380 | 215870 | 1567000 | 530140 | 1786900 | 9911200 | 1713000 | 0 | 1505100 | 1520300 | 530290 | 1354300 |
| VNTQDFR | TJP2; TJP2; TJP2 | Q9UDY2; A0 | ZO2_HUMAN T | Unmodified | 310210 | 403790 | 376590 | 353390 | 0 | 0 | 0 | 0 | 2493300 | 741640 | 48037 | 1216300 | 0 | 0 | 166610 | 199590 | 0 | 0 |
| VNVEAPDV | AHNAK | Q09666 | AHNK_HUMAN | Unmodified | 0 | 0 | 0 | 0 | 8320300 | 1846900 | 0 | 0 | 0 | 330370 | 0 | 0 | 0 | 0 | 0 | 0 | 1125700 | 3771200 |
| VNVEHMTE | PFKL | P17858 | PFKAL_HUMAN | Unmodified | 0 | 0 | 0 | 0 | 0 | 0 | 0 | 0 | 371920 | 0 | 0 | 319190 | 0 | 0 | 0 | 56609 | 0 | 0 |
| VNVHGIFS | HSPA4; HSPA4 | P34932; A0A | HSP74_HUMAN | Oxidation (M) | 0 | 0 | 0 | 0 | 0 | 0 | 0 | 0 | 0 | 0 | 0 | 2240400 | 0 | 0 | 0 | 0 | 0 | 0 |
| VNVHVLGV | TTC37 | Q6PGP7 | TTC37_HUMAN | Unmodified | 0 | 0 | 57495 | 0 | 0 | 0 | 0 | 22361 | 0 | 0 | 0 | 541290 | 0 | 0 | 26513 | 40425 | 0 | 0 |

| | | | | | | | | | | | | | | | | | | | | | | |
|---|---|---|---|---|---|---|---|---|---|---|---|---|---|---|---|---|---|---|---|---|---|---|
| VNVIHAR | AP2M1; AP2M1; | Q96CW1; C9 | AP2M1_HUMAI | Unmodified | 0 | 0 | 0 | 0 | 0 | 0 | 0 | 0 | 0 | 0 | 0 | 0 | 0 | 0 | 0 | 0 | 0 | 0 |
| VNVPGSQ/ | NUCB1; NUCB | Q02818; H7I | NUCB1_HUMA | Unmodified | 0 | 0 | 301850 | 226460 | 664410 | 669020 | 0 | 0 | 6371600 | 891410 | 3344600 | 809630 | 242700 | 226750 | 173690 | 264340 | 0 | 1070200 |
| VNVPGSQ/ | NUCB1; NUCB | Q02818; H7I | NUCB1_HUMA | Unmodified | 0 | 0 | 0 | 0 | 0 | 0 | 4623300 | 0 | 0 | 3234900 | 0 | 0 | 717820 | 0 | 0 | 0 | 0 | 2438200 |
| VNVTVDYII | SND1 | Q7KZF4 | SND1_HUMAN | Unmodified | 342410 | 0 | 0 | 0 | 0 | 2534700 | 474020 | 212150 | 0 | 0 | 672250 | 5206400 | 0 | 0 | 1225700 | 1365200 | 0 | 0 |
| VNVVTCTK | ST14 | Q9Y5Y6 | ST14_HUMAN S | Unmodified | 0 | 312060 | 795960 | 2727800 | 1848700 | 522980 | 23102 | 0 | 3603000 | 1373000 | 0 | 0 | 3211900 | 1978800 | 0 | 3891300 | 376700 | 406760 |
| VNWATTPS | TIA1; TIAL1; TIA | P31483; Q01 | TIA1_HUMAN N | Unmodified | 0 | 0 | 0 | 0 | 0 | 0 | 0 | 0 | 0 | 0 | 0 | 391120 | 14211 | 0 | 0 | 0 | 0 | 0 |
| VNWATTPS | TIA1; TIAL1; TIA | P31483; Q01 | TIA1_HUMAN N | Unmodified | 519110 | 371810 | 0 | 0 | 694380 | 1653100 | 404590 | 170230 | 455170 | 228400 | 283910 | 1648600 | 0 | 609160 | 336200 | 0 | 132010 | 430110 |
| VNWTPEIN | SMCHD1; SMC | A6NHR9; J3 | SMHD1_HUMA | Unmodified | 0 | 0 | 0 | 0 | 0 | 0 | 579940 | 0 | 0 | 0 | 0 | 0 | 0 | 0 | 0 | 0 | 0 | 0 |
| VNYQANIV | MTAP; MTAP; N | Q13126; A0A | MTAP_HUMAN | Unmodified | 0 | 0 | 0 | 0 | 964960 | 0 | 0 | 0 | 485510 | 0 | 558450 | 2929200 | 0 | 0 | 0 | 0 | 0 | 0 |
| VPADLGAE | SPR | P35270 | SPRE_HUMAN | Unmodified | 0 | 37222 | 0 | 0 | 0 | 0 | 0 | 0 | 0 | 0 | 0 | 0 | 0 | 0 | 78714 | 0 | 0 | 0 |
| VPADTEVV | TPI1 | P60174 | TPIS_HUMAN T | Unmodified | 200150 | 0 | 194620 | 73962 | 0 | 884750 | 0 | 0 | 1854100 | 443960 | 2887700 | 19338000 | 0 | 1275600 | 0 | 0 | 0 | 1382100 |
| VPADTEVV | TPI1 | P60174 | TPIS_HUMAN T | Unmodified | 0 | 0 | 0 | 0 | 2654300 | 0 | 928830 | 0 | 0 | 0 | 0 | 0 | 0 | 0 | 0 | 0 | 2209100 | 1706100 |
| VPAEGLEE | CORO7; CORO | P57737; A0A | CORO7_HUMA | Unmodified | 0 | 0 | 0 | 129100 | 0 | 0 | 0 | 0 | 0 | 0 | 507600 | 2351500 | 0 | 0 | 0 | 0 | 0 | 0 |
| VPAGSAAV | HSPG2 | P98160 | PGBM_HUMAN | Unmodified | 0 | 0 | 211740 | 153480 | 0 | 0 | 0 | 0 | 0 | 0 | 0 | 0 | 0 | 0 | 0 | 281070 | 0 | 0 |
| VPAHAVVV | MELTF | P08582 | TRFM_HUMAN | Unmodified | 0 | 0 | 0 | 0 | 0 | 0 | 0 | 0 | 1037800 | 0 | 0 | 185850 | 0 | 0 | 0 | 0 | 0 | 0 |
| VPAILYYVK | PAN2 | Q504Q3 | PAN2_HUMAN | Unmodified | 384550 | 791800 | 254710 | 131640 | 0 | 0 | 0 | 0 | 0 | 0 | 0 | 0 | 0 | 0 | 0 | 0 | 0 | 0 |
| VPAILYYVK | PAN2 | Q504Q3 | PAN2_HUMAN | Unmodified | 154950 | 143580 | 107250 | 0 | 0 | 0 | 0 | 0 | 0 | 0 | 0 | 0 | 0 | 0 | 76041 | 91093 | 0 | 0 |
| VPAINVND | AHCY | P23526 | SAHH_HUMAN | Unmodified | 1111400 | 1131000 | 3805400 | 3677700 | 5452100 | 1323900 | 533100 | 364620 | 5771100 | 1381500 | 11790000 | 58880000 | 3272700 | 2254200 | 3917100 | 4924800 | 1184400 | 4239900 |
| VPAPEAFR | MYORG | Q6NSJ0 | MYORG_HUMA | Unmodified | 0 | 0 | 0 | 0 | 0 | 0 | 1986200 | 0 | 128560 | 0 | 0 | 100060 | 0 | 0 | 203530 | 0 | 0 | 0 |
| VPAPFSPP | KIAA1522 | Q9P206 | K1522_HUMAN | Unmodified | 0 | 0 | 514620 | 1329500 | 0 | 0 | 0 | 0 | 618900 | 0 | 0 | 683210 | 0 | 0 | 1624700 | 1392500 | 0 | 0 |
| VPDEAPKV | STEAP3; STEAP | Q658P3; B8 | STEA3_HUMAN | Unmodified | 0 | 0 | 0 | 289460 | 0 | 0 | 0 | 0 | 210040 | 0 | 0 | 0 | 0 | 0 | 0 | 0 | 0 | 0 |
| VPDGKPEN | PPA1 | Q15181 | IPYR_HUMAN I | Unmodified | 0 | 0 | 0 | 0 | 0 | 0 | 0 | 0 | 0 | 0 | 0 | 5989700 | 162590 | 0 | 0 | 0 | 0 | 0 |
| VPDGKPEN | PPA1 | Q15181 | IPYR_HUMAN I | Unmodified | 0 | 0 | 0 | 0 | 308450 | 0 | 0 | 0 | 0 | 0 | 374440 | 4063800 | 0 | 0 | 0 | 0 | 0 | 0 |
| VPDGKPEN | PPA2; PPA2 | Q9H2U2; D6 | IPYR2_HUMAN | Unmodified | 0 | 0 | 0 | 0 | 0 | 0 | 0 | 0 | 645640 | 0 | 146050 | 2207000 | 0 | 0 | 0 | 0 | 0 | 0 |
| VPDSVSLG | BMX | P51813 | BMX_HUMAN ( | Unmodified | 0 | 0 | 0 | 0 | 0 | 0 | 0 | 0 | 0 | 0 | 187180 | 453780 | 0 | 0 | 0 | 0 | 0 | 0 |
| VPDVLFTTI | C2CD5 | Q86YS7 | C2CD5_HUMA | Unmodified | 0 | 0 | 18747 | 60279 | 0 | 10593 | 0 | 0 | 90980 | 29856 | 40318 | 39582 | 0 | 0 | 88755 | 68682 | 0 | 0 |
| VPDVLVAD | GANAB; GANAE | Q14697; F5 | GANAB_HUMA | Unmodified | 0 | 0 | 0 | 0 | 0 | 0 | 0 | 0 | 3457000 | 571370 | 430780 | 0 | 0 | 0 | 0 | 0 | 0 | 0 |
| VPEALGWV | PIK3C2B; PIK3C | O00750; F5C | P3C2B_HUMAI | Unmodified | 35433 | 37149 | 0 | 0 | 0 | 0 | 0 | 0 | 0 | 0 | 41916 | 0 | 0 | 0 | 103750 | 0 | 0 | 0 |
| VPEANSSV | DLAT; DLAT; DL | P10515; A0A | ODP2_HUMAN | Unmodified | 0 | 0 | 0 | 0 | 908770 | 0 | 189260 | 0 | 1091600 | 0 | 0 | 0 | 0 | 0 | 0 | 90298 | 0 | 0 |
| VPEEVLRP | HELZ2 | Q9BYK8 | HELZ2_HUMAN | Unmodified | 0 | 0 | 149880 | 357100 | 0 | 0 | 0 | 0 | 0 | 0 | 0 | 0 | 0 | 0 | 866830 | 449870 | 0 | 0 |
| VPEGPIPPS | GLB1; GLB1 | P16278; E7E | BGAL_HUMAN | Unmodified | 0 | 0 | 0 | 0 | 0 | 0 | 0 | 0 | 480430 | 224750 | 0 | 0 | 0 | 0 | 0 | 0 | 0 | 0 |
| VPEILQLSD | CARS1; CARS1 | P49589; B4D | SYCC_HUMAN | Unmodified | 0 | 0 | 0 | 0 | 0 | 0 | 0 | 0 | 0 | 0 | 0 | 728170 | 0 | 0 | 0 | 0 | 0 | 0 |
| VPELNVQN | SF3B1; SF3B1 | O75533; H7 | SF3B1_HUMAN | Unmodified | 0 | 0 | 0 | 0 | 0 | 0 | 0 | 0 | 0 | 0 | 0 | 680190 | 0 | 0 | 0 | 0 | 0 | 0 |
| VPEPWVPP | JSRP1 | Q96MG2 | JSPR1_HUMAN | Unmodified | 0 | 0 | 0 | 0 | 0 | 0 | 0 | 0 | 1414000 | 0 | 0 | 0 | 0 | 0 | 0 | 0 | 0 | 0 |
| VPERAPLP | VGF; VGF | O15240; Q0 | VGF_HUMAN N | Unmodified | 247610 | 221700 | 0 | 0 | 0 | 0 | 0 | 0 | 1227600 | 0 | 0 | 0 | 0 | 0 | 0 | 0 | 0 | 0 |
| VPEWCLDI | NDUFB9; NDUF | Q9Y6M9; E7 | NDUB9_HUMA | Unmodified | 0 | 0 | 0 | 0 | 0 | 0 | 0 | 0 | 716720 | 0 | 38033 | 0 | 0 | 0 | 0 | 0 | 0 | 0 |
| VPEWVDTV | RPS19; RPS19; | P39019; M0 | RS19_HUMAN | Unmodified | 2012300 | 1370800 | 2437300 | 1078700 | 117870 | 187440 | 520830 | 0 | 0 | 0 | 0 | 0 | 0 | 0 | 2163600 | 1663100 | 0 | 504040 |
| VPFCPMVG | RUVBL1; RUVBI | Q9Y265; E7E | RUVB1_HUMAI | Oxidation (M) | 0 | 162090 | 1362000 | 0 | 0 | 0 | 0 | 0 | 0 | 0 | 192940 | 1344700 | 0 | 0 | 0 | 0 | 0 | 0 |
| VPFDLQNN | FARP2 | H7C3M7 | H7C3M7_HUM | Unmodified | 0 | 0 | 0 | 0 | 411360 | 0 | 0 | 38555 | 830200 | 0 | 0 | 0 | 0 | 0 | 1292700 | 913470 | 0 | 0 |
| VPFILTYDF | PIK3CB; PIK3CE | P42338; H0 | PK3CB_HUMAI | Unmodified | 0 | 0 | 0 | 0 | 0 | 0 | 0 | 0 | 0 | 0 | 0 | 78536 | 0 | 0 | 0 | 0 | 0 | 0 |
| VPFLSSLEC | ANLN; ANLN; A | Q9NQW6; H | ANLN_HUMAN | Unmodified | 0 | 0 | 0 | 0 | 2442700 | 0 | 0 | 0 | 0 | 0 | 0 | 0 | 0 | 0 | 0 | 0 | 0 | 0 |
| VPFSLLR | HSPB1; HSPB1; | P04792; A0A | HSPB1_HUMAI | Unmodified | 0 | 0 | 801050 | 298630 | 0 | 0 | 0 | 0 | 365430 | 0 | 0 | 0 | 0 | 0 | 1214100 | 0 | 0 | 0 |
| VPGFADDF | PTGFRN | Q9P2B2 | FPRP_HUMAN | Unmodified | 0 | 0 | 6841900 | 100140 | 0 | 0 | 0 | 0 | 0 | 0 | 0 | 2881000 | 228830 | 0 | 4966600 | 3563700 | 0 | 0 |
| VPGFADDF | PTGFRN | Q9P2B2 | FPRP_HUMAN | Unmodified | 0 | 0 | 1273600 | 542640 | 0 | 0 | 0 | 0 | 0 | 0 | 0 | 0 | 0 | 0 | 0 | 0 | 0 | 0 |
| VPGIDATTK | AHNAK | Q09666 | AHNK_HUMAN | Unmodified | 0 | 0 | 0 | 0 | 0 | 0 | 0 | 0 | 0 | 0 | 0 | 0 | 0 | 0 | 0 | 0 | 0 | 237850 |
| VPGNFHVS | ERGIC1 | Q969X5 | ERGI1_HUMAN | Unmodified | 0 | 0 | 0 | 0 | 0 | 0 | 0 | 0 | 6432700 | 385920 | 0 | 0 | 0 | 0 | 594770 | 0 | 0 | 0 |
| VPGQEETV | USP9X | Q93008 | USP9X_HUMAI | Unmodified | 11836000 | 11645000 | 7875600 | 8848700 | 0 | 0 | 0 | 0 | 0 | 0 | 0 | 0 | 961100 | 625660 | 8821900 | 10513000 | 0 | 0 |
| VPGQGSSE | SCG2 | P13521 | SCG2_HUMAN | Unmodified | 0 | 0 | 692930 | 0 | 0 | 0 | 427810 | 140240 | 4271400 | 772910 | 0 | 0 | 0 | 676190 | 0 | 0 | 90915 | 319570 |
| VPGTLLPR | AIDA | Q96BJ3 | AIDA_HUMAN | Unmodified | 204370 | 112360 | 440990 | 234530 | 0 | 0 | 0 | 0 | 0 | 0 | 0 | 737140 | 0 | 0 | 2140900 | 2099100 | 0 | 0 |
| VPGTSTSA | FN1; FN1 | P02751; H0 | FINC_HUMAN | Unmodified | 0 | 0 | 3461500 | 419600 | 0 | 0 | 0 | 0 | 0 | 0 | 0 | 0 | 0 | 0 | 0 | 1202000 | 0 | 0 |
| VPHNAAVC | MVP | Q14764 | MVP_HUMAN N | Unmodified | 1043700 | 609710 | 0 | 0 | 0 | 0 | 0 | 0 | 0 | 0 | 0 | 10492000 | 0 | 0 | 0 | 0 | 0 | 0 |
| VPHNAAVC | MVP | Q14764 | MVP_HUMAN N | Unmodified | 0 | 0 | 0 | 0 | 0 | 0 | 55915 | 0 | 0 | 0 | 60601 | 1230800 | 0 | 0 | 0 | 0 | 0 | 353850 |
| VPIDGPPID | IGF2R | P11717 | MPRI_HUMAN | Unmodified | 0 | 0 | 1951200 | 909510 | 373400 | 0 | 0 | 90420 | 49362000 | 2114900 | 4289000 | 11357000 | 0 | 0 | 0 | 2322000 | 0 | 139530 |
| VPIISNDVC | HPN; HPN | P05981; M0 | HEPS_HUMAN | Unmodified | 0 | 372700 | 520680 | 1413600 | 0 | 0 | 0 | 0 | 0 | 0 | 0 | 0 | 0 | 0 | 1576200 | 1040600 | 0 | 0 |
| VPINESFISI | WASHC5; WAS | Q12768; E7E | WASC5_HUMA | Oxidation (M) | 12417 | 0 | 1138300 | 0 | 42310 | 0 | 32831 | 0 | 47837 | 25120 | 0 | 288540 | 0 | 0 | 61499 | 15304 | 0 | 0 |
| VPLALFALN | PEPD; PEPD; PE | P12955; A0A | PEPD_HUMAN | Unmodified | 0 | 0 | 382040 | 0 | 0 | 0 | 0 | 0 | 1959600 | 163130 | 1260300 | 8446800 | 0 | 0 | 0 | 249530 | 0 | 0 |
| VPLAPITDP | RRM2; RRM2 | P31350; A0A | RIR2_HUMAN F | Unmodified | 29050 | 0 | 0 | 31957 | 0 | 0 | 0 | 0 | 0 | 0 | 100360 | 97656 | 85822 | 0 | 0 | 26546 | 0 | 0 |
| VPLEVQEAI | VARS1; VARS1 | P26640; A0A | SYVC_HUMAN | Unmodified | 0 | 0 | 386640 | 76956 | 0 | 0 | 0 | 48739 | 441500 | 222960 | 980850 | 8918800 | 0 | 0 | 240860 | 215810 | 0 | 0 |
| VPLEVQEAI | VARS1; VARS1 | P26640; A0A | SYVC_HUMAN | Unmodified | 0 | 0 | 0 | 0 | 0 | 0 | 0 | 0 | 281530 | 0 | 280160 | 2911500 | 0 | 0 | 0 | 0 | 0 | 0 |
| VPLILVGNK | RAP2C; RAP2C | Q9Y3L5; A0A | RAP2C_HUMAI | Unmodified | 0 | 0 | 0 | 0 | 0 | 0 | 0 | 0 | 0 | 0 | 143540 | 214630 | 0 | 0 | 0 | 0 | 25205 | 20308 |
| VPLILVGNK | RAP2C; RAP2C | Q9Y3L5; A0A | RAP2C_HUMAI | Oxidation (M) | 2257000 | 2807700 | 31694 | 0 | 656380 | 212190 | 0 | 0 | 1552600 | 0 | 4724700 | 10239000 | 0 | 984160 | 91641 | 3441100 | 488430 | 911120 |
| VPLILVGNK | RAP2C; RAP2C | Q9Y3L5; A0A | RAP2C_HUMAI | Unmodified | 0 | 0 | 0 | 611610 | 671040 | 91419 | 0 | 0 | 4325100 | 273490 | 709570 | 1657200 | 720010 | 370780 | 0 | 0 | 201610 | 709150 |
| VPLKVEQAI | MYO6; MYO6; N | Q9UM54; A0 | MYO6_HUMAN | Unmodified | 341660 | 144590 | 0 | 429100 | 0 | 0 | 0 | 0 | 380240 | 0 | 0 | 1291500 | 0 | 0 | 782130 | 1217800 | 0 | 0 |
| VPLQQNFC | LCN2; LCN2 | P80188; X6R | NGAL_HUMAN | Unmodified | 0 | 0 | 0 | 0 | 0 | 0 | 0 | 0 | 355960 | 0 | 0 | 0 | 0 | 0 | 0 | 0 | 409590 | 0 |
| VPLSVQLKI | RBCK1; RBCK1; | Q9BYM8; A6 | HOIL1_HUMAN | Unmodified | 182430 | 104130 | 0 | 0 | 0 | 0 | 0 | 0 | 0 | 0 | 0 | 0 | 0 | 0 | 0 | 0 | 0 | 0 |
| VPLVAEKPI | GARS1; GARS1; | P41250; A0A | GARS_HUMAN | Unmodified | 0 | 0 | 0 | 0 | 0 | 0 | 0 | 0 | 366290 | 0 | 366040 | 3478800 | 0 | 0 | 300210 | 237610 | 0 | 0 |
| VPLVAPEDI | PRMT5; PRMT5; | O14744; H0 | ANM5_HUMAN | Unmodified | 0 | 0 | 0 | 0 | 499770 | 0 | 568340 | 963380 | 522600 | 438890 | 0 | 6206800 | 640250 | 0 | 0 | 0 | 379410 | 489010 |
| VPLVAPEDI | PRMT5; PRMT5; | O14744; H0 | ANM5_HUMAN | Unmodified | 0 | 0 | 0 | 1200200 | 0 | 0 | 0 | 0 | 1591700 | 0 | 2004300 | 28152000 | 0 | 0 | 769360 | 1363300 | 0 | 0 |
| VPLVVRLEC | SUCLG2; SUCL | Q96I99; H0Y | SUCB2_HUMAI | Unmodified | 0 | 0 | 0 | 0 | 0 | 0 | 0 | 0 | 7682100 | 595460 | 0 | 0 | 0 | 0 | 0 | 0 | 283680 | 254520 |
| VPMGSGQI | PTPN23; PTPN2 | Q9H3S7; C9 | PTN23_HUMAN | Oxidation (M) | 1583000 | 1634800 | 0 | 0 | 0 | 0 | 0 | 0 | 0 | 0 | 0 | 0 | 0 | 0 | 0 | 195960 | 0 | 0 |
| VPMILVGNI | RAP2B | P61225 | RAP2B_HUMAN | Unmodified | 0 | 0 | 231840 | 0 | 0 | 0 | 0 | 0 | 542410 | 0 | 0 | 385910 | 0 | 0 | 254980 | 0 | 0 | 0 |
| VPNQYTHL | RAB3GAP1 | Q15042 | RB3GP_HUMAI | Unmodified | 0 | 0 | 27409 | 0 | 0 | 0 | 0 | 0 | 44938 | 29625 | 54988 | 397050 | 0 | 0 | 99913 | 71983 | 0 | 0 |
| VPNSNPPE | MAGED2; MAGI | Q9UNF1; Q5 | MAGD2_HUMA | Unmodified | 0 | 0 | 0 | 0 | 0 | 234960 | 0 | 0 | 133460 | 0 | 28049 | 238730 | 0 | 0 | 25374 | 0 | 20638 | 0 |
| VPPAINQFI | RPL7A; RPL7A | P62424; Q5 | RL7A_HUMAN | Unmodified | 6918800 | 15601000 | 2407500 | 1757700 | 90826 | 74822 | 0 | 0 | 300760 | 51278 | 2418900 | 55876000 | 0 | 0 | 6559000 | 5179000 | 0 | 67735 |
| VPPAINQFI | RPL7A; RPL7A | P62424; Q5 | RL7A_HUMAN | Unmodified | 7398700 | 8366700 | 4079300 | 0 | 0 | 0 | 0 | 0 | 1903100 | 0 | 5544600 | 31460000 | 0 | 0 | 7324800 | 0 | 0 | 0 |
| VPPEFLVQI | MOGS; MOGS; | Q13724; A0A | MOGS_HUMAN | Unmodified | 0 | 0 | 0 | 0 | 0 | 0 | 0 | 0 | 8610500 | 799870 | 1323800 | 261220 | 0 | 0 | 0 | 0 | 0 | 0 |
| VPPLSLGY | KLRG2 | A4D1S0 | KLRG2_HUMAN | Unmodified | 0 | 0 | 649830 | 0 | 0 | 0 | 0 | 0 | 37036 | 0 | 0 | 0 | 0 | 0 | 381270 | 203840 | 391600 | 0 |
| VPPRLQAV | BLVRB; BLVRB | P30043; A0A | BLVRB_HUMAN | Unmodified | 0 | 0 | 1004800 | 0 | 0 | 0 | 0 | 0 | 0 | 0 | 0 | 0 | 0 | 0 | 617880 | 0 | 0 | 0 |
| VPPVQVSP | ATP5ME | P56385 | ATP5I_HUMAN | Unmodified | 0 | 0 | 0 | 0 | 0 | 0 | 1334400 | 510730 | 9698600 | 4544000 | 251240 | 0 | 0 | 0 | 0 | 0 | 0 | 0 |
| VPQIEVETH | DYNC1H1; DYN | Q14204; A0A | DYHC1_HUMA | Unmodified | 0 | 0 | 0 | 0 | 0 | 0 | 0 | 0 | 0 | 0 | 0 | 627950 | 0 | 0 | 43289 | 43846 | 0 | 0 |
| VPSEGAYD | GPRC5C; GPRC | Q9NQ84; A8 | GPC5C_HUMA | Unmodified | 0 | 0 | 5224200 | 5384700 | 332960 | 0 | 0 | 0 | 6302900 | 876200 | 0 | 48335 | 2980900 | 912460 | 5669300 | 4293900 | 0 | 1066600 |
| VPSGGYSS | MAN1B1 | Q9UKM7 | MA1B1_HUMAI | Unmodified | 0 | 0 | 0 | 0 | 0 | 0 | 0 | 0 | 0 | 455060 | 0 | 0 | 0 | 0 | 0 | 0 | 0 | 0 |
| VPSLVGSFI | GSTO1; GSTO1 | P78417; Q5 | GSTO1_HUMAI | Unmodified | 0 | 0 | 0 | 0 | 0 | 0 | 0 | 0 | 0 | 0 | 1714500 | 20254000 | 0 | 0 | 0 | 0 | 0 | 0 |
| VPSNITDV/ | LRRC8D | Q7L1W4 | LRC8D_HUMAI | Unmodified | 0 | 0 | 0 | 307570 | 0 | 931900 | 269200 | 222610 | 0 | 86734 | 0 | 2694800 | 47243 | 0 | 465350 | 631720 | 86553 | 0 |
| VPSPLTDEI | CC2D1B; CC2D | Q5T0F9; H0 | C2D1B_HUMAI | Unmodified | 0 | 0 | 0 | 0 | 0 | 0 | 107840 | 0 | 1125000 | 0 | 0 | 662740 | 0 | 0 | 0 | 0 | 0 | 0 |
| VPSTEAEAI | GDI2; GDI2; GD | P50395; Q5 | GDIB_HUMAN | Oxidation (M) | 0 | 0 | 0 | 0 | 0 | 0 | 0 | 0 | 0 | 0 | 0 | 149660 | 0 | 0 | 0 | 0 | 0 | 0 |
| VPSYLNYH | FZD1 | Q9UP38 | FZD1_HUMAN | Unmodified | 434960 | 329920 | 848420 | 464900 | 227400 | 0 | 0 | 0 | 2019100 | 0 | 0 | 458690 | 0 | 0 | 775320 | 555210 | 0 | 0 |
| VPTANVSV | GAPDH; GAPDI | P04406; E7E | G3P_HUMAN G | Unmodified | 900680 | 1447900 | 283860 | 585820 | 723720 | 0 | 380810 | 149690 | 1281700 | 639890 | 3915200 | 19875000 | 0 | 0 | 1740600 | 1376400 | 586180 | 33457 |
| VPTANVSV | GAPDH; GAPDI | P04406; E7E | G3P_HUMAN G | Unmodified | 3320100 | 1304900 | 7070100 | 1051900 | 822910 | 0 | 285420 | 14553 | 5281400 | 252320 | 5543400 | 96664000 | 770470 | 239850 | 7409500 | 7668100 | 345310 | 724330 |
| VPTATLVR | PTGFRN | Q9P2B2 | FPRP_HUMAN | Unmodified | 9256700 | 3873800 | 13754000 | 5271100 | 736170 | 0 | 1648500 | 0 | 1624500 | 760990 | 423150 | 12152000 | 817130 | 233610 | 22443000 | 19506000 | 0 | 36700 |
| VPTHVPVC | RABL6; RABL6; | Q3YEC7; F2 | RABL6_HUMAN | Unmodified | 0 | 0 | 0 | 0 | 0 | 0 | 0 | 0 | 0 | 0 | 0 | 2770300 | 0 | 0 | 0 | 0 | 0 | 0 |
| VPTISINK | CAP2; CAP1; C/ | P40123; Q01 | CAP2_HUMAN | Unmodified | 967410 | 473610 | 1521600 | 459740 | 0 | 0 | 88395 | 24402 | 121100 | 0 | 939180 | 5577000 | 58469 | 0 | 2424200 | 1088500 | 0 | 0 |
| VPTWGPLR | ILF3; ILF3; ILF3 | Q12906; K7E | ILF3_HUMAN I | Unmodified | 1151800 | 323410 | 0 | 43442 | 0 | 0 | 0 | 0 | 0 | 272460 | 0 | 1901100 | 64784 | 0 | 275650 | 195530 | 0 | 0 |
| VPVAIRVLD | CDH11; CDH11 | P55287; H3E | CAD11_HUMAI | Unmodified | 0 | 0 | 0 | 0 | 0 | 0 | 0 | 0 | 0 | 0 | 0 | 0 | 0 | 0 | 0 | 0 | 0 | 0 |
| VPVGDQPK | DNM1L; DNM1I | O00429; F8V | DNM1L_HUMA | Unmodified | 0 | 0 | 0 | 0 | 0 | 0 | 85614 | 23200 | 0 | 0 | 96790 | 973820 | 0 | 0 | 0 | 0 | 0 | 0 |
| VPVILVGNK | RAP2A | P10114 | RAP2A_HUMAN | Unmodified | 0 | 0 | 0 | 0 | 0 | 0 | 0 | 0 | 0 | 0 | 0 | 426440 | 0 | 0 | 0 | 0 | 0 | 0 |
| VPVILVGNK | RAP2A | P10114 | RAP2A_HUMAN | Unmodified | 130400 | 0 | 39815 | 0 | 0 | 0 | 0 | 0 | 0 | 154010 | 288700 | 907300 | 307700 | 0 | 48685 | 30915 | 0 | 0 |
| VPVILVGNK | RAP2A | P10114 | RAP2A_HUMAN | Unmodified | 2511800 | 2053500 | 0 | 831990 | 484430 | 0 | 0 | 0 | 19328000 | 1667100 | 6739100 | 21731000 | 0 | 0 | 1300800 | 1394500 | 0 | 0 |
| VPVLQLDS | MARS1; MARS1 | P56192; F8V | SYMC_HUMAN | Unmodified | 0 | 0 | 0 | 0 | 0 | 0 | 0 | 0 | 0 | 0 | 0 | 650970 | 0 | 0 | 0 | 0 | 0 | 0 |
| VPVPDERFI | OLA1; OLA1; OL | Q9NTK5; J3 | OLA1_HUMAN | Unmodified | 212500 | 0 | 0 | 0 | 0 | 0 | 0 | 0 | 0 | 0 | 0 | 2045000 | 0 | 0 | 342150 | 0 | 0 | 0 |
| VPVPEDKY | ATP5PD | O75947 | ATP5H_HUMAN | Unmodified | 0 | 0 | 0 | 0 | 136500 | 0 | 30280 | 0 | 2248900 | 0 | 0 | 0 | 65771 | 86919 | 0 | 0 | 0 | 137110 |
| VPVPEDKY | ATP5PD | O75947 | ATP5H_HUMAN | Unmodified | 0 | 141930 | 0 | 0 | 0 | 0 | 0 | 0 | 10785000 | 413620 | 0 | 0 | 0 | 0 | 0 | 0 | 0 | 0 |
| VPVSVNLL | RSL1D1; RSL1D | O76021; J3Q | RL1D1_HUMAN | Unmodified | 54552 | 26923 | 0 | 0 | 0 | 0 | 0 | 0 | 0 | 0 | 0 | 718290 | 0 | 0 | 0 | 0 | 0 | 0 |
| VPVVHVDE | NUP210 | Q8TEM1 | PO210_HUMAN | Unmodified | 0 | 0 | 0 | 0 | 387130 | 0 | 0 | 0 | 1166800 | 288770 | 0 | 79510 | 0 | 0 | 0 | 0 | 373960 | 0 |
| VPVVQAYA | NT5E; NT5E | P21589; H0 | 5NTD_HUMAN | Unmodified | 0 | 0 | 0 | 0 | 3126100 | 1550300 | 0 | 0 | 0 | 0 | 0 | 0 | 0 | 0 | 0 | 0 | 767190 | 782430 |
| VPYYINHQ | UTRN; UTRN | P46939; Q5 | UTRN_HUMAN | Unmodified | 0 | 0 | 0 | 346780 | 0 | 0 | 0 | 0 | 219400 | 0 | 0 | 0 | 0 | 0 | 0 | 291460 | 0 | 0 |
| VQADQVD | AHNAK2 | Q8IVF2 | AHNK2_HUMAI | Unmodified | 0 | 0 | 0 | 0 | 6587000 | 1127100 | 0 | 0 | 0 | 0 | 0 | 0 | 0 | 0 | 0 | 0 | 0 | 7088000 |
| VQAEAERLI | BECN1; BECN1 | Q14457; E7E | BECN1_HUMAI | Unmodified | 0 | 0 | 0 | 0 | 0 | 0 | 0 | 0 | 0 | 0 | 0 | 979330 | 0 | 0 | 0 | 0 | 0 | 0 |
| VQAELDETI | YKT6; YKT6; YK | O15498; A0A | YKT6_HUMAN S | Unmodified | 3308500 | 2970200 | 745550 | 1983000 | 6017400 | 0 | 0 | 0 | 8574400 | 969450 | 79066 | 7067200 | 3779500 | 716900 | 3118100 | 3139600 | 267580 | 7478500 |

| | | | | | | | | | | | | | | | | | | | | | | |
|---|---|---|---|---|---|---|---|---|---|---|---|---|---|---|---|---|---|---|---|---|---|---|
| VQAGDVITI | RUVBL2; RUVBI | Q9Y230; M0I | RUVB2_HUMAI | Unmodified | 435810 | 165170 | 697370 | 190220 | 0 | 0 | 0 | 0 | 0 | 0 | 1664100 | 3321500 | 444230 | 1151300 | 612530 | 366590 | 0 | 0 |
| VQAGQMD | AHNAK2 | Q8IVF2 | AHNK2_HUMAI | Unmodified | 0 | 0 | 0 | 0 | 2766800 | 0 | 0 | 0 | 0 | 0 | 0 | 0 | 0 | 0 | 0 | 0 | 63668 | 2616500 |
| VQAGQMD | AHNAK2 | Q8IVF2 | AHNK2_HUMAI | Oxidation (M) | 0 | 0 | 0 | 0 | 2240500 | 413890 | 0 | 0 | 0 | 0 | 0 | 0 | 0 | 402600 | 0 | 0 | 169830 | 904510 |
| VQAGQVD\ | AHNAK2 | Q8IVF2 | AHNK2_HUMAI | Unmodified | 0 | 0 | 0 | 0 | 949570 | 0 | 0 | 0 | 0 | 0 | 0 | 0 | 0 | 0 | 0 | 0 | 0 | 1499900 |
| VQAHAAAA | IPO5; IPO5 | O00410; H0Y | IPO5_HUMAN I | Unmodified | 0 | 0 | 0 | 0 | 0 | 0 | 0 | 0 | 612610 | 0 | 0 | 4713700 | 0 | 0 | 0 | 0 | 271380 | 0 |
| VQAINVSSI | EFHD2 | Q96C19 | EFHD2_HUMAI | Unmodified | 0 | 0 | 0 | 0 | 0 | 0 | 164100 | 0 | 1254900 | 0 | 0 | 0 | 0 | 0 | 0 | 0 | 154850 | 0 |
| VQAMQISS | CDV3; CDV3 | Q9UKY7; H0 | CDV3_HUMAN | Unmodified | 0 | 0 | 73484 | 43029 | 215890 | 0 | 0 | 0 | 193370 | 0 | 83244 | 841270 | 32976 | 0 | 116480 | 87003 | 0 | 0 |
| VQANLGAF | AHNAK | Q09666 | AHNK_HUMAN | Unmodified | 0 | 0 | 0 | 0 | 9835100 | 1831400 | 0 | 0 | 0 | 281090 | 0 | 1350700 | 257540 | 92860 | 175360 | 0 | 1778900 | 5203100 |
| VQANLGAF | AHNAK | Q09666 | AHNK_HUMAN | Unmodified | 0 | 0 | 0 | 0 | 1953500 | 0 | 0 | 0 | 0 | 0 | 0 | 0 | 0 | 0 | 0 | 0 | 547320 | 2222700 |
| VQAQGHTI | ALDH16A1; ALD | Q8IZ83; F5H | A16A1_HUMAN | Unmodified | 0 | 135410 | 0 | 0 | 0 | 0 | 0 | 0 | 0 | 0 | 0 | 2986300 | 0 | 0 | 0 | 233400 | 0 | 0 |
| VQAQGPGI | FLNB; FLNB | O75369; E7E | FLNB_HUMAN | Unmodified | 0 | 0 | 99503 | 0 | 217000 | 144410 | 0 | 0 | 0 | 0 | 0 | 2658900 | 349150 | 374590 | 334040 | 164760 | 0 | 0 |
| VQAQHDY1 | BIN1 | O00499 | BIN1_HUMAN I | Unmodified | 0 | 0 | 0 | 0 | 0 | 0 | 0 | 0 | 0 | 0 | 0 | 0 | 672810 | 0 | 0 | 96639 | 0 | 0 |
| VQASLAAN | BTF3; BTF3; BTF | P20290; H0Y | BTF3_HUMAN ` | Unmodified | 2205800 | 1760700 | 698560 | 1362600 | 5513500 | 566610 | 1398800 | 595930 | 669950 | 410930 | 3938200 | 29737000 | 4683200 | 2284200 | 893070 | 972610 | 527850 | 868020 |
| VQAVQPTL | MBLAC2 | Q68D91 | MBLC2_HUMAI | Unmodified | 200720 | 0 | 697130 | 1222900 | 0 | 0 | 0 | 0 | 0 | 0 | 197350 | 2580500 | 0 | 0 | 2556000 | 918320 | 0 | 0 |
| VQCPALIVI | BPHL | Q86WA6 | BPHL_HUMAN | Unmodified | 0 | 0 | 0 | 0 | 0 | 0 | 0 | 0 | 702310 | 230750 | 0 | 0 | 0 | 0 | 0 | 0 | 120620 | 238110 |
| VQDAFAAA | SLC7A5 | Q01650 | LAT1_HUMAN I | Unmodified | 0 | 0 | 0 | 0 | 0 | 230150 | 490100 | 231650 | 85464 | 104240 | 163080 | 574290 | 0 | 0 | 196570 | 68497 | 0 | 0 |
| VQDDEVGI | CCT2; CCT2; C( | P78371; F5G | TCPB_HUMAN | Unmodified | 0 | 0 | 0 | 276720 | 0 | 0 | 0 | 0 | 377590 | 0 | 0 | 1085000 | 0 | 0 | 0 | 0 | 0 | 0 |
| VQDFQEDI | PHYH; PHYH; P | O14832; B1/ | PAHX_HUMAN | Unmodified | 26191 | 0 | 0 | 100500 | 0 | 0 | 17811 | 0 | 7993700 | 607090 | 166330 | 0 | 248040 | 0 | 0 | 15661 | 0 | 0 |
| VQDFQEDI | PHYH; PHYH; P | O14832; B1/ | PAHX_HUMAN | Unmodified | 0 | 0 | 0 | 0 | 0 | 0 | 0 | 0 | 1813600 | 191580 | 0 | 0 | 0 | 0 | 0 | 0 | 0 | 0 |
| VQDIKYEP( | C1orf43; C1orf4 | Q9BWL3; X6 | CA043_HUMAN | Unmodified | 0 | 0 | 0 | 0 | 0 | 0 | 0 | 0 | 0 | 0 | 0 | 1778900 | 0 | 0 | 0 | 0 | 0 | 0 |
| VQDLENQL | GULP1; GULP1 | Q9UBP9; H0 | GULP1_HUMAI | Unmodified | 306020 | 0 | 514030 | 0 | 0 | 0 | 0 | 0 | 0 | 0 | 79794 | 1431100 | 0 | 0 | 249850 | 568310 | 0 | 0 |
| VQDLGRD! | ADGRE5 | P48960 | AGRE5_HUMAI | Unmodified | 0 | 0 | 0 | 0 | 3755600 | 0 | 0 | 0 | 411490 | 0 | 0 | 0 | 0 | 0 | 0 | 0 | 598890 | 1303800 |
| VQDNFDKI | SWAP70 | Q9UH65 | SWP70_HUMAI | Unmodified | 0 | 0 | 0 | 0 | 256450 | 0 | 0 | 0 | 0 | 0 | 0 | 820930 | 0 | 0 | 0 | 0 | 0 | 0 |
| VQDSNAEI | UACA; UACA; U | Q9BZF9; H0Y | UACA_HUMAN | Unmodified | 0 | 0 | 0 | 152180 | 1107800 | 103680 | 0 | 0 | 0 | 115800 | 0 | 457250 | 282240 | 119290 | 117720 | 102280 | 258160 | 1278400 |
| VQDYIQSK | PFKFB2 | O60825 | F262_HUMAN ( | Unmodified | 0 | 0 | 0 | 0 | 0 | 0 | 0 | 0 | 0 | 0 | 0 | 820710 | 0 | 0 | 0 | 0 | 0 | 0 |
| VQEAESPV | SLC20A2 | Q08357 | S20A2_HUMAN | Unmodified | 0 | 0 | 0 | 0 | 0 | 0 | 0 | 0 | 0 | 0 | 0 | 687270 | 0 | 465720 | 78864 | 0 | 0 | 0 |
| VQEAVESM | FAM136A; FAM1 | Q96C01; C9J | F136A_HUMAN | Unmodified | 10382 | 0 | 0 | 0 | 257160 | 3443400 | 0 | 0 | 95625 | 84418 | 0 | 514010 | 0 | 0 | 0 | 0 | 0 | 64686000 |
| VQEDEFR | PTGFRN | Q9P2B2 | FPRP_HUMAN | Unmodified | 5370900 | 4358500 | 18385000 | 9600600 | 0 | 234570 | 0 | 0 | 0 | 0 | 0 | 20434000 | 0 | 0 | 14323000 | 17139000 | 0 | 0 |
| VQEDEFRY | PTGFRN | Q9P2B2 | FPRP_HUMAN | Unmodified | 557380 | 0 | 505130 | 6016500 | 0 | 626340 | 0 | 0 | 0 | 0 | 0 | 0 | 3493600 | 0 | 169870 | 0 | 0 | 0 |
| VQEFTVK | ANK3; ANK3 | D6RF31; H0Y | D6RF31_HUMA | Unmodified | 0 | 0 | 436290 | 578150 | 0 | 0 | 0 | 0 | 88926 | 0 | 0 | 191940 | 0 | 294970 | 1556800 | 1313300 | 0 | 0 |
| VQEFTVKSI | ANK3; ANK3 | D6RF31; H0Y | D6RF31_HUMA | Unmodified | 0 | 0 | 0 | 235670 | 0 | 0 | 0 | 0 | 0 | 0 | 0 | 0 | 804370 | 668030 | 0 | 0 | 0 | 0 |
| VQEFTVKSI | ANK3; ANK3 | D6RF31; H0Y | D6RF31_HUMA | Oxidation (M) | 0 | 0 | 192970 | 0 | 0 | 0 | 0 | 0 | 0 | 0 | 0 | 0 | 278600 | 149790 | 0 | 0 | 0 | 0 |
| VQELGLSA | CBS; CBSL; CBS | P35520; P0D | CBS_HUMAN C | Unmodified | 0 | 0 | 0 | 0 | 0 | 0 | 0 | 0 | 0 | 0 | 0 | 1428100 | 0 | 0 | 0 | 0 | 0 | 0 |
| VQELQNLL | KTN1; KTN1; KT | Q86UP2; G3 | KTN1_HUMAN | Unmodified | 0 | 0 | 33142 | 0 | 0 | 12693 | 0 | 404080 | 8716400 | 1385000 | 71060 | 1583400 | 0 | 0 | 106140 | 77091 | 0 | 0 |
| VQEMNYIQ | ACSL4 | O60488 | ACSL4_HUMAI | Oxidation (M) | 0 | 0 | 0 | 0 | 127300 | 221930 | 0 | 0 | 0 | 0 | 193800 | 1539300 | 0 | 0 | 0 | 0 | 140710 | 57322 |
| VQEMNYIQ | ACSL4 | O60488 | ACSL4_HUMAI | Unmodified | 0 | 0 | 0 | 148890 | 867300 | 615380 | 0 | 0 | 0 | 0 | 776650 | 6800200 | 0 | 0 | 0 | 0 | 600250 | 376580 |
| VQEQELKS | IMMT; IMMT; IM | Q16891; B9/ | MIC60_HUMAN | Unmodified | 0 | 0 | 0 | 0 | 0 | 0 | 288670 | 0 | 1153900 | 0 | 0 | 0 | 0 | 0 | 0 | 0 | 188130 | 0 |
| VQEQELTR | DSP | P15924 | DESP_HUMAN | Unmodified | 211590 | 0 | 0 | 137540 | 1368100 | 387900 | 0 | 42734 | 322470 | 45606 | 14024 | 184030 | 684270 | 158010 | 0 | 0 | 79372 | 142040 |
| VQEQLGNI | PMM2; PMM2; F | O15305; H3I | PMM2_HUMAN | Unmodified | 0 | 0 | 0 | 0 | 0 | 0 | 0 | 0 | 177900 | 0 | 217420 | 1123800 | 0 | 0 | 121750 | 0 | 0 | 0 |
| VQEQVHTL | CKAP4 | Q07065 | CKAP4_HUMAI | Unmodified | 0 | 0 | 0 | 0 | 1971500 | 0 | 0 | 0 | 8324700 | 346940 | 700260 | 0 | 0 | 0 | 0 | 0 | 0 | 0 |
| VQEQVHTL | CKAP4 | Q07065 | CKAP4_HUMAI | Unmodified | 0 | 0 | 0 | 0 | 1225600 | 0 | 2823100 | 0 | 0 | 10594000 | 0 | 0 | 3234800 | 1558200 | 0 | 0 | 0 | 4504300 |
| VQESADEL | ARF5; ARF5 | P84085; Q5I | ARF5_HUMAN . | Unmodified | 1317100 | 1629300 | 1184200 | 1536000 | 0 | 0 | 0 | 0 | 1115600 | 257570 | 0 | 8516100 | 0 | 56280000 | 1693800 | 1396000 | 0 | 0 |
| VQESLKKQ | PDCD6IP | Q8WUM4 | PDC6I_HUMAN | Unmodified | 539830 | 290700 | 0 | 912630 | 14727000 | 1087000 | 1654200 | 666620 | 493560 | 6641300 | 0 | 0 | 20364000 | 15459000 | 0 | 0 | 2488000 | 7408500 |
| VQETGDIVI | HSD17B4; HSD | P51659; E7E | DHB4_HUMAN | Unmodified | 0 | 0 | 0 | 0 | 0 | 0 | 0 | 0 | 6803700 | 874790 | 1811600 | 0 | 0 | 0 | 0 | 0 | 0 | 0 |
| VQEVLLK | HYOU1; HYOU: | Q9Y4L1; E9F | HYOU1_HUMA | Unmodified | 0 | 902850 | 1063200 | 885790 | 319460 | 0 | 122990 | 47711 | 40382000 | 8621000 | 3823100 | 2351000 | 932200 | 0 | 1120300 | 899130 | 0 | 364430 |
| VQEYPTLK\ | MAEA; MAEA; M | Q7L5Y9; D6I | MAEA_HUMAN | Unmodified | 0 | 0 | 0 | 0 | 364770 | 0 | 219990 | 183330 | 0 | 299350 | 0 | 0 | 306500 | 215340 | 0 | 0 | 0 | 198510 |
| VQFAPEKP | ARRB2; ARRB2; | P32121; K7E | ARRB2_HUMAI | Unmodified | 0 | 0 | 146550 | 0 | 0 | 0 | 0 | 0 | 98123 | 0 | 0 | 410280 | 0 | 0 | 0 | 0 | 0 | 0 |
| VQFRDLNF | ATP6V0A1; ATP( | Q93050; B7Z | VPP1_HUMAN ` | Unmodified | 341590 | 354290 | 0 | 0 | 112310 | 0 | 0 | 26859 | 0 | 366520 | 96216 | 525040 | 92812 | 0 | 245950 | 0 | 0 | 0 |
| VQFRDLNF | ATP6V0A1; ATP( | Q93050; B7Z | VPP1_HUMAN ` | Unmodified | 0 | 0 | 0 | 0 | 0 | 887770 | 1683700 | 0 | 0 | 2261400 | 0 | 0 | 659090 | 0 | 0 | 0 | 0 | 0 |
| VQFRNQG! | ITGA6 | P23229 | ITA6_HUMAN I | Unmodified | 0 | 0 | 0 | 0 | 1454800 | 0 | 0 | 0 | 0 | 0 | 0 | 0 | 1593800 | 0 | 0 | 0 | 0 | 0 |
| VQFRNQG! | ITGA6 | P23229 | ITA6_HUMAN I | Unmodified | 0 | 0 | 0 | 0 | 7080300 | 1496400 | 0 | 0 | 0 | 0 | 0 | 0 | 5006700 | 2665900 | 0 | 0 | 9321900 | 31526000 |
| VQFVPFHS | ABHD12; ABHD | Q8N2K0; A0 | ABD12_HUMAI | Unmodified | 0 | 0 | 0 | 0 | 0 | 0 | 227520 | 0 | 4146600 | 0 | 0 | 0 | 0 | 103570 | 0 | 0 | 0 | 0 |
| VQFVPFHS | ABHD12; ABHD | Q8N2K0; A0 | ABD12_HUMAI | Unmodified | 0 | 0 | 0 | 0 | 0 | 0 | 0 | 0 | 1237300 | 426450 | 0 | 0 | 0 | 0 | 0 | 0 | 0 | 0 |
| VQGEAVSN | NACA; NACA; N | E9PAV3; Q1: | NACAM_HUMA | Unmodified | 0 | 0 | 0 | 0 | 0 | 0 | 0 | 0 | 0 | 0 | 1045800 | 1343600 | 0 | 0 | 0 | 0 | 0 | 0 |
| VQGGALEL | CCT7 | Q99832 | TCPH_HUMAN | Unmodified | 0 | 0 | 0 | 0 | 433170 | 0 | 0 | 26965 | 177730 | 0 | 583460 | 1132200 | 0 | 0 | 0 | 139800 | 162730 | 0 |
| VQGGALEL | CCT7 | Q99832 | TCPH_HUMAN | Unmodified | 448190 | 490300 | 312550 | 520730 | 2501900 | 377590 | 1953900 | 1168400 | 1432200 | 2153200 | 5111700 | 21085000 | 4204900 | 2756000 | 668040 | 880970 | 1836500 | 0 |
| VQGILAMS | PPM1L | Q5SGD2 | PPM1L_HUMAN | Unmodified | 0 | 0 | 0 | 0 | 0 | 0 | 0 | 0 | 1331800 | 0 | 0 | 0 | 0 | 0 | 0 | 0 | 0 | 0 |
| VQGSDSDI | VARS1; VARS1 | P26640; A0/ | SYVC_HUMAN | Unmodified | 0 | 0 | 0 | 0 | 0 | 0 | 0 | 0 | 0 | 0 | 349590 | 4541600 | 0 | 0 | 0 | 0 | 0 | 0 |
| VQHIQLLQ | FGA; FGA | P02671; A0/ | FIBA_HUMAN F | Unmodified | 0 | 1236900 | 0 | 0 | 0 | 0 | 0 | 0 | 0 | 0 | 0 | 0 | 0 | 0 | 0 | 0 | 0 | 0 |
| VQHQTSST | RALGAPB; RAL( | Q86X10; A2/ | RLGPB_HUMAN | Unmodified | 0 | 0 | 0 | 0 | 0 | 0 | 0 | 0 | 0 | 0 | 235550 | 728140 | 0 | 0 | 0 | 230320 | 0 | 0 |
| VQIAVANA( | TIMM13 | Q9Y5L4 | TIM13_HUMAN | Unmodified | 0 | 522350 | 0 | 0 | 34884 | 0 | 0 | 0 | 411320 | 300340 | 705710 | 3280100 | 511570 | 133960 | 0 | 107340 | 0 | 0 |
| VQIAVANA( | TIMM13 | Q9Y5L4 | TIM13_HUMAN | Unmodified | 0 | 0 | 0 | 0 | 0 | 1226800 | 1718100 | 0 | 0 | 797890 | 0 | 0 | 5035400 | 2361100 | 0 | 0 | 172410 | 1318800 |
| VQIAVANA( | TIMM13 | Q9Y5L4 | TIM13_HUMAN | Oxidation (M) | 0 | 0 | 0 | 0 | 0 | 0 | 139150 | 0 | 0 | 54728 | 0 | 0 | 728480 | 156480 | 0 | 0 | 0 | 0 |
| VQIDVYEK | PCYOX1L; PCY( | Q8NBM8; D6 | PCYXL_HUMAI | Unmodified | 0 | 0 | 0 | 0 | 0 | 0 | 0 | 0 | 3974100 | 0 | 0 | 0 | 0 | 0 | 0 | 0 | 0 | 0 |
| VQIEHISSL | PSMD11 | O00231 | PSD11_HUMAN | Unmodified | 0 | 0 | 0 | 0 | 0 | 0 | 0 | 0 | 0 | 0 | 0 | 2135400 | 0 | 0 | 0 | 0 | 0 | 0 |
| VQIFEYNEI | SEH1L; SEH1L | Q96EE3; K7I | SEH1_HUMAN | Unmodified | 0 | 0 | 0 | 0 | 0 | 0 | 0 | 0 | 0 | 0 | 0 | 315550 | 0 | 0 | 0 | 0 | 0 | 0 |
| VQIPIMLVG | RHEB; RHEB; RI | Q15382; C9J | RHEB_HUMAN | Oxidation (M) | 1950800 | 1645600 | 1912800 | 1644100 | 864310 | 0 | 0 | 200560 | 606390 | 336130 | 0 | 1708000 | 299580 | 0 | 2225800 | 3206900 | 264310 | 0 |
| VQIPIMLVG | RHEB; RHEB; RI | Q15382; C9J | RHEB_HUMAN | Unmodified | 0 | 0 | 0 | 0 | 915530 | 431350 | 0 | 0 | 542480 | 0 | 0 | 0 | 0 | 190120 | 0 | 0 | 0 | 999510 |
| VQIQESVR | PRPSAP2; PRPS | O60256; E7E | KPRB_HUMAN | Unmodified | 0 | 0 | 0 | 0 | 0 | 0 | 0 | 0 | 0 | 0 | 0 | 3250600 | 0 | 0 | 0 | 0 | 0 | 0 |
| VQIQSNRE | MACF1; MACF1 | Q9UPN3; H3 | MACF1_HUMAN | Unmodified | 0 | 0 | 0 | 0 | 0 | 0 | 0 | 0 | 0 | 0 | 0 | 0 | 0 | 0 | 0 | 0 | 0 | 0 |
| VQISPDSG | KHSRP; KHSRP | Q92945; M0I | FUBP2_HUMAI | Unmodified | 0 | 0 | 0 | 0 | 0 | 0 | 0 | 0 | 791960 | 0 | 536080 | 1192200 | 0 | 0 | 0 | 0 | 0 | 0 |
| VQISPDSG | KHSRP; KHSRP | Q92945; M0I | FUBP2_HUMAI | Unmodified | 0 | 0 | 0 | 0 | 3787900 | 0 | 0 | 750690 | 0 | 1139400 | 0 | 0 | 0 | 0 | 0 | 0 | 0 | 1841200 |
| VQIVSIKDY | GNA13 | Q14344 | GNA13_HUMAI | Unmodified | 137050 | 116560 | 0 | 0 | 468030 | 0 | 0 | 0 | 310560 | 240140 | 71347 | 82143 | 0 | 0 | 0 | 0 | 580740 | 0 |
| VQIYHNPT/ | VASP; VASP; VA | P50552; K7E | VASP_HUMAN | Unmodified | 291890 | 547290 | 0 | 2120100 | 0 | 0 | 0 | 0 | 449940 | 0 | 0 | 9718900 | 0 | 232750 | 311690 | 301410 | 0 | 36568 |
| VQKEDIEV\ | RELA; RELA; RE | Q04206; A0/ | TF65_HUMAN ` | Unmodified | 0 | 0 | 0 | 0 | 0 | 0 | 35596 | 97057 | 0 | 119100 | 0 | 462440 | 50997 | 62722 | 0 | 62465 | 286370 | 236670 |
| VQKLEESLI | KIF15; KIF15 | Q9NS87; C9 | KIF15_HUMAN | Unmodified | 0 | 0 | 0 | 0 | 241050 | 23376 | 3125500 | 68270 | 85986 | 147660 | 2138800 | 329020 | 94968 | 75749 | 0 | 0 | 11577000 | 23365000 |
| VQKLLQDF | HSPA1A; HSPA1 | P0DMV8; P0 | HS71A_HUMAN | Unmodified | 0 | 0 | 0 | 0 | 193840 | 0 | 1495100 | 0 | 0 | 0 | 0 | 0 | 0 | 0 | 0 | 0 | 523970 | 0 |
| VQLEHLEFI | PLCB4; PLCB4 | Q15147; A0/ | PLCB4_HUMAI | Unmodified | 0 | 0 | 0 | 0 | 0 | 0 | 0 | 0 | 0 | 0 | 0 | 0 | 0 | 0 | 0 | 0 | 0 | 983070 |
| VQLETIIK | EPS15L1; EPS1 | Q9UBC2; M( | EP15R_HUMAN | Unmodified | 0 | 0 | 166340 | 0 | 0 | 0 | 0 | 0 | 1642600 | 71541 | 241260 | 0 | 0 | 0 | 125920 | 403150 | 0 | 0 |
| VQLFEISLS | ARRDC1; ARRD | Q8N5I2; Q5I | ARRD1_HUMAN | Unmodified | 0 | 0 | 809420 | 489320 | 0 | 0 | 0 | 0 | 0 | 0 | 0 | 0 | 0 | 0 | 0 | 0 | 0 | 0 |
| VQLLQDLQ | SRGAP2; SRGA | O75044; Q7. | SRGP2_HUMAI | Unmodified | 511550 | 440720 | 319890 | 568530 | 5900800 | 1987900 | 484830 | 0 | 1782900 | 355480 | 650960 | 6030900 | 1537600 | 163270 | 1046800 | 711930 | 348410 | 245160 |
| VQLNVFDE | DIAPH1; DIAPH | O60610; A0/ | DIAP1_HUMAN | Unmodified | 0 | 0 | 0 | 0 | 0 | 0 | 0 | 0 | 0 | 0 | 0 | 2794100 | 0 | 0 | 320070 | 269050 | 0 | 0 |
| VQLPAYLK | CISD2; CISD2; ( | Q8N5K1; I3L | CISD2_HUMAN | Unmodified | 0 | 0 | 0 | 0 | 0 | 0 | 0 | 64893 | 4360900 | 886280 | 0 | 742750 | 0 | 0 | 0 | 0 | 0 | 0 |
| VQLSGPQE | COPS3; COPS3 | Q9UNS2; H7 | CSN3_HUMAN | Unmodified | 0 | 0 | 0 | 0 | 0 | 0 | 0 | 0 | 0 | 0 | 0 | 3329200 | 0 | 0 | 0 | 0 | 0 | 0 |
| VQLTGEPV | PVR; PVR; PVR | P15151; A0/ | PVR_HUMAN P | Unmodified | 728120 | 630720 | 0 | 0 | 1093600 | 0 | 0 | 0 | 764360 | 0 | 1143500 | 5057200 | 0 | 0 | 890470 | 477710 | 0 | 0 |
| VQLTGEPV | PVR; PVR; PVR | P15151; A0/ | PVR_HUMAN P | Oxidation (M) | 1386800 | 987220 | 582530 | 1145400 | 670720 | 439850 | 0 | 0 | 281730 | 0 | 815010 | 7379800 | 0 | 0 | 16152000 | 786300 | 0 | 0 |
| VQLVTDRD | P4HTM; P4HTM | Q9NXG6; A0 | P4HTM_HUMAI | Unmodified | 0 | 0 | 0 | 0 | 0 | 0 | 0 | 0 | 1648200 | 0 | 0 | 0 | 0 | 0 | 0 | 0 | 0 | 0 |
| VQMPSLK | AHNAK2 | Q8IVF2 | AHNK2_HUMAI | Unmodified | 0 | 0 | 0 | 0 | 363420 | 0 | 0 | 0 | 0 | 0 | 0 | 0 | 0 | 0 | 0 | 0 | 215390 | 405410 |
| VQNATLAV | STOM; STOM | P27105; F8V | STOM_HUMAN | Unmodified | 1061600 | 2.94E+08 | 3184600 | 614650 | 0 | 0 | 0 | 11469 | 0 | 0 | 0 | 0 | 0 | 0 | 1541500 | 1390400 | 0 | 0 |
| VQNGGIVT | SLC19A2; SLC1 | O60779; A0/ | S19A2_HUMAN | Unmodified | 284480 | 261090 | 329410 | 0 | 0 | 0 | 0 | 0 | 0 | 0 | 0 | 2438400 | 0 | 0 | 0 | 381850 | 0 | 0 |
| VQNHEIEL | ATP11C; ATP11 | Q8NB49; A0. | AT11C_HUMAN | Unmodified | 0 | 0 | 0 | 670860 | 0 | 0 | 114680 | 0 | 574000 | 209070 | 371660 | 1145000 | 69718 | 66431 | 0 | 0 | 0 | 0 |
| VQNIHPVE | NUCB2; NUCB: | P80303; E9P | NUCB2_HUMA | Unmodified | 0 | 0 | 184000 | 0 | 0 | 0 | 0 | 0 | 1150700 | 0 | 0 | 43391 | 0 | 0 | 0 | 0 | 0 | 0 |
| VQNIHPVE | NUCB2; NUCB: | P80303; E9P | NUCB2_HUMA | Unmodified | 0 | 0 | 0 | 0 | 0 | 0 | 664770 | 0 | 90368000 | 12439000 | 0 | 0 | 0 | 0 | 1598400 | 0 | 0 | 0 |
| VQPFHQGI | ACTL8 | Q9H568 | ACTL8_HUMAN | Unmodified | 0 | 0 | 0 | 0 | 0 | 0 | 0 | 0 | 0 | 0 | 320730 | 5110600 | 0 | 0 | 0 | 0 | 0 | 0 |
| VQPGDRVI | MCM4; MCM4; | P33991; A0/ | MCM4_HUMAN | Unmodified | 0 | 0 | 0 | 0 | 0 | 0 | 0 | 0 | 816690 | 0 | 758110 | 1916700 | 87881 | 0 | 0 | 0 | 124290 | 63236 |
| VQPNEAVY | G6PD | P11413 | G6PD_HUMAN | Unmodified | 0 | 0 | 0 | 1525300 | 4264000 | 1119400 | 52326 | 46032 | 1897800 | 163720 | 0 | 4514400 | 0 | 0 | 821140 | 624960 | 352870 | 659770 |
| VQPQEPLV | SPINT1; SPINT1 | O43278; H3I | SPIT1_HUMAN | Unmodified | 0 | 1796000 | 1322400 | 3130100 | 1365500 | 936730 | 0 | 0 | 3673600 | 155970 | 0 | 0 | 0 | 641670 | 2245500 | 143140 | 0 | 0 |
| VQPSDQV( | CBS; CBSL; CB! | P35520; P0D | CBS_HUMAN C | Unmodified | 0 | 0 | 0 | 0 | 864040 | 0 | 0 | 0 | 0 | 0 | 64473 | 1430200 | 0 | 0 | 0 | 0 | 0 | 0 |
| VQQAELHT | ACACA | Q13085 | ACACA_HUMAI | Unmodified | 0 | 0 | 578960 | 0 | 0 | 0 | 0 | 41825 | 0 | 0 | 194420 | 1984900 | 0 | 0 | 361080 | 371760 | 0 | 0 |
| VQQEAREK | N/A | N/A | N/A | Unmodified | 0 | 0 | 0 | 0 | 0 | 0 | 3474600 | 122370 | 0 | 0 | 0 | 0 | 1567700 | 0 | 0 | 0 | 2284100 | 0 |
| VQQLERSI( | TRAF2 | Q12933 | TRAF2_HUMAN | Unmodified | 0 | 0 | 0 | 0 | 5491100 | 1284100 | 52503 | 30672 | 0 | 0 | 0 | 0 | 0 | 0 | 0 | 0 | 0 | 0 |
| VQQLLHIC: | PSMD2; PSMD2 | Q13200; H7( | PSMD2_HUMA | Unmodified | 0 | 0 | 0 | 0 | 0 | 0 | 0 | 0 | 0 | 0 | 0 | 1970400 | 117280 | 0 | 0 | 0 | 0 | 0 |
| VQQLVPK | ACTN4; ACTN4; | O43707; F5( | ACTN4_HUMAI | Unmodified | 377650 | 0 | 1262400 | 0 | 1790000 | 1529300 | 142050 | 0 | 1836300 | 938730 | 0 | 11287000 | 139760 | 60942 | 1061700 | 871530 | 708620 | 0 |
| VQQLVPKR | ACTN4; ACTN4; | O43707; F5( | ACTN4_HUMAI | Unmodified | 0 | 0 | 0 | 283000 | 23917000 | 1308500 | 1063400 | 106760 | 2504200 | 3188300 | 458200 | 3100500 | 1242400 | 0 | 151610 | 0 | 7847400 | 267950 |
| VQQTVQDI | HSPA9; HSPA9; | P38646; A0/ | GRP75_HUMAI | Unmodified | 0 | 0 | 0 | 0 | 0 | 0 | 0 | 0 | 5328800 | 0 | 0 | 188170 | 0 | 0 | 0 | 0 | 0 | 0 |
| VQQTVQDI | HSPA9; HSPA9; | P38646; A0/ | GRP75_HUMAI | Unmodified | 23935 | 0 | 0 | 0 | 2879600 | 621660 | 3382500 | 946940 | 23176000 | 8014100 | 277840 | 705050 | 4652400 | 2604500 | 0 | 0 | 517580 | 1373100 |
| VQQVQPAI | NIBAN2 | Q96TA1 | NIBA2_HUMAN | Unmodified | 361920 | 105200 | 0 | 175690 | 0 | 0 | 0 | 0 | 903270 | 88892 | 0 | 504310 | 0 | 0 | 225060 | 113720 | 0 | 0 |
| VQQVQPAI | NIBAN2 | Q96TA1 | NIBA2_HUMAN | Oxidation (M) | 0 | 0 | 0 | 0 | 2883200 | 589490 | 0 | 0 | 0 | 326350 | 0 | 0 | 517060 | 0 | 0 | 0 | 0 | 0 |
| VQQVQPAI | NIBAN2 | Q96TA1 | NIBA2_HUMAN | Unmodified | 61190 | 0 | 0 | 0 | 11127000 | 1849700 | 0 | 0 | 1754300 | 787740 | 0 | 0 | 0 | 0 | 0 | 0 | 0 | 0 |

| | | | | | | | | | | | | | | | | | | | | | | |
|---|---|---|---|---|---|---|---|---|---|---|---|---|---|---|---|---|---|---|---|---|---|---|
| VQQWDSQ | ABR; ABR; ABR; | Q12979; A0/ | ABR_HUMAN A | Unmodified | 0 | 0 | 0 | 0 | 648710 | 0 | 0 | 0 | 0 | 0 | 0 | 0 | 0 | 0 | 0 | 0 | 464940 | 0 |
| VQRAVCMI | TUBA3C; TUBA: | P0DPH7; Q7 | TBA3C_HUMAN | Oxidation (M) | 0 | 0 | 0 | 0 | 0 | 0 | 1381600 | 0 | 0 | 738230 | 0 | 0 | 0 | 0 | 0 | 0 | 223680 | 0 |
| VQREPLLTI | PSMD2 | Q13200 | PSMD2_HUMA | Unmodified | 85892 | 0 | 178940 | 109580 | 983790 | 0 | 0 | 0 | 168840 | 105390 | 172180 | 2260700 | 666130 | 165560 | 85834 | 119660 | 321870 | 0 |
| VQRQLLTL! | EZR; EZR | P15311; E7E | EZRI_HUMAN E | Unmodified | 0 | 0 | 0 | 0 | 2384600 | 0 | 0 | 0 | 0 | 758910 | 0 | 0 | 821110 | 0 | 0 | 0 | 0 | 626840 |
| VQSFQMK | VPS35; VPS35; | Q96QK1; A0, | VPS35_HUMAN | Unmodified | 111290 | 0 | 0 | 0 | 0 | 0 | 0 | 0 | 802540 | 130160 | 176990 | 1774400 | 0 | 0 | 0 | 0 | 253450 | 386380 |
| VQSGNINA | CCT3; CCT3; C( | P49368; Q5S | TCPG_HUMAN | Unmodified | 0 | 0 | 0 | 0 | 150350 | 0 | 425650 | 0 | 0 | 0 | 0 | 0 | 273810 | 0 | 0 | 0 | 837610 | 206460 |
| VQSGSESV | PLEC; PLEC | Q15149; H0' | PLEC_HUMAN | Unmodified | 0 | 0 | 0 | 0 | 249420 | 0 | 0 | 0 | 0 | 0 | 0 | 0 | 0 | 0 | 0 | 0 | 697540 | 1667100 |
| VQSLQATF( | CKAP4 | Q07065 | CKAP4_HUMAI | Unmodified | 0 | 0 | 0 | 280300 | 0 | 0 | 0 | 0 | 3274400 | 636900 | 0 | 0 | 0 | 0 | 0 | 0 | 0 | 0 |
| VQSLQATF( | CKAP4 | Q07065 | CKAP4_HUMAI | Unmodified | 0 | 0 | 0 | 0 | 564510 | 0 | 315370 | 0 | 0 | 898930 | 0 | 0 | 814710 | 714760 | 0 | 0 | 0 | 676760 |
| VQSLVDTIF | SRXN1 | Q9BYN0 | SRXN1_HUMAI | Unmodified | 0 | 0 | 0 | 0 | 2041500 | 190220 | 0 | 0 | 0 | 0 | 0 | 0 | 198500 | 0 | 0 | 252930 | 0 | 0 |
| VQSPEPPA | SCRIB; SCRIB; S | Q14160; A0/ | SCRIB_HUMAN | Unmodified | 0 | 0 | 0 | 0 | 1621900 | 0 | 0 | 0 | 0 | 0 | 0 | 0 | 0 | 0 | 0 | 0 | 1666800 | 3836600 |
| VQSSIAFLC | IGF2R; IGF2R; I | P11717; A0A | MPRI_HUMAN ( | Unmodified | 0 | 0 | 0 | 0 | 684850 | 1083800 | 485730 | 0 | 7449500 | 966570 | 0 | 2273800 | 0 | 0 | 0 | 0 | 0 | 607880 |
| VQSVEQIR | CTBP2; CTBP1; | P56545; Q1: | CTBP2_HUMAI | Unmodified | 0 | 336550 | 136700 | 0 | 0 | 24502 | 83935 | 0 | 18244 | 12728 | 0 | 2093700 | 0 | 0 | 210760 | 188220 | 0 | 272140 |
| VQTANEVK | KIF5B | P33176 | KINH_HUMAN | Unmodified | 0 | 0 | 0 | 0 | 0 | 0 | 692470 | 86071 | 0 | 0 | 0 | 0 | 370970 | 727040 | 0 | 0 | 873870 | 0 |
| VQTAVTMG | CHMP1B | Q7LBR1 | CHM1B_HUMA | Unmodified | 1553400 | 1397000 | 2020900 | 2478300 | 4975900 | 0 | 0 | 0 | 24303000 | 2603200 | 6124000 | 34131000 | 2112800 | 617600 | 1828900 | 1312000 | 0 | 0 |
| VQTAVTMG | CHMP1B | Q7LBR1 | CHM1B_HUMA | Unmodified | 0 | 0 | 0 | 0 | 5838900 | 0 | 176580 | 97169 | 1123500 | 4760400 | 350130 | 91771 | 5350000 | 0 | 0 | 0 | 388240 | 1320700 |
| VQTAVTMK | CHMP1A; CHM | Q9HD42; A0 | CHM1A_HUMA | Unmodified | 0 | 0 | 0 | 0 | 0 | 1220700 | 387810 | 246940 | 0 | 172190 | 0 | 0 | 131180 | 483790 | 0 | 0 | 0 | 0 |
| VQTDPPSV | METAP2; META | P50579; F8V | MAP2_HUMAN | Unmodified | 0 | 0 | 0 | 0 | 0 | 0 | 0 | 0 | 0 | 0 | 0 | 1702400 | 0 | 0 | 0 | 0 | 0 | 0 |
| VQTERPQP | DCTN6 | O00399 | DCTN6_HUMAI | Oxidation (M) | 45318 | 0 | 0 | 0 | 1226700 | 0 | 0 | 0 | 0 | 0 | 1075600 | 969520 | 446470 | 0 | 0 | 0 | 2937900 | 2147400 |
| VQTLQFHP | PWP1; PWP1 | Q13610; B4I | PWP1_HUMAN | Unmodified | 0 | 0 | 0 | 0 | 0 | 0 | 0 | 0 | 0 | 0 | 0 | 2825000 | 0 | 0 | 0 | 0 | 0 | 0 |
| VQTPEVDV | AHNAK | Q09666 | AHNK_HUMAN | Unmodified | 129020 | 124540 | 97297 | 111920 | 13083000 | 1552900 | 47400 | 0 | 0 | 0 | 0 | 0 | 0 | 220510 | 123370 | 55643 | 3939400 | 6659100 |
| VQTRLEET( | DCTN1; DCTN1 | Q14203; A0/ | DCTN1_HUMAI | Unmodified | 0 | 0 | 0 | 0 | 1011900 | 199320 | 61025 | 26806 | 902310 | 234870 | 593230 | 2815100 | 2159300 | 602110 | 0 | 263470 | 181670 | 600520 |
| VQVALEEL( | DYNC1H1; DYN | Q14204; A0/ | DYHC1_HUMA | Unmodified | 0 | 0 | 0 | 0 | 0 | 0 | 0 | 0 | 0 | 0 | 766560 | 1536000 | 0 | 0 | 0 | 0 | 0 | 0 |
| VQVALGNI! | DCTN3; DCTN3 | O75935; X6I | DCTN3_HUMAI | Unmodified | 0 | 0 | 0 | 232000 | 660140 | 0 | 0 | 0 | 718770 | 0 | 0 | 2749000 | 0 | 170250 | 0 | 142260 | 0 | 0 |
| VQVEAIEG( | SIL1; SIL1; SIL1 | Q9H173; A0I | SIL1_HUMAN N | Unmodified | 0 | 0 | 0 | 0 | 0 | 0 | 0 | 0 | 0 | 647000 | 0 | 0 | 0 | 0 | 0 | 0 | 0 | 0 |
| VQVEYKGE | HSPA8; HSPA2; | P11142; P54 | HSP7C_HUMAI | Unmodified | 7072300 | 8047400 | 8098900 | 14072000 | 46501000 | 0 | 11263000 | 3907000 | 14614000 | 9471300 | 2584700 | 28533000 | 48097000 | 60001000 | 6497100 | 5086000 | 6335100 | 14004000 |
| VQVEYKGE | HSPA8; HSPA8; | P11142; E9P | HSP7C_HUMAI | Unmodified | 0 | 0 | 0 | 0 | 506050 | 0 | 2799800 | 49795 | 0 | 153370 | 0 | 0 | 839720 | 1023900 | 0 | 0 | 1645400 | 0 |
| VQVEYKGE | HSPA8; HSPA8; | P11142; E9P | HSP7C_HUMAI | Oxidation (M) | 0 | 0 | 0 | 0 | 572330 | 0 | 2742900 | 299810 | 0 | 0 | 0 | 0 | 0 | 0 | 0 | 0 | 332110 | 0 |
| VQVLAAQL | MEAK7; MEAK7 | Q6P9B6; H3 | MEAK7_HUMAI | Unmodified | 0 | 0 | 0 | 0 | 318010 | 0 | 0 | 20245 | 358930 | 16388 | 0 | 0 | 0 | 298760 | 0 | 0 | 0 | 0 |
| VQVMGKVI | CAP1 | Q01518 | CAP1_HUMAN | Unmodified | 76283 | 0 | 0 | 315520 | 1695000 | 0 | 0 | 0 | 1568700 | 147520 | 915750 | 3091900 | 579200 | 0 | 144120 | 0 | 133090 | 0 |
| VQVMGKVI | CAP1 | Q01518 | CAP1_HUMAN | Oxidation (M) | 0 | 0 | 499120 | 443740 | 661230 | 367890 | 100470 | 0 | 594300 | 0 | 1437200 | 3018700 | 427970 | 181820 | 363780 | 323040 | 0 | 374920 |
| VQVMGKVI | CAP1 | Q01518 | CAP1_HUMAN | Oxidation (M) | 0 | 0 | 0 | 168430 | 5436500 | 1188300 | 799620 | 112950 | 441790 | 1324200 | 0 | 979080 | 972330 | 1289700 | 0 | 0 | 1959900 | 2597600 |
| VQVMGKVI | CAP1 | Q01518 | CAP1_HUMAN | Unmodified | 0 | 0 | 0 | 0 | 16840000 | 937480 | 993360 | 0 | 0 | 2315000 | 0 | 593640 | 5023900 | 2362800 | 0 | 0 | 2580400 | 6307800 |
| VQVMVPEA | NOMO1; NOMC | Q15155; P69 | NOMO1_HUM/ | Unmodified | 0 | 0 | 0 | 0 | 0 | 0 | 0 | 0 | 15818000 | 0 | 382640 | 0 | 0 | 0 | 0 | 0 | 0 | 0 |
| VQVMVPEA | NOMO1; NOMC | Q15155; P69 | NOMO1_HUM/ | Oxidation (M) | 0 | 0 | 0 | 0 | 0 | 0 | 0 | 0 | 2586500 | 0 | 0 | 0 | 0 | 0 | 0 | 0 | 185330 | 170220 |
| VQVPNCDI | COPA; COPA; C | P53621; A0A | COPA_HUMAN | Unmodified | 1133800 | 2273500 | 0 | 4494500 | 0 | 221060 | 0 | 0 | 0 | 0 | 0 | 1041800 | 0 | 0 | 0 | 0 | 0 | 0 |
| VQVPNMQI | SF3A1 | Q15459 | SF3A1_HUMAI | Unmodified | 1815200 | 91748 | 0 | 0 | 0 | 0 | 0 | 0 | 0 | 0 | 0 | 0 | 0 | 0 | 0 | 0 | 0 | 0 |
| VQVQCLSK | SORL1; SORL1; | Q92673; E9I | SORL_HUMAN | Unmodified | 0 | 0 | 0 | 0 | 863770 | 0 | 89283 | 42976 | 729720 | 357240 | 0 | 0 | 65466 | 0 | 0 | 0 | 72319 | 0 |
| VQVQVVER | FLOT1; FLOT1; I | O75955; A0/ | FLOT1_HUMAN | Unmodified | 175700 | 161250 | 2692800 | 900090 | 0 | 0 | 236670 | 213450 | 6162300 | 955880 | 525170 | 6205000 | 0 | 0 | 3392300 | 1796500 | 0 | 0 |
| VQVSLDEH | UBR4 | Q5T4S7 | UBR4_HUMAN | Unmodified | 8484 | 0 | 0 | 0 | 0 | 0 | 0 | 0 | 0 | 0 | 0 | 1086500 | 997830 | 0 | 0 | 0 | 0 | 0 |
| VQVSPERT( | HSPG2 | P98160 | PGBM_HUMAN | Unmodified | 0 | 0 | 727520 | 742620 | 55011 | 0 | 0 | 0 | 274540 | 120550 | 0 | 0 | 0 | 0 | 123480 | 240120 | 0 | 0 |
| VQVSYKGE | HSPA1A; HSPA: | P0DMV8; P0 | HS71A_HUMAI | Unmodified | 1865400 | 2158500 | 1584900 | 2239100 | 4489100 | 1221000 | 737690 | 235670 | 4373500 | 1723800 | 0 | 6110500 | 4050900 | 2721000 | 1553500 | 1167500 | 619970 | 2134500 |
| VQVSYVIRI | WDR48; WDR4 | Q8TAF3; F8V | WDR48_HUMA | Unmodified | 0 | 0 | 0 | 0 | 1501800 | 0 | 0 | 98962 | 562470 | 438650 | 0 | 0 | 296000 | 0 | 0 | 0 | 472800 | 704830 |
| VQVTEAKAI | ABCC3; ABCC3 | O15438; H0' | MRP3_HUMAN | Unmodified | 0 | 0 | 0 | 0 | 656680 | 0 | 0 | 0 | 0 | 0 | 0 | 0 | 759140 | 0 | 0 | 0 | 0 | 0 |
| VQVVGTYR | MCM3; MCM3; | P25205; A0A | MCM3_HUMAN | Unmodified | 0 | 0 | 0 | 0 | 387730 | 0 | 88407 | 0 | 45287 | 0 | 0 | 1146600 | 0 | 0 | 0 | 62479 | 79457 | 0 |
| VQVVVTVLI | SYT1; SYT1 | P21579; J3K | SYT1_HUMAN ! | Unmodified | 0 | 0 | 0 | 0 | 0 | 0 | 0 | 0 | 0 | 559080 | 0 | 0 | 0 | 0 | 0 | 0 | 0 | 0 |
| VQVVVTVLI | SYT2 | Q8N9I0 | SYT2_HUMAN ! | Unmodified | 0 | 0 | 0 | 0 | 0 | 0 | 1924500 | 0 | 0 | 0 | 0 | 0 | 0 | 0 | 0 | 0 | 0 | 0 |
| VQVYLHES | CUL1; CUL1; CI | Q13616; A0/ | CUL1_HUMAN | Unmodified | 98333 | 53587 | 0 | 235270 | 121530 | 0 | 0 | 0 | 0 | 0 | 488320 | 2099800 | 0 | 0 | 113610 | 141340 | 45672 | 160240 |
| VQYAPERP | ARRB1; ARRB1 | P49407; E9P | ARRB1_HUMAI | Unmodified | 80675 | 84542 | 0 | 0 | 0 | 21157 | 37929 | 23913 | 644000 | 0 | 0 | 463480 | 301380 | 0 | 515390 | 267990 | 37732 | 134450 |
| VQYLSAQN | EVI5; EVI5 | O60447; A0/ | EVI5_HUMAN E | Unmodified | 0 | 0 | 0 | 0 | 226840 | 0 | 0 | 0 | 0 | 0 | 0 | 0 | 100150 | 0 | 0 | 0 | 59881 | 0 |
| VQYPQSQ/ | DYNC1H1; DYN | Q14204; A0/ | DYHC1_HUMA | Unmodified | 0 | 0 | 0 | 0 | 0 | 0 | 0 | 0 | 0 | 0 | 0 | 732920 | 0 | 0 | 0 | 0 | 0 | 0 |
| VQYRPEEF | DLG5; DLG5 | Q8TDM6; A0 | DLG5_HUMAN | Unmodified | 0 | 0 | 0 | 0 | 0 | 0 | 0 | 0 | 0 | 0 | 0 | 0 | 0 | 0 | 0 | 0 | 1449600 | 0 |
| VRAAMFPE | CTNNB1; CTNN | P35222; A0A | CTNB1_HUMAI | Oxidation (M) | 0 | 0 | 0 | 623020 | 3826300 | 0 | 1931000 | 40215 | 0 | 0 | 0 | 0 | 1866500 | 0 | 208130 | 176060 | 689300 | 4185600 |
| VRAAMFPE | CTNNB1; CTNN | P35222; A0A | CTNB1_HUMAI | 3 Oxidation (M) | 0 | 0 | 0 | 0 | 0 | 946390 | 0 | 0 | 0 | 0 | 0 | 0 | 1553800 | 4118400 | 0 | 0 | 3128000 | 0 |
| VRAEMEVLI | PLEC | Q15149 | PLEC_HUMAN | Unmodified | 0 | 0 | 0 | 0 | 258140 | 0 | 0 | 0 | 145740 | 91816 | 0 | 0 | 0 | 0 | 0 | 0 | 391800 | 471210 |
| VRAGGPGI | FLNA; FLNA; FL | P21333; A0A | FLNA_HUMAN | Unmodified | 0 | 963980 | 0 | 0 | 3223700 | 817490 | 708080 | 0 | 709570 | 317280 | 0 | 0 | 1696200 | 1471800 | 0 | 0 | 0 | 558950 |
| VRAGGPGI | FLNB; FLNB | O75369; E7E | FLNB_HUMAN | Unmodified | 0 | 0 | 977050 | 4637400 | 1862900 | 881780 | 0 | 0 | 646930 | 0 | 290600 | 800580 | 985880 | 0 | 0 | 0 | 541820 | 0 |
| VRALLVEP\ | TGM2 | P21980 | TGM2_HUMAN | Unmodified | 0 | 0 | 0 | 0 | 15375000 | 582730 | 0 | 0 | 0 | 0 | 0 | 0 | 0 | 0 | 0 | 0 | 656810 | 638450 |
| VRALYDYA( | PACSIN3; PACS | Q9UKS6; A0, | PACN3_HUMAI | Unmodified | 5237000 | 5673700 | 1365700 | 1513900 | 568850 | 36730 | 391930 | 0 | 0 | 200790 | 0 | 0 | 1727600 | 0 | 3035300 | 2196300 | 1166000 | 2540600 |
| VRALYDYA( | PACSIN3; PACS | Q9UKS6; A0, | PACN3_HUMAI | Unmodified | 0 | 0 | 777190 | 0 | 0 | 0 | 0 | 0 | 0 | 0 | 0 | 0 | 13447000 | 0 | 0 | 0 | 0 | 0 |
| VRALYDYD | PACSIN1; PACS | Q9BY11; F6I | PACN1_HUMAI | Unmodified | 1560500 | 1726900 | 0 | 291970 | 389490 | 0 | 0 | 0 | 439920 | 0 | 531950 | 1709300 | 156240 | 0 | 914810 | 0 | 0 | 0 |
| VRALYDYE( | PACSIN2; PACS | Q9UNF0; H0 | PACN2_HUMAI | Unmodified | 3412300 | 3187400 | 1407300 | 1464600 | 0 | 0 | 0 | 0 | 64522 | 0 | 0 | 239570 | 548280 | 0 | 4067800 | 2559000 | 0 | 0 |
| VRALYDYE( | PACSIN2 | Q9UNF0 | PACN2_HUMAI | Unmodified | 370880 | 290120 | 0 | 0 | 0 | 0 | 0 | 0 | 0 | 0 | 0 | 0 | 0 | 0 | 506590 | 181410 | 0 | 0 |
| VRAPMVNI | NNT; NNT; NNT | Q13423; A0/ | NNTM_HUMAN | Oxidation (M) | 0 | 0 | 0 | 0 | 0 | 0 | 385500 | 0 | 4868600 | 1381800 | 0 | 0 | 0 | 0 | 0 | 0 | 0 | 0 |
| VRAPMVNI | NNT; NNT; NNT | Q13423; A0/ | NNTM_HUMAN | Unmodified | 0 | 0 | 0 | 0 | 0 | 0 | 205480 | 609290 | 6189600 | 992510 | 0 | 0 | 0 | 0 | 0 | 0 | 0 | 0 |
| VRATLAENI | PRR15L | Q9BU68 | PR15L_HUMAN | Unmodified | 224960 | 0 | 438060 | 1179900 | 0 | 0 | 245790 | 0 | 0 | 363370 | 0 | 0 | 2871200 | 2716900 | 679640 | 548170 | 0 | 0 |
| VRAVFDNL | NRBP1; NRBP1 | Q9UHY1; F8' | NRBP_HUMAN | Unmodified | 0 | 0 | 0 | 29251 | 330830 | 0 | 119640 | 0 | 0 | 92631 | 0 | 0 | 92514 | 0 | 0 | 0 | 142940 | 117460 |
| VRAVIGSTY | MVP | Q14764 | MVP_HUMAN N | Oxidation (M) | 222310 | 679410 | 0 | 0 | 0 | 319450 | 0 | 0 | 409680 | 895540 | 0 | 1192000 | 0 | 0 | 0 | 0 | 0 | 911070 |
| VRAWGPGI | FLNA; FLNA; FL | P21333; A0A | FLNA_HUMAN | Unmodified | 1523100 | 2305300 | 2463800 | 8298300 | 62642000 | 10431000 | 55889000 | 25680000 | 3203200 | 3489500 | 801170 | 4429900 | 35278000 | 44233000 | 3159700 | 2187400 | 33005000 | 19938000 |
| VRAWGPGI | FLNC | Q14315 | FLNC_HUMAN | Unmodified | 0 | 0 | 0 | 0 | 12705000 | 2254800 | 0 | 0 | 0 | 71805 | 0 | 0 | 0 | 0 | 0 | 0 | 5319800 | 1595900 |
| VRDGPQW | IGF2R | P11717 | MPRI_HUMAN ( | Unmodified | 0 | 0 | 0 | 85304 | 512030 | 0 | 208400 | 0 | 15065000 | 18104000 | 5927400 | 0 | 2172300 | 510380 | 0 | 0 | 0 | 99925 |
| VRDKLNNI | RPS25 | P62851 | RS25_HUMAN | Unmodified | 0 | 71347 | 0 | 53265 | 313610 | 344360 | 568000 | 137310 | 0 | 90748 | 0 | 0 | 984450 | 372910 | 0 | 0 | 30195 | 224820 |
| VRDLTIQK/ | DARS1; DARS1; | P14868; H7( | SYDC_HUMAN | Unmodified | 0 | 0 | 0 | 82881 | 212240 | 0 | 0 | 0 | 250600 | 146800 | 0 | 660240 | 349210 | 0 | 0 | 0 | 0 | 176310 |
| VREAFQPQ | VCL | P18206 | VINC_HUMAN ' | Unmodified | 0 | 0 | 0 | 0 | 0 | 0 | 0 | 0 | 0 | 320960 | 0 | 1918100 | 0 | 0 | 283300 | 523340 | 0 | 0 |
| VREEYPDRI | TUBB3; N/A; TU | Q13509; A0/ | TBB3_HUMAN ' | Oxidation (M) | 0 | 0 | 0 | 0 | 1577500 | 0 | 1531300 | 0 | 0 | 1494600 | 0 | 0 | 0 | 0 | 0 | 0 | 0 | 1309000 |
| VREEYPDRI | TUBB3; N/A; TU | Q13509; A0/ | TBB3_HUMAN ' | Unmodified | 0 | 0 | 0 | 0 | 6540200 | 0 | 2301500 | 108130 | 2841400 | 1077000 | 0 | 274040 | 0 | 0 | 0 | 0 | 0 | 1300000 |
| VREFSITDL | HSPA4L; HSPA | O95757; E9I | HS74L_HUMAI | Unmodified | 0 | 0 | 0 | 0 | 0 | 0 | 0 | 0 | 0 | 0 | 315650 | 464330 | 0 | 0 | 0 | 0 | 0 | 0 |
| VREFSITDV | HSPA4; HSPA4; | P34932; A0A | HSP74_HUMAI | Unmodified | 305630 | 375110 | 0 | 1399300 | 0 | 1485700 | 530730 | 0 | 4505000 | 1107500 | 3478700 | 11772000 | 4294400 | 1462300 | 2101700 | 780470 | 692610 | 1536100 |
| VREFSITDV | HSPA4; HSPA4; | P34932; A0A | HSP74_HUMAI | Unmodified | 0 | 0 | 0 | 0 | 1401700 | 0 | 0 | 0 | 0 | 0 | 0 | 0 | 2984100 | 2196900 | 0 | 0 | 996680 | 788480 |
| VREFTMAEI | GARS1; GARS1 | P41250; A0A | GARS_HUMAN | Unmodified | 0 | 0 | 0 | 0 | 0 | 0 | 0 | 47501 | 1187300 | 0 | 498550 | 2594400 | 0 | 0 | 0 | 0 | 0 | 0 |
| VREFVGQL | VWA1; VWA1; V | Q6PCB0; J3( | VWA1_HUMAN | Unmodified | 0 | 0 | 407720 | 476470 | 0 | 0 | 0 | 0 | 348460 | 35914 | 0 | 0 | 0 | 0 | 214010 | 298360 | 0 | 0 |
| VRELELVY/ | MVP | Q14764 | MVP_HUMAN N | Unmodified | 255550 | 937380 | 0 | 0 | 0 | 245990 | 280440 | 272000 | 1697400 | 221350 | 0 | 16712000 | 0 | 0 | 0 | 0 | 0 | 0 |
| VRELFVQSI | GMPS | P49915 | GUAA_HUMAN | Unmodified | 0 | 0 | 0 | 222470 | 1676600 | 326690 | 233480 | 0 | 158820 | 623840 | 112200 | 545000 | 1669000 | 965120 | 0 | 41302 | 654420 | 883340 |
| VRESGSLSI | PTPN1; PTPN1 | P18031; B4I | PTN1_HUMAN | Unmodified | 0 | 0 | 0 | 0 | 0 | 0 | 0 | 0 | 0 | 37254 | 0 | 732350 | 560660 | 385860 | 0 | 0 | 0 | 0 |
| VREVSQPD | PDIA4; PDIA4 | P13667; A0A | PDIA4_HUMAN | Unmodified | 645510 | 785210 | 732570 | 684540 | 2569800 | 511630 | 5606200 | 1747300 | 1.07E+08 | 60185000 | 15175000 | 6691800 | 2820400 | 2730200 | 2483600 | 1021000 | 642190 | 919660 |
| VRFLEQQN | VIM; VIM; VIM | P08670; B0Y | VIME_HUMAN ' | Unmodified | 7920500 | 0 | 0 | 0 | 556650 | 0 | 0 | 0 | 0 | 0 | 0 | 0 | 0 | 0 | 0 | 0 | 1813600 | 2521800 |
| VRFQEAEEI | HSPH1; HSPH1 | Q92598; A0/ | HS105_HUMAI | Unmodified | 0 | 0 | 0 | 0 | 337890 | 0 | 362530 | 250720 | 0 | 238870 | 0 | 0 | 540330 | 0 | 0 | 0 | 574150 | 660260 |
| VRGAVASV | MVP | Q14764 | MVP_HUMAN N | Unmodified | 0 | 0 | 0 | 0 | 2228800 | 473300 | 0 | 0 | 1628700 | 702760 | 0 | 20706000 | 0 | 214140 | 0 | 790420 | 347200 | 316720 |
| VRGEVAPD | LGALS1 | P09382 | LEG1_HUMAN | Unmodified | 0 | 0 | 0 | 0 | 0 | 0 | 0 | 0 | 0 | 0 | 0 | 0 | 0 | 0 | 0 | 0 | 2147700 | 2423800 |
| VRGLPWS( | HNRNPH1; HNI | P31943; D6I | HNRH1_HUMA | Unmodified | 275110 | 89496 | 0 | 0 | 0 | 0 | 88718 | 54607 | 474440 | 0 | 0 | 832540 | 130930 | 0 | 85386 | 320740 | 0 | 101740 |
| VRGLPWS( | HNRNPH1; HNI | P31943; E9P | HNRH1_HUMA | Unmodified | 0 | 0 | 0 | 0 | 0 | 0 | 1135000 | 0 | 0 | 0 | 0 | 0 | 0 | 0 | 0 | 0 | 0 | 0 |
| VRGRLYPW | SEPTIN2; SEPTI | Q15019; H7( | SEPT2_HUMAN | Unmodified | 0 | 0 | 0 | 0 | 0 | 436550 | 0 | 0 | 0 | 0 | 0 | 0 | 1209500 | 550070 | 0 | 0 | 1407800 | 620710 |
| VRGVDVVV | NT5E | P21589 | 5NTD_HUMAN | Unmodified | 682800 | 0 | 0 | 0 | 83873000 | 25117000 | 0 | 0 | 0 | 0 | 386170 | 0 | 0 | 0 | 0 | 0 | 7677700 | 19974000 |
| VRHVDTKG | EHBP1L1 | Q8N3D4 | EH1L1_HUMAI | Unmodified | 0 | 0 | 0 | 0 | 1593300 | 554440 | 0 | 0 | 157700 | 0 | 0 | 0 | 0 | 0 | 0 | 0 | 538360 | 1255800 |
| VRIDEYDYS | SDHA; SDHA; S | P31040; D6I | SDHA_HUMAN | Unmodified | 0 | 0 | 0 | 0 | 0 | 0 | 0 | 5613.4 | 1112500 | 0 | 0 | 0 | 0 | 0 | 0 | 0 | 0 | 0 |
| VRIDEYDYS | SDHA; SDHA; S | P31040; D6I | SDHA_HUMAN | Unmodified | 0 | 0 | 0 | 0 | 39483 | 0 | 0 | 0 | 9634200 | 963600 | 167620 | 0 | 0 | 0 | 0 | 0 | 0 | 0 |
| VRIDQYQG | TXNL1; TXNL1; | O43396; K7I | TXNL1_HUMAN | Unmodified | 470670 | 295440 | 1145500 | 1267100 | 1555400 | 453610 | 829860 | 652170 | 2629800 | 1424400 | 1455800 | 4392000 | 3921700 | 3028200 | 1667700 | 799230 | 6002600 | 0 |
| VRIDQYQG | TXNL1; TXNL1; | O43396; K7I | TXNL1_HUMAN | Unmodified | 0 | 0 | 0 | 0 | 0 | 0 | 472420 | 0 | 409280 | 0 | 0 | 0 | 0 | 199030 | 0 | 0 | 411790 | 0 |
| VRIPAEVLIL | GMPPA | Q96IJ6 | GMPPA_HUMA | Unmodified | 0 | 0 | 0 | 0 | 0 | 0 | 0 | 0 | 0 | 0 | 0 | 316370 | 0 | 0 | 0 | 0 | 0 | 0 |
| VRIQETQAI | MCM6 | Q14566 | MCM6_HUMAN | Unmodified | 79174 | 38887 | 0 | 0 | 92618 | 0 | 0 | 222810 | 446390 | 0 | 152000 | 1228500 | 72905 | 0 | 0 | 0 | 14416 | 0 |
| VRIQETQAI | MCM6 | Q14566 | MCM6_HUMAN | Unmodified | 0 | 0 | 0 | 0 | 866240 | 0 | 374080 | 0 | 0 | 905920 | 0 | 0 | 916320 | 0 | 0 | 0 | 0 | 0 |
| VRIQTQPG' | SLC25A3; SLC2 | Q00325; F8V | MPCP_HUMAN | Unmodified | 0 | 0 | 238580 | 557210 | 1092600 | 0 | 0 | 0 | 13756000 | 387400 | 145400 | 127180 | 0 | 0 | 0 | 261560 | 0 | 0 |
| VRIQTQPG' | SLC25A3; SLC2 | Q00325; F8V | MPCP_HUMAN | Unmodified | 0 | 0 | 0 | 0 | 0 | 0 | 181580 | 197150 | 26455000 | 17924000 | 0 | 0 | 0 | 0 | 0 | 0 | 0 | 0 |
| VRKPALTAI | SEL1L | Q9UBV2 | SE1L1_HUMAN | Unmodified | 0 | 0 | 0 | 0 | 254330 | 0 | 0 | 0 | 11433000 | 1030900 | 0 | 0 | 0 | 0 | 0 | 0 | 0 | 0 |
| VRKPVVSTI | SLC7A11 | Q9UPY5 | XCT_HUMAN C | Unmodified | 78144 | 96753 | 338310 | 397560 | 2131600 | 0 | 0 | 0 | 463650 | 278580 | 94700 | 339240 | 0 | 320650 | 0 | 269130 | 555230 | 0 |
| VRKSSTPEI | CFL1; CFL1; CF | P23528; G3\ | COF1_HUMAN | Unmodified | 0 | 0 | 0 | 0 | 0 | 0 | 0 | 1232300 | 0 | 0 | 0 | 0 | 0 | 0 | 0 | 0 | 5445600 | 414120 |
| VRLDDDDE | CDK5 | Q00535 | CDK5_HUMAN | Unmodified | 0 | 0 | 0 | 455260 | 0 | 0 | 0 | 0 | 0 | 0 | 0 | 0 | 0 | 0 | 0 | 254890 | 0 | 0 |
| VRLGDVISI | VCP; VCP; VCP; | P55072; C9J | TERA_HUMAN ' | Unmodified | 748160 | 448630 | 0 | 670210 | 2929500 | 662070 | 0 | 0 | 3521100 | 1134800 | 1842400 | 6572700 | 3337900 | 1911400 | 0 | 805440 | 372990 | 0 |
| VRLGDVISI | VCP; VCP; VCP; | P55072; C9J | TERA_HUMAN ' | Unmodified | 0 | 0 | 0 | 0 | 4290100 | 0 | 0 | 0 | 0 | 2223000 | 0 | 0 | 0 | 8189700 | 0 | 0 | 1716000 | 1219300 |

| | | | | | | | | | | | | | | | | | | | | | | |
|---|---|---|---|---|---|---|---|---|---|---|---|---|---|---|---|---|---|---|---|---|---|---|
| VRLGSEIRD | SEMA4B | Q9NPR2 | SEM4B_HUMAI | Unmodified | 0 | 0 | 0 | 0 | 2878400 | 466500 | 1505600 | 0 | 51972 | 0 | 0 | 0 | 77803 | 4229500 | 0 | 0 | 60998 | 0 |
| VRLHSHDII | SDF2L1 | Q9HCN8 | SDF2L_HUMAN | Unmodified | 0 | 0 | 0 | 0 | 0 | 0 | 0 | 0 | 0 | 771840 | 0 | 0 | 0 | 0 | 0 | 0 | 0 | 0 |
| VRLLADPT( | PRDX5 | P30044 | PRDX5_HUMAI | Unmodified | 0 | 0 | 0 | 0 | 0 | 0 | 0 | 0 | 1013900 | 0 | 438140 | 358770 | 215740 | 157050 | 160830 | 94418 | 0 | 0 |
| VRLLIDEAIL | SARS1; SARS1 | P49591; Q5T | SYSC_HUMAN | Unmodified | 0 | 0 | 0 | 0 | 396670 | 0 | 344540 | 27081 | 369380 | 368930 | 265030 | 463470 | 710700 | 565850 | 0 | 0 | 186110 | 229710 |
| VRLPDGCL | UBXN6; UBXN6 | Q9BZV1; K7E | UBXN6_HUMAI | Unmodified | 284110 | 488740 | 0 | 0 | 0 | 0 | 0 | 0 | 0 | 0 | 0 | 0 | 0 | 0 | 469730 | 290930 | 0 | 0 |
| VRLSNKVE | MCM4; MCM4; | P33991; A0A | MCM4_HUMAN | Unmodified | 0 | 0 | 0 | 0 | 469560 | 0 | 80217 | 0 | 0 | 132030 | 117800 | 343300 | 181110 | 75035 | 0 | 0 | 70515 | 0 |
| VRLSPEPW | ACSL4 | O60488 | ACSL4_HUMAN | Unmodified | 65237 | 131050 | 0 | 98787 | 14151000 | 1654400 | 974910 | 0 | 0 | 373090 | 3891900 | 6347700 | 8707800 | 2767300 | 223230 | 43587 | 11200000 | 14689000 |
| VRLSPEPW | ACSL4 | O60488 | ACSL4_HUMAN | Unmodified | 0 | 0 | 0 | 0 | 5327300 | 0 | 2880900 | 0 | 0 | 0 | 0 | 0 | 0 | 0 | 0 | 0 | 13511000 | 10393000 |
| VRNTVQDF | OPRK1; OPRK1 | P41145; A0A | OPRK_HUMAN | Unmodified | 0 | 0 | 0 | 0 | 0 | 0 | 0 | 0 | 935360 | 193330 | 0 | 618830 | 0 | 0 | 0 | 0 | 0 | 0 |
| VRPAELEQI | EPB41L2; EPB4 | O43491; A0A | E41L2_HUMAN | Unmodified | 96306 | 0 | 327770 | 502040 | 0 | 0 | 0 | 0 | 0 | 0 | 0 | 1305000 | 0 | 0 | 1404000 | 401900 | 0 | 0 |
| VRPCVVYG | DDX3X; DDX3Y; | O00571; O15 | DDX3X_HUMAI | Unmodified | 0 | 0 | 0 | 661350 | 0 | 0 | 0 | 0 | 0 | 0 | 0 | 0 | 0 | 0 | 566270 | 284120 | 0 | 0 |
| VRPGSALA | GPRIN1 | Q7Z2K8 | GRIN1_HUMAN | Unmodified | 0 | 0 | 0 | 0 | 0 | 0 | 0 | 0 | 2952800 | 762780 | 0 | 0 | 0 | 899970 | 0 | 0 | 0 | 1878400 |
| VRPLNNEG | CDC42BPA; CD | Q5VT25; A0A | MRCKA_HUMA | Unmodified | 0 | 0 | 193250 | 592900 | 0 | 0 | 0 | 0 | 0 | 0 | 0 | 0 | 0 | 0 | 871850 | 562020 | 0 | 0 |
| VRPSEEML | AKAP2; PALM2A | Q9Y2D5; C9 | AKAP2_HUMAN | Unmodified | 0 | 0 | 0 | 0 | 3391400 | 0 | 0 | 0 | 416880 | 189390 | 0 | 0 | 0 | 0 | 0 | 0 | 859850 | 881790 |
| VRPSSDLN | MARK3; MARK3 | P27448; A0A | MARK3_HUMAI | Unmodified | 0 | 0 | 195530 | 0 | 1290700 | 297150 | 0 | 29996 | 278030 | 247920 | 87067 | 296310 | 461640 | 466190 | 405330 | 268540 | 0 | 163460 |
| VRQFMIFT( | DCUN1D1; DCI | Q96GG9; F8 | DCNL1_HUMAI | Oxidation (M) | 0 | 0 | 0 | 0 | 232290 | 1249500 | 0 | 0 | 0 | 0 | 0 | 0 | 409310 | 0 | 0 | 0 | 108610 | 0 |
| VRQLVEQV | TMED9 | Q9BVK6 | TMED9_HUMAI | Unmodified | 0 | 0 | 0 | 1369000 | 0 | 0 | 0 | 23244 | 2.73E+08 | 2622800 | 3542200 | 1441200 | 0 | 137280 | 0 | 0 | 0 | 0 |
| VRQLVEQV | TMED9 | Q9BVK6 | TMED9_HUMAI | Unmodified | 0 | 0 | 0 | 0 | 0 | 0 | 0 | 0 | 14496000 | 2480100 | 0 | 0 | 0 | 0 | 0 | 0 | 0 | 0 |
| VRQMQNT( | IFIT3; IFIT3; IFIT | O14879; A0A | IFIT3_HUMAN I | Unmodified | 0 | 0 | 0 | 0 | 209440 | 0 | 0 | 0 | 22933 | 0 | 0 | 107760 | 0 | 0 | 0 | 0 | 205110 | 0 |
| VRQPEGPE | KLRG2 | A4D1S0 | KLRG2_HUMAN | Unmodified | 49271 | 0 | 906550 | 1200300 | 541790 | 150310 | 0 | 0 | 784540 | 593020 | 0 | 43394 | 1145300 | 717650 | 1213800 | 669070 | 0 | 50840 |
| VRQQAADL | SF3B1 | O75533 | SF3B1_HUMAN | Unmodified | 0 | 170680 | 0 | 0 | 4088800 | 1204100 | 0 | 443470 | 414940 | 972210 | 393530 | 2381900 | 1254700 | 1534400 | 0 | 168120 | 0 | 1345800 |
| VRSDVKPV | FGFRL1 | Q8N441 | FGRL1_HUMAN | Unmodified | 165820 | 0 | 495730 | 445590 | 0 | 0 | 0 | 0 | 0 | 0 | 0 | 0 | 1168700 | 998990 | 379940 | 441770 | 0 | 0 |
| VRSLETENS | LMNB1; LMNB1 | P20700; A0A | LMNB1_HUMAI | Unmodified | 0 | 0 | 0 | 0 | 0 | 0 | 0 | 0 | 525500 | 0 | 0 | 1421500 | 0 | 0 | 0 | 0 | 0 | 0 |
| VRSLKTDIE | HOMER2 | Q9NSB8 | HOME2_HUMA | Unmodified | 0 | 0 | 0 | 0 | 0 | 0 | 0 | 0 | 0 | 0 | 0 | 0 | 550960 | 776820 | 0 | 0 | 0 | 0 |
| VRTDITYPA | RPS4X | P62701 | RS4X_HUMAN | Oxidation (M) | 1424300 | 2432800 | 0 | 0 | 1134000 | 153980 | 0 | 1126900 | 1062800 | 0 | 0 | 12171000 | 0 | 0 | 245250 | 0 | 8780400 | 18688000 |
| VRVADEGS | CD276; CD276; | Q5ZPR3; H0 | CD276_HUMAI | Unmodified | 0 | 0 | 168980 | 0 | 0 | 0 | 0 | 0 | 0 | 0 | 0 | 0 | 0 | 0 | 0 | 740110 | 0 | 0 |
| VRVELSNG | SRSF3 | P84103 | SRSF3_HUMAN | Unmodified | 1480100 | 1907700 | 302190 | 591960 | 0 | 0 | 0 | 71020 | 247870 | 193220 | 97133 | 0 | 262280 | 0 | 489730 | 534540 | 212470 | 76595 |
| VRVESGYF | MPRIP; MPRIP; | Q6WCQ1; J3 | MPRIP_HUMAN | Unmodified | 0 | 0 | 0 | 0 | 0 | 0 | 0 | 0 | 0 | 0 | 0 | 0 | 0 | 0 | 0 | 0 | 1476000 | 0 |
| VRVGEAQE | CCNDBP1; CCI | O95273; H3I | CCDB1_HUMA | Unmodified | 0 | 0 | 0 | 0 | 0 | 0 | 0 | 0 | 0 | 0 | 0 | 0 | 649420 | 0 | 0 | 671600 | 84148 | 0 |
| VRVLEEQL | FUT8; FUT8 | Q9BYC5; A0A | FUT8_HUMAN | Unmodified | 0 | 0 | 0 | 0 | 0 | 0 | 0 | 0 | 8059800 | 0 | 469920 | 1987000 | 0 | 0 | 0 | 0 | 0 | 0 |
| VRVPTTGII | GNAQ | P50148 | GNAQ_HUMAN | Unmodified | 139440 | 406940 | 0 | 417920 | 116810 | 0 | 50957 | 0 | 1199700 | 876110 | 337790 | 1327300 | 0 | 0 | 331930 | 0 | 0 | 0 |
| VRVSPQDA | CTNND1; CTNN | O60716; A0A | CTND1_HUMAI | Unmodified | 0 | 0 | 0 | 0 | 0 | 0 | 0 | 0 | 0 | 0 | 0 | 0 | 0 | 0 | 0 | 0 | 307820 | 584870 |
| VRVSPQDA | CTNND1; CTNN | O60716; A0A | CTND1_HUMAI | Oxidation (M) | 0 | 0 | 0 | 0 | 0 | 0 | 0 | 0 | 0 | 0 | 0 | 0 | 0 | 0 | 211290 | 0 | 153140 | 227550 |
| VRWPDFN( | GALNT2 | Q10471 | GALT2_HUMAN | Oxidation (M) | 0 | 0 | 339620 | 0 | 656650 | 387030 | 0 | 0 | 28996000 | 4209900 | 1141000 | 1219600 | 534760 | 189880 | 0 | 0 | 0 | 1555700 |
| VRWPDFN( | GALNT2 | Q10471 | GALT2_HUMAN | Unmodified | 0 | 0 | 0 | 0 | 0 | 0 | 0 | 0 | 7372400 | 508710 | 0 | 0 | 108070 | 67748 | 0 | 0 | 0 | 70742 |
| VRYEGSYA | CAVIN2 | O95810 | CAVN2_HUMAI | Unmodified | 0 | 0 | 0 | 0 | 1648000 | 0 | 0 | 0 | 0 | 0 | 0 | 0 | 0 | 0 | 0 | 0 | 0 | 0 |
| VRYGLIPEE | ATP5PB | P24539 | AT5F1_HUMAN | Unmodified | 0 | 0 | 0 | 0 | 0 | 0 | 0 | 0 | 209500 | 211810 | 0 | 0 | 0 | 0 | 0 | 0 | 0 | 0 |
| VRYMVADK | PPP2R1A; PPP2 | P30153; B3K | 2AAA_HUMAN | Oxidation (M) | 0 | 0 | 0 | 385170 | 1627500 | 150710 | 269450 | 328400 | 68277 | 37162 | 0 | 447520 | 3436500 | 972860 | 0 | 0 | 1021100 | 1555000 |
| VRYMVADK | PPP2R1A; PPP2 | P30153; B3K | 2AAA_HUMAN | Unmodified | 0 | 0 | 0 | 0 | 0 | 0 | 0 | 0 | 0 | 196820 | 0 | 129810 | 756520 | 284750 | 0 | 0 | 225040 | 533840 |
| VRYSLDPEI | RPL17; RPL17; I | P18621; A0A | RL17_HUMAN | Unmodified | 5363700 | 6379600 | 2258900 | 2142900 | 5134200 | 869770 | 3985300 | 3499000 | 1372900 | 645300 | 1192100 | 22160000 | 4968300 | 2858000 | 3405100 | 1815600 | 1089700 | 2354300 |
| VRYSLDPEI | RPL17; RPL17; I | P18621; A0A | RL17_HUMAN | Unmodified | 0 | 0 | 0 | 0 | 0 | 0 | 203800 | 197090 | 0 | 0 | 0 | 0 | 312890 | 123050 | 0 | 0 | 0 | 0 |
| VSADEDLK | SNX6; SNX6 | Q9UNH7; A0 | SNX6_HUMAN | Unmodified | 0 | 36857 | 0 | 78865 | 602270 | 159460 | 0 | 0 | 255490 | 0 | 0 | 1315900 | 0 | 0 | 0 | 92071 | 0 | 227920 |
| VSADEDLK | SNX6; SNX6 | Q9UNH7; A0 | SNX6_HUMAN | Unmodified | 0 | 0 | 0 | 0 | 5407800 | 0 | 181960 | 36504 | 0 | 0 | 0 | 0 | 10181 | 0 | 0 | 0 | 230710 | 0 |
| VSAEIKNDI | SH3BP4 | Q9P0V3 | SH3B4_HUMAI | Unmodified | 0 | 0 | 0 | 0 | 1566900 | 0 | 0 | 0 | 612180 | 976470 | 0 | 0 | 0 | 0 | 0 | 0 | 740150 | 0 |
| VSAENLQK | DIAPH1; DIAPH | O60610; A0A | DIAP1_HUMAN | Unmodified | 0 | 0 | 0 | 0 | 0 | 0 | 0 | 0 | 0 | 0 | 0 | 831000 | 0 | 0 | 0 | 0 | 0 | 0 |
| VSAEPTAN( | ERICH5 | Q6P6B1 | ERIC5_HUMAN | Unmodified | 0 | 0 | 0 | 0 | 0 | 0 | 0 | 0 | 0 | 0 | 0 | 0 | 0 | 0 | 0 | 0 | 523120 | 1891200 |
| VSAFENPY | GNAQ; GNAQ | P50148; B1A | GNAQ_HUMAN | Unmodified | 972910 | 891520 | 412990 | 1094100 | 4130500 | 1272600 | 2172500 | 189040 | 6620600 | 2444000 | 3879800 | 13715000 | 0 | 0 | 706150 | 0 | 0 | 0 |
| VSAGEAVV | PSMB9; PSMB9 | P28065; A0A | PSB9_HUMAN | Unmodified | 0 | 0 | 0 | 0 | 0 | 0 | 0 | 0 | 0 | 0 | 0 | 978100 | 0 | 0 | 0 | 0 | 0 | 0 |
| VSAGEPGS | FARP1; FARP1; | Q9Y4F1; A0A | FARP1_HUMAN | Unmodified | 0 | 0 | 0 | 0 | 0 | 0 | 0 | 0 | 5314900 | 2636700 | 5891200 | 8414300 | 0 | 0 | 0 | 0 | 0 | 0 |
| VSAGNGG | MUC1; MUC1; I | P15941; A6Z | MUC1_HUMAN | Unmodified | 0 | 0 | 0 | 0 | 28396000 | 2509200 | 0 | 0 | 0 | 225930 | 0 | 0 | 0 | 985210 | 0 | 0 | 0 | 0 |
| VSALDLAVI | DCTN2; DCTN2 | Q13561; A8K | DCTN2_HUMAI | Unmodified | 12602 | 45666 | 0 | 0 | 289660 | 0 | 0 | 0 | 0 | 132860 | 546990 | 2447700 | 978290 | 0 | 0 | 112430 | 80125 | 0 |
| VSALEEELA | PPFIA2; PPFIA2 | O75334; G3 | LIPA2_HUMAN | Unmodified | 0 | 0 | 194410 | 5164800 | 0 | 0 | 0 | 0 | 0 | 0 | 0 | 0 | 0 | 0 | 0 | 345100 | 0 | 0 |
| VSALSVVR | SLC25A13 | Q9UJS0 | CMC2_HUMAN | Unmodified | 68683 | 79057 | 0 | 0 | 310880 | 0 | 312630 | 536140 | 2356800 | 0 | 0 | 78784 | 0 | 0 | 0 | 0 | 0 | 0 |
| VSAMEDEN | L1RE1 | Q9UN81 | LORF1_HUMAN | Oxidation (M) | 569210 | 553670 | 0 | 0 | 0 | 0 | 0 | 0 | 0 | 0 | 0 | 0 | 0 | 0 | 57781 | 57720 | 0 | 0 |
| VSAMEDEN | L1RE1 | Q9UN81 | LORF1_HUMAN | Unmodified | 401400 | 0 | 0 | 0 | 0 | 0 | 0 | 0 | 0 | 0 | 0 | 0 | 0 | 0 | 0 | 69428 | 0 | 0 |
| VSAPEEQF | POGLUT2 | Q6UW63 | PLGT2_HUMAN | Unmodified | 0 | 0 | 0 | 0 | 0 | 0 | 0 | 0 | 1059000 | 0 | 0 | 142510 | 0 | 0 | 0 | 0 | 0 | 0 |
| VSAPGVQC | AHNAK | Q09666 | AHNK_HUMAN | Unmodified | 0 | 0 | 0 | 0 | 0 | 0 | 0 | 0 | 0 | 0 | 0 | 0 | 0 | 0 | 0 | 0 | 0 | 0 |
| VSAPGVQC | AHNAK | Q09666 | AHNK_HUMAN | Unmodified | 0 | 0 | 0 | 0 | 0 | 0 | 0 | 0 | 0 | 0 | 0 | 0 | 0 | 0 | 0 | 0 | 0 | 0 |
| VSAQEVRK | EZR; EZR | P15311; E7E | EZRI_HUMAN E | Unmodified | 0 | 0 | 0 | 0 | 7948200 | 3421800 | 478640 | 0 | 4196800 | 2173700 | 1647900 | 0 | 1302800 | 0 | 0 | 0 | 1614000 | 0 |
| VSAQGITLT | TNS2; TNS1; TN | Q63HR2; Q9 | TNS2_HUMAN | Unmodified | 0 | 0 | 0 | 0 | 0 | 0 | 0 | 0 | 112550 | 580230 | 0 | 0 | 0 | 0 | 0 | 0 | 468560 | 0 |
| VSAQYLSEI | FLOT1; FLOT1; I | O75955; A2A | FLOT1_HUMAN | Oxidation (M) | 134700 | 0 | 1151100 | 963230 | 6770900 | 1068100 | 965710 | 1952100 | 3535000 | 1723800 | 925940 | 2894400 | 1923800 | 3044300 | 2007500 | 873300 | 830530 | 670180 |
| VSAQYLSEI | FLOT1; FLOT1; I | O75955; A2A | FLOT1_HUMAN | Unmodified | 0 | 0 | 0 | 0 | 2747200 | 0 | 1124800 | 0 | 1961600 | 32348 | 134220 | 113140 | 0 | 909710 | 0 | 0 | 361130 | 1225700 |
| VSAQYLSEI | FLOT1; FLOT1; I | O75955; A2A | FLOT1_HUMAN | Unmodified | 0 | 0 | 0 | 0 | 0 | 0 | 1247500 | 0 | 0 | 0 | 0 | 0 | 0 | 0 | 0 | 0 | 1635500 | 0 |
| VSASPLLYT | TFRC; TFRC | P02786; G3 | TFR1_HUMAN | Unmodified | 8174300 | 11534000 | 18995000 | 6707000 | 3179000 | 2188900 | 1860400 | 0 | 38045000 | 7860100 | 1769700 | 18446000 | 0 | 0 | 12205000 | 11028000 | 0 | 2031900 |
| VSASVAEV( | INF2; INF2; INF | Q27J81; A0A | INF2_HUMAN I | Unmodified | 0 | 0 | 0 | 0 | 314170 | 0 | 0 | 0 | 515540 | 0 | 0 | 1643600 | 0 | 0 | 0 | 0 | 0 | 0 |
| VSAVEEVR | GGA2 | Q9UJY4 | GGA2_HUMAN | Unmodified | 0 | 20913 | 15043 | 0 | 529050 | 30746 | 120720 | 0 | 1168400 | 179390 | 632850 | 578870 | 0 | 0 | 0 | 0 | 180440 | 64516 |
| VSAYVLR | PNMA2 | Q9UL42 | PNMA2_HUMA | Unmodified | 5663200 | 6699000 | 0 | 0 | 29807 | 0 | 0 | 0 | 0 | 0 | 0 | 0 | 0 | 0 | 0 | 0 | 0 | 0 |
| VSAYVLRLE | PNMA2 | Q9UL42 | PNMA2_HUMA | Unmodified | 2501200 | 3777400 | 0 | 0 | 1064800 | 331070 | 0 | 0 | 0 | 15956 | 0 | 0 | 26983 | 0 | 0 | 0 | 0 | 0 |
| VSCLGVTD | GNB2; GNB1; G | P62879; P62 | GBB2_HUMAN | Unmodified | 0 | 0 | 0 | 0 | 0 | 0 | 0 | 0 | 467280 | 0 | 0 | 0 | 0 | 0 | 0 | 0 | 0 | 0 |
| VSCVTPNF | WFDC2 | Q14508 | WFDC2_HUMA | Unmodified | 0 | 0 | 0 | 0 | 0 | 31539 | 94204 | 0 | 1523900 | 854040 | 0 | 0 | 0 | 0 | 0 | 0 | 0 | 211870 |
| VSDATGQN | CAPG | P40121 | CAPG_HUMAN | Oxidation (M) | 0 | 0 | 0 | 0 | 0 | 87393 | 0 | 0 | 0 | 0 | 0 | 0 | 0 | 0 | 0 | 0 | 368650 | 265450 |
| VSDATGQN | CAPG | P40121 | CAPG_HUMAN | Unmodified | 0 | 0 | 0 | 0 | 2445200 | 726210 | 0 | 0 | 0 | 0 | 107570 | 280030 | 0 | 0 | 0 | 0 | 2334400 | 1342400 |
| VSDFGLSR | EPHB3; EPHB1; | P54753; P54 | EPHB3_HUMAI | Unmodified | 4020000 | 3429600 | 946310 | 2220400 | 332130 | 0 | 0 | 0 | 2198300 | 308370 | 2474800 | 8380000 | 1011500 | 0 | 1623700 | 1910500 | 669050 | 1961300 |
| VSDFGLTK | CSK | P41240 | CSK_HUMAN T | Unmodified | 0 | 342440 | 160460 | 593760 | 0 | 0 | 491740 | 264860 | 0 | 37454 | 1245900 | 4180100 | 1478600 | 795240 | 430300 | 176650 | 0 | 0 |
| VSDFREWI | HPN; HPN | P05981; M0 | HEPS_HUMAN | Unmodified | 1340400 | 885300 | 765410 | 2593200 | 0 | 0 | 0 | 0 | 14411 | 28433 | 0 | 0 | 2895300 | 0 | 1963900 | 1973100 | 0 | 30063 |
| VSDGGSSS | HUWE1; HUWE | Q7Z6Z7; A0A | HUWE1_HUMA | Unmodified | 0 | 0 | 0 | 0 | 0 | 0 | 0 | 0 | 0 | 125550 | 0 | 537660 | 0 | 6789 | 0 | 0 | 0 | 698500 |
| VSDKEKID( | OPA1; OPA1; O | O60313; E5K | OPA1_HUMAN | Unmodified | 0 | 0 | 0 | 0 | 2859300 | 0 | 0 | 0 | 0 | 74667 | 0 | 0 | 174330 | 70121 | 0 | 0 | 0 | 144240 |
| VSDLAGLD | EHHADH | Q08426 | ECHP_HUMAN | Unmodified | 0 | 0 | 22442 | 0 | 0 | 0 | 0 | 0 | 1678100 | 81884 | 0 | 0 | 0 | 0 | 38408 | 12058 | 0 | 0 |
| VSDLENEA | LAMC1 | P11047 | LAMC1_HUMAI | Unmodified | 21863 | 0 | 0 | 0 | 91132 | 172440 | 0 | 60491 | 514370 | 0 | 0 | 279250 | 0 | 0 | 0 | 175210 | 71145 | 302730 |
| VSDSISAQ | SDHA; SDHA; S | P31040; D6F | SDHA_HUMAN | Unmodified | 0 | 0 | 0 | 244320 | 48828 | 41388 | 0 | 0 | 7102100 | 821580 | 0 | 0 | 62949 | 0 | 0 | 0 | 0 | 0 |
| VSDYISPLL | ENTPD4; ENTPI | Q9Y227; Q8I | ENTP4_HUMAN | Unmodified | 0 | 0 | 0 | 0 | 0 | 296080 | 0 | 0 | 794040 | 0 | 503940 | 0 | 0 | 0 | 0 | 0 | 0 | 0 |
| VSDYISPLL | ENTPD4; ENTPI | Q9Y227; Q8I | ENTP4_HUMAN | Unmodified | 0 | 0 | 0 | 0 | 0 | 0 | 0 | 0 | 164520 | 818510 | 0 | 0 | 0 | 0 | 0 | 0 | 0 | 0 |
| VSEAIVSIR | ABCC4; ABCC4 | O15439; A0A | MRP4_HUMAN | Unmodified | 0 | 0 | 0 | 0 | 0 | 0 | 108390 | 0 | 0 | 0 | 0 | 0 | 140210 | 203590 | 2708400 | 1136000 | 70652 | 0 |
| VSEDLNQA | CUL5 | Q93034 | CUL5_HUMAN | Unmodified | 0 | 0 | 0 | 0 | 0 | 0 | 0 | 0 | 0 | 0 | 0 | 1374200 | 0 | 0 | 0 | 0 | 0 | 0 |
| VSEDLVSIH | UBR1; UBR1 | Q8IWV7; A0A | UBR1_HUMAN | Unmodified | 21780 | 17458 | 0 | 0 | 53448 | 0 | 0 | 0 | 0 | 0 | 15298 | 1159000 | 0 | 0 | 78362 | 91419 | 0 | 0 |
| VSEDVYR | CRAT; CRAT; CF | P43155; A0A | CACP_HUMAN | Unmodified | 0 | 0 | 0 | 0 | 0 | 1188300 | 177370 | 0 | 1392100 | 43873 | 0 | 577920 | 153710 | 0 | 65364 | 0 | 0 | 0 |
| VSEEIEDIIK | GGCT; N/A | O75223; M0 | GGCT_HUMAN | Unmodified | 0 | 0 | 0 | 0 | 0 | 0 | 0 | 0 | 2078800 | 214140 | 1541700 | 373210 | 0 | 0 | 0 | 0 | 0 | 0 |
| VSEEIEDIIK | GGCT; N/A | O75223; M0 | GGCT_HUMAN | Unmodified | 287430 | 431910 | 0 | 746500 | 0 | 0 | 0 | 0 | 2388700 | 0 | 0 | 1945700 | 3383200 | 0 | 0 | 432440 | 0 | 0 |
| VSEEIFFGR | AFG3L2 | Q9Y4W6 | AFG32_HUMAN | Unmodified | 0 | 0 | 0 | 0 | 0 | 0 | 0 | 143380 | 2204200 | 0 | 0 | 386790 | 0 | 0 | 0 | 0 | 0 | 0 |
| VSEGDRLR | MYH10 | P35580 | MYH10_HUMAI | Unmodified | 0 | 0 | 0 | 505300 | 13389000 | 1476600 | 0 | 0 | 0 | 442510 | 0 | 0 | 2618000 | 3087800 | 0 | 0 | 998730 | 892450 |
| VSEGGPAE | TAX1BP3 | O14907 | TX1B3_HUMAN | Unmodified | 608350 | 190690 | 709780 | 976040 | 0 | 0 | 0 | 0 | 0 | 0 | 290390 | 1954500 | 51021 | 0 | 632860 | 1641500 | 0 | 0 |
| VSEHSGGF | TXNDC5 | Q8NBS9 | TXND5_HUMAI | Unmodified | 0 | 0 | 0 | 0 | 0 | 0 | 0 | 0 | 5315100 | 86579 | 0 | 0 | 0 | 0 | 0 | 0 | 0 | 0 |
| VSEIPLTRD | BAZ2A; BAZ2A; | Q9UIF9; F8V | BAZ2A_HUMAN | Unmodified | 608900 | 311920 | 0 | 0 | 517590 | 0 | 0 | 0 | 0 | 324540 | 0 | 0 | 0 | 0 | 0 | 0 | 0 | 119520 |
| VSELFDK | SNX6; SNX6 | Q9UNH7; A0 | SNX6_HUMAN | Unmodified | 0 | 0 | 0 | 632830 | 6856300 | 198270 | 0 | 0 | 0 | 0 | 0 | 0 | 0 | 2726600 | 0 | 0 | 653610 | 2877600 |
| VSELHLEV | PACSIN2; PACS | Q9UNF0; B0 | PACN2_HUMAI | Unmodified | 661520 | 699780 | 674960 | 462750 | 0 | 0 | 0 | 0 | 0 | 0 | 0 | 0 | 0 | 0 | 0 | 1238400 | 0 | 0 |
| VSELHQEV | PACSIN1; PACS | Q9BY11; F6 | PACN1_HUMAI | Unmodified | 713440 | 0 | 0 | 0 | 157740 | 0 | 0 | 0 | 3566600 | 1297600 | 71067 | 0 | 209150 | 97965 | 797420 | 117160 | 0 | 0 |
| VSELKEELK | HNRNPU; HNR | Q00839; A0A | HNRPU_HUMA | Unmodified | 0 | 0 | 0 | 0 | 0 | 0 | 0 | 0 | 284350 | 143010 | 409460 | 1457900 | 80786 | 0 | 0 | 0 | 0 | 0 |
| VSELKEELK | HNRNPU; HNR | Q00839; A0A | HNRPU_HUMA | Unmodified | 182330 | 0 | 0 | 0 | 0 | 0 | 0 | 67623 | 886300 | 561310 | 2304400 | 5139000 | 589460 | 0 | 0 | 0 | 0 | 0 |
| VSEMDSER | PPFIBP1; PPFIB | Q86W92; F5 | LIPB1_HUMAN | Unmodified | 0 | 0 | 0 | 661230 | 3902000 | 768580 | 0 | 0 | 0 | 0 | 0 | 492320 | 0 | 1291100 | 241390 | 185690 | 0 | 1466400 |
| VSEMDSER | PPFIBP1; PPFIB | Q86W92; F5 | LIPB1_HUMAN | Unmodified | 0 | 0 | 14017 | 0 | 470700 | 135640 | 0 | 0 | 0 | 0 | 43676 | 70696 | 300970 | 107270 | 24200 | 0 | 231630 | 246560 |
| VSEPVLSLF | ALKBH5 | Q6P6C2 | ALKB5_HUMAN | Unmodified | 0 | 0 | 0 | 0 | 0 | 0 | 0 | 0 | 0 | 0 | 0 | 930620 | 0 | 0 | 0 | 0 | 0 | 0 |
| VSEQGLIEI | PDCD5; PDCD5 | O14737; K7E | PDCD5_HUMA | Unmodified | 1129200 | 0 | 1552900 | 2600600 | 382330 | 350650 | 0 | 66939 | 3203200 | 858290 | 1777500 | 5035800 | 2072100 | 565810 | 805250 | 1774200 | 449200 | 0 |
| VSEQGLIEI | PDCD5; PDCD5 | O14737; K7E | PDCD5_HUMA | Unmodified | 0 | 827790 | 1762700 | 2600400 | 1206400 | 0 | 647440 | 0 | 1873000 | 599970 | 3278100 | 4272700 | 1333100 | 723370 | 1578600 | 1586100 | 380750 | 0 |
| VSEQGLIEI | PDCD5; PDCD5 | O14737; K7E | PDCD5_HUMA | Unmodified | 0 | 0 | 0 | 0 | 272290 | 0 | 530240 | 0 | 0 | 333740 | 0 | 0 | 978470 | 0 | 0 | 0 | 104130 | 142250 |
| VSEVEAVLS | CIT; CIT; CIT; CI | O14578; A0A | CTRO_HUMAN | Unmodified | 0 | 0 | 0 | 0 | 827780 | 741730 | 0 | 0 | 0 | 0 | 0 | 0 | 0 | 0 | 0 | 0 | 0 | 612310 |
| VSEVEAVLS | CIT; CIT; CIT; CI | O14578; A0A | CTRO_HUMAN | Unmodified | 0 | 0 | 0 | 0 | 1679000 | 0 | 0 | 0 | 0 | 0 | 0 | 0 | 0 | 0 | 0 | 0 | 0 | 0 |
| VSEVKPTYF | PURA | Q00577 | PURA_HUMAN | Unmodified | 0 | 0 | 0 | 0 | 0 | 0 | 0 | 0 | 0 | 41091 | 0 | 0 | 716010 | 83267 | 0 | 0 | 0 | 0 |
| VSEVLQNR | ENTPD6; ENTPI | O75354; H0 | ENTP6_HUMAN | Unmodified | 2502600 | 10005000 | 234510 | 192340 | 0 | 355730 | 45207 | 17604 | 473100 | 0 | 0 | 0 | 0 | 39504 | 2428400 | 2711800 | 0 | 0 |
| VSEVYPEK | SLC2A5 | P22732 | GTR5_HUMAN | Unmodified | 0 | 0 | 0 | 0 | 0 | 0 | 416050 | 0 | 0 | 0 | 526120 | 0 | 0 | 0 | 0 | 120850 | 0 | 0 |

| | | | | | | | | | | | | | | | | | | | | | | |
|---|---|---|---|---|---|---|---|---|---|---|---|---|---|---|---|---|---|---|---|---|---|---|
| VSEVYPEKI | SLC2A5 | P22732 | GTR5_HUMAN | Unmodified | 0 | 0 | 0 | 0 | 0 | 987960 | 0 | 0 | 0 | 0 | 0 | 0 | 0 | 124240 | 0 | 2836500 | 0 | 0 |
| VSEYARRF( | IMPDH2; IMPDH | P12268; A0A | IMDH2_HUMAN | Unmodified | 0 | 0 | 0 | 0 | 0 | 0 | 19438000 | 0 | 0 | 0 | 0 | 0 | 0 | 0 | 0 | 0 | 9127100 | 0 |
| VSEYVPEIV | CARS1; CARS1 | P49589; B4D | SYCC_HUMAN | Unmodified | 0 | 0 | 0 | 0 | 245890 | 0 | 0 | 0 | 655320 | 0 | 684170 | 7410100 | 0 | 0 | 0 | 0 | 152700 | 203970 |
| VSFCPLSL\ | CLPTM1 | O96005 | CLPT1_HUMAN | Unmodified | 0 | 0 | 0 | 0 | 0 | 0 | 0 | 0 | 1600800 | 0 | 160440 | 281670 | 0 | 0 | 0 | 0 | 0 | 0 |
| VSFFTSGSI | OGFOD3; OGF( | Q6PK18; J3Q | OGFD3_HUMA | Unmodified | 168500 | 0 | 0 | 0 | 0 | 0 | 48559 | 0 | 9165400 | 41660 | 0 | 0 | 265350 | 0 | 0 | 0 | 0 | 0 |
| VSFNKPDS | NCAM2; NCAM | O15394; H9I | NCAM2_HUMA | Unmodified | 376630 | 268640 | 7602600 | 4583600 | 0 | 0 | 0 | 0 | 0 | 0 | 0 | 0 | 0 | 0 | 25878000 | 16962000 | 0 | 0 |
| VSFSYAGL | TMEM43 | Q9BTV4 | TMM43_HUMA | Unmodified | 0 | 0 | 0 | 0 | 0 | 0 | 0 | 0 | 1E+08 | 9396100 | 487110 | 223650 | 0 | 0 | 0 | 0 | 28231 | 0 |
| VSFTGSVP | ALDH9A1 | P49189 | AL9A1_HUMAN | Unmodified | 0 | 0 | 0 | 0 | 803040 | 0 | 497500 | 0 | 899870 | 0 | 0 | 557590 | 616640 | 0 | 0 | 0 | 484710 | 575220 |
| VSFYQLSH | ATP2A2; ATP2A | P16615; H7C | AT2A2_HUMAN | Unmodified | 28447 | 0 | 0 | 0 | 0 | 0 | 0 | 0 | 939720 | 148100 | 425730 | 484850 | 0 | 0 | 0 | 0 | 0 | 0 |
| VSGAGFSP | TNKS1BP1 | Q9C0C2 | TB182_HUMAN | Unmodified | 20454000 | 19085000 | 20445000 | 15712000 | 14667000 | 11194000 | 6944500 | 9065000 | 21660000 | 5230800 | 533870 | 17170000 | 15725000 | 14785000 | 19750000 | 19548000 | 10476000 | 15184000 |
| VSGALIRPL | TMX3 | Q96JJ7 | TMX3_HUMAN | Unmodified | 0 | 0 | 0 | 0 | 0 | 0 | 0 | 0 | 3557200 | 0 | 0 | 0 | 0 | 0 | 0 | 0 | 0 | 0 |
| VSGAQEM\ | PLIN3; PLIN3; P | O60664; K7E | PLIN3_HUMAN | Unmodified | 0 | 0 | 0 | 0 | 492170 | 0 | 0 | 0 | 0 | 0 | 0 | 0 | 0 | 0 | 0 | 0 | 82809 | 1033000 |
| VSGDYGHI | EFNB3 | Q15768 | EFNB3_HUMAN | Unmodified | 0 | 0 | 0 | 0 | 0 | 0 | 0 | 0 | 0 | 0 | 0 | 0 | 0 | 0 | 0 | 0 | 0 | 3175700 |
| VSGDYGHI | EFNB1; EFNB2 | P98172; P52 | EFNB1_HUMAN | Oxidation (M) | 0 | 0 | 0 | 875300 | 1352800 | 0 | 4803800 | 503380 | 0 | 649510 | 0 | 4288100 | 548720 | 0 | 341520 | 2152000 | 20221000 | 46204000 |
| VSGDYGHI | EFNB1; EFNB2 | P98172; P52 | EFNB1_HUMAN | Unmodified | 0 | 0 | 0 | 0 | 0 | 0 | 0 | 0 | 1027200 | 0 | 0 | 0 | 0 | 0 | 0 | 0 | 3474800 | 12775000 |
| VSGGLEVL | ANP32A; ANP32 | P39687; O43 | AN32A_HUMAN | Unmodified | 0 | 306760 | 0 | 0 | 0 | 0 | 0 | 161950 | 0 | 0 | 0 | 15055000 | 0 | 0 | 0 | 402440 | 0 | 0 |
| VSGGPSLE | EIF3L; EIF3L; EI | Q9Y262; B0( | EIF3L_HUMAN | Unmodified | 167960 | 64163 | 0 | 0 | 41137 | 0 | 0 | 0 | 0 | 37012 | 308440 | 4100800 | 0 | 0 | 0 | 140120 | 0 | 0 |
| VSGGTPIKI | ADD3 | Q9UEY8 | ADDG_HUMAN | Unmodified | 0 | 0 | 0 | 1756500 | 0 | 0 | 0 | 0 | 0 | 0 | 0 | 0 | 3451000 | 3733500 | 0 | 0 | 0 | 0 |
| VSGLGEKV | FLNA; FLNA; FL | P21333; A0A | FLNA_HUMAN | Unmodified | 1799300 | 2853800 | 0 | 6772600 | 6771700 | 6475200 | 27457000 | 23169000 | 79565 | 2179200 | 0 | 936470 | 21387000 | 29829000 | 1053600 | 1639600 | 15606000 | 9366200 |
| VSGPDLDL | AHNAK | Q09666 | AHNK_HUMAN | Unmodified | 0 | 0 | 0 | 0 | 6199700 | 1130700 | 309530 | 381680 | 0 | 0 | 0 | 523470 | 0 | 0 | 0 | 0 | 1247100 | 3283800 |
| VSGPDLDL | AHNAK | Q09666 | AHNK_HUMAN | Unmodified | 0 | 0 | 0 | 0 | 508560 | 0 | 0 | 81064 | 0 | 0 | 0 | 0 | 684060 | 517480 | 0 | 0 | 370240 | 0 |
| VSGPDLDL | AHNAK | Q09666 | AHNK_HUMAN | Oxidation (M) | 0 | 0 | 0 | 0 | 4786200 | 508750 | 0 | 0 | 0 | 0 | 0 | 0 | 2228800 | 0 | 0 | 0 | 4873600 | 3107600 |
| VSGPDLDL | AHNAK | Q09666 | AHNK_HUMAN | Unmodified | 0 | 0 | 0 | 0 | 1102300 | 106280 | 0 | 0 | 0 | 0 | 0 | 0 | 0 | 0 | 0 | 0 | 694940 | 1094700 |
| VSGPYVEH | B4GALNT1; B4( | Q00973; B4D | B4GN1_HUMA | Unmodified | 0 | 0 | 0 | 0 | 0 | 0 | 364780 | 0 | 800490 | 0 | 0 | 817130 | 0 | 0 | 298050 | 0 | 0 | 0 |
| VSGSELVQ | PSMC5; PSMC5 | P62195; J3Q | PRS8_HUMAN | Unmodified | 0 | 0 | 0 | 0 | 0 | 0 | 263990 | 0 | 0 | 0 | 0 | 0 | 0 | 0 | 0 | 0 | 1464500 | 0 |
| VSGSQIVDI | GABARAPL2; G/ | P60520; H3E | GBRL2_HUMAN | Unmodified | 1067700 | 0 | 384040 | 232990 | 0 | 0 | 50709 | 0 | 410950 | 0 | 82750 | 2981900 | 0 | 43816 | 757560 | 521910 | 23016 | 54620 |
| VSGSQVEV | SRGAP2; SRGA | O75044; Q7 | SRGP2_HUMAN | Unmodified | 172210 | 0 | 0 | 0 | 0 | 0 | 0 | 0 | 0 | 206530 | 0 | 3058400 | 0 | 0 | 277440 | 337700 | 0 | 0 |
| VSGWGTIS | KLK14; KLK14 | Q9P0G3; A0 | KLK14_HUMAN | Unmodified | 0 | 0 | 0 | 0 | 0 | 0 | 0 | 0 | 8710500 | 0 | 166510 | 0 | 0 | 0 | 0 | 0 | 0 | 0 |
| VSHGSSPS | TRIM56; TRIM5 | Q9BRZ2; C9. | TRI56_HUMAN | Acetyl (Protein N | 0 | 0 | 0 | 0 | 603330 | 0 | 0 | 0 | 0 | 0 | 0 | 0 | 0 | 0 | 0 | 0 | 0 | 0 |
| VSHITFAHE | ADAM10; ADAM | O14672; H0 | ADA10_HUMAN | Unmodified | 12610000 | 5720700 | 13351000 | 20159000 | 16118000 | 577820 | 1405100 | 27596 | 2420800 | 974380 | 1032200 | 12939000 | 21081000 | 13644000 | 21268000 | 20570000 | 1409900 | 6703200 |
| VSHKPSGL | MAP2K1 | Q02750 | MP2K1_HUMAN | Unmodified | 0 | 0 | 0 | 0 | 249050 | 52563 | 0 | 31051 | 193130 | 0 | 76832 | 724900 | 0 | 0 | 0 | 342810 | 199490 | 115450 |
| VSHLLGIN\ | MYH9 | P35579 | MYH9_HUMAN | Unmodified | 1483500 | 1859600 | 5494400 | 8277000 | 13701000 | 3851300 | 1244900 | 161510 | 4993800 | 5803500 | 4850000 | 44139000 | 4300100 | 3552700 | 7989500 | 5410700 | 1815900 | 3562800 |
| VSHLLGIN\ | MYH9 | P35579 | MYH9_HUMAN | Unmodified | 0 | 0 | 16466 | 112200 | 11552000 | 2838000 | 1269000 | 0 | 0 | 3194800 | 82670 | 0 | 2251000 | 4544100 | 0 | 0 | 5355200 | 4202000 |
| VSHLLGIN\ | MYH9 | P35579 | MYH9_HUMAN | Unmodified | 0 | 0 | 0 | 0 | 476380 | 0 | 0 | 0 | 0 | 0 | 0 | 0 | 297940 | 0 | 0 | 0 | 437570 | 352580 |
| VSHPLLQQ | PARP10; PARP1 | Q53GL7; E9 | PAR10_HUMAN | Unmodified | 0 | 0 | 269570 | 99140 | 0 | 0 | 0 | 0 | 0 | 0 | 0 | 0 | 0 | 0 | 0 | 0 | 0 | 0 |
| VSHPQEPN | NOS1AP; NOS1 | O75052; E9 | CAPON_HUMA | Unmodified | 0 | 0 | 0 | 217690 | 565720 | 118950 | 0 | 0 | 459700 | 0 | 0 | 0 | 844730 | 611270 | 170160 | 0 | 0 | 421000 |
| VSHQAGDI | MVP | Q14764 | MVP_HUMAN N | Unmodified | 268070 | 0 | 0 | 0 | 0 | 0 | 0 | 0 | 0 | 0 | 0 | 3804300 | 0 | 0 | 0 | 0 | 0 | 0 |
| VSHQGYST | LRPAP1 | P30533 | AMRP_HUMAN | Unmodified | 0 | 0 | 0 | 0 | 0 | 0 | 0 | 0 | 2156300 | 0 | 0 | 0 | 0 | 0 | 0 | 0 | 0 | 0 |
| VSHSHVG\ | HSH2D; HSH2D | Q96JZ2; K7E | HSH2D_HUMA | Unmodified | 0 | 0 | 392400 | 243620 | 0 | 0 | 0 | 0 | 0 | 0 | 0 | 55742 | 0 | 0 | 1215700 | 444180 | 0 | 0 |
| VSHVLAAL( | TLN1 | Q9Y490 | TLN1_HUMAN | Unmodified | 344780 | 0 | 0 | 0 | 0 | 0 | 0 | 0 | 0 | 0 | 0 | 2539000 | 0 | 0 | 650890 | 855320 | 0 | 0 |
| VSHVSTGG | PGK1; PGK2 | P00558; P07 | PGK1_HUMAN | Unmodified | 0 | 0 | 326060 | 144020 | 0 | 0 | 0 | 0 | 1102900 | 62515 | 0 | 8507200 | 667390 | 0 | 378620 | 751170 | 380430 | 2117700 |
| VSHVSTGG | PGK1 | P00558 | PGK1_HUMAN | Unmodified | 0 | 0 | 0 | 0 | 751150 | 0 | 2037300 | 0 | 0 | 0 | 0 | 0 | 1016100 | 0 | 0 | 0 | 25669000 | 6847200 |
| VSHYIINSS | CRK; CRK | P46108; I3L | CRK_HUMAN A | Unmodified | 0 | 363620 | 0 | 0 | 0 | 0 | 0 | 0 | 0 | 0 | 0 | 2697600 | 0 | 0 | 0 | 836560 | 0 | 0 |
| VSIFFDYAR | ATP6V1H; ATP6 | Q9UI12; G3\ | VATH_HUMAN | Unmodified | 0 | 0 | 0 | 0 | 0 | 0 | 181690 | 0 | 0 | 0 | 0 | 485970 | 0 | 0 | 0 | 118880 | 0 | 0 |
| VSILESLDK | PRKAR1A; PRK/ | P10644; K7E | KAP0_HUMAN | Unmodified | 227640 | 142920 | 291840 | 573040 | 1150800 | 0 | 0 | 0 | 0 | 656670 | 0 | 3684900 | 128800 | 0 | 558710 | 384650 | 94801 | 303580 |
| VSIRLPSTS | EPHA2 | P29317 | EPHA2_HUMAN | Unmodified | 801910 | 988690 | 785850 | 239900 | 0 | 0 | 0 | 0 | 402920 | 0 | 205520 | 3837600 | 0 | 0 | 1157800 | 664380 | 0 | 0 |
| VSISEGDDI | FBL; FBL; FBL; F | P22087; M0 | FBRL_HUMAN | Unmodified | 98239 | 0 | 51840 | 82816 | 0 | 0 | 0 | 0 | 0 | 0 | 133030 | 407770 | 65132 | 0 | 0 | 0 | 48062 | 0 |
| VSITSEVES | ATP7A | Q04656 | ATP7A_HUMAN | Unmodified | 32142 | 0 | 0 | 1581300 | 0 | 0 | 0 | 0 | 176560 | 0 | 0 | 2505100 | 0 | 0 | 1602300 | 1837900 | 0 | 0 |
| VSIVTPEDII | USP14; USP14 | P54578; A6N | UBP14_HUMAN | Unmodified | 0 | 0 | 782690 | 0 | 348580 | 0 | 0 | 0 | 1208500 | 0 | 7137500 | 6046700 | 0 | 0 | 1886100 | 972850 | 89061 | 0 |
| VSKNEEKP | SERPINB6; SER | P35237; A0A | SPB6_HUMAN | Unmodified | 0 | 0 | 0 | 0 | 584390 | 190370 | 0 | 0 | 0 | 118790 | 778280 | 822910 | 0 | 208900 | 0 | 0 | 224520 | 572240 |
| VSKNEEKP | SERPINB6; SER | P35237; A0A | SPB6_HUMAN | Oxidation (M) | 0 | 0 | 0 | 0 | 795400 | 0 | 104120 | 0 | 411030 | 276240 | 294680 | 727030 | 0 | 0 | 0 | 0 | 0 | 0 |
| VSKNSETFI | DIAPH1; DIAPH | O60610; A0/ | DIAP1_HUMAN | Unmodified | 0 | 0 | 0 | 0 | 0 | 0 | 0 | 0 | 0 | 0 | 0 | 0 | 379320 | 555010 | 811260 | 555840 | 690700 | 344680 |
| VSKTGAEG | KIF5B; KIF5C; K | P33176; O6( | KINH_HUMAN | Unmodified | 0 | 0 | 0 | 0 | 0 | 0 | 1198800 | 751930 | 0 | 0 | 0 | 0 | 0 | 0 | 0 | 0 | 965910 | 0 |
| VSLAEDDD | SEZ6; SEZ6; SE | Q53EL9; H0 | SEZ6_HUMAN | Unmodified | 0 | 87040 | 0 | 0 | 0 | 0 | 0 | 0 | 782750 | 0 | 684190 | 0 | 0 | 0 | 0 | 0 | 0 | 0 |
| VSLDEDNE | SEZ6L2; SEZ6L | Q6UXD5; H3 | SE6L2_HUMAN | Unmodified | 0 | 0 | 57508 | 0 | 0 | 0 | 216640 | 0 | 7127900 | 82057 | 0 | 0 | 0 | 53891 | 0 | 0 | 0 | 0 |
| VSLDVNHF | HSPB1; HSPB1 | P04792; A0A | HSPB1_HUMAN | Unmodified | 6118500 | 2916000 | 25059000 | 35805000 | 3194200 | 955460 | 936660 | 473960 | 1.81E+08 | 10008000 | 921220 | 0 | 11584000 | 0 | 27112000 | 17692000 | 2468300 | 2413200 |
| VSLEDLYN( | DNAJA2; DNAJ/ | O60884; A0/ | DNJA2_HUMAN | Unmodified | 0 | 0 | 0 | 0 | 12739000 | 1049700 | 0 | 0 | 5020500 | 2717100 | 557440 | 6263700 | 3322600 | 6225400 | 0 | 0 | 373070 | 1563400 |
| VSLEDLYN( | DNAJA2; DNAJ/ | O60884; A0/ | DNJA2_HUMAN | Unmodified | 0 | 60065 | 0 | 0 | 2149300 | 568510 | 216280 | 155110 | 0 | 769010 | 4466600 | 0 | 1173000 | 1974000 | 0 | 0 | 0 | 744850 |
| VSLEDLYN( | DNAJA2; DNAJ/ | O60884; A0/ | DNJA2_HUMAN | Unmodified | 0 | 0 | 0 | 0 | 1098900 | 0 | 650260 | 75157 | 0 | 0 | 0 | 0 | 846120 | 0 | 0 | 0 | 367640 | 0 |
| VSLEETAQI | PYM1 | Q9BRP8 | PYM1_HUMAN | Unmodified | 551560 | 610670 | 0 | 0 | 0 | 0 | 0 | 0 | 0 | 0 | 0 | 0 | 0 | 0 | 481160 | 306290 | 0 | 0 |
| VSLFSHLP( | EIF2B4; EIF2B4 | Q9UI10; A0A | EI2BD_HUMAN | Unmodified | 0 | 0 | 0 | 101100 | 0 | 0 | 0 | 0 | 0 | 0 | 0 | 1977800 | 0 | 0 | 0 | 0 | 0 | 0 |
| VSLGIDICH | ACTL8 | Q9H568 | ACTL8_HUMAN | Unmodified | 0 | 0 | 0 | 0 | 0 | 0 | 2859600 | 0 | 0 | 0 | 0 | 1340200 | 0 | 7754500 | 0 | 0 | 0 | 9521900 |
| VSLGSQIDI | COBL; COBL; C | O75128; J3K | COBL_HUMAN | Unmodified | 660500 | 675370 | 0 | 227930 | 152520 | 5443500 | 0 | 0 | 355150 | 1650000 | 18609000 | 753910 | 743690 | 165910 | 0 | 163760 | 356120 | 0 |
| VSLLDDTV | EPB41; EPB41; | P11171; A0A | EPB41_HUMAN | Unmodified | 600630 | 0 | 174770 | 1092700 | 913670 | 0 | 3699300 | 815890 | 1466500 | 461530 | 0 | 678440 | 0 | 3422400 | 0 | 0 | 540180 | 859820 |
| VSLLDDTV | EPB41; EPB41; | P11171; A0A | EPB41_HUMAN | Unmodified | 0 | 0 | 0 | 0 | 0 | 0 | 557160 | 68094 | 0 | 0 | 0 | 0 | 0 | 0 | 0 | 0 | 0 | 0 |
| VSLLDGTD | EPB41L5 | Q9HCM4 | E41L5_HUMAN | Unmodified | 0 | 0 | 1018200 | 2153600 | 673620 | 0 | 0 | 117730 | 2499300 | 721550 | 0 | 817330 | 0 | 0 | 3054500 | 2459100 | 0 | 0 |
| VSLLGPVTI | RACGAP1 | Q9H0H5 | RGAP1_HUMAN | Unmodified | 1270600 | 294900 | 4207400 | 4625500 | 50851000 | 2806400 | 161810 | 45080 | 24728000 | 5575200 | 4542300 | 11414000 | 2330800 | 1186400 | 1044600 | 1296400 | 1688200 | 990140 |
| VSLLGPVTI | RACGAP1 | Q9H0H5 | RGAP1_HUMAN | Unmodified | 0 | 0 | 0 | 0 | 1349500 | 112040 | 0 | 0 | 0 | 146580 | 0 | 0 | 249120 | 0 | 0 | 0 | 650640 | 0 |
| VSLLQGDL | ERC1; ERC1; EF | Q8IUD2; G8 | RB6I2_HUMAN | Unmodified | 0 | 0 | 0 | 0 | 795700 | 0 | 0 | 0 | 0 | 0 | 0 | 0 | 0 | 0 | 0 | 0 | 0 | 0 |
| VSLLQNES | TRAF3; TRAF3; | Q13114; A6N | TRAF3_HUMAN | Unmodified | 0 | 0 | 0 | 0 | 0 | 0 | 0 | 0 | 0 | 0 | 0 | 0 | 0 | 0 | 0 | 0 | 0 | 0 |
| VSLMPATL/ | TAPBP; TAPBP; | O15533; A0/ | TPSN_HUMAN | Oxidation (M) | 0 | 0 | 314210 | 0 | 0 | 468140 | 493050 | 70083 | 933830 | 119890 | 163640 | 64545 | 0 | 0 | 0 | 0 | 0 | 0 |
| VSLMPATL/ | TAPBP; TAPBP; | O15533; A0/ | TPSN_HUMAN | Unmodified | 0 | 0 | 0 | 0 | 0 | 0 | 0 | 0 | 5957800 | 252960 | 245520 | 362180 | 0 | 0 | 0 | 0 | 0 | 0 |
| VSLPLFDFK | PIK3C2A | O00443 | P3C2A_HUMAN | Unmodified | 0 | 0 | 0 | 0 | 0 | 0 | 0 | 0 | 0 | 0 | 0 | 448530 | 0 | 0 | 0 | 0 | 0 | 0 |
| VSLPLRTN | NPTX2 | P47972 | NPTX2_HUMAN | Unmodified | 0 | 0 | 0 | 0 | 0 | 0 | 0 | 62427 | 5988800 | 5472400 | 0 | 0 | 0 | 0 | 0 | 0 | 0 | 0 |
| VSLPPIPAV | UBAP1; UBAP1 | Q9NZ09; A0 | UBAP1_HUMAN | Unmodified | 5878800 | 3934300 | 1006800 | 1704900 | 3783400 | 1424300 | 0 | 0 | 0 | 0 | 174500 | 0 | 0 | 0 | 1988800 | 1134100 | 0 | 594300 |
| VSLQVHPD | DNAJC9 | Q8WXX5 | DNJC9_HUMAN | Unmodified | 0 | 0 | 0 | 0 | 196890 | 0 | 0 | 0 | 575330 | 0 | 316140 | 1042900 | 0 | 0 | 0 | 0 | 91444 | 0 |
| VSLRAPQEI | ADPGK | Q9BRR6 | ADPGK_HUMA | Unmodified | 0 | 0 | 0 | 0 | 0 | 0 | 0 | 0 | 2976600 | 0 | 0 | 0 | 0 | 0 | 0 | 0 | 0 | 0 |
| VSLSGSSD | ENTPD2 | Q9Y5L3 | ENTP2_HUMAN | Unmodified | 0 | 353110 | 0 | 0 | 0 | 0 | 0 | 0 | 1216300 | 0 | 0 | 0 | 0 | 14271 | 902830 | 891120 | 19677 | 64602 |
| VSLSHGNL | N/A | N/A | N/A | Oxidation (M) | 1837100 | 1367300 | 0 | 0 | 0 | 0 | 0 | 0 | 0 | 0 | 0 | 0 | 0 | 0 | 1137600 | 1666800 | 0 | 0 |
| VSMDNRLN | BZW1; BZW1 | Q7L1Q6; C9 | BZW1_HUMAN | Unmodified | 0 | 0 | 0 | 0 | 0 | 0 | 0 | 0 | 0 | 0 | 0 | 0 | 0 | 0 | 0 | 0 | 0 | 0 |
| VSMEAVQK | CRYZL1; CRYZL | O95825; A6N | QORL1_HUMAN | Unmodified | 0 | 0 | 0 | 0 | 580680 | 95663 | 101660 | 0 | 0 | 100480 | 55436 | 106490 | 246840 | 0 | 212910 | 157040 | 66193 | 246870 |
| VSMFVLGT | MYOF | Q9NZM1 | MYOF_HUMAN | Unmodified | 0 | 0 | 0 | 0 | 1699200 | 371220 | 0 | 29029 | 0 | 0 | 0 | 0 | 0 | 0 | 0 | 0 | 74490 | 100220 |
| VSMKPEFLI | UMPS | P11172 | UMPS_HUMAN | Oxidation (M) | 0 | 0 | 0 | 0 | 0 | 0 | 0 | 0 | 737070 | 0 | 363300 | 830880 | 0 | 0 | 0 | 0 | 0 | 0 |
| VSMPDVDL | AHNAK | Q09666 | AHNK_HUMAN | Unmodified | 0 | 0 | 0 | 0 | 2867100 | 0 | 0 | 0 | 0 | 0 | 0 | 0 | 0 | 0 | 0 | 0 | 58993 | 1345600 |
| VSMPDVDL | AHNAK | Q09666 | AHNK_HUMAN | Unmodified | 0 | 0 | 0 | 0 | 1304600 | 0 | 0 | 0 | 0 | 0 | 0 | 0 | 0 | 0 | 0 | 0 | 668480 | 443620 |
| VSMPDVEL | AHNAK | Q09666 | AHNK_HUMAN | Oxidation (M) | 0 | 0 | 0 | 0 | 2397000 | 267690 | 0 | 0 | 0 | 0 | 0 | 0 | 0 | 0 | 0 | 0 | 818260 | 872380 |
| VSMPDVEL | AHNAK | Q09666 | AHNK_HUMAN | Unmodified | 0 | 0 | 0 | 0 | 5628300 | 0 | 0 | 0 | 0 | 0 | 0 | 0 | 0 | 0 | 0 | 0 | 1032200 | 2836800 |
| VSMPDVEL | AHNAK | Q09666 | AHNK_HUMAN | Unmodified | 0 | 0 | 0 | 0 | 0 | 0 | 0 | 0 | 0 | 0 | 0 | 0 | 0 | 0 | 0 | 0 | 1066300 | 0 |
| VSMSPVLH | NUP210 | Q8TEM1 | PO210_HUMAN | Unmodified | 0 | 0 | 0 | 0 | 0 | 0 | 0 | 0 | 548660 | 0 | 0 | 0 | 0 | 0 | 0 | 0 | 0 | 0 |
| VSMSSQNI | MYO1B; MYO1E | O43795; H7( | MYO1B_HUMA | Unmodified | 0 | 0 | 15396 | 0 | 9687800 | 752750 | 0 | 0 | 1975800 | 0 | 967900 | 5266300 | 0 | 0 | 1143900 | 0 | 1110200 | 1427400 |
| VSMSSQNI | MYO1B; MYO1E | O43795; H7( | MYO1B_HUMA | Oxidation (M) | 0 | 0 | 0 | 417230 | 5450900 | 0 | 0 | 359580 | 1135700 | 292240 | 3655400 | 12561000 | 0 | 0 | 0 | 261230 | 942620 | 2586100 |
| VSMVGQVI | PPME1 | Q9Y570 | PPME1_HUMAN | Unmodified | 0 | 0 | 0 | 0 | 193000 | 0 | 0 | 0 | 0 | 80454 | 299290 | 1309300 | 414280 | 0 | 181650 | 0 | 0 | 0 |
| VSNAQSVT | SLC16A7 | O60669 | MOT2_HUMAN | Unmodified | 152420 | 336980 | 288010 | 278130 | 0 | 0 | 2765700 | 764920 | 2179300 | 1355400 | 2572900 | 2323800 | 0 | 0 | 0 | 57370 | 740230 | 1073900 |
| VSNDIIQSK | NCAM2; NCAM | O15394; H9I | NCAM2_HUMA | Unmodified | 0 | 0 | 4511200 | 884530 | 0 | 0 | 0 | 0 | 0 | 0 | 0 | 0 | 0 | 0 | 4340500 | 3433200 | 0 | 0 |
| VSNDIIQSK | NCAM2; NCAM | O15394; H9I | NCAM2_HUMA | Unmodified | 0 | 0 | 90557 | 152270 | 0 | 0 | 27119 | 0 | 0 | 19263 | 0 | 0 | 0 | 0 | 135440 | 105330 | 0 | 0 |
| VSNDIIQSK | NCAM2; NCAM | O15394; H9I | NCAM2_HUMA | Unmodified | 2456900 | 1247700 | 3084800 | 8527200 | 0 | 0 | 156590 | 71423 | 0 | 0 | 0 | 0 | 0 | 0 | 14549000 | 8561000 | 66086 | 182290 |
| VSNEEVTEI | TOM1L2; TOM1 | Q6ZVM7; F5I | TM1L2_HUMAN | Unmodified | 1106100 | 5277600 | 44963 | 1416800 | 0 | 0 | 0 | 0 | 0 | 0 | 0 | 0 | 0 | 0 | 5521200 | 378150 | 0 | 0 |
| VSNLQYLH | SRP68 | Q9UHB9 | SRP68_HUMAN | Unmodified | 0 | 0 | 147470 | 0 | 0 | 0 | 0 | 0 | 0 | 0 | 1356500 | 7855800 | 0 | 0 | 0 | 0 | 35657 | 0 |
| VSNTQDAS | CD109 | Q6YHK3 | CD109_HUMAN | Unmodified | 0 | 0 | 0 | 0 | 0 | 0 | 0 | 0 | 4092500 | 230310 | 0 | 83847 | 0 | 0 | 0 | 0 | 0 | 427070 |
| VSNTQDAS | CD109 | Q6YHK3 | CD109_HUMAN | Unmodified | 0 | 455180 | 0 | 0 | 0 | 0 | 0 | 0 | 3809000 | 366440 | 1630900 | 0 | 0 | 0 | 0 | 0 | 0 | 3842700 |
| VSPEEAIK | EIF2S1; EIF2S1; | P05198; H0Y | IF2A_HUMAN E | Unmodified | 345890 | 0 | 0 | 0 | 0 | 0 | 0 | 0 | 0 | 0 | 0 | 1020100 | 0 | 0 | 0 | 0 | 0 | 0 |
| VSPEEAIKC | EIF2S1; EIF2S1; | P05198; H0Y | IF2A_HUMAN E | Unmodified | 0 | 0 | 0 | 0 | 504280 | 0 | 0 | 0 | 44952 | 0 | 100790 | 635020 | 0 | 0 | 0 | 70064 | 255160 | 61832 |
| VSPEEVTIY | NUP210 | Q8TEM1 | PO210_HUMAN | Unmodified | 0 | 0 | 0 | 0 | 0 | 0 | 0 | 21594 | 15479000 | 1813400 | 0 | 595790 | 0 | 0 | 0 | 0 | 0 | 0 |
| VSPGAFTPI | DCN; DCN; DC | P07585; F8V | PGS2_HUMAN | Unmodified | 85981 | 0 | 0 | 0 | 0 | 0 | 2193900 | 0 | 0 | 0 | 0 | 0 | 0 | 0 | 0 | 0 | 0 | 0 |
| VSPHASLQ | COBLL1; COBL | Q53SF7; A0/ | COBL1_HUMAN | Unmodified | 2992100 | 1152300 | 694410 | 685430 | 0 | 0 | 0 | 0 | 0 | 0 | 0 | 238220 | 0 | 0 | 1476100 | 1006200 | 0 | 0 |
| VSPHSGVV | FBLN1 | B1AHL2 | B1AHL2_HUMA | Unmodified | 0 | 0 | 0 | 0 | 0 | 0 | 0 | 0 | 5357300 | 1047400 | 0 | 0 | 0 | 0 | 0 | 0 | 378280 | 0 |
| VSPHSGVV | FBLN1 | B1AHL2 | B1AHL2_HUMA | Unmodified | 0 | 0 | 0 | 0 | 0 | 0 | 0 | 0 | 0 | 2552200 | 0 | 0 | 0 | 0 | 0 | 0 | 0 | 0 |
| VSPIQIDGA | TOR1AIP2 | Q8NFQ8 | TOIP2_HUMAN | Unmodified | 0 | 0 | 0 | 0 | 0 | 0 | 0 | 0 | 13859000 | 0 | 0 | 0 | 0 | 0 | 0 | 0 | 0 | 0 |
| VSPIQIDGA | TOR1AIP2 | Q8NFQ8 | TOIP2_HUMAN | Unmodified | 0 | 0 | 0 | 0 | 0 | 0 | 0 | 402470 | 6953500 | 5566600 | 967460 | 0 | 0 | 0 | 0 | 0 | 0 | 0 |

| | | | | | | | | | | | | | | | | | | | | | | |
|---|---|---|---|---|---|---|---|---|---|---|---|---|---|---|---|---|---|---|---|---|---|---|
| VSPPLMAP | TMED8 | Q6PL24 | TMED8_HUMAI | Oxidation (M) | 0 | 0 | 38235 | 0 | 0 | 0 | 0 | 0 | 0 | 0 | 145170 | 804120 | 0 | 0 | 0 | 0 | 0 | 0 |
| VSPSGALEI | HMCN2 | Q8NDA2 | HMCN2_HUMA | Unmodified | 0 | 0 | 0 | 0 | 0 | 0 | 22100 | 0 | 197640 | 0 | 0 | 0 | 0 | 0 | 0 | 0 | 0 | 0 |
| VSPSKSPSI | MAP1B | P46821 | MAP1B_HUMAI | Unmodified | 0 | 0 | 0 | 0 | 0 | 0 | 264810 | 0 | 0 | 171640 | 0 | 0 | 638590 | 775340 | 0 | 0 | 144530 | 201410 |
| VSPVLRDFI | PAK6 | Q9NQU5 | PAK6_HUMAN | Unmodified | 56978 | 233800 | 0 | 0 | 0 | 0 | 0 | 0 | 0 | 0 | 349560 | 471990 | 0 | 0 | 0 | 0 | 396850 | 1490300 |
| VSPVSYLR | NUP210 | Q8TEM1 | PO210_HUMAN | Unmodified | 0 | 0 | 0 | 0 | 0 | 0 | 0 | 0 | 8666800 | 0 | 0 | 0 | 0 | 0 | 0 | 0 | 0 | 0 |
| VSPVSYLR\ | NUP210 | Q8TEM1 | PO210_HUMAN | Unmodified | 0 | 0 | 0 | 0 | 0 | 0 | 0 | 0 | 0 | 275180 | 0 | 0 | 0 | 0 | 0 | 0 | 0 | 0 |
| VSQAAADL | GNG5 | P63218 | GBG5_HUMAN | Unmodified | 24551000 | 21929000 | 5033600 | 8782700 | 31351000 | 7070500 | 7740600 | 0 | 32740000 | 11789000 | 5432800 | 1.18E+08 | 42685000 | 7011000 | 10525000 | 10332000 | 22272000 | 84544000 |
| VSQAAADL | GNG5 | P63218 | GBG5_HUMAN | Unmodified | 0 | 0 | 0 | 0 | 0 | 0 | 1119500 | 0 | 0 | 0 | 0 | 0 | 0 | 0 | 0 | 0 | 0 | 0 |
| VSQAAADL | GNG4 | P50150 | GBG4_HUMAN | Unmodified | 0 | 0 | 1602000 | 0 | 0 | 0 | 772880 | 161710 | 0 | 0 | 244560 | 1022600 | 0 | 0 | 0 | 0 | 1999500 | 0 |
| VSQAAADL | GNG4 | P50150 | GBG4_HUMAN | Unmodified | 0 | 0 | 0 | 0 | 0 | 0 | 0 | 0 | 907190 | 0 | 0 | 0 | 0 | 0 | 0 | 0 | 0 | 0 |
| VSQAAADL | GNG4 | P50150 | GBG4_HUMAN | Unmodified | 0 | 0 | 0 | 0 | 5056200 | 0 | 13324000 | 0 | 24055000 | 29861000 | 0 | 0 | 0 | 0 | 0 | 0 | 0 | 0 |
| VSQAAAEL( | GNG10; DNAJC | P50151; A0A | GBG10_HUMAI | Unmodified | 124300 | 0 | 0 | 0 | 2893400 | 0 | 0 | 0 | 0 | 0 | 0 | 1967200 | 0 | 211070 | 0 | 0 | 841140 | 972620 |
| VSQAAAEL( | GNG10; DNAJC | P50151; A0A | GBG10_HUMAI | Oxidation (M) | 0 | 0 | 0 | 0 | 1292100 | 734730 | 148500 | 0 | 0 | 319170 | 187270 | 1031500 | 0 | 0 | 0 | 0 | 232670 | 540660 |
| VSQGKEPV | SCIN | Q9Y6U3 | ADSV_HUMAN | Unmodified | 45548 | 0 | 0 | 0 | 0 | 0 | 0 | 0 | 486350 | 0 | 0 | 412660 | 0 | 0 | 0 | 0 | 0 | 0 |
| VSQGQLVV | PDIA4; PDIA4 | P13667; A0A | PDIA4_HUMAN | Oxidation (M) | 0 | 0 | 0 | 0 | 0 | 0 | 213750 | 0 | 5447300 | 2126100 | 1099400 | 538720 | 0 | 0 | 0 | 0 | 0 | 0 |
| VSQGQLVV | PDIA4; PDIA4 | P13667; A0A | PDIA4_HUMAN | Unmodified | 114580 | 0 | 0 | 0 | 0 | 0 | 492610 | 195610 | 32476000 | 5844500 | 2682400 | 970940 | 0 | 0 | 0 | 0 | 0 | 0 |
| VSQGQLVV | PDIA4; PDIA4 | P13667; A0A | PDIA4_HUMAN | Unmodified | 0 | 74868 | 0 | 0 | 0 | 0 | 0 | 0 | 1218600 | 891980 | 0 | 0 | 0 | 0 | 0 | 0 | 0 | 0 |
| VSQGQLVV | PDIA4; PDIA4 | P13667; A0A | PDIA4_HUMAN | Unmodified | 0 | 0 | 0 | 0 | 0 | 0 | 208490 | 0 | 1559800 | 1812300 | 0 | 0 | 0 | 0 | 0 | 0 | 0 | 0 |
| VSQGQLVV | PDIA4; PDIA4 | P13667; A0A | PDIA4_HUMAN | Oxidation (M) | 0 | 0 | 0 | 0 | 285070 | 0 | 0 | 0 | 0 | 1595500 | 0 | 0 | 0 | 0 | 0 | 0 | 0 | 0 |
| VSQGSKDF | GANAB; GANAE | Q14697; F5I | GANAB_HUMA | Unmodified | 81544 | 0 | 155260 | 0 | 0 | 0 | 0 | 0 | 13669000 | 1707100 | 3641300 | 1208000 | 0 | 0 | 0 | 259460 | 0 | 0 |
| VSQGVEDC | SAE1; SAE1; SA | Q9UBE0; B3 | SAE1_HUMAN | Unmodified | 0 | 0 | 0 | 0 | 0 | 0 | 0 | 0 | 500050 | 23835 | 60059 | 97468 | 0 | 0 | 0 | 0 | 0 | 0 |
| VSQLDIQIK | EEA1 | Q15075 | EEA1_HUMAN | Unmodified | 0 | 0 | 0 | 0 | 539810 | 0 | 0 | 0 | 0 | 0 | 0 | 0 | 0 | 0 | 0 | 0 | 0 | 0 |
| VSQLEAQL | PDE4DIP; PDE4 | Q5VU43; A0A | MYOME_HUMA | Unmodified | 0 | 0 | 0 | 210800 | 0 | 0 | 1051700 | 223830 | 0 | 218060 | 0 | 911610 | 1447200 | 0 | 299480 | 0 | 0 | 0 |
| VSQLMEW | MAGT1; MAGT1 | Q9H0U3; A0 | MAGT1_HUMAI | Unmodified | 0 | 0 | 0 | 0 | 0 | 0 | 0 | 158720 | 2353100 | 0 | 0 | 0 | 0 | 0 | 0 | 0 | 0 | 0 |
| VSQLMEW | MAGT1; MAGT1 | Q9H0U3; A0 | MAGT1_HUMAI | Oxidation (M) | 0 | 0 | 0 | 0 | 0 | 0 | 32475 | 0 | 1110500 | 0 | 0 | 0 | 0 | 0 | 0 | 0 | 0 | 0 |
| VSQLMEW | MAGT1; MAGT1 | Q9H0U3; A0 | MAGT1_HUMAI | Oxidation (M) | 0 | 0 | 0 | 0 | 0 | 0 | 0 | 0 | 5707000 | 0 | 0 | 0 | 0 | 0 | 0 | 0 | 0 | 0 |
| VSQLYMNF | PYGL | P06737 | PYGL_HUMAN | Unmodified | 0 | 0 | 0 | 0 | 1690900 | 0 | 414460 | 0 | 513090 | 0 | 0 | 0 | 243090 | 0 | 0 | 0 | 501870 | 2956700 |
| VSQLYMNF | PYGL | P06737 | PYGL_HUMAN | Oxidation (M) | 0 | 0 | 0 | 0 | 912740 | 0 | 370170 | 197550 | 0 | 0 | 0 | 0 | 0 | 0 | 0 | 0 | 0 | 0 |
| VSQMLILTF | HAT1 | O14929 | HAT1_HUMAN | Oxidation (M) | 0 | 0 | 0 | 0 | 0 | 0 | 0 | 0 | 0 | 0 | 0 | 4806100 | 0 | 0 | 0 | 0 | 0 | 0 |
| VSQNSLGE | KIDINS220; KID | Q9ULH0; E9 | KDIS_HUMAN | Unmodified | 0 | 442010 | 0 | 0 | 0 | 0 | 105270 | 0 | 0 | 0 | 0 | 906670 | 0 | 0 | 0 | 0 | 0 | 0 |
| VSQPIEGH | CLTC; CLTC | Q00610; A0A | CLH1_HUMAN | Unmodified | 3371600 | 3694000 | 2807500 | 1770400 | 308150 | 125780 | 137480 | 29215 | 437770 | 174500 | 165380 | 8826500 | 232420 | 102300 | 3076000 | 3634600 | 0 | 0 |
| VSQQILEK | ACAD9; ACAD9 | Q9H845; H0 | ACAD9_HUMA | Unmodified | 0 | 0 | 0 | 0 | 0 | 0 | 675630 | 28720 | 949970 | 165300 | 0 | 0 | 0 | 187750 | 48818 | 0 | 280910 | 0 |
| VSQTTWD | CORO1A; COR | P31146; H3E | COR1A_HUMA | Unmodified | 0 | 0 | 0 | 0 | 0 | 0 | 0 | 0 | 0 | 0 | 0 | 3598100 | 0 | 0 | 0 | 0 | 0 | 0 |
| VSRDTLYE | RPL10A | P62906 | RL10A_HUMAN | Unmodified | 2364600 | 3025100 | 1145100 | 537980 | 0 | 0 | 0 | 0 | 335700 | 0 | 260320 | 41245000 | 0 | 0 | 2815900 | 1734300 | 0 | 0 |
| VSRDTLYE | RPL10A | P62906 | RL10A_HUMAN | Unmodified | 1085700 | 1613100 | 106180 | 1001900 | 177600 | 0 | 998260 | 0 | 0 | 117640 | 227520 | 2931400 | 2169400 | 656390 | 667130 | 298830 | 293820 | 113950 |
| VSREELSQI | EVPL; EVPL | Q92817; K7E | EVPL_HUMAN | Unmodified | 0 | 0 | 0 | 0 | 2017800 | 217470 | 0 | 0 | 0 | 0 | 0 | 0 | 167750 | 0 | 0 | 0 | 0 | 0 |
| VSRELAGH | GNB1; GNB1; C | P62873; B3K | GBB1_HUMAN | Unmodified | 0 | 0 | 0 | 0 | 2512200 | 551220 | 0 | 0 | 0 | 0 | 0 | 0 | 32827 | 1336800 | 0 | 0 | 785910 | 2049200 |
| VSRELPGH | GNB2; GNB4; C | P62879; Q9I | GBB2_HUMAN | Unmodified | 0 | 0 | 0 | 0 | 0 | 0 | 0 | 0 | 0 | 0 | 285920 | 0 | 0 | 0 | 0 | 0 | 278040 | 598530 |
| VSREMDRE | CCT6A | P40227 | TCPZ_HUMAN | Unmodified | 0 | 0 | 0 | 214840 | 769350 | 149520 | 0 | 0 | 0 | 0 | 0 | 0 | 1237000 | 466080 | 0 | 0 | 144540 | 0 |
| VSRLETELC | CGN | Q9P2M7 | CING_HUMAN | Unmodified | 19638 | 0 | 0 | 0 | 485590 | 0 | 49735 | 0 | 0 | 502940 | 475370 | 1128000 | 0 | 0 | 0 | 0 | 0 | 260890 |
| VSRVEFLP | SEC23IP | Q9Y6Y8 | S23IP_HUMAN | Unmodified | 0 | 0 | 0 | 0 | 0 | 0 | 0 | 0 | 0 | 0 | 0 | 495230 | 0 | 0 | 0 | 0 | 0 | 0 |
| VSSAEGAA | HMGN1; HMGN | P05114; H7E | HMGN1_HUMA | Unmodified | 0 | 0 | 0 | 0 | 0 | 0 | 0 | 32725 | 0 | 0 | 96976 | 863300 | 0 | 0 | 32948 | 258930 | 0 | 0 |
| VSSDEDLK | SNX5; SNX5; SN | Q9Y5X3; Q5( | SNX5_HUMAN | Unmodified | 0 | 0 | 0 | 150770 | 0 | 0 | 0 | 0 | 0 | 0 | 0 | 0 | 0 | 0 | 0 | 44842 | 0 | 0 |
| VSSDNVAL | PSMB5 | P28074 | PSB5_HUMAN | Unmodified | 0 | 83712 | 0 | 1089600 | 0 | 0 | 0 | 48572 | 3170800 | 323520 | 1326500 | 9798400 | 40237 | 0 | 794880 | 719320 | 46398 | 0 |
| VSSDVIDQ | EIF3L; EIF3L; EI | Q9Y262; B0( | EIF3L_HUMAN | Unmodified | 901580 | 0 | 606920 | 444460 | 0 | 0 | 0 | 0 | 0 | 0 | 0 | 4835300 | 0 | 0 | 1349300 | 1877300 | 0 | 0 |
| VSSFEEK | MDH2; MDH2 | P40926; G3) | MDHM_HUMAI | Unmodified | 0 | 48343 | 191760 | 191320 | 0 | 0 | 1359200 | 0 | 12506000 | 1711400 | 0 | 403160 | 1130400 | 439930 | 214200 | 113520 | 0 | 0 |
| VSSINDKSI | MYO1C; MYO1( | O00159; F5I | MYO1C_HUMA | Unmodified | 4574400 | 3093900 | 739420 | 7895300 | 10279000 | 6241300 | 456250 | 14773 | 583320 | 237050 | 133630 | 922330 | 1519200 | 744510 | 4794300 | 4577400 | 3257200 | 879280 |
| VSSKNSLE | HSPA8; HSPA8 | P11142; E9P | HSP7C_HUMA | Unmodified | 0 | 0 | 0 | 160170 | 703640 | 0 | 0 | 34564 | 83405 | 429970 | 151040 | 0 | 304410 | 343230 | 0 | 0 | 519620 | 150100 |
| VSSLGKDV | CRIP2 | P52943 | CRIP2_HUMAN | Unmodified | 0 | 0 | 0 | 0 | 0 | 0 | 453960 | 85225 | 0 | 31891 | 0 | 0 | 0 | 0 | 0 | 0 | 309970 | 0 |
| VSSLGKDV | CRIP2 | P52943 | CRIP2_HUMAN | Unmodified | 0 | 0 | 0 | 0 | 0 | 124540 | 393600 | 0 | 0 | 0 | 0 | 0 | 0 | 0 | 0 | 0 | 488660 | 979290 |
| VSSLIER | PDIA5 | Q14554 | PDIA5_HUMAN | Unmodified | 0 | 1155900 | 1678000 | 0 | 0 | 277990 | 251930 | 599410 | 11879000 | 0 | 3092000 | 341130 | 515810 | 0 | 0 | 0 | 0 | 0 |
| VSSLPLLLP | IAH1; IAH1 | Q2TAA2; H7( | IAH1_HUMAN | Unmodified | 0 | 0 | 0 | 0 | 0 | 0 | 0 | 0 | 0 | 0 | 0 | 300000 | 0 | 0 | 0 | 0 | 0 | 0 |
| VSSQGNLII | ADAM9; ADAMS | Q13443; A0A | ADAM9_HUMA | Unmodified | 0 | 7399400 | 7840600 | 0 | 58918000 | 7834900 | 263440 | 305590 | 1203000 | 792000 | 332180 | 0 | 33594000 | 7883900 | 0 | 18876000 | 4307700 | 4991500 |
| VSSQNLVA | AP3M1 | Q9Y2T2 | AP3M1_HUMAI | Unmodified | 0 | 0 | 0 | 0 | 0 | 0 | 0 | 0 | 0 | 0 | 0 | 2697800 | 0 | 0 | 0 | 0 | 0 | 0 |
| VSSSPAPA | NCAM2; NCAM | O15394; H9I | NCAM2_HUMA | Unmodified | 0 | 0 | 0 | 0 | 0 | 0 | 0 | 0 | 0 | 0 | 0 | 0 | 0 | 0 | 7011000 | 5110600 | 0 | 0 |
| VSSTATTQ | AFDN; AFDN; A | P55196; Q51 | AFAD_HUMAN | Unmodified | 458660 | 556740 | 138260 | 155980 | 0 | 0 | 0 | 0 | 1090800 | 64369 | 1734400 | 4060600 | 0 | 0 | 487010 | 278680 | 468510 | 0 |
| VSSTEDWS | MANBA; MANB/ | O00462; A0A | MANBA_HUMA | Unmodified | 0 | 0 | 0 | 0 | 0 | 0 | 0 | 0 | 881120 | 274680 | 0 | 0 | 0 | 0 | 0 | 0 | 0 | 0 |
| VSSVFKDE | RANGAP1 | P46060 | RAGP1_HUMAI | Unmodified | 0 | 45884 | 0 | 361730 | 3715300 | 12949000 | 14276000 | 6938200 | 333570 | 396510 | 577660 | 7230800 | 1821100 | 10899000 | 0 | 137060 | 1550100 | 20014000 |
| VSSVPNTS | ABRAXAS2 | Q15018 | ABRX2_HUMAN | Unmodified | 0 | 0 | 0 | 0 | 146570 | 0 | 0 | 0 | 0 | 0 | 0 | 673450 | 0 | 0 | 0 | 0 | 0 | 0 |
| VSTAVLSIT | PSMD1; PSMD1 | Q99460; A0A | PSMD1_HUMA | Unmodified | 116030 | 0 | 270060 | 516400 | 3203700 | 0 | 1038000 | 676900 | 1174900 | 764380 | 320850 | 5541200 | 4411900 | 791650 | 0 | 118150 | 2102800 | 3744800 |
| VSTCVPGE | DNER | Q8NFT8 | DNER_HUMAN | Unmodified | 0 | 0 | 0 | 0 | 0 | 0 | 0 | 0 | 333660 | 98122 | 0 | 0 | 0 | 0 | 0 | 0 | 0 | 0 |
| VSTDLLR | PGAM5; PGAM5 | Q96HS1; F5( | PGAM5_HUMA | Unmodified | 28469 | 0 | 36605 | 29104 | 0 | 0 | 0 | 0 | 973220 | 181630 | 0 | 542900 | 0 | 0 | 0 | 0 | 0 | 0 |
| VSTEANRSI | RP2 | O75695 | XRP2_HUMAN | Unmodified | 0 | 0 | 0 | 0 | 104350 | 0 | 0 | 0 | 0 | 568850 | 0 | 0 | 684060 | 0 | 0 | 0 | 19232 | 405880 |
| VSTKIQQLL | DIP2B | Q9P265 | DIP2B_HUMAN | Unmodified | 0 | 0 | 0 | 0 | 0 | 0 | 0 | 0 | 0 | 0 | 0 | 0 | 0 | 0 | 0 | 0 | 0 | 0 |
| VSTLKDIIPI | PRSS8 | Q16651 | PRSS8_HUMAN | Unmodified | 0 | 0 | 0 | 0 | 0 | 0 | 0 | 0 | 0 | 5168900 | 0 | 0 | 0 | 717170 | 0 | 0 | 0 | 0 |
| VSTNGSDL | ITCH; ITCH; ITC | Q96J02; A0A | ITCH_HUMAN | Unmodified | 968660 | 644790 | 554430 | 421100 | 0 | 0 | 0 | 0 | 0 | 0 | 0 | 0 | 0 | 0 | 747750 | 791780 | 0 | 0 |
| VSTNGSDL | ITCH; ITCH; ITC | Q96J02; A0A | ITCH_HUMAN | Unmodified | 0 | 3047100 | 0 | 0 | 0 | 0 | 0 | 0 | 0 | 0 | 0 | 0 | 0 | 0 | 0 | 0 | 0 | 0 |
| VSTSDQYD | ITGB1BP1; ITGE | O14713; C9. | ITBP1_HUMAN | Unmodified | 0 | 0 | 0 | 275600 | 0 | 37246 | 0 | 45891 | 412180 | 0 | 68454 | 0 | 144160 | 118140 | 0 | 593610 | 0 | 0 |
| VSTTLNVA( | MYO1B; MYO1E | O43795; E9I | MYO1B_HUMA | Unmodified | 197350 | 449020 | 117520 | 0 | 0 | 150390 | 103520 | 35608 | 1297500 | 82240 | 1575200 | 21636000 | 0 | 0 | 880030 | 212330 | 145140 | 42475 |
| VSTTLNVA( | MYO1B; MYO1E | O43795; E9I | MYO1B_HUMA | Unmodified | 0 | 0 | 0 | 0 | 1022800 | 4662500 | 2100600 | 1945900 | 0 | 408750 | 930910 | 0 | 0 | 0 | 0 | 0 | 7188700 | 0 |
| VSTVFWK | EIF3A | Q14152 | EIF3A_HUMAN | Unmodified | 164110 | 636530 | 234200 | 169180 | 0 | 0 | 0 | 0 | 0 | 0 | 0 | 3737800 | 0 | 0 | 702350 | 1021000 | 0 | 0 |
| VSTWNRLN | FAM20B | O75063 | XYLK_HUMAN | Unmodified | 0 | 0 | 0 | 0 | 0 | 0 | 0 | 0 | 6866700 | 0 | 0 | 0 | 0 | 0 | 0 | 0 | 0 | 0 |
| VSVADHSL | PRDX4; PRDX4 | Q13162; A6I | PRDX4_HUMAI | Unmodified | 125010 | 0 | 77017 | 0 | 1629400 | 146130 | 2344500 | 788190 | 82813000 | 16249000 | 8276600 | 6345300 | 0 | 60594 | 0 | 0 | 2034200 | 254200 |
| VSVELTNSI | JUP | P14923 | PLAK_HUMAN | Unmodified | 0 | 0 | 0 | 0 | 503110 | 0 | 0 | 0 | 397980 | 0 | 419790 | 896250 | 0 | 0 | 0 | 0 | 0 | 0 |
| VSVFFGGL | DDX39A; DDX3 | O00148; A0A | DX39A_HUMAN | Unmodified | 91602 | 0 | 0 | 0 | 0 | 0 | 1716700 | 199340 | 0 | 108370 | 0 | 418730 | 0 | 0 | 0 | 0 | 0 | 0 |
| VSVFQTQL | SEC24B | O95487 | SC24B_HUMAI | Unmodified | 0 | 0 | 0 | 0 | 0 | 0 | 0 | 0 | 0 | 0 | 0 | 980580 | 0 | 0 | 0 | 0 | 0 | 0 |
| VSVGAPDL | AHNAK | Q09666 | AHNK_HUMAN | Unmodified | 0 | 0 | 0 | 143060 | 27594000 | 0 | 0 | 0 | 0 | 0 | 0 | 0 | 0 | 0 | 0 | 0 | 16882000 | 22837000 |
| VSVGEFVG | STAU1; STAU1 | O95793; Q5. | STAU1_HUMAN | Unmodified | 304160 | 0 | 0 | 0 | 0 | 847850 | 0 | 0 | 0 | 0 | 0 | 828480 | 0 | 0 | 0 | 0 | 0 | 0 |
| VSVHVIEGI | FASN; FASN | P49327; A0A | FAS_HUMAN F | Unmodified | 0 | 336830 | 1805400 | 0 | 0 | 0 | 0 | 0 | 51188 | 0 | 3611100 | 50118000 | 0 | 0 | 1960800 | 2288200 | 0 | 0 |
| VSVHVLAR | IDE; IDE; IDE; ID | P14735; A0A | IDE_HUMAN In | Unmodified | 0 | 0 | 0 | 0 | 37207 | 0 | 0 | 0 | 0 | 0 | 0 | 1071700 | 0 | 0 | 0 | 0 | 18403 | 0 |
| VSVIQFLEA | SCRIB; SCRIB; S | Q14160; A0A | SCRIB_HUMAN | Unmodified | 0 | 0 | 309020 | 930490 | 0 | 0 | 0 | 0 | 4678200 | 533390 | 137790 | 0 | 571990 | 0 | 0 | 0 | 0 | 527300 |
| VSVIQFLEA | SCRIB; SCRIB; S | Q14160; A0A | SCRIB_HUMAN | Unmodified | 0 | 0 | 0 | 0 | 0 | 0 | 0 | 0 | 0 | 0 | 388890 | 0 | 0 | 0 | 0 | 0 | 0 | 0 |
| VSVPLIAQ( | LAMC1 | P11047 | LAMC1_HUMAI | Unmodified | 44666 | 0 | 1111200 | 0 | 1097000 | 150160 | 0 | 0 | 1674700 | 0 | 0 | 0 | 0 | 0 | 0 | 0 | 0 | 0 |
| VSVPQTDN | ITGB4; ITGB4 | P16144; J3Q | ITB4_HUMAN I | Unmodified | 0 | 65945 | 0 | 0 | 6958200 | 1484600 | 0 | 0 | 0 | 0 | 0 | 0 | 4273900 | 1405800 | 0 | 0 | 12210000 | 29607000 |
| VSVPQTDN | ITGB4; ITGB4 | P16144; J3Q | ITB4_HUMAN I | Oxidation (M) | 0 | 0 | 0 | 0 | 1944500 | 333950 | 0 | 0 | 0 | 0 | 0 | 78149 | 748290 | 190350 | 96300 | 102840 | 1213000 | 3031400 |
| VSVPSNLS | KTN1; KTN1; KT | Q86UP2; G3 | KTN1_HUMAN | Unmodified | 0 | 0 | 0 | 0 | 0 | 0 | 0 | 0 | 0 | 0 | 0 | 1936900 | 0 | 0 | 281820 | 326320 | 0 | 0 |
| VSVPTHPE | VCAN; VCAN; V | P13611; Q86 | CSPG2_HUMAI | Unmodified | 0 | 0 | 0 | 0 | 0 | 0 | 0 | 0 | 717460 | 236500 | 0 | 0 | 0 | 0 | 0 | 0 | 0 | 0 |
| VSVVNPTA | GPC4 | O75487 | GPC4_HUMAN | Unmodified | 1408200 | 1831100 | 0 | 0 | 108800 | 0 | 146650 | 40012 | 704270 | 471220 | 0 | 0 | 1589700 | 137110 | 0 | 491650 | 0 | 44813 |
| VSVYLQNV | GPS1; GPS1; GI | Q13098; A8I | CSN1_HUMAN | Unmodified | 0 | 24402 | 0 | 0 | 0 | 0 | 130820 | 37761 | 0 | 96743 | 197920 | 133050 | 0 | 91421 | 316800 | 689330 | 50487 | 151630 |
| VSWSPAEE | L1CAM | P32004 | L1CAM_HUMAI | Unmodified | 0 | 0 | 0 | 0 | 0 | 0 | 0 | 0 | 3380700 | 0 | 0 | 0 | 0 | 0 | 0 | 0 | 0 | 0 |
| VSYALLFGI | SEL1L | Q9UBV2 | SE1L1_HUMAN | Unmodified | 0 | 0 | 0 | 0 | 0 | 0 | 0 | 0 | 7507400 | 933650 | 32974 | 144160 | 0 | 0 | 0 | 0 | 0 | 0 |
| VSYARPSSI | ELAVL1; ELAVL | Q15717; M0I | ELAV1_HUMAN | Unmodified | 119960 | 0 | 0 | 0 | 0 | 0 | 0 | 0 | 443170 | 189950 | 0 | 1045000 | 0 | 0 | 0 | 0 | 0 | 0 |
| VSYASLAD | ALDH1L2; ALDH | Q3SY69; A0A | AL1L2_HUMAN | Unmodified | 0 | 0 | 0 | 0 | 0 | 0 | 0 | 26379 | 1114200 | 0 | 0 | 0 | 0 | 0 | 0 | 0 | 0 | 0 |
| VSYFPTVP( | FLNB; FLNB; FL | O75369; H7( | FLNB_HUMAN | Unmodified | 0 | 0 | 0 | 0 | 4311100 | 572600 | 0 | 0 | 0 | 0 | 337500 | 2321800 | 0 | 0 | 0 | 0 | 0 | 0 |
| VSYFPTVP( | FLNB; FLNB | O75369; E7E | FLNB_HUMAN | Unmodified | 0 | 0 | 0 | 383670 | 4998100 | 700990 | 209060 | 0 | 0 | 360780 | 0 | 0 | 889980 | 0 | 0 | 0 | 2454900 | 773880 |
| VSYGIGDEI | APEX1; APEX1; | P27695; G3\ | APEX1_HUMAN | Unmodified | 0 | 0 | 0 | 0 | 0 | 0 | 0 | 0 | 2482700 | 80776 | 467640 | 5823900 | 0 | 0 | 0 | 0 | 0 | 0 |
| VSYHHGNI | PIPOX | Q9P0Z9 | SOX_HUMAN P | Unmodified | 0 | 0 | 0 | 0 | 0 | 0 | 0 | 0 | 2311300 | 60372 | 0 | 0 | 0 | 0 | 0 | 0 | 0 | 0 |
| VSYLPFTEA | SELENON; SELI | Q9NZV5; H9 | SELN_HUMAN | Unmodified | 0 | 0 | 0 | 0 | 0 | 0 | 0 | 0 | 2322800 | 0 | 0 | 29330 | 0 | 0 | 0 | 0 | 0 | 0 |
| VSYLRPLDI | KIFBP; KIFBP; K | Q96EK5; A0A | KBP_HUMAN K | Unmodified | 44690 | 54001 | 0 | 33728 | 0 | 0 | 0 | 0 | 0 | 0 | 0 | 791600 | 0 | 0 | 131150 | 102520 | 0 | 0 |
| VSYLTLVTD | ATG5; ATG5; AT | Q9H1Y0; A0A | ATG5_HUMAN | Unmodified | 0 | 0 | 0 | 0 | 0 | 0 | 0 | 0 | 235480 | 0 | 0 | 1516100 | 0 | 0 | 0 | 0 | 1034900 | 0 |
| VTAAIASNI | LAMTOR2 | Q9Y2Q5 | LTOR2_HUMAN | Unmodified | 0 | 0 | 0 | 0 | 0 | 0 | 0 | 0 | 0 | 0 | 0 | 2050800 | 0 | 0 | 0 | 0 | 0 | 0 |
| VTADVINA/ | PHGDH; PHGD | O43175; A0A | SERA_HUMAN | Unmodified | 928950 | 454140 | 3645300 | 5408100 | 2737400 | 510210 | 2315400 | 1595100 | 6651600 | 2168500 | 3839700 | 41654000 | 26277000 | 206050 | 6350700 | 8028900 | 897630 | 2810200 |
| VTADVINA/ | PHGDH; PHGD | O43175; A0A | SERA_HUMAN | Unmodified | 0 | 0 | 0 | 0 | 0 | 0 | 373590 | 0 | 0 | 358120 | 0 | 0 | 3265100 | 1231300 | 0 | 0 | 0 | 0 |
| VTAEEFK | FKBP9; FKBP9P | O95302; Q75 | FKBP9_HUMAN | Unmodified | 0 | 0 | 0 | 0 | 822710 | 114340 | 85473 | 0 | 4071000 | 656570 | 741400 | 248270 | 0 | 0 | 12110 | 0 | 134570 | 687310 |
| VTAEVVLAI | RPN1; RPN1 | P04843; F8V | RPN1_HUMAN | Unmodified | 0 | 0 | 1099700 | 756210 | 0 | 0 | 343040 | 96633 | 5.5E+08 | 79877000 | 17721000 | 7342900 | 655750 | 0 | 1130900 | 928190 | 0 | 0 |
| VTAFLDDL | NLN; NLN; NLN | Q9BYT8; F5I | NEUL_HUMAN | Unmodified | 0 | 0 | 0 | 106180 | 0 | 0 | 0 | 0 | 1324500 | 0 | 1402700 | 1881000 | 0 | 0 | 0 | 0 | 0 | 611530 |
| VTAFPATR | EMC1 | Q8N766 | EMC1_HUMAN | Unmodified | 0 | 0 | 0 | 0 | 164070 | 704820 | 11456 | 0 | 377490 | 39805 | 0 | 1105800 | 0 | 0 | 0 | 0 | 167450 | 0 |
| VTAGISFAII | HTRA1; HTRA1 | Q92743; A0A | HTRA1_HUMAN | Unmodified | 0 | 0 | 0 | 0 | 935380 | 0 | 0 | 0 | 0 | 747630 | 0 | 0 | 0 | 0 | 0 | 0 | 685270 | 4002700 |
| VTAGISFAII | HTRA1; HTRA1 | Q92743; A0A | HTRA1_HUMAN | Unmodified | 0 | 0 | 0 | 0 | 0 | 0 | 0 | 0 | 0 | 134200 | 0 | 0 | 0 | 0 | 144650 | 0 | 539290 | 1660400 |

| | | | | | | | | | | | | | | | | | | | | | | |
|---|---|---|---|---|---|---|---|---|---|---|---|---|---|---|---|---|---|---|---|---|---|---|
| VTAGISFAII | HTRA1; HTRA1; | Q92743; A0A | HTRA1_HUMAN | Unmodified | 0 | 0 | 0 | 0 | 0 | 0 | 0 | 0 | 0 | 0 | 0 | 0 | 0 | 0 | 0 | 0 | 558640 | 1458200 |
| VTAGISFAII | HTRA2; HTRA2 | O43464; A0A | HTRA2_HUMAN | Unmodified | 0 | 0 | 0 | 239370 | 0 | 0 | 0 | 0 | 942760 | 153490 | 0 | 0 | 0 | 0 | 0 | 0 | 0 | 0 |
| VTAIHIDPA | FASN; FASN | P49327; A0A | FAS_HUMAN F | Unmodified | 385130 | 419100 | 890970 | 879910 | 0 | 0 | 0 | 0 | 765560 | 0 | 653650 | 85497000 | 0 | 0 | 2480600 | 2602100 | 0 | 0 |
| VTAIWSGW | SERGEF; SERGI | Q9UGK8; E9 | SRGEF_HUMAN | Unmodified | 0 | 0 | 0 | 0 | 0 | 716710 | 0 | 0 | 154420 | 0 | 128060 | 614840 | 404620 | 112530 | 0 | 0 | 0 | 0 |
| VTALCWHF | GEMIN5 | Q8TEQ6 | GEMI5_HUMAN | Unmodified | 0 | 0 | 0 | 0 | 0 | 0 | 0 | 0 | 0 | 0 | 0 | 847640 | 0 | 0 | 0 | 0 | 0 | 0 |
| VTALTQME | TMED3 | Q9Y3Q3 | TMED3_HUMAI | Unmodified | 0 | 684840 | 0 | 0 | 72309 | 0 | 0 | 0 | 1990200 | 776320 | 0 | 0 | 0 | 0 | 0 | 0 | 0 | 0 |
| VTALVGPR | LIFR | P42702 | LIFR_HUMAN L | Unmodified | 0 | 0 | 0 | 0 | 0 | 0 | 0 | 20431 | 694750 | 0 | 155840 | 0 | 0 | 0 | 0 | 0 | 0 | 0 |
| VTANQMEN | DIPK1A; DIPK1A | Q5T7M9; A0 | DIK1A_HUMAN | Oxidation (M) | 0 | 0 | 0 | 0 | 0 | 0 | 0 | 0 | 732010 | 0 | 0 | 0 | 0 | 0 | 0 | 0 | 0 | 0 |
| VTANQMEN | DIPK1A; DIPK1A | Q5T7M9; A0 | DIK1A_HUMAN | Unmodified | 2980700 | 0 | 0 | 0 | 0 | 0 | 0 | 0 | 945670 | 0 | 213610 | 121720 | 0 | 0 | 0 | 0 | 52979 | 0 |
| VTAPDVDLI | AHNAK | Q09666 | AHNK_HUMAN | Unmodified | 0 | 237050 | 0 | 0 | 0 | 1042500 | 451580 | 0 | 0 | 261510 | 0 | 871440 | 0 | 0 | 0 | 0 | 954040 | 2176100 |
| VTAPQPAA | ATP2C1; ATP2C | P98194; B4E | AT2C1_HUMAN | Unmodified | 0 | 0 | 0 | 81588 | 0 | 0 | 0 | 0 | 628160 | 135940 | 0 | 599200 | 0 | 296810 | 0 | 0 | 44529 | 0 |
| VTAQGPGL | FLNA; FLNA; FL | P21333; A0A | FLNA_HUMAN | Unmodified | 547630 | 723040 | 650510 | 1628200 | 548760 | 247350 | 293930 | 162990 | 360340 | 0 | 0 | 1947100 | 346180 | 1131400 | 874820 | 1026500 | 0 | 198140 |
| VTASGPGL | FLNB; FLNB | O75369; E7E | FLNB_HUMAN | Unmodified | 0 | 0 | 0 | 469970 | 0 | 0 | 0 | 0 | 0 | 0 | 0 | 1717200 | 0 | 288290 | 680540 | 0 | 0 | 0 |
| VTASPLQR | DECR2; DECR2 | Q9NUI1; Q4 | DECR2_HUMAI | Unmodified | 0 | 0 | 913530 | 0 | 0 | 0 | 102520 | 145260 | 3869400 | 289740 | 203080 | 327400 | 1096700 | 815000 | 0 | 381720 | 0 | 0 |
| VTASQLSA | CDC42BPA; CD | Q5VT25; A0A | MRCKA_HUMA | Unmodified | 0 | 0 | 0 | 0 | 0 | 0 | 0 | 0 | 200260 | 0 | 0 | 376010 | 0 | 0 | 0 | 0 | 0 | 0 |
| VTAVIPCFP | PRPS2; PRPS1; | P11908; P60 | PRPS2_HUMAN | Unmodified | 0 | 0 | 0 | 0 | 0 | 0 | 0 | 0 | 920580 | 0 | 315590 | 1845500 | 0 | 9881900 | 0 | 462220 | 408040 | 0 |
| VTAVPTLLK | TXNDC17; TXNI | Q9BRA2; I3L | TXD17_HUMAN | Unmodified | 0 | 44237 | 2787400 | 7319600 | 28158000 | 6504800 | 3869300 | 2498000 | 6902300 | 7708900 | 6152800 | 11606000 | 21933000 | 12711000 | 45928 | 7552900 | 16277000 | 13757000 |
| VTAYTVDVI | AHNAK | Q09666 | AHNK_HUMAN | Unmodified | 0 | 193070 | 0 | 189820 | 1792400 | 519940 | 0 | 0 | 0 | 0 | 0 | 0 | 0 | 0 | 0 | 0 | 101080 | 767720 |
| VTAYTVDVI | AHNAK | Q09666 | AHNK_HUMAN | Unmodified | 0 | 0 | 0 | 0 | 5603100 | 0 | 0 | 0 | 0 | 0 | 0 | 0 | 0 | 0 | 0 | 0 | 2027500 | 2922200 |
| VTCIGSCG | ARRDC1; ARRD | Q8N5I2; Q51 | ARRD1_HUMAI | Unmodified | 1847400 | 679190 | 0 | 0 | 0 | 0 | 0 | 0 | 0 | 0 | 0 | 493740 | 0 | 0 | 0 | 3235900 | 0 | 0 |
| VTCLSCELI | DPP4; DPP4; DI | P27487; A0A | DPP4_HUMAN | Unmodified | 0 | 0 | 0 | 1923500 | 0 | 0 | 0 | 0 | 0 | 0 | 0 | 0 | 0 | 0 | 3380600 | 2631400 | 0 | 0 |
| VTCLSCELI | DPP4; DPP4; DI | P27487; A0A | DPP4_HUMAN | Unmodified | 0 | 0 | 0 | 2382500 | 0 | 0 | 0 | 0 | 0 | 0 | 0 | 0 | 0 | 0 | 609430 | 0 | 1856700 | 322100 |
| VTCQSSEL | MGAT5 | Q09328 | MGT5A_HUMAI | Unmodified | 0 | 0 | 0 | 0 | 232980 | 0 | 0 | 0 | 1536800 | 0 | 80080 | 64629 | 0 | 0 | 0 | 0 | 36234 | 33851 |
| VTDAIVLLD | UMPS; UMPS | P11172; E9P | UMPS_HUMAN | Unmodified | 0 | 0 | 0 | 0 | 0 | 0 | 0 | 0 | 0 | 0 | 0 | 7301400 | 0 | 0 | 0 | 0 | 0 | 0 |
| VTDAIVLLD | UMPS; UMPS | P11172; E9P | UMPS_HUMAN | Unmodified | 0 | 0 | 0 | 0 | 0 | 0 | 0 | 0 | 517710 | 0 | 0 | 836780 | 0 | 0 | 0 | 56865 | 81799 | 0 |
| VTDALNATI | HSPD1; HSPD1 | P10809; A0A | CH60_HUMAN | Unmodified | 0 | 1947900 | 1008200 | 0 | 0 | 0 | 0 | 820860 | 3118900 | 0 | 1064200 | 295340 | 96869 | 0 | 0 | 86794 | 0 | 23010 |
| VTDDLVCL | SRP9 | P49458 | SRP09_HUMAN | Unmodified | 2346800 | 0 | 2509900 | 961380 | 0 | 0 | 0 | 0 | 0 | 0 | 0 | 1462100 | 0 | 0 | 7917700 | 4223900 | 0 | 0 |
| VTDDLVCL | SRP9 | P49458 | SRP09_HUMAN | Unmodified | 28554000 | 4294300 | 0 | 12618000 | 0 | 0 | 0 | 0 | 0 | 474990 | 881470 | 4657200 | 0 | 0 | 5089700 | 9368400 | 0 | 57362 |
| VTDDLVCL | SRP9 | P49458 | SRP09_HUMAN | Unmodified | 0 | 303670 | 0 | 469800 | 0 | 0 | 0 | 0 | 0 | 0 | 0 | 0 | 0 | 0 | 670330 | 883880 | 0 | 0 |
| VTDEILHLV | PAPOLA; PAPOI | P51003; Q9 | PAPOA_HUMAI | Unmodified | 0 | 107000 | 0 | 0 | 0 | 0 | 0 | 0 | 0 | 0 | 0 | 289830 | 0 | 0 | 0 | 0 | 57894 | 0 |
| VTDEMFLE | MYO1D; MYO1[ | O94832; J3Q | MYO1D_HUMA | Oxidation (M) | 506760 | 662880 | 1798100 | 0 | 0 | 0 | 0 | 0 | 0 | 0 | 0 | 242280 | 0 | 0 | 2677200 | 1322700 | 0 | 0 |
| VTDFGDKV | DYNC1H1; DYN | Q14204; A0A | DYHC1_HUMA | Unmodified | 0 | 0 | 0 | 0 | 0 | 0 | 0 | 0 | 1519400 | 991900 | 944010 | 10934000 | 0 | 0 | 0 | 432020 | 0 | 0 |
| VTDRFGFLI | USP6NL; USP6I | Q92738; X6F | US6NL_HUMAI | Unmodified | 257980 | 0 | 0 | 0 | 0 | 0 | 0 | 0 | 2060800 | 0 | 0 | 2710300 | 0 | 0 | 409480 | 660660 | 0 | 0 |
| VTDTDFDG | NCEH1; NCEH1 | Q6PIU2; H7 | NCEH1_HUMA | Unmodified | 0 | 0 | 305760 | 135070 | 0 | 0 | 0 | 0 | 1492900 | 144210 | 0 | 0 | 0 | 0 | 0 | 0 | 0 | 0 |
| VTDYIAEK | EXOC7; EXOC7 | Q9UPT5; B4I | EXOC7_HUMAI | Unmodified | 0 | 0 | 0 | 0 | 2407700 | 0 | 0 | 0 | 0 | 0 | 0 | 0 | 0 | 0 | 0 | 0 | 0 | 0 |
| VTEDTSSVI | PCCA; PCCA; P | P05165; A0A | PCCA_HUMAN | Unmodified | 0 | 0 | 0 | 0 | 0 | 0 | 0 | 0 | 319890 | 0 | 0 | 70831 | 148890 | 0 | 0 | 0 | 0 | 0 |
| VTEEDFYK | CEL | P19835 | CEL_HUMAN B | Unmodified | 0 | 0 | 0 | 0 | 0 | 0 | 0 | 0 | 2430500 | 285330 | 0 | 197450 | 0 | 0 | 0 | 0 | 0 | 0 |
| VTEEDFYKL | CEL | P19835 | CEL_HUMAN B | Unmodified | 0 | 0 | 0 | 0 | 35389 | 0 | 0 | 0 | 138840 | 403860 | 0 | 0 | 0 | 0 | 0 | 0 | 0 | 0 |
| VTEGGEPY | GANAB; GANAE | Q14697; F5F | GANAB_HUMA | Unmodified | 0 | 0 | 566290 | 0 | 127420 | 0 | 184950 | 16731 | 41053000 | 829900 | 357790 | 2406600 | 0 | 0 | 642190 | 590830 | 0 | 0 |
| VTEGLTDVI | SYNCRIP; SYN( | O60506; B7Z | HNRPQ_HUMA | Unmodified | 633790 | 354480 | 940590 | 586730 | 334270 | 91216 | 0 | 79205 | 3791500 | 634470 | 3792300 | 40942000 | 1787600 | 233330 | 628990 | 1255000 | 0 | 492890 |
| VTEGLTDVI | SYNCRIP; SYN( | O60506; B7Z | HNRPQ_HUMA | Unmodified | 0 | 0 | 0 | 217620 | 0 | 0 | 0 | 0 | 319830 | 128740 | 613170 | 8935600 | 0 | 0 | 0 | 58872 | 0 | 0 |
| VTEGLTDVI | SYNCRIP; SYN( | O60506; B7Z | HNRPQ_HUMA | Unmodified | 0 | 0 | 0 | 0 | 2007900 | 0 | 472790 | 104010 | 530480 | 655390 | 475080 | 2114800 | 2466900 | 503460 | 0 | 0 | 131100 | 258250 |
| VTEGLVDVI | HNRNPR; HNRI | O43390; B4I | HNRPR_HUMA | Unmodified | 0 | 0 | 0 | 0 | 0 | 0 | 0 | 155560 | 0 | 0 | 0 | 9868000 | 0 | 0 | 0 | 0 | 0 | 0 |
| VTEGLVDVI | HNRNPR; HNRI | O43390; B4I | HNRPR_HUMA | Unmodified | 0 | 0 | 0 | 0 | 0 | 0 | 0 | 0 | 0 | 0 | 0 | 2294700 | 0 | 0 | 0 | 0 | 0 | 0 |
| VTEGSFVYI | GDI2; GDI2; GD | P50395; V9G | GDIB_HUMAN | Unmodified | 1252800 | 813470 | 0 | 333370 | 84194 | 194760 | 0 | 0 | 1491200 | 0 | 1088500 | 16874000 | 0 | 0 | 3450200 | 3434400 | 0 | 0 |
| VTEGSFVYI | GDI2; GDI2; GD | P50395; V9G | GDIB_HUMAN | Unmodified | 146660 | 0 | 0 | 321980 | 565060 | 242490 | 160680 | 0 | 475340 | 1108000 | 0 | 1284700 | 392240 | 203020 | 0 | 162920 | 181300 | 144060 |
| VTEGTIREE | EPB41L2; EPB4 | O43491; E9F | E41L2_HUMAN | Unmodified | 0 | 0 | 929350 | 0 | 0 | 0 | 0 | 0 | 0 | 0 | 0 | 2098600 | 0 | 0 | 1309600 | 1701200 | 181910 | 0 |
| VTEIWQEVI | SOD3; SOD3 | P08294; M0I | SODE_HUMAN | Unmodified | 0 | 0 | 0 | 0 | 0 | 0 | 0 | 0 | 1704700 | 20008 | 0 | 0 | 0 | 0 | 0 | 0 | 0 | 0 |
| VTEIWQEVI | SOD3; SOD3 | P08294; M0I | SODE_HUMAN | Unmodified | 0 | 0 | 0 | 0 | 0 | 389210 | 0 | 0 | 5257000 | 0 | 0 | 0 | 0 | 0 | 0 | 0 | 0 | 0 |
| VTEIWQEVI | SOD3 | P08294 | SODE_HUMAN | Oxidation (M) | 0 | 0 | 0 | 0 | 0 | 0 | 0 | 0 | 0 | 628880 | 0 | 0 | 0 | 0 | 0 | 0 | 0 | 0 |
| VTELALTAS | PRKDC | P78527 | PRKDC_HUMAI | Unmodified | 14026 | 0 | 0 | 0 | 0 | 0 | 0 | 0 | 0 | 394000 | 0 | 1726300 | 0 | 0 | 0 | 0 | 0 | 71724 |
| VTELERGN | NPTX2 | P47972 | NPTX2_HUMAN | Unmodified | 0 | 0 | 0 | 0 | 0 | 0 | 0 | 0 | 2572900 | 162470 | 0 | 0 | 0 | 0 | 0 | 0 | 0 | 0 |
| VTELLRTFC | STXBP2; STXBP | Q15833; M0I | STXB2_HUMAN | Unmodified | 0 | 0 | 0 | 0 | 584560 | 0 | 0 | 0 | 0 | 126580 | 0 | 0 | 0 | 250390 | 0 | 0 | 575070 | 525110 |
| VTELQPR | PRSS8 | Q16651 | PRSS8_HUMAN | Unmodified | 72572 | 294320 | 0 | 1421100 | 0 | 0 | 16095 | 0 | 11654000 | 628250 | 265880 | 1384900 | 337100 | 195180 | 1488100 | 1535800 | 0 | 0 |
| VTELQQQP | LRRC59 | Q96AG4 | LRC59_HUMAN | Unmodified | 0 | 0 | 0 | 0 | 0 | 0 | 0 | 0 | 24623000 | 0 | 0 | 0 | 802810 | 0 | 0 | 0 | 0 | 0 |
| VTELQQQP | LRRC59 | Q96AG4 | LRC59_HUMAN | Unmodified | 0 | 0 | 0 | 0 | 0 | 0 | 0 | 0 | 2328100 | 0 | 0 | 0 | 0 | 0 | 0 | 0 | 0 | 0 |
| VTENNLQT | PAN3 | Q58A45 | PAN3_HUMAN | Unmodified | 308450 | 286010 | 0 | 0 | 0 | 0 | 0 | 0 | 0 | 0 | 0 | 0 | 0 | 0 | 0 | 133400 | 0 | 0 |
| VTENQIPAI | ANLN | Q9NQW6 | ANLN_HUMAN | Unmodified | 0 | 0 | 259850 | 321570 | 14068000 | 1294800 | 179050 | 0 | 0 | 0 | 0 | 133330 | 0 | 236070 | 0 | 0 | 1656100 | 2178100 |
| VTENQIPAI | ANLN | Q9NQW6 | ANLN_HUMAN | Unmodified | 0 | 0 | 0 | 0 | 1037000 | 0 | 66685 | 72824 | 148050 | 0 | 0 | 0 | 128100 | 0 | 0 | 0 | 497120 | 304780 |
| VTENQIPAI | ANLN | Q9NQW6 | ANLN_HUMAN | Unmodified | 0 | 0 | 0 | 0 | 0 | 0 | 0 | 0 | 0 | 0 | 0 | 0 | 0 | 0 | 0 | 0 | 817160 | 0 |
| VTEQEWR | CLTB | P09497 | CLCB_HUMAN | Unmodified | 3792400 | 2372500 | 1639700 | 1123900 | 0 | 0 | 36703 | 13113 | 0 | 229620 | 38299 | 4098100 | 0 | 0 | 2614000 | 2467500 | 0 | 0 |
| VTEQEWRE | CLTB | P09497 | CLCB_HUMAN | Unmodified | 4396000 | 2324800 | 986230 | 4608300 | 1568200 | 107300 | 502940 | 0 | 224360 | 46407 | 129710 | 3821500 | 5012300 | 2222300 | 1522200 | 1322800 | 396160 | 0 |
| VTEQLIEAI | CAMK2D; CAMI | Q13557; E9F | KCC2D_HUMA | Unmodified | 643110 | 0 | 0 | 0 | 0 | 0 | 0 | 0 | 0 | 730800 | 0 | 0 | 0 | 0 | 0 | 0 | 0 | 0 |
| VTETLGEQI | CCER2 | I3L3R5 | CCER2_HUMAI | Unmodified | 0 | 0 | 0 | 0 | 0 | 0 | 0 | 279510 | 0 | 0 | 0 | 0 | 0 | 0 | 0 | 0 | 0 | 0 |
| VTEWQQTY | JUP; JUP; JUP; J | P14923; C9J | PLAK_HUMAN . | Unmodified | 0 | 0 | 1795700 | 3243400 | 4256400 | 0 | 0 | 0 | 1948400 | 0 | 1039900 | 4441000 | 0 | 0 | 3778200 | 1154800 | 0 | 1216200 |
| VTEYLYANI | ME2; ME2; ME2 | P23368; A0A | MAOM_HUMAN | Unmodified | 0 | 0 | 0 | 0 | 0 | 0 | 137680 | 0 | 1492000 | 111910 | 0 | 0 | 0 | 0 | 0 | 0 | 0 | 0 |
| VTFEIYDPN | PLCG2; PLCG2 | P16885; A0A | PLCG2_HUMAI | Unmodified | 0 | 0 | 0 | 0 | 0 | 0 | 0 | 0 | 0 | 57137 | 486920 | 1733900 | 0 | 0 | 0 | 0 | 0 | 0 |
| VTFHIEPYS | MANEA | Q5SRI9 | MANEA_HUMA | Oxidation (M) | 0 | 0 | 0 | 0 | 0 | 0 | 0 | 0 | 3707400 | 1281100 | 0 | 0 | 0 | 0 | 0 | 0 | 0 | 0 |
| VTFHIEPYS | MANEA | Q5SRI9 | MANEA_HUMA | Unmodified | 0 | 0 | 0 | 0 | 0 | 0 | 0 | 37481 | 9013700 | 0 | 0 | 467870 | 0 | 0 | 0 | 0 | 0 | 0 |
| VTFLGLQH | MFGE8; MFGE8 | Q08431; H0 | MFGM_HUMAN | Unmodified | 15173000 | 4813900 | 11385000 | 29040000 | 12545000 | 6037600 | 58972 | 0 | 32250000 | 1970300 | 933560 | 16470000 | 1453500 | 328660 | 30803000 | 21760000 | 1204400 | 1160900 |
| VTFLGLQH | MFGE8; MFGE8 | Q08431; H0 | MFGM_HUMAN | Unmodified | 0 | 0 | 0 | 639410 | 3078700 | 1082600 | 0 | 0 | 0 | 235340 | 0 | 0 | 940290 | 0 | 0 | 0 | 343200 | 204600 |
| VTFLPTGITI | F11R; F11R | Q9Y624; A0A | JAM1_HUMAN . | Unmodified | 1842300 | 2038400 | 1599300 | 517820 | 0 | 0 | 0 | 345740 | 2260100 | 0 | 0 | 5090200 | 0 | 0 | 3359000 | 3575500 | 0 | 0 |
| VTFNSALA( | ITM2B; ITM2B; I | Q9Y287; A0A | ITM2B_HUMAN | Unmodified | 155360 | 100660 | 841970 | 429510 | 0 | 234050 | 0 | 0 | 14098000 | 832960 | 0 | 135820 | 336610 | 0 | 349780 | 308600 | 0 | 0 |
| VTFQFSYG | SAMM50 | Q9Y512 | SAM50_HUMAI | Unmodified | 0 | 0 | 0 | 0 | 0 | 0 | 0 | 0 | 1529100 | 930060 | 0 | 0 | 0 | 0 | 0 | 0 | 0 | 0 |
| VTFSCAAG | DCTN1; DCTN1 | Q14203; A0A | DCTN1_HUMAI | Unmodified | 0 | 0 | 0 | 0 | 0 | 0 | 0 | 0 | 0 | 117280 | 99117 | 894800 | 833250 | 0 | 0 | 0 | 0 | 0 |
| VTFSGLRD | EPPK1; EPPK1 | P58107; A0A | EPIPL_HUMAN | Unmodified | 0 | 0 | 0 | 0 | 0 | 0 | 0 | 0 | 147550 | 0 | 0 | 0 | 0 | 0 | 0 | 0 | 0 | 0 |
| VTFVNFTVI | DYNC1H1; DYN | Q14204; A0A | DYHC1_HUMA | Unmodified | 143240 | 0 | 0 | 100990 | 0 | 0 | 0 | 0 | 669180 | 165430 | 484920 | 2953600 | 0 | 0 | 253670 | 325630 | 0 | 0 |
| VTGADVPM | PDHB | P11177 | ODPB_HUMAN | Unmodified | 0 | 0 | 0 | 0 | 0 | 0 | 0 | 0 | 0 | 0 | 0 | 0 | 0 | 0 | 0 | 0 | 0 | 0 |
| VTGDLFYTI | SLC43A1; SLC4 | O75387; E9F | LAT3_HUMAN I | Unmodified | 0 | 0 | 540440 | 402690 | 0 | 0 | 0 | 0 | 1097500 | 67451 | 191940 | 0 | 0 | 0 | 1112200 | 505780 | 0 | 0 |
| VTGDLFYTI | SLC43A1; SLC4 | O75387; E9F | LAT3_HUMAN I | Oxidation (M) | 349820 | 0 | 2640700 | 0 | 0 | 0 | 0 | 0 | 0 | 0 | 0 | 1600900 | 0 | 0 | 2566200 | 2094000 | 0 | 0 |
| VTGDRKYQ | MAN1B1; MAN1 | Q9UKM7; H( | MA1B1_HUMAI | Unmodified | 0 | 0 | 0 | 0 | 0 | 0 | 0 | 0 | 1197900 | 59173 | 0 | 0 | 0 | 0 | 0 | 0 | 0 | 0 |
| VTGESHIG( | FERMT3; FERM1 | Q86UX7; F5I | URP2_HUMAN | Unmodified | 406960 | 572010 | 255620 | 726350 | 0 | 0 | 0 | 0 | 0 | 0 | 0 | 0 | 0 | 0 | 561940 | 1050200 | 0 | 0 |
| VTGEVLDIL | FLOT1; FLOT1; I | O75955; A0A | FLOT1_HUMAN | Unmodified | 0 | 0 | 0 | 0 | 0 | 0 | 380490 | 118360 | 1771300 | 887350 | 2872800 | 2641300 | 0 | 0 | 0 | 114160 | 0 | 0 |
| VTGEVLDIL | FLOT1; FLOT1; I | O75955; A0A | FLOT1_HUMAN | Unmodified | 0 | 0 | 2360000 | 0 | 0 | 0 | 0 | 0 | 40914000 | 4423700 | 0 | 0 | 0 | 0 | 11848000 | 0 | 0 | 0 |
| VTGEVTDP | OSBP | P22059 | OSBP1_HUMAI | Unmodified | 357630 | 486150 | 0 | 0 | 0 | 0 | 247160 | 0 | 0 | 31537 | 0 | 762570 | 0 | 0 | 0 | 0 | 0 | 0 |
| VTGIITQGA | EDIL3 | O43854 | EDIL3_HUMAN | Unmodified | 1212100 | 1060800 | 182250 | 0 | 1.97E+08 | 65322000 | 3909900 | 1778700 | 29966000 | 4544100 | 104210 | 0 | 0 | 103970 | 0 | 0 | 4696100 | 82889000 |
| VTGIITQGA | EDIL3 | O43854 | EDIL3_HUMAN | Unmodified | 0 | 0 | 0 | 0 | 44362000 | 13790000 | 3815000 | 0 | 366060 | 1306700 | 0 | 0 | 0 | 0 | 0 | 0 | 3677100 | 21722000 |
| VTGIITQGA | EDIL3 | O43854 | EDIL3_HUMAN | Unmodified | 0 | 0 | 0 | 0 | 23670000 | 0 | 0 | 0 | 0 | 0 | 0 | 0 | 0 | 0 | 0 | 0 | 576810 | 7465100 |
| VTGLHKVT | FLNB; FLNB | O75369; E7E | FLNB_HUMAN | Unmodified | 0 | 0 | 0 | 0 | 2291200 | 0 | 0 | 0 | 0 | 338160 | 806710 | 0 | 0 | 0 | 0 | 0 | 1131400 | 0 |
| VTGLNCTTI | ITGA5 | P08648 | ITA5_HUMAN I | Unmodified | 0 | 0 | 45354 | 0 | 0 | 0 | 37263 | 0 | 5464100 | 1375000 | 24724 | 0 | 0 | 0 | 221590 | 0 | 0 | 0 |
| VTGPRLVSI | FLNA; FLNA; FL | P21333; A0A | FLNA_HUMAN | Unmodified | 0 | 0 | 0 | 0 | 3837000 | 0 | 0 | 0 | 0 | 0 | 0 | 0 | 332520 | 1685500 | 0 | 0 | 12904000 | 1359300 |
| VTGSGGPF | FAM234A; FAM2 | Q9H0X4; B4I | F234A_HUMAN | Unmodified | 413000 | 0 | 541320 | 0 | 630460 | 65265 | 42148 | 0 | 164430 | 34828 | 0 | 826780 | 81535 | 0 | 993460 | 694530 | 544460 | 354800 |
| VTGSLETK | VDAC1; VDAC1 | P21796; C9J | VDAC1_HUMAI | Unmodified | 30647 | 0 | 0 | 0 | 0 | 0 | 0 | 0 | 10631000 | 465540 | 0 | 0 | 0 | 0 | 0 | 0 | 0 | 0 |
| VTGSLETKY | VDAC1; VDAC1 | P21796; C9J | VDAC1_HUMAI | Unmodified | 0 | 0 | 0 | 0 | 56856 | 0 | 0 | 0 | 789050 | 501350 | 0 | 0 | 0 | 0 | 114450 | 0 | 0 | 0 |
| VTGTLDAN | FSCN1; FSCN1 | Q16658; A0A | FSCN1_HUMAI | Unmodified | 0 | 0 | 110320 | 0 | 0 | 0 | 30097 | 14544 | 2949000 | 217940 | 696760 | 13987000 | 47190 | 0 | 54243 | 54761 | 0 | 0 |
| VTGTLDAN | FSCN1 | Q16658 | FSCN1_HUMAI | Unmodified | 0 | 0 | 0 | 0 | 7592800 | 0 | 0 | 0 | 0 | 326240 | 1076800 | 0 | 0 | 0 | 0 | 0 | 556360 | 4251700 |
| VTGTLETK | VDAC2; VDAC2 | P45880; Q5J | VDAC2_HUMAI | Unmodified | 0 | 0 | 60531 | 0 | 0 | 779490 | 0 | 0 | 63234000 | 1425300 | 0 | 0 | 2199200 | 0 | 54530 | 18336 | 0 | 0 |
| VTGTLETKY | VDAC2; VDAC2 | P45880; Q5J | VDAC2_HUMAI | Unmodified | 0 | 0 | 0 | 0 | 0 | 0 | 0 | 0 | 0 | 148600 | 0 | 0 | 0 | 0 | 0 | 0 | 0 | 0 |
| VTGVITQG/ | EDIL3 | O43854 | EDIL3_HUMAN | Unmodified | 0 | 0 | 0 | 0 | 24203000 | 41472000 | 333700 | 72781 | 14571000 | 1671000 | 0 | 0 | 0 | 0 | 0 | 0 | 2737900 | 14754000 |
| VTGVITQG/ | EDIL3 | O43854 | EDIL3_HUMAN | Unmodified | 0 | 0 | 0 | 0 | 4156700 | 1907900 | 930110 | 0 | 0 | 318640 | 0 | 0 | 0 | 0 | 0 | 0 | 993050 | 2200100 |
| VTGVPGTE | MAPK13 | O15264 | MK13_HUMAN | Unmodified | 0 | 0 | 0 | 0 | 0 | 0 | 0 | 0 | 0 | 0 | 0 | 1904300 | 0 | 0 | 0 | 0 | 0 | 0 |
| VTGVVLFR | SOD3 | P08294 | SODE_HUMAN | Unmodified | 0 | 0 | 0 | 0 | 0 | 0 | 0 | 0 | 12795000 | 1506000 | 21517 | 145780 | 0 | 0 | 0 | 51548 | 0 | 0 |
| VTHAVVTVI | HSPA5; HSPA5; | P11021; A0A | BIP_HUMAN Er | Unmodified | 370240 | 0 | 837160 | 702410 | 0 | 0 | 594920 | 770770 | 84946000 | 11013000 | 3862700 | 2319500 | 0 | 0 | 1289900 | 450630 | 0 | 0 |
| VTHAVVTVI | HSPA5; HSPA5; | P11021; A0A | BIP_HUMAN Er | Unmodified | 0 | 0 | 0 | 0 | 1569000 | 0 | 9863000 | 2283000 | 1354500 | 10114000 | 0 | 0 | 1134700 | 1982700 | 0 | 0 | 0 | 622540 |
| VTHAVVTVI | HSPA5; HSPA5; | P11021; A0A | BIP_HUMAN Er | Oxidation (M) | 0 | 0 | 0 | 0 | 0 | 0 | 0 | 0 | 0 | 1595400 | 0 | 0 | 0 | 1050300 | 0 | 0 | 0 | 0 |
| VTHELQAM | PSMC3; PSMC3 | P17980; E9P | PRS6A_HUMAN | Unmodified | 0 | 0 | 0 | 0 | 0 | 0 | 0 | 19683 | 172490 | 0 | 0 | 905350 | 0 | 0 | 0 | 0 | 0 | 0 |
| VTHETSAH | UCHL3; UCHL3 | P15374; A0A | UCHL3_HUMAI | Unmodified | 120200 | 0 | 26822 | 0 | 0 | 0 | 0 | 0 | 0 | 0 | 563130 | 4678100 | 0 | 0 | 48878 | 322470 | 0 | 0 |
| VTIAQGGVI | H2AX; H2AC6; I | P16104; Q9 | H2AX_HUMAN | Unmodified | 1.33E+09 | 8.1E+08 | 5605200 | 4863700 | 93432000 | 47222000 | 56278000 | 15512000 | 39710000 | 9721900 | 25291000 | 3.08E+09 | 10137000 | 5543800 | 6211400 | 4317100 | 3041300 | 30160000 |
| VTIAQGGVI | H2AX; H2AC6; I | P16104; Q9 | H2AX_HUMAN | Unmodified | 4305100 | 3374300 | 107100 | 0 | 830610 | 0 | 696540 | 159490 | 161380 | 0 | 0 | 22711000 | 0 | 0 | 0 | 0 | 0 | 0 |

| | | | | | | | | | | | | | | | | | | | | | | |
|---|---|---|---|---|---|---|---|---|---|---|---|---|---|---|---|---|---|---|---|---|---|---|
| VTIAQGGVI | H2AC6; H2AW; | Q93077; Q7I | H2A1C_HUMAI | Unmodified | 359280 | 0 | 0 | 0 | 6168500 | 1250900 | 24464000 | 2490600 | 0 | 559770 | 0 | 213140 | 2982900 | 384910 | 0 | 0 | 0 | 2329300 |
| VTIAQGGVI | H2AC21; H2AC | Q8IUE6; Q16 | H2A2B_HUMAI | Unmodified | 0 | 0 | 0 | 0 | 457120 | 0 | 2331900 | 665980 | 0 | 0 | 0 | 0 | 153940 | 0 | 0 | 0 | 0 | 0 |
| VTIAQGYDA | NNT; NNT; NNT | Q13423; E9I | NNTM_HUMAN | Unmodified | 0 | 17065 | 0 | 0 | 0 | 0 | 125510 | 0 | 5632800 | 0 | 0 | 0 | 0 | 60420 | 0 | 0 | 0 | 0 |
| VTIEYYSQL | ALDH9A1 | P49189 | AL9A1_HUMAN | Unmodified | 0 | 0 | 0 | 0 | 0 | 0 | 0 | 0 | 814910 | 0 | 0 | 1153200 | 0 | 0 | 0 | 0 | 0 | 0 |
| VTILELFR | SLC2A1; SLC2A | P11166; A0A | GTR1_HUMAN | Unmodified | 3008100 | 3009500 | 6617300 | 1414800 | 47011 | 0 | 58416 | 0 | 1661900 | 765430 | 0 | 24163000 | 0 | 0 | 5517900 | 3844100 | 0 | 0 |
| VTIMPKDIQ | H3-4; H3-3A; H | Q16695; P84 | H31T_HUMAN | Unmodified | 0 | 0 | 0 | 0 | 427820 | 0 | 371290 | 734550 | 0 | 0 | 0 | 0 | 0 | 0 | 0 | 0 | 0 | 557110 |
| VTIMPKDIQ | H3-4; H3-3A; H | Q16695; P84 | H31T_HUMAN | Oxidation (M) | 0 | 0 | 0 | 0 | 405280 | 1217000 | 0 | 1513000 | 0 | 0 | 0 | 773130 | 0 | 570110 | 0 | 0 | 0 | 0 |
| VTIPTDLIAS | DAG1; DAG1; D | Q14118; C9J | DAG1_HUMAN | Unmodified | 0 | 0 | 0 | 0 | 0 | 0 | 0 | 0 | 1647200 | 0 | 0 | 0 | 0 | 0 | 0 | 0 | 0 | 0 |
| VTISNRLVS | HSP90AB1 | P08238 | HS90B_HUMAI | 2 Oxidation (M) | 0 | 0 | 0 | 0 | 942580 | 0 | 0 | 0 | 486810 | 0 | 0 | 0 | 1019700 | 0 | 0 | 0 | 0 | 0 |
| VTIVTHQD\ | GNPTAB | Q3T906 | GNPTA_HUMAI | Unmodified | 0 | 0 | 289690 | 279690 | 0 | 0 | 0 | 0 | 1126300 | 0 | 165440 | 0 | 0 | 0 | 0 | 101350 | 0 | 0 |
| VTKEQWDT | DECR1 | Q16698 | DECR_HUMAN | Unmodified | 0 | 0 | 0 | 0 | 0 | 0 | 0 | 0 | 511990 | 460560 | 0 | 0 | 0 | 0 | 0 | 0 | 0 | 0 |
| VTKEQWDT | DECR1 | Q16698 | DECR_HUMAN | Unmodified | 8088500 | 2412700 | 2834700 | 4443900 | 0 | 0 | 0 | 9656.5 | 0 | 599850 | 0 | 0 | 5937600 | 2283200 | 232660 | 1970100 | 0 | 0 |
| VTKLQVELI | MYH9 | P35579 | MYH9_HUMAN | Unmodified | 2368200 | 12439000 | 0 | 0 | 1666000 | 0 | 843720 | 0 | 0 | 594920 | 0 | 0 | 0 | 0 | 0 | 508050 | 0 | 797270 |
| VTKPCTPK | MDK; MDK; MDI | P21741; E9P | MK_HUMAN Mi | Unmodified | 0 | 0 | 13892 | 0 | 0 | 0 | 0 | 0 | 1489300 | 58215 | 0 | 0 | 0 | 0 | 0 | 0 | 0 | 0 |
| VTKPNIPEA | ERLIN2; ERLIN2 | O94905; E5F | ERLN2_HUMAN | Unmodified | 0 | 0 | 0 | 0 | 129450 | 0 | 0 | 293870 | 1958900 | 198440 | 0 | 0 | 0 | 0 | 0 | 0 | 0 | 0 |
| VTKPNIPEA | ERLIN2; ERLIN2 | O94905; E5F | ERLN2_HUMAN | Unmodified | 0 | 0 | 0 | 0 | 0 | 0 | 0 | 0 | 11294000 | 1278200 | 0 | 0 | 0 | 0 | 0 | 0 | 0 | 0 |
| VTLAVSDL( | GLOD4; GLOD4 | Q9HC38; F6 | GLOD4_HUMA | Unmodified | 230420 | 0 | 241770 | 448690 | 4004500 | 473610 | 1275300 | 834930 | 1500800 | 1048700 | 2191400 | 7645800 | 2712100 | 1740200 | 1094300 | 489850 | 776890 | 1301700 |
| VTLDLQNS | HTT; HTT; HTT; | P42858; A0A | HD_HUMAN Hu | Unmodified | 0 | 0 | 0 | 364220 | 0 | 0 | 0 | 0 | 985350 | 182190 | 998420 | 0 | 203270 | 0 | 0 | 0 | 0 | 0 |
| VTLELGGK | ALDH2; ALDH1A | P05091; P00 | ALDH2_HUMAI | Unmodified | 0 | 0 | 0 | 0 | 0 | 0 | 1218500 | 1197900 | 200360 | 82829 | 7789700 | 52686000 | 0 | 0 | 0 | 0 | 64112 | 0 |
| VTLELKPLS | TTC5 | Q8N0Z6 | TTC5_HUMAN ' | Unmodified | 0 | 0 | 0 | 0 | 0 | 0 | 0 | 0 | 0 | 0 | 0 | 675870 | 0 | 0 | 0 | 0 | 0 | 0 |
| VTLEYRPVI | SDHA; SDHA; S | P31040; D6F | SDHA_HUMAN | Unmodified | 0 | 0 | 0 | 0 | 0 | 0 | 0 | 0 | 13370000 | 0 | 0 | 0 | 0 | 0 | 71463 | 33704 | 0 | 0 |
| VTLGTQPT\ | DDB1; DDB1; D | Q16531; A0A | DDB1_HUMAN | Unmodified | 0 | 0 | 0 | 0 | 310680 | 0 | 0 | 0 | 0 | 0 | 274990 | 3479200 | 1165200 | 0 | 0 | 0 | 0 | 0 |
| VTLLDASEY | EPB41L1; EPB4 | Q9H4G0; Q4 | E41L1_HUMAN | Unmodified | 0 | 0 | 0 | 0 | 136860 | 0 | 0 | 0 | 0 | 0 | 1151700 | 2467200 | 0 | 0 | 0 | 0 | 0 | 0 |
| VTLLDGTEY | EPB41L2; EPB4 | O43491; A0A | E41L2_HUMAN | Unmodified | 0 | 0 | 0 | 0 | 1749700 | 0 | 0 | 0 | 0 | 0 | 0 | 256780 | 137390 | 0 | 0 | 0 | 0 | 0 |
| VTLLDLMIA | ATXN10; ATXN1 | Q9UBB4; A0 | ATX10_HUMAN | Unmodified | 0 | 0 | 0 | 0 | 0 | 0 | 0 | 0 | 24989 | 0 | 10423 | 281200 | 0 | 0 | 62202 | 0 | 0 | 0 |
| VTLLEANR | FMNL2; FMNL3 | Q96PY5; Q8I | FMNL2_HUMAI | Unmodified | 0 | 0 | 0 | 4452800 | 1409500 | 1690300 | 387110 | 30025 | 832020 | 1626200 | 1421300 | 795620 | 0 | 0 | 0 | 0 | 0 | 0 |
| VTLLHHVLI | INF2; INF2; INF | Q27J81; A0A | INF2_HUMAN I | Unmodified | 0 | 0 | 0 | 0 | 742710 | 308930 | 0 | 0 | 1645000 | 118300 | 0 | 3555600 | 220160 | 0 | 0 | 0 | 229200 | 0 |
| VTLMQLPTI | EIF3B | P55884 | EIF3B_HUMAN | Unmodified | 0 | 0 | 0 | 0 | 0 | 0 | 0 | 0 | 0 | 0 | 0 | 3341100 | 0 | 0 | 0 | 71563 | 0 | 0 |
| VTLMQLPTI | EIF3B | P55884 | EIF3B_HUMAN | Oxidation (M) | 190480 | 283240 | 369520 | 446220 | 0 | 0 | 0 | 0 | 208140 | 0 | 228040 | 3524600 | 0 | 0 | 224240 | 294510 | 0 | 0 |
| VTLMQLPTI | EIF3B | P55884 | EIF3B_HUMAN | Unmodified | 231390 | 212920 | 325360 | 330750 | 0 | 0 | 0 | 0 | 1012800 | 0 | 0 | 4742400 | 0 | 0 | 234580 | 154810 | 0 | 0 |
| VTLNGVPA | ICAM1; ICAM1 | P05362; E7E | ICAM1_HUMAN | Unmodified | 0 | 0 | 745400 | 892940 | 23032000 | 6806900 | 0 | 0 | 0 | 0 | 0 | 0 | 0 | 0 | 893300 | 988080 | 1127700 | 4201000 |
| VTLNGVPA | ICAM1; ICAM1 | P05362; E7E | ICAM1_HUMAN | Unmodified | 0 | 0 | 0 | 0 | 1.51E+08 | 22874000 | 20216 | 0 | 0 | 331900 | 0 | 0 | 0 | 368880 | 0 | 0 | 18185000 | 20849000 |
| VTLNPPGTI | NDUFA11; NDU | Q86Y39; K7I | NDUAB_HUMA | Unmodified | 0 | 0 | 0 | 0 | 0 | 0 | 0 | 0 | 923060 | 0 | 0 | 0 | 0 | 0 | 0 | 0 | 0 | 0 |
| VTLNRLNTI | LAMC1 | P11047 | LAMC1_HUMAI | Unmodified | 0 | 0 | 0 | 0 | 0 | 0 | 0 | 104100 | 2306500 | 54317000 | 0 | 0 | 0 | 0 | 0 | 0 | 413890 | 112410 |
| VTLPAGPD | MAL2 | Q969L2 | MAL2_HUMAN | Unmodified | 5344100 | 6544200 | 3176700 | 727670 | 0 | 0 | 0 | 0 | 1.14E+08 | 24751000 | 356620 | 2543300 | 0 | 0 | 5364800 | 5001700 | 0 | 0 |
| VTLPCHHQ | CLMP | Q9H6B4 | CLMP_HUMAN | Unmodified | 0 | 0 | 0 | 0 | 1207900 | 0 | 547490 | 0 | 0 | 0 | 0 | 0 | 0 | 9411900 | 0 | 0 | 0 | 1252600 |
| VTLPGLKEI | SMCHD1; SMCI | A6NHR9; J3I | SMHD1_HUMA | Unmodified | 0 | 132220 | 0 | 0 | 0 | 0 | 1410000 | 636970 | 0 | 0 | 0 | 0 | 0 | 0 | 0 | 0 | 0 | 0 |
| VTLRPYLTP | ANPEP | P15144 | AMPN_HUMAN | Unmodified | 0 | 0 | 0 | 0 | 0 | 0 | 0 | 0 | 0 | 0 | 0 | 0 | 0 | 0 | 0 | 0 | 2586800 | 7897800 |
| VTLSEGPH | OPA1; OPA1; OI | O60313; E5I | OPA1_HUMAN | Unmodified | 0 | 0 | 0 | 0 | 0 | 0 | 0 | 15486 | 0 | 0 | 0 | 643330 | 0 | 0 | 0 | 0 | 0 | 0 |
| VTLSFPSTL | NPEPPSL1; NPI | A6NEC2; P5 | PSAL_HUMAN | Unmodified | 97429 | 149580 | 537590 | 268060 | 194850 | 61797 | 0 | 0 | 1230600 | 70081 | 1221600 | 8686200 | 0 | 0 | 1018500 | 669110 | 0 | 131080 |
| VTLSVLSR | PIR | O00625 | PIR_HUMAN Pi | Unmodified | 0 | 0 | 0 | 0 | 0 | 0 | 0 | 0 | 0 | 0 | 101370 | 3198200 | 0 | 0 | 0 | 84155 | 0 | 0 |
| VTLSVLSRE | PIR | O00625 | PIR_HUMAN Pi | Unmodified | 0 | 0 | 0 | 0 | 282930 | 0 | 0 | 25005 | 0 | 0 | 298000 | 190350 | 789390 | 200540 | 0 | 0 | 439260 | 177240 |
| VTLTLPVLN | PGLS; PGLS | O95336; M0 | 6PGL_HUMAN | Unmodified | 0 | 0 | 0 | 0 | 0 | 0 | 0 | 0 | 3217600 | 243860 | 389520 | 3951000 | 0 | 0 | 0 | 307230 | 0 | 0 |
| VTLTSEEEA | LDHA | P00338 | LDHA_HUMAN | Unmodified | 0 | 1504900 | 2061800 | 0 | 3933400 | 0 | 0 | 225390 | 8366100 | 1296400 | 29355000 | 1.62E+08 | 0 | 0 | 4804700 | 0 | 0 | 0 |
| VTLTSEEEA | LDHA | P00338 | LDHA_HUMAN | Unmodified | 1287400 | 399630 | 733490 | 2297800 | 10548000 | 0 | 780860 | 207350 | 2934600 | 2227800 | 4049200 | 40800000 | 3342500 | 1117600 | 2094400 | 2367200 | 8160100 | 19307000 |
| VTLTSEEEA | LDHA | P00338 | LDHA_HUMAN | Unmodified | 0 | 0 | 281820 | 678970 | 6180100 | 0 | 416580 | 0 | 526420 | 3807200 | 3642200 | 1769800 | 5762100 | 1433200 | 0 | 287810 | 4438200 | 18198000 |
| VTLVVGAS( | COMT; COMT; ( | P21964; E7E | COMT_HUMAN | Unmodified | 0 | 0 | 0 | 0 | 0 | 0 | 0 | 0 | 0 | 0 | 0 | 0 | 0 | 0 | 0 | 0 | 0 | 0 |
| VTMESALT/ | MYO1C; MYO1( | O00159; I3L! | MYO1C_HUMA | Unmodified | 887060 | 1298000 | 1353300 | 1438500 | 0 | 0 | 82015 | 0 | 331190 | 0 | 0 | 0 | 0 | 0 | 3293100 | 1925600 | 0 | 0 |
| VTMESALT/ | MYO1C; MYO1( | O00159; I3L! | MYO1C_HUMA | Oxidation (M) | 0 | 0 | 510480 | 300010 | 0 | 0 | 0 | 0 | 0 | 0 | 0 | 265640 | 0 | 0 | 888150 | 247670 | 0 | 0 |
| VTMLFLGLI | DARS1; DARS1 | P14868; H7E | SYDC_HUMAN | Oxidation (M) | 0 | 0 | 123800 | 143360 | 0 | 0 | 0 | 0 | 0 | 0 | 220890 | 2605900 | 0 | 0 | 0 | 139100 | 0 | 0 |
| VTMLFLGLI | DARS1; DARS1 | P14868; H7E | SYDC_HUMAN | Unmodified | 0 | 0 | 0 | 0 | 0 | 0 | 0 | 0 | 438860 | 0 | 0 | 4459500 | 0 | 0 | 0 | 0 | 0 | 0 |
| VTMNFEPN | ITGA1 | P56199 | ITA1_HUMAN I | Unmodified | 830100 | 483920 | 0 | 0 | 470250 | 72092 | 0 | 0 | 0 | 0 | 0 | 0 | 0 | 0 | 0 | 0 | 820100 | 1887300 |
| VTMNFEPN | ITGA1 | P56199 | ITA1_HUMAN I | Unmodified | 0 | 0 | 0 | 0 | 548890 | 0 | 987740 | 0 | 0 | 398750 | 0 | 0 | 57254 | 0 | 0 | 641260 | 4629300 | 4874300 |
| VTMNFEPN | ITGA1 | P56199 | ITA1_HUMAN I | Unmodified | 0 | 0 | 0 | 0 | 363900 | 0 | 0 | 0 | 0 | 156340 | 0 | 0 | 0 | 0 | 0 | 0 | 3140200 | 6440900 |
| VTMNFEPN | ITGA1 | P56199 | ITA1_HUMAN I | Oxidation (M) | 0 | 0 | 0 | 0 | 80326 | 0 | 0 | 0 | 0 | 0 | 0 | 0 | 96464 | 0 | 0 | 0 | 425090 | 773220 |
| VTNDGSLT | PTPRB; PTPRB; | P23467; F8V | PTPRB_HUMAN | Unmodified | 370700 | 756990 | 0 | 0 | 0 | 0 | 0 | 0 | 0 | 0 | 0 | 0 | 0 | 0 | 0 | 0 | 0 | 0 |
| VTNEFVHIN | PPP6C | O00743 | PPP6_HUMAN | Unmodified | 0 | 0 | 0 | 0 | 416850 | 126650 | 529020 | 828010 | 481280 | 415050 | 407790 | 4085800 | 0 | 0 | 294300 | 0 | 510550 | 255980 |
| VTNGAFTG | TPI1 | P60174 | TPIS_HUMAN T | Unmodified | 0 | 0 | 0 | 0 | 17129000 | 2579000 | 0 | 0 | 4692800 | 2460500 | 1195000 | 5066800 | 14037000 | 4200700 | 0 | 0 | 3255800 | 7905100 |
| VTNGAFTG | TPI1 | P60174 | TPIS_HUMAN T | Oxidation (M) | 691350 | 533010 | 938330 | 1264400 | 4655700 | 0 | 0 | 69583 | 0 | 615550 | 2429900 | 9213700 | 5027400 | 2098600 | 472730 | 1888600 | 1975300 | 841630 |
| VTNHEDGS | NCAM2; NCAM | O15394; H9I | NCAM2_HUMA | Unmodified | 0 | 0 | 418990 | 0 | 0 | 0 | 0 | 0 | 0 | 0 | 0 | 0 | 0 | 0 | 491120 | 269880 | 0 | 0 |
| VTNHEDGS | NCAM2; NCAM | O15394; H9I | NCAM2_HUMA | Unmodified | 174730 | 0 | 1131600 | 447180 | 0 | 0 | 0 | 0 | 0 | 0 | 0 | 0 | 202340 | 0 | 6030500 | 4496000 | 0 | 0 |
| VTNINDRS/ | WIPF2; WIPF2 | Q8TF74; J3Q | WIPF2_HUMAN | Unmodified | 0 | 0 | 0 | 198200 | 0 | 0 | 0 | 0 | 454820 | 0 | 0 | 0 | 1128200 | 0 | 0 | 0 | 0 | 376670 |
| VTNLSEDTI | EIF3G; EIF3G | O75821; K7E | EIF3G_HUMAN | Unmodified | 4091700 | 3449600 | 1200000 | 5421000 | 103080 | 32230 | 81082 | 0 | 2532900 | 107910 | 1527700 | 6436900 | 261990 | 40405 | 4237300 | 2372900 | 0 | 98797 |
| VTNRDIICC | RPL5; RPL5; RP | P46777; Q5I | RL5_HUMAN 6 | Unmodified | 2330100 | 1394400 | 516960 | 585210 | 0 | 0 | 0 | 0 | 0 | 70843 | 0 | 6219800 | 0 | 0 | 960120 | 446150 | 0 | 0 |
| VTNSTELQI | EEA1 | Q15075 | EEA1_HUMAN | Unmodified | 0 | 0 | 0 | 0 | 1123700 | 370270 | 0 | 0 | 0 | 0 | 0 | 433300 | 541650 | 0 | 0 | 0 | 186360 | 287210 |
| VTPDTDW/ | ACLY; ACLY | P53396; K7E | ACLY_HUMAN | Unmodified | 0 | 0 | 0 | 0 | 0 | 0 | 0 | 0 | 2086900 | 0 | 1512200 | 0 | 0 | 0 | 0 | 0 | 0 | 0 |
| VTPDTDW/ | ACLY; ACLY | P53396; K7E | ACLY_HUMAN | Unmodified | 0 | 0 | 0 | 0 | 1764900 | 0 | 0 | 0 | 0 | 0 | 0 | 0 | 1870800 | 1315400 | 0 | 0 | 488880 | 1031400 |
| VTPERNRTI | SCEL | O95171 | SCEL_HUMAN | Unmodified | 0 | 0 | 0 | 0 | 1755800 | 0 | 0 | 0 | 0 | 0 | 0 | 0 | 0 | 0 | 0 | 0 | 0 | 0 |
| VTPMAAIQI | SCG3 | Q8WXD2 | SCG3_HUMAN | Unmodified | 0 | 0 | 0 | 0 | 0 | 0 | 0 | 0 | 1979400 | 0 | 0 | 0 | 0 | 0 | 0 | 0 | 0 | 0 |
| VTPMAAIQI | SCG3 | Q8WXD2 | SCG3_HUMAN | Oxidation (M) | 0 | 0 | 17282000 | 9161700 | 13798000 | 62479000 | 19742000 | 3456900 | 2921100 | 2037000 | 24767000 | 1731200 | 3176600 | 22313000 | 0 | 0 | 0 | 29202000 |
| VTPPEGYE\ | PDLIM1 | O00151 | PDLI1_HUMAN | Unmodified | 28024 | 0 | 0 | 830180 | 3053600 | 0 | 0 | 0 | 508630 | 234540 | 0 | 0 | 1304700 | 0 | 0 | 0 | 913130 | 4574800 |
| VTPQREEG | DCTN4 | Q9UJW0 | DCTN4_HUMAI | Unmodified | 0 | 0 | 0 | 0 | 395270 | 130020 | 157730 | 80678 | 0 | 0 | 199650 | 1630100 | 732960 | 407800 | 142880 | 234890 | 344190 | 269970 |
| VTPQSDGS | HSPA4; HSPA4 | P34932; A0A | HSP74_HUMAI | Unmodified | 0 | 0 | 0 | 0 | 0 | 0 | 207910 | 0 | 0 | 0 | 0 | 16200 | 13223 | 0 | 0 | 0 | 452380 | 180370 |
| VTPSTVLL( | ASPSCR1; ASPS | Q9BZE9; C9. | ASPC1_HUMAI | Unmodified | 0 | 0 | 0 | 0 | 36897 | 141290 | 0 | 0 | 444130 | 0 | 0 | 74902 | 0 | 0 | 0 | 0 | 44212 | 10811 |
| VTPTRTEIIII | RPS3; RPS3; RP | P23396; H0\ | RS3_HUMAN 4 | Unmodified | 3371800 | 4049500 | 2314900 | 2971600 | 114770 | 0 | 53892 | 0 | 1901500 | 0 | 4185700 | 91925000 | 0 | 0 | 2476100 | 2544000 | 0 | 62613 |
| VTPTRTEIII | RPS3; RPS3; RP | P23396; H0\ | RS3_HUMAN 4 | Unmodified | 0 | 0 | 0 | 976410 | 1169500 | 0 | 1161000 | 32786 | 314460 | 0 | 1199000 | 0 | 0 | 0 | 0 | 0 | 203670 | 2776100 |
| VTQDATPG | HMGCS1 | Q01581 | HMCS1_HUMA | Unmodified | 0 | 0 | 0 | 0 | 1929700 | 425570 | 364570 | 209540 | 1533700 | 434290 | 1827600 | 10623000 | 0 | 0 | 0 | 0 | 79505 | 376270 |
| VTQDATPG | HMGCS1 | Q01581 | HMCS1_HUMA | Unmodified | 987190 | 0 | 567040 | 0 | 2640500 | 0 | 1929300 | 0 | 1393300 | 0 | 38732 | 2208600 | 0 | 0 | 2442300 | 1417800 | 590910 | 0 |
| VTQEIVTER | DSG2 | Q14126 | DSG2_HUMAN | Unmodified | 363090 | 0 | 0 | 0 | 10269000 | 1928000 | 0 | 32429 | 3431700 | 0 | 3382800 | 4397200 | 2752700 | 1165700 | 0 | 0 | 337520 | 1279900 |
| VTQEIVTER | DSG2 | Q14126 | DSG2_HUMAN | Unmodified | 0 | 0 | 0 | 0 | 43385000 | 8071900 | 0 | 0 | 0 | 1653400 | 171800 | 0 | 10344000 | 7293400 | 0 | 0 | 1830700 | 2733400 |
| VTQEIVTER | DSG2 | Q14126 | DSG2_HUMAN | Unmodified | 0 | 0 | 0 | 0 | 4965100 | 394600 | 0 | 0 | 0 | 83506 | 0 | 0 | 713100 | 818370 | 0 | 0 | 344590 | 724760 |
| VTQHIHGL | MAN1B1; MAN1 | Q9UKM7; H( | MA1B1_HUMA | Unmodified | 0 | 0 | 25141 | 0 | 225880 | 0 | 0 | 0 | 2575400 | 536450 | 42249 | 240370 | 0 | 0 | 88089 | 0 | 0 | 0 |
| VTQLDPKEI | WASF2 | Q9Y6W5 | WASF2_HUMAI | Unmodified | 761010 | 436320 | 1471600 | 1845300 | 9861100 | 1293100 | 1721400 | 0 | 3697600 | 2586800 | 0 | 11045000 | 5421300 | 5432300 | 2438000 | 3699900 | 5227500 | 5458400 |
| VTQLTDRW | DSP | P15924 | DESP_HUMAN | Unmodified | 0 | 0 | 0 | 0 | 0 | 0 | 0 | 0 | 396530 | 0 | 0 | 1369300 | 0 | 0 | 133130 | 190080 | 0 | 0 |
| VTQLYMQL | ANGPTL2; ANG | Q9UKU9; OS | ANGL2_HUMAI | Unmodified | 0 | 0 | 0 | 0 | 0 | 0 | 0 | 0 | 331980 | 0 | 0 | 92296 | 0 | 0 | 0 | 0 | 0 | 0 |
| VTQSAYAQ | AFG3L2 | Q9Y4W6 | AFG32_HUMAN | Unmodified | 0 | 0 | 0 | 0 | 0 | 0 | 0 | 0 | 2042400 | 17649 | 0 | 0 | 0 | 0 | 0 | 0 | 0 | 0 |
| VTQSFLPP( | NEDD4L; NEDD | Q96PU5; A0A | NED4L_HUMAI | Oxidation (M) | 0 | 66280 | 218760 | 179220 | 0 | 0 | 0 | 0 | 0 | 25596 | 0 | 0 | 0 | 0 | 514430 | 388850 | 0 | 0 |
| VTQSNFAV | VDAC1; VDAC1 | P21796; C9J | VDAC1_HUMAI | Unmodified | 0 | 0 | 0 | 0 | 0 | 0 | 0 | 0 | 6991600 | 358830 | 0 | 224990 | 0 | 0 | 0 | 0 | 0 | 0 |
| VTQSNFAV | VDAC1 | P21796 | VDAC1_HUMAI | Unmodified | 0 | 0 | 0 | 0 | 0 | 0 | 0 | 0 | 4442800 | 0 | 0 | 0 | 0 | 0 | 0 | 0 | 0 | 0 |
| VTQSTEGH | PRKG2 | Q13237 | KGP2_HUMAN | Unmodified | 0 | 0 | 0 | 0 | 0 | 0 | 1570600 | 687240 | 0 | 0 | 0 | 0 | 0 | 0 | 0 | 0 | 0 | 0 |
| VTQVDGNS | ARCN1; ARCN1 | P48444; B0Y | COPD_HUMAN | Unmodified | 0 | 0 | 112520 | 67090 | 0 | 0 | 0 | 0 | 350900 | 0 | 0 | 2729900 | 0 | 0 | 134090 | 0 | 55549 | 0 |
| VTRADVDV | GTPBP4 | Q9BZE4 | GTPB4_HUMAI | Unmodified | 0 | 0 | 0 | 0 | 0 | 0 | 0 | 933800 | 0 | 0 | 0 | 0 | 0 | 0 | 0 | 0 | 0 | 0 |
| VTRDSRTY | IGF2R | P11717 | MPRI_HUMAN | Unmodified | 0 | 0 | 0 | 0 | 0 | 0 | 0 | 0 | 0 | 507600 | 0 | 0 | 0 | 0 | 0 | 0 | 0 | 0 |
| VTRSNMSF | SLC4A7 | H7C3C4 | H7C3C4_HUM | Unmodified | 0 | 0 | 0 | 0 | 3306400 | 0 | 0 | 0 | 0 | 0 | 0 | 0 | 1297400 | 0 | 0 | 0 | 602120 | 8070200 |
| VTSAHKGP | PAICS; PAICS; I | P22234; D6I | PUR6_HUMAN | Unmodified | 0 | 0 | 0 | 0 | 0 | 0 | 0 | 0 | 0 | 0 | 0 | 0 | 0 | 0 | 0 | 0 | 0 | 0 |
| VTSAHKGP | PAICS; PAICS; I | P22234; D6I | PUR6_HUMAN | Unmodified | 0 | 0 | 0 | 116500 | 0 | 0 | 0 | 0 | 0 | 0 | 386580 | 0 | 0 | 0 | 0 | 0 | 0 | 0 |
| VTSALSRD | BCAM; BCAM | P50895; A0A | BCAM_HUMAN | Unmodified | 610900 | 158400 | 0 | 0 | 555180 | 236140 | 0 | 0 | 0 | 48371 | 0 | 0 | 23269 | 0 | 60034 | 0 | 0 | 117600 |
| VTSAVEALL | USP5 | P45974 | UBP5_HUMAN | Unmodified | 0 | 0 | 0 | 0 | 0 | 0 | 0 | 0 | 0 | 0 | 1207400 | 878650 | 0 | 0 | 0 | 0 | 0 | 0 |
| VTSCHDPC | SEZ6; SEZ6; SEZ | Q53EL9; C9J | SEZ6_HUMAN | Unmodified | 0 | 0 | 0 | 0 | 0 | 96335 | 0 | 0 | 1904000 | 39309 | 0 | 0 | 0 | 0 | 0 | 0 | 0 | 0 |
| VTSEELHYI | UQCRC2; UQCI | P22695; H3I | QCR2_HUMAN | Unmodified | 0 | 0 | 0 | 0 | 0 | 0 | 0 | 0 | 71732000 | 4485900 | 478360 | 0 | 0 | 0 | 0 | 0 | 23247 | 0 |
| VTSEGAGL | GAA | P10253 | LYAG_HUMAN | Unmodified | 257300 | 0 | 1272600 | 1838500 | 0 | 0 | 0 | 0 | 10264000 | 1539300 | 0 | 591590 | 733710 | 381640 | 1855300 | 1236800 | 0 | 0 |
| VTSELLR | XPNPEP1; MAN | Q9NQW7; O | XPP1_HUMAN | Unmodified | 84010 | 0 | 143330 | 279610 | 226580 | 69164 | 68859 | 77330 | 521180 | 126310 | 473500 | 3081200 | 591060 | 152330 | 431400 | 316970 | 109260 | 191120 |
| VTSELVIPS | MKRN3; MKRN | Q13064; A0A | MKRN3_HUMA | Unmodified | 496370 | 0 | 0 | 216680 | 0 | 0 | 0 | 0 | 0 | 0 | 0 | 0 | 0 | 0 | 0 | 76675 | 0 | 0 |
| VTSEVNRLI | FLOT2; FLOT2; | Q14254; J3Q | FLOT2_HUMAN | Unmodified | 0 | 0 | 0 | 0 | 1710300 | 0 | 171890 | 0 | 0 | 162230 | 0 | 0 | 455260 | 777180 | 0 | 0 | 0 | 0 |
| VTSFASNLI | COBL; COBL; C | O75128; J3K | COBL_HUMAN | Unmodified | 373710 | 326920 | 0 | 0 | 0 | 0 | 0 | 0 | 0 | 0 | 0 | 0 | 87695 | 0 | 60782 | 71649 | 0 | 0 |
| VTSLRSPG | KIAA1522 | Q9P206 | K1522_HUMAN | Unmodified | 0 | 0 | 0 | 0 | 0 | 0 | 0 | 0 | 0 | 0 | 0 | 0 | 0 | 0 | 450620 | 269680 | 0 | 0 |
| VTSLTACLV | THY1; THY1; TH | P04216; E9P | THY1_HUMAN | Unmodified | 0 | 274060 | 0 | 0 | 0 | 0 | 0 | 142270 | 0 | 115990 | 0 | 847890 | 0 | 0 | 0 | 0 | 132320 | 0 |
| VTSMSLAR | PLBD2 | Q8NHP8 | PLBL2_HUMAN | Unmodified | 0 | 0 | 0 | 0 | 0 | 0 | 0 | 0 | 486120 | 177790 | 0 | 0 | 0 | 0 | 0 | 0 | 0 | 0 |
| VTSRGPDE | SLC4A7 | E9PFN4 | E9PFN4_HUMA | Unmodified | 0 | 0 | 279870 | 384320 | 1229600 | 132860 | 473830 | 184240 | 333480 | 157550 | 287120 | 580720 | 0 | 211180 | 0 | 140830 | 0 | 688360 |

| | | | | | | | | | | | | | | | | | | | | | | |
|---|---|---|---|---|---|---|---|---|---|---|---|---|---|---|---|---|---|---|---|---|---|---|
| VTSSETDEI | SLC9A2 | Q9UBY0 | SL9A2_HUMAN | Unmodified | 0 | 0 | 1510800 | 424860 | 0 | 0 | 0 | 0 | 39006 | 0 | 0 | 132530 | 851840 | 0 | 1059800 | 630530 | 0 | 4400000 |
| VTSSGPVIN | GAS6 | Q14393 | GAS6_HUMAN | Oxidation (M) | 0 | 0 | 0 | 0 | 0 | 0 | 0 | 0 | 2038000 | 0 | 0 | 0 | 0 | 0 | 0 | 0 | 354390 | 522640 |
| VTSTNNMA | PDLIM5; PDLIM | Q96HC4; A0 | PDLI5_HUMAN | Unmodified | 0 | 23839 | 0 | 263390 | 420230 | 0 | 0 | 0 | 0 | 0 | 0 | 357970 | 0 | 0 | 0 | 96586 | 81366 | 462460 |
| VTSVVFHP | PRPF19 | Q9UMS4 | PRP19_HUMAN | Unmodified | 0 | 0 | 0 | 0 | 0 | 0 | 0 | 0 | 0 | 0 | 0 | 2178800 | 0 | 0 | 0 | 0 | 0 | 0 |
| VTSVVTGFI | B3GNT2 | Q9NY97 | B3GN2_HUMAN | Unmodified | 0 | 0 | 0 | 0 | 0 | 0 | 0 | 0 | 6048000 | 0 | 241350 | 782950 | 0 | 0 | 0 | 0 | 0 | 0 |
| VTSVVDV | APEH; APEH; AI | P13798; C9J | ACPH_HUMAN | Unmodified | 699430 | 613150 | 0 | 0 | 0 | 0 | 0 | 0 | 0 | 0 | 0 | 1928200 | 0 | 0 | 0 | 291330 | 0 | 0 |
| VTSYEVQLI | PTPRB; PTPRB; | P23467; F8V | PTPRB_HUMAN | Unmodified | 434230 | 584900 | 0 | 0 | 144200 | 0 | 144300 | 0 | 0 | 170490 | 0 | 0 | 0 | 0 | 0 | 0 | 0 | 307620 |
| VTTFEHQY | GNA11 | P29992 | GNA11_HUMAN | Unmodified | 0 | 0 | 0 | 0 | 6352000 | 682890 | 0 | 661140 | 0 | 575000 | 2289800 | 9163800 | 0 | 0 | 0 | 0 | 1309300 | 3476300 |
| VTTGSVPV | ITGA2 | P17301 | ITA2_HUMAN | Oxidation (M) | 0 | 954910 | 0 | 400800 | 665190 | 0 | 0 | 0 | 0 | 0 | 0 | 0 | 0 | 0 | 1498400 | 870950 | 905970 | 0 |
| VTTGTSNT | RBBP5 | Q15291 | RBBP5_HUMAN | Unmodified | 0 | 0 | 0 | 0 | 0 | 0 | 0 | 0 | 0 | 648150 | 0 | 295200 | 0 | 930130 | 0 | 0 | 0 | 0 |
| VTTHPLAK | PARK7; PARK7; | Q99497; K7E | PARK7_HUMAN | Unmodified | 0 | 0 | 0 | 0 | 0 | 0 | 0 | 86123 | 0 | 534240 | 0 | 2547500 | 0 | 0 | 0 | 0 | 1016200 | 2110000 |
| VTTMDAEL | RDX; EZR; MSN | P35241; P15 | RADI_HUMAN | Oxidation (M) | 833620 | 0 | 1229200 | 1084200 | 0 | 846630 | 0 | 88384 | 2773800 | 0 | 873290 | 5412300 | 0 | 0 | 1820800 | 1800000 | 695550 | 0 |
| VTTNPNLR | SMPDL3A | Q92484 | ASM3A_HUMAN | Unmodified | 6149900 | 9057400 | 72081 | 0 | 0 | 0 | 0 | 481440 | 1472300 | 358090 | 1243600 | 386930 | 0 | 0 | 100080 | 42534 | 0 | 0 |
| VTTQTSLPI | SNX1; SNX1 | Q13596; H0 | SNX1_HUMAN | Unmodified | 0 | 0 | 0 | 0 | 403290 | 0 | 0 | 0 | 332770 | 0 | 0 | 1667100 | 0 | 0 | 0 | 0 | 0 | 279890 |
| VTTQTSLPI | SNX1 | Q13596 | SNX1_HUMAN | Unmodified | 0 | 0 | 0 | 0 | 609810 | 73366 | 176570 | 132800 | 0 | 0 | 0 | 0 | 161570 | 259930 | 0 | 0 | 0 | 192930 |
| VTTRMGNT | SUMF2; SUMF2 | Q8NBJ7; J3C | SUMF2_HUMAN | Oxidation (M) | 0 | 0 | 0 | 0 | 0 | 0 | 0 | 0 | 0 | 754270 | 0 | 0 | 0 | 0 | 0 | 274620 | 0 | 0 |
| VTTRMGNT | SUMF2; SUMF2 | Q8NBJ7; J3C | SUMF2_HUMAN | Unmodified | 0 | 0 | 0 | 0 | 0 | 0 | 0 | 0 | 0 | 2163100 | 0 | 0 | 0 | 0 | 0 | 0 | 0 | 0 |
| VTTVASHTS | CLU; CLU; CLU | P10909; H0Y | CLUS_HUMAN | Unmodified | 283770 | 191870 | 139260 | 0 | 0 | 0 | 0 | 0 | 1029500 | 0 | 0 | 0 | 0 | 0 | 0 | 0 | 0 | 0 |
| VTVTEIGK | COPA; COPA; C | P53621; A0A | COPA_HUMAN | Unmodified | 0 | 0 | 0 | 0 | 0 | 0 | 0 | 0 | 807780 | 0 | 592260 | 3823100 | 0 | 0 | 446860 | 168320 | 0 | 0 |
| VTVTEIGK | COPA; COPA; C | P53621; A0A | COPA_HUMAN | Unmodified | 0 | 0 | 0 | 0 | 0 | 0 | 39131 | 0 | 1037200 | 0 | 514390 | 987100 | 563020 | 0 | 0 | 0 | 0 | 0 |
| VTVVATPC | GSK3B; GSK3B | P49841; A0A | GSK3B_HUMAN | Unmodified | 0 | 586090 | 0 | 1137200 | 2792100 | 0 | 0 | 0 | 3698200 | 978000 | 0 | 4923100 | 1481400 | 0 | 0 | 0 | 0 | 5064000 |
| VTVAGGVH | FASN; FASN | P49327; A0A | FAS_HUMAN F | Unmodified | 0 | 0 | 1304900 | 5295800 | 586850 | 227710 | 0 | 51693 | 956190 | 178710 | 5477500 | 89643000 | 629890 | 0 | 4145200 | 5473900 | 167530 | 1192600 |
| VTVAGGVH | FASN; FASN | P49327; A0A | FAS_HUMAN F | Unmodified | 0 | 0 | 0 | 0 | 0 | 0 | 0 | 0 | 0 | 0 | 541830 | 1536000 | 0 | 0 | 0 | 0 | 0 | 0 |
| VTVAGLAG | PARK7; PARK7 | Q99497; K7E | PARK7_HUMAN | Unmodified | 98184 | 0 | 632200 | 94230 | 756910 | 0 | 484970 | 743820 | 489590 | 252670 | 650760 | 8180300 | 190770 | 28039 | 0 | 259220 | 365650 | 1128500 |
| VTVAGLAG | PARK7; PARK7 | Q99497; K7E | PARK7_HUMAN | Unmodified | 4544500 | 1960800 | 5018600 | 6458100 | 1639700 | 646900 | 1016900 | 0 | 24377000 | 1521700 | 4336400 | 27270000 | 4243600 | 1548400 | 8435800 | 5185500 | 333640 | 1547000 |
| VTVDQVTD | NHLRC2 | Q8NBF2 | NHLC2_HUMA | Unmodified | 0 | 0 | 0 | 94144 | 0 | 0 | 0 | 124140 | 0 | 0 | 234000 | 2160500 | 0 | 0 | 0 | 0 | 0 | 0 |
| VTVDQVTD | NHLRC2 | Q8NBF2 | NHLC2_HUMA | Unmodified | 0 | 0 | 0 | 0 | 0 | 0 | 0 | 0 | 0 | 0 | 841380 | 3289000 | 0 | 0 | 0 | 0 | 0 | 0 |
| VTVGDITC | PRKRA; PRKRA; | O75569; A0 | PRKRA_HUMAN | Unmodified | 155650 | 0 | 0 | 144710 | 0 | 0 | 0 | 0 | 0 | 0 | 0 | 763850 | 0 | 0 | 77555 | 0 | 0 | 0 |
| VTVGVIGS | STEAP2; STEAP | Q8NFT2; C9 | STEA2_HUMAN | Unmodified | 2709600 | 10590000 | 12960000 | 3086900 | 0 | 0 | 33384 | 0 | 0 | 0 | 0 | 0 | 881400 | 902740 | 37619000 | 37132000 | 0 | 0 |
| VTVLFAGQ | FLNA; FLNA; FL | P21333; A0A | FLNA_HUMAN | Unmodified | 0 | 0 | 0 | 829160 | 3830800 | 1717700 | 2361900 | 110980 | 541620 | 0 | 0 | 2948100 | 298870 | 0 | 0 | 0 | 1111600 | 363310 |
| VTVLFAGQ | FLNB; FLNB | O75369; E7E | FLNB_HUMAN | Unmodified | 240860 | 0 | 1759700 | 1306500 | 6480000 | 2701700 | 0 | 0 | 771190 | 0 | 1197200 | 7926400 | 1017200 | 0 | 1036900 | 1087300 | 1624800 | 985500 |
| VTVLFAGQ | FLNB; FLNB | O75369; E7E | FLNB_HUMAN | Unmodified | 0 | 0 | 0 | 0 | 2426600 | 0 | 0 | 0 | 0 | 0 | 0 | 0 | 0 | 0 | 0 | 0 | 0 | 0 |
| VTVLGVAT | GAA | P10253 | LYAG_HUMAN | Unmodified | 0 | 0 | 0 | 0 | 0 | 0 | 0 | 0 | 1752200 | 0 | 0 | 0 | 0 | 0 | 0 | 0 | 0 | 0 |
| VTVLTLQDI | UBA2 | Q9UBT2 | SAE2_HUMAN | Unmodified | 0 | 0 | 0 | 0 | 475680 | 0 | 164240 | 68916 | 352010 | 419010 | 1181800 | 2303400 | 347760 | 0 | 0 | 0 | 0 | 126070 |
| VTVNTNMV | COPB1 | P53618 | COPB_HUMAN | Oxidation (M) | 0 | 0 | 0 | 0 | 0 | 0 | 0 | 15179 | 0 | 0 | 222720 | 1394000 | 87901 | 85536 | 0 | 0 | 0 | 0 |
| VTVNYPPTI | LSAMP; LSAMP | Q13449; F5H | LSAMP_HUMAN | Unmodified | 0 | 0 | 0 | 0 | 0 | 0 | 0 | 0 | 1754100 | 405920 | 0 | 0 | 0 | 0 | 0 | 0 | 0 | 0 |
| VTVQAACG | SORCS2; SORC | Q96PQ0; B5 | SORC2_HUMAN | Unmodified | 0 | 0 | 0 | 0 | 0 | 0 | 0 | 0 | 1097300 | 0 | 0 | 0 | 0 | 0 | 0 | 0 | 0 | 0 |
| VTVRFPPE | EPRS1; EPRS1 | P07814; V9C | SYEP_HUMAN | Unmodified | 246280 | 0 | 352490 | 167830 | 414960 | 0 | 0 | 0 | 0 | 0 | 696400 | 4428900 | 1078600 | 0 | 216860 | 261980 | 0 | 0 |
| VTVTNPDT | CNTN1 | Q12860 | CNTN1_HUMA | Unmodified | 420200 | 292730 | 0 | 1487600 | 254090 | 0 | 119610 | 83379 | 413860 | 0 | 0 | 28616 | 206160 | 0 | 0 | 0 | 0 | 0 |
| VTVTSEGG | HSPG2 | P98160 | PGBM_HUMAN | Unmodified | 0 | 0 | 821220 | 654180 | 0 | 0 | 0 | 0 | 657590 | 0 | 0 | 0 | 0 | 0 | 0 | 177440 | 0 | 0 |
| VTVVDVNE | UGDH; UGDH; | O60701; E7E | UGDH_HUMAN | Unmodified | 0 | 0 | 0 | 0 | 0 | 0 | 0 | 0 | 0 | 0 | 0 | 1495100 | 0 | 0 | 0 | 0 | 0 | 0 |
| VTVVDVNE | UGDH; UGDH; | O60701; E7E | UGDH_HUMAN | Unmodified | 0 | 0 | 0 | 0 | 11211000 | 0 | 0 | 0 | 0 | 0 | 0 | 0 | 431970 | 0 | 0 | 0 | 0 | 0 |
| VTWAPPPS | FN1; FN1 | P02751; H0Y | FINC_HUMAN | Unmodified | 0 | 0 | 2272000 | 2025400 | 0 | 0 | 0 | 0 | 462490 | 412800 | 73362 | 132450 | 0 | 0 | 0 | 930740 | 0 | 0 |
| VTWDGHS | SF3A1; SF3A1 | Q15459; H7C | SF3A1_HUMAN | Unmodified | 588090 | 617300 | 0 | 0 | 52077 | 0 | 75522 | 0 | 0 | 50479 | 0 | 408880 | 140100 | 0 | 0 | 218980 | 0 | 0 |
| VTWDSSFC | CORO1C; COR | Q9ULV4; F8 | COR1C_HUMAN | Unmodified | 0 | 151270 | 0 | 0 | 130140 | 0 | 42500 | 14517 | 218130 | 0 | 416850 | 6294300 | 0 | 0 | 0 | 0 | 0 | 0 |
| VTWDSTFC | CORO1B; COR | Q9BR76; F5H | COR1B_HUMA | Unmodified | 0 | 0 | 109800 | 134910 | 0 | 0 | 0 | 0 | 0 | 0 | 0 | 3236100 | 0 | 63676 | 508690 | 1019000 | 39166 | 0 |
| VTWTMNG | DCDC1 | M0R2J8 | DCDC1_HUMA | Oxidation (M) | 0 | 0 | 0 | 0 | 0 | 0 | 0 | 0 | 123440 | 0 | 1299200 | 0 | 0 | 0 | 0 | 0 | 0 | 0 |
| VTYHPDGP | KARS1 | Q15046 | SYK_HUMAN L | Unmodified | 0 | 0 | 0 | 0 | 0 | 0 | 0 | 0 | 741360 | 0 | 0 | 6098400 | 0 | 0 | 0 | 215520 | 0 | 0 |
| VTYHPDGP | KARS1 | Q15046 | SYK_HUMAN L | Unmodified | 0 | 0 | 0 | 0 | 0 | 0 | 0 | 0 | 0 | 0 | 229110 | 7075000 | 0 | 0 | 0 | 0 | 0 | 359930 |
| VTYKAPVP | CANX; CANX; C | P27824; A0A | CALX_HUMAN | Unmodified | 0 | 0 | 0 | 0 | 0 | 0 | 0 | 0 | 11061000 | 967460 | 0 | 0 | 0 | 0 | 0 | 0 | 0 | 0 |
| VTYKAPVP | CANX; CANX; C | P27824; A0A | CALX_HUMAN | Unmodified | 0 | 0 | 0 | 0 | 0 | 0 | 0 | 0 | 0 | 4798500 | 0 | 0 | 0 | 0 | 0 | 0 | 0 | 0 |
| VTYLVHNF | IDH1 | O75874 | IDHC_HUMAN | 2 Oxidation (M) | 261950 | 281160 | 217290 | 332040 | 0 | 0 | 0 | 0 | 0 | 0 | 0 | 2676900 | 1018100 | 0 | 1106200 | 522060 | 0 | 0 |
| VTYMEASA | RRAS2; RRAS2 | P62070; E9P | RRAS2_HUMAN | Unmodified | 1544000 | 1285200 | 2390300 | 1915000 | 1451000 | 474960 | 44517 | 30478 | 2005700 | 400560 | 0 | 582350 | 832010 | 724370 | 2957400 | 2102500 | 410400 | 409230 |
| VTYMEASA | RRAS2; RRAS2 | P62070; E9P | RRAS2_HUMAN | Oxidation (M) | 550620 | 0 | 0 | 968920 | 511120 | 0 | 0 | 0 | 0 | 0 | 0 | 0 | 0 | 90027 | 1369500 | 812910 | 18775 | 0 |
| VTYMEEER | HSPA4; HSPA4; | P34932; A0A | HSP74_HUMAN | Unmodified | 0 | 0 | 0 | 0 | 0 | 0 | 0 | 0 | 0 | 0 | 0 | 1037600 | 0 | 0 | 0 | 0 | 0 | 0 |
| VTYRPLDP | PFKFB2 | O60825 | F262_HUMAN | Unmodified | 0 | 0 | 0 | 93644 | 0 | 0 | 0 | 0 | 60807 | 0 | 109780 | 1165200 | 0 | 0 | 0 | 0 | 0 | 0 |
| VVACNLYP | ATIC; ATIC | P31939; C9J | PUR9_HUMAN | Unmodified | 0 | 0 | 0 | 912970 | 853100 | 0 | 0 | 0 | 832830 | 221820 | 0 | 3860400 | 1125900 | 0 | 0 | 529000 | 0 | 0 |
| VVADGAGL | GFUS; GFUS; G | Q13630; E9P | FCL_HUMAN G | Unmodified | 90818 | 140180 | 119760 | 163700 | 317520 | 42986 | 26835 | 10713 | 733490 | 92786 | 171810 | 859030 | 0 | 65444 | 650560 | 272850 | 228110 | 458300 |
| VVADGAGL | GFUS; GFUS; G | Q13630; E9P | FCL_HUMAN G | Unmodified | 0 | 0 | 0 | 0 | 0 | 0 | 0 | 0 | 0 | 0 | 0 | 0 | 0 | 0 | 0 | 0 | 0 | 3801500 |
| VVAEEAWE | USP15 | Q9Y4E8 | UBP15_HUMAN | Unmodified | 0 | 0 | 0 | 0 | 0 | 0 | 0 | 0 | 0 | 0 | 0 | 1059100 | 75118 | 0 | 0 | 0 | 0 | 0 |
| VVAEGFDS | PON2; PON2; P | Q15165; A0A | PON2_HUMAN | Unmodified | 0 | 0 | 0 | 0 | 0 | 0 | 0 | 0 | 1810700 | 0 | 0 | 866020 | 0 | 0 | 0 | 0 | 0 | 0 |
| VVAEPQTQ | LARP4B; LARP4 | Q92615; X6F | LAR4B_HUMAN | Unmodified | 98772 | 0 | 0 | 0 | 0 | 0 | 0 | 0 | 0 | 0 | 0 | 0 | 0 | 0 | 0 | 0 | 0 | 0 |
| VVAGDEWI | MVP; MVP | Q14764; H3I | MVP_HUMAN | Unmodified | 73902 | 28006 | 140590 | 0 | 0 | 0 | 0 | 0 | 67720 | 354310 | 0 | 706250 | 0 | 0 | 0 | 0 | 0 | 0 |
| VVAGDEWI | MVP; MVP | Q14764; H3I | MVP_HUMAN | Unmodified | 0 | 0 | 0 | 0 | 0 | 0 | 0 | 0 | 0 | 0 | 0 | 2173800 | 0 | 0 | 0 | 0 | 0 | 0 |
| VVAGEVQV | IGSF8; IGSF8 | Q969P0; C9 | IGSF8_HUMAN | Unmodified | 3748300 | 2680400 | 1962900 | 1357900 | 0 | 0 | 0 | 0 | 0 | 0 | 0 | 80679 | 0 | 0 | 1555700 | 2056100 | 0 | 0 |
| VVAGEVQV | IGSF8; IGSF8 | Q969P0; C9 | IGSF8_HUMAN | Unmodified | 2428400 | 499130 | 310050 | 1436300 | 535690 | 141640 | 0 | 34572 | 289150 | 0 | 0 | 0 | 1872700 | 554660 | 166570 | 160120 | 362630 | 804670 |
| VVAIYDYTK | ABI1; ABI2; ABI | Q8IZP0; Q9 | ABI1_HUMAN | Unmodified | 861850 | 0 | 2350000 | 1592000 | 1937400 | 599380 | 325910 | 254270 | 3491900 | 968400 | 912740 | 5989200 | 0 | 585580 | 4228700 | 5489700 | 549700 | 559150 |
| VVAKLPFTF | TMCO1; TMCO | Q9UM00; J3I | TMCO1_HUMA | Unmodified | 0 | 0 | 0 | 0 | 0 | 0 | 0 | 0 | 1404000 | 0 | 0 | 0 | 0 | 0 | 0 | 0 | 0 | 0 |
| VVALIGVAT | ABHD10 | Q9NUJ1 | ABHDA_HUMA | Unmodified | 0 | 0 | 0 | 0 | 0 | 0 | 0 | 0 | 1766300 | 849420 | 0 | 95425 | 0 | 0 | 0 | 0 | 0 | 0 |
| VVANSKES | RPL22; RPL22; | P35268; K7E | RL22_HUMAN | Unmodified | 1087000 | 923890 | 0 | 0 | 168380 | 0 | 0 | 0 | 187470 | 48174 | 0 | 0 | 0 | 240610 | 0 | 0 | 0 | 55235 |
| VVAPTISSP | TLN1 | Q9Y490 | TLN1_HUMAN | Unmodified | 1173800 | 1047400 | 0 | 1145700 | 0 | 0 | 0 | 0 | 0 | 0 | 341690 | 3992300 | 0 | 0 | 1318800 | 1072200 | 0 | 0 |
| VVAPTISSP | TLN1 | Q9Y490 | TLN1_HUMAN | Unmodified | 0 | 0 | 2233700 | 0 | 19023000 | 1131500 | 32794000 | 16083000 | 885420 | 2383600 | 0 | 0 | 19214000 | 26911000 | 0 | 0 | 11380000 | 14781000 |
| VVASDKET | RPL22L1; RPL2 | Q6P5R6; H0 | RL22L_HUMAN | Unmodified | 4454600 | 4056500 | 2444300 | 3919000 | 0 | 0 | 0 | 26437 | 0 | 0 | 0 | 4577800 | 368630 | 124870 | 6508700 | 4653200 | 0 | 101570 |
| VVATNTLG | CNTN1 | Q12860 | CNTN1_HUMA | Unmodified | 277240 | 369450 | 0 | 0 | 0 | 0 | 0 | 0 | 515940 | 0 | 0 | 0 | 0 | 0 | 0 | 0 | 0 | 0 |
| VVATTQMQ | ANPEP | P15144 | AMPN_HUMAN | Unmodified | 0 | 0 | 0 | 0 | 0 | 0 | 0 | 0 | 0 | 584050 | 0 | 0 | 0 | 0 | 0 | 0 | 1774500 | 1784500 |
| VVATTQMQ | ANPEP | P15144 | AMPN_HUMAN | Unmodified | 0 | 0 | 0 | 0 | 355040 | 0 | 0 | 0 | 0 | 0 | 330680 | 0 | 0 | 0 | 0 | 0 | 45885000 | 1.46E+08 |
| VVATTQMQ | ANPEP | P15144 | AMPN_HUMAN | Oxidation (M) | 0 | 0 | 0 | 0 | 115280 | 0 | 17940 | 0 | 325660 | 107360 | 0 | 0 | 128140 | 134820 | 0 | 0 | 7264200 | 43506000 |
| VVAVDGDS | PCDHB8; PCDH | Q9UN66; Q9 | PCDB8_HUMAI | Unmodified | 0 | 0 | 0 | 38643 | 0 | 0 | 0 | 0 | 3663800 | 1177500 | 0 | 0 | 0 | 0 | 0 | 0 | 0 | 819770 |
| VVCDENGS | PABPC1; PABP | P11940; A0A | PABP1_HUMAN | Unmodified | 2075500 | 857240 | 0 | 1234400 | 0 | 0 | 0 | 0 | 0 | 0 | 0 | 0 | 0 | 0 | 0 | 0 | 0 | 0 |
| VVCDENGS | PABPC1; PABP | P11940; A0A | PABP1_HUMAN | Unmodified | 0 | 0 | 0 | 1561600 | 0 | 0 | 0 | 0 | 0 | 0 | 0 | 0 | 319130 | 231220 | 0 | 0 | 0 | 0 |
| VVCLIQNG | SRP72; SRP72; | O76094; D6I | SRP72_HUMAN | Unmodified | 0 | 0 | 0 | 0 | 0 | 0 | 448820 | 0 | 0 | 649110 | 0 | 0 | 0 | 296790 | 0 | 0 | 1203400 | 220400 |
| VVCVDQQI | ACVR1 | Q04771 | ACVR1_HUMAN | Unmodified | 0 | 0 | 0 | 0 | 2861300 | 0 | 0 | 0 | 0 | 0 | 0 | 448880 | 545670 | 357220 | 0 | 0 | 2194100 | 6438500 |
| VVDALGNA | ATP5F1A; ATP5 | P25705; K7E | ATPA_HUMAN | Unmodified | 0 | 0 | 0 | 0 | 182230 | 0 | 0 | 0 | 2060200 | 231980 | 1515300 | 0 | 0 | 0 | 0 | 0 | 0 | 0 |
| VVDALGNA | ATP5F1A; ATP5 | P25705; K7E | ATPA_HUMAN | Unmodified | 0 | 0 | 0 | 0 | 876360 | 148080 | 93248 | 19749 | 16995000 | 840440 | 1023800 | 614090 | 0 | 0 | 0 | 0 | 2334500 | 884460 |
| VVDELNLLI | APPL2; APPL2; | Q8NEU8; F8 | DP13B_HUMAN | Unmodified | 210360 | 0 | 0 | 553400 | 2501300 | 1819900 | 0 | 36658 | 1027900 | 804150 | 0 | 196670 | 2350500 | 3032900 | 458910 | 498730 | 0 | 3504600 |
| VVDEREQN | SNAP25 | P60880 | SNP25_HUMAN | Oxidation (M) | 287160 | 308880 | 0 | 0 | 0 | 0 | 0 | 42261 | 800080 | 104820 | 0 | 0 | 0 | 0 | 0 | 0 | 0 | 0 |
| VVDEREQN | SNAP25 | P60880 | SNP25_HUMAN | Unmodified | 0 | 0 | 0 | 0 | 0 | 0 | 0 | 0 | 1809100 | 0 | 0 | 0 | 0 | 0 | 0 | 0 | 0 | 0 |
| VVDGAVGG | DLAT; DLAT; DL | P10515; A0A | ODP2_HUMAN | Unmodified | 0 | 0 | 0 | 228930 | 0 | 0 | 0 | 0 | 7046200 | 0 | 0 | 0 | 0 | 0 | 0 | 0 | 0 | 0 |
| VVDHFGRL | HPGD; HPGD | P15428; E9P | PGDH_HUMAN | Unmodified | 113610 | 80805 | 0 | 1189800 | 4236500 | 750390 | 2742300 | 0 | 0 | 0 | 0 | 0 | 2992600 | 4780200 | 379980 | 0 | 0 | 10282000 |
| VVDLEGSL | LRRC47 | Q8N1G4 | LRC47_HUMAN | Unmodified | 0 | 355090 | 405970 | 349560 | 0 | 0 | 0 | 0 | 904310 | 0 | 655040 | 4958600 | 188450 | 0 | 552780 | 334700 | 0 | 0 |
| VVDLLAPY | ATP5F1B; ATP5 | P06576; H0 | ATPB_HUMAN | Unmodified | 0 | 0 | 0 | 0 | 378250 | 0 | 0 | 13869 | 46278000 | 4204400 | 422000 | 1212400 | 0 | 24041 | 198150 | 0 | 126370 | 587430 |
| VVDLLAQD | ADSS2 | P30520 | PURA2_HUMAN | Unmodified | 37136 | 0 | 0 | 0 | 0 | 0 | 0 | 0 | 0 | 27801 | 843250 | 6191300 | 0 | 0 | 108280 | 60209 | 0 | 0 |
| VVDLMAHN | GAPDH; GAPDH | P04406; E7E | G3P_HUMAN | Oxidation (M) | 0 | 0 | 91541 | 0 | 0 | 0 | 0 | 0 | 163920 | 0 | 2774400 | 850980 | 0 | 0 | 0 | 0 | 0 | 0 |
| VVDLMAHN | GAPDH; GAPDH | P04406; E7E | G3P_HUMAN | Unmodified | 3749700 | 3570200 | 2636300 | 4416900 | 11973000 | 1560200 | 542200 | 0 | 23159000 | 1205700 | 26940000 | 2.97E+08 | 3661800 | 840150 | 9575800 | 5631200 | 3677900 | 6323100 |
| VVDLMAHN | GAPDH; GAPDH | P04406; E7E | G3P_HUMAN | Oxidation (M) | 2602200 | 1193300 | 3036500 | 1439100 | 984280 | 265760 | 70904 | 101130 | 4513500 | 599930 | 10417000 | 75552000 | 1032800 | 299780 | 4093400 | 3965200 | 433460 | 830960 |
| VVDLMAHN | GAPDH; GAPDH | P04406; E7E | G3P_HUMAN | 2 Oxidation (M) | 1648200 | 1436200 | 1744500 | 933890 | 665820 | 8466.1 | 204760 | 39421 | 1954100 | 0 | 3052300 | 33796000 | 2757400 | 57282 | 3145200 | 3764100 | 247070 | 321920 |
| VVDNTTAK | RAB1C; RAB1B | Q92928; Q9I | RAB1C_HUMAI | Unmodified | 0 | 0 | 0 | 0 | 1943900 | 0 | 1691900 | 0 | 0 | 998810 | 0 | 0 | 1676900 | 0 | 0 | 0 | 934340 | 0 |
| VVDPFSKK | RPS3A; RPS3A | P61247; H0 | RS3A_HUMAN | Unmodified | 0 | 0 | 0 | 0 | 0 | 0 | 0 | 0 | 0 | 0 | 517140 | 2670100 | 0 | 0 | 0 | 0 | 0 | 0 |
| VVDRDSEE | PARP1; PARP1 | P09874; A0A | PARP1_HUMAN | Unmodified | 0 | 262250 | 0 | 0 | 0 | 0 | 382870 | 526530 | 4076400 | 2083900 | 852170 | 8379500 | 0 | 0 | 0 | 0 | 0 | 0 |
| VVDSLYNK | PSMD11 | O00231 | PSD11_HUMAN | Unmodified | 0 | 0 | 0 | 0 | 0 | 0 | 0 | 2585700 | 0 | 0 | 0 | 0 | 0 | 0 | 0 | 0 | 0 | 0 |
| VVDSMEDE | FARSA; FARSA; | Q9Y285; K7E | SYFA_HUMAN | Unmodified | 0 | 0 | 0 | 0 | 0 | 0 | 0 | 0 | 0 | 0 | 0 | 565280 | 0 | 0 | 0 | 0 | 0 | 0 |
| VVDSMEDE | FARSA; FARSA; | Q9Y285; K7E | SYFA_HUMAN | Unmodified | 0 | 0 | 0 | 0 | 0 | 0 | 0 | 0 | 392720 | 0 | 52329 | 2023600 | 633180 | 0 | 0 | 0 | 0 | 0 |
| VVDTLYDG | TJP2; TJP2; TJP2 | Q9UDY2; A0 | ZO2_HUMAN T | Unmodified | 0 | 0 | 105370 | 0 | 0 | 0 | 0 | 0 | 4640100 | 200990 | 0 | 3860500 | 0 | 0 | 0 | 0 | 249770 | 158640 |
| VVDTLYNG | TJP1; TJP1; TJP1 | Q07157; A0A | ZO1_HUMAN T | Unmodified | 0 | 0 | 538320 | 431010 | 0 | 28096 | 0 | 0 | 0 | 75740 | 0 | 74717 | 0 | 0 | 0 | 619550 | 0 | 0 |
| VVDYSQFQ | NUCKS1 | Q9H1E3 | NUCKS_HUMA | Unmodified | 263080 | 0 | 0 | 0 | 1070900 | 0 | 0 | 0 | 644640 | 1278900 | 0 | 0 | 595490 | 0 | 0 | 0 | 715140 | 1037200 |
| VVDYSQFQ | NUCKS1 | Q9H1E3 | NUCKS_HUMA | Unmodified | 0 | 0 | 0 | 0 | 858430 | 0 | 0 | 0 | 1171000 | 992510 | 0 | 0 | 348830 | 0 | 0 | 0 | 540360 | 718850 |
| VVDYTTAK | RAB1A; RAB1A | P62820; E7E | RAB1A_HUMAN | Unmodified | 0 | 0 | 561030 | 659260 | 0 | 0 | 654140 | 407060 | 0 | 0 | 255360 | 4781400 | 77317 | 853330 | 0 | 257860 | 0 | 0 |
| VVEATAYGI | KIT | P10721 | KIT_HUMAN M | Unmodified | 0 | 0 | 0 | 0 | 0 | 0 | 0 | 0 | 1874900 | 351950 | 822150 | 4360700 | 0 | 0 | 0 | 0 | 0 | 390350 |
| VVEDGILK | CTNNA1; CTNN | P35221; E5R | CTNA1_HUMAN | Unmodified | 482280 | 745830 | 2507800 | 541290 | 1914400 | 904360 | 30855 | 0 | 450420 | 250150 | 767950 | 8293300 | 421850 | 725420 | 4610300 | 4759300 | 2131500 | 0 |

| | | | | | | | | | | | | | | | | | | | | | | |
|---|---|---|---|---|---|---|---|---|---|---|---|---|---|---|---|---|---|---|---|---|---|---|
| VVEEAPSIF | PCCA; PCCA; P | P05165; A0A | PCCA_HUMAN | Unmodified | 0 | 0 | 0 | 0 | 0 | 0 | 0 | 0 | 1371300 | 41260 | 0 | 30858 | 1617300 | 0 | 0 | 0 | 0 | 0 |
| VVEEEKQEI | MACF1; MACF1 | Q9UPN3; H3 | MACF1_HUMAI | Unmodified | 0 | 0 | 0 | 0 | 0 | 0 | 0 | 0 | 0 | 0 | 0 | 0 | 0 | 0 | 0 | 0 | 0 | 0 |
| VVEELTR | DDB1; NCS1; D | Q16531; P62 | DDB1_HUMAN | Unmodified | 0 | 0 | 0 | 0 | 0 | 0 | 0 | 0 | 0 | 0 | 0 | 3570500 | 0 | 0 | 0 | 0 | 0 | 0 |
| VVEEPNAF | GNPTG; GNPTG | Q9UJJ9; E9P | GNPTG_HUMA | Unmodified | 0 | 0 | 0 | 0 | 0 | 129170 | 0 | 0 | 3471400 | 262640 | 0 | 0 | 300000 | 0 | 0 | 0 | 0 | 2400100 |
| VVEEPYTVF | CLUH; CLUH; C | O75153; I3L | CLU_HUMAN C | Unmodified | 0 | 0 | 66039 | 176350 | 0 | 0 | 2263700 | 2180300 | 0 | 0 | 121040 | 2662300 | 0 | 1237600 | 0 | 71381 | 0 | 0 |
| VVEFLVK | CDKN2C | P42773 | CDN2C_HUMA | Unmodified | 0 | 0 | 0 | 0 | 0 | 147600 | 41649 | 0 | 623390 | 79287 | 0 | 0 | 0 | 0 | 0 | 0 | 0 | 0 |
| VVEGFTDEI | UBE3C | Q15386 | UBE3C_HUMAI | Unmodified | 0 | 0 | 0 | 0 | 0 | 0 | 0 | 0 | 446390 | 45845 | 0 | 1010800 | 0 | 0 | 0 | 0 | 0 | 0 |
| VVEGSFVYI | GDI1; GDI1 | P31150; G5E | GDIA_HUMAN | Unmodified | 0 | 535330 | 426090 | 343330 | 0 | 0 | 0 | 27035 | 1122600 | 275790 | 0 | 2449300 | 98692 | 0 | 1032100 | 1879300 | 0 | 0 |
| VVEGSFVYI | GDI1; GDI1 | P31150; G5E | GDIA_HUMAN | Unmodified | 0 | 0 | 0 | 0 | 0 | 0 | 0 | 0 | 489210 | 543010 | 0 | 0 | 452020 | 0 | 0 | 0 | 330030 | 1218200 |
| VVEIAPAAH | PC; PC; PC; PC; | P11498; E9P | PYC_HUMAN P | Unmodified | 0 | 0 | 0 | 0 | 0 | 0 | 0 | 0 | 1127100 | 0 | 0 | 816770 | 0 | 0 | 0 | 0 | 289250 | 0 |
| VVEINHEDI | SLC39A10; SLC | Q9ULF5; E7E | S39AA_HUMAN | Unmodified | 2068100 | 1429200 | 2225100 | 2766200 | 9586400 | 1566900 | 0 | 0 | 951960 | 0 | 0 | 39804000 | 3811900 | 574840 | 3327600 | 3479300 | 1331900 | 11899000 |
| VVEINPGLI | PIPOX; PIPOX; F | Q9P0Z9; K7E | SOX_HUMAN P | Unmodified | 0 | 0 | 0 | 0 | 0 | 0 | 0 | 0 | 1636000 | 676140 | 0 | 0 | 0 | 0 | 0 | 0 | 0 | 0 |
| VVEISRLQE | STX18; STX18 | Q9P2W9; D6 | STX18_HUMAN | Unmodified | 19334 | 0 | 0 | 0 | 160660 | 0 | 93595 | 0 | 523260 | 131290 | 284670 | 81462 | 0 | 0 | 0 | 24159 | 0 | 0 |
| VVEIVDEK | LARP1; LARP1; | Q6PKG0; H0 | LARP1_HUMAN | Unmodified | 694000 | 0 | 0 | 0 | 0 | 0 | 0 | 0 | 0 | 0 | 0 | 0 | 0 | 0 | 751850 | 878980 | 0 | 0 |
| VVELPKTDI | LIN7A; LIN7A; L | O14910; H0Y | LIN7A_HUMAN | Unmodified | 0 | 0 | 0 | 0 | 0 | 0 | 0 | 0 | 925320 | 0 | 0 | 0 | 0 | 0 | 0 | 0 | 0 | 0 |
| VVENLQDD | STAT3; STAT3; S | P40763; A0A | STAT3_HUMAN | Unmodified | 0 | 0 | 0 | 0 | 823900 | 0 | 0 | 0 | 0 | 0 | 171870 | 1205700 | 549370 | 0 | 0 | 0 | 0 | 1684200 |
| VVEQLEK | DPP3; DPP3; DI | Q9NY33; G3 | DPP3_HUMAN | Unmodified | 91977 | 0 | 0 | 0 | 902780 | 147910 | 292540 | 0 | 0 | 0 | 372180 | 2813000 | 491540 | 449730 | 0 | 0 | 821760 | 948870 |
| VVEQMCIT | PRNP; PRNP | P04156; A2A | PRIO_HUMAN I | Oxidation (M) | 269040 | 152870 | 141160 | 0 | 0 | 0 | 0 | 0 | 0 | 0 | 0 | 0 | 0 | 0 | 117900 | 130820 | 0 | 0 |
| VVEREEAVI | EPB41L2; EPB4 | O43491; E9F | E41L2_HUMAN | Unmodified | 0 | 0 | 0 | 0 | 14528000 | 2311000 | 0 | 0 | 0 | 0 | 0 | 0 | 0 | 0 | 0 | 0 | 6761700 | 11655000 |
| VVESPDFSI | SEC23IP | Q9Y6Y8 | S23IP_HUMAN | Unmodified | 0 | 0 | 0 | 0 | 522510 | 0 | 139410 | 0 | 0 | 0 | 0 | 1366800 | 0 | 0 | 0 | 0 | 0 | 0 |
| VVETEDAYI | RAB35; RAB35 | Q15286; F5H | RAB35_HUMAN | Unmodified | 4579300 | 6080800 | 11564000 | 3902800 | 0 | 0 | 0 | 0 | 775190 | 136750 | 1500300 | 11927000 | 595350 | 285210 | 11156000 | 9567400 | 96810 | 0 |
| VVEVLAGH | FASN | P49327 | FAS_HUMAN F | Unmodified | 1763200 | 2816800 | 967470 | 6904600 | 0 | 0 | 0 | 0 | 165760 | 0 | 288020 | 44253000 | 1569700 | 1325000 | 2154100 | 1837900 | 0 | 0 |
| VVEVWWD | GALNT7; GALN | Q86SF2; H0Y | GALT7_HUMAN | Unmodified | 0 | 0 | 0 | 0 | 0 | 0 | 0 | 0 | 1964000 | 144110 | 0 | 475960 | 0 | 0 | 0 | 0 | 0 | 0 |
| VVFAADGII | PLOD2; PLOD2; | O00469; C9J | PLOD2_HUMAI | Unmodified | 0 | 0 | 0 | 251220 | 0 | 0 | 0 | 0 | 2431100 | 612750 | 0 | 0 | 0 | 0 | 0 | 0 | 0 | 0 |
| VVFAADGII | PLOD2; PLOD2; | O00469; C9J | PLOD2_HUMAI | Unmodified | 0 | 0 | 0 | 0 | 0 | 0 | 0 | 0 | 1861300 | 593010 | 0 | 0 | 37157 | 0 | 0 | 0 | 0 | 0 |
| VVFDSAQR | APEH; APEH; AI | P13798; H0Y | ACPH_HUMAN | Unmodified | 89905 | 63062 | 149880 | 778890 | 0 | 0 | 0 | 367560 | 0 | 0 | 0 | 12581000 | 0 | 0 | 0 | 1116900 | 0 | 0 |
| VVFEQTK | TPI1; TPI1; TPI1 | P60174; U3K | TPIS_HUMAN T | Unmodified | 7902100 | 0 | 5098500 | 12345000 | 77463000 | 12816000 | 4955800 | 3028000 | 742910 | 7097100 | 12165000 | 48705000 | 69427000 | 30786000 | 11458000 | 7846800 | 25953000 | 39829000 |
| VVFEQTKVI | TPI1; TPI1; TPI1 | P60174; U3K | TPIS_HUMAN T | Unmodified | 0 | 0 | 0 | 0 | 0 | 0 | 3001300 | 1369800 | 0 | 0 | 0 | 0 | 0 | 5344400 | 0 | 0 | 16635000 | 3233300 |
| VVFEQTKVI | TPI1; TPI1; TPI1 | P60174; U3K | TPIS_HUMAN T | Unmodified | 0 | 0 | 0 | 0 | 2452800 | 0 | 0 | 0 | 0 | 463720 | 0 | 0 | 1670300 | 0 | 0 | 0 | 19897000 | 1490300 |
| VVFLTGAR | CHPF; CHPF | Q8IZ52; F8W | CHSS2_HUMAI | Unmodified | 0 | 0 | 0 | 0 | 0 | 0 | 0 | 0 | 712090 | 0 | 156140 | 0 | 0 | 0 | 0 | 0 | 0 | 0 |
| VVFMGDD | PIGO | Q8TEQ8 | PIGO_HUMAN | Oxidation (M) | 0 | 0 | 0 | 21077 | 20583 | 1449100 | 0 | 0 | 3307700 | 399470 | 383710 | 0 | 0 | 0 | 0 | 0 | 0 | 0 |
| VVFNLNTYI | ITGA2; ITGA2; I | P17301; E9P | ITA2_HUMAN I | Unmodified | 0 | 3501400 | 5458000 | 5167700 | 4811900 | 2590300 | 171670 | 0 | 0 | 0 | 0 | 981970 | 418960 | 0 | 9034400 | 8360600 | 2169100 | 2464200 |
| VVFQEFR | MYH9 | P35579 | MYH9_HUMAN | Unmodified | 652560 | 0 | 0 | 0 | 0 | 0 | 0 | 53073 | 0 | 0 | 0 | 10613000 | 0 | 0 | 1577500 | 1574900 | 0 | 0 |
| VVFTPSICK | LTBP1; LTBP1; L | Q14766; E7E | LTBP1_HUMAN | Unmodified | 0 | 0 | 0 | 0 | 0 | 0 | 0 | 510140 | 0 | 0 | 0 | 0 | 0 | 0 | 0 | 0 | 431390 | 1219100 |
| VVFVFGPD | PRDX6 | P30041 | PRDX6_HUMAI | Unmodified | 0 | 0 | 0 | 0 | 1208800 | 0 | 0 | 0 | 1162200 | 0 | 0 | 22543000 | 0 | 0 | 0 | 1947700 | 1537400 | 1348800 |
| VVGAGEFG | EPHA3; EPHA3 | P29320; C9J | EPHA3_HUMAI | Unmodified | 1837100 | 1221300 | 202050 | 0 | 0 | 0 | 0 | 0 | 0 | 0 | 0 | 0 | 0 | 0 | 0 | 0 | 0 | 0 |
| VVGAMQLY | CLTCL1; CLTC; | P53675; Q00 | CLH2_HUMAN | Oxidation (M) | 1090500 | 857400 | 671960 | 527380 | 0 | 0 | 0 | 0 | 0 | 45076 | 0 | 1549400 | 0 | 0 | 615630 | 803260 | 0 | 0 |
| VVGAMQLY | CLTCL1; CLTC; | P53675; Q00 | CLH2_HUMAN | Unmodified | 855330 | 1326300 | 0 | 363280 | 0 | 0 | 0 | 0 | 24369 | 0 | 0 | 495050 | 0 | 0 | 113590 | 425870 | 121190 | 0 |
| VVGAMQLY | CLTCL1; CLTC; | P53675; Q00 | CLH2_HUMAN | Unmodified | 6361900 | 7805100 | 0 | 4202600 | 0 | 0 | 0 | 0 | 1438800 | 0 | 0 | 13910000 | 0 | 0 | 10247000 | 3747100 | 0 | 0 |
| VVGAMQLY | CLTCL1; CLTC; | P53675; Q00 | CLH2_HUMAN | Oxidation (M) | 7160100 | 5305600 | 2294300 | 1572700 | 0 | 0 | 0 | 0 | 289580 | 0 | 0 | 6009800 | 0 | 0 | 4032600 | 5932400 | 74161 | 0 |
| VVGAQSLK | PGD; PGD; PGD | P52209; K7E | 6PGD_HUMAN | Unmodified | 197930 | 0 | 638270 | 391590 | 18553000 | 1171400 | 3061700 | 2139400 | 4636900 | 3194500 | 1794400 | 10147000 | 1950700 | 4877000 | 762770 | 714000 | 2382000 | 4309300 |
| VVGAQSLK | PGD; PGD; PGD | P52209; K7E | 6PGD_HUMAN | Unmodified | 0 | 0 | 0 | 0 | 1237300 | 187990 | 0 | 0 | 0 | 0 | 0 | 0 | 731630 | 0 | 0 | 0 | 540480 | 407040 |
| VVGAVGSD | PTGR1; PTGR1 | Q14914; F22 | PTGR1_HUMAN | Unmodified | 107500 | 0 | 0 | 0 | 8056000 | 668070 | 0 | 34315 | 475100 | 242600 | 0 | 0 | 0 | 0 | 258070 | 0 | 1821100 | 1070900 |
| VVGAVGSD | PTGR1; PTGR1 | Q14914; F22 | PTGR1_HUMAN | Unmodified | 0 | 0 | 0 | 0 | 1070600 | 0 | 54557 | 80150 | 0 | 214970 | 0 | 0 | 0 | 0 | 0 | 0 | 587030 | 0 |
| VVGCSCVV | RPS12 | P25398 | RS12_HUMAN | Unmodified | 2419000 | 1116800 | 403180 | 1249500 | 836400 | 0 | 0 | 0 | 679110 | 0 | 844790 | 17170000 | 1613300 | 195660 | 1757000 | 1480300 | 128170 | 274590 |
| VVGDREN | TPD52L2; TPD5 | O43399; A0A | TPD54_HUMAN | Unmodified | 0 | 0 | 0 | 0 | 1653400 | 0 | 0 | 117400 | 0 | 0 | 349290 | 491060 | 0 | 0 | 0 | 0 | 790920 | 0 |
| VVGDVAYD | MTHFD1; MTHF | P11586; V9G | C1TC_HUMAN | Unmodified | 191440 | 141880 | 487500 | 866260 | 2590400 | 539240 | 659210 | 620920 | 4301500 | 1281400 | 966480 | 7016600 | 1526000 | 1057200 | 767700 | 499070 | 1939100 | 1620000 |
| VVGDVAYD | MTHFD1; MTHF | P11586; V9G | C1TC_HUMAN | Unmodified | 0 | 31027 | 0 | 0 | 0 | 0 | 490070 | 0 | 56499 | 322160 | 0 | 283270 | 36279 | 128890 | 0 | 0 | 185450 | 0 |
| VVGEHVEV | HSPB6; HSPB6; | O14558; A0A | HSPB6_HUMAI | Unmodified | 522390 | 0 | 0 | 0 | 0 | 0 | 0 | 0 | 0 | 0 | 0 | 0 | 0 | 0 | 0 | 0 | 0 | 0 |
| VVGESIWK | SORL1; SORL1; | Q92673; E9F | SORL_HUMAN | Unmodified | 0 | 0 | 0 | 38421 | 26531 | 0 | 0 | 0 | 1978900 | 178900 | 0 | 0 | 0 | 0 | 147150 | 0 | 19260 | 0 |
| VVGFALQA | SNX33; SNX33 | Q8WV41; H3 | SNX33_HUMAN | Unmodified | 199740 | 152520 | 905510 | 413360 | 0 | 0 | 0 | 0 | 0 | 0 | 0 | 0 | 0 | 0 | 3519600 | 2100400 | 56009 | 0 |
| VVGFALQA | SNX33; SNX33 | Q8WV41; H3 | SNX33_HUMAN | Unmodified | 0 | 0 | 254120 | 676450 | 0 | 0 | 0 | 0 | 0 | 0 | 0 | 0 | 0 | 0 | 365030 | 190290 | 0 | 0 |
| VVGFHVLG | TXNRD1; TXNRI | Q16881; A0A | TRXR1_HUMAN | Unmodified | 0 | 0 | 287490 | 1228900 | 4015000 | 804500 | 0 | 0 | 24498000 | 1214800 | 5031100 | 24167000 | 1478100 | 0 | 1332900 | 890740 | 1962500 | 1533100 |
| VVGKEQG | RAP1B; RAP1A; | P61224; P62 | RAP1B_HUMAN | Unmodified | 984690 | 1524500 | 0 | 0 | 160860 | 0 | 0 | 0 | 0 | 10582 | 137850 | 418060 | 0 | 0 | 0 | 236480 | 25673 | 0 |
| VVGMEESC | SSH3 | Q8TE77 | SSH3_HUMAN | Unmodified | 0 | 0 | 0 | 0 | 376770 | 0 | 0 | 0 | 435130 | 0 | 150270 | 437970 | 0 | 0 | 0 | 0 | 0 | 444750 |
| VVGQTTPE | AKAP12 | Q02952 | AKA12_HUMAN | Unmodified | 0 | 0 | 0 | 0 | 2000800 | 380900 | 0 | 0 | 0 | 0 | 0 | 0 | 0 | 0 | 0 | 0 | 0 | 3653100 |
| VVGSEFVQ | PSMC4 | P43686 | PRS6B_HUMAN | Unmodified | 258190 | 268110 | 287610 | 0 | 2053700 | 52601 | 272990 | 0 | 0 | 262360 | 496230 | 7949700 | 926810 | 0 | 283920 | 576010 | 325900 | 782790 |
| VVGSELIQK | PSMC1 | P62191 | PRS4_HUMAN | Unmodified | 101910 | 0 | 1452800 | 0 | 1090800 | 188480 | 131760 | 148860 | 2632200 | 461560 | 2826800 | 2192400 | 0 | 0 | 0 | 0 | 283350 | 412190 |
| VVGSSGTQ | HSPG2 | P98160 | PGBM_HUMAN | Unmodified | 0 | 0 | 0 | 0 | 0 | 0 | 0 | 0 | 0 | 0 | 0 | 0 | 373100 | 0 | 0 | 654600 | 0 | 0 |
| VVGVLLGS | PSMD7; PSMD7 | P51665; H3E | PSMD7_HUMA | Unmodified | 0 | 0 | 0 | 0 | 0 | 0 | 0 | 0 | 0 | 1118700 | 0 | 2092600 | 0 | 0 | 0 | 0 | 0 | 0 |
| VVGVLLGS | PSMD7; PSMD7 | P51665; H3E | PSMD7_HUMA | Unmodified | 0 | 0 | 0 | 0 | 0 | 0 | 0 | 0 | 0 | 0 | 0 | 1729100 | 0 | 0 | 0 | 0 | 0 | 0 |
| VVGWRSG | SBDS; SBDS | Q9Y3A5; F8W | SBDS_HUMAN | Unmodified | 0 | 0 | 0 | 0 | 486720 | 0 | 322110 | 0 | 0 | 0 | 0 | 0 | 0 | 469000 | 0 | 0 | 223140 | 669510 |
| VVGYKGSV | CANT1 | Q8WVQ1 | CANT1_HUMAI | Unmodified | 0 | 0 | 0 | 0 | 0 | 0 | 0 | 0 | 6797300 | 1729200 | 0 | 0 | 0 | 0 | 0 | 0 | 0 | 0 |
| VVHASGDA | EPB41L3; EPB4 | Q9Y2J2; A0A | E41L3_HUMAN | Unmodified | 0 | 0 | 0 | 0 | 0 | 0 | 0 | 0 | 0 | 0 | 0 | 0 | 0 | 0 | 0 | 0 | 252740 | 4119600 |
| VVHLSQEIN | MYO5C | Q9NQX4 | MYO5C_HUMA | Unmodified | 0 | 0 | 0 | 0 | 601040 | 0 | 0 | 0 | 0 | 0 | 0 | 0 | 592740 | 207570 | 0 | 0 | 0 | 0 |
| VVHVNGY | DLD; DLD | P09622; E9P | DLDH_HUMAN | Unmodified | 126580 | 0 | 0 | 0 | 0 | 0 | 0 | 0 | 505160 | 45715 | 0 | 0 | 0 | 0 | 0 | 0 | 0 | 55356 |
| VVHWYFS | SMOC1 | Q9H4F8 | SMOC1_HUMA | Unmodified | 0 | 0 | 0 | 0 | 0 | 0 | 0 | 0 | 6568800 | 283520 | 0 | 0 | 256460 | 696510 | 0 | 0 | 0 | 0 |
| VVIDAFR | PSMD14 | O00487 | PSDE_HUMAN | Unmodified | 0 | 0 | 133850 | 0 | 0 | 0 | 0 | 0 | 0 | 0 | 0 | 3087200 | 32455 | 0 | 99194 | 0 | 0 | 0 |
| VVIEDEQLV | TWF2; TWF2; TV | Q6IBS0; D6F | TWF2_HUMAN | Unmodified | 0 | 0 | 0 | 0 | 0 | 0 | 0 | 0 | 0 | 0 | 0 | 1399600 | 159360 | 0 | 0 | 0 | 0 | 0 |
| VVIFQQEQI | PPP2R2A; PPP2 | P63151; E5R | 2ABA_HUMAN | Unmodified | 452360 | 203920 | 632470 | 393620 | 0 | 0 | 0 | 0 | 715720 | 0 | 1312000 | 5722700 | 0 | 0 | 724250 | 740450 | 0 | 0 |
| VVIGMDVA | ENO1; ENO1 | P06733; A0A | ENOA_HUMAN | Oxidation (M) | 0 | 0 | 0 | 147870 | 0 | 259620 | 304080 | 0 | 16108000 | 4060800 | 440890 | 19453000 | 1475800 | 45854 | 990530 | 200260 | 33439 | 177070 |
| VVIGMDVA | ENO1; ENO1 | P06733; A0A | ENOA_HUMAN | Unmodified | 0 | 0 | 0 | 0 | 0 | 0 | 0 | 0 | 85385 | 117980 | 0 | 0 | 150690 | 0 | 0 | 0 | 0 | 0 |
| VVIIESVLCI | ACTR10; ACTR1 | Q9NZ32; F6 | ARP10_HUMAN | Unmodified | 0 | 0 | 0 | 0 | 0 | 0 | 0 | 0 | 0 | 0 | 0 | 926930 | 0 | 0 | 0 | 0 | 0 | 0 |
| VVIIGAGKP | GANAB; GANAE | Q14697; F5H | GANAB_HUMA | Unmodified | 598040 | 0 | 1186200 | 1590800 | 0 | 0 | 0 | 0 | 7.27E+08 | 35560000 | 9259100 | 6447300 | 0 | 0 | 1680700 | 1391400 | 0 | 0 |
| VVILRYNEN | CIT; CIT; CIT; CI | O14578; A0A | CTRO_HUMAN | Unmodified | 0 | 0 | 0 | 0 | 0 | 0 | 0 | 0 | 0 | 0 | 0 | 0 | 0 | 0 | 0 | 0 | 2324700 | 0 |
| VVINGQEV | LTA4H; LTA4H | P09960; B4D | LKHA4_HUMAN | Unmodified | 3694000 | 16208000 | 37855 | 0 | 1584300 | 450190 | 0 | 0 | 1011000 | 10042 | 740080 | 7006900 | 0 | 0 | 1643800 | 898050 | 1239300 | 771280 |
| VVINVPIFK | GCLC; GCLC; G | P48506; A0A | GSH1_HUMAN | Unmodified | 312020 | 0 | 234120 | 0 | 9958900 | 2867300 | 439480 | 103620 | 761180 | 1357300 | 900980 | 5737000 | 1313800 | 893180 | 0 | 142740 | 513080 | 878570 |
| VVIQSNDD | USP9X | Q93008 | USP9X_HUMAN | Unmodified | 7887400 | 0 | 0 | 0 | 0 | 0 | 0 | 0 | 0 | 0 | 0 | 2146800 | 0 | 0 | 2667700 | 58679 | 0 | 0 |
| VVIRGAGN | GLT8D1 | Q68CQ7 | GL8D1_HUMAI | Unmodified | 0 | 0 | 0 | 0 | 0 | 0 | 0 | 0 | 3327700 | 2086200 | 577020 | 0 | 0 | 0 | 0 | 0 | 0 | 0 |
| VVISGFGDI | AGRN | O00468 | AGRIN_HUMAN | Unmodified | 2774300 | 366230 | 2119100 | 2307100 | 0 | 0 | 0 | 0 | 2557900 | 118430 | 473990 | 0 | 1309300 | 245530 | 0 | 1465900 | 0 | 430670 |
| VVIVNPETK | DIP2B | Q9P265 | DIP2B_HUMAN | Unmodified | 0 | 0 | 0 | 0 | 0 | 0 | 0 | 0 | 383820 | 199780 | 0 | 8933500 | 0 | 144470 | 280730 | 0 | 0 | 0 |
| VVKEEEAIA | RAB27A; RAB27 | P51159; H3E | RB27A_HUMAN | Unmodified | 0 | 0 | 2654000 | 1008700 | 0 | 0 | 0 | 0 | 0 | 0 | 0 | 0 | 0 | 0 | 1185700 | 823300 | 0 | 0 |
| VVKLENGE | APMAP; APMAP | Q9HDC9; H0 | APMAP_HUMAI | Unmodified | 0 | 0 | 0 | 1239000 | 0 | 0 | 0 | 0 | 3.09E+08 | 7336700 | 2806800 | 0 | 0 | 0 | 543270 | 470490 | 0 | 0 |
| VVKQASEG | GAPDH; GAPDH | P04406; E7E | G3P_HUMAN G | Unmodified | 0 | 0 | 0 | 0 | 975070 | 0 | 2559600 | 2584600 | 0 | 219470 | 0 | 0 | 2198600 | 2061200 | 0 | 0 | 6932000 | 1132400 |
| VVLAASPLF | PARP12; PARP1 | Q9H0J9; G5I | PAR12_HUMAN | Unmodified | 0 | 321590 | 990030 | 499890 | 0 | 0 | 0 | 0 | 0 | 0 | 0 | 0 | 0 | 0 | 743930 | 916710 | 0 | 0 |
| VVLAEVIQA | C12orf57; C12c | Q99622; F5C | C10_HUMAN P | Unmodified | 664400 | 719430 | 137300 | 838370 | 101330 | 18355 | 36615 | 0 | 690950 | 90322 | 274810 | 842360 | 188760 | 80908 | 1877000 | 804570 | 0 | 0 |
| VVLAPQDV | PTK7; PTK7 | Q13308; Q8 | PTK7_HUMAN I | Unmodified | 0 | 0 | 268420 | 131250 | 0 | 0 | 0 | 0 | 3555900 | 0 | 0 | 0 | 0 | 0 | 580310 | 487740 | 0 | 0 |
| VVLAYEPV | TPI1; TPI1; TPI1 | P60174; U3K | TPIS_HUMAN T | Unmodified | 19501000 | 16066000 | 8176000 | 21848000 | 2.18E+08 | 34946000 | 29674000 | 8739600 | 36937000 | 30892000 | 38368000 | 2.43E+08 | 1.72E+08 | 80365000 | 25102000 | 13889000 | 2.02E+08 | 1.29E+08 |
| VVLAYEPV | TPI1; TPI1 | P60174; U3K | TPIS_HUMAN T | Unmodified | 0 | 0 | 0 | 0 | 2382300 | 0 | 4721100 | 1533600 | 0 | 0 | 0 | 0 | 0 | 2539000 | 0 | 0 | 36268000 | 1839300 |
| VVLCELGN | ITGA2B; ITGA2E | P08514; A0A | ITA2B_HUMAN | Unmodified | 0 | 0 | 0 | 0 | 1427500 | 437020 | 2582300 | 792280 | 0 | 0 | 0 | 0 | 1540400 | 1424400 | 0 | 0 | 255030 | 798330 |
| VVLCELGN | ITGA2B; ITGA2E | P08514; A0A | ITA2B_HUMAN | Oxidation (M) | 1388000 | 1185800 | 1979200 | 3421800 | 0 | 0 | 1208500 | 970980 | 0 | 0 | 0 | 0 | 1656600 | 1809500 | 2405900 | 3261400 | 0 | 0 |
| VVLDDKDY | HSPE1; HSPE1 | P61604; B8Z | CH10_HUMAN | Unmodified | 108020 | 0 | 0 | 510140 | 291450 | 0 | 66044 | 20585 | 27386000 | 0 | 446430 | 1445000 | 125410 | 0 | 0 | 95612 | 0 | 0 |
| VVLDDKDY | HSPE1; HSPE1 | P61604; B8Z | CH10_HUMAN | Unmodified | 0 | 0 | 83806 | 0 | 0 | 0 | 0 | 59195 | 20465000 | 2290800 | 799180 | 0 | 1976400 | 739430 | 0 | 0 | 0 | 0 |
| VVLEAPDE | PTRHD1 | Q6GMV3 | PTRD1_HUMAN | Unmodified | 0 | 0 | 0 | 0 | 2932400 | 0 | 0 | 0 | 0 | 213920 | 0 | 10881000 | 2840000 | 0 | 0 | 920410 | 843860 | 1401100 |
| VVLEAPDE | PTRHD1 | Q6GMV3 | PTRD1_HUMAN | Unmodified | 0 | 0 | 0 | 0 | 6065500 | 62594 | 323360 | 0 | 0 | 763880 | 0 | 0 | 650910 | 4848600 | 0 | 0 | 5490600 | 1471000 |
| VVLFEMEAI | CPNE3; CPNE3 | O75131; H0Y | CPNE3_HUMAI | Oxidation (M) | 0 | 0 | 0 | 0 | 0 | 0 | 0 | 0 | 1315200 | 0 | 0 | 0 | 0 | 0 | 0 | 0 | 0 | 0 |
| VVLFEMEAI | CPNE3; CPNE3 | O75131; H0Y | CPNE3_HUMAI | Unmodified | 0 | 0 | 0 | 0 | 0 | 0 | 0 | 0 | 4991100 | 0 | 0 | 0 | 0 | 297080 | 0 | 0 | 0 | 0 |
| VVLGDQDI | HABP2 | Q14520 | HABP2_HUMAI | Unmodified | 0 | 0 | 0 | 0 | 124350 | 0 | 0 | 609250 | 0 | 0 | 24199 | 1891300 | 269680 | 0 | 0 | 0 | 0 | 0 |
| VVLGYLDD | PLD1 | Q13393 | PLD1_HUMAN | Unmodified | 0 | 0 | 0 | 0 | 0 | 0 | 0 | 0 | 0 | 0 | 0 | 2599700 | 0 | 0 | 766910 | 0 | 0 | 0 |
| VVLIGDSG | RAB11A; RAB11 | P62491; Q15 | RB11A_HUMAN | Unmodified | 8096100 | 8115600 | 17283000 | 30563000 | 56623000 | 15441000 | 14962000 | 9089100 | 11463000 | 6211400 | 15139000 | 45830000 | 14547000 | 16489000 | 14096000 | 18591000 | 2567100 | 11766000 |
| VVLIGDSG | RAB11A; RAB11 | P62491; Q15 | RB11A_HUMAN | Unmodified | 0 | 0 | 0 | 0 | 14592000 | 376280 | 1626400 | 754570 | 0 | 544970 | 0 | 0 | 7985800 | 5536000 | 0 | 0 | 205380 | 2674800 |
| VVLIGESG | RAB25 | P57735 | RAB25_HUMAN | Unmodified | 0 | 0 | 1653500 | 2499500 | 0 | 0 | 0 | 84033 | 0 | 0 | 1036400 | 5899600 | 0 | 0 | 1777100 | 2377600 | 0 | 228340 |
| VVLIGESG | RAB25 | P57735 | RAB25_HUMAN | Unmodified | 0 | 0 | 0 | 397920 | 0 | 0 | 0 | 0 | 0 | 0 | 0 | 0 | 301250 | 0 | 0 | 0 | 0 | 0 |
| VVLIGGKPI | HNRNPK; HNRI | P61978; Q5T | HNRPK_HUMA | Unmodified | 643000 | 0 | 0 | 0 | 0 | 0 | 308930 | 142150 | 0 | 132070 | 0 | 5176500 | 0 | 0 | 0 | 0 | 0 | 476000 |
| VVLLGEFLH | GAR1 | Q9NY12 | GAR1_HUMAN | Unmodified | 767340 | 0 | 0 | 29240 | 0 | 0 | 0 | 0 | 0 | 0 | 0 | 168100 | 0 | 0 | 0 | 0 | 0 | 0 |
| VVLLGEGC | RAB21 | Q9UL25 | RAB21_HUMAN | Unmodified | 927160 | 666810 | 1584000 | 1926500 | 1341800 | 0 | 415340 | 0 | 1798000 | 0 | 1458500 | 1591100 | 1404300 | 822280 | 2516000 | 1449500 | 0 | 0 |
| VVLLTGETS | SNRNP200 | O75643 | U520_HUMAN | Unmodified | 120770 | 0 | 0 | 0 | 0 | 0 | 0 | 0 | 0 | 0 | 0 | 2539500 | 0 | 0 | 0 | 0 | 0 | 0 |
| VVLNKADQ | EHD1; EHD3; EI | Q9H4M9; Q9 | EHD1_HUMAN | Oxidation (M) | 0 | 525750 | 0 | 0 | 0 | 0 | 0 | 0 | 0 | 0 | 0 | 3656400 | 0 | 0 | 642700 | 803370 | 0 | 0 |
| VVLNKADQ | EHD1; EHD3; EI | Q9H4M9; Q9 | EHD1_HUMAN | Unmodified | 171390 | 255750 | 0 | 0 | 0 | 0 | 0 | 0 | 0 | 0 | 0 | 1831200 | 0 | 0 | 390970 | 187720 | 0 | 0 |

| | | | | | | | | | | | | | | | | | | | | | | |
|---|---|---|---|---|---|---|---|---|---|---|---|---|---|---|---|---|---|---|---|---|---|---|
| VVLNKADQ | EHD4 | Q9H223 | EHD4_HUMAN | Oxidation (M) | 264360 | 357500 | 0 | 523780 | 400750 | 0 | 0 | 0 | 0 | 0 | 0 | 470580 | 0 | 0 | 589770 | 505220 | 33388 | 0 |
| VVLNKADQ | EHD4 | Q9H223 | EHD4_HUMAN | Unmodified | 0 | 439800 | 0 | 415490 | 0 | 0 | 0 | 0 | 0 | 170650 | 0 | 871680 | 0 | 0 | 826770 | 414060 | 0 | 0 |
| VVLPIEAPIF | MT-CO2 | P00403 | COX2_HUMAN | Unmodified | 37023 | 0 | 148020 | 0 | 0 | 0 | 0 | 28478 | 6588800 | 885090 | 620570 | 423860 | 0 | 0 | 0 | 0 | 0 | 0 |
| VVLPTFILEI | OSBPL8; OSBPI | Q9BZF1; F8V | OSBL8_HUMAN | Unmodified | 115770 | 0 | 0 | 0 | 0 | 0 | 0 | 0 | 0 | 0 | 39743 | 880410 | 0 | 0 | 0 | 0 | 0 | 0 |
| VVLPTLASL | PLCB3 | Q01970 | PLCB3_HUMAN | Unmodified | 218420 | 0 | 0 | 143740 | 0 | 0 | 0 | 10285 | 0 | 0 | 832610 | 4448600 | 31387 | 0 | 0 | 145110 | 0 | 0 |
| VVLPYLVPK | GCN1 | Q92616 | GCN1_HUMAN | Unmodified | 0 | 0 | 0 | 0 | 0 | 0 | 0 | 0 | 0 | 0 | 0 | 1006700 | 0 | 0 | 0 | 0 | 0 | 0 |
| VVLQDEDI | PLXNA2 | O75051 | PLXA2_HUMAN | Unmodified | 0 | 0 | 0 | 0 | 0 | 0 | 0 | 0 | 744960 | 197980 | 0 | 612910 | 0 | 0 | 0 | 0 | 0 | 0 |
| VVLQDEDI | PLXNA2 | O75051 | PLXA2_HUMAN | Unmodified | 0 | 0 | 0 | 0 | 0 | 0 | 0 | 0 | 8995400 | 1584200 | 0 | 0 | 0 | 0 | 0 | 0 | 144890 | 0 |
| VVLQELR | OR56B2P; UBE2 | Q8NGI1; Q1 | O56B2_HUMAN | Unmodified | 732160 | 754130 | 610770 | 409680 | 1093800 | 0 | 0 | 0 | 797600 | 953870 | 432060 | 3943000 | 1933800 | 1621800 | 1495500 | 1101400 | 451140 | 829210 |
| VVLRGVGG | ANPEP | P15144 | AMPN_HUMAN | Unmodified | 0 | 0 | 0 | 0 | 0 | 0 | 0 | 0 | 0 | 0 | 0 | 0 | 0 | 0 | 0 | 0 | 2507400 | 6188700 |
| VVLSKLPIG | CCT7 | Q99832 | TCPH_HUMAN | Unmodified | 0 | 0 | 0 | 0 | 0 | 0 | 0 | 41114 | 0 | 65211 | 34998 | 497260 | 0 | 0 | 0 | 0 | 0 | 0 |
| VVLSSQDK | NOMO1; NOMC | Q15155; P69 | NOMO1_HUMA | Unmodified | 0 | 0 | 0 | 0 | 40474 | 236950 | 66636 | 48837 | 4224400 | 219910 | 332630 | 438530 | 158960 | 72438 | 43615 | 0 | 255360 | 0 |
| VVLTAEVS( | SORT1; SORT1; | Q99523; A0A | SORT_HUMAN | Unmodified | 31630 | 33521 | 0 | 0 | 0 | 0 | 0 | 0 | 9793000 | 948990 | 1296600 | 4673200 | 0 | 0 | 0 | 0 | 0 | 0 |
| VVLTAEVS( | SORT1; SORT1; | Q99523; A0A | SORT_HUMAN | Unmodified | 0 | 0 | 0 | 0 | 0 | 0 | 0 | 0 | 8043600 | 826860 | 2236100 | 255690 | 0 | 0 | 0 | 0 | 0 | 0 |
| VVLTDPEA | GBA; GBA; GBA | P04062; A0A | GLCM_HUMAN | Unmodified | 0 | 0 | 0 | 110300 | 0 | 0 | 506030 | 417180 | 4182100 | 866350 | 160810 | 0 | 0 | 0 | 574140 | 162620 | 289930 | 0 |
| VVLTQANK | SEC31A; SEC31 | O94979; H0 | SC31A_HUMAN | Unmodified | 0 | 0 | 0 | 0 | 0 | 0 | 628090 | 788920 | 546880 | 971410 | 0 | 0 | 2093200 | 0 | 0 | 0 | 465350 | 0 |
| VVLVLAGR | RPL27; RPL27; | P61353; K7E | RL27_HUMAN ( | Unmodified | 14517000 | 11630000 | 8240300 | 5966600 | 0 | 0 | 0 | 0 | 1321800 | 0 | 1639900 | 50033000 | 479660 | 0 | 8883000 | 7714600 | 0 | 0 |
| VVLVNNIL( | SNAPIN | O95295 | SNAPN_HUMA | Unmodified | 37967 | 35781 | 34320 | 46162 | 0 | 0 | 0 | 0 | 170210 | 0 | 122400 | 964350 | 0 | 0 | 65830 | 29606 | 14349 | 0 |
| VVLVSSASI | UTRN | P46939 | UTRN_HUMAN | Unmodified | 0 | 0 | 2890700 | 2284100 | 382820 | 0 | 0 | 0 | 5419400 | 362800 | 0 | 302230 | 273540 | 0 | 2725700 | 2542400 | 201800 | 0 |
| VVLVSSASI | UTRN | P46939 | UTRN_HUMAN | Unmodified | 0 | 0 | 0 | 0 | 6934200 | 0 | 0 | 0 | 0 | 1454800 | 0 | 0 | 657180 | 549980 | 0 | 0 | 1038600 | 2516700 |
| VVLVTGAG | HSD17B4; HSD | P51659; A0A | DHB4_HUMAN | Unmodified | 0 | 0 | 0 | 0 | 201070 | 0 | 0 | 0 | 26139000 | 718010 | 1437400 | 0 | 0 | 0 | 116330 | 106670 | 559260 | 0 |
| VVMDGVIS | P3H1; P3H1 | Q32P28; H7 | P3H1_HUMAN | Oxidation (M) | 23593 | 0 | 0 | 0 | 0 | 0 | 0 | 0 | 683990 | 0 | 0 | 0 | 0 | 0 | 0 | 0 | 0 | 0 |
| VVMDGVIS | P3H1; P3H1 | Q32P28; H7 | P3H1_HUMAN | Unmodified | 0 | 0 | 0 | 0 | 0 | 0 | 127650 | 234190 | 4349400 | 61974 | 0 | 0 | 0 | 0 | 0 | 0 | 0 | 0 |
| VVMITGDN | ATP2A3 | Q93084 | AT2A3_HUMAN | Unmodified | 0 | 0 | 0 | 0 | 717850 | 0 | 0 | 0 | 0 | 0 | 0 | 642070 | 0 | 0 | 0 | 0 | 453430 | 2170100 |
| VVMRVDFN | PGK1 | P00558 | PGK1_HUMAN | Oxidation (M) | 0 | 0 | 0 | 0 | 329300 | 0 | 133290 | 0 | 0 | 0 | 0 | 0 | 1579000 | 0 | 0 | 0 | 3088400 | 810860 |
| VVMRVDFN | PGK1 | P00558 | PGK1_HUMAN | Oxidation (M) | 0 | 0 | 0 | 0 | 0 | 0 | 0 | 0 | 0 | 0 | 0 | 0 | 0 | 0 | 0 | 0 | 609410 | 242580 |
| VVMRVDFN | PGK1 | P00558 | PGK1_HUMAN | Unmodified | 0 | 0 | 0 | 0 | 0 | 0 | 0 | 0 | 0 | 0 | 0 | 0 | 0 | 0 | 0 | 0 | 540580 | 208810 |
| VVNAFRSS | ATP2B1; ATP2B | P20020; E7E | AT2B1_HUMAN | Unmodified | 0 | 0 | 0 | 0 | 0 | 0 | 40693 | 0 | 0 | 245550 | 0 | 0 | 1488000 | 0 | 0 | 0 | 0 | 1365500 |
| VVNAPIFH | OGDH; OGDH; | Q02218; E9F | ODO1_HUMAN | Unmodified | 0 | 0 | 0 | 0 | 1354700 | 0 | 0 | 0 | 974030 | 0 | 0 | 0 | 0 | 0 | 0 | 0 | 0 | 0 |
| VVNCARG( | PHGDH; PHGD | O43175; A0A | SERA_HUMAN | Unmodified | 0 | 0 | 0 | 0 | 0 | 0 | 0 | 0 | 0 | 0 | 0 | 0 | 1486200 | 485960 | 0 | 0 | 0 | 297350 |
| VVNDDCTI | DPYSL5; DPYSL | Q9BPU6; E9 | DPYL5_HUMAN | Unmodified | 0 | 0 | 0 | 0 | 0 | 0 | 0 | 0 | 0 | 0 | 316060 | 2288700 | 0 | 0 | 0 | 0 | 0 | 0 |
| VVNEINIED | CISD2; CISD2 | Q8N5K1; D6 | CISD2_HUMAN | Unmodified | 0 | 0 | 0 | 0 | 0 | 0 | 301570 | 0 | 4073000 | 1106700 | 812400 | 0 | 0 | 0 | 0 | 0 | 683430 | 0 |
| VVNGPEILI | NSF; NSF | P46459; I3L( | NSF_HUMAN V | Unmodified | 0 | 0 | 450900 | 560710 | 976910 | 0 | 0 | 0 | 841290 | 115420 | 0 | 1198300 | 4197400 | 1341400 | 0 | 472200 | 595370 | 821200 |
| VVNIIPVIAK | SEPTIN3; SEPTI | Q9UH03; A0 | SEPT3_HUMAN | Oxidation (M) | 0 | 0 | 0 | 0 | 0 | 0 | 1008400 | 0 | 0 | 0 | 0 | 0 | 478870 | 0 | 0 | 0 | 0 | 0 |
| VVNIVPVIA | SEPTIN9; SEPTI | Q9UHD8; K7 | SEPT9_HUMAN | Unmodified | 0 | 184180 | 0 | 1738900 | 15185000 | 0 | 1928000 | 1391300 | 2267100 | 1421500 | 3738500 | 18081000 | 2640200 | 2009000 | 1461800 | 692280 | 6714300 | 8568600 |
| VVNIVPVIA | SEPTIN9; SEPTI | Q9UHD8; K7 | SEPT9_HUMAN | Unmodified | 0 | 0 | 0 | 0 | 2558900 | 0 | 563690 | 0 | 0 | 0 | 0 | 0 | 0 | 0 | 0 | 0 | 3707400 | 3543900 |
| VVNKESLVI | INSR | P06213 | INSR_HUMAN I | Unmodified | 0 | 0 | 0 | 34587 | 0 | 0 | 0 | 0 | 761680 | 0 | 0 | 0 | 137100 | 0 | 0 | 0 | 67677 | 0 |
| VVNLLQER | SEC24A | O95486 | SC24A_HUMAN | Unmodified | 0 | 0 | 0 | 0 | 0 | 0 | 0 | 0 | 0 | 0 | 0 | 736780 | 0 | 0 | 0 | 0 | 0 | 0 |
| VVNPYYLR | PRKAA1 | Q13131 | AAPK1_HUMAN | Unmodified | 217420 | 448700 | 0 | 280600 | 0 | 0 | 0 | 0 | 679620 | 420790 | 0 | 361300 | 0 | 574590 | 190370 | 189870 | 295690 | 720160 |
| VVNSDPVE | EHHADH | Q08426 | ECHP_HUMAN | Unmodified | 0 | 0 | 0 | 0 | 152280 | 0 | 0 | 0 | 2587800 | 0 | 0 | 0 | 0 | 0 | 0 | 0 | 0 | 0 |
| VVNSILAFR | ACY1; ABHD14 | Q03154; A0A | ACY1_HUMAN | Unmodified | 0 | 0 | 0 | 0 | 1062700 | 0 | 0 | 0 | 0 | 1064300 | 0 | 809520 | 881490 | 564310 | 0 | 0 | 0 | 262070 |
| VVNTQCG | TSPAN14; TSPA | Q8NG11; H7 | TSN14_HUMAN | Unmodified | 2423300 | 1063800 | 3480400 | 3666200 | 0 | 0 | 28917 | 21139 | 996730 | 225500 | 764450 | 4765600 | 76253 | 68287 | 3528600 | 2856700 | 0 | 0 |
| VVNVANVG | AP3B1; AP3B1 | O00203; H0 | AP3B1_HUMAN | Unmodified | 0 | 0 | 0 | 0 | 0 | 0 | 0 | 0 | 0 | 0 | 244610 | 3592400 | 0 | 0 | 0 | 0 | 0 | 0 |
| VVNVANVG | AP3B1 | O00203 | AP3B1_HUMAN | Unmodified | 0 | 0 | 0 | 0 | 1902700 | 0 | 644130 | 514450 | 0 | 476060 | 451430 | 0 | 1096200 | 806150 | 0 | 0 | 1159800 | 544800 |
| VVNVELPIE | HSPH1; HSPH1 | Q92598; A0A | HS105_HUMAN | Unmodified | 0 | 0 | 0 | 0 | 0 | 0 | 0 | 0 | 0 | 0 | 0 | 0 | 94194 | 20739 | 38012 | 0 | 0 | 0 |
| VVNVSSIM | CBR1 | P16152 | CBR1_HUMAN | Unmodified | 0 | 0 | 0 | 438910 | 3185800 | 793310 | 0 | 0 | 1311200 | 533020 | 96216 | 2762800 | 548550 | 0 | 401600 | 375120 | 1350600 | 1501400 |
| VVNVSSIM | CBR1 | P16152 | CBR1_HUMAN | Unmodified | 0 | 0 | 0 | 0 | 0 | 0 | 327750 | 247670 | 0 | 0 | 0 | 0 | 42822 | 116770 | 0 | 0 | 721910 | 73851 |
| VVNYLPR | LRBA; LRBA; LR | P50851; A0A | LRBA_HUMAN | Unmodified | 0 | 0 | 0 | 0 | 0 | 0 | 0 | 0 | 0 | 72355 | 0 | 1025000 | 0 | 0 | 0 | 27689 | 0 | 0 |
| VVPCLVTP | FLNB; FLNB | O75369; E7E | FLNB_HUMAN | Unmodified | 0 | 0 | 0 | 0 | 9125100 | 1648800 | 0 | 72369 | 0 | 0 | 0 | 943500 | 0 | 0 | 0 | 0 | 1636900 | 1511900 |
| VVPEMTEIL | ETFA; ETFA; ETF | P13804; H0 | ETFA_HUMAN I | Unmodified | 0 | 0 | 95999 | 36632 | 0 | 51715 | 0 | 28632 | 3198100 | 331340 | 0 | 0 | 41607 | 0 | 23970 | 0 | 0 | 73178 |
| VVPGFIVQ( | CWC27; CWC2 | Q6UX04; D6 | CWC27_HUMA | Unmodified | 0 | 0 | 0 | 0 | 0 | 0 | 0 | 0 | 0 | 0 | 0 | 1367900 | 0 | 0 | 0 | 0 | 0 | 0 |
| VVPGYGHA | CS; CS; CS; CS | O75390; A0A | CISY_HUMAN ( | Unmodified | 0 | 0 | 0 | 0 | 0 | 0 | 0 | 0 | 1569700 | 0 | 0 | 0 | 0 | 0 | 0 | 0 | 0 | 0 |
| VVPGYGHA | CS; CS; CS; CS | O75390; A0A | CISY_HUMAN ( | Unmodified | 0 | 0 | 0 | 0 | 0 | 0 | 0 | 0 | 519540 | 0 | 0 | 0 | 0 | 0 | 0 | 0 | 0 | 0 |
| VVPIASLTP | RPA1 | P27694 | RFA1_HUMAN | Unmodified | 248280 | 0 | 126060 | 0 | 0 | 0 | 0 | 0 | 0 | 0 | 225400 | 3968000 | 0 | 0 | 0 | 0 | 0 | 0 |
| VVPLADIITI | PDXK; PDXK | O00764; F2Z | PDXK_HUMAN | Unmodified | 201180 | 0 | 283870 | 242420 | 6263900 | 1036000 | 0 | 0 | 0 | 403600 | 266040 | 2234900 | 2375200 | 0 | 21048 | 1829000 | 2006700 | 2685400 |
| VVPPLDED | CDK2; CDK2; C | P24941; G3 | CDK2_HUMAN | Unmodified | 0 | 0 | 0 | 0 | 654560 | 331410 | 0 | 0 | 1357600 | 309170 | 0 | 0 | 0 | 0 | 0 | 0 | 159480 | 0 |
| VVPQQSLV | PROM2 | Q8N271 | PROM2_HUMA | Unmodified | 1288400 | 792960 | 0 | 1016500 | 0 | 0 | 0 | 0 | 0 | 0 | 0 | 1651500 | 0 | 0 | 1335900 | 1041000 | 0 | 0 |
| VVPSFLPVI | DSG2 | Q14126 | DSG2_HUMAN | Unmodified | 234480 | 6480500 | 3173800 | 2725400 | 1383800 | 415480 | 0 | 0 | 311010 | 295070 | 5682100 | 35908000 | 548960 | 0 | 1494100 | 11039000 | 89780 | 36095 |
| VVPTTDHII | TIPRL | O75663 | TIPRL_HUMAN | Unmodified | 364240 | 198520 | 292280 | 877020 | 0 | 144760 | 119840 | 84408 | 800090 | 0 | 1109000 | 4021900 | 601770 | 261400 | 474270 | 323130 | 244490 | 0 |
| VVPTVYMK | ERGIC3; ERGIC | Q9Y282; A0A | ERGI3_HUMAN | Unmodified | 0 | 0 | 0 | 0 | 120360 | 0 | 0 | 0 | 2517100 | 0 | 0 | 0 | 0 | 0 | 0 | 0 | 0 | 0 |
| VVPVRLDL | GGH; GGH; GG | Q92820; A0A | GGH_HUMAN ( | Unmodified | 0 | 0 | 589130 | 0 | 0 | 0 | 0 | 0 | 51620000 | 3225200 | 4209000 | 1873900 | 62494 | 0 | 0 | 0 | 0 | 0 |
| VVPVRLDL | GGH; GGH; GG | Q92820; A0A | GGH_HUMAN ( | Unmodified | 13367 | 0 | 0 | 0 | 0 | 0 | 0 | 0 | 8784400 | 8761400 | 50424 | 0 | 135200 | 0 | 0 | 0 | 0 | 0 |
| VVPYFTQTI | HSPG2; HSPG2 | P98160; A0A | PGBM_HUMAN | Unmodified | 0 | 0 | 166590 | 0 | 0 | 0 | 0 | 0 | 184910 | 0 | 0 | 0 | 0 | 0 | 0 | 99188 | 0 | 0 |
| VVQDAQTF | NAGA | P17050 | NAGAB_HUMA | Unmodified | 0 | 0 | 0 | 0 | 0 | 0 | 0 | 86678 | 825990 | 0 | 123520 | 0 | 0 | 0 | 0 | 0 | 0 | 0 |
| VVQDFIDIF | PTPRE | P23469 | PTPRE_HUMAN | Unmodified | 0 | 0 | 0 | 0 | 292810 | 15587 | 0 | 0 | 0 | 0 | 0 | 0 | 0 | 0 | 0 | 0 | 0 | 0 |
| VVQDTYQII | HGS; HGS; HG | O14964; A0A | HGS_HUMAN I | Oxidation (M) | 240420 | 205700 | 161570 | 0 | 0 | 0 | 0 | 0 | 290320 | 0 | 0 | 0 | 0 | 0 | 94105 | 131230 | 0 | 0 |
| VVQETVLVI | EPB41L3; EPB4 | Q9Y2J2; A0A | E41L3_HUMAN | Unmodified | 0 | 0 | 0 | 0 | 0 | 0 | 0 | 0 | 151050 | 0 | 0 | 0 | 0 | 153530 | 0 | 0 | 565230 | 3316600 |
| VVQETVLVI | EPB41L3; EPB4 | Q9Y2J2; A0A | E41L3_HUMAN | Unmodified | 0 | 0 | 0 | 0 | 0 | 0 | 0 | 0 | 0 | 0 | 0 | 0 | 0 | 0 | 0 | 0 | 524310 | 4537100 |
| VVQETVLVI | EPB41L3; EPB4 | Q9Y2J2; A0A | E41L3_HUMAN | Unmodified | 0 | 0 | 0 | 0 | 0 | 0 | 0 | 0 | 0 | 0 | 0 | 0 | 583220 | 0 | 0 | 0 | 6185700 | 31967000 |
| VVQEYIDAI | PTPRA | P18433 | PTPRA_HUMAN | Unmodified | 0 | 0 | 0 | 185920 | 0 | 0 | 0 | 0 | 0 | 118880 | 0 | 387870 | 0 | 473830 | 898540 | 0 | 767160 | 1231700 |
| VVQLHEYS | SMC1A; SMC1A | Q14683; G8 | SMC1A_HUMA | Unmodified | 0 | 0 | 0 | 0 | 0 | 0 | 0 | 0 | 0 | 54312 | 0 | 733560 | 0 | 0 | 0 | 0 | 0 | 0 |
| VVQMTNDI | EPHA2 | P29317 | EPHA2_HUMAN | Oxidation (M) | 0 | 0 | 0 | 0 | 0 | 149840 | 0 | 0 | 0 | 0 | 0 | 0 | 0 | 0 | 0 | 0 | 377340 | 365300 |
| VVQMTNDI | EPHA2 | P29317 | EPHA2_HUMAN | Unmodified | 0 | 0 | 0 | 0 | 1395100 | 562540 | 0 | 206690 | 0 | 0 | 0 | 573820 | 0 | 196080 | 0 | 0 | 1896200 | 2785200 |
| VVQMTNDI | EPHA2 | P29317 | EPHA2_HUMAN | Unmodified | 863350 | 394900 | 192110 | 31306 | 676690 | 285770 | 0 | 0 | 198930 | 0 | 244500 | 2684700 | 0 | 143090 | 307230 | 200560 | 1930300 | 1567700 |
| VVQPTLEQ | AOC1 | P19801 | AOC1_HUMAN | Unmodified | 0 | 0 | 0 | 228750 | 0 | 0 | 0 | 0 | 1575600 | 100660 | 0 | 0 | 0 | 0 | 0 | 18189 | 76204 | 0 |
| VVQQLRLE | GNG5 | P63218 | GBG5_HUMAN | Unmodified | 0 | 0 | 0 | 294160 | 948260 | 131530 | 373120 | 169000 | 1061600 | 985480 | 1266900 | 0 | 1544200 | 458300 | 0 | 0 | 863650 | 1108600 |
| VVQQLRLE | GNG5 | P63218 | GBG5_HUMAN | Unmodified | 0 | 0 | 0 | 0 | 26675000 | 4386500 | 20888000 | 4492200 | 1695200 | 18516000 | 2541700 | 509790 | 54891000 | 33529000 | 0 | 0 | 22350000 | 32835000 |
| VVQSLEQT | TSN; TSN; TSN; | Q15631; H7 | TSN_HUMAN T | Unmodified | 0 | 0 | 0 | 445510 | 0 | 0 | 0 | 0 | 0 | 0 | 447170 | 3658900 | 0 | 0 | 0 | 0 | 0 | 0 |
| VVQSLVIAK | PTPRB; PTPRB; | P23467; F8V | PTPRB_HUMAN | Unmodified | 396600 | 0 | 0 | 0 | 0 | 0 | 0 | 0 | 0 | 0 | 0 | 0 | 0 | 0 | 0 | 0 | 0 | 0 |
| VVQSMTQL | UBE2O; UBE2O | Q9C0C9; K7 | UBE2O_HUMA | Unmodified | 0 | 0 | 0 | 0 | 0 | 0 | 0 | 0 | 0 | 0 | 0 | 1718900 | 0 | 0 | 0 | 0 | 0 | 0 |
| VVQTDETA | UBA6 | A0AVT1 | UBA6_HUMAN | Unmodified | 0 | 0 | 0 | 0 | 0 | 0 | 0 | 0 | 0 | 0 | 134670 | 1634200 | 0 | 0 | 31095 | 53810 | 0 | 0 |
| VVQTKYVE | PFDN5; PFDN5 | Q99471; A0A | PFD5_HUMAN | Unmodified | 0 | 0 | 0 | 0 | 0 | 0 | 0 | 0 | 0 | 0 | 0 | 0 | 0 | 0 | 0 | 0 | 1463600 | 0 |
| VVQVSAGD | RCC1; RCC1; R | P18754; C9J | RCC1_HUMAN | Unmodified | 0 | 0 | 0 | 0 | 0 | 0 | 0 | 0 | 0 | 0 | 0 | 559660 | 0 | 0 | 0 | 0 | 350720 | 528650 |
| VVQYDGYI | GLCE; GLCE | O94923; H0 | GLCE_HUMAN | Unmodified | 0 | 0 | 0 | 0 | 0 | 0 | 90458 | 30315 | 2738300 | 834430 | 589730 | 1606000 | 0 | 0 | 0 | 0 | 0 | 0 |
| VVRELQAQ | MYH10 | P35580 | MYH10_HUMA | Unmodified | 0 | 0 | 0 | 0 | 6091200 | 0 | 0 | 0 | 27648 | 272790 | 115610 | 0 | 2612400 | 0 | 0 | 0 | 0 | 0 |
| VVRFEVIPC | TM9SF4; TM9SF | Q92544; A0A | TM9S4_HUMAN | Unmodified | 0 | 0 | 0 | 0 | 0 | 0 | 0 | 0 | 10917000 | 342260 | 0 | 577030 | 0 | 0 | 0 | 0 | 0 | 0 |
| VVRFGDED | C1orf198; C1or | Q9H425; E5 | CA198_HUMAN | Unmodified | 0 | 0 | 0 | 0 | 3203300 | 877070 | 2170000 | 70849 | 628540 | 810150 | 551810 | 813100 | 0 | 1498000 | 0 | 0 | 602010 | 1859300 |
| VVRGEGMI | DNAJA2 | O60884 | DNJA2_HUMAN | Unmodified | 393170 | 392040 | 297590 | 822560 | 1828300 | 0 | 0 | 0 | 3145500 | 611380 | 0 | 1531700 | 642480 | 266750 | 353180 | 290030 | 158330 | 0 |
| VVSDRWLS | SNRNP200; SN | O75643; A0A | U520_HUMAN | Unmodified | 129240 | 0 | 0 | 0 | 0 | 0 | 0 | 0 | 0 | 0 | 0 | 979520 | 0 | 0 | 0 | 0 | 0 | 0 |
| VVSEDFLQ | PARP1; PARP1 | P09874; A0A | PARP1_HUMAN | Unmodified | 1195900 | 689330 | 182430 | 132000 | 0 | 0 | 0 | 127610 | 696740 | 0 | 906840 | 0 | 0 | 0 | 0 | 0 | 46269 | 0 |
| VVSEGHTL | ADAM15 | Q13444 | ADA15_HUMAN | Unmodified | 0 | 0 | 86865 | 0 | 0 | 0 | 0 | 0 | 251810 | 0 | 26141 | 0 | 0 | 0 | 0 | 0 | 0 | 0 |
| VVSEIPVLK | UFC1 | Q9Y3C8 | UFC1_HUMAN | Unmodified | 54548 | 0 | 0 | 16514 | 386560 | 138380 | 0 | 0 | 510930 | 124050 | 1293500 | 4639000 | 0 | 0 | 29624 | 0 | 140220 | 144610 |
| VVSEQKDT | FGFBP1 | Q14512 | FGFP1_HUMAN | Unmodified | 0 | 0 | 0 | 0 | 0 | 0 | 0 | 0 | 0 | 0 | 0 | 0 | 0 | 0 | 0 | 0 | 669840 | 254320 |
| VVSGAANV | FAM3A; FAM3A; | P98173; D3 | FAM3A_HUMAN | Unmodified | 0 | 0 | 0 | 0 | 0 | 0 | 0 | 0 | 7442500 | 0 | 0 | 0 | 0 | 0 | 0 | 0 | 0 | 0 |
| VVSHLIEK | PKP3 | Q9Y446 | PKP3_HUMAN | Unmodified | 0 | 0 | 0 | 0 | 0 | 0 | 0 | 0 | 0 | 0 | 125420 | 587910 | 0 | 0 | 0 | 0 | 0 | 0 |
| VVSHLIEKL | PKP3 | Q9Y446 | PKP3_HUMAN | Unmodified | 237830 | 0 | 0 | 729170 | 0 | 0 | 0 | 0 | 0 | 165030 | 317060 | 1920600 | 303800 | 0 | 425850 | 521960 | 123870 | 0 |
| VVSHPECR | RACGAP1 | Q9H0H5 | RGAP1_HUMAN | Unmodified | 0 | 0 | 0 | 0 | 494680 | 355080 | 890380 | 0 | 0 | 0 | 0 | 0 | 0 | 0 | 0 | 0 | 444690 | 123480 |
| VVSHPTLP | STRN3 | Q13033 | STRN3_HUMAN | Unmodified | 0 | 0 | 0 | 0 | 0 | 0 | 0 | 0 | 0 | 0 | 0 | 2949600 | 0 | 0 | 0 | 113920 | 0 | 0 |
| VVSIIAELLS | MYO1D; MYO1D | O94832; J3Q | MYO1D_HUMA | Unmodified | 1174500 | 1653600 | 789210 | 14895 | 0 | 0 | 0 | 0 | 0 | 0 | 0 | 136890 | 0 | 0 | 2460400 | 1220100 | 181560 | 0 |
| VVSIIAELLS | MYO1D; MYO1D | O94832; J3Q | MYO1D_HUMA | Oxidation (M) | 5920300 | 9755600 | 0 | 0 | 0 | 0 | 0 | 0 | 120130 | 331880 | 0 | 499280 | 0 | 0 | 3831300 | 1661200 | 0 | 0 |
| VVSKEIPVE | OGA; OGA | O60502; H7 | OGA_HUMAN I | Unmodified | 0 | 0 | 0 | 0 | 81528 | 0 | 342990 | 0 | 0 | 38404 | 0 | 0 | 253790 | 140690 | 0 | 0 | 132840 | 152100 |
| VVSLSEYR | NOP56; NOP56 | O00567; A0A | NOP56_HUMA | Unmodified | 72073 | 0 | 65181 | 0 | 0 | 0 | 0 | 0 | 0 | 0 | 0 | 1503900 | 0 | 0 | 0 | 0 | 0 | 0 |
| VVSMQVQ | NAALADL2 | Q58DX5 | NADL2_HUMA | Unmodified | 0 | 0 | 0 | 0 | 0 | 0 | 0 | 0 | 0 | 0 | 0 | 0 | 0 | 0 | 399650 | 0 | 0 | 0 |
| VVSNFILQ | UXS1; UXS1; U | Q8NBZ7; C9 | UXS1_HUMAN | Unmodified | 0 | 0 | 0 | 0 | 0 | 0 | 0 | 0 | 906420 | 50176 | 0 | 0 | 0 | 0 | 0 | 0 | 0 | 0 |
| VVSPWNSI | PDHB; PDHB; P | P11177; F8W | ODPB_HUMAN | Unmodified | 0 | 0 | 0 | 0 | 0 | 0 | 123480 | 0 | 1612000 | 426640 | 0 | 0 | 0 | 0 | 0 | 0 | 0 | 0 |
| VVSQYHEL | IMMT; IMMT; IM | Q16891; B9 | MIC60_HUMAN | Unmodified | 0 | 0 | 0 | 0 | 138750 | 0 | 0 | 105660 | 1533800 | 0 | 0 | 0 | 0 | 0 | 0 | 0 | 0 | 229690 |
| VVSQYSSLI | CCT4 | P50991 | TCPD_HUMAN | 2 Oxidation (M) | 0 | 0 | 0 | 0 | 0 | 0 | 0 | 0 | 0 | 681010 | 447780 | 3597900 | 1290900 | 0 | 0 | 0 | 0 | 0 |
| VVSSIEQK | YWHAZ; YWHA | P63104; E5R | 1433Z_HUMAN | Unmodified | 2877800 | 2080700 | 4313100 | 2676400 | 2444200 | 622870 | 319800 | 13320 | 628010 | 543430 | 615860 | 20066000 | 1059000 | 414330 | 5672400 | 3899100 | 1082800 | 3022500 |
| VVSSIEQKT | YWHAZ; YWHA | P63104; E5R | 1433Z_HUMAN | Unmodified | 10371000 | 8130100 | 3355600 | 43477000 | 92185000 | 34320000 | 13600000 | 5735300 | 1.42E+08 | 53999000 | 28717000 | 80848000 | 54125000 | 41411000 | 12910000 | 17630000 | 75519000 | 1.64E+08 |

| | | | | | | | | | | | | | | | | | | | | | | |
|---|---|---|---|---|---|---|---|---|---|---|---|---|---|---|---|---|---|---|---|---|---|---|
| VVSSIEQKT | YWHAZ; YWHAZ | P63104; E5R | 1433Z_HUMAN | Unmodified | 3265300 | 1131900 | 237260 | 8080800 | 18715000 | 10445000 | 1883500 | 2276000 | 24315000 | 9168600 | 6724300 | 19734000 | 10511000 | 8418500 | 2652600 | 2122900 | 10765000 | 25810000 |
| VVSSSIVDK | PSMC6; PSMC6 | P62333; A0A | PRS10_HUMAN | Unmodified | 0 | 0 | 272520 | 0 | 1786700 | 0 | 0 | 0 | 2202100 | 0 | 628640 | 9858600 | 634700 | 0 | 265810 | 262990 | 1303000 | 1296100 |
| VVSSSIVDK | PSMC6; PSMC6 | P62333; A0A | PRS10_HUMAN | Unmodified | 0 | 0 | 0 | 0 | 0 | 0 | 672140 | 463630 | 336580 | 676040 | 0 | 0 | 0 | 661300 | 0 | 0 | 731020 | 0 |
| VVSSTSEEE | OXR1 | Q8N573 | OXR1_HUMAN | Unmodified | 0 | 0 | 0 | 0 | 0 | 0 | 0 | 0 | 214790 | 0 | 0 | 646760 | 0 | 0 | 0 | 0 | 0 | 0 |
| VVSSVLQF | MYH10 | P35580 | MYH10_HUMAN | Unmodified | 0 | 0 | 0 | 0 | 0 | 0 | 0 | 0 | 266010 | 0 | 0 | 1420800 | 0 | 0 | 38643 | 18048 | 0 | 0 |
| VVSSVLQF | MYH10 | P35580 | MYH10_HUMAN | Unmodified | 0 | 0 | 1172100 | 186490 | 1378800 | 0 | 0 | 278590 | 0 | 251720 | 0 | 3284700 | 1208000 | 342530 | 2196800 | 472490 | 1424400 | 3882500 |
| VVSVLTVLH | IGHG4; IGHG3; | P01861; P01 | IGHG4_HUMAN | Unmodified | 0 | 381680 | 0 | 372940 | 0 | 0 | 0 | 0 | 0 | 46831 | 59026 | 81344 | 0 | 0 | 0 | 0 | 14819 | 0 |
| VVSYRVPH | MVP | Q14764 | MVP_HUMAN | Unmodified | 0 | 0 | 0 | 0 | 0 | 0 | 0 | 0 | 0 | 0 | 0 | 7504900 | 0 | 0 | 0 | 0 | 0 | 0 |
| VVSYRVPH | MVP | Q14764 | MVP_HUMAN | Unmodified | 0 | 0 | 0 | 0 | 0 | 0 | 0 | 0 | 0 | 0 | 0 | 1926300 | 0 | 0 | 0 | 0 | 0 | 0 |
| VVTATLWG | RPA1; RPA1 | P27694; I3L | RFA1_HUMAN | Unmodified | 467450 | 0 | 185370 | 0 | 991750 | 0 | 0 | 0 | 418280 | 797010 | 0 | 7941800 | 804050 | 0 | 177670 | 234490 | 0 | 0 |
| VVTCAHRY | ITGA6; ITGA6 | P23229; C9J | ITA6_HUMAN | Unmodified | 0 | 0 | 2953000 | 1683400 | 8323800 | 2549600 | 1235900 | 894470 | 129610 | 0 | 249620 | 1663600 | 5566500 | 3322000 | 1684000 | 4684000 | 12940000 | 38319000 |
| VVTDTDETI | EIF2S1 | P05198 | IF2A_HUMAN | Unmodified | 669040 | 421700 | 1196100 | 856250 | 0 | 0 | 0 | 0 | 1785700 | 141520 | 1586200 | 10790000 | 191010 | 0 | 842790 | 899910 | 0 | 0 |
| VVTHVQSG | TANC1 | Q9C0D5 | TANC1_HUMAN | Unmodified | 0 | 0 | 0 | 156360 | 0 | 0 | 0 | 323400 | 30532 | 138100 | 44729 | 63618 | 0 | 72259 | 126690 | 132180 | 0 | 0 |
| VVTLLYDLV | SIL1; SIL1 | Q9H173; D6 | SIL1_HUMAN | Unmodified | 0 | 0 | 200350 | 302220 | 0 | 0 | 0 | 0 | 2065100 | 0 | 0 | 0 | 0 | 0 | 316890 | 242710 | 0 | 0 |
| VVTMEVEA | CPNE1; CPNE1 | Q99829; Q5 | CPNE1_HUMAN | Oxidation (M) | 0 | 0 | 0 | 0 | 0 | 0 | 0 | 0 | 0 | 0 | 0 | 436710 | 0 | 0 | 0 | 0 | 0 | 0 |
| VVTMEVEA | CPNE1; CPNE1 | Q99829; Q5 | CPNE1_HUMAN | Unmodified | 0 | 159400 | 162390 | 323330 | 0 | 0 | 0 | 0 | 0 | 0 | 363740 | 1929100 | 0 | 0 | 196960 | 176960 | 0 | 0 |
| VVTPLHINL | DLG5; DLG5 | Q8TDM6; A0 | DLG5_HUMAN | Unmodified | 0 | 0 | 0 | 0 | 3521300 | 0 | 0 | 0 | 0 | 0 | 0 | 0 | 0 | 0 | 0 | 0 | 613230 | 1959600 |
| VVTRGPGL | FLNC | Q14315 | FLNC_HUMAN | Unmodified | 0 | 0 | 0 | 0 | 7728400 | 1794000 | 0 | 0 | 0 | 304230 | 0 | 0 | 0 | 0 | 0 | 0 | 4473700 | 1412700 |
| VVTRGPGL | FLNC | Q14315 | FLNC_HUMAN | Unmodified | 0 | 0 | 0 | 0 | 0 | 0 | 0 | 0 | 0 | 0 | 0 | 0 | 0 | 0 | 0 | 0 | 942410 | 0 |
| VVTRHPDL | CAPNS1; CAPN | P04632; A0A | CPNS1_HUMAN | Unmodified | 0 | 170730 | 0 | 385920 | 126560 | 0 | 0 | 0 | 1141900 | 104190 | 2179600 | 1379200 | 205370 | 0 | 181940 | 0 | 0 | 0 |
| VVTTAIKEE | DST; DST; DST; | Q03001; A0A | DYST_HUMAN | Unmodified | 0 | 0 | 0 | 0 | 731580 | 0 | 0 | 0 | 0 | 0 | 0 | 0 | 0 | 0 | 59749 | 0 | 0 | 0 |
| VVTTERFDI | ANKRD13A; AN | Q8IZ07; H0Y | AN13A_HUMAN | Oxidation (M) | 0 | 0 | 179940 | 0 | 0 | 0 | 0 | 0 | 0 | 0 | 0 | 0 | 0 | 0 | 270680 | 229030 | 0 | 0 |
| VVTTPAFM | GATD3B; GATD | A0A0B4J2D5 | GAL3B_HUMAN | Unmodified | 0 | 0 | 0 | 0 | 0 | 0 | 0 | 0 | 956440 | 0 | 0 | 0 | 0 | 0 | 0 | 0 | 0 | 0 |
| VVTVFSVA | WDR1 | O75083 | WDR1_HUMAN | Unmodified | 4807900 | 5729400 | 10080000 | 1565900 | 14375000 | 7685700 | 4000300 | 0 | 3996700 | 2166400 | 6340200 | 44816000 | 11151000 | 8399600 | 4501600 | 13292000 | 3722500 | 2728000 |
| VVTVGQFK | VPS29; VPS29 | Q9UBQ0; F8 | VPS29_HUMAN | Unmodified | 0 | 0 | 83364 | 813660 | 2052600 | 587010 | 365250 | 262820 | 136960 | 66792 | 802080 | 9929400 | 0 | 1150100 | 943500 | 0 | 346200 | 677990 |
| VVTVVTTK | DDX46; DDX46 | Q7L014; A0A | DDX46_HUMAN | Unmodified | 0 | 0 | 0 | 0 | 184160 | 22970 | 86361 | 131060 | 72231 | 43389 | 0 | 679800 | 446580 | 131150 | 0 | 0 | 50340 | 72998 |
| VVTWETVQ | TRPM5; TRPM5 | Q9NZQ8; A0 | TRPM5_HUMAN | Unmodified | 0 | 0 | 0 | 0 | 0 | 0 | 0 | 0 | 0 | 0 | 0 | 1921000 | 0 | 0 | 0 | 0 | 0 | 0 |
| VVTYDHIGI | INPP5A; INPP5 | Q14642; B1 | I5P1_HUMAN | Oxidation (M) | 0 | 0 | 505550 | 770910 | 0 | 0 | 0 | 0 | 0 | 0 | 248950 | 1638400 | 0 | 198050 | 582100 | 1038700 | 0 | 0 |
| VVTYGMAN | VAT1; VAT1 | Q99536; K7E | VAT1_HUMAN | Oxidation (M) | 0 | 0 | 0 | 0 | 0 | 0 | 0 | 0 | 0 | 0 | 0 | 1916300 | 0 | 0 | 0 | 0 | 0 | 0 |
| VVTYGMAN | VAT1; VAT1 | Q99536; K7E | VAT1_HUMAN | Unmodified | 0 | 0 | 0 | 0 | 530080 | 125470 | 0 | 0 | 335070 | 64281 | 0 | 202280 | 0 | 0 | 0 | 0 | 0 | 0 |
| VVTYGMAN | VAT1; VAT1 | Q99536; K7E | VAT1_HUMAN | Unmodified | 151750 | 0 | 0 | 0 | 0 | 0 | 0 | 0 | 648740 | 0 | 125050 | 1691400 | 0 | 0 | 109530 | 0 | 0 | 0 |
| VVVAENFD | PDIA3 | P30101 | PDIA3_HUMAN | Unmodified | 1457100 | 980360 | 1750200 | 1003300 | 1324100 | 0 | 580350 | 678690 | 1.47E+08 | 11120000 | 20363000 | 8274500 | 1105200 | 694910 | 2468200 | 1442900 | 211160 | 172430 |
| VVVCDNG1 | ACTR2; ACTR2 | P61160; A0A | ARP2_HUMAN | Unmodified | 672020 | 402540 | 0 | 501240 | 3525800 | 544980 | 960340 | 517950 | 571770 | 320800 | 776320 | 9480900 | 5538100 | 1803400 | 585650 | 431880 | 1778600 | 1977400 |
| VVVDALSG | CKMT1A; CKMT | P12532; C9J | KCRU_HUMAN | Unmodified | 0 | 0 | 2241400 | 0 | 0 | 0 | 0 | 0 | 0 | 0 | 3390000 | 8518700 | 0 | 0 | 548460 | 657340 | 0 | 0 |
| VVVDALSG | CKMT1A; CKMT | P12532; C9J | KCRU_HUMAN | Unmodified | 1374500 | 484300 | 632990 | 1134900 | 0 | 0 | 0 | 0 | 1079100 | 0 | 1416000 | 15250000 | 0 | 0 | 822560 | 1475500 | 0 | 0 |
| VVVDEGQI | SLC38A10 | Q9HBR0 | S38AA_HUMAN | Unmodified | 0 | 0 | 0 | 0 | 0 | 0 | 0 | 0 | 9660400 | 106970 | 0 | 343770 | 0 | 0 | 0 | 0 | 0 | 0 |
| VVVDGSG | SMURF2; SMUF | Q9HAU4; J3 | SMUF2_HUMAN | Unmodified | 178970 | 0 | 130370 | 0 | 163710 | 0 | 0 | 0 | 200070 | 52308 | 0 | 523310 | 0 | 0 | 306320 | 223890 | 0 | 0 |
| VVVFIKPTC | GLRX | P35754 | GLRX1_HUMAN | Unmodified | 414560 | 503040 | 0 | 0 | 0 | 0 | 0 | 0 | 0 | 0 | 959270 | 6897700 | 0 | 0 | 0 | 0 | 1760400 | 2404700 |
| VVVGREPG | TMEM132A; TM | Q24JP5; H0Y | T132A_HUMAN | Unmodified | 0 | 0 | 0 | 0 | 0 | 0 | 0 | 0 | 4078000 | 32436 | 0 | 0 | 0 | 0 | 0 | 0 | 0 | 0 |
| VVVHKETEI | EPB41L3; EPB4 | Q9Y2J2; A0A | E41L3_HUMAN | Unmodified | 0 | 12393 | 0 | 0 | 0 | 0 | 0 | 0 | 0 | 210950 | 0 | 0 | 62805 | 1171300 | 0 | 0 | 9962100 | 33097000 |
| VVVHKETEI | EPB41L2; EPB4 | O43491; Q6 | E41L2_HUMAN | Unmodified | 0 | 0 | 0 | 0 | 6857700 | 1091200 | 20891 | 194970 | 0 | 0 | 138110 | 1339100 | 2250300 | 214850 | 0 | 0 | 1984400 | 1777800 |
| VVVHPLVLI | PSMD7; PSMD7 | P51665; H3E | PSMD7_HUMAN | Unmodified | 288040 | 0 | 363250 | 535810 | 0 | 0 | 0 | 0 | 452670 | 63156 | 708400 | 12716000 | 139990 | 0 | 547890 | 384780 | 0 | 0 |
| VVVHPLVLI | PSMD7; PSMD7 | P51665; H3E | PSMD7_HUMAN | Unmodified | 0 | 0 | 0 | 0 | 3102100 | 0 | 427640 | 0 | 357190 | 297840 | 1401400 | 234920 | 7674500 | 0 | 0 | 0 | 722740 | 3320600 |
| VVVHQETE | EPB41; EPB41; | P11171; A0A | EPB41_HUMAN | Unmodified | 349220 | 148160 | 0 | 0 | 2215900 | 315020 | 620980 | 1207900 | 278730 | 1860300 | 2888700 | 1901000 | 8820900 | 4131600 | 0 | 0 | 2297200 | 2350400 |
| VVVLESRP | MAP4K5 | Q9Y4K4 | M4K5_HUMAN | Unmodified | 0 | 0 | 0 | 0 | 0 | 0 | 0 | 0 | 0 | 0 | 0 | 1401700 | 0 | 0 | 0 | 0 | 0 | 0 |
| VVVLGLLPI | PAFAH1B3; PAF | Q15102; M0 | PA1B3_HUMAN | Unmodified | 0 | 0 | 0 | 0 | 0 | 0 | 0 | 0 | 4223200 | 840190 | 3189500 | 39943000 | 0 | 0 | 0 | 0 | 0 | 0 |
| VVVLGSGG | RAP2C; RAP2A | Q9Y3L5; P10 | RAP2C_HUMAN | Unmodified | 2548000 | 3566500 | 10945000 | 16434000 | 14991000 | 4994400 | 3025400 | 2602200 | 6094600 | 4252000 | 7485600 | 39063000 | 8431100 | 8450100 | 11052000 | 8241700 | 8457000 | 13374000 |
| VVVLMGST | PAICS; PAICS; F | P22234; D6F | PUR6_HUMAN | Unmodified | 0 | 0 | 0 | 639450 | 2531700 | 0 | 195660 | 0 | 3267300 | 505970 | 2227600 | 15766000 | 0 | 719750 | 455130 | 589950 | 0 | 1102800 |
| VVVLMGST | PAICS; PAICS; F | P22234; D6F | PUR6_HUMAN | Oxidation (M) | 195440 | 0 | 0 | 868790 | 0 | 0 | 0 | 0 | 0 | 0 | 3253200 | 7857600 | 1174200 | 330430 | 557150 | 485730 | 0 | 0 |
| VVVPPLPG | SNX3 | O60493 | SNX3_HUMAN | Unmodified | 0 | 0 | 112040 | 85434 | 946960 | 0 | 850420 | 117370 | 691390 | 0 | 43245 | 0 | 979190 | 149840 | 0 | 0 | 0 | 374340 |
| VVVQEVVV | EVPL; EVPL | Q92817; K7E | EVPL_HUMAN | Unmodified | 0 | 0 | 0 | 0 | 1272500 | 0 | 57295 | 0 | 0 | 0 | 0 | 0 | 0 | 0 | 0 | 0 | 0 | 0 |
| VVVQVLAEI | FASN; FASN | P49327; A0A | FAS_HUMAN | Unmodified | 0 | 0 | 205220 | 877010 | 274490 | 197640 | 0 | 97907 | 0 | 0 | 1788300 | 9182900 | 921940 | 300360 | 1416500 | 769470 | 0 | 0 |
| VVVSNRLV | HSP90AA1 | P07900 | HS90A_HUMAN | Oxidation (M) | 0 | 0 | 0 | 0 | 0 | 0 | 0 | 0 | 0 | 0 | 0 | 0 | 0 | 0 | 0 | 0 | 2088600 | 2439800 |
| VVVAGET | DHX29; DHX29 | Q7Z478; A0A | DHX29_HUMAN | Unmodified | 0 | 0 | 0 | 0 | 0 | 0 | 94386 | 0 | 0 | 0 | 117710 | 840410 | 0 | 0 | 0 | 0 | 0 | 0 |
| VVVDSRP | EIF2B4; EIF2B4 | Q9UI10; A0A | EI2BD_HUMAN | Unmodified | 0 | 0 | 0 | 0 | 0 | 0 | 0 | 0 | 0 | 0 | 0 | 1761700 | 88965 | 0 | 0 | 0 | 0 | 0 |
| VVVGDQS | OPA1; OPA1; O | O60313; E5K | OPA1_HUMAN | Unmodified | 0 | 0 | 406120 | 532730 | 0 | 0 | 90069 | 0 | 861750 | 0 | 1034500 | 5179000 | 0 | 0 | 0 | 0 | 0 | 0 |
| VVVTGAN | RDH11; RDH11 | Q8TC12; G3 | RDH11_HUMAN | Unmodified | 0 | 0 | 0 | 0 | 0 | 0 | 0 | 0 | 0 | 0 | 0 | 0 | 0 | 0 | 0 | 0 | 0 | 0 |
| VVVTGAN | RDH11; RDH11 | Q8TC12; G3 | RDH11_HUMAN | Unmodified | 0 | 0 | 0 | 454460 | 279410 | 0 | 497430 | 0 | 547650 | 129820 | 70391 | 215310 | 0 | 0 | 0 | 0 | 0 | 439580 |
| VVVVPNE | HNRNPUL2; HN | Q1KMD3; H3 | HNRL2_HUMAN | Unmodified | 0 | 0 | 0 | 0 | 0 | 0 | 0 | 0 | 0 | 404970 | 0 | 204920 | 1146200 | 0 | 0 | 0 | 130990 | 0 |
| VVYLQK | CTSC; CTSC; C | P53634; A0A | CATC_HUMAN | Unmodified | 0 | 0 | 25243 | 22521 | 81211 | 0 | 292950 | 353480 | 184110 | 60091 | 4109500 | 9082400 | 144020 | 109590 | 0 | 38759 | 37520 | 0 |
| VVYSNTIC | GNAI3 | P08754 | GNAI3_HUMAN | Unmodified | 0 | 295710 | 0 | 0 | 0 | 0 | 103370 | 0 | 29825 | 227200 | 54655 | 0 | 0 | 86859 | 1413400 | 0 | 0 | 0 |
| VVWEESD | DDX3X; DDX3X | O00571; A0A | DDX3X_HUMAN | Unmodified | 2403700 | 105010 | 3594600 | 2025900 | 0 | 0 | 0 | 0 | 0 | 0 | 0 | 0 | 0 | 0 | 3594500 | 3005000 | 0 | 0 |
| VVYENAYG | TXNRD1; TXNRI | Q16881; A0A | TRXR1_HUMAN | Unmodified | 253030 | 0 | 259080 | 563030 | 0 | 0 | 0 | 0 | 0 | 0 | 825690 | 7079400 | 0 | 0 | 388090 | 386260 | 0 | 0 |
| VVYENAYG | TXNRD1; TXNRI | Q16881; A0A | TRXR1_HUMAN | Unmodified | 0 | 0 | 0 | 0 | 3389100 | 323550 | 196810 | 0 | 0 | 423880 | 324590 | 0 | 1437000 | 504220 | 0 | 0 | 604400 | 1794000 |
| VVYGGGA | CCT5; CCT5; C | P48643; B7Z | TCPE_HUMAN | Unmodified | 0 | 0 | 177890 | 0 | 0 | 0 | 0 | 0 | 0 | 0 | 0 | 3062100 | 0 | 0 | 0 | 114000 | 0 | 0 |
| VVYIFGPP | GMPS | P49915 | GUAA_HUMAN | Unmodified | 0 | 0 | 0 | 0 | 0 | 0 | 0 | 0 | 0 | 0 | 0 | 1610800 | 0 | 0 | 0 | 0 | 0 | 0 |
| VVYIFGPP | GMPS | P49915 | GUAA_HUMAN | Unmodified | 0 | 0 | 0 | 0 | 0 | 0 | 0 | 0 | 170800 | 0 | 0 | 237650 | 0 | 0 | 0 | 0 | 0 | 0 |
| VVYLASETF | PI4K2A; N/A | Q9BTU6; E9 | P4K2A_HUMAN | Unmodified | 0 | 27653 | 0 | 124760 | 0 | 0 | 0 | 0 | 405990 | 233270 | 0 | 798850 | 0 | 0 | 0 | 135730 | 0 | 0 |
| VVYLRCTG | RPL12 | P30050 | RL12_HUMAN | Unmodified | 0 | 0 | 0 | 0 | 594630 | 0 | 524880 | 263760 | 0 | 0 | 0 | 0 | 0 | 0 | 0 | 0 | 0 | 709030 |
| VVYPSKAD | LRRC47 | Q8N1G4 | LRC47_HUMAN | Unmodified | 0 | 0 | 0 | 0 | 0 | 0 | 901940 | 874360 | 0 | 683890 | 0 | 0 | 2663000 | 1265800 | 0 | 0 | 1071700 | 0 |
| VVYSAPRS | CLDN3 | O15551 | CLD3_HUMAN | Unmodified | 0 | 0 | 0 | 0 | 2103200 | 0 | 0 | 0 | 0 | 252670 | 0 | 0 | 213910 | 0 | 0 | 0 | 0 | 1113000 |
| VVYSPGEP | ARRDC1; ARRD | Q8N5I2; Q5 | ARRD1_HUMAN | Unmodified | 5213300 | 8894600 | 4308900 | 5263400 | 0 | 10401000 | 535090 | 635710 | 2791700 | 209330 | 821250 | 6870800 | 0 | 437770 | 9087200 | 7388800 | 782150 | 16673000 |
| VVYSSKDP | ABCC1; ABCC1 | P33527; I3L | MRP1_HUMAN | Unmodified | 0 | 59368 | 208960 | 707320 | 7524100 | 0 | 0 | 0 | 0 | 0 | 736700 | 4502700 | 1444600 | 314740 | 599220 | 1084800 | 0 | 261690 |
| VWAHYEE | SND1; SND1 | Q7KZF4; H7 | SND1_HUMAN | Unmodified | 0 | 0 | 0 | 0 | 1694100 | 0 | 0 | 0 | 0 | 0 | 0 | 1202900 | 0 | 0 | 0 | 0 | 232070 | 1053100 |
| VWAHYEE | SND1; SND1 | Q7KZF4; H7 | SND1_HUMAN | Oxidation (M) | 0 | 0 | 0 | 537770 | 0 | 0 | 0 | 0 | 0 | 0 | 0 | 3143600 | 639910 | 422510 | 0 | 0 | 0 | 0 |
| VWDAVSG | STRAP | Q9Y3F4 | STRAP_HUMAN | Unmodified | 0 | 406470 | 0 | 125520 | 2039800 | 0 | 160740 | 0 | 1531900 | 104530 | 0 | 1821000 | 2031600 | 675500 | 236140 | 156320 | 545060 | 2516600 |
| VWDAVSG | STRAP | Q9Y3F4 | STRAP_HUMAN | Oxidation (M) | 461610 | 0 | 362580 | 492090 | 1288300 | 428770 | 572100 | 394120 | 431760 | 627450 | 1819300 | 9673900 | 2645300 | 1046900 | 464620 | 395740 | 0 | 718070 |
| VWDISGLR | COPA; COPA; C | P53621; A0A | COPA_HUMAN | Unmodified | 7082700 | 172820 | 0 | 0 | 0 | 0 | 18598 | 176310 | 310210 | 0 | 0 | 5999800 | 0 | 0 | 0 | 0 | 0 | 0 |
| VWDLAQQ | WDR77; WDR7 | Q9BQA1; H0 | MEP50_HUMAN | Unmodified | 0 | 0 | 0 | 0 | 0 | 0 | 0 | 0 | 0 | 0 | 0 | 2478900 | 0 | 0 | 0 | 0 | 0 | 0 |
| VWDLFPEA | EIF3CL; EIF3C | B5ME19; Q9 | EIFCL_HUMAN | Unmodified | 219920 | 0 | 746120 | 0 | 0 | 0 | 0 | 0 | 191290 | 0 | 1654000 | 8833400 | 1426000 | 0 | 0 | 536190 | 0 | 0 |
| VWDLFPEA | EIF3CL; EIF3C | B5ME19; Q9 | EIFCL_HUMAN | Unmodified | 20430 | 0 | 0 | 0 | 0 | 67604 | 0 | 25637 | 212820 | 164290 | 412700 | 842670 | 0 | 0 | 0 | 0 | 0 | 0 |
| VWDVIGR | CCS; CCS; CCS | O14618; J3K | CCS_HUMAN | Unmodified | 0 | 0 | 0 | 0 | 0 | 0 | 0 | 0 | 0 | 0 | 0 | 0 | 0 | 0 | 0 | 0 | 0 | 0 |
| VWDYETGI | PAFAH1B1; PAF | P43034; A0A | LIS1_HUMAN | Unmodified | 110700 | 0 | 178790 | 197550 | 0 | 0 | 0 | 0 | 586990 | 0 | 744630 | 2157000 | 0 | 0 | 264860 | 0 | 0 | 0 |
| VWEQATPL | MMP15 | P51511 | MMP15_HUMAN | Unmodified | 1725900 | 1134300 | 286220 | 693170 | 0 | 0 | 0 | 0 | 436410 | 0 | 0 | 0 | 0 | 0 | 3995500 | 1971200 | 0 | 0 |
| VWEQIDQ | DYNC1H1; DYN | Q14204; A0A | DYHC1_HUMAN | Unmodified | 0 | 0 | 0 | 0 | 319520 | 0 | 0 | 0 | 337400 | 0 | 115690 | 1496400 | 0 | 0 | 0 | 0 | 287920 | 0 |
| VWFRIPNQ | HEBP1; HEBP1 | Q9NRV9; F5 | HEBP1_HUMAN | Unmodified | 296080 | 349920 | 0 | 1162700 | 7680000 | 0 | 0 | 78818 | 824660 | 0 | 0 | 0 | 8458600 | 7899500 | 515600 | 565400 | 4645900 | 4151700 |
| VWGNVGT | SYNCRIP; SYNC | O60506; B7 | HNRPQ_HUMAN | Oxidation (M) | 225270 | 0 | 0 | 0 | 1647000 | 216550 | 1514900 | 133300 | 0 | 784430 | 1008000 | 3096700 | 6305200 | 1579600 | 0 | 0 | 0 | 0 |
| VWGSPGG | OS9; OS9 | Q13438; B4 | OS9_HUMAN | Unmodified | 0 | 0 | 0 | 0 | 0 | 0 | 0 | 0 | 800360 | 0 | 0 | 0 | 0 | 0 | 0 | 0 | 0 | 0 |
| VWHLVRPT | PLXNB2; PLXNE | O15031; A6 | PLXB2_HUMAN | Unmodified | 0 | 0 | 125150 | 0 | 0 | 0 | 0 | 0 | 0 | 0 | 0 | 208780 | 0 | 0 | 214520 | 164950 | 0 | 0 |
| VWINTSDI | EIF1AY; EIF1AX | O14602; P4 | IF1AY_HUMAN | Unmodified | 0 | 0 | 0 | 0 | 0 | 0 | 0 | 0 | 0 | 0 | 0 | 0 | 4391000 | 0 | 0 | 0 | 0 | 0 |
| VWKYEVTY | EPHA2 | P29317 | EPHA2_HUMAN | Unmodified | 0 | 0 | 0 | 0 | 0 | 0 | 0 | 0 | 0 | 0 | 0 | 1682700 | 0 | 0 | 0 | 0 | 0 | 0 |
| VWLDPNE | RPL19; RPL19; | P84098; J3K | RL19_HUMAN | Unmodified | 5247500 | 4798000 | 2468300 | 2825500 | 0 | 0 | 0 | 115130 | 0 | 0 | 4297600 | 16057000 | 1922900 | 0 | 3012700 | 2397500 | 0 | 0 |
| VWLDPNE | RPL19; RPL19; | P84098; J3K | RL19_HUMAN | Unmodified | 0 | 0 | 0 | 0 | 1895700 | 0 | 0 | 508210 | 0 | 0 | 0 | 0 | 4345700 | 1403800 | 0 | 0 | 0 | 1400200 |
| VWLDPNE | RPL19; RPL19; | P84098; J3K | RL19_HUMAN | Unmodified | 0 | 0 | 0 | 0 | 1567800 | 0 | 1085300 | 0 | 0 | 0 | 0 | 0 | 1978600 | 0 | 0 | 0 | 0 | 410030 |
| VWLEVEPV | MCAM | P43121 | MUC18_HUMAN | Oxidation (M) | 0 | 0 | 0 | 0 | 0 | 0 | 0 | 0 | 0 | 0 | 0 | 0 | 386010 | 0 | 0 | 0 | 300360 | 427190 |
| VWLEVEPV | MCAM | P43121 | MUC18_HUMAN | Unmodified | 0 | 0 | 0 | 0 | 593850 | 0 | 0 | 0 | 0 | 0 | 0 | 0 | 0 | 0 | 0 | 0 | 203550 | 325780 |
| VWNLANC | RACK1; RACK1 | P63244; D6 | RACK1_HUMAN | Unmodified | 0 | 0 | 717530 | 0 | 589640 | 450240 | 1851800 | 0 | 0 | 0 | 1250400 | 18160000 | 0 | 596030 | 613330 | 817990 | 0 | 433190 |
| VWNTLSIK | GEMIN5 | Q8TEQ6 | GEMI5_HUMAN | Unmodified | 0 | 0 | 0 | 0 | 0 | 0 | 0 | 0 | 0 | 0 | 0 | 1677600 | 33710 | 0 | 0 | 0 | 0 | 0 |
| VWPDLPN | MGAM2 | Q2M2H8 | MGAL_HUMAN | Unmodified | 0 | 0 | 0 | 0 | 0 | 0 | 0 | 0 | 0 | 0 | 0 | 1447400 | 0 | 0 | 0 | 0 | 0 | 0 |
| VWQCGGS | GALNT2 | Q10471 | GALT2_HUMAN | Unmodified | 28230 | 198850 | 0 | 0 | 0 | 0 | 0 | 0 | 1746400 | 0 | 0 | 0 | 0 | 0 | 35431 | 0 | 0 | 0 |
| VWQLGSS | COPB2; COPB2 | P35606; H0 | COPB2_HUMAN | Unmodified | 0 | 0 | 397790 | 307230 | 476180 | 165780 | 283920 | 0 | 0 | 1758600 | 1039600 | 7389700 | 0 | 0 | 559980 | 265190 | 370690 | 448690 |
| VWQLQDL | UGGT1 | Q9NYU2 | UGGG1_HUMAN | Unmodified | 0 | 0 | 26665 | 0 | 297660 | 0 | 0 | 0 | 26244000 | 10703000 | 1959700 | 622870 | 66526 | 64035 | 65850 | 0 | 0 | 0 |
| VWQVTIGT | RACK1; RACK1 | P63244; D6 | RACK1_HUMAN | Unmodified | 3792700 | 4071100 | 1615500 | 1072900 | 0 | 0 | 118430 | 0 | 1281300 | 352470 | 2903900 | 23447000 | 276210 | 0 | 2688100 | 2401000 | 0 | 396130 |
| VWRDQHF | TFRC; TFRC | P02786; G3 | TFR1_HUMAN | Unmodified | 0 | 0 | 2405700 | 2601800 | 0 | 0 | 871680 | 161250 | 10691000 | 3160400 | 755240 | 1988800 | 0 | 0 | 1417000 | 1554800 | 343610 | 1571700 |
| VWRDQHF | TFRC; TFRC | P02786; G3 | TFR1_HUMAN | Unmodified | 0 | 0 | 0 | 0 | 708090 | 0 | 0 | 0 | 0 | 0 | 0 | 0 | 0 | 0 | 0 | 166910 | 227930 | 0 |
| VWSPLVTE | PDLIM1 | O00151 | PDLI1_HUMAN | Unmodified | 212190 | 0 | 0 | 0 | 3709000 | 595750 | 0 | 0 | 0 | 0 | 0 | 0 | 877180 | 549350 | 0 | 0 | 584080 | 3925200 |
| VWSRNEDI | GSTK1; GSTK1 | Q9Y2Q3; E9 | GSTK1_HUMAN | Unmodified | 0 | 0 | 0 | 0 | 0 | 0 | 0 | 0 | 2180000 | 522620 | 0 | 0 | 0 | 0 | 0 | 0 | 0 | 0 |
| VWSVASTV | CADM4 | Q8NFZ8 | CADM4_HUMAN | Unmodified | 0 | 0 | 0 | 0 | 0 | 0 | 130430 | 0 | 3686400 | 415380 | 0 | 296020 | 0 | 0 | 0 | 0 | 81823 | 0 |
| VWTSGQV | AQP1; AQP1; A | P29972; A0A | AQP1_HUMAN | Unmodified | 0 | 0 | 0 | 0 | 0 | 0 | 0 | 0 | 970280 | 1013300 | 0 | 0 | 0 | 0 | 0 | 0 | 0 | 0 |

| | | | | | | | | | | | | | | | | | | | | | | |
|---|---|---|---|---|---|---|---|---|---|---|---|---|---|---|---|---|---|---|---|---|---|---|
| VWTSGQVI | AQP1; AQP1; N | P29972; A0A | AQP1_HUMAN | Oxidation (M) | 0 | 0 | 0 | 0 | 0 | 0 | 0 | 0 | 4047000 | 1593000 | 0 | 0 | 0 | 0 | 0 | 0 | 0 | 1333200 |
| VWTSGQVI | AQP1; AQP1; N | P29972; A0A | AQP1_HUMAN | Unmodified | 0 | 0 | 0 | 0 | 0 | 0 | 0 | 0 | 12814000 | 0 | 0 | 0 | 0 | 0 | 0 | 0 | 0 | 0 |
| VWVLTGDK | ATP11C; ATP11 | Q8NB49; P9 | AT11C_HUMAN | Unmodified | 0 | 0 | 0 | 0 | 0 | 0 | 0 | 0 | 0 | 0 | 0 | 682110 | 0 | 0 | 0 | 0 | 0 | 0 |
| VWVVATK | PAFAH1B1; PAF | P43034; I3L3 | LIS1_HUMAN P | Unmodified | 0 | 199530 | 0 | 121860 | 0 | 0 | 0 | 0 | 14990000 | 5016900 | 289460 | 2054700 | 0 | 0 | 269850 | 243030 | 0 | 0 |
| VWYVSNID | GPI; GPI; GPI; G | P06744; A0A | G6PI_HUMAN ( | Unmodified | 2031400 | 2314100 | 2718800 | 2371500 | 856960 | 0 | 0 | 0 | 6614200 | 1727200 | 3182700 | 58839000 | 420460 | 0 | 4652700 | 7125100 | 42793 | 0 |
| VYAALGNH | SMPDL3B; SMP | Q92485; Q5 | ASM3B_HUMAI | Unmodified | 1819300 | 1600400 | 8027800 | 6434600 | 1788800 | 8045.9 | 0 | 0 | 2907800 | 621330 | 722910 | 0 | 6729900 | 3456200 | 8272100 | 5638700 | 373770 | 0 |
| VYAENAIR | CHMP1A; CHM | Q9HD42; F8 | CHM1A_HUMA | Unmodified | 1210300 | 1491600 | 621760 | 196060 | 874360 | 409100 | 0 | 0 | 2139400 | 0 | 180290 | 674790 | 0 | 322200 | 1436800 | 678340 | 57047 | 157590 |
| VYAILTHGI | PRPS2; PRPS1; | P11908; P60 | PRPS2_HUMAN | Unmodified | 0 | 570910 | 1706000 | 3033000 | 457420 | 295230 | 1186300 | 85368 | 7115200 | 3312200 | 3439500 | 19419000 | 6033100 | 1050100 | 2541300 | 1473500 | 328670 | 470470 |
| VYALPEDL | PLS3; PLS3 | P13797; A0A | PLST_HUMAN I | Unmodified | 0 | 0 | 0 | 0 | 19543000 | 6396900 | 1456400 | 919820 | 794320 | 1161500 | 0 | 0 | 0 | 5164000 | 0 | 870440 | 19849000 | 13699000 |
| VYAPASTL | DSG2 | Q14126 | DSG2_HUMAN | Unmodified | 0 | 1988200 | 2381800 | 5526400 | 0 | 0 | 0 | 0 | 7693700 | 0 | 4116700 | 7791400 | 0 | 0 | 4245800 | 6258700 | 0 | 0 |
| VYATILNAG | FASN; FASN | P49327; A0A | FAS_HUMAN F | Unmodified | 0 | 0 | 237710 | 0 | 0 | 0 | 0 | 0 | 0 | 0 | 277290 | 3998600 | 71711 | 0 | 276520 | 262750 | 0 | 0 |
| VYCDMNTE | FGB; FGB | P02675; D6F | FIBB_HUMAN F | Oxidation (M) | 0 | 1232600 | 0 | 0 | 0 | 0 | 0 | 0 | 0 | 0 | 0 | 0 | 0 | 0 | 47378 | 0 | 0 | 0 |
| VYCENQPN | CSNK2B; CSNK | P67870; Q5S | CSK2B_HUMAI | 2 Oxidation (M) | 0 | 0 | 0 | 0 | 0 | 0 | 0 | 0 | 0 | 0 | 761880 | 902740 | 0 | 0 | 0 | 0 | 0 | 0 |
| VYCGHEYT | HAGH; HAGH; I | Q16775; H3I | GLO2_HUMAN | Unmodified | 56671 | 0 | 0 | 0 | 150680 | 0 | 0 | 0 | 383910 | 0 | 0 | 554890 | 0 | 0 | 0 | 98139 | 73530 | 67838 |
| VYDESIQLI | LRP1 | Q07954 | LRP1_HUMAN I | Unmodified | 0 | 100570 | 0 | 0 | 0 | 0 | 388390 | 0 | 1535000 | 79340 | 0 | 0 | 0 | 0 | 0 | 0 | 0 | 0 |
| VYDPKNEE | NUCB2; NUCB2 | P80303; Q2I | NUCB2_HUMA | Unmodified | 0 | 0 | 0 | 0 | 0 | 0 | 0 | 0 | 894680 | 0 | 0 | 0 | 0 | 0 | 0 | 0 | 0 | 0 |
| VYDPKNEE | NUCB2; NUCB2 | P80303; Q2I | NUCB2_HUMA | 2 Oxidation (M) | 0 | 0 | 0 | 242060 | 0 | 0 | 0 | 0 | 6310800 | 1147100 | 0 | 0 | 0 | 0 | 887300 | 0 | 0 | 0 |
| VYDPKNEE | NUCB2; NUCB2 | P80303; Q2I | NUCB2_HUMA | Oxidation (M) | 0 | 0 | 0 | 849200 | 0 | 0 | 0 | 0 | 58708000 | 2133700 | 0 | 0 | 0 | 186410 | 717150 | 0 | 0 | 0 |
| VYDPKNEE | NUCB2; NUCB2 | P80303; Q2I | NUCB2_HUMA | Unmodified | 0 | 0 | 0 | 0 | 0 | 0 | 0 | 0 | 56480000 | 2593100 | 0 | 0 | 0 | 0 | 0 | 0 | 0 | 0 |
| VYDSLLALF | CLDN9; CLDN3 | O95484; O1 | CLD9_HUMAN | Unmodified | 4936800 | 3325600 | 144550 | 170720 | 612800 | 0 | 0 | 0 | 171040 | 0 | 75787 | 1851000 | 0 | 0 | 4558700 | 2292800 | 0 | 0 |
| VYEDPALS | EXOC7; EXOC7 | Q9UPT5; B4I | EXOC7_HUMAI | Unmodified | 150050 | 0 | 154250 | 0 | 2039300 | 358900 | 518610 | 0 | 1866200 | 1572900 | 150600 | 299770 | 660610 | 0 | 709970 | 229500 | 531530 | 387270 |
| VYEELRATC | ABCF1; ABCF1; | Q8NE71; H0 | ABCF1_HUMAN | Unmodified | 66647 | 84943 | 0 | 194990 | 301650 | 160650 | 213610 | 189460 | 977180 | 796980 | 0 | 4732000 | 0 | 122720 | 0 | 0 | 1578200 | 149480 |
| VYEEVLSVT | ASPH | Q12797 | ASPH_HUMAN | Unmodified | 0 | 0 | 0 | 0 | 0 | 0 | 0 | 0 | 2860900 | 663340 | 0 | 0 | 0 | 381200 | 0 | 0 | 0 | 0 |
| VYEFLDK | PSMD3; PSMD3 | O43242; H0 | PSMD3_HUMA | Unmodified | 0 | 0 | 0 | 0 | 0 | 0 | 0 | 0 | 0 | 0 | 0 | 851260 | 0 | 0 | 0 | 0 | 0 | 0 |
| VYEFLDKLI | PSMD3; PSMD3 | O43242; H0 | PSMD3_HUMA | Unmodified | 135680 | 152980 | 0 | 0 | 0 | 0 | 0 | 0 | 179210 | 237820 | 0 | 5559900 | 0 | 0 | 0 | 384480 | 0 | 0 |
| VYEGERPL | HSPA5; HSPA5 | P11021; A0A | BIP_HUMAN Er | Unmodified | 0 | 0 | 0 | 0 | 0 | 0 | 2704600 | 596820 | 17787000 | 14448000 | 1648100 | 857870 | 300360 | 239640 | 854060 | 0 | 0 | 0 |
| VYEGERPL | HSPA5; HSPA5 | P11021; A0A | BIP_HUMAN Er | Unmodified | 0 | 0 | 0 | 0 | 0 | 0 | 1960700 | 0 | 2261400 | 11372000 | 0 | 0 | 0 | 0 | 0 | 0 | 0 | 0 |
| VYEILPTFD | PANX1 | Q96RD7 | PANX1_HUMAN | Unmodified | 0 | 0 | 0 | 0 | 0 | 0 | 0 | 0 | 381860 | 169890 | 359300 | 2856100 | 463390 | 301080 | 0 | 362010 | 0 | 0 |
| VYEIQDIYE | EIF3L; EIF3L; EI | Q9Y262; B0 | EIF3L_HUMAN | Unmodified | 0 | 0 | 0 | 0 | 0 | 0 | 0 | 0 | 0 | 0 | 627340 | 3641700 | 0 | 0 | 0 | 0 | 0 | 0 |
| VYELPFLVA | DAGLB; DAGLB | Q8NCG7; E7 | DGLB_HUMAN | Unmodified | 0 | 0 | 0 | 0 | 0 | 0 | 0 | 0 | 0 | 0 | 0 | 570600 | 0 | 0 | 0 | 0 | 0 | 0 |
| VYELQASR | EIF3L; EIF3L; EI | Q9Y262; B0 | EIF3L_HUMAN | Unmodified | 346280 | 82181 | 0 | 0 | 0 | 0 | 0 | 0 | 0 | 0 | 864450 | 6484000 | 0 | 0 | 418180 | 582990 | 0 | 0 |
| VYELSNVQ | ICAM1; ICAM1 | P05362; K7E | ICAM1_HUMAN | Oxidation (M) | 0 | 0 | 0 | 276430 | 6628200 | 2691500 | 0 | 170030 | 0 | 0 | 0 | 0 | 0 | 400500 | 0 | 0 | 2691000 | 3036200 |
| VYELSNVQ | ICAM1; ICAM1 | P05362; K7E | ICAM1_HUMAN | Unmodified | 0 | 0 | 0 | 0 | 19829000 | 2394700 | 0 | 0 | 0 | 0 | 0 | 0 | 0 | 0 | 0 | 0 | 4447100 | 5162600 |
| VYENYPTYI | DEK; DEK; DEK; | P35659; B4D | DEK_HUMAN P | Unmodified | 159100 | 0 | 0 | 0 | 124620 | 0 | 0 | 0 | 0 | 546940 | 0 | 0 | 0 | 0 | 0 | 0 | 685510 | 0 |
| VYFAEKVTS | CRIP2; CRIP2; ( | P52943; H0 | CRIP2_HUMAN | Unmodified | 0 | 0 | 0 | 0 | 0 | 0 | 1130800 | 473960 | 70780 | 0 | 0 | 0 | 0 | 0 | 0 | 0 | 0 | 0 |
| VYFLPITPH | CAD; CAD | P27708; F8V | PYR1_HUMAN | Unmodified | 0 | 0 | 0 | 0 | 0 | 0 | 0 | 0 | 329200 | 0 | 0 | 5004500 | 0 | 0 | 0 | 0 | 0 | 0 |
| VYFQSPPG | PPP1R14B | Q96C90 | PP14B_HUMAN | Unmodified | 409810 | 87363 | 0 | 0 | 0 | 0 | 0 | 0 | 0 | 380570 | 139120 | 651740 | 0 | 0 | 366320 | 140910 | 0 | 136900 |
| VYFQSPPG | PPP1R14B | Q96C90 | PP14B_HUMAN | Unmodified | 1346300 | 1005200 | 0 | 2553900 | 0 | 0 | 0 | 0 | 599670 | 0 | 941910 | 7220100 | 0 | 0 | 0 | 0 | 0 | 0 |
| VYFQSPPG | PPP1R14B | Q96C90 | PP14B_HUMAN | Unmodified | 197480 | 0 | 0 | 408630 | 921430 | 0 | 0 | 0 | 0 | 0 | 449300 | 0 | 1197600 | 0 | 0 | 0 | 521210 | 724360 |
| VYGALMWS | EHD4; EHD1; E | Q9H223; Q9 | EHD4_HUMAN | Oxidation (M) | 683150 | 495810 | 0 | 0 | 0 | 0 | 0 | 0 | 0 | 0 | 0 | 2412300 | 0 | 0 | 1594600 | 1006200 | 0 | 0 |
| VYGAPQLQ | MGAT1 | P26572 | MGAT1_HUMAI | Unmodified | 0 | 0 | 0 | 0 | 0 | 0 | 0 | 0 | 7030100 | 756050 | 0 | 0 | 0 | 0 | 0 | 0 | 0 | 0 |
| VYGGDDRI | HAGH; HAGH; I | Q16775; H3I | GLO2_HUMAN | Unmodified | 0 | 0 | 0 | 0 | 0 | 0 | 0 | 0 | 0 | 481850 | 0 | 0 | 0 | 0 | 0 | 0 | 2605900 | 0 |
| VYGGLTTK | PDCD6IP | Q8WUM4 | PDC6I_HUMAN | Unmodified | 0 | 0 | 171420 | 0 | 0 | 105750 | 80111 | 0 | 780540 | 470850 | 0 | 1432700 | 0 | 0 | 0 | 0 | 34269 | 124350 |
| VYGTIKPAF | SHANK2; SHAN | Q9UPX8; A6I | SHAN2_HUMAI | Unmodified | 0 | 0 | 0 | 64554 | 0 | 0 | 0 | 0 | 0 | 0 | 0 | 0 | 231300 | 180140 | 47154 | 69572 | 0 | 0 |
| VYGVESDL | MVD; MVD | P53602; H3E | MVD1_HUMAN | Unmodified | 105830 | 0 | 0 | 0 | 0 | 0 | 0 | 0 | 0 | 0 | 0 | 1370100 | 0 | 0 | 0 | 0 | 0 | 0 |
| VYGVESDL | MVD; MVD | P53602; H3E | MVD1_HUMAN | Unmodified | 0 | 0 | 0 | 0 | 0 | 0 | 0 | 0 | 0 | 0 | 0 | 709260 | 0 | 0 | 0 | 0 | 0 | 0 |
| VYIASSSGS | SH3BGRL | O75368 | SH3L1_HUMAN | Unmodified | 0 | 0 | 0 | 0 | 175780 | 0 | 0 | 0 | 601550 | 0 | 1083200 | 3752000 | 0 | 0 | 0 | 502970 | 92869 | 0 |
| VYIDPFTYE | EPHB3; EPHB4; | P54753; P54 | EPHB3_HUMAN | Unmodified | 0 | 286560 | 0 | 0 | 0 | 0 | 0 | 0 | 0 | 0 | 0 | 1648100 | 0 | 0 | 0 | 0 | 0 | 0 |
| VYIDPFTYE | EPHB3; EPHB4; | P54753; P54 | EPHB3_HUMAN | Unmodified | 1192400 | 1064300 | 0 | 556520 | 9221900 | 730190 | 0 | 0 | 2721200 | 0 | 1868800 | 3400000 | 0 | 0 | 0 | 379920 | 2681300 | 8898200 |
| VYIGEEEK | PIP4K2C; PIP4K | Q8TBX8; H0 | PI42C_HUMAN | Unmodified | 0 | 0 | 0 | 0 | 0 | 57706 | 0 | 0 | 117800 | 813330 | 273590 | 0 | 0 | 0 | 20202 | 0 | 0 | 1237900 |
| VYIGEEEKK | PIP4K2C; PIP4K | Q8TBX8; H0 | PI42C_HUMAN | Unmodified | 0 | 0 | 0 | 0 | 689970 | 0 | 414370 | 30929 | 0 | 325850 | 0 | 0 | 273600 | 1600700 | 0 | 0 | 828950 | 1682600 |
| VYIGSFWA | EHD4 | Q9H223 | EHD4_HUMAN | Unmodified | 1562600 | 975880 | 1210900 | 293610 | 0 | 0 | 0 | 0 | 53359 | 10967 | 23726 | 1411300 | 90560 | 0 | 3953000 | 2331400 | 0 | 0 |
| VYIGSFWA | EHD4 | Q9H223 | EHD4_HUMAN | Unmodified | 2879400 | 2483200 | 3027200 | 1842400 | 0 | 0 | 0 | 0 | 0 | 0 | 0 | 2603400 | 0 | 0 | 6115800 | 3755100 | 190630 | 0 |
| VYIGSFWS | EHD1; EHD3; E | Q9H4M9; Q | EHD1_HUMAN | Unmodified | 1374000 | 1503800 | 2121100 | 642480 | 263390 | 243780 | 0 | 0 | 0 | 0 | 0 | 13647000 | 859740 | 0 | 4106200 | 2663700 | 459790 | 0 |
| VYIYHSSSK | ITGA3; ITGA3; I | P26006; K7E | ITA3_HUMAN I | Unmodified | 0 | 0 | 0 | 0 | 23645000 | 7982200 | 0 | 0 | 283840 | 203390 | 123350 | 1174700 | 0 | 0 | 118530 | 53642 | 14657000 | 11298000 |
| VYIYHSSSK | ITGA3; ITGA3; I | P26006; K7E | ITA3_HUMAN I | Unmodified | 0 | 0 | 0 | 0 | 4076500 | 333300 | 0 | 0 | 0 | 0 | 0 | 0 | 0 | 0 | 0 | 0 | 4052500 | 1736800 |
| VYIYSSGSV | ENOPH1; ENOF | Q9UHY7; D6 | ENOPH_HUMA | Unmodified | 0 | 0 | 0 | 0 | 343570 | 0 | 0 | 0 | 0 | 335090 | 537120 | 0 | 777930 | 517220 | 116960 | 239160 | 0 | 224940 |
| VYKAQNKE | SLK | Q9H2G2 | SLK_HUMAN S | Unmodified | 0 | 0 | 0 | 0 | 66360 | 0 | 87612 | 0 | 0 | 76973 | 0 | 0 | 361330 | 350930 | 0 | 0 | 113850 | 0 |
| VYKGEFQL | TPST1 | O60507 | TPST1_HUMAN | Unmodified | 0 | 0 | 0 | 0 | 470660 | 0 | 0 | 0 | 2574300 | 472490 | 0 | 0 | 0 | 0 | 0 | 0 | 194470 | 0 |
| VYKGHLFG | ROR2; ROR2 | Q01974; B1 | ROR2_HUMAN | Unmodified | 0 | 0 | 0 | 0 | 0 | 0 | 144190 | 65150 | 0 | 1296800 | 0 | 0 | 0 | 0 | 0 | 0 | 0 | 154790 |
| VYKPCGPI | MUC5B | Q9HC84 | MUC5B_HUMA | Unmodified | 0 | 0 | 0 | 0 | 0 | 0 | 0 | 0 | 0 | 0 | 0 | 0 | 0 | 0 | 0 | 0 | 0 | 860400 |
| VYLACRDV | RDH11 | Q8TC12 | RDH11_HUMAI | Unmodified | 0 | 0 | 311740 | 145000 | 0 | 0 | 0 | 0 | 0 | 0 | 0 | 287890 | 0 | 0 | 0 | 0 | 0 | 0 |
| VYLATDDP | FUT8; FUT8; FU | Q9BYC5; A0 | FUT8_HUMAN | Unmodified | 0 | 0 | 0 | 0 | 0 | 0 | 0 | 0 | 4142900 | 587190 | 0 | 0 | 0 | 0 | 0 | 0 | 0 | 0 |
| VYLCNQDE | DSC2; DSC2 | Q02487; A0 | DSC2_HUMAN | Unmodified | 0 | 0 | 0 | 0 | 1706700 | 0 | 0 | 0 | 702960 | 127560 | 0 | 0 | 1754100 | 1750600 | 0 | 439780 | 816840 | 2241900 |
| VYLFDFPE | SEMA7A | O75326 | SEM7A_HUMAI | Unmodified | 0 | 0 | 0 | 0 | 0 | 0 | 0 | 0 | 0 | 0 | 545350 | 0 | 0 | 0 | 0 | 0 | 0 | 0 |
| VYLFTIK | ITGA2; ITGA2; I | P17301; E9P | ITA2_HUMAN I | Unmodified | 3291800 | 2715400 | 2536900 | 885660 | 385040 | 69083 | 0 | 0 | 0 | 0 | 0 | 1735000 | 18411 | 0 | 4223500 | 3163700 | 69631 | 12955 |
| VYLGPGSD | PLXNB1 | O43157 | PLXB1_HUMAN | Unmodified | 0 | 0 | 0 | 0 | 0 | 0 | 0 | 0 | 3391000 | 0 | 0 | 0 | 0 | 0 | 0 | 0 | 0 | 0 |
| VYLHCTNT | HPSE; HPSE | Q9Y251; A0 | HPSE_HUMAN | Unmodified | 129830 | 143390 | 0 | 0 | 0 | 0 | 0 | 0 | 299380 | 0 | 0 | 0 | 0 | 0 | 570600 | 182790 | 0 | 0 |
| VYLLYRPGI | OTUB1; OTUB1 | Q96FW1; J3 | OTUB1_HUMAI | Unmodified | 1039900 | 1099100 | 199300 | 273690 | 4865200 | 2739500 | 890900 | 0 | 2580900 | 1941500 | 3698900 | 27931000 | 7171300 | 2364400 | 1792400 | 1455200 | 5009000 | 9077800 |
| VYLPEPPFF | INTS9; INTS9 | Q9NV88; G3 | INT9_HUMAN I | Unmodified | 0 | 0 | 0 | 0 | 115980 | 0 | 0 | 0 | 102630 | 0 | 79338 | 888220 | 0 | 0 | 0 | 0 | 0 | 0 |
| VYLQNEYK | LRPPRC; LRPPF | P42704; B8Z | LPPRC_HUMAN | Unmodified | 0 | 0 | 54415 | 0 | 0 | 0 | 0 | 0 | 1281700 | 0 | 68043 | 0 | 0 | 0 | 0 | 0 | 0 | 0 |
| VYLQNEYK | LRPPRC; LRPPF | P42704; B8Z | LPPRC_HUMAN | Unmodified | 0 | 0 | 0 | 0 | 0 | 0 | 105040 | 0 | 989230 | 271990 | 0 | 0 | 0 | 0 | 0 | 0 | 0 | 0 |
| VYLTFDELF | SEMA3C | Q99985 | SEM3C_HUMAI | Unmodified | 0 | 0 | 1735800 | 1324100 | 0 | 721780 | 0 | 0 | 0 | 0 | 0 | 0 | 0 | 0 | 986380 | 568510 | 0 | 0 |
| VYLTPDGT | PLXNB2; PLXNE | O15031; E2F | PLXB2_HUMAN | Unmodified | 0 | 0 | 0 | 0 | 3249100 | 0 | 0 | 0 | 0 | 655610 | 0 | 0 | 4770500 | 1306300 | 0 | 0 | 1186600 | 0 |
| VYLTWPVP | GNPTAB | Q3T906 | GNPTA_HUMAI | Unmodified | 0 | 0 | 0 | 0 | 0 | 0 | 0 | 0 | 642390 | 0 | 0 | 0 | 0 | 0 | 0 | 0 | 0 | 29594 |
| VYMGHGG | CNDP2 | Q96KP4 | CNDP2_HUMA | Unmodified | 0 | 0 | 0 | 0 | 0 | 0 | 0 | 0 | 281810 | 0 | 0 | 1428500 | 1169100 | 542240 | 0 | 384740 | 0 | 0 |
| VYMGHGG | CNDP2 | Q96KP4 | CNDP2_HUMA | Oxidation (M) | 0 | 0 | 0 | 0 | 0 | 0 | 0 | 0 | 0 | 0 | 0 | 1621200 | 941770 | 999620 | 1152700 | 608610 | 0 | 0 |
| VYMGHGG | CNDP2 | Q96KP4 | CNDP2_HUMA | Unmodified | 0 | 0 | 0 | 129220 | 0 | 0 | 0 | 0 | 0 | 82612 | 0 | 0 | 1807300 | 1151800 | 0 | 0 | 0 | 401360 |
| VYNAVANL | SMPDL3A | Q92484 | ASM3A_HUMAI | Unmodified | 88226 | 0 | 0 | 0 | 0 | 0 | 0 | 0 | 1938400 | 2130300 | 0 | 0 | 0 | 0 | 0 | 0 | 0 | 0 |
| VYNENLVH | BCAS2 | O75934 | SPF27_HUMAN | Unmodified | 0 | 0 | 0 | 0 | 579550 | 0 | 95846 | 21544 | 160890 | 212060 | 127250 | 435330 | 544520 | 124780 | 0 | 0 | 125650 | 483570 |
| VYNIEFNPF | AK3 | Q9UIJ7 | KAD3_HUMAN | Unmodified | 0 | 0 | 0 | 0 | 0 | 0 | 0 | 0 | 1960400 | 161340 | 0 | 0 | 0 | 0 | 0 | 0 | 20854 | 0 |
| VYNVDGPS | VANGL1 | Q8TAA9 | VANG1_HUMAI | Unmodified | 0 | 0 | 177230 | 267540 | 1359300 | 257480 | 0 | 28683 | 615600 | 264500 | 25328 | 0 | 0 | 0 | 165010 | 944640 | 0 | 219400 |
| VYNVTQHA | RPL21; RPL21; I | P46778; G3 | RL21_HUMAN ( | Unmodified | 17519000 | 17060000 | 8562400 | 6383400 | 2923200 | 128870 | 1999900 | 1426900 | 2593600 | 528100 | 7758500 | 2.06E+08 | 2826600 | 1087500 | 8154800 | 8476000 | 160300 | 1510900 |
| VYNVTQHA | RPL21; RPL21 | P46778; M0I | RL21_HUMAN ( | Unmodified | 0 | 0 | 0 | 0 | 0 | 0 | 544470 | 384450 | 0 | 0 | 0 | 0 | 827690 | 0 | 0 | 0 | 0 | 0 |
| VYNYNHLN | RPL27; RPL27 | P61353; K7E | RL27_HUMAN ( | Oxidation (M) | 1994500 | 1532200 | 328860 | 492000 | 0 | 0 | 0 | 0 | 0 | 0 | 0 | 7758200 | 0 | 0 | 1726000 | 1315900 | 0 | 0 |
| VYNYNHLN | RPL27; RPL27 | P61353; K7E | RL27_HUMAN ( | Unmodified | 5159500 | 6952900 | 1633200 | 1140900 | 158540 | 0 | 0 | 0 | 0 | 0 | 0 | 23581000 | 0 | 0 | 8357700 | 11640000 | 0 | 0 |
| VYNYNHLN | RPL27; RPL27 | P61353; K7E | RL27_HUMAN ( | Oxidation (M) | 0 | 0 | 0 | 36989000 | 2335800 | 0 | 1343200 | 0 | 0 | 188000 | 0 | 0 | 974020 | 0 | 0 | 0 | 628980 | 1088400 |
| VYNYNHLN | RPL27; RPL27 | P61353; K7E | RL27_HUMAN ( | Unmodified | 2054700 | 0 | 680820 | 2007800 | 2787400 | 0 | 1548100 | 0 | 0 | 404470 | 0 | 0 | 0 | 0 | 1787700 | 0 | 0 | 0 |
| VYPAEPEL | ACAD10; ACAD | Q6JQN1; F8 | ACD10_HUMAI | Unmodified | 0 | 0 | 0 | 0 | 0 | 0 | 0 | 0 | 289410 | 0 | 0 | 0 | 0 | 0 | 0 | 0 | 0 | 0 |
| VYQETSEM | MLF2; MLF2 | Q15773; F5 | MLF2_HUMAN | Unmodified | 5053600 | 6789500 | 2714100 | 3911200 | 0 | 0 | 0 | 0 | 220970 | 0 | 0 | 0 | 446990 | 138360 | 0 | 0 | 31475 | 0 |
| VYQHQVV | SHMT1 | P34896 | GLYC_HUMAN | Unmodified | 0 | 0 | 81153 | 0 | 0 | 0 | 0 | 0 | 0 | 0 | 0 | 991070 | 0 | 0 | 75853 | 43094 | 0 | 0 |
| VYQWDDP | FASN; FASN | P49327; A0A | FAS_HUMAN F | Unmodified | 0 | 0 | 0 | 0 | 0 | 0 | 0 | 0 | 738150 | 101090 | 983530 | 28921000 | 0 | 0 | 0 | 145700 | 0 | 0 |
| VYRDDTFV | KIAA1522 | Q9P206 | K1522_HUMAN | Unmodified | 0 | 0 | 338950 | 771630 | 0 | 0 | 0 | 0 | 384990 | 56620 | 0 | 1060200 | 0 | 0 | 753460 | 557030 | 0 | 0 |
| VYSATNVEI | ANKRD13D | Q6ZTN6 | AN13D_HUMAI | Unmodified | 859870 | 871610 | 0 | 0 | 0 | 0 | 0 | 0 | 0 | 0 | 0 | 0 | 0 | 0 | 320780 | 306550 | 0 | 0 |
| VYSDLHAF | MANEAL; MANE | Q5VSG8; H0 | MANEL_HUMAI | Unmodified | 0 | 0 | 0 | 109490 | 0 | 0 | 0 | 0 | 3347000 | 0 | 0 | 0 | 0 | 0 | 0 | 0 | 0 | 2320400 |
| VYSLFLDES | RUVBL2 | Q9Y230 | RUVB2_HUMAI | Unmodified | 0 | 0 | 0 | 0 | 0 | 0 | 0 | 0 | 227670 | 0 | 0 | 3210300 | 0 | 0 | 0 | 0 | 353150 | 0 |
| VYSPHVLN | DNM3; DNM2; I | Q9UQ16; P5 | DYN3_HUMAN | Unmodified | 1130500 | 793720 | 987080 | 395700 | 0 | 0 | 0 | 0 | 1058900 | 0 | 171000 | 1256800 | 0 | 0 | 2198400 | 758030 | 0 | 0 |
| VYSPHVLN | DNM2 | P50570 | DYN2_HUMAN | Unmodified | 0 | 0 | 0 | 459780 | 1424400 | 0 | 0 | 0 | 0 | 0 | 0 | 0 | 2167300 | 0 | 1314600 | 0 | 0 | 955690 |
| VYSPHVLN | DNM1; DNM1; I | Q05193; A0 | DYN1_HUMAN | Unmodified | 0 | 0 | 0 | 0 | 0 | 0 | 0 | 0 | 885540 | 0 | 0 | 0 | 0 | 0 | 0 | 0 | 0 | 0 |
| VYSSHTGY | MRPL13; MRPL | Q9BYD1; E5 | RM13_HUMAN | Unmodified | 0 | 0 | 0 | 0 | 0 | 0 | 0 | 0 | 867260 | 0 | 133790 | 0 | 0 | 0 | 0 | 0 | 0 | 0 |
| VYSSSTTG | UNC5B | Q8IZJ1 | UNC5B_HUMA | Unmodified | 0 | 0 | 0 | 0 | 0 | 0 | 0 | 0 | 985390 | 0 | 0 | 0 | 0 | 0 | 0 | 0 | 0 | 0 |
| VYSTSVTG | SH3BGRL3; SH | Q9H299; Q5 | SH3L3_HUMAN | Unmodified | 6083900 | 4453800 | 0 | 1805700 | 492820 | 0 | 60367 | 0 | 4476500 | 31705 | 960790 | 878150 | 0 | 0 | 5089100 | 2516500 | 188860 | 926290 |
| VYSTSVTG | SH3BGRL3; SH | Q9H299; Q5 | SH3L3_HUMAN | Unmodified | 0 | 0 | 0 | 0 | 2308800 | 0 | 313740 | 0 | 0 | 0 | 510810 | 0 | 0 | 1245800 | 0 | 0 | 1668500 | 7707400 |
| VYSYALALK | NUDT5; NUDT5 | Q9UKK9; A6 | NUDT5_HUMAI | Unmodified | 0 | 232610 | 0 | 546130 | 0 | 0 | 0 | 0 | 303610 | 1874100 | 4951700 | 11930000 | 125860 | 0 | 407110 | 0 | 0 | 0 |
| VYSYFECR | NAMPT; NAMPT | P43490; A0A | NAMPT_HUMAI | Unmodified | 0 | 0 | 0 | 0 | 0 | 0 | 0 | 0 | 0 | 0 | 0 | 2529100 | 0 | 0 | 0 | 0 | 0 | 0 |
| VYSYFECRI | NAMPT; NAMPT | P43490; A0A | NAMPT_HUMAI | Unmodified | 0 | 0 | 0 | 0 | 6492800 | 268430 | 0 | 0 | 0 | 150230 | 0 | 362720 | 981970 | 576820 | 0 | 0 | 625960 | 703460 |
| VYTAQNPS | PLAT; PLAT; PL | P00750; E7E | TPA_HUMAN Ti | Unmodified | 0 | 0 | 0 | 0 | 0 | 0 | 0 | 0 | 0 | 0 | 0 | 0 | 0 | 0 | 0 | 0 | 635650 | 6849500 |
| VYTDVQQV | TTLL12 | Q14166 | TTL12_HUMAN | Unmodified | 0 | 0 | 0 | 0 | 0 | 0 | 0 | 0 | 0 | 0 | 50907 | 3781900 | 0 | 0 | 1814800 | 340760 | 0 | 0 |
| VYTDVQQV | TTLL12 | Q14166 | TTL12_HUMAN | Unmodified | 0 | 0 | 0 | 0 | 0 | 0 | 0 | 0 | 0 | 0 | 0 | 0 | 1803100 | 0 | 0 | 0 | 0 | 0 |

| | | | | | | | | | | | | | | | | | | | | | | |
|---|---|---|---|---|---|---|---|---|---|---|---|---|---|---|---|---|---|---|---|---|---|---|
| VYTFNSVR | ATP2B2; ATP2B | Q01814; Q1( | AT2B2_HUMAN | Unmodified | 402400 | 0 | 676590 | 746430 | 0 | 0 | 0 | 80085 | 448280 | 0 | 0 | 8496700 | 151530 | 0 | 2180200 | 1489200 | 0 | 0 |
| VYTFNSVRI | ATP2B2; ATP2B | Q01814; Q1( | AT2B2_HUMAN | Unmodified | 128510 | 70417 | 494680 | 447380 | 1651500 | 326070 | 1107700 | 465410 | 340730 | 204930 | 0 | 3887700 | 576080 | 0 | 440170 | 0 | 245060 | 262370 |
| VYTHEVVTL | CDK1; CDK1; C | P06493; A0A | CDK1_HUMAN | Unmodified | 254750 | 0 | 0 | 206830 | 0 | 0 | 0 | 0 | 703370 | 0 | 0 | 1582500 | 0 | 0 | 1048300 | 712090 | 0 | 0 |
| VYTLSVSGI | BUB3; BUB3 | O43684; J3C | BUB3_HUMAN | Unmodified | 0 | 130700 | 0 | 273430 | 2077900 | 0 | 757550 | 988500 | 187890 | 0 | 0 | 3901500 | 0 | 2147400 | 0 | 0 | 0 | 1429700 |
| VYTLSVSGI | BUB3; BUB3 | O43684; J3C | BUB3_HUMAN | Unmodified | 0 | 0 | 0 | 303700 | 0 | 0 | 0 | 62696 | 0 | 0 | 0 | 2928200 | 0 | 0 | 0 | 0 | 0 | 0 |
| VYTLTPFLA | ARRB1; ARRB1 | P49407; H0Y | ARRB1_HUMAN | Unmodified | 0 | 923410 | 0 | 656850 | 580970 | 3366500 | 1435800 | 260900 | 0 | 338150 | 0 | 0 | 1617700 | 800510 | 1179000 | 0 | 380990 | 1314200 |
| VYTPVGKAI | NPEPPSL1; NPE | A6NEC2; P5! | PSAL_HUMAN | Unmodified | 0 | 0 | 0 | 173700 | 606410 | 219850 | 0 | 0 | 448810 | 493440 | 362070 | 1100700 | 312330 | 0 | 73756 | 73713 | 206060 | 657150 |
| VYTPVGKAI | NPEPPSL1; NPE | A6NEC2; P5! | PSAL_HUMAN | Unmodified | 0 | 0 | 0 | 0 | 662540 | 145590 | 0 | 0 | 0 | 221250 | 0 | 0 | 0 | 0 | 0 | 0 | 739180 | 814330 |
| VYTRHGVP | FLOT1; FLOT1; | O75955; A2A | FLOT1_HUMAN | Unmodified | 0 | 0 | 0 | 0 | 6448000 | 1296800 | 0 | 516780 | 0 | 1844200 | 0 | 0 | 2829100 | 1288000 | 0 | 0 | 2472800 | 2230500 |
| VYTRHGVP | FLOT1; FLOT1; | O75955; A2A | FLOT1_HUMAN | Unmodified | 0 | 0 | 0 | 0 | 0 | 0 | 0 | 0 | 0 | 0 | 0 | 0 | 0 | 0 | 0 | 0 | 692950 | 0 |
| VYTRTPTLY | PIR | O00625 | PIR_HUMAN Pi | Unmodified | 0 | 0 | 0 | 0 | 0 | 0 | 0 | 0 | 171390 | 60085 | 739580 | 0 | 148520 | 108630 | 0 | 0 | 0 | 0 |
| VYTVNNVN | ANKRD13A; AN | Q8IZ07; H0Y | AN13A_HUMAN | Unmodified | 2911700 | 2173900 | 548510 | 986540 | 4015900 | 438530 | 277920 | 70384 | 0 | 186130 | 0 | 0 | 0 | 378000 | 1010400 | 1076500 | 850110 | 1114400 |
| VYVAKVDC | TXNDC5 | Q8NBS9 | TXND5_HUMAN | Unmodified | 0 | 0 | 0 | 0 | 0 | 0 | 0 | 0 | 1263500 | 391540 | 0 | 265340 | 0 | 0 | 0 | 0 | 0 | 0 |
| VYVAKVDC | TXNDC5 | Q8NBS9 | TXND5_HUMAN | Unmodified | 0 | 0 | 0 | 0 | 0 | 0 | 0 | 0 | 0 | 4762900 | 0 | 0 | 0 | 458630 | 0 | 0 | 0 | 0 |
| VYVAVDGT | POMGNT1; POI | Q8WZA1; Q6 | PMGT1_HUMAN | Unmodified | 0 | 0 | 0 | 0 | 0 | 0 | 0 | 0 | 3283500 | 0 | 0 | 0 | 68774 | 0 | 0 | 0 | 0 | 0 |
| VYVGNLGN | SRSF3; SRSF3 | P84103; A0A | SRSF3_HUMAN | Unmodified | 83060 | 58638 | 295280 | 0 | 0 | 0 | 0 | 0 | 0 | 0 | 0 | 0 | 0 | 0 | 0 | 0 | 0 | 0 |
| VYVGNLGN | SRSF3; SRSF3 | P84103; A0A | SRSF3_HUMAN | Unmodified | 7959700 | 6705000 | 3546400 | 2429000 | 511090 | 0 | 68601 | 60904 | 597290 | 102790 | 920020 | 8334900 | 780180 | 216840 | 3129100 | 3596900 | 0 | 0 |
| VYVGNLGT | SRSF7; SRSF7; | Q16629; A0A | SRSF7_HUMAN | Unmodified | 6630900 | 4846200 | 1501300 | 493870 | 0 | 0 | 0 | 0 | 0 | 0 | 0 | 2872100 | 0 | 0 | 2112800 | 1907700 | 0 | 0 |
| VYVIEPHSN | NUP210 | Q8TEM1 | PO210_HUMAN | Oxidation (M) | 422450 | 0 | 0 | 161430 | 440390 | 0 | 0 | 0 | 16391000 | 2262700 | 451510 | 0 | 0 | 0 | 0 | 0 | 0 | 0 |
| VYVIEPHSN | NUP210 | Q8TEM1 | PO210_HUMAN | Unmodified | 0 | 0 | 0 | 0 | 0 | 0 | 32276 | 0 | 26800000 | 1014500 | 0 | 0 | 0 | 0 | 0 | 0 | 0 | 0 |
| VYVIGSFVL | TRMT10C | Q7L0Y3 | TM10C_HUMAN | Unmodified | 0 | 0 | 0 | 0 | 404950 | 0 | 0 | 0 | 2626200 | 0 | 0 | 0 | 0 | 0 | 0 | 0 | 0 | 0 |
| VYVPTGFS | EPHX1 | P07099 | HYEP_HUMAN | Unmodified | 73773 | 0 | 2285100 | 6753200 | 918000 | 0 | 1493400 | 0 | 8451300 | 3062700 | 0 | 241260 | 1243800 | 2395400 | 5220100 | 1017400 | 0 | 0 |
| VYVSDNIR | NUP210 | Q8TEM1 | PO210_HUMAN | Unmodified | 0 | 0 | 0 | 0 | 0 | 0 | 0 | 0 | 1434300 | 0 | 0 | 0 | 0 | 0 | 0 | 0 | 0 | 0 |
| VYVSGLMK | APMAP; APMAP | Q9HDC9; H( | APMAP_HUMAN | Unmodified | 0 | 0 | 0 | 0 | 0 | 0 | 0 | 0 | 18908000 | 2938400 | 0 | 0 | 0 | 0 | 0 | 0 | 0 | 0 |
| VYVVEGSK | ZMPSTE24 | O75844 | FACE1_HUMAN | Unmodified | 216410 | 0 | 177270 | 0 | 334300 | 361790 | 0 | 0 | 2023700 | 0 | 0 | 0 | 461570 | 0 | 0 | 0 | 470960 | 0 |
| VYVYLQHP | ITGA5 | P08648 | ITA5_HUMAN In | Unmodified | 0 | 18436000 | 0 | 0 | 0 | 0 | 0 | 0 | 0 | 0 | 0 | 0 | 0 | 0 | 5857700 | 4253400 | 0 | 0 |
| VYVYPQQK | EXT1 | Q16394 | EXT1_HUMAN I | Unmodified | 0 | 0 | 0 | 0 | 0 | 0 | 516030 | 309020 | 837620 | 196950 | 0 | 0 | 0 | 0 | 0 | 0 | 0 | 0 |
| VYWDNGA | PGM2 | Q96G03 | PGM2_HUMAN | Unmodified | 0 | 0 | 0 | 0 | 0 | 0 | 0 | 0 | 0 | 0 | 362640 | 971340 | 0 | 0 | 0 | 277370 | 0 | 0 |
| VYWELTDL | ITFG1; ITFG1 | Q8TB96; F5( | TIP_HUMAN T-( | Unmodified | 0 | 0 | 132660 | 48994 | 0 | 0 | 0 | 0 | 2568800 | 611540 | 0 | 0 | 0 | 0 | 231480 | 155100 | 0 | 0 |
| VYWEVEVE | TRIM26 | Q12899 | TRI26_HUMAN | Unmodified | 0 | 1830400 | 0 | 0 | 0 | 0 | 0 | 0 | 0 | 0 | 0 | 141510 | 0 | 0 | 1983200 | 0 | 237190 | 285090 |
| VYYDGTVE | C6orf120 | Q7Z4R8 | CF120_HUMAN | Unmodified | 0 | 0 | 0 | 0 | 0 | 0 | 0 | 0 | 6172200 | 443800 | 0 | 0 | 0 | 0 | 0 | 0 | 0 | 0 |
| VYYFNHIT | PIN1; PIN1 | Q13526; K7E | PIN1_HUMAN I | Unmodified | 16623000 | 12907000 | 11553000 | 6566000 | 0 | 0 | 0 | 179960 | 1520600 | 0 | 0 | 5821500 | 495130 | 0 | 11817000 | 9832800 | 0 | 0 |
| VYYLSWLR | TNIK; MINK1; M | Q9UKE5; Q8 | TNIK_HUMAN T | Unmodified | 0 | 0 | 0 | 0 | 0 | 0 | 0 | 0 | 0 | 0 | 0 | 1845200 | 0 | 0 | 0 | 0 | 0 | 0 |
| VYYPADPP | CAPN7 | Q9Y6W3 | CAN7_HUMAN | Unmodified | 0 | 0 | 0 | 0 | 0 | 0 | 0 | 0 | 0 | 0 | 0 | 1332300 | 0 | 0 | 0 | 0 | 0 | 0 |
| VYYPADPP | CAPN7 | Q9Y6W3 | CAN7_HUMAN | Unmodified | 0 | 0 | 0 | 0 | 0 | 0 | 0 | 0 | 0 | 1881100 | 434730 | 0 | 0 | 0 | 0 | 0 | 0 | 0 |
| VYYTAGYN | HEBP2 | Q9Y5Z4 | HEBP2_HUMAN | Unmodified | 0 | 0 | 0 | 287470 | 0 | 0 | 0 | 0 | 1703300 | 1190900 | 589370 | 0 | 395810 | 708890 | 0 | 599930 | 535000 | 301630 |
| VYYVDHVE | ITCH; ITCH; ITC | Q96J02; A0A | ITCH_HUMAN I | Unmodified | 2583600 | 2196800 | 0 | 0 | 0 | 15904 | 0 | 0 | 0 | 0 | 0 | 135870 | 0 | 0 | 1331000 | 1335700 | 228050 | 0 |
| WAAQTIR | MYO1C; MYO1( | O00159; F5H | MYO1C_HUMA | Unmodified | 0 | 769590 | 4826000 | 875590 | 0 | 0 | 0 | 0 | 0 | 0 | 0 | 0 | 0 | 0 | 4389400 | 2576300 | 0 | 0 |
| WAASLENL | RGS10 | O43665 | RGS10_HUMAN | Unmodified | 0 | 0 | 0 | 0 | 196120 | 0 | 6939.3 | 0 | 0 | 150570 | 0 | 0 | 0 | 0 | 0 | 0 | 107640 | 86906 |
| WAAVVVPS | HLA-E; HLA-B; | P13747; P01 | HLAE_HUMAN | Unmodified | 857820 | 1613100 | 594250 | 631280 | 1039500 | 255470 | 0 | 0 | 28932000 | 1927300 | 1829200 | 1.24E+08 | 0 | 0 | 1008500 | 1027600 | 355050 | 1254500 |
| WAAVVVPS | HLA-B; HLA-A; | P01889; P04 | HLAB_HUMAN | Unmodified | 0 | 0 | 0 | 0 | 0 | 0 | 0 | 0 | 839580 | 0 | 0 | 2494400 | 0 | 0 | 0 | 0 | 0 | 924300 |
| WAAVVVPS | HLA-B; HLA-A; | P01889; P04 | HLAB_HUMAN | Unmodified | 0 | 0 | 0 | 0 | 0 | 0 | 0 | 124470 | 2039000 | 0 | 0 | 3302500 | 0 | 0 | 0 | 0 | 0 | 0 |
| WAAVVVPS | HLA-C; HLA-A; | P10321; Q5! | HLAC_HUMAN | Unmodified | 483580 | 281660 | 1397200 | 802320 | 0 | 0 | 0 | 0 | 28792000 | 1415700 | 12217000 | 83158000 | 0 | 0 | 1019900 | 704280 | 0 | 0 |
| WACLVAG( | ADGRE5 | P48960 | AGRE5_HUMAN | Unmodified | 177610 | 0 | 0 | 0 | 0 | 117380 | 0 | 0 | 0 | 0 | 0 | 0 | 0 | 0 | 0 | 0 | 0 | 0 |
| WADNPYV | SORCS2; SORC | Q96PQ0; B5 | SORC2_HUMAN | Unmodified | 0 | 0 | 0 | 0 | 0 | 0 | 0 | 0 | 8704700 | 403570 | 0 | 262730 | 0 | 0 | 0 | 437370 | 0 | 243790 |
| WAELLPLL | RNH1; RNH1; R | P13489; E9P | RINI_HUMAN R | Unmodified | 116820 | 0 | 28150 | 65789 | 0 | 0 | 0 | 0 | 0 | 0 | 106410 | 2087300 | 0 | 0 | 399130 | 121580 | 0 | 0 |
| WAEQYLK | ERP29; ERP29 | P30040; F8V | ERP29_HUMAN | Unmodified | 267760 | 0 | 95626 | 0 | 0 | 0 | 0 | 0 | 76918000 | 20307000 | 0 | 0 | 0 | 0 | 329230 | 148440 | 0 | 0 |
| WAEQYLKI | ERP29; ERP29 | P30040; F8V | ERP29_HUMAN | Unmodified | 0 | 0 | 0 | 0 | 0 | 0 | 0 | 0 | 0 | 0 | 0 | 0 | 0 | 0 | 0 | 0 | 0 | 0 |
| WAEYHADI | PHPT1 | Q9NRX4 | PHP14_HUMAN | Unmodified | 1102700 | 724120 | 918680 | 823050 | 1892700 | 0 | 373240 | 0 | 0 | 361760 | 952980 | 6159300 | 827570 | 0 | 2048400 | 2052300 | 0 | 0 |
| WAFSCGTV | AASDHPPT | Q9NRN7 | ADPPT_HUMAN | Unmodified | 0 | 0 | 0 | 0 | 0 | 0 | 0 | 0 | 0 | 0 | 0 | 1098600 | 0 | 0 | 0 | 0 | 0 | 0 |
| WAHLDIAG | LAP3 | P28838 | AMPL_HUMAN | Unmodified | 0 | 0 | 0 | 0 | 1755800 | 0 | 462350 | 0 | 1098500 | 416990 | 161440 | 0 | 1863100 | 0 | 0 | 0 | 0 | 1252800 |
| WAIAQLNY | DYNC1H1; DYN | Q14204; A0A | DYHC1_HUMA | Oxidation (M) | 0 | 0 | 0 | 30608 | 0 | 24675 | 0 | 0 | 109090 | 79091 | 72966 | 448800 | 82146 | 0 | 0 | 29559 | 46923 | 105940 |
| WALSQSNI | VCP; VCP; VCP; | P55072; C9J | TERA_HUMAN | Unmodified | 1190500 | 1378900 | 2550500 | 1243300 | 0 | 0 | 0 | 0 | 4084300 | 0 | 1359400 | 49104000 | 0 | 0 | 1899400 | 2811500 | 0 | 0 |
| WALWFFK | EIF4E1B; EIF4E | A6NMX2; P0 | I4E1B_HUMAN | Unmodified | 118870 | 150930 | 36224 | 0 | 215000 | 54831 | 0 | 0 | 852010 | 0 | 243420 | 2902800 | 148660 | 52276 | 108650 | 79994 | 0 | 0 |
| WANFHLEN | PLS3; PLS3 | P13797; A0A | PLST_HUMAN I | Unmodified | 0 | 221060 | 0 | 0 | 318210 | 78986 | 0 | 0 | 0 | 0 | 0 | 0 | 0 | 0 | 37430 | 47155 | 503540 | 0 |
| WANLLTTS | MANEA | Q5SRI9 | MANEA_HUMA | Unmodified | 0 | 0 | 0 | 0 | 0 | 0 | 92165 | 0 | 3627500 | 243570 | 630770 | 0 | 121880 | 0 | 0 | 89370 | 0 | 0 |
| WAPNENK | ARPC1B; ARPC | O15143; C9J | ARC1B_HUMAN | Unmodified | 0 | 0 | 0 | 0 | 0 | 43937 | 0 | 0 | 0 | 0 | 105000 | 0 | 0 | 0 | 540080 | 0 | 0 | 0 |
| WAQALQE | PROM2 | Q8N271 | PROM2_HUMA | Unmodified | 558530 | 0 | 0 | 0 | 0 | 0 | 0 | 0 | 0 | 0 | 0 | 0 | 0 | 0 | 398180 | 256780 | 0 | 0 |
| WARPPISM | AP2M1; AP2M1 | Q96CW1; E! | AP2M1_HUMA | Oxidation (M) | 1223800 | 2582400 | 814270 | 0 | 3768200 | 651480 | 2917100 | 46915 | 2581300 | 1714200 | 2333300 | 9645900 | 2601200 | 1122800 | 2807500 | 1589000 | 2223900 | 3391900 |
| WARPPISM | AP2M1; AP2M1 | Q96CW1; E! | AP2M1_HUMA | Unmodified | 153140 | 290060 | 35706 | 152270 | 1309600 | 164040 | 307090 | 0 | 3292600 | 227350 | 159350 | 646450 | 505020 | 236170 | 220410 | 125160 | 340420 | 848350 |
| WARVEGTF | GMPPA; GMPP/ | Q96IJ6; A0A | GMPPA_HUMA | Unmodified | 0 | 0 | 0 | 0 | 0 | 0 | 0 | 0 | 0 | 0 | 0 | 458040 | 0 | 0 | 0 | 0 | 0 | 0 |
| WASEPEHI | CYC1 | P08574 | CY1_HUMAN C | Unmodified | 0 | 0 | 0 | 0 | 0 | 0 | 0 | 0 | 806250 | 175830 | 0 | 0 | 0 | 0 | 0 | 57957 | 0 | 0 |
| WASGLTPA | TAPBP; TAPBP; | O15533; A0A | TPSN_HUMAN | Unmodified | 0 | 0 | 0 | 0 | 0 | 0 | 0 | 0 | 3370700 | 646830 | 0 | 51831 | 0 | 0 | 0 | 0 | 0 | 0 |
| WASVVVPS | HLA-A; HLA-A; | Q9GJ45; A0/ | Q9GJ45_HUMA | Unmodified | 0 | 0 | 0 | 31794 | 10121000 | 3942100 | 622420 | 0 | 0 | 0 | 0 | 0 | 0 | 0 | 0 | 87303 | 0 | 0 |
| WASVVVPS | HLA-A; HLA-A; | Q9GJ45; A0/ | Q9GJ45_HUMA | Unmodified | 0 | 0 | 0 | 0 | 925160 | 250130 | 0 | 0 | 0 | 0 | 0 | 0 | 0 | 0 | 0 | 0 | 0 | 139380 |
| WCATSDPI | MELTF; MELTF | P08582; C9J | TRFM_HUMAN | Unmodified | 0 | 0 | 0 | 0 | 495640 | 0 | 0 | 0 | 1655800 | 136610 | 33456 | 0 | 0 | 0 | 0 | 0 | 38623 | 0 |
| WCAVQGQ | RAET1L; ULBP2 | Q5VY80; Q9I | ULBP6_HUMAN | Unmodified | 0 | 0 | 0 | 202880 | 0 | 0 | 0 | 0 | 366430 | 0 | 0 | 0 | 315400 | 0 | 184810 | 0 | 0 | 0 |
| WCEYGLTF | VDAC2; VDAC2 | P45880; Q5J | VDAC2_HUMAN | Unmodified | 0 | 0 | 0 | 0 | 0 | 0 | 0 | 0 | 63018000 | 7649100 | 878090 | 0 | 0 | 0 | 0 | 0 | 0 | 0 |
| WCGTTQN | FN1 | P02751 | FINC_HUMAN I | Unmodified | 307150 | 455880 | 5793800 | 2345000 | 298250 | 2586100 | 0 | 0 | 677210 | 0 | 699680 | 0 | 2048900 | 3262100 | 432800 | 1381500 | 237850 | 2205200 |
| WCHDNGV | FN1; FN1 | P02751; H0Y | FINC_HUMAN I | Unmodified | 101140 | 534810 | 5000000 | 161560 | 112890 | 1038500 | 0 | 0 | 121690 | 46391 | 29453 | 112330 | 348320 | 586020 | 295150 | 1171800 | 0 | 299430 |
| WCSWSLS | SLC3A2; SLC3A | P08195; A0A | 4F2_HUMAN 4I | Unmodified | 2047500 | 686200 | 509210 | 0 | 0 | 0 | 0 | 0 | 674980 | 0 | 0 | 11108000 | 0 | 0 | 4202700 | 1699600 | 0 | 0 |
| WCVLSTPE | MELTF | P08582 | TRFM_HUMAN | Unmodified | 0 | 0 | 0 | 0 | 1205700 | 496920 | 0 | 0 | 1955700 | 309840 | 1403400 | 0 | 0 | 0 | 0 | 0 | 0 | 0 |
| WDICAGN/ | BPNT2 | Q9NX62 | IMPA3_HUMAN | Unmodified | 0 | 0 | 0 | 0 | 656390 | 0 | 0 | 0 | 948690 | 0 | 0 | 0 | 0 | 0 | 0 | 0 | 0 | 0 |
| WDLGLNK | UPF1 | Q92900 | RENT1_HUMAN | Unmodified | 2552000 | 1887700 | 1663600 | 278080 | 0 | 0 | 134990 | 12843 | 0 | 0 | 0 | 1354400 | 48329 | 0 | 898130 | 1194600 | 87292 | 59456 |
| WDLSPEQI | NLN | Q9BYT8 | NEUL_HUMAN | Unmodified | 0 | 0 | 0 | 0 | 0 | 0 | 0 | 0 | 0 | 197670 | 0 | 1176200 | 0 | 0 | 0 | 0 | 89167 | 0 |
| WDPTANEI | RCN2; RCN2 | Q14257; A8I | RCN2_HUMAN | Unmodified | 0 | 0 | 0 | 0 | 0 | 0 | 0 | 0 | 4383000 | 271410 | 1079600 | 361790 | 0 | 0 | 112160 | 0 | 0 | 0 |
| WDRELISEI | HPX; HPX | P02790; Q9I | HEMO_HUMAN | Unmodified | 0 | 0 | 0 | 0 | 0 | 0 | 0 | 0 | 0 | 0 | 0 | 962700 | 0 | 0 | 0 | 0 | 0 | 0 |
| WDRQGEN | FN1; FN1 | P02751; H0Y | FINC_HUMAN I | 2 Oxidation (M) | 0 | 0 | 615260 | 0 | 0 | 0 | 0 | 0 | 0 | 0 | 0 | 0 | 0 | 0 | 0 | 0 | 0 | 0 |
| WDTANNPI | ITGB3 | P05106 | ITB3_HUMAN In | Unmodified | 997770 | 1169700 | 1136800 | 1474900 | 0 | 0 | 0 | 0 | 0 | 0 | 0 | 0 | 0 | 0 | 1297800 | 1313200 | 0 | 0 |
| WDTERVFE | LARS1; LARS1; | Q9P2J5; B4E | SYLC_HUMAN | Unmodified | 0 | 0 | 256570 | 259560 | 0 | 0 | 0 | 20644 | 370190 | 0 | 0 | 3859000 | 0 | 0 | 216450 | 115950 | 0 | 0 |
| WDTGENPI | ITGB1; ITGB1; I | P05556; H7( | ITB1_HUMAN In | Unmodified | 2524000 | 3166900 | 5092700 | 3405800 | 6411100 | 3064800 | 169150 | 86462 | 2944800 | 895810 | 1426400 | 9575900 | 0 | 0 | 7955800 | 4985800 | 1216700 | 10522000 |
| WDTGENPI | ITGB1; ITGB1; I | P05556; H7( | ITB1_HUMAN In | Unmodified | 0 | 0 | 0 | 0 | 0 | 0 | 3339900 | 0 | 28832000 | 66161000 | 2215200 | 0 | 726300 | 1290100 | 0 | 0 | 1225600 | 0 |
| WDVSTTK | EPRS1; EPRS1; | P07814; V9( | SYEP_HUMAN | Unmodified | 0 | 0 | 0 | 0 | 0 | 0 | 0 | 0 | 0 | 0 | 0 | 807990 | 0 | 0 | 412250 | 0 | 0 | 0 |
| WEAAEGLI | COPE; COPE | O14579; M0 | COPE_HUMAN | Unmodified | 0 | 0 | 0 | 0 | 0 | 0 | 0 | 0 | 0 | 0 | 0 | 1991500 | 0 | 0 | 0 | 0 | 0 | 0 |
| WEAAHVAI | HLA-A; HLA-A; | Q53Z42; A0/ | Q53Z42_HUMA | Unmodified | 0 | 0 | 0 | 0 | 0 | 0 | 0 | 0 | 0 | 0 | 2890500 | 0 | 0 | 0 | 0 | 0 | 0 | 0 |
| WEATQQVI | KLK14; KLK14 | Q9P0G3; A0. | KLK14_HUMAN | Unmodified | 0 | 0 | 0 | 0 | 0 | 0 | 0 | 0 | 2142400 | 0 | 0 | 0 | 0 | 0 | 0 | 0 | 0 | 0 |
| WEDAIPLAI | LTA4H | P09960 | LKHA4_HUMAN | Unmodified | 24761 | 26662 | 28462 | 0 | 0 | 29418 | 0 | 0 | 91638 | 22680 | 485230 | 5672900 | 0 | 0 | 68351 | 49533 | 37237 | 0 |
| WEDRGPS( | MOGS; MOGS; | Q13724; A0/ | MOGS_HUMAN | Unmodified | 0 | 0 | 0 | 0 | 0 | 0 | 0 | 0 | 3038700 | 0 | 412650 | 615210 | 0 | 1597000 | 0 | 0 | 0 | 0 |
| WEFKHPQI | EPRS1 | P07814 | SYEP_HUMAN | Unmodified | 528060 | 1677100 | 0 | 15399000 | 214390 | 2199500 | 0 | 110120 | 0 | 0 | 421240 | 991960 | 0 | 0 | 19587000 | 1559000 | 0 | 877840 |
| WELEYQLT | SMPDL3B; SMP | Q92485; F8V | ASM3B_HUMA | Oxidation (M) | 0 | 1081300 | 0 | 0 | 0 | 0 | 0 | 0 | 0 | 0 | 0 | 189190 | 0 | 0 | 3482700 | 1022400 | 0 | 0 |
| WENIATILE | KIF4A; KIF4B | O95239; Q2( | KIF4A_HUMAN | Unmodified | 121730 | 0 | 0 | 0 | 0 | 1921400 | 0 | 0 | 0 | 0 | 0 | 0 | 0 | 0 | 0 | 0 | 0 | 332680 |
| WENTLENC | NOSTRIN; NOS | Q8IVI9; E9PI | NOSTN_HUMA | Unmodified | 261760 | 2396400 | 413500 | 268020 | 0 | 0 | 0 | 0 | 97470 | 0 | 0 | 0 | 0 | 0 | 3683000 | 376850 | 0 | 0 |
| WEPDPTG | APLP2; APLP2; | Q06481; E9F | APLP2_HUMAN | Unmodified | 0 | 0 | 0 | 0 | 0 | 0 | 0 | 0 | 0 | 0 | 152120 | 364940 | 0 | 0 | 0 | 0 | 0 | 0 |
| WEQDYHL | ANO6; ANO6; A | Q4KMQ2; A0 | ANO6_HUMAN | Unmodified | 0 | 0 | 0 | 0 | 0 | 0 | 0 | 0 | 0 | 0 | 0 | 578350 | 0 | 0 | 0 | 0 | 0 | 0 |
| WEQIEGNS | GALNT2 | Q10471 | GALT2_HUMAN | Unmodified | 0 | 0 | 0 | 0 | 0 | 0 | 0 | 0 | 2721100 | 138600 | 0 | 431760 | 0 | 0 | 0 | 0 | 0 | 0 |
| WEVEEMK | CANX; CANX; C | P27824; A0A | CALX_HUMAN | Unmodified | 0 | 0 | 0 | 0 | 0 | 0 | 0 | 0 | 2361900 | 216380 | 0 | 0 | 0 | 0 | 0 | 0 | 0 | 0 |
| WEYDESYC | FAM20B | O75063 | XYLK_HUMAN ( | Unmodified | 0 | 0 | 0 | 0 | 0 | 0 | 0 | 0 | 1295300 | 0 | 0 | 0 | 0 | 0 | 0 | 0 | 0 | 0 |
| WEYMHSP | CPOX | P36551 | HEM6_HUMAN | Unmodified | 0 | 0 | 0 | 0 | 0 | 0 | 0 | 0 | 161690 | 0 | 0 | 412850 | 0 | 0 | 0 | 0 | 0 | 0 |
| WEYYDSVY | DPP4; DPP4; DI | P27487; A0A | DPP4_HUMAN | Unmodified | 150570 | 170190 | 1698500 | 0 | 0 | 0 | 0 | 0 | 0 | 0 | 0 | 0 | 0 | 0 | 0 | 1315700 | 0 | 0 |
| WFATTDWI | PRKACA; PRKA( | P17612; P22 | KAPCA_HUMAN | Unmodified | 0 | 0 | 0 | 0 | 1235500 | 0 | 1132900 | 0 | 0 | 289190 | 0 | 0 | 0 | 250050 | 0 | 0 | 0 | 816520 |
| WFCHIQH | EEF1E1 | O43324 | MCA3_HUMAN | Unmodified | 0 | 0 | 0 | 0 | 0 | 0 | 0 | 0 | 0 | 0 | 0 | 3255000 | 0 | 0 | 0 | 0 | 0 | 0 |
| WFDFPFTR | UBR4; UBR4 | Q5T4S7; A0/ | UBR4_HUMAN | Unmodified | 0 | 0 | 0 | 43806 | 0 | 0 | 0 | 0 | 137070 | 55569 | 53640 | 844500 | 0 | 0 | 0 | 63957 | 0 | 0 |
| WFDVDFPS | MUC5AC | P98088 | MUC5A_HUMA | Unmodified | 0 | 0 | 0 | 0 | 0 | 0 | 0 | 0 | 0 | 23867 | 0 | 0 | 0 | 0 | 0 | 0 | 0 | 736740 |
| WFDVDFPS | MUC5AC | P98088 | MUC5A_HUMA | Unmodified | 0 | 0 | 0 | 0 | 0 | 0 | 0 | 0 | 0 | 0 | 0 | 0 | 0 | 0 | 0 | 0 | 0 | 4037100 |
| WFDVDFPS | MUC5AC | P98088 | MUC5A_HUMA | Unmodified | 0 | 0 | 0 | 0 | 0 | 0 | 0 | 0 | 0 | 0 | 0 | 0 | 0 | 0 | 0 | 0 | 0 | 38363000 |
| WFEENASC | ILF2; ILF2 | Q12905; B4I | ILF2_HUMAN Ir | Unmodified | 0 | 798970 | 0 | 0 | 0 | 0 | 0 | 0 | 0 | 80855 | 303830 | 6335500 | 0 | 0 | 0 | 118750 | 0 | 0 |
| WFGFLEAC | EPRS1; EPRS1; | P07814; V9( | SYEP_HUMAN | Unmodified | 38118 | 0 | 0 | 0 | 0 | 0 | 106890 | 0 | 0 | 0 | 0 | 241180 | 0 | 0 | 92267 | 36918 | 0 | 0 |
| WFHPNITG | PTPN11; PTPN1 | Q06124; A0/ | PTN11_HUMAN | Unmodified | 0 | 0 | 0 | 0 | 0 | 0 | 0 | 0 | 658360 | 535390 | 0 | 0 | 459190 | 0 | 0 | 0 | 0 | 0 |
| WFKFDDHI | USP8 | P40818 | UBP8_HUMAN | Unmodified | 0 | 0 | 0 | 0 | 457740 | 0 | 0 | 156800 | 0 | 84490 | 0 | 0 | 72928 | 0 | 0 | 0 | 0 | 442040 |

| | | | | | | | | | | | | | | | | | | | | | | |
|---|---|---|---|---|---|---|---|---|---|---|---|---|---|---|---|---|---|---|---|---|---|---|
| WFKNDAP\ | ROR1; ROR1 | Q01973; A0/ | ROR1_HUMAN | Unmodified | 0 | 0 | 0 | 108910 | 937550 | 277570 | 0 | 0 | 0 | 0 | 0 | 0 | 0 | 0 | 0 | 0 | 569950 | 190280 |
| WFLTCINQ | EEF1G | P26641 | EF1G_HUMAN | Unmodified | 165160 | 110050 | 201220 | 219120 | 62323 | 22266 | 50992 | 0 | 444480 | 0 | 0 | 38199000 | 0 | 0 | 501970 | 342920 | 36120 | 0 |
| WFNGQPII | U2AF1; U2AF1L | Q01081; P0l | U2AF1_HUMAN | Unmodified | 339260 | 289900 | 104040 | 0 | 0 | 0 | 0 | 37928 | 90383 | 0 | 0 | 516460 | 3821800 | 3326800 | 94964 | 148970 | 0 | 0 |
| WFPEVTHF | RHOF; RHOF; R | Q9HBH0; V9 | RHOF_HUMAN | Unmodified | 1345300 | 691540 | 0 | 0 | 0 | 0 | 0 | 0 | 0 | 0 | 0 | 0 | 0 | 0 | 0 | 705880 | 0 | 0 |
| WFQQKYD | RPL11; RPL11 | P62913; Q5\ | RL11_HUMAN ( | Unmodified | 4520700 | 6884600 | 0 | 1751300 | 0 | 0 | 0 | 0 | 0 | 0 | 0 | 4882100 | 2241900 | 0 | 351970 | 0 | 0 | 0 |
| WFVLGVG | FKBP7; FKBP7 | Q9Y680; H7l | FKBP7_HUMAN | Unmodified | 0 | 0 | 0 | 0 | 0 | 0 | 0 | 0 | 2485600 | 1554800 | 564820 | 218340 | 0 | 0 | 0 | 0 | 0 | 0 |
| WFVPWGP | WLS; WLS; WLS | Q5T9L3; H0\ | WLS_HUMAN F | Unmodified | 0 | 0 | 0 | 0 | 0 | 0 | 0 | 0 | 1748800 | 0 | 0 | 0 | 0 | 0 | 0 | 0 | 0 | 0 |
| WFYIDFGG | SLC25A13 | Q9UJS0 | CMC2_HUMAN | Oxidation (M) | 0 | 0 | 0 | 0 | 0 | 0 | 0 | 0 | 485540 | 122910 | 0 | 0 | 378890 | 0 | 0 | 0 | 0 | 0 |
| WFYIDFGG | SLC25A13 | Q9UJS0 | CMC2_HUMAN | Unmodified | 0 | 0 | 0 | 0 | 0 | 0 | 0 | 0 | 354760 | 272740 | 0 | 0 | 0 | 0 | 0 | 0 | 0 | 0 |
| WGDAGAE | GAPDH; GAPDH | P04406; E7E | G3P_HUMAN ( | Oxidation (M) | 5415300 | 5376700 | 5338300 | 6951700 | 34225000 | 4464200 | 11049000 | 889570 | 0 | 2761300 | 10747000 | 20965000 | 22319000 | 5336300 | 13576000 | 6441600 | 11717000 | 20503000 |
| WGDAGAE | GAPDH; GAPDH | P04406; E7E | G3P_HUMAN ( | Unmodified | 576640 | 0 | 0 | 0 | 7646200 | 941990 | 0 | 0 | 2823000 | 0 | 401570 | 1188200 | 0 | 0 | 1129800 | 209190 | 0 | 4676500 |
| WGDAGAE | GAPDH; GAPDH | P04406; E7E | G3P_HUMAN ( | Unmodified | 0 | 586340 | 0 | 0 | 0 | 64187 | 649030 | 0 | 0 | 0 | 0 | 0 | 0 | 0 | 2852700 | 0 | 2150100 | 0 |
| WGDEHIP( | FLNA; FLNA; FL | P21333; A0/ | FLNA_HUMAN | Unmodified | 0 | 0 | 0 | 0 | 8065800 | 3748800 | 296910 | 11404 | 1101500 | 572470 | 1030800 | 49702 | 1030000 | 906090 | 0 | 0 | 3909400 | 2921400 |
| WGDESVP( | FLNC | Q14315 | FLNC_HUMAN | Unmodified | 0 | 0 | 0 | 148030 | 1579400 | 0 | 0 | 0 | 0 | 0 | 0 | 0 | 0 | 0 | 0 | 0 | 436700 | 0 |
| WGDIEFPP | ACLY | P53396 | ACLY_HUMAN | Unmodified | 0 | 0 | 0 | 0 | 0 | 0 | 0 | 0 | 0 | 0 | 0 | 2131800 | 0 | 0 | 0 | 0 | 0 | 0 |
| WGDKGSTI | ANTXR2; ANTX | P58335; J3K | ANTR2_HUMAN | Unmodified | 0 | 0 | 0 | 0 | 1299300 | 0 | 0 | 96669 | 233450 | 54307 | 0 | 0 | 0 | 0 | 0 | 0 | 1082200 | 3153500 |
| WGDPVTR\ | NSF; NSF | P46459; I3L( | NSF_HUMAN V | Unmodified | 0 | 0 | 0 | 0 | 0 | 0 | 0 | 0 | 0 | 0 | 0 | 0 | 2646100 | 0 | 0 | 0 | 165810 | 0 |
| WGDYDSH | PRPF8 | Q6P2Q9 | PRP8_HUMAN | Unmodified | 0 | 45780 | 0 | 0 | 0 | 0 | 0 | 0 | 0 | 0 | 0 | 1145900 | 0 | 0 | 0 | 0 | 0 | 0 |
| WGEEEAEL | CARS1; CARS1 | P49589; B4D | SYCC_HUMAN | Unmodified | 0 | 0 | 0 | 0 | 0 | 0 | 0 | 0 | 0 | 0 | 0 | 2767700 | 0 | 0 | 0 | 0 | 0 | 0 |
| WGEEHIPG | FLNB; FLNB | O75369; E7E | FLNB_HUMAN | Unmodified | 265220 | 202480 | 1124900 | 1918500 | 5552200 | 1117400 | 101470 | 0 | 1342500 | 141150 | 1104200 | 6926800 | 594930 | 0 | 1262100 | 813910 | 962020 | 1464800 |
| WGFCETEE | SEL1L; SEL1L | Q9UBV2; G3 | SE1L1_HUMAN | Unmodified | 0 | 0 | 0 | 0 | 0 | 0 | 0 | 0 | 4737700 | 0 | 0 | 0 | 0 | 0 | 0 | 0 | 0 | 0 |
| WGGIMVYI | PIGS | Q96S52 | PIGS_HUMAN ( | Oxidation (M) | 0 | 348070 | 0 | 0 | 0 | 219000 | 0 | 0 | 1138200 | 183600 | 0 | 152810 | 0 | 0 | 0 | 0 | 98422 | 0 |
| WGGIMVYI | PIGS | Q96S52 | PIGS_HUMAN ( | Unmodified | 0 | 0 | 0 | 0 | 0 | 0 | 0 | 0 | 999660 | 0 | 261030 | 0 | 0 | 0 | 0 | 0 | 0 | 0 |
| WGLGGTC | TXNRD3; TXNR | Q86VQ6; Q9 | TRXR3_HUMAN | Unmodified | 0 | 0 | 333730 | 0 | 0 | 0 | 0 | 0 | 0 | 0 | 0 | 1833500 | 0 | 0 | 0 | 0 | 0 | 0 |
| WGLLSMP\ | DNAJC13; DNA | O75165; A0/ | DJC13_HUMAN | Oxidation (M) | 0 | 0 | 102930 | 0 | 0 | 0 | 0 | 0 | 224600 | 129640 | 0 | 719110 | 0 | 0 | 0 | 0 | 0 | 0 |
| WGLQDND | RP1L1 | Q8IWN7 | RP1L1_HUMAN | Oxidation (M) | 0 | 0 | 0 | 0 | 0 | 0 | 5744000 | 570900 | 0 | 0 | 0 | 0 | 638580 | 1658700 | 0 | 0 | 0 | 0 |
| WGQALPV/ | IL17C | Q9P0M4 | IL17C_HUMAN | Unmodified | 0 | 0 | 0 | 0 | 8198.5 | 0 | 0 | 0 | 947330 | 0 | 0 | 0 | 0 | 0 | 0 | 0 | 0 | 0 |
| WGQASAA/ | FARP1 | A0A1B0GV6 | A0A1B0GV68_I | Unmodified | 0 | 0 | 156920 | 0 | 0 | 0 | 0 | 0 | 942110 | 132780 | 0 | 0 | 0 | 0 | 425610 | 229360 | 88004 | 239340 |
| WGQYHTS | MAN1A2; MAN1 | O60476; H0\ | MA1A2_HUMAI | Unmodified | 0 | 0 | 0 | 0 | 0 | 0 | 0 | 0 | 3072500 | 295650 | 0 | 0 | 0 | 0 | 0 | 0 | 0 | 0 |
| WGRDDIEI | SMARCA5 | O60264 | SMCA5_HUMA | Unmodified | 1166200 | 0 | 0 | 0 | 0 | 0 | 0 | 0 | 0 | 0 | 0 | 0 | 0 | 0 | 0 | 0 | 0 | 0 |
| WGSKPSN | CEP55 | Q53EZ4 | CEP55_HUMAN | Unmodified | 0 | 0 | 0 | 0 | 5615200 | 325830 | 0 | 0 | 269570 | 313260 | 0 | 0 | 0 | 0 | 0 | 0 | 0 | 20928 |
| WGTDEAQI | ANXA6 | P08133 | ANXA6_HUMAI | Unmodified | 241400 | 877170 | 644390 | 786660 | 0 | 20228 | 0 | 0 | 12606000 | 529060 | 0 | 0 | 0 | 0 | 5276200 | 961580 | 0 | 0 |
| WGTDEAQI | ANXA6 | P08133 | ANXA6_HUMAI | Unmodified | 0 | 0 | 0 | 0 | 854230 | 1187000 | 928250 | 0 | 0 | 2321600 | 0 | 0 | 0 | 0 | 0 | 0 | 0 | 810730 |
| WGTFSNL\ | ECE1; ECE1; EC | P42892; B4D | ECE1_HUMAN | Unmodified | 470350 | 322540 | 488980 | 732650 | 0 | 202820 | 0 | 0 | 31582000 | 1602400 | 0 | 882070 | 0 | 0 | 962780 | 518460 | 0 | 0 |
| WGTIEVEN | SEPTIN9 | Q9UHD8 | SEPT9_HUMAN | Unmodified | 0 | 169320 | 0 | 250660 | 0 | 0 | 0 | 0 | 101960 | 0 | 89014 | 2342000 | 0 | 0 | 442360 | 0 | 0 | 0 |
| WGTLTDC\ | HNRNPA3; HNI | P51991; A0/ | ROA3_HUMAN | Unmodified | 0 | 0 | 0 | 0 | 1207800 | 0 | 1455400 | 314850 | 790030 | 954930 | 0 | 0 | 894630 | 377870 | 0 | 0 | 188260 | 1674000 |
| WGTLTDC\ | HNRNPA3; HNI | P51991; A0/ | ROA3_HUMAN | Oxidation (M) | 0 | 0 | 0 | 0 | 379470 | 0 | 1853600 | 687080 | 0 | 331020 | 0 | 1294700 | 670090 | 1199800 | 0 | 78986 | 0 | 843330 |
| WGTPVPLE | MARS1 | P56192 | SYMC_HUMAN | Unmodified | 0 | 0 | 0 | 0 | 0 | 0 | 0 | 0 | 0 | 0 | 0 | 3877100 | 0 | 0 | 0 | 0 | 703270 | 0 |
| WHLGVGP | ARSA; ARSA | P15289; A0/ | ARSA_HUMAN | Unmodified | 0 | 0 | 0 | 0 | 0 | 0 | 0 | 0 | 1332600 | 0 | 0 | 0 | 0 | 0 | 0 | 0 | 0 | 0 |
| WHNLPAH | LRBA; LRBA; LR | P50851; A0/ | LRBA_HUMAN | Oxidation (M) | 0 | 0 | 0 | 0 | 0 | 0 | 0 | 0 | 0 | 0 | 0 | 1726900 | 0 | 0 | 0 | 0 | 0 | 0 |
| WIADRTDV | MAGT1; MAGT1 | Q9H0U3; A0 | MAGT1_HUMAI | Unmodified | 0 | 0 | 0 | 0 | 0 | 0 | 114530 | 0 | 25310000 | 1625400 | 0 | 1918500 | 0 | 0 | 0 | 0 | 0 | 0 |
| WIAIESLAD | AXL; MERTK; ME | P30530; Q12 | UFO_HUMAN T | Unmodified | 0 | 0 | 0 | 0 | 2150500 | 1362700 | 0 | 0 | 0 | 0 | 0 | 0 | 0 | 0 | 0 | 149130 | 0 | 0 |
| WIANNWIN | P4HTM; P4HTM | Q9NXG6; V9 | P4HTM_HUMAI | Unmodified | 0 | 0 | 0 | 0 | 0 | 0 | 0 | 0 | 0 | 0 | 0 | 0 | 0 | 0 | 0 | 0 | 0 | 0 |
| WIDFDNDY | IARS1; IARS1; I/ | P41252; A0/ | SYIC_HUMAN I | Unmodified | 0 | 0 | 0 | 0 | 0 | 0 | 0 | 0 | 0 | 0 | 0 | 4035000 | 0 | 0 | 0 | 0 | 0 | 0 |
| WIDIHNPA | ALDH6A1 | Q02252 | MMSA_HUMAN | Unmodified | 0 | 0 | 0 | 0 | 0 | 0 | 0 | 0 | 464020 | 0 | 0 | 0 | 0 | 0 | 0 | 0 | 0 | 0 |
| WIDNPTVD | VCL | P18206 | VINC_HUMAN ' | Unmodified | 0 | 0 | 0 | 0 | 0 | 0 | 0 | 0 | 0 | 0 | 104480 | 1914100 | 0 | 0 | 0 | 0 | 0 | 0 |
| WIDNPTVD | VCL | P18206 | VINC_HUMAN ' | Unmodified | 2506100 | 4592700 | 994830 | 2966800 | 1596900 | 0 | 0 | 0 | 2797300 | 0 | 238040 | 12585000 | 0 | 0 | 3020400 | 2574900 | 1340800 | 0 |
| WIEAGPVV | PTK7; PTK7; PTI | Q13308; Q8( | PTK7_HUMAN I | Unmodified | 81940 | 50592 | 218900 | 356020 | 0 | 0 | 82350 | 0 | 4329200 | 0 | 287210 | 200570 | 309500 | 76434 | 1213500 | 807840 | 0 | 340220 |
| WIEAGPVV | PTK7; PTK7; PTI | Q13308; Q8( | PTK7_HUMAN I | Unmodified | 0 | 0 | 0 | 0 | 1441600 | 0 | 0 | 0 | 1407300 | 1488500 | 0 | 0 | 1405600 | 0 | 0 | 0 | 0 | 1182700 |
| WIEDGIPKI | GNG13 | Q9P2W3 | GBG13_HUMAI | Oxidation (M) | 0 | 0 | 0 | 0 | 0 | 0 | 0 | 0 | 0 | 0 | 38009000 | 1.04E+08 | 0 | 0 | 0 | 0 | 456090 | 0 |
| WIEDGIPKI | GNG13 | Q9P2W3 | GBG13_HUMAI | Unmodified | 0 | 0 | 0 | 0 | 0 | 133930 | 44411 | 44368 | 0 | 0 | 1534700 | 3686000 | 0 | 0 | 0 | 0 | 0 | 329670 |
| WIEYDRTE | CPNE2; CPNE2 | Q96FN4; H3 | CPNE2_HUMAN | Unmodified | 170660 | 0 | 77814 | 174660 | 4834900 | 0 | 0 | 0 | 7369900 | 549240 | 0 | 111130 | 91255 | 0 | 398720 | 162240 | 2049300 | 0 |
| WIEYDRTE | CPNE2; CPNE2 | Q96FN4; H3 | CPNE2_HUMAI | Unmodified | 0 | 0 | 0 | 0 | 2033600 | 227840 | 324150 | 93859 | 1004900 | 130710 | 0 | 786760 | 245010 | 0 | 0 | 0 | 0 | 475810 |
| WIFVGGKG | GET3 | O43681 | GET3_HUMAN | Unmodified | 0 | 0 | 0 | 0 | 0 | 0 | 635010 | 0 | 0 | 0 | 0 | 0 | 740600 | 0 | 0 | 0 | 677090 | 2652500 |
| WIGLDLSN | TCP1; TCP1; TC | P17987; F5H | TCPA_HUMAN | Unmodified | 0 | 0 | 0 | 1037300 | 255530 | 57598 | 0 | 0 | 888390 | 120070 | 1118500 | 25263000 | 611420 | 0 | 0 | 0 | 86459 | 121740 |
| WIGLDLSN | TCP1; TCP1; TC | P17987; F5H | TCPA_HUMAN | Unmodified | 0 | 0 | 210930 | 145880 | 1112600 | 0 | 0 | 0 | 322250 | 300420 | 0 | 220410 | 1621500 | 0 | 0 | 0 | 0 | 0 |
| WIHDVRTI/ | ARRDC1; ARRD | Q8N5I2; Q5l | ARRD1_HUMAI | Unmodified | 0 | 0 | 0 | 75955 | 0 | 0 | 0 | 0 | 0 | 232610 | 0 | 0 | 370970 | 309040 | 0 | 0 | 229860 | 454470 |
| WILSQTHN | SLC30A7 | Q8NEW0 | ZNT7_HUMAN . | Unmodified | 0 | 0 | 0 | 0 | 0 | 0 | 0 | 0 | 621640 | 0 | 0 | 0 | 0 | 0 | 0 | 0 | 0 | 0 |
| WINATDPS | ASNS | P08243 | ASNS_HUMAN | Unmodified | 679610 | 0 | 820950 | 2118500 | 2108700 | 0 | 0 | 0 | 1072300 | 0 | 1608500 | 6418100 | 3858000 | 1100600 | 1253500 | 2105100 | 103630 | 974520 |
| WINATDPS | ASNS | P08243 | ASNS_HUMAN | Unmodified | 0 | 0 | 0 | 0 | 1975300 | 0 | 934790 | 800780 | 0 | 292680 | 0 | 0 | 2177700 | 3529900 | 0 | 0 | 864130 | 773600 |
| WINDVEDS | ATP1A1; ATP1A | P05023; Q5l | AT1A1_HUMAN | Unmodified | 0 | 0 | 1130100 | 0 | 0 | 0 | 0 | 0 | 0 | 0 | 0 | 15649000 | 0 | 0 | 4237600 | 3307600 | 0 | 0 |
| WINDVEDS | ATP1A1; ATP1A | P05023; Q5l | AT1A1_HUMAN | Unmodified | 0 | 0 | 0 | 0 | 0 | 0 | 0 | 0 | 0 | 0 | 0 | 1735800 | 0 | 0 | 0 | 0 | 0 | 0 |
| WIQEAETT\ | UTRN | P46939 | UTRN_HUMAN | Unmodified | 0 | 0 | 0 | 661610 | 0 | 0 | 0 | 0 | 0 | 0 | 0 | 0 | 0 | 0 | 1089800 | 0 | 0 | 0 |
| WIQGLQET | ECSIT | Q9BQ95 | ECSIT_HUMAN | Unmodified | 0 | 0 | 0 | 0 | 0 | 0 | 0 | 0 | 565800 | 0 | 0 | 0 | 0 | 0 | 0 | 0 | 0 | 0 |
| WIRENISE\ | PRPS1; PRPS1; | P60891; A0/ | PRPS1_HUMAN | Unmodified | 74570 | 0 | 426280 | 493400 | 319350 | 28518 | 0 | 15726 | 1039300 | 0 | 776730 | 4956400 | 800040 | 0 | 0 | 518020 | 102110 | 0 |
| WIRENISE\ | PRPS1; PRPS1; | P60891; A0/ | PRPS1_HUMAN | Unmodified | 0 | 0 | 0 | 0 | 645890 | 0 | 32029 | 0 | 0 | 0 | 0 | 0 | 1320300 | 879140 | 0 | 477350 | 0 | 0 |
| WISDHEYL | DPP4; DPP4; DI | P27487; A0/ | DPP4_HUMAN | Unmodified | 526790 | 471630 | 6520500 | 1963400 | 0 | 0 | 0 | 0 | 0 | 0 | 0 | 0 | 0 | 0 | 16490000 | 5355800 | 0 | 0 |
| WISDWNL | EIF3M; EIF3M; E | Q7L2H7; E9l | EIF3M_HUMAN | Unmodified | 0 | 0 | 326440 | 0 | 0 | 0 | 0 | 0 | 0 | 0 | 0 | 1756600 | 0 | 0 | 0 | 0 | 0 | 0 |
| WISIMTER | ANXA2P2; ANX/ | A6NMY6; P0 | AXA2L_HUMAN | Unmodified | 300810 | 117130 | 0 | 0 | 26891000 | 10718000 | 0 | 0 | 8348000 | 3108000 | 0 | 607190 | 0 | 0 | 0 | 0 | 2508600 | 6719200 |
| WISIMTER | ANXA2P2; ANX/ | A6NMY6; P0 | AXA2L_HUMAN | Oxidation (M) | 54900 | 0 | 0 | 0 | 11071000 | 1218500 | 0 | 0 | 0 | 1599700 | 0 | 0 | 0 | 0 | 0 | 0 | 1462900 | 1635200 |
| WISIMTERS | ANXA2P2; ANX/ | A6NMY6; P0 | AXA2L_HUMAN | Unmodified | 0 | 0 | 0 | 0 | 27177000 | 2612200 | 252670 | 0 | 0 | 770700 | 0 | 0 | 0 | 0 | 0 | 0 | 7609000 | 5533600 |
| WISIMTERS | ANXA2P2; ANX/ | A6NMY6; P0 | AXA2L_HUMAN | Oxidation (M) | 0 | 0 | 0 | 0 | 6142500 | 148060 | 237030 | 0 | 0 | 0 | 0 | 0 | 0 | 0 | 0 | 0 | 1757100 | 3152900 |
| WISLNTVAI | CLTC; CLTC | Q00610; A0/ | CLH1_HUMAN | Oxidation (M) | 0 | 468750 | 0 | 0 | 0 | 0 | 0 | 0 | 0 | 0 | 0 | 0 | 0 | 0 | 487380 | 0 | 0 | 0 |
| WITAKEDD | LTA4H | P09960 | LKHA4_HUMAI | Unmodified | 409860 | 0 | 78287 | 0 | 609000 | 0 | 0 | 0 | 954890 | 0 | 1243100 | 9908300 | 0 | 0 | 175230 | 0 | 182800 | 0 |
| WIYEDVER | CALU | O43852 | CALU_HUMAN | Unmodified | 0 | 0 | 0 | 0 | 0 | 0 | 0 | 0 | 2367400 | 0 | 0 | 127930 | 0 | 0 | 0 | 0 | 0 | 0 |
| WKAYDATH | ESD; ESD; ESD; | P10768; X6R | ESTD_HUMAN | Unmodified | 0 | 92505 | 0 | 0 | 644370 | 0 | 374930 | 314580 | 0 | 166830 | 0 | 0 | 346530 | 284050 | 0 | 0 | 736980 | 0 |
| WKDSDEAI | CBX3 | Q13185 | CBX3_HUMAN | Unmodified | 0 | 0 | 0 | 0 | 0 | 0 | 0 | 207020 | 0 | 0 | 510640 | 2299700 | 600830 | 0 | 0 | 0 | 83752 | 0 |
| WKEATIPG | FN1 | P02751 | FINC_HUMAN I | Unmodified | 0 | 113520 | 73651 | 55689 | 0 | 0 | 0 | 0 | 0 | 227860 | 0 | 0 | 0 | 0 | 0 | 1322500 | 0 | 0 |
| WKEYRGQ | EPB41L5 | Q9HCM4 | E41L5_HUMAN | Unmodified | 0 | 0 | 0 | 0 | 0 | 0 | 0 | 0 | 0 | 809240 | 0 | 0 | 0 | 213440 | 0 | 0 | 317140 | 39576 |
| WKGEIQEF | RND3 | P61587 | RND3_HUMAN | Unmodified | 0 | 60486 | 0 | 0 | 386880 | 81438 | 0 | 0 | 0 | 0 | 0 | 403880 | 0 | 0 | 39954 | 0 | 692150 | 979530 |
| WKGQYDV | BMX | P51813 | BMX_HUMAN ( | Unmodified | 0 | 0 | 0 | 0 | 0 | 0 | 0 | 0 | 0 | 0 | 258370 | 1223800 | 367980 | 0 | 0 | 0 | 0 | 0 |
| WKIARQES | DSG2 | Q14126 | DSG2_HUMAN | Unmodified | 0 | 0 | 0 | 0 | 1986200 | 413200 | 0 | 0 | 0 | 1164100 | 762690 | 0 | 344870 | 596040 | 0 | 0 | 0 | 160920 |
| WKPDYDS/ | NAPG; NAPG | Q99747; J3K | SNAG_HUMAN | Unmodified | 0 | 0 | 0 | 0 | 0 | 0 | 0 | 0 | 0 | 75303 | 104560 | 234240 | 0 | 0 | 67417 | 0 | 520780 | 621790 |
| WKPGSLAS | CDK1; CDK1 | P06493; A0/ | CDK1_HUMAN | Unmodified | 0 | 0 | 0 | 0 | 0 | 0 | 0 | 0 | 0 | 0 | 0 | 1033600 | 0 | 0 | 0 | 0 | 0 | 1079900 |
| WKPLIHSP | EIF2AK3 | Q9NZJ5 | E2AK3_HUMAN | Unmodified | 0 | 0 | 0 | 0 | 0 | 0 | 0 | 0 | 3125500 | 356530 | 0 | 0 | 0 | 0 | 0 | 0 | 0 | 0 |
| WKPPMIDI | CANX; CANX; C | P27824; A0/ | CALX_HUMAN | Oxidation (M) | 0 | 0 | 0 | 0 | 0 | 40647 | 0 | 0 | 2866800 | 0 | 321550 | 0 | 0 | 0 | 0 | 0 | 0 | 4687700 |
| WKPPMIDI | CANX; CANX; C | P27824; A0/ | CALX_HUMAN | Unmodified | 0 | 0 | 0 | 0 | 0 | 0 | 0 | 0 | 2166100 | 0 | 0 | 0 | 0 | 0 | 0 | 0 | 0 | 0 |
| WKPPMIDI | CANX; CANX; C | P27824; A0/ | CALX_HUMAN | Oxidation (M) | 525710 | 0 | 0 | 0 | 0 | 0 | 0 | 0 | 1.64E+08 | 32617000 | 15842000 | 6323900 | 0 | 0 | 1170000 | 570030 | 0 | 0 |
| WKPPMIDI | CANX; CANX; C | P27824; A0/ | CALX_HUMAN | Unmodified | 0 | 0 | 0 | 0 | 0 | 0 | 0 | 0 | 1.97E+08 | 5687400 | 489680 | 0 | 426370 | 0 | 0 | 0 | 0 | 0 |
| WKQPLTVI | ITCH; ITCH; ITC | Q96J02; A0/ | ITCH_HUMAN I | Unmodified | 6432200 | 11564000 | 4597800 | 0 | 3023800 | 1218000 | 787790 | 103980 | 0 | 762690 | 0 | 0 | 1263000 | 1275000 | 6494300 | 5123100 | 691560 | 2022300 |
| WKRPPENI | PIGT; PIGT; PIG | Q969N2; A0 | PIGT_HUMAN ( | Unmodified | 0 | 0 | 0 | 0 | 0 | 0 | 0 | 0 | 16123000 | 0 | 0 | 0 | 0 | 0 | 0 | 0 | 0 | 0 |
| WKRPTELL | CAPN1; CAPN1 | P07384; E9P | CAN1_HUMAN | Unmodified | 103300 | 0 | 0 | 0 | 0 | 0 | 0 | 0 | 0 | 0 | 0 | 10450000 | 0 | 0 | 490210 | 161170 | 101120 | 0 |
| WKSDEVDI | AKAP12 | Q02952 | AKA12_HUMAN | Unmodified | 0 | 122570 | 0 | 0 | 985310 | 223410 | 0 | 0 | 0 | 0 | 0 | 0 | 0 | 0 | 0 | 0 | 563770 | 1402200 |
| WKSDYPM | UBTD1; UBTD2 | Q9HAC8; Q8 | UBTD1_HUMAI | Oxidation (M) | 0 | 0 | 40943 | 0 | 0 | 0 | 0 | 21431 | 110240 | 0 | 0 | 254660 | 0 | 0 | 0 | 0 | 123050 | 561660 |
| WKSDYPM | UBTD1; UBTD2 | Q9HAC8; Q8 | UBTD1_HUMAI | Unmodified | 0 | 0 | 122960 | 173080 | 0 | 155210 | 0 | 0 | 1184000 | 0 | 0 | 0 | 367460 | 183040 | 217160 | 122200 | 0 | 1702600 |
| WKYEKPD( | ALCAM; ALCAM | Q13740; F5( | CD166_HUMAI | Unmodified | 9042700 | 10917000 | 2525200 | 281520 | 4279800 | 1142700 | 0 | 0 | 9443300 | 2019300 | 286580 | 1650200 | 2239800 | 0 | 6345600 | 2689000 | 685140 | 4578400 |
| WLAIDANA | KPNB1; KPNB1 | Q14974; J3( | IMB1_HUMAN I | Unmodified | 115740 | 159460 | 214620 | 191830 | 0 | 0 | 146540 | 18010 | 584330 | 324670 | 0 | 1406400 | 0 | 0 | 431140 | 582710 | 0 | 0 |
| WLAPDGLI | PRMT1; PRMT1; | Q99873; F5H | ANM1_HUMAN | Unmodified | 0 | 0 | 0 | 0 | 0 | 0 | 0 | 0 | 201000 | 189900 | 483380 | 7778100 | 0 | 0 | 0 | 0 | 0 | 0 |
| WLAVDYEI | OAT | P04181 | OAT_HUMAN C | Unmodified | 0 | 0 | 0 | 0 | 0 | 0 | 0 | 0 | 8442300 | 1598300 | 214020 | 0 | 0 | 0 | 0 | 0 | 0 | 0 |
| WLCPLSGI | SRRT; SRRT | Q9BXP5; H7 | SRRT_HUMAN | Unmodified | 0 | 0 | 0 | 0 | 0 | 0 | 0 | 99215 | 149320 | 67290 | 0 | 253680 | 72502 | 0 | 0 | 0 | 0 | 0 |
| WLDAIIR | PREX1 | Q8TCU6 | PREX1_HUMAN | Unmodified | 0 | 0 | 0 | 0 | 0 | 0 | 0 | 0 | 677790 | 0 | 0 | 517340 | 0 | 0 | 0 | 0 | 0 | 0 |
| WLEMYGVI | EPB41L5 | Q9HCM4 | E41L5_HUMAN | 2 Oxidation (M) | 0 | 0 | 0 | 460330 | 0 | 0 | 0 | 0 | 0 | 0 | 0 | 0 | 183310 | 0 | 800000 | 0 | 0 | 0 |
| WLEMYGVI | EPB41L4B | Q9H329 | E41LB_HUMAN | 2 Oxidation (M) | 541070 | 0 | 322630 | 0 | 0 | 0 | 0 | 13387 | 147060 | 60763 | 0 | 310380 | 0 | 0 | 2621400 | 0 | 0 | 0 |
| WLEQAAR | KNTC1; KNTC1 | P50748; E7E | KNTC1_HUMAI | Unmodified | 0 | 0 | 0 | 0 | 1042300 | 209620 | 0 | 0 | 226410 | 0 | 61060 | 302150 | 0 | 0 | 0 | 34299 | 183770 | 689730 |
| WLGANAFI | GUSB | P08236 | BGLR_HUMAN | Unmodified | 0 | 0 | 0 | 0 | 0 | 0 | 0 | 0 | 22199000 | 2294300 | 0 | 0 | 0 | 0 | 225140 | 0 | 0 | 0 |
| WLGESEK | VPS4B; VPS4A; | O75351; Q9l | VPS4B_HUMAN | Unmodified | 0 | 0 | 1818600 | 0 | 5105100 | 630440 | 822780 | 322270 | 4522800 | 3064500 | 1116000 | 3509900 | 4214400 | 5776000 | 1262100 | 1586000 | 675890 | 1554600 |
| WLGLEEAC | NUDT2 | P50583 | AP4A_HUMAN | Unmodified | 56158 | 0 | 0 | 118750 | 0 | 0 | 0 | 0 | 0 | 45647 | 0 | 95817 | 0 | 0 | 213650 | 65324 | 0 | 0 |
| WLHEINQI | RAB35; RAB35 | Q15286; F5H | RAB35_HUMAI | Unmodified | 4105500 | 2537200 | 0 | 1807500 | 41549 | 0 | 0 | 0 | 758980 | 0 | 0 | 1852600 | 210850 | 0 | 2207100 | 1845300 | 244180 | 80109 |
| WLHGLES( | SPTBN1; SPTBN | Q01082; A0/ | SPTB2_HUMAN | Unmodified | 0 | 0 | 0 | 0 | 1152300 | 250410 | 92179 | 0 | 911800 | 209660 | 401660 | 1669600 | 50449 | 0 | 0 | 0 | 177240 | 549960 |

| | | | | | | | | | | | | | | | | | | | | | | |
|---|---|---|---|---|---|---|---|---|---|---|---|---|---|---|---|---|---|---|---|---|---|---|
| WLHNEDQ | TALDO1 | P37837 | TALDO_HUMAN | Unmodified | 82450 | 0 | 0 | 0 | 1489500 | 0 | 0 | 0 | 3245600 | 366700 | 6737700 | 16958000 | 1259600 | 172310 | 0 | 218520 | 414870 | 0 |
| WLHNEDQ | TALDO1 | P37837 | TALDO_HUMAN | Oxidation (M) | 0 | 0 | 0 | 243940 | 572730 | 0 | 0 | 0 | 772370 | 0 | 1655700 | 7549800 | 308220 | 0 | 0 | 0 | 0 | 0 |
| WLIPLLEGI | ACAD10 | Q6JQN1 | ACD10_HUMAI | Unmodified | 0 | 0 | 0 | 0 | 0 | 0 | 0 | 0 | 471520 | 25381 | 0 | 0 | 0 | 0 | 0 | 0 | 0 | 0 |
| WLISTDLD( | LRP2; LRP2 | P98164; A0A | LRP2_HUMAN | Unmodified | 0 | 0 | 0 | 0 | 0 | 0 | 0 | 0 | 835810 | 424140 | 0 | 0 | 0 | 0 | 0 | 0 | 0 | 0 |
| WLITLNK | EIF4E; EIF4E; EI | P06730; D6F | IF4E_HUMAN E | Unmodified | 258770 | 2460400 | 0 | 0 | 421400 | 0 | 0 | 0 | 567040 | 0 | 1748600 | 8753400 | 0 | 0 | 588810 | 0 | 0 | 258250 |
| WLKEDKLE | CLTCL1; CLTC; | P53675; Q0( | CLH2_HUMAN | Unmodified | 14575000 | 8141900 | 0 | 8766100 | 13848000 | 3859700 | 9087700 | 111130 | 0 | 2381500 | 189290 | 19574000 | 12696000 | 2734300 | 1722800 | 3232900 | 5741100 | 7138000 |
| WLLAAGA( | TPP1; TPP1; TPI | O14773; E7E | TPP1_HUMAN | Unmodified | 0 | 0 | 0 | 0 | 0 | 0 | 0 | 0 | 1024500 | 628170 | 0 | 170410 | 0 | 0 | 0 | 0 | 0 | 0 |
| WLLDLNQI | TRPM5; TRPM5; | Q9NZQ8; A0 | TRPM5_HUMAI | Unmodified | 0 | 0 | 484450 | 532850 | 0 | 0 | 0 | 0 | 0 | 0 | 0 | 2008900 | 261310 | 0 | 758160 | 673840 | 0 | 0 |
| WLLLCNP( | PYGB | P11216 | PYGB_HUMAN | Unmodified | 0 | 0 | 0 | 0 | 0 | 0 | 0 | 0 | 0 | 54528 | 40868 | 0 | 0 | 0 | 0 | 0 | 0 | 252620 |
| WLLLNDPE | MYOF | Q9NZM1 | MYOF_HUMAN | Unmodified | 0 | 0 | 0 | 0 | 1080300 | 375580 | 0 | 0 | 0 | 0 | 0 | 0 | 0 | 0 | 0 | 0 | 57641 | 81545 |
| WLLLTGIS/ | CLTC; CLTC | Q00610; A0/ | CLH1_HUMAN | Unmodified | 4422300 | 18890000 | 0 | 4713900 | 236730 | 92020 | 0 | 0 | 1487600 | 0 | 360950 | 22245000 | 0 | 0 | 25368000 | 14122000 | 0 | 0 |
| WLLSKYDP | C1GALT1 | Q9NS00 | C1GLT_HUMAI | Unmodified | 0 | 0 | 0 | 0 | 0 | 0 | 0 | 0 | 2523500 | 0 | 0 | 0 | 0 | 0 | 0 | 0 | 0 | 0 |
| WLLSKYDP | C1GALT1 | Q9NS00 | C1GLT_HUMAI | Unmodified | 0 | 0 | 0 | 0 | 870940 | 0 | 0 | 0 | 5716900 | 0 | 0 | 0 | 0 | 0 | 0 | 0 | 405780 | 0 |
| WLLTAAHC | KLK11; KLK11; I | Q9UBX7; M0 | KLK11_HUMAN | Unmodified | 0 | 0 | 0 | 0 | 0 | 199740 | 0 | 0 | 2388000 | 966710 | 0 | 0 | 0 | 0 | 0 | 0 | 0 | 0 |
| WLLYLVKVI | ITGA6 | P23229 | ITA6_HUMAN I | Unmodified | 0 | 0 | 0 | 0 | 2498700 | 0 | 1267200 | 0 | 0 | 148360 | 0 | 0 | 1534300 | 2528400 | 0 | 0 | 3793700 | 6631100 |
| WLNENAVE | AHCY | P23526 | SAHH_HUMAN | Unmodified | 0 | 2101500 | 0 | 2327000 | 1767500 | 780200 | 934830 | 238790 | 1798300 | 688180 | 3943300 | 23849000 | 1620000 | 1281700 | 0 | 1814000 | 1063500 | 1536900 |
| WLNVHYH | CAMK2G; CAMI | Q13555; A0/ | KCC2G_HUMA | Unmodified | 0 | 0 | 0 | 0 | 571940 | 0 | 0 | 0 | 0 | 0 | 0 | 0 | 0 | 0 | 0 | 0 | 1874200 | 1847400 |
| WLPAGDAI | EEF2 | P13639 | EF2_HUMAN El | Oxidation (M) | 0 | 586930 | 0 | 159040 | 2694700 | 264830 | 1308900 | 0 | 0 | 847320 | 0 | 553570 | 1927000 | 1500600 | 678120 | 0 | 691670 | 1797300 |
| WLPAGTGI | VASP; VASP; VA | P50552; K7E | VASP_HUMAN | Unmodified | 0 | 0 | 460190 | 976480 | 0 | 0 | 0 | 0 | 429310 | 241810 | 243340 | 3251900 | 1725400 | 470880 | 567970 | 0 | 0 | 0 |
| WLPLEANP | UCHL3 | P15374 | UCHL3_HUMAI | Unmodified | 573980 | 0 | 0 | 466860 | 0 | 0 | 250570 | 0 | 758850 | 0 | 2077300 | 8157700 | 0 | 767510 | 0 | 407980 | 0 | 0 |
| WLPLGLGL | PSMD2; PSMD2 | Q13200; H7( | PSMD2_HUMA | Unmodified | 0 | 0 | 0 | 0 | 0 | 0 | 0 | 0 | 0 | 0 | 0 | 1394400 | 0 | 0 | 0 | 0 | 0 | 0 |
| WLPSLVLAI | SLC29A1 | Q99808 | S29A1_HUMAN | Unmodified | 1120700 | 656490 | 495360 | 0 | 0 | 0 | 0 | 0 | 0 | 0 | 0 | 4941400 | 0 | 0 | 2923200 | 1506300 | 0 | 0 |
| WLPSSSPV | FN1; FN1 | P02751; H0Y | FINC_HUMAN I | Unmodified | 977870 | 4995100 | 29588000 | 9466500 | 0 | 376360 | 0 | 0 | 547690 | 0 | 558640 | 1912700 | 502930 | 0 | 3725000 | 10022000 | 0 | 0 |
| WLPSSSPV | FN1; FN1 | P02751; H0Y | FINC_HUMAN I | Unmodified | 0 | 0 | 1214600 | 3644700 | 546330 | 604090 | 0 | 0 | 0 | 0 | 95240 | 0 | 4622600 | 7886600 | 0 | 0 | 1862100 | 12342000 |
| WLPVQESS | SPCS1; SPCS1; | Q9Y6A9; C9J | SPCS1_HUMAN | Unmodified | 0 | 0 | 0 | 0 | 0 | 0 | 0 | 60683 | 8550500 | 1362800 | 1784900 | 527470 | 0 | 0 | 0 | 0 | 0 | 0 |
| WLQDLQEI | KTN1; KTN1 | Q86UP2; G3 | KTN1_HUMAN | Unmodified | 0 | 0 | 0 | 0 | 0 | 0 | 0 | 0 | 0 | 0 | 0 | 1417800 | 0 | 0 | 0 | 0 | 341870 | 0 |
| WLSAEIED\ | NDUFA9; NDUI | Q16795; F5H | NDUA9_HUMA | Unmodified | 0 | 246420 | 0 | 0 | 0 | 0 | 0 | 0 | 1655100 | 0 | 0 | 0 | 0 | 0 | 48375 | 0 | 0 | 0 |
| WLSALRHI | TAPBP; TAPBP; | O15533; A0/ | TPSN_HUMAN | Unmodified | 0 | 0 | 0 | 0 | 0 | 0 | 0 | 0 | 1445300 | 0 | 382960 | 0 | 0 | 0 | 0 | 0 | 0 | 0 |
| WLSDECTN | ECI2; ECI2 | O75521; A0/ | ECI2_HUMAN E | Unmodified | 0 | 0 | 0 | 333990 | 652780 | 0 | 0 | 0 | 444640 | 0 | 0 | 0 | 0 | 0 | 0 | 0 | 0 | 0 |
| WLSDECTN | ECI2; ECI2 | O75521; A0/ | ECI2_HUMAN E | Unmodified | 0 | 0 | 0 | 661890 | 0 | 0 | 0 | 0 | 13453000 | 1378000 | 0 | 0 | 0 | 0 | 0 | 0 | 0 | 0 |
| WLSEPMFV | SEMA3C | Q99985 | SEM3C_HUMAI | Oxidation (M) | 166080 | 0 | 716810 | 1791300 | 0 | 338310 | 0 | 0 | 664790 | 0 | 0 | 0 | 0 | 0 | 1035100 | 569520 | 0 | 0 |
| WLSNVDG | MTMR1; MTMR1 | Q13613; F8W | MTMR1_HUMA | Unmodified | 0 | 0 | 0 | 0 | 0 | 0 | 0 | 0 | 0 | 0 | 0 | 178940 | 0 | 0 | 113530 | 64357 | 0 | 0 |
| WLSTRPEV | NUP210 | Q8TEM1 | PO210_HUMAI | Unmodified | 0 | 0 | 0 | 0 | 0 | 0 | 0 | 0 | 76700000 | 3251100 | 1343600 | 0 | 0 | 0 | 0 | 0 | 0 | 0 |
| WLSTSIPE/ | FASN; FASN | P49327; A0A | FAS_HUMAN F | Unmodified | 2682000 | 0 | 4689500 | 11845000 | 3374400 | 1407200 | 750010 | 0 | 3999400 | 1347800 | 8729200 | 1.21E+08 | 5757700 | 886390 | 9004800 | 8093200 | 1970200 | 1753300 |
| WLTDEDTR | OS9; OS9 | Q13438; B4E | OS9_HUMAN P | Unmodified | 0 | 0 | 0 | 0 | 0 | 0 | 0 | 0 | 239430 | 162340 | 1857400 | 0 | 0 | 0 | 0 | 0 | 0 | 0 |
| WLVEPRCL | LLGL2; LLGL2 | Q6P1M3; J3( | L2GL2_HUMAN | Unmodified | 0 | 0 | 0 | 0 | 0 | 0 | 0 | 0 | 0 | 0 | 0 | 0 | 0 | 0 | 0 | 0 | 561740 | 0 |
| WLVIDPVP/ | CC2D1A | Q6P1N0 | C2D1A_HUMAI | Unmodified | 1398300 | 0 | 724440 | 793970 | 0 | 0 | 136700 | 0 | 1504400 | 0 | 243180 | 1375500 | 0 | 0 | 1611900 | 722680 | 0 | 0 |
| WMDEAQA | NAA15; NAA15 | Q9BXJ9; A0A | NAA15_HUMAI | Oxidation (M) | 0 | 0 | 0 | 0 | 0 | 0 | 0 | 0 | 0 | 0 | 0 | 1639800 | 0 | 0 | 0 | 0 | 0 | 0 |
| WMEEQAQ | SNRPC; SNRPC | P09234; A0A | RU1C_HUMAN | Oxidation (M) | 182530 | 169710 | 373830 | 382930 | 333320 | 0 | 0 | 0 | 0 | 0 | 0 | 0 | 769370 | 0 | 377270 | 346800 | 0 | 0 |
| WMEPAIPV | POGLUT1; POG | Q8NBL1; B4I | PGLT1_HUMAN | Oxidation (M) | 0 | 0 | 0 | 0 | 0 | 0 | 0 | 0 | 2527600 | 1063500 | 228460 | 0 | 0 | 0 | 0 | 0 | 0 | 0 |
| WMEWNP( | ALOX5 | P09917 | LOX5_HUMAN | Oxidation (M) | 0 | 0 | 0 | 0 | 0 | 0 | 0 | 0 | 322320 | 0 | 115860 | 0 | 0 | 0 | 0 | 422320 | 0 | 0 |
| WMHPQVT | ACY1; ABHD14/ | Q03154; A0/ | ACY1_HUMAN | Oxidation (M) | 0 | 0 | 0 | 557760 | 0 | 0 | 0 | 0 | 0 | 0 | 2333900 | 1528700 | 0 | 0 | 774650 | 390290 | 0 | 0 |
| WMIDSRDI | SDHB | P21912 | SDHB_HUMAN | Unmodified | 0 | 0 | 0 | 0 | 0 | 0 | 0 | 0 | 1160700 | 0 | 0 | 0 | 0 | 0 | 0 | 0 | 0 | 0 |
| WMNYLEVI | ACSL4; ACSL4 | O60488; D6I | ACSL4_HUMAN | Oxidation (M) | 0 | 0 | 0 | 0 | 0 | 113350 | 0 | 0 | 0 | 0 | 0 | 2008600 | 0 | 0 | 0 | 0 | 0 | 0 |
| WMNYLEVI | ACSL4; ACSL4 | O60488; D6I | ACSL4_HUMAN | Unmodified | 0 | 0 | 0 | 0 | 0 | 0 | 0 | 0 | 0 | 0 | 0 | 1647700 | 0 | 0 | 0 | 0 | 0 | 0 |
| WMQLGAF | MGAM2 | Q2M2H8 | MGAL_HUMAN | Oxidation (M) | 0 | 0 | 0 | 0 | 0 | 0 | 0 | 0 | 0 | 0 | 0 | 1285500 | 3677700 | 0 | 138120 | 0 | 0 | 0 |
| WMQSELN | MIX23; MIX23; I | Q4VC31; C9 | MIX23_HUMAN | Oxidation (M) | 0 | 0 | 0 | 0 | 0 | 0 | 0 | 0 | 0 | 0 | 735080 | 0 | 730030 | 0 | 0 | 0 | 0 | 0 |
| WNATNKDI | PPFIBP2; PPFIB | Q8ND30; H( | LIPB2_HUMAN | Unmodified | 0 | 0 | 299140 | 328340 | 0 | 0 | 0 | 0 | 0 | 0 | 0 | 0 | 78411 | 0 | 356660 | 330960 | 0 | 0 |
| WNDNCPS | CAPN2 | P17655 | CAN2_HUMAN | Unmodified | 0 | 0 | 0 | 0 | 0 | 0 | 0 | 0 | 460570 | 0 | 0 | 157860 | 0 | 0 | 87556 | 0 | 0 | 0 |
| WNDVAGLI | VPS4A; N/A | Q9UN37; I3L | VPS4A_HUMAN | Unmodified | 917560 | 1009200 | 1524900 | 1481100 | 655200 | 0 | 0 | 0 | 3149000 | 69846 | 1221800 | 8060800 | 767500 | 0 | 829140 | 1074100 | 0 | 0 |
| WNEEFYFR | NEDD4L; NEDD | Q96PU5; A0/ | NED4L_HUMAI | Unmodified | 0 | 0 | 0 | 0 | 0 | 0 | 0 | 0 | 410650 | 0 | 125240 | 2083200 | 0 | 0 | 0 | 0 | 0 | 0 |
| WNFDDLLE | DRG1 | Q9Y295 | DRG1_HUMAN | Unmodified | 0 | 0 | 0 | 9371.8 | 0 | 0 | 0 | 0 | 0 | 0 | 0 | 1235200 | 0 | 0 | 51536 | 0 | 0 | 0 |
| WNFTTPRD | CKAP5 | Q14008 | CKAP5_HUMAI | Unmodified | 93535 | 127480 | 0 | 235810 | 971570 | 0 | 251270 | 23860 | 602380 | 405600 | 398750 | 1163600 | 719980 | 507720 | 0 | 271970 | 371620 | 329420 |
| WNGHTDM | PCYOX1 | Q9UHG3 | PCYOX_HUMAI | Unmodified | 0 | 0 | 0 | 0 | 0 | 0 | 0 | 0 | 5978200 | 0 | 0 | 0 | 0 | 0 | 0 | 0 | 0 | 0 |
| WNGWGYI | AGPS; AGPS; A( | O00116; A0/ | ADAS_HUMAN | Unmodified | 0 | 0 | 0 | 0 | 0 | 0 | 0 | 0 | 1544500 | 217660 | 0 | 0 | 0 | 0 | 0 | 0 | 0 | 0 |
| WNKPSDV! | USP6NL; USP6I | Q92738; X6F | US6NL_HUMAI | Unmodified | 0 | 0 | 0 | 0 | 147370 | 0 | 0 | 0 | 0 | 48816 | 0 | 0 | 83928 | 10582 | 0 | 0 | 0 | 715520 |
| WNNHHLT | MMP15 | P51511 | MMP15_HUMA | Unmodified | 0 | 0 | 77884 | 84766 | 0 | 0 | 0 | 0 | 0 | 0 | 0 | 0 | 0 | 0 | 165790 | 72467 | 0 | 0 |
| WNNVKPIF | ITGA6 | P23229 | ITA6_HUMAN I | Unmodified | 38561 | 0 | 268110 | 0 | 0 | 0 | 0 | 0 | 0 | 0 | 0 | 693410 | 0 | 0 | 357130 | 357850 | 0 | 0 |
| WNPFIDNII | CORO2B | Q9UQ03 | COR2B_HUMA | Unmodified | 0 | 0 | 0 | 0 | 0 | 0 | 0 | 0 | 0 | 0 | 0 | 371080 | 0 | 0 | 0 | 0 | 0 | 0 |
| WNQDTME | SF1; SF1; SF1 | Q15637; F8V | SF01_HUMAN ! | Unmodified | 0 | 0 | 0 | 0 | 0 | 0 | 0 | 0 | 0 | 0 | 0 | 508300 | 0 | 0 | 0 | 0 | 0 | 0 |
| WNQQQLD | DCPS; DCPS | Q96C86; A0/ | DCPS_HUMAN | Unmodified | 0 | 0 | 0 | 0 | 0 | 0 | 0 | 0 | 663250 | 370420 | 0 | 1241200 | 0 | 0 | 0 | 0 | 0 | 0 |
| WNRLLMSI | UTRN; UTRN | P46939; Q5T | UTRN_HUMAN | Oxidation (M) | 0 | 0 | 13028 | 139390 | 153900 | 27285 | 0 | 0 | 20415 | 72494 | 0 | 0 | 83901 | 49088 | 79543 | 0 | 69477 | 74507 |
| WNSCSTPI | KIF23; KIF23; KI | A0A2U3TZL8 | A0A2U3TZL8_H | Unmodified | 0 | 0 | 0 | 0 | 1419800 | 0 | 0 | 0 | 0 | 0 | 2146500 | 0 | 0 | 0 | 0 | 0 | 0 | 0 |
| WNSPAEE( | HSPA4; HSPA4; | P34932; A0A | HSP74_HUMAI | Unmodified | 162910 | 91514 | 0 | 0 | 0 | 0 | 0 | 0 | 527080 | 0 | 843180 | 4584800 | 438410 | 0 | 199820 | 155550 | 0 | 209190 |
| WNTDNTL( | VDAC2; VDAC2 | P45880; Q5J | VDAC2_HUMAI | Unmodified | 0 | 0 | 0 | 0 | 0 | 0 | 0 | 0 | 3340900 | 2374100 | 0 | 0 | 0 | 0 | 0 | 0 | 0 | 0 |
| WNTDNTL( | VDAC3; VDAC3 | Q9Y277; E5F | VDAC3_HUMAI | Unmodified | 0 | 0 | 0 | 0 | 0 | 0 | 0 | 0 | 10727000 | 785490 | 178170 | 0 | 0 | 0 | 0 | 0 | 0 | 0 |
| WNTDNTL( | VDAC3; VDAC3 | Q9Y277; E5F | VDAC3_HUMAI | Unmodified | 0 | 9777300 | 0 | 0 | 0 | 0 | 0 | 72001 | 2075300 | 3137300 | 0 | 796700 | 0 | 0 | 385790 | 1600500 | 1146100 | 0 |
| WNTDNTL( | VDAC1; VDAC1 | P21796; C9J | VDAC1_HUMAI | Unmodified | 0 | 0 | 0 | 0 | 0 | 0 | 0 | 0 | 18712000 | 4397500 | 0 | 0 | 0 | 0 | 0 | 0 | 0 | 0 |
| WNTDNTL( | VDAC1; VDAC1 | P21796; C9J | VDAC1_HUMAI | Unmodified | 0 | 0 | 0 | 0 | 0 | 0 | 1147800 | 0 | 1039200 | 22115000 | 0 | 725750 | 0 | 0 | 0 | 0 | 0 | 0 |
| WNTDSVEI | SELENOF; SELE | O60613; A0/ | SEP15_HUMAN | Unmodified | 0 | 0 | 0 | 0 | 0 | 0 | 0 | 0 | 2106900 | 0 | 0 | 0 | 0 | 0 | 0 | 0 | 0 | 0 |
| WNTDSVEI | SELENOF; SELE | O60613; A0/ | SEP15_HUMAN | Unmodified | 0 | 0 | 0 | 0 | 0 | 0 | 0 | 0 | 266620 | 2421100 | 0 | 0 | 0 | 0 | 0 | 0 | 0 | 0 |
| WNTEDKVS | PGK1; PGK2 | P00558; P07 | PGK1_HUMAN | Unmodified | 0 | 0 | 0 | 204890 | 21917000 | 3007800 | 6933900 | 1108300 | 0 | 10143000 | 0 | 0 | 20205000 | 14546000 | 0 | 0 | 40065000 | 58326000 |
| WNTEDKVS | PGK1 | P00558 | PGK1_HUMAN | Unmodified | 0 | 0 | 0 | 0 | 0 | 0 | 1644600 | 0 | 0 | 0 | 0 | 0 | 0 | 0 | 0 | 0 | 11636000 | 4892600 |
| WPDFNQE/ | GALNT2 | Q10471 | GALT2_HUMAN | Unmodified | 0 | 0 | 0 | 0 | 0 | 0 | 0 | 0 | 1179200 | 0 | 0 | 0 | 0 | 0 | 0 | 0 | 0 | 0 |
| WPDVARPI | MMP15 | P51511 | MMP15_HUMA | Unmodified | 827210 | 1161100 | 1834300 | 1282700 | 0 | 0 | 0 | 0 | 0 | 0 | 0 | 0 | 0 | 0 | 4847900 | 2392500 | 0 | 0 |
| WPFWLSPI | TARS1; TARS3; | P26639; A2F | SYTC_HUMAN | Unmodified | 0 | 0 | 0 | 0 | 0 | 0 | 0 | 0 | 0 | 0 | 19695 | 463780 | 0 | 0 | 0 | 0 | 0 | 0 |
| WPYQNGF | LRBA; LRBA; LR | P50851; A0A | LRBA_HUMAN | Unmodified | 21386 | 0 | 0 | 0 | 215060 | 0 | 0 | 0 | 0 | 0 | 0 | 644010 | 0 | 0 | 75563 | 0 | 348650 | 743260 |
| WQALLVQI | DYNC1H1; DYN | Q14204; A0/ | DYHC1_HUMA | Unmodified | 0 | 0 | 0 | 0 | 0 | 0 | 0 | 0 | 0 | 0 | 0 | 2034300 | 0 | 0 | 0 | 0 | 0 | 0 |
| WQASSLP/ | DYNC1H1; DYN | Q14204; A0/ | DYHC1_HUMA | Oxidation (M) | 0 | 0 | 0 | 0 | 0 | 0 | 0 | 0 | 176360 | 0 | 178620 | 2075500 | 0 | 0 | 0 | 255960 | 0 | 0 |
| WQCVEDA | SULF2 | Q8IWU5 | SULF2_HUMAN | Unmodified | 0 | 0 | 0 | 0 | 0 | 0 | 0 | 0 | 0 | 0 | 0 | 0 | 0 | 0 | 0 | 0 | 0 | 405940 |
| WQDAQATI | SNX1; SNX1 | Q13596; H0Y | SNX1_HUMAN | Unmodified | 0 | 0 | 0 | 0 | 0 | 0 | 0 | 0 | 560540 | 0 | 0 | 1100700 | 11295 | 0 | 35794 | 79308 | 0 | 0 |
| WQDIQNDI | CHD4; CHD5; ( | Q14839; Q8 | CHD4_HUMAN | Unmodified | 0 | 0 | 0 | 0 | 0 | 0 | 0 | 0 | 115840 | 0 | 0 | 805300 | 0 | 0 | 0 | 0 | 0 | 0 |
| WQDNIKEE | CACNA2D2; CA | Q9NY47; C9 | CA2D2_HUMAI | Unmodified | 90062 | 69176 | 0 | 0 | 0 | 0 | 0 | 0 | 877250 | 0 | 0 | 697600 | 0 | 0 | 0 | 0 | 0 | 0 |
| WQETCVD/ | BAIAP2L1 | Q9UHR4 | BI2L1_HUMAN | Unmodified | 0 | 0 | 0 | 724540 | 1188400 | 1017600 | 0 | 0 | 1370100 | 0 | 234150 | 580020 | 833590 | 1040600 | 519230 | 693720 | 0 | 0 |
| WQFLTLR | LARS1; LARS1; | Q9P2J5; A0A | SYLC_HUMAN | Unmodified | 74938 | 38723 | 325530 | 123540 | 0 | 0 | 0 | 0 | 366170 | 0 | 828140 | 13517000 | 0 | 0 | 467660 | 354190 | 0 | 0 |
| WQLYAAQ! | CLPTM1 | O96005 | CLPT1_HUMAN | Unmodified | 0 | 0 | 0 | 0 | 0 | 0 | 0 | 0 | 2468300 | 0 | 586110 | 0 | 0 | 0 | 0 | 0 | 0 | 0 |
| WQMNPYD | DDX1; DDX1; D | Q92499; A0/ | DDX1_HUMAN | Unmodified | 0 | 0 | 0 | 0 | 0 | 0 | 0 | 0 | 0 | 0 | 0 | 553430 | 0 | 0 | 0 | 0 | 0 | 0 |
| WQNADLN | IDE; IDE; IDE; ID | P14735; A0A | IDE_HUMAN In | Unmodified | 0 | 0 | 0 | 30184 | 61422 | 0 | 0 | 0 | 0 | 0 | 0 | 701620 | 70012 | 0 | 0 | 0 | 52951 | 0 |
| WQNNLLP! | RPS15A; RPS15 | P62244; H3E | RS15A_HUMAN | Unmodified | 2343500 | 591960 | 1185900 | 901380 | 0 | 0 | 0 | 0 | 1031200 | 75346 | 1315300 | 54212000 | 267460 | 0 | 1726700 | 1830300 | 0 | 0 |
| WQNSYSIK | UBE2V2; UBE2\ | Q15819; Q1 | UB2V2_HUMAI | Unmodified | 1347600 | 1562600 | 1148300 | 719390 | 1614100 | 0 | 715410 | 341440 | 1260400 | 1052800 | 762020 | 3246200 | 1745000 | 0 | 2221900 | 1603200 | 1192800 | 1052400 |
| WQPDTEEE | SF3A3 | Q12874 | SF3A3_HUMAN | Unmodified | 665540 | 615190 | 0 | 0 | 0 | 0 | 0 | 0 | 0 | 0 | 688440 | 1018600 | 0 | 0 | 0 | 300570 | 0 | 0 |
| WQPGEEP/ | UCHL5; UCHL5 | Q9Y5K5; Q5I | UCHL5_HUMAI | Unmodified | 0 | 0 | 0 | 0 | 0 | 0 | 0 | 0 | 2813000 | 1111900 | 0 | 1720200 | 0 | 0 | 0 | 0 | 0 | 0 |
| WQQGNVF | IGHG2; IGHG1; | P01859; P01 | IGHG2_HUMAN | Unmodified | 0 | 0 | 0 | 0 | 0 | 0 | 0 | 0 | 1079700 | 0 | 0 | 0 | 0 | 0 | 0 | 0 | 0 | 0 |
| WQQQGDI | CHGB | P05060 | SCG1_HUMAN | Unmodified | 2607200 | 504400 | 0 | 0 | 0 | 0 | 0 | 0 | 549370 | 0 | 0 | 0 | 0 | 0 | 2634800 | 1569600 | 0 | 86292 |
| WQTLLSVD | GNS; GNS; GN! | P15586; H7( | GNS_HUMAN I | Unmodified | 0 | 0 | 0 | 0 | 0 | 0 | 0 | 0 | 1865600 | 450250 | 0 | 0 | 0 | 0 | 0 | 0 | 0 | 0 |
| WREEQMEI | CLTA; CLTA | P09496; F8V | CLCA_HUMAN | Unmodified | 9458300 | 7408600 | 2042400 | 4705900 | 1080100 | 0 | 0 | 0 | 4937700 | 0 | 167930 | 42789000 | 363670 | 0 | 3242600 | 3404100 | 0 | 108410 |
| WREEQMEI | CLTA; CLTA | P09496; F8V | CLCA_HUMAN | Oxidation (M) | 538150 | 372910 | 206370 | 0 | 0 | 0 | 0 | 0 | 0 | 0 | 0 | 9431100 | 0 | 0 | 199400 | 1017600 | 0 | 0 |
| WRGTVEK( | PACSIN3; PACS | Q9UKS6; E9I | PACN3_HUMAI | Unmodified | 38710 | 0 | 0 | 130500 | 1648600 | 0 | 0 | 289150 | 0 | 25226 | 0 | 0 | 7334100 | 2236600 | 0 | 0 | 688740 | 706940 |
| WRLVDSK( | FABP5 | Q01469 | FABP5_HUMAN | Oxidation (M) | 0 | 0 | 0 | 0 | 109890 | 0 | 0 | 0 | 360390 | 0 | 0 | 0 | 0 | 0 | 0 | 0 | 1876900 | 0 |
| WRLVDSK( | FABP5 | Q01469 | FABP5_HUMAN | Unmodified | 0 | 0 | 0 | 0 | 883390 | 245180 | 0 | 0 | 113720 | 0 | 0 | 0 | 0 | 0 | 0 | 0 | 3076300 | 3940800 |
| WRQLVEK( | PACSIN2; PACS | Q9UNF0; B0 | PACN2_HUMAI | Unmodified | 0 | 897350 | 0 | 246680 | 1479600 | 99941 | 723070 | 178710 | 0 | 296650 | 0 | 0 | 22130000 | 8120000 | 0 | 0 | 291440 | 1144100 |
| WRSEWKF | CAPZA1 | P52907 | CAZA1_HUMAN | Unmodified | 0 | 0 | 0 | 0 | 0 | 0 | 0 | 0 | 0 | 0 | 0 | 0 | 1609300 | 0 | 0 | 0 | 0 | 0 |
| WRVENQEI | CAP1 | Q01518 | CAP1_HUMAN | Unmodified | 0 | 0 | 0 | 0 | 9886000 | 0 | 0 | 0 | 0 | 559720 | 0 | 0 | 3061500 | 0 | 0 | 0 | 1846700 | 3054500 |
| WRVENSW | BLMH; BLMH | Q13867; K7E | BLMH_HUMAN | Unmodified | 0 | 0 | 0 | 0 | 537400 | 0 | 206630 | 84289 | 1069000 | 602700 | 708410 | 1483800 | 0 | 30936 | 0 | 0 | 98424 | 0 |
| WRVEYQEI | CAP2; CAP2; C/ | P40123; B7Z | CAP2_HUMAN | Unmodified | 0 | 0 | 0 | 0 | 573470 | 0 | 114430 | 0 | 0 | 0 | 0 | 0 | 295300 | 149810 | 0 | 0 | 233240 | 233930 |
| WRYPSLSL | CNDP2 | Q96KP4 | CNDP2_HUMA | Unmodified | 800970 | 0 | 829590 | 5354400 | 1212500 | 158280 | 160500 | 0 | 1400600 | 845140 | 2588600 | 8333700 | 10619000 | 3805700 | 3327000 | 6799500 | 715450 | 559760 |
| WSAPEVFI | BMX | P51813 | BMX_HUMAN ( | Unmodified | 0 | 0 | 0 | 0 | 0 | 0 | 0 | 0 | 0 | 0 | 707740 | 2929800 | 0 | 0 | 0 | 0 | 0 | 0 |
| WSDQGPA | TNKS1BP1 | Q9C0C2 | TB182_HUMAN | Unmodified | 0 | 0 | 0 | 0 | 0 | 0 | 0 | 0 | 0 | 0 | 0 | 437860 | 0 | 0 | 213440 | 0 | 0 | 0 |
| WSDVAGLI | VPS4B | O75351 | VPS4B_HUMAN | Unmodified | 1317200 | 0 | 0 | 1725000 | 437500 | 0 | 0 | 209770 | 5304500 | 963260 | 1750700 | 6166400 | 105470 | 0 | 1830800 | 1353700 | 71922 | 0 |

| | | | | | | | | | | | | | | | | | | | | | | |
|---|---|---|---|---|---|---|---|---|---|---|---|---|---|---|---|---|---|---|---|---|---|---|
| WSEPNEEE | FEN1; FEN1 | P39748; I3L3 | FEN1_HUMAN | Unmodified | 0 | 0 | 0 | 0 | 0 | 0 | 0 | 0 | 0 | 0 | 401960 | 2018700 | 0 | 0 | 0 | 0 | 0 | 0 |
| WSETSVER | TM9SF1; TM9SF | O15321; E9F | TM9S1_HUMAN | Unmodified | 0 | 0 | 0 | 103310 | 0 | 0 | 0 | 0 | 2350300 | 750170 | 50568 | 255340 | 0 | 0 | 0 | 56912 | 122270 | 0 |
| WSEVPEAD | SDK2; SDK2 | Q58EX2; H7( | SDK2_HUMAN | Unmodified | 0 | 0 | 0 | 0 | 0 | 0 | 0 | 0 | 1194800 | 0 | 0 | 0 | 0 | 0 | 0 | 0 | 80392 | 0 |
| WSFVESLE | MRPL45; MRPL | Q9BRJ2; A0A | RM45_HUMAN | Unmodified | 0 | 0 | 0 | 0 | 0 | 0 | 0 | 0 | 2987700 | 0 | 0 | 0 | 0 | 0 | 0 | 0 | 0 | 0 |
| WSGPLSLC | PEBP1 | P30086 | PEBP1_HUMAN | Unmodified | 5204700 | 5439900 | 9150900 | 20324000 | 91037000 | 10580000 | 5703800 | 60230 | 58769000 | 54008000 | 50604000 | 54029000 | 1.39E+08 | 70695000 | 46132000 | 26320000 | 1E+08 | 1.01E+08 |
| WSGPLSLC | PEBP1 | P30086 | PEBP1_HUMAN | Unmodified | 0 | 0 | 0 | 0 | 0 | 0 | 0 | 0 | 0 | 0 | 0 | 0 | 0 | 0 | 0 | 0 | 20538000 | 6672600 |
| WSGQSAG | CAPN7 | Q9Y6W3 | CAN7_HUMAN | Unmodified | 133700 | 119980 | 84864 | 135060 | 0 | 0 | 0 | 0 | 630870 | 107190 | 336840 | 3859400 | 0 | 0 | 30917 | 53559 | 0 | 0 |
| WSPESPLC | COLGALT1 | Q8NBJ5 | GT251_HUMAN | Unmodified | 0 | 45053 | 27420 | 0 | 0 | 0 | 0 | 0 | 539820 | 0 | 0 | 0 | 0 | 0 | 66136 | 50421 | 0 | 0 |
| WSPIASTLF | DCTN2; DCTN2 | Q13561; A8I | DCTN2_HUMAI | Unmodified | 0 | 0 | 0 | 795470 | 127490 | 0 | 170310 | 0 | 0 | 117450 | 780690 | 7656000 | 0 | 0 | 287390 | 98384 | 70659 | 18740 |
| WSPLENK | ARPC1A; ARPC | Q92747; E9F | ARC1A_HUMAN | Unmodified | 465710 | 499150 | 0 | 241620 | 0 | 0 | 0 | 0 | 131930 | 0 | 0 | 11705000 | 0 | 0 | 0 | 670250 | 0 | 0 |
| WSQLLANS | SPTAN1; SPTAN | Q13813; A0A | SPTN1_HUMAN | Unmodified | 0 | 0 | 0 | 0 | 0 | 0 | 0 | 0 | 0 | 0 | 0 | 1520400 | 0 | 0 | 81854 | 0 | 0 | 0 |
| WSQTAFPK | SPON2; SPON2 | Q9BUD6; DE | SPON2_HUMAI | Unmodified | 0 | 0 | 20429000 | 7580200 | 0 | 0 | 134040 | 0 | 0 | 0 | 0 | 0 | 1500700 | 3799900 | 10766000 | 5172000 | 0 | 0 |
| WSQVAPP\ | EMC1 | Q8N766 | EMC1_HUMAN | Unmodified | 0 | 0 | 0 | 0 | 0 | 0 | 0 | 0 | 2121100 | 0 | 266790 | 0 | 0 | 0 | 0 | 0 | 0 | 0 |
| WSRPQAPI | FN1 | P02751 | FINC_HUMAN I | Unmodified | 0 | 0 | 1127700 | 299100 | 0 | 0 | 0 | 0 | 0 | 0 | 0 | 0 | 0 | 0 | 69829 | 498400 | 0 | 0 |
| WSSLACNI | CCT3; CCT3; C( | P49368; Q5S | TCPG_HUMAN | Unmodified | 0 | 0 | 0 | 0 | 0 | 0 | 0 | 0 | 2144600 | 539690 | 1779800 | 9819900 | 0 | 0 | 0 | 0 | 0 | 0 |
| WSSQPEGI | AHNAK2 | Q8IVF2 | AHNK2_HUMAI | Unmodified | 0 | 0 | 0 | 0 | 2218700 | 159940 | 0 | 0 | 0 | 0 | 0 | 0 | 0 | 0 | 0 | 0 | 210970 | 1022500 |
| WSTAVEFC | CD55; CD55; C | P08174; H3E | DAF_HUMAN C | Unmodified | 1528800 | 0 | 737640 | 933960 | 3710200 | 164770 | 0 | 0 | 679610 | 241680 | 1445500 | 7916400 | 425330 | 0 | 787090 | 543690 | 12625000 | 14927000 |
| WSTAVEFC | CD55; CD55; C | P08174; H3E | DAF_HUMAN C | Unmodified | 0 | 0 | 0 | 0 | 1102400 | 211840 | 0 | 0 | 160910 | 0 | 0 | 1120200 | 0 | 0 | 0 | 0 | 2022000 | 1856300 |
| WSTLVEDY | PSMD12 | O00232 | PSD12_HUMAN | Oxidation (M) | 0 | 0 | 255760 | 0 | 0 | 0 | 0 | 0 | 0 | 0 | 0 | 2356900 | 108720 | 0 | 410920 | 0 | 0 | 0 |
| WSTLVEDY | PSMD12 | O00232 | PSD12_HUMAN | Unmodified | 0 | 0 | 0 | 79992 | 0 | 0 | 0 | 0 | 0 | 0 | 469330 | 560670 | 0 | 0 | 0 | 0 | 0 | 0 |
| WSTPSGAS | EHHADH | Q08426 | ECHP_HUMAN | Unmodified | 247160 | 220660 | 0 | 0 | 0 | 0 | 0 | 94342 | 1328300 | 0 | 0 | 0 | 150980 | 0 | 0 | 139190 | 0 | 0 |
| WSTRGSES | CARMIL1; CARN | Q5VZK9; A0A | CARL1_HUMAN | Unmodified | 0 | 0 | 0 | 0 | 173030 | 0 | 0 | 0 | 0 | 0 | 0 | 0 | 0 | 0 | 0 | 0 | 35074 | 59609 |
| WSVHIAMH | FSCN1 | Q16658 | FSCN1_HUMAI | Oxidation (M) | 0 | 0 | 0 | 0 | 0 | 0 | 0 | 0 | 0 | 0 | 0 | 2640200 | 432950 | 0 | 0 | 0 | 0 | 0 |
| WSVHIAMH | FSCN1 | Q16658 | FSCN1_HUMAI | Oxidation (M) | 0 | 0 | 0 | 0 | 329460 | 121640 | 0 | 0 | 0 | 300660 | 2113900 | 0 | 576840 | 785260 | 0 | 0 | 704300 | 998640 |
| WSVLTSKT | NRP1; NRP1; N | O14786; E9F | NRP1_HUMAN | Unmodified | 0 | 0 | 0 | 0 | 9010900 | 2598500 | 0 | 42524 | 0 | 0 | 0 | 0 | 0 | 0 | 0 | 0 | 4104900 | 23518000 |
| WSYLYPLA | SUSD2 | Q9UGT4 | SUSD2_HUMAI | Unmodified | 2122900 | 1804900 | 95118 | 49507 | 657120 | 554450 | 0 | 0 | 12706000 | 710250 | 0 | 0 | 0 | 559580 | 1398200 | 350300 | 0 | 0 |
| WSYLYPLA | SUSD2 | Q9UGT4 | SUSD2_HUMAI | Unmodified | 5328700 | 3403600 | 0 | 321680 | 2708900 | 1241500 | 0 | 0 | 41594000 | 2816500 | 0 | 0 | 0 | 0 | 789950 | 0 | 0 | 0 |
| WTAPEAAL | YES1; SRC; FYN | P07947; P12 | YES_HUMAN Ty | Unmodified | 441180 | 828810 | 1503200 | 909750 | 0 | 0 | 0 | 0 | 5265000 | 678530 | 811380 | 0 | 0 | 479230 | 2895800 | 2129200 | 1079200 | 0 |
| WTAPEAAL | YES1; SRC; FYN | P07947; P12 | YES_HUMAN Ty | Unmodified | 0 | 0 | 0 | 0 | 821750 | 0 | 0 | 0 | 0 | 0 | 0 | 0 | 0 | 0 | 0 | 0 | 749800 | 1059100 |
| WTAPEAIQ | EPHB2; EPHA7; | P29323; Q15 | EPHB2_HUMAN | Unmodified | 0 | 0 | 0 | 0 | 0 | 0 | 0 | 50417 | 634520 | 0 | 566660 | 0 | 0 | 0 | 0 | 0 | 0 | 0 |
| WTAPEAIS | EPHA2 | P29317 | EPHA2_HUMAN | Unmodified | 0 | 0 | 0 | 0 | 0 | 0 | 0 | 0 | 0 | 0 | 0 | 1715100 | 0 | 0 | 0 | 0 | 0 | 0 |
| WTAPEALR | CSK | P41240 | CSK_HUMAN T | Unmodified | 75721 | 291430 | 0 | 0 | 0 | 0 | 0 | 0 | 0 | 0 | 0 | 1917700 | 0 | 0 | 37126 | 48021 | 0 | 0 |
| WTDENIDT | DYNC1H1; DYN | Q14204; A0A | DYHC1_HUMA | Unmodified | 0 | 0 | 0 | 0 | 0 | 0 | 0 | 0 | 0 | 0 | 0 | 4575200 | 0 | 0 | 417360 | 0 | 0 | 0 |
| WTELAGCT | LMAN2; LMAN2 | Q12907; D6I | LMAN2_HUMAI | Unmodified | 437760 | 0 | 0 | 0 | 0 | 0 | 0 | 0 | 33925000 | 1768700 | 0 | 1079800 | 0 | 0 | 0 | 0 | 0 | 0 |
| WTELAGCT | LMAN2; LMAN2 | Q12907; D6I | LMAN2_HUMAI | Unmodified | 0 | 0 | 0 | 0 | 0 | 0 | 0 | 0 | 11714000 | 250480 | 0 | 0 | 0 | 0 | 0 | 0 | 0 | 0 |
| WTETYVR | EIF3B; EIF3B; EI | P55884; C9J | EIF3B_HUMAN | Unmodified | 287160 | 0 | 0 | 0 | 0 | 0 | 0 | 0 | 359410 | 19788 | 243860 | 5749500 | 0 | 0 | 278300 | 527570 | 0 | 0 |
| WTEYGLTF | VDAC1; VDAC1 | P21796; C9J | VDAC1_HUMAI | Unmodified | 0 | 298360 | 0 | 0 | 0 | 0 | 0 | 0 | 42970000 | 1907900 | 132320 | 1134800 | 0 | 0 | 0 | 0 | 0 | 0 |
| WTFEACR | NPTX1 | Q15818 | NPTX1_HUMAN | Unmodified | 0 | 0 | 0 | 0 | 0 | 0 | 0 | 0 | 16279000 | 769050 | 0 | 0 | 0 | 0 | 0 | 0 | 0 | 0 |
| WTFEACRC | NPTX1 | Q15818 | NPTX1_HUMAN | Unmodified | 0 | 0 | 0 | 0 | 0 | 0 | 0 | 0 | 0 | 1811200 | 0 | 0 | 0 | 0 | 0 | 0 | 0 | 0 |
| WTFEILDET | KIT | P10721 | KIT_HUMAN Ma | Unmodified | 0 | 0 | 0 | 0 | 0 | 0 | 0 | 0 | 886080 | 0 | 0 | 0 | 0 | 0 | 0 | 0 | 0 | 0 |
| WTFGGRDI | SEMA4C | Q9C0C4 | SEM4C_HUMAI | Unmodified | 0 | 57236 | 0 | 87626 | 0 | 0 | 0 | 0 | 9444700 | 0 | 70705 | 0 | 0 | 0 | 84535 | 50555 | 0 | 0 |
| WTGMIIGP | UBE2V2; UBE2\ | Q15819; Q1 | UB2V2_HUMAN | Oxidation (M) | 662650 | 357280 | 1111800 | 148640 | 0 | 0 | 0 | 0 | 427170 | 135950 | 93989 | 3803400 | 0 | 0 | 1006500 | 1717000 | 0 | 0 |
| WTHLRKPE | SORT1 | Q99523 | SORT_HUMAN | Unmodified | 0 | 0 | 0 | 173140 | 0 | 0 | 0 | 19996 | 2064800 | 0 | 159680 | 581700 | 0 | 0 | 0 | 0 | 0 | 0 |
| WTLKEDPV | ERN1; ERN1 | O75460; A0A | ERN1_HUMAN | Unmodified | 0 | 0 | 0 | 0 | 0 | 0 | 0 | 0 | 704300 | 0 | 0 | 0 | 0 | 0 | 0 | 0 | 0 | 0 |
| WTNIPLLVI | IPO9 | Q96P70 | IPO9_HUMAN I | Unmodified | 0 | 0 | 180400 | 0 | 0 | 0 | 0 | 0 | 0 | 0 | 222380 | 1234100 | 0 | 0 | 0 | 0 | 0 | 0 |
| WTNQLNSI | COPS2; COPS2 | P61201; B4D | CSN2_HUMAN | Unmodified | 0 | 0 | 104940 | 0 | 95471 | 0 | 0 | 0 | 0 | 0 | 272060 | 0 | 162230 | 116760 | 0 | 0 | 0 | 169140 |
| WTPIIKDIM | STXBP1; STXBP | P61764; A0A | STXB1_HUMAN | Unmodified | 0 | 0 | 0 | 0 | 0 | 0 | 252570 | 0 | 0 | 505390 | 0 | 0 | 0 | 0 | 0 | 0 | 0 | 0 |
| WTSPEAIA | EPHA3; EPHA3 | P29320; C9J | EPHA3_HUMAN | Unmodified | 945390 | 1265400 | 0 | 0 | 0 | 0 | 281420 | 86320 | 0 | 0 | 169370 | 0 | 0 | 0 | 0 | 0 | 546870 | 0 |
| WTSQDSLL | FASN; FASN | P49327; A0A | FAS_HUMAN F | Oxidation (M) | 0 | 0 | 625160 | 140710 | 0 | 0 | 0 | 0 | 0 | 0 | 0 | 20507000 | 0 | 0 | 0 | 397820 | 0 | 0 |
| WTSQDSLL | FASN; FASN | P49327; A0A | FAS_HUMAN F | Unmodified | 0 | 0 | 0 | 0 | 0 | 0 | 0 | 0 | 0 | 0 | 0 | 1988500 | 0 | 0 | 0 | 0 | 0 | 0 |
| WTTEQQQI | ARL6IP1; N/A; A | Q15041; H3I | AR6P1_HUMAN | Unmodified | 0 | 0 | 0 | 0 | 0 | 0 | 0 | 0 | 0 | 0 | 0 | 1046300 | 0 | 0 | 0 | 212880 | 0 | 0 |
| WTTLNSLQ | NSUN2 | Q08J23 | NSUN2_HUMA | Unmodified | 95996 | 0 | 32423 | 390600 | 4072300 | 171560 | 1687200 | 31970 | 373580 | 971570 | 949230 | 2499200 | 2929600 | 0 | 102310 | 259230 | 2236400 | 3108300 |
| WTTSQLLQ | SLK | Q9H2G2 | SLK_HUMAN S | Unmodified | 0 | 0 | 0 | 0 | 0 | 0 | 0 | 0 | 1193100 | 0 | 663780 | 4019300 | 0 | 0 | 0 | 572710 | 0 | 0 |
| WTVGEGAI | WDR20; WDR2( | Q8TBZ3; A0A | WDR20_HUMA | Unmodified | 65683 | 0 | 0 | 0 | 353560 | 0 | 0 | 0 | 0 | 564360 | 0 | 611270 | 254780 | 380960 | 0 | 305650 | 107630 | 689140 |
| WVAIQSVS | CNDP2; CNDP2 | Q96KP4; J3Q | CNDP2_HUMA | Unmodified | 0 | 0 | 745060 | 1171800 | 2931800 | 329000 | 622990 | 321590 | 2303100 | 1237900 | 6478700 | 14578000 | 12805000 | 9678900 | 2890400 | 2029300 | 1652100 | 1351600 |
| WVASAPEA | MKRN3; MKRN3 | Q13064; A0A | MKRN3_HUMA | Unmodified | 1968400 | 1787100 | 643690 | 660500 | 0 | 0 | 0 | 0 | 0 | 0 | 0 | 0 | 1728600 | 0 | 566680 | 507020 | 0 | 0 |
| WVASSTAE | EXOC1 | Q9NV70 | EXOC1_HUMAI | Unmodified | 0 | 58676 | 82930 | 158360 | 0 | 0 | 263280 | 0 | 523260 | 236810 | 288210 | 1304900 | 387270 | 195860 | 353680 | 435310 | 0 | 0 |
| WVCSSLR | FASN; FASN | P49327; A0A | FAS_HUMAN F | Unmodified | 481740 | 0 | 293760 | 743470 | 104110 | 0 | 61608 | 0 | 278660 | 58179 | 924080 | 22624000 | 0 | 0 | 2071800 | 0 | 0 | 0 |
| WVDPNSP\ | AKR1C4; AKR1( | P17516; P52 | AK1C4_HUMAN | Unmodified | 293380 | 108120 | 0 | 0 | 233940 | 0 | 0 | 0 | 0 | 0 | 0 | 0 | 106810 | 0 | 0 | 0 | 0 | 0 |
| WVFKEEGV | DDOST; DDOST | P39656; A0A | OST48_HUMAN | Unmodified | 0 | 0 | 0 | 0 | 0 | 0 | 0 | 0 | 16400000 | 3793400 | 0 | 435920 | 0 | 0 | 0 | 0 | 0 | 0 |
| WVGADHA | CEL | P19835 | CEL_HUMAN B | Unmodified | 0 | 0 | 0 | 0 | 0 | 0 | 0 | 0 | 2163000 | 233350 | 1571500 | 210750 | 0 | 0 | 0 | 0 | 0 | 0 |
| WVGADHA | CEL | P19835 | CEL_HUMAN B | Unmodified | 0 | 0 | 0 | 0 | 0 | 0 | 0 | 0 | 78306000 | 4596700 | 58143000 | 3719600 | 0 | 0 | 0 | 0 | 0 | 0 |
| WVGGPEIE | CCT5; CCT5; C( | P48643; B7Z | TCPE_HUMAN | Unmodified | 0 | 56918 | 0 | 330320 | 223160 | 0 | 0 | 0 | 363910 | 291550 | 197510 | 1519200 | 973040 | 0 | 365440 | 25393 | 61736 | 0 |
| WVGGPEIE | CCT5; CCT5; C( | P48643; B7Z | TCPE_HUMAN | Unmodified | 0 | 0 | 0 | 0 | 2487200 | 0 | 0 | 0 | 0 | 734900 | 0 | 0 | 4332700 | 1508300 | 0 | 0 | 0 | 2794500 |
| WVHSAEPT | IGSF21 | Q96ID5 | IGS21_HUMAN | Unmodified | 0 | 0 | 0 | 0 | 0 | 0 | 0 | 0 | 952140 | 0 | 0 | 0 | 0 | 0 | 0 | 0 | 0 | 0 |
| WVIEAAEG | SEZ6L2; SEZ6L2 | Q6UXD5; H3 | SE6L2_HUMAN | Unmodified | 0 | 0 | 0 | 176270 | 0 | 0 | 274990 | 0 | 5086100 | 0 | 0 | 400430 | 194010 | 0 | 0 | 58568 | 0 | 0 |
| WVINPSGG | SCP2; SCP2 | P22307; E9P | SCP2_HUMAN | Unmodified | 0 | 0 | 0 | 0 | 0 | 0 | 0 | 0 | 9231700 | 465030 | 0 | 0 | 0 | 0 | 0 | 0 | 0 | 0 |
| WVIYNDQK | USP5 | P45974 | UBP5_HUMAN | Unmodified | 953350 | 1168100 | 0 | 261020 | 375040 | 0 | 153480 | 0 | 610070 | 72412 | 467470 | 1799000 | 674250 | 451590 | 428140 | 298650 | 177450 | 165380 |
| WVKENKEV | ANPEP | P15144 | AMPN_HUMAN | Unmodified | 0 | 0 | 0 | 0 | 0 | 0 | 0 | 0 | 58023 | 0 | 0 | 0 | 0 | 0 | 0 | 0 | 5686000 | 3979100 |
| WVNQTLK | PLS1 | Q14651 | PLSI_HUMAN F | Unmodified | 121170 | 0 | 0 | 0 | 467720 | 492670 | 0 | 0 | 0 | 0 | 389060 | 4150800 | 816030 | 0 | 0 | 0 | 191100 | 194490 |
| WVPAGGS | ENAH; ENAH; E | Q8N8S7; A0A | ENAH_HUMAN | Unmodified | 0 | 0 | 0 | 71075 | 0 | 0 | 0 | 0 | 0 | 0 | 0 | 3753300 | 0 | 0 | 246390 | 0 | 0 | 0 |
| WVPEITHH | CDC42; CDC42 | P60953; Q5J | CDC42_HUMA | Unmodified | 5577900 | 3352100 | 16014000 | 3681300 | 3362400 | 0 | 1377500 | 0 | 1440200 | 637240 | 2308500 | 6968300 | 1951600 | 2154900 | 0 | 3342100 | 873760 | 2864900 |
| WVPETPVP | GPC3 | P51654 | GPC3_HUMAN | Unmodified | 0 | 0 | 0 | 0 | 0 | 0 | 0 | 0 | 6295800 | 575430 | 369660 | 0 | 0 | 0 | 0 | 0 | 0 | 0 |
| WVPFDGDI | DDOST; DDOST | P39656; A0A | OST48_HUMAN | Unmodified | 0 | 0 | 0 | 0 | 0 | 0 | 0 | 0 | 0 | 199920 | 0 | 0 | 0 | 0 | 0 | 0 | 0 | 0 |
| WVPQMGD | HMGCL | P35914 | HMGCL_HUMA | Unmodified | 63883 | 0 | 0 | 0 | 0 | 0 | 0 | 0 | 5082100 | 0 | 0 | 1068400 | 0 | 0 | 0 | 0 | 0 | 0 |
| WVTALELAI | OSBP | P22059 | OSBP1_HUMAI | Unmodified | 156490 | 0 | 0 | 579770 | 235090 | 134580 | 0 | 43612 | 665860 | 0 | 634670 | 5303400 | 358190 | 0 | 399130 | 487560 | 126310 | 234970 |
| WVTTASLLI | SF3B3 | Q15393 | SF3B3_HUMAN | Unmodified | 0 | 934700 | 93554 | 0 | 0 | 0 | 0 | 0 | 0 | 0 | 0 | 2185400 | 0 | 0 | 0 | 0 | 0 | 0 |
| WVTYFNKP | COX5A; COX5A | P20674; H3E | COX5A_HUMAI | Unmodified | 24766 | 0 | 0 | 0 | 0 | 0 | 0 | 37774 | 757910 | 33086 | 0 | 0 | 0 | 0 | 0 | 0 | 0 | 0 |
| WVTYFNKP | COX5A; COX5A | P20674; H3E | COX5A_HUMAI | Unmodified | 133510 | 0 | 88611 | 0 | 128750 | 0 | 0 | 0 | 30429000 | 1632700 | 1576400 | 729760 | 0 | 0 | 144680 | 0 | 0 | 0 |
| WVVIGDEN | ACO2; ACO2; A | Q99798; A2A | ACON_HUMAN | Unmodified | 0 | 0 | 0 | 0 | 0 | 0 | 0 | 0 | 3371400 | 71682 | 198600 | 0 | 0 | 0 | 0 | 0 | 0 | 0 |
| WVVTDSK | DIP2B; DIP2B | Q9P265; H0 | DIP2B_HUMAN | Unmodified | 25922 | 197360 | 0 | 518080 | 215340 | 94444 | 94574 | 34567 | 986510 | 384400 | 546430 | 4455800 | 213420 | 273710 | 959910 | 641180 | 434840 | 270560 |
| WWDIIDIQ | MAN2A1 | Q16706 | MA2A1_HUMAI | Unmodified | 0 | 0 | 0 | 0 | 0 | 0 | 0 | 0 | 1369600 | 261070 | 0 | 856060 | 0 | 0 | 0 | 0 | 0 | 0 |
| WWEEERYI | TOP1 | P11387 | TOP1_HUMAN | Unmodified | 253180 | 131320 | 105480 | 0 | 202310 | 0 | 0 | 0 | 0 | 0 | 112570 | 2254300 | 0 | 0 | 129490 | 36330 | 0 | 0 |
| WWHTGAL | SLC3A2; SLC3A | P08195; A0A | 4F2_HUMAN 4I | Unmodified | 1273200 | 1457000 | 0 | 0 | 266690 | 0 | 0 | 0 | 104270 | 0 | 1510300 | 13684000 | 0 | 0 | 0 | 1456000 | 0 | 0 |
| WWHTGAL | SLC3A2; SLC3A | P08195; A0A | 4F2_HUMAN 4I | Unmodified | 0 | 0 | 0 | 203000 | 35343000 | 7577600 | 3074100 | 0 | 195650 | 2970800 | 3325700 | 0 | 2253000 | 0 | 0 | 0 | 2180500 | 5433200 |
| WWNHIDE | TVP23B; TVP23 | Q9NYZ1; Q9 | TV23B_HUMAN | Unmodified | 0 | 0 | 0 | 0 | 0 | 0 | 0 | 0 | 0 | 0 | 0 | 790620 | 0 | 0 | 0 | 426950 | 0 | 0 |
| WWNVLPG | ELAPOR2; ELAP | A8MWY0; H7 | ELAP2_HUMAN | Oxidation (M) | 0 | 0 | 0 | 0 | 0 | 0 | 0 | 0 | 1180400 | 0 | 0 | 0 | 0 | 0 | 0 | 0 | 0 | 0 |
| WWNVLPG | ELAPOR2; ELAP | A8MWY0; H7 | ELAP2_HUMAN | Unmodified | 0 | 0 | 0 | 0 | 379970 | 0 | 0 | 0 | 1081400 | 0 | 0 | 0 | 0 | 0 | 0 | 0 | 0 | 0 |
| WWTCFVK | ARPC3 | O15145 | ARPC3_HUMAI | Unmodified | 1647500 | 0 | 433520 | 978950 | 48229 | 0 | 0 | 19378 | 0 | 0 | 840260 | 3583900 | 2654200 | 1748400 | 1007400 | 655050 | 422220 | 332560 |
| WYEEALAIS | TTC7B; TTC7B; | Q86TV6; A0A | TTC7B_HUMAN | Unmodified | 0 | 0 | 0 | 0 | 0 | 0 | 0 | 0 | 0 | 0 | 0 | 259950 | 0 | 0 | 535150 | 0 | 0 | 0 |
| WYEIEKIPT | APOD; APOD; A | P05090; C9J | APOD_HUMAN | Unmodified | 0 | 0 | 0 | 0 | 0 | 0 | 0 | 0 | 45464000 | 2000700 | 0 | 0 | 0 | 0 | 0 | 0 | 0 | 0 |
| WYFDVERN | SPINT2; SPINT2 | O43291; K7E | SPIT2_HUMAN | Unmodified | 0 | 0 | 0 | 0 | 0 | 0 | 0 | 0 | 6535300 | 279230 | 0 | 0 | 0 | 0 | 0 | 0 | 0 | 0 |
| WYFDVERN | SPINT2; SPINT2 | O43291; K7E | SPIT2_HUMAN | Unmodified | 0 | 0 | 0 | 0 | 0 | 0 | 0 | 0 | 5754100 | 1031900 | 0 | 0 | 0 | 0 | 0 | 0 | 0 | 0 |
| WYFDVTEG | APP; APP; APP | P05067; H7C | A4_HUMAN Am | Unmodified | 0 | 0 | 0 | 0 | 735920 | 323140 | 0 | 0 | 11271000 | 1183700 | 1745500 | 0 | 0 | 0 | 0 | 0 | 84870 | 1200100 |
| WYHSLTWA | ARHGEF39 | Q8N4T4 | ARG39_HUMAN | Unmodified | 183670 | 174190 | 70955 | 190750 | 529670 | 0 | 0 | 0 | 144550 | 0 | 0 | 1620600 | 0 | 0 | 273520 | 162120 | 0 | 0 |
| WYIPDPTG | GLT8D1 | Q68CQ7 | GL8D1_HUMAN | Unmodified | 0 | 0 | 0 | 0 | 0 | 0 | 0 | 0 | 12130000 | 869010 | 134810 | 815110 | 0 | 0 | 0 | 0 | 0 | 0 |
| WYIPDPTG | GLT8D1 | Q68CQ7 | GL8D1_HUMAN | Unmodified | 0 | 0 | 0 | 0 | 0 | 0 | 29971 | 0 | 7682200 | 2029200 | 47650 | 111650 | 0 | 0 | 0 | 0 | 0 | 0 |
| WYIPDPTG | GLT8D1 | Q68CQ7 | GL8D1_HUMAN | Unmodified | 0 | 0 | 0 | 0 | 0 | 248910 | 0 | 0 | 3174400 | 1587400 | 0 | 0 | 0 | 0 | 0 | 0 | 0 | 320230 |
| WYLENVYP | GALNT2 | Q10471 | GALT2_HUMAN | Unmodified | 0 | 0 | 0 | 0 | 0 | 0 | 0 | 0 | 3160300 | 0 | 0 | 0 | 0 | 0 | 0 | 0 | 0 | 0 |
| WYNGTNN | GOT1 | P17174 | AATC_HUMAN | Unmodified | 95684 | 0 | 122180 | 0 | 0 | 0 | 0 | 0 | 272920 | 185550 | 115030 | 2613000 | 0 | 0 | 0 | 0 | 0 | 0 |
| WYPEEYEF | REXO2; REXO2; | Q9Y3B8; F5( | ORN_HUMAN ( | Unmodified | 0 | 0 | 0 | 63660 | 0 | 0 | 0 | 0 | 176700 | 0 | 0 | 919340 | 1099200 | 0 | 93006 | 113840 | 155580 | 0 |
| WYPYYARL | EDIL3 | O43854 | EDIL3_HUMAN | Unmodified | 0 | 0 | 0 | 0 | 79198000 | 11976000 | 5124100 | 637500 | 0 | 1683000 | 0 | 0 | 0 | 0 | 0 | 0 | 9832900 | 22727000 |
| WYPYYARL | EDIL3 | O43854 | EDIL3_HUMAN | Unmodified | 0 | 0 | 0 | 0 | 30458000 | 0 | 3039800 | 0 | 0 | 574260 | 0 | 0 | 363110 | 0 | 0 | 0 | 201250 | 0 |
| WYQDLDKI | PCYOX1L; PCY( | Q8NBM8; E7 | PCYXL_HUMAN | Unmodified | 0 | 0 | 0 | 0 | 0 | 0 | 0 | 0 | 1396100 | 0 | 0 | 0 | 0 | 0 | 0 | 0 | 0 | 0 |
| WYQMGAY | GANAB; GANAE | Q14697; F5H | GANAB_HUMA | Oxidation (M) | 0 | 0 | 0 | 0 | 257400 | 123930 | 0 | 0 | 4754700 | 616330 | 0 | 0 | 0 | 0 | 297150 | 211420 | 113660 | 0 |
| WYQMGAY | GANAB; GANAE | Q14697; F5H | GANAB_HUMA | Unmodified | 0 | 0 | 0 | 0 | 0 | 0 | 0 | 0 | 7810000 | 170040 | 0 | 0 | 0 | 0 | 0 | 0 | 0 | 0 |
| WYRAPEIM | MAPK3; MAPK1 | P27361; P28 | MK03_HUMAN | Unmodified | 0 | 99025 | 0 | 111200 | 0 | 0 | 0 | 0 | 429500 | 0 | 0 | 496420 | 0 | 0 | 74862 | 71355 | 0 | 0 |
| WYRAPEIM | MAPK3; MAPK1 | P27361; P28 | MK03_HUMAN | Oxidation (M) | 436830 | 384570 | 354720 | 421090 | 0 | 0 | 64569 | 39805 | 357200 | 129180 | 0 | 903350 | 0 | 96107 | 696290 | 543920 | 0 | 97074 |
| WYVDGPAS | LSS; LSS | P48449; A0A | LSS_HUMAN La | Unmodified | 0 | 0 | 0 | 0 | 0 | 0 | 0 | 0 | 422440 | 97343 | 0 | 1823200 | 0 | 0 | 108230 | 0 | 0 | 0 |

| | | | | | | | | | | | | | | | | | | | | | | |
|---|---|---|---|---|---|---|---|---|---|---|---|---|---|---|---|---|---|---|---|---|---|---|
| WYVNLHSI | ENDOD1 | O94919 | ENDD1_HUMA | Oxidation (M) | 0 | 0 | 531420 | 625490 | 0 | 0 | 0 | 0 | 2176400 | 0 | 0 | 0 | 0 | 0 | 0 | 339140 | 0 | 0 |
| WYVNLHSI | ENDOD1 | O94919 | ENDD1_HUMA | Unmodified | 0 | 0 | 0 | 0 | 0 | 0 | 0 | 0 | 8632600 | 115820 | 0 | 0 | 0 | 0 | 205390 | 85130 | 0 | 0 |
| WYVVQTNY | ASAH1; ASAH1; | Q13510; A0A | ASAH1_HUMAN | Unmodified | 0 | 0 | 0 | 0 | 1604700 | 0 | 0 | 1319800 | 7284100 | 216470 | 0 | 0 | 0 | 868430 | 689450 | 0 | 0 | 0 |
| WYYDPTEQ | SPINT1; SPINT1 | O43278; H3I | SPIT1_HUMAN | Unmodified | 1123800 | 0 | 1883800 | 551390 | 0 | 0 | 0 | 0 | 1380300 | 0 | 0 | 0 | 0 | 0 | 0 | 1034900 | 0 | 0 |
| WYYNAAGI | UQCRB; UQCRI | P14927; B7Z | QCR7_HUMAN | Unmodified | 0 | 0 | 0 | 0 | 0 | 0 | 0 | 0 | 11436000 | 2590600 | 0 | 0 | 0 | 0 | 0 | 0 | 0 | 0 |
| YAAELAENI | PTPRJ; PTPRJ; P | Q12913; A0A | PTPRJ_HUMAN | Unmodified | 0 | 0 | 0 | 0 | 0 | 0 | 0 | 0 | 0 | 0 | 0 | 2485300 | 131440 | 0 | 182760 | 72306 | 0 | 0 |
| YAAELAENI | PTPRJ; PTPRJ; P | Q12913; A0A | PTPRJ_HUMAN | Unmodified | 0 | 0 | 519150 | 267280 | 5196800 | 116600 | 681100 | 217930 | 511650 | 0 | 360200 | 5140900 | 1801500 | 179300 | 0 | 0 | 55443 | 2157500 |
| YAALYQPLI | NAP1L4; NAP1L | Q99733; C9J | NP1L4_HUMAN | Unmodified | 0 | 0 | 0 | 0 | 0 | 0 | 0 | 0 | 0 | 614480 | 1394500 | 11386000 | 0 | 0 | 0 | 0 | 495830 | 1445800 |
| YAAPELEIA | SERINC5; SERII | Q86VE9; A0A | SERC5_HUMAN | Unmodified | 11195000 | 0 | 34991000 | 19648000 | 0 | 0 | 160710 | 0 | 2921600 | 0 | 203520 | 0 | 0 | 0 | 50472000 | 38489000 | 0 | 0 |
| YAASLPALL | TXNDC16 | Q9P2K2 | TXD16_HUMAN | Unmodified | 0 | 0 | 0 | 0 | 0 | 0 | 117910 | 0 | 918770 | 428510 | 197300 | 431760 | 0 | 0 | 0 | 0 | 0 | 0 |
| YAAVTQFEA | NPEPPSL1; NPE | A6NEC2; P5 | PSAL_HUMAN | Unmodified | 101990 | 0 | 0 | 0 | 0 | 0 | 0 | 0 | 787700 | 0 | 627190 | 5197900 | 0 | 0 | 168490 | 0 | 0 | 0 |
| YAAVTQFEA | NPEPPSL1; NPE | A6NEC2; P5 | PSAL_HUMAN | Unmodified | 90193 | 109130 | 851450 | 804250 | 339910 | 0 | 0 | 0 | 3397000 | 0 | 1907800 | 15816000 | 276780 | 156960 | 1030000 | 769260 | 168440 | 366440 |
| YAAYQQLQ | SRXN1 | Q9BYN0 | SRXN1_HUMAN | Unmodified | 0 | 210760 | 0 | 1331800 | 1048800 | 137420 | 185760 | 0 | 0 | 0 | 0 | 0 | 404030 | 351110 | 0 | 0 | 0 | 0 |
| YACGGWLI | MME; MME; MM | P08473; C9I | NEP_HUMAN N | Unmodified | 0 | 0 | 10811000 | 18871000 | 0 | 0 | 0 | 0 | 8343500 | 2523200 | 0 | 177110 | 20670000 | 7659200 | 23307000 | 14363000 | 0 | 0 |
| YADALQEII | SQOR; N/A | Q9Y6N5; H3 | SQOR_HUMAN | Unmodified | 0 | 0 | 0 | 187410 | 0 | 39512 | 0 | 0 | 2099700 | 0 | 0 | 0 | 0 | 0 | 0 | 0 | 0 | 0 |
| YADEEIPRS | FLNB; FLNB | O75369; E7E | FLNB_HUMAN | Unmodified | 0 | 0 | 0 | 0 | 8284800 | 2204200 | 0 | 183940 | 0 | 190400 | 0 | 6227800 | 0 | 104130 | 0 | 46476 | 1990200 | 1150800 |
| YADEEIPRS | FLNB; FLNB | O75369; E7E | FLNB_HUMAN | Unmodified | 0 | 0 | 0 | 0 | 571390 | 0 | 0 | 0 | 0 | 0 | 0 | 0 | 0 | 0 | 0 | 0 | 744730 | 179680 |
| YADGTIDIE | IGF1R; IGF1R | P08069; C9J | IGF1R_HUMAN | Unmodified | 0 | 0 | 0 | 0 | 1953900 | 0 | 0 | 0 | 0 | 0 | 0 | 0 | 0 | 0 | 0 | 0 | 0 | 0 |
| YADIDIRGC | KIDINS220; KID | Q9ULH0; E9 | KDIS_HUMAN | Unmodified | 0 | 0 | 0 | 0 | 2898600 | 0 | 353090 | 0 | 357610 | 157740 | 0 | 518700 | 0 | 31367 | 229610 | 0 | 0 | 611870 |
| YADLSHNF | RARS1 | P54136 | SYRC_HUMAN | Unmodified | 0 | 0 | 223030 | 225600 | 243740 | 0 | 416600 | 0 | 0 | 517890 | 770600 | 6373300 | 583460 | 1376100 | 172150 | 100490 | 0 | 471040 |
| YADLTEDQ | PGAM1 | P18669 | PGAM1_HUMA | Unmodified | 646990 | 223730 | 1050700 | 1310700 | 2193600 | 496280 | 0 | 0 | 944940 | 0 | 6802000 | 23190000 | 2581700 | 575050 | 2024100 | 1508800 | 228920 | 602880 |
| YADLTEDQ | PGAM1 | P18669 | PGAM1_HUMA | Unmodified | 1362700 | 2221200 | 0 | 7810100 | 15548000 | 2289100 | 0 | 262100 | 1006200 | 1334600 | 10789000 | 16992000 | 12752000 | 3658400 | 3707800 | 1533100 | 4032100 | 7398000 |
| YADPVADL | CTDSP1; CTDSI | Q9GZU7; H7 | CTDS1_HUMAN | Unmodified | 1363500 | 244270 | 1053900 | 510900 | 0 | 0 | 0 | 0 | 0 | 0 | 0 | 183560 | 0 | 0 | 1900400 | 451280 | 0 | 0 |
| YADPVADL | CTDSP1; CTDSI | Q9GZU7; H7 | CTDS1_HUMAN | Unmodified | 1036700 | 1007200 | 0 | 457040 | 132980 | 0 | 71835 | 0 | 251870 | 0 | 39084 | 117800 | 0 | 289680 | 410070 | 235670 | 227960 | 199700 |
| YADPVADL | CTDSP1; CTDSI | Q9GZU7; H7 | CTDS1_HUMAN | Unmodified | 0 | 0 | 0 | 0 | 382180 | 115530 | 176010 | 0 | 0 | 120070 | 0 | 0 | 552220 | 0 | 0 | 0 | 423850 | 384570 |
| YADPVADL | CTDSPL; CTDSI | O15194; H7 | CTDSL_HUMAN | Unmodified | 137280 | 335860 | 0 | 0 | 0 | 13254 | 0 | 0 | 22979 | 67696 | 11176 | 11243 | 87071 | 0 | 0 | 0 | 0 | 0 |
| YADQEVPR | FLNC | Q14315 | FLNC_HUMAN | Unmodified | 0 | 51108 | 0 | 0 | 5271700 | 1637200 | 0 | 0 | 0 | 0 | 0 | 0 | 0 | 0 | 0 | 0 | 2604500 | 300790 |
| YADQEVPR | FLNC | Q14315 | FLNC_HUMAN | Unmodified | 150840 | 0 | 0 | 0 | 0 | 0 | 0 | 0 | 248700 | 0 | 0 | 0 | 0 | 0 | 0 | 0 | 588810 | 0 |
| YADQRLSS | GLG1 | Q92896 | GSLG1_HUMAN | Unmodified | 0 | 0 | 0 | 0 | 0 | 0 | 0 | 0 | 421600 | 29065 | 0 | 0 | 0 | 0 | 0 | 0 | 0 | 0 |
| YAEAVTR | PGK1 | P00558 | PGK1_HUMAN | Unmodified | 33060 | 0 | 503450 | 806420 | 0 | 343490 | 0 | 0 | 2377200 | 0 | 0 | 597530 | 0 | 607500 | 0 | 35380 | 1688000 | 3809300 |
| YAEAVTRAI | PGK1 | P00558 | PGK1_HUMAN | Unmodified | 0 | 0 | 0 | 0 | 27152 | 0 | 0 | 20445 | 258530 | 502620 | 0 | 0 | 0 | 0 | 0 | 0 | 7703900 | 3422500 |
| YAECGLR | GPX4; GPX4; GI | P36969; A0A | GPX4_HUMAN | Unmodified | 652860 | 482570 | 0 | 0 | 0 | 0 | 0 | 0 | 0 | 0 | 48916 | 942370 | 298600 | 221270 | 334130 | 552690 | 0 | 88830 |
| YAEDIFGEL | WASF2 | Q9Y6W5 | WASF2_HUMAN | Unmodified | 0 | 0 | 26823 | 174380 | 250020 | 28416 | 81983 | 0 | 0 | 0 | 0 | 27403 | 422860 | 346020 | 385130 | 27112 | 100500 | 205750 |
| YAEIVHLTL | ATP6V1B2; ATP | P21281; H0Y | VATB2_HUMAN | Unmodified | 0 | 118500 | 594470 | 0 | 1076300 | 0 | 591170 | 297870 | 1014000 | 292460 | 670150 | 5318600 | 2193600 | 617760 | 589960 | 573090 | 345620 | 1724400 |
| YAEIVHLTL | ATP6V1B2; ATP | P21281; H0Y | VATB2_HUMAN | Unmodified | 0 | 0 | 0 | 0 | 0 | 0 | 0 | 0 | 0 | 0 | 541870 | 0 | 0 | 0 | 0 | 0 | 0 | 0 |
| YAELLVSQ | OGDH; OGDH; | Q02218; E9F | ODO1_HUMAN | Unmodified | 0 | 0 | 0 | 0 | 0 | 0 | 0 | 0 | 1178500 | 0 | 247050 | 0 | 0 | 0 | 0 | 0 | 0 | 126090 |
| YAELSEEDI | NADK2 | Q4G0N4 | NAKD2_HUMAN | Unmodified | 0 | 0 | 0 | 0 | 0 | 0 | 0 | 0 | 664960 | 69667 | 0 | 921510 | 0 | 0 | 0 | 0 | 74112 | 401790 |
| YAEQLPVA | PIGK; PIGK | Q92643; B1A | GPI8_HUMAN | Unmodified | 0 | 0 | 0 | 0 | 0 | 0 | 0 | 0 | 2722300 | 0 | 0 | 0 | 0 | 0 | 0 | 0 | 0 | 0 |
| YAEQYSAT | DST; DST; DST; | Q03001; A0A | DYST_HUMAN | Unmodified | 0 | 0 | 0 | 0 | 275130 | 51301 | 0 | 8049.6 | 53070 | 0 | 0 | 128060 | 0 | 0 | 0 | 0 | 0 | 0 |
| YAETKDSG | DIPK2A | Q8NDZ4 | DIK2A_HUMAN | Unmodified | 0 | 0 | 0 | 0 | 0 | 0 | 0 | 0 | 0 | 0 | 0 | 0 | 0 | 0 | 0 | 0 | 0 | 0 |
| YAEVLRYVI | PPID | Q08752 | PPID_HUMAN | Unmodified | 0 | 0 | 0 | 0 | 230650 | 0 | 65231 | 0 | 0 | 390830 | 0 | 0 | 612870 | 512030 | 0 | 0 | 611310 | 343110 |
| YAFGQETN | NDUFA2 | O43678 | NDUA2_HUMA | Unmodified | 72954 | 52398 | 0 | 0 | 0 | 0 | 0 | 0 | 2399000 | 0 | 0 | 60275 | 0 | 0 | 0 | 42447 | 0 | 0 |
| YAFVNWIN | LCP1; PLS3; PL | P13796; P13 | PLSL_HUMAN | Unmodified | 0 | 0 | 0 | 0 | 6930200 | 2583400 | 777530 | 117340 | 513040 | 709560 | 0 | 0 | 1813200 | 945880 | 0 | 0 | 4372600 | 2069700 |
| YAGEPVPFI | LRPPRC | P42704 | LPPRC_HUMAN | Unmodified | 0 | 0 | 0 | 0 | 0 | 0 | 0 | 0 | 5186400 | 1279600 | 0 | 0 | 0 | 0 | 0 | 0 | 0 | 0 |
| YAGESFPG | FOLH1B; FOLH | Q9HBA9; Q0 | FOH1B_HUMAN | Unmodified | 65248 | 334580 | 740750 | 500370 | 0 | 0 | 0 | 0 | 0 | 0 | 0 | 0 | 0 | 0 | 2463900 | 535010 | 0 | 0 |
| YAGESFPG | FOLH1B; FOLH | Q9HBA9; Q0 | FOH1B_HUMAN | Unmodified | 0 | 3556200 | 508550 | 6651800 | 0 | 0 | 0 | 0 | 0 | 0 | 0 | 0 | 927540 | 580140 | 9439800 | 340050 | 0 | 0 |
| YAGKDGYN | S100A11 | P31949 | S10AB_HUMAN | Unmodified | 0 | 0 | 0 | 0 | 766230 | 0 | 0 | 0 | 0 | 169030 | 0 | 0 | 161850 | 0 | 0 | 0 | 102150 | 351990 |
| YAGKDGYN | S100A11 | P31949 | S10AB_HUMAN | Unmodified | 0 | 0 | 0 | 0 | 4567000 | 0 | 0 | 0 | 0 | 781120 | 0 | 0 | 1807200 | 0 | 0 | 0 | 1387100 | 5145100 |
| YAGLSTCFI | SARS1; SARS1 | P49591; Q5T | SYSC_HUMAN | Unmodified | 106770 | 0 | 0 | 0 | 0 | 0 | 0 | 25095 | 284010 | 0 | 0 | 407230 | 0 | 0 | 81927 | 140260 | 0 | 0 |
| YAGSALQY | VTA1; VTA1; VTA | Q9NP79; Q5 | VTA1_HUMAN | Unmodified | 3457400 | 3826200 | 2889300 | 8223500 | 43661000 | 7421300 | 6881600 | 2673300 | 24912000 | 19174000 | 13548000 | 13761000 | 29051000 | 17294000 | 3572000 | 2479100 | 8439500 | 15258000 |
| YAGVFAEN | C1GALT1C1 | Q96EU7 | C1GLC_HUMAN | Unmodified | 0 | 0 | 0 | 0 | 0 | 0 | 0 | 0 | 989280 | 0 | 0 | 0 | 0 | 0 | 0 | 0 | 130940 | 0 |
| YAGVFAEN | C1GALT1C1 | Q96EU7 | C1GLC_HUMAN | Unmodified | 0 | 0 | 0 | 0 | 0 | 0 | 0 | 220880 | 8276500 | 2983300 | 513770 | 433410 | 0 | 0 | 0 | 0 | 0 | 0 |
| YAHVVLR | RPS18 | P62269 | RS18_HUMAN | Unmodified | 547800 | 368740 | 708600 | 0 | 0 | 0 | 58976 | 20225 | 0 | 40193 | 77072 | 4255000 | 0 | 0 | 0 | 0 | 0 | 74296 |
| YAIAVNDL | GLS; GLS | O94925; H7I | GLSK_HUMAN | Unmodified | 219100 | 729700 | 0 | 0 | 0 | 0 | 0 | 0 | 2670000 | 0 | 337870 | 0 | 112140 | 0 | 0 | 0 | 0 | 0 |
| YAICSALAA | RPL4; RPL4; RP | P36578; H3E | RL4_HUMAN 6 | Oxidation (M) | 797640 | 3002300 | 0 | 673970 | 0 | 0 | 0 | 0 | 380240 | 227980 | 259500 | 666520 | 1329500 | 0 | 0 | 0 | 0 | 0 |
| YAILNEPFK | CHD4; CHD4; C | Q14839; A0A | CHD4_HUMAN | Unmodified | 0 | 0 | 0 | 0 | 0 | 0 | 0 | 0 | 320500 | 0 | 0 | 2261400 | 0 | 0 | 0 | 0 | 0 | 0 |
| YAIQLITAA | RUVBL2 | Q9Y230 | RUVB2_HUMAN | Unmodified | 0 | 0 | 0 | 0 | 0 | 0 | 0 | 0 | 0 | 0 | 0 | 0 | 0 | 0 | 0 | 0 | 0 | 0 |
| YALELQK | ALPL | P05186 | PPBT_HUMAN | Unmodified | 326650 | 541280 | 599410 | 382530 | 462000 | 180110 | 76843 | 138150 | 1590100 | 1554700 | 0 | 0 | 0 | 0 | 0 | 570810 | 157620 | 0 |
| YALGERQS | LTA4H; LTA4H | P09960; B4D | LKHA4_HUMAN | Unmodified | 0 | 0 | 0 | 0 | 294210 | 0 | 0 | 13920 | 98229 | 59802 | 73609 | 0 | 0 | 0 | 0 | 0 | 49866 | 236900 |
| YALPLVGH | PDIA4; PDIA4 | P13667; A0A | PDIA4_HUMAN | Unmodified | 0 | 0 | 0 | 0 | 0 | 0 | 0 | 0 | 3934300 | 406180 | 293960 | 0 | 0 | 0 | 0 | 0 | 0 | 0 |
| YALQSQQF | ACAA2; ACAA2; | P42765; A0A | THIM_HUMAN | Unmodified | 2709200 | 0 | 0 | 0 | 229640 | 50161 | 23023 | 0 | 1673000 | 75270 | 76839 | 539730 | 0 | 77228 | 0 | 76528 | 83432 | 0 |
| YALSVGYR | AKR1A1; AKR1A | P14550; V9C | AK1A1_HUMAN | Unmodified | 27499 | 0 | 79042 | 0 | 687050 | 0 | 0 | 0 | 3360900 | 484030 | 526870 | 547550 | 1231300 | 860500 | 0 | 0 | 227360 | 0 |
| YALTGDEVI | RPS4Y2; RPS4X | Q8TD47; P6 | RS4Y2_HUMAN | Unmodified | 0 | 908460 | 0 | 0 | 3225100 | 34566 | 1328500 | 1193200 | 647440 | 791440 | 1597600 | 15716000 | 637180 | 450730 | 0 | 0 | 0 | 442730 |
| YALTGDEVI | RPS4Y2; RPS4X | Q8TD47; P6 | RS4Y2_HUMAN | Unmodified | 0 | 0 | 0 | 0 | 101790 | 0 | 264630 | 285340 | 0 | 40480 | 165990 | 4593100 | 0 | 0 | 0 | 0 | 0 | 0 |
| YALTTTLSA | MYG1; MYG1 | Q9HB07; F8 | MYG1_HUMAN | Unmodified | 0 | 58795 | 97163 | 0 | 2214400 | 201670 | 0 | 0 | 832300 | 74786 | 150810 | 2154300 | 0 | 0 | 124020 | 192460 | 0 | 0 |
| YALVLQDV | SORCS2; SORC | Q96PQ0; B5 | SORC2_HUMAN | Unmodified | 0 | 0 | 0 | 97394 | 0 | 0 | 0 | 0 | 4060800 | 76360 | 279470 | 0 | 0 | 0 | 0 | 0 | 0 | 0 |
| YALYDASFI | DSTN; DSTN | P60981; F6R | DEST_HUMAN | Unmodified | 6152700 | 0 | 3532400 | 3894300 | 5832700 | 1699200 | 303620 | 176970 | 5767000 | 734370 | 5652200 | 41758000 | 7497100 | 2525600 | 5472500 | 4746000 | 1010900 | 2316600 |
| YALYDATYI | CFL1; CFL2; CF | P23528; Q9Y | COF1_HUMAN | Unmodified | 46608000 | 644770 | 21812000 | 34997000 | 2.13E+08 | 23734000 | 4112900 | 2226800 | 47507000 | 7213500 | 25025000 | 2.94E+08 | 82676000 | 15667000 | 42317000 | 42023000 | 9125400 | 26723000 |
| YANEVNSD | SRP68 | Q9UHB9 | SRP68_HUMAN | Unmodified | 0 | 0 | 0 | 0 | 0 | 0 | 0 | 0 | 0 | 0 | 0 | 1674600 | 0 | 0 | 0 | 0 | 0 | 0 |
| YANNSNYK | MAGOH; MAGC | P61326; Q96 | MGN_HUMAN | Unmodified | 206210 | 116950 | 0 | 0 | 0 | 0 | 0 | 0 | 0 | 0 | 0 | 0 | 0 | 0 | 0 | 0 | 0 | 0 |
| YANVIAYDI | PTPRD; PTPRF; | P23468; P10 | PTPRD_HUMAN | Unmodified | 1308100 | 0 | 1521300 | 702290 | 0 | 0 | 0 | 0 | 0 | 310230 | 0 | 2081200 | 0 | 0 | 4695800 | 3037100 | 0 | 0 |
| YAPDDIPNI | AFDN; AFDN; A | P55196; H0Y | AFAD_HUMAN | Unmodified | 0 | 0 | 20736 | 0 | 0 | 128370 | 0 | 535480 | 158540 | 0 | 81491 | 276990 | 117940 | 121160 | 64483 | 46676 | 0 | 0 |
| YAPENLDE | ACAD9; ACAD9 | Q9H845; H0 | ACAD9_HUMAN | Unmodified | 0 | 0 | 0 | 0 | 0 | 0 | 0 | 0 | 628850 | 0 | 0 | 0 | 0 | 0 | 0 | 0 | 0 | 0 |
| YAPISGGDI | TNC; TNC; TNC | P24821; F5H | TENA_HUMAN | Unmodified | 0 | 0 | 0 | 0 | 0 | 0 | 354200 | 156190 | 0 | 153570 | 0 | 0 | 0 | 0 | 0 | 0 | 447480 | 0 |
| YAPLHLVPI | CYFIP1; CYFIP2 | Q7L576; Q96 | CYFP1_HUMAN | Unmodified | 4278100 | 0 | 4334000 | 3384000 | 0 | 0 | 0 | 0 | 4974900 | 0 | 0 | 15222000 | 0 | 0 | 6013600 | 4523400 | 0 | 0 |
| YAPRPIFIS | MTR; MTR; MTR | Q99707; A0A | METH_HUMAN | Unmodified | 0 | 0 | 0 | 0 | 89738 | 0 | 0 | 0 | 468090 | 173950 | 80025 | 852440 | 0 | 0 | 0 | 22164 | 20640 | 0 |
| YAPSEAGLI | FLNA; FLNA; FL | P21333; H0Y | FLNA_HUMAN | Oxidation (M) | 249760 | 1022800 | 0 | 0 | 0 | 0 | 0 | 0 | 0 | 0 | 106620 | 191630 | 0 | 0 | 518320 | 893300 | 0 | 0 |
| YAPSEAGLI | FLNA; FLNA; FL | P21333; H0Y | FLNA_HUMAN | Unmodified | 298600 | 472190 | 123410 | 0 | 0 | 0 | 0 | 37210 | 104930 | 0 | 0 | 0 | 0 | 0 | 488340 | 394540 | 79221 | 0 |
| YAPSGFYIA | WDR1; WDR1 | O75083; D6I | WDR1_HUMAN | Unmodified | 4411100 | 2960200 | 4169100 | 3702600 | 44033000 | 10130000 | 5195800 | 992010 | 7641900 | 4901500 | 6639700 | 20099000 | 17318000 | 14254000 | 5334500 | 4745600 | 8858300 | 10468000 |
| YAPSGFYIA | WDR1; WDR1 | O75083; D6I | WDR1_HUMAN | Unmodified | 0 | 0 | 0 | 0 | 3077900 | 0 | 273390 | 245890 | 0 | 400240 | 0 | 0 | 0 | 0 | 0 | 0 | 1129600 | 693560 |
| YAPSGFYIA | WDR1; WDR1 | O75083; D6I | WDR1_HUMAN | Unmodified | 0 | 0 | 0 | 0 | 0 | 0 | 4621200 | 0 | 0 | 741210 | 0 | 0 | 2797000 | 7058400 | 0 | 0 | 3215900 | 1210200 |
| YAPTEVGLI | FLNB; FLNB | O75369; E7E | FLNB_HUMAN | Unmodified | 0 | 0 | 0 | 0 | 0 | 0 | 0 | 0 | 0 | 0 | 0 | 2107200 | 0 | 0 | 0 | 0 | 87824 | 0 |
| YAPTIAPSP | SLC6A9; SLC6A | P48067; B7Z | SC6A9_HUMAN | Unmodified | 0 | 0 | 3390900 | 3255100 | 0 | 0 | 0 | 0 | 2972300 | 0 | 1209300 | 3443800 | 0 | 0 | 3808300 | 2524700 | 0 | 0 |
| YAQDFGLV | CTSC; CTSC; C | P53634; A0A | CATC_HUMAN | Unmodified | 0 | 0 | 0 | 0 | 0 | 0 | 0 | 0 | 803700 | 0 | 1016200 | 649120 | 0 | 0 | 0 | 0 | 0 | 0 |
| YAQEVAGL | CLIP2 | Q9UDT6 | CLIP2_HUMAN | Unmodified | 0 | 0 | 0 | 0 | 0 | 0 | 0 | 0 | 0 | 0 | 0 | 0 | 0 | 0 | 0 | 0 | 0 | 147280 |
| YAQQDLK | TSPAN4; TSPAN | O14817; A8I | TSN4_HUMAN | Unmodified | 566100 | 126250 | 198010 | 124530 | 0 | 0 | 0 | 334250 | 41263 | 224070 | 148360 | 171780 | 127160 | 93639 | 1278400 | 820300 | 73200 | 338040 |
| YARTDGKV | RPL24; RPL24; | P83731; C9J | RL24_HUMAN | Unmodified | 0 | 0 | 0 | 0 | 0 | 0 | 425100 | 0 | 0 | 0 | 0 | 0 | 1528800 | 0 | 0 | 0 | 0 | 0 |
| YARVTPDTI | ACLY; ACLY | P53396; K7E | ACLY_HUMAN | Unmodified | 0 | 0 | 0 | 0 | 1378900 | 0 | 454220 | 0 | 0 | 256330 | 0 | 0 | 0 | 0 | 0 | 0 | 0 | 0 |
| YASACQMI | IGF2R; IGF2R | P11717; A0A | MPRI_HUMAN | Unmodified | 0 | 0 | 0 | 0 | 0 | 0 | 0 | 0 | 4661200 | 0 | 0 | 1212200 | 0 | 0 | 0 | 0 | 0 | 0 |
| YASACQMI | IGF2R; IGF2R | P11717; A0A | MPRI_HUMAN | Unmodified | 0 | 0 | 0 | 0 | 0 | 0 | 0 | 0 | 3876900 | 374130 | 135370 | 0 | 0 | 0 | 0 | 0 | 0 | 0 |
| YASFQVEN | SEC24C; SEC24 | P53992; G5E | SC24C_HUMAN | Unmodified | 0 | 0 | 0 | 0 | 0 | 0 | 0 | 0 | 214070 | 125560 | 0 | 900770 | 0 | 229650 | 0 | 0 | 0 | 0 |
| YASINAHL | CSNK1A1L; CSI | Q8N752; P4 | KC1AL_HUMAN | Unmodified | 336530 | 264650 | 546660 | 405110 | 293100 | 0 | 235890 | 60601 | 0 | 0 | 0 | 1212300 | 0 | 0 | 834830 | 791630 | 0 | 149790 |
| YASNAVSE | DNPEP; DNPEP | Q9ULA0; E5I | DNPEP_HUMAN | Unmodified | 0 | 403300 | 0 | 0 | 0 | 0 | 0 | 0 | 0 | 0 | 0 | 1512000 | 0 | 0 | 0 | 0 | 0 | 0 |
| YASRLAGKI | BROX; BROX; BI | Q5VW32; F5 | BROX_HUMAN | Unmodified | 0 | 0 | 0 | 94681 | 333800 | 84489 | 54405 | 34767 | 0 | 312370 | 0 | 0 | 167910 | 150030 | 0 | 0 | 0 | 0 |
| YATALYSA | ATP5PO; ATP5P | P48047; H7C | ATPO_HUMAN | Unmodified | 0 | 0 | 0 | 57110 | 0 | 0 | 118010 | 0 | 10689000 | 1088800 | 0 | 0 | 92829 | 61773 | 0 | 0 | 0 | 0 |
| YATATRFSY | KIRREL1; KIRRE | Q96J84; Q5V | KIRR1_HUMAN | Unmodified | 0 | 0 | 0 | 0 | 7340600 | 2273600 | 2081600 | 851680 | 0 | 0 | 0 | 0 | 284350 | 5362500 | 0 | 0 | 364680 | 1435700 |
| YATDTFAGI | COPS3; COPS3 | Q9UNS2; C9 | CSN3_HUMAN | Unmodified | 2569500 | 1411000 | 0 | 0 | 0 | 0 | 0 | 0 | 76934 | 0 | 32816 | 1093100 | 0 | 0 | 153220 | 495380 | 0 | 0 |
| YATGENTV | OSBPL9; OSBPI | Q96SU4; A0 | OSBL9_HUMAN | Unmodified | 0 | 0 | 0 | 0 | 500770 | 174580 | 0 | 0 | 0 | 214060 | 0 | 296740 | 221750 | 123460 | 0 | 0 | 207950 | 293240 |
| YATGGWEI | FUT8; FUT8; FU | Q9BYC5; A0 | FUT8_HUMAN | Unmodified | 0 | 0 | 0 | 0 | 0 | 0 | 0 | 0 | 1307900 | 238950 | 0 | 0 | 0 | 0 | 0 | 0 | 0 | 0 |
| YATLATVSI | DYNC1H1; DYN | Q14204; A0A | DYHC1_HUMA | Unmodified | 0 | 0 | 2359300 | 0 | 1264300 | 0 | 1317600 | 439840 | 0 | 144040 | 642390 | 3479400 | 0 | 3503800 | 211800 | 0 | 0 | 2132600 |
| YAVLYQPLI | NAP1L1; NAP1L | P55209; F8V | NP1L1_HUMAN | Unmodified | 3748000 | 3976300 | 5970800 | 4754800 | 6275700 | 1317300 | 1559000 | 1048300 | 5722800 | 0 | 2211700 | 31497000 | 5512700 | 6157000 | 4633000 | 5258600 | 1501400 | 2233500 |
| YAVPDEILV | ACSS1 | Q9NUB1 | ACS2L_HUMAN | Unmodified | 0 | 0 | 0 | 0 | 0 | 0 | 0 | 0 | 1689100 | 388070 | 0 | 0 | 0 | 0 | 0 | 0 | 0 | 0 |
| YAVPVLR | PDIA5 | Q14554 | PDIA5_HUMAN | Unmodified | 0 | 0 | 0 | 0 | 118860 | 0 | 0 | 0 | 16955000 | 5487400 | 292570 | 0 | 0 | 0 | 0 | 0 | 146450 | 0 |
| YAVTTGDH | COPA; COPA; C | P53621; A0A | COPA_HUMAN | Unmodified | 139310 | 100710 | 0 | 0 | 0 | 0 | 0 | 0 | 557820 | 346850 | 0 | 79835 | 0 | 0 | 0 | 447310 | 0 | 0 |
| YAVWFGG | ACTR3; ACTR3 | P61158; B4D | ARP3_HUMAN | Oxidation (M) | 37914 | 45583 | 0 | 92184 | 0 | 0 | 0 | 0 | 0 | 0 | 0 | 1574700 | 0 | 0 | 0 | 50048 | 0 | 0 |
| YAYIQVTH | DOCK9; DOCK | Q9BZ29; A0 | DOCK9_HUMA | Unmodified | 154570 | 139300 | 122310 | 287260 | 313080 | 0 | 46057 | 0 | 1054700 | 191840 | 97513 | 863460 | 130350 | 42260 | 595980 | 452290 | 0 | 55124 |
| YAYSAASG | ITGA2; ITGA2; I | P17301; E9P | ITA2_HUMAN | Unmodified | 445520 | 311500 | 559480 | 263870 | 1589100 | 63291 | 0 | 0 | 0 | 0 | 0 | 0 | 88262 | 0 | 986060 | 0 | 190000 | 573810 |
| YCAGWAD | ALDH1A1; ALDI | P00352; Q5 | AL1A1_HUMAN | Unmodified | 0 | 0 | 0 | 0 | 0 | 0 | 0 | 0 | 395890 | 0 | 4247800 | 1.94E+08 | 0 | 0 | 0 | 0 | 0 | 0 |
| YCALLLPLL | EIF2B5; EIF2B5 | Q13144; A0A | EI2BE_HUMAN | Unmodified | 0 | 0 | 0 | 0 | 0 | 0 | 0 | 0 | 15232 | 0 | 0 | 273510 | 0 | 0 | 59695 | 0 | 112910 | 0 |

| | | | | | | | | | | | | | | | | | | | | | | |
|---|---|---|---|---|---|---|---|---|---|---|---|---|---|---|---|---|---|---|---|---|---|---|
| YCAQDAFF | ECH1; ECH1 | Q13011; M0 | ECH1_HUMAN | Unmodified | 0 | 0 | 0 | 0 | 0 | 0 | 0 | 0 | 4415700 | 0 | 0 | 0 | 0 | 0 | 0 | 0 | 0 | 0 |
| YCDHENA/ | TOR1AIP2; TOR | Q8NFQ8; Q5 | TOIP2_HUMAN | Unmodified | 0 | 0 | 0 | 0 | 0 | 0 | 0 | 0 | 6778000 | 0 | 444540 | 0 | 0 | 0 | 0 | 0 | 0 | 0 |
| YCECDDFS | ITGB3; N/A | P05106; H3E | ITB3_HUMAN I | Unmodified | 750700 | 643850 | 0 | 1229300 | 0 | 0 | 0 | 0 | 0 | 0 | 0 | 0 | 0 | 0 | 610720 | 573250 | 0 | 0 |
| YCFEIVPK | PTPRJ; PTPRJ; P | Q12913; A0/ | PTPRJ_HUMAN | Unmodified | 0 | 67171 | 0 | 0 | 0 | 0 | 0 | 0 | 305390 | 0 | 0 | 1160000 | 0 | 0 | 58235 | 0 | 0 | 0 |
| YCFGVEDT | ETF1; ETF1 | P62495; B7Z | ERF1_HUMAN | Unmodified | 90826 | 0 | 0 | 0 | 0 | 0 | 0 | 0 | 0 | 0 | 0 | 1333700 | 0 | 0 | 108600 | 97179 | 0 | 0 |
| YCGERSQF | ST14 | Q9Y5Y6 | ST14_HUMAN | Unmodified | 0 | 0 | 0 | 0 | 325600 | 0 | 0 | 0 | 0 | 0 | 0 | 0 | 1879300 | 0 | 0 | 0 | 81829 | 147360 |
| YCPIPDVL( | EHHADH | Q08426 | ECHP_HUMAN | Unmodified | 0 | 0 | 0 | 0 | 0 | 0 | 0 | 0 | 139200 | 0 | 0 | 0 | 0 | 0 | 0 | 0 | 0 | 0 |
| YCRSISPEV | MYO1C; MYO1( | O00159; F5H | MYO1C_HUMA | Unmodified | 0 | 0 | 0 | 0 | 1130000 | 0 | 0 | 0 | 0 | 0 | 0 | 0 | 0 | 0 | 0 | 0 | 1278800 | 0 |
| YCVEEEEK | GTF2I; GTF2I; G | P78347; A0A | GTF2I_HUMAN | Unmodified | 31421 | 0 | 0 | 0 | 0 | 0 | 0 | 0 | 0 | 38315 | 0 | 1619200 | 0 | 0 | 0 | 0 | 0 | 0 |
| YDAFLASES | RPL10A | P62906 | RL10A_HUMAN | Unmodified | 0 | 0 | 0 | 0 | 0 | 0 | 0 | 0 | 0 | 1276600 | 1148100 | 2115600 | 0 | 0 | 0 | 0 | 0 | 0 |
| YDAIRINQL | FERMT2; FERM1 | Q96AC1; A0, | FERM2_HUMAI | Unmodified | 0 | 0 | 0 | 0 | 1261200 | 0 | 0 | 0 | 0 | 0 | 0 | 0 | 0 | 0 | 0 | 0 | 52722 | 434430 |
| YDDMAAAN | YWHAB; YWHA | P31946; P61 | 1433B_HUMAN | Unmodified | 0 | 75994 | 0 | 177330 | 116680 | 0 | 0 | 0 | 328250 | 32206 | 1387900 | 2922900 | 0 | 0 | 359230 | 0 | 35388 | 63550 |
| YDDMAAAN | YWHAB; YWHA | P31946; P61 | 1433B_HUMAN | Oxidation (M) | 0 | 0 | 0 | 0 | 0 | 0 | 0 | 0 | 58399 | 0 | 395800 | 1253900 | 0 | 0 | 385740 | 267840 | 0 | 0 |
| YDDMAACI | YWHAZ; YWHA; | P63104; E5R | 1433Z_HUMAN | Unmodified | 0 | 0 | 0 | 0 | 0 | 0 | 0 | 0 | 496140 | 0 | 591410 | 1387000 | 0 | 0 | 0 | 99830 | 0 | 0 |
| YDDMAACI | YWHAZ; YWHA; | P63104; E5R | 1433Z_HUMAN | Oxidation (M) | 0 | 0 | 0 | 0 | 0 | 0 | 0 | 0 | 89627 | 0 | 314030 | 489320 | 0 | 0 | 60534 | 44998 | 0 | 0 |
| YDEAASYIC | GNAI2 | P04899 | GNAI2_HUMAN | Unmodified | 0 | 0 | 545370 | 0 | 0 | 0 | 0 | 0 | 0 | 0 | 0 | 11855000 | 0 | 0 | 0 | 117490 | 0 | 0 |
| YDEAASYIC | GNAI2 | P04899 | GNAI2_HUMAN | Unmodified | 294920 | 0 | 296640 | 0 | 789580 | 0 | 261620 | 0 | 0 | 154440 | 877900 | 7377900 | 0 | 0 | 417310 | 194760 | 666800 | 877740 |
| YDEDAKRP | NECTIN1 | Q15223 | NECT1_HUMAI | Unmodified | 508120 | 0 | 0 | 0 | 0 | 0 | 0 | 0 | 0 | 0 | 0 | 1472200 | 0 | 0 | 0 | 233800 | 0 | 0 |
| YDEIFLK | EPS15; EPS15 | P42566; B1A | EPS15_HUMAN | Unmodified | 0 | 0 | 0 | 0 | 0 | 0 | 0 | 0 | 0 | 41078 | 1122100 | 0 | 0 | 0 | 114420 | 71882 | 0 | 0 |
| YDEIVKEVS | EEF1A2; EEF1A: | Q05639; A0/ | EF1A2_HUMAN | Unmodified | 83385 | 63858 | 0 | 204300 | 0 | 0 | 37822 | 0 | 0 | 0 | 0 | 0 | 0 | 0 | 0 | 0 | 116610 | 0 |
| YDEMVESM | YWHAE; YWHAI | P62258; I3L: | 1433E_HUMAN | Unmodified | 0 | 0 | 0 | 0 | 0 | 0 | 0 | 0 | 0 | 0 | 0 | 3977800 | 0 | 0 | 0 | 0 | 0 | 0 |
| YDEYVNVK | PDCD6IP | Q8WUM4 | PDC6I_HUMAN | Unmodified | 5770400 | 0 | 2707100 | 1632400 | 0 | 0 | 0 | 0 | 3008900 | 249120 | 611000 | 0 | 186820 | 0 | 3412500 | 4018000 | 0 | 0 |
| YDEYVNVK | PDCD6IP | Q8WUM4 | PDC6I_HUMAN | Unmodified | 158140 | 156950 | 0 | 0 | 0 | 0 | 0 | 0 | 0 | 0 | 0 | 0 | 0 | 0 | 0 | 0 | 0 | 0 |
| YDFGIYDD | DNAJC10; DNA | Q8IXB1; A0A | DJC10_HUMAN | Unmodified | 0 | 0 | 0 | 0 | 0 | 0 | 38755 | 0 | 2468100 | 0 | 99826 | 0 | 0 | 0 | 2938000 | 0 | 0 | 0 |
| YDFKATAD | GRB2; GRB2; G | P62993; J3Q | GRB2_HUMAN | Unmodified | 907510 | 896200 | 107950 | 304140 | 0 | 0 | 0 | 0 | 0 | 215340 | 475720 | 445780 | 633390 | 253460 | 619400 | 720110 | 111970 | 0 |
| YDGIILPGK | RPL11; RPL11 | P62913; Q5\ | RL11_HUMAN ( | Unmodified | 6009300 | 5607500 | 4280200 | 1227700 | 0 | 0 | 196920 | 349480 | 522070 | 349720 | 2088200 | 36611000 | 865310 | 543900 | 8488600 | 4636800 | 0 | 0 |
| YDGKWEVI | CANX; CANX; C | P27824; A0A | CALX_HUMAN | Oxidation (M) | 0 | 0 | 0 | 0 | 0 | 280220 | 0 | 0 | 7784800 | 213590 | 0 | 49892 | 0 | 0 | 0 | 0 | 273510 | 0 |
| YDGKWEVI | CANX; CANX; C | P27824; A0A | CALX_HUMAN | Unmodified | 0 | 0 | 0 | 0 | 134110 | 0 | 140990 | 0 | 36414000 | 634520 | 917430 | 1219500 | 0 | 0 | 0 | 0 | 0 | 0 |
| YDGVDAGH | HMGCS1; HMG | Q01581; D6I | HMCS1_HUMA | Unmodified | 0 | 0 | 0 | 0 | 245590 | 0 | 0 | 0 | 94026 | 0 | 0 | 1501700 | 0 | 0 | 0 | 0 | 0 | 0 |
| YDKSLHQA | ANXA6; ANXA6; | P08133; E5R | ANXA6_HUMAI | Unmodified | 0 | 0 | 0 | 0 | 0 | 0 | 349120 | 39141 | 0 | 0 | 0 | 0 | 0 | 0 | 0 | 0 | 0 | 66768 |
| YDLASGATI | SORL1 | Q92673 | SORL_HUMAN | Unmodified | 0 | 0 | 0 | 0 | 0 | 0 | 0 | 0 | 450330 | 0 | 0 | 0 | 0 | 0 | 0 | 0 | 0 | 0 |
| YDLDFKSP | ENO1; ENO1 | P06733; A0A | ENOA_HUMAN | Unmodified | 39369 | 114410 | 299200 | 0 | 0 | 0 | 0 | 7949.7 | 863280 | 0 | 2439600 | 3787600 | 0 | 0 | 383090 | 225080 | 0 | 0 |
| YDLDFKSP | ENO1; ENO1 | P06733; A0A | ENOA_HUMAN | Unmodified | 0 | 0 | 0 | 0 | 5180100 | 3956200 | 0 | 0 | 0 | 0 | 0 | 0 | 0 | 0 | 0 | 0 | 4163000 | 1970200 |
| YDLSALVR | IGF2R; IGF2R; I | P11717; A0A | MPRI_HUMAN ( | Unmodified | 0 | 75746 | 0 | 89194 | 0 | 0 | 0 | 0 | 2857900 | 1951600 | 121820 | 0 | 74896 | 485410 | 0 | 0 | 0 | 0 |
| YDLSPITVK | ERGIC1 | Q969X5 | ERGI1_HUMAN | Unmodified | 592650 | 87675 | 815730 | 279960 | 0 | 0 | 79437 | 66763 | 26991000 | 2226700 | 0 | 921280 | 43208 | 0 | 1032200 | 758900 | 0 | 272680 |
| YDNMAELF | VPS28; VPS28; | Q9UK41; E9I | VPS28_HUMAN | Unmodified | 4834100 | 3549000 | 0 | 0 | 0 | 0 | 0 | 0 | 2213200 | 0 | 0 | 663810 | 0 | 0 | 4990200 | 2524200 | 0 | 0 |
| YDNMAELF | VPS28; VPS28; | Q9UK41; E9I | VPS28_HUMAN | Oxidation (M) | 9212900 | 9351100 | 10787000 | 8369500 | 0 | 0 | 0 | 0 | 1423200 | 0 | 0 | 1492700 | 2008500 | 1118200 | 19028000 | 12268000 | 0 | 0 |
| YDNQIHIID | EIPR1; EIPR1; E | Q53HC9; D3 | EIPR1_HUMAN | Unmodified | 0 | 0 | 0 | 0 | 0 | 0 | 0 | 0 | 0 | 0 | 0 | 2226400 | 0 | 0 | 0 | 0 | 0 | 0 |
| YDNSLKIIS | GAPDH; GAPDH | P04406; E7E | G3P_HUMAN G | Unmodified | 0 | 0 | 0 | 0 | 0 | 0 | 230590 | 232990 | 363530 | 0 | 0 | 0 | 0 | 0 | 0 | 0 | 1185300 | 0 |
| YDPEEPIYF | C1GALT1 | Q9NS00 | C1GLT_HUMAN | Unmodified | 0 | 0 | 0 | 0 | 0 | 0 | 0 | 0 | 1060200 | 103920 | 0 | 0 | 0 | 0 | 0 | 0 | 20733 | 0 |
| YDPMPPES | PLCG2; PLCG2 | P16885; A0A | PLCG2_HUMAI | Unmodified | 0 | 0 | 208820 | 0 | 0 | 61524 | 0 | 0 | 124900 | 0 | 298470 | 1719900 | 0 | 0 | 0 | 0 | 159550 | 169800 |
| YDPNVYSII | ITGAV | P06756 | ITAV_HUMAN I | Unmodified | 362240 | 0 | 1333400 | 243850 | 0 | 0 | 0 | 0 | 884360 | 0 | 0 | 2444500 | 0 | 0 | 1083300 | 811760 | 0 | 0 |
| YDPTIEDFY | RAP2C; RAP2A; | Q9Y3L5; P1C | RAP2C_HUMAN | Unmodified | 51902 | 0 | 635650 | 0 | 0 | 0 | 0 | 0 | 0 | 0 | 401320 | 871950 | 0 | 0 | 205740 | 144080 | 929550 | 0 |
| YDPTIEDFY | RAP2C; RAP2A; | Q9Y3L5; P1C | RAP2C_HUMAI | Unmodified | 0 | 136890 | 2885000 | 1703100 | 0 | 0 | 0 | 0 | 1922400 | 0 | 3742300 | 6745900 | 192230 | 0 | 3082800 | 1189300 | 0 | 566900 |
| YDPTIEDSY | RAP1B; RAP1A; | P61224; P62 | RAP1B_HUMAN | Unmodified | 318910 | 299920 | 1114900 | 709590 | 0 | 0 | 0 | 0 | 0 | 0 | 0 | 2741200 | 0 | 0 | 916940 | 796470 | 0 | 0 |
| YDPTIEDSY | RAP1B; RAP1A; | P61224; P62 | RAP1B_HUMAN | Unmodified | 4774600 | 4554000 | 6192800 | 9198700 | 6296800 | 1230900 | 1411800 | 462700 | 3468300 | 415210 | 6811300 | 23197000 | 3307000 | 1405900 | 9357000 | 9251400 | 429830 | 3026800 |
| YDSDRTEL( | PDIA5 | Q14554 | PDIA5_HUMAN | Unmodified | 268290 | 0 | 195740 | 0 | 0 | 0 | 0 | 0 | 16552000 | 2419100 | 403500 | 0 | 0 | 0 | 0 | 0 | 0 | 0 |
| YDSRTTIFS | PSMA4; PSMA4 | P25789; H0\ | PSA4_HUMAN | Unmodified | 0 | 0 | 0 | 0 | 604520 | 21357 | 24340 | 0 | 0 | 148660 | 0 | 0 | 502670 | 211980 | 0 | 0 | 83438 | 0 |
| YDTADRILT | ERMP1; ERMP1 | Q7Z2K6; A0/ | ERMP1_HUMAI | Unmodified | 0 | 0 | 20097 | 0 | 0 | 0 | 0 | 0 | 810730 | 64275 | 0 | 152100 | 0 | 0 | 0 | 19712 | 0 | 0 |
| YDVGGGEI | PTPN11; PTPN1 | Q06124; H0\ | PTN11_HUMAN | Unmodified | 0 | 0 | 111810 | 0 | 0 | 0 | 0 | 0 | 0 | 0 | 1261400 | 3754600 | 0 | 0 | 333450 | 0 | 0 | 0 |
| YDYNSGEE | STRAP; STRAP | Q9Y3F4; H0\ | STRAP_HUMAN | Unmodified | 0 | 0 | 0 | 0 | 0 | 0 | 0 | 0 | 0 | 0 | 191210 | 2294400 | 0 | 0 | 0 | 0 | 0 | 0 |
| YDYQRQPI | RPN1; RPN1 | P04843; B7Z | RPN1_HUMAN | Unmodified | 0 | 0 | 0 | 0 | 0 | 0 | 0 | 0 | 1679400 | 233990 | 0 | 0 | 0 | 0 | 0 | 0 | 0 | 0 |
| YEAFQQGH | SEZ6; SEZ6; SEZ | Q53EL9; C9J | SEZ6_HUMAN | Unmodified | 0 | 0 | 0 | 0 | 0 | 0 | 0 | 0 | 2855500 | 0 | 0 | 0 | 0 | 30190 | 0 | 0 | 0 | 503170 |
| YEALKEPM | SPTAN1; SPTAN | Q13813; A0/ | SPTN1_HUMAN | Unmodified | 0 | 0 | 0 | 0 | 0 | 0 | 0 | 0 | 0 | 0 | 0 | 839820 | 0 | 0 | 0 | 0 | 0 | 0 |
| YEAPQATD | TOM1 | O60784 | TOM1_HUMAN | Unmodified | 28247000 | 245320 | 183260 | 0 | 0 | 0 | 0 | 0 | 0 | 0 | 0 | 0 | 0 | 0 | 3102600 | 498280 | 0 | 0 |
| YEAVGSVH | MRI1 | Q9BV20 | MTNA_HUMAN | Unmodified | 0 | 0 | 0 | 0 | 0 | 0 | 0 | 0 | 0 | 0 | 0 | 1474800 | 0 | 0 | 0 | 0 | 0 | 0 |
| YEDFKEEG | OLA1; OLA1 | Q9NTK5; J3K | OLA1_HUMAN | Unmodified | 0 | 0 | 0 | 0 | 0 | 0 | 0 | 0 | 114750 | 0 | 750630 | 3069800 | 0 | 0 | 0 | 0 | 18806 | 0 |
| YEDFSNLG | UQCRC2; UQC | P22695; H3E | QCR2_HUMAN | Unmodified | 0 | 0 | 0 | 0 | 0 | 0 | 0 | 0 | 2373100 | 0 | 0 | 0 | 0 | 0 | 0 | 0 | 0 | 0 |
| YEDLAPCIT | TPP2; TPP2; TPI | P29144; A0A | TPP2_HUMAN | Unmodified | 0 | 0 | 0 | 0 | 0 | 0 | 0 | 0 | 0 | 0 | 0 | 1207200 | 0 | 0 | 0 | 0 | 0 | 0 |
| YEDLLTR | GDF15; GDF15 | Q99988; A0/ | GDF15_HUMAI | Unmodified | 0 | 158790 | 0 | 0 | 0 | 0 | 0 | 0 | 175750 | 304470 | 14045 | 0 | 0 | 0 | 352420 | 0 | 0 | 0 |
| YEEIVKEVS | EEF1A1P5; EEF | Q5VTE0; P68 | EF1A3_HUMAN | Unmodified | 0 | 1128700 | 0 | 2106800 | 0 | 0 | 0 | 0 | 0 | 0 | 0 | 0 | 601400 | 0 | 0 | 0 | 2810700 | 0 |
| YEEIVKEVS | EEF1A1P5; EEF | Q5VTE0; P68 | EF1A3_HUMAN | Unmodified | 392350 | 221080 | 0 | 0 | 238090 | 0 | 0 | 0 | 0 | 0 | 0 | 0 | 0 | 0 | 62460 | 88555 | 421290 | 0 |
| YEELFPAFS | NAPA; NAPA; N. | P54920; M0F | SNAA_HUMAN | Unmodified | 0 | 0 | 495770 | 0 | 0 | 0 | 0 | 0 | 0 | 0 | 0 | 1254500 | 0 | 0 | 423510 | 310080 | 0 | 0 |
| YEEVSVSG | TXNDC17; TXNI | Q9BRA2; I3L | TXD17_HUMAN | Unmodified | 293830 | 210700 | 0 | 0 | 0 | 0 | 0 | 0 | 1490900 | 0 | 258610 | 1954700 | 0 | 0 | 344470 | 608870 | 0 | 0 |
| YEGGRELS | PDIA3 | P30101 | PDIA3_HUMAN | Unmodified | 0 | 0 | 0 | 0 | 0 | 0 | 0 | 0 | 4056100 | 0 | 428020 | 0 | 0 | 0 | 28764 | 0 | 0 | 0 |
| YEGGYPAL | NIPSNAP2; NIP | O75323; H7( | NIPS2_HUMAN | Unmodified | 0 | 0 | 0 | 0 | 0 | 0 | 0 | 0 | 1562600 | 0 | 0 | 0 | 0 | 0 | 0 | 0 | 0 | 0 |
| YEHKEIEYV | BAIAP2L1 | Q9UHR4 | BI2L1_HUMAN | Unmodified | 0 | 0 | 0 | 0 | 2323000 | 0 | 0 | 0 | 0 | 0 | 0 | 0 | 450980 | 404480 | 0 | 0 | 0 | 0 |
| YEIEWITEY/ | IGF2R; IGF2R; I | P11717; A0A | MPRI_HUMAN ( | Unmodified | 0 | 0 | 0 | 0 | 0 | 0 | 0 | 0 | 0 | 454370 | 0 | 689720 | 0 | 0 | 0 | 0 | 0 | 0 |
| YEIEWITEY/ | IGF2R; IGF2R; I | P11717; A0A | MPRI_HUMAN ( | Unmodified | 0 | 0 | 0 | 0 | 0 | 0 | 0 | 0 | 3342400 | 141500 | 0 | 340820 | 0 | 0 | 0 | 0 | 0 | 0 |
| YEILTPNSII | MYH9 | P35579 | MYH9_HUMAN | Unmodified | 126040 | 77368 | 68980 | 622460 | 0 | 0 | 0 | 0 | 0 | 0 | 0 | 3243100 | 627180 | 0 | 0 | 495940 | 0 | 0 |
| YEIMDGAP' | VPS26A | O75436 | VP26A_HUMAN | Unmodified | 0 | 0 | 0 | 0 | 0 | 0 | 0 | 0 | 0 | 0 | 0 | 1991300 | 0 | 0 | 0 | 0 | 0 | 0 |
| YEIMDGAP' | VPS26A | O75436 | VP26A_HUMAN | Oxidation (M) | 0 | 0 | 0 | 0 | 0 | 0 | 0 | 0 | 0 | 0 | 0 | 0 | 0 | 0 | 0 | 0 | 0 | 0 |
| YEIMDGAP' | VPS26B | Q4G0F5 | VP26B_HUMAN | Unmodified | 0 | 0 | 0 | 0 | 0 | 0 | 0 | 0 | 0 | 0 | 0 | 656570 | 0 | 0 | 0 | 0 | 0 | 0 |
| YEISSVPTF | GLRX3 | O76003 | GLRX3_HUMAN | Unmodified | 0 | 0 | 183030 | 0 | 251940 | 0 | 0 | 0 | 0 | 0 | 0 | 993740 | 0 | 0 | 0 | 0 | 0 | 0 |
| YEKPDGSP | ALCAM; ALCAM | Q13740; F5C | CD166_HUMAN | Unmodified | 18654 | 0 | 730220 | 0 | 0 | 0 | 0 | 0 | 300720 | 14316 | 0 | 0 | 0 | 0 | 0 | 0 | 0 | 0 |
| YELGRPAAI | RPS8; RPS8 | P62241; Q5J | RS8_HUMAN 4 | Unmodified | 194830 | 0 | 9948100 | 0 | 3591200 | 145750 | 94974 | 612140 | 4747300 | 4146800 | 1712900 | 47183000 | 0 | 0 | 0 | 736780 | 349190 | 429740 |
| YELISETGG | STRBP; ILF3 | Q96SI9; Q12 | STRBP_HUMAN | Unmodified | 213180 | 221410 | 695670 | 0 | 0 | 0 | 0 | 0 | 74963 | 0 | 448690 | 532040 | 0 | 0 | 0 | 73854 | 0 | 0 |
| YELTGKFEF | ANXA6; ANXA6; | P08133; E5R | ANXA6_HUMAI | Unmodified | 0 | 0 | 1355600 | 0 | 0 | 0 | 0 | 0 | 2266600 | 0 | 0 | 0 | 0 | 0 | 0 | 0 | 0 | 0 |
| YENGCAYF | PDXDC1; PDXD | Q6P996; Q6I | PDXD1_HUMAI | Unmodified | 0 | 0 | 0 | 1330800 | 2057700 | 0 | 1251700 | 508480 | 237090 | 0 | 1931400 | 139980 | 564560 | 5276600 | 348460 | 0 | 0 | 7357300 |
| YENNVMNI | PPP3CA; PPP3( | Q08209; P16 | PP2BA_HUMAN | Oxidation (M) | 0 | 0 | 0 | 0 | 0 | 0 | 0 | 0 | 0 | 0 | 100130 | 853250 | 0 | 0 | 0 | 0 | 0 | 0 |
| YENVPLIGF | TTYH2; TTYH2 | Q9BSA4; B4I | TTYH2_HUMAN | Unmodified | 0 | 0 | 632470 | 0 | 1765800 | 676900 | 0 | 0 | 1802700 | 384570 | 0 | 0 | 0 | 0 | 0 | 595860 | 1432700 | 0 |
| YENYELTLK | APOB | P04114 | APOB_HUMAN | Unmodified | 0 | 928600 | 2092100 | 0 | 0 | 0 | 570180 | 0 | 0 | 0 | 0 | 0 | 0 | 0 | 1444800 | 651090 | 0 | 0 |
| YEPAAVSE( | ATP1A1 | P05023 | AT1A1_HUMAN | Unmodified | 0 | 0 | 1183500 | 173020 | 0 | 0 | 0 | 0 | 0 | 82370 | 0 | 1050300 | 0 | 0 | 843030 | 0 | 0 | 0 |
| YEPAAVSE( | ATP1A1 | P05023 | AT1A1_HUMAN | Unmodified | 0 | 0 | 10564000 | 1013700 | 463470 | 0 | 306030 | 0 | 937160 | 14428 | 1889900 | 10001000 | 0 | 0 | 7077600 | 5190800 | 19860 | 0 |
| YEQGTGCV | PRKCSH; PRKC | P14314; K7E | GLU2B_HUMAI | Unmodified | 0 | 0 | 0 | 0 | 0 | 0 | 0 | 0 | 38021000 | 0 | 754770 | 688850 | 0 | 0 | 0 | 0 | 0 | 0 |
| YEQGTGCV | PRKCSH; PRKC | P14314; K7E | GLU2B_HUMAI | Unmodified | 0 | 0 | 0 | 0 | 76008 | 31637 | 0 | 0 | 9727800 | 1740200 | 1274100 | 0 | 0 | 0 | 0 | 0 | 0 | 71197 |
| YESLELCRI | CLTC; CLTC | Q00610; A0/ | CLH1_HUMAN | Unmodified | 1104000 | 265050 | 970200 | 122040 | 0 | 0 | 0 | 0 | 0 | 0 | 0 | 2941300 | 0 | 0 | 614450 | 754690 | 0 | 0 |
| YESLELCRI | CLTC; CLTC | Q00610; A0/ | CLH1_HUMAN | Unmodified | 277460 | 138880 | 812840 | 424210 | 0 | 0 | 0 | 0 | 0 | 0 | 0 | 7399600 | 0 | 0 | 253540 | 1262800 | 0 | 0 |
| YESLTDPSI | HSP90AB4P; HS | Q58FF6; Q14 | H90B4_HUMAN | Unmodified | 0 | 0 | 0 | 0 | 0 | 0 | 0 | 0 | 0 | 0 | 0 | 6007300 | 0 | 0 | 0 | 0 | 0 | 0 |
| YESLTDPSI | HSP90AA2P; HS | Q14568; P08 | HS902_HUMAN | Unmodified | 1348900 | 406160 | 553140 | 530410 | 191480 | 274810 | 111900 | 36900 | 7649800 | 758080 | 9326900 | 62491000 | 87775 | 0 | 1938200 | 1946900 | 0 | 598600 |
| YESQLSTNI | SLC39A6; SLC3 | Q13433; K7E | S39A6_HUMAN | Unmodified | 0 | 0 | 1452400 | 0 | 0 | 0 | 0 | 0 | 0 | 0 | 262510 | 845810 | 0 | 0 | 0 | 0 | 0 | 0 |
| YESQLSTNI | SLC39A6; SLC3 | Q13433; K7E | S39A6_HUMAN | Unmodified | 164620 | 110020 | 0 | 0 | 0 | 0 | 0 | 0 | 0 | 31177 | 0 | 2806300 | 0 | 0 | 722290 | 679660 | 0 | 0 |
| YESSALPS( | TRAP1; TRAP1 | Q12931; I3L | TRAP1_HUMAN | Unmodified | 253490 | 349590 | 259480 | 159710 | 0 | 0 | 35195 | 0 | 1068000 | 36035 | 316020 | 133300 | 67280 | 40340 | 642590 | 221860 | 0 | 0 |
| YETELAMR | KRT18; KRT18 | P05783; F8V | K1C18_HUMAI | Unmodified | 0 | 0 | 194660 | 0 | 0 | 137410 | 0 | 0 | 630770 | 0 | 1309800 | 2423000 | 0 | 0 | 0 | 0 | 0 | 474920 |
| YETELAMR | KRT18; KRT18 | P05783; F8V | K1C18_HUMAI | Oxidation (M) | 0 | 0 | 0 | 0 | 17788 | 0 | 0 | 0 | 0 | 0 | 0 | 812020 | 0 | 27411 | 0 | 0 | 0 | 157690 |
| YETVIMPVF | SUPT16H | Q9Y5B9 | SP16H_HUMAN | Oxidation (M) | 121770 | 0 | 0 | 0 | 0 | 0 | 0 | 0 | 0 | 0 | 0 | 1109100 | 0 | 0 | 0 | 0 | 0 | 0 |
| YEVFRTEEE | UCHL3; UCHL3 | P15374; A0A | UCHL3_HUMAI | Unmodified | 0 | 0 | 0 | 0 | 0 | 0 | 0 | 0 | 0 | 0 | 0 | 4385600 | 0 | 0 | 791150 | 40899 | 0 | 0 |
| YEVSVYALK | FN1; FN1 | P02751; H0\ | FINC_HUMAN | Unmodified | 89925 | 531110 | 16266000 | 0 | 180670 | 201170 | 0 | 0 | 0 | 148350 | 0 | 310470 | 0 | 0 | 457050 | 1317300 | 0 | 123580 |
| YEWDVAEA | EEF2 | P13639 | EF2_HUMAN El | Unmodified | 1019000 | 696600 | 3560200 | 954000 | 0 | 0 | 313650 | 0 | 1135100 | 0 | 4435200 | 46193000 | 0 | 0 | 1447000 | 236030 | 0 | 0 |
| YEYHWAD( | MOB1B; MOB1/ | Q7L9L4; Q9I | MOB1B_HUMA | Unmodified | 0 | 0 | 0 | 0 | 0 | 0 | 0 | 0 | 164280 | 0 | 0 | 2111100 | 0 | 0 | 616580 | 1097300 | 0 | 0 |
| YEYQPFAG | WDR1; WDR1 | O75083; D6I | WDR1_HUMAN | Unmodified | 2536400 | 1584000 | 2492300 | 1881000 | 16973000 | 3260700 | 0 | 332420 | 0 | 2359000 | 3810500 | 15495000 | 4117700 | 2548500 | 0 | 3086900 | 803090 | 3695500 |
| YEYQPFAG | WDR1; WDR1 | O75083; D6I | WDR1_HUMAN | Unmodified | 0 | 0 | 0 | 0 | 551190 | 0 | 0 | 0 | 0 | 0 | 0 | 0 | 165020 | 170690 | 0 | 0 | 225310 | 0 |
| YFAATQFEI | LNPEP | Q9UIQ6 | LCAP_HUMAN | Unmodified | 0 | 0 | 155730 | 81080 | 0 | 0 | 0 | 0 | 0 | 0 | 0 | 198570 | 0 | 0 | 34201 | 39343 | 0 | 0 |
| YFADTQPA | LRRC8A | Q8IWT6 | LRC8A_HUMAN | Unmodified | 0 | 0 | 0 | 0 | 0 | 0 | 0 | 0 | 293220 | 0 | 0 | 1011700 | 0 | 0 | 0 | 48651 | 0 | 0 |
| YFAEALGPI | SULT1A3; SULT | P0DMM9; PC | ST1A3_HUMAN | Unmodified | 0 | 0 | 0 | 0 | 2321900 | 0 | 0 | 0 | 0 | 0 | 0 | 0 | 0 | 0 | 0 | 0 | 0 | 0 |
| YFDEGRGS | MKRN1 | Q9UHC7 | MKRN1_HUMA | Unmodified | 7093900 | 2624200 | 841450 | 2205600 | 0 | 0 | 0 | 0 | 0 | 0 | 0 | 0 | 1017400 | 229180 | 769160 | 568760 | 0 | 0 |
| YFDEHYEY( | CKS2 | P33552 | CKS2_HUMAN | Unmodified | 988730 | 693270 | 1063500 | 2335200 | 97087 | 0 | 0 | 0 | 444340 | 0 | 95057 | 558730 | 0 | 0 | 1746600 | 1355300 | 0 | 0 |
| YFDEISQD1 | ETF1; ETF1 | P62495; B7Z | ERF1_HUMAN | Unmodified | 0 | 0 | 214340 | 0 | 325790 | 93066 | 0 | 0 | 755560 | 371100 | 791050 | 2588700 | 0 | 0 | 374790 | 262850 | 0 | 0 |
| YFDGSGGI | USP5 | P45974 | UBP5_HUMAN | Unmodified | 639240 | 542750 | 265410 | 0 | 0 | 0 | 0 | 0 | 0 | 0 | 74848 | 1253300 | 0 | 0 | 390690 | 342680 | 0 | 0 |

| | | | | | | | | | | | | | | | | | | | | | | |
|---|---|---|---|---|---|---|---|---|---|---|---|---|---|---|---|---|---|---|---|---|---|---|
| YFDGSGGI | USP5 | P45974 | UBP5_HUMAN | Unmodified | 2280200 | 925690 | 778930 | 1058100 | 2459200 | 307110 | 0 | 0 | 522340 | 123420 | 666030 | 2450700 | 1749400 | 639750 | 598400 | 417310 | 153340 | 995650 |
| YFDLGLPN | IDH2 | P48735 | IDHP_HUMAN | Unmodified | 0 | 0 | 0 | 0 | 883220 | 0 | 2464100 | 0 | 5347100 | 8871500 | 0 | 341280 | 7051600 | 0 | 0 | 0 | 426950 | 0 |
| YFDPANGK | EEF2 | P13639 | EF2_HUMAN EI | Unmodified | 405480 | 0 | 1011100 | 261810 | 0 | 0 | 0 | 0 | 0 | 0 | 48941 | 22705000 | 0 | 0 | 1006100 | 0 | 0 | 0 |
| YFDSGDYN | ARPP19; ENSA; | P56211; O43 | ARP19_HUMAN | Oxidation (M) | 203610 | 0 | 298530 | 0 | 1201800 | 0 | 195690 | 0 | 0 | 82134 | 0 | 0 | 831780 | 0 | 192850 | 0 | 662660 | 1197600 |
| YFDSGDYN | ARPP19; ENSA; | P56211; O43 | ARP19_HUMAN | Unmodified | 0 | 0 | 0 | 0 | 2133900 | 207140 | 353010 | 0 | 0 | 0 | 0 | 0 | 2072800 | 1475700 | 0 | 0 | 1207600 | 1639200 |
| YFDTAVSRI | HLA-C; HLA-C; | P10321; Q5( | HLAC_HUMAN | Unmodified | 0 | 0 | 0 | 0 | 85644 | 12981 | 425970 | 0 | 9409400 | 0 | 0 | 0 | 0 | 0 | 0 | 0 | 0 | 0 |
| YFEFLLPSS | CACNA2D2 | C9JE82 | C9JE82_HUMA | Unmodified | 0 | 0 | 0 | 0 | 0 | 17624000 | 1126300 | 319130 | 0 | 1955200 | 693340 | 0 | 0 | 6601500 | 0 | 0 | 0 | 0 |
| YFEITDESP | CORO1C; COR( | Q9ULV4; B4I | COR1C_HUMA | Unmodified | 829910 | 713330 | 135170 | 3118000 | 852410 | 276230 | 0 | 0 | 4989900 | 0 | 1745400 | 9712000 | 0 | 0 | 2505600 | 560860 | 0 | 0 |
| YFEITDESP | CORO1C; COR( | Q9ULV4; B4I | COR1C_HUMA | Unmodified | 2367400 | 0 | 0 | 734080 | 0 | 0 | 1401800 | 0 | 4763900 | 0 | 1522100 | 12360000 | 0 | 0 | 0 | 0 | 0 | 0 |
| YFEITEEPP | CORO1B | Q9BR76 | COR1B_HUMA | Unmodified | 217550 | 228200 | 114940 | 418280 | 0 | 0 | 0 | 0 | 73021 | 0 | 0 | 717610 | 0 | 0 | 1479300 | 155610 | 0 | 0 |
| YFEITEEPP | CORO1B | Q9BR76 | COR1B_HUMA | Unmodified | 0 | 1388800 | 0 | 423100 | 0 | 0 | 0 | 0 | 2129800 | 0 | 176580 | 0 | 0 | 0 | 0 | 174390 | 0 | 0 |
| YFELEVRVF | SCRIB; SCRIB; | Q14160; A0/ | SCRIB_HUMAN | Unmodified | 0 | 0 | 201640 | 618310 | 0 | 0 | 0 | 0 | 2791400 | 2318500 | 214980 | 0 | 2969900 | 1543700 | 317700 | 586870 | 1171000 | 0 |
| YFEQLLEEE | P4HA2 | O15460 | P4HA2_HUMAN | Unmodified | 0 | 0 | 0 | 0 | 0 | 0 | 0 | 0 | 1203900 | 156250 | 0 | 0 | 0 | 0 | 0 | 0 | 0 | 0 |
| YFEYIMAK | P4HA1 | P13674 | P4HA1_HUMAN | Unmodified | 0 | 0 | 0 | 0 | 0 | 0 | 0 | 0 | 2210900 | 762840 | 0 | 0 | 0 | 0 | 0 | 0 | 0 | 0 |
| YFEYIMAK | P4HA1 | P13674 | P4HA1_HUMAN | Oxidation (M) | 0 | 0 | 0 | 0 | 0 | 0 | 0 | 0 | 848300 | 1147800 | 0 | 0 | 0 | 0 | 0 | 0 | 0 | 0 |
| YFFDFLDE( | PLXNB2; PLXNE | O15031; A6( | PLXB2_HUMAN | Unmodified | 1715500 | 643350 | 709840 | 291100 | 0 | 0 | 0 | 0 | 547480 | 0 | 239620 | 3889500 | 0 | 0 | 3316900 | 1712900 | 0 | 0 |
| YFFFDDGN | SPCS3 | P61009 | SPCS3_HUMAN | Unmodified | 0 | 0 | 0 | 0 | 0 | 0 | 0 | 0 | 28456000 | 1634200 | 3507100 | 3333500 | 0 | 0 | 0 | 0 | 0 | 0 |
| YFFTSVSRF | HLA-A; HLA-A; I | P04439; A0A | HLAA_HUMAN | Unmodified | 0 | 0 | 0 | 0 | 0 | 0 | 0 | 0 | 0 | 0 | 0 | 986740 | 0 | 0 | 0 | 0 | 0 | 0 |
| YFGDIISVG | ISOC1; ISOC1 | Q96CN7; D6 | ISOC1_HUMAN | Unmodified | 0 | 0 | 0 | 0 | 0 | 0 | 0 | 0 | 0 | 0 | 376200 | 3339800 | 0 | 0 | 0 | 0 | 0 | 0 |
| YFGDTDMY | DNAJB12; DNA. | Q9NXW2; V9 | DJB12_HUMAN | Oxidation (M) | 0 | 0 | 0 | 0 | 0 | 0 | 0 | 0 | 889590 | 51139 | 0 | 0 | 0 | 0 | 0 | 0 | 0 | 0 |
| YFGDTDMY | DNAJB12; DNA. | Q9NXW2; V9 | DJB12_HUMAN | Unmodified | 0 | 0 | 0 | 0 | 0 | 0 | 0 | 0 | 655140 | 31454 | 0 | 0 | 0 | 0 | 0 | 0 | 0 | 0 |
| YFGDTDMY | DNAJB12; DNA. | Q9NXW2; V9 | DJB12_HUMAN | Unmodified | 0 | 0 | 0 | 0 | 0 | 0 | 0 | 0 | 0 | 251890 | 0 | 0 | 0 | 0 | 0 | 0 | 0 | 0 |
| YFGENPKT | FMNL2; FMNL2 | Q96PY5; C9I | FMNL2_HUMAI | Unmodified | 0 | 0 | 0 | 0 | 0 | 0 | 0 | 0 | 0 | 0 | 0 | 0 | 0 | 0 | 0 | 0 | 0 | 0 |
| YFGENPKT | FMNL2; FMNL2 | Q96PY5; C9I | FMNL2_HUMAI | Unmodified | 0 | 0 | 0 | 0 | 1123400 | 0 | 0 | 0 | 0 | 0 | 0 | 0 | 0 | 0 | 0 | 0 | 531550 | 968070 |
| YFGGTEDR | FSCN1 | Q16658 | FSCN1_HUMAI | Unmodified | 0 | 0 | 0 | 0 | 3062700 | 993010 | 0 | 45969 | 2496200 | 884670 | 4704300 | 10670000 | 0 | 0 | 0 | 0 | 1761600 | 1626500 |
| YFGKGTYTI | COL6A1; COL6/ | P12109; A0A | CO6A1_HUMAI | Unmodified | 0 | 0 | 0 | 0 | 0 | 0 | 754230 | 309760 | 0 | 319830 | 0 | 110570 | 0 | 0 | 0 | 0 | 0 | 1114700 |
| YFGRSLEL | RNASET2; N/A; | O00584; H0 | RNT2_HUMAN | Unmodified | 0 | 0 | 0 | 0 | 0 | 0 | 0 | 0 | 0 | 260440 | 0 | 0 | 709550 | 516840 | 0 | 0 | 0 | 0 |
| YFHAQCNI | ASAP2; ASAP2 | O43150; A0/ | ASAP2_HUMAN | Unmodified | 0 | 0 | 0 | 0 | 233560 | 0 | 53330 | 0 | 0 | 0 | 0 | 331810 | 241320 | 277640 | 0 | 0 | 338300 | 662670 |
| YFHPPAHL | MBNL1; MBNL2 | Q9NR56; Q5 | MBNL1_HUMAI | Unmodified | 553980 | 491230 | 233810 | 0 | 518010 | 118240 | 0 | 0 | 0 | 0 | 0 | 264710 | 991100 | 278520 | 0 | 0 | 0 | 0 |
| YFIRDEFLR | GNAS; GNAS; C | Q5JWF2; P6 | GNAS1_HUMAI | Unmodified | 875340 | 758030 | 2384700 | 0 | 5534100 | 3541700 | 1138400 | 0 | 1750800 | 375160 | 4279400 | 6331900 | 6303700 | 4532500 | 2755000 | 4191200 | 2027200 | 8619300 |
| YFIRDEFLR | GNAS; GNAS; C | Q5JWF2; P6 | GNAS1_HUMAI | Unmodified | 0 | 0 | 0 | 0 | 919110 | 121950 | 0 | 0 | 0 | 0 | 0 | 0 | 0 | 429930 | 0 | 0 | 0 | 1257300 |
| YFKLEPLQ/ | POFUT1 | Q9H488 | OFUT1_HUMAN | Unmodified | 0 | 0 | 0 | 0 | 0 | 0 | 0 | 0 | 2553100 | 820460 | 0 | 0 | 0 | 0 | 0 | 0 | 0 | 0 |
| YFKLEPLQ/ | POFUT1 | Q9H488 | OFUT1_HUMAN | Oxidation (M) | 0 | 0 | 0 | 0 | 0 | 0 | 0 | 0 | 0 | 4260600 | 0 | 0 | 251220 | 0 | 0 | 0 | 0 | 0 |
| YFKLEPLQ/ | POFUT1 | Q9H488 | OFUT1_HUMAN | Unmodified | 0 | 0 | 0 | 0 | 0 | 0 | 93227 | 0 | 0 | 2605700 | 0 | 0 | 0 | 0 | 0 | 0 | 0 | 0 |
| YFLDNLDK | GNA13 | Q14344 | GNA13_HUMAI | Unmodified | 0 | 0 | 0 | 259450 | 0 | 0 | 0 | 0 | 1666000 | 0 | 2426200 | 5035900 | 0 | 0 | 0 | 0 | 0 | 0 |
| YFLDNLDK | GNA13 | Q14344 | GNA13_HUMAI | Unmodified | 0 | 0 | 0 | 0 | 0 | 0 | 0 | 0 | 822930 | 0 | 0 | 248430 | 0 | 0 | 0 | 0 | 0 | 0 |
| YFLFYDGE | VPS26B; VPS26 | Q4G0F5; H0 | VP26B_HUMAN | Unmodified | 0 | 0 | 0 | 0 | 0 | 0 | 0 | 0 | 0 | 0 | 0 | 2077700 | 0 | 0 | 0 | 0 | 0 | 0 |
| YFLHQSHE | FTH1; FTH1; FTI | P02794; G3\ | FRIH_HUMAN I | Unmodified | 65049 | 0 | 0 | 0 | 0 | 0 | 0 | 0 | 1942700 | 0 | 0 | 0 | 0 | 0 | 107950 | 0 | 0 | 0 |
| YFLHQSHE | FTH1; FTH1; FTI | P02794; G3\ | FRIH_HUMAN I | Unmodified | 0 | 0 | 0 | 0 | 0 | 0 | 2550200 | 0 | 37119000 | 39441000 | 0 | 0 | 0 | 0 | 0 | 0 | 2497900 | 0 |
| YFLHQSHE | FTH1; FTH1; FTI | P02794; G3\ | FRIH_HUMAN I | Unmodified | 0 | 0 | 0 | 0 | 0 | 0 | 664320 | 0 | 0 | 529450 | 0 | 0 | 258700 | 389570 | 0 | 0 | 611410 | 0 |
| YFLKPEVLC | NCEH1; NCEH1 | Q6PIU2; H7( | NCEH1_HUMA | Unmodified | 0 | 0 | 0 | 0 | 815040 | 0 | 0 | 0 | 2834300 | 454540 | 241650 | 205190 | 0 | 0 | 0 | 0 | 0 | 0 |
| YFLLKNDG | AKT1 | P31749 | AKT1_HUMAN I | Unmodified | 0 | 0 | 0 | 75485 | 0 | 0 | 0 | 0 | 0 | 0 | 0 | 0 | 462500 | 561440 | 0 | 0 | 0 | 0 |
| YFLQATHV | TMTC3 | Q6ZXV5 | TMTC3_HUMAI | Unmodified | 0 | 0 | 0 | 0 | 0 | 0 | 0 | 249840 | 6626400 | 0 | 0 | 0 | 0 | 0 | 0 | 0 | 0 | 0 |
| YFLQGMG\ | NDRG3; NDRG3 | Q9UGV2; F8 | NDRG3_HUMA | 2 Oxidation (M) | 0 | 0 | 0 | 0 | 1055300 | 0 | 0 | 0 | 0 | 0 | 504800 | 4083600 | 626590 | 0 | 0 | 283490 | 0 | 0 |
| YFLQGMG\ | NDRG3; NDRG3 | Q9UGV2; F8 | NDRG3_HUMA | Oxidation (M) | 0 | 0 | 0 | 0 | 988070 | 0 | 0 | 0 | 0 | 0 | 0 | 1126300 | 87499 | 170770 | 0 | 0 | 0 | 406480 |
| YFLTFAVER | JAK2; JAK2 | O60674; A0/ | JAK2_HUMAN 1 | Unmodified | 0 | 0 | 0 | 206330 | 282210 | 0 | 0 | 0 | 0 | 7792300 | 56497 | 0 | 103430 | 0 | 198750 | 0 | 139030 | 453720 |
| YFLVGAGA | UBA1 | P22314 | UBA1_HUMAN | Unmodified | 0 | 0 | 0 | 0 | 1924400 | 0 | 3659800 | 0 | 2130600 | 1556200 | 823970 | 4641600 | 898330 | 708270 | 367940 | 0 | 4598200 | 4074700 |
| YFLWVVK | GRB2 | P62993 | GRB2_HUMAN | Unmodified | 5876500 | 9695800 | 5458500 | 754360 | 268470 | 277700 | 186680 | 0 | 453710 | 420750 | 235090 | 5460200 | 130270 | 26755 | 14274000 | 6495000 | 473140 | 50599 |
| YFNLASQG | SEL1L | Q9UBV2 | SE1L1_HUMAN | 2 Oxidation (M) | 0 | 0 | 0 | 0 | 0 | 0 | 0 | 0 | 2380800 | 1203800 | 0 | 0 | 0 | 0 | 0 | 0 | 0 | 0 |
| YFNVHELE. | MIA3; MIA3 | Q5JRA6; A0A | TGO1_HUMAN | Unmodified | 0 | 0 | 0 | 0 | 9774 | 0 | 0 | 0 | 579090 | 396850 | 0 | 0 | 0 | 0 | 0 | 0 | 0 | 0 |
| YFNVHELE. | MIA3; MIA3 | Q5JRA6; A0A | TGO1_HUMAN | Unmodified | 0 | 0 | 33665 | 0 | 0 | 0 | 337170 | 0 | 0 | 90974 | 0 | 0 | 0 | 0 | 0 | 0 | 0 | 0 |
| YFPAGNVV | GLIPR2; GLIPR2 | Q9H4G4; Q5 | GAPR1_HUMAI | Unmodified | 4368800 | 4846500 | 1188100 | 3230700 | 4643100 | 1326900 | 0 | 0 | 3294800 | 1098200 | 0 | 0 | 12078000 | 2299500 | 8655100 | 4806100 | 0 | 7785000 |
| YFPEYAEKI | NIT2; NIT2; NIT2 | Q9NQR4; F8 | NIT2_HUMAN ( | Unmodified | 0 | 0 | 0 | 0 | 736730 | 0 | 0 | 0 | 0 | 0 | 0 | 67874 | 0 | 1029900 | 0 | 0 | 1689400 | 2125900 |
| YFPIPVEHL | PTPRE; PTPRE; | P23469; S4F | PTPRE_HUMAN | Unmodified | 0 | 0 | 0 | 0 | 1202000 | 150030 | 0 | 0 | 0 | 0 | 0 | 0 | 0 | 0 | 0 | 0 | 0 | 0 |
| YFPTQALNI | SLC25A4; SLC2 | P12235; P12 | ADT1_HUMAN | Unmodified | 523250 | 522770 | 315120 | 190960 | 0 | 0 | 0 | 0 | 10223000 | 2609200 | 270880 | 2342100 | 0 | 0 | 765450 | 336200 | 0 | 0 |
| YFPTQALNI | SLC25A4; SLC2 | P12235; P12 | ADT1_HUMAN | Unmodified | 0 | 0 | 0 | 0 | 0 | 0 | 0 | 0 | 0 | 1051100 | 0 | 0 | 0 | 0 | 0 | 0 | 0 | 0 |
| YFPYYGK | ATP1B2; ATP1B | P14415; P54 | AT1B2_HUMAN | Unmodified | 0 | 233230 | 2479800 | 2549500 | 7955900 | 1147100 | 487800 | 171230 | 2957900 | 552400 | 3465200 | 15458000 | 4328100 | 4523200 | 3250400 | 3527800 | 1728900 | 1527700 |
| YFQFQEEG | ARPC2 | O15144 | ARPC2_HUMAI | Unmodified | 0 | 295800 | 611880 | 0 | 0 | 0 | 0 | 0 | 0 | 0 | 0 | 2387900 | 0 | 0 | 971340 | 448890 | 0 | 0 |
| YFQFQEEG | ARPC2 | O15144 | ARPC2_HUMAI | Unmodified | 2272000 | 2012800 | 3143600 | 2982100 | 1014700 | 0 | 347760 | 202140 | 174870 | 0 | 1926400 | 22336000 | 920900 | 223990 | 3404400 | 2660000 | 205250 | 292250 |
| YFQHLLGK | HYOU1; HYOU1 | Q9Y4L1; J3Q | HYOU1_HUMA | Unmodified | 0 | 0 | 270140 | 122040 | 0 | 0 | 0 | 0 | 8730000 | 2974200 | 1262100 | 579340 | 0 | 199600 | 190740 | 101740 | 0 | 0 |
| YFQINQDE | RPL22; RPL22 | P35268; K7E | RL22_HUMAN ( | Unmodified | 1062100 | 1125400 | 0 | 0 | 0 | 0 | 0 | 93887 | 0 | 0 | 959110 | 6975100 | 0 | 0 | 0 | 0 | 0 | 0 |
| YFQPPGTV | NMD3; NMD3; I | Q96D46; C9 | NMD3_HUMAN | Unmodified | 92716 | 0 | 0 | 0 | 0 | 0 | 0 | 9095.3 | 0 | 31080 | 0 | 329710 | 0 | 0 | 0 | 0 | 0 | 0 |
| YFSASGID( | N/A | N/A | N/A | Unmodified | 0 | 98222 | 0 | 0 | 91174 | 4606900 | 143210 | 0 | 0 | 0 | 0 | 0 | 0 | 0 | 0 | 0 | 157300 | 0 |
| YFSERGDA | PSME2; PSME2 | Q9UL46; A0/ | PSME2_HUMAI | Unmodified | 0 | 0 | 0 | 35428 | 0 | 0 | 0 | 0 | 0 | 0 | 0 | 1095000 | 0 | 0 | 0 | 0 | 0 | 0 |
| YFSERGDA | PSME1; PSME1 | Q06323; H0 | PSME1_HUMAI | Unmodified | 0 | 0 | 0 | 0 | 0 | 0 | 68026 | 31008 | 0 | 79751 | 0 | 628780 | 54329 | 49925 | 0 | 0 | 111240 | 0 |
| YFSLDLTHI | FRMD6 | Q96NE9 | FRMD6_HUMAI | Unmodified | 0 | 0 | 0 | 0 | 0 | 38561 | 0 | 0 | 0 | 0 | 0 | 0 | 0 | 28593 | 0 | 0 | 0 | 818620 |
| YFSLIPTGF. | ETNK1; ETNK1; | Q9HBU6; H( | EKI1_HUMAN E | Unmodified | 0 | 0 | 40546 | 0 | 0 | 0 | 0 | 0 | 0 | 0 | 0 | 1166800 | 0 | 0 | 0 | 15818 | 0 | 0 |
| YFSLKPEEK | CD2AP | Q9Y5K6 | CD2AP_HUMAI | Unmodified | 2215500 | 2618700 | 852730 | 1648200 | 0 | 0 | 0 | 0 | 0 | 0 | 47553 | 0 | 2032800 | 765270 | 1890100 | 1282500 | 0 | 0 |
| YFSLPFCV( | TM9SF3; TM9SF | Q9HD45; Q5 | TM9S3_HUMAN | Unmodified | 0 | 0 | 0 | 53518 | 0 | 0 | 0 | 0 | 18151000 | 4071500 | 373660 | 0 | 0 | 0 | 0 | 0 | 0 | 0 |
| YFSLPFCV( | TM9SF3; TM9SF | Q9HD45; Q5 | TM9S3_HUMAN | Unmodified | 0 | 0 | 264960 | 244180 | 540470 | 0 | 843020 | 132330 | 61429000 | 9571700 | 2702000 | 3267000 | 298550 | 450010 | 0 | 0 | 0 | 0 |
| YFSNERIPF | CNKSR3; CNKS | Q6P9H4; A0. | CNKR3_HUMAI | Unmodified | 0 | 0 | 0 | 0 | 0 | 0 | 0 | 0 | 760970 | 0 | 207800 | 571350 | 0 | 0 | 0 | 0 | 0 | 0 |
| YFSNIHWC | PLAU; PLAU; PL | P00749; E7E | UROK_HUMAN | Unmodified | 0 | 0 | 0 | 0 | 1488800 | 1546500 | 0 | 0 | 0 | 66059 | 0 | 0 | 0 | 0 | 0 | 0 | 1620300 | 3034300 |
| YFSSKWSF | WDR45B; WDR | Q5MNZ6; I3I | WIPI3_HUMAN | Unmodified | 0 | 0 | 166620 | 102850 | 287810 | 0 | 197800 | 0 | 0 | 137820 | 0 | 247480 | 103000 | 111330 | 0 | 203390 | 135520 | 0 |
| YFSTLQPGI | TXNDC16 | Q9P2K2 | TXD16_HUMAN | Unmodified | 0 | 0 | 0 | 0 | 358510 | 0 | 0 | 0 | 1242100 | 378120 | 0 | 0 | 0 | 0 | 0 | 0 | 0 | 0 |
| YFTDAGLK | BZW2; BZW2; B | Q9Y6E2; B5I | BZW2_HUMAN | Unmodified | 258580 | 0 | 0 | 0 | 817620 | 0 | 268840 | 179950 | 225810 | 37862 | 845040 | 3025300 | 198790 | 104260 | 0 | 0 | 0 | 0 |
| YFTEAGLK | BZW1; BZW1 | Q7L1Q6; C9I | BZW1_HUMAN | Unmodified | 0 | 1195200 | 781640 | 710490 | 3245800 | 461520 | 608190 | 194790 | 836160 | 330100 | 600070 | 7538800 | 1060500 | 812260 | 0 | 559610 | 1022800 | 4997500 |
| YFTLGESW | MSMP | Q1L6U9 | MSMP_HUMAN | Unmodified | 0 | 0 | 0 | 0 | 0 | 0 | 0 | 0 | 0 | 0 | 0 | 0 | 259420 | 0 | 0 | 0 | 5851800 | 19730000 |
| YFTLGLPT( | GNPDA2; GNPI | Q8TDQ7; P4 | GNPI2_HUMAN | Unmodified | 0 | 0 | 20548 | 0 | 187040 | 0 | 0 | 0 | 0 | 411110 | 111670 | 391170 | 199580 | 191710 | 122600 | 82061 | 0 | 0 |
| YFTVVVR | PTPRB; PTPRB | P23467; F8V | PTPRB_HUMAN | Unmodified | 283590 | 366390 | 0 | 0 | 0 | 0 | 0 | 0 | 0 | 0 | 0 | 0 | 0 | 0 | 0 | 65065 | 0 | 0 |
| YFTWDPSR | GANAB; GANAE | Q14697; F5I | GANAB_HUMA | Unmodified | 287820 | 158550 | 787370 | 0 | 0 | 0 | 0 | 0 | 3.18E+08 | 15468000 | 1397400 | 0 | 135140 | 0 | 546960 | 585150 | 68409 | 156790 |
| YFVEAGAM | TCP1; TCP1; TC | P17987; F5G | TCPA_HUMAN | Oxidation (M) | 0 | 0 | 0 | 0 | 0 | 212960 | 0 | 0 | 0 | 0 | 0 | 4638700 | 0 | 0 | 0 | 0 | 0 | 0 |
| YFVEAGAM | TCP1; TCP1; TC | P17987; F5G | TCPA_HUMAN | Unmodified | 0 | 0 | 0 | 0 | 0 | 0 | 259750 | 0 | 603600 | 0 | 0 | 7991000 | 0 | 0 | 0 | 0 | 0 | 0 |
| YFVEAGAM | TCP1; TCP1; TC | P17987; F5G | TCPA_HUMAN | Unmodified | 0 | 0 | 0 | 0 | 0 | 0 | 0 | 0 | 0 | 0 | 0 | 3689300 | 0 | 0 | 0 | 0 | 0 | 0 |
| YFVQGMG\ | NDRG1; NDRG1 | Q92597; E7E | NDRG1_HUMA | Oxidation (M) | 0 | 0 | 0 | 0 | 245720 | 0 | 0 | 0 | 797730 | 93034 | 0 | 0 | 59172 | 0 | 0 | 0 | 288000 | 904850 |
| YFVQGMG\ | NDRG1; NDRG1 | Q92597; E7E | NDRG1_HUMA | 3 Oxidation (M) | 0 | 0 | 0 | 0 | 0 | 0 | 0 | 0 | 0 | 170010 | 0 | 0 | 0 | 0 | 0 | 50119 | 628650 | 879990 |
| YFVQGMG\ | NDRG1; NDRG1 | Q92597; E7E | NDRG1_HUMA | Unmodified | 0 | 0 | 0 | 0 | 204310 | 0 | 0 | 0 | 1157000 | 0 | 0 | 0 | 0 | 0 | 0 | 0 | 53061 | 438410 |
| YFVTFPYP | LARS1; LARS1; | Q9P2J5; B4D | SYLC_HUMAN | Oxidation (M) | 0 | 0 | 0 | 0 | 0 | 0 | 0 | 0 | 0 | 0 | 0 | 3756300 | 0 | 511470 | 0 | 0 | 0 | 0 |
| YFWNDTIH | ATP13A3; ATP1 | Q9H7F0; A0, | AT133_HUMAN | Unmodified | 0 | 0 | 17783 | 128080 | 0 | 0 | 0 | 24320 | 472910 | 126380 | 0 | 107710 | 0 | 0 | 0 | 0 | 0 | 0 |
| YFYLFPNRI | GRK3; GRK2 | P35626; P25 | ARBK2_HUMAN | Unmodified | 44362 | 0 | 0 | 0 | 0 | 0 | 0 | 0 | 0 | 0 | 0 | 838550 | 0 | 0 | 286330 | 216550 | 0 | 0 |
| YFYTAMSRI | HLA-B; HLA-B; I | D5H3J5; A0/ | D5H3J5_HUMA | Unmodified | 77169 | 0 | 0 | 0 | 0 | 0 | 0 | 0 | 0 | 0 | 0 | 5825400 | 0 | 0 | 40897 | 0 | 0 | 0 |
| YFYTAMSRI | HLA-B; HLA-B; I | D5H3J5; A0/ | D5H3J5_HUMA | Oxidation (M) | 0 | 0 | 0 | 0 | 0 | 0 | 0 | 0 | 0 | 0 | 0 | 2641900 | 0 | 0 | 0 | 0 | 0 | 0 |
| YFYTSVSRF | HLA-B; HLA-A; I | P01889; A0A | HLAB_HUMAN | Unmodified | 0 | 0 | 0 | 0 | 0 | 0 | 0 | 0 | 3676200 | 378310 | 0 | 133600 | 0 | 0 | 0 | 0 | 38442 | 0 |
| YFYVSAEQ | TXNDC12 | O95881 | TXD12_HUMAN | Unmodified | 0 | 0 | 0 | 23793 | 0 | 0 | 0 | 0 | 4476500 | 1013800 | 0 | 0 | 0 | 0 | 0 | 0 | 0 | 0 |
| YFYVSAEQ | TXNDC12 | O95881 | TXD12_HUMAN | Oxidation (M) | 0 | 0 | 0 | 0 | 0 | 0 | 0 | 43632 | 1507600 | 937170 | 385600 | 0 | 0 | 0 | 0 | 0 | 0 | 0 |
| YGAAPAAP | FKRP | Q9H9S5 | FKRP_HUMAN | Unmodified | 0 | 0 | 0 | 0 | 0 | 0 | 0 | 0 | 1855600 | 58582 | 0 | 0 | 0 | 0 | 0 | 0 | 0 | 0 |
| YGAATANY | GPD2; GPD2 | P43304; F5G | GPDM_HUMAN | Unmodified | 0 | 0 | 0 | 0 | 79412 | 161300 | 0 | 0 | 3126400 | 186200 | 0 | 0 | 0 | 0 | 0 | 0 | 0 | 0 |
| YGAATANY | GPD2; GPD2 | P43304; F5G | GPDM_HUMAN | Oxidation (M) | 0 | 0 | 0 | 0 | 0 | 0 | 0 | 0 | 1525400 | 742300 | 0 | 0 | 0 | 0 | 0 | 0 | 0 | 0 |
| YGAFVKPA | TBCB; TBCB | Q99426; A0/ | TBCB_HUMAN | Unmodified | 0 | 0 | 0 | 0 | 2261300 | 0 | 1635200 | 0 | 0 | 4187200 | 0 | 0 | 2675400 | 2038400 | 0 | 0 | 1738400 | 2535300 |
| YGAHNYHI | OAT | P04181 | OAT_HUMAN C | Unmodified | 0 | 0 | 0 | 0 | 0 | 0 | 0 | 0 | 3126400 | 74335 | 0 | 0 | 0 | 0 | 0 | 0 | 0 | 0 |
| YGAITFGN\ | ABCA2; ABCA2; | Q9BZC7; A0, | ABCA2_HUMAI | Unmodified | 0 | 0 | 0 | 0 | 0 | 0 | 0 | 0 | 2799200 | 0 | 0 | 0 | 0 | 0 | 0 | 0 | 0 | 0 |
| YGALALQEI | CSE1L | P55060 | XPO2_HUMAN | Unmodified | 0 | 0 | 0 | 0 | 30163 | 33277 | 104960 | 17718 | 50022 | 93587 | 0 | 120810 | 0 | 0 | 0 | 0 | 42577 | 23299 |
| YGAPEDWI | TBCE; TBCE; TB | Q15813; A0/ | TBCE_HUMAN | Oxidation (M) | 0 | 0 | 0 | 0 | 519800 | 0 | 0 | 319400 | 0 | 0 | 0 | 1155200 | 0 | 0 | 0 | 0 | 212380 | 0 |
| YGASWTAE | MME; MME; MM | P08473; A0A | NEP_HUMAN N | Unmodified | 0 | 0 | 24008000 | 33950000 | 0 | 0 | 756510 | 663800 | 8479800 | 2067900 | 0 | 760940 | 7323300 | 3134400 | 35607000 | 22335000 | 0 | 0 |
| YGCLAGVR | RCC2 | Q9P258 | RCC2_HUMAN | Unmodified | 0 | 209970 | 0 | 0 | 0 | 0 | 0 | 0 | 0 | 0 | 178010 | 7960800 | 0 | 0 | 0 | 0 | 0 | 0 |
| YGDFFIR | EIF3L; EIF3L; EII | Q9Y262; B0( | EIF3L_HUMAN | Unmodified | 0 | 220300 | 0 | 476530 | 0 | 0 | 0 | 0 | 0 | 0 | 0 | 3391800 | 0 | 0 | 0 | 153270 | 0 | 0 |
| YGDFNKEV | MSN | P26038 | MOES_HUMAN | Unmodified | 0 | 0 | 0 | 0 | 2862700 | 940090 | 0 | 0 | 0 | 0 | 0 | 0 | 0 | 0 | 0 | 0 | 0 | 764120 |
| YGDFNKEV | MSN | P26038 | MOES_HUMAN | Unmodified | 0 | 0 | 0 | 0 | 2077200 | 0 | 0 | 0 | 0 | 0 | 0 | 0 | 0 | 78004 | 0 | 0 | 4126300 | 932200 |
| YGDFRADI | SND1 | Q7KZF4 | SND1_HUMAN | Unmodified | 1012300 | 1002900 | 842940 | 1647300 | 1239600 | 455980 | 183240 | 66617 | 513420 | 324610 | 335150 | 1244400 | 2231600 | 862010 | 1400100 | 1051100 | 793100 | 1116900 |
| YGDGGSTI | HNRNPH1; HNI | P31943; H0Y | HNRH1_HUMA | Unmodified | 0 | 0 | 0 | 0 | 0 | 0 | 0 | 0 | 0 | 0 | 0 | 2126500 | 0 | 0 | 0 | 0 | 0 | 0 |

| | | | | | | | | | | | | | | | | | | | | | | |
|---|---|---|---|---|---|---|---|---|---|---|---|---|---|---|---|---|---|---|---|---|---|---|
| YGDLLPAD | ATP2B4; ATP2B | P23634; P20 | AT2B4_HUMAN | Unmodified | 0 | 0 | 5890900 | 0 | 0 | 0 | 0 | 0 | 1971500 | 607180 | 3803500 | 13987000 | 0 | 0 | 0 | 3790600 | 0 | 0 |
| YGDLLPAD | ATP2B4; ATP2B | P23634; P20 | AT2B4_HUMAN | Unmodified | 2564200 | 1160900 | 1365000 | 12366000 | 12951000 | 1652400 | 0 | 0 | 4520800 | 1012100 | 8537000 | 18771000 | 9069900 | 0 | 16091000 | 5231500 | 1288800 | 0 |
| YGDPGSLF | ITGA6 | P23229 | ITA6_HUMAN | Oxidation (M) | 0 | 0 | 0 | 0 | 176660 | 0 | 0 | 0 | 0 | 0 | 0 | 0 | 499210 | 0 | 0 | 0 | 0 | 1379300 |
| YGDPGSLF | ITGA6 | P23229 | ITA6_HUMAN | Unmodified | 0 | 0 | 0 | 0 | 0 | 0 | 0 | 0 | 0 | 0 | 0 | 0 | 0 | 0 | 0 | 0 | 0 | 354430 |
| YGDSEFTV | HNRNPF | P52597 | HNRPF_HUMAN | Unmodified | 0 | 0 | 0 | 0 | 0 | 0 | 0 | 0 | 0 | 0 | 0 | 1190100 | 0 | 0 | 0 | 0 | 0 | 0 |
| YGDSGEQI | CTSA; CTSA; CT | P10619; X6R | PPGB_HUMAN | Unmodified | 0 | 0 | 0 | 0 | 0 | 0 | 0 | 149310 | 828840 | 390790 | 0 | 0 | 0 | 0 | 0 | 0 | 0 | 0 |
| YGEAGEGF | VARS1; VARS1; | P26640; A2A | SYVC_HUMAN | Unmodified | 129470 | 40321 | 458180 | 0 | 315260 | 0 | 126830 | 0 | 0 | 0 | 1900400 | 9773100 | 588370 | 0 | 0 | 152850 | 0 | 181910 |
| YGEDSEQF | ATP6AP2; ATP6 | O75787; A0A | RENR_HUMAN | Unmodified | 0 | 0 | 0 | 0 | 0 | 0 | 0 | 0 | 4050000 | 326410 | 440000 | 0 | 0 | 0 | 0 | 0 | 0 | 0 |
| YGEGDSLT | SLC39A14; SLC | Q15043; E5F | S39AE_HUMAN | Unmodified | 1009200 | 662770 | 990900 | 601950 | 0 | 0 | 0 | 0 | 593360 | 0 | 0 | 2608400 | 0 | 0 | 731230 | 1506600 | 0 | 0 |
| YGEGGHYI | P4HTM | Q9NXG6 | P4HTM_HUMAN | Unmodified | 0 | 0 | 0 | 0 | 0 | 0 | 0 | 0 | 6853500 | 0 | 0 | 0 | 0 | 0 | 0 | 0 | 0 | 0 |
| YGEGHQA | SEPHS1 | P49903 | SPS1_HUMAN | Unmodified | 78761 | 62195 | 78183 | 209100 | 748200 | 0 | 249770 | 64659 | 644330 | 376330 | 1005400 | 2599900 | 1549900 | 304930 | 116310 | 75297 | 297820 | 118400 |
| YGEGHQA | SEPHS2 | Q99611 | SPS2_HUMAN | Unmodified | 0 | 0 | 0 | 0 | 141470 | 0 | 0 | 0 | 106120 | 0 | 406960 | 684800 | 219230 | 0 | 0 | 0 | 0 | 0 |
| YGEHEASP | UTRN; UTRN | P46939; Q5 | UTRN_HUMAN | Unmodified | 0 | 0 | 777810 | 954630 | 1510500 | 550490 | 0 | 0 | 1225200 | 199210 | 0 | 0 | 494890 | 223380 | 1312200 | 1070300 | 583880 | 1363500 |
| YGENGRLS | SLC39A10; SLC | Q9ULF5; E7 | S39AA_HUMAN | Unmodified | 0 | 0 | 0 | 0 | 1476700 | 0 | 0 | 0 | 15429000 | 761250 | 259510 | 0 | 849480 | 0 | 109680 | 0 | 321290 | 1243200 |
| YGEPGEVF | SFPQ | P23246 | SFPQ_HUMAN | Unmodified | 0 | 0 | 0 | 0 | 0 | 0 | 887530 | 635470 | 1074300 | 1079700 | 1480200 | 9250900 | 0 | 464610 | 0 | 0 | 0 | 0 |
| YGEPGEVF | SFPQ | P23246 | SFPQ_HUMAN | Unmodified | 0 | 0 | 0 | 0 | 0 | 0 | 0 | 189340 | 0 | 0 | 0 | 0 | 0 | 0 | 0 | 0 | 0 | 0 |
| YGEPSEVF | PSPC1; PSPC1 | Q8WXF1; X6 | PSPC1_HUMAN | Unmodified | 0 | 0 | 0 | 0 | 0 | 0 | 0 | 0 | 0 | 0 | 0 | 854050 | 0 | 0 | 0 | 0 | 0 | 0 |
| YGEPSEVF | PSPC1; PSPC1 | Q8WXF1; X6 | PSPC1_HUMAN | Unmodified | 0 | 0 | 0 | 0 | 0 | 0 | 0 | 0 | 0 | 0 | 0 | 2875800 | 0 | 0 | 0 | 0 | 0 | 28594 |
| YGEPTLR | PSMD2; PSMD2 | Q13200; H7 | PSMD2_HUMA | Unmodified | 69237 | 44252 | 226450 | 137410 | 84820 | 0 | 0 | 36742 | 0 | 0 | 0 | 1876800 | 0 | 0 | 0 | 0 | 0 | 2168800 |
| YGEQGLR | DNAJA2 | O60884 | DNJA2_HUMAN | Unmodified | 236680 | 210960 | 176540 | 0 | 52361 | 19167 | 0 | 48265 | 65033 | 278990 | 0 | 414230 | 0 | 0 | 271980 | 185640 | 10893 | 0 |
| YGEYFPGT | HDAC2; HDAC | Q92769; Q1 | HDAC2_HUMA | Unmodified | 333350 | 0 | 290750 | 0 | 0 | 0 | 0 | 0 | 0 | 0 | 397490 | 9861100 | 0 | 0 | 0 | 260300 | 0 | 0 |
| YGFIEGHV | CD44; CD44; C | P16070; H0 | CD44_HUMAN | Unmodified | 13868000 | 13126000 | 3865600 | 3074000 | 1165100 | 0 | 0 | 0 | 0 | 0 | 0 | 0 | 0 | 0 | 2482900 | 2005000 | 424370 | 1438500 |
| YGGAPQAL | AP2A1 | O95782 | AP2A1_HUMAN | Unmodified | 400900 | 493100 | 0 | 305780 | 1012500 | 0 | 437290 | 119510 | 557880 | 977190 | 701960 | 1474400 | 0 | 0 | 0 | 203080 | 1841800 | 1725900 |
| YGGCLGNI | TFPI; TFPI | P10646; C9 | TFPI1_HUMAN | Oxidation (M) | 0 | 0 | 776330 | 1074200 | 0 | 0 | 0 | 0 | 0 | 0 | 0 | 0 | 0 | 0 | 0 | 0 | 0 | 0 |
| YGGDEIPF | FLNA; FLNA | P21333; Q6 | FLNA_HUMAN | Unmodified | 902010 | 1414900 | 357210 | 1009700 | 0 | 0 | 0 | 69194 | 843250 | 0 | 0 | 1721900 | 0 | 0 | 1276100 | 0 | 0 | 0 |
| YGGDEIPF | FLNA; FLNA | P21333; Q6 | FLNA_HUMAN | Unmodified | 634450 | 702030 | 853410 | 3493100 | 7509600 | 3473200 | 3470200 | 814420 | 1775900 | 1756600 | 0 | 567610 | 5313500 | 7347400 | 1182300 | 0 | 3134300 | 1919600 |
| YGGDEIPF | FLNA; FLNA | P21333; Q6 | FLNA_HUMAN | Unmodified | 0 | 0 | 0 | 0 | 766010 | 0 | 762960 | 229080 | 0 | 0 | 0 | 0 | 112150 | 329220 | 0 | 0 | 437180 | 0 |
| YGGDEIPY | FLNC | Q14315 | FLNC_HUMAN | Unmodified | 0 | 0 | 0 | 0 | 0 | 0 | 0 | 0 | 0 | 0 | 0 | 0 | 0 | 0 | 0 | 1666700 | 0 | 0 |
| YGGEEEDC | NENF | Q9UMX5 | NENF_HUMAN | Unmodified | 0 | 0 | 0 | 0 | 0 | 0 | 0 | 0 | 4868000 | 271760 | 89696 | 388410 | 0 | 0 | 0 | 0 | 0 | 0 |
| YGGELVPH | FLNB; FLNB | O75369; E7 | FLNB_HUMAN | Unmodified | 0 | 0 | 412380 | 552510 | 2775200 | 687280 | 0 | 0 | 0 | 0 | 204870 | 1856000 | 437570 | 165140 | 0 | 431000 | 568250 | 895950 |
| YGGPPPDS | SYNCRIP; SYN | O60506; B7 | HNRPQ_HUMA | Unmodified | 0 | 0 | 0 | 0 | 0 | 0 | 0 | 0 | 0 | 0 | 0 | 1276600 | 0 | 0 | 0 | 0 | 0 | 0 |
| YGGPPPDS | SYNCRIP; SYN | O60506; B7 | HNRPQ_HUMA | Unmodified | 0 | 0 | 0 | 0 | 0 | 0 | 0 | 0 | 0 | 305540 | 0 | 7905300 | 0 | 0 | 0 | 0 | 0 | 0 |
| YGGPPPDS | HNRNPR; HNR | O43390; A0 | HNRPR_HUMA | Unmodified | 0 | 232410 | 0 | 0 | 0 | 0 | 0 | 0 | 911710 | 0 | 297780 | 2742500 | 0 | 0 | 0 | 0 | 0 | 0 |
| YGGPPPDS | HNRNPR; HNR | O43390; A0 | HNRPR_HUMA | Unmodified | 0 | 0 | 0 | 0 | 0 | 0 | 0 | 0 | 0 | 0 | 0 | 5171600 | 0 | 0 | 0 | 0 | 0 | 0 |
| YGGPQHIV | FLNC | Q14315 | FLNC_HUMAN | Unmodified | 66504 | 332920 | 0 | 0 | 0 | 936900 | 0 | 0 | 0 | 0 | 0 | 0 | 0 | 0 | 0 | 0 | 2642100 | 436480 |
| YGGPYHIG | FLNA; FLNA; FL | P21333; A0 | FLNA_HUMAN | Unmodified | 141250 | 0 | 0 | 0 | 6647500 | 2473600 | 301820 | 77575 | 1457900 | 2674200 | 0 | 4347000 | 2875600 | 1250400 | 0 | 510760 | 10101000 | 2733900 |
| YGGQPVP | FLNA; FLNA | P21333; Q6 | FLNA_HUMAN | Unmodified | 2569200 | 2762300 | 0 | 3507100 | 6128000 | 1460800 | 2329700 | 1018100 | 0 | 377390 | 0 | 1429900 | 2768700 | 2413300 | 2301000 | 0 | 2279900 | 1392000 |
| YGGRDYSI | MYOF; MYOF | Q9NZM1; H | MYOF_HUMAN | Unmodified | 0 | 0 | 0 | 0 | 3206300 | 715770 | 0 | 0 | 255210 | 0 | 0 | 0 | 0 | 0 | 0 | 0 | 109550 | 288000 |
| YGGTFQN | AP2A2; AP2A2 | O94973; A0 | AP2A2_HUMAN | Unmodified | 216990 | 288010 | 287670 | 557760 | 784660 | 168090 | 77273 | 0 | 1042800 | 1311900 | 1486900 | 4730700 | 2635600 | 417590 | 2001900 | 1071100 | 432540 | 0 |
| YGGWSFG | ABCA12 | Q86UK0 | ABCAC_HUMA | Unmodified | 0 | 0 | 0 | 0 | 0 | 0 | 0 | 0 | 2967000 | 0 | 0 | 38106 | 0 | 0 | 0 | 0 | 0 | 0 |
| YGIENVK | GSR | P00390 | GSHR_HUMAN | Unmodified | 0 | 866110 | 0 | 0 | 0 | 479400 | 664480 | 0 | 590250 | 114950 | 440940 | 1494200 | 836000 | 0 | 47388 | 948420 | 0 | 0 |
| YGIEPTMV | UBA6 | A0AVT1 | UBA6_HUMAN | Oxidation (M) | 0 | 0 | 0 | 0 | 0 | 0 | 0 | 0 | 0 | 0 | 0 | 957330 | 0 | 0 | 0 | 0 | 0 | 0 |
| YGIICMEDL | RPL7; RPL7 | P18124; A8 | RL7_HUMAN 6 | Unmodified | 0 | 66058 | 0 | 0 | 0 | 0 | 231160 | 0 | 0 | 0 | 0 | 45369 | 0 | 0 | 0 | 0 | 0 | 0 |
| YGIICMEDL | RPL7; RPL7 | P18124; A8 | RL7_HUMAN 6 | Oxidation (M) | 509530 | 0 | 0 | 0 | 0 | 0 | 0 | 0 | 0 | 0 | 0 | 1689600 | 0 | 0 | 101650 | 0 | 0 | 0 |
| YGIICMEDL | RPL7; RPL7 | P18124; A8 | RL7_HUMAN 6 | Unmodified | 1026500 | 1497000 | 66053 | 169250 | 0 | 0 | 0 | 0 | 0 | 0 | 0 | 2281700 | 0 | 0 | 1136700 | 155720 | 0 | 0 |
| YGINNIR | FARSA; FARSA | Q9Y285; K7 | SYFA_HUMAN | Unmodified | 0 | 0 | 0 | 0 | 0 | 0 | 0 | 0 | 0 | 0 | 0 | 2047900 | 0 | 0 | 0 | 0 | 0 | 0 |
| YGINTTDIF | TAGLN2; TAGL | P37802; X6 | TAGL2_HUMAN | Unmodified | 0 | 0 | 118860 | 266500 | 0 | 0 | 0 | 0 | 298790 | 166310 | 155950 | 0 | 17377 | 0 | 803880 | 0 | 0 | 287460 |
| YGIQMPAF | IQGAP1; IQGA | P46940; A0 | IQGA1_HUMAN | Unmodified | 0 | 0 | 0 | 0 | 4732700 | 0 | 0 | 0 | 2155900 | 0 | 0 | 5060200 | 0 | 0 | 0 | 0 | 0 | 0 |
| YGISSMIQ | DARS1; DARS1 | P14868; H7 | SYDC_HUMAN | Unmodified | 0 | 0 | 0 | 0 | 0 | 0 | 0 | 0 | 0 | 0 | 402680 | 3490100 | 0 | 0 | 0 | 0 | 0 | 0 |
| YGIVDYMIE | PDIA4; PDIA4 | P13667; A0 | PDIA4_HUMAN | Unmodified | 66703 | 0 | 67548 | 67520 | 160070 | 59825 | 567630 | 519190 | 45811000 | 10792000 | 4759000 | 2023500 | 399190 | 231660 | 252240 | 97990 | 58169 | 52417 |
| YGIVDYMIE | PDIA4; PDIA4 | P13667; A0 | PDIA4_HUMAN | Oxidation (M) | 0 | 0 | 0 | 0 | 0 | 0 | 0 | 47126 | 2877900 | 1705100 | 1484500 | 0 | 130190 | 0 | 171110 | 99345 | 0 | 0 |
| YGIVDYMIE | PDIA4; PDIA4 | P13667; A0 | PDIA4_HUMAN | Unmodified | 4024100 | 0 | 0 | 0 | 601730 | 0 | 1297000 | 0 | 13522000 | 17449000 | 228860 | 0 | 1582400 | 612780 | 0 | 0 | 520130 | 0 |
| YGIVDYMIE | PDIA4; PDIA4 | P13667; A0 | PDIA4_HUMAN | Oxidation (M) | 0 | 0 | 0 | 0 | 0 | 0 | 324270 | 0 | 805260 | 5682000 | 43084 | 0 | 95103 | 117010 | 0 | 0 | 0 | 0 |
| YGKDATNV | ENO1; ENO1 | P06733; A0 | ENOA_HUMAN | Unmodified | 0 | 0 | 0 | 0 | 9302800 | 730180 | 2109200 | 1346100 | 356860 | 2403000 | 0 | 0 | 4824900 | 5500600 | 0 | 0 | 4185900 | 5674600 |
| YGKDATNV | ENO1; ENO1 | P06733; A0 | ENOA_HUMAN | Unmodified | 0 | 0 | 0 | 0 | 1.63E+08 | 14393000 | 0 | 0 | 2457200 | 44194000 | 0 | 0 | 62226000 | 43219000 | 0 | 0 | 1.72E+08 | 1.17E+08 |
| YGKIETIEV | HNRNPA3; HN | P51991; A0 | ROA3_HUMAN | Unmodified | 10406 | 0 | 0 | 0 | 207830 | 0 | 317380 | 113010 | 0 | 0 | 0 | 5210000 | 0 | 0 | 0 | 0 | 0 | 0 |
| YGKIETIEV | HNRNPA3; HN | P51991; A0 | ROA3_HUMAN | Unmodified | 0 | 0 | 0 | 0 | 2754200 | 0 | 0 | 966730 | 2196600 | 1106000 | 1494100 | 3723400 | 5586200 | 909970 | 0 | 0 | 0 | 0 |
| YGKLEVAN | ANK3 | Q12955 | ANK3_HUMAN | Unmodified | 34909 | 17547 | 854270 | 2468500 | 0 | 0 | 21109 | 0 | 0 | 0 | 0 | 335200 | 10995000 | 8858300 | 4402200 | 3518000 | 107220 | 128060 |
| YGKSPYLY | GDI1; GDI2; GD | P31150; P5 | GDIA_HUMAN | Unmodified | 0 | 2613200 | 0 | 1798000 | 5865900 | 1824600 | 1227200 | 0 | 2145300 | 6300500 | 253360 | 0 | 6783500 | 1843400 | 0 | 6631500 | 2547200 | 4441300 |
| YGLDSDLS | SERPINF1; SER | P36955; I3L | PEDF_HUMAN | Unmodified | 0 | 0 | 0 | 0 | 0 | 0 | 0 | 0 | 538200 | 0 | 1989000 | 956200 | 0 | 0 | 0 | 0 | 0 | 0 |
| YGLEECTC | ADAM10 | O14672 | ADA10_HUMAN | Unmodified | 2928200 | 0 | 2949100 | 3193500 | 1270800 | 443310 | 0 | 0 | 927910 | 93891 | 610740 | 7315800 | 5782800 | 2813300 | 6730200 | 4584400 | 100990 | 582350 |
| YGLFKEEN | PTTG1IP; PTTG | P53801; A8 | PTTG_HUMAN | Unmodified | 1383500 | 1291100 | 1069200 | 10765000 | 2406000 | 1429800 | 0 | 0 | 12938000 | 3661600 | 0 | 2558800 | 347900 | 0 | 4390500 | 4896400 | 36233 | 278080 |
| YGLFKEEN | PTTG1IP; PTTG | P53801; A8 | PTTG_HUMAN | Unmodified | 1086100 | 1270200 | 0 | 19139000 | 98118000 | 24611000 | 11801000 | 833890 | 9582200 | 13636000 | 0 | 793250 | 16525000 | 14752000 | 1864600 | 2642400 | 2017700 | 15382000 |
| YGLFPANY | CTTN | Q14247 | SRC8_HUMAN | Unmodified | 352830 | 505430 | 143400 | 389920 | 273490 | 136650 | 0 | 0 | 119800 | 0 | 0 | 500040 | 0 | 148580 | 650370 | 286700 | 135440 | 117770 |
| YGLIFHSTF | NOP56; NOP56 | O00567; A0 | NOP56_HUMA | Unmodified | 0 | 0 | 0 | 0 | 0 | 0 | 1080500 | 22521 | 0 | 0 | 0 | 0 | 0 | 0 | 0 | 0 | 0 | 0 |
| YGLIYHASL | NOP58 | Q9Y2X3 | NOP58_HUMA | Unmodified | 345440 | 0 | 430630 | 199130 | 172350 | 0 | 770740 | 642150 | 824650 | 0 | 0 | 1610300 | 131000 | 0 | 319740 | 205520 | 0 | 32828 |
| YGLLPSHA | YRDC | Q86U90 | YRDC_HUMAN | Unmodified | 0 | 0 | 0 | 0 | 0 | 0 | 0 | 0 | 0 | 0 | 0 | 0 | 0 | 0 | 0 | 0 | 0 | 0 |
| YGLLVGGA | QPRT | Q15274 | NADC_HUMAN | Oxidation (M) | 0 | 0 | 0 | 0 | 0 | 0 | 243100 | 0 | 0 | 0 | 571000 | 2397600 | 0 | 0 | 0 | 0 | 0 | 0 |
| YGLQLPAF | IQGAP3; IQGA | Q86VI3; F2Z | IQGA3_HUMAN | Unmodified | 381960 | 0 | 216200 | 163600 | 554050 | 146880 | 0 | 0 | 604590 | 273450 | 70326 | 661360 | 0 | 0 | 0 | 0 | 0 | 0 |
| YGMGTSVE | PDHA1 | P08559 | ODPA_HUMAN | Unmodified | 0 | 0 | 0 | 0 | 0 | 0 | 0 | 0 | 275770 | 0 | 0 | 46781 | 0 | 0 | 0 | 0 | 0 | 0 |
| YGMNPHQ | ATIC; ATIC | P31939; C9 | PUR9_HUMAN | Oxidation (M) | 261010 | 0 | 931730 | 1079700 | 0 | 0 | 0 | 0 | 0 | 0 | 536100 | 6464800 | 304270 | 0 | 1475800 | 976610 | 0 | 0 |
| YGMNPHQ | ATIC; ATIC | P31939; C9 | PUR9_HUMAN | Unmodified | 200620 | 0 | 0 | 472780 | 0 | 0 | 0 | 0 | 0 | 0 | 0 | 6934500 | 0 | 0 | 1301100 | 317970 | 0 | 0 |
| YGMPPQD | CHD4; CHD4; | Q14839; A0 | CHD4_HUMAN | Oxidation (M) | 0 | 0 | 0 | 0 | 0 | 0 | 0 | 0 | 0 | 0 | 65579 | 284380 | 0 | 0 | 0 | 0 | 0 | 0 |
| YGMRFNVI | DECR1; DECR1 | Q16698; H0 | DECR_HUMAN | Unmodified | 0 | 0 | 0 | 0 | 0 | 30172 | 127430 | 0 | 0 | 447630 | 0 | 0 | 0 | 0 | 0 | 0 | 0 | 256510 |
| YGNFDILRI | MME; MME; MM | P08473; C9 | NEP_HUMAN | Unmodified | 0 | 0 | 0 | 2821700 | 0 | 0 | 0 | 0 | 0 | 0 | 0 | 0 | 1278200 | 111980 | 5342100 | 0 | 0 | 0 |
| YGNFQLYC | IFIT2; IFIT2; IFIT | P09913; A0 | IFIT2_HUMAN | Oxidation (M) | 0 | 0 | 0 | 0 | 0 | 168120 | 0 | 0 | 0 | 0 | 437820 | 0 | 0 | 0 | 0 | 0 | 0 | 0 |
| YGNIIAYDH | PTPRM; PTPRK; | P28827; Q1 | PTPRM_HUMA | Unmodified | 158040 | 102340 | 0 | 0 | 0 | 0 | 0 | 0 | 2009300 | 186320 | 0 | 0 | 0 | 0 | 132710 | 89589 | 0 | 0 |
| YGNQLVR | MCM7 | P33993 | MCM7_HUMAN | Unmodified | 0 | 0 | 0 | 0 | 529840 | 0 | 379800 | 169750 | 270710 | 0 | 130760 | 577820 | 0 | 0 | 0 | 0 | 163890 | 150040 |
| YGPADVED | EEF1B2; EEF1B | P24534; C9 | EF1B_HUMAN | Unmodified | 0 | 0 | 0 | 0 | 0 | 0 | 0 | 0 | 0 | 0 | 2766000 | 10393000 | 0 | 0 | 0 | 0 | 0 | 0 |
| YGPDAFFN | LRBA; LRBA; LR | P50851; A0 | LRBA_HUMAN | Unmodified | 0 | 0 | 0 | 0 | 0 | 0 | 0 | 0 | 0 | 0 | 0 | 7393500 | 0 | 0 | 0 | 0 | 0 | 0 |
| YGPFVADF | ATP5PB; ATP5P | P24539; Q5 | AT5F1_HUMAN | Unmodified | 0 | 0 | 0 | 0 | 0 | 0 | 23227 | 0 | 0 | 920460 | 0 | 0 | 0 | 0 | 0 | 0 | 0 | 0 |
| YGPIADVSI | TRA2B | P62995 | TRA2B_HUMAN | Unmodified | 704220 | 0 | 0 | 0 | 849460 | 0 | 0 | 0 | 0 | 231440 | 570570 | 1133400 | 0 | 0 | 0 | 223430 | 0 | 349710 |
| YGPMEEPL | GPX4; GPX4; GI | P36969; A0 | GPX4_HUMAN | Unmodified | 0 | 0 | 0 | 0 | 0 | 0 | 0 | 0 | 730450 | 0 | 0 | 0 | 1226300 | 0 | 0 | 0 | 0 | 0 |
| YGPMEEPL | GPX4; GPX4; GI | P36969; A0 | GPX4_HUMAN | Oxidation (M) | 0 | 0 | 0 | 0 | 0 | 0 | 0 | 0 | 0 | 0 | 0 | 0 | 0 | 0 | 0 | 0 | 0 | 0 |
| YGPQYGHI | HNRNPL; HNRN | P14866; M0 | HNRPL_HUMAI | Unmodified | 0 | 0 | 0 | 0 | 0 | 0 | 2342600 | 0 | 689500 | 0 | 1660800 | 5891300 | 0 | 0 | 0 | 0 | 0 | 0 |
| YGPSLMPC | PGD; PGD; PGD | P52209; K7E | 6PGD_HUMAN | Oxidation (M) | 78458 | 42114 | 156560 | 0 | 0 | 0 | 0 | 0 | 0 | 0 | 305200 | 1979000 | 0 | 0 | 256820 | 146600 | 0 | 0 |
| YGPSLMPC | PGD; PGD; PGD | P52209; K7E | 6PGD_HUMAN | Unmodified | 0 | 0 | 0 | 0 | 0 | 0 | 0 | 0 | 0 | 0 | 0 | 7196000 | 0 | 0 | 0 | 0 | 0 | 0 |
| YGPVVYAA | RGS13 | O14921 | RGS13_HUMAI | Unmodified | 0 | 0 | 0 | 0 | 0 | 0 | 0 | 0 | 0 | 0 | 430140 | 1095900 | 0 | 0 | 0 | 0 | 0 | 0 |
| YGQDYSTH | TNFRSF10B | O14763 | TR10B_HUMAN | Unmodified | 169830 | 144920 | 92213 | 890980 | 1363600 | 294890 | 0 | 0 | 0 | 0 | 447090 | 847600 | 802150 | 316430 | 675960 | 373990 | 144740 | 475730 |
| YGQFSGLN | PPP1CA; PPP1 | P62136; E9 | PP1A_HUMAN | Unmodified | 0 | 0 | 0 | 472320 | 0 | 0 | 0 | 0 | 795520 | 0 | 842780 | 4280700 | 0 | 0 | 0 | 0 | 31633 | 0 |
| YGQGAGE | TNKS1BP1; TN | Q9C0C2; A0 | TB182_HUMAN | Unmodified | 0 | 44009 | 125940 | 0 | 313090 | 38943 | 0 | 0 | 48974 | 0 | 0 | 191710 | 0 | 0 | 136950 | 330340 | 36600 | 175590 |
| YGQGFYLI | LLGL2; LLGL2 | Q6P1M3; J3 | L2GL2_HUMAN | Unmodified | 0 | 0 | 0 | 0 | 0 | 0 | 0 | 0 | 0 | 0 | 0 | 795650 | 0 | 0 | 0 | 352560 | 0 | 0 |
| YGQGFYLI | LLGL2; LLGL2 | Q6P1M3; J3 | L2GL2_HUMAN | Unmodified | 0 | 0 | 0 | 450640 | 0 | 0 | 0 | 0 | 999480 | 1600800 | 0 | 0 | 595200 | 0 | 625730 | 0 | 1198900 | 0 |
| YGQHPNII | RPS6KA6; RPS6 | Q9UK32; Q1 | KS6A6_HUMAN | Unmodified | 0 | 0 | 0 | 0 | 0 | 0 | 0 | 28984 | 0 | 109280 | 258570 | 1465900 | 0 | 351960 | 362090 | 645940 | 0 | 0 |
| YGQISEVV | CIRBP; CIRBP; | Q14011; K7 | CIRBP_HUMAN | Unmodified | 4098900 | 3693100 | 3281000 | 3372700 | 479420 | 0 | 0 | 0 | 1490000 | 511730 | 890930 | 3024800 | 1576700 | 0 | 4855700 | 4819700 | 0 | 0 |
| YGQISEVV | CIRBP; CIRBP; | Q14011; K7 | CIRBP_HUMAN | Unmodified | 69707 | 127510 | 0 | 171500 | 0 | 0 | 29535 | 0 | 0 | 115370 | 0 | 0 | 315640 | 87659 | 0 | 0 | 0 | 0 |
| YGQLSEK | PDCD5; PDCD5 | O14737; K7 | PDCD5_HUMA | Unmodified | 277030 | 0 | 487440 | 974430 | 0 | 234850 | 0 | 21437 | 338700 | 99333 | 0 | 1070400 | 311360 | 0 | 302090 | 224400 | 0 | 0 |
| YGQPIQMK | HSPA4L; HSPA | O95757; E9 | HS74L_HUMAN | Unmodified | 0 | 0 | 0 | 0 | 737160 | 0 | 0 | 0 | 0 | 0 | 235230 | 983640 | 469770 | 0 | 0 | 0 | 226870 | 0 |
| YGSDIVPF | XRCC5 | P13010 | XRCC5_HUMAI | Unmodified | 0 | 0 | 0 | 0 | 0 | 0 | 0 | 0 | 0 | 0 | 0 | 5204200 | 0 | 0 | 51793 | 0 | 0 | 0 |
| YGSDIVPF | XRCC5 | P13010 | XRCC5_HUMAI | Unmodified | 0 | 0 | 0 | 0 | 0 | 0 | 0 | 0 | 651200 | 0 | 0 | 3207500 | 0 | 0 | 0 | 0 | 0 | 0 |
| YGSDIVPF | XRCC5 | P13010 | XRCC5_HUMAI | Oxidation (M) | 594760 | 335610 | 0 | 0 | 0 | 0 | 0 | 0 | 0 | 0 | 0 | 8704000 | 0 | 0 | 0 | 123590 | 0 | 0 |
| YGSGSGQ | SDF2L1 | Q9HCN8 | SDF2L_HUMAN | Unmodified | 0 | 0 | 95695 | 0 | 0 | 0 | 0 | 0 | 10815000 | 0 | 1641400 | 1209300 | 0 | 0 | 148310 | 0 | 0 | 0 |
| YGSGSGQ | SDF2; SDF2 | Q99470; K7 | SDF2_HUMAN | Unmodified | 0 | 0 | 0 | 0 | 0 | 0 | 0 | 0 | 14607000 | 774800 | 0 | 472890 | 0 | 0 | 0 | 0 | 0 | 0 |
| YGSGSGQ | SDF2; SDF2 | Q99470; K7 | SDF2_HUMAN | Unmodified | 0 | 0 | 0 | 0 | 0 | 0 | 0 | 0 | 0 | 945530 | 0 | 0 | 0 | 0 | 0 | 0 | 0 | 0 |
| YGSIVDDE | IQGAP2 | Q13576 | IQGA2_HUMAN | Oxidation (M) | 0 | 0 | 900410 | 241280 | 0 | 349760 | 347900 | 0 | 0 | 0 | 0 | 0 | 0 | 0 | 725510 | 727450 | 0 | 0 |
| YGSIVDDE | IQGAP2 | Q13576 | IQGA2_HUMAN | Unmodified | 0 | 0 | 373720 | 306470 | 0 | 0 | 0 | 0 | 0 | 0 | 0 | 0 | 0 | 0 | 463650 | 241680 | 0 | 0 |
| YGSPYTK | GAS6 | Q14393 | GAS6_HUMAN | Unmodified | 0 | 0 | 0 | 0 | 0 | 0 | 0 | 0 | 3437800 | 333600 | 27610 | 0 | 25031 | 20118 | 10787 | 0 | 0 | 0 |
| YGSQEHPI | RASA3 | Q14644 | RASA3_HUMAN | Unmodified | 247460 | 0 | 322980 | 0 | 0 | 0 | 0 | 1668000 | 0 | 108880 | 0 | 0 | 0 | 0 | 0 | 0 | 0 | 0 |
| YGSSLEFTE | CASP6 | P55212 | CASP6_HUMAN | Unmodified | 0 | 0 | 0 | 0 | 0 | 0 | 0 | 0 | 0 | 54850 | 238460 | 259970 | 355670 | 0 | 0 | 0 | 0 | 0 |

| | | | | | | | | | | | | | | | | | | | | | | |
|---|---|---|---|---|---|---|---|---|---|---|---|---|---|---|---|---|---|---|---|---|---|---|
| YGSVAFPN | AGPS; AGPS; A | O00116; A0A | ADAS_HUMAN | Unmodified | 145170 | 0 | 0 | 408520 | 0 | 0 | 0 | 0 | 11849000 | 119120 | 0 | 0 | 70883 | 0 | 0 | 18234 | 0 | 0 |
| YGSVAFPN | AGPS; AGPS; A | O00116; A0A | ADAS_HUMAN | Unmodified | 0 | 0 | 0 | 0 | 0 | 299990 | 2498400 | 0 | 7233700 | 7004500 | 0 | 0 | 0 | 0 | 0 | 0 | 0 | 0 |
| YGSYNVFW | MOGS; MOGS; | Q13724; A0A | MOGS_HUMAN | Oxidation (M) | 0 | 0 | 0 | 0 | 0 | 0 | 0 | 0 | 437220 | 0 | 0 | 0 | 0 | 0 | 0 | 0 | 0 | 0 |
| YGTDLSR | ATP1A1; ATP1A | P05023; Q5T | AT1A1_HUMAN | Unmodified | 698850 | 186700 | 11719000 | 8959300 | 2822900 | 882090 | 144790 | 238430 | 14283000 | 1607400 | 7274200 | 1.03E+08 | 1405400 | 666570 | 11793000 | 13419000 | 712530 | 920470 |
| YGTDLSRG | ATP1A1; ATP1A | P05023; Q5T | AT1A1_HUMAN | Unmodified | 0 | 0 | 0 | 285910 | 3811800 | 1551800 | 589890 | 144170 | 56910 | 544550 | 1628400 | 485230 | 5708500 | 1186700 | 0 | 0 | 0 | 924300 |
| YGVAARPG | ELN; ELN | B3KRT8; E7E | B3KRT8_HUMA | Unmodified | 0 | 0 | 0 | 0 | 0 | 0 | 1474900 | 0 | 0 | 0 | 0 | 0 | 0 | 0 | 0 | 0 | 0 | 0 |
| YGVGTPAA | ELN; ELN | E7EN65; E7E | E7EN65_HUMA | Unmodified | 0 | 358000 | 0 | 0 | 2132100 | 0 | 546020 | 708510 | 0 | 0 | 0 | 50390 | 0 | 0 | 0 | 0 | 263410 | 1910500 |
| YGVIILDEAI | DHX15 | O43143 | DHX15_HUMAI | Unmodified | 229750 | 289530 | 0 | 111500 | 0 | 0 | 0 | 0 | 0 | 0 | 1163900 | 13206000 | 0 | 0 | 67096 | 107520 | 0 | 0 |
| YGVIIVGNF | UPF1 | Q92900 | RENT1_HUMAN | Unmodified | 1263600 | 1214700 | 993920 | 791730 | 143160 | 0 | 172570 | 352160 | 0 | 0 | 0 | 1791400 | 0 | 0 | 729610 | 617460 | 0 | 0 |
| YGVNPGPI | TMPO; TMPO; T | P42167; P42 | LAP2B_HUMAN | Unmodified | 234360 | 222210 | 0 | 0 | 0 | 0 | 0 | 0 | 1855400 | 1139900 | 0 | 1476900 | 0 | 0 | 0 | 0 | 0 | 0 |
| YGVQADR\ | CTTN | Q14247 | SRC8_HUMAN | Unmodified | 245180 | 178570 | 0 | 142570 | 0 | 0 | 0 | 0 | 0 | 0 | 0 | 0 | 422240 | 0 | 269890 | 264400 | 0 | 0 |
| YGVSGYPT | PDIA3; PDIA3 | P30101; H7E | PDIA3_HUMAN | Unmodified | 374340 | 247230 | 1283000 | 1644400 | 2013000 | 0 | 1876200 | 1237000 | 1.56E+08 | 22278000 | 40540000 | 14420000 | 296870 | 1815200 | 1277800 | 970340 | 311800 | 1702800 |
| YGVSSLQR | BTBD17 | A6NE02 | BTBDH_HUMAI | Unmodified | 0 | 76251 | 0 | 0 | 43445 | 330640 | 159500 | 83052 | 750490 | 54028 | 0 | 0 | 0 | 0 | 0 | 0 | 33065 | 0 |
| YGVTTLVR | PTP4A2; PTP4A | Q12974; E9F | TP4A2_HUMAN | Unmodified | 653500 | 743410 | 334040 | 826240 | 0 | 0 | 0 | 0 | 2185600 | 0 | 394890 | 6456700 | 0 | 0 | 1298900 | 994380 | 0 | 0 |
| YGVTTLVR\ | PTP4A2; PTP4A | Q12974; E9F | TP4A2_HUMAN | Unmodified | 0 | 0 | 0 | 0 | 0 | 0 | 130070 | 0 | 0 | 301210 | 0 | 0 | 255670 | 227210 | 0 | 0 | 587360 | 911210 |
| YGWSADES | EIF3K; EIF3K; EI | Q9UBQ5; B4 | EIF3K_HUMAN | Unmodified | 0 | 0 | 0 | 0 | 0 | 0 | 0 | 0 | 0 | 0 | 0 | 1509200 | 0 | 0 | 0 | 0 | 0 | 0 |
| YGYEIPVDI | PSMA6; PSMA6 | P60900; G3\ | PSA6_HUMAN | Oxidation (M) | 0 | 0 | 0 | 0 | 389000 | 0 | 0 | 0 | 0 | 246520 | 0 | 2420000 | 0 | 0 | 0 | 0 | 257280 | 253960 |
| YGYILPDITI | MRPL15; MRPL | Q9P015; E5F | RM15_HUMAN | Unmodified | 0 | 0 | 0 | 0 | 104190 | 22013 | 0 | 0 | 2770600 | 0 | 0 | 0 | 0 | 0 | 0 | 0 | 0 | 0 |
| YGYTHLSA | CMPK1; CMPK1 | P30085; Q5T | KCY_HUMAN U | Unmodified | 65996 | 0 | 149620 | 0 | 0 | 0 | 0 | 0 | 303440 | 0 | 0 | 732170 | 0 | 0 | 207070 | 232170 | 0 | 0 |
| YGYTHLSA | CMPK1; CMPK1 | P30085; Q5T | KCY_HUMAN U | Unmodified | 0 | 200840 | 1186700 | 743920 | 1996500 | 0 | 0 | 0 | 0 | 0 | 0 | 2859900 | 1338300 | 0 | 1173300 | 1293900 | 496800 | 1223000 |
| YGYTHLST | AK1; AK1 | P00568; Q5T | KAD1_HUMAN | Unmodified | 528070 | 510830 | 1029300 | 1032900 | 470270 | 0 | 183200 | 103130 | 3909300 | 747990 | 840870 | 9765100 | 558320 | 0 | 2562500 | 1770800 | 171620 | 374840 |
| YGYTHLST | AK1; AK1 | P00568; Q5T | KAD1_HUMAN | Unmodified | 324430 | 145330 | 288330 | 970050 | 0 | 0 | 0 | 0 | 2246900 | 1794300 | 621120 | 710870 | 1747000 | 1390200 | 556750 | 430550 | 1379800 | 1012500 |
| YGYTHLST | AK1; AK1 | P00568; Q5T | KAD1_HUMAN | Unmodified | 0 | 0 | 0 | 0 | 5271500 | 0 | 3961800 | 543000 | 0 | 2548100 | 0 | 0 | 5214500 | 0 | 0 | 0 | 3998300 | 0 |
| YHAASAEE | MRPS7; MRPS7 | Q9Y2R9; J3C | RT07_HUMAN : | Unmodified | 0 | 0 | 0 | 0 | 0 | 0 | 0 | 0 | 3547100 | 0 | 0 | 0 | 0 | 0 | 0 | 0 | 0 | 0 |
| YHEEFEK | LASP1; LASP1 | Q14847; C9J | LASP1_HUMAN | Unmodified | 1839900 | 679200 | 484400 | 837080 | 1486600 | 808730 | 0 | 0 | 0 | 0 | 0 | 0 | 0 | 486950 | 950450 | 501910 | 211050 | 0 |
| YHEQLSTQ | CLTC; CLTC | Q00610; A0A | CLH1_HUMAN | Unmodified | 0 | 615110 | 0 | 46048 | 0 | 0 | 0 | 0 | 0 | 84759 | 0 | 0 | 55922 | 82107 | 684280 | 24579 | 0 | 0 |
| YHGNVMLI | FASN; FASN | P49327; A0A | FAS_HUMAN F | Unmodified | 0 | 0 | 0 | 0 | 598670 | 114210 | 150770 | 38452 | 88552 | 98807 | 0 | 242390 | 282900 | 0 | 0 | 0 | 0 | 330060 |
| YHIDLDPH | CSNK2A2; CSN | P19784; H3E | CSK22_HUMAN | Unmodified | 0 | 0 | 0 | 0 | 0 | 0 | 0 | 0 | 0 | 0 | 0 | 259090 | 0 | 0 | 0 | 0 | 0 | 0 |
| YHIIHADIYI | MARS1; MARS1 | P56192; H0Y | SYMC_HUMAN | Unmodified | 113490 | 0 | 0 | 0 | 0 | 0 | 0 | 0 | 0 | 0 | 0 | 1976300 | 0 | 0 | 0 | 0 | 0 | 0 |
| YHISALYVV | UGGT2; UGGT1 | Q9NYU1; Q9 | UGGG2_HUMA | Unmodified | 0 | 0 | 0 | 0 | 0 | 0 | 0 | 0 | 0 | 1730500 | 0 | 0 | 0 | 0 | 0 | 0 | 0 | 0 |
| YHLTVAQV | FOLH1B; FOLH | Q9HBA9; Q0 | FOH1B_HUMAI | Unmodified | 2273600 | 2420800 | 31164000 | 5037000 | 0 | 0 | 93632 | 0 | 0 | 0 | 0 | 0 | 0 | 0 | 14140000 | 10391000 | 0 | 0 |
| YHNRDTWI | RASA3 | Q14644 | RASA3_HUMAN | Unmodified | 293030 | 0 | 0 | 0 | 0 | 0 | 976670 | 0 | 0 | 0 | 0 | 0 | 602490 | 0 | 0 | 303510 | 404770 | 0 |
| YHNVGLSK | HNRNPD; HNR | Q14103; H0Y | HNRPD_HUMA | Unmodified | 283990 | 0 | 0 | 0 | 0 | 0 | 0 | 0 | 0 | 0 | 394710 | 3270100 | 0 | 32142 | 0 | 0 | 0 | 0 |
| YHNVGLSK | HNRNPD; HNR | Q14103; H0Y | HNRPD_HUMA | Unmodified | 0 | 0 | 0 | 0 | 0 | 0 | 151950 | 169940 | 0 | 157350 | 0 | 0 | 0 | 0 | 0 | 0 | 0 | 0 |
| YHPDKGGI | SEC63; SEC63 | Q9UGP8; A6 | SEC63_HUMAN | Oxidation (M) | 0 | 0 | 0 | 0 | 0 | 0 | 0 | 0 | 2465800 | 347760 | 0 | 0 | 0 | 0 | 0 | 0 | 0 | 0 |
| YHPDKGGI | SEC63; SEC63 | Q9UGP8; A6 | SEC63_HUMAN | Unmodified | 0 | 0 | 0 | 0 | 0 | 0 | 0 | 0 | 9587400 | 705780 | 179340 | 0 | 0 | 0 | 273160 | 0 | 142310 | 0 |
| YHPDKNPI | DNAJC5 | Q9H3Z4 | DNJC5_HUMAI | Unmodified | 0 | 250040 | 0 | 0 | 0 | 1177700 | 254610 | 0 | 0 | 0 | 0 | 11742000 | 0 | 0 | 0 | 2972200 | 0 | 0 |
| YHPDKNPI | DNAJC5 | Q9H3Z4 | DNJC5_HUMAI | Unmodified | 1420300 | 1228800 | 0 | 65977 | 0 | 0 | 0 | 22618 | 477540 | 232330 | 101810 | 10847000 | 97868 | 77752 | 0 | 2000500 | 0 | 0 |
| YHPDKNPI | DNAJA1 | P31689 | DNJA1_HUMAN | Unmodified | 0 | 0 | 0 | 0 | 0 | 0 | 0 | 0 | 211780 | 0 | 0 | 0 | 0 | 0 | 1929200 | 0 | 0 | 0 |
| YHPGYFGK | RPL27A; RPL27 | P46776; E9P | RL27A_HUMAN | Unmodified | 1619700 | 1484100 | 661460 | 515840 | 0 | 51190 | 176750 | 148340 | 0 | 0 | 0 | 4696200 | 289150 | 111090 | 733070 | 451380 | 0 | 578200 |
| YHPGYFGK | RPL27A; RPL27 | P46776; E9P | RL27A_HUMAN | Unmodified | 0 | 0 | 0 | 0 | 143030 | 44024 | 152260 | 34458 | 0 | 0 | 0 | 0 | 313240 | 0 | 0 | 0 | 0 | 0 |
| YHPIDIETS | DIP2A; DIP2C; I | Q14689; Q9 | DIP2A_HUMAN | Unmodified | 130230 | 0 | 0 | 0 | 0 | 0 | 0 | 53090 | 439790 | 317770 | 0 | 0 | 0 | 0 | 784230 | 438630 | 21052 | 0 |
| YHPIDIETS | DIP2B | Q9P265 | DIP2B_HUMAN | Unmodified | 0 | 0 | 0 | 0 | 0 | 0 | 0 | 0 | 0 | 0 | 0 | 0 | 0 | 0 | 0 | 0 | 0 | 0 |
| YHQVSLFY | FMNL2; FMNL2 | Q96PY5; C9I | FMNL2_HUMAI | Unmodified | 0 | 0 | 0 | 0 | 734350 | 0 | 259200 | 0 | 0 | 0 | 157020 | 1077000 | 0 | 0 | 0 | 0 | 0 | 0 |
| YHSDEYIK | HDAC2; HDAC2 | Q92769; E5F | HDAC2_HUMA | Unmodified | 0 | 0 | 0 | 0 | 163470 | 148130 | 116280 | 24502 | 0 | 0 | 164260 | 794190 | 0 | 0 | 0 | 0 | 5656.9 | 263710 |
| YHSGDVRL | PTGFRN | Q9P2B2 | FPRP_HUMAN | Unmodified | 4160600 | 3477500 | 8870200 | 11275000 | 0 | 0 | 0 | 0 | 0 | 0 | 0 | 1995600 | 381060 | 0 | 11112000 | 9932500 | 0 | 0 |
| YHSGDVRL | PTGFRN | Q9P2B2 | FPRP_HUMAN | Unmodified | 0 | 0 | 0 | 2288200 | 455640 | 0 | 0 | 0 | 0 | 0 | 0 | 0 | 1509900 | 1044000 | 0 | 0 | 0 | 0 |
| YHSGFFVD | BMX | P51813 | BMX_HUMAN ( | Unmodified | 0 | 0 | 0 | 0 | 0 | 86032 | 0 | 0 | 0 | 0 | 163410 | 1031000 | 0 | 0 | 0 | 0 | 0 | 0 |
| YHSLAPMY | RAB5B; RAB5C; | P61020; P51 | RAB5B_HUMAN | Oxidation (M) | 2772200 | 2255800 | 3983000 | 1101300 | 0 | 0 | 0 | 0 | 776370 | 113040 | 0 | 6574500 | 0 | 0 | 3339300 | 3099800 | 0 | 209680 |
| YHSLAPMY | RAB5B; RAB5C; | P61020; P51 | RAB5B_HUMAN | Unmodified | 956330 | 1741000 | 688180 | 0 | 0 | 0 | 0 | 0 | 1851700 | 0 | 223870 | 7217900 | 0 | 0 | 2109700 | 1547300 | 0 | 0 |
| YHSYKDLS | CPD; CPD | O75976; J3C | CBPD_HUMAN | Unmodified | 0 | 0 | 0 | 250810 | 0 | 0 | 0 | 0 | 146280 | 398560 | 0 | 0 | 0 | 37859 | 0 | 0 | 0 | 0 |
| YHTEIVFAR | ATP1A1 | P05023 | AT1A1_HUMAN | Unmodified | 1022500 | 76257 | 7858500 | 4947900 | 3281400 | 0 | 0 | 0 | 4827900 | 898040 | 8211200 | 27863000 | 4210000 | 2190700 | 9402400 | 9365100 | 134520 | 283540 |
| YHTEIVFAR | ATP1A1 | P05023 | AT1A1_HUMAN | Unmodified | 0 | 0 | 0 | 0 | 25473000 | 2906300 | 6757900 | 647590 | 0 | 2771200 | 777080 | 0 | 17100000 | 15710000 | 0 | 0 | 1552200 | 4666400 |
| YHTINGHN | HNRNPA2B1; H | P22626; A0A | ROA2_HUMAN | Unmodified | 104140 | 85709 | 51766 | 38022 | 487900 | 0 | 0 | 0 | 311640 | 523170 | 413920 | 1279700 | 1098100 | 595950 | 43838 | 29312 | 96460 | 227930 |
| YHTINGHN | HNRNPA3; HNI | P51991; A0A | ROA3_HUMAN | Unmodified | 0 | 0 | 0 | 0 | 92938 | 0 | 0 | 183050 | 35606 | 0 | 0 | 388600 | 273070 | 160490 | 0 | 0 | 67248 | 149210 |
| YHTSQSGD | HSP90AB1 | P08238 | HS90B_HUMAI | Unmodified | 3583500 | 3800600 | 3862000 | 14864000 | 7069400 | 3758400 | 121780 | 58332 | 51185000 | 1781700 | 18780000 | 1.73E+08 | 7190900 | 2026300 | 9395400 | 6394300 | 1402800 | 1942000 |
| YHTSQSGD | HSP90AB1 | P08238 | HS90B_HUMAI | Oxidation (M) | 3261700 | 2312600 | 6228700 | 6500500 | 1695700 | 536410 | 164840 | 103900 | 10632000 | 1775200 | 13841000 | 1.08E+08 | 2300600 | 435010 | 5777000 | 5768100 | 710010 | 472280 |
| YHTSQSGD | HSP90AB1 | P08238 | HS90B_HUMAI | Unmodified | 0 | 0 | 0 | 0 | 1709600 | 0 | 0 | 0 | 0 | 0 | 0 | 0 | 755610 | 0 | 0 | 0 | 312460 | 1392500 |
| YHTSQSGD | HSP90AB1 | P08238 | HS90B_HUMAI | Oxidation (M) | 0 | 0 | 0 | 0 | 1501700 | 0 | 0 | 0 | 0 | 0 | 0 | 0 | 0 | 225810 | 0 | 0 | 0 | 731930 |
| YHTSQSGD | HSP90AB1 | P08238 | HS90B_HUMAI | Unmodified | 0 | 0 | 0 | 0 | 2136500 | 474390 | 0 | 0 | 0 | 0 | 0 | 0 | 0 | 0 | 0 | 0 | 1449700 | 842850 |
| YHTVNGHI | HNRNPA1; HNI | P09651; F8V | ROA1_HUMAN | Unmodified | 108130 | 33307 | 0 | 86379 | 0 | 35680 | 0 | 0 | 223300 | 0 | 134340 | 2743700 | 0 | 0 | 65570 | 0 | 0 | 0 |
| YHTVNGHI | HNRNPA1; HNI | P09651; F8V | ROA1_HUMAN | Unmodified | 0 | 68038 | 45301 | 0 | 1956800 | 588660 | 0 | 753340 | 1397200 | 2778400 | 2391200 | 7321700 | 2649100 | 1513600 | 17796 | 0 | 333870 | 216980 |
| YHVPVVVV | RPN2; RPN2; RI | P04844; Q5J | RPN2_HUMAN | Unmodified | 0 | 0 | 0 | 0 | 0 | 263850 | 0 | 0 | 43007000 | 4748400 | 3985500 | 4560600 | 0 | 0 | 0 | 0 | 0 | 0 |
| YHYVGTFP | FKBP9 | O95302 | FKBP9_HUMAN | Unmodified | 0 | 0 | 0 | 0 | 0 | 0 | 0 | 0 | 2515600 | 100590 | 441530 | 285470 | 0 | 0 | 0 | 0 | 0 | 0 |
| YHYVGTFP | FKBP9 | O95302 | FKBP9_HUMAN | Unmodified | 0 | 0 | 0 | 0 | 0 | 0 | 0 | 0 | 40918000 | 1419500 | 3198800 | 0 | 0 | 0 | 0 | 0 | 0 | 0 |
| YIAAQGPRI | PTPRC; PTPRC; | P08575; A0A | PTPRC_HUMAN | Unmodified | 247670 | 696270 | 344320 | 1002300 | 89769 | 0 | 0 | 0 | 0 | 0 | 0 | 0 | 132770 | 49540 | 660500 | 733140 | 0 | 0 |
| YIAENGTDI | PRPF19; PRPF1 | Q9UMS4; F5 | PRP19_HUMAN | Unmodified | 0 | 0 | 0 | 0 | 1810600 | 0 | 0 | 272050 | 2626200 | 767210 | 2814500 | 6966500 | 0 | 0 | 0 | 0 | 841980 | 3831100 |
| YIAIVSTTVE | GDI2 | P50395 | GDIB_HUMAN | Unmodified | 2915400 | 2140100 | 4227700 | 6505000 | 0 | 2899400 | 1311800 | 627680 | 14068000 | 6103800 | 10501000 | 40388000 | 6349400 | 4746300 | 9052700 | 7193700 | 2347900 | 3745100 |
| YIANTVELR | RPN2 | P04844 | RPN2_HUMAN | Unmodified | 0 | 0 | 0 | 0 | 201060 | 239060 | 0 | 142910 | 3405000 | 360480 | 742820 | 0 | 0 | 0 | 0 | 0 | 0 | 0 |
| YIAPTVLTD | ALDH3A2; ALDI | P51648; A0A | AL3A2_HUMAN | Unmodified | 0 | 0 | 0 | 0 | 0 | 0 | 0 | 0 | 2595700 | 425380 | 0 | 0 | 0 | 0 | 0 | 0 | 0 | 0 |
| YIAVSFIDP | PLOD2; PLOD2 | O00469; E7E | PLOD2_HUMAI | Unmodified | 0 | 0 | 0 | 0 | 0 | 0 | 0 | 0 | 657280 | 195300 | 27179 | 63844 | 0 | 0 | 0 | 0 | 0 | 0 |
| YIAVSFVDP | PLOD1 | Q02809 | PLOD1_HUMAI | Unmodified | 360500 | 0 | 12328 | 18854 | 0 | 0 | 0 | 0 | 2278800 | 547330 | 62974 | 361200 | 0 | 0 | 0 | 14871 | 0 | 133020 |
| YICPDKINS | DTX3L | Q8TDB6 | DTX3L_HUMAN | Unmodified | 0 | 82445 | 0 | 0 | 269540 | 0 | 0 | 0 | 0 | 0 | 0 | 0 | 507660 | 476790 | 192210 | 0 | 533050 | 638900 |
| YIDDKDVF | CUL2; CUL2; CI | Q13617; Q5T | CUL2_HUMAN | Unmodified | 20953 | 0 | 0 | 0 | 0 | 0 | 0 | 51945 | 226690 | 0 | 0 | 660520 | 0 | 0 | 42624 | 59597 | 236020 | 467500 |
| YIDENQDR | CNDP2; CNDP2 | Q96KP4; J3C | CNDP2_HUMA | Unmodified | 370170 | 39910 | 0 | 486520 | 1063700 | 0 | 0 | 169000 | 0 | 57346 | 0 | 1730900 | 494420 | 0 | 808160 | 470570 | 370490 | 0 |
| YIDENQDR | CNDP2; CNDP2 | Q96KP4; J3C | CNDP2_HUMA | Unmodified | 151340 | 117720 | 1000600 | 1330200 | 3209000 | 394000 | 1175700 | 659680 | 2835300 | 2345800 | 421960 | 7652900 | 27102000 | 10561000 | 1598700 | 1110000 | 1527200 | 1505400 |
| YIDENQDR | CNDP2; CNDP2 | Q96KP4; J3C | CNDP2_HUMA | Unmodified | 0 | 0 | 0 | 0 | 1435600 | 119590 | 152280 | 110980 | 403360 | 273990 | 0 | 861210 | 0 | 1919200 | 0 | 0 | 292310 | 713820 |
| YIDEVTR | CACNA2D2; CA | Q9NY47; C9 | CA2D2_HUMAI | Unmodified | 0 | 269240 | 0 | 0 | 114930 | 100430 | 0 | 0 | 803160 | 593780 | 671210 | 431920 | 24027 | 170210 | 0 | 0 | 245130 | 0 |
| YIDQEELNI | HSP90AA2P; H | Q14568; P08 | HS902_HUMAN | Unmodified | 5405100 | 4067700 | 10336000 | 15399000 | 89936000 | 17931000 | 9672000 | 4318400 | 52671000 | 29231000 | 55114000 | 2.94E+08 | 40476000 | 24058000 | 14558000 | 13683000 | 25362000 | 34317000 |
| YIDQEELNI | HSP90AA2P; H | Q14568; P08 | HS902_HUMAN | Unmodified | 0 | 0 | 0 | 0 | 504210 | 0 | 0 | 100970 | 0 | 218120 | 0 | 0 | 0 | 0 | 0 | 0 | 508000 | 332090 |
| YIDSADLEF | CTPS1; CTPS1; | P17812; B4E | PYRG1_HUMAI | Unmodified | 0 | 0 | 0 | 0 | 0 | 0 | 0 | 226510 | 0 | 0 | 1312600 | 7764700 | 0 | 0 | 0 | 0 | 0 | 0 |
| YIEACAR | UGDH; UGDH; | O60701; E7E | UGDH_HUMAN | Unmodified | 0 | 0 | 161830 | 278290 | 7619100 | 805140 | 177700 | 113820 | 0 | 19053 | 0 | 496330 | 1587100 | 1719500 | 65981 | 14827 | 0 | 470200 |
| YIEEAIEK | GLUL; GLUL | P15104; A0A | GLNA_HUMAN | Unmodified | 0 | 0 | 0 | 284740 | 627520 | 122730 | 222220 | 123040 | 548760 | 712780 | 533910 | 5750500 | 754150 | 211710 | 0 | 0 | 593940 | 327250 |
| YIEELAR | MFAP3L; MFAP | O75121; P5 | MFA3L_HUMAN | Unmodified | 0 | 0 | 0 | 0 | 0 | 0 | 0 | 0 | 0 | 0 | 0 | 0 | 0 | 0 | 364350 | 0 | 0 | 0 |
| YIEERLHTL | SMC3 | Q9UQE7 | SMC3_HUMAN | Unmodified | 0 | 0 | 0 | 0 | 785710 | 223300 | 0 | 0 | 0 | 0 | 0 | 0 | 0 | 0 | 0 | 0 | 0 | 0 |
| YIEIYVQK | CLTCL1; CLTC; | P53675; Q00 | CLH2_HUMAN | Unmodified | 4683400 | 3927400 | 1478000 | 790170 | 1059600 | 952240 | 1198100 | 173570 | 0 | 408460 | 0 | 8345600 | 540290 | 493790 | 3844100 | 3173700 | 474910 | 0 |
| YIELLTR | DSP | P15924 | DESP_HUMAN | Unmodified | 0 | 0 | 304920 | 69468 | 224010 | 0 | 45431 | 228180 | 1877900 | 292560 | 0 | 158580 | 0 | 328750 | 189270 | 197300 | 46161 | 0 |
| YIESPVLFLI | TTLL12 | Q14166 | TTL12_HUMAN | Unmodified | 0 | 0 | 227200 | 0 | 0 | 0 | 0 | 0 | 0 | 0 | 0 | 5205200 | 0 | 0 | 451360 | 307580 | 0 | 0 |
| YIESPVLFLI | TTLL12 | Q14166 | TTL12_HUMAN | Unmodified | 0 | 0 | 0 | 0 | 0 | 0 | 0 | 0 | 0 | 0 | 0 | 4217600 | 0 | 0 | 311750 | 0 | 0 | 0 |
| YIFPLDDK | PARP4 | Q9UKK3 | PARP4_HUMAN | Unmodified | 0 | 0 | 0 | 0 | 0 | 0 | 0 | 0 | 0 | 0 | 0 | 2113800 | 0 | 0 | 0 | 0 | 0 | 0 |
| YIFPTLDKP | ARHGEF2; ARH | Q92974; V9C | ARHG2_HUMA | Unmodified | 0 | 0 | 0 | 0 | 0 | 0 | 0 | 0 | 0 | 0 | 0 | 611680 | 0 | 0 | 0 | 0 | 0 | 0 |
| YIGNMHGI | CPE | P16870 | CBPE_HUMAN | Oxidation (M) | 0 | 0 | 0 | 0 | 0 | 473830 | 0 | 0 | 4256200 | 62030 | 0 | 0 | 0 | 328480 | 0 | 0 | 17003 | 0 |
| YIGNMHGI | CPE | P16870 | CBPE_HUMAN | Unmodified | 0 | 0 | 0 | 0 | 0 | 35667 | 29226 | 0 | 37263000 | 761610 | 109090 | 0 | 0 | 0 | 0 | 0 | 0 | 111940 |
| YIGNMHGI | CPD | O75976 | CBPD_HUMAN | Oxidation (M) | 0 | 0 | 0 | 0 | 0 | 0 | 0 | 0 | 8104300 | 258600 | 0 | 0 | 0 | 0 | 0 | 0 | 0 | 0 |
| YIGNMHGI | CPD | O75976 | CBPD_HUMAN | Unmodified | 0 | 0 | 616980 | 464230 | 133350 | 43237 | 19013 | 66588 | 41448000 | 1180000 | 254020 | 0 | 0 | 0 | 0 | 0 | 135410 | 0 |
| YIGTGHAD | SF3B5 | Q9BWJ5 | SF3B5_HUMAN | Unmodified | 575090 | 893220 | 247650 | 0 | 0 | 0 | 0 | 0 | 0 | 0 | 0 | 3053500 | 0 | 0 | 0 | 468380 | 0 | 0 |
| YIIDTYGSH | MANEAL; MANE | Q5VSG8; E9I | MANEL_HUMAI | Unmodified | 0 | 0 | 0 | 0 | 0 | 0 | 0 | 0 | 2743800 | 0 | 0 | 0 | 0 | 0 | 0 | 0 | 0 | 0 |
| YIIEELNVR | IARS1; IARS1; I | P41252; A0A | SYIC_HUMAN I | Unmodified | 85303 | 0 | 0 | 165270 | 0 | 0 | 0 | 0 | 0 | 243380 | 506220 | 1493700 | 347970 | 0 | 0 | 78454 | 0 | 0 |
| YIIEELNVR | IARS1; IARS1; I | P41252; A0A | SYIC_HUMAN I | Unmodified | 0 | 0 | 0 | 0 | 239120 | 0 | 0 | 0 | 512970 | 0 | 116200 | 8325300 | 0 | 0 | 0 | 0 | 0 | 0 |
| YIIHKYPSLI | HP1BP3; HP1BI | Q5SSJ5; B0C | HP1B3_HUMAI | Unmodified | 1606000 | 0 | 0 | 0 | 0 | 0 | 310190 | 0 | 0 | 0 | 0 | 0 | 0 | 0 | 0 | 0 | 0 | 224710 |
| YIIIGDMGV | RAB14; RAB14 | P61106; X6R | RAB14_HUMAN | Oxidation (M) | 0 | 647070 | 0 | 1185100 | 3393600 | 654240 | 1659600 | 1760600 | 1121900 | 896600 | 213110 | 4348100 | 0 | 0 | 0 | 1039000 | 384760 | 1039400 |
| YIIIGDMGV | RAB14; RAB14 | P61106; X6R | RAB14_HUMAN | Unmodified | 0 | 0 | 0 | 0 | 6341600 | 0 | 0 | 425420 | 1962000 | 1573400 | 0 | 2535900 | 949010 | 0 | 109090 | 91607 | 0 | 1906700 |
| YIIIGDTGV | RAB2A; RAB2B; | P61019; Q8 | RAB2A_HUMAN | Unmodified | 1049300 | 939230 | 658070 | 944800 | 1661100 | 0 | 1563800 | 1648900 | 7340400 | 11271000 | 2363600 | 10583000 | 0 | 0 | 812470 | 1466800 | 0 | 388910 |
| YIISLDQDS | PTGFRN | Q9P2B2 | FPRP_HUMAN | Unmodified | 0 | 0 | 6641300 | 140670 | 0 | 0 | 0 | 0 | 0 | 0 | 0 | 374270 | 0 | 0 | 0 | 0 | 0 | 0 |
| YIKDKYPNI | IMPDH2; IMPDI | P12268; A0A | IMDH2_HUMAN | Unmodified | 0 | 0 | 0 | 0 | 0 | 0 | 1152300 | 417570 | 0 | 0 | 0 | 0 | 0 | 0 | 0 | 0 | 1863300 | 0 |
| YILDGSGYI | ADI1; ADI1 | Q9BV57; A0A | MTND_HUMAN | Unmodified | 61525 | 85176 | 0 | 0 | 0 | 0 | 0 | 0 | 0 | 0 | 0 | 806320 | 0 | 0 | 0 | 0 | 0 | 84769 |
| YILGNPLTP | ALDH1A1 | P00352 | AL1A1_HUMAN | Unmodified | 0 | 0 | 0 | 0 | 351270 | 0 | 155300 | 0 | 0 | 571040 | 4856500 | 93467000 | 806480 | 635460 | 0 | 0 | 0 | 0 |
| YILGNPLTP | ALDH1A1 | P00352 | AL1A1_HUMAN | Unmodified | 0 | 0 | 0 | 0 | 0 | 0 | 0 | 0 | 3006800 | 1900700 | 27182000 | 5.77E+08 | 0 | 0 | 0 | 0 | 0 | 0 |
| YILRGDETF | NT5DC2; NT5D | Q9H857; H7 | NT5D2_HUMAI | Unmodified | 0 | 0 | 0 | 0 | 137910 | 0 | 0 | 0 | 1199600 | 194960 | 0 | 0 | 0 | 57133 | 0 | 0 | 49115 | 0 |

| | | | | | | | | | | | | | | | | | | | | | | |
|---|---|---|---|---|---|---|---|---|---|---|---|---|---|---|---|---|---|---|---|---|---|---|
| YILVTGGVI | CTPS2; CTPS1; | Q9NRF8; P1 | PYRG2_HUMAN | Unmodified | 0 | 0 | 0 | 0 | 618700 | 0 | 0 | 155840 | 1012500 | 779100 | 0 | 3401500 | 0 | 0 | 0 | 0 | 708500 | 0 |
| YILYDFSVT | MGAM2 | Q2M2H8 | MGAL_HUMAN | Unmodified | 0 | 0 | 0 | 0 | 0 | 0 | 0 | 0 | 0 | 111040 | 337110 | 1090400 | 0 | 0 | 0 | 0 | 0 | 0 |
| YIMQYIAAI | MYO1D; MYO1 | O94832; J3Q | MYO1D_HUMA | Oxidation (M) | 3962700 | 2473000 | 336930 | 673230 | 0 | 0 | 121590 | 0 | 150940 | 0 | 0 | 0 | 0 | 0 | 1602200 | 1195300 | 0 | 0 |
| YIMQYIAAI | MYO1D; MYO1 | O94832; J3Q | MYO1D_HUMA | Unmodified | 1612600 | 1475900 | 0 | 372640 | 0 | 67749 | 0 | 0 | 0 | 168540 | 0 | 99864 | 0 | 0 | 855420 | 409360 | 0 | 0 |
| YIMQYIAAI | MYO1D; MYO1 | O94832; J3Q | MYO1D_HUMA | Oxidation (M) | 726290 | 350880 | 0 | 0 | 0 | 0 | 0 | 0 | 0 | 0 | 0 | 0 | 0 | 0 | 0 | 0 | 0 | 0 |
| YINENLIVN | TCP1; TCP1; TC | P17987; F5H | TCPA_HUMAN | Unmodified | 134260 | 0 | 0 | 99564 | 355220 | 0 | 0 | 64284 | 0 | 145960 | 208590 | 865550 | 0 | 0 | 0 | 0 | 229460 | 0 |
| YINENLIVN | TCP1; TCP1; TC | P17987; F5H | TCPA_HUMAN | Unmodified | 0 | 0 | 0 | 0 | 0 | 0 | 0 | 0 | 0 | 0 | 0 | 2221500 | 0 | 0 | 0 | 0 | 0 | 0 |
| YINEQYEK | SEPTIN3; SEPTI | Q9UH03; B1 | SEPT3_HUMAN | Unmodified | 0 | 0 | 0 | 0 | 79957 | 0 | 388250 | 166740 | 0 | 0 | 0 | 114970 | 146310 | 222080 | 0 | 0 | 0 | 0 |
| YINILAYDH | PTPRG | P23470 | PTPRG_HUMAN | Unmodified | 105720 | 0 | 0 | 0 | 0 | 0 | 0 | 0 | 0 | 0 | 0 | 0 | 0 | 0 | 0 | 659730 | 0 | 0 |
| YINTEHGG | SEC23B; SEC23 | Q15437; A0A | SC23B_HUMAN | Unmodified | 0 | 0 | 33198 | 0 | 0 | 0 | 0 | 0 | 382980 | 15550 | 82433 | 1019500 | 206630 | 56028 | 0 | 41500 | 0 | 0 |
| YINTEHGG | SEC23B; SEC23 | Q15437; A0A | SC23B_HUMAN | Unmodified | 0 | 0 | 0 | 0 | 111490 | 0 | 139250 | 0 | 337630 | 209840 | 0 | 0 | 0 | 124370 | 0 | 0 | 383180 | 893890 |
| YIPLIQGIDI | DIP2A; DIP2A | Q14689; A0A | DIP2A_HUMAN | Unmodified | 0 | 0 | 0 | 0 | 2762500 | 0 | 0 | 0 | 5731900 | 3177000 | 324730 | 2795500 | 0 | 0 | 0 | 0 | 0 | 0 |
| YIPQQDALI | FARP1; FARP1; | Q9Y4F1; A0A | FARP1_HUMAN | Unmodified | 120640 | 0 | 597920 | 0 | 0 | 0 | 0 | 0 | 2529500 | 182850 | 1163100 | 2089000 | 0 | 0 | 1027300 | 539870 | 0 | 0 |
| YIQAACKT( | CLTCL1; CLTC; | P53675; Q00 | CLH2_HUMAN | Unmodified | 0 | 0 | 0 | 0 | 0 | 36468 | 278650 | 0 | 0 | 0 | 0 | 0 | 0 | 468250 | 0 | 0 | 241830 | 270130 |
| YIQELWR | RPL15; RPL15; | P61313; E7E | RL15_HUMAN ( | Unmodified | 4134200 | 3345800 | 2045200 | 0 | 52142 | 0 | 0 | 34841 | 0 | 0 | 252470 | 31494000 | 153780 | 0 | 1703900 | 1420100 | 17607 | 0 |
| YIQKPDNSI | ABI3BP; ABI3BF | Q7Z7G0; D3 | TARSH_HUMAN | Unmodified | 0 | 0 | 0 | 0 | 0 | 0 | 0 | 0 | 0 | 289590 | 0 | 0 | 0 | 0 | 0 | 0 | 0 | 222020 |
| YIQLPVTHL | PGAP1 | Q75T13 | PGAP1_HUMAN | Unmodified | 0 | 0 | 0 | 0 | 0 | 0 | 0 | 0 | 466570 | 174680 | 0 | 0 | 0 | 0 | 0 | 0 | 0 | 0 |
| YIQQTKPLT | NUDT21 | O43809 | CPSF5_HUMAN | Unmodified | 19276000 | 17238000 | 3754200 | 8649700 | 2050400 | 0 | 107340 | 192910 | 0 | 0 | 0 | 6698900 | 3281000 | 0 | 2129900 | 3900300 | 423190 | 0 |
| YIRCTSYNI | SEC24C; SEC24 | P53992; G5E | SC24C_HUMAN | Oxidation (M) | 0 | 0 | 143010 | 0 | 0 | 0 | 32620 | 23651 | 546320 | 0 | 0 | 0 | 0 | 0 | 0 | 0 | 247570 | 235840 |
| YIRCTSYNI | SEC24C; SEC24 | P53992; G5E | SC24C_HUMAN | Unmodified | 0 | 0 | 0 | 0 | 0 | 0 | 0 | 0 | 0 | 91759 | 0 | 368780 | 0 | 90215 | 0 | 0 | 548940 | 898250 |
| YIRHYEGLS | ADAM10; ADAM | O14672; C9J | ADA10_HUMAN | Unmodified | 0 | 0 | 0 | 0 | 259170 | 0 | 0 | 106380 | 0 | 458790 | 0 | 0 | 0 | 244520 | 0 | 0 | 0 | 1892800 |
| YIRKPIDYTI | ABI2; ABI2; ABI2 | Q9NYB9; H7 | ABI2_HUMAN A | Unmodified | 0 | 0 | 0 | 0 | 987900 | 0 | 0 | 0 | 0 | 0 | 0 | 1452100 | 0 | 0 | 0 | 0 | 0 | 2342900 |
| YIRKPIDYTI | ABI1; ABI1; ABI1 | Q8IZP0; A0A | ABI1_HUMAN A | Unmodified | 504740 | 0 | 699370 | 2276900 | 6595000 | 2734100 | 0 | 0 | 597010 | 276840 | 0 | 0 | 6760000 | 2005900 | 1212100 | 818720 | 5513700 | 22834000 |
| YISIGSEAEI | LNPEP | Q9UIQ6 | LCAP_HUMAN | Unmodified | 0 | 0 | 293410 | 47705 | 0 | 0 | 0 | 0 | 0 | 0 | 252000 | 926560 | 0 | 0 | 412450 | 224480 | 0 | 0 |
| YISPDQLAD | ENO1; ENO1 | P06733; A0A | ENOA_HUMAN | Unmodified | 3459700 | 0 | 5666100 | 6059900 | 11972000 | 3100300 | 0 | 496410 | 68202000 | 5481000 | 29459000 | 2.28E+08 | 4758300 | 1924900 | 13879000 | 11480000 | 3910400 | 3815000 |
| YISPDQLAD | ENO1; ENO1 | P06733; A0A | ENOA_HUMAN | Unmodified | 0 | 0 | 0 | 0 | 223450 | 0 | 0 | 0 | 0 | 65637 | 0 | 0 | 50846 | 229920 | 0 | 0 | 562910 | 0 |
| YISSQIERPI | SNRNP200 | O75643 | U520_HUMAN | Unmodified | 0 | 0 | 0 | 0 | 0 | 0 | 0 | 0 | 371300 | 0 | 127940 | 9076100 | 0 | 0 | 0 | 0 | 0 | 0 |
| YITDWQNV | PDCD6; PDCD6 | O75340; A0A | PDCD6_HUMA | Unmodified | 985160 | 29591000 | 323860 | 409580 | 0 | 379250 | 0 | 0 | 23498000 | 1598300 | 0 | 5719800 | 257780 | 0 | 1087900 | 613790 | 0 | 0 |
| YITLFIEK | STAMBP; STAMI | O95630; C9J | STABP_HUMAN | Unmodified | 974950 | 782120 | 140240 | 360130 | 0 | 0 | 0 | 55181 | 930090 | 0 | 215030 | 2694500 | 664060 | 386220 | 380710 | 293020 | 0 | 0 |
| YITLFIEKLP | STAMBP; STAMI | O95630; C9J | STABP_HUMAN | Unmodified | 0 | 0 | 0 | 0 | 1869400 | 584480 | 848430 | 0 | 0 | 942550 | 0 | 0 | 0 | 1415800 | 0 | 0 | 322020 | 583010 |
| YITQNGDY( | AP1M1; AP1M1 | Q9BXS5; K7I | AP1M1_HUMAN | Unmodified | 0 | 0 | 0 | 0 | 0 | 0 | 0 | 0 | 222400 | 0 | 0 | 929980 | 0 | 0 | 0 | 0 | 0 | 0 |
| YITQSGDY( | AP1M2; AP1M2 | Q9Y6Q5; K7I | AP1M2_HUMAN | Unmodified | 853800 | 36327 | 0 | 654860 | 0 | 0 | 0 | 0 | 0 | 341080 | 0 | 1567900 | 0 | 0 | 467020 | 0 | 0 | 0 |
| YIVERTMLC | NLN | Q9BYT8 | NEUL_HUMAN | Unmodified | 0 | 0 | 0 | 0 | 419550 | 0 | 0 | 0 | 0 | 0 | 0 | 0 | 97598 | 0 | 0 | 0 | 0 | 98811 |
| YIVHKEFDI | PLAT; PLAT; PLA | P00750; E7E | TPA_HUMAN Ti | Unmodified | 0 | 0 | 0 | 0 | 0 | 275840 | 0 | 0 | 0 | 0 | 0 | 0 | 0 | 0 | 0 | 0 | 330190 | 2068400 |
| YIVHLGQH | KLK11; KLK11 | Q9UBX7; M0 | KLK11_HUMAN | Unmodified | 0 | 0 | 0 | 0 | 0 | 0 | 0 | 0 | 2853200 | 665190 | 74480 | 367180 | 0 | 0 | 0 | 0 | 0 | 0 |
| YIVHLGQH | KLK11; KLK11 | Q9UBX7; M0 | KLK11_HUMAN | Unmodified | 0 | 0 | 0 | 0 | 0 | 0 | 0 | 0 | 11405000 | 484910 | 0 | 0 | 0 | 0 | 0 | 0 | 0 | 0 |
| YIVLEGSK | CIT; CIT; CIT; CI | O14578; A0A | CTRO_HUMAN | Unmodified | 0 | 0 | 0 | 0 | 616010 | 268300 | 0 | 0 | 0 | 0 | 0 | 0 | 0 | 0 | 83904 | 0 | 312090 | 0 |
| YIVQVDGK | MTCH1; MTCH1 | Q9NZJ7; H0 | MTCH1_HUMA | Unmodified | 2158900 | 550750 | 0 | 0 | 0 | 250620 | 0 | 110780 | 366440 | 122850 | 39233 | 0 | 230340 | 0 | 256030 | 0 | 104420 | 56487 |
| YIVQVSPLC | CPSF1 | Q10570 | CPSF1_HUMAN | Unmodified | 0 | 0 | 0 | 0 | 0 | 0 | 0 | 0 | 0 | 0 | 0 | 771170 | 0 | 0 | 0 | 0 | 0 | 0 |
| YIVSGTPTF | NRCAM; NRCAI | Q92823; C9J | NRCAM_HUMA | Unmodified | 0 | 0 | 0 | 0 | 0 | 0 | 0 | 0 | 848720 | 0 | 0 | 0 | 0 | 0 | 0 | 0 | 0 | 0 |
| YIYDQCPA\ | UQCRC1 | P31930 | QCR1_HUMAN | Unmodified | 0 | 1094300 | 0 | 0 | 0 | 0 | 0 | 0 | 7188200 | 0 | 0 | 0 | 0 | 0 | 0 | 0 | 331350 | 0 |
| YIYDQCPA\ | UQCRC1 | P31930 | QCR1_HUMAN | Unmodified | 0 | 0 | 0 | 0 | 0 | 0 | 0 | 0 | 20263000 | 5174300 | 0 | 0 | 0 | 0 | 0 | 0 | 894900 | 0 |
| YIYDSAFHI | SFXN1; SFXN1; | Q9H9B4; D6 | SFXN1_HUMAN | Unmodified | 0 | 0 | 0 | 56115 | 0 | 0 | 120030 | 60635 | 5622600 | 948120 | 0 | 822260 | 0 | 0 | 0 | 47201 | 0 | 0 |
| YIYKGFEKP | ARPC5L | Q9BPX5 | ARP5L_HUMAN | Unmodified | 0 | 0 | 0 | 0 | 0 | 0 | 1452500 | 0 | 0 | 0 | 0 | 0 | 0 | 991090 | 0 | 0 | 946970 | 0 |
| YIYNREEYA | HLA-DQB1; HL/ | Q5SU54; A0 | Q5SU54_HUM/ | Unmodified | 0 | 0 | 0 | 0 | 0 | 0 | 421140 | 396200 | 7104500 | 184550 | 0 | 0 | 0 | 0 | 0 | 0 | 0 | 0 |
| YIYNREEYC | HLA-DQB2; HL/ | P05538; Q5S | DQB2_HUMAN | Unmodified | 0 | 0 | 0 | 0 | 0 | 1673600 | 308220 | 0 | 3611400 | 463430 | 0 | 0 | 0 | 241040 | 0 | 0 | 0 | 0 |
| YIYNRSPAI | PMPCB | O75439 | MPPB_HUMAN | Unmodified | 0 | 0 | 0 | 0 | 233010 | 0 | 0 | 0 | 2423400 | 886980 | 0 | 0 | 0 | 0 | 0 | 0 | 0 | 0 |
| YIYVADILAI | PON2; PON2; P | Q15165; A0A | PON2_HUMAN | Unmodified | 0 | 0 | 0 | 0 | 0 | 0 | 0 | 0 | 757210 | 101160 | 114590 | 0 | 0 | 0 | 0 | 0 | 0 | 0 |
| YKAEDEVQ | HSPA1A; HSPA | P0DMV8; P0 | HS71A_HUMAI | Unmodified | 0 | 7698200 | 2850500 | 0 | 0 | 0 | 63084 | 0 | 0 | 0 | 0 | 2005000 | 0 | 0 | 10818 | 5860500 | 0 | 0 |
| YKAEWRAV | NCAM1; NCAM | P13591; A0A | NCAM1_HUMAN | Unmodified | 0 | 0 | 0 | 0 | 0 | 0 | 2009400 | 0 | 0 | 335540 | 0 | 0 | 0 | 0 | 0 | 0 | 0 | 0 |
| YKAGGVAS | SCIN | Q9Y6U3 | ADSV_HUMAN | Unmodified | 0 | 0 | 0 | 0 | 0 | 0 | 0 | 0 | 0 | 0 | 0 | 0 | 0 | 0 | 0 | 0 | 0 | 0 |
| YKASITALE/ | MYH9 | P35579 | MYH9_HUMAN | Unmodified | 0 | 0 | 0 | 0 | 1075300 | 0 | 0 | 0 | 0 | 0 | 0 | 0 | 0 | 0 | 0 | 0 | 1703800 | 364770 |
| YKASYDVS | PFKP; PFKP; PF | Q01813; A0A | PFKAP_HUMAN | Unmodified | 0 | 0 | 0 | 0 | 0 | 0 | 0 | 0 | 0 | 0 | 0 | 0 | 372910 | 0 | 0 | 0 | 2122600 | 1871000 |
| YKDAANLLI | KLC1; KLC1; KL | Q07866; G3\ | KLC1_HUMAN | Unmodified | 0 | 0 | 0 | 0 | 3343100 | 393110 | 401490 | 0 | 315610 | 558120 | 0 | 0 | 725250 | 740240 | 0 | 0 | 838380 | 1939100 |
| YKDAIHFYI | STIP1 | P31948 | STIP1_HUMAN | Unmodified | 0 | 0 | 0 | 453760 | 4409300 | 0 | 619870 | 235560 | 1703000 | 2651300 | 0 | 582120 | 5832300 | 4528800 | 0 | 0 | 2863500 | 2286200 |
| YKDDDDD( | PLOD3; PLOD3 | O60568; H7( | PLOD3_HUMAI | Unmodified | 0 | 0 | 0 | 67608 | 0 | 0 | 0 | 0 | 7248000 | 361510 | 0 | 0 | 0 | 33485 | 0 | 0 | 0 | 0 |
| YKDIIRQPS | PEA15; PEA15 | Q15121; B1/ | PEA15_HUMAN | Unmodified | 0 | 0 | 0 | 469610 | 1303500 | 243950 | 67972 | 117850 | 0 | 0 | 0 | 0 | 1988900 | 2250400 | 68825 | 95636 | 872330 | 2145600 |
| YKDLDEDE | TMOD3; TMOD3 | Q9NYL9; H0 | TMOD3_HUMA | Unmodified | 0 | 0 | 0 | 652780 | 9101500 | 1193300 | 0 | 0 | 738720 | 320020 | 0 | 1947000 | 1883100 | 1356400 | 0 | 848460 | 0 | 1073900 |
| YKDNFTAA( | EPHA7; EPHA7 | Q15375; A0A | EPHA7_HUMAI | Unmodified | 238880 | 544810 | 0 | 77043 | 0 | 0 | 0 | 0 | 141190 | 0 | 0 | 749470 | 0 | 0 | 0 | 0 | 0 | 0 |
| YKDSAQRD | ATP1B1; ATP1B | P05026; V9C | AT1B1_HUMAN | Oxidation (M) | 711150 | 254420 | 0 | 2526800 | 1101000 | 324760 | 572420 | 179690 | 4906900 | 1312400 | 2717500 | 17295000 | 2863800 | 0 | 3135200 | 2277900 | 349100 | 0 |
| YKDSAQRD | ATP1B1; ATP1B | P05026; V9C | AT1B1_HUMAN | Unmodified | 0 | 0 | 0 | 3075700 | 2544200 | 0 | 0 | 0 | 8918900 | 1119000 | 0 | 8467100 | 1392900 | 0 | 3012300 | 681640 | 556040 | 0 |
| YKEAAHLLI | KLC2; KLC2 | Q9H0B6; A8 | KLC2_HUMAN | Unmodified | 0 | 0 | 0 | 0 | 452310 | 0 | 0 | 0 | 0 | 85803 | 0 | 0 | 756360 | 0 | 0 | 0 | 615990 | 0 |
| YKEEFEK | LASP1; LASP1 | Q14847; C9J | LASP1_HUMAN | Unmodified | 1517300 | 549750 | 0 | 0 | 0 | 0 | 0 | 0 | 0 | 0 | 0 | 419580 | 0 | 233580 | 0 | 398790 | 0 | 0 |
| YKEETIEK | RPL26 | P61254 | RL26_HUMAN ( | Unmodified | 1842800 | 0 | 0 | 0 | 0 | 0 | 0 | 0 | 28856 | 0 | 0 | 4830700 | 0 | 0 | 0 | 1150200 | 0 | 0 |
| YKENPDIVI | ARIH2 | O95376 | ARI2_HUMAN E | Unmodified | 0 | 0 | 0 | 0 | 0 | 0 | 0 | 0 | 101210 | 0 | 302150 | 671810 | 0 | 0 | 0 | 0 | 0 | 0 |
| YKEQVETLV | DST; DST; DST; | Q03001; A0A | DYST_HUMAN | Unmodified | 0 | 0 | 0 | 0 | 2178100 | 0 | 0 | 0 | 93809 | 0 | 0 | 180960 | 0 | 0 | 225350 | 0 | 0 | 0 |
| YKERGTVL/ | SNF8; SNF8; SN | Q96H20; D6 | SNF8_HUMAN | Unmodified | 0 | 0 | 0 | 0 | 245750 | 0 | 0 | 83802 | 0 | 0 | 0 | 0 | 461550 | 480620 | 0 | 0 | 0 | 0 |
| YKESFDVG | CLIC2; CLIC2 | O15247; A6I | CLIC2_HUMAN | Unmodified | 0 | 0 | 0 | 0 | 0 | 0 | 0 | 0 | 0 | 0 | 0 | 0 | 0 | 3665900 | 0 | 0 | 0 | 0 |
| YKETDLLILI | SSB | P05455 | LA_HUMAN Lu | Unmodified | 0 | 0 | 0 | 0 | 0 | 0 | 0 | 0 | 0 | 402650 | 2717600 | 1283200 | 0 | 0 | 0 | 0 | 0 | 0 |
| YKETFSAPS | KIF14 | Q15058 | KIF14_HUMAN | Unmodified | 0 | 0 | 0 | 0 | 1078400 | 233360 | 218000 | 0 | 0 | 0 | 0 | 0 | 0 | 0 | 0 | 0 | 491730 | 348320 |
| YKFENWG/ | MDK; MDK; MD | P21741; E9P | MK_HUMAN Mi | Unmodified | 0 | 0 | 0 | 0 | 0 | 0 | 0 | 0 | 4919500 | 1442100 | 0 | 0 | 0 | 0 | 0 | 0 | 0 | 0 |
| YKFEVYEKI | DHFR | P00374 | DYR_HUMAN D | Unmodified | 0 | 0 | 0 | 129040 | 1081700 | 159470 | 378280 | 242280 | 0 | 145680 | 0 | 0 | 1089700 | 731680 | 0 | 0 | 201530 | 238840 |
| YKFMELNL | VBP1 | P61758 | PFD3_HUMAN | Unmodified | 0 | 0 | 0 | 0 | 564270 | 0 | 0 | 0 | 0 | 0 | 0 | 0 | 0 | 0 | 0 | 0 | 1106900 | 0 |
| YKGESVQC | TMEM87A; TME | Q8NBN3; H3 | TM87A_HUMAN | Unmodified | 0 | 0 | 0 | 0 | 0 | 0 | 0 | 0 | 0 | 0 | 0 | 240590 | 0 | 0 | 0 | 0 | 0 | 0 |
| YKGICTED( | JAK1; JAK1; JAK | P23458; A0A | JAK1_HUMAN T | Unmodified | 0 | 0 | 0 | 0 | 200870 | 0 | 0 | 0 | 0 | 0 | 0 | 0 | 0 | 0 | 0 | 0 | 64859 | 101660 |
| YKGIFNGF | SLC25A3; SLC2 | Q00325; F8\ | MPCP_HUMAN | Unmodified | 0 | 0 | 0 | 0 | 0 | 0 | 0 | 0 | 1341000 | 133860 | 0 | 0 | 0 | 0 | 0 | 0 | 45262 | 0 |
| YKGQEAFV | CA12; CA12 | O43570; B3I | CAH12_HUMAI | Unmodified | 1421000 | 464520 | 0 | 0 | 0 | 0 | 0 | 0 | 0 | 0 | 0 | 188540 | 0 | 0 | 327790 | 180310 | 0 | 0 |
| YKGQEAFV | CA12; CA12 | O43570; B3I | CAH12_HUMAI | Unmodified | 954120 | 1001400 | 0 | 336830 | 0 | 0 | 0 | 0 | 0 | 0 | 0 | 0 | 0 | 0 | 343010 | 0 | 0 | 0 |
| YKGQHVP( | FLNA; FLNA; FL | P21333; A0A | FLNA_HUMAN | Unmodified | 0 | 0 | 0 | 1209000 | 2300200 | 853660 | 6644300 | 0 | 0 | 424390 | 0 | 0 | 737190 | 0 | 0 | 0 | 2262700 | 0 |
| YKGVWRPI | CLGN | O14967 | CLGN_HUMAN | Unmodified | 0 | 0 | 0 | 0 | 0 | 0 | 0 | 0 | 3545600 | 2324500 | 0 | 0 | 0 | 0 | 0 | 0 | 0 | 0 |
| YKHFDWLY | SNX9; SNX9; SN | Q9Y5X1; A0/ | SNX9_HUMAN | Unmodified | 1193800 | 993650 | 0 | 0 | 0 | 0 | 0 | 0 | 0 | 0 | 0 | 0 | 0 | 0 | 0 | 0 | 0 | 0 |
| YKIQILTVS( | PTPRB; PTPRB; | P23467; F8V | PTPRB_HUMAN | Unmodified | 84261 | 59655 | 0 | 0 | 0 | 0 | 69180 | 0 | 0 | 0 | 0 | 0 | 80836 | 0 | 0 | 0 | 0 | 0 |
| YKKVENLIA | LAMB1; LAMB1 | P07942; G3\ | LAMB1_HUMAN | Unmodified | 0 | 0 | 0 | 0 | 74154 | 0 | 38752 | 19489 | 0 | 0 | 0 | 0 | 39158 | 88794 | 0 | 0 | 43818 | 797980 |
| YKLDKESYI | ERP29; ERP29 | P30040; F8V | ERP29_HUMAN | Unmodified | 0 | 0 | 0 | 0 | 0 | 0 | 620670 | 0 | 3220700 | 1059100 | 0 | 0 | 0 | 216440 | 0 | 0 | 0 | 0 |
| YKLLPEYP( | DHFR | P00374 | DYR_HUMAN D | Unmodified | 105680 | 0 | 0 | 0 | 1087500 | 0 | 225640 | 0 | 0 | 0 | 0 | 0 | 1605300 | 0 | 0 | 0 | 0 | 0 |
| YKLPVHTEI | GEMIN5 | Q8TEQ6 | GEMI5_HUMAN | Unmodified | 0 | 0 | 0 | 0 | 0 | 0 | 0 | 0 | 0 | 108840 | 0 | 691890 | 0 | 0 | 0 | 0 | 0 | 0 |
| YKLPVHTEI | GEMIN5 | Q8TEQ6 | GEMI5_HUMAN | Unmodified | 0 | 0 | 0 | 0 | 0 | 0 | 0 | 0 | 0 | 0 | 305520 | 1421100 | 0 | 0 | 0 | 0 | 0 | 0 |
| YKLSEIHHL | ZC3HAV1; ZC3I | Q7Z2W4; C9 | ZCCHV_HUMA | Unmodified | 0 | 0 | 0 | 0 | 1187300 | 149910 | 0 | 0 | 0 | 0 | 0 | 0 | 0 | 504450 | 0 | 0 | 144120 | 121220 |
| YKPAVNQII | AKR1B1; AKR1E | P15121; E9P | ALDR_HUMAN | Unmodified | 0 | 0 | 0 | 0 | 9558300 | 1270900 | 1242500 | 428410 | 0 | 0 | 2626000 | 22407000 | 0 | 232320 | 0 | 0 | 0 | 393960 |
| YKPESEELT | P4HB; P4HB; P4 | P07237; A0A | PDIA1_HUMAN | Unmodified | 564480 | 115790 | 496990 | 198030 | 0 | 0 | 110530 | 141880 | 94553000 | 3317700 | 7270300 | 7215500 | 124920 | 136830 | 765410 | 768990 | 0 | 0 |
| YKPESEELT | P4HB; P4HB; P4 | P07237; A0A | PDIA1_HUMAN | Unmodified | 0 | 0 | 0 | 0 | 0 | 193140 | 0 | 0 | 1.14E+08 | 4065200 | 6385700 | 0 | 0 | 0 | 0 | 0 | 0 | 0 |
| YKPLSSSA\ | LRATD2 | Q96KN1 | LRAT2_HUMAN | Unmodified | 41038 | 0 | 0 | 0 | 0 | 0 | 216860 | 64385 | 1049100 | 0 | 0 | 0 | 21730 | 0 | 130240 | 203310 | 0 | 0 |
| YKPNSSHC | KIF14 | Q15058 | KIF14_HUMAN | Unmodified | 0 | 0 | 0 | 0 | 2608200 | 255550 | 0 | 0 | 0 | 0 | 0 | 0 | 0 | 0 | 0 | 0 | 1729700 | 336360 |
| YKPVTNQV | AKR1B15; AKR1 | C9JRZ8; O6C | AK1BF_HUMAN | Unmodified | 0 | 0 | 0 | 0 | 333400 | 0 | 580440 | 1548300 | 0 | 0 | 0 | 0 | 0 | 0 | 0 | 0 | 0 | 0 |
| YKQDVYLLI | AHCYL1; AHCY | O43865; Q9I | SAHH2_HUMAI | Unmodified | 1064300 | 0 | 2111800 | 1437000 | 320560 | 1289800 | 0 | 0 | 756010 | 0 | 0 | 534270 | 486790 | 0 | 2516000 | 1813700 | 846410 | 707870 |
| YKRPGYGA | EIF2S1; EIF2S1 | P05198; H0\ | IF2A_HUMAN E | Unmodified | 0 | 149650 | 0 | 277630 | 294360 | 0 | 28984 | 16262 | 281230 | 0 | 154320 | 1165900 | 185720 | 0 | 0 | 0 | 0 | 115010 |
| YKTPQPIGE | CLGN; CLGN | O14967; D6I | CLGN_HUMAN | Unmodified | 0 | 0 | 0 | 0 | 0 | 0 | 0 | 384050 | 7881700 | 0 | 0 | 0 | 0 | 0 | 0 | 0 | 0 | 0 |
| YKTTNQSTI | SLITRK6 | Q9H5Y7 | SLIK6_HUMAN | Unmodified | 0 | 0 | 0 | 0 | 0 | 0 | 808360 | 216690 | 0 | 625070 | 0 | 0 | 0 | 1246800 | 0 | 0 | 0 | 4055100 |
| YKTTNQSTI | SLITRK6 | Q9H5Y7 | SLIK6_HUMAN | Unmodified | 0 | 0 | 0 | 0 | 1140200 | 0 | 0 | 0 | 0 | 2931700 | 0 | 0 | 0 | 0 | 0 | 0 | 0 | 0 |
| YKVEGFPTI | PDIA4; PDIA4 | P13667; A0A | PDIA4_HUMAN | Unmodified | 26955 | 29331 | 0 | 0 | 0 | 0 | 0 | 0 | 4329900 | 116810 | 0 | 0 | 0 | 0 | 115760 | 50456 | 0 | 0 |
| YKVPDGKP | PPA1 | Q15181 | IPYR_HUMAN I | Unmodified | 69569 | 0 | 0 | 0 | 567170 | 0 | 0 | 60621 | 295630 | 0 | 3302700 | 6526500 | 769790 | 0 | 0 | 0 | 0 | 218100 |
| YKVTCQSS | MGAT5 | Q09328 | MGT5A_HUMAI | Unmodified | 0 | 0 | 0 | 0 | 0 | 0 | 0 | 0 | 682860 | 114700 | 97595 | 69793 | 0 | 0 | 0 | 0 | 0 | 0 |
| YKVVLSSQI | NOMO1; NOMC | Q15155; P6 | NOMO1_HUM/ | Unmodified | 0 | 0 | 0 | 0 | 0 | 0 | 51479 | 30849 | 449600 | 1156400 | 0 | 52100 | 0 | 0 | 0 | 0 | 0 | 26314 |
| YKWCEYGI | VDAC2; VDAC2 | P45880; Q5J | VDAC2_HUMAN | Unmodified | 0 | 0 | 416680 | 0 | 393270 | 0 | 1174900 | 30336 | 67245000 | 48825000 | 476650 | 0 | 247190 | 399300 | 0 | 12809 | 186400 | 597690 |
| YKYWFSQF | LMF2 | Q9BU23 | LMF2_HUMAN | Unmodified | 0 | 0 | 0 | 0 | 0 | 0 | 0 | 0 | 0 | 239260 | 0 | 0 | 0 | 0 | 0 | 0 | 0 | 0 |
| YLAADKDG | FSCN1 | Q16658 | FSCN1_HUMAI | Unmodified | 0 | 0 | 0 | 0 | 702570 | 62621 | 0 | 0 | 0 | 24632 | 726660 | 16372000 | 0 | 0 | 0 | 0 | 183120 | 650710 |
| YLAADKDG | FSCN1 | Q16658 | FSCN1_HUMAI | Unmodified | 0 | 0 | 0 | 0 | 764000 | 0 | 0 | 0 | 813950 | 0 | 991790 | 3495400 | 0 | 0 | 0 | 0 | 0 | 0 |
| YLAEFATGI | YWHAE; YWHAI | P62258; K7E | 1433E_HUMAN | Unmodified | 0 | 0 | 288860 | 169970 | 0 | 0 | 0 | 0 | 0 | 0 | 0 | 8875400 | 0 | 0 | 1152300 | 1567800 | 0 | 0 |
| YLAEFATGI | YWHAE; YWHAI | P62258; K7E | 1433E_HUMAN | Unmodified | 3736900 | 3634000 | 13775000 | 6980800 | 3444800 | 614980 | 205250 | 91033 | 9132800 | 1309300 | 8463200 | 1.54E+08 | 657320 | 0 | 49534000 | 37546000 | 343140 | 308830 |

| | | | | | | | | | | | | | | | | | | | | | | |
|---|---|---|---|---|---|---|---|---|---|---|---|---|---|---|---|---|---|---|---|---|---|---|
| YLAEFATGI | YWHAE; YWHAI | P62258; K7E | 1433E_HUMAN | Unmodified | 2073400 | 0 | 0 | 21129000 | 3362900 | 2728400 | 0 | 97178 | 12986000 | 3341500 | 6073500 | 18559000 | 4993100 | 3604000 | 7522700 | 6259000 | 3506100 | 469980 |
| YLAEKYEW | EEF2 | P13639 | EF2_HUMAN EI | Unmodified | 1282300 | 904000 | 183580 | 2208900 | 6082000 | 997230 | 648270 | 0 | 2381000 | 868190 | 2085000 | 17746000 | 2701300 | 1252100 | 1304100 | 1344100 | 3200800 | 2320200 |
| YLAEKYEW | EEF2 | P13639 | EF2_HUMAN EI | Unmodified | 3202500 | 5429800 | 252930 | 4633000 | 44735000 | 8368400 | 5221300 | 899980 | 5226900 | 5082500 | 16216000 | 16513000 | 18158000 | 7226800 | 1553100 | 3186300 | 8756800 | 18576000 |
| YLAEVAAGI | YWHAZ; YWHA | P63104; B7Z | 1433Z_HUMAN | Unmodified | 1496100 | 1284700 | 2305200 | 1162000 | 428500 | 514510 | 0 | 0 | 7210000 | 711380 | 2212400 | 15436000 | 0 | 0 | 5391700 | 2624800 | 180790 | 122740 |
| YLAEVAAGI | YWHAZ; YWHA | P63104; B7Z | 1433Z_HUMAN | Unmodified | 3426500 | 2017900 | 6599000 | 5870500 | 2769100 | 0 | 288370 | 343720 | 26431000 | 4027300 | 8498600 | 50373000 | 921990 | 0 | 9841000 | 6440000 | 1468600 | 1968500 |
| YLAEVACG | YWHAQ; YWHA | P27348; E9P | 1433T_HUMAN | Unmodified | 0 | 0 | 0 | 0 | 0 | 0 | 0 | 40272 | 448800 | 0 | 221260 | 4332100 | 0 | 303910 | 0 | 103230 | 0 | 0 |
| YLAEVACG | YWHAQ; YWHA | P27348; E9P | 1433T_HUMAN | Unmodified | 0 | 0 | 0 | 381390 | 0 | 263520 | 59966 | 0 | 712100 | 0 | 1816500 | 19818000 | 0 | 0 | 0 | 0 | 207780 | 0 |
| YLAEVASGI | YWHAH; YWHA | Q04917; A2I | 1433F_HUMAN | Unmodified | 0 | 0 | 0 | 1145900 | 3588700 | 1094400 | 0 | 0 | 0 | 0 | 2794700 | 9859600 | 1080400 | 901600 | 1483700 | 1647900 | 0 | 0 |
| YLAEVASGI | YWHAH; YWHA | Q04917; A2I | 1433F_HUMAN | Unmodified | 95718 | 0 | 154980 | 0 | 0 | 0 | 0 | 0 | 241730 | 51784 | 245160 | 1309700 | 69433 | 0 | 174030 | 192480 | 0 | 0 |
| YLAEVATGI | SFN | P31947 | 1433S_HUMAN | Unmodified | 953280 | 633850 | 0 | 0 | 0 | 83096 | 156700 | 245190 | 1803800 | 0 | 0 | 0 | 0 | 0 | 0 | 0 | 0 | 0 |
| YLAEVATGI | YWHAG | P61981 | 1433G_HUMAN | Unmodified | 423220 | 287110 | 5725100 | 6884000 | 5581100 | 4105200 | 877830 | 663450 | 5114400 | 2266000 | 7421300 | 33066000 | 7608500 | 159360 | 7932200 | 6682600 | 6503100 | 911970 |
| YLAEVATGI | YWHAG | P61981 | 1433G_HUMAN | Unmodified | 0 | 0 | 0 | 0 | 149050 | 0 | 0 | 0 | 1381700 | 0 | 2451400 | 27531000 | 0 | 0 | 0 | 0 | 355690 | 0 |
| YLAEVATGI | YWHAG | P61981 | 1433G_HUMAN | Unmodified | 0 | 0 | 0 | 0 | 551330 | 0 | 0 | 89686 | 0 | 0 | 0 | 0 | 1362800 | 0 | 0 | 0 | 1616100 | 266090 |
| YLAIAPPIIK | DDB1; DDB1; D | Q16531; A0A | DDB1_HUMAN | Unmodified | 0 | 0 | 160290 | 0 | 120740 | 0 | 0 | 0 | 920090 | 466500 | 1219900 | 4008500 | 0 | 0 | 96811 | 157430 | 0 | 0 |
| YLALESMC | AP2A2; AP2A1; | O94973; O9 | AP2A2_HUMAN | Unmodified | 0 | 0 | 0 | 0 | 0 | 0 | 0 | 0 | 0 | 0 | 0 | 1202500 | 0 | 0 | 0 | 0 | 0 | 0 |
| YLANIEQQI | NUP50 | Q9UKX7 | NUP50_HUMAI | Unmodified | 0 | 0 | 0 | 0 | 0 | 303890 | 0 | 0 | 89742 | 0 | 307500 | 1573100 | 0 | 0 | 0 | 0 | 195090 | 0 |
| YLANMNYV | EPHA2 | P29317 | EPHA2_HUMAI | Unmodified | 0 | 0 | 0 | 0 | 0 | 0 | 0 | 0 | 0 | 0 | 0 | 2615100 | 0 | 0 | 0 | 0 | 0 | 0 |
| YLANMNYV | EPHA2 | P29317 | EPHA2_HUMAI | Oxidation (M) | 0 | 0 | 0 | 0 | 0 | 0 | 0 | 0 | 0 | 0 | 34724 | 552160 | 0 | 0 | 0 | 0 | 55364 | 45371 |
| YLANMNYV | EPHA2 | P29317 | EPHA2_HUMAI | Unmodified | 238360 | 0 | 0 | 0 | 2198000 | 713140 | 0 | 0 | 0 | 0 | 0 | 0 | 0 | 0 | 0 | 152790 | 2142600 | 1474000 |
| YLANMNYV | EPHA2 | P29317 | EPHA2_HUMAI | Unmodified | 0 | 0 | 0 | 0 | 0 | 0 | 0 | 0 | 0 | 0 | 0 | 0 | 0 | 0 | 2020500 | 0 | 1481600 | 0 |
| YLAPSGPS | FSCN1 | Q16658 | FSCN1_HUMAI | Unmodified | 0 | 0 | 0 | 0 | 3536600 | 802000 | 71266 | 0 | 5612300 | 507440 | 10908000 | 46951000 | 0 | 0 | 0 | 0 | 1016200 | 2570200 |
| YLATASTMI | MBP; MBP; MBF | P02686; H7E | MBP_HUMAN N | Oxidation (M) | 113280 | 78354 | 0 | 286460 | 386900 | 113460 | 0 | 0 | 633980 | 15157 | 142770 | 61186 | 335270 | 0 | 141070 | 37076 | 128180 | 231140 |
| YLATLNFVI | DDR1; DDR1; D | Q08345; A0A | DDR1_HUMAN | Unmodified | 0 | 0 | 0 | 0 | 0 | 0 | 199140 | 0 | 338050 | 752120 | 0 | 1289500 | 0 | 0 | 0 | 0 | 0 | 0 |
| YLATSQPRI | TTYH3 | Q9C0H2 | TTYH3_HUMAN | Unmodified | 464530 | 339630 | 0 | 798300 | 209830 | 599160 | 309780 | 364200 | 1944200 | 230100 | 764170 | 1739400 | 1916600 | 922020 | 0 | 582830 | 1937200 | 4678500 |
| YLAYKQPG | FUT11 | Q495W5 | FUT11_HUMAN | Unmodified | 0 | 0 | 0 | 0 | 0 | 0 | 0 | 0 | 3173900 | 2617100 | 0 | 0 | 0 | 0 | 0 | 0 | 0 | 0 |
| YLCQSNAL | WWP1; WWP1 | Q9H0M0; H0 | WWP1_HUMAN | Unmodified | 540040 | 257870 | 503150 | 1230400 | 763580 | 0 | 0 | 0 | 0 | 0 | 0 | 0 | 0 | 0 | 1529600 | 1140300 | 0 | 0 |
| YLCSACR | TRAF2; TRAF2; T | Q12933; B1A | TRAF2_HUMAN | Unmodified | 290050 | 0 | 0 | 0 | 1125700 | 743930 | 0 | 0 | 0 | 0 | 0 | 404600 | 0 | 0 | 0 | 0 | 0 | 0 |
| YLDEDTIYI | MAT2A | P31153 | METK2_HUMAN | Unmodified | 228960 | 0 | 596130 | 648860 | 0 | 0 | 0 | 0 | 1198700 | 0 | 1056500 | 9153600 | 0 | 0 | 764840 | 1030300 | 321010 | 0 |
| YLDELMK | IQGAP1; IQGAP | P46940; A0A | IQGA1_HUMAN | Unmodified | 0 | 0 | 0 | 0 | 0 | 0 | 0 | 0 | 0 | 27489 | 0 | 247360 | 0 | 0 | 0 | 0 | 36103 | 0 |
| YLDESSGG | DNASE2; DNAS | O00115; K7E | DNS2A_HUMAI | Unmodified | 0 | 0 | 0 | 0 | 0 | 0 | 0 | 0 | 709630 | 0 | 0 | 0 | 0 | 0 | 0 | 0 | 0 | 0 |
| YLDESSGG | DNASE2; DNAS | O00115; K7E | DNS2A_HUMAI | Unmodified | 0 | 0 | 0 | 0 | 1019800 | 0 | 263410 | 0 | 0 | 13558000 | 0 | 0 | 869880 | 0 | 0 | 0 | 0 | 506810 |
| YLDFKVTE | GDI2; GDI2; GD | P50395; V9G | GDIB_HUMAN | Unmodified | 332200 | 283770 | 0 | 418910 | 257650 | 0 | 40972 | 0 | 1055000 | 304970 | 0 | 0 | 0 | 0 | 414050 | 254800 | 0 | 0 |
| YLDFVFAVI | LAMP2; LAMP2 | P13473; H0Y | LAMP2_HUMAI | Unmodified | 20593 | 194650 | 301020 | 491030 | 0 | 0 | 0 | 0 | 1705900 | 3118000 | 1281500 | 223670 | 0 | 0 | 368270 | 167200 | 0 | 10122 |
| YLDFVFAVI | LAMP2; LAMP2 | P13473; H0Y | LAMP2_HUMAI | Unmodified | 0 | 0 | 0 | 0 | 0 | 0 | 0 | 0 | 1452100 | 0 | 0 | 0 | 0 | 0 | 0 | 0 | 0 | 0 |
| YLDFVFAVI | LAMP2; LAMP2 | P13473; H0Y | LAMP2_HUMAI | Unmodified | 0 | 0 | 0 | 0 | 0 | 0 | 608080 | 612710 | 0 | 2885100 | 0 | 0 | 804140 | 0 | 0 | 0 | 604850 | 0 |
| YLDIPKMLI | ACTN1; ACTN1 | P12814; G3V | ACTN1_HUMAI | Oxidation (M) | 0 | 0 | 0 | 0 | 394790 | 0 | 389700 | 42177 | 0 | 0 | 0 | 0 | 0 | 0 | 0 | 0 | 1476000 | 0 |
| YLDIPKMLI | ACTN1; ACTN1 | P12814; G3V | ACTN1_HUMAI | Unmodified | 0 | 0 | 0 | 0 | 492850 | 0 | 446980 | 0 | 0 | 155280 | 0 | 0 | 0 | 0 | 0 | 0 | 1574100 | 1029700 |
| YLDIPKMLI | ACTN4 | O43707 | ACTN4_HUMAI | Oxidation (M) | 0 | 465350 | 0 | 224820 | 1898800 | 0 | 467370 | 0 | 0 | 0 | 0 | 1033500 | 0 | 0 | 0 | 0 | 3128700 | 0 |
| YLDIPKMLI | ACTN4 | O43707 | ACTN4_HUMAI | Unmodified | 0 | 0 | 0 | 0 | 2504900 | 0 | 0 | 0 | 0 | 298910 | 0 | 0 | 0 | 0 | 0 | 0 | 1166200 | 0 |
| YLDLEEEAI | BAG5 | Q9UL15 | BAG5_HUMAN | Unmodified | 380960 | 361630 | 280940 | 389780 | 563050 | 0 | 0 | 0 | 1905600 | 1268100 | 220530 | 831540 | 274250 | 0 | 346310 | 282020 | 0 | 0 |
| YLDNIHLHI | UBXN6; UBXN6 | Q9BZV1; K7E | UBXN6_HUMAI | Unmodified | 605900 | 814960 | 648580 | 0 | 3077300 | 0 | 229220 | 142240 | 1274400 | 465860 | 112590 | 578630 | 3272600 | 2023900 | 1046600 | 1160400 | 878560 | 1653300 |
| YLDNIHLHI | UBXN6; UBXN6 | Q9BZV1; K7E | UBXN6_HUMAI | Unmodified | 0 | 0 | 0 | 0 | 0 | 0 | 691570 | 0 | 0 | 146300 | 0 | 0 | 427390 | 0 | 0 | 0 | 0 | 0 |
| YLDPEYIDF | DCTN3; DCTN3 | O75935; X6F | DCTN3_HUMAI | Unmodified | 0 | 0 | 0 | 0 | 0 | 0 | 0 | 0 | 0 | 0 | 0 | 2954800 | 0 | 0 | 0 | 0 | 0 | 0 |
| YLDPSFFQI | TM9SF3; TM9SF | Q9HD45; Q5 | TM9S3_HUMAN | Unmodified | 0 | 0 | 0 | 0 | 0 | 505710 | 0 | 0 | 956640 | 101880 | 0 | 582790 | 0 | 0 | 0 | 0 | 0 | 0 |
| YLDQIGNM | LMO7; LMO7; LI | Q8WWI1; A0 | LMO7_HUMAN | Unmodified | 0 | 0 | 0 | 0 | 1134600 | 0 | 0 | 0 | 350530 | 0 | 0 | 0 | 0 | 0 | 0 | 0 | 0 | 0 |
| YLDSAMQE | CLIC3 | O95833 | CLIC3_HUMAN | Unmodified | 0 | 0 | 0 | 0 | 615810 | 0 | 0 | 0 | 0 | 0 | 0 | 0 | 0 | 0 | 0 | 0 | 0 | 0 |
| YLEANMTQ | OLA1; OLA1 | Q9NTK5; J3K | OLA1_HUMAN | Unmodified | 0 | 0 | 0 | 0 | 0 | 0 | 244230 | 151910 | 606700 | 0 | 764500 | 6561100 | 171910 | 253650 | 275600 | 232030 | 0 | 138510 |
| YLEANMTQ | OLA1; OLA1 | Q9NTK5; J3K | OLA1_HUMAN | Oxidation (M) | 0 | 0 | 0 | 0 | 0 | 0 | 0 | 0 | 0 | 0 | 466240 | 2823800 | 0 | 0 | 0 | 405630 | 0 | 0 |
| YLEATGQLI | GFM1 | Q96RP9 | EFGM_HUMAN | Unmodified | 0 | 0 | 0 | 0 | 233550 | 0 | 0 | 0 | 2757100 | 1009600 | 0 | 0 | 84605 | 0 | 0 | 0 | 0 | 0 |
| YLEAVEK | RO60 | P10155 | RO60_HUMAN | Unmodified | 14483000 | 16034000 | 4690900 | 4204400 | 137990 | 268960 | 301650 | 201800 | 3688600 | 0 | 2538000 | 2048500 | 1658500 | 928670 | 6530800 | 4401100 | 139400 | 113640 |
| YLECRVEEI | PPM1A | P35813 | PPM1A_HUMAI | Unmodified | 217480 | 87174 | 0 | 0 | 0 | 0 | 0 | 0 | 344640 | 428750 | 0 | 187080 | 294870 | 0 | 205300 | 0 | 0 | 0 |
| YLECSALQ | RHOG | P84095 | RHOG_HUMAN | Unmodified | 722830 | 605650 | 300160 | 1225700 | 122860 | 0 | 0 | 0 | 0 | 265410 | 832360 | 5950100 | 84307 | 56549 | 320960 | 1136100 | 18539 | 0 |
| YLECSALT | RAC2; RAC3; R | P15153; P60 | RAC2_HUMAN | Unmodified | 11213000 | 9121600 | 7160900 | 10032000 | 8152100 | 2329700 | 3479500 | 763340 | 3405800 | 982620 | 8412800 | 20120000 | 9179400 | 6038500 | 11560000 | 11224000 | 2763900 | 9862800 |
| YLECSALT | RAC2; RAC3; R | P15153; P60 | RAC2_HUMAN | Unmodified | 0 | 0 | 0 | 420920 | 17892000 | 2979000 | 5196800 | 3323700 | 236900 | 2809900 | 442650 | 0 | 15042000 | 18194000 | 0 | 0 | 7009300 | 19482000 |
| YLECSALT | RAC2; RAC3; R | P15153; P60 | RAC2_HUMAN | Unmodified | 0 | 0 | 0 | 0 | 747810 | 125900 | 684310 | 0 | 0 | 0 | 0 | 0 | 1588700 | 2082700 | 0 | 0 | 762910 | 969490 |
| YLEDRGEV | GRK2; GRK2 | P25098; E9P | ARBK1_HUMAN | Unmodified | 0 | 120040 | 0 | 389220 | 831010 | 0 | 367400 | 237780 | 0 | 154020 | 0 | 1081800 | 1558400 | 1157800 | 510130 | 363050 | 0 | 0 |
| YLEEIATQM | PFKP; PFKP; PF | Q01813; A0A | PFKAP_HUMAN | Unmodified | 0 | 0 | 0 | 0 | 0 | 0 | 0 | 0 | 0 | 0 | 302260 | 2314500 | 0 | 0 | 201270 | 0 | 0 | 0 |
| YLEESNFVI | SYK | P43405 | KSYK_HUMAN | Unmodified | 34232 | 62227 | 0 | 0 | 0 | 0 | 0 | 0 | 162100 | 0 | 59345 | 1227900 | 0 | 0 | 0 | 0 | 0 | 0 |
| YLEGYVAN | PRKG2 | Q13237 | KGP2_HUMAN | Unmodified | 0 | 0 | 0 | 0 | 0 | 0 | 656490 | 395990 | 0 | 0 | 0 | 0 | 0 | 0 | 0 | 0 | 0 | 0 |
| YLEHGEYE | PALS2; PALS2 | Q9NZW5; B8 | PALS2_HUMAN | Unmodified | 54522 | 0 | 771780 | 273470 | 536430 | 237480 | 0 | 0 | 905410 | 63567 | 1630300 | 3182900 | 49885 | 40589 | 992810 | 622820 | 81117 | 201730 |
| YLEHGEYE | MPP2; MPP2; M | Q14168; A0A | MPP2_HUMAN | Unmodified | 0 | 0 | 0 | 0 | 0 | 0 | 0 | 0 | 414140 | 0 | 0 | 0 | 0 | 0 | 850100 | 395420 | 0 | 0 |
| YLEKPITML | DLAT; DLAT; DL | P10515; A0A | ODP2_HUMAN | Unmodified | 0 | 0 | 0 | 0 | 0 | 0 | 0 | 0 | 1447600 | 356780 | 0 | 0 | 0 | 0 | 0 | 0 | 0 | 0 |
| YLELFQR | EXTL2; EXTL2 | Q9UBQ6; C9 | EXTL2_HUMAN | Unmodified | 0 | 0 | 0 | 0 | 0 | 0 | 0 | 0 | 1831300 | 0 | 0 | 0 | 0 | 0 | 0 | 0 | 0 | 0 |
| YLENGKETI | HLA-A; HLA-H; I | P04439; P01 | HLAA_HUMAN | Unmodified | 460520 | 1095000 | 0 | 728420 | 113050 | 601600 | 0 | 183910 | 18695000 | 1110800 | 3304900 | 24506000 | 0 | 72074 | 1774900 | 281950 | 1044200 | 2962000 |
| YLENGKETI | HLA-C; HLA-C; | Q5QT33; A0A | Q5QT33_HUMA | Unmodified | 0 | 0 | 0 | 0 | 1583800 | 1203200 | 0 | 0 | 506270 | 159180 | 0 | 0 | 0 | 0 | 0 | 0 | 112080 | 826480 |
| YLENGKETI | HLA-A; HLA-A; I | Q9GJ45; Q5 | Q9GJ45_HUMA | Unmodified | 0 | 0 | 0 | 0 | 0 | 0 | 65756 | 21062 | 700210 | 1167100 | 55357 | 138010 | 0 | 0 | 0 | 0 | 0 | 0 |
| YLEQMQI | APLP2 | Q06481 | APLP2_HUMAN | Oxidation (M) | 16487 | 0 | 0 | 224680 | 1350300 | 0 | 284100 | 80900 | 0 | 0 | 0 | 0 | 0 | 0 | 0 | 0 | 0 | 479230 |
| YLERGESIE | ERBB3; ERBB3; | P21860; B4D | ERBB3_HUMAN | Unmodified | 98680 | 48939 | 0 | 210880 | 539790 | 179250 | 0 | 0 | 0 | 0 | 247070 | 551230 | 180990 | 143330 | 0 | 201030 | 0 | 0 |
| YLESAGAR | GGH; GGH; GG | Q92820; A0A | GGH_HUMAN ( | Unmodified | 274860 | 72006 | 200430 | 149840 | 0 | 0 | 0 | 0 | 25758000 | 4108400 | 805680 | 562790 | 119290 | 43070 | 210710 | 209960 | 0 | 0 |
| YLESAGAR | GGH; GGH; GG | Q92820; A0A | GGH_HUMAN ( | Unmodified | 0 | 0 | 0 | 0 | 0 | 0 | 0 | 0 | 0 | 4520600 | 0 | 0 | 0 | 0 | 0 | 0 | 0 | 0 |
| YLESLGEE | NUCB1; NUCB | Q02818; H7I | NUCB1_HUMA | Unmodified | 0 | 0 | 0 | 552150 | 741650 | 349850 | 15501 | 0 | 42820000 | 2989300 | 4275100 | 11455000 | 0 | 0 | 76715 | 104570 | 487080 | 2678800 |
| YLESLVQT | SNX2 | O60749 | SNX2_HUMAN | Unmodified | 0 | 0 | 0 | 221420 | 3847200 | 0 | 621480 | 359070 | 193810 | 274430 | 1037800 | 2530000 | 1882500 | 1400500 | 372260 | 293420 | 913020 | 0 |
| YLESNVLD | N/A | N/A | N/A | Unmodified | 0 | 0 | 1244900 | 232070 | 0 | 832740 | 0 | 34869 | 0 | 0 | 0 | 0 | 0 | 130350 | 0 | 0 | 0 | 161150 |
| YLETLLYSQ | SNX1; SNX1 | Q13596; H0Y | SNX1_HUMAN | Unmodified | 0 | 0 | 0 | 0 | 2490000 | 151700 | 259880 | 0 | 2180700 | 559770 | 0 | 0 | 963030 | 0 | 0 | 0 | 0 | 309150 |
| YLETPGDEI | APP; APP; APP; | P05067; H7C | A4_HUMAN Am | Unmodified | 0 | 0 | 0 | 0 | 1350900 | 0 | 0 | 0 | 12224000 | 1485700 | 1217900 | 543910 | 0 | 0 | 0 | 0 | 0 | 353870 |
| YLFTGLK | USP14; USP14 | P54578; A6N | UBP14_HUMAI | Unmodified | 426050 | 306470 | 966460 | 1216600 | 4349400 | 516300 | 2737600 | 954910 | 1389700 | 1842800 | 2957300 | 13694000 | 6245600 | 3464700 | 1184300 | 819940 | 2175100 | 4364200 |
| YLGHVEVD | NUMB; NUMB; | P49757; G3V | NUMB_HUMAN | Unmodified | 207160 | 303130 | 254970 | 582040 | 1001400 | 92092 | 0 | 23279 | 1595700 | 49247 | 0 | 896790 | 188560 | 42411 | 515100 | 377260 | 55101 | 768910 |
| YLGHVEVD | NUMB; NUMB; | P49757; G3V | NUMB_HUMAN | Unmodified | 0 | 0 | 0 | 0 | 3683300 | 600700 | 0 | 0 | 0 | 0 | 0 | 0 | 0 | 0 | 0 | 0 | 241100 | 1592600 |
| YLGHVEVD | NUMB; NUMB; | P49757; G3V | NUMB_HUMAN | Oxidation (M) | 0 | 0 | 0 | 0 | 1038500 | 331040 | 0 | 0 | 0 | 0 | 0 | 0 | 0 | 294730 | 0 | 0 | 0 | 0 |
| YLGHVEVE | NUMBL; NUMB | Q9Y6R0; A0A | NUMBL_HUMA | Unmodified | 0 | 0 | 0 | 120970 | 0 | 0 | 0 | 0 | 120250 | 39419 | 14046 | 269900 | 298530 | 0 | 0 | 0 | 0 | 0 |
| YLGLLENLI | MYO1C; MYO1 | O00159; F5I | MYO1C_HUMA | Unmodified | 0 | 0 | 356540 | 1387700 | 1861800 | 428280 | 160790 | 0 | 0 | 179110 | 0 | 0 | 226630 | 125230 | 0 | 0 | 196150 | 92735 |
| YLGLLENVI | MYO1D; MYO1 | O94832; B0I | MYO1D_HUMA | Unmodified | 0 | 0 | 0 | 0 | 903640 | 939350 | 0 | 0 | 0 | 146730 | 306870 | 0 | 47724 | 0 | 0 | 0 | 150510 | 0 |
| YLGLPGSLI | MOGS; MOGS; | Q13724; A0A | MOGS_HUMAN | Unmodified | 0 | 0 | 0 | 0 | 0 | 0 | 0 | 0 | 3415400 | 0 | 0 | 0 | 0 | 0 | 0 | 0 | 0 | 0 |
| YLGPAVLM | SDHB | P21912 | SDHB_HUMAN | Unmodified | 0 | 0 | 0 | 0 | 0 | 0 | 0 | 0 | 4878500 | 0 | 0 | 0 | 0 | 0 | 0 | 0 | 18097 | 0 |
| YLGQDYEC | CAPN1; CAPN1 | P07384; E9P | CAN1_HUMAN | Unmodified | 0 | 0 | 383680 | 650230 | 0 | 0 | 0 | 0 | 0 | 0 | 3243900 | 5446100 | 627710 | 290980 | 0 | 356130 | 0 | 0 |
| YLGQDYEC | CAPN1; CAPN1 | P07384; E9P | CAN1_HUMAN | Unmodified | 0 | 100760 | 0 | 183010 | 4604400 | 974530 | 0 | 51867 | 687310 | 1810600 | 2567100 | 0 | 2944100 | 0 | 341860 | 195980 | 1958700 | 4683300 |
| YLGQLHYL | PTPN23; PTPN2 | Q9H3S7; C9 | PTN23_HUMAN | Unmodified | 1367000 | 0 | 0 | 0 | 0 | 0 | 0 | 0 | 0 | 0 | 0 | 0 | 0 | 0 | 0 | 0 | 0 | 0 |
| YLGRWYEII | APOD; APOD; A | P05090; C9J | APOD_HUMAN | Unmodified | 0 | 185630 | 44673 | 42704 | 0 | 0 | 0 | 0 | 0 | 1262900 | 0 | 0 | 0 | 0 | 53250 | 62337 | 0 | 0 |
| YLGRWYEII | APOD; APOD; A | P05090; C9J | APOD_HUMAN | Unmodified | 0 | 0 | 0 | 0 | 0 | 0 | 0 | 0 | 536800 | 4478800 | 0 | 0 | 0 | 0 | 0 | 0 | 0 | 0 |
| YLGSYSGK | IGSF8; IGSF8 | Q969P0; C9J | IGSF8_HUMAN | Unmodified | 22793000 | 21503000 | 0 | 0 | 0 | 0 | 0 | 0 | 1556300 | 0 | 639510 | 0 | 866440 | 152430 | 9255300 | 13555000 | 0 | 0 |
| YLHDESGL | SEPTIN2; SEPTI | Q15019; C9J | SEPT2_HUMAN | Unmodified | 0 | 193630 | 0 | 0 | 0 | 0 | 0 | 0 | 196290 | 167470 | 386160 | 1653600 | 0 | 0 | 115280 | 85105 | 121950 | 265270 |
| YLHDESGL | SEPTIN2; SEPTI | Q15019; C9J | SEPT2_HUMAN | Unmodified | 0 | 0 | 0 | 0 | 0 | 0 | 0 | 0 | 0 | 0 | 0 | 4591000 | 0 | 0 | 0 | 0 | 0 | 0 |
| YLHVGYIVI | SNX2; SNX2 | O60749; D6I | SNX2_HUMAN | Unmodified | 0 | 0 | 0 | 0 | 1812600 | 0 | 1928500 | 930310 | 0 | 0 | 0 | 1705400 | 1846200 | 1182400 | 0 | 0 | 1162900 | 646580 |
| YLHVRPQP | RNF31; RNF31; | Q96EP0; H0Y | RNF31_HUMAN | Unmodified | 418210 | 491620 | 0 | 0 | 0 | 0 | 0 | 0 | 0 | 0 | 0 | 0 | 0 | 0 | 0 | 0 | 0 | 0 |
| YLIANATNI | YWHAQ; YWHA | P27348; E9P | 1433T_HUMAN | Unmodified | 1861900 | 1672700 | 4296100 | 6072100 | 13168000 | 4116700 | 1861600 | 1536000 | 6275400 | 3630900 | 11419000 | 51158000 | 6781000 | 4140300 | 5674700 | 6678700 | 7361400 | 5767100 |
| YLIANATNI | YWHAQ; YWHA | P27348; E9P | 1433T_HUMAN | Unmodified | 0 | 0 | 0 | 0 | 7060700 | 686810 | 4470300 | 1435300 | 128850 | 1841900 | 170490 | 0 | 4240900 | 7367100 | 0 | 0 | 16354000 | 5186200 |
| YLIATSEQP | SARS1; SARS1 | P49591; Q5T | SYSC_HUMAN | Unmodified | 0 | 0 | 0 | 0 | 2873700 | 671750 | 2587400 | 0 | 0 | 0 | 1374600 | 3801800 | 2341000 | 1817200 | 0 | 0 | 503770 | 1530800 |
| YLIEIAK | IST1; IST1; IST1 | P53990; H3E | IST1_HUMAN IS | Unmodified | 0 | 0 | 0 | 0 | 1899400 | 471450 | 0 | 0 | 3351200 | 924230 | 2049400 | 6450800 | 0 | 0 | 0 | 0 | 313730 | 0 |
| YLIFPNPVC | LAMB2 | P55268 | LAMB2_HUMAI | Unmodified | 0 | 0 | 22256 | 0 | 0 | 0 | 0 | 0 | 298050 | 124710 | 0 | 0 | 0 | 0 | 0 | 0 | 0 | 0 |
| YLIGELVSS | KIF4A | O95239 | KIF4A_HUMAN | Unmodified | 0 | 0 | 0 | 0 | 0 | 0 | 1695300 | 0 | 0 | 254060 | 0 | 0 | 1508400 | 1638300 | 0 | 0 | 5110600 | 0 |
| YLIIMNPGI | MGAM2 | Q2M2H8 | MGAL_HUMAN | Oxidation (M) | 0 | 0 | 0 | 0 | 0 | 0 | 0 | 0 | 0 | 0 | 591100 | 1385800 | 0 | 0 | 0 | 0 | 0 | 0 |
| YLIPNATQF | YWHAB | P31946 | 1433B_HUMAN | Unmodified | 1933200 | 2047000 | 7024200 | 8935000 | 21741000 | 7250300 | 4080300 | 2671700 | 14283000 | 5759700 | 14856000 | 38261000 | 10666000 | 8700600 | 7334500 | 7083400 | 13511000 | 26250000 |
| YLIPNATQF | YWHAB | P31946 | 1433B_HUMAN | Unmodified | 0 | 0 | 0 | 0 | 3112200 | 365780 | 871270 | 0 | 0 | 490070 | 0 | 0 | 2757100 | 3599800 | 0 | 0 | 10386000 | 2095800 |
| YLIRPQPWI | FUT8; FUT8; FU | Q9BYC5; A0A | FUT8_HUMAN | Unmodified | 0 | 0 | 0 | 0 | 0 | 0 | 0 | 0 | 4029900 | 0 | 0 | 0 | 0 | 0 | 0 | 0 | 0 | 0 |
| YLIVVTDGI | COL6A1; COL6 | P12109; A0A | CO6A1_HUMAI | Unmodified | 0 | 0 | 0 | 0 | 0 | 0 | 0 | 0 | 2259500 | 0 | 0 | 0 | 0 | 0 | 0 | 0 | 0 | 3822600 |
| YLIYRGLVG | N/A | N/A | N/A | Oxidation (M) | 0 | 0 | 0 | 0 | 0 | 0 | 4878200 | 0 | 0 | 0 | 0 | 0 | 0 | 14849000 | 0 | 0 | 0 | 0 |
| YLKGDHAG | FSCN1 | Q16658 | FSCN1_HUMAI | Unmodified | 0 | 0 | 0 | 0 | 909780 | 193690 | 29376 | 0 | 112720 | 78599 | 201630 | 495240 | 0 | 0 | 0 | 0 | 336510 | 402330 |
| YLKGDHAG | FSCN1 | Q16658 | FSCN1_HUMAI | Unmodified | 0 | 0 | 0 | 0 | 473160 | 0 | 0 | 0 | 0 | 0 | 0 | 0 | 0 | 0 | 0 | 0 | 1662200 | 0 |
| YLKSEPIPE | PDIA3 | P30101 | PDIA3_HUMAN | Unmodified | 0 | 0 | 0 | 0 | 0 | 0 | 0 | 72301 | 737890 | 669120 | 36457 | 0 | 0 | 0 | 0 | 0 | 0 | 0 |

| | | | | | | | | | | | | | | | | | | | | | | |
|---|---|---|---|---|---|---|---|---|---|---|---|---|---|---|---|---|---|---|---|---|---|---|
| YLLEGTAET | VPS13C; VPS13 | Q709C8; A0 | VP13C_HUMAN | Unmodified | 0 | 0 | 0 | 0 | 0 | 0 | 0 | 0 | 0 | 0 | 0 | 1270700 | 0 | 0 | 0 | 0 | 0 | 0 |
| YLLEQDFP | EHD4; EHD3 | Q9H223; Q9 | EHD4_HUMAN | Oxidation (M) | 1135800 | 874250 | 0 | 3928300 | 0 | 0 | 0 | 0 | 0 | 0 | 0 | 1185600 | 0 | 0 | 2678900 | 1291400 | 0 | 0 |
| YLLETSGNI | NAMPT; NAMPT | P43490; A0A | NAMPT_HUMA | Unmodified | 0 | 0 | 0 | 67456 | 1209900 | 0 | 0 | 0 | 0 | 42187 | 0 | 2085200 | 153640 | 66948 | 436300 | 115130 | 0 | 0 |
| YLLFTSPQE | AOC1 | P19801 | AOC1_HUMAN | Unmodified | 0 | 0 | 0 | 0 | 0 | 0 | 43945 | 0 | 1874000 | 1146700 | 0 | 0 | 0 | 0 | 0 | 0 | 41155 | 0 |
| YLLFTSPQE | AOC1 | P19801 | AOC1_HUMAN | Unmodified | 145630 | 0 | 0 | 0 | 0 | 0 | 0 | 0 | 1640300 | 372940 | 0 | 0 | 0 | 0 | 0 | 0 | 0 | 0 |
| YLLLAGAPI | ITGA3; ITGA3 | P26006; D6F | ITA3_HUMAN I | Unmodified | 691870 | 586020 | 0 | 0 | 20758000 | 16558000 | 0 | 0 | 2153500 | 95945 | 460060 | 12731000 | 0 | 0 | 721320 | 0 | 6499000 | 20587000 |
| YLLLEETEK | MTSS2; MTSS1; | Q765P7; O4 | MTSS2_HUMAN | Unmodified | 0 | 0 | 0 | 0 | 0 | 0 | 0 | 0 | 926190 | 0 | 0 | 2652000 | 0 | 0 | 0 | 0 | 0 | 0 |
| YLLLLAAGI | ECPAS; ECPAS; | Q5VYK3; R4 | ECM29_HUMA | Unmodified | 0 | 0 | 0 | 0 | 0 | 0 | 0 | 0 | 0 | 576670 | 0 | 1725300 | 2030500 | 0 | 0 | 0 | 783210 | 0 |
| YLLLSDLPC | DMAC2L; DMAC | Q99766; A0A | ATP5S_HUMAN | Unmodified | 0 | 0 | 0 | 0 | 2634100 | 0 | 35445 | 89585 | 1177300 | 440760 | 0 | 0 | 684180 | 21938 | 0 | 0 | 0 | 0 |
| YLLNLNLEI | TGM2 | P21980 | TGM2_HUMAN | Unmodified | 0 | 0 | 0 | 0 | 6600200 | 1292400 | 0 | 0 | 0 | 0 | 0 | 0 | 0 | 0 | 0 | 0 | 659590 | 316950 |
| YLLPEVTVL | CTDSPL; CTDS | O15194; H7 | CTDSL_HUMAN | Unmodified | 0 | 756640 | 0 | 0 | 0 | 0 | 0 | 0 | 0 | 0 | 456490 | 0 | 2007700 | 1328900 | 0 | 0 | 0 | 0 |
| YLLQEQLK | LONP1; LONP1 | P36776; K7E | LONM_HUMAN | Unmodified | 1295300 | 0 | 0 | 0 | 0 | 0 | 0 | 149190 | 2577100 | 1351900 | 0 | 0 | 0 | 0 | 104710 | 187450 | 0 | 0 |
| YLLQETWL | ERO1A | Q96HE7 | ERO1A_HUMA | Unmodified | 0 | 0 | 0 | 0 | 0 | 4102400 | 0 | 94905 | 0 | 761730 | 0 | 0 | 0 | 0 | 0 | 0 | 0 | 0 |
| YLLSEEEDI | GNPAT | O15228 | GNPAT_HUMA | Unmodified | 0 | 8890700 | 0 | 0 | 0 | 0 | 0 | 0 | 783800 | 249900 | 0 | 0 | 0 | 0 | 0 | 0 | 0 | 0 |
| YLLSQSSP | TWF1; TWF1 | Q12792; F8V | TWF1_HUMAN | Unmodified | 0 | 1236600 | 0 | 0 | 3947800 | 968110 | 0 | 0 | 520000 | 806690 | 246930 | 436340 | 3114300 | 1986000 | 0 | 0 | 1245700 | 573530 |
| YLLTPREPN | SLC6A6; SLC6A | P31641; A0A | SC6A6_HUMAN | Unmodified | 0 | 0 | 0 | 0 | 0 | 0 | 0 | 0 | 0 | 0 | 0 | 1026700 | 0 | 0 | 0 | 0 | 0 | 0 |
| YLMASDPN | PKP3; PKP3 | Q9Y446; E9F | PKP3_HUMAN | Oxidation (M) | 535230 | 458460 | 1029200 | 1680900 | 3294700 | 0 | 0 | 324270 | 1015900 | 0 | 2153200 | 2026200 | 4472500 | 4742000 | 1859600 | 1559500 | 1756700 | 630640 |
| YLMDPDTF | APOBEC3B; AP | Q9UH17; B0 | ABC3B_HUMA | Oxidation (M) | 0 | 0 | 28386 | 0 | 0 | 0 | 0 | 0 | 0 | 0 | 0 | 0 | 0 | 0 | 234420 | 35338 | 0 | 0 |
| YLMEEDED | RPL5; RPL5 | P46777; A0A | RL5_HUMAN 6 | Unmodified | 4099000 | 2393800 | 545590 | 875160 | 1060600 | 127780 | 59074 | 42296 | 1848700 | 572320 | 4900600 | 24874000 | 1045300 | 122750 | 2285500 | 1356500 | 67104 | 107050 |
| YLMEEDED | RPL5; RPL5 | P46777; A0A | RL5_HUMAN 6 | Oxidation (M) | 2668800 | 1450800 | 590130 | 535670 | 0 | 0 | 0 | 0 | 182980 | 75394 | 1081700 | 6160300 | 0 | 0 | 803420 | 961890 | 42204 | 0 |
| YLNEYGAP | GNS; GNS; GN | P15586; H7 | GNS_HUMAN | Unmodified | 0 | 0 | 0 | 0 | 0 | 0 | 0 | 0 | 875610 | 782570 | 0 | 0 | 0 | 0 | 0 | 0 | 0 | 0 |
| YLNFFTK | PCNA | P12004 | PCNA_HUMAN | Unmodified | 0 | 0 | 0 | 1121800 | 3038700 | 848000 | 680640 | 444490 | 2976300 | 3423200 | 2032200 | 12049000 | 7963000 | 1025100 | 0 | 0 | 1328000 | 0 |
| YLNFVKGV | HSP90B1; HSP | P14625; A0A | ENPL_HUMAN | Unmodified | 0 | 0 | 0 | 0 | 0 | 0 | 142230 | 0 | 8555700 | 3308000 | 57515 | 0 | 0 | 0 | 0 | 0 | 0 | 0 |
| YLNFVKGV | HSP90B1; HSP | P14625; A0A | ENPL_HUMAN | Unmodified | 0 | 0 | 0 | 0 | 204840 | 0 | 9444200 | 0 | 0 | 63113000 | 0 | 0 | 0 | 0 | 0 | 0 | 0 | 0 |
| YLNRNYWI | HGS; HGS; HG | O14964; A0 | HGS_HUMAN | Unmodified | 188300 | 0 | 0 | 147580 | 1601900 | 0 | 499750 | 81301 | 0 | 618570 | 63526 | 0 | 679780 | 204950 | 0 | 0 | 305870 | 1417200 |
| YLNWIGVP | PFKFB2; PFKFB | O60825; Q1 | F262_HUMAN | Unmodified | 0 | 0 | 0 | 0 | 0 | 0 | 0 | 0 | 0 | 0 | 0 | 1523700 | 0 | 0 | 0 | 0 | 0 | 0 |
| YLPDTLLLE | COMT; COMT | P21964; E7E | COMT_HUMAN | Unmodified | 0 | 0 | 0 | 0 | 0 | 0 | 0 | 239270 | 0 | 0 | 127380 | 327250 | 0 | 0 | 0 | 4230100 | 0 | 0 |
| YLPHSAGR | RPS5; RPS5; RP | P46782; M0 | RS5_HUMAN 4 | Unmodified | 0 | 0 | 0 | 0 | 3265600 | 0 | 3223300 | 1569700 | 0 | 602850 | 0 | 0 | 0 | 0 | 0 | 0 | 0 | 0 |
| YLPRPANP | CDH1 | P12830 | CADH1_HUMA | Unmodified | 7770000 | 6571900 | 14553000 | 37315000 | 98487000 | 14656000 | 0 | 0 | 8030700 | 2088700 | 24349000 | 30250000 | 99415000 | 71233000 | 53531000 | 50693000 | 3870500 | 493190 |
| YLQDSTFA | NUP85; NUP85 | Q9BW27; J3 | NUP85_HUMA | Unmodified | 0 | 0 | 0 | 0 | 0 | 0 | 0 | 0 | 0 | 0 | 0 | 1190500 | 0 | 282910 | 0 | 0 | 0 | 0 |
| YLQEEVNI | SEPTIN9; SEPTI | Q9UHD8; K7 | SEPT9_HUMAN | Unmodified | 0 | 0 | 0 | 263700 | 425520 | 0 | 25249 | 0 | 0 | 169910 | 0 | 3898900 | 0 | 0 | 105940 | 0 | 319650 | 0 |
| YLQEEVNI | SEPTIN9; SEPTI | Q9UHD8; K7 | SEPT9_HUMAN | Unmodified | 0 | 0 | 0 | 1481100 | 12944000 | 195780 | 1045800 | 521640 | 740350 | 792930 | 126150 | 6692700 | 250030 | 0 | 519300 | 0 | 5874400 | 1903900 |
| YLQEEVNI | SEPTIN9; SEPTI | Q9UHD8; K7 | SEPT9_HUMAN | Unmodified | 0 | 0 | 0 | 0 | 3254000 | 391660 | 323430 | 258890 | 0 | 198470 | 0 | 443390 | 0 | 1190900 | 0 | 0 | 1861600 | 1569900 |
| YLQEIYNS | FGG; FGG; FG | P02679; C9J | FIBG_HUMAN | Unmodified | 0 | 4502900 | 0 | 0 | 0 | 0 | 0 | 0 | 0 | 0 | 0 | 0 | 0 | 0 | 0 | 0 | 0 | 0 |
| YLQEIYNS | FGG; FGG; FG | P02679; C9J | FIBG_HUMAN | Unmodified | 0 | 232220 | 0 | 0 | 0 | 0 | 0 | 0 | 0 | 0 | 0 | 0 | 0 | 0 | 0 | 0 | 0 | 0 |
| YLQEVIDVL | NUCB1; NUCB | Q02818; H7 | NUCB1_HUMA | Unmodified | 0 | 0 | 0 | 37923 | 0 | 0 | 81845 | 0 | 740590 | 0 | 575090 | 0 | 0 | 0 | 0 | 0 | 76713 | 0 |
| YLQEVIDVL | NUCB1; NUCB | Q02818; H7 | NUCB1_HUMA | Unmodified | 110450 | 0 | 0 | 0 | 0 | 0 | 0 | 0 | 8217900 | 386470 | 1626400 | 0 | 0 | 0 | 0 | 311280 | 0 | 0 |
| YLQLAEELI | MCM6 | Q14566 | MCM6_HUMAN | Unmodified | 0 | 0 | 0 | 0 | 0 | 0 | 0 | 25894 | 404060 | 0 | 269270 | 3321400 | 0 | 0 | 0 | 0 | 0 | 51091 |
| YLQLQQEK | GOLIM4; GOLI | O00461; F8 | GOLI4_HUMAN | Unmodified | 0 | 0 | 0 | 0 | 199750 | 0 | 134910 | 120800 | 22592000 | 2414300 | 776950 | 980850 | 116120 | 0 | 99978 | 0 | 0 | 294230 |
| YLQLQQEK | GOLIM4; GOLI | O00461; F8 | GOLI4_HUMAN | Unmodified | 0 | 0 | 0 | 0 | 404770 | 160370 | 671810 | 637020 | 9203000 | 11825000 | 148680 | 0 | 541930 | 622950 | 0 | 0 | 95587 | 809110 |
| YLQLQQEK | GOLIM4; GOLI | O00461; F8 | GOLI4_HUMAN | Unmodified | 0 | 0 | 0 | 0 | 0 | 0 | 182420 | 358150 | 0 | 1338900 | 0 | 0 | 415630 | 0 | 0 | 0 | 0 | 215130 |
| YLQSNSNN | HUWE1 | Q7Z6Z7 | HUWE1_HUMA | Unmodified | 0 | 67176 | 0 | 0 | 0 | 0 | 0 | 0 | 0 | 0 | 0 | 950240 | 0 | 122230 | 0 | 0 | 0 | 0 |
| YLQSTQAI | SETD3; SETD3 | Q86TU7; Q6 | SETD3_HUMAN | Unmodified | 0 | 0 | 0 | 0 | 0 | 0 | 0 | 0 | 0 | 600870 | 0 | 1083200 | 0 | 0 | 0 | 0 | 0 | 0 |
| YLQTLTTIA | STOM | P27105 | STOM_HUMAN | Unmodified | 6035200 | 5158700 | 2891500 | 0 | 25597000 | 4581700 | 1718200 | 0 | 0 | 0 | 0 | 872170 | 2425400 | 1895300 | 4390100 | 6302100 | 0 | 1218600 |
| YLQWEIGN | TRMT10C | Q7L0Y3 | TM10C_HUMA | Unmodified | 0 | 0 | 0 | 0 | 0 | 0 | 0 | 0 | 1661100 | 463030 | 0 | 190080 | 0 | 0 | 0 | 0 | 629770 | 0 |
| YLRAEPEDI | ACTR3; ACTR3; | P61158; B4D | ARP3_HUMAN | Unmodified | 2011900 | 2353400 | 410820 | 3264100 | 0 | 0 | 0 | 0 | 1317000 | 92668 | 0 | 38811000 | 0 | 0 | 6112200 | 272770 | 0 | 0 |
| YLRIPDEIID | LSM4; LSM4; LS | Q9Y4Z0; V9 | LSM4_HUMAN | Oxidation (M) | 0 | 0 | 0 | 0 | 0 | 0 | 0 | 0 | 0 | 0 | 0 | 143310 | 0 | 0 | 0 | 0 | 0 | 0 |
| YLRIPDEIID | LSM4; LSM4; LS | Q9Y4Z0; V9 | LSM4_HUMAN | Oxidation (M) | 0 | 0 | 0 | 0 | 923870 | 0 | 670400 | 0 | 0 | 349460 | 0 | 0 | 1532000 | 231240 | 0 | 0 | 286970 | 367240 |
| YLSAGPTL | VANGL1 | Q8TAA9 | VANG1_HUMA | Unmodified | 0 | 0 | 533220 | 236880 | 0 | 0 | 0 | 0 | 0 | 0 | 0 | 0 | 0 | 0 | 519280 | 654660 | 0 | 0 |
| YLSAQKPLI | CPT2; CPT2; CF | P23786; A0A | CPT2_HUMAN | Unmodified | 0 | 0 | 0 | 0 | 0 | 0 | 0 | 0 | 2313900 | 0 | 0 | 0 | 0 | 0 | 0 | 0 | 0 | 0 |
| YLSEMNYV | EPHB3 | P54753 | EPHB3_HUMA | Oxidation (M) | 101840 | 114500 | 127490 | 31546 | 0 | 0 | 0 | 0 | 0 | 0 | 0 | 1010300 | 0 | 0 | 136730 | 0 | 0 | 0 |
| YLSEMNYV | EPHB3 | P54753 | EPHB3_HUMA | Unmodified | 0 | 0 | 0 | 0 | 0 | 0 | 0 | 0 | 0 | 0 | 0 | 1089000 | 0 | 0 | 300120 | 0 | 0 | 0 |
| YLSEVASGI | YWHAB | P31946 | 1433B_HUMAN | Unmodified | 3006900 | 3959300 | 5633000 | 4804200 | 0 | 492380 | 0 | 0 | 4333600 | 323910 | 7159500 | 21794000 | 613300 | 397620 | 12017000 | 5560400 | 377960 | 1049600 |
| YLSEVASGI | YWHAB | P31946 | 1433B_HUMAN | Unmodified | 2465800 | 1186700 | 0 | 12292000 | 14458000 | 288500 | 1373400 | 0 | 9994300 | 3333500 | 7194200 | 13881000 | 16280000 | 9306100 | 2713400 | 2400900 | 9785500 | 14568000 |
| YLSFTPPEK | PAK2 | Q13177 | PAK2_HUMAN | Unmodified | 110730 | 70412 | 219870 | 57762 | 0 | 0 | 0 | 0 | 0 | 0 | 783490 | 0 | 1057700 | 0 | 0 | 278930 | 837530 | 1182600 |
| YLSFTPPEK | PAK2 | Q13177 | PAK2_HUMAN | Unmodified | 0 | 0 | 0 | 0 | 1801700 | 0 | 75744 | 49793 | 0 | 219100 | 0 | 0 | 279340 | 407790 | 0 | 0 | 1531400 | 290270 |
| YLSGIAHFL | PMPCA | Q10713 | MPPA_HUMAN | Unmodified | 0 | 0 | 0 | 0 | 0 | 0 | 316960 | 0 | 2827000 | 893210 | 0 | 217420 | 0 | 0 | 0 | 0 | 0 | 524060 |
| YLSLEQKTE | EEA1 | Q15075 | EEA1_HUMAN | Unmodified | 1075900 | 0 | 217130 | 547120 | 2730900 | 690730 | 0 | 0 | 248790 | 173240 | 0 | 686700 | 1153200 | 628780 | 374910 | 533460 | 275410 | 0 |
| YLSLLYPAR | PARP4 | Q9UKK3 | PARP4_HUMAN | Unmodified | 0 | 0 | 0 | 0 | 230890 | 0 | 0 | 0 | 0 | 0 | 0 | 2447600 | 0 | 0 | 0 | 0 | 0 | 0 |
| YLSNAYAR | CLIC1 | O00299 | CLIC1_HUMAN | Unmodified | 0 | 0 | 0 | 0 | 0 | 0 | 0 | 0 | 0 | 0 | 0 | 7730200 | 0 | 0 | 0 | 0 | 0 | 0 |
| YLSNAYARI | CLIC1 | O00299 | CLIC1_HUMAN | Unmodified | 0 | 0 | 0 | 0 | 12719000 | 17238000 | 0 | 0 | 2325300 | 823120 | 0 | 0 | 6476700 | 3444500 | 0 | 0 | 4884200 | 5828900 |
| YLSSANPN | ROCK1; ROCK1 | Q13464; A0 | ROCK1_HUMA | Unmodified | 0 | 0 | 0 | 0 | 0 | 0 | 0 | 0 | 0 | 0 | 42626 | 407720 | 0 | 0 | 0 | 32562 | 0 | 0 |
| YLSSANPN | ROCK1; ROCK1 | Q13464; A0 | ROCK1_HUMA | Unmodified | 0 | 0 | 0 | 0 | 3250800 | 561310 | 0 | 0 | 588090 | 350000 | 0 | 0 | 855830 | 523140 | 0 | 0 | 1267900 | 1753400 |
| YLSSANPN | ROCK1; ROCK1 | Q13464; A0 | ROCK1_HUMA | Unmodified | 0 | 0 | 0 | 0 | 0 | 0 | 0 | 0 | 0 | 0 | 0 | 0 | 123860 | 0 | 0 | 0 | 423890 | 0 |
| YLTAEAFG | FSCN1 | Q16658 | FSCN1_HUMA | Unmodified | 0 | 0 | 0 | 0 | 31150000 | 8736400 | 2959500 | 1779700 | 7383000 | 3497400 | 13372000 | 29557000 | 0 | 0 | 0 | 0 | 9686900 | 22798000 |
| YLTAEAFG | FSCN1 | Q16658 | FSCN1_HUMA | Unmodified | 0 | 0 | 0 | 0 | 3818800 | 220770 | 732920 | 40384 | 0 | 687210 | 0 | 0 | 0 | 0 | 0 | 0 | 5070400 | 1582000 |
| YLTAEAFG | FSCN1 | Q16658 | FSCN1_HUMA | Unmodified | 0 | 0 | 0 | 0 | 420660 | 0 | 0 | 0 | 1730300 | 0 | 0 | 0 | 0 | 0 | 0 | 0 | 1795800 | 0 |
| YLTEHPDP | PRPF8 | Q6P2Q9 | PRP8_HUMAN | Unmodified | 0 | 0 | 0 | 0 | 0 | 0 | 0 | 0 | 0 | 0 | 0 | 1021800 | 0 | 441640 | 0 | 0 | 0 | 0 |
| YLTESYGT | MYO6; MYO6; M | Q9UM54; A0 | MYO6_HUMAN | Unmodified | 0 | 0 | 1322600 | 260170 | 0 | 0 | 0 | 0 | 0 | 0 | 0 | 0 | 0 | 0 | 473780 | 319760 | 0 | 0 |
| YLTESYGT | MYO6; MYO6; M | Q9UM54; A0 | MYO6_HUMAN | Unmodified | 0 | 0 | 425730 | 2452400 | 670370 | 0 | 0 | 0 | 1723100 | 490650 | 200710 | 238660 | 2080200 | 0 | 3566800 | 1032100 | 0 | 0 |
| YLTETEDTV | DOCK10; DOCK | Q96BY6; A0 | DOC10_HUMA | Unmodified | 0 | 0 | 0 | 0 | 0 | 412000 | 0 | 0 | 0 | 0 | 0 | 0 | 0 | 0 | 0 | 0 | 237630 | 240610 |
| YLTETEDTV | DOCK10; DOCK | Q96BY6; A0 | DOC10_HUMA | Unmodified | 381850 | 0 | 0 | 0 | 753130 | 231010 | 36875 | 31572 | 210760 | 0 | 0 | 0 | 0 | 0 | 471730 | 146500 | 189330 | 0 |
| YLTGIGSD | EXT1; EXT1 | Q16394; H7 | EXT1_HUMAN | Unmodified | 0 | 0 | 0 | 0 | 0 | 848700 | 0 | 0 | 598420 | 0 | 0 | 0 | 0 | 0 | 0 | 0 | 83296 | 0 |
| YLTHEEK | PROSER2; PRO | Q86WR7; X6 | PRSR2_HUMAN | Unmodified | 0 | 0 | 0 | 0 | 326710 | 340050 | 0 | 0 | 20514 | 0 | 48869 | 534480 | 0 | 0 | 12159 | 0 | 0 | 0 |
| YLTLENVAI | TPP1; TPP1; TP | O14773; E7E | TPP1_HUMAN | Unmodified | 99496 | 0 | 0 | 0 | 0 | 892480 | 1277100 | 0 | 6912700 | 3216400 | 662190 | 0 | 0 | 0 | 328450 | 0 | 0 | 782090 |
| YLTMDDLT | BCAT1 | P54687 | BCAT1_HUMAN | Unmodified | 0 | 0 | 0 | 0 | 0 | 0 | 0 | 0 | 0 | 0 | 597560 | 2455100 | 0 | 0 | 0 | 0 | 0 | 0 |
| YLTMDDLT | BCAT1 | P54687 | BCAT1_HUMAN | Oxidation (M) | 0 | 0 | 0 | 0 | 0 | 0 | 0 | 0 | 0 | 0 | 456890 | 2127800 | 0 | 0 | 0 | 0 | 0 | 0 |
| YLTNAYSRI | CLIC4 | Q9Y696 | CLIC4_HUMAN | Unmodified | 0 | 0 | 1345000 | 0 | 222980 | 0 | 0 | 0 | 1187400 | 0 | 0 | 1807500 | 0 | 0 | 0 | 0 | 0 | 298140 |
| YLTNAYSRI | CLIC4 | Q9Y696 | CLIC4_HUMAN | Unmodified | 223950 | 0 | 0 | 0 | 10616000 | 3985900 | 0 | 0 | 5919600 | 2333700 | 0 | 3119100 | 5384600 | 4107000 | 0 | 0 | 8049900 | 7402900 |
| YLTTAVITNI | EIF3E; EIF3E; EI | P60228; B3K | EIF3E_HUMAN | Unmodified | 388370 | 268060 | 525370 | 128320 | 0 | 0 | 0 | 0 | 0 | 0 | 281870 | 14620000 | 0 | 0 | 314670 | 612380 | 0 | 0 |
| YLTTAVITNI | EIF3E; EIF3E; EI | P60228; B3K | EIF3E_HUMAN | Unmodified | 71543 | 99766 | 0 | 56561 | 0 | 0 | 60676 | 0 | 0 | 0 | 0 | 0 | 0 | 0 | 0 | 10901 | 0 | 0 |
| YLTVAAIFR | TUBB2A; TUBB | Q13885; A6 | TBB2A_HUMAN | Unmodified | 0 | 0 | 112590 | 181330 | 0 | 0 | 0 | 0 | 14074000 | 1579200 | 6621100 | 53473000 | 0 | 0 | 148610 | 0 | 0 | 0 |
| YLTVAAVFF | TUBB4A; TUBB; | P04350; P07 | TBB4A_HUMAN | Unmodified | 9691900 | 8735200 | 6455000 | 9398000 | 3508700 | 882750 | 1609500 | 294550 | 44106000 | 4553200 | 18092000 | 1.45E+08 | 6993800 | 1254400 | 11798000 | 10112000 | 651350 | 2783800 |
| YLTVAAVFF | TUBB4A; TUBB; | P04350; P07 | TBB4A_HUMAN | Unmodified | 0 | 0 | 0 | 0 | 521810 | 0 | 294960 | 0 | 0 | 364760 | 0 | 0 | 0 | 0 | 0 | 0 | 124310 | 653630 |
| YLTVATVFR | TUBB6; TUBB3; | Q9BUF5; Q1 | TBB6_HUMAN | Unmodified | 337450 | 0 | 0 | 0 | 484930 | 0 | 148340 | 51252 | 4637900 | 0 | 420280 | 6132000 | 0 | 0 | 0 | 0 | 0 | 0 |
| YLTYLNAR | SLC44A2; SLC4 | Q8IWA5; K7 | CTL2_HUMAN | Unmodified | 73266 | 0 | 0 | 179470 | 0 | 0 | 0 | 0 | 333990 | 0 | 0 | 4479000 | 0 | 0 | 336290 | 57708 | 0 | 0 |
| YLVIQGDE | EGFR; EGFR; E | P00533; Q5 | EGFR_HUMAN | Unmodified | 0 | 0 | 2125900 | 0 | 0 | 0 | 0 | 0 | 812430 | 823510 | 0 | 412850 | 0 | 0 | 1770600 | 2401900 | 0 | 0 |
| YLVLDEADI | DDX5; DDX17; | P17844; Q9 | DDX5_HUMAN | Unmodified | 399180 | 438470 | 309140 | 103630 | 0 | 0 | 0 | 85894 | 286030 | 169910 | 232890 | 922900 | 0 | 218970 | 318280 | 339500 | 0 | 0 |
| YLVNPEGQ | GPX8; GPX8 | Q8TED1; E7 | GPX8_HUMAN | Unmodified | 0 | 0 | 0 | 0 | 249840 | 0 | 0 | 0 | 1790800 | 267280 | 0 | 0 | 0 | 0 | 0 | 0 | 0 | 0 |
| YLVPSDLT | GABARAP; GAB | O95166; Q9 | GBRAP_HUMA | Unmodified | 0 | 0 | 0 | 0 | 0 | 0 | 0 | 0 | 0 | 304390 | 0 | 0 | 0 | 0 | 0 | 0 | 0 | 0 |
| YLVQDGAC | ANK3 | Q12955 | ANK3_HUMAN | Unmodified | 0 | 0 | 1075500 | 987250 | 0 | 0 | 38752 | 0 | 1199400 | 0 | 0 | 0 | 860720 | 0 | 2964600 | 2962400 | 0 | 0 |
| YLVVYNPL | MAN2A1 | Q16706 | MA2A1_HUMA | Unmodified | 0 | 0 | 0 | 0 | 0 | 0 | 0 | 0 | 0 | 0 | 0 | 0 | 0 | 0 | 0 | 0 | 0 | 0 |
| YLYGRHM | RSU1 | Q15404 | RSU1_HUMAN | Unmodified | 0 | 0 | 0 | 0 | 741350 | 177560 | 279340 | 616210 | 0 | 61142 | 0 | 0 | 0 | 343690 | 279270 | 0 | 154560 | 257320 |
| YLYHGQTL | TGFBI; TGFBI; T | Q15582; H0 | BGH3_HUMAN | Unmodified | 0 | 0 | 0 | 0 | 61557 | 0 | 1400600 | 0 | 675750 | 0 | 0 | 206790 | 0 | 0 | 34269 | 0 | 0 | 0 |
| YLYHGQTL | TGFBI; TGFBI; T | Q15582; H0 | BGH3_HUMAN | Unmodified | 0 | 0 | 0 | 0 | 0 | 0 | 215550 | 162560 | 0 | 0 | 0 | 0 | 0 | 0 | 0 | 0 | 0 | 0 |
| YLYPNIDK | STAT1; STAT1; | P42224; A0A | STAT1_HUMAN | Unmodified | 0 | 0 | 0 | 0 | 1678000 | 562750 | 0 | 59781 | 0 | 0 | 0 | 0 | 137250 | 258760 | 420480 | 73258 | 297980 | 3640800 |
| YLYSASML | WWP1 | Q9H0M0 | WWP1_HUMAN | Oxidation (M) | 0 | 0 | 0 | 860080 | 0 | 0 | 0 | 0 | 276880 | 0 | 0 | 0 | 0 | 0 | 1079800 | 412680 | 0 | 0 |
| YLYTLVITDI | RPL38; RPL38; | P63173; J3K | RL38_HUMAN | Unmodified | 87385 | 0 | 0 | 0 | 663700 | 0 | 0 | 0 | 0 | 0 | 2774400 | 13609000 | 1107400 | 322130 | 0 | 0 | 0 | 1218700 |
| YLYVDKNF | MYH9; MYH9; M | P35579; Q5 | MYH9_HUMAN | Unmodified | 25669000 | 0 | 0 | 10564000 | 0 | 0 | 595310 | 0 | 0 | 0 | 0 | 0 | 0 | 2930400 | 0 | 0 | 2580700 | 0 |
| YMACCLLY | TUBA1A; TUBA | Q71U36; P6 | TBA1A_HUMAN | Oxidation (M) | 907000 | 0 | 0 | 176920 | 0 | 0 | 216760 | 254070 | 997290 | 365310 | 268910 | 22986000 | 0 | 0 | 248300 | 216700 | 0 | 0 |
| YMACCLLY | TUBA1A; TUBA | Q71U36; P6 | TBA1A_HUMAN | Unmodified | 0 | 0 | 0 | 65126 | 0 | 0 | 0 | 0 | 1804000 | 143970 | 134710 | 8478000 | 0 | 0 | 0 | 0 | 0 | 303580 |
| YMACCLLY | TUBA1A; TUBA | Q71U36; P6 | TBA1A_HUMAN | Unmodified | 0 | 73189 | 0 | 0 | 19618000 | 3021000 | 17050000 | 1157800 | 18867000 | 6925900 | 1245100 | 8994000 | 8451800 | 8989100 | 0 | 0 | 5648600 | 23249000 |
| YMACCLLY | TUBA1A; TUBA | Q71U36; P6 | TBA1A_HUMAN | Oxidation (M) | 3415000 | 825100 | 1935000 | 5179000 | 16544000 | 4517100 | 32899000 | 6603500 | 11808000 | 14203000 | 9841000 | 40174000 | 19955000 | 19194000 | 2436600 | 2499400 | 13243000 | 23573000 |
| YMACCLLY | TUBA1A; TUBA1 | Q71U36; Q9 | TBA1A_HUMAN | Oxidation (M) | 0 | 0 | 0 | 0 | 3423700 | 0 | 19787000 | 73397 | 0 | 2116100 | 0 | 0 | 2807900 | 0 | 0 | 0 | 7878800 | 0 |
| YMAFAHDL | ARSA; ARSA | P15289; A0A | ARSA_HUMAN | Oxidation (M) | 0 | 0 | 0 | 0 | 0 | 0 | 0 | 0 | 662610 | 0 | 0 | 0 | 0 | 0 | 38519 | 0 | 0 | 0 |
| YMAFAHDL | ARSA; ARSA | P15289; A0A | ARSA_HUMAN | Unmodified | 0 | 0 | 0 | 823480 | 0 | 0 | 0 | 0 | 1296400 | 26537 | 0 | 0 | 0 | 0 | 0 | 0 | 0 | 0 |
| YMDINFDF | MYO1D; MYO1 | O94832; B0 | MYO1D_HUMA | Oxidation (M) | 140360 | 0 | 189790 | 56335 | 0 | 0 | 0 | 0 | 0 | 0 | 0 | 49293 | 0 | 0 | 287370 | 155000 | 0 | 0 |
| YMDNDYA | NAPG | Q99747 | SNAG_HUMAN | Unmodified | 40543 | 0 | 127300 | 239570 | 0 | 0 | 183300 | 0 | 0 | 0 | 127270 | 512960 | 514900 | 0 | 225730 | 0 | 424160 | 1054300 |
| YMDVQFDI | MYO1C; MYO1 | O00159; F5 | MYO1C_HUMA | Oxidation (M) | 0 | 1488800 | 1676900 | 2198900 | 1629300 | 0 | 0 | 0 | 532310 | 0 | 0 | 0 | 0 | 0 | 3850800 | 2643200 | 528050 | 0 |

| | | | | | | | | | | | | | | | | | | | | | | |
|---|---|---|---|---|---|---|---|---|---|---|---|---|---|---|---|---|---|---|---|---|---|---|
| YMDVQFDI | MYO1C; MYO1( | O00159; F5H | MYO1C_HUMA | Oxidation (M) | 0 | 0 | 0 | 0 | 506070 | 14060000 | 0 | 0 | 84187 | 0 | 0 | 109780 | 205300 | 0 | 38890 | 0 | 419060 | 1623000 |
| YMEAFKPFI | KPNB1; KPNB1 | Q14974; J3K | IMB1_HUMAN I | Oxidation (M) | 15453 | 31734 | 273100 | 0 | 0 | 0 | 0 | 0 | 0 | 388940 | 51605 | 4497000 | 190490 | 0 | 70204 | 0 | 176330 | 0 |
| YMEDMEQ/ | PACSIN3; PACS | Q9UKS6; E9I | PACN3_HUMAI | 2 Oxidation (M) | 1124000 | 179450 | 81150 | 0 | 0 | 0 | 0 | 0 | 0 | 0 | 0 | 0 | 0 | 0 | 273450 | 360280 | 0 | 0 |
| YMEDMEQ/ | PACSIN3; PACS | Q9UKS6; E9I | PACN3_HUMAI | Oxidation (M) | 2055500 | 1557900 | 0 | 0 | 0 | 0 | 0 | 0 | 0 | 0 | 0 | 0 | 0 | 0 | 1188400 | 706130 | 0 | 0 |
| YMEDMEQ/ | PACSIN3; PACS | Q9UKS6; E9I | PACN3_HUMAI | Unmodified | 182160 | 0 | 0 | 474030 | 0 | 0 | 0 | 0 | 0 | 0 | 0 | 0 | 968910 | 0 | 0 | 0 | 0 | 0 |
| YMEHEERP | HSPA4L; HSPA4 | O95757; E9F | HS74L_HUMAN | Unmodified | 0 | 0 | 0 | 0 | 136460 | 39213 | 30020 | 0 | 216430 | 98934 | 350190 | 1278300 | 0 | 420650 | 39126 | 0 | 142760 | 59332 |
| YMGDLSG( | HMOX2; HMOX | P30519; I3L1 | HMOX2_HUMA | Unmodified | 0 | 0 | 0 | 0 | 0 | 0 | 0 | 0 | 0 | 0 | 0 | 511040 | 0 | 0 | 0 | 0 | 0 | 0 |
| YMGLPTPE | DPP4; DPP4; DI | P27487; A0A | DPP4_HUMAN | Oxidation (M) | 0 | 0 | 673760 | 0 | 0 | 0 | 0 | 0 | 0 | 0 | 0 | 0 | 0 | 0 | 276740 | 277460 | 0 | 0 |
| YMHSGPVI | NME1; NME1-N | P15531; J3KI | NDKA_HUMAN | 2 Oxidation (M) | 0 | 0 | 962970 | 454720 | 14322000 | 0 | 5317700 | 0 | 0 | 4161400 | 26325000 | 13912000 | 18999000 | 9710200 | 566040 | 2735600 | 9064800 | 9257200 |
| YMHSGPVI | NME1; NME1-N | P15531; J3KI | NDKA_HUMAN | Oxidation (M) | 0 | 0 | 0 | 106860 | 2273200 | 0 | 165550 | 0 | 249490 | 230610 | 392030 | 692250 | 3002500 | 478110 | 0 | 0 | 1224900 | 4288600 |
| YMIGNSVQ | ERBIN | Q96RT1 | ERBIN_HUMAN | Unmodified | 0 | 0 | 0 | 0 | 10544000 | 1127800 | 154540 | 120890 | 174930 | 163790 | 0 | 486500 | 595650 | 1003600 | 197260 | 89074 | 487010 | 1056100 |
| YMLDCRTS | VDAC3 | Q9Y277 | VDAC3_HUMAI | Unmodified | 0 | 49087 | 0 | 0 | 0 | 0 | 75601 | 13523 | 482790 | 1625700 | 0 | 0 | 0 | 0 | 0 | 0 | 0 | 0 |
| YMLDCRTS | VDAC3 | Q9Y277 | VDAC3_HUMAI | Oxidation (M) | 586180 | 0 | 0 | 0 | 0 | 0 | 0 | 23092 | 0 | 550060 | 0 | 0 | 400200 | 0 | 0 | 0 | 0 | 0 |
| YMLEPDPD | AAK1; AAK1; AA | Q2M2I8; A0A | AAK1_HUMAN | Oxidation (M) | 0 | 0 | 77080 | 194010 | 0 | 0 | 0 | 0 | 0 | 0 | 0 | 0 | 169870 | 732660 | 531830 | 399030 | 0 | 0 |
| YMLGTNH( | DOP1B | Q9Y3R5 | DOP2_HUMAN | Unmodified | 0 | 0 | 0 | 0 | 0 | 0 | 0 | 0 | 588320 | 0 | 122300 | 502740 | 0 | 0 | 0 | 0 | 0 | 0 |
| YMLTHQEL | KIF23; KIF23; KI | Q02241; A0A | KIF23_HUMAN | Oxidation (M) | 0 | 0 | 211350 | 333180 | 4442400 | 3205500 | 0 | 67886 | 0 | 449630 | 362950 | 475720 | 0 | 0 | 263490 | 0 | 868390 | 548910 |
| YMLTHQEL | KIF23; KIF23; KI | Q02241; A0A | KIF23_HUMAN | Unmodified | 99383 | 0 | 273090 | 722240 | 35547000 | 3161900 | 411440 | 96656 | 2927500 | 1554800 | 0 | 1047500 | 1177900 | 821340 | 285340 | 81475 | 2920900 | 2218000 |
| YMLTHQEL | KIF23; KIF23; KI | Q02241; A0A | KIF23_HUMAN | Unmodified | 0 | 0 | 0 | 0 | 7534100 | 0 | 560500 | 0 | 0 | 1240500 | 0 | 0 | 0 | 0 | 0 | 0 | 0 | 0 |
| YMMIVDPA | GAA | P10253 | LYAG_HUMAN | 2 Oxidation (M) | 0 | 0 | 243730 | 0 | 0 | 0 | 0 | 0 | 1547100 | 0 | 0 | 0 | 0 | 0 | 0 | 0 | 2337700 | 0 |
| YMMIVDPA | GAA | P10253 | LYAG_HUMAN | Oxidation (M) | 0 | 0 | 0 | 0 | 1117900 | 0 | 1128300 | 0 | 2227000 | 0 | 0 | 0 | 0 | 0 | 0 | 0 | 0 | 0 |
| YMNATLK | BAIAP2L1 | Q9UHR4 | BI2L1_HUMAN | Unmodified | 0 | 0 | 0 | 0 | 0 | 760520 | 61338 | 0 | 580810 | 369190 | 0 | 0 | 0 | 0 | 0 | 0 | 0 | 0 |
| YMNSGPVI | NME2P1; NME2 | O60361; P22 | NDK8_HUMAN | Oxidation (M) | 0 | 0 | 0 | 0 | 1676000 | 0 | 0 | 0 | 131220 | 206600 | 0 | 0 | 664440 | 0 | 0 | 0 | 1340200 | 0 |
| YMNVLFSC | MYO1D; MYO1I | O94832; J3Q | MYO1D_HUMA | Oxidation (M) | 1298000 | 708510 | 259400 | 410920 | 0 | 0 | 0 | 0 | 0 | 0 | 0 | 174570 | 0 | 0 | 853380 | 489850 | 0 | 0 |
| YMQMNHA | PLCG2; PLCG2 | P16885; A0A | PLCG2_HUMAI | 2 Oxidation (M) | 0 | 0 | 71327 | 0 | 0 | 0 | 0 | 0 | 0 | 0 | 0 | 2695100 | 0 | 0 | 0 | 78720 | 0 | 0 |
| YMQMNHA | PLCG2; PLCG2 | P16885; A0A | PLCG2_HUMAI | Oxidation (M) | 0 | 0 | 0 | 0 | 0 | 0 | 0 | 0 | 0 | 0 | 0 | 0 | 0 | 0 | 0 | 0 | 0 | 0 |
| YMRALDEY | ATP6V1A | P38606 | VATA_HUMAN | Unmodified | 0 | 0 | 0 | 0 | 0 | 0 | 408540 | 0 | 0 | 0 | 0 | 0 | 0 | 0 | 0 | 0 | 0 | 0 |
| YMTISDEW | EIF3B; EIF3B | P55884; C9J | EIF3B_HUMAN | Oxidation (M) | 531240 | 0 | 0 | 629400 | 0 | 0 | 0 | 0 | 0 | 0 | 413110 | 2484500 | 138960 | 0 | 227540 | 0 | 0 | 0 |
| YMVADKFT | PPP2R1A; PPP2 | P30153; B3K | 2AAA_HUMAN | Oxidation (M) | 1338100 | 0 | 0 | 1112000 | 2860600 | 637050 | 434390 | 473040 | 0 | 678250 | 999680 | 5443200 | 6168500 | 2130300 | 0 | 696450 | 1184200 | 2199300 |
| YMVADKFT | PPP2R1A; PPP2 | P30153; B3K | 2AAA_HUMAN | Unmodified | 0 | 0 | 0 | 0 | 3468300 | 1633600 | 819460 | 154040 | 2190500 | 2041700 | 0 | 0 | 2095600 | 0 | 542710 | 0 | 1554700 | 5026900 |
| YMVSGTN\ | GPAA1; GPAA1; | O43292; E9I | GPAA1_HUMAI | Unmodified | 0 | 0 | 0 | 0 | 0 | 0 | 0 | 0 | 860070 | 0 | 0 | 0 | 98230 | 0 | 0 | 0 | 0 | 0 |
| YMVSGTN\ | GPAA1; GPAA1; | O43292; E9I | GPAA1_HUMAI | Oxidation (M) | 0 | 0 | 0 | 0 | 308110 | 0 | 0 | 0 | 0 | 701240 | 0 | 0 | 0 | 0 | 0 | 0 | 0 | 0 |
| YMVSGTN\ | GPAA1; GPAA1; | O43292; E9I | GPAA1_HUMAI | Oxidation (M) | 0 | 0 | 0 | 0 | 1245700 | 0 | 0 | 0 | 3391500 | 0 | 0 | 0 | 0 | 0 | 0 | 0 | 90018 | 0 |
| YMVSGTN\ | GPAA1; GPAA1; | O43292; E9I | GPAA1_HUMAI | Unmodified | 0 | 0 | 0 | 0 | 0 | 0 | 0 | 0 | 5398000 | 0 | 0 | 0 | 0 | 0 | 0 | 0 | 0 | 0 |
| YMVWSDEI | POFUT1 | Q9H488 | OFUT1_HUMAI | Oxidation (M) | 0 | 0 | 0 | 0 | 0 | 0 | 0 | 0 | 4067500 | 530090 | 0 | 0 | 0 | 0 | 0 | 0 | 0 | 0 |
| YNADEARS | EIF1AY; EIF1AX; | O14602; P47 | IF1AY_HUMAN | Unmodified | 0 | 0 | 0 | 0 | 176360 | 0 | 0 | 156990 | 0 | 31556 | 0 | 0 | 1690600 | 319650 | 0 | 0 | 102790 | 224590 |
| YNADISTFF | TYRP1; TYRP1 | P17643; E7E | TYRP1_HUMAN | Unmodified | 0 | 0 | 0 | 0 | 0 | 0 | 0 | 0 | 610300 | 0 | 0 | 0 | 0 | 0 | 0 | 0 | 0 | 0 |
| YNAESTERI | PSEN1; PSEN1 | P49768; E7E | PSN1_HUMAN | Unmodified | 185390 | 0 | 278460 | 0 | 4038300 | 94790 | 0 | 0 | 1983400 | 0 | 0 | 521650 | 0 | 0 | 0 | 305990 | 0 | 0 |
| YNALLAVQ | ATP6V1H; ATP6 | Q9UI12; G3\ | VATH_HUMAN | Unmodified | 0 | 82537 | 72641 | 130750 | 0 | 0 | 0 | 0 | 446720 | 0 | 260360 | 2416900 | 0 | 0 | 147970 | 133390 | 0 | 0 |
| YNAPTSHV | SPAG9; SPAG9 | O60271; A0A | JIP4_HUMAN C | Unmodified | 0 | 0 | 0 | 0 | 349320 | 58715 | 347930 | 293470 | 0 | 91106 | 175960 | 425300 | 335030 | 168800 | 0 | 0 | 444480 | 1930000 |
| YNAPTSHV | SPAG9; SPAG9 | O60271; A0A | JIP4_HUMAN C | Unmodified | 0 | 0 | 0 | 0 | 0 | 0 | 0 | 36850 | 0 | 0 | 0 | 0 | 0 | 0 | 0 | 0 | 0 | 239530 |
| YNAQCQE1 | MDK; MDK; MD | P21741; E9P | MK_HUMAN Mi | Unmodified | 0 | 14343 | 0 | 67873 | 40089 | 0 | 0 | 0 | 16117000 | 557800 | 123790 | 82076 | 0 | 0 | 0 | 0 | 0 | 0 |
| YNAQCQE1 | MDK; MDK; MD | P21741; E9P | MK_HUMAN Mi | Unmodified | 0 | 0 | 0 | 0 | 1262300 | 0 | 134860 | 0 | 4397800 | 4791600 | 0 | 166350 | 0 | 0 | 0 | 0 | 300270 | 842350 |
| YNDFLHDF | PLBD2 | Q8NHP8 | PLBL2_HUMAN | Unmodified | 0 | 28401 | 0 | 66400 | 1317400 | 24044 | 0 | 0 | 664840 | 136220 | 126700 | 0 | 0 | 0 | 0 | 0 | 393760 | 0 |
| YNDTFWK | HSP90B2P; HSI | Q58FF3; P14 | ENPLL_HUMAN | Unmodified | 0 | 0 | 2410300 | 0 | 0 | 0 | 0 | 0 | 24818000 | 4346600 | 11841000 | 9366300 | 0 | 0 | 995170 | 863440 | 0 | 73123 |
| YNDTFWKE | HSP90B2P; HSI | Q58FF3; P14 | ENPLL_HUMAN | Unmodified | 0 | 0 | 0 | 0 | 386950 | 37095 | 0 | 17034 | 334610 | 1051500 | 0 | 138060 | 0 | 0 | 0 | 0 | 0 | 77751 |
| YNEQHVPC | FLNA; FLNA | P21333; Q60 | FLNA_HUMAN | Unmodified | 0 | 0 | 290930 | 0 | 1290800 | 706070 | 0 | 0 | 287660 | 0 | 0 | 1444900 | 804310 | 332980 | 635990 | 323580 | 580620 | 0 |
| YNFFTGCP | CCT7 | Q99832 | TCPH_HUMAN | Unmodified | 0 | 0 | 0 | 0 | 0 | 0 | 0 | 0 | 0 | 0 | 0 | 4183500 | 104200 | 0 | 0 | 0 | 0 | 0 |
| YNFHGTAE | CSK; CSK; CSK | P41240; H3E | CSK_HUMAN T | Unmodified | 59197 | 0 | 138580 | 35343 | 124900 | 0 | 0 | 0 | 0 | 0 | 286480 | 1477200 | 73992 | 39671 | 132390 | 134650 | 0 | 0 |
| YNFHPSLL | PGAP4 | Q9BRR3 | PGAP4_HUMAI | Unmodified | 0 | 0 | 0 | 0 | 0 | 0 | 0 | 0 | 2178600 | 0 | 0 | 0 | 0 | 0 | 0 | 0 | 0 | 0 |
| YNFIADVVE | HTRA1; HTRA2; | Q92743; O43 | HTRA1_HUMAN | Unmodified | 0 | 0 | 0 | 0 | 1271000 | 0 | 0 | 58966 | 3700500 | 1263000 | 0 | 0 | 0 | 0 | 0 | 0 | 1171400 | 7072600 |
| YNFNNRG | XPNPEP1; XPNI | Q9NQW7; Q | XPP1_HUMAN | Unmodified | 0 | 0 | 0 | 135390 | 1331800 | 0 | 1568100 | 48489 | 0 | 397060 | 292130 | 0 | 1228200 | 1515100 | 0 | 0 | 1635600 | 0 |
| YNGEPVQI | VIL1; VIL1; VIL1 | P09327; H7C | VILI_HUMAN Vi | Unmodified | 0 | 0 | 0 | 0 | 0 | 0 | 0 | 0 | 0 | 0 | 0 | 3195300 | 0 | 0 | 0 | 0 | 0 | 0 |
| YNHIDESEI | HSPH1; HSPH1 | Q92598; Q5T | HS105_HUMAI | Unmodified | 0 | 0 | 0 | 0 | 0 | 0 | 0 | 0 | 1632600 | 0 | 1263700 | 900370 | 136330 | 0 | 0 | 30856 | 0 | 0 |
| YNHIDESEI | HSPH1; HSPH1 | Q92598; Q5T | HS105_HUMAI | Unmodified | 0 | 0 | 0 | 0 | 378930 | 65460 | 286150 | 143330 | 0 | 148550 | 0 | 0 | 558400 | 430980 | 0 | 0 | 510310 | 582670 |
| YNHSHDQ | EIPR1; EIPR1; E | Q53HC9; A8 | EIPR1_HUMAN | Unmodified | 0 | 0 | 0 | 0 | 0 | 0 | 0 | 0 | 262180 | 0 | 83012 | 443110 | 0 | 0 | 0 | 43085 | 0 | 0 |
| YNIAITAVS | PTPRB; PTPRB; | P23467; F8V | PTPRB_HUMAN | Unmodified | 386530 | 0 | 0 | 0 | 0 | 0 | 0 | 198680 | 0 | 0 | 0 | 0 | 0 | 0 | 0 | 0 | 0 | 0 |
| YNIEVVCE | EIF2S3; EIF2S3I | P41091; Q2I | IF2G_HUMAN E | Unmodified | 0 | 0 | 0 | 0 | 0 | 0 | 0 | 0 | 0 | 0 | 0 | 1897300 | 0 | 0 | 0 | 0 | 335190 | 0 |
| YNIMLVR | CCT8; CCT8 | P50990; H7C | TCPQ_HUMAN | Unmodified | 0 | 0 | 0 | 0 | 0 | 234710 | 0 | 0 | 1344100 | 0 | 941670 | 5486100 | 1370000 | 0 | 0 | 0 | 0 | 696580 |
| YNIMLVR | CCT8; CCT8 | P50990; H7C | TCPQ_HUMAN | Oxidation (M) | 0 | 0 | 0 | 0 | 0 | 214060 | 0 | 0 | 2288100 | 800050 | 1388900 | 5414000 | 0 | 0 | 0 | 0 | 209210 | 482810 |
| YNIWCSTE | YTHDF2 | Q9Y5A9 | YTHD2_HUMAI | Unmodified | 753470 | 304570 | 318740 | 240380 | 0 | 0 | 0 | 0 | 0 | 0 | 0 | 1089300 | 0 | 0 | 261700 | 320310 | 0 | 0 |
| YNLDLTNL | DICER1; DICER | Q9UPY3; A0 | DICER_HUMAN | Unmodified | 0 | 0 | 0 | 0 | 0 | 0 | 0 | 0 | 0 | 0 | 0 | 0 | 0 | 0 | 0 | 0 | 0 | 0 |
| YNLIAVSNI | USP15; USP15 | Q9Y4E8; H0 | UBP15_HUMAN | Oxidation (M) | 0 | 0 | 0 | 0 | 0 | 0 | 0 | 0 | 0 | 0 | 0 | 1531700 | 0 | 0 | 0 | 172900 | 0 | 0 |
| YNLPLDIQ | TOM1L1; TOM1 | O75674; J3K | TM1L1_HUMAN | Unmodified | 5257800 | 5641800 | 3890200 | 1610100 | 0 | 0 | 0 | 0 | 0 | 0 | 0 | 12082000 | 873710 | 233000 | 7396400 | 4800200 | 0 | 0 |
| YNLPLDIQ | TOM1L1; TOM1 | O75674; J3K | TM1L1_HUMAN | Unmodified | 0 | 137660 | 0 | 95172 | 526480 | 0 | 93990 | 0 | 0 | 745970 | 0 | 0 | 1894900 | 1988700 | 294440 | 0 | 147380 | 154470 |
| YNLPSWCI | INPP5D; INPP5I | Q92835; H0 | SHIP1_HUMAN | Unmodified | 0 | 0 | 0 | 0 | 0 | 0 | 47144 | 0 | 0 | 0 | 130740 | 243480 | 0 | 0 | 0 | 0 | 0 | 23542 |
| YNLSPSIFF | SORD; SORD | Q00796; A0A | DHSO_HUMAN | Unmodified | 0 | 0 | 0 | 0 | 0 | 0 | 0 | 0 | 172940 | 0 | 180360 | 676200 | 0 | 0 | 1763100 | 0 | 0 | 0 |
| YNMERDIN | PLCG2; PLCG2 | P16885; A0A | PLCG2_HUMAI | Unmodified | 155570 | 86319000 | 34401000 | 0 | 0 | 0 | 0 | 152970 | 488310 | 0 | 0 | 736350 | 81229 | 0 | 1.05E+08 | 41529000 | 0 | 0 |
| YNNCWLAI | PCK2; PCK2 | Q16822; H0 | PCKGM_HUMA | Unmodified | 0 | 0 | 0 | 0 | 0 | 0 | 0 | 0 | 761840 | 0 | 0 | 0 | 0 | 0 | 0 | 0 | 0 | 0 |
| YNNGSTEL | PHKB | Q93100 | KPBB_HUMAN | Unmodified | 0 | 0 | 0 | 0 | 0 | 0 | 0 | 0 | 0 | 0 | 0 | 496650 | 0 | 0 | 0 | 0 | 0 | 0 |
| YNNPAMLI | COPS3; COPS3 | Q9UNS2; H7 | CSN3_HUMAN | Unmodified | 0 | 0 | 0 | 0 | 586560 | 0 | 0 | 0 | 394890 | 332050 | 0 | 813820 | 337800 | 131310 | 0 | 0 | 168290 | 439370 |
| YNNPAMLI | COPS3; COPS3 | Q9UNS2; H7 | CSN3_HUMAN | 2 Oxidation (M) | 0 | 0 | 0 | 101870 | 0 | 0 | 0 | 0 | 0 | 0 | 0 | 477300 | 90780 | 0 | 0 | 0 | 0 | 0 |
| YNNPAMLI | COPS3; COPS3 | Q9UNS2; H7 | CSN3_HUMAN | Oxidation (M) | 0 | 0 | 542470 | 0 | 132850 | 0 | 63137 | 20651 | 0 | 0 | 240360 | 813930 | 353220 | 190080 | 0 | 0 | 0 | 0 |
| YNNQLATR | ITGAV | P06756 | ITAV_HUMAN Ii | Unmodified | 0 | 0 | 0 | 0 | 0 | 0 | 0 | 0 | 2391000 | 126210 | 0 | 5392200 | 0 | 0 | 0 | 0 | 0 | 0 |
| YNPNVLPV | ATP1B1 | P05026 | AT1B1_HUMAN | Unmodified | 195410 | 0 | 1336500 | 995850 | 0 | 0 | 119420 | 0 | 2301600 | 562970 | 654390 | 9131600 | 0 | 0 | 1197200 | 1132900 | 0 | 0 |
| YNPNVLPV | ATP1B1 | P05026 | AT1B1_HUMAN | Unmodified | 0 | 0 | 0 | 0 | 0 | 0 | 0 | 0 | 3769300 | 476910 | 182720 | 11197000 | 0 | 0 | 2725500 | 1851600 | 139880 | 0 |
| YNPNVLPV | ATP1B1 | P05026 | AT1B1_HUMAN | Unmodified | 700190 | 371210 | 9244100 | 5448400 | 5197100 | 1569400 | 1404500 | 132240 | 14865000 | 1438100 | 5860500 | 40831000 | 4926000 | 1855800 | 7319900 | 6622900 | 347030 | 2693400 |
| YNPTWHCI | DYNLL2; DYNLI | Q96FJ2; P63 | DYL2_HUMAN | Unmodified | 1865200 | 1271400 | 0 | 0 | 0 | 305010 | 0 | 0 | 0 | 0 | 0 | 8654600 | 0 | 0 | 0 | 965040 | 97790 | 0 |
| YNQATPNF | VASP | P50552 | VASP_HUMAN | Unmodified | 0 | 0 | 0 | 0 | 0 | 0 | 0 | 0 | 0 | 0 | 0 | 457670 | 0 | 0 | 0 | 0 | 0 | 0 |
| YNQATPNF | VASP | P50552 | VASP_HUMAN | Unmodified | 32935 | 0 | 98773 | 58667 | 1249100 | 121150 | 279470 | 0 | 0 | 203410 | 0 | 0 | 1632700 | 895810 | 0 | 0 | 179100 | 965060 |
| YNQATPTF | EVL; EVL; EVL; E | Q9UI08; G3\ | EVL_HUMAN Ei | Unmodified | 29754 | 62365 | 0 | 0 | 0 | 0 | 0 | 0 | 0 | 0 | 0 | 451160 | 0 | 0 | 241930 | 331710 | 0 | 0 |
| YNQATQTF | ENAH; ENAH; E | Q8N8S7; A0 | ENAH_HUMAN | Unmodified | 0 | 0 | 480130 | 255410 | 0 | 0 | 0 | 0 | 1413500 | 0 | 134280 | 23403000 | 0 | 0 | 933430 | 1953000 | 0 | 0 |
| YNQLLRIEE | ENO1; ENO1 | P06733; A0A | ENOA_HUMAN | Unmodified | 3622000 | 2307900 | 5074000 | 32434000 | 1.01E+09 | 1.01E+08 | 36878000 | 3365900 | 87511000 | 2.78E+08 | 18552000 | 46414000 | 4.35E+08 | 1.66E+08 | 13204000 | 6705700 | 4.25E+08 | 5.57E+08 |
| YNQLLRIEE | ENO1; ENO1 | P06733; A0A | ENOA_HUMAN | Unmodified | 0 | 0 | 0 | 0 | 25426000 | 3810900 | 10984000 | 629530 | 0 | 4340400 | 0 | 0 | 6185200 | 7404000 | 0 | 0 | 1.6E+08 | 14422000 |
| YNQLMRIEI | ENO3 | P13929 | ENOB_HUMAN | Oxidation (M) | 0 | 0 | 0 | 0 | 0 | 0 | 0 | 0 | 0 | 732040 | 0 | 0 | 0 | 0 | 0 | 0 | 0 | 0 |
| YNQLMRIEI | ENO2; ENO2 | P09104; F5H | ENOG_HUMAN | Oxidation (M) | 0 | 0 | 0 | 0 | 79858 | 0 | 0 | 0 | 0 | 500390 | 0 | 0 | 0 | 0 | 0 | 0 | 0 | 0 |
| YNQMDSTI | TM9SF2 | Q99805 | TM9S2_HUMAI | Oxidation (M) | 759560 | 0 | 0 | 0 | 0 | 0 | 0 | 0 | 0 | 0 | 0 | 2416600 | 0 | 0 | 223270 | 0 | 0 | 0 |
| YNQPFFVT | TPP2; TPP2; TPI | P29144; A0A | TPP2_HUMAN | Unmodified | 0 | 0 | 0 | 0 | 0 | 235770 | 0 | 0 | 1617400 | 443550 | 1758100 | 12980000 | 833690 | 0 | 371580 | 0 | 0 | 560210 |
| YNRDFESH | EIF3CL; EIF3C; | B5ME19; Q9 | EIFCL_HUMAN | Unmodified | 0 | 0 | 355020 | 0 | 455290 | 0 | 0 | 0 | 201040 | 18104 | 409890 | 4782800 | 472740 | 0 | 0 | 463050 | 0 | 68587 |
| YNREGDLL | EIF3I; EIF3I; EIF | Q13347; A0A | EIF3I_HUMAN | Unmodified | 0 | 0 | 252370 | 103260 | 0 | 0 | 0 | 0 | 0 | 0 | 0 | 0 | 41678 | 0 | 181070 | 124880 | 0 | 0 |
| YNRLDIIETI | ELP1; ELP1; ELF | O95163; A0A | ELP1_HUMAN | Unmodified | 0 | 0 | 0 | 0 | 0 | 0 | 101350 | 0 | 0 | 219850 | 332820 | 1912300 | 0 | 0 | 0 | 0 | 108290 | 0 |
| YNSMEDAI | TXNDC5 | Q8NBS9 | TXND5_HUMAI | Unmodified | 0 | 0 | 0 | 0 | 0 | 0 | 0 | 0 | 0 | 0 | 0 | 0 | 0 | 0 | 0 | 0 | 0 | 0 |
| YNSMPVPC | SLC31A1 | O15431 | COPT1_HUMAI | 2 Oxidation (M) | 503610 | 666900 | 1121200 | 529720 | 6001400 | 329920 | 0 | 0 | 0 | 0 | 0 | 12380000 | 0 | 0 | 2825600 | 2913600 | 2863800 | 0 |
| YNSMPVPC | SLC31A1 | O15431 | COPT1_HUMAI | Oxidation (M) | 177550 | 454780 | 1241000 | 1510700 | 1326900 | 0 | 0 | 0 | 0 | 0 | 0 | 28447000 | 745650 | 301700 | 4077500 | 2726200 | 0 | 0 |
| YNSQLLSP | TFRC; TFRC; TF | P02786; H7C | TFR1_HUMAN | Unmodified | 699040 | 0 | 495680 | 0 | 0 | 0 | 0 | 676690 | 0 | 0 | 0 | 985290 | 0 | 0 | 0 | 499190 | 0 | 0 |
| YNSWPSSL | UGGT1 | Q9NYU2 | UGGG1_HUMA | Unmodified | 0 | 0 | 820680 | 0 | 691290 | 141540 | 0 | 0 | 11815000 | 341540 | 96246 | 0 | 680370 | 0 | 0 | 0 | 367740 | 2386800 |
| YNTDCVQ( | ATP1A3; ATP1A | P13637; M0 | AT1A3_HUMAN | Unmodified | 0 | 0 | 0 | 0 | 0 | 0 | 262050 | 0 | 493060 | 0 | 0 | 0 | 0 | 0 | 0 | 0 | 0 | 0 |
| YNVPFLTFI | GFM1; GFM1; G | Q96RP9; C9I | EFGM_HUMAN | Unmodified | 0 | 0 | 0 | 0 | 0 | 0 | 0 | 0 | 2991700 | 0 | 0 | 0 | 0 | 0 | 0 | 0 | 0 | 0 |
| YNVPLNR | LIMA1; LIMA1; L | Q9UHB6; F8 | LIMA1_HUMAN | Unmodified | 0 | 0 | 0 | 494870 | 464600 | 0 | 0 | 0 | 68512 | 0 | 0 | 476570 | 0 | 215920 | 859940 | 0 | 287950 | 0 |
| YNYHLDSS | AARS1; AARS1; | P49588; A0A | SYAC_HUMAN | Oxidation (M) | 0 | 0 | 0 | 0 | 805960 | 0 | 0 | 0 | 711380 | 267170 | 707020 | 1081900 | 531430 | 0 | 9055800 | 0 | 0 | 0 |
| YNYLSLDS, | MYO1B; MYO1E | O43795; E9I | MYO1B_HUMA | Unmodified | 146900 | 232110 | 0 | 0 | 0 | 170920 | 0 | 0 | 609250 | 0 | 1000100 | 21566000 | 0 | 0 | 568590 | 358700 | 0 | 0 |
| YNYTLLR | UBXN6; UBXN6 | Q9BZV1; K7I | UBXN6_HUMAI | Unmodified | 155180 | 0 | 0 | 0 | 0 | 0 | 36927 | 0 | 96218 | 133380 | 79978 | 0 | 92268 | 90607 | 0 | 179580 | 0 | 0 |
| YPASLQCV | KLK14; KLK14 | Q9P0G3; A0 | KLK14_HUMAN | Unmodified | 357020 | 0 | 0 | 0 | 0 | 0 | 0 | 0 | 2385200 | 0 | 0 | 0 | 0 | 455240 | 0 | 0 | 0 | 0 |
| YPAVDYNL | DLL1 | O00548 | DLL1_HUMAN | Unmodified | 0 | 0 | 0 | 0 | 0 | 0 | 0 | 0 | 106220 | 628870 | 0 | 447980 | 0 | 0 | 0 | 0 | 0 | 0 |
| YPDVIQELK | CNNM4 | Q6P4Q7 | CNNM4_HUMA | Unmodified | 0 | 0 | 485380 | 0 | 0 | 0 | 110110 | 0 | 0 | 0 | 0 | 916610 | 798260 | 0 | 0 | 0 | 0 | 0 |
| YPDVIQELK | CNNM4 | Q6P4Q7 | CNNM4_HUMA | Unmodified | 0 | 0 | 179010 | 0 | 0 | 0 | 0 | 0 | 0 | 0 | 572950 | 1135700 | 0 | 0 | 140560 | 228800 | 0 | 0 |
| YPENFFLLF | PPP1CA; PPP1( | P62136; P62 | PP1A_HUMAN | Unmodified | 0 | 0 | 612330 | 0 | 870850 | 0 | 1777200 | 28040 | 0 | 88783 | 156450 | 3985200 | 0 | 0 | 233020 | 0 | 0 | 0 |
| YPEVGDLR | TMEM43 | Q9BTV4 | TMM43_HUMAI | Unmodified | 0 | 0 | 0 | 0 | 0 | 0 | 0 | 0 | 1340800 | 204020 | 177530 | 507240 | 0 | 0 | 0 | 0 | 0 | 0 |
| YPEWLQSF | PEX19; PEX19; | P40855; Q5( | PEX19_HUMAN | Unmodified | 222880 | 243950 | 0 | 19554 | 0 | 0 | 0 | 0 | 0 | 0 | 0 | 981260 | 0 | 0 | 0 | 66982 | 0 | 0 |
| YPFESAEA( | RRM1; RRM1 | P23921; E9P | RIR1_HUMAN F | Unmodified | 0 | 0 | 0 | 0 | 0 | 0 | 0 | 0 | 0 | 0 | 0 | 2880200 | 0 | 0 | 0 | 0 | 0 | 0 |

| | | | | | | | | | | | | | | | | | | | | | | |
|---|---|---|---|---|---|---|---|---|---|---|---|---|---|---|---|---|---|---|---|---|---|---|
| YPFILPHQ( | MCTS1; MCTS2 | Q9ULC4; A0. | MCTS1_HUMAI | Unmodified | 327940 | 0 | 1340700 | 576550 | 661750 | 0 | 997170 | 777640 | 2046300 | 268540 | 978260 | 4053600 | 526100 | 0 | 1301000 | 752290 | 323410 | 473180 |
| YPFIVTSDD | NT5E; NT5E | P21589; H0' | 5NTD_HUMAN | Unmodified | 0 | 0 | 163780 | 154640 | 1327000 | 2374200 | 0 | 61960 | 0 | 0 | 0 | 0 | 0 | 479400 | 0 | 0 | 0 | 0 |
| YPGPQAEG | CHGA | P10645 | CMGA_HUMAN | Unmodified | 0 | 0 | 0 | 0 | 0 | 0 | 0 | 0 | 2210900 | 2987900 | 0 | 0 | 0 | 0 | 0 | 0 | 0 | 0 |
| YPGPQAEG | CHGA | P10645 | CMGA_HUMAN | Unmodified | 0 | 0 | 0 | 0 | 0 | 0 | 0 | 0 | 1.09E+08 | 6121800 | 0 | 0 | 0 | 235920 | 0 | 0 | 0 | 0 |
| YPGQGSP( | PRNP; PRNP; PI | P04156; A2A | PRIO_HUMAN I | Unmodified | 2063100 | 593070 | 2236600 | 0 | 0 | 0 | 0 | 0 | 131110 | 0 | 132060 | 0 | 0 | 0 | 138800 | 378570 | 0 | 0 |
| YPHLPCLQ | AGO3; AGO1; A | Q9H9G7; Q9 | AGO3_HUMAN | Unmodified | 0 | 0 | 179990 | 106470 | 0 | 0 | 0 | 0 | 0 | 179230 | 0 | 689270 | 0 | 0 | 0 | 0 | 0 | 0 |
| YPHVEDYR | PSME3; PSME3 | P61289; A0A | PSME3_HUMAI | Unmodified | 0 | 0 | 0 | 55658 | 0 | 0 | 0 | 0 | 0 | 0 | 0 | 979720 | 0 | 0 | 0 | 0 | 0 | 0 |
| YPIEHGIITN | ACTC1; ACTA1; | P68032; P68 | ACTC_HUMAN | Oxidation (M) | 2.57E+08 | 1.67E+08 | 3.31E+08 | 2.61E+08 | 4.04E+08 | 1.63E+08 | 37906000 | 29377000 | 1.61E+08 | 26092000 | 3.19E+08 | 1.07E+09 | 1.41E+08 | 51113000 | 4.24E+08 | 3.71E+08 | 41137000 | 96966000 |
| YPIEHGIITN | ACTC1; ACTA1; | P68032; P68 | ACTC_HUMAN | Unmodified | 30709000 | 30236000 | 29398000 | 99726000 | 4.41E+08 | 96201000 | 19135000 | 1471400 | 3.27E+08 | 10596000 | 46642000 | 3.53E+08 | 79721000 | 37924000 | 70020000 | 33753000 | 28940000 | 1.28E+08 |
| YPIEHGIVT | ACTG1; ACTB; / | P63261; P60 | ACTG_HUMAN | Oxidation (M) | 568490 | 601670 | 1414600 | 1394300 | 3101200 | 1185600 | 0 | 0 | 1158100 | 0 | 1081800 | 9888800 | 0 | 0 | 1817000 | 845110 | 557650 | 661190 |
| YPIEHGIVT | ACTG1; ACTB; / | P63261; P60 | ACTG_HUMAN | Unmodified | 0 | 0 | 106320 | 378440 | 3618300 | 0 | 0 | 0 | 2088100 | 0 | 0 | 2096600 | 248260 | 139730 | 573830 | 97858 | 890930 | 0 |
| YPLAPGVPI | PLCB3 | Q01970 | PLCB3_HUMAN | Oxidation (M) | 30625 | 0 | 0 | 0 | 0 | 0 | 0 | 0 | 0 | 0 | 41813 | 488580 | 0 | 0 | 32007 | 26938 | 0 | 0 |
| YPLAVRPF\ | DARS1 | P14868 | SYDC_HUMAN | Oxidation (M) | 0 | 0 | 180710 | 0 | 0 | 0 | 0 | 0 | 0 | 0 | 0 | 2883200 | 0 | 0 | 0 | 0 | 0 | 0 |
| YPLFEGQE | GARS1; GARS1 | P41250; A0A | GARS_HUMAN | Unmodified | 0 | 0 | 0 | 0 | 0 | 0 | 0 | 0 | 0 | 0 | 910610 | 389280 | 0 | 0 | 0 | 0 | 0 | 0 |
| YPLFEGQE | GARS1; GARS1 | P41250; A0A | GARS_HUMAN | Unmodified | 495470 | 0 | 0 | 0 | 0 | 0 | 0 | 0 | 2800000 | 0 | 0 | 19041000 | 0 | 0 | 0 | 1105700 | 0 | 0 |
| YPLLISR | ARHGEF2; ARH | Q92974; V9( | ARHG2_HUMA | Unmodified | 324620 | 0 | 473300 | 0 | 164740 | 0 | 1152200 | 978960 | 0 | 0 | 0 | 412870 | 0 | 521980 | 146200 | 10329 | 55703 | 1495500 |
| YPLLLDVY/ | DPP4; DPP4; DI | P27487; A0A | DPP4_HUMAN | Unmodified | 0 | 0 | 103220 | 144790 | 0 | 0 | 0 | 0 | 0 | 0 | 0 | 0 | 0 | 0 | 357450 | 182820 | 0 | 0 |
| YPLLLQSIG | ARHGEF1; ARH | Q92888; M0 | ARHG1_HUMA | Unmodified | 0 | 0 | 0 | 0 | 0 | 398690 | 0 | 0 | 0 | 0 | 0 | 376120 | 0 | 94188 | 0 | 0 | 0 | 0 |
| YPLRNLQP | FN1 | P02751 | FINC_HUMAN I | Unmodified | 0 | 0 | 276520 | 0 | 0 | 0 | 0 | 22196 | 0 | 106610 | 0 | 0 | 0 | 0 | 0 | 0 | 0 | 0 |
| YPNAELAW | OGDH; OGDH; | Q02218; E9F | ODO1_HUMAN | Unmodified | 0 | 0 | 0 | 0 | 0 | 0 | 0 | 0 | 495670 | 0 | 0 | 0 | 0 | 0 | 0 | 0 | 0 | 0 |
| YPPASQLR | CPD | O75976 | CBPD_HUMAN | Unmodified | 0 | 0 | 0 | 0 | 0 | 0 | 0 | 0 | 2314400 | 0 | 0 | 0 | 0 | 0 | 0 | 0 | 0 | 0 |
| YPPPTELLD | SNRNP200; SN | O75643; A0/ | U520_HUMAN | Unmodified | 0 | 0 | 44031 | 0 | 0 | 0 | 0 | 0 | 0 | 0 | 0 | 452490 | 0 | 0 | 0 | 0 | 0 | 0 |
| YPQLLPGIF | AHCY | P23526 | SAHH_HUMAN | Unmodified | 173870 | 128110 | 0 | 210520 | 55211 | 23177 | 0 | 13732 | 0 | 57590 | 3297900 | 26640000 | 0 | 0 | 1924100 | 894140 | 0 | 0 |
| YPSATNNT | GEMIN5 | Q8TEQ6 | GEMI5_HUMAN | Unmodified | 0 | 438200 | 0 | 0 | 0 | 0 | 0 | 0 | 0 | 0 | 0 | 802920 | 0 | 0 | 167380 | 0 | 0 | 0 |
| YPSHSFIGE | IMPA1; IMPA1; I | P29218; E5F | IMPA1_HUMAN | Unmodified | 0 | 0 | 0 | 0 | 0 | 0 | 52564 | 0 | 621080 | 0 | 0 | 0 | 0 | 0 | 95606 | 59964 | 0 | 0 |
| YPSLSLHG | CNDP2 | Q96KP4 | CNDP2_HUMA | Unmodified | 16842 | 57089 | 0 | 0 | 0 | 0 | 542170 | 0 | 0 | 0 | 0 | 462560 | 368590 | 0 | 0 | 0 | 0 | 0 |
| YPSPFFVF( | DHX9 | Q08211 | DHX9_HUMAN | Unmodified | 0 | 0 | 283010 | 0 | 0 | 0 | 0 | 0 | 0 | 0 | 396560 | 4499200 | 0 | 0 | 0 | 0 | 0 | 0 |
| YPSPPMGS | CRKL | P46109 | CRKL_HUMAN | Oxidation (M) | 0 | 0 | 0 | 0 | 0 | 0 | 0 | 0 | 0 | 0 | 0 | 2304900 | 0 | 0 | 0 | 0 | 0 | 0 |
| YPTPYPDEI | ERBIN | Q96RT1 | ERBIN_HUMAN | Unmodified | 0 | 0 | 0 | 0 | 661400 | 29353 | 0 | 0 | 0 | 0 | 0 | 0 | 267790 | 0 | 0 | 0 | 0 | 0 |
| YPTYPEAEK | FUT8; FUT8 | Q9BYC5; G3 | FUT8_HUMAN | Unmodified | 0 | 0 | 0 | 0 | 0 | 0 | 0 | 0 | 805160 | 0 | 0 | 0 | 0 | 0 | 0 | 0 | 0 | 0 |
| YPVIQTDDI | MARVELD2; MA | Q8N4S9; A1 | MALD2_HUMAI | Unmodified | 0 | 0 | 0 | 226720 | 0 | 0 | 0 | 0 | 124700 | 0 | 265790 | 589500 | 0 | 0 | 2939100 | 1510200 | 0 | 0 |
| YPVNSVNII | TCP1; TCP1; TC | P17987; E7E | TCPA_HUMAN | Unmodified | 965800 | 660270 | 927570 | 691970 | 0 | 0 | 226500 | 302830 | 2508800 | 216150 | 4865400 | 51135000 | 898110 | 0 | 1141700 | 957070 | 168360 | 0 |
| YPVTPELLE | PLCG2; PLCG2 | P16885; A0A | PLCG2_HUMAI | Unmodified | 0 | 116150 | 0 | 0 | 0 | 0 | 0 | 0 | 224050 | 0 | 691900 | 3848100 | 0 | 0 | 0 | 0 | 0 | 0 |
| YPVYGVQV | GGH; GGH | Q92820; A0A | GGH_HUMAN ( | Unmodified | 0 | 0 | 1659600 | 0 | 0 | 0 | 0 | 0 | 7135900 | 678460 | 411490 | 1007800 | 0 | 0 | 0 | 0 | 0 | 0 |
| YQAEIPDRI | MTA2 | O94776 | MTA2_HUMAN | Unmodified | 287330 | 234630 | 0 | 0 | 175360 | 0 | 0 | 0 | 0 | 0 | 0 | 2159500 | 0 | 248150 | 0 | 0 | 0 | 0 |
| YQAIGAYYI | PKP1 | Q13835 | PKP1_HUMAN | Unmodified | 0 | 0 | 0 | 0 | 0 | 0 | 0 | 0 | 0 | 0 | 0 | 0 | 0 | 0 | 0 | 0 | 205070 | 0 |
| YQAMVHC( | GTPBP1; GTPBI | O00178; B0( | GTPB1_HUMAN | Unmodified | 112510 | 0 | 54003 | 0 | 37505 | 0 | 0 | 0 | 0 | 0 | 0 | 1862300 | 0 | 0 | 135800 | 89374 | 0 | 0 |
| YQAVQEAV | UTRN | P46939 | UTRN_HUMAN | Unmodified | 0 | 0 | 227400 | 0 | 0 | 0 | 0 | 0 | 0 | 0 | 0 | 0 | 0 | 0 | 368350 | 80501 | 0 | 0 |
| YQAVTATLE | RPL13AP3; RPL | Q6NVV1; P4( | R13P3_HUMAN | Unmodified | 6893300 | 5117900 | 3479800 | 2614000 | 1282300 | 467120 | 484950 | 1620300 | 1067800 | 0 | 3531300 | 33744000 | 363060 | 309430 | 4260100 | 2967900 | 0 | 906270 |
| YQCYCYGF | FN1 | P02751 | FINC_HUMAN I | Unmodified | 0 | 516530 | 0 | 1132400 | 0 | 198240 | 0 | 0 | 368080 | 108740 | 0 | 514780 | 0 | 0 | 0 | 854690 | 0 | 0 |
| YQDAETLYI | KLC2; KLC2 | Q9H0B6; A8 | KLC2_HUMAN | Unmodified | 144770 | 0 | 196080 | 0 | 33560 | 0 | 0 | 0 | 345560 | 0 | 0 | 354860 | 0 | 0 | 0 | 0 | 0 | 0 |
| YQDEVFG( | G3BP1; G3BP1; | Q13283; A0/ | G3BP1_HUMAI | Unmodified | 0 | 0 | 0 | 0 | 0 | 0 | 0 | 0 | 0 | 0 | 1942800 | 3981900 | 0 | 0 | 0 | 0 | 0 | 0 |
| YQDILNEIA | IQGAP2; IQGAP | Q13576; E7E | IQGA2_HUMAN | Unmodified | 0 | 0 | 1236500 | 104950 | 0 | 0 | 0 | 0 | 0 | 0 | 0 | 0 | 0 | 37518 | 1064000 | 747780 | 0 | 0 |
| YQDILVFR | SRM; SRM | P19623; K7E | SPEE_HUMAN | Unmodified | 155520 | 143290 | 201550 | 307080 | 158770 | 62842 | 36330 | 0 | 2007300 | 522330 | 484640 | 3268700 | 448770 | 124010 | 362830 | 397960 | 0 | 211520 |
| YQDILVFRS | SRM | P19623 | SPEE_HUMAN | Unmodified | 0 | 0 | 0 | 0 | 254290 | 0 | 0 | 147150 | 0 | 383510 | 0 | 0 | 873120 | 0 | 0 | 0 | 604160 | 368320 |
| YQDLYTVEI | CACNA2D1; CA | P54289; E7E | CA2D1_HUMAI | Unmodified | 819650 | 697030 | 0 | 0 | 0 | 0 | 0 | 0 | 1748400 | 404900 | 0 | 0 | 0 | 0 | 0 | 0 | 0 | 0 |
| YQDLYTVEI | CACNA2D1; CA | P54289; E7E | CA2D1_HUMAI | Unmodified | 0 | 0 | 0 | 0 | 0 | 0 | 10787000 | 0 | 0 | 1813000 | 0 | 0 | 0 | 0 | 0 | 0 | 0 | 0 |
| YQDTPGVK | ACLY | P53396 | ACLY_HUMAN | Unmodified | 266140 | 143040 | 522080 | 788070 | 354470 | 259990 | 1568500 | 881040 | 0 | 443560 | 981370 | 20360000 | 364680 | 0 | 1115100 | 937060 | 0 | 1029800 |
| YQDWGWI | MAN1B1; MAN1 | Q9UKM7; H( | MA1B1_HUMAI | Unmodified | 0 | 0 | 0 | 0 | 0 | 0 | 0 | 0 | 202680 | 42932 | 0 | 0 | 0 | 0 | 0 | 0 | 0 | 0 |
| YQEAAPNV | CTNND1; CTNI | O60716; A0/ | CTND1_HUMAI | Unmodified | 565550 | 292870 | 894250 | 1046100 | 1440500 | 0 | 0 | 0 | 0 | 0 | 776050 | 4814500 | 785600 | 494330 | 1327800 | 1803900 | 607620 | 904930 |
| YQEAAPNV | CTNND1; CTNI | O60716; A0/ | CTND1_HUMAI | Unmodified | 0 | 0 | 0 | 0 | 0 | 0 | 0 | 0 | 0 | 0 | 134090 | 1027600 | 0 | 0 | 366560 | 413270 | 400750 | 0 |
| YQEALAK | SOD2; SOD2; S | P04179; G8J | SODM_HUMAN | Unmodified | 68587 | 15937 | 0 | 0 | 942170 | 213000 | 477350 | 193280 | 3324200 | 107750 | 0 | 0 | 0 | 0 | 0 | 0 | 20342 | 536360 |
| YQEAQEEII | RAI14 | Q9P0K7 | RAI14_HUMAN | Unmodified | 0 | 0 | 0 | 0 | 508900 | 0 | 0 | 0 | 87284 | 0 | 0 | 433760 | 0 | 0 | 0 | 74438 | 191390 | 347480 |
| YQEEFEHF | LMAN1 | P49257 | LMAN1_HUMAI | Unmodified | 0 | 0 | 1773100 | 854880 | 101930 | 973370 | 0 | 35792 | 2.03E+08 | 3713900 | 5227600 | 2787600 | 0 | 0 | 2140600 | 726100 | 0 | 870310 |
| YQEGGVES | CAPG; CAPG; C | P40121; E7E | CAPG_HUMAN | Unmodified | 79761 | 0 | 0 | 0 | 2017900 | 329420 | 0 | 0 | 0 | 0 | 114760 | 348450 | 0 | 0 | 0 | 0 | 805480 | 650980 |
| YQEIPFYHI | UNC5B | Q8IZJ1 | UNC5B_HUMA | Unmodified | 0 | 0 | 0 | 0 | 0 | 0 | 256550 | 0 | 1168600 | 170530 | 0 | 954540 | 0 | 0 | 0 | 0 | 0 | 0 |
| YQELINDIA | IQGAP1; IQGAP | P46940; H0 | IQGA1_HUMAN | Unmodified | 282440 | 0 | 1339700 | 0 | 0 | 0 | 0 | 0 | 0 | 0 | 248590 | 2502800 | 0 | 0 | 602800 | 368980 | 0 | 0 |
| YQELLER | PAIP1; PAIP1 | Q9H074; H0 | PAIP1_HUMAN | Unmodified | 675960 | 0 | 0 | 0 | 174050 | 0 | 53670 | 55412 | 2909500 | 2214100 | 367430 | 0 | 0 | 0 | 0 | 34636 | 0 | 217950 |
| YQELQVLFI | VPS37B; VPS37 | Q9H9H4; F5 | VP37B_HUMAN | Unmodified | 0 | 693360 | 0 | 759570 | 816140 | 0 | 112600 | 0 | 0 | 0 | 0 | 0 | 1360600 | 0 | 1778700 | 0 | 0 | 0 |
| YQELQVLFI | VPS37B; VPS37 | Q9H9H4; F5 | VP37B_HUMAN | Unmodified | 8606500 | 16831000 | 3308900 | 19183000 | 5621900 | 1225400 | 0 | 0 | 35787 | 1298000 | 0 | 3037700 | 0 | 13609000 | 28433000 | 14193 | 703780 | 0 |
| YQETFNVIE | ECH1; ECH1; E | Q13011; M0 | ECH1_HUMAN | Unmodified | 0 | 0 | 0 | 0 | 0 | 0 | 0 | 0 | 4687000 | 0 | 309130 | 0 | 0 | 0 | 0 | 106400 | 0 | 0 |
| YQEVTNNL | CAPRIN1; CAPF | Q14444; G3 | CAPR1_HUMAN | Unmodified | 0 | 0 | 0 | 0 | 0 | 162570 | 0 | 0 | 1169400 | 0 | 1402300 | 4196800 | 434650 | 0 | 0 | 184760 | 0 | 0 |
| YQFTPAFLF | UNC119B; UNC | A6NIH7; Q1 | U119B_HUMAN | Unmodified | 101460 | 0 | 0 | 148110 | 0 | 0 | 0 | 0 | 0 | 0 | 0 | 1473900 | 0 | 0 | 726860 | 0 | 0 | 0 |
| YQGRDDE/ | ARHGAP18 | Q8N392 | RHG18_HUMAI | Unmodified | 45842 | 0 | 0 | 0 | 288430 | 0 | 0 | 15941 | 0 | 14703 | 189910 | 805350 | 102630 | 66147 | 0 | 48293 | 0 | 0 |
| YQGVNLYII | PABPC4; PABP( | Q13310; B1/ | PABP4_HUMAN | Unmodified | 15018000 | 8144800 | 3915600 | 2501700 | 0 | 0 | 0 | 0 | 0 | 0 | 0 | 786100 | 290770 | 0 | 3065600 | 4248500 | 0 | 0 |
| YQGVNLYV | PABPC1; PABP( | P11940; Q4 | PABP1_HUMAN | Unmodified | 79747000 | 59160000 | 22628000 | 10569000 | 0 | 0 | 0 | 62743 | 928370 | 256520 | 653800 | 8349500 | 2696700 | 890030 | 13347000 | 21156000 | 0 | 0 |
| YQHLITNSI | ARIH1; ARIH1 | Q9Y4X5; H3 | ARI1_HUMAN E | Unmodified | 0 | 51922 | 0 | 0 | 0 | 0 | 0 | 0 | 0 | 0 | 0 | 826300 | 0 | 0 | 0 | 0 | 993870 | 0 |
| YQHTGAVL | BUB3; BUB3; BI | O43684; J3( | BUB3_HUMAN | Unmodified | 0 | 0 | 0 | 0 | 852320 | 0 | 0 | 0 | 0 | 483540 | 0 | 6754000 | 0 | 0 | 0 | 0 | 0 | 0 |
| YQIDPDAC | VDAC1 | P21796 | VDAC1_HUMAI | Unmodified | 0 | 0 | 0 | 0 | 0 | 105030 | 0 | 0 | 54635000 | 12687000 | 1928400 | 1013100 | 492660 | 475590 | 0 | 0 | 0 | 0 |
| YQILNNEVI | CYFIP2; CYFIP2 | Q96F07; H7( | CYFP2_HUMAN | Unmodified | 0 | 0 | 0 | 0 | 0 | 0 | 0 | 0 | 1201300 | 0 | 121700 | 1966900 | 124090 | 0 | 605130 | 238710 | 0 | 0 |
| YQISVNK | FGB; FGB | P02675; D6F | FIBB_HUMAN F | Unmodified | 0 | 3999300 | 90851 | 144140 | 0 | 0 | 0 | 148440 | 0 | 0 | 621450 | 216430 | 0 | 0 | 104070 | 0 | 0 | 0 |
| YQKPIIQSE | GUSB | P08236 | BGLR_HUMAN | Oxidation (M) | 0 | 1530100 | 0 | 0 | 0 | 0 | 0 | 0 | 9674100 | 5714000 | 0 | 0 | 0 | 0 | 0 | 0 | 0 | 0 |
| YQKPIIQSE | GUSB | P08236 | BGLR_HUMAN | Unmodified | 0 | 0 | 0 | 0 | 0 | 0 | 0 | 0 | 14346000 | 1029300 | 0 | 0 | 0 | 0 | 0 | 0 | 0 | 0 |
| YQKSTELLI | H3-2; H3-4; H3 | Q5TEC6; Q1 | H3PS2_HUMAN | Unmodified | 0 | 0 | 0 | 0 | 0 | 0 | 0 | 379450 | 0 | 0 | 0 | 631060 | 0 | 0 | 0 | 0 | 0 | 0 |
| YQLAFQR | GNG13 | Q9P2W3 | GBG13_HUMAI | Unmodified | 0 | 63989 | 0 | 0 | 0 | 0 | 0 | 0 | 0 | 0 | 5035800 | 73976000 | 0 | 0 | 16417 | 0 | 0 | 0 |
| YQLAFQRE | GNG13 | Q9P2W3 | GBG13_HUMAI | Oxidation (M) | 0 | 0 | 0 | 0 | 0 | 0 | 0 | 0 | 0 | 0 | 410820 | 0 | 0 | 0 | 0 | 0 | 0 | 0 |
| YQLDKDG\ | P4HB; P4HB; P4 | P07237; A0/ | PDIA1_HUMAN | Unmodified | 94101 | 55740 | 31576 | 163550 | 444950 | 302230 | 0 | 273420 | 22778000 | 40367000 | 6344100 | 450390 | 470190 | 545980 | 137420 | 116410 | 0 | 151940 |
| YQLDPTASI | VDAC2; VDAC2 | P45880; A0A | VDAC2_HUMAI | Unmodified | 0 | 0 | 0 | 441260 | 776600 | 0 | 1386300 | 1250900 | 8.38E+08 | 1.84E+08 | 5285700 | 2452000 | 463880 | 3961600 | 0 | 0 | 0 | 1271300 |
| YQLEIKIPE | UBE2K; UBE2K | P61086; D6F | UBE2K_HUMAN | Unmodified | 373880 | 0 | 0 | 0 | 0 | 0 | 38976 | 0 | 0 | 0 | 619870 | 1307700 | 0 | 0 | 451280 | 539140 | 0 | 180820 |
| YQLSIHKNI | ATP1A1 | P05023 | AT1A1_HUMAN | Unmodified | 218280 | 0 | 0 | 4779000 | 34123000 | 2143400 | 2356300 | 0 | 9779700 | 5331300 | 7145000 | 19722000 | 15065000 | 12829000 | 0 | 532350 | 2069300 | 4326300 |
| YQLSIHKNI | ATP1A1 | P05023 | AT1A1_HUMAN | Oxidation (M) | 0 | 0 | 0 | 7706000 | 7681500 | 852180 | 813120 | 0 | 1216400 | 1850300 | 6244600 | 10187000 | 9576100 | 5337700 | 0 | 0 | 446540 | 0 |
| YQMGDQN | USP15; USP15; | Q9Y4E8; H0 | UBP15_HUMAI | Oxidation (M) | 0 | 0 | 0 | 0 | 0 | 0 | 0 | 0 | 0 | 0 | 73256 | 684300 | 0 | 0 | 0 | 0 | 0 | 0 |
| YQNLENDE | TMEM134; TME | Q9H6X4; G3 | TM134_HUMAN | Unmodified | 0 | 0 | 132710 | 0 | 0 | 0 | 0 | 0 | 0 | 0 | 0 | 0 | 0 | 0 | 880510 | 580130 | 0 | 0 |
| YQNYIMEG | RNF114; RNF1 | Q9Y508; A0/ | RN114_HUMAN | Unmodified | 0 | 221240 | 0 | 0 | 885380 | 0 | 0 | 0 | 0 | 0 | 0 | 0 | 1702600 | 463330 | 0 | 0 | 177700 | 602120 |
| YQPCLPST | GFM1 | Q96RP9 | EFGM_HUMAN | Unmodified | 0 | 0 | 0 | 0 | 0 | 0 | 0 | 0 | 343450 | 0 | 51330 | 0 | 0 | 0 | 0 | 0 | 0 | 0 |
| YQPFKPLSI | PSMD1; PSMD1 | Q99460; A0/ | PSMD1_HUMA | Unmodified | 0 | 0 | 200900 | 456120 | 0 | 0 | 0 | 0 | 385170 | 69373 | 473250 | 1509700 | 281010 | 0 | 334540 | 206830 | 0 | 0 |
| YQPNIDVQ | CD109 | Q6YHK3 | CD109_HUMAI | Unmodified | 0 | 0 | 0 | 0 | 0 | 0 | 0 | 0 | 174660 | 180960 | 0 | 0 | 0 | 0 | 0 | 0 | 0 | 0 |
| YQPTLPER | C1GALT1 | Q9NS00 | C1GLT_HUMAN | Unmodified | 0 | 0 | 0 | 0 | 0 | 0 | 0 | 0 | 2973200 | 0 | 0 | 0 | 0 | 0 | 0 | 0 | 0 | 0 |
| YQQAGLPL | ACO1 | P21399 | ACOC_HUMAN | Unmodified | 0 | 0 | 0 | 0 | 0 | 0 | 0 | 0 | 0 | 0 | 0 | 992880 | 0 | 0 | 0 | 0 | 0 | 0 |
| YQQDQVVI | PIK3R1; PIK3R1 | P27986; H0 | P85A_HUMAN | Unmodified | 0 | 0 | 121970 | 155260 | 0 | 0 | 0 | 200690 | 369510 | 0 | 0 | 687820 | 0 | 59488 | 217240 | 242610 | 0 | 0 |
| YQQGDFG | CSNK2B; CSNK | P67870; Q5( | CSK2B_HUMAN | Unmodified | 0 | 0 | 0 | 0 | 0 | 0 | 0 | 0 | 0 | 0 | 282130 | 3480500 | 0 | 0 | 0 | 0 | 0 | 0 |
| YQQHQESI | GOLIM4; GOLIN | O00461; F8V | GOLI4_HUMAN | Unmodified | 0 | 0 | 0 | 0 | 0 | 0 | 0 | 0 | 24148000 | 2131100 | 0 | 1186800 | 0 | 0 | 0 | 102750 | 0 | 0 |
| YQQLFEDIF | MCM4; MCM4; | P33991; A0/ | MCM4_HUMAN | Unmodified | 155730 | 0 | 0 | 0 | 0 | 0 | 0 | 0 | 959140 | 262480 | 566990 | 819890 | 903730 | 0 | 0 | 0 | 0 | 0 |
| YQRLLHEV | DCTN2; DCTN2 | Q13561; A8( | DCTN2_HUMAI | Unmodified | 0 | 0 | 0 | 0 | 2311400 | 0 | 331560 | 0 | 0 | 267300 | 0 | 0 | 2360800 | 1265400 | 0 | 0 | 529190 | 3465200 |
| YQSHPDA/ | PDPK1; PDPK1 | O15530; E9F | PDPK1_HUMAN | Unmodified | 0 | 0 | 0 | 0 | 0 | 0 | 0 | 0 | 80630 | 0 | 0 | 0 | 0 | 0 | 476140 | 440540 | 0 | 64343 |
| YQTDCAPN | NOMO1; NOMC | Q15155; P6( | NOMO1_HUMA | Oxidation (M) | 0 | 0 | 0 | 0 | 0 | 0 | 0 | 0 | 723300 | 0 | 205270 | 0 | 0 | 0 | 0 | 0 | 0 | 0 |
| YQTDCAPN | NOMO1; NOMC | Q15155; P6( | NOMO1_HUMA | Unmodified | 0 | 0 | 0 | 0 | 0 | 0 | 0 | 0 | 607320 | 0 | 0 | 0 | 0 | 0 | 0 | 0 | 0 | 0 |
| YQTNPCIG | ITGB5; ITGB5; I | P18084; V9( | ITB5_HUMAN I | Unmodified | 222400 | 0 | 0 | 306850 | 0 | 0 | 0 | 0 | 253110 | 0 | 0 | 1753900 | 235870 | 0 | 956150 | 616530 | 0 | 0 |
| YQTPDHW | GSTT2; GSTT2B | P0CG29; P0 | GST2_HUMAN | Unmodified | 0 | 0 | 0 | 0 | 0 | 0 | 0 | 0 | 0 | 0 | 0 | 1126700 | 366410 | 0 | 120140 | 0 | 0 | 0 |
| YQVAAAQA | CC2D1A; CC2D | Q6P1N0; K7 | C2D1A_HUMAN | Unmodified | 14297000 | 12160000 | 10038000 | 88325 | 32105000 | 21158000 | 3244500 | 1113900 | 22029000 | 12625000 | 9308700 | 18842000 | 12294000 | 0 | 14241000 | 9518100 | 0 | 0 |
| YQVDLGD( | SORCS2; SORC | Q96PQ0; B5 | SORC2_HUMAI | Unmodified | 0 | 0 | 368580 | 0 | 204010 | 0 | 0 | 0 | 835810 | 65848 | 0 | 0 | 0 | 0 | 0 | 202910 | 0 | 124300 |
| YQVFFFGTI | HDGF | P51858 | HDGF_HUMAN | Unmodified | 68450 | 214910 | 0 | 253760 | 3072500 | 0 | 0 | 16362 | 3736000 | 758410 | 6290900 | 12394000 | 1521300 | 257360 | 0 | 0 | 0 | 0 |
| YQVGVHYE | DYNC1H1; DYN | Q14204; A0/ | DYHC1_HUMAN | Unmodified | 0 | 0 | 0 | 0 | 0 | 0 | 0 | 0 | 1071600 | 0 | 1047400 | 6341400 | 0 | 0 | 0 | 0 | 0 | 0 |
| YQVLDGEN | PTGS1; PTGS1; | P23219; A0A | PGH1_HUMAN 2 | Oxidation (M) | 0 | 0 | 0 | 0 | 0 | 0 | 0 | 0 | 0 | 0 | 0 | 0 | 0 | 0 | 0 | 0 | 0 | 0 |
| YQVPGSVI | PLD1 | Q13393 | PLD1_HUMAN | Unmodified | 0 | 0 | 0 | 0 | 0 | 0 | 0 | 0 | 0 | 0 | 0 | 856890 | 0 | 0 | 180940 | 60587 | 0 | 0 |
| YQVSWSLI | SSR4 | P51571 | SSRD_HUMAN | Unmodified | 0 | 0 | 0 | 0 | 0 | 0 | 0 | 0 | 4645100 | 244720 | 0 | 0 | 0 | 0 | 0 | 0 | 0 | 0 |
| YQWGIMK | LARS1; LARS1 | Q9P2J5; A0A | SYLC_HUMAN | Oxidation (M) | 0 | 0 | 0 | 0 | 127860 | 0 | 103230 | 0 | 485450 | 0 | 0 | 1373000 | 0 | 0 | 0 | 0 | 0 | 0 |
| YQYAIDEY | FAM177A1; FAI | Q8N128; G3 | F177A_HUMAN | Unmodified | 178010 | 244430 | 0 | 1003600 | 0 | 0 | 0 | 0 | 0 | 0 | 0 | 0 | 345640 | 0 | 1019100 | 0 | 0 | 0 |
| YQYGGLNS | PPP1CB | P62140 | PP1B_HUMAN | Unmodified | 0 | 0 | 1058300 | 0 | 0 | 0 | 0 | 0 | 2096900 | 409000 | 912040 | 5114200 | 0 | 0 | 1208100 | 1620700 | 0 | 0 |

| | | | | | | | | | | | | | | | | | | | | | | |
|---|---|---|---|---|---|---|---|---|---|---|---|---|---|---|---|---|---|---|---|---|---|---|
| YRAAHVFF | STXBP1; STXBP | P61764; A0A | STXB1_HUMAN | Unmodified | 0 | 0 | 0 | 0 | 87825 | 111700 | 11973000 | 0 | 1042500 | 4306700 | 0 | 0 | 0 | 6798600 | 0 | 0 | 2503100 | 2291300 |
| YRAFITNIPI | HNRNPM; HNR | P52272; M0 | HNRPM_HUMA | Unmodified | 0 | 0 | 0 | 0 | 66992 | 0 | 1938400 | 0 | 0 | 161650 | 0 | 0 | 507430 | 584360 | 0 | 0 | 0 | 0 |
| YRAQLLEPS | DRG2; DRG2; D | P55039; J3Q | DRG2_HUMAN | Unmodified | 0 | 0 | 0 | 0 | 0 | 0 | 0 | 0 | 261490 | 235160 | 113790 | 568430 | 0 | 0 | 0 | 323800 | 0 | 550050 |
| YRCELLYE( | EEF2 | P13639 | EF2_HUMAN El | Oxidation (M) | 2603400 | 974480 | 1519800 | 2209400 | 5884700 | 1714200 | 2985900 | 1884400 | 1685600 | 1469700 | 5751200 | 45215000 | 2580400 | 2884100 | 3230600 | 2082900 | 2792100 | 3083500 |
| YRCELLYE( | EEF2 | P13639 | EF2_HUMAN El | Unmodified | 726070 | 487020 | 224530 | 824900 | 16137000 | 0 | 3165400 | 243640 | 6224600 | 1107500 | 2307700 | 12200000 | 3584200 | 2543300 | 507710 | 328660 | 2594300 | 8652500 |
| YRCNVENL | GLG1; GLG1 | Q92896; H3I | GSLG1_HUMA | Unmodified | 0 | 86898 | 0 | 0 | 0 | 0 | 0 | 0 | 2435000 | 0 | 676660 | 383390 | 0 | 0 | 0 | 0 | 0 | 0 |
| YRDCLTESI | TGM2 | P21980 | TGM2_HUMAN | Unmodified | 0 | 0 | 0 | 0 | 10683000 | 1405700 | 0 | 0 | 0 | 0 | 0 | 0 | 0 | 0 | 0 | 0 | 1407800 | 457980 |
| YRDLTVRE | TSG101; TSG10 | Q99816; F5H | TS101_HUMAN | Unmodified | 270780 | 0 | 340020 | 19325000 | 2496800 | 711800 | 296090 | 0 | 1591500 | 4324900 | 0 | 0 | 26569000 | 7215500 | 0 | 0 | 572310 | 2183200 |
| YRDQFTDQ | MYO1B; MYO1E | O43795; H7 | MYO1B_HUMA | Unmodified | 280230 | 172290 | 0 | 88662 | 1972000 | 942300 | 233090 | 250840 | 655200 | 266380 | 942230 | 15447000 | 0 | 0 | 672030 | 379530 | 847370 | 546490 |
| YRDQIPSP( | ATP1B3; ATP1B | P54709; C9J | AT1B3_HUMAN | Oxidation (M) | 0 | 0 | 1939400 | 243100 | 0 | 0 | 130660 | 0 | 0 | 0 | 124140 | 1463600 | 0 | 0 | 2652100 | 1818700 | 0 | 0 |
| YRDQIPSP( | ATP1B3; ATP1B | P54709; C9J | AT1B3_HUMAN | Oxidation (M) | 0 | 0 | 1277200 | 2952400 | 0 | 0 | 0 | 0 | 2390400 | 0 | 1348900 | 8837700 | 0 | 0 | 14927000 | 14184000 | 0 | 0 |
| YRDVSPFD | PTPN1 | P18031 | PTN1_HUMAN | Unmodified | 0 | 0 | 0 | 0 | 0 | 0 | 0 | 0 | 0 | 0 | 0 | 458740 | 0 | 0 | 0 | 0 | 0 | 0 |
| YREDIINVR | RABEP1 | Q15276 | RABE1_HUMAN | Unmodified | 0 | 0 | 146930 | 0 | 669040 | 120820 | 55751 | 0 | 0 | 0 | 0 | 0 | 173880 | 127560 | 0 | 0 | 69474 | 108820 |
| YREDLDFV | ABCC1; ABCC1 | P33527; I3L | MRP1_HUMAN | Unmodified | 0 | 0 | 0 | 158190 | 0 | 0 | 0 | 60170 | 0 | 409010 | 363920 | 997170 | 0 | 0 | 524660 | 266670 | 0 | 0 |
| YRELGNIIQ | METAP1; METAI | P53582; H0Y | MAP11_HUMA | Unmodified | 270520 | 0 | 0 | 295210 | 656110 | 0 | 140460 | 63687 | 424100 | 0 | 0 | 1814500 | 966550 | 409530 | 344440 | 371200 | 426430 | 0 |
| YRENLPLVI | ABCC5 | O15440 | MRP5_HUMAN | Unmodified | 0 | 0 | 0 | 0 | 0 | 0 | 0 | 0 | 136460 | 0 | 0 | 2916800 | 0 | 0 | 0 | 0 | 0 | 0 |
| YREREQQII | DBN1; DBN1 | Q16643; D6I | DREB_HUMAN | Unmodified | 0 | 0 | 34287 | 119670 | 0 | 158250 | 0 | 0 | 102300 | 0 | 41690 | 206210 | 0 | 0 | 93953 | 35109 | 0 | 55876 |
| YRESNTAG | CLIC3 | O95833 | CLIC3_HUMAN | Unmodified | 0 | 0 | 35132 | 0 | 862910 | 0 | 0 | 0 | 0 | 0 | 0 | 0 | 0 | 0 | 0 | 0 | 0 | 0 |
| YREVQLPD | IDE; IDE; IDE; ID | P14735; A0A | IDE_HUMAN In | Unmodified | 0 | 0 | 0 | 0 | 0 | 0 | 0 | 0 | 0 | 0 | 0 | 744580 | 0 | 0 | 0 | 0 | 0 | 0 |
| YREWHHFI | PEBP1 | P30086 | PEBP1_HUMAN | Unmodified | 1149700 | 1220600 | 123880 | 1291000 | 3524900 | 481650 | 258120 | 0 | 4615800 | 1532600 | 3023200 | 8683600 | 6001100 | 2384900 | 1532200 | 0 | 1317000 | 2871600 |
| YREWHHFI | PEBP1 | P30086 | PEBP1_HUMAN | Oxidation (M) | 3644700 | 725290 | 1841000 | 2176900 | 1685200 | 391540 | 172330 | 0 | 2203000 | 1115700 | 2753900 | 4407000 | 2130400 | 0 | 2837800 | 1497600 | 1457200 | 1156000 |
| YRGGPGDI | CYB5D2 | Q8WUJ1 | NEUFC_HUMA | Unmodified | 0 | 0 | 0 | 0 | 0 | 0 | 0 | 0 | 409350 | 0 | 31622 | 0 | 0 | 0 | 0 | 0 | 0 | 0 |
| YRGMGSLI | IMPDH2; IMPDI | P12268; A0A | IMDH2_HUMA | Unmodified | 0 | 0 | 0 | 0 | 443350 | 0 | 246090 | 71243 | 126900 | 81039 | 0 | 151910 | 74398 | 0 | 0 | 0 | 302350 | 218270 |
| YRGPGIFEI | TMX4 | Q9H1E5 | TMX4_HUMAN | Unmodified | 0 | 18330 | 0 | 0 | 0 | 0 | 163010 | 0 | 4229100 | 2254100 | 617950 | 42792 | 0 | 0 | 0 | 0 | 0 | 0 |
| YRGQHVP( | FLNC | Q14315 | FLNC_HUMAN | Unmodified | 0 | 0 | 0 | 0 | 6227700 | 0 | 0 | 0 | 1261800 | 142300 | 0 | 0 | 0 | 0 | 0 | 0 | 2838400 | 691840 |
| YRGQHVT( | FLNB; FLNB; FL | O75369; H7 | FLNB_HUMAN | Unmodified | 0 | 0 | 574580 | 0 | 5416900 | 1708500 | 0 | 0 | 0 | 0 | 0 | 796530 | 0 | 0 | 1237500 | 0 | 1486400 | 0 |
| YRGSIHDFI | ANXA6; ANXA6; | P08133; E5R | ANXA6_HUMAI | Oxidation (M) | 0 | 0 | 0 | 3969400 | 12177000 | 4189400 | 12573000 | 487710 | 16448000 | 27657000 | 0 | 0 | 10421000 | 9042100 | 4503500 | 0 | 8555000 | 29744000 |
| YRGSIHDFI | ANXA6; ANXA6; | P08133; E5R | ANXA6_HUMAI | Unmodified | 0 | 0 | 0 | 0 | 20938000 | 2885400 | 24960000 | 0 | 1.42E+08 | 48367000 | 0 | 0 | 18982000 | 12803000 | 3132200 | 0 | 8807400 | 49187000 |
| YRGVSFYE | MYO1C; MYO1( | O00159; I3L | MYO1C_HUMA | Unmodified | 749560 | 575100 | 328230 | 776540 | 0 | 0 | 0 | 0 | 0 | 0 | 0 | 0 | 0 | 84167 | 2420200 | 1129900 | 0 | 0 |
| YRHPDMSI | ZC3H10; ZC3H | Q96K80; F8V | ZC3HA_HUMAI | Unmodified | 0 | 0 | 403280 | 353390 | 0 | 0 | 0 | 0 | 0 | 0 | 0 | 0 | 0 | 0 | 429850 | 562480 | 0 | 0 |
| YRHPDMSI | ZC3H10; ZC3H | Q96K80; F8V | ZC3HA_HUMAI | Oxidation (M) | 1082000 | 737000 | 0 | 1854300 | 0 | 0 | 0 | 0 | 0 | 0 | 0 | 0 | 1266300 | 784800 | 0 | 0 | 0 | 0 |
| YRHPDMSI | ZC3H10; ZC3H | Q96K80; F8V | ZC3HA_HUMAI | Unmodified | 0 | 0 | 0 | 0 | 0 | 0 | 0 | 0 | 0 | 0 | 0 | 0 | 1580200 | 0 | 0 | 0 | 0 | 0 |
| YRHSDGNI | SRP9; SRP9; SR | P49458; Q6 | SRP09_HUMAN | Unmodified | 0 | 0 | 0 | 2429500 | 0 | 0 | 76433 | 0 | 340520 | 0 | 0 | 0 | 0 | 0 | 0 | 1207800 | 0 | 0 |
| YRIPFYMST | MARK3; MARK2 | P27448; Q7 | MARK3_HUMAI | Oxidation (M) | 0 | 85700 | 0 | 208330 | 306170 | 164210 | 133430 | 34656 | 0 | 211280 | 0 | 309260 | 0 | 0 | 325580 | 1038400 | 61774 | 0 |
| YRIQEQES | PPP1R2B; PPP1 | Q6NXS1; P4 | IPP2B_HUMAN | Unmodified | 0 | 0 | 0 | 0 | 0 | 0 | 0 | 0 | 419450 | 0 | 0 | 1150700 | 0 | 0 | 0 | 0 | 0 | 0 |
| YRISKYPTLI | ERP44 | Q9BS26 | ERP44_HUMAN | Unmodified | 0 | 0 | 0 | 0 | 0 | 0 | 0 | 0 | 2653800 | 1548700 | 0 | 0 | 0 | 0 | 0 | 0 | 0 | 0 |
| YRKGPEDT | STXBP2; STXBP | Q15833; M0 | STXB2_HUMAN | Unmodified | 221520 | 305580 | 80471 | 967060 | 1524200 | 577770 | 0 | 0 | 347730 | 216980 | 0 | 996600 | 1231800 | 1079800 | 0 | 202320 | 1080100 | 0 |
| YRLPNTLKI | ANPEP; ANPEP | P15144; H0Y | AMPN_HUMAN | Unmodified | 0 | 0 | 0 | 0 | 0 | 0 | 0 | 0 | 0 | 0 | 0 | 0 | 0 | 65315 | 0 | 0 | 390460 | 564760 |
| YRLVGSQE | PSMD2; PSMD2 | Q13200; H7 | PSMD2_HUMA | Unmodified | 0 | 106790 | 0 | 0 | 721610 | 0 | 0 | 0 | 906310 | 0 | 922450 | 3392000 | 941690 | 0 | 0 | 0 | 0 | 0 |
| YRLVGSQE | PSMD2 | Q13200 | PSMD2_HUMA | Unmodified | 0 | 0 | 0 | 0 | 2586800 | 0 | 0 | 0 | 0 | 0 | 0 | 0 | 1180500 | 1564200 | 0 | 0 | 632590 | 2032100 |
| YRLVVVGG | RRAS2; RRAS2; | P62070; E9P | RRAS2_HUMAI | Unmodified | 1241000 | 2096600 | 1288500 | 2689400 | 9955300 | 2020300 | 272500 | 196330 | 1688800 | 1494400 | 0 | 0 | 4889500 | 2879900 | 3017900 | 2105500 | 1727200 | 2934700 |
| YRMYQTQ\ | PTGFRN | Q9P2B2 | FPRP_HUMAN | Oxidation (M) | 0 | 0 | 0 | 3291600 | 0 | 0 | 0 | 0 | 0 | 0 | 0 | 0 | 0 | 0 | 0 | 0 | 0 | 0 |
| YRMYQTQ\ | PTGFRN | Q9P2B2 | FPRP_HUMAN | Unmodified | 87657 | 64686 | 142710 | 616680 | 0 | 58406 | 0 | 0 | 37660 | 0 | 0 | 129330 | 229620 | 49926 | 226790 | 84429 | 0 | 0 |
| YRNWNPPI | CUEDC1; CUEI | Q9NWM3; J3 | CUED1_HUMA | Unmodified | 42694 | 0 | 0 | 0 | 396390 | 27572 | 0 | 53088 | 877270 | 279160 | 0 | 0 | 0 | 69601 | 0 | 0 | 44996 | 390580 |
| YRPDEQPS | PLOD3; PLOD3 | O60568; H7 | PLOD3_HUMAI | Unmodified | 0 | 0 | 0 | 0 | 50630 | 0 | 0 | 0 | 12405000 | 4937300 | 0 | 0 | 0 | 0 | 0 | 0 | 0 | 0 |
| YRPENTPEI | YES1; YES1 | P07947; J3Q | YES_HUMAN Ty | Unmodified | 1779700 | 1348600 | 2152100 | 3894000 | 6989700 | 0 | 482450 | 0 | 0 | 0 | 0 | 3749400 | 4274200 | 3763900 | 3816400 | 3543300 | 2848200 | 15075000 |
| YRPGYSSS | ZDHHC5; ZDHI | Q9C0B5; H0 | ZDHC5_HUMA | Unmodified | 307040 | 0 | 248590 | 1788700 | 6008900 | 553440 | 0 | 69077 | 1234900 | 831410 | 730980 | 1543700 | 3655700 | 4190200 | 2586300 | 728120 | 6005100 | 10219000 |
| YRPLFDYFI | IARS1; IARS1; I/ | P41252; A0A | SYIC_HUMAN I | Unmodified | 15235 | 0 | 0 | 0 | 0 | 0 | 1301600 | 0 | 438780 | 259140 | 942050 | 5535800 | 0 | 0 | 0 | 0 | 0 | 0 |
| YRPLILDQV | CNTFR | P26992 | CNTFR_HUMAI | Unmodified | 0 | 0 | 0 | 0 | 0 | 0 | 0 | 0 | 4379400 | 1150100 | 0 | 0 | 0 | 0 | 0 | 0 | 0 | 0 |
| YRPQTLND | RFC5; RFC5; RI | P40937; C9J | RFC5_HUMAN | Unmodified | 0 | 521200 | 0 | 0 | 2322600 | 404300 | 1638600 | 0 | 1078800 | 147820 | 42106 | 164600 | 0 | 322240 | 258460 | 194180 | 103440 | 0 |
| YRQDDDQI | SND1; SND1 | Q7KZF4; H7 | SND1_HUMAN | Unmodified | 129120 | 0 | 0 | 871210 | 644260 | 0 | 0 | 0 | 0 | 0 | 0 | 0 | 0 | 486510 | 234750 | 0 | 0 | 428300 |
| YRQENQEA | NIPSNAP1 | Q9BPW8 | NIPS1_HUMAN | Unmodified | 0 | 0 | 0 | 0 | 0 | 0 | 26180 | 0 | 0 | 434160 | 0 | 0 | 0 | 0 | 0 | 0 | 0 | 0 |
| YRQQGRN' | OLA1; OLA1 | Q9NTK5; J3K | OLA1_HUMAN | Unmodified | 0 | 0 | 0 | 0 | 3142100 | 0 | 2447000 | 0 | 0 | 508140 | 0 | 0 | 2338600 | 0 | 0 | 0 | 1863500 | 1932100 |
| YRSDGALLI | WDR77 | Q9BQA1 | MEP50_HUMAI | Unmodified | 0 | 0 | 0 | 0 | 0 | 0 | 0 | 0 | 0 | 0 | 0 | 0 | 0 | 0 | 0 | 0 | 0 | 0 |
| YRSDIHTE/ | DIP2B | Q9P265 | DIP2B_HUMAN | Unmodified | 153100 | 125880 | 491200 | 235080 | 0 | 0 | 0 | 0 | 260290 | 0 | 0 | 1056900 | 0 | 0 | 246380 | 194750 | 86151 | 0 |
| YRSEALPTI | PRKCSH; PRKC | P14314; K7E | GLU2B_HUMAI | Unmodified | 58332 | 0 | 0 | 0 | 0 | 225350 | 0 | 0 | 2250500 | 0 | 0 | 0 | 0 | 0 | 0 | 54238 | 0 | 0 |
| YRSLYGDLI | SOGA1; SOGA1 | O94964; X6F | SOGA1_HUMAI | Unmodified | 0 | 0 | 0 | 0 | 0 | 0 | 0 | 0 | 0 | 420220 | 0 | 0 | 1233500 | 1548400 | 0 | 0 | 0 | 0 |
| YRSWIEETI | KLK14; KLK14 | Q9P0G3; A0 | KLK14_HUMAN | Unmodified | 0 | 0 | 0 | 0 | 0 | 0 | 0 | 0 | 14191000 | 3184900 | 0 | 0 | 0 | 0 | 0 | 0 | 0 | 0 |
| YRSWIEETI | KLK14; KLK14 | Q9P0G3; A0 | KLK14_HUMAN | Oxidation (M) | 0 | 0 | 0 | 0 | 0 | 0 | 0 | 0 | 966170 | 541680 | 0 | 0 | 0 | 0 | 0 | 0 | 0 | 0 |
| YRVEDSVD | CRLF1; CRLF1 | O75462; M0 | CRLF1_HUMAN | Unmodified | 0 | 0 | 0 | 0 | 0 | 0 | 0 | 0 | 884510 | 0 | 0 | 112180 | 0 | 0 | 0 | 0 | 0 | 0 |
| YRVPDVLV | GANAB; GANAE | Q14697; F5H | GANAB_HUMA | Unmodified | 0 | 0 | 0 | 0 | 0 | 0 | 0 | 0 | 54094000 | 10199000 | 0 | 672590 | 0 | 0 | 0 | 0 | 0 | 0 |
| YRWTEYGL | VDAC1; VDAC1 | P21796; C9J | VDAC1_HUMAI | Unmodified | 0 | 0 | 0 | 246940 | 1066400 | 0 | 3395100 | 0 | 1.87E+08 | 89470000 | 4119000 | 416290 | 1821700 | 4876000 | 183450 | 98254 | 62705 | 2302200 |
| YRYPGSLD | B4GALNT1 | Q00973 | B4GN1_HUMAI | Unmodified | 0 | 0 | 0 | 0 | 0 | 0 | 0 | 0 | 809580 | 0 | 0 | 0 | 0 | 0 | 0 | 0 | 0 | 0 |
| YRYQDEDT | DLG1; DLG1; D | Q12959; A0A | DLG1_HUMAN | Unmodified | 0 | 0 | 0 | 1578100 | 32064000 | 5380600 | 0 | 0 | 8179400 | 9367400 | 402160 | 1534900 | 15749000 | 17078000 | 1024100 | 3711700 | 15861000 | 38197000 |
| YSAARSAA | CLDN4 | O14493 | CLD4_HUMAN | Unmodified | 286130 | 47527 | 0 | 0 | 0 | 0 | 0 | 0 | 0 | 2650800 | 0 | 0 | 7328400 | 2897300 | 0 | 0 | 5848300 | 15819000 |
| YSAEAQAM | MUC5B | Q9HC84 | MUC5B_HUMA | Unmodified | 0 | 0 | 0 | 0 | 0 | 0 | 0 | 0 | 0 | 0 | 0 | 0 | 0 | 0 | 0 | 0 | 0 | 537490 |
| YSAFQLAPI | USP6NL; USP6I | Q92738; X6F | US6NL_HUMAI | Unmodified | 0 | 0 | 0 | 0 | 0 | 0 | 0 | 0 | 0 | 1461400 | 0 | 0 | 0 | 0 | 0 | 0 | 1033900 | 24717000 |
| YSALNVQH | GOLIM4; GOLIN | O00461; F8V | GOLI4_HUMAN | Unmodified | 111160 | 0 | 0 | 0 | 0 | 0 | 0 | 0 | 30161000 | 0 | 1502000 | 0 | 0 | 0 | 292790 | 0 | 0 | 0 |
| YSALNVQH | GOLIM4; GOLIN | O00461; F8V | GOLI4_HUMAN | Oxidation (M) | 0 | 0 | 0 | 0 | 0 | 0 | 0 | 0 | 1674800 | 549080 | 0 | 205440 | 0 | 0 | 0 | 0 | 0 | 0 |
| YSAPANLY | PTPRD; PTPRF; | P23468; P10 | PTPRD_HUMAI | Unmodified | 1064500 | 591590 | 0 | 0 | 0 | 0 | 0 | 0 | 1345500 | 0 | 0 | 0 | 42464 | 0 | 0 | 340760 | 0 | 0 |
| YSDEENLPI | AIF1L; AIF1L | Q9BQI0; Q5J | AIF1L_HUMAN | Unmodified | 0 | 0 | 393900 | 0 | 0 | 0 | 0 | 0 | 0 | 0 | 0 | 0 | 0 | 0 | 874140 | 303640 | 0 | 0 |
| YSDFEWLR | SNX3 | O60493 | SNX3_HUMAN | Unmodified | 153030 | 247030 | 0 | 701640 | 7639200 | 0 | 816810 | 0 | 1487900 | 497680 | 0 | 122130 | 2021100 | 1293500 | 313640 | 207030 | 0 | 946410 |
| YSDGDLTLI | IGF2R; IGF2R; I | P11717; A0A | MPRI_HUMAN ( | Unmodified | 0 | 0 | 0 | 0 | 0 | 0 | 0 | 0 | 7686900 | 802080 | 740760 | 0 | 0 | 0 | 0 | 0 | 0 | 0 |
| YSDHAGPA | MARK3; MARK3 | P27448; A0A | MARK3_HUMAI | Unmodified | 0 | 0 | 399320 | 1197000 | 0 | 0 | 0 | 0 | 0 | 0 | 0 | 0 | 0 | 0 | 0 | 695260 | 0 | 0 |
| YSDIDIILLK | TNPO1; TNPO1 | Q92973; A0/ | TNPO1_HUMAI | Unmodified | 0 | 187250 | 125960 | 396390 | 0 | 0 | 1652700 | 65294 | 0 | 0 | 0 | 365380 | 938680 | 131480 | 538290 | 252300 | 0 | 0 |
| YSDMIVAAI | H1-0 | P07305 | H10_HUMAN H | Oxidation (M) | 931110 | 523180 | 0 | 0 | 0 | 0 | 0 | 404810 | 0 | 0 | 0 | 0 | 0 | 0 | 0 | 0 | 0 | 0 |
| YSDNWEAI | IGF2R; IGF2R | P11717; A0A | MPRI_HUMAN ( | Unmodified | 0 | 0 | 0 | 0 | 0 | 0 | 0 | 0 | 60360000 | 8588800 | 3401600 | 2683500 | 0 | 0 | 0 | 0 | 0 | 0 |
| YSDPTQW | ST14 | Q9Y5Y6 | ST14_HUMAN S | Unmodified | 5981800 | 4519400 | 0 | 8929700 | 0 | 0 | 0 | 0 | 5053500 | 0 | 422900 | 0 | 0 | 0 | 15568000 | 10014000 | 0 | 0 |
| YSEAAELR | AKAP2; PALM2/ | Q9Y2D5; C9 | AKAP2_HUMAN | Unmodified | 0 | 0 | 163280 | 0 | 1360900 | 129530 | 0 | 0 | 0 | 31656 | 0 | 0 | 0 | 206130 | 0 | 235390 | 0 | 244930 |
| YSEAVFYY | CAPN7; CAPN7 | Q9Y6W3; F8 | CAN7_HUMAN | Unmodified | 294350 | 0 | 0 | 0 | 0 | 0 | 0 | 0 | 714450 | 0 | 0 | 3108500 | 0 | 0 | 0 | 0 | 71084 | 0 |
| YSEDRIGL/ | SUN2; SUN2 | Q9UH99; B0 | SUN2_HUMAN | Unmodified | 0 | 0 | 0 | 0 | 0 | 0 | 0 | 0 | 40325000 | 0 | 200810 | 0 | 0 | 0 | 0 | 0 | 0 | 0 |
| YSEEANNL | ERO1A; ERO1A; | Q96HE7; G3 | ERO1A_HUMAI | Unmodified | 557550 | 583800 | 174220 | 289550 | 0 | 0 | 0 | 0 | 1694300 | 106450 | 0 | 0 | 391450 | 0 | 1085500 | 489640 | 0 | 0 |
| YSEEANNL | ERO1A; ERO1A; | Q96HE7; G3 | ERO1A_HUMAI | Unmodified | 0 | 0 | 0 | 0 | 0 | 0 | 0 | 0 | 7587700 | 904590 | 0 | 0 | 0 | 0 | 0 | 0 | 0 | 0 |
| YSEELIGAL | PLXNA2 | O75051 | PLXA2_HUMAN | Unmodified | 0 | 0 | 167670 | 0 | 0 | 0 | 0 | 0 | 4186100 | 137520 | 129950 | 2318600 | 0 | 0 | 0 | 310830 | 0 | 0 |
| YSEFFTGS | SDF4; SDF4 | Q9BRK5; A0 | CAB45_HUMAI | Unmodified | 2997600 | 3706100 | 301370 | 301670 | 140300 | 0 | 0 | 0 | 17370000 | 148950 | 567850 | 1836800 | 663230 | 400540 | 2215100 | 219140 | 0 | 0 |
| YSEFFTGS | SDF4; SDF4 | Q9BRK5; A0 | CAB45_HUMAI | Unmodified | 0 | 0 | 0 | 0 | 76683 | 0 | 141620 | 0 | 0 | 649160 | 0 | 0 | 0 | 0 | 0 | 0 | 296260 | 0 |
| YSEFTSTTS | ADGRE5 | P48960 | AGRE5_HUMAI | Unmodified | 0 | 0 | 133980 | 108750 | 184440 | 0 | 0 | 0 | 0 | 0 | 0 | 28205 | 0 | 0 | 89754 | 109660 | 28115 | 34116 |
| YSEFTSTTS | ADGRE5 | P48960 | AGRE5_HUMAI | Unmodified | 0 | 0 | 0 | 0 | 522330 | 0 | 0 | 0 | 0 | 0 | 0 | 0 | 0 | 0 | 0 | 0 | 230190 | 514190 |
| YSEGYPGC | SHMT1; SHMT1 | P34896; J3K | GLYC_HUMAN | Unmodified | 0 | 0 | 185270 | 52114 | 0 | 0 | 0 | 0 | 0 | 0 | 0 | 1449100 | 0 | 0 | 99168 | 117150 | 0 | 0 |
| YSEIEPSTE | LAMB2; LAMB2 | P55268; F5H | LAMB2_HUMAI | Unmodified | 0 | 0 | 0 | 0 | 0 | 0 | 0 | 0 | 844890 | 0 | 0 | 0 | 0 | 0 | 0 | 0 | 0 | 0 |
| YSEKEDKYI | TPM4; TPM4; TF | P67936; A0A | TPM4_HUMAN | Unmodified | 0 | 0 | 0 | 202590 | 5037600 | 1164400 | 0 | 0 | 0 | 0 | 262180 | 585500 | 847490 | 336400 | 0 | 0 | 570950 | 338680 |
| YSEKQERN | CNN2; CNN2; ( | Q99439; H3I | CNN2_HUMAN | Unmodified | 0 | 8780600 | 0 | 0 | 1996800 | 500060 | 33117 | 0 | 0 | 0 | 0 | 0 | 13470000 | 10118000 | 0 | 0 | 202920 | 0 |
| YSELTTVQ( | XPOT; XPOT | O43592; F8V | XPOT_HUMAN | Unmodified | 0 | 0 | 0 | 0 | 0 | 0 | 0 | 0 | 0 | 0 | 0 | 912560 | 0 | 0 | 42290 | 0 | 0 | 0 |
| YSELVQNH | HIP1 | O00291 | HIP1_HUMAN I | Unmodified | 0 | 0 | 0 | 0 | 0 | 0 | 0 | 0 | 0 | 0 | 0 | 0 | 554510 | 263700 | 0 | 0 | 0 | 0 |
| YSESLLCSI | TOMM34 | Q15785 | TOM34_HUMAI | Unmodified | 0 | 0 | 0 | 0 | 196590 | 0 | 0 | 28465 | 0 | 0 | 0 | 925630 | 333100 | 0 | 0 | 181840 | 0 | 471770 |
| YSESLLCSI | TOMM34 | Q15785 | TOM34_HUMAI | Unmodified | 0 | 0 | 0 | 0 | 0 | 0 | 0 | 0 | 0 | 0 | 0 | 0 | 0 | 0 | 0 | 0 | 1909400 | 0 |
| YSETLKPDI | CACNA2D1; CA | P54289; H0Y | CA2D1_HUMAI | Unmodified | 2080800 | 1736600 | 0 | 0 | 0 | 0 | 0 | 0 | 5868300 | 0 | 0 | 0 | 0 | 0 | 0 | 0 | 0 | 0 |
| YSEVFEAIN | CSNK2A3; CSN | Q8NEV1; P6 | CSK23_HUMAN | Unmodified | 0 | 0 | 0 | 0 | 0 | 0 | 0 | 0 | 0 | 0 | 0 | 1619600 | 0 | 0 | 0 | 0 | 0 | 0 |
| YSEVVLDSI | SIT1; SIT1 | Q9Y3P8; A0/ | SIT1_HUMAN S | Unmodified | 0 | 0 | 0 | 0 | 0 | 0 | 0 | 0 | 3704100 | 0 | 0 | 0 | 0 | 0 | 0 | 0 | 0 | 0 |
| YSFCTDHT | FN1 | P02751 | FINC_HUMAN I | Unmodified | 1365200 | 3810100 | 19748000 | 14130000 | 0 | 0 | 40432 | 0 | 2709900 | 1790300 | 2244100 | 1496700 | 2906900 | 3286200 | 1611000 | 7030300 | 0 | 78348 |
| YSFGATCV | EGFR; EGFR; E( | P00533; Q5( | EGFR_HUMAN | Unmodified | 0 | 0 | 0 | 0 | 1653400 | 661910 | 0 | 0 | 0 | 0 | 0 | 0 | 0 | 0 | 0 | 0 | 0 | 0 |
| YSFHPVED | WIPF2; WIPF2 | Q8TF74; A8N | WIPF2_HUMAN | Unmodified | 4540900 | 3665500 | 3118000 | 2298600 | 507550 | 110400 | 0 | 0 | 0 | 0 | 81461 | 233510 | 1010300 | 509180 | 9117300 | 5206700 | 40968 | 282580 |
| YSFKGPHL | LMAN1 | P49257 | LMAN1_HUMAI | Unmodified | 0 | 0 | 0 | 580500 | 0 | 0 | 0 | 0 | 1.33E+08 | 28780000 | 1793400 | 0 | 0 | 0 | 0 | 0 | 0 | 0 |
| YSFKGPHL | LMAN1 | P49257 | LMAN1_HUMAI | Unmodified | 0 | 0 | 0 | 0 | 0 | 0 | 10609000 | 0 | 0 | 1.23E+08 | 0 | 0 | 4510800 | 9332800 | 0 | 0 | 0 | 17340000 |
| YSFLQFDP/ | PPP2CB; PPP2( | P62714; P67 | PP2AB_HUMAN | Unmodified | 0 | 0 | 2553600 | 0 | 0 | 0 | 0 | 168140 | 3461800 | 756950 | 3029300 | 17457000 | 0 | 0 | 2731300 | 2946400 | 0 | 0 |
| YSGGNYRI | RBM3 | P98179 | RBM3_HUMAN | Unmodified | 758280 | 673880 | 296470 | 238960 | 0 | 0 | 0 | 0 | 0 | 0 | 0 | 0 | 0 | 0 | 299380 | 584170 | 0 | 0 |
| YSGKEETV( | COL21A1; COL | Q96P44; F5( | COLA1_HUMAI | Unmodified | 0 | 0 | 0 | 0 | 225090 | 0 | 0 | 0 | 8168400 | 958820 | 0 | 0 | 0 | 0 | 0 | 0 | 71909 | 0 |
| YSGLCPHV | MTHFD1; MTHF | P11586; V9( | C1TC_HUMAN | Unmodified | 0 | 0 | 0 | 0 | 0 | 0 | 0 | 0 | 699070 | 0 | 0 | 1863000 | 0 | 0 | 0 | 0 | 287220 | 0 |
| YSGPEDDA | TOP2A | P11388 | TOP2A_HUMAN | Unmodified | 0 | 0 | 3712600 | 0 | 0 | 485030 | 0 | 0 | 9784600 | 802240 | 14543000 | 0 | 0 | 491560 | 1016400 | 113750 | 0 | 536250 |

| | | | | | | | | | | | | | | | | | | | | | | |
|---|---|---|---|---|---|---|---|---|---|---|---|---|---|---|---|---|---|---|---|---|---|---|
| YSGRDSLIF | XRCC6 | P12956 | XRCC6_HUMAI | Unmodified | 474970 | 0 | 0 | 0 | 1342400 | 430300 | 620300 | 0 | 890700 | 414290 | 1205500 | 7556200 | 0 | 0 | 0 | 0 | 0 | 550920 |
| YSGSEGST | S100P | P25815 | S100P_HUMAN | Unmodified | 172100 | 117200 | 0 | 0 | 0 | 121990 | 0 | 0 | 0 | 0 | 0 | 0 | 0 | 0 | 0 | 93147 | 0 | 0 |
| YSGSEGST | S100P | P25815 | S100P_HUMAN | Unmodified | 832780 | 1047700 | 0 | 27757 | 0 | 0 | 0 | 15481 | 139090 | 0 | 0 | 0 | 124150 | 0 | 0 | 28383 | 0 | 0 |
| YSGTLNLD | FASN; FASN | P49327; A0A | FAS_HUMAN F | Unmodified | 0 | 0 | 0 | 0 | 0 | 0 | 0 | 372220 | 0 | 0 | 405780 | 57924000 | 0 | 0 | 0 | 0 | 0 | 0 |
| YSGTLNLD | FASN; FASN | P49327; A0A | FAS_HUMAN F | Unmodified | 709900 | 0 | 3497700 | 0 | 0 | 0 | 0 | 0 | 0 | 1198300 | 3332500 | 47873000 | 0 | 0 | 0 | 0 | 0 | 0 |
| YSGYFGGV | B4GALT4; B4GA | O60513; C9J | B4GT4_HUMAN | Unmodified | 0 | 0 | 0 | 0 | 0 | 0 | 0 | 0 | 3599000 | 0 | 0 | 0 | 0 | 0 | 0 | 0 | 0 | 2038800 |
| YSGYFGGV | B4GALT4; B4GA | O60513; C9J | B4GT4_HUMAN | Unmodified | 0 | 0 | 0 | 0 | 91931 | 0 | 0 | 0 | 2345200 | 840490 | 0 | 374180 | 0 | 0 | 0 | 0 | 43441 | 0 |
| YSHDFNFH | HYOU1; HYOU1 | Q9Y4L1; E9F | HYOU1_HUMA | Unmodified | 0 | 0 | 0 | 0 | 0 | 0 | 0 | 0 | 993660 | 0 | 0 | 0 | 0 | 0 | 0 | 0 | 0 | 0 |
| YSHDLSDS | MKRN1; MKRN1 | Q9UHC7; C9 | MKRN1_HUMA | Unmodified | 279580 | 0 | 133640 | 133210 | 0 | 0 | 0 | 0 | 0 | 0 | 0 | 0 | 0 | 0 | 75672 | 0 | 0 | 0 |
| YSHEVLSEI | ZC3HAV1; ZC3I | Q7Z2W4; C9 | ZCCHV_HUMA | Unmodified | 1250400 | 1219700 | 3021800 | 3969400 | 33461000 | 9961800 | 0 | 2076600 | 1137300 | 852170 | 471440 | 928280 | 8338600 | 3542600 | 3014800 | 2236800 | 2481900 | 3327100 |
| YSHEVLSEI | ZC3HAV1; ZC3I | Q7Z2W4; C9 | ZCCHV_HUMA | Unmodified | 0 | 0 | 0 | 0 | 774480 | 0 | 258850 | 0 | 0 | 0 | 0 | 0 | 299620 | 382460 | 0 | 0 | 489790 | 193450 |
| YSHISMLD | OLFM1 | Q99784 | NOE1_HUMAN | Unmodified | 0 | 0 | 0 | 0 | 0 | 0 | 0 | 0 | 5127000 | 0 | 0 | 0 | 0 | 0 | 0 | 0 | 170150 | 0 |
| YSHLQPGD | KARS1; KARS1 | Q15046; H3I | SYK_HUMAN Ly | Unmodified | 0 | 0 | 218190 | 447390 | 1202000 | 448450 | 367180 | 0 | 870940 | 0 | 1193000 | 14203000 | 1548000 | 475030 | 261570 | 582610 | 312730 | 268450 |
| YSIGGLSPF | PTPRF; PTPRF; | P10586; H0Y | PTPRF_HUMAN | Unmodified | 181080 | 186040 | 0 | 0 | 0 | 0 | 0 | 0 | 175800 | 40163 | 0 | 0 | 0 | 0 | 0 | 0 | 0 | 6032.1 |
| YSIMHPGT | ASPH | Q12797 | ASPH_HUMAN | Unmodified | 0 | 0 | 77978 | 0 | 0 | 0 | 0 | 0 | 2411500 | 151050 | 0 | 0 | 0 | 0 | 0 | 0 | 0 | 0 |
| YSIMHPGT | ASPH | Q12797 | ASPH_HUMAN | Oxidation (M) | 0 | 0 | 0 | 0 | 0 | 0 | 0 | 0 | 626970 | 157850 | 0 | 0 | 0 | 0 | 0 | 0 | 0 | 0 |
| YSISRTEAA | CD44; CD44; C | P16070; Q8 | CD44_HUMAN | Unmodified | 6046100 | 2103100 | 0 | 0 | 1285300 | 80325 | 0 | 0 | 0 | 0 | 0 | 0 | 0 | 0 | 0 | 0 | 329730 | 1535200 |
| YSITFTGK | SPON2; SPON2 | Q9BUD6; D6 | SPON2_HUMA | Unmodified | 0 | 0 | 35685000 | 18973000 | 0 | 0 | 0 | 0 | 0 | 21908 | 0 | 0 | 3365400 | 4502100 | 12888000 | 8519100 | 0 | 0 |
| YSITFTGKV | SPON2; SPON2 | Q9BUD6; D6 | SPON2_HUMA | Unmodified | 33224 | 0 | 0 | 0 | 0 | 0 | 0 | 0 | 0 | 0 | 0 | 0 | 9797700 | 0 | 0 | 0 | 0 | 0 |
| YSIWCSTEI | YTHDF1; YTHDI | Q9BYJ9; Q7 | YTHD1_HUMAI | Unmodified | 653050 | 309410 | 0 | 0 | 0 | 0 | 0 | 0 | 76049 | 31541 | 0 | 366190 | 0 | 43054 | 58434 | 71548 | 0 | 0 |
| YSKAIAVTA | TLN2; TLN2; TLI | Q9Y4G6; H0 | TLN2_HUMAN | Oxidation (M) | 422610 | 0 | 0 | 0 | 0 | 0 | 0 | 0 | 0 | 0 | 0 | 2009500 | 0 | 0 | 0 | 0 | 122950 | 0 |
| YSLDPENP | RPL17; RPL17; | P18621; A0A | RL17_HUMAN | Unmodified | 340290 | 921680 | 0 | 0 | 0 | 0 | 0 | 0 | 0 | 71838 | 1777800 | 6457600 | 0 | 0 | 0 | 0 | 0 | 0 |
| YSLEPVAVE | PGK1 | P00558 | PGK1_HUMAN | Unmodified | 960170 | 749460 | 5622200 | 3457200 | 5320000 | 0 | 1568100 | 732720 | 11806000 | 1561600 | 12325000 | 61825000 | 0 | 2406400 | 7672600 | 6952000 | 5531600 | 17617000 |
| YSLEQDIRE | EPS15L1; EPS1 | Q9UBC2; M0 | EP15R_HUMAN | Unmodified | 252110 | 0 | 595640 | 148590 | 0 | 0 | 0 | 0 | 0 | 0 | 127460 | 832060 | 251240 | 0 | 135720 | 209220 | 0 | 0 |
| YSLEQRLN | PKN2; PKN2; PI | Q16513; B1A | PKN2_HUMAN | Unmodified | 0 | 0 | 0 | 0 | 1875400 | 596420 | 173470 | 74754 | 0 | 168040 | 0 | 0 | 0 | 0 | 0 | 224830 | 0 | 0 |
| YSLKPNDQ | ADK; ADK; ADK; | P55263; A0A | ADK_HUMAN A | Unmodified | 184910 | 129420 | 565940 | 449610 | 1214100 | 0 | 0 | 0 | 2176600 | 101830 | 2078600 | 18122000 | 0 | 0 | 932830 | 649550 | 0 | 0 |
| YSLLPFWY | GANAB; GANAE | Q14697; F5H | GANAB_HUMA | Unmodified | 0 | 0 | 77092 | 0 | 0 | 0 | 0 | 0 | 1517900 | 370560 | 0 | 0 | 0 | 0 | 459510 | 82795 | 0 | 0 |
| YSLSEDKLI | PLXNA1; PLXNA | Q9UIW2; O7 | PLXA1_HUMAN | Unmodified | 0 | 0 | 0 | 571700 | 0 | 0 | 0 | 0 | 0 | 328010 | 0 | 0 | 0 | 0 | 0 | 0 | 0 | 0 |
| YSLSGGGT | STK24; STK24; | Q9Y6E0; Q5 | STK24_HUMAN | Unmodified | 283920 | 249480 | 111350 | 218750 | 2161500 | 354590 | 108370 | 0 | 656460 | 311590 | 290290 | 2391900 | 520760 | 305870 | 188980 | 260160 | 493750 | 1138000 |
| YSLTYIYTG | AZGP1; AZGP1 | P25311; C9J | ZA2G_HUMAN | Unmodified | 0 | 0 | 184830 | 0 | 0 | 0 | 0 | 0 | 0 | 0 | 1707400 | 883680 | 0 | 0 | 524000 | 0 | 0 | 771980 |
| YSLVAKQE | INCENP | Q9NQS7 | INCE_HUMAN | Unmodified | 0 | 0 | 0 | 251620 | 1733900 | 0 | 0 | 0 | 0 | 210710 | 0 | 0 | 1159100 | 672020 | 0 | 0 | 0 | 0 |
| YSLVLELSD | APMAP; APMAP | Q9HDC9; H0 | APMAP_HUMA | Unmodified | 0 | 0 | 0 | 0 | 0 | 0 | 47261 | 0 | 3039100 | 1063600 | 0 | 0 | 0 | 0 | 0 | 0 | 0 | 0 |
| YSLYEVHVS | AFDN; AFDN; A | A8MQ02; F8 | A8MQ02_HUM | Unmodified | 0 | 0 | 0 | 0 | 0 | 0 | 0 | 0 | 1422900 | 0 | 917320 | 4912100 | 0 | 0 | 0 | 0 | 0 | 0 |
| YSLYEVHVS | AFDN; AFDN; A | P55196; Q5 | AFAD_HUMAN | Unmodified | 134640 | 0 | 725750 | 0 | 0 | 0 | 140100 | 49338 | 0 | 0 | 925230 | 1060200 | 0 | 0 | 1263800 | 955580 | 0 | 0 |
| YSNIMISQF | EXTL2 | Q9UBQ6 | EXTL2_HUMAN | Oxidation (M) | 0 | 63427 | 0 | 111400 | 296040 | 0 | 984540 | 20298 | 1857000 | 120300 | 599830 | 0 | 0 | 1234600 | 75414 | 39635 | 0 | 49693 |
| YSNIMISQF | EXTL2 | Q9UBQ6 | EXTL2_HUMAN | Unmodified | 0 | 0 | 0 | 0 | 0 | 0 | 0 | 0 | 408790 | 0 | 0 | 0 | 0 | 0 | 0 | 0 | 0 | 0 |
| YSNIMISQF | EXTL2 | Q9UBQ6 | EXTL2_HUMAN | Oxidation (M) | 0 | 0 | 0 | 0 | 811940 | 0 | 0 | 0 | 915350 | 0 | 0 | 0 | 0 | 0 | 0 | 0 | 541980 | 2703000 |
| YSNSALGH | RTN4; RTN4; RT | Q9NQC3; F8 | RTN4_HUMAN | Unmodified | 199990 | 0 | 295740 | 101350 | 98022 | 0 | 69349 | 0 | 978900 | 390570 | 275980 | 360680 | 56446 | 53380 | 258280 | 232650 | 35909 | 142710 |
| YSNSEVVT | CYFIP2; CYFIP2 | Q96F07; H7 | CYFP2_HUMAN | Unmodified | 0 | 0 | 0 | 0 | 0 | 0 | 0 | 0 | 0 | 0 | 0 | 525260 | 126480 | 0 | 0 | 0 | 17273 | 0 |
| YSNSEVVT | CYFIP1; CYFIP1 | Q7L576; A0A | CYFP1_HUMAN | Unmodified | 289900 | 369510 | 178670 | 53675 | 0 | 0 | 0 | 30322 | 0 | 0 | 245160 | 1303600 | 0 | 0 | 601950 | 732030 | 0 | 0 |
| YSPAPGGP | EVPL; EVPL | Q92817; K7E | EVPL_HUMAN | Unmodified | 0 | 0 | 0 | 0 | 1406800 | 389590 | 0 | 0 | 725560 | 0 | 0 | 0 | 432090 | 0 | 0 | 0 | 0 | 0 |
| YSPTSPTYS | POLR2A; POLR2 | P24928; A0A | RPB1_HUMAN | Unmodified | 0 | 0 | 0 | 0 | 0 | 0 | 0 | 0 | 420540 | 293440 | 0 | 0 | 0 | 0 | 0 | 0 | 0 | 0 |
| YSQFINFPI | HSP90B1; HSP | P14625; A0A | ENPL_HUMAN | Unmodified | 189310 | 99679 | 178850 | 85548 | 0 | 0 | 98822 | 0 | 25356000 | 1718400 | 1057000 | 1327900 | 0 | 0 | 624620 | 215470 | 0 | 0 |
| YSQKEDKY | TPM3; TPM3; TF | P06753; J3K | TPM3_HUMAN | Unmodified | 1402700 | 0 | 422860 | 5821200 | 85190000 | 17024000 | 1186200 | 1281500 | 4922300 | 3321600 | 8356700 | 13391000 | 22856000 | 9264200 | 627710 | 2638400 | 8021400 | 7495500 |
| YSQKEDRY | TPM1; TPM1; TF | P09493; H0 | TPM1_HUMAN | Unmodified | 113160 | 0 | 0 | 0 | 3504500 | 625660 | 0 | 0 | 0 | 0 | 140680 | 2189300 | 0 | 0 | 0 | 226270 | 0 | 0 |
| YSQNIYIQN | PKP2 | Q99959 | PKP2_HUMAN | Unmodified | 0 | 0 | 0 | 0 | 0 | 0 | 0 | 0 | 0 | 0 | 383070 | 2089100 | 0 | 0 | 0 | 0 | 290930 | 436260 |
| YSQNIYIQN | PKP2 | Q99959 | PKP2_HUMAN | Unmodified | 0 | 0 | 0 | 0 | 0 | 0 | 0 | 0 | 0 | 0 | 0 | 0 | 0 | 0 | 0 | 0 | 1143800 | 1454800 |
| YSQSDLEQ | SMC1A; SMC1A | Q14683; G8 | SMC1A_HUMA | Unmodified | 0 | 0 | 0 | 0 | 0 | 0 | 103250 | 80410 | 195620 | 0 | 78522 | 163460 | 0 | 0 | 0 | 0 | 0 | 0 |
| YSQTGNYE | DDOST; DDOST | P39656; A0A | OST48_HUMAN | Unmodified | 0 | 0 | 0 | 0 | 0 | 0 | 0 | 0 | 37448000 | 3585200 | 0 | 527290 | 0 | 61919 | 0 | 0 | 0 | 0 |
| YSQVLANG | RPS18; RPS18 | P62269; J3J | RS18_HUMAN | Unmodified | 177230 | 173070 | 1036600 | 0 | 0 | 0 | 0 | 0 | 0 | 0 | 208510 | 15462000 | 0 | 0 | 207550 | 248880 | 229460 | 0 |
| YSQVLANG | RPS18; RPS18 | P62269; J3J | RS18_HUMAN | Unmodified | 1269800 | 876680 | 0 | 283570 | 0 | 0 | 0 | 0 | 131500 | 0 | 0 | 0 | 0 | 0 | 0 | 0 | 0 | 0 |
| YSQYQQAI | PRMT5; PRMT5 | O14744; H0 | ANM5_HUMAN | Unmodified | 0 | 0 | 0 | 0 | 0 | 0 | 0 | 0 | 0 | 0 | 0 | 5262500 | 0 | 0 | 0 | 262140 | 0 | 0 |
| YSRHYSPE | N/A; AAK1 | Q6ZSR9; A0 | YJ005_HUMAN | Unmodified | 0 | 0 | 0 | 0 | 0 | 0 | 0 | 0 | 0 | 0 | 0 | 0 | 1283900 | 1622300 | 0 | 0 | 595410 | 1130400 |
| YSRKPNVG | UBA6 | A0AVT1 | UBA6_HUMAN | Unmodified | 0 | 0 | 0 | 18541 | 630680 | 45488 | 52741 | 32183 | 0 | 0 | 170230 | 434580 | 149720 | 88779 | 0 | 0 | 577700 | 1018800 |
| YSRLSLLMI | N/A | N/A | | Oxidation (M) | 0 | 0 | 0 | 0 | 0 | 0 | 0 | 0 | 0 | 0 | 0 | 1249000 | 0 | 0 | 0 | 0 | 0 | 0 |
| YSSAFCNK | TUSC3; TUSC3 | Q13454; D6 | TUSC3_HUMAI | Unmodified | 1734800 | 2060000 | 0 | 0 | 105890 | 0 | 0 | 64970 | 1335900 | 170290 | 35563 | 0 | 0 | 0 | 2636500 | 2364600 | 0 | 0 |
| YSSAGTVEI | PCCA; PCCA; P | P05165; A0A | PCCA_HUMAN | Unmodified | 0 | 0 | 0 | 0 | 203370 | 0 | 0 | 0 | 666100 | 459580 | 0 | 0 | 0 | 0 | 0 | 0 | 41014 | 0 |
| YSSDPTGA | EGFR; EGFR | P00533; E9P | EGFR_HUMAN | Unmodified | 0 | 0 | 0 | 862190 | 0 | 0 | 0 | 0 | 0 | 0 | 0 | 0 | 0 | 0 | 0 | 0 | 0 | 0 |
| YSSDYFQA | LGALS3BP | Q08380 | LG3BP_HUMAI | Unmodified | 0 | 0 | 0 | 0 | 0 | 0 | 0 | 0 | 1603900 | 0 | 406310 | 800630 | 0 | 0 | 0 | 0 | 0 | 64373 |
| YSSDYFQA | LGALS3BP | Q08380 | LG3BP_HUMAI | Unmodified | 0 | 0 | 0 | 0 | 1326500 | 789170 | 0 | 0 | 4019100 | 6620000 | 1472400 | 0 | 17138000 | 14913000 | 0 | 0 | 0 | 78522000 |
| YSSGGNFE | OCLN; OCLN | Q16625; A0 | OCLN_HUMAN | Unmodified | 1145300 | 1297100 | 3057200 | 3226400 | 2991100 | 0 | 0 | 0 | 2064800 | 776250 | 42371 | 132290 | 5769800 | 1320600 | 3732800 | 4180100 | 148080 | 445830 |
| YSSNKEEW | ALG9; N/A; ALG | Q9H6U8; A0 | ALG9_HUMAN | Unmodified | 0 | 0 | 0 | 0 | 139870 | 0 | 16397 | 0 | 14778000 | 0 | 0 | 0 | 79576 | 213060 | 0 | 0 | 0 | 237350 |
| YSSNRPPI | KIF14 | Q15058 | KIF14_HUMAN | Unmodified | 0 | 0 | 0 | 1263500 | 1773000 | 0 | 0 | 23013 | 0 | 0 | 0 | 582380 | 758150 | 181050 | 0 | 0 | 393450 | 264500 |
| YSSQFGG | EIF3D; EIF3D; E | O15371; B0 | EIF3D_HUMAN | Unmodified | 0 | 0 | 0 | 0 | 2046600 | 0 | 0 | 0 | 0 | 701360 | 0 | 0 | 1687600 | 1133000 | 0 | 0 | 0 | 1093600 |
| YSSSSLSHI | PHLDB2; PHLD | Q86SQ0; G5 | PHLB2_HUMAN | Unmodified | 0 | 0 | 0 | 0 | 245850 | 0 | 0 | 0 | 0 | 0 | 0 | 0 | 0 | 0 | 0 | 0 | 0 | 419240 |
| YSSTEEVLV | ITGA1; ITGA1 | P56199; A0A | ITA1_HUMAN I | Unmodified | 0 | 262250 | 0 | 0 | 0 | 0 | 0 | 0 | 0 | 106370 | 0 | 0 | 0 | 0 | 0 | 0 | 803010 | 2328700 |
| YSSTEEVLV | ITGA1; ITGA1 | P56199; A0A | ITA1_HUMAN I | Unmodified | 0 | 0 | 0 | 0 | 0 | 0 | 17761 | 0 | 0 | 0 | 0 | 0 | 0 | 0 | 0 | 61726 | 640770 | 1903800 |
| YSTDESITS | DNAJC13; DNA | O75165; A0 | DJC13_HUMAN | Unmodified | 26428 | 106010 | 0 | 36185 | 1073200 | 0 | 0 | 0 | 30178 | 0 | 0 | 184070 | 0 | 0 | 0 | 13025 | 0 | 0 |
| YSTHLHLAI | STXBP2; STXBP | Q15833; M0 | STXB2_HUMAN | Unmodified | 0 | 0 | 110850 | 0 | 849470 | 651540 | 0 | 0 | 0 | 0 | 0 | 0 | 943570 | 569100 | 0 | 0 | 464140 | 1370300 |
| YSTHLHLAI | STXBP2; STXBP | Q15833; M0 | STXB2_HUMAN | Oxidation (M) | 0 | 0 | 0 | 127720 | 553220 | 153820 | 171330 | 549480 | 88465 | 0 | 0 | 682850 | 990700 | 430850 | 0 | 271520 | 365330 | 568040 |
| YSTHLHLAI | STXBP1; STXBP | P61764; A0A | STXB1_HUMAN | Unmodified | 66564 | 0 | 0 | 83600 | 1065600 | 81417 | 1653500 | 455860 | 1567200 | 688810 | 0 | 897070 | 684500 | 724700 | 0 | 0 | 379370 | 1246700 |
| YSTIPKFGI | HGS; HGS; HGS | O14964; A0 | HGS_HUMAN H | Unmodified | 0 | 0 | 0 | 0 | 0 | 0 | 962980 | 431100 | 0 | 275550 | 0 | 0 | 0 | 0 | 0 | 0 | 461920 | 300260 |
| YSTKEDKY | TPM2; TPM2; TF | P07951; A7X | TPM2_HUMAN | Unmodified | 0 | 0 | 0 | 0 | 0 | 0 | 0 | 0 | 0 | 0 | 0 | 483090 | 0 | 0 | 0 | 0 | 0 | 0 |
| YSTLHTQS | CD99L2; CD99 | Q8TCZ2; A0 | C99L2_HUMAN | Unmodified | 1312100 | 1624900 | 0 | 0 | 15297000 | 0 | 3881900 | 1082900 | 1391800 | 0 | 0 | 0 | 0 | 0 | 318660 | 1076200 | 6337100 | 18810000 |
| YSTLTAR | LMCD1; LMCD1 | Q9NZU5; H7 | LMCD1_HUMA | Unmodified | 0 | 0 | 0 | 0 | 339230 | 38941 | 972190 | 37847 | 78391 | 0 | 81284 | 2296500 | 98084 | 38302 | 14421 | 17402 | 152850 | 361770 |
| YSTNEGET | SORL1 | Q92673 | SORL_HUMAN | Unmodified | 0 | 0 | 0 | 0 | 0 | 0 | 0 | 0 | 473140 | 0 | 0 | 0 | 0 | 0 | 0 | 0 | 0 | 0 |
| YSTPRLPA | NBR1; NBR1 | Q14596; B7 | NBR1_HUMAN | Unmodified | 0 | 0 | 39566 | 0 | 0 | 0 | 88105 | 0 | 676240 | 25247 | 0 | 0 | 0 | 0 | 0 | 0 | 0 | 87616 |
| YSTPRLPA | NBR1; NBR1 | Q14596; B7 | NBR1_HUMAN | Unmodified | 0 | 0 | 0 | 0 | 0 | 0 | 1934000 | 0 | 0 | 0 | 0 | 0 | 309130 | 0 | 0 | 0 | 0 | 0 |
| YSTSGSSG | IMMT; IMMT; IM | Q16891; D6 | MIC60_HUMAN | Unmodified | 0 | 0 | 0 | 0 | 0 | 0 | 0 | 0 | 1589100 | 518420 | 0 | 0 | 0 | 0 | 0 | 0 | 0 | 0 |
| YSTSLYASF | ERN1; ERN1 | O75460; A0 | ERN1_HUMAN | Oxidation (M) | 0 | 0 | 0 | 0 | 0 | 1065400 | 0 | 88199 | 508990 | 189620 | 0 | 0 | 0 | 0 | 39189 | 0 | 0 | 0 |
| YSTSLYASF | ERN1; ERN1 | O75460; A0 | ERN1_HUMAN | Unmodified | 0 | 0 | 0 | 0 | 0 | 0 | 0 | 0 | 1594600 | 0 | 0 | 0 | 0 | 0 | 0 | 0 | 0 | 0 |
| YSTSVVEK | RAB11FIP5; RAI | Q9BXF6; A0 | RFIP5_HUMAN | Unmodified | 0 | 0 | 0 | 0 | 984350 | 0 | 0 | 34660 | 379000 | 0 | 0 | 0 | 399230 | 213590 | 0 | 0 | 0 | 161970 |
| YSVDIPLDK | RPL27; RPL27 | P61353; K7E | RL27_HUMAN | Unmodified | 14971000 | 8736600 | 5979100 | 3491400 | 93004 | 0 | 0 | 0 | 718320 | 480980 | 2598900 | 28024000 | 0 | 0 | 7604100 | 6348100 | 0 | 0 |
| YSVDPSIVN | SLC4A7 | H7C3C4 | H7C3C4_HUM | Unmodified | 0 | 0 | 0 | 0 | 1771200 | 0 | 0 | 0 | 0 | 0 | 0 | 186910 | 0 | 187940 | 0 | 0 | 0 | 0 |
| YSVGIHNLI | IFIT1 | P09914 | IFIT1_HUMAN | Unmodified | 0 | 0 | 0 | 0 | 502400 | 118690 | 0 | 0 | 0 | 0 | 0 | 0 | 0 | 0 | 0 | 0 | 0 | 1094500 |
| YSVGSLVD | PPFIA2; PPFIA2 | O75334; F8 | LIPA2_HUMAN | Unmodified | 0 | 0 | 211530 | 307630 | 543660 | 0 | 0 | 25849 | 0 | 0 | 0 | 0 | 159790 | 222660 | 272080 | 0 | 132800 | 41559 |
| YSVLIMAEK | PTGFRN | Q9P2B2 | FPRP_HUMAN | Oxidation (M) | 439000 | 0 | 20359000 | 8588700 | 0 | 0 | 0 | 0 | 0 | 0 | 0 | 434670 | 0 | 0 | 8991400 | 0 | 0 | 0 |
| YSVLIMAEK | PTGFRN | Q9P2B2 | FPRP_HUMAN | Unmodified | 0 | 0 | 1761800 | 0 | 0 | 0 | 0 | 0 | 0 | 0 | 0 | 0 | 0 | 0 | 0 | 66655 | 0 | 0 |
| YSVQLLTP | RUVBL1; RUVBI | Q9Y265; H7 | RUVB1_HUMAI | Unmodified | 158780 | 116480 | 1033900 | 1124300 | 13364 | 143750 | 0 | 0 | 1117300 | 60982 | 1765900 | 20182000 | 3630500 | 16233 | 2086300 | 197340 | 67362 | 16714 |
| YSVQTADH | FSCN1 | Q16658 | FSCN1_HUMAI | Unmodified | 0 | 0 | 0 | 0 | 0 | 0 | 0 | 0 | 1396900 | 0 | 233990 | 9669700 | 0 | 0 | 0 | 0 | 0 | 0 |
| YSVWIGGS | ACTG1; ACTB; | P63261; P60 | ACTG_HUMAN | Oxidation (M) | 0 | 803310 | 0 | 0 | 9490600 | 0 | 2862500 | 0 | 0 | 521160 | 0 | 0 | 17742000 | 8792500 | 0 | 0 | 0 | 6349100 |
| YSVWIGGS | ACTG1; ACTB; | P63261; P60 | ACTG_HUMAN | Unmodified | 0 | 0 | 0 | 0 | 655460 | 0 | 0 | 0 | 0 | 0 | 0 | 0 | 376690 | 0 | 0 | 0 | 0 | 406000 |
| YSYNTEWR | TMEM43 | Q9BTV4 | TMM43_HUMAI | Unmodified | 0 | 0 | 0 | 0 | 0 | 0 | 0 | 0 | 6253800 | 292040 | 0 | 0 | 72010 | 0 | 0 | 0 | 0 | 0 |
| YSYNTEWR | TMEM43 | Q9BTV4 | TMM43_HUMAI | Unmodified | 0 | 0 | 0 | 0 | 0 | 0 | 0 | 0 | 1660400 | 0 | 0 | 0 | 0 | 0 | 0 | 0 | 0 | 0 |
| YSYQYTVAI | ERGIC1 | Q969X5 | ERGI1_HUMAN | Unmodified | 0 | 0 | 248370 | 0 | 0 | 0 | 0 | 0 | 10562000 | 800730 | 0 | 0 | 0 | 0 | 345040 | 0 | 0 | 0 |
| YSYQYTVAI | ERGIC1 | Q969X5 | ERGI1_HUMAN | Unmodified | 0 | 0 | 0 | 0 | 99349 | 0 | 0 | 0 | 73616 | 1033100 | 0 | 0 | 0 | 0 | 0 | 0 | 0 | 0 |
| YSYVCPDL | ACTR3; ACTR3 | P61158; B4D | ARP3_HUMAN | Unmodified | 212640 | 0 | 170380 | 0 | 0 | 0 | 139630 | 74663 | 0 | 0 | 554990 | 838290 | 523240 | 0 | 141790 | 154880 | 0 | 0 |
| YSYYDESQ | MEMO1 | Q9Y316 | MEMO1_HUMA | Unmodified | 123700 | 132160 | 0 | 0 | 0 | 0 | 0 | 0 | 0 | 0 | 0 | 289780 | 767230 | 0 | 158470 | 0 | 0 | 0 |
| YTAEEIEVLI | CAPN7 | Q9Y6W3 | CAN7_HUMAN | Unmodified | 124430 | 125700 | 679230 | 522090 | 0 | 0 | 0 | 0 | 0 | 0 | 0 | 4916800 | 0 | 0 | 0 | 0 | 0 | 0 |
| YTAESSDTI | IARS2 | Q9NSE4 | SYIM_HUMAN I | Unmodified | 0 | 0 | 0 | 0 | 0 | 0 | 0 | 0 | 742580 | 0 | 0 | 0 | 0 | 0 | 0 | 0 | 0 | 0 |
| YTALDKWT | COPS2; COPS2 | P61201; B4D | CSN2_HUMAN | Unmodified | 0 | 0 | 0 | 0 | 1340900 | 0 | 0 | 0 | 0 | 481880 | 0 | 0 | 1992400 | 646130 | 0 | 0 | 0 | 1444900 |
| YTAQVDAE | ATP5PD | O75947 | ATP5H_HUMAN | Unmodified | 0 | 0 | 0 | 0 | 0 | 0 | 0 | 0 | 2812200 | 262810 | 307820 | 0 | 0 | 0 | 0 | 0 | 0 | 0 |
| YTASVTVG | RHOF; RHOF | Q9HBH0; F5 | RHOF_HUMAN | Unmodified | 0 | 0 | 0 | 0 | 0 | 0 | 0 | 0 | 0 | 0 | 0 | 0 | 0 | 0 | 0 | 0 | 0 | 0 |
| YTCHVQHE | HLA-H; HLA-C; | P01893; Q5 | HLAH_HUMAN | Unmodified | 0 | 0 | 0 | 0 | 0 | 0 | 0 | 0 | 846990 | 294610 | 0 | 10143000 | 0 | 0 | 0 | 45457 | 0 | 0 |
| YTCHVQHE | HLA-B; HLA-A; | P01889; P04 | HLAB_HUMAN | Unmodified | 0 | 0 | 0 | 0 | 0 | 0 | 0 | 0 | 0 | 0 | 0 | 9590000 | 0 | 0 | 0 | 0 | 0 | 0 |
| YTCHVQHE | HLA-B; HLA-A; | P01889; P04 | HLAB_HUMAN | Unmodified | 684440 | 532000 | 0 | 0 | 0 | 0 | 0 | 0 | 5430400 | 0 | 0 | 1.35E+08 | 0 | 0 | 0 | 827840 | 0 | 0 |
| YTCSFCGK | RPL37AP8; RPL | A6NKH3; P6 | RL37L_HUMAN | Unmodified | 1587100 | 710610 | 192040 | 224520 | 0 | 0 | 0 | 0 | 0 | 0 | 0 | 8745100 | 0 | 0 | 0 | 0 | 0 | 0 |
| YTDIDYQV | PIGM | Q9H3S5 | PIGM_HUMAN | Unmodified | 0 | 0 | 0 | 34973 | 0 | 0 | 0 | 0 | 2669900 | 104390 | 0 | 0 | 0 | 0 | 63543 | 0 | 0 | 0 |
| YTDIRGQPI | TCP1; TCP1; TC | P17987; E7E | TCPA_HUMAN | Unmodified | 0 | 0 | 0 | 386670 | 1937700 | 311140 | 638170 | 165580 | 0 | 979230 | 0 | 0 | 2263700 | 0 | 0 | 0 | 822150 | 0 |

| | | | | | | | | | | | | | | | | | | | | | | |
|---|---|---|---|---|---|---|---|---|---|---|---|---|---|---|---|---|---|---|---|---|---|---|
| YTDQLKPF( | GPC6 | Q9Y625 | GPC6_HUMAN | Unmodified | 3629900 | 0 | 0 | 21924 | 0 | 0 | 0 | 0 | 0 | 0 | 0 | 0 | 0 | 0 | 0 | 0 | 0 | 0 |
| YTDTLGSIC | VAV1; VAV1; VA | P15498; Q9( | VAV_HUMAN P | Unmodified | 0 | 0 | 0 | 0 | 0 | 0 | 0 | 0 | 0 | 0 | 0 | 715330 | 0 | 0 | 0 | 0 | 0 | 0 |
| YTEGVQSLI | SUPT16H | Q9Y5B9 | SP16H_HUMAI | Unmodified | 0 | 81467 | 0 | 0 | 149710 | 0 | 415290 | 101230 | 0 | 0 | 0 | 1796400 | 0 | 0 | 0 | 0 | 0 | 0 |
| YTEISNIK | GLT8D1 | Q68CQ7 | GL8D1_HUMAI | Unmodified | 207820 | 0 | 0 | 0 | 0 | 0 | 0 | 0 | 2927800 | 156080 | 0 | 475360 | 0 | 0 | 174510 | 151050 | 0 | 0 |
| YTELEYIHA | EXOC4 | Q96A65 | EXOC4_HUMAI | Unmodified | 49240 | 0 | 125870 | 107790 | 0 | 0 | 0 | 0 | 333220 | 233720 | 142120 | 1070500 | 0 | 0 | 426100 | 221900 | 0 | 0 |
| YTELPYGRI | CTSS; CTSS; CT | P25774; U3k | CATS_HUMAN | Unmodified | 0 | 0 | 0 | 0 | 0 | 0 | 0 | 27537 | 4323700 | 1007400 | 0 | 0 | 0 | 0 | 0 | 0 | 0 | 0 |
| YTEQLKPF( | GPC4 | O75487 | GPC4_HUMAN | Unmodified | 1066200 | 724010 | 216390 | 0 | 0 | 0 | 0 | 0 | 0 | 0 | 0 | 0 | 0 | 0 | 0 | 305110 | 0 | 0 |
| YTFAAHMD | TMED2; TMED2 | Q15363; E7E | TMED2_HUMAI | Oxidation (M) | 16598 | 0 | 27053 | 0 | 0 | 0 | 0 | 0 | 5575500 | 19057 | 0 | 467550 | 0 | 0 | 207570 | 84689 | 0 | 0 |
| YTFAAHMD | TMED2; TMED2 | Q15363; E7E | TMED2_HUMAI | Unmodified | 0 | 0 | 0 | 0 | 0 | 0 | 0 | 0 | 6880900 | 460910 | 0 | 0 | 0 | 0 | 0 | 0 | 0 | 0 |
| YTFEVQAVI | EPHB3 | P54753 | EPHB3_HUMAI | Unmodified | 154810 | 809460 | 0 | 0 | 0 | 0 | 0 | 0 | 0 | 0 | 0 | 0 | 0 | 0 | 32518 | 15710 | 0 | 675320 |
| YTFHEVITA | NDST1 | P52848 | NDST1_HUMAI | Unmodified | 0 | 0 | 0 | 0 | 0 | 0 | 0 | 0 | 1213300 | 168580 | 0 | 0 | 0 | 0 | 0 | 0 | 0 | 131360 |
| YTGEDFDE | DYNC1H1; DYN | Q14204; A0/ | DYHC1_HUMA | Unmodified | 0 | 0 | 0 | 0 | 0 | 0 | 0 | 0 | 164410 | 0 | 0 | 1840900 | 0 | 0 | 0 | 0 | 0 | 0 |
| YTGLRTHL( | ZDHHC5; ZDHH | Q9C0B5; H0 | ZDHC5_HUMA | Unmodified | 0 | 0 | 0 | 0 | 240660 | 0 | 0 | 0 | 0 | 0 | 0 | 0 | 237660 | 0 | 0 | 0 | 45479 | 172310 |
| YTGNTYRV | FN1 | P02751 | FINC_HUMAN I | Unmodified | 0 | 548690 | 5788800 | 8898100 | 1199000 | 0 | 143730 | 0 | 1509100 | 512750 | 970050 | 0 | 2050900 | 2126000 | 0 | 802450 | 200100 | 3567400 |
| YTGYKGPD | ERO1A; ERO1A; | Q96HE7; G3 | ERO1A_HUMAI | Unmodified | 0 | 0 | 0 | 0 | 0 | 0 | 0 | 0 | 3236700 | 0 | 0 | 645840 | 0 | 0 | 0 | 0 | 0 | 0 |
| YTHAANTV | WDR82 | Q6UXN9 | WDR82_HUMA | Unmodified | 543300 | 447290 | 0 | 1094500 | 0 | 91842 | 0 | 0 | 0 | 0 | 0 | 5448900 | 0 | 0 | 0 | 638520 | 0 | 0 |
| YTIGHAWT | PREP; PREP | P48147; A0A | PPCE_HUMAN | Unmodified | 25593 | 0 | 0 | 0 | 0 | 0 | 0 | 0 | 0 | 0 | 0 | 846250 | 0 | 0 | 86087 | 139700 | 0 | 0 |
| YTIGLGQAI | HMGCS1; HMG | Q01581; D6I | HMCS1_HUMA | Unmodified | 0 | 0 | 0 | 0 | 268360 | 0 | 0 | 45559 | 674850 | 0 | 344040 | 5591300 | 0 | 0 | 0 | 0 | 127590 | 47856 |
| YTIIIPENLK | ACO1 | P21399 | ACOC_HUMAN | Oxidation (M) | 0 | 0 | 0 | 0 | 0 | 0 | 0 | 0 | 0 | 0 | 0 | 1443700 | 0 | 0 | 0 | 165870 | 0 | 0 |
| YTISQEAYD | TBCB; TBCB; TE | Q99426; K7E | TBCB_HUMAN | Unmodified | 0 | 0 | 0 | 0 | 6214300 | 3901300 | 659480 | 519730 | 0 | 0 | 0 | 805130 | 0 | 0 | 0 | 0 | 3151300 | 1139000 |
| YTISQEAYD | TBCB; TBCB; TE | Q99426; K7E | TBCB_HUMAN | Unmodified | 885400 | 411500 | 1380700 | 1823100 | 1391500 | 584230 | 0 | 0 | 8619500 | 1080500 | 357370 | 5530000 | 1932700 | 600850 | 959720 | 1148000 | 0 | 831320 |
| YTISQEAYD | TBCB; TBCB; TE | Q99426; K7E | TBCB_HUMAN | Unmodified | 0 | 0 | 0 | 0 | 3683100 | 177120 | 1207200 | 0 | 0 | 1278700 | 0 | 0 | 6323400 | 3149200 | 0 | 0 | 297150 | 2792500 |
| YTIVQQQIP | TUBGCP2; TUB | Q9BSJ2; F2Z | GCP2_HUMAN | Unmodified | 0 | 0 | 0 | 0 | 0 | 0 | 0 | 0 | 0 | 0 | 0 | 703780 | 0 | 0 | 0 | 0 | 0 | 0 |
| YTIYSPK | LARS1; LARS1; | Q9P2J5; A0A | SYLC_HUMAN | Unmodified | 0 | 0 | 0 | 0 | 0 | 0 | 0 | 0 | 0 | 0 | 0 | 1121100 | 0 | 0 | 0 | 91476 | 0 | 0 |
| YTLETRPN( | IDE; IDE; IDE; ID | P14735; A0A | IDE_HUMAN In | Unmodified | 0 | 0 | 0 | 0 | 0 | 0 | 0 | 0 | 0 | 0 | 532570 | 2578200 | 0 | 0 | 0 | 0 | 0 | 0 |
| YTLFSASSE | GEMIN5 | Q8TEQ6 | GEMI5_HUMAI | Unmodified | 0 | 0 | 0 | 0 | 0 | 0 | 0 | 0 | 446380 | 0 | 0 | 1515600 | 0 | 0 | 0 | 0 | 0 | 0 |
| YTLRSHFD( | STRN3 | Q13033 | STRN3_HUMAI | Unmodified | 0 | 0 | 0 | 0 | 1891800 | 0 | 478680 | 0 | 0 | 0 | 0 | 0 | 0 | 0 | 0 | 0 | 0 | 0 |
| YTPEEIAMA | ALDOC; ALDOC | P09972; A8N | ALDOC_HUMA | Oxidation (M) | 0 | 0 | 0 | 0 | 0 | 0 | 0 | 0 | 0 | 0 | 0 | 2676300 | 0 | 0 | 0 | 0 | 0 | 0 |
| YTPSGQAG | ALDOA; ALDOA | P04075; J3KI | ALDOA_HUMAI | Unmodified | 0 | 0 | 5734600 | 0 | 20966000 | 3752600 | 1060800 | 549560 | 22263000 | 1246400 | 17261000 | 69878000 | 2687000 | 1022000 | 6899000 | 6673900 | 5907000 | 8468900 |
| YTPTQQGN | FLNB; FLNB | O75369; E7E | FLNB_HUMAN | Oxidation (M) | 0 | 424780 | 0 | 0 | 928880 | 0 | 0 | 0 | 0 | 2051400 | 0 | 766870 | 0 | 0 | 213530 | 0 | 0 | 0 |
| YTPVQQGF | FLNA; FLNA; FL | P21333; A0A | FLNA_HUMAN | Unmodified | 0 | 0 | 0 | 0 | 0 | 1296800 | 0 | 0 | 0 | 449690 | 544360 | 0 | 0 | 0 | 0 | 0 | 1576500 | 0 |
| YTQELTLK | ITGA6 | P23229 | ITA6_HUMAN I | Unmodified | 170640 | 443840 | 1725500 | 804740 | 10777000 | 2645700 | 0 | 0 | 312670 | 0 | 626380 | 6518700 | 10033000 | 6007900 | 3528300 | 3467700 | 6724100 | 35862000 |
| YTQELTLKF | ITGA6 | P23229 | ITA6_HUMAN I | Unmodified | 0 | 0 | 0 | 0 | 139130 | 0 | 0 | 87076 | 0 | 0 | 0 | 0 | 133950 | 176230 | 0 | 0 | 2008500 | 1202300 |
| YTQQIIQGI | PNPT1 | Q8TCS8 | PNPT1_HUMAI | Unmodified | 0 | 0 | 0 | 0 | 0 | 0 | 0 | 0 | 3300700 | 250380 | 0 | 0 | 0 | 0 | 0 | 0 | 0 | 0 |
| YTQSNSVC | ATIC; ATIC | P31939; H7( | PUR9_HUMAN | Unmodified | 0 | 0 | 612900 | 523910 | 995360 | 202530 | 0 | 162400 | 282920 | 161400 | 0 | 2220400 | 1202900 | 904650 | 0 | 0 | 406380 | 752030 |
| YTQVLWSG | ITGA3 | P26006 | ITA3_HUMAN I | Unmodified | 0 | 0 | 0 | 0 | 6747800 | 5150700 | 0 | 0 | 1832500 | 0 | 567430 | 7598600 | 0 | 0 | 0 | 0 | 2476900 | 3747700 |
| YTQVLWSG | ITGA3 | P26006 | ITA3_HUMAN I | Oxidation (M) | 0 | 0 | 0 | 0 | 4291800 | 1034100 | 0 | 0 | 0 | 0 | 0 | 0 | 0 | 0 | 0 | 0 | 1784300 | 2899600 |
| YTQVLWSG | ITGA3 | P26006 | ITA3_HUMAN I | Unmodified | 0 | 0 | 0 | 0 | 11271000 | 2618600 | 0 | 0 | 0 | 0 | 0 | 0 | 0 | 0 | 0 | 0 | 3996300 | 9040700 |
| YTSGFDEL( | SPINT1; SPINT1 | O43278; H3I | SPIT1_HUMAN | Unmodified | 0 | 5905000 | 750720 | 688270 | 0 | 0 | 0 | 0 | 2315700 | 0 | 427060 | 0 | 0 | 0 | 809350 | 772570 | 0 | 0 |
| YTSLMLR | WDR1 | O75083 | WDR1_HUMAN | Unmodified | 355600 | 397840 | 254320 | 348360 | 0 | 0 | 0 | 0 | 1881000 | 0 | 264430 | 4573400 | 0 | 0 | 517970 | 213380 | 0 | 0 |
| YTSQLPPLT | MMAB; MMAB | Q96EY8; S4F | MMAB_HUMAN | Unmodified | 0 | 0 | 1532800 | 0 | 0 | 0 | 0 | 0 | 566250 | 0 | 0 | 0 | 0 | 0 | 0 | 0 | 0 | 0 |
| YTTPEDATF | GNAS; GNAS; G | Q5JWF2; P6: | GNAS1_HUMAI | Unmodified | 0 | 508310 | 4774200 | 1169600 | 0 | 0 | 164470 | 234410 | 936800 | 238240 | 2774900 | 12455000 | 37381 | 101660 | 2428500 | 3585200 | 0 | 0 |
| YTTPEDATF | GNAS; GNAS; G | Q5JWF2; P6: | GNAS1_HUMAI | Unmodified | 2111900 | 1985200 | 6809900 | 14276000 | 2962300 | 906820 | 380450 | 214640 | 6969600 | 483950 | 8501300 | 27836000 | 3159400 | 1017700 | 8824700 | 11665000 | 1873500 | 2261000 |
| YTTPEDATF | GNAS; GNAS; G | Q5JWF2; P6: | GNAS1_HUMAI | Unmodified | 0 | 0 | 0 | 0 | 5323700 | 1368100 | 1676100 | 0 | 0 | 191390 | 148560 | 0 | 2405500 | 1666400 | 0 | 0 | 1657200 | 3937200 |
| YTTPSGEVI | NPEPPSL1; NPI | A6NEC2; P5! | PSAL_HUMAN I | Unmodified | 0 | 0 | 0 | 0 | 0 | 0 | 0 | 0 | 0 | 0 | 0 | 3672700 | 0 | 0 | 0 | 0 | 0 | 0 |
| YTVQDESH | RACK1; RACK1; | P63244; D6F | RACK1_HUMAI | Unmodified | 1089200 | 499280 | 506630 | 884490 | 0 | 49872 | 0 | 43983 | 501290 | 87780 | 1346300 | 11031000 | 0 | 141570 | 551380 | 1209200 | 0 | 302020 |
| YTYGKPVK( | CD109 | Q6YHK3 | CD109_HUMAI | Unmodified | 0 | 0 | 0 | 0 | 0 | 0 | 0 | 0 | 3922700 | 3015800 | 0 | 0 | 0 | 0 | 0 | 0 | 1409500 | 2706800 |
| YTYSEWHS | DSC3; DSC2; D | Q14574; Q0: | DSC3_HUMAN | Unmodified | 0 | 152860 | 376200 | 0 | 0 | 0 | 0 | 0 | 1215700 | 0 | 0 | 0 | 0 | 0 | 163600 | 0 | 0 | 0 |
| YVAAAFPS/ | PCK2; PCK2; P( | Q16822; H0 | PCKGM_HUMA | Unmodified | 0 | 0 | 0 | 0 | 0 | 0 | 0 | 0 | 1808500 | 0 | 0 | 0 | 0 | 0 | 0 | 0 | 31352 | 0 |
| YVAADTLYS | DST; DST; DST; | Q03001; A0/ | DYST_HUMAN | Unmodified | 0 | 0 | 0 | 70494 | 0 | 665520 | 0 | 0 | 340000 | 367140 | 32011 | 0 | 246680 | 0 | 0 | 0 | 0 | 328390 |
| YVAAQLHL | CA14 | Q9ULX7 | CAH14_HUMAI | Unmodified | 0 | 0 | 0 | 0 | 0 | 0 | 0 | 0 | 0 | 0 | 0 | 1205600 | 0 | 0 | 0 | 0 | 0 | 0 |
| YVAEIEKEK | PSMA7; PSMA7 | O14818; H0 | PSA7_HUMAN | Unmodified | 63735 | 0 | 0 | 993500 | 2515700 | 0 | 57947 | 40793 | 1026100 | 1036900 | 527760 | 812870 | 5992700 | 697460 | 0 | 21519 | 228880 | 711140 |
| YVALTSLLK | AP1G1 | O43747 | AP1G1_HUMAI | Unmodified | 0 | 0 | 0 | 0 | 1616000 | 0 | 198000 | 0 | 564120 | 0 | 0 | 4015400 | 0 | 0 | 0 | 0 | 178150 | 0 |
| YVALVQEK | COPB1 | P53618 | COPB_HUMAN | Unmodified | 0 | 0 | 0 | 0 | 602560 | 0 | 245500 | 202880 | 141850 | 131220 | 0 | 1835800 | 0 | 0 | 0 | 0 | 299320 | 346250 |
| YVANLFPYI | PPA1; PPA1 | Q15181; Q5! | IPYR_HUMAN I | Unmodified | 541010 | 635680 | 1611200 | 0 | 4333600 | 0 | 620920 | 441950 | 1235200 | 397670 | 2863500 | 51547000 | 3326800 | 1400100 | 1800600 | 2162100 | 2688900 | 2571600 |
| YVANMHGI | CPM; CPM; CPI | P14384; F8V | CBPM_HUMAN | Oxidation (M) | 0 | 0 | 0 | 0 | 0 | 0 | 173310 | 0 | 952290 | 232260 | 0 | 0 | 0 | 0 | 0 | 0 | 0 | 0 |
| YVANMHGI | CPM; CPM; CPI | P14384; F8V | CBPM_HUMAN | Unmodified | 0 | 0 | 0 | 0 | 0 | 0 | 0 | 0 | 1148700 | 57435 | 0 | 0 | 0 | 0 | 0 | 40734 | 0 | 0 |
| YVASYLLA/ | RPLP2 | P05387 | RLA2_HUMAN ( | Unmodified | 5026800 | 31926000 | 14259000 | 15723000 | 1831700 | 0 | 0 | 105780 | 2718500 | 0 | 2213900 | 46251000 | 3969200 | 0 | 44112000 | 31508000 | 0 | 0 |
| YVASYLLA/ | RPLP2 | P05387 | RLA2_HUMAN ( | Unmodified | 0 | 0 | 0 | 0 | 0 | 0 | 0 | 0 | 0 | 361720 | 0 | 247820 | 2649100 | 235920 | 0 | 0 | 0 | 0 |
| YVATLGVE\ | RAN; RAN; RAN | P62826; H0 | RAN_HUMAN G | Unmodified | 0 | 0 | 0 | 77022 | 0 | 0 | 0 | 0 | 0 | 0 | 0 | 22413000 | 0 | 0 | 0 | 0 | 0 | 0 |
| YVATLGVE\ | RAN; RAN; RAN | P62826; H0 | RAN_HUMAN G | Unmodified | 0 | 0 | 0 | 0 | 5065500 | 0 | 1265700 | 0 | 3566000 | 391290 | 0 | 41408000 | 2574500 | 1483600 | 0 | 0 | 279800 | 2192900 |
| YVAVELK | CNPY3; CNPY3 | Q9BT09; A0/ | CNPY3_HUMAI | Unmodified | 0 | 0 | 0 | 0 | 0 | 0 | 0 | 0 | 22056000 | 176850 | 0 | 669440 | 0 | 0 | 0 | 35552 | 0 | 0 |
| YVAVELKS/ | CNPY3; CNPY3 | Q9BT09; A0/ | CNPY3_HUMAI | Unmodified | 0 | 0 | 0 | 0 | 0 | 0 | 0 | 0 | 0 | 215050 | 0 | 0 | 0 | 0 | 0 | 0 | 0 | 461590 |
| YVAVMPPH | BLVRB; BLVRB | P30043; A0A | BLVRB_HUMAN | Oxidation (M) | 3792600 | 1676800 | 0 | 6745900 | 0 | 0 | 0 | 0 | 420140 | 0 | 0 | 0 | 2126300 | 0 | 7219800 | 2970600 | 0 | 0 |
| YVAVPACEI | WDR3 | Q9UNX4 | WDR3_HUMAN | Unmodified | 34964 | 118290 | 0 | 0 | 1178300 | 2379200 | 3207400 | 3810600 | 0 | 293300 | 0 | 193840 | 0 | 0 | 0 | 0 | 216760 | 491440 |
| YVCEADGF | MASP2 | O00187 | MASP2_HUMAI | Unmodified | 0 | 0 | 466780 | 151270 | 0 | 0 | 0 | 0 | 0 | 0 | 0 | 0 | 0 | 0 | 578310 | 181260 | 0 | 0 |
| YVDGDLCF | IGF2R | P11717 | MPRI_HUMAN ( | Unmodified | 0 | 0 | 0 | 0 | 0 | 0 | 0 | 0 | 10015000 | 2212200 | 966520 | 2433300 | 0 | 0 | 0 | 402600 | 0 | 0 |
| YVDGDLCF | IGF2R | P11717 | MPRI_HUMAN ( | Unmodified | 0 | 0 | 0 | 0 | 0 | 0 | 0 | 0 | 0 | 910990 | 544360 | 1149200 | 0 | 0 | 0 | 244430 | 0 | 0 |
| YVDGDLCF | IGF2R | P11717 | MPRI_HUMAN ( | Unmodified | 0 | 0 | 0 | 0 | 0 | 0 | 127580 | 0 | 5947500 | 4605700 | 808000 | 27155 | 508150 | 475530 | 0 | 0 | 0 | 0 |
| YVDIAIPCN | RPSA; RPSA; RF | P08865; A0A | RSSA_HUMAN | Unmodified | 2672400 | 1090800 | 0 | 1456700 | 465330 | 0 | 226350 | 0 | 594820 | 115710 | 3386800 | 18554000 | 433370 | 0 | 1268400 | 952430 | 0 | 221680 |
| YVDLSDNR | TLR9; N/A | Q9NR96; H0 | TLR9_HUMAN T | Unmodified | 0 | 0 | 0 | 0 | 0 | 0 | 0 | 10361 | 92983 | 0 | 376070 | 258350 | 0 | 0 | 0 | 0 | 0 | 0 |
| YVDQVLQL | IGF2R | P11717 | MPRI_HUMAN ( | Unmodified | 0 | 0 | 766520 | 515440 | 0 | 0 | 0 | 0 | 45369000 | 5451000 | 1036100 | 7766100 | 0 | 0 | 984160 | 1346100 | 0 | 0 |
| YVDQVLQL | IGF2R | P11717 | MPRI_HUMAN ( | Unmodified | 0 | 0 | 0 | 0 | 0 | 0 | 0 | 50955 | 5107600 | 5462600 | 0 | 0 | 0 | 0 | 0 | 0 | 0 | 0 |
| YVDQVLQL | IGF2R | P11717 | MPRI_HUMAN ( | Unmodified | 0 | 0 | 0 | 0 | 130650 | 78508 | 0 | 0 | 0 | 754830 | 0 | 0 | 1187100 | 731070 | 0 | 0 | 0 | 0 |
| YVDSEGHL | CAV1 | Q03135 | CAV1_HUMAN | Unmodified | 0 | 0 | 0 | 0 | 10965000 | 1602700 | 0 | 0 | 0 | 0 | 0 | 0 | 0 | 0 | 0 | 0 | 600000 | 1864700 |
| YVDTPFGK | MTAP; N/A; MT/ | Q13126; F2Z | MTAP_HUMAN | Unmodified | 449350 | 265340 | 0 | 1470900 | 8873000 | 0 | 0 | 704560 | 6366500 | 2696300 | 6582700 | 19292000 | 4051500 | 0 | 569850 | 644340 | 3924800 | 6949700 |
| YVDTPFGK | MTAP; N/A; MT/ | Q13126; F2Z | MTAP_HUMAN | Unmodified | 0 | 0 | 0 | 0 | 0 | 0 | 0 | 0 | 0 | 0 | 0 | 0 | 0 | 0 | 0 | 0 | 637740 | 0 |
| YVDWIQET | KLK11; KLK11; I | Q9UBX7; M0 | KLK11_HUMAN | Oxidation (M) | 0 | 0 | 0 | 0 | 0 | 0 | 0 | 0 | 2508500 | 1341500 | 0 | 0 | 0 | 0 | 0 | 0 | 0 | 0 |
| YVDWIQET | KLK11; KLK11; I | Q9UBX7; M0 | KLK11_HUMAN | Unmodified | 0 | 0 | 0 | 0 | 0 | 0 | 0 | 0 | 7219900 | 1330200 | 0 | 0 | 0 | 304230 | 0 | 0 | 0 | 73009 |
| YVEAAWYP | VARS1; VARS1; | P26640; A0/ | SYVC_HUMAN | Unmodified | 0 | 0 | 0 | 750270 | 0 | 0 | 0 | 0 | 1887800 | 0 | 0 | 2241900 | 0 | 0 | 1286100 | 372260 | 0 | 0 |
| YVEAKDCL | PFDN5; PFDN5 | Q99471; A0/ | PFD5_HUMAN | Unmodified | 0 | 0 | 0 | 0 | 1192000 | 262080 | 853360 | 155660 | 0 | 801670 | 0 | 0 | 1768400 | 1292600 | 0 | 0 | 2087500 | 1258800 |
| YVECSALT( | RHOQ; CDC42 | P17081; P60 | RHOQ_HUMAN | Unmodified | 8154300 | 5901500 | 4753100 | 6293200 | 46513000 | 7351600 | 7543100 | 5930700 | 5632900 | 5257800 | 6290200 | 17478000 | 30659000 | 22711000 | 6777600 | 5277700 | 12003000 | 37345000 |
| YVECSALT( | RHOQ; CDC42 | P17081; P60 | RHOQ_HUMAN | Unmodified | 0 | 14804 | 0 | 0 | 51192 | 0 | 865440 | 668770 | 0 | 0 | 0 | 0 | 217670 | 442220 | 0 | 0 | 1136800 | 0 |
| YVELFLNST | HNRNPH1; HNI | P31943; G8J | HNRH1_HUMA | Unmodified | 204970 | 0 | 0 | 0 | 0 | 0 | 0 | 0 | 579110 | 0 | 0 | 3444800 | 0 | 0 | 0 | 92513 | 0 | 0 |
| YVELINQA/ | ACE; ACE; ACE; | P12821; J3K | ACE_HUMAN A | Unmodified | 1266600 | 611230 | 0 | 0 | 0 | 0 | 0 | 0 | 1021800 | 0 | 0 | 0 | 0 | 0 | 0 | 0 | 0 | 0 |
| YVEQLLTLF | CUL5 | Q93034 | CUL5_HUMAN | Unmodified | 0 | 0 | 0 | 0 | 0 | 0 | 0 | 0 | 0 | 0 | 0 | 241270 | 0 | 0 | 0 | 21301 | 0 | 0 |
| YVESALNN | N/A | N/A | N/A | Unmodified | 0 | 0 | 0 | 0 | 0 | 0 | 0 | 0 | 0 | 0 | 0 | 0 | 0 | 0 | 0 | 0 | 1492300 | 121010 |
| YVESLWK | PLA2G4A | P47712 | PA24A_HUMAN | Unmodified | 0 | 0 | 0 | 0 | 0 | 0 | 0 | 0 | 0 | 0 | 126560 | 954520 | 0 | 0 | 0 | 0 | 0 | 0 |
| YVEVSPATE | RBM12 | Q9NTZ6 | RBM12_HUMAI | Unmodified | 0 | 0 | 0 | 0 | 0 | 0 | 0 | 0 | 0 | 0 | 0 | 1403500 | 0 | 0 | 0 | 0 | 0 | 0 |
| YVFFLDPCI | SLC44A1 | Q8WWI5 | CTL1_HUMAN ( | Unmodified | 78932 | 15393 | 0 | 0 | 32196 | 0 | 0 | 0 | 41730 | 76937 | 19641 | 27335 | 0 | 0 | 60503 | 0 | 0 | 0 |
| YVFFLDPCI | SLC44A1 | Q8WWI5 | CTL1_HUMAN ( | Unmodified | 3055300 | 0 | 2500100 | 4722600 | 3506700 | 760320 | 1027300 | 0 | 4916900 | 2227500 | 1086300 | 2766300 | 4538100 | 1813100 | 3645700 | 3654500 | 143080 | 754460 |
| YVFRLHEAT | LAMC1 | P11047 | LAMC1_HUMAI | Unmodified | 0 | 0 | 0 | 0 | 446820 | 0 | 104280 | 0 | 0 | 648700 | 0 | 347370 | 597980 | 0 | 0 | 1372500 | 0 | 2134500 |
| YVGATTNIH | PLCB4; PLCB4 | Q15147; A0/ | PLCB4_HUMAN | Oxidation (M) | 0 | 0 | 201800 | 893920 | 0 | 0 | 0 | 0 | 0 | 0 | 0 | 0 | 0 | 0 | 564110 | 686600 | 0 | 0 |
| YVGESEAN | NSF; NSF | P46459; I3L( | NSF_HUMAN V | Unmodified | 299620 | 655970 | 626040 | 1087700 | 1321700 | 222270 | 569320 | 449180 | 1487500 | 521880 | 303460 | 3652100 | 6862900 | 3792500 | 961560 | 989020 | 694830 | 2315900 |
| YVGTGGID | CEMIP2 | Q9UHN6 | CEIP2_HUMAN | Unmodified | 0 | 32191 | 217260 | 383500 | 1271500 | 286240 | 8461.2 | 0 | 1225800 | 248650 | 44055 | 96053 | 0 | 0 | 284340 | 0 | 539020 | 0 |
| YVHAMNNI | TMTC4 | Q5T4D3 | TMTC4_HUMAI | Unmodified | 0 | 0 | 0 | 0 | 0 | 0 | 0 | 0 | 817460 | 92163 | 0 | 0 | 0 | 0 | 0 | 0 | 0 | 0 |
| YVIHTVGPI | MACROD1; MA( | Q9BQ69; A0. | MACD1_HUMA | Unmodified | 667050 | 301330 | 39782 | 0 | 0 | 0 | 0 | 0 | 5571400 | 258250 | 0 | 716710 | 0 | 0 | 0 | 0 | 0 | 0 |
| YVIIPTTFEP | CAPN5; CAPN5 | O15484; E7E | CAN5_HUMAN | Unmodified | 2440500 | 6081600 | 2621200 | 5046200 | 439560 | 108790 | 320940 | 0 | 9609300 | 635000 | 903520 | 6308000 | 1211600 | 373680 | 8453200 | 6372800 | 0 | 180140 |
| YVIWSADM | COPA; COPA; C | P53621; A0A | COPA_HUMAN | Oxidation (M) | 0 | 0 | 0 | 0 | 0 | 0 | 0 | 0 | 0 | 0 | 0 | 0 | 0 | 0 | 0 | 0 | 0 | 0 |
| YVIWSADM | COPA; COPA; C | P53621; A0A | COPA_HUMAN | Unmodified | 0 | 0 | 0 | 0 | 0 | 0 | 0 | 0 | 0 | 0 | 0 | 0 | 0 | 0 | 0 | 0 | 0 | 0 |
| YVIYIER | RPL26; RPL26L | P61254; Q9U | RL26_HUMAN ( | Unmodified | 2915500 | 2788700 | 1327000 | 1749300 | 138750 | 0 | 0 | 81916 | 1005300 | 0 | 0 | 30844000 | 163160 | 0 | 1855900 | 1352400 | 0 | 0 |
| YVKLEDRD | DSG2 | Q14126 | DSG2_HUMAN | Unmodified | 205000 | 0 | 0 | 0 | 2762500 | 1087600 | 0 | 0 | 9506200 | 2427900 | 0 | 273160 | 0 | 0 | 0 | 0 | 151010 | 0 |
| YVLCPSTTF | AXL; AXL | P30530; M0I | UFO_HUMAN T | Unmodified | 0 | 0 | 0 | 0 | 1077700 | 0 | 0 | 0 | 0 | 0 | 0 | 0 | 0 | 0 | 0 | 0 | 199840 | 0 |
| YVLDDQYV | BMX | P51813 | BMX_HUMAN ( | Unmodified | 0 | 0 | 0 | 0 | 0 | 0 | 277080 | 0 | 0 | 0 | 0 | 1460800 | 958930 | 1325200 | 0 | 0 | 0 | 0 |
| YVLDDQYV | BMX | P51813 | BMX_HUMAN ( | Unmodified | 0 | 0 | 0 | 0 | 0 | 0 | 0 | 0 | 0 | 0 | 0 | 11163000 | 0 | 0 | 0 | 0 | 0 | 0 |
| YVLEDGPE | TBCE; TBCE; TB | Q15813; A0/ | TBCE_HUMAN | Unmodified | 0 | 0 | 0 | 198710 | 0 | 0 | 0 | 54691 | 0 | 0 | 0 | 456750 | 0 | 0 | 164420 | 0 | 0 | 0 |
| YVLEPEISF | UGGT1 | Q9NYU2 | UGGG1_HUMA | Unmodified | 24413 | 93447 | 497360 | 0 | 0 | 295580 | 30368 | 27229 | 10142000 | 15661 | 892220 | 0 | 0 | 0 | 519130 | 67185 | 0 | 0 |
| YVLEPEISF | UGGT1 | Q9NYU2 | UGGG1_HUMA | Unmodified | 0 | 0 | 0 | 0 | 0 | 0 | 621510 | 0 | 0 | 666090 | 0 | 0 | 0 | 0 | 0 | 0 | 0 | 0 |
| YVLGYKQTI | RPL30; RPL30; I | P62888; A0A | RL30_HUMAN ( | Unmodified | 0 | 290140 | 0 | 1059700 | 648920 | 0 | 760390 | 797530 | 0 | 0 | 0 | 0 | 1297600 | 687310 | 0 | 0 | 0 | 0 |

| | | | | | | | | | | | | | | | | | | | | | | |
|---|---|---|---|---|---|---|---|---|---|---|---|---|---|---|---|---|---|---|---|---|---|---|
| YVLHCQGT | ETF1; ETF1 | P62495; B7Z | ERF1_HUMAN I | Unmodified | 236830 | 100700 | 94555 | 153290 | 0 | 0 | 0 | 0 | 0 | 0 | 0 | 2992300 | 0 | 0 | 269850 | 256530 | 0 | 0 |
| YVLINWVG | DBN1; DBN1; D | Q16643; D6I | DREB_HUMAN | Unmodified | 0 | 0 | 1259000 | 3172400 | 21065000 | 3085200 | 129610 | 267860 | 3635900 | 4704300 | 4053500 | 20615000 | 11074000 | 8910800 | 2980700 | 2597700 | 19243000 | 13113000 |
| YVLINWVG | DBN1; DBN1; D | Q16643; D6I | DREB_HUMAN | Unmodified | 0 | 0 | 0 | 0 | 0 | 3133600 | 0 | 0 | 0 | 0 | 0 | 0 | 0 | 0 | 0 | 0 | 1547000 | 0 |
| YVLSNQYR | AFDN; AFDN; A | P55196; Q5T | AFAD_HUMAN | Unmodified | 277980 | 0 | 276250 | 47079 | 0 | 0 | 0 | 86101 | 3694100 | 0 | 521070 | 4304400 | 0 | 0 | 587160 | 0 | 0 | 0 |
| YVLWVVTH | PTPRB; PTPRB; | P23467; F8V | PTPRB_HUMAN | Unmodified | 0 | 0 | 0 | 0 | 0 | 0 | 0 | 0 | 0 | 0 | 0 | 0 | 0 | 0 | 0 | 0 | 0 | 0 |
| YVMPSCES | SMOC1 | Q9H4F8 | SMOC1_HUMA | Unmodified | 0 | 22433 | 0 | 0 | 0 | 0 | 0 | 0 | 465870 | 0 | 0 | 0 | 0 | 0 | 0 | 0 | 0 | 0 |
| YVMTTTTLE | EIF2S1; EIF2S1; | P05198; H0Y | IF2A_HUMAN E | Unmodified | 0 | 0 | 0 | 0 | 0 | 0 | 0 | 0 | 0 | 0 | 191520 | 1834400 | 0 | 0 | 0 | 0 | 0 | 0 |
| YVMYPIAVS | ADAM17 | P78536 | ADA17_HUMAN | Unmodified | 0 | 0 | 214230 | 0 | 484350 | 0 | 0 | 0 | 1007400 | 170920 | 0 | 0 | 0 | 0 | 432820 | 0 | 0 | 492820 |
| YVNILPYDH | PTPRA | P18433 | PTPRA_HUMAN | Unmodified | 0 | 0 | 0 | 0 | 0 | 0 | 0 | 0 | 0 | 0 | 0 | 3096100 | 0 | 0 | 0 | 0 | 0 | 0 |
| YVPGSSGS | PLAA; PLAA; PL | Q9Y263; H0Y | PLAP_HUMAN I | Unmodified | 337960 | 0 | 0 | 0 | 0 | 0 | 0 | 0 | 0 | 0 | 0 | 1142200 | 0 | 0 | 0 | 0 | 0 | 0 |
| YVPPAAGR | FLNB; FLNB | O75369; E7E | FLNB_HUMAN | Unmodified | 0 | 0 | 0 | 0 | 4473800 | 1094100 | 0 | 201340 | 0 | 216250 | 0 | 136850 | 715130 | 0 | 0 | 0 | 2773700 | 1283800 |
| YVPPSSTDI | MUC1; MUC1; I | P15941; A6Z | MUC1_HUMAN | Unmodified | 0 | 0 | 0 | 0 | 9056900 | 1888500 | 0 | 0 | 4342300 | 877220 | 160920 | 492590 | 0 | 0 | 0 | 0 | 19416 | 0 |
| YVPRAILVD | TUBB2A; TUBB; | Q13885; P07 | TBB2A_HUMAN | Oxidation (M) | 0 | 0 | 0 | 0 | 2963200 | 0 | 488450 | 0 | 0 | 923640 | 0 | 0 | 428030 | 163530 | 0 | 0 | 220290 | 1465500 |
| YVPRAILVD | TUBB2A; TUBB; | Q13885; P07 | TBB2A_HUMAN | Unmodified | 0 | 0 | 0 | 0 | 466780 | 0 | 150040 | 0 | 1119000 | 189840 | 0 | 0 | 975820 | 0 | 0 | 0 | 24165 | 1251400 |
| YVPTAGKLT | SYT1; SYT1 | P21579; J3K | SYT1_HUMAN S | Unmodified | 0 | 0 | 0 | 0 | 321200 | 0 | 139130 | 0 | 0 | 534040 | 0 | 0 | 0 | 0 | 0 | 0 | 0 | 0 |
| YVQLPADE | LSM2 | Q9Y333 | LSM2_HUMAN | Unmodified | 0 | 0 | 0 | 0 | 0 | 0 | 0 | 0 | 0 | 0 | 0 | 0 | 0 | 0 | 0 | 0 | 0 | 0 |
| YVQQENFV | RHBDF1 | Q96CC6 | RHDF1_HUMAI | Unmodified | 46615 | 78658 | 57525 | 612730 | 0 | 0 | 0 | 0 | 4675900 | 3245200 | 0 | 0 | 0 | 0 | 0 | 116570 | 0 | 0 |
| YVRELEQAI | NDEL1; NDEL1; | Q9GZM8; J3I | NDEL1_HUMAI | Unmodified | 0 | 0 | 0 | 0 | 0 | 0 | 0 | 0 | 0 | 0 | 0 | 0 | 0 | 0 | 0 | 0 | 0 | 0 |
| YVRPFLNSI | PSMD11; PSMD | O00231; J3Q | PSD11_HUMAN | Unmodified | 0 | 0 | 0 | 0 | 241740 | 0 | 192720 | 0 | 0 | 0 | 0 | 4393600 | 0 | 0 | 222030 | 247630 | 0 | 158240 |
| YVRPGGGF | GPX1; GPX1 | P07203; A0A | GPX1_HUMAN | Oxidation (M) | 0 | 0 | 0 | 869600 | 2346900 | 0 | 2662600 | 317750 | 3464900 | 1926400 | 1298100 | 4646500 | 3725600 | 3277000 | 75099 | 0 | 754140 | 4683100 |
| YVRPGGGF | GPX1; GPX1 | P07203; A0A | GPX1_HUMAN | Unmodified | 0 | 0 | 0 | 0 | 0 | 0 | 0 | 0 | 2207400 | 245100 | 0 | 0 | 0 | 214030 | 0 | 0 | 0 | 1708400 |
| YVRPGGGF | GPX3; GPX3; GI | P22352; H0Y | GPX3_HUMAN | Unmodified | 0 | 0 | 0 | 0 | 0 | 0 | 0 | 0 | 3855000 | 897200 | 0 | 0 | 0 | 0 | 0 | 0 | 0 | 0 |
| YVSGSSPD | PTPN14 | Q15678 | PTN14_HUMAN | Unmodified | 0 | 0 | 0 | 0 | 1302200 | 176390 | 0 | 0 | 0 | 0 | 0 | 0 | 0 | 0 | 0 | 0 | 0 | 237950 |
| YVSGYGLQ | PIGT; PIGT; PIG | Q969N2; A0 | PIGT_HUMAN ( | Unmodified | 0 | 0 | 0 | 0 | 0 | 0 | 0 | 0 | 3729600 | 275740 | 0 | 282280 | 0 | 0 | 0 | 0 | 0 | 0 |
| YVSHFETE( | C4A; C4B; C4B_ | P0C0L4; P0C | CO4A_HUMAN | Unmodified | 614200 | 0 | 6089100 | 10394000 | 0 | 0 | 0 | 0 | 0 | 0 | 0 | 0 | 0 | 4936600 | 27748000 | 15243000 | 0 | 0 |
| YVSHGATG | ASS1; ASS1 | P00966; Q5T | ASSY_HUMAN | Unmodified | 32067 | 0 | 0 | 0 | 0 | 0 | 2748600 | 92242 | 0 | 183870 | 0 | 0 | 349110 | 0 | 0 | 0 | 0 | 0 |
| YVSILYNGF | N/A | N/A | N/A | Oxidation (M) | 4895100 | 3772200 | 2991600 | 3711500 | 4242000 | 1488900 | 612000 | 903740 | 1998600 | 613290 | 0 | 2071200 | 1477200 | 1538000 | 3719000 | 2483200 | 1270200 | 0 |
| YVSLITSYQ | IFNGR1; IFNGR | P15260; A0A | INGR1_HUMAN | Unmodified | 0 | 0 | 13673 | 0 | 153890 | 0 | 278900 | 0 | 1212000 | 258140 | 134710 | 472200 | 0 | 0 | 26728 | 0 | 0 | 17825 |
| YVSQYYPK | HP1BP3; HP1BI | Q5SSJ5; B0C | HP1B3_HUMAI | Unmodified | 685550 | 0 | 0 | 0 | 0 | 0 | 1263800 | 1515600 | 0 | 41156 | 0 | 0 | 0 | 0 | 0 | 0 | 0 | 214510 |
| YVSSLTEEI | LMAN1 | P49257 | LMAN1_HUMAI | Unmodified | 0 | 0 | 731760 | 300770 | 192500 | 0 | 0 | 0 | 85104000 | 7458700 | 2189400 | 2098800 | 116580 | 0 | 609350 | 615490 | 0 | 0 |
| YVSSLTEEI | LMAN1 | P49257 | LMAN1_HUMAI | Unmodified | 0 | 0 | 0 | 0 | 0 | 0 | 0 | 0 | 34166000 | 2771000 | 2445400 | 0 | 0 | 0 | 0 | 0 | 0 | 0 |
| YVTFNSK | SLC38A1; SLC3 | Q9H2H9; F8 | S38A1_HUMAN | Unmodified | 842110 | 422550 | 637780 | 463370 | 80303 | 0 | 0 | 0 | 6317700 | 847950 | 659130 | 16499000 | 92264 | 0 | 795220 | 908270 | 32195 | 215220 |
| YVTGFDSV | AMPD3; AMPD3 | Q01432; E9F | AMPD3_HUMA | Unmodified | 0 | 0 | 0 | 0 | 0 | 0 | 0 | 0 | 0 | 0 | 0 | 5660700 | 0 | 0 | 0 | 0 | 0 | 0 |
| YVTVYNLIK | USP8; USP8; US | P40818; H0Y | UBP8_HUMAN | Unmodified | 0 | 0 | 0 | 0 | 4041300 | 0 | 250560 | 0 | 0 | 0 | 0 | 0 | 0 | 0 | 0 | 0 | 174540 | 0 |
| YVVGELIWI | GUSB | P08236 | BGLR_HUMAN | Unmodified | 0 | 0 | 0 | 0 | 0 | 0 | 0 | 0 | 0 | 1626600 | 0 | 0 | 32358 | 0 | 0 | 0 | 0 | 0 |
| YVVGLIIK | XPO1; XPO1; XF | O14980; A0A | XPO1_HUMAN | Unmodified | 62927 | 0 | 41320 | 45752 | 133570 | 19894 | 196350 | 0 | 223570 | 178820 | 281760 | 3068300 | 58637 | 21885 | 130210 | 115750 | 0 | 48089 |
| YVVISQGLI | LRP1 | Q07954 | LRP1_HUMAN I | Unmodified | 0 | 0 | 0 | 207910 | 0 | 0 | 0 | 0 | 2571700 | 106950 | 0 | 0 | 0 | 0 | 0 | 0 | 0 | 0 |
| YVVLTTK | FUCA1 | P04066 | FUCO_HUMAN | Unmodified | 42818 | 0 | 93010 | 74867 | 0 | 0 | 0 | 25615 | 1089100 | 154750 | 162440 | 182700 | 0 | 0 | 130940 | 85169 | 0 | 0 |
| YVVVKNDY | NIBAN1 | Q9BZQ8 | NIBA1_HUMAN | Unmodified | 0 | 0 | 0 | 0 | 1223300 | 663550 | 0 | 82759 | 969340 | 987400 | 0 | 0 | 0 | 0 | 0 | 0 | 0 | 259750 |
| YVVVTGITP | MTHFD1; MTHF | P11586; V9C | C1TC_HUMAN | Unmodified | 68292 | 0 | 291660 | 273960 | 562280 | 0 | 0 | 271430 | 3202900 | 227070 | 1287500 | 14013000 | 461450 | 0 | 772240 | 332280 | 179070 | 308390 |
| YVWLVYEQ | PEBP1 | P30086 | PEBP1_HUMAN | Unmodified | 4457500 | 2151200 | 3939200 | 3151700 | 706610 | 165750 | 0 | 0 | 5942300 | 323880 | 5569000 | 13628000 | 0 | 0 | 7449900 | 4700000 | 0 | 0 |
| YVWLVYEQ | PEBP1 | P30086 | PEBP1_HUMAN | Unmodified | 2401000 | 0 | 0 | 0 | 602850 | 0 | 0 | 0 | 1760200 | 0 | 4296500 | 1738400 | 0 | 0 | 1086300 | 633800 | 0 | 0 |
| YVYQPMEK | ARCN1; ARCN1 | P48444; Q6F | COPD_HUMAN | Oxidation (M) | 0 | 0 | 0 | 0 | 0 | 0 | 0 | 0 | 0 | 0 | 0 | 794800 | 0 | 0 | 0 | 0 | 0 | 0 |
| YWAAEKVT | SCRN1; SCRN1 | Q12765; C9I | SCRN1_HUMAI | Unmodified | 0 | 0 | 0 | 0 | 138200 | 0 | 0 | 0 | 0 | 0 | 0 | 0 | 0 | 0 | 0 | 0 | 138340 | 0 |
| YWAAHDKI | CNTN1 | Q12860 | CNTN1_HUMA | Unmodified | 360100 | 348740 | 0 | 0 | 0 | 0 | 0 | 0 | 512960 | 0 | 0 | 0 | 0 | 0 | 0 | 0 | 0 | 0 |
| YWEAFLPE | SNX1; SNX2; SN | Q13596; O6 | SNX1_HUMAN | Unmodified | 0 | 0 | 0 | 0 | 4961500 | 0 | 0 | 0 | 1074900 | 622790 | 255160 | 2352100 | 2590400 | 0 | 0 | 0 | 681530 | 3429700 |
| YWFSQPGI | LMF2 | Q9BU23 | LMF2_HUMAN | Unmodified | 0 | 0 | 0 | 0 | 0 | 0 | 0 | 0 | 1415100 | 0 | 0 | 0 | 0 | 0 | 0 | 0 | 0 | 0 |
| YWGEIISQ( | IDE; IDE; IDE; ID | P14735; A0A | IDE_HUMAN In | Unmodified | 0 | 0 | 0 | 2720100 | 3052000 | 0 | 0 | 0 | 0 | 0 | 342640 | 1043400 | 0 | 806460 | 0 | 0 | 605200 | 871290 |
| YWIQGDSE | ITGB4 | P16144 | ITB4_HUMAN I | Oxidation (M) | 0 | 0 | 0 | 0 | 0 | 0 | 0 | 0 | 0 | 0 | 0 | 0 | 0 | 0 | 0 | 0 | 4532500 | 6351100 |
| YWLAQQA | MRPL9; MRPL9 | Q9BYD2; Q5 | RM09_HUMAN | Unmodified | 184730 | 0 | 0 | 0 | 0 | 0 | 0 | 109490 | 960610 | 0 | 0 | 508660 | 0 | 0 | 0 | 0 | 0 | 0 |
| YWLCAATG | RACK1; RACK1; | P63244; D6F | RACK1_HUMAI | Unmodified | 578000 | 263790 | 179870 | 0 | 0 | 0 | 0 | 0 | 0 | 0 | 0 | 6652200 | 0 | 0 | 0 | 0 | 0 | 0 |
| YWLYYLHP | B4GALT1; B4G | P15291; Q8 | B4GT1_HUMAN | Unmodified | 0 | 0 | 0 | 0 | 0 | 0 | 0 | 0 | 2234200 | 338740 | 0 | 0 | 0 | 0 | 0 | 0 | 0 | 0 |
| YWNIGPQ | GLB1L3; GLB1L | Q8NCI6; Q8I | GLBL3_HUMAN | Unmodified | 0 | 0 | 0 | 0 | 0 | 0 | 80463 | 0 | 701640 | 0 | 0 | 0 | 0 | 0 | 0 | 0 | 0 | 0 |
| YWQMDIFT | N/A | N/A | N/A | Unmodified | 0 | 0 | 97345 | 0 | 7324400 | 413700 | 394230 | 76019 | 0 | 870960 | 0 | 4928600 | 3261900 | 1639700 | 0 | 0 | 2476000 | 0 |
| YWRDQAQ | ALPL | P05186 | PPBT_HUMAN | Unmodified | 323210 | 0 | 0 | 0 | 0 | 0 | 0 | 0 | 0 | 320160 | 0 | 0 | 0 | 0 | 0 | 0 | 0 | 0 |
| YWTLTATG | FSCN1; FSCN1 | Q16658; C9J | FSCN1_HUMAI | Unmodified | 0 | 0 | 0 | 0 | 5369700 | 0 | 442630 | 492340 | 1388800 | 798050 | 3884000 | 5357200 | 0 | 0 | 0 | 0 | 1117000 | 1964800 |
| YYADGEDA | NAA11; NAA10; | Q9BSU3; P4 | NAA11_HUMAI | Unmodified | 0 | 0 | 0 | 0 | 0 | 0 | 0 | 0 | 0 | 0 | 0 | 1877600 | 336810 | 0 | 0 | 25917 | 0 | 0 |
| YYAELCAPI | PGM2 | Q96G03 | PGM2_HUMAN | Unmodified | 0 | 0 | 0 | 411850 | 0 | 0 | 1129700 | 615690 | 0 | 5579000 | 0 | 540570 | 380010 | 203670 | 0 | 0 | 0 | 0 |
| YYALCGFG | SLC25A3; SLC2 | F8VWQ0; F8 | F8VWQ0_HUM | Unmodified | 0 | 0 | 0 | 0 | 0 | 0 | 0 | 0 | 1555800 | 5440700 | 0 | 0 | 0 | 0 | 0 | 0 | 0 | 0 |
| YYAVNFPM | HDAC2 | Q92769 | HDAC2_HUMA | Oxidation (M) | 0 | 0 | 0 | 0 | 0 | 0 | 833380 | 0 | 0 | 1041800 | 0 | 3518100 | 0 | 0 | 0 | 0 | 0 | 0 |
| YYCLNVPLI | HDAC3; HDAC3 | O15379; E7E | HDAC3_HUMA | Unmodified | 0 | 0 | 0 | 0 | 0 | 0 | 0 | 0 | 0 | 0 | 0 | 1091000 | 0 | 0 | 0 | 0 | 0 | 0 |
| YYDAIACLK | AIP; AIP; AIP | O00170; E9F | AIP_HUMAN AI | Unmodified | 0 | 0 | 0 | 0 | 0 | 0 | 0 | 0 | 0 | 0 | 0 | 0 | 974070 | 0 | 0 | 74262 | 779400 | 1020000 |
| YYDDTYPS | COPZ1; COPZ1 | P61923; F8V | COPZ1_HUMAI | Unmodified | 113850 | 147310 | 0 | 0 | 409140 | 0 | 0 | 0 | 495710 | 111470 | 0 | 3604600 | 214700 | 0 | 324330 | 0 | 0 | 0 |
| YYDEIINALI | PLXNB2 | O15031 | PLXB2_HUMAN | Unmodified | 1085000 | 947190 | 860100 | 3241500 | 6438300 | 675810 | 390130 | 59853 | 5620300 | 652010 | 1625900 | 3828900 | 1761900 | 620630 | 6124500 | 3956500 | 409740 | 0 |
| YYDQICSIE | PDCD6IP; PDCI | Q8WUM4; C | PDC6I_HUMAN | Unmodified | 9811100 | 13155000 | 3268700 | 1914700 | 0 | 0 | 0 | 0 | 0 | 0 | 0 | 4246400 | 0 | 0 | 6621200 | 5364600 | 0 | 0 |
| YYDSRPGG | RBM3 | P98179 | RBM3_HUMAN | Unmodified | 6161500 | 2705600 | 2416000 | 2841500 | 0 | 0 | 44932 | 0 | 123030 | 0 | 0 | 121010 | 393050 | 0 | 2051800 | 2366700 | 0 | 0 |
| YYDVMSDE | P4HA2 | O15460 | P4HA2_HUMAI | Unmodified | 0 | 0 | 0 | 0 | 0 | 0 | 0 | 0 | 2062500 | 0 | 0 | 0 | 0 | 0 | 0 | 0 | 0 | 0 |
| YYECDVLPI | WLS; WLS; WLS | Q5T9L3; H0Y | WLS_HUMAN F | Unmodified | 0 | 0 | 38412 | 0 | 0 | 0 | 0 | 0 | 856660 | 64830 | 0 | 0 | 0 | 0 | 0 | 0 | 0 | 0 |
| YYEKQEQE | EPHA3; EPHA3 | P29320; C9J | EPHA3_HUMAI | Unmodified | 218840 | 524560 | 0 | 0 | 0 | 0 | 0 | 0 | 45562 | 0 | 0 | 0 | 0 | 0 | 0 | 0 | 0 | 130330 |
| YYEVLGAA | BAG6; BAG6; B | P46379; F6S | BAG6_HUMAN | Unmodified | 0 | 0 | 0 | 0 | 0 | 0 | 0 | 0 | 0 | 0 | 0 | 4136700 | 0 | 0 | 0 | 0 | 0 | 0 |
| YYEVSADKI | CORO2A | Q92828 | COR2A_HUMAI | Unmodified | 263500 | 0 | 264360 | 0 | 0 | 0 | 0 | 0 | 0 | 0 | 0 | 1182500 | 0 | 0 | 718540 | 421770 | 0 | 0 |
| YYFEGIK | AHSA1; AHSA1; | O95433; H0Y | AHSA1_HUMAI | Unmodified | 0 | 0 | 0 | 0 | 93305 | 0 | 0 | 0 | 881290 | 103670 | 659770 | 7122600 | 137310 | 50685 | 508140 | 492600 | 0 | 0 |
| YYGAQTVR | FH | P07954 | FUMH_HUMAN | Unmodified | 109100 | 59622 | 0 | 0 | 759120 | 0 | 363380 | 28372 | 3953100 | 202450 | 364010 | 76419 | 0 | 0 | 97705 | 141830 | 181250 | 826000 |
| YYGAQTVR | FH | P07954 | FUMH_HUMAN | Unmodified | 0 | 0 | 0 | 0 | 0 | 0 | 760030 | 0 | 0 | 0 | 0 | 0 | 0 | 0 | 0 | 0 | 1235900 | 2277500 |
| YYGGAEVV | SHMT2; SHMT2 | P34897; G3V | GLYM_HUMAN | Unmodified | 0 | 0 | 0 | 0 | 0 | 0 | 0 | 0 | 5185700 | 0 | 0 | 0 | 0 | 0 | 0 | 0 | 0 | 0 |
| YYGGGYGS | C3 | P01024 | CO3_HUMAN ( | Oxidation (M) | 0 | 0 | 531930 | 1252100 | 0 | 0 | 0 | 0 | 0 | 0 | 0 | 0 | 0 | 0 | 0 | 0 | 0 | 0 |
| YYGGTEFID | SHMT1; SHMT1 | P34896; J3K | GLYC_HUMAN | Unmodified | 0 | 0 | 0 | 0 | 0 | 0 | 0 | 0 | 0 | 0 | 0 | 569230 | 0 | 0 | 0 | 84580 | 0 | 0 |
| YYGLQILEN | XPO1; XPO1; XF | O14980; A0A | XPO1_HUMAN | Unmodified | 0 | 0 | 0 | 0 | 0 | 0 | 141040 | 0 | 512360 | 0 | 178080 | 2031800 | 0 | 0 | 0 | 0 | 0 | 0 |
| YYGTKDPV | NAMPT; NAMPT | P43490; A0A | NAMPT_HUMAI | Unmodified | 0 | 0 | 0 | 144000 | 1553900 | 0 | 0 | 0 | 0 | 0 | 228840 | 971880 | 0 | 0 | 0 | 96460 | 0 | 0 |
| YYGTKDPV | NAMPT; NAMPT | P43490; A0A | NAMPT_HUMAI | Unmodified | 0 | 0 | 0 | 0 | 5397000 | 0 | 0 | 0 | 0 | 0 | 0 | 0 | 0 | 262340 | 0 | 0 | 0 | 0 |
| YYGTREAVI | ADAM10 | O14672 | ADA10_HUMAN | Unmodified | 0 | 0 | 0 | 390140 | 15412000 | 3632100 | 4086300 | 319490 | 0 | 1774500 | 0 | 95064 | 19681000 | 13524000 | 0 | 0 | 3490100 | 3581800 |
| YYHDFPVY | VANGL2 | Q9ULK5 | VANG2_HUMAI | Unmodified | 0 | 0 | 0 | 0 | 0 | 0 | 0 | 0 | 0 | 0 | 0 | 1339600 | 0 | 0 | 0 | 105510 | 0 | 0 |
| YYIAASYVK | GGH; GGH; GG | Q92820; A0A | GGH_HUMAN ( | Unmodified | 223720 | 0 | 109920 | 0 | 0 | 0 | 0 | 0 | 32875000 | 178720 | 0 | 0 | 0 | 0 | 0 | 215390 | 0 | 0 |
| YYINVCRPI | IGF2R; IGF2R | P11717; A0A | MPRI_HUMAN ( | Unmodified | 0 | 0 | 0 | 0 | 0 | 32710 | 0 | 0 | 1347600 | 0 | 0 | 0 | 0 | 0 | 0 | 0 | 62838 | 0 |
| YYITIIDAPG | EEF1A2; EEF1A1 | Q05639; A0A | EF1A2_HUMAN | Unmodified | 0 | 23431000 | 21535000 | 0 | 0 | 156170 | 742260 | 358060 | 650070 | 101220 | 0 | 0 | 0 | 0 | 52597000 | 44346000 | 65432 | 0 |
| YYITIIDAPG | EEF1A2; EEF1A1 | Q05639; A0A | EF1A2_HUMAN | Unmodified | 18405000 | 14649000 | 31295000 | 30491000 | 16338000 | 2373400 | 9856700 | 401840 | 505630 | 2339000 | 0 | 0 | 30470000 | 13715000 | 37427000 | 27580000 | 3811400 | 7469400 |
| YYKVEPLDI | HEXB; HEXB; HI | P07686; H0Y | HEXB_HUMAN | Unmodified | 0 | 303210 | 892420 | 1207300 | 806400 | 734630 | 0 | 0 | 6223300 | 3155700 | 604410 | 0 | 4386500 | 3363500 | 1095200 | 0 | 0 | 277650 |
| YYLAPKIED | PCNA | P12004 | PCNA_HUMAN | Unmodified | 371390 | 412060 | 327050 | 642840 | 5696000 | 1155900 | 4816900 | 4225600 | 0 | 4531800 | 1563600 | 8797500 | 13304000 | 4031000 | 910230 | 609770 | 5234200 | 0 |
| YYLDSLDRI | GNAO1; GNAO1 | P09471; H3E | GNAO_HUMAN | Unmodified | 0 | 0 | 0 | 0 | 0 | 0 | 0 | 0 | 4457800 | 0 | 0 | 0 | 0 | 0 | 0 | 0 | 0 | 0 |
| YYLNDLDR | GNAQ; GNAQ | P50148; B1A | GNAQ_HUMAN | Unmodified | 0 | 0 | 0 | 0 | 0 | 0 | 0 | 0 | 1573600 | 0 | 2798000 | 10787000 | 0 | 0 | 0 | 0 | 0 | 0 |
| YYLQDGDL | IGF2R | P11717 | MPRI_HUMAN ( | Unmodified | 0 | 29690 | 0 | 0 | 0 | 0 | 0 | 0 | 4377600 | 1291600 | 1073800 | 463150 | 0 | 0 | 0 | 0 | 0 | 0 |
| YYLTDVDRI | GNA11; GNA11 | P29992; K7E | GNA11_HUMAI | Unmodified | 0 | 0 | 0 | 0 | 0 | 616980 | 0 | 0 | 3543200 | 849690 | 367040 | 803210 | 0 | 0 | 0 | 0 | 0 | 1199200 |
| YYNSDVHF | SRM; SRM | P19623; K7E | SPEE_HUMAN S | Unmodified | 0 | 0 | 0 | 0 | 2731000 | 0 | 1176000 | 0 | 0 | 939210 | 0 | 0 | 5601500 | 3902600 | 0 | 0 | 5292300 | 3401000 |
| YYPASPWV | CACNA2D1; CA | P54289; E7E | CA2D1_HUMAI | Unmodified | 865470 | 462860 | 0 | 0 | 0 | 0 | 241840 | 137780 | 988610 | 0 | 500020 | 0 | 0 | 0 | 0 | 0 | 234290 | 0 |
| YYPASPWV | CACNA2D1; CA | P54289; E7E | CA2D1_HUMAI | Unmodified | 76340 | 0 | 0 | 0 | 0 | 0 | 0 | 0 | 0 | 0 | 0 | 0 | 0 | 0 | 0 | 0 | 222930 | 0 |
| YYPASPWV | CACNA2D1; CA | P54289; E7E | CA2D1_HUMAI | Unmodified | 593470 | 834950 | 0 | 0 | 0 | 0 | 0 | 0 | 18667000 | 200710 | 0 | 54942 | 0 | 0 | 0 | 0 | 0 | 0 |
| YYPATPWR | CACNA2D2; CA | Q9NY47; C9 | CA2D2_HUMAI | Unmodified | 0 | 0 | 0 | 79157 | 0 | 0 | 0 | 39907 | 5851900 | 530390 | 0 | 0 | 0 | 0 | 0 | 0 | 0 | 0 |
| YYPGGSQH | KIDINS220; KID | Q9ULH0; A0 | KDIS_HUMAN I | Unmodified | 0 | 0 | 445030 | 869220 | 2553200 | 0 | 0 | 0 | 0 | 0 | 0 | 1474300 | 0 | 0 | 531570 | 634940 | 0 | 0 |
| YYPIAVLLK | AVIL | O75366 | AVIL_HUMAN A | Unmodified | 0 | 0 | 0 | 0 | 0 | 0 | 116700 | 0 | 0 | 0 | 11925000 | 61943000 | 0 | 0 | 0 | 0 | 0 | 0 |
| YYPTEDVPI | RPL6; RPL6; RP | Q02878; U3I | RL6_HUMAN 60 | Unmodified | 1631900 | 1606000 | 414470 | 927340 | 0 | 0 | 58499 | 98629 | 0 | 0 | 0 | 0 | 0 | 0 | 0 | 395160 | 0 | 0 |
| YYPTEDVPI | RPL6; RPL6; RP | Q02878; U3I | RL6_HUMAN 60 | Unmodified | 3527300 | 2548100 | 0 | 1583000 | 256060 | 0 | 499900 | 0 | 0 | 189080 | 15227000 | 860180 | 0 | 0 | 1581600 | 0 | 0 | 0 |
| YYPYQSFQ | LGALS3BP | Q08380 | LG3BP_HUMAN | Unmodified | 0 | 0 | 0 | 513960 | 0 | 0 | 0 | 261360 | 0 | 225560 | 46678000 | 11516000 | 0 | 0 | 0 | 0 | 0 | 3724000 |
| YYPYQSFQ | LGALS3BP | Q08380 | LG3BP_HUMAN | Unmodified | 0 | 0 | 0 | 0 | 0 | 0 | 0 | 0 | 3187100 | 427690 | 2166000 | 3204700 | 0 | 0 | 0 | 0 | 0 | 0 |
| YYQLFLDSI | KIFBP; KIFBP; K | Q96EK5; A0 | KBP_HUMAN K | Unmodified | 0 | 0 | 0 | 0 | 0 | 0 | 0 | 0 | 0 | 0 | 0 | 1231400 | 0 | 0 | 0 | 0 | 0 | 0 |
| YYQLRCSG | DPP4; DPP4; DI | P27487; A0A | DPP4_HUMAN | Unmodified | 0 | 0 | 0 | 0 | 0 | 0 | 0 | 0 | 0 | 0 | 0 | 0 | 0 | 391980 | 0 | 0 | 4082300 | 0 |
| YYQQLKLE( | SLC29A1 | Q99808 | S29A1_HUMAN | Unmodified | 1617400 | 459530 | 0 | 0 | 0 | 0 | 0 | 0 | 0 | 0 | 0 | 4447200 | 0 | 0 | 340390 | 713150 | 0 | 0 |
| YYQTIGNH | PSMD13; PSMD | Q9UNM6; A0 | PSD13_HUMAN | Unmodified | 0 | 0 | 0 | 1159600 | 489870 | 263410 | 504160 | 197990 | 426090 | 99357 | 0 | 5284600 | 583380 | 1024700 | 0 | 0 | 337510 | 305270 |
| YYRESADPI | PLEC; PLEC | Q15149; H0Y | PLEC_HUMAN | Unmodified | 0 | 0 | 0 | 0 | 117510 | 123700 | 0 | 0 | 501480 | 160780 | 0 | 201240 | 0 | 0 | 0 | 0 | 287430 | 0 |
| YYREVLPGI | PPAT | Q06203 | PUR1_HUMAN | Unmodified | 0 | 0 | 0 | 0 | 0 | 0 | 0 | 11224 | 233590 | 0 | 170650 | 3265400 | 0 | 0 | 166870 | 115380 | 0 | 0 |

| | | | | | | | | | | | | | | | | | | | | | | |
|---|---|---|---|---|---|---|---|---|---|---|---|---|---|---|---|---|---|---|---|---|---|---|
| YYREVLPGI | PPAT | Q06203 | PUR1_HUMAN | Unmodified | 0 | 0 | 0 | 0 | 0 | 0 | 0 | 0 | 193180 | 0 | 219200 | 886940 | 0 | 0 | 0 | 0 | 0 | 0 |
| YYRFNEELI | MMP14; MMP14 | P50281; A0A | MMP14_HUMA | Unmodified | 0 | 0 | 0 | 0 | 7148400 | 11003000 | 0 | 0 | 0 | 0 | 0 | 0 | 0 | 0 | 0 | 38822 | 1200700 | 2582000 |
| YYRPTEVDI | GMDS | O60547 | GMDS_HUMAN | Unmodified | 0 | 0 | 0 | 0 | 713970 | 0 | 0 | 0 | 255890 | 0 | 0 | 660350 | 0 | 0 | 0 | 0 | 225840 | 509300 |
| YYRQNPDII | EHHADH | Q08426 | ECHP_HUMAN | Unmodified | 0 | 0 | 0 | 0 | 0 | 0 | 0 | 0 | 3119100 | 1016400 | 107850 | 799220 | 0 | 0 | 0 | 0 | 0 | 0 |
| YYRVCTLAI | RPL30; RPL30; I | P62888; A0A | RL30_HUMAN ( | Unmodified | 65394 | 0 | 0 | 504220 | 230940 | 0 | 1699400 | 19290 | 0 | 319670 | 0 | 0 | 0 | 186820 | 38127 | 0 | 109460 | 205250 |
| YYSFFDLDI | FERMT3; FERMT | Q86UX7; F5I | URP2_HUMAN | Unmodified | 0 | 0 | 0 | 517400 | 0 | 0 | 0 | 0 | 0 | 0 | 0 | 0 | 0 | 0 | 1384600 | 912420 | 0 | 0 |
| YYSFFDLDI | FERMT3; FERMT | Q86UX7; F5I | URP2_HUMAN | Unmodified | 196020 | 678490 | 316890 | 0 | 0 | 0 | 0 | 0 | 0 | 228040 | 0 | 0 | 0 | 0 | 753300 | 407230 | 0 | 0 |
| YYSFFDLNI | FERMT2; FERMT | Q96AC1; A0A | FERM2_HUMAN | Unmodified | 0 | 0 | 0 | 0 | 8556000 | 2037000 | 506540 | 0 | 0 | 0 | 0 | 0 | 0 | 365360 | 0 | 0 | 1648100 | 2780500 |
| YYSFNYEG | ENPP1 | P22413 | ENPP1_HUMAN | Unmodified | 0 | 0 | 0 | 0 | 0 | 0 | 0 | 0 | 0 | 0 | 0 | 0 | 0 | 0 | 0 | 0 | 0 | 0 |
| YYSSEPNN | EPB41L4B | Q9H329 | E41LB_HUMAN | Unmodified | 1417300 | 1458200 | 1302400 | 825080 | 0 | 0 | 0 | 0 | 0 | 0 | 0 | 0 | 0 | 0 | 5188000 | 3699900 | 0 | 0 |
| YYSSEYHY | CTSC; CTSC; CT | P53634; A0A | CATC_HUMAN | Oxidation (M) | 0 | 0 | 0 | 0 | 0 | 0 | 0 | 0 | 0 | 0 | 2799700 | 2622900 | 0 | 0 | 0 | 0 | 0 | 0 |
| YYSSEYHY | CTSC; CTSC; CT | P53634; A0A | CATC_HUMAN | Unmodified | 0 | 0 | 0 | 0 | 0 | 0 | 0 | 0 | 0 | 0 | 0 | 1882100 | 0 | 0 | 0 | 0 | 0 | 0 |
| YYTEFPTVL | HAT1 | O14929 | HAT1_HUMAN | Unmodified | 0 | 0 | 0 | 0 | 0 | 0 | 0 | 0 | 25323 | 0 | 0 | 530240 | 0 | 0 | 0 | 0 | 0 | 0 |
| YYTLFGR | EPRS1 | P07814 | SYEP_HUMAN | Unmodified | 166520 | 66043 | 0 | 272460 | 717790 | 0 | 121430 | 104100 | 1332300 | 663230 | 1613400 | 9787500 | 2984900 | 500100 | 854260 | 222290 | 372520 | 794540 |
| YYTLNGSK | ACADVL | P49748 | ACADV_HUMAN | Unmodified | 0 | 0 | 0 | 0 | 0 | 0 | 0 | 0 | 3975500 | 164350 | 84247 | 0 | 0 | 0 | 0 | 0 | 0 | 0 |
| YYTPTISR | PSMB2; PSMB2 | P49721; A0A | PSB2_HUMAN | Unmodified | 0 | 300190 | 1822000 | 1008600 | 0 | 0 | 0 | 0 | 0 | 0 | 1490000 | 2210100 | 0 | 0 | 782000 | 0 | 0 | 0 |
| YYTRLGND | RPS17; RPS17; | P08708; H0Y | RS17_HUMAN | Unmodified | 0 | 0 | 0 | 0 | 0 | 0 | 784260 | 351620 | 0 | 0 | 0 | 0 | 0 | 0 | 0 | 0 | 0 | 0 |
| YYTSASGD | HSP90AA1 | P07900 | HS90A_HUMAN | Oxidation (M) | 407080 | 410960 | 1277800 | 1570100 | 2578700 | 899350 | 316660 | 257050 | 3477700 | 1432700 | 5172300 | 27423000 | 793740 | 354450 | 1325300 | 1462500 | 585080 | 651010 |
| YYTSASGD | HSP90AA1 | P07900 | HS90A_HUMAN | Unmodified | 424780 | 896370 | 1155500 | 927730 | 5151700 | 1319100 | 502960 | 153740 | 17497000 | 3669100 | 6623300 | 30898000 | 1189200 | 582510 | 1143100 | 838460 | 1036100 | 1804800 |
| YYVNTTTNI | GAS7; GAS7; GA | O60861; A0A | GAS7_HUMAN | Unmodified | 0 | 0 | 0 | 0 | 0 | 0 | 0 | 0 | 0 | 0 | 1001000 | 2801000 | 0 | 0 | 0 | 0 | 0 | 0 |
| YYVTIIDAP( | EEF1A1P5; EEF1 | Q5VTE0; P68 | EF1A3_HUMAN | Unmodified | 2.51E+08 | 2.04E+08 | 1.02E+08 | 44707000 | 4724700 | 1235800 | 2871400 | 1464600 | 18118000 | 4378400 | 15070000 | 2.46E+08 | 2050300 | 851260 | 1.84E+08 | 1.5E+08 | 701880 | 2329700 |
| YYVTIIDAP( | EEF1A1P5; EEF1 | Q5VTE0; P68 | EF1A3_HUMAN | Unmodified | 2.9E+08 | 2.75E+08 | 1.36E+08 | 2.69E+08 | 2.12E+08 | 59388000 | 58648000 | 5061900 | 52527000 | 41303000 | 50005000 | 83053000 | 3.08E+08 | 1.04E+08 | 2E+08 | 1.09E+08 | 73974000 | 1.89E+08 |
| YYYDGKDY | AZGP1; AZGP1 | P25311; C9J | ZA2G_HUMAN | Unmodified | 0 | 0 | 0 | 0 | 0 | 0 | 0 | 0 | 0 | 0 | 1068000 | 0 | 0 | 0 | 25466 | 0 | 0 | 0 |
| YYYVQNVY | PEG10; PEG10; | Q86TG7; A0A | PEG10_HUMAN | Unmodified | 304850 | 247050 | 0 | 0 | 0 | 202470 | 0 | 0 | 255730 | 0 | 0 | 0 | 0 | 0 | 0 | 0 | 0 | 0 |

| Gene | 22RV1 #1 | 22RV1 #2 | C4-2 #1 | C4-2 #2 | DU145 #1 | DU145 #2 | EF1 #1 | EF1 #2 | NCI-H660 #1 | NCI-H660#2 | LASCPC-01 #1 | LASCPC-01 #2 | LNCaP 95 #1 | LNCaP 95 #2 | LNCaP #1 | LNCaP #2 | PC3 #1 | PC3 #2 |
|---|---|---|---|---|---|---|---|---|---|---|---|---|---|---|---|---|---|---|
| AAK1 | 16.76471 | 14.59233 | 17.40863 | 16.79212 | 18.00065 | 17.36609 | 16.36175 | 12.2946 | 14.04118 | 17.12007 | 14.67227 | 13.79651 | 18.87813 | 20.37992 | 17.83411 | 17.5999 | 17.35673 | 18.13672 |
| AAMDC | 16.74161 | 17.21935 | 16.31683 | 17.73042 | 16.15001 | 18.51921 | 15.98044 | 16.33516 | 14.3146 | 15.18544 | 15.65392 | 14.47851 | 17.57518 | 16.13262 | 17.57283 | 17.27866 | 17.60205 | 19.0392 |
| AAMP | 13.91158 | 15.88854 | 12.54818 | 15.37559 | 18.18886 | 13.13826 | 14.68497 | 16.72377 | 13.03286 | 15.80335 | 17.21472 | 16.61048 | 15.26092 | 13.30393 | 12.74038 | 14.62086 | 17.72341 | 17.12519 |
| AARS1 | 17.4565 | 17.43788 | 17.98986 | 17.70581 | 20.41726 | 19.81968 | 18.3626 | 17.69752 | 18.55086 | 19.21803 | 19.59834 | 20.09564 | 20.76942 | 20.81769 | 19.26656 | 18.89872 | 19.24885 | 19.31262 |
| AARSD1 | 15.45183 | 13.43352 | 16.12439 | 16.98271 | 13.69145 | 14.79106 | 15.02542 | 16.87268 | 15.31598 | 17.68869 | 18.9995 | 19.82319 | 14.84817 | 15.26935 | 14.84953 | 17.02056 | 15.44397 | 12.39389 |
| AASDHPPT | 15.8072 | 15.90526 | 14.97793 | 19.38766 | 16.16937 | 14.95434 | 15.04191 | 14.7619 | 13.96252 | 15.12409 | 17.28678 | 18.94537 | 16.9929 | 14.75826 | 16.72746 | 15.54193 | 15.56749 | 16.4147 |
| ABAT | 15.83915 | 16.08891 | 15.09935 | 14.85597 | 15.03426 | 15.33833 | 16.51643 | 17.77493 | 19.37433 | 18.00227 | 15.74624 | 13.21654 | 14.56909 | 15.12954 | 15.93579 | 15.56107 | 15.22632 | 14.98252 |
| ABCA12 | 15.682 | 16.31929 | 16.76198 | 14.93831 | 14.49917 | 16.75124 | 16.4529 | 16.05265 | 18.38485 | 16.74328 | 16.24554 | 14.28617 | 14.47538 | 15.22286 | 13.59988 | 14.7977 | 15.70341 | 14.46478 |
| ABCA2 | 18.76532 | 14.15292 | 14.78387 | 14.16816 | 15.42402 | 17.30991 | 16.12901 | 16.24436 | 19.27442 | 16.84331 | 14.91659 | 14.00038 | 13.72382 | 15.39993 | 14.57736 | 16.07044 | 15.43568 | 14.38822 |
| ABCA3 | 16.34577 | 17.11573 | 15.71213 | 17.13835 | 13.70176 | 17.39025 | 16.20451 | 16.30143 | 19.15422 | 14.38123 | 16.03474 | 13.20445 | 15.98483 | 13.66594 | 13.8816 | 14.20609 | 16.16718 | 13.53025 |
| ABCB6 | 19.18609 | 16.01088 | 19.26848 | 19.91506 | 17.65225 | 18.66005 | 18.62817 | 17.51792 | 18.22808 | 16.59052 | 18.43307 | 18.33256 | 18.48447 | 19.40373 | 20.56695 | 19.54526 | 16.26288 | 17.58277 |
| ABCC1 | 18.0954 | 15.84949 | 18.80662 | 16.97003 | 18.26957 | 16.74027 | 15.84725 | 16.64335 | 14.43919 | 15.89206 | 18.41014 | 19.74034 | 16.47082 | 15.84105 | 18.49453 | 19.34993 | 16.64653 | 17.01036 |
| ABCC3 | 15.76973 | 14.32489 | 16.77129 | 16.78625 | 19.91111 | 17.82774 | 15.71905 | 15.87601 | 13.30982 | 15.80163 | 15.05358 | 16.84981 | 16.86885 | 15.03537 | 15.9645 | 15.28023 | 17.37395 | 16.36917 |
| ABCC4 | 16.42816 | 16.9136 | 21.42092 | 19.99458 | 19.49828 | 16.37778 | 18.00296 | 16.68367 | 15.25214 | 15.59136 | 15.83488 | 16.85552 | 16.80055 | 16.87618 | 20.767 | 20.58508 | 15.91213 | 15.49589 |
| ABCC5 | 15.68653 | 16.25057 | 15.19687 | 15.79118 | 14.63112 | 16.69576 | 15.1385 | 16.11069 | 13.59729 | 14.48266 | 17.68752 | 19.42559 | 14.37318 | 15.58095 | 15.48384 | 15.1174 | 15.77854 | 15.43319 |
| ABCD3 | 16.24115 | 14.80776 | 16.34032 | 15.1833 | 11.39075 | 15.87615 | 15.48219 | 15.27583 | 19.27851 | 11.39748 | 14.57278 | 10.88242 | 14.92233 | 18.11651 | 16.88639 | 14.54995 | 15.59598 | 16.47128 |
| ABCE1 | 18.96808 | 18.8786 | 18.61589 | 18.59706 | 19.08213 | 17.69959 | 17.08853 | 17.15098 | 15.89646 | 16.33181 | 18.91888 | 19.98336 | 18.3742 | 17.21296 | 18.58061 | 18.43369 | 18.35251 | 17.39571 |
| ABCF1 | 16.58024 | 16.98104 | 19.26297 | 18.7759 | 18.45522 | 20.82889 | 17.49948 | 17.44305 | 15.99858 | 16.29437 | 18.16454 | 19.48842 | 16.23664 | 18.05796 | 17.63164 | 16.86709 | 17.78542 | 17.277 |
| ABCF2 | 16.76735 | 16.08154 | 15.96766 | 17.20337 | 18.05785 | 15.55017 | 16.22233 | 16.30798 | 12.62272 | 15.04597 | 14.33873 | 17.27491 | 13.09382 | 17.1365 | 17.54046 | 17.95941 | 17.78802 | 16.93594 |
| ABHD10 | 16.47565 | 15.29444 | 14.20949 | 16.3493 | 12.11857 | 13.46165 | 15.30879 | 15.42253 | 20.15522 | 20.31484 | 16.01584 | 14.052 | 17.29627 | 17.84978 | 15.30062 | 14.47402 | 16.43621 | 16.1408 |
| ABHD11 | 17.9325 | 18.95126 | 15.14162 | 18.41039 | 17.63114 | 14.96916 | 16.13265 | 17.07583 | 19.07828 | 16.27748 | 16.75862 | 15.64255 | 15.13269 | 15.29578 | 18.1415 | 16.95521 | 16.71661 | 16.73207 |
| ABHD12 | 16.34894 | 19.32092 | 16.5254 | 15.99106 | 16.18018 | 16.1971 | 16.63204 | 16.89821 | 22.52186 | 20.69142 | 16.61118 | 16.07702 | 16.27113 | 16.59817 | 17.32037 | 17.31931 | 17.33197 | 15.88405 |
| ABHD14B | 17.64701 | 17.50253 | 17.99076 | 16.67583 | 19.63921 | 18.57483 | 16.5772 | 16.14725 | 16.74424 | 17.09066 | 18.81027 | 18.93227 | 18.50378 | 16.73697 | 17.67127 | 18.01507 | 19.1963 | 18.97642 |
| ABHD17B | 14.65083 | 17.42138 | 16.79848 | 15.98457 | 16.87002 | 16.50462 | 18.3649 | 16.96993 | 16.09944 | 15.77805 | 18.14499 | 13.09519 | 13.95127 | 14.35526 | 20.40151 | 13.35544 | 16.36868 | 15.09241 |
| ABI1 | 19.13782 | 18.13724 | 19.68718 | 20.06142 | 20.41661 | 19.51032 | 17.1426 | 17.37348 | 18.63503 | 18.07541 | 18.88961 | 19.76291 | 20.52866 | 19.63778 | 20.30697 | 20.44926 | 20.10126 | 21.04324 |
| ABI2 | 18.22824 | 16.80095 | 16.56031 | 16.71484 | 17.94762 | 17.25127 | 16.75928 | 15.17754 | 15.74658 | 15.47167 | 18.70433 | 18.96423 | 16.09628 | 16.42158 | 16.01221 | 16.09835 | 17.86103 | 19.27749 |
| ABI3BP | 14.72709 | 14.88849 | 13.81395 | 16.02743 | 14.0078 | 17.17341 | 16.39833 | 18.31004 | 18.02624 | 17.24957 | 15.97323 | 12.98409 | 15.89707 | 15.30329 | 14.86562 | 13.17295 | 13.54668 | 16.73535 |
| ABR | 15.31721 | 19.59338 | 17.68244 | 15.0773 | 18.1925 | 15.00412 | 16.47942 | 15.58634 | 13.42328 | 14.84484 | 14.12118 | 14.78252 | 15.88176 | 16.85553 | 17.26404 | 16.80556 | 18.70947 | 18.15355 |
| ABRACL | 21.33416 | 21.41626 | 16.48062 | 19.99073 | 19.60427 | 19.49077 | 19.20541 | 17.48858 | 19.48286 | 20.95292 | 15.16228 | 13.62522 | 19.51263 | 19.25493 | 19.19444 | 15.8887 | 19.99182 | 19.16176 |
| ABRAXAS2 | 15.76742 | 15.52703 | 15.54556 | 15.64086 | 16.81085 | 15.09602 | 16.12728 | 15.74312 | 14.67379 | 15.1235 | 15.62569 | 17.26562 | 15.28834 | 15.11098 | 14.47479 | 14.44872 | 16.70965 | 16.57072 |
| ACAA1 | 17.16566 | 18.90622 | 18.89301 | 22.40239 | 17.36194 | 19.92295 | 17.79377 | 16.4981 | 21.81343 | 20.91251 | 17.6902 | 14.66075 | 15.8514 | 15.83819 | 15.87667 | 16.78331 | 17.38248 | 17.51129 |
| ACAA2 | 18.404 | 18.34547 | 15.17597 | 15.57389 | 15.59037 | 16.40197 | 17.0044 | 16.80377 | 20.19067 | 18.90268 | 14.88128 | 14.3637 | 15.15716 | 15.24893 | 14.7513 | 15.68749 | 16.43654 | 15.17739 |
| ACACA | 14.83336 | 21.30606 | 21.40678 | 20.59166 | 15.27548 | 15.96016 | 16.76903 | 16.35284 | 16.38795 | 14.51865 | 16.20682 | 17.75271 | 15.96288 | 16.7818 | 21.8294 | 21.30258 | 15.10322 | 15.40693 |
| ACACB | 13.24633 | 14.44679 | 18.34062 | 17.54817 | 15.66612 | 15.86653 | 16.56896 | 16.45109 | 11.80108 | 14.4733 | 14.02604 | 18.96203 | 13.49185 | 15.1549 | 18.22182 | 18.38765 | 15.08432 | 14.82553 |
| ACAD10 | 15.99862 | 15.29437 | 13.8363 | 14.21627 | 14.89832 | 14.70029 | 15.84737 | 16.54333 | 16.92287 | 14.25351 | 14.25902 | 14.08118 | 13.22577 | 14.69069 | 16.01378 | 14.75389 | 15.26691 | 14.63694 |
| ACAD11 | 14.29787 | 17.95118 | 13.73106 | 14.34705 | 16.34601 | 13.72935 | 15.19138 | 18.54281 | 18.94571 | 17.40809 | 15.96227 | 15.51858 | 15.32185 | 14.64419 | 15.83503 | 15.07624 | 13.74047 | 14.0698 |
| ACAD9 | 15.28314 | 15.57672 | 15.26492 | 17.25627 | 17.06213 | 16.01475 | 17.07481 | 16.37182 | 19.32742 | 17.20245 | 14.99558 | 16.74983 | 16.60811 | 17.47608 | 16.21042 | 15.411 | 16.83052 | 16.55542 |
| ACADM | 15.45386 | 13.47837 | 15.57841 | 16.22432 | 15.0049 | 15.36982 | 15.56003 | 17.15178 | 19.37325 | 17.21031 | 16.50628 | 12.77281 | 13.96476 | 16.16765 | 16.04459 | 14.67178 | 16.24358 | 12.98075 |
| ACADVL | 16.87868 | 17.64291 | 16.73107 | 15.28234 | 16.10061 | 17.24067 | 16.78883 | 16.28644 | 20.58232 | 19.87828 | 16.74136 | 15.2943 | 15.0414 | 16.05958 | 16.11602 | 15.64664 | 18.56274 | 15.96612 |
| ACAT1 | 16.46778 | 15.88552 | 16.13646 | 15.62955 | 16.08072 | 16.79684 | 17.42805 | 17.07653 | 19.67832 | 18.4634 | 15.83106 | 14.18839 | 19.18484 | 17.46039 | 16.3193 | 15.86971 | 16.71342 | 16.61759 |
| ACAT2 | 18.89092 | 18.06204 | 17.70456 | 17.67134 | 20.88215 | 19.03818 | 18.58834 | 18.15064 | 19.02983 | 19.80146 | 19.29946 | 20.5558 | 20.2642 | 16.32157 | 17.0593 | 16.94409 | 18.09712 | 17.90103 |
| ACBD3 | 15.6864 | 16.00646 | 15.29691 | 16.17151 | 15.15446 | 16.22055 | 17.35186 | 16.27768 | 15.0994 | 14.77311 | 17.79372 | 18.3141 | 18.36406 | 16.24563 | 14.33166 | 15.62864 | 13.58467 | 16.06758 |
| ACBD7 | 17.4294 | 13.92508 | 15.77295 | 14.17615 | 13.74059 | 15.55479 | 13.06435 | 18.7321 | 13.53847 | 13.81105 | 14.01481 | 16.02274 | 13.58677 | 15.99407 | 14.31163 | 13.90488 | 15.23595 | 13.87134 |
| ACE | 20.59991 | 19.35372 | 14.92154 | 15.39699 | 14.3837 | 15.78639 | 15.15636 | 15.2355 | 18.3455 | 16.57779 | 15.99368 | 14.10326 | 14.99246 | 15.23617 | 16.47508 | 15.77179 | 15.5549 | 17.604 |
| ACIN1 | 15.68384 | 14.64439 | 16.82166 | 14.8121 | 13.26402 | 14.33734 | 16.34993 | 17.3033 | 12.56845 | 17.56042 | 15.09538 | 17.27965 | 12.33689 | 16.18614 | 14.01446 | 15.95236 | 14.55895 | 14.00828 |
| ACLY | 17.29139 | 17.27315 | 18.02934 | 18.40994 | 19.579 | 17.7689 | 21.02926 | 19.39404 | 17.7484 | 17.52359 | 18.62494 | 21.02956 | 18.49421 | 17.07921 | 18.38718 | 18.15805 | 17.77828 | 17.33535 |
| ACO1 | 17.19828 | 16.98924 | 17.13562 | 17.36527 | 18.20343 | 17.32659 | 18.78394 | 17.35697 | 15.74725 | 16.28939 | 18.37164 | 19.65292 | 17.60481 | 17.49547 | 17.79104 | 17.57336 | 17.7214 | 17.61037 |
| ACO2 | 15.58271 | 16.69689 | 15.89248 | 15.80482 | 16.29972 | 16.91866 | 18.79778 | 16.80825 | 20.34146 | 18.71809 | 16.82563 | 14.94266 | 16.23249 | 16.20197 | 16.44509 | 16.64007 | 16.31781 | 15.59893 |
| ACOT13 | 16.76348 | 16.29176 | 16.2351 | 17.14514 | 19.11846 | 17.28333 | 17.21068 | 17.60935 | 18.954 | 18.8082 | 16.89982 | 12.88441 | 23.32873 | 22.75919 | 17.56314 | 15.69305 | 18.23191 | 16.77844 |
| ACOT2 | 15.82487 | 18.6584 | 14.71295 | 15.07028 | 14.00043 | 18.02798 | 14.95356 | 16.50532 | 18.54723 | 16.55894 | 15.35331 | 14.23136 | 13.18736 | 15.56039 | 16.91816 | 18.00718 | 15.96487 | 14.90555 |
| ACOT7 | 17.04837 | 16.55612 | 17.8736 | 17.71954 | 20.44726 | 18.7598 | 20.3683 | 19.6337 | 20.25204 | 20.95423 | 19.8624 | 20.78267 | 19.77693 | 19.70193 | 17.38434 | 17.26324 | 21.69165 | 21.70762 |
| ACOT8 | 17.08916 | 17.2499 | 16.52836 | 16.98677 | 14.56447 | 17.7335 | 15.58363 | 16.74598 | 21.675 | 18.1912 | 16.312 | 15.59232 | 15.45282 | 15.14635 | 16.39491 | 16.47048 | 15.26029 | 16.84147 |
| ACOX1 | 15.73159 | 16.62153 | 15.76278 | 15.75125 | 15.64366 | 15.47316 | 17.1432 | 16.73056 | 20.42857 | 19.10087 | 19.78724 | 14.77137 | 16.39487 | 15.72853 | 15.91551 | 15.12192 | 16.28083 | 16.18897 |
| ACOX3 | 14.8333 | 15.1912 | 15.36187 | 14.50851 | 17.49185 | 16.29278 | 17.31153 | 18.29408 | 18.26499 | 15.84688 | 14.75132 | 13.07827 | 14.58046 | 14.62482 | 13.81669 | 15.50415 | 14.59331 | 13.38762 |
| ACP1 | 20.31222 | 19.77264 | 21.54387 | 19.2788 | 21.05175 | 18.64824 | 19.13739 | 19.91929 | 17.98359 | 17.90023 | 19.54983 | 20.38255 | 21.04067 | 20.61446 | 19.33633 | 19.60136 | 21.24466 | 20.84578 |
| ACP2 | 14.96285 | 17.4159 | 15.84977 | 15.21339 | 17.04268 | 16.36913 | 16.19047 | 15.76624 | 20.58473 | 19.29537 | 18.04342 | 16.16951 | 15.99673 | 16.15781 | 17.20436 | 14.29526 | 15.15694 | 15.90473 |
| ACP3 | 18.34802 | 18.278 | 18.09864 | 16.58325 | 15.43597 | 14.82102 | 16.39176 | 16.03898 | 20.70061 | 20.42981 | 15.73278 | 14.51765 | 15.61874 | 16.36658 | 17.49719 | 18.90419 | 16.57151 | 15.17334 |
| ACSF2 | 14.62371 | 15.41429 | 15.28854 | 15.53761 | 14.5907 | 14.0636 | 19.0312 | 16.01655 | 18.66259 | 15.27309 | 13.43445 | 12.5601 | 13.34223 | 13.86547 | 14.5908 | 14.86266 | 14.63225 | 13.01249 |
| ACSL3 | 17.53959 | 16.04608 | 17.98549 | 17.3249 | 19.11321 | 15.08159 | 19.82146 | 17.42188 | 14.32686 | 16.97087 | 16.93667 | 18.51908 | 20.6259 | 19.2738 | 13.89028 | 18.37506 | 17.16972 | 19.06019 |
| ACSL4 | 16.00998 | 22.29778 | 18.69496 | 19.49142 | 20.22666 | 20.60114 | 19.54596 | 19.41662 | 19.62147 | 18.92162 | 20.02358 | 20.33768 | 21.45413 | 20.92465 | 18.50591 | 18.12923 | 22.07359 | 21.96625 |
| ACSS1 | 15.69965 | 14.65733 | 13.5134 | 14.6077 | 11.3965 | 15.036 | 19.00564 | 15.40855 | 18.29848 | 18.37975 | 13.71441 | 12.28406 | 15.76463 | 15.19474 | 14.26109 | 15.03949 | 15.91079 | 17.26464 |
| ACSS2 | 21.0346 | 17.5121 | 17.37025 | 22.29665 | 20.25143 | 21.29578 | 20.4782 | 22.0464 | 12.83601 | 15.61421 | 16.27636 | 11.82716 | 20.13179 | 19.30431 | 17.7783 | 21.38502 | 22.01374 | 12.08874 |
| ACTBL2 | 28.47778 | 27.76335 | 28.90124 | 28.91061 | 27.70382 | 27.48138 | 21.90637 | 21.42699 | 10.88144 | 19.63661 | 23.45033 | 24.39439 | 22.93511 | 22.47163 | 29.13978 | 24.30163 | 21.41874 | 26.44643 |
| ACTC1 | 26.81431 | 26.77829 | 26.92396 | 27.42389 | 30.09953 | 29.05991 | 27.57044 | 23.96794 | 25.71909 | 26.30878 | 26.70028 | 26.64411 | 28.75123 | 28.45449 | 27.03852 | 26.93028 | 28.10244 | 28.4121 |
| ACTG1 | 26.1527 | 25.96891 | 26.38401 | 26.49669 | 27.70406 | 26.86913 | 25.86065 | 25.9469 | 24.95042 | 24.15128 | 25.73487 | 26.59071 | 26.7029 | 26.37008 | 26.87487 | 26.6112 | 26.5195 | 26.25958 |
| ACTL6A | 18.5093 | 15.9868 | 17.75348 | 17.2285 | 16.95728 | 16.40337 | 15.83735 | 16.72588 | 18.83848 | 18.54665 | 19.71967 | 20.08839 | 16.41749 | 15.59538 | 16.17176 | 17.98369 | 17.10161 | 14.60739 |
| ACTL8 | 14.63599 | 15.49912 | 15.28624 | 14.99188 | 17.82979 | 15.88777 | 19.35196 | 15.92729 | 15.18139 | 15.49787 | 19.77656 | 19.47711 | 16.3911 | 20.37946 | 15.81316 | 14.90985 | 14.63088 | 20.493 |
| ACTN1 | 18.80553 | 18.01502 | 17.92147 | 19.48524 | 21.7562 | 21.54518 | 19.85887 | 18.71747 | 17.57791 | 17.46499 | 19.78741 | 21.12662 | 19.72443 | 20.02275 | 19.42838 | 19.45538 | 21.49757 | 21.20714 |
| ACTN2 | 18.6261 | 18.09662 | 17.52453 | 19.13936 | 20.99453 | 19.24879 | 18.34372 | 17.34777 | 18.29747 | 18.51335 | 18.89385 | 21.62135 | 19.84503 | 20.57258 | 20.32433 | 19.93434 | 21.20771 | 19.80324 |
| ACTN4 | 19.32458 | 18.9966 | 20.16484 | 20.05165 | 22.70859 | 22.46632 | 19.62245 | 19.03045 | 19.98416 | 20.52187 | 20.59983 | 21.90776 | 21.01397 | 21.19846 | 20.71206 | 20.74849 | 21.91782 | 20.16337 |
| ACTR10 | 16.9396 | 15.98517 | 15.78066 | 14.38563 | 12.26398 | 14.03296 | 15.73186 | 16.64878 | 12.21425 | 15.11706 | 15.53239 | 18.60542 | 18.33373 | 17.73698 | 12.38017 | 16.12227 | 15.00944 | 13.67601 |
| ACTR1A | 16.74776 | 17.38413 | 17.63227 | 17.82258 | 19.2255 | 18.4255 | 19.5567 | 19.89201 | 17.1144 | 18.33504 | 17.45001 | 20.95025 | 17.71536 | 18.08981 | 17.44344 | 17.96521 | 19.29265 | 19.7353 |
| ACTR2 | 19.65293 | 19.58018 | 18.29281 | 18.83767 | 20.53628 | 20.99955 | 19.97444 | 19.50132 | 17.39359 | 17.83965 | 18.73157 | 20.7775 | 20.85541 | 20.40036 | 19.76927 | 20.17137 | 20.65003 | 20.12127 |
| ACTR3 | 20.0213 | 20.47704 | 18.68058 | 20.51762 | 20.34761 | 19.65351 | 19.54801 | 18.88274 | 18.36106 | 18.62589 | 19.11051 | 21.25336 | 20.32232 | 20.71603 | 20.48804 | 20.08152 | 20.26084 | 19.83067 |
| ACTR3B | 21.14798 | 20.78656 | 20.44464 | 19.69543 | 18.44287 | 17.01468 | 16.99316 | 16.84971 | 18.53171 | 17.06748 | 19.06484 | 22.03671 | 19.05906 | 17.80488 | 21.02779 | 21.63192 | 16.71247 | 16.89852 |
| ACTRT1 | 14.75705 | 15.01062 | 15.29074 | 15.1573 | 20.10462 | 25.33988 | 23.33369 | 20.55458 | 12.68728 | 13.80746 | 13.52375 | 12.75337 | 21.45962 | 24.53483 | 13.97648 | 13.64054 | 20.63161 | 25.57327 |
| ACVR1 | 15.09499 | 15.19378 | 16.16114 | 16.70505 | 20.34526 | 17.64648 | 15.89884 | 15.40066 | 13.81162 | 13.61269 | 15.9724 | 16.03884 | 18.7451 | 18.14311 | 16.89974 | 16.95894 | 20.50519 | 21.60942 |
| ACY1 | 15.43224 | 15.9292 | 16.91015 | 17.80781 | 16.79503 | 17.55464 | 16.49964 | 16.20566 | 17.18667 | 17.88919 | 19.25663 | 19.68472 | 16.58868 | 16.30535 | 17.50374 | 16.98306 | 16.67395 | 17.0276 |
| ADA | 15.68924 | 17.16929 | 18.27467 | 18.94169 | 13.35942 | 13.86802 | 13.72088 | 16.37801 | 12.83747 | 14.08658 | 18.03999 | 18.47041 | 14.90636 | 15.9668 | 17.5846 | 17.34683 | 15.52699 | 15.63737 |
| ADAM10 | 21.26159 | 20.69215 | 22.24443 | 22.01613 | 20.60883 | 19.73357 | 19.4909 | 17.43753 | 18.23598 | 17.90166 | 17.73467 | 20.07436 | 21.52515 | 21.29893 | 22.58064 | 22.36512 | 19.23957 | 19.89664 |
| ADAM15 | 20.61298 | 19.48242 | 18.50209 | 18.63109 | 15.14988 | 15.96979 | 15.79121 | 15.9953 | 18.20244 | 15.22137 | 16.31978 | 17.96272 | 15.36116 | 16.59972 | 19.18109 | 19.37607 | 15.56454 | 15.92466 |
| ADAM17 | 22.82331 | 22.82477 | 22.80452 | 17.2767 | 19.5628 | 18.01404 | 15.98675 | 17.70715 | 18.19876 | 17.56727 | 17.0056 | 15.10644 | 18.46237 | 18.81484 | 21.83511 | 22.34839 | 19.79134 | 20.57855 |
| ADAM9 | 20.34673 | 20.32235 | 20.72314 | 21.00734 | 21.55787 | 20.21895 | 16.5642 | 17.21651 | 17.46267 | 17.0792 | 17.44998 | 15.90331 | 20.9536 | 19.76689 | 20.99193 | 21.42674 | 18.62151 | 18.62619 |
| ADAMTS1 | 14.9182 | 14.27622 | 19.58677 | 16.85283 | 14.09959 | 16.24882 | 15.12222 | 16.98367 | 13.5377 | 15.70292 | 15.29179 | 11.75923 | 16.73543 | 16.11503 | 16.19961 | 16.84809 | 18.32318 | 18.3683 |

| | | | | | | | | | | | | | | | | | | |
|---|---|---|---|---|---|---|---|---|---|---|---|---|---|---|---|---|---|---|
| ADAMTS13 | 15.86008 | 17.34672 | 22.09143 | 19.29673 | 13.42517 | 20.76433 | 15.00061 | 15.59851 | 13.99552 | 16.4263 | 18.94956 | 11.86083 | 13.39608 | 16.65389 | 19.25221 | 17.51667 | 15.59963 | 13.96724 |
| ADAMTS4 | 16.5764 | 16.0618 | 14.44478 | 14.92992 | 12.02675 | 12.97926 | 15.96622 | 13.07149 | 13.93231 | 14.66784 | 15.46148 | 16.59996 | 20.25146 | 13.35311 | 14.07613 | 15.58745 | 15.2448 | 15.18222 |
| ADAR | 17.17869 | 17.85387 | 17.89655 | 15.1922 | 12.90415 | 17.45257 | 16.816 | 17.41308 | 15.98569 | 15.1778 | 16.10052 | 18.64831 | 16.66763 | 15.57027 | 17.00127 | 16.88351 | 16.0594 | 18.31521 |
| ADCY6 | 16.91962 | 18.45898 | 16.85399 | 18.66111 | 16.89226 | 17.20846 | 15.09017 | 16.42524 | 14.79787 | 16.18004 | 15.30264 | 15.057 | 12.924 | 13.78408 | 16.87863 | 17.93343 | 15.64106 | 15.83948 |
| ADD1 | 16.60918 | 14.84819 | 17.19592 | 18.81299 | 20.69044 | 18.65912 | 17.06144 | 17.04495 | 16.58327 | 17.13228 | 15.27948 | 13.94467 | 17.02077 | 17.65028 | 19.32627 | 18.79373 | 18.36067 | 20.09859 |
| ADD2 | 15.58998 | 14.52202 | 14.08553 | 15.50516 | 19.7986 | 18.84254 | 16.24731 | 17.5075 | 13.27644 | 15.01859 | 14.57189 | 12.41436 | 14.36346 | 14.13752 | 14.60406 | 15.11544 | 14.82739 | 14.70835 |
| ADD3 | 15.42961 | 15.40089 | 18.17294 | 19.00355 | 15.90068 | 17.73615 | 15.52586 | 18.46984 | 14.58367 | 16.54909 | 16.79689 | 13.99074 | 19.32724 | 19.9714 | 18.50263 | 18.5477 | 17.90123 | 17.37794 |
| ADGRE2 | 14.84593 | 15.36344 | 14.80578 | 13.81292 | 16.82359 | 15.28287 | 17.19081 | 15.95797 | 12.8437 | 13.28605 | 20.44894 | 19.29607 | 15.54443 | 16.02363 | 16.08479 | 14.71868 | 14.17938 | 15.10308 |
| ADGRE5 | 19.18736 | 18.48065 | 20.11248 | 19.66523 | 22.56755 | 21.63025 | 16.67372 | 16.93199 | 16.37943 | 15.2902 | 17.4782 | 16.53189 | 18.73401 | 16.56745 | 19.25814 | 19.33621 | 20.3882 | 22.99623 |
| ADGRG1 | 16.91055 | 15.57325 | 18.2641 | 18.68078 | 15.18504 | 16.35172 | 16.73292 | 16.75901 | 19.73554 | 17.80488 | 18.38027 | 14.36894 | 16.06388 | 16.07489 | 18.0954 | 18.40746 | 17.08196 | 17.95821 |
| ADGRG2 | 20.14949 | 19.76291 | 19.87163 | 19.26356 | 20.35912 | 19.59122 | 16.99876 | 15.93788 | 20.39369 | 20.92834 | 15.64299 | 17.45117 | 16.03015 | 18.17829 | 19.73763 | 19.92705 | 17.01268 | 15.39201 |
| ADGRG6 | 17.29413 | 16.5379 | 19.00431 | 20.10278 | 21.47067 | 20.64059 | 16.05222 | 16.50792 | 14.38384 | 15.01209 | 16.90464 | 15.19559 | 17.43844 | 16.74908 | 19.67584 | 20.43284 | 15.20403 | 17.30263 |
| ADH1B | 16.77292 | 18.0204 | 17.31898 | 16.37501 | 16.07845 | 17.12999 | 16.45114 | 15.18327 | 17.4439 | 15.50468 | 21.205 | 18.22484 | 15.95653 | 15.83405 | 16.0832 | 16.84244 | 16.30728 | 15.60893 |
| ADH1C | 16.13013 | 19.40411 | 13.78446 | 13.19225 | 14.5249 | 15.20398 | 15.8943 | 14.69246 | 12.68933 | 15.83003 | 21.59084 | 17.76891 | 15.45058 | 14.10518 | 16.4318 | 16.74729 | 15.45739 | 16.20086 |
| ADH5 | 17.78572 | 16.70655 | 17.74054 | 17.38182 | 20.30168 | 18.82659 | 18.66236 | 18.17833 | 16.87037 | 17.01029 | 21.02779 | 21.57787 | 18.94994 | 19.03984 | 17.76082 | 16.93728 | 19.76042 | 20.12684 |
| ADI1 | 15.30058 | 17.68444 | 17.97085 | 17.80269 | 19.45753 | 18.74814 | 17.35277 | 15.519 | 16.13341 | 17.52902 | 19.38096 | 19.46498 | 19.51801 | 18.67484 | 17.74752 | 17.47872 | 19.34673 | 18.31637 |
| ADIRF | 20.22855 | 19.63493 | 16.6581 | 16.97469 | 14.4049 | 16.27783 | 19.34514 | 19.64987 | 16.18761 | 17.64042 | 17.37092 | 14.79118 | 16.05044 | 16.95706 | 16.4935 | 16.20853 | 21.96005 | 22.84206 |
| ADK | 16.69903 | 17.00564 | 17.71319 | 17.74653 | 20.42379 | 19.71704 | 19.74308 | 18.64603 | 17.79626 | 17.60544 | 18.89497 | 20.29612 | 18.80291 | 19.01091 | 18.09076 | 18.32118 | 20.66218 | 20.93015 |
| ADPGK | 14.95845 | 15.78209 | 15.088 | 16.43824 | 14.77967 | 15.6072 | 16.96003 | 15.96563 | 20.12066 | 18.12091 | 15.85295 | 15.44077 | 14.74988 | 15.49657 | 15.15914 | 15.69604 | 14.62952 | 14.72229 |
| ADRA2A | 16.18516 | 16.64768 | 19.3836 | 17.37454 | 15.76023 | 15.08294 | 15.85475 | 17.77971 | 18.41066 | 18.10268 | 15.31286 | 13.19892 | 16.79526 | 17.67323 | 16.65885 | 20.85321 | 17.55687 | 16.42979 |
| ADRM1 | 16.54951 | 14.95747 | 16.88547 | 17.18911 | 18.76028 | 17.18511 | 17.25535 | 19.03019 | 17.5089 | 15.89518 | 17.29679 | 16.93683 | 20.25606 | 17.92932 | 17.09175 | 15.89056 | 18.89603 | 19.01508 |
| ADSL | 16.80379 | 16.5223 | 17.02457 | 17.99951 | 17.59447 | 19.26631 | 18.21594 | 17.74568 | 15.5903 | 15.58898 | 19.89138 | 20.95737 | 17.94015 | 17.64938 | 19.03614 | 18.31133 | 16.8161 | 18.30474 |
| ADSS2 | 15.63663 | 15.52833 | 16.4674 | 17.27135 | 18.18495 | 16.86243 | 17.96858 | 17.48775 | 16.67069 | 16.04525 | 20.01887 | 20.94579 | 17.74122 | 16.25733 | 15.79954 | 17.54357 | 18.24991 | 17.74644 |
| AFDN | 17.243 | 17.72956 | 16.92058 | 16.4361 | 18.91378 | 18.24227 | 17.70956 | 17.60854 | 16.89669 | 16.1729 | 19.14226 | 18.51269 | 19.58847 | 20.06619 | 16.79519 | 16.98317 | 17.47968 | 17.36893 |
| AFG3L2 | 18.2136 | 16.46855 | 16.07473 | 16.26771 | 17.8148 | 16.28463 | 18.91887 | 18.06892 | 19.04052 | 17.54293 | 15.03446 | 14.75076 | 15.73467 | 15.28118 | 16.03719 | 15.9988 | 17.83563 | 15.9737 |
| AGA | 16.85701 | 16.04389 | 15.76649 | 16.70575 | 13.66577 | 16.60374 | 16.67934 | 16.37419 | 19.99899 | 19.31694 | 15.15589 | 15.49603 | 14.16945 | 14.68456 | 14.5187 | 17.92711 | 15.63595 | 14.89257 |
| AGAP3 | 16.75228 | 15.87403 | 16.31172 | 17.12611 | 12.04963 | 14.7498 | 15.8344 | 14.65855 | 15.98307 | 15.08233 | 17.10498 | 18.12163 | 14.32098 | 14.46469 | 12.70622 | 15.06843 | 13.77495 | 15.63441 |
| AGFG1 | 18.35692 | 15.61358 | 16.22366 | 13.73928 | 20.62739 | 18.72226 | 17.60258 | 17.33887 | 16.34527 | 16.76951 | 14.7915 | 15.6991 | 18.11929 | 18.09395 | 15.56041 | 16.55714 | 18.67385 | 18.99937 |
| AGL | 16.20188 | 16.04807 | 16.85491 | 16.03377 | 15.12382 | 17.05642 | 16.30421 | 16.71484 | 15.04532 | 16.29885 | 17.64935 | 18.7749 | 16.32123 | 15.63257 | 15.59823 | 16.18458 | 15.00785 | 14.67316 |
| AGO3 | 13.73068 | 14.25914 | 17.3825 | 18.65559 | 12.01345 | 14.27123 | 14.80588 | 13.43123 | 15.21334 | 17.59477 | 14.77066 | 18.09133 | 13.47761 | 16.2324 | 15.24144 | 15.99429 | 14.86452 | 14.81729 |
| AGPS | 16.87872 | 16.88866 | 16.5576 | 16.93942 | 16.0415 | 16.45133 | 19.64964 | 17.77663 | 21.63766 | 20.62565 | 18.11101 | 14.05303 | 16.17684 | 15.29417 | 16.09549 | 15.37787 | 16.12974 | 15.77753 |
| AGRN | 20.0953 | 17.63043 | 20.86623 | 19.416 | 14.93009 | 17.88316 | 16.19506 | 16.86305 | 17.82609 | 16.357 | 17.02877 | 15.88438 | 18.22602 | 17.53036 | 19.32347 | 19.417 | 15.89831 | 17.3271 |
| AGTRAP | 17.65787 | 15.57472 | 20.03262 | 20.84642 | 14.78144 | 16.9755 | 17.41556 | 21.5416 | 12.99332 | 15.81895 | 14.46394 | 17.02384 | 14.25631 | 13.82187 | 18.28886 | 18.70555 | 14.10646 | 15.89855 |
| AHCY | 19.21245 | 19.59844 | 20.86605 | 20.62034 | 21.40763 | 20.22915 | 19.23181 | 19.60349 | 19.41318 | 19.97728 | 21.86746 | 22.53851 | 20.80205 | 20.62679 | 20.86591 | 20.79696 | 21.77374 | 21.44539 |
| AHCYL1 | 18.587 | 18.08379 | 18.9403 | 19.41541 | 21.22909 | 20.36626 | 18.83718 | 17.74177 | 16.34503 | 17.69826 | 17.19794 | 17.76342 | 19.66426 | 19.31808 | 20.45757 | 19.5318 | 21.01972 | 20.25494 |
| AHCYL2 | 16.14205 | 18.38953 | 16.43932 | 17.23761 | 20.08829 | 18.51822 | 20.15537 | 15.28765 | 13.78325 | 14.8553 | 16.49038 | 12.13561 | 17.23593 | 17.7157 | 18.89004 | 14.8471 | 19.68156 | 17.73362 |
| AHNAK | 16.90935 | 16.94564 | 16.64646 | 16.67932 | 21.91082 | 20.14649 | 17.02821 | 17.13848 | 15.412 | 16.4067 | 16.19518 | 15.31308 | 16.83426 | 17.07253 | 16.92066 | 16.76893 | 21.1592 | 21.51491 |
| AHNAK2 | 20.95675 | 16.0346 | 15.50516 | 15.88025 | 20.13849 | 18.64453 | 16.48173 | 16.52843 | 15.09733 | 16.60267 | 15.52783 | 14.40376 | 20.67267 | 15.60075 | 15.40521 | 17.51626 | 18.45702 | 20.88051 |
| AHSA1 | 16.22709 | 16.5164 | 16.36601 | 16.47377 | 16.30826 | 16.41468 | 17.59042 | 16.64166 | 17.26002 | 16.80077 | 19.65796 | 19.95925 | 16.86267 | 16.28863 | 18.11554 | 19.47293 | 17.12336 | 16.44682 |
| AIDA | 18.77634 | 18.95345 | 20.48137 | 21.62157 | 18.90429 | 18.65517 | 18.34199 | 18.80227 | 17.41399 | 17.61275 | 18.68942 | 18.07774 | 21.62521 | 21.65665 | 21.76967 | 21.62227 | 20.42897 | 21.6175 |
| AIF1L | 16.49979 | 16.0891 | 18.23167 | 17.80424 | 14.31171 | 13.81155 | 15.62107 | 15.30054 | 15.4007 | 17.97277 | 15.76569 | 12.95635 | 20.6534 | 18.47316 | 18.80336 | 18.14651 | 16.24222 | 15.90658 |
| AIFM1 | 15.33293 | 15.55774 | 15.28837 | 16.89669 | 14.06785 | 16.16807 | 17.05566 | 17.49604 | 17.67334 | 17.62612 | 17.74715 | 18.16107 | 18.44367 | 17.9181 | 14.72091 | 16.06639 | 14.77089 | 16.61561 |
| AIMP1 | 20.07201 | 17.31018 | 16.32376 | 17.70061 | 17.91713 | 17.06821 | 17.05169 | 17.53027 | 15.93346 | 16.92371 | 18.94357 | 19.51544 | 19.83602 | 19.06727 | 15.54648 | 16.77722 | 17.47161 | 17.29401 |
| AIMP2 | 16.45762 | 15.97197 | 16.10208 | 16.31931 | 17.67731 | 16.48332 | 16.16155 | 16.11033 | 15.77848 | 18.07938 | 17.73967 | 20.64117 | 19.46789 | 18.04907 | 16.11616 | 16.21598 | 15.11993 | 17.98082 |
| AIP | 16.0702 | 16.36064 | 18.30191 | 15.93503 | 19.64164 | 18.77561 | 18.24718 | 17.24888 | 20.09731 | 18.24566 | 17.39546 | 18.14051 | 20.67227 | 19.49263 | 16.7968 | 18.8343 | 20.70789 | 21.19369 |
| AK1 | 22.77426 | 22.36649 | 21.14021 | 22.36451 | 21.05392 | 19.27003 | 21.00099 | 20.16156 | 19.61235 | 20.90678 | 19.10499 | 19.52902 | 20.75632 | 21.16467 | 20.13509 | 22.05797 | 21.19864 | 20.11068 |
| AK2 | 18.2404 | 17.01673 | 16.08364 | 16.35176 | 17.41685 | 16.0125 | 18.25372 | 17.30875 | 18.97512 | 19.12632 | 19.19911 | 19.82658 | 19.38576 | 18.35344 | 17.14232 | 16.72529 | 16.44094 | 16.35769 |
| AK3 | 14.94943 | 14.59602 | 14.9159 | 16.40219 | 16.38697 | 16.39712 | 17.54869 | 16.81819 | 19.9216 | 19.08246 | 15.38347 | 13.38648 | 14.89938 | 15.81822 | 16.04576 | 15.10664 | 15.16533 | 14.20431 |
| AKAP12 | 15.8137 | 16.16642 | 18.36407 | 15.79449 | 19.48547 | 17.91998 | 15.80247 | 15.85553 | 14.82225 | 15.56116 | 16.12027 | 14.64714 | 15.20843 | 16.28319 | 15.91086 | 16.96733 | 19.99892 | 20.86078 |
| AKAP2 | 15.94556 | 16.50935 | 16.25381 | 15.7621 | 20.29874 | 17.93332 | 16.78531 | 17.62489 | 15.30376 | 15.22223 | 15.5296 | 14.90088 | 15.13772 | 16.07927 | 16.21213 | 16.15018 | 19.18222 | 19.30317 |
| AKAP5 | 15.56464 | 16.0008 | 17.12643 | 16.1823 | 18.22458 | 15.31026 | 14.53859 | 16.41313 | 16.20703 | 14.11597 | 14.31788 | 13.36505 | 20.16123 | 20.48404 | 17.20295 | 17.04128 | 16.13449 | 19.51503 |
| AKR1A1 | 16.46268 | 16.37586 | 18.09764 | 18.76221 | 19.36982 | 17.65991 | 17.80551 | 17.51205 | 19.43765 | 19.66159 | 18.24189 | 19.49711 | 19.84623 | 19.75206 | 19.53899 | 19.19662 | 19.65335 | 18.8724 |
| AKR1B1 | 16.10904 | 17.01204 | 16.32837 | 16.78675 | 21.94914 | 20.38679 | 17.89201 | 16.95875 | 13.86023 | 18.13071 | 20.96432 | 21.72425 | 16.58341 | 15.84665 | 15.51929 | 16.58453 | 18.35354 | 17.13827 |
| AKR1B10 | 15.50511 | 15.93603 | 15.76778 | 15.06099 | 19.24267 | 16.15901 | 21.90398 | 21.53818 | 12.94467 | 15.65687 | 16.1934 | 13.34916 | 16.5236 | 14.7408 | 15.81951 | 15.16685 | 15.04932 | 13.89119 |
| AKR1B15 | 16.29604 | 14.9872 | 14.01554 | 16.6214 | 16.32811 | 17.61117 | 20.14287 | 21.04408 | 17.10952 | 17.13226 | 16.37243 | 18.82242 | 15.15992 | 15.40916 | 16.32681 | 15.19792 | 15.01204 | 15.95838 |
| AKR1C2 | 17.92682 | 17.32939 | 21.72913 | 15.39418 | 21.18115 | 16.87554 | 16.91668 | 17.23672 | 14.13212 | 15.35975 | 15.3398 | 13.38833 | 15.45634 | 15.55765 | 16.28691 | 16.52021 | 18.26922 | 15.45749 |
| AKR1C3 | 17.06427 | 15.51884 | 14.61161 | 14.56382 | 21.04075 | 18.14142 | 16.90183 | 15.90471 | 14.21131 | 16.29775 | 13.13313 | 12.68386 | 15.64259 | 16.15495 | 16.97409 | 15.5128 | 19.75773 | 16.23832 |
| AKR1C4 | 19.51697 | 17.72136 | 14.66556 | 15.23839 | 22.83959 | 20.3744 | 15.66252 | 17.18029 | 14.97359 | 16.94466 | 15.27169 | 15.62262 | 16.75376 | 16.5775 | 16.35974 | 18.15784 | 19.29163 | 18.46054 |
| AKR7A2 | 17.57324 | 12.79576 | 17.15532 | 17.23539 | 15.83119 | 17.46744 | 16.16923 | 15.03847 | 17.76432 | 15.8599 | 15.42569 | 18.05994 | 15.69024 | 15.21921 | 17.52656 | 17.38141 | 16.03369 | 15.58972 |
| AKR7L | 16.47256 | 16.64834 | 16.7703 | 17.0366 | 18.91733 | 17.72324 | 20.91226 | 19.48324 | 19.24337 | 20.2594 | 16.08378 | 18.58521 | 19.30789 | 20.47076 | 16.26591 | 16.55889 | 21.02917 | 18.79909 |
| AKT1 | 17.293 | 16.22685 | 20.10277 | 20.24195 | 17.21262 | 16.30577 | 15.9297 | 16.40378 | 16.21566 | 15.18008 | 19.69755 | 19.48254 | 18.60828 | 18.28036 | 21.55971 | 20.60637 | 16.55082 | 15.77912 |
| AKT2 | 14.64375 | 16.03007 | 18.69713 | 18.55858 | 14.08618 | 16.50372 | 15.45111 | 15.95683 | 14.00169 | 15.12025 | 16.89106 | 14.21837 | 17.15553 | 14.46022 | 19.22491 | 19.62661 | 14.78375 | 15.63228 |
| AKT3 | 13.77732 | 14.32552 | 17.36503 | 13.76749 | 13.69574 | 15.96711 | 14.98697 | 15.24134 | 13.37008 | 14.29672 | 15.06899 | 18.20806 | 13.58548 | 15.31751 | 15.91576 | 18.68694 | 15.44143 | 15.36688 |
| ALCAM | 22.39796 | 22.2545 | 20.74022 | 20.52577 | 21.11402 | 19.10778 | 19.5164 | 18.56901 | 20.10369 | 19.6702 | 16.078 | 17.72467 | 19.32922 | 19.1919 | 20.38721 | 20.12424 | 20.65012 | 21.76436 |
| ALDH16A1 | 19.88861 | 17.26781 | 16.33179 | 15.72829 | 14.42874 | 17.17328 | 16.90825 | 17.0649 | 14.15404 | 15.47887 | 17.22237 | 18.38735 | 14.1479 | 14.51064 | 19.36611 | 19.68932 | 15.24413 | 16.3593 |
| ALDH18A1 | 14.6593 | 16.27086 | 16.78852 | 14.22062 | 16.88152 | 15.97747 | 16.68514 | 15.61818 | 18.44021 | 16.41972 | 14.66526 | 16.63743 | 14.86367 | 15.75532 | 14.7218 | 16.01461 | 14.74688 | 16.55804 |
| ALDH1A1 | 16.47101 | 16.88195 | 17.1746 | 16.93692 | 16.10932 | 17.12872 | 17.43002 | 17.75566 | 18.57606 | 19.01525 | 23.38877 | 26.29807 | 16.10154 | 16.40624 | 16.82229 | 16.4997 | 16.6021 | 16.44274 |
| ALDH1A3 | 16.03505 | 16.63061 | 17.15291 | 15.00596 | 18.64969 | 17.45609 | 16.50548 | 17.78269 | 16.33638 | 16.52926 | 17.59032 | 16.27789 | 17.52764 | 16.51523 | 16.28024 | 15.89237 | 23.25243 | 22.33064 |
| ALDH1B1 | 14.50743 | 14.38555 | 15.23524 | 15.65921 | 15.61821 | 15.05538 | 16.73452 | 15.344 | 17.52389 | 15.85769 | 16.61052 | 12.50672 | 14.24549 | 16.65104 | 15.59814 | 15.99393 | 16.29785 | 15.92272 |
| ALDH1L2 | 15.54126 | 15.57926 | 17.00758 | 16.46501 | 16.42824 | 18.17157 | 14.97871 | 16.39593 | 18.53771 | 17.43784 | 15.16895 | 13.71741 | 15.58548 | 16.34093 | 15.45286 | 15.62974 | 14.33192 | 17.32522 |
| ALDH2 | 16.61483 | 18.05188 | 16.87716 | 17.28722 | 16.30446 | 17.53208 | 19.59799 | 19.90792 | 19.92926 | 17.93379 | 22.86702 | 25.0678 | 17.61589 | 16.16489 | 15.95392 | 16.90417 | 16.88328 | 18.01278 |
| ALDH3A2 | 18.7704 | 16.92906 | 20.1209 | 16.90884 | 16.86941 | 17.40661 | 16.82063 | 16.28848 | 21.3595 | 20.54189 | 16.58087 | 18.31871 | 16.09254 | 15.53711 | 17.80108 | 18.03169 | 16.34554 | 17.77696 |
| ALDH3B1 | 15.67017 | 16.25065 | 15.51553 | 16.7486 | 20.98281 | 20.48967 | 17.90464 | 18.30927 | 20.25203 | 16.14263 | 15.29172 | 13.78247 | 14.79566 | 15.70086 | 15.96896 | 16.25848 | 14.66056 | 15.96718 |
| ALDH4A1 | 17.44933 | 14.02998 | 14.06839 | 14.91174 | 13.52597 | 18.45048 | 17.06317 | 17.8259 | 17.73382 | 17.25481 | 14.83086 | 13.918 | 18.41512 | 16.84644 | 16.51113 | 16.62096 | 15.05107 | 16.60684 |
| ALDH6A1 | 16.58527 | 15.77299 | 13.4382 | 16.04435 | 14.32047 | 15.36211 | 16.55581 | 16.09518 | 18.82245 | 17.02161 | 14.421 | 12.0619 | 15.16892 | 17.05571 | 15.34371 | 15.73716 | 16.51633 | 14.39482 |
| ALDH7A1 | 15.21179 | 16.42331 | 15.44141 | 15.12711 | 19.1819 | 17.40292 | 17.39668 | 16.78509 | 16.58605 | 15.95801 | 17.3683 | 18.89806 | 15.23791 | 17.46093 | 16.51669 | 15.40072 | 19.86567 | 21.30743 |
| ALDH9A1 | 17.47798 | 15.87514 | 17.88906 | 21.20256 | 19.76004 | 17.59786 | 19.18038 | 18.94321 | 18.01903 | 19.0069 | 17.58439 | 18.13124 | 20.95288 | 20.9863 | 17.83235 | 17.58393 | 20.85622 | 19.86227 |
| ALDOA | 19.8501 | 19.22344 | 20.94042 | 21.2453 | 24.92661 | 23.58222 | 21.94159 | 21.22184 | 21.94768 | 22.18828 | 23.62362 | 23.79491 | 22.24944 | 21.98493 | 21.40147 | 21.53281 | 24.05826 | 23.43827 |
| ALDOC | 18.89111 | 16.87634 | 19.73806 | 19.16861 | 21.01965 | 19.94677 | 18.58809 | 18.64425 | 18.81103 | 18.53747 | 21.23429 | 21.36486 | 19.60573 | 18.98821 | 18.40745 | 17.86884 | 20.76619 | 20.66374 |
| ALG9 | 16.25637 | 16.52347 | 16.72566 | 16.6284 | 18.4318 | 15.14609 | 17.35455 | 17.11971 | 21.55013 | 17.68294 | 17.27665 | 14.93753 | 15.56066 | 17.56356 | 15.90041 | 15.68829 | 14.94571 | 17.32453 |
| ALKBH5 | 15.07001 | 16.96167 | 15.81605 | 15.78468 | 15.00835 | 15.42849 | 16.72714 | 16.73803 | 13.59148 | 16.34243 | 16.2709 | 17.82787 | 13.61717 | 14.79718 | 17.42548 | 14.80648 | 15.28876 | 15.70139 |
| ALOX5 | 16.28672 | 16.07099 | 15.72651 | 15.23891 | 14.20486 | 15.31581 | 14.75202 | 15.60854 | 17.83428 | 14.09209 | 17.52441 | 17.91649 | 14.74033 | 16.24596 | 14.4613 | 16.8359 | 13.8882 | 15.30197 |
| ALPL | 19.23664 | 18.61855 | 17.32635 | 16.87943 | 16.39213 | 17.68184 | 15.3438 | 17.6028 | 19.08308 | 18.6647 | 14.24159 | 13.45262 | 14.52721 | 15.26821 | 14.36671 | 16.95284 | 15.69416 | 16.60195 |
| AMPD2 | 18.11748 | 15.21114 | 14.68262 | 12.9814 | 15.34796 | 15.19999 | 15.41324 | 15.87945 | 14.17488 | 16.56473 | 15.2451 | 17.57788 | 16.12518 | 14.66797 | 15.23876 | 15.54191 | 14.44281 | 14.2104 |
| AMPD3 | 16.37108 | 15.57602 | 18.65347 | 16.67977 | 13.94189 | 19.25988 | 15.9093 | 16.6734 | 16.82534 | 14.99895 | 15.04935 | 18.88011 | 15.68444 | 15.2361 | 16.6378 | 13.88598 | 15.51238 | 16.98881 |
| ANAPC5 | 13.83188 | 14.78059 | 15.47349 | 13.90155 | 15.58855 | 14.22336 | 14.20682 | 17.4121 | 11.93707 | 14.9215 | 19.82046 | 17.75362 | 14.80992 | 12.96149 | 19.20717 | 18.44437 | 15.01887 | 14.83879 |

| | | | | | | | | | | | | | | | | | | |
|---|---|---|---|---|---|---|---|---|---|---|---|---|---|---|---|---|---|---|
| ANAPC7 | 16.97672 | 16.32415 | 15.9404 | 16.79877 | 14.09299 | 16.37066 | 16.90211 | 14.84939 | 12.51401 | 15.2035 | 15.31541 | 17.72258 | 14.92861 | 15.56846 | 15.09283 | 15.75088 | 14.68663 | 15.30484 |
| ANGPTL2 | 17.99885 | 13.77215 | 14.44905 | 13.73683 | 15.45042 | 15.62434 | 16.7176 | 14.38205 | 17.709 | 17.4517 | 15.46846 | 13.84235 | 17.251 | 12.86263 | 15.3818 | 13.09701 | 13.80813 | 15.8367 |
| ANK1 | 17.85412 | 18.69711 | 17.23315 | 18.20584 | 15.10644 | 15.75262 | 19.98679 | 18.00661 | 13.75604 | 15.69385 | 15.97813 | 13.98317 | 16.77538 | 15.47935 | 17.93054 | 17.99722 | 14.20613 | 15.25312 |
| ANK2 | 17.53078 | 16.30196 | 15.04509 | 19.46075 | 18.93959 | 19.06619 | 15.50035 | 15.37481 | 16.77101 | 17.56523 | 13.91151 | 16.95801 | 19.04464 | 20.4992 | 17.68421 | 20.63882 | 15.587 | 18.04 |
| ANK3 | 15.75858 | 15.92079 | 19.69617 | 20.54592 | 16.90118 | 17.18999 | 16.24456 | 17.13361 | 15.87412 | 16.01769 | 17.41552 | 16.53895 | 21.2028 | 21.41278 | 21.30707 | 21.12801 | 16.49248 | 16.37525 |
| ANKFY1 | 16.26096 | 15.46135 | 16.21131 | 17.75107 | 19.91833 | 18.77151 | 18.63416 | 17.52087 | 16.56389 | 16.19632 | 16.13753 | 16.94856 | 16.49043 | 18.64226 | 16.37583 | 16.86195 | 16.54815 | 16.59351 |
| ANKH | 17.04851 | 14.99722 | 17.73848 | 15.00632 | 14.4864 | 14.32529 | 16.81933 | 16.43914 | 18.83554 | 12.22622 | 14.68837 | 14.83041 | 16.87609 | 14.54961 | 16.38495 | 16.35813 | 13.04631 | 16.20472 |
| ANKRD13A | 20.19511 | 19.95552 | 18.79567 | 19.24965 | 19.11896 | 16.77751 | 16.76538 | 15.79575 | 16.25119 | 15.88806 | 16.19664 | 15.00541 | 15.17927 | 16.26297 | 19.05063 | 18.90616 | 17.41524 | 17.55116 |
| ANKRD13D | 19.87971 | 19.93595 | 17.66057 | 17.45837 | 13.21142 | 16.61436 | 16.03319 | 17.7368 | 12.96624 | 14.41889 | 16.10358 | 15.4401 | 15.25698 | 15.28202 | 17.93481 | 18.6872 | 14.53321 | 14.4361 |
| ANKRD26 | 14.15051 | 17.73435 | 14.46401 | 16.40406 | 18.89406 | 15.88017 | 15.79418 | 14.11146 | 13.67384 | 17.0097 | 15.7252 | 12.84872 | 19.42758 | 18.36827 | 13.89898 | 18.45772 | 19.05488 | 20.48515 |
| ANKRD28 | 16.84149 | 15.94817 | 16.73377 | 17.97625 | 19.79547 | 16.08481 | 15.58994 | 16.11445 | 16.94977 | 17.59726 | 14.52335 | 18.20669 | 15.89345 | 16.0217 | 16.24249 | 17.59662 | 15.09785 | 17.58977 |
| ANLN | 16.53249 | 16.48543 | 16.80118 | 17.7497 | 21.41099 | 20.01142 | 17.67184 | 17.62637 | 16.25357 | 15.29112 | 16.13622 | 15.16215 | 17.19174 | 15.55967 | 16.17946 | 16.14844 | 21.17973 | 19.81375 |
| ANO6 | 16.61739 | 16.65989 | 17.92834 | 18.1099 | 16.77499 | 19.3153 | 17.95016 | 15.23236 | 16.72784 | 17.1379 | 18.04055 | 18.79703 | 16.44729 | 16.22371 | 18.57318 | 18.57695 | 16.25407 | 16.25384 |
| ANP32A | 20.14545 | 22.37889 | 16.16449 | 15.97918 | 21.10631 | 19.0674 | 19.51176 | 18.33472 | 18.02521 | 19.38979 | 22.52743 | 21.4579 | 21.86041 | 19.79794 | 16.86819 | 17.11609 | 19.3385 | 18.10143 |
| ANP32B | 16.61684 | 17.62913 | 17.86722 | 17.18676 | 18.96192 | 19.2413 | 18.24268 | 18.17747 | 16.40739 | 18.73522 | 20.27716 | 20.42856 | 18.69364 | 18.0901 | 15.89359 | 16.78016 | 18.05127 | 15.55141 |
| ANP32E | 17.02249 | 17.18466 | 14.33267 | 14.92765 | 20.65366 | 15.46575 | 16.46782 | 17.68464 | 16.35929 | 19.5273 | 19.43992 | 18.38696 | 19.84465 | 16.5795 | 15.16037 | 16.40706 | 18.76858 | 17.30218 |
| ANPEP | 18.50062 | 19.01384 | 18.73001 | 18.13439 | 16.39956 | 17.90964 | 17.5353 | 16.70614 | 15.34279 | 16.80635 | 17.3037 | 14.9395 | 17.49531 | 19.46579 | 18.64755 | 18.39388 | 23.60655 | 24.01833 |
| ANTXR1 | 14.63876 | 14.45824 | 14.88036 | 15.50815 | 20.57412 | 19.26367 | 18.05753 | 16.64599 | 17.03825 | 14.46336 | 15.92456 | 12.55449 | 13.44431 | 15.37241 | 14.55048 | 16.68726 | 18.08762 | 19.06167 |
| ANTXR2 | 17.03719 | 18.20105 | 17.03987 | 18.11723 | 18.56578 | 15.59874 | 17.58401 | 17.58991 | 16.40376 | 15.55071 | 16.6746 | 14.60553 | 17.37986 | 17.43894 | 15.41606 | 17.81147 | 18.78143 | 21.34706 |
| ANXA1 | 18.84455 | 19.70153 | 18.28942 | 19.89922 | 23.86929 | 23.31736 | 23.64085 | 20.35856 | 16.93509 | 18.41129 | 26.70216 | 27.76982 | 20.55683 | 22.02998 | 18.75037 | 18.93766 | 21.44588 | 22.51312 |
| ANXA11 | 19.87956 | 19.7626 | 19.17095 | 20.29993 | 22.77085 | 21.81392 | 21.46192 | 20.65217 | 24.55618 | 23.61252 | 21.86668 | 20.71938 | 22.3918 | 23.066 | 19.52149 | 19.72731 | 22.2633 | 24.41586 |
| ANXA2 | 20.19222 | 20.04432 | 17.94929 | 19.09467 | 22.68329 | 22.53902 | 19.56839 | 18.82118 | 22.93392 | 21.1887 | 18.37703 | 15.24606 | 15.69313 | 16.70329 | 18.16955 | 17.85291 | 20.65883 | 21.65531 |
| ANXA2P2 | 20.52021 | 19.59992 | 17.25949 | 19.99134 | 26.466 | 25.43621 | 22.35115 | 20.82183 | 22.6933 | 23.33572 | 18.76576 | 16.31876 | 18.77039 | 20.22256 | 18.53232 | 17.93016 | 26.46143 | 25.2641 |
| ANXA3 | 16.50114 | 17.58015 | 16.28375 | 15.67658 | 23.46492 | 22.32692 | 16.63166 | 17.31125 | 15.11463 | 16.33011 | 15.47314 | 16.86233 | 16.19954 | 15.96957 | 16.71652 | 16.90576 | 21.72869 | 21.30635 |
| ANXA4 | 20.08534 | 19.28842 | 19.26019 | 19.77999 | 22.09342 | 21.5665 | 20.52469 | 19.74233 | 23.02804 | 22.73802 | 23.61492 | 20.60822 | 20.80011 | 21.25364 | 19.47124 | 19.12674 | 20.04667 | 21.0902 |
| ANXA6 | 25.43737 | 19.20327 | 21.43774 | 21.46656 | 21.68726 | 20.73517 | 22.17472 | 20.54659 | 23.02656 | 23.094 | 17.10963 | 16.51061 | 21.63469 | 21.45756 | 21.56403 | 20.27189 | 21.32073 | 22.72351 |
| ANXA7 | 19.7766 | 19.68848 | 20.09218 | 20.32957 | 20.17699 | 20.12435 | 19.98464 | 18.30784 | 24.62011 | 23.24391 | 20.36986 | 19.22103 | 19.61703 | 20.42154 | 20.74626 | 20.40057 | 21.29848 | 22.2151 |
| AOC1 | 16.97468 | 15.45779 | 17.69863 | 16.77118 | 18.86948 | 18.69437 | 16.3107 | 16.66954 | 19.43648 | 18.15664 | 15.79754 | 14.56989 | 15.74254 | 18.02967 | 15.39796 | 16.36477 | 18.99404 | 19.32189 |
| AP1B1 | 17.4398 | 17.45929 | 18.67781 | 18.20866 | 17.02494 | 17.72549 | 16.03093 | 16.64594 | 16.57593 | 17.70602 | 18.34557 | 19.6172 | 18.1882 | 18.0287 | 19.05349 | 18.68358 | 17.9264 | 18.11345 |
| AP1G1 | 15.80758 | 15.26045 | 17.32708 | 18.9473 | 18.38093 | 13.87319 | 16.58083 | 13.70347 | 16.84962 | 14.49597 | 16.80396 | 19.37832 | 12.07695 | 14.23394 | 17.92929 | 18.53789 | 17.36813 | 14.93215 |
| AP1M1 | 16.66249 | 15.77243 | 18.79847 | 18.24384 | 14.88755 | 15.37053 | 15.14375 | 14.568 | 16.65753 | 15.1589 | 15.54921 | 17.86503 | 17.99724 | 16.73205 | 16.27111 | 17.79862 | 16.41539 | 15.65787 |
| AP1M2 | 17.66022 | 17.34526 | 17.32055 | 17.88923 | 14.75666 | 16.33862 | 15.03914 | 15.88344 | 12.85241 | 16.38524 | 15.96886 | 18.43087 | 14.29593 | 15.82062 | 17.88421 | 18.59747 | 14.08914 | 15.07819 |
| AP1S1 | 13.63481 | 16.35486 | 18.21497 | 15.32322 | 15.69163 | 14.3524 | 14.1628 | 16.84168 | 16.09617 | 15.48163 | 18.02032 | 18.482 | 15.74182 | 12.30262 | 16.68631 | 18.17138 | 13.15888 | 13.93281 |
| AP1S2 | 14.041 | 16.19665 | 16.07197 | 16.24042 | 13.68303 | 15.41006 | 14.98 | 15.37709 | 15.55283 | 13.98162 | 16.01488 | 18.38238 | 15.82989 | 14.6466 | 16.46884 | 15.26549 | 13.71634 | 11.65039 |
| AP2A1 | 18.09902 | 17.89185 | 18.43967 | 18.96064 | 17.91101 | 16.68738 | 17.73155 | 16.81094 | 16.25677 | 17.90582 | 17.38859 | 18.86478 | 17.22115 | 18.75861 | 17.87497 | 18.6441 | 19.81073 | 19.37839 |
| AP2A2 | 18.66517 | 19.05223 | 19.02686 | 19.17299 | 18.17624 | 17.5955 | 16.83421 | 16.35492 | 16.93187 | 17.9024 | 17.54534 | 19.20097 | 18.7344 | 17.50043 | 20.25855 | 19.98969 | 18.11582 | 17.77947 |
| AP2B1 | 18.01354 | 18.81586 | 18.23678 | 17.95085 | 18.82515 | 18.49112 | 17.77255 | 16.8032 | 17.90762 | 17.98088 | 18.77593 | 19.25651 | 18.28704 | 19.04794 | 18.71466 | 18.08335 | 19.32903 | 18.97332 |
| AP2M1 | 17.90426 | 19.47397 | 18.88448 | 18.93496 | 19.06463 | 18.67866 | 19.05103 | 16.87093 | 18.37326 | 18.44566 | 18.71984 | 19.72946 | 18.42078 | 18.41029 | 19.32592 | 19.17948 | 18.99113 | 19.19945 |
| AP2S1 | 18.95749 | 18.89563 | 19.12731 | 18.25258 | 13.74068 | 15.73714 | 17.04307 | 18.27359 | 17.20784 | 15.76293 | 18.03261 | 20.01052 | 14.43577 | 18.90691 | 18.89046 | 19.2927 | 16.36274 | 18.13909 |
| AP3B1 | 15.97342 | 15.88635 | 15.53032 | 15.65021 | 17.27733 | 15.90516 | 17.93639 | 17.09493 | 17.89477 | 18.87474 | 17.49232 | 19.25517 | 17.1486 | 16.67487 | 16.15073 | 15.67229 | 21.11445 | 18.24469 |
| AP3B2 | 12.3547 | 13.87387 | 13.881 | 14.88216 | 12.81736 | 16.96638 | 14.84502 | 16.58988 | 14.11105 | 16.20145 | 12.85003 | 18.56485 | 12.99796 | 13.17423 | 14.24237 | 14.91652 | 15.18521 | 13.4239 |
| AP3D1 | 17.53007 | 17.71302 | 16.16693 | 15.99501 | 14.2984 | 16.90436 | 15.56279 | 16.39112 | 16.78445 | 15.04745 | 19.19098 | 18.69634 | 15.39728 | 16.28754 | 15.41251 | 16.01278 | 14.70896 | 15.21812 |
| AP3M1 | 17.75109 | 18.22295 | 17.59072 | 15.15166 | 19.54921 | 14.77693 | 18.35503 | 19.32029 | 21.58103 | 21.85232 | 16.8657 | 19.56243 | 18.02384 | 16.80455 | 15.7094 | 15.89225 | 16.12735 | 16.86897 |
| AP3M2 | 17.51988 | 17.81352 | 15.02282 | 17.34195 | 19.30767 | 17.90417 | 18.2632 | 18.43017 | 15.77497 | 17.53059 | 16.99989 | 19.32178 | 17.98901 | 18.63966 | 14.85909 | 16.65254 | 17.3485 | 18.75945 |
| AP3S1 | 16.89283 | 16.17933 | 17.10097 | 17.10509 | 17.73667 | 15.01935 | 17.502 | 18.71724 | 15.84183 | 14.63404 | 16.46166 | 18.52804 | 18.08874 | 15.51301 | 14.57288 | 16.11624 | 17.37377 | 17.44316 |
| APBB2 | 14.06313 | 15.61082 | 13.198 | 15.96626 | 17.82144 | 19.1677 | 15.01256 | 13.89243 | 14.52576 | 14.22582 | 14.15256 | 12.08411 | 14.56262 | 13.92838 | 17.10881 | 13.78775 | 15.54076 | 12.98037 |
| APEH | 20.10969 | 21.01749 | 19.16392 | 19.99905 | 18.72659 | 18.3296 | 17.13927 | 17.63241 | 16.68124 | 17.11238 | 19.15414 | 20.38782 | 18.49379 | 18.91784 | 19.18464 | 19.56841 | 19.33559 | 19.34746 |
| APEX1 | 15.14614 | 15.92261 | 17.40554 | 16.80215 | 18.17776 | 17.14477 | 18.22764 | 17.92836 | 17.40237 | 18.14439 | 19.37305 | 20.0711 | 18.12295 | 17.52716 | 16.44028 | 16.96403 | 17.03344 | 16.67449 |
| API5 | 17.06879 | 16.16855 | 15.90274 | 16.47229 | 17.21503 | 17.72017 | 18.29911 | 16.97884 | 15.57903 | 16.75355 | 19.37774 | 19.76262 | 17.85602 | 17.29009 | 15.31347 | 18.07629 | 15.50622 | 17.75847 |
| APLP1 | 15.87233 | 17.04821 | 15.54146 | 17.04816 | 15.32939 | 17.73428 | 15.9934 | 17.06663 | 20.03915 | 17.92837 | 16.39527 | 13.76814 | 15.26201 | 16.8376 | 16.07402 | 16.81824 | 16.17979 | 15.27252 |
| APLP2 | 15.49219 | 15.97448 | 16.56828 | 16.05757 | 18.77383 | 17.72491 | 17.43926 | 16.55381 | 19.49646 | 18.39625 | 17.23169 | 15.17516 | 17.47207 | 17.38273 | 15.22391 | 15.66894 | 15.88888 | 17.82797 |
| APMAP | 16.58559 | 16.78359 | 17.05569 | 17.70849 | 15.21514 | 16.41004 | 16.17799 | 16.68296 | 24.3251 | 21.65914 | 18.73181 | 16.36198 | 15.87991 | 17.82408 | 17.39036 | 17.00215 | 16.20473 | 16.64626 |
| APOB | 16.9378 | 18.963 | 20.35947 | 17.34003 | 13.19299 | 19.6566 | 20.13193 | 16.39713 | 11.8509 | 14.44839 | 15.89084 | 14.70175 | 17.14308 | 15.56469 | 18.60661 | 18.1801 | 15.03952 | 14.9711 |
| APOBEC3B | 19.86147 | 20.20787 | 19.72726 | 18.73748 | 14.79499 | 15.80262 | 15.46568 | 16.54333 | 14.17988 | 15.56187 | 14.9714 | 14.73847 | 16.02234 | 16.28265 | 20.78563 | 20.07315 | 15.30642 | 14.79784 |
| APOD | 17.40642 | 30.00438 | 30.02198 | 30.4128 | 17.51813 | 20.194 | 25.55669 | 17.88931 | 23.9835 | 21.67316 | 19.17631 | 14.39084 | 28.8662 | 28.3525 | 16.59943 | 17.27223 | 18.10737 | 18.24069 |
| APOM | 21.66838 | 23.24813 | 21.75585 | 21.63362 | 18.36163 | 19.58006 | 18.19074 | 17.40296 | 15.02508 | 14.20288 | 15.86104 | 13.6782 | 20.15993 | 20.9469 | 20.86826 | 20.6991 | 16.65228 | 19.14387 |
| APP | 17.36609 | 16.56177 | 16.31512 | 16.31798 | 18.78973 | 18.87255 | 16.49022 | 17.10946 | 20.5438 | 19.32218 | 18.32483 | 14.52961 | 16.55243 | 15.94417 | 16.52592 | 16.13318 | 16.33091 | 18.59935 |
| APPL1 | 18.90987 | 19.38418 | 17.86568 | 18.03184 | 20.11822 | 19.47337 | 17.73899 | 17.39584 | 19.01851 | 19.06636 | 19.10863 | 18.86602 | 19.64127 | 18.92237 | 18.20716 | 18.20763 | 18.74487 | 18.87232 |
| APPL2 | 16.70612 | 15.67014 | 15.98874 | 20.1675 | 19.39713 | 19.23576 | 16.8963 | 16.74945 | 17.05453 | 17.53381 | 17.60189 | 15.24657 | 20.23679 | 21.0076 | 17.49547 | 19.55367 | 18.50284 | 19.67204 |
| APRT | 18.1446 | 18.16942 | 19.10435 | 18.38581 | 18.98193 | 17.35258 | 16.62491 | 16.31481 | 16.07765 | 16.04398 | 18.15829 | 20.4901 | 18.55435 | 20.9482 | 19.98731 | 20.49541 | 18.22917 | 17.67926 |
| ARAP1 | 17.06739 | 14.3022 | 17.3055 | 14.91434 | 13.27193 | 15.99793 | 16.90221 | 16.76018 | 12.96921 | 15.95682 | 15.66563 | 16.6377 | 15.93503 | 16.22281 | 16.8633 | 16.54493 | 16.20266 | 15.15746 |
| ARCN1 | 16.05047 | 16.05705 | 16.46929 | 15.46135 | 16.97943 | 16.65829 | 17.25936 | 17.02983 | 16.14705 | 17.11598 | 18.1052 | 19.2375 | 18.01961 | 15.33259 | 16.09668 | 16.29798 | 17.68555 | 17.44157 |
| ARF3 | 21.5221 | 21.83026 | 22.41159 | 20.28955 | 14.99374 | 17.06835 | 17.61518 | 16.34034 | 19.24767 | 18.677 | 20.59637 | 22.65656 | 16.54521 | 15.0478 | 22.04457 | 21.82231 | 17.38804 | 16.06723 |
| ARF4 | 21.49351 | 21.47553 | 21.33254 | 20.41142 | 17.12048 | 17.21679 | 16.48415 | 15.98738 | 19.41983 | 17.55291 | 18.8656 | 21.76541 | 15.71239 | 16.18782 | 22.09465 | 21.76011 | 16.74939 | 20.37107 |
| ARF5 | 19.70617 | 19.92084 | 19.42967 | 20.21056 | 15.14734 | 17.27467 | 15.98361 | 15.25766 | 17.49287 | 17.05224 | 16.8677 | 19.72716 | 15.33194 | 23.96042 | 19.53613 | 21.02124 | 15.99788 | 15.14691 |
| ARF6 | 20.72045 | 20.97588 | 21.27055 | 20.72866 | 19.9705 | 19.38121 | 17.33058 | 17.35433 | 20.04682 | 19.36131 | 19.24186 | 20.09297 | 15.40774 | 16.63482 | 21.92589 | 22.06965 | 17.62846 | 18.3474 |
| ARFGEF1 | 14.42345 | 16.36178 | 16.78026 | 14.1584 | 14.21745 | 14.85924 | 16.25264 | 15.50559 | 17.06868 | 16.01399 | 16.36002 | 12.87414 | 15.59651 | 14.07935 | 14.95199 | 15.65309 | 13.35143 | 15.32885 |
| ARFIP1 | 16.73036 | 16.1032 | 18.09752 | 19.56683 | 15.23401 | 15.23175 | 18.31323 | 17.9836 | 14.70167 | 16.17547 | 16.00512 | 17.00043 | 18.42318 | 17.61015 | 16.10328 | 17.23735 | 16.1559 | 16.08164 |
| ARG2 | 16.71406 | 15.00217 | 15.56032 | 15.74322 | 13.46461 | 16.66247 | 16.51853 | 13.38387 | 18.53957 | 14.36757 | 15.54954 | 13.08762 | 14.20412 | 16.02584 | 13.88078 | 14.61157 | 14.42095 | 16.05643 |
| ARHGAP1 | 19.5456 | 17.31226 | 16.34666 | 17.34424 | 19.36542 | 18.60044 | 19.02918 | 18.16531 | 17.0712 | 17.85314 | 17.67626 | 16.77945 | 18.17303 | 18.72099 | 17.07554 | 16.74758 | 17.32512 | 19.03392 |
| ARHGAP10 | 15.51083 | 18.35973 | 17.22695 | 18.8421 | 17.24484 | 16.37322 | 19.2161 | 17.65339 | 13.41023 | 17.69918 | 14.51893 | 17.40899 | 16.96482 | 14.81893 | 18.21036 | 17.7559 | 18.594 | 15.03774 |
| ARHGAP17 | 17.84574 | 17.58336 | 17.22967 | 17.88845 | 18.65513 | 17.10529 | 15.74729 | 15.88364 | 16.55608 | 16.63153 | 17.39883 | 17.51479 | 19.056 | 18.87222 | 17.39561 | 17.92356 | 16.79333 | 17.12641 |
| ARHGAP18 | 15.10123 | 15.79382 | 14.79147 | 16.58985 | 17.92185 | 15.87987 | 17.87339 | 16.12078 | 11.91956 | 13.93635 | 16.66883 | 16.82854 | 16.29601 | 17.44226 | 13.65704 | 15.32604 | 17.70121 | 14.43598 |
| ARHGAP29 | 16.35791 | 16.93578 | 16.50525 | 15.76611 | 19.39462 | 16.93626 | 17.77443 | 17.10192 | 14.20128 | 16.03904 | 15.76152 | 15.72868 | 17.46167 | 17.12991 | 15.64345 | 16.11575 | 20.15365 | 20.15199 |
| ARHGAP39 | 16.52931 | 16.39058 | 15.63477 | 16.55426 | 14.26535 | 14.92409 | 16.29826 | 16.33419 | 16.90926 | 16.03107 | 14.47908 | 13.9278 | 17.10486 | 14.61084 | 15.15948 | 17.41609 | 18.63605 | 16.37348 |
| ARHGAP4 | 21.60103 | 21.71514 | 19.4305 | 16.47789 | 15.1878 | 16.88071 | 15.46154 | 14.9386 | 13.68934 | 15.46976 | 14.87407 | 17.1334 | 17.26404 | 15.44095 | 20.05218 | 19.94619 | 15.35117 | 14.61553 |
| ARHGAP5 | 15.47841 | 17.12493 | 17.23164 | 16.45871 | 18.96877 | 17.44318 | 14.69743 | 17.87398 | 13.47834 | 15.59416 | 13.06686 | 14.5317 | 13.98535 | 17.08608 | 15.50132 | 15.03218 | 16.48339 | 15.68851 |
| ARHGDIA | 22.14984 | 20.22312 | 21.83651 | 22.50511 | 23.67253 | 22.26454 | 23.0614 | 23.72536 | 20.22394 | 19.78116 | 23.24774 | 23.15159 | 23.73299 | 23.22946 | 22.79898 | 22.47403 | 24.65462 | 24.37681 |
| ARHGEF1 | 20.39941 | 15.1788 | 18.88049 | 13.64026 | 19.02014 | 18.72719 | 15.69488 | 15.73871 | 14.62975 | 16.89882 | 15.55746 | 17.7549 | 15.38397 | 15.62076 | 20.19737 | 19.98291 | 19.18808 | 19.81549 |
| ARHGEF16 | 16.44742 | 15.7135 | 15.79668 | 16.85742 | 16.90332 | 17.28046 | 15.52491 | 15.65618 | 18.51004 | 17.70268 | 15.84916 | 15.35671 | 17.77087 | 19.04001 | 16.15405 | 17.8573 | 15.46991 | 13.90515 |
| ARHGEF2 | 15.97973 | 17.16139 | 17.35524 | 15.83106 | 15.92541 | 18.2107 | 17.31503 | 17.99721 | 16.59796 | 16.45793 | 17.47874 | 18.55363 | 16.66087 | 16.22983 | 16.62594 | 16.61976 | 16.5078 | 17.73153 |
| ARHGEF26 | 19.0961 | 15.98678 | 16.27076 | 13.31927 | 19.97607 | 12.1558 | 14.91136 | 17.27999 | 11.815 | 16.9988 | 11.25863 | 15.80644 | 17.57563 | 13.74776 | 14.12932 | 15.90087 | 16.78905 | 15.32941 |
| ARHGEF39 | 19.91896 | 18.77118 | 19.31095 | 20.53161 | 17.70588 | 16.4986 | 18.78485 | 16.50523 | 17.76833 | 16.9991 | 16.32141 | 16.94847 | 15.24122 | 16.70992 | 20.08407 | 18.6595 | 15.50985 | 16.38658 |
| ARHGEF7 | 22.46442 | 20.06405 | 18.71117 | 18.36636 | 20.43203 | 18.22227 | 20.0746 | 18.3 | 18.72101 | 19.59529 | 17.70662 | 17.88373 | 19.0044 | 18.9486 | 18.97656 | 18.76151 | 20.42668 | 20.4971 |
| ARIH1 | 17.1234 | 15.4002 | 14.19156 | 17.35287 | 15.87331 | 14.16491 | 13.4884 | 13.39237 | 12.08265 | 16.27692 | 15.89441 | 18.14086 | 13.7254 | 14.6022 | 14.89605 | 15.85772 | 19.08747 | 16.97622 |
| ARIH2 | 17.01375 | 14.82502 | 15.87539 | 16.51196 | 14.74578 | 16.4719 | 16.26316 | 20.53158 | 14.71248 | 14.88209 | 17.27531 | 17.71221 | 14.62089 | 14.12763 | 14.47318 | 14.98322 | 16.68392 | 15.96111 |

| | | | | | | | | | | | | | | | | | | |
|---|---|---|---|---|---|---|---|---|---|---|---|---|---|---|---|---|---|---|
| ARL1 | 19.00046 | 16.54804 | 13.70842 | 17.34847 | 17.10847 | 17.58197 | 16.0766 | 15.49796 | 17.96154 | 18.24191 | 21.32826 | 17.64723 | 16.49219 | 16.32857 | 19.82654 | 18.19921 | 13.3983 | 17.59316 |
| ARL15 | 16.06395 | 14.94693 | 16.97292 | 16.25182 | 15.24656 | 16.37925 | 16.56925 | 15.69961 | 16.41849 | 16.18632 | 15.0239 | 18.45902 | 15.10881 | 14.70751 | 14.94693 | 15.91536 | 14.66403 | 14.92567 |
| ARL2 | 14.26518 | 18.15666 | 19.46253 | 17.42925 | 13.84887 | 13.46895 | 15.6151 | 18.34146 | 12.14629 | 15.9096 | 17.89153 | 19.43826 | 14.17885 | 17.18555 | 16.92265 | 16.87902 | 14.81278 | 16.83824 |
| ARL3 | 18.00327 | 17.66942 | 19.41976 | 20.09801 | 19.61328 | 18.31139 | 21.79345 | 21.02781 | 17.91316 | 19.12615 | 19.36238 | 20.6901 | 22.21024 | 22.00198 | 20.01037 | 19.99493 | 20.66846 | 20.8861 |
| ARL6 | 17.2545 | 17.31934 | 15.59824 | 16.84789 | 17.23729 | 14.6511 | 14.53041 | 15.40131 | 15.76269 | 17.05657 | 14.41025 | 17.58362 | 17.75339 | 18.51398 | 18.13796 | 17.92507 | 17.6683 | 18.72967 |
| ARL6IP1 | 14.12331 | 15.27713 | 16.44704 | 17.75085 | 14.92762 | 13.56253 | 16.18814 | 14.64171 | 17.00941 | 16.95412 | 15.76162 | 18.37209 | 14.50788 | 14.96541 | 14.16335 | 16.93103 | 13.52933 | 14.68388 |
| ARL8A | 15.98212 | 16.13233 | 14.25577 | 13.59241 | 18.5227 | 19.68071 | 18.17288 | 18.80193 | 17.01583 | 17.38223 | 16.54952 | 18.47089 | 17.15564 | 16.26486 | 17.1738 | 16.52471 | 18.81795 | 16.3935 |
| ARL8B | 15.40478 | 17.21097 | 16.87756 | 15.82846 | 17.94352 | 16.8193 | 18.06854 | 17.20537 | 16.58064 | 18.04774 | 17.14779 | 17.88135 | 17.5218 | 19.3256 | 14.93923 | 15.47393 | 16.97411 | 18.72636 |
| ARMT1 | 16.10683 | 16.19147 | 15.04494 | 15.83014 | 13.5601 | 14.51268 | 15.87912 | 15.2435 | 11.39849 | 17.1917 | 15.97864 | 19.12039 | 13.44266 | 14.1106 | 15.86847 | 18.14488 | 15.12208 | 14.98769 |
| ARPC1A | 20.17959 | 19.5568 | 20.41052 | 19.23065 | 15.06267 | 15.56517 | 16.37218 | 17.39281 | 16.94371 | 14.27198 | 18.02065 | 21.03087 | 16.68074 | 15.69064 | 19.72829 | 20.13447 | 16.72622 | 15.57493 |
| ARPC1B | 20.20641 | 19.64236 | 17.1578 | 18.48288 | 21.19869 | 19.97806 | 20.52124 | 20.18105 | 18.06425 | 18.73327 | 17.47237 | 20.82493 | 19.84443 | 20.11532 | 19.123 | 18.92097 | 21.52793 | 20.89353 |
| ARPC2 | 20.44399 | 20.33425 | 20.37812 | 19.92541 | 18.20796 | 18.04583 | 18.57982 | 18.49546 | 17.71114 | 17.99656 | 19.63642 | 21.60779 | 18.74129 | 17.32972 | 20.93736 | 20.69432 | 18.23601 | 17.56215 |
| ARPC3 | 20.96962 | 20.91267 | 20.91162 | 21.06928 | 20.38574 | 19.50979 | 19.45565 | 18.77683 | 18.60413 | 17.98768 | 19.31501 | 21.37198 | 21.05952 | 20.23714 | 21.28355 | 21.24265 | 20.33285 | 19.65875 |
| ARPC4 | 20.21026 | 20.61395 | 18.21854 | 20.33455 | 20.17384 | 19.40424 | 18.57903 | 18.8357 | 17.03125 | 17.99458 | 19.52649 | 20.54506 | 22.64689 | 18.88046 | 19.81241 | 19.71573 | 18.59654 | 19.93369 |
| ARPC5 | 19.5218 | 18.63326 | 19.01553 | 18.38592 | 19.9106 | 18.834 | 19.73735 | 19.96483 | 17.35893 | 17.67318 | 18.52101 | 20.49054 | 20.96597 | 20.50581 | 18.18214 | 18.29321 | 19.84417 | 18.96852 |
| ARPC5L | 18.50556 | 17.469 | 19.06627 | 19.28317 | 20.39341 | 18.66213 | 19.98989 | 18.82991 | 18.98016 | 18.70827 | 17.65578 | 18.70816 | 20.37019 | 19.92795 | 19.59322 | 19.07602 | 20.02778 | 19.62866 |
| ARPP19 | 17.31242 | 16.43761 | 18.81038 | 17.12243 | 19.94605 | 17.72545 | 18.47578 | 17.9143 | 13.87945 | 15.62068 | 17.33188 | 11.58397 | 20.34775 | 19.76876 | 17.2167 | 17.72521 | 20.00333 | 20.23889 |
| ARRB1 | 17.22905 | 18.63566 | 14.37234 | 18.7713 | 16.61779 | 20.4616 | 18.94196 | 17.33673 | 16.29523 | 16.50974 | 14.37478 | 15.50632 | 18.64363 | 17.98125 | 19.26905 | 17.88444 | 17.15347 | 18.3349 |
| ARRB2 | 17.26288 | 15.15561 | 17.57744 | 13.89021 | 18.57913 | 15.51572 | 17.75221 | 17.76011 | 16.87656 | 15.58379 | 17.2416 | 17.61114 | 13.44083 | 15.45136 | 17.22308 | 17.5065 | 15.78631 | 15.02518 |
| ARRDC1 | 21.96594 | 21.86664 | 21.43479 | 21.64873 | 18.14081 | 20.54714 | 17.37613 | 17.75878 | 19.10121 | 18.03345 | 18.42809 | 19.14468 | 19.12382 | 18.0314 | 22.60623 | 22.23046 | 18.36306 | 20.56415 |
| ARRDC3 | 14.63222 | 14.46486 | 16.74899 | 18.33279 | 19.40787 | 17.63394 | 19.07825 | 16.71975 | 16.09798 | 14.83568 | 13.8712 | 13.23581 | 16.80105 | 16.50651 | 13.68992 | 13.72202 | 13.81103 | 14.97653 |
| ARSA | 16.38742 | 16.31756 | 16.43756 | 17.95424 | 15.85017 | 16.79046 | 17.09108 | 16.47667 | 19.29611 | 18.10205 | 16.3977 | 16.59308 | 15.76104 | 17.94443 | 16.0907 | 16.50392 | 17.67838 | 17.2517 |
| ARVCF | 16.98927 | 17.03493 | 19.18397 | 19.84173 | 16.17437 | 16.77722 | 15.89378 | 16.46338 | 15.70285 | 15.37718 | 16.23603 | 17.24468 | 18.08601 | 18.72123 | 20.45915 | 20.53085 | 16.28267 | 17.99305 |
| ASAH1 | 16.17033 | 15.11548 | 17.87779 | 17.93197 | 17.26197 | 15.57646 | 15.47957 | 18.77744 | 20.58163 | 18.55323 | 18.87761 | 16.0492 | 15.64716 | 17.82679 | 17.96591 | 17.15371 | 15.56522 | 16.77063 |
| ASAP1 | 16.95761 | 17.328 | 18.23762 | 17.31764 | 17.15612 | 15.67119 | 13.75222 | 16.28059 | 15.60542 | 16.30746 | 15.35282 | 14.79516 | 16.6773 | 17.5513 | 15.75213 | 17.11807 | 18.22877 | 19.05616 |
| ASAP2 | 15.28392 | 16.86969 | 14.72145 | 16.2212 | 18.96039 | 18.19666 | 15.23919 | 16.8369 | 16.05478 | 15.58754 | 16.44688 | 16.61265 | 18.8527 | 18.12265 | 16.68403 | 15.51263 | 19.2789 | 19.77908 |
| ASDURF | 15.56668 | 14.06878 | 13.61711 | 12.21883 | 16.20083 | 16.5537 | 13.48877 | 17.61775 | 14.21514 | 15.31407 | 16.38727 | 18.2916 | 16.47155 | 16.5656 | 12.82634 | 12.1335 | 12.41486 | 17.28911 |
| ASL | 16.23099 | 15.50558 | 15.4328 | 16.44573 | 18.12494 | 16.61548 | 16.29251 | 17.03795 | 16.21481 | 15.45481 | 16.97426 | 16.21601 | 15.0827 | 16.76501 | 14.47203 | 16.33466 | 16.41226 | 17.37399 |
| ASNS | 17.20199 | 16.63939 | 18.36387 | 26.20178 | 24.81915 | 17.69341 | 24.06671 | 18.323 | 17.43118 | 17.41282 | 20.2227 | 20.93998 | 25.14253 | 24.96194 | 19.61675 | 19.39616 | 17.66405 | 17.81733 |
| ASPH | 17.05621 | 17.28423 | 15.83872 | 16.32788 | 16.97419 | 16.67645 | 16.50899 | 16.78508 | 20.88087 | 18.99426 | 17.40635 | 14.79303 | 16.91735 | 17.56549 | 15.61544 | 15.87984 | 16.26748 | 17.11347 |
| ASRGL1 | 15.67477 | 16.48189 | 15.53636 | 17.46147 | 16.08754 | 21.69337 | 18.6776 | 16.76202 | 18.53687 | 19.84495 | 17.47518 | 17.43898 | 16.79372 | 16.4673 | 16.70457 | 16.136 | 16.90581 | 16.23813 |
| ASS1 | 17.80257 | 17.457 | 17.2765 | 17.71644 | 17.35694 | 16.55706 | 21.39612 | 20.12709 | 17.88621 | 18.51715 | 16.12662 | 15.76643 | 19.15186 | 19.83237 | 19.59852 | 19.1693 | 16.76279 | 17.74933 |
| ATAD3A | 16.00752 | 15.44607 | 15.74273 | 15.23945 | 19.80455 | 20.14617 | 22.86384 | 16.04098 | 19.59484 | 17.21852 | 16.06776 | 13.66135 | 16.5583 | 14.72276 | 14.20554 | 16.02969 | 18.41021 | 20.32057 |
| ATF6 | 14.46392 | 15.36808 | 18.70132 | 13.74725 | 14.43149 | 16.66895 | 14.09725 | 17.99101 | 17.61989 | 14.78405 | 15.69563 | 12.30536 | 14.20909 | 14.05465 | 15.24716 | 15.25853 | 13.02916 | 15.7942 |
| ATG3 | 17.09734 | 14.75277 | 16.56445 | 16.94811 | 16.85306 | 17.67038 | 17.53347 | 17.9379 | 16.54638 | 17.39856 | 17.19139 | 17.73245 | 20.33949 | 19.2258 | 17.622 | 18.10793 | 19.84268 | 18.02014 |
| ATG5 | 16.63639 | 18.09387 | 14.93903 | 16.52074 | 15.81672 | 16.04647 | 18.51259 | 16.87065 | 17.22526 | 17.73221 | 16.94407 | 19.04116 | 17.81491 | 17.35246 | 17.15253 | 16.78004 | 20.03716 | 16.40035 |
| ATG7 | 13.59529 | 12.97778 | 14.1277 | 16.34374 | 18.03081 | 16.57489 | 15.83256 | 17.42147 | 11.98554 | 16.83091 | 16.12843 | 16.5179 | 18.59152 | 18.5622 | 16.37691 | 15.25702 | 19.29964 | 20.04269 |
| ATIC | 17.60916 | 17.09879 | 18.98224 | 19.56775 | 20.89516 | 18.75485 | 18.79145 | 17.71424 | 18.12515 | 18.83822 | 19.73423 | 20.54902 | 21.69177 | 20.9558 | 19.70843 | 19.39095 | 20.80709 | 20.06816 |
| ATL3 | 17.11056 | 14.56045 | 19.02588 | 17.43975 | 19.32428 | 17.41986 | 19.64164 | 18.81188 | 17.78399 | 18.98374 | 16.70671 | 15.49158 | 18.67982 | 17.87797 | 18.6587 | 18.13646 | 17.5455 | 19.18218 |
| ATOX1 | 19.34434 | 18.67734 | 18.87679 | 19.90545 | 19.12116 | 19.48868 | 16.16313 | 17.1905 | 16.5542 | 16.47414 | 18.21005 | 16.72559 | 21.5381 | 21.1105 | 19.96459 | 20.66877 | 20.23364 | 20.12976 |
| ATP11A | 15.12387 | 16.20796 | 14.36798 | 17.13454 | 18.51741 | 17.82063 | 19.93705 | 18.44391 | 13.75839 | 16.47433 | 17.86081 | 17.32696 | 15.40612 | 16.00756 | 18.5099 | 17.25973 | 14.62825 | 18.06319 |
| ATP11C | 16.33708 | 16.81236 | 18.45969 | 18.57962 | 15.55274 | 15.69315 | 15.98911 | 16.70454 | 17.05054 | 16.55478 | 17.80293 | 18.72655 | 15.69416 | 15.55569 | 19.21388 | 18.55389 | 17.31824 | 14.37344 |
| ATP13A3 | 15.61631 | 15.44292 | 16.18088 | 16.72107 | 17.65429 | 16.80258 | 14.02359 | 15.68171 | 19.63296 | 18.6243 | 16.00486 | 15.91995 | 15.23048 | 15.26611 | 15.18634 | 15.72027 | 14.56163 | 15.64025 |
| ATP1A1 | 20.30803 | 19.78772 | 24.22916 | 23.90121 | 23.4745 | 21.08244 | 21.90687 | 19.72839 | 21.78384 | 21.80813 | 23.45891 | 24.23025 | 24.03299 | 23.89709 | 24.25238 | 24.23354 | 21.80337 | 21.19839 |
| ATP1A3 | 18.27996 | 15.95388 | 16.33611 | 15.83322 | 16.86943 | 16.79116 | 19.09204 | 16.46663 | 17.74575 | 17.25413 | 15.17476 | 13.46246 | 15.95846 | 16.22572 | 13.61107 | 14.97489 | 15.77088 | 14.93861 |
| ATP1B1 | 19.96511 | 19.57295 | 22.90253 | 22.90328 | 22.00096 | 18.93327 | 21.22571 | 20.30199 | 22.20546 | 20.80423 | 21.82235 | 22.71919 | 21.64661 | 21.48706 | 22.93662 | 22.73717 | 20.51145 | 20.86232 |
| ATP1B3 | 17.59688 | 17.99657 | 21.83138 | 20.25498 | 19.02273 | 18.09131 | 18.36053 | 18.40261 | 19.37904 | 17.62886 | 20.4916 | 22.43976 | 18.76481 | 19.15378 | 22.66507 | 21.87901 | 16.80893 | 17.42843 |
| ATP2A1 | 14.96856 | 16.51541 | 14.32172 | 15.57973 | 17.95749 | 14.95465 | 19.51736 | 17.70669 | 14.58307 | 15.39638 | 16.53978 | 17.55543 | 13.93719 | 15.78382 | 15.54875 | 15.58606 | 16.981 | 14.17613 |
| ATP2A2 | 16.29278 | 16.49094 | 16.82852 | 17.59249 | 16.25657 | 16.08518 | 18.99575 | 19.10885 | 16.93651 | 16.57932 | 16.94906 | 17.18493 | 16.25388 | 16.27097 | 16.90084 | 16.40535 | 15.72341 | 16.69999 |
| ATP2A3 | 17.90444 | 17.4202 | 15.74912 | 16.74144 | 16.3122 | 15.25985 | 14.72081 | 17.91055 | 14.25521 | 15.49041 | 17.39588 | 18.51417 | 16.01322 | 15.12658 | 15.15861 | 15.9378 | 17.4038 | 17.92727 |
| ATP2B1 | 19.32032 | 18.66657 | 20.27814 | 20.97009 | 19.18623 | 17.6815 | 18.55171 | 17.50844 | 18.31617 | 18.36007 | 19.73037 | 20.68966 | 21.47739 | 21.63443 | 21.86991 | 21.23174 | 18.89956 | 19.17843 |
| ATP2B2 | 19.22617 | 18.84018 | 21.05288 | 21.2917 | 22.29962 | 19.56182 | 21.32988 | 20.02699 | 19.03345 | 19.16462 | 20.67048 | 21.6549 | 22.79883 | 23.29559 | 21.40959 | 21.87513 | 21.07923 | 20.89344 |
| ATP2B3 | 19.48701 | 13.41372 | 17.96431 | 18.30203 | 16.93458 | 17.79403 | 18.14433 | 16.43037 | 15.74703 | 16.70537 | 17.30817 | 19.7694 | 17.17587 | 17.40743 | 18.63886 | 18.05788 | 16.20877 | 15.13777 |
| ATP2B4 | 17.78815 | 17.42494 | 19.92497 | 19.87206 | 21.04541 | 19.22142 | 20.12648 | 18.5873 | 17.67568 | 18.06078 | 20.55273 | 20.93383 | 18.57411 | 16.31199 | 20.22528 | 20.08894 | 19.44314 | 19.81878 |
| ATP2C1 | 16.1043 | 16.93894 | 16.31075 | 15.71402 | 15.56767 | 17.58805 | 15.31465 | 16.446 | 18.74309 | 17.02607 | 14.77545 | 16.52282 | 14.22652 | 17.1269 | 15.20626 | 16.48317 | 16.81169 | 16.21678 |
| ATP5F1A | 16.23887 | 16.86574 | 16.42744 | 16.96128 | 17.7148 | 17.59175 | 18.36393 | 17.1279 | 21.85862 | 20.02942 | 18.77362 | 16.11972 | 16.74589 | 17.00908 | 16.48779 | 16.24652 | 17.4774 | 16.8161 |
| ATP5F1B | 17.27322 | 27.49449 | 17.86196 | 16.78326 | 17.1207 | 18.38314 | 19.52187 | 17.88002 | 23.31695 | 21.8892 | 18.90246 | 17.38102 | 17.56986 | 18.73799 | 17.08054 | 17.0645 | 17.69477 | 18.86818 |
| ATP5F1C | 19.06673 | 16.99771 | 18.88109 | 17.03807 | 18.18776 | 16.91466 | 17.35135 | 17.20847 | 21.03593 | 19.68634 | 16.4989 | 15.14007 | 16.63434 | 16.34849 | 16.30645 | 18.28334 | 16.93042 | 16.66155 |
| ATP5F1D | 16.15793 | 17.1557 | 17.41821 | 16.20586 | 17.58427 | 15.56348 | 17.96673 | 17.18311 | 21.39579 | 16.77004 | 17.83756 | 18.43651 | 18.39928 | 18.21151 | 17.60968 | 14.04642 | 17.18797 | 13.43778 |
| ATP5ME | 15.28876 | 15.72551 | 17.22581 | 17.37423 | 15.17518 | 16.48481 | 19.35848 | 18.31928 | 20.42484 | 20.44967 | 18.3627 | 11.95741 | 15.92512 | 16.76575 | 14.76211 | 15.76381 | 15.5109 | 14.94248 |
| ATP5MG | 17.64613 | 15.3132 | 16.31129 | 15.91278 | 16.14879 | 16.22437 | 16.66408 | 17.19854 | 19.95243 | 19.49068 | 14.8154 | 13.94057 | 15.9563 | 15.14004 | 15.409 | 15.56087 | 16.49308 | 14.82945 |
| ATP5PB | 17.05824 | 17.05859 | 15.33491 | 17.2064 | 18.05826 | 17.959 | 17.76687 | 16.73119 | 21.58531 | 20.76526 | 16.43566 | 14.52254 | 17.23885 | 17.45457 | 15.70735 | 16.84362 | 17.20236 | 16.05585 |
| ATP5PD | 17.14367 | 16.57872 | 17.39209 | 16.58198 | 16.4098 | 16.51083 | 18.91244 | 17.79468 | 20.87879 | 20.11336 | 16.58827 | 15.01459 | 16.60108 | 17.72962 | 15.36486 | 15.45805 | 17.23617 | 15.86758 |
| ATP5PF | 14.79877 | 13.81392 | 14.1836 | 16.47227 | 14.88956 | 18.04993 | 18.79876 | 15.43389 | 21.24568 | 20.11079 | 15.33687 | 14.91759 | 13.42032 | 17.68283 | 14.419 | 15.23919 | 15.45357 | 16.28204 |
| ATP5PO | 16.3619 | 16.56096 | 16.04718 | 18.05921 | 17.67623 | 17.59024 | 18.64698 | 17.08921 | 21.31477 | 20.10679 | 19.92024 | 15.91189 | 17.08107 | 17.60648 | 16.07064 | 16.79286 | 17.60858 | 15.91348 |
| ATP6AP1 | 17.6449 | 17.84162 | 17.22959 | 15.61306 | 15.41192 | 15.5769 | 16.06133 | 17.36655 | 20.57707 | 19.30096 | 18.44879 | 16.98937 | 14.69985 | 13.79107 | 15.84233 | 17.07831 | 15.65805 | 15.16258 |
| ATP6AP2 | 16.70732 | 15.83417 | 17.19004 | 17.62499 | 21.43787 | 20.50557 | 17.97736 | 17.279 | 20.77344 | 19.20761 | 18.87985 | 17.26185 | 15.03747 | 17.2714 | 17.18101 | 16.62624 | 20.27774 | 21.31955 |
| ATP6V0A1 | 18.04527 | 17.93101 | 18.92523 | 18.03675 | 15.03946 | 17.70995 | 18.46168 | 17.68683 | 20.11721 | 19.71631 | 19.35128 | 18.66227 | 16.78002 | 16.01199 | 18.1908 | 17.72535 | 16.31905 | 15.62337 |
| ATP6V0A2 | 16.65053 | 13.8409 | 16.85836 | 15.17283 | 15.44317 | 15.86146 | 14.68461 | 15.59172 | 17.81157 | 17.8082 | 15.76924 | 17.45284 | 16.00148 | 15.30388 | 15.19833 | 17.17113 | 15.14718 | 13.56655 |
| ATP6V0A4 | 17.12275 | 17.26465 | 14.39532 | 15.81595 | 16.23467 | 16.51426 | 14.99367 | 14.92788 | 13.49573 | 15.34475 | 15.3732 | 18.21765 | 14.81452 | 15.01426 | 17.35168 | 13.10398 | 13.35852 | 15.27825 |
| ATP6V0D1 | 17.27395 | 18.13285 | 18.43501 | 18.72787 | 17.4706 | 16.76112 | 19.39158 | 17.30959 | 20.23459 | 19.15956 | 19.97094 | 19.45079 | 17.22387 | 18.58936 | 19.37436 | 18.30491 | 15.51137 | 16.72223 |
| ATP6V1A | 17.52226 | 17.40171 | 17.20991 | 17.29458 | 17.9242 | 16.72408 | 19.06327 | 17.84037 | 17.25443 | 17.33226 | 17.98113 | 19.0865 | 19.0344 | 19.2063 | 17.81363 | 18.02815 | 18.1081 | 17.48534 |
| ATP6V1B2 | 16.98981 | 16.22517 | 17.47257 | 17.25471 | 18.50986 | 17.41857 | 19.06269 | 18.01744 | 17.82134 | 17.29395 | 18.86137 | 19.60246 | 18.53347 | 18.54567 | 17.2394 | 17.59485 | 16.53269 | 18.64354 |
| ATP6V1C1 | 15.74605 | 16.46107 | 15.86679 | 15.88245 | 16.43766 | 15.18177 | 16.96855 | 16.40078 | 17.28512 | 17.69078 | 15.00923 | 17.35326 | 15.04567 | 16.03796 | 16.0074 | 15.68187 | 17.3609 | 15.19394 |
| ATP6V1D | 16.78791 | 16.30912 | 16.14216 | 17.22224 | 13.69294 | 15.35507 | 17.60513 | 18.4064 | 15.52914 | 14.173 | 18.41511 | 18.80447 | 14.39531 | 17.84073 | 16.67114 | 18.70111 | 18.2112 | 15.96172 |
| ATP6V1E1 | 17.13137 | 14.29632 | 16.01956 | 14.22003 | 19.74194 | 16.67865 | 22.18673 | 15.73184 | 15.8538 | 18.08171 | 19.17691 | 19.28792 | 20.38707 | 20.85489 | 14.85421 | 14.5391 | 18.6156 | 19.13194 |
| ATP6V1E2 | 17.43703 | 15.40335 | 18.38745 | 16.74111 | 17.47421 | 15.01685 | 18.51008 | 19.21093 | 14.49453 | 17.11957 | 16.21352 | 17.77563 | 15.52724 | 17.51455 | 17.18217 | 17.43041 | 14.42787 | 13.57968 |
| ATP6V1F | 16.41955 | 14.50642 | 16.04452 | 16.13647 | 16.58436 | 14.37204 | 16.85163 | 15.90678 | 17.72298 | 14.94192 | 18.75767 | 19.33141 | 17.80102 | 17.99569 | 16.57798 | 16.33304 | 16.08871 | 19.68781 |
| ATP6V1G1 | 16.69226 | 16.07134 | 19.16243 | 16.45001 | 18.66954 | 18.49577 | 21.34575 | 20.46021 | 14.81415 | 17.23544 | 17.20888 | 18.62212 | 20.72542 | 20.596 | 15.46232 | 17.57308 | 18.20375 | 19.0593 |
| ATP6V1H | 15.25001 | 15.48172 | 16.10653 | 16.75573 | 14.48615 | 15.20329 | 16.76835 | 16.55578 | 16.28815 | 14.97156 | 15.79405 | 19.13379 | 16.3655 | 15.76863 | 15.57931 | 16.26041 | 15.96063 | 14.79414 |
| ATP7A | 15.94709 | 16.29697 | 18.66476 | 19.86861 | 15.91197 | 15.65248 | 17.86028 | 18.10411 | 15.25367 | 16.52489 | 17.23694 | 17.79609 | 18.6823 | 19.14849 | 19.362 | 19.86767 | 15.74044 | 16.7479 |
| ATP8A1 | 15.47332 | 15.56541 | 15.53521 | 14.85662 | 13.36976 | 14.73796 | 16.64084 | 14.53418 | 17.12564 | 15.79379 | 14.37238 | 15.16876 | 13.78044 | 15.51021 | 15.57746 | 14.98969 | 15.04627 | 14.52264 |
| ATP8B1 | 16.11345 | 15.08999 | 18.01521 | 16.03113 | 14.48644 | 16.44844 | 16.4718 | 15.3225 | 15.12771 | 14.71928 | 14.67875 | 18.90412 | 14.82282 | 15.52451 | 16.8139 | 16.78494 | 15.23621 | 14.69004 |
| ATP9A | 16.03384 | 16.18198 | 16.2203 | 15.03339 | 14.78256 | 17.73885 | 14.79302 | 16.08746 | 18.36701 | 17.24686 | 17.96903 | 12.51698 | 14.43823 | 15.90409 | 16.72105 | 16.93378 | 15.57475 | 14.19196 |
| ATRN | 15.30574 | 17.17565 | 15.93906 | 17.29807 | 15.1711 | 16.60593 | 16.43954 | 15.18409 | 18.96546 | 18.39007 | 17.31933 | 16.46244 | 15.03637 | 16.88331 | 16.50267 | 15.92511 | 14.65068 | 16.23922 |
| ATXN10 | 17.08466 | 17.20371 | 18.33453 | 17.01038 | 14.88705 | 16.06828 | 16.31768 | 16.76502 | 17.8194 | 17.37069 | 17.55452 | 19.01026 | 16.79086 | 17.015 | 18.77893 | 18.96701 | 15.40633 | 15.57866 |

| | | | | | | | | | | | | | | | | | | |
|---|---|---|---|---|---|---|---|---|---|---|---|---|---|---|---|---|---|---|
| ATXN2L | 19.3381 | 18.09684 | 15.61138 | 17.21209 | 16.98035 | 17.72025 | 16.43464 | 16.43603 | 15.53563 | 16.36989 | 17.06791 | 16.81153 | 17.47876 | 16.38572 | 17.62788 | 16.93774 | 15.90727 | 16.06434 |
| ATXN3 | 19.0592 | 19.70834 | 18.63899 | 17.96308 | 14.11252 | 16.37996 | 18.21821 | 17.49599 | 14.94147 | 16.48501 | 15.12721 | 12.78361 | 18.18363 | 14.68416 | 18.53387 | 18.30903 | 14.96075 | 14.2235 |
| AURKA | 17.14844 | 17.85171 | 17.17677 | 19.02105 | 14.75453 | 19.07284 | 14.82446 | 16.85785 | 17.00751 | 16.968 | 16.81477 | 12.82076 | 18.60255 | 16.29689 | 18.17886 | 17.45993 | 18.20539 | 14.23095 |
| AURKB | 17.2357 | 18.10095 | 18.88537 | 20.19433 | 18.95566 | 17.71942 | 16.64057 | 17.84576 | 15.49568 | 16.36868 | 15.05359 | 14.80182 | 17.628 | 16.85932 | 18.81695 | 17.60493 | 17.52702 | 16.30118 |
| AVIL | 16.35416 | 16.98336 | 16.31913 | 15.90714 | 16.8876 | 16.67938 | 16.73978 | 16.83933 | 14.54288 | 15.77185 | 21.60903 | 22.66283 | 15.85601 | 16.9601 | 16.81387 | 17.12527 | 18.25667 | 16.37992 |
| AXL | 15.48051 | 15.61811 | 14.51426 | 15.78875 | 19.75238 | 18.76692 | 16.5528 | 16.30397 | 15.32277 | 15.00871 | 15.10247 | 12.87678 | 16.45109 | 17.07891 | 14.31547 | 15.91194 | 15.91889 | 15.07118 |
| AZGP1 | 16.12268 | 17.16702 | 19.99437 | 19.10957 | 16.53035 | 17.24958 | 15.48752 | 16.09385 | 14.96909 | 14.70395 | 22.85675 | 17.53255 | 17.04348 | 16.98627 | 19.90518 | 17.72293 | 17.17701 | 16.6689 |
| B3GALT6 | 16.09688 | 15.32098 | 16.31216 | 14.99733 | 17.33792 | 14.87047 | 15.36305 | 18.58838 | 20.09965 | 16.62332 | 18.26718 | 17.22603 | 17.34365 | 14.30464 | 15.323 | 16.28018 | 17.33767 | 14.4574 |
| B3GAT3 | 16.06525 | 16.65041 | 16.55009 | 17.2006 | 16.31637 | 16.21464 | 16.42592 | 16.83932 | 21.29265 | 18.63357 | 17.52533 | 17.75272 | 16.38266 | 16.20363 | 16.28065 | 15.66092 | 18.25701 | 17.52836 |
| B3GLCT | 15.67724 | 15.14312 | 15.7172 | 15.51151 | 16.7003 | 16.37293 | 14.15566 | 15.82423 | 18.926 | 18.00383 | 14.95726 | 13.6633 | 15.05068 | 15.3219 | 16.71429 | 16.44597 | 16.41451 | 16.5035 |
| B3GNT2 | 15.97369 | 16.15317 | 15.54657 | 15.3921 | 14.56511 | 15.65606 | 18.21658 | 15.16953 | 19.65281 | 18.49792 | 17.11345 | 15.75771 | 14.09792 | 14.33624 | 16.08078 | 14.6178 | 15.13799 | 17.50696 |
| B3GNT8 | 15.16553 | 15.80266 | 16.08069 | 15.49506 | 14.59232 | 15.59841 | 16.19365 | 15.79271 | 18.58236 | 17.04269 | 14.12805 | 11.85505 | 14.66149 | 16.61011 | 14.38847 | 15.69485 | 14.5178 | 14.60829 |
| B4GALNT1 | 16.48999 | 17.3667 | 17.17944 | 17.61955 | 16.8094 | 14.50036 | 16.63771 | 15.80732 | 19.89165 | 17.06777 | 17.60991 | 15.60758 | 14.14573 | 15.65672 | 17.33005 | 17.71867 | 14.54998 | 14.87464 |
| B4GALT1 | 14.70119 | 16.26325 | 14.99582 | 14.28306 | 13.25926 | 17.23548 | 15.5971 | 15.87877 | 19.61094 | 18.34207 | 15.02345 | 16.16064 | 13.98482 | 15.70725 | 17.11764 | 13.37666 | 14.41129 | 14.49889 |
| B4GALT3 | 14.40639 | 15.3636 | 16.58959 | 14.84543 | 13.94238 | 15.4593 | 15.49348 | 14.92491 | 19.098 | 15.35331 | 16.9369 | 16.1458 | 18.5823 | 15.67899 | 15.70888 | 14.95809 | 16.24 | 15.14578 |
| B4GALT4 | 16.59599 | 15.6954 | 16.63772 | 15.59403 | 15.34923 | 16.32824 | 16.76036 | 18.72827 | 21.28636 | 20.0598 | 15.35083 | 15.59557 | 15.84434 | 17.02962 | 16.40011 | 17.32364 | 16.24418 | 18.89144 |
| B4GALT5 | 15.06791 | 14.51911 | 19.24694 | 17.14229 | 15.60194 | 16.37215 | 13.27217 | 14.77282 | 20.56674 | 18.33994 | 18.97686 | 19.71473 | 14.97597 | 14.69014 | 14.16612 | 12.75157 | 14.75816 | 13.09281 |
| BABAM2 | 16.40161 | 16.65856 | 15.62492 | 16.29636 | 15.31619 | 16.12749 | 14.91897 | 17.48449 | 14.14681 | 15.545 | 15.38503 | 17.61413 | 15.2752 | 15.80583 | 14.6964 | 15.69987 | 15.86303 | 16.20694 |
| BAG2 | 15.0656 | 17.2616 | 17.67442 | 17.56073 | 19.84221 | 19.7653 | 18.67326 | 17.17874 | 17.33956 | 18.56339 | 19.50965 | 16.81637 | 19.20158 | 19.01436 | 17.68205 | 17.02042 | 20.02191 | 18.46323 |
| BAG3 | 15.31181 | 15.70402 | 15.70904 | 17.56428 | 19.48794 | 17.73579 | 17.03159 | 16.92404 | 15.36108 | 14.54622 | 11.74751 | 11.95952 | 19.05474 | 19.719 | 15.04991 | 11.37342 | 19.9474 | 19.83083 |
| BAG5 | 18.08326 | 18.9441 | 17.84641 | 19.72831 | 19.44911 | 17.93672 | 19.17144 | 16.95058 | 20.14714 | 20.56174 | 16.76057 | 17.27496 | 19.63411 | 19.27833 | 19.04979 | 18.3355 | 16.64705 | 17.17588 |
| BAG6 | 16.93085 | 14.62958 | 15.59582 | 16.88603 | 14.7075 | 16.13521 | 15.87284 | 15.51073 | 15.42646 | 15.5252 | 16.89374 | 19.37583 | 15.21039 | 16.45789 | 16.14901 | 16.67499 | 18.6055 | 17.88366 |
| BAIAP2 | 17.72105 | 19.4452 | 19.57248 | 20.76108 | 21.09407 | 20.9491 | 17.50489 | 18.05102 | 18.4575 | 17.93949 | 17.39548 | 18.29804 | 20.77804 | 21.112 | 20.73409 | 20.87406 | 19.52119 | 19.72509 |
| BAIAP2L1 | 17.79116 | 16.46294 | 18.79703 | 19.17151 | 21.58105 | 20.62533 | 15.64008 | 18.09324 | 18.30121 | 17.96779 | 16.92428 | 16.55318 | 20.88302 | 21.08586 | 19.00748 | 18.23352 | 18.27938 | 17.12879 |
| BANF1 | 21.34919 | 21.38936 | 19.98111 | 19.85294 | 22.39007 | 21.71505 | 21.34997 | 20.53402 | 19.807 | 20.41189 | 20.72964 | 23.45392 | 21.78867 | 21.89087 | 20.03281 | 20.05091 | 21.00399 | 22.71598 |
| BASP1 | 16.72005 | 17.89613 | 16.7156 | 16.49349 | 24.92249 | 23.72044 | 22.21655 | 23.92438 | 23.03803 | 22.7595 | 16.0068 | 13.976 | 19.50777 | 20.33462 | 16.89343 | 15.80217 | 24.16889 | 24.39139 |
| BAX | 25.05121 | 21.93189 | 18.31763 | 17.43762 | 18.5786 | 16.93353 | 18.35688 | 18.19087 | 16.68759 | 16.59191 | 17.02922 | 18.49471 | 15.10395 | 17.19735 | 15.77268 | 19.46165 | 17.03585 | 17.99843 |
| BAZ1B | 18.40594 | 18.99202 | 15.50618 | 16.11317 | 15.25076 | 16.89825 | 17.35544 | 18.77717 | 21.08585 | 22.43436 | 25.60154 | 15.78447 | 15.08914 | 15.88406 | 15.73727 | 15.42833 | 15.17069 | 13.87233 |
| BAZ2A | 18.56939 | 17.35527 | 16.7534 | 15.87918 | 16.95309 | 15.12612 | 17.80352 | 15.6983 | 14.22082 | 16.59986 | 15.98547 | 13.90878 | 16.25812 | 15.84208 | 12.84686 | 14.49707 | 14.9457 | 15.21913 |
| BCAM | 21.13533 | 20.91807 | 15.56819 | 15.58421 | 17.5028 | 17.8144 | 15.69816 | 16.74283 | 17.88746 | 15.63748 | 16.61537 | 16.27873 | 15.31128 | 16.11812 | 16.1813 | 17.4479 | 15.86344 | 17.05824 |
| BCAP31 | 15.9285 | 15.56643 | 18.57202 | 17.24925 | 17.53576 | 16.33205 | 19.55092 | 17.85262 | 18.87685 | 17.38321 | 16.54649 | 16.57274 | 17.07789 | 19.14264 | 18.59173 | 18.09663 | 16.71442 | 18.26373 |
| BCAT1 | 15.18295 | 15.37414 | 16.72953 | 16.60367 | 18.04596 | 16.97187 | 16.34013 | 15.82924 | 14.37409 | 15.88917 | 21.10549 | 21.53562 | 14.82976 | 14.18585 | 16.20647 | 15.34133 | 15.9805 | 15.30606 |
| BCCIP | 17.98158 | 16.76037 | 16.5154 | 17.42871 | 16.41188 | 16.93004 | 16.08628 | 15.367 | 15.57075 | 15.59763 | 19.70919 | 19.57313 | 17.25271 | 15.45391 | 16.88885 | 18.16471 | 15.86158 | 14.67627 |
| BCKDHA | 15.24742 | 14.32139 | 15.26111 | 14.11256 | 13.59688 | 14.23239 | 15.34624 | 14.40179 | 18.81768 | 13.13194 | 18.82288 | 13.92135 | 13.49537 | 15.74937 | 14.21312 | 15.61942 | 15.38444 | 16.54654 |
| BCLAF1 | 15.07357 | 20.75996 | 17.55036 | 17.55115 | 16.225 | 18.80167 | 19.44405 | 16.83569 | 15.48806 | 16.08451 | 19.67146 | 17.9336 | 19.67789 | 17.51233 | 19.30183 | 18.33867 | 15.98283 | 16.19663 |
| BCS1L | 16.1921 | 15.12826 | 16.32841 | 14.49313 | 15.5477 | 14.52983 | 16.59443 | 16.18818 | 16.36981 | 14.59174 | 14.181 | 12.98396 | 15.32145 | 15.38326 | 14.16814 | 14.61589 | 14.95385 | 14.46137 |
| BDH2 | 15.95479 | 14.49067 | 16.75096 | 15.16662 | 16.68197 | 14.77408 | 14.35662 | 14.41458 | 15.99716 | 15.09117 | 12.62783 | 12.44235 | 16.41054 | 14.53844 | 14.56685 | 15.16192 | 14.36943 | 14.36044 |
| BECN1 | 15.29716 | 17.5033 | 15.8388 | 16.11237 | 16.9793 | 16.28508 | 16.38678 | 15.68999 | 16.32472 | 15.35675 | 16.7401 | 17.20891 | 17.34239 | 15.13255 | 16.08289 | 17.0355 | 16.45438 | 17.36273 |
| BICD2 | 17.01997 | 16.93055 | 15.07826 | 16.6319 | 18.46075 | 16.78812 | 14.07704 | 17.01064 | 14.36863 | 14.88065 | 19.41334 | 13.68435 | 19.4379 | 15.27568 | 13.95292 | 13.20531 | 17.19451 | 16.49761 |
| BIN1 | 16.22117 | 17.50134 | 17.01066 | 18.67154 | 16.83794 | 17.91072 | 17.10622 | 15.96132 | 15.49225 | 16.90486 | 19.4988 | 13.08575 | 19.36602 | 17.39171 | 17.92251 | 17.85684 | 18.8736 | 17.93199 |
| BIN3 | 16.69695 | 17.34432 | 17.22629 | 17.54404 | 17.85183 | 16.66452 | 12.9603 | 17.11256 | 13.26379 | 14.71788 | 13.30535 | 11.21011 | 16.01491 | 14.38338 | 16.43545 | 17.0183 | 17.22791 | 13.39786 |
| BIRC2 | 16.13995 | 16.91469 | 16.06179 | 15.51095 | 20.69869 | 19.98272 | 16.12666 | 17.16646 | 16.15699 | 16.91167 | 15.26791 | 14.21763 | 17.41588 | 15.41814 | 15.7444 | 14.44489 | 16.83663 | 17.38152 |
| BIRC5 | 15.20949 | 14.58273 | 17.72658 | 19.59128 | 20.74024 | 18.67519 | 17.23401 | 17.42221 | 16.74263 | 17.66455 | 14.28632 | 14.74746 | 19.77447 | 19.71432 | 17.48628 | 16.96978 | 20.37531 | 18.73191 |
| BIRC6 | 15.8541 | 15.86692 | 16.21068 | 15.62973 | 15.96273 | 17.0843 | 16.13476 | 15.13392 | 13.0568 | 14.93527 | 15.82898 | 17.19655 | 15.76896 | 15.16088 | 16.22148 | 14.94374 | 16.33678 | 14.92847 |
| BLMH | 16.68573 | 15.7649 | 18.81868 | 15.74976 | 18.26735 | 15.80296 | 19.68983 | 19.44714 | 19.22737 | 19.46816 | 18.65329 | 19.81345 | 18.93152 | 19.29892 | 16.9092 | 16.08326 | 17.64014 | 18.47925 |
| BLOC1S1 | 15.36049 | 14.86931 | 15.35628 | 14.26716 | 17.94167 | 15.97131 | 13.97217 | 17.35587 | 15.66018 | 16.0606 | 15.07848 | 18.65098 | 18.14386 | 17.63246 | 14.97342 | 14.10775 | 12.77714 | 16.73582 |
| BLOC1S5 | 16.56686 | 12.52466 | 15.77699 | 15.72971 | 17.48417 | 15.35227 | 14.88409 | 16.13517 | 14.05573 | 16.40068 | 15.05507 | 16.46101 | 17.66297 | 15.96961 | 14.89423 | 11.94376 | 13.91677 | 16.84215 |
| BLVRA | 18.42125 | 18.93816 | 17.78366 | 18.97764 | 19.98636 | 16.57947 | 19.19231 | 19.29589 | 16.54573 | 16.80853 | 16.88867 | 16.80232 | 20.17914 | 20.29051 | 18.7875 | 18.81015 | 20.28206 | 19.60423 |
| BLVRB | 21.06087 | 19.898 | 19.93346 | 21.55257 | 17.85151 | 14.81186 | 16.11578 | 15.95419 | 16.88887 | 15.30551 | 14.32763 | 14.26141 | 19.37861 | 18.50028 | 21.84572 | 21.19955 | 18.90668 | 17.59585 |
| BMP3 | 16.11676 | 15.20967 | 14.60806 | 14.96865 | 18.82915 | 19.33365 | 19.98974 | 20.96901 | 19.0432 | 18.24925 | 17.93958 | 19.61408 | 18.62609 | 15.49921 | 18.00066 | 16.49997 | 17.06493 | 18.67438 |
| BMP8B | 15.70368 | 17.27883 | 14.12708 | 14.06771 | 14.52595 | 15.3948 | 15.47195 | 16.24459 | 18.21415 | 17.36717 | 14.31772 | 14.49561 | 15.86155 | 15.64091 | 15.62878 | 14.57602 | 14.03351 | 14.56484 |
| BMPR2 | 15.39088 | 16.30504 | 16.40218 | 16.82214 | 19.89251 | 17.14295 | 20.17583 | 16.95642 | 17.20696 | 18.00138 | 14.76544 | 15.33545 | 16.33336 | 18.35211 | 16.94748 | 17.41182 | 18.06809 | 19.59263 |
| BMX | 15.88152 | 16.42895 | 17.52243 | 16.30162 | 14.34702 | 16.23906 | 15.89037 | 17.11469 | 14.94984 | 15.08744 | 19.7585 | 20.34763 | 16.38287 | 17.48591 | 16.00771 | 16.04628 | 15.3411 | 15.10277 |
| BOLA2 | 20.37244 | 20.53289 | 19.44479 | 19.89511 | 20.6847 | 18.44137 | 18.17735 | 17.72222 | 18.298 | 17.22658 | 20.99668 | 21.17575 | 20.38614 | 19.51546 | 19.96129 | 19.69577 | 19.29318 | 19.06473 |
| BORCS7 | 16.09232 | 17.92234 | 16.61925 | 12.54349 | 18.96695 | 20.97221 | 14.4986 | 16.74855 | 15.8735 | 15.83237 | 18.87048 | 17.98609 | 17.79185 | 18.53549 | 17.02165 | 14.26989 | 17.59441 | 12.89262 |
| BPGM | 15.43791 | 16.15592 | 16.74823 | 16.33118 | 14.20388 | 14.7254 | 14.59594 | 15.42371 | 11.17851 | 14.98634 | 17.76959 | 16.03468 | 15.11814 | 14.23879 | 14.94009 | 14.32997 | 15.79225 | 15.52841 |
| BPHL | 15.09051 | 15.90548 | 15.25782 | 14.38781 | 15.68553 | 15.86047 | 14.71564 | 15.43414 | 18.45901 | 16.33002 | 14.75722 | 11.78316 | 15.74319 | 15.19922 | 14.33839 | 13.82629 | 17.43517 | 17.07672 |
| BPNT1 | 16.67619 | 16.70556 | 17.97074 | 18.20031 | 18.52888 | 17.64782 | 18.54045 | 17.86851 | 17.43055 | 18.13263 | 19.68549 | 19.70938 | 19.87314 | 18.08266 | 18.49219 | 18.12678 | 18.41938 | 19.3366 |
| BPNT2 | 16.35219 | 16.41408 | 17.44033 | 15.91213 | 16.11263 | 17.21608 | 16.77674 | 17.39493 | 22.98482 | 22.21968 | 18.74038 | 18.11423 | 15.94885 | 16.67422 | 16.26607 | 17.53605 | 16.54961 | 16.49496 |
| BRCC3 | 15.46728 | 15.95422 | 15.81785 | 15.60278 | 15.44337 | 14.64977 | 14.87169 | 15.71029 | 15.41187 | 14.06722 | 15.54348 | 17.56942 | 15.97626 | 14.89226 | 16.14679 | 14.64207 | 14.48889 | 16.57326 |
| BRI3BP | 17.31968 | 16.42024 | 13.31414 | 13.28928 | 15.99349 | 15.42021 | 19.72746 | 16.92316 | 20.31438 | 19.22334 | 15.4521 | 14.63871 | 14.23964 | 13.2064 | 16.94201 | 14.55771 | 15.48445 | 14.29759 |
| BRIX1 | 15.77199 | 15.29721 | 15.62615 | 15.53874 | 15.17572 | 18.56435 | 18.09154 | 17.77668 | 13.70791 | 15.59353 | 17.80285 | 19.01884 | 14.78354 | 14.16165 | 14.98423 | 14.8855 | 13.83254 | 15.54031 |
| BRK1 | 16.72454 | 17.33206 | 17.92762 | 18.82237 | 21.43013 | 19.27587 | 20.23564 | 17.62491 | 16.94756 | 18.23371 | 16.4835 | 17.94967 | 20.75951 | 20.39427 | 18.66543 | 18.87711 | 22.55173 | 20.88792 |
| BROX | 22.18493 | 22.68387 | 20.95932 | 21.76485 | 20.51821 | 20.1212 | 18.004 | 18.30696 | 19.76116 | 20.46977 | 18.57901 | 19.99777 | 20.03365 | 19.67726 | 21.88182 | 21.3642 | 18.34013 | 18.49824 |
| BSCL2 | 14.58174 | 15.48221 | 16.4555 | 13.7452 | 18.36534 | 16.19017 | 16.74433 | 14.21449 | 18.03642 | 14.7013 | 14.47517 | 15.82997 | 13.2919 | 12.77078 | 14.5565 | 16.95372 | 16.55946 | 14.3274 |
| BSG | 20.23949 | 20.05919 | 23.03288 | 22.44262 | 22.36399 | 21.55512 | 20.72537 | 19.6592 | 21.11797 | 19.04291 | 19.35661 | 22.0962 | 22.27854 | 22.1178 | 23.34511 | 23.45446 | 22.24843 | 23.57458 |
| BST2 | 18.49444 | 15.96071 | 17.14156 | 18.83505 | 14.87624 | 16.29285 | 17.75099 | 14.24449 | 16.84726 | 15.1304 | 17.05489 | 16.99191 | 20.17689 | 21.09454 | 22.50102 | 22.71858 | 15.71456 | 16.82933 |
| BTBD17 | 16.49347 | 16.49006 | 16.78602 | 18.17103 | 18.19903 | 16.59333 | 19.65107 | 19.09523 | 19.43273 | 17.6968 | 14.39064 | 12.92096 | 18.08529 | 19.29006 | 20.20034 | 19.84549 | 17.89153 | 17.70369 |
| BTF3 | 19.42969 | 19.0589 | 18.06145 | 18.60452 | 20.21968 | 20.74813 | 19.3734 | 18.35123 | 15.73527 | 17.68927 | 20.21773 | 21.3404 | 20.93524 | 19.78145 | 17.76553 | 18.27982 | 17.58197 | 19.68249 |
| BTF3L4 | 18.70866 | 17.74275 | 19.07617 | 19.80132 | 20.23509 | 15.29894 | 14.39416 | 16.98133 | 17.36087 | 18.23981 | 15.43361 | 19.30061 | 20.20228 | 14.71361 | 18.84539 | 18.99709 | 14.5774 | 16.87644 |
| BTN2A1 | 16.69867 | 15.3874 | 17.88169 | 18.04593 | 17.37026 | 17.42937 | 16.2194 | 16.08917 | 16.62005 | 15.08367 | 17.73624 | 19.14425 | 18.05231 | 15.50155 | 18.19636 | 18.45751 | 17.2218 | 17.91842 |
| BTN2A2 | 15.68322 | 15.89203 | 14.91299 | 14.92205 | 17.45473 | 15.24588 | 14.65777 | 13.7548 | 17.91282 | 14.57864 | 14.0003 | 18.53973 | 19.61684 | 14.59232 | 16.55297 | 17.75939 | 16.27085 | 15.39901 |
| BTN3A1 | 16.29165 | 15.06273 | 17.96399 | 15.60841 | 16.11385 | 17.31597 | 15.43824 | 17.26715 | 17.81403 | 15.13632 | 16.66895 | 18.08916 | 14.1825 | 14.3533 | 16.04089 | 16.08585 | 16.21719 | 17.32821 |
| BTN3A2 | 16.4464 | 16.84269 | 14.72009 | 14.85951 | 17.38434 | 17.04122 | 16.40435 | 18.20893 | 16.36189 | 17.18406 | 16.57686 | 17.81322 | 16.83128 | 16.61462 | 15.55459 | 16.23902 | 17.85643 | 18.76559 |
| BTN3A3 | 15.66599 | 15.15319 | 15.67347 | 14.76599 | 16.48039 | 15.67207 | 16.34553 | 17.45681 | 14.33187 | 14.9555 | 16.87721 | 17.18213 | 15.14514 | 15.28288 | 14.67825 | 13.98673 | 16.10149 | 17.53134 |
| BUB3 | 17.16366 | 16.95914 | 16.51387 | 15.97418 | 18.03688 | 17.37143 | 17.71219 | 19.2427 | 15.21057 | 17.09345 | 17.72127 | 20.67094 | 16.68394 | 17.86129 | 15.81146 | 15.55556 | 17.09555 | 17.42858 |
| BUD31 | 15.95559 | 16.25022 | 17.4867 | 17.87519 | 14.06674 | 17.71422 | 19.27185 | 15.42504 | 14.06076 | 15.00577 | 18.41364 | 14.03572 | 17.20155 | 17.6791 | 13.70793 | 15.81588 | 16.61489 | 15.71278 |
| BYSL | 14.49003 | 14.15379 | 15.93139 | 15.26437 | 13.11954 | 15.29207 | 14.65325 | 15.30291 | 12.34189 | 16.58421 | 13.62496 | 17.2539 | 15.2399 | 13.4574 | 14.20596 | 14.8817 | 12.73079 | 14.72544 |
| BZW1 | 16.55169 | 18.63343 | 17.96934 | 17.51186 | 19.33032 | 17.76398 | 18.12799 | 17.30183 | 15.7153 | 15.86076 | 18.31321 | 19.73874 | 18.44224 | 18.36007 | 16.96898 | 18.0524 | 18.71116 | 20.20208 |
| BZW2 | 16.86608 | 17.60063 | 16.72461 | 16.4206 | 17.35271 | 17.40808 | 17.43209 | 17.02495 | 15.59647 | 17.35403 | 18.3145 | 19.44047 | 17.84545 | 15.50545 | 16.07104 | 14.61044 | 14.92001 | 16.70898 |
| C11orf52 | 22.4063 | 22.11835 | 20.58805 | 20.42938 | 15.85058 | 15.76194 | 17.76172 | 15.34792 | 19.90863 | 19.26694 | 14.82696 | 16.21131 | 21.35374 | 20.70773 | 21.18987 | 21.05504 | 15.80019 | 19.24159 |
| C11orf54 | 18.38349 | 16.22259 | 16.19681 | 16.08293 | 16.88762 | 18.01489 | 16.29204 | 17.56082 | 16.4175 | 19.78316 | 16.46979 | 17.02023 | 19.10275 | 17.80035 | 15.44691 | 15.64169 | 20.64078 | 20.81197 |
| C12orf57 | 19.05661 | 19.37901 | 16.63928 | 19.51639 | 19.61313 | 18.71055 | 19.98544 | 20.01314 | 17.33214 | 18.0018 | 17.22934 | 17.76983 | 20.34214 | 20.63589 | 19.68183 | 18.74163 | 20.4136 | 19.24516 |
| C1GALT1 | 15.99159 | 16.39304 | 15.9309 | 15.51643 | 19.75499 | 19.60495 | 16.80437 | 18.03694 | 20.27391 | 17.35894 | 16.29201 | 18.43635 | 15.85913 | 16.86368 | 16.06758 | 17.0822 | 18.30727 | 18.16205 |
| C1GALT1C | 15.38595 | 16.42856 | 16.29377 | 17.27433 | 18.77778 | 17.11637 | 17.39763 | 18.24272 | 19.88452 | 19.506 | 18.32658 | 16.63251 | 18.03663 | 17.70138 | 15.45199 | 14.48772 | 19.09408 | 16.23195 |
| C1orf198 | 16.56837 | 18.38443 | 17.67705 | 17.0087 | 19.56427 | 18.72176 | 20.39784 | 19.11539 | 17.75808 | 18.47144 | 19.86068 | 19.15329 | 18.6368 | 19.10239 | 18.65666 | 17.10336 | 18.16057 | 19.3955 |

| | | | | | | | | | | | | | | | | | | |
|---|---|---|---|---|---|---|---|---|---|---|---|---|---|---|---|---|---|---|
| C1orf21 | 16.26006 | 15.85121 | 15.61456 | 18.83909 | 16.32179 | 15.58335 | 16.53239 | 16.88104 | 12.94052 | 15.51558 | 15.75207 | 14.05978 | 20.2284 | 20.02832 | 17.92598 | 17.48656 | 15.65836 | 16.71856 |
| C1orf226 | 15.68419 | 17.04343 | 14.91063 | 18.16784 | 19.5578 | 21.65821 | 18.94298 | 16.00069 | 15.18677 | 15.67536 | 14.97894 | 13.57091 | 20.68669 | 21.10955 | 13.23354 | 15.21831 | 13.9688 | 18.28096 |
| C1orf43 | 15.93263 | 14.89744 | 15.88427 | 13.40338 | 10.72488 | 17.05958 | 14.2236 | 15.55276 | 12.45594 | 14.31963 | 16.55202 | 18.63685 | 15.58188 | 14.43853 | 15.64634 | 14.73706 | 13.95529 | 14.60276 |
| C1QBP | 19.16196 | 20.5304 | 16.28073 | 16.73412 | 18.89249 | 15.77621 | 19.08608 | 17.09776 | 21.3261 | 18.48154 | 20.14505 | 17.18005 | 19.00904 | 18.81395 | 18.25087 | 18.38954 | 15.41748 | 17.32052 |
| C1QL1 | 14.20215 | 18.64042 | 14.21827 | 15.16501 | 14.52194 | 15.9915 | 14.84446 | 16.49389 | 20.53883 | 19.07238 | 14.5329 | 13.7515 | 13.93808 | 16.03875 | 16.16875 | 14.36859 | 13.05303 | 13.85672 |
| C1R | 16.1745 | 13.75555 | 14.12242 | 14.81189 | 17.85573 | 16.39754 | 14.99109 | 15.59541 | 18.24745 | 16.88027 | 15.22225 | 13.64813 | 15.61319 | 16.34297 | 14.21654 | 17.14063 | 14.9216 | 16.88142 |
| C20orf27 | 18.79366 | 14.85939 | 18.93107 | 17.71436 | 18.68344 | 17.27009 | 16.02883 | 15.6944 | 13.68023 | 15.67984 | 16.6521 | 18.38723 | 17.5343 | 15.53271 | 17.62531 | 17.48917 | 18.41009 | 18.80688 |
| C2CD5 | 16.24343 | 16.63332 | 16.44969 | 16.72674 | 17.66943 | 16.3047 | 17.13213 | 17.3816 | 17.46864 | 16.6426 | 16.25874 | 16.84073 | 16.84276 | 16.52753 | 16.38209 | 16.97493 | 18.31113 | 18.05839 |
| C2orf72 | 17.64713 | 17.36224 | 16.89161 | 15.98767 | 16.45658 | 16.35044 | 14.78874 | 16.4661 | 14.24727 | 14.8094 | 16.76898 | 14.74208 | 13.51017 | 14.28499 | 19.02099 | 18.83355 | 14.94574 | 16.41149 |
| C3 | 14.81418 | 16.54114 | 18.25662 | 19.27571 | 14.58617 | 21.14426 | 17.14632 | 17.06296 | 15.13731 | 16.91116 | 17.89337 | 11.73609 | 14.66806 | 19.49602 | 16.46985 | 14.05119 | 15.78215 | 18.86712 |
| C4A | 19.01813 | 17.98732 | 22.891 | 25.89551 | 20.98341 | 21.27 | 18.49575 | 14.81332 | 11.59039 | 12.55723 | 14.0662 | 11.10968 | 22.64441 | 22.83835 | 25.69591 | 24.16772 | 19.45056 | 21.96688 |
| C6orf89 | 16.50421 | 15.24009 | 16.64815 | 16.23575 | 14.54312 | 13.49191 | 13.68282 | 13.89336 | 19.39901 | 17.65616 | 14.09749 | 12.72646 | 14.41014 | 14.93678 | 14.70869 | 13.3301 | 15.13647 | 13.5455 |
| C8orf82 | 16.49318 | 17.58336 | 16.36359 | 15.05539 | 13.62188 | 17.13547 | 16.07667 | 15.16783 | 19.02234 | 18.30422 | 14.83458 | 12.86002 | 14.78829 | 18.89424 | 14.15244 | 15.34878 | 14.87331 | 17.8939 |
| C9orf64 | 17.03504 | 17.5946 | 14.86804 | 16.79742 | 21.01281 | 13.80345 | 18.3591 | 19.12932 | 16.971 | 18.20294 | 18.15435 | 19.71896 | 19.27075 | 15.6715 | 17.5289 | 16.07099 | 17.55912 | 15.12467 |
| CA12 | 20.33983 | 19.58903 | 15.12256 | 17.51622 | 15.65787 | 18.07283 | 15.35836 | 15.46748 | 14.97547 | 14.48888 | 15.63193 | 16.63981 | 14.53983 | 15.24104 | 18.29928 | 17.40102 | 15.91901 | 15.62823 |
| CA14 | 17.09414 | 15.74036 | 15.54043 | 15.70522 | 15.69966 | 16.6041 | 15.61868 | 16.39734 | 13.66259 | 14.03316 | 18.15244 | 20.32521 | 13.22325 | 15.12285 | 15.71108 | 15.03255 | 16.16183 | 14.85483 |
| CAB39 | 19.01573 | 19.49443 | 19.99355 | 19.78181 | 18.81891 | 18.78888 | 17.43385 | 16.89851 | 17.68085 | 17.10316 | 19.48057 | 20.5752 | 16.83907 | 17.91433 | 20.44363 | 20.60229 | 18.90879 | 18.4296 |
| CAB39L | 15.58558 | 15.59686 | 17.67519 | 18.1939 | 14.0509 | 17.5983 | 15.06585 | 15.54227 | 15.19601 | 14.2084 | 17.92647 | 18.7786 | 17.68591 | 16.97522 | 18.09987 | 18.80042 | 14.53254 | 14.17789 |
| CACNA2D1 | 20.43786 | 20.22382 | 16.04549 | 16.65764 | 15.40208 | 17.88431 | 21.69454 | 20.35394 | 20.17702 | 19.06067 | 16.35406 | 14.36899 | 18.29014 | 18.12543 | 16.26914 | 15.87987 | 16.7473 | 15.73943 |
| CACNA2D2 | 17.56925 | 16.76841 | 15.61462 | 16.58079 | 17.30412 | 20.29731 | 19.33179 | 17.61662 | 20.91332 | 20.23346 | 18.38884 | 17.47732 | 13.35699 | 18.58332 | 16.32735 | 16.20558 | 16.2289 | 16.08374 |
| CACYBP | 17.42312 | 16.68826 | 18.31394 | 18.27532 | 20.17246 | 19.13805 | 20.54978 | 20.34455 | 19.46836 | 20.0533 | 21.33523 | 21.69268 | 21.78566 | 21.1805 | 17.08283 | 18.43383 | 20.03097 | 18.56333 |
| CAD | 16.59442 | 16.96371 | 17.38772 | 19.15863 | 15.6159 | 15.33622 | 15.82721 | 16.0114 | 16.0678 | 14.98075 | 19.71273 | 18.82593 | 16.74724 | 19.13693 | 16.76357 | 16.80978 | 15.57499 | 16.24983 |
| CADM1 | 14.30362 | 15.51451 | 15.67336 | 16.22069 | 20.86698 | 18.36838 | 20.78074 | 19.25986 | 19.12065 | 17.7804 | 20.41546 | 17.69578 | 15.69321 | 14.54899 | 15.51664 | 17.53439 | 14.06357 | 14.30813 |
| CADM4 | 15.77115 | 16.1825 | 16.29677 | 17.32922 | 14.6746 | 14.51506 | 16.04702 | 17.56824 | 19.09885 | 16.90419 | 14.9398 | 19.34069 | 14.6335 | 15.60799 | 19.22106 | 18.12978 | 15.98612 | 15.89959 |
| CALB2 | 16.58244 | 16.69907 | 16.79614 | 17.80029 | 14.77454 | 17.11391 | 20.24833 | 19.83877 | 14.60861 | 15.89875 | 15.84758 | 16.61785 | 16.22443 | 16.49022 | 17.65675 | 17.49938 | 16.22031 | 14.89593 |
| CALCA | 16.97173 | 16.37464 | 15.21257 | 17.96919 | 15.16264 | 17.72942 | 16.55488 | 16.12898 | 22.00636 | 20.55994 | 15.80503 | 15.57175 | 17.14996 | 15.6025 | 16.04294 | 16.71343 | 17.74982 | 16.15976 |
| CALCB | 16.75974 | 16.30836 | 16.38447 | 16.58993 | 17.43464 | 17.58498 | 16.86609 | 17.8979 | 18.91089 | 19.22325 | 15.57161 | 14.5331 | 16.12104 | 16.38746 | 14.31394 | 16.84678 | 18.0555 | 17.92971 |
| CALCOCO1 | 17.97041 | 18.39562 | 15.75915 | 17.03483 | 18.66268 | 17.62192 | 20.93303 | 18.09069 | 18.10123 | 17.8872 | 15.71163 | 14.66853 | 18.44676 | 17.93155 | 16.77999 | 16.64841 | 19.88346 | 21.14434 |
| CALD1 | 17.60819 | 16.70584 | 18.96518 | 19.53203 | 18.13515 | 17.71134 | 18.0546 | 16.10446 | 13.79721 | 16.9022 | 18.36172 | 16.98798 | 20.07427 | 20.07863 | 19.42467 | 18.6402 | 20.31352 | 18.31163 |
| CALM2 | 22.70562 | 22.55806 | 21.98183 | 22.08682 | 22.19738 | 22.25367 | 20.12348 | 19.75632 | 20.92885 | 20.67386 | 22.32387 | 22.95654 | 21.16803 | 20.82039 | 23.40675 | 22.85464 | 21.91772 | 21.65151 |
| CALML3 | 15.47041 | 14.81466 | 16.01242 | 15.28488 | 13.38927 | 16.29867 | 16.86603 | 17.71647 | 18.34194 | 15.98249 | 18.29707 | 13.2748 | 13.19446 | 14.18295 | 15.48073 | 15.78539 | 14.2765 | 14.2715 |
| CALR | 18.14489 | 17.65862 | 18.45391 | 17.27554 | 18.37949 | 18.07771 | 22.25817 | 21.09086 | 25.32607 | 25.18958 | 23.09926 | 19.51136 | 18.4671 | 19.12509 | 19.46765 | 18.58879 | 17.79134 | 19.49543 |
| CALU | 16.31352 | 16.54459 | 19.39249 | 17.70482 | 19.06541 | 19.14541 | 18.7344 | 18.0039 | 22.42411 | 20.90771 | 21.51123 | 17.76989 | 17.51597 | 18.45047 | 17.69097 | 17.03716 | 16.63146 | 18.51138 |
| CAMK2D | 18.66249 | 17.61359 | 16.93442 | 17.5413 | 18.14499 | 17.32348 | 16.65725 | 17.25275 | 17.46406 | 17.82975 | 18.38153 | 19.44438 | 18.68396 | 17.98342 | 17.41726 | 17.5137 | 19.00841 | 17.72226 |
| CAMK2G | 15.58497 | 14.61613 | 15.57904 | 16.32048 | 18.06606 | 16.00513 | 15.97214 | 15.98764 | 17.66914 | 15.63055 | 16.34682 | 15.76949 | 12.97561 | 16.24059 | 14.53177 | 15.4604 | 21.02927 | 20.79612 |
| CAMP | 14.92807 | 15.14087 | 16.75755 | 17.69587 | 15.23037 | 16.44639 | 14.43807 | 16.70549 | 20.28475 | 17.79466 | 15.5258 | 14.87383 | 18.01034 | 15.0644 | 17.12494 | 16.99609 | 16.09337 | 15.60197 |
| CAMSAP3 | 16.66262 | 15.70472 | 16.38146 | 17.75757 | 17.92281 | 15.72021 | 17.91208 | 18.62454 | 15.01405 | 16.77562 | 15.23122 | 14.32226 | 19.56273 | 18.25481 | 16.61359 | 16.23307 | 17.5932 | 17.09729 |
| CAND1 | 17.21255 | 16.83935 | 17.88355 | 17.99431 | 17.89143 | 17.81153 | 18.13089 | 17.49725 | 18.5417 | 18.31705 | 18.96676 | 20.36387 | 18.15348 | 18.04364 | 18.33266 | 18.88412 | 18.30412 | 18.40068 |
| CAND2 | 12.86494 | 16.99471 | 15.55258 | 17.82936 | 20.82185 | 19.324 | 20.74097 | 20.62175 | 17.51329 | 19.59951 | 18.90865 | 18.92553 | 21.50452 | 21.42729 | 15.17771 | 14.5523 | 21.72867 | 21.17229 |
| CANT1 | 15.73381 | 16.51206 | 19.35686 | 17.11811 | 15.13231 | 18.56348 | 16.6106 | 17.12648 | 22.05481 | 20.37745 | 18.31658 | 15.32835 | 16.74488 | 17.38398 | 16.61322 | 16.58253 | 16.79421 | 17.61781 |
| CANX | 19.48937 | 18.75208 | 18.23661 | 18.93871 | 18.54644 | 18.04915 | 19.53668 | 17.8861 | 25.80146 | 23.77592 | 22.14601 | 19.35161 | 16.5608 | 17.53946 | 19.1748 | 19.12223 | 16.68741 | 17.81409 |
| CAP1 | 18.38003 | 16.51174 | 17.62963 | 19.05383 | 22.41945 | 21.46114 | 21.99635 | 20.73807 | 19.25079 | 20.2149 | 19.63391 | 20.31288 | 21.30082 | 21.73336 | 19.05572 | 18.53241 | 21.73585 | 21.90089 |
| CAP2 | 18.1175 | 17.2153 | 18.92591 | 17.14848 | 18.54454 | 16.93428 | 17.90692 | 18.47895 | 14.558 | 15.90749 | 17.72143 | 18.3531 | 17.52692 | 16.22424 | 19.19011 | 18.14877 | 17.93805 | 18.25044 |
| CAPG | 18.11726 | 15.91506 | 16.91426 | 16.09637 | 19.97201 | 18.6604 | 16.11201 | 16.32966 | 14.12209 | 15.2514 | 16.46944 | 15.49557 | 17.60751 | 14.77276 | 16.25263 | 14.73468 | 20.75701 | 19.36971 |
| CAPN1 | 24.99442 | 24.91785 | 22.42561 | 24.02539 | 17.91861 | 17.58138 | 19.17384 | 17.10131 | 17.67376 | 17.20336 | 19.30914 | 19.73941 | 21.48484 | 20.69462 | 22.9648 | 23.45806 | 18.13078 | 18.31276 |
| CAPN15 | 19.4655 | 19.30342 | 15.63956 | 14.56285 | 16.11346 | 13.35964 | 17.02173 | 15.04824 | 13.05333 | 16.02459 | 14.3442 | 12.86153 | 13.63395 | 16.48735 | 14.13666 | 16.33555 | 13.91523 | 15.88197 |
| CAPN2 | 17.90669 | 15.70442 | 15.84063 | 16.53631 | 20.33787 | 16.57662 | 20.51613 | 16.11485 | 17.56695 | 17.51285 | 15.6303 | 15.06019 | 16.04028 | 17.62296 | 17.12873 | 15.23612 | 20.4694 | 19.9826 |
| CAPN5 | 18.22312 | 19.2919 | 18.7516 | 19.31816 | 18.16811 | 17.71514 | 17.11624 | 16.62516 | 18.34132 | 17.34372 | 16.3165 | 17.72218 | 19.26861 | 20.24297 | 19.81413 | 19.75442 | 18.3622 | 19.28326 |
| CAPN7 | 17.05658 | 16.42858 | 17.05518 | 16.95602 | 16.69817 | 15.58577 | 16.0726 | 16.91411 | 17.62353 | 18.76266 | 16.97424 | 19.55229 | 16.13442 | 16.43316 | 15.39895 | 16.00343 | 16.87651 | 15.64179 |
| CAPNS1 | 19.07536 | 17.56446 | 15.99731 | 17.45864 | 17.42155 | 15.28807 | 15.12101 | 16.28755 | 17.92859 | 16.52121 | 19.04729 | 18.59382 | 16.07267 | 15.37748 | 18.19571 | 17.79925 | 18.6348 | 17.03784 |
| CAPRIN1 | 16.71228 | 17.517 | 16.09118 | 15.44363 | 18.45583 | 16.43417 | 16.01467 | 17.03596 | 16.79922 | 16.16765 | 19.84296 | 20.5077 | 17.74066 | 15.59919 | 15.7919 | 16.86677 | 16.97124 | 15.75818 |
| CAPZA1 | 20.37648 | 20.35712 | 19.95119 | 19.33912 | 21.75205 | 19.90149 | 19.20205 | 18.58551 | 18.88839 | 19.74222 | 20.22063 | 20.82612 | 21.63444 | 20.97983 | 20.51287 | 20.45494 | 20.91819 | 20.60705 |
| CAPZA2 | 21.66528 | 21.38706 | 19.71437 | 19.99009 | 20.16737 | 17.886 | 19.21941 | 17.874 | 19.05868 | 17.51381 | 20.60679 | 21.43282 | 20.00638 | 19.96483 | 21.22047 | 20.93106 | 20.15365 | 19.73554 |
| CAPZB | 20.7571 | 21.15799 | 19.18342 | 19.58743 | 21.28028 | 19.63953 | 19.10125 | 18.61715 | 18.94445 | 19.09218 | 19.15831 | 21.18014 | 20.6818 | 20.48202 | 20.45694 | 19.91543 | 20.3585 | 20.55397 |
| CARHSP1 | 17.93648 | 17.12076 | 17.26763 | 18.23562 | 20.06883 | 17.10869 | 21.00002 | 20.51219 | 19.34191 | 20.67667 | 18.43993 | 18.3362 | 18.83896 | 20.58095 | 17.35331 | 15.10424 | 22.08898 | 21.30599 |
| CARM1 | 18.893 | 19.19094 | 16.95465 | 19.11248 | 16.03273 | 18.40292 | 14.8288 | 17.19823 | 13.44932 | 13.34867 | 14.97933 | 19.48298 | 16.04462 | 15.5405 | 17.21065 | 17.56506 | 16.17002 | 15.16384 |
| CARMIL1 | 16.32437 | 16.28966 | 17.53409 | 16.44408 | 19.50139 | 16.82479 | 17.38585 | 16.98846 | 14.04224 | 15.18807 | 17.14308 | 18.63075 | 17.91351 | 17.02151 | 16.29976 | 16.15451 | 18.14105 | 18.12398 |
| CARS1 | 15.41762 | 16.21975 | 15.70258 | 15.12881 | 16.00667 | 16.029 | 16.41312 | 16.1641 | 15.73648 | 16.43019 | 17.76884 | 19.75851 | 18.17088 | 16.12516 | 17.04335 | 16.37309 | 16.43047 | 16.04034 |
| CASC3 | 19.96583 | 20.02259 | 18.79391 | 18.62493 | 15.36995 | 16.93257 | 15.83062 | 15.2744 | 14.04424 | 14.8331 | 15.20499 | 14.33022 | 16.85305 | 17.46861 | 17.60583 | 19.352 | 15.89295 | 14.65284 |
| CASD1 | 15.33873 | 15.71128 | 18.36685 | 14.48484 | 13.66982 | 16.38967 | 15.63188 | 16.14124 | 18.44172 | 17.81788 | 14.42879 | 14.08783 | 15.21978 | 16.37278 | 15.98763 | 18.92307 | 13.12121 | 15.16963 |
| CASK | 17.56034 | 17.32131 | 19.77443 | 19.98252 | 19.95458 | 19.12605 | 18.42871 | 17.9966 | 19.24567 | 19.79596 | 17.0761 | 17.82662 | 20.05984 | 20.50216 | 20.23797 | 19.92645 | 19.29149 | 19.83151 |
| CASKIN2 | 16.1206 | 15.35853 | 16.46903 | 17.90997 | 16.16508 | 14.41368 | 15.16779 | 17.01375 | 12.33578 | 14.41709 | 16.30003 | 17.07782 | 14.14749 | 14.64804 | 18.06063 | 17.55377 | 17.31731 | 16.64068 |
| CASP3 | 16.39672 | 15.20745 | 17.19079 | 16.49704 | 15.36211 | 16.96147 | 15.50542 | 16.6978 | 16.47256 | 15.50901 | 19.09808 | 19.67321 | 14.29997 | 14.36002 | 15.17483 | 15.85314 | 17.09032 | 16.81261 |
| CASP6 | 16.68897 | 14.84074 | 14.669 | 15.8248 | 13.74107 | 16.58749 | 14.91047 | 17.21704 | 15.5007 | 15.00132 | 17.94999 | 19.76101 | 17.01286 | 15.41642 | 14.29212 | 16.3048 | 16.62519 | 14.08095 |
| CASP8AP2 | 21.88019 | 20.41189 | 20.29239 | 19.84293 | 19.16543 | 14.93898 | 18.35476 | 18.9088 | 15.83605 | 16.8053 | 14.87074 | 17.45183 | 19.19891 | 16.73119 | 20.80539 | 21.92771 | 19.27397 | 19.97235 |
| CAST | 17.69589 | 15.9187 | 16.21416 | 15.91602 | 18.32157 | 17.51109 | 15.53937 | 15.41029 | 16.0151 | 17.97754 | 17.79467 | 17.74497 | 16.83826 | 16.8402 | 16.80975 | 16.19438 | 18.48338 | 18.79515 |
| CAV1 | 16.38158 | 15.61462 | 17.3372 | 17.94591 | 22.38197 | 21.21655 | 19.72774 | 18.93551 | 15.97632 | 16.77362 | 17.03921 | 17.49894 | 16.21209 | 15.48779 | 15.78621 | 16.44834 | 19.74551 | 20.22758 |
| CAVIN1 | 15.63841 | 16.00026 | 15.49524 | 15.02266 | 22.8515 | 20.79735 | 15.37389 | 17.37157 | 14.18586 | 16.30802 | 15.75712 | 14.05781 | 13.40231 | 17.52764 | 15.68028 | 14.9826 | 16.00793 | 14.60044 |
| CAVIN2 | 17.27329 | 15.52163 | 15.85524 | 16.161 | 20.84842 | 18.20743 | 18.23477 | 16.53149 | 17.4573 | 17.71227 | 16.69542 | 16.15754 | 16.73371 | 16.97061 | 15.92378 | 17.16188 | 18.14202 | 14.97273 |
| CBFB | 14.03124 | 16.37568 | 16.8653 | 17.32249 | 15.73058 | 14.32118 | 15.91613 | 17.30758 | 13.86705 | 13.50951 | 17.35363 | 18.2639 | 17.70218 | 17.31503 | 17.47425 | 16.56847 | 15.56541 | 17.64495 |
| CBR1 | 17.15836 | 16.55226 | 18.90972 | 18.84554 | 21.38535 | 20.28414 | 18.63357 | 17.6418 | 18.59375 | 18.61841 | 17.32355 | 19.31366 | 19.34569 | 19.23565 | 19.18831 | 18.97343 | 21.19616 | 21.6098 |
| CBR3 | 18.38119 | 18.75515 | 20.07521 | 20.56623 | 21.18806 | 21.17273 | 18.84439 | 19.08639 | 18.68522 | 18.39074 | 17.39688 | 20.53156 | 19.73957 | 19.2273 | 21.00107 | 20.71112 | 20.76751 | 21.51882 |
| CBR4 | 19.15462 | 13.22381 | 14.97238 | 15.82165 | 14.88932 | 15.71698 | 14.79956 | 16.77292 | 18.75176 | 14.08613 | 13.73708 | 12.72781 | 14.09675 | 14.79751 | 13.95675 | 18.53234 | 14.46706 | 17.37517 |
| CBS | 16.5156 | 19.26774 | 15.57403 | 16.19722 | 17.10604 | 15.60996 | 17.32876 | 17.20778 | 16.72969 | 16.9606 | 18.72775 | 20.26375 | 16.73157 | 16.03284 | 16.00006 | 16.45768 | 16.35385 | 16.04503 |
| CBWD2 | 13.66868 | 13.51344 | 13.25095 | 14.0033 | 14.80261 | 15.75868 | 15.30931 | 15.53253 | 14.38605 | 16.9725 | 14.66829 | 17.55744 | 19.39835 | 18.87446 | 14.90205 | 17.4614 | 16.61966 | 15.08625 |
| CBX3 | 17.97416 | 16.61122 | 15.27855 | 14.30877 | 16.44774 | 15.47644 | 15.78709 | 18.8727 | 16.41099 | 16.94782 | 18.42635 | 19.57887 | 17.01946 | 16.59778 | 14.5063 | 16.48727 | 17.25253 | 14.96117 |
| CBX5 | 18.03852 | 17.97489 | 16.08056 | 16.22308 | 13.56524 | 15.3745 | 16.74196 | 18.46881 | 17.28208 | 17.41446 | 18.63492 | 17.65752 | 16.08079 | 16.30564 | 14.46002 | 14.87625 | 15.15909 | 16.20655 |
| CC2D1A | 21.23001 | 20.76344 | 20.25766 | 18.78921 | 20.37885 | 20.97157 | 18.2016 | 17.62812 | 19.24871 | 19.43478 | 19.25574 | 18.67671 | 19.52287 | 17.0181 | 20.51563 | 20.23211 | 16.43453 | 16.52184 |
| CC2D1B | 17.88999 | 18.0219 | 16.19128 | 16.51127 | 16.30427 | 16.61222 | 16.51646 | 15.84679 | 17.8871 | 16.68845 | 14.77985 | 15.92231 | 17.70445 | 16.17526 | 17.23146 | 16.91965 | 17.08293 | 15.35439 |
| CCAR1 | 15.44686 | 15.7765 | 14.457 | 15.42655 | 15.91275 | 15.82483 | 17.28939 | 17.70674 | 14.29328 | 14.57827 | 18.0089 | 18.42939 | 14.35863 | 15.16961 | 16.76996 | 13.69272 | 16.68843 | 15.69858 |
| CCAR2 | 16.30383 | 15.69308 | 15.63682 | 16.50315 | 14.01003 | 16.27377 | 18.93694 | 17.76122 | 14.53648 | 14.57369 | 16.43642 | 16.50097 | 18.16165 | 17.06173 | 16.4972 | 15.66239 | 16.39553 | 16.49172 |
| CCDC124 | 19.30379 | 20.72271 | 20.73115 | 19.80286 | 15.18805 | 16.30436 | 15.36953 | 14.91274 | 16.84164 | 16.47321 | 17.306 | 15.76589 | 19.39638 | 18.81536 | 19.21935 | 20.40459 | 14.45369 | 15.32348 |
| CCDC126 | 16.74044 | 14.41443 | 13.68994 | 16.80259 | 15.13513 | 13.95638 | 16.31844 | 16.81589 | 19.71162 | 15.86624 | 16.34229 | 16.42494 | 13.62923 | 16.02733 | 15.73221 | 14.44394 | 15.05622 | 15.51517 |
| CCDC134 | 15.78702 | 15.00677 | 15.92341 | 15.70415 | 14.06019 | 17.82519 | 20.23227 | 15.36926 | 20.44892 | 20.95984 | 18.65361 | 15.51022 | 14.4382 | 15.21938 | 15.17978 | 14.88726 | 14.9193 | 14.10317 |
| CCDC22 | 16.89244 | 18.50134 | 15.72917 | 16.87246 | 18.93952 | 16.88301 | 17.124 | 16.33336 | 15.96774 | 15.76648 | 16.40373 | 17.52347 | 17.2441 | 17.13451 | 14.41574 | 16.73757 | 18.72396 | 17.98084 |
| CCDC28A | 18.97347 | 19.21016 | 14.13272 | 17.10707 | 16.08813 | 16.30304 | 18.09992 | 16.6013 | 13.97196 | 13.19635 | 15.75057 | 17.02465 | 15.43204 | 16.78997 | 12.31628 | 14.17985 | 17.54148 | 16.74822 |

| | | | | | | | | | | | | | | | | | | |
|---|---|---|---|---|---|---|---|---|---|---|---|---|---|---|---|---|---|---|
| CCDC47 | 14.44659 | 13.52501 | 17.12385 | 17.18664 | 14.98358 | 17.37956 | 18.89857 | 15.0054 | 16.14938 | 16.41957 | 15.5069 | 17.49767 | 18.25066 | 19.28424 | 16.09023 | 16.49433 | 15.32837 | 14.28412 |
| CCDC50 | 22.57647 | 22.89155 | 22.95065 | 22.9635 | 15.92586 | 17.54295 | 16.62689 | 16.73133 | 17.23365 | 16.83262 | 19.28678 | 15.76731 | 21.61531 | 19.91734 | 23.44622 | 23.08184 | 17.17403 | 17.5021 |
| CCDC6 | 17.33937 | 16.05314 | 17.37728 | 16.642 | 16.30701 | 18.01608 | 16.43766 | 16.89812 | 19.04659 | 17.09198 | 18.54721 | 19.65454 | 19.19935 | 16.61909 | 17.11069 | 16.42844 | 17.65595 | 18.36206 |
| CCDC88A | 14.48306 | 16.02775 | 15.57227 | 14.93221 | 17.82719 | 17.46447 | 16.38329 | 17.80852 | 19.60254 | 18.86743 | 21.00319 | 16.5397 | 16.42722 | 15.26015 | 16.96939 | 16.69948 | 18.22137 | 17.54383 |
| CCER2 | 14.90051 | 15.12699 | 15.61536 | 14.95374 | 16.37252 | 15.31108 | 20.07783 | 22.59195 | 15.74034 | 15.42571 | 15.61548 | 12.06632 | 15.25909 | 15.81564 | 15.69791 | 15.4956 | 15.94441 | 16.31861 |
| CCM2 | 14.14632 | 16.82163 | 16.59718 | 17.41441 | 14.46893 | 16.14888 | 15.60189 | 14.36708 | 17.12485 | 17.42929 | 16.19532 | 14.45874 | 18.60307 | 19.95958 | 16.95776 | 18.18698 | 20.05369 | 14.94502 |
| CCNB1 | 18.53304 | 14.31903 | 16.59857 | 18.55253 | 18.33821 | 18.09745 | 17.5066 | 17.14465 | 18.0752 | 18.25959 | 18.24347 | 18.14156 | 18.63451 | 19.06717 | 17.04296 | 17.76325 | 17.88595 | 17.5734 |
| CCNB2 | 15.46747 | 16.14789 | 13.68865 | 14.25288 | 19.3432 | 18.26584 | 15.87452 | 18.39951 | 14.61276 | 18.22231 | 16.48673 | 15.06751 | 16.53875 | 14.55986 | 17.38647 | 15.90523 | 12.47935 | 17.84687 |
| CCNDBP1 | 19.98838 | 20.28064 | 19.12628 | 19.18414 | 15.80209 | 15.8845 | 15.53329 | 15.3722 | 13.74767 | 14.85733 | 14.49223 | 14.53427 | 17.84831 | 16.55569 | 20.23575 | 21.01085 | 15.21547 | 16.12771 |
| CCNY | 17.8664 | 17.65349 | 17.54825 | 18.16005 | 16.4704 | 16.0323 | 15.86603 | 16.4297 | 12.20387 | 14.49697 | 17.6558 | 19.54 | 13.87365 | 13.79149 | 20.71207 | 20.30764 | 17.74755 | 18.50812 |
| CCPG1 | 15.40193 | 15.70085 | 16.87395 | 15.92433 | 16.09653 | 17.49663 | 14.76424 | 17.16158 | 18.28248 | 18.04443 | 13.17578 | 15.24911 | 15.12473 | 15.81934 | 13.77956 | 15.32344 | 14.408 | 15.21734 |
| CCS | 17.84208 | 18.10817 | 16.69738 | 17.96042 | 18.98733 | 17.75574 | 14.21627 | 17.03967 | 17.47185 | 17.08344 | 18.84544 | 19.18794 | 16.88237 | 15.95377 | 18.58203 | 15.46189 | 18.24712 | 18.14027 |
| CCT2 | 18.61407 | 17.94559 | 18.58765 | 18.83977 | 20.63775 | 19.21011 | 20.90592 | 19.95871 | 19.35964 | 19.83018 | 21.05499 | 21.63291 | 22.17497 | 21.63628 | 19.03987 | 19.22244 | 21.00587 | 20.96131 |
| CCT3 | 17.84579 | 18.46591 | 18.68622 | 18.75169 | 19.48202 | 19.30872 | 19.84419 | 18.29592 | 19.09696 | 19.50963 | 21.62727 | 22.0772 | 21.22715 | 20.54314 | 18.89716 | 19.07392 | 19.79092 | 19.87557 |
| CCT4 | 17.07006 | 16.70014 | 18.1333 | 18.05776 | 20.17074 | 18.66223 | 20.16515 | 19.88587 | 17.83389 | 19.68798 | 20.3362 | 21.03069 | 21.60785 | 21.34 | 17.47998 | 17.94985 | 20.26894 | 20.30738 |
| CCT5 | 17.11991 | 17.20142 | 17.80907 | 18.6838 | 19.7908 | 18.20643 | 19.50699 | 18.48732 | 18.23534 | 19.1599 | 20.29449 | 21.02674 | 21.37308 | 20.61312 | 17.94084 | 18.11488 | 19.84669 | 20.40542 |
| CCT6A | 17.46767 | 18.22199 | 18.77055 | 19.20969 | 20.88725 | 20.9646 | 20.7177 | 20.00515 | 19.35546 | 20.16026 | 21.19826 | 21.77092 | 22.40644 | 21.85336 | 18.86088 | 18.96294 | 21.13552 | 21.19136 |
| CCT7 | 17.58968 | 17.64226 | 17.94293 | 18.24343 | 19.2764 | 17.86614 | 19.62129 | 18.85133 | 17.9896 | 19.15523 | 20.17042 | 21.10911 | 21.46102 | 21.08454 | 18.21468 | 18.14707 | 19.85992 | 19.08143 |
| CCT8 | 18.18844 | 18.00479 | 18.84954 | 19.62167 | 21.02285 | 19.56563 | 20.81212 | 19.43884 | 19.08603 | 19.8936 | 20.99436 | 21.5972 | 22.38665 | 21.54123 | 19.28568 | 19.30033 | 21.34622 | 21.14445 |
| CCZ1 | 15.07707 | 16.02394 | 14.2801 | 16.56082 | 18.00735 | 16.77779 | 16.14855 | 16.7759 | 13.46442 | 15.54299 | 16.60708 | 16.59109 | 16.22825 | 16.18291 | 16.07769 | 14.92317 | 14.9823 | 17.22582 |
| CD109 | 18.15134 | 18.41409 | 17.33451 | 16.23287 | 16.13374 | 16.17121 | 15.70566 | 17.69464 | 20.13182 | 20.22969 | 16.94039 | 14.76187 | 15.34437 | 16.04869 | 16.90738 | 16.95864 | 20.05382 | 19.06037 |
| CD151 | 23.41788 | 22.6194 | 21.01404 | 21.03888 | 18.3669 | 18.34781 | 16.88573 | 18.18942 | 18.14496 | 15.18849 | 15.97639 | 19.51703 | 16.46956 | 16.63121 | 22.25056 | 22.00353 | 18.4258 | 19.05859 |
| CD276 | 19.30585 | 18.91286 | 21.52197 | 20.51085 | 19.79178 | 18.626 | 16.30461 | 17.53198 | 17.16138 | 17.93017 | 15.1345 | 18.46012 | 19.49348 | 20.4103 | 20.74665 | 21.21004 | 18.29261 | 18.33509 |
| CD2AP | 22.7419 | 22.84126 | 22.68608 | 22.35555 | 19.24585 | 18.67159 | 18.00504 | 17.25339 | 16.78623 | 16.78791 | 17.78962 | 18.12482 | 21.68133 | 20.76325 | 23.18866 | 22.54342 | 20.30226 | 18.42607 |
| CD320 | 17.464 | 17.53961 | 19.45185 | 19.05167 | 18.05056 | 15.42267 | 15.73094 | 14.89416 | 19.79401 | 17.89689 | 17.05094 | 15.98093 | 15.67562 | 17.55814 | 18.5093 | 16.73073 | 15.98478 | 18.68318 |
| CD44 | 22.20508 | 21.82313 | 19.51622 | 18.90877 | 21.76664 | 18.49564 | 16.84208 | 18.39527 | 15.8112 | 17.52738 | 15.67098 | 15.60574 | 17.26148 | 16.27499 | 18.82931 | 18.5304 | 19.52086 | 22.19166 |
| CD46 | 22.41321 | 21.98427 | 21.56685 | 21.10415 | 19.92181 | 18.24848 | 16.76154 | 16.79273 | 19.47978 | 18.99573 | 18.93311 | 20.96813 | 19.46089 | 19.32465 | 21.58643 | 21.17313 | 17.95524 | 19.47111 |
| CD47 | 21.66966 | 21.18699 | 22.62024 | 22.4097 | 21.17265 | 19.63205 | 19.3772 | 20.63425 | 23.82918 | 22.63826 | 24.32937 | 24.61296 | 21.68005 | 20.52895 | 22.34232 | 22.87076 | 18.85398 | 21.55165 |
| CD55 | 21.26106 | 19.73626 | 18.38062 | 19.44666 | 19.7259 | 19.9941 | 17.21769 | 17.85924 | 18.09933 | 16.6864 | 19.67207 | 20.94623 | 17.38415 | 18.26102 | 19.43645 | 18.73377 | 21.60656 | 22.2456 |
| CD58 | 14.27159 | 15.33297 | 18.28002 | 13.84373 | 18.8577 | 16.99934 | 14.61812 | 16.66463 | 15.88168 | 16.90981 | 16.96385 | 12.86664 | 20.11174 | 15.45318 | 15.14551 | 14.34989 | 18.04157 | 20.65871 |
| CD59 | 24.03713 | 21.96466 | 23.74405 | 23.74082 | 24.59558 | 23.36365 | 21.96986 | 21.73398 | 21.4494 | 21.72038 | 19.09901 | 21.45528 | 24.43026 | 24.59827 | 23.992 | 23.74999 | 25.76374 | 27.16068 |
| CD63 | 22.70977 | 21.60729 | 21.12438 | 20.46712 | 14.67083 | 16.97099 | 17.94344 | 16.59496 | 22.18989 | 21.65979 | 17.93739 | 19.74373 | 16.57171 | 16.84049 | 22.84995 | 22.08095 | 16.95191 | 16.367 |
| CD74 | 15.56474 | 16.01175 | 14.38531 | 13.72213 | 15.28475 | 14.4786 | 15.84763 | 16.6948 | 20.72394 | 17.7075 | 14.53559 | 14.38701 | 15.17937 | 15.40242 | 15.43026 | 14.0435 | 14.43497 | 15.23475 |
| CD81 | 27.08378 | 25.38701 | 26.51996 | 25.23323 | 22.41446 | 21.88853 | 19.88884 | 17.14131 | 21.34255 | 21.18393 | 22.27787 | 24.07433 | 25.15834 | 24.93309 | 27.31343 | 27.05519 | 20.49056 | 22.73765 |
| CD82 | 19.40346 | 18.54296 | 20.70473 | 16.09547 | 15.92824 | 15.07183 | 16.61235 | 17.31675 | 14.04683 | 17.12654 | 14.80166 | 18.81546 | 14.67909 | 14.75456 | 20.79754 | 18.51772 | 16.91665 | 14.8127 |
| CD9 | 27.30663 | 27.16075 | 28.35134 | 27.55653 | 24.0468 | 23.14546 | 20.44376 | 19.09919 | 24.77417 | 23.57503 | 23.02246 | 23.7164 | 25.3961 | 24.54877 | 28.77251 | 28.38891 | 21.59182 | 23.11325 |
| CD99 | 21.00382 | 21.2878 | 23.13591 | 22.40423 | 22.85974 | 21.76271 | 19.48404 | 19.56484 | 16.35997 | 16.59291 | 16.92352 | 15.50384 | 21.67626 | 20.94698 | 23.74583 | 23.62585 | 22.63044 | 24.31848 |
| CD99L2 | 19.74322 | 19.66537 | 18.47048 | 17.21091 | 21.64256 | 14.62716 | 20.9002 | 19.5202 | 17.28462 | 16.89956 | 16.07682 | 15.05111 | 13.93245 | 14.78388 | 17.23112 | 18.88686 | 21.25075 | 22.43317 |
| CDA | 14.08024 | 14.31838 | 15.79646 | 18.24045 | 20.82662 | 15.94438 | 15.12447 | 16.6021 | 14.10602 | 13.65322 | 17.46199 | 13.72988 | 14.8609 | 16.13109 | 15.3451 | 16.12651 | 20.26841 | 15.7642 |
| CDC16 | 16.55793 | 15.23406 | 15.19122 | 13.96467 | 13.54673 | 17.76416 | 15.44937 | 17.04032 | 18.10226 | 14.23749 | 15.03225 | 16.20842 | 15.24201 | 14.94436 | 13.08174 | 16.5179 | 18.62157 | 16.5894 |
| CDC23 | 15.8345 | 14.60744 | 15.74708 | 15.399 | 14.55243 | 15.49607 | 15.29466 | 15.46525 | 14.90163 | 15.8536 | 12.62394 | 17.9515 | 13.06835 | 13.17715 | 14.73952 | 16.5265 | 14.52025 | 15.62875 |
| CDC37 | 16.35892 | 17.35278 | 18.07607 | 18.62388 | 20.13893 | 18.66472 | 18.45657 | 18.56632 | 17.96966 | 17.99577 | 18.83072 | 19.09855 | 19.11638 | 19.1008 | 19.6936 | 18.42311 | 20.17592 | 20.83929 |
| CDC42 | 22.34245 | 22.83961 | 22.5138 | 21.90554 | 23.21582 | 21.52684 | 22.72706 | 22.21502 | 19.95433 | 21.34606 | 21.22915 | 21.28244 | 22.73921 | 23.37052 | 22.27776 | 21.83391 | 23.3512 | 23.55028 |
| CDC42BPA | 16.38865 | 16.74967 | 16.4569 | 18.161 | 18.01636 | 17.81777 | 16.80203 | 16.79614 | 17.34477 | 17.52753 | 16.88115 | 17.18837 | 18.66633 | 19.03964 | 18.04634 | 18.42661 | 18.92289 | 20.10362 |
| CDC42BPE | 14.8593 | 14.86693 | 15.70394 | 16.15664 | 17.99079 | 15.72354 | 14.63899 | 17.60031 | 16.30334 | 16.3471 | 18.00239 | 16.13863 | 14.29053 | 16.85114 | 17.15012 | 16.9151 | 17.50789 | 19.13683 |
| CDC42EP2 | 18.30697 | 17.30627 | 15.90776 | 14.20966 | 16.58072 | 15.91759 | 11.80331 | 16.6941 | 13.26218 | 15.98524 | 16.63268 | 10.88341 | 12.19017 | 13.62179 | 15.90936 | 15.56619 | 20.65166 | 20.00051 |
| CDC42SE2 | 17.38871 | 17.41985 | 17.21085 | 20.58631 | 21.33655 | 17.43462 | 16.40077 | 17.69655 | 15.3104 | 18.45657 | 16.62839 | 16.2312 | 20.1173 | 18.83925 | 22.38214 | 21.43508 | 18.76875 | 20.4064 |
| CDCA3 | 19.20461 | 18.69779 | 19.57619 | 19.10921 | 21.03144 | 19.14743 | 17.53933 | 17.50758 | 17.01745 | 18.21682 | 17.12409 | 16.93648 | 20.66128 | 19.88151 | 18.22557 | 18.05939 | 18.78608 | 19.47048 |
| CDCA8 | 16.68917 | 18.97894 | 19.50421 | 19.96273 | 20.30754 | 18.18155 | 16.25619 | 17.81228 | 16.75761 | 17.46626 | 16.6826 | 15.77485 | 19.64401 | 18.88773 | 17.71648 | 16.86973 | 17.61054 | 14.99355 |
| CDCP1 | 18.71125 | 17.65683 | 20.67387 | 20.01047 | 21.07224 | 19.52084 | 16.50923 | 18.8265 | 15.79541 | 17.74004 | 19.43746 | 20.77092 | 19.76364 | 20.00366 | 21.15022 | 20.64804 | 22.29916 | 21.89207 |
| CDH1 | 20.17021 | 19.69395 | 21.09595 | 22.05065 | 22.13684 | 21.03004 | 16.02931 | 15.53826 | 19.23401 | 17.28553 | 20.78676 | 19.53204 | 22.557 | 22.60493 | 22.57728 | 22.38732 | 18.62393 | 16.64299 |
| CDH11 | 14.65043 | 15.02407 | 14.34091 | 13.73309 | 14.41647 | 15.24724 | 12.45808 | 15.29944 | 14.08813 | 14.21804 | 15.67516 | 14.66581 | 12.75691 | 16.53362 | 13.87336 | 16.71189 | 22.42794 | 22.01635 |
| CDH2 | 16.19869 | 16.62718 | 15.77104 | 15.91025 | 15.24735 | 15.88154 | 16.32236 | 17.06578 | 18.99509 | 17.82806 | 19.84249 | 18.48349 | 15.20624 | 16.40075 | 16.09152 | 16.23129 | 21.85058 | 21.35647 |
| CDH3 | 15.31439 | 20.98754 | 13.22052 | 15.41204 | 23.05414 | 21.90742 | 15.52863 | 15.67791 | 15.85207 | 15.79379 | 12.58677 | 10.77041 | 14.78927 | 16.62543 | 14.94606 | 16.17068 | 17.6022 | 18.04727 |
| CDK1 | 18.62156 | 19.36379 | 19.49943 | 20.31275 | 20.56651 | 19.43913 | 19.34048 | 19.43257 | 18.55632 | 19.32926 | 17.75532 | 20.11351 | 21.06592 | 20.49269 | 20.2485 | 19.84829 | 20.29038 | 20.63028 |
| CDK2 | 18.59174 | 18.45564 | 15.8347 | 16.27242 | 17.27195 | 18.72812 | 16.33164 | 15.15263 | 17.53643 | 16.50245 | 14.59031 | 16.52818 | 15.49702 | 15.34672 | 15.73161 | 16.56666 | 16.50703 | 13.90915 |
| CDK4 | 17.93139 | 14.94541 | 16.47448 | 14.47243 | 13.6607 | 16.56754 | 16.59187 | 16.26085 | 17.30698 | 16.91326 | 12.69469 | 14.77209 | 16.1133 | 17.22264 | 16.77822 | 17.34621 | 17.46722 | 17.71148 |
| CDK5 | 18.20279 | 16.77729 | 18.40026 | 17.41302 | 18.36148 | 17.81095 | 18.88923 | 18.65198 | 16.1102 | 16.79366 | 17.05411 | 17.61141 | 18.40209 | 18.70946 | 18.45071 | 18.5086 | 19.61214 | 19.33791 |
| CDKN2C | 13.60596 | 13.39922 | 16.56743 | 15.43144 | 13.89827 | 16.95363 | 16.35764 | 15.07214 | 17.38282 | 17.03821 | 14.88027 | 12.81976 | 14.79095 | 16.01713 | 15.29818 | 17.02487 | 16.6337 | 13.41023 |
| CDS2 | 15.64882 | 14.04839 | 13.52972 | 15.48263 | 13.03483 | 13.99559 | 15.5683 | 14.0528 | 19.03599 | 17.95372 | 13.37385 | 11.24364 | 13.41948 | 16.14329 | 14.67305 | 14.73436 | 14.20409 | 15.59161 |
| CDV3 | 17.87858 | 17.19209 | 17.07862 | 17.66738 | 19.03663 | 15.12606 | 15.48577 | 16.01789 | 16.43388 | 15.87992 | 18.386 | 18.03883 | 19.66847 | 18.35906 | 17.05908 | 17.6285 | 17.77613 | 18.07222 |
| CEACAM1 | 15.83405 | 15.27728 | 15.92677 | 15.96848 | 16.60921 | 16.96437 | 16.7052 | 16.33521 | 18.38584 | 18.87658 | 21.341 | 20.75153 | 15.31496 | 14.34242 | 16.34764 | 14.24266 | 18.74017 | 18.83179 |
| CEACAM5 | 16.82271 | 15.99007 | 16.41181 | 15.45646 | 15.30503 | 16.2821 | 15.4088 | 16.5592 | 23.93847 | 21.13482 | 15.28422 | 14.81253 | 16.89597 | 15.36369 | 16.21266 | 16.14858 | 15.76226 | 15.07912 |
| CEACAM6 | 14.85063 | 16.16684 | 14.41355 | 14.89298 | 14.61811 | 15.52389 | 15.38964 | 15.98005 | 21.59041 | 20.17753 | 15.82551 | 13.6949 | 14.49934 | 14.28863 | 13.61064 | 16.14932 | 15.86362 | 18.13657 |
| CEL | 17.41005 | 16.96691 | 16.41803 | 15.79611 | 16.79729 | 17.11346 | 16.46768 | 17.10621 | 22.46582 | 21.4113 | 23.54695 | 18.00764 | 16.35882 | 16.96142 | 16.32088 | 16.0998 | 16.40692 | 15.97776 |
| CELF1 | 20.64363 | 20.02437 | 19.36236 | 19.58909 | 15.34535 | 16.53979 | 19.08939 | 17.89484 | 14.91044 | 16.25237 | 15.90366 | 17.09614 | 19.48348 | 18.1918 | 18.84942 | 19.03074 | 15.74871 | 16.3317 |
| CELSR3 | 14.43976 | 15.4359 | 15.6416 | 17.91059 | 14.52418 | 14.94953 | 16.32945 | 15.96119 | 17.09383 | 18.00978 | 15.02148 | 13.07466 | 15.21666 | 15.61586 | 17.01431 | 14.42191 | 17.47303 | 14.73329 |
| CEMIP2 | 18.54442 | 18.08034 | 19.08167 | 18.49684 | 20.08409 | 19.02778 | 17.16361 | 17.03103 | 20.08418 | 19.11732 | 16.74685 | 16.0189 | 17.3141 | 17.24908 | 19.57284 | 19.1526 | 16.28846 | 15.87845 |
| CENPF | 14.60712 | 15.67592 | 15.66213 | 15.68211 | 20.81481 | 22.17856 | 20.7104 | 19.77166 | 16.70665 | 16.118 | 21.43071 | 13.1528 | 19.39325 | 22.33922 | 15.28151 | 15.80186 | 16.80466 | 21.99368 |
| CENPV | 17.26449 | 14.23804 | 14.87249 | 16.73607 | 15.61156 | 17.09529 | 21.12536 | 19.53618 | 13.33515 | 15.09548 | 16.71975 | 19.05295 | 14.44401 | 14.66466 | 13.23332 | 12.66851 | 13.86598 | 13.34487 |
| CEP55 | 18.5022 | 18.79849 | 19.41436 | 20.88349 | 22.62257 | 21.24763 | 18.80753 | 17.89841 | 19.75221 | 21.15083 | 18.04285 | 17.74535 | 19.26872 | 19.14795 | 19.92472 | 18.27498 | 19.48309 | 19.2177 |
| CES2 | 15.65979 | 15.54247 | 15.98412 | 14.97112 | 13.34044 | 14.91108 | 14.61875 | 15.69875 | 18.79312 | 17.57581 | 14.35559 | 12.94864 | 15.28063 | 14.39516 | 16.31541 | 15.06192 | 14.93796 | 13.95734 |
| CETN2 | 15.43353 | 17.66466 | 13.56548 | 12.62369 | 18.42123 | 15.10322 | 19.38622 | 17.42736 | 15.23089 | 15.61862 | 14.47046 | 12.84776 | 15.81874 | 14.65105 | 14.19263 | 14.9648 | 19.4278 | 19.50708 |
| CFAP298 | 13.17855 | 17.36695 | 15.07511 | 16.29831 | 17.96734 | 14.78807 | 14.78487 | 14.98562 | 13.69857 | 16.47724 | 14.81265 | 16.11305 | 18.7483 | 18.14692 | 16.536 | 16.98106 | 18.11976 | 17.93966 |
| CFAP36 | 18.77765 | 16.73813 | 15.94171 | 13.81995 | 16.48693 | 16.09315 | 17.13996 | 15.31972 | 13.21531 | 15.83493 | 13.92391 | 9.945668 | 17.56156 | 15.82987 | 14.61601 | 15.68581 | 16.54166 | 15.79172 |
| CFC1 | 20.38881 | 20.10149 | 14.81205 | 15.6228 | 16.48252 | 14.35144 | 15.4256 | 14.84123 | 15.198 | 14.77659 | 15.02228 | 12.71369 | 14.8405 | 15.77887 | 16.97557 | 14.32508 | 14.14899 | 17.2585 |
| CFL1 | 23.79637 | 23.74642 | 22.93577 | 23.5056 | 27.06044 | 25.6133 | 24.59024 | 23.42429 | 22.13151 | 23.23408 | 23.08306 | 23.6885 | 25.89191 | 25.26137 | 23.70435 | 23.58272 | 26.07529 | 25.61507 |
| CFL2 | 19.87938 | 19.59161 | 18.25756 | 19.09994 | 24.5912 | 23.43949 | 20.21161 | 19.3203 | 16.84498 | 17.32649 | 17.15953 | 17.97307 | 19.83359 | 20.67749 | 19.29507 | 18.64797 | 22.43085 | 21.4883 |
| CGN | 14.70658 | 14.90203 | 16.17242 | 15.89542 | 16.70857 | 15.05574 | 15.08001 | 17.1363 | 15.88131 | 16.41091 | 16.21329 | 17.71545 | 15.1973 | 18.73749 | 17.67601 | 15.49816 | 14.82765 | 15.16254 |
| CHAMP1 | 15.78469 | 15.3596 | 15.86834 | 14.03098 | 14.09406 | 16.85334 | 17.5679 | 17.01016 | 14.58807 | 16.63114 | 15.1775 | 15.83548 | 16.74224 | 15.81664 | 14.97993 | 15.22698 | 18.7041 | 14.32683 |
| CHAT | 16.31806 | 19.42372 | 14.68669 | 16.71298 | 16.4814 | 15.25692 | 16.1118 | 17.10152 | 15.47649 | 14.95107 | 17.40621 | 19.04585 | 18.14959 | 17.09446 | 13.278 | 15.39959 | 16.59556 | 15.30674 |
| CHCHD3 | 15.94624 | 13.76601 | 16.44364 | 15.57158 | 13.79192 | 14.62822 | 16.93195 | 15.92565 | 16.86101 | 17.97457 | 14.98513 | 14.4872 | 13.94956 | 17.05355 | 15.37742 | 14.62612 | 16.98806 | 14.74199 |
| CHD4 | 16.22971 | 15.02902 | 16.3126 | 15.58003 | 13.84341 | 15.42577 | 15.89888 | 16.30067 | 14.70293 | 15.2513 | 17.24565 | 18.08148 | 16.23161 | 14.48014 | 13.90915 | 16.1278 | 14.25498 | 14.57837 |
| CHDH | 17.29672 | 16.03964 | 16.62493 | 15.81362 | 15.09373 | 16.5127 | 15.34229 | 15.71897 | 19.35111 | 18.08507 | 15.17162 | 18.20942 | 15.39109 | 15.87336 | 18.32716 | 15.30067 | 16.49402 | 15.49543 |
| CHGA | 16.72926 | 16.56101 | 16.44207 | 16.45774 | 15.6959 | 17.16216 | 17.13412 | 17.34611 | 23.6368 | 23.26425 | 16.64189 | 14.38811 | 15.62426 | 16.47353 | 15.89036 | 16.19598 | 16.23532 | 16.06257 |
| CHGB | 18.5287 | 17.10618 | 18.95072 | 14.88538 | 15.78385 | 15.40406 | 16.17512 | 21.47934 | 18.72443 | 17.68141 | 15.17208 | 12.99001 | 14.8474 | 25.75473 | 18.18247 | 17.94164 | 16.99405 | 15.71961 |

| | | | | | | | | | | | | | | | | | | |
|---|---|---|---|---|---|---|---|---|---|---|---|---|---|---|---|---|---|---|
| CHID1 | 15.81233 | 16.14644 | 16.45138 | 16.14469 | 14.68959 | 15.88376 | 17.71026 | 18.28773 | 21.69672 | 20.04026 | 18.68063 | 17.77345 | 16.1492 | 17.08699 | 16.36392 | 15.78735 | 16.26744 | 16.16646 |
| CHMP1A | 19.92157 | 19.63264 | 18.25675 | 18.00869 | 20.91065 | 19.8861 | 19.45106 | 19.35707 | 18.72256 | 19.32628 | 17.90985 | 16.88531 | 20.73089 | 20.26746 | 18.53028 | 18.23013 | 18.51339 | 19.70776 |
| CHMP1B | 18.58636 | 19.66285 | 18.61016 | 21.17965 | 23.64264 | 22.05295 | 19.47071 | 18.57372 | 21.39411 | 22.72244 | 21.69473 | 21.73899 | 21.89648 | 21.69701 | 18.7748 | 18.4682 | 19.02253 | 20.76435 |
| CHMP2A | 21.47583 | 21.83984 | 20.96295 | 22.03332 | 22.40121 | 21.47115 | 18.98588 | 19.01058 | 21.71411 | 21.4919 | 20.17255 | 21.30627 | 21.18347 | 21.43148 | 21.29282 | 21.06823 | 19.37308 | 20.31158 |
| CHMP2B | 20.75344 | 20.28545 | 18.4212 | 20.04607 | 21.57059 | 19.97608 | 18.43741 | 19.0075 | 19.47591 | 20.58638 | 18.56145 | 18.67965 | 21.99026 | 21.95193 | 19.32605 | 18.6737 | 19.74391 | 21.13838 |
| CHMP3 | 19.81202 | 19.97877 | 19.10371 | 19.35765 | 19.80272 | 18.81738 | 19.13122 | 18.79203 | 19.34954 | 20.17327 | 17.65942 | 17.1379 | 19.36811 | 20.48971 | 19.29619 | 19.50724 | 18.81449 | 18.2747 |
| CHMP4A | 20.69779 | 20.84159 | 17.53696 | 17.77774 | 19.29127 | 17.59056 | 15.96225 | 16.37133 | 18.9093 | 19.41671 | 17.78668 | 18.37133 | 17.64718 | 18.73469 | 18.93852 | 19.09918 | 18.22531 | 17.72268 |
| CHMP4B | 23.34358 | 23.4659 | 21.87324 | 22.83089 | 23.05418 | 21.56774 | 20.3696 | 17.78228 | 22.07482 | 21.32806 | 21.09955 | 19.73671 | 22.65094 | 22.02226 | 22.61055 | 22.36955 | 18.94552 | 21.64066 |
| CHMP4C | 21.57514 | 21.29652 | 18.59304 | 19.06796 | 18.67054 | 16.58302 | 17.4501 | 17.65503 | 19.78305 | 19.60419 | 16.57796 | 15.26799 | 20.07432 | 18.7689 | 19.52477 | 20.05989 | 16.11446 | 16.99088 |
| CHMP5 | 20.12795 | 20.67072 | 19.36945 | 21.46837 | 23.60548 | 22.17524 | 20.09227 | 19.31351 | 20.97577 | 22.45583 | 21.15042 | 20.04429 | 22.5819 | 22.73941 | 19.27434 | 19.61304 | 20.48789 | 21.28732 |
| CHMP6 | 18.2771 | 18.4761 | 16.81404 | 15.35703 | 16.01213 | 15.51473 | 15.36775 | 15.51058 | 16.7446 | 15.70128 | 16.49618 | 18.02959 | 13.68048 | 17.1675 | 19.48486 | 19.70928 | 15.55848 | 15.09137 |
| CHORDC1 | 14.60553 | 18.60091 | 16.69252 | 15.48154 | 18.38938 | 18.37682 | 17.9237 | 17.16956 | 17.04172 | 18.94834 | 19.07571 | 19.27163 | 20.29307 | 19.79337 | 15.33334 | 16.33656 | 21.12021 | 19.26267 |
| CHP1 | 18.94612 | 19.99652 | 20.17114 | 19.97936 | 18.2458 | 18.53514 | 16.38212 | 16.973 | 17.23493 | 16.12975 | 17.84108 | 19.66266 | 19.65371 | 18.49478 | 21.02297 | 20.63048 | 16.26362 | 18.07503 |
| CHPF | 15.38931 | 15.80415 | 15.30393 | 14.25716 | 14.04549 | 15.63392 | 13.81961 | 16.34757 | 17.68165 | 14.20242 | 17.68789 | 11.46747 | 13.29007 | 14.98612 | 12.93951 | 14.53125 | 16.03331 | 16.00253 |
| CHPF2 | 16.33041 | 15.37433 | 16.06752 | 16.26235 | 14.64339 | 17.71407 | 15.96443 | 15.80111 | 18.86629 | 14.37268 | 15.90574 | 17.08062 | 15.16127 | 15.05653 | 15.41517 | 15.76862 | 15.75191 | 16.40525 |
| CHST1 | 16.38752 | 15.6316 | 16.48575 | 15.69178 | 15.53387 | 15.56135 | 16.48808 | 16.79359 | 20.34917 | 17.92663 | 16.65408 | 14.37025 | 15.02995 | 15.30178 | 16.41152 | 15.18887 | 16.08304 | 14.53059 |
| CHSY1 | 16.59679 | 17.02908 | 14.34543 | 14.26297 | 17.90851 | 16.70382 | 14.66942 | 14.95629 | 18.8126 | 16.13579 | 15.94339 | 11.42526 | 12.10082 | 14.24471 | 15.04791 | 16.73635 | 13.51707 | 14.7228 |
| CIAO1 | 18.02858 | 20.28341 | 14.84917 | 16.4768 | 16.18956 | 15.08036 | 15.88471 | 15.62819 | 15.1331 | 15.67754 | 18.07805 | 17.82983 | 18.0931 | 15.97061 | 16.8328 | 13.89901 | 17.80088 | 16.14099 |
| CIAPIN1 | 16.01923 | 14.52726 | 15.44832 | 16.12435 | 16.84293 | 16.13648 | 15.27503 | 17.72225 | 13.92755 | 16.6 | 18.09748 | 18.45526 | 17.04449 | 16.84981 | 18.38155 | 16.64559 | 17.30175 | 17.24481 |
| CIB1 | 17.89177 | 18.44748 | 16.14544 | 17.01392 | 16.35352 | 17.12735 | 17.68727 | 15.95928 | 16.48197 | 13.66799 | 16.74624 | 16.76123 | 17.78728 | 15.6924 | 17.89348 | 17.28227 | 15.9168 | 15.18352 |
| CIRBP | 21.03221 | 21.01146 | 20.6159 | 20.33542 | 16.08501 | 15.17084 | 17.09126 | 16.99277 | 16.71843 | 17.7509 | 17.28906 | 18.86635 | 18.42046 | 15.47086 | 20.87391 | 20.8387 | 15.38166 | 15.87488 |
| CISD2 | 21.18097 | 21.90472 | 22.15947 | 20.72538 | 20.54109 | 18.43476 | 18.0928 | 20.71095 | 21.07445 | 19.77908 | 19.88522 | 16.64769 | 16.17206 | 18.9773 | 18.42125 | 19.09395 | 17.68562 | 19.58358 |
| CIT | 16.01742 | 15.90531 | 18.13909 | 17.423 | 21.1382 | 19.524 | 18.08854 | 17.76659 | 17.51255 | 16.66544 | 16.40098 | 16.51538 | 16.89652 | 16.84224 | 16.73702 | 16.3137 | 19.68725 | 18.0435 |
| CKAP4 | 17.17003 | 16.0757 | 17.08924 | 16.94918 | 17.92248 | 18.96983 | 18.74917 | 18.11787 | 20.58362 | 19.55898 | 17.93077 | 14.68251 | 17.90887 | 19.14753 | 16.58886 | 17.14693 | 16.49333 | 19.33011 |
| CKAP5 | 17.28412 | 17.66716 | 16.22412 | 17.20905 | 17.29557 | 17.66375 | 17.47558 | 18.01058 | 16.71017 | 17.6154 | 17.40891 | 18.39966 | 18.45814 | 17.84375 | 16.79247 | 17.04218 | 17.12105 | 17.85351 |
| CKB | 20.05603 | 20.24454 | 21.59822 | 23.31734 | 23.32738 | 21.48268 | 25.29975 | 23.18117 | 21.13251 | 21.52619 | 20.04473 | 22.50686 | 24.02969 | 23.01792 | 23.09698 | 22.62235 | 17.40519 | 16.52958 |
| CKMT1A | 18.31311 | 17.1257 | 20.14819 | 18.0095 | 14.00406 | 16.18166 | 18.83758 | 16.34954 | 16.46546 | 15.30796 | 19.11662 | 20.06309 | 16.16534 | 15.47945 | 17.61454 | 18.15398 | 16.3269 | 14.25845 |
| CKS1B | 20.64157 | 20.97103 | 19.31329 | 17.61174 | 16.46898 | 16.76201 | 16.49743 | 16.4361 | 17.5923 | 14.81106 | 14.93972 | 17.04058 | 17.86383 | 14.96564 | 19.95648 | 19.48601 | 15.9543 | 16.0403 |
| CLASP1 | 15.48085 | 16.83878 | 14.51661 | 16.39705 | 16.04607 | 15.61773 | 14.79189 | 16.5397 | 14.05087 | 15.93327 | 15.89026 | 17.04184 | 16.21937 | 16.51916 | 17.37938 | 15.31423 | 16.34746 | 15.08496 |
| CLASP2 | 16.43618 | 16.09993 | 15.85473 | 15.9713 | 17.31414 | 18.93543 | 17.26356 | 17.51512 | 14.9312 | 15.2867 | 15.93576 | 18.37103 | 15.47815 | 17.57763 | 13.95329 | 16.16338 | 16.62937 | 16.23898 |
| CLCN3 | 14.45472 | 14.17781 | 19.36772 | 15.65688 | 16.76876 | 14.6776 | 17.34989 | 17.9551 | 18.55398 | 14.90643 | 17.39736 | 19.09732 | 14.34999 | 15.8428 | 17.62993 | 19.1139 | 15.64975 | 13.26558 |
| CLDN12 | 21.47611 | 21.36323 | 21.38212 | 22.40569 | 20.67773 | 20.0006 | 17.53067 | 15.92742 | 20.30075 | 21.00431 | 15.34703 | 17.19468 | 21.81624 | 21.51849 | 22.89923 | 22.49909 | 19.04642 | 18.31711 |
| CLDN3 | 22.78522 | 22.42049 | 23.80885 | 23.41623 | 22.48247 | 21.92234 | 15.46997 | 16.87929 | 22.05914 | 22.65808 | 18.36482 | 18.17118 | 21.90487 | 21.19743 | 24.06345 | 23.61033 | 22.34106 | 23.37 |
| CLDN4 | 18.44068 | 16.2066 | 15.1111 | 14.84384 | 12.33041 | 16.25314 | 15.71213 | 13.89536 | 14.88474 | 20.11059 | 18.47768 | 13.17593 | 21.5098 | 20.69236 | 15.8985 | 16.51969 | 21.6427 | 22.71441 |
| CLDND1 | 21.00857 | 20.16176 | 18.65074 | 17.39271 | 17.95374 | 16.82296 | 17.1326 | 15.72679 | 18.57227 | 18.21373 | 19.18382 | 20.43986 | 16.53228 | 17.12626 | 20.4033 | 19.74051 | 16.00668 | 14.97342 |
| CLGN | 15.57286 | 16.7147 | 16.15245 | 16.85371 | 15.0513 | 16.46424 | 16.94131 | 17.21311 | 21.6944 | 20.00077 | 19.52382 | 15.86706 | 16.3831 | 16.71165 | 15.90692 | 15.31512 | 15.68688 | 14.73172 |
| CLIC1 | 21.85168 | 21.77386 | 20.70212 | 22.02367 | 24.86022 | 24.06272 | 22.41416 | 22.29235 | 21.66026 | 22.00924 | 21.2332 | 21.88947 | 24.09727 | 24.58196 | 22.39889 | 21.94532 | 25.21706 | 23.98421 |
| CLIC2 | 14.81317 | 15.52491 | 14.50842 | 16.00845 | 18.32299 | 15.06266 | 17.12671 | 15.70542 | 13.96339 | 14.43577 | 15.85189 | 13.08141 | 19.88379 | 21.99344 | 16.55963 | 16.16051 | 15.90022 | 17.46341 |
| CLIC3 | 15.15428 | 13.92664 | 14.96033 | 14.71914 | 19.4091 | 17.95797 | 14.64554 | 15.23389 | 15.83417 | 16.62855 | 14.10506 | 15.18725 | 13.71439 | 14.08634 | 15.13942 | 13.76643 | 14.37039 | 14.4025 |
| CLIC4 | 18.49855 | 18.17147 | 18.93892 | 19.3598 | 22.85994 | 21.98691 | 21.68542 | 21.94949 | 19.5429 | 20.66699 | 19.64994 | 19.94246 | 21.51618 | 22.7749 | 19.50495 | 19.56463 | 23.17448 | 22.57227 |
| CLIC5 | 16.18753 | 14.86721 | 14.22754 | 15.03845 | 18.8582 | 15.44551 | 17.47918 | 17.29671 | 16.19449 | 16.96908 | 16.05125 | 18.23956 | 17.63466 | 14.93263 | 18.37912 | 15.0841 | 17.45599 | 15.70077 |
| CLINT1 | 16.88368 | 17.23653 | 18.27261 | 18.07553 | 16.63436 | 13.72318 | 15.19342 | 16.11909 | 17.18452 | 13.67484 | 19.22274 | 19.27487 | 18.00671 | 18.2466 | 17.67969 | 17.84975 | 14.86731 | 16.39154 |
| CLIP1 | 16.37924 | 16.31243 | 15.68181 | 15.4273 | 17.96693 | 15.88138 | 16.98583 | 16.8966 | 14.99639 | 15.9172 | 15.85675 | 17.65565 | 17.83549 | 17.28952 | 14.49257 | 15.06558 | 16.53795 | 17.36054 |
| CLIP2 | 14.8361 | 14.7423 | 20.45108 | 20.31463 | 19.03389 | 15.4196 | 18.0991 | 14.76795 | 14.68058 | 14.35091 | 14.78649 | 14.06914 | 13.83501 | 17.75517 | 15.45532 | 15.79339 | 14.27832 | 17.9142 |
| CLMP | 16.25518 | 16.13168 | 15.46019 | 17.15981 | 18.0077 | 14.67835 | 18.09248 | 16.27784 | 14.10138 | 15.39296 | 15.93116 | 14.32237 | 15.35248 | 21.82828 | 17.45641 | 13.6414 | 19.16539 | 21.72664 |
| CLNS1A | 14.82152 | 15.33497 | 16.86078 | 16.47468 | 18.13209 | 19.631 | 14.80112 | 17.88795 | 15.87523 | 15.79515 | 19.77623 | 20.38596 | 17.60456 | 16.01627 | 15.01086 | 13.97879 | 18.5987 | 18.79882 |
| CLPB | 13.50407 | 15.56376 | 16.68851 | 16.59698 | 13.41273 | 13.47548 | 13.64917 | 17.95948 | 17.7919 | 12.09672 | 16.77346 | 18.57171 | 13.07796 | 14.56414 | 14.36894 | 14.47706 | 15.16039 | 13.84266 |
| CLPP | 17.38464 | 17.10168 | 17.39942 | 15.05281 | 13.79852 | 18.40728 | 14.76591 | 17.93037 | 19.79581 | 18.36352 | 15.58769 | 13.07054 | 14.36287 | 14.69719 | 13.72708 | 15.66859 | 16.64327 | 17.05679 |
| CLPTM1 | 17.10068 | 16.46511 | 14.05185 | 15.76391 | 14.90806 | 17.80192 | 16.20655 | 16.91884 | 20.34388 | 14.12557 | 17.65367 | 15.6962 | 15.26675 | 16.61626 | 14.97626 | 16.22113 | 15.56403 | 15.77923 |
| CLPTM1L | 15.99955 | 15.91628 | 15.77162 | 16.17282 | 15.63536 | 17.03193 | 16.08232 | 16.84457 | 21.32783 | 18.76801 | 19.70486 | 14.92481 | 18.79139 | 20.34789 | 20.93332 | 16.26569 | 16.50698 | 14.8454 |
| CLSTN1 | 16.47718 | 16.53238 | 18.72082 | 16.66916 | 16.32441 | 17.04231 | 18.3022 | 16.56657 | 20.47879 | 19.07389 | 18.42274 | 15.64579 | 17.66322 | 16.13918 | 16.95903 | 15.92781 | 16.22553 | 15.76878 |
| CLTA | 22.58586 | 22.1649 | 20.42448 | 21.38564 | 22.96184 | 21.74634 | 21.58732 | 18.83796 | 19.28623 | 20.49236 | 18.56461 | 22.3946 | 22.74135 | 22.2544 | 20.95132 | 21.15086 | 21.42421 | 22.31652 |
| CLTB | 22.62216 | 21.85175 | 21.58039 | 21.79936 | 21.13449 | 19.64914 | 20.20845 | 20.16741 | 16.82691 | 17.38611 | 17.12581 | 20.63107 | 22.90642 | 22.04508 | 21.09896 | 21.45569 | 20.17054 | 20.56474 |
| CLTC | 23.60727 | 23.27492 | 22.25996 | 22.23661 | 19.57525 | 19.90876 | 20.00094 | 18.67719 | 18.72912 | 18.39578 | 19.22416 | 23.11702 | 20.69743 | 20.04894 | 22.50354 | 22.77266 | 19.56 | 19.81096 |
| CLTCL1 | 23.79456 | 23.07845 | 21.36116 | 21.43416 | 20.30012 | 19.75313 | 20.68496 | 19.06712 | 17.40706 | 18.5023 | 17.48544 | 22.64509 | 20.63541 | 19.56167 | 22.02693 | 22.44288 | 19.58662 | 19.50984 |
| CLU | 17.01778 | 18.82766 | 17.14315 | 15.64768 | 20.7576 | 21.74239 | 17.16434 | 18.18936 | 20.09222 | 20.09338 | 17.83579 | 14.72118 | 16.84409 | 18.29813 | 16.89858 | 16.84374 | 18.32196 | 21.75734 |
| CLUH | 15.92103 | 15.59516 | 15.67635 | 15.95438 | 17.4362 | 16.78157 | 17.65429 | 18.09602 | 16.45778 | 16.60545 | 17.38045 | 19.46497 | 16.22327 | 16.72398 | 15.17972 | 14.88192 | 16.30627 | 16.30609 |
| CMBL | 16.93828 | 15.87553 | 18.32509 | 18.09423 | 16.28585 | 16.94668 | 15.10234 | 17.43467 | 15.24643 | 13.88264 | 15.79205 | 17.71551 | 15.82364 | 16.92099 | 18.50208 | 17.28059 | 18.67373 | 16.98634 |
| CMIP | 16.43732 | 16.92823 | 13.72912 | 14.27974 | 19.14286 | 17.32929 | 19.47109 | 20.356 | 17.21037 | 15.19222 | 16.40064 | 18.49555 | 14.75997 | 16.49475 | 16.56965 | 15.42568 | 17.50694 | 18.33053 |
| CMPK1 | 15.80731 | 16.9869 | 18.24758 | 18.39886 | 20.76375 | 19.48126 | 19.66407 | 17.75102 | 17.59723 | 19.17628 | 16.72292 | 18.01837 | 21.64816 | 20.78976 | 18.76933 | 17.894 | 21.49412 | 21.11561 |
| CMTM4 | 19.20596 | 20.61847 | 19.17244 | 19.29146 | 18.28966 | 15.27539 | 15.90192 | 17.33844 | 13.50491 | 14.42728 | 17.50023 | 18.31477 | 16.28527 | 15.08297 | 20.26234 | 20.22935 | 16.62958 | 16.28775 |
| CMTM6 | 18.63784 | 20.03951 | 18.49148 | 18.77064 | 16.89431 | 16.74206 | 15.09338 | 17.76584 | 19.39233 | 17.88977 | 19.99642 | 20.24701 | 14.91302 | 15.86655 | 18.00293 | 18.65998 | 17.30019 | 16.22715 |
| CMTM7 | 15.10137 | 14.25798 | 17.21358 | 16.6821 | 12.5427 | 15.82788 | 15.84654 | 15.98402 | 17.46645 | 17.15726 | 15.12031 | 13.93007 | 13.13036 | 16.0173 | 17.16177 | 17.82297 | 18.78713 | 18.69811 |
| CNBP | 19.14612 | 17.67961 | 17.95621 | 17.37412 | 17.6565 | 16.92021 | 17.64701 | 16.67743 | 14.82662 | 16.78263 | 16.83024 | 18.37255 | 19.04899 | 19.56433 | 18.253 | 18.23329 | 19.42802 | 18.0694 |
| CNDP2 | 17.64147 | 16.81636 | 19.18355 | 20.04071 | 19.08832 | 18.14935 | 18.39689 | 18.40076 | 18.37489 | 19.0246 | 21.16926 | 21.23374 | 22.60026 | 22.3004 | 20.62163 | 20.92408 | 19.05464 | 18.94607 |
| CNK3/IPCE | 16.00956 | 15.02857 | 16.33363 | 13.50613 | 15.38613 | 15.79472 | 16.52497 | 16.66966 | 16.39099 | 16.64884 | 14.34847 | 15.79309 | 15.79792 | 14.34464 | 14.62008 | 15.17682 | 16.98799 | 14.64159 |
| CNKSR3 | 16.93164 | 16.33067 | 14.77117 | 14.16783 | 18.25518 | 16.87014 | 15.38433 | 14.88249 | 17.60908 | 18.95996 | 17.59216 | 16.67351 | 17.86896 | 18.2596 | 16.99692 | 14.13357 | 17.41885 | 18.07007 |
| CNN2 | 16.07879 | 20.1723 | 17.64196 | 16.97329 | 21.53555 | 19.19849 | 18.1807 | 17.85313 | 15.7906 | 16.23435 | 16.3432 | 13.94302 | 20.07047 | 20.37664 | 16.40472 | 16.97154 | 18.99919 | 18.05221 |
| CNN3 | 16.87431 | 16.85871 | 15.442 | 16.2835 | 20.38932 | 19.59779 | 16.57957 | 16.62696 | 16.27704 | 19.03584 | 18.92363 | 18.52505 | 14.85789 | 14.86206 | 15.45522 | 16.32527 | 20.13614 | 18.86737 |
| CNNM3 | 16.01252 | 15.51855 | 16.4636 | 15.80989 | 14.27278 | 16.02192 | 15.78278 | 16.09426 | 17.21016 | 16.94708 | 16.19024 | 17.42508 | 16.16588 | 14.46282 | 17.36168 | 17.19782 | 16.98499 | 13.85951 |
| CNNM4 | 15.60839 | 16.46544 | 17.26248 | 17.13835 | 15.13352 | 16.21391 | 15.92554 | 16.15049 | 15.84249 | 16.2336 | 18.89932 | 19.91108 | 17.72775 | 16.47563 | 17.2452 | 18.11111 | 14.81508 | 15.04887 |
| CNOT1 | 15.83033 | 15.87251 | 16.03734 | 16.19629 | 16.08539 | 16.72823 | 16.30504 | 16.00778 | 13.55491 | 16.05124 | 15.03362 | 17.97584 | 15.56845 | 15.73284 | 16.20117 | 15.63024 | 15.70186 | 14.29501 |
| CNOT10 | 16.02558 | 16.33577 | 14.46178 | 15.41705 | 17.80539 | 17.19289 | 15.10812 | 16.027 | 13.91105 | 16.12509 | 15.46214 | 15.52702 | 16.48699 | 15.09748 | 16.41715 | 16.51697 | 16.36424 | 14.31072 |
| CNOT11 | 16.68121 | 16.17255 | 14.95934 | 13.13036 | 18.853 | 15.2544 | 16.25374 | 18.19979 | 15.44886 | 13.68558 | 17.78985 | 17.89808 | 15.64882 | 16.04097 | 14.27445 | 18.23585 | 16.15317 | 17.23596 |
| CNP | 18.92636 | 18.52977 | 20.39554 | 20.80209 | 21.79483 | 20.68087 | 20.41685 | 20.55008 | 19.4046 | 19.73821 | 23.10197 | 22.49801 | 21.45089 | 21.62601 | 21.16125 | 20.83156 | 21.36786 | 22.12268 |
| CNPY1 | 14.06199 | 16.18935 | 17.2878 | 17.53703 | 14.50534 | 16.41019 | 15.06602 | 15.76454 | 14.10989 | 15.17405 | 20.55324 | 17.37513 | 14.50377 | 14.24849 | 14.69734 | 14.73386 | 17.16349 | 15.78212 |
| CNPY2 | 16.92771 | 16.49892 | 16.87971 | 17.05866 | 15.66372 | 16.85243 | 16.76891 | 17.2685 | 23.89887 | 22.99101 | 20.86541 | 16.97207 | 17.18453 | 17.18503 | 16.50717 | 15.98019 | 17.40378 | 16.44631 |
| CNPY3 | 16.09755 | 15.04062 | 15.76139 | 15.35455 | 14.3772 | 16.85489 | 19.39386 | 18.22148 | 22.23588 | 21.84142 | 18.79071 | 15.47145 | 16.62628 | 15.18937 | 16.2426 | 16.15975 | 15.71249 | 16.75643 |
| CNPY4 | 16.88113 | 16.73221 | 15.71103 | 14.888 | 14.33443 | 18.13671 | 16.16056 | 17.02567 | 20.49947 | 20.14094 | 18.09053 | 13.78141 | 15.60792 | 15.39142 | 15.09442 | 17.55543 | 16.09151 | 15.72831 |
| CNRIP1 | 13.09285 | 12.39269 | 15.46193 | 16.68149 | 12.71592 | 14.6685 | 19.1862 | 16.55469 | 13.47425 | 15.97833 | 15.0111 | 13.41376 | 16.45893 | 18.20272 | 17.72134 | 15.36648 | 18.93107 | 15.40146 |
| CNTFR | 19.66902 | 20.20421 | 16.07029 | 15.16464 | 16.64622 | 14.68752 | 16.08591 | 15.73189 | 19.23963 | 18.39852 | 13.79818 | 15.95975 | 14.91068 | 19.38411 | 15.11787 | 15.00812 | 14.64565 | 14.44262 |
| CNTN1 | 18.97539 | 19.55106 | 15.79559 | 17.37448 | 14.89116 | 15.97599 | 16.01645 | 16.62368 | 18.87343 | 16.75806 | 15.52976 | 15.23524 | 15.00216 | 15.78141 | 16.37399 | 15.03452 | 15.1823 | 15.6832 |
| CNTN2 | 16.5665 | 13.97693 | 14.33952 | 21.81782 | 12.09447 | 15.38581 | 21.44064 | 16.33294 | 15.05673 | 12.73509 | 14.98172 | 12.99789 | 14.73152 | 15.97381 | 16.23217 | 15.59276 | 15.47881 | 14.94915 |
| COASY | 17.35044 | 16.40566 | 18.01046 | 18.05731 | 17.98558 | 14.74897 | 14.40119 | 15.43227 | 16.65458 | 15.07042 | 17.09403 | 16.75783 | 14.15444 | 14.62524 | 17.77372 | 18.47634 | 14.18877 | 19.04037 |
| COBL | 20.60045 | 20.6563 | 16.92328 | 16.6443 | 19.34917 | 19.77629 | 15.88615 | 16.69734 | 14.30738 | 17.14423 | 20.52747 | 15.60275 | 17.49787 | 16.04328 | 16.71729 | 16.7481 | 16.3956 | 17.43136 |
| COBLL1 | 20.62049 | 20.50175 | 18.94492 | 18.78354 | 15.55804 | 16.20806 | 15.87825 | 16.61647 | 14.81682 | 15.38684 | 15.73174 | 14.42661 | 19.54339 | 19.30557 | 18.57361 | 19.11358 | 16.30708 | 16.82204 |

| | | | | | | | | | | | | | | | | | | |
|---|---|---|---|---|---|---|---|---|---|---|---|---|---|---|---|---|---|---|
| COG3 | 16.74097 | 15.05548 | 14.71883 | 18.1192 | 18.41502 | 19.07243 | 21.29845 | 15.89414 | 14.05561 | 14.65662 | 14.59734 | 16.40348 | 18.79408 | 20.90402 | 15.64554 | 14.46392 | 15.50268 | 19.6154 |
| COG4 | 14.75222 | 14.83886 | 15.85679 | 15.27221 | 13.73725 | 14.63962 | 14.19853 | 15.88813 | 12.1989 | 17.17687 | 15.17019 | 16.57468 | 16.11081 | 14.98942 | 15.28333 | 15.26492 | 13.37452 | 13.81948 |
| COG8 | 14.16696 | 15.36074 | 17.49274 | 16.22931 | 13.02392 | 14.6512 | 17.62243 | 16.7432 | 12.69197 | 14.85791 | 14.83659 | 17.42778 | 13.02067 | 16.49455 | 14.94447 | 14.61954 | 15.04258 | 15.7569 |
| COL12A1 | 16.13012 | 17.15559 | 17.2996 | 15.95059 | 17.89763 | 16.51652 | 17.12954 | 16.08301 | 18.16929 | 17.25176 | 15.54942 | 16.15838 | 14.9141 | 17.04939 | 16.45062 | 17.7583 | 15.68207 | 16.38012 |
| COL18A1 | 16.50106 | 16.5387 | 18.12886 | 15.68157 | 15.12641 | 21.89524 | 18.73333 | 17.55585 | 18.93467 | 18.7353 | 16.98725 | 14.32386 | 15.10688 | 15.73918 | 15.29083 | 18.15026 | 17.14228 | 19.99052 |
| COL1A1 | 15.66836 | 15.51714 | 19.01519 | 19.16548 | 16.06923 | 19.90036 | 16.93854 | 16.25939 | 13.23898 | 15.34711 | 15.88728 | 14.3559 | 13.61602 | 18.2343 | 17.48709 | 15.04178 | 16.35578 | 17.88267 |
| COL1A2 | 14.16914 | 16.62137 | 19.4072 | 14.904 | 16.14152 | 18.95681 | 15.23838 | 17.28095 | 17.74404 | 16.23789 | 17.25737 | 12.36437 | 15.03653 | 15.17028 | 16.62624 | 15.65584 | 15.56159 | 17.89407 |
| COL21A1 | 16.19644 | 16.83359 | 16.2242 | 15.09101 | 16.72822 | 20.40081 | 17.9949 | 17.67119 | 20.60173 | 19.2126 | 16.17849 | 14.24491 | 14.65413 | 16.49652 | 15.31717 | 15.44012 | 16.55827 | 18.04762 |
| COL2A1 | 15.62809 | 16.25002 | 17.6027 | 16.50699 | 14.10496 | 17.68749 | 16.72476 | 17.76549 | 14.93081 | 16.19143 | 22.32031 | 15.252 | 16.05118 | 17.61911 | 16.73703 | 15.73995 | 15.97659 | 16.63866 |
| COL4A2 | 14.89874 | 14.72945 | 16.86247 | 18.88597 | 12.47224 | 18.78489 | 14.99489 | 15.45937 | 17.41822 | 17.15179 | 15.90592 | 13.75652 | 18.3629 | 14.39487 | 14.89799 | 13.82532 | 16.16799 | 16.35226 |
| COL5A1 | 15.06543 | 16.11287 | 19.71483 | 19.34329 | 17.75652 | 20.14357 | 18.52783 | 18.22676 | 13.41998 | 16.66007 | 14.69488 | 16.09474 | 13.90535 | 19.20973 | 17.19379 | 15.77772 | 17.08876 | 19.09273 |
| COL6A1 | 16.50862 | 17.4737 | 20.37085 | 18.69234 | 16.89102 | 18.88055 | 18.81058 | 18.30826 | 19.53765 | 19.26427 | 14.8338 | 17.04114 | 17.29042 | 17.8166 | 18.17275 | 18.14892 | 17.51469 | 20.75462 |
| COL6A3 | 17.4587 | 16.6389 | 16.86211 | 16.81616 | 13.72453 | 15.3953 | 18.08687 | 15.32962 | 15.58863 | 16.92845 | 16.3632 | 12.67059 | 15.85748 | 17.91123 | 13.82322 | 15.63835 | 14.56078 | 16.33758 |
| COLEC12 | 17.34049 | 16.65652 | 19.11801 | 19.92678 | 14.36709 | 14.61666 | 17.43619 | 17.5896 | 15.28672 | 15.34037 | 15.06226 | 12.16859 | 14.22982 | 14.42807 | 17.80085 | 17.98689 | 16.20872 | 16.99845 |
| COLGALT1 | 18.47823 | 15.63436 | 16.33216 | 16.48327 | 15.68396 | 16.901 | 16.85849 | 16.28381 | 21.55042 | 21.14558 | 17.18251 | 15.3675 | 14.67664 | 16.59198 | 15.52224 | 16.09191 | 16.19575 | 17.47813 |
| COMMD10 | 14.34675 | 13.18696 | 15.44479 | 14.84985 | 19.4514 | 15.49542 | 15.77235 | 14.77313 | 13.15956 | 11.07023 | 17.38153 | 13.91952 | 15.85845 | 14.75808 | 15.99866 | 15.72619 | 17.53248 | 13.02682 |
| COMT | 19.4428 | 17.9749 | 17.10584 | 18.01827 | 16.34958 | 17.58575 | 17.17238 | 17.01146 | 15.79956 | 14.52829 | 16.15375 | 16.93876 | 20.01849 | 19.77566 | 18.85593 | 20.42404 | 14.24721 | 17.65645 |
| COPA | 18.79511 | 17.63745 | 17.61846 | 18.01523 | 17.1138 | 17.53642 | 18.34651 | 17.46898 | 17.35619 | 18.13964 | 18.175 | 20.30883 | 17.97237 | 17.84498 | 17.14851 | 17.69753 | 17.49975 | 17.74345 |
| COPB1 | 15.05795 | 16.78247 | 17.03368 | 16.58396 | 16.82038 | 16.80906 | 16.15908 | 16.14569 | 16.22191 | 16.57 | 18.23728 | 19.71974 | 18.15761 | 16.32402 | 16.03307 | 16.8339 | 17.21379 | 17.34317 |
| COPB2 | 17.00517 | 16.33729 | 16.44827 | 16.2653 | 17.91362 | 17.47734 | 18.07753 | 16.59541 | 16.34557 | 18.42758 | 18.44305 | 19.30489 | 18.04833 | 17.98241 | 17.59172 | 16.6261 | 18.87262 | 17.62724 |
| COPE | 17.38823 | 16.27053 | 16.4478 | 17.21741 | 17.79503 | 16.48662 | 17.10521 | 15.59613 | 17.79015 | 17.85718 | 19.20922 | 20.13982 | 17.76201 | 17.27185 | 17.77625 | 17.02667 | 17.91848 | 18.21029 |
| COPG1 | 19.5472 | 16.50125 | 16.37196 | 18.72007 | 17.52882 | 16.92483 | 17.34009 | 16.75584 | 15.66031 | 17.21259 | 17.06418 | 19.22468 | 18.82921 | 17.29116 | 17.80351 | 18.0968 | 17.95727 | 19.17921 |
| COPS2 | 15.54924 | 15.31991 | 15.22268 | 15.37867 | 17.1227 | 16.28545 | 17.39885 | 16.88474 | 15.34989 | 16.3887 | 16.24026 | 19.0811 | 17.68625 | 16.92734 | 16.63485 | 17.69538 | 17.69525 | 18.01833 |
| COPS3 | 19.36014 | 18.41382 | 17.50268 | 17.15692 | 17.00912 | 17.49134 | 15.62379 | 18.08189 | 16.0927 | 15.95019 | 16.72521 | 18.74958 | 17.79396 | 17.46347 | 15.88037 | 17.35419 | 17.01008 | 17.44716 |
| COPS4 | 16.62185 | 16.8765 | 15.72693 | 16.08758 | 17.06758 | 16.26685 | 17.65965 | 17.57324 | 15.62567 | 15.85735 | 17.06883 | 18.91204 | 17.83931 | 17.97783 | 16.554 | 16.99523 | 18.60886 | 17.20635 |
| COPS5 | 16.2679 | 16.15488 | 17.5877 | 17.027 | 17.24715 | 17.09008 | 18.3997 | 15.64992 | 17.44966 | 15.8541 | 17.76443 | 18.89936 | 17.6669 | 16.32801 | 16.57459 | 17.40358 | 18.78098 | 19.72604 |
| COPS6 | 17.20848 | 17.41804 | 17.3909 | 17.15571 | 15.43044 | 16.87576 | 15.49981 | 17.99058 | 17.31892 | 17.18056 | 18.94732 | 19.6563 | 17.17647 | 16.07049 | 17.83082 | 17.8202 | 16.55746 | 16.30977 |
| COPS7B | 14.96022 | 18.09769 | 17.26687 | 14.54976 | 17.2098 | 14.77906 | 17.83653 | 17.53518 | 14.73488 | 16.34892 | 17.84793 | 17.66799 | 17.53551 | 16.28813 | 15.09435 | 14.38637 | 18.63807 | 17.7078 |
| COPS8 | 14.58337 | 16.61029 | 19.04907 | 15.47999 | 16.55174 | 17.7442 | 15.01986 | 15.61844 | 16.10354 | 16.14452 | 19.13611 | 19.75119 | 18.6957 | 18.41657 | 16.43673 | 17.93798 | 16.3914 | 17.43561 |
| COPZ1 | 18.16378 | 19.89294 | 18.52598 | 18.80184 | 18.139 | 17.91982 | 18.57835 | 16.1031 | 18.65537 | 19.17836 | 20.55071 | 21.09775 | 20.04076 | 18.58708 | 18.6938 | 18.75841 | 18.39027 | 19.37329 |
| CORO1A | 17.29539 | 16.4833 | 18.00701 | 17.04064 | 17.28046 | 18.19098 | 19.10011 | 18.82965 | 17.31653 | 17.69839 | 21.11394 | 21.34955 | 16.4917 | 18.69116 | 16.94675 | 17.29712 | 17.18521 | 16.80384 |
| CORO1B | 19.11943 | 19.34825 | 18.51531 | 19.37183 | 18.66705 | 18.1627 | 16.73729 | 16.64928 | 17.89647 | 16.88809 | 18.86697 | 19.61339 | 19.77798 | 19.34873 | 20.27847 | 19.90906 | 18.01871 | 18.29135 |
| CORO1C | 18.33293 | 18.62271 | 17.40569 | 19.15088 | 22.9268 | 21.7428 | 19.38251 | 19.09827 | 19.16521 | 19.54415 | 20.57283 | 21.20305 | 20.28295 | 19.83404 | 18.25303 | 17.64627 | 20.94408 | 20.46094 |
| CORO2A | 16.89267 | 14.36738 | 16.98654 | 15.18748 | 14.33243 | 16.37475 | 14.68738 | 17.88639 | 12.04127 | 14.76679 | 13.15186 | 17.19072 | 14.61064 | 14.79256 | 18.81454 | 18.24075 | 16.25628 | 11.85897 |
| CORO2B | 16.19656 | 16.28602 | 15.43793 | 15.86606 | 14.31505 | 15.81431 | 17.05318 | 16.0994 | 13.24052 | 14.09129 | 17.50856 | 19.59195 | 16.01218 | 15.11943 | 15.03897 | 15.0241 | 16.01741 | 15.53144 |
| CORO7 | 17.89305 | 19.18763 | 18.95896 | 18.6927 | 15.44619 | 15.86444 | 18.18463 | 18.33584 | 15.19687 | 15.77392 | 19.32224 | 19.03114 | 17.67109 | 19.69369 | 19.36631 | 17.37844 | 14.97568 | 18.02451 |
| COTL1 | 17.78781 | 19.83931 | 17.93181 | 18.18464 | 21.65343 | 21.00102 | 20.68426 | 18.90055 | 20.4889 | 21.5534 | 16.02303 | 18.58603 | 19.32054 | 20.3917 | 18.14806 | 17.19796 | 22.87275 | 21.11893 |
| COX4I1 | 17.3245 | 16.09821 | 15.24509 | 14.86254 | 14.3677 | 19.04282 | 16.49642 | 15.76454 | 20.64226 | 19.72004 | 17.09037 | 15.69141 | 15.69225 | 15.82414 | 14.3645 | 16.1066 | 16.09528 | 15.3867 |
| COX5A | 16.9515 | 15.91219 | 16.64521 | 18.88945 | 17.98721 | 17.42708 | 17.99336 | 17.94981 | 21.65075 | 20.17641 | 19.51956 | 16.02046 | 17.94661 | 19.0908 | 17.22225 | 16.41211 | 16.31692 | 18.92985 |
| COX5B | 14.98947 | 16.5498 | 19.26804 | 14.54339 | 15.14076 | 15.50376 | 18.98431 | 18.35123 | 20.15396 | 20.63357 | 19.10191 | 15.70638 | 16.31611 | 14.06185 | 17.35279 | 14.46865 | 15.48268 | 14.38563 |
| CPD | 18.92659 | 21.57096 | 18.43735 | 18.18353 | 18.97773 | 18.69391 | 16.51206 | 17.33089 | 22.79303 | 21.67111 | 18.80369 | 16.30323 | 16.53596 | 16.75074 | 17.14936 | 17.4605 | 16.31359 | 15.93588 |
| CPE | 16.31236 | 16.60017 | 16.41295 | 16.42975 | 16.03469 | 17.38133 | 18.27829 | 19.49101 | 23.8278 | 22.35268 | 16.9756 | 14.5714 | 16.99963 | 18.05654 | 16.79722 | 16.42773 | 16.05158 | 16.68718 |
| CPM | 16.92505 | 16.67081 | 16.07839 | 15.98704 | 19.94756 | 20.74274 | 16.91333 | 17.07051 | 21.66405 | 19.55514 | 17.43261 | 14.77269 | 15.7231 | 15.56634 | 16.07173 | 15.70119 | 16.77523 | 17.11515 |
| CPNE1 | 17.73782 | 18.64177 | 19.03262 | 19.63875 | 18.48263 | 20.17799 | 17.97349 | 16.63869 | 19.75441 | 18.15778 | 19.41442 | 19.94103 | 17.64972 | 18.56619 | 19.65017 | 19.63325 | 19.34579 | 19.61099 |
| CPNE2 | 17.54265 | 15.83947 | 16.77431 | 17.81666 | 20.46097 | 19.194 | 20.67902 | 17.92986 | 19.97357 | 18.95151 | 15.18462 | 15.66181 | 18.4791 | 18.33418 | 17.03534 | 17.36173 | 18.30595 | 19.19602 |
| CPNE3 | 19.19218 | 19.01409 | 21.10868 | 21.27806 | 20.13768 | 18.65356 | 21.01339 | 20.53908 | 22.34174 | 21.77407 | 18.17888 | 18.58671 | 20.58655 | 21.13232 | 20.75706 | 20.63885 | 20.89718 | 21.84463 |
| CPNE8 | 14.51172 | 15.91692 | 16.26756 | 16.61773 | 20.27631 | 19.28126 | 17.48708 | 17.13082 | 15.04001 | 16.88082 | 16.20985 | 13.70338 | 15.77444 | 16.19052 | 16.53719 | 15.93784 | 15.97379 | 17.45914 |
| CPOX | 14.61736 | 14.66475 | 16.32995 | 14.9225 | 15.00968 | 16.60755 | 14.08963 | 16.95836 | 18.30264 | 18.88814 | 16.23492 | 16.85182 | 14.32112 | 15.9976 | 15.35697 | 15.4136 | 15.43582 | 15.60635 |
| CPSF1 | 19.67371 | 21.86224 | 16.77729 | 18.96574 | 14.43488 | 15.76237 | 14.10253 | 15.77552 | 15.48478 | 15.96844 | 14.39833 | 17.55051 | 15.16998 | 15.39007 | 20.1156 | 20.27766 | 15.28723 | 12.93379 |
| CPSF3 | 14.86462 | 15.38446 | 14.6685 | 14.62033 | 15.85373 | 15.67527 | 16.64209 | 17.25303 | 12.40819 | 15.71167 | 15.27583 | 18.18678 | 14.60671 | 14.5674 | 11.80003 | 12.72863 | 16.77094 | 14.34724 |
| CPSF6 | 20.96312 | 20.59214 | 18.8417 | 19.81442 | 15.05998 | 23.25121 | 16.32165 | 15.85342 | 14.53636 | 15.81107 | 16.52261 | 16.92168 | 18.7161 | 18.06499 | 17.69172 | 18.68339 | 15.89469 | 15.41344 |
| CPSF7 | 21.09966 | 20.58643 | 21.5439 | 20.96794 | 17.62068 | 15.77131 | 18.47623 | 18.05148 | 18.55243 | 17.33423 | 17.48587 | 17.7239 | 18.9396 | 18.3882 | 20.06979 | 19.76754 | 15.16131 | 16.34757 |
| CPT2 | 15.57615 | 15.39344 | 16.03047 | 16.08018 | 16.29584 | 16.70101 | 16.03971 | 16.30707 | 18.80722 | 17.03946 | 15.63982 | 15.02831 | 14.05928 | 17.27365 | 16.30433 | 15.9695 | 15.24601 | 15.84178 |
| CRABP1 | 18.98438 | 18.20813 | 19.2469 | 18.52416 | 16.64669 | 17.03217 | 15.57245 | 16.97203 | 19.03966 | 19.42911 | 18.27863 | 13.86994 | 16.11163 | 18.19412 | 18.19808 | 18.86147 | 20.83347 | 21.46502 |
| CRABP2 | 17.58214 | 16.33236 | 17.55939 | 16.65297 | 21.26537 | 19.4891 | 21.9474 | 20.74129 | 19.44695 | 21.29079 | 15.08479 | 15.41481 | 15.70778 | 17.25291 | 16.24652 | 15.60699 | 25.09295 | 24.06662 |
| CRAT | 15.40701 | 17.00303 | 16.69562 | 16.67774 | 15.88752 | 18.60276 | 17.1297 | 17.95577 | 18.93597 | 18.36747 | 14.47877 | 14.83066 | 16.24987 | 15.74035 | 16.11333 | 16.93932 | 15.38704 | 16.23624 |
| CREG1 | 15.70327 | 15.1109 | 15.56124 | 16.0771 | 14.78029 | 15.87201 | 15.24875 | 15.19478 | 20.44064 | 19.69573 | 19.05565 | 13.98011 | 17.22965 | 15.92631 | 16.69767 | 13.60459 | 15.63023 | 13.28725 |
| CRELD1 | 15.56153 | 14.78114 | 15.10629 | 15.53949 | 13.85442 | 16.68614 | 15.47609 | 15.17023 | 20.04372 | 18.58698 | 19.74981 | 13.34494 | 15.31977 | 15.06448 | 15.18104 | 15.38596 | 14.94075 | 14.41563 |
| CRELD2 | 15.72056 | 15.31593 | 15.91285 | 15.56764 | 12.45586 | 16.59722 | 17.80919 | 18.11881 | 19.6426 | 18.63146 | 17.77331 | 15.62952 | 19.06359 | 19.382 | 15.5264 | 15.46908 | 15.79584 | 14.81838 |
| CRIM1 | 15.48423 | 15.88421 | 14.14395 | 16.35367 | 21.47845 | 20.21009 | 19.12322 | 18.02691 | 14.21235 | 15.42337 | 14.92574 | 15.26885 | 15.01268 | 14.31782 | 16.07053 | 18.61954 | 16.12724 | 18.93923 |
| CRIP1 | 18.25751 | 17.5258 | 15.91915 | 15.34148 | 23.44045 | 22.62435 | 25.42362 | 24.00926 | 19.38318 | 22.01007 | 18.40434 | 13.95822 | 17.51666 | 18.76595 | 15.5164 | 16.22191 | 20.31027 | 22.02605 |
| CRIP2 | 18.50917 | 18.36643 | 16.90904 | 16.12035 | 18.71967 | 16.54582 | 21.65377 | 21.62551 | 20.6718 | 20.93401 | 16.52638 | 14.31635 | 15.92518 | 16.79701 | 17.04349 | 15.9004 | 22.86781 | 22.75713 |
| CRISPLD1 | 17.05913 | 14.13562 | 15.76416 | 16.87846 | 17.9031 | 14.66565 | 15.71209 | 18.55485 | 18.20004 | 16.30641 | 13.99573 | 14.79803 | 14.51766 | 18.92872 | 14.34637 | 13.49005 | 14.63671 | 16.81276 |
| CRK | 19.48761 | 20.17547 | 17.92263 | 17.91141 | 15.67186 | 17.62915 | 15.88204 | 16.36905 | 16.63645 | 16.71399 | 14.58969 | 18.24808 | 16.92906 | 15.52729 | 18.36727 | 19.31623 | 16.57684 | 14.9135 |
| CRKL | 19.27817 | 18.83814 | 17.24455 | 17.85013 | 19.38513 | 18.20638 | 16.42156 | 17.02494 | 14.55383 | 15.68148 | 17.22859 | 18.55507 | 16.76089 | 17.69304 | 19.11756 | 17.5912 | 17.54782 | 17.55404 |
| CRLF1 | 19.57929 | 16.89375 | 16.53237 | 15.93325 | 13.65753 | 14.68555 | 16.00592 | 17.82746 | 20.29889 | 17.10446 | 14.74678 | 13.84424 | 16.41586 | 14.9628 | 14.29742 | 15.38855 | 18.21122 | 14.60951 |
| CRLF3 | 13.97584 | 15.70869 | 16.29107 | 16.18142 | 17.43934 | 16.7079 | 14.84744 | 17.50418 | 13.85003 | 15.6457 | 16.15916 | 17.49728 | 15.40094 | 14.33634 | 15.49747 | 15.48552 | 15.47416 | 13.10093 |
| CRMP1 | 22.92874 | 22.47775 | 22.67801 | 20.23587 | 19.49789 | 18.728 | 21.15251 | 21.04514 | 18.59955 | 19.36262 | 20.53591 | 21.35036 | 17.27439 | 17.49131 | 23.07443 | 22.74363 | 17.61407 | 17.79736 |
| CRTAP | 16.3126 | 18.45988 | 16.04811 | 14.77649 | 14.13258 | 15.86202 | 15.59376 | 15.83151 | 20.51183 | 19.92593 | 14.29998 | 14.86344 | 15.2655 | 16.13749 | 15.00937 | 15.75056 | 15.22172 | 16.49923 |
| CRYZ | 15.69439 | 15.13757 | 16.17498 | 15.37506 | 18.41716 | 15.13253 | 16.35346 | 17.77889 | 18.61406 | 18.27707 | 15.56636 | 17.5097 | 18.31245 | 17.59209 | 15.30818 | 14.84601 | 17.77619 | 17.21402 |
| CRYZL1 | 16.84986 | 17.30582 | 16.65362 | 17.69694 | 20.50305 | 19.35283 | 18.7765 | 16.68637 | 17.70158 | 18.56448 | 18.74671 | 18.67901 | 19.23482 | 18.45071 | 17.95888 | 17.5017 | 20.32765 | 19.54746 |
| CS | 16.55623 | 17.23196 | 17.11739 | 16.6553 | 16.88468 | 18.23457 | 17.52748 | 16.98467 | 22.53229 | 20.92716 | 17.39638 | 17.71587 | 16.88728 | 18.26586 | 16.0183 | 16.58217 | 16.28853 | 18.71288 |
| CSDE1 | 16.70543 | 17.21474 | 17.05083 | 17.55376 | 16.76549 | 16.23973 | 16.60006 | 16.75381 | 14.23702 | 16.57351 | 17.3872 | 18.5556 | 18.67937 | 17.6208 | 17.28751 | 18.029 | 16.87821 | 15.43719 |
| CSE1L | 16.58041 | 16.24145 | 17.17817 | 17.52602 | 17.40918 | 17.19186 | 18.45859 | 17.56015 | 17.43809 | 17.89101 | 18.82431 | 20.79348 | 16.36077 | 16.42704 | 17.13884 | 16.87323 | 17.47231 | 17.08263 |
| CSGALNAC | 16.65285 | 19.71087 | 17.88583 | 17.37213 | 17.74175 | 16.55144 | 15.53374 | 16.01708 | 18.23546 | 16.45807 | 16.46599 | 14.69065 | 15.29626 | 15.84467 | 17.00174 | 16.43694 | 18.34396 | 15.05967 |
| CSK | 16.83684 | 17.21762 | 17.37784 | 17.2128 | 15.08937 | 16.93278 | 17.902 | 16.47986 | 16.63443 | 16.66044 | 20.06312 | 19.68801 | 18.01891 | 18.36932 | 17.51938 | 17.23415 | 17.08923 | 15.47041 |
| CSNK1A1 | 18.86523 | 18.30389 | 19.26039 | 19.49291 | 16.61901 | 15.63737 | 16.30648 | 16.10769 | 17.44763 | 18.25575 | 15.7793 | 18.71638 | 16.15273 | 15.00809 | 20.54029 | 19.52828 | 16.25477 | 16.25111 |
| CSNK1A1L | 18.9513 | 17.06826 | 18.51406 | 20.51509 | 15.61053 | 15.77764 | 17.73972 | 15.71039 | 16.39552 | 16.65894 | 15.40154 | 18.06808 | 14.8078 | 14.61571 | 20.00892 | 19.59582 | 13.80224 | 15.906 |
| CSNK1D | 17.94563 | 18.24314 | 18.78014 | 18.89171 | 13.07369 | 15.84621 | 15.27625 | 15.03823 | 11.92025 | 12.58803 | 16.02001 | 16.22441 | 15.20466 | 14.30376 | 18.18216 | 18.4901 | 14.27442 | 14.40442 |
| CSNK1G1 | 17.72637 | 18.59089 | 18.45419 | 17.5773 | 17.00272 | 17.3139 | 17.25445 | 16.38184 | 17.13652 | 15.30639 | 18.67603 | 18.28441 | 15.84341 | 16.50995 | 18.99581 | 18.92246 | 16.69468 | 16.74568 |
| CSNK1G2 | 16.45168 | 16.84262 | 18.42915 | 17.04592 | 20.77186 | 17.37436 | 14.94372 | 17.7796 | 16.4826 | 14.39448 | 14.8675 | 16.34062 | 13.92346 | 14.22914 | 19.27468 | 18.3962 | 19.89508 | 21.49044 |
| CSNK1G3 | 16.87794 | 21.12498 | 18.97122 | 16.47596 | 16.62194 | 15.40863 | 14.85576 | 14.29646 | 13.74966 | 17.76821 | 16.11913 | 18.97635 | 17.73655 | 19.63684 | 18.86795 | 19.34196 | 15.09643 | 14.42242 |
| CSNK2A1 | 15.90765 | 14.59216 | 17.9824 | 17.28714 | 15.84497 | 18.13908 | 15.79355 | 16.12084 | 12.83479 | 16.86666 | 16.10599 | 19.12146 | 13.65108 | 19.61485 | 17.28965 | 15.64656 | 15.12443 | 15.95932 |
| CSNK2A2 | 16.94184 | 17.59067 | 14.51681 | 16.07876 | 14.66443 | 15.38699 | 16.37528 | 15.31466 | 15.96094 | 15.57229 | 16.56204 | 18.66169 | 14.12144 | 14.08302 | 16.00438 | 15.77013 | 16.5061 | 14.82845 |
| CSNK2A3 | 19.79046 | 18.99471 | 19.20268 | 19.90032 | 16.09005 | 19.0171 | 15.59091 | 16.18868 | 18.05124 | 15.92971 | 19.31775 | 21.18214 | 16.25978 | 15.45185 | 19.38401 | 19.10747 | 15.03936 | 15.58879 |
| CSNK2B | 19.08895 | 16.95778 | 17.25951 | 17.18317 | 18.97011 | 17.30416 | 18.05364 | 17.29132 | 15.87903 | 16.60441 | 18.3419 | 19.89108 | 18.27754 | 17.69292 | 15.31646 | 18.06225 | 18.84348 | 17.22488 |

| | | | | | | | | | | | | | | | | | | |
|---|---|---|---|---|---|---|---|---|---|---|---|---|---|---|---|---|---|---|
| CSRP1 | 20.42646 | 19.86537 | 20.0989 | 20.03292 | 22.14986 | 21.33477 | 21.01899 | 21.47986 | 20.11001 | 21.01764 | 18.58904 | 18.49519 | 22.03939 | 22.10995 | 20.49109 | 20.15414 | 21.89649 | 20.50702 |
| CSRP2 | 18.73046 | 17.68369 | 16.28175 | 15.85097 | 17.33531 | 15.86358 | 18.44687 | 18.98249 | 14.65858 | 16.79836 | 18.57995 | 19.82844 | 14.90384 | 17.72887 | 15.49002 | 17.05381 | 19.70417 | 18.13874 |
| CST3 | 15.80046 | 15.21881 | 19.95728 | 17.85387 | 15.98988 | 18.36363 | 16.29073 | 16.29462 | 22.33604 | 22.03245 | 15.80558 | 13.6278 | 17.45577 | 17.68474 | 18.13426 | 16.55717 | 16.80801 | 17.40973 |
| CST4 | 15.77187 | 16.04138 | 18.00311 | 14.61235 | 13.49193 | 17.68609 | 14.51187 | 17.00666 | 20.17452 | 16.03056 | 16.60031 | 13.31139 | 16.24724 | 16.00555 | 16.45992 | 13.89802 | 18.09891 | 14.64359 |
| CSTB | 17.16758 | 18.15876 | 16.15346 | 18.13293 | 19.32914 | 17.32288 | 18.73583 | 15.04146 | 17.53717 | 19.56827 | 13.79454 | 19.53021 | 14.68832 | 17.56663 | 22.01577 | 21.28688 | 20.92664 | 14.98664 |
| CSTF1 | 16.9601 | 16.36585 | 16.88986 | 16.30178 | 14.09568 | 17.36589 | 14.84931 | 16.27327 | 13.7284 | 14.1369 | 14.94457 | 18.96495 | 19.48942 | 13.92371 | 17.47009 | 17.61696 | 12.74981 | 12.59646 |
| CSTF2 | 15.29734 | 15.90063 | 15.5222 | 17.92783 | 15.42667 | 15.34399 | 15.31794 | 18.37396 | 15.06933 | 16.57839 | 15.92079 | 18.53294 | 18.82131 | 16.28283 | 14.7093 | 14.46999 | 16.46197 | 15.30535 |
| CTBP1 | 24.23876 | 23.63494 | 23.24813 | 22.6747 | 15.18505 | 15.30971 | 16.59148 | 17.20082 | 16.06805 | 16.38432 | 15.45065 | 18.59939 | 20.45442 | 18.82143 | 22.80531 | 23.16108 | 15.48041 | 14.60821 |
| CTBP2 | 15.37864 | 17.12088 | 16.1087 | 15.13884 | 16.55687 | 15.19653 | 16.3025 | 18.42559 | 15.69704 | 15.4912 | 14.8881 | 18.36237 | 15.67448 | 16.31241 | 16.09432 | 15.98208 | 15.39018 | 17.76767 |
| CTDSP1 | 20.27079 | 19.91148 | 21.34723 | 18.63216 | 17.80118 | 18.16045 | 16.83599 | 15.63797 | 15.78032 | 15.45267 | 15.8712 | 17.62918 | 21.00816 | 17.17447 | 20.27295 | 19.7065 | 17.81512 | 17.96831 |
| CTDSPL | 19.27056 | 19.97208 | 14.57565 | 15.95551 | 16.93943 | 14.38853 | 20.17682 | 18.33139 | 12.49097 | 16.86672 | 17.98728 | 12.13771 | 20.47687 | 19.15644 | 15.05576 | 16.67608 | 13.3964 | 15.57139 |
| CTH | 14.03786 | 15.01908 | 15.34491 | 16.82063 | 14.40695 | 14.77453 | 15.89081 | 17.01064 | 13.96352 | 16.06001 | 16.05828 | 18.01349 | 20.06522 | 19.93107 | 17.08184 | 15.84658 | 17.77732 | 18.94633 |
| CTHRC1 | 15.18689 | 15.19069 | 13.90391 | 17.80268 | 14.86302 | 14.75182 | 15.84376 | 16.01689 | 20.80242 | 20.26128 | 15.01971 | 15.07378 | 15.53197 | 15.56893 | 14.40908 | 17.53255 | 17.53381 | 16.52567 |
| CTNNA1 | 20.41702 | 20.30495 | 21.55602 | 22.498 | 22.87219 | 22.33443 | 16.99051 | 17.26901 | 17.78128 | 18.25957 | 20.60382 | 20.81659 | 22.74546 | 22.93767 | 22.72197 | 22.66813 | 17.4402 | 17.14474 |
| CTNNB1 | 20.4353 | 19.66172 | 21.33467 | 21.88116 | 22.16345 | 21.53499 | 17.65413 | 17.47087 | 17.64583 | 17.74079 | 19.6499 | 20.67225 | 22.05199 | 22.25805 | 22.38745 | 22.48172 | 22.87154 | 22.48292 |
| CTNND1 | 19.39855 | 19.48402 | 20.94831 | 21.64157 | 21.90738 | 20.83365 | 17.86795 | 17.3841 | 15.1299 | 16.5673 | 20.20699 | 20.26239 | 22.03293 | 22.56002 | 22.17578 | 21.98083 | 22.62869 | 22.9163 |
| CTPS1 | 18.47635 | 18.06185 | 18.92462 | 17.36599 | 16.01226 | 16.03355 | 17.93822 | 17.89548 | 16.68389 | 17.11467 | 18.07785 | 19.63958 | 17.45981 | 16.96533 | 17.97931 | 18.62213 | 17.597 | 17.31694 |
| CTPS2 | 15.55401 | 15.4959 | 16.60371 | 14.75973 | 16.80817 | 16.32799 | 15.40933 | 16.56665 | 15.91924 | 17.39405 | 16.8768 | 19.16054 | 14.78611 | 15.86183 | 16.44671 | 16.70865 | 17.2427 | 14.01122 |
| CTSA | 15.53925 | 16.03982 | 16.09274 | 17.24003 | 17.49054 | 18.99368 | 16.96926 | 17.14951 | 18.34555 | 18.47839 | 16.60018 | 15.33602 | 16.30904 | 16.94591 | 16.15244 | 14.71477 | 14.97505 | 18.73976 |
| CTSB | 14.81928 | 14.7094 | 16.68459 | 16.38131 | 19.13707 | 19.56265 | 15.81237 | 16.45888 | 16.41279 | 14.8294 | 22.35719 | 15.37167 | 15.61541 | 17.79469 | 14.11307 | 14.26817 | 16.98375 | 15.38518 |
| CTSC | 16.55126 | 14.91759 | 17.77047 | 15.93922 | 16.04997 | 17.2751 | 16.62156 | 17.49845 | 18.1591 | 16.03081 | 22.00047 | 20.52887 | 17.80308 | 16.09724 | 15.33686 | 17.12639 | 16.4646 | 15.1228 |
| CTSF | 15.75013 | 16.30022 | 14.46173 | 17.89468 | 12.28405 | 13.64612 | 15.64817 | 15.57706 | 18.89504 | 16.09303 | 13.33442 | 14.3044 | 14.77191 | 14.27683 | 14.77956 | 14.10614 | 12.84254 | 15.64057 |
| CTSS | 15.87674 | 15.89709 | 16.32911 | 15.7743 | 13.68631 | 16.57433 | 16.19434 | 15.40696 | 19.50304 | 19.11837 | 15.41421 | 13.15409 | 17.63904 | 14.31484 | 14.76089 | 15.91141 | 16.07756 | 16.40275 |
| CTSZ | 15.87748 | 14.91438 | 18.44663 | 18.50842 | 17.08302 | 18.21736 | 16.30074 | 16.68958 | 19.36023 | 19.06677 | 15.8605 | 13.5404 | 18.45926 | 16.44271 | 18.59603 | 18.55214 | 16.79592 | 15.34084 |
| CTTN | 20.16237 | 19.89469 | 19.04283 | 19.94062 | 21.75044 | 20.36216 | 17.50551 | 17.5836 | 16.68792 | 17.60936 | 16.73885 | 18.67423 | 21.21273 | 21.52084 | 20.01422 | 19.71735 | 20.99505 | 21.06207 |
| CUEDC1 | 17.77815 | 15.92048 | 14.86976 | 14.53471 | 17.61188 | 14.62653 | 14.76462 | 15.76602 | 17.27282 | 17.21809 | 15.13234 | 12.49275 | 14.14865 | 15.66402 | 15.32299 | 14.75326 | 15.17528 | 17.47853 |
| CUL1 | 16.12714 | 15.97091 | 14.99876 | 16.26425 | 18.53899 | 18.60839 | 16.05924 | 16.19723 | 14.6634 | 15.97888 | 17.41687 | 18.18648 | 17.32273 | 18.24988 | 15.82295 | 15.80818 | 17.60754 | 17.77263 |
| CUL2 | 15.90053 | 15.38541 | 16.25853 | 16.73717 | 16.18375 | 16.75385 | 17.36377 | 16.68405 | 14.82172 | 15.6044 | 16.07207 | 18.1151 | 13.16229 | 15.79575 | 16.46585 | 16.4519 | 18.24326 | 17.9603 |
| CUL3 | 16.17804 | 15.66379 | 16.48748 | 16.06035 | 18.49241 | 16.70294 | 15.50466 | 16.66608 | 15.00686 | 16.12916 | 16.13621 | 19.1159 | 16.53916 | 15.06216 | 15.62542 | 17.27215 | 15.887 | 17.01687 |
| CUL4A | 15.58404 | 17.90687 | 16.11908 | 14.69402 | 14.55202 | 16.76386 | 15.6893 | 17.43435 | 18.5956 | 16.77906 | 16.43561 | 18.40859 | 15.56722 | 14.49607 | 15.74186 | 16.31651 | 14.89895 | 14.53914 |
| CUL4B | 16.8233 | 17.44549 | 15.8479 | 16.20717 | 17.86263 | 21.45604 | 18.25702 | 18.51329 | 15.55578 | 15.21645 | 17.18858 | 18.87546 | 15.38784 | 18.65897 | 16.7339 | 16.18925 | 16.73885 | 18.8037 |
| CUL5 | 16.0053 | 15.74017 | 15.48117 | 17.84732 | 17.47378 | 16.75204 | 17.3315 | 17.44848 | 15.64285 | 16.06578 | 15.47779 | 17.97496 | 17.12862 | 17.15606 | 16.39587 | 17.33582 | 17.57296 | 17.45386 |
| CUTA | 16.02368 | 15.93756 | 16.79364 | 21.16686 | 20.50304 | 16.78782 | 18.17418 | 18.56699 | 20.27398 | 20.14018 | 19.55994 | 19.97294 | 20.34685 | 17.81168 | 19.13165 | 15.4103 | 20.88082 | 20.93756 |
| CXADR | 19.74358 | 19.18528 | 18.65222 | 19.04673 | 22.94736 | 20.89729 | 19.68974 | 18.79469 | 18.89391 | 18.10283 | 18.71488 | 20.50889 | 18.56032 | 18.2824 | 18.85485 | 18.88022 | 19.27905 | 17.75481 |
| CXCR4 | 18.25768 | 19.78547 | 16.19872 | 16.75365 | 15.66865 | 14.23275 | 17.62522 | 14.73153 | 19.07197 | 19.82816 | 14.32088 | 13.64428 | 15.17626 | 15.19663 | 14.77295 | 15.22501 | 16.97066 | 15.58822 |
| CYB5D2 | 15.41856 | 15.45744 | 14.57185 | 14.49421 | 14.70902 | 15.59753 | 13.87879 | 15.39553 | 17.5746 | 14.49224 | 15.34053 | 12.77062 | 15.11989 | 15.06233 | 14.95527 | 15.17906 | 13.15624 | 16.05783 |
| CYB5R3 | 16.5521 | 16.37473 | 17.70188 | 16.62845 | 16.8558 | 17.25079 | 17.92337 | 18.26304 | 15.56846 | 16.88775 | 17.00633 | 17.92978 | 16.53316 | 15.85624 | 16.3722 | 17.07075 | 16.15224 | 16.57441 |
| CYC1 | 18.00457 | 16.1949 | 17.06813 | 15.56403 | 14.86667 | 17.03428 | 15.68008 | 16.01777 | 22.33633 | 18.99307 | 17.2711 | 15.32151 | 15.29931 | 15.45062 | 16.51053 | 15.99969 | 15.38353 | 15.28736 |
| CYFIP1 | 19.01327 | 17.62662 | 19.29254 | 19.08538 | 16.85284 | 18.06608 | 17.74297 | 17.2296 | 17.42675 | 16.76701 | 17.45842 | 19.81395 | 15.88739 | 17.80139 | 20.02279 | 19.79045 | 17.01501 | 17.39633 |
| CYFIP2 | 16.27614 | 16.81041 | 17.25795 | 18.28114 | 16.32288 | 15.70247 | 17.15696 | 16.96364 | 17.62414 | 16.78437 | 17.29696 | 18.82353 | 17.36832 | 17.37793 | 19.36792 | 18.75932 | 15.95262 | 15.03967 |
| CYLD | 17.77044 | 18.2101 | 16.30224 | 17.49511 | 16.86947 | 14.35321 | 17.75048 | 17.08941 | 14.68692 | 12.17874 | 17.08353 | 13.89422 | 17.60049 | 12.92958 | 16.58499 | 16.54022 | 15.26146 | 15.01375 |
| CYP51A1 | 15.35709 | 16.34007 | 15.16283 | 15.21565 | 15.48484 | 20.42589 | 17.71733 | 17.81488 | 14.57913 | 15.62596 | 16.33271 | 16.70754 | 15.10038 | 15.45927 | 14.10376 | 16.06338 | 14.83798 | 15.3849 |
| CYRIA | 15.7103 | 14.91906 | 15.34239 | 16.77017 | 14.38761 | 16.55779 | 15.88832 | 16.85553 | 17.20954 | 17.39715 | 16.65644 | 19.81164 | 19.57758 | 17.26585 | 15.24213 | 14.55733 | 15.2735 | 14.53025 |
| CYRIB | 16.85579 | 18.57558 | 17.68713 | 18.56918 | 20.41881 | 19.05657 | 17.11696 | 17.5593 | 19.3459 | 18.73864 | 19.06325 | 20.29133 | 17.30478 | 17.61276 | 18.11182 | 17.7798 | 20.51254 | 20.99477 |
| CZIB | 14.75226 | 15.32082 | 14.96284 | 19.20767 | 21.13788 | 21.006 | 21.80178 | 16.99526 | 18.54846 | 19.18644 | 17.97344 | 19.27766 | 21.02441 | 20.72404 | 18.56791 | 17.58762 | 21.82118 | 21.79203 |
| DAAM1 | 17.1664 | 16.90714 | 16.64388 | 18.10323 | 18.94228 | 17.04413 | 19.48998 | 17.8489 | 17.73609 | 17.94617 | 17.83172 | 19.20061 | 18.70979 | 18.26223 | 18.23609 | 18.21272 | 18.16073 | 17.26554 |
| DAD1 | 16.8726 | 14.4858 | 13.76418 | 16.25828 | 14.24863 | 17.40045 | 17.48609 | 15.91912 | 19.72445 | 16.38662 | 13.94027 | 14.83208 | 15.53573 | 15.04172 | 14.43989 | 15.3101 | 15.76955 | 15.20701 |
| DAG1 | 21.09908 | 20.69866 | 19.14334 | 18.849 | 18.94739 | 18.28467 | 16.49467 | 16.77069 | 19.20986 | 17.30855 | 16.67587 | 17.94176 | 19.19332 | 18.88781 | 19.23849 | 19.20889 | 18.74906 | 17.93645 |
| DAGLB | 16.68482 | 18.18494 | 15.73621 | 14.67139 | 16.12853 | 15.27384 | 15.16547 | 16.23441 | 15.54018 | 16.37077 | 17.15716 | 16.63315 | 15.65637 | 14.93211 | 14.59129 | 14.60441 | 16.79694 | 15.81412 |
| DAP3 | 14.36428 | 14.47506 | 14.3517 | 15.30875 | 15.23599 | 15.07078 | 14.2972 | 16.41691 | 18.47798 | 14.55193 | 18.28965 | 11.03467 | 15.8014 | 15.53857 | 13.91609 | 16.32407 | 15.8755 | 15.04424 |
| DARS1 | 15.57985 | 15.84397 | 17.93392 | 17.59586 | 17.61564 | 16.82352 | 17.16049 | 17.33985 | 17.05334 | 17.36632 | 18.76734 | 20.52398 | 18.41221 | 18.21271 | 16.80031 | 17.48717 | 17.75674 | 17.4412 |
| DARS2 | 15.15853 | 16.30587 | 15.38798 | 16.03983 | 16.87485 | 18.07236 | 15.76801 | 16.63416 | 19.49855 | 18.35774 | 16.50466 | 13.35961 | 15.25028 | 15.58304 | 15.60424 | 15.50718 | 16.50484 | 14.59891 |
| DAZAP1 | 17.40452 | 18.05728 | 15.97232 | 16.15398 | 16.97714 | 15.59831 | 19.06946 | 16.43545 | 17.8411 | 17.75942 | 19.12441 | 20.5756 | 18.85364 | 18.59564 | 16.7908 | 16.15068 | 16.53007 | 17.56775 |
| DBI | 20.94375 | 21.45983 | 22.43767 | 19.99975 | 18.24235 | 17.98842 | 19.06432 | 19.45632 | 19.78299 | 20.40888 | 20.66016 | 20.25873 | 18.41956 | 17.93722 | 22.83116 | 22.46158 | 18.65424 | 19.40741 |
| DBN1 | 16.85173 | 16.64687 | 19.58563 | 19.75153 | 20.71286 | 19.90869 | 17.76504 | 18.41748 | 18.24172 | 18.41134 | 19.87677 | 20.95239 | 19.9311 | 20.10605 | 19.88416 | 19.5425 | 21.15902 | 20.27318 |
| DBNL | 18.98014 | 18.92961 | 18.86118 | 19.38638 | 20.02053 | 18.90592 | 19.17239 | 19.12706 | 17.53371 | 16.38985 | 18.53073 | 19.42895 | 20.7177 | 20.58811 | 19.62073 | 18.8177 | 20.10343 | 20.25045 |
| DBT | 15.5703 | 16.81314 | 15.54487 | 14.74952 | 15.65061 | 15.54153 | 16.3465 | 15.94012 | 18.43388 | 16.53429 | 14.91314 | 13.62913 | 14.1584 | 15.16284 | 15.84329 | 15.27655 | 17.36214 | 14.7613 |
| DCAF11 | 14.55171 | 14.904 | 16.00551 | 17.03208 | 19.46076 | 12.66005 | 14.88662 | 14.81123 | 13.36178 | 14.75953 | 14.87935 | 18.08673 | 18.94633 | 13.1082 | 14.38353 | 12.95154 | 18.80143 | 18.67247 |
| DCAF7 | 16.21765 | 15.24788 | 14.62957 | 15.58384 | 16.0725 | 14.76214 | 14.56094 | 14.89971 | 13.60783 | 15.52375 | 16.41602 | 18.52733 | 16.18481 | 17.54465 | 16.53681 | 14.68704 | 13.17927 | 13.87756 |
| DCD | 16.43235 | 16.39356 | 17.81541 | 14.25475 | 20.23703 | 17.30014 | 17.84688 | 14.39204 | 16.66889 | 14.52026 | 16.3439 | 13.04677 | 15.56685 | 15.15443 | 16.70164 | 14.43162 | 14.9276 | 13.85346 |
| DCDC1 | 23.82664 | 23.65566 | 22.64252 | 24.10049 | 14.0956 | 14.69679 | 17.79127 | 17.6405 | 14.51823 | 13.91934 | 19.35415 | 11.17691 | 21.13242 | 19.43105 | 21.94837 | 22.02318 | 15.00796 | 15.39976 |
| DCHS1 | 15.95385 | 15.40174 | 13.8658 | 16.09079 | 19.71589 | 19.68749 | 22.15695 | 14.7996 | 18.2201 | 17.1119 | 16.48128 | 13.88752 | 16.02465 | 15.70092 | 14.44605 | 14.62328 | 13.96592 | 18.93253 |
| DCLK1 | 16.06903 | 14.6177 | 15.30538 | 15.3014 | 16.04981 | 14.37795 | 14.33491 | 16.70124 | 15.94511 | 14.02889 | 17.33459 | 17.3137 | 14.64614 | 15.22109 | 14.64576 | 14.9609 | 14.78112 | 15.44242 |
| DCN | 16.48497 | 14.49908 | 16.78002 | 17.45845 | 14.89108 | 16.62778 | 20.17511 | 17.2271 | 12.61095 | 15.55325 | 16.14451 | 15.0999 | 15.17564 | 14.7352 | 13.43406 | 16.57051 | 15.29117 | 13.44403 |
| DCP1A | 17.3025 | 14.1202 | 14.77855 | 15.11262 | 14.26852 | 15.73926 | 14.40401 | 15.80017 | 12.38956 | 14.29212 | 15.70811 | 16.64186 | 15.77122 | 16.63023 | 16.18839 | 15.876 | 15.64754 | 15.24524 |
| DCPS | 14.37762 | 15.28549 | 16.15216 | 14.57242 | 14.2823 | 16.26912 | 15.54246 | 16.19149 | 16.42928 | 16.61002 | 18.58795 | 19.09051 | 16.35074 | 15.49648 | 15.3083 | 14.79479 | 16.02449 | 15.52895 |
| DCTD | 14.32408 | 15.64294 | 18.56937 | 16.49497 | 18.96148 | 17.77144 | 18.20883 | 19.17631 | 14.10485 | 17.72459 | 16.8125 | 18.31955 | 20.02514 | 18.77788 | 17.84526 | 17.89842 | 20.38002 | 19.55058 |
| DCTN1 | 15.578 | 16.43192 | 16.60087 | 16.43031 | 18.09029 | 16.59157 | 18.22453 | 17.29993 | 17.26763 | 17.49597 | 17.01612 | 18.96026 | 19.02986 | 18.16344 | 16.84724 | 17.02616 | 18.10621 | 18.37268 |
| DCTN2 | 16.53508 | 15.85504 | 16.14054 | 18.00653 | 19.2118 | 17.50071 | 19.42485 | 18.1956 | 16.1117 | 17.33935 | 18.17975 | 18.87179 | 19.6698 | 19.45954 | 17.34363 | 17.77584 | 17.70639 | 20.35568 |
| DCTN3 | 14.43574 | 14.66304 | 14.06433 | 16.75849 | 17.07744 | 13.93771 | 14.44353 | 14.17791 | 16.56405 | 16.46696 | 18.23931 | 19.35128 | 13.30013 | 16.28187 | 17.10273 | 16.30555 | 15.05168 | 17.98561 |
| DCTN4 | 15.17958 | 15.4439 | 17.47955 | 15.78396 | 16.44098 | 15.91532 | 16.03659 | 16.05661 | 15.5832 | 17.44181 | 16.98725 | 18.40228 | 18.64499 | 18.84108 | 16.8579 | 16.8771 | 16.90697 | 16.56646 |
| DCTPP1 | 15.66696 | 16.16032 | 17.30142 | 16.9184 | 15.15056 | 16.12809 | 15.93935 | 15.73719 | 15.3384 | 14.87505 | 18.86105 | 19.58521 | 17.75169 | 15.38284 | 14.44591 | 16.41893 | 18.3382 | 16.69936 |
| DCUN1D1 | 16.73677 | 15.17538 | 15.40586 | 18.24251 | 16.22598 | 19.95784 | 17.32889 | 15.61438 | 14.39844 | 15.32815 | 17.4974 | 14.98188 | 17.37516 | 16.28522 | 15.75174 | 15.56741 | 16.54703 | 14.78399 |
| DCUN1D3 | 16.08067 | 15.70636 | 17.13816 | 17.55282 | 16.9579 | 18.28114 | 14.92832 | 17.50121 | 15.91274 | 16.36748 | 16.00689 | 15.21501 | 15.28267 | 17.35077 | 17.72029 | 17.44986 | 18.59565 | 19.81234 |
| DCXR | 16.26619 | 16.03442 | 15.31752 | 18.06345 | 14.35753 | 17.80836 | 13.45275 | 15.81276 | 17.90135 | 16.40319 | 14.63642 | 18.18572 | 16.89963 | 14.92605 | 18.63899 | 18.11621 | 16.01568 | 15.26312 |
| DDAH1 | 18.01181 | 17.95295 | 18.90909 | 18.33601 | 19.02818 | 18.28043 | 16.39994 | 16.4555 | 18.39544 | 18.64531 | 17.8005 | 17.87013 | 20.40371 | 19.75491 | 18.75989 | 18.97753 | 20.02879 | 20.35122 |
| DDAH2 | 18.36142 | 17.93457 | 18.04909 | 18.35161 | 20.14455 | 17.39742 | 16.04306 | 16.41159 | 19.1958 | 18.00275 | 18.83813 | 17.30904 | 16.75279 | 16.98532 | 18.08762 | 17.97598 | 22.82734 | 21.55678 |
| DDB1 | 16.46133 | 17.95985 | 16.52328 | 16.21435 | 16.74902 | 16.98057 | 16.21589 | 16.42033 | 16.61617 | 17.2887 | 17.36613 | 19.47551 | 16.95902 | 16.27695 | 16.21312 | 17.19122 | 17.3493 | 16.43751 |
| DDC | 18.64957 | 18.98987 | 21.47872 | 19.02196 | 14.4525 | 16.17835 | 15.53372 | 15.93186 | 19.87956 | 20.94763 | 16.06952 | 16.04729 | 22.73624 | 16.46333 | 16.61429 | 18.51693 | 16.09089 | 16.28423 |
| DDO | 15.46401 | 14.24525 | 15.01174 | 16.00867 | 15.84267 | 18.31108 | 19.58949 | 19.06115 | 18.96566 | 18.30017 | 14.59768 | 15.50632 | 17.49001 | 15.70616 | 14.78243 | 15.91194 | 16.19407 | 18.72374 |
| DDOST | 15.4995 | 16.02552 | 17.59744 | 16.10503 | 14.57555 | 16.37443 | 16.84724 | 17.27761 | 24.57526 | 23.0284 | 21.39113 | 18.26637 | 15.16191 | 17.38834 | 17.31916 | 17.56794 | 15.30456 | 15.94035 |
| DDR1 | 19.12193 | 18.69327 | 17.15388 | 16.23592 | 15.5782 | 17.93414 | 16.22965 | 16.98277 | 18.58609 | 17.76805 | 17.52409 | 17.11696 | 16.04209 | 15.66466 | 16.87697 | 16.49005 | 16.98066 | 16.21545 |
| DDT | 20.19669 | 20.41775 | 20.53925 | 20.09417 | 19.78473 | 19.86685 | 19.88539 | 17.27625 | 16.31732 | 19.25514 | 20.27388 | 21.41989 | 20.79961 | 20.97046 | 22.81477 | 21.62689 | 19.99928 | 18.90306 |
| DDX1 | 18.01111 | 17.25631 | 18.09888 | 19.19734 | 18.76386 | 17.69032 | 18.51101 | 17.23402 | 16.23292 | 16.32878 | 18.91388 | 19.73839 | 19.86988 | 19.17465 | 18.11756 | 17.98024 | 19.37887 | 18.55257 |
| DDX17 | 15.64334 | 17.16388 | 17.84292 | 15.53844 | 18.35536 | 16.76603 | 18.35718 | 17.92236 | 16.82511 | 17.0489 | 20.4287 | 20.61102 | 18.88871 | 16.05349 | 16.52895 | 16.21758 | 15.44435 | 16.48011 |
| DDX19B | 18.26234 | 17.64757 | 16.5012 | 16.85721 | 20.09107 | 16.26191 | 16.61543 | 17.96386 | 16.02726 | 16.39182 | 17.02602 | 17.95084 | 18.40654 | 19.13819 | 15.55053 | 16.90576 | 16.52861 | 20.14614 |

| | | | | | | | | | | | | | | | | | | |
|---|---|---|---|---|---|---|---|---|---|---|---|---|---|---|---|---|---|---|
| DDX21 | 18.80089 | 15.8774 | 15.18336 | 15.24471 | 18.07395 | 15.73426 | 18.22309 | 18.64332 | 13.66399 | 16.6538 | 17.66885 | 17.57303 | 16.43492 | 17.74512 | 16.41249 | 16.07106 | 19.76923 | 19.78347 |
| DDX39A | 16.29123 | 15.3805 | 14.78422 | 15.06228 | 14.72049 | 15.47529 | 19.98753 | 19.13676 | 13.66935 | 15.70791 | 16.38144 | 16.33119 | 17.08075 | 13.72975 | 16.8287 | 15.78977 | 15.91885 | 15.69463 |
| DDX39B | 17.93249 | 17.64653 | 17.23322 | 16.72609 | 17.33464 | 17.50901 | 18.45943 | 17.58794 | 17.91601 | 18.39539 | 19.55007 | 21.12223 | 16.29663 | 16.01562 | 16.41073 | 17.57755 | 15.81225 | 15.78551 |
| DDX3X | 20.09409 | 19.43006 | 19.8223 | 19.61958 | 16.20644 | 16.41734 | 16.85371 | 16.5061 | 15.86427 | 16.54751 | 19.30673 | 17.88225 | 17.08661 | 17.0062 | 20.35772 | 19.63083 | 16.24782 | 14.57498 |
| DDX42 | 15.28769 | 14.62962 | 14.77369 | 16.1851 | 15.64822 | 16.27608 | 15.90668 | 14.76485 | 14.18489 | 15.86521 | 19.05436 | 17.79366 | 17.23622 | 15.22621 | 14.83486 | 15.92921 | 15.04599 | 14.66489 |
| DDX46 | 15.24865 | 15.04014 | 15.88654 | 14.28972 | 15.91175 | 15.6251 | 19.18056 | 17.12804 | 13.37098 | 15.88433 | 15.85362 | 18.13612 | 18.4204 | 17.36753 | 16.28716 | 15.38783 | 14.69197 | 15.89988 |
| DDX5 | 17.1054 | 17.25502 | 16.36782 | 15.8896 | 17.32454 | 16.70036 | 19.55936 | 18.8473 | 17.05763 | 17.41437 | 18.34295 | 20.11222 | 17.58715 | 17.71542 | 16.575 | 16.98211 | 15.64985 | 16.79486 |
| DDX50 | 16.38912 | 19.19212 | 16.18663 | 14.34671 | 13.956 | 14.88175 | 19.28741 | 20.33085 | 16.16854 | 15.26141 | 14.70642 | 13.89256 | 13.86499 | 14.71091 | 15.61375 | 15.10943 | 14.99834 | 13.92663 |
| DDX6 | 19.94124 | 20.95636 | 19.31965 | 20.11862 | 19.1442 | 18.29827 | 19.44635 | 18.00505 | 17.39552 | 18.65516 | 19.66584 | 20.2212 | 20.19211 | 19.24421 | 19.96255 | 19.81119 | 18.59254 | 19.21826 |
| DECR1 | 19.95018 | 18.41914 | 18.51063 | 19.08451 | 15.53922 | 16.57637 | 17.78761 | 17.17248 | 21.12552 | 19.74942 | 15.558 | 16.72452 | 18.66744 | 17.95357 | 16.08492 | 17.8876 | 16.99999 | 18.80885 |
| DECR2 | 18.13896 | 18.89431 | 17.7905 | 15.54617 | 16.28122 | 14.70159 | 16.46553 | 17.01389 | 19.7263 | 17.18921 | 15.7616 | 14.13087 | 17.46113 | 17.22457 | 16.29171 | 16.61889 | 16.79611 | 14.01932 |
| DEGS1 | 16.61405 | 16.58432 | 14.57849 | 17.0008 | 17.18079 | 17.50901 | 14.31818 | 14.03196 | 15.10457 | 19.81135 | 12.64237 | 13.45011 | 13.47183 | 13.8417 | 15.46917 | 11.15824 | 16.86863 | 17.83678 | 15.45203 |
| DEK | 15.9343 | 14.88661 | 14.82427 | 14.20076 | 15.34453 | 16.8318 | 18.04055 | 19.25577 | 12.40302 | 17.42107 | 15.22747 | 15.64599 | 20.17346 | 19.1474 | 14.23042 | 15.79712 | 17.81812 | 14.21908 |
| DENR | 17.88859 | 17.92292 | 17.29637 | 18.58226 | 19.19524 | 18.33449 | 16.89599 | 14.34099 | 17.43876 | 15.59221 | 18.62161 | 18.95246 | 20.15552 | 17.33998 | 17.09916 | 17.91776 | 18.67767 | 20.67369 |
| DERA | 16.78296 | 14.43952 | 15.4375 | 14.52855 | 16.06542 | 14.50083 | 14.40021 | 17.05607 | 16.55804 | 16.31524 | 14.89348 | 18.93764 | 15.37248 | 16.56769 | 14.89214 | 16.94245 | 16.9288 | 15.33915 |
| DFFA | 14.15046 | 12.97247 | 13.19342 | 16.12703 | 17.67399 | 13.05278 | 16.99633 | 15.48756 | 12.71548 | 13.7669 | 15.90265 | 13.64446 | 19.77423 | 19.98097 | 15.66766 | 14.53525 | 18.25629 | 16.51283 |
| DGKH | 16.09545 | 15.14762 | 15.40673 | 15.81926 | 12.81269 | 15.52227 | 16.06275 | 14.2885 | 15.99012 | 15.68456 | 15.55709 | 12.54392 | 16.50981 | 17.97276 | 14.76136 | 15.31951 | 18.26062 | 17.11889 |
| DGLUCY | 13.83774 | 16.89189 | 16.62544 | 17.22385 | 14.88555 | 15.38357 | 15.05666 | 15.43415 | 17.33084 | 17.83865 | 13.80363 | 12.41935 | 12.18763 | 16.10691 | 14.04391 | 16.34676 | 15.36289 | 13.45659 |
| DHCR24 | 15.51534 | 14.32733 | 15.68642 | 16.18935 | 15.7423 | 13.88091 | 15.17179 | 16.07407 | 15.73087 | 14.95438 | 15.46879 | 18.73896 | 15.92179 | 18.00254 | 17.57171 | 16.44469 | 14.61196 | 14.04112 |
| DHFR | 16.21596 | 15.91856 | 16.51124 | 18.45165 | 19.70444 | 18.00274 | 17.71598 | 17.70227 | 16.22974 | 16.5628 | 17.08121 | 16.11565 | 19.36565 | 18.60499 | 17.37564 | 16.40105 | 16.00153 | 18.28014 |
| DHFR2 | 17.76428 | 18.0374 | 18.39218 | 17.92571 | 16.57881 | 16.74131 | 13.69873 | 13.98612 | 15.7877 | 16.14063 | 18.94227 | 15.6826 | 19.38645 | 13.66831 | 18.38992 | 18.90632 | 12.88791 | 14.72937 |
| DHRS4 | 14.96316 | 15.96255 | 16.73878 | 15.6055 | 15.7142 | 18.44109 | 20.14603 | 19.47011 | 21.79397 | 21.34402 | 16.97028 | 12.78239 | 15.68328 | 13.9448 | 13.77518 | 16.80306 | 15.88759 | 16.03295 |
| DHRS4L1 | 15.50118 | 15.42713 | 16.21847 | 14.86442 | 15.10133 | 16.0921 | 18.44852 | 17.94212 | 21.22089 | 20.50946 | 15.87982 | 11.3928 | 14.60986 | 14.81978 | 13.48354 | 15.02158 | 15.48637 | 16.17108 |
| DHRS4L2 | 17.69758 | 18.22773 | 18.109 | 19.42818 | 16.44426 | 17.34015 | 20.0447 | 17.26952 | 21.5002 | 21.27538 | 15.48024 | 13.91938 | 15.11287 | 16.83881 | 17.43829 | 18.53882 | 16.50306 | 15.68458 |
| DHRS7 | 18.94045 | 18.69298 | 20.65364 | 14.94381 | 15.58505 | 14.99449 | 15.58328 | 15.90049 | 19.32177 | 16.4686 | 20.29907 | 19.22661 | 12.59915 | 13.22241 | 19.0934 | 19.27203 | 15.39765 | 15.00304 |
| DHTKD1 | 18.76743 | 19.15817 | 20.44429 | 19.97321 | 14.97629 | 16.77069 | 16.095 | 17.18482 | 18.69468 | 16.66114 | 14.53003 | 13.6618 | 17.64161 | 16.58878 | 19.96898 | 20.10422 | 15.28827 | 15.96085 |
| DHX15 | 17.14672 | 18.61292 | 16.66696 | 15.70513 | 15.9457 | 16.17406 | 16.77267 | 16.10456 | 16.47366 | 16.14973 | 18.44386 | 20.08187 | 16.10202 | 16.41966 | 16.51901 | 16.54662 | 15.46396 | 15.27073 |
| DHX16 | 14.7382 | 14.4628 | 14.54097 | 12.57504 | 14.28039 | 13.78347 | 14.63442 | 16.92076 | 14.67872 | 14.93551 | 15.23762 | 16.99158 | 13.42717 | 14.80957 | 14.51589 | 14.09586 | 12.68202 | 15.48606 |
| DHX29 | 15.94983 | 15.45611 | 15.50163 | 15.17216 | 15.663 | 15.58491 | 16.19172 | 16.86311 | 14.90198 | 14.48719 | 17.5094 | 18.19011 | 16.53993 | 16.15795 | 16.44909 | 16.33258 | 16.65446 | 17.36762 |
| DHX30 | 16.42712 | 16.33392 | 16.10129 | 16.26343 | 14.09242 | 13.86202 | 16.1842 | 18.5036 | 17.34202 | 16.13208 | 16.64935 | 17.58422 | 15.41009 | 14.66079 | 14.00136 | 14.98517 | 14.50459 | 14.03484 |
| DHX36 | 18.46809 | 18.64228 | 17.85888 | 16.94481 | 14.95642 | 16.17298 | 14.31335 | 16.27171 | 13.13744 | 14.98438 | 15.86087 | 18.54248 | 15.7842 | 15.73414 | 18.80375 | 18.14526 | 16.49812 | 15.83913 |
| DHX9 | 17.42595 | 25.36602 | 18.11325 | 26.12663 | 16.65575 | 19.16652 | 18.41788 | 17.75537 | 16.05049 | 17.03861 | 18.03678 | 19.85095 | 18.21347 | 17.68879 | 19.80475 | 25.23897 | 15.52673 | 15.9889 |
| DIABLO | 18.33703 | 17.30295 | 14.327 | 14.92075 | 14.55179 | 14.49997 | 15.89153 | 16.38292 | 20.07182 | 18.77304 | 17.36949 | 18.08541 | 15.39995 | 15.46492 | 15.99384 | 16.14071 | 14.5507 | 14.83419 |
| DIAPH1 | 16.30934 | 16.15165 | 16.22294 | 16.74356 | 18.3821 | 16.51558 | 17.39816 | 16.43735 | 16.20594 | 16.85362 | 17.29202 | 18.56591 | 18.05326 | 18.44904 | 17.7371 | 17.18898 | 18.97442 | 18.39028 |
| DICER1 | 16.44924 | 13.78767 | 14.3895 | 14.59605 | 13.15363 | 15.78212 | 16.35472 | 16.51789 | 13.30247 | 14.66786 | 12.02792 | 15.87802 | 14.34007 | 11.96072 | 13.52701 | 16.05773 | 16.5697 | 13.99397 |
| DIMT1 | 15.61917 | 15.50942 | 15.25181 | 14.23576 | 14.3572 | 16.87882 | 15.35477 | 14.99728 | 13.57972 | 15.37292 | 18.01608 | 18.2012 | 16.99669 | 15.36616 | 12.96746 | 15.20672 | 14.27857 | 14.35638 |
| DIP2A | 16.8081 | 19.178 | 18.62016 | 19.00195 | 18.05247 | 18.67709 | 16.52774 | 16.44673 | 18.44952 | 18.65178 | 17.79269 | 19.02099 | 16.04537 | 17.20314 | 19.1747 | 19.82485 | 16.19766 | 18.66262 |
| DIP2B | 19.76235 | 18.00692 | 17.39327 | 18.57689 | 17.42021 | 17.35017 | 17.43993 | 17.17728 | 17.8402 | 17.57433 | 19.16514 | 19.58522 | 18.56075 | 17.93131 | 19.00168 | 18.7756 | 19.28839 | 20.50446 |
| DIP2C | 14.84452 | 16.80995 | 18.44028 | 14.94214 | 13.60888 | 17.68612 | 16.92205 | 14.81184 | 17.93196 | 18.0108 | 15.78244 | 13.80783 | 17.81695 | 16.51962 | 19.20351 | 18.62874 | 17.97028 | 14.97922 |
| DIPK1A | 21.05616 | 16.00927 | 16.2522 | 14.86427 | 15.14102 | 15.79698 | 15.66672 | 15.47641 | 19.14813 | 15.99909 | 16.88803 | 14.51801 | 16.10334 | 15.88725 | 14.32424 | 16.60834 | 16.78231 | 15.71203 |
| DIPK1B | 13.39031 | 14.36025 | 14.88292 | 16.31466 | 11.8122 | 16.42528 | 17.60591 | 16.64095 | 20.74891 | 16.19057 | 14.53031 | 18.51474 | 14.56557 | 16.52345 | 15.34094 | 14.05013 | 14.66352 | 14.34106 |
| DIPK2A | 17.24545 | 15.72662 | 17.15763 | 16.81097 | 15.52117 | 19.55889 | 17.19988 | 16.45978 | 22.1929 | 20.08874 | 16.42383 | 16.53522 | 16.045 | 16.72734 | 16.29804 | 15.68364 | 16.99711 | 15.94775 |
| DIS3L2 | 16.83588 | 16.34514 | 16.69693 | 17.21961 | 14.29956 | 15.7633 | 17.62727 | 16.01372 | 13.89304 | 17.36845 | 16.02694 | 18.18952 | 14.51185 | 14.92022 | 17.22601 | 16.61632 | 15.64438 | 15.41016 |
| DKC1 | 19.26617 | 18.28194 | 18.07343 | 18.17437 | 13.72066 | 15.58181 | 15.92162 | 17.36187 | 15.65562 | 16.31503 | 16.28476 | 18.032 | 17.22364 | 15.75426 | 15.56174 | 18.30138 | 15.84879 | 15.24003 |
| DKK1 | 16.19577 | 14.83922 | 14.98722 | 14.88979 | 14.22708 | 16.07467 | 15.85138 | 15.46593 | 16.55887 | 13.83351 | 15.6197 | 13.32158 | 15.6843 | 15.90946 | 15.28873 | 15.99303 | 19.56432 | 17.60329 |
| DKK3 | 14.75723 | 17.80262 | 15.08786 | 16.54906 | 14.85915 | 15.75239 | 18.45318 | 14.8463 | 16.13218 | 15.02899 | 21.31988 | 14.92956 | 14.71421 | 14.22158 | 17.71798 | 15.28044 | 15.31021 | 14.0759 |
| DLAT | 17.82387 | 18.18483 | 15.01594 | 16.80565 | 16.17434 | 16.36025 | 16.69183 | 17.5204 | 19.96126 | 18.88117 | 15.95083 | 18.58555 | 16.01832 | 16.2492 | 15.49983 | 15.76994 | 17.41423 | 15.81516 |
| DLD | 17.63676 | 17.77916 | 16.88804 | 17.13414 | 16.76049 | 16.40156 | 17.97588 | 16.87571 | 20.03513 | 18.60991 | 17.13485 | 13.15857 | 17.65528 | 17.60269 | 16.85795 | 16.98532 | 16.3333 | 16.99089 |
| DLG1 | 17.30397 | 18.20378 | 18.60559 | 18.36539 | 21.31335 | 20.62209 | 16.02879 | 16.67514 | 19.33836 | 20.32497 | 18.76057 | 17.53327 | 19.96656 | 20.61576 | 18.79629 | 19.06594 | 20.9169 | 21.40703 |
| DLG2 | 15.2286 | 17.22151 | 18.54184 | 17.98476 | 19.11868 | 19.4378 | 16.28716 | 16.6252 | 17.57084 | 19.10774 | 18.09746 | 17.04795 | 22.77606 | 21.30763 | 18.64672 | 17.68892 | 19.67368 | 17.42055 |
| DLG3 | 16.35689 | 15.5846 | 20.12928 | 19.20127 | 16.77914 | 15.17071 | 16.16134 | 16.57879 | 18.59741 | 18.57038 | 18.19892 | 19.17615 | 18.45385 | 17.80539 | 19.31185 | 18.90019 | 19.35664 | 17.18689 |
| DLG5 | 16.93525 | 16.32998 | 16.79778 | 16.19892 | 18.95281 | 17.0894 | 15.28858 | 16.62507 | 16.73728 | 15.97755 | 15.6209 | 14.53979 | 16.82409 | 16.4668 | 15.5961 | 15.59854 | 19.99247 | 19.81892 |
| DLGAP4 | 15.89446 | 17.62055 | 17.09112 | 19.5203 | 16.09843 | 13.81612 | 15.40302 | 16.97176 | 14.92463 | 15.09972 | 16.83759 | 16.9257 | 16.86996 | 17.77084 | 16.16007 | 15.87409 | 18.99263 | 18.44062 |
| DLL1 | 14.53241 | 14.50343 | 16.61387 | 16.97857 | 13.61444 | 13.88188 | 12.34024 | 14.4712 | 14.2439 | 18.2255 | 13.95256 | 18.38536 | 13.11261 | 16.70741 | 14.92261 | 15.88479 | 14.65409 | 14.51185 |
| DLL3 | 16.42945 | 16.32067 | 14.69903 | 15.42108 | 14.38709 | 15.07434 | 15.91115 | 16.69228 | 19.28866 | 15.38073 | 15.60162 | 16.41609 | 15.73014 | 15.2416 | 14.61043 | 15.05866 | 16.86729 | 15.40158 |
| DLST | 20.00195 | 20.52132 | 17.00674 | 16.41976 | 15.60601 | 15.52646 | 20.95231 | 18.00426 | 20.99949 | 20.25905 | 16.16093 | 15.40914 | 18.59848 | 18.27135 | 17.97626 | 17.0709 | 15.12924 | 15.62445 |
| DMAC2L | 14.3648 | 14.63118 | 15.31862 | 14.8109 | 19.64003 | 15.37009 | 17.82936 | 17.00967 | 18.4798 | 18.14496 | 15.75144 | 13.37352 | 18.14956 | 15.86549 | 15.74134 | 13.37725 | 14.86902 | 13.2268 |
| DMXL1 | 14.77712 | 17.10035 | 13.47146 | 13.34973 | 16.76039 | 14.45853 | 19.09857 | 16.43273 | 12.56215 | 12.10438 | 15.46127 | 16.19161 | 12.86345 | 13.17293 | 13.83064 | 15.13055 | 12.79481 | 14.44732 |
| DNAH17 | 15.66179 | 15.70676 | 15.7808 | 13.75582 | 13.26767 | 13.52741 | 16.04919 | 15.38209 | 18.59477 | 18.84042 | 19.67442 | 12.53672 | 13.91529 | 15.26934 | 13.96475 | 15.19855 | 17.73332 | 14.09301 |
| DNAJA1 | 20.28879 | 20.96747 | 19.95288 | 21.39366 | 21.28455 | 21.29703 | 19.37585 | 18.04755 | 20.9622 | 21.29838 | 18.25155 | 18.7305 | 22.08488 | 21.9084 | 21.1124 | 20.28737 | 18.63202 | 19.12546 |
| DNAJA2 | 19.74342 | 19.49741 | 18.98183 | 18.58709 | 20.21717 | 18.97667 | 18.01428 | 19.18808 | 18.17747 | 18.59154 | 18.50682 | 18.39756 | 19.71937 | 20.16963 | 19.43855 | 19.26048 | 17.92662 | 18.02701 |
| DNAJA3 | 16.4944 | 16.30703 | 14.91072 | 16.94323 | 12.40936 | 15.67702 | 15.73356 | 15.88985 | 19.97972 | 16.61814 | 14.34264 | 13.53685 | 11.70468 | 14.27437 | 17.2173 | 14.7229 | 14.14989 | 14.86198 |
| DNAJB1 | 18.9099 | 18.88941 | 17.81782 | 18.98159 | 20.31176 | 18.76201 | 19.37206 | 17.37773 | 18.50381 | 19.09341 | 17.8319 | 16.54246 | 20.27476 | 19.48851 | 17.54212 | 17.3427 | 20.24874 | 20.14454 |
| DNAJB11 | 21.10964 | 17.66912 | 17.61456 | 16.92508 | 17.08618 | 17.32534 | 18.46658 | 17.45606 | 22.26058 | 20.89587 | 20.69751 | 18.06531 | 16.83599 | 19.46685 | 16.8773 | 16.60692 | 16.32774 | 16.04723 |
| DNAJB12 | 18.27776 | 17.53213 | 16.46927 | 15.53611 | 14.81787 | 16.63852 | 17.19268 | 16.12107 | 20.1941 | 18.99302 | 15.54993 | 14.97643 | 15.48116 | 15.6052 | 15.64322 | 16.0886 | 16.06705 | 15.77813 |
| DNAJB2 | 19.40075 | 19.89719 | 17.87111 | 18.67085 | 15.89761 | 17.09889 | 15.21352 | 16.86267 | 14.53553 | 15.42067 | 14.86621 | 14.28531 | 16.54491 | 15.82996 | 18.46971 | 18.61118 | 16.73245 | 15.81328 |
| DNAJB6 | 21.72228 | 22.57039 | 21.66801 | 19.43496 | 17.06044 | 15.51543 | 15.81279 | 17.62953 | 16.36558 | 15.83632 | 15.26283 | 16.5317 | 19.75381 | 17.67872 | 21.38653 | 20.81416 | 16.93894 | 15.53351 |
| DNAJC1 | 14.56387 | 14.78918 | 16.34969 | 16.07567 | 15.29839 | 14.10968 | 12.81094 | 16.49139 | 19.45575 | 16.51637 | 14.47051 | 14.78694 | 12.99798 | 15.82038 | 17.83456 | 15.20026 | 14.52574 | 13.02821 |
| DNAJC10 | 15.73906 | 17.19152 | 16.58796 | 17.05637 | 18.63289 | 19.38349 | 17.01414 | 16.97788 | 21.343 | 20.75412 | 18.38696 | 17.22484 | 15.33474 | 16.53188 | 18.08907 | 16.60446 | 17.46516 | 17.79086 |
| DNAJC11 | 14.85803 | 15.61427 | 16.78438 | 16.7 | 15.58384 | 16.07078 | 15.06917 | 14.72896 | 19.60569 | 18.59045 | 14.80235 | 13.26804 | 15.0164 | 15.09714 | 15.30674 | 15.59599 | 14.44156 | 14.54084 |
| DNAJC13 | 16.2351 | 16.00614 | 18.20147 | 15.95975 | 18.46165 | 17.6664 | 16.31892 | 16.47351 | 14.79812 | 15.97943 | 14.82332 | 19.01647 | 17.85564 | 17.4767 | 15.79197 | 18.04654 | 17.17837 | 16.43611 |
| DNAJC16 | 15.46268 | 15.32856 | 15.31633 | 16.1228 | 16.72289 | 16.31654 | 13.50242 | 16.01708 | 18.00893 | 17.37663 | 14.60971 | 13.40264 | 16.86946 | 17.12134 | 15.19924 | 15.92434 | 15.85636 | 16.98086 |
| DNAJC2 | 15.16069 | 13.14065 | 16.80989 | 14.86875 | 16.73532 | 13.53695 | 17.11136 | 17.49061 | 13.019 | 14.86558 | 14.70415 | 15.11789 | 18.29868 | 17.67882 | 15.24608 | 16.21458 | 16.8596 | 16.23559 |
| DNAJC3 | 17.04031 | 15.54744 | 16.15482 | 15.5411 | 15.59548 | 15.81277 | 17.0451 | 16.85307 | 20.60442 | 19.65653 | 17.20935 | 14.40366 | 15.2386 | 16.08549 | 15.74968 | 15.30298 | 16.36829 | 16.32036 |
| DNAJC5 | 19.2943 | 19.13635 | 21.19425 | 18.74646 | 15.68533 | 19.02858 | 16.6184 | 15.93356 | 15.40235 | 16.79217 | 15.71632 | 21.33592 | 16.88881 | 16.16661 | 20.21769 | 21.09321 | 15.70538 | 13.63499 |
| DNAJC7 | 16.45817 | 16.48446 | 17.1754 | 17.94467 | 18.47232 | 17.0649 | 18.7569 | 17.7457 | 17.32257 | 17.80301 | 17.22159 | 17.7205 | 19.74114 | 18.70122 | 17.33009 | 16.38227 | 16.84505 | 16.65184 |
| DNAJC8 | 15.13538 | 16.17506 | 18.0209 | 15.63676 | 14.13528 | 14.06499 | 15.54464 | 19.06731 | 17.24261 | 17.60414 | 13.75387 | 19.63346 | 13.65261 | 15.32359 | 14.65343 | 13.19883 | 14.58121 | 15.46631 |
| DNAJC9 | 16.8548 | 15.51731 | 15.92842 | 15.89092 | 15.16399 | 14.77436 | 15.89433 | 17.38109 | 17.00643 | 19.14981 | 18.13033 | 17.13304 | 16.7784 | 15.39237 | 15.04296 | 14.52966 | 15.75461 | 16.36326 |
| DNASE1L1 | 19.84802 | 21.04417 | 19.29525 | 19.00749 | 13.52573 | 14.12661 | 15.30302 | 16.56715 | 17.48521 | 15.59905 | 15.89411 | 17.85941 | 15.44652 | 17.57899 | 19.6348 | 19.08865 | 12.94267 | 13.6947 |
| DNASE2 | 16.96297 | 18.86512 | 16.52681 | 17.40746 | 17.22724 | 17.51982 | 17.26901 | 16.42559 | 21.31199 | 21.68985 | 19.32793 | 16.72428 | 17.99631 | 17.65541 | 15.64237 | 17.0917 | 15.33584 | 18.23582 |
| DNER | 14.43026 | 14.00662 | 17.39683 | 14.18578 | 13.97238 | 15.89821 | 14.80858 | 17.41722 | 19.2084 | 17.05857 | 14.82287 | 12.46854 | 14.37566 | 18.17708 | 15.62206 | 14.72904 | 13.20677 | 14.86784 |
| DNM1 | 15.26853 | 14.55885 | 16.94964 | 15.72457 | 15.52137 | 16.86169 | 15.06083 | 16.39987 | 18.2184 | 17.33951 | 15.43693 | 14.5981 | 14.62877 | 16.22579 | 15.43033 | 13.76114 | 15.66852 | 15.72256 |
| DNM1L | 16.10993 | 16.29686 | 16.44481 | 16.08369 | 15.51566 | 15.95439 | 19.0438 | 16.77062 | 16.45822 | 16.27993 | 17.28396 | 19.33874 | 18.10878 | 17.62642 | 16.50752 | 15.93334 | 15.92496 | 16.16264 |
| DNM2 | 19.97052 | 20.1782 | 19.76233 | 19.20089 | 18.68083 | 17.31258 | 17.63062 | 17.43813 | 17.48185 | 17.75268 | 17.36855 | 17.93965 | 20.06183 | 19.38717 | 18.91913 | 19.29885 | 18.0409 | 18.18102 |
| DNM3 | 21.38026 | 21.66027 | 20.42053 | 20.9587 | 20.29755 | 19.0853 | 19.23475 | 16.17285 | 20.33141 | 20.59703 | 18.28127 | 18.99191 | 20.52722 | 19.57673 | 21.26955 | 20.17344 | 19.07604 | 19.34954 |
| DNMT1 | 15.96517 | 14.44371 | 14.57258 | 14.70846 | 14.281 | 14.31711 | 15.04415 | 15.69612 | 15.06233 | 16.54842 | 16.85061 | 15.65241 | 13.60443 | 15.68679 | 14.18046 | 16.89144 | 13.33488 | 16.53573 |

| | | | | | | | | | | | | | | | | | | |
|---|---|---|---|---|---|---|---|---|---|---|---|---|---|---|---|---|---|---|
| DNPEP | 15.94156 | 17.19271 | 15.56493 | 15.98776 | 15.08226 | 16.99496 | 15.36575 | 16.84894 | 16.89678 | 16.77174 | 15.56715 | 18.75758 | 17.1343 | 16.76591 | 15.47297 | 16.33143 | 17.66314 | 16.84953 |
| DNPH1 | 18.39348 | 18.32424 | 18.87264 | 17.4488 | 14.73147 | 15.95564 | 16.72713 | 16.18871 | 15.1143 | 14.31105 | 19.29404 | 20.13706 | 16.07781 | 13.04823 | 19.57489 | 19.0953 | 15.00897 | 15.05363 |
| DOCK1 | 15.25745 | 15.22987 | 17.41074 | 17.90267 | 16.41297 | 15.49403 | 16.76034 | 16.7014 | 13.9511 | 13.90917 | 15.72905 | 14.95687 | 12.02928 | 16.43712 | 19.52747 | 16.84177 | 15.89816 | 14.74969 |
| DOCK10 | 16.25411 | 15.70091 | 16.8303 | 16.35039 | 19.88199 | 17.70144 | 16.72074 | 16.36528 | 17.20701 | 15.91712 | 15.93596 | 16.51052 | 13.90658 | 15.37647 | 16.64967 | 16.13476 | 17.58698 | 17.8037 |
| DOCK7 | 16.55246 | 16.38206 | 17.69881 | 20.13346 | 19.48614 | 20.93968 | 19.36441 | 17.1608 | 14.50607 | 18.17409 | 20.15581 | 18.15027 | 17.8742 | 21.16259 | 17.77956 | 17.87387 | 18.65807 | 20.78804 |
| DOCK9 | 17.47377 | 19.46791 | 17.22001 | 16.53343 | 17.75646 | 16.39282 | 15.81362 | 15.71094 | 16.68491 | 16.88311 | 14.90249 | 15.76639 | 18.17732 | 17.10927 | 19.04085 | 18.41764 | 15.57285 | 15.24569 |
| DOP1B | 15.46776 | 16.2573 | 14.17593 | 15.50939 | 14.18095 | 17.01765 | 15.55201 | 16.39893 | 17.52376 | 16.16297 | 16.19725 | 17.07277 | 14.74554 | 15.45005 | 15.86805 | 16.06121 | 15.69236 | 15.51617 |
| DPP3 | 16.71394 | 16.23209 | 17.3009 | 17.50691 | 16.61952 | 16.92344 | 16.30333 | 16.60906 | 17.80043 | 17.54711 | 19.39935 | 19.95007 | 17.5226 | 16.5669 | 17.5996 | 17.68452 | 18.24635 | 17.63151 |
| DPP4 | 19.11783 | 18.88588 | 22.71633 | 22.80028 | 16.07535 | 16.35117 | 15.43195 | 16.6768 | 15.2513 | 16.02804 | 17.30438 | 15.3665 | 16.23866 | 16.81278 | 22.89363 | 22.36723 | 20.09387 | 19.28453 |
| DPP7 | 17.7235 | 17.19142 | 16.63921 | 15.88576 | 15.31234 | 18.26036 | 17.17521 | 16.31393 | 19.12876 | 15.64858 | 16.9629 | 15.2946 | 16.16121 | 15.73821 | 18.12667 | 16.08662 | 17.09286 | 15.5992 |
| DPP9 | 15.64657 | 17.53554 | 17.00317 | 14.90561 | 18.96488 | 17.25294 | 15.47977 | 16.04789 | 14.60629 | 14.9337 | 16.028 | 16.855 | 17.15608 | 15.62234 | 15.09734 | 16.01168 | 19.37127 | 18.85156 |
| DPY19L1 | 13.0915 | 16.87645 | 15.83578 | 13.93314 | 16.6453 | 15.96385 | 16.0643 | 16.86702 | 19.96202 | 18.19874 | 15.6735 | 11.38876 | 14.53786 | 14.61941 | 16.95938 | 15.51389 | 15.60199 | 16.50057 |
| DPY19L4 | 15.06044 | 15.47396 | 15.5873 | 15.4886 | 17.03754 | 15.83718 | 18.89345 | 17.92197 | 18.24473 | 16.32845 | 15.50954 | 14.7453 | 15.05859 | 16.04974 | 15.11451 | 15.2429 | 16.54389 | 15.83767 |
| DPYSL2 | 18.08652 | 17.63207 | 18.4332 | 17.61448 | 21.44487 | 20.16546 | 19.48418 | 18.70385 | 19.19545 | 18.86441 | 18.30572 | 20.11632 | 16.38156 | 17.48765 | 18.18884 | 17.56743 | 18.50897 | 18.868 |
| DPYSL3 | 16.49257 | 17.23416 | 17.4208 | 17.02775 | 19.22065 | 16.81479 | 20.06304 | 18.72521 | 17.03357 | 15.92514 | 23.48759 | 23.34854 | 16.75455 | 16.91327 | 16.36599 | 16.94034 | 17.73498 | 16.58741 |
| DPYSL4 | 13.679 | 16.35307 | 16.62747 | 17.08175 | 18.6177 | 15.48976 | 17.08103 | 16.11964 | 13.81704 | 17.22573 | 15.71894 | 13.75617 | 15.10127 | 14.43896 | 14.00589 | 15.72967 | 17.58212 | 18.19722 |
| DPYSL5 | 16.07807 | 15.98284 | 16.97326 | 15.71294 | 16.3504 | 15.93425 | 18.75054 | 16.45115 | 15.55707 | 16.98232 | 18.61544 | 19.52235 | 15.34442 | 15.17634 | 15.85383 | 15.69642 | 16.26426 | 14.90734 |
| DRG1 | 15.8384 | 16.45681 | 16.49237 | 17.18256 | 16.6724 | 18.37467 | 15.87871 | 17.95656 | 15.43231 | 15.81756 | 17.65851 | 18.67818 | 17.1553 | 17.44435 | 16.44785 | 16.1913 | 16.3552 | 17.17077 |
| DRG2 | 18.35733 | 18.40359 | 17.81672 | 17.07214 | 13.79513 | 17.03043 | 15.63832 | 16.46267 | 18.4193 | 16.84658 | 15.95153 | 17.7486 | 16.63988 | 13.71021 | 16.93494 | 18.36301 | 14.91626 | 16.98214 |
| DSC2 | 14.9347 | 15.25265 | 17.00523 | 15.42966 | 18.38695 | 16.56776 | 17.09136 | 17.32243 | 15.91768 | 18.62443 | 17.05436 | 16.56056 | 18.43351 | 18.47005 | 17.0243 | 16.31761 | 18.64257 | 19.33929 |
| DSG2 | 17.46172 | 18.90368 | 19.69989 | 19.98917 | 22.01489 | 20.6766 | 18.22348 | 17.04123 | 20.07755 | 19.46734 | 20.05592 | 19.76184 | 19.83594 | 20.11382 | 19.53963 | 20.36525 | 18.91932 | 19.09366 |
| DSP | 16.51496 | 16.12792 | 16.62246 | 16.05562 | 16.19182 | 16.40783 | 16.38324 | 16.45921 | 17.59783 | 17.52669 | 14.90537 | 15.42897 | 16.7146 | 16.87865 | 15.37154 | 16.7757 | 18.14847 | 16.1856 |
| DST | 16.60947 | 17.0734 | 18.00013 | 17.91166 | 18.95926 | 17.56398 | 16.11119 | 16.36706 | 16.21615 | 17.10858 | 16.30401 | 16.07612 | 16.25995 | 17.78451 | 16.93282 | 16.82357 | 17.14749 | 17.1414 |
| DSTN | 20.51501 | 19.2094 | 18.95744 | 19.59355 | 21.70248 | 19.89552 | 20.22181 | 18.78227 | 17.93263 | 18.94408 | 19.88861 | 21.076 | 22.23615 | 22.2761 | 19.55376 | 19.99827 | 21.84723 | 21.05865 |
| DTD1 | 15.69986 | 13.62524 | 14.56576 | 12.45295 | 12.35141 | 17.00889 | 13.94554 | 15.99727 | 17.40814 | 15.774 | 17.96991 | 19.20989 | 13.9059 | 15.19428 | 15.56018 | 13.57097 | 17.12503 | 16.3029 |
| DTNA | 17.1374 | 17.37984 | 17.91154 | 18.71789 | 18.35057 | 16.75244 | 18.22906 | 16.80885 | 19.29103 | 18.8997 | 16.36167 | 16.61577 | 19.05698 | 16.60499 | 19.94893 | 19.7375 | 16.8406 | 17.40555 |
| DTNB | 15.70808 | 17.59208 | 17.17217 | 19.36822 | 16.29495 | 15.24116 | 15.6125 | 16.21318 | 18.72697 | 18.68728 | 14.80448 | 14.80545 | 17.78896 | 17.16819 | 19.51557 | 19.39012 | 15.82232 | 16.48141 |
| DTX3L | 14.43731 | 15.5399 | 16.08798 | 14.70355 | 16.33371 | 16.4841 | 16.31007 | 16.21379 | 16.78186 | 15.79662 | 14.59409 | 12.98848 | 17.87448 | 18.10776 | 16.21227 | 16.12156 | 17.5895 | 18.85511 |
| DTYMK | 17.21073 | 15.97452 | 15.31659 | 17.46971 | 21.30856 | 19.77948 | 20.61932 | 19.27713 | 17.72587 | 18.70357 | 17.45898 | 17.52293 | 20.61966 | 19.55335 | 16.49668 | 16.6848 | 20.52936 | 20.79907 |
| DUSP23 | 18.18303 | 17.76292 | 15.13333 | 16.18234 | 15.22106 | 14.61122 | 13.63184 | 17.05109 | 14.18683 | 14.90769 | 13.33841 | 18.23311 | 13.97709 | 16.58192 | 17.10791 | 15.60705 | 17.28047 | 13.01156 |
| DUSP3 | 18.46754 | 15.09502 | 16.54379 | 15.14684 | 13.20252 | 15.90252 | 15.11092 | 14.17331 | 11.71724 | 15.80801 | 15.82864 | 16.85711 | 17.43596 | 17.19296 | 14.71125 | 15.10085 | 12.15419 | 12.94304 |
| DUT | 15.37332 | 16.91955 | 19.29491 | 15.07815 | 19.51347 | 16.30767 | 15.56116 | 16.81481 | 18.4059 | 15.58264 | 20.14492 | 20.40382 | 16.85436 | 16.67208 | 15.11544 | 16.74387 | 16.58055 | 16.73253 |
| DVL2 | 16.88642 | 15.42873 | 17.41992 | 16.54945 | 17.24685 | 16.79274 | 14.76281 | 16.13264 | 15.23099 | 16.56592 | 15.01511 | 12.79052 | 19.5078 | 19.93184 | 14.476 | 15.62582 | 16.17305 | 17.34872 |
| DYNC1H1 | 16.80579 | 16.76468 | 17.76535 | 17.16012 | 17.68049 | 19.84183 | 17.68933 | 17.31426 | 17.34415 | 17.61931 | 18.11162 | 19.82251 | 17.40304 | 17.39362 | 17.68176 | 17.6738 | 16.73247 | 18.21848 |
| DYNC1I2 | 16.2544 | 15.95983 | 16.44701 | 17.38004 | 18.59452 | 16.64336 | 17.51869 | 17.73643 | 17.91489 | 18.27597 | 20.61516 | 19.32279 | 19.28516 | 19.24117 | 17.27306 | 17.99902 | 19.36961 | 19.84868 |
| DYNC1LI1 | 16.0636 | 16.42207 | 17.10442 | 16.19562 | 16.16059 | 16.53051 | 17.40844 | 17.2152 | 16.75547 | 17.19487 | 18.53318 | 19.30285 | 17.82399 | 17.18975 | 16.71062 | 17.24157 | 16.42062 | 17.91791 |
| DYNC1LI2 | 16.90743 | 20.42632 | 17.55365 | 18.05047 | 18.98485 | 18.60915 | 19.1203 | 16.49875 | 16.76604 | 18.30754 | 19.52808 | 19.646 | 19.17152 | 19.67406 | 17.88408 | 17.26541 | 19.3254 | 18.45573 |
| DYNLL2 | 21.50735 | 20.59839 | 20.01837 | 19.39671 | 17.06993 | 18.0725 | 20.33577 | 19.27331 | 17.63528 | 16.47199 | 16.82271 | 20.7929 | 21.17403 | 21.51798 | 18.58682 | 19.75185 | 18.91283 | 17.76248 |
| DYNLRB1 | 17.81114 | 14.37113 | 15.04005 | 15.30243 | 19.96774 | 16.94832 | 19.10752 | 16.46705 | 18.19133 | 19.92238 | 16.64713 | 17.71596 | 20.88061 | 20.05864 | 13.31951 | 13.67011 | 19.94357 | 21.51099 |
| DYNLRB2 | 19.22908 | 14.59353 | 17.03139 | 18.00208 | 19.12047 | 16.26221 | 20.1804 | 18.91153 | 17.81279 | 18.81352 | 19.29563 | 19.59568 | 19.50632 | 19.86229 | 17.97721 | 16.08818 | 21.6394 | 20.11513 |
| DYNLT1 | 15.16427 | 14.58845 | 16.81646 | 14.8121 | 16.78111 | 16.60687 | 18.27229 | 16.36014 | 15.33466 | 18.66238 | 18.70769 | 20.39115 | 15.44691 | 18.19854 | 16.43234 | 16.54897 | 16.34005 | 14.46432 |
| DYSF | 13.8125 | 16.55824 | 16.14618 | 15.99533 | 22.84001 | 20.94201 | 15.29034 | 14.81954 | 15.64788 | 16.02565 | 14.68108 | 13.27478 | 15.68947 | 14.59418 | 16.11807 | 16.38762 | 17.55809 | 20.04873 |
| EBP | 17.57609 | 17.87267 | 18.37499 | 16.6116 | 20.16091 | 18.49358 | 14.76619 | 16.54429 | 23.15813 | 21.80748 | 22.63131 | 22.69719 | 15.2752 | 16.36249 | 17.36868 | 17.62456 | 14.10152 | 17.91963 |
| ECE1 | 17.89221 | 18.31847 | 20.38614 | 17.83885 | 19.74868 | 20.55274 | 16.8893 | 16.83627 | 21.61047 | 19.8165 | 15.9736 | 15.81949 | 16.19416 | 16.52298 | 17.98756 | 17.89771 | 17.15066 | 16.75592 |
| ECH1 | 15.18392 | 17.69018 | 16.21377 | 15.91811 | 14.70398 | 16.99461 | 19.4003 | 17.0066 | 20.87055 | 20.44807 | 17.00654 | 16.86651 | 17.07482 | 17.25942 | 16.39977 | 16.19526 | 15.20678 | 15.24027 |
| ECHDC1 | 17.168 | 15.702 | 17.10791 | 16.08472 | 18.64734 | 16.25557 | 16.94765 | 17.40575 | 15.06889 | 14.82592 | 19.64686 | 19.96719 | 17.47868 | 17.70297 | 15.73047 | 15.82898 | 17.69173 | 13.95314 |
| ECHS1 | 20.73601 | 20.43936 | 16.41514 | 16.60489 | 15.74453 | 16.56618 | 21.28656 | 18.17271 | 18.2985 | 17.35769 | 15.64131 | 13.05658 | 18.27194 | 19.18311 | 16.13443 | 17.11282 | 16.78862 | 17.8185 |
| ECI1 | 17.14059 | 16.40813 | 17.28289 | 16.58873 | 16.55558 | 16.86515 | 17.64874 | 16.37834 | 20.83363 | 17.87725 | 15.03767 | 14.76739 | 15.56085 | 17.15932 | 17.00208 | 15.96136 | 16.83462 | 15.32863 |
| ECI2 | 16.25206 | 16.09308 | 16.60373 | 17.62022 | 19.53286 | 16.9229 | 17.85718 | 17.94363 | 21.4066 | 20.06203 | 16.76526 | 14.50699 | 15.8785 | 15.52266 | 17.28459 | 17.14324 | 20.52368 | 17.05813 |
| ECPAS | 16.54395 | 16.25926 | 16.09484 | 15.76961 | 16.65584 | 15.05026 | 17.73715 | 16.66038 | 15.21902 | 16.19727 | 16.90637 | 18.24049 | 17.2286 | 17.34027 | 16.11984 | 16.40531 | 18.18511 | 16.35929 |
| ECSIT | 14.63162 | 16.20266 | 16.97811 | 11.8685 | 12.33836 | 15.51653 | 17.10493 | 15.31739 | 17.94973 | 13.60137 | 17.35692 | 13.07157 | 12.53089 | 13.39725 | 13.09473 | 14.37145 | 14.90966 | 14.45561 |
| ECT2 | 15.08868 | 14.93047 | 17.34667 | 19.50994 | 17.99542 | 16.77872 | 14.82054 | 15.17047 | 19.8597 | 17.18684 | 13.21035 | 11.97551 | 13.87092 | 19.53627 | 15.71308 | 15.60927 | 13.69104 | 15.67112 |
| EDEM3 | 15.12599 | 14.49896 | 15.10181 | 15.10088 | 15.08235 | 14.18746 | 15.53167 | 15.10985 | 18.68991 | 12.54635 | 16.4563 | 11.69723 | 14.07765 | 15.02354 | 13.49432 | 13.17062 | 12.66098 | 14.99492 |
| EDF1 | 20.85411 | 21.34364 | 21.69505 | 21.20622 | 16.08154 | 17.35242 | 17.21605 | 17.46489 | 19.04145 | 15.606 | 19.82697 | 19.70086 | 17.96652 | 17.69865 | 21.77245 | 21.84743 | 18.89777 | 17.82004 |
| EDIL3 | 19.39805 | 18.40574 | 16.71069 | 16.75267 | 26.42835 | 25.84953 | 22.65344 | 21.25958 | 22.34698 | 21.86554 | 17.21631 | 15.27726 | 16.30477 | 16.34233 | 17.08659 | 16.06769 | 22.85951 | 25.50664 |
| EEA1 | 16.69139 | 16.81905 | 16.41939 | 17.36497 | 19.97981 | 17.96004 | 17.43397 | 16.66632 | 14.876 | 16.08686 | 16.32218 | 15.67791 | 18.245 | 16.92298 | 16.05869 | 15.88235 | 16.61938 | 17.51711 |
| EEF1A1P5 | 26.51453 | 26.48148 | 25.16315 | 25.53477 | 23.9744 | 23.24866 | 23.00734 | 20.775 | 21.70292 | 22.06155 | 23.35511 | 23.47257 | 24.72966 | 24.01011 | 25.92503 | 25.53418 | 23.75637 | 24.33984 |
| EEF1A2 | 25.97091 | 26.17036 | 25.319 | 25.44115 | 24.26836 | 22.45301 | 23.53941 | 21.00629 | 21.23048 | 21.63261 | 22.71007 | 23.53489 | 24.32787 | 23.67225 | 26.14232 | 25.75638 | 23.67974 | 23.9277 |
| EEF1AKNM | 14.06892 | 15.46479 | 15.42509 | 16.54224 | 16.76805 | 16.05687 | 17.68857 | 15.96261 | 12.28056 | 15.69266 | 16.32123 | 17.07207 | 16.80521 | 18.24393 | 16.0137 | 15.51753 | 15.81316 | 16.83636 |
| EEF1B2 | 17.65326 | 17.16204 | 18.20808 | 19.38944 | 21.89595 | 20.95394 | 19.97998 | 18.01298 | 19.2681 | 20.66079 | 22.67344 | 23.26356 | 22.73493 | 21.24855 | 20.17896 | 17.80036 | 21.90297 | 21.23941 |
| EEF1D | 17.87752 | 17.49529 | 18.59358 | 18.74694 | 20.24365 | 19.61173 | 18.02317 | 17.77344 | 19.01948 | 18.11152 | 20.84059 | 22.28739 | 19.66409 | 19.28633 | 18.82273 | 19.14606 | 21.04406 | 20.20804 |
| EEF1E1 | 17.77558 | 14.64466 | 17.60654 | 15.43992 | 18.00079 | 15.9262 | 15.27487 | 18.48963 | 16.63139 | 17.60907 | 18.51627 | 20.39181 | 18.99212 | 18.52705 | 14.59966 | 16.61277 | 17.79517 | 17.83156 |
| EEF1G | 19.20112 | 18.41838 | 20.12787 | 18.85675 | 19.95164 | 19.4222 | 16.90281 | 17.92104 | 18.42142 | 18.25974 | 20.37347 | 22.60159 | 18.68729 | 18.05492 | 19.48496 | 19.48731 | 19.67121 | 19.87964 |
| EEF2 | 20.32148 | 20.27366 | 20.49222 | 20.92023 | 22.43196 | 21.31916 | 21.31925 | 19.99712 | 19.21389 | 19.77835 | 21.91435 | 22.70501 | 21.18233 | 20.9063 | 20.65115 | 20.52271 | 21.75861 | 21.55428 |
| EEPD1 | 15.51506 | 16.13995 | 17.91832 | 17.93336 | 17.7931 | 16.04626 | 17.16756 | 17.23464 | 17.30411 | 17.78103 | 15.88891 | 17.32393 | 18.74804 | 19.75449 | 19.0576 | 18.13135 | 21.55614 | 21.8412 |
| EFCAB14 | 16.80061 | 16.36994 | 17.1425 | 15.62124 | 16.39245 | 16.3456 | 17.35076 | 16.17597 | 21.68629 | 21.39897 | 17.4236 | 14.69645 | 16.02205 | 16.47742 | 16.45711 | 16.18489 | 16.88945 | 16.65928 |
| EFHD2 | 16.00542 | 16.41552 | 15.9213 | 17.23956 | 21.49254 | 21.88706 | 16.95371 | 16.8759 | 18.85316 | 18.32278 | 18.41974 | 17.66738 | 17.49108 | 18.27067 | 16.46863 | 15.66271 | 18.53128 | 19.39572 |
| EFL1 | 14.55128 | 14.53334 | 16.10729 | 15.20705 | 15.6771 | 16.98681 | 16.2276 | 14.89671 | 16.99916 | 15.3692 | 16.09975 | 17.20327 | 13.0997 | 16.61008 | 15.70207 | 14.56864 | 15.366 | 15.44273 |
| EFNA1 | 20.39781 | 20.357 | 18.94884 | 19.82484 | 14.11919 | 16.52647 | 15.96228 | 18.53443 | 18.18566 | 15.44766 | 16.08256 | 12.17079 | 14.98984 | 16.01444 | 18.31268 | 18.47641 | 16.1713 | 17.09273 |
| EFNB1 | 16.92957 | 15.96441 | 15.63767 | 18.12134 | 18.56596 | 18.88313 | 20.02094 | 18.21485 | 16.29205 | 17.28075 | 15.14913 | 18.11717 | 16.80736 | 16.7348 | 17.68515 | 19.16018 | 22.44106 | 23.52346 |
| EFNB2 | 17.37617 | 16.8559 | 16.21091 | 16.46813 | 14.17537 | 16.53973 | 16.97885 | 16.19339 | 14.77031 | 14.90604 | 16.94349 | 18.01411 | 14.33674 | 16.90062 | 15.51166 | 20.71817 | 14.13588 | 16.93226 |
| EFNB3 | 18.2135 | 17.80078 | 15.056 | 15.37529 | 14.48031 | 15.73997 | 13.99987 | 17.19128 | 11.86607 | 13.72819 | 15.08444 | 13.68547 | 14.39528 | 14.49657 | 13.14188 | 14.15117 | 14.83602 | 20.39501 |
| EFR3A | 16.73431 | 16.59141 | 18.09542 | 18.65065 | 18.73004 | 17.37216 | 16.91735 | 16.87554 | 17.89737 | 16.75756 | 16.01701 | 17.45757 | 17.62289 | 16.28324 | 18.74319 | 18.48593 | 17.57158 | 17.42794 |
| EFR3B | 14.00823 | 16.08073 | 15.18563 | 15.12118 | 17.05182 | 16.86252 | 16.07923 | 15.17845 | 17.37217 | 17.11427 | 16.92759 | 14.7351 | 16.47196 | 15.91483 | 18.39102 | 18.27167 | 16.68008 | 17.30709 |
| EFTUD2 | 19.46173 | 19.14215 | 25.80907 | 17.51715 | 17.88628 | 16.48156 | 17.84692 | 17.82323 | 16.44802 | 15.44263 | 18.88648 | 20.82243 | 18.86238 | 17.27197 | 18.22077 | 17.92422 | 17.97735 | 17.89178 |
| EGFR | 17.46265 | 17.80787 | 19.88849 | 19.70847 | 21.9704 | 20.91359 | 16.26144 | 17.18605 | 16.27065 | 16.08784 | 16.65168 | 16.00077 | 17.79495 | 19.64783 | 20.00954 | 20.52289 | 19.6478 | 20.31863 |
| EHBP1 | 13.6337 | 16.21464 | 14.0297 | 15.52717 | 18.61222 | 17.35966 | 18.31509 | 16.27488 | 16.68963 | 16.99707 | 15.47476 | 15.22746 | 15.67722 | 16.30463 | 13.67012 | 15.43516 | 17.71964 | 18.04442 |
| EHBP1L1 | 15.44359 | 16.29725 | 16.8861 | 15.46754 | 19.18146 | 17.384 | 16.17452 | 16.726 | 16.16712 | 15.4412 | 16.13549 | 13.78461 | 13.68981 | 14.82738 | 15.31454 | 16.44887 | 17.90576 | 17.53927 |
| EHD1 | 19.97581 | 20.47718 | 19.73407 | 20.32688 | 22.59197 | 21.38732 | 19.2442 | 18.01579 | 17.63146 | 18.37182 | 20.77225 | 21.08273 | 21.22604 | 20.79593 | 21.13956 | 20.65138 | 22.77219 | 23.11012 |
| EHD2 | 16.31975 | 16.41877 | 15.92236 | 15.71891 | 21.57938 | 20.67083 | 16.39171 | 17.7066 | 15.87046 | 16.42947 | 16.06081 | 13.23272 | 16.48324 | 16.75091 | 15.72885 | 16.23812 | 16.88884 | 16.96334 |
| EHD3 | 18.20309 | 20.49905 | 20.64428 | 21.14302 | 19.67879 | 18.00873 | 21.00472 | 21.2621 | 14.0252 | 16.82152 | 17.04084 | 14.74687 | 20.06181 | 20.9524 | 20.89156 | 20.11455 | 18.9596 | 20.04392 |
| EHD4 | 20.53973 | 20.76375 | 20.55018 | 20.87888 | 21.1414 | 20.36914 | 17.63061 | 17.02282 | 16.85138 | 17.06088 | 16.71643 | 19.12498 | 20.04216 | 20.58272 | 21.7032 | 21.22664 | 21.23672 | 21.73273 |
| EHHADH | 15.94539 | 16.37142 | 15.61188 | 15.67965 | 14.31908 | 16.70446 | 16.14049 | 16.7944 | 19.73055 | 18.07427 | 15.81564 | 15.14316 | 15.68884 | 15.35984 | 16.3598 | 16.13165 | 15.60749 | 15.07882 |
| EIF1 | 19.10729 | 19.58744 | 16.68155 | 19.96439 | 17.34892 | 16.52288 | 17.88817 | 16.75385 | 16.55455 | 16.39564 | 16.67378 | 14.43531 | 22.50456 | 21.70774 | 20.09093 | 18.91516 | 20.87099 | 19.76461 |
| EIF1AD | 17.14905 | 16.75477 | 16.42185 | 16.49309 | 15.85557 | 15.47579 | 15.26307 | 16.9674 | 15.36886 | 16.54671 | 18.04658 | 17.66628 | 13.73126 | 13.48186 | 15.97067 | 16.12615 | 14.08034 | 14.27743 |
| EIF1AY | 21.21263 | 20.6308 | 20.67909 | 19.74061 | 16.61837 | 15.87032 | 17.30221 | 16.67917 | 17.12358 | 16.36386 | 19.34766 | 20.43786 | 20.17809 | 16.73387 | 20.59606 | 21.23714 | 16.58751 | 17.26473 |

| | | | | | | | | | | | | | | | | | | |
|---|---|---|---|---|---|---|---|---|---|---|---|---|---|---|---|---|---|---|
| EIF1B | 19.78975 | 20.11314 | 19.79204 | 20.73917 | 20.17008 | 18.0335 | 19.54901 | 19.37098 | 18.76783 | 17.76755 | 17.2503 | 17.94716 | 21.83403 | 21.80336 | 20.62571 | 20.59252 | 20.81548 | 20.67735 |
| EIF2A | 17.27424 | 17.02274 | 17.4758 | 17.51038 | 16.14127 | 15.72654 | 16.584 | 18.17247 | 15.65786 | 15.27323 | 18.28482 | 19.89473 | 17.14677 | 16.59286 | 17.35011 | 17.95009 | 17.16299 | 16.91328 |
| EIF2AK2 | 17.09171 | 17.63198 | 14.13682 | 13.16492 | 17.65083 | 15.63017 | 16.43708 | 17.31363 | 14.21653 | 17.5545 | 17.91229 | 18.64646 | 17.61072 | 17.98311 | 13.72282 | 14.99699 | 16.45239 | 17.49168 |
| EIF2AK3 | 15.06363 | 14.85651 | 15.463 | 15.21214 | 13.0554 | 15.63826 | 14.73969 | 14.53916 | 18.79607 | 17.4899 | 15.21226 | 14.58421 | 15.73831 | 17.63926 | 15.7505 | 14.37431 | 15.23127 | 15.67721 |
| EIF2B1 | 17.81887 | 17.97249 | 15.74228 | 13.84521 | 12.96072 | 19.00643 | 14.40153 | 15.84298 | 16.47665 | 14.71215 | 18.48053 | 17.41904 | 15.21075 | 14.93211 | 13.48854 | 13.53293 | 15.80263 | 14.29829 |
| EIF2B2 | 16.32608 | 18.00474 | 16.41572 | 15.4597 | 17.61005 | 19.30428 | 19.37595 | 18.98518 | 14.90859 | 15.93848 | 18.57762 | 18.88698 | 17.56095 | 16.59952 | 17.27443 | 14.92632 | 17.99118 | 18.62938 |
| EIF2B3 | 21.05177 | 14.96599 | 15.86293 | 15.44287 | 17.11334 | 16.6254 | 17.82113 | 17.92536 | 15.13671 | 15.92064 | 16.17114 | 18.34946 | 17.29859 | 17.55492 | 13.12246 | 15.31724 | 18.86545 | 18.16543 |
| EIF2B4 | 16.46262 | 17.58551 | 15.7016 | 15.908 | 14.6346 | 15.02806 | 16.72367 | 15.76917 | 13.71019 | 15.24088 | 15.77945 | 18.67321 | 16.43511 | 14.75876 | 16.84392 | 15.54035 | 16.3583 | 15.98637 |
| EIF2B5 | 16.66689 | 14.07451 | 14.52903 | 15.70212 | 12.91952 | 17.67209 | 15.17334 | 16.73075 | 16.75466 | 15.99903 | 15.32366 | 19.17796 | 13.88122 | 12.86794 | 14.99563 | 14.8758 | 15.6572 | 15.63514 |
| EIF2S1 | 18.98332 | 17.32086 | 18.55195 | 18.47256 | 16.80847 | 17.23339 | 14.9949 | 18.18625 | 17.0273 | 15.5766 | 18.64345 | 19.92395 | 18.77379 | 18.38819 | 19.15684 | 19.20372 | 16.35178 | 15.85685 |
| EIF2S2 | 17.08927 | 16.9627 | 17.00795 | 17.4206 | 18.57894 | 16.75203 | 17.767 | 17.09958 | 17.54146 | 16.53228 | 18.49789 | 19.32247 | 17.80037 | 17.91744 | 17.37629 | 18.51898 | 18.21372 | 19.90319 |
| EIF2S3 | 17.30571 | 17.30581 | 18.54344 | 19.89693 | 19.38605 | 17.57831 | 17.40864 | 18.2761 | 18.63727 | 19.22323 | 19.66941 | 20.27627 | 20.04625 | 19.00163 | 17.87457 | 18.32171 | 19.16717 | 19.19516 |
| EIF3A | 18.48222 | 18.22491 | 18.71324 | 18.33542 | 17.01397 | 17.20645 | 17.08684 | 17.94802 | 15.17021 | 16.06895 | 17.77186 | 20.47522 | 17.29403 | 17.01409 | 18.51255 | 18.72898 | 15.98225 | 15.7417 |
| EIF3B | 18.45854 | 18.82343 | 18.6891 | 19.0992 | 17.81498 | 18.79921 | 16.04699 | 15.99516 | 16.87955 | 17.03683 | 19.14627 | 20.67926 | 17.47874 | 17.22902 | 18.5872 | 18.83059 | 16.97104 | 16.08113 |
| EIF3CL | 17.97519 | 17.68016 | 18.63434 | 17.47801 | 15.93761 | 15.83806 | 16.57191 | 16.3477 | 16.33794 | 16.05977 | 19.13862 | 19.99431 | 17.66537 | 16.06036 | 17.67679 | 18.27727 | 15.47367 | 14.78518 |
| EIF3D | 17.65658 | 20.6007 | 20.23245 | 18.40338 | 18.89046 | 17.10003 | 16.99343 | 16.55793 | 17.22655 | 17.89502 | 17.86571 | 19.54467 | 19.13158 | 17.96057 | 18.1125 | 18.60604 | 17.19787 | 18.27115 |
| EIF3E | 18.43989 | 18.4716 | 18.57357 | 18.41434 | 17.18069 | 16.08342 | 17.10181 | 17.37186 | 16.74682 | 16.6963 | 17.84599 | 20.68852 | 18.46034 | 16.82151 | 18.6975 | 18.17847 | 16.20774 | 17.33745 |
| EIF3F | 17.87814 | 17.72427 | 17.73505 | 18.04314 | 15.99079 | 16.88795 | 15.89329 | 16.21754 | 15.15663 | 15.97485 | 19.0118 | 21.40371 | 14.55389 | 15.49322 | 17.93702 | 18.08934 | 16.56013 | 16.22887 |
| EIF3G | 20.16844 | 20.17141 | 19.40966 | 20.59722 | 19.04114 | 17.64008 | 17.70792 | 17.86953 | 18.06116 | 16.87853 | 19.79296 | 20.18293 | 20.29512 | 19.75715 | 19.74734 | 19.82893 | 19.29106 | 19.7133 |
| EIF3H | 18.86875 | 16.5109 | 16.532 | 18.50106 | 19.08681 | 18.11212 | 17.77336 | 18.8421 | 15.81253 | 17.2513 | 17.65451 | 19.99655 | 20.14222 | 19.12772 | 17.89211 | 16.77298 | 18.20274 | 19.29782 |
| EIF3I | 20.13382 | 18.56282 | 19.2955 | 18.46231 | 19.1436 | 18.03796 | 18.15656 | 18.14895 | 17.97877 | 17.25628 | 20.51582 | 21.67844 | 20.62073 | 20.17444 | 18.6648 | 19.06153 | 19.86754 | 19.2733 |
| EIF3J | 15.23913 | 16.13228 | 16.27383 | 16.90386 | 18.3898 | 16.47085 | 18.29213 | 16.12267 | 16.50466 | 16.48319 | 19.47669 | 19.71879 | 18.90008 | 16.21382 | 16.54319 | 16.15545 | 16.2981 | 16.73284 |
| EIF3K | 14.73947 | 16.82038 | 15.36186 | 18.52986 | 18.59178 | 18.9689 | 18.48158 | 17.3372 | 14.27298 | 15.21622 | 18.61754 | 18.36404 | 19.75633 | 18.6854 | 18.45783 | 17.17913 | 15.30662 | 15.03934 |
| EIF3L | 17.48345 | 17.78678 | 18.57358 | 18.49931 | 14.74509 | 17.05993 | 17.84041 | 16.35197 | 15.31185 | 16.05801 | 18.83523 | 20.50803 | 17.69475 | 17.02388 | 18.15861 | 18.45907 | 15.21928 | 14.93924 |
| EIF3M | 18.2371 | 17.76274 | 18.56267 | 17.37781 | 16.49864 | 15.84702 | 16.03142 | 15.45067 | 14.80581 | 15.16478 | 17.68096 | 19.85216 | 13.28789 | 13.9788 | 18.08054 | 19.02834 | 15.46908 | 14.4353 |
| EIF4A1 | 20.8419 | 20.39309 | 21.61337 | 21.03753 | 19.39314 | 19.36748 | 19.62308 | 19.89707 | 18.85041 | 18.86544 | 20.46248 | 21.18293 | 19.82001 | 18.51675 | 21.33267 | 21.25003 | 18.72375 | 19.01706 |
| EIF4A2 | 16.97474 | 17.9847 | 17.92846 | 16.96738 | 14.04251 | 15.48329 | 15.40704 | 16.55874 | 15.50775 | 15.99723 | 16.92823 | 19.33358 | 14.09718 | 14.78211 | 17.40056 | 17.7125 | 16.1995 | 14.64344 |
| EIF4A3 | 20.55874 | 20.24929 | 19.41498 | 20.27991 | 17.83911 | 17.99608 | 17.94451 | 17.50345 | 18.14492 | 17.29757 | 19.07158 | 20.52313 | 19.34979 | 17.79043 | 19.94644 | 20.16658 | 18.01835 | 18.03341 |
| EIF4B | 18.23115 | 16.30568 | 17.96694 | 17.54331 | 19.30898 | 18.48399 | 16.41439 | 15.9187 | 16.34164 | 17.12276 | 18.33879 | 18.69766 | 18.39343 | 17.92663 | 17.9599 | 18.23305 | 18.96936 | 18.54407 |
| EIF4E | 19.19881 | 19.93884 | 17.89928 | 19.62466 | 19.74585 | 20.85573 | 18.35885 | 19.10682 | 19.03623 | 17.98129 | 20.89297 | 21.27817 | 20.82726 | 20.56661 | 18.34313 | 18.04531 | 18.31594 | 18.66472 |
| EIF4G1 | 17.5371 | 16.92128 | 20.93839 | 17.78963 | 20.97986 | 18.39031 | 16.95536 | 17.82446 | 17.70164 | 18.68326 | 18.25678 | 20.32771 | 18.39148 | 16.90817 | 16.11486 | 21.31512 | 18.8617 | 19.82612 |
| EIF4G2 | 16.82657 | 16.12155 | 17.66383 | 18.92646 | 18.03517 | 16.34241 | 17.09422 | 15.33337 | 14.9436 | 15.4725 | 16.06396 | 18.06951 | 18.39074 | 18.83096 | 16.67799 | 17.85439 | 16.66258 | 17.25112 |
| EIF4H | 22.12507 | 21.7006 | 21.6796 | 22.84639 | 19.01598 | 19.39197 | 16.48041 | 15.68696 | 18.7037 | 16.61293 | 17.92334 | 18.57128 | 19.71208 | 17.72954 | 22.71605 | 22.00495 | 15.54633 | 13.63033 |
| EIF5 | 18.7397 | 16.85721 | 17.91986 | 17.79642 | 19.72977 | 20.64671 | 17.10904 | 18.3833 | 16.37959 | 16.06908 | 17.41091 | 19.31425 | 17.99717 | 17.72185 | 17.00268 | 16.51766 | 19.59183 | 19.55043 |
| EIF5A | 22.87887 | 22.31667 | 21.23737 | 21.96676 | 24.00604 | 22.79902 | 22.44598 | 21.40355 | 20.85302 | 21.18947 | 22.29981 | 22.49518 | 23.61736 | 23.10184 | 22.48495 | 22.34774 | 23.40648 | 23.41542 |
| EIF5B | 17.10778 | 17.411 | 17.97587 | 17.93508 | 18.03063 | 16.00928 | 16.08822 | 16.08858 | 15.65202 | 16.33989 | 18.24005 | 19.09296 | 20.09394 | 18.90639 | 17.24677 | 17.58755 | 17.26943 | 18.41072 |
| EIF6 | 20.4927 | 21.29816 | 19.07668 | 19.05107 | 18.7189 | 17.88703 | 16.85486 | 17.15939 | 16.6214 | 17.13608 | 17.03332 | 19.40714 | 19.05532 | 18.67762 | 20.71251 | 19.28336 | 19.93583 | 19.627 |
| EIPR1 | 14.70724 | 15.50852 | 15.1598 | 16.07369 | 19.55116 | 17.03273 | 15.41575 | 16.58898 | 15.00223 | 15.32962 | 14.96196 | 17.99706 | 17.74888 | 16.14694 | 19.79779 | 15.10807 | 14.92192 | 17.80619 |
| ELAC2 | 14.82761 | 16.39753 | 15.00552 | 14.99459 | 15.25051 | 16.69021 | 17.63846 | 16.22747 | 14.44627 | 14.33265 | 13.12674 | 16.89474 | 13.93159 | 13.36051 | 13.13408 | 17.13872 | 16.99647 | 15.50425 |
| ELAPOR1 | 18.75301 | 17.0971 | 15.69295 | 15.34147 | 13.20094 | 15.24118 | 16.55952 | 15.03124 | 17.77468 | 17.0974 | 14.66727 | 13.33438 | 13.67341 | 14.80606 | 14.01022 | 14.39119 | 14.52265 | 14.12633 |
| ELAPOR2 | 15.61653 | 16.65883 | 16.30476 | 15.27387 | 16.7728 | 16.8618 | 17.03266 | 16.67743 | 19.88238 | 19.01342 | 17.01785 | 15.34394 | 16.59252 | 16.00582 | 15.20105 | 15.48231 | 15.24911 | 14.9875 |
| ELAVL1 | 17.26053 | 17.60383 | 19.06772 | 18.2977 | 17.08246 | 15.46003 | 17.23144 | 17.1339 | 16.50203 | 17.2317 | 17.76587 | 20.38514 | 17.0193 | 16.89318 | 18.21005 | 18.69178 | 17.38806 | 17.92654 |
| ELFN2 | 16.59232 | 17.3802 | 15.16992 | 17.13844 | 20.01077 | 19.58116 | 18.33269 | 16.07649 | 15.88562 | 17.51646 | 17.5881 | 14.80461 | 15.38195 | 14.76712 | 16.50743 | 16.53688 | 18.3011 | 16.31698 |
| ELN | 15.00042 | 17.54292 | 19.69146 | 15.04194 | 18.72635 | 14.97632 | 20.01689 | 19.27238 | 11.54853 | 15.47899 | 14.28682 | 12.88699 | 14.24533 | 15.63694 | 15.33261 | 15.79078 | 16.85786 | 19.13006 |
| ELOB | 18.09313 | 18.2128 | 19.16607 | 18.53697 | 21.21619 | 20.86695 | 17.65187 | 19.29501 | 18.6345 | 18.58609 | 19.52341 | 20.52609 | 21.18804 | 20.94763 | 18.7788 | 19.09834 | 21.05477 | 20.89526 |
| ELOC | 18.89728 | 17.87997 | 18.13412 | 18.35586 | 22.34756 | 19.88465 | 21.22938 | 18.07951 | 18.34827 | 18.78633 | 19.17065 | 20.55337 | 21.36932 | 22.25428 | 18.8924 | 18.52266 | 22.48571 | 22.46071 |
| ELOVL7 | 14.8062 | 14.15191 | 16.2067 | 15.37459 | 14.49231 | 16.37751 | 12.92662 | 14.55185 | 19.64861 | 18.53454 | 16.2247 | 12.33823 | 14.2745 | 15.22501 | 12.77358 | 16.57178 | 14.65192 | 15.54935 |
| ELP1 | 15.02167 | 16.55983 | 15.27507 | 15.53354 | 16.06116 | 15.87333 | 16.74619 | 18.17785 | 14.93601 | 16.2337 | 18.03774 | 18.74509 | 14.91399 | 15.25011 | 16.41117 | 15.97381 | 16.19505 | 16.21931 |
| ELP2 | 14.46381 | 15.60297 | 15.42585 | 15.29155 | 14.39809 | 19.78991 | 15.58306 | 17.39043 | 14.46629 | 13.1302 | 14.68965 | 19.16101 | 14.32876 | 14.79412 | 17.0518 | 16.17776 | 16.79082 | 13.81016 |
| ELP3 | 15.88956 | 15.0801 | 18.61775 | 16.00488 | 13.83997 | 14.64456 | 15.0128 | 16.48217 | 15.25613 | 15.8942 | 14.47294 | 18.73748 | 14.98018 | 14.89115 | 18.04951 | 16.01908 | 17.26857 | 13.9016 |
| EMB | 15.4293 | 14.9604 | 18.29395 | 14.47552 | 19.83225 | 19.03493 | 16.28734 | 16.34816 | 17.58254 | 16.38586 | 15.10405 | 13.25723 | 15.36939 | 17.75551 | 18.61065 | 18.38698 | 15.5306 | 15.15711 |
| EMC1 | 16.42073 | 17.42561 | 16.34729 | 15.66274 | 18.9932 | 17.75244 | 17.10363 | 17.61768 | 21.35752 | 19.50854 | 18.45099 | 16.14982 | 19.12127 | 17.1883 | 16.07294 | 15.32719 | 19.11649 | 19.3523 |
| EMC10 | 18.67166 | 14.71824 | 16.36662 | 15.53714 | 16.08017 | 16.15393 | 17.58435 | 18.26161 | 22.47235 | 21.20132 | 19.06772 | 15.45083 | 15.22894 | 14.12988 | 14.82111 | 14.3656 | 14.19036 | 16.09591 |
| EMC7 | 15.49528 | 16.46641 | 17.98096 | 14.06817 | 16.37904 | 16.2198 | 14.94834 | 15.3356 | 21.12642 | 18.84277 | 20.67187 | 17.11159 | 14.76637 | 15.88596 | 15.0896 | 16.80855 | 16.04084 | 14.93903 |
| EMG1 | 14.69557 | 15.32446 | 12.49273 | 14.4895 | 14.43751 | 14.78037 | 13.72017 | 16.02081 | 12.63949 | 15.42831 | 14.22741 | 19.21106 | 13.33857 | 15.66612 | 15.81591 | 14.42837 | 13.67012 | 14.53544 |
| EML2 | 17.36876 | 17.80199 | 19.24315 | 17.42746 | 14.01932 | 16.4783 | 17.7672 | 17.84821 | 15.62171 | 14.72716 | 15.58661 | 17.89581 | 17.59697 | 17.8219 | 14.97929 | 16.99998 | 16.4928 | 18.65463 |
| ENAH | 16.79339 | 16.78815 | 18.55249 | 19.25139 | 20.95085 | 19.82946 | 19.38939 | 18.88897 | 19.02728 | 19.46619 | 20.04872 | 20.98156 | 21.24555 | 20.8451 | 19.56442 | 19.63093 | 20.3463 | 19.68587 |
| ENDOD1 | 16.69581 | 16.45039 | 18.34804 | 18.92139 | 17.647 | 16.91667 | 16.55192 | 16.90658 | 20.99156 | 19.01666 | 16.36591 | 15.72163 | 18.5025 | 17.99987 | 17.77499 | 17.90921 | 18.14422 | 18.89645 |
| ENO1 | 21.27696 | 21.05332 | 21.80134 | 23.19135 | 26.35232 | 24.7957 | 22.70889 | 22.02177 | 23.65141 | 24.76744 | 24.0569 | 24.12223 | 25.50499 | 24.8108 | 22.6261 | 22.74908 | 26.11835 | 25.83807 |
| ENO2 | 20.87277 | 17.45106 | 21.56209 | 18.51285 | 18.77117 | 18.07256 | 19.78641 | 19.06022 | 19.21671 | 19.54661 | 16.12234 | 17.96081 | 17.22973 | 17.39978 | 15.32043 | 21.04137 | 20.8078 | 20.98707 |
| ENO3 | 15.20265 | 14.58891 | 14.5322 | 16.99126 | 13.963 | 15.56977 | 16.02366 | 16.48235 | 17.21692 | 18.76016 | 16.71679 | 17.59013 | 15.47809 | 15.84723 | 15.6485 | 17.37787 | 15.53655 | 17.92631 |
| ENOPH1 | 16.66042 | 16.44107 | 15.47209 | 15.00405 | 16.90507 | 16.71517 | 15.0812 | 16.20182 | 16.435 | 17.98595 | 18.97373 | 19.08558 | 18.82829 | 17.41534 | 16.66721 | 18.02597 | 17.78359 | 16.71078 |
| ENPP1 | 20.5432 | 17.8944 | 18.46978 | 16.53882 | 14.85612 | 16.8011 | 15.691 | 15.38621 | 17.06995 | 16.49812 | 15.50153 | 17.14534 | 18.41823 | 17.78398 | 19.90931 | 19.83556 | 19.36719 | 20.18477 |
| ENPP4 | 18.81525 | 18.12581 | 18.32912 | 19.1471 | 18.33172 | 17.50165 | 18.77469 | 18.42415 | 22.4132 | 22.24263 | 21.49869 | 18.69198 | 18.57643 | 18.71311 | 18.21453 | 18.03561 | 17.31383 | 17.32126 |
| ENSA | 19.0979 | 18.79075 | 18.36226 | 19.37653 | 20.76885 | 20.58671 | 19.87245 | 20.08712 | 18.73276 | 18.81361 | 18.81567 | 17.33793 | 21.05605 | 21.13366 | 18.47764 | 18.96393 | 20.95131 | 21.83256 |
| ENTPD2 | 17.28572 | 19.34427 | 17.29533 | 16.46061 | 16.61646 | 14.86291 | 14.41383 | 15.84783 | 17.12085 | 14.64626 | 14.75009 | 15.98864 | 15.63153 | 15.95215 | 20.17479 | 19.40994 | 13.69021 | 14.55969 |
| ENTPD3 | 15.99666 | 18.20886 | 15.61616 | 16.02855 | 14.99551 | 17.2957 | 14.10936 | 14.718 | 18.33197 | 17.87073 | 15.13605 | 13.16321 | 15.69332 | 14.21308 | 15.35258 | 16.60363 | 13.94047 | 16.66057 |
| ENTPD4 | 16.43737 | 15.35172 | 14.44766 | 16.86881 | 15.95106 | 18.54438 | 15.68736 | 16.22554 | 19.25822 | 18.0514 | 16.88211 | 14.77139 | 15.62014 | 15.30755 | 15.49388 | 16.28426 | 16.43745 | 17.20916 |
| ENTPD6 | 20.28559 | 22.25003 | 16.82776 | 17.02884 | 12.55375 | 17.68601 | 14.90905 | 14.55498 | 18.48893 | 14.40591 | 15.83866 | 16.28998 | 15.25225 | 14.93878 | 19.90235 | 20.10177 | 15.03666 | 14.26701 |
| ENTPD8 | 16.36825 | 15.18589 | 13.98723 | 12.37806 | 12.55171 | 15.36941 | 12.39532 | 13.39554 | 19.58612 | 15.88718 | 14.46356 | 11.09342 | 14.6509 | 13.22107 | 16.21216 | 15.2503 | 16.9376 | 15.19623 |
| ENTR1 | 16.75252 | 18.15109 | 17.23652 | 18.0197 | 18.1678 | 16.10524 | 14.04983 | 18.72018 | 12.88326 | 13.5363 | 15.40451 | 12.23714 | 17.01451 | 16.05103 | 17.57362 | 16.8816 | 14.58302 | 16.40924 |
| ENY2 | 13.76734 | 15.44118 | 14.7879 | 14.82761 | 20.0808 | 14.31441 | 18.49102 | 15.25929 | 18.51791 | 18.88955 | 19.79159 | 14.09864 | 20.16548 | 17.6946 | 18.47436 | 15.56949 | 20.60703 | 20.29298 |
| EPB41 | 18.16367 | 17.33446 | 19.02315 | 19.55905 | 18.22975 | 16.94514 | 20.11321 | 19.90825 | 17.85396 | 18.67491 | 18.61057 | 18.64843 | 19.76257 | 20.44222 | 18.76324 | 19.21637 | 19.35682 | 20.01527 |
| EPB41L1 | 17.86558 | 17.65171 | 18.01789 | 18.87396 | 19.24471 | 18.79172 | 17.5473 | 16.18001 | 16.15292 | 16.4816 | 19.38393 | 18.14247 | 18.27638 | 19.40021 | 20.02391 | 19.34805 | 18.92549 | 19.30308 |
| EPB41L2 | 16.84239 | 15.91205 | 17.64057 | 17.19917 | 21.48295 | 19.76039 | 17.83598 | 18.88819 | 14.81706 | 15.6087 | 16.90834 | 16.15762 | 18.11058 | 18.21392 | 17.82006 | 17.93681 | 20.89364 | 21.07376 |
| EPB41L3 | 15.9403 | 16.39945 | 17.10929 | 16.84698 | 15.1388 | 16.00904 | 16.46027 | 16.63263 | 15.45099 | 15.97605 | 16.25122 | 13.8983 | 17.1279 | 17.99127 | 16.01919 | 15.6718 | 20.87352 | 22.7231 |
| EPB41L4B | 19.94434 | 20.09971 | 20.32468 | 20.3149 | 15.05689 | 16.19775 | 17.70679 | 15.29977 | 16.8885 | 16.59955 | 15.79461 | 15.79984 | 15.96108 | 18.07362 | 21.92761 | 21.08595 | 15.61826 | 16.72542 |
| EPB41L5 | 19.73719 | 18.26965 | 19.71429 | 20.15931 | 17.46772 | 18.14997 | 15.39106 | 18.99655 | 20.3117 | 22.27808 | 18.52094 | 16.80415 | 19.11632 | 19.66692 | 21.65172 | 20.74518 | 17.79399 | 19.23115 |
| EPCAM | 23.86817 | 23.08503 | 23.55898 | 23.64503 | 21.6013 | 20.59942 | 16.66428 | 17.23561 | 23.22277 | 23.47512 | 22.30443 | 21.85989 | 23.99114 | 23.86219 | 23.8052 | 24.03533 | 20.22624 | 20.17558 |
| EPDR1 | 15.79571 | 14.56986 | 14.03173 | 14.49823 | 12.36762 | 15.82155 | 16.74311 | 16.89135 | 18.413 | 12.83558 | 18.36617 | 16.79537 | 13.86985 | 14.69203 | 14.16535 | 15.31105 | 15.39213 | 14.75009 |
| EPHA1 | 21.46314 | 20.83344 | 19.22826 | 18.17234 | 16.78196 | 16.70311 | 19.26457 | 19.01272 | 14.16575 | 15.44171 | 16.00985 | 15.04593 | 16.25584 | 16.99568 | 20.45462 | 19.42572 | 15.51811 | 16.78991 |
| EPHA2 | 18.34242 | 18.14021 | 17.86907 | 17.92911 | 21.56618 | 20.81221 | 17.57913 | 17.07431 | 15.63224 | 15.93611 | 19.40403 | 19.35086 | 18.12437 | 18.15669 | 18.23128 | 17.73222 | 23.42366 | 23.5538 |
| EPHA3 | 20.37236 | 20.7299 | 17.43111 | 17.6464 | 15.47321 | 16.8198 | 17.6371 | 15.89031 | 14.23067 | 14.52655 | 16.44452 | 16.39501 | 14.90836 | 16.76296 | 15.44034 | 16.28289 | 16.81823 | 16.43234 |
| EPHA7 | 17.55703 | 17.89702 | 17.6144 | 16.08428 | 13.90733 | 15.81941 | 17.06998 | 16.64516 | 18.70294 | 17.39977 | 18.09599 | 16.40952 | 14.81849 | 15.62345 | 15.57732 | 16.36555 | 16.67863 | 16.90373 |
| EPHB2 | 14.77515 | 15.85312 | 13.97683 | 16.46053 | 16.08474 | 14.34931 | 19.38813 | 18.78549 | 18.03584 | 18.43821 | 18.16438 | 14.66565 | 15.95992 | 17.50672 | 14.40533 | 14.7495 | 20.01842 | 19.45346 |
| EPHB3 | 19.95811 | 19.99222 | 17.81588 | 18.05936 | 18.77454 | 18.11982 | 16.50606 | 17.18583 | 16.9419 | 16.90189 | 18.52702 | 19.54659 | 16.69812 | 16.4718 | 18.13331 | 18.06354 | 18.43923 | 19.57677 |

| | | | | | | | | | | | | | | | | | | |
|---|---|---|---|---|---|---|---|---|---|---|---|---|---|---|---|---|---|---|
| EPHB4 | 18.44745 | 18.29517 | 17.00957 | 15.22134 | 16.7699 | 17.33313 | 16.58661 | 15.83829 | 13.48006 | 16.7683 | 14.9927 | 14.05005 | 16.44183 | 14.98163 | 17.55897 | 16.89834 | 16.71455 | 17.70954 |
| EPHX1 | 15.40393 | 15.65389 | 20.15341 | 19.51171 | 16.51873 | 16.22276 | 17.62347 | 17.42413 | 19.70063 | 18.66177 | 15.58237 | 15.7627 | 16.91011 | 18.09956 | 19.91678 | 18.27795 | 15.58703 | 16.22733 |
| EPN1 | 21.67857 | 21.622 | 20.28476 | 20.0747 | 18.27912 | 16.54147 | 17.11751 | 17.15363 | 17.1836 | 16.98326 | 17.36294 | 15.27791 | 19.89175 | 18.9322 | 19.44743 | 20.61647 | 19.29987 | 19.99304 |
| EPN2 | 19.38841 | 19.45542 | 18.44505 | 18.62777 | 15.0962 | 15.35072 | 14.75646 | 15.69184 | 16.55504 | 15.48642 | 14.50115 | 15.12625 | 18.27808 | 17.8541 | 19.92184 | 19.6626 | 15.10373 | 17.09314 |
| EPN3 | 19.83859 | 19.63306 | 17.54687 | 17.06446 | 14.9503 | 15.75837 | 17.80483 | 17.79674 | 13.98427 | 14.91345 | 16.05556 | 14.73924 | 16.62269 | 14.04415 | 18.84718 | 19.47153 | 15.88631 | 14.99556 |
| EPPK1 | 17.04781 | 13.94652 | 15.18599 | 14.50099 | 16.76001 | 16.09496 | 14.76691 | 19.04441 | 18.13306 | 14.16084 | 12.45541 | 13.70893 | 19.28348 | 19.67187 | 16.04952 | 14.64134 | 14.78869 | 14.39981 |
| EPRS1 | 17.32791 | 16.96371 | 19.38115 | 19.16742 | 18.23821 | 18.8078 | 18.50212 | 17.88084 | 17.35324 | 17.84978 | 19.60042 | 20.47652 | 19.91545 | 19.29619 | 19.40215 | 18.19954 | 18.52797 | 18.61255 |
| EPS15 | 18.85217 | 19.89291 | 17.50827 | 17.24429 | 17.45894 | 17.50714 | 17.39004 | 17.23657 | 16.35604 | 16.64828 | 17.49755 | 15.57216 | 19.40427 | 18.24811 | 16.71804 | 17.49891 | 17.46684 | 18.36529 |
| EPS15L1 | 18.90795 | 18.88224 | 17.44096 | 16.81356 | 16.58348 | 14.94819 | 17.20565 | 16.097 | 16.37779 | 16.19866 | 17.48572 | 17.9028 | 18.42197 | 17.91418 | 16.1547 | 18.26646 | 16.66182 | 17.13231 |
| EPS8 | 17.4135 | 17.03318 | 16.02579 | 16.85053 | 19.70287 | 18.13345 | 15.70023 | 17.39721 | 14.39498 | 15.12069 | 16.00277 | 15.90098 | 16.10227 | 17.86792 | 15.81665 | 15.62369 | 17.91469 | 16.94087 |
| EPS8L2 | 15.43556 | 17.09161 | 17.31046 | 18.25272 | 19.09193 | 19.77429 | 16.27993 | 17.55243 | 17.641 | 16.90885 | 17.07091 | 16.04144 | 19.79008 | 19.11047 | 19.67901 | 18.58353 | 16.72366 | 16.71371 |
| ERAP1 | 15.02109 | 14.84129 | 14.87888 | 14.28744 | 13.99311 | 14.43915 | 12.55472 | 15.39016 | 18.81372 | 18.86894 | 20.34849 | 15.34658 | 14.39875 | 18.29198 | 17.7687 | 14.83064 | 13.78937 | 13.81435 |
| ERBB2 | 18.32281 | 18.56501 | 20.22182 | 20.51855 | 21.30746 | 19.94728 | 17.60326 | 15.55441 | 17.42827 | 15.72292 | 18.13045 | 11.8021 | 18.44586 | 18.57083 | 20.82 | 20.58417 | 16.43843 | 18.99647 |
| ERBB3 | 18.33701 | 16.89525 | 15.96539 | 17.88323 | 16.83924 | 16.9091 | 15.98606 | 15.78536 | 15.57206 | 15.77999 | 17.99477 | 16.54601 | 16.51251 | 16.68367 | 16.06501 | 17.29066 | 15.31805 | 14.35351 |
| ERBB4 | 15.05554 | 15.38928 | 15.61137 | 15.63137 | 19.39186 | 17.70269 | 15.56329 | 15.3091 | 16.0689 | 14.90407 | 15.64615 | 17.79227 | 16.84992 | 17.59922 | 15.15198 | 15.79609 | 16.73588 | 18.81706 |
| ERBIN | 17.1477 | 16.8816 | 17.99107 | 18.83738 | 22.51486 | 21.17714 | 17.77394 | 18.27996 | 17.39456 | 17.31756 | 18.19068 | 18.15208 | 19.48253 | 19.37119 | 17.84438 | 17.89689 | 19.16249 | 19.87767 |
| ERC1 | 14.37526 | 14.65533 | 14.4859 | 15.02337 | 19.07323 | 18.01436 | 16.66585 | 16.90798 | 13.28272 | 15.72318 | 14.71509 | 11.50411 | 14.91565 | 21.05832 | 16.56742 | 14.77801 | 16.69496 | 15.61779 |
| ERC2 | 15.37359 | 16.43144 | 15.19459 | 16.11833 | 18.03142 | 17.30909 | 16.43216 | 15.20114 | 16.50593 | 15.53661 | 14.51495 | 16.07336 | 15.9895 | 15.53829 | 16.52711 | 13.92518 | 16.12651 | 15.61454 |
| ERGIC1 | 17.43781 | 16.92371 | 18.95853 | 17.55821 | 15.4718 | 16.78778 | 17.37933 | 16.8586 | 22.32731 | 20.65409 | 16.8901 | 16.33334 | 17.42353 | 15.50128 | 19.20369 | 18.69638 | 16.51208 | 16.37375 |
| ERGIC2 | 15.16625 | 16.42607 | 15.54736 | 17.46331 | 15.95884 | 15.32321 | 17.4804 | 16.0028 | 20.67188 | 17.60168 | 17.10464 | 14.74454 | 14.89138 | 16.19897 | 16.05166 | 15.59224 | 15.10193 | 15.63233 |
| ERGIC3 | 16.80138 | 15.37625 | 16.98886 | 15.65867 | 18.53704 | 15.91557 | 16.26667 | 17.00598 | 19.63663 | 17.20149 | 18.70139 | 16.6428 | 18.79995 | 18.23129 | 14.40272 | 13.8658 | 16.69132 | 18.88963 |
| ERH | 20.04001 | 20.33566 | 20.56582 | 20.29194 | 17.21693 | 20.18558 | 19.25161 | 19.28711 | 19.50959 | 19.90612 | 22.31678 | 21.31654 | 20.86778 | 19.60269 | 18.51329 | 20.22217 | 17.48893 | 15.92254 |
| ERI3 | 20.85037 | 19.28295 | 18.27882 | 19.1578 | 15.74391 | 15.49415 | 14.53094 | 15.99136 | 14.35929 | 16.59533 | 14.69433 | 15.39931 | 18.89207 | 18.60211 | 19.05497 | 20.15144 | 14.55155 | 17.01259 |
| ERICH5 | 16.40494 | 13.53573 | 15.81159 | 17.34743 | 13.25817 | 15.59675 | 17.05361 | 16.66568 | 14.76254 | 15.29671 | 15.00788 | 14.61708 | 14.81238 | 17.14868 | 15.83783 | 15.24175 | 19.84889 | 20.90436 |
| ERLEC1 | 16.107 | 16.43556 | 15.58967 | 14.71371 | 14.38826 | 15.81654 | 17.01003 | 15.83482 | 21.01145 | 19.71424 | 17.25601 | 14.96701 | 15.43778 | 16.42513 | 14.95027 | 16.45464 | 16.27132 | 15.03489 |
| ERLIN1 | 17.35022 | 17.96581 | 16.68311 | 16.82234 | 18.37621 | 17.75649 | 18.54582 | 18.59462 | 20.93961 | 20.03061 | 18.76561 | 14.49049 | 16.74862 | 17.05615 | 17.00794 | 16.39121 | 15.1142 | 15.58007 |
| ERLIN2 | 16.26285 | 18.15813 | 16.21606 | 17.0208 | 17.26515 | 20.88686 | 19.30641 | 17.99331 | 21.98659 | 21.27584 | 17.93444 | 16.13382 | 17.34937 | 19.03703 | 17.36107 | 16.89483 | 17.17044 | 18.76452 |
| ERMP1 | 15.96232 | 17.2816 | 16.90406 | 16.54816 | 15.1449 | 15.11466 | 16.04049 | 16.32529 | 20.03359 | 16.5603 | 15.68241 | 15.31647 | 15.81691 | 14.93213 | 16.78783 | 17.18665 | 16.04258 | 14.23147 |
| ERN1 | 16.00062 | 16.03523 | 16.53294 | 17.35574 | 22.22916 | 26.484 | 23.58852 | 22.38063 | 18.99396 | 16.28435 | 14.3123 | 13.02384 | 22.30379 | 25.72414 | 15.23888 | 15.21498 | 15.07792 | 25.17916 |
| ERO1A | 16.68603 | 16.89663 | 16.66105 | 15.77409 | 16.56782 | 18.9802 | 17.71697 | 17.09369 | 20.99689 | 21.46017 | 19.25236 | 16.2778 | 15.68989 | 16.45309 | 17.13945 | 17.47736 | 16.6274 | 17.1973 |
| ERO1B | 15.70737 | 15.4308 | 16.74009 | 17.27371 | 14.74066 | 17.59336 | 16.31264 | 17.17274 | 19.26775 | 19.60453 | 18.35841 | 12.99017 | 14.90411 | 18.58397 | 16.43647 | 14.75858 | 15.97888 | 14.96166 |
| ERP29 | 17.82287 | 16.35704 | 18.25921 | 17.14294 | 17.54758 | 17.68748 | 20.61924 | 20.03653 | 24.96387 | 25.09184 | 21.05501 | 18.024 | 17.18245 | 18.55928 | 17.3121 | 17.77996 | 17.00853 | 17.14156 |
| ERP44 | 17.17682 | 17.15638 | 17.21011 | 17.41613 | 17.30082 | 17.26039 | 19.86451 | 18.10801 | 23.73494 | 22.99779 | 19.87006 | 17.37459 | 17.57883 | 17.48503 | 17.12179 | 17.10005 | 16.05661 | 17.12159 |
| ESD | 17.02674 | 17.44288 | 18.86475 | 19.79972 | 19.57863 | 18.62659 | 19.67773 | 19.95787 | 18.40006 | 18.54488 | 19.68521 | 20.21113 | 19.72254 | 19.87324 | 20.26137 | 19.12554 | 20.97723 | 19.65923 |
| ESPN | 15.53704 | 14.28655 | 15.12579 | 16.75432 | 14.6663 | 15.7689 | 16.18522 | 16.29876 | 12.818 | 15.79418 | 16.70467 | 18.47118 | 15.05888 | 14.16828 | 15.10115 | 15.99631 | 14.47 | 14.31041 |
| ESYT1 | 18.05086 | 16.95525 | 17.71476 | 16.97607 | 18.87085 | 17.43131 | 19.67192 | 16.2756 | 17.49759 | 17.76981 | 16.54842 | 16.91519 | 14.71725 | 15.73141 | 17.58838 | 16.59882 | 16.32641 | 16.96808 |
| ESYT2 | 16.23272 | 17.37308 | 16.73561 | 14.38367 | 19.33828 | 16.78807 | 16.28396 | 17.31663 | 14.35541 | 16.52486 | 16.46616 | 14.33576 | 15.51701 | 16.68666 | 15.16568 | 16.69775 | 15.49572 | 15.59843 |
| ETF1 | 18.84701 | 17.9147 | 18.68877 | 18.71555 | 18.39997 | 17.86593 | 17.1397 | 17.45188 | 17.07751 | 16.65514 | 19.78128 | 20.09203 | 18.51006 | 18.30883 | 18.95989 | 18.98674 | 17.90848 | 17.80864 |
| ETFA | 15.24586 | 15.61502 | 16.22801 | 15.48687 | 16.68698 | 15.98519 | 19.2841 | 15.82478 | 19.53241 | 18.44811 | 14.72634 | 13.33679 | 15.69192 | 15.06887 | 15.60298 | 15.94498 | 15.3255 | 14.17408 |
| ETFB | 19.35713 | 21.29706 | 15.27938 | 17.37306 | 15.38776 | 15.33265 | 15.95109 | 16.04603 | 19.3628 | 16.21973 | 16.67106 | 17.23652 | 16.25456 | 16.57089 | 18.82831 | 19.06741 | 15.21481 | 14.7624 |
| ETFDH | 15.07003 | 14.46048 | 16.96369 | 14.04653 | 15.29467 | 15.07851 | 16.8845 | 13.25942 | 17.83785 | 17.10082 | 16.00332 | 10.89825 | 15.62996 | 17.6639 | 15.44409 | 16.47496 | 16.59661 | 14.70598 |
| ETNK1 | 15.10681 | 19.04661 | 16.50288 | 15.14326 | 14.91175 | 16.07592 | 16.5106 | 17.7816 | 13.09093 | 15.48583 | 15.9341 | 17.38672 | 14.51326 | 14.3909 | 14.19484 | 13.6813 | 15.40121 | 16.95705 |
| EVA1B | 18.66866 | 19.10109 | 20.36343 | 20.29081 | 13.70682 | 16.28033 | 16.24393 | 15.80688 | 15.07384 | 13.92165 | 14.72777 | 13.40589 | 15.85729 | 14.31208 | 20.82189 | 20.04644 | 19.2883 | 19.25955 |
| EVI5 | 16.2387 | 14.88572 | 15.3382 | 14.73842 | 17.10749 | 16.43322 | 16.63822 | 17.37123 | 13.81147 | 14.97251 | 17.18456 | 17.23411 | 15.41962 | 16.31683 | 14.24213 | 17.24397 | 15.24279 | 17.22358 |
| EVL | 16.63419 | 15.84198 | 16.73773 | 15.9131 | 14.25516 | 17.15786 | 15.14952 | 19.84981 | 16.30729 | 14.61922 | 14.05561 | 17.1954 | 17.08739 | 16.46002 | 16.60593 | 18.00176 | 16.85264 | 16.51425 |
| EVPL | 15.06398 | 15.87649 | 15.20739 | 15.91709 | 20.38527 | 21.38566 | 17.97472 | 17.63146 | 16.86617 | 14.15391 | 16.52806 | 13.76472 | 17.50864 | 19.46831 | 13.95209 | 15.06087 | 17.27618 | 20.03585 |
| EWSR1 | 16.5097 | 15.56631 | 16.19073 | 16.60713 | 18.16957 | 17.73844 | 16.95932 | 16.04268 | 15.89518 | 15.71908 | 18.62217 | 19.80933 | 16.25774 | 14.97793 | 15.05899 | 16.08342 | 15.13129 | 17.28214 |
| EXOC1 | 16.44937 | 15.69308 | 16.24725 | 15.60345 | 19.21424 | 17.00571 | 19.17349 | 16.92395 | 19.179 | 16.95633 | 16.62093 | 17.15221 | 18.05426 | 17.70258 | 16.53745 | 16.53312 | 17.47607 | 18.01709 |
| EXOC2 | 16.25264 | 16.39904 | 17.94181 | 18.31115 | 19.50265 | 18.23652 | 17.78032 | 16.96178 | 17.88807 | 18.39853 | 17.98302 | 18.61291 | 17.6891 | 17.0452 | 17.93194 | 18.2126 | 17.81257 | 17.77774 |
| EXOC3 | 16.11684 | 16.4752 | 20.119 | 18.35676 | 19.14512 | 18.0216 | 16.73252 | 16.00928 | 17.34497 | 17.84996 | 17.49104 | 18.01291 | 17.75333 | 17.84469 | 17.52519 | 17.31861 | 17.67639 | 17.56514 |
| EXOC4 | 15.89797 | 16.03881 | 16.94332 | 17.25138 | 18.30617 | 17.63672 | 17.83915 | 16.69651 | 17.06812 | 17.15973 | 17.25644 | 18.01773 | 17.39338 | 16.95431 | 17.41745 | 17.33135 | 17.36416 | 17.1919 |
| EXOC5 | 16.53877 | 15.9743 | 17.22483 | 17.62512 | 20.78031 | 19.75186 | 18.33164 | 16.10371 | 17.78035 | 18.69634 | 17.07416 | 17.79467 | 18.27015 | 18.6367 | 17.69707 | 17.47369 | 19.33652 | 19.17473 |
| EXOC6 | 15.79943 | 16.17987 | 15.60905 | 15.50639 | 18.18231 | 17.51565 | 15.95975 | 15.65697 | 14.41349 | 15.79049 | 13.79454 | 17.1106 | 16.5365 | 16.92625 | 16.56363 | 14.97622 | 15.4048 | 15.17115 |
| EXOC6B | 20.85574 | 21.24113 | 15.64752 | 15.68122 | 18.58205 | 16.42665 | 16.92821 | 17.7562 | 15.82202 | 16.06566 | 14.11393 | 15.09588 | 18.21511 | 16.6102 | 14.72217 | 15.50725 | 17.88648 | 17.42697 |
| EXOC7 | 16.15867 | 17.38274 | 16.45337 | 16.76771 | 20.51524 | 19.05633 | 18.07075 | 17.49499 | 17.58137 | 18.18153 | 17.95955 | 17.29464 | 18.16557 | 17.62068 | 17.41435 | 16.68234 | 18.58607 | 18.48377 |
| EXOC8 | 14.91867 | 14.76503 | 16.04139 | 16.90216 | 19.9745 | 18.35265 | 17.56385 | 17.29156 | 16.41156 | 18.0255 | 16.16375 | 16.46881 | 17.78181 | 18.16366 | 15.20194 | 15.25208 | 18.87662 | 18.39358 |
| EXOSC2 | 18.0699 | 16.36973 | 16.40488 | 18.30075 | 18.19227 | 16.48576 | 13.66545 | 15.37001 | 13.60028 | 14.92489 | 19.15386 | 17.34034 | 18.01781 | 17.1879 | 14.15357 | 17.54014 | 13.65609 | 13.90198 |
| EXOSC4 | 15.6881 | 17.14937 | 15.67856 | 17.17104 | 17.80157 | 15.98274 | 15.53612 | 16.60848 | 14.6856 | 15.55263 | 14.76696 | 18.6756 | 18.78971 | 17.56641 | 14.44823 | 15.988 | 18.48101 | 14.69048 |
| EXT1 | 15.2574 | 16.09896 | 18.45636 | 16.27465 | 16.89644 | 20.19571 | 20.08895 | 17.31792 | 20.60386 | 21.23119 | 14.6938 | 14.31381 | 20.82722 | 20.22903 | 16.96955 | 16.24906 | 21.82182 | 22.67182 |
| EXT2 | 15.43682 | 15.42141 | 14.84849 | 15.22564 | 14.97386 | 17.04581 | 16.0759 | 17.36972 | 19.6558 | 16.63563 | 16.39125 | 15.86275 | 15.69254 | 15.3339 | 15.55743 | 15.43488 | 15.86712 | 17.8502 |
| EXTL2 | 16.4889 | 16.6026 | 16.47625 | 15.64485 | 16.70978 | 15.87627 | 17.69031 | 16.6989 | 20.13264 | 17.2717 | 18.78102 | 15.98068 | 15.51591 | 17.96999 | 15.91229 | 14.75945 | 16.82156 | 18.53075 |
| EXTL3 | 14.51357 | 18.9502 | 16.16879 | 15.23846 | 14.95456 | 19.49289 | 16.64845 | 16.20328 | 18.56699 | 17.58323 | 17.18043 | 13.66332 | 16.83689 | 18.70527 | 16.43443 | 14.56942 | 14.32261 | 15.88855 |
| EZR | 20.04413 | 20.04551 | 19.53992 | 20.23127 | 22.28579 | 21.52426 | 18.09907 | 18.62953 | 21.53266 | 20.86087 | 20.7668 | 21.75986 | 20.83256 | 20.88356 | 20.16644 | 20.50384 | 20.30307 | 19.79399 |
| F10 | 18.26679 | 20.89631 | 20.42376 | 21.7264 | 20.52761 | 19.5373 | 22.63888 | 22.60286 | 15.34304 | 14.02673 | 17.00772 | 13.51052 | 21.61746 | 22.09658 | 20.19781 | 19.32925 | 18.78422 | 20.23385 |
| F11R | 22.3193 | 22.34435 | 22.15101 | 22.34412 | 21.22143 | 19.63137 | 16.92369 | 17.85326 | 20.55819 | 18.86888 | 19.58661 | 19.69773 | 21.89808 | 20.90474 | 22.26186 | 21.85592 | 19.70103 | 20.44969 |
| F2RL1 | 12.4414 | 14.68549 | 17.06444 | 18.85275 | 12.53484 | 17.82307 | 12.80444 | 15.64517 | 12.44485 | 14.3452 | 13.01142 | 11.1437 | 16.07629 | 15.67359 | 15.07145 | 14.40481 | 17.54457 | 18.35968 |
| F3 | 21.6039 | 21.26574 | 16.95631 | 13.59776 | 22.91863 | 21.47024 | 14.41768 | 17.0099 | 13.27591 | 13.32028 | 13.38566 | 13.19906 | 15.31307 | 15.04044 | 13.93152 | 14.37651 | 14.71527 | 13.88149 |
| FABP5 | 19.26396 | 17.09625 | 16.90083 | 16.73473 | 21.83639 | 21.08277 | 19.535 | 18.10768 | 16.37131 | 18.03574 | 21.91875 | 21.31642 | 18.28411 | 17.39288 | 15.90664 | 18.87099 | 23.76498 | 24.02832 |
| FABP5P3 | 20.41386 | 19.59973 | 17.78886 | 17.47395 | 23.58475 | 21.49896 | 19.75594 | 18.34413 | 18.28719 | 17.39217 | 22.61906 | 22.03349 | 16.68889 | 18.01601 | 14.35896 | 15.88122 | 25.35381 | 25.63544 |
| FAF1 | 19.43816 | 17.82603 | 17.88946 | 17.26062 | 14.46932 | 16.99447 | 16.30576 | 13.18537 | 16.69665 | 14.16902 | 16.53528 | 18.59256 | 12.79234 | 12.74449 | 16.41124 | 15.87971 | 16.68882 | 13.88094 |
| FAH | 14.79517 | 17.08425 | 16.45752 | 14.50375 | 14.39088 | 15.65006 | 14.84417 | 16.73952 | 14.60404 | 15.08009 | 16.48473 | 18.5762 | 16.01402 | 15.79882 | 18.84454 | 16.82498 | 15.81113 | 16.07843 |
| FAHD1 | 20.53321 | 15.91012 | 15.85125 | 15.91047 | 15.27151 | 16.90818 | 14.72446 | 14.67394 | 20.24941 | 18.45345 | 15.82151 | 13.7292 | 14.16413 | 16.36919 | 14.88985 | 15.89281 | 16.39855 | 14.2647 |
| FAM111B | 19.15884 | 15.9446 | 17.09406 | 14.79732 | 17.5762 | 15.36236 | 16.19237 | 14.87688 | 15.25993 | 14.35183 | 14.33365 | 12.97243 | 15.3384 | 15.40024 | 14.83566 | 15.06613 | 14.44883 | 13.64627 |
| FAM114A1 | 15.67818 | 16.02541 | 14.66566 | 15.86323 | 17.33031 | 16.84059 | 18.31526 | 17.59392 | 16.47614 | 16.90745 | 15.79414 | 13.54849 | 19.36991 | 20.194 | 16.51939 | 16.97648 | 19.1608 | 20.27022 |
| FAM120A | 18.58116 | 17.38072 | 16.96909 | 18.21066 | 15.56393 | 17.94554 | 17.62959 | 16.55632 | 17.54818 | 16.69813 | 17.11981 | 17.92181 | 17.56831 | 15.98389 | 14.87954 | 17.77777 | 15.10515 | 16.97989 |
| FAM126A | 16.17588 | 14.71368 | 17.12938 | 16.55703 | 19.71669 | 16.73836 | 16.5896 | 17.93013 | 15.03316 | 15.64854 | 15.34251 | 14.64373 | 18.06916 | 16.10985 | 18.15347 | 17.75201 | 16.80479 | 20.83817 |
| FAM131B | 14.6499 | 15.53149 | 16.15189 | 20.07119 | 14.55845 | 15.79149 | 15.18036 | 15.39419 | 17.48305 | 13.07343 | 18.18123 | 13.19753 | 14.99245 | 15.14195 | 17.12458 | 17.36855 | 17.17798 | 18.2882 |
| FAM136A | 17.21147 | 15.64102 | 14.5519 | 15.51328 | 16.55205 | 21.42946 | 17.72435 | 16.92224 | 15.09858 | 16.14815 | 13.05294 | 17.34814 | 16.59136 | 17.90758 | 13.54525 | 12.09925 | 16.83995 | 24.74081 |
| FAM168A | 16.5962 | 17.3468 | 17.09507 | 15.85369 | 19.10259 | 16.55475 | 18.69121 | 18.24736 | 17.63961 | 18.61134 | 15.86942 | 13.24908 | 17.4901 | 17.07351 | 16.98336 | 17.82968 | 13.85009 | 20.58959 |
| FAM171A1 | 14.55539 | 15.57683 | 15.69123 | 16.19577 | 17.92785 | 13.6849 | 15.93424 | 16.0475 | 14.626 | 14.97 | 17.85953 | 17.62725 | 16.99773 | 14.80646 | 15.97754 | 16.70059 | 16.26395 | 16.33668 |
| FAM171A2 | 15.50652 | 15.99806 | 17.1881 | 16.34843 | 16.78661 | 19.41679 | 17.13638 | 16.96496 | 17.85717 | 17.87365 | 16.11344 | 14.99645 | 16.38022 | 15.95895 | 15.32884 | 15.8831 | 19.7923 | 20.79279 |
| FAM177A1 | 16.48147 | 17.82963 | 18.82976 | 19.09575 | 13.24107 | 15.95532 | 15.35543 | 18.80542 | 12.00725 | 14.81234 | 14.7647 | 13.90403 | 16.68779 | 17.55736 | 19.62726 | 20.14874 | 14.56578 | 16.6959 |
| FAM20B | 16.14163 | 18.63281 | 19.99635 | 19.39753 | 15.51168 | 21.80684 | 18.55146 | 17.2563 | 21.9702 | 20.56334 | 19.03723 | 16.9327 | 15.58526 | 15.56898 | 18.69059 | 16.2433 | 15.77561 | 20.09806 |
| FAM234A | 19.01272 | 18.44551 | 17.78299 | 18.14013 | 16.59453 | 19.93469 | 18.61085 | 16.77378 | 16.95533 | 17.13896 | 17.29201 | 18.88022 | 15.15483 | 17.32277 | 19.07299 | 19.57684 | 17.97384 | 15.96175 |
| FAM234B | 16.109 | 15.15767 | 15.30881 | 16.14354 | 14.66386 | 17.22978 | 14.60949 | 15.39186 | 19.94323 | 18.15778 | 16.97034 | 16.5858 | 14.11413 | 15.01173 | 16.14311 | 16.34077 | 12.89998 | 13.5753 |
| FAM3A | 17.23196 | 15.32207 | 14.57475 | 17.04199 | 15.37993 | 15.81029 | 16.77276 | 16.27998 | 20.47089 | 18.81832 | 18.2036 | 17.20144 | 14.80679 | 15.32956 | 16.84261 | 16.11606 | 14.49966 | 14.4543 |

| | | | | | | | | | | | | | | | | | | |
|---|---|---|---|---|---|---|---|---|---|---|---|---|---|---|---|---|---|---|
| FAM3B | 22.14076 | 14.33372 | 16.91711 | 14.06831 | 16.03482 | 16.03966 | 14.34732 | 16.38435 | 19.36904 | 18.2711 | 19.32954 | 17.36525 | 14.00912 | 15.11082 | 16.80187 | 14.49751 | 14.13917 | 16.39607 |
| FAM3C | 17.47791 | 18.14275 | 17.48862 | 16.58327 | 17.34011 | 17.25559 | 17.94697 | 17.64385 | 22.23333 | 21.03847 | 19.29516 | 17.73089 | 17.9336 | 18.01624 | 16.83212 | 18.97181 | 16.58091 | 18.81457 |
| FAM83F | 15.73208 | 17.9505 | 18.29514 | 19.24951 | 13.69735 | 17.79885 | 16.43482 | 16.02813 | 17.2193 | 18.02639 | 18.11487 | 15.99423 | 17.11034 | 16.97326 | 19.98414 | 19.78132 | 15.03762 | 17.50386 |
| FAM91A1 | 16.03105 | 15.73854 | 15.31663 | 15.18721 | 16.67341 | 16.42377 | 17.15159 | 16.51881 | 16.87182 | 16.55122 | 16.18558 | 18.30362 | 14.89035 | 15.99647 | 16.78461 | 17.06834 | 17.59088 | 16.06115 |
| FAM98A | 16.6364 | 15.67939 | 17.34605 | 16.82746 | 19.44222 | 16.77163 | 19.80768 | 17.14424 | 14.82686 | 14.89571 | 18.39201 | 19.38311 | 20.19817 | 18.46151 | 16.89824 | 17.8204 | 17.32459 | 18.66347 |
| FAM98B | 14.16224 | 16.26772 | 14.49084 | 15.22435 | 18.3504 | 17.04344 | 18.63991 | 17.04942 | 12.26517 | 16.56507 | 16.51049 | 16.79664 | 18.83055 | 18.39375 | 14.25544 | 15.41178 | 17.65653 | 18.49118 |
| FAR1 | 15.3068 | 16.19416 | 15.38319 | 14.69802 | 16.36645 | 15.27315 | 17.78297 | 16.61974 | 13.16764 | 13.86121 | 12.66818 | 13.92895 | 14.92597 | 14.92652 | 14.54606 | 15.49272 | 12.66836 | 15.1951 |
| FARP1 | 16.80347 | 16.78219 | 17.79331 | 18.03486 | 18.49363 | 17.88334 | 17.15554 | 17.20261 | 19.52608 | 19.41951 | 19.8698 | 19.14944 | 18.53812 | 17.95194 | 19.44222 | 18.83879 | 18.49328 | 18.52975 |
| FARP2 | 16.52306 | 15.56815 | 18.16308 | 19.24405 | 17.64291 | 16.73026 | 15.73661 | 16.38464 | 16.89445 | 16.27506 | 15.62077 | 14.37056 | 17.57304 | 18.21684 | 19.95564 | 19.24285 | 17.10741 | 16.3018 |
| FARSA | 15.64833 | 17.11259 | 16.92327 | 16.04821 | 17.20832 | 17.27851 | 17.98658 | 16.61708 | 16.21867 | 18.14098 | 17.76915 | 19.50434 | 19.27123 | 18.30248 | 17.10804 | 16.88516 | 17.226 | 18.04875 |
| FARSB | 15.5771 | 17.15993 | 17.36297 | 16.4581 | 18.04359 | 18.9855 | 19.22636 | 17.90777 | 16.66024 | 17.58318 | 18.33446 | 19.89293 | 19.14109 | 18.31804 | 17.73308 | 16.7973 | 17.70166 | 17.30171 |
| FAS | 17.4604 | 18.24366 | 17.76393 | 17.36806 | 18.95946 | 17.75929 | 16.90703 | 19.35287 | 14.68718 | 14.38978 | 13.99368 | 16.10386 | 18.33308 | 17.25785 | 18.64423 | 17.79752 | 17.48348 | 15.14259 |
| FASN | 19.1473 | 18.1897 | 20.58075 | 20.85128 | 18.20414 | 18.89992 | 18.77805 | 17.49238 | 18.17542 | 17.66538 | 21.5209 | 23.53809 | 19.04602 | 18.1222 | 21.39772 | 20.9743 | 18.4756 | 18.08721 |
| FAT1 | 17.30109 | 16.69069 | 17.63426 | 17.27393 | 14.92404 | 15.6429 | 15.61191 | 17.07655 | 13.8442 | 15.47649 | 17.08059 | 17.01318 | 17.38766 | 17.79474 | 17.35447 | 17.84502 | 17.01922 | 18.48469 |
| FAT3 | 15.01515 | 15.99508 | 14.11055 | 18.94328 | 17.84792 | 14.96732 | 11.69994 | 14.85897 | 13.42843 | 15.62463 | 13.33335 | 17.86973 | 15.92203 | 13.83489 | 14.58612 | 14.84191 | 15.33826 | 16.45706 |
| FBL | 17.76955 | 17.65203 | 18.25077 | 16.93297 | 17.06402 | 17.57688 | 20.63878 | 19.68492 | 14.67404 | 16.30565 | 17.17464 | 18.40646 | 18.68531 | 18.60867 | 17.1399 | 16.96053 | 17.89705 | 17.85988 |
| FBLN1 | 15.79707 | 16.08546 | 16.20901 | 17.40582 | 13.97314 | 16.4723 | 15.818 | 17.16085 | 20.99191 | 20.0816 | 16.80118 | 14.64847 | 16.05117 | 15.49403 | 14.81682 | 15.76155 | 15.88016 | 15.37558 |
| FBP1 | 14.73323 | 15.76246 | 15.1741 | 13.59717 | 16.20685 | 15.33011 | 22.76844 | 18.80045 | 18.0055 | 18.76721 | 17.74596 | 16.09161 | 16.62969 | 18.47183 | 16.20707 | 16.01755 | 18.00367 | 15.97409 |
| FDPS | 19.25982 | 18.9559 | 17.5617 | 18.0043 | 19.22939 | 22.97513 | 19.17273 | 18.31912 | 19.3562 | 18.9969 | 20.15668 | 21.52754 | 18.50989 | 17.41423 | 18.52745 | 17.81071 | 19.82044 | 17.13705 |
| FDX1 | 13.91873 | 14.9232 | 13.59293 | 15.47269 | 14.13526 | 14.10155 | 16.97148 | 15.77954 | 18.17381 | 15.10357 | 18.53877 | 15.22857 | 16.29904 | 14.72913 | 15.66427 | 16.00775 | 15.20143 | 15.66547 |
| FDXR | 12.58421 | 15.91676 | 15.99192 | 13.42071 | 16.85218 | 16.92394 | 18.88835 | 15.42171 | 17.61345 | 17.99756 | 16.16946 | 14.54222 | 16.68545 | 14.14162 | 17.56233 | 15.30933 | 17.47088 | 16.27085 |
| FECH | 16.59134 | 15.3113 | 14.61295 | 14.56863 | 14.61634 | 17.72524 | 14.70945 | 16.94059 | 18.75916 | 17.04438 | 16.64142 | 13.45376 | 15.61231 | 17.07317 | 17.4381 | 15.82666 | 16.52293 | 16.08497 |
| FEN1 | 17.79282 | 16.27067 | 16.96497 | 17.39008 | 18.7857 | 17.87732 | 19.79804 | 19.88998 | 15.75001 | 18.53823 | 19.83677 | 20.2086 | 18.71566 | 17.61308 | 16.27078 | 15.73741 | 18.34398 | 16.77525 |
| FERMT1 | 15.6165 | 15.10042 | 14.76029 | 15.61824 | 17.36491 | 16.19803 | 15.74614 | 16.32554 | 12.99407 | 16.57186 | 14.6246 | 13.81261 | 16.24964 | 15.99286 | 13.77398 | 14.65927 | 17.83288 | 19.79594 |
| FERMT2 | 16.63466 | 17.03736 | 16.48735 | 17.30678 | 21.58653 | 20.50643 | 16.69171 | 16.58475 | 15.24744 | 15.64129 | 15.75112 | 13.84315 | 15.24169 | 16.06395 | 16.27094 | 16.54052 | 20.77583 | 21.87144 |
| FERMT3 | 18.58748 | 19.16952 | 19.05728 | 20.19292 | 17.74295 | 15.54362 | 20.06278 | 19.77599 | 14.01107 | 16.00145 | 15.55361 | 13.58988 | 18.58216 | 19.36524 | 19.33385 | 19.53534 | 17.6449 | 17.12532 |
| FGA | 15.44128 | 22.39431 | 16.22182 | 15.76173 | 14.74665 | 15.88376 | 14.51297 | 17.17339 | 12.91065 | 14.73279 | 16.04321 | 13.85058 | 15.97296 | 19.17916 | 16.09526 | 16.49712 | 16.02699 | 16.34986 |
| FGB | 16.58747 | 21.98912 | 15.69019 | 15.95601 | 15.13631 | 18.33389 | 15.64287 | 16.1772 | 13.11264 | 16.40591 | 16.15761 | 14.60732 | 15.3977 | 19.21485 | 16.11924 | 15.2697 | 15.76395 | 14.90866 |
| FGD4 | 14.57839 | 20.81319 | 15.71532 | 14.56594 | 15.09277 | 13.79936 | 16.38167 | 15.69643 | 12.07588 | 13.7234 | 14.32379 | 17.64254 | 16.79992 | 15.51655 | 16.19476 | 15.23551 | 16.56182 | 13.68389 |
| FGD6 | 15.32819 | 15.03123 | 15.83913 | 14.702 | 19.07097 | 18.31016 | 13.47012 | 13.79465 | 13.3203 | 16.71946 | 14.70791 | 11.92108 | 16.92554 | 16.53014 | 16.4912 | 14.41799 | 14.04125 | 16.07577 |
| FGFBP1 | 16.00346 | 16.70515 | 14.62005 | 13.76949 | 15.36261 | 18.00138 | 18.62845 | 19.38888 | 13.51068 | 13.94689 | 13.83468 | 13.31243 | 14.99466 | 13.23492 | 15.97828 | 14.50736 | 18.85118 | 18.31841 |
| FGFRL1 | 20.0909 | 19.98359 | 20.38882 | 19.97873 | 17.02278 | 16.36821 | 14.05978 | 17.91954 | 14.84125 | 14.36549 | 15.7606 | 13.29149 | 19.00585 | 19.21712 | 20.06003 | 20.03657 | 14.99892 | 15.83981 |
| FGG | 15.25333 | 22.83169 | 15.57999 | 14.62698 | 15.96804 | 16.06146 | 15.06924 | 16.08214 | 14.65933 | 15.00336 | 15.6383 | 14.08078 | 14.98233 | 20.46607 | 16.10805 | 14.98552 | 16.13721 | 16.17595 |
| FH | 16.36487 | 16.50593 | 17.0076 | 15.88537 | 17.61437 | 17.45009 | 19.9014 | 18.13542 | 19.80047 | 19.79913 | 18.31241 | 18.74372 | 19.62517 | 19.24356 | 16.30128 | 17.13801 | 19.62514 | 18.98316 |
| FHL1 | 15.94061 | 14.71604 | 15.33495 | 15.46871 | 18.64906 | 17.26345 | 20.69957 | 20.04412 | 12.22323 | 17.08383 | 16.85005 | 11.76446 | 13.07032 | 18.70581 | 16.2704 | 15.07711 | 19.31087 | 18.68669 |
| FHL2 | 16.74252 | 15.77377 | 15.13206 | 16.5578 | 18.07746 | 17.10478 | 15.44579 | 15.96518 | 14.41629 | 15.22992 | 16.17796 | 15.2926 | 14.98656 | 15.42182 | 15.502 | 15.87783 | 19.76427 | 19.3976 |
| FKBP10 | 16.7951 | 16.77301 | 14.58861 | 16.20116 | 18.28933 | 20.23268 | 16.76986 | 16.29984 | 15.76191 | 16.67485 | 20.38669 | 18.43168 | 15.28285 | 14.81866 | 15.21248 | 16.17696 | 15.28039 | 16.1328 |
| FKBP15 | 15.8912 | 15.88164 | 17.6468 | 15.68073 | 19.03137 | 17.55694 | 15.44817 | 18.41251 | 15.07416 | 16.9372 | 15.92213 | 16.57694 | 15.67223 | 14.6627 | 15.23562 | 15.97875 | 15.88521 | 16.15619 |
| FKBP1A | 23.65461 | 23.28335 | 22.6543 | 22.81198 | 22.59396 | 22.45671 | 22.40023 | 21.85691 | 20.21339 | 20.97732 | 20.66657 | 20.61762 | 23.69234 | 23.59034 | 23.23108 | 22.7773 | 23.7175 | 23.44778 |
| FKBP2 | 16.36854 | 16.6725 | 16.34498 | 17.29497 | 15.41001 | 17.64616 | 16.70162 | 16.35556 | 23.06773 | 20.41754 | 19.80174 | 13.86785 | 15.38982 | 16.32516 | 15.26335 | 16.12379 | 15.40228 | 15.2275 |
| FKBP3 | 20.4536 | 20.7019 | 19.76435 | 19.39877 | 20.40542 | 18.8224 | 18.05256 | 17.88338 | 19.20723 | 19.44498 | 19.91741 | 20.2154 | 19.95551 | 18.7266 | 20.41191 | 19.99692 | 18.06176 | 19.78941 |
| FKBP4 | 18.57568 | 18.21543 | 19.44276 | 20.07897 | 18.81804 | 18.10582 | 19.56083 | 19.36929 | 17.55671 | 17.78234 | 20.95362 | 21.59516 | 22.12454 | 21.54862 | 19.6997 | 19.70934 | 19.85743 | 19.07023 |
| FKBP5 | 16.9893 | 16.64745 | 16.77402 | 16.45618 | 19.0444 | 18.24897 | 16.73758 | 17.21788 | 16.90863 | 15.44267 | 16.35433 | 17.82373 | 20.18597 | 18.79315 | 15.93298 | 16.65682 | 19.58293 | 17.46264 |
| FKBP7 | 17.62584 | 14.74144 | 14.53873 | 16.44844 | 16.57511 | 15.07019 | 15.55074 | 16.61399 | 19.50372 | 19.63246 | 18.9186 | 15.64055 | 16.30835 | 17.35007 | 16.07758 | 15.90251 | 14.9585 | 16.09021 |
| FKBP9 | 16.0862 | 22.3365 | 16.4681 | 17.07318 | 15.83455 | 17.03346 | 16.87639 | 17.29354 | 21.25265 | 19.80336 | 19.98387 | 16.51172 | 15.36548 | 16.1284 | 16.79452 | 15.56618 | 16.67705 | 16.65022 |
| FKRP | 16.19622 | 15.751 | 15.45971 | 16.74693 | 14.3069 | 16.12839 | 15.74063 | 17.23217 | 19.64832 | 17.16883 | 16.63672 | 13.46837 | 15.2132 | 15.22873 | 15.67302 | 15.07282 | 16.04861 | 15.83527 |
| FLII | 15.79915 | 17.67659 | 16.17037 | 16.12098 | 17.06683 | 17.17444 | 17.05414 | 17.13856 | 15.30948 | 16.32867 | 15.7099 | 17.26325 | 16.88103 | 16.02353 | 16.22154 | 16.66531 | 18.15891 | 17.62214 |
| FLNA | 19.38155 | 20.20592 | 19.6856 | 20.99933 | 21.42944 | 20.67712 | 21.80739 | 21.2267 | 17.22395 | 18.24776 | 17.43667 | 18.8209 | 20.84549 | 21.4607 | 20.22231 | 20.05781 | 21.83279 | 20.25359 |
| FLNB | 17.79826 | 17.1626 | 19.10884 | 19.07709 | 21.86243 | 20.79143 | 17.34915 | 17.33773 | 16.66716 | 17.1017 | 18.04539 | 19.71392 | 18.53032 | 18.53354 | 19.31394 | 18.61247 | 21.19992 | 19.67744 |
| FLNC | 17.83579 | 18.30753 | 18.89549 | 19.85834 | 20.90625 | 20.10881 | 18.8717 | 18.12012 | 15.91813 | 16.36975 | 16.55897 | 17.55825 | 18.35422 | 18.88818 | 19.2478 | 19.4757 | 20.94763 | 19.06083 |
| FLOT1 | 17.14008 | 17.38308 | 21.05401 | 20.67142 | 21.29736 | 20.56596 | 19.55714 | 18.78234 | 21.2803 | 20.42417 | 20.16795 | 20.42569 | 19.70694 | 19.20684 | 21.78017 | 20.55078 | 21.15421 | 19.66706 |
| FLOT2 | 19.27843 | 18.78399 | 20.90464 | 21.51288 | 21.5971 | 20.09188 | 20.79324 | 19.67281 | 21.30099 | 21.28212 | 20.48632 | 20.36109 | 20.56658 | 21.00419 | 21.75427 | 21.09753 | 17.92792 | 19.06919 |
| FLRT1 | 15.55678 | 15.02971 | 21.64264 | 20.92395 | 16.77187 | 13.57108 | 16.31064 | 16.25876 | 14.11366 | 20.68171 | 14.50732 | 12.42039 | 13.36688 | 14.08127 | 22.09466 | 22.59824 | 16.01498 | 13.38723 |
| FLVCR1 | 16.98663 | 17.89196 | 20.34968 | 19.69968 | 20.38789 | 18.47476 | 19.44899 | 19.62393 | 18.91854 | 19.47231 | 19.92241 | 19.56585 | 20.32398 | 20.20544 | 19.18884 | 19.74731 | 19.91111 | 19.49329 |
| FMNL1 | 15.98938 | 15.87671 | 15.44284 | 16.81291 | 20.72737 | 20.25449 | 16.89244 | 16.13554 | 13.87058 | 16.52078 | 15.56263 | 12.82751 | 16.44058 | 16.02839 | 15.87507 | 15.54595 | 16.79468 | 16.93954 |
| FMNL2 | 18.89965 | 18.30259 | 15.60982 | 19.55164 | 19.3489 | 18.62804 | 17.44138 | 16.16022 | 16.78299 | 17.71487 | 18.21513 | 18.27279 | 16.86799 | 15.73121 | 15.85124 | 15.94933 | 17.37036 | 18.9907 |
| FN1 | 17.90467 | 20.16923 | 23.31333 | 21.78462 | 16.78985 | 19.89627 | 17.00182 | 16.74746 | 17.69786 | 18.50794 | 18.23267 | 17.12929 | 19.33185 | 20.24804 | 19.31254 | 21.42742 | 17.66032 | 19.75386 |
| FN3KRP | 16.08001 | 17.6231 | 17.64253 | 17.63746 | 16.70023 | 17.55532 | 17.3838 | 16.09668 | 16.52787 | 17.53272 | 16.80676 | 17.26515 | 17.71023 | 17.98349 | 18.19127 | 16.43805 | 18.86129 | 19.71258 |
| FNBP1 | 18.54073 | 18.6351 | 16.28441 | 16.04901 | 20.50833 | 20.30157 | 17.92837 | 19.35896 | 13.7607 | 15.55165 | 16.29457 | 17.64123 | 18.18684 | 17.4689 | 15.69944 | 16.31033 | 18.47204 | 20.04943 |
| FNBP1L | 20.66064 | 21.14822 | 17.09564 | 17.62068 | 17.38789 | 16.2049 | 19.13874 | 17.11598 | 16.58042 | 16.79086 | 16.19408 | 15.20788 | 18.78382 | 18.25898 | 19.36167 | 17.80962 | 17.76275 | 18.08034 |
| FOCAD | 16.1012 | 15.61109 | 15.40739 | 14.98888 | 13.50692 | 16.70554 | 12.6462 | 16.15326 | 18.39648 | 14.37659 | 16.38603 | 17.80907 | 14.88245 | 16.84281 | 15.61609 | 16.27358 | 15.66907 | 14.43465 |
| FOLH1 | 23.12974 | 23.03839 | 24.63405 | 25.50846 | 15.19165 | 16.87008 | 16.93976 | 16.83397 | 16.0664 | 16.09855 | 15.77602 | 14.80182 | 21.02759 | 20.34228 | 24.98172 | 24.6533 | 16.57927 | 16.06059 |
| FOLH1B | 22.96731 | 23.25976 | 25.12782 | 25.7928 | 16.10807 | 16.64958 | 16.42558 | 16.95607 | 16.02153 | 16.05046 | 17.09568 | 14.18514 | 19.94977 | 19.11685 | 25.23236 | 24.85836 | 15.6959 | 16.49143 |
| FOXRED2 | 12.8293 | 15.16961 | 15.02011 | 16.55767 | 15.99618 | 16.41881 | 15.24592 | 16.08878 | 20.06987 | 16.5753 | 14.77801 | 12.94191 | 14.42024 | 15.66034 | 16.13091 | 16.26679 | 15.35267 | 13.84548 |
| FPGS | 15.87302 | 15.33589 | 16.53873 | 17.54853 | 13.76262 | 15.92505 | 14.73788 | 13.78871 | 17.48356 | 15.42654 | 14.42653 | 12.33571 | 15.6828 | 15.9397 | 17.61067 | 17.27418 | 14.47573 | 15.94826 |
| FREM1 | 12.98918 | 13.73624 | 13.00246 | 17.49424 | 21.92913 | 15.17211 | 14.37927 | 16.12003 | 16.59785 | 17.20908 | 13.3597 | 16.55921 | 13.65821 | 12.89743 | 14.63969 | 14.85686 | 14.60371 | 13.0545 |
| FRMD6 | 17.20767 | 15.49495 | 15.43853 | 15.08141 | 18.59156 | 17.37612 | 14.0263 | 15.70348 | 14.67345 | 12.13197 | 13.85669 | 11.04879 | 12.77563 | 14.8603 | 16.7051 | 15.16068 | 19.06903 | 20.34773 |
| FRMD8 | 16.64242 | 15.53828 | 18.11737 | 16.35232 | 17.71327 | 17.22163 | 14.92714 | 18.09631 | 16.62673 | 16.69458 | 16.23164 | 16.04816 | 16.61785 | 16.69715 | 17.61599 | 17.67483 | 18.57722 | 19.20417 |
| FRYL | 15.60497 | 15.3585 | 16.06702 | 15.63088 | 17.33014 | 17.81672 | 16.76895 | 17.17282 | 17.37022 | 17.01691 | 18.27617 | 17.84889 | 17.71012 | 15.39573 | 14.84642 | 15.45287 | 18.1438 | 17.29979 |
| FSCN1 | 16.69173 | 16.9322 | 16.41049 | 17.36414 | 21.51659 | 21.37357 | 19.24897 | 19.38811 | 18.89445 | 19.00735 | 21.52527 | 21.08385 | 16.68739 | 18.99769 | 16.01993 | 16.22837 | 21.34529 | 21.39166 |
| FSD1L | 15.36444 | 17.62267 | 16.74382 | 14.51663 | 14.59478 | 14.87814 | 19.10942 | 18.79938 | 16.90058 | 12.80606 | 16.48025 | 17.59396 | 13.34327 | 14.90126 | 14.59068 | 14.43912 | 15.6075 | 13.78661 |
| FTH1 | 16.44047 | 17.84956 | 18.31653 | 17.20375 | 18.61174 | 18.37483 | 21.48361 | 19.97115 | 23.98242 | 24.46729 | 18.7522 | 17.63037 | 20.17339 | 20.831 | 18.51752 | 17.28789 | 20.91696 | 20.18524 |
| FTL | 14.79775 | 16.06 | 16.67563 | 14.68248 | 16.36739 | 16.54849 | 18.06378 | 15.80852 | 21.69518 | 19.86098 | 17.54069 | 15.3918 | 15.60476 | 17.85928 | 17.73006 | 17.70066 | 17.11486 | 16.01774 |
| FUBP1 | 15.81752 | 15.89142 | 15.64345 | 15.76341 | 16.38059 | 16.4918 | 16.1609 | 16.48828 | 17.2866 | 17.59727 | 19.00893 | 18.98302 | 16.68564 | 16.84034 | 16.02058 | 15.37046 | 15.87789 | 15.9086 |
| FUBP3 | 18.34109 | 17.9583 | 16.41853 | 14.47211 | 14.94593 | 17.90435 | 15.16036 | 17.83985 | 15.71252 | 12.01957 | 17.93304 | 18.21361 | 14.10994 | 15.38794 | 14.56757 | 16.01619 | 17.34038 | 16.29785 |
| FUCA1 | 15.27621 | 17.03883 | 15.41585 | 14.75675 | 16.836 | 16.24228 | 15.09796 | 16.50234 | 18.60362 | 17.21084 | 15.98616 | 16.4425 | 14.80166 | 15.32404 | 15.90876 | 15.41044 | 15.28305 | 15.69508 |
| FUCA2 | 14.55462 | 15.27002 | 16.60302 | 15.37753 | 15.8843 | 15.71956 | 17.62922 | 15.69738 | 18.18669 | 18.36251 | 17.61187 | 18.51274 | 14.20987 | 15.89651 | 17.46454 | 15.98636 | 17.3617 | 16.67212 |
| FURIN | 15.84532 | 15.10116 | 15.93255 | 15.78321 | 16.48134 | 17.60684 | 16.34445 | 16.04971 | 18.16539 | 16.46147 | 20.75504 | 15.75992 | 15.27141 | 15.90405 | 15.80785 | 15.42132 | 16.91278 | 15.09652 |
| FUS | 19.28562 | 16.19494 | 17.07596 | 15.90008 | 19.39631 | 16.03509 | 19.14162 | 16.26146 | 18.98458 | 19.27277 | 21.37375 | 20.05631 | 18.40195 | 15.70077 | 15.67203 | 16.47996 | 19.67321 | 17.2318 |
| FUT11 | 14.87338 | 14.45751 | 17.70014 | 15.01201 | 15.33993 | 16.07719 | 15.96387 | 16.35075 | 19.64467 | 18.52642 | 15.76784 | 13.47821 | 15.69531 | 16.19391 | 14.28892 | 16.72307 | 15.87933 | 16.12812 |
| FUT8 | 19.01014 | 16.10845 | 16.44521 | 15.9083 | 15.70173 | 15.66419 | 17.47812 | 16.66541 | 21.23649 | 18.86809 | 16.64779 | 15.42019 | 15.15601 | 16.31137 | 16.719 | 17.05492 | 16.29541 | 16.60442 |
| FYB1 | 18.04777 | 18.35044 | 19.31904 | 17.60734 | 15.18918 | 14.1238 | 14.97877 | 15.56919 | 13.06167 | 15.15672 | 15.16421 | 18.66243 | 13.16664 | 13.79888 | 16.68073 | 17.24212 | 16.13925 | 15.30949 |
| FYN | 16.22487 | 15.35772 | 16.83564 | 15.38257 | 13.74467 | 14.72863 | 18.62281 | 16.10236 | 19.43355 | 18.9378 | 18.04547 | 13.37011 | 15.64363 | 14.56036 | 14.13783 | 13.86371 | 16.91738 | 17.23932 |
| FZD6 | 15.02807 | 15.54821 | 15.73913 | 16.238 | 20.11508 | 17.48439 | 15.07651 | 16.88793 | 18.4151 | 16.64517 | 13.59038 | 12.41922 | 14.71555 | 14.87816 | 16.45131 | 17.42674 | 14.60156 | 17.52076 |
| G3BP1 | 17.54434 | 16.9381 | 16.76182 | 15.90089 | 17.08462 | 17.24688 | 16.40118 | 17.10981 | 16.93686 | 17.03799 | 19.48736 | 20.50269 | 18.11268 | 17.60303 | 15.99382 | 15.57108 | 18.23702 | 16.02274 |
| G3BP2 | 16.11072 | 15.66132 | 16.79778 | 14.96758 | 15.96639 | 16.58925 | 16.87253 | 16.98308 | 13.41455 | 15.73294 | 17.78158 | 19.37508 | 18.8148 | 17.48391 | 15.88257 | 16.24959 | 16.39036 | 14.44367 |

| | | | | | | | | | | | | | | | | | | |
|---|---|---|---|---|---|---|---|---|---|---|---|---|---|---|---|---|---|---|
| G6PD | 15.77596 | 15.10607 | 15.99459 | 18.28404 | 21.92004 | 20.73682 | 18.18073 | 18.18161 | 17.84417 | 16.15693 | 17.20798 | 18.89799 | 19.24195 | 19.17061 | 17.12591 | 16.86727 | 19.74467 | 20.16406 |
| GAA | 16.20144 | 17.29784 | 17.61544 | 17.98707 | 18.71128 | 16.20543 | 17.75137 | 17.56403 | 20.60115 | 17.71606 | 15.2394 | 15.95385 | 16.12506 | 16.82515 | 17.67269 | 17.18637 | 15.9447 | 16.61041 |
| GABARAP | 15.76244 | 16.75434 | 17.21788 | 14.56174 | 15.95396 | 14.24648 | 20.28802 | 16.60584 | 17.25308 | 19.10511 | 15.6665 | 17.03488 | 17.91748 | 17.41522 | 17.09825 | 16.49343 | 18.63936 | 19.33753 |
| GAK | 16.30471 | 16.44424 | 15.68775 | 14.98555 | 13.94349 | 15.24802 | 16.05984 | 17.44542 | 13.80583 | 15.58015 | 14.68686 | 17.65211 | 16.80393 | 15.55302 | 16.70359 | 16.5675 | 15.35845 | 16.2716 |
| GALE | 15.15061 | 16.10748 | 17.8945 | 17.24022 | 18.77871 | 18.80385 | 14.76996 | 16.0595 | 13.75801 | 14.80475 | 16.36085 | 15.34074 | 17.7506 | 16.49135 | 17.99317 | 18.41265 | 19.20633 | 17.21017 |
| GALK1 | 16.31224 | 18.41161 | 18.29105 | 18.57628 | 18.38445 | 17.57643 | 17.52192 | 18.2087 | 15.66501 | 16.19272 | 15.75389 | 17.69216 | 18.11135 | 17.99277 | 19.46935 | 17.91069 | 18.50803 | 17.95399 |
| GALNS | 14.35262 | 14.24371 | 16.81452 | 13.93159 | 13.96045 | 15.24768 | 15.63511 | 14.42262 | 18.49701 | 17.45341 | 14.94527 | 13.39284 | 13.91777 | 19.9177 | 16.29488 | 13.73222 | 16.37597 | 13.64136 |
| GALNT1 | 17.25655 | 17.44281 | 18.01114 | 18.56591 | 16.19785 | 16.21419 | 15.58198 | 16.83565 | 20.36646 | 17.59774 | 19.20029 | 18.67329 | 16.7358 | 15.56683 | 17.40001 | 17.71402 | 15.70402 | 15.0663 |
| GALNT10 | 15.99997 | 16.41455 | 18.23291 | 14.81386 | 15.27418 | 16.90313 | 16.46849 | 16.53403 | 19.24206 | 14.53754 | 15.07625 | 13.90076 | 14.77076 | 15.77088 | 14.10884 | 13.65115 | 16.39485 | 15.44079 |
| GALNT12 | 13.65909 | 14.63157 | 15.67742 | 16.92181 | 12.28409 | 15.3057 | 16.40215 | 15.36229 | 18.5556 | 15.65601 | 16.63651 | 10.49169 | 14.70297 | 14.90389 | 15.15779 | 15.05264 | 13.26367 | 14.09212 |
| GALNT13 | 14.99253 | 16.55688 | 13.9534 | 14.27476 | 13.95064 | 14.15294 | 15.32408 | 16.27149 | 17.00413 | 15.56405 | 15.44095 | 15.41183 | 14.20828 | 13.6823 | 18.97012 | 15.96909 | 15.58401 | 15.82468 |
| GALNT14 | 15.98721 | 15.0746 | 16.62229 | 15.50422 | 14.98921 | 15.89831 | 13.77174 | 16.13043 | 18.69523 | 16.24467 | 15.82208 | 11.361 | 14.67142 | 16.91057 | 13.25111 | 15.53331 | 14.33366 | 14.59081 |
| GALNT16 | 15.5793 | 15.88769 | 15.28582 | 15.33845 | 16.60448 | 15.60246 | 15.32998 | 16.11083 | 20.60656 | 19.66041 | 15.3076 | 14.9332 | 15.54765 | 15.41769 | 16.76927 | 17.11355 | 15.92658 | 14.8815 |
| GALNT2 | 17.5132 | 17.67017 | 17.12016 | 16.81509 | 17.22859 | 17.35342 | 16.61834 | 16.61279 | 22.19597 | 20.85589 | 18.26501 | 16.91913 | 16.54265 | 16.79873 | 17.32911 | 16.73391 | 16.39748 | 16.52719 |
| GALNT3 | 16.4304 | 17.36267 | 16.582 | 15.96226 | 14.30587 | 14.96604 | 16.65462 | 16.90682 | 18.81795 | 17.99842 | 18.06788 | 15.85558 | 17.27109 | 17.969 | 15.70823 | 15.67934 | 16.93468 | 15.15065 |
| GALNT4 | 16.1199 | 19.28267 | 18.87289 | 19.33324 | 18.58972 | 22.56843 | 19.31505 | 17.57101 | 20.53642 | 19.09055 | 20.99826 | 14.36088 | 17.15153 | 19.54551 | 17.92871 | 14.88251 | 17.68946 | 20.6907 |
| GALNT6 | 17.16516 | 16.28841 | 15.18781 | 14.20323 | 16.65677 | 16.31416 | 16.00926 | 18.03758 | 17.82906 | 14.04326 | 14.3664 | 16.40643 | 13.78533 | 18.98963 | 14.93998 | 15.89351 | 18.31025 | 19.18728 |
| GALNT7 | 18.61557 | 18.60395 | 18.55947 | 18.48707 | 15.72306 | 17.65769 | 16.82359 | 17.53317 | 19.32576 | 18.31162 | 16.96206 | 16.66158 | 17.01501 | 16.48541 | 17.4271 | 17.33525 | 16.469 | 17.33027 |
| GANAB | 18.45287 | 16.85606 | 18.47764 | 18.03353 | 16.07654 | 16.86296 | 16.89101 | 17.35213 | 24.49808 | 22.73105 | 20.13492 | 17.98779 | 16.60113 | 16.73852 | 17.94156 | 17.87777 | 16.37241 | 16.7804 |
| GAP43 | 17.08526 | 16.11965 | 14.73715 | 14.86185 | 14.36355 | 14.91855 | 21.80879 | 23.16943 | 16.03976 | 15.49197 | 15.77034 | 13.43206 | 14.22181 | 13.86617 | 16.57747 | 15.52294 | 16.84682 | 12.76654 |
| GAPDH | 23.15245 | 22.54279 | 22.9886 | 23.5452 | 25.69577 | 23.72778 | 24.83313 | 22.87464 | 21.97035 | 22.67098 | 24.51877 | 24.92961 | 24.96913 | 24.46294 | 23.66576 | 23.24411 | 26.03232 | 25.93934 |
| GAPVD1 | 16.54221 | 15.07049 | 16.28719 | 16.01338 | 14.47865 | 16.7678 | 16.38643 | 14.48288 | 15.78646 | 14.70968 | 15.61748 | 19.04883 | 14.94779 | 15.6766 | 17.18158 | 18.9036 | 16.28117 | 18.46158 |
| GAR1 | 18.12358 | 13.83342 | 16.66433 | 15.80153 | 14.61886 | 13.90356 | 18.21652 | 18.97057 | 12.98191 | 15.71552 | 15.48424 | 17.36408 | 15.63816 | 16.88699 | 15.15573 | 17.0846 | 14.09827 | 14.37665 |
| GARS1 | 16.97152 | 16.61996 | 18.77009 | 18.42623 | 18.89545 | 18.50817 | 19.77725 | 18.45761 | 19.12852 | 19.31923 | 20.29196 | 20.84991 | 19.5761 | 19.62134 | 19.05231 | 18.94053 | 19.26472 | 18.62239 |
| GART | 17.16152 | 17.41017 | 16.89378 | 18.38085 | 19.36931 | 18.83399 | 19.15584 | 18.2952 | 16.69796 | 18.37938 | 18.24648 | 19.47206 | 19.8284 | 19.61713 | 18.11727 | 17.78721 | 19.7576 | 19.28947 |
| GAS6 | 15.657 | 15.58783 | 16.07479 | 15.39807 | 14.81833 | 16.99979 | 16.10087 | 18.03735 | 20.59811 | 16.73691 | 15.11967 | 14.00459 | 15.74768 | 17.66332 | 16.44815 | 15.72705 | 16.45506 | 16.748 |
| GAS7 | 15.32934 | 17.03954 | 15.58243 | 15.6638 | 13.34375 | 17.44328 | 15.94294 | 15.11637 | 14.47605 | 15.48008 | 19.09716 | 18.65277 | 15.95909 | 14.67026 | 17.45731 | 15.90669 | 15.78536 | 14.19401 |
| GATD1 | 14.72474 | 13.68356 | 14.59802 | 16.34135 | 17.08008 | 17.21388 | 17.775 | 18.23679 | 17.50784 | 16.09382 | 16.80399 | 17.56755 | 15.34059 | 17.88737 | 17.62831 | 17.18643 | 15.25092 | 17.10952 |
| GATD3B | 16.98925 | 16.96403 | 17.90685 | 15.34011 | 16.31962 | 16.86701 | 19.15092 | 17.37779 | 20.51513 | 18.81914 | 15.91909 | 15.36671 | 18.57448 | 16.30951 | 15.89624 | 16.21213 | 16.08334 | 17.14296 |
| GBA | 16.3579 | 15.47416 | 15.6771 | 17.3211 | 15.22425 | 16.55476 | 17.0145 | 17.29235 | 19.81833 | 18.88728 | 16.35931 | 14.63016 | 15.71829 | 15.87413 | 17.16732 | 16.42602 | 16.58034 | 14.91098 |
| GBE1 | 15.94402 | 16.13537 | 16.37123 | 16.62965 | 16.83262 | 15.77323 | 16.15122 | 17.56993 | 17.89068 | 18.89032 | 16.12674 | 17.35572 | 15.64555 | 17.02589 | 16.00895 | 16.2543 | 19.02743 | 18.36547 |
| GCA | 16.85856 | 16.73977 | 17.7628 | 17.33276 | 15.87134 | 15.02082 | 16.70156 | 17.03233 | 18.77242 | 17.92507 | 17.77657 | 18.49744 | 15.56944 | 16.4871 | 17.94971 | 17.45219 | 17.2295 | 15.96336 |
| GCDH | 17.62726 | 18.87903 | 19.7409 | 19.35621 | 18.3487 | 15.44688 | 18.87053 | 16.12032 | 20.227 | 17.72074 | 13.36597 | 17.09509 | 15.56904 | 17.63068 | 18.92591 | 18.27151 | 16.89814 | 14.85454 |
| GCLC | 17.05063 | 17.07708 | 17.17892 | 16.74054 | 20.977 | 19.68697 | 16.43438 | 17.52076 | 15.90867 | 17.16509 | 18.216 | 19.3278 | 17.34286 | 18.20916 | 16.93884 | 16.18234 | 17.09661 | 17.09046 |
| GCLM | 17.01639 | 16.49703 | 17.71482 | 16.93683 | 20.1773 | 18.18366 | 18.3364 | 17.23275 | 17.14238 | 17.51863 | 15.90757 | 16.71048 | 20.28871 | 19.17898 | 17.77025 | 17.58951 | 19.01155 | 18.28705 |
| GCN1 | 16.19456 | 16.65123 | 17.18828 | 15.89329 | 16.163 | 15.87462 | 17.48407 | 17.03288 | 15.85519 | 15.0188 | 15.83269 | 18.27935 | 16.90324 | 15.92566 | 17.32343 | 16.89071 | 16.08034 | 16.89386 |
| GDE1 | 13.53156 | 15.31286 | 14.67954 | 13.3071 | 16.37211 | 16.22275 | 18.97856 | 16.724 | 18.90088 | 16.63672 | 17.2886 | 17.66059 | 19.14875 | 19.0429 | 16.91725 | 14.31418 | 18.9013 | 17.1591 |
| GDF15 | 15.73366 | 15.69572 | 18.75977 | 17.68988 | 15.83491 | 16.36828 | 15.30496 | 17.14155 | 19.50826 | 18.12813 | 13.98622 | 13.03725 | 15.08785 | 15.70074 | 19.55479 | 18.1858 | 17.45821 | 16.37134 |
| GDI1 | 19.49211 | 19.67458 | 19.75608 | 20.28787 | 19.93892 | 19.33324 | 19.51524 | 19.07606 | 20.49363 | 20.79822 | 20.35739 | 20.71633 | 20.63978 | 20.33753 | 20.77175 | 20.74442 | 20.40054 | 20.38782 |
| GDI2 | 20.51068 | 19.84591 | 20.92021 | 21.33457 | 21.77284 | 21.0155 | 20.08768 | 18.84592 | 21.26097 | 20.813 | 22.03462 | 21.9421 | 21.24825 | 21.41169 | 21.38505 | 21.57184 | 21.19971 | 21.54128 |
| GEMIN5 | 15.55166 | 16.51015 | 15.12624 | 15.39201 | 16.10098 | 15.92944 | 15.00434 | 17.90794 | 15.20658 | 15.80889 | 17.12537 | 19.02137 | 15.68996 | 16.59045 | 16.1199 | 14.97605 | 15.20639 | 15.17697 |
| GEMIN6 | 17.15067 | 15.68285 | 13.8759 | 13.31261 | 16.37365 | 16.56086 | 15.95969 | 14.46257 | 13.90185 | 16.1462 | 15.15673 | 16.62479 | 17.12392 | 16.11372 | 15.29471 | 15.51158 | 15.44333 | 13.63653 |
| GET3 | 15.90499 | 15.96577 | 15.90328 | 17.82576 | 18.27058 | 16.65021 | 18.12381 | 15.70498 | 15.47762 | 15.44334 | 17.89013 | 19.43494 | 18.59298 | 17.58034 | 17.09039 | 16.0693 | 19.8927 | 21.28587 |
| GFM1 | 16.47024 | 15.05735 | 16.88559 | 15.34303 | 15.07109 | 16.4162 | 14.85277 | 16.81141 | 18.63735 | 16.68118 | 16.80124 | 13.79826 | 14.64737 | 15.7643 | 15.93491 | 14.94607 | 16.73468 | 15.89789 |
| GFM2 | 16.3939 | 16.52454 | 15.06478 | 16.80712 | 17.28476 | 14.06152 | 15.46437 | 13.66621 | 14.85393 | 16.68717 | 17.70609 | 16.62234 | 15.21366 | 15.55102 | 13.8813 | 14.86026 | 17.05705 | 18.49707 |
| GFPT1 | 17.51093 | 16.88368 | 18.73641 | 18.3927 | 17.22728 | 17.27725 | 18.0271 | 17.74953 | 17.92724 | 17.77498 | 17.21512 | 18.10486 | 20.37527 | 19.75177 | 18.21371 | 17.65972 | 16.97851 | 18.11869 |
| GFUS | 15.95895 | 16.06771 | 16.61332 | 17.17139 | 19.04316 | 17.01068 | 16.3121 | 16.8993 | 17.43761 | 18.29969 | 15.727 | 17.58912 | 18.76632 | 16.36231 | 17.67264 | 17.85694 | 20.75713 | 20.64736 |
| GGA1 | 18.42571 | 19.20296 | 12.85033 | 16.61494 | 16.64441 | 15.66165 | 13.67046 | 17.36389 | 10.68989 | 17.03566 | 18.33378 | 17.78896 | 17.38872 | 16.76556 | 17.50638 | 17.27704 | 16.95736 | 17.01313 |
| GGA3 | 16.14904 | 16.79299 | 21.28329 | 18.54704 | 16.20044 | 15.51182 | 17.57188 | 16.67917 | 14.7803 | 13.93852 | 14.56346 | 14.17151 | 15.86205 | 14.39212 | 19.84886 | 18.18818 | 18.58588 | 18.51026 |
| GGCT | 18.03195 | 17.90503 | 17.91443 | 17.77343 | 17.06983 | 15.75732 | 17.31076 | 15.28416 | 18.80942 | 19.46794 | 19.53407 | 19.20697 | 20.24333 | 19.24442 | 18.45788 | 18.85503 | 17.94298 | 15.98746 |
| GGCX | 16.12997 | 13.90941 | 15.79534 | 16.63545 | 13.45032 | 16.7457 | 15.36168 | 15.72789 | 19.17836 | 14.48488 | 14.01377 | 15.03402 | 13.90725 | 14.83852 | 14.33248 | 13.5823 | 16.99182 | 15.02074 |
| GGH | 17.71116 | 15.85917 | 18.64511 | 16.75776 | 17.41595 | 17.64421 | 15.50917 | 16.92286 | 23.5574 | 23.42927 | 20.66873 | 17.96033 | 18.45696 | 17.85667 | 16.93782 | 18.07266 | 16.81678 | 16.48377 |
| GGT1 | 21.08635 | 21.34161 | 22.09493 | 22.12886 | 16.75758 | 15.9082 | 15.64512 | 16.90334 | 20.47559 | 19.37011 | 19.88297 | 19.86846 | 17.76817 | 17.96705 | 22.54236 | 21.98827 | 16.11836 | 15.83223 |
| GGT7 | 16.85321 | 17.392 | 17.09803 | 16.22295 | 15.71743 | 15.99005 | 15.7889 | 16.22943 | 20.69268 | 18.70013 | 15.65097 | 13.59228 | 16.45992 | 15.90435 | 16.21021 | 15.74041 | 16.2554 | 15.88114 |
| GIPC1 | 18.05261 | 16.2335 | 19.03063 | 19.21568 | 21.97325 | 20.77373 | 18.71786 | 17.24482 | 17.83759 | 19.18128 | 18.17097 | 18.03072 | 21.02516 | 21.18763 | 19.04197 | 19.35089 | 22.12527 | 22.19727 |
| GIT1 | 15.87492 | 16.5142 | 16.64119 | 16.84093 | 14.60647 | 19.20719 | 14.35309 | 15.66295 | 16.66218 | 14.738 | 14.28667 | 18.4313 | 16.51114 | 17.16196 | 14.27199 | 16.05223 | 15.81088 | 15.77553 |
| GIT2 | 14.00999 | 14.57079 | 13.87846 | 16.31211 | 17.64851 | 15.18608 | 16.383 | 18.06421 | 16.95003 | 17.32794 | 17.00393 | 18.0993 | 15.49331 | 15.90246 | 17.19102 | 16.77084 | 17.42917 | 17.04988 |
| GJC1 | 16.84626 | 16.35313 | 13.81559 | 14.43325 | 22.37517 | 21.29606 | 15.86462 | 15.23725 | 12.81109 | 12.73055 | 15.75411 | 12.31304 | 15.46631 | 16.05399 | 15.18694 | 18.35848 | 14.31768 | 16.30348 |
| GLB1 | 15.16142 | 15.48873 | 15.75666 | 16.2976 | 14.22947 | 15.48181 | 15.45493 | 16.71585 | 19.25645 | 19.00026 | 16.8475 | 15.17665 | 14.57295 | 15.7051 | 16.79355 | 14.80357 | 15.4179 | 16.79387 |
| GLB1L2 | 14.30896 | 15.68969 | 16.17094 | 15.55473 | 13.80854 | 14.30083 | 14.93561 | 16.28391 | 19.33303 | 18.36557 | 15.10375 | 15.41203 | 13.29826 | 12.73068 | 13.66612 | 17.26439 | 14.85509 | 16.14979 |
| GLCE | 15.02675 | 16.5978 | 17.65769 | 15.27669 | 16.15459 | 16.59107 | 15.47215 | 15.72139 | 18.80057 | 17.37588 | 18.115 | 17.39386 | 15.84909 | 14.63673 | 15.49114 | 14.52707 | 15.67726 | 15.46599 |
| GLG1 | 22.41812 | 22.69215 | 17.06364 | 16.7136 | 16.94335 | 17.70443 | 16.58932 | 17.32065 | 21.11738 | 19.81116 | 20.31842 | 18.23894 | 16.39992 | 17.69062 | 16.92118 | 16.26621 | 15.6352 | 15.88018 |
| GLIPR2 | 20.71668 | 21.21576 | 20.74285 | 21.48791 | 22.1 | 20.18806 | 20.8746 | 19.23592 | 18.33247 | 19.32642 | 15.99414 | 14.89584 | 21.72658 | 21.63187 | 21.89643 | 21.47514 | 21.04908 | 22.02851 |
| GLMN | 17.26111 | 13.00424 | 16.27364 | 15.93018 | 17.26221 | 13.39023 | 17.40206 | 15.38957 | 14.66498 | 16.05978 | 16.53526 | 17.39707 | 17.4899 | 13.99049 | 14.12484 | 15.35239 | 16.99014 | 11.79483 |
| GLO1 | 18.53574 | 18.84875 | 18.16803 | 18.55119 | 19.71992 | 18.14375 | 16.55887 | 16.76144 | 19.11965 | 18.33529 | 19.4423 | 20.21014 | 21.42982 | 20.31441 | 20.10498 | 19.75248 | 18.3601 | 18.97908 |
| GLOD4 | 17.91437 | 16.92566 | 18.14417 | 18.81235 | 19.92938 | 18.96958 | 18.80701 | 18.98828 | 17.90861 | 18.81283 | 20.53323 | 20.12719 | 20.76665 | 20.62785 | 19.04342 | 19.00756 | 18.53313 | 18.99309 |
| GLRX | 17.57386 | 17.25835 | 17.32801 | 16.94721 | 16.40422 | 20.01871 | 15.93896 | 15.66805 | 17.02748 | 15.37593 | 19.31354 | 19.96926 | 17.21971 | 16.24103 | 17.14731 | 16.74281 | 20.8455 | 18.93598 |
| GLRX3 | 16.22049 | 15.44836 | 16.2633 | 17.25106 | 20.16489 | 18.96957 | 19.39427 | 18.28169 | 15.93689 | 17.77055 | 19.3713 | 19.75465 | 19.30692 | 18.73091 | 17.16472 | 17.20528 | 19.37495 | 19.3339 |
| GLRX5 | 16.31645 | 16.67923 | 18.20658 | 15.09452 | 13.98566 | 18.33085 | 18.25408 | 16.12881 | 20.96901 | 17.95593 | 16.07091 | 13.36678 | 15.09668 | 16.96008 | 15.61032 | 15.80152 | 18.8571 | 17.6461 |
| GLS | 17.37907 | 18.56263 | 15.59462 | 14.52384 | 17.4533 | 18.63185 | 16.16109 | 16.63104 | 19.9521 | 13.94963 | 17.21394 | 12.11398 | 16.39115 | 16.4023 | 15.5618 | 18.78365 | 15.81758 | 13.61408 |
| GLT8D1 | 17.0039 | 16.42009 | 16.02223 | 17.49946 | 15.32992 | 17.20262 | 16.96273 | 17.7594 | 22.55345 | 20.52048 | 17.63577 | 16.76808 | 15.84656 | 16.21147 | 16.40723 | 16.51019 | 18.98539 | 16.14386 |
| GLUL | 16.60987 | 14.78725 | 15.86098 | 17.00531 | 17.14282 | 15.99337 | 17.9024 | 17.68318 | 17.3147 | 17.26171 | 17.28544 | 19.60081 | 16.95057 | 16.1708 | 15.75314 | 14.09347 | 17.81674 | 16.9951 |
| GM2A | 15.08438 | 17.62557 | 18.24448 | 16.91705 | 14.15189 | 13.75331 | 15.6845 | 13.92887 | 19.16806 | 16.95038 | 18.57367 | 15.09343 | 18.21697 | 17.60814 | 18.60473 | 18.13404 | 15.69473 | 17.16728 |
| GMDS | 14.90767 | 15.69877 | 15.93307 | 16.61542 | 17.16508 | 16.56262 | 16.69347 | 17.42845 | 16.35426 | 14.68077 | 15.95493 | 18.48344 | 15.63234 | 15.55632 | 16.97065 | 16.39206 | 16.42548 | 17.31284 |
| GMFB | 19.81267 | 18.40382 | 19.13199 | 18.83461 | 20.2971 | 19.60527 | 19.87372 | 17.27089 | 17.28581 | 19.52638 | 18.42037 | 19.1845 | 20.63463 | 21.26861 | 19.59977 | 19.54847 | 21.07275 | 20.48786 |
| GMPPA | 16.04923 | 16.10175 | 16.3838 | 16.97711 | 17.87254 | 19.5175 | 19.60867 | 18.25237 | 16.38665 | 17.6583 | 16.10047 | 18.29184 | 17.60031 | 16.96378 | 16.00269 | 17.13802 | 19.55743 | 18.23648 |
| GMPPB | 17.47698 | 18.179 | 15.78957 | 15.40444 | 14.24815 | 16.67127 | 15.04463 | 15.77318 | 16.504 | 17.356 | 13.33427 | 19.02616 | 15.02476 | 14.05395 | 15.29175 | 18.35901 | 15.18349 | 14.15627 |
| GMPR2 | 15.59368 | 14.01609 | 15.10612 | 13.64352 | 16.08636 | 16.99754 | 17.32354 | 16.86568 | 16.46129 | 15.73826 | 17.51337 | 19.13873 | 17.46251 | 15.4148 | 13.54258 | 15.9682 | 18.38159 | 17.49942 |
| GMPS | 16.2468 | 16.72848 | 16.654 | 17.7393 | 19.09607 | 18.00554 | 18.42475 | 17.65675 | 17.1355 | 18.37079 | 19.34999 | 20.11542 | 19.4477 | 17.8364 | 18.30617 | 18.25186 | 18.01667 | 18.62244 |
| GNA11 | 17.98835 | 18.25712 | 15.9647 | 15.62177 | 20.26203 | 19.26785 | 18.5669 | 18.47109 | 18.10243 | 18.57564 | 18.97714 | 20.00878 | 14.84749 | 15.38737 | 16.02796 | 16.01897 | 19.69962 | 20.29137 |
| GNA13 | 18.12873 | 17.93382 | 16.86727 | 18.1538 | 19.8907 | 19.08212 | 20.13374 | 19.38112 | 19.26386 | 18.80002 | 20.07558 | 20.86344 | 16.16373 | 17.56244 | 16.87257 | 16.6367 | 20.12202 | 20.26613 |
| GNAI1 | 15.48327 | 15.33913 | 16.63297 | 17.18805 | 16.4791 | 17.36968 | 17.15873 | 17.05494 | 17.02706 | 16.5995 | 20.3736 | 21.13739 | 20.12477 | 20.13438 | 16.46291 | 17.90824 | 17.20412 | 16.74602 |
| GNAI2 | 21.11786 | 21.50898 | 19.01072 | 21.01504 | 20.12318 | 19.65708 | 19.42177 | 18.45507 | 20.4967 | 19.77482 | 22.31078 | 23.14052 | 17.51241 | 18.0305 | 20.61649 | 20.47214 | 21.063 | 21.50193 |
| GNAI3 | 20.89471 | 20.92597 | 20.2825 | 21.22473 | 21.18292 | 20.03566 | 21.14563 | 20.5757 | 20.18962 | 20.34301 | 22.0227 | 22.51851 | 21.65558 | 21.87316 | 21.06275 | 20.89414 | 21.94965 | 22.49783 |
| GNAO1 | 17.34908 | 16.18833 | 15.49776 | 15.54993 | 18.63203 | 17.67265 | 19.70141 | 19.01353 | 19.39201 | 19.31151 | 15.76461 | 13.86115 | 16.7188 | 16.43827 | 16.14074 | 15.50368 | 16.23951 | 16.36222 |
| GNAQ | 19.27848 | 19.80344 | 18.12307 | 18.63932 | 19.44627 | 19.22205 | 20.57582 | 19.67086 | 18.76125 | 19.11082 | 19.84131 | 21.74389 | 15.20309 | 18.15459 | 17.82478 | 17.56084 | 18.25448 | 18.70605 |

| | | | | | | | | | | | | | | | | | | |
|---|---|---|---|---|---|---|---|---|---|---|---|---|---|---|---|---|---|---|
| GNAS | 20.73369 | 20.9613 | 21.65773 | 22.07611 | 22.21441 | 21.43155 | 22.005 | 22.2757 | 20.09923 | 20.45644 | 22.28259 | 23.02471 | 22.39893 | 22.46769 | 22.23139 | 22.20112 | 22.76689 | 23.30404 |
| GNAT3 | 15.89293 | 17.94543 | 14.27405 | 17.70949 | 14.97622 | 18.83994 | 14.85222 | 15.65054 | 15.97466 | 13.85927 | 15.89419 | 20.13436 | 14.51627 | 16.46225 | 14.75612 | 14.62636 | 15.92799 | 14.02778 |
| GNAZ | 17.22099 | 17.01027 | 15.21568 | 16.73776 | 15.36177 | 14.22879 | 18.34528 | 17.67556 | 16.471 | 13.84614 | 15.66001 | 15.21375 | 15.0899 | 14.85382 | 15.46944 | 16.77633 | 14.56837 | 13.36116 |
| GNB1 | 21.56631 | 21.18946 | 20.39695 | 21.8053 | 23.88749 | 23.03152 | 23.11337 | 21.93678 | 22.93018 | 22.70888 | 23.01825 | 23.56123 | 22.88678 | 23.57366 | 21.0756 | 21.29498 | 24.48252 | 24.94683 |
| GNB2 | 22.79764 | 22.70777 | 21.66786 | 22.07346 | 20.95109 | 20.56389 | 21.24136 | 19.61274 | 21.31054 | 20.46242 | 23.07999 | 24.73493 | 19.13855 | 21.58532 | 23.1082 | 22.75721 | 21.80237 | 21.57364 |
| GNB4 | 15.04119 | 15.22303 | 14.22621 | 19.21928 | 18.54136 | 19.22073 | 16.07317 | 16.64163 | 16.42536 | 15.39469 | 18.14284 | 18.17393 | 15.28308 | 14.06351 | 14.93426 | 17.95653 | 19.08628 | 17.66479 |
| GNG10 | 19.10116 | 19.2638 | 17.57025 | 18.35868 | 22.19259 | 19.79358 | 20.10388 | 20.35032 | 19.33657 | 20.39572 | 16.72167 | 21.05223 | 19.76535 | 20.16252 | 18.05096 | 18.27922 | 20.93411 | 21.53844 |
| GNG11 | 19.50319 | 18.79092 | 18.56149 | 19.58495 | 18.99629 | 18.40361 | 22.12696 | 22.41407 | 14.22073 | 16.11942 | 16.01944 | 14.65792 | 20.2005 | 21.37557 | 16.39769 | 17.11406 | 18.81992 | 20.73897 |
| GNG12 | 20.13354 | 18.81715 | 19.44276 | 19.83911 | 22.89822 | 20.82917 | 18.64695 | 18.55089 | 19.13226 | 19.84732 | 15.89373 | 15.82305 | 22.64786 | 22.89444 | 20.70535 | 20.72767 | 22.19558 | 23.88161 |
| GNG13 | 17.02988 | 18.39194 | 18.43324 | 16.24499 | 17.76989 | 17.23443 | 16.79255 | 17.1618 | 13.875 | 16.47396 | 23.38404 | 23.6219 | 15.33028 | 15.6796 | 15.98836 | 15.28099 | 17.34975 | 18.21929 |
| GNG2 | 16.65122 | 18.36265 | 16.44584 | 16.10323 | 19.76474 | 20.66727 | 20.41435 | 19.28002 | 15.05233 | 16.38198 | 16.28049 | 15.55387 | 17.51977 | 19.90552 | 16.63416 | 16.41485 | 22.34648 | 23.0707 |
| GNG4 | 17.6606 | 17.26954 | 19.28637 | 16.77621 | 19.96728 | 16.8906 | 22.55466 | 20.69659 | 21.14993 | 22.78313 | 17.59318 | 16.5394 | 15.09859 | 17.29762 | 17.25654 | 15.78267 | 16.16464 | 15.75368 |
| GNG5 | 22.86696 | 22.65897 | 21.83215 | 21.22829 | 23.17104 | 22.0487 | 23.17483 | 21.66302 | 21.84811 | 22.81777 | 21.89334 | 23.36446 | 24.13369 | 23.42758 | 21.64847 | 21.67034 | 23.53199 | 24.58446 |
| GNPAT | 18.59898 | 21.15181 | 15.4283 | 18.60248 | 19.90667 | 17.46721 | 18.28395 | 15.40195 | 19.31334 | 18.31201 | 17.58386 | 16.25163 | 16.11073 | 20.30795 | 15.35735 | 15.83109 | 21.31577 | 22.08894 |
| GNPDA1 | 15.8554 | 16.84425 | 17.77963 | 16.83831 | 16.74231 | 17.65074 | 18.03384 | 16.93184 | 15.74187 | 16.54402 | 19.05796 | 17.68423 | 17.62034 | 18.99508 | 17.22299 | 16.95348 | 17.94061 | 16.59812 |
| GNPDA2 | 14.69466 | 14.63802 | 16.67638 | 14.29486 | 16.61695 | 14.01228 | 14.8313 | 16.06985 | 11.74296 | 17.48472 | 18.86786 | 18.45005 | 17.08653 | 17.05052 | 16.18523 | 18.65201 | 14.02146 | 14.68724 |
| GNPNAT1 | 18.24568 | 15.85489 | 19.16113 | 18.82497 | 19.48268 | 17.35837 | 16.48131 | 16.43288 | 14.69502 | 18.17606 | 17.16579 | 17.51589 | 19.38543 | 19.99138 | 19.56635 | 18.31927 | 20.38714 | 18.43056 |
| GNPTAB | 15.9264 | 16.03128 | 16.85573 | 15.82774 | 15.40171 | 17.27211 | 15.73508 | 15.92766 | 18.67402 | 17.07107 | 17.41997 | 13.17118 | 17.15931 | 14.39642 | 16.01424 | 16.02881 | 16.06178 | 15.13347 |
| GNPTG | 15.14674 | 15.52455 | 14.99172 | 16.28401 | 14.64549 | 15.86681 | 14.13752 | 16.97829 | 20.0339 | 15.76917 | 15.21409 | 14.01092 | 15.911 | 15.09463 | 17.49156 | 16.52519 | 12.89878 | 18.81713 |
| GNS | 15.91609 | 16.66897 | 17.36494 | 17.52557 | 17.78495 | 17.01644 | 17.13977 | 16.9048 | 20.81194 | 20.79504 | 18.76813 | 16.59896 | 19.26791 | 18.83302 | 17.07633 | 16.2654 | 16.70678 | 17.91091 |
| GOLGA3 | 14.74408 | 14.88135 | 16.09402 | 15.09765 | 17.0226 | 15.57941 | 16.27109 | 16.13165 | 13.45892 | 14.55397 | 14.92613 | 15.18357 | 17.10711 | 17.89656 | 15.15337 | 15.53755 | 15.70758 | 18.28409 |
| GOLGA7 | 18.9399 | 19.24181 | 20.04594 | 19.72515 | 19.87598 | 17.1747 | 17.87673 | 15.74402 | 16.71216 | 18.01045 | 18.93276 | 20.52014 | 18.77645 | 18.35189 | 20.39703 | 20.34997 | 18.5299 | 18.41439 |
| GOLGA7B | 17.03718 | 17.60553 | 15.93383 | 17.09072 | 14.4514 | 16.93999 | 16.29842 | 17.97711 | 19.5744 | 18.79637 | 20.72538 | 20.0333 | 15.86999 | 15.3486 | 13.64349 | 15.59146 | 14.08314 | 15.97127 |
| GOLGB1 | 15.34318 | 15.69051 | 18.8325 | 14.85788 | 15.80739 | 16.53126 | 15.05568 | 16.76284 | 14.0002 | 14.5837 | 16.59473 | 16.61494 | 16.99783 | 17.06511 | 15.91397 | 16.1792 | 15.20839 | 13.63257 |
| GOLIM4 | 16.38833 | 16.53131 | 16.17968 | 16.15612 | 18.32567 | 17.70997 | 17.76345 | 18.54426 | 22.3185 | 21.18483 | 18.51893 | 16.96952 | 16.30926 | 16.71074 | 15.91144 | 16.30793 | 16.20981 | 17.56568 |
| GOLM1 | 17.14129 | 17.57889 | 17.09821 | 17.9911 | 17.34399 | 17.63056 | 16.55341 | 17.05378 | 21.99248 | 19.37185 | 16.92403 | 15.28629 | 16.3201 | 15.98349 | 18.109 | 17.57828 | 15.99606 | 17.22042 |
| GOLM2 | 15.76819 | 15.40583 | 15.29142 | 16.8635 | 16.54566 | 16.20353 | 15.82042 | 16.74563 | 20.17118 | 18.8782 | 17.00857 | 16.83875 | 17.01841 | 17.30832 | 15.52736 | 15.85719 | 15.84423 | 17.38734 |
| GOLPH3 | 18.12503 | 20.39215 | 16.59619 | 19.5911 | 18.54703 | 17.78609 | 15.83022 | 16.42714 | 15.04347 | 14.53702 | 17.87793 | 18.32804 | 19.09937 | 17.71973 | 17.0371 | 20.05805 | 16.42711 | 18.22347 |
| GOT1 | 18.64557 | 18.44513 | 19.13346 | 19.84348 | 20.17703 | 19.50369 | 18.99864 | 16.89593 | 18.79179 | 19.5466 | 21.51384 | 21.92988 | 21.08732 | 20.80206 | 20.13193 | 20.23433 | 19.18348 | 18.63858 |
| GOT2 | 19.30576 | 17.76952 | 16.18619 | 16.92513 | 16.024 | 15.80916 | 16.00969 | 16.29646 | 21.18309 | 21.01973 | 17.52527 | 16.929 | 16.16871 | 20.73708 | 15.81993 | 17.51695 | 16.73422 | 15.66349 |
| GP1BB | 21.48017 | 22.71523 | 21.94192 | 22.58663 | 13.74347 | 15.71211 | 13.57593 | 17.77038 | 11.76734 | 15.84279 | 15.79674 | 12.04126 | 16.75087 | 15.04724 | 22.46684 | 22.41214 | 15.15021 | 14.21576 |
| GPAA1 | 17.07685 | 17.1449 | 16.09885 | 16.27123 | 17.95255 | 18.03504 | 16.63757 | 15.63662 | 21.16365 | 19.84031 | 16.5377 | 13.99904 | 16.26653 | 16.03173 | 15.76937 | 16.26561 | 16.40014 | 15.01128 |
| GPC1 | 19.27851 | 19.55148 | 19.61555 | 20.67859 | 20.19253 | 20.16427 | 18.47536 | 19.48734 | 17.86087 | 16.14492 | 20.49913 | 15.29272 | 18.2344 | 18.94129 | 19.42073 | 19.40915 | 16.20497 | 16.0554 |
| GPC2 | 13.62971 | 15.90989 | 14.61256 | 14.50348 | 17.78178 | 16.30122 | 16.22823 | 15.73789 | 18.15729 | 15.82132 | 16.50229 | 15.24597 | 15.62713 | 17.2231 | 14.37731 | 15.4154 | 15.91039 | 16.0673 |
| GPC3 | 16.4578 | 15.41432 | 15.93705 | 19.70755 | 15.47735 | 18.08613 | 16.31373 | 16.48499 | 20.33901 | 20.26945 | 17.50788 | 14.5677 | 15.88201 | 16.27875 | 17.90583 | 15.43657 | 16.5872 | 16.18787 |
| GPC4 | 20.40089 | 20.31423 | 16.82292 | 17.3944 | 15.80293 | 15.66739 | 16.16021 | 15.87725 | 18.47191 | 17.37916 | 15.55316 | 14.64585 | 20.17701 | 19.9529 | 15.96232 | 17.59274 | 18.00709 | 19.00514 |
| GPC6 | 21.09703 | 21.3512 | 19.93866 | 21.23346 | 16.24627 | 17.23471 | 16.00911 | 15.76844 | 16.69146 | 16.18819 | 15.0354 | 17.43803 | 19.07939 | 18.42508 | 15.79449 | 19.04078 | 16.46192 | 16.86642 |
| GPD1L | 16.78315 | 15.8711 | 16.57123 | 16.7131 | 18.11351 | 19.86331 | 18.3795 | 19.06475 | 16.98076 | 17.291 | 16.73757 | 18.42937 | 17.9569 | 19.80228 | 14.95029 | 15.21903 | 18.14539 | 19.58588 |
| GPD2 | 15.47536 | 15.84507 | 16.62401 | 16.27925 | 18.95884 | 18.37724 | 17.75063 | 18.54665 | 20.74112 | 19.86909 | 16.04864 | 14.85791 | 16.93482 | 17.15256 | 15.47324 | 15.47815 | 18.67962 | 19.13632 |
| GPHN | 16.3793 | 16.83032 | 15.90361 | 14.03322 | 16.48727 | 15.53553 | 18.04296 | 17.95327 | 15.83704 | 14.72855 | 17.90453 | 18.23635 | 17.25825 | 18.22529 | 16.07314 | 15.93181 | 14.21997 | 17.59482 |
| GPI | 20.09844 | 20.24884 | 20.82698 | 21.87616 | 23.18511 | 22.33673 | 21.7237 | 20.39987 | 21.92407 | 22.17328 | 22.92646 | 23.70252 | 22.22686 | 21.86963 | 21.61799 | 21.69123 | 22.17733 | 21.85767 |
| GPM6A | 23.89518 | 23.12325 | 17.57776 | 14.92021 | 16.78826 | 17.45351 | 17.50079 | 18.47624 | 15.13252 | 14.30335 | 14.87116 | 17.42544 | 17.43201 | 14.00453 | 18.68744 | 17.74366 | 15.68148 | 13.86796 |
| GPN1 | 16.24578 | 19.8047 | 16.59446 | 16.85631 | 16.41582 | 14.46162 | 15.99924 | 14.93113 | 14.36208 | 13.26757 | 14.83275 | 14.09032 | 16.75191 | 14.95312 | 16.31649 | 17.49421 | 16.25334 | 14.79996 |
| GPR107 | 16.21735 | 13.3122 | 16.22922 | 15.76856 | 18.23821 | 18.95643 | 16.97877 | 17.31401 | 21.22673 | 19.67798 | 17.35414 | 16.11014 | 17.60699 | 16.30274 | 15.49285 | 16.57181 | 15.17277 | 14.85275 |
| GPR108 | 17.22834 | 15.30382 | 15.37955 | 15.76536 | 15.84445 | 16.09659 | 16.00698 | 15.2718 | 21.78464 | 19.48921 | 18.04269 | 15.27754 | 15.06778 | 14.4118 | 16.14548 | 15.86083 | 14.52584 | 15.59314 |
| GPR180 | 15.58513 | 15.56661 | 14.62909 | 16.59621 | 17.25513 | 15.56158 | 15.3078 | 16.1145 | 20.6986 | 19.33215 | 16.56193 | 13.86005 | 13.9316 | 15.22356 | 16.2834 | 15.93905 | 14.65309 | 14.01589 |
| GPR39 | 13.79455 | 15.1947 | 16.73249 | 15.29266 | 18.77859 | 17.24029 | 16.1327 | 17.70503 | 11.81501 | 14.00063 | 15.94628 | 14.33614 | 15.89279 | 15.60948 | 14.86557 | 15.40784 | 18.92599 | 20.70046 |
| GPRASP2 | 18.59259 | 18.81785 | 19.1429 | 18.00973 | 16.42134 | 13.9575 | 14.41202 | 15.5282 | 13.74041 | 13.11248 | 14.11352 | 13.69099 | 17.39064 | 16.38422 | 18.97904 | 19.07973 | 15.85883 | 12.70504 |
| GPRC5A | 16.16421 | 17.09007 | 16.74077 | 16.32454 | 23.82927 | 23.5329 | 15.66641 | 16.74956 | 15.86096 | 14.79354 | 15.51276 | 14.95255 | 16.82124 | 15.98605 | 20.88676 | 16.61305 | 16.57694 | 15.12723 |
| GPRC5B | 14.62468 | 16.88623 | 20.10172 | 19.91508 | 16.41485 | 16.92127 | 17.00644 | 16.04248 | 16.56648 | 17.03643 | 19.5206 | 17.80575 | 16.84577 | 16.32572 | 18.51728 | 18.65328 | 14.68763 | 16.12037 |
| GPRC5C | 16.17858 | 17.5152 | 21.4334 | 22.54496 | 19.15881 | 18.63386 | 17.03513 | 17.2196 | 20.44255 | 19.56499 | 17.37856 | 16.05312 | 21.40656 | 20.1169 | 21.47106 | 21.6274 | 18.40409 | 20.34358 |
| GPRIN1 | 17.77468 | 17.06817 | 17.06795 | 16.21472 | 16.62345 | 16.27206 | 17.96841 | 18.46691 | 18.97726 | 18.43444 | 17.24313 | 14.70412 | 17.65348 | 17.65567 | 16.52931 | 16.01683 | 19.22354 | 19.89163 |
| GPS1 | 15.89332 | 16.36295 | 16.54342 | 14.63533 | 16.25176 | 16.22639 | 17.25807 | 17.05728 | 13.37891 | 14.67673 | 16.95255 | 18.17962 | 16.34139 | 16.48871 | 16.68324 | 17.76479 | 16.34092 | 16.92908 |
| GPSM1 | 16.65063 | 16.6578 | 15.83459 | 15.02253 | 18.3619 | 14.98717 | 16.56876 | 18.11053 | 16.66881 | 14.43319 | 16.6691 | 14.133 | 15.24821 | 16.43565 | 15.52672 | 15.11116 | 17.59614 | 17.015 |
| GPX1 | 15.36629 | 15.2217 | 16.31165 | 17.32331 | 17.58863 | 15.32691 | 19.22784 | 17.45506 | 19.13772 | 19.59317 | 18.14949 | 18.75201 | 18.60001 | 19.10874 | 16.48694 | 16.04416 | 18.77448 | 20.05717 |
| GPX4 | 20.12695 | 19.7968 | 18.10165 | 18.52229 | 15.27432 | 16.94592 | 17.96017 | 19.2537 | 19.30109 | 19.06924 | 16.98799 | 17.59374 | 19.19281 | 19.44422 | 19.87078 | 19.357 | 18.58854 | 17.29423 |
| GPX8 | 17.72681 | 19.01432 | 15.43195 | 15.79125 | 16.93159 | 15.20506 | 15.48535 | 16.68583 | 18.62642 | 16.38314 | 18.8841 | 12.93338 | 14.03863 | 13.45917 | 14.69313 | 14.56001 | 17.20691 | 18.34573 |
| GRB2 | 22.25574 | 22.22311 | 21.61665 | 19.95273 | 17.78154 | 19.29883 | 18.13757 | 17.56974 | 17.76943 | 17.50549 | 17.83654 | 19.66254 | 19.41805 | 17.24733 | 22.17004 | 21.67513 | 19.00477 | 19.09724 |
| GRHPR | 18.90819 | 13.87425 | 19.35124 | 18.29175 | 21.54498 | 20.12967 | 18.88225 | 17.25274 | 19.30069 | 19.713 | 19.86475 | 19.91312 | 20.97205 | 20.59047 | 20.80859 | 20.0988 | 19.19765 | 19.62099 |
| GRIK3 | 13.90143 | 17.87667 | 18.60164 | 18.24726 | 14.73501 | 15.39882 | 16.82602 | 15.8187 | 17.81844 | 16.37746 | 14.73211 | 16.93394 | 16.22789 | 15.71877 | 18.18573 | 16.17649 | 14.84315 | 14.36738 |
| GRK2 | 14.0811 | 16.52877 | 12.98229 | 18.07289 | 17.97857 | 13.96049 | 17.89883 | 17.91046 | 12.18753 | 16.71889 | 14.96159 | 18.34687 | 19.29463 | 19.36641 | 18.09621 | 18.01199 | 14.37879 | 13.78153 |
| GRK5 | 16.19419 | 17.26182 | 16.41423 | 15.8898 | 16.07924 | 16.44109 | 16.32274 | 16.43553 | 14.09584 | 16.43334 | 20.42547 | 19.42674 | 15.59134 | 16.2114 | 17.57168 | 15.65399 | 17.54294 | 17.79625 |
| GRK6 | 17.81697 | 17.43351 | 19.35181 | 19.29408 | 14.97199 | 16.64408 | 16.49096 | 17.73115 | 17.86727 | 15.21855 | 18.56435 | 18.13428 | 17.69169 | 17.47193 | 19.21977 | 19.31266 | 15.98228 | 16.3047 |
| GRN | 16.54898 | 14.25419 | 17.1018 | 17.81604 | 15.10627 | 17.02899 | 14.79526 | 16.9418 | 19.58432 | 18.07299 | 15.63969 | 15.98939 | 14.03324 | 17.25091 | 16.86534 | 16.42077 | 14.87134 | 18.30938 |
| GRP | 17.2421 | 16.5248 | 13.86884 | 15.58937 | 14.80331 | 15.9652 | 14.78773 | 15.46299 | 18.75638 | 17.93373 | 13.80517 | 12.6826 | 14.00575 | 14.44333 | 14.66114 | 14.88373 | 15.93475 | 14.74186 |
| GRPEL1 | 16.44338 | 16.08307 | 16.06784 | 14.42401 | 19.81532 | 17.46843 | 20.17264 | 15.4253 | 20.23623 | 19.68751 | 19.44402 | 15.61977 | 18.45732 | 18.7094 | 13.47133 | 14.86721 | 18.14017 | 18.49707 |
| GRSF1 | 16.95324 | 15.39924 | 22.44772 | 18.1638 | 13.70008 | 16.63159 | 15.02832 | 13.49977 | 17.99895 | 14.23842 | 13.35402 | 13.83333 | 16.57789 | 14.71439 | 19.15987 | 18.63594 | 15.48319 | 13.66874 |
| GRTP1 | 17.54899 | 17.46119 | 19.24831 | 21.58088 | 15.03209 | 16.49817 | 15.27875 | 14.74874 | 14.43815 | 15.42497 | 16.86929 | 13.85937 | 14.16224 | 15.92316 | 19.96815 | 19.96688 | 16.01861 | 13.60041 |
| GSDMD | 15.6919 | 14.91117 | 14.94301 | 14.70691 | 17.14543 | 20.22141 | 17.79988 | 16.56771 | 17.3701 | 17.6813 | 15.78422 | 15.23663 | 14.5442 | 18.88679 | 14.29157 | 15.72015 | 19.73384 | 21.16321 |
| GSK3A | 16.92277 | 17.20994 | 15.37482 | 17.29964 | 18.21457 | 17.58674 | 17.74716 | 17.16108 | 15.41404 | 14.08399 | 17.22436 | 17.42634 | 17.88277 | 14.06086 | 17.74968 | 16.75193 | 15.94916 | 16.87098 |
| GSK3B | 16.07053 | 18.00914 | 15.30597 | 18.07724 | 18.39379 | 16.17695 | 16.99753 | 17.24241 | 18.2803 | 17.40134 | 15.85312 | 18.26374 | 18.12122 | 16.96189 | 16.97599 | 15.29017 | 16.96683 | 19.71765 |
| GSPT1 | 17.07958 | 16.21146 | 18.25372 | 17.93399 | 18.98421 | 17.494 | 19.2579 | 18.61804 | 17.0627 | 17.55504 | 18.41021 | 19.19637 | 20.23054 | 19.5575 | 16.86984 | 17.69695 | 19.57889 | 19.81289 |
| GSPT2 | 21.76012 | 22.66615 | 20.39193 | 21.84076 | 14.19897 | 14.7595 | 20.58713 | 16.41182 | 16.85017 | 14.41216 | 18.3253 | 19.04594 | 22.36797 | 21.37513 | 21.18522 | 20.72961 | 14.70303 | 16.09206 |
| GSR | 18.00374 | 17.7227 | 17.53071 | 18.9902 | 22.62158 | 21.2859 | 19.14522 | 18.4791 | 18.36652 | 19.39288 | 19.3911 | 19.77674 | 21.73156 | 21.22844 | 19.33534 | 19.62188 | 20.15237 | 19.41918 |
| GSS | 16.50401 | 17.72616 | 17.29083 | 17.79902 | 18.65739 | 17.32872 | 17.40043 | 17.22556 | 17.43693 | 17.61503 | 18.85233 | 19.802 | 17.14282 | 15.42249 | 18.23259 | 18.72416 | 18.71744 | 17.76237 |
| GSTK1 | 16.35609 | 15.82636 | 15.49805 | 17.80733 | 14.55579 | 15.19155 | 15.64882 | 18.04509 | 19.81604 | 18.24239 | 16.48407 | 13.32754 | 14.95965 | 15.01244 | 15.68705 | 16.88516 | 15.03481 | 17.07663 |
| GSTM3 | 14.91478 | 16.96309 | 16.99703 | 16.67415 | 15.73401 | 15.4219 | 17.08747 | 17.18573 | 14.91996 | 16.05807 | 14.69754 | 11.73709 | 19.95753 | 20.63012 | 19.6656 | 18.7796 | 18.57602 | 18.18745 |
| GSTO1 | 17.24435 | 17.30883 | 16.27164 | 17.06562 | 20.10689 | 18.74975 | 18.84824 | 18.28769 | 14.22865 | 15.9304 | 19.85856 | 20.59397 | 18.15565 | 17.57468 | 18.23738 | 17.09403 | 19.11015 | 18.11418 |
| GSTT2 | 15.03643 | 13.46317 | 14.59937 | 15.51747 | 12.03865 | 15.31333 | 13.93349 | 15.94578 | 13.77309 | 13.2122 | 15.56811 | 18.36173 | 17.21266 | 13.74834 | 16.20355 | 15.39885 | 14.66776 | 12.73156 |
| GSTZ1 | 15.73487 | 16.2993 | 15.72086 | 14.86439 | 15.79031 | 15.90361 | 15.69616 | 16.69619 | 19.21912 | 18.3219 | 15.10293 | 15.57021 | 16.84029 | 14.90335 | 16.8729 | 15.32531 | 16.57768 | 14.77396 |
| GTF2I | 15.61726 | 14.92415 | 14.81331 | 17.81327 | 15.35438 | 16.61721 | 16.6535 | 16.25135 | 13.91115 | 14.76936 | 17.31136 | 18.64183 | 15.4546 | 15.49368 | 15.85492 | 15.7202 | 16.44566 | 14.79472 |
| GTPBP1 | 18.78906 | 18.80906 | 17.57813 | 19.21919 | 16.88244 | 15.32379 | 16.9485 | 16.91064 | 16.37366 | 15.80275 | 18.14715 | 21.52598 | 18.61059 | 17.71394 | 19.0611 | 18.7752 | 16.9327 | 16.98774 |
| GTPBP4 | 14.57737 | 15.97209 | 16.3637 | 16.00897 | 16.44353 | 14.32719 | 17.28368 | 19.62614 | 14.19788 | 14.34043 | 15.50664 | 14.49872 | 14.8701 | 15.61199 | 16.95086 | 16.05292 | 17.67119 | 14.98868 |
| GUF1 | 13.2414 | 15.47943 | 11.52129 | 13.83624 | 15.33027 | 14.24065 | 16.4661 | 15.9256 | 16.51806 | 18.90732 | 13.65975 | 13.31206 | 14.50281 | 16.58007 | 15.62194 | 17.57547 | 15.36629 | 16.18297 |
| GUK1 | 14.53586 | 15.0878 | 15.72114 | 15.87476 | 18.76594 | 15.00791 | 19.17405 | 15.58444 | 16.83117 | 16.523 | 13.19765 | 14.17851 | 19.3171 | 19.68607 | 15.27642 | 14.778 | 19.8986 | 20.41286 |
| GULP1 | 17.30983 | 16.813 | 18.59213 | 18.8028 | 16.77637 | 16.91263 | 18.44353 | 15.81735 | 14.7482 | 15.76595 | 16.24831 | 15.92699 | 18.6929 | 18.91095 | 15.97938 | 17.36919 | 17.18196 | 17.04882 |

| | | | | | | | | | | | | | | | | | | |
|---|---|---|---|---|---|---|---|---|---|---|---|---|---|---|---|---|---|---|
| GUSB | 16.05237 | 17.5317 | 17.42957 | 16.95357 | 17.97289 | 19.46321 | 17.9635 | 17.33185 | 22.39119 | 22.17314 | 18.08232 | 17.17949 | 18.42672 | 18.32213 | 18.80764 | 16.09842 | 15.86535 | 16.02308 |
| GXYLT1 | 15.70096 | 14.94696 | 15.01974 | 14.28207 | 14.73249 | 16.9086 | 14.38848 | 14.39263 | 18.70475 | 17.95303 | 16.22328 | 15.4174 | 13.68896 | 14.60261 | 15.22595 | 13.79998 | 12.87177 | 12.99096 |
| GYG1 | 14.67457 | 14.33351 | 16.73775 | 16.62688 | 17.59195 | 17.49834 | 17.43225 | 17.58178 | 16.18047 | 16.70161 | 18.22872 | 20.03789 | 18.45963 | 17.96404 | 15.70985 | 16.06177 | 16.48378 | 16.62637 |
| GYS1 | 16.29065 | 16.78269 | 15.66673 | 16.98606 | 16.88189 | 19.22396 | 16.65804 | 16.5281 | 13.6461 | 15.08847 | 18.55314 | 19.8511 | 15.49689 | 16.15893 | 15.48243 | 15.87446 | 17.26927 | 17.43151 |
| H1-0 | 21.22149 | 20.99819 | 15.36115 | 16.71752 | 17.35167 | 16.65826 | 15.98046 | 21.88537 | 14.6561 | 16.40023 | 16.64575 | 15.09771 | 17.71019 | 15.56585 | 15.98026 | 16.00675 | 16.74273 | 18.05673 |
| H1-10 | 14.66355 | 14.46009 | 16.75912 | 17.897 | 21.97738 | 20.55445 | 18.25362 | 20.66599 | 17.98268 | 15.11336 | 18.14253 | 17.57528 | 16.29215 | 20.57906 | 15.58316 | 15.17577 | 21.12328 | 21.97034 |
| H1-2 | 24.38948 | 20.39017 | 16.71522 | 17.67294 | 18.55767 | 18.28885 | 22.21883 | 28.06425 | 17.77574 | 16.22634 | 18.15629 | 18.23343 | 18.25581 | 16.99466 | 17.57369 | 17.30977 | 15.32548 | 18.18402 |
| H1-3 | 23.72892 | 23.1979 | 17.40485 | 17.01163 | 20.99509 | 20.09952 | 22.99312 | 28.85544 | 16.62327 | 17.95234 | 18.50186 | 16.97444 | 18.13474 | 17.07668 | 17.03704 | 17.19122 | 17.5575 | 20.13837 |
| H1-5 | 24.0704 | 23.41959 | 16.20402 | 17.34345 | 19.03332 | 18.4153 | 20.23776 | 25.49823 | 15.80163 | 17.83988 | 19.36096 | 17.78627 | 17.56618 | 20.42736 | 16.21664 | 16.8654 | 16.67705 | 15.56043 |
| H2AX | 28.84444 | 28.26952 | 21.63095 | 20.5321 | 24.02313 | 24.14269 | 24.81771 | 25.71508 | 21.35905 | 20.94433 | 22.3961 | 28.6677 | 21.52734 | 21.03778 | 20.25605 | 20.41092 | 19.77693 | 23.36358 |
| H2AZ2 | 23.14362 | 22.06212 | 14.20728 | 17.00237 | 18.74586 | 17.94115 | 21.80522 | 26.31976 | 16.15555 | 16.95557 | 17.19849 | 23.58881 | 14.00113 | 16.8659 | 14.72148 | 16.9096 | 16.19121 | 19.01466 |
| H2BC19P | 25.86848 | 22.7428 | 16.67684 | 15.73874 | 13.97471 | 16.61144 | 21.83693 | 24.72692 | 13.10255 | 14.22663 | 17.24687 | 25.79696 | 17.73105 | 12.45084 | 16.70255 | 16.31998 | 15.4926 | 15.87359 |
| H2BC4 | 26.24409 | 25.60284 | 19.0694 | 18.1413 | 22.39058 | 22.56659 | 23.70943 | 26.09377 | 18.22234 | 18.55434 | 20.32722 | 26.83123 | 19.58014 | 19.04557 | 18.05753 | 18.38789 | 18.06715 | 21.69223 |
| H2BE1 | 28.47029 | 27.81481 | 21.70831 | 20.52011 | 22.75172 | 24.17641 | 24.79581 | 28.11657 | 20.39487 | 20.34589 | 23.15839 | 28.66068 | 22.30911 | 20.89695 | 20.02957 | 20.89164 | 19.58379 | 23.25831 |
| H2BU1 | 14.84225 | 15.40483 | 13.70734 | 15.07013 | 14.2899 | 17.96465 | 16.70537 | 19.43349 | 11.75388 | 14.98573 | 12.21599 | 16.23788 | 15.87749 | 14.39303 | 14.55704 | 18.34253 | 13.7888 | 13.65988 |
| H3-2 | 27.59632 | 26.65731 | 21.07793 | 20.15778 | 22.78758 | 22.68824 | 24.0144 | 26.3496 | 21.83923 | 22.96809 | 23.30703 | 27.95673 | 20.73857 | 21.0343 | 19.23443 | 20.34592 | 18.10224 | 21.10386 |
| H3-4 | 26.39868 | 25.40607 | 19.59603 | 18.08993 | 18.58503 | 19.72743 | 21.19945 | 22.35464 | 20.32945 | 18.62829 | 21.09822 | 27.18349 | 15.98779 | 18.2353 | 17.0966 | 18.07148 | 16.92116 | 19.8829 |
| H4C1 | 29.22039 | 28.90498 | 21.82567 | 20.48842 | 23.31742 | 22.38136 | 26.50056 | 25.16222 | 21.83035 | 23.84014 | 22.74535 | 29.26836 | 20.75212 | 21.86014 | 18.73036 | 20.69717 | 19.92523 | 22.47567 |
| H6PD | 15.4544 | 15.60996 | 15.47235 | 16.38329 | 16.35001 | 16.82624 | 17.72299 | 17.30314 | 19.20663 | 17.59487 | 15.59136 | 14.876 | 15.46864 | 14.81243 | 15.26749 | 15.1967 | 16.68377 | 14.8277 |
| HABP2 | 15.65475 | 15.11456 | 16.28464 | 17.6192 | 15.09746 | 16.65602 | 18.27553 | 19.55519 | 20.03022 | 20.24752 | 14.38706 | 17.54125 | 16.79039 | 17.55566 | 14.37875 | 15.20052 | 16.38044 | 14.73096 |
| HACL1 | 16.37424 | 15.68128 | 16.03229 | 14.87548 | 17.04604 | 16.97276 | 17.06568 | 17.01342 | 19.47574 | 18.00403 | 18.89803 | 15.24415 | 16.7847 | 15.9825 | 15.44644 | 15.73805 | 15.10871 | 15.59128 |
| HADH | 17.52283 | 16.28784 | 15.98393 | 17.69504 | 15.69556 | 18.92629 | 20.07764 | 18.94585 | 20.74002 | 19.33674 | 16.88646 | 17.13897 | 17.09885 | 16.99617 | 17.03718 | 16.56923 | 15.66257 | 15.37995 |
| HADHA | 21.00464 | 20.73643 | 20.74673 | 20.13292 | 16.39726 | 17.71702 | 18.71703 | 17.12975 | 19.87296 | 18.85051 | 17.14526 | 14.40354 | 18.35735 | 17.3751 | 20.49242 | 20.47298 | 15.72883 | 17.28397 |
| HADHB | 14.83571 | 15.36164 | 16.75938 | 16.29925 | 15.71046 | 15.15398 | 16.03978 | 18.75965 | 19.98021 | 17.70605 | 16.81908 | 13.30148 | 15.18409 | 15.60731 | 14.26576 | 15.89119 | 16.93134 | 15.85569 |
| HAGH | 15.72868 | 18.1674 | 17.27185 | 15.11533 | 18.53389 | 15.81405 | 16.9046 | 18.27868 | 18.29858 | 18.54278 | 17.94833 | 16.91047 | 19.11568 | 18.90695 | 16.75441 | 17.89514 | 20.28158 | 18.91845 |
| HAPLN1 | 14.58993 | 15.25982 | 16.68941 | 16.62593 | 16.85258 | 16.57793 | 16.44859 | 17.4 | 20.15965 | 19.30386 | 15.15397 | 13.13565 | 14.43372 | 16.40965 | 17.35848 | 16.26104 | 16.07484 | 16.47168 |
| HAT1 | 15.89371 | 14.33791 | 15.19737 | 14.1605 | 15.58252 | 16.67014 | 17.17808 | 16.02589 | 12.29984 | 13.42937 | 16.56791 | 19.54966 | 11.16824 | 12.73823 | 15.6302 | 14.5572 | 12.87802 | 17.17068 |
| HAUS3 | 14.19598 | 16.59383 | 13.95259 | 13.35286 | 15.99056 | 17.51528 | 14.16178 | 17.83509 | 15.48775 | 18.82363 | 16.24487 | 16.13921 | 16.41339 | 15.80724 | 15.85445 | 16.03354 | 14.88869 | 12.97083 |
| HBS1L | 19.34141 | 19.04628 | 17.96473 | 18.39053 | 18.07479 | 18.18574 | 17.21404 | 15.48501 | 16.90342 | 17.72615 | 17.51872 | 17.91396 | 17.52732 | 17.23938 | 18.42384 | 18.23176 | 15.72823 | 17.79575 |
| HCFC1 | 14.77612 | 15.19303 | 14.57367 | 15.86194 | 17.23557 | 16.04276 | 15.56561 | 14.10777 | 16.04111 | 15.98604 | 16.47091 | 17.6003 | 13.86582 | 15.06425 | 13.40268 | 15.07437 | 18.21681 | 16.12433 |
| HCK | 18.21963 | 18.85815 | 18.86105 | 18.19795 | 19.16082 | 18.69658 | 17.94774 | 16.576 | 18.68914 | 18.95109 | 20.16817 | 20.09929 | 17.67964 | 17.92077 | 18.89492 | 19.22701 | 19.6627 | 17.9596 |
| HDAC2 | 17.27045 | 16.16703 | 16.91858 | 16.02408 | 14.83554 | 16.71561 | 17.59753 | 16.85213 | 16.90377 | 17.42905 | 17.29185 | 20.00817 | 15.53908 | 16.37349 | 16.00197 | 16.67248 | 16.05313 | 16.70896 |
| HDAC3 | 16.08401 | 16.20531 | 15.53652 | 15.56323 | 15.67471 | 14.63398 | 16.04685 | 15.76842 | 15.70428 | 14.06493 | 15.7078 | 17.41811 | 15.91139 | 16.14243 | 14.91398 | 14.70449 | 16.19546 | 14.48725 |
| HDDC3 | 13.55624 | 15.98359 | 16.95045 | 15.70126 | 16.82897 | 18.08346 | 14.8999 | 15.80478 | 14.34645 | 16.8171 | 17.16897 | 17.29409 | 12.81683 | 17.29433 | 15.51775 | 14.40245 | 16.45034 | 14.01077 |
| HDGF | 16.09778 | 16.48214 | 16.16536 | 16.97364 | 19.42289 | 18.17459 | 16.09372 | 16.65329 | 17.60975 | 18.05406 | 20.3311 | 19.77387 | 19.41438 | 16.62542 | 16.58778 | 16.71883 | 18.49897 | 17.7908 |
| HDHD2 | 15.83017 | 15.0358 | 14.53241 | 15.85433 | 16.27629 | 13.80551 | 15.48909 | 15.94307 | 17.43226 | 16.26565 | 19.13386 | 19.17985 | 14.27161 | 15.48098 | 18.50996 | 14.89153 | 15.26803 | 14.34376 |
| HDLBP | 18.21143 | 18.09397 | 17.69198 | 18.17317 | 17.58144 | 16.65859 | 18.53407 | 18.17032 | 16.16563 | 15.99568 | 18.28752 | 18.47853 | 18.89967 | 18.91355 | 17.66671 | 18.22618 | 18.26179 | 17.53193 |
| HEBP1 | 18.7935 | 17.10759 | 17.12545 | 18.66075 | 20.59904 | 15.54352 | 18.20606 | 16.9903 | 16.60362 | 16.89376 | 16.8069 | 16.61252 | 21.19817 | 21.3156 | 18.25101 | 18.18868 | 20.17753 | 20.39295 |
| HEBP2 | 18.14748 | 18.44049 | 17.66448 | 18.91044 | 17.19338 | 16.75521 | 16.98693 | 14.39305 | 19.35913 | 20.61326 | 19.84802 | 18.72686 | 19.869 | 19.84314 | 18.2133 | 19.61418 | 18.81393 | 17.91198 |
| HECTD1 | 15.29516 | 13.75097 | 15.6748 | 15.5098 | 19.50634 | 15.77866 | 14.20839 | 14.70533 | 13.92812 | 16.45701 | 16.90877 | 15.69581 | 18.82739 | 12.9018 | 15.25533 | 16.79613 | 17.91989 | 16.56943 |
| HECW2 | 15.10715 | 16.1325 | 17.30136 | 11.63358 | 11.35225 | 14.21234 | 15.11617 | 16.7287 | 17.59397 | 16.8702 | 13.56406 | 13.14616 | 14.89246 | 13.23411 | 14.68552 | 13.91451 | 15.71909 | 13.65348 |
| HEG1 | 18.04593 | 17.27106 | 17.88427 | 19.13372 | 19.92058 | 15.92327 | 20.09372 | 15.89295 | 16.78214 | 15.93505 | 16.36379 | 12.78885 | 18.34528 | 19.87403 | 18.10686 | 18.17585 | 16.21203 | 18.57445 |
| HELZ2 | 16.26132 | 16.04219 | 16.64779 | 18.0233 | 14.09346 | 14.08983 | 13.46619 | 18.25339 | 14.01329 | 15.77641 | 19.10628 | 13.0593 | 12.74171 | 14.93717 | 18.59167 | 17.9417 | 18.59839 | 15.0273 |
| HERC1 | 19.64476 | 13.00131 | 14.37176 | 14.43098 | 14.58458 | 15.42458 | 15.94697 | 16.11425 | 13.69285 | 15.24606 | 14.90978 | 15.23286 | 18.22841 | 17.85111 | 16.54032 | 13.41144 | 13.77387 | 14.08318 |
| HEXA | 17.679 | 18.92525 | 18.38015 | 17.51619 | 17.39872 | 15.43461 | 15.28248 | 16.87859 | 18.53464 | 17.16427 | 17.48431 | 14.29449 | 14.90842 | 15.44115 | 17.47155 | 17.01405 | 17.36034 | 13.56337 |
| HEXB | 17.93145 | 18.58836 | 19.12646 | 19.34628 | 17.93427 | 18.09733 | 16.62895 | 16.61031 | 20.2141 | 19.14917 | 18.03248 | 15.30158 | 19.67799 | 19.54212 | 19.28067 | 17.06058 | 16.65188 | 15.86292 |
| HGD | 16.12381 | 15.30491 | 13.94063 | 18.09181 | 13.76378 | 15.37739 | 17.03206 | 16.51564 | 13.86455 | 14.84843 | 15.85736 | 13.09067 | 20.02053 | 20.57867 | 14.20524 | 14.00736 | 14.00478 | 17.27403 |
| HGS | 19.09557 | 18.99466 | 18.34228 | 16.90602 | 19.92693 | 17.32302 | 20.07434 | 17.09589 | 18.0528 | 19.40008 | 17.37358 | 16.72935 | 16.83418 | 17.98359 | 17.09449 | 17.07252 | 18.12608 | 18.27686 |
| HIBADH | 15.05981 | 14.06862 | 16.08169 | 16.54157 | 16.84261 | 17.14027 | 17.17882 | 17.22928 | 19.66852 | 19.20826 | 16.80284 | 13.34329 | 15.73014 | 16.79045 | 15.97157 | 16.63684 | 16.48575 | 16.05501 |
| HIKESHI | 16.92476 | 17.08949 | 15.41456 | 17.37653 | 12.26044 | 15.09994 | 16.12178 | 15.662 | 12.66778 | 15.58414 | 14.41173 | 17.73568 | 13.43188 | 17.25063 | 13.05491 | 17.20548 | 16.0759 | 12.37128 |
| HINT1 | 19.96877 | 19.59899 | 19.59414 | 21.03796 | 23.08468 | 21.26002 | 23.52364 | 22.98318 | 21.39157 | 22.67064 | 20.81251 | 22.26532 | 24.40449 | 23.54981 | 21.09477 | 20.682 | 24.03691 | 22.52797 |
| HINT2 | 16.54171 | 16.55282 | 15.40385 | 16.2821 | 14.21727 | 17.02656 | 18.48216 | 17.28455 | 21.64205 | 20.95121 | 15.29017 | 14.01712 | 15.55866 | 15.13806 | 16.03527 | 17.5726 | 15.2665 | 13.82679 |
| HINT3 | 16.35341 | 16.62743 | 17.02101 | 18.52542 | 18.74767 | 16.71664 | 17.3923 | 16.53331 | 17.73838 | 17.81162 | 18.01161 | 19.9717 | 18.94024 | 19.80906 | 17.3592 | 18.30606 | 17.46039 | 19.69275 |
| HIP1 | 21.08278 | 23.97343 | 22.02141 | 22.36023 | 21.11838 | 20.69448 | 17.5856 | 17.07143 | 19.5808 | 20.38075 | 17.83647 | 18.08361 | 22.57816 | 20.16491 | 21.73688 | 21.6086 | 19.2166 | 20.45532 |
| HIP1R | 15.05053 | 16.11125 | 17.48044 | 16.39947 | 16.2232 | 15.51643 | 15.26914 | 16.65666 | 15.99932 | 15.85013 | 16.63588 | 17.35173 | 18.32994 | 18.81398 | 16.68722 | 16.92944 | 15.41446 | 15.43743 |
| HK2 | 17.9184 | 19.11056 | 21.22948 | 18.15504 | 15.81591 | 14.74441 | 14.86664 | 16.14854 | 13.79884 | 16.00946 | 15.68794 | 19.09928 | 16.81245 | 17.41103 | 19.62335 | 18.55092 | 16.40742 | 19.51697 |
| HLA-A | 16.79222 | 17.1213 | 17.23141 | 19.2238 | 19.61298 | 19.91643 | 17.02266 | 17.11643 | 21.45331 | 19.24273 | 20.76841 | 22.06671 | 16.53041 | 17.35931 | 17.34459 | 17.38115 | 18.40288 | 20.7024 |
| HLA-B | 18.34928 | 18.63119 | 16.84937 | 17.34992 | 22.16992 | 21.56236 | 16.67924 | 17.48081 | 22.25277 | 20.15589 | 21.1014 | 23.67449 | 15.54327 | 16.35474 | 16.85724 | 18.90716 | 16.76384 | 20.51608 |
| HLA-C | 16.38369 | 19.10304 | 18.90247 | 16.80318 | 19.00248 | 17.59992 | 16.3649 | 16.64873 | 22.09512 | 18.86074 | 21.75774 | 22.63034 | 15.6079 | 15.47171 | 17.74367 | 17.80736 | 16.60385 | 18.9033 |
| HLA-DQB1 | 15.20484 | 14.88623 | 15.34986 | 14.21288 | 16.03271 | 16.32764 | 17.16077 | 17.45008 | 19.81303 | 16.13934 | 15.11655 | 14.771 | 14.16243 | 13.87253 | 16.47899 | 15.61782 | 16.04613 | 15.96534 |
| HLA-E | 18.98736 | 20.29442 | 18.85388 | 19.26521 | 18.35068 | 17.90425 | 16.96496 | 17.18277 | 22.28646 | 19.39304 | 19.34574 | 23.5846 | 15.16363 | 15.79966 | 19.19983 | 18.91262 | 17.95792 | 19.38381 |
| HLA-G | 18.07383 | 18.88613 | 18.55472 | 17.87273 | 12.0813 | 17.75369 | 13.65739 | 17.36623 | 21.95143 | 13.86216 | 19.10943 | 20.3771 | 13.1482 | 17.34485 | 18.6796 | 18.68983 | 16.4896 | 16.69288 |
| HM13 | 15.9308 | 16.43809 | 15.84128 | 14.56097 | 17.10226 | 17.31998 | 18.42593 | 18.59374 | 19.2477 | 18.84796 | 17.51646 | 17.63918 | 16.77008 | 16.17216 | 15.57391 | 16.38866 | 17.75291 | 18.42263 |
| HMBS | 14.73026 | 18.26643 | 17.44562 | 15.84366 | 18.47974 | 18.05114 | 18.96943 | 17.55305 | 15.07892 | 16.54823 | 18.70288 | 18.78778 | 18.89207 | 18.2965 | 15.89657 | 16.41067 | 18.11742 | 17.99532 |
| HMCN2 | 15.61786 | 15.79068 | 13.87211 | 15.76334 | 15.16774 | 16.07542 | 17.24655 | 16.40047 | 16.00116 | 13.85459 | 17.58325 | 16.96334 | 17.30637 | 15.4642 | 13.07094 | 14.46794 | 14.18929 | 14.20143 |
| HMGA1 | 14.72553 | 14.20969 | 14.53659 | 15.36601 | 20.12306 | 19.80733 | 17.41869 | 17.39793 | 14.95297 | 15.99094 | 21.9594 | 20.7678 | 17.82051 | 17.44906 | 14.25396 | 14.19352 | 19.50891 | 20.75982 |
| HMGB1 | 16.81925 | 15.57542 | 18.20244 | 17.2193 | 19.09251 | 17.39706 | 19.80668 | 19.19542 | 15.8336 | 18.77415 | 20.8391 | 21.27191 | 19.6261 | 16.84212 | 17.84643 | 17.8415 | 17.35522 | 16.0112 |
| HMGB1P1 | 16.78733 | 16.57996 | 19.57239 | 18.31714 | 14.51953 | 17.74067 | 19.04758 | 18.91313 | 17.23712 | 17.35576 | 20.09767 | 20.16848 | 17.95726 | 16.61956 | 18.49958 | 18.14787 | 15.85503 | 15.19914 |
| HMGB2 | 16.34387 | 15.75881 | 15.97712 | 17.05853 | 14.91545 | 16.77304 | 17.76182 | 17.46568 | 16.9602 | 18.03452 | 18.84897 | 18.86079 | 18.04302 | 16.95979 | 17.10619 | 16.89032 | 17.19736 | 16.5566 |
| HMGB3 | 15.16197 | 16.23576 | 15.56761 | 15.6415 | 17.70184 | 17.15958 | 15.48235 | 15.55426 | 18.72343 | 19.86718 | 18.6312 | 17.62992 | 16.23325 | 16.44561 | 16.28939 | 15.21233 | 15.69506 | 16.89441 |
| HMGCL | 17.50073 | 17.82277 | 16.03895 | 15.18811 | 16.31264 | 15.50526 | 15.5061 | 18.3181 | 20.38074 | 19.74589 | 15.32525 | 15.75454 | 14.92848 | 15.85413 | 14.18864 | 16.30038 | 17.17177 | 14.81973 |
| HMGCS1 | 16.98092 | 16.50872 | 17.37889 | 16.40427 | 19.18694 | 17.65657 | 18.53387 | 17.94682 | 17.71049 | 17.84134 | 19.03562 | 20.346 | 16.95887 | 16.66064 | 17.80594 | 17.48397 | 16.62208 | 16.43775 |
| HMGN1 | 14.3381 | 14.39277 | 14.59535 | 16.74724 | 13.89586 | 13.97633 | 15.12554 | 15.798 | 12.7411 | 14.53252 | 15.98816 | 18.22567 | 17.35003 | 15.03769 | 14.24075 | 17.55494 | 13.40103 | 14.41878 |
| HMOX2 | 14.63232 | 16.76847 | 15.51874 | 14.76828 | 17.02066 | 18.63315 | 16.00599 | 14.24397 | 10.66627 | 17.10583 | 14.26518 | 19.60262 | 16.78432 | 14.64545 | 14.2325 | 13.35538 | 16.50586 | 16.3678 |
| HNRNPA0 | 16.07971 | 15.54201 | 16.72424 | 16.60394 | 16.63101 | 14.37248 | 17.90263 | 17.71588 | 16.43774 | 18.9609 | 15.25615 | 17.16514 | 18.00797 | 16.47253 | 14.79464 | 15.89147 | 16.05846 | 16.16691 |
| HNRNPA1 | 17.27238 | 17.01344 | 16.63083 | 18.92157 | 18.94663 | 18.60034 | 17.00362 | 20.12757 | 19.23519 | 20.22943 | 21.45632 | 20.78245 | 19.92529 | 18.88854 | 16.1876 | 15.86418 | 18.11458 | 16.58285 |
| HNRNPA1L | 18.33723 | 17.56758 | 17.94905 | 17.12847 | 19.61975 | 18.51098 | 19.1914 | 18.23365 | 19.12029 | 20.11604 | 21.08173 | 21.36893 | 19.80948 | 19.45745 | 17.60803 | 17.24564 | 18.11761 | 19.38135 |
| HNRNPA2E | 18.60095 | 17.67003 | 17.32746 | 16.99928 | 19.1042 | 19.20671 | 18.87711 | 19.34386 | 19.2439 | 19.67688 | 20.62472 | 20.78092 | 20.43865 | 19.09214 | 16.936 | 16.60377 | 18.14151 | 17.85073 |
| HNRNPA3 | 16.38256 | 16.48355 | 16.08047 | 17.25906 | 19.52197 | 17.08801 | 22.14291 | 20.07215 | 17.92413 | 19.55636 | 20.04939 | 19.72316 | 20.93134 | 20.1978 | 16.51206 | 16.59507 | 18.01511 | 20.50671 |
| HNRNPAB | 17.93001 | 19.3309 | 17.27424 | 15.35493 | 18.47308 | 16.46863 | 18.14715 | 18.15234 | 18.76919 | 19.55057 | 20.39821 | 21.659 | 18.36408 | 16.00813 | 17.218 | 18.89643 | 15.84524 | 16.31432 |
| HNRNPC | 19.52925 | 18.48195 | 17.46622 | 17.02437 | 20.75788 | 19.66218 | 21.24595 | 20.78578 | 18.80629 | 20.5611 | 20.18167 | 20.03439 | 21.07286 | 19.76024 | 16.5854 | 17.42423 | 18.57579 | 19.95433 |
| HNRNPCL: | 19.57505 | 19.01836 | 18.93208 | 18.21323 | 20.29819 | 19.79693 | 22.35318 | 21.30486 | 19.31509 | 21.23177 | 21.23297 | 20.98887 | 20.60423 | 19.08277 | 17.02881 | 16.4505 | 16.53289 | 19.71473 |
| HNRNPD | 18.06969 | 16.44506 | 16.97192 | 17.30041 | 18.36753 | 18.33644 | 18.774 | 18.59239 | 18.49447 | 19.14127 | 21.25585 | 21.4606 | 19.50462 | 18.06596 | 15.92034 | 16.805 | 16.36936 | 17.64173 |
| HNRNPDL | 16.99074 | 15.59217 | 16.32278 | 16.07734 | 17.6416 | 19.04014 | 17.91695 | 17.87114 | 17.83418 | 18.46234 | 20.14976 | 21.08122 | 17.11968 | 16.69954 | 16.55798 | 16.18097 | 16.80316 | 17.33266 |
| HNRNPF | 17.16511 | 16.48826 | 17.37286 | 16.71973 | 18.22217 | 17.62724 | 19.3334 | 19.96695 | 17.86408 | 18.43444 | 19.49965 | 19.53383 | 19.93645 | 18.5518 | 16.91932 | 17.51825 | 18.63448 | 18.04027 |
| HNRNPH1 | 19.12966 | 18.34103 | 17.70487 | 15.92594 | 16.98062 | 17.6552 | 19.49078 | 17.32385 | 17.60257 | 17.6716 | 18.7174 | 20.04273 | 17.91093 | 15.15285 | 18.83332 | 17.89697 | 17.30231 | 18.73184 |

| | | | | | | | | | | | | | | | | | | |
|---|---|---|---|---|---|---|---|---|---|---|---|---|---|---|---|---|---|---|
| HNRNPH3 | 19.49331 | 19.00481 | 17.49298 | 17.76532 | 16.17525 | 16.78072 | 18.20986 | 18.83993 | 14.78506 | 16.55425 | 16.70318 | 19.03206 | 17.31645 | 17.20392 | 18.38033 | 17.70337 | 15.6087 | 17.20847 |
| HNRNPK | 20.37301 | 20.07293 | 19.12876 | 19.70033 | 19.91181 | 19.01979 | 20.87311 | 19.83653 | 19.02897 | 19.70162 | 21.27409 | 22.07658 | 20.63293 | 19.96591 | 18.99044 | 18.60661 | 18.57637 | 19.20029 |
| HNRNPL | 17.63615 | 16.8015 | 16.03312 | 15.93535 | 17.41361 | 16.26339 | 20.08916 | 19.68768 | 16.98105 | 17.84922 | 19.51331 | 20.53084 | 19.06736 | 17.63331 | 15.91578 | 15.71309 | 16.06962 | 17.30921 |
| HNRNPLL | 15.50591 | 17.67545 | 15.86655 | 13.57338 | 13.06622 | 14.22571 | 14.38819 | 17.26833 | 14.05999 | 15.72799 | 15.16545 | 17.10093 | 15.34945 | 14.62765 | 15.11266 | 15.7864 | 14.70138 | 15.82071 |
| HNRNPM | 20.27222 | 16.88182 | 18.09937 | 19.72773 | 18.50703 | 17.6463 | 20.95535 | 19.0534 | 17.24655 | 18.52822 | 17.5832 | 18.52603 | 19.91289 | 19.718 | 16.814 | 16.15463 | 18.58846 | 19.22373 |
| HNRNPR | 16.08424 | 16.65866 | 16.10542 | 16.72265 | 17.61952 | 16.83306 | 19.29535 | 18.32786 | 16.72428 | 18.08903 | 19.06174 | 20.19481 | 18.52138 | 18.4599 | 15.60503 | 16.83536 | 16.99284 | 17.71272 |
| HNRNPU | 19.59658 | 17.45359 | 16.14015 | 16.51776 | 18.26335 | 17.96519 | 22.05798 | 21.52064 | 17.48493 | 18.6019 | 19.50624 | 19.77308 | 19.86264 | 19.33312 | 16.38948 | 16.31868 | 17.44085 | 18.38314 |
| HNRNPUL2 | 15.92225 | 16.14854 | 15.77206 | 16.07664 | 14.84345 | 16.10039 | 18.12245 | 17.53286 | 15.08672 | 16.61908 | 17.20213 | 18.00731 | 18.87377 | 17.47129 | 15.61945 | 16.82419 | 15.76684 | 16.69338 |
| HOMER1 | 15.90938 | 16.34113 | 17.65109 | 17.02357 | 15.40902 | 15.59397 | 16.39879 | 18.24311 | 14.11483 | 15.85427 | 15.7904 | 16.70733 | 20.50744 | 20.24337 | 15.99034 | 17.27148 | 15.57031 | 15.21825 |
| HOMER2 | 16.27915 | 16.49594 | 16.0555 | 15.99676 | 14.3775 | 17.68137 | 17.33898 | 15.77546 | 14.30034 | 16.23073 | 14.18393 | 12.67272 | 21.09896 | 21.17631 | 15.99594 | 16.25695 | 14.2614 | 14.83777 |
| HOOK1 | 16.00392 | 17.03089 | 13.9549 | 18.42093 | 16.38754 | 15.5873 | 15.09276 | 17.64603 | 18.52178 | 16.79228 | 14.90094 | 16.2936 | 18.3267 | 18.77852 | 17.49664 | 16.92947 | 15.80737 | 16.37839 |
| HP1BP3 | 21.31732 | 20.3708 | 17.00045 | 14.4667 | 18.08794 | 15.63761 | 16.86989 | 19.34359 | 16.29201 | 16.23566 | 15.96513 | 15.41864 | 16.29315 | 15.07802 | 16.85614 | 17.03166 | 16.10726 | 17.84739 |
| HPCA | 14.90334 | 16.56104 | 16.14623 | 16.22547 | 18.84896 | 17.06254 | 16.19529 | 17.57432 | 16.11242 | 15.77272 | 16.17994 | 17.07665 | 13.19124 | 15.95485 | 15.00552 | 16.19545 | 19.21884 | 19.31368 |
| HPGD | 15.5898 | 17.23893 | 15.58386 | 18.40436 | 18.99004 | 18.22121 | 19.66532 | 16.63588 | 14.68711 | 15.20357 | 13.2101 | 14.87404 | 19.49291 | 20.18416 | 17.05927 | 15.95264 | 16.92646 | 21.82986 |
| HPN | 18.58798 | 18.64412 | 18.91632 | 20.09959 | 16.39335 | 16.11276 | 15.77304 | 17.49747 | 12.77946 | 15.24403 | 16.23962 | 14.19727 | 18.63038 | 14.82394 | 19.88678 | 20.04028 | 15.57691 | 14.23374 |
| HPRT1 | 17.88028 | 17.98165 | 14.85164 | 16.40879 | 18.50955 | 17.28816 | 18.46995 | 17.66403 | 18.79702 | 18.47784 | 19.89461 | 19.89678 | 16.45 | 15.6516 | 15.85306 | 15.49226 | 18.58686 | 18.00876 |
| HPSE | 16.30698 | 16.35634 | 15.1321 | 14.49486 | 11.99613 | 15.50633 | 14.68807 | 17.55017 | 17.85831 | 14.99168 | 13.70479 | 12.49858 | 15.32277 | 16.16653 | 17.87625 | 16.42499 | 15.62867 | 15.29011 |
| HPX | 14.29037 | 16.57046 | 15.05108 | 17.15447 | 14.23335 | 16.24747 | 14.80467 | 15.81659 | 14.21612 | 13.61589 | 15.23719 | 20.15277 | 14.18197 | 16.84003 | 16.08646 | 15.29803 | 17.31162 | 14.5751 |
| HS2ST1 | 15.598 | 16.22738 | 15.59898 | 16.63074 | 15.43821 | 17.22998 | 17.00535 | 16.92705 | 21.1064 | 19.72443 | 16.9957 | 16.65635 | 16.31355 | 15.16992 | 16.25432 | 15.66701 | 17.25147 | 16.31984 |
| HSD17B10 | 16.19563 | 16.73348 | 16.62755 | 16.05278 | 15.15522 | 17.27355 | 19.06697 | 18.09121 | 21.92377 | 19.19679 | 18.35564 | 15.57522 | 16.5942 | 15.63515 | 16.7962 | 15.22606 | 15.54071 | 16.7707 |
| HSD17B11 | 16.21121 | 15.65968 | 17.16875 | 15.85793 | 16.55915 | 16.54665 | 14.88437 | 16.38951 | 17.93222 | 16.84601 | 14.45056 | 14.01792 | 16.38008 | 15.33768 | 14.55819 | 14.03281 | 17.46638 | 14.74817 |
| HSD17B12 | 16.53611 | 14.89609 | 15.5282 | 17.03731 | 18.21571 | 18.39507 | 16.49556 | 16.38398 | 18.84133 | 13.94915 | 15.55002 | 17.99108 | 13.81328 | 17.2903 | 13.46273 | 17.31305 | 18.86397 | 18.66244 |
| HSD17B4 | 17.19603 | 17.33021 | 16.98913 | 17.61314 | 16.91815 | 17.69613 | 18.69849 | 17.06157 | 22.65494 | 20.73649 | 19.60069 | 15.91124 | 16.2748 | 16.24668 | 16.90623 | 16.60715 | 16.94867 | 16.98115 |
| HSD17B8 | 14.35455 | 17.02786 | 16.73472 | 13.33161 | 15.5459 | 16.87549 | 14.73049 | 14.08926 | 17.42252 | 15.45885 | 14.05555 | 16.05073 | 13.29828 | 15.60965 | 15.26267 | 13.66093 | 16.98956 | 16.2925 |
| HSDL2 | 16.06865 | 17.39207 | 16.25128 | 16.251 | 16.52545 | 16.68134 | 19.00051 | 17.78483 | 21.73401 | 21.71625 | 15.38757 | 13.99569 | 17.66482 | 16.8278 | 16.46023 | 16.52552 | 16.77173 | 17.94789 |
| HSH2D | 17.92799 | 18.0326 | 20.04349 | 19.83361 | 14.54513 | 15.75893 | 16.42908 | 16.86829 | 14.00987 | 14.31704 | 15.56621 | 12.58939 | 14.97196 | 14.86811 | 20.73595 | 20.39314 | 15.22926 | 14.96966 |
| HSP90AA1 | 20.81336 | 18.95967 | 21.20769 | 21.59382 | 22.59904 | 21.86988 | 20.61757 | 19.85131 | 21.63292 | 22.108 | 22.85684 | 23.45391 | 21.11125 | 20.20098 | 20.65903 | 20.95805 | 22.14555 | 22.46666 |
| HSP90AA2 | 21.62684 | 21.70898 | 22.71921 | 23.02638 | 24.21231 | 23.75948 | 22.09638 | 21.09326 | 22.84034 | 23.45121 | 24.51517 | 25.12477 | 23.69164 | 22.7467 | 22.83007 | 22.73653 | 23.5427 | 23.67169 |
| HSP90AA5 | 20.99913 | 21.64455 | 20.42826 | 21.33055 | 22.58532 | 22.42482 | 20.91103 | 17.89575 | 22.49144 | 22.70367 | 23.13631 | 23.52432 | 21.78294 | 20.96672 | 21.39146 | 20.57502 | 22.28094 | 22.46344 |
| HSP90AB1 | 21.08983 | 20.78506 | 21.99073 | 22.48256 | 23.26099 | 22.42296 | 20.91292 | 20.0642 | 21.92346 | 22.16544 | 23.48959 | 24.36701 | 23.13799 | 22.22563 | 22.27207 | 22.0901 | 22.60818 | 22.70074 |
| HSP90AB2 | 23.18019 | 22.62827 | 22.21451 | 23.46125 | 24.07367 | 23.47781 | 21.67928 | 14.5356 | 22.52004 | 22.71311 | 23.83464 | 24.36965 | 23.29736 | 22.99965 | 22.35931 | 23.1239 | 23.40664 | 24.0312 |
| HSP90AB4 | 22.80225 | 22.73624 | 22.83522 | 23.65491 | 23.19492 | 23.20931 | 20.74798 | 20.31163 | 24.91936 | 23.37545 | 25.71505 | 25.9833 | 22.95896 | 21.56722 | 23.52796 | 23.58487 | 22.54421 | 23.07385 |
| HSP90B1 | 18.79572 | 17.99169 | 20.2129 | 19.11114 | 19.49497 | 19.0643 | 22.778 | 20.89052 | 25.43287 | 25.80487 | 24.07476 | 21.7678 | 20.25172 | 20.78816 | 19.74958 | 19.29301 | 18.6569 | 19.74071 |
| HSP90B2P | 17.57479 | 18.55873 | 20.50975 | 17.80321 | 18.48648 | 18.06478 | 20.68172 | 20.24834 | 24.81955 | 24.93427 | 23.6989 | 21.60104 | 18.51656 | 19.15566 | 18.86001 | 19.30081 | 17.31167 | 18.13718 |
| HSPA13 | 18.01594 | 15.43704 | 16.31665 | 16.93744 | 16.70718 | 17.11026 | 16.2616 | 15.95456 | 20.95928 | 19.60644 | 17.44467 | 15.37614 | 15.25923 | 16.35051 | 16.70005 | 15.65367 | 15.49249 | 16.34992 |
| HSPA14 | 16.05488 | 16.26074 | 15.58928 | 15.53093 | 20.35406 | 24.34015 | 21.10777 | 16.48906 | 14.52304 | 15.48829 | 15.10454 | 17.40022 | 16.38805 | 22.07155 | 15.23132 | 15.79352 | 17.8141 | 22.23343 |
| HSPA1A | 23.51057 | 23.65312 | 22.30927 | 22.94007 | 23.75637 | 22.3606 | 22.69123 | 21.29826 | 22.64603 | 23.3801 | 20.74993 | 21.08348 | 24.19404 | 23.68356 | 22.09241 | 21.98339 | 22.88886 | 23.18086 |
| HSPA1L | 24.67384 | 24.82427 | 24.47495 | 24.19719 | 23.51863 | 22.57871 | 22.60641 | 21.5055 | 25.07206 | 25.33104 | 23.56377 | 23.36086 | 24.36775 | 22.47411 | 24.11322 | 23.93755 | 21.9769 | 22.69197 |
| HSPA2 | 19.87275 | 19.87959 | 16.24764 | 16.77041 | 20.43294 | 18.7038 | 16.16041 | 16.68595 | 13.50663 | 17.55244 | 17.2757 | 17.89514 | 15.51927 | 17.59978 | 16.74497 | 16.31413 | 19.07396 | 18.55933 |
| HSPA4 | 23.10592 | 20.88199 | 19.41543 | 20.4472 | 19.33144 | 18.13153 | 17.99875 | 17.80525 | 17.95914 | 20.68048 | 19.84793 | 20.76716 | 19.68297 | 19.57072 | 20.24587 | 19.37415 | 19.0057 | 18.64838 |
| HSPA4L | 17.10336 | 17.49722 | 16.05735 | 16.64612 | 18.37072 | 17.75871 | 17.67181 | 16.5841 | 16.32783 | 16.97107 | 18.61757 | 19.07894 | 19.74855 | 18.64902 | 17.87256 | 16.16307 | 18.04697 | 17.07902 |
| HSPA5 | 18.26871 | 17.79637 | 21.0266 | 19.48351 | 20.31099 | 19.2361 | 23.35854 | 22.50901 | 25.37696 | 25.27136 | 23.24247 | 20.05072 | 19.75949 | 20.95566 | 21.29676 | 19.60978 | 18.42001 | 20.49184 |
| HSPA6 | 23.81267 | 24.08116 | 24.38491 | 24.53397 | 24.91269 | 23.48873 | 23.9257 | 22.53011 | 22.9844 | 23.7994 | 22.50113 | 23.3684 | 25.55554 | 25.03834 | 23.4836 | 23.56524 | 23.37173 | 23.54678 |
| HSPA8 | 24.32118 | 24.61809 | 24.42313 | 24.74034 | 25.44446 | 24.32039 | 24.85498 | 23.44273 | 23.52808 | 24.50124 | 23.33567 | 23.75259 | 25.9021 | 25.5034 | 23.93769 | 23.70376 | 23.79446 | 24.40952 |
| HSPA9 | 17.41985 | 19.62183 | 21.75447 | 17.84534 | 19.00701 | 20.5118 | 19.38093 | 18.27494 | 21.72776 | 20.56426 | 18.53401 | 16.16693 | 18.57467 | 17.75914 | 19.84229 | 17.40357 | 17.49313 | 18.91322 |
| HSPB1 | 21.01328 | 20.31653 | 22.80465 | 23.04201 | 20.09251 | 19.34578 | 19.96183 | 18.61682 | 22.95131 | 21.55088 | 16.97406 | 14.704 | 22.16684 | 20.93475 | 22.88731 | 22.26058 | 21.57635 | 20.20248 |
| HSPB11 | 17.83396 | 14.76298 | 17.99998 | 19.04927 | 17.8283 | 17.87861 | 18.53003 | 14.77782 | 18.20209 | 19.092 | 17.24838 | 17.26175 | 19.4326 | 18.10575 | 18.60675 | 20.16343 | 19.38854 | 18.09544 |
| HSPB6 | 19.86518 | 18.06687 | 13.74807 | 16.52624 | 20.54668 | 20.13447 | 18.80675 | 16.34284 | 18.30106 | 18.38035 | 16.28348 | 12.47145 | 16.37904 | 13.50688 | 15.42031 | 14.76208 | 14.15505 | 13.99552 |
| HSPBP1 | 15.13469 | 16.40615 | 16.83058 | 13.91663 | 15.4311 | 16.45517 | 18.1419 | 17.09173 | 15.8883 | 15.17468 | 18.57246 | 18.04886 | 15.09637 | 18.38232 | 16.2714 | 15.45179 | 18.46592 | 17.16984 |
| HSPD1 | 17.35622 | 17.85884 | 18.18352 | 17.40691 | 18.62359 | 18.17436 | 21.25667 | 19.89935 | 22.41492 | 21.75564 | 19.5686 | 16.8011 | 20.50177 | 21.19407 | 17.55349 | 17.5351 | 18.71647 | 19.13145 |
| HSPE1 | 18.20845 | 17.22603 | 18.80711 | 18.12923 | 18.46233 | 17.90152 | 21.09937 | 18.84744 | 23.83059 | 22.89402 | 20.93148 | 18.11142 | 21.23981 | 21.9289 | 17.53887 | 17.77922 | 18.05921 | 18.55917 |
| HSPG2 | 16.19383 | 15.98181 | 21.20949 | 19.78622 | 16.20925 | 18.16378 | 16.19598 | 16.46421 | 17.97769 | 16.64798 | 16.66858 | 15.74206 | 17.49085 | 18.61618 | 17.39203 | 19.42419 | 16.94982 | 19.44981 |
| HSPH1 | 16.35671 | 18.36422 | 18.67403 | 19.12047 | 17.77752 | 16.51341 | 19.9504 | 19.25407 | 17.76714 | 17.69661 | 19.41342 | 19.7446 | 19.46512 | 18.56319 | 18.05452 | 18.13328 | 19.6113 | 19.20351 |
| HTATIP2 | 18.00465 | 17.36541 | 16.29698 | 17.58334 | 19.4503 | 18.70442 | 19.35515 | 17.68362 | 20.53091 | 20.37507 | 17.70027 | 18.74665 | 18.41128 | 18.23364 | 15.53829 | 15.87363 | 18.89718 | 18.82984 |
| HTATSF1 | 18.81797 | 16.45648 | 16.4831 | 13.25914 | 14.76792 | 14.33415 | 16.13398 | 14.96065 | 14.53009 | 12.63008 | 17.89293 | 17.2485 | 14.50998 | 13.43744 | 15.43332 | 15.34079 | 15.69819 | 12.36792 |
| HTRA1 | 15.5216 | 17.495 | 16.9995 | 16.71741 | 17.94571 | 16.07665 | 15.5662 | 16.19649 | 18.83371 | 18.40699 | 15.86282 | 14.08911 | 15.7405 | 17.32867 | 16.03987 | 16.94589 | 19.12385 | 21.29801 |
| HTRA2 | 16.25569 | 16.40676 | 15.39273 | 17.30071 | 14.60285 | 13.308 | 15.05948 | 15.3339 | 18.19174 | 16.25338 | 12.53556 | 16.68525 | 12.76993 | 13.1222 | 16.63822 | 15.14531 | 14.30491 | 15.18275 |
| HTT | 15.52916 | 15.19375 | 14.8806 | 16.73788 | 13.28292 | 15.31068 | 16.37429 | 16.34269 | 16.43497 | 16.42566 | 17.81123 | 17.87983 | 16.12011 | 18.39504 | 15.22969 | 15.55725 | 16.72923 | 16.09591 |
| HUWE1 | 16.10758 | 16.2364 | 15.93594 | 17.41858 | 14.98732 | 15.91612 | 16.04827 | 17.22115 | 15.71934 | 16.01827 | 15.74236 | 19.00713 | 16.04443 | 15.92822 | 16.44206 | 16.33455 | 17.8686 | 16.48557 |
| HYAL2 | 21.10088 | 19.47868 | 19.71783 | 18.12541 | 15.2564 | 14.96869 | 15.00687 | 15.90598 | 16.73242 | 16.6354 | 13.91215 | 18.385 | 14.60424 | 15.46393 | 22.06335 | 17.95631 | 14.54778 | 13.72306 |
| HYOU1 | 17.44931 | 16.58077 | 19.30913 | 18.14409 | 17.2525 | 18.39637 | 19.03595 | 18.39798 | 22.58496 | 22.60449 | 20.91847 | 18.14163 | 18.24462 | 19.29995 | 19.16512 | 18.65242 | 17.02415 | 18.36639 |
| IAH1 | 15.98548 | 16.30354 | 15.41402 | 13.88233 | 14.30008 | 16.36562 | 17.78709 | 13.02193 | 14.49636 | 16.59424 | 14.99997 | 19.17045 | 14.22718 | 16.25686 | 13.57128 | 16.55352 | 16.25327 | 14.24578 |
| IARS1 | 16.28078 | 16.24719 | 17.10392 | 17.09615 | 18.19671 | 17.22134 | 18.30412 | 17.13838 | 16.71764 | 17.40891 | 19.18578 | 20.668 | 17.30818 | 16.39126 | 16.98051 | 16.15344 | 16.44388 | 16.85777 |
| IARS2 | 15.99065 | 16.49093 | 15.46576 | 16.95334 | 14.99536 | 17.25011 | 16.99624 | 16.59535 | 19.33555 | 18.02574 | 15.91481 | 15.03864 | 15.84049 | 15.70359 | 15.46207 | 15.98286 | 16.40285 | 16.05288 |
| ICAM1 | 16.47141 | 19.11621 | 19.04231 | 19.26012 | 24.2114 | 23.01656 | 16.45811 | 17.43427 | 15.3091 | 16.24289 | 15.02757 | 13.86032 | 16.43515 | 16.99613 | 18.83456 | 19.01419 | 22.18998 | 22.1817 |
| IDE | 16.11214 | 15.92414 | 15.7436 | 17.5679 | 18.05006 | 16.90842 | 16.20071 | 16.26776 | 15.33417 | 16.72648 | 18.00458 | 19.56909 | 16.39853 | 16.61583 | 16.88817 | 16.5726 | 16.42249 | 16.18407 |
| IDH1 | 19.54913 | 19.14379 | 19.7495 | 20.29306 | 20.32829 | 18.98749 | 19.7458 | 17.44358 | 20.36739 | 19.76234 | 20.40019 | 20.713 | 22.80195 | 22.1352 | 21.05653 | 21.0433 | 19.3591 | 18.97102 |
| IDH2 | 18.88346 | 18.07542 | 19.02887 | 17.43925 | 16.35407 | 16.01627 | 19.43964 | 17.35904 | 22.63089 | 21.26508 | 18.28748 | 15.99154 | 19.08386 | 18.61558 | 17.39891 | 17.10312 | 16.3508 | 16.3461 |
| IDH3A | 16.29606 | 19.67623 | 16.34221 | 16.34864 | 14.87505 | 15.97665 | 20.80876 | 18.84879 | 18.83097 | 16.94629 | 15.33362 | 13.18641 | 19.31629 | 14.95653 | 14.99896 | 16.76462 | 15.21999 | 16.15734 |
| IDH3B | 16.34508 | 14.51461 | 16.12655 | 15.6309 | 16.74845 | 15.62611 | 15.19112 | 15.88228 | 18.61994 | 15.58232 | 13.32699 | 14.68335 | 13.99399 | 15.65322 | 17.8249 | 14.38495 | 15.52633 | 14.21824 |
| IDI1 | 18.68072 | 15.15482 | 14.99092 | 15.67441 | 18.00806 | 17.63768 | 19.10287 | 17.75501 | 18.04366 | 18.12188 | 19.2654 | 19.40951 | 17.92594 | 14.6186 | 14.65051 | 18.72205 | 16.37479 | 16.62219 |
| IFIT1 | 13.94804 | 16.32972 | 15.33278 | 14.28667 | 17.96185 | 16.93419 | 14.46218 | 17.2627 | 14.4784 | 13.36356 | 14.71664 | 15.37435 | 15.19944 | 15.71968 | 11.76638 | 14.12449 | 15.33113 | 15.4896 |
| IFIT2 | 16.78794 | 15.61461 | 13.18956 | 15.19458 | 18.22157 | 17.01318 | 14.57893 | 15.63205 | 13.00518 | 14.95852 | 17.24105 | 13.0196 | 14.71122 | 15.0321 | 14.56651 | 13.76506 | 14.80614 | 16.53968 |
| IFIT3 | 16.51458 | 15.28333 | 15.5289 | 14.91076 | 17.19523 | 16.52302 | 17.60219 | 14.13525 | 13.25416 | 14.39513 | 15.59708 | 13.52757 | 14.75893 | 16.95757 | 13.38398 | 13.45768 | 16.43044 | 20.56115 |
| IFITM3 | 16.59413 | 12.43073 | 23.60676 | 21.2948 | 15.43053 | 19.02547 | 14.26736 | 15.45774 | 15.13038 | 13.51909 | 14.66575 | 15.62515 | 12.68828 | 19.0021 | 22.0896 | 22.06744 | 15.65163 | 13.24532 |
| IFNGR1 | 13.46163 | 15.23633 | 14.36776 | 13.78952 | 16.84558 | 17.06794 | 17.0128 | 15.79315 | 17.97989 | 16.57967 | 16.79417 | 16.12473 | 13.3782 | 15.57653 | 14.77124 | 15.12013 | 14.99843 | 15.72778 |
| IGF1R | 15.03979 | 15.29206 | 16.31963 | 15.76179 | 19.5758 | 18.21592 | 17.64977 | 15.1821 | 14.44363 | 13.6773 | 14.33732 | 14.96013 | 14.05406 | 18.23171 | 15.10725 | 14.74977 | 15.5404 | 17.22457 |
| IGF2BP1 | 19.19565 | 18.87834 | 14.33635 | 13.7967 | 14.37707 | 16.01691 | 19.55215 | 19.2018 | 14.89084 | 13.7387 | 19.10812 | 12.94542 | 15.47865 | 18.69802 | 13.57477 | 15.91523 | 16.14366 | 17.48739 |
| IGF2BP3 | 20.14113 | 19.68608 | 17.1248 | 18.30214 | 14.74591 | 15.87104 | 15.74334 | 17.94473 | 16.0971 | 15.10412 | 17.25563 | 18.74339 | 19.02799 | 17.37013 | 16.52017 | 15.76763 | 15.42688 | 15.63803 |
| IGF2R | 17.04441 | 17.09689 | 18.91876 | 18.59717 | 16.89943 | 18.78623 | 17.54553 | 17.56259 | 22.58752 | 21.36591 | 20.62127 | 19.156 | 18.38908 | 18.79613 | 17.79549 | 18.07462 | 16.47175 | 17.7121 |
| IGFBP2 | 16.26954 | 15.83671 | 17.52946 | 16.95145 | 16.33841 | 17.75994 | 17.93115 | 17.77862 | 23.52766 | 21.73367 | 15.86674 | 13.34798 | 15.65486 | 16.32293 | 16.70845 | 15.72898 | 18.20322 | 16.20549 |
| IGFBP5 | 17.12567 | 17.34272 | 16.30857 | 16.30896 | 15.61117 | 15.09035 | 15.52658 | 16.98939 | 20.78538 | 19.85558 | 14.81049 | 13.82454 | 15.54095 | 16.03318 | 16.95794 | 15.09116 | 16.54243 | 16.11407 |
| IGFBP6 | 15.30831 | 14.3805 | 14.97425 | 16.98627 | 13.44684 | 18.73805 | 10.87172 | 14.95001 | 18.4886 | 18.38484 | 15.21075 | 15.38791 | 18.00022 | 15.30947 | 14.94941 | 15.55172 | 15.08068 | 18.23468 |
| IGSF21 | 16.02808 | 15.99804 | 15.85086 | 15.68326 | 15.01408 | 15.26473 | 18.13239 | 16.25631 | 19.12461 | 15.87063 | 15.79312 | 17.22096 | 15.36306 | 18.15628 | 16.78971 | 16.10276 | 16.43147 | 14.68573 |
| IGSF3 | 20.23934 | 20.16191 | 19.06274 | 18.99711 | 18.3006 | 18.4584 | 18.26168 | 17.62412 | 16.9338 | 15.70757 | 16.03263 | 16.68459 | 19.20343 | 17.47656 | 20.42665 | 19.83981 | 15.87742 | 15.94992 |
| IGSF8 | 24.07742 | 23.93537 | 22.55921 | 23.01804 | 19.89844 | 19.07478 | 18.73808 | 17.4194 | 18.10774 | 17.35866 | 18.1552 | 18.76092 | 22.58955 | 21.20713 | 22.84303 | 22.99164 | 19.22242 | 20.3754 |

| | | | | | | | | | | | | | | | | | | |
|---|---|---|---|---|---|---|---|---|---|---|---|---|---|---|---|---|---|---|
| IGSF9 | 16.72186 | 16.56643 | 13.63651 | 16.01051 | 14.24243 | 16.81583 | 14.78931 | 17.26589 | 13.11211 | 15.50755 | 16.63467 | 17.34061 | 15.27898 | 14.19469 | 13.96282 | 15.18629 | 16.80812 | 15.41625 |
| IKBIP | 14.66002 | 14.2426 | 14.98858 | 14.91571 | 16.56072 | 17.19645 | 18.82282 | 13.87033 | 19.08628 | 18.35353 | 15.70338 | 14.53762 | 15.63663 | 17.23549 | 14.98978 | 14.41219 | 14.82088 | 15.89672 |
| IKBKG | 19.02126 | 15.84589 | 15.12354 | 15.55665 | 21.34796 | 19.93163 | 13.23306 | 17.06239 | 15.48839 | 17.47396 | 13.72363 | 15.94143 | 20.3479 | 19.21155 | 14.40918 | 16.48956 | 12.9901 | 15.41958 |
| IL17C | 14.57438 | 17.92666 | 15.0095 | 24.94982 | 12.05726 | 15.98224 | 15.82373 | 16.19921 | 18.04945 | 12.27189 | 17.85138 | 14.16323 | 15.77123 | 11.0603 | 23.8773 | 24.83688 | 16.14214 | 16.16797 |
| IL1RAP | 15.51725 | 16.61978 | 18.54547 | 18.41442 | 18.51236 | 18.30167 | 17.93234 | 16.26911 | 17.79667 | 16.58885 | 17.12502 | 16.0701 | 17.82644 | 17.62285 | 18.96744 | 18.61182 | 15.06174 | 17.80569 |
| IL1RAPL1 | 17.12036 | 14.11456 | 18.27905 | 18.85023 | 20.72808 | 19.88156 | 17.19625 | 15.67125 | 12.59066 | 15.50299 | 14.17017 | 17.22068 | 11.68917 | 19.10783 | 18.82225 | 19.28627 | 15.08268 | 19.85118 |
| IL6ST | 16.27985 | 16.62342 | 17.22873 | 17.48668 | 19.17607 | 17.51672 | 17.40646 | 17.2293 | 19.69169 | 16.54255 | 14.90147 | 15.84846 | 15.87229 | 16.82134 | 16.19448 | 17.25843 | 16.61401 | 18.23971 |
| ILF2 | 21.32016 | 21.2702 | 19.6739 | 20.2541 | 16.66035 | 16.7325 | 16.56588 | 17.78329 | 17.71795 | 17.49903 | 19.38327 | 21.02646 | 18.63767 | 17.56677 | 18.63495 | 19.44857 | 16.12823 | 17.36163 |
| ILF3 | 21.5532 | 21.05242 | 19.74678 | 20.25577 | 16.23119 | 16.48123 | 17.77116 | 17.62917 | 16.39918 | 17.30016 | 18.40172 | 19.36581 | 19.1512 | 18.37284 | 18.8509 | 19.36583 | 16.35183 | 16.36223 |
| ILK | 18.02128 | 18.31535 | 19.12575 | 19.51636 | 19.57195 | 17.70639 | 20.66512 | 18.19628 | 15.94981 | 15.86783 | 16.20069 | 15.28438 | 18.77415 | 20.39795 | 18.49899 | 18.3401 | 18.40989 | 19.83279 |
| ILKAP | 13.96879 | 17.28246 | 13.79514 | 16.80369 | 16.87346 | 16.47146 | 19.43403 | 19.2237 | 10.68792 | 14.63792 | 16.47041 | 18.4794 | 17.49529 | 14.176 | 14.26752 | 22.22406 | 15.58356 | 16.98629 |
| ILVBL | 15.13465 | 16.39346 | 16.14632 | 17.028 | 17.62641 | 16.39876 | 20.74625 | 18.88056 | 15.40123 | 14.37367 | 16.34122 | 16.92488 | 16.54393 | 17.27738 | 15.03326 | 17.03985 | 14.78626 | 17.3218 |
| IMMT | 16.11984 | 15.39302 | 15.53402 | 15.92276 | 15.60346 | 15.3289 | 17.09337 | 17.23237 | 18.26345 | 17.06466 | 15.27267 | 13.25764 | 14.67472 | 14.59835 | 15.28557 | 15.95439 | 16.0946 | 15.14545 |
| IMPA1 | 16.88664 | 16.54992 | 15.68793 | 16.07534 | 18.85139 | 17.56898 | 18.70191 | 19.17719 | 18.86227 | 19.71239 | 17.44477 | 18.41568 | 19.71689 | 18.62567 | 15.44414 | 17.63896 | 19.88941 | 20.1675 |
| IMPDH1 | 15.61737 | 17.03441 | 16.81267 | 19.03582 | 17.94718 | 18.06367 | 17.0784 | 17.28683 | 13.91489 | 14.52404 | 16.72223 | 15.4687 | 20.10863 | 17.78715 | 17.23217 | 16.12878 | 16.11638 | 17.43576 |
| IMPDH2 | 17.46075 | 16.88928 | 18.76804 | 18.53353 | 19.91179 | 18.28661 | 21.75288 | 20.37025 | 18.56729 | 18.92986 | 19.51498 | 18.83282 | 19.56783 | 18.02334 | 18.11644 | 18.27308 | 21.33516 | 20.28672 |
| IMUP | 15.31054 | 17.05578 | 16.21658 | 14.69125 | 20.67443 | 14.70093 | 14.17848 | 16.94843 | 13.37092 | 16.84985 | 15.3147 | 10.21423 | 15.27087 | 15.686 | 15.34649 | 14.66933 | 14.57411 | 19.08828 |
| INCENP | 15.41632 | 15.95739 | 17.17606 | 19.58169 | 19.33079 | 19.12811 | 16.6321 | 16.74925 | 16.20118 | 16.16955 | 14.81879 | 14.69591 | 19.04962 | 18.24931 | 17.34504 | 17.73107 | 18.04533 | 16.20565 |
| INF2 | 15.10401 | 15.31587 | 16.87229 | 15.70831 | 16.48978 | 16.70273 | 15.84677 | 17.6147 | 17.88912 | 17.18381 | 18.65534 | 18.6081 | 15.50518 | 17.94001 | 17.19901 | 15.23685 | 16.83235 | 16.4394 |
| ING3 | 15.89053 | 24.94619 | 13.04107 | 14.37592 | 13.31675 | 21.12204 | 21.33417 | 19.728 | 12.26756 | 18.60598 | 15.70502 | 14.21276 | 15.93858 | 15.26971 | 15.86511 | 14.65599 | 21.27001 | 14.28553 |
| INHBE | 17.60981 | 15.07096 | 14.99604 | 19.63291 | 11.81044 | 15.18936 | 15.94213 | 15.02992 | 19.03083 | 17.83719 | 15.80275 | 13.39164 | 15.11499 | 15.00888 | 13.68091 | 18.68273 | 13.44264 | 14.5256 |
| INPP5A | 16.12962 | 16.20704 | 18.24283 | 19.43254 | 17.5559 | 16.02761 | 17.00113 | 15.7506 | 16.45789 | 16.89776 | 15.7751 | 17.21874 | 17.01087 | 16.96366 | 18.18389 | 19.96152 | 16.37163 | 17.23545 |
| INPP5D | 15.60859 | 16.36045 | 17.13824 | 14.90795 | 16.43904 | 15.95235 | 14.83893 | 16.80047 | 14.83162 | 15.65587 | 18.2769 | 18.72505 | 15.91147 | 16.43856 | 14.79559 | 15.03167 | 14.91009 | 14.31792 |
| INPPL1 | 16.70715 | 17.0001 | 15.43217 | 16.2137 | 15.85422 | 16.45407 | 15.64817 | 15.90124 | 15.38372 | 16.01046 | 18.7261 | 18.94618 | 14.96414 | 16.49946 | 15.51528 | 16.67572 | 15.79608 | 17.76442 |
| INSR | 25.11952 | 24.94801 | 24.584 | 23.46297 | 17.83248 | 16.44391 | 17.25642 | 16.93732 | 18.77275 | 16.5045 | 15.77544 | 14.06383 | 17.13061 | 16.95777 | 25.06259 | 24.65612 | 15.87856 | 16.78563 |
| IPO11 | 13.97289 | 19.26158 | 15.48198 | 15.57638 | 14.52684 | 16.16889 | 14.8827 | 17.88861 | 15.44353 | 16.00504 | 14.52706 | 19.02057 | 16.14675 | 15.38158 | 15.95079 | 15.20805 | 16.45083 | 16.09325 |
| IPO4 | 15.74874 | 15.71025 | 15.85481 | 14.48721 | 18.12858 | 14.71822 | 17.6658 | 14.87908 | 15.4195 | 15.23801 | 14.73481 | 17.8898 | 15.03298 | 15.53046 | 15.91932 | 16.36271 | 16.03633 | 14.86952 |
| IPO5 | 18.86143 | 19.94898 | 17.69629 | 16.95469 | 20.68798 | 19.85795 | 21.11538 | 17.5046 | 17.54638 | 18.62641 | 17.99655 | 20.48869 | 16.8443 | 14.48892 | 18.54315 | 17.18213 | 17.67927 | 19.78519 |
| IPO7 | 16.46545 | 17.23013 | 17.44957 | 19.46123 | 18.41103 | 16.8907 | 17.4122 | 16.36332 | 16.52102 | 16.88375 | 18.88174 | 19.83858 | 19.53436 | 18.67843 | 16.71412 | 16.78866 | 18.24049 | 18.11183 |
| IPO8 | 16.32986 | 15.75703 | 18.38841 | 17.10289 | 19.63236 | 19.14827 | 16.5701 | 17.19132 | 16.17422 | 17.83343 | 17.57231 | 21.59408 | 17.66409 | 18.63087 | 16.78814 | 17.15063 | 18.36497 | 17.96339 |
| IPO9 | 15.40185 | 15.05219 | 16.55168 | 15.50676 | 17.38617 | 14.37749 | 17.90758 | 16.68365 | 17.05907 | 18.41644 | 18.55741 | 19.39286 | 16.05494 | 14.95775 | 16.88311 | 15.82672 | 16.92913 | 16.46636 |
| IQGAP1 | 19.16796 | 19.27753 | 18.90396 | 19.20287 | 21.95856 | 20.89481 | 18.93972 | 19.01996 | 18.66514 | 18.90348 | 18.06517 | 19.21177 | 19.21174 | 19.11605 | 19.25901 | 19.24953 | 19.1549 | 18.59767 |
| IQGAP2 | 16.50834 | 16.73018 | 20.59185 | 19.44259 | 17.98326 | 16.90817 | 16.72363 | 15.89835 | 16.38984 | 16.48618 | 16.02677 | 17.72377 | 17.97731 | 17.67999 | 19.96177 | 19.78723 | 15.91742 | 15.72127 |
| IQGAP3 | 21.55882 | 17.5207 | 17.41687 | 17.97346 | 19.37951 | 18.39034 | 16.70988 | 17.11491 | 17.09929 | 17.7304 | 17.27534 | 17.83042 | 21.3531 | 15.83679 | 17.04924 | 17.1267 | 17.54422 | 16.03765 |
| IQSEC2 | 17.54234 | 16.77909 | 14.52754 | 15.86643 | 18.7251 | 16.66114 | 16.41299 | 16.56217 | 14.54739 | 17.28454 | 14.73343 | 13.42012 | 14.62613 | 13.82319 | 16.68072 | 17.5157 | 15.89601 | 14.28181 |
| IRAG2 | 14.90667 | 15.50069 | 14.35952 | 15.40868 | 14.64167 | 15.32098 | 15.67155 | 15.34486 | 14.70039 | 14.81298 | 18.29301 | 18.8651 | 14.06645 | 16.62755 | 16.24735 | 14.20335 | 16.43476 | 15.11596 |
| IREB2 | 15.39738 | 17.48024 | 15.42243 | 15.16122 | 15.5476 | 16.52311 | 15.49713 | 16.48436 | 13.3399 | 13.95123 | 16.01728 | 18.44559 | 16.17667 | 15.06786 | 16.26871 | 15.44905 | 14.92468 | 16.39774 |
| IRGQ | 15.74606 | 16.27844 | 15.46711 | 15.96629 | 19.21371 | 16.76543 | 18.95398 | 19.03239 | 16.66228 | 17.46558 | 18.67928 | 18.87206 | 18.43198 | 13.49961 | 15.84497 | 15.66328 | 18.94423 | 18.56494 |
| ISG15 | 20.02013 | 19.50118 | 18.74079 | 15.09704 | 16.9823 | 17.20385 | 15.84399 | 14.5662 | 12.55585 | 15.87422 | 17.36059 | 18.30089 | 18.05038 | 17.61281 | 19.29295 | 18.33612 | 18.77554 | 19.01433 |
| ISOC1 | 15.69898 | 16.6589 | 14.86415 | 16.48539 | 17.24048 | 16.94344 | 17.8675 | 19.12755 | 17.5057 | 18.27869 | 19.68466 | 20.38156 | 17.75022 | 17.94114 | 14.54647 | 17.24537 | 17.93964 | 14.74099 |
| ISOC2 | 16.78385 | 17.37445 | 17.27191 | 16.48373 | 14.20125 | 15.38681 | 18.46573 | 17.25194 | 19.4081 | 20.11434 | 14.65421 | 13.80696 | 16.71262 | 14.69591 | 17.25871 | 14.8697 | 16.00828 | 16.97652 |
| IST1 | 19.3869 | 19.32342 | 19.93352 | 20.73893 | 22.88834 | 21.45166 | 19.79799 | 17.29984 | 21.15032 | 21.54669 | 21.59957 | 22.00549 | 21.38941 | 21.61368 | 19.33412 | 19.53724 | 19.47241 | 20.42501 |
| ITCH | 21.72831 | 22.19058 | 20.89421 | 20.23433 | 18.76036 | 17.13399 | 16.71534 | 16.81456 | 17.23318 | 16.67341 | 16.47673 | 16.4196 | 17.1978 | 16.85565 | 21.32497 | 21.09753 | 18.00749 | 19.0832 |
| ITFG1 | 16.27872 | 17.54823 | 18.91199 | 18.55721 | 17.40678 | 16.91907 | 16.26424 | 16.23792 | 21.86484 | 20.65368 | 16.73337 | 16.05603 | 16.82745 | 16.39205 | 18.39192 | 17.64349 | 14.48195 | 15.3914 |
| ITGA1 | 19.55038 | 19.54041 | 15.97013 | 17.8942 | 17.37858 | 16.92554 | 17.29743 | 16.89029 | 16.10668 | 16.8785 | 16.99158 | 15.24022 | 16.38305 | 16.14706 | 15.89984 | 16.95256 | 21.18364 | 22.05173 |
| ITGA2 | 21.13596 | 20.69565 | 21.26307 | 20.92551 | 22.09388 | 21.21408 | 18.74627 | 17.30155 | 15.23717 | 16.34013 | 16.03741 | 17.00746 | 19.69651 | 19.02583 | 21.6775 | 21.55305 | 22.1763 | 22.47555 |
| ITGA2B | 20.65531 | 19.77227 | 21.34267 | 21.05738 | 18.76914 | 18.64484 | 21.27837 | 20.79228 | 16.42379 | 15.24723 | 17.15608 | 12.9807 | 20.31671 | 20.83881 | 22.36733 | 21.88274 | 17.53499 | 18.48141 |
| ITGA3 | 18.17941 | 17.73196 | 18.24914 | 17.25621 | 24.36658 | 24.25639 | 16.99997 | 17.79622 | 17.26586 | 17.57933 | 18.39818 | 19.5071 | 16.93277 | 17.14731 | 17.99904 | 17.20936 | 23.82977 | 24.50102 |
| ITGA5 | 19.95746 | 20.90262 | 19.68757 | 19.86162 | 21.35432 | 20.87046 | 16.44159 | 20.84898 | 16.68975 | 16.50317 | 15.12722 | 14.55502 | 16.62678 | 21.33186 | 20.32626 | 20.24782 | 19.60273 | 20.79966 |
| ITGA6 | 18.47367 | 17.88817 | 20.23862 | 20.88101 | 22.42299 | 21.19541 | 19.5385 | 18.45739 | 16.15638 | 16.64233 | 18.54829 | 19.95036 | 21.98982 | 21.85076 | 21.02132 | 21.16248 | 23.72586 | 25.1631 |
| ITGA7 | 15.12824 | 15.36471 | 16.0943 | 15.79504 | 16.58413 | 18.43473 | 20.44296 | 18.51077 | 18.01034 | 18.75205 | 15.75199 | 15.28902 | 16.26744 | 14.96794 | 15.9178 | 15.43618 | 15.33898 | 16.37622 |
| ITGAV | 19.51612 | 18.41031 | 19.73174 | 19.57763 | 19.50369 | 18.7042 | 16.8327 | 16.10768 | 19.01252 | 17.95362 | 17.2854 | 19.44748 | 18.11162 | 17.1581 | 20.37021 | 19.99661 | 18.63331 | 18.07423 |
| ITGB1 | 21.78348 | 21.59838 | 21.72247 | 22.23639 | 26.65068 | 25.25486 | 21.83215 | 20.61746 | 22.64062 | 22.38441 | 20.48834 | 20.81711 | 22.26566 | 22.26841 | 22.58446 | 22.37843 | 25.37955 | 26.90549 |
| ITGB1BP1 | 13.76698 | 15.49354 | 14.36399 | 17.93435 | 13.19944 | 15.15208 | 14.65474 | 17.44691 | 16.1542 | 15.85179 | 16.36584 | 10.66328 | 17.74709 | 17.96861 | 15.07798 | 18.50401 | 17.85477 | 18.68339 |
| ITGB2 | 17.2202 | 19.05497 | 16.92001 | 19.45974 | 17.3621 | 13.59604 | 19.7905 | 19.71642 | 15.98773 | 15.83148 | 16.25337 | 14.66565 | 17.69591 | 18.77062 | 16.95699 | 17.91687 | 15.38623 | 17.12258 |
| ITGB3 | 21.65174 | 21.99173 | 21.68197 | 23.07291 | 21.63989 | 19.86819 | 23.58242 | 20.9633 | 14.7494 | 15.50135 | 15.85579 | 13.0775 | 22.08632 | 22.72739 | 21.75034 | 21.69498 | 18.11701 | 20.47539 |
| ITGB4 | 16.21576 | 17.38267 | 16.14766 | 16.59749 | 20.18557 | 19.27151 | 15.95066 | 17.44419 | 15.64913 | 16.31651 | 15.57588 | 14.92438 | 18.00405 | 17.20191 | 16.84208 | 16.4903 | 22.25263 | 22.3431 |
| ITGB5 | 19.5296 | 18.16437 | 19.54901 | 19.5376 | 17.71542 | 17.97966 | 15.52487 | 16.78434 | 17.29096 | 16.83605 | 17.93853 | 19.86054 | 18.02515 | 18.26003 | 20.4352 | 19.47629 | 18.18702 | 17.50575 |
| ITGB6 | 18.10234 | 16.55891 | 21.50692 | 21.69781 | 22.92929 | 21.83068 | 16.45219 | 17.83699 | 21.20807 | 15.28316 | 19.98381 | 20.52307 | 19.83497 | 19.05733 | 21.49692 | 21.32942 | 21.20262 | 23.25345 |
| ITGB8 | 13.5958 | 16.96879 | 16.35582 | 14.94905 | 14.97594 | 14.53707 | 18.56589 | 16.02073 | 15.73335 | 15.74254 | 14.81367 | 11.95816 | 13.79481 | 15.61597 | 17.9485 | 14.68014 | 17.8639 | 16.70402 |
| ITM2B | 16.87747 | 17.18086 | 17.94193 | 17.99295 | 20.75911 | 19.36722 | 21.04268 | 18.71538 | 20.42943 | 19.93432 | 15.55385 | 13.71447 | 18.93855 | 18.51344 | 16.74603 | 16.81245 | 15.9557 | 19.10682 |
| ITM2C | 15.52677 | 16.42322 | 16.87415 | 16.40712 | 19.67372 | 19.48201 | 16.96298 | 17.42703 | 21.7248 | 18.46866 | 17.75938 | 15.83347 | 15.99033 | 16.60017 | 16.34308 | 15.04538 | 14.75324 | 18.10332 |
| ITPA | 16.85819 | 16.42709 | 16.96765 | 16.24229 | 19.0489 | 20.42781 | 20.60101 | 19.09866 | 15.49658 | 16.85521 | 19.19015 | 20.4341 | 17.7205 | 19.96908 | 16.13233 | 15.49369 | 19.67391 | 20.20704 |
| ITSN2 | 14.55918 | 13.8356 | 16.09458 | 14.26652 | 18.33282 | 15.13311 | 14.98213 | 14.33907 | 12.41819 | 13.90205 | 16.03539 | 13.30616 | 16.50568 | 16.86821 | 15.98231 | 16.30172 | 16.31074 | 12.65662 |
| JAG1 | 15.11849 | 16.0697 | 16.58472 | 15.64638 | 15.3326 | 16.31853 | 16.80375 | 16.11914 | 15.19223 | 16.99871 | 17.77231 | 15.69377 | 20.43175 | 21.01031 | 15.17836 | 15.95923 | 19.36598 | 20.11885 |
| JAG2 | 16.45751 | 15.70043 | 16.15169 | 16.78562 | 13.67338 | 15.70156 | 14.07643 | 14.32277 | 17.77788 | 15.63993 | 17.89513 | 19.42476 | 15.33046 | 15.50329 | 16.05612 | 17.36415 | 14.56763 | 14.62001 |
| JAK1 | 16.96451 | 15.64911 | 16.30869 | 15.51227 | 19.6286 | 18.66333 | 17.95186 | 15.7386 | 16.33894 | 16.67332 | 15.44902 | 15.23527 | 15.09817 | 15.43496 | 15.19704 | 14.93175 | 18.22943 | 18.27678 |
| JAK2 | 15.07111 | 13.47779 | 14.85565 | 17.27365 | 15.97648 | 16.41328 | 15.15056 | 18.44895 | 14.8993 | 21.15512 | 15.29218 | 13.86182 | 16.79893 | 15.29804 | 16.26127 | 16.38015 | 17.78088 | 19.37772 |
| JAM3 | 18.83664 | 17.87559 | 17.98638 | 18.6854 | 16.66406 | 16.30312 | 17.41067 | 16.12061 | 20.81502 | 19.12861 | 18.22757 | 19.95088 | 15.55789 | 15.64738 | 17.57617 | 18.58644 | 17.5128 | 17.0692 |
| JMJD1C | 14.65573 | 22.86424 | 16.62823 | 14.36871 | 14.62652 | 15.41684 | 14.67049 | 24.19035 | 13.01298 | 15.09287 | 13.31697 | 17.07534 | 15.09429 | 15.64209 | 13.74461 | 14.1394 | 17.46079 | 12.70322 |
| JPT1 | 16.80717 | 16.8478 | 19.26892 | 21.07983 | 21.03163 | 19.41027 | 17.91882 | 18.77459 | 16.39561 | 17.37902 | 18.18938 | 17.89695 | 21.45754 | 21.46151 | 21.71687 | 20.92792 | 21.09763 | 21.08517 |
| JPT2 | 17.94454 | 18.35843 | 18.45209 | 18.79408 | 17.87859 | 20.07744 | 15.26378 | 17.15462 | 18.19717 | 15.44477 | 18.07139 | 17.43212 | 18.16659 | 17.44536 | 18.41997 | 18.6046 | 16.15445 | 17.52293 |
| JTB | 16.37516 | 16.7071 | 13.66999 | 13.69361 | 18.5461 | 17.29491 | 17.29463 | 17.42292 | 20.99753 | 19.16987 | 17.23039 | 17.17627 | 16.81254 | 15.89137 | 13.2993 | 14.93631 | 15.97095 | 17.41375 |
| JUP | 19.49883 | 19.05069 | 22.11359 | 22.43814 | 22.72314 | 21.72809 | 17.17922 | 17.1533 | 19.51611 | 19.49614 | 20.42741 | 21.07974 | 22.2456 | 22.22966 | 22.79419 | 22.80863 | 21.2787 | 21.45237 |
| KANK2 | 15.41936 | 17.01682 | 17.05204 | 15.4936 | 18.61284 | 15.80617 | 16.37737 | 16.80556 | 13.39413 | 15.83873 | 13.41195 | 11.89428 | 14.79039 | 14.82275 | 15.82422 | 13.48184 | 17.97353 | 16.3209 |
| KARS1 | 16.83792 | 16.62982 | 18.0781 | 17.63791 | 17.42399 | 18.61223 | 17.76261 | 17.44297 | 16.91379 | 15.67351 | 18.81849 | 20.88469 | 17.89778 | 17.06056 | 16.43948 | 17.38597 | 16.94165 | 16.78474 |
| KCTD12 | 15.10599 | 17.26301 | 15.54388 | 14.82775 | 13.87968 | 16.50022 | 16.86712 | 15.98995 | 18.95266 | 17.83809 | 18.35229 | 17.54759 | 16.08801 | 14.798 | 15.31261 | 14.37798 | 14.40155 | 15.58545 |
| KCTD3 | 15.86011 | 15.95479 | 14.31714 | 13.56401 | 16.09272 | 15.86257 | 15.83613 | 16.71735 | 13.70184 | 17.05413 | 16.83671 | 12.30922 | 17.56517 | 18.53605 | 17.70156 | 16.1749 | 18.95137 | 19.76743 |
| KCTD9 | 13.4233 | 16.36042 | 14.96705 | 16.90507 | 16.80049 | 15.80232 | 14.86074 | 14.90351 | 18.91862 | 15.99979 | 15.44534 | 15.47085 | 15.44371 | 15.10034 | 13.83977 | 15.23337 | 18.02272 | 18.84006 |
| KDF1 | 15.8793 | 16.03159 | 20.10345 | 18.98976 | 14.67341 | 16.6242 | 16.50785 | 15.77354 | 18.51063 | 18.05183 | 18.54031 | 17.01134 | 18.28958 | 18.50822 | 19.79966 | 19.17537 | 16.78521 | 15.35087 |
| KDM1A | 16.09127 | 14.89101 | 16.4612 | 16.29667 | 14.31955 | 14.03856 | 15.38319 | 16.53439 | 15.66859 | 15.33644 | 16.57239 | 18.90356 | 13.6954 | 13.77965 | 15.65533 | 14.89121 | 15.149 | 14.05008 |
| KDSR | 16.53262 | 16.06458 | 15.12782 | 16.29715 | 13.25527 | 18.25281 | 15.20657 | 17.85828 | 18.17218 | 17.33588 | 15.97026 | 12.8452 | 14.93952 | 15.69755 | 16.64819 | 15.93497 | 16.53864 | 16.27029 |
| KHDRBS1 | 16.35972 | 14.93427 | 21.51848 | 17.4972 | 17.65263 | 18.73802 | 15.9825 | 16.22596 | 17.25688 | 19.32602 | 19.25584 | 19.43579 | 17.05697 | 17.26723 | 15.31409 | 14.89965 | 16.37626 | 16.73251 |
| KHNYN | 18.49041 | 17.84841 | 16.58823 | 16.3407 | 17.58519 | 17.43079 | 15.57397 | 16.04097 | 15.68492 | 14.85088 | 15.07892 | 14.61392 | 15.86999 | 14.57659 | 16.71971 | 17.48828 | 15.77093 | 15.49413 |
| KHSRP | 17.12988 | 15.64867 | 16.39425 | 16.6869 | 18.59618 | 15.91768 | 18.29235 | 18.85419 | 18.4369 | 19.17927 | 19.66479 | 19.26843 | 17.19149 | 16.68209 | 15.86908 | 15.39681 | 17.81395 | 18.68936 |
| KIAA0319L | 17.03312 | 17.05329 | 18.46557 | 17.33293 | 16.64721 | 17.44974 | 16.12374 | 16.41517 | 20.43745 | 19.48415 | 17.27113 | 15.79863 | 14.57431 | 17.90625 | 18.06166 | 18.52164 | 15.50927 | 16.01167 |

| | | | | | | | | | | | | | | | | | | |
|---|---|---|---|---|---|---|---|---|---|---|---|---|---|---|---|---|---|---|
| KIAA1109 | 15.23257 | 17.85027 | 15.19096 | 15.40855 | 17.21932 | 14.43304 | 15.0092 | 14.64872 | 12.17968 | 15.58185 | 17.50197 | 18.68574 | 14.47561 | 15.37349 | 15.69561 | 16.17542 | 14.05511 | 16.21777 |
| KIAA1217 | 17.18962 | 14.79742 | 16.1926 | 15.46738 | 14.19178 | 15.39178 | 17.38174 | 17.49452 | 15.11989 | 16.82272 | 15.91027 | 13.91127 | 16.89546 | 19.085 | 17.45805 | 14.91043 | 18.52401 | 16.90168 |
| KIAA1522 | 17.22533 | 16.14415 | 18.71828 | 20.07073 | 18.91936 | 17.57247 | 17.06396 | 22.42019 | 17.28932 | 15.86793 | 17.02814 | 17.71583 | 19.99359 | 19.96273 | 20.21536 | 19.81934 | 19.01818 | 18.98556 |
| KIAA1549 | 16.03182 | 14.77336 | 16.75667 | 15.06169 | 20.39705 | 17.63202 | 14.9202 | 15.6836 | 16.31736 | 14.52746 | 13.6876 | 14.23285 | 18.13382 | 14.93366 | 17.22988 | 16.03546 | 15.15232 | 18.07398 |
| KIAA2013 | 16.77895 | 17.71755 | 15.71665 | 16.22494 | 14.84945 | 17.79398 | 18.23273 | 16.48202 | 22.22414 | 20.65067 | 16.75131 | 17.82964 | 15.35152 | 15.15613 | 15.78506 | 15.72738 | 17.20193 | 15.84415 |
| KIDINS220 | 15.56947 | 16.46987 | 17.76984 | 18.14066 | 19.6764 | 16.65232 | 17.42221 | 17.03059 | 18.01823 | 16.39564 | 16.3771 | 17.76228 | 18.23004 | 17.84751 | 17.35404 | 18.03724 | 15.94012 | 16.32276 |
| KIF11 | 14.34941 | 14.24073 | 15.11624 | 13.10544 | 17.808 | 15.90527 | 16.06317 | 15.41804 | 14.22219 | 14.63766 | 17.62357 | 12.74539 | 16.57341 | 15.58455 | 12.6649 | 15.24598 | 17.8832 | 18.50375 |
| KIF13A | 15.96981 | 14.60229 | 15.58122 | 16.69985 | 20.19599 | 15.84248 | 18.62062 | 17.84904 | 15.60381 | 15.90988 | 15.92867 | 14.72119 | 25.3689 | 16.18013 | 15.32346 | 15.17427 | 17.99532 | 17.98837 |
| KIF14 | 16.06705 | 16.62284 | 16.13385 | 17.32145 | 19.92645 | 17.89358 | 17.52247 | 16.5643 | 14.78581 | 15.15871 | 17.59174 | 15.73957 | 16.53432 | 16.91969 | 16.41499 | 16.33104 | 19.13763 | 17.37085 |
| KIF15 | 13.72459 | 14.11769 | 16.59455 | 15.24802 | 16.5329 | 14.71626 | 21.00156 | 16.33475 | 16.00037 | 16.66943 | 20.16281 | 16.17005 | 15.94798 | 15.67469 | 14.55206 | 16.21418 | 22.63096 | 23.27491 |
| KIF1A | 15.39196 | 17.24984 | 13.48438 | 13.30586 | 14.87359 | 15.70877 | 17.09994 | 14.00948 | 15.14095 | 14.92691 | 13.96522 | 16.70153 | 15.22733 | 14.18212 | 15.65296 | 16.02675 | 11.56535 | 16.14418 |
| KIF20A | 17.4119 | 16.47849 | 18.72371 | 20.13389 | 19.80304 | 16.9661 | 16.52083 | 16.39332 | 17.36984 | 17.57041 | 16.17699 | 14.06012 | 19.27692 | 19.72264 | 17.09976 | 16.59857 | 19.57974 | 17.78087 |
| KIF20B | 15.60857 | 16.14983 | 16.21856 | 16.41101 | 19.59542 | 18.89784 | 16.69477 | 17.77353 | 14.28141 | 15.18349 | 16.66028 | 15.56924 | 17.62953 | 16.78161 | 14.75866 | 15.71595 | 18.95771 | 17.5784 |
| KIF23 | 17.63425 | 17.8013 | 19.19647 | 20.10601 | 23.0258 | 21.80094 | 19.21983 | 17.75489 | 19.46296 | 20.3985 | 19.89211 | 18.6456 | 18.52342 | 18.80956 | 18.94005 | 17.72315 | 21.05307 | 18.78283 |
| KIF2A | 16.72072 | 15.9726 | 15.98315 | 16.45422 | 17.4252 | 16.0722 | 19.53764 | 18.7076 | 15.03291 | 16.28387 | 15.75908 | 18.14066 | 17.32848 | 18.18269 | 16.20394 | 14.53046 | 15.09179 | 14.77186 |
| KIF4A | 15.93428 | 16.41662 | 17.31946 | 16.15093 | 18.52662 | 18.26132 | 18.79464 | 19.4391 | 14.84691 | 16.22371 | 15.43935 | 13.8161 | 19.24159 | 19.0361 | 16.0118 | 15.90462 | 20.17732 | 18.58859 |
| KIF5B | 16.8876 | 24.58572 | 17.18671 | 17.44774 | 20.26989 | 18.81831 | 19.63078 | 19.07482 | 18.18877 | 18.65483 | 18.61178 | 19.49433 | 19.36514 | 19.06314 | 17.52588 | 17.34074 | 19.63544 | 19.61914 |
| KIF5C | 13.72182 | 15.06295 | 15.26925 | 14.9752 | 14.93281 | 15.15625 | 17.06163 | 15.84292 | 15.71167 | 15.4401 | 15.10277 | 14.74321 | 18.54645 | 17.71765 | 15.68537 | 14.74058 | 15.52102 | 14.0879 |
| KIFBP | 15.17078 | 15.99881 | 14.2754 | 15.15687 | 14.69354 | 16.99126 | 16.61493 | 15.06304 | 16.09076 | 16.52048 | 19.84315 | 17.95265 | 15.81495 | 17.54913 | 16.09151 | 16.11226 | 16.28294 | 15.66574 |
| KIRREL1 | 15.13743 | 16.65914 | 16.57419 | 16.13104 | 21.35396 | 20.54567 | 19.18183 | 18.96962 | 14.38279 | 17.00945 | 14.55593 | 15.64295 | 16.18511 | 20.33606 | 14.9882 | 15.7403 | 18.45079 | 19.65589 |
| KIT | 15.16513 | 16.91292 | 20.58612 | 17.36578 | 16.26392 | 17.74718 | 17.27734 | 16.68828 | 18.90905 | 17.83532 | 18.62536 | 19.11873 | 16.00529 | 14.89401 | 18.03849 | 17.29984 | 17.11523 | 16.18523 |
| KLC1 | 19.90456 | 16.25057 | 17.13538 | 17.33496 | 20.15771 | 18.11076 | 18.97775 | 18.12333 | 17.6747 | 18.40986 | 17.97608 | 18.85961 | 19.13411 | 18.94047 | 17.39726 | 16.95101 | 19.97899 | 20.30587 |
| KLC2 | 16.12955 | 18.13351 | 16.53888 | 15.50935 | 16.35734 | 15.79362 | 16.77063 | 16.92899 | 14.96573 | 15.28818 | 13.65745 | 14.6513 | 18.07491 | 17.86581 | 15.2433 | 15.1964 | 17.73333 | 15.92071 |
| KLK11 | 17.03406 | 16.53875 | 16.39776 | 16.0856 | 15.73515 | 16.5162 | 17.25854 | 18.12739 | 20.82056 | 19.90314 | 15.91157 | 15.02408 | 15.64709 | 16.48025 | 16.98503 | 15.65355 | 15.60268 | 15.15788 |
| KLK12 | 16.24638 | 14.23504 | 21.29484 | 14.64487 | 14.66515 | 16.13263 | 13.72558 | 16.67572 | 21.53406 | 16.8455 | 14.64362 | 13.89756 | 12.11598 | 13.10385 | 18.11496 | 13.9389 | 14.8185 | 14.39238 |
| KLK14 | 19.10225 | 16.10401 | 15.77805 | 15.88628 | 15.7129 | 16.11305 | 17.74053 | 16.6435 | 20.70653 | 18.56375 | 15.89012 | 14.00234 | 16.46385 | 19.06829 | 15.32374 | 15.77443 | 16.15675 | 16.73489 |
| KLK3 | 18.03795 | 16.28879 | 21.28416 | 20.96976 | 14.14873 | 14.49542 | 14.65539 | 16.22561 | 14.22903 | 15.48959 | 15.03461 | 13.21274 | 17.39344 | 14.57793 | 22.12469 | 20.72879 | 14.08166 | 14.67649 |
| KLRG2 | 14.94713 | 15.55204 | 19.21919 | 18.86647 | 16.95701 | 16.56636 | 15.40914 | 15.52439 | 16.33212 | 17.43438 | 14.87463 | 13.6045 | 18.01958 | 17.97703 | 18.84182 | 18.16923 | 16.98119 | 14.04868 |
| KNTC1 | 16.07123 | 16.30181 | 15.12129 | 15.78579 | 16.82293 | 17.27171 | 18.55358 | 16.62235 | 16.92223 | 19.12313 | 15.25661 | 20.07145 | 13.91864 | 19.09052 | 16.45813 | 15.59459 | 17.27801 | 18.45667 |
| KPNA1 | 16.89499 | 18.6369 | 16.1873 | 18.7368 | 17.872 | 18.6739 | 18.48733 | 18.74464 | 15.75876 | 16.9729 | 16.89971 | 16.84437 | 19.77122 | 19.90854 | 16.45341 | 16.65246 | 20.28633 | 20.03101 |
| KPNA2 | 18.33502 | 18.16991 | 18.92528 | 19.00568 | 19.72972 | 18.90214 | 18.09943 | 17.79156 | 18.64733 | 18.64423 | 18.72124 | 19.06655 | 19.30912 | 18.84745 | 18.60193 | 18.63774 | 20.0682 | 18.76537 |
| KPNA3 | 18.35248 | 17.29675 | 18.17283 | 18.03655 | 19.80934 | 18.13419 | 18.95024 | 17.80948 | 17.39713 | 17.498 | 19.33079 | 18.73997 | 18.88948 | 17.37709 | 17.81573 | 18.11962 | 19.63158 | 19.40902 |
| KPNA4 | 21.25928 | 20.6453 | 18.60942 | 20.25111 | 19.62533 | 19.61473 | 18.19266 | 17.53875 | 17.84228 | 16.24357 | 17.64654 | 18.45588 | 19.64058 | 20.12992 | 20.68239 | 21.39198 | 21.50325 | 20.00717 |
| KPNA6 | 18.68029 | 18.65059 | 18.21034 | 17.27769 | 14.64062 | 16.85872 | 16.87649 | 17.27136 | 16.81567 | 16.56505 | 16.28899 | 15.9688 | 15.60067 | 16.97779 | 17.651 | 17.99926 | 18.34365 | 16.13599 |
| KPNB1 | 19.51499 | 19.4817 | 20.2255 | 19.88127 | 20.89589 | 19.53824 | 20.34775 | 19.28603 | 18.8132 | 19.98106 | 20.13674 | 20.61607 | 20.77058 | 19.42115 | 20.10649 | 20.00168 | 21.13644 | 20.39912 |
| KRAS | 19.86694 | 20.62337 | 20.08692 | 18.27058 | 20.20212 | 18.41163 | 18.39072 | 17.90669 | 19.26463 | 18.76317 | 19.43258 | 19.06847 | 20.77147 | 20.4511 | 19.59027 | 20.10211 | 19.21116 | 19.66606 |
| KRIT1 | 16.09399 | 14.46695 | 16.58477 | 17.9408 | 18.92655 | 18.09689 | 19.11396 | 16.94289 | 15.55697 | 17.46939 | 12.55082 | 15.29731 | 17.9302 | 19.82113 | 17.27381 | 16.60428 | 18.56029 | 19.1782 |
| KRT1 | 21.80467 | 21.71792 | 20.77597 | 20.72471 | 20.42177 | 14.77568 | 15.52897 | 19.89696 | 20.61141 | 18.14996 | 19.04555 | 19.58282 | 18.95826 | 18.99764 | 21.07094 | 15.13617 | 19.74485 | 19.96634 |
| KRT18 | 17.78011 | 17.71312 | 17.82881 | 17.66659 | 18.32271 | 18.22372 | 16.68073 | 17.04303 | 19.0119 | 16.66507 | 21.37208 | 21.22126 | 18.94323 | 17.88848 | 17.14257 | 17.90076 | 18.19138 | 17.07375 |
| KRTCAP3 | 20.46061 | 20.05182 | 20.94139 | 20.66128 | 17.30692 | 18.13059 | 16.11905 | 17.74764 | 15.91118 | 14.95207 | 19.40079 | 19.02319 | 17.70478 | 17.68045 | 20.96121 | 21.00385 | 15.02297 | 17.18266 |
| KTN1 | 17.77194 | 15.74116 | 15.22134 | 16.66107 | 15.41837 | 15.84301 | 16.25199 | 16.79235 | 17.08747 | 16.65517 | 17.28803 | 18.75443 | 16.64896 | 16.48254 | 16.11804 | 15.93964 | 16.33162 | 15.57445 |
| KYAT3 | 14.62161 | 16.78895 | 14.14308 | 16.93499 | 15.55344 | 12.98258 | 13.89004 | 15.9546 | 16.28983 | 14.59102 | 16.81479 | 16.61712 | 14.56738 | 14.83886 | 16.15729 | 16.32554 | 13.25159 | 16.23961 |
| KYNU | 14.33212 | 15.95504 | 18.0721 | 14.73966 | 19.56376 | 16.15472 | 16.64171 | 17.21322 | 14.34756 | 16.32832 | 16.25387 | 14.66019 | 14.67839 | 18.44172 | 13.4202 | 13.90712 | 16.52653 | 17.81816 |
| L1CAM | 19.0081 | 18.54009 | 23.39693 | 15.73839 | 16.11004 | 23.85761 | 19.47024 | 15.7688 | 19.20003 | 19.08786 | 19.54704 | 18.06297 | 14.92514 | 15.76203 | 21.2521 | 15.54855 | 16.66506 | 16.9812 |
| L1RE1 | 22.00438 | 21.68502 | 19.72403 | 19.76991 | 15.19794 | 16.96339 | 17.08475 | 17.26523 | 15.874 | 17.51059 | 15.01814 | 14.75567 | 18.02245 | 18.46888 | 18.64128 | 19.84327 | 16.1267 | 15.54617 |
| LACTB2 | 17.11062 | 16.29482 | 17.28032 | 17.71076 | 19.17957 | 18.77118 | 16.2593 | 16.98382 | 21.59139 | 20.18868 | 16.33406 | 16.84247 | 17.05742 | 16.87033 | 17.63464 | 16.73024 | 18.6657 | 18.86992 |
| LAD1 | 15.68713 | 16.38955 | 17.43682 | 16.71722 | 16.58414 | 17.05333 | 15.92928 | 16.67185 | 14.32954 | 16.43128 | 17.94535 | 17.9212 | 20.15887 | 19.54661 | 16.15004 | 16.54106 | 14.08187 | 16.74622 |
| LAMA4 | 16.27413 | 14.00921 | 15.80031 | 14.80146 | 12.60158 | 13.24387 | 15.37355 | 14.11011 | 17.34073 | 13.52961 | 14.38452 | 11.60278 | 14.85378 | 13.01494 | 16.62109 | 13.11678 | 15.29831 | 16.14968 |
| LAMA5 | 18.43066 | 16.45234 | 18.24438 | 16.78712 | 16.21771 | 16.60746 | 16.99331 | 16.26535 | 15.67138 | 18.93806 | 14.7193 | 13.6756 | 15.7597 | 16.55407 | 16.84884 | 16.04289 | 17.01262 | 18.6127 |
| LAMB1 | 16.79267 | 16.18089 | 18.30044 | 16.63002 | 15.11381 | 16.49771 | 17.76105 | 17.86256 | 15.15459 | 15.64757 | 17.97243 | 15.15312 | 15.83711 | 17.16528 | 15.57781 | 15.75338 | 17.17969 | 19.61567 |
| LAMB2 | 16.41868 | 16.72475 | 17.18635 | 16.04489 | 14.94856 | 16.37187 | 16.12115 | 16.64231 | 18.74897 | 16.99456 | 15.23966 | 13.80319 | 17.32432 | 17.06348 | 16.23642 | 16.89549 | 15.38694 | 16.20721 |
| LAMC1 | 18.13405 | 16.79094 | 18.13512 | 17.91177 | 17.77361 | 18.59357 | 17.07799 | 17.69578 | 18.42308 | 21.38095 | 16.73828 | 15.5066 | 17.9331 | 18.68427 | 16.98084 | 17.96685 | 16.9319 | 20.30772 |
| LAMP1 | 20.13008 | 19.67293 | 21.15806 | 21.06338 | 21.62322 | 20.5309 | 21.27066 | 20.27965 | 25.16871 | 24.11972 | 22.66494 | 19.44914 | 21.19488 | 21.91349 | 20.53957 | 19.9595 | 18.13602 | 21.15305 |
| LAMP2 | 17.16601 | 19.11444 | 20.74205 | 20.7095 | 14.36859 | 16.29372 | 18.26239 | 18.81127 | 21.79522 | 20.99858 | 19.4101 | 17.47691 | 17.77607 | 17.07581 | 20.62972 | 19.8218 | 17.77934 | 16.17748 |
| LAMP3 | 14.8961 | 15.97027 | 15.75244 | 17.64894 | 17.40087 | 16.48608 | 17.2531 | 16.63274 | 20.86467 | 19.54026 | 14.62972 | 14.48384 | 17.49381 | 15.71765 | 16.51502 | 15.73089 | 15.63478 | 16.33283 |
| LAMTOR1 | 15.21348 | 16.8082 | 20.11672 | 19.46739 | 19.46671 | 15.75712 | 18.06586 | 17.61024 | 19.51339 | 19.0642 | 19.92652 | 19.59304 | 18.48985 | 19.58454 | 19.84958 | 18.6313 | 15.3343 | 18.77553 |
| LAMTOR2 | 15.91292 | 16.43498 | 16.09642 | 16.5003 | 19.30069 | 18.66195 | 20.62101 | 17.0492 | 17.13946 | 19.23361 | 17.29424 | 17.63545 | 20.23728 | 20.3349 | 14.35225 | 15.02177 | 19.99095 | 19.49888 |
| LAMTOR3 | 19.26426 | 19.58023 | 19.95924 | 21.0605 | 22.78156 | 20.8604 | 21.53878 | 16.59396 | 21.4538 | 21.98208 | 22.4584 | 21.40313 | 23.37317 | 23.1324 | 21.38944 | 20.03967 | 19.47155 | 22.46839 |
| LANCL1 | 18.7943 | 19.21141 | 18.06531 | 17.28831 | 16.43532 | 17.89365 | 15.81415 | 17.16662 | 17.01711 | 17.04044 | 15.82427 | 19.33665 | 16.58068 | 16.84347 | 19.15418 | 18.79808 | 16.89028 | 16.92419 |
| LAP3 | 16.02584 | 15.516 | 16.61208 | 17.21305 | 20.18564 | 17.63257 | 19.69915 | 18.82221 | 18.27122 | 18.51939 | 18.81074 | 18.893 | 20.73946 | 20.19802 | 16.84485 | 17.26544 | 19.31764 | 19.92316 |
| LAPTM4A | 18.18714 | 16.76375 | 19.73514 | 13.41283 | 14.60984 | 18.14927 | 17.38363 | 15.69797 | 19.06149 | 16.44666 | 15.4838 | 12.50648 | 15.47845 | 14.19299 | 15.00446 | 17.58406 | 14.12446 | 15.50765 |
| LARGE2 | 14.98555 | 15.54388 | 19.03471 | 17.80268 | 14.94375 | 14.40088 | 18.81857 | 17.41885 | 12.98316 | 16.29465 | 16.63721 | 18.19243 | 14.90426 | 14.41791 | 14.24813 | 15.80787 | 12.41266 | 17.5096 |
| LARP1 | 19.08034 | 19.20153 | 18.49707 | 19.45969 | 14.98159 | 17.14259 | 16.91324 | 15.95912 | 14.38068 | 14.55148 | 16.67626 | 17.11126 | 18.10394 | 17.6549 | 18.62166 | 18.91704 | 16.54682 | 16.41767 |
| LARP4 | 18.35611 | 18.50082 | 17.10553 | 17.92485 | 13.67549 | 16.36894 | 15.89931 | 17.29351 | 14.53095 | 15.18199 | 16.2156 | 15.425 | 16.29226 | 14.73271 | 16.8458 | 16.72201 | 14.92252 | 14.54619 |
| LARP4B | 18.59827 | 13.24687 | 14.60959 | 16.85181 | 14.84758 | 13.30627 | 15.55849 | 14.80438 | 14.87465 | 13.31398 | 15.06577 | 14.49192 | 12.94085 | 15.04301 | 13.88972 | 15.93311 | 15.07883 | 14.29132 |
| LARS1 | 16.23377 | 16.28825 | 18.36309 | 16.39495 | 16.39566 | 16.30372 | 18.46137 | 17.93698 | 16.19056 | 15.94476 | 18.28073 | 20.43896 | 17.53825 | 16.48709 | 18.4743 | 18.16835 | 16.20707 | 16.19225 |
| LASP1 | 20.69707 | 20.2146 | 19.16653 | 19.26521 | 21.62985 | 20.62612 | 19.22375 | 21.04452 | 17.29746 | 18.19761 | 17.38338 | 17.74048 | 20.92813 | 21.11139 | 19.48569 | 19.27431 | 21.14294 | 21.57697 |
| LCK | 18.2134 | 17.40173 | 14.64426 | 19.19019 | 20.70199 | 17.14892 | 16.21515 | 18.29585 | 18.84022 | 19.62278 | 14.42234 | 17.61255 | 18.30542 | 17.48675 | 18.83235 | 19.23211 | 15.62452 | 15.78191 |
| LCP1 | 17.19051 | 17.17009 | 17.45578 | 16.96962 | 21.5145 | 20.97269 | 18.7945 | 18.61581 | 16.87905 | 17.99233 | 15.3338 | 15.40326 | 18.77646 | 19.84099 | 17.63616 | 16.95352 | 22.39212 | 20.69187 |
| LDHA | 19.26858 | 19.48144 | 20.33923 | 20.90087 | 22.62427 | 21.29357 | 20.15522 | 19.77096 | 19.79402 | 20.8553 | 22.46998 | 24.04286 | 21.35073 | 20.62998 | 21.02121 | 21.23394 | 23.208 | 23.22636 |
| LDHAL6A | 21.86655 | 21.82584 | 21.78282 | 22.95848 | 20.3502 | 19.23041 | 18.03561 | 17.77076 | 21.55537 | 20.38964 | 23.63573 | 25.82877 | 19.35724 | 18.7207 | 22.8314 | 23.00916 | 20.92153 | 20.9526 |
| LDHB | 20.48075 | 20.18157 | 20.6926 | 20.64608 | 21.96023 | 20.33655 | 17.84427 | 17.87004 | 15.35976 | 16.72021 | 23.12955 | 24.17114 | 21.32878 | 20.73561 | 20.73982 | 20.50949 | 21.75575 | 21.11127 |
| LDLR | 16.54719 | 15.98705 | 19.30954 | 16.44028 | 17.58634 | 19.05846 | 19.61335 | 18.04663 | 19.03488 | 17.13832 | 17.21931 | 15.43191 | 14.47668 | 16.78112 | 16.7136 | 15.69478 | 17.49836 | 16.39363 |
| LDOC1 | 20.74144 | 20.59212 | 21.6466 | 20.46362 | 15.1533 | 15.7852 | 16.92556 | 17.20597 | 13.70749 | 14.02852 | 16.95962 | 16.39239 | 13.9434 | 15.16432 | 22.22373 | 21.32858 | 15.93561 | 16.3739 |
| LEMD2 | 17.71041 | 16.66775 | 16.82135 | 13.31153 | 13.48371 | 13.68137 | 15.71758 | 17.71841 | 18.57591 | 17.3355 | 16.14993 | 11.88927 | 13.67626 | 14.05316 | 16.2098 | 14.34609 | 15.13495 | 13.72679 |
| LETM1 | 15.74839 | 16.71906 | 14.21754 | 15.13661 | 13.65803 | 14.90882 | 16.23632 | 15.57212 | 18.79678 | 18.84324 | 16.09844 | 14.17393 | 16.29581 | 14.86023 | 15.0435 | 14.26787 | 17.12585 | 15.16027 |
| LFNG | 16.39545 | 16.22598 | 18.47527 | 18.30957 | 16.68599 | 15.06307 | 15.84593 | 16.32492 | 18.32672 | 14.69602 | 14.67136 | 12.34851 | 18.36117 | 17.83952 | 16.47649 | 17.36432 | 16.41596 | 14.72056 |
| LGALS1 | 19.87994 | 16.83748 | 16.84167 | 16.61965 | 22.158 | 21.71381 | 17.27972 | 16.76735 | 17.11858 | 15.12598 | 17.91318 | 14.69406 | 15.0922 | 15.74179 | 16.37545 | 16.00644 | 24.26139 | 23.61011 |
| LGALS3 | 15.47501 | 16.46336 | 16.28617 | 14.32179 | 17.3988 | 18.36165 | 16.09604 | 16.89344 | 12.85035 | 14.37765 | 16.32471 | 19.99545 | 17.94032 | 14.92989 | 18.8273 | 15.9661 | 17.88265 | 16.7864 |
| LGALS3BP | 17.88687 | 16.45167 | 17.6953 | 18.38774 | 17.64712 | 19.54767 | 20.17162 | 20.20385 | 22.77992 | 22.266 | 24.1355 | 21.05083 | 21.77404 | 22.26222 | 17.57428 | 17.77688 | 18.86604 | 23.94321 |
| LGMN | 15.92458 | 15.27492 | 14.87741 | 12.67612 | 16.94029 | 15.0195 | 14.28837 | 14.92849 | 19.40371 | 19.03536 | 18.23456 | 14.08467 | 14.98645 | 16.22985 | 17.46152 | 13.56609 | 16.00112 | 14.6068 |
| LGR4 | 13.94273 | 14.80665 | 13.5711 | 15.23635 | 20.46102 | 18.70504 | 14.70717 | 16.08396 | 16.89238 | 15.75989 | 16.43217 | 14.68325 | 17.37338 | 14.65118 | 13.68314 | 15.34609 | 12.24808 | 17.09321 |
| LIFR | 18.2417 | 14.17622 | 16.72582 | 13.35576 | 16.56776 | 14.96946 | 13.27143 | 19.44232 | 17.77418 | 16.96436 | 15.74953 | 12.59106 | 16.51519 | 14.74593 | 15.0509 | 14.76887 | 16.16538 | 14.6499 |
| LIMA1 | 19.45816 | 18.86604 | 19.34661 | 19.47364 | 19.63754 | 17.88128 | 16.02676 | 16.76373 | 15.29939 | 15.36661 | 15.24764 | 17.65796 | 19.0556 | 19.08717 | 19.80798 | 18.99385 | 17.62819 | 16.91439 |
| LIMCH1 | 14.54494 | 15.57261 | 16.27629 | 17.13369 | 14.83077 | 15.83475 | 18.0076 | 18.93541 | 15.11697 | 16.54911 | 16.87995 | 15.71817 | 18.75531 | 19.56907 | 17.44339 | 17.62417 | 15.29774 | 15.37981 |
| LIMS1 | 16.57199 | 15.81422 | 16.21898 | 14.91909 | 15.59479 | 19.39671 | 20.32939 | 14.42326 | 16.40213 | 13.94599 | 12.37336 | 12.93771 | 16.5606 | 18.99742 | 16.08351 | 17.69562 | 15.24965 | 15.04837 |

| | | | | | | | | | | | | | | | | | | |
|---|---|---|---|---|---|---|---|---|---|---|---|---|---|---|---|---|---|---|
| LIMS2 | 17.85326 | 18.75008 | 17.29166 | 14.85637 | 17.20285 | 19.44461 | 19.86504 | 19.45311 | 15.03953 | 17.38974 | 16.07618 | 16.67921 | 15.85685 | 17.05845 | 13.27898 | 16.9546 | 15.07212 | 17.12893 |
| LIN7A | 18.46477 | 18.06643 | 20.64741 | 19.2779 | 18.60104 | 18.49474 | 18.28062 | 18.50835 | 19.62325 | 19.59748 | 18.80125 | 18.59875 | 18.40355 | 18.90119 | 20.91334 | 20.80344 | 18.77558 | 18.75801 |
| LIN7C | 17.30663 | 18.54961 | 21.37302 | 21.48671 | 20.56845 | 20.92114 | 19.7082 | 19.82722 | 20.26715 | 18.99213 | 20.01371 | 19.41664 | 20.19012 | 18.78306 | 21.27037 | 21.11922 | 21.09355 | 21.51659 |
| LLGL1 | 18.27111 | 18.10446 | 17.9902 | 18.66812 | 17.19059 | 17.73811 | 16.46202 | 17.09128 | 17.52433 | 16.3661 | 16.10653 | 18.58938 | 16.11798 | 15.98631 | 19.06565 | 19.2289 | 17.99541 | 17.82271 |
| LLGL2 | 18.68203 | 18.76304 | 17.93148 | 18.8423 | 17.01622 | 17.15776 | 15.46638 | 17.25555 | 19.17741 | 18.69566 | 17.97362 | 18.44697 | 16.55495 | 15.87332 | 19.46267 | 18.98778 | 18.32032 | 17.70099 |
| LMAN1 | 16.83796 | 17.67986 | 19.54836 | 19.27062 | 16.27103 | 17.59816 | 19.62369 | 17.53276 | 24.98113 | 23.21257 | 21.44133 | 18.59845 | 18.12716 | 19.29628 | 19.30653 | 18.27044 | 16.01676 | 20.02208 |
| LMAN2 | 17.66606 | 17.89999 | 18.79188 | 17.99012 | 16.37835 | 18.31837 | 18.01307 | 17.0266 | 24.25466 | 22.2198 | 19.67215 | 18.02004 | 17.1436 | 16.9478 | 18.65284 | 17.78653 | 16.58666 | 16.79196 |
| LMAN2L | 16.0157 | 16.24387 | 16.14806 | 15.96228 | 14.50886 | 18.29493 | 16.63523 | 16.35812 | 22.20088 | 19.50938 | 14.9772 | 13.82199 | 18.01678 | 14.8582 | 15.04843 | 15.25056 | 14.34937 | 13.94752 |
| LMBRD1 | 14.91737 | 16.31535 | 16.5359 | 19.70955 | 13.41262 | 14.17632 | 15.62859 | 13.71807 | 22.32043 | 21.12133 | 15.72541 | 14.00654 | 14.47906 | 13.72199 | 14.83874 | 13.85716 | 18.80711 | 12.01606 |
| LMCD1 | 15.28828 | 16.87795 | 17.41323 | 16.19067 | 16.93599 | 19.24122 | 17.87474 | 17.00834 | 15.05843 | 15.73459 | 18.2241 | 19.2796 | 16.80267 | 16.61382 | 16.58024 | 14.63519 | 15.50838 | 16.50276 |
| LMF2 | 18.3566 | 19.84758 | 15.53076 | 17.73362 | 15.38494 | 16.10715 | 18.48086 | 16.96904 | 19.75651 | 17.77137 | 15.85321 | 14.97145 | 16.97357 | 15.5171 | 16.224 | 17.16314 | 15.51585 | 15.24787 |
| LMNA | 16.97889 | 17.20732 | 16.30207 | 16.91828 | 20.62169 | 18.99383 | 17.2101 | 18.47503 | 16.51123 | 17.97843 | 17.44432 | 17.07624 | 18.98527 | 17.12001 | 15.38205 | 16.23662 | 19.77975 | 19.22317 |
| LMNB1 | 16.49187 | 16.34977 | 16.73519 | 16.98105 | 18.14518 | 17.64775 | 17.15868 | 17.4749 | 17.25101 | 17.67234 | 18.86677 | 18.69339 | 18.5016 | 16.22661 | 15.49711 | 16.17234 | 16.6198 | 16.4632 |
| LMNB2 | 16.27212 | 15.63425 | 15.79508 | 17.22754 | 19.51875 | 18.07133 | 15.07976 | 16.35453 | 17.36997 | 18.41601 | 17.53193 | 17.89689 | 16.99034 | 15.18737 | 15.18927 | 15.61286 | 15.73574 | 15.44339 |
| LMO7 | 16.04681 | 15.60225 | 16.87541 | 16.47752 | 18.28309 | 16.94773 | 18.04315 | 15.28266 | 15.417 | 16.57454 | 15.70954 | 14.04483 | 14.39634 | 14.58383 | 15.231 | 15.20319 | 19.93078 | 19.5067 |
| LNPEP | 16.30864 | 15.738 | 18.46079 | 17.82703 | 17.33378 | 16.22911 | 16.48044 | 15.94792 | 17.80576 | 16.73994 | 17.26383 | 17.49514 | 16.23694 | 16.05891 | 18.39123 | 18.06242 | 15.77922 | 15.36708 |
| LONP1 | 18.59955 | 18.34152 | 17.38721 | 17.03297 | 14.95939 | 16.24055 | 18.08137 | 16.94131 | 20.01352 | 18.88792 | 17.73292 | 14.04647 | 15.60394 | 16.78038 | 16.86861 | 17.40352 | 17.16637 | 17.2592 |
| LONP2 | 15.89705 | 16.59587 | 16.87248 | 17.72875 | 15.12616 | 17.23264 | 15.35513 | 15.86289 | 19.32135 | 19.56676 | 17.89352 | 17.06156 | 16.13809 | 15.31772 | 16.15462 | 14.81511 | 15.35556 | 14.26197 |
| LOXL2 | 15.30241 | 16.32553 | 15.90317 | 15.94578 | 17.24992 | 19.3548 | 17.25994 | 16.46683 | 19.92667 | 19.18722 | 15.47727 | 12.8357 | 14.28066 | 16.33013 | 16.11897 | 15.45792 | 16.75821 | 18.73096 |
| LPAR2 | 14.21379 | 15.75141 | 14.15097 | 17.18508 | 12.69737 | 15.35978 | 16.718 | 17.50865 | 15.45551 | 12.41215 | 13.53199 | 14.41393 | 16.73763 | 15.04726 | 17.90818 | 17.39522 | 15.71359 | 14.85547 |
| LPIN1 | 15.89151 | 16.39421 | 14.74981 | 15.03734 | 15.78611 | 17.23937 | 16.04106 | 15.31245 | 12.64462 | 16.24201 | 16.29786 | 16.16572 | 13.06808 | 14.25502 | 14.04431 | 16.58919 | 14.42941 | 15.06813 |
| LPL | 15.87048 | 14.43025 | 16.23114 | 14.30935 | 14.80254 | 16.2693 | 14.95933 | 16.56882 | 18.80057 | 17.00231 | 15.84579 | 13.25657 | 18.98415 | 19.13693 | 16.85474 | 13.9128 | 16.46467 | 17.24766 |
| LPP | 17.92237 | 15.58368 | 16.69185 | 18.42646 | 21.15368 | 16.5973 | 17.53893 | 16.80501 | 12.25642 | 16.64408 | 15.305 | 15.72617 | 21.02742 | 21.0266 | 16.45392 | 18.50208 | 19.60507 | 19.16551 |
| LPXN | 15.81438 | 15.19833 | 15.78217 | 14.57354 | 16.34164 | 17.6515 | 14.27924 | 13.41499 | 13.69375 | 12.30632 | 15.70079 | 11.78088 | 16.59905 | 20.95711 | 14.26792 | 14.22926 | 18.51031 | 18.11442 |
| LRATD2 | 17.32555 | 14.45678 | 20.92957 | 17.35531 | 17.31958 | 16.67451 | 17.21586 | 15.53535 | 18.99402 | 18.58035 | 19.29909 | 15.08431 | 19.05669 | 16.78838 | 16.66595 | 16.37494 | 20.19622 | 17.80503 |
| LRBA | 16.22859 | 16.38111 | 15.38034 | 16.3804 | 15.755 | 16.45131 | 15.25609 | 16.40739 | 15.16757 | 14.90145 | 15.16867 | 19.03403 | 16.18283 | 15.80991 | 14.88432 | 15.85618 | 16.30085 | 16.59496 |
| LRIG1 | 14.93174 | 15.41422 | 17.73163 | 17.0208 | 13.23899 | 17.93647 | 15.7297 | 16.50914 | 17.97095 | 15.51586 | 14.30349 | 11.54957 | 14.99381 | 16.83247 | 14.18874 | 12.9489 | 15.9858 | 16.41421 |
| LRP1 | 15.82104 | 15.71491 | 17.35376 | 16.35194 | 17.47878 | 18.03986 | 17.39006 | 17.30597 | 19.03059 | 17.05808 | 15.48659 | 13.74808 | 15.2184 | 16.06602 | 15.68936 | 15.88033 | 16.71388 | 17.51239 |
| LRP11 | 15.9107 | 16.4974 | 15.14942 | 15.37127 | 15.4761 | 15.87653 | 16.72537 | 15.76887 | 20.38846 | 18.38517 | 15.69193 | 13.47307 | 16.24955 | 15.48292 | 15.75661 | 17.1544 | 16.45549 | 15.54717 |
| LRP2 | 13.20803 | 16.15465 | 16.21489 | 15.72746 | 13.24064 | 14.90249 | 16.28309 | 17.68267 | 18.8132 | 18.18501 | 13.67977 | 11.77414 | 15.345 | 14.85429 | 15.48059 | 13.79975 | 14.93342 | 14.0165 |
| LRP8 | 15.6236 | 16.04648 | 17.12973 | 14.87773 | 16.18168 | 18.85084 | 16.56531 | 16.87927 | 19.16248 | 18.57185 | 17.79285 | 14.50066 | 15.10294 | 14.36134 | 15.10903 | 14.77598 | 18.99085 | 19.56923 |
| LRPAP1 | 17.26331 | 17.62734 | 18.0191 | 15.37297 | 17.54097 | 16.72192 | 15.55316 | 17.0705 | 19.89491 | 19.81635 | 16.77321 | 14.55934 | 15.79131 | 17.13667 | 14.61468 | 16.27782 | 15.36547 | 17.01506 |
| LRPPRC | 16.39361 | 16.16794 | 16.7032 | 15.6287 | 15.67516 | 16.46993 | 17.35757 | 17.75599 | 19.92767 | 19.02886 | 16.72063 | 14.92886 | 15.62272 | 16.03822 | 17.52606 | 16.04 | 17.34864 | 16.13349 |
| LRRC1 | 16.71561 | 16.64227 | 17.47771 | 17.13559 | 18.3811 | 17.30286 | 17.91429 | 15.60011 | 18.566 | 16.90667 | 18.51497 | 19.13721 | 18.05628 | 17.65334 | 17.79857 | 18.58728 | 17.66862 | 18.03517 |
| LRRC20 | 14.30181 | 16.2591 | 17.1157 | 14.20927 | 20.56477 | 18.19874 | 18.56247 | 17.73794 | 14.97875 | 18.50142 | 16.19608 | 17.58151 | 17.39107 | 20.49175 | 14.47044 | 15.94621 | 20.73281 | 16.21408 |
| LRRC40 | 16.93818 | 15.68924 | 15.88 | 15.56354 | 13.82741 | 15.62612 | 17.18817 | 16.13715 | 16.16377 | 15.9108 | 16.48018 | 18.72234 | 15.86889 | 15.03707 | 16.79499 | 15.51124 | 15.80286 | 16.03913 |
| LRRC47 | 16.87712 | 17.06822 | 17.49648 | 17.05666 | 15.76881 | 16.8409 | 17.29298 | 17.961 | 17.53553 | 17.49819 | 17.71302 | 18.85398 | 18.6616 | 18.41196 | 17.6351 | 17.85946 | 17.55955 | 16.43394 |
| LRRC57 | 19.79137 | 19.84079 | 17.85928 | 20.42368 | 21.45418 | 19.94955 | 19.3868 | 18.1624 | 16.05045 | 19.21512 | 19.23808 | 19.46214 | 19.54397 | 19.29514 | 19.57376 | 19.1018 | 21.04599 | 21.96677 |
| LRRC59 | 16.74829 | 16.65091 | 17.28683 | 16.44231 | 19.7081 | 17.60586 | 19.45238 | 17.65591 | 21.39583 | 18.61401 | 18.62826 | 17.88982 | 19.22313 | 19.8492 | 17.06654 | 16.70196 | 18.6306 | 19.98642 |
| LRRC7 | 18.16533 | 15.89001 | 18.94796 | 19.68415 | 23.04702 | 20.36708 | 16.90945 | 17.82917 | 13.57622 | 14.76344 | 14.6838 | 19.94811 | 17.73569 | 14.25193 | 18.64876 | 15.21374 | 19.09722 | 17.85324 |
| LRRC8A | 17.71729 | 15.6235 | 15.52408 | 15.67612 | 14.10048 | 16.68262 | 15.05557 | 16.65711 | 14.28588 | 15.45176 | 15.54566 | 18.73806 | 14.08865 | 14.90601 | 15.22836 | 15.56898 | 14.91188 | 15.67103 |
| LRRC8C | 15.73439 | 15.58422 | 14.01752 | 14.9332 | 16.89026 | 16.47353 | 15.74313 | 14.7135 | 16.12522 | 13.96063 | 15.78514 | 16.04613 | 13.25047 | 15.59844 | 13.65037 | 16.88547 | 18.13991 | 15.65308 |
| LRRC8D | 15.67471 | 15.6231 | 14.61379 | 17.39423 | 17.02099 | 18.97062 | 17.05466 | 17.57812 | 16.24523 | 16.25769 | 15.1912 | 18.32119 | 17.09072 | 16.62173 | 17.48501 | 18.17896 | 15.62694 | 17.4449 |
| LRRFIP1 | 17.20648 | 16.81929 | 16.60359 | 16.97761 | 17.74018 | 20.11081 | 17.17455 | 17.04319 | 17.36557 | 17.51861 | 14.30452 | 17.14008 | 18.94772 | 17.18384 | 14.4172 | 15.12507 | 16.70146 | 19.09338 |
| LRRFIP2 | 13.99336 | 16.804 | 17.32124 | 16.02618 | 19.70264 | 17.35642 | 15.94373 | 17.84204 | 15.58092 | 16.93146 | 15.99134 | 12.59596 | 19.19148 | 19.25971 | 14.38273 | 14.23467 | 19.28885 | 19.39803 |
| LRSAM1 | 14.77512 | 16.72688 | 16.44564 | 16.37245 | 18.02027 | 15.74419 | 16.09071 | 14.62627 | 15.42849 | 14.18498 | 16.03506 | 12.15931 | 16.77458 | 16.6661 | 16.65308 | 17.07988 | 14.40924 | 16.97698 |
| LSAMP | 15.09226 | 17.91982 | 17.21132 | 15.03303 | 14.8604 | 15.80721 | 16.02433 | 16.43446 | 19.15164 | 17.25106 | 16.04632 | 13.99448 | 15.50801 | 14.89016 | 17.67183 | 15.58336 | 15.74313 | 14.83962 |
| LSM12 | 16.09388 | 19.28181 | 14.10316 | 13.28465 | 16.25236 | 15.58629 | 15.45381 | 18.08633 | 14.36131 | 15.25798 | 17.01588 | 19.2894 | 17.23693 | 16.95934 | 14.7123 | 15.88625 | 18.44182 | 16.53178 |
| LSM14B | 19.36935 | 18.3972 | 19.98863 | 20.19033 | 13.92811 | 15.0592 | 14.88712 | 16.05511 | 14.01754 | 16.66207 | 14.22417 | 15.31959 | 19.59689 | 18.50682 | 20.13474 | 20.16483 | 16.52206 | 13.83117 |
| LSM2 | 17.2242 | 17.59535 | 17.5289 | 14.98556 | 18.17933 | 17.35541 | 18.83705 | 20.29282 | 15.3875 | 18.63407 | 18.43824 | 18.75163 | 18.47468 | 17.17812 | 11.52454 | 17.23721 | 19.37482 | 17.74325 |
| LSM3 | 14.12026 | 13.75902 | 15.68576 | 13.62789 | 14.02638 | 13.56597 | 16.60382 | 17.0394 | 17.08576 | 13.93278 | 17.03646 | 20.60355 | 14.93773 | 15.40803 | 14.64933 | 17.87582 | 15.01761 | 14.22149 |
| LSM4 | 17.42814 | 17.87814 | 15.24104 | 17.59538 | 17.79045 | 14.30684 | 18.58248 | 17.43557 | 14.75292 | 18.24132 | 17.70003 | 17.74388 | 19.65235 | 18.43409 | 15.36393 | 16.8839 | 17.45875 | 17.46151 |
| LSM6 | 15.63036 | 15.91355 | 16.34037 | 16.86304 | 19.88354 | 15.51726 | 19.35492 | 18.48618 | 15.16145 | 17.99102 | 17.18842 | 18.30068 | 20.08761 | 19.44359 | 16.85208 | 14.34254 | 19.63163 | 18.40554 |
| LSM8 | 15.64294 | 15.35905 | 15.23141 | 15.63307 | 13.51876 | 15.07318 | 17.39623 | 16.03726 | 12.86342 | 16.01148 | 17.71572 | 18.38528 | 14.8267 | 13.78186 | 15.21705 | 16.60555 | 16.21163 | 13.98775 |
| LSR | 21.12545 | 20.64285 | 22.13815 | 22.15239 | 20.76394 | 19.60558 | 17.21036 | 17.14965 | 22.4373 | 22.12999 | 20.33254 | 20.51881 | 20.18719 | 19.72208 | 22.15534 | 22.25727 | 18.45155 | 18.10585 |
| LSS | 15.78681 | 16.00637 | 15.76709 | 15.20503 | 16.01849 | 16.87514 | 18.16936 | 16.72929 | 17.0089 | 17.10413 | 15.73183 | 17.82033 | 15.59096 | 15.43428 | 15.63626 | 16.21417 | 16.16895 | 15.50422 |
| LTA4H | 18.10854 | 19.93799 | 16.29505 | 16.63006 | 19.44884 | 18.16574 | 16.83247 | 16.61805 | 17.27296 | 17.34183 | 19.64872 | 20.3252 | 17.57899 | 16.98198 | 17.47928 | 17.31484 | 18.74548 | 18.17147 |
| LTBP1 | 16.52268 | 16.43373 | 14.84492 | 15.62195 | 14.48811 | 15.56996 | 17.23314 | 17.48057 | 14.15146 | 15.7267 | 15.63412 | 12.17405 | 13.94277 | 15.78069 | 14.83054 | 15.02841 | 19.28647 | 20.80957 |
| LTBP2 | 16.45274 | 15.98678 | 15.309 | 17.48629 | 16.29803 | 13.5353 | 16.77043 | 14.82386 | 16.73508 | 15.41178 | 15.02614 | 12.77167 | 14.59002 | 13.3614 | 14.95109 | 17.3823 | 15.07229 | 12.38843 |
| LTV1 | 18.41309 | 18.25192 | 18.18096 | 18.42007 | 13.11465 | 16.88443 | 15.9354 | 16.50652 | 12.97548 | 15.90294 | 15.83999 | 13.25122 | 17.24101 | 13.44162 | 16.8252 | 18.10125 | 16.29224 | 15.26434 |
| LUC7L2 | 16.92226 | 17.73172 | 16.35441 | 15.77672 | 17.91419 | 15.67427 | 17.25024 | 16.63157 | 17.86457 | 18.55906 | 20.08052 | 17.47379 | 18.46075 | 17.09898 | 15.23387 | 16.8446 | 18.92122 | 17.2286 |
| LUZP1 | 15.97113 | 15.40683 | 15.26874 | 13.96056 | 18.44503 | 16.47152 | 15.4418 | 17.05977 | 16.89056 | 14.67085 | 15.9721 | 12.89218 | 16.86076 | 15.96673 | 15.13765 | 15.80723 | 17.87398 | 17.09134 |
| LXN | 17.45759 | 15.83439 | 14.9543 | 15.86207 | 17.89276 | 18.56836 | 18.50612 | 18.48623 | 12.87061 | 14.18749 | 17.28373 | 14.32046 | 15.51305 | 15.20806 | 18.32628 | 15.67612 | 20.97469 | 21.82795 |
| LY6D | 16.5606 | 16.03651 | 16.15149 | 16.26305 | 12.96889 | 18.22146 | 22.57711 | 22.26809 | 13.43402 | 15.04325 | 15.46639 | 12.60144 | 17.56016 | 16.36263 | 14.93858 | 15.38524 | 15.22229 | 14.29301 |
| LYN | 18.70695 | 18.69042 | 18.41419 | 18.75787 | 20.27517 | 21.12273 | 19.63352 | 18.30473 | 19.35663 | 18.80597 | 16.71871 | 15.65933 | 18.35619 | 20.07399 | 19.57338 | 19.79179 | 17.22116 | 19.77613 |
| LYPLA1 | 18.25462 | 18.25683 | 18.57794 | 17.48816 | 18.887 | 19.18663 | 17.80819 | 18.19335 | 18.7301 | 18.93599 | 19.62666 | 20.87951 | 18.6753 | 19.19777 | 18.38122 | 18.14159 | 17.02811 | 16.35193 |
| LYPLA2 | 18.45548 | 17.81583 | 17.12068 | 17.27753 | 18.37139 | 17.0798 | 20.14171 | 17.21498 | 16.28367 | 16.46597 | 18.4069 | 20.08157 | 16.2084 | 18.1122 | 18.67215 | 17.77737 | 19.8143 | 19.73788 |
| LYPLAL1 | 15.96042 | 13.96173 | 17.48122 | 18.26594 | 19.72449 | 17.6954 | 13.52571 | 15.53905 | 16.84715 | 15.8664 | 16.6283 | 11.62442 | 19.83319 | 19.52162 | 18.86319 | 18.37293 | 18.15175 | 18.10342 |
| LYRM7 | 15.05371 | 14.53649 | 15.43648 | 14.98563 | 13.18493 | 15.11817 | 14.65733 | 15.70538 | 19.04738 | 17.55949 | 16.89546 | 14.29994 | 15.99271 | 17.08205 | 16.76987 | 16.26666 | 16.953 | 14.86859 |
| LZTFL1 | 17.48323 | 14.91663 | 15.59913 | 15.84287 | 17.32385 | 17.39173 | 16.12161 | 16.51347 | 17.53687 | 17.86439 | 14.43002 | 16.61017 | 15.68225 | 17.78991 | 14.98332 | 14.43483 | 16.45376 | 18.39282 |
| LZTS1 | 14.70238 | 16.67451 | 15.95401 | 17.12877 | 14.05298 | 15.53582 | 14.95431 | 14.84169 | 16.76736 | 14.73605 | 15.94915 | 17.08682 | 16.25116 | 14.67464 | 16.5767 | 13.67859 | 16.6351 | 15.14073 |
| M6PR | 19.59947 | 19.12441 | 19.78865 | 19.18591 | 16.97641 | 17.09339 | 19.65867 | 19.94393 | 22.73295 | 21.17428 | 20.26266 | 21.51082 | 20.24015 | 20.25988 | 19.08584 | 19.29381 | 16.78205 | 15.76504 |
| MACF1 | 15.98337 | 15.37639 | 16.33585 | 16.87684 | 17.7667 | 17.99267 | 16.68272 | 17.38485 | 16.67141 | 17.45279 | 16.57078 | 16.96214 | 16.35981 | 17.98185 | 15.84358 | 15.86219 | 17.98252 | 17.30216 |
| MACROD1 | 19.03294 | 25.32082 | 19.14178 | 18.46354 | 20.59653 | 20.20978 | 21.33082 | 15.69182 | 20.51401 | 18.43561 | 18.99503 | 16.72445 | 17.65884 | 17.74715 | 14.70226 | 17.08326 | 12.52007 | 19.69315 |
| MACROH2. | 22.66949 | 21.18836 | 19.28268 | 18.21311 | 15.14616 | 19.64671 | 19.87518 | 21.92197 | 14.94365 | 16.29641 | 16.30968 | 20.17196 | 17.22188 | 19.04048 | 17.41777 | 16.86905 | 17.0034 | 19.43301 |
| MACROH2. | 22.28414 | 21.80431 | 17.44367 | 16.54632 | 17.85099 | 18.32232 | 16.08166 | 19.58397 | 14.25028 | 15.03324 | 16.69801 | 22.43244 | 17.19187 | 15.55373 | 14.91337 | 15.17116 | 14.51221 | 19.03687 |
| MAGEA4 | 19.32232 | 19.86417 | 15.61685 | 15.59364 | 11.69692 | 15.10473 | 14.88477 | 16.51856 | 15.35514 | 15.1693 | 15.06944 | 15.37261 | 15.07432 | 15.34748 | 16.15067 | 18.01832 | 15.6525 | 16.34527 |
| MAGED2 | 17.34527 | 16.32263 | 16.38321 | 17.253 | 17.48189 | 17.60982 | 18.43719 | 18.67503 | 17.15943 | 17.96861 | 17.50606 | 15.18239 | 18.1674 | 18.27282 | 16.00237 | 16.19812 | 18.09358 | 19.20111 |
| MAGI1 | 14.74492 | 14.67088 | 16.14432 | 17.12893 | 15.25774 | 15.82657 | 15.40371 | 14.56659 | 16.27395 | 18.66798 | 17.00248 | 16.2703 | 17.97545 | 18.10089 | 16.89923 | 17.86889 | 17.07082 | 16.61201 |
| MAGOH | 19.35386 | 19.73996 | 17.83454 | 19.29034 | 19.97026 | 19.32315 | 17.47544 | 16.68744 | 14.39507 | 19.27501 | 16.99359 | 18.57177 | 19.67838 | 19.60203 | 18.46927 | 18.00642 | 18.33333 | 19.24273 |
| MAGT1 | 15.75187 | 16.19872 | 15.69267 | 17.45145 | 16.87697 | 16.41861 | 18.12925 | 17.52377 | 22.14635 | 19.47346 | 17.23486 | 17.64275 | 16.82587 | 14.84327 | 15.90449 | 16.32755 | 16.67336 | 15.07263 |
| MAIP1 | 16.28125 | 12.60241 | 14.40681 | 15.79436 | 14.12005 | 14.47563 | 17.65929 | 12.56168 | 18.26025 | 12.96989 | 12.93418 | 12.57357 | 15.49476 | 20.38703 | 11.81783 | 13.13633 | 14.89033 | 14.43179 |
| MAL2 | 22.27204 | 22.74894 | 21.09724 | 20.25676 | 15.6636 | 17.58567 | 17.71606 | 15.14852 | 24.21746 | 23.33245 | 17.49158 | 18.51479 | 16.38694 | 15.60366 | 22.01112 | 21.79758 | 15.37842 | 15.03772 |
| MAN1A1 | 24.10683 | 24.25724 | 16.34537 | 23.329 | 15.14233 | 16.46606 | 15.97905 | 17.32954 | 19.3662 | 18.0242 | 19.6513 | 19.00724 | 15.11576 | 17.10687 | 17.9471 | 16.32393 | 15.85275 | 15.49507 |
| MAN1A2 | 17.95143 | 16.30991 | 15.65392 | 20.64412 | 17.31514 | 18.61491 | 17.37942 | 16.70049 | 20.53762 | 17.95894 | 16.00946 | 16.8052 | 17.43774 | 16.57219 | 18.34415 | 16.14938 | 17.08933 | 17.96857 |
| MAN1B1 | 15.40028 | 16.87925 | 16.96034 | 15.05707 | 14.59241 | 15.51734 | 19.21174 | 16.77992 | 20.39522 | 18.65265 | 15.49848 | 15.03961 | 16.50107 | 16.5499 | 15.83081 | 16.06672 | 17.81652 | 15.30909 |

| | | | | | | | | | | | | | | | | | | |
|---|---|---|---|---|---|---|---|---|---|---|---|---|---|---|---|---|---|---|
| MAN1C1 | 15.65889 | 15.4118 | 15.90122 | 15.83656 | 14.85568 | 17.09992 | 16.21343 | 15.93885 | 18.54551 | 15.9255 | 14.76445 | 13.0061 | 14.66786 | 14.68538 | 15.44483 | 16.72236 | 15.87423 | 16.86647 |
| MAN2A1 | 16.03626 | 16.51491 | 16.58307 | 16.03718 | 16.04145 | 17.44171 | 16.92332 | 16.20754 | 19.19631 | 18.7115 | 17.84653 | 18.14018 | 15.81509 | 15.98544 | 16.67691 | 16.22041 | 16.33387 | 17.22241 |
| MAN2B1 | 16.84504 | 19.71258 | 17.55702 | 17.13873 | 15.70628 | 14.88955 | 15.23321 | 15.51012 | 19.34658 | 18.09107 | 18.56985 | 15.27602 | 15.78789 | 16.65388 | 17.15771 | 16.61331 | 13.94353 | 16.94583 |
| MANBA | 15.94303 | 15.98106 | 17.82904 | 16.9556 | 14.42308 | 16.76853 | 15.34926 | 15.8311 | 17.55312 | 18.65984 | 15.84764 | 13.94317 | 17.21257 | 16.30582 | 14.78948 | 14.01193 | 15.489 | 15.31979 |
| MANEA | 16.31922 | 17.053 | 16.42422 | 19.07623 | 15.00225 | 18.2397 | 17.91536 | 16.93418 | 21.41439 | 18.41274 | 18.09151 | 16.52038 | 16.02578 | 16.66061 | 15.31093 | 19.84541 | 16.18051 | 16.43947 |
| MANEAL | 15.13407 | 16.95514 | 16.05888 | 16.1354 | 15.55388 | 15.34455 | 16.34616 | 17.18037 | 19.42858 | 14.37162 | 15.01163 | 13.82878 | 14.22531 | 15.30966 | 15.87921 | 16.17046 | 16.21352 | 18.19804 |
| MANF | 17.69963 | 16.50236 | 17.4426 | 17.18274 | 18.62731 | 18.25221 | 19.6403 | 19.17589 | 23.85471 | 23.48576 | 22.28061 | 18.91488 | 19.07463 | 19.34584 | 17.24583 | 17.67049 | 17.46498 | 19.54495 |
| MANSC1 | 17.68268 | 14.13991 | 16.9058 | 15.72416 | 14.82096 | 13.7746 | 15.19265 | 15.73621 | 17.33237 | 18.63618 | 15.42759 | 14.18355 | 15.04069 | 16.39414 | 14.46266 | 15.39588 | 16.21226 | 13.96147 |
| MAOA | 15.33776 | 15.7794 | 20.2038 | 15.2502 | 15.33969 | 14.53412 | 15.52752 | 15.88988 | 14.32485 | 15.06992 | 14.384 | 12.67017 | 15.68167 | 17.91436 | 20.53766 | 18.85356 | 14.87517 | 14.69671 |
| MAP1B | 15.32945 | 16.51626 | 17.94283 | 15.50734 | 15.7616 | 16.46344 | 20.1705 | 18.74049 | 16.79354 | 18.3297 | 18.59655 | 17.98006 | 18.84233 | 18.88095 | 16.3984 | 15.60541 | 17.90596 | 18.38173 |
| MAP2K1 | 17.56215 | 15.45919 | 17.85513 | 18.09161 | 18.3853 | 17.03792 | 15.39604 | 15.98489 | 16.46459 | 16.28999 | 18.05583 | 18.43857 | 18.26523 | 17.84904 | 18.63307 | 18.56367 | 16.67875 | 17.55174 |
| MAP2K2 | 14.99532 | 15.87702 | 16.0833 | 17.60943 | 18.043 | 16.52902 | 19.41349 | 17.21007 | 16.33362 | 16.94427 | 16.82402 | 15.89389 | 16.75961 | 15.83151 | 17.65956 | 16.54543 | 18.047 | 17.4707 |
| MAP2K7 | 17.90478 | 18.3029 | 14.90861 | 15.77749 | 16.69182 | 15.42178 | 12.76774 | 16.04036 | 16.04423 | 17.89428 | 13.61785 | 15.79285 | 16.86643 | 18.54897 | 15.9133 | 17.00012 | 19.17748 | 15.85384 |
| MAP4 | 16.18597 | 16.55267 | 16.15206 | 14.98831 | 17.41087 | 16.16469 | 15.82753 | 17.42267 | 17.0076 | 16.14019 | 17.6974 | 18.25695 | 17.84878 | 16.96088 | 15.9465 | 15.40625 | 17.83573 | 18.5542 |
| MAP4K3 | 17.2634 | 22.33906 | 14.38892 | 16.3552 | 15.637 | 15.83574 | 16.07939 | 15.76466 | 11.58349 | 13.53955 | 13.62131 | 17.80287 | 12.65042 | 15.77134 | 15.5107 | 12.03204 | 16.3447 | 14.17975 |
| MAP4K4 | 16.19834 | 16.35144 | 17.34377 | 17.78836 | 17.46696 | 17.59442 | 16.5546 | 16.33957 | 17.75598 | 17.15149 | 18.99553 | 18.79387 | 17.01525 | 17.10852 | 17.86026 | 16.87526 | 18.97358 | 20.09137 |
| MAP4K5 | 18.0745 | 15.74695 | 16.58021 | 17.7704 | 18.12114 | 17.05428 | 16.37638 | 15.56166 | 16.41551 | 16.33088 | 17.81157 | 19.57196 | 14.8161 | 17.94669 | 15.81064 | 14.85355 | 15.66773 | 16.07569 |
| MAPK1 | 18.60104 | 18.14367 | 18.72936 | 19.18556 | 19.75532 | 17.62967 | 20.01779 | 17.27879 | 17.78504 | 18.6032 | 17.82043 | 18.57163 | 20.23765 | 19.76124 | 19.27193 | 19.03468 | 19.55929 | 19.79757 |
| MAPK13 | 17.28537 | 20.05503 | 15.46163 | 16.4684 | 16.60844 | 16.75839 | 17.28427 | 17.52205 | 18.14665 | 18.41916 | 16.82619 | 18.89108 | 16.62723 | 15.47758 | 15.563 | 15.35556 | 16.60761 | 16.83132 |
| MAPK14 | 18.65338 | 19.4597 | 17.45849 | 17.99832 | 18.77187 | 15.73182 | 16.46772 | 16.54022 | 16.89592 | 14.93687 | 15.58709 | 17.16985 | 18.26898 | 18.77921 | 18.26268 | 17.27693 | 19.3245 | 18.84668 |
| MAPK3 | 18.3002 | 18.45796 | 18.1165 | 18.92653 | 15.2153 | 17.49766 | 16.25375 | 17.30727 | 17.07643 | 16.79617 | 15.10203 | 18.08586 | 14.80618 | 16.25799 | 19.02985 | 18.75812 | 14.02471 | 16.21981 |
| MAPKAP1 | 15.73851 | 14.52336 | 16.44977 | 17.13803 | 17.61891 | 14.67967 | 15.92699 | 15.98718 | 12.09146 | 13.99471 | 16.7674 | 18.00551 | 17.39025 | 15.98257 | 18.17446 | 17.94286 | 15.16969 | 14.22092 |
| MAPKAPK2 | 16.89082 | 16.03452 | 14.40275 | 14.82465 | 13.28659 | 14.72642 | 14.94602 | 15.14303 | 15.94923 | 16.90653 | 15.09209 | 18.97825 | 14.71744 | 13.2806 | 14.46086 | 16.55871 | 15.41079 | 13.46062 |
| MAPRE1 | 19.16542 | 18.43596 | 18.7318 | 18.11172 | 18.94694 | 17.48463 | 18.69787 | 16.18084 | 17.38763 | 17.87139 | 20.1906 | 20.47627 | 19.11913 | 17.91752 | 18.35898 | 18.38864 | 19.42696 | 19.13283 |
| MAPRE2 | 17.7503 | 19.12782 | 21.89267 | 20.68015 | 17.51044 | 20.60088 | 20.82006 | 20.11774 | 20.07592 | 18.78809 | 16.89621 | 19.35502 | 18.75153 | 20.8326 | 19.56782 | 20.93027 | 16.9493 | 19.48871 |
| MAPRE3 | 15.04503 | 15.14652 | 15.11704 | 14.32662 | 16.62152 | 17.27494 | 20.09816 | 16.78401 | 15.35461 | 17.20116 | 17.08667 | 17.95013 | 16.47938 | 18.44262 | 13.17826 | 15.80763 | 13.01685 | 17.56914 |
| MARCKS | 20.67823 | 20.31533 | 21.45645 | 22.16945 | 18.23827 | 18.27452 | 21.04815 | 21.73927 | 24.03463 | 22.98338 | 22.86643 | 22.57607 | 21.43703 | 20.72643 | 21.82644 | 21.64775 | 22.90576 | 23.16976 |
| MARCKSL1 | 21.3864 | 20.23789 | 23.14192 | 23.85272 | 21.35155 | 19.96489 | 21.30294 | 22.88986 | 22.9684 | 23.5684 | 24.67114 | 24.13553 | 24.39333 | 24.79661 | 23.5711 | 23.67405 | 25.11626 | 26.22023 |
| MARK2 | 18.44454 | 16.41741 | 18.21927 | 17.14303 | 16.30045 | 16.6489 | 16.41782 | 16.38857 | 15.26533 | 16.84241 | 17.98275 | 18.17277 | 18.64498 | 17.30202 | 18.68739 | 19.11425 | 17.51854 | 16.6946 |
| MARK3 | 18.25133 | 19.35909 | 17.19899 | 17.91404 | 17.58425 | 16.79199 | 17.16969 | 18.28266 | 16.13469 | 16.83325 | 16.35803 | 17.01471 | 17.74189 | 17.49064 | 17.99368 | 18.80809 | 15.9439 | 15.48506 |
| MARS1 | 16.82985 | 16.71004 | 18.12896 | 17.22502 | 15.83665 | 15.87754 | 16.15157 | 16.55738 | 16.59548 | 15.179 | 17.58349 | 19.84414 | 16.30475 | 17.09379 | 16.80873 | 16.72139 | 16.24503 | 15.69109 |
| MARVELD2 | 19.09461 | 17.60729 | 17.70894 | 19.05575 | 16.27541 | 15.57047 | 15.00816 | 17.21121 | 15.42273 | 14.31831 | 18.2063 | 17.2636 | 16.3481 | 15.76385 | 20.37576 | 20.02869 | 16.03356 | 16.79151 |
| MARVELD3 | 22.04147 | 18.80181 | 20.72318 | 18.65154 | 15.10956 | 15.42588 | 14.65545 | 17.36829 | 13.90102 | 15.63103 | 20.96463 | 19.04623 | 16.18576 | 17.13371 | 20.72588 | 20.89778 | 16.82227 | 14.2305 |
| MASP1 | 18.27029 | 18.27032 | 21.09163 | 15.08714 | 15.18976 | 16.33328 | 16.09561 | 16.28769 | 15.78625 | 12.94893 | 17.92777 | 17.75549 | 14.69142 | 15.59967 | 20.09639 | 19.38239 | 13.98275 | 13.65937 |
| MAT2A | 17.82465 | 18.29951 | 17.90199 | 17.5555 | 17.70197 | 15.80413 | 18.57661 | 16.79136 | 17.51929 | 15.69897 | 17.82003 | 20.23166 | 17.13363 | 15.95939 | 17.5703 | 18.57452 | 18.19966 | 18.92171 |
| MAT2B | 15.55647 | 15.42511 | 17.05066 | 15.88934 | 17.65908 | 17.57037 | 15.31336 | 17.65966 | 15.87901 | 16.6014 | 18.3619 | 18.99661 | 18.13878 | 16.77598 | 16.6086 | 15.50544 | 17.65647 | 18.36001 |
| MATN2 | 15.19943 | 16.19028 | 16.46063 | 16.57178 | 14.44393 | 16.56376 | 17.74304 | 17.30297 | 19.0873 | 18.2877 | 16.30794 | 14.09594 | 16.09934 | 16.12511 | 15.53317 | 15.43496 | 16.36322 | 16.62627 |
| MATR3 | 15.44505 | 15.91984 | 16.94262 | 16.37374 | 14.06306 | 15.95179 | 15.45189 | 18.95678 | 16.71806 | 17.12576 | 18.55227 | 20.15058 | 19.64416 | 18.5201 | 14.63768 | 16.34095 | 16.40432 | 17.89384 |
| MBLAC2 | 17.61581 | 16.44589 | 20.00183 | 19.30357 | 16.60105 | 17.15022 | 16.0077 | 17.54607 | 16.12801 | 15.98316 | 19.16736 | 19.66007 | 16.64668 | 17.22698 | 19.51324 | 19.69994 | 17.33761 | 16.46611 |
| MBNL1 | 21.51103 | 20.84917 | 18.8615 | 20.77459 | 18.59844 | 17.48283 | 17.52282 | 17.51092 | 14.87819 | 15.54557 | 16.79565 | 16.98998 | 20.66425 | 18.75571 | 18.85616 | 19.45017 | 15.95953 | 18.21084 |
| MBP | 16.35711 | 18.08843 | 17.10395 | 19.07313 | 19.84701 | 21.62161 | 18.32055 | 15.50227 | 15.94476 | 14.98066 | 16.97756 | 16.31232 | 20.51107 | 19.96695 | 18.78702 | 18.46259 | 18.89126 | 19.88092 |
| MBTPS1 | 18.23897 | 15.59284 | 15.23071 | 18.31617 | 15.48789 | 20.02074 | 15.61037 | 15.76794 | 18.10872 | 16.29589 | 16.36291 | 13.6959 | 14.03389 | 15.43078 | 14.99688 | 15.50154 | 14.63187 | 16.58866 |
| MCAM | 16.70074 | 15.99438 | 16.12469 | 18.6265 | 18.85424 | 17.24337 | 16.90095 | 17.59624 | 17.6943 | 17.58526 | 16.05638 | 17.23161 | 16.3299 | 16.52984 | 16.2244 | 16.23698 | 19.53237 | 21.13986 |
| MCCC1 | 15.92747 | 16.39645 | 16.61599 | 16.78532 | 15.60374 | 16.82101 | 16.99778 | 16.14531 | 18.54906 | 15.48499 | 15.98744 | 12.342 | 14.66814 | 15.21647 | 15.28089 | 15.002 | 13.99703 | 14.7205 |
| MCEE | 15.8092 | 15.75293 | 16.42538 | 14.10444 | 13.06502 | 16.59071 | 13.57209 | 16.21604 | 12.6835 | 16.06795 | 15.86437 | 16.90348 | 14.74147 | 13.43411 | 19.73017 | 14.16171 | 12.46327 | 15.54288 |
| MCFD2 | 19.19751 | 15.86649 | 16.34436 | 16.13557 | 14.54724 | 16.87359 | 15.95819 | 17.6585 | 20.82301 | 17.37322 | 16.81615 | 14.98327 | 14.76027 | 14.89538 | 16.96077 | 16.68707 | 16.19443 | 15.3512 |
| MCM2 | 14.60129 | 17.2211 | 15.29671 | 14.8907 | 17.23016 | 16.57059 | 17.32953 | 16.1236 | 15.81778 | 17.04092 | 17.64579 | 18.47237 | 16.95959 | 16.4403 | 15.85291 | 17.18212 | 16.233 | 16.03042 |
| MCM3 | 17.19619 | 17.23361 | 16.39629 | 17.80492 | 18.57138 | 17.07303 | 18.91212 | 18.30938 | 16.31309 | 17.36749 | 17.13113 | 18.02321 | 16.98037 | 14.85075 | 15.45197 | 15.11007 | 15.5862 | 15.10918 |
| MCM4 | 16.67049 | 16.01175 | 16.19825 | 16.58829 | 18.04583 | 16.23376 | 17.86938 | 16.72244 | 16.7891 | 16.4523 | 17.91711 | 18.19086 | 17.59997 | 15.52373 | 15.20724 | 16.24473 | 16.59485 | 15.49147 |
| MCM5 | 15.95869 | 17.89298 | 16.97452 | 16.13352 | 16.70026 | 15.98509 | 16.40968 | 17.05841 | 16.34432 | 15.69116 | 17.59613 | 17.9824 | 15.69462 | 15.10063 | 17.01692 | 15.42701 | 16.80853 | 15.65613 |
| MCM6 | 16.85154 | 15.91521 | 16.99536 | 16.50558 | 17.72214 | 17.26596 | 17.72706 | 17.81324 | 16.46299 | 17.46881 | 18.05025 | 18.94696 | 18.11815 | 15.42272 | 15.10005 | 15.22958 | 17.1471 | 16.39324 |
| MCM7 | 18.71372 | 15.59143 | 16.40949 | 16.31794 | 18.09645 | 17.22686 | 19.449 | 16.92666 | 17.42095 | 17.35172 | 17.96998 | 18.14544 | 15.25949 | 15.02592 | 15.13283 | 17.86894 | 16.98475 | 17.10075 |
| MCRIP2 | 19.53583 | 16.66869 | 15.95004 | 12.03163 | 11.8592 | 16.18823 | 16.49922 | 14.20231 | 12.7629 | 13.90203 | 15.75603 | 13.36701 | 13.40258 | 13.95407 | 18.10754 | 15.41456 | 13.81247 | 14.77144 |
| MCTS1 | 18.03222 | 17.39648 | 18.97894 | 19.07143 | 17.848 | 16.91819 | 18.46673 | 18.11126 | 17.58749 | 17.80898 | 17.6823 | 18.93123 | 19.37316 | 19.36671 | 18.76691 | 18.51577 | 19.14286 | 19.50243 |
| MCU | 14.59778 | 15.28596 | 13.68933 | 15.81966 | 14.17037 | 15.7736 | 15.86097 | 14.91438 | 18.15557 | 14.97552 | 12.9205 | 11.80721 | 13.11831 | 16.25251 | 14.6681 | 13.4376 | 14.75621 | 15.12735 |
| MDH1 | 19.4005 | 19.2433 | 18.41688 | 19.0371 | 22.5846 | 20.9978 | 21.84775 | 21.04212 | 19.98346 | 20.71064 | 21.85398 | 22.33395 | 21.78093 | 21.93942 | 19.11938 | 19.03185 | 22.693 | 22.40373 |
| MDH2 | 20.93185 | 21.67849 | 18.21504 | 18.0117 | 19.91015 | 18.71635 | 22.51016 | 19.37808 | 23.08482 | 22.94273 | 19.68062 | 18.02694 | 20.48085 | 21.53832 | 21.94654 | 21.95767 | 19.06883 | 19.84196 |
| MDK | 17.06863 | 16.96071 | 18.55713 | 17.29658 | 17.28492 | 20.54819 | 16.11958 | 16.11412 | 21.60991 | 20.59288 | 15.95562 | 14.431 | 16.06451 | 15.11949 | 17.49311 | 16.93607 | 17.10174 | 16.99 |
| ME1 | 16.36428 | 16.20888 | 15.7472 | 17.74031 | 18.42417 | 19.50605 | 16.97479 | 17.11825 | 15.06879 | 16.11267 | 17.34365 | 19.24287 | 17.13791 | 17.22868 | 17.26172 | 17.44724 | 18.45692 | 18.02601 |
| ME2 | 17.27322 | 15.93088 | 15.67787 | 15.33431 | 15.27098 | 14.8787 | 16.55997 | 16.72584 | 19.12802 | 15.94397 | 13.90259 | 14.57867 | 15.58356 | 15.93547 | 14.54079 | 15.54068 | 14.51738 | 15.27059 |
| MEAK7 | 17.18069 | 17.05128 | 17.02958 | 17.41523 | 22.42751 | 19.77598 | 15.54 | 16.56365 | 19.58333 | 19.33817 | 16.70544 | 18.02324 | 17.48927 | 18.78327 | 18.31046 | 18.22611 | 22.08153 | 21.92593 |
| MEGF8 | 15.43887 | 14.32592 | 17.1339 | 15.39732 | 16.12049 | 15.58049 | 14.89884 | 15.55039 | 18.11421 | 16.49526 | 15.91758 | 15.42788 | 14.35362 | 14.12845 | 16.34344 | 16.20092 | 16.14228 | 15.93146 |
| MEGF9 | 15.1945 | 15.10302 | 15.60093 | 13.38937 | 17.36607 | 17.01184 | 17.44128 | 16.56825 | 19.13678 | 18.21483 | 16.59584 | 16.13579 | 14.40522 | 15.19139 | 15.40583 | 15.41173 | 15.30751 | 14.53727 |
| MELTF | 17.21336 | 16.95471 | 16.42638 | 15.59809 | 20.56201 | 20.27587 | 16.67133 | 17.19973 | 19.92215 | 18.86491 | 17.89414 | 18.18559 | 16.57632 | 15.7695 | 15.21084 | 17.12756 | 18.7136 | 18.15437 |
| MEMO1 | 18.09831 | 18.59787 | 16.42633 | 18.13809 | 16.16131 | 17.6517 | 16.34833 | 16.8055 | 14.02754 | 16.49192 | 16.0282 | 17.37558 | 17.16339 | 16.08674 | 17.80322 | 17.12571 | 17.8534 | 17.76995 |
| MESD | 18.76525 | 18.15848 | 17.88159 | 18.5737 | 14.67258 | 15.542 | 14.68592 | 15.90899 | 20.2384 | 15.62219 | 16.51382 | 15.07848 | 13.98175 | 14.72644 | 16.58246 | 18.78839 | 15.15302 | 16.30041 |
| MET | 17.74971 | 16.88317 | 16.1306 | 19.31815 | 18.66551 | 17.10095 | 16.69965 | 17.00153 | 18.17066 | 16.75155 | 15.81385 | 15.27575 | 14.33709 | 15.05704 | 16.11407 | 16.37186 | 18.8719 | 18.99013 |
| METAP1 | 17.82422 | 15.61291 | 16.31004 | 17.6913 | 19.13682 | 13.68984 | 17.08426 | 16.77292 | 16.21786 | 11.96424 | 15.41054 | 18.56001 | 18.94481 | 18.90176 | 17.53332 | 17.84284 | 17.89534 | 17.15257 |
| METAP2 | 17.73838 | 16.42278 | 19.95055 | 16.02201 | 12.51437 | 17.63281 | 17.30256 | 16.14417 | 14.6203 | 13.66591 | 17.94029 | 19.07732 | 15.07594 | 14.39795 | 15.28443 | 15.8755 | 14.63554 | 14.33202 |
| METTL26 | 17.87362 | 18.5625 | 20.09597 | 19.17403 | 12.61134 | 17.13365 | 14.56206 | 16.90536 | 15.32024 | 16.53057 | 15.14414 | 17.84568 | 14.3589 | 14.61369 | 20.28526 | 19.55888 | 15.75537 | 15.61946 |
| MFAP3L | 17.21797 | 16.35389 | 14.29986 | 16.04451 | 16.39519 | 17.84954 | 14.45512 | 16.14175 | 15.78856 | 14.9595 | 12.87789 | 19.572 | 15.80862 | 15.70055 | 19.24017 | 18.78329 | 14.5709 | 15.52257 |
| MFAP4 | 18.06679 | 17.51621 | 18.87159 | 18.9211 | 17.07567 | 20.73155 | 19.2221 | 16.41534 | 13.34928 | 18.68767 | 16.60063 | 15.15483 | 20.10748 | 19.66041 | 20.88522 | 20.01347 | 18.23485 | 20.86002 |
| MFGE8 | 21.67309 | 20.86239 | 22.65352 | 23.51061 | 24.65763 | 24.17381 | 20.53252 | 18.17236 | 21.40302 | 20.70145 | 19.10325 | 20.73088 | 21.91487 | 20.73662 | 23.30521 | 22.68611 | 22.06579 | 22.47464 |
| MFSD6 | 15.88573 | 15.84555 | 15.45291 | 15.62148 | 14.78419 | 15.40048 | 15.18308 | 15.03691 | 18.35019 | 15.55998 | 15.50636 | 17.46462 | 14.31836 | 14.78794 | 17.09744 | 16.92905 | 16.72966 | 14.19104 |
| MGAM | 15.16807 | 17.92279 | 16.81556 | 20.53216 | 14.38168 | 21.30462 | 15.19789 | 19.02056 | 19.88432 | 13.49008 | 14.2157 | 12.70769 | 15.26366 | 15.68708 | 15.76386 | 18.50727 | 14.25964 | 13.32123 |
| MGAM2 | 15.79115 | 15.19468 | 16.72894 | 17.43624 | 15.4351 | 16.17646 | 14.73692 | 15.65894 | 14.54681 | 15.1658 | 17.88532 | 19.15733 | 17.57049 | 18.23412 | 17.5872 | 17.47647 | 15.40398 | 13.67007 |
| MGAT1 | 15.15901 | 16.32211 | 15.88283 | 15.49744 | 16.14667 | 15.63599 | 15.51906 | 16.9697 | 20.97839 | 18.47185 | 16.05457 | 14.03055 | 14.96995 | 15.09493 | 14.48129 | 16.20024 | 14.73223 | 16.30855 |
| MGAT2 | 17.0176 | 16.3562 | 16.70179 | 16.86099 | 16.42018 | 16.93034 | 17.32637 | 17.39278 | 21.43017 | 19.49598 | 17.01426 | 16.30623 | 16.62038 | 16.09405 | 16.30091 | 16.03943 | 16.63535 | 17.33945 |
| MGAT3 | 17.68813 | 16.96201 | 15.06556 | 13.07322 | 13.48618 | 15.88074 | 14.95574 | 15.90623 | 18.35141 | 14.24396 | 16.73003 | 14.3036 | 15.29554 | 13.51041 | 17.78194 | 15.34414 | 15.32982 | 14.64686 |
| MGAT4A | 16.25218 | 17.41893 | 15.49618 | 16.80128 | 15.51573 | 16.76667 | 17.44506 | 17.382 | 19.64592 | 16.95472 | 17.25746 | 16.73786 | 14.25566 | 15.8406 | 14.53922 | 17.75209 | 15.39905 | 15.42651 |
| MGAT4B | 15.61247 | 15.65684 | 17.36147 | 18.28758 | 17.10455 | 16.83123 | 17.35756 | 16.58895 | 19.54042 | 18.4886 | 15.35324 | 15.04549 | 14.54243 | 17.21962 | 17.58998 | 18.37777 | 14.961 | 19.24915 |
| MGAT5 | 15.57448 | 18.1695 | 15.98539 | 15.0229 | 19.23809 | 15.93846 | 15.0488 | 16.80205 | 19.94036 | 18.60372 | 18.26481 | 17.13241 | 15.16075 | 16.28167 | 15.39657 | 15.57027 | 14.97039 | 17.87846 |
| MGLL | 15.3843 | 14.56488 | 15.92623 | 14.49733 | 19.69678 | 15.56897 | 14.71156 | 20.756 | 13.71619 | 13.16263 | 18.24081 | 18.09451 | 19.7085 | 15.19894 | 15.58272 | 16.09081 | 19.27958 | 20.06229 |
| MGRN1 | 16.44862 | 15.77673 | 17.15222 | 16.63096 | 14.76652 | 15.68057 | 14.47207 | 15.55679 | 16.31365 | 13.1348 | 16.03168 | 15.58501 | 14.95532 | 15.94083 | 17.02215 | 17.37418 | 14.57542 | 14.83618 |
| MIA3 | 15.83442 | 16.15398 | 16.74421 | 14.80366 | 15.03106 | 16.03257 | 16.44163 | 16.39419 | 20.14763 | 19.37729 | 18.53493 | 14.27363 | 17.48525 | 16.5127 | 15.11259 | 16.04152 | 15.56916 | 15.23254 |
| MIB1 | 15.57114 | 17.34328 | 16.40972 | 14.42227 | 18.89413 | 19.5862 | 18.71369 | 15.95956 | 18.44503 | 13.61888 | 14.9051 | 15.78446 | 15.56531 | 15.8668 | 15.34624 | 16.02123 | 17.5722 | 16.90746 |

| | | | | | | | | | | | | | | | | | | |
|---|---|---|---|---|---|---|---|---|---|---|---|---|---|---|---|---|---|---|
| MICAL1 | 14.75786 | 16.73021 | 17.48832 | 18.05716 | 15.50623 | 16.15008 | 15.6785 | 17.32747 | 17.09666 | 14.92826 | 14.57974 | 13.38963 | 18.24687 | 16.42362 | 17.01892 | 16.28526 | 13.13575 | 14.80109 |
| MICAL3 | 16.33956 | 16.57816 | 18.08807 | 17.621 | 19.91704 | 19.78901 | 15.4235 | 17.53417 | 16.50033 | 14.5286 | 15.47065 | 14.39085 | 15.34542 | 16.42203 | 16.8547 | 16.50216 | 16.55791 | 15.07314 |
| MICALL1 | 17.82619 | 18.33632 | 17.31428 | 18.39247 | 19.3706 | 20.81953 | 18.34636 | 18.02822 | 15.02581 | 14.82702 | 16.03214 | 15.76361 | 19.22125 | 20.31288 | 17.26979 | 16.76997 | 17.8384 | 20.54544 |
| MICU2 | 15.35368 | 14.09782 | 16.92949 | 15.63361 | 15.22556 | 13.28718 | 13.36 | 14.01357 | 17.81411 | 17.99671 | 16.21119 | 13.81267 | 13.61483 | 17.29292 | 12.88571 | 14.69783 | 13.8719 | 14.74998 |
| MID1 | 15.72316 | 15.04501 | 16.37333 | 15.66991 | 17.5639 | 15.64806 | 16.56359 | 14.76322 | 14.13616 | 15.35256 | 17.26694 | 11.22257 | 16.88996 | 17.71596 | 15.5428 | 15.75077 | 19.77051 | 18.38586 |
| MIF | 22.93036 | 22.10599 | 22.61382 | 23.07446 | 23.10492 | 22.63793 | 20.86599 | 20.37436 | 21.57335 | 22.53826 | 23.55348 | 23.56775 | 24.31798 | 23.93373 | 23.31568 | 23.33815 | 24.60238 | 23.80729 |
| MINDY3 | 20.52515 | 20.50925 | 17.87799 | 18.6702 | 15.64317 | 15.37627 | 16.27797 | 18.08124 | 14.78969 | 15.82885 | 14.01017 | 15.68961 | 15.61903 | 15.63525 | 19.03183 | 18.76238 | 16.80299 | 15.54369 |
| MINK1 | 16.62409 | 16.78567 | 18.99889 | 19.72842 | 19.29061 | 17.03497 | 15.75058 | 16.34089 | 19.2387 | 18.53077 | 18.57321 | 19.00812 | 19.44908 | 19.76071 | 19.13281 | 19.17929 | 17.88365 | 19.46493 |
| MISP | 14.48284 | 15.32787 | 13.4241 | 15.69974 | 19.42751 | 18.3868 | 14.40524 | 15.54105 | 11.99752 | 16.09776 | 15.13639 | 15.7014 | 17.44603 | 13.61236 | 15.04356 | 15.78031 | 15.10921 | 12.95342 |
| MITD1 | 19.26638 | 19.25456 | 17.49126 | 18.03266 | 19.56913 | 19.62113 | 16.66016 | 16.72941 | 19.25927 | 18.58297 | 19.77608 | 20.70633 | 16.0927 | 15.52792 | 16.73464 | 16.85074 | 15.97451 | 15.96447 |
| MIX23 | 15.19903 | 15.45508 | 16.08059 | 14.91886 | 15.07251 | 16.51917 | 16.74476 | 16.34521 | 12.25588 | 15.87161 | 19.78052 | 18.48919 | 17.69919 | 16.73142 | 14.45474 | 14.87372 | 14.63426 | 14.80159 |
| MKRN1 | 22.16987 | 21.66766 | 18.94347 | 20.21258 | 15.25424 | 16.5453 | 16.23709 | 16.56333 | 13.99041 | 15.3168 | 15.9381 | 14.69382 | 18.18489 | 16.53889 | 19.74954 | 19.54805 | 16.38892 | 15.68434 |
| MKRN2 | 19.00239 | 18.43816 | 17.16058 | 17.96549 | 14.68552 | 14.11413 | 15.55803 | 17.21363 | 14.82352 | 15.67434 | 14.11653 | 15.36789 | 18.59815 | 17.2867 | 16.61924 | 17.29947 | 15.80608 | 14.53468 |
| MKRN3 | 20.37214 | 19.76874 | 18.69515 | 18.53633 | 13.73921 | 16.48292 | 14.89236 | 15.62357 | 13.719 | 15.01228 | 14.56958 | 13.53397 | 18.87813 | 13.83654 | 19.34009 | 17.89859 | 15.63662 | 15.32605 |
| MLEC | 17.09915 | 16.76959 | 15.80118 | 16.53066 | 16.43181 | 17.29116 | 15.55347 | 17.11141 | 21.61877 | 19.74025 | 17.67062 | 15.4961 | 15.44958 | 15.88436 | 17.37388 | 16.63315 | 15.29605 | 15.69917 |
| MLF1 | 18.16746 | 18.248 | 18.1035 | 18.26296 | 14.48066 | 13.43464 | 14.72443 | 16.70155 | 14.01543 | 14.739 | 15.52932 | 12.1322 | 18.25218 | 14.33273 | 17.64304 | 17.32051 | 14.12814 | 19.35899 |
| MLF2 | 23.12092 | 23.66524 | 21.57393 | 22.0638 | 15.61392 | 15.95606 | 16.20426 | 17.10659 | 14.7636 | 14.86614 | 14.87153 | 14.96027 | 20.25109 | 17.621 | 20.35663 | 20.67239 | 16.30328 | 15.82626 |
| MLPH | 15.72784 | 14.99065 | 14.17377 | 18.68581 | 16.73193 | 16.15179 | 13.4088 | 17.4784 | 13.96594 | 13.45764 | 15.32463 | 13.19766 | 18.21919 | 15.95255 | 13.39889 | 16.6992 | 17.7377 | 18.04997 |
| MMAB | 18.38481 | 18.13231 | 19.61649 | 18.30764 | 15.03686 | 16.08653 | 14.22631 | 16.95006 | 18.70075 | 16.3749 | 16.92092 | 15.15175 | 17.51105 | 16.261 | 15.6342 | 17.14867 | 15.95036 | 14.51639 |
| MME | 17.58049 | 17.61898 | 23.6507 | 24.37426 | 17.4571 | 18.42564 | 18.02273 | 17.39613 | 21.00847 | 19.81277 | 18.09307 | 17.77455 | 22.36203 | 21.24862 | 24.49594 | 24.02041 | 17.21102 | 18.06442 |
| MMP14 | 15.45525 | 15.58722 | 16.80469 | 16.04397 | 21.3104 | 22.15253 | 16.42378 | 15.9126 | 15.00175 | 16.07586 | 17.50608 | 13.86863 | 16.50811 | 15.266 | 16.0726 | 16.13549 | 20.22775 | 20.50171 |
| MMP15 | 19.45524 | 19.81627 | 20.25781 | 19.61524 | 14.31821 | 15.05586 | 17.28566 | 16.54964 | 15.39571 | 13.94339 | 13.61049 | 13.40675 | 17.0528 | 16.03687 | 21.3241 | 20.77868 | 14.94074 | 14.7505 |
| MMS19 | 15.72033 | 15.41566 | 15.34814 | 14.54285 | 14.92778 | 16.39083 | 15.49988 | 16.44368 | 14.81558 | 16.63952 | 15.31032 | 18.53866 | 17.16807 | 15.48241 | 14.28417 | 16.52455 | 15.6966 | 16.5072 |
| MMUT | 15.61236 | 14.74058 | 17.20692 | 15.6271 | 16.1461 | 16.43556 | 16.07858 | 16.78573 | 19.19282 | 18.384 | 15.08323 | 13.42708 | 16.56985 | 16.84514 | 16.57572 | 14.89487 | 17.25322 | 14.54953 |
| MOB1B | 18.27343 | 17.93044 | 15.87125 | 16.99417 | 20.30721 | 19.53347 | 17.08411 | 17.81314 | 16.73068 | 17.81128 | 19.31355 | 19.48596 | 19.67572 | 19.88704 | 18.27156 | 19.46795 | 20.18821 | 21.26777 |
| MOB4 | 14.56008 | 14.30975 | 17.13157 | 15.63464 | 17.11739 | 15.41224 | 14.91218 | 16.62805 | 15.70982 | 13.55602 | 15.17638 | 18.17941 | 14.01265 | 17.80239 | 16.24003 | 17.879 | 17.70546 | 16.05928 |
| MOGS | 16.68458 | 15.70945 | 17.31748 | 17.90611 | 17.20386 | 16.50113 | 19.2468 | 18.03287 | 22.51358 | 21.36231 | 18.83131 | 16.351 | 17.76023 | 19.04345 | 17.2464 | 15.757 | 16.9265 | 16.90945 |
| MON2 | 17.86652 | 16.55876 | 15.98568 | 15.5392 | 16.03796 | 17.39786 | 16.18426 | 16.6666 | 16.08474 | 17.74647 | 16.78538 | 18.09477 | 14.49823 | 14.97864 | 14.79966 | 15.52727 | 17.08493 | 14.77803 |
| MOV10 | 22.44698 | 22.44864 | 20.31022 | 20.47956 | 16.39594 | 16.37244 | 18.18018 | 17.07246 | 16.66925 | 16.99333 | 16.5261 | 16.26398 | 18.93116 | 18.20403 | 21.13641 | 21.64033 | 16.44994 | 16.05959 |
| MPC2 | 13.19439 | 16.30412 | 14.5653 | 14.15879 | 13.37239 | 17.29949 | 11.58303 | 14.28165 | 18.15372 | 13.95083 | 15.10177 | 12.27866 | 15.96991 | 11.95444 | 15.05443 | 15.44337 | 16.18746 | 14.5885 |
| MPDU1 | 18.21451 | 18.59665 | 15.75082 | 15.56776 | 14.52411 | 16.69352 | 16.51377 | 16.17754 | 20.89972 | 20.2968 | 19.74067 | 15.98737 | 17.53983 | 14.11114 | 15.59616 | 16.79782 | 15.929 | 14.54394 |
| MPDZ | 16.06845 | 16.20154 | 14.57416 | 16.07466 | 18.56905 | 16.08504 | 16.29806 | 17.97464 | 14.70037 | 15.06928 | 15.45298 | 14.98111 | 13.19763 | 13.49038 | 16.19928 | 15.69286 | 16.78361 | 17.50519 |
| MPI | 18.48701 | 20.18572 | 17.52179 | 17.67059 | 19.51861 | 17.88916 | 16.65059 | 17.40829 | 14.86021 | 17.7122 | 18.0344 | 18.448 | 19.40225 | 19.07043 | 18.12355 | 17.79224 | 18.73279 | 17.96357 |
| MPP1 | 15.90352 | 16.0575 | 15.88452 | 15.61034 | 20.13263 | 18.61393 | 16.06527 | 16.51308 | 15.91596 | 15.23794 | 15.72065 | 14.38427 | 14.40058 | 16.53096 | 14.51636 | 15.10136 | 17.53368 | 20.30397 |
| MPP2 | 16.05145 | 15.3134 | 18.19117 | 16.33803 | 15.82371 | 15.00531 | 15.45028 | 16.6617 | 15.88117 | 16.77929 | 14.33581 | 12.80792 | 14.87563 | 14.42345 | 19.449 | 18.49187 | 15.34052 | 15.70231 |
| MPP7 | 15.5639 | 15.71519 | 17.38812 | 17.34807 | 20.81548 | 18.60711 | 16.34065 | 16.72133 | 17.53216 | 18.06839 | 16.64365 | 16.14121 | 19.14416 | 19.42692 | 17.53399 | 17.14124 | 18.0976 | 18.84805 |
| MPPE1 | 15.8633 | 15.90852 | 14.90929 | 14.52907 | 14.49365 | 15.955 | 14.26302 | 15.93194 | 17.95696 | 16.43529 | 14.46402 | 13.03809 | 15.34331 | 13.8456 | 14.28734 | 16.78979 | 14.35346 | 14.47183 |
| MPRIP | 14.56744 | 14.20306 | 16.50103 | 15.58834 | 14.16514 | 15.58789 | 16.94617 | 16.55021 | 14.4777 | 15.25803 | 16.07809 | 17.42154 | 15.7347 | 16.36524 | 17.28348 | 17.28547 | 18.14637 | 15.71104 |
| MPST | 19.61526 | 18.70783 | 16.95914 | 16.92561 | 15.32187 | 16.07064 | 16.8512 | 17.11758 | 18.64756 | 17.40479 | 18.57099 | 18.9368 | 18.23877 | 17.4922 | 17.88767 | 16.77843 | 17.69644 | 17.65524 |
| MPZL1 | 20.79598 | 19.81172 | 21.5666 | 20.35548 | 22.01095 | 20.65448 | 19.74304 | 18.24997 | 19.00227 | 18.38977 | 18.26379 | 21.28054 | 20.02873 | 20.2296 | 20.67591 | 20.48264 | 20.30268 | 21.96318 |
| MPZL2 | 18.36244 | 17.13116 | 19.81293 | 18.50488 | 17.28218 | 16.56813 | 17.51816 | 16.20651 | 11.71161 | 14.92031 | 17.21239 | 18.07143 | 19.37725 | 16.55775 | 19.63513 | 19.67952 | 15.97383 | 15.1914 |
| MRE11 | 14.79968 | 17.34076 | 16.06385 | 16.07631 | 17.40549 | 18.3091 | 14.95454 | 16.27939 | 13.87272 | 16.03259 | 15.64389 | 17.985 | 12.95043 | 15.83222 | 15.64576 | 16.68105 | 16.92613 | 15.57782 |
| MRI1 | 14.76163 | 15.99342 | 15.62155 | 15.8961 | 14.22499 | 16.99051 | 16.43469 | 16.45468 | 16.44138 | 14.79528 | 17.14131 | 18.08332 | 14.47839 | 12.7275 | 15.95335 | 13.70328 | 16.1607 | 15.60071 |
| MRPL1 | 15.63792 | 15.05807 | 16.80704 | 13.21058 | 14.42925 | 14.73158 | 15.93923 | 16.71548 | 19.61352 | 17.67634 | 16.19799 | 15.86706 | 13.83463 | 13.22908 | 17.90702 | 14.04283 | 15.47872 | 14.30099 |
| MRPL11 | 14.745 | 16.53266 | 14.76739 | 12.97619 | 19.10279 | 17.41284 | 15.64141 | 15.58933 | 19.49759 | 18.77519 | 14.37422 | 15.32066 | 17.79281 | 16.82761 | 14.32138 | 14.95217 | 16.20002 | 12.72957 |
| MRPL12 | 19.8565 | 17.02368 | 13.45994 | 13.42125 | 14.40614 | 16.63714 | 18.2534 | 17.19319 | 18.9578 | 19.16332 | 16.05182 | 14.78619 | 14.48768 | 15.10237 | 16.06094 | 10.34075 | 15.70244 | 15.7868 |
| MRPL13 | 13.66419 | 15.6175 | 14.46264 | 15.16262 | 13.80211 | 15.1327 | 16.90728 | 15.99953 | 17.98243 | 16.30495 | 16.18139 | 13.07856 | 19.08687 | 16.56638 | 16.37897 | 14.92771 | 16.76518 | 15.8284 |
| MRPL15 | 15.55466 | 14.06515 | 15.15342 | 15.15105 | 15.02266 | 14.06836 | 14.65724 | 15.32171 | 19.31957 | 15.79844 | 14.72717 | 13.99162 | 13.99547 | 15.15827 | 16.11134 | 15.28565 | 15.34188 | 14.15459 |
| MRPL21 | 15.09976 | 15.6442 | 16.2184 | 16.02581 | 17.47305 | 15.68134 | 14.29716 | 14.53007 | 17.94272 | 12.22847 | 16.14188 | 11.15255 | 17.10762 | 17.98622 | 13.87247 | 16.00228 | 17.66493 | 13.45554 |
| MRPL24 | 20.3062 | 18.16548 | 14.99059 | 13.30097 | 14.18785 | 13.94633 | 13.43102 | 19.21105 | 19.15114 | 14.72703 | 15.08586 | 20.84188 | 13.37809 | 14.25495 | 13.7929 | 13.64041 | 15.47554 | 13.33497 |
| MRPL37 | 15.13802 | 18.2807 | 21.18282 | 14.92392 | 13.9171 | 14.26573 | 16.06963 | 15.63973 | 17.66666 | 13.10836 | 15.69252 | 13.32908 | 19.27285 | 15.2115 | 16.38199 | 16.75353 | 13.52989 | 14.58773 |
| MRPL39 | 13.3098 | 15.96474 | 13.24385 | 15.37509 | 18.52959 | 17.17737 | 17.97227 | 16.26711 | 18.98297 | 19.07259 | 15.77265 | 15.67524 | 18.44172 | 21.15783 | 14.16143 | 12.17955 | 18.73618 | 17.07421 |
| MRPL40 | 14.49649 | 16.37574 | 16.6313 | 14.13441 | 14.05926 | 15.071 | 13.94919 | 15.10534 | 17.24657 | 17.0233 | 15.58165 | 12.33161 | 13.79061 | 13.6287 | 14.48527 | 12.6945 | 13.53972 | 13.87098 |
| MRPL41 | 14.65711 | 14.85722 | 17.95423 | 14.5466 | 10.79501 | 15.87282 | 14.89965 | 15.90449 | 16.7233 | 17.55456 | 18.30074 | 12.04276 | 12.78125 | 16.14011 | 15.9072 | 14.84829 | 12.07075 | 14.3496 |
| MRPL43 | 14.27302 | 14.27311 | 15.50134 | 15.61537 | 14.38337 | 15.75042 | 15.34718 | 16.06998 | 17.70948 | 15.924 | 16.4068 | 13.76803 | 14.88569 | 15.71174 | 13.3473 | 13.77037 | 13.95542 | 16.03286 |
| MRPL44 | 17.24749 | 16.88885 | 14.97673 | 17.38423 | 18.05272 | 14.89808 | 19.14795 | 17.66358 | 16.78111 | 16.82635 | 16.75257 | 16.06888 | 17.82885 | 14.39581 | 17.30452 | 17.69247 | 16.04387 | 15.78575 |
| MRPL45 | 16.30206 | 13.5493 | 15.0499 | 16.11675 | 17.96828 | 19.66772 | 16.27638 | 15.19738 | 19.10535 | 14.11143 | 14.11425 | 11.59582 | 13.98838 | 14.31274 | 14.4667 | 15.35344 | 14.34586 | 13.70949 |
| MRPL46 | 14.91781 | 14.73048 | 15.22742 | 16.55688 | 15.3931 | 16.40914 | 15.17564 | 14.98223 | 17.55696 | 15.33538 | 16.01579 | 13.76663 | 13.67544 | 15.87403 | 15.2419 | 16.0653 | 16.51281 | 14.88082 |
| MRPL53 | 15.99986 | 15.6896 | 13.88019 | 13.57747 | 14.85218 | 13.52361 | 15.06354 | 15.90528 | 17.75738 | 17.41995 | 17.74268 | 14.12725 | 13.65714 | 15.55548 | 14.1278 | 15.60221 | 16.0108 | 17.07594 |
| MRPL9 | 18.47101 | 14.67576 | 13.93449 | 15.56578 | 14.03831 | 14.43509 | 12.55113 | 17.04646 | 18.49998 | 17.66877 | 14.36987 | 16.20113 | 13.75103 | 14.04005 | 14.17168 | 16.3703 | 13.29637 | 14.54811 |
| MRPS22 | 16.5823 | 15.76626 | 17.41629 | 14.91769 | 15.80678 | 15.89311 | 17.99655 | 15.58466 | 19.318 | 16.80405 | 16.29089 | 14.06437 | 17.69968 | 17.58375 | 16.01923 | 15.23862 | 14.9112 | 14.5179 |
| MRPS23 | 14.84467 | 14.18198 | 15.82426 | 14.88554 | 14.36241 | 14.82363 | 15.04396 | 15.98327 | 18.04767 | 14.35693 | 15.2585 | 14.1091 | 13.93732 | 15.19845 | 15.15505 | 15.57703 | 12.32837 | 14.56942 |
| MRPS25 | 16.29282 | 16.48965 | 14.99322 | 14.88156 | 14.81221 | 13.46704 | 15.0685 | 13.52801 | 17.6145 | 17.36142 | 13.8425 | 12.9912 | 13.36534 | 16.55669 | 14.13763 | 16.67 | 13.46207 | 12.58299 |
| MRPS26 | 15.16414 | 16.41117 | 17.00794 | 13.97558 | 14.79856 | 14.3034 | 15.12834 | 19.4183 | 17.65593 | 15.87276 | 15.87999 | 12.56951 | 15.57053 | 14.5962 | 15.7073 | 15.41315 | 15.84289 | 15.29372 |
| MRPS27 | 17.38327 | 14.99423 | 15.74847 | 15.98459 | 16.29181 | 14.75848 | 16.12772 | 15.66346 | 18.5142 | 17.01186 | 16.98622 | 15.42156 | 15.22623 | 15.46104 | 14.98195 | 15.34492 | 15.82141 | 14.60166 |
| MRPS30 | 14.75679 | 12.5014 | 17.02635 | 13.50332 | 18.55204 | 18.17215 | 14.63362 | 14.35079 | 17.67935 | 17.35312 | 14.58784 | 12.50078 | 16.9783 | 12.8667 | 14.43445 | 14.93541 | 14.00261 | 14.86042 |
| MRPS35 | 15.0362 | 15.9674 | 14.22061 | 15.37196 | 14.46945 | 15.35785 | 15.10388 | 17.36332 | 17.50606 | 12.90855 | 14.91154 | 13.80004 | 13.94344 | 15.5765 | 15.11485 | 14.95706 | 15.9013 | 13.55157 |
| MRPS7 | 16.23391 | 14.91052 | 15.87518 | 14.76657 | 13.82289 | 14.98351 | 13.12678 | 15.13414 | 19.14834 | 15.50909 | 15.97585 | 14.31613 | 15.89901 | 14.82153 | 14.24827 | 15.11222 | 14.83844 | 13.99523 |
| MSH2 | 19.57588 | 22.14957 | 14.98469 | 14.51348 | 16.97375 | 16.91475 | 20.33012 | 19.44337 | 14.89246 | 17.44435 | 15.81731 | 16.0224 | 16.82675 | 19.23597 | 17.40766 | 17.07605 | 16.74667 | 18.69315 |
| MSH6 | 14.66279 | 13.25589 | 15.9547 | 13.40003 | 12.93526 | 15.51219 | 21.5862 | 18.32406 | 13.49529 | 16.8147 | 10.43858 | 16.51518 | 12.53241 | 15.68606 | 13.7117 | 14.78187 | 13.99864 | 13.76406 |
| MSMP | 15.80863 | 16.66229 | 16.1105 | 13.88575 | 12.126 | 16.06198 | 14.83486 | 13.24759 | 14.57903 | 13.12269 | 12.24194 | 14.27025 | 16.71855 | 16.43432 | 15.46165 | 17.71995 | 21.69909 | 23.12123 |
| MSN | 18.45933 | 19.50396 | 19.05393 | 20.31996 | 24.24839 | 23.30432 | 20.51463 | 19.85866 | 16.28067 | 16.53961 | 17.01534 | 15.215 | 21.56225 | 21.88671 | 19.39546 | 19.31474 | 23.7806 | 23.65715 |
| MST1R | 15.49504 | 15.63009 | 16.60316 | 15.89746 | 18.19402 | 17.59985 | 16.99684 | 16.50116 | 17.18617 | 16.40443 | 16.22718 | 14.33412 | 16.13037 | 16.6122 | 15.23371 | 14.54318 | 16.53457 | 18.10842 |
| MT-CO2 | 15.59457 | 16.3574 | 17.05893 | 15.53942 | 15.37126 | 18.10843 | 16.86967 | 16.09876 | 20.41087 | 19.69705 | 17.76884 | 15.54477 | 14.67315 | 16.25274 | 13.37928 | 15.88103 | 15.99782 | 14.54725 |
| MT1X | 15.00395 | 15.08177 | 16.2751 | 14.5045 | 15.41791 | 16.47575 | 14.81093 | 15.59693 | 15.12161 | 14.96917 | 14.49487 | 14.31926 | 12.4448 | 16.61859 | 15.39163 | 12.32017 | 18.18924 | 15.26785 |
| MT2A | 15.56826 | 16.77669 | 17.52102 | 16.0888 | 14.65276 | 18.04666 | 15.51667 | 16.68516 | 15.28419 | 15.53313 | 15.18855 | 16.27187 | 15.90188 | 14.94125 | 16.30451 | 16.13978 | 22.44349 | 14.63577 |
| MTA2 | 17.13882 | 17.06655 | 12.64786 | 14.98589 | 15.48211 | 16.33057 | 15.30914 | 15.50817 | 19.62695 | 14.30642 | 17.81835 | 18.37586 | 15.13196 | 16.63866 | 16.54 | 14.3186 | 15.15722 | 14.64945 |
| MTAP | 16.98955 | 16.85459 | 19.33609 | 17.74912 | 19.95198 | 18.89803 | 17.66395 | 18.15733 | 17.88098 | 18.60382 | 20.14457 | 19.83169 | 19.54729 | 18.68735 | 16.47489 | 16.83981 | 19.99076 | 20.03649 |
| MTCH1 | 20.52627 | 18.64314 | 14.51943 | 15.79331 | 14.45079 | 17.67492 | 14.15894 | 16.9644 | 18.23158 | 15.90273 | 14.52712 | 13.54728 | 16.5573 | 12.89988 | 17.18636 | 15.24192 | 16.35947 | 15.16494 |
| MTCH2 | 14.16546 | 17.01296 | 14.54659 | 15.1347 | 17.66799 | 16.73546 | 16.5954 | 15.84919 | 20.01584 | 19.30705 | 17.77848 | 14.13173 | 14.92817 | 15.87708 | 15.15173 | 16.09127 | 15.45963 | 14.95123 |
| MTDH | 19.64027 | 19.21025 | 19.76216 | 17.59499 | 15.17528 | 14.59692 | 17.20649 | 17.87853 | 19.48039 | 17.86367 | 18.05009 | 16.42769 | 15.99502 | 15.46304 | 19.99326 | 19.85833 | 15.99594 | 17.10856 |
| MTHFD1 | 20.01783 | 18.4675 | 18.94543 | 19.0103 | 20.67237 | 21.77084 | 20.98763 | 19.85003 | 19.5279 | 19.77809 | 21.5353 | 20.94476 | 19.77818 | 19.18545 | 18.67683 | 18.79599 | 21.42103 | 20.98305 |
| MTHFD1L | 16.99959 | 17.9765 | 16.64696 | 16.81239 | 17.45767 | 17.21759 | 18.92756 | 18.9862 | 18.68426 | 16.28245 | 16.84939 | 17.46464 | 17.92873 | 18.26242 | 19.58032 | 16.2712 | 19.66022 | 19.06405 |
| MTHFD2 | 18.4676 | 17.88502 | 15.82484 | 16.6554 | 17.12802 | 16.13811 | 16.92785 | 17.61541 | 19.63155 | 17.83433 | 16.5851 | 16.12382 | 18.76699 | 17.64209 | 17.22074 | 17.78611 | 16.03367 | 16.22465 |
| MTMR1 | 13.76552 | 15.00725 | 15.60706 | 14.05137 | 17.4247 | 15.74153 | 14.69732 | 17.39513 | 12.27742 | 13.82164 | 15.04165 | 14.84919 | 13.14065 | 13.97334 | 16.3699 | 15.35703 | 16.59819 | 10.54603 |
| MTMR12 | 15.3047 | 15.96038 | 14.96885 | 16.44866 | 20.57301 | 18.66996 | 16.17232 | 16.40024 | 14.36031 | 14.99436 | 14.91617 | 16.63263 | 18.64093 | 16.84848 | 15.605 | 15.46471 | 17.5837 | 18.15496 |

| | | | | | | | | | | | | | | | | | | |
|---|---|---|---|---|---|---|---|---|---|---|---|---|---|---|---|---|---|---|
| MTOR | 16.58022 | 16.22486 | 14.86799 | 16.2823 | 15.93017 | 18.61532 | 15.46614 | 15.22414 | 16.653 | 15.87815 | 15.43979 | 17.05612 | 15.63168 | 18.18657 | 16.02656 | 18.46174 | 17.43062 | 15.41921 |
| MTPAP | 17.15409 | 15.74962 | 14.51312 | 14.92071 | 10.30382 | 15.98597 | 14.85647 | 18.01805 | 17.46113 | 16.63522 | 18.07404 | 13.64442 | 15.01903 | 14.99661 | 16.98462 | 15.94601 | 15.90702 | 15.6392 |
| MTPN | 17.92922 | 18.49729 | 18.75964 | 20.09665 | 20.28203 | 19.74657 | 21.51053 | 21.04183 | 17.25248 | 19.05846 | 20.58261 | 20.70956 | 20.91713 | 21.24803 | 20.19468 | 19.67788 | 21.48906 | 21.07998 |
| MTR | 16.32953 | 14.49227 | 16.13189 | 13.40882 | 14.84769 | 14.45231 | 17.07983 | 15.37952 | 16.69734 | 15.87701 | 15.33324 | 16.41261 | 13.84317 | 15.77362 | 14.95542 | 15.58458 | 15.71265 | 13.00531 |
| MTREX | 14.62258 | 15.6918 | 13.41532 | 13.09779 | 16.45359 | 15.48176 | 15.70413 | 16.41233 | 15.41095 | 15.27936 | 17.57185 | 18.31453 | 14.83001 | 17.75304 | 15.25651 | 15.25596 | 17.56713 | 13.21634 |
| MTSS1 | 16.13374 | 14.68708 | 14.63806 | 13.16315 | 14.74685 | 15.45917 | 17.17279 | 17.82013 | 16.97473 | 15.76758 | 13.36978 | 15.37449 | 16.48935 | 15.74765 | 14.43549 | 13.93293 | 14.92437 | 16.25045 |
| MTSS2 | 16.91911 | 16.17036 | 14.94024 | 15.12709 | 16.36823 | 15.77074 | 16.31956 | 17.46786 | 16.15879 | 14.9143 | 15.75905 | 18.2667 | 13.63423 | 14.45953 | 15.02751 | 14.25802 | 16.68331 | 14.43101 |
| MTTP | 13.05747 | 14.63958 | 17.67798 | 17.92625 | 15.15147 | 17.73751 | 16.35801 | 19.0687 | 18.50624 | 18.48412 | 16.2633 | 18.58846 | 18.5904 | 18.10875 | 18.59371 | 17.65537 | 14.82951 | 15.7972 |
| MUC1 | 15.56837 | 16.40375 | 17.45509 | 16.77491 | 22.78901 | 21.19073 | 15.4616 | 17.19692 | 22.03011 | 20.10985 | 15.88223 | 15.99955 | 15.60694 | 18.00688 | 14.98911 | 15.32845 | 16.36618 | 14.7766 |
| MUC13 | 21.89049 | 19.13718 | 15.64867 | 15.66081 | 15.91131 | 17.42622 | 16.73693 | 15.61133 | 19.3341 | 17.97972 | 15.23803 | 13.29256 | 14.76804 | 15.17856 | 15.0857 | 15.34013 | 16.2529 | 16.25907 |
| MUC5AC | 17.99078 | 15.72475 | 17.84973 | 16.87539 | 15.04108 | 17.18106 | 16.23569 | 16.49837 | 14.69311 | 14.66732 | 14.79475 | 13.26391 | 15.14018 | 16.11233 | 15.85008 | 17.89649 | 15.89743 | 22.55519 |
| MUC5B | 15.97357 | 16.41156 | 15.85598 | 16.72551 | 16.6142 | 17.69021 | 16.55512 | 17.77975 | 14.65912 | 15.33028 | 15.97907 | 13.66012 | 15.70221 | 17.20097 | 15.1884 | 16.03388 | 15.87941 | 21.08063 |
| MUL1 | 17.00118 | 14.54324 | 14.51434 | 18.29581 | 15.77665 | 14.15226 | 15.95297 | 14.5937 | 19.78506 | 17.14063 | 14.82537 | 13.27131 | 14.26943 | 15.35495 | 15.1909 | 16.53816 | 16.84471 | 14.97549 |
| MVB12A | 23.65503 | 23.57291 | 22.18526 | 23.04835 | 19.34765 | 19.91477 | 18.16658 | 17.95271 | 17.97489 | 20.30747 | 17.82169 | 18.59731 | 22.75411 | 21.77851 | 23.48384 | 23.23754 | 18.90854 | 20.24648 |
| MVB12B | 20.76872 | 20.57858 | 21.16599 | 21.15199 | 16.99664 | 17.78763 | 17.27392 | 17.8663 | 19.5791 | 18.95277 | 17.83197 | 16.31924 | 19.0679 | 18.10045 | 21.83802 | 21.23168 | 15.33486 | 18.0826 |
| MVD | 16.37293 | 20.43672 | 16.45563 | 16.01619 | 19.00379 | 14.82951 | 17.8421 | 18.28675 | 16.18443 | 17.52607 | 16.64749 | 18.79353 | 16.12732 | 15.24121 | 15.56862 | 14.58006 | 19.8044 | 15.58957 |
| MVK | 15.94674 | 14.62796 | 15.80174 | 14.25014 | 13.3766 | 15.0361 | 15.56 | 16.80645 | 17.86133 | 14.99817 | 15.73916 | 16.16995 | 19.75826 | 15.55134 | 14.12762 | 16.63723 | 13.6135 | 13.91872 |
| MVP | 20.57163 | 20.97116 | 18.1216 | 16.72599 | 20.62881 | 20.20688 | 18.19913 | 18.37723 | 18.7656 | 18.13312 | 17.17718 | 21.98009 | 15.968 | 16.53822 | 16.48222 | 17.36217 | 16.70153 | 18.09902 |
| MX1 | 15.93363 | 15.53764 | 15.3722 | 15.22624 | 16.96029 | 16.49944 | 16.49146 | 15.05869 | 19.90464 | 18.14323 | 15.09725 | 16.97184 | 14.53918 | 15.53712 | 16.63214 | 14.91946 | 14.85775 | 19.19815 |
| MYADM | 22.46292 | 23.09214 | 22.87702 | 22.4981 | 13.80311 | 13.03646 | 15.94853 | 18.6979 | 17.0437 | 17.23044 | 14.46432 | 15.4365 | 17.64311 | 14.81331 | 22.74907 | 23.28957 | 17.12932 | 16.29163 |
| MYCBP | 19.32363 | 19.23992 | 16.62671 | 20.38489 | 21.61032 | 19.78331 | 21.54416 | 19.91843 | 19.32273 | 19.61074 | 19.13785 | 18.94314 | 21.65258 | 20.86006 | 17.54059 | 17.85875 | 20.63426 | 20.91534 |
| MYDGF | 16.41801 | 17.02254 | 17.01314 | 17.06315 | 19.49674 | 18.12379 | 19.98733 | 20.27386 | 24.10331 | 24.38748 | 21.86642 | 18.28287 | 18.80494 | 19.9765 | 15.52816 | 16.25989 | 20.0279 | 19.48247 |
| MYG1 | 17.08358 | 18.11919 | 17.7374 | 16.23592 | 17.79325 | 16.75737 | 16.33365 | 16.94168 | 18.61184 | 17.40104 | 18.04351 | 19.31916 | 16.47567 | 15.30929 | 17.11631 | 16.87503 | 16.55163 | 15.73663 |
| MYH10 | 15.86032 | 16.56263 | 18.70423 | 18.22714 | 20.95069 | 19.40593 | 18.7552 | 17.71025 | 16.374 | 16.34049 | 17.10843 | 19.35638 | 19.61331 | 19.81221 | 18.91226 | 18.32688 | 18.20811 | 18.14918 |
| MYH14 | 17.28703 | 16.97593 | 19.71365 | 18.64601 | 20.27071 | 18.576 | 19.91618 | 18.83859 | 17.58345 | 19.10084 | 18.01493 | 19.75889 | 19.46478 | 20.34108 | 19.57269 | 19.00747 | 20.26684 | 18.55532 |
| MYH9 | 23.28924 | 18.98544 | 22.0568 | 22.59392 | 22.48039 | 21.40194 | 20.56411 | 19.27797 | 18.12338 | 19.77409 | 19.34067 | 20.35455 | 21.58433 | 21.53213 | 22.91456 | 22.6456 | 21.99472 | 21.48997 |
| MYL12A | 17.51732 | 16.827 | 17.99393 | 19.24394 | 20.88969 | 19.26383 | 17.83966 | 17.23964 | 17.21211 | 17.1015 | 18.95901 | 20.73387 | 19.89908 | 19.08027 | 20.28608 | 19.30828 | 20.2279 | 19.84089 |
| MYL6 | 20.91205 | 20.81822 | 20.57047 | 21.83016 | 24.97656 | 24.36972 | 23.19223 | 20.98784 | 20.93538 | 22.04977 | 21.26098 | 21.6492 | 23.93074 | 23.73612 | 21.63478 | 21.17187 | 24.59072 | 23.66381 |
| MYL6B | 19.54671 | 20.30731 | 20.32454 | 20.80537 | 23.51988 | 22.5016 | 18.84174 | 18.65153 | 20.1697 | 20.07678 | 20.56121 | 21.40717 | 21.81731 | 21.87738 | 20.63613 | 20.05932 | 21.95701 | 21.78672 |
| MYO10 | 15.33333 | 14.90291 | 22.58774 | 17.04955 | 16.39526 | 19.23529 | 18.69176 | 16.23539 | 15.63952 | 17.91229 | 15.39316 | 13.25157 | 14.72717 | 17.34556 | 18.14438 | 16.83742 | 16.38565 | 15.51592 |
| MYO18A | 16.64601 | 17.08092 | 15.16858 | 14.87614 | 16.62893 | 19.45745 | 14.61797 | 15.30111 | 12.29162 | 16.35183 | 15.06988 | 17.33682 | 16.58236 | 14.66873 | 16.2591 | 14.54384 | 14.53117 | 13.82336 |
| MYO1A | 16.58312 | 16.85942 | 15.35899 | 17.61432 | 17.69012 | 13.58871 | 19.84956 | 14.94351 | 20.21233 | 22.86322 | 14.58294 | 12.22621 | 13.86653 | 12.1809 | 12.64406 | 14.7679 | 13.40685 | 12.2353 |
| MYO1B | 17.53112 | 17.90934 | 17.37231 | 17.40091 | 22.01858 | 21.06285 | 19.90184 | 19.61344 | 18.12649 | 18.63239 | 21.07977 | 21.59666 | 18.19303 | 17.7795 | 18.91474 | 17.99913 | 21.06636 | 20.22748 |
| MYO1C | 21.25613 | 21.04058 | 21.46451 | 22.08425 | 21.65646 | 21.57412 | 18.27926 | 16.85004 | 16.96129 | 17.1195 | 16.52175 | 17.38426 | 20.16723 | 20.37567 | 22.88561 | 22.26274 | 20.69978 | 20.2122 |
| MYO1D | 22.57303 | 22.70412 | 20.87169 | 21.08394 | 17.9367 | 18.24859 | 16.28815 | 17.14034 | 17.87374 | 17.8337 | 17.8269 | 17.85097 | 18.6819 | 19.10788 | 21.88957 | 21.31587 | 18.8379 | 20.69307 |
| MYO1E | 16.3728 | 18.2347 | 16.53727 | 17.31511 | 14.75021 | 16.17506 | 15.43268 | 16.97553 | 14.44255 | 15.68345 | 15.53421 | 16.16195 | 14.86584 | 14.18539 | 17.1903 | 16.2927 | 18.20931 | 19.14118 |
| MYO5A | 17.12627 | 16.0707 | 15.67949 | 17.57217 | 18.08326 | 12.57024 | 13.71718 | 13.39005 | 15.4132 | 15.04656 | 15.57867 | 12.8214 | 13.45618 | 14.83708 | 15.06332 | 13.8523 | 16.79083 | 15.53988 |
| MYO5C | 18.91974 | 17.1599 | 15.6948 | 20.16345 | 18.13965 | 17.53671 | 18.57983 | 18.06188 | 17.32167 | 17.06703 | 15.99697 | 15.09128 | 17.60068 | 16.35665 | 19.05701 | 19.4861 | 16.82742 | 15.22463 |
| MYO6 | 17.62142 | 17.59265 | 20.63142 | 21.06103 | 17.41921 | 18.11208 | 17.34794 | 16.61331 | 17.92276 | 17.97507 | 17.73976 | 18.41383 | 19.36814 | 19.09012 | 21.53271 | 21.3059 | 16.74983 | 16.08238 |
| MYO9B | 15.65654 | 16.52555 | 15.09043 | 15.86873 | 18.26755 | 15.91465 | 14.51735 | 15.98015 | 14.15802 | 16.39952 | 16.35638 | 15.89665 | 18.01086 | 20.40839 | 14.20921 | 14.62511 | 16.68156 | 15.26772 |
| MYOF | 17.62495 | 18.08859 | 16.55289 | 17.05527 | 22.59437 | 21.28066 | 17.48494 | 16.60084 | 15.97866 | 16.66215 | 16.69505 | 16.21875 | 15.55281 | 16.22868 | 16.08607 | 17.65206 | 17.59859 | 19.53613 |
| MYORG | 16.05079 | 16.29158 | 16.09523 | 13.37198 | 13.98485 | 15.66541 | 19.23253 | 16.60176 | 16.56725 | 14.90587 | 13.90486 | 16.25399 | 14.52086 | 16.46389 | 16.56818 | 14.72966 | 14.7271 | 14.61783 |
| N/A | 25.46143 | 26.05951 | 25.03087 | 25.93959 | 23.01497 | 23.09086 | 22.15798 | 21.73766 | 23.87122 | 23.24553 | 24.18957 | 21.83986 | 23.49083 | 23.22037 | 25.09183 | 25.17749 | 22.09275 | 22.74232 |
| N4BP1 | 18.69199 | 19.1869 | 17.26204 | 17.16537 | 21.22602 | 19.84034 | 16.22459 | 19.34228 | 16.96914 | 17.75486 | 16.58877 | 16.85796 | 19.56748 | 19.37049 | 17.68427 | 17.42894 | 19.4856 | 20.36099 |
| NAA10 | 15.5323 | 19.93112 | 19.46165 | 16.87762 | 17.76576 | 16.90109 | 20.07259 | 17.3543 | 15.12779 | 14.82084 | 18.46342 | 19.83402 | 16.73469 | 16.30246 | 19.52941 | 19.50722 | 15.83602 | 15.97885 |
| NAA11 | 15.3047 | 15.78294 | 17.16758 | 16.91693 | 17.33105 | 14.95546 | 17.27542 | 17.48779 | 14.7164 | 18.06251 | 18.04407 | 20.23339 | 17.19869 | 15.77626 | 18.13549 | 16.22925 | 15.87361 | 17.51287 |
| NAA15 | 18.22553 | 16.51582 | 18.16657 | 17.77834 | 15.61054 | 17.19776 | 16.36447 | 16.63295 | 14.94861 | 14.44512 | 16.55929 | 19.23993 | 15.80207 | 16.85607 | 17.61427 | 18.04541 | 17.2413 | 15.02618 |
| NAA16 | 14.64334 | 18.77813 | 18.50531 | 19.98213 | 16.46409 | 15.36525 | 16.07046 | 17.04453 | 14.19116 | 15.20579 | 16.02609 | 18.93947 | 17.29505 | 18.20509 | 18.97288 | 18.40296 | 15.87338 | 15.2918 |
| NAA25 | 16.57195 | 15.72676 | 15.80638 | 17.31164 | 13.77759 | 14.97036 | 15.85054 | 14.8827 | 15.10321 | 15.22683 | 17.10806 | 18.13879 | 15.20526 | 16.65884 | 16.16431 | 15.17678 | 15.60597 | 17.41928 |
| NAA50 | 17.08036 | 16.82017 | 15.50148 | 16.04924 | 18.33474 | 16.85102 | 18.70924 | 18.79495 | 16.15754 | 17.43021 | 17.5752 | 20.00688 | 18.70771 | 18.23907 | 15.86004 | 16.36479 | 19.41199 | 17.88304 |
| NAALADL2 | 15.31549 | 18.26845 | 19.14078 | 19.48137 | 16.43638 | 14.62983 | 16.01373 | 16.95527 | 13.13778 | 16.39347 | 17.10957 | 15.40043 | 18.86364 | 16.73898 | 20.03766 | 18.59852 | 15.71029 | 15.7076 |
| NACA | 19.55849 | 18.95988 | 18.37595 | 18.18962 | 16.65456 | 15.43914 | 16.27339 | 16.42715 | 15.19491 | 14.98823 | 18.64071 | 20.19379 | 16.0548 | 15.76347 | 17.8391 | 17.25774 | 14.5562 | 15.52592 |
| NADK2 | 15.60331 | 16.45579 | 16.86774 | 15.30503 | 13.38796 | 17.48692 | 15.86774 | 16.57487 | 17.78826 | 16.62205 | 15.43475 | 15.58914 | 15.18077 | 15.67334 | 17.15357 | 16.65568 | 14.87761 | 16.34677 |
| NAE1 | 15.75524 | 14.89876 | 14.25593 | 15.9932 | 17.66172 | 19.01782 | 16.56923 | 17.81767 | 16.3236 | 15.05537 | 18.6626 | 18.65847 | 18.87581 | 16.54798 | 17.19208 | 14.88109 | 16.75017 | 16.81144 |
| NAGA | 15.88667 | 15.29472 | 17.41196 | 14.2987 | 14.56333 | 16.3553 | 15.74651 | 16.71421 | 18.51042 | 17.76354 | 17.36877 | 13.10789 | 16.34142 | 16.77427 | 16.17351 | 14.81489 | 14.77769 | 13.01464 |
| NAGK | 16.90019 | 16.73779 | 16.75832 | 16.23056 | 17.44321 | 16.32644 | 14.7921 | 15.51581 | 17.77835 | 16.51142 | 17.54258 | 17.83209 | 17.64402 | 17.28144 | 16.9103 | 15.87553 | 16.44353 | 18.48241 |
| NAGLU | 16.63221 | 15.31233 | 16.11685 | 15.35577 | 15.07742 | 18.11117 | 19.01238 | 18.01879 | 18.60769 | 19.40991 | 15.57645 | 12.40667 | 14.10556 | 15.43542 | 14.77697 | 14.38234 | 17.53154 | 16.11047 |
| NAGPA | 13.6669 | 15.44063 | 15.72351 | 14.35287 | 16.48593 | 14.80386 | 17.52923 | 17.09844 | 18.18345 | 18.3134 | 17.25461 | 13.83169 | 15.24243 | 15.85256 | 14.45067 | 14.73682 | 15.00594 | 15.0222 |
| NAMPT | 16.23653 | 16.25941 | 18.4266 | 18.61163 | 22.04385 | 20.37692 | 18.40139 | 16.85285 | 17.47691 | 17.89179 | 18.71084 | 19.62173 | 20.00112 | 19.40429 | 18.54862 | 18.61864 | 18.46251 | 18.84132 |
| NANS | 16.07375 | 18.2108 | 19.46018 | 18.41211 | 19.98614 | 15.91529 | 18.82651 | 17.6288 | 15.42438 | 16.91571 | 18.09819 | 18.70285 | 20.55206 | 20.0308 | 18.80935 | 18.26058 | 18.57322 | 18.54487 |
| NAP1L1 | 22.16345 | 22.33971 | 21.9083 | 22.64911 | 22.12818 | 21.04483 | 21.89011 | 20.00452 | 20.90217 | 21.35356 | 21.40058 | 22.25754 | 21.80457 | 21.93737 | 21.26914 | 21.44641 | 22.00527 | 21.59519 |
| NAP1L4 | 19.65825 | 20.11719 | 20.05846 | 19.88566 | 20.21043 | 19.46331 | 19.71574 | 18.56695 | 18.55411 | 18.69946 | 20.03507 | 21.47778 | 20.32746 | 19.98443 | 20.0115 | 19.62992 | 19.69453 | 20.48289 |
| NAPA | 20.68839 | 20.06619 | 18.6477 | 19.00638 | 19.42953 | 17.56555 | 20.82034 | 20.20781 | 18.85454 | 18.86117 | 19.02302 | 19.54508 | 20.27029 | 20.03166 | 19.55837 | 19.58625 | 20.00472 | 20.01642 |
| NAPB | 17.48713 | 17.76573 | 14.98797 | 14.92278 | 14.41256 | 16.50698 | 18.15327 | 17.94914 | 17.17827 | 17.79581 | 14.98507 | 16.00362 | 15.7775 | 15.31061 | 17.45597 | 16.8522 | 14.38365 | 17.61296 |
| NAPG | 17.02195 | 17.48715 | 19.685 | 18.74409 | 21.01658 | 18.90627 | 19.35188 | 18.88815 | 17.90497 | 18.37293 | 20.60364 | 18.76251 | 20.13368 | 20.1203 | 18.02302 | 18.18839 | 20.97331 | 21.39083 |
| NAPRT | 17.91226 | 14.73699 | 15.02337 | 16.19007 | 17.72476 | 17.21453 | 15.58687 | 17.37787 | 14.60981 | 16.88981 | 14.67562 | 14.92212 | 18.82929 | 17.8674 | 16.25687 | 16.52476 | 14.22618 | 14.91118 |
| NARS1 | 17.86111 | 17.82969 | 18.14117 | 18.81475 | 17.76292 | 19.0802 | 17.32006 | 17.78993 | 18.2936 | 17.9171 | 18.9913 | 20.85224 | 18.55432 | 18.97959 | 19.65421 | 19.40765 | 17.3275 | 17.71919 |
| NASP | 16.81254 | 17.19716 | 15.26942 | 16.96473 | 18.92878 | 17.33368 | 18.89815 | 18.76814 | 18.53232 | 19.71192 | 18.91407 | 18.73067 | 20.21233 | 17.67229 | 17.54395 | 17.04528 | 18.07808 | 18.75589 |
| NAXE | 18.79208 | 17.43639 | 18.10799 | 17.98923 | 18.6954 | 16.53506 | 18.22631 | 17.34781 | 18.60635 | 19.62277 | 19.92763 | 19.37504 | 19.06825 | 18.943 | 17.80268 | 18.65092 | 19.96231 | 19.54678 |
| NBEA | 14.73851 | 15.97684 | 15.25058 | 15.26011 | 20.01961 | 17.43361 | 19.45319 | 20.17879 | 14.0206 | 14.96543 | 15.57531 | 18.70821 | 15.47811 | 20.68162 | 19.59614 | 15.1211 | 17.52482 | 20.65521 |
| NBEAL2 | 17.32927 | 16.3385 | 16.36888 | 17.31099 | 15.12095 | 15.78031 | 15.36796 | 16.12277 | 14.03916 | 14.54096 | 16.29387 | 18.10421 | 16.49153 | 18.09921 | 17.38745 | 16.98587 | 13.83297 | 14.7927 |
| NBR1 | 14.23344 | 15.41916 | 16.13349 | 23.92156 | 15.27095 | 16.61467 | 21.22285 | 16.50661 | 17.31435 | 16.35994 | 16.41988 | 12.80399 | 16.49267 | 18.11085 | 15.39028 | 15.84301 | 16.99899 | 17.27552 |
| NCALD | 18.03493 | 17.49222 | 18.41024 | 17.29557 | 18.99077 | 18.1693 | 15.52492 | 16.21765 | 19.00171 | 16.04733 | 18.45728 | 19.09282 | 16.3197 | 16.43161 | 18.14608 | 18.48045 | 16.86621 | 18.54178 |
| NCAM1 | 16.09134 | 15.70362 | 16.49937 | 16.56321 | 14.81551 | 16.48921 | 18.33187 | 18.92821 | 19.44063 | 19.72872 | 16.0898 | 18.10901 | 15.83647 | 16.59973 | 15.54057 | 16.11493 | 15.789 | 15.42531 |
| NCAM2 | 18.24257 | 18.84578 | 21.73518 | 21.29439 | 16.84694 | 19.12287 | 17.75557 | 17.37387 | 14.18747 | 15.97174 | 16.3217 | 14.86931 | 15.69341 | 18.59572 | 22.63252 | 22.09583 | 17.07025 | 17.74458 |
| NCAPD2 | 13.20788 | 17.88846 | 15.14563 | 13.95622 | 13.77035 | 13.87407 | 13.31482 | 16.72586 | 14.01449 | 11.98719 | 12.96381 | 15.52233 | 12.85263 | 15.15074 | 14.02347 | 14.12434 | 12.23943 | 13.09403 |
| NCAPH | 15.53709 | 16.30683 | 13.82609 | 16.96871 | 16.88381 | 14.18809 | 20.116 | 19.34774 | 14.72048 | 17.86735 | 15.83913 | 17.00612 | 15.60533 | 16.76719 | 17.11499 | 16.6595 | 17.22693 | 18.24221 |
| NCBP1 | 14.98798 | 16.33673 | 18.88684 | 16.94821 | 15.65852 | 15.65932 | 16.2575 | 17.11911 | 13.59848 | 14.81949 | 15.82528 | 18.80107 | 15.08904 | 15.78613 | 17.80237 | 15.9792 | 15.81677 | 15.09147 |
| NCDN | 16.38254 | 16.92745 | 17.60262 | 17.22883 | 14.43346 | 17.07369 | 16.22044 | 17.54114 | 16.49654 | 16.22984 | 16.27243 | 18.92534 | 17.39037 | 16.84715 | 17.53926 | 18.25531 | 18.9804 | 17.40376 |
| NCEH1 | 15.24481 | 17.25917 | 16.57948 | 16.04076 | 18.45242 | 17.2329 | 16.77602 | 15.52655 | 19.76215 | 18.89615 | 16.81367 | 15.78224 | 16.31314 | 15.70979 | 16.18394 | 16.59866 | 15.13038 | 16.3904 |
| NCK1 | 18.35212 | 18.46609 | 19.7282 | 17.41262 | 17.19737 | 13.55996 | 16.95723 | 14.21149 | 15.7815 | 14.99496 | 15.40726 | 15.59136 | 16.44391 | 17.39458 | 19.18135 | 18.80203 | 14.63777 | 16.44851 |
| NCKAP1 | 17.42982 | 16.72761 | 18.13771 | 18.45801 | 17.96704 | 17.46294 | 16.89901 | 17.64173 | 17.02129 | 17.09373 | 17.07524 | 19.24911 | 18.18581 | 17.75334 | 19.28476 | 19.21868 | 18.00284 | 17.81137 |
| NCL | 17.43502 | 16.86559 | 16.93338 | 20.69582 | 18.15631 | 19.50516 | 19.00891 | 18.28774 | 18.30866 | 19.58688 | 20.91625 | 20.13078 | 19.70039 | 19.52192 | 16.26277 | 16.56767 | 18.45456 | 17.80172 |
| NCLN | 16.78185 | 17.50178 | 18.32287 | 17.39856 | 16.28473 | 16.47082 | 17.55426 | 17.4273 | 21.63771 | 19.48247 | 17.36842 | 16.02973 | 16.88773 | 14.43801 | 16.31565 | 16.58575 | 16.16015 | 14.94332 |
| NCOA4 | 15.485 | 15.87124 | 15.61392 | 15.01906 | 15.91177 | 16.56132 | 20.26566 | 16.84931 | 18.52484 | 17.03496 | 16.28945 | 15.04783 | 18.58667 | 18.42065 | 15.92965 | 15.87349 | 18.87373 | 19.80433 |
| NCR3LG1 | 18.36233 | 18.08785 | 16.9511 | 16.56802 | 13.88194 | 16.11208 | 12.11739 | 14.79363 | 18.90736 | 14.06242 | 15.39729 | 13.84349 | 15.34286 | 15.14662 | 14.86374 | 15.97488 | 15.63247 | 14.53688 |

| | | | | | | | | | | | | | | | | | | |
|---|---|---|---|---|---|---|---|---|---|---|---|---|---|---|---|---|---|---|
| NCS1 | 13.71276 | 16.43288 | 14.3597 | 18.2278 | 15.27369 | 14.55804 | 16.18079 | 17.23906 | 15.39258 | 16.60041 | 14.57026 | 17.04298 | 18.35221 | 17.79423 | 18.0632 | 18.12665 | 15.80512 | 14.79728 |
| NCSTN | 17.80384 | 17.87184 | 19.16617 | 18.76907 | 18.32923 | 18.11882 | 16.7044 | 17.31035 | 22.04075 | 20.55264 | 18.47302 | 20.29482 | 15.49238 | 15.34295 | 19.66819 | 19.40016 | 15.72133 | 15.31196 |
| NDC80 | 15.27417 | 15.21036 | 16.1948 | 16.20988 | 18.71489 | 16.25482 | 16.84464 | 14.99102 | 14.90026 | 17.50399 | 14.57529 | 14.10059 | 17.93308 | 18.53903 | 16.43902 | 15.75642 | 18.01477 | 17.2665 |
| NDEL1 | 15.35368 | 14.85302 | 13.99292 | 13.1761 | 18.04522 | 16.69962 | 15.59309 | 16.32063 | 16.23812 | 17.90428 | 16.50193 | 11.42534 | 18.70142 | 15.51113 | 16.17625 | 15.0817 | 17.3268 | 17.82 |
| NDRG1 | 19.82776 | 19.98413 | 20.82972 | 20.72564 | 21.54555 | 20.51159 | 16.79841 | 17.15519 | 20.3232 | 20.74268 | 17.11688 | 18.66438 | 18.98039 | 18.90418 | 19.90294 | 20.21157 | 22.9112 | 22.75655 |
| NDRG2 | 17.46925 | 18.95105 | 17.91966 | 20.09027 | 18.96055 | 17.3857 | 19.16942 | 18.29872 | 14.36002 | 15.74967 | 19.98988 | 19.12542 | 20.84991 | 19.67337 | 19.77522 | 20.04653 | 16.09019 | 17.12136 |
| NDRG3 | 17.56181 | 17.20486 | 18.40297 | 19.27827 | 20.78839 | 18.49839 | 17.56999 | 16.82659 | 17.78474 | 17.20793 | 21.56451 | 22.61768 | 20.4772 | 20.43196 | 19.83728 | 19.04356 | 19.18994 | 19.13762 |
| NDRG4 | 14.86397 | 15.15363 | 14.93843 | 15.16124 | 18.73793 | 16.24161 | 19.15635 | 18.65642 | 18.81749 | 19.0592 | 18.38553 | 18.45821 | 14.79176 | 15.02046 | 16.69479 | 13.40524 | 17.87842 | 15.24494 |
| NDST1 | 16.57426 | 16.20215 | 15.89612 | 14.50742 | 14.34524 | 15.817 | 16.0236 | 16.2479 | 18.76383 | 16.12654 | 14.58841 | 15.48898 | 13.75262 | 15.63016 | 15.53513 | 13.57431 | 14.37233 | 15.51555 |
| NDST2 | 15.73663 | 15.89024 | 15.17241 | 15.63424 | 17.27454 | 19.44177 | 17.37228 | 16.35632 | 20.23747 | 18.37989 | 16.6813 | 15.10258 | 14.27673 | 18.1074 | 15.8955 | 15.25676 | 16.63631 | 14.8753 |
| NDUFA10 | 15.99779 | 13.76838 | 13.2756 | 15.42941 | 12.79743 | 14.40771 | 16.11542 | 18.08502 | 18.60062 | 18.44884 | 17.11428 | 12.34345 | 14.05402 | 18.23825 | 14.71934 | 15.36917 | 13.68829 | 16.41875 |
| NDUFA11 | 16.3671 | 12.97546 | 16.86766 | 15.05821 | 11.9024 | 19.84006 | 17.56738 | 14.7894 | 17.57895 | 15.84598 | 15.14429 | 12.52748 | 13.55484 | 14.90159 | 14.03133 | 14.55335 | 14.36039 | 13.42249 |
| NDUFA2 | 18.03085 | 18.02838 | 16.93914 | 17.54305 | 17.08457 | 17.60454 | 16.9169 | 16.46385 | 18.92727 | 17.6317 | 14.72693 | 13.18678 | 16.02467 | 14.63179 | 18.64175 | 16.93579 | 16.22275 | 15.35998 |
| NDUFA9 | 16.31978 | 17.08973 | 15.11074 | 16.01958 | 14.67171 | 16.19831 | 17.52133 | 15.20842 | 19.15139 | 16.596 | 17.17409 | 17.99458 | 14.75591 | 13.10929 | 14.32806 | 14.89157 | 16.29197 | 14.78419 |
| NDUFAF4 | 15.37975 | 15.13702 | 16.0453 | 15.9318 | 15.29609 | 15.04415 | 16.69439 | 20.82377 | 19.40773 | 18.37098 | 15.42101 | 12.71616 | 15.41849 | 13.41662 | 15.40891 | 13.18399 | 18.71031 | 15.83318 |
| NDUFB10 | 14.87885 | 15.81603 | 15.8603 | 14.66085 | 16.19429 | 15.51474 | 18.00092 | 17.52258 | 19.86681 | 18.70475 | 14.02104 | 16.79955 | 18.92524 | 17.63769 | 14.27683 | 16.43845 | 14.89508 | 16.0153 |
| NDUFB4 | 15.54812 | 14.19278 | 15.71251 | 13.89145 | 17.01692 | 15.97026 | 17.51218 | 17.83667 | 19.38249 | 16.01526 | 13.85512 | 15.90277 | 14.14171 | 17.08668 | 15.73269 | 13.03001 | 16.00956 | 14.80283 |
| NDUFB9 | 15.82126 | 15.84369 | 14.93961 | 14.37727 | 16.14666 | 14.90631 | 14.85583 | 14.99136 | 19.12609 | 15.00672 | 15.24154 | 13.93207 | 16.48054 | 14.70404 | 15.38732 | 13.24509 | 14.97967 | 15.29515 |
| NDUFS1 | 15.93375 | 13.6671 | 16.07845 | 14.74849 | 15.29449 | 17.24513 | 15.49529 | 16.38444 | 19.47474 | 18.13406 | 16.92806 | 15.28255 | 15.25094 | 14.91211 | 14.18463 | 15.06966 | 16.11752 | 15.47643 |
| NDUFS2 | 14.66887 | 15.82911 | 15.32075 | 16.83594 | 14.56495 | 15.68001 | 17.84689 | 16.24182 | 19.27179 | 18.46848 | 15.60563 | 14.44084 | 15.17787 | 17.1231 | 16.43495 | 16.91908 | 15.37712 | 16.0771 |
| NDUFS3 | 16.60787 | 14.88418 | 16.67843 | 16.29946 | 14.89195 | 16.45486 | 15.82314 | 16.29352 | 18.28938 | 16.23617 | 14.43571 | 15.10402 | 14.86764 | 16.09461 | 17.73684 | 15.96408 | 14.14133 | 16.2311 |
| NDUFS8 | 15.96503 | 15.2803 | 14.96785 | 11.6847 | 12.91514 | 17.4117 | 16.73355 | 15.18446 | 18.73572 | 16.72412 | 14.78139 | 14.42537 | 14.77231 | 14.34441 | 14.22241 | 15.52161 | 16.05694 | 13.81648 |
| NDUFV1 | 14.62488 | 16.37814 | 16.09037 | 16.29715 | 17.45314 | 14.69322 | 16.67287 | 15.19135 | 15.76481 | 17.7006 | 17.51373 | 17.51806 | 19.36115 | 15.47721 | 15.7166 | 12.47913 | 18.60186 | 14.10546 |
| NECAP2 | 17.99923 | 17.37665 | 17.65323 | 16.82211 | 19.96406 | 16.64566 | 17.67199 | 16.73279 | 16.69714 | 16.64969 | 18.6015 | 15.26407 | 16.88057 | 17.02531 | 15.39031 | 17.94401 | 19.45336 | 20.68273 |
| NECTIN1 | 19.51703 | 19.16037 | 17.8535 | 17.34479 | 17.29049 | 18.89481 | 16.72997 | 16.06373 | 16.40812 | 14.79814 | 17.89269 | 19.25638 | 17.23877 | 16.65077 | 17.65071 | 18.21279 | 17.07712 | 16.30751 |
| NECTIN2 | 17.21501 | 16.63965 | 15.76627 | 16.38487 | 17.64642 | 16.33052 | 15.11197 | 15.82505 | 17.25656 | 16.64321 | 17.75782 | 15.65367 | 14.15933 | 16.55529 | 14.77667 | 17.8505 | 18.51822 | 19.10372 |
| NECTIN3 | 18.94035 | 17.25758 | 18.95605 | 19.71399 | 20.31242 | 14.63088 | 18.27357 | 16.87059 | 19.57125 | 18.95709 | 14.52874 | 13.72612 | 20.4592 | 19.54917 | 19.04881 | 18.90609 | 19.47348 | 21.50197 |
| NEDD1 | 16.69056 | 16.48336 | 15.15892 | 16.5682 | 15.77256 | 18.33569 | 19.15819 | 17.59869 | 18.48084 | 13.03897 | 16.18837 | 15.97603 | 15.75279 | 19.17856 | 17.00746 | 17.91974 | 14.22516 | 19.59092 |
| NEDD4 | 20.00514 | 21.11928 | 18.89486 | 18.27556 | 16.63446 | 17.92066 | 17.05981 | 15.7916 | 17.7894 | 16.73453 | 18.51012 | 17.83547 | 16.09221 | 17.73032 | 20.99854 | 20.80491 | 16.38773 | 15.82836 |
| NEDD4L | 20.60893 | 20.81378 | 20.73718 | 20.26479 | 18.18585 | 17.62874 | 16.39118 | 17.18633 | 16.00147 | 18.43971 | 16.74258 | 17.72485 | 19.3605 | 18.36586 | 20.54806 | 20.45815 | 18.7127 | 18.87309 |
| NEK9 | 13.647 | 13.96861 | 16.56181 | 14.43955 | 17.64694 | 18.449 | 17.31541 | 16.75347 | 12.88281 | 17.14854 | 14.29191 | 17.48171 | 17.56116 | 17.70628 | 15.73197 | 12.69469 | 12.15908 | 13.81351 |
| NELL1 | 14.95788 | 16.73779 | 16.29838 | 16.75282 | 15.69034 | 15.9422 | 14.57731 | 16.74136 | 18.61519 | 19.15497 | 16.72561 | 13.9859 | 14.51711 | 13.38164 | 14.30724 | 15.09569 | 14.95804 | 15.19699 |
| NENF | 17.61542 | 17.91291 | 16.63582 | 17.21527 | 15.88837 | 18.47792 | 16.95744 | 18.57547 | 23.0547 | 23.44888 | 18.51788 | 16.75979 | 16.98264 | 16.87027 | 16.1516 | 16.01702 | 16.54034 | 16.72394 |
| NEO1 | 17.91802 | 18.57958 | 18.55346 | 19.13386 | 19.03342 | 14.92049 | 17.80974 | 17.24689 | 16.16561 | 15.55474 | 15.69497 | 13.46425 | 17.34852 | 17.79091 | 19.22213 | 19.24589 | 13.51069 | 14.57424 |
| NF2 | 16.02392 | 16.30139 | 17.59306 | 18.20963 | 19.90273 | 18.26844 | 16.44073 | 17.64455 | 15.21209 | 18.1909 | 16.89065 | 14.97578 | 18.88616 | 19.42123 | 16.59587 | 17.39227 | 20.25393 | 21.72733 |
| NFKBIB | 15.89862 | 14.1268 | 16.17313 | 18.76062 | 14.84958 | 14.58513 | 15.59943 | 15.0047 | 13.93202 | 14.63578 | 15.53702 | 16.10663 | 13.95859 | 16.66781 | 13.76086 | 15.45717 | 17.46828 | 16.3872 |
| NHLRC2 | 16.38794 | 17.06382 | 16.08352 | 15.82408 | 17.93617 | 17.80493 | 19.14564 | 17.06287 | 15.28849 | 17.87213 | 19.14553 | 19.11111 | 19.11995 | 18.49973 | 13.89576 | 17.16495 | 18.46977 | 18.48753 |
| NHS | 16.09173 | 16.38323 | 17.77786 | 14.83017 | 19.08671 | 17.46292 | 16.16237 | 17.30794 | 14.92487 | 16.08213 | 14.59604 | 14.27564 | 14.93182 | 13.81295 | 14.27141 | 16.72753 | 18.50948 | 18.81282 |
| NIBAN1 | 16.96635 | 18.25194 | 17.57706 | 19.24104 | 19.66664 | 20.09792 | 19.64189 | 18.15353 | 20.74584 | 20.78132 | 15.5453 | 13.42557 | 16.57833 | 16.99783 | 17.28688 | 17.52707 | 17.06457 | 18.44717 |
| NIBAN2 | 20.1511 | 20.03389 | 20.43282 | 20.17518 | 22.35809 | 21.13834 | 17.80164 | 17.01022 | 19.81924 | 19.79807 | 17.76505 | 18.71621 | 20.55401 | 20.39512 | 20.55573 | 20.10384 | 21.18332 | 20.88101 |
| NID1 | 16.57667 | 15.19819 | 17.3495 | 17.30183 | 15.23442 | 16.41374 | 19.9925 | 18.06013 | 20.26818 | 19.29582 | 17.64911 | 13.83353 | 17.83847 | 19.99839 | 14.68236 | 16.33815 | 18.76201 | 19.07308 |
| NIF3L1 | 13.78051 | 15.60022 | 16.83577 | 17.12225 | 17.31031 | 21.21704 | 15.98706 | 17.32674 | 20.20123 | 16.48571 | 18.93734 | 20.05998 | 16.80126 | 15.0366 | 16.66619 | 16.77575 | 16.28208 | 16.43775 |
| NINJ1 | 17.48881 | 14.79451 | 20.11451 | 19.2468 | 16.81189 | 15.75411 | 14.11687 | 16.9234 | 16.20167 | 17.58626 | 13.71629 | 16.04463 | 16.82369 | 16.17987 | 21.10771 | 21.40529 | 15.85452 | 12.77452 |
| NIPSNAP1 | 14.23462 | 14.56654 | 16.3288 | 14.60196 | 14.00604 | 16.03989 | 15.13418 | 14.92473 | 18.12085 | 17.47023 | 15.14302 | 17.19008 | 14.77759 | 17.06186 | 14.79782 | 16.87468 | 15.59834 | 14.9878 |
| NIPSNAP2 | 16.43489 | 17.15123 | 17.442 | 15.2334 | 16.27881 | 19.76696 | 13.94602 | 15.96853 | 19.91214 | 17.98943 | 17.85911 | 16.46416 | 16.45674 | 15.91822 | 15.59811 | 14.54556 | 16.23672 | 15.82049 |
| NIPSNAP3/ | 15.41552 | 17.67538 | 16.22612 | 15.47756 | 15.16692 | 15.94318 | 15.68578 | 18.61638 | 20.09751 | 18.34951 | 15.74597 | 13.67549 | 16.06855 | 15.29508 | 15.75606 | 15.80328 | 16.27884 | 15.00615 |
| NISCH | 14.55898 | 15.42282 | 16.01983 | 16.08845 | 18.39706 | 18.0678 | 17.791 | 17.25744 | 16.91503 | 18.5638 | 16.44354 | 17.69552 | 17.16473 | 16.31229 | 14.7538 | 14.82423 | 17.2515 | 16.34974 |
| NIT1 | 15.87329 | 14.30803 | 16.03003 | 15.45061 | 12.8981 | 16.19042 | 17.59693 | 16.5258 | 14.04584 | 15.78836 | 16.94439 | 18.80378 | 13.14807 | 12.18651 | 16.64814 | 14.18319 | 12.66206 | 13.7207 |
| NIT2 | 16.20892 | 14.37889 | 18.67499 | 16.3253 | 19.02078 | 17.26896 | 16.5285 | 16.38217 | 17.28448 | 17.55742 | 18.49038 | 18.48663 | 19.48691 | 19.45535 | 17.49451 | 17.85891 | 19.95267 | 19.26644 |
| NLE1 | 15.49906 | 16.56954 | 15.47937 | 15.17005 | 16.54563 | 15.90171 | 15.03709 | 15.9797 | 14.69379 | 15.58097 | 15.68321 | 17.78047 | 17.07453 | 16.55865 | 17.51647 | 14.4704 | 17.70389 | 15.33223 |
| NLGN1 | 18.35944 | 17.79391 | 15.91132 | 16.64599 | 13.8335 | 15.95922 | 15.13623 | 16.43985 | 13.58497 | 14.66568 | 15.46602 | 13.48019 | 13.45892 | 14.61441 | 12.25428 | 15.55108 | 13.73262 | 16.27635 |
| NLN | 16.60222 | 15.4215 | 14.39001 | 15.863 | 17.79122 | 16.52024 | 15.61506 | 16.54127 | 17.77512 | 17.40399 | 17.7764 | 18.57925 | 16.16203 | 15.4969 | 15.80907 | 15.2804 | 15.37532 | 15.9616 |
| NMD3 | 17.6636 | 17.71636 | 16.56685 | 17.71709 | 13.54644 | 13.75535 | 16.50057 | 18.34027 | 14.29949 | 13.96207 | 16.27901 | 18.10101 | 13.68416 | 14.67578 | 16.81502 | 16.84926 | 16.22567 | 15.09766 |
| NME1 | 19.80962 | 19.18231 | 21.49669 | 22.15718 | 24.25541 | 22.62848 | 22.25169 | 19.41457 | 20.41352 | 21.35378 | 24.58738 | 24.28921 | 24.61908 | 23.53185 | 21.47568 | 21.5251 | 23.55523 | 24.22667 |
| NME2 | 17.48651 | 17.3772 | 17.83804 | 19.59498 | 24.92528 | 23.68332 | 22.36258 | 21.92265 | 16.75787 | 22.02355 | 21.72037 | 18.10247 | 25.0774 | 24.12842 | 18.42834 | 19.18793 | 25.49522 | 25.03549 |
| NME2P1 | 20.6293 | 19.08281 | 20.65737 | 21.13893 | 22.44365 | 20.76291 | 20.91313 | 18.24656 | 19.82307 | 19.06665 | 22.05211 | 24.21803 | 22.46766 | 20.12053 | 21.26276 | 21.274 | 23.16533 | 22.28623 |
| NMI | 13.73526 | 14.72476 | 15.17388 | 14.39712 | 19.20825 | 15.69941 | 15.21731 | 16.52768 | 12.68927 | 15.16784 | 14.15918 | 11.99735 | 18.14606 | 16.34676 | 16.89413 | 15.32132 | 18.21696 | 19.37939 |
| NMT1 | 15.67441 | 17.52493 | 19.43594 | 16.37016 | 17.63563 | 16.6452 | 16.94756 | 17.73787 | 15.81664 | 16.13438 | 16.40644 | 19.31261 | 15.55409 | 14.62942 | 15.782 | 14.2651 | 14.73641 | 16.65116 |
| NNMT | 14.78545 | 15.18463 | 14.96387 | 14.98172 | 21.93422 | 20.10163 | 15.29243 | 16.30655 | 16.53547 | 14.95273 | 15.06997 | 14.65265 | 14.2825 | 14.68608 | 14.09337 | 14.72127 | 16.35915 | 17.56106 |
| NNT | 16.7175 | 18.67029 | 16.05965 | 15.35503 | 14.96685 | 18.62393 | 18.09651 | 17.7059 | 20.8791 | 19.75741 | 16.29585 | 14.08926 | 14.84745 | 17.25322 | 15.77169 | 16.69839 | 16.52646 | 15.18078 |
| NOL6 | 14.90234 | 15.21152 | 16.29518 | 14.12376 | 13.61692 | 14.19483 | 14.68885 | 14.72087 | 12.23685 | 14.75852 | 17.65873 | 16.65679 | 15.92473 | 13.41081 | 14.81893 | 14.94494 | 14.69554 | 13.3719 |
| NOLC1 | 16.49425 | 18.33186 | 14.61021 | 17.00585 | 15.90878 | 15.01011 | 14.49763 | 15.35012 | 15.60995 | 14.20519 | 15.69855 | 16.12823 | 17.23955 | 16.47197 | 16.34423 | 16.22057 | 16.12769 | 16.81603 |
| NOMO1 | 15.71116 | 16.34029 | 15.59494 | 15.879 | 15.88724 | 16.75579 | 16.97458 | 17.5304 | 21.10154 | 19.94368 | 19.06578 | 15.65273 | 16.10685 | 17.01976 | 15.43998 | 16.60012 | 16.30869 | 15.32329 |
| NONO | 15.62216 | 15.46726 | 18.02062 | 18.09239 | 17.32562 | 18.55019 | 19.55228 | 19.48177 | 16.52073 | 19.03165 | 18.00571 | 18.38514 | 18.36049 | 17.68189 | 18.88371 | 18.50548 | 17.38875 | 18.60516 |
| NOP56 | 17.16871 | 16.93584 | 14.87062 | 16.49521 | 16.983 | 14.91351 | 18.98476 | 16.70476 | 14.98163 | 15.92142 | 17.17077 | 16.99294 | 16.37256 | 15.84077 | 16.02436 | 16.01322 | 17.50161 | 17.73841 |
| NOP58 | 16.68811 | 15.72962 | 17.3709 | 16.11781 | 14.73158 | 16.07469 | 18.6901 | 18.88728 | 16.49219 | 15.45045 | 16.04665 | 17.40354 | 15.32118 | 15.47246 | 16.96744 | 17.07115 | 15.98357 | 14.73569 |
| NOS1AP | 16.14536 | 15.15434 | 17.30948 | 17.55393 | 17.79259 | 17.64875 | 15.23984 | 15.91221 | 17.64042 | 17.27738 | 16.04993 | 14.81627 | 19.35972 | 19.42118 | 16.77465 | 17.23598 | 17.30329 | 16.52129 |
| NOSTRIN | 21.54082 | 21.44033 | 22.05153 | 22.38138 | 14.88645 | 16.40817 | 16.1524 | 16.38242 | 15.57083 | 15.55046 | 16.54874 | 16.89581 | 19.81612 | 20.00375 | 22.98072 | 21.98524 | 16.21061 | 15.72623 |
| NOTCH1 | 17.6602 | 17.20089 | 17.14024 | 18.20035 | 14.51424 | 17.28497 | 16.42594 | 17.96667 | 17.83501 | 14.24441 | 18.4196 | 19.25999 | 15.61433 | 15.75579 | 16.89373 | 16.85047 | 16.1359 | 15.27131 |
| NOTCH2 | 16.96939 | 16.53167 | 16.90161 | 17.72666 | 20.16823 | 18.02326 | 18.51643 | 15.34824 | 16.66418 | 17.68793 | 14.51687 | 12.59632 | 19.37544 | 20.38278 | 17.52075 | 17.29141 | 20.50065 | 22.51996 |
| NOXO1 | 15.68386 | 16.43334 | 16.01808 | 14.47885 | 14.40644 | 17.98888 | 15.52377 | 17.66111 | 14.64637 | 15.6768 | 19.53861 | 18.36866 | 15.17221 | 17.1314 | 16.50955 | 16.31937 | 15.49418 | 14.26559 |
| NPC1 | 16.21517 | 18.88432 | 16.07194 | 16.30125 | 18.90599 | 18.09349 | 19.3761 | 18.30876 | 19.46553 | 17.52664 | 17.52159 | 16.51616 | 17.23309 | 16.97576 | 19.03609 | 16.9795 | 15.58791 | 17.95964 |
| NPC2 | 16.87415 | 16.59109 | 17.70417 | 16.68989 | 19.07199 | 20.32827 | 19.21707 | 19.00014 | 22.04459 | 21.90851 | 16.85605 | 15.14287 | 18.96477 | 19.17087 | 17.40354 | 17.32628 | 17.14574 | 16.23739 |
| NPDC1 | 18.96217 | 19.98733 | 20.65488 | 21.32388 | 15.22376 | 17.16409 | 14.85134 | 16.38867 | 20.74319 | 16.86232 | 17.17446 | 13.5151 | 14.66745 | 16.84099 | 21.17154 | 19.91352 | 15.10881 | 13.81134 |
| NPEPPS | 16.17032 | 16.78115 | 18.13748 | 17.71161 | 18.96469 | 18.51616 | 16.74961 | 16.91715 | 17.70968 | 17.60622 | 18.69686 | 21.32479 | 19.36394 | 18.55259 | 18.54964 | 18.71652 | 19.34453 | 19.61578 |
| NPEPPSL1 | 16.19633 | 17.85732 | 18.61808 | 18.04513 | 17.71402 | 17.20171 | 15.40195 | 15.78591 | 17.81447 | 17.16996 | 19.15722 | 20.34387 | 16.78372 | 17.63764 | 18.45388 | 18.07268 | 18.58938 | 18.50288 |
| NPLOC4 | 22.08313 | 21.76992 | 22.54506 | 23.51528 | 20.53209 | 20.11333 | 15.41732 | 16.99715 | 18.25931 | 19.76394 | 18.28199 | 20.13339 | 22.75437 | 19.58263 | 23.17387 | 23.83844 | 18.74891 | 18.4241 |
| NPM1 | 18.79094 | 18.76564 | 18.98415 | 19.12696 | 19.94137 | 18.28892 | 19.41995 | 18.05985 | 20.34892 | 20.44375 | 22.82173 | 20.9154 | 21.45448 | 20.46417 | 17.68618 | 17.15639 | 19.55944 | 20.37131 |
| NPRL2 | 16.9803 | 16.13389 | 15.31719 | 15.5211 | 12.7644 | 15.572 | 15.05304 | 16.18836 | 15.13312 | 15.20758 | 12.09474 | 18.28623 | 16.16918 | 14.55421 | 21.82003 | 13.82113 | 17.68804 | 14.55162 |
| NPTN | 18.6316 | 18.26847 | 19.77852 | 19.53782 | 20.74732 | 19.08137 | 18.38142 | 18.25528 | 19.52322 | 18.03262 | 21.00274 | 21.34196 | 20.26256 | 21.42598 | 20.4028 | 20.02984 | 19.94319 | 21.64226 |
| NPTX1 | 17.12324 | 17.36099 | 17.43996 | 17.22827 | 16.73868 | 17.91825 | 16.34304 | 17.29092 | 24.12212 | 21.99662 | 16.87993 | 16.52503 | 20.9201 | 18.5554 | 16.98812 | 18.26285 | 15.71235 | 16.01271 |
| NPTX2 | 15.20552 | 15.7706 | 15.65941 | 15.86012 | 17.80944 | 16.77262 | 15.86135 | 16.92968 | 19.81422 | 19.54252 | 16.2376 | 14.37985 | 14.59711 | 15.68799 | 14.97022 | 15.56567 | 14.33292 | 14.39593 |
| NPTXR | 16.75387 | 16.90156 | 16.37105 | 16.00023 | 16.67731 | 16.04427 | 18.09191 | 16.34279 | 21.3373 | 20.3429 | 18.22934 | 15.35655 | 13.94492 | 17.90325 | 18.42923 | 15.85712 | 16.77673 | 16.51424 |
| NRAS | 19.75647 | 22.06397 | 21.15686 | 22.30953 | 22.00822 | 21.83015 | 18.1528 | 21.99935 | 21.05565 | 21.27454 | 22.45633 | 21.0458 | 22.92142 | 23.05718 | 21.93375 | 22.09465 | 21.31935 | 21.27772 |
| NRBP1 | 15.92191 | 16.20852 | 15.52066 | 17.11697 | 18.37848 | 14.41832 | 18.09643 | 17.20545 | 16.01412 | 17.37219 | 16.2526 | 19.54239 | 17.67501 | 17.47185 | 16.44162 | 17.16178 | 15.96858 | 18.25889 |
| NRCAM | 15.67534 | 15.07278 | 15.48521 | 15.26954 | 16.81742 | 17.27324 | 14.94945 | 14.70868 | 18.25463 | 15.70649 | 16.52227 | 14.05967 | 14.40002 | 15.24578 | 15.155 | 15.74151 | 14.92467 | 13.89865 |

| | | | | | | | | | | | | | | | | | | |
|---|---|---|---|---|---|---|---|---|---|---|---|---|---|---|---|---|---|---|
| NRDC | 14.26086 | 15.34113 | 16.71856 | 16.42555 | 18.21611 | 13.2117 | 14.04787 | 16.03286 | 16.99339 | 17.46908 | 12.77076 | 17.94467 | 14.15166 | 14.29905 | 17.5829 | 14.43338 | 18.8717 | 15.28087 |
| NRP1 | 17.4137 | 17.48804 | 18.54823 | 18.44608 | 21.13004 | 20.59363 | 15.93853 | 16.32087 | 16.50373 | 15.88468 | 14.8782 | 14.17778 | 18.20251 | 17.50916 | 17.91948 | 19.64299 | 20.65289 | 22.9102 |
| NRP2 | 16.51205 | 15.40776 | 15.7763 | 15.90849 | 14.18202 | 16.75834 | 18.19802 | 16.03197 | 18.73294 | 16.74923 | 17.17871 | 13.88769 | 16.83979 | 16.23385 | 16.9791 | 16.81993 | 16.76974 | 16.483 |
| NRXN1 | 16.12772 | 14.54778 | 13.32805 | 14.09127 | 16.10941 | 15.50338 | 15.5394 | 14.76265 | 16.82421 | 15.16243 | 16.16953 | 13.87919 | 20.95055 | 15.15063 | 15.17061 | 14.23614 | 14.05645 | 15.61554 |
| NSDHL | 16.84985 | 14.75066 | 15.05477 | 15.5614 | 14.7783 | 14.1601 | 18.26694 | 17.49434 | 18.92293 | 15.84662 | 16.99629 | 16.86519 | 15.16917 | 15.92201 | 15.85422 | 15.83602 | 14.57012 | 15.23612 |
| NSF | 17.60751 | 16.93422 | 18.30546 | 18.49985 | 17.98899 | 19.60145 | 19.50941 | 18.81467 | 17.54108 | 17.2457 | 17.55189 | 17.97456 | 21.10862 | 20.89438 | 18.29446 | 18.51963 | 19.48153 | 20.03677 |
| NSFL1C | 18.72537 | 17.89458 | 17.256 | 17.23893 | 15.68235 | 15.05964 | 15.36596 | 15.43635 | 17.05209 | 14.16698 | 17.81195 | 18.62439 | 17.55307 | 16.18734 | 15.72026 | 17.88688 | 16.90439 | 15.36062 |
| NSG1 | 13.92458 | 16.02668 | 17.03849 | 15.00281 | 13.98951 | 16.83744 | 21.12105 | 18.25756 | 14.85296 | 16.18273 | 15.17127 | 12.63335 | 14.5894 | 15.25505 | 14.59823 | 15.4122 | 14.12463 | 14.05372 |
| NSUN2 | 16.09878 | 15.22983 | 17.39827 | 18.00002 | 19.39533 | 17.32311 | 19.54761 | 18.43695 | 15.92287 | 17.77833 | 18.50575 | 18.77242 | 19.42748 | 16.6997 | 17.45562 | 17.29931 | 19.50207 | 19.51926 |
| NT5DC2 | 15.65881 | 15.49339 | 16.04317 | 15.3404 | 14.68824 | 16.09268 | 15.32888 | 14.94385 | 18.18799 | 16.31158 | 14.89542 | 14.59375 | 15.23397 | 15.03136 | 17.05998 | 16.0997 | 15.97453 | 15.08066 |
| NT5E | 17.93447 | 16.05409 | 17.22535 | 15.74627 | 23.72602 | 23.14879 | 16.99223 | 16.97729 | 15.5558 | 16.12235 | 16.08432 | 14.96182 | 17.8925 | 16.31532 | 15.87136 | 16.73402 | 22.04819 | 22.70757 |
| NTPCR | 17.65815 | 17.33005 | 17.90368 | 18.0836 | 14.25915 | 16.11389 | 19.56526 | 17.44354 | 16.383 | 13.42873 | 16.9374 | 17.27659 | 20.54744 | 19.11984 | 18.10457 | 18.47545 | 17.04244 | 16.48004 |
| NUBP1 | 21.93752 | 17.29078 | 17.23461 | 14.27295 | 17.13793 | 14.93798 | 17.1518 | 16.58175 | 15.50681 | 15.26702 | 16.10815 | 17.28041 | 17.34514 | 18.83793 | 14.28472 | 18.4258 | 19.46453 | 20.11037 |
| NUBP2 | 19.16241 | 21.79894 | 16.87027 | 17.82073 | 18.67228 | 18.28688 | 21.20683 | 16.74429 | 16.62349 | 17.75644 | 15.78977 | 12.52913 | 19.42323 | 19.97485 | 18.69257 | 16.89647 | 19.81022 | 19.21045 |
| NUCB1 | 16.31614 | 16.53244 | 17.27062 | 16.65409 | 18.19399 | 19.09663 | 19.63047 | 18.4826 | 21.30781 | 20.18375 | 20.88374 | 18.34571 | 16.9938 | 16.912 | 16.52028 | 17.05899 | 17.17541 | 19.67133 |
| NUCB2 | 17.99307 | 17.23519 | 19.06722 | 19.67172 | 17.43508 | 17.68888 | 19.48546 | 18.40319 | 23.9176 | 22.79068 | 18.88505 | 16.4323 | 19.7289 | 20.55167 | 20.6795 | 18.50251 | 16.55871 | 18.09071 |
| NUCKS1 | 16.56701 | 14.8463 | 15.46734 | 16.27094 | 19.30152 | 18.4681 | 17.53742 | 17.67857 | 18.40267 | 19.80298 | 18.47312 | 18.44938 | 18.8458 | 17.85523 | 17.05082 | 16.97586 | 19.42943 | 19.6772 |
| NUDC | 16.47791 | 16.41079 | 18.3633 | 18.59306 | 18.66838 | 17.2286 | 17.74319 | 18.6997 | 19.63274 | 19.1926 | 19.9501 | 20.69149 | 20.42745 | 19.47889 | 19.18116 | 18.349 | 19.62821 | 19.19111 |
| NUDCD1 | 14.99136 | 15.6599 | 14.48572 | 15.96749 | 16.60561 | 17.38637 | 16.94498 | 16.29052 | 15.65947 | 16.15669 | 17.20226 | 18.05433 | 16.8283 | 16.30626 | 15.54385 | 16.27002 | 17.7447 | 16.04556 |
| NUDCD2 | 17.47154 | 17.96343 | 18.07648 | 18.44682 | 20.18649 | 21.57624 | 18.0655 | 17.95481 | 17.56763 | 18.22584 | 18.82732 | 18.05891 | 19.54558 | 20.25704 | 17.61037 | 18.12882 | 19.66463 | 19.23316 |
| NUDT1 | 17.31229 | 20.16094 | 14.18388 | 16.56841 | 16.57684 | 15.54608 | 18.52571 | 17.70017 | 16.85533 | 16.51196 | 13.53899 | 17.75865 | 15.22211 | 17.44313 | 15.64067 | 17.49188 | 17.33737 | 16.35327 |
| NUDT16L1 | 16.6421 | 16.79739 | 16.22068 | 15.70236 | 14.0908 | 16.18609 | 15.85808 | 16.68145 | 13.66914 | 16.18489 | 13.20367 | 17.76554 | 14.3892 | 15.16964 | 17.94689 | 17.92399 | 15.73869 | 15.55373 |
| NUDT19 | 17.03058 | 15.25269 | 16.17764 | 15.08598 | 14.9145 | 18.26413 | 15.65733 | 16.69454 | 20.78933 | 16.56632 | 18.02007 | 15.27917 | 15.56662 | 16.50687 | 15.68721 | 15.13427 | 14.45152 | 18.81041 |
| NUDT21 | 22.87366 | 22.5554 | 20.76127 | 21.53912 | 18.56291 | 17.75735 | 17.04306 | 17.58495 | 17.7202 | 17.99867 | 18.116 | 20.08203 | 19.6628 | 17.32552 | 19.60321 | 20.65521 | 17.12371 | 15.43128 |
| NUDT3 | 15.33247 | 14.24538 | 16.92087 | 16.65329 | 16.15686 | 18.78049 | 13.74025 | 15.23618 | 16.14728 | 15.61553 | 17.92851 | 18.07933 | 13.55782 | 15.78069 | 17.79763 | 13.99621 | 12.35052 | 12.91689 |
| NUDT4B | 14.31484 | 14.35434 | 13.36737 | 16.08672 | 15.97362 | 14.88218 | 15.63844 | 15.47774 | 12.12363 | 13.89623 | 15.0139 | 16.0481 | 14.16092 | 18.09287 | 14.23706 | 14.20023 | 16.49366 | 14.23329 |
| NUDT5 | 19.29169 | 17.14868 | 17.29551 | 18.41806 | 20.97434 | 19.91418 | 20.43913 | 17.6142 | 19.15113 | 20.07315 | 21.30159 | 20.88644 | 20.01997 | 19.58502 | 17.55151 | 17.05292 | 20.04328 | 17.41571 |
| NUDT9 | 15.7321 | 14.28287 | 15.45868 | 16.87279 | 18.76544 | 14.32465 | 13.25463 | 16.80484 | 17.76896 | 18.75694 | 17.02895 | 17.20196 | 18.80667 | 17.02456 | 17.59462 | 14.91162 | 17.51703 | 15.79632 |
| NUMA1 | 16.27439 | 16.18379 | 15.66792 | 15.67135 | 15.19214 | 15.46899 | 15.33601 | 16.85051 | 16.16809 | 17.47135 | 16.96645 | 16.94478 | 18.33789 | 16.70769 | 16.22917 | 16.33469 | 17.02607 | 16.02712 |
| NUMB | 17.02047 | 17.82814 | 17.84882 | 18.90606 | 22.78111 | 20.94928 | 17.04765 | 17.41646 | 18.52159 | 18.31833 | 16.423 | 16.26736 | 20.54949 | 18.88879 | 19.22838 | 18.38629 | 19.20652 | 22.03072 |
| NUMBL | 15.15361 | 16.22224 | 16.60763 | 16.91001 | 19.99271 | 18.1718 | 15.92377 | 16.26363 | 15.93068 | 17.09677 | 14.21114 | 14.9722 | 18.27982 | 18.22429 | 16.57283 | 16.7003 | 14.16517 | 17.9189 |
| NUP210 | 17.57313 | 17.89626 | 16.77613 | 17.38663 | 15.99645 | 16.68568 | 16.95938 | 16.46808 | 22.60295 | 20.33442 | 18.81466 | 15.16676 | 16.15253 | 17.27829 | 16.88336 | 17.42367 | 17.01851 | 16.07762 |
| NUP50 | 14.66682 | 16.05109 | 13.74782 | 15.46775 | 14.61131 | 17.44207 | 15.70066 | 16.52553 | 13.60869 | 15.69897 | 16.81882 | 17.72962 | 14.96038 | 18.60604 | 14.11682 | 16.93264 | 17.11857 | 14.41999 |
| NUP62 | 14.09296 | 15.22064 | 16.14657 | 14.10331 | 18.74071 | 17.43677 | 16.17003 | 16.29495 | 14.90391 | 14.98882 | 14.33552 | 16.7973 | 17.56044 | 16.4763 | 15.30853 | 15.81745 | 17.19117 | 17.80559 |
| NUP85 | 14.45942 | 16.73257 | 15.51886 | 15.27015 | 13.79021 | 15.15994 | 16.21786 | 14.78367 | 12.44967 | 15.30347 | 14.86859 | 17.65447 | 11.97934 | 17.36159 | 15.87673 | 13.3646 | 14.84409 | 13.81708 |
| NUP88 | 16.98707 | 12.86784 | 15.15499 | 15.24049 | 16.70622 | 15.8597 | 15.56421 | 16.97516 | 13.81209 | 16.09274 | 14.92541 | 18.58608 | 14.92985 | 15.5345 | 14.43997 | 17.90314 | 15.80322 | 14.9582 |
| NUP93 | 15.92727 | 15.73612 | 16.74542 | 16.00077 | 14.82555 | 16.95179 | 17.18043 | 16.29465 | 13.06523 | 14.37577 | 16.38637 | 18.55293 | 15.16871 | 14.41843 | 17.04453 | 17.03969 | 15.39816 | 15.5975 |
| NUTF2 | 17.73006 | 19.80599 | 20.47406 | 19.75402 | 20.90966 | 19.67841 | 20.79522 | 14.91902 | 18.6407 | 13.96985 | 20.20564 | 20.62809 | 21.49811 | 21.36184 | 14.85369 | 20.46171 | 21.39226 | 20.44289 |
| NXF1 | 16.94722 | 17.02569 | 15.41028 | 18.62402 | 21.41601 | 20.38639 | 20.46583 | 20.84142 | 16.61341 | 18.39585 | 17.73779 | 17.55555 | 20.18944 | 20.61382 | 15.93409 | 17.82798 | 22.68611 | 22.49122 |
| NXN | 14.09124 | 15.75406 | 14.86624 | 15.67491 | 17.09155 | 16.90264 | 16.9376 | 15.65505 | 14.33321 | 15.79854 | 16.6722 | 19.22441 | 16.44944 | 16.67707 | 16.36412 | 16.01724 | 14.59905 | 14.93214 |
| OAT | 16.81211 | 16.12674 | 15.53379 | 15.53981 | 15.85573 | 15.83385 | 16.17197 | 16.50401 | 20.3085 | 19.33853 | 16.65094 | 15.00595 | 15.08913 | 16.25234 | 15.91246 | 15.59683 | 15.56245 | 15.26761 |
| OCC1 | 22.09958 | 22.76622 | 20.50112 | 20.10475 | 18.39732 | 16.66847 | 17.50671 | 17.54574 | 18.32268 | 16.41837 | 16.32489 | 16.82104 | 17.46055 | 17.43973 | 18.58414 | 20.10148 | 19.98388 | 21.4858 |
| OCIAD1 | 15.77034 | 18.05228 | 16.04106 | 16.14059 | 14.60668 | 16.25082 | 16.03234 | 15.95189 | 18.54547 | 17.60279 | 15.14426 | 16.16907 | 12.62753 | 16.2916 | 15.4675 | 15.56914 | 14.11131 | 13.52541 |
| OCLN | 20.14661 | 20.26352 | 21.41007 | 21.98264 | 21.75466 | 20.27128 | 16.8434 | 16.50723 | 20.35968 | 20.4643 | 16.27415 | 17.89388 | 22.27911 | 22.74692 | 22.86128 | 21.94768 | 16.88956 | 19.70602 |
| OCRL | 18.20798 | 17.16301 | 16.52836 | 16.7371 | 17.96082 | 15.7609 | 16.53919 | 16.63082 | 14.89302 | 15.89401 | 15.95315 | 16.74039 | 17.48563 | 17.80738 | 16.12189 | 16.76602 | 18.26623 | 18.12264 |
| OGA | 14.46832 | 16.97559 | 15.80965 | 16.8531 | 15.2232 | 17.31632 | 17.21212 | 17.39263 | 16.05684 | 16.28198 | 15.83939 | 15.37538 | 17.90824 | 17.6483 | 18.92425 | 17.99959 | 17.32184 | 17.43173 |
| OGDH | 16.87401 | 15.78548 | 15.20099 | 15.77898 | 20.41918 | 23.935 | 20.48181 | 16.43657 | 18.86639 | 18.03669 | 16.59699 | 14.27044 | 15.02551 | 22.50695 | 16.12985 | 14.94974 | 19.07913 | 21.98063 |
| OGDHL | 15.30306 | 18.55991 | 19.33448 | 14.70613 | 15.7322 | 13.13472 | 16.82197 | 14.74843 | 19.4934 | 13.28319 | 17.95301 | 19.68717 | 17.3566 | 14.22989 | 19.02147 | 15.74329 | 16.29488 | 16.63935 |
| OGFOD3 | 16.81572 | 16.94204 | 15.59961 | 14.04166 | 15.7605 | 17.71587 | 15.55871 | 15.75032 | 20.09045 | 17.22388 | 15.5028 | 13.46778 | 16.45024 | 16.28085 | 15.04716 | 14.39343 | 17.16829 | 17.83586 |
| OLA1 | 17.94126 | 17.72931 | 18.30085 | 18.6965 | 18.83615 | 17.59913 | 19.527 | 18.95111 | 17.42162 | 17.64199 | 20.00621 | 20.62836 | 19.75575 | 19.05568 | 19.04454 | 19.15052 | 19.50983 | 18.82757 |
| OLFM1 | 15.59579 | 17.3075 | 14.92372 | 14.7378 | 16.11989 | 15.60629 | 16.51386 | 16.33245 | 19.72803 | 17.4483 | 13.33665 | 13.97829 | 13.97054 | 17.04887 | 15.06076 | 14.39123 | 16.70332 | 16.1243 |
| OPA1 | 15.82303 | 15.36301 | 17.07234 | 15.88471 | 16.87743 | 16.21019 | 16.76307 | 15.63368 | 17.22548 | 16.45176 | 18.50969 | 19.72907 | 16.37953 | 15.51911 | 15.25904 | 15.70079 | 15.61536 | 15.87898 |
| OPLAH | 14.42949 | 16.2944 | 15.62722 | 15.26136 | 15.79037 | 17.39857 | 14.49372 | 15.6827 | 14.67731 | 16.27749 | 16.11565 | 17.6832 | 14.47433 | 15.15231 | 16.30653 | 15.44897 | 16.80166 | 14.89003 |
| OPRK1 | 16.49192 | 16.67689 | 13.92896 | 16.79824 | 14.04792 | 15.30099 | 14.08946 | 16.33596 | 19.56259 | 18.09371 | 14.23427 | 16.85596 | 18.15892 | 14.59732 | 16.97947 | 19.10651 | 16.34689 | 14.92809 |
| OPTN | 13.94697 | 16.31135 | 12.54382 | 15.30568 | 12.92903 | 12.46969 | 18.86488 | 17.13608 | 15.00484 | 13.27552 | 16.11462 | 14.76838 | 15.83469 | 13.1581 | 18.24433 | 16.82707 | 15.51023 | 16.42516 |
| OS9 | 17.66161 | 16.61844 | 16.32255 | 16.58399 | 16.34058 | 16.54622 | 16.19467 | 16.5439 | 20.77356 | 18.02857 | 18.30971 | 15.31723 | 16.18302 | 15.06616 | 15.41021 | 16.51469 | 16.30636 | 15.27689 |
| OSBP | 17.50432 | 17.10158 | 17.4521 | 18.46828 | 18.57719 | 17.70233 | 17.77192 | 15.72999 | 17.1865 | 17.00072 | 18.7381 | 19.54661 | 19.08335 | 18.75178 | 17.99886 | 18.38089 | 16.6428 | 17.88434 |
| OSBPL1A | 14.47647 | 14.19999 | 17.27309 | 22.07823 | 14.6537 | 14.98749 | 14.95938 | 16.52882 | 12.78962 | 15.77441 | 14.4964 | 15.22276 | 15.18045 | 20.43003 | 15.10789 | 21.59141 | 12.57785 | 13.95869 |
| OSBPL8 | 16.42647 | 15.18465 | 15.6395 | 15.26874 | 13.1327 | 13.74742 | 15.57931 | 13.83071 | 16.29946 | 14.47804 | 14.95161 | 17.81082 | 13.05196 | 14.18931 | 12.65489 | 16.09133 | 15.38642 | 17.37698 |
| OSBPL9 | 14.07347 | 15.15375 | 18.69705 | 17.62431 | 17.24337 | 19.23868 | 15.4244 | 14.73584 | 13.13465 | 17.0512 | 20.10606 | 15.60806 | 16.78977 | 16.76953 | 17.20485 | 15.25002 | 16.86235 | 17.3866 |
| OSGEP | 13.39421 | 14.937 | 15.5909 | 15.76802 | 15.60197 | 14.71527 | 14.38973 | 16.50897 | 13.06261 | 15.04126 | 17.07537 | 18.20845 | 15.26102 | 15.92578 | 16.84469 | 15.01657 | 13.85162 | 14.49466 |
| OSMR | 15.45591 | 14.70853 | 15.91086 | 17.35551 | 20.46207 | 18.56083 | 15.86887 | 15.68596 | 16.41928 | 14.90969 | 16.43961 | 13.01211 | 15.44003 | 14.01876 | 14.87423 | 14.86596 | 15.48141 | 14.17785 |
| OSTF1 | 22.54547 | 26.53152 | 23.52822 | 24.21113 | 23.69547 | 22.91686 | 21.05648 | 19.33086 | 23.96959 | 21.52194 | 23.81787 | 23.08398 | 21.82616 | 22.74788 | 23.62837 | 21.89662 | 22.54115 | 23.29116 |
| OSTM1 | 14.2835 | 13.75538 | 15.02283 | 11.21725 | 17.17806 | 13.82155 | 16.97749 | 18.69321 | 19.80277 | 15.28734 | 16.97246 | 12.12608 | 16.87491 | 17.67625 | 17.21151 | 11.79125 | 13.85358 | 15.03557 |
| OTUB1 | 18.67266 | 18.56639 | 17.55406 | 18.3904 | 19.71619 | 20.1321 | 17.89221 | 14.79689 | 17.91176 | 18.58333 | 20.13414 | 21.08041 | 20.34807 | 19.49432 | 19.38047 | 19.14012 | 20.47941 | 20.70094 |
| OTUD6B | 20.76654 | 20.2442 | 18.95692 | 20.73629 | 12.44485 | 15.88289 | 14.43821 | 15.57327 | 16.27699 | 16.17094 | 17.4979 | 18.41753 | 12.9467 | 13.85231 | 21.2568 | 19.78217 | 16.81747 | 12.67621 |
| OTUD7B | 15.62628 | 17.61464 | 18.41872 | 17.61208 | 19.4671 | 17.72386 | 14.09862 | 17.15333 | 16.53335 | 16.12113 | 15.34723 | 15.71324 | 19.72968 | 19.27636 | 16.80572 | 17.69154 | 17.73238 | 19.32876 |
| OVCA2 | 18.98953 | 17.26841 | 17.23613 | 16.58517 | 13.78278 | 15.14189 | 15.66731 | 16.38304 | 12.05369 | 15.34185 | 14.89163 | 18.31448 | 14.80207 | 14.16411 | 18.13673 | 17.7166 | 14.97329 | 16.00981 |
| OXCT1 | 15.69068 | 16.53209 | 16.04397 | 16.16462 | 16.98225 | 15.47892 | 18.97393 | 16.03647 | 19.52088 | 18.3334 | 15.76587 | 16.29745 | 15.11909 | 16.46007 | 16.44645 | 17.11949 | 16.14423 | 15.54313 |
| OXR1 | 15.6564 | 17.97801 | 15.52647 | 15.78319 | 13.26034 | 16.27996 | 16.26918 | 15.52139 | 17.28366 | 16.34115 | 15.78684 | 18.11629 | 15.01844 | 15.52261 | 13.8804 | 15.84034 | 13.51373 | 15.04097 |
| OXSR1 | 15.57697 | 15.85058 | 17.29669 | 16.636 | 17.23791 | 17.36763 | 13.70242 | 16.45048 | 15.9398 | 16.57917 | 17.35409 | 17.97798 | 18.45749 | 18.04468 | 15.83483 | 15.75576 | 15.97734 | 17.16782 |
| P2RX4 | 16.04765 | 15.11793 | 14.32012 | 13.49178 | 15.73167 | 15.09604 | 15.59832 | 17.57332 | 19.93832 | 17.69061 | 15.07735 | 15.80095 | 15.80792 | 16.93457 | 13.89179 | 15.8242 | 13.02926 | 14.07364 |
| P3H1 | 15.07154 | 15.68839 | 15.56449 | 15.80205 | 16.36421 | 18.1598 | 16.31404 | 17.95122 | 20.03352 | 18.96537 | 16.32359 | 14.8014 | 16.41057 | 17.05598 | 15.98085 | 15.88208 | 16.557 | 17.22773 |
| P4HA1 | 16.80974 | 15.81518 | 16.23578 | 16.21429 | 15.45212 | 16.56816 | 15.54583 | 16.3191 | 20.51765 | 20.37794 | 19.23066 | 15.47423 | 16.01231 | 16.31178 | 15.4283 | 15.88336 | 17.01677 | 18.43655 |
| P4HA2 | 15.56675 | 16.03662 | 16.75784 | 16.10948 | 15.98161 | 16.85709 | 16.44233 | 17.37163 | 19.88354 | 19.77205 | 15.39384 | 13.07712 | 14.62675 | 16.67513 | 14.78034 | 16.44957 | 16.4612 | 15.88714 |
| P4HB | 17.30852 | 17.28526 | 18.88344 | 18.58952 | 19.57761 | 18.86058 | 21.71125 | 20.62425 | 24.20129 | 24.23665 | 22.10101 | 18.81424 | 19.9044 | 20.51891 | 19.25137 | 18.70605 | 18.58839 | 19.52561 |
| P4HTM | 17.11472 | 15.85782 | 16.58975 | 15.4223 | 15.54891 | 15.76163 | 15.45639 | 16.59606 | 20.36424 | 17.11801 | 14.57901 | 14.0248 | 14.9872 | 14.75119 | 15.90204 | 15.87186 | 14.83742 | 14.7605 |
| PA2G4 | 21.00988 | 21.10133 | 19.83127 | 19.96099 | 19.22816 | 18.96966 | 19.10832 | 18.77664 | 18.66787 | 19.4303 | 20.14463 | 21.74141 | 19.93254 | 19.60467 | 20.18699 | 20.11148 | 19.04363 | 18.37808 |
| PAAF1 | 23.49905 | 22.96143 | 19.59667 | 15.05157 | 15.70827 | 16.16595 | 17.07488 | 18.25783 | 15.83107 | 14.29763 | 14.87031 | 17.49939 | 14.90873 | 16.95885 | 17.39591 | 22.15808 | 17.26103 | 18.3639 |
| PAAT | 20.38656 | 19.34033 | 15.2744 | 15.07809 | 13.11252 | 14.69736 | 18.18503 | 16.66118 | 12.45045 | 13.8064 | 15.13071 | 14.23614 | 16.07828 | 15.46736 | 16.13586 | 16.53603 | 16.36631 | 14.09405 |
| PABPC1 | 25.79008 | 25.57085 | 24.16922 | 24.50347 | 18.71672 | 17.58039 | 18.92654 | 18.37042 | 17.90019 | 18.50113 | 19.2067 | 19.99584 | 21.71671 | 20.62701 | 23.16588 | 24.09444 | 17.96325 | 17.63065 |
| PABPC4 | 24.58941 | 24.03491 | 22.91075 | 22.8896 | 17.17375 | 17.34432 | 18.3827 | 17.99293 | 16.59053 | 17.2032 | 17.14153 | 18.46141 | 19.76515 | 19.04717 | 21.90678 | 22.39038 | 18.0784 | 16.17911 |
| PABPC4L | 26.23457 | 25.83157 | 24.83106 | 24.51834 | 20.38172 | 20.26711 | 21.97391 | 20.73357 | 15.32011 | 19.18286 | 17.16507 | 17.08125 | 22.34386 | 21.48653 | 23.78105 | 24.59739 | 21.75107 | 19.78136 |
| PABPC5 | 25.67211 | 25.4405 | 23.79861 | 23.91935 | 20.04024 | 15.4545 | 21.49904 | 20.84319 | 16.74522 | 14.98013 | 19.13612 | 20.57122 | 22.76766 | 19.57649 | 22.46914 | 23.7596 | 20.68967 | 16.6504 |
| PABPN1 | 14.45818 | 15.60426 | 16.48778 | 17.98706 | 16.86058 | 15.11111 | 15.11652 | 19.01559 | 16.5381 | 17.18098 | 18.14939 | 19.3992 | 14.81417 | 19.06025 | 14.44495 | 16.73008 | 16.82754 | 16.08498 |
| PACC1 | 17.35978 | 17.3779 | 17.40669 | 14.49472 | 14.58444 | 17.93267 | 15.78781 | 16.59704 | 20.29249 | 17.13007 | 17.3641 | 19.56574 | 18.23368 | 15.79909 | 17.43162 | 17.24451 | 16.29098 | 15.58546 |

| | | | | | | | | | | | | | | | | | | |
|---|---|---|---|---|---|---|---|---|---|---|---|---|---|---|---|---|---|---|
| PACS1 | 16.42755 | 13.51882 | 16.54733 | 15.39897 | 18.89894 | 16.98314 | 16.72722 | 15.64613 | 13.36806 | 16.57212 | 16.84976 | 11.89636 | 17.64037 | 17.76616 | 17.52209 | 17.84445 | 17.51344 | 16.98168 |
| PACS2 | 13.55064 | 13.74322 | 14.21439 | 16.20988 | 13.64187 | 13.88225 | 16.10195 | 14.95424 | 12.8078 | 12.74745 | 15.35048 | 17.94261 | 14.99967 | 13.00426 | 14.71117 | 17.04429 | 14.04949 | 13.76934 |
| PACSIN1 | 23.04047 | 22.73782 | 19.46795 | 19.75633 | 16.17148 | 15.86826 | 15.82783 | 17.19035 | 16.60194 | 17.2367 | 16.49572 | 16.60201 | 19.3089 | 18.27322 | 20.86448 | 19.56632 | 16.15138 | 16.83684 |
| PACSIN2 | 24.51583 | 24.37428 | 23.77738 | 24.03086 | 20.55417 | 19.24138 | 19.20108 | 19.15032 | 17.82271 | 18.77303 | 19.10578 | 19.04636 | 24.57949 | 23.45604 | 24.61156 | 24.09091 | 19.26136 | 20.00738 |
| PACSIN3 | 23.69038 | 23.71488 | 22.03061 | 22.15035 | 18.40657 | 17.41185 | 17.59981 | 16.70283 | 16.25803 | 16.38619 | 17.66769 | 14.44602 | 21.85217 | 20.10107 | 22.80016 | 22.09761 | 18.5223 | 18.62082 |
| PAFAH1B1 | 16.22893 | 16.61862 | 18.06766 | 18.27914 | 19.3804 | 18.54223 | 19.00885 | 18.3259 | 19.27972 | 19.1776 | 19.18196 | 19.44854 | 18.72814 | 19.33113 | 18.27115 | 17.71675 | 18.32398 | 17.99518 |
| PAFAH1B2 | 18.53496 | 17.87903 | 19.5641 | 17.30492 | 17.55441 | 16.33176 | 17.53087 | 15.90698 | 17.67993 | 15.61655 | 19.70892 | 22.0248 | 18.39184 | 18.33588 | 20.07722 | 19.86492 | 18.49658 | 17.4613 |
| PAFAH1B3 | 18.31147 | 17.60668 | 17.68225 | 17.71919 | 17.42866 | 16.17756 | 18.57165 | 17.37864 | 19.72815 | 19.60889 | 21.05223 | 21.75551 | 17.42951 | 17.43077 | 17.39057 | 17.99416 | 15.67235 | 16.01323 |
| PAFAH2 | 16.45849 | 16.76979 | 15.68662 | 14.81915 | 14.20928 | 16.00217 | 15.40554 | 14.31596 | 18.06957 | 15.33327 | 16.90496 | 13.44582 | 15.75897 | 14.64214 | 14.07508 | 16.17847 | 16.22568 | 13.6861 |
| PAICS | 16.54122 | 16.98311 | 17.9674 | 18.49221 | 19.29521 | 18.50868 | 19.04079 | 18.59178 | 17.81914 | 18.57794 | 20.15113 | 21.08435 | 20.68635 | 20.43363 | 17.82282 | 20.91975 | 20.66787 | 20.02022 |
| PAIP1 | 20.81354 | 19.81892 | 19.11716 | 19.09984 | 15.51342 | 18.6212 | 16.11576 | 16.23756 | 18.10456 | 19.06472 | 17.18433 | 17.93393 | 17.63885 | 15.20183 | 18.30632 | 18.85001 | 16.29773 | 15.93734 |
| PAIP2B | 19.52 | 19.90193 | 17.8515 | 14.79224 | 19.27262 | 18.49985 | 16.37231 | 14.01209 | 13.19978 | 14.59462 | 15.35376 | 15.38011 | 16.84397 | 20.8115 | 18.00873 | 17.84013 | 14.58886 | 12.30789 |
| PAK1 | 14.16188 | 17.24115 | 19.00939 | 17.909 | 16.72307 | 16.59785 | 14.84143 | 15.54486 | 15.42933 | 15.62512 | 15.76963 | 16.36851 | 16.76619 | 16.58909 | 14.23567 | 15.72565 | 15.42872 | 15.09301 |
| PAK2 | 17.41126 | 16.71307 | 16.11917 | 18.16075 | 20.03147 | 18.65889 | 17.24246 | 17.99671 | 17.725 | 17.80147 | 19.22271 | 19.41448 | 20.12929 | 19.48052 | 17.22916 | 17.36059 | 20.5002 | 19.90035 |
| PAK4 | 18.30665 | 18.80355 | 18.37958 | 18.87762 | 17.38749 | 15.50816 | 15.60621 | 15.65831 | 14.20377 | 14.29601 | 16.91389 | 17.72907 | 17.05025 | 16.16191 | 18.79722 | 18.93705 | 18.36737 | 17.05297 |
| PAK6 | 16.4069 | 16.57544 | 15.18814 | 17.67842 | 18.17917 | 17.11859 | 18.73434 | 17.40558 | 15.17197 | 15.66827 | 17.10788 | 15.81271 | 14.63654 | 17.4674 | 14.3874 | 13.89969 | 19.14989 | 19.16225 |
| PALD1 | 15.87068 | 14.91231 | 15.55416 | 16.59116 | 13.62848 | 18.2964 | 15.63644 | 15.68784 | 15.25404 | 15.69419 | 17.82278 | 18.7422 | 16.83859 | 15.95695 | 16.0172 | 15.47112 | 16.83207 | 17.19791 |
| PALM | 16.69036 | 18.85161 | 18.61334 | 15.70257 | 18.94692 | 17.90832 | 14.8346 | 16.31911 | 17.29846 | 14.69009 | 15.21065 | 15.46213 | 13.91301 | 14.75835 | 17.20173 | 16.37531 | 17.10506 | 18.71599 |
| PALM2 | 15.19835 | 16.16311 | 17.64276 | 14.99614 | 18.72767 | 16.46856 | 17.69167 | 18.4165 | 17.16332 | 16.88203 | 14.06384 | 11.89754 | 13.84593 | 14.17595 | 15.1265 | 15.26172 | 16.55103 | 17.58283 |
| PALS1 | 16.15097 | 15.92145 | 16.4437 | 18.47271 | 20.56863 | 19.69832 | 17.26997 | 15.53077 | 17.14384 | 17.87989 | 18.24573 | 15.63351 | 19.78724 | 19.31063 | 17.31167 | 16.1666 | 19.77428 | 20.30499 |
| PALS2 | 15.68687 | 15.86228 | 18.775 | 18.81608 | 20.49722 | 19.10378 | 17.15519 | 18.74946 | 17.79435 | 17.42362 | 20.63026 | 18.35984 | 17.76676 | 18.09801 | 18.896 | 18.45377 | 20.02293 | 20.54438 |
| PAM | 14.83148 | 15.90115 | 15.47805 | 15.94576 | 15.50803 | 15.28628 | 16.53793 | 16.80233 | 19.24205 | 13.93306 | 20.46307 | 17.43102 | 13.46595 | 13.5345 | 15.46988 | 15.84185 | 14.10691 | 14.18589 |
| PAM16 | 12.7974 | 15.14727 | 15.54006 | 16.45327 | 15.24087 | 15.80523 | 15.81274 | 17.58716 | 19.7612 | 13.04636 | 13.85108 | 14.66012 | 15.31112 | 15.80051 | 15.03122 | 15.13254 | 13.25499 | 13.94109 |
| PAN2 | 19.59842 | 19.64451 | 18.20756 | 18.9878 | 16.42347 | 15.29773 | 16.3669 | 16.23749 | 15.64112 | 15.4836 | 16.01839 | 15.02485 | 15.524 | 16.1432 | 17.38286 | 17.20136 | 15.53806 | 15.6826 |
| PAN3 | 19.93625 | 18.63548 | 15.69804 | 18.68239 | 13.58496 | 15.4972 | 15.31349 | 15.31012 | 13.89398 | 14.34733 | 16.69649 | 14.28881 | 13.92562 | 16.20704 | 15.78504 | 16.38866 | 16.27661 | 14.73038 |
| PANK4 | 20.20965 | 15.90676 | 15.30844 | 20.11438 | 20.79711 | 20.52821 | 20.33963 | 21.78161 | 18.93507 | 19.56157 | 19.74036 | 17.49854 | 16.09623 | 14.72521 | 19.018 | 19.05001 | 21.04325 | 20.86353 |
| PANX1 | 13.54766 | 17.24868 | 20.09011 | 18.22362 | 16.65615 | 18.67389 | 14.95009 | 15.26301 | 16.006 | 16.96181 | 19.61379 | 18.79701 | 17.57707 | 17.50229 | 13.68598 | 18.48102 | 17.53841 | 15.82255 |
| PAPOLA | 15.59167 | 15.90457 | 15.99107 | 14.12055 | 12.28926 | 15.57288 | 16.07456 | 16.79532 | 13.33448 | 14.33361 | 14.87921 | 17.65755 | 14.81807 | 15.84188 | 17.08917 | 16.41013 | 15.05272 | 15.04464 |
| PAPSS1 | 17.36305 | 18.9013 | 16.58689 | 18.20561 | 18.86349 | 19.61676 | 17.51756 | 16.43131 | 13.66957 | 15.28078 | 17.52708 | 18.70844 | 17.95524 | 17.73464 | 17.40042 | 17.62247 | 14.63367 | 15.05485 |
| PAPSS2 | 16.27674 | 16.84394 | 16.05326 | 15.1631 | 18.60446 | 18.56505 | 18.33334 | 16.70487 | 14.34935 | 15.38027 | 15.47419 | 16.59486 | 14.74734 | 15.82361 | 13.58496 | 15.54496 | 20.41307 | 20.77151 |
| PARD3 | 15.85562 | 15.19092 | 19.33284 | 16.20313 | 19.30431 | 17.94434 | 17.12362 | 16.33607 | 18.10618 | 16.68137 | 16.40366 | 16.91035 | 15.98433 | 16.59922 | 14.69165 | 16.26046 | 20.97257 | 20.03237 |
| PARK7 | 20.10392 | 19.04383 | 20.4829 | 20.75302 | 21.94349 | 19.19712 | 19.85053 | 18.98901 | 21.366 | 20.92578 | 21.01736 | 21.3079 | 23.13794 | 22.83126 | 20.95438 | 21.30404 | 21.67524 | 22.05466 |
| PARP1 | 19.71254 | 19.1247 | 17.12454 | 16.7642 | 17.57447 | 17.92644 | 19.31918 | 20.62693 | 17.98642 | 19.14588 | 19.28909 | 19.30444 | 18.73834 | 18.11828 | 16.06628 | 16.2576 | 17.98756 | 18.30959 |
| PARP12 | 16.4308 | 18.92515 | 19.57825 | 19.40537 | 14.03746 | 15.74415 | 15.92048 | 16.91288 | 14.05796 | 15.20602 | 14.94465 | 14.15177 | 17.11033 | 14.97818 | 19.48176 | 19.50162 | 14.96683 | 15.40563 |
| PARP4 | 21.08193 | 21.29714 | 15.92086 | 20.89849 | 15.9639 | 16.57441 | 16.04058 | 16.25675 | 15.06633 | 16.60634 | 14.93095 | 18.84937 | 16.14904 | 15.62748 | 20.28636 | 20.09697 | 15.43096 | 16.55453 |
| PARVA | 15.4436 | 15.35089 | 15.40956 | 19.5521 | 18.90681 | 15.61339 | 15.8093 | 15.5004 | 16.93394 | 16.34662 | 15.39337 | 12.51029 | 14.21369 | 13.9398 | 13.9184 | 15.47118 | 18.65815 | 19.44492 |
| PARVB | 18.60535 | 19.16518 | 20.04218 | 19.97101 | 15.69748 | 16.38635 | 17.99495 | 18.75773 | 15.44918 | 12.7717 | 16.15715 | 17.2165 | 17.27374 | 15.84868 | 19.41423 | 19.37555 | 16.33316 | 16.71515 |
| PATJ | 15.90626 | 14.88864 | 17.26158 | 17.91729 | 20.42159 | 18.58511 | 16.16821 | 15.67778 | 17.11021 | 16.65954 | 15.17742 | 14.9298 | 19.55698 | 20.51953 | 18.26152 | 16.88928 | 20.19086 | 20.10564 |
| PAWR | 18.11517 | 18.66777 | 16.4672 | 17.40432 | 18.34398 | 15.99876 | 16.12688 | 16.19217 | 15.74532 | 14.10917 | 15.80197 | 15.87847 | 15.28777 | 17.64113 | 14.74618 | 18.63538 | 18.34157 | 14.48516 |
| PBDC1 | 16.7541 | 16.23539 | 16.86392 | 16.60353 | 17.06074 | 17.45583 | 17.09443 | 17.10633 | 15.27096 | 17.86012 | 16.7828 | 18.64596 | 17.94503 | 17.80745 | 16.25407 | 16.23038 | 19.95255 | 17.59256 |
| PBK | 17.451 | 17.71795 | 18.14814 | 17.43958 | 18.684 | 16.64021 | 17.71379 | 18.63025 | 17.39506 | 17.2299 | 16.80584 | 15.31058 | 19.3277 | 18.36695 | 17.15864 | 17.63096 | 19.47441 | 16.19296 |
| PBXIP1 | 18.5413 | 16.91339 | 17.53934 | 18.22951 | 16.20363 | 16.2132 | 16.8794 | 17.17587 | 20.55208 | 18.63157 | 20.05035 | 17.14456 | 16.57269 | 16.50143 | 16.9126 | 17.79757 | 17.25514 | 16.60526 |
| PCBD1 | 17.11674 | 19.60386 | 19.84124 | 20.05702 | 19.88561 | 18.79541 | 17.4229 | 15.97823 | 21.12522 | 21.32422 | 19.58616 | 20.06004 | 21.3368 | 20.86986 | 20.21276 | 19.26055 | 20.5627 | 20.64488 |
| PCBD2 | 15.40878 | 15.76797 | 16.73797 | 17.47311 | 16.48143 | 16.97823 | 16.43137 | 16.6557 | 16.95394 | 17.28399 | 17.65387 | 15.9748 | 18.32804 | 17.22088 | 17.7865 | 17.35881 | 15.50916 | 13.69859 |
| PCBP1 | 23.70114 | 23.23955 | 22.13336 | 22.4766 | 19.68951 | 19.06484 | 20.56028 | 19.44394 | 18.80753 | 19.16682 | 21.12154 | 21.82111 | 22.38818 | 21.02447 | 21.58248 | 22.30068 | 19.27399 | 19.45127 |
| PCBP2 | 24.71675 | 24.45682 | 23.12829 | 24.28633 | 19.41265 | 17.15624 | 21.35421 | 18.58124 | 18.89614 | 18.39223 | 19.63111 | 20.91602 | 23.38324 | 22.27637 | 22.79121 | 23.30994 | 18.50639 | 19.32991 |
| PCBP3 | 24.84549 | 23.16084 | 22.65336 | 23.5183 | 18.70028 | 18.57212 | 20.64287 | 20.62604 | 18.68318 | 18.63033 | 19.78312 | 20.64211 | 22.7929 | 21.54077 | 21.92056 | 22.59808 | 19.81027 | 18.7347 |
| PCCA | 16.47635 | 15.06237 | 16.14102 | 15.79421 | 15.66887 | 16.11695 | 14.89119 | 18.87565 | 18.55967 | 17.5097 | 15.75142 | 13.7717 | 18.62231 | 13.73863 | 15.76736 | 15.79858 | 14.85347 | 15.51858 |
| PCCB | 17.61938 | 19.13358 | 15.28739 | 14.23957 | 17.97533 | 16.48358 | 16.23507 | 18.86765 | 18.49965 | 16.3561 | 16.00178 | 16.26164 | 16.53716 | 14.93034 | 18.36271 | 16.31519 | 18.68217 | 15.12878 |
| PCDH1 | 18.24153 | 18.44897 | 18.24803 | 18.14695 | 14.93494 | 16.84443 | 16.28144 | 17.05631 | 17.71228 | 15.77101 | 21.19581 | 20.14153 | 17.90306 | 17.7194 | 18.7412 | 19.12812 | 17.10568 | 18.25443 |
| PCDHA4 | 14.18243 | 13.51586 | 16.07113 | 16.07688 | 18.73556 | 18.04288 | 19.74386 | 13.86216 | 19.46928 | 18.26887 | 14.45499 | 11.85014 | 13.97469 | 13.51616 | 15.83559 | 13.98342 | 18.34358 | 18.65803 |
| PCDHB5 | 16.27714 | 16.11287 | 15.63825 | 15.29376 | 17.5497 | 14.14928 | 15.42229 | 17.26109 | 18.92962 | 18.1324 | 13.37038 | 18.88483 | 17.82772 | 17.24164 | 12.15547 | 18.90092 | 14.86138 | 12.70912 |
| PCDHB8 | 16.87765 | 17.61525 | 16.05201 | 15.78215 | 15.12238 | 16.78059 | 14.87128 | 16.12808 | 19.06772 | 18.17047 | 18.57849 | 20.90726 | 15.37324 | 16.24339 | 14.04321 | 15.35312 | 13.90412 | 17.52735 |
| PCID2 | 15.63634 | 17.03304 | 13.57377 | 14.83667 | 14.49646 | 17.09651 | 13.95647 | 16.25958 | 12.556 | 15.53261 | 15.29129 | 18.61022 | 11.92236 | 13.27737 | 16.67402 | 15.81975 | 14.83547 | 13.72176 |
| PCK2 | 20.17024 | 22.08642 | 15.74779 | 17.73538 | 15.65733 | 16.43813 | 16.98026 | 16.6751 | 20.3412 | 18.4851 | 15.52757 | 14.72833 | 15.97086 | 15.33599 | 16.73441 | 19.02241 | 18.08301 | 15.55771 |
| PCMT1 | 18.83558 | 18.65043 | 18.2842 | 16.45424 | 19.18096 | 20.21282 | 18.86656 | 18.26322 | 20.19154 | 20.66232 | 19.71307 | 19.80892 | 18.64484 | 20.0962 | 17.78909 | 18.60693 | 19.46526 | 18.88468 |
| PCNA | 16.10926 | 18.4473 | 17.85558 | 18.26191 | 19.53597 | 18.80717 | 20.35196 | 20.59304 | 19.84776 | 21.75257 | 19.94935 | 20.01573 | 21.40676 | 19.45711 | 18.52902 | 17.2128 | 19.70423 | 18.26802 |
| PCNP | 14.10103 | 15.25594 | 16.36047 | 14.44709 | 15.88703 | 16.27593 | 14.21094 | 16.68813 | 17.05643 | 16.71118 | 19.04541 | 15.10322 | 15.80303 | 14.92257 | 15.23401 | 14.41306 | 16.61063 | 16.59275 |
| PCSK1 | 16.94567 | 15.00872 | 16.33226 | 16.05107 | 13.78081 | 16.0445 | 16.63579 | 16.57323 | 18.75628 | 18.98682 | 17.42603 | 13.66515 | 15.57593 | 15.32369 | 15.10411 | 14.0401 | 16.22425 | 13.23185 |
| PCSK1N | 17.58813 | 15.95882 | 16.79451 | 15.37259 | 15.10123 | 18.11604 | 16.3921 | 16.47812 | 21.48066 | 21.66122 | 15.24029 | 13.9166 | 15.11476 | 14.95229 | 14.2194 | 17.86159 | 17.00217 | 17.00826 |
| PCYOX1 | 16.62779 | 16.48123 | 17.55953 | 16.10346 | 16.50757 | 21.50775 | 18.88796 | 16.42447 | 21.62651 | 21.13706 | 17.69219 | 14.13162 | 15.68841 | 16.27338 | 16.897 | 15.64525 | 18.09964 | 16.7519 |
| PCYOX1L | 14.96415 | 16.5194 | 15.59774 | 15.77896 | 15.11228 | 16.01576 | 15.66505 | 17.65346 | 20.78177 | 18.37662 | 17.33272 | 14.39695 | 15.50927 | 15.2365 | 15.97634 | 15.7193 | 15.12516 | 16.55131 |
| PCYT2 | 15.62082 | 16.23648 | 17.44287 | 18.78086 | 18.35806 | 20.65803 | 19.30068 | 19.30742 | 16.33228 | 16.4636 | 17.10467 | 18.61783 | 16.90628 | 19.57589 | 19.93517 | 16.64863 | 15.82187 | 19.21798 |
| PDAP1 | 19.45991 | 19.37206 | 19.69568 | 19.90509 | 19.45614 | 18.05751 | 19.95499 | 18.60287 | 16.48908 | 16.97864 | 17.43963 | 17.30738 | 18.86366 | 17.89433 | 19.07836 | 19.09465 | 19.09968 | 20.40345 |
| PDCD10 | 16.30658 | 16.62423 | 15.51368 | 18.40321 | 19.95051 | 16.52477 | 17.23659 | 17.53998 | 17.46522 | 17.53515 | 20.14728 | 18.98518 | 19.92942 | 19.88805 | 15.74362 | 18.58503 | 18.85922 | 19.71769 |
| PDCD11 | 13.94589 | 15.21573 | 14.62083 | 13.33457 | 14.09837 | 16.11385 | 17.62656 | 16.99445 | 13.45094 | 17.48529 | 14.12944 | 13.5533 | 15.69282 | 15.22817 | 15.15488 | 15.55908 | 16.00562 | 14.89833 |
| PDCD2L | 15.99914 | 13.2345 | 14.83998 | 17.96992 | 20.69692 | 19.13006 | 16.3653 | 16.39942 | 15.7669 | 18.18061 | 15.99228 | 16.46783 | 21.55195 | 20.88643 | 15.55292 | 17.11173 | 20.42858 | 20.37852 |
| PDCD4 | 16.11307 | 17.94315 | 15.68324 | 16.83194 | 14.13007 | 17.65613 | 16.1663 | 16.87687 | 16.17318 | 18.35666 | 18.12706 | 19.2908 | 17.08508 | 15.61256 | 18.08744 | 16.40758 | 18.04576 | 13.39771 |
| PDCD5 | 19.32328 | 17.78613 | 20.33003 | 20.80016 | 17.99008 | 18.15091 | 18.30324 | 15.93906 | 19.19858 | 18.17972 | 20.29624 | 19.27671 | 19.65771 | 18.4723 | 20.31483 | 20.37544 | 17.90604 | 16.54797 |
| PDCD6 | 22.97785 | 23.46063 | 21.48093 | 21.60727 | 19.23069 | 20.21969 | 17.27061 | 16.49077 | 22.76998 | 21.43923 | 17.36167 | 20.51151 | 18.59684 | 18.36841 | 22.97045 | 22.25277 | 18.23444 | 19.18772 |
| PDCD6IP | 25.54504 | 25.55739 | 24.10805 | 24.20457 | 23.70558 | 22.62856 | 21.36466 | 20.21226 | 22.19536 | 23.15612 | 21.10317 | 20.97986 | 24.35683 | 23.92683 | 24.25229 | 24.23132 | 21.8239 | 23.16191 |
| PDE2A | 15.45534 | 15.63458 | 16.06059 | 15.5499 | 17.97807 | 18.27299 | 16.28186 | 16.55647 | 13.10711 | 15.4067 | 17.91184 | 20.0136 | 16.37485 | 16.38013 | 16.60768 | 15.51118 | 16.85382 | 17.38958 |
| PDE4DIP | 14.74516 | 14.04259 | 17.08358 | 17.04334 | 15.59734 | 12.18394 | 19.41693 | 17.83677 | 19.11384 | 17.90366 | 12.94816 | 17.00843 | 19.43245 | 14.69101 | 17.41674 | 16.60946 | 15.19246 | 13.97972 |
| PDGFD | 14.87449 | 15.76537 | 15.22614 | 14.85651 | 18.91193 | 19.10086 | 14.19229 | 15.23836 | 13.09147 | 13.33702 | 17.79505 | 11.68019 | 15.03753 | 15.53129 | 15.1447 | 15.45717 | 17.42425 | 18.55015 |
| PDHA1 | 17.25351 | 18.54336 | 19.18175 | 16.11812 | 15.83937 | 15.22556 | 16.72703 | 16.91736 | 18.58282 | 17.41289 | 14.46726 | 14.19232 | 17.63544 | 15.6464 | 18.02262 | 17.33059 | 16.10756 | 16.20513 |
| PDHB | 15.46127 | 15.99421 | 14.72178 | 15.2035 | 13.79827 | 21.68387 | 18.57353 | 15.3975 | 20.20336 | 19.00745 | 16.63745 | 14.67447 | 16.87833 | 16.26501 | 15.5308 | 16.58753 | 16.34355 | 15.99169 |
| PDIA3 | 18.38254 | 18.96622 | 19.21313 | 18.77649 | 20.5113 | 19.52149 | 21.61966 | 20.46705 | 24.9159 | 24.76843 | 23.86279 | 20.54498 | 20.51459 | 21.16024 | 20.33904 | 19.52691 | 19.5424 | 20.92354 |
| PDIA4 | 18.21078 | 17.68 | 17.94017 | 17.5157 | 18.73443 | 19.24757 | 21.1236 | 20.98482 | 23.03268 | 22.79683 | 21.73433 | 18.2725 | 18.63144 | 19.27322 | 18.54499 | 17.73174 | 17.48365 | 18.83335 |
| PDIA5 | 16.5897 | 17.13672 | 17.34999 | 15.94111 | 21.94866 | 19.40611 | 18.26883 | 17.70178 | 21.46355 | 20.96244 | 17.68415 | 14.4087 | 20.21274 | 20.4522 | 16.20429 | 16.3835 | 21.67623 | 21.28378 |
| PDIA6 | 17.12921 | 17.19027 | 18.0875 | 16.55824 | 18.86383 | 18.45819 | 19.92929 | 19.03336 | 24.38392 | 24.12746 | 22.60932 | 19.08042 | 18.2306 | 18.5841 | 18.72977 | 18.20464 | 17.88428 | 18.62011 |
| PDK3 | 17.15975 | 17.24089 | 13.96518 | 16.62667 | 14.68584 | 18.86762 | 17.41953 | 16.38951 | 18.95373 | 18.03452 | 16.84189 | 15.73404 | 14.268 | 13.96901 | 16.6809 | 16.31984 | 16.28288 | 17.49165 |
| PDLIM1 | 16.20211 | 15.77884 | 16.27542 | 17.29007 | 20.30306 | 19.00206 | 18.28629 | 18.00947 | 16.08699 | 16.79121 | 16.37271 | 14.4076 | 17.87364 | 18.42245 | 15.83791 | 15.51648 | 21.43471 | 21.52363 |
| PDLIM5 | 16.19951 | 16.6245 | 17.29622 | 17.2558 | 17.60074 | 16.32061 | 16.02811 | 16.12483 | 13.75875 | 16.399 | 18.00732 | 18.71346 | 15.47464 | 15.63412 | 17.63684 | 17.88391 | 18.75608 | 18.619 |
| PDLIM7 | 16.79083 | 17.42001 | 16.90514 | 17.21953 | 20.41596 | 19.48058 | 19.13729 | 16.21298 | 13.39078 | 16.26864 | 14.75225 | 13.88774 | 18.60064 | 17.10003 | 16.78529 | 18.14557 | 19.03554 | 18.92666 |
| PDPK1 | 16.80364 | 17.25197 | 19.67651 | 20.20895 | 17.18432 | 15.25576 | 17.68238 | 17.4494 | 15.9712 | 17.83876 | 14.4447 | 17.90842 | 20.31505 | 20.6152 | 20.33281 | 20.26876 | 17.99768 | 18.63462 |
| PDPK2P | 16.26073 | 16.72468 | 19.9574 | 19.22087 | 16.99518 | 16.1059 | 16.62261 | 17.19992 | 16.18823 | 16.62182 | 15.34262 | 14.48544 | 20.49394 | 20.34595 | 19.679 | 19.92718 | 15.64272 | 17.87075 |

| | | | | | | | | | | | | | | | | | | |
|---|---|---|---|---|---|---|---|---|---|---|---|---|---|---|---|---|---|---|
| PDS5A | 17.42949 | 16.84977 | 15.64496 | 15.9803 | 14.91159 | 17.70405 | 19.69042 | 17.92279 | 14.57534 | 14.64056 | 16.6735 | 16.94688 | 15.44533 | 21.38571 | 15.15348 | 15.15756 | 13.56021 | 16.54916 |
| PDS5B | 15.31971 | 16.36679 | 16.46039 | 16.1317 | 16.69465 | 14.65349 | 14.72417 | 18.88543 | 13.76432 | 14.28905 | 15.97281 | 17.32699 | 15.0804 | 14.79274 | 16.47788 | 17.2743 | 16.05489 | 14.97748 |
| PDXDC1 | 15.91736 | 15.25723 | 17.66357 | 18.42649 | 18.02314 | 15.44354 | 18.44736 | 18.09081 | 14.79306 | 15.22692 | 18.79054 | 17.84994 | 18.28976 | 20.25203 | 16.87307 | 16.175 | 16.96517 | 20.34797 |
| PDXK | 18.45611 | 16.63829 | 17.65415 | 18.49586 | 20.87244 | 20.02245 | 15.2558 | 16.96599 | 15.98693 | 17.97763 | 17.16304 | 18.70842 | 19.93704 | 18.74035 | 15.38992 | 20.84761 | 20.90107 | 20.63341 |
| PDXP | 13.44072 | 15.22636 | 16.50193 | 14.04261 | 15.68513 | 16.52715 | 13.74504 | 15.931 | 14.5284 | 15.45085 | 16.1571 | 18.3581 | 14.5484 | 15.23445 | 14.43388 | 15.85123 | 15.62612 | 17.16172 |
| PEA15 | 19.26262 | 20.87666 | 18.84882 | 18.3443 | 18.57852 | 17.59018 | 16.7189 | 17.01395 | 16.5709 | 16.56307 | 15.31342 | 14.03154 | 19.73931 | 20.11827 | 17.68385 | 18.44748 | 18.15865 | 18.99918 |
| PEAK1 | 16.60102 | 13.56907 | 16.1259 | 14.10399 | 17.84209 | 16.44981 | 14.54294 | 16.79001 | 12.96507 | 14.03633 | 16.57655 | 15.53221 | 15.11711 | 16.11129 | 14.57826 | 15.81154 | 17.49015 | 15.18058 |
| PEBP1 | 22.39057 | 21.8649 | 22.09656 | 22.53379 | 22.93109 | 21.57804 | 20.68865 | 19.08533 | 21.92452 | 22.86873 | 23.46476 | 22.52315 | 24.23462 | 23.88684 | 23.14773 | 22.91156 | 24.36524 | 23.81379 |
| PECR | 15.5649 | 17.39274 | 15.64394 | 15.55911 | 17.64006 | 20.14129 | 16.88742 | 16.21205 | 19.21647 | 18.57053 | 15.59949 | 14.9938 | 15.11609 | 15.90079 | 15.58531 | 14.42983 | 19.85654 | 20.08745 |
| PEF1 | 18.73541 | 19.64084 | 19.09055 | 19.82686 | 17.81723 | 18.06737 | 15.87951 | 16.36169 | 21.27287 | 17.65011 | 17.31188 | 18.07225 | 17.59573 | 17.74083 | 20.61332 | 19.93966 | 15.46341 | 16.52809 |
| PEG10 | 17.83379 | 18.06502 | 19.87515 | 16.55151 | 13.56354 | 18.15532 | 14.29442 | 14.97619 | 15.44876 | 15.26126 | 14.92954 | 12.32957 | 14.19256 | 15.87331 | 15.52986 | 14.35054 | 13.19163 | 15.78502 |
| PELI2 | 17.71346 | 15.53559 | 15.55823 | 16.40578 | 14.66036 | 14.61812 | 14.9989 | 15.01902 | 18.75953 | 18.66592 | 16.57208 | 17.52213 | 13.88827 | 14.94675 | 15.61225 | 14.69704 | 14.53721 | 14.83801 |
| PELO | 18.99361 | 17.84908 | 15.5999 | 16.65773 | 17.11369 | 19.11002 | 15.84373 | 16.52515 | 12.42661 | 14.84754 | 13.42367 | 16.30127 | 17.25638 | 17.24868 | 17.64351 | 17.7025 | 14.77809 | 17.89207 |
| PEPD | 16.87219 | 15.91493 | 17.53426 | 14.82764 | 17.91875 | 15.48747 | 18.71241 | 18.64598 | 17.04137 | 17.40669 | 17.75911 | 19.00703 | 15.58123 | 17.76241 | 16.3439 | 16.35845 | 18.86637 | 18.28503 |
| PEX11B | 16.77234 | 16.0904 | 15.19494 | 16.51407 | 15.05286 | 15.17552 | 16.01461 | 24.1159 | 20.32079 | 17.8966 | 18.28388 | 13.747 | 14.39611 | 17.81922 | 16.6375 | 16.44391 | 14.14523 | 15.47607 |
| PEX14 | 16.7237 | 16.39669 | 15.87282 | 16.60112 | 14.20018 | 15.53736 | 16.2469 | 16.68331 | 21.9112 | 20.90236 | 14.94929 | 15.75161 | 15.58756 | 14.98184 | 14.76902 | 17.17751 | 15.45693 | 16.88242 |
| PEX19 | 17.13726 | 17.64909 | 17.06931 | 16.76643 | 17.24235 | 14.42581 | 15.09452 | 15.63964 | 16.70196 | 16.50059 | 16.20612 | 17.0102 | 18.85291 | 19.03446 | 16.27314 | 17.07862 | 17.51316 | 18.60967 |
| PFAS | 15.48316 | 16.12076 | 16.23852 | 16.73906 | 17.3236 | 15.98225 | 16.52668 | 16.77545 | 14.84022 | 14.2767 | 17.67807 | 18.90872 | 16.13745 | 15.52565 | 15.94527 | 16.31024 | 16.28218 | 16.57601 |
| PFDN1 | 17.36803 | 19.42723 | 18.03444 | 17.64741 | 20.09135 | 17.94331 | 17.76238 | 18.77879 | 18.15116 | 18.67862 | 18.98845 | 19.61175 | 19.90016 | 19.5852 | 15.79923 | 17.78316 | 21.36994 | 20.31608 |
| PFDN2 | 17.98051 | 17.23333 | 17.31828 | 18.40032 | 20.17923 | 18.80023 | 19.37489 | 18.54605 | 18.0135 | 19.55152 | 21.17964 | 20.76689 | 21.31036 | 20.58993 | 18.57005 | 17.93976 | 21.07487 | 20.96248 |
| PFDN4 | 17.40595 | 17.30415 | 17.39699 | 17.62124 | 19.4276 | 17.86137 | 16.5486 | 18.08653 | 18.97618 | 18.98115 | 20.30961 | 20.23148 | 19.84891 | 19.23011 | 17.41312 | 17.34337 | 20.79626 | 19.28391 |
| PFDN5 | 16.8308 | 16.82815 | 16.12007 | 17.00811 | 19.93011 | 17.43966 | 18.77818 | 17.99672 | 17.64642 | 19.59616 | 17.87621 | 19.6287 | 20.83144 | 19.58953 | 16.74383 | 16.19039 | 20.8216 | 19.86232 |
| PFDN6 | 16.43047 | 16.19826 | 16.46005 | 16.0627 | 19.86081 | 17.38444 | 19.14802 | 18.42247 | 17.57779 | 18.58151 | 18.25372 | 19.38008 | 20.73109 | 19.74531 | 17.00822 | 16.82826 | 21.1085 | 19.75326 |
| PFKFB2 | 14.97669 | 17.04311 | 15.34439 | 16.29973 | 14.88555 | 15.71308 | 15.1419 | 16.39971 | 14.79845 | 13.94561 | 16.96266 | 18.32215 | 16.47263 | 13.03815 | 14.5325 | 14.87112 | 15.59435 | 14.84237 |
| PFKL | 17.66855 | 18.2965 | 17.84718 | 19.29162 | 18.44967 | 16.29547 | 18.90602 | 19.09601 | 16.99 | 16.97544 | 17.71884 | 17.89194 | 18.3943 | 17.93447 | 18.49506 | 18.72349 | 19.43595 | 19.83848 |
| PFKM | 17.36952 | 16.67183 | 17.73451 | 17.92812 | 16.61368 | 17.39752 | 18.38544 | 17.83683 | 14.00017 | 14.82718 | 17.8764 | 19.77879 | 19.2593 | 18.93037 | 18.63311 | 18.40008 | 18.00755 | 17.88823 |
| PFKP | 16.31919 | 16.86907 | 16.22218 | 15.3601 | 20.19662 | 19.67779 | 17.30523 | 17.58291 | 16.97967 | 17.04535 | 18.89408 | 19.90417 | 16.79178 | 16.73869 | 16.71924 | 17.08156 | 19.92307 | 19.98662 |
| PFN1 | 24.00411 | 24.58092 | 23.20638 | 24.28144 | 26.17407 | 24.91314 | 25.27077 | 24.63018 | 23.35307 | 24.66054 | 24.31804 | 23.66058 | 25.80522 | 25.22503 | 24.06885 | 24.05092 | 26.02147 | 25.3653 |
| PFN2 | 20.88963 | 20.98508 | 22.59044 | 19.01778 | 21.91237 | 20.80902 | 18.76711 | 18.75125 | 19.1858 | 20.36479 | 19.64778 | 20.9013 | 21.02598 | 20.98965 | 19.06417 | 19.45408 | 22.35666 | 21.47807 |
| PGAM1 | 19.12387 | 18.95187 | 20.32687 | 20.85678 | 22.52733 | 22.21045 | 19.24505 | 19.04755 | 18.36231 | 18.97487 | 21.64295 | 22.09254 | 22.27635 | 21.26749 | 20.50134 | 20.42663 | 21.96667 | 22.64852 |
| PGAM5 | 13.94974 | 16.15081 | 14.70486 | 14.12406 | 11.9288 | 16.0183 | 14.94382 | 16.83037 | 18.74691 | 18.19002 | 16.6196 | 17.56395 | 15.85602 | 17.17418 | 15.89759 | 13.84918 | 16.95549 | 14.71537 |
| PGAP1 | 15.96538 | 14.9639 | 15.02295 | 14.38765 | 14.39536 | 18.92249 | 16.26845 | 15.36445 | 19.63458 | 17.30017 | 15.00009 | 12.46791 | 16.72628 | 15.81921 | 15.72888 | 16.33819 | 15.01703 | 14.48313 |
| PGAP4 | 16.0829 | 16.25025 | 15.01963 | 14.27054 | 15.95366 | 16.00656 | 14.89173 | 16.48687 | 19.52105 | 15.75926 | 14.69963 | 14.66976 | 15.14418 | 15.14977 | 15.11815 | 15.49507 | 16.35412 | 16.12193 |
| PGAP6 | 18.39125 | 18.19199 | 18.7826 | 17.73445 | 12.7687 | 15.76216 | 14.31779 | 13.9055 | 17.1021 | 14.25952 | 17.70992 | 18.73296 | 14.90697 | 14.79324 | 19.56198 | 18.56428 | 13.33674 | 15.55597 |
| PGD | 18.84695 | 17.63367 | 19.00359 | 19.92019 | 21.92158 | 20.34014 | 20.11137 | 19.45896 | 20.29666 | 20.9943 | 21.41944 | 21.60235 | 21.14769 | 20.93646 | 20.31929 | 19.54697 | 20.07918 | 20.29164 |
| PGK1 | 19.70266 | 19.69928 | 21.47746 | 21.98138 | 23.61501 | 22.36559 | 23.49071 | 22.46267 | 21.62423 | 22.8267 | 22.98319 | 23.29354 | 24.24304 | 23.91178 | 21.71329 | 21.59293 | 26.2788 | 25.94945 |
| PGLS | 18.55017 | 17.61325 | 15.93434 | 16.1789 | 18.93365 | 17.81342 | 19.21023 | 18.1597 | 19.66925 | 20.43798 | 18.48518 | 19.64497 | 18.67569 | 19.73641 | 15.73651 | 18.31948 | 20.2194 | 21.00139 |
| PGM1 | 14.50732 | 17.11928 | 16.45949 | 17.47071 | 17.01229 | 17.75965 | 17.59019 | 17.25593 | 16.45832 | 16.91038 | 15.28435 | 17.07076 | 17.61904 | 18.30641 | 15.84825 | 16.29453 | 18.9491 | 20.486 |
| PGM2 | 15.55269 | 16.73749 | 15.38461 | 18.01059 | 18.17646 | 17.2969 | 18.15514 | 17.91125 | 15.0493 | 19.52003 | 18.72736 | 18.61385 | 20.13177 | 19.03825 | 17.49331 | 17.52293 | 17.24673 | 18.2598 |
| PGM3 | 16.68718 | 15.78421 | 16.07646 | 16.85419 | 18.0538 | 18.87525 | 16.05944 | 17.10632 | 16.15514 | 17.04931 | 17.2656 | 17.58524 | 20.06016 | 20.27297 | 16.33265 | 17.41815 | 18.20154 | 18.98855 |
| PGP | 16.53169 | 16.71554 | 16.70966 | 17.38214 | 14.4207 | 16.07398 | 16.43739 | 15.22504 | 16.40727 | 15.53637 | 18.90275 | 19.07854 | 16.53698 | 14.87771 | 17.10237 | 17.19751 | 16.90091 | 15.09694 |
| PGRMC1 | 16.81374 | 16.83175 | 15.94843 | 16.82334 | 13.68638 | 16.17986 | 19.13706 | 16.43905 | 17.99566 | 18.45794 | 14.67783 | 18.07943 | 15.44832 | 14.38964 | 14.34527 | 15.3681 | 14.89765 | 13.96593 |
| PGRMC2 | 15.31661 | 15.5944 | 17.03818 | 15.15648 | 16.55997 | 17.7078 | 18.38198 | 16.69199 | 15.36486 | 17.91701 | 16.4245 | 18.83605 | 16.9896 | 15.51008 | 15.50512 | 13.85336 | 14.75243 | 13.85187 |
| PHACTR4 | 16.29788 | 16.24313 | 16.21377 | 16.38655 | 19.09422 | 16.73872 | 15.15293 | 16.71881 | 18.48297 | 17.93372 | 15.62798 | 14.8019 | 18.428 | 18.25329 | 15.81118 | 16.72794 | 17.58274 | 19.97401 |
| PHB | 15.67921 | 15.90177 | 16.67933 | 16.64762 | 16.12169 | 17.43777 | 16.69944 | 18.98203 | 19.20989 | 18.78482 | 16.14376 | 15.0241 | 17.14609 | 17.31111 | 16.34863 | 15.19248 | 15.90526 | 14.82662 |
| PHB2 | 14.93929 | 15.78651 | 15.84219 | 16.1569 | 16.7777 | 16.23612 | 18.96668 | 18.46954 | 18.47276 | 19.22598 | 16.69101 | 16.13835 | 16.77597 | 17.58783 | 16.24974 | 15.18944 | 16.73435 | 15.75562 |
| PHGDH | 18.91726 | 18.55604 | 21.33188 | 21.70156 | 19.54093 | 19.40526 | 21.91724 | 21.47653 | 20.04911 | 20.24762 | 20.74724 | 21.74191 | 23.56612 | 23.15115 | 21.38278 | 21.46634 | 20.57222 | 20.469 |
| PHKB | 15.30256 | 13.75728 | 16.67614 | 12.9764 | 14.7317 | 14.761 | 15.0548 | 16.90263 | 12.61455 | 14.17601 | 16.84027 | 16.55727 | 13.44147 | 13.7223 | 13.891 | 14.29321 | 15.36837 | 16.23376 |
| PHLDA1 | 16.24733 | 16.42909 | 16.70613 | 16.01173 | 20.25086 | 19.09142 | 16.51832 | 17.38571 | 15.97264 | 14.7865 | 15.62158 | 15.51346 | 15.30439 | 16.57934 | 14.33887 | 15.77695 | 22.61344 | 21.38259 |
| PHLDA2 | 18.34374 | 20.03162 | 19.0563 | 19.85667 | 20.43607 | 19.67433 | 18.4536 | 18.3507 | 15.17328 | 15.90614 | 16.46364 | 16.20455 | 20.87931 | 19.64207 | 19.7906 | 19.2291 | 18.76023 | 20.91506 |
| PHLDA3 | 17.28802 | 15.43662 | 16.20029 | 15.20474 | 17.50805 | 17.52591 | 14.42256 | 17.99716 | 12.15485 | 12.71301 | 14.6431 | 12.37836 | 18.8615 | 17.54978 | 18.03519 | 18.43826 | 16.2349 | 15.64081 |
| PHLDB2 | 14.58236 | 15.88768 | 13.12986 | 13.36999 | 17.91128 | 14.38022 | 13.95142 | 15.95685 | 13.38448 | 15.25589 | 14.94882 | 15.13709 | 15.10318 | 15.70945 | 14.07102 | 14.24122 | 16.32969 | 18.21869 |
| PHPT1 | 19.0872 | 17.82093 | 18.19688 | 18.93922 | 20.03616 | 16.89018 | 19.48047 | 17.87033 | 18.01806 | 19.298 | 18.46864 | 19.63138 | 20.69268 | 20.25217 | 20.22386 | 19.82555 | 19.76751 | 20.05907 |
| PHYH | 17.76023 | 16.15567 | 16.64475 | 16.22282 | 14.47357 | 16.75016 | 17.01556 | 16.57147 | 21.68285 | 20.72585 | 15.38557 | 16.30777 | 15.58595 | 15.26053 | 16.04744 | 16.10444 | 15.71908 | 15.65343 |
| PHYHIPL | 13.22514 | 16.35859 | 13.29244 | 14.92233 | 11.91995 | 17.39739 | 17.55056 | 18.97476 | 16.63478 | 16.95182 | 19.08379 | 13.96319 | 14.70188 | 14.86154 | 17.30165 | 12.85838 | 14.37872 | 16.64341 |
| PI4K2A | 14.95261 | 15.24004 | 17.0972 | 16.36744 | 13.26195 | 15.20375 | 16.66035 | 16.20624 | 16.86871 | 16.57654 | 16.36549 | 17.14259 | 14.76776 | 15.66619 | 16.47128 | 17.0308 | 15.89242 | 14.23095 |
| PI4KA | 15.43353 | 16.33315 | 18.72074 | 17.29162 | 15.60936 | 15.55447 | 16.73067 | 16.22284 | 16.53814 | 15.69619 | 18.8059 | 17.50967 | 15.09877 | 16.00867 | 19.05411 | 18.88179 | 17.48592 | 16.78283 |
| PICALM | 20.39798 | 20.70241 | 18.36452 | 17.76908 | 20.35205 | 19.25172 | 18.08354 | 17.95272 | 18.63204 | 19.52461 | 18.14156 | 18.22192 | 21.18874 | 21.05649 | 18.80929 | 19.26409 | 19.68721 | 19.52505 |
| PICK1 | 17.76287 | 14.75059 | 16.31284 | 19.19591 | 13.14758 | 15.28407 | 15.62067 | 16.35355 | 14.17825 | 15.15242 | 16.6097 | 16.54673 | 15.40315 | 14.46394 | 16.47142 | 17.14706 | 12.13747 | 14.96288 |
| PIGG | 17.1544 | 16.03367 | 15.15205 | 15.51152 | 13.2845 | 16.38366 | 16.27747 | 17.332 | 19.88825 | 15.62071 | 15.94319 | 14.17175 | 15.71846 | 17.45188 | 16.0672 | 17.73796 | 15.69957 | 16.33928 |
| PIGK | 15.8357 | 15.83201 | 15.25355 | 15.19545 | 16.96299 | 16.24273 | 15.3392 | 15.91481 | 21.12219 | 18.63729 | 17.0412 | 13.76375 | 16.41385 | 15.94082 | 15.40052 | 15.85524 | 17.58092 | 15.05217 |
| PIGO | 16.05993 | 17.67592 | 16.46052 | 15.50914 | 13.86134 | 20.01598 | 14.21182 | 15.48078 | 20.21914 | 18.78181 | 16.86544 | 15.18573 | 15.41758 | 13.76292 | 15.17752 | 14.44344 | 17.4058 | 14.76142 |
| PIGS | 16.56601 | 16.77225 | 17.18801 | 16.71956 | 14.98132 | 17.58308 | 16.01679 | 17.0723 | 20.46883 | 18.15396 | 15.83134 | 15.28395 | 14.5242 | 16.4237 | 16.11792 | 15.35037 | 16.31927 | 15.73493 |
| PIGT | 15.05234 | 14.55828 | 16.12651 | 16.69566 | 15.63058 | 16.876 | 13.94229 | 16.18433 | 21.08938 | 18.39631 | 15.30936 | 16.9087 | 14.95461 | 14.7616 | 17.12959 | 17.0146 | 15.19264 | 13.80144 |
| PIGX | 16.24559 | 16.42919 | 16.87659 | 17.70214 | 14.93766 | 16.56724 | 17.61954 | 16.58173 | 20.57798 | 19.20544 | 14.59043 | 15.22647 | 15.49401 | 15.47268 | 16.03256 | 16.29143 | 14.81436 | 15.80206 |
| PIK3C2A | 16.06827 | 15.40871 | 16.38415 | 16.16107 | 15.69491 | 16.29857 | 15.2999 | 16.2197 | 15.66906 | 16.4666 | 16.24243 | 18.35142 | 14.306 | 15.42633 | 15.94154 | 15.45002 | 15.18789 | 14.6054 |
| PIK3C2B | 17.90295 | 17.25319 | 17.78285 | 17.56159 | 15.03073 | 15.74982 | 18.55983 | 19.10201 | 13.65778 | 18.32318 | 14.52528 | 15.99923 | 17.00391 | 17.22963 | 19.19877 | 19.33916 | 19.38226 | 19.06038 |
| PIK3CB | 15.88231 | 16.01891 | 14.28148 | 14.20092 | 10.33659 | 15.42902 | 13.68547 | 15.87803 | 9.59809 | 14.56699 | 14.91145 | 16.08368 | 14.26591 | 20.05943 | 16.9401 | 16.71723 | 13.0683 | 14.63689 |
| PIK3R1 | 14.0828 | 16.44876 | 16.74237 | 17.08387 | 15.25307 | 18.19021 | 16.2462 | 17.10357 | 15.48036 | 15.82746 | 15.90042 | 16.01822 | 19.07617 | 17.46688 | 16.91172 | 17.23568 | 16.71326 | 14.23526 |
| PIK3R2 | 15.68756 | 16.03838 | 16.22733 | 15.68382 | 16.31997 | 16.24698 | 15.91268 | 17.86132 | 17.17624 | 15.31808 | 13.99358 | 16.22671 | 16.02089 | 14.45672 | 15.91692 | 16.46969 | 15.91779 | 14.94102 |
| PIMREG | 15.01164 | 16.54274 | 17.13077 | 14.42034 | 16.11787 | 13.36521 | 15.99148 | 17.46571 | 15.66656 | 17.28239 | 16.625 | 12.279 | 16.12158 | 16.33279 | 14.32255 | 16.77678 | 16.30016 | 16.8672 |
| PIN1 | 23.31714 | 23.1086 | 21.80961 | 22.20589 | 19.30392 | 19.07913 | 20.95455 | 20.47417 | 17.74255 | 18.80487 | 17.81462 | 18.0132 | 21.64484 | 21.12728 | 22.0563 | 22.13829 | 20.6895 | 20.06427 |
| PIN4 | 15.07915 | 16.84155 | 15.77173 | 19.7545 | 17.33577 | 16.63729 | 16.1218 | 16.50244 | 12.95387 | 16.25107 | 16.87861 | 17.61534 | 17.50528 | 17.60042 | 19.24044 | 16.58702 | 18.53904 | 18.22075 |
| PIP4K2A | 16.48983 | 16.39186 | 15.65036 | 17.57207 | 18.42988 | 17.37396 | 19.79149 | 19.1237 | 15.02724 | 15.55418 | 17.29303 | 17.65964 | 18.12188 | 18.68853 | 15.55092 | 16.06388 | 18.67337 | 19.21832 |
| PIP4K2B | 16.75342 | 17.77685 | 17.31362 | 18.44191 | 19.05632 | 17.66331 | 20.54379 | 21.54009 | 16.17283 | 16.47062 | 17.05715 | 18.21102 | 19.8239 | 20.40209 | 17.21337 | 17.76374 | 19.47751 | 20.03561 |
| PIP4K2C | 16.10875 | 16.84451 | 15.98312 | 17.62554 | 19.29072 | 18.50049 | 18.40316 | 18.43526 | 18.63213 | 18.80166 | 17.05708 | 16.8151 | 18.98535 | 20.04249 | 17.27472 | 16.98232 | 19.99354 | 20.38508 |
| PIP4P1 | 17.62981 | 17.52613 | 17.37565 | 16.52064 | 17.05806 | 19.69982 | 14.88494 | 16.40988 | 20.19302 | 19.34249 | 15.97531 | 17.96034 | 14.03009 | 17.02664 | 15.14388 | 15.91952 | 14.79303 | 16.81457 |
| PIP5K1A | 17.64721 | 16.79089 | 18.42314 | 17.50928 | 17.61167 | 16.82001 | 16.31171 | 16.31797 | 16.34193 | 15.99818 | 17.0506 | 19.27692 | 18.08995 | 24.91593 | 18.40366 | 18.43299 | 17.67985 | 18.48032 |
| PIPOX | 16.04806 | 15.93016 | 16.20757 | 15.42978 | 14.23786 | 16.42127 | 14.51155 | 15.8444 | 19.81939 | 18.59949 | 14.67571 | 13.65525 | 16.73978 | 15.98273 | 15.0229 | 15.48642 | 15.70402 | 13.68955 |
| PIR | 17.48935 | 16.51083 | 16.9941 | 16.55032 | 18.85171 | 17.32272 | 17.66807 | 16.84205 | 16.36967 | 17.55893 | 20.03328 | 20.37253 | 18.49933 | 18.93502 | 16.62997 | 17.70927 | 18.60797 | 18.58963 |
| PITPNA | 17.53285 | 16.58593 | 17.71463 | 18.36697 | 18.83942 | 16.80314 | 17.93992 | 17.54008 | 15.57468 | 15.47218 | 16.4983 | 17.97249 | 18.61662 | 17.6152 | 18.72071 | 17.5914 | 18.57541 | 19.65426 |
| PITPNB | 18.68352 | 18.03277 | 18.06543 | 18.62752 | 19.14669 | 16.96651 | 17.48839 | 17.38501 | 17.23714 | 16.70125 | 19.2553 | 19.05436 | 19.42319 | 18.49619 | 19.55844 | 18.76577 | 18.70176 | 19.06542 |
| PITRM1 | 15.79204 | 13.97729 | 14.42286 | 15.47786 | 16.61244 | 14.93592 | 14.95479 | 17.00124 | 17.80626 | 16.88037 | 15.58903 | 14.87058 | 15.32161 | 15.50391 | 14.42446 | 15.94033 | 15.87263 | 15.61825 |
| PKHD1L1 | 16.31322 | 17.51951 | 18.7724 | 17.49514 | 12.30407 | 15.14586 | 15.44503 | 16.82583 | 19.0302 | 16.92096 | 12.5151 | 16.57329 | 14.19061 | 14.48194 | 19.09132 | 17.9243 | 15.97698 | 15.15449 |
| PKM | 20.64612 | 20.54762 | 21.73743 | 22.21722 | 24.5328 | 23.18124 | 22.00194 | 21.04733 | 21.24939 | 20.63166 | 23.55025 | 24.08632 | 22.28595 | 21.68139 | 22.46257 | 21.91921 | 23.80689 | 23.88685 |
| PKN1 | 16.34291 | 16.70479 | 16.96462 | 18.21438 | 18.55799 | 18.26656 | 18.18259 | 16.65241 | 18.15274 | 18.08816 | 17.31315 | 17.05279 | 18.13272 | 18.01141 | 17.65129 | 17.8957 | 19.00238 | 18.61681 |

| | | | | | | | | | | | | | | | | | | |
|---|---|---|---|---|---|---|---|---|---|---|---|---|---|---|---|---|---|---|
| PKN2 | 21.89183 | 21.89083 | 17.22057 | 20.4123 | 19.31416 | 18.18375 | 18.78628 | 16.57146 | 19.93814 | 19.34894 | 20.3619 | 18.99819 | 19.48654 | 19.45022 | 16.7877 | 17.52621 | 18.22147 | 18.1149 |
| PKP1 | 15.85289 | 15.95092 | 15.40507 | 15.43257 | 13.59471 | 17.23765 | 14.42219 | 16.73713 | 14.00027 | 14.7642 | 15.65087 | 13.00994 | 16.81232 | 16.30602 | 15.93698 | 15.28249 | 19.55475 | 18.42043 |
| PKP2 | 15.52093 | 17.30893 | 17.03596 | 16.37007 | 16.65679 | 16.12923 | 16.70141 | 16.11203 | 14.62706 | 16.52991 | 17.87229 | 18.14323 | 15.84823 | 16.65116 | 16.20781 | 16.04311 | 19.62945 | 19.19402 |
| PKP3 | 19.21175 | 20.00801 | 20.45007 | 21.2379 | 18.40018 | 17.24578 | 17.19677 | 16.84277 | 17.50037 | 16.74249 | 19.59034 | 19.80344 | 19.72889 | 18.92599 | 21.26531 | 21.18796 | 18.5256 | 17.36286 |
| PKP4 | 23.74351 | 23.13191 | 30.27789 | 22.00685 | 16.72833 | 17.71024 | 22.51487 | 21.64449 | 17.83052 | 17.33158 | 19.32007 | 16.83762 | 30.04812 | 21.84285 | 29.82056 | 30.03372 | 18.25854 | 18.04267 |
| PLA2G3 | 18.67361 | 17.63049 | 16.02918 | 14.86445 | 12.27942 | 14.79543 | 15.47897 | 14.67511 | 17.3053 | 16.12927 | 15.36742 | 14.91091 | 14.37535 | 15.21574 | 14.50585 | 13.31435 | 15.79851 | 13.37311 |
| PLA2G4A | 15.53998 | 14.70933 | 16.70245 | 16.68012 | 14.25507 | 17.97051 | 16.01277 | 16.69762 | 12.70196 | 14.47945 | 18.33669 | 19.03524 | 14.42311 | 15.63683 | 16.01782 | 14.82457 | 15.26563 | 16.48314 |
| PLAA | 16.22031 | 16.37366 | 16.38485 | 17.0024 | 15.14548 | 15.87741 | 16.54128 | 17.30402 | 15.99141 | 16.2328 | 17.4177 | 18.79907 | 16.26965 | 16.2007 | 16.88997 | 16.87895 | 15.96928 | 15.17471 |
| PLAT | 16.65276 | 17.15233 | 16.23873 | 15.83432 | 13.48101 | 16.67915 | 14.27129 | 14.80345 | 14.56046 | 13.93701 | 13.87125 | 12.73497 | 15.9571 | 14.83727 | 15.47253 | 15.21573 | 18.02306 | 21.39998 |
| PLAU | 14.82566 | 19.09265 | 15.82071 | 17.59595 | 19.05754 | 20.24003 | 14.78304 | 16.92435 | 15.39953 | 15.2312 | 14.70411 | 14.88804 | 15.62907 | 16.28713 | 16.0233 | 14.69129 | 19.72783 | 21.21709 |
| PLBD2 | 15.14924 | 15.55808 | 17.1581 | 16.55165 | 17.33137 | 15.74951 | 15.8811 | 15.19694 | 17.78 | 16.36868 | 15.22896 | 13.76695 | 14.01193 | 15.21043 | 15.50387 | 15.68626 | 16.85083 | 14.84877 |
| PLCB1 | 18.27754 | 16.90097 | 14.55808 | 14.33084 | 15.06359 | 15.95823 | 15.41991 | 17.58117 | 17.71291 | 16.38654 | 15.95641 | 12.95906 | 16.06954 | 15.50422 | 15.63932 | 15.5602 | 17.13871 | 14.95515 |
| PLCB2 | 16.73327 | 16.10383 | 16.0247 | 16.59815 | 14.89918 | 16.38618 | 16.84121 | 16.16922 | 16.09593 | 15.83941 | 18.83551 | 19.96809 | 15.26573 | 17.30315 | 15.067 | 16.20076 | 15.96431 | 16.8959 |
| PLCB3 | 17.50765 | 16.44393 | 16.25945 | 15.52126 | 18.58535 | 17.27786 | 16.79539 | 16.69518 | 15.80175 | 15.04505 | 17.63015 | 18.01058 | 14.8669 | 16.62093 | 15.60396 | 16.05787 | 17.35311 | 17.42572 |
| PLCB4 | 17.25362 | 16.19154 | 18.52217 | 19.28809 | 15.95171 | 16.49446 | 15.99838 | 16.99781 | 15.10263 | 16.56283 | 16.25358 | 14.38794 | 18.01843 | 18.09239 | 18.99995 | 19.47009 | 19.60528 | 20.24709 |
| PLCD3 | 16.49839 | 17.45898 | 16.9149 | 16.6681 | 18.71664 | 15.83454 | 17.33838 | 16.10977 | 14.15296 | 15.70206 | 15.03749 | 15.66326 | 18.60252 | 18.12587 | 15.42001 | 16.05768 | 20.42584 | 20.45475 |
| PLCG1 | 17.88807 | 19.42172 | 16.09145 | 18.87103 | 14.71843 | 16.26472 | 14.88935 | 16.0007 | 15.19181 | 16.45208 | 16.8272 | 18.61591 | 17.4137 | 15.45722 | 18.60041 | 16.82778 | 19.34055 | 18.10962 |
| PLCG2 | 17.86105 | 21.20517 | 19.94199 | 17.55528 | 16.06313 | 17.05107 | 16.63155 | 16.34999 | 15.77943 | 15.93274 | 20.40967 | 21.47639 | 16.99077 | 16.73418 | 21.12766 | 19.97244 | 16.0171 | 16.55609 |
| PLCH1 | 15.80907 | 14.55003 | 14.89999 | 17.53994 | 18.69175 | 16.48643 | 18.88475 | 18.60671 | 20.01271 | 20.5368 | 16.40943 | 17.18423 | 17.91107 | 17.22546 | 16.19704 | 15.53035 | 20.71553 | 15.96848 |
| PLD1 | 15.7028 | 15.14809 | 15.75177 | 15.79726 | 13.18511 | 14.21057 | 14.83468 | 15.33536 | 15.71017 | 14.02492 | 15.42434 | 18.21803 | 13.13618 | 13.68865 | 17.77011 | 15.87493 | 15.25761 | 14.45553 |
| PLD3 | 16.31345 | 16.01448 | 16.47523 | 15.49224 | 18.7323 | 17.70531 | 16.99222 | 16.47305 | 20.05498 | 19.47061 | 20.80812 | 17.86406 | 18.1229 | 15.55889 | 17.62047 | 17.22548 | 17.83297 | 16.24726 |
| PLEC | 18.38586 | 18.72978 | 16.91955 | 16.63152 | 18.50202 | 19.39119 | 16.76476 | 16.68875 | 15.45271 | 15.95824 | 16.47358 | 16.16293 | 17.2255 | 16.68071 | 17.80697 | 16.69409 | 21.217 | 21.16126 |
| PLEK2 | 16.81016 | 15.0787 | 18.10877 | 16.14689 | 14.63481 | 16.0314 | 15.447 | 17.12333 | 18.84928 | 16.54681 | 16.48907 | 14.16299 | 15.0019 | 14.22546 | 17.43282 | 15.82825 | 17.41427 | 15.17387 |
| PLEKHA2 | 15.6127 | 17.42713 | 16.33874 | 16.07018 | 18.85889 | 18.72943 | 22.46819 | 23.3105 | 11.93244 | 16.23687 | 15.6613 | 14.39704 | 14.39814 | 21.40239 | 16.3533 | 14.96316 | 20.16395 | 21.17939 |
| PLEKHA6 | 16.6488 | 17.21185 | 16.31736 | 17.3633 | 16.43106 | 17.00535 | 13.72474 | 16.44251 | 16.15871 | 15.77396 | 16.46199 | 17.18439 | 16.17408 | 15.55024 | 18.0849 | 18.01619 | 17.13659 | 14.69128 |
| PLEKHA7 | 15.37937 | 15.29765 | 15.33499 | 17.22842 | 15.23157 | 15.14461 | 14.68951 | 15.7282 | 13.19624 | 12.87179 | 14.06138 | 14.17655 | 18.84807 | 18.10653 | 17.13164 | 17.84998 | 14.9638 | 14.23716 |
| PLEKHB1 | 16.13518 | 16.62258 | 18.20014 | 14.26338 | 15.48396 | 16.69693 | 13.27425 | 15.14401 | 14.79083 | 15.6966 | 15.43598 | 17.13114 | 15.39305 | 13.44305 | 17.84159 | 19.2425 | 15.2842 | 13.45808 |
| PLEKHB2 | 19.44307 | 19.53696 | 20.06368 | 17.50192 | 14.83158 | 14.37076 | 15.92069 | 15.28239 | 17.27423 | 15.57911 | 15.78496 | 13.205 | 16.65532 | 17.671 | 19.75645 | 19.65691 | 15.39304 | 15.46271 |
| PLEKHG3 | 14.00106 | 14.74831 | 14.12403 | 13.75654 | 18.40847 | 17.21955 | 15.57895 | 17.60887 | 11.99173 | 14.67868 | 16.41683 | 13.15044 | 15.61438 | 13.72942 | 14.57507 | 15.34857 | 18.11864 | 17.36908 |
| PLEKHN1 | 16.604 | 17.23116 | 16.9625 | 16.84953 | 13.28421 | 15.19708 | 16.33752 | 14.57384 | 12.9066 | 14.62358 | 13.78423 | 12.66514 | 14.39316 | 14.80079 | 18.24252 | 18.67347 | 12.40291 | 14.85423 |
| PLIN3 | 17.30455 | 17.94602 | 16.87687 | 15.99613 | 18.76203 | 18.09933 | 15.65692 | 15.03095 | 14.68517 | 15.06005 | 15.60047 | 14.93796 | 16.39911 | 15.62047 | 16.86414 | 16.38099 | 18.72005 | 19.464 |
| PLIN4 | 15.18627 | 15.37333 | 17.81949 | 17.24162 | 12.8723 | 15.0582 | 15.21203 | 17.18936 | 13.41525 | 16.16333 | 13.79341 | 15.146 | 16.62155 | 17.10471 | 14.68293 | 14.12016 | 17.55179 | 18.29122 |
| PLK1 | 15.64819 | 15.36735 | 15.70133 | 16.14489 | 15.95673 | 15.02173 | 16.08598 | 17.26243 | 18.02534 | 18.52682 | 17.30907 | 16.72013 | 15.49959 | 14.48206 | 17.71554 | 14.94853 | 13.26921 | 13.106 |
| PLOD1 | 17.94685 | 16.9895 | 16.00323 | 15.97294 | 16.97089 | 16.28108 | 16.13555 | 15.90392 | 20.56868 | 19.59019 | 16.96517 | 15.03129 | 15.60634 | 15.66643 | 16.88565 | 16.69966 | 15.52878 | 18.17552 |
| PLOD2 | 15.52616 | 16.23178 | 16.92698 | 16.04102 | 16.77941 | 17.51835 | 16.35703 | 16.54404 | 20.67716 | 20.25546 | 16.82494 | 14.54201 | 14.43166 | 16.30099 | 15.08659 | 15.4634 | 15.84606 | 16.06026 |
| PLOD3 | 17.12709 | 17.13213 | 16.03546 | 15.49783 | 16.24589 | 16.2835 | 16.13978 | 16.64329 | 21.24595 | 19.57898 | 18.63052 | 14.63188 | 15.59744 | 15.61311 | 15.06747 | 15.83055 | 17.21908 | 15.65036 |
| PLPBP | 17.7113 | 16.91053 | 16.95964 | 16.57975 | 19.64809 | 15.82045 | 17.8518 | 15.71852 | 16.32848 | 16.88786 | 16.37063 | 16.70187 | 16.24779 | 15.47795 | 16.30134 | 17.86066 | 19.30209 | 17.43722 |
| PLPP1 | 14.49054 | 16.16791 | 22.28049 | 22.14833 | 15.98089 | 16.2636 | 16.64215 | 17.43558 | 13.76547 | 15.08373 | 13.58234 | 19.02463 | 15.23776 | 15.40189 | 22.65081 | 21.80437 | 15.67118 | 15.14323 |
| PLPP2 | 14.48672 | 16.26955 | 14.28329 | 15.10346 | 14.87358 | 17.63661 | 15.09464 | 15.94162 | 20.51267 | 18.24761 | 17.89446 | 20.39751 | 15.28168 | 15.14802 | 16.00034 | 16.76892 | 16.64738 | 14.77713 |
| PLPP3 | 16.76051 | 16.18195 | 16.17944 | 17.19331 | 21.97503 | 18.09204 | 17.22264 | 16.64333 | 14.51529 | 16.97057 | 15.23866 | 15.88678 | 18.5584 | 18.07528 | 14.36852 | 15.6468 | 19.09338 | 21.03568 |
| PLS1 | 18.17619 | 17.08028 | 16.17183 | 16.00059 | 19.92401 | 19.56736 | 16.81493 | 17.08272 | 17.712 | 17.01463 | 18.90561 | 19.22188 | 19.22827 | 19.43807 | 16.10642 | 16.69959 | 18.32027 | 17.3955 |
| PLS3 | 19.51747 | 17.44442 | 23.5135 | 22.62797 | 21.0139 | 21.71597 | 17.45627 | 17.69659 | 15.63485 | 17.25641 | 16.04113 | 13.59387 | 18.21308 | 19.49939 | 21.36612 | 19.53001 | 21.93638 | 20.09645 |
| PLSCR1 | 19.33741 | 16.0269 | 16.94469 | 14.10139 | 12.64112 | 14.03711 | 16.93251 | 15.51889 | 19.03462 | 18.66741 | 15.81409 | 16.54312 | 14.156 | 15.66757 | 18.81522 | 16.41193 | 14.55715 | 15.02279 |
| PLSCR3 | 22.72678 | 22.05798 | 19.71859 | 19.05978 | 16.86191 | 15.85561 | 15.29818 | 16.64332 | 18.44926 | 18.24397 | 16.71157 | 19.8828 | 16.11947 | 16.09642 | 21.16908 | 21.08646 | 16.27592 | 15.4824 |
| PLXNA1 | 16.65928 | 16.77231 | 16.0391 | 17.06218 | 19.22774 | 17.39632 | 15.7729 | 16.41237 | 16.53136 | 16.28919 | 16.24054 | 17.40563 | 18.62593 | 19.07685 | 17.44796 | 17.15573 | 19.34443 | 19.56703 |
| PLXNA2 | 16.87058 | 16.29414 | 16.69429 | 16.72989 | 15.26552 | 16.85435 | 17.43961 | 15.88557 | 19.35369 | 18.36456 | 15.96753 | 17.36613 | 16.09425 | 16.35614 | 17.68678 | 16.51473 | 16.85657 | 16.15702 |
| PLXNA4 | 17.07881 | 19.92876 | 16.73283 | 15.49413 | 13.74683 | 14.00715 | 19.16331 | 17.73268 | 14.29016 | 16.09508 | 16.02372 | 15.35668 | 14.90374 | 18.86256 | 18.75638 | 19.42336 | 16.35556 | 17.45678 |
| PLXNB1 | 16.88501 | 14.09424 | 14.80746 | 15.02867 | 14.26775 | 15.33804 | 17.80364 | 18.89471 | 18.75751 | 15.7047 | 15.28113 | 15.71928 | 13.86471 | 16.73304 | 16.39059 | 14.17291 | 16.73968 | 15.35786 |
| PLXNB2 | 19.20213 | 18.38636 | 20.27873 | 20.39949 | 19.89025 | 18.84668 | 17.50993 | 17.37057 | 18.47165 | 17.23746 | 17.87923 | 19.40466 | 19.4616 | 19.14523 | 21.45206 | 21.08678 | 19.53723 | 19.97946 |
| PLXNB3 | 16.76521 | 14.7117 | 17.31334 | 15.95869 | 12.65789 | 17.39644 | 15.5144 | 12.76624 | 13.22961 | 14.53276 | 14.39821 | 14.1172 | 14.01931 | 15.92214 | 18.0863 | 17.7252 | 16.54501 | 16.16442 |
| PMM2 | 15.45213 | 14.97233 | 15.35108 | 13.22592 | 17.43172 | 15.27163 | 14.8116 | 16.9635 | 16.4089 | 15.79824 | 17.24094 | 17.54251 | 18.49817 | 19.31737 | 16.10497 | 15.27157 | 17.86898 | 17.41689 |
| PMPCA | 15.68197 | 16.74724 | 15.40925 | 15.43796 | 14.77205 | 15.1921 | 17.70237 | 18.83489 | 19.1463 | 21.9437 | 16.90496 | 14.92707 | 14.80043 | 15.34756 | 14.88762 | 14.53208 | 16.53634 | 17.06941 |
| PMPCB | 15.65913 | 19.48358 | 17.17319 | 16.73221 | 15.41012 | 16.29028 | 15.66768 | 17.02937 | 18.95607 | 17.33481 | 18.70463 | 13.35914 | 14.80668 | 14.17243 | 16.98633 | 17.74249 | 16.76147 | 16.09769 |
| PMVK | 18.3541 | 18.48199 | 16.87675 | 17.20442 | 18.94239 | 16.88973 | 19.05512 | 17.30723 | 17.48235 | 18.24146 | 16.59003 | 17.96427 | 19.69048 | 18.72976 | 16.28674 | 17.48706 | 18.58498 | 20.02205 |
| PNMA2 | 23.04775 | 23.40844 | 16.40693 | 16.14923 | 17.65965 | 17.01642 | 16.26102 | 16.01472 | 18.0134 | 17.41853 | 15.09942 | 18.14488 | 15.57795 | 15.47226 | 15.28312 | 16.36703 | 17.52658 | 17.64345 |
| PNP | 16.58129 | 17.58302 | 16.38928 | 17.63945 | 18.15194 | 17.60554 | 16.2283 | 16.04225 | 17.4132 | 17.29985 | 19.28422 | 21.09152 | 17.28463 | 16.62333 | 17.15843 | 17.98984 | 16.40929 | 17.05224 |
| PNPLA8 | 17.07168 | 18.48441 | 15.55165 | 12.69837 | 17.73679 | 16.93128 | 15.27974 | 16.49443 | 19.27742 | 18.93298 | 14.0091 | 17.87337 | 15.69801 | 17.31291 | 18.30126 | 15.73364 | 17.52973 | 13.27429 |
| PNPO | 17.8966 | 17.67323 | 19.31603 | 20.03009 | 20.29241 | 19.24547 | 18.19548 | 16.08561 | 17.18697 | 16.7198 | 18.85614 | 19.31082 | 21.73486 | 21.20943 | 20.00611 | 19.36756 | 20.46749 | 21.52031 |
| PNPT1 | 15.89572 | 14.45159 | 14.79914 | 14.75102 | 15.74438 | 15.08231 | 17.81126 | 16.04129 | 18.5363 | 16.06989 | 15.51323 | 12.31793 | 13.75155 | 15.98789 | 14.94324 | 14.80322 | 15.58553 | 15.92068 |
| PODXL | 17.89405 | 18.27652 | 18.20223 | 19.71995 | 19.17809 | 15.58492 | 19.20679 | 18.72445 | 22.77496 | 20.84601 | 21.9063 | 19.05769 | 21.11111 | 20.69805 | 18.54613 | 18.098 | 18.8616 | 19.9357 |
| PODXL2 | 13.99018 | 15.65639 | 18.72147 | 18.81882 | 16.04152 | 15.30626 | 16.82632 | 17.68041 | 19.38978 | 17.68747 | 17.0551 | 12.88924 | 15.32585 | 17.85542 | 13.73306 | 17.17374 | 15.97906 | 16.00547 |
| POFUT1 | 15.91498 | 16.11003 | 17.04919 | 17.14075 | 15.06861 | 17.00567 | 16.50871 | 16.4698 | 21.54466 | 20.8652 | 18.52836 | 15.12694 | 15.59417 | 14.8436 | 15.35525 | 17.0447 | 15.20951 | 13.91845 |
| POFUT2 | 15.54047 | 14.4075 | 16.53459 | 14.89024 | 14.64976 | 14.80125 | 15.43821 | 17.94612 | 19.56571 | 17.81672 | 16.12532 | 12.06441 | 14.18402 | 17.57621 | 13.34527 | 13.89959 | 14.78355 | 15.11033 |
| POGLUT1 | 15.92935 | 15.88212 | 18.75081 | 18.32507 | 13.76255 | 18.95217 | 15.6087 | 17.43018 | 19.50334 | 18.5778 | 17.51164 | 14.81615 | 14.8534 | 17.4834 | 18.74804 | 16.61234 | 17.31844 | 17.72333 |
| POGLUT2 | 15.49422 | 16.43264 | 14.70011 | 16.05844 | 15.64578 | 15.19716 | 16.95588 | 17.07808 | 17.96718 | 13.75367 | 13.52095 | 14.03203 | 13.95967 | 16.09716 | 14.9405 | 15.66806 | 15.24943 | 13.77942 |
| POGLUT3 | 15.88585 | 16.41361 | 15.91169 | 15.52451 | 15.79645 | 16.74934 | 15.85475 | 16.23167 | 20.24449 | 19.54026 | 16.38993 | 16.22631 | 16.72564 | 18.57179 | 16.20798 | 16.41695 | 15.90353 | 15.27182 |
| POLDIP2 | 16.17238 | 16.89936 | 15.24869 | 14.88315 | 13.79644 | 14.45882 | 14.57812 | 16.4568 | 18.18132 | 16.11138 | 15.8673 | 12.40361 | 12.28619 | 14.68282 | 15.25032 | 16.55036 | 15.29313 | 16.56206 |
| POLDIP3 | 18.66429 | 11.95784 | 17.22095 | 15.0167 | 13.72233 | 14.81132 | 17.25237 | 16.71544 | 15.8416 | 15.14928 | 15.88972 | 17.18547 | 13.69748 | 14.66526 | 15.44134 | 15.13349 | 14.03913 | 14.59044 |
| POLR1A | 15.89881 | 16.92636 | 15.50989 | 14.71128 | 18.9171 | 15.59255 | 18.69637 | 15.61754 | 21.4756 | 22.61162 | 15.85488 | 16.79218 | 17.27188 | 14.74031 | 14.37473 | 13.89271 | 16.09318 | 16.82246 |
| POLR1C | 17.60508 | 17.43716 | 17.41684 | 17.77061 | 13.58465 | 17.15887 | 13.74109 | 14.11525 | 16.24871 | 13.02079 | 17.02194 | 18.23638 | 16.66201 | 14.30072 | 17.79853 | 18.21261 | 14.8091 | 14.93968 |
| POLR2A | 17.64232 | 17.13821 | 17.88361 | 17.66035 | 12.94598 | 14.89703 | 17.49859 | 17.00397 | 16.22467 | 18.08535 | 17.38893 | 18.02217 | 15.19416 | 15.9114 | 15.14272 | 15.45198 | 16.79552 | 15.76961 |
| POLR2B | 16.65338 | 16.89816 | 15.69919 | 15.56954 | 15.98778 | 12.52691 | 17.82969 | 14.2738 | 16.55542 | 16.36418 | 16.08293 | 17.29399 | 15.18189 | 14.51401 | 15.15178 | 14.2985 | 16.78796 | 16.5415 |
| POLR2C | 24.2284 | 24.16747 | 22.37034 | 22.13408 | 12.46693 | 18.04565 | 18.98652 | 17.94167 | 18.02571 | 14.18247 | 15.27781 | 17.4439 | 21.37418 | 17.85468 | 21.75562 | 22.53183 | 17.17061 | 14.77031 |
| POMGNT1 | 16.13133 | 17.12036 | 15.85951 | 15.21277 | 14.04278 | 15.57566 | 15.83167 | 17.46225 | 19.95467 | 17.33535 | 15.6221 | 15.70052 | 16.6265 | 14.51643 | 15.21958 | 16.91865 | 15.66168 | 15.49219 |
| POMGNT2 | 16.33168 | 15.7757 | 16.62455 | 15.4343 | 15.03071 | 16.55611 | 14.54471 | 14.7043 | 18.72322 | 16.71783 | 15.23485 | 13.30066 | 14.84936 | 20.16477 | 14.64431 | 17.14287 | 15.34396 | 15.30071 |
| POMK | 16.84427 | 18.55213 | 15.87595 | 17.60123 | 15.5332 | 17.66066 | 17.21408 | 17.28451 | 20.95654 | 19.72118 | 17.64032 | 15.77325 | 14.81306 | 16.71957 | 16.66248 | 15.63468 | 14.82577 | 16.35304 |
| POMT1 | 16.56717 | 15.54698 | 16.47756 | 13.64539 | 14.05437 | 14.86517 | 15.54989 | 15.68444 | 19.29208 | 15.14257 | 14.89424 | 12.86102 | 15.29722 | 15.77595 | 15.68264 | 15.76838 | 14.70646 | 14.65493 |
| POMT2 | 15.33544 | 16.59266 | 16.44666 | 15.3477 | 14.51822 | 17.59814 | 12.5962 | 18.55906 | 19.37631 | 15.83121 | 14.43603 | 12.76268 | 13.99216 | 14.78948 | 13.57502 | 13.43322 | 18.21827 | 16.33064 |
| PON2 | 16.68896 | 16.14603 | 15.78324 | 14.14552 | 14.98919 | 16.14232 | 15.54148 | 17.69323 | 20.97558 | 18.52209 | 17.28522 | 15.33222 | 13.97869 | 14.98959 | 15.30093 | 14.7028 | 14.71475 | 17.47337 |
| POR | 15.34974 | 14.80489 | 15.97768 | 16.96133 | 14.80456 | 16.91964 | 17.66124 | 18.43723 | 17.52972 | 15.88574 | 15.26831 | 18.03299 | 15.70587 | 17.60606 | 16.32771 | 15.80916 | 17.30397 | 16.49836 |
| POTEE | 25.10529 | 24.84613 | 25.07341 | 26.09738 | 26.47492 | 26.48875 | 23.88886 | 23.2208 | 24.44831 | 23.7124 | 25.28543 | 25.83942 | 25.84544 | 24.97684 | 25.68845 | 25.58676 | 25.3447 | 25.28742 |
| POTEJ | 26.40764 | 26.69006 | 26.41987 | 26.3239 | 30.17169 | 29.01699 | 28.00459 | 24.86983 | 24.98583 | 26.05368 | 25.84987 | 26.81399 | 28.94149 | 28.98602 | 27.19672 | 27.16951 | 28.26755 | 28.64959 |
| PPA1 | 19.04636 | 17.83877 | 18.68092 | 19.25986 | 20.8762 | 18.79199 | 18.21711 | 18.20656 | 17.62352 | 17.42114 | 21.24431 | 21.95453 | 21.00313 | 20.37417 | 19.94288 | 19.6311 | 20.54052 | 20.70101 |
| PPA2 | 17.14702 | 17.9982 | 17.20758 | 16.73463 | 17.65952 | 15.86098 | 17.50119 | 17.8405 | 19.669 | 18.63569 | 17.33084 | 18.98403 | 17.63496 | 16.64555 | 16.96695 | 16.80624 | 16.46872 | 16.0377 |
| PPAT | 16.82465 | 16.46046 | 16.29468 | 15.96317 | 15.39409 | 17.39152 | 16.38608 | 16.57901 | 15.85262 | 16.90497 | 18.76459 | 20.19253 | 18.50272 | 17.34154 | 17.25597 | 16.58804 | 17.22161 | 17.93198 |
| PPFIA1 | 16.36113 | 15.50356 | 17.02768 | 18.05808 | 19.86254 | 19.21784 | 17.55668 | 17.03862 | 17.6956 | 17.43003 | 17.00811 | 15.90958 | 18.42119 | 16.05739 | 17.85472 | 17.7718 | 18.93284 | 19.30548 |

| | | | | | | | | | | | | | | | | | | |
|---|---|---|---|---|---|---|---|---|---|---|---|---|---|---|---|---|---|---|
| PPFIA2 | 14.10701 | 16.82704 | 17.50322 | 19.97183 | 15.83215 | 16.25155 | 15.70227 | 16.75989 | 15.64542 | 15.72337 | 15.46556 | 13.89926 | 18.12547 | 15.94391 | 18.53019 | 17.97242 | 15.73247 | 14.49937 |
| PPFIA3 | 14.33286 | 16.76496 | 13.35057 | 14.74364 | 18.20714 | 15.67757 | 17.53457 | 15.8121 | 16.32965 | 14.78142 | 14.74994 | 18.64057 | 17.39512 | 15.29542 | 18.84041 | 18.11456 | 15.55744 | 14.50667 |
| PPFIBP1 | 16.39481 | 16.53532 | 16.99885 | 18.0995 | 19.94429 | 18.85144 | 16.38677 | 15.13274 | 14.63372 | 15.77254 | 14.89587 | 16.18609 | 18.3003 | 19.48915 | 17.1712 | 16.88766 | 20.04414 | 19.58165 |
| PPFIBP2 | 15.55841 | 15.32574 | 17.75367 | 19.12108 | 16.33618 | 16.17861 | 17.42644 | 15.95719 | 14.72176 | 14.89252 | 15.8348 | 16.19937 | 17.14158 | 17.12835 | 18.08166 | 18.23302 | 17.58955 | 17.96235 |
| PPIB | 18.78156 | 17.80173 | 20.56516 | 18.97814 | 17.83994 | 18.72546 | 21.40757 | 19.9143 | 24.7972 | 24.22588 | 23.57342 | 21.16128 | 19.28358 | 21.43064 | 20.4329 | 20.08261 | 18.44068 | 18.06744 |
| PPIC | 14.20586 | 16.05715 | 16.80907 | 15.12975 | 12.37988 | 13.79834 | 19.80202 | 19.10861 | 17.82298 | 19.06663 | 13.54512 | 14.33638 | 13.2048 | 17.35797 | 17.92807 | 13.04709 | 15.97405 | 15.64355 |
| PPID | 17.75605 | 16.74869 | 14.62616 | 16.45176 | 16.5704 | 16.35948 | 17.05606 | 16.63744 | 16.22908 | 17.06181 | 19.28632 | 19.63606 | 17.10685 | 16.87416 | 16.45066 | 15.83891 | 17.92928 | 16.47276 |
| PPIF | 15.81686 | 15.36856 | 15.9798 | 14.473 | 16.15544 | 15.29789 | 18.95278 | 17.05877 | 19.66692 | 19.81163 | 16.16953 | 15.55803 | 16.88834 | 16.40599 | 17.72851 | 15.94071 | 18.15321 | 17.36716 |
| PPIH | 16.53489 | 16.96501 | 17.08182 | 14.97142 | 16.9771 | 14.89575 | 17.65039 | 15.87419 | 13.71585 | 16.48659 | 15.65162 | 15.36508 | 20.65249 | 18.7477 | 17.79797 | 16.60411 | 17.55034 | 17.39363 |
| PPIL1 | 21.04099 | 20.64904 | 20.24004 | 19.90137 | 18.56443 | 16.48626 | 16.32359 | 17.61823 | 17.07179 | 16.0168 | 18.50263 | 19.74676 | 18.9581 | 17.18272 | 20.70364 | 20.55404 | 17.46935 | 16.86837 |
| PPIL3 | 18.83386 | 19.69197 | 17.10189 | 18.80622 | 21.18807 | 19.07853 | 18.29283 | 18.81344 | 17.25538 | 18.93774 | 17.95201 | 16.59884 | 20.12585 | 20.44886 | 18.24596 | 18.11211 | 19.19672 | 19.45012 |
| PPL | 15.14239 | 15.41547 | 16.3545 | 15.81482 | 20.44769 | 20.21122 | 16.0461 | 15.74442 | 14.90329 | 14.32203 | 15.44821 | 14.59454 | 18.07617 | 18.11161 | 15.75759 | 15.81038 | 17.4949 | 17.08262 |
| PPM1A | 19.45337 | 18.26713 | 18.60639 | 18.54696 | 17.50382 | 17.5604 | 17.14199 | 16.66819 | 16.47921 | 16.66767 | 17.07907 | 18.69156 | 17.61711 | 17.33429 | 20.54415 | 20.04353 | 18.11351 | 18.36299 |
| PPM1G | 15.56237 | 15.29092 | 14.1906 | 16.38838 | 16.52278 | 15.44967 | 16.65885 | 16.53778 | 14.8187 | 15.13018 | 17.40125 | 18.65891 | 15.86995 | 15.00917 | 15.77149 | 14.79415 | 15.86036 | 15.61702 |
| PPM1H | 16.39792 | 17.19068 | 16.10647 | 15.69841 | 15.41601 | 17.35608 | 17.12111 | 16.83783 | 17.28004 | 16.83694 | 17.81775 | 19.17639 | 16.78842 | 16.45094 | 15.43925 | 15.87636 | 16.84987 | 16.80968 |
| PPM1L | 15.4726 | 16.61061 | 15.08023 | 16.01008 | 18.20698 | 18.28781 | 16.73212 | 16.87367 | 20.86267 | 18.6301 | 15.40847 | 14.26207 | 15.97752 | 15.88978 | 14.8707 | 14.68699 | 16.30934 | 15.68896 |
| PPME1 | 16.47145 | 17.54309 | 17.7916 | 16.19114 | 18.50551 | 15.80397 | 16.79031 | 16.71323 | 18.24489 | 17.52696 | 18.30427 | 18.87669 | 19.66272 | 18.78153 | 16.83534 | 17.4735 | 18.48306 | 18.21546 |
| PPP1CA | 19.33659 | 19.11601 | 19.33357 | 19.36969 | 20.89557 | 21.099 | 20.1733 | 18.4094 | 19.50434 | 20.08196 | 19.56517 | 21.0561 | 19.26227 | 20.77374 | 20.57557 | 19.92134 | 20.53076 | 20.08718 |
| PPP1CB | 17.42012 | 18.33106 | 19.02404 | 18.7831 | 19.25992 | 18.83485 | 18.73062 | 18.87199 | 20.73745 | 19.18172 | 19.43399 | 20.72552 | 21.08923 | 18.25743 | 19.83256 | 19.89758 | 19.68639 | 17.59402 |
| PPP1CC | 15.56151 | 17.25032 | 18.66021 | 19.34245 | 18.39422 | 17.8312 | 16.11884 | 18.98066 | 19.30085 | 16.78697 | 19.47594 | 20.01204 | 14.23164 | 15.57208 | 18.70273 | 18.15047 | 17.9539 | 15.17499 |
| PPP1R12A | 16.46139 | 17.50579 | 15.27363 | 16.7881 | 19.79312 | 18.02263 | 15.8544 | 16.23121 | 15.09959 | 16.25151 | 18.15348 | 16.07286 | 18.65255 | 18.66257 | 15.55566 | 16.33036 | 18.98357 | 17.95865 |
| PPP1R14B | 20.00732 | 20.17399 | 18.76002 | 20.81084 | 19.30534 | 15.08345 | 15.62858 | 17.58137 | 17.05935 | 17.39534 | 21.20619 | 20.59819 | 20.75564 | 19.29224 | 19.86392 | 18.46669 | 20.22893 | 19.86999 |
| PPP1R16A | 15.63536 | 17.74398 | 15.91641 | 14.63361 | 15.66205 | 15.9579 | 16.21187 | 16.02139 | 16.89736 | 15.36565 | 18.30173 | 13.69622 | 16.3027 | 18.86556 | 16.31823 | 16.39317 | 19.48037 | 18.96375 |
| PPP1R18 | 17.08733 | 15.14902 | 15.688 | 17.45173 | 18.87752 | 17.16074 | 15.39228 | 16.97678 | 14.68333 | 13.78884 | 14.75286 | 11.91098 | 15.06925 | 14.65606 | 14.00533 | 16.23476 | 18.76564 | 17.6097 |
| PPP1R1B | 15.07289 | 15.7009 | 13.68866 | 16.7619 | 15.48108 | 15.54752 | 16.62096 | 15.65275 | 14.73644 | 14.62828 | 19.57432 | 19.99135 | 13.97737 | 13.252 | 15.51436 | 14.88536 | 14.65481 | 14.84986 |
| PPP1R2 | 15.69765 | 14.87314 | 14.74451 | 14.01016 | 17.47666 | 15.39059 | 15.85955 | 15.1729 | 13.78549 | 16.41949 | 12.6912 | 12.56918 | 18.0701 | 16.88428 | 15.36954 | 16.0804 | 18.30938 | 18.02075 |
| PPP1R2B | 17.68034 | 13.33218 | 15.25789 | 17.33166 | 14.34943 | 14.19731 | 15.17476 | 15.42903 | 16.30177 | 14.19249 | 14.84225 | 18.55081 | 12.71661 | 12.79305 | 15.75979 | 14.55063 | 14.4198 | 14.24136 |
| PPP1R7 | 15.86867 | 15.57244 | 19.76472 | 17.7176 | 17.65676 | 16.34856 | 16.28228 | 17.25643 | 16.66063 | 18.40299 | 18.30814 | 19.70631 | 17.27981 | 16.73078 | 17.22969 | 18.05032 | 16.52093 | 16.06394 |
| PPP1R8 | 16.65635 | 14.91503 | 14.07926 | 16.43587 | 17.1507 | 14.96692 | 16.02551 | 13.94931 | 14.01604 | 16.91514 | 14.8784 | 16.05069 | 17.701 | 15.02504 | 14.68655 | 14.97809 | 17.05845 | 18.03731 |
| PPP2CA | 16.83708 | 15.88427 | 16.62528 | 16.55014 | 19.9122 | 17.77863 | 19.97141 | 19.9732 | 18.66274 | 18.45728 | 14.57066 | 18.44039 | 19.3558 | 20.13275 | 17.20941 | 17.60291 | 21.30563 | 19.42905 |
| PPP2CB | 15.5925 | 16.93614 | 19.05781 | 15.30881 | 17.20158 | 16.8132 | 16.01672 | 17.07325 | 18.58089 | 17.10574 | 19.21477 | 20.05098 | 17.08774 | 18.28345 | 18.48848 | 19.75649 | 16.70482 | 17.14688 |
| PPP2R1A | 18.8699 | 18.08492 | 20.14749 | 20.51263 | 21.29911 | 20.22671 | 20.47851 | 19.11935 | 19.59988 | 19.99419 | 20.63012 | 21.48344 | 21.59138 | 21.63149 | 20.15554 | 20.0239 | 21.3197 | 21.72887 |
| PPP2R1B | 16.08805 | 16.05287 | 17.40777 | 14.41305 | 19.7111 | 18.23203 | 18.04244 | 15.15933 | 16.60903 | 15.33802 | 18.12182 | 18.05437 | 20.52059 | 20.32646 | 17.68437 | 17.24967 | 18.51568 | 19.45098 |
| PPP2R2A | 19.1055 | 16.6355 | 18.41238 | 17.32486 | 17.14378 | 17.00376 | 16.66686 | 17.88384 | 18.28072 | 16.82033 | 18.30586 | 19.45913 | 18.54 | 18.10051 | 19.22933 | 18.78943 | 18.92044 | 17.61916 |
| PPP2R2C | 16.22859 | 14.30137 | 14.54213 | 15.84763 | 19.47784 | 19.00491 | 17.7599 | 17.45085 | 19.57254 | 14.2614 | 13.94874 | 18.76538 | 18.82044 | 14.7138 | 13.53045 | 18.87086 | 17.88055 | 18.28065 |
| PPP2R5D | 15.40018 | 17.13287 | 15.70035 | 17.03247 | 17.47027 | 17.8537 | 15.47159 | 18.35631 | 15.08233 | 16.04976 | 15.0731 | 20.22892 | 17.74302 | 16.32319 | 18.14987 | 17.5392 | 16.50377 | 15.22026 |
| PPP2R5E | 17.15052 | 16.5536 | 19.42453 | 18.43257 | 16.85448 | 14.56229 | 19.30666 | 20.10086 | 17.89312 | 15.79382 | 14.80275 | 20.3347 | 14.64658 | 15.09789 | 19.20273 | 18.11027 | 20.55804 | 17.37278 |
| PPP3CA | 16.96217 | 16.0137 | 17.33425 | 18.68387 | 17.08876 | 16.61035 | 17.69315 | 17.33391 | 17.31552 | 16.0019 | 17.52361 | 18.69549 | 19.57264 | 19.39554 | 18.0724 | 17.99692 | 18.31623 | 18.41773 |
| PPP3CB | 19.31709 | 19.06079 | 16.3879 | 15.22026 | 17.80767 | 16.09582 | 16.25678 | 16.71075 | 17.0008 | 16.30254 | 14.80904 | 18.56512 | 14.55054 | 15.68199 | 16.04829 | 19.77436 | 18.46522 | 19.07822 |
| PPP3R1 | 17.48272 | 19.37451 | 17.31513 | 18.58388 | 17.08702 | 16.90221 | 16.85306 | 16.02257 | 15.30993 | 17.15558 | 18.59519 | 19.49051 | 19.01471 | 18.62115 | 18.30684 | 18.45621 | 17.22311 | 14.64375 |
| PPP4C | 16.35791 | 14.32891 | 15.74915 | 15.85482 | 13.5752 | 16.91755 | 14.94549 | 16.44268 | 13.80432 | 15.62619 | 16.8329 | 18.54818 | 13.00278 | 14.58483 | 15.46586 | 15.06302 | 13.57511 | 15.38649 |
| PPP4R3A | 15.44104 | 16.72178 | 13.68755 | 16.96104 | 19.34612 | 18.59065 | 16.05485 | 18.33525 | 15.97805 | 17.50741 | 16.20747 | 17.90029 | 16.99511 | 16.45755 | 17.44188 | 15.91902 | 19.10379 | 18.64109 |
| PPP5C | 16.33401 | 17.30865 | 16.98443 | 15.60009 | 18.42249 | 18.02124 | 17.98324 | 17.6655 | 16.9173 | 17.02009 | 18.75335 | 19.56269 | 18.16658 | 16.59178 | 16.56742 | 15.97377 | 17.98542 | 17.98773 |
| PPP6C | 15.72713 | 14.26577 | 14.8842 | 15.49549 | 17.25195 | 16.20467 | 17.98566 | 19.14833 | 15.97243 | 17.10006 | 17.24576 | 19.38371 | 14.4263 | 14.90659 | 16.83752 | 15.33754 | 17.68691 | 16.41895 |
| PPP6R3 | 15.89801 | 14.56 | 14.56587 | 14.67021 | 14.8725 | 17.03349 | 15.79396 | 16.17857 | 13.87687 | 16.93183 | 16.5754 | 19.52507 | 13.99899 | 14.82239 | 17.06869 | 16.10058 | 14.83334 | 17.14516 |
| PPT1 | 16.52061 | 14.66159 | 13.43065 | 15.44372 | 13.85092 | 13.53868 | 18.53742 | 18.14098 | 19.01327 | 14.45786 | 14.3842 | 14.32243 | 18.01371 | 13.88084 | 15.14463 | 14.18102 | 17.40039 | 18.82019 |
| PPWD1 | 13.61144 | 14.96941 | 19.08141 | 18.5714 | 16.55982 | 21.3738 | 16.97216 | 16.1915 | 14.15561 | 17.12538 | 16.08311 | 17.70397 | 15.2548 | 14.94258 | 15.30173 | 15.79841 | 16.19581 | 14.92556 |
| PRCP | 16.92512 | 16.21241 | 14.50306 | 15.40178 | 15.68258 | 16.40539 | 14.46888 | 15.71974 | 18.82476 | 17.31927 | 15.79446 | 15.24962 | 14.08692 | 15.94219 | 16.21517 | 16.2162 | 13.98033 | 13.64673 |
| PRDX2 | 22.06026 | 21.15392 | 21.59453 | 22.31535 | 19.04836 | 19.66374 | 19.8802 | 20.08366 | 22.31625 | 21.14229 | 23.50914 | 24.9462 | 20.50377 | 19.19314 | 21.8431 | 21.91479 | 20.72955 | 20.00307 |
| PRDX3 | 16.76564 | 16.61622 | 16.78304 | 17.46748 | 15.30333 | 16.99309 | 18.07668 | 17.79085 | 22.66272 | 21.17687 | 18.30559 | 17.40695 | 16.8406 | 18.59327 | 15.51661 | 16.72048 | 16.44579 | 15.01678 |
| PRDX4 | 16.44383 | 16.01285 | 16.43055 | 17.01127 | 16.31919 | 16.22154 | 18.26831 | 17.37122 | 23.99081 | 22.77974 | 23.0999 | 19.96898 | 15.60786 | 15.6315 | 17.50186 | 17.17316 | 17.33455 | 17.66624 |
| PRDX5 | 19.44259 | 17.93298 | 20.39718 | 20.50028 | 22.0877 | 20.50636 | 21.53932 | 19.8976 | 22.05757 | 22.18475 | 20.54955 | 20.87688 | 22.72697 | 22.46148 | 21.48642 | 20.82657 | 23.12704 | 22.5322 |
| PRDX6 | 20.05181 | 20.11767 | 20.66713 | 21.68861 | 22.13964 | 20.56085 | 19.07472 | 17.9107 | 19.76138 | 20.31623 | 20.69288 | 21.50423 | 24.25394 | 23.27985 | 21.32474 | 21.2926 | 24.11017 | 24.63655 |
| PREP | 15.3052 | 16.99194 | 17.39757 | 16.4138 | 16.49011 | 16.23612 | 18.96642 | 16.41754 | 15.84616 | 18.07143 | 18.68092 | 18.46594 | 17.02809 | 18.6064 | 16.72334 | 16.05402 | 17.12325 | 16.93527 |
| PREX1 | 15.75037 | 15.65195 | 13.75419 | 15.99928 | 15.04251 | 16.33718 | 15.78324 | 14.95928 | 15.58241 | 14.19325 | 14.0362 | 17.01461 | 14.01531 | 15.79227 | 16.74342 | 15.67884 | 17.01039 | 15.46189 |
| PRKAA1 | 18.93625 | 18.69615 | 17.46365 | 17.7896 | 18.27804 | 17.70399 | 17.37876 | 17.02373 | 18.15877 | 18.08365 | 16.78939 | 16.35329 | 18.42579 | 18.65507 | 19.36801 | 18.49939 | 18.22916 | 19.83503 |
| PRKAB1 | 19.40696 | 16.4841 | 15.13963 | 15.37121 | 15.13424 | 14.23003 | 15.42202 | 18.78637 | 16.48908 | 16.70781 | 15.01614 | 15.30654 | 16.45048 | 16.71144 | 16.04919 | 17.53834 | 17.13352 | 17.00651 |
| PRKACA | 16.2424 | 15.89586 | 15.60674 | 16.80108 | 20.40305 | 18.0728 | 21.37564 | 18.60514 | 18.63928 | 20.12094 | 15.29374 | 16.77727 | 19.81454 | 19.8744 | 15.78215 | 15.68325 | 17.91572 | 19.62427 |
| PRKAG1 | 17.18667 | 16.50296 | 17.22694 | 15.44056 | 17.99489 | 12.34186 | 17.30201 | 17.49934 | 17.06962 | 13.89471 | 13.71934 | 16.12496 | 13.23817 | 19.26705 | 16.3509 | 20.90993 | 17.5698 | 19.69803 |
| PRKAR1A | 16.72394 | 16.53513 | 16.60329 | 16.96744 | 19.44357 | 17.41907 | 19.1994 | 18.1055 | 18.74913 | 18.38948 | 16.10818 | 17.91372 | 17.0013 | 17.0159 | 17.6681 | 16.4164 | 20.54653 | 20.28151 |
| PRKAR2A | 16.57378 | 16.9963 | 18.90609 | 18.94449 | 19.1751 | 18.33645 | 17.94517 | 17.60824 | 17.94277 | 17.73506 | 18.2649 | 18.36245 | 21.2225 | 21.20337 | 19.98266 | 19.35709 | 18.52409 | 19.47604 |
| PRKCA | 15.95596 | 16.35046 | 16.20842 | 16.18738 | 19.93604 | 16.66154 | 21.3721 | 19.70979 | 13.50955 | 15.85827 | 16.7919 | 15.36594 | 18.69917 | 16.44581 | 16.12002 | 14.41376 | 20.13813 | 20.16847 |
| PRKCB | 17.55599 | 17.05234 | 17.21627 | 18.48443 | 17.82388 | 16.30982 | 21.45174 | 21.08358 | 14.5874 | 16.24208 | 15.81658 | 15.65639 | 19.04499 | 20.20291 | 17.66215 | 17.59233 | 16.33676 | 16.66381 |
| PRKCD | 16.55049 | 16.87226 | 16.70361 | 17.01403 | 17.52226 | 16.64777 | 16.92031 | 16.65786 | 17.41626 | 17.41634 | 18.35996 | 18.63829 | 18.36616 | 17.28462 | 17.40611 | 16.29796 | 19.62738 | 20.22519 |
| PRKCSH | 17.60027 | 17.60906 | 18.14557 | 16.95527 | 16.7622 | 20.18018 | 17.32845 | 17.74679 | 24.78589 | 23.01033 | 21.55412 | 18.16254 | 16.34039 | 16.45892 | 17.47426 | 17.63422 | 17.04223 | 17.21679 |
| PRKD1 | 16.38691 | 14.13271 | 16.28617 | 15.00451 | 12.90872 | 17.12802 | 16.24764 | 15.69645 | 16.51765 | 15.35687 | 14.52263 | 12.68999 | 18.16395 | 18.11591 | 17.69287 | 16.41119 | 16.1601 | 14.90669 |
| PRKDC | 16.3842 | 16.77715 | 17.49921 | 15.88383 | 17.18734 | 16.53558 | 18.6451 | 18.36885 | 16.02885 | 16.75041 | 16.45467 | 18.50961 | 15.93166 | 16.82131 | 15.60602 | 15.78575 | 16.14064 | 17.93877 |
| PRKG2 | 15.81396 | 15.68519 | 14.62681 | 14.34207 | 14.48063 | 15.84885 | 21.12738 | 20.9615 | 14.42194 | 15.08375 | 13.90703 | 13.68154 | 14.41344 | 14.91008 | 15.90173 | 15.07298 | 13.96126 | 14.51992 |
| PRKRA | 17.36993 | 17.5464 | 17.41502 | 18.3299 | 14.4495 | 15.61656 | 19.49054 | 18.77867 | 13.33669 | 15.02475 | 15.70351 | 16.16323 | 17.86399 | 16.50419 | 17.75828 | 17.15804 | 15.21767 | 16.20921 |
| PRMT1 | 16.85666 | 16.35488 | 18.47802 | 15.94061 | 13.7191 | 17.41534 | 17.08034 | 16.30459 | 13.76169 | 16.15214 | 20.70152 | 21.58387 | 16.42323 | 16.38639 | 15.83503 | 16.92088 | 16.76184 | 16.6861 |
| PRMT5 | 15.92224 | 16.43616 | 15.4716 | 17.52019 | 16.30335 | 16.20442 | 16.76568 | 17.85662 | 16.70384 | 16.32219 | 18.65058 | 20.80862 | 17.47912 | 15.9903 | 17.53157 | 17.82814 | 16.8532 | 17.37665 |
| PRNP | 19.8841 | 18.87537 | 20.04024 | 17.01064 | 15.03552 | 17.54151 | 15.23669 | 16.50369 | 14.01728 | 15.44888 | 16.12005 | 13.75354 | 16.36941 | 14.36412 | 18.66147 | 19.18929 | 14.12993 | 17.06891 |
| PROM1 | 17.12375 | 15.63976 | 17.25127 | 15.65082 | 16.71909 | 16.17137 | 16.92974 | 17.82856 | 15.22034 | 15.41365 | 20.53656 | 21.35506 | 15.17636 | 15.87795 | 15.70341 | 16.42672 | 16.41422 | 16.48509 |
| PROM2 | 21.03559 | 20.43382 | 20.39483 | 20.66729 | 16.98285 | 16.94784 | 16.43854 | 16.74345 | 15.37109 | 15.19863 | 17.52631 | 18.34901 | 16.88404 | 16.02616 | 20.80049 | 20.69306 | 16.34795 | 16.32296 |
| PROSER2 | 17.04224 | 15.1287 | 14.87246 | 16.09876 | 20.69499 | 17.94749 | 16.11459 | 16.7927 | 15.2004 | 16.57019 | 16.61187 | 15.29217 | 14.98043 | 14.7817 | 16.2953 | 15.41088 | 16.69913 | 16.93581 |
| PRPF19 | 16.19876 | 15.49851 | 17.7343 | 15.18378 | 18.02901 | 17.61684 | 18.67183 | 18.65445 | 17.39774 | 18.2443 | 19.11226 | 20.04562 | 18.50767 | 16.80408 | 16.3133 | 16.91905 | 19.33507 | 19.00186 |
| PRPF4 | 14.66826 | 17.10219 | 16.18825 | 16.52621 | 16.0878 | 16.46214 | 15.53134 | 17.17989 | 12.56347 | 14.84062 | 15.05965 | 18.09194 | 16.9032 | 14.36157 | 15.34564 | 14.82398 | 17.01612 | 17.62331 |
| PRPF40A | 14.77377 | 18.01742 | 16.01614 | 15.94648 | 13.31323 | 16.62835 | 15.30683 | 16.04883 | 14.01186 | 13.46728 | 17.95226 | 18.2842 | 15.20342 | 15.8255 | 14.27755 | 15.28406 | 15.1246 | 14.40676 |
| PRPF8 | 17.09983 | 16.72198 | 16.93936 | 16.32247 | 16.9343 | 16.47495 | 15.25492 | 16.48771 | 14.46475 | 16.17583 | 16.23886 | 18.6129 | 15.68286 | 16.70726 | 15.68356 | 16.6779 | 15.97845 | 15.62533 |
| PRPH | 15.79986 | 16.121 | 16.65879 | 15.30667 | 19.27964 | 18.07095 | 20.86588 | 20.09267 | 13.22371 | 16.11278 | 15.56999 | 12.94729 | 13.77859 | 15.89194 | 14.32223 | 15.23814 | 20.42715 | 20.37117 |
| PRPS1 | 16.95005 | 16.76941 | 18.72721 | 19.21719 | 20.04383 | 18.3496 | 21.87926 | 20.02022 | 17.0455 | 18.93402 | 19.31345 | 19.95428 | 21.85278 | 21.09045 | 18.02677 | 18.98458 | 20.97137 | 20.34048 |
| PRPS2 | 17.98977 | 18.49889 | 18.49948 | 19.43709 | 19.93402 | 18.11185 | 20.74408 | 17.48975 | 19.30238 | 19.75286 | 19.588 | 20.39635 | 21.88738 | 20.85538 | 18.62285 | 19.15803 | 18.53626 | 19.82122 |
| PRPSAP1 | 14.95078 | 16.4611 | 17.87854 | 16.47043 | 16.53649 | 17.85805 | 17.90628 | 16.71688 | 19.6582 | 17.33832 | 18.49249 | 18.93343 | 17.34 | 17.37179 | 16.8966 | 16.92899 | 17.82677 | 17.87991 |
| PRPSAP2 | 15.53772 | 16.00476 | 15.94059 | 17.22594 | 17.49973 | 17.0917 | 17.87442 | 17.48186 | 17.32983 | 17.30533 | 18.7802 | 19.94561 | 17.58081 | 17.3108 | 17.58458 | 18.04336 | 18.12127 | 16.81525 |
| PRR15L | 19.43231 | 14.01181 | 17.78781 | 19.04698 | 13.97911 | 14.46611 | 17.33196 | 15.00853 | 13.53429 | 18.66707 | 16.10433 | 12.97052 | 21.95496 | 22.24977 | 20.13264 | 20.64989 | 13.98382 | 13.93568 |
| PRRC2C | 19.07907 | 18.55122 | 17.3691 | 17.74356 | 15.4715 | 15.93746 | 15.77186 | 16.32421 | 14.88123 | 15.12691 | 16.96544 | 16.74435 | 16.74736 | 15.65616 | 15.63034 | 16.64524 | 15.96029 | 16.27875 |

| | | | | | | | | | | | | | | | | | | |
|---|---|---|---|---|---|---|---|---|---|---|---|---|---|---|---|---|---|---|
| PRSS23 | 18.36699 | 16.82668 | 15.79779 | 15.70799 | 20.23675 | 19.67209 | 15.93923 | 15.68203 | 13.87433 | 14.82101 | 15.3732 | 14.60627 | 16.66291 | 14.94777 | 16.45603 | 16.25504 | 14.70897 | 18.58732 |
| PRSS8 | 18.06723 | 18.6523 | 18.81291 | 22.34995 | 16.70352 | 14.91189 | 15.54967 | 16.09722 | 21.23963 | 21.1152 | 18.4526 | 16.68487 | 20.15128 | 19.68218 | 21.00017 | 20.17941 | 17.23312 | 16.44088 |
| PRUNE1 | 12.76334 | 17.72131 | 14.98695 | 15.8407 | 16.76786 | 14.06376 | 14.68588 | 16.8619 | 16.48871 | 13.25428 | 16.59952 | 18.87689 | 15.65519 | 13.83951 | 16.1822 | 16.7625 | 15.58602 | 16.73663 |
| PRXL2A | 16.09841 | 16.66747 | 16.15539 | 16.27925 | 15.15063 | 16.09255 | 19.42645 | 17.98806 | 21.94618 | 20.44086 | 17.45949 | 16.40404 | 15.43894 | 16.82844 | 15.72185 | 15.58734 | 16.44828 | 16.81603 |
| PSAP | 17.22769 | 17.05348 | 19.38171 | 18.93843 | 19.28776 | 20.62579 | 20.17601 | 20.04673 | 23.51477 | 23.08479 | 20.57306 | 17.11217 | 20.88883 | 21.38962 | 18.66107 | 17.70306 | 17.35407 | 19.61109 |
| PSAT1 | 17.65981 | 16.71474 | 18.69458 | 19.58038 | 20.61019 | 19.02152 | 19.08311 | 17.74381 | 17.48298 | 18.838 | 20.29149 | 20.47975 | 21.64444 | 21.60179 | 19.40742 | 19.71932 | 19.51429 | 18.60731 |
| PSD2 | 16.64295 | 16.90839 | 16.45728 | 16.00267 | 14.09873 | 16.37345 | 16.2375 | 16.12644 | 15.24185 | 15.68171 | 20.85743 | 18.01867 | 14.90171 | 16.39876 | 15.95769 | 16.39716 | 14.95998 | 15.48745 |
| PSD3 | 15.36637 | 18.39511 | 19.09264 | 19.40598 | 19.24312 | 18.10543 | 16.97207 | 17.00722 | 15.63398 | 15.63022 | 15.34982 | 16.52898 | 17.97792 | 18.35019 | 18.00727 | 17.68348 | 19.42632 | 18.45703 |
| PSEN1 | 16.5753 | 16.11271 | 17.68652 | 15.27162 | 18.85811 | 15.9083 | 15.05628 | 16.81766 | 17.86964 | 17.70455 | 15.37748 | 19.8337 | 15.99532 | 14.41423 | 18.15049 | 18.65198 | 14.56101 | 16.33491 |
| PSIP1 | 16.62074 | 16.64748 | 16.14817 | 16.8151 | 16.09562 | 15.89251 | 18.02898 | 18.62312 | 15.24631 | 16.55844 | 16.81731 | 18.25586 | 15.62421 | 15.88544 | 16.74426 | 16.98398 | 16.12353 | 15.52191 |
| PSMA1 | 18.57595 | 16.89476 | 19.01104 | 18.93918 | 20.23833 | 19.06487 | 20.08493 | 19.29389 | 19.48674 | 19.70898 | 19.64104 | 21.08152 | 19.80569 | 19.91596 | 18.98087 | 18.87405 | 20.07251 | 19.67969 |
| PSMA2 | 16.20204 | 18.57922 | 17.722 | 17.46854 | 20.54856 | 18.00376 | 18.79682 | 19.06905 | 18.44933 | 19.6277 | 19.57133 | 20.35927 | 20.60998 | 19.39116 | 17.49455 | 18.32484 | 19.80174 | 19.64805 |
| PSMA3 | 17.53331 | 17.13047 | 18.73306 | 18.06723 | 19.29604 | 19.13938 | 17.84937 | 18.01873 | 19.18423 | 19.01575 | 19.37962 | 21.1444 | 19.29251 | 19.09647 | 17.19389 | 19.03323 | 19.38649 | 20.08705 |
| PSMA4 | 18.16345 | 18.35323 | 19.48358 | 19.40795 | 19.79602 | 18.35885 | 18.32514 | 18.46408 | 19.12061 | 19.7482 | 20.19262 | 20.73978 | 19.83222 | 19.20029 | 18.55352 | 19.01492 | 19.59374 | 18.97258 |
| PSMA5 | 17.75125 | 18.3426 | 18.60034 | 19.08555 | 20.23991 | 19.45009 | 17.91516 | 18.38539 | 19.3705 | 19.6941 | 20.42934 | 20.98828 | 20.96531 | 19.3798 | 19.06471 | 19.15898 | 20.33422 | 20.44368 |
| PSMA6 | 17.37366 | 20.13584 | 19.13418 | 20.27382 | 20.99936 | 19.94685 | 19.90248 | 19.60556 | 19.41205 | 20.58708 | 20.0087 | 21.25012 | 20.99153 | 21.02445 | 19.0539 | 18.94135 | 21.05482 | 20.83693 |
| PSMA7 | 18.56562 | 16.85856 | 19.10893 | 20.06824 | 21.3176 | 19.63956 | 19.23383 | 17.80253 | 20.2038 | 20.70123 | 21.11913 | 21.42241 | 21.94324 | 21.61492 | 19.38308 | 19.41889 | 20.85799 | 20.93105 |
| PSMB1 | 19.02063 | 17.83712 | 19.06884 | 19.92135 | 20.28965 | 19.32024 | 17.13409 | 18.48879 | 20.40596 | 19.56471 | 20.4758 | 21.79532 | 20.50484 | 18.70582 | 19.10941 | 19.85814 | 18.51276 | 19.87807 |
| PSMB2 | 17.0742 | 17.55418 | 18.94296 | 17.8822 | 15.63813 | 17.09265 | 15.25681 | 15.86494 | 17.47644 | 16.40686 | 18.81228 | 20.08082 | 14.7408 | 15.11587 | 17.80473 | 16.9368 | 16.5705 | 14.66732 |
| PSMB3 | 21.42228 | 21.09102 | 18.9151 | 17.75056 | 15.52777 | 14.45266 | 17.07062 | 19.80587 | 17.22434 | 18.03201 | 18.80937 | 19.37999 | 20.33314 | 18.16965 | 18.6732 | 19.71805 | 17.38188 | 15.9809 |
| PSMB4 | 14.94 | 16.00543 | 16.62242 | 17.93603 | 18.1607 | 18.39344 | 16.63645 | 16.46871 | 19.29706 | 18.83841 | 18.84962 | 20.70634 | 19.30325 | 16.1805 | 16.86021 | 19.60203 | 17.41867 | 17.23769 |
| PSMB5 | 17.65885 | 17.6532 | 17.09781 | 18.1643 | 15.26638 | 16.18984 | 15.99058 | 15.88781 | 18.06144 | 16.69779 | 18.12793 | 20.37272 | 14.86492 | 19.83729 | 17.37787 | 18.27535 | 16.59353 | 15.65785 |
| PSMB6 | 16.89546 | 17.47776 | 19.46073 | 19.68278 | 18.38392 | 17.26566 | 17.73781 | 18.22386 | 21.61381 | 21.41214 | 19.7528 | 21.55382 | 20.24916 | 19.12472 | 19.272 | 19.59333 | 18.27341 | 15.75365 |
| PSMB7 | 26.54631 | 26.33092 | 26.42061 | 27.4724 | 19.67285 | 18.51643 | 20.45318 | 17.48474 | 18.36365 | 18.48867 | 19.6525 | 20.3256 | 24.80156 | 23.66251 | 26.63992 | 25.58701 | 19.02853 | 19.16442 |
| PSMB8 | 16.19623 | 16.83227 | 14.78255 | 15.89572 | 15.58437 | 16.9288 | 15.22881 | 17.27448 | 17.30564 | 16.25598 | 18.64014 | 20.26766 | 14.62854 | 16.42789 | 14.88195 | 15.52326 | 15.08614 | 17.30051 |
| PSMB9 | 17.05124 | 18.17061 | 15.61643 | 15.67292 | 11.56981 | 15.68454 | 15.85276 | 17.81213 | 14.10757 | 15.15423 | 14.51281 | 17.74614 | 14.91378 | 15.12212 | 12.14368 | 13.24671 | 15.90435 | 13.67964 |
| PSMC1 | 17.15189 | 16.92198 | 19.22863 | 18.2653 | 20.34643 | 19.51818 | 20.25892 | 17.57114 | 18.00005 | 17.90978 | 20.24318 | 18.72167 | 19.72824 | 19.36192 | 16.99582 | 17.63053 | 20.18338 | 19.68024 |
| PSMC2 | 16.23098 | 17.13414 | 17.89383 | 17.50116 | 18.66842 | 18.15406 | 18.70817 | 17.88885 | 17.13446 | 17.12815 | 17.87624 | 19.58697 | 19.19035 | 18.78711 | 17.82528 | 17.24087 | 19.39954 | 18.84841 |
| PSMC3 | 18.87322 | 17.76602 | 18.48903 | 18.46744 | 19.97704 | 18.8226 | 19.72097 | 18.27526 | 18.68172 | 18.98636 | 18.38103 | 19.14358 | 20.07707 | 19.82787 | 17.87966 | 17.32207 | 20.87453 | 20.64406 |
| PSMC4 | 17.25519 | 17.43744 | 18.18292 | 18.29061 | 19.82868 | 18.6786 | 20.17806 | 18.40597 | 18.14881 | 19.04684 | 19.11114 | 19.5914 | 21.17983 | 20.8534 | 16.76526 | 17.74965 | 21.00776 | 20.59692 |
| PSMC5 | 17.21944 | 16.2907 | 17.7876 | 17.33768 | 18.52031 | 16.07197 | 17.80358 | 18.63998 | 16.82526 | 17.35793 | 17.06551 | 18.82014 | 19.67689 | 19.56569 | 16.80727 | 16.98535 | 19.45476 | 18.43301 |
| PSMC6 | 15.83991 | 16.63923 | 16.52611 | 17.52979 | 18.74888 | 16.75216 | 19.28789 | 17.95195 | 17.3796 | 17.99483 | 17.26765 | 19.0374 | 19.28579 | 18.70505 | 15.93337 | 17.21464 | 20.0456 | 18.33192 |
| PSMD1 | 17.05606 | 16.12556 | 17.46284 | 17.115 | 17.17479 | 18.20469 | 18.13712 | 18.23235 | 16.17682 | 16.90179 | 17.13051 | 19.19588 | 18.44122 | 17.20465 | 16.7798 | 16.55814 | 17.61954 | 17.69412 |
| PSMD10 | 17.12015 | 16.96176 | 18.10603 | 17.98118 | 18.31331 | 17.94195 | 19.57122 | 19.06116 | 18.2435 | 19.07094 | 17.06301 | 18.01444 | 20.784 | 21.06071 | 18.66092 | 19.08298 | 21.73361 | 20.67624 |
| PSMD11 | 16.45967 | 15.65689 | 17.20411 | 17.84717 | 17.94533 | 17.44388 | 18.17685 | 18.31416 | 16.59277 | 17.05523 | 18.94495 | 20.34591 | 18.68392 | 18.59473 | 17.29633 | 17.48779 | 17.63485 | 16.87467 |
| PSMD12 | 16.14634 | 16.25549 | 17.22979 | 15.86064 | 17.8063 | 18.94349 | 18.14097 | 17.37864 | 14.50758 | 14.93587 | 16.89424 | 19.52216 | 16.09265 | 16.93168 | 17.05543 | 17.75113 | 16.96504 | 16.87291 |
| PSMD13 | 16.17952 | 15.86 | 17.6396 | 18.15529 | 18.43422 | 17.57636 | 19.22394 | 18.30164 | 16.1031 | 17.43082 | 17.38686 | 19.77022 | 19.46615 | 19.32645 | 16.8978 | 17.82433 | 17.36124 | 19.62951 |
| PSMD14 | 16.27842 | 15.23219 | 18.16952 | 17.29519 | 18.79269 | 17.12526 | 17.82687 | 18.80131 | 15.77338 | 16.49834 | 18.70294 | 20.39355 | 19.32752 | 19.16752 | 16.09123 | 18.18401 | 18.17432 | 19.27074 |
| PSMD2 | 18.17228 | 16.71015 | 17.36624 | 17.4683 | 17.87833 | 17.79014 | 17.23172 | 17.44179 | 17.07558 | 16.03795 | 18.25339 | 19.26341 | 18.30605 | 17.83456 | 17.02837 | 17.3808 | 18.13251 | 18.63076 |
| PSMD3 | 16.37775 | 16.43683 | 18.0219 | 17.63911 | 17.0694 | 17.46592 | 17.47106 | 16.98038 | 17.16643 | 16.49272 | 17.54968 | 20.21785 | 17.70161 | 17.61918 | 17.57079 | 18.24591 | 17.05125 | 17.60334 |
| PSMD4 | 15.38627 | 15.6025 | 15.87823 | 16.693 | 18.70989 | 15.04667 | 18.71183 | 17.4819 | 13.78246 | 16.31935 | 15.46556 | 19.58538 | 19.45887 | 18.86007 | 16.45296 | 14.43046 | 18.56868 | 18.65014 |
| PSMD5 | 14.611 | 15.93282 | 17.0105 | 16.32206 | 16.94823 | 17.22327 | 17.10126 | 15.26969 | 14.60881 | 17.07625 | 17.49712 | 17.90317 | 15.47827 | 14.93621 | 16.03578 | 15.38837 | 16.68877 | 16.58239 |
| PSMD6 | 16.90371 | 15.98671 | 17.81197 | 16.97543 | 16.31152 | 15.64754 | 17.09135 | 16.74792 | 17.15627 | 16.91292 | 19.3697 | 20.39248 | 18.41218 | 18.4595 | 18.25102 | 19.32702 | 17.3348 | 17.26835 |
| PSMD7 | 16.75309 | 19.19799 | 17.21967 | 18.40253 | 18.37292 | 16.23753 | 17.58072 | 16.31967 | 16.37251 | 17.71128 | 19.21664 | 20.22625 | 19.87426 | 16.94804 | 18.08807 | 17.01351 | 18.28972 | 18.88228 |
| PSMD8 | 15.73385 | 13.78525 | 16.03492 | 15.96301 | 14.98175 | 15.39031 | 17.87666 | 15.46599 | 13.63196 | 15.36799 | 17.91415 | 19.69582 | 16.77303 | 16.67749 | 14.22476 | 15.00826 | 15.50281 | 16.90678 |
| PSMD9 | 15.69013 | 17.45286 | 15.88028 | 16.00142 | 18.63809 | 17.35005 | 17.99793 | 17.4838 | 14.67556 | 16.14561 | 17.59922 | 18.51536 | 18.9482 | 18.33891 | 17.51195 | 17.84316 | 19.95428 | 19.39956 |
| PSME1 | 15.08937 | 16.41501 | 16.81271 | 17.43486 | 17.30312 | 17.28315 | 17.12501 | 18.35143 | 18.79337 | 19.1315 | 20.34827 | 19.9742 | 18.60997 | 17.27781 | 14.84495 | 18.33746 | 19.17241 | 20.43488 |
| PSME2 | 16.85776 | 17.17344 | 17.1644 | 16.73305 | 18.18332 | 16.60082 | 19.20404 | 16.8997 | 17.14635 | 18.23556 | 17.44454 | 18.59636 | 18.98136 | 15.72934 | 15.83099 | 15.96202 | 19.28657 | 17.65616 |
| PSME3 | 16.16396 | 15.44789 | 15.56341 | 16.0232 | 16.22602 | 15.85773 | 16.58331 | 17.05624 | 14.30979 | 15.41482 | 19.09885 | 18.98736 | 17.26474 | 14.97024 | 14.46096 | 15.53513 | 17.03122 | 14.45472 |
| PSME4 | 14.56316 | 15.55034 | 14.99991 | 16.05405 | 14.72276 | 14.1321 | 15.43746 | 15.32939 | 14.07942 | 14.79677 | 14.21462 | 18.08116 | 15.20414 | 15.48274 | 17.16389 | 13.59774 | 14.20848 | 15.39239 |
| PSMF1 | 16.15945 | 15.32888 | 16.04551 | 15.50189 | 12.8907 | 14.69229 | 17.23107 | 15.5412 | 14.18797 | 18.98222 | 17.76601 | 16.45986 | 16.3221 | 13.38663 | 13.90441 | 15.60937 | 19.3961 | 15.8014 |
| PSMG2 | 14.5287 | 14.47937 | 16.49175 | 15.37181 | 14.7777 | 16.65951 | 17.50082 | 18.02277 | 12.62031 | 15.33225 | 16.92637 | 18.65647 | 15.76938 | 13.83963 | 15.78881 | 15.84191 | 14.66201 | 12.88886 |
| PSMG3 | 16.7865 | 15.40654 | 16.19907 | 15.76473 | 18.78843 | 19.02093 | 19.30192 | 17.30307 | 12.18049 | 18.02249 | 17.96696 | 19.41931 | 19.25487 | 19.36855 | 17.25483 | 14.46461 | 19.09792 | 18.47652 |
| PSPC1 | 16.48321 | 15.86455 | 16.22492 | 15.38164 | 18.23818 | 18.82425 | 18.96603 | 16.9241 | 15.87909 | 17.19055 | 19.38256 | 19.19763 | 17.36206 | 16.89424 | 13.5787 | 16.06008 | 17.54734 | 17.6666 |
| PSPH | 16.64122 | 17.1139 | 17.72952 | 17.02798 | 17.95222 | 16.2275 | 17.04181 | 16.67479 | 17.54545 | 18.34533 | 18.68985 | 18.91169 | 20.41552 | 19.11607 | 17.97713 | 18.78084 | 17.72833 | 16.96378 |
| PTBP1 | 17.22376 | 18.40211 | 17.23171 | 16.35959 | 17.01326 | 16.98664 | 16.06651 | 17.14475 | 17.85932 | 18.66891 | 19.66455 | 20.06118 | 15.82407 | 17.99978 | 15.72801 | 17.77902 | 15.7807 | 13.62615 |
| PTBP2 | 16.95867 | 16.16222 | 14.91512 | 15.2068 | 13.84007 | 15.57009 | 13.35264 | 16.9916 | 12.20489 | 14.9268 | 16.40655 | 18.06881 | 15.21516 | 17.09461 | 14.12289 | 14.87304 | 16.17151 | 14.41089 |
| PTGES2 | 16.20373 | 15.27289 | 15.54554 | 14.7517 | 13.93533 | 17.53975 | 16.69435 | 17.47451 | 16.92963 | 15.65859 | 17.48268 | 17.00073 | 15.31099 | 14.99742 | 16.91082 | 16.01789 | 15.47027 | 15.00592 |
| PTGES3 | 19.9491 | 19.43758 | 18.94826 | 20.13382 | 22.08536 | 20.65168 | 21.31689 | 21.20994 | 20.89904 | 21.51956 | 20.93036 | 21.58925 | 22.49111 | 22.25432 | 19.66108 | 19.58607 | 22.41507 | 21.46244 |
| PTGFRN | 22.44024 | 22.04281 | 24.04235 | 23.6045 | 20.18016 | 19.75639 | 19.6809 | 17.9068 | 16.18062 | 16.82633 | 17.22039 | 21.44058 | 22.54048 | 22.38217 | 23.67858 | 23.75051 | 17.52989 | 18.05308 |
| PTGR1 | 17.16516 | 16.69502 | 16.27448 | 18.26842 | 21.88419 | 20.60885 | 17.41861 | 16.98546 | 17.89367 | 19.21656 | 14.72919 | 15.01956 | 18.04913 | 17.83491 | 17.59399 | 17.24713 | 18.96715 | 18.25362 |
| PTGR2 | 15.62653 | 17.10369 | 16.40514 | 17.05848 | 14.77668 | 16.7233 | 14.97218 | 15.57338 | 13.62948 | 16.50012 | 14.07269 | 17.98717 | 16.83215 | 17.41434 | 18.31815 | 19.12294 | 16.62699 | 15.12413 |
| PTGS1 | 15.29867 | 16.50346 | 15.90125 | 15.56084 | 14.66705 | 14.89134 | 15.62486 | 17.51825 | 15.14357 | 15.79938 | 21.52406 | 18.83046 | 14.71897 | 16.0653 | 15.81209 | 14.64221 | 15.40526 | 14.69319 |
| PTK7 | 16.18986 | 16.24685 | 18.21553 | 19.05814 | 20.67953 | 20.79665 | 19.86177 | 18.65198 | 20.11487 | 20.03196 | 19.93142 | 19.15225 | 20.7287 | 20.64565 | 20.38585 | 19.71658 | 20.35391 | 21.3425 |
| PTMA | 17.7166 | 16.49632 | 18.36589 | 17.27843 | 18.18821 | 18.5546 | 20.58718 | 20.73576 | 19.02194 | 17.45103 | 19.2009 | 20.3246 | 15.71982 | 14.91495 | 18.22339 | 18.55997 | 21.09971 | 16.95072 |
| PTMS | 19.47274 | 19.62103 | 19.0939 | 19.3167 | 20.43264 | 19.64772 | 20.07432 | 21.6562 | 18.84999 | 21.14111 | 21.57626 | 21.65361 | 22.40074 | 21.78708 | 15.14329 | 18.35524 | 20.17089 | 19.14451 |
| PTP4A1 | 20.82296 | 21.15191 | 21.24943 | 21.36627 | 18.62517 | 18.36685 | 18.51644 | 17.44821 | 19.84743 | 19.63328 | 20.98325 | 21.72947 | 20.39465 | 19.98529 | 21.35358 | 21.70882 | 19.54139 | 19.1669 |
| PTP4A2 | 18.98376 | 21.63497 | 17.90508 | 19.05949 | 12.32993 | 16.95093 | 16.61183 | 16.79782 | 18.58903 | 17.24635 | 17.66301 | 19.97703 | 22.29047 | 17.32286 | 19.50346 | 19.4306 | 18.39593 | 18.66132 |
| PTP4A3 | 15.38415 | 15.47465 | 15.60791 | 17.0039 | 14.99673 | 16.11544 | 14.04605 | 17.11686 | 19.0294 | 15.31455 | 19.38709 | 13.64201 | 14.19136 | 14.50347 | 14.63667 | 15.97708 | 14.19614 | 16.58051 |
| PTPA | 17.36687 | 17.44817 | 17.04795 | 18.0086 | 18.50552 | 18.55959 | 17.29142 | 16.59158 | 16.28927 | 17.05153 | 17.11034 | 19.31339 | 17.89509 | 17.4435 | 18.78344 | 16.86669 | 19.66135 | 17.31632 |
| PTPN1 | 14.61118 | 17.65697 | 14.64677 | 16.05274 | 16.52287 | 14.57852 | 16.10714 | 16.66586 | 15.1032 | 14.10616 | 16.77923 | 17.96672 | 16.88466 | 17.42622 | 14.82524 | 15.01448 | 13.5855 | 13.60573 |
| PTPN11 | 17.62081 | 17.6094 | 21.31538 | 15.82006 | 16.21458 | 16.07267 | 16.46406 | 18.10675 | 16.24275 | 16.45525 | 17.34924 | 18.70754 | 16.20566 | 15.67266 | 17.90431 | 17.02683 | 16.73419 | 18.24937 |
| PTPN12 | 16.91558 | 17.182 | 15.87488 | 16.14619 | 17.44222 | 15.63301 | 16.74159 | 12.48824 | 12.51658 | 13.24892 | 15.04841 | 11.50688 | 13.65571 | 17.76415 | 12.26969 | 13.13949 | 15.16213 | 19.06208 |
| PTPN14 | 13.77715 | 17.72104 | 16.6295 | 13.87988 | 20.06202 | 18.73045 | 14.64125 | 17.20619 | 13.24642 | 14.39295 | 13.98323 | 12.82905 | 14.76163 | 13.76259 | 13.5691 | 15.80363 | 19.14198 | 19.25086 |
| PTPN23 | 19.59127 | 19.82667 | 16.87953 | 15.73347 | 19.56911 | 18.58434 | 17.74696 | 18.10107 | 18.30736 | 18.08138 | 16.37577 | 16.01698 | 16.40164 | 16.52826 | 16.81183 | 15.99835 | 17.06527 | 18.74455 |
| PTPN6 | 19.08926 | 14.05948 | 14.88833 | 14.73643 | 16.65116 | 17.25853 | 14.64543 | 17.02701 | 16.21513 | 15.79173 | 15.37372 | 17.45354 | 16.36184 | 16.53259 | 17.77622 | 16.06651 | 15.64865 | 15.39666 |
| PTPRA | 17.26905 | 16.29115 | 17.73539 | 17.33614 | 18.21647 | 17.59938 | 15.63675 | 16.85314 | 16.35977 | 15.58099 | 18.35533 | 18.96336 | 17.25838 | 16.87596 | 18.70883 | 18.71658 | 18.10109 | 19.86099 |
| PTPRB | 19.3616 | 19.15682 | 16.17384 | 16.18595 | 14.59328 | 15.25563 | 16.63055 | 16.90511 | 13.39913 | 16.71639 | 16.32018 | 13.34355 | 15.59517 | 15.09268 | 14.68758 | 14.88333 | 14.99561 | 15.59345 |
| PTPRC | 17.63695 | 19.08566 | 18.16393 | 19.2875 | 16.73886 | 17.66078 | 14.55937 | 14.82848 | 13.97238 | 17.15802 | 14.83515 | 10.70282 | 17.66659 | 18.14054 | 18.49008 | 18.91825 | 15.54931 | 16.42509 |
| PTPRD | 20.21367 | 19.09159 | 19.6822 | 19.4025 | 18.53249 | 18.15128 | 15.07006 | 16.56278 | 18.57522 | 17.92958 | 18.80924 | 17.72498 | 17.56646 | 16.92253 | 20.90451 | 20.63282 | 15.96109 | 15.73834 |
| PTPRE | 15.04511 | 16.12867 | 17.28262 | 15.2167 | 19.61109 | 18.9377 | 14.92866 | 16.62457 | 12.8435 | 15.27288 | 16.10823 | 14.13528 | 14.03213 | 14.45038 | 15.45303 | 14.84296 | 15.75892 | 12.45857 |
| PTPRF | 20.04373 | 19.86707 | 18.30106 | 18.57596 | 20.01415 | 21.18664 | 16.0697 | 17.16154 | 18.93062 | 17.63991 | 17.48469 | 16.37092 | 17.82416 | 17.12222 | 20.54112 | 19.82512 | 17.49606 | 20.61132 |
| PTPRG | 16.5069 | 17.32863 | 17.6002 | 17.94296 | 20.02401 | 18.7593 | 16.71592 | 16.714 | 14.29813 | 15.31039 | 16.47872 | 16.74142 | 16.67668 | 16.72404 | 17.0912 | 18.6932 | 17.22251 | 14.41993 |
| PTPRJ | 18.67092 | 18.85193 | 17.27089 | 18.06483 | 20.17402 | 19.6697 | 18.18309 | 17.76118 | 19.32178 | 18.44963 | 21.07367 | 20.51624 | 18.98399 | 17.95641 | 17.85736 | 18.30934 | 18.59353 | 18.93822 |
| PTPRK | 18.45774 | 18.72247 | 17.06388 | 16.69463 | 16.25701 | 14.97397 | 15.05418 | 15.79743 | 18.93466 | 17.91927 | 18.78161 | 17.11426 | 15.96732 | 15.35996 | 16.39293 | 16.40817 | 18.43217 | 16.91478 |
| PTPRM | 16.19359 | 15.54324 | 15.84222 | 15.75645 | 18.81298 | 15.98941 | 13.93673 | 14.77736 | 17.63258 | 17.32125 | 15.81492 | 14.80825 | 16.14772 | 15.77251 | 16.71066 | 17.44864 | 16.54404 | 18.0016 |

| | | | | | | | | | | | | | | | | | | |
|---|---|---|---|---|---|---|---|---|---|---|---|---|---|---|---|---|---|---|
| PTPRS | 16.14439 | 14.77478 | 16.4018 | 18.13399 | 19.36315 | 19.81138 | 19.10549 | 16.70003 | 18.56828 | 17.86329 | 14.68886 | 17.8454 | 17.53 | 17.19479 | 15.64627 | 17.02465 | 18.20079 | 19.07987 |
| PTRHD1 | 18.94486 | 18.65151 | 18.12652 | 16.10635 | 20.90592 | 17.72591 | 17.63315 | 16.70915 | 14.43538 | 18.45234 | 19.02782 | 20.49055 | 19.55945 | 20.4355 | 17.73604 | 19.40942 | 21.07603 | 19.64817 |
| PTTG1IP | 19.98837 | 19.99664 | 21.90041 | 23.30654 | 23.9617 | 23.44907 | 21.97212 | 19.32705 | 21.58818 | 21.99806 | 16.77359 | 18.41914 | 21.7657 | 22.08432 | 21.55323 | 21.72601 | 19.16956 | 21.8449 |
| PUF60 | 15.82659 | 16.09239 | 15.40154 | 15.7261 | 17.2143 | 18.6074 | 17.21573 | 18.79233 | 19.09422 | 19.2278 | 19.06032 | 19.56095 | 17.58702 | 17.8291 | 16.15188 | 15.19365 | 18.77888 | 19.23449 |
| PURA | 18.26894 | 18.16588 | 16.01977 | 14.22936 | 14.61857 | 15.549 | 15.30426 | 16.18568 | 15.98894 | 15.07616 | 16.13856 | 19.87471 | 17.04147 | 15.00675 | 16.10636 | 18.48088 | 17.12883 | 15.03821 |
| PURB | 18.55573 | 18.85604 | 16.89246 | 17.32389 | 14.16553 | 14.74532 | 15.8586 | 16.90149 | 12.75916 | 15.09042 | 15.53746 | 16.53778 | 16.86582 | 15.90407 | 16.24757 | 17.7509 | 16.03249 | 14.3417 |
| PUS1 | 15.66218 | 14.94912 | 15.66781 | 16.32907 | 12.16201 | 15.63226 | 14.67616 | 14.28838 | 16.8845 | 18.02228 | 15.0073 | 17.44434 | 11.95852 | 14.67853 | 14.93289 | 12.56237 | 17.2327 | 14.11694 |
| PVR | 21.68939 | 21.89141 | 20.78466 | 20.47681 | 18.90611 | 18.73449 | 17.11645 | 16.94634 | 17.8183 | 16.00195 | 21.32933 | 21.95307 | 17.09785 | 16.79221 | 22.09831 | 20.7565 | 16.89148 | 17.63007 |
| PWP1 | 14.67821 | 15.88518 | 13.80528 | 15.38515 | 13.08842 | 15.48733 | 16.62704 | 18.28563 | 12.59522 | 15.72247 | 16.34028 | 18.24831 | 15.07396 | 14.59481 | 16.66406 | 15.45407 | 16.01799 | 12.80952 |
| PXDN | 16.13302 | 15.04192 | 15.20073 | 14.00642 | 14.92606 | 16.73875 | 15.14017 | 16.29928 | 17.97787 | 16.63411 | 16.02046 | 14.31501 | 13.92042 | 15.33188 | 15.23108 | 14.64714 | 15.2727 | 16.29306 |
| PXN | 15.78092 | 16.35061 | 17.564 | 15.8643 | 18.01176 | 14.77356 | 15.09196 | 15.45952 | 14.45852 | 14.69164 | 14.64195 | 15.04736 | 16.6239 | 17.41764 | 14.19072 | 14.85348 | 21.0048 | 20.27306 |
| PYCR1 | 15.70007 | 16.68894 | 16.3705 | 16.27085 | 14.18663 | 15.0639 | 16.63417 | 16.82496 | 19.4682 | 17.69431 | 15.30981 | 13.3031 | 14.96706 | 14.5898 | 16.60417 | 16.99812 | 14.8171 | 16.18866 |
| PYCR2 | 14.05073 | 16.49877 | 15.83214 | 14.91215 | 15.41891 | 17.01718 | 17.82012 | 17.04169 | 19.52822 | 18.50655 | 14.46831 | 13.38426 | 14.26617 | 14.88231 | 15.51326 | 17.29542 | 15.42995 | 16.29047 |
| PYCR3 | 14.02494 | 14.07848 | 14.60849 | 14.2189 | 17.14497 | 15.30489 | 15.78586 | 16.38575 | 12.90484 | 18.03743 | 17.52316 | 19.05216 | 19.50374 | 15.2278 | 14.62491 | 16.5444 | 17.53147 | 17.51162 |
| PYGB | 15.69799 | 16.25788 | 17.28208 | 17.41898 | 18.10852 | 16.56978 | 17.84557 | 17.64756 | 17.53587 | 18.49652 | 17.18443 | 18.28673 | 18.21566 | 18.10587 | 17.53339 | 17.58393 | 19.84936 | 21.65884 |
| PYGL | 15.59196 | 17.98297 | 20.56678 | 22.14833 | 20.87571 | 24.81162 | 20.26706 | 22.06905 | 16.9727 | 18.70975 | 22.89034 | 16.35721 | 19.00022 | 23.44341 | 20.1373 | 17.37873 | 19.37488 | 23.83835 |
| PYM1 | 19.64462 | 19.25495 | 16.55654 | 16.49943 | 14.81874 | 15.84968 | 14.45659 | 18.87975 | 14.81508 | 17.41247 | 15.59936 | 15.98539 | 19.27114 | 16.18532 | 17.56192 | 18.82925 | 15.24196 | 14.67276 |
| QARS1 | 15.80009 | 16.26073 | 15.96155 | 16.23387 | 16.60755 | 16.63352 | 17.13484 | 17.3195 | 16.03761 | 16.58664 | 17.97092 | 19.75312 | 17.34021 | 15.79388 | 17.01871 | 16.2929 | 17.46656 | 17.43613 |
| QDPR | 15.58735 | 14.50251 | 15.58666 | 15.48215 | 20.54015 | 17.79951 | 20.12392 | 18.9696 | 16.40264 | 16.63037 | 18.45558 | 18.12863 | 21.1566 | 19.27164 | 15.91819 | 16.80066 | 20.03575 | 19.84827 |
| QPRT | 15.47087 | 15.13623 | 14.70718 | 12.95364 | 13.98131 | 14.63787 | 17.29825 | 16.607 | 15.69869 | 14.85046 | 17.48529 | 19.82579 | 14.92556 | 15.61084 | 15.2717 | 14.79675 | 15.54094 | 16.2801 |
| QSOX1 | 16.48389 | 16.13767 | 18.06169 | 17.37558 | 15.58323 | 17.9131 | 17.25545 | 16.94666 | 19.67283 | 18.14513 | 17.7057 | 14.26793 | 15.56593 | 17.76491 | 16.57311 | 16.57835 | 15.45821 | 16.46769 |
| QSOX2 | 15.91187 | 15.57117 | 16.01765 | 16.50255 | 15.12362 | 18.2062 | 16.44797 | 17.01947 | 22.17548 | 19.90098 | 18.11251 | 17.01396 | 14.54618 | 15.15765 | 15.15643 | 16.26339 | 17.02648 | 17.43588 |
| QTRT1 | 14.96158 | 14.83802 | 14.25889 | 16.74033 | 13.65054 | 14.91956 | 14.49206 | 15.32415 | 14.76759 | 11.91431 | 14.45983 | 17.49052 | 15.73487 | 15.76521 | 14.96348 | 16.04394 | 14.37384 | 14.5968 |
| QTRT2 | 18.35318 | 18.16172 | 15.76419 | 14.47722 | 16.04679 | 15.79281 | 15.38428 | 16.51712 | 16.42067 | 15.71398 | 16.59312 | 19.60275 | 12.13767 | 13.63495 | 15.57631 | 14.08159 | 14.88151 | 15.19381 |
| RAB10 | 22.04111 | 22.30229 | 22.38709 | 22.22345 | 19.49182 | 17.93019 | 18.22073 | 17.97708 | 19.51433 | 19.43701 | 20.17554 | 21.17808 | 19.19416 | 19.57104 | 22.3861 | 22.38304 | 17.85557 | 19.23745 |
| RAB11A | 22.03282 | 22.19371 | 22.71165 | 23.3568 | 23.75918 | 22.85823 | 22.50591 | 21.85648 | 20.17288 | 20.67079 | 22.40798 | 21.80546 | 22.42468 | 22.47756 | 22.28617 | 22.82407 | 20.73431 | 21.43718 |
| RAB11B | 21.29797 | 21.68014 | 20.58552 | 21.97929 | 19.01922 | 17.83192 | 18.27653 | 16.6437 | 20.17149 | 18.54116 | 19.95705 | 21.46038 | 16.90951 | 17.032 | 19.27586 | 20.74399 | 16.0073 | 14.22559 |
| RAB11FIP1 | 16.42717 | 16.24156 | 16.20487 | 17.20386 | 20.48398 | 17.39296 | 16.22171 | 16.12471 | 13.67413 | 16.01626 | 16.63319 | 15.69789 | 16.867 | 17.03335 | 15.82743 | 15.68693 | 16.92075 | 16.01952 |
| RAB11FIP5 | 14.80373 | 16.04481 | 15.78249 | 17.08587 | 21.00282 | 19.4841 | 16.59254 | 16.86806 | 14.68929 | 16.68202 | 16.45845 | 13.94184 | 20.03929 | 18.28058 | 14.90713 | 14.88844 | 18.0127 | 19.47099 |
| RAB12 | 15.1672 | 16.12761 | 17.07346 | 17.50305 | 18.49334 | 14.27239 | 15.31921 | 14.35437 | 14.28184 | 14.42607 | 15.60058 | 16.94647 | 16.65426 | 14.67702 | 14.83067 | 18.03476 | 14.95748 | 17.60276 |
| RAB13 | 21.83689 | 22.09489 | 21.17329 | 20.97795 | 22.54416 | 20.5 | 19.09069 | 17.32749 | 18.82354 | 19.95277 | 17.6 | 17.7953 | 22.01235 | 22.09996 | 20.96338 | 20.40679 | 20.63829 | 21.49664 |
| RAB14 | 20.71562 | 20.57836 | 21.0663 | 21.88585 | 20.84943 | 18.87534 | 20.26681 | 20.07935 | 20.71999 | 20.33052 | 21.58777 | 22.04727 | 20.12296 | 18.3034 | 20.5895 | 21.04421 | 17.94302 | 19.38656 |
| RAB18 | 18.54709 | 18.82445 | 16.73719 | 18.90038 | 18.12642 | 18.29928 | 18.8704 | 18.76078 | 17.67145 | 17.94171 | 19.99198 | 20.49091 | 18.68892 | 19.12064 | 17.50651 | 17.47846 | 16.39681 | 15.98614 |
| RAB1A | 21.5337 | 20.71156 | 20.11213 | 20.40652 | 20.65043 | 19.83528 | 21.0807 | 20.33128 | 19.47422 | 20.10912 | 20.87763 | 21.01 | 21.10712 | 21.18302 | 20.5606 | 20.51252 | 20.07343 | 20.50405 |
| RAB1B | 21.539 | 21.59518 | 21.08875 | 22.27678 | 21.45976 | 14.71069 | 14.10834 | 14.36158 | 19.29292 | 17.99855 | 21.16795 | 20.48335 | 18.21177 | 17.5797 | 22.70325 | 22.39651 | 19.02389 | 18.71802 |
| RAB1C | 23.54019 | 23.50012 | 23.16235 | 23.44326 | 22.11718 | 21.09142 | 22.18746 | 21.10979 | 21.71068 | 21.86364 | 22.70955 | 23.18387 | 22.37054 | 21.98906 | 23.529 | 23.47835 | 21.40089 | 21.99307 |
| RAB20 | 15.88932 | 17.32024 | 16.88575 | 16.43359 | 15.22341 | 18.07291 | 15.26332 | 15.98099 | 18.40523 | 17.7771 | 12.29971 | 17.47884 | 16.65937 | 15.94097 | 17.84073 | 17.88817 | 15.64891 | 15.88361 |
| RAB21 | 19.96443 | 20.45039 | 19.89066 | 20.11474 | 20.27056 | 18.82858 | 19.21714 | 17.95704 | 18.86066 | 18.83504 | 19.53261 | 19.93668 | 19.41307 | 19.3734 | 19.59916 | 19.95004 | 17.85479 | 19.29485 |
| RAB22A | 20.23936 | 21.07158 | 19.63293 | 20.12162 | 19.81558 | 18.11016 | 20.63167 | 17.56619 | 18.90911 | 18.72389 | 17.00169 | 18.23417 | 19.54431 | 20.16087 | 20.9474 | 20.4963 | 18.56243 | 18.88297 |
| RAB23 | 18.10257 | 20.2201 | 19.15276 | 19.81624 | 18.50346 | 18.40949 | 16.49258 | 17.93874 | 18.20433 | 18.32376 | 16.62355 | 17.07518 | 18.49995 | 17.40991 | 19.77555 | 19.87972 | 17.92961 | 18.91671 |
| RAB24 | 19.56125 | 18.64011 | 18.58374 | 18.75383 | 17.16689 | 16.92366 | 15.92487 | 21.15058 | 11.97444 | 14.73527 | 14.95214 | 15.85132 | 18.13159 | 15.52196 | 18.18332 | 17.31823 | 15.63184 | 16.3713 |
| RAB25 | 18.79827 | 18.61578 | 20.34392 | 20.33989 | 17.14212 | 17.32805 | 15.94887 | 16.08382 | 15.64431 | 15.89299 | 19.29071 | 20.42081 | 16.66479 | 15.63474 | 20.12803 | 20.63352 | 15.65574 | 16.0846 |
| RAB27A | 17.67101 | 18.13769 | 20.56388 | 20.19639 | 16.33587 | 16.53132 | 19.35781 | 17.69471 | 15.00551 | 15.8062 | 13.80587 | 19.35225 | 16.99564 | 18.00385 | 19.86106 | 20.56299 | 15.8526 | 13.59447 |
| RAB27B | 19.22188 | 18.64113 | 18.54601 | 19.50657 | 19.22962 | 16.59725 | 20.07587 | 20.04876 | 13.77029 | 17.22597 | 17.05094 | 15.27004 | 18.39657 | 19.95999 | 19.18001 | 19.08455 | 16.38791 | 18.61183 |
| RAB2A | 19.16026 | 19.71622 | 19.88179 | 20.06194 | 18.63045 | 16.97904 | 20.18515 | 19.39752 | 21.49118 | 21.73851 | 19.74689 | 20.35025 | 18.27067 | 20.08674 | 19.46238 | 20.10005 | 15.6885 | 16.90712 |
| RAB2B | 16.9039 | 16.49339 | 15.21542 | 13.14847 | 15.61387 | 16.07135 | 17.1312 | 18.83902 | 17.79835 | 16.51795 | 17.90833 | 19.42376 | 14.40594 | 15.07082 | 16.81828 | 18.14261 | 15.17991 | 14.37171 |
| RAB31 | 17.62846 | 16.57 | 15.57102 | 15.83814 | 21.0227 | 19.78613 | 15.56368 | 16.53976 | 12.71308 | 15.47343 | 15.42364 | 13.48592 | 14.29311 | 15.20717 | 16.53668 | 16.8898 | 14.97382 | 12.65768 |
| RAB32 | 16.27084 | 15.1224 | 18.24965 | 18.5982 | 22.00162 | 19.40044 | 16.88922 | 14.99647 | 15.37011 | 16.31883 | 17.51788 | 12.21779 | 16.49194 | 15.79427 | 15.94428 | 15.90966 | 21.55868 | 20.95705 |
| RAB34 | 16.22724 | 16.52265 | 16.71805 | 16.03984 | 18.68538 | 18.56788 | 17.67735 | 17.13669 | 17.75862 | 17.77039 | 15.90524 | 17.08058 | 14.63979 | 15.77055 | 15.90289 | 14.7623 | 17.20689 | 14.12773 |
| RAB35 | 22.01239 | 22.24054 | 21.13825 | 21.8661 | 19.27025 | 17.69445 | 18.74255 | 18.14011 | 17.66568 | 16.83977 | 19.55315 | 19.46365 | 19.91375 | 19.06379 | 22.16429 | 22.06431 | 18.53486 | 19.04726 |
| RAB3A | 17.92432 | 15.31064 | 20.596 | 20.17917 | 16.1188 | 14.74738 | 16.85409 | 15.5845 | 12.77937 | 16.77764 | 18.04187 | 18.43082 | 16.11209 | 16.62845 | 19.85263 | 19.46792 | 18.09826 | 16.13892 |
| RAB3B | 17.50262 | 15.68231 | 13.69957 | 15.82972 | 17.89792 | 15.72565 | 15.18958 | 15.66483 | 14.06231 | 16.72478 | 17.39639 | 15.4749 | 18.14613 | 15.05604 | 15.35802 | 13.454 | 14.20597 | 16.14495 |
| RAB3D | 21.16442 | 21.04388 | 20.39307 | 21.83377 | 17.25599 | 15.98619 | 15.20201 | 16.49329 | 16.63484 | 17.49021 | 18.26176 | 18.94977 | 16.48161 | 15.97152 | 21.55571 | 21.33056 | 15.75769 | 15.6754 |
| RAB3GAP1 | 14.7047 | 16.22382 | 15.56691 | 14.32957 | 17.18721 | 16.00577 | 16.09061 | 16.86001 | 15.62617 | 17.88084 | 17.08726 | 18.36404 | 16.3938 | 16.31254 | 16.27775 | 15.45637 | 16.03009 | 16.78029 |
| RAB3GAP2 | 20.31406 | 14.08391 | 15.7312 | 17.44772 | 21.73763 | 19.94989 | 20.31255 | 16.99632 | 20.82838 | 22.20821 | 20.9066 | 18.19021 | 23.47708 | 22.38179 | 15.23543 | 15.30655 | 20.08049 | 20.78116 |
| RAB4A | 18.02747 | 17.43207 | 18.71194 | 19.68575 | 17.01279 | 16.54705 | 18.06097 | 19.15214 | 16.38014 | 16.35914 | 16.70834 | 17.96547 | 18.80793 | 16.97059 | 18.9511 | 19.45904 | 15.54744 | 13.61957 |
| RAB4B | 17.4605 | 16.93892 | 14.84158 | 13.99022 | 13.71875 | 13.80127 | 17.45616 | 17.16856 | 16.39493 | 17.05768 | 15.46606 | 17.36856 | 13.58088 | 15.37856 | 18.16109 | 17.70267 | 16.05206 | 14.6056 |
| RAB5A | 20.5938 | 20.99984 | 19.15867 | 19.8805 | 19.74773 | 17.22577 | 18.30349 | 18.19848 | 19.85006 | 19.5426 | 19.00141 | 19.75062 | 18.94634 | 18.69467 | 19.75685 | 19.79805 | 17.66577 | 18.867 |
| RAB5B | 22.87539 | 22.90352 | 21.51864 | 21.98252 | 20.72133 | 19.46015 | 21.29213 | 20.49856 | 20.49292 | 21.13061 | 21.38609 | 21.19148 | 20.27844 | 20.59871 | 22.58326 | 22.48163 | 19.12395 | 18.66426 |
| RAB5C | 22.09206 | 22.28992 | 21.72172 | 22.6154 | 20.39519 | 18.27074 | 21.03992 | 19.67722 | 20.95883 | 20.90866 | 20.82607 | 21.51582 | 21.86427 | 21.0434 | 22.87661 | 21.9995 | 16.42762 | 15.8036 |
| RAB6A | 20.59942 | 21.93185 | 21.61009 | 21.77589 | 18.95932 | 18.35932 | 19.64798 | 18.6093 | 20.33034 | 19.28782 | 21.96113 | 21.81607 | 18.89817 | 18.53932 | 22.03773 | 21.97524 | 17.61125 | 18.38211 |
| RAB6B | 16.56693 | 16.66242 | 17.10686 | 18.09463 | 17.18122 | 14.79844 | 17.2984 | 17.37566 | 17.39078 | 15.25413 | 18.79022 | 19.03262 | 17.24443 | 15.40256 | 15.95445 | 16.26058 | 15.97243 | 15.79753 |
| RAB7A | 22.4376 | 22.8931 | 22.18945 | 21.96937 | 22.74375 | 21.74349 | 21.664 | 21.55632 | 21.63174 | 21.95889 | 22.96269 | 22.85138 | 22.63121 | 22.1641 | 22.61785 | 22.28873 | 20.97536 | 22.15807 |
| RAB8A | 21.50988 | 21.56826 | 21.98501 | 22.20092 | 22.78294 | 20.49916 | 21.3737 | 18.36935 | 19.77374 | 20.38159 | 20.99292 | 21.12837 | 21.31254 | 21.73955 | 21.91394 | 21.95734 | 20.60708 | 21.73557 |
| RAB8B | 16.98002 | 16.01166 | 20.89146 | 19.44594 | 18.76207 | 17.64074 | 17.29605 | 17.06714 | 17.24027 | 15.777 | 17.60696 | 18.70776 | 18.95886 | 16.80795 | 20.31846 | 20.2216 | 17.55661 | 18.45682 |
| RAB9A | 15.78482 | 17.0269 | 16.77319 | 16.42721 | 11.55395 | 16.84388 | 16.90681 | 16.42911 | 14.59963 | 16.09769 | 17.22041 | 13.62237 | 18.94534 | 18.23797 | 17.23103 | 16.89184 | 14.71477 | 14.49737 |
| RABEP1 | 14.73706 | 15.83638 | 16.19853 | 16.30031 | 19.0471 | 17.74973 | 15.89543 | 16.42757 | 14.84483 | 15.9895 | 14.84164 | 18.05899 | 17.54317 | 16.68402 | 16.82415 | 17.89974 | 16.29679 | 16.29149 |
| RABGAP1 | 15.7437 | 16.16681 | 15.34433 | 16.27709 | 18.82155 | 16.68444 | 17.15987 | 17.1674 | 14.06821 | 16.47808 | 16.6807 | 16.98882 | 17.40476 | 16.07922 | 15.12874 | 15.70034 | 16.49872 | 16.48441 |
| RABGAP1L | 16.82493 | 16.80662 | 14.32719 | 16.81586 | 16.53421 | 18.67093 | 16.2863 | 16.57474 | 16.63279 | 16.48844 | 18.25007 | 18.03148 | 16.99735 | 16.18119 | 16.46696 | 18.12184 | 15.44738 | 14.74431 |
| RABGGTA | 16.72357 | 14.85953 | 15.49119 | 15.99602 | 14.09222 | 16.42015 | 14.94088 | 15.92069 | 15.1696 | 17.51182 | 15.58326 | 18.21486 | 16.46429 | 15.07927 | 15.57021 | 21.16829 | 15.10849 | 14.64511 |
| RABL6 | 16.09172 | 17.01326 | 15.362 | 14.8036 | 15.12921 | 14.71051 | 18.31606 | 16.96404 | 16.24655 | 15.7305 | 16.8272 | 18.52923 | 15.36863 | 16.98677 | 15.95506 | 14.04444 | 16.44872 | 13.92883 |
| RAC1 | 21.55482 | 21.52036 | 21.81187 | 20.65729 | 21.3873 | 17.13537 | 20.86334 | 20.24673 | 17.40109 | 17.37416 | 18.72291 | 20.44552 | 20.79291 | 21.32811 | 21.47187 | 21.40892 | 21.4202 | 21.51285 |
| RAC2 | 23.93169 | 24.19909 | 23.80268 | 23.55448 | 22.307 | 21.61048 | 21.55613 | 20.74175 | 19.69492 | 19.78879 | 22.25637 | 22.22769 | 22.60425 | 22.88512 | 23.67721 | 23.87656 | 22.00394 | 22.99092 |
| RAC3 | 22.10666 | 19.25142 | 14.16318 | 22.2035 | 18.75348 | 17.56343 | 14.6712 | 18.12968 | 18.04544 | 16.92456 | 18.77798 | 19.56498 | 18.8097 | 18.73782 | 21.96726 | 21.71584 | 18.63016 | 20.69513 |
| RACGAP1 | 18.09888 | 17.34064 | 18.9693 | 19.88468 | 22.70502 | 21.08926 | 17.92016 | 17.61391 | 19.91938 | 19.92987 | 19.77026 | 18.88318 | 19.16085 | 18.64599 | 18.52238 | 17.50226 | 21.09025 | 19.81894 |
| RACK1 | 20.90396 | 20.8309 | 21.5249 | 20.51545 | 19.01448 | 17.64123 | 20.07178 | 19.49931 | 17.29473 | 18.22585 | 21.69852 | 22.40991 | 18.92928 | 19.56286 | 20.39669 | 20.46769 | 18.26485 | 19.99849 |
| RAD23A | 22.2547 | 22.25148 | 19.83053 | 20.409 | 16.80893 | 15.64402 | 18.53837 | 17.59635 | 14.01198 | 15.52855 | 16.13142 | 15.32075 | 19.26109 | 18.40305 | 19.35744 | 20.16976 | 16.38321 | 14.27543 |
| RAD23B | 22.00521 | 22.08354 | 20.24049 | 19.31192 | 18.83463 | 17.5195 | 18.99263 | 19.05324 | 18.17844 | 19.5504 | 19.0063 | 17.78046 | 19.48235 | 17.0377 | 21.08139 | 21.41841 | 19.44999 | 16.16618 |
| RAD50 | 13.54737 | 14.46011 | 16.05575 | 14.98621 | 14.98591 | 15.61258 | 18.84511 | 16.28105 | 14.14129 | 17.02619 | 16.42679 | 18.20884 | 15.82459 | 15.2794 | 14.64369 | 15.57648 | 14.48028 | 17.41088 |
| RAE1 | 18.92425 | 17.15691 | 16.81458 | 19.81814 | 17.44422 | 16.49129 | 16.79611 | 18.40055 | 12.90008 | 18.33289 | 17.36797 | 18.81151 | 14.99444 | 15.32607 | 16.67957 | 17.20508 | 19.25873 | 16.50726 |
| RAET1L | 16.82707 | 17.77376 | 18.43108 | 19.31105 | 14.53491 | 15.5033 | 15.05689 | 16.55956 | 18.56842 | 16.24764 | 14.32493 | 12.85577 | 16.55843 | 17.57089 | 19.22604 | 18.70291 | 14.67472 | 16.58882 |
| RAI14 | 15.896 | 15.62896 | 16.11171 | 17.15449 | 20.21972 | 17.35082 | 15.73389 | 16.87114 | 14.71502 | 15.69034 | 16.02899 | 17.62988 | 15.89994 | 16.86593 | 16.09677 | 15.91692 | 19.25985 | 18.55943 |
| RALA | 22.53651 | 22.25652 | 22.44904 | 22.41881 | 22.17287 | 21.09421 | 21.83856 | 22.1085 | 22.4282 | 21.95707 | 23.40031 | 23.11474 | 22.74828 | 23.308 | 22.8198 | 22.93437 | 22.01681 | 22.46522 |
| RALB | 18.8936 | 18.70011 | 19.45488 | 18.98012 | 19.53201 | 18.88419 | 16.06963 | 17.0038 | 14.69145 | 16.16607 | 20.40942 | 21.01 | 18.6191 | 18.52892 | 20.19296 | 20.01439 | 19.36159 | 18.91939 |
| RALBP1 | 16.00286 | 14.9185 | 17.65047 | 17.79325 | 13.93916 | 15.47879 | 18.40611 | 17.99741 | 16.12906 | 14.77665 | 16.85359 | 16.34916 | 17.6167 | 18.02307 | 18.08634 | 18.85782 | 17.5593 | 17.22582 |
| RALGAPB | 15.29829 | 15.1982 | 14.97572 | 14.7437 | 15.1739 | 12.87415 | 16.93884 | 13.62142 | 14.30136 | 14.69011 | 17.35671 | 17.854 | 13.06554 | 15.10795 | 16.03438 | 17.60641 | 15.81487 | 14.92767 |

| | | | | | | | | | | | | | | | | | | |
|---|---|---|---|---|---|---|---|---|---|---|---|---|---|---|---|---|---|---|
| RALY | 15.95473 | 15.39844 | 16.58399 | 17.0824 | 14.41781 | 15.54453 | 20.59945 | 18.89012 | 15.29214 | 18.97011 | 15.65555 | 16.6502 | 18.11459 | 18.44194 | 17.39473 | 15.07951 | 18.56708 | 17.77277 |
| RAN | 21.2176 | 21.3028 | 21.65459 | 20.7117 | 22.93758 | 21.89384 | 22.57579 | 21.83822 | 20.42308 | 21.19903 | 22.602 | 23.95294 | 22.50413 | 22.70387 | 21.46394 | 21.09572 | 21.85581 | 22.70581 |
| RANBP1 | 20.91657 | 18.19036 | 21.09384 | 20.06962 | 21.72879 | 19.05709 | 19.49368 | 19.39662 | 18.9272 | 17.73035 | 21.6264 | 20.43401 | 19.95034 | 19.28397 | 20.30769 | 21.2708 | 19.38797 | 20.34241 |
| RANBP2 | 16.73484 | 16.13779 | 16.12123 | 16.32063 | 16.10325 | 18.50457 | 15.84061 | 16.74106 | 14.40483 | 15.04697 | 15.59199 | 18.2491 | 17.64717 | 16.4381 | 15.71409 | 15.94914 | 15.52483 | 16.42831 |
| RANBP3 | 16.97286 | 16.39902 | 16.61383 | 18.33393 | 16.99354 | 14.35458 | 18.42875 | 16.26373 | 14.48018 | 16.03656 | 17.25063 | 17.82537 | 16.62441 | 16.81439 | 16.80281 | 15.72711 | 14.72796 | 17.10527 |
| RANGAP1 | 16.43693 | 15.5816 | 16.70442 | 16.42553 | 18.79088 | 20.43097 | 20.34634 | 20.10459 | 16.09478 | 17.19873 | 16.87762 | 18.78643 | 19.73792 | 20.27928 | 15.81936 | 15.90635 | 18.98916 | 20.46347 |
| RAP1A | 19.07713 | 19.10077 | 20.76528 | 20.87733 | 15.09712 | 17.98193 | 17.06271 | 15.93334 | 16.79567 | 16.11662 | 18.91215 | 19.71939 | 19.19177 | 19.05195 | 20.35737 | 20.08544 | 17.03665 | 17.99963 |
| RAP1B | 23.35569 | 24.06812 | 23.03903 | 24.66152 | 23.41361 | 22.51824 | 24.42855 | 23.11919 | 19.93372 | 20.88257 | 22.68902 | 22.79673 | 24.12507 | 24.18202 | 23.52447 | 23.35401 | 22.48654 | 23.31699 |
| RAP1GDS1 | 18.94216 | 17.74109 | 18.25091 | 16.64534 | 16.02777 | 16.70667 | 17.52352 | 15.97553 | 16.69568 | 15.70692 | 15.68848 | 19.0204 | 15.88395 | 17.12376 | 16.28188 | 17.40139 | 17.89989 | 14.1925 |
| RAP2A | 20.34073 | 19.9745 | 16.86561 | 18.85756 | 17.12743 | 16.58635 | 16.50984 | 17.40458 | 21.10558 | 19.02422 | 21.25282 | 21.16525 | 16.75427 | 15.61352 | 18.98798 | 19.24814 | 17.8541 | 18.04755 |
| RAP2B | 16.98426 | 16.80178 | 20.88174 | 21.10576 | 17.33205 | 17.59479 | 18.05109 | 17.54193 | 17.72781 | 16.42285 | 18.2517 | 18.44108 | 17.61394 | 19.09706 | 20.99951 | 19.47249 | 20.26043 | 19.82679 |
| RAP2C | 20.24543 | 20.63973 | 21.38422 | 21.61462 | 20.93161 | 20.15484 | 20.21005 | 19.9281 | 19.24813 | 19.27974 | 21.04354 | 21.05365 | 20.4791 | 21.10105 | 21.36832 | 20.91873 | 21.10489 | 21.21699 |
| RARS1 | 17.0675 | 15.24019 | 17.52523 | 16.58585 | 17.88018 | 16.74931 | 17.57605 | 17.4708 | 16.44634 | 17.67165 | 18.93774 | 20.37701 | 19.20885 | 18.39339 | 16.03418 | 17.08326 | 16.54845 | 18.02798 |
| RARS2 | 15.71254 | 15.09247 | 15.35768 | 13.47881 | 20.23597 | 19.81042 | 14.42616 | 14.83007 | 17.75324 | 14.67989 | 15.70484 | 15.31246 | 14.37585 | 13.43261 | 14.59537 | 12.23312 | 19.90426 | 20.32017 |
| RASA3 | 17.71214 | 18.39375 | 17.42961 | 16.80352 | 16.76744 | 16.1515 | 17.99824 | 18.47564 | 15.68249 | 16.6755 | 14.94731 | 14.72975 | 15.94013 | 15.91846 | 18.10146 | 18.42652 | 19.75952 | 20.01003 |
| RASAL1 | 15.30517 | 19.02935 | 16.28318 | 15.80401 | 19.14688 | 17.32806 | 18.35413 | 16.16068 | 19.53183 | 18.68498 | 15.91375 | 16.78258 | 19.73129 | 15.68993 | 17.04015 | 15.19825 | 16.97193 | 17.4892 |
| RASAL2 | 14.06695 | 16.69805 | 14.95672 | 17.0543 | 19.25043 | 17.99416 | 15.97589 | 16.23492 | 14.75344 | 16.8623 | 15.75928 | 15.62974 | 14.77556 | 16.85947 | 15.91496 | 15.40773 | 18.07936 | 17.18329 |
| RASSF8 | 12.46631 | 14.07278 | 18.90243 | 15.18709 | 15.35566 | 14.92849 | 15.51473 | 15.53119 | 10.84172 | 13.57425 | 13.50005 | 11.72132 | 14.7676 | 15.47073 | 13.29017 | 18.33261 | 18.61859 | 18.50157 |
| RBBP4 | 15.15584 | 15.99731 | 15.29154 | 15.8464 | 18.96425 | 18.00527 | 18.93617 | 18.9111 | 18.01721 | 20.48156 | 20.01038 | 19.76529 | 19.54124 | 19.87587 | 15.77275 | 13.75294 | 19.60769 | 18.2984 |
| RBBP7 | 17.63423 | 16.1333 | 17.73115 | 17.76683 | 17.67696 | 16.15027 | 18.20409 | 16.06487 | 18.37484 | 18.08883 | 18.8706 | 20.23901 | 19.57136 | 18.21716 | 17.36684 | 17.84735 | 18.42683 | 17.07225 |
| RBCK1 | 18.36259 | 18.15836 | 14.64222 | 14.70594 | 16.6639 | 14.82188 | 16.29134 | 15.56228 | 13.56524 | 15.57553 | 15.04243 | 13.85177 | 14.92204 | 16.46519 | 16.03141 | 16.46136 | 14.58041 | 16.25466 |
| RBM12 | 16.48488 | 14.15128 | 16.42756 | 15.96133 | 13.96253 | 15.65092 | 16.79797 | 15.97942 | 12.68117 | 15.39581 | 18.48485 | 19.56293 | 15.17218 | 14.80268 | 15.14108 | 15.21326 | 14.25271 | 12.87 |
| RBM17 | 16.29543 | 16.35193 | 16.21302 | 14.7969 | 16.32833 | 16.05891 | 17.10402 | 15.94132 | 13.25968 | 13.98413 | 16.37371 | 19.24387 | 17.19787 | 13.74116 | 13.51344 | 15.27191 | 12.89036 | 16.53889 |
| RBM25 | 14.76204 | 14.30335 | 15.53079 | 15.07149 | 15.16216 | 15.14575 | 14.53984 | 16.57789 | 12.5826 | 15.30987 | 16.33049 | 18.19963 | 14.92394 | 14.64612 | 16.3922 | 15.18863 | 14.33338 | 17.39668 |
| RBM3 | 23.9072 | 23.74073 | 22.53013 | 22.57902 | 15.77465 | 16.19469 | 15.72903 | 16.92802 | 16.05861 | 17.11628 | 16.39677 | 15.82471 | 19.98662 | 18.37441 | 22.35713 | 22.92617 | 15.33561 | 15.96832 |
| RBM39 | 18.99654 | 14.9619 | 20.23492 | 15.58325 | 18.04814 | 17.22368 | 17.11667 | 17.08401 | 17.68889 | 17.60232 | 19.53031 | 17.99308 | 20.01942 | 17.53084 | 17.42407 | 17.93861 | 20.15925 | 15.55238 |
| RBM4 | 17.35319 | 18.41643 | 19.45772 | 18.33571 | 13.11341 | 15.22968 | 15.19231 | 16.23016 | 13.03935 | 15.01132 | 18.66512 | 17.21812 | 16.21614 | 20.86685 | 18.13322 | 18.27113 | 14.88174 | 17.27661 |
| RBM8A | 21.31406 | 20.5842 | 20.85996 | 19.43222 | 18.04031 | 16.95737 | 18.65942 | 19.27892 | 18.46573 | 18.26722 | 19.25994 | 19.03359 | 18.99472 | 18.25109 | 18.40076 | 19.3473 | 14.50036 | 15.34893 |
| RBMX | 16.87818 | 17.80857 | 16.69697 | 17.09877 | 18.0421 | 15.7763 | 18.77393 | 19.61023 | 18.40562 | 19.9232 | 21.0056 | 20.61823 | 19.1315 | 18.11813 | 18.42694 | 16.96747 | 16.61395 | 15.52709 |
| RBMXL3 | 15.65279 | 16.29131 | 15.78372 | 15.75665 | 14.39598 | 14.5134 | 18.73384 | 17.00652 | 17.42265 | 18.69155 | 20.36038 | 20.44574 | 21.935 | 15.15936 | 15.0935 | 19.52465 | 14.83412 | 12.75283 |
| RBX1 | 16.1262 | 15.74204 | 15.69308 | 18.57944 | 18.88055 | 16.38688 | 19.40319 | 19.0888 | 13.17379 | 15.84517 | 14.93655 | 18.34331 | 19.06501 | 20.50745 | 16.3019 | 16.37437 | 21.08398 | 18.91514 |
| RCC1 | 19.98965 | 18.42753 | 16.64726 | 20.84456 | 18.31673 | 17.12303 | 16.84658 | 14.38688 | 17.29593 | 17.07541 | 17.97756 | 17.93153 | 19.28074 | 17.96185 | 16.83388 | 17.04649 | 19.04324 | 19.03479 |
| RCC2 | 19.71801 | 16.34559 | 15.89647 | 16.38102 | 17.77815 | 16.94804 | 17.81036 | 17.18542 | 17.60566 | 18.04904 | 19.8065 | 21.09744 | 16.85251 | 16.16339 | 15.87449 | 16.11387 | 16.41398 | 18.26609 |
| RCN1 | 16.66163 | 16.3574 | 18.63729 | 21.02339 | 18.23103 | 18.05395 | 18.49762 | 17.79911 | 22.18179 | 21.36263 | 20.69505 | 18.2402 | 18.65635 | 18.62806 | 17.96888 | 17.39424 | 15.97608 | 18.27299 |
| RCN2 | 16.53556 | 16.28006 | 15.9672 | 15.44687 | 15.62392 | 15.30468 | 18.09059 | 16.90121 | 21.47737 | 20.39858 | 19.58897 | 16.10328 | 14.71829 | 15.74873 | 16.21242 | 16.28903 | 15.5434 | 16.63282 |
| RDH11 | 18.74918 | 18.4187 | 19.52995 | 20.08639 | 17.46797 | 16.44973 | 19.10622 | 18.25331 | 16.91813 | 16.67458 | 16.77799 | 17.26352 | 17.09867 | 20.45038 | 19.89641 | 18.85612 | 15.51788 | 17.90775 |
| RDX | 20.43771 | 19.94228 | 21.08746 | 21.7851 | 23.40776 | 22.71583 | 20.90312 | 20.46335 | 20.94391 | 21.47483 | 20.2875 | 21.67288 | 22.51368 | 23.22024 | 21.69194 | 20.93258 | 21.94064 | 21.34515 |
| RECQL | 16.10434 | 14.61924 | 16.731 | 16.81639 | 16.57374 | 16.52184 | 19.4212 | 18.62879 | 17.82914 | 17.00964 | 16.0762 | 15.92753 | 16.24115 | 16.41825 | 15.63568 | 15.47781 | 16.55214 | 17.82326 |
| REEP5 | 15.69169 | 17.56505 | 17.97083 | 16.41132 | 18.18021 | 16.60447 | 20.04034 | 17.81918 | 16.11432 | 15.13883 | 18.62787 | 14.84768 | 16.28544 | 14.30221 | 17.16572 | 17.52436 | 16.79869 | 15.73949 |
| RELA | 22.10029 | 21.07584 | 21.15002 | 21.69482 | 13.19549 | 16.38657 | 18.92727 | 17.61838 | 17.27178 | 15.58911 | 16.21155 | 17.83577 | 20.14332 | 19.64119 | 20.78861 | 20.82948 | 17.36561 | 17.8933 |
| RELCH | 15.16165 | 16.93307 | 15.5009 | 19.28989 | 19.16382 | 15.62605 | 17.04413 | 16.38436 | 13.70382 | 15.82351 | 15.73071 | 15.62351 | 20.29431 | 20.7897 | 14.66147 | 14.63668 | 17.53858 | 17.69482 |
| RELL1 | 17.23698 | 19.46161 | 18.07116 | 18.34946 | 20.46557 | 18.79276 | 18.48897 | 18.97924 | 18.12428 | 18.17514 | 16.90439 | 13.67801 | 19.69049 | 18.84537 | 17.24562 | 18.37217 | 15.99275 | 17.03596 |
| RELT | 14.24941 | 15.21416 | 16.5742 | 15.57969 | 16.163 | 16.68695 | 17.30914 | 15.7096 | 15.92088 | 14.05915 | 16.86652 | 13.70066 | 16.82008 | 17.15404 | 16.71794 | 16.26921 | 16.21108 | 15.66682 |
| REPS1 | 16.08542 | 16.1926 | 17.6978 | 16.82726 | 16.79761 | 15.69143 | 17.6516 | 17.17031 | 14.84353 | 15.09752 | 14.95134 | 14.39717 | 16.18379 | 15.84235 | 17.82235 | 17.78755 | 16.46472 | 15.59412 |
| REPS2 | 17.87386 | 17.83726 | 16.44737 | 18.72075 | 13.40028 | 15.63459 | 15.17928 | 16.39647 | 16.80247 | 14.79002 | 14.27744 | 16.19843 | 17.36291 | 16.33471 | 18.56001 | 18.53645 | 14.69147 | 17.35046 |
| RET | 16.61102 | 15.73243 | 16.10985 | 16.65639 | 17.22018 | 16.63968 | 16.33328 | 17.16149 | 20.50738 | 16.59472 | 15.23761 | 13.64876 | 16.37837 | 15.12534 | 15.72138 | 15.67808 | 17.28499 | 16.21189 |
| REXO2 | 16.48196 | 17.14435 | 17.26016 | 18.1665 | 19.13942 | 18.61002 | 16.2146 | 16.26382 | 17.50806 | 17.76136 | 18.90745 | 19.04064 | 20.90124 | 19.47647 | 18.36466 | 18.36053 | 18.33232 | 19.22464 |
| RFC4 | 16.97458 | 16.31137 | 17.91881 | 16.76431 | 18.54717 | 17.20497 | 17.69665 | 17.5869 | 17.48943 | 17.28987 | 15.66093 | 14.86334 | 17.82813 | 17.69181 | 14.42413 | 15.907 | 16.73279 | 16.00293 |
| RFC5 | 14.92528 | 18.29642 | 14.48228 | 14.13646 | 18.96716 | 18.05223 | 19.51336 | 16.23211 | 17.63304 | 17.16753 | 14.85633 | 15.74741 | 15.32019 | 17.5643 | 16.88104 | 16.78878 | 15.84466 | 15.76342 |
| RFFL | 14.97684 | 15.8664 | 16.9692 | 18.00248 | 20.13814 | 19.95322 | 23.65304 | 20.25617 | 13.64678 | 14.24919 | 15.38637 | 12.41492 | 20.04541 | 15.43729 | 13.97482 | 16.72218 | 15.45547 | 17.52298 |
| RFTN1 | 19.20808 | 19.1705 | 18.38388 | 17.92432 | 20.60647 | 19.7342 | 16.40811 | 17.28539 | 16.2265 | 16.01282 | 15.71208 | 17.57975 | 16.84341 | 16.34588 | 17.94083 | 18.24291 | 17.04346 | 17.9155 |
| RGPD3 | 16.27863 | 17.43846 | 13.94451 | 16.43263 | 14.25733 | 17.41327 | 15.68873 | 15.89834 | 13.3291 | 16.47894 | 16.17081 | 17.96422 | 16.31376 | 16.54991 | 14.66819 | 15.69024 | 15.09268 | 15.33973 |
| RGS10 | 15.73036 | 15.32166 | 14.76488 | 14.89325 | 19.04196 | 17.5298 | 18.16793 | 18.46068 | 13.64416 | 17.01812 | 15.45857 | 11.95658 | 18.74428 | 17.30188 | 15.19816 | 15.53762 | 17.97529 | 17.68076 |
| RGS13 | 16.77035 | 15.50215 | 14.86659 | 15.3013 | 14.26294 | 15.12632 | 16.25965 | 15.98307 | 14.2395 | 13.96543 | 18.83387 | 18.26967 | 14.7253 | 14.33123 | 15.1502 | 14.54306 | 15.62767 | 14.84264 |
| RGS14 | 12.50499 | 15.08657 | 14.04959 | 14.49621 | 16.29277 | 15.91013 | 14.68159 | 15.0356 | 13.20291 | 14.87951 | 16.93528 | 18.51044 | 11.58743 | 15.28394 | 15.64399 | 14.61121 | 15.57131 | 16.87037 |
| RGS19 | 15.71304 | 17.09116 | 17.49994 | 17.33024 | 17.9213 | 15.19117 | 16.16462 | 16.74177 | 14.82843 | 17.12265 | 14.78886 | 12.58895 | 13.71603 | 15.481 | 15.90443 | 14.45757 | 20.36305 | 20.29116 |
| RHBDD2 | 14.19902 | 14.94371 | 14.74032 | 13.94633 | 12.98468 | 17.58074 | 14.9395 | 16.38021 | 17.66029 | 16.4013 | 13.71299 | 15.54407 | 15.76483 | 14.94432 | 14.94609 | 15.35963 | 14.52305 | 15.54011 |
| RHBDF1 | 17.85845 | 18.02135 | 18.19908 | 19.0929 | 15.24093 | 16.55616 | 14.01201 | 14.25731 | 19.84566 | 19.80952 | 16.98237 | 14.2736 | 16.62254 | 13.97053 | 18.14003 | 18.8585 | 14.00869 | 17.69349 |
| RHEB | 20.91832 | 20.62952 | 20.48672 | 20.51414 | 20.82402 | 20.5329 | 18.73274 | 19.0312 | 18.89466 | 18.76782 | 17.91466 | 18.58978 | 19.49875 | 18.59471 | 20.47613 | 20.45766 | 18.47294 | 20.25523 |
| RHOA | 22.34409 | 22.18029 | 21.41498 | 22.48681 | 23.14711 | 22.36327 | 21.28043 | 21.05961 | 21.77583 | 22.08655 | 22.76324 | 22.41856 | 22.96201 | 22.72396 | 22.49414 | 22.21811 | 22.79968 | 23.47657 |
| RHOB | 17.56577 | 17.81894 | 18.80054 | 19.76492 | 18.45263 | 14.87565 | 18.21716 | 17.12302 | 17.31406 | 15.77322 | 18.24767 | 19.47193 | 19.07748 | 18.11358 | 19.6717 | 19.89712 | 16.01454 | 14.04973 |
| RHOBTB3 | 17.75095 | 15.07583 | 14.64558 | 16.11064 | 18.15749 | 11.16499 | 15.16078 | 15.09838 | 18.16983 | 18.53692 | 17.65641 | 17.04008 | 10.90961 | 14.89291 | 15.82397 | 15.2157 | 15.01293 | 17.30928 |
| RHOC | 20.28583 | 20.2875 | 16.79634 | 20.83323 | 19.68392 | 16.9308 | 15.20622 | 16.90736 | 14.1606 | 15.20773 | 13.60639 | 17.16563 | 20.78201 | 18.3575 | 21.35251 | 21.00007 | 19.52495 | 21.89235 |
| RHOF | 20.25126 | 19.15455 | 16.49868 | 15.9388 | 20.06736 | 18.92568 | 15.28674 | 16.94707 | 15.75744 | 15.46455 | 15.1995 | 15.30608 | 15.2591 | 14.77758 | 17.79799 | 18.70334 | 19.92346 | 19.73695 |
| RHOG | 21.17491 | 20.66158 | 20.26685 | 21.29302 | 20.14233 | 19.02646 | 18.56308 | 18.71208 | 15.85458 | 16.59745 | 20.48941 | 20.40504 | 19.98277 | 19.4825 | 20.5999 | 20.63311 | 19.44717 | 20.09287 |
| RHOQ | 22.90681 | 23.25737 | 20.9167 | 22.73815 | 23.31412 | 22.69445 | 22.63828 | 22.21402 | 19.45811 | 20.4877 | 21.66773 | 21.74219 | 23.12215 | 23.70145 | 23.08775 | 22.95301 | 23.23058 | 23.615 |
| RHOU | 20.15654 | 20.04652 | 19.46327 | 19.48788 | 17.86882 | 15.81578 | 18.35308 | 17.30477 | 17.33233 | 16.56499 | 18.24034 | 16.55186 | 19.35514 | 19.6347 | 19.01831 | 18.51297 | 13.16252 | 15.87466 |
| RICTOR | 15.96892 | 15.26738 | 14.23416 | 18.24805 | 17.45528 | 16.34975 | 15.33491 | 16.10166 | 14.99787 | 14.51934 | 13.87384 | 17.74549 | 15.35464 | 18.15729 | 17.06379 | 16.42228 | 18.11131 | 15.96001 |
| RIDA | 14.96375 | 16.2556 | 15.63735 | 15.08869 | 18.44455 | 16.90104 | 19.00938 | 17.88721 | 18.06018 | 19.61751 | 15.68697 | 15.66874 | 16.05264 | 16.86794 | 17.14027 | 16.25543 | 19.31996 | 18.15186 |
| RIPK1 | 13.99266 | 15.52838 | 14.12181 | 16.06011 | 17.88788 | 15.70746 | 14.91346 | 16.98371 | 15.75548 | 17.79643 | 14.39284 | 13.64245 | 18.28337 | 18.25119 | 14.24697 | 15.7233 | 19.5383 | 18.00386 |
| RIPOR1 | 16.32485 | 17.96526 | 15.58279 | 17.06166 | 14.67341 | 15.02245 | 15.20935 | 15.9199 | 15.70811 | 13.43929 | 13.65045 | 17.04569 | 15.75595 | 15.61432 | 17.4133 | 17.64273 | 14.5995 | 14.55519 |
| RIPOR2 | 16.47429 | 16.88665 | 16.6861 | 15.75816 | 15.18441 | 16.76364 | 17.32058 | 16.95194 | 15.5261 | 16.1162 | 17.55618 | 18.36102 | 18.83344 | 19.02867 | 14.59969 | 15.83274 | 16.04008 | 15.39489 |
| RIT1 | 14.66175 | 14.99252 | 17.17618 | 16.05161 | 16.8038 | 15.63034 | 16.82644 | 16.72285 | 15.28661 | 15.10503 | 18.77986 | 11.48582 | 14.60267 | 16.7218 | 13.40192 | 14.72264 | 18.9889 | 18.80738 |
| RMC1 | 29.25661 | 29.51925 | 29.24627 | 29.31303 | 17.38263 | 13.51972 | 15.25982 | 15.72636 | 12.39416 | 12.55932 | 14.6743 | 15.35797 | 29.36119 | 28.5086 | 26.07731 | 28.877 | 13.82246 | 13.69754 |
| RMDN1 | 14.12403 | 16.42155 | 17.43341 | 15.71236 | 16.94111 | 15.33476 | 17.1252 | 18.38799 | 16.61573 | 17.37906 | 15.80736 | 12.26735 | 15.85493 | 13.32348 | 15.94828 | 13.36814 | 14.36749 | 12.4791 |
| RMDN3 | 13.57847 | 16.61446 | 15.28197 | 16.06129 | 13.21313 | 14.97809 | 16.42013 | 16.20337 | 14.66375 | 12.97702 | 16.89738 | 16.73989 | 16.6906 | 17.38163 | 15.58309 | 15.70063 | 15.14838 | 14.33758 |
| RNASET2 | 15.91767 | 16.61552 | 17.33924 | 16.9268 | 15.42886 | 16.28396 | 15.8619 | 15.99415 | 19.24285 | 19.82913 | 19.62132 | 17.01406 | 20.72428 | 20.28918 | 16.38023 | 16.8595 | 15.71239 | 16.85552 |
| RND3 | 16.64339 | 16.30772 | 17.19519 | 16.32706 | 17.52522 | 17.10781 | 18.17283 | 18.18216 | 13.9553 | 15.71343 | 13.53677 | 15.21326 | 14.18153 | 15.00848 | 15.46982 | 16.57865 | 19.72343 | 19.24435 |
| RNF114 | 18.86199 | 17.23531 | 16.08474 | 18.55562 | 17.81832 | 17.31549 | 17.17123 | 17.1904 | 14.76889 | 13.83047 | 16.24233 | 15.12827 | 20.53568 | 19.98848 | 18.12939 | 15.63037 | 18.79285 | 18.84676 |
| RNF141 | 18.7533 | 19.67 | 12.98506 | 15.50287 | 15.818 | 15.98273 | 14.17809 | 16.4973 | 14.0934 | 15.72763 | 15.27467 | 11.95825 | 15.33378 | 16.09188 | 17.58597 | 17.71814 | 15.36251 | 13.79431 |
| RNF31 | 19.45654 | 19.63853 | 15.08189 | 16.72673 | 15.63713 | 16.0956 | 15.52594 | 16.88359 | 14.88178 | 13.34497 | 15.64537 | 13.31968 | 15.72143 | 17.05048 | 15.4765 | 16.59438 | 15.18129 | 15.02436 |
| RNF41 | 15.33427 | 13.64529 | 15.10728 | 16.86868 | 19.81611 | 18.98279 | 15.94386 | 15.51962 | 12.63576 | 13.96679 | 13.63172 | 12.25729 | 16.77808 | 18.37213 | 14.96611 | 14.16855 | 16.21832 | 13.93174 |
| RNF7 | 16.75407 | 14.9488 | 17.29088 | 16.28172 | 16.98437 | 15.64365 | 17.92033 | 14.56971 | 12.53777 | 12.66207 | 15.97892 | 12.58103 | 17.76957 | 16.09138 | 14.48787 | 16.61567 | 20.23464 | 19.17057 |
| RNH1 | 19.29387 | 17.54218 | 17.14885 | 18.74467 | 17.84174 | 16.8362 | 17.40378 | 18.00012 | 16.74475 | 15.98663 | 18.13864 | 19.99615 | 17.8473 | 17.09384 | 18.66911 | 18.41263 | 18.57675 | 18.80417 |
| RNMT | 13.48731 | 17.02673 | 13.88236 | 12.95215 | 13.74967 | 16.47621 | 14.37653 | 17.31354 | 15.76776 | 15.29686 | 18.47411 | 18.55932 | 13.6682 | 16.36572 | 15.48167 | 16.65501 | 14.74418 | 15.29459 |
| RNPEP | 16.4525 | 16.20528 | 17.38649 | 16.25356 | 18.94282 | 18.63836 | 16.48744 | 17.08542 | 16.77005 | 17.12184 | 19.12071 | 20.13444 | 18.0144 | 15.28764 | 16.47501 | 17.13057 | 17.98686 | 16.23996 |

| | | | | | | | | | | | | | | | | | | |
|---|---|---|---|---|---|---|---|---|---|---|---|---|---|---|---|---|---|---|
| RO60 | 24.09286 | 23.83911 | 21.65468 | 22.23653 | 15.66477 | 16.30812 | 17.33632 | 17.45674 | 16.87614 | 15.72362 | 18.04829 | 18.002 | 19.06862 | 18.50561 | 22.8625 | 22.12458 | 16.12868 | 16.88087 |
| ROBO1 | 15.08686 | 16.00164 | 15.89072 | 15.65088 | 14.29924 | 18.29051 | 16.14429 | 17.04296 | 18.05949 | 17.31688 | 16.63961 | 16.10019 | 16.38227 | 15.82905 | 14.8511 | 14.36636 | 17.37017 | 20.20473 |
| ROCK1 | 16.38066 | 17.17942 | 17.10908 | 17.60137 | 19.75588 | 18.17146 | 17.32199 | 17.7268 | 17.16195 | 17.92285 | 16.13371 | 16.72749 | 18.37293 | 17.55622 | 17.10363 | 16.56033 | 20.2308 | 19.67189 |
| ROCK2 | 15.46391 | 16.60601 | 16.3058 | 17.14043 | 19.1623 | 19.67405 | 16.58511 | 17.73721 | 19.88611 | 20.17673 | 20.74382 | 17.22807 | 18.36394 | 18.47454 | 15.78413 | 15.99719 | 19.91755 | 20.16372 |
| ROR1 | 15.34565 | 15.76329 | 17.52438 | 18.25168 | 19.45017 | 20.08055 | 17.5138 | 16.51769 | 14.28888 | 15.18949 | 15.63629 | 17.16011 | 16.02081 | 15.50555 | 17.9341 | 17.78803 | 17.67664 | 16.61989 |
| ROR2 | 16.25556 | 15.05591 | 15.55727 | 15.09801 | 16.31626 | 16.94356 | 17.15676 | 15.88478 | 18.50685 | 18.50676 | 16.95152 | 15.19422 | 14.38485 | 14.68892 | 14.71523 | 15.59024 | 14.90625 | 15.67296 |
| RP2 | 19.34036 | 19.01356 | 19.64872 | 19.84453 | 19.15224 | 19.19865 | 17.93647 | 18.10742 | 19.35985 | 19.44567 | 20.42023 | 21.41018 | 19.92661 | 19.94504 | 20.34536 | 20.42483 | 20.25447 | 21.01043 |
| RPA1 | 17.73202 | 16.58273 | 16.77435 | 17.63816 | 18.24114 | 16.00855 | 18.39383 | 16.89894 | 16.07946 | 17.38898 | 16.8587 | 19.77957 | 17.39644 | 15.64975 | 15.81604 | 16.05821 | 18.21276 | 13.04835 |
| RPA2 | 15.27098 | 14.02257 | 15.032 | 15.14933 | 12.5168 | 14.01006 | 16.85856 | 16.62741 | 15.2719 | 16.55147 | 15.12919 | 19.62445 | 15.51582 | 16.84503 | 16.15706 | 14.13075 | 14.80919 | 15.46356 |
| RPA3 | 17.96318 | 18.05057 | 16.8383 | 16.51976 | 20.11241 | 18.50618 | 20.97186 | 16.32808 | 16.4298 | 19.44204 | 19.06565 | 19.38963 | 21.83626 | 19.80705 | 16.7803 | 16.65104 | 21.00599 | 20.22344 |
| RPAP1 | 16.78967 | 21.53218 | 18.70968 | 19.16935 | 12.44108 | 16.00506 | 14.10686 | 14.43592 | 13.60553 | 14.89723 | 13.80285 | 15.983 | 16.26194 | 16.15536 | 20.21624 | 20.85425 | 17.27513 | 13.57939 |
| RPAP3 | 14.41555 | 14.98498 | 14.68297 | 14.64688 | 12.48037 | 15.87544 | 14.14209 | 16.39527 | 11.89742 | 14.89427 | 16.24483 | 20.94596 | 14.56219 | 14.96387 | 15.20355 | 14.69385 | 17.37858 | 14.48454 |
| RPF2 | 16.87116 | 15.96138 | 16.98671 | 16.24651 | 13.9106 | 16.16946 | 18.42516 | 16.42575 | 12.78146 | 17.39857 | 17.34142 | 13.962 | 13.1162 | 17.76712 | 15.36763 | 15.56296 | 13.76946 | 13.8808 |
| RPIA | 14.5198 | 15.785 | 13.72871 | 14.53965 | 16.81857 | 15.95121 | 17.47373 | 16.55586 | 16.57498 | 17.53427 | 19.35711 | 19.99572 | 17.41057 | 16.67834 | 15.60809 | 16.21528 | 15.35223 | 14.95834 |
| RPL10 | 20.78885 | 21.08876 | 19.5662 | 19.40351 | 16.76901 | 18.26838 | 17.18087 | 17.15005 | 17.12329 | 15.22916 | 19.72312 | 21.7694 | 15.64377 | 17.19109 | 20.2034 | 19.66082 | 15.97906 | 16.26405 |
| RPL10A | 21.75221 | 21.92577 | 20.31857 | 20.27493 | 19.57232 | 17.83366 | 20.27864 | 19.81557 | 17.6836 | 18.90451 | 21.16579 | 23.32154 | 20.45961 | 20.29562 | 21.084 | 20.33581 | 17.90309 | 18.75334 |
| RPL11 | 21.64347 | 21.74799 | 20.76175 | 20.03313 | 19.64458 | 18.11323 | 21.68255 | 21.17449 | 17.48725 | 18.26232 | 20.46385 | 21.76947 | 19.90687 | 20.36977 | 21.15639 | 20.33703 | 17.93179 | 19.40912 |
| RPL12 | 22.57765 | 22.94961 | 20.48327 | 21.41872 | 19.52826 | 17.3694 | 19.02443 | 18.94112 | 17.54213 | 18.70699 | 21.73636 | 23.20419 | 20.64214 | 19.17872 | 21.91747 | 21.60676 | 17.73576 | 19.58928 |
| RPL13 | 21.60365 | 21.8665 | 21.09178 | 19.94239 | 19.45485 | 16.98361 | 20.46512 | 20.80809 | 18.03715 | 17.89932 | 21.18235 | 22.06337 | 19.80178 | 19.73534 | 21.04137 | 20.26533 | 18.48199 | 19.48541 |
| RPL13A | 20.99611 | 20.24662 | 17.88531 | 18.75769 | 18.64711 | 17.74826 | 20.41837 | 17.58918 | 15.63709 | 17.08329 | 18.27917 | 19.59248 | 19.14324 | 18.63906 | 18.43528 | 18.1318 | 16.68631 | 17.98876 |
| RPL13AP3 | 22.6563 | 22.35691 | 21.59515 | 21.0948 | 19.43058 | 18.66112 | 19.19668 | 21.02092 | 18.62118 | 17.02085 | 21.61559 | 22.76768 | 18.53105 | 18.69898 | 21.17285 | 21.30083 | 16.75125 | 19.22351 |
| RPL14 | 22.0923 | 20.41643 | 21.04033 | 20.38604 | 18.63036 | 16.81284 | 20.36564 | 18.16938 | 17.85227 | 17.98195 | 20.48596 | 22.45287 | 18.4496 | 18.11272 | 20.66687 | 20.75175 | 16.09832 | 21.02002 |
| RPL15 | 21.86919 | 21.7524 | 21.2566 | 20.85179 | 17.97216 | 16.6629 | 19.6967 | 17.28506 | 18.11641 | 17.56521 | 19.30727 | 22.73156 | 17.8999 | 16.53214 | 21.04015 | 20.72349 | 15.61682 | 16.16041 |
| RPL17 | 22.4529 | 22.59875 | 20.89593 | 20.90762 | 20.94743 | 19.74605 | 21.80466 | 20.58429 | 18.18339 | 19.04707 | 21.83425 | 22.56093 | 21.64671 | 21.02162 | 20.88189 | 20.82258 | 19.26761 | 20.41362 |
| RPL18 | 22.15165 | 22.19381 | 21.44675 | 19.8837 | 18.79985 | 18.24259 | 21.77514 | 20.75339 | 16.79828 | 17.44337 | 20.54086 | 22.69161 | 19.9577 | 18.13903 | 21.56081 | 20.82493 | 17.57162 | 19.10578 |
| RPL18A | 21.51375 | 19.63889 | 18.61438 | 19.04249 | 15.80393 | 16.90007 | 18.40863 | 18.6526 | 16.49417 | 15.92544 | 18.77859 | 21.57566 | 16.71693 | 17.28692 | 18.50343 | 19.01231 | 15.78835 | 15.77498 |
| RPL19 | 20.36284 | 20.44937 | 19.21004 | 19.47515 | 18.3639 | 18.57592 | 18.55929 | 18.60147 | 15.67117 | 16.05014 | 19.70288 | 20.10753 | 20.3077 | 18.50894 | 18.57363 | 18.89129 | 16.49998 | 18.33721 |
| RPL21 | 22.39622 | 22.46354 | 21.41388 | 21.72294 | 20.77466 | 18.43747 | 22.38907 | 20.86096 | 17.47771 | 18.17676 | 21.00753 | 23.37439 | 20.86491 | 21.36966 | 20.8707 | 21.1936 | 18.33455 | 21.61175 |
| RPL22 | 21.46875 | 21.80908 | 16.34577 | 16.92672 | 19.46604 | 18.07002 | 21.12391 | 21.02799 | 17.46983 | 18.90435 | 20.9027 | 22.41101 | 15.29341 | 16.49835 | 16.25432 | 16.2248 | 16.70292 | 18.43171 |
| RPL22L1 | 21.75367 | 21.96709 | 21.65763 | 21.37691 | 15.21791 | 15.07159 | 18.82063 | 16.93765 | 14.03756 | 15.99562 | 17.16491 | 19.4835 | 17.50129 | 17.06598 | 22.17499 | 21.67818 | 14.87592 | 16.12658 |
| RPL23 | 21.61611 | 21.25769 | 19.77718 | 18.34649 | 20.30123 | 20.29429 | 21.38016 | 20.97195 | 16.77484 | 17.65686 | 20.24313 | 21.62431 | 19.75972 | 21.59928 | 19.39262 | 20.39402 | 17.83866 | 19.80567 |
| RPL23A | 20.79718 | 20.32421 | 19.40922 | 19.28504 | 17.71582 | 15.73182 | 18.11875 | 17.76838 | 16.4162 | 17.18384 | 19.4537 | 20.49543 | 18.85044 | 18.15565 | 19.44127 | 18.94405 | 17.50272 | 15.96542 |
| RPL24 | 21.88283 | 21.94395 | 20.74502 | 19.9827 | 18.91835 | 17.76661 | 20.52927 | 20.3147 | 18.12775 | 17.83259 | 20.19007 | 22.05784 | 20.50692 | 19.68769 | 20.72834 | 20.5736 | 19.06373 | 19.04459 |
| RPL26 | 21.14978 | 21.18886 | 20.19888 | 19.38424 | 17.52837 | 18.40036 | 17.2983 | 17.75211 | 16.92912 | 16.56187 | 18.95102 | 21.51482 | 18.27947 | 16.0357 | 20.50358 | 20.06049 | 17.05697 | 17.522 |
| RPL27 | 23.68197 | 23.78916 | 22.59016 | 23.90276 | 20.19638 | 18.24466 | 20.07149 | 18.04322 | 17.9788 | 17.78275 | 19.78941 | 22.99778 | 20.15609 | 18.99495 | 22.86292 | 22.88248 | 17.6247 | 19.2809 |
| RPL27A | 21.73295 | 22.22593 | 21.38531 | 21.86826 | 19.15453 | 19.78044 | 21.59728 | 20.88255 | 17.28948 | 17.86798 | 20.18016 | 22.13801 | 20.97846 | 21.03813 | 20.74572 | 20.88858 | 17.81702 | 19.27268 |
| RPL28 | 21.51263 | 21.53486 | 20.87186 | 20.2219 | 19.67018 | 18.01175 | 20.52798 | 20.39214 | 17.13643 | 17.50374 | 18.84565 | 21.70443 | 19.95783 | 18.72436 | 20.50995 | 19.97216 | 17.43552 | 18.89395 |
| RPL29 | 22.79841 | 22.19662 | 21.0742 | 21.16486 | 20.84649 | 16.39713 | 19.94757 | 21.4864 | 18.45594 | 18.4319 | 18.66476 | 21.82492 | 22.40427 | 20.60186 | 21.64101 | 21.55287 | 18.5863 | 20.22769 |
| RPL3 | 22.62525 | 22.93282 | 21.29226 | 21.22781 | 19.4153 | 18.8785 | 22.08789 | 19.32628 | 17.68196 | 19.00323 | 21.30414 | 22.77313 | 19.41439 | 19.07341 | 22.17799 | 21.05661 | 17.68924 | 19.87559 |
| RPL30 | 22.40955 | 21.98101 | 21.73876 | 21.35605 | 18.18543 | 16.71463 | 20.16243 | 19.77056 | 16.65103 | 17.15225 | 19.40442 | 21.98305 | 19.21403 | 19.32783 | 22.07775 | 22.07285 | 16.08917 | 17.10321 |
| RPL31 | 21.59128 | 21.6494 | 18.7964 | 18.19049 | 19.34976 | 18.78704 | 20.83727 | 19.3718 | 13.40937 | 17.12193 | 19.71722 | 21.88865 | 19.44152 | 19.0963 | 20.04085 | 19.89717 | 17.44165 | 18.9615 |
| RPL32 | 20.43804 | 19.92828 | 18.76242 | 18.84377 | 18.27945 | 17.14293 | 20.33291 | 20.21884 | 16.38654 | 17.56248 | 18.26779 | 20.67791 | 19.7268 | 18.6146 | 18.6862 | 18.11706 | 17.71377 | 18.14282 |
| RPL34 | 21.52727 | 21.24977 | 21.27469 | 20.98497 | 18.62764 | 17.29539 | 19.207 | 19.07301 | 16.66205 | 15.98276 | 20.49658 | 20.82822 | 20.51036 | 19.20203 | 20.89945 | 20.41876 | 16.63283 | 18.40612 |
| RPL35 | 21.31922 | 21.77288 | 21.49897 | 20.58544 | 19.52941 | 18.34957 | 17.89648 | 18.77735 | 18.31311 | 17.46991 | 21.72859 | 23.52556 | 19.85049 | 19.05982 | 20.59056 | 20.77503 | 15.90003 | 18.83894 |
| RPL35A | 22.58729 | 22.40801 | 21.94907 | 21.3401 | 19.34605 | 17.84434 | 21.18136 | 20.79553 | 18.06303 | 17.92143 | 20.36085 | 22.80346 | 18.91233 | 18.55312 | 21.42015 | 21.27087 | 17.67477 | 19.47108 |
| RPL36 | 21.33491 | 21.74003 | 20.11541 | 20.54955 | 19.68086 | 18.33251 | 18.55686 | 17.59066 | 16.53758 | 15.6813 | 18.15647 | 20.89608 | 20.36214 | 18.95584 | 20.27552 | 20.33119 | 17.42731 | 18.86721 |
| RPL36A | 18.2992 | 19.98159 | 17.96516 | 18.87394 | 17.52157 | 17.24997 | 18.43045 | 18.5141 | 15.5344 | 15.66954 | 18.22126 | 20.07642 | 18.52012 | 19.06744 | 16.38985 | 16.84038 | 16.19054 | 18.21191 |
| RPL37A | 22.21631 | 21.99009 | 20.33631 | 20.18466 | 18.43441 | 15.91476 | 20.62986 | 21.09976 | 17.53602 | 17.61803 | 20.25775 | 21.91698 | 17.95638 | 18.68331 | 20.88746 | 20.60141 | 17.7931 | 19.55134 |
| RPL38 | 19.95145 | 20.29606 | 19.64666 | 19.17313 | 18.47565 | 16.05568 | 17.78389 | 18.98656 | 16.44377 | 16.69649 | 20.39026 | 21.54976 | 19.77377 | 18.07735 | 19.65812 | 19.14982 | 16.29368 | 18.43835 |
| RPL3L | 21.23939 | 20.5944 | 20.34136 | 19.75659 | 18.61447 | 17.71258 | 18.71425 | 18.9699 | 16.93923 | 16.85753 | 19.74407 | 21.94535 | 18.58237 | 18.50029 | 19.31895 | 20.20555 | 17.4251 | 18.56928 |
| RPL4 | 22.09736 | 21.8728 | 20.82473 | 20.29807 | 18.4268 | 22.80307 | 19.69357 | 18.7957 | 18.08116 | 17.36099 | 19.09336 | 22.09213 | 20.28757 | 18.54126 | 20.42256 | 20.46095 | 17.18118 | 18.26862 |
| RPL5 | 22.05097 | 21.43402 | 20.13368 | 20.03975 | 19.59083 | 19.37215 | 19.8497 | 19.48755 | 17.21225 | 18.21306 | 20.39077 | 21.76843 | 19.87901 | 19.19775 | 20.39799 | 20.52103 | 18.67629 | 19.19277 |
| RPL6 | 21.49288 | 21.41345 | 20.64676 | 20.31129 | 17.83143 | 18.51072 | 17.50154 | 17.4974 | 16.56013 | 17.28377 | 19.8964 | 22.31073 | 19.59484 | 17.60031 | 20.30394 | 20.45843 | 17.35423 | 16.96178 |
| RPL7 | 21.80848 | 21.94141 | 20.83689 | 20.59875 | 17.83171 | 17.4115 | 18.81043 | 18.33902 | 17.31216 | 17.52732 | 20.00475 | 22.63556 | 19.51066 | 20.85886 | 20.79717 | 20.53873 | 16.84424 | 16.64438 |
| RPL7A | 22.58888 | 22.72442 | 21.36948 | 20.84837 | 18.39927 | 17.38105 | 19.89608 | 20.70226 | 17.76712 | 17.59534 | 20.92961 | 22.78981 | 20.74038 | 20.04738 | 21.55424 | 21.29455 | 17.75614 | 18.03649 |
| RPL8 | 21.52457 | 21.60715 | 21.1191 | 19.97781 | 20.86704 | 18.52615 | 21.33092 | 19.47189 | 17.39351 | 18.38909 | 20.07939 | 22.0381 | 21.5011 | 20.54868 | 20.35835 | 20.59261 | 18.49503 | 20.11282 |
| RPL9 | 21.31176 | 22.26799 | 19.61974 | 20.64083 | 19.81083 | 18.90077 | 20.68726 | 20.30937 | 16.77286 | 18.74645 | 19.54399 | 21.91862 | 20.65299 | 19.59909 | 20.48774 | 20.30661 | 17.97358 | 20.05606 |
| RPLP0 | 22.85202 | 23.35603 | 22.44567 | 22.51318 | 19.22461 | 18.0321 | 19.78002 | 19.75662 | 17.53791 | 18.45938 | 21.04881 | 22.93107 | 19.73249 | 20.3291 | 23.67754 | 23.0935 | 18.24111 | 19.86259 |
| RPLP1 | 23.98096 | 24.17204 | 22.04606 | 23.13531 | 18.03287 | 12.66335 | 21.94622 | 17.47476 | 19.32571 | 20.23458 | 19.13494 | 20.29466 | 23.2595 | 22.66102 | 24.17715 | 23.86185 | 21.30346 | 20.89306 |
| RPLP2 | 25.29393 | 25.64131 | 24.94402 | 25.05098 | 21.81652 | 20.21947 | 20.3455 | 19.6549 | 20.5123 | 20.44005 | 23.4945 | 24.68141 | 24.41694 | 23.32485 | 25.71808 | 25.66739 | 20.61746 | 21.44306 |
| RPN1 | 17.64454 | 18.77945 | 18.0362 | 17.80161 | 17.20157 | 18.48861 | 19.36436 | 17.00724 | 25.19995 | 23.70207 | 21.568 | 18.94306 | 20.72761 | 18.41487 | 17.65776 | 17.81168 | 16.68725 | 18.44325 |
| RPN2 | 16.05606 | 17.32714 | 16.89108 | 16.18059 | 15.68116 | 17.77058 | 17.04229 | 16.90062 | 22.96541 | 22.16434 | 19.93284 | 17.26229 | 16.37385 | 16.58236 | 17.69897 | 16.17253 | 17.41933 | 16.46065 |
| RPRD1A | 14.4557 | 18.7405 | 12.7393 | 15.15536 | 14.52117 | 18.35487 | 20.79468 | 15.82006 | 13.56454 | 17.13662 | 16.95795 | 17.36333 | 15.46792 | 17.75245 | 15.66687 | 16.15758 | 18.58693 | 16.29851 |
| RPRD1B | 16.34268 | 15.92479 | 15.683 | 15.38438 | 18.41048 | 15.50318 | 15.84629 | 16.15976 | 17.24957 | 19.36621 | 14.58926 | 14.39536 | 18.54746 | 15.68394 | 16.33396 | 15.14898 | 17.04574 | 16.79948 |
| RPS10 | 22.424 | 22.36211 | 21.08904 | 21.56115 | 20.25569 | 18.77019 | 19.81903 | 16.25993 | 19.30551 | 19.35678 | 21.31116 | 22.26312 | 19.94903 | 19.57919 | 21.55564 | 20.90013 | 15.29742 | 20.44245 |
| RPS11 | 21.21113 | 20.33369 | 19.88921 | 19.69985 | 19.91045 | 18.77202 | 19.68582 | 20.62162 | 16.14015 | 17.8969 | 19.92388 | 22.25375 | 21.03404 | 19.58885 | 20.06228 | 19.67783 | 18.73307 | 19.58928 |
| RPS12 | 20.19477 | 19.54585 | 18.33152 | 19.87509 | 19.42853 | 18.21155 | 20.01516 | 18.95838 | 17.47858 | 18.36697 | 20.32767 | 21.43048 | 21.48506 | 19.64353 | 19.04069 | 19.01063 | 20.26343 | 20.0637 |
| RPS13 | 21.74716 | 22.11269 | 20.95014 | 21.21549 | 17.78651 | 16.94051 | 18.56002 | 18.15607 | 17.55651 | 17.88063 | 19.85845 | 23.26451 | 19.34652 | 17.81133 | 21.14342 | 21.31942 | 16.35608 | 15.54367 |
| RPS14 | 21.97919 | 21.98407 | 21.02899 | 21.37237 | 20.09177 | 18.46431 | 19.14855 | 19.62085 | 18.23686 | 17.97893 | 21.2012 | 22.2443 | 20.87151 | 19.8838 | 20.54775 | 20.80072 | 16.36685 | 18.27181 |
| RPS15 | 18.29344 | 18.09282 | 17.51397 | 21.20963 | 16.70534 | 17.34194 | 15.50046 | 14.64065 | 16.55295 | 12.85989 | 17.93524 | 21.33638 | 19.82196 | 12.45781 | 17.44109 | 18.00766 | 16.42377 | 19.5075 |
| RPS15A | 21.05428 | 20.50339 | 19.90823 | 19.54251 | 17.3669 | 17.32106 | 20.14768 | 19.67136 | 17.10406 | 16.93531 | 19.90147 | 23.27178 | 18.03076 | 18.06902 | 20.41668 | 20.53392 | 15.97671 | 17.19466 |
| RPS16 | 22.02286 | 22.28201 | 21.27254 | 20.81149 | 20.09691 | 18.22125 | 22.81531 | 23.17484 | 18.20332 | 18.89991 | 21.1316 | 23.61256 | 19.99179 | 20.22697 | 21.30784 | 21.43079 | 18.42681 | 19.97648 |
| RPS17 | 20.4106 | 20.63883 | 17.9665 | 19.41229 | 19.97089 | 18.57266 | 17.69105 | 19.2509 | 12.98335 | 16.58629 | 17.96868 | 20.22117 | 20.82001 | 20.20314 | 19.44981 | 18.22237 | 18.41058 | 19.48139 |
| RPS18 | 20.84963 | 20.38039 | 19.89632 | 19.3647 | 18.16731 | 16.69701 | 19.17275 | 19.56358 | 17.38424 | 17.47014 | 19.55063 | 22.69412 | 18.27813 | 17.13816 | 19.73274 | 19.78513 | 17.65317 | 17.70838 |
| RPS19 | 21.34931 | 21.48101 | 20.38798 | 20.60343 | 19.77063 | 17.80778 | 20.6726 | 17.83023 | 17.8054 | 18.97381 | 19.81254 | 21.51423 | 21.14213 | 20.65035 | 20.14906 | 20.4193 | 19.52623 | 19.52161 |
| RPS2 | 21.13002 | 21.3403 | 20.97401 | 20.63043 | 18.93386 | 19.07549 | 22.26036 | 20.77725 | 18.28943 | 19.06295 | 20.99692 | 23.07281 | 20.54159 | 20.17656 | 20.60163 | 20.44072 | 18.71186 | 20.55327 |
| RPS20 | 21.21614 | 21.04062 | 20.51716 | 20.55843 | 19.7011 | 18.80797 | 19.83312 | 19.1416 | 18.14975 | 19.10282 | 21.74064 | 22.03566 | 21.24112 | 18.98077 | 20.34605 | 20.3835 | 17.6388 | 20.25353 |
| RPS21 | 21.30315 | 21.55447 | 19.6842 | 20.63774 | 19.14926 | 19.08838 | 20.33774 | 19.36214 | 17.95009 | 18.32174 | 20.09838 | 21.57161 | 20.69793 | 20.62843 | 19.31532 | 19.89683 | 19.91267 | 20.64899 |
| RPS23 | 20.64993 | 20.9514 | 20.48988 | 20.44375 | 20.60512 | 18.99561 | 22.18485 | 22.28444 | 17.94993 | 20.24421 | 20.5896 | 21.29549 | 21.45579 | 20.83831 | 19.93236 | 19.62118 | 17.7905 | 20.8984 |
| RPS24 | 19.89932 | 19.54456 | 19.30936 | 19.05404 | 18.27833 | 16.2732 | 19.86323 | 19.11579 | 16.74167 | 15.9521 | 19.26162 | 21.0084 | 18.87629 | 18.73243 | 18.71493 | 19.45044 | 16.70035 | 17.87424 |
| RPS25 | 22.46471 | 22.20012 | 21.65798 | 20.73951 | 19.03557 | 17.7345 | 21.13176 | 21.02427 | 17.48981 | 17.76981 | 21.39739 | 22.74545 | 21.4348 | 20.67585 | 21.32686 | 21.15335 | 17.68505 | 19.35654 |
| RPS26 | 19.2662 | 19.77913 | 18.58007 | 17.75808 | 12.45109 | 15.48624 | 19.87011 | 20.0224 | 15.96175 | 16.65499 | 18.49404 | 20.29875 | 12.89477 | 18.43401 | 17.89269 | 18.65683 | 14.01878 | 17.96594 |
| RPS27L | 20.20742 | 19.89518 | 19.74123 | 19.02823 | 19.06945 | 16.0588 | 19.67336 | 19.34711 | 14.88481 | 17.73365 | 19.72508 | 20.89193 | 19.52778 | 19.08268 | 18.98359 | 19.0206 | 16.04449 | 18.11542 |
| RPS28 | 22.12531 | 21.7512 | 21.13562 | 21.91603 | 20.11509 | 19.01363 | 18.86425 | 19.489 | 18.99487 | 18.24518 | 20.83658 | 21.28572 | 21.23268 | 20.06188 | 20.92261 | 20.74736 | 20.26972 | 20.27932 |
| RPS3 | 21.23925 | 21.01128 | 20.75498 | 20.51773 | 18.06936 | 16.71244 | 19.83674 | 18.89062 | 17.635 | 17.11027 | 20.76804 | 22.75113 | 18.00426 | 17.19753 | 20.41546 | 20.70119 | 16.80885 | 18.20705 |
| RPS3A | 21.53777 | 21.0908 | 20.41275 | 20.56109 | 19.68065 | 17.98448 | 21.10357 | 20.14457 | 18.18008 | 19.34332 | 20.67882 | 21.92449 | 19.86507 | 19.64215 | 20.28926 | 20.52383 | 17.62797 | 19.69227 |

| | | | | | | | | | | | | | | | | | | |
|---|---|---|---|---|---|---|---|---|---|---|---|---|---|---|---|---|---|---|
| RPS4X | 19.77892 | 19.76379 | 20.10167 | 19.73364 | 17.75819 | 17.73925 | 20.05116 | 20.15737 | 17.49161 | 17.8386 | 19.68746 | 22.05781 | 17.64563 | 17.44037 | 19.5738 | 19.28183 | 19.82708 | 20.64179 |
| RPS4Y1 | 19.28404 | 19.99526 | 13.47616 | 15.49017 | 12.31848 | 15.48638 | 15.82692 | 15.32213 | 13.26922 | 14.13857 | 19.04427 | 20.02114 | 14.35742 | 16.48867 | 15.08276 | 16.30262 | 15.558 | 14.39231 |
| RPS4Y2 | 19.07152 | 18.47375 | 18.58221 | 18.27605 | 18.43616 | 16.79855 | 19.79084 | 19.27321 | 17.54097 | 19.56163 | 19.03332 | 21.60942 | 17.49569 | 17.55661 | 18.9264 | 17.35767 | 16.62589 | 17.64062 |
| RPS5 | 19.17261 | 19.16263 | 17.53406 | 18.37835 | 18.67293 | 17.1968 | 19.77973 | 19.21868 | 16.55305 | 17.78126 | 19.38854 | 19.76658 | 17.53746 | 17.95857 | 18.07174 | 17.21611 | 15.86035 | 17.18517 |
| RPS6 | 20.63012 | 20.07189 | 19.92258 | 19.82461 | 18.18093 | 20.22221 | 19.72201 | 19.84134 | 16.83491 | 17.14586 | 18.33161 | 21.01109 | 19.37605 | 18.18909 | 19.28977 | 19.20608 | 17.37467 | 18.27408 |
| RPS6KA1 | 15.77885 | 19.10142 | 18.54611 | 19.62117 | 18.10491 | 16.80689 | 17.55047 | 17.4002 | 17.39069 | 17.61239 | 16.46187 | 17.502 | 18.07106 | 18.26262 | 20.07295 | 19.76956 | 18.43408 | 18.57516 |
| RPS6KA3 | 14.09447 | 14.51964 | 14.03904 | 17.81007 | 12.66697 | 15.79791 | 15.42672 | 15.92931 | 16.23577 | 14.45149 | 15.75704 | 16.03739 | 19.61818 | 15.59125 | 13.60441 | 16.00574 | 15.77127 | 19.88245 |
| RPS6KA6 | 14.42802 | 16.50853 | 14.81396 | 16.31552 | 15.46611 | 13.89679 | 14.17024 | 16.98508 | 13.36702 | 17.96137 | 17.63091 | 18.56927 | 18.50103 | 17.68959 | 17.72071 | 19.07611 | 14.64293 | 12.25987 |
| RPS7 | 21.04184 | 21.75662 | 19.3235 | 21.10061 | 20.41779 | 18.71336 | 21.06044 | 19.67863 | 17.25816 | 19.07754 | 20.3633 | 20.60191 | 22.01403 | 20.7943 | 20.70628 | 20.04177 | 19.17097 | 20.62561 |
| RPS8 | 21.76079 | 21.23639 | 21.80025 | 20.74235 | 19.94918 | 19.83417 | 22.05153 | 21.68401 | 18.44857 | 19.19808 | 20.61517 | 22.54 | 20.53975 | 19.73779 | 20.04962 | 19.94558 | 18.77202 | 20.14914 |
| RPS9 | 21.0145 | 21.25795 | 20.40379 | 19.50029 | 17.43957 | 17.92801 | 19.92611 | 17.41453 | 17.35909 | 18.47176 | 20.11885 | 22.87393 | 17.88375 | 17.52932 | 19.94334 | 19.51294 | 17.5061 | 17.55202 |
| RPSA | 22.0633 | 21.88757 | 20.69498 | 20.77715 | 19.88489 | 17.70858 | 20.69547 | 19.53632 | 17.99436 | 19.68833 | 21.68494 | 23.00326 | 20.99351 | 20.6234 | 21.16743 | 21.27397 | 19.01244 | 19.0344 |
| RRAS | 19.98491 | 20.50143 | 20.66506 | 19.59095 | 19.75674 | 19.68385 | 17.63896 | 17.19936 | 17.32474 | 16.30615 | 14.82823 | 16.12259 | 18.32454 | 17.32029 | 21.247 | 21.12053 | 19.35415 | 20.34919 |
| RRAS2 | 21.0914 | 21.15878 | 20.96237 | 21.24483 | 20.0686 | 19.05395 | 16.36865 | 16.90688 | 18.62791 | 17.86835 | 15.45467 | 14.44356 | 19.36906 | 19.28016 | 21.6626 | 21.4041 | 18.36694 | 18.48957 |
| RRBP1 | 16.71969 | 15.53904 | 15.10807 | 15.46635 | 14.34316 | 15.537 | 16.01577 | 16.08202 | 13.69925 | 15.28381 | 14.92859 | 15.3523 | 19.7297 | 18.06989 | 14.04026 | 15.24119 | 15.83295 | 15.57424 |
| RRM1 | 16.59559 | 17.19372 | 17.23309 | 17.58668 | 20.07512 | 17.69678 | 18.25742 | 17.6085 | 15.90544 | 16.52666 | 18.22979 | 19.60966 | 18.50513 | 17.31917 | 18.84953 | 18.1226 | 20.39996 | 20.47133 |
| RRM2 | 14.20923 | 15.93016 | 15.46726 | 16.59755 | 17.03965 | 16.38584 | 14.97132 | 17.69576 | 15.37643 | 16.09764 | 15.71592 | 15.97132 | 18.50828 | 17.98569 | 15.0857 | 16.07467 | 15.15223 | 18.86334 |
| RRP9 | 15.7118 | 16.13525 | 15.60711 | 15.68955 | 14.35502 | 14.49549 | 16.93387 | 16.62964 | 11.17431 | 13.41485 | 18.56925 | 18.43466 | 14.56109 | 13.55899 | 14.24581 | 16.60577 | 15.2328 | 14.93157 |
| RSL1D1 | 17.22714 | 16.8458 | 16.07739 | 15.24693 | 12.66891 | 14.29494 | 17.62832 | 17.92541 | 13.84113 | 15.85055 | 15.79771 | 19.61148 | 16.31396 | 15.15067 | 15.16037 | 15.31493 | 14.53679 | 14.20877 |
| RSU1 | 19.23555 | 20.64131 | 19.34482 | 21.27253 | 20.45884 | 19.20375 | 22.27916 | 20.50026 | 17.84923 | 18.39199 | 17.95997 | 16.97429 | 20.39167 | 21.80575 | 20.81905 | 19.39272 | 19.74512 | 19.88146 |
| RTBDN | 15.5448 | 15.46103 | 14.30796 | 12.89148 | 13.18392 | 15.47 | 14.19926 | 17.76975 | 18.99307 | 18.21034 | 14.91294 | 16.24832 | 13.87817 | 14.50926 | 13.27386 | 15.38656 | 14.2412 | 14.2085 |
| RTCA | 16.15427 | 18.00252 | 17.29038 | 17.72392 | 19.03856 | 18.69458 | 19.25923 | 16.35276 | 17.09002 | 16.80428 | 16.97314 | 12.55364 | 21.50679 | 21.10424 | 18.12081 | 19.32184 | 18.66412 | 19.99374 |
| RTCB | 16.8211 | 17.18634 | 18.85656 | 18.50647 | 18.40473 | 16.91094 | 19.04648 | 18.06976 | 16.62554 | 16.88814 | 19.40929 | 20.16725 | 19.70181 | 19.24167 | 17.7872 | 17.3356 | 19.24031 | 18.96914 |
| RTKN | 16.09568 | 15.06858 | 12.07715 | 16.79677 | 18.97514 | 18.88954 | 14.31884 | 14.73317 | 15.80522 | 16.69576 | 15.9706 | 19.07319 | 14.50606 | 13.43853 | 14.54397 | 14.80385 | 18.7274 | 18.09335 |
| RTN3 | 18.32563 | 15.40384 | 21.17461 | 19.49768 | 16.89836 | 15.12873 | 19.93737 | 19.10776 | 18.17084 | 18.47631 | 18.98807 | 18.87816 | 19.86075 | 16.46844 | 19.21592 | 19.65538 | 15.42756 | 21.32051 |
| RTN4 | 17.05587 | 13.12757 | 17.98033 | 17.04991 | 18.49423 | 14.84229 | 17.17407 | 17.04166 | 16.58694 | 16.31051 | 18.69238 | 17.35 | 15.44816 | 16.87128 | 17.56594 | 18.25226 | 17.50519 | 17.93241 |
| RTN4IP1 | 16.28685 | 16.11846 | 15.52638 | 16.81425 | 14.30881 | 21.69198 | 14.15799 | 16.12592 | 18.3232 | 16.38251 | 14.98016 | 11.51523 | 14.88045 | 16.10977 | 17.13263 | 12.92419 | 16.09473 | 15.26655 |
| RTN4R | 18.54799 | 17.85288 | 18.55242 | 17.84457 | 16.82087 | 15.24642 | 12.04822 | 16.57335 | 17.52605 | 14.44839 | 16.05934 | 12.06195 | 16.28668 | 14.57647 | 17.43593 | 16.81124 | 12.84608 | 15.07669 |
| RTRAF | 17.7543 | 17.49575 | 18.37519 | 19.6977 | 19.68326 | 19.61333 | 17.60723 | 16.72954 | 15.85385 | 17.57115 | 22.42602 | 19.43618 | 20.79197 | 19.13973 | 17.91347 | 18.30682 | 18.57736 | 19.72091 |
| RUFY2 | 16.10377 | 14.02133 | 15.14593 | 16.83843 | 18.25057 | 17.07541 | 19.91963 | 15.53242 | 12.81614 | 18.64666 | 13.84507 | 13.77157 | 17.8736 | 19.98328 | 15.69571 | 14.02425 | 15.77742 | 18.0608 |
| RUVBL1 | 18.35885 | 17.48021 | 18.86182 | 18.51588 | 19.81573 | 18.87746 | 19.14598 | 18.89216 | 17.18872 | 17.38143 | 19.86057 | 21.15132 | 20.40666 | 19.73045 | 18.54696 | 18.05125 | 18.85697 | 19.06361 |
| RUVBL2 | 17.55032 | 16.44544 | 17.46113 | 17.18381 | 18.26485 | 18.23111 | 17.22164 | 17.63035 | 15.52919 | 16.11011 | 18.61994 | 20.21649 | 18.40275 | 18.18557 | 17.04397 | 17.44791 | 17.2402 | 17.83268 |
| RXYLT1 | 16.04201 | 15.98219 | 14.86877 | 15.68875 | 15.19734 | 16.59061 | 14.56546 | 15.76883 | 18.14952 | 15.76035 | 13.48479 | 15.47343 | 15.77081 | 15.53385 | 14.75943 | 14.45493 | 14.94906 | 14.62757 |
| S100A10 | 19.59869 | 19.33799 | 17.62089 | 18.21304 | 24.33787 | 23.25102 | 16.41737 | 18.39955 | 18.17038 | 17.06294 | 20.78203 | 15.06027 | 17.4179 | 17.66916 | 18.1732 | 17.12607 | 22.32472 | 22.6923 |
| S100A11 | 18.79535 | 19.51825 | 18.91574 | 19.90166 | 24.01378 | 22.91864 | 18.97084 | 18.81051 | 20.83979 | 21.60724 | 17.48006 | 14.21486 | 22.31755 | 22.70145 | 19.8608 | 18.88919 | 23.13486 | 23.80488 |
| S100A13 | 17.46494 | 18.49136 | 16.79053 | 17.51853 | 23.93319 | 22.20944 | 16.65845 | 16.70873 | 19.28514 | 19.77743 | 17.93989 | 17.68808 | 18.08888 | 18.08977 | 17.14989 | 17.81328 | 23.76667 | 24.59931 |
| S100A14 | 13.75806 | 17.14807 | 16.32883 | 16.17512 | 13.85023 | 15.53809 | 14.85211 | 15.72879 | 19.37019 | 19.06524 | 15.01015 | 14.59932 | 16.24183 | 15.7084 | 15.19328 | 14.66176 | 17.18169 | 17.03716 |
| S100A16 | 17.51705 | 18.08016 | 17.72129 | 16.27386 | 22.7804 | 20.82386 | 18.21403 | 16.85767 | 18.77648 | 18.80663 | 16.66934 | 14.10109 | 16.11015 | 17.29237 | 16.0494 | 16.45461 | 23.48166 | 24.29095 |
| S100A4 | 18.58917 | 18.70913 | 18.41757 | 18.49097 | 20.33742 | 18.65989 | 20.58298 | 19.65626 | 12.45687 | 17.49001 | 16.38258 | 16.81394 | 17.93672 | 20.04359 | 18.72595 | 18.00638 | 19.38773 | 19.92255 |
| S100A6 | 18.64475 | 19.9655 | 18.3149 | 14.43951 | 19.15217 | 18.95809 | 14.61525 | 17.57025 | 19.59209 | 17.91915 | 15.46923 | 16.11138 | 14.32614 | 13.95815 | 15.29336 | 16.84665 | 19.59813 | 19.77461 |
| S100A8 | 13.3223 | 15.05145 | 16.18145 | 20.62383 | 14.40912 | 17.18255 | 16.509 | 15.0288 | 15.02748 | 14.73884 | 16.80046 | 13.67228 | 15.9539 | 15.23381 | 14.6589 | 15.58676 | 12.53128 | 17.62846 |
| S100A9 | 15.37131 | 17.00827 | 16.75734 | 22.11672 | 19.59333 | 17.57049 | 16.00797 | 17.92645 | 16.46127 | 15.15583 | 15.43898 | 13.61463 | 16.93522 | 15.87535 | 16.35394 | 16.06276 | 15.86086 | 17.34624 |
| S100P | 19.80044 | 19.92577 | 14.17053 | 15.44972 | 14.76533 | 18.92396 | 15.84291 | 18.01297 | 14.78955 | 14.63043 | 15.02991 | 12.43994 | 16.33568 | 13.18901 | 13.78712 | 16.32538 | 13.72853 | 14.65762 |
| SAAL1 | 14.8965 | 21.58034 | 15.59656 | 18.53663 | 12.59268 | 14.7251 | 15.89766 | 15.22756 | 13.77664 | 14.66953 | 15.71275 | 18.32615 | 17.01581 | 15.92702 | 15.27734 | 16.39098 | 13.47613 | 13.98591 |
| SAE1 | 15.71545 | 15.50684 | 16.52576 | 15.6848 | 16.03376 | 17.01026 | 15.65239 | 15.87776 | 16.44371 | 16.97145 | 17.56348 | 17.56442 | 14.56247 | 14.88043 | 15.67552 | 15.97083 | 15.15985 | 13.98289 |
| SAFB | 15.11115 | 15.40136 | 15.46374 | 16.65027 | 13.89203 | 19.35002 | 15.58507 | 16.88155 | 15.92976 | 16.1696 | 19.53419 | 18.63451 | 17.35565 | 14.73695 | 16.48701 | 14.80938 | 17.23468 | 15.56269 |
| SAMHD1 | 15.21696 | 16.64014 | 14.82352 | 17.2233 | 16.90034 | 14.86195 | 20.00938 | 19.19082 | 12.99294 | 13.47195 | 14.91739 | 16.89248 | 14.73124 | 13.27554 | 16.79075 | 14.92227 | 17.00652 | 18.42007 |
| SAMM50 | 16.52836 | 15.95226 | 16.27781 | 16.10335 | 18.72233 | 17.09669 | 18.87352 | 17.32561 | 20.38971 | 19.11659 | 15.98734 | 14.33032 | 15.39579 | 16.67285 | 14.48058 | 14.98179 | 15.86014 | 17.59136 |
| SAR1A | 17.14049 | 17.76655 | 15.41207 | 16.98148 | 13.25583 | 16.4916 | 19.3365 | 16.56283 | 17.7295 | 17.51086 | 17.08363 | 18.66009 | 16.73328 | 16.72639 | 15.8606 | 15.62813 | 17.21483 | 19.32184 |
| SAR1B | 17.11121 | 14.7738 | 17.78431 | 16.88234 | 18.84819 | 23.22186 | 18.60644 | 16.69477 | 17.54295 | 19.35269 | 18.13214 | 19.28786 | 16.07957 | 14.74793 | 18.66053 | 18.10639 | 19.43398 | 15.97683 |
| SARG | 18.3884 | 17.76994 | 18.81787 | 19.40748 | 17.2405 | 17.19163 | 16.12077 | 16.0951 | 15.44371 | 16.13091 | 16.17913 | 14.72552 | 22.12646 | 22.43762 | 18.41132 | 18.95309 | 16.91655 | 15.59526 |
| SARNP | 12.96362 | 16.26063 | 14.57663 | 15.59249 | 18.49096 | 15.96328 | 16.05601 | 18.24598 | 16.85162 | 18.66915 | 16.96831 | 18.84201 | 12.71241 | 17.24482 | 14.49198 | 15.63926 | 14.77562 | 12.42101 |
| SARS1 | 16.30338 | 16.50524 | 18.15133 | 18.27601 | 19.30834 | 17.85749 | 18.84794 | 17.14195 | 18.49034 | 18.08368 | 18.46561 | 19.78822 | 20.1468 | 20.13531 | 18.55342 | 18.70896 | 18.35846 | 19.04799 |
| SBDS | 20.45182 | 19.88186 | 18.7997 | 18.46704 | 18.26652 | 17.60849 | 17.49817 | 17.62941 | 17.25278 | 17.60949 | 17.68085 | 18.42714 | 19.03032 | 18.91553 | 18.55769 | 19.59222 | 17.51183 | 18.67827 |
| SBF1 | 16.1115 | 15.54027 | 15.22374 | 16.54514 | 16.56353 | 17.13089 | 16.47232 | 16.5663 | 14.21721 | 15.50786 | 16.37924 | 17.56844 | 15.80265 | 16.10612 | 15.17336 | 16.4739 | 20.04891 | 21.80304 |
| SCAMP1 | 19.84759 | 21.11339 | 21.75649 | 22.43909 | 21.99739 | 19.35366 | 22.18011 | 18.11886 | 19.00339 | 19.51045 | 19.12525 | 18.53654 | 21.58832 | 23.50532 | 21.04388 | 21.7644 | 17.724 | 19.20608 |
| SCAMP2 | 19.5011 | 18.26122 | 21.33314 | 21.29621 | 18.12958 | 17.52649 | 16.2192 | 16.2421 | 17.91086 | 15.2851 | 17.0409 | 16.70544 | 16.6044 | 14.55509 | 19.23526 | 21.00032 | 13.71151 | 16.59519 |
| SCAMP3 | 18.90824 | 18.71547 | 19.76245 | 19.0848 | 18.45165 | 16.30423 | 15.24847 | 16.6544 | 17.87456 | 16.5643 | 16.49981 | 18.43227 | 17.98368 | 16.84677 | 18.43933 | 18.55211 | 16.63797 | 17.37181 |
| SCAMP4 | 21.55083 | 21.87867 | 23.02936 | 23.05074 | 17.24717 | 17.09038 | 15.99222 | 14.70238 | 18.18804 | 16.16591 | 16.73295 | 17.82442 | 18.37358 | 18.02894 | 23.22895 | 22.96051 | 15.21671 | 14.85608 |
| SCARB1 | 20.56997 | 19.93124 | 19.92221 | 19.14039 | 20.73774 | 19.15236 | 16.08846 | 17.37594 | 16.62926 | 16.58248 | 20.00352 | 20.56058 | 18.00055 | 16.15745 | 20.80096 | 19.79175 | 17.54026 | 17.8186 |
| SCARB2 | 18.57771 | 17.1243 | 18.16928 | 18.33942 | 15.39637 | 16.46352 | 17.00436 | 15.82645 | 20.97398 | 20.08873 | 17.43024 | 18.03406 | 16.24465 | 16.1326 | 20.14944 | 19.22 | 15.96307 | 14.32811 |
| SCEL | 15.01403 | 14.86887 | 15.00864 | 14.26076 | 20.1944 | 16.86271 | 14.58707 | 15.79976 | 16.09941 | 14.19289 | 16.02771 | 13.62611 | 15.99102 | 15.12208 | 14.53568 | 14.93213 | 15.90648 | 14.39892 |
| SCFD1 | 16.26559 | 15.47821 | 16.03906 | 17.40438 | 19.67045 | 18.56636 | 19.25378 | 15.39169 | 15.95125 | 16.64343 | 17.41353 | 17.36945 | 16.29732 | 15.79613 | 16.65862 | 16.69772 | 15.81907 | 16.48437 |
| SCG2 | 16.70269 | 15.48232 | 17.55339 | 16.43813 | 16.93614 | 16.89626 | 18.08949 | 19.00838 | 19.36802 | 19.65477 | 17.1528 | 16.31592 | 17.01444 | 17.67285 | 16.05065 | 16.64117 | 18.31897 | 17.41455 |
| SCG3 | 19.28781 | 19.63489 | 19.9393 | 19.48165 | 18.58545 | 21.93488 | 20.03751 | 18.89656 | 23.00931 | 22.51146 | 19.99682 | 15.52298 | 17.05047 | 20.03144 | 18.22071 | 15.57693 | 16.25933 | 19.97401 |
| SCIN | 17.96141 | 16.21543 | 16.08068 | 16.4634 | 15.49927 | 16.52703 | 14.99489 | 17.05113 | 18.89177 | 19.58396 | 15.043 | 16.79181 | 12.8867 | 15.51135 | 15.96061 | 16.6897 | 15.66889 | 16.95946 |
| SCNN1A | 14.11945 | 13.90818 | 16.04253 | 15.2116 | 15.90615 | 16.05116 | 16.38615 | 15.49321 | 18.59167 | 16.99556 | 13.28274 | 15.7147 | 15.6257 | 12.764 | 14.42491 | 14.25124 | 14.38313 | 14.00941 |
| SCP2 | 17.17441 | 16.19769 | 16.14651 | 16.87635 | 16.25927 | 16.66306 | 18.19651 | 17.33331 | 22.32122 | 21.4819 | 18.27125 | 15.92385 | 16.51224 | 17.48667 | 16.63118 | 15.4703 | 16.53981 | 18.97517 |
| SCPEP1 | 17.60459 | 18.91633 | 15.79825 | 17.44421 | 15.20777 | 16.7122 | 16.42926 | 16.96753 | 18.64379 | 19.19668 | 15.88238 | 16.83078 | 15.89069 | 16.57431 | 16.32283 | 15.91105 | 16.02471 | 17.52971 |
| SCRIB | 17.49052 | 17.56615 | 19.64066 | 19.40105 | 19.14835 | 18.21976 | 17.85298 | 17.34888 | 20.73085 | 20.34685 | 18.73058 | 18.88235 | 19.72638 | 18.76785 | 19.40944 | 18.69919 | 21.1938 | 20.91973 |
| SCRN1 | 16.05775 | 17.22726 | 15.46023 | 15.04318 | 19.02296 | 16.80812 | 16.36624 | 18.15656 | 18.56211 | 16.55128 | 14.51553 | 14.1048 | 13.93985 | 16.44708 | 14.6486 | 14.55497 | 18.29694 | 18.71956 |
| SCRN3 | 13.5928 | 14.11795 | 16.70424 | 16.0482 | 18.22192 | 15.71298 | 19.06562 | 17.62462 | 17.31349 | 14.47254 | 15.44737 | 14.3819 | 18.83368 | 18.70473 | 15.62255 | 14.55836 | 18.21683 | 17.68711 |
| SCYL1 | 14.69846 | 15.64851 | 16.6107 | 17.88907 | 16.61919 | 18.94879 | 17.17028 | 15.9715 | 15.23187 | 16.12166 | 16.45352 | 17.98792 | 15.96893 | 15.73346 | 16.58125 | 16.96958 | 17.62561 | 16.52797 |
| SCYL2 | 16.97206 | 16.11564 | 16.74894 | 17.50606 | 16.41333 | 14.73936 | 15.07608 | 16.19876 | 16.46554 | 15.98021 | 17.19289 | 18.47156 | 17.59907 | 17.21593 | 15.38138 | 19.37311 | 16.61048 | 16.62059 |
| SDC1 | 20.7845 | 20.94779 | 19.68313 | 18.55544 | 22.04353 | 20.91298 | 15.40867 | 17.12904 | 19.10786 | 16.21948 | 17.79375 | 13.28579 | 13.49017 | 17.80497 | 17.68173 | 18.8897 | 20.17563 | 21.40264 |
| SDC2 | 18.5467 | 19.30631 | 15.37134 | 15.99793 | 15.01146 | 14.13801 | 16.6775 | 15.21364 | 12.94508 | 14.10721 | 15.92839 | 15.35246 | 17.96332 | 13.82666 | 14.64335 | 17.37141 | 15.59674 | 15.68358 |
| SDC4 | 20.0566 | 20.05121 | 22.0047 | 21.34732 | 17.73761 | 18.8593 | 15.65123 | 17.6133 | 18.25714 | 15.23966 | 15.851 | 14.53966 | 16.41304 | 16.28221 | 20.40608 | 21.2166 | 15.30438 | 16.44669 |
| SDCBP | 24.10426 | 24.71888 | 23.13889 | 24.66695 | 23.30351 | 22.55244 | 21.07517 | 19.44987 | 24.63897 | 23.83419 | 21.6372 | 21.9986 | 23.12993 | 23.1119 | 24.51537 | 24.02952 | 18.98136 | 21.15146 |
| SDCBP2 | 22.56016 | 22.05516 | 23.84514 | 22.53945 | 23.07634 | 21.81378 | 19.39122 | 19.22027 | 22.55849 | 22.13616 | 20.1389 | 20.67799 | 22.5605 | 21.81316 | 23.24607 | 22.91852 | 18.67888 | 20.70871 |
| SDF2 | 17.07377 | 16.44192 | 17.05869 | 16.3703 | 18.34335 | 17.81977 | 19.75899 | 19.20472 | 22.69452 | 22.31244 | 20.16583 | 16.8169 | 15.18228 | 18.55769 | 15.58359 | 15.20617 | 16.00914 | 19.64771 |
| SDF2L1 | 15.46708 | 16.74408 | 17.6983 | 15.99481 | 16.92014 | 15.26454 | 19.22331 | 19.03153 | 21.84879 | 20.58311 | 20.15536 | 17.29642 | 17.33449 | 17.15739 | 16.73184 | 16.34339 | 16.32345 | 17.15443 |
| SDF4 | 18.79698 | 19.03774 | 18.65895 | 18.12538 | 17.35215 | 17.57594 | 17.44485 | 17.57062 | 22.8442 | 21.60704 | 20.04531 | 17.8442 | 19.48249 | 19.95246 | 18.71025 | 17.96878 | 17.03137 | 18.4262 |
| SDHA | 15.85096 | 17.63486 | 15.79839 | 15.76974 | 14.87269 | 16.42299 | 16.93665 | 16.22505 | 20.15515 | 18.44346 | 16.22905 | 13.99428 | 14.6004 | 16.52314 | 15.68259 | 16.6604 | 15.68629 | 17.08189 |
| SDHB | 17.40517 | 18.75982 | 18.33877 | 17.92161 | 17.97815 | 18.80293 | 18.65612 | 15.53053 | 19.9894 | 18.78072 | 15.89929 | 18.7793 | 17.27152 | 20.14298 | 17.49356 | 17.20102 | 18.96099 | 15.33713 |
| SDK2 | 14.91172 | 17.34466 | 16.13111 | 15.31437 | 12.33784 | 15.87129 | 15.71671 | 16.19667 | 17.65275 | 16.5645 | 14.86327 | 13.64159 | 15.99106 | 18.05408 | 13.88053 | 15.58962 | 15.79678 | 15.06927 |
| SDSL | 18.55698 | 16.39309 | 17.32586 | 14.38076 | 13.12033 | 16.46749 | 14.14268 | 15.69032 | 17.21482 | 16.39996 | 16.12632 | 16.0729 | 18.49049 | 14.41227 | 18.86448 | 18.40899 | 17.04729 | 16.71565 |

| | | | | | | | | | | | | | | | | | | |
|---|---|---|---|---|---|---|---|---|---|---|---|---|---|---|---|---|---|---|
| SEC11A | 16.67243 | 15.64388 | 16.62472 | 15.14309 | 15.79641 | 16.60747 | 16.21422 | 17.19796 | 21.91478 | 19.67236 | 20.00495 | 17.93939 | 15.05434 | 16.41837 | 16.62388 | 15.67583 | 16.43206 | 18.44283 |
| SEC11C | 16.34668 | 15.45664 | 14.77876 | 15.05866 | 15.53775 | 16.29918 | 16.14497 | 16.59471 | 21.60725 | 18.53045 | 20.13265 | 17.40336 | 16.27114 | 15.36017 | 15.58386 | 15.46679 | 16.09478 | 16.33932 |
| SEC13 | 16.27288 | 14.99765 | 14.89136 | 16.42066 | 18.13493 | 17.84144 | 17.23552 | 17.39421 | 15.81913 | 16.51733 | 16.47768 | 18.52059 | 16.6288 | 18.29225 | 14.46365 | 15.61075 | 17.78516 | 18.23396 |
| SEC14L1 | 14.48837 | 16.41961 | 17.43051 | 16.16795 | 14.91818 | 15.78711 | 13.41166 | 13.80266 | 16.92943 | 14.65438 | 15.96225 | 17.52234 | 13.76545 | 14.20151 | 14.17689 | 14.0588 | 16.49128 | 16.94063 |
| SEC14L2 | 12.05317 | 14.32321 | 15.11097 | 15.06799 | 14.55003 | 13.91829 | 16.20917 | 14.48298 | 14.86003 | 14.19613 | 16.43254 | 16.10046 | 15.65092 | 15.31116 | 13.82107 | 13.19329 | 16.37175 | 14.23741 |
| SEC16A | 17.81091 | 15.75004 | 20.20887 | 18.09938 | 20.02088 | 20.38361 | 20.34256 | 19.47821 | 14.31865 | 14.65919 | 15.17163 | 19.19272 | 18.31951 | 22.17281 | 20.16597 | 17.46144 | 19.43493 | 19.70733 |
| SEC22B | 16.09062 | 15.02683 | 17.11012 | 18.6235 | 14.46625 | 16.29637 | 17.05506 | 15.83664 | 17.85568 | 17.53674 | 17.73665 | 18.11783 | 19.2343 | 19.22711 | 17.66071 | 16.84603 | 17.01295 | 15.55376 |
| SEC23A | 17.46282 | 20.16638 | 15.83137 | 15.81495 | 16.10085 | 17.23482 | 15.51957 | 16.07563 | 14.2975 | 16.53253 | 15.98722 | 18.65443 | 15.98252 | 15.76732 | 16.75105 | 16.40881 | 16.52295 | 17.29324 |
| SEC23B | 16.79419 | 16.81502 | 15.94957 | 15.37987 | 16.16975 | 17.71567 | 16.55539 | 16.60977 | 16.9351 | 16.29505 | 16.46864 | 19.42593 | 15.89195 | 17.06743 | 16.45174 | 16.7836 | 18.00079 | 17.7492 |
| SEC23IP | 16.21934 | 15.80189 | 14.88925 | 15.03496 | 16.57457 | 16.83975 | 16.75557 | 15.22903 | 14.39413 | 15.28467 | 14.13076 | 18.02306 | 17.07256 | 14.89211 | 14.42362 | 14.39095 | 15.84868 | 16.96343 |
| SEC24A | 17.82722 | 15.14996 | 15.70873 | 16.85168 | 15.03276 | 16.90011 | 14.9378 | 16.2553 | 12.98673 | 14.59672 | 15.11855 | 18.23943 | 15.53219 | 16.28382 | 15.02363 | 15.72678 | 15.37828 | 16.01117 |
| SEC24B | 15.29898 | 14.30648 | 15.48134 | 14.23877 | 12.54036 | 14.9923 | 15.88539 | 15.68441 | 13.04388 | 15.89599 | 14.7873 | 17.7455 | 14.74498 | 16.69031 | 13.09798 | 15.53844 | 14.45173 | 14.68082 |
| SEC24C | 16.35094 | 17.03623 | 16.13396 | 15.66761 | 15.91482 | 16.26905 | 16.74042 | 18.2603 | 17.22154 | 16.9704 | 15.39198 | 18.05526 | 15.64011 | 16.82537 | 15.62807 | 15.15521 | 18.51086 | 19.4919 |
| SEC24D | 15.31858 | 17.093 | 14.27071 | 16.20939 | 18.99923 | 16.04847 | 19.08079 | 16.21112 | 14.01823 | 16.98458 | 16.07272 | 17.68147 | 14.11148 | 15.29262 | 14.10724 | 14.58006 | 15.18132 | 15.35405 |
| SEC31A | 16.15686 | 16.08719 | 16.5665 | 16.26458 | 16.63781 | 16.22865 | 17.43666 | 17.51103 | 16.12386 | 17.30772 | 16.8063 | 18.67033 | 18.62118 | 18.2978 | 15.79178 | 15.12399 | 17.5303 | 17.85558 |
| SEC61A1 | 16.27189 | 16.572 | 14.36597 | 17.54276 | 15.48297 | 16.83891 | 17.14471 | 16.41084 | 19.99484 | 19.24025 | 18.08206 | 17.18022 | 13.43844 | 15.49652 | 14.63697 | 15.36424 | 16.78474 | 15.25083 |
| SEC62 | 16.56745 | 19.56972 | 18.51471 | 20.23584 | 16.62094 | 18.20823 | 18.78709 | 19.39902 | 18.58922 | 20.22961 | 19.18619 | 23.92356 | 14.82713 | 15.77903 | 19.42889 | 19.43616 | 17.67149 | 17.00941 |
| SEC63 | 16.35353 | 16.07179 | 16.5786 | 16.53504 | 15.18743 | 18.22333 | 16.5421 | 17.39493 | 20.93003 | 18.92073 | 16.5465 | 16.3532 | 15.48018 | 15.59508 | 17.16117 | 16.12181 | 17.13992 | 15.30351 |
| SEH1L | 21.16773 | 13.55005 | 15.45171 | 16.17986 | 13.7595 | 14.85485 | 15.12368 | 16.58162 | 15.57041 | 16.15853 | 13.73572 | 17.83216 | 14.07873 | 18.6958 | 15.59603 | 15.40495 | 15.61503 | 14.49933 |
| SEL1L | 15.75954 | 16.00035 | 17.65096 | 16.31705 | 15.03393 | 17.69062 | 16.13109 | 16.61059 | 20.81564 | 18.45921 | 16.54776 | 14.36691 | 15.84299 | 15.40721 | 16.48793 | 17.14639 | 15.81737 | 16.58697 |
| SEL1L3 | 12.00131 | 15.80741 | 15.64683 | 17.93845 | 14.25529 | 16.79416 | 15.44551 | 17.92501 | 18.30529 | 16.97509 | 12.64022 | 15.0606 | 12.37385 | 13.30754 | 15.60644 | 13.6621 | 19.03931 | 13.0336 |
| SELENBP1 | 16.58154 | 16.61074 | 19.90457 | 19.73198 | 15.03129 | 17.59396 | 16.15919 | 16.83569 | 16.19368 | 16.37147 | 16.16767 | 13.85118 | 20.6506 | 20.27993 | 20.54089 | 20.68385 | 18.0762 | 20.85816 |
| SELENOF | 15.98089 | 19.09525 | 16.45518 | 15.80875 | 18.00973 | 16.61156 | 16.66198 | 16.41005 | 19.83271 | 20.05693 | 19.11644 | 14.15106 | 16.72979 | 16.80229 | 16.71338 | 13.2242 | 17.42881 | 15.00239 |
| SELENON | 16.08848 | 15.57372 | 16.62563 | 14.5627 | 14.64353 | 15.1245 | 16.19089 | 16.00519 | 19.04116 | 16.12962 | 15.52764 | 14.32576 | 15.19127 | 14.66944 | 14.03308 | 15.45695 | 14.11141 | 15.32323 |
| SELENOT | 16.40254 | 17.59849 | 17.41853 | 15.90782 | 13.08987 | 13.37926 | 16.1602 | 15.90653 | 19.31202 | 13.58074 | 17.40748 | 10.14837 | 15.32988 | 16.246 | 14.88408 | 16.42983 | 15.59398 | 14.49948 |
| SELPLG | 15.40748 | 18.96788 | 14.87 | 14.5182 | 13.16941 | 15.84881 | 13.99075 | 15.51178 | 12.78915 | 13.95016 | 18.83332 | 18.75218 | 17.57281 | 14.46901 | 15.03753 | 15.01155 | 16.18862 | 13.11549 |
| SEMA3C | 20.64531 | 17.5989 | 19.82159 | 19.67696 | 16.13803 | 17.75332 | 15.95692 | 16.03252 | 15.31311 | 15.49251 | 18.12026 | 14.03747 | 15.56057 | 15.54856 | 20.89606 | 18.94529 | 15.39932 | 15.94711 |
| SEMA4B | 15.48332 | 16.79221 | 16.09372 | 14.44884 | 19.36426 | 18.19528 | 18.15089 | 15.41779 | 17.65904 | 15.94581 | 17.21914 | 13.64997 | 15.43827 | 19.25514 | 14.87372 | 14.42774 | 15.88135 | 15.61081 |
| SEMA4C | 16.79374 | 17.3145 | 16.02106 | 15.95622 | 13.77728 | 18.18274 | 15.35273 | 15.35 | 20.21456 | 16.01197 | 14.31799 | 14.12069 | 14.80279 | 14.99681 | 16.30546 | 16.01003 | 16.45021 | 15.42701 |
| SEMA7A | 15.86843 | 14.42458 | 16.36459 | 13.85588 | 19.43752 | 17.79991 | 16.59545 | 17.36571 | 14.37557 | 11.82335 | 19.67638 | 14.31161 | 18.03224 | 13.33673 | 14.10681 | 15.24964 | 18.00806 | 18.5077 |
| SEPHS1 | 15.8639 | 15.82577 | 16.09987 | 17.05545 | 16.99646 | 14.99928 | 16.69484 | 16.59578 | 17.33175 | 16.64906 | 19.11441 | 18.87192 | 20.12913 | 17.28981 | 16.47544 | 16.28143 | 16.97195 | 15.86775 |
| SEPHS2 | 14.8319 | 17.40898 | 14.64095 | 16.87558 | 15.95048 | 16.37341 | 14.55537 | 14.21391 | 15.88221 | 14.68907 | 18.02548 | 17.79197 | 16.48144 | 14.95501 | 16.78409 | 15.26673 | 15.16958 | 15.79849 |
| SEPTIN10 | 14.95023 | 16.14727 | 14.79999 | 15.35239 | 21.20792 | 19.71756 | 16.84296 | 15.7508 | 14.07523 | 15.4738 | 15.26306 | 14.12968 | 17.12976 | 16.88378 | 15.85854 | 15.62407 | 20.14907 | 18.75452 |
| SEPTIN11 | 18.36072 | 18.31816 | 17.93389 | 18.73087 | 22.73545 | 21.02575 | 20.77351 | 18.2801 | 17.25901 | 19.8855 | 18.75093 | 19.284 | 20.20085 | 20.12742 | 17.46806 | 17.53185 | 22.92399 | 21.23428 |
| SEPTIN2 | 17.12233 | 17.78228 | 18.01548 | 19.31454 | 22.12046 | 21.46706 | 19.92891 | 19.53692 | 17.46519 | 18.73489 | 19.42611 | 19.87677 | 20.3636 | 19.74932 | 18.76547 | 18.12143 | 22.07082 | 21.26671 |
| SEPTIN3 | 14.11246 | 15.95743 | 15.33765 | 16.26628 | 15.81769 | 15.32413 | 19.76495 | 19.44582 | 14.18867 | 13.83844 | 21.0191 | 15.04041 | 17.71765 | 16.71963 | 16.00574 | 16.76698 | 15.58001 | 16.21324 |
| SEPTIN5 | 15.07403 | 16.86551 | 17.41686 | 18.69887 | 18.98439 | 19.46432 | 20.45713 | 13.09875 | 15.50308 | 16.92873 | 13.96915 | 15.50095 | 20.31296 | 20.17887 | 18.0971 | 17.90167 | 20.16626 | 19.01859 |
| SEPTIN6 | 16.07612 | 17.50089 | 17.90523 | 16.55716 | 19.87887 | 18.07901 | 18.632 | 16.69315 | 15.68302 | 14.9102 | 14.70028 | 18.92751 | 18.45103 | 18.50963 | 17.24108 | 16.96721 | 21.47814 | 20.36047 |
| SEPTIN7 | 17.33472 | 17.91819 | 18.2927 | 19.41581 | 22.64444 | 21.51849 | 20.38708 | 19.21939 | 19.68042 | 19.29083 | 19.30874 | 19.38889 | 20.44089 | 20.72539 | 18.87244 | 18.45129 | 21.82312 | 21.24818 |
| SEPTIN8 | 15.68616 | 16.3171 | 16.78349 | 17.48196 | 21.54199 | 21.09937 | 18.80315 | 16.88656 | 17.67587 | 18.2973 | 16.18402 | 19.49726 | 19.4738 | 16.46904 | 16.79181 | 16.65858 | 20.32058 | 20.49321 |
| SEPTIN9 | 16.78319 | 16.46131 | 17.10781 | 18.83442 | 21.33056 | 19.73968 | 19.17104 | 18.95929 | 18.34325 | 18.72893 | 19.10476 | 19.66494 | 19.16075 | 19.24423 | 17.7266 | 17.77169 | 21.74947 | 21.44112 |
| SERBP1 | 17.61603 | 17.45715 | 16.79879 | 17.44392 | 16.35814 | 15.69314 | 15.85708 | 15.34688 | 15.99772 | 16.07415 | 18.90097 | 18.19945 | 17.3749 | 15.82197 | 18.19384 | 17.19285 | 17.03243 | 16.76369 |
| SERINC1 | 17.85851 | 20.83755 | 20.42142 | 19.02366 | 13.08689 | 15.49413 | 14.28476 | 16.93561 | 18.63929 | 18.24006 | 16.72976 | 19.03484 | 13.82876 | 15.79828 | 19.42628 | 19.41555 | 14.59714 | 17.08744 |
| SERINC3 | 19.77534 | 18.45008 | 19.65959 | 18.31687 | 15.22502 | 15.41505 | 14.80535 | 17.78957 | 16.1826 | 16.7092 | 16.44031 | 17.31266 | 15.90884 | 13.09925 | 16.78458 | 18.50279 | 14.70336 | 16.06404 |
| SERINC5 | 22.0655 | 17.43555 | 23.76604 | 22.57701 | 13.78261 | 15.85938 | 16.16531 | 16.66879 | 17.97105 | 15.72552 | 16.21877 | 14.01254 | 14.26446 | 15.09158 | 24.1397 | 23.77111 | 14.05123 | 16.5578 |
| SERPINB1 | 17.86886 | 17.06957 | 16.34867 | 17.3831 | 17.4211 | 18.03501 | 18.95878 | 18.62788 | 18.68405 | 19.6115 | 18.94661 | 18.89093 | 15.82862 | 18.35082 | 14.73316 | 15.86913 | 17.22068 | 19.51415 |
| SERPINB5 | 16.75613 | 18.08648 | 15.98533 | 16.56788 | 18.31467 | 18.84175 | 16.73612 | 17.65906 | 17.28869 | 17.01248 | 16.02297 | 14.55621 | 19.93718 | 19.8916 | 15.98627 | 16.24173 | 21.70886 | 21.59137 |
| SERPINB6 | 17.54369 | 17.40163 | 17.33468 | 18.85986 | 18.7165 | 18.21014 | 19.51399 | 17.0714 | 15.484 | 17.86118 | 18.4795 | 20.70612 | 18.8065 | 18.18644 | 19.1868 | 18.44159 | 19.32653 | 17.27529 |
| SERPINE2 | 15.32605 | 16.88055 | 15.79552 | 19.32125 | 14.55921 | 17.11772 | 14.53509 | 15.47794 | 15.63797 | 14.28143 | 17.67697 | 18.02415 | 18.33463 | 18.75747 | 17.73101 | 18.24081 | 16.94486 | 19.04972 |
| SERPINF1 | 16.97303 | 15.19704 | 14.93748 | 15.19809 | 15.77942 | 15.79549 | 16.23343 | 16.78798 | 18.58136 | 18.60243 | 22.89562 | 17.74419 | 15.14957 | 15.46039 | 14.45328 | 15.43077 | 15.92256 | 13.86098 |
| SERPINH1 | 16.74214 | 16.77948 | 16.63491 | 16.26733 | 18.3105 | 17.25979 | 18.73612 | 18.37454 | 23.49711 | 22.50481 | 16.87135 | 15.35219 | 16.64054 | 16.72786 | 16.51609 | 16.63073 | 16.58063 | 17.2529 |
| SERPINI1 | 15.5583 | 17.2921 | 14.67078 | 16.15185 | 13.80532 | 17.675 | 17.91933 | 17.2217 | 18.64854 | 17.29468 | 16.83905 | 12.93393 | 14.87103 | 13.57743 | 16.11042 | 13.23107 | 13.93352 | 12.81654 |
| SETD2 | 16.05218 | 15.66726 | 14.52272 | 15.49728 | 22.18213 | 20.51779 | 18.78088 | 19.00161 | 11.32059 | 19.11291 | 14.61561 | 15.70541 | 18.09476 | 18.48675 | 14.55647 | 13.98503 | 21.97405 | 23.72346 |
| SETD3 | 13.28806 | 14.81304 | 13.62543 | 16.13952 | 13.18715 | 14.82956 | 13.5858 | 15.10913 | 15.25049 | 17.97981 | 14.19431 | 17.49254 | 12.92188 | 16.68629 | 16.00124 | 14.70409 | 15.02553 | 13.87169 |
| SETSIP | 17.15349 | 16.39944 | 18.12881 | 18.4377 | 20.22411 | 19.07366 | 17.76456 | 18.50944 | 19.82491 | 21.20422 | 22.7895 | 21.19868 | 21.02542 | 18.70587 | 17.01802 | 17.49284 | 18.41094 | 17.47665 |
| SEZ6 | 16.18726 | 15.84399 | 16.0448 | 15.24174 | 14.92443 | 15.76539 | 15.51285 | 16.13886 | 19.81401 | 17.1759 | 16.68935 | 13.31126 | 15.30062 | 14.87692 | 16.47485 | 16.0031 | 15.38874 | 16.29268 |
| SEZ6L | 16.53651 | 14.9437 | 15.14109 | 16.07616 | 15.22389 | 16.09581 | 16.19544 | 16.39275 | 19.57041 | 17.24151 | 14.36882 | 14.07972 | 14.97904 | 15.25392 | 14.54213 | 15.29822 | 15.57088 | 13.45047 |
| SEZ6L2 | 18.92808 | 19.1983 | 18.55245 | 17.69246 | 14.35605 | 15.62377 | 17.17846 | 15.85161 | 20.55769 | 16.6595 | 13.72979 | 15.92752 | 15.79084 | 15.34953 | 18.83503 | 18.86181 | 14.2216 | 15.20198 |
| SF1 | 18.14353 | 20.08926 | 17.2722 | 17.68127 | 17.56984 | 17.17599 | 15.31958 | 16.5167 | 16.1202 | 17.02547 | 17.59587 | 16.85198 | 18.27739 | 16.16854 | 16.55009 | 16.87556 | 17.02486 | 16.11553 |
| SF3A1 | 19.55334 | 19.73697 | 17.31694 | 17.9823 | 14.98069 | 16.68619 | 17.73093 | 16.52279 | 15.00681 | 17.15226 | 17.85083 | 16.72852 | 18.29784 | 17.43643 | 16.38123 | 17.62231 | 16.2823 | 15.46955 |
| SF3A3 | 19.86195 | 18.37637 | 15.94893 | 17.85546 | 16.7604 | 15.7707 | 17.29618 | 16.21672 | 16.9838 | 16.95443 | 17.03888 | 18.03601 | 18.80819 | 17.65187 | 16.48166 | 17.41739 | 16.55084 | 14.41226 |
| SF3B1 | 18.00468 | 17.92157 | 17.28893 | 16.4982 | 17.66142 | 19.56576 | 16.2337 | 17.08546 | 16.72565 | 16.67668 | 17.45978 | 20.53792 | 16.92149 | 16.98664 | 16.72702 | 16.61151 | 17.04513 | 16.39884 |
| SF3B2 | 19.80309 | 19.0317 | 17.88045 | 17.69198 | 17.81542 | 15.95478 | 18.75912 | 17.51431 | 15.1903 | 15.17054 | 17.72913 | 19.27262 | 18.04405 | 17.3678 | 16.88032 | 17.89462 | 16.69989 | 17.75775 |
| SF3B3 | 20.5992 | 20.37468 | 19.17449 | 19.20457 | 14.71478 | 15.94919 | 15.98533 | 16.28935 | 14.87501 | 14.64214 | 17.05913 | 20.04971 | 15.49545 | 16.07676 | 19.05654 | 18.50344 | 15.98358 | 15.84201 |
| SF3B4 | 19.15447 | 19.54587 | 17.11068 | 16.50453 | 13.55384 | 16.04976 | 16.0745 | 19.69586 | 13.16937 | 12.166 | 17.78834 | 13.02426 | 15.94279 | 15.34574 | 14.93708 | 16.06615 | 17.23552 | 15.18113 |
| SF3B5 | 18.71121 | 19.37496 | 17.54744 | 14.59247 | 17.95173 | 16.94699 | 16.72651 | 17.46561 | 14.25195 | 16.53119 | 16.75694 | 19.28306 | 19.89564 | 18.73091 | 17.08009 | 18.36167 | 17.46606 | 17.18712 |
| SF3B6 | 19.33778 | 18.50218 | 17.92166 | 18.17443 | 13.07275 | 17.02469 | 16.00186 | 18.8924 | 16.829 | 12.95613 | 17.09248 | 18.33575 | 13.46177 | 13.51275 | 13.65505 | 17.27863 | 15.94348 | 11.66359 |
| SFN | 21.52928 | 24.8891 | 22.52385 | 21.82271 | 23.7128 | 22.81627 | 18.55676 | 18.64447 | 21.6178 | 21.52642 | 21.4739 | 23.44828 | 20.36778 | 20.02314 | 22.86733 | 23.00248 | 23.49718 | 22.74587 |
| SFPQ | 16.82035 | 17.10677 | 19.07133 | 16.92437 | 16.69415 | 18.27377 | 17.94753 | 18.41564 | 18.0291 | 18.15796 | 19.16177 | 19.98703 | 19.17767 | 17.74373 | 16.32015 | 16.37942 | 16.13318 | 17.8543 |
| SFRP1 | 15.88387 | 16.63119 | 16.74152 | 15.67336 | 13.38851 | 15.47275 | 13.00244 | 15.41982 | 18.23022 | 19.02379 | 15.27454 | 11.56047 | 16.91611 | 14.80976 | 13.44866 | 17.10742 | 15.89484 | 14.16771 |
| SFRP5 | 21.36465 | 19.84184 | 14.11034 | 13.87786 | 13.59762 | 13.17611 | 15.98753 | 14.97624 | 20.63381 | 18.59461 | 14.4495 | 12.55976 | 14.8735 | 14.65805 | 14.23014 | 14.16991 | 15.73185 | 12.48288 |
| SFTPA2 | 15.12731 | 15.20014 | 14.01769 | 14.66784 | 14.15106 | 16.03174 | 17.84481 | 16.03623 | 19.55492 | 19.83711 | 16.28937 | 12.70088 | 17.24702 | 15.72336 | 15.14153 | 15.6464 | 17.68574 | 17.37317 |
| SFTPB | 14.86564 | 15.3079 | 13.16994 | 15.05621 | 18.5639 | 16.41253 | 15.86781 | 15.64644 | 19.54521 | 19.49611 | 15.38403 | 12.83316 | 15.49576 | 16.73203 | 14.98755 | 16.33931 | 14.40866 | 15.84647 |
| SFXN1 | 16.16622 | 16.26227 | 16.6424 | 16.53521 | 14.37898 | 17.3518 | 16.19684 | 17.27131 | 20.40522 | 18.42298 | 16.24925 | 16.29305 | 14.68258 | 15.16162 | 16.19 | 15.64687 | 15.86968 | 15.04551 |
| SGCE | 19.71032 | 20.36168 | 14.89213 | 14.99833 | 18.10537 | 15.11846 | 13.59456 | 16.10259 | 16.56583 | 17.43143 | 15.5481 | 13.1856 | 14.07844 | 13.25087 | 12.50352 | 14.07012 | 14.70476 | 15.39046 |
| SGPL1 | 18.80755 | 16.45199 | 14.99173 | 16.63452 | 18.18175 | 15.92504 | 15.54876 | 16.87556 | 21.88983 | 20.87244 | 17.38864 | 16.16154 | 18.27461 | 15.72052 | 17.91796 | 15.26561 | 16.49288 | 15.61712 |
| SGTA | 15.8776 | 17.72234 | 17.29346 | 17.83706 | 18.62451 | 19.39177 | 20.47274 | 19.19796 | 16.9639 | 18.20732 | 19.93408 | 19.65025 | 20.469 | 18.99207 | 16.04905 | 17.19853 | 19.93139 | 18.57027 |
| SH2D4A | 15.89992 | 16.67078 | 15.39876 | 16.89644 | 19.66074 | 17.39983 | 15.12691 | 16.38442 | 16.23608 | 16.70455 | 14.89604 | 15.94561 | 19.12565 | 18.97417 | 18.01543 | 17.0778 | 19.35382 | 18.13672 |
| SH3BGRL | 17.09709 | 15.97546 | 14.993 | 15.33665 | 18.01002 | 14.12268 | 16.78922 | 16.19582 | 15.81974 | 18.02105 | 18.47713 | 18.76186 | 13.21293 | 14.26624 | 17.2301 | 17.64564 | 17.40491 | 18.39445 |
| SH3BGRL3 | 21.83197 | 21.61924 | 17.23038 | 20.25744 | 23.24355 | 22.10902 | 23.27615 | 23.04285 | 20.21889 | 21.30502 | 19.16172 | 16.66365 | 21.73428 | 22.21475 | 20.89483 | 20.21781 | 25.84076 | 25.99545 |
| SH3BP1 | 18.5077 | 18.15477 | 13.89796 | 13.08727 | 12.25004 | 15.4004 | 15.78759 | 14.22978 | 11.01552 | 16.42354 | 14.63437 | 10.64448 | 16.01627 | 15.7737 | 15.05926 | 16.03142 | 14.51033 | 14.81817 |
| SH3BP4 | 16.40792 | 15.56929 | 18.17349 | 17.49484 | 18.19027 | 16.23697 | 17.32988 | 15.5796 | 18.52643 | 18.34057 | 16.87878 | 16.95114 | 15.22571 | 16.50488 | 17.91635 | 17.27225 | 19.02617 | 17.54011 |
| SH3D19 | 15.73203 | 14.22464 | 14.35998 | 19.56459 | 23.19707 | 23.24679 | 19.04147 | 19.20397 | 13.54094 | 22.08748 | 19.49894 | 15.99897 | 17.80921 | 21.37104 | 14.89833 | 14.54647 | 19.18765 | 25.55298 |
| SH3GL1 | 24.06421 | 23.73934 | 23.41669 | 23.12179 | 19.54882 | 17.49219 | 20.00018 | 19.32231 | 18.02568 | 18.33493 | 17.20848 | 16.49201 | 22.96982 | 21.94168 | 23.54284 | 23.52155 | 17.87422 | 18.80938 |
| SH3GL2 | 23.67731 | 23.88233 | 23.48894 | 23.01616 | 19.86758 | 18.78186 | 19.4004 | 19.52964 | 19.49301 | 19.95406 | 17.0088 | 16.65841 | 23.12459 | 21.97647 | 23.44702 | 23.61728 | 18.65298 | 18.82885 |
| SH3GL3 | 15.05579 | 20.34331 | 20.13305 | 19.48735 | 13.44033 | 15.49105 | 15.71725 | 14.98052 | 13.86047 | 14.46795 | 15.48385 | 14.25921 | 19.44749 | 19.7878 | 18.2776 | 16.66556 | 15.88343 | 16.4798 |

| | | | | | | | | | | | | | | | | | | |
|---|---|---|---|---|---|---|---|---|---|---|---|---|---|---|---|---|---|---|
| SH3GLB1 | 15.93427 | 16.30125 | 16.0333 | 17.71656 | 17.76148 | 16.30212 | 16.86164 | 17.4674 | 17.62292 | 17.51191 | 16.97861 | 16.4481 | 19.52813 | 14.69387 | 17.80597 | 18.62462 | 17.53492 | 14.86553 |
| SH3GLB2 | 15.60936 | 16.54648 | 14.40791 | 16.0563 | 19.10674 | 19.26028 | 17.89606 | 17.41984 | 17.53479 | 18.32237 | 15.72181 | 16.94909 | 17.85031 | 17.97374 | 13.75774 | 16.56916 | 17.78643 | 18.15607 |
| SH3KBP1 | 16.84496 | 20.03408 | 19.53387 | 18.97706 | 18.77054 | 16.9605 | 14.86524 | 15.34347 | 13.41785 | 15.88046 | 15.40174 | 13.44287 | 19.01963 | 17.08313 | 18.46798 | 18.39979 | 15.85134 | 14.17437 |
| SHANK2 | 15.12374 | 17.13198 | 22.65944 | 16.55731 | 14.28535 | 16.55832 | 15.52001 | 16.54545 | 14.04153 | 13.71697 | 15.30559 | 12.19074 | 17.98144 | 18.32025 | 17.63158 | 16.22678 | 15.44225 | 15.52019 |
| SHB | 15.31301 | 14.28035 | 16.09411 | 15.98517 | 17.50505 | 14.24212 | 14.22437 | 15.97716 | 13.3474 | 15.61401 | 19.99233 | 20.10193 | 19.36614 | 13.26077 | 15.33319 | 15.39944 | 19.39382 | 19.06416 |
| SHC1 | 17.79345 | 15.31199 | 15.13237 | 15.55499 | 18.71111 | 15.70365 | 17.62069 | 16.74164 | 17.06976 | 15.78993 | 17.23851 | 18.84986 | 18.21783 | 17.2311 | 16.05273 | 16.97407 | 17.90621 | 18.201 |
| SHCBP1 | 16.63039 | 17.86584 | 15.63617 | 16.57882 | 21.17506 | 18.97541 | 15.86593 | 16.25123 | 15.25486 | 15.70144 | 16.43763 | 15.47191 | 15.01045 | 15.13544 | 16.1651 | 15.9666 | 16.30541 | 16.24115 |
| SHMT1 | 16.08309 | 15.19411 | 17.34573 | 17.54468 | 18.75665 | 16.51918 | 15.3024 | 15.76559 | 15.47374 | 17.18707 | 17.51424 | 19.45156 | 18.39514 | 16.65041 | 17.39056 | 17.08602 | 15.7814 | 15.19371 |
| SHMT2 | 18.51394 | 19.40886 | 18.46069 | 18.05329 | 17.42477 | 17.96675 | 17.22547 | 16.90043 | 22.36911 | 20.9044 | 17.89351 | 17.04872 | 17.26651 | 17.66661 | 17.92275 | 17.85324 | 17.24255 | 17.84768 |
| SHPK | 15.80432 | 14.80988 | 16.36279 | 16.09231 | 15.63262 | 15.18508 | 13.72684 | 15.82759 | 13.82558 | 16.71674 | 13.39317 | 18.82147 | 14.82405 | 17.06037 | 17.40804 | 13.9244 | 17.58615 | 15.44598 |
| SHPRH | 20.76253 | 22.51301 | 20.47705 | 21.38354 | 21.32784 | 20.56485 | 21.97087 | 21.66018 | 26.25939 | 25.12926 | 18.42603 | 18.2536 | 22.21267 | 19.67668 | 20.90373 | 14.28684 | 20.47766 | 22.50985 |
| SHTN1 | 17.19077 | 15.78569 | 18.34745 | 17.14605 | 19.02434 | 16.92037 | 18.5939 | 18.72172 | 17.73831 | 18.70126 | 17.46459 | 17.73935 | 18.67221 | 19.66432 | 16.66523 | 17.03147 | 18.26529 | 17.62721 |
| SIAE | 14.13038 | 16.35869 | 16.05841 | 14.89326 | 14.58475 | 16.34498 | 16.21905 | 16.19134 | 17.87935 | 17.13137 | 18.17536 | 15.02964 | 14.39941 | 16.77089 | 16.28195 | 14.12774 | 14.34015 | 16.05776 |
| SIGIRR | 17.61689 | 17.90492 | 19.79234 | 19.94968 | 16.38773 | 16.77672 | 15.68302 | 15.83566 | 18.28652 | 17.23563 | 13.53346 | 17.99327 | 15.0518 | 15.10205 | 20.40097 | 20.09434 | 17.40547 | 15.46058 |
| SIGMAR1 | 13.11538 | 13.76722 | 14.64415 | 13.81716 | 14.32529 | 14.88162 | 13.04017 | 15.59155 | 19.80908 | 18.04214 | 15.08053 | 16.92133 | 12.52125 | 15.52459 | 15.25555 | 15.65331 | 14.35135 | 14.42521 |
| SIL1 | 15.53953 | 17.02303 | 18.17505 | 18.5018 | 16.22563 | 20.20761 | 16.42601 | 17.71196 | 19.09779 | 18.81425 | 16.63487 | 13.96554 | 16.01624 | 17.67764 | 17.83377 | 16.08988 | 15.69247 | 19.12426 |
| SIPA1L1 | 15.86474 | 16.33827 | 16.27014 | 17.43299 | 18.84781 | 17.99418 | 13.28431 | 15.46839 | 15.41275 | 14.14555 | 16.31612 | 15.7785 | 17.37143 | 18.04715 | 17.00552 | 16.82329 | 17.20183 | 18.32084 |
| SIRPA | 17.11554 | 14.51982 | 16.26402 | 15.84714 | 19.64168 | 17.13733 | 15.6073 | 17.01468 | 14.9433 | 18.44679 | 17.20162 | 16.66465 | 18.01808 | 15.96189 | 14.97085 | 15.06867 | 19.76983 | 20.83783 |
| SIT1 | 16.39497 | 16.21137 | 16.11367 | 15.50716 | 13.53023 | 16.24253 | 15.16975 | 16.76408 | 20.44241 | 19.96909 | 15.86801 | 13.70206 | 14.53924 | 14.81771 | 17.40847 | 15.40982 | 15.07559 | 14.15067 |
| SKAP2 | 16.01252 | 14.89739 | 16.43685 | 14.85526 | 14.78957 | 13.69775 | 17.38129 | 17.16594 | 15.12549 | 14.79389 | 16.93085 | 17.38723 | 16.67502 | 17.08348 | 15.49953 | 14.88563 | 14.75715 | 14.15398 |
| SKIV2L | 15.14124 | 15.28122 | 16.23561 | 17.24803 | 13.56662 | 16.23559 | 15.01757 | 16.19654 | 13.95981 | 12.54837 | 15.77442 | 17.86098 | 15.82846 | 13.86438 | 16.26012 | 19.35304 | 16.43985 | 14.97674 |
| SKP1 | 18.77613 | 16.0062 | 19.14391 | 18.02226 | 19.5553 | 17.91562 | 16.98254 | 15.458 | 17.35737 | 16.49171 | 18.84568 | 20.18379 | 19.19705 | 17.94637 | 19.45406 | 19.24945 | 15.90111 | 17.50537 |
| SLC12A2 | 21.55385 | 21.7544 | 19.13095 | 19.15264 | 17.97457 | 17.01411 | 17.64339 | 17.15743 | 16.11663 | 15.59209 | 18.87634 | 17.62985 | 18.49377 | 17.4599 | 21.32931 | 21.36032 | 19.01811 | 21.14277 |
| SLC12A6 | 16.89875 | 17.14849 | 17.00916 | 13.81132 | 18.91768 | 17.69899 | 15.6405 | 15.54869 | 16.19858 | 15.39818 | 18.4132 | 18.57407 | 17.66076 | 13.54031 | 16.34448 | 17.35705 | 15.82956 | 18.00915 |
| SLC12A7 | 18.00949 | 17.89416 | 18.54346 | 17.98665 | 16.74801 | 16.60928 | 18.24858 | 18.4785 | 18.47249 | 16.66946 | 16.71806 | 19.16377 | 18.27576 | 17.97269 | 19.8557 | 19.81634 | 16.72889 | 17.20902 |
| SLC16A1 | 22.75931 | 19.54393 | 22.92461 | 22.84511 | 20.97256 | 19.22145 | 21.3717 | 20.10657 | 19.51344 | 18.96158 | 21.72122 | 21.6933 | 22.51113 | 23.61684 | 23.06961 | 23.37315 | 22.9852 | 23.15574 |
| SLC16A10 | 16.76008 | 16.8893 | 19.65628 | 17.91987 | 15.34925 | 16.90972 | 15.43851 | 15.81382 | 14.63805 | 14.55687 | 17.19287 | 19.05705 | 14.29569 | 15.39437 | 19.64537 | 19.03049 | 15.38404 | 14.82296 |
| SLC16A3 | 17.56183 | 16.65653 | 20.56918 | 19.92916 | 21.74874 | 21.6443 | 19.75759 | 15.74274 | 17.15527 | 17.22037 | 17.0005 | 19.9331 | 14.5071 | 19.10016 | 20.26317 | 20.23138 | 21.75773 | 21.80041 |
| SLC19A1 | 17.41155 | 16.3631 | 19.58767 | 19.39981 | 19.36862 | 17.5188 | 16.60999 | 16.70055 | 16.67349 | 15.9916 | 17.73024 | 16.56454 | 18.52056 | 17.7424 | 19.65131 | 19.46642 | 17.60396 | 18.53787 |
| SLC19A2 | 19.31999 | 18.51027 | 18.02784 | 17.69155 | 12.87225 | 16.04607 | 14.9704 | 13.78687 | 16.35323 | 16.39667 | 14.3643 | 18.83262 | 16.37392 | 13.49482 | 17.51162 | 17.93671 | 15.60894 | 12.26177 |
| SLC1A4 | 19.58061 | 19.26116 | 19.55105 | 16.35991 | 17.26724 | 15.85879 | 18.28131 | 17.80349 | 21.78508 | 21.58045 | 19.17796 | 21.93964 | 17.84783 | 15.65328 | 19.70578 | 19.22315 | 16.54092 | 15.21167 |
| SLC1A5 | 21.13334 | 20.79534 | 21.62418 | 21.86498 | 23.32947 | 22.21328 | 21.32151 | 19.89678 | 22.37866 | 21.82003 | 23.68769 | 24.52168 | 22.76526 | 22.17684 | 22.55924 | 22.76226 | 22.25306 | 21.09402 |
| SLC20A1 | 17.47743 | 17.52187 | 17.42188 | 16.63359 | 15.15497 | 17.02606 | 15.43909 | 17.59337 | 17.59958 | 15.99814 | 19.861 | 19.53522 | 15.09194 | 15.52371 | 17.06112 | 18.21058 | 16.65617 | 16.7402 |
| SLC20A2 | 14.88536 | 15.1536 | 18.76324 | 14.54289 | 18.60219 | 15.69335 | 14.85211 | 14.03841 | 14.62794 | 12.92215 | 19.19293 | 17.88995 | 14.43539 | 18.04602 | 18.1383 | 17.85878 | 15.21144 | 14.62662 |
| SLC22A3 | 14.70453 | 15.47248 | 17.40029 | 16.20297 | 21.58096 | 17.54184 | 15.37538 | 15.03087 | 14.82989 | 14.51818 | 21.37389 | 12.35441 | 14.8023 | 16.18693 | 15.95024 | 20.87123 | 18.09596 | 17.08354 |
| SLC23A2 | 16.00605 | 15.78486 | 15.38642 | 16.93398 | 16.37986 | 15.5291 | 14.72203 | 16.22926 | 14.5636 | 18.04536 | 17.2302 | 18.57849 | 13.03112 | 14.72339 | 15.02678 | 16.48005 | 16.92083 | 16.90502 |
| SLC25A1 | 14.5773 | 17.11346 | 15.29747 | 15.65513 | 13.632 | 17.02996 | 16.17177 | 18.26391 | 18.42568 | 17.96524 | 16.76378 | 14.37047 | 14.73629 | 15.23712 | 14.90287 | 14.86457 | 16.59229 | 14.79603 |
| SLC25A10 | 14.48622 | 16.97312 | 17.46265 | 16.24773 | 14.34891 | 16.13292 | 14.6284 | 16.80594 | 19.39843 | 17.78544 | 15.88925 | 13.27599 | 14.66433 | 14.78462 | 14.90823 | 14.52276 | 15.22571 | 14.47708 |
| SLC25A11 | 17.43327 | 17.60276 | 18.1731 | 16.60325 | 15.26308 | 15.83284 | 17.27279 | 16.38085 | 19.54529 | 18.39328 | 16.56882 | 16.28655 | 16.35587 | 15.57988 | 16.11022 | 16.6979 | 16.01489 | 17.5251 |
| SLC25A12 | 16.10437 | 12.73844 | 15.52611 | 15.41634 | 16.26476 | 15.71195 | 14.9125 | 15.19982 | 19.72648 | 18.71527 | 15.58334 | 13.07678 | 14.25441 | 13.63315 | 15.01312 | 16.36668 | 13.66664 | 13.83273 |
| SLC25A13 | 15.7144 | 23.03305 | 14.8066 | 17.88719 | 16.06165 | 17.26281 | 16.73695 | 17.43306 | 19.60494 | 17.40028 | 15.83044 | 15.55842 | 16.55589 | 15.82223 | 19.92972 | 19.32183 | 17.30521 | 15.109 |
| SLC25A24 | 15.54572 | 15.61148 | 15.90157 | 16.47737 | 19.3374 | 17.81876 | 18.45474 | 17.44721 | 20.08306 | 19.03848 | 15.83095 | 15.54748 | 15.24649 | 15.75336 | 16.86482 | 16.92234 | 18.29845 | 17.68954 |
| SLC25A3 | 16.47106 | 16.04027 | 17.10377 | 16.88853 | 16.65757 | 15.87417 | 17.85611 | 16.63124 | 22.67544 | 22.01031 | 17.17529 | 16.7877 | 15.80329 | 15.57656 | 17.09201 | 15.96226 | 16.56974 | 15.42776 |
| SLC25A4 | 17.55564 | 17.14218 | 17.5506 | 16.18714 | 17.66407 | 16.6896 | 18.10358 | 17.49172 | 21.47033 | 20.45609 | 18.82298 | 17.83983 | 18.07118 | 17.04238 | 17.81623 | 17.44103 | 16.27238 | 17.75195 |
| SLC25A5 | 17.14467 | 16.30643 | 16.68243 | 17.23539 | 16.86067 | 16.11887 | 21.68439 | 20.9774 | 21.3603 | 21.49895 | 20.17515 | 15.70468 | 16.87416 | 18.16334 | 15.66247 | 15.55065 | 16.50324 | 17.05852 |
| SLC25A6 | 14.58082 | 17.11822 | 17.04427 | 15.04299 | 17.95244 | 16.12322 | 19.14548 | 18.00787 | 21.74431 | 20.7431 | 19.52248 | 17.44027 | 14.76612 | 15.31521 | 15.49234 | 17.81001 | 15.92954 | 14.67511 |
| SLC26A6 | 17.39248 | 17.31929 | 15.66021 | 16.11959 | 19.06581 | 16.91595 | 16.23307 | 16.64153 | 14.08665 | 16.0781 | 16.39056 | 18.18596 | 17.88503 | 17.54059 | 18.03041 | 16.44676 | 17.26184 | 16.90572 |
| SLC29A1 | 22.24895 | 21.97803 | 22.56323 | 20.95693 | 15.23229 | 17.10205 | 16.72994 | 16.42971 | 18.48496 | 18.98074 | 20.19198 | 22.65474 | 16.08779 | 15.51181 | 22.78425 | 22.73215 | 16.94343 | 16.0406 |
| SLC29A2 | 18.63995 | 18.26682 | 19.89988 | 16.91891 | 14.6999 | 19.46331 | 14.57835 | 15.43496 | 15.44284 | 16.10485 | 17.8399 | 19.62142 | 18.03704 | 14.68811 | 19.24269 | 19.40708 | 14.67656 | 15.05758 |
| SLC2A1 | 21.18846 | 21.51753 | 22.89171 | 22.33511 | 20.70899 | 21.10338 | 18.28291 | 19.25296 | 20.48591 | 19.37924 | 19.74369 | 22.65226 | 16.12549 | 17.23017 | 22.52122 | 22.49988 | 20.72135 | 17.58538 |
| SLC2A5 | 13.94857 | 14.59373 | 14.64662 | 15.9365 | 13.78455 | 19.29996 | 17.5753 | 16.28271 | 13.83593 | 15.57126 | 17.97774 | 13.28494 | 15.22451 | 16.91452 | 15.6343 | 21.10353 | 15.89963 | 14.92598 |
| SLC30A1 | 18.51362 | 16.9649 | 20.23689 | 20.66033 | 20.96201 | 18.82743 | 18.68488 | 17.12529 | 17.59826 | 17.71691 | 18.6949 | 20.19193 | 19.69961 | 18.71467 | 20.44815 | 20.03111 | 19.81069 | 20.12935 |
| SLC30A7 | 16.7138 | 14.23201 | 15.77838 | 15.11281 | 14.31374 | 15.98925 | 16.31085 | 15.61289 | 18.51297 | 17.10577 | 13.82408 | 14.37608 | 15.43517 | 18.23996 | 15.67586 | 14.66254 | 17.38229 | 15.04479 |
| SLC31A1 | 18.84559 | 19.68453 | 20.74752 | 20.31204 | 21.0831 | 18.4973 | 16.58543 | 18.29578 | 16.32694 | 17.70924 | 15.86509 | 22.48581 | 18.22699 | 17.57192 | 21.93797 | 21.78943 | 20.63723 | 14.916 |
| SLC35A4 | 21.5046 | 19.0271 | 14.85393 | 17.51801 | 13.52962 | 16.34503 | 19.91942 | 18.61885 | 18.79645 | 19.29847 | 18.05859 | 16.06315 | 12.61667 | 14.39646 | 14.26267 | 14.36579 | 15.65725 | 14.69048 |
| SLC35B2 | 14.98189 | 16.27151 | 14.9142 | 13.89002 | 13.24315 | 15.45291 | 15.56668 | 19.09438 | 19.01189 | 18.03562 | 13.7747 | 18.91776 | 15.94925 | 14.59938 | 15.35497 | 14.75943 | 15.20007 | 13.72658 |
| SLC36A1 | 15.56297 | 16.03801 | 13.54432 | 15.80477 | 15.56188 | 13.71137 | 16.21345 | 16.10489 | 19.53749 | 14.87061 | 16.13626 | 16.54919 | 15.49501 | 15.43281 | 16.93154 | 16.83398 | 16.62513 | 15.60695 |
| SLC38A1 | 19.00216 | 18.39407 | 18.54303 | 18.13333 | 15.01626 | 19.20413 | 15.30208 | 15.48978 | 19.57206 | 17.93759 | 17.8606 | 21.3677 | 14.7384 | 14.72574 | 19.11416 | 19.36133 | 15.90261 | 17.6564 |
| SLC38A10 | 18.65964 | 18.16354 | 17.49672 | 16.48056 | 15.66342 | 17.09064 | 16.56207 | 16.78083 | 20.70472 | 17.2545 | 15.7782 | 17.56157 | 15.94849 | 16.16012 | 16.95273 | 17.00226 | 15.42732 | 15.33396 |
| SLC38A2 | 22.21662 | 22.35248 | 21.63512 | 21.7623 | 22.6186 | 20.87127 | 22.03871 | 18.63762 | 20.9845 | 21.09529 | 21.60086 | 21.71765 | 21.63521 | 21.23945 | 20.95522 | 21.03343 | 21.63902 | 23.18717 |
| SLC38A3 | 15.80617 | 17.56915 | 14.80436 | 16.85923 | 14.38361 | 15.58505 | 16.4009 | 19.30505 | 18.96375 | 16.34963 | 16.45926 | 16.53856 | 15.78145 | 16.61481 | 16.3719 | 14.68121 | 16.17058 | 15.67377 |
| SLC38A9 | 15.73554 | 17.07628 | 16.19798 | 14.07546 | 12.57369 | 17.05998 | 13.86925 | 16.1288 | 16.49934 | 13.23897 | 14.92765 | 12.49949 | 19.07586 | 22.88605 | 15.67735 | 15.53873 | 13.00654 | 17.07144 |
| SLC39A10 | 18.32697 | 17.92708 | 19.00486 | 18.67543 | 21.5553 | 19.60249 | 18.57191 | 16.833 | 18.96725 | 16.83083 | 17.0269 | 20.80995 | 20.16635 | 19.3505 | 19.34516 | 19.31756 | 20.12667 | 21.51502 |
| SLC39A14 | 20.24865 | 19.77874 | 19.96099 | 19.72066 | 15.80041 | 14.84391 | 14.16816 | 16.34435 | 17.81534 | 14.29556 | 16.74708 | 18.7267 | 14.88622 | 18.49162 | 20.05083 | 20.26593 | 17.08804 | 17.33472 |
| SLC39A6 | 16.49984 | 17.60533 | 19.15486 | 17.81712 | 19.82445 | 18.92146 | 17.13604 | 17.65939 | 16.82439 | 15.8421 | 18.12922 | 19.43616 | 18.92131 | 18.3325 | 18.722 | 19.10541 | 19.5073 | 20.48854 |
| SLC39A8 | 19.19275 | 12.91005 | 20.9202 | 21.37658 | 16.64652 | 17.20431 | 14.92498 | 15.00847 | 17.78098 | 13.24612 | 14.17894 | 16.6687 | 13.79929 | 13.09207 | 20.94106 | 19.5614 | 15.95705 | 15.02103 |
| SLC3A2 | 23.36158 | 22.9724 | 22.75423 | 22.8987 | 23.06278 | 22.32208 | 20.28431 | 18.22774 | 22.52907 | 21.29015 | 23.89848 | 24.28697 | 20.50506 | 19.1607 | 23.80563 | 23.00878 | 20.00192 | 20.84724 |
| SLC43A1 | 17.49138 | 17.81805 | 21.62353 | 21.13612 | 18.72574 | 22.24744 | 20.56953 | 20.49995 | 17.02685 | 16.11208 | 17.22733 | 17.55284 | 17.19236 | 20.58527 | 21.93996 | 21.82238 | 18.82558 | 21.58752 |
| SLC44A1 | 21.80706 | 20.38328 | 21.75494 | 21.43383 | 19.89331 | 18.5227 | 18.55361 | 18.04932 | 19.66788 | 19.88417 | 19.32042 | 20.31879 | 21.02148 | 19.85952 | 21.99116 | 21.76054 | 17.88174 | 18.66828 |
| SLC44A2 | 22.05658 | 21.49281 | 21.79845 | 20.53594 | 16.4035 | 16.96008 | 15.68928 | 16.38232 | 19.86006 | 19.46321 | 18.59072 | 20.1749 | 17.75598 | 16.80901 | 22.01164 | 21.85848 | 16.91789 | 17.44945 |
| SLC44A4 | 17.4907 | 15.40758 | 21.81224 | 15.73203 | 15.96019 | 16.67876 | 15.18099 | 17.78927 | 14.58459 | 15.19025 | 16.0931 | 15.94874 | 16.68656 | 17.17902 | 21.25733 | 20.94902 | 14.59316 | 15.86279 |
| SLC4A2 | 18.69469 | 18.19192 | 20.43052 | 20.28275 | 16.97216 | 16.32464 | 17.46597 | 16.11057 | 15.8306 | 14.9461 | 16.7839 | 17.75419 | 17.26852 | 16.8718 | 21.20548 | 20.44867 | 15.22352 | 16.34175 |
| SLC4A4 | 15.94972 | 15.32954 | 17.38138 | 16.34094 | 19.91373 | 18.14313 | 16.56573 | 16.71812 | 12.64726 | 14.27019 | 14.8853 | 13.34568 | 18.4264 | 18.09182 | 17.6803 | 18.80548 | 15.95231 | 17.09293 |
| SLC4A7 | 16.10023 | 16.35295 | 17.43223 | 17.72604 | 19.04044 | 16.21679 | 17.80233 | 17.66394 | 16.68451 | 16.75983 | 18.26309 | 18.62547 | 16.84692 | 16.21312 | 17.20462 | 17.77993 | 18.82624 | 19.78834 |
| SLC5A3 | 13.88615 | 14.12642 | 14.52792 | 14.37105 | 13.22527 | 15.91167 | 14.9506 | 16.62401 | 13.49283 | 14.93293 | 14.49458 | 15.25779 | 13.60216 | 13.5181 | 17.42948 | 18.359 | 18.01243 | 19.26119 |
| SLC5A6 | 17.69563 | 16.46292 | 19.45189 | 18.1356 | 12.93775 | 17.26303 | 16.72147 | 15.70539 | 16.54489 | 14.89665 | 20.40418 | 20.62886 | 13.32553 | 15.7922 | 17.15341 | 18.80318 | 14.62136 | 13.97434 |
| SLC6A6 | 18.85935 | 15.44843 | 14.91856 | 14.33619 | 15.96944 | 15.21183 | 15.00821 | 16.04228 | 11.63913 | 12.97674 | 13.69618 | 19.62562 | 14.23648 | 15.7367 | 15.78422 | 15.69453 | 14.92837 | 14.24171 |
| SLC6A8 | 15.56567 | 14.78096 | 19.56925 | 19.7924 | 14.44389 | 15.74123 | 17.07426 | 15.30652 | 15.12549 | 16.99776 | 19.54558 | 20.32459 | 16.63796 | 14.96076 | 19.44383 | 20.60743 | 15.30793 | 17.32882 |
| SLC7A1 | 16.83031 | 17.58981 | 20.65666 | 20.75997 | 16.32783 | 16.86755 | 16.88877 | 17.72819 | 18.1757 | 18.41512 | 16.93947 | 20.85461 | 17.64471 | 16.52471 | 22.16058 | 20.44964 | 18.04123 | 15.59232 |
| SLC7A11 | 16.29487 | 16.57534 | 17.75472 | 18.3934 | 19.3502 | 16.40863 | 14.7905 | 15.03903 | 15.64011 | 16.42739 | 15.84986 | 17.66428 | 18.0564 | 18.38407 | 18.3268 | 19.0522 | 17.46506 | 16.27216 |
| SLC7A2 | 14.91636 | 16.46352 | 13.88802 | 18.06261 | 13.41211 | 15.90087 | 14.79226 | 13.23882 | 13.43247 | 17.14028 | 15.70541 | 18.32246 | 17.52998 | 15.77121 | 17.66759 | 16.64231 | 14.98945 | 14.36117 |
| SLC7A5 | 22.71423 | 22.59556 | 21.96601 | 21.25692 | 21.33921 | 20.76676 | 19.75537 | 19.46019 | 19.96911 | 19.76304 | 22.21644 | 23.36767 | 21.25226 | 19.64926 | 22.15627 | 22.09804 | 19.87183 | 18.53345 |
| SLC7A6 | 16.14218 | 17.58596 | 15.18693 | 19.07856 | 14.58048 | 14.36958 | 15.75767 | 16.10739 | 13.14377 | 15.86001 | 14.01726 | 20.42926 | 15.99338 | 16.12787 | 16.71656 | 18.70612 | 16.09062 | 13.54911 |
| SLC9A1 | 17.92333 | 16.95166 | 19.43277 | 18.44442 | 18.30939 | 18.25869 | 16.23553 | 17.52506 | 16.03793 | 16.17827 | 17.15363 | 17.26991 | 20.51884 | 21.20938 | 20.2216 | 19.46847 | 17.39544 | 18.56489 |
| SLC9A3R1 | 17.83812 | 17.86975 | 20.44327 | 21.17904 | 21.62855 | 20.54325 | 20.53266 | 20.16438 | 18.35496 | 18.73106 | 19.75443 | 20.33169 | 23.62444 | 24.1112 | 21.20732 | 21.25952 | 21.53116 | 20.34383 |
| SLC9A3R2 | 18.89565 | 18.70202 | 18.73865 | 19.74109 | 19.93037 | 18.37489 | 16.03081 | 16.44604 | 17.52773 | 17.20388 | 17.22948 | 18.77746 | 20.32967 | 19.41587 | 19.8062 | 19.57804 | 19.98533 | 19.97456 |

| | | | | | | | | | | | | | | | | | | |
|---|---|---|---|---|---|---|---|---|---|---|---|---|---|---|---|---|---|---|
| SLC9A6 | 14.57528 | 15.04074 | 16.90028 | 16.93049 | 15.17847 | 16.39922 | 16.21406 | 16.17258 | 20.24557 | 18.51861 | 13.79542 | 15.99471 | 16.19746 | 16.06908 | 16.89378 | 15.22765 | 15.60615 | 15.19252 |
| SLIRP | 19.51054 | 19.2799 | 16.81495 | 19.77371 | 15.0412 | 18.58759 | 17.48333 | 19.94563 | 19.91692 | 19.36056 | 17.5126 | 15.65276 | 17.60182 | 18.45016 | 16.36416 | 19.24422 | 17.5615 | 16.70465 |
| SLIT2 | 16.18604 | 17.4076 | 16.06745 | 15.52957 | 14.87114 | 14.80948 | 17.52723 | 18.85593 | 15.99105 | 16.19406 | 17.57765 | 16.87943 | 14.90562 | 15.32574 | 16.45589 | 15.60851 | 18.06992 | 21.02934 |
| SLITRK5 | 19.1235 | 19.06179 | 15.73832 | 13.51842 | 13.48899 | 14.42825 | 15.74484 | 15.69565 | 17.56823 | 16.88173 | 13.64201 | 12.44097 | 14.29345 | 16.5074 | 15.08172 | 15.53685 | 15.16835 | 14.44562 |
| SLITRK6 | 21.09394 | 20.97455 | 15.44694 | 16.08561 | 15.97517 | 15.8478 | 16.70123 | 16.96198 | 19.89979 | 20.87112 | 17.13069 | 14.02668 | 16.06934 | 17.37361 | 17.26167 | 16.67901 | 17.89251 | 19.45304 |
| SLK | 17.9887 | 17.05666 | 17.0903 | 17.12131 | 18.0786 | 16.82721 | 17.6804 | 18.46343 | 17.95921 | 17.33269 | 18.84332 | 18.81316 | 19.69214 | 19.97131 | 17.48517 | 17.96695 | 17.29791 | 16.85218 |
| SMAD1 | 17.05092 | 16.01759 | 17.52516 | 18.35613 | 17.30655 | 18.0122 | 13.88423 | 16.66298 | 16.28142 | 16.77429 | 18.26622 | 18.2776 | 18.29655 | 14.02034 | 18.33237 | 18.46644 | 17.20389 | 17.92677 |
| SMAD2 | 14.95568 | 17.69818 | 15.31364 | 17.61725 | 13.29015 | 15.71995 | 15.57679 | 13.82852 | 13.74211 | 16.23191 | 15.49233 | 18.12978 | 14.84938 | 17.77835 | 15.32835 | 15.92494 | 14.71318 | 12.69791 |
| SMAD3 | 15.54262 | 17.0776 | 16.58186 | 18.84203 | 16.28069 | 16.12585 | 15.64113 | 18.01348 | 15.12306 | 17.1556 | 16.34898 | 15.57248 | 16.03873 | 16.25515 | 17.45048 | 19.119 | 18.40759 | 19.05704 |
| SMAP | 14.57464 | 13.74359 | 14.93015 | 15.63464 | 13.11487 | 14.79039 | 15.17325 | 14.46331 | 16.70586 | 14.89268 | 17.33784 | 17.38284 | 12.68019 | 15.26677 | 16.66457 | 15.11675 | 14.36987 | 16.51967 |
| SMARCA1 | 18.90365 | 18.89748 | 16.44714 | 16.27098 | 16.68971 | 16.75211 | 17.17481 | 15.99909 | 15.31012 | 16.90859 | 15.66767 | 17.38354 | 17.1754 | 16.62698 | 15.66552 | 16.65101 | 16.93689 | 16.61292 |
| SMARCA5 | 19.37006 | 18.42367 | 16.21265 | 16.10115 | 15.72353 | 18.60444 | 17.80491 | 17.51784 | 14.43436 | 14.34092 | 16.36029 | 16.87899 | 16.88263 | 16.79014 | 16.79343 | 16.69261 | 17.22195 | 16.33758 |
| SMARCC1 | 14.77243 | 13.28912 | 15.50375 | 14.41754 | 14.86208 | 14.29547 | 15.17778 | 14.62452 | 15.42246 | 16.42357 | 18.72422 | 16.98711 | 14.97428 | 11.89904 | 15.87403 | 15.10547 | 12.4634 | 15.68946 |
| SMC1A | 17.99875 | 15.36586 | 16.22151 | 14.70626 | 17.81797 | 18.22527 | 17.44303 | 17.45035 | 14.42942 | 16.54359 | 15.45728 | 16.83406 | 15.9674 | 17.70729 | 15.71488 | 16.40562 | 16.28437 | 18.30782 |
| SMC2 | 14.97023 | 15.89257 | 16.86592 | 16.12994 | 16.91564 | 16.03296 | 17.66824 | 17.24869 | 17.51909 | 17.0596 | 16.63842 | 17.68356 | 15.86179 | 15.84635 | 14.83151 | 16.76109 | 16.64539 | 15.8063 |
| SMC3 | 16.68399 | 16.49932 | 16.3935 | 14.96882 | 18.11202 | 18.09418 | 18.8015 | 16.49677 | 14.59831 | 16.64562 | 16.65546 | 17.33904 | 16.67682 | 15.75884 | 15.69794 | 14.95314 | 15.7253 | 15.26712 |
| SMC4 | 19.98272 | 18.74594 | 20.27902 | 16.93174 | 18.32472 | 18.16161 | 19.29409 | 18.07678 | 16.5113 | 17.38523 | 17.69315 | 16.96541 | 18.39155 | 17.70257 | 19.10551 | 18.40072 | 17.48195 | 17.38304 |
| SMCHD1 | 16.63468 | 15.77219 | 15.97039 | 16.73604 | 15.79463 | 15.55353 | 19.65419 | 19.29752 | 13.52135 | 15.26972 | 14.09857 | 14.68607 | 17.18188 | 15.14816 | 15.29853 | 15.52999 | 14.85699 | 15.07536 |
| SMG9 | 14.73612 | 14.63407 | 13.65805 | 14.91846 | 17.44691 | 16.32042 | 15.63968 | 15.03542 | 13.827 | 14.61143 | 15.64318 | 14.88253 | 13.47621 | 13.83108 | 12.82296 | 17.99302 | 15.86107 | 15.38052 |
| SMOC1 | 16.70132 | 16.58856 | 15.21439 | 14.59294 | 14.24441 | 16.18854 | 16.11581 | 15.41448 | 19.40381 | 18.5196 | 15.33698 | 13.52735 | 15.17119 | 17.04453 | 16.64085 | 16.05196 | 15.09973 | 15.31867 |
| SMPDL3A | 22.03602 | 22.66948 | 16.20445 | 12.42576 | 15.34792 | 15.76759 | 15.40093 | 19.06223 | 19.14531 | 20.01533 | 19.29722 | 15.77603 | 13.84063 | 14.4889 | 16.32478 | 14.92948 | 17.07919 | 13.77628 |
| SMPDL3B | 21.38218 | 21.0635 | 23.14367 | 22.8276 | 19.02908 | 18.38557 | 15.90912 | 16.99948 | 20.37406 | 18.56073 | 19.74191 | 20.37391 | 21.12457 | 20.38497 | 23.31894 | 22.80336 | 20.06437 | 17.49183 |
| SMS | 16.99295 | 16.87389 | 17.81027 | 17.76375 | 19.55884 | 17.27291 | 19.56034 | 16.57456 | 16.31476 | 16.70496 | 17.97103 | 19.26103 | 20.98602 | 20.43952 | 18.01309 | 15.85421 | 20.68189 | 19.86846 |
| SMURF1 | 19.30309 | 20.11062 | 14.28922 | 17.62609 | 17.39897 | 16.81797 | 16.34763 | 15.49147 | 15.11593 | 16.88529 | 15.98452 | 15.52856 | 17.12667 | 14.52609 | 18.98671 | 19.17139 | 17.2412 | 17.64017 |
| SNAP23 | 19.4654 | 20.07676 | 20.07748 | 20.74003 | 22.20618 | 20.85702 | 16.8267 | 17.25135 | 17.21318 | 17.01563 | 20.29091 | 20.49114 | 21.6081 | 21.04327 | 21.1262 | 20.57627 | 21.4667 | 22.06977 |
| SNAP25 | 17.74541 | 18.06562 | 16.91087 | 15.62819 | 17.1791 | 15.3794 | 17.99851 | 18.50819 | 19.17288 | 17.06135 | 15.2143 | 12.89354 | 17.45019 | 16.19755 | 14.84664 | 14.73644 | 16.13074 | 13.88174 |
| SNAP29 | 15.69496 | 15.86508 | 15.78055 | 15.78317 | 17.15526 | 19.44615 | 15.35961 | 16.65349 | 17.77218 | 17.7822 | 17.08604 | 16.27108 | 17.23347 | 16.22694 | 16.49049 | 17.44093 | 15.31487 | 18.96353 |
| SNCA | 15.53154 | 14.55679 | 15.40792 | 17.26114 | 18.37927 | 15.76422 | 18.14639 | 18.51516 | 11.86007 | 15.09091 | 17.77767 | 19.03459 | 13.45332 | 13.12351 | 14.90247 | 15.57459 | 13.72698 | 18.56586 |
| SND1 | 18.67605 | 18.50011 | 19.7042 | 20.12842 | 18.00782 | 18.19886 | 18.88742 | 17.93982 | 17.09311 | 17.72222 | 19.21741 | 20.37762 | 19.70679 | 19.27355 | 20.05705 | 20.00736 | 18.97432 | 18.67045 |
| SNF8 | 21.2732 | 21.49492 | 21.01382 | 20.40515 | 17.71838 | 17.3383 | 17.52325 | 17.0981 | 15.90214 | 15.56746 | 16.42821 | 17.50756 | 20.03319 | 19.39223 | 21.67487 | 21.38343 | 15.72107 | 17.46615 |
| SNRNP200 | 17.66123 | 16.74297 | 20.36654 | 16.96921 | 14.70563 | 16.51161 | 16.14979 | 17.55941 | 15.98871 | 15.70786 | 15.62633 | 19.49073 | 15.89997 | 16.19361 | 16.43539 | 16.36296 | 15.96508 | 15.70177 |
| SNRNP40 | 16.92302 | 16.71899 | 16.51843 | 16.39083 | 14.56697 | 16.91027 | 16.7107 | 16.61673 | 13.30328 | 18.72517 | 15.60574 | 18.68574 | 16.31464 | 15.52642 | 16.18249 | 16.73166 | 15.57459 | 13.91584 |
| SNRNP70 | 19.10899 | 17.10206 | 16.77936 | 16.34914 | 18.02874 | 16.55793 | 16.51773 | 18.41897 | 17.70813 | 17.9846 | 18.0657 | 18.37103 | 16.90249 | 16.90605 | 17.68589 | 18.26548 | 17.09505 | 18.57539 |
| SNRPA | 19.02496 | 18.24705 | 17.94772 | 15.64505 | 16.20043 | 15.47293 | 16.10357 | 17.61784 | 16.70964 | 16.8899 | 19.26396 | 20.86151 | 16.71805 | 15.95319 | 17.87383 | 17.8139 | 13.64867 | 14.91829 |
| SNRPA1 | 19.21101 | 18.80289 | 17.74222 | 17.48848 | 17.60679 | 17.93859 | 16.7609 | 16.51504 | 14.61299 | 15.99387 | 18.00874 | 19.11529 | 15.02144 | 15.73015 | 17.16821 | 17.51686 | 14.96665 | 16.09791 |
| SNRPB | 19.89846 | 19.77683 | 19.06545 | 19.43725 | 19.1829 | 17.49477 | 18.87973 | 16.9491 | 17.63414 | 19.65256 | 20.98477 | 20.1002 | 20.67404 | 19.51582 | 18.21257 | 18.62413 | 18.88906 | 18.90094 |
| SNRPB2 | 19.45979 | 19.78551 | 15.01989 | 15.07369 | 18.51533 | 15.04054 | 14.34568 | 16.45519 | 13.18303 | 14.5208 | 14.29173 | 15.59981 | 16.63534 | 15.37373 | 18.41083 | 17.20418 | 13.97906 | 16.38697 |
| SNRPD1 | 21.12929 | 20.84397 | 19.18775 | 19.63878 | 18.67308 | 19.27073 | 20.8194 | 20.65041 | 17.93171 | 19.73088 | 21.31979 | 21.53731 | 19.25411 | 17.26221 | 18.99792 | 19.70078 | 19.13017 | 17.80322 |
| SNRPD2 | 19.97445 | 20.37113 | 16.50509 | 18.63024 | 19.94818 | 17.63208 | 21.15814 | 19.29706 | 17.55637 | 19.09333 | 20.03838 | 19.42487 | 21.00832 | 19.89761 | 16.56092 | 17.64087 | 17.79844 | 18.55237 |
| SNRPD3 | 21.26071 | 20.53587 | 20.13211 | 21.01899 | 20.39977 | 18.67947 | 19.77126 | 23.86006 | 19.79099 | 20.14907 | 22.00937 | 21.48172 | 22.66638 | 21.02609 | 19.45386 | 19.69119 | 20.03673 | 17.65739 |
| SNRPE | 21.39579 | 21.16047 | 19.47474 | 18.62779 | 17.7268 | 17.55975 | 16.90292 | 16.09022 | 16.68564 | 18.49715 | 17.27235 | 20.55044 | 18.92846 | 16.22379 | 19.62418 | 18.96303 | 19.89686 | 17.12574 |
| SNRPF | 22.61743 | 22.75212 | 19.46651 | 13.78621 | 19.46841 | 17.29276 | 19.53705 | 18.44945 | 15.66298 | 15.37045 | 21.32098 | 20.52067 | 21.49196 | 20.61314 | 18.8177 | 19.33464 | 15.32578 | 14.25428 |
| SNTA1 | 16.37633 | 16.25112 | 18.05944 | 17.18925 | 16.99194 | 17.43026 | 15.31267 | 16.68098 | 16.73932 | 16.59291 | 16.92446 | 18.58685 | 14.67659 | 17.02219 | 20.3358 | 18.92278 | 16.60355 | 15.88736 |
| SNTB1 | 17.0207 | 16.1989 | 17.65957 | 15.4384 | 17.47967 | 15.81931 | 16.78696 | 17.06048 | 19.70209 | 19.33837 | 15.76408 | 14.185 | 15.84326 | 16.04152 | 17.15825 | 17.69377 | 17.81767 | 18.33532 |
| SNTB2 | 16.74611 | 17.67186 | 20.46901 | 19.20541 | 19.20728 | 17.92305 | 15.15387 | 16.37964 | 16.87264 | 17.68732 | 20.4888 | 16.6613 | 19.59862 | 18.62861 | 20.154 | 20.06519 | 18.34284 | 19.21709 |
| SNU13 | 17.30016 | 16.01771 | 15.43141 | 15.3925 | 17.31384 | 17.89781 | 19.99327 | 19.2448 | 13.20079 | 17.71371 | 16.40357 | 16.36811 | 19.52694 | 19.37732 | 17.68688 | 17.40167 | 18.61701 | 18.9503 |
| SNW1 | 16.1538 | 16.68751 | 14.36364 | 16.63787 | 18.96892 | 17.56186 | 15.56221 | 16.18019 | 16.72434 | 18.58074 | 18.80243 | 15.80465 | 17.35304 | 16.88997 | 15.80763 | 15.30612 | 17.013 | 17.83732 |
| SNX1 | 15.98783 | 15.69792 | 15.75941 | 15.81241 | 19.44323 | 17.78212 | 17.00833 | 17.1794 | 17.65557 | 17.16426 | 15.92831 | 17.92509 | 18.57554 | 17.7501 | 16.194 | 16.28821 | 16.94621 | 18.78145 |
| SNX11 | 17.06545 | 14.90176 | 15.72483 | 14.04291 | 17.5435 | 16.85916 | 16.79761 | 14.47818 | 16.07198 | 16.03703 | 17.29245 | 14.64999 | 18.58367 | 18.17729 | 16.62164 | 14.87569 | 20.50072 | 20.35374 |
| SNX12 | 16.9242 | 17.62387 | 18.41755 | 17.53092 | 21.33271 | 15.13722 | 21.77113 | 16.14285 | 18.77661 | 19.14391 | 19.69486 | 19.33716 | 20.5304 | 20.70123 | 18.60606 | 17.91555 | 20.27664 | 20.15505 |
| SNX16 | 16.41183 | 15.45141 | 15.02753 | 15.47552 | 17.15597 | 16.39763 | 15.27993 | 17.75706 | 14.75855 | 14.34992 | 13.45528 | 14.4306 | 15.41013 | 16.63697 | 16.91808 | 17.23203 | 14.40206 | 15.44919 |
| SNX17 | 16.96644 | 16.43443 | 15.69426 | 17.58456 | 20.95719 | 18.97317 | 18.7048 | 14.97956 | 15.65486 | 14.16711 | 14.51114 | 11.62346 | 16.55303 | 19.7522 | 13.86169 | 17.93842 | 16.34467 | 18.40831 |
| SNX18 | 19.99671 | 20.05798 | 21.85247 | 21.66167 | 17.12019 | 16.4281 | 16.30698 | 16.3469 | 16.24909 | 17.16032 | 15.30559 | 14.42113 | 21.12231 | 20.53452 | 21.93697 | 21.86565 | 16.89843 | 15.96788 |
| SNX2 | 14.95938 | 16.29519 | 17.19118 | 17.08318 | 20.21138 | 17.61815 | 19.93685 | 19.83297 | 16.96264 | 16.44781 | 18.24217 | 18.91702 | 20.43059 | 19.85416 | 17.10921 | 18.43458 | 19.20052 | 18.8808 |
| SNX27 | 18.15115 | 18.14773 | 17.0194 | 16.68248 | 22.29882 | 17.82466 | 16.93208 | 16.1143 | 16.03269 | 15.7387 | 18.90777 | 17.48155 | 15.61253 | 15.59786 | 17.04629 | 18.17807 | 20.50446 | 20.75536 |
| SNX3 | 17.43285 | 18.05338 | 18.30193 | 19.20532 | 20.41082 | 16.15592 | 20.19087 | 16.83501 | 19.11325 | 17.11084 | 16.61718 | 16.89799 | 19.64178 | 19.50323 | 18.87786 | 18.96444 | 16.98138 | 18.11336 |
| SNX33 | 20.21126 | 20.82524 | 22.79442 | 22.64686 | 15.31693 | 16.30147 | 16.66218 | 16.93766 | 16.59483 | 15.91252 | 16.33605 | 14.79424 | 17.4165 | 17.21377 | 23.90135 | 22.91212 | 16.29029 | 15.58824 |
| SNX4 | 14.82814 | 16.35841 | 15.98887 | 14.179 | 20.12858 | 21.06839 | 18.71754 | 18.00829 | 17.55856 | 18.29234 | 13.61494 | 13.65378 | 18.29098 | 17.32363 | 16.79902 | 16.27128 | 15.29409 | 18.53472 |
| SNX5 | 16.95493 | 16.06396 | 16.45791 | 17.51733 | 14.88847 | 16.7918 | 15.87703 | 16.98337 | 16.50925 | 16.70088 | 17.26789 | 18.26147 | 18.10575 | 18.36632 | 18.06616 | 17.02909 | 15.97521 | 16.94619 |
| SNX6 | 17.3807 | 16.8934 | 16.77543 | 17.12131 | 20.60529 | 18.69723 | 18.76606 | 18.36313 | 16.37968 | 16.70603 | 18.79872 | 18.402 | 19.18879 | 19.41925 | 17.53803 | 16.67822 | 18.59852 | 19.03171 |
| SNX9 | 23.47249 | 22.97222 | 19.1201 | 19.55008 | 17.92551 | 17.00881 | 15.92382 | 16.45345 | 16.6128 | 16.61706 | 18.25032 | 19.16052 | 20.09298 | 17.80295 | 19.5824 | 20.40196 | 15.91111 | 15.86725 |
| SOAT1 | 16.72308 | 14.20917 | 15.48182 | 14.08049 | 17.93575 | 15.09289 | 15.39963 | 16.70275 | 19.97147 | 15.96991 | 17.61129 | 10.16324 | 17.84595 | 17.9571 | 15.09282 | 15.75478 | 17.17687 | 18.84752 |
| SOD2 | 16.61277 | 16.94275 | 16.57036 | 16.05873 | 17.0391 | 16.45776 | 17.40451 | 17.1199 | 20.91654 | 19.46924 | 17.18697 | 13.05701 | 15.4617 | 16.792 | 15.5317 | 15.27352 | 17.00849 | 17.25496 |
| SOD3 | 17.82117 | 17.56971 | 16.49905 | 15.39566 | 16.42711 | 18.64452 | 18.67755 | 17.70286 | 21.45012 | 19.84511 | 15.66111 | 15.05073 | 15.56157 | 15.4445 | 17.47723 | 16.07712 | 15.12645 | 17.57889 |
| SOGA1 | 15.03895 | 16.91212 | 15.51651 | 15.82979 | 15.03667 | 15.7 | 14.64133 | 15.79835 | 12.28731 | 16.29283 | 15.26375 | 14.28236 | 19.3906 | 19.26796 | 15.58311 | 16.58063 | 16.32802 | 16.22783 |
| SORBS3 | 20.25307 | 20.39188 | 15.80919 | 13.99803 | 15.422 | 15.91832 | 16.43534 | 16.60505 | 13.45413 | 15.69814 | 15.75476 | 13.57896 | 15.01775 | 16.09428 | 16.06277 | 16.63473 | 18.13286 | 13.54937 |
| SORCS2 | 16.47472 | 19.54173 | 16.49714 | 18.68044 | 16.00994 | 15.92844 | 17.75436 | 16.89223 | 20.06688 | 18.72406 | 15.90709 | 14.45188 | 15.88992 | 18.11569 | 15.72421 | 16.84071 | 17.01559 | 17.68355 |
| SORD | 16.96583 | 17.85878 | 20.1271 | 21.41297 | 19.50264 | 17.08014 | 17.06705 | 17.35299 | 16.89156 | 17.74629 | 20.16512 | 20.34215 | 21.69462 | 21.05927 | 20.97666 | 20.75625 | 18.71463 | 18.02505 |
| SORL1 | 17.76404 | 16.31567 | 16.12683 | 15.47497 | 15.81776 | 16.22847 | 16.18397 | 16.50809 | 19.26543 | 17.31729 | 15.74924 | 13.8879 | 15.66057 | 15.03706 | 16.78137 | 15.51334 | 17.4646 | 14.3139 |
| SORT1 | 15.73746 | 16.50098 | 17.51758 | 16.34181 | 14.1284 | 17.67248 | 16.7842 | 17.70562 | 20.54333 | 19.58315 | 19.20065 | 18.07001 | 16.11297 | 16.2889 | 16.9178 | 15.54462 | 16.5501 | 15.72084 |
| SPAG1 | 17.5648 | 17.29873 | 16.76384 | 16.73988 | 20.15199 | 18.92592 | 16.90947 | 16.78111 | 15.40101 | 16.33537 | 14.56475 | 12.10111 | 18.70448 | 18.79645 | 15.62906 | 16.2761 | 20.06722 | 20.22066 |
| SPAG9 | 16.45231 | 16.00944 | 15.94604 | 15.71026 | 17.3222 | 16.41652 | 17.41373 | 17.73411 | 15.61903 | 15.56819 | 17.24552 | 17.50027 | 17.09623 | 17.17597 | 15.47002 | 16.01399 | 18.27391 | 15.59981 |
| SPART | 13.30753 | 17.22233 | 17.52164 | 18.11702 | 13.87861 | 14.82439 | 16.14495 | 15.06088 | 12.73608 | 14.84749 | 16.316 | 16.8835 | 14.12611 | 16.70547 | 20.60991 | 17.61659 | 13.50322 | 14.51427 |
| SPAST | 16.91665 | 17.09801 | 18.10008 | 17.88758 | 20.24637 | 19.01266 | 16.59619 | 16.21388 | 15.62218 | 17.80615 | 17.17341 | 15.78862 | 17.46956 | 19.22502 | 17.52573 | 15.63509 | 16.57687 | 17.60589 |
| SPATS2L | 15.2211 | 16.55302 | 15.59023 | 14.88421 | 14.43642 | 14.75119 | 16.01427 | 18.02943 | 15.71081 | 16.87533 | 13.92504 | 18.41104 | 16.02117 | 14.22051 | 15.29742 | 16.00607 | 15.50632 | 14.31619 |
| SPC24 | 17.76276 | 14.96224 | 16.06964 | 18.87703 | 18.55758 | 17.1742 | 18.17876 | 16.29285 | 15.94472 | 17.0264 | 15.21766 | 17.9198 | 20.14433 | 17.70516 | 17.83542 | 16.93684 | 16.87306 | 18.18918 |
| SPCS1 | 17.18309 | 17.15072 | 17.76445 | 15.49337 | 17.60875 | 20.01722 | 16.0484 | 17.19015 | 21.16482 | 19.64018 | 19.95872 | 17.17139 | 16.29409 | 15.01549 | 16.80664 | 14.60564 | 16.23294 | 18.94025 |
| SPCS2 | 17.02598 | 17.15768 | 15.9915 | 18.354 | 16.92615 | 17.99104 | 18.6101 | 18.28403 | 21.19857 | 21.64335 | 16.88598 | 16.89216 | 18.44481 | 19.97382 | 16.23821 | 16.53114 | 16.20521 | 16.63255 |
| SPCS3 | 15.73879 | 16.86545 | 14.61014 | 15.7158 | 16.43956 | 15.54912 | 16.12634 | 15.90339 | 22.17205 | 20.19039 | 20.23481 | 18.4631 | 15.79238 | 16.07901 | 15.11591 | 15.16436 | 16.03815 | 15.14178 |
| SPECC1 | 15.12625 | 15.73127 | 14.40239 | 17.31793 | 17.27451 | 15.78492 | 14.02872 | 16.19151 | 17.71509 | 17.16245 | 15.40192 | 14.36329 | 17.45695 | 18.2873 | 16.61972 | 15.09316 | 15.99884 | 17.42157 |
| SPINT1 | 18.91632 | 20.29352 | 19.66184 | 20.09855 | 16.73766 | 17.20506 | 16.01975 | 15.58157 | 18.82731 | 16.92027 | 17.54529 | 15.51689 | 17.90663 | 18.13052 | 19.72476 | 19.35254 | 16.73405 | 14.64196 |
| SPINT2 | 18.53379 | 17.88545 | 18.56243 | 19.26546 | 19.39696 | 19.15311 | 17.48034 | 16.20335 | 23.69122 | 22.214 | 17.25458 | 14.92078 | 18.33563 | 17.63644 | 17.53158 | 17.78887 | 16.15359 | 18.14678 |
| SPN | 17.17675 | 17.63315 | 14.37955 | 18.32949 | 18.23063 | 14.85339 | 20.61881 | 16.37762 | 15.90037 | 12.50345 | 16.70602 | 21.02236 | 19.29535 | 19.3809 | 16.88156 | 17.24832 | 18.25658 | 17.12771 |
| SPNS1 | 16.98405 | 14.82768 | 15.72197 | 22.13882 | 19.75113 | 24.962 | 20.24655 | 15.66283 | 15.7297 | 12.85956 | 14.4198 | 13.5978 | 18.9826 | 21.186 | 16.5687 | 17.14338 | 20.08076 | 19.9901 |
| SPON2 | 16.49233 | 16.34595 | 23.5249 | 22.81386 | 14.99337 | 16.30918 | 17.17363 | 16.09459 | 13.83221 | 15.45138 | 16.18242 | 13.71335 | 20.62419 | 21.43366 | 22.62415 | 21.87842 | 16.56862 | 15.5364 |
| SPPL2B | 14.55153 | 17.32711 | 15.25215 | 15.72721 | 12.41027 | 15.12726 | 15.60454 | 15.22136 | 20.13594 | 16.62278 | 15.22657 | 17.91768 | 13.88687 | 15.83168 | 16.85899 | 16.55706 | 15.2388 | 14.13328 |

| | | | | | | | | | | | | | | | | | | |
|---|---|---|---|---|---|---|---|---|---|---|---|---|---|---|---|---|---|---|
| SPR | 15.6104 | 15.16877 | 14.6176 | 15.57576 | 18.52895 | 16.41974 | 18.61788 | 17.27796 | 16.73744 | 16.93217 | 15.59573 | 16.52847 | 18.87208 | 18.6306 | 17.00041 | 17.47372 | 18.31461 | 18.46402 |
| SPRED1 | 16.81609 | 14.62899 | 16.31134 | 18.83678 | 15.3285 | 15.64894 | 15.79747 | 14.75146 | 13.34112 | 15.26186 | 16.45086 | 15.32458 | 15.46683 | 15.52011 | 16.10456 | 15.14928 | 18.65719 | 18.86366 |
| SPRED2 | 15.85853 | 15.30458 | 16.31524 | 16.9863 | 17.33875 | 16.02358 | 16.2109 | 17.07832 | 13.97243 | 14.44887 | 15.0022 | 12.05225 | 12.87398 | 18.43306 | 17.49038 | 17.65739 | 20.29283 | 19.85923 |
| SPRY4 | 15.19276 | 15.696 | 16.46286 | 14.84821 | 13.30623 | 15.3566 | 17.49927 | 15.97434 | 15.82651 | 14.68392 | 15.24509 | 13.98855 | 14.75158 | 13.17934 | 13.44193 | 13.40231 | 15.81026 | 20.87156 |
| SPTAN1 | 15.88898 | 16.66688 | 17.09669 | 15.79676 | 17.45212 | 17.54903 | 16.24536 | 16.79997 | 16.67126 | 16.94725 | 18.20819 | 19.49945 | 16.52728 | 15.85122 | 17.36574 | 16.84271 | 17.86359 | 16.46815 |
| SPTBN1 | 16.93592 | 16.53985 | 16.22912 | 15.62248 | 17.87027 | 17.52512 | 16.95876 | 16.71881 | 16.59079 | 16.35233 | 17.6095 | 19.41353 | 15.5278 | 16.00656 | 15.44523 | 16.58105 | 17.25882 | 16.93699 |
| SPTBN2 | 16.42685 | 16.28656 | 18.28575 | 17.10136 | 15.5429 | 16.65077 | 15.84194 | 17.6369 | 14.35813 | 16.02337 | 15.72653 | 17.87196 | 17.29247 | 17.94131 | 19.02004 | 18.14173 | 16.56535 | 15.7879 |
| SPTLC2 | 13.89598 | 15.30025 | 17.40659 | 16.06729 | 15.89708 | 18.00068 | 16.25113 | 15.37789 | 14.87518 | 16.24623 | 15.36197 | 16.99155 | 14.88033 | 15.82904 | 13.9095 | 14.11923 | 15.51105 | 15.72689 |
| SQOR | 18.64722 | 16.05659 | 15.53503 | 16.73152 | 14.5377 | 15.92214 | 16.35811 | 17.18165 | 20.61491 | 19.20086 | 14.91252 | 14.17829 | 17.73675 | 17.01247 | 15.50527 | 15.49389 | 15.50239 | 16.71321 |
| SQSTM1 | 19.52463 | 19.34202 | 20.32979 | 20.38872 | 21.24232 | 19.85852 | 19.67003 | 17.87731 | 20.58128 | 19.84233 | 16.30722 | 16.38905 | 20.80999 | 20.28042 | 18.81738 | 19.30345 | 20.77956 | 21.10385 |
| SRA1 | 15.95575 | 14.53868 | 14.32886 | 17.9394 | 17.5667 | 13.62818 | 15.76827 | 15.3595 | 16.97499 | 14.6037 | 17.18911 | 16.5306 | 17.93741 | 18.77212 | 13.77244 | 15.56134 | 17.52734 | 20.06541 |
| SRC | 19.29743 | 29.61972 | 20.77558 | 20.53156 | 16.7617 | 18.54249 | 22.01659 | 20.65049 | 18.03462 | 17.64586 | 17.30343 | 16.76873 | 15.57673 | 17.09001 | 28.94157 | 29.26051 | 20.90775 | 21.02639 |
| SRCIN1 | 16.49122 | 16.23483 | 16.06252 | 16.49343 | 18.44788 | 13.70675 | 16.21573 | 15.69092 | 13.3984 | 13.56764 | 16.60528 | 18.00207 | 14.35789 | 14.21646 | 12.60223 | 13.65394 | 14.88541 | 17.70908 |
| SRGAP1 | 16.2786 | 17.53385 | 15.33609 | 16.22436 | 18.95747 | 20.86889 | 18.96773 | 17.76625 | 15.84157 | 17.4127 | 21.12789 | 17.97443 | 17.16827 | 16.91763 | 15.0642 | 15.52428 | 18.3159 | 17.65479 |
| SRGAP2 | 16.6561 | 16.95364 | 16.15075 | 17.48453 | 20.04509 | 18.47993 | 17.61816 | 17.51117 | 16.5371 | 17.21405 | 17.10927 | 18.49998 | 19.50307 | 18.78983 | 17.46874 | 17.16009 | 18.37326 | 18.79331 |
| SRI | 20.57781 | 20.8279 | 19.70669 | 20.52835 | 18.70656 | 17.3196 | 19.72799 | 18.28758 | 22.2671 | 19.89589 | 21.11956 | 20.37489 | 18.12176 | 19.27251 | 20.59391 | 20.31969 | 18.06406 | 19.10046 |
| SRM | 19.80072 | 17.98008 | 18.8742 | 19.01087 | 18.52618 | 17.19153 | 18.04208 | 24.45962 | 19.81999 | 18.83192 | 18.68732 | 19.76129 | 20.15042 | 26.58304 | 18.50575 | 19.45789 | 20.3824 | 19.14194 |
| SRP14 | 22.47956 | 22.46745 | 21.90115 | 23.28561 | 19.76976 | 21.01656 | 20.31252 | 19.9381 | 17.11601 | 17.4612 | 20.0056 | 20.23857 | 19.35298 | 19.6067 | 24.13512 | 23.76589 | 18.7042 | 20.55657 |
| SRP19 | 17.53064 | 17.00806 | 16.48446 | 21.27464 | 22.24238 | 21.02438 | 21.5676 | 21.30224 | 15.11195 | 16.26586 | 16.25383 | 18.8249 | 23.00484 | 22.99344 | 19.67081 | 18.82897 | 19.27368 | 19.5996 |
| SRP54 | 15.69858 | 14.80082 | 15.95528 | 16.70324 | 17.90865 | 19.98557 | 18.38385 | 17.88222 | 15.14977 | 16.75461 | 17.38125 | 18.44536 | 18.37841 | 17.92468 | 16.83375 | 17.66291 | 16.56853 | 18.25766 |
| SRP68 | 17.01256 | 16.08212 | 17.68537 | 16.10638 | 15.31097 | 16.32836 | 17.78104 | 16.45212 | 14.63079 | 16.30024 | 18.3005 | 19.33606 | 16.89526 | 15.37853 | 16.17452 | 15.0984 | 17.58825 | 16.16499 |
| SRP72 | 16.68277 | 15.54053 | 17.13039 | 16.06099 | 17.51008 | 17.63995 | 19.06247 | 18.52424 | 14.00205 | 16.59845 | 17.11576 | 18.82654 | 18.05832 | 17.47487 | 15.86569 | 15.58755 | 18.10471 | 17.78293 |
| SRP9 | 23.39568 | 21.88654 | 21.77393 | 22.70616 | 19.84952 | 18.59975 | 21.24413 | 20.1196 | 16.74602 | 18.89887 | 19.28459 | 20.08135 | 20.26114 | 19.81777 | 23.08797 | 23.23725 | 20.08734 | 19.34399 |
| SRPK1 | 16.82806 | 16.40445 | 16.61125 | 17.37955 | 16.21084 | 16.68456 | 14.59566 | 15.46067 | 15.32736 | 16.33731 | 16.47715 | 19.13039 | 17.00462 | 16.58875 | 14.96236 | 15.31969 | 16.2385 | 16.0523 |
| SRPRB | 19.43339 | 21.20539 | 16.90748 | 16.77831 | 15.96973 | 16.77118 | 17.80981 | 15.54872 | 18.0865 | 16.18188 | 17.7016 | 16.93113 | 18.08647 | 19.3116 | 17.86396 | 17.22536 | 16.546 | 15.12965 |
| SRRM2 | 15.00574 | 14.93899 | 13.98952 | 16.43137 | 12.71251 | 15.22492 | 14.0045 | 15.98347 | 15.00292 | 15.21015 | 18.34785 | 17.86893 | 17.27006 | 14.96121 | 15.72885 | 14.59601 | 17.28108 | 13.33304 |
| SRRT | 16.15791 | 14.41298 | 14.8565 | 15.1611 | 18.72377 | 15.8397 | 15.87019 | 16.94351 | 15.09817 | 16.46005 | 17.73712 | 18.42666 | 17.05768 | 14.33037 | 15.40337 | 16.06775 | 15.23676 | 17.67036 |
| SRSF1 | 20.38924 | 20.38857 | 18.96974 | 18.2821 | 15.40741 | 16.29578 | 16.58927 | 16.54272 | 13.19318 | 15.82804 | 15.68285 | 16.90509 | 17.77022 | 16.79216 | 19.04072 | 19.0015 | 16.35017 | 15.32138 |
| SRSF3 | 22.38637 | 22.8945 | 21.2011 | 20.82002 | 16.98764 | 19.42604 | 17.71062 | 17.97552 | 16.77082 | 16.37426 | 19.17791 | 19.89291 | 18.49121 | 18.55368 | 21.21755 | 20.95632 | 16.55572 | 16.40292 |
| SRSF6 | 19.37313 | 18.90668 | 17.80495 | 17.82635 | 15.53379 | 15.43112 | 20.6294 | 16.53276 | 16.65076 | 16.41159 | 17.81696 | 18.57907 | 18.55589 | 17.29558 | 17.86579 | 18.15931 | 14.33531 | 15.09845 |
| SRSF7 | 22.45208 | 22.47592 | 20.21234 | 18.42052 | 13.84974 | 16.58586 | 15.58756 | 16.64934 | 13.41637 | 13.17199 | 17.10413 | 19.08181 | 14.86539 | 14.39597 | 20.98603 | 19.99855 | 14.54983 | 14.80984 |
| SRSF9 | 19.96236 | 20.39929 | 17.45113 | 18.79405 | 15.27356 | 16.01007 | 17.03809 | 16.92957 | 14.34516 | 16.00466 | 17.8513 | 14.64701 | 16.19546 | 14.74633 | 19.10645 | 18.83706 | 18.34487 | 16.69317 |
| SRXN1 | 17.23716 | 17.48013 | 17.26735 | 19.607 | 19.36135 | 17.92098 | 16.41411 | 19.03684 | 12.1226 | 15.72858 | 14.27912 | 13.12938 | 17.37314 | 17.10775 | 17.66691 | 18.41094 | 15.68973 | 14.09762 |
| SSB | 16.83142 | 16.28858 | 16.25521 | 17.30528 | 15.95003 | 17.2496 | 15.75317 | 16.71355 | 15.44146 | 16.83915 | 18.861 | 19.10592 | 16.10269 | 17.40725 | 17.62857 | 15.63772 | 15.84369 | 15.73429 |
| SSBP1 | 16.24346 | 15.93064 | 15.84987 | 16.50101 | 15.42069 | 15.84557 | 15.15455 | 17.59991 | 20.68271 | 20.05711 | 17.52348 | 19.03944 | 16.10219 | 16.10819 | 15.69454 | 14.96799 | 16.05653 | 15.74839 |
| SSR1 | 13.9108 | 14.2955 | 19.13156 | 15.92968 | 14.69101 | 14.76214 | 13.99496 | 17.10171 | 22.74243 | 20.68023 | 19.05268 | 16.27757 | 15.37406 | 12.97429 | 18.09972 | 17.45391 | 15.48835 | 13.27942 |
| SSR4 | 17.3048 | 15.56852 | 17.43843 | 16.54349 | 15.91582 | 16.99353 | 15.61476 | 17.12019 | 23.19152 | 21.88425 | 20.51214 | 18.49399 | 15.27396 | 15.27813 | 16.19074 | 16.23334 | 16.79945 | 15.23993 |
| SSRP1 | 22.30014 | 15.19788 | 16.12893 | 18.6322 | 17.73001 | 16.75304 | 18.90187 | 18.08661 | 16.14556 | 16.58635 | 15.7456 | 18.44394 | 14.9007 | 18.17551 | 15.04981 | 16.94395 | 16.33372 | 19.50405 |
| ST13 | 16.63022 | 16.64655 | 17.32173 | 15.41314 | 15.97263 | 16.88149 | 18.35673 | 17.50114 | 17.71878 | 17.63683 | 17.58032 | 18.61145 | 19.92531 | 20.62157 | 16.72591 | 17.93871 | 18.25713 | 18.9002 |
| ST13P4 | 19.9659 | 18.8227 | 20.36862 | 20.68517 | 19.64159 | 18.05424 | 21.49611 | 18.59004 | 20.2119 | 20.87428 | 21.49892 | 21.32579 | 22.32583 | 22.31454 | 19.74299 | 19.76048 | 20.33556 | 19.42513 |
| ST14 | 20.4935 | 20.81932 | 22.05661 | 21.88567 | 16.19306 | 16.53944 | 14.97327 | 17.32816 | 18.81217 | 16.61849 | 17.71293 | 16.32174 | 19.46555 | 18.31939 | 22.28172 | 21.95186 | 16.1619 | 15.56166 |
| ST3GAL1 | 16.0128 | 15.63006 | 17.8851 | 18.53496 | 17.87502 | 16.48825 | 17.43156 | 17.18126 | 14.55977 | 15.05514 | 15.30053 | 12.10267 | 15.97149 | 14.29775 | 12.73793 | 17.81997 | 15.51934 | 19.22425 |
| ST3GAL4 | 18.58281 | 15.85543 | 15.97427 | 16.19023 | 14.79338 | 16.58218 | 14.79757 | 16.54961 | 18.05331 | 15.71025 | 16.79306 | 16.6213 | 15.23776 | 16.14655 | 16.22061 | 14.05432 | 16.28529 | 14.95215 |
| ST6GALNA | 16.95142 | 17.69892 | 15.33467 | 15.34928 | 15.42334 | 17.2686 | 16.32998 | 16.29627 | 19.59516 | 16.32265 | 14.7129 | 15.46936 | 16.73035 | 18.48975 | 15.80545 | 17.06888 | 15.64231 | 16.11005 |
| STAM | 20.08353 | 19.42419 | 17.12103 | 18.6662 | 20.82632 | 19.01998 | 20.13379 | 19.89911 | 19.50344 | 20.14633 | 16.18597 | 15.94814 | 20.32337 | 18.64445 | 17.69334 | 17.99988 | 18.4155 | 19.69818 |
| STAM2 | 20.04111 | 19.49894 | 20.87545 | 19.20609 | 19.93571 | 23.58467 | 20.07308 | 19.1286 | 17.87156 | 18.24053 | 16.03799 | 16.48627 | 19.10636 | 19.87707 | 20.10822 | 18.03799 | 19.50339 | 21.95581 |
| STAMBP | 20.3643 | 19.77834 | 18.31642 | 18.92972 | 19.69692 | 18.83443 | 17.8005 | 17.43577 | 18.51602 | 18.52833 | 17.84961 | 18.00918 | 19.79695 | 19.42866 | 18.54452 | 18.66677 | 17.64593 | 17.90522 |
| STAT1 | 16.14173 | 16.56941 | 16.9278 | 15.17824 | 19.77307 | 19.02719 | 16.96605 | 17.1182 | 15.43639 | 16.829 | 18.94182 | 18.74644 | 17.42526 | 18.29933 | 16.63982 | 16.74623 | 18.71847 | 20.78844 |
| STAT3 | 16.39179 | 16.60937 | 17.8155 | 15.6035 | 18.26192 | 16.57426 | 16.57481 | 17.29274 | 16.66732 | 15.03558 | 15.68999 | 18.23846 | 16.93972 | 14.81661 | 18.66202 | 18.244 | 15.4395 | 18.01505 |
| STAU1 | 18.44667 | 17.44967 | 17.50434 | 17.29358 | 16.1264 | 18.36912 | 16.50794 | 16.02316 | 14.38939 | 14.54684 | 16.03956 | 17.65795 | 14.96184 | 14.81728 | 13.58701 | 17.42423 | 14.95108 | 14.80224 |
| STC2 | 15.70168 | 15.94395 | 14.48521 | 15.06442 | 12.9945 | 16.39862 | 15.57792 | 16.45755 | 18.34144 | 16.00853 | 14.50366 | 16.94258 | 14.76532 | 15.12039 | 16.07687 | 15.50316 | 14.90228 | 15.75113 |
| STEAP1 | 19.84056 | 19.15928 | 22.94264 | 23.13615 | 15.3591 | 17.89726 | 15.34565 | 16.40279 | 15.09366 | 15.25595 | 16.02382 | 14.31411 | 21.66165 | 21.38416 | 23.29593 | 22.96509 | 15.76517 | 16.61194 |
| STEAP1B | 17.65482 | 16.37171 | 21.46172 | 16.39101 | 18.94521 | 16.12463 | 14.57462 | 16.80902 | 12.33051 | 15.39832 | 14.93438 | 15.54821 | 14.32637 | 16.7072 | 22.24431 | 21.7651 | 17.73694 | 16.77186 |
| STEAP2 | 20.23953 | 20.90319 | 23.1418 | 22.3769 | 16.2164 | 15.89058 | 16.78577 | 17.72643 | 15.0206 | 15.81426 | 15.92974 | 16.08203 | 18.31974 | 18.4927 | 24.02706 | 23.63144 | 16.89031 | 16.3683 |
| STEAP3 | 17.89723 | 21.51833 | 19.7992 | 19.17702 | 16.43341 | 15.69798 | 14.76945 | 17.09265 | 17.84629 | 16.96348 | 19.46544 | 19.09773 | 17.05659 | 16.98624 | 19.91859 | 19.55344 | 14.57708 | 14.89727 |
| STIM1 | 17.63547 | 15.82186 | 15.84787 | 15.00541 | 14.83963 | 15.2743 | 16.16189 | 16.08567 | 21.40841 | 17.75282 | 18.22222 | 15.12827 | 14.87952 | 15.55465 | 16.23192 | 15.45634 | 16.27672 | 14.70492 |
| STIP1 | 17.36027 | 17.80489 | 18.96299 | 19.06578 | 20.49658 | 19.14113 | 18.92207 | 18.47839 | 19.5517 | 19.94543 | 20.63845 | 21.21696 | 21.54692 | 21.24443 | 19.27509 | 18.68493 | 21.88533 | 21.22556 |
| STK10 | 15.63313 | 15.86881 | 18.21471 | 17.53367 | 20.23685 | 18.33886 | 16.8883 | 18.48459 | 16.29336 | 17.58448 | 19.00166 | 17.95887 | 18.90042 | 19.3711 | 16.78079 | 15.94416 | 20.49139 | 20.41665 |
| STK11 | 16.67865 | 14.98004 | 16.07477 | 17.07676 | 15.44127 | 17.27377 | 15.89543 | 15.30987 | 15.72119 | 13.86607 | 14.79044 | 17.76876 | 14.55716 | 14.18309 | 17.44336 | 16.96956 | 15.94088 | 14.28627 |
| STK24 | 17.56136 | 17.764 | 18.73402 | 18.19155 | 21.55276 | 20.7255 | 18.08238 | 19.33697 | 18.46361 | 19.09226 | 20.16509 | 18.83297 | 21.04651 | 20.99167 | 18.19597 | 18.36354 | 20.18114 | 21.15859 |
| STK25 | 16.46825 | 14.68955 | 17.85633 | 18.46686 | 20.86403 | 19.70735 | 17.26247 | 14.36095 | 16.08301 | 17.24351 | 19.07249 | 18.3409 | 20.83229 | 20.56434 | 18.64935 | 18.96505 | 19.88475 | 21.16391 |
| STK26 | 15.94738 | 15.44433 | 17.11077 | 17.74279 | 20.3268 | 20.25228 | 17.29785 | 18.37119 | 15.27496 | 18.48152 | 18.72323 | 19.16587 | 21.08358 | 21.36924 | 18.33537 | 18.04564 | 20.55753 | 20.69914 |
| STK3 | 16.67342 | 16.73051 | 16.32632 | 15.84779 | 19.40699 | 16.59515 | 16.70284 | 16.28642 | 17.18937 | 17.87477 | 16.55749 | 16.59091 | 17.33305 | 16.93444 | 16.32708 | 16.84328 | 17.53496 | 17.72956 |
| STK38 | 14.11429 | 17.05077 | 16.71741 | 15.49981 | 19.1757 | 16.0354 | 15.81928 | 16.46352 | 14.7284 | 15.67845 | 15.53091 | 18.10647 | 15.50698 | 16.32425 | 15.46052 | 16.22526 | 16.63423 | 19.02976 |
| STK39 | 15.29014 | 15.43812 | 17.6524 | 17.87782 | 17.09359 | 15.2439 | 17.81768 | 17.40842 | 16.57593 | 15.58688 | 16.87589 | 16.76244 | 20.07432 | 19.12123 | 18.02092 | 18.36498 | 15.78984 | 17.6133 |
| STK4 | 15.9871 | 16.17541 | 15.76566 | 16.06286 | 16.17692 | 15.36497 | 16.48958 | 15.69418 | 15.31875 | 13.00203 | 15.94188 | 18.29855 | 15.05851 | 15.74455 | 15.31667 | 16.66511 | 16.67652 | 17.24567 |
| STMN1 | 18.04059 | 17.48831 | 18.11435 | 18.52125 | 20.73794 | 19.70757 | 21.86914 | 22.07301 | 21.87586 | 22.71541 | 21.78672 | 20.18636 | 20.55442 | 19.00242 | 17.77594 | 17.90264 | 20.77747 | 21.13515 |
| STMN2 | 18.88098 | 19.05019 | 18.50522 | 18.81588 | 21.21579 | 19.16006 | 22.41558 | 22.90762 | 20.75149 | 22.29004 | 21.54315 | 21.34942 | 19.90576 | 19.77477 | 17.92749 | 18.17468 | 21.11475 | 21.30202 |
| STMN3 | 19.13923 | 18.97164 | 18.38374 | 17.60065 | 17.55015 | 15.99533 | 21.37351 | 19.97175 | 17.40546 | 17.71786 | 19.23161 | 17.92146 | 16.89882 | 16.19305 | 19.62464 | 17.69057 | 17.8063 | 15.91201 |
| STOM | 21.55634 | 25.76663 | 21.30156 | 21.43837 | 24.02528 | 22.36632 | 22.20197 | 21.3168 | 16.03416 | 18.24294 | 17.18921 | 15.89573 | 20.75963 | 20.95894 | 21.01242 | 21.17862 | 17.98149 | 19.01003 |
| STOML2 | 16.38467 | 16.65468 | 17.13396 | 16.32281 | 16.46783 | 17.03657 | 19.11394 | 16.55315 | 20.42373 | 18.3516 | 17.32864 | 14.50778 | 15.06672 | 15.05701 | 16.42816 | 15.5222 | 14.80567 | 15.14487 |
| STON1 | 17.98583 | 18.15757 | 14.23215 | 14.52575 | 15.82186 | 16.00449 | 16.05611 | 17.65411 | 16.48734 | 14.21033 | 17.17468 | 17.21212 | 17.40618 | 16.09757 | 18.99958 | 19.7475 | 14.39486 | 16.49302 |
| STRAP | 18.61349 | 18.48295 | 18.39943 | 18.94992 | 19.82117 | 19.95494 | 19.40553 | 19.37705 | 18.28783 | 18.22734 | 19.50649 | 20.79954 | 20.81463 | 20.04536 | 18.59628 | 18.52938 | 19.38286 | 19.87998 |
| STRBP | 21.41843 | 19.66899 | 18.92518 | 16.50265 | 13.06558 | 14.90788 | 15.05303 | 17.03102 | 13.96332 | 15.03558 | 17.82335 | 17.90758 | 14.98714 | 16.64241 | 13.08259 | 16.18733 | 15.18939 | 12.0737 |
| STRN | 15.14129 | 16.05491 | 15.68805 | 15.37222 | 20.65633 | 20.1014 | 18.9626 | 16.94846 | 15.80402 | 17.18995 | 18.54223 | 18.53109 | 19.8637 | 20.65392 | 16.93541 | 15.42162 | 20.52296 | 20.40671 |
| STRN3 | 13.97939 | 15.45865 | 15.7028 | 14.17967 | 17.79242 | 17.12726 | 17.00391 | 15.16016 | 16.50645 | 15.76247 | 16.56117 | 19.05939 | 13.9221 | 15.32762 | 15.03132 | 16.05457 | 16.82402 | 14.8645 |
| STRN4 | 15.20559 | 15.91383 | 14.57147 | 15.73994 | 16.16768 | 15.7392 | 17.48064 | 19.10162 | 10.84533 | 16.54072 | 14.27377 | 18.03288 | 14.42005 | 17.47539 | 15.30636 | 14.4889 | 17.11275 | 14.55161 |
| STS | 14.78429 | 17.16352 | 16.09288 | 15.86241 | 16.10608 | 15.57304 | 16.14992 | 16.55144 | 19.77366 | 18.5137 | 14.67052 | 13.16435 | 14.42492 | 16.69028 | 13.64221 | 17.07978 | 15.97643 | 16.6285 |
| STT3A | 16.54324 | 17.10941 | 16.58852 | 15.31829 | 15.24571 | 16.81278 | 16.34601 | 16.65586 | 21.62209 | 19.58826 | 18.50106 | 17.09211 | 17.27654 | 17.17666 | 15.88394 | 16.09556 | 16.52457 | 16.08644 |
| STT3B | 16.29938 | 17.95049 | 17.51706 | 17.04756 | 15.16148 | 19.10961 | 17.14959 | 17.42612 | 21.9856 | 21.15913 | 21.6594 | 17.8767 | 15.8141 | 17.26671 | 18.33821 | 17.42563 | 16.46281 | 15.55578 |
| STUB1 | 18.01741 | 18.01003 | 18.13466 | 19.0415 | 19.82456 | 18.20842 | 20.13106 | 17.05678 | 16.98504 | 16.89121 | 18.02875 | 18.73413 | 20.92145 | 21.0762 | 18.16724 | 18.49951 | 20.17614 | 20.49572 |
| STX12 | 18.38457 | 18.61738 | 20.77776 | 20.24463 | 17.22824 | 16.00927 | 17.54892 | 16.50488 | 19.45102 | 17.40558 | 17.51579 | 17.33308 | 18.32234 | 18.30074 | 20.20989 | 20.41484 | 16.50901 | 16.4841 |
| STX1A | 17.90407 | 17.74757 | 14.32501 | 14.98021 | 14.25529 | 14.94181 | 16.40839 | 17.45876 | 18.05376 | 18.82649 | 19.27561 | 13.84405 | 13.2655 | 15.06519 | 15.37327 | 14.25082 | 16.13981 | 15.89888 |
| STX2 | 16.37676 | 16.82895 | 17.99689 | 17.29769 | 15.65896 | 17.04762 | 16.37919 | 17.14169 | 16.60285 | 15.74262 | 17.29452 | 20.47269 | 16.76519 | 16.77993 | 17.96571 | 17.84325 | 17.71946 | 15.94797 |
| STX3 | 19.42944 | 17.36737 | 20.47298 | 17.87959 | 16.07031 | 14.78981 | 16.07081 | 16.20121 | 19.1182 | 18.29947 | 17.2919 | 17.83281 | 13.60452 | 17.28992 | 20.31192 | 19.61866 | 16.65621 | 16.7944 |
| STX4 | 19.37458 | 19.18816 | 19.5937 | 19.25511 | 20.7499 | 19.94372 | 18.56158 | 17.16846 | 19.10959 | 19.06759 | 19.72383 | 20.3774 | 19.86054 | 19.33549 | 19.89828 | 19.94423 | 19.96682 | 20.44266 |

| | | | | | | | | | | | | | | | | | | |
|---|---|---|---|---|---|---|---|---|---|---|---|---|---|---|---|---|---|---|
| STX6 | 16.89195 | 19.40528 | 17.19359 | 17.35449 | 16.13379 | 16.84363 | 17.57349 | 17.93506 | 18.15368 | 15.58586 | 14.75096 | 17.62968 | 15.82648 | 15.39507 | 16.64431 | 15.76115 | 16.09069 | 19.74392 |
| STX7 | 17.29558 | 17.27653 | 18.50357 | 19.18041 | 20.01253 | 18.88035 | 17.44805 | 17.43924 | 18.10888 | 17.09368 | 18.70089 | 17.52604 | 19.92337 | 17.31495 | 16.94028 | 18.08721 | 16.50765 | 18.1593 |
| STX8 | 13.44843 | 15.22057 | 16.38603 | 17.09109 | 13.22027 | 14.63847 | 13.20706 | 17.95965 | 18.03645 | 17.15641 | 18.4743 | 18.38905 | 14.07857 | 13.29705 | 18.78069 | 17.37518 | 16.39403 | 14.46583 |
| STXBP1 | 18.55837 | 18.54942 | 17.02477 | 18.42289 | 19.15079 | 18.45169 | 21.7546 | 20.74905 | 19.21922 | 19.4625 | 17.91472 | 19.35615 | 18.68087 | 19.66438 | 17.53417 | 17.86391 | 18.75965 | 19.46179 |
| STXBP2 | 18.54255 | 18.68979 | 18.61119 | 19.83743 | 19.80619 | 18.88591 | 17.67841 | 17.37434 | 17.38377 | 17.22969 | 18.14499 | 18.69988 | 19.79957 | 19.87225 | 18.66534 | 19.11841 | 20.12338 | 19.99534 |
| STXBP3 | 18.97263 | 18.13298 | 18.9264 | 19.30839 | 20.65517 | 19.56529 | 19.53427 | 19.74766 | 18.20765 | 18.99856 | 19.32784 | 19.91178 | 20.94483 | 21.70494 | 19.74729 | 20.13745 | 21.20842 | 21.60304 |
| SUB1 | 16.44541 | 14.74012 | 16.42569 | 15.55724 | 19.43875 | 18.81623 | 17.15363 | 15.92723 | 18.10906 | 18.34646 | 20.0175 | 21.05242 | 19.69772 | 15.80288 | 16.60993 | 16.90232 | 16.07264 | 17.22983 |
| SUCLA2 | 14.8849 | 20.36195 | 15.7376 | 19.01646 | 16.64612 | 15.29826 | 16.44099 | 16.18452 | 18.4637 | 17.4105 | 15.89653 | 16.40927 | 15.32824 | 14.85038 | 19.1301 | 19.95599 | 15.57835 | 16.33926 |
| SUCLG1 | 16.17618 | 15.76711 | 16.66214 | 16.84265 | 15.64901 | 14.99914 | 20.38743 | 17.77737 | 19.81349 | 20.32749 | 15.59749 | 12.1672 | 17.56006 | 17.72439 | 14.91922 | 16.26173 | 16.80667 | 18.94323 |
| SUCLG2 | 20.89014 | 20.88635 | 20.31316 | 19.79016 | 16.51046 | 16.9079 | 17.18911 | 16.77091 | 19.70037 | 18.08696 | 16.94127 | 14.90054 | 15.56614 | 16.4173 | 20.42176 | 20.28786 | 16.18727 | 17.40055 |
| SUCO | 16.03519 | 14.13041 | 16.02145 | 15.24826 | 15.29478 | 17.95574 | 14.73901 | 16.60743 | 17.87023 | 16.81399 | 16.16152 | 15.8669 | 13.66908 | 13.48073 | 13.32335 | 15.11353 | 16.12312 | 13.4887 |
| SUGT1 | 16.78382 | 15.86326 | 21.01882 | 15.79564 | 19.12628 | 17.67208 | 15.80921 | 17.17451 | 16.32102 | 17.154 | 18.22921 | 18.31554 | 16.6139 | 16.11393 | 17.00503 | 21.36631 | 18.28769 | 17.18421 |
| SULF2 | 16.27257 | 17.29212 | 16.66404 | 15.71587 | 15.92624 | 17.21703 | 15.65817 | 17.35202 | 13.9597 | 15.0766 | 15.86702 | 14.40577 | 17.43163 | 15.21347 | 15.60173 | 15.26127 | 16.03338 | 20.47093 |
| SULT1A3 | 17.64681 | 17.93415 | 14.73769 | 18.53467 | 20.06645 | 15.0493 | 15.02499 | 14.84236 | 13.32441 | 14.4057 | 15.87637 | 14.74176 | 17.1846 | 17.57279 | 17.36673 | 16.85108 | 17.61125 | 13.29666 |
| SUMF1 | 15.77578 | 15.77825 | 14.97117 | 14.94679 | 15.06239 | 14.36465 | 16.86802 | 18.72484 | 18.89373 | 18.03431 | 17.14953 | 16.04222 | 11.32631 | 14.92382 | 15.45272 | 12.11348 | 13.09452 | 15.76667 |
| SUMF2 | 17.24076 | 16.22953 | 16.77786 | 15.75087 | 16.63462 | 16.30337 | 17.99748 | 16.93171 | 22.90175 | 21.98442 | 18.06857 | 15.78323 | 16.74915 | 16.75653 | 15.4928 | 16.75453 | 15.17538 | 16.21964 |
| SUMO2 | 19.09221 | 19.29774 | 17.71814 | 18.49595 | 15.78556 | 15.73297 | 15.23557 | 17.98227 | 16.22684 | 14.08424 | 18.35436 | 19.32429 | 17.79924 | 16.89959 | 19.7923 | 18.97021 | 19.09357 | 13.75806 |
| SUMO3 | 20.54364 | 20.69285 | 20.71068 | 20.23988 | 22.25439 | 21.14184 | 21.33401 | 20.59176 | 21.39564 | 23.24717 | 17.09838 | 21.37097 | 20.6722 | 20.28763 | 20.92561 | 20.23634 | 23.24867 | 22.38145 |
| SUN1 | 17.89348 | 17.73536 | 17.01105 | 15.2136 | 16.83936 | 15.80522 | 16.64734 | 17.16716 | 21.14385 | 17.67892 | 17.78883 | 15.40245 | 17.10229 | 14.91315 | 17.15974 | 16.7976 | 15.90624 | 14.85814 |
| SUN2 | 19.28305 | 19.5147 | 16.38538 | 19.04906 | 16.25814 | 16.80917 | 17.54851 | 17.28037 | 22.48573 | 18.66452 | 15.94806 | 15.7232 | 17.33679 | 18.03711 | 19.24541 | 18.93099 | 14.97136 | 15.33187 |
| SUPT16H | 16.24185 | 16.52457 | 15.53707 | 15.93349 | 17.00002 | 16.74476 | 19.57907 | 16.93646 | 15.30306 | 17.46986 | 17.18779 | 18.73007 | 15.65597 | 15.54735 | 15.42625 | 15.3462 | 15.58548 | 15.84843 |
| SUSD2 | 20.97015 | 20.87154 | 16.71259 | 16.53847 | 20.58647 | 19.5536 | 15.94452 | 16.74215 | 22.07638 | 20.13421 | 15.91179 | 14.08989 | 15.50365 | 16.72425 | 20.1721 | 17.24666 | 15.42324 | 16.07801 |
| SV2A | 14.41529 | 17.09657 | 17.75212 | 20.599 | 15.78324 | 16.3434 | 17.39733 | 16.68525 | 19.38093 | 18.62255 | 14.64915 | 18.38483 | 19.39548 | 18.85135 | 16.42484 | 15.53765 | 16.90394 | 18.32936 |
| SVIP | 21.59224 | 21.4439 | 20.57939 | 20.32918 | 19.83973 | 17.25194 | 20.54537 | 20.66832 | 20.27003 | 21.80332 | 19.56357 | 19.09625 | 22.06166 | 22.49067 | 20.10429 | 20.51431 | 21.17792 | 22.32261 |
| SWAP70 | 15.56048 | 17.37537 | 15.24551 | 14.41968 | 19.02859 | 17.99878 | 17.37818 | 16.22622 | 14.8611 | 16.99923 | 16.20545 | 15.57022 | 16.00605 | 15.86802 | 15.6065 | 16.43925 | 15.69743 | 18.21925 |
| SYAP1 | 15.71449 | 15.90731 | 15.41535 | 15.6965 | 17.6197 | 15.80171 | 18.01295 | 17.3016 | 14.94135 | 16.40765 | 16.07988 | 17.2689 | 19.05599 | 18.97536 | 19.49721 | 17.17135 | 19.22861 | 18.68381 |
| SYK | 16.16482 | 15.47547 | 13.57497 | 15.6244 | 16.38985 | 18.04126 | 16.88223 | 16.01372 | 16.16173 | 15.81018 | 17.32862 | 17.85556 | 15.43669 | 15.75551 | 14.29087 | 14.4645 | 15.79779 | 13.18824 |
| SYNCRIP | 17.55741 | 16.70964 | 17.6432 | 17.43729 | 17.30139 | 16.83694 | 18.83794 | 17.00037 | 17.46354 | 18.1957 | 20.09025 | 21.16136 | 19.61688 | 18.15521 | 17.46236 | 17.70805 | 16.37006 | 16.57951 |
| SYP | 20.31915 | 19.28864 | 16.69854 | 14.89613 | 12.53662 | 15.64271 | 16.67412 | 15.66036 | 22.70861 | 20.78495 | 14.21808 | 17.19052 | 13.22141 | 17.11051 | 15.87088 | 16.33389 | 17.14952 | 17.08236 |
| SYPL1 | 17.55623 | 16.74198 | 19.1961 | 17.41091 | 15.34665 | 16.30185 | 15.96436 | 16.92825 | 18.6705 | 18.66105 | 17.77755 | 20.61169 | 15.06643 | 16.34836 | 22.37939 | 21.79199 | 16.46392 | 15.29766 |
| SYT1 | 15.42662 | 16.44313 | 15.23518 | 14.70554 | 18.55773 | 14.35636 | 17.69085 | 17.73032 | 19.19909 | 20.06958 | 15.92295 | 12.38135 | 14.00374 | 16.73484 | 15.13658 | 15.04759 | 16.17903 | 16.48534 |
| SYT13 | 16.78503 | 13.94349 | 14.14589 | 16.98159 | 14.90957 | 16.96529 | 19.98091 | 14.41299 | 18.25524 | 15.41869 | 14.33559 | 13.75737 | 16.37314 | 15.38019 | 13.70818 | 13.9271 | 15.51623 | 15.07812 |
| SYT2 | 17.78559 | 15.36494 | 16.57126 | 16.58068 | 16.04546 | 15.63452 | 21.97979 | 17.04388 | 17.9223 | 17.80033 | 13.72904 | 12.00593 | 14.10403 | 16.44209 | 16.48825 | 17.21921 | 15.64088 | 14.01958 |
| SYT4 | 17.58486 | 16.79584 | 21.30869 | 21.26419 | 16.06145 | 16.70316 | 19.27806 | 17.12345 | 16.17407 | 17.17941 | 16.72561 | 16.19673 | 16.56469 | 15.70992 | 22.77842 | 22.19898 | 15.52738 | 16.42329 |
| SYT7 | 16.47552 | 16.34293 | 17.11787 | 17.31457 | 15.79881 | 14.24166 | 17.94203 | 18.20381 | 18.99373 | 18.53791 | 15.14651 | 17.79656 | 16.42455 | 14.74679 | 17.7146 | 16.93035 | 16.05844 | 14.96167 |
| SYTL2 | 17.14512 | 20.9682 | 19.6336 | 25.2666 | 22.88211 | 25.84647 | 25.41972 | 25.32135 | 11.52882 | 16.42726 | 17.60638 | 13.18529 | 22.56676 | 24.57841 | 17.94212 | 15.85859 | 22.48379 | 25.12459 |
| SYTL4 | 15.00227 | 18.95474 | 13.76813 | 15.10833 | 18.94227 | 16.47837 | 17.0192 | 16.74502 | 16.03563 | 17.1395 | 14.78328 | 13.47798 | 15.83739 | 15.31308 | 14.24347 | 16.63258 | 18.07454 | 17.56308 |
| TACC1 | 15.04039 | 14.64399 | 15.19254 | 14.6357 | 17.69834 | 20.22067 | 15.70703 | 16.70566 | 14.09875 | 14.4326 | 14.18162 | 11.56015 | 14.42494 | 15.89726 | 17.50939 | 16.0183 | 16.84373 | 19.66781 |
| TACC2 | 18.4515 | 16.05271 | 15.66795 | 15.12933 | 17.94139 | 16.67562 | 14.40835 | 16.30559 | 15.24512 | 16.16356 | 15.32103 | 15.98132 | 17.86979 | 17.79146 | 15.54569 | 14.2381 | 15.01177 | 17.10907 |
| TACSTD2 | 14.07061 | 14.76944 | 15.00516 | 17.72031 | 22.95301 | 22.24197 | 16.37597 | 16.17298 | 15.89919 | 15.69414 | 15.24175 | 13.41095 | 14.85458 | 13.49174 | 16.47549 | 14.79273 | 17.2507 | 18.47059 |
| TAGLN2 | 21.98797 | 21.70342 | 19.75354 | 21.43349 | 23.65147 | 22.9941 | 21.44619 | 20.06232 | 19.98708 | 20.99204 | 20.96192 | 20.8474 | 21.89359 | 20.99784 | 20.90339 | 20.06458 | 22.6658 | 22.71924 |
| TAGLN3 | 20.06646 | 19.65328 | 18.31939 | 18.53466 | 18.10891 | 17.73529 | 16.42638 | 15.69107 | 17.58641 | 15.60175 | 15.86737 | 18.35359 | 16.71356 | 16.5515 | 17.58552 | 17.58946 | 16.47512 | 17.50317 |
| TALDO1 | 18.40002 | 18.01559 | 18.45717 | 18.45114 | 21.0952 | 19.28739 | 18.42006 | 17.6622 | 17.89437 | 19.22439 | 21.12152 | 21.17572 | 20.72687 | 20.10467 | 19.00846 | 17.65892 | 20.44716 | 19.46614 |
| TANC1 | 15.02872 | 15.31158 | 17.65158 | 18.49777 | 16.34189 | 16.29815 | 15.33413 | 16.90067 | 13.61732 | 15.83178 | 15.74991 | 12.96612 | 18.16363 | 18.33382 | 18.33203 | 18.21104 | 16.27725 | 17.17417 |
| TAOK1 | 18.32185 | 17.95603 | 17.96378 | 19.00129 | 18.54572 | 17.74877 | 18.24422 | 18.25555 | 17.80938 | 17.46741 | 17.94124 | 18.57064 | 17.3974 | 18.04994 | 18.3972 | 18.7796 | 19.25257 | 20.36173 |
| TAOK2 | 17.04112 | 16.38402 | 18.18223 | 18.00732 | 14.45107 | 17.00865 | 15.81402 | 16.32677 | 17.80158 | 15.38224 | 15.63481 | 17.85059 | 13.26552 | 16.56487 | 18.4624 | 18.89472 | 15.22587 | 16.37601 |
| TAOK3 | 17.95791 | 18.37966 | 18.98571 | 19.46454 | 20.38043 | 16.95529 | 19.63282 | 18.94967 | 15.14364 | 17.09689 | 16.42556 | 15.98809 | 19.71533 | 20.50699 | 18.5684 | 18.77685 | 18.74866 | 18.86303 |
| TAPBP | 14.72825 | 15.08068 | 17.40408 | 16.3249 | 14.49704 | 17.78587 | 17.32266 | 17.31177 | 21.55117 | 19.00286 | 19.33895 | 16.55953 | 17.09416 | 15.98448 | 15.68819 | 15.05983 | 16.03908 | 14.86973 |
| TARDBP | 19.20727 | 18.30585 | 18.73507 | 17.73993 | 18.39052 | 16.80723 | 21.84696 | 19.75388 | 16.89958 | 18.16254 | 16.2361 | 18.69687 | 20.18209 | 19.1887 | 17.46479 | 17.50769 | 16.70439 | 20.18077 |
| TARS1 | 17.25193 | 17.11024 | 18.39465 | 18.71003 | 19.56603 | 19.31826 | 17.13705 | 16.79018 | 17.90419 | 17.4244 | 19.50418 | 20.56541 | 18.8611 | 18.69476 | 18.93047 | 19.28942 | 17.4036 | 18.37715 |
| TARS2 | 14.995 | 13.87168 | 15.79919 | 14.79786 | 13.65152 | 15.51781 | 15.06214 | 15.32097 | 16.98759 | 15.22772 | 13.37992 | 12.45639 | 14.30036 | 18.45459 | 13.41666 | 14.25977 | 15.99565 | 19.23721 |
| TAX1BP1 | 16.81275 | 16.80072 | 15.85705 | 16.63219 | 18.66852 | 19.39622 | 20.74464 | 17.72173 | 18.06094 | 18.23687 | 15.98601 | 16.10728 | 17.47763 | 16.79153 | 15.59815 | 15.74756 | 19.53927 | 20.15035 |
| TAX1BP3 | 18.79888 | 20.03657 | 20.45138 | 21.58154 | 22.94726 | 23.09959 | 19.71262 | 17.21222 | 15.3013 | 17.79809 | 20.73278 | 19.59394 | 21.34156 | 22.43864 | 21.4473 | 21.26937 | 22.38545 | 21.88014 |
| TBC1D10A | 18.50898 | 12.84449 | 18.74066 | 20.03079 | 21.05439 | 15.66512 | 18.46519 | 18.14434 | 18.13035 | 18.91709 | 13.83891 | 18.26345 | 21.88935 | 21.81204 | 17.15086 | 17.00776 | 21.18163 | 21.44463 |
| TBC1D10B | 17.52771 | 17.39166 | 17.43366 | 15.97189 | 19.51776 | 16.95204 | 16.17345 | 18.15412 | 18.67473 | 16.76371 | 18.20945 | 17.90342 | 17.36598 | 17.00924 | 19.43449 | 18.76876 | 14.62753 | 16.32196 |
| TBC1D13 | 15.372 | 14.40453 | 15.42745 | 15.40199 | 14.76498 | 15.31506 | 13.18879 | 15.89834 | 12.30499 | 13.84382 | 17.12982 | 17.35794 | 14.88699 | 13.61603 | 16.08336 | 15.86789 | 13.33047 | 14.8206 |
| TBC1D15 | 15.56392 | 16.68769 | 13.92397 | 15.94045 | 16.2777 | 16.6974 | 18.53874 | 17.76215 | 12.80655 | 16.66372 | 14.29401 | 15.52942 | 16.94167 | 15.95318 | 14.0332 | 15.66264 | 15.23973 | 14.39426 |
| TBC1D2 | 14.45258 | 17.27009 | 13.42017 | 16.05887 | 18.65339 | 18.1348 | 14.54193 | 16.21683 | 13.01349 | 14.12309 | 14.7369 | 12.30905 | 14.20541 | 14.75022 | 15.38701 | 12.56667 | 15.76307 | 14.47058 |
| TBC1D24 | 14.7029 | 15.94197 | 15.5784 | 15.86216 | 15.5136 | 18.99496 | 14.77599 | 17.29033 | 16.78515 | 14.67764 | 16.69613 | 18.06327 | 17.56277 | 16.62312 | 17.46315 | 16.72828 | 16.66755 | 17.21192 |
| TBC1D4 | 13.9885 | 15.21863 | 13.70603 | 15.35148 | 15.84379 | 15.83344 | 14.89558 | 16.23313 | 13.10425 | 14.94881 | 13.82886 | 16.73865 | 14.70523 | 13.91996 | 17.79901 | 14.19038 | 14.28242 | 13.10567 |
| TBCA | 20.72972 | 19.75978 | 20.31025 | 20.67153 | 19.55425 | 18.37402 | 15.90469 | 17.63959 | 20.28335 | 19.03286 | 20.54359 | 21.40769 | 21.19295 | 19.82269 | 20.65993 | 21.43419 | 18.36258 | 18.37628 |
| TBCB | 17.95428 | 17.08017 | 19.53061 | 18.72658 | 20.65161 | 20.43967 | 19.92326 | 18.12033 | 20.31419 | 20.35366 | 18.58729 | 19.86119 | 20.56243 | 20.18527 | 18.03781 | 18.28395 | 20.19267 | 20.18344 |
| TBCD | 18.57943 | 14.69651 | 18.1782 | 17.66273 | 14.21648 | 14.79634 | 16.61723 | 16.86555 | 15.6534 | 14.53927 | 15.88966 | 19.18412 | 15.43445 | 15.4848 | 18.74128 | 18.62772 | 16.48701 | 15.82917 |
| TBCE | 16.35271 | 15.32091 | 16.52256 | 15.90939 | 17.42759 | 16.57015 | 16.74335 | 17.3701 | 14.27576 | 16.02747 | 17.94784 | 19.14002 | 16.75669 | 16.48695 | 16.17063 | 15.5747 | 18.74055 | 17.82063 |
| TBK1 | 14.803 | 14.70268 | 14.47343 | 15.61339 | 16.83757 | 15.57565 | 16.20658 | 17.0165 | 17.15806 | 14.87085 | 16.9443 | 12.80196 | 16.49038 | 12.70488 | 14.15947 | 15.79041 | 16.21528 | 14.77969 |
| TBL1XR1 | 16.0852 | 15.71557 | 14.9189 | 16.52499 | 16.8846 | 15.93762 | 16.0214 | 16.57977 | 16.85236 | 12.58528 | 18.20586 | 18.39252 | 16.99831 | 14.7135 | 14.95898 | 15.87564 | 17.9811 | 16.38512 |
| TBL1Y | 16.50514 | 16.33703 | 17.27573 | 18.39555 | 17.07602 | 15.462 | 15.20061 | 18.51 | 18.34379 | 16.46323 | 17.56537 | 19.62461 | 18.53855 | 17.53315 | 17.72891 | 17.74064 | 16.87954 | 16.97852 |
| TC2N | 16.21387 | 15.52515 | 18.02151 | 19.05967 | 14.52425 | 18.88995 | 14.99852 | 17.06565 | 12.86977 | 14.8159 | 15.1101 | 12.69044 | 13.16294 | 15.29102 | 18.16604 | 19.18355 | 15.76201 | 14.14145 |
| TCAF1 | 18.69595 | 19.3632 | 16.02698 | 16.02998 | 14.13879 | 13.29052 | 14.25205 | 17.5671 | 11.98952 | 14.35897 | 14.5689 | 15.92079 | 15.01332 | 15.13775 | 16.55243 | 18.20048 | 16.66659 | 14.97565 |
| TCEA1 | 18.3702 | 14.76637 | 16.98173 | 17.0111 | 19.24956 | 17.23401 | 17.04914 | 17.78147 | 17.29545 | 18.62067 | 17.03084 | 17.19468 | 17.28263 | 17.97165 | 18.10929 | 17.42147 | 17.1892 | 14.82931 |
| TCF25 | 18.2639 | 19.66345 | 18.3105 | 17.972 | 14.99915 | 16.57162 | 15.94913 | 17.0343 | 14.36794 | 14.32907 | 15.14141 | 13.94943 | 14.70954 | 14.64604 | 19.40706 | 18.79699 | 15.92125 | 14.54673 |
| TCP1 | 17.85278 | 17.94432 | 18.37131 | 18.36065 | 18.84138 | 17.68078 | 19.27931 | 18.265 | 18.54636 | 18.95399 | 20.36371 | 21.5883 | 20.18552 | 20.14282 | 18.90898 | 18.54222 | 19.63643 | 19.62709 |
| TECR | 15.07615 | 14.40573 | 15.38014 | 17.65222 | 18.49544 | 17.41583 | 20.25311 | 17.73129 | 17.76808 | 17.8833 | 15.30869 | 14.29362 | 19.1102 | 18.93846 | 14.75869 | 15.71528 | 14.95977 | 15.2373 |
| TENM4 | 14.73658 | 14.76354 | 14.61528 | 15.2212 | 13.70759 | 14.47282 | 14.40628 | 15.46287 | 14.60826 | 12.99914 | 15.48226 | 12.59857 | 14.27784 | 13.95511 | 15.34992 | 14.92276 | 17.67765 | 17.39569 |
| TEP1 | 17.42971 | 17.05391 | 15.63811 | 15.7407 | 15.84937 | 15.51173 | 15.0751 | 16.06515 | 15.43591 | 14.91048 | 17.62757 | 18.57037 | 13.9712 | 15.7659 | 15.88136 | 16.51759 | 14.23311 | 16.53653 |
| TES | 16.92565 | 16.18056 | 16.4905 | 18.67836 | 20.07088 | 18.55869 | 17.31922 | 17.48128 | 16.43648 | 18.24446 | 16.60109 | 18.33922 | 20.16949 | 19.72702 | 18.0385 | 18.04907 | 20.26546 | 21.24103 |
| TEX10 | 16.05269 | 15.92885 | 14.8499 | 15.00735 | 15.02903 | 17.29132 | 14.2701 | 15.62916 | 10.82176 | 13.5895 | 14.57132 | 16.08387 | 14.15912 | 14.67015 | 13.79222 | 14.53858 | 14.98017 | 14.18336 |
| TFAM | 17.52471 | 16.70118 | 19.54693 | 19.23261 | 16.86997 | 16.84612 | 15.45694 | 17.72168 | 19.24336 | 16.72997 | 17.98673 | 15.90026 | 12.98566 | 12.40232 | 18.92831 | 18.43706 | 17.87423 | 16.69866 |
| TFAP2A | 20.18409 | 21.05184 | 14.10322 | 16.26327 | 13.82171 | 14.66895 | 15.57892 | 18.89523 | 12.73032 | 14.41666 | 13.30114 | 12.43769 | 15.81125 | 15.34339 | 16.78654 | 17.11014 | 17.13855 | 16.91204 |
| TFCP2 | 15.48084 | 16.59712 | 16.71162 | 15.16591 | 14.78785 | 16.25464 | 14.517 | 14.73964 | 14.43818 | 15.05188 | 16.43226 | 17.27217 | 15.4948 | 16.2455 | 15.74653 | 16.92999 | 15.01006 | 14.16578 |
| TFF3 | 13.22588 | 15.12906 | 15.47511 | 15.06333 | 13.97663 | 14.8999 | 14.23801 | 14.20608 | 21.28506 | 20.01754 | 15.50881 | 11.61182 | 17.0739 | 14.72425 | 13.71066 | 14.84096 | 13.47311 | 14.30432 |
| TFG | 17.75926 | 21.0773 | 17.61472 | 15.80887 | 21.01469 | 17.97675 | 18.1282 | 17.22031 | 19.49878 | 19.08176 | 17.48702 | 19.31329 | 19.50032 | 19.23353 | 18.40844 | 19.19562 | 18.98445 | 20.8065 |
| TFPI | 14.93126 | 16.88558 | 20.45138 | 20.95378 | 17.10954 | 16.88763 | 16.76672 | 15.77883 | 15.5194 | 13.5636 | 16.15179 | 13.56358 | 15.48631 | 16.35491 | 14.90649 | 18.65266 | 18.38757 | 18.40136 |
| TFRC | 21.27658 | 21.5641 | 22.26321 | 22.21347 | 20.44753 | 19.92106 | 19.55513 | 17.72921 | 22.05033 | 20.98873 | 19.18615 | 19.50982 | 19.981 | 19.01602 | 21.85288 | 21.64465 | 18.65471 | 20.77053 |
| TGFBI | 16.92255 | 14.35668 | 15.97433 | 15.2369 | 18.64301 | 18.85898 | 20.48003 | 19.20722 | 16.82583 | 14.27577 | 16.83663 | 15.23889 | 15.00595 | 14.7383 | 15.55442 | 14.08039 | 16.89127 | 17.73715 |
| TGM2 | 15.48588 | 17.82548 | 16.45401 | 16.57479 | 22.39776 | 20.93776 | 18.02094 | 17.08475 | 17.72207 | 17.28386 | 17.32767 | 14.28175 | 16.9381 | 16.7221 | 16.31478 | 15.7094 | 19.10139 | 18.45293 |

| | | | | | | | | | | | | | | | | | | |
|---|---|---|---|---|---|---|---|---|---|---|---|---|---|---|---|---|---|---|
| TGOLN2 | 16.02664 | 16.78482 | 18.36138 | 15.50027 | 16.25689 | 15.93028 | 15.67778 | 16.46324 | 22.05239 | 19.66265 | 19.11888 | 16.9334 | 16.88028 | 16.31192 | 15.95264 | 16.11279 | 15.99961 | 16.52169 |
| THBS1 | 15.62494 | 14.84417 | 18.88505 | 13.79177 | 16.42903 | 16.11999 | 14.31261 | 15.81352 | 10.63562 | 12.85087 | 14.7834 | 12.689 | 14.28764 | 15.08433 | 18.00299 | 16.85307 | 15.89434 | 13.42531 |
| THBS4 | 15.30523 | 18.10628 | 20.17402 | 17.92611 | 12.48467 | 14.91154 | 14.42512 | 15.62827 | 14.88515 | 14.32978 | 14.20885 | 12.47367 | 14.09934 | 17.2067 | 17.50425 | 16.99507 | 14.76372 | 13.22312 |
| THEM6 | 26.99187 | 26.98135 | 26.49997 | 26.30867 | 18.09722 | 16.51949 | 18.71021 | 16.56004 | 20.19854 | 18.66458 | 15.85182 | 17.50983 | 20.06935 | 18.42065 | 26.64865 | 26.47697 | 18.61195 | 17.55157 |
| THNSL1 | 18.5715 | 18.79676 | 16.07151 | 19.73152 | 13.67742 | 16.11836 | 13.11857 | 15.27802 | 18.2549 | 16.62218 | 14.71223 | 15.58167 | 13.47982 | 13.37653 | 18.77419 | 15.56905 | 16.38625 | 15.1804 |
| THOC1 | 16.41013 | 13.7385 | 15.96831 | 14.66121 | 13.92089 | 14.87954 | 13.66297 | 17.30656 | 11.58044 | 13.35617 | 15.92943 | 15.20295 | 17.03243 | 16.63618 | 15.64403 | 15.301 | 14.62331 | 13.7882 |
| THOC2 | 14.48813 | 15.89298 | 16.39858 | 14.93105 | 15.01202 | 16.2625 | 16.68669 | 15.02373 | 15.20631 | 14.13517 | 16.5902 | 17.6184 | 13.08037 | 13.52045 | 14.36243 | 17.01534 | 14.51157 | 14.33742 |
| THOC3 | 17.59749 | 17.02822 | 16.08826 | 16.17156 | 18.0913 | 15.72332 | 18.24721 | 19.68764 | 17.01205 | 18.14309 | 18.56434 | 18.81256 | 21.20383 | 20.03575 | 15.24919 | 16.88873 | 20.13336 | 20.55421 |
| THOC6 | 15.59991 | 17.72068 | 17.54946 | 13.92536 | 14.80774 | 14.67191 | 14.75917 | 16.91512 | 15.4022 | 14.86258 | 16.20168 | 17.46511 | 17.94063 | 16.31993 | 15.44595 | 15.50031 | 16.33774 | 15.58472 |
| THOC7 | 14.23295 | 16.22352 | 14.94896 | 14.83111 | 11.54829 | 14.0004 | 14.61045 | 17.92022 | 16.2522 | 16.1341 | 15.37543 | 16.72342 | 13.59595 | 14.90179 | 14.87445 | 17.12313 | 17.10559 | 14.83617 |
| THOP1 | 17.01382 | 16.67663 | 17.74098 | 16.82219 | 16.48425 | 17.69369 | 15.33231 | 15.28468 | 15.61332 | 18.50145 | 19.33294 | 19.5446 | 15.75221 | 16.35991 | 17.74035 | 17.62457 | 16.86911 | 15.39977 |
| THRAP3 | 16.34626 | 14.02484 | 16.83588 | 15.15519 | 14.64167 | 15.63833 | 21.44411 | 20.42557 | 16.33961 | 16.25487 | 14.93956 | 19.02072 | 19.08702 | 15.11935 | 15.6805 | 16.18956 | 14.54824 | 14.83188 |
| THUMPD1 | 17.03302 | 15.89492 | 15.85556 | 15.9769 | 15.99374 | 14.56136 | 14.38169 | 14.60203 | 14.81834 | 13.52434 | 13.40113 | 16.82022 | 13.4936 | 12.93256 | 15.25199 | 17.8701 | 14.29443 | 15.58981 |
| THUMPD3 | 17.69111 | 18.23951 | 17.45493 | 17.11642 | 16.38207 | 15.2189 | 18.0912 | 17.49193 | 15.28896 | 18.4714 | 15.86782 | 20.79157 | 16.88869 | 16.42492 | 18.5587 | 18.32846 | 16.94383 | 15.21844 |
| THY1 | 15.46988 | 17.80398 | 15.50362 | 16.89275 | 13.21686 | 16.96404 | 16.47569 | 17.38831 | 12.51186 | 16.25151 | 12.79639 | 18.3052 | 13.98945 | 16.06242 | 15.18847 | 15.96457 | 16.49402 | 11.62517 |
| THYN1 | 14.16704 | 12.20505 | 14.60547 | 14.55845 | 12.51635 | 16.174 | 13.61833 | 13.8733 | 13.38053 | 15.20391 | 16.79955 | 17.98844 | 15.22738 | 15.78239 | 13.52237 | 15.83015 | 14.73063 | 16.75325 |
| TIA1 | 18.02222 | 17.19774 | 15.84213 | 15.65627 | 17.06251 | 19.42117 | 17.7463 | 17.3717 | 16.40605 | 16.00168 | 16.65406 | 18.17854 | 13.93921 | 17.71995 | 16.6724 | 16.06965 | 16.36049 | 17.02926 |
| TIAL1 | 17.53955 | 17.77049 | 18.61287 | 18.25786 | 18.04112 | 17.61783 | 18.19591 | 17.10288 | 16.11507 | 16.78168 | 16.9571 | 18.48172 | 19.46477 | 18.839 | 19.70503 | 18.86026 | 17.95983 | 17.26361 |
| TIGAR | 16.14326 | 15.58999 | 14.87671 | 15.65097 | 17.37864 | 16.08349 | 16.31769 | 16.75013 | 13.59222 | 15.4725 | 17.28487 | 18.36318 | 18.63713 | 16.18087 | 15.90633 | 15.95743 | 18.17234 | 15.81216 |
| TIMM13 | 14.57751 | 18.74597 | 15.71704 | 16.07461 | 14.86952 | 20.13028 | 20.21803 | 16.08767 | 16.44844 | 19.10169 | 18.64119 | 19.69569 | 21.63334 | 20.61724 | 16.45479 | 17.73954 | 17.53142 | 19.15778 |
| TIMM9 | 16.85603 | 16.42901 | 17.32871 | 15.56591 | 12.70854 | 13.7926 | 14.14022 | 16.11744 | 18.06804 | 13.89558 | 15.04963 | 16.69732 | 17.75479 | 14.16757 | 14.07723 | 16.36908 | 13.52114 | 16.206 |
| TIMP2 | 14.87704 | 15.76356 | 13.15431 | 15.42591 | 20.22739 | 21.52096 | 16.17177 | 17.88651 | 18.66531 | 19.38282 | 17.64076 | 16.06825 | 14.60958 | 16.01188 | 16.89892 | 13.60763 | 19.77005 | 20.90077 |
| TIPRL | 19.7487 | 17.33252 | 16.90229 | 20.08721 | 19.5269 | 18.56857 | 19.23473 | 18.89497 | 17.86305 | 19.76409 | 19.65281 | 20.12319 | 20.29738 | 19.33288 | 17.68732 | 18.30669 | 19.86605 | 19.15434 |
| TJP1 | 15.54593 | 16.82312 | 18.79643 | 19.54332 | 21.26933 | 19.87066 | 17.11675 | 17.53044 | 16.56185 | 17.50918 | 16.64604 | 17.00917 | 21.13146 | 20.92478 | 18.62744 | 19.17531 | 18.77989 | 18.85417 |
| TJP2 | 16.84939 | 16.42849 | 16.90133 | 17.47515 | 19.96998 | 19.5216 | 16.43413 | 16.60732 | 19.87398 | 19.86922 | 17.64905 | 17.88334 | 17.20643 | 17.51783 | 16.35008 | 16.90728 | 17.57923 | 16.16089 |
| TK1 | 16.97757 | 17.10174 | 16.91895 | 16.90072 | 17.81142 | 16.37344 | 13.5699 | 15.38267 | 14.85723 | 15.31437 | 16.22238 | 15.85222 | 16.89251 | 15.80505 | 17.84813 | 16.17989 | 18.89689 | 16.40442 |
| TKFC | 16.63862 | 16.07171 | 17.1779 | 16.96955 | 15.79022 | 17.03339 | 16.86949 | 15.79488 | 14.13788 | 15.88304 | 17.82927 | 18.93249 | 18.58161 | 17.74761 | 18.59549 | 18.32797 | 16.97097 | 16.5431 |
| TKT | 17.09864 | 17.46447 | 17.34249 | 16.98626 | 22.34985 | 20.98582 | 19.06133 | 18.48322 | 19.56254 | 20.8127 | 21.00872 | 21.07808 | 19.78466 | 18.15037 | 17.18632 | 17.87349 | 21.10323 | 21.1052 |
| TLL2 | 15.56925 | 16.68225 | 17.03431 | 15.09095 | 15.12797 | 16.02884 | 15.80518 | 17.53851 | 18.31656 | 16.02711 | 16.74905 | 14.54842 | 15.63354 | 15.29821 | 15.34555 | 15.25091 | 14.69254 | 16.24514 |
| TLN1 | 19.80624 | 20.06264 | 19.65295 | 20.26195 | 21.42938 | 20.42579 | 21.99751 | 21.05961 | 17.53014 | 18.63846 | 18.1464 | 19.26984 | 20.74522 | 21.51824 | 19.63161 | 19.61463 | 20.72693 | 20.69431 |
| TLN2 | 18.52984 | 19.36266 | 19.20612 | 20.09855 | 20.33987 | 19.79713 | 21.47229 | 20.99485 | 17.02331 | 17.43878 | 17.17665 | 18.50115 | 20.4439 | 21.29196 | 19.06038 | 19.28055 | 20.33884 | 20.32238 |
| TLR9 | 15.17124 | 14.59571 | 15.23817 | 15.45954 | 15.82007 | 14.87003 | 16.09703 | 15.33669 | 15.34831 | 16.29991 | 18.23169 | 18.0428 | 14.11711 | 16.70787 | 13.94342 | 16.01589 | 16.05799 | 13.96617 |
| TM9SF1 | 15.68578 | 18.12232 | 17.45492 | 17.39487 | 17.7911 | 16.58498 | 17.50083 | 17.28459 | 22.43161 | 20.84601 | 18.63072 | 17.61917 | 16.24106 | 15.53486 | 15.73657 | 15.81275 | 17.02096 | 15.02953 |
| TM9SF2 | 18.40758 | 17.50495 | 17.82183 | 17.3088 | 17.11559 | 16.70761 | 17.30131 | 17.01647 | 22.35476 | 20.28146 | 19.20762 | 20.22621 | 14.82832 | 16.94528 | 17.3802 | 17.41925 | 15.94286 | 16.15589 |
| TM9SF3 | 16.14619 | 16.93048 | 16.70696 | 16.19656 | 15.55963 | 17.11322 | 17.65718 | 16.77879 | 22.32218 | 20.8633 | 18.96265 | 18.03061 | 16.14531 | 16.56274 | 16.21106 | 15.37019 | 15.44118 | 14.54717 |
| TM9SF4 | 15.88778 | 15.71177 | 16.82851 | 16.22997 | 14.66446 | 17.15342 | 16.23711 | 17.3812 | 22.47996 | 20.36707 | 18.41905 | 19.39687 | 15.91314 | 17.0073 | 16.50442 | 15.70473 | 16.89215 | 15.94457 |
| TMA7 | 19.18125 | 19.18972 | 18.37589 | 19.10666 | 19.90843 | 15.29277 | 17.86749 | 18.06489 | 17.31636 | 19.39728 | 16.6933 | 16.17327 | 20.4782 | 20.48959 | 19.0283 | 19.24294 | 19.38206 | 21.02496 |
| TMBIM1 | 23.14081 | 22.94007 | 23.10869 | 20.44632 | 16.18611 | 15.2443 | 15.99263 | 15.40334 | 14.32704 | 15.87688 | 14.14916 | 14.50092 | 13.6606 | 17.07995 | 23.97053 | 23.36081 | 15.87525 | 16.65701 |
| TMCO1 | 15.31953 | 14.67167 | 15.88055 | 17.60048 | 12.66108 | 16.02661 | 16.3601 | 14.73614 | 20.10443 | 19.04053 | 14.89091 | 17.93619 | 15.55964 | 15.55827 | 17.21229 | 14.77411 | 16.20722 | 15.20432 |
| TMCO3 | 14.88851 | 14.25792 | 15.82636 | 16.20047 | 14.11742 | 16.6784 | 15.96158 | 18.52586 | 20.8331 | 19.31191 | 17.81106 | 15.67758 | 13.96352 | 16.18953 | 15.36514 | 15.64106 | 15.35799 | 16.02153 |
| TMED1 | 15.73104 | 17.1788 | 15.66414 | 16.19409 | 17.28936 | 18.28348 | 15.99703 | 17.39577 | 21.9882 | 20.78744 | 16.35416 | 15.12014 | 16.1594 | 19.20306 | 16.02155 | 16.94078 | 16.47453 | 16.30162 |
| TMED10 | 17.68997 | 17.31368 | 18.9754 | 19.2248 | 20.19681 | 18.7794 | 20.62152 | 19.98829 | 25.59995 | 24.35692 | 22.31429 | 19.43117 | 20.29622 | 21.19104 | 19.17996 | 19.25606 | 18.59764 | 21.32816 |
| TMED2 | 16.77258 | 15.79438 | 15.83237 | 15.93216 | 15.25824 | 16.90032 | 16.87653 | 16.87285 | 20.5167 | 17.74744 | 16.57968 | 15.91608 | 16.08719 | 15.00745 | 16.69181 | 16.29432 | 15.57678 | 15.02405 |
| TMED3 | 14.90405 | 18.07205 | 14.1682 | 16.04419 | 15.19782 | 15.92455 | 15.60931 | 16.51141 | 20.53796 | 19.7784 | 17.50478 | 16.62726 | 15.3651 | 15.39046 | 15.02673 | 16.01805 | 15.00405 | 14.67336 |
| TMED4 | 19.79647 | 16.03543 | 17.39638 | 20.27709 | 18.45187 | 18.49658 | 20.71611 | 18.56311 | 23.33 | 22.37869 | 19.10918 | 16.87725 | 17.95021 | 16.62011 | 18.62355 | 17.00808 | 18.61362 | 18.46248 |
| TMED5 | 17.38903 | 15.22122 | 16.31073 | 15.81627 | 14.5863 | 16.85928 | 15.17606 | 18.22395 | 23.62967 | 20.70475 | 16.44548 | 12.73211 | 15.91294 | 15.47537 | 15.24901 | 16.92082 | 14.88365 | 16.5516 |
| TMED7 | 16.60531 | 15.62323 | 16.192 | 19.14211 | 13.6953 | 14.32127 | 18.03473 | 17.20023 | 22.02768 | 20.59277 | 19.49282 | 17.42145 | 16.87887 | 15.4193 | 19.5076 | 18.76224 | 14.41103 | 15.97556 |
| TMED8 | 15.58162 | 15.16981 | 15.16474 | 13.93944 | 13.45007 | 14.85242 | 13.96452 | 15.15453 | 13.45806 | 13.69892 | 18.65387 | 18.40069 | 14.45741 | 14.93661 | 16.53585 | 13.66695 | 13.85774 | 16.38089 |
| TMED9 | 17.50121 | 18.81316 | 17.25925 | 18.34725 | 17.15416 | 16.29192 | 17.93615 | 18.49151 | 24.44391 | 21.79501 | 20.62845 | 18.49751 | 18.156 | 18.31544 | 19.081 | 18.25791 | 16.08925 | 17.88457 |
| TMEFF2 | 19.24211 | 14.80914 | 16.37509 | 16.61662 | 13.13711 | 16.03639 | 14.72441 | 17.73223 | 13.93613 | 14.28389 | 16.53856 | 12.65393 | 16.412 | 16.90608 | 14.19943 | 17.00603 | 13.52529 | 15.64439 |
| TMEM106B | 13.8549 | 15.99726 | 17.41668 | 16.08055 | 15.50419 | 15.6005 | 16.97628 | 16.59768 | 17.70626 | 18.78401 | 15.45065 | 18.39749 | 15.9361 | 17.26214 | 14.26766 | 14.99561 | 14.78074 | 16.94424 |
| TMEM109 | 15.99279 | 15.67114 | 18.81495 | 17.64129 | 14.60787 | 14.20278 | 14.67066 | 15.00788 | 22.41221 | 20.61162 | 19.56001 | 17.79344 | 12.08246 | 16.95532 | 17.39682 | 14.41959 | 16.38615 | 14.62598 |
| TMEM131 | 13.46658 | 15.61182 | 15.34552 | 14.05232 | 13.66949 | 16.99768 | 14.01251 | 16.59352 | 18.71498 | 17.22245 | 16.94912 | 12.60373 | 18.08199 | 16.85875 | 15.40812 | 15.23895 | 15.88231 | 13.9923 |
| TMEM132A | 16.70875 | 16.87412 | 16.69591 | 15.98267 | 13.80382 | 15.84145 | 15.03434 | 15.71946 | 18.93412 | 15.56379 | 14.97002 | 14.66219 | 16.15656 | 15.00736 | 16.83732 | 15.83487 | 15.75888 | 15.0147 |
| TMEM165 | 17.04293 | 19.88363 | 15.11051 | 15.74374 | 15.68641 | 17.60739 | 15.59734 | 15.7311 | 24.46329 | 21.40412 | 21.10413 | 21.34219 | 15.47631 | 16.52209 | 16.40619 | 14.9785 | 16.50982 | 16.22704 |
| TMEM181 | 17.21433 | 16.27523 | 15.56738 | 16.4841 | 18.68678 | 18.53552 | 18.16922 | 15.44465 | 21.59388 | 19.52537 | 16.91986 | 18.09254 | 15.34927 | 15.64237 | 16.88455 | 16.81873 | 18.93332 | 18.53282 |
| TMEM219 | 13.1787 | 16.53297 | 14.65721 | 15.774 | 13.61304 | 15.80385 | 16.46553 | 16.32678 | 17.50244 | 16.13057 | 14.99308 | 13.05699 | 17.03035 | 15.85508 | 13.96899 | 15.02308 | 17.02404 | 13.92063 |
| TMEM222 | 14.73301 | 16.87384 | 13.58698 | 19.45062 | 13.73933 | 15.55731 | 15.69645 | 14.35803 | 15.14463 | 13.87048 | 13.08409 | 17.3798 | 12.50382 | 13.96558 | 17.63227 | 20.5564 | 16.46159 | 14.56481 |
| TMEM237 | 15.15763 | 16.44841 | 17.9272 | 17.17713 | 21.45575 | 19.2595 | 17.7348 | 18.45291 | 17.48985 | 16.33458 | 18.47105 | 17.32945 | 18.54792 | 19.62554 | 16.3682 | 18.15541 | 20.39359 | 20.11803 |
| TMEM256 | 15.78635 | 17.29223 | 15.94066 | 15.7246 | 14.73648 | 17.76285 | 14.66861 | 16.36663 | 20.97859 | 20.36199 | 15.25613 | 12.42214 | 13.74828 | 15.34831 | 15.1535 | 15.73837 | 13.40909 | 14.3631 |
| TMEM260 | 15.38342 | 17.58925 | 15.47814 | 15.52441 | 14.20456 | 15.09592 | 16.52399 | 16.27803 | 18.93213 | 17.06288 | 13.98777 | 16.66221 | 14.58986 | 15.58994 | 15.81874 | 15.48849 | 14.5612 | 15.16346 |
| TMEM30A | 17.97787 | 17.85661 | 19.45779 | 19.31261 | 19.87996 | 18.24974 | 18.79818 | 16.3597 | 21.23883 | 20.0563 | 20.89153 | 19.79922 | 18.99966 | 16.91716 | 19.67997 | 19.02499 | 17.59358 | 17.13236 |
| TMEM33 | 16.3377 | 17.12272 | 16.09643 | 15.99073 | 14.94758 | 14.25327 | 13.45851 | 16.67305 | 19.52352 | 18.35707 | 16.88259 | 18.61929 | 13.45248 | 15.52773 | 15.30033 | 15.6137 | 14.48644 | 15.39349 |
| TMEM43 | 16.50458 | 16.49084 | 16.47239 | 15.74678 | 18.13344 | 17.45501 | 15.81608 | 16.50898 | 22.92991 | 20.66516 | 18.5065 | 16.22406 | 15.7703 | 16.09752 | 16.29063 | 16.4228 | 17.7468 | 16.74576 |
| TMEM59 | 16.35482 | 17.34865 | 15.90291 | 15.2737 | 20.20532 | 17.92981 | 21.715 | 20.03219 | 20.72287 | 20.27539 | 14.72174 | 14.06425 | 14.57891 | 17.77688 | 17.14336 | 17.11419 | 15.55458 | 18.43115 |
| TMEM63B | 16.68623 | 16.91574 | 17.46755 | 16.74598 | 13.38569 | 13.66195 | 13.32644 | 14.28599 | 16.69865 | 13.99554 | 17.73243 | 19.01031 | 14.55145 | 13.99809 | 18.37561 | 17.73126 | 15.19546 | 13.75962 |
| TMEM68 | 14.87794 | 16.95168 | 15.60461 | 15.66046 | 16.17588 | 15.77757 | 13.71512 | 18.27037 | 17.62751 | 14.95279 | 14.27307 | 15.64247 | 15.36243 | 17.50874 | 15.79038 | 13.7663 | 16.18834 | 17.19126 |
| TMEM87A | 15.86579 | 15.00636 | 16.40045 | 15.26302 | 17.06786 | 16.66371 | 15.30721 | 16.47056 | 17.63594 | 18.08757 | 17.64291 | 18.15578 | 16.83727 | 16.11782 | 16.12306 | 16.00109 | 15.53628 | 15.85934 |
| TMEM87B | 15.79296 | 16.07732 | 14.09521 | 14.76517 | 14.60774 | 15.30737 | 15.37404 | 15.72276 | 19.28813 | 13.45423 | 16.19542 | 15.79448 | 14.48836 | 12.87528 | 14.2405 | 14.86687 | 16.37733 | 14.66743 |
| TMEM9 | 16.93037 | 14.41308 | 19.03531 | 19.02032 | 11.31892 | 12.8395 | 14.28706 | 15.92339 | 19.59702 | 17.82213 | 17.68638 | 16.16844 | 15.30642 | 15.34484 | 18.38613 | 18.20995 | 15.27376 | 16.38727 |
| TMEM94 | 16.70417 | 14.9053 | 17.43678 | 15.53686 | 18.65026 | 14.30153 | 17.95657 | 18.07253 | 14.52791 | 16.06781 | 14.69055 | 17.82812 | 13.13514 | 18.35976 | 16.37473 | 16.59869 | 19.49626 | 18.58071 |
| TMOD3 | 18.07637 | 16.8294 | 17.33085 | 17.9433 | 21.4676 | 20.51966 | 17.76183 | 17.44058 | 17.19712 | 17.86909 | 17.87033 | 18.45677 | 20.22013 | 19.74284 | 17.00963 | 18.83996 | 18.6182 | 18.18027 |
| TMPO | 18.5581 | 17.19883 | 17.04365 | 15.84917 | 16.98981 | 15.14315 | 20.01181 | 17.7666 | 18.00115 | 18.29857 | 18.20786 | 18.30986 | 16.58485 | 17.13789 | 14.87263 | 14.70022 | 18.15788 | 16.40111 |
| TMPPE | 15.23595 | 17.27113 | 15.22532 | 16.89998 | 16.8176 | 16.7732 | 16.13664 | 15.33996 | 19.18134 | 17.98738 | 15.07869 | 14.00864 | 17.42481 | 16.9668 | 14.22656 | 16.03234 | 14.31479 | 14.31118 |
| TMPRSS2 | 22.07622 | 23.22145 | 20.98229 | 21.83685 | 15.61775 | 15.653 | 16.52015 | 16.07403 | 13.6078 | 17.20552 | 17.87076 | 13.42132 | 15.73337 | 15.42611 | 20.86026 | 20.79237 | 16.10466 | 15.61286 |
| TMSB10 | 23.73363 | 23.13855 | 18.72847 | 19.65856 | 24.89355 | 23.91446 | 22.05594 | 23.07621 | 22.2332 | 23.06206 | 20.19238 | 17.78044 | 20.1703 | 21.32835 | 19.15654 | 17.71973 | 26.32179 | 25.69723 |
| TMSB15B | 17.61935 | 16.70529 | 16.57707 | 16.64918 | 15.74073 | 17.89702 | 18.62613 | 20.6298 | 19.88892 | 21.03836 | 17.00451 | 15.83617 | 19.58168 | 18.26962 | 14.12723 | 16.29866 | 17.56592 | 16.83184 |
| TMSB4X | 18.07976 | 20.60768 | 19.55821 | 21.78835 | 25.10572 | 23.96351 | 25.30276 | 25.71334 | 24.32657 | 25.50976 | 20.86031 | 17.95428 | 23.34313 | 24.22424 | 18.97172 | 19.07906 | 26.54241 | 27.31358 |
| TMTC3 | 16.07026 | 16.4123 | 14.98502 | 15.17429 | 14.37751 | 18.00545 | 16.58011 | 17.49489 | 19.84135 | 17.88123 | 17.04483 | 15.26555 | 15.74979 | 15.8759 | 15.00754 | 15.98373 | 16.10181 | 14.83159 |
| TMTC4 | 17.09273 | 15.73011 | 15.31181 | 15.8301 | 14.08803 | 17.7274 | 16.2491 | 14.6964 | 18.36364 | 15.18936 | 14.07353 | 11.29373 | 14.18806 | 15.25653 | 14.53009 | 14.47488 | 15.04153 | 15.55741 |
| TMX1 | 16.05282 | 16.97405 | 19.80624 | 20.05242 | 16.58572 | 17.11771 | 16.77857 | 17.65079 | 23.77751 | 21.40268 | 20.61699 | 18.16264 | 21.06406 | 18.65336 | 16.04151 | 19.45609 | 16.54559 | 14.79684 |
| TMX3 | 16.59542 | 17.05268 | 16.86275 | 15.7438 | 16.15716 | 16.5436 | 16.70441 | 17.70965 | 21.09736 | 20.00284 | 18.64278 | 17.53963 | 18.91608 | 18.84605 | 16.83157 | 17.33849 | 18.66077 | 17.45081 |
| TMX4 | 18.56436 | 17.37346 | 15.47181 | 18.3443 | 14.57196 | 14.53838 | 19.20344 | 16.55558 | 20.65445 | 19.01471 | 17.42856 | 14.48259 | 14.07029 | 17.36723 | 17.8166 | 17.83672 | 17.14264 | 16.55567 |
| TNC | 15.7053 | 18.8208 | 15.86693 | 13.11266 | 12.97441 | 18.53782 | 17.82219 | 17.36417 | 17.12786 | 16.26658 | 15.39056 | 14.00436 | 14.68028 | 15.74423 | 17.37762 | 14.09387 | 19.71299 | 13.96537 |
| TNFAIP2 | 17.07606 | 15.12471 | 15.67265 | 15.45626 | 18.98735 | 18.97336 | 17.11523 | 15.82869 | 14.7943 | 15.5686 | 15.5288 | 18.11209 | 16.02368 | 16.60755 | 14.12512 | 14.37742 | 18.79325 | 16.76594 |
| TNFRSF10B | 17.24296 | 21.56905 | 17.2672 | 19.40537 | 20.09574 | 18.54539 | 14.7125 | 15.58262 | 17.03795 | 16.11172 | 18.35749 | 17.06995 | 18.88401 | 18.03111 | 19.97252 | 18.70887 | 17.83975 | 18.65467 |

| | | | | | | | | | | | | | | | | | | |
|---|---|---|---|---|---|---|---|---|---|---|---|---|---|---|---|---|---|---|
| TNFSF11 | 14.87568 | 15.89603 | 14.29302 | 16.54693 | 15.6905 | 15.28604 | 16.57548 | 15.86016 | 17.18934 | 13.34659 | 14.7892 | 15.07033 | 14.37458 | 15.29252 | 14.95519 | 14.13148 | 14.19783 | 14.00407 |
| TNIK | 18.27664 | 16.91922 | 17.94423 | 18.22146 | 18.23666 | 17.99531 | 17.35964 | 18.31435 | 18.11772 | 18.38168 | 18.27519 | 19.04226 | 16.89885 | 17.74758 | 18.74973 | 18.49908 | 17.47805 | 18.25338 |
| TNIP1 | 16.12908 | 15.80017 | 17.21369 | 20.99891 | 17.19779 | 16.35186 | 16.33058 | 16.51576 | 14.98119 | 17.58775 | 15.23024 | 15.02259 | 15.91643 | 15.25255 | 15.93599 | 15.14031 | 18.1689 | 18.67431 |
| TNK2 | 15.35791 | 16.51989 | 17.8719 | 17.3464 | 15.77761 | 14.58471 | 14.05435 | 14.35202 | 13.62236 | 12.04874 | 14.28649 | 15.87326 | 17.92682 | 17.52439 | 17.34537 | 17.05869 | 15.81794 | 15.69502 |
| TNKS1BP1 | 20.40314 | 20.24413 | 20.34871 | 20.01651 | 18.85838 | 19.68151 | 18.62129 | 19.72368 | 18.29221 | 17.72635 | 16.34047 | 18.51186 | 19.89546 | 20.56622 | 20.14228 | 20.31367 | 19.32329 | 19.35398 |
| TNPO1 | 16.64061 | 16.58205 | 19.1314 | 17.7448 | 15.16608 | 15.95829 | 18.77416 | 16.28571 | 12.7511 | 16.35686 | 16.8494 | 18.63913 | 17.82343 | 15.01342 | 18.30905 | 17.67436 | 16.41221 | 15.85756 |
| TNPO2 | 18.23994 | 14.63848 | 15.07756 | 16.83893 | 19.66437 | 16.46195 | 18.65336 | 19.60326 | 15.73544 | 17.24542 | 17.86629 | 18.68039 | 18.87402 | 18.18763 | 16.52777 | 16.01236 | 18.77922 | 18.11179 |
| TNPO3 | 17.49754 | 16.68668 | 16.73189 | 17.0963 | 15.22371 | 16.78942 | 15.05616 | 15.48424 | 16.8219 | 16.64742 | 14.8364 | 17.56447 | 15.6035 | 15.95122 | 15.95965 | 14.72584 | 16.90414 | 15.45055 |
| TNS3 | 15.58143 | 19.23397 | 14.9776 | 16.36354 | 19.83044 | 17.09519 | 16.99722 | 16.23403 | 12.82309 | 16.00494 | 16.97577 | 15.03156 | 14.48806 | 16.30482 | 14.94455 | 16.22429 | 20.65147 | 19.52179 |
| TOLLIP | 23.38077 | 22.9739 | 22.03771 | 22.24547 | 19.32409 | 19.54255 | 19.44523 | 17.76818 | 19.00607 | 19.51679 | 18.77315 | 18.28715 | 22.25004 | 21.89884 | 23.30818 | 23.44022 | 20.18431 | 20.47299 |
| TOM1 | 22.4753 | 21.26444 | 19.95404 | 20.83056 | 18.21473 | 17.56654 | 17.71692 | 16.17279 | 17.91208 | 17.25252 | 15.40775 | 16.02496 | 20.22096 | 19.64724 | 21.3184 | 20.64087 | 17.01996 | 18.61362 |
| TOM1L1 | 23.00014 | 23.04489 | 23.04336 | 22.28253 | 19.63571 | 19.05346 | 19.14112 | 17.78705 | 18.51295 | 19.48483 | 17.81375 | 19.38144 | 22.10858 | 21.79618 | 23.32249 | 22.65603 | 20.44089 | 20.87055 |
| TOM1L2 | 21.4202 | 21.20224 | 20.30706 | 20.58475 | 18.82288 | 18.36293 | 20.59196 | 20.52813 | 16.88819 | 16.90415 | 15.1338 | 16.40332 | 19.90074 | 19.39872 | 21.4309 | 20.97583 | 17.37387 | 18.55085 |
| TOMM34 | 16.53614 | 16.5573 | 16.35705 | 17.6445 | 17.43977 | 16.26036 | 16.48973 | 16.23854 | 16.87524 | 16.91633 | 17.81157 | 17.7642 | 18.23958 | 18.01922 | 17.52472 | 17.31623 | 19.09779 | 18.34123 |
| TOMM40 | 16.33779 | 15.60806 | 15.91055 | 17.31524 | 15.64068 | 16.3503 | 15.74717 | 16.30329 | 19.71921 | 19.20471 | 14.75424 | 13.27681 | 16.35115 | 15.40022 | 14.94844 | 15.3795 | 15.58114 | 14.95848 |
| TOP1 | 18.26653 | 17.45006 | 15.07816 | 15.14978 | 17.21637 | 17.02108 | 17.86676 | 17.95494 | 14.24102 | 14.94035 | 15.87905 | 20.00112 | 15.57617 | 15.34857 | 16.65376 | 16.43693 | 15.07748 | 16.67952 |
| TOP2A | 15.32698 | 15.91242 | 20.36982 | 14.41508 | 18.7507 | 18.2609 | 19.60281 | 19.31585 | 19.70216 | 17.8676 | 21.84019 | 16.20893 | 17.5675 | 17.90958 | 18.31203 | 15.84225 | 17.0462 | 18.49772 |
| TOP2B | 18.06164 | 16.48212 | 15.63689 | 15.88414 | 15.83206 | 16.41807 | 18.21763 | 17.70293 | 15.19853 | 16.25529 | 15.74303 | 18.07496 | 14.75713 | 16.06649 | 15.60045 | 16.61515 | 16.10208 | 15.65858 |
| TOR1A | 16.92174 | 16.44618 | 17.21389 | 15.98813 | 17.08225 | 19.89448 | 18.69459 | 18.94179 | 20.82302 | 18.83114 | 17.04093 | 14.84 | 16.25584 | 18.44061 | 16.87004 | 16.77431 | 17.24152 | 18.49035 |
| TOR1AIP1 | 16.95412 | 15.51443 | 16.34797 | 15.2296 | 13.92504 | 14.55732 | 16.09772 | 17.36356 | 21.37662 | 17.15313 | 19.04208 | 14.68431 | 14.5113 | 16.4733 | 15.6718 | 15.95693 | 18.91576 | 15.26021 |
| TOR1AIP2 | 15.88129 | 16.1877 | 16.22111 | 15.12936 | 16.8147 | 17.24553 | 17.85918 | 17.51422 | 21.46926 | 20.76969 | 18.29239 | 15.58955 | 15.29684 | 16.58482 | 15.86503 | 16.37844 | 16.16614 | 16.10825 |
| TOR1B | 15.53038 | 16.02439 | 15.71395 | 14.7964 | 16.41687 | 15.29618 | 15.17546 | 15.31449 | 19.46905 | 18.90697 | 15.64102 | 13.49272 | 15.87393 | 14.7741 | 16.13223 | 16.01919 | 15.62037 | 14.93392 |
| TOR2A | 16.0095 | 16.2867 | 14.96168 | 16.72417 | 14.34676 | 15.90441 | 16.4837 | 16.80743 | 20.90448 | 18.75588 | 16.77518 | 12.85776 | 15.02539 | 14.6705 | 15.03804 | 14.5892 | 16.62757 | 15.42953 |
| TOR3A | 14.50155 | 15.40879 | 16.3657 | 15.93835 | 16.88961 | 15.56155 | 16.48769 | 17.89516 | 18.91642 | 16.48716 | 14.95154 | 12.40899 | 14.5319 | 13.90308 | 15.74501 | 14.4949 | 15.9505 | 18.48079 |
| TP53 | 13.43891 | 16.0008 | 14.12358 | 13.99423 | 14.36726 | 14.2887 | 20.39601 | 18.53907 | 14.46304 | 13.57297 | 15.37326 | 11.30373 | 17.54654 | 13.81222 | 15.71586 | 13.52781 | 15.17979 | 13.6957 |
| TP53BP2 | 15.21871 | 15.06457 | 16.35318 | 15.33851 | 18.12537 | 16.32158 | 17.65757 | 17.15726 | 13.60414 | 14.33522 | 16.19385 | 12.6079 | 14.63078 | 17.34872 | 14.00826 | 15.16739 | 18.53619 | 18.81671 |
| TP53I3 | 13.53921 | 15.89518 | 14.36636 | 15.01821 | 20.22097 | 18.60101 | 14.50237 | 18.05915 | 17.50207 | 19.49499 | 18.33153 | 18.59591 | 19.92097 | 15.4594 | 14.96638 | 14.45176 | 20.43361 | 20.25693 |
| TPBG | 17.76437 | 18.10123 | 15.5937 | 15.46743 | 18.25924 | 18.56196 | 16.2662 | 16.05454 | 18.83421 | 17.66707 | 18.80542 | 21.84764 | 14.74537 | 16.20707 | 14.77537 | 15.58263 | 15.53661 | 15.32119 |
| TPBGL | 16.34635 | 15.06965 | 14.12453 | 14.2779 | 14.59954 | 16.84704 | 15.84464 | 15.4806 | 18.81405 | 16.1652 | 14.6533 | 15.7615 | 18.02454 | 15.5852 | 14.619 | 15.44301 | 15.16344 | 13.54804 |
| TPD52 | 16.50098 | 17.15594 | 16.78069 | 17.52718 | 20.00645 | 18.23429 | 19.27104 | 19.19966 | 19.88503 | 20.97367 | 18.37475 | 19.34877 | 19.63318 | 20.02499 | 17.23281 | 17.47478 | 19.61875 | 19.04739 |
| TPD52L1 | 15.15125 | 15.63678 | 17.39073 | 16.12172 | 16.83362 | 17.78285 | 18.05699 | 15.09114 | 13.92716 | 15.93129 | 15.03566 | 12.18356 | 19.7857 | 19.49275 | 16.81739 | 16.20439 | 17.20301 | 18.70593 |
| TPD52L2 | 15.22656 | 15.00066 | 15.85134 | 16.32912 | 19.23589 | 16.93431 | 16.11848 | 17.49428 | 16.5787 | 16.92983 | 17.03638 | 16.44643 | 18.28027 | 17.25726 | 17.66761 | 17.64526 | 18.5071 | 18.2623 |
| TPI1 | 22.44598 | 21.65767 | 21.71112 | 22.54196 | 24.87062 | 23.2401 | 23.04637 | 21.21251 | 20.74766 | 22.42674 | 23.08132 | 23.35889 | 24.87712 | 24.5428 | 22.71749 | 22.42644 | 25.47022 | 24.97482 |
| TPM1 | 16.20068 | 15.99791 | 16.40388 | 16.66027 | 20.37491 | 18.49842 | 16.08521 | 16.67178 | 15.02679 | 16.21824 | 17.37812 | 17.96324 | 18.35226 | 17.31808 | 17.88925 | 17.42982 | 16.83297 | 16.74299 |
| TPM2 | 16.57112 | 16.72806 | 17.23982 | 15.25101 | 15.76058 | 17.07737 | 14.73661 | 15.03144 | 15.7181 | 13.14589 | 17.18404 | 19.51231 | 14.75037 | 16.2554 | 16.09198 | 16.74714 | 15.47548 | 20.89425 |
| TPM3 | 19.15681 | 18.23244 | 20.00056 | 20.42909 | 23.09829 | 22.18483 | 19.18095 | 20.41146 | 19.74056 | 19.03353 | 20.84812 | 21.43001 | 21.80789 | 20.67991 | 20.19316 | 20.11184 | 21.24177 | 19.97997 |
| TPM4 | 19.11468 | 20.74484 | 17.4332 | 19.69928 | 23.1031 | 22.65877 | 18.82815 | 18.57307 | 16.7831 | 18.05564 | 19.47034 | 19.03721 | 21.57031 | 20.40068 | 18.82915 | 17.99037 | 19.98133 | 19.25161 |
| TPMT | 15.65864 | 15.99179 | 14.6593 | 16.53691 | 19.54025 | 17.33506 | 14.48778 | 16.99419 | 16.10846 | 17.67 | 13.41271 | 17.19774 | 16.96746 | 16.14055 | 14.63338 | 16.25931 | 18.95981 | 17.13008 |
| TPP1 | 16.07611 | 15.69195 | 17.78616 | 17.20189 | 13.00055 | 19.75181 | 18.49346 | 15.73086 | 19.92576 | 19.64907 | 17.29059 | 14.44364 | 16.23835 | 14.46392 | 17.73818 | 16.67354 | 15.74856 | 17.19126 |
| TPP2 | 16.85988 | 16.69831 | 16.29315 | 18.05093 | 16.66239 | 20.01073 | 18.08262 | 16.96828 | 17.01801 | 16.91046 | 18.32195 | 20.38798 | 16.89227 | 17.37819 | 17.36291 | 16.23942 | 16.37676 | 17.80561 |
| TPPP | 15.20029 | 13.51728 | 13.89716 | 20.76357 | 15.30846 | 17.39561 | 19.80289 | 16.22037 | 14.64489 | 16.26929 | 15.07104 | 12.57536 | 15.54214 | 15.09895 | 14.45767 | 14.78924 | 14.67857 | 13.67633 |
| TPPP3 | 17.26106 | 16.8885 | 15.89858 | 15.24136 | 15.13368 | 16.76073 | 21.79212 | 21.03531 | 14.21091 | 14.85641 | 15.21383 | 13.57981 | 14.17857 | 15.87959 | 15.6062 | 15.21784 | 15.93678 | 15.56201 |
| TPR | 16.91301 | 15.19776 | 16.12661 | 14.42883 | 17.6513 | 16.62141 | 16.92092 | 14.43704 | 15.30515 | 17.05488 | 17.37112 | 16.10488 | 18.55013 | 16.407 | 14.76991 | 15.3328 | 17.78123 | 14.45575 |
| TPRKB | 14.22873 | 14.49639 | 14.02715 | 15.95238 | 14.69077 | 15.87962 | 17.39786 | 15.46604 | 14.48341 | 14.87035 | 12.99717 | 18.11292 | 16.12716 | 16.31418 | 15.59273 | 15.33771 | 14.06298 | 13.09487 |
| TPRN | 14.73987 | 14.23749 | 14.59023 | 16.95933 | 19.76797 | 17.81635 | 19.31966 | 16.13099 | 12.39656 | 17.47803 | 14.802 | 10.95349 | 18.65976 | 18.0631 | 17.34418 | 17.0114 | 20.1668 | 18.25707 |
| TPST1 | 17.83647 | 15.91305 | 15.70006 | 16.41512 | 16.73723 | 15.25981 | 15.69925 | 17.26201 | 19.18859 | 18.30555 | 18.23081 | 13.2183 | 14.44025 | 15.45095 | 13.82089 | 18.36112 | 17.71411 | 15.08889 |
| TPT1 | 20.53422 | 21.00538 | 19.89561 | 22.11706 | 21.37032 | 19.80084 | 19.91515 | 16.03554 | 18.18772 | 19.44757 | 20.06111 | 20.38945 | 21.94347 | 22.02297 | 21.8758 | 21.72838 | 21.79139 | 20.21136 |
| TRA2B | 18.962 | 13.38612 | 13.45383 | 14.51733 | 17.98225 | 15.15462 | 17.72546 | 16.03646 | 12.25619 | 16.62123 | 18.19593 | 18.13828 | 14.79779 | 16.39211 | 15.11866 | 17.31812 | 16.76112 | 17.43109 |
| TRAF2 | 17.42404 | 17.53328 | 15.41731 | 15.57118 | 21.54275 | 21.15159 | 15.58799 | 17.13539 | 15.65514 | 16.64701 | 16.14264 | 17.84497 | 16.88393 | 16.6062 | 15.88657 | 14.70931 | 16.57437 | 16.71428 |
| TRAF3 | 17.13984 | 16.53243 | 15.69128 | 15.57134 | 22.1124 | 20.935 | 17.07054 | 18.41662 | 15.59016 | 18.38705 | 14.92977 | 14.74978 | 15.58138 | 15.50005 | 16.37904 | 15.49049 | 18.11674 | 17.15476 |
| TRAF4 | 20.30892 | 18.47634 | 16.54298 | 18.31315 | 13.31557 | 15.24141 | 15.66783 | 15.96586 | 14.74028 | 15.15822 | 17.02498 | 18.87018 | 17.61104 | 16.67821 | 19.51778 | 19.21718 | 15.58611 | 14.64152 |
| TRAF5 | 16.89435 | 15.50955 | 14.5138 | 15.95582 | 14.84413 | 14.59708 | 16.53076 | 16.4429 | 14.20172 | 13.38686 | 16.23712 | 17.54128 | 13.87084 | 13.96729 | 16.44698 | 16.52405 | 15.93426 | 13.78558 |
| TRAFD1 | 19.61595 | 19.82399 | 16.82606 | 15.36897 | 14.71507 | 15.93544 | 14.71334 | 15.70826 | 13.9984 | 14.95213 | 14.99977 | 13.02839 | 16.18203 | 15.94464 | 15.7058 | 15.97589 | 16.95787 | 15.2782 |
| TRAP1 | 15.87401 | 16.71811 | 17.14844 | 15.21327 | 16.70221 | 17.20285 | 16.92649 | 16.43891 | 19.43045 | 18.13205 | 16.70318 | 15.552 | 16.54448 | 16.28162 | 17.21637 | 16.21132 | 16.28902 | 15.93361 |
| TRAPPC3 | 15.86875 | 16.94463 | 17.9948 | 19.059 | 13.89966 | 16.98077 | 15.78796 | 16.94036 | 17.10829 | 16.47643 | 15.92528 | 19.40109 | 16.58694 | 17.17777 | 17.44488 | 18.75391 | 16.69622 | 16.13722 |
| TRIM14 | 15.31966 | 16.12072 | 17.33562 | 19.0989 | 18.09978 | 15.77107 | 18.03299 | 18.60448 | 14.84279 | 14.64412 | 16.12828 | 15.82469 | 16.16234 | 17.00561 | 17.38502 | 16.68046 | 16.91214 | 17.16928 |
| TRIM16 | 19.59653 | 15.69224 | 16.43638 | 17.1769 | 19.03535 | 17.69755 | 17.38474 | 17.81343 | 15.25684 | 12.88596 | 11.4527 | 13.3673 | 17.45947 | 16.82907 | 15.92292 | 15.66444 | 12.83773 | 13.92366 |
| TRIM25 | 16.5574 | 16.57587 | 16.45271 | 17.50128 | 21.49798 | 20.78966 | 17.2579 | 16.41315 | 14.96204 | 15.91248 | 17.4186 | 18.11852 | 18.85533 | 18.84975 | 16.90968 | 17.14293 | 19.1643 | 19.60554 |
| TRIM26 | 22.29434 | 22.57783 | 22.10337 | 21.7544 | 16.61756 | 18.02068 | 15.92106 | 16.80363 | 14.27578 | 15.17328 | 15.85736 | 15.129 | 18.48336 | 17.02466 | 22.4798 | 21.96072 | 16.51834 | 16.61197 |
| TRIM28 | 17.91674 | 16.74955 | 16.97686 | 16.58645 | 17.33561 | 17.02939 | 19.97297 | 19.54383 | 17.98471 | 18.95433 | 17.86658 | 19.09662 | 18.18295 | 16.91177 | 16.03414 | 16.91069 | 18.68802 | 18.73337 |
| TRIM29 | 15.45383 | 16.45518 | 17.62479 | 13.62544 | 14.55998 | 16.0535 | 15.53358 | 15.99875 | 12.58302 | 16.72592 | 17.54711 | 18.80558 | 14.67563 | 15.58072 | 15.20067 | 16.73055 | 16.33357 | 13.57815 |
| TRIM32 | 15.80748 | 16.3056 | 18.05375 | 16.36062 | 16.86024 | 16.79787 | 19.04694 | 16.47647 | 12.39066 | 15.98594 | 14.41742 | 14.93063 | 13.01756 | 14.76399 | 14.92473 | 15.84029 | 16.61364 | 15.43108 |
| TRIM41 | 23.13022 | 19.57935 | 17.72008 | 15.7468 | 14.23819 | 14.89929 | 14.73846 | 16.0517 | 12.26299 | 16.72078 | 15.40358 | 13.43537 | 15.49325 | 14.36668 | 18.25577 | 17.14843 | 17.22825 | 14.52082 |
| TRIM56 | 14.39576 | 15.63151 | 13.97893 | 14.90936 | 18.13558 | 15.95526 | 15.62266 | 15.52339 | 13.75835 | 14.21461 | 16.95635 | 16.30885 | 15.81161 | 14.90229 | 15.25591 | 13.97927 | 15.31566 | 14.68758 |
| TRIO | 16.57123 | 18.74091 | 16.29746 | 16.01955 | 17.30547 | 17.31231 | 17.18048 | 16.20068 | 18.27526 | 17.45684 | 23.97326 | 15.5571 | 15.09233 | 17.05971 | 15.70385 | 15.89455 | 15.88616 | 15.91109 |
| TRIP10 | 18.42126 | 16.169 | 16.61352 | 15.51161 | 21.24813 | 18.64895 | 14.39825 | 15.92473 | 15.08886 | 17.04837 | 14.36386 | 12.89197 | 17.7725 | 13.7987 | 15.2864 | 16.12566 | 18.87107 | 19.75575 |
| TRIP11 | 15.65872 | 15.91226 | 14.39089 | 17.76236 | 14.56229 | 13.57634 | 19.54672 | 16.5848 | 15.03581 | 12.83219 | 13.76579 | 15.76983 | 14.92462 | 16.58114 | 20.3763 | 16.48753 | 16.05829 | 13.55546 |
| TRMT10C | 16.24318 | 17.10502 | 16.09868 | 16.65571 | 15.28948 | 15.74538 | 15.52049 | 16.31386 | 18.44042 | 16.94693 | 14.46638 | 14.37497 | 16.46879 | 14.0964 | 16.10259 | 15.34177 | 16.82685 | 14.58227 |
| TRMT112 | 17.08929 | 14.76166 | 17.66677 | 16.01541 | 13.91155 | 15.61136 | 14.18864 | 15.30229 | 12.48966 | 16.14625 | 15.59851 | 18.70086 | 17.63474 | 14.33818 | 17.79057 | 17.26367 | 15.82455 | 15.71749 |
| TRNT1 | 16.68611 | 19.80303 | 17.1045 | 14.9865 | 15.95222 | 16.47655 | 17.18945 | 16.01392 | 14.21492 | 14.80673 | 17.92712 | 18.76265 | 16.08184 | 16.18325 | 15.76327 | 14.79792 | 14.46598 | 14.04472 |
| TRPM4 | 18.56892 | 19.00712 | 18.51856 | 17.20639 | 12.85065 | 15.35675 | 15.02076 | 15.26982 | 13.84506 | 14.52972 | 15.73572 | 11.90487 | 15.01032 | 14.57717 | 19.67023 | 19.48125 | 16.57583 | 14.09231 |
| TRPM5 | 19.22211 | 16.01285 | 18.57385 | 18.48832 | 15.04152 | 15.71236 | 14.41686 | 16.25451 | 17.16054 | 14.13822 | 16.52429 | 18.71189 | 15.72799 | 15.85136 | 18.78504 | 17.03732 | 15.04773 | 13.60416 |
| TRRAP | 16.54725 | 17.81491 | 14.43882 | 15.46747 | 15.32925 | 16.47437 | 16.49304 | 19.76528 | 14.89364 | 14.402 | 18.79987 | 17.47296 | 13.16487 | 14.76486 | 15.13531 | 15.48457 | 17.10488 | 15.25449 |
| TSC22D1 | 16.31402 | 15.67654 | 14.94873 | 16.38231 | 17.767 | 16.22848 | 16.18611 | 14.5842 | 13.35342 | 17.75081 | 13.90063 | 15.79552 | 19.33891 | 17.7848 | 17.17588 | 14.1952 | 18.1276 | 18.71061 |
| TSFM | 18.13323 | 18.42297 | 16.00069 | 16.36751 | 14.36115 | 16.1689 | 18.17732 | 17.78165 | 19.19977 | 17.50883 | 17.30981 | 16.77583 | 17.33691 | 17.70596 | 17.93131 | 17.34371 | 17.72957 | 18.22124 |
| TSG101 | 24.56565 | 24.68886 | 24.72542 | 24.1972 | 19.38948 | 22.07559 | 21.76738 | 21.60407 | 20.56292 | 20.3556 | 19.63803 | 20.22617 | 21.63833 | 21.46184 | 25.38655 | 24.94028 | 17.90246 | 21.10127 |
| TSN | 17.49945 | 15.69924 | 16.36157 | 16.97689 | 16.38731 | 15.66581 | 16.49927 | 17.42696 | 17.53914 | 15.4786 | 18.78999 | 21.17603 | 16.78226 | 15.99913 | 15.82705 | 17.0628 | 16.27829 | 16.22675 |
| TSNAX | 16.75666 | 18.61666 | 17.96784 | 15.67283 | 17.1292 | 17.97059 | 17.63434 | 20.41576 | 18.72092 | 17.56658 | 18.06696 | 18.83697 | 16.73437 | 16.81847 | 18.96712 | 15.26941 | 18.73151 | 17.67079 |
| TSPAN1 | 21.90412 | 20.90187 | 18.13481 | 21.1849 | 15.78966 | 17.34865 | 15.72601 | 14.99545 | 20.79532 | 19.43897 | 14.75168 | 14.72897 | 15.22712 | 13.93787 | 20.97549 | 13.87078 | 18.12165 | 12.89558 |
| TSPAN14 | 21.53488 | 21.35364 | 21.0306 | 21.5082 | 20.69995 | 17.52661 | 21.43518 | 21.40889 | 19.55773 | 22.05216 | 25.00535 | 19.79683 | 19.55136 | 22.59423 | 22.88517 | 21.45789 | 19.34907 | 22.26031 |
| TSPAN15 | 17.51701 | 16.73222 | 19.40064 | 17.64833 | 18.29957 | 17.10738 | 17.13008 | 16.90524 | 16.47356 | 16.73844 | 14.79191 | 18.18241 | 15.00516 | 18.75786 | 20.87014 | 20.24379 | 15.02935 | 17.4911 |
| TSPAN4 | 18.69073 | 19.56031 | 17.19823 | 18.64807 | 12.49575 | 14.69933 | 16.24694 | 18.48908 | 13.40089 | 16.64292 | 19.9757 | 15.17584 | 15.67955 | 15.8598 | 21.8745 | 18.9827 | 15.34463 | 18.93999 |
| TSPAN6 | 19.58072 | 17.92294 | 20.30282 | 20.28392 | 17.86873 | 16.12802 | 14.54757 | 17.29627 | 20.74239 | 19.14618 | 15.93671 | 20.07943 | 17.92461 | 15.56415 | 20.64747 | 20.14942 | 12.66972 | 13.51304 |
| TSPAN8 | 26.861 | 26.24474 | 16.88096 | 18.00174 | 14.36487 | 17.642 | 15.33117 | 15.71522 | 20.05758 | 19.63201 | 16.65636 | 14.13892 | 16.72004 | 16.01238 | 18.20067 | 18.25411 | 17.52487 | 21.38389 |
| TSPAN9 | 16.21773 | 15.20241 | 20.31706 | 19.86683 | 16.56604 | 19.3953 | 16.26168 | 13.81504 | 13.88983 | 13.23245 | 14.0198 | 12.68149 | 15.22361 | 14.31974 | 19.56699 | 18.93494 | 14.22991 | 15.39516 |
| TST | 21.60019 | 21.53741 | 19.46859 | 18.08612 | 14.59516 | 17.0137 | 14.59402 | 16.7397 | 19.44517 | 18.02337 | 17.20077 | 13.88964 | 15.17233 | 15.55995 | 14.71594 | 20.37949 | 15.58809 | 15.1144 |

| | | | | | | | | | | | | | | | | | | |
|---|---|---|---|---|---|---|---|---|---|---|---|---|---|---|---|---|---|---|
| TSTD1 | 19.51059 | 18.99677 | 16.55384 | 18.17938 | 12.4649 | 15.58993 | 15.60836 | 15.43225 | 19.07576 | 17.87542 | 15.12574 | 15.8141 | 17.04192 | 17.88169 | 18.78646 | 16.9159 | 15.26816 | 14.2124 |
| TTC13 | 15.48155 | 16.51613 | 14.77625 | 17.16573 | 16.08474 | 17.796 | 15.72624 | 15.07793 | 19.17292 | 15.74893 | 16.28481 | 14.37019 | 18.67281 | 14.60179 | 16.68505 | 16.94599 | 15.60185 | 14.41936 |
| TTC17 | 18.68985 | 16.03764 | 16.61573 | 17.77277 | 15.57443 | 15.83254 | 16.40081 | 16.48829 | 18.93952 | 17.38752 | 15.2682 | 13.35793 | 14.38032 | 15.39159 | 15.93659 | 17.90041 | 16.87 | 15.43134 |
| TTC4 | 15.98856 | 16.58027 | 16.41113 | 14.97779 | 13.99789 | 13.51913 | 16.57228 | 13.59751 | 12.72981 | 14.4581 | 14.79519 | 13.38534 | 17.93332 | 16.88866 | 14.02306 | 17.38155 | 17.77851 | 16.78469 |
| TTC5 | 15.18167 | 14.8829 | 15.64371 | 13.75105 | 18.06998 | 17.163 | 18.64982 | 19.35451 | 14.48989 | 12.62289 | 13.6752 | 16.6014 | 17.39972 | 17.29089 | 15.45201 | 13.48364 | 17.71832 | 12.49964 |
| TTC7B | 19.34208 | 18.57888 | 18.4468 | 18.05408 | 17.94488 | 16.9428 | 17.85615 | 16.46169 | 15.50352 | 15.88675 | 15.94271 | 16.20245 | 17.60772 | 16.79151 | 19.22106 | 18.94894 | 18.50084 | 19.32799 |
| TTLL12 | 15.91863 | 20.72791 | 18.31604 | 17.44133 | 15.46303 | 16.81793 | 16.33821 | 17.1892 | 14.63849 | 14.93935 | 17.62845 | 19.91476 | 17.45348 | 16.343 | 21.21886 | 18.44728 | 15.54113 | 16.24919 |
| TTYH2 | 15.80679 | 16.89744 | 18.78164 | 18.94871 | 17.58984 | 17.85958 | 15.38485 | 16.25432 | 20.22451 | 19.59862 | 15.39434 | 17.16411 | 16.09174 | 15.17982 | 19.19344 | 19.42973 | 18.34378 | 13.74254 |
| TTYH3 | 19.21915 | 18.93026 | 19.35186 | 19.45705 | 15.42271 | 17.839 | 17.47959 | 18.17941 | 19.54353 | 19.98386 | 17.92612 | 19.69318 | 18.09808 | 17.58901 | 19.50488 | 20.04618 | 18.57356 | 19.45942 |
| TUBA1A | 21.68042 | 21.40622 | 21.38035 | 22.51398 | 22.65964 | 21.74298 | 23.87471 | 22.44045 | 23.04983 | 22.65246 | 23.12957 | 23.80972 | 22.24133 | 22.30834 | 21.89386 | 21.93922 | 22.47745 | 22.81316 |
| TUBA1C | 18.48594 | 16.7955 | 15.85151 | 18.59472 | 21.6352 | 20.55041 | 20.18854 | 15.61914 | 19.37046 | 19.99246 | 18.76821 | 19.2085 | 19.44936 | 19.29266 | 17.16479 | 15.71942 | 20.59635 | 21.72354 |
| TUBA3C | 22.65488 | 22.28229 | 23.33738 | 22.88162 | 23.90824 | 23.33156 | 25.22971 | 25.04382 | 23.69193 | 23.94915 | 23.67016 | 25.59501 | 23.76854 | 23.53014 | 22.85377 | 22.94475 | 23.13681 | 24.08995 |
| TUBA4A | 20.80551 | 21.00173 | 21.61213 | 22.13186 | 21.54021 | 20.65486 | 23.07754 | 21.29654 | 20.22348 | 21.79616 | 20.33807 | 21.74647 | 22.22662 | 22.75098 | 21.69211 | 21.39038 | 21.77709 | 22.1656 |
| TUBB | 21.74749 | 21.26473 | 22.45367 | 22.66689 | 24.13254 | 23.02747 | 25.46692 | 23.93515 | 23.50697 | 23.18032 | 23.2411 | 25.13123 | 24.07044 | 23.34552 | 22.90723 | 22.57945 | 22.91406 | 24.17342 |
| TUBB2A | 21.87567 | 21.59369 | 21.58117 | 22.29484 | 23.04584 | 21.78363 | 24.09522 | 21.1366 | 22.98755 | 22.48613 | 23.38239 | 24.34667 | 23.5232 | 22.5657 | 22.86342 | 22.44535 | 21.74876 | 23.1748 |
| TUBB3 | 16.84403 | 16.84956 | 16.57167 | 16.59355 | 21.25372 | 19.50752 | 20.54198 | 18.00552 | 20.45432 | 19.91807 | 18.0904 | 19.72454 | 16.81281 | 16.88813 | 16.58224 | 16.78237 | 17.68077 | 19.62725 |
| TUBB4A | 21.99445 | 21.52007 | 21.6284 | 21.76135 | 22.98129 | 21.86851 | 23.94678 | 22.47517 | 22.52951 | 22.34499 | 22.89167 | 23.93257 | 22.74913 | 22.36393 | 22.0853 | 21.78252 | 21.87491 | 22.87867 |
| TUBB6 | 16.5969 | 16.34032 | 16.02747 | 16.14659 | 19.31916 | 17.26249 | 17.88709 | 17.67224 | 18.38617 | 16.62386 | 16.98799 | 20.21085 | 15.43416 | 16.61944 | 15.55526 | 16.62268 | 16.5967 | 17.29462 |
| TUBG1 | 17.57532 | 17.12837 | 17.34178 | 18.27085 | 17.98284 | 17.93069 | 15.67046 | 17.67948 | 15.75107 | 16.28657 | 18.10073 | 19.65437 | 17.21507 | 16.30961 | 18.47832 | 17.12747 | 16.28632 | 14.65096 |
| TUBGCP2 | 14.62322 | 14.96158 | 16.25376 | 15.37435 | 17.37298 | 16.47224 | 17.38152 | 16.83306 | 13.61051 | 15.8278 | 16.47299 | 17.67488 | 15.77706 | 16.33484 | 16.21547 | 15.66945 | 15.85888 | 17.22656 |
| TUBGCP3 | 17.12721 | 19.0961 | 17.59918 | 14.58217 | 15.34133 | 14.42574 | 15.53491 | 14.20675 | 15.40528 | 19.32385 | 17.5153 | 18.47219 | 17.0046 | 16.53998 | 16.83011 | 16.87098 | 15.02625 | 14.54128 |
| TUBGCP4 | 14.42557 | 14.72157 | 15.99155 | 14.87907 | 13.21578 | 16.30874 | 14.8954 | 15.61077 | 12.54296 | 15.07835 | 13.88385 | 18.62508 | 14.6754 | 13.41712 | 15.09212 | 17.08546 | 15.79208 | 13.40329 |
| TUFM | 16.91028 | 16.62337 | 16.86399 | 17.03562 | 18.00937 | 17.05971 | 19.68336 | 18.27604 | 21.33582 | 19.44912 | 17.26228 | 14.73021 | 17.76816 | 18.01535 | 16.48686 | 14.9008 | 17.15732 | 17.54124 |
| TUSC3 | 19.09277 | 19.44871 | 15.87579 | 14.98187 | 15.29247 | 15.25961 | 15.06558 | 15.71783 | 19.42754 | 17.41866 | 15.10864 | 16.10947 | 15.11305 | 14.15879 | 19.20909 | 19.29049 | 13.59168 | 14.51585 |
| TVP23B | 14.72488 | 15.80909 | 16.42597 | 16.29179 | 16.8081 | 14.9645 | 16.71713 | 17.14676 | 12.28983 | 16.4477 | 15.19749 | 16.84343 | 17.65096 | 18.89915 | 14.56169 | 18.09199 | 14.86725 | 17.568 |
| TWF1 | 19.58911 | 20.12494 | 18.88765 | 18.27562 | 19.69583 | 18.32539 | 16.10076 | 18.08679 | 16.03358 | 17.24606 | 17.22269 | 18.14756 | 19.71473 | 20.38497 | 19.56599 | 19.31631 | 19.13989 | 18.71749 |
| TWF2 | 17.18884 | 17.24814 | 16.99862 | 16.02599 | 18.39753 | 17.7811 | 17.07988 | 16.85946 | 17.6995 | 18.06608 | 19.40332 | 20.22477 | 15.79511 | 16.09792 | 17.11652 | 16.32733 | 20.22954 | 19.32672 |
| TXLNA | 15.26327 | 16.62465 | 17.20375 | 15.58297 | 17.18835 | 16.61501 | 16.16426 | 16.48202 | 15.18434 | 15.66183 | 14.93171 | 15.99532 | 19.05699 | 15.92222 | 15.96023 | 15.87741 | 18.84696 | 19.63004 |
| TXNDC11 | 14.89545 | 17.18322 | 14.95437 | 13.80678 | 14.82698 | 14.28986 | 14.43476 | 15.55237 | 18.73161 | 15.11792 | 15.7555 | 12.03059 | 14.84237 | 15.15346 | 15.79137 | 14.86222 | 16.06258 | 16.10747 |
| TXNDC12 | 16.1673 | 15.39743 | 17.55162 | 17.03087 | 17.10945 | 17.33344 | 17.59657 | 18.04076 | 22.31714 | 21.6442 | 19.64985 | 15.63686 | 18.2843 | 18.84047 | 16.11895 | 17.0719 | 17.01274 | 16.29435 |
| TXNDC15 | 14.64996 | 15.16693 | 14.93926 | 16.60544 | 15.97987 | 15.41998 | 14.6968 | 15.95397 | 19.9519 | 17.34058 | 15.30599 | 13.09382 | 15.06847 | 15.05458 | 16.23198 | 17.17408 | 15.8925 | 15.4928 |
| TXNDC16 | 17.031 | 14.18525 | 14.88783 | 16.59605 | 16.67285 | 14.38937 | 15.51191 | 15.50513 | 18.50744 | 18.20751 | 15.83365 | 14.95889 | 13.53484 | 13.60257 | 13.91382 | 14.345 | 15.43268 | 14.99848 |
| TXNDC17 | 19.87123 | 18.76989 | 20.68589 | 21.58406 | 23.03204 | 21.8749 | 20.78529 | 20.20228 | 20.51437 | 21.49167 | 21.40348 | 20.1527 | 23.13966 | 22.88673 | 20.65016 | 21.38285 | 23.07203 | 22.77255 |
| TXNDC5 | 23.65067 | 16.71495 | 16.66863 | 16.76251 | 18.55933 | 18.92296 | 19.29932 | 24.9809 | 23.49275 | 22.69195 | 19.94485 | 16.99743 | 17.92413 | 19.13646 | 17.09135 | 17.70099 | 17.98049 | 18.29537 |
| TXNDC9 | 17.4493 | 15.42902 | 16.4619 | 16.93908 | 17.78979 | 16.77493 | 16.10169 | 16.86427 | 14.5337 | 17.84994 | 17.66129 | 17.64927 | 16.63241 | 15.95136 | 16.05603 | 15.58619 | 18.26083 | 17.94371 |
| TXNIP | 17.17397 | 18.88937 | 16.58966 | 16.86178 | 18.04315 | 15.46245 | 16.84534 | 17.13855 | 15.66981 | 11.59311 | 17.7555 | 17.47624 | 17.02864 | 15.56609 | 16.54158 | 18.27809 | 16.63965 | 18.09498 |
| TXNL1 | 19.52784 | 18.16801 | 20.0276 | 20.44514 | 20.41057 | 18.21037 | 18.12689 | 18.85857 | 19.15509 | 19.93764 | 19.776 | 19.76878 | 21.01714 | 19.85786 | 20.34552 | 20.12927 | 22.14779 | 19.09487 |
| TXNRD1 | 18.30618 | 17.47479 | 17.25631 | 17.97856 | 22.49342 | 20.11653 | 19.07731 | 17.76069 | 19.71339 | 20.1928 | 20.77005 | 20.43538 | 21.20474 | 20.01682 | 17.78162 | 18.41 | 21.52575 | 21.57392 |
| TXNRD2 | 15.74839 | 13.91498 | 15.70579 | 16.12901 | 14.27491 | 14.74455 | 15.83178 | 15.53046 | 17.40917 | 16.75154 | 15.23141 | 11.54684 | 14.29521 | 15.31437 | 14.89249 | 16.97725 | 16.2438 | 14.44851 |
| TYK2 | 16.48256 | 17.24564 | 17.13803 | 17.90511 | 18.3064 | 17.91799 | 15.41592 | 15.79841 | 15.89552 | 14.35927 | 15.80942 | 16.74402 | 14.78799 | 15.94734 | 17.25806 | 17.29784 | 18.23164 | 19.04983 |
| TYMP | 15.33626 | 16.1388 | 17.88661 | 17.58665 | 18.23394 | 16.92436 | 15.44686 | 17.3824 | 13.62263 | 16.08647 | 16.16554 | 14.09714 | 17.24252 | 16.18794 | 18.07642 | 17.49632 | 16.31523 | 15.93838 |
| TYRP1 | 16.63141 | 14.79666 | 15.4026 | 15.46937 | 14.10565 | 15.75115 | 14.81353 | 16.81031 | 19.88963 | 16.38446 | 15.25126 | 10.92481 | 13.57312 | 15.2288 | 14.88431 | 13.9567 | 15.97637 | 13.31532 |
| TYSND1 | 15.1398 | 15.72942 | 17.12022 | 16.98076 | 12.56578 | 15.86342 | 15.70516 | 16.4776 | 18.92753 | 15.49532 | 15.75773 | 12.71895 | 14.2425 | 14.39126 | 15.74485 | 15.16793 | 13.27337 | 16.25885 |
| U2AF1 | 20.39828 | 19.88312 | 18.92801 | 18.7681 | 14.94692 | 15.73585 | 16.53936 | 15.59466 | 16.24223 | 15.82124 | 17.18388 | 18.27362 | 19.9975 | 20.31132 | 17.82498 | 18.2418 | 14.85105 | 14.34288 |
| U2AF2 | 21.60214 | 21.33424 | 19.06985 | 19.91546 | 19.29409 | 18.6187 | 19.94737 | 20.11255 | 17.27197 | 18.89922 | 19.45742 | 20.63388 | 20.42004 | 19.82618 | 18.79838 | 19.86073 | 18.88212 | 19.28567 |
| U2SURP | 14.84596 | 12.34054 | 15.59532 | 14.32878 | 14.76674 | 14.36779 | 13.53497 | 14.66007 | 12.82046 | 15.32724 | 14.24733 | 18.3753 | 16.16657 | 15.74224 | 14.69224 | 13.81105 | 15.58616 | 13.77492 |
| UACA | 16.01461 | 16.91322 | 15.88377 | 16.69793 | 17.99065 | 16.91267 | 17.08303 | 15.62654 | 14.58592 | 15.45772 | 15.96722 | 15.62151 | 17.27935 | 17.35749 | 16.42974 | 15.21798 | 19.41935 | 18.58818 |
| UAP1 | 17.1658 | 15.98062 | 18.41746 | 18.24442 | 20.98373 | 18.94973 | 18.01141 | 17.34642 | 16.33507 | 16.84493 | 16.96174 | 17.02038 | 20.85349 | 20.20865 | 17.53505 | 16.98097 | 19.36253 | 19.78164 |
| UAP1L1 | 16.44179 | 17.48034 | 14.4109 | 19.19576 | 13.12146 | 14.01555 | 20.09954 | 18.92716 | 13.34636 | 16.71778 | 18.86767 | 15.5507 | 15.25324 | 18.30167 | 17.59315 | 16.61852 | 17.96154 | 15.00084 |
| UBA1 | 18.29593 | 18.04647 | 19.17205 | 18.99467 | 19.23604 | 18.81047 | 20.33329 | 18.88839 | 19.69543 | 19.79617 | 20.36604 | 20.92461 | 20.01555 | 19.28622 | 19.52939 | 18.8772 | 20.26867 | 19.98279 |
| UBA2 | 14.20284 | 15.29186 | 14.27005 | 15.48223 | 16.81726 | 16.21652 | 16.18924 | 17.8162 | 16.33477 | 16.43904 | 18.95192 | 18.6369 | 16.00034 | 14.36581 | 15.48612 | 15.87255 | 17.06125 | 15.44573 |
| UBA3 | 15.27301 | 15.87836 | 18.20656 | 16.12294 | 14.94931 | 14.94836 | 15.74174 | 17.16959 | 16.76466 | 16.00526 | 18.13529 | 19.12021 | 17.37055 | 13.65745 | 16.49649 | 17.06089 | 16.75847 | 16.72757 |
| UBA5 | 15.72965 | 15.31505 | 13.7771 | 13.33377 | 14.92063 | 14.15637 | 16.67742 | 15.39856 | 15.41855 | 14.67336 | 16.54662 | 18.60209 | 15.44491 | 16.42824 | 14.20422 | 14.40054 | 11.4114 | 15.957 |
| UBA6 | 16.08277 | 17.8028 | 16.13606 | 15.5831 | 17.38378 | 16.59011 | 17.08178 | 17.02213 | 15.8153 | 16.23712 | 17.13961 | 19.04044 | 17.96412 | 17.7018 | 17.43167 | 15.86033 | 19.08181 | 19.55069 |
| UBAP1 | 21.48337 | 21.41736 | 17.62391 | 18.52686 | 20.34511 | 19.90523 | 16.54642 | 15.82502 | 14.44122 | 16.22895 | 17.99031 | 15.0763 | 18.08656 | 19.74224 | 18.5387 | 18.45678 | 16.20043 | 17.94382 |
| UBAP2L | 16.61655 | 15.27882 | 15.65024 | 13.15092 | 16.64263 | 16.37441 | 15.29933 | 15.3216 | 15.97099 | 15.20821 | 17.50107 | 18.47851 | 15.19 | 16.79621 | 17.27995 | 15.85469 | 16.375 | 15.84794 |
| UBE2D3 | 21.44698 | 21.96706 | 16.20309 | 19.65581 | 14.83537 | 14.93192 | 14.99988 | 17.70047 | 18.94635 | 15.02824 | 15.83257 | 20.67216 | 15.10404 | 13.81369 | 22.37112 | 21.88112 | 15.9151 | 12.86905 |
| UBE2K | 18.42477 | 15.69795 | 16.05391 | 14.42006 | 15.54182 | 15.85953 | 15.27528 | 16.08659 | 16.03368 | 16.20488 | 18.86852 | 18.72164 | 14.53254 | 15.92189 | 17.88894 | 18.51472 | 15.32448 | 15.86611 |
| UBE2L3 | 17.10451 | 17.53892 | 18.33066 | 15.31398 | 18.33021 | 15.34851 | 14.89814 | 16.26706 | 14.58509 | 17.44393 | 19.15867 | 19.62378 | 18.52727 | 20.33631 | 19.46969 | 19.19519 | 16.76527 | 12.91903 |
| UBE2L5 | 17.63369 | 17.78457 | 18.38597 | 17.77107 | 17.67835 | 17.6096 | 17.46982 | 18.38038 | 15.07567 | 16.12743 | 18.43333 | 19.45841 | 17.66396 | 18.29982 | 20.86604 | 18.52406 | 18.55622 | 17.51454 |
| UBE2M | 19.02241 | 19.33378 | 19.42128 | 18.83805 | 17.06999 | 16.04666 | 18.18094 | 17.9447 | 18.97751 | 18.68791 | 19.10783 | 19.57943 | 18.77318 | 18.04731 | 19.04275 | 19.12632 | 19.71755 | 18.29825 |
| UBE2NL | 21.35606 | 21.93594 | 20.77956 | 18.85831 | 15.70555 | 16.95177 | 16.46652 | 17.11928 | 18.86491 | 18.18968 | 21.42254 | 21.8469 | 18.24783 | 15.70655 | 22.36729 | 21.56585 | 17.28479 | 15.01268 |
| UBE2O | 15.63934 | 16.45444 | 17.82938 | 19.23475 | 18.79311 | 21.48028 | 18.64203 | 17.61678 | 14.69982 | 13.77907 | 16.94855 | 19.2439 | 18.22154 | 18.32212 | 17.75899 | 16.26754 | 18.87043 | 20.26384 |
| UBE2T | 14.95474 | 15.15946 | 13.96431 | 17.01696 | 13.34577 | 15.27743 | 14.94747 | 12.65207 | 11.77465 | 14.32244 | 13.84993 | 19.33857 | 13.10004 | 15.06121 | 13.11342 | 14.25867 | 14.63474 | 15.49668 |
| UBE2V1 | 19.19646 | 18.0158 | 16.17897 | 19.63751 | 19.42496 | 19.42442 | 17.80085 | 19.8812 | 18.22693 | 21.51243 | 19.59353 | 18.96422 | 20.11802 | 19.39371 | 17.95223 | 19.64847 | 20.69151 | 20.47671 |
| UBE2V2 | 20.77227 | 20.78683 | 20.85895 | 19.6689 | 19.35503 | 17.93975 | 18.32928 | 17.88038 | 18.56422 | 19.77746 | 20.44382 | 20.51395 | 20.44937 | 18.99151 | 20.84494 | 20.93496 | 19.93958 | 21.11612 |
| UBE2Z | 16.04533 | 15.92808 | 15.34872 | 15.30858 | 16.48743 | 16.79329 | 16.29482 | 16.59395 | 15.13695 | 13.97775 | 15.60997 | 16.63103 | 16.31858 | 15.00037 | 16.1469 | 16.06894 | 19.70585 | 18.77663 |
| UBE3A | 17.10632 | 17.82781 | 15.67512 | 18.05486 | 17.60476 | 18.3115 | 17.41663 | 16.81641 | 16.51712 | 17.19016 | 17.19269 | 18.10016 | 17.19781 | 16.26521 | 16.1752 | 18.1009 | 19.91577 | 16.36077 |
| UBE3C | 22.44251 | 21.72398 | 14.799 | 19.14314 | 12.9191 | 14.48463 | 16.5004 | 16.64869 | 15.98127 | 14.6386 | 15.58484 | 16.58282 | 19.20408 | 16.17247 | 21.8813 | 18.82073 | 14.38387 | 13.8374 |
| UBE4A | 17.56386 | 17.65875 | 16.58611 | 16.14134 | 13.24447 | 17.61268 | 13.37129 | 16.31849 | 12.42862 | 14.76608 | 14.5839 | 18.83526 | 15.45257 | 16.11661 | 15.1527 | 17.19079 | 14.4404 | 14.26915 |
| UBFD1 | 16.68549 | 18.37973 | 18.34961 | 14.89155 | 15.51853 | 15.7787 | 16.96502 | 16.34314 | 13.99567 | 17.43835 | 18.32337 | 17.61697 | 19.90944 | 18.90901 | 15.86663 | 14.13463 | 19.41512 | 20.2135 |
| UBL5 | 18.97636 | 18.61394 | 19.7974 | 19.95425 | 18.22475 | 17.052 | 18.33667 | 18.57069 | 15.65111 | 16.81849 | 17.09273 | 15.86034 | 19.51836 | 19.35261 | 19.48448 | 19.84749 | 20.16978 | 19.64344 |
| UBLCP1 | 16.12676 | 16.85855 | 15.86144 | 13.6392 | 15.21502 | 15.63053 | 15.14005 | 16.57072 | 13.75075 | 16.69195 | 14.60782 | 18.29529 | 17.86872 | 15.51621 | 13.99168 | 15.24147 | 13.83639 | 13.30066 |
| UBQLN2 | 19.58203 | 18.84026 | 18.85169 | 16.04615 | 14.97028 | 14.87131 | 15.37707 | 16.25502 | 15.65684 | 16.8953 | 15.54403 | 18.07764 | 15.49305 | 14.54124 | 18.24317 | 19.17804 | 17.85316 | 15.32889 |
| UBR1 | 15.35061 | 15.27104 | 14.33296 | 14.69154 | 13.66186 | 13.89268 | 15.9589 | 14.92205 | 13.49322 | 14.56411 | 14.98756 | 17.01247 | 15.92633 | 16.1111 | 15.59324 | 16.41384 | 15.928 | 13.35027 |
| UBR4 | 15.46785 | 16.5288 | 15.47276 | 15.35653 | 16.83225 | 15.87849 | 16.30504 | 16.09975 | 15.16661 | 16.56469 | 16.66034 | 18.59688 | 17.00255 | 15.22382 | 15.1164 | 16.04791 | 15.78911 | 15.27825 |
| UBTD1 | 18.43534 | 17.27655 | 17.77393 | 17.82242 | 19.52448 | 16.8992 | 14.77072 | 15.57327 | 18.12797 | 17.22855 | 17.31447 | 19.05143 | 17.98086 | 18.34624 | 18.34533 | 18.24389 | 20.02341 | 20.6949 |
| UBTF | 20.02576 | 20.27249 | 19.8025 | 22.31159 | 24.64935 | 17.12218 | 17.4158 | 17.81031 | 22.28164 | 22.43587 | 22.48729 | 21.6498 | 23.05244 | 22.18182 | 21.42172 | 21.08257 | 21.88918 | 23.30973 |
| UBXN1 | 18.16298 | 18.99989 | 17.94541 | 19.75592 | 16.03884 | 15.1361 | 16.93636 | 16.73889 | 17.25234 | 12.81624 | 19.19622 | 20.11757 | 12.81571 | 12.06407 | 17.63211 | 18.68685 | 15.2716 | 14.84553 |
| UBXN6 | 18.2211 | 19.1052 | 17.67141 | 15.00747 | 18.1765 | 16.46346 | 17.68459 | 16.99427 | 16.85431 | 17.04888 | 16.04542 | 16.29623 | 19.09836 | 19.68666 | 18.44212 | 18.79585 | 17.34272 | 18.49858 |
| UCHL1 | 18.03237 | 16.97135 | 18.15399 | 17.38575 | 24.55133 | 23.35288 | 22.47035 | 21.32677 | 21.75802 | 22.56066 | 23.03004 | 23.46031 | 16.05161 | 17.27794 | 16.843 | 17.19142 | 16.80419 | 15.97741 |
| UCHL3 | 17.04229 | 15.3162 | 15.07106 | 16.78762 | 17.40457 | 15.7355 | 18.69027 | 15.73301 | 15.19242 | 16.71185 | 18.63848 | 19.46472 | 19.45251 | 20.04248 | 17.65741 | 18.0865 | 19.54062 | 18.13272 |
| UCHL5 | 16.50218 | 16.19838 | 16.77681 | 16.40018 | 16.10654 | 15.98624 | 16.42383 | 17.21234 | 17.75447 | 17.99006 | 17.44173 | 19.22285 | 17.68896 | 16.93402 | 15.08949 | 15.67973 | 16.25284 | 16.16507 |
| UCK2 | 16.07692 | 18.47109 | 14.31138 | 16.15007 | 19.08267 | 17.06204 | 18.8458 | 19.04327 | 13.72803 | 13.75936 | 16.35472 | 15.94749 | 20.01958 | 19.70663 | 14.82144 | 15.96752 | 19.73832 | 17.56214 |
| UEVLD | 16.6705 | 15.59142 | 16.99952 | 18.47193 | 21.0774 | 20.19972 | 16.53127 | 16.1521 | 18.81891 | 19.21096 | 18.54375 | 18.55753 | 19.58043 | 19.64192 | 15.27708 | 16.67977 | 18.54903 | 18.85649 |
| UFC1 | 15.54774 | 16.65986 | 16.37733 | 14.38497 | 20.69352 | 18.00651 | 19.37208 | 16.88993 | 15.92317 | 16.39749 | 19.00578 | 19.29366 | 19.72809 | 20.03706 | 15.48495 | 15.35081 | 21.10329 | 19.99145 |

| | | | | | | | | | | | | | | | | | | |
|---|---|---|---|---|---|---|---|---|---|---|---|---|---|---|---|---|---|---|
| UFD1 | 16.53474 | 14.63343 | 16.33564 | 16.7227 | 18.02727 | 16.89297 | 14.94516 | 17.52072 | 14.20019 | 15.82339 | 12.4221 | 16.40577 | 17.90728 | 14.27133 | 14.57125 | 15.87493 | 18.23722 | 12.66346 |
| UFL1 | 15.46357 | 15.30132 | 16.43248 | 16.96218 | 16.60188 | 14.87768 | 16.87417 | 16.95095 | 15.23566 | 17.10123 | 15.26634 | 15.85852 | 15.99688 | 17.29607 | 14.76738 | 14.66407 | 15.46133 | 13.56551 |
| UFM1 | 20.49852 | 20.23205 | 22.64703 | 21.32227 | 19.18345 | 16.53514 | 18.64303 | 18.28496 | 18.32489 | 19.15166 | 20.87087 | 19.8042 | 19.60371 | 19.36123 | 23.20628 | 22.60513 | 20.25886 | 20.08623 |
| UGDH | 17.51533 | 17.21937 | 16.88471 | 17.18301 | 22.23051 | 20.69639 | 17.79785 | 17.31234 | 15.75567 | 16.51102 | 17.92457 | 17.34009 | 19.99778 | 20.60503 | 16.26023 | 17.39739 | 18.4794 | 18.90665 |
| UGGT1 | 16.46002 | 16.81298 | 17.09173 | 16.1709 | 16.96548 | 16.88303 | 16.78329 | 16.83854 | 21.61298 | 21.04135 | 19.02982 | 16.11389 | 16.75394 | 16.47641 | 17.0105 | 16.55311 | 16.43219 | 16.80298 |
| UGGT2 | 14.58999 | 14.78875 | 15.89134 | 15.72056 | 13.29778 | 14.90789 | 17.08272 | 18.32375 | 19.52922 | 21.0783 | 17.06991 | 16.15279 | 16.20679 | 16.06585 | 15.94616 | 15.72229 | 14.36385 | 13.5918 |
| UGP2 | 16.83251 | 17.49685 | 19.88941 | 19.09072 | 20.58517 | 21.63782 | 19.45734 | 18.77741 | 18.76739 | 20.18312 | 20.60541 | 19.15783 | 19.62364 | 20.1418 | 18.0827 | 16.52191 | 19.89561 | 20.47202 |
| UGT8 | 19.79083 | 19.43416 | 16.28425 | 15.1871 | 16.84974 | 16.77599 | 15.50099 | 13.93939 | 19.04516 | 15.26198 | 17.72551 | 17.41857 | 16.37257 | 14.8451 | 16.92981 | 16.69936 | 16.67555 | 14.77453 |
| ULK3 | 15.85601 | 15.15582 | 16.84647 | 17.40045 | 15.14267 | 15.94972 | 16.05603 | 16.8813 | 12.43381 | 16.80527 | 17.72864 | 17.67148 | 13.77662 | 16.59166 | 16.95347 | 17.07823 | 16.11968 | 15.80605 |
| UMAD1 | 22.32069 | 22.29331 | 22.24015 | 21.76388 | 15.17093 | 16.47004 | 16.80301 | 16.90568 | 16.94931 | 16.68893 | 18.21673 | 14.971 | 19.10836 | 15.88834 | 22.24014 | 21.90181 | 17.73862 | 15.74934 |
| UMPS | 16.4883 | 15.85412 | 16.42576 | 16.90934 | 16.78508 | 17.32875 | 17.83037 | 15.66788 | 17.67516 | 17.47736 | 18.21108 | 19.69206 | 15.41213 | 16.75267 | 16.02119 | 16.45723 | 16.94907 | 16.50724 |
| UNC119B | 19.60554 | 19.08286 | 16.35535 | 16.00711 | 17.33455 | 16.14754 | 15.44601 | 16.28868 | 12.12677 | 16.12984 | 15.78258 | 17.12936 | 18.92584 | 17.98302 | 17.98063 | 16.12848 | 17.87242 | 18.45777 |
| UNC13D | 15.98801 | 16.68766 | 13.03388 | 14.59362 | 17.20616 | 14.75565 | 19.74844 | 15.21023 | 16.92976 | 19.29974 | 14.07408 | 13.36146 | 20.34695 | 15.04136 | 17.20981 | 16.92368 | 17.78136 | 15.45207 |
| UNC45A | 17.61774 | 15.76607 | 15.97945 | 17.35598 | 18.51814 | 16.53981 | 17.80609 | 18.10861 | 15.05719 | 16.12858 | 17.00383 | 18.41068 | 19.1473 | 18.5742 | 17.51413 | 16.47446 | 18.06651 | 18.25427 |
| UNC5B | 16.58694 | 14.98776 | 14.27276 | 13.83074 | 14.15887 | 17.85221 | 18.10585 | 15.85456 | 18.06182 | 15.95317 | 14.52227 | 18.17323 | 14.12846 | 14.632 | 14.35856 | 14.6042 | 15.40399 | 16.72016 |
| UPF1 | 21.03468 | 21.44155 | 20.63614 | 20.53355 | 17.2845 | 16.64186 | 18.679 | 17.90799 | 16.58124 | 17.03252 | 17.29701 | 18.3251 | 18.76774 | 18.1101 | 19.8363 | 20.23408 | 17.5906 | 17.33527 |
| UPF2 | 15.24852 | 14.77245 | 15.22816 | 15.93586 | 12.5684 | 15.68268 | 14.8487 | 15.10874 | 12.89746 | 14.52033 | 15.38586 | 16.78146 | 11.41242 | 16.14776 | 13.43957 | 16.04821 | 15.60805 | 17.15177 |
| UQCRB | 19.62704 | 17.47425 | 16.80204 | 17.04941 | 15.92037 | 16.70342 | 20.56726 | 20.71477 | 21.62892 | 21.11015 | 17.76475 | 16.03161 | 15.20414 | 15.69199 | 16.34703 | 16.17918 | 19.20297 | 17.13624 |
| UQCRC1 | 15.7881 | 17.38301 | 17.11773 | 15.72148 | 15.61644 | 16.36454 | 16.68151 | 17.35228 | 22.46561 | 20.95817 | 18.02097 | 15.88662 | 15.30777 | 16.10544 | 15.66818 | 16.03409 | 17.49786 | 17.2048 |
| UQCRC2 | 15.7066 | 16.41999 | 16.23394 | 16.29711 | 15.84911 | 16.89119 | 17.83521 | 17.58962 | 22.32594 | 21.48124 | 17.96511 | 15.35852 | 16.34265 | 17.64398 | 15.6634 | 15.97424 | 15.05935 | 16.10929 |
| UQCRFS1P | 15.04925 | 15.57799 | 18.07948 | 16.95374 | 14.13858 | 15.0936 | 17.73996 | 16.53883 | 20.9325 | 18.89976 | 17.97159 | 14.49467 | 16.58288 | 17.10626 | 16.15632 | 15.4275 | 15.42226 | 14.5158 |
| UQCRH | 16.09534 | 16.54354 | 17.5026 | 16.87346 | 14.74465 | 16.05686 | 17.05517 | 15.8147 | 23.38579 | 20.84761 | 17.47445 | 15.97105 | 17.02942 | 14.83714 | 16.62685 | 15.74333 | 15.78224 | 16.01676 |
| UQCRQ | 15.99347 | 16.5597 | 15.10814 | 14.12724 | 15.06513 | 15.80164 | 14.64624 | 18.24886 | 20.00152 | 18.34991 | 16.79886 | 13.43681 | 12.44235 | 15.76742 | 15.53154 | 15.1751 | 13.69307 | 13.61996 |
| UROD | 15.09086 | 13.08425 | 14.75373 | 15.21671 | 13.3553 | 16.20808 | 16.07646 | 15.54275 | 11.70107 | 13.88237 | 16.22308 | 18.41973 | 14.79093 | 13.99435 | 14.9505 | 16.59653 | 17.23396 | 12.8639 |
| USO1 | 16.07805 | 16.32236 | 15.82136 | 15.67138 | 17.40762 | 17.24406 | 17.76538 | 17.30094 | 15.48599 | 17.81788 | 17.25912 | 19.33524 | 18.67397 | 18.119 | 15.87786 | 15.66537 | 17.94186 | 17.90795 |
| USP10 | 14.85686 | 15.83297 | 15.63295 | 15.06433 | 14.46499 | 15.41285 | 14.7668 | 16.433 | 14.57014 | 15.1032 | 17.59753 | 19.52217 | 15.43746 | 15.19371 | 16.164 | 14.39465 | 14.28956 | 13.25996 |
| USP13 | 19.74181 | 19.32004 | 17.52739 | 18.70865 | 16.17421 | 15.86018 | 16.83564 | 15.00479 | 14.30193 | 15.27568 | 16.49424 | 16.54392 | 17.55105 | 15.37953 | 17.52683 | 18.61193 | 18.81127 | 15.4425 |
| USP14 | 17.49126 | 16.85022 | 18.1335 | 18.27255 | 18.86909 | 17.74856 | 18.41537 | 18.26309 | 17.13609 | 17.61447 | 20.06686 | 20.41725 | 19.52703 | 19.07698 | 18.65458 | 18.12216 | 18.84185 | 18.65292 |
| USP15 | 16.76141 | 15.49536 | 15.72461 | 15.1716 | 14.03846 | 15.4714 | 14.05958 | 17.71028 | 14.13628 | 15.21652 | 16.869 | 18.51645 | 16.82637 | 15.74054 | 14.8458 | 16.19687 | 14.73232 | 15.48235 |
| USP24 | 16.39395 | 17.92664 | 15.49874 | 15.41607 | 14.71631 | 14.81995 | 15.2376 | 15.92494 | 15.30918 | 13.92268 | 15.28405 | 17.87693 | 15.61488 | 17.22501 | 16.30982 | 16.45401 | 14.44052 | 14.54134 |
| USP25 | 17.51164 | 16.57315 | 14.71942 | 15.36281 | 18.39702 | 16.75353 | 15.50721 | 17.34487 | 17.69552 | 15.61805 | 16.69337 | 12.26947 | 15.43166 | 15.24288 | 14.19521 | 14.38182 | 16.49895 | 16.3203 |
| USP43 | 15.77963 | 16.75664 | 16.71906 | 17.7558 | 14.5536 | 17.61922 | 15.4893 | 16.60144 | 13.89035 | 16.07514 | 15.54836 | 13.63917 | 17.91728 | 16.82709 | 17.63081 | 17.81759 | 15.69018 | 14.92686 |
| USP47 | 15.37519 | 14.31959 | 16.10014 | 15.34071 | 14.99993 | 15.38358 | 15.5952 | 14.99527 | 15.51638 | 16.34625 | 17.22359 | 17.58055 | 14.44809 | 15.33776 | 15.95377 | 16.509 | 16.114 | 15.91772 |
| USP5 | 20.9341 | 21.04435 | 18.7236 | 19.45143 | 18.79888 | 17.25625 | 18.63535 | 17.22746 | 17.17929 | 17.66656 | 18.51099 | 18.84361 | 20.52237 | 19.8625 | 19.1423 | 18.83558 | 19.23894 | 19.5077 |
| USP54 | 16.09008 | 17.06912 | 17.36935 | 17.17952 | 14.728 | 14.79559 | 17.47745 | 16.02202 | 14.221 | 15.5615 | 14.68671 | 13.77605 | 18.95261 | 18.90076 | 16.19199 | 16.19591 | 15.05461 | 14.23817 |
| USP6NL | 16.27068 | 15.10357 | 14.66577 | 14.52393 | 15.5965 | 15.652 | 17.03126 | 15.61347 | 16.71194 | 17.37035 | 15.16713 | 16.664 | 16.11307 | 15.5439 | 17.66077 | 16.74198 | 17.7429 | 21.59038 |
| USP7 | 16.13137 | 16.39989 | 16.23298 | 15.96228 | 15.57954 | 16.54186 | 15.8006 | 15.66195 | 15.80933 | 17.14721 | 18.3813 | 18.09045 | 15.82006 | 15.41994 | 16.24404 | 16.00553 | 15.96695 | 16.01334 |
| USP8 | 15.93734 | 15.48222 | 15.73053 | 15.58208 | 19.54839 | 17.30479 | 16.69805 | 16.45635 | 15.73663 | 15.91062 | 14.55299 | 15.01567 | 16.54712 | 16.10985 | 14.83524 | 15.24639 | 17.00501 | 17.31562 |
| USP9X | 20.83071 | 20.30178 | 19.65404 | 19.62315 | 14.44505 | 14.84312 | 15.34723 | 15.88562 | 14.68929 | 15.53759 | 15.97784 | 18.28913 | 16.43904 | 16.7671 | 19.75638 | 19.82818 | 15.34139 | 14.49365 |
| UTRN | 17.25963 | 16.69074 | 19.10287 | 19.66707 | 19.18503 | 18.04466 | 16.14612 | 16.7529 | 17.68214 | 18.44201 | 16.38342 | 15.12398 | 18.28978 | 18.09565 | 19.82815 | 19.85007 | 19.13757 | 18.96586 |
| UXS1 | 16.52738 | 16.60195 | 16.33192 | 17.0154 | 16.05153 | 17.28459 | 17.86426 | 17.05634 | 20.29795 | 19.03851 | 15.59729 | 14.91928 | 15.43973 | 16.32319 | 19.0002 | 16.70704 | 17.97023 | 17.41069 |
| UXT | 17.14722 | 16.67815 | 14.64554 | 15.72904 | 16.61257 | 15.38359 | 15.72718 | 13.3275 | 16.69911 | 14.71664 | 15.40241 | 16.90834 | 14.97835 | 14.05281 | 14.53452 | 14.98336 | 15.55986 | 15.23504 |
| VAC14 | 14.75382 | 15.69693 | 16.15997 | 15.95971 | 13.83029 | 17.00774 | 16.35632 | 15.22844 | 12.85699 | 14.31108 | 13.60034 | 17.1799 | 13.84873 | 15.05234 | 15.89374 | 15.9286 | 17.90955 | 15.8782 |
| VAMP3 | 18.94167 | 18.93892 | 22.54994 | 21.21095 | 20.18171 | 15.23003 | 16.58961 | 17.95359 | 20.22452 | 19.31536 | 17.40042 | 20.11623 | 19.89784 | 18.7272 | 20.16307 | 18.9634 | 15.70454 | 20.76201 |
| VAMP7 | 18.13932 | 16.6742 | 19.17993 | 17.89487 | 16.7527 | 17.3736 | 18.06581 | 15.94146 | 17.85326 | 16.93746 | 17.26981 | 17.876 | 16.92265 | 16.09585 | 18.39594 | 17.91811 | 15.94223 | 16.37087 |
| VAMP8 | 19.38966 | 18.94449 | 19.50502 | 20.11283 | 20.07459 | 19.02133 | 15.6801 | 17.61316 | 19.96814 | 19.41131 | 21.14168 | 20.23594 | 20.21186 | 19.38746 | 19.13404 | 19.09412 | 19.3506 | 18.94441 |
| VANGL1 | 16.26993 | 14.90446 | 18.31321 | 17.7666 | 19.30843 | 17.56051 | 17.22253 | 15.73482 | 17.0047 | 17.49313 | 15.00439 | 14.42019 | 16.06995 | 17.89245 | 17.95461 | 18.89971 | 18.34902 | 17.77317 |
| VANGL2 | 16.10527 | 15.26143 | 15.38057 | 15.95439 | 17.00692 | 16.18203 | 15.31795 | 16.89015 | 15.02045 | 16.44173 | 17.86351 | 18.73294 | 14.17741 | 17.19521 | 16.80347 | 15.55177 | 16.64049 | 12.94556 |
| VAPA | 16.40468 | 16.8052 | 17.12298 | 15.38214 | 21.69848 | 19.20428 | 20.43828 | 19.73309 | 17.3238 | 19.02751 | 19.43901 | 18.93212 | 20.79356 | 19.67061 | 16.04356 | 16.06217 | 17.92854 | 19.94423 |
| VAPB | 15.94259 | 16.47133 | 17.72803 | 16.27618 | 20.38738 | 18.7965 | 19.68229 | 19.48233 | 18.31013 | 19.52917 | 17.70574 | 18.95459 | 19.37788 | 19.91697 | 14.89578 | 15.87558 | 18.39407 | 19.2433 |
| VARS1 | 16.00825 | 16.31872 | 16.95417 | 16.974 | 16.13463 | 16.41501 | 16.87577 | 17.25756 | 15.78116 | 16.00387 | 18.0282 | 20.28973 | 16.12136 | 15.99288 | 17.55189 | 17.17027 | 17.06933 | 15.80456 |
| VASP | 17.01039 | 17.26368 | 17.21761 | 18.7855 | 19.11386 | 15.93438 | 18.16267 | 17.32427 | 16.35299 | 17.27959 | 18.61188 | 19.51382 | 18.81819 | 19.83691 | 17.38906 | 16.76443 | 18.06995 | 17.73467 |
| VAT1 | 17.91159 | 18.18016 | 16.34908 | 17.72191 | 23.00336 | 22.00841 | 19.18093 | 19.92726 | 19.93931 | 19.2137 | 18.92705 | 19.18923 | 19.28571 | 20.45639 | 17.80228 | 17.12094 | 21.38118 | 21.85527 |
| VAV1 | 15.10072 | 15.97871 | 16.86457 | 15.34319 | 15.36388 | 15.41297 | 15.65759 | 15.78311 | 13.66643 | 16.94969 | 16.94864 | 18.84731 | 14.19288 | 14.8409 | 16.23768 | 16.39858 | 16.16009 | 14.87496 |
| VBP1 | 16.40578 | 16.69314 | 16.84361 | 16.77091 | 19.75243 | 16.4383 | 18.84005 | 17.63381 | 18.16997 | 19.45579 | 18.35653 | 19.85238 | 20.65536 | 19.65979 | 17.59663 | 17.6893 | 20.68394 | 20.0185 |
| VCAN | 17.03889 | 15.73954 | 16.35768 | 17.01509 | 16.18841 | 14.37525 | 16.00752 | 17.0572 | 19.42141 | 17.9847 | 16.70418 | 17.01292 | 15.9877 | 15.38113 | 14.92765 | 16.63935 | 15.50826 | 16.5478 |
| VCL | 18.44072 | 19.13965 | 18.24138 | 19.2475 | 20.31806 | 19.82411 | 21.17563 | 19.5076 | 17.89066 | 19.21659 | 18.07334 | 19.75303 | 19.58604 | 20.11557 | 19.13558 | 19.13716 | 23.51104 | 20.29222 |
| VCP | 19.32776 | 19.70165 | 18.57442 | 20.07612 | 21.46236 | 20.40901 | 19.95514 | 19.81533 | 19.72189 | 20.55295 | 20.52062 | 21.17547 | 22.08206 | 21.72759 | 18.94311 | 19.87962 | 20.26707 | 19.56332 |
| VDAC1 | 17.01179 | 17.11282 | 17.03851 | 16.67705 | 17.59801 | 17.17995 | 20.67372 | 18.31214 | 24.75135 | 24.52353 | 20.81519 | 17.39915 | 18.83622 | 20.47556 | 17.26696 | 17.18682 | 17.08089 | 18.62478 |
| VDAC2 | 16.5545 | 17.22931 | 17.73275 | 17.89689 | 18.24887 | 20.86624 | 20.50743 | 20.1067 | 26.49702 | 25.51574 | 20.52904 | 17.30099 | 18.03999 | 19.31173 | 16.52739 | 16.82154 | 17.6577 | 19.26218 |
| VDAC3 | 17.16583 | 20.27679 | 15.89193 | 16.09903 | 15.91822 | 18.26112 | 16.87872 | 17.72611 | 22.04315 | 21.1668 | 21.13819 | 15.17331 | 16.37658 | 16.48869 | 16.73166 | 17.65165 | 18.28087 | 16.68552 |
| VGF | 19.30348 | 19.6699 | 15.74945 | 16.65368 | 16.31056 | 20.35576 | 18.31316 | 18.92751 | 21.17228 | 20.22012 | 16.9395 | 15.51608 | 15.24627 | 15.91068 | 17.51088 | 16.65319 | 15.94078 | 16.40344 |
| VIL1 | 18.27649 | 17.99496 | 19.55838 | 19.02618 | 16.29897 | 19.11391 | 18.11615 | 17.24034 | 20.182 | 19.8004 | 21.40008 | 21.23236 | 15.3884 | 16.47623 | 16.57138 | 17.89489 | 16.27214 | 16.862 |
| VIM | 19.66466 | 15.93166 | 15.33974 | 16.73281 | 22.08054 | 21.38213 | 19.57022 | 19.11388 | 15.91748 | 15.3612 | 15.30753 | 15.15582 | 17.24856 | 17.16116 | 14.74731 | 16.65962 | 21.88637 | 22.61587 |
| VIPAS39 | 14.54231 | 12.90129 | 15.24689 | 17.49866 | 18.95552 | 16.90495 | 17.33879 | 16.0728 | 14.47927 | 15.08197 | 14.58997 | 16.2838 | 16.063 | 16.23321 | 15.029 | 15.55094 | 16.11253 | 15.56841 |
| VLDLR | 15.38899 | 14.29347 | 14.90237 | 14.47112 | 17.42709 | 18.42661 | 17.17959 | 15.33212 | 18.42819 | 17.7522 | 14.60006 | 13.12573 | 16.04647 | 15.43615 | 14.93432 | 14.84821 | 15.28849 | 15.253 |
| VPS11 | 15.75949 | 14.93957 | 17.02595 | 15.32584 | 18.14984 | 16.8666 | 17.03822 | 18.05609 | 13.14194 | 14.51764 | 16.13161 | 18.33496 | 16.07178 | 16.6842 | 17.61273 | 18.21414 | 17.67321 | 17.00621 |
| VPS13A | 15.54218 | 15.063 | 14.08505 | 14.78051 | 16.6231 | 16.48319 | 16.86731 | 17.20571 | 15.46379 | 16.0227 | 17.33738 | 18.85982 | 19.08333 | 15.73737 | 20.83426 | 14.14793 | 17.44716 | 15.09268 |
| VPS13C | 17.63333 | 15.96151 | 15.81525 | 15.76084 | 15.8738 | 17.82547 | 16.20226 | 16.2834 | 14.36739 | 16.86685 | 15.68511 | 18.62811 | 17.19376 | 16.32645 | 17.59776 | 17.45007 | 16.39589 | 15.15836 |
| VPS25 | 22.50042 | 22.77702 | 21.63457 | 21.07458 | 15.6185 | 16.18677 | 18.33884 | 16.90796 | 16.75491 | 15.36699 | 17.28386 | 19.13839 | 17.4347 | 15.88423 | 23.21381 | 22.51301 | 17.01421 | 15.35193 |
| VPS26A | 14.182 | 15.78856 | 16.8109 | 16.57903 | 19.38241 | 18.29211 | 16.63576 | 16.27908 | 17.68431 | 16.70963 | 18.02762 | 19.26445 | 18.11981 | 16.6262 | 18.24991 | 18.05985 | 18.10328 | 17.80514 |
| VPS26B | 15.99021 | 15.91715 | 16.44012 | 15.72816 | 18.47613 | 22.65459 | 22.07586 | 21.53926 | 14.31132 | 16.04577 | 14.64157 | 18.18889 | 18.10511 | 22.83914 | 15.71199 | 15.60799 | 21.90802 | 21.84406 |
| VPS28 | 24.56906 | 24.38602 | 23.71804 | 23.88202 | 21.61624 | 20.76216 | 20.61834 | 19.29064 | 20.27355 | 21.28188 | 19.68366 | 18.35071 | 23.91764 | 23.31908 | 24.62514 | 24.02967 | 21.16993 | 22.04071 |
| VPS29 | 18.21445 | 18.11087 | 18.81946 | 18.95695 | 20.1861 | 19.50078 | 18.94967 | 17.066 | 18.53157 | 18.57709 | 19.21104 | 19.97098 | 19.3927 | 20.68435 | 19.54813 | 19.33185 | 20.13338 | 18.76135 |
| VPS33B | 17.09905 | 16.18636 | 16.14051 | 17.23985 | 19.32276 | 17.71056 | 18.80122 | 18.86005 | 14.9053 | 16.32902 | 16.57869 | 17.55835 | 16.56207 | 17.9832 | 17.13634 | 16.74658 | 17.5478 | 16.84133 |
| VPS35 | 17.19022 | 18.44918 | 18.21133 | 17.56458 | 17.64534 | 16.20798 | 16.13321 | 17.71046 | 17.59378 | 16.35909 | 18.97776 | 20.03645 | 17.15318 | 15.03885 | 18.27377 | 17.9451 | 17.50846 | 16.7274 |
| VPS36 | 22.18187 | 22.01453 | 21.69136 | 20.78775 | 16.30787 | 17.36234 | 17.29442 | 19.12126 | 16.18876 | 16.67196 | 17.32681 | 17.45165 | 20.1635 | 18.44256 | 22.25369 | 21.80421 | 17.59457 | 18.17492 |
| VPS37A | 22.12242 | 22.23743 | 19.56361 | 19.04949 | 19.73709 | 18.73132 | 16.03424 | 16.94704 | 17.51221 | 16.29647 | 17.16902 | 16.54418 | 18.33485 | 17.57637 | 19.54314 | 19.77126 | 17.25955 | 16.60293 |
| VPS37B | 24.15399 | 23.8628 | 23.51207 | 23.71532 | 20.67179 | 19.33075 | 18.66204 | 17.29902 | 18.40126 | 19.16908 | 19.86284 | 19.51654 | 22.91876 | 22.61728 | 24.66416 | 23.85093 | 19.10575 | 21.71128 |
| VPS37C | 22.75957 | 22.70985 | 22.2791 | 21.68362 | 16.51996 | 18.14861 | 17.40554 | 17.68267 | 19.37154 | 17.44682 | 15.40612 | 15.13713 | 19.48212 | 18.4208 | 21.97127 | 22.42348 | 16.71985 | 17.23515 |
| VPS37D | 20.2805 | 19.26913 | 18.54293 | 18.52427 | 13.36209 | 15.70327 | 17.2005 | 16.6517 | 13.25954 | 16.62746 | 14.85253 | 13.5585 | 17.36792 | 15.92694 | 21.17249 | 19.83477 | 16.56219 | 15.83555 |
| VPS39 | 17.15557 | 16.95077 | 18.29619 | 13.26377 | 14.64749 | 19.67167 | 19.51993 | 18.81049 | 15.26571 | 12.92727 | 15.33824 | 18.10314 | 13.61313 | 15.08923 | 14.2297 | 14.84018 | 16.20482 | 21.28713 |
| VPS45 | 19.25856 | 19.71982 | 18.43592 | 15.78473 | 17.39654 | 16.13495 | 17.56638 | 17.81605 | 17.45551 | 17.91573 | 17.52037 | 18.64179 | 16.80727 | 16.98438 | 16.99418 | 18.66648 | 16.04592 | 16.23128 |
| VPS4A | 20.0511 | 20.05561 | 19.79754 | 20.42069 | 20.21334 | 20.89097 | 18.61823 | 17.46437 | 19.48935 | 19.8048 | 20.06227 | 20.3057 | 20.82393 | 20.85969 | 19.29191 | 19.53319 | 17.7946 | 18.4045 |
| VPS4B | 20.32722 | 20.6225 | 20.70727 | 21.00936 | 20.53439 | 19.59234 | 17.57412 | 17.20894 | 20.86248 | 20.74662 | 20.92251 | 20.18357 | 20.85712 | 20.63122 | 20.27465 | 20.70562 | 18.07109 | 19.31214 |
| VPS9D1 | 15.27246 | 12.71134 | 16.54553 | 15.89691 | 18.24172 | 16.55709 | 17.29665 | 18.58297 | 11.49455 | 15.5494 | 15.90565 | 13.40986 | 16.48643 | 14.48817 | 14.41078 | 15.98014 | 14.93095 | 15.14173 |

| | | | | | | | | | | | | | | | | | | |
|---|---|---|---|---|---|---|---|---|---|---|---|---|---|---|---|---|---|---|
| VTA1 | 21.31001 | 20.63074 | 19.97527 | 21.34642 | 21.76614 | 21.06977 | 20.26217 | 19.72489 | 21.232 | 21.79658 | 21.12399 | 20.89724 | 21.7415 | 21.62531 | 20.46419 | 20.03752 | 20.48791 | 20.91901 |
| VWA1 | 16.98031 | 15.24661 | 19.41744 | 18.29988 | 14.82826 | 16.13633 | 15.94053 | 16.17799 | 15.83199 | 14.9672 | 16.045 | 13.91495 | 15.21333 | 16.29074 | 17.913 | 17.9147 | 15.5565 | 15.27291 |
| VWDE | 14.40631 | 14.25779 | 13.84794 | 15.16857 | 20.9845 | 14.55663 | 18.77917 | 15.5303 | 16.39774 | 12.85707 | 13.84218 | 12.15409 | 19.77877 | 15.24472 | 15.29488 | 12.37499 | 13.89123 | 15.96811 |
| WARS1 | 17.424 | 16.40651 | 17.24241 | 17.90526 | 20.22517 | 19.00954 | 17.98842 | 16.83482 | 19.47672 | 19.43042 | 17.74526 | 18.60661 | 19.81875 | 19.13969 | 17.56378 | 18.15319 | 18.74088 | 20.35482 |
| WASF1 | 15.56726 | 16.49853 | 16.18028 | 15.77185 | 18.25155 | 21.54625 | 17.91609 | 16.43784 | 17.3004 | 16.8792 | 18.26187 | 18.98931 | 15.82179 | 14.54858 | 16.49488 | 15.9117 | 17.33385 | 16.84309 |
| WASF2 | 17.95246 | 17.12086 | 18.19662 | 18.33594 | 20.23065 | 18.16147 | 19.07183 | 17.95424 | 17.90955 | 18.64558 | 16.23553 | 18.72158 | 19.48051 | 20.31057 | 18.57645 | 19.12128 | 20.49047 | 20.0358 |
| WASF3 | 17.61511 | 16.43432 | 16.73635 | 17.58386 | 16.88535 | 17.23616 | 15.71669 | 17.55168 | 13.82007 | 15.64614 | 15.26244 | 16.51979 | 16.79859 | 17.48713 | 17.30625 | 18.08639 | 16.45355 | 17.98343 |
| WASHC2A | 16.22708 | 16.52323 | 18.86958 | 18.32582 | 19.64817 | 17.92088 | 17.58494 | 18.01091 | 17.09255 | 16.15307 | 15.87303 | 16.15195 | 17.7478 | 17.27674 | 17.88455 | 16.31946 | 18.31724 | 20.51468 |
| WASHC5 | 16.70303 | 15.73292 | 18.72626 | 15.63671 | 14.95073 | 14.96943 | 15.73193 | 15.33136 | 14.05974 | 15.71501 | 14.40175 | 17.57568 | 13.76027 | 15.76781 | 15.15491 | 15.527 | 15.07689 | 15.83461 |
| WASL | 20.28942 | 19.94013 | 19.24709 | 20.56596 | 18.38007 | 19.37104 | 16.74021 | 17.23276 | 17.60523 | 16.80758 | 15.9447 | 15.69281 | 19.0161 | 19.21552 | 20.79272 | 20.07064 | 17.74272 | 17.85918 |
| WBP2 | 16.91466 | 17.14232 | 17.64379 | 14.59533 | 18.91927 | 14.84994 | 19.27065 | 19.39568 | 14.79254 | 17.55384 | 18.02395 | 14.06208 | 17.9267 | 19.84727 | 17.82657 | 17.74076 | 20.14323 | 14.49782 |
| WDFY1 | 21.75033 | 21.52791 | 22.49208 | 21.97643 | 18.87654 | 16.70737 | 12.75399 | 15.74917 | 13.50317 | 18.4671 | 15.22334 | 19.78709 | 15.56327 | 18.49835 | 22.97097 | 22.50633 | 12.843 | 14.54182 |
| WDR1 | 20.93815 | 20.65111 | 20.58371 | 20.72762 | 23.46894 | 22.39576 | 21.18048 | 20.32749 | 19.80227 | 20.38102 | 20.79246 | 21.43674 | 22.74197 | 22.63867 | 20.99469 | 21.10257 | 22.07733 | 21.85401 |
| WDR11 | 15.70826 | 16.12114 | 13.86912 | 15.84557 | 16.14927 | 16.09563 | 13.58931 | 14.75563 | 11.57735 | 15.55882 | 16.00556 | 17.37195 | 13.80282 | 15.96226 | 16.57857 | 15.17087 | 13.99993 | 16.61904 |
| WDR20 | 20.5155 | 20.52371 | 16.12105 | 16.70305 | 19.20116 | 17.88671 | 17.43366 | 19.01616 | 15.66641 | 17.67515 | 14.82382 | 15.89243 | 18.40941 | 18.53254 | 15.63968 | 16.61557 | 18.94496 | 19.62264 |
| WDR26 | 14.78914 | 18.15659 | 17.75997 | 18.44163 | 14.35725 | 17.01575 | 16.6843 | 16.08469 | 16.0635 | 16.27053 | 16.23529 | 18.04927 | 16.98533 | 14.67969 | 18.06195 | 15.50006 | 15.5848 | 15.98905 |
| WDR33 | 14.99748 | 14.84505 | 16.68443 | 17.7718 | 14.71926 | 15.15372 | 17.91521 | 14.42143 | 13.89743 | 15.59078 | 16.44361 | 16.82018 | 17.21173 | 17.04146 | 14.27464 | 15.47421 | 15.40251 | 17.48519 |
| WDR36 | 14.73716 | 15.70299 | 16.31565 | 16.67422 | 14.52297 | 13.93188 | 16.06029 | 14.6383 | 13.1397 | 12.53663 | 17.35194 | 16.96138 | 17.69941 | 13.41074 | 15.77957 | 15.61448 | 16.96986 | 15.02113 |
| WDR43 | 15.68289 | 13.45713 | 16.03366 | 14.85759 | 16.67246 | 16.5061 | 15.8346 | 16.47644 | 14.78482 | 12.23569 | 12.92577 | 17.50131 | 18.04737 | 14.47865 | 15.57955 | 15.55851 | 16.57289 | 13.60546 |
| WDR44 | 16.05729 | 15.28987 | 15.12582 | 14.93045 | 18.01103 | 15.74509 | 16.1531 | 14.67837 | 14.08159 | 13.59444 | 15.24703 | 14.54618 | 18.40164 | 15.85473 | 17.03148 | 15.96238 | 17.91569 | 20.22591 |
| WDR45B | 18.1242 | 17.90631 | 17.50859 | 17.97293 | 16.61215 | 16.28658 | 18.15648 | 15.32952 | 16.45045 | 16.1517 | 17.29889 | 17.7591 | 15.7995 | 17.07185 | 19.22561 | 19.23286 | 16.45561 | 14.55189 |
| WDR48 | 15.66173 | 15.61131 | 17.92679 | 16.13599 | 19.26631 | 18.17722 | 17.76074 | 18.1071 | 15.90687 | 17.49177 | 15.34353 | 17.56787 | 18.18594 | 18.0116 | 16.84791 | 16.96124 | 19.86934 | 19.07979 |
| WDR59 | 14.32627 | 17.30191 | 17.10195 | 18.02038 | 14.14224 | 15.55915 | 15.74174 | 17.27967 | 17.17071 | 15.98292 | 18.06789 | 17.96597 | 17.64528 | 14.99675 | 18.03815 | 18.11065 | 14.08841 | 16.83926 |
| WDR6 | 18.46383 | 19.66404 | 17.03001 | 16.06842 | 15.79751 | 16.13587 | 17.39731 | 15.61695 | 13.25478 | 14.44849 | 16.27088 | 18.07762 | 14.34698 | 14.53562 | 15.58396 | 18.44607 | 16.50103 | 15.96616 |
| WDR77 | 15.39674 | 15.8087 | 17.49875 | 15.41027 | 14.627 | 17.04412 | 14.61971 | 15.45642 | 14.65896 | 15.50341 | 15.90779 | 19.86313 | 14.8622 | 16.22661 | 14.77654 | 15.94794 | 13.88119 | 13.45537 |
| WDR82 | 17.90541 | 17.50665 | 20.36039 | 18.25395 | 13.20516 | 15.26104 | 16.23377 | 18.36868 | 13.85405 | 15.59028 | 14.4341 | 19.73981 | 15.16859 | 15.04122 | 15.46593 | 17.5174 | 14.33237 | 15.71093 |
| WFDC2 | 14.91427 | 13.42804 | 15.35776 | 21.34239 | 12.64388 | 14.82599 | 15.81212 | 16.57972 | 19.24643 | 18.28952 | 15.02834 | 12.98707 | 16.0301 | 15.6866 | 21.33032 | 14.57882 | 15.7262 | 17.66104 |
| WIPF2 | 20.9061 | 20.80224 | 20.57481 | 20.74531 | 16.77567 | 16.53536 | 15.50366 | 16.1526 | 14.23465 | 16.02439 | 15.92164 | 16.01051 | 18.99757 | 17.82075 | 21.62487 | 20.74505 | 16.36158 | 17.02326 |
| WLS | 15.12464 | 14.79732 | 15.627 | 16.57268 | 15.30481 | 15.03192 | 15.59614 | 15.40751 | 20.31037 | 17.55976 | 15.95696 | 14.6063 | 16.16459 | 16.79857 | 15.86854 | 17.43964 | 15.07691 | 15.18195 |
| WNK1 | 15.31701 | 18.54108 | 15.55492 | 14.72872 | 14.41547 | 16.13602 | 14.55061 | 19.88799 | 13.43278 | 13.06978 | 13.07319 | 16.47429 | 14.63396 | 15.11852 | 13.03177 | 15.5602 | 13.97658 | 16.99526 |
| WWP1 | 19.90337 | 20.21937 | 18.86427 | 19.23804 | 18.01324 | 16.65879 | 16.53212 | 16.87358 | 15.11363 | 15.64656 | 16.45667 | 15.01174 | 18.07772 | 18.75375 | 20.79778 | 19.89553 | 17.81817 | 17.97667 |
| WWP2 | 20.72627 | 21.08637 | 19.79973 | 19.70631 | 14.5309 | 16.87171 | 15.10392 | 15.60692 | 15.37973 | 15.45631 | 15.39952 | 15.15426 | 15.40139 | 16.47837 | 20.8262 | 20.30187 | 15.58383 | 15.37774 |
| XPNPEP1 | 16.34416 | 16.56479 | 16.78798 | 16.74564 | 16.9618 | 16.8472 | 18.25259 | 17.02763 | 16.15179 | 16.23803 | 18.01789 | 19.39353 | 18.12992 | 18.40543 | 17.80938 | 17.59973 | 18.27721 | 17.19985 |
| XPO1 | 16.95525 | 17.92191 | 16.59847 | 15.76485 | 16.43367 | 15.8714 | 17.14195 | 16.2703 | 16.37601 | 16.7092 | 17.51217 | 19.94651 | 15.64791 | 18.37144 | 16.17445 | 16.30005 | 16.67497 | 16.00781 |
| XPO4 | 16.96234 | 14.91508 | 16.00985 | 14.39475 | 13.89977 | 15.69792 | 17.89238 | 17.13546 | 15.26596 | 16.00077 | 15.2734 | 17.74112 | 15.62054 | 17.51068 | 16.21532 | 16.76911 | 17.35784 | 14.88594 |
| XPO7 | 15.68718 | 15.9239 | 15.65771 | 14.88993 | 17.18424 | 16.59408 | 16.24035 | 16.50128 | 14.1848 | 15.62145 | 16.17093 | 18.54843 | 15.80837 | 14.77255 | 15.30313 | 15.69609 | 16.92388 | 15.99631 |
| XPOT | 17.94375 | 16.32846 | 17.76499 | 14.51714 | 14.77852 | 14.90401 | 16.34188 | 15.4895 | 13.73371 | 16.39367 | 15.80438 | 18.69047 | 15.39984 | 15.09245 | 17.17401 | 17.07865 | 16.39259 | 14.57202 |
| XPR1 | 17.48678 | 15.33727 | 17.1903 | 17.00269 | 18.85341 | 15.97011 | 17.26482 | 16.88867 | 15.06088 | 16.70436 | 17.42397 | 17.6697 | 17.30984 | 16.84597 | 15.8967 | 16.39226 | 16.09892 | 15.97262 |
| XRCC5 | 18.01702 | 17.27433 | 17.53734 | 16.09439 | 18.15888 | 16.92926 | 19.18074 | 19.67749 | 16.2254 | 17.56782 | 19.47153 | 20.75749 | 17.21102 | 16.55508 | 17.30348 | 16.57149 | 16.83624 | 17.68978 |
| XRCC6 | 18.57638 | 17.44639 | 16.72836 | 16.98479 | 18.03073 | 17.71957 | 18.6435 | 18.81458 | 17.30628 | 17.73399 | 19.80982 | 20.76192 | 17.87848 | 17.21834 | 17.07236 | 16.41281 | 16.97424 | 17.62207 |
| XRN2 | 16.60068 | 16.31605 | 15.8305 | 16.70377 | 15.68085 | 19.37639 | 14.87888 | 16.51366 | 15.01804 | 14.05599 | 17.63303 | 18.52474 | 14.39285 | 14.38699 | 16.87697 | 16.05994 | 16.99872 | 16.58405 |
| XXYLT1 | 18.17644 | 16.16219 | 15.75169 | 16.55834 | 15.1986 | 16.44438 | 16.20403 | 16.29603 | 19.29248 | 16.3969 | 15.90303 | 13.84832 | 15.39083 | 15.01612 | 15.95568 | 15.84447 | 16.2907 | 15.645 |
| XYLT2 | 17.37476 | 18.3962 | 14.84606 | 16.9431 | 12.75096 | 15.61009 | 15.1875 | 15.85729 | 17.76514 | 15.86657 | 15.76685 | 14.90253 | 15.62534 | 16.45102 | 17.74467 | 16.12613 | 14.59172 | 13.5169 |
| YARS1 | 17.22992 | 17.84714 | 17.88203 | 18.22933 | 19.39822 | 17.84904 | 19.06509 | 18.4134 | 17.88662 | 18.39244 | 18.42586 | 19.25311 | 20.88602 | 20.79885 | 18.50371 | 18.67822 | 19.26596 | 19.55916 |
| YBX1 | 21.42656 | 20.93871 | 21.30007 | 21.55857 | 17.41517 | 16.8734 | 16.76808 | 18.21473 | 16.40557 | 16.06814 | 20.43585 | 21.25269 | 18.42274 | 16.30974 | 19.93157 | 20.74489 | 16.7898 | 16.897 |
| YES1 | 19.86607 | 19.79068 | 20.1829 | 20.01414 | 19.60329 | 19.06954 | 16.76035 | 17.23927 | 19.05146 | 18.49797 | 18.36616 | 18.94017 | 19.10082 | 19.0712 | 20.74409 | 20.44643 | 20.5341 | 20.94023 |
| YIPF6 | 15.92983 | 14.6039 | 21.30817 | 18.86809 | 14.32971 | 16.10377 | 15.21038 | 15.48075 | 14.27976 | 13.50195 | 14.78069 | 18.60085 | 15.8588 | 14.19187 | 18.13799 | 18.04181 | 16.6261 | 15.27992 |
| YKT6 | 20.15793 | 20.29405 | 18.78622 | 18.4703 | 18.32674 | 15.44636 | 16.28915 | 15.87032 | 19.14209 | 17.78661 | 17.43361 | 19.06344 | 18.3224 | 17.92964 | 19.85332 | 19.72759 | 17.00442 | 19.11572 |
| YOD1 | 20.07057 | 20.43591 | 17.58048 | 18.0577 | 16.74789 | 16.83161 | 16.32614 | 14.85614 | 13.1155 | 16.11638 | 14.04291 | 15.50659 | 18.75393 | 18.27166 | 18.39665 | 18.6789 | 17.73914 | 16.76234 |
| YRDC | 18.26508 | 14.54797 | 16.02932 | 14.53734 | 14.1839 | 15.22871 | 15.36071 | 15.42039 | 17.68519 | 16.44357 | 17.35569 | 18.16008 | 13.37592 | 16.99546 | 16.29061 | 16.23192 | 13.59759 | 14.98592 |
| YTHDF1 | 19.0552 | 18.06263 | 17.39577 | 16.27585 | 14.34098 | 15.51179 | 19.04319 | 15.95152 | 16.28015 | 14.23512 | 15.79957 | 16.87641 | 18.05814 | 16.39415 | 17.026 | 18.51301 | 15.82548 | 15.84992 |
| YTHDF2 | 19.38487 | 18.76916 | 18.84318 | 17.42872 | 15.47327 | 17.50065 | 16.04175 | 16.71084 | 14.63454 | 15.57682 | 16.23254 | 17.06601 | 17.97214 | 17.92373 | 18.85773 | 17.68964 | 15.14899 | 15.49947 |
| YTHDF3 | 19.55229 | 19.34675 | 16.95204 | 17.43202 | 18.26665 | 17.66889 | 17.36422 | 21.04733 | 16.07373 | 16.29629 | 15.47113 | 15.30557 | 17.65363 | 14.7512 | 15.89652 | 17.70536 | 15.67772 | 13.35821 |
| YWHAB | 21.35101 | 21.25664 | 22.35971 | 23.22171 | 24.63642 | 23.55953 | 23.02911 | 21.6129 | 21.61679 | 22.14087 | 22.79035 | 22.54424 | 24.06073 | 24.13605 | 22.93841 | 22.73142 | 25.13229 | 25.40924 |
| YWHAE | 21.29346 | 20.62812 | 23.31811 | 23.266 | 22.4655 | 22.63048 | 21.62345 | 21.09755 | 21.19013 | 21.93166 | 22.90752 | 23.91313 | 22.99063 | 23.0872 | 23.90208 | 23.95848 | 22.96875 | 21.30149 |
| YWHAG | 20.34406 | 20.40809 | 21.7204 | 22.72153 | 23.28509 | 22.73842 | 22.83 | 21.98589 | 20.25143 | 21.63244 | 22.28237 | 22.46078 | 23.40299 | 24.38593 | 23.19979 | 22.40805 | 24.59312 | 23.62407 |
| YWHAH | 19.27905 | 19.23308 | 19.54582 | 20.65277 | 22.72286 | 21.93859 | 21.0461 | 20.53468 | 19.05122 | 20.42885 | 20.25524 | 21.00329 | 21.65179 | 21.83625 | 20.04618 | 20.54742 | 22.41567 | 22.33123 |
| YWHAQ | 19.97177 | 19.03294 | 20.86238 | 21.49116 | 23.41154 | 22.82212 | 22.12848 | 21.80011 | 20.69437 | 21.25261 | 22.51296 | 22.54425 | 22.47716 | 23.06547 | 21.82817 | 21.28886 | 24.48928 | 23.66832 |
| YWHAZ | 22.36852 | 22.43927 | 22.55844 | 23.75399 | 24.98654 | 24.29362 | 24.82136 | 24.25455 | 23.83829 | 24.5505 | 23.62468 | 23.54192 | 24.13346 | 24.79229 | 23.0818 | 23.24528 | 26.70258 | 26.14263 |
| ZC2HC1A | 15.0102 | 16.29723 | 17.19211 | 14.56162 | 16.71631 | 14.08956 | 15.59826 | 16.14398 | 16.30992 | 16.8838 | 14.52246 | 14.16608 | 17.85905 | 17.47124 | 15.24812 | 15.30531 | 16.45329 | 17.52393 |
| ZC3H10 | 21.08477 | 19.95656 | 19.31734 | 21.0727 | 14.60448 | 16.66488 | 18.37014 | 16.95188 | 14.49822 | 15.16218 | 15.84668 | 14.30898 | 20.26468 | 18.58203 | 18.26414 | 19.70317 | 18.03939 | 15.69871 |
| ZC3H15 | 17.99809 | 16.04398 | 18.63429 | 16.52753 | 17.78404 | 15.59772 | 15.51004 | 17.10183 | 12.90591 | 13.53818 | 18.19683 | 18.1087 | 17.95058 | 16.79398 | 15.1628 | 14.71898 | 16.7812 | 18.31025 |
| ZC3HAV1 | 19.34216 | 18.81518 | 19.97702 | 20.92469 | 21.59269 | 20.873 | 17.89385 | 18.36405 | 17.20421 | 17.07735 | 16.86109 | 16.47067 | 19.91552 | 19.66054 | 19.50293 | 19.597 | 18.51069 | 18.56163 |
| ZC3HAV1L | 17.02237 | 16.60861 | 18.04753 | 18.89383 | 18.39064 | 17.53261 | 16.37318 | 16.17863 | 15.39747 | 15.38445 | 16.60149 | 17.79143 | 15.66606 | 16.21547 | 18.73293 | 18.14773 | 17.45219 | 16.76816 |
| ZCCHC3 | 18.26137 | 18.26694 | 15.72798 | 15.12519 | 12.44154 | 16.00198 | 16.76132 | 17.3911 | 16.25427 | 16.42168 | 15.40064 | 16.61492 | 18.00507 | 17.22552 | 16.51029 | 14.86607 | 14.27293 | 14.13495 |
| ZDHHC20 | 17.5773 | 17.59492 | 20.51113 | 20.95815 | 20.24838 | 13.97445 | 17.37873 | 13.97127 | 17.93505 | 17.09549 | 18.70632 | 16.89109 | 18.34291 | 16.12215 | 20.10102 | 20.56363 | 15.12171 | 17.32881 |
| ZDHHC5 | 18.50397 | 18.21067 | 20.12475 | 20.50368 | 21.51523 | 19.49962 | 18.68056 | 16.63166 | 18.45395 | 19.61706 | 19.82032 | 19.45008 | 21.79903 | 22.28927 | 20.49007 | 20.59794 | 21.97608 | 22.56913 |
| ZFAND2B | 20.73046 | 20.42305 | 16.67011 | 17.87174 | 13.06034 | 16.30718 | 14.65919 | 15.99127 | 18.56511 | 14.75507 | 16.00035 | 14.02085 | 15.46842 | 16.37165 | 18.75067 | 18.68594 | 15.41872 | 18.10297 |
| ZFP36L1 | 20.28066 | 19.0937 | 15.63708 | 13.24097 | 13.58002 | 15.67677 | 13.17427 | 15.50633 | 14.10885 | 14.64421 | 16.95178 | 12.8105 | 14.1129 | 16.72434 | 13.42604 | 15.87587 | 17.1656 | 14.67381 |
| ZFP91 | 14.70737 | 16.79023 | 13.78691 | 17.62247 | 15.4188 | 12.74866 | 17.35998 | 16.38365 | 13.0732 | 15.48964 | 16.23425 | 12.57136 | 18.66749 | 17.13852 | 16.60945 | 27.02831 | 14.80253 | 15.93701 |
| ZFYVE16 | 14.96626 | 16.30699 | 13.63464 | 13.14902 | 18.91128 | 14.53539 | 15.47512 | 16.17504 | 15.26087 | 14.99079 | 15.72079 | 14.96684 | 17.72647 | 16.61264 | 15.02896 | 15.4254 | 14.30747 | 14.13826 |
| ZMPSTE24 | 15.99197 | 17.18057 | 16.6735 | 15.34144 | 15.04425 | 17.33281 | 16.56774 | 16.99719 | 19.89046 | 16.41935 | 16.77575 | 16.44559 | 15.92476 | 15.34809 | 15.15375 | 15.59928 | 16.77941 | 18.8663 |
| ZNF185 | 16.82513 | 16.64369 | 16.41595 | 16.07206 | 18.59043 | 17.46352 | 16.16996 | 17.09743 | 16.683 | 16.12261 | 16.40021 | 16.28554 | 14.51187 | 15.72326 | 16.2656 | 16.31572 | 20.98634 | 17.9408 |
| ZNF330 | 16.04892 | 14.5572 | 14.72281 | 12.71463 | 14.62007 | 13.8366 | 15.29318 | 17.48361 | 13.84991 | 14.32268 | 14.45504 | 15.38091 | 16.60097 | 17.54085 | 16.25928 | 13.54534 | 16.9551 | 17.36543 |
| ZNF512 | 19.22348 | 17.49955 | 15.65209 | 13.72855 | 13.16916 | 15.49078 | 14.42915 | 15.8513 | 15.03773 | 17.66746 | 15.9444 | 12.87463 | 14.78793 | 14.92334 | 14.41175 | 15.35804 | 15.17 | 13.44088 |
| ZNF622 | 17.38295 | 19.65557 | 15.81755 | 17.15161 | 13.2699 | 17.00483 | 15.21372 | 15.45768 | 13.96442 | 14.48532 | 19.04258 | 13.67468 | 17.44067 | 16.79835 | 18.28423 | 18.03772 | 15.47562 | 17.38073 |
| ZNF638 | 14.63112 | 14.65522 | 12.41277 | 14.99244 | 14.58234 | 13.80145 | 16.21565 | 13.3479 | 14.80266 | 14.77668 | 19.9106 | 16.19418 | 19.51694 | 17.09977 | 13.5022 | 13.42783 | 14.40718 | 14.80903 |
| ZNF706 | 19.38218 | 16.52149 | 11.99666 | 16.07152 | 12.37634 | 13.82834 | 13.95965 | 14.78098 | 17.1883 | 12.03789 | 18.27266 | 15.25806 | 15.33221 | 15.08625 | 13.40097 | 14.10977 | 18.13796 | 20.20526 |
| ZNRF2 | 16.24975 | 18.19504 | 14.89344 | 15.41605 | 16.25445 | 17.39324 | 17.65653 | 15.13119 | 17.89398 | 13.94882 | 14.16555 | 12.87598 | 13.84301 | 12.74412 | 15.54493 | 16.20511 | 15.38355 | 14.59189 |
| ZPR1 | 16.09243 | 15.60229 | 18.55441 | 17.79542 | 15.17293 | 16.68443 | 15.75983 | 15.99639 | 15.14269 | 15.62627 | 18.06165 | 18.46056 | 17.13314 | 16.70295 | 18.15469 | 17.05503 | 16.2529 | 16.26658 |
| ZRANB2 | 12.05183 | 16.26892 | 15.05468 | 15.80403 | 16.19407 | 14.1199 | 13.82514 | 15.96146 | 14.22311 | 14.2989 | 16.26676 | 15.74915 | 15.66283 | 16.04988 | 13.54771 | 16.00491 | 16.47645 | 14.31058 |
| ZW10 | 16.43174 | 16.99374 | 16.55364 | 16.02067 | 17.97806 | 16.44357 | 16.72348 | 17.12551 | 16.01602 | 16.65619 | 17.08412 | 16.98333 | 20.36313 | 16.42103 | 15.15132 | 15.19142 | 17.80233 | 16.3365 |
| ZYG11B | 14.11242 | 14.28612 | 15.53462 | 15.13927 | 17.85404 | 16.17 | 15.06669 | 14.16264 | 13.79477 | 18.58754 | 15.28392 | 16.4426 | 15.78273 | 14.92285 | 15.36675 | 15.46847 | 15.16426 | 17.23828 |
| ZYX | 16.1846 | 16.16613 | 16.77651 | 17.21202 | 19.23911 | 17.6386 | 14.68294 | 16.97967 | 14.86858 | 15.49155 | 16.75172 | 14.54566 | 17.28743 | 17.13429 | 17.73062 | 17.25984 | 20.40171 | 21.81945 |

| NAME | GS<br> follow link to MSigDB | GS DETAILS | SIZE | ES | NES | NOM p-val | FDR q-val | FWER p-val | RANK AT M/ | LEADING EDGE |
|---|---|---|---|---|---|---|---|---|---|---|
| KEGG_RIBOSOME | KEGG_RIBOSOME | Details ... | 75 | 0.686165 | 2.564688 | 0 | 0 | 0 | 1052 | tags=84%, list=27%, signal=112% |
| KEGG_ENDOCYTOSIS | KEGG_ENDOCYTOSIS | Details ... | 115 | 0.528894 | 2.068208 | 0 | 0 | 0 | 736 | tags=42%, list=19%, signal=50% |
| KEGG_ENDOMETRIAL_CANCER | KEGG_ENDOMETRIAL_CANCER | Details ... | 22 | 0.639747 | 2.032773 | 0 | 0 | 0 | 1018 | tags=73%, list=26%, signal=97% |
| KEGG_MTOR_SIGNALING_PATHWAY | KEGG_MTOR_SIGNALING_PATHWA | Details ... | 23 | 0.63665 | 1.99321 | 0 | 0 | 0 | 1080 | tags=61%, list=27%, signal=83% |
| KEGG_PANCREATIC_CANCER | KEGG_PANCREATIC_CANCER | Details ... | 29 | 0.561472 | 1.865829 | 0 | 0.014549 | 0.08 | 1018 | tags=62%, list=26%, signal=83% |
| KEGG_B_CELL_RECEPTOR_SIGNALING_PATHWAY | KEGG_B_CELL_RECEPTOR_SIGNAL | Details ... | 32 | 0.588921 | 1.844488 | 0 | 0.013613 | 0.09 | 1018 | tags=59%, list=26%, signal=79% |
| KEGG_ECM_RECEPTOR_INTERACTION | KEGG_ECM_RECEPTOR_INTERACT | Details ... | 40 | 0.525001 | 1.810747 | 0 | 0.014273 | 0.11 | 567 | tags=33%, list=14%, signal=38% |
| KEGG_ACUTE_MYELOID_LEUKEMIA | KEGG_ACUTE_MYELOID_LEUKEMI/ | Details ... | 19 | 0.613962 | 1.794644 | 0 | 0.014799 | 0.13 | 1018 | tags=63%, list=26%, signal=85% |
| KEGG_CHRONIC_MYELOID_LEUKEMIA | KEGG_CHRONIC_MYELOID_LEUKE | Details ... | 25 | 0.570729 | 1.74836 | 0.013889 | 0.026767 | 0.26 | 1018 | tags=56%, list=26%, signal=75% |
| KEGG_T_CELL_RECEPTOR_SIGNALING_PATHWAY | KEGG_T_CELL_RECEPTOR_SIGNAL | Details ... | 40 | 0.516182 | 1.729357 | 0.013699 | 0.02947 | 0.31 | 1018 | tags=52%, list=26%, signal=70% |
| KEGG_NON_SMALL_CELL_LUNG_CANCER | KEGG_NON_SMALL_CELL_LUNG_C | Details ... | 23 | 0.571775 | 1.72675 | 0 | 0.026791 | 0.31 | 1018 | tags=65%, list=26%, signal=87% |
| KEGG_VEGF_SIGNALING_PATHWAY | KEGG_VEGF_SIGNALING_PATHWA\ | Details ... | 32 | 0.543178 | 1.679382 | 0 | 0.037828 | 0.43 | 1018 | tags=59%, list=26%, signal=79% |
| KEGG_HEMATOPOIETIC_CELL_LINEAGE | KEGG_HEMATOPOIETIC_CELL_LINI | Details ... | 16 | 0.60605 | 1.67008 | 0 | 0.041015 | 0.48 | 134 | tags=31%, list=3%, signal=32% |
| KEGG_COMPLEMENT_AND_COAGULATION_CASCAI | KEGG_COMPLEMENT_AND_COAGU | Details ... | 15 | 0.617759 | 1.637152 | 0.046154 | 0.056781 | 0.56 | 721 | tags=53%, list=18%, signal=65% |
| KEGG_COLORECTAL_CANCER | KEGG_COLORECTAL_CANCER | Details ... | 25 | 0.533576 | 1.632484 | 0 | 0.053591 | 0.57 | 1018 | tags=56%, list=26%, signal=75% |
| KEGG_ALDOSTERONE_REGULATED_SODIUM_REAB: | KEGG_ALDOSTERONE_REGULATEI | Details ... | 18 | 0.558984 | 1.612752 | 0.015625 | 0.061207 | 0.64 | 967 | tags=61%, list=25%, signal=81% |
| KEGG_ADHERENS_JUNCTION | KEGG_ADHERENS_JUNCTION | Details ... | 50 | 0.446405 | 1.611108 | 0 | 0.058087 | 0.64 | 909 | tags=42%, list=23%, signal=54% |
| KEGG_PROSTATE_CANCER | KEGG_PROSTATE_CANCER | Details ... | 29 | 0.528024 | 1.607819 | 0 | 0.055357 | 0.64 | 1018 | tags=55%, list=26%, signal=74% |
| KEGG_FC_GAMMA_R_MEDIATED_PHAGOCYTOSIS | KEGG_FC_GAMMA_R_MEDIATED_P | Details ... | 58 | 0.444276 | 1.586389 | 0 | 0.066903 | 0.71 | 1018 | tags=48%, list=26%, signal=64% |
| KEGG_MAPK_SIGNALING_PATHWAY | KEGG_MAPK_SIGNALING_PATHWA' | Details ... | 72 | 0.415286 | 1.562225 | 0 | 0.07889 | 0.77 | 1118 | tags=54%, list=28%, signal=74% |
| KEGG_SMALL_CELL_LUNG_CANCER | KEGG_SMALL_CELL_LUNG_CANCER | | 30 | 0.486243 | 1.541413 | 0 | 0.088638 | 0.81 | 477 | tags=30%, list=12%, signal=34% |
| KEGG_ERBB_SIGNALING_PATHWAY | KEGG_ERBB_SIGNALING_PATHWAY | | 35 | 0.488069 | 1.533175 | 0.042857 | 0.091282 | 0.85 | 1018 | tags=63%, list=26%, signal=84% |
| KEGG_LEUKOCYTE_TRANSENDOTHELIAL_MIGRATIC | KEGG_LEUKOCYTE_TRANSENDOTHELIAL_MIGI | | 48 | 0.445793 | 1.519272 | 0.054054 | 0.09398 | 0.88 | 944 | tags=40%, list=24%, signal=51% |
| KEGG_FC_EPSILON_RI_SIGNALING_PATHWAY | KEGG_FC_EPSILON_RI_SIGNALING_PATHWAY | | 32 | 0.455066 | 1.502668 | 0.058824 | 0.098481 | 0.88 | 1018 | tags=56%, list=26%, signal=75% |
| KEGG_NEUROTROPHIN_SIGNALING_PATHWAY | KEGG_NEUROTROPHIN_SIGNALING_PATHWAY | | 53 | 0.420082 | 1.492032 | 0.042857 | 0.102645 | 0.92 | 1018 | tags=43%, list=26%, signal=58% |
| KEGG_PATHWAYS_IN_CANCER | KEGG_PATHWAYS_IN_CANCER | | 97 | 0.3823 | 1.491812 | 0 | 0.099031 | 0.92 | 1039 | tags=41%, list=26%, signal=55% |
| KEGG_RENAL_CELL_CARCINOMA | KEGG_RENAL_CELL_CARCINOMA | | 31 | 0.428703 | 1.472401 | 0.067568 | 0.109857 | 0.97 | 1073 | tags=55%, list=27%, signal=75% |
| KEGG_MELANOMA | KEGG_MELANOMA | | 19 | 0.494086 | 1.442442 | 0.081967 | 0.127814 | 0.98 | 1018 | tags=53%, list=26%, signal=71% |
| KEGG_FOCAL_ADHESION | KEGG_FOCAL_ADHESION | | 96 | 0.351827 | 1.43757 | 0.040541 | 0.125797 | 0.99 | 1039 | tags=39%, list=26%, signal=51% |
| KEGG_REGULATION_OF_ACTIN_CYTOSKELETON | KEGG_REGULATION_OF_ACTIN_CYTOSKELETO | | 99 | 0.336629 | 1.357412 | 0.026316 | 0.206108 | 1 | 1018 | tags=37%, list=26%, signal=49% |
| KEGG_NOTCH_SIGNALING_PATHWAY | KEGG_NOTCH_SIGNALING_PATHWAY | | 18 | 0.50219 | 1.351149 | 0.092308 | 0.206529 | 1 | 355 | tags=17%, list=9%, signal=18% |
| KEGG_UBIQUITIN_MEDIATED_PROTEOLYSIS | KEGG_UBIQUITIN_MEDIATED_PROTEOLYSIS | | 46 | 0.3938 | 1.34874 | 0.071429 | 0.203055 | 1 | 716 | tags=28%, list=18%, signal=34% |
| KEGG_INSULIN_SIGNALING_PATHWAY | KEGG_INSULIN_SIGNALING_PATHWAY | | 52 | 0.374164 | 1.321927 | 0.071429 | 0.230896 | 1 | 1192 | tags=54%, list=30%, signal=76% |
| KEGG_CELL_ADHESION_MOLECULES_CAMS | KEGG_CELL_ADHESION_MOLECULES_CAMS | | 49 | 0.354368 | 1.313281 | 0.0625 | 0.239045 | 1 | 1079 | tags=47%, list=27%, signal=64% |
| KEGG_SPLICEOSOME | KEGG_SPLICEOSOME | | 75 | 0.33018 | 1.296125 | 0.102564 | 0.254559 | 1 | 690 | tags=27%, list=17%, signal=32% |
| KEGG_ADIPOCYTOKINE_SIGNALING_PATHWAY | KEGG_ADIPOCYTOKINE_SIGNALING_PATHWAY | | 20 | 0.449733 | 1.285795 | 0.101695 | 0.262679 | 1 | 1181 | tags=55%, list=30%, signal=78% |
| KEGG_TOLL_LIKE_RECEPTOR_SIGNALING_PATHWA' | KEGG_TOLL_LIKE_RECEPTOR_SIGNALING_PATI | | 22 | 0.464225 | 1.270607 | 0.109375 | 0.274258 | 1 | 1018 | tags=59%, list=26%, signal=79% |
| KEGG_SYSTEMIC_LUPUS_ERYTHEMATOSUS | KEGG_SYSTEMIC_LUPUS_ERYTHEMATOSUS | | 20 | 0.442196 | 1.266141 | 0.142857 | 0.274765 | 1 | 30 | tags=10%, list=1%, signal=10% |
| KEGG_EPITHELIAL_CELL_SIGNALING_IN_HELICOBA | KEGG_EPITHELIAL_CELL_SIGNALING_IN_HELIC | | 36 | 0.381606 | 1.256256 | 0.185714 | 0.278615 | 1 | 1140 | tags=39%, list=29%, signal=54% |
| KEGG_APOPTOSIS | KEGG_APOPTOSIS | | 30 | 0.40513 | 1.250552 | 0.142857 | 0.277545 | 1 | 1021 | tags=47%, list=26%, signal=62% |
| KEGG_GLIOMA | KEGG_GLIOMA | | 27 | 0.38695 | 1.203469 | 0.231884 | 0.344936 | 1 | 1018 | tags=48%, list=26%, signal=64% |
| KEGG_TYROSINE_METABOLISM | KEGG_TYROSINE_METABOLISM | | 15 | 0.404592 | 1.170905 | 0.349206 | 0.388276 | 1 | 542 | tags=27%, list=14%, signal=31% |
| KEGG_NATURAL_KILLER_CELL_MEDIATED_CYTOTO: | KEGG_NATURAL_KILLER_CELL_MEDIATED_CYT | | 41 | 0.317761 | 1.133986 | 0.227273 | 0.441402 | 1 | 1040 | tags=49%, list=26%, signal=66% |
| KEGG_RIG_I_LIKE_RECEPTOR_SIGNALING_PATHWA | KEGG_RIG_I_LIKE_RECEPTOR_SIGNALING_PAT | | 16 | 0.41569 | 1.122166 | 0.355932 | 0.454118 | 1 | 847 | tags=44%, list=21%, signal=55% |
| KEGG_TIGHT_JUNCTION | KEGG_TIGHT_JUNCTION | | 65 | 0.289391 | 1.079734 | 0.315068 | 0.525944 | 1 | 1302 | tags=42%, list=33%, signal=61% |
| KEGG_PROGESTERONE_MEDIATED_OOCYTE_MATU | KEGG_PROGESTERONE_MEDIATED_OOCYTE_I | | 32 | 0.341434 | 1.067003 | 0.369863 | 0.539036 | 1 | 1063 | tags=41%, list=27%, signal=55% |
| KEGG_SNARE_INTERACTIONS_IN_VESICULAR_TRAN | KEGG_SNARE_INTERACTIONS_IN_VESICULAR_ | | 16 | 0.417437 | 1.060597 | 0.35 | 0.538543 | 1 | 1123 | tags=44%, list=28%, signal=61% |
| KEGG_WNT_SIGNALING_PATHWAY | KEGG_WNT_SIGNALING_PATHWAY | | 53 | 0.294057 | 1.060423 | 0.375 | 0.527323 | 1 | 673 | tags=23%, list=17%, signal=27% |
| KEGG_CHEMOKINE_SIGNALING_PATHWAY | KEGG_CHEMOKINE_SIGNALING_PATHWAY | | 67 | 0.286135 | 1.056159 | 0.345679 | 0.524311 | 1 | 1018 | tags=37%, list=26%, signal=49% |
| KEGG_P53_SIGNALING_PATHWAY | KEGG_P53_SIGNALING_PATHWAY | | 17 | 0.35222 | 1.000264 | 0.485714 | 0.61514 | 1 | 731 | tags=35%, list=19%, signal=43% |
| KEGG_JAK_STAT_SIGNALING_PATHWAY | KEGG_JAK_STAT_SIGNALING_PATHWAY | | 24 | 0.314752 | 0.966723 | 0.557377 | 0.669481 | 1 | 308 | tags=17%, list=8%, signal=18% |
| KEGG_ARRHYTHMOGENIC_RIGHT_VENTRICULAR_C | KEGG_ARRHYTHMOGENIC_RIGHT_VENTRICUL | | 31 | 0.310253 | 0.95912 | 0.542857 | 0.673342 | 1 | 698 | tags=29%, list=18%, signal=35% |
| KEGG_INOSITOL_PHOSPHATE_METABOLISM | KEGG_INOSITOL_PHOSPHATE_METABOLISM | | 21 | 0.341139 | 0.957413 | 0.5 | 0.66429 | 1 | 850 | tags=33%, list=22%, signal=42% |
| KEGG_TGF_BETA_SIGNALING_PATHWAY | KEGG_TGF_BETA_SIGNALING_PATHWAY | | 25 | 0.310888 | 0.955292 | 0.542857 | 0.65593 | 1 | 737 | tags=24%, list=19%, signal=29% |
| KEGG_DILATED_CARDIOMYOPATHY | KEGG_DILATED_CARDIOMYOPATHY | | 28 | 0.294927 | 0.909592 | 0.5 | 0.726878 | 1 | 567 | tags=21%, list=14%, signal=25% |
| KEGG_HYPERTROPHIC_CARDIOMYOPATHY_HCM | KEGG_HYPERTROPHIC_CARDIOMYOPATHY_HC | | 29 | 0.292397 | 0.861056 | 0.65625 | 0.804271 | 1 | 567 | tags=21%, list=14%, signal=24% |
| KEGG_AXON_GUIDANCE | KEGG_AXON_GUIDANCE | | 60 | 0.237552 | 0.824919 | 0.744186 | 0.858321 | 1 | 589 | tags=22%, list=15%, signal=25% |
| KEGG_PHOSPHATIDYLINOSITOL_SIGNALING_SYSTE | KEGG_PHOSPHATIDYLINOSITOL_SIGNALING_S | | 28 | 0.258495 | 0.819858 | 0.6875 | 0.852721 | 1 | 937 | tags=32%, list=24%, signal=42% |
| KEGG_VASOPRESSIN_REGULATED_WATER_REABSC | KEGG_VASOPRESSIN_REGULATED_WATER_RE/ | | 19 | 0.285249 | 0.807206 | 0.647059 | 0.859569 | 1 | 1011 | tags=32%, list=26%, signal=42% |
| KEGG_LONG_TERM_POTENTIATION | KEGG_LONG_TERM_POTENTIATION | | 31 | 0.246937 | 0.781733 | 0.802817 | 0.885756 | 1 | 1228 | tags=45%, list=31%, signal=65% |
| KEGG_AMYOTROPHIC_LATERAL_SCLEROSIS_ALS | KEGG_AMYOTROPHIC_LATERAL_SCLEROSIS_A | | 15 | 0.270195 | 0.774203 | 0.777778 | 0.882534 | 1 | 731 | tags=33%, list=19%, signal=41% |
| KEGG_PATHOGENIC_ESCHERICHIA_COLI_INFECTIC | KEGG_PATHOGENIC_ESCHERICHIA_COLI_INFI | | 36 | 0.231366 | 0.767756 | 0.794521 | 0.879225 | 1 | 740 | tags=25%, list=19%, signal=30% |
| KEGG_PROPANOATE_METABOLISM | KEGG_PROPANOATE_METABOLISM | | 27 | 0.248787 | 0.751886 | 0.823529 | 0.884236 | 1 | 415 | tags=15%, list=11%, signal=16% |
| KEGG_PRION_DISEASES | KEGG_PRION_DISEASES | | 15 | 0.269398 | 0.708226 | 0.863636 | 0.928663 | 1 | 731 | tags=27%, list=19%, signal=33% |
| KEGG_THYROID_CANCER | KEGG_THYROID_CANCER | | 15 | 0.271862 | 0.695049 | 0.890909 | 0.927277 | 1 | 1301 | tags=47%, list=33%, signal=69% |
| KEGG_AMINO_SUGAR_AND_NUCLEOTIDE_SUGAR_I | KEGG_AMINO_SUGAR_AND_NUCLEOTIDE_SUC | | 26 | 0.234116 | 0.69496 | 0.846154 | 0.913472 | 1 | 1212 | tags=42%, list=31%, signal=61% |
| KEGG_CYTOKINE_CYTOKINE_RECEPTOR_INTERACT | KEGG_CYTOKINE_CYTOKINE_RECEPTOR_INTEI | | 17 | 0.226869 | 0.594847 | 1 | 0.970538 | 1 | 789 | tags=29%, list=20%, signal=37% |

| NAME | GS<br> foll | GS DETAILS | SIZE | ES | NES | NOM p-val | FDR q-val | FWER p-val | RANK AT M/ | LEADING EDGE |
|---|---|---|---|---|---|---|---|---|---|---|
| HALLMARK_ | HALLMARK_ | Details ... | 66 | 0.434157 | 1.608342 | 0 | 0.121962 | 0.18 | 922 | tags=44%, list=23%, signal=56% |
| HALLMARK_ | HALLMARK_ | Details ... | 55 | 0.418099 | 1.485793 | 0.042254 | 0.248369 | 0.51 | 1073 | tags=44%, list=27%, signal=59% |
| HALLMARK_ | HALLMARK_ | Details ... | 34 | 0.421615 | 1.483721 | 0.086957 | 0.170236 | 0.52 | 469 | tags=26%, list=12%, signal=30% |
| HALLMARK_ | HALLMARK_ | Details ... | 41 | 0.431538 | 1.448415 | 0.070423 | 0.191213 | 0.7 | 589 | tags=39%, list=15%, signal=45% |
| HALLMARK_ | HALLMARK_ | Details ... | 41 | 0.429565 | 1.412187 | 0.058824 | 0.206019 | 0.85 | 615 | tags=32%, list=16%, signal=37% |
| HALLMARK_ | HALLMARK_ | Details ... | 19 | 0.454304 | 1.307453 | 0.106061 | 0.36146 | 0.96 | 877 | tags=42%, list=22%, signal=54% |
| HALLMARK_ | HALLMARK_ | Details ... | 57 | 0.348641 | 1.28149 | 0.125 | 0.369085 | 0.97 | 718 | tags=30%, list=18%, signal=36% |
| HALLMARK_ | HALLMARK_ | Details ... | 95 | 0.318363 | 1.251813 | 0.1625 | 0.380173 | 0.98 | 1085 | tags=41%, list=28%, signal=55% |
| HALLMARK_ | HALLMARK_ | Details ... | 16 | 0.448927 | 1.220454 | 0.1875 | 0.404567 | 0.99 | 876 | tags=44%, list=22%, signal=56% |
| HALLMARK_ | HALLMARK_ | Details ... | 50 | 0.309132 | 1.133263 | 0.263158 | 0.591863 | 1 | 927 | tags=32%, list=23%, signal=41% |
| HALLMARK_ | HALLMARK_ | Details ... | 72 | 0.311194 | 1.1202 | 0.28 | 0.571493 | 1 | 967 | tags=33%, list=25%, signal=43% |
| HALLMARK_ | HALLMARK_ | Details ... | 39 | 0.324481 | 1.091849 | 0.375 | 0.596018 | 1 | 566 | tags=21%, list=14%, signal=24% |
| HALLMARK_ | HALLMARK_ | Details ... | 70 | 0.273602 | 1.004271 | 0.413333 | 0.78369 | 1 | 604 | tags=20%, list=15%, signal=23% |
| HALLMARK_ | HALLMARK_ | Details ... | 47 | 0.27705 | 0.99196 | 0.476923 | 0.761631 | 1 | 790 | tags=28%, list=20%, signal=34% |
| HALLMARK_ | HALLMARK_ | Details ... | 67 | 0.254828 | 0.958995 | 0.526316 | 0.77782 | 1 | 893 | tags=25%, list=23%, signal=32% |
| HALLMARK_ | HALLMARK_ | Details ... | 17 | 0.331096 | 0.869854 | 0.651515 | 0.938455 | 1 | 804 | tags=35%, list=20%, signal=44% |
| HALLMARK_ | HALLMARK_ | Details ... | 71 | 0.235479 | 0.86745 | 0.676471 | 0.888074 | 1 | 876 | tags=27%, list=22%, signal=34% |
| HALLMARK_ | HALLMARK_ | Details ... | 55 | 0.24034 | 0.859736 | 0.621212 | 0.853131 | 1 | 1036 | tags=36%, list=26%, signal=49% |
| HALLMARK_ | HALLMARK_ | Details ... | 33 | 0.25113 | 0.813248 | 0.693548 | 0.91256 | 1 | 499 | tags=15%, list=13%, signal=17% |
| HALLMARK_ | HALLMARK_ | Details ... | 51 | 0.223472 | 0.809815 | 0.80303 | 0.873104 | 1 | 1040 | tags=35%, list=26%, signal=47% |
| HALLMARK_ | HALLMARK_UV_RESPOI | | 74 | 0.196962 | 0.76754 | 0.833333 | 0.893602 | 1 | 776 | tags=22%, list=20%, signal=26% |
| HALLMARK_ | HALLMARK_MYC_TARG | | 26 | 0.194381 | 0.626019 | 0.905405 | 0.979241 | 1 | 733 | tags=19%, list=19%, signal=23% |

| NAME | GS<br> foll | GS DETAILS | SIZE | ES | NES | NOM p-val | FDR q-val | FWER p-val | RANK AT M | LEADING EDGE |
|---|---|---|---|---|---|---|---|---|---|---|
| HALLMARK_EPITHELIAL_MESENCHYMAL_TRANSITION | HALLMARK | Details ... | 84 | 0.549916 | 2.170175 | 0 | 0 | 0 | 504 | tags=32%, list=13%, signal=36% |
| HALLMARK_APICAL_JUNCTION | HALLMARK | Details ... | 95 | 0.475958 | 2.140142 | 0 | 0 | 0 | 877 | tags=42%, list=22%, signal=53% |
| HALLMARK_HYPOXIA | HALLMARK | Details ... | 83 | 0.494998 | 2.105294 | 0 | 0 | 0 | 737 | tags=37%, list=19%, signal=45% |
| HALLMARK_MITOTIC_SPINDLE | HALLMARK | Details ... | 103 | 0.475103 | 2.078243 | 0 | 0 | 0 | 1145 | tags=56%, list=29%, signal=77% |
| HALLMARK_UV_RESPONSE_DN | HALLMARK | Details ... | 51 | 0.552017 | 2.034813 | 0 | 0 | 0 | 747 | tags=47%, list=19%, signal=57% |
| HALLMARK_COAGULATION | HALLMARK | Details ... | 41 | 0.497068 | 1.839792 | 0 | 0 | 0 | 784 | tags=44%, list=20%, signal=54% |
| HALLMARK_TNFA_SIGNALING_VIA_NFKB | HALLMARK | Details ... | 39 | 0.518015 | 1.809001 | 0 | 0.001703 | 0.01 | 846 | tags=56%, list=21%, signal=71% |
| HALLMARK_COMPLEMENT | HALLMARK | Details ... | 74 | 0.428747 | 1.767145 | 0 | 0.003175 | 0.02 | 1002 | tags=47%, list=25%, signal=62% |
| HALLMARK_ESTROGEN_RESPONSE_EARLY | HALLMARK | Details ... | 54 | 0.452796 | 1.731849 | 0 | 0.004258 | 0.03 | 610 | tags=33%, list=15%, signal=39% |
| HALLMARK_GLYCOLYSIS | HALLMARK | Details ... | 115 | 0.380998 | 1.728286 | 0 | 0.003832 | 0.03 | 827 | tags=34%, list=21%, signal=42% |
| HALLMARK_REACTIVE_OXYGEN_SPECIES_PATHWAY | HALLMARK | Details ... | 26 | 0.529958 | 1.721041 | 0 | 0.004567 | 0.04 | 894 | tags=54%, list=23%, signal=69% |
| HALLMARK_IL2_STAT5_SIGNALING | HALLMARK | Details ... | 70 | 0.427432 | 1.663414 | 0 | 0.008825 | 0.08 | 466 | tags=27%, list=12%, signal=30% |
| HALLMARK_KRAS_SIGNALING_UP | HALLMARK | Details ... | 47 | 0.388001 | 1.421755 | 0.071429 | 0.076869 | 0.56 | 282 | tags=21%, list=7%, signal=23% |
| HALLMARK_INFLAMMATORY_RESPONSE | HALLMARK | Details ... | 41 | 0.374493 | 1.361117 | 0.06 | 0.121522 | 0.73 | 536 | tags=29%, list=14%, signal=34% |
| HALLMARK_INTERFERON_GAMMA_RESPONSE | HALLMARK | Details ... | 62 | 0.338728 | 1.304372 | 0.076923 | 0.174908 | 0.87 | 819 | tags=31%, list=21%, signal=38% |
| HALLMARK_G2M_CHECKPOINT | HALLMARK | Details ... | 87 | 0.292474 | 1.283342 | 0.102941 | 0.196088 | 0.91 | 669 | tags=22%, list=17%, signal=26% |
| HALLMARK_ESTROGEN_RESPONSE_LATE | HALLMARK | Details ... | 71 | 0.311754 | 1.183352 | 0.236364 | 0.356181 | 0.99 | 818 | tags=32%, list=21%, signal=40% |
| HALLMARK_APOPTOSIS | HALLMARK | Details ... | 61 | 0.290905 | 1.166288 | 0.192982 | 0.368395 | 1 | 821 | tags=28%, list=21%, signal=35% |
| HALLMARK_XENOBIOTIC_METABOLISM | HALLMARK | Details ... | 78 | 0.288792 | 1.144234 | 0.183333 | 0.391983 | 1 | 763 | tags=28%, list=19%, signal=34% |
| HALLMARK_IL6_JAK_STAT3_SIGNALING | HALLMARK | Details ... | 16 | 0.382946 | 1.135546 | 0.317073 | 0.390954 | 1 | 378 | tags=31%, list=10%, signal=34% |
| HALLMARK_ANDROGEN_RESPONSE | HALLMARK_ANDROGEI | | 50 | 0.298399 | 1.071473 | 0.306452 | 0.49362 | 1 | 779 | tags=36%, list=20%, signal=44% |
| HALLMARK_UV_RESPONSE_UP | HALLMARK_UV_RESPO | | 74 | 0.24382 | 1.051331 | 0.355932 | 0.511356 | 1 | 835 | tags=24%, list=21%, signal=30% |
| HALLMARK_TGF_BETA_SIGNALING | HALLMARK_TGF_BETA_ | | 19 | 0.35558 | 1.050366 | 0.456522 | 0.491327 | 1 | 1259 | tags=63%, list=32%, signal=92% |
| HALLMARK_MYOGENESIS | HALLMARK_MYOGENES | | 55 | 0.276981 | 1.049901 | 0.366667 | 0.47147 | 1 | 817 | tags=31%, list=21%, signal=38% |
| HALLMARK_FATTY_ACID_METABOLISM | HALLMARK_FATTY_ACII | | 92 | 0.20107 | 0.84429 | 0.789474 | 0.945293 | 1 | 706 | tags=20%, list=18%, signal=23% |
| HALLMARK_SPERMATOGENESIS | HALLMARK_SPERMATC | | 33 | 0.237066 | 0.826966 | 0.786885 | 0.946465 | 1 | 354 | tags=12%, list=9%, signal=13% |
| HALLMARK_HEME_METABOLISM | HALLMARK_HEME_MET | | 67 | 0.210656 | 0.817736 | 0.789474 | 0.926445 | 1 | 1041 | tags=36%, list=26%, signal=48% |
| HALLMARK_CHOLESTEROL_HOMEOSTASIS | HALLMARK_CHOLESTE | | 38 | 0.224741 | 0.802999 | 0.763636 | 0.919589 | 1 | 1353 | tags=55%, list=34%, signal=83% |
| HALLMARK_ALLOGRAFT_REJECTION | HALLMARK_ALLOGRAF | | 51 | 0.19843 | 0.75269 | 0.836066 | 0.956947 | 1 | 252 | tags=10%, list=6%, signal=10% |
| HALLMARK_MTORC1_SIGNALING | HALLMARK_MTORC1_S | | 139 | 0.148191 | 0.686324 | 1 | 0.984791 | 1 | 1220 | tags=35%, list=31%, signal=49% |
| HALLMARK_E2F_TARGETS | HALLMARK_E2F_TARGE | | 93 | 0.1125 | 0.469883 | 1 | 0.999583 | 1 | 801 | tags=15%, list=20%, signal=18% |

| NAME | GS<br> foll | GS DETAILS | SIZE | ES | NES | NOM p-val | FDR q-val | FWER p-val | RANK AT M | LEADING EDGE |
|---|---|---|---|---|---|---|---|---|---|---|
| HALLMARK_OXIDATIVE_PHOSPHORYLATION | HALLMARK | Details ... | 110 | 0.616721 | 3.01734 | 0 | 0 | 0 | 936 | tags=61%, list=24%, signal=78% |
| HALLMARK_UNFOLDED_PROTEIN_RESPONSE | HALLMARK | Details ... | 61 | 0.460994 | 1.938361 | 0 | 0.005333 | 0.01 | 573 | tags=28%, list=15%, signal=32% |
| HALLMARK_ADIPOGENESIS | HALLMARK | Details ... | 91 | 0.421298 | 1.811373 | 0 | 0.015569 | 0.05 | 943 | tags=43%, list=24%, signal=55% |
| HALLMARK_PEROXISOME | HALLMARK | Details ... | 52 | 0.460463 | 1.803631 | 0 | 0.011677 | 0.05 | 454 | tags=35%, list=12%, signal=39% |
| HALLMARK_FATTY_ACID_METABOLISM | HALLMARK | Details ... | 92 | 0.39156 | 1.794015 | 0 | 0.009341 | 0.05 | 991 | tags=42%, list=25%, signal=55% |
| HALLMARK_MYC_TARGETS_V1 | HALLMARK | Details ... | 164 | 0.353671 | 1.785653 | 0 | 0.007784 | 0.05 | 1290 | tags=45%, list=33%, signal=64% |
| HALLMARK_BILE_ACID_METABOLISM | HALLMARK | Details ... | 33 | 0.46989 | 1.693801 | 0.071429 | 0.010853 | 0.08 | 1242 | tags=58%, list=31%, signal=83% |
| HALLMARK_MTORC1_SIGNALING | HALLMARK | Details ... | 139 | 0.329976 | 1.590233 | 0 | 0.02037 | 0.16 | 570 | tags=21%, list=14%, signal=24% |
| HALLMARK_E2F_TARGETS | HALLMARK | Details ... | 93 | 0.296862 | 1.397634 | 0.085714 | 0.076812 | 0.49 | 1022 | tags=37%, list=26%, signal=48% |
| HALLMARK_CHOLESTEROL_HOMEOSTASIS | HALLMARK | Details ... | 38 | 0.315821 | 1.223706 | 0.191489 | 0.213187 | 0.87 | 1123 | tags=50%, list=28%, signal=69% |
| HALLMARK_G2M_CHECKPOINT | HALLMARK | Details ... | 87 | 0.222278 | 1.041085 | 0.393939 | 0.531678 | 0.99 | 999 | tags=36%, list=25%, signal=47% |
| HALLMARK_MYC_TARGETS_V2 | HALLMARK | Details ... | 26 | 0.303768 | 1.039207 | 0.446809 | 0.49412 | 0.99 | 406 | tags=23%, list=10%, signal=26% |
| HALLMARK_GLYCOLYSIS | HALLMARK | Details ... | 115 | 0.216372 | 1.036444 | 0.424242 | 0.459922 | 0.99 | 898 | tags=30%, list=23%, signal=37% |
| HALLMARK_DNA_REPAIR | HALLMARK | Details ... | 57 | 0.208861 | 0.944936 | 0.69697 | 0.629034 | 1 | 814 | tags=28%, list=21%, signal=35% |
| HALLMARK_UV_RESPONSE_UP | HALLMARK | Details ... | 74 | 0.201061 | 0.921544 | 0.657143 | 0.651863 | 1 | 755 | tags=26%, list=19%, signal=31% |
| HALLMARK_PROTEIN_SECRETION | HALLMARK | Details ... | 72 | 0.205191 | 0.911134 | 0.675676 | 0.63476 | 1 | 904 | tags=29%, list=23%, signal=37% |

Highlighted cell indicates overlap with cell line EV proteome

| AR Activity | | | Neuroendocrine Activity (Che et al) | Luminal Activity (Che et al) | Basal Activity (Che et al) | NE Score: 70 genes (Beltran et al) |
|---|---|---|---|---|---|---|
| Spratt et al | Che et al | Nelson et al | CHGA | MSMB | KRT14 | CYLD |
| KLK3 | ABCC4 | HPGD | GRP | KLK3 | DST | CIITA |
| KLK2 | ACSL3 | PPAP2A | CALCA | ACPP | KRT15 | CREBBP |
| FKBP5 | ADAM7 | SCD | SCG2 | PLA2G2A | TAGLN | HERPUD1 |
| STEAP1 | C1orf116 | SORD | TPH1 | KLK2 | CYR61 | MMP2 |
| STEAP2 | CENPN | DHCR24 | ASCL1 | TFF1 | KRT5 | NUP93 |
| PPAP2A | EAF2 | SMS | SCG5 | AZGP1 | RGCC | PAX8 |
| RAB3B | ELL2 | LOC51171 | PHGR1 | MT1G | TIMP3 | CCND1 |
| ACSL3 | FKBP5 | ALDH1A3 | MIAT | TSPAN8 | CAV1 | CSDE1 |
| NKX3-1 | GNMT | SAT | MAOB | CLDN3 | MYC | DICER1 |
| | HERC3 | HMGCS1 | ZACN | NEAT1 | KRT17 | FHIT |
| | KLK2 | DBI | FGF14 | SEC11C | PHLDA1 | MYH9 |
| | KLK3 | B4GALT1 | ST18 | CPE | S100A2 | RBBP6 |
| | MAF | PHYH | MS4A8 | RDH11 | MYL9 | TRIM33 |
| | MED28 | IDI1 | STMN2 | AGR2 | PALLD | FOXP1 |
| | NKX3-1 | HELO1 | DDC | TMPRSS2 | ZFP36L2 | RB1 |
| | NNMT | UGDH | SCG3 | ZG16B | HSPA1A | NKX3-1 |
| | PMEPA1 | FACL3 | ELAVL4 | NPDC1 | SERPINF1 | ETV5 |
| | PTGER4 | DDC | SLC35D3 | DHRS7 | MEG3 | MYCN |
| | ZBTB10 | MAF | MIR7-3HG | SLC12A2 | CLU | AURKA |
| | | NDRG1 | HTR3C | CD24 | FHL2 | EZH2 |
| | | ELL2 | SYT4 | KLK4 | CAV2 | GATA2 |
| | | GG2-1 | TAAR1 | SMIM14 | STK17A | KLK3 |
| | | CDC14 | INSM1 | ADIRF | HBEGF | GNAO1 |
| | | B | CRMP1 | TMC5 | EDN1 | ARHGAP8 |
| | | ID2 | PPFIA2 | PPP3CA | F3 | PRR5-ARHGAP8 |
| | | HES6 | RGS7 | TPD52 | HSPA1B | MAPKAPK3 |
| | | CTBP1 | SLC18A1 | PART1 | DEFB1 | EPN3 |
| | | CDK8 | NDST4 | DSC2 | DKK1 | SCG3 |
| | | FKBP5 | PCSK2 | CIB1 | CRYAB | PIEZO1 |
| | | UNC13 | C11orf16 | SLC44A4 | | CAND2 |
| | | ABCC4 | CELF3 | RASEF | | EFNA4 |
| | | ANKH | KCNJ6 | INSIG1 | | HOXB13 |
| | | KIAA1382 | NELL2 | SLC39A6 | | PSCA |
| | | SNAP25 | PTPRN | UQCRQ | | SLC25A37 |
| | | APPBP2 | TMEM176A | STEAP2 | | SYT11 |
| | | AZGP1 | KLK12 | SERF2 | | SVOP |
| | | LMAN1 | LINGO2 | RNF144B | | PCSK1 |
| | | RAB4 | NPPA | ABHD2 | | LMAN1L |
| | | SRP19 | LINC00299 | LINC00844 | | JAKMIP2 |
| | | SLC26A2 | RP11-1C8.7 | GOLM1 | | ASXL3 |
| | | SEC24D | KCNB2 | RAB27B | | BRINP1 |
| | | VAPA | PKD2L1 | BACE2 | | DNMT1 |
| | | SCNN1A | LINC01497 | MT1F | | TRIM9 |
| | | SLC29A | NOL4 | PSCA | | TC2N |
| | | NKX3A | NR0B2 | ELOVL5 | | SPDEF |
| | | PDEF | NAT8L | FLJ20021 | | RGS10 |
| | | RINZF | SLC8A1 | B2M | | CATSPERB |
| | | DDXBP1 | TMEM176B | PMEPA1 | | EVPL |
| | | TSC22 | LCN15 | HMGCS1 | | GPX2 |
| | | MRF2-hom | INA | LMO7 | | KCND2 |
| | | KLF4 | ST8SIA5 | SORD | | KIAA0408 |
| | | MYC | SNAP25 | FXYD3 | | KLK4 |
| | | TMPRSS2 | SYT13 | ALDH1A3 | | LRRC16B |
| | | KLK2 | NELL1 | IL6ST | | MAP10 |
| | | KLK3 | AOC1 | TSC22D1 | | NRSN1 |
| | | KLK4 | SCGN | DBI | | OPHN1 |
| | | CPD | LY6H | PLPP1 | | PROX1 |
| | | SERPIN1 | TAGLN3 | H2AFJ | | RAB27B |
| | | LIFR | SRRM4 | ELF3 | | RGS7 |
| | | IQGAP2 | EPHB1 | SPINK1 | | SEC11C |
| | | AGM1 | SLC38A11 | NDRG1 | | SLC44A4 |
| | | HOMER-2B | NRXN1 | RPS27L | | SOGA3 |
| | | AKAP12 | SYT11 | SAT1 | | ST8SIA3 |
| | | TPD52 | HEPACAM2 | NKX3-1 | | UPK2 |
| | | LIM | CDK5R2 | TXNIP | | AR |
| | | CAMKK2 | CEND1 | MT1E | | C7orf76 |
| | | PTPN21 | GRIN3A | FDPS | | KCNB2 |
| | | GUCY1A3 | FLJ46066 | MT2A | | RIPK2 |
| | | INPP4B | RBFOX1 | JUN | | SEZ6 |
| | | MERTK | GPR55 | GADD45B | | |
| | | AKAP2 | RP11-867O8.5 | | | |
| | | BCHE | CCDC187 | | | |
| | | PRKCM | IQSEC3 | | | |
| | | SDC4 | AC015849.2 | | | |
| | | PEG3 | GPC5-AS1 | | | |
| | | TUSP | TDRG1 | | | |
| | | PIK3R3 | RP11-768G7.2 | | | |
| | | RDC1 | NKAIN2 | | | |
| | | KRT8 | PEX5L | | | |
| | | ADRM1 | RP11-675F6.4 | | | |
| | | ACTN1 | ANK2 | | | |
| | | ITGAV | FOXA2 | | | |
| | | KRT19 | CADPS | | | |

| | |
|---|---|
| H1F0 | FIBCD1 |
| DCTN3 | LINC00261 |
| MLCB | SYT1 |
| MMP16 | PCSK1 |
| FN1 | UCHL1 |
| ORM1 | AC010746.3 |
| SEPP1 | KIAA1644 |
| GSR | ASGR2 |
| SGK | RP11-386M24.9 |
| SPAK | AC009410.1 |
| DNAJB9 | RP11-317N12.1 |
| TMEPAI | RP11-482M8.3 |
| B2M | NRSN1 |
| TCRG/TARP | ISX |
| HERC3 | RIIAD1 |
| FLJ23153 | DAB1 |
| ADAMTS | MAP1A |
| HS1-2 | DUSP26 |
| RBM10 | RAB39B |
| UAP1 | KIF1A |
| ST7 | FXYD6 |
| DPH2L1 | SORCS1 |
| | AC053503.4 |
| | KCNJ12 |
| | CXCR5 |
| | RP11-310E22.4 |
| | ARHGDIG |
| | RP11-977B10.2 |
| | B3GALT2 |
| | RP11-632L2.2 |
| | JAKMIP2 |
| | BSN |
| | SPX |
| | SYN3 |
| | HCN2 |
| | SCN3A |
| | SYP |

| | Peptides | Proteins | Protein Overlap |
|---|---|---|---|
| **22RV1** | 10,562 | 2,499 | 2,104 (73%) |
| | 10,433 | 2,480 | |
| **C4-2** | 10,567 | 2,481 | 2,073 (72%) |
| | 10,653 | 2,465 | |
| **LNCaP** | 10,693 | 2,509 | 2,211 (78%) |
| | 10,742 | 2,549 | |
| **LNCaP 95** | 10,815 | 2,498 | 2,095 (73%) |
| | 10,574 | 2,466 | |
| **EF1** | 10,577 | 2,601 | 2,179 (71%) |
| | 10,402 | 2,637 | |
| **NCI-H660** | 11,391 | 2,809 | 2,370 (76%) |
| | 11,228 | 2,700 | |
| **LASCPC-01** | 10,819 | 2,598 | 2,246 (72%) |
| | 11,476 | 2,778 | |
| **DU145** | 10,977 | 2,567 | 2,032 (67%) |
| | 10,416 | 2,477 | |
| **PC3** | 10,805 | 2,510 | 2,122 (74%) |
| | 10,746 | 2,483 | |

| | mCRPC Patient | | | | | | | | | | | | | | | | | | | | | | | | | | |
|---|---|---|---|---|---|---|---|---|---|---|---|---|---|---|---|---|---|---|---|---|---|---|---|---|---|---|---|
| Gene | 1 | 2 | 3 | 4 | 5 | 6 | 7 | 8 | 9 | 10 | 11 | 12 | 13 | 14 | 15 | 16 | 17 | 18 | 19 | 20 | 21 | 22 | 23 | 24 | 25 | 26 | 27 |
| AGRN | 16.69479 | 16.48617 | 17.9618 | 15.66444 | 16.98859 | 17.96679 | 17.0218 | 16.82759 | 15.90978 | 17.63477 | 16.68679 | 17.42026 | 16.35974 | 17.16656 | 16.57967 | 18.16606 | 16.5357 | 16.86997 | 17.52498 | 17.09607 | 16.90016 | 15.20639 | 16.62628 | 16.98766 | 17.10956 | 17.56729 | 17.22606 |
| ATXN3 | 17.54731 | 16.33855 | 17.43198 | 16.40482 | 16.65248 | 16.657 | 17.31849 | 15.69819 | 17.16587 | 15.65499 | 16.21207 | 16.66561 | 16.66781 | 17.55208 | 17.10296 | 16.3207 | 17.64923 | 17.12425 | 18.20512 | 17.53388 | 16.51635 | 16.63156 | 16.80753 | 16.97973 | 18.19611 | 16.75574 | 18.45494 |
| CCDC124 | 18.70845 | 19.31824 | 16.11427 | 17.41339 | 16.60678 | 18.24055 | 16.26479 | 18.35607 | 17.4412 | 17.23467 | 15.45421 | 17.90864 | 17.73231 | 15.79233 | 19.24149 | 16.44376 | 17.36244 | 19.13217 | 17.38874 | 17.92222 | 17.04416 | 19.71693 | 16.85674 | 16.21399 | 19.77126 | 20.57311 | 19.72618 |
| CD276 | 16.86815 | 20.29894 | 18.79484 | 16.31507 | 16.16416 | 16.30091 | 17.10522 | 18.28672 | 20.23023 | 17.25226 | 17.92807 | 19.30862 | 16.89886 | 19.69892 | 20.17036 | 16.76305 | 19.40328 | 16.13364 | 19.62986 | 16.49147 | 16.47208 | 17.35245 | 19.43868 | 19.24606 | 16.62743 | 19.70699 | 14.56986 |
| CLTCL1 | 16.95228 | 16.27751 | 19.01064 | 16.51046 | 17.49089 | 16.96425 | 17.54345 | 17.6524 | 15.95524 | 15.61751 | 16.38952 | 16.64418 | 16.21983 | 17.13814 | 16.46476 | 17.61904 | 15.80012 | 17.61306 | 17.2956 | 17.64825 | 16.85456 | 17.26824 | 17.40513 | 16.59063 | 17.31046 | 17.46188 | 17.41239 |
| CSNK1G3 | 22.03926 | 22.13946 | 21.42726 | 22.1519 | 21.78756 | 21.2764 | 17.55756 | 21.65403 | 22.27923 | 22.5285 | 21.96014 | 22.10149 | 22.22435 | 22.09184 | 22.40665 | 21.3538 | 21.83461 | 21.77957 | 22.10149 | 21.77555 | 21.98468 | 22.44253 | 22.39362 | 22.03591 | 22.18556 | 22.23608 | 21.8423 |
| CTBP1 | 15.60566 | 20.18253 | 19.66004 | 16.39125 | 16.90366 | 16.1427 | 17.67396 | 19.74525 | 20.40665 | 19.05021 | 20.75532 | 20.0559 | 17.31454 | 19.31005 | 19.40471 | 19.75531 | 19.85977 | 17.25276 | 20.1332 | 20.34299 | 16.97656 | 19.44403 | 20.09507 | 17.01443 | 19.80549 | 20.73066 | 18.62114 |
| EDF1 | 17.17622 | 20.01563 | 19.58469 | 19.97421 | 20.89504 | 18.82105 | 16.4074 | 18.72743 | 19.83439 | 18.32144 | 19.10596 | 19.83162 | 19.86162 | 19.49366 | 19.62691 | 16.45408 | 19.667 | 19.78896 | 20.5451 | 18.97423 | 19.07057 | 19.57233 | 19.29852 | 19.46154 | 19.51135 | 19.68338 | 19.66006 |
| EEF1A2 | 16.96748 | 17.65163 | 17.15136 | 17.70989 | 16.65357 | 16.47801 | 17.11147 | 17.22264 | 17.16539 | 16.71134 | 17.18468 | 19.48395 | 16.90143 | 16.5324 | 17.88238 | 15.61517 | 15.86839 | 17.07649 | 19.8037 | 17.30353 | 18.64665 | 16.58665 | 17.71388 | 16.35805 | 16.95749 | 15.38805 | 16.65529 |
| EIF3G | 21.2764 | 21.0625 | 20.53564 | 21.22435 | 20.73892 | 22.00881 | 21.91342 | 21.24771 | 20.51653 | 22.63423 | 21.06907 | 21.66275 | 21.31008 | 21.92071 | 21.7348 | 21.10789 | 21.83461 | 20.32114 | 21.84613 | 21.10149 | 22.04926 | 21.50203 | 21.21252 | 20.5545 | 21.67573 | 21.49228 | 22.50446 |
| EPCAM | 16.16655 | 19.37372 | 17.15846 | 17.2422 | 17.28045 | 16.62114 | 16.85834 | 16.71544 | 17.49054 | 18.16194 | 17.36858 | 19.20827 | 15.7712 | 18.43641 | 19.93157 | 17.79186 | 17.60296 | 16.17304 | 21.33211 | 16.76766 | 16.97401 | 18.9329 | 20.1332 | 16.16404 | 15.74109 | 16.61602 | 16.20708 |
| FOLH1 | 18.11796 | 20.77153 | 19.66375 | 19.5198 | 20.39624 | 23.417 | 26.0296 | 21.81127 | 19.71861 | 21.48738 | 21.51653 | 20.45764 | 23.33756 | 21.50688 | 21.79947 | 16.84865 | 18.99581 | 20.79551 | 23.82881 | 15.30704 | 17.20194 | 19.98815 | 21.01563 | 20.78756 | 20.71398 | 20.02918 | 20.1206 |
| HIP1 | 18.50671 | 17.34313 | 15.93983 | 16.95091 | 17.71989 | 18.31971 | 17.9569 | 18.64538 | 16.13382 | 17.36837 | 18.3105 | 18.72256 | 17.1688 | 17.68286 | 19.58304 | 16.78255 | 19.81914 | 17.08231 | 20.29894 | 18.83733 | 16.97373 | 18.78109 | 17.37143 | 16.96609 | 17.62628 | 17.00778 | 16.98706 |
| LARP1 | 19.09928 | 19.8774 | 19.58937 | 19.66895 | 19.60363 | 20.14569 | 19.78322 | 19.14093 | 19.78801 | 19.87372 | 18.96607 | 19.64928 | 19.96014 | 19.79497 | 19.74962 | 19.92377 | 19.47417 | 16.58493 | 20.23023 | 20.17036 | 19.94592 | 19.65044 | 18.85374 | 16.25458 | 19.75632 | 20.40665 | 20.38574 |
| MBNL1 | 19.60373 | 20.10789 | 16.90012 | 16.4129 | 20.0559 | 20.37518 | 15.31963 | 19.92004 | 21.48247 | 20.61863 | 21.63644 | 21.04926 | 16.43193 | 20.77957 | 20.64526 | 21.12692 | 21.22435 | 20.10789 | 22.80341 | 21.50688 | 21.5451 | 20.66275 | 19.66594 | 20.48738 | 21.08861 | 20.36453 | 21.12692 |
| MKRN2 | 19.10736 | 20.29894 | 16.10155 | 21.6006 | 19.34178 | 15.92319 | 16.6048 | 19.78736 | 17.31253 | 16.52598 | 19.25487 | 19.33002 | 17.45146 | 17.1142 | 15.7824 | 19.49047 | 16.71158 | 15.73857 | 19.56317 | 19.3055 | 16.21544 | 19.01154 | 15.46903 | 19.97421 | 16.74149 | 17.39441 | 16.62164 |
| MVB12A | 18.8877 | 20.0559 | 19.75642 | 20.2764 | 19.17731 | 19.8097 | 20.85757 | 19.20195 | 18.15129 | 19.82464 | 18.9568 | 19.03409 | 20.18253 | 19.27231 | 19.81557 | 16.46084 | 19.92262 | 20.90976 | 20.73892 | 18.84252 | 20.01563 | 19.7024 | 20.39624 | 21.33756 | 20.48738 | 20.66275 | 20.62756 |
| MYO6 | 18.22603 | 19.26907 | 18.99235 | 17.72395 | 18.60405 | 20.52612 | 16.24282 | 18.37073 | 17.88347 | 18.0386 | 18.04219 | 18.85449 | 17.88255 | 18.63111 | 18.82824 | 18.81775 | 18.46107 | 19.52249 | 19.07423 | 18.29283 | 18.20737 | 18.51783 | 18.75608 | 19.32213 | 18.48711 | 18.56025 | 19.36775 |
| NEO1 | 17.70927 | 19.29184 | 17.16528 | 15.6365 | 17.72363 | 16.61226 | 18.52547 | 18.21833 | 16.7473 | 16.2916 | 17.04521 | 16.68285 | 18.02215 | 16.32098 | 17.31634 | 15.87804 | 17.39285 | 18.91181 | 17.08886 | 17.63794 | 17.22665 | 17.19347 | 18.0617 | 17.2364 | 16.96687 | 16.34324 | 17.60518 |
| OCLN | 17.22181 | 19.54144 | 20.47754 | 19.87322 | 16.56714 | 17.79949 | 16.69113 | 19.6085 | 16.35305 | 16.91922 | 18.45753 | 19.14627 | 21.00196 | 20.61863 | 18.70379 | 16.62836 | 18.68916 | 21.07561 | 21.06907 | 19.62401 | 16.91731 | 19.54501 | 20.64526 | 20.2535 | 19.40286 | 22.56151 | 15.90974 |
| PABPC4 | 18.75247 | 19.77348 | 16.3208 | 16.60941 | 16.35897 | 18.98543 | 15.84159 | 18.86155 | 18.74545 | 19.22161 | 18.5299 | 18.76371 | 19.67635 | 16.99506 | 19.59363 | 17.70715 | 19.61753 | 17.72377 | 22.19761 | 19.0615 | 17.51956 | 16.45303 | 20.86514 | 17.17708 | 20.17036 | 20.28771 | 19.42097 |
| PACSIN3 | 17.54544 | 17.60902 | 16.15918 | 17.47446 | 18.39712 | 16.64984 | 16.29871 | 16.16986 | 17.30635 | 18.71215 | 16.96508 | 18.20302 | 20.00196 | 17.2527 | 18.63908 | 17.56272 | 17.57051 | 16.15122 | 18.70468 | 15.61857 | 16.28011 | 15.62497 | 17.02096 | 16.29719 | 15.46201 | 16.79896 | 20.09507 |
| PDPK1 | 18.52693 | 19.19965 | 16.01024 | 19.12043 | 18.6082 | 18.69341 | 18.09593 | 18.66891 | 18.92911 | 18.71293 | 19.02286 | 18.02911 | 18.60525 | 18.74726 | 19.09897 | 18.73102 | 19.52438 | 19.5215 | 19.9214 | 19.07771 | 17.24179 | 19.31919 | 19.52436 | 19.98815 | 19.2394 | 16.87268 | 19.3245 |
| PKP4 | 16.774 | 21.0559 | 16.61298 | 17.35985 | 17.38298 | 17.0286 | 16.56819 | 18.08305 | 15.63245 | 16.32802 | 15.31768 | 16.86086 | 17.569 | 17.27267 | 19.33907 | 16.19205 | 16.84047 | 15.6257 | 16.42379 | 16.67219 | 18.06783 | 17.08719 | 16.95497 | 17.45708 | 17.02867 | 17.70574 | 21.47259 |
| PPIL1 | 16.80032 | 17.69142 | 17.42716 | 16.77018 | 16.06914 | 17.02877 | 16.1333 | 18.32453 | 20.37518 | 15.90546 | 20.06907 | 18.86906 | 16.97728 | 19.01486 | 15.16749 | 17.14768 | 20.0426 | 17.14445 | 19.33466 | 19.13712 | 17.36791 | 18.6139 | 16.61614 | 16.34609 | 19.55724 | 17.46801 | 16.40202 |
| PROM2 | 17.61732 | 16.80846 | 17.06815 | 17.78733 | 17.0908 | 16.34684 | 16.21679 | 16.43302 | 15.87859 | 18.11779 | 18.26177 | 18.75941 | 17.32778 | 16.63055 | 17.1649 | 17.22064 | 17.10079 | 17.17696 | 20.43746 | 16.96745 | 16.675 | 16.59473 | 16.42321 | 16.50276 | 16.88805 | 15.79126 | 15.86418 |
| PTP4A1 | 20.36453 | 21.40665 | 20.47754 | 20.71398 | 20.99507 | 19.39281 | 20.88763 | 20.45764 | 22.8839 | 20.60964 | 22.42982 | 21.31008 | 20.33211 | 21.00881 | 20.6006 | 20.10789 | 20.67142 | 21.38047 | 21.11426 | 22.03255 | 21.14569 | 20.68859 | 21.28207 | 17.12865 | 21.9812 | 20.28771 | 20.1206 |
| RBM3 | 16.62962 | 20.23023 | 16.83505 | 17.10737 | 18.32874 | 16.43627 | 15.50645 | 19.12102 | 20.48738 | 17.13716 | 19.97421 | 19.81716 | 17.58735 | 17.12938 | 19.74061 | 19.02383 | 19.51408 | 17.43482 | 19.69157 | 17.00379 | 19.77056 | 16.07364 | 16.38576 | 15.53023 | 16.71239 | 19.31293 | 17.22799 |
| REPS2 | 19.73113 | 19.53813 | 17.27927 | 19.71959 | 19.14467 | 20.46762 | 20.20658 | 19.48063 | 19.47893 | 19.78903 | 20.38574 | 18.84562 | 19.09505 | 16.86376 | 19.49416 | 20.28771 | 19.92311 | 20.5451 | 20.58233 | 20.29894 | 20.26499 | 20.15808 | 19.81978 | 20.29894 | 19.97421 | 19.89277 | 20.50688 |
| RPLP1 | 16.73419 | 21.21845 | 20.56384 | 20.96719 | 20.14569 | 16.68374 | 17.61009 | 20.3538 | 21.1332 | 20.59149 | 20.93876 | 21.45262 | 16.68265 | 20.10789 | 20.72234 | 20.56384 | 21.25926 | 17.83029 | 17.04175 | 21.08861 | 16.03774 | 20.5451 | 21.24191 | 15.73848 | 17.19721 | 21.51171 | 17.24315 |
| SPON2 | 17.47816 | 17.76799 | 18.93183 | 16.47845 | 16.99247 | 16.38673 | 17.43919 | 16.06024 | 17.02794 | 16.79237 | 16.74217 | 17.44361 | 16.76021 | 16.82085 | 17.93327 | 17.15688 | 15.53035 | 16.38255 | 17.5209 | 16.43651 | 16.44607 | 16.33866 | 17.75545 | 17.06038 | 18.30084 | 16.27784 | 17.41555 |
| SRSF3 | 16.3552 | 17.1191 | 17.70926 | 15.14948 | 18.63452 | 15.55601 | 16.70806 | 16.75958 | 19.91416 | 16.81914 | 16.92022 | 17.26258 | 16.69055 | 16.76339 | 18.1258 | 16.88261 | 18.6772 | 17.47788 | 17.1944 | 16.47141 | 19.80576 | 17.07209 | 17.11844 | 17.1935 | 21.70556 | 17.86925 | 16.24484 |
| STEAP1 | 16.77836 | 16.71399 | 17.53761 | 15.56976 | 16.77063 | 17.91433 | 16.25511 | 16.01886 | 17.00937 | 16.18236 | 16.70686 | 16.8445 | 16.41762 | 16.88706 | 16.87047 | 17.81782 | 17.47363 | 16.28832 | 18.74723 | 16.98123 | 16.63168 | 17.13838 | 16.45093 | 16.9264 | 17.03934 | 17.33665 | 16.14549 |
| STEAP2 | 16.94023 | 16.81334 | 15.90429 | 16.23101 | 16.923 | 18.47427 | 21.08861 | 16.60452 | 17.68374 | 16.98483 | 16.0034 | 17.43607 | 17.58009 | 16.21575 | 16.22179 | 16.11038 | 17.01961 | 17.22082 | 20.43746 | 16.97343 | 16.92493 | 16.40918 | 16.79277 | 15.6812 | 16.55205 | 16.29767 | 16.7654 |
| THEM6 | 19.30502 | 18.79881 | 19.73446 | 16.95987 | 18.79419 | 15.91161 | 18.80527 | 18.79231 | 15.90064 | 17.54791 | 19.78946 | 18.50606 | 18.53254 | 19.1326 | 18.00986 | 19.36504 | 19.401 | 17.67666 | 18.98201 | 19.77965 | 16.39902 | 16.57281 | 17.34957 | 16.5224 | 17.19221 | 18.30447 | 18.74364 |
| VPS37A | 19.16719 | 20.02918 | 20.01563 | 19.85231 | 19.98815 | 19.72099 | 16.81184 | 20.32114 | 19.79576 | 19.4299 | 20.08213 | 19.74395 | 19.408 | 20.10789 | 19.66305 | 20.0559 | 20.2764 | 19.83721 | 20.18253 | 20.44758 | 20.10789 | 19.88428 | 19.8108 | 19.61104 | 19.83664 | 20.0559 | 20.36453 |

| Gene | mCRPC Patient 1 | 2 | 3 | 4 | 5 | 6 | 7 | 8 | 9 | 10 | 11 | 12 | 13 | 14 | 15 | 16 | 17 | 18 | 19 | 20 | 21 | 22 | 23 | 24 | 25 | 26 | 27 |
|---|---|---|---|---|---|---|---|---|---|---|---|---|---|---|---|---|---|---|---|---|---|---|---|---|---|---|---|
| ADGRE5 | 21.36453 | 21.80341 | 21.09507 | 21.75939 | 21.10149 | 17.39664 | 21.86891 | 20.98815 | 20.2764 | 21.61414 | 20.6006 | 20.39624 | 21.0426 | 20.88017 | 21.89134 | 19.6679 | 22.35917 | 21.58692 | 22.68432 | 21.31008 | 21.36453 | 21.75124 | 21.55918 | 22.45513 | 22.1673 | 21.53564 | 21.84995 |
| ADK | 20.29894 | 20.90242 | 20.09507 | 19.6562 | 20.28771 | 19.42612 | 19.92047 | 20.70556 | 20.38574 | 19.73795 | 20.50688 | 20.52612 | 19.88875 | 20.2764 | 20.49717 | 20.00196 | 20.2535 | 20.15808 | 19.96014 | 20.5451 | 19.82134 | 20.62756 | 20.81127 | 19.34835 | 19.93157 | 20.29894 | 20.38574 |
| ATAD3A | 21.07561 | 21.27071 | 19.62553 | 19.1604 | 20.74714 | 21.8839 | 20.38574 | 21.34841 | 16.65946 | 18.48308 | 20.5451 | 20.24191 | 17.1439 | 20.57311 | 19.58731 | 21.1332 | 20.56384 | 16.98326 | 20.02918 | 21.10789 | 20.06907 | 15.92938 | 20.0559 | 21.08213 | 18.20755 | 20.09507 | 20.70556 |
| CFL2 | 16.81843 | 16.68138 | 19.40318 | 16.98033 | 20.28771 | 19.48291 | 19.60792 | 18.20577 | 18.87722 | 19.53476 | 19.86153 | 19.5289 | 19.76923 | 20.0426 | 19.37966 | 16.64389 | 19.90863 | 20.1332 | 20.1946 | 19.77093 | 19.63126 | 19.63125 | 19.62558 | 19.49633 | 17.20175 | 16.5173 | 19.93157 |
| CLNS1A | 16.22844 | 20.14569 | 21.46762 | 20.10789 | 16.28427 | 17.04339 | 17.55174 | 19.2744 | 18.17985 | 19.94592 | 19.96014 | 20.70556 | 17.11151 | 16.55052 | 20.20658 | 17.69074 | 21.09507 | 16.34248 | 16.36199 | 19.50938 | 17.10114 | 16.72144 | 16.57493 | 16.64258 | 16.17749 | 22.29334 | 15.84757 |
| EGFR | 18.46378 | 18.21329 | 16.2738 | 16.25403 | 15.74673 | 16.29071 | 16.64382 | 16.35746 | 17.28985 | 16.45484 | 16.10592 | 16.77285 | 20.2535 | 17.8364 | 17.78661 | 16.89709 | 15.80884 | 18.86594 | 19.14303 | 18.26089 | 18.10343 | 16.29938 | 21.56384 | 16.17606 | 16.94121 | 16.8075 | 18.52456 |
| EHBP1 | 17.81106 | 19.20274 | 17.75803 | 16.60501 | 18.14055 | 19.01308 | 16.59474 | 18.33945 | 19.07138 | 18.84416 | 18.95285 | 19.10265 | 18.78899 | 20.46762 | 19.63176 | 18.38422 | 18.991 | 18.71159 | 18.90313 | 18.44779 | 19.3433 | 19.36092 | 19.10292 | 19.07395 | 19.00514 | 19.25021 | 18.73573 |
| GIPC1 | 20.08213 | 19.97421 | 20.76345 | 20.46762 | 20.3538 | 20.08213 | 17.56038 | 19.84936 | 16.42331 | 19.63371 | 19.28794 | 19.98815 | 21.06907 | 20.62756 | 19.88104 | 20.17036 | 19.69248 | 21.24191 | 20.88017 | 20.45764 | 21.85377 | 20.28771 | 20.88763 | 21.28207 | 20.44758 | 21.0625 | 20.45764 |
| ITPA | 18.28738 | 20.68003 | 14.97046 | 16.05451 | 17.26808 | 18.21704 | 17.4358 | 19.92857 | 20.68003 | 17.28382 | 20.0559 | 20.70556 | 18.56556 | 18.23146 | 19.84733 | 18.60551 | 20.58233 | 16.64595 | 18.49882 | 20.17036 | 20.01563 | 19.18265 | 20.0559 | 18.95626 | 19.23851 | 15.95587 | 17.03354 |
| PALS1 | 18.99577 | 16.26829 | 16.28586 | 17.43304 | 16.7124 | 16.81106 | 21.417 | 16.69215 | 17.16857 | 17.09419 | 17.65797 | 17.67527 | 20.42726 | 16.02294 | 17.20896 | 16.33918 | 17.35281 | 16.87961 | 16.56353 | 18.0562 | 17.31204 | 17.52081 | 16.72022 | 16.89036 | 17.8074 | 16.44229 | 16.56175 |
| PALS2 | 17.24542 | 20.44758 | 20.28771 | 16.27509 | 16.12049 | 20.0559 | 17.01912 | 20.89504 | 20.34299 | 16.82413 | 16.44467 | 18.90479 | 20.53564 | 17.3467 | 19.46888 | 16.23505 | 19.06801 | 17.05659 | 20.2764 | 18.31378 | 17.53237 | 17.89989 | 17.81514 | 19.18185 | 16.60221 | 20.93876 | 19.52389 |
| PANK4 | 19.83434 | 19.54636 | 17.18598 | 20.80341 | 19.90425 | 19.33258 | 15.94243 | 19.76684 | 19.44888 | 20.01563 | 19.80809 | 19.64498 | 16.67576 | 20.01563 | 19.65889 | 19.46166 | 19.91772 | 19.93157 | 20.5545 | 20.23023 | 16.08694 | 19.62611 | 20.00196 | 19.67001 | 18.75922 | 19.69658 | 19.97421 |
| PATJ | 17.84155 | 16.41034 | 16.15601 | 16.89254 | 16.64802 | 16.27851 | 16.1571 | 16.88435 | 17.72562 | 18.79956 | 17.06897 | 16.86881 | 17.77311 | 16.80109 | 18.41281 | 15.81406 | 17.4886 | 18.78162 | 17.69872 | 18.10933 | 17.90032 | 16.41746 | 18.06407 | 17.25546 | 16.96983 | 18.10649 | 18.99173 |
| PDXK | 19.61812 | 20.32114 | 19.21914 | 20.52612 | 19.96014 | 16.11549 | 20.28771 | 19.94592 | 18.69302 | 20.73066 | 19.91077 | 19.60562 | 17.64573 | 19.48479 | 20.49717 | 18.98263 | 20.2764 | 16.64327 | 20.44758 | 19.83953 | 20.66275 | 20.26499 | 20.1206 | 21.15808 | 21.86514 | 15.7683 | 19.60766 |
| PLAU | 17.56483 | 16.65352 | 16.94071 | 15.36019 | 17.09381 | 17.61907 | 17.94442 | 13.28914 | 17.8079 | 17.70257 | 16.90545 | 16.75542 | 17.28682 | 16.81311 | 18.17171 | 17.34199 | 17.77063 | 15.48283 | 17.0801 | 16.66578 | 17.00345 | 16.26575 | 16.83364 | 16.33546 | 17.13673 | 17.32761 | 15.99138 |
| PLEKHG3 | 15.58467 | 16.43201 | 15.9518 | 17.46763 | 17.41503 | 15.71992 | 16.8082 | 16.75727 | 16.86527 | 16.33083 | 17.07179 | 16.89812 | 17.19545 | 16.5395 | 18.60995 | 17.07779 | 17.97112 | 18.36784 | 17.95401 | 16.64886 | 17.00609 | 17.04742 | 15.9766 | 16.51829 | 15.8976 | 16.27097 | 18.23355 |
| PPFIA1 | 18.63196 | 20.23023 | 19.33859 | 19.26798 | 16.35075 | 17.10953 | 20.68859 | 19.88046 | 18.77237 | 19.29669 | 19.77022 | 20.21845 | 19.30752 | 16.63984 | 20.17036 | 19.31313 | 20.00196 | 19.84559 | 20.59149 | 19.33752 | 19.87424 | 19.55328 | 19.11893 | 20.64526 | 19.66152 | 20.0426 | 19.86514 |
| PPFIBP1 | 17.97582 | 18.79729 | 17.3875 | 17.27359 | 16.68041 | 19.02508 | 17.78296 | 18.01666 | 18.16014 | 16.31677 | 17.69351 | 18.50598 | 16.64961 | 16.99115 | 18.49393 | 16.82392 | 16.08933 | 16.00326 | 16.30019 | 16.32256 | 16.59036 | 17.49401 | 16.59328 | 17.63636 | 15.7482 | 18.4145 | 17.38794 |
| SDC1 | 16.51942 | 19.4347 | 19.72166 | 19.50566 | 16.16951 | 17.25806 | 20.10789 | 17.89559 | 19.10338 | 18.99215 | 17.11159 | 19.28503 | 22.09828 | 15.95672 | 15.9441 | 17.22079 | 16.09134 | 20.94592 | 20.79551 | 17.81858 | 20.2535 | 16.79345 | 21.32663 | 21.10789 | 16.06217 | 20.21845 | 20.58233 |
| SPAG1 | 16.81717 | 18.51858 | 17.11721 | 18.42163 | 18.10064 | 16.85384 | 17.621 | 17.13426 | 16.47285 | 19.31582 | 17.23982 | 15.94608 | 19.81109 | 16.51055 | 19.14925 | 16.69169 | 19.45297 | 17.85503 | 18.96307 | 18.53229 | 16.71336 | 18.49002 | 19.18005 | 17.73788 | 18.94292 | 19.40384 | 19.80238 |
| STRN | 21.43237 | 21.53564 | 20.89504 | 20.78756 | 21.40665 | 22.05258 | 20.57311 | 21.70978 | 21.61414 | 20.45764 | 21.417 | 21.3538 | 20.32114 | 21.56384 | 21.52133 | 21.86136 | 21.69285 | 20.52612 | 21.23608 | 21.70556 | 20.36453 | 21.02242 | 21.21252 | 20.68003 | 21.21845 | 21.1946 | 21.91707 |
| TAX1BP1 | 16.47419 | 16.56688 | 16.72671 | 17.1808 | 17.90422 | 15.30514 | 18.19104 | 16.31796 | 16.81649 | 16.08979 | 16.74511 | 16.44942 | 16.07227 | 16.21153 | 17.41239 | 16.93545 | 16.32057 | 19.13041 | 16.97764 | 17.49808 | 17.01311 | 18.20356 | 16.72096 | 16.37948 | 18.05232 | 16.75925 | 18.93431 |
| TAX1BP3 | 16.35465 | 15.93585 | 15.89156 | 16.43495 | 16.40929 | 16.21759 | 15.59846 | 15.82379 | 20.15808 | 16.52603 | 19.42263 | 17.57935 | 17.37484 | 17.7271 | 17.29264 | 17.84336 | 16.23671 | 16.89894 | 16.64446 | 19.33059 | 17.63396 | 16.85465 | 17.66508 | 17.2217 | 18.39981 | 16.42174 | 17.57453 |
| TMSB10 | 21.51653 | 21.84613 | 20.70556 | 21.75532 | 20.96719 | 21.70556 | 21.83461 | 20.46762 | 22.37252 | 21.77957 | 22.42982 | 20.02918 | 22.23023 | 22.72234 | 21.24771 | 16.72653 | 20.70556 | 22.38047 | 22.74509 | 23.23316 | 21.31562 | 22.45764 | 22.239 | 22.83074 | 22.30452 | 21.89874 | 22.72025 |
| TMSB4X | 26.12534 | 25.54628 | 24.32389 | 24.83364 | 26.35615 | 25.2571 | 25.94503 | 25.56384 | 26.87408 | 27.16039 | 28.35363 | 26.07521 | 25.08942 | 26.07725 | 25.89041 | 24.53445 | 25.87174 | 24.75939 | 25.71293 | 28.00838 | 26.239 | 26.41538 | 25.72598 | 25.53445 | 27.78332 | 25.59548 | 26.64581 |
| TP53I3 | 21.28207 | 20.47754 | 16.99024 | 21.1519 | 20.65403 | 19.6711 | 21.33211 | 18.38225 | 21.30452 | 21.36986 | 20.89504 | 20.23023 | 20.26499 | 20.01563 | 19.87539 | 21.75124 | 18.28202 | 16.7492 | 20.74714 | 20.29894 | 17.64109 | 16.23911 | 20.02918 | 20.51653 | 19.72291 | 16.5476 | 21.33756 |
| TUBA1C | 27.67304 | 27.2535 | 26.68646 | 27.0296 | 27.68646 | 27.70636 | 27.44727 | 27.22619 | 27.509 | 27.98142 | 27.56093 | 27.48616 | 26.33926 | 27.62506 | 27.3662 | 28.42342 | 27.3662 | 26.87408 | 27.42342 | 28.07623 | 28.00303 | 27.42342 | 26.67304 | 27.07112 | 27.13124 | 27.90885 | 28.258 |

| | mCRPC Patient | | | | | | | | | | | | | | | | | | | | | | | | | | |
|---|---|---|---|---|---|---|---|---|---|---|---|---|---|---|---|---|---|---|---|---|---|---|---|---|---|---|---|
| **Gene** | **1** | **2** | **3** | **4** | **5** | **6** | **7** | **8** | **9** | **10** | **11** | **12** | **13** | **14** | **15** | **16** | **17** | **18** | **19** | **20** | **21** | **22** | **23** | **24** | **25** | **26** | **27** |
| AIFM1 | 21.95305 | 22.3538 | 19.3402 | 20.0559 | 21.76749 | 22.32114 | 21.2764 | 22.48001 | 17.53006 | 19.61868 | 21.87267 | 21.21845 | 19.82942 | 21.40665 | 20.5451 | 22.02242 | 22.00881 | 20.10789 | 20.97421 | 22.29053 | 20.77153 | 19.50018 | 21.25926 | 20.42726 | 16.62853 | 20.88763 | 21.81127 |
| IDI1 | 17.38225 | 20.09507 | 17.68207 | 15.59325 | 18.4309 | 17.44189 | 16.04591 | 18.73722 | 16.45492 | 17.16805 | 19.09053 | 19.14904 | 16.5734 | 18.79013 | 15.27728 | 18.85376 | 18.72524 | 18.05124 | 16.31568 | 18.97132 | 15.19556 | 18.61258 | 17.28738 | 17.34524 | 17.42284 | 16.51767 | 16.91054 |
| SMC2 | 16.48525 | 18.28501 | 18.72768 | 17.50586 | 18.20641 | 18.9312 | 16.18514 | 17.00393 | 16.30415 | 18.41899 | 16.77976 | 18.05501 | 16.83021 | 17.64088 | 18.15818 | 17.52059 | 18.48134 | 17.60503 | 16.94426 | 17.3022 | 16.22795 | 18.30418 | 16.7193 | 18.44205 | 17.84936 | 18.4301 | 17.58992 |
| SUMF1 | 18.71631 | 18.77105 | 16.02804 | 16.20584 | 19.14633 | 19.69998 | 16.62528 | 18.93748 | 17.6374 | 17.60164 | 18.45294 | 18.07144 | 19.50818 | 16.78473 | 18.13325 | 19.16846 | 19.0566 | 17.18908 | 17.14542 | 19.1182 | 17.67256 | 16.31137 | 16.93738 | 17.21361 | 16.21522 | 19.03828 | 19.62893 |
